# GLYATL1 is associated with metabolic and epigenetic changes and with endocrine resistance in luminal breast cancer

**DOI:** 10.1101/2025.10.17.682994

**Authors:** Janina Müller, Emre Sofyali, Luisa Schwarzmüller, Yael Aylon, Eviatar Weizman, Lisa Schlicker, Katherine Kelly, Simone Borgoni, Simin Oz, Cole Stocker, Sara Burmester, Angelika Wörner, Sabine Karolus, Birgitta E Michels, Daniela Heiss, Rainer Will, Veronica Rodrigues de Melo Costa, Pavlo Lutsik, Dieter Weichenhan, Ilse Hofmann, Nishanth Belugali Nataraj, Yosef Yarden, Luca Magnani, Christoph Plass, Almut Schulze, Cindy Körner, Moshe Oren, Stefan Wiemann

## Abstract

Estrogen receptor alpha (ERα)-positive luminal breast cancer is commonly treated with aromatase inhibitors (AI) to block estrogen signaling; however, resistance frequently develops, limiting therapy success.

We observed that *GLYATL1* (Glycine-N-Acyltransferase Like 1) expression is upregulated in AI-resistant breast cancer cell models and in patients undergoing AI therapy, correlating with poorer survival. Here we demonstrate that GLYATL1 promotes resistance to estrogen deprivation by elevating succinate levels and altering epigenetic histone marks associated with active transcription. Knockdown or knockout of *GLYATL1* reverses these effects and reduces proliferation under estrogen-deprived conditions. Notably, *GLYATL1* expression is positively regulated by estrogen receptor alpha signaling independently of estrogen.

These findings reveal GLYATL1 as a metabolic and epigenetic mediator of endocrine therapy resistance, suggesting it as a potential target to overcome AI resistance in luminal breast cancer.

## Introduction

Breast cancer remains the most prevalent malignancy and the second leading cause of cancer-related mortality among women globally (1, 2). It is a heterogeneous disease, requiring molecular subtyping in clinical practice. This classification is based on the expression of estrogen receptor alpha (ERα), progesterone receptor (PR), human epidermal growth factor receptor 2 (HER2), and the proliferation marker protein Ki-67 (3). Approximately two-thirds of breast tumors are classified as ER-positive (ER+) (4). Patients with such luminal-like disease typically receive endocrine therapies designed to interfere with estrogen signaling. These therapy approaches either target ERα directly, or prevent estrogen synthesis by inhibiting the essential enzyme aromatase (CYP19A1) with aromatase inhibitors (AI), thereby depriving the cancer cells of estrogen (5). Despite the initial efficacy of endocrine therapies, up to 40% of patients presenting with locally advanced disease at the time of diagnosis relapse during or after endocrine therapy (6). However, resistance acquisition is a highly individual process that may involve various mechanisms, including the activation of alternative signaling pathways (7, 8), acquisition of mutations in *ESR1* (9, 10), i.e., the gene encoding ERα, metabolic rewiring (11), and epigenetic reprogramming (12, 13). These mechanisms contribute to inter- and intratumor heterogeneity and allow tumors to sustain growth independently of estrogen supply (14, 15). Uncovering further resistance mechanisms is of key relevance to facilitate the development of personalized therapeutic approaches.

Along these lines, we characterized long-term estrogen deprived (LTED) sublines of two ER+ breast cancer cell line models (MCF7 and T47D), mirroring resistance to aromatase inhibition. Multi-omics profiling of cells resistant to estrogen-deprivation included RNA sequencing, ATAC-sequencing, methylation array analysis, metabolomics, and proteomic profiling. We identified significant alterations in gene expression and chromatin accessibility as well alterations at metabolite, proteomic, and epigenomic levels. Among the top deregulated genes was Glycine-N-Acyltransferase Like 1 (*GLYATL1*), which was upregulated in both *in vitro* models. While GLYATL1 has been implicated in tumor growth and disease progression across various cancer types including breast cancer (16–18), its potential role in endocrine resistance in luminal breast cancer remains poorly understood.

Here, we present a comprehensive investigation of GLYATL1, delineating its functional role in sustaining a resistance phenotype, and its critical impact on cell proliferation and tumor cell survival, on mediating succinate accumulation and on epigenetic reprogramming under estrogen-deprived conditions. We further explore the regulatory mechanisms underlying *GLYATL1* expression, suggesting that this is regulated by estrogen-independent ERα activity specifically in LTED cells. These findings highlight GLYATL1 as a potential biomarker and therapeutic target for addressing endocrine resistance to aromatase-inhibition in ER+ breast cancer.

## Methods

### Cell lines

MCF7 (RRID:CVCL_0031) was obtained from ATCC (Manassas, VA, USA). T47D (RRID:CVCL_0553) and HEK293FT (RRID:CVCL_6911) were obtained from LGC Standard GmbH (Wesel, Germany). These parental cell lines were cultivated in Dulbecco’s Modified Eagle Medium (DMEM #11574486, Gibco, Thermo Fisher Scientific, Waltham, MA, USA) supplemented with 10% FCS, 10 nM 17-β-estradiol (E8875, Sigma-Aldrich, Saint-Louis, MI, USA), 2 mM L-glutamine (#25030081, Gibco, Thermo Fisher Scientific, Waltham, MA, USA), 1 mM sodium pyruvate (#11360070, Gibco, Thermo Fisher Scientific, Waltham, MA, USA), and 50 units/mL penicillin / 50 µg/mL streptomycin sulfate (#15140122, Gibco, Thermo Fisher Scientific, Waltham, MA, USA), and incubated at 37°C with 5% CO_2_ in a humidified incubator. Long-term estrogen-deprived (LTED) cells were cultured in DMEM (w/o phenolred, #31053028) supplemented with 10% charcoal-stripped FBS (#12676029), 2 mM L-glutamine (#25030081), 1 mM sodium pyruvate (#11360070), and 50 units/mL penicillin / 50 µg/mL 50 µg/mL streptomycin sulfate (#15140122, all Gibco, Thermo Fisher Scientific, Waltham, MA, USA), until the cells had regained proliferative capacities similar to parental MCF7 and T47D cells respectively (19, 20). All cell lines were regularly authenticated by STR profiling (Multiplexion GmbH Heidelberg, Germany) and free of mycoplasma contamination.

### Generation of stable *GLYATL1* knockout clones using the CRISPR/cas9 approach

*GLYATL1* was knocked out in the MCF7 LTED cell line using CRISPR/Cas9 technology with a sgRNA targeting exon 3 (Reference transcript: NM_001389711.2) of the *GLYATL1* gene (sgRNA: 5’-GUUUCUUCUUCAAGGUCUCA-3’) (Synthego, Redwood City, CA, USA). Transfection of cells was performed using Lipofectamin CRISPRMAX Cas9 transfection reagent (#CMAX00001, Thermo Fisher Scientific, Waltham, MA, USA) according to the manufacturer’s instructions. Knockout cultures were cultivated in DMEM (w/o phenol red #31053028) supplemented with 10% charcoal-stripped FBS (#12676029), 2 mM L-glutamine (#25030081), 1 mM sodium pyruvate (#11360070), and 50 units/mL penicillin as well as 50 µg/mL streptomycin sulfate (#15140122, all Gibco, Thermo Fisher Scientific, Waltham, MA, USA), and incubated at 37°C with 5% CO_2_ in a humidified incubator. Single clones were isolated in 96well plates utilizing an F.SIGHT single-cell dispenser (Cytena, Freiburg, Germany). PCR primers were designed upstream of exon 3 (forward primer: 5’- AAAATCTATGCCTACTCCTGCTCC-3’) and downstream of exon 4 (reverse primer: 5’- ACTCCCATACCCTGACCTCC-3’) within the *GLYATL1* gene to amplify the region of the CRISPR knockout, using Phusion Hot Start II DNA-Polymerase (2 U/µL) (#F549L, Thermo Fisher Scientific, Waltham, MA, USA). The thermocycling protocol included: 98 °C – 2 min, 35x (98 °C – 10 s, 64 °C – 20 s, 72 °C – 60 s), and 72 °C – 10 min. PCR-products were Sanger-sequenced (Eurofins Genomics Germany GmbH, Ebersberg, Germany), and sequences analyzed using SnapGene software (Dotmatics, Bishop’s Stortford, UK).

Two clones (KO1 and KO2) were selected based on a predicted knockout by SnapGene. Gel analysis of PCR products from these clones revealed bands of about 1100 and 1550 bp, respectively. The bands were gel-purified using the Wizard SV Gel and PCR Clean-up System (#A9281, Promega, Madison, WI, USA), according to the manufacturer’s instructions, Sanger-sequenced (Eurofins Genomics Germany GmbH, Ebersberg, Germany), and sequences mapped to hg38 using the BLAT-tool in the UCSC genome browser (21). A 454bp deletion affecting most of exon 3 and ranging into downstream intron 3 was observed in one allele and in both knockout clones. The resulting transcripts lacked the start-codon for translation of the GLYATL1 protein, which maps to exon 3, and extended into intronic sequence between exons 3 and 4 (compare Supplementary Figure 2A). These alterations rendered this allele non-functional. A deletion of 20bp was present in the second allele of the *GLYATL1* gene and again in both knockout clones. This deletion was associated with aberrant splice patterns of *GLYATL1* transcripts. The fractions of reads supporting retention of the *GLYATL1* start codon differed between *GLYATL1* knockout clones KO1 and KO2 as some canonical splicing of exon 3 to exon 4 was observed only in clone KO2 but not in KO1 (Supplementary Figure 2B/C).

### RNA-sequencing

Total RNA was extracted using the RNeasy Midi kit (#75144, Qiagen, Hilden, Germany) according to the manufacturer’s instructions and RNA-sequencing was performed at the DKFZ NGS Core Facility, using different protocols for MCF7 parental and LTED, vs. MCF7 LTED and *GLYATL1* knockout clones KO1 and KO2. MCF7 parental and LTED conditions: RNA-seq libraries were prepared using the Illumina TruSeq RNA Library Preparation Kit (Illumina, San Diego, CA, USA) and sequenced on an Illumina HiSeq 4000 platform (Illumina, San Diego, CA, USA), generating on average 45 million paired-end reads with a length of 2 × 100 base pairs. Sequencing reads were pre-processed using a combination of open-source tools and *in-house* scripts. Quality filtering and artifact removal were carried out using the FastX Toolkit v0.0.13 (https://github.com/agordon/fastx_toolkit), specifically employing *fastq_quality_filter* with parameters -q 20 -p 90 and *fastx_artifacts_filter*. Poly-A tails were trimmed using HOMER tools v4.7 (22) with the trim -3 AAAAAAAAA command. Reads containing ambiguous bases (‘N’) or characters outside the canonical nucleotide set (A, C, G, T, U) were removed using custom in-house Perl scripts. Additionally, reads shorter than 17 bases or longer than 10,000 bases were excluded. To ensure proper pairing after these filtering steps, orphaned reads were discarded using an in-house script. Ribosomal RNA (rRNA) contamination was removed if the alignment rate against rRNA sequences exceeded 1%. For this purpose, reads were first aligned to an rRNA reference using Bowtie2 v2.2.4 (23). Where necessary, rRNA-aligned reads were filtered out using samtools *view* v0.1.18 (24) with flags -f 12 -F 256, and the remaining reads were converted back to FASTQ format using *SamToFastq* from Picard tools v1.78 (https://broadinstitute.github.io/picard/). Duplicate reads were removed using Sambamba v0.6.5 (25) with the *sort* and *markdup* commands executed consecutively. Reads were aligned to the human reference genome GRCh38.p13 using STAR aligner v2.3 (26), and gene-level quantification was performed using HTSeq-count v0.6.0 (27). GENCODE release 34 was used for annotation during both alignment and read counting. Differential gene expression analysis was performed with DESeq2 v1.28.1 (28) with differentially expressed genes identified at a false discovery rate (FDR) threshold of 0.05.

MCF7 LTED and LTED *GLYATL1* knockout KO1 and KO2: RNA libraries were prepared using the TruSeq Stranded RNA Library Prep Kit (Illumina, San Diego, CA, USA) according to the manufacturer’s protocol. Sequencing was conducted on the Illumina NovaSeq 6000 platform (Illumina, San Diego, CA, USA), generating paired-end reads with a length of 2 × 100 base pairs (S1 flow cell). Each sample yielded an average of approximately 59 million read pairs. Quality control of the raw sequencing data was performed using FastQC v0.11.9 (https://www.bioinformatics.babraham.ac.uk/projects/fastqc/). Strand-specific reads were aligned to the human reference genome GRCh38.p13 using STAR v2.5.2a (26), with the genome index generated from GENCODE v34 annotations. Genome-wide similarity between sequencing replicates was evaluated using deepTools v3.5.2 (29). Gene-level read counts were obtained using the htseq-count tool from HTSeq v0.11.1 (27). Differential gene expression analysis was performed with DESeq2 v1.40.2 (28), comparing the indicated cell lines. Genes with fewer than 10 total reads across all samples were excluded from the analysis. Differentially expressed genes were identified at an FDR threshold of 0.05.

### ATAC-Sequencing

Libraries for ATAC-sequencing were prepared with modifications based on published protocols (30). Cells were lysed using 1% NP40, followed by tagmentation at 55°C for 8 minutes in a reaction mixture consisting of 2.5 µL of TDE1 (Nextera Illumina DNA Kit, Illumina, San Diego, CA, USA), 25 µL of tagmentation buffer (Nextera Illumina DNA Kit), and 25 µL of the lysed cells. The tagmentation reaction was terminated by the addition of 10 µL of 5M guanidine thiocyanate, and samples were subsequently purified with AMpure Beads (Beckman Coulter, Brea, CA, USA). Library construction was performed using NEBNext High Fidelity PCR Mix, and sequencing (> 5 million reads/sample) was conducted on an Illumina HiSeq 2500 platform at the DKFZ NGS Core Facility. Sequencing reads were processed to remove adapter sequences using TrimGalore (v.0.6.7). Trimmed reads were aligned to the hg38 canonical reference using Bowtie2 (v.2.4.2) with very sensitive end-to-end presets (23). A maximum fragment length of 1000 was selected for paired alignments and mate dovetailing was allowed. Alignments were filtered to remove mitochondrial reads, non-proper pairs and reads with mapping quality score < 30. Duplicates were removed using Picard MarkDuplicates (v.2.18.2.2, http://broadinstitute.github.io/picard). Peak calling was performed using MACS2 (v.2.2.9.1) (31) using the parameters “--qvalue ‘0.05’ --nomodel --extsize ‘200’ --shift ‘-100’”. All identified peaks were merged to create a common bed file containing read counts, which was then utilized for differential analysis using edgeR (v3.36.0) (32). Heatmaps of differential accessibility were created using ChIPpeakAnno (v3.0.0) (23, 33) and ComplexHeatmap (v2.8.0).

### EPIC 850k methylome array

Total DNA was extracted from MCF7 parental and LTED cells using the DNeasy Blood and Tissue Kit (Qiagen, Hilden, Germany), following the manufacturer’s protocol. Methylation profiling was performed by the DKFZ Microarray Core Facility using the Illumina MethylationEPIC BeadChip platform (Illumina, San Diego, CA, USA). Quality control, filtering and normalization were performed using Rnbeads (v.2.21.3) (34). Probes overlapping with single nucleotide polymorphisms (SNPs), cross-reactive probes, those outside of CpG context and those mapping to sex chromosomes were filtered out. Probes with detection p-value > 0.05 and sites covered by fewer than three beads were further excluded. Normalization and background subtraction were carried out using the “scaling.internal” and “sesame.noobsb” methods of Rnbeads. Methylation beta-values were summarised by mean across each pair of replicates and the difference in mean methylation was calculated per-CpG as a measure of differential methylation between parental and LTED cells.

### Immunofluorescence imaging

For recombinant expression of the GLYATL1 protein, the *GLYATL1* open reading frame was cloned from a Gateway entry clone (pENTR223-GLYATL1, IMAGE:100071537) (35) into a Gateway-compatible expression vector (pFLAG-C) to obtain plasmid pFLAG-C-GLYATL1, which encodes a protein with a FLAG-tag fused to the C-terminus of GLYATL1. MCF7, T47D and HEK293cells transiently transfected with pFLAG-C-GLYATL1 plasmid and grown on 12 mm coverslips until reaching 70-80% confluency. Mitochondria were stained with 500 nM abberior LIVE ORANGE mito (LVORANGE, Abberior, Göttingen, Germany) for 1 hour at 37°C in 5% CO_2_. Subsequently, cells were fixed with 4% formaldehyde for 15 minutes at room temperature and permeabilized with 0.1% Triton X-100 in PBS for 10 minutes. To prevent non-specific binding, cells were incubated with 3% BSA in PBS for 1 hour at room temperature. Mouse anti FLAG antibodies (1:1000, #F3165, Sigma Aldrich, St. Louis, MI, USA) in 3% BSA were then added and incubated overnight. Following three washing steps with PBS, secondary goat anti mouse antibodies conjugated to alexa-488 (1:400, #ab150113, Abcam, Cambridge, UK) were added and incubated for 1 hour at room temperature in the dark. After washing off excess secondary antibodies with PBS, cells were mounted onto glass slides using ProLong Diamond antifade mounting media with DAPI (#P36966, Thermo Fisher Scientific, Waltham, MA, USA). Samples were examined using a fluorescence microscope (LSM-900, Zeiss, Oberkochen, Germany). Negative controls, which lacked primary or secondary antibodies, were included to verify specificity and antibody reactivity. Analysis of images was performed using Zen Blue software (Carl Zeiss, Jena, Germany) and ImageJ (https://imagej.net/ij/). Manders’ coefficient (36) was calculated using the JACoP in ImageJ to quantify the degree of colocalization between fluorophores.

### Western Blot

Cell were lysed using RIPA buffer (#89900, Thermo Fisher Scientific, Waltham, MA, USA) supplemented with cOmplete EDTA-free protease inhibitor (#11697498001; Merck, Darmstadt, Germany) and PhosSTOP phosphatase inhibitor (#4906845001; Merck, Darmstadt, Germany). Following incubation on ice for 30 minutes, lysates were centrifuged for 30 minutes at 15,000 xg and 4°C. Protein concentration of the supernatant was measured utilizing Pierce™ BCA Protein Assay Kit (#23250, Thermo Fisher Scientific, Waltham, MA, USA). Proteins from cell lysates and PageRuler™ Prestained Protein Ladder (#26616, Thermo Fisher, Waltham, MA, USA) as size maker were separated by SDS-PAGE (37). Proteins were blotted using Trans-Blot Turbo™ Mini PVDF Transfer Packs and the Trans-Blot Turbo™ Transfer System (Bio-Rad Laboratories, Hercules, CA, US) according to the manufacturer’s instructions. Membranes were blocked in Blocking Buffer for Fluorescent Western Blotting (MB-070, Rockland, Philadelphia, PA, USA): TBS supplemented with 10 mM NaF and 1 mM Na_3_VO_4_. Subsequently, the membranes were incubated with a rabbit anti GLYATL1 antibody (1:1,000 dilution, #PA039501, Sigma-Aldrich, Taufkirchen, Germany) in blocking buffer at 4 °C overnight. The next day, membranes were washed three times for 5 min each with TBS containing 0.1% Tween 20 (TBST #91414, Merck, Taufkirchen, Germany). Then, blots were incubated with a goat anti-rabbit (Alexa Fluor™ 680 conjugated, 1:10,000 dilution, #A-21077, Thermo Fisher Scientific, Waltham, MA, USA) secondary antibody, and washed again. Proteins were visualized with a LI-COR Odyssey 9120 scanner (LI-COR, Lincoln, NE, USA) scanning in the 700 and 800 nm channels for Alexa Fluor™ 680 and DyLight™ 800 fluorophores, respectively. Then, gels were incubated with a mouse anti ß-Actin (1:10,000 dilution, #0869100-CF, MP Biologicals, Irvine, CA, USA, 1:1500 dilution) over night, and washed three times for 5 min each with TBS containing 0.1% Tween 20. A goat anti-mouse (DyLight™ 800 4X PEG conjugated, 1:10,000 dilution, #SA5-35521, Thermo Fisher Scientific, Waltham, MA, USA) was used to visualize actin bands with a LI-COR Odyssey 9120 scanner (LI-COR, Lincoln, NE, USA) scanning in both, 700 and 800 nm channels for Alexa Fluor™ 680 and DyLight™ 800 fluorophores, respectively.

### Single-pot solid-phase sample preparation (SP3) for proteome measurements

Single-pot solid-phase sample preparation (SP3) was performed using 20 µg of protein in a total volume of 60 µL of RIPA buffer (#89900, Thermo Fisher Scientific, Waltham, MA, USA), supplemented with 100 mM triethylammonium bicarbonate (TEAB). To facilitate protein reduction and alkylation, chloracetamide (CAA) and tris(2-carboxyethyl)phosphine (TCEP) were added to final concentrations of 40 mM and 10 mM, respectively, and the mixture was incubated at 95°C for 5 minutes. Subsequently, 1 µL of each type of paramagnetic bead (#45152105050250 and #65152105050250; Sigma-Aldrich, St. Louis, MO, USA) was mixed in a 1:1 ratio. Ethanol was added to achieve a final concentration of 50%, and the mixture was incubated for 15 minutes at 650 rpm. To enhance bead-protein binding, the sample was shaken at room temperature for an additional 15 minutes at the same speed. Bound proteins were washed twice with 80% ethanol, followed by a single wash with 100% acetonitrile. After aspirating any residual acetonitrile, the beads were resuspended in 100 mM TEAB containing trypsin at a protease-to-protein ratio of 1:25. Digestion was facilitated by sonication in a water bath for 30 seconds, followed by incubation at 37°C while shaking at 800 rpm for 16 hours. The next day, the supernatant containing the digested peptides was collected and vacuum-dried in a speed-vac at 30°C and 1300 rpm. The resulting peptides were stored at −20°C until further analysis.

### Liquid chromatography (LC) separation of peptides

Peptides were solubilized in mass spectrometry (MS)-grade water supplemented with 0.1% trifluoroacetic acid (TFA) and 2.5% hexafluoroisopropanol (HFIP) prior to analysis. Using an UltiMate 3000 liquid chromatography (LC) system (Thermo Fisher Scientific, Waltham, MA, USA) 1 µg of peptides was separated on a 25 cm column (#186008795, Waters nanoEase™ BEH C18, 130 Å, 1.7 µm, 75 µm x 250 mm, Waters, Milford, MA, USA). A linear gradient of acetonitrile from 4% to 30% over 100 minutes at a flow rate of 300 nL/min was employed, culminating in a total method duration of 120 minutes for peptide separation.

### Parallel Reaction Monitoring (PRM) mass spectrometry (MS) measurements

Mass spectrometric analysis was performed using an Orbitrap Exploris 480 (Thermo Fisher Scientific, Waltham, MA, USA), with MS1 scans acquired at a resolution of 120,000 and an automatic gain control (AGC) target set to 3e6 ions. This approach ensured high sensitivity and resolution during the analysis of the peptide samples. The MS2 spectra were collected for pre-selected peptides of GLYATL1 at a resolution of 120,000. Data analysis was conducted using Skyline (version 3.1.0.7312). The GLYATL1 protein FASTA files were sourced from UniProt (Q969I3) and subjected to *in silico* digestion with trypsin, permitting one missed cleavage site. The identification of Y and b fragment ions facilitated the selection of filtered peptides corresponding to three pre-measured GLYATL1 peptides. The retention time of the peptide R.ALLLVTEDILK.L was determined using spectra from an MCF7 cell line stably overexpressing *GLYATL1*. This retention time was subsequently applied across all samples. Peak areas were computed by defining peak boundaries and summing the areas of the fragment ions.

### Data-independent acquisition (DIA) for total proteome analysis

MS1 spectra were acquired using an Orbitrap Exploris 480 (Thermo Fisher Scientific, Waltham, MA, USA) at a resolution of 120,000 and a maximum of 3e6 ions were collected for each spectrum. MS2 spectra were acquired in 47 isolation windows of variable width covering 400 m/z to 1000 m/z. The peptides were fragmented with a collision energy of 28%, the MS2 resolution was 60,000 and up to 1e6 ions were collected for each spectrum. Using Spectronaut (version 15.6; Biognosys, Schlieren, Switzerland) proteins were identified by searching the raw data against the proteome-wide human fasta file (79,052 entries, downloaded from Uniprot 15.03.2022). Up to two missed cleavage sites by trypsin were allowed, carbamidomethylation (C) was set as a fixed modification, oxidation (M) and acetylation (N-terminus) as variable modifications. Peptides were quantified on the MS2 level by summing up the signal area under the curve of the 3-6 highest abundant fragment ions. Proteome data were further analyzed using R (version 4.3.1). Protein intensities were log2 transformed and principal component analysis was performed using the prcomp() function. For statistical analysis, p-values were computed using unpaired, two-sided t-tests for proteins that were identified and quantified in all six samples of a given comparison. Benjamini-Hochberg adjustment (38) was applied to correct p-values for multiple hypothesis testing.

### Metabolite extraction

For extraction of polar metabolites, cells (∼1x10e^6^) were washed with cold ammonium acetate (154 mM) and scraped off in 0.5 mL ice-cold MeOH/H_2_O/Acetonitrile (50/20/30 v/v) containing internal standards (stable isotope labeled amino acids and ^13^C_3_-Malonyl-CoA as well as deuterium labeled TCA cycle intermediates, Cambridge Isotope laboratories, Tewsbury, MA, USA). After vortexing and sonication, samples were centrifuged at 13,000 x g for 5 minutes and the supernatant was applied onto a C18 8B-S001-DAK solid phase column (Phenomenex, Torrance, CA, USA), previously activated using acetonitrile and equilibrated using MeOH/H_2_O/Acetonitrile (50/20/30 v/v). The remaining pellet was extracted one more time using 0.5 mL ice-cold MeOH/H_2_O/Acetonitrile (50/20/30 v/v). The eluate was dried in a refrigerated vacuum concentrator (Labconco, Kansas City, MO, USA) at 10°C overnight. The residual pellet was used for protein quantification by BCA assay according to the instructions from the manufacturer. For this, the pellet was dissolved in 0.2 M NaOH and incubated for 20 min at 95°C.

### LC-MS measurement of metabolites

LC-MS measurement of metabolites was performed on an Ultimate 3000 coupled to a Q Exactive Plus mass spectrometer (Thermo Fisher Scientific, Waltham, MA, USA). Dried metabolite extracts were dissolved in 100 μL 5 mM NH_4_OAc in CH_3_CN/H_2_O (75/25, v/v), and 3 µL were applied onto an amide-HILIC column (16726–012105, 2.6 μm, 2.1x100 mm, Thermo Fisher Scientific, Waltham, MA, USA), where the temperature was kept at 30°C. The following solvents were used: solvent A consisting of 5 mM NH_4_OAc in CH_3_CN/H_2_O (5:95, v/v) and solvent B consisting of 5 mM NH_4_OAc in CH_3_CN/H_2_O (95:5, v/v). The following gradient was applied: 98% solvent B for 2 min, followed by a linear decrease to 40% solvent B within 5 min, then maintaining 40% solvent B for 13 min, then returning to 98% solvent B within 1 min and maintaining 98% solvent B for 5 min for column equilibration before each injection. The flow rate was maintained at 350 μL/min. The eluent was directed to the hESI source from 1.5 min to 21.0 min after sample injection and the following HESI source parameters were applied: Sheath gas flow rate – 30, Auxiliary gas flow rate – 10, Spray voltage – 3.6 kV (pos)/ 2.5 kV (neg), Capillary temperature – 320°C, S-lens RF level – 55.0. For the detection of acyl-CoAs, acquisition was done in positive mode with a scan range from 760 to 1100 m/z, a resolution of 70,000, AGC target of 1E^6,^ and max. injection time of 50 ms. For data-dependent MS2 (ddMS2), the resolution was set to 17,500, AGC target to 5E^4^, max. injection time of 50 ms and stepped collision energies of 20, 50 and 80. For broad-spectrum detection of water-soluble metabolites, the acquisition was performed in polarity switching mode within the scan range of 69-1000 m/z, with otherwise similar settings and ddMS2. Peaks corresponding to the calculated metabolite masses taken from an in-house metabolite library were integrated using the El-MAVEN software (https://docs.polly.elucidata.io/Apps/Metabolomic Data/El-MAVEN.html) and metabolite identification was supported by fragmentation patterns (39). Peak intensities were normalized by their respective internal standard levels.

### Epigenetic-focused cytometry time-of-flight (EpiTOF)

MCF7 (parental, LTED, LTED *GLYATL1* KO1 and KO2) single-cell suspensions were generated and washed with Maxpar PBS (#201058, Fluidigm, San Francisco, CA. USA) and subsequently stained with 1.25 µM Cisplatin for one minute to identify dead cells. The staining reaction was quenched using DMEM supplemented with 10% FBS. Cells were washed with Maxpar Cell Staining Buffer (#201068, Fluidigm, San Francisco, CA, USA), and about 3 × 10^6^ cells per sample were fixed, permeabilized using the Maxpar Nuclear Antigen Staining Buffer Set (#201063, Fluidigm, San Francisco, CA, USA), and barcoded with the Cell-ID 20-Plex Pd Barcoding Kit (#201060, Fluidigm, San Francisco, CA, USA) according to the manufacturer’s instructions. Following two washing steps with Maxpar Nuclear Antigen Staining Buffer Set permeabilization buffer, equal amounts of barcoded samples of the different MCF7 cell lines (i.e., parental, LTED, LTED *GLYATL1* KO1 and KO2) were combined and incubated with an antibody cocktail (Supplementary Table 11) for 30 minutes at room temperature. Following antibody incubation, the cells were washed twice with Maxpar Cell Staining Buffer and fixed at 4 °C with fresh 4% formaldehyde (#28908, Thermo Fisher Scientific, Waltham, MA, USA), ensuring gentle rocking to prevent clumping. Following overnight incubation, 125 nM Cell-ID Intercalator-Ir (#201192A, Fluidigm, San Francisco, CA, USA) was added and incubated for 45 minutes at room temperature to label DNA. Afterward, cells were washed twice with Maxpar Cell Staining Buffer and once with Maxpar Water (#201069, Fluidigm, San Francisco, CA, USA). Cells were resuspended in a dilution of EQ Four Element Calibration Beads (#28908, Fluidigm, San Francisco, CA, USA) in Maxpar Water, achieving a concentration of approximately 250K cells/ml, and subsequently filtered through a 35 μm mesh. Data was acquired on a CyTOF Helios platform (Fluidigm, San Francisco, CA, USA). Normalization and data cleanup to isolate live single cells were conducted as described by Bagwell et al. (40). Metal-conjugated antibodies (Supplementary Table 11) were either obtained from Fluidigm or conjugated utilizing appropriate Maxpar X8 Antibody Labeling Kits (Fluidigm, San Francisco, CA, USA). The mass cytometry antibody panel was designed to minimize signal spillover utilizing the Maxpar Panel Designer (Fluidigm, San Francisco, CA, USA). Analysis of EpiTOF data was executed through an R-based pipeline (41). Data were imported into R (version 4.0.2) and transformed using arcsine transformation with a cofactor of 5 and regressed to H3 and H3.3 expression. For statistical comparison, the median arcsine-transformed intensities were used (n=2). Using a two-sided, unpaired T-test, LTED was compared against the parental cells and cells from both GLYATL1 KO clones simultaneously.

### Cell proliferation assay

Cells were plated into black clear F-bottom 96-well plates (#655090, Greiner Bio-One International GmbH, Kremsmünster, Austria) and cultivated for seven days. Then, cells were labeled using Hoechst-33342 (#62249, Thermo Fisher Scientific, Waltham, MA, USA). Imaging of the plates was conducted with an IXM XLS microscope (Molecular Devices, San Jose, CA, USA). Image analysis was performed using the Molecular Devices Analysis Software, MetaXpress. Here, nuclei were detected based on the size and intensity of fluorescence in the DAPI channel and automatically counted. The median of six technical replicates was determined for each biological replicate.

### siRNA and plasmid transfection

RNAi knockdown was accomplished with pools of siRNAs (*GLYATL1* (#si-G020-92292), *ESR1* (#si-G020-2099), both from siTOOLs Biotech, Planegg, Germany). Untarget Control siPools negC-120 (#si-C002; siTOOLs Biotech, Planegg, Germany) served as a negative control. Cells were transfected using RNAiMAX® (Thermo Fisher Scientific, Waltham, MA, USA) according to the manufacturer’s instructions and with 3 nM of siRNA. Transfected cells were analyzed by RT-qPCR 72 hours post-transfection to assess knockdown efficiency (see below). Plasmids were transiently transfected into MCF7 and T47D cell lines with 2,500 ng plasmid DNA and using 6 µL Lipofectamin^TM^ 2000 (Thermo Fisher Scientific, Waltham, MA, USA), according to the manufacturer’s instructions. HEK293FT cells were transiently transfected using plasmid DNA and linear polyethylenimine MW 25000 (PEI, #23966, Polysciences, Warrington, PA, USA) as transfection reagent. HEK293FT cells were first trypsinized and subsequently seeded at a density of 1.5 × 10^6^ cells in 9 mL of fresh growth medium. 4 µg plasmid DNA with 20 µL PEI (1 mg/mL) were mixed and incubated for 15 minutes at room temperature to allow the formation of DNA-PEI complexes. Following the incubation period, the transfection mixture was added dropwise to the prediluted cell suspension and plated on a 100 mm dish. After 24 hours of transfection, the culture medium was replaced with fresh growth medium.

### RT-qPCR

RNA was isolated using the RNeasy Mini Kit (#74104, Qiagen, Hilden, Germany) with on-column DNA digestion, according to the manufacturer’s instructions. Purified RNA was reverse transcribed utilizing the RevertAid RT Reverse Transcription Kit (#K1691, Thermo Fisher Scientific, Waltham, MA, USA) according to the manufacturer’s instructions. Quantitative PCR was conducted using the Power SYBR Green PCR Master Mix (#4368706, Thermo Fisher Scientific, Waltham, MA, USA), along with the QuantStudio™ 5 Real-Time PCR System (Thermo Fisher Scientific, Waltham, MA, USA). Data was analyzed utilizing the QuantStudio™ Design & Analysis Software v1.5.0. Relative changes in gene expression were determined using the comparative Ct (ΔΔCt) method (42) and visualized as 2^−ΔΔCt^. The median Ct values of technical replicates and the mean Ct values of biological replicates were used as basis for the ΔΔCt method and expression of the genes of interest was normalized to *ACTB* and *PUM1* housekeeping genes. The following primer pairs were used: *ACTB* forward 5’-ATTGGCAATGAGCGGTTC-3’, reverse 5’-GGATGCCACAGGACTCCA-3’; *PUM1* forward 5’-TCACATGGATCCTCTTCAAGC-3’, reverse 5’-CCTGGAGCAGCAGAGATGTAT-3’; *ESR1* forward 5’-GATGGGCTTACTGACCAACC-3’, reverse 5’-AAAGCCTGGCACCCTCTT-3’; *GLYATL1* forward 5’-CACATCAATCACGGGAACC-3’, reverse 5’-CCATGTCATCAGTCATCTCCTG-3’.

### Transcription factor activity and pathway analysis

Differential gene expression analysis based on RNA-sequencing data was performed using the R package limma (43). The resulting t-statistics values were used as input for the decoupleR R package (44) (version 2.8.0) to estimate transcription factor activities, which were based on the DoRothEA (v1.16.0) (45, 46) database containing signed and confidence-weighted TF–target gene-interactions.

Gene Set Enrichment Analysis (GSEA) (47) was performed with RNA-sequencing data using R package GSEA v4.1.0 (Build 27), while ReactomePA (version 1.46.0) (48) was used for gene set enrichment analysis in proteomic data, based on Reactome molecular pathways (49).

### Patient data analysis

GEO datasets (GSE55374 (50), GSE10281 (51)) were used to assess *GLYATL1* expression in ER+ breast cancer patients pretreated and after neoadjuvant therapy with 2.5 mg/day of the aromatase inhibitor letrozole. Patient samples of dataset GSE55374 had been profiled on Illumina HT-12v4 (*GLYATL1* probe: ILMN_1769032) and patients in the GSE10281 cohort on Affymetrix Human Genome U133 Plus 2.0 Array (*GLYATL1* probe: 1562089_at). Data from The Cancer Genome Atlas (TCGA) (52) were utilized for patient survival analysis. Clinical and survival data were extracted from the Cancer Genome Atlas (TCGA) for Kaplan-Meier survival analysis as described previously (53). Patients were filtered for PAM50 subtype ‘luminal A’ and sorted by *GLYATL1* expression levels. Patients with an expression level below the median expression were assigned to the ‘low’ group, patients with above-median expression to the ‘high’ group. Overall survival rates were visualized as Kaplan-Meier plots using GraphPad Prism (v. 10), and statistical significance between patient groups was analyzed using the log-rank (Mantel-Cox) test.

## Results

### GLYATL1 is upregulated in two luminal endocrine therapy-resistant cell line models

Endocrine therapy-resistant MCF7 and T47D cell lines were generated by continuously depriving cells of estrogen for one year, thereby mimicking the effects of aromatase inhibition. This resulted in long-term estrogen-deprived (LTED) MCF7 and T47D cell lines that had regained proliferative capacities in estrogen-depleted media (19, 20). To elucidate transcriptomic and epigenomic alterations associated with resistance to estrogen deprivation, we characterized MCF7 parental and LTED cell lines using RNA-as well as ATAC-sequencing. ATAC-sequencing peaks within ± 2 kbp of the transcription start sites (TSS) which were gained in resistant MCF7 LTED cells compared to their sensitive counterpart (i.e., parental), were associated with differentially expressed genes (Figure 1A). *GLYATL1* was among the top ten upregulated genes in MCF7 LTED cells (log2FC 11.79; padj 4.79E-18, Supplementary Table 1) and the *GLYATL1*-promoter was among the top 1% of genomic loci with gains in chromatin accessibility (chr11:58926244-58926663: log2FC 2.4; FDR 7.59E-06, Supplementary Table 2) compared to parental cells (Figure 1A). The upregulation of GLYATL1 in MCF7 LTED cells was validated at both RNA and protein levels (Figure 1B), and a similar upregulation of GLYATL1 was observed also in T47D LTED cells (Supplementary Figure 1A). Clinical relevance of these *in vitro* findings was supported by two patient datasets comparing transcriptomic profiles of matching pre-and post-treatment samples from luminal breast cancer patients. This analysis revealed significant increases in *GLYATL1* expression upon neoadjuvant administration of the AI letrozole for 2 weeks (GSE55374, (50) Figure 1B) and 3 months (GSE10281, (51) Figure 1C). Survival analysis of luminal A patient samples from the TCGA cohort (52) further showed that high *GLYATL1* expression in these treatment-naïve patients was significantly associated with a worse overall patient survival (Supplementary Figure 1B).

**Figure 1:**
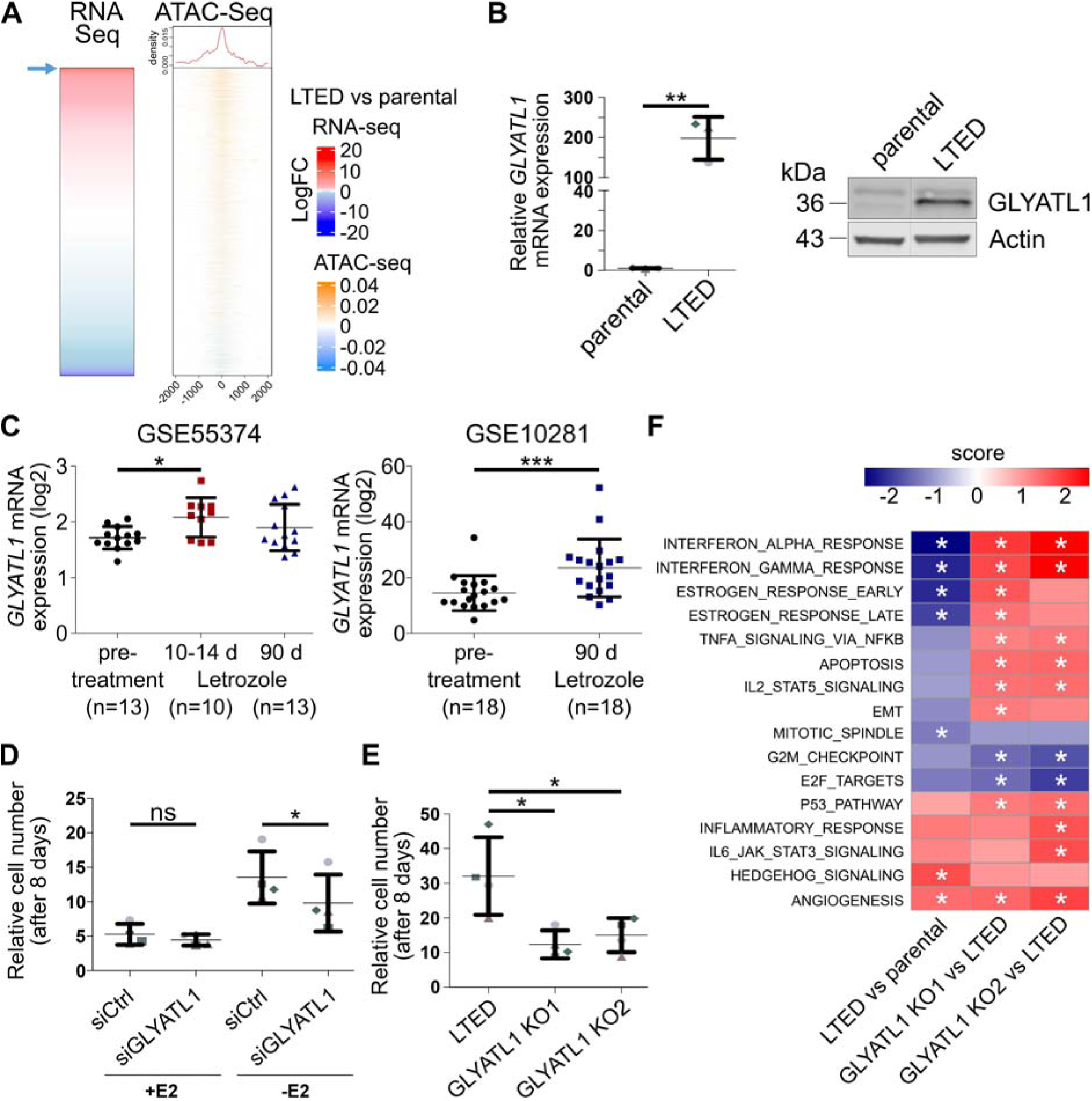
*GLYATL1* expression is associated with endocrine therapy *in vitro* and in patients, and with cell viability as well as estrogen response. (A) Gene expression of MCF7 long-term estrogen-deprived (LTED) and parental cells was assessed by RNA-sequencing (n=3) and differentially expressed genes were ranked by log2FC. The rank position of *GLYATL1* in the heatmap is indicated by a blue arrow. Open chromatin was analyzed by ATAC-sequencing (n=2) in MCF7 parental and LTED cell lines, probing TSS ± 2kb for long term estrogen deprived (LTED) compared to sensitive (i.e., parental) cells in the figure. The ATAC-seq heatmap is sorted in the same order of loci/genes as the heatmap from RNA-sequencing. (B) Expression of *GLYATL1* at mRNA and protein levels in parental and LTED MCF7 cell lines as detected by RT-qPCR and Western blot, respectively. mRNA expression was normalized to *ACTB* and *PUM1* levels and relative changes to parental were calculated (left). Statistical significance was assessed using unpaired Student’s t-test, ** indicates p<0.01. Data are represented as the mean ± SEM (n=3). Western Blot (right): β-actin was used as a loading control. Uncropped images of Western blots, data replication and validation of anti-GLYATL1 antibody are presented in the Supplementary File. (C) GLYATL1 expression levels in two GEO datasets (GSE55374 (50), GSE10281 (51)) of matched tumor biopsies from ER+ breast cancer patients before and after letrozole administration. Gene expression values are represented as log2 expression ± SD. Statistical significance was assessed using one-way ANOVA with Bonferroni post-test (GSE55374) or paired Student’s t-test (GSE10281). * indicates p<0.05, *** indicates p<0.001. (D) Equal numbers of MCF7 LTED and LTED *GLYATL1* knockdown cells were cultivated in media with (+E2) and without (-E2) estrogen for 8 days. Cell numbers were then quantified via microscopy-based nuclear count and normalized to the seeding control. Statistical significance was assessed using paired Student’s t-test. “ns” indicates a non-significant effect, * indicates p<0.05, *** indicates p<0.001. Data are presented as mean ± SEM (n=4). (E) *GLYATL1* was knocked out in MCF7 LTED cells, and proliferation rates of LTED cells and LTED *GLYATL1* knockout clones KO1 and KO2 were measured in media without estrogen via nuclear count, and normalized to the respective seeding control. Statistical significance was assessed using one-way ANOVA with Bonferroni post-test. “ns” indicates a non-significant effect, * indicates p<0.05, ** indicates p<0.01, *** indicates p<0.001. Data are presented as mean ± SEM (n=4). (F) Geneset enrichment analysis of HALLMARK genes based on RNA-sequencing data from MCF7 parental and LTED cells, and the two LTED *GLYATL1* knockout clones KO1 and KO2 was performed using R package GSEA v4.1.0 (Build 27). Heatmap shows comparisons for LTED vs. parental cells and for LTED *GLYATL1* knockout clones KO1 and KO2 vs. LTED cells, respectively. The cut-off for statistical significance was set to a False Discovery Rate (FDR) q-value of < 0.25), and * indicates an FDR q<0.25.

Hypothesizing that GLYATL1 contributes to the resistance of the LTED cells, we then examined whether perturbation of *GLYATL1* expression in resistant LTED cells would affect their growth under anti-estrogen treatment. To this end, we first tested whether reducing *GLYATL1* gene expression through siRNA-mediated knockdown influenced cell growth rates. Transient knockdown of *GLYATL1* in MCF7 LTED (Supplementary Figure 1C) as well as in T47D LTED (Supplementary Figure 1D) cells resulted in a significant reduction in proliferation under estrogen-deprived conditions in both cell line models, but had no effect in the presence of estrogen (Figure 1D, Supplementary Figure 1E), suggesting a functional role of GLYATL1 in endocrine resistance. To validate these findings, we knocked out *GLYATL1* in MCF7 LTED cells and isolated two knockout (KO) clones. Of note, no viable *GLYATL1* KO cells could be obtained in the T47D cell line. Proliferation was significantly reduced in both MCF7 LTED *GLYATL1* knockout clones compared to the parental LTED cells (Figure 1E), thereby corroborating our results from *GLYATL1* knockdown. RNA-sequencing followed by Gene Set Enrichment Analysis (GSEA) (47) also of the two LTED *GLYATL1* KO clones (Supplementary Table 3) revealed that several Hallmark capabilities were altered in LTED compared to parental MCF7 cells; importantly, some of those were reversed when *GLYATL1* was knocked out in the LTED cells (Figure 1F). As expected, expression of genes associated with ESTROGEN RESPONSE was strongly reduced in the LTED condition, which lacked estrogen in the media. Unexpectedly, this Hallmark increased again in both LTED *GLYATL1* KO clones (Figure 1F), even though also these cells were cultivated in media lacking estrogen. Assessment of *ESR1* transcript sequence reads in MCF7 parental and LTED cells, and the two LTED *GLYATL1* knockout clones did not reveal any pathogenic mutation in the *ESR1* gene that could be associated with resistance. Only a minor allele (c.1965G>A; p.T593= relative to the reference sequence NM_001291241.2) having a low variant allele frequency in the parental MCF7 cell line, was enriched in the LTED cells and in the two LTED GLYATL1 knockout clones (Supplementary Figure 1F). The wildtype and mutant *ESR1* alleles were similarly expressed in all LTED conditions, collectively suggesting that alterations in *ESR1* were not associated with endocrine resistance in our MCF7 LTED cell line model and that a subclone from the parental cell line had potentially been selected in the course of resistance development (15). Our findings thus indicate that while GLYATL1 might affect ERα signaling, it likely does not directly impact *ESR1* gene expression. Other Hallmark capabilities showing consistent differences across conditions included INTERFERON-ALPHA- and INTEFERON-GAMMA.RESPONSE, while cell cycle-related hallmarks were negatively regulated in LTED cells, and even further downregulated in the two LTED *GLYATL1* KO clones. The latter is in agreement with the proliferation data shown in Figure 1E.

Based on the RNA sequencing data, we next assessed transcription factor activities in MCF7 parental and LTED cells, and the two LTED *GLYATL1* knockout clones using the DoRothEA-decoupleR pipeline (44, 54). Consistent with reduced proliferation of both MCF7 LTED *GLYATL1* knockout clones in estrogen-deprivation (Figure 1E), activities of E2F1 and E2F4 were significantly repressed in the knockout clones (Supplementary Figure 1F). Further, several transcription factors associated with stem cell-like activities (e.g., SOX2, PAX6, SNAI1,) were activated in MCF7 LTED cells, and this regulation was reversed in the LTED *GLYATL1* KO clones (Supplementary Figure 1G). Concordant with GSEA, *ESR1*/ERα activity was reduced in the LTED condition compared to parental cells (Supplementary Figure 1F, Supplementary Table 4). Yet, *ESR1* expression was upregulated in MCF7 LTED (RNA-seq: log2FC 1.14, adj. p-value 0.0006; MS-proteomics: log2FC 3.40, adj. p-value 0.0295) and in T47D LTED cells (Supplementary Figure 1H), relative to their respective parental cell lines, probably owing to compensatory feedback. Additionally, the activity of *ESR1*/ERα was increased in LTED *GLYATL1* knockout clone KO1 relative to control LTED, although this was not evident in LTED *GLYATL1* clone KO2 (Supplementary Figure 1G).

The observed differences between the two LTED *GLYATL1* knockout clones in some Hallmark capabilities as well as in transcription factor activities prompted us to investigate whether the GLYATL1 protein was similarly depleted in the two LTED GLYATL1 knockout clones. Parallel reaction monitoring-based mass spectrometry of the MCF7 LTED *GLYATL1* KO clones showed different knockout levels of GLYATL1 at the protein level (Supplementary Figure 1I, Supplementary Table 5). GLYATL1 protein levels were ∼1% in LTED *GLYATL1* knockout clone KO1 but ∼25% in clone KO2, compared to the parental LTED cells. No GLYATL1-peptides were reliably detected in the parental MCF7 cells, which was in line with a very high log2 foldchange difference between LTED vs. parental cells (Supplementary Table 1). Notably, these protein levels were consistent with the CRISPR-induced aberrations: while both knockout clones shared the same genomic alterations resulting in two different KO alleles (Supplementary Figure 2A) they had different extents of reduced read-coverage in the exon covering the start of the *GLYATL1* open reading frame (Supplementary Figure 2B). Furthermore, aberrant splicing was particularly prominent in the KO1 clone (Supplementary Figure 2C,D). Combined, this resulted in either reduced levels (clone KO2) or an almost complete loss (clone KO1) of GLYATL1-protein and of functional mRNA (Supplementary Figures 1I, 2C). In summary, we found that *GLYATL1* expression increased upon aromatase inhibitor therapy in patients and upon estrogen-deprivation in two independent *in vitro* models. *GLYATL1* expression was required for the proliferative capacity of the endocrine resistant cells *in vitro* and its expression correlated with patient survival. Our findings suggest that GLYATL1 is associated with proliferation under estrogen-deprived conditions and might affect ERα activity.

### GLYATL1 expression is associated with succinate levels

We then performed quantitative proteomic analysis in the MCF7 parental, LTED and the two LTED *GLYATL1* knockout clones, to assess whether the effects of GLYATL1-expression were reflected also at the proteome level. While GLYATL1 was not detected in parental cells and in LTED *GLYATL1* knockout clone KO1, it was within the top 1% upregulated proteins in LTED compared to parental MCF7 cells (Supplementary Table 6), consistent with the results from PRM-analysis (Supplementary Figure 1H). Principal component analysis of proteomic data revealed that the data obtained from the two knockout clones clustered away from both parental and LTED conditions (Supplementary Figure 3A). A number of proteins were significantly upregulated in LTED compared to parental cells and downregulated in either one or both LTED *GLYATL1* knockout clones compared to LTED cells (Supplementary Figure 3B, Supplementary Table 6). Thus, ACSL3 and LEO1 were upregulated in LTED compared to parental cells and were strongly downregulated again in both LTED *GYLATL1* knockout clones (Supplementary Figure 3C). ACSL3 activates long-chain fatty acids for synthesis of cellular lipids as well as for degradation via beta-oxidation (55), while LEO1 is a component of the PAF complex (PAF1C) and has been linked to epigenetic regulation of gene expression through histone modifications such as monoubiquitination of histone H2B and tri-methylation of H3K4 (56). Congruent with the observed regulation of ACSL3, Reactome pathway analysis of the proteomic data revealed induction of lipid as well as of steroid metabolism and cholesterol biosynthesis in the LTED cells (Figure 2A). Regulation of the latter is consistent with a previous report showing that 27-hydoxycholesterol can compensate for the lack of estrogen in MCF7 LTED cells (19). Reduced protein levels of DNA polymerase epsilon (POLE) in the two knockout clones (Supplementary Figure 3C) were in accordance with the lower proliferation rate (Figure 1E) and with the downregulation of cell cycle activities in both LTED *GLYATL1* KO clones compared to their parental LTED cells (Figure 1F). These results collectively demonstrate that GLYATL1 is critical for maintaining proliferation under endocrine therapy conditions in the two LTED cell line models, and its loss compromises fitness upon estrogen deprivation.

**Figure 2.**
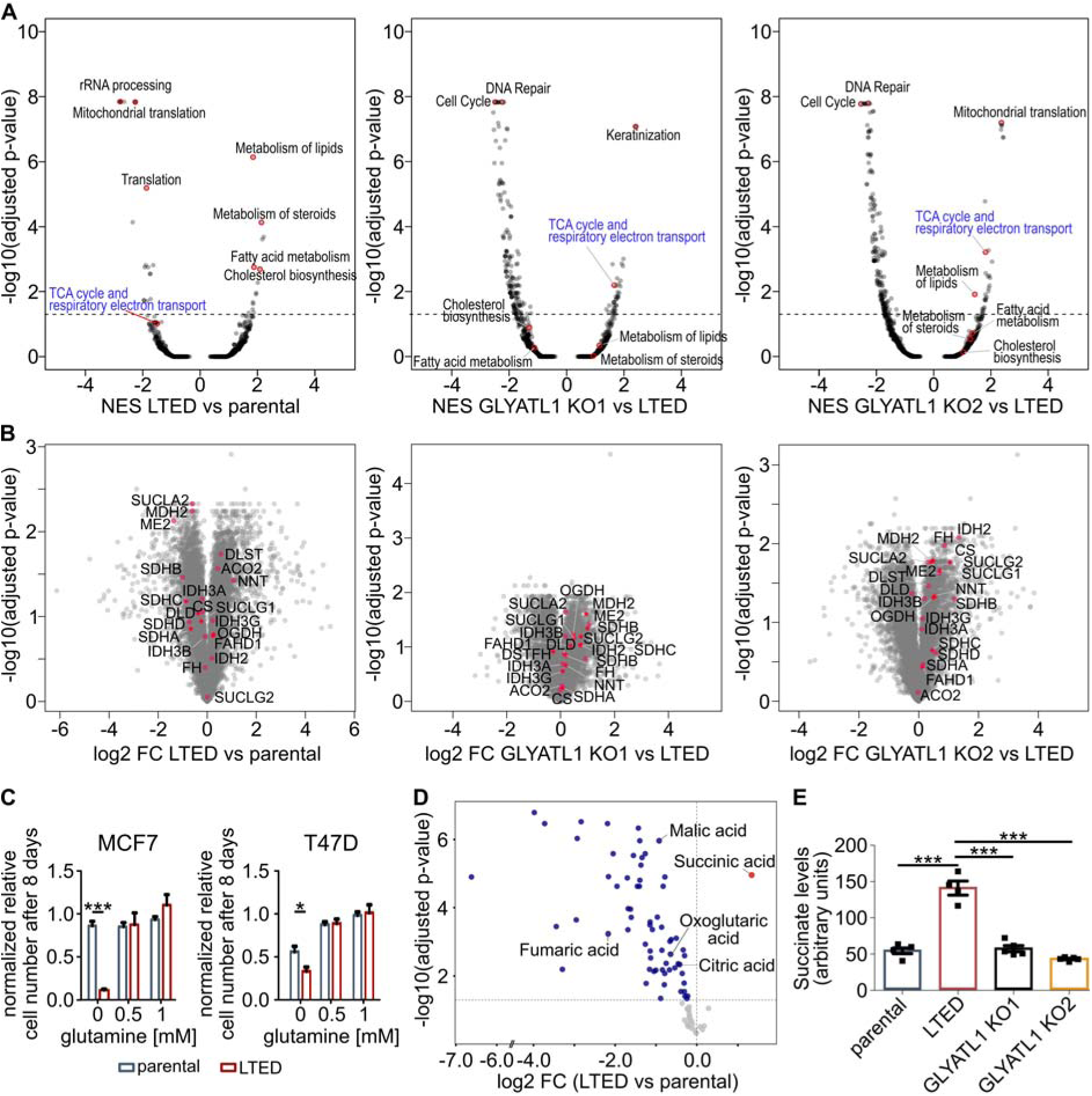
GLYATL1-expression is associated with metabolic pathways and TCA cycle metabolites. (A) Log2 fold-changes of protein groups were calculated between the respective comparisons (n=3) and sorted in decreasing order. Protein groups with duplicated gene names were removed and gene names were mapped to Entrez gene IDs based on the org.Hs.eg.db package (version 3.18.0) using the mapIds function of the AnnotationDbi package (version 1.65.1). The gsePathway function of the ReactomePA package (version 1.46.0) (48) was then applied to the sorted log2 fold-changes to perform gene set enrichment analysis based on Reactome molecular pathways (49). Shown are the GSEA statistics (normalized enrichment scores (NES) and Benjamini-Hochberg adjusted p-values) for all Reactome pathways for the indicated comparisons. The term <TCA cycle and respiratory electron chain> is highlighted in the comparisons. (B) Volcano plot showing differential abundance (log2 fold changes (log2 FC) of members of the Reactome pathway “TCA-cycle and respiratory electron chain” measured in a proteomic workflow between MCF7 LTED vs. parental cells and the two LTED *GLYATL1* knockout clones KO1 and KO2 vs. parental LTED cells. Differential protein expression was tested using unpaired Student’s t-test followed by Benjamini-Hochberg adjustment of p-values. (C) MCF7 as well as T47D parental and LTED cell lines were cultivated in media with varying glutamine concentrations for eight days. The proliferation rate was measured via microscopy-based nuclear counting. Proliferation was normalized to respective cells grown in 2 mM glutamine. Statistical significance was assessed compared to the parental condition using two-way ANOVA with Bonferroni post-test, *** indicates p<0.001, * indicates p<0.05. Data are presented as mean ± SEM (n=3 with 6 technical replicates each). (D) Steady-state levels of soluble metabolites were quantified in MCF7 parental and LTED by mass spectrometry (n≥4). Colored dots represent significantly higher (red) or lower (blue) steady-state abundance levels metabolites. Quantified TCA intermediates are indicated. (E) Steady-state levels of succinate in MCF7 parental and LTED, and the two LTED *GLYATL1* knockout clones KO1 and KO2. Peak intensities were normalized by their respective internal standard and by protein levels. Statistical significance was assessed using one-way ANOVA with Bonferroni post-test, *** indicates p<0.001. Data are presented as mean ± SEM of the biological replicates (n≥4).

Reactome analysis of the proteomic data further suggested that the term ‘TCA cycle and respiratory electron transport chain’ was downregulated in LTED as compared to parental MCF7 cells, whereas it was significantly enriched in both LTED *GLYATL1* KO clones (Figure 2A, Supplementary Table 7). While the log2 foldchange differences appeared to be minor for this Reactome term, down- and upregulation was consistent for the vast majority of enzymes mapping to the TCA cycle in the LTED and LTED *GLYATL1* KO clones, respectively (Figure 2B).

Since Reactome pathway analysis suggested metabolic effects upon GLYATL1 perturbation, and because GLYATL1 is annotated also as a glutamine-*N*-acyltransferase (57), we hypothesized that its relevance in endocrine therapy-resistance is associated with cellular metabolic homeostasis. Therefore, we first tested the effect glutamine limitation had on the viability of the MCF7 as well as T47 parental and LTED cell lines. Parental MCF7 cells were not affected by glutamine-restriction, while growth was moderately slowed down in the parental T47D cells (Figure 2C). In contrast, both LTED cell lines were severely impacted by the lack of glutamine (Figure 2C) suggesting that those cells were addicted glutamine. GLYATL1 and glutamine thus seemed to be essential in both endocrine resistant cell models. To uncover metabolic effects of GLYATL1 more comprehensively, we next quantified steady-state levels of intracellular soluble metabolites as well as N-acyl-CoA species in our set of MCF7 cell lines. Intracellular steady-state levels of glutamine and of glutathione were reduced in MCF7 LTED compared to the parental cells (Supplementary Figure 4A, Supplementary Table 8) even though the genes encoding SLC38A1 and SLC38A2 glutamine transporters were upregulated in LTED cells (Supplementary Table 1). Furthermore, and in agreement with the proteomic data, the steady-state levels of tricarboxylic acid cycle (TCA) intermediates, including the direct oxidation product of succinate (i.e., fumarate), the direct precursor of succinate (succinyl-CoA), and other TCA-metabolites were significantly downregulated in the LTED compared to parental MCF7 cells (Figure 2D, Supplementary Figure 4A,B, Supplementary Tables 8,9). This was in line with a general downregulation of TCA cycle-associated genes in the LTED condition, which was partially reverted in the LTED *GLYALT1* KO clones (Supplementary Figure 4C). The proteomic data matched these findings as the TCA-cycle proteins ME2, SDHB, SUCLA2 and MDH2 were among the most strongly downregulated proteins in the LTED vs. parental cells (Figure 2B). Congruent with a potential role GLYATL1 might have in these alterations, TCA cycle-associated genes and proteins were elevated in LTED *GLYATL1* knockout cells, thus reversing the direction we had observed in the LTED cells (Figure 2B, Supplementary Figure 4C).

Succinate was the only TCA metabolite that had significantly higher steady-state levels in LTED cells while its levels were reversed back to the baseline in the two LTED *GLYATL1* knockout clones (Figure 2 D,E, Supplementary Figure 4A, Supplementary Table 8). This might be related, at least in part, to the reduced expression of *SDHB* (58) that we observed in LTED cells, and which was reverted in the two LTED *GLYATL1* knockout clones (Supplementary Figure 4C). Of note, neither any of the *SDH* or *IDH* genes nor *FH* were mutated in the MCF7 LTED cells or LTED *GLYATL1* knockout clones, as determined based on the RNA-seq data (not shown), while recurrent mutations have been found within these genes in several cancer entities (59). Furthermore, there was also no indication of aberrant splicing of these genes in the RNA-sequencing data that could have explained our observations (not shown). However, components of the 2-oxoglutarate dehydrogenase complex, *OGDH* and *DLD,* were strongly downregulated in the LTED cells, while *OGDH* was increased again in the two LTED *GLYATL1* KO clones (Supplementary Figure 4C, Supplementary Tables 1,3). The 2-oxoglutarate dehydrogenase complex catalyzes the conversion of 2-oxoglutarate to succinyl-CoA, a precursor of succinate. *OGDH*, as well as succinate, have previously been associated with cancer (60, 61). Yet, the steady-state levels of 2-oxoglutarate and of succinyl-CoA were rather downregulated in the LTED and the LTED GLYATL1 knockout conditions (Supplementary Figure 4A,B). Combined, these results suggest that GLYATL1 specifically increases the steady state levels specifically of succinate in the MCF7 LTED cells, while the levels of the other TCA-cycle metabolites are likely not directly affected by GLYATL1.

### GLYATL1 is associated with epigenetic reprogramming

Succinate has been described as a competitive inhibitor of oxoglutarate-dependent dioxygenases, including lysine demethylases (KDMs) (62). We thus hypothesized that GLYATL1 might impact the epigenetic landscape by modulating histone modifications via alteration of succinate levels. Along these lines, we next profiled the epigenetic landscape using Epigenetic-focused cytometry by Time-of-Flight (EpiTOF) (63, 64). There, we measured signal intensities for target proteins and modifications in on average approximately 40,500 single cells for every condition (spread: 22,261 to 57,873 cells). Analysis of the resulting data indeed revealed consistent differences in histone mark abundance between parental, LTED, and LTED *GLYATL1* knockout cells (Figure 3A, Supplementary Table 10). Specifically, H3K64ac and total histone H4 were significantly less abundant in parental MCF7 cells and the two LTED *GLYATL1* knockout clones as compared to LTED cells. A similar trend was observed for H3K4me3, which marks open chromatin (65), similarly to H3K64ac (66). For H3K4me3, the EpiTOF pattern was consistent with the increased expression of LEO1 protein observed in our proteomic data (Supplementary Figure 3B), as LEO1 is involved in the tri-methylation of histone H3K4 (56). The H3K4me3-pattern matched also the increased steady-state level of succinate in LTED cells and the expression of *KDM5B* and *KDM5C* (65, 67), which was increased in the LTED *GLYATL1* knockout clone KO1 compared to the LTED cells (Supplementary Figure 5). H3K36me3, which has been described as a tumor suppressor mark (68), showed the reverse order, with lowest levels in the LTED cells. This pattern was in line with the expression of *KDM4D* (Supplementary Figure 5), which catalyzes H3K36me3 demethylation and which was most highly expressed in the LTED condition. These findings indicate that GLYATL1 may impact histone modifications via altered methylation and acetylation. However, further studies are warranted to fully uncover the effects of GLYATL1 on epigenetic regulation of chromatin. Of note, inclusion of Ki67 in the EpiTOF panel confirmed that Ki67 levels were congruent with the proliferation characteristics as well as with the results from GSEA of parental, LTED, and the two LTED *GLYATL1* knockout clones (compare Figure 1 E,F), thus validating our approach.

**Figure 3:**
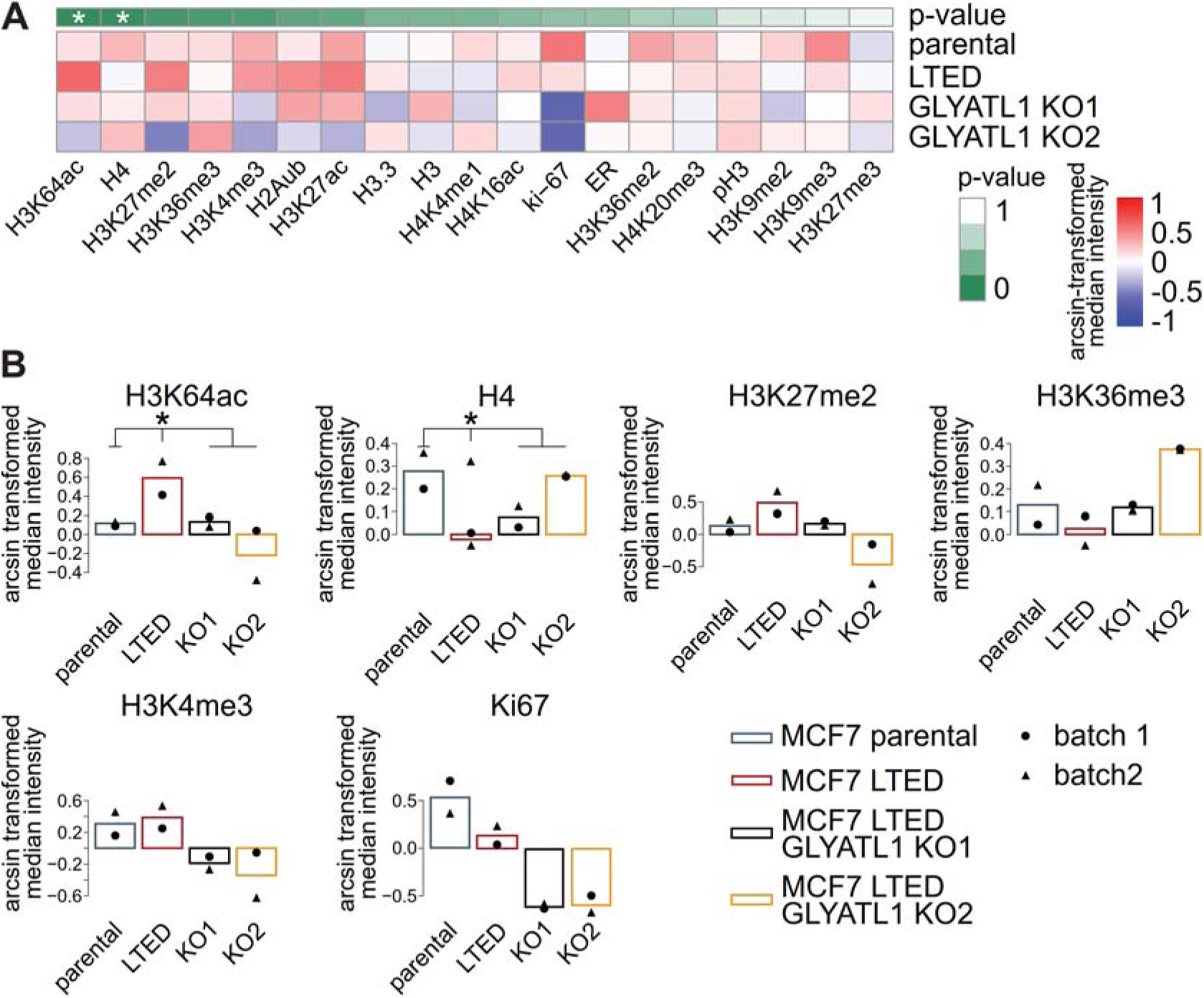
GLYATL1 affects histone modifications. EpiTOF technology (63, 64) was applied to analyze epigenetic histone marks in MCF7 parental and long-term estrogen deprived (LTED) cells, and the two LTED *GLYATL1* knockout clones KO1 and KO2. (A) Data was arcsine transformed and normalized on core histones H3 and H3.3. For statistical comparison, median values of signal intensities were calculated for every condition. Using a two-sided, unpaired t-test, LTED was compared against the parental cells and both KO cells simultaneously. (B) Two replicate experiments (filled triangles and circles, respectively) were performed. The means of the two replicates are indicated for modifications that reached significance or showed a consistent trend (e.g., upregulated in LTED compared to the parental cells and downregulated in the LTED *GLYATL1* KO clones compared to the LTED cells).

As we had found GLYATL1 to be related to the mitochondrial TCA-cycle as well as to nuclear histone modifications, we next investigated the subcellular localization of GLYATL1 protein. Immunofluorescence staining indicated that recombinantly overexpressed GLYATL1 co-localizes predominantly with mitochondria in MCF7 (Manders’ coefficient 0.861), T47D (Manders coefficient 0.467), as well as HEK293FT (Manders’ coefficient 0.607) cells (Supplementary Figure 6). These findings suggest that the epigenetic regulation that is associated with GLYATL1 is likely not direct, but potentially (also) via succinate. In conclusion, our findings suggest that GLYATL1 mediates alterations in cellular metabolism and epigenetic reprogramming, and these mechanisms could be related to endocrine drug resistance in luminal breast cancer.

### Expression of *GLYATL1* is regulated by estrogen-independent estrogen receptor activity

Having uncovered metabolic and epigenetic alterations that are associated with *GLYATL1*, we next aimed to understand how expression of this gene is regulated in the context of resistance to endocrine therapy. Gene set enrichment analysis had highlighted ERα signaling as a strongly downregulated pathway in LTED compared to parental cells (Figure 1F). While loss of ERα activity was expected upon estrogen deprivation, ERα signaling appeared to be elevated in the LTED *GLYATL1* knockout cells (Figure 1F). This observation indicated that ERα signaling and *GLYATL1* expression might be related. Hence, we next investigated this potential connection.

Consistent with previous findings (69), *ESR1* gene expression as well as ERα protein levels were upregulated in LTED compared to parental MCF7 cells (Supplementary Figure 1G, Supplementary Tables 1,6). However, this upregulation seemed to be independent of GLYATL1 as *ESR1*/ERα gene and protein expression was not affected by the knockout of *GLYATL1* at either RNA or protein levels (Supplementary Tables 3,6). To still uncover a potential relationship between ERα activity and *GLYATL1* expression, we next investigated how expression of the latter was regulated in the MCF7 parental and LTED cells.

In line with the findings described above (Figure 1 A,B), genome-mapping of individual RNA-seq reads from MCF7 parental and LTED cells validated that *GLYATL1* was expressed only in the LTED but not the parental cells (Figure 4A). *GLYATL1* expression was induced mostly from the second annotated promoter of the gene, while there was no expression from either of the more distal and proximal promoters. The sequence reads thus matched RefSeq isoform 2, which is encoded by transcript variant 6 (NM_001389711.2) and is equivalent to ENSEMBL transcript ENST00000689150.1. Only in the LTED and not in parental MCF7, two ERα binding regions were identified in ChIP-seq data (MACS_peak 3254). The ChIP-seq data had been previously generated using the same LTED cells we used in our study (19). These peaks were <1 kb upstream and downstream of the transcription start site, respectively, the upstream region coinciding with open chromatin (ATAC-seq) as well as reduced DNA methylation specifically in the LTED cells (Figure 4A). A third ERα ChIP-seq peak further upstream (MACS_peak_3253) was also detected only in LTED cells; however, this was not associated with altered chromatin accessibility. These findings suggested that *GLYATL1* might potentially be regulated by ERα. However, this was in conflict with our results from Reactome and TF-activity analysis, which suggested that ERα signaling was rather downregulated in the LTED cells and was upregulated again in LTED *GLYATL1* knockout conditions (compare Figure 1F, Supplementary Figure 1F).

**Figure 4:**
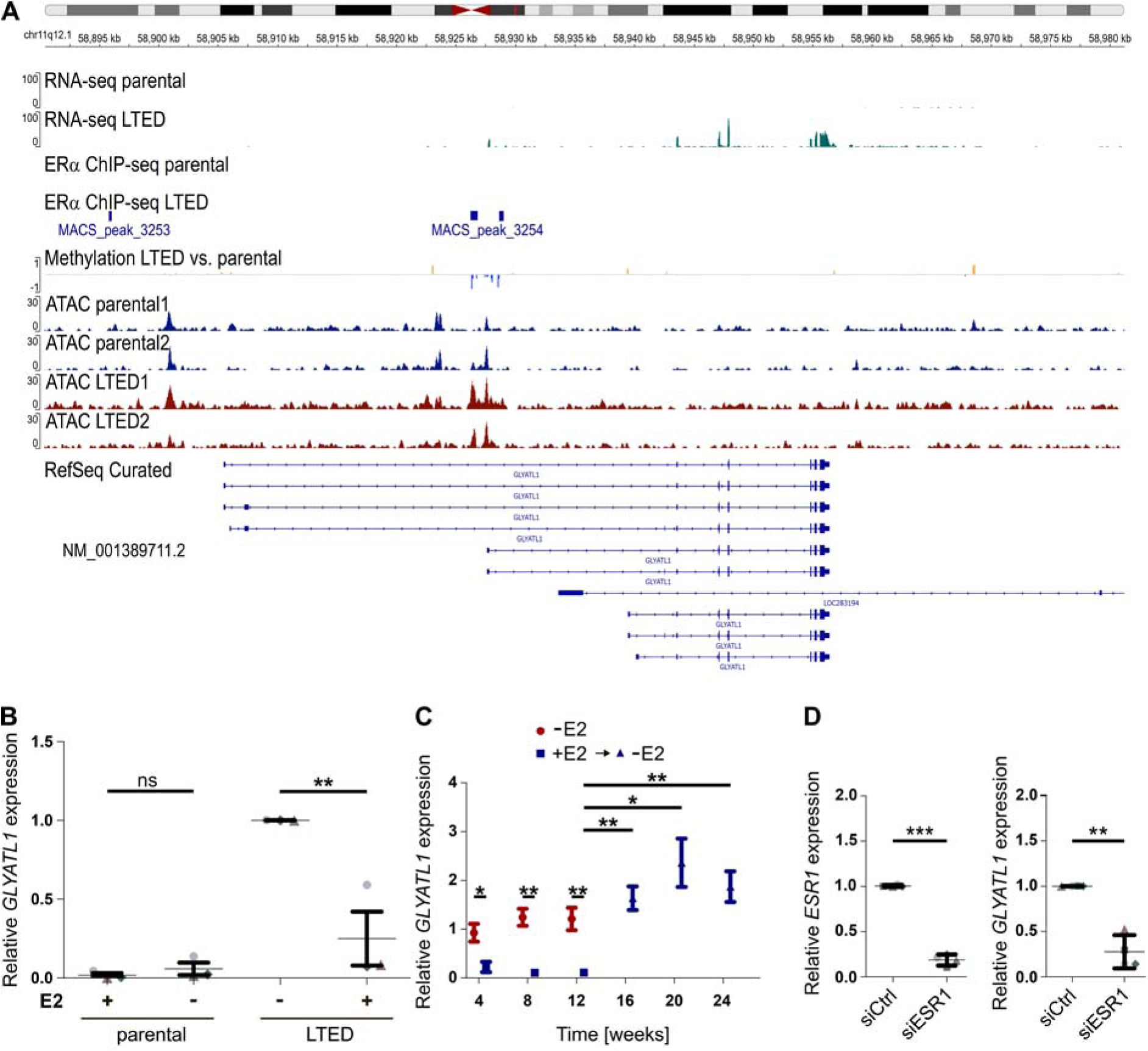
*GLYATL1* expression is regulated by non-canonical estrogen-signaling in long term estrogen deprived MCF7 cells. (A) RNA-seq, ERα ChIP-seq (from GSE60517), methylation changes (EPIC-array), as well as ATAC-seq peaks acquired from MCF7 parental and long-term estrogen deprived (LTED) cell lines were mapped to the human genome (GRCh38/hg38). The RefSeq curated gene structure of *GLYATL1* and a RefSeq ID matching the RNA-seq peaks are indicated. Red and blue peaks in the <Methylation LTED vs. parental> track depict hypermethylated and hypomethylated CpG positions in the LTED cell line, respectively. (B) *GLYATL1* mRNA levels were assessed via RT-qPCR after 48 hours of estrogen treatment or estrogen deprivation in parental MCF7 cells (left) and MCF7 LTED cells (right). Data from all conditions were normalized to LTED cells cultivated in estrogen-depleted media (n=3, each with 3 technical replicates). Statistical significance was assessed using one-way ANOVA with Bonferroni post-test. ** indicates p<0.01, ns: not significant. (C) MCF7 LTED cells were cultured in the presence (+E2, blue squares) or absence (-E2, red circles) of estrogen for 12 weeks. Then, cells were deprived of estrogen again and cultivation was continued for another 12 weeks (+E2 -> -E2, black triangles). mRNA levels were determined by RT-qPCR from cultures harvested at the indicated time points. Relative changes to LTED cells cultivated in estrogen-deprived media were calculated (n≥4, with 3 technical replicates each). Statistical significance was assessed using unpaired Student’s t-tests. * indicates p<0.05, ** indicates p<0.01. (D) MCF7 LTED cells were transfected with non-targeting siRNA (siCtrl) or a pool of siRNAs targeting *ESR1* (siESR1) for 72h. Knockdown efficiency (left) and effect on *GLYATL1* expression (right) were validated by RT-qPCR and relative levels with respect to control transfected cells were calculated. Values for mRNA expression were normalized to *ACTB* and *PUM1* expression levels. Statistical significance was assessed using paired Student’s t-test, ** indicates p<0.01, *** indicates p<0.001. Data are represented by mean ± SEM (n=3).

To resolve these contradicting observations, we next tested the effect estrogen has on *GLYATL1* expression in parental and LTED MCF7 cells. Parental MCF7 cells showed marginally increased *GLYATL1* expression when these cells were kept under estrogen deprivation for 48 hours compared to cells grown in medium including estrogen, while the expression of *GLYATL1* was strongly reduced in LTED cells upon short time estrogen treatment (Figure 4B). These findings were corroborated in T47D, where the same trend and significant regulation were observed, respectively (Supplementary Figure 7A). Hence, negative regulation of *GLYATL1* expression appeared to be a rapid response to estrogen supplementation. We then tested if this negative effect of estrogen on *GLYATL1* expression would persist also for a longer time. To that end, we cultivated MCF7 and T47D LTED cells for three months in media either with or without estrogen. *GLYATL1* expression was indeed consistently reduced in MCF7 LTED cells grown in full medium with continuous presence of estrogen (Figure 4C). After three months, we deprived the LTED cells of estrogen again and observed a significant increase in *GLYATL1* expression back to similar levels to those present in the original LTED cells (Figure 4C). Consistent results were observed also in the T47D LTED cell line (Supplementary Figure 7B), demonstrating that this dependency was not cell line-specific. Taken together, these results imply a strong and direct association between estrogen deprivation and *GLYATL1* expression, suggesting that *GLYATL1* is negatively regulated by estrogen and thus, putatively, also by estrogen-dependent ERα activity.

In light of these observations and the results from the analysis of ERα ChIP-seq data, we then knocked down *ESR1*/ERα in MCF7 and T47D LTED cells that were kept in estrogen-deprived media. This revealed strongly reduced expression of *GLYATL1* when ERα was depleted in MCF7 and in T47D LTED cells (Figure 4D, Supplementary Figure 7C), suggesting that *GLYATL1* expression was positively regulated by ERα, but only in the absence of estrogen. Accordingly, binding of ERα at the promoter of GLYATL1 (proximal peak of MACS_peak_3254) coincided with open chromatin specifically in the MCF7 LTED cells (Figure 4A). Induction of ERα activity by ligands other than estrogen has been described before. In line with a report that had identified 27-hydroxycholesterol as a *bona fide* alternative regulator of ERα activity (19), expression of *CYP27A1*, which encodes the monooxygenase that converts cholesterol to 27-hydroxycholesterol, was upregulated in the MCF7 LTED cells compared to parental cells (log2FC 2.46, adjusted p-value 0.002). While CYP27A1 was not detected at the protein level, transcription factor activity analysis predicted activation of SREBF1 and SREBF2 in LTED cells (Supplementary Figure 1F). These transcription factors, which regulate genes that are involved in cholesterol biosynthesis and lipid homeostasis (70), were the strongest downregulated TFs in the LTED *GLYATL1* KO1 clone (Supplementary Figure 1F). Furthermore, cholesterol biosynthesis was significantly upregulated in the MCF7 LTED vs. parental cells, while this Reactome geneset was downregulated in LTED *GLYATL1* KO1 cells, supporting a potential involvement of GLYATL1 in the regulation of cholesterol biosynthesis (Figure 2A). Collectively, these observations indicate that *GLYALT1* expression can be induced by the estrogen receptor, but only in the absence of estrogen.

Taken together, our observations suggest that GLYATL1 and its activity support the ability of luminal breast cancer cells to overcome their dependency on estrogen (71).

## Discussion

In the present study, we identify GLYATL1 as a novel player in the crossroads of metabolic and epigenetic mechanisms of resistance to estrogen deprivation, and provide insights into its regulation. GLYATL1 was strongly upregulated in two long term estrogen deprived (LTED) breast cancer cell line models mimicking aromatase inhibition, and perturbation of *GLYATL1* expression resulted in a significant proliferation disadvantage. This suggests that GLYATL1 contributes to the growth and survival of therapy-resistant breast cancer cells, particularly under conditions of estrogen deprivation. GLYATL1 is annotated as an N-acyl-transferase family member that conjugates acyl groups to glutamine (57). While we did not identify the particular substrate or product of the GLYATL1 enzymatic reaction, GLYATL1 was significantly associated with glutamine, as the viability of LTED cells was dependent on the supply of this amino acid. Furthermore, the steady-state levels of succinate were elevated in the MCF7 LTED model, while the levels of all other TCA cycle metabolites were reduced. GLYATL1 seemed to be relevant in this regulation as the levels specifically of succinate reverted back to baseline levels upon knockout of *GLYATL1* in the LTED cells. Succinate has been implicated in tumor aggressiveness (61) and the LTED cells we employed in our study indeed had elevated expression of stem-like marker genes and had shown increased metastatic potential *in vitro* and *in vivo* (19). In contrast to succinate, several other metabolites (e.g., nucleosides and nucleotides, and a number of amino acids) were downregulated in the LTED condition. However, this seemed to be independent of GLYATL1, as their levels were unchanged in the LTED *GLYATL1* knockout clones. Yet, these observations are in agreement with the reduced proliferation potential we observed in LTED cells, which was even more prominent in the *GLYATL1* knockout clones. GLYATL1 thus seems to be an essential survival factor in our endocrine resistance model.

Beyond its metabolic implications in the TCA cycle, succinate has been recognized as an onco-metabolite due to its ability to inhibit the activity of various oxoglutarate-dependent dioxygenases, key regulators of epigenetic modifications such as DNA methylation and histone modifications (62, 72). Specifically, succinate can inhibit members of the histone demethylase family KDM, thereby influencing the methylation status of diverse histone residues (73). In line with these findings, we observed increased levels of H3K4me3 in LTED cells. Elevated levels of this epigenetic mark have been reported to correlate with reduced progression-free survival and increased metastasis in breast cancer (74, 75). Along the same lines, downregulation of H3K4me3 has been shown to enhance the response to the AI fulvestrant and reduce tumor growth and invasiveness by modulating cancer stemness (76). These data align with increased activities of several transcription factors associated with stemness in the long-term estrogen deprived MCF7 cells, and with decreased H3K4me3 in the LTED *GLYATL1* knockout clones. H3K4me3 is targeted by KDM5 (65, 67), and we indeed observed reduced expression of *KDM5B* and *KDM5C* in the LTED compared to the parental cells, while expression of those demethylases was elevated in response to *GLYATL1* knockout.

In addition to these methylation marks, we observed alterations in acetylated histone residues, significantly for H3K64ac, which were elevated in LTED cells and diminished in response to *GLYATL1* perturbation. This mark defines transcriptionally active chromatin via regulation of nucleosome positioning at transcription start sites (66), and its downregulation in the *GLYATL1* knockout clones was in accordance with the reduced activity of E2F1 and E2F4 transcription factors and the attenuated proliferative potential of these cells. Increased acetylation of H3K27, which did not reach significance in our EpiTOF analysis, was previously observed in the MCF7 LTED cells that we used in our study (19) and has been associated with endocrine therapy resistance in tamoxifen- and fulvestrant-resistant cells (77). Elevated H3K27ac levels support resistance by promoting transcriptional activity, while inhibition of EP300/CREBBP, which catalyzes acetylation of H3K27, suppresses breast cancer growth *in vitro* and *in vivo* (74, 78).

Our findings further demonstrate that *GLYATL1* expression was negatively associated with estrogen availability, despite a positive association with ERα transcriptional activity. Analysis of chromatin immunoprecipitation sequencing (ChIP-seq) data (19) revealed direct ERα binding upstream of the *GLYATL1* transcription start site, exclusively in LTED cells. A few mechanisms have been reported for estrogen-independent activation of estrogen receptor signaling (79, 80). For example, activation of ERα by IKKE (*IKBKE*) and DDX3X with an associated anti-viral response has been implicated with a potential link to endocrine resistance in luminal breast cancer (81). However, while DDX3X was marginally upregulated in the LTED condition, IKKE was not changed and the associated interferon alpha and gamma responses were even reduced in LTED as compared to parental MCF7 cells (compare Figure 1F). This suggests that this link is likely not relevant in our model. Upregulation of the estrogen receptor and estrogen-independent activity has been previously observed in the same LTED cells that we employed in our study (19). There, this activation along with chromatin association and gene regulation had been attributed to the binding of 27-hydroxycholesterol to ERα and to epigenetic reprogramming (19). In line with these findings, we observed ‘cholesterol biosynthesis’ and ‘metabolism of steroids’ as strongly upregulated gene sets in GSEA of transcriptomic data from LTED compared to parental cells. While canonical estrogen signaling was significantly downregulated in LTED compared to parental cells (compare Figure 1F), our data suggests that estrogen receptor signaling that is independent of estrogen activates a distinct set of target genes, including *GLYATL1*.

In our initial prioritization of candidate factors for endocrine resistance, we had focused on GLYATL1 also because we observed a correlation between *GLYATL1* expression in clinical samples and the prognosis of breast cancer patients. The significant upregulation of *GLYATL1* expression in patient samples following letrozole treatment, coupled with its correlation with poor overall survival in ER-positive patients, supports its role in AI therapy resistance. Furthermore, endocrine therapy-resistance was associated with a dependence of tumor cells on glutamine in both cell line models and this could, at least in part (82), be related to the activity of GLYALT1. The dependency of LTED cells on glutamine could thus suggest that GLYTL1 might be central in a mechanism that requires glutamine to overcome the negative impact of estrogen depletion on tumor cell viability upon aromatase inhibition. Glutamine addiction has been described as a therapeutic opportunity in cancer (83). Targeting of glutaminase (84) might thus be a vulnerability also in progressing GLYATL1-high endocrine breast cancer.

In conclusion, we have identified GLYATL1 as a potential regulator of endocrine therapy resistance, particularly to aromatase inhibition, in ER+ breast cancer. We have linked its activity to glutamine dependence, metabolic rewiring, and epigenetic reprogramming. There, GLYATL1 may contribute to transcriptional programs associated with endocrine therapy resistance by promoting succinate accumulation and influencing histone modifications. Targeting GLYATL1 and associated mechanisms might eventually be established as a treatment for AI-resistant luminal breast cancer.

## Abbreviations

AI: aromatase inhibitor
ATAC-seq: Assay for Transposase-Accessible Chromatin using sequencing
ChIP-seq: Chromatin immunoprecipitation sequencing
E2: estrogen, 17-β-estradiol
EpiTOF: Epigenetic-focused cytometry by Time-of-Flight
ERα: estrogen receptor alpha
ER+: estrogen receptor (α) positive (breast cancer)
GLYATL1: Glycine-N-Acyltransferase Like 1
GSEA: Gene Set Enrichment Analysis
KDM: Lysine demethylase
KO (CRISPR/Cas9): knock out
LTED: long term estrogen deprived
TCA: tricarboxylic acid cycle/citrate cycle

## Declarations

### Ethics approval and consent to participate

Not applicable

### Consent for publication

Not applicable

### Availability of data and materials

OMICs datasets generated in the course of the current study are available in respective repositories: RNA-sequencing data of MCF7 parental, LTED, and LTED *GLYTL1* knockout clones KO1 and KO2 as well as ATAC-sequencing data of MCF7 parental and LTED cell lines are available at the GHGA Data Portal (https://data.ghga.de/) in the study GHGAS95686857321950 with accession numbers GHGAD23940263267179 (RNA-seq) and GHGAD24495022399035 (ATAC seq). EPIC 850k methylome data of MCF7 parental and LTED cell lines are available at EMBL-EBI ArrayExpress database (https://www.ebi.ac.uk/biostudies/arrayexpress) with accession number E-MTAB-15519. Total DIA mass spectrometry proteomic data of MCF7 parental and LTED cells, and LTED *GLYTL1* knockout clones KO1 and KO2 have been deposited to the ProteomeXchange Consortium via the PRIDE (85) partner repository with the dataset identifier PXD067936. Raw EpiTOF data (i.e., z-scores for every mark measured in every cell) from MCF7 parental and LTED cells, and LTED *GLYTL1* knockout clones KO1 and KO2 are available at Zenodo (https://zenodo.org/) with record number 16947455 (DOI:10.5281/zenodo.16947455). We re-analyzed the following datasets, which are available at public repositories: GEO datasets GSE55374 (50) and GSE10281 (51) are available from the Gene Expression Omnibus (https://www.ncbi.nlm.nih.gov/geo/). Breast cancer RNA-sequencing data (52) were downloaded from cBioPortal (https://www.cbioportal.org/). Other supporting data (Tables, Figures) are provided as Supplementary Information with the manuscript.

### Author’ Contributions

J.M., E.S. and S.W. conceived the study. J.M, E.S., Lu.S., Y.A., Li.S., S.O., C.S., S.B., A.W., S.K., D.H., R.W., I.H., and N.B.N, designed and performed *in vitro* experiments. Lu.S., E.W., K.K., S.O., B.E.M, V.R.d.M.C., C.S., P.L, and D.W. performed bioinformatic data analysis. J.M., E.S., Lu.S, and K.K. generated figures and tables. Y.Y., L.M., C.P., A.S., C.K, M.O., and S.W. supervised the study. J.M., Lu.S., C.K., and S.W. wrote the manuscript draft. All authors participated in discussions and interpretation of the data and results as well as read and approved the final manuscript.

## Acknowledgements

We thank the DKFZ High-throughput DNA sequencing, OMICS Data IT, Microarray, Cellular Tools, Antibody, and Proteomics Core Facilities for performing excellent services. We thank Anke Heit-Mondrzyk for bioinformatics and Thomas Hielscher for statistical support. The results shown are in part, based upon data generated by the TCGA Research Network: (http://www.cancer.gov/tcga).

## Supplementary Tables

**Supplementary Table 1: Analysis of differentially expressed genes in RNA-sequencing data of MCF7 parental and long time estrogen deprived (LTED) cells.** Gene expression levels (raw read counts from RNA-sequencing of MCF7 parental and LTED cells were used to identify differential gene expression employing DESeq2 v1.28.1 (28). Indicated are log2 foldchanges and non-adjusted as well as FDR-adjusted p-values for all identified ENSEMBL/HGNC genes. Rawdata are available at GHGA (dataset ID: GHGAD23940263267179).

**Supplementary Table 2: ATAC-sequencing of MCF7 parental and long-term estrogen deprived (LTED) cells.** ATAC libraries were generated using a tagmentation protocol (30). Sequencing reads were aligned to the hg38 reference genome and peaks were identified using MACS2 (v.2.2.9.1) (31). Chromosomal coordinates of differentially accessible regions are annotated with log2 fold changes, non-adjusted P-values, and FDR adjusted p-values, each in comparisons of LTED and parental cells. Rawdata are available at GHGA (accession number: GHGAD24495022399035)

**Supplementary Table 3: Analysis of differentially expressed genes in RNA-sequencing data of long-time estrogen deprived (LTED) cells and LTED *GLYATL1* knockout clones KO1 and KO2.** Gene expression levels (CPM) from RNA-sequencing of MCF7 LTED cells and LTED *GLYATL1* knockout clones KO1 and KO2 were used to identify differential gene expression employing DESeq2 v1.28.1 (28). Indicated are log2 foldchanges and non-adjusted as well as FDR-adjusted p-values for all identified ENSEMBL/HGNC genes. Results for comparison of *GLYATL1* knockout clone KO1 vs. LTED are presented in sheet 1 of the table, while results for comparison of LTED *GLYATL1* knockout clone KO2 vs. LTED are shown in sheet 2. Rawdata are available at GHGA (dataset ID: GHGAD23940263267179).

**Supplementary Table 4: DoRothEA prediction of transcription factor activities in comparisons between MCF7 parental and long-term estrogen deprived (LTED) cells, and in LTED *GLYATL1* knockout clones KO1 and KO2.** Differential gene expression analysis was performed using RNA-sequencing data and the R package limma (43). The resulting t-statistics values were used as input for the decoupleR R package (44) to estimate transcription factor activities, which were based on the DoRothEA (v1.16.0) (45, 46) database containing signed and confidence-weighted TF–target gene-interactions. Shown are DoRothEA scores and p-values for every comparison.

**Supplementary Table 5: PRM based mass spectrometric analysis of GLYATL1 protein expression in MCF7 parental and long-term estrogen deprived (LTED) cells, and in LTED *GLYATL1* knockout clones KO1 and KO2.** The areas of the GLYATL1 fragment ions R.ALLLVTEDILK.L as measured in MS2 spectra from PRM mass spectrometry are indicated. Peak areas were computed by defining peak boundaries and summing up the areas of the fragment ions (n=4).

**Supplementary Table 6: Proteomic analysis of in MCF7 parental and long-term estrogen deprived (LTED) cells, and in LTED *GLYATL1* knockout clones KO1 and KO2.** DIA mass spectrometry was applied to identify and quantify proteins in the indicated cell lines. Peptides and protein groups were identified using Spectronaut and quantified by summing up precursor signal areas under the curve. Shown are log2 transformed protein intensities for the different conditions and log2 fold-changes (FC) as well as Benjamini-Hochberg adjusted p-values (padj) for indicated comparisons (n=3). NA not detected. Rawdata are available via ProteomeXchange with identifier PXD067936.

**Supplementary Table 7: Reactome analysis of proteomic data from MCF7 parental and long-term estrogen deprived (LTED) cells, and in LTED *GLYATL1* knockout clones KO1 and KO2.** Log2 fold-changes of protein groups were calculated between the respective comparisons (n=3) and sorted in decreasing order. Protein groups with duplicated gene names were removed and gene names were mapped to Entrez gene IDs based on the org.Hs.eg.db package (version 3.18.0) using the mapIds function of the AnnotationDbi package (version 1.65.1). The gsePathway function of the ReactomePA package (version 1.46.0) (48) was then applied to the sorted log2 fold-changes to perform gene set enrichment analysis based on Reactome molecular pathways (49). Shown are the GSEA statistics, including Benjamini-Hochberg adjusted p-values, for all Reactome pathways for the indicated comparisons.

**Supplementary Table 8: Steady state levels of soluble metabolites measured in MCF7 parental and long-term estrogen deprived (LTED) cells, and in LTED *GLYATL1* knockout clones KO1 and KO2.** Indicated metabolites (with corresponding identifiers in the Human Metabolome Database – HMDB_ID) were quantified using mass spectrometry. Peaks corresponding to the calculated metabolite masses taken from an in-house metabolite library were integrated using the El-MAVEN software (https://docs.polly.elucidata.io/Apps/Metabolomic Data/El-MAVEN.html) and metabolite identification was supported by fragmentation patterns (40). Peak intensities were normalized by their respective internal standard levels. Shown are peak intensities (C) of indicated conditions (n=5), mean intensities (MEAN) used for calculation of foldchanges (FC) as well as log2 fold changes (log2FC), and the p-values (pcal) and adjusted p-values (padj) of indicated comparisons.

**Supplementary Table 9: Steady state levels of acyl-CoA species measured in MCF7 parental and long-term estrogen deprived (LTED) cells, and in LTED *GLYATL1* knockout clones KO1 and KO2.** Acyl-CoA species (Metabolite) were quantified using mass spectrometry. Peaks corresponding to the calculated metabolite masses taken from an in-house metabolite library were integrated using the El-MAVEN software (https://docs.polly.elucidata.io/Apps/Metabolomic Data/El-MAVEN.html) and acyl-CoA identification was supported by fragmentation patterns (39). Peak intensities (C) were normalized by their respective internal standard levels. Shown are peak intensities of indicated conditions (n=5).

**Supplementary Table 10: Median values from EpiTOF data measured in MCF7 parental, long-term estrogen deprived (LTED), and in LTED *GLYATL1* knockout KO1 and KO2 cell lines.** Median values from two replicate EpiTOF experiments of MCF7 parental, LTED, LTED *GLYATL1* KO1 and KO2 conditions are shown for the indicated marks. Rawdata, i.e., z-scores for individual cells having been measured in the two replicates are available at Zenodo (https://doi.org/10.5281/zenodo.16947455).

**Supplementary Table 11: List of primary antibodies used in EpiTOF analysis.** Target specificities, conjugated elements and their isotopes, vendors, and order numbers (cat#) are indicated for antibodies that were used in EpiTOF profiling.

## Supplementary Figures

**Supplementary Figure 1:**
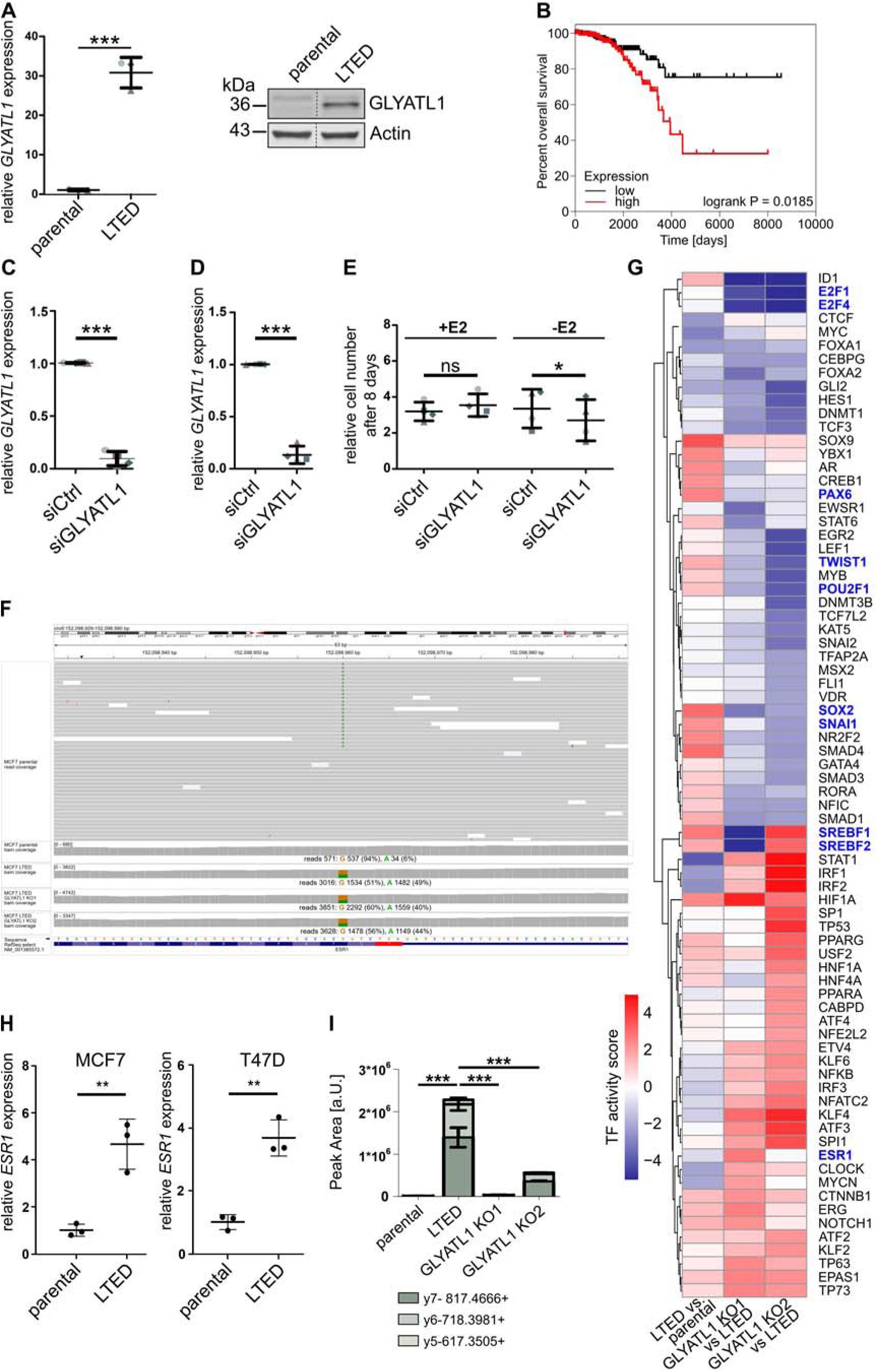
*GLYATL1* is upregulated in long time estrogen deprived (LTED) T47D cell line, and GLYALT1 expression correlates with patient survival as well as affects transcription factor activities. (A) Relative *GLYATL1* mRNA expression in T47D parental and long-term estrogen-deprived (LTED) cell lines analyzed by RT-qPCR (left). mRNA expression was normalized to *ACTB* and *PUM1* expression and relative changes to the parental cells were calculated. Data are represented as the mean ± SEM (n=3). Statistical significance was assessed using unpaired Student’s t-test, *** indicates p<0.001. GLYATL1 protein levels in T47D parental and LTED cells was analyzed by Western Blot (right). β-actin was used as a loading control. Uncropped images of blots are presented in the Supplementary File. (B) Median-based overall survival analysis of luminal A breast cancer patients in the TCGA dataset (52) of *GLYATL1* low (black) versus high (red) gene expression (n=271 per group). Statistical significance was assessed using log-rank (Mantel-Cox) test using GraphPad Prism (v. 10). (C) *GLYATL1* was knocked down via RNA interference in MCF7 LTED cells and knockdown verified via RT-qPCR, compared to MCF7 LTED cells transfected with a non-targeting control siRNA (siCtrl). Data are represented as the mean ± SEM (n=3). Statistical significance was assessed using unpaired Student’s t-test, *** indicates p<0.001. (D) *GLYATL1* was knocked down via RNA interference in T47D LTED cells and knockdown verified via RT-qPCR, compared to T47D LTED cells transfected with a non-targeting control siRNA (siCtrl). Data are represented as the mean ± SEM (n=3). Statistical significance was assessed using unpaired Student’s t-test, *** indicates p<0.001. (E) Equal numbers of MCF7 LTED *GLYATL1* knockdown and control LTED cells were cultivated in media with (+E2) and without (-E2) estrogen for 8 days and cell numbers were then quantified via microscopy-based nuclear count and normalized to the respective seeding control. Statistical significance was assessed using paired Student’s t-test. ns indicates non-significant p-value, * indicates p<0.05. (F) Read-coverage in a window covering 63bp in the terminal exon of the *ESR1* gene in MCF7 parental and LTED cells, and in LTED *GLYATL1* knockout clones KO1 and KO2. A synonymous mutation (A-allele) relative to the G-allele in RefSeq reference sequence (NM_001385572.1) is highlighted. Bam coverage at this position is shown for all conditions. Graphics adapted from Integrative Genomics Viewer (IGV) (86). (G) Relative transcription factor (TF) activities were estimated using DoRothEA (45, 46) R package. The heatmap visualizes TF activity scores in comparisons between MCF7 LTED vs. parental, and between the respective LTED *GLYATL1* knockout clones (KO1 and KO2) vs. the LTED condition. TFs were filtered for an absolute(Score)>2 observed in any comparison. (H) RT-qPCR analysis of *ESR1* expression in MCF7 and in T47D parental and long-term estrogen deprived (LTED) cell lines. Data are presented as mean ± SEM (n=3). Statistical significance was assessed using unpaired Student’s t-test, ** indicates p<0.01. (I) Peak area of all GLYATL1 peptide (ALLLVTEDILK) fragment ions (indicated in color) in MCF7 parental and LTED, and LTED *GLYATL1* knockout clones KO1 and KO2, determined by PRM-based targeted mass spectrometry. Data are represented as mean ± SEM (n=4). Statistical significance was assessed using unpaired Student’s t-test based on the sum of all fragment ion intensities per samples, *** indicates p<0.001.

**Supplementary Figure 2:**
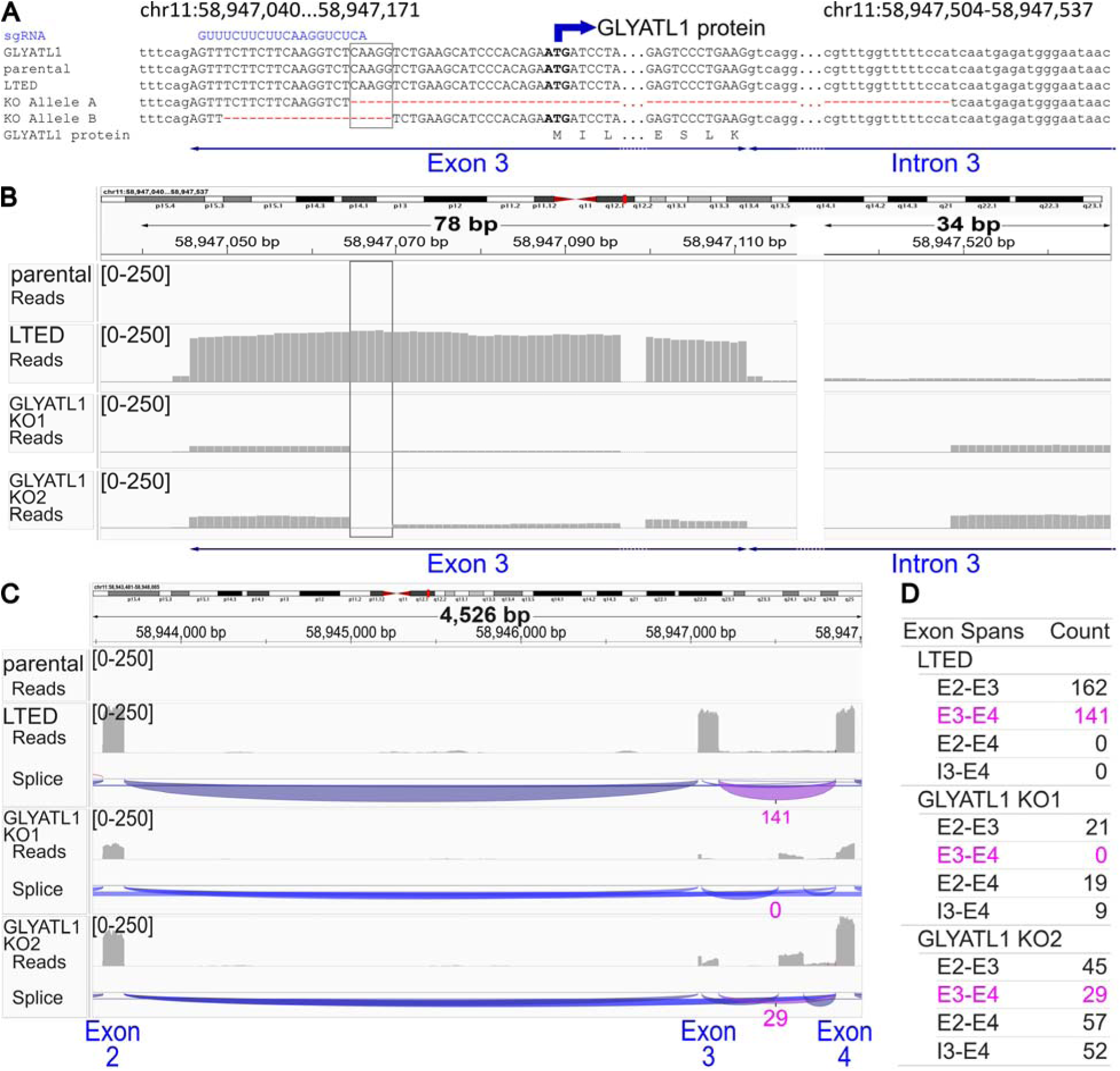
Genotyping and assessment of coding potential of LTED *GLYATL1* knockout clones. (A) Sequence of GLYATL1 gene at the 5’-end of exon 3 and a breakpoint in intron 3 (present in KO allele A) is shown. Sanger sequences of PCR amplified genomic sequence from MCF7 parental, long-term estrogen-deprived (LTED), and two LTED *GLYATL1* knockout clones (KO1, KO2) are shown. A single PCR product was amplified in parental and LTED cells, while two products were amplified in the knockout clones. These were sequenced individually. The sequence and position of the sgRNA binding site is indicated (in blue). Deleted sequence caused by CRISPR/Cas9 editing are indicated by red dashes. A five bp overlap of deleted sequence in alleles A and B is indicated by a box. Exonic sequence is indicated in upper case letters of nucleic acid sequence while intronic sequences are in lower case. The translation start of the GLYATL1 open reading frame is indicated above and the encoded amino acid sequence is indicated below the nucleotide sequences. Dots indicate sequence (black dots) and deleted sequence (red dots) that is not shown. (B) Read-coverage of spliced transcripts in MCF7 parental, LTED and in LTED *GLYATL1* knockout clones KO1 and KO2 is indicated for sequence within exon 3 and around the breakpoint within intron 3. No sequence reads were detected in the parental condition. A five bp overlap of deleted sequence in alleles A and B is indicated by a box and this sequence does not have read coverage in either knockout clone. Graphics adapted from Integrative Genomics Viewer (IGV) (86). (C) Inferred splice junction patterns supported by reads from RNA-sequencing of MCF7 parental, LTED, and LTED *GLYATL1* knockout clones KO1 and KO2. Read counts supporting splicing from exon 3 to exon 4 are indicated in pink for LTED, *GLATLY1* KO1 and KO2 clones. No sequence reads were detected in the parental condition. Graphics adapted from IGV. (D) Read coverage and splice junctions in RNA-sequencing data obtained from parental, LTED, and LTED *GLYATL1* knockout clones KO1 and KO2. Read counts supporting respective splice forms are indicated. Splicing of exon 2 to exon 3 (E2-E3) and of exon3 to exon 4 (E3-E4) indicates canonical splicing events of functional *GLYATL1* transcripts, where splicing E3-E4 (indicated in pink) is critical for the coding potential of GLYATL1 mRNA. Splicing from exon 2 to exon 4 (E2-E4) indicates skipping of exon 3 and of the translational start of the GLYATL1 protein. Splicing from intron 3 to exon 4 (I3-E4) is indicative of transcription from allele A (545bp deletion), which lacks GLYATL1 coding potential.

**Supplementary Figure 3:**
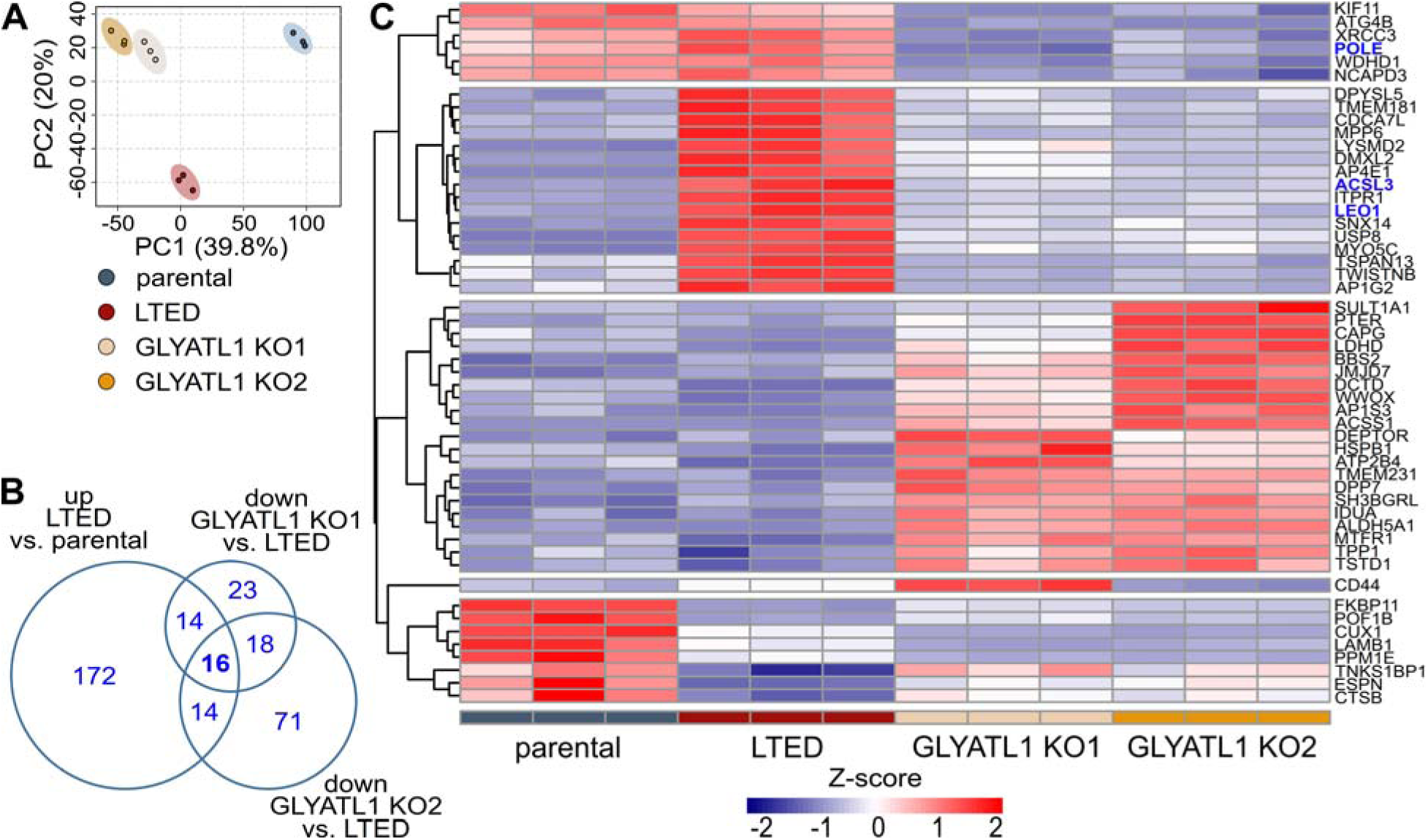
Proteomic changes associated with estrogen deprivation and with GLYATL1 protein expression. (A) Total proteins in MCF7 parental, long term estrogen deprived (LTED), and LTED *GLYATL1* KO1 and KO2 cell lines were analyzed by mass spectrometry and protein intensities were used for principal component analysis. (B) VENN diagram depicting numbers of differentially expressed proteins, filtered for proteins significantly upregulated in MCF7 LTED vs. parental cells, and proteins significantly downregulated in MCF7 LTED *GLYATL1* knockout clones KO1 or KO2 vs. LTED cells. Note: GLYATL1 is not listed as this protein was detected neither in parental nor in LTED *GLYATL1* KO1 cells. Statistical significance was determined by Student’s unpaired t-test and p-values were adjusted using the Benjamini-Hochberg method. Thresholds: log2FC >1 (LTED vs. parental) and <-1 (KO vs. LTED); significance: adjusted p-value < 0.05. (C) Hierarchically clustered heatmap showing z-scaled intensities of proteins with significant changes (adjusted p-values < 0.05) in LTED *GLYATL1* KO1 and KO2 cell lines compared to parental or long-term estrogen deprived (LTED) MCF7 cell lines, with an absolute log2 fold-change of at least 1 in any comparison. Biological replicates (n=3) are displayed separately. Statistical significance was determined by Student’s unpaired t-test and p-values were adjusted using the Benjamini-Hochberg method.

**Supplementary Figure 4:**
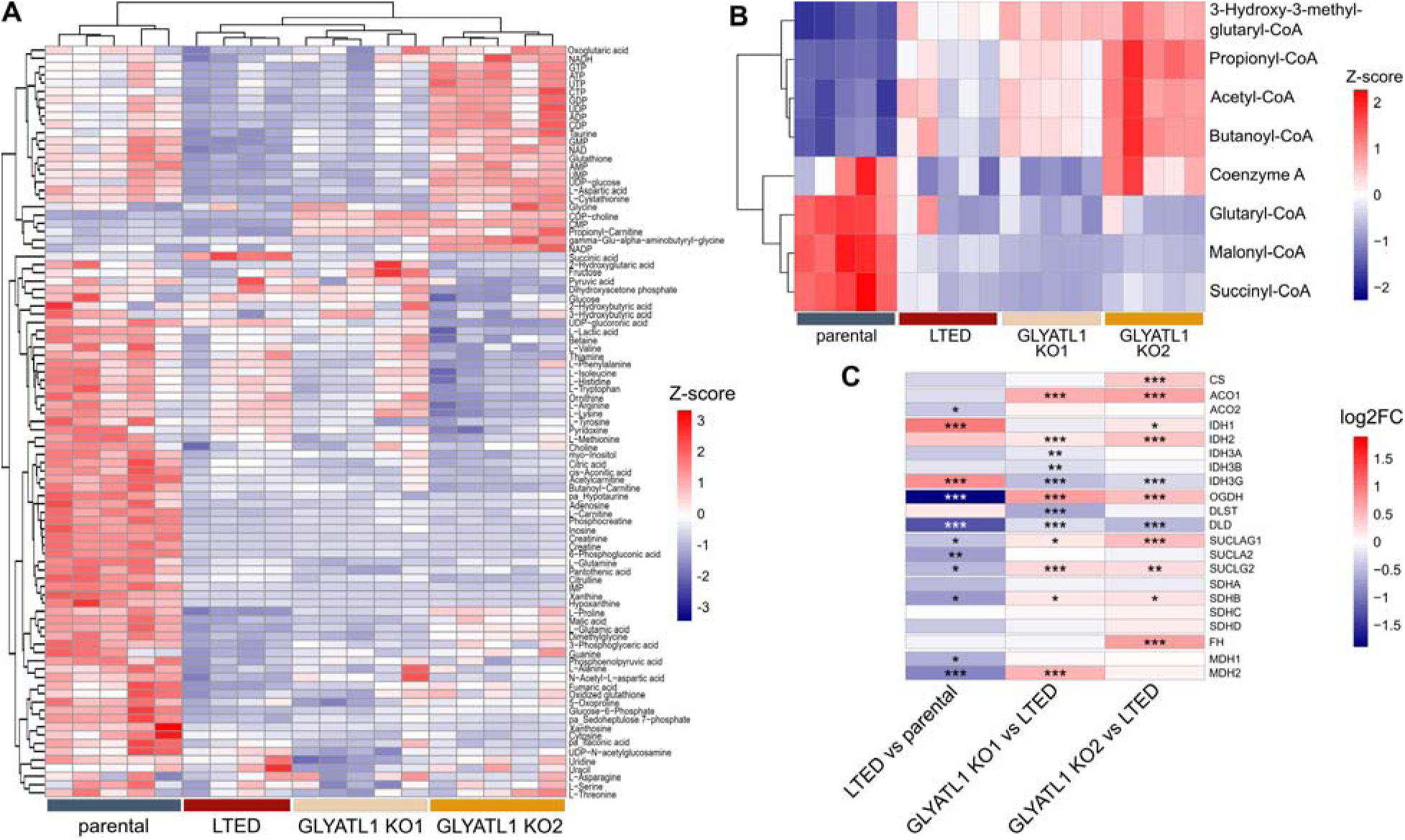
Metabolomic analysis of MCF7 parental and long-term estrogen deprived (LTED) cells, and in LTED *GLYATL1* knockout clones KO1 and KO2. (A) Hierarchical clustering showing z-scaled intensities, normalized by their respective internal standard levels, of soluble metabolites that were measured by mass spectrometry in MCF7 parental, LTED, and the LTED *GLYATL1* knockout clones KO1 and KO2. Biological replicates (n≥4) are displayed individually. (B) Heatmap showing z-scaled values of acyl-CoA species measured via LC-MS in MCF7 parental and LTED cells, and two *GLYATL1* knockout clones KO1 and KO2 (n=5). (C) Differential gene expression (RNA-sequencing) of indicated genes in the TCA-cycle as determined by DESeq2 analysis, sorted by their respective position in that cycle. Data extracted from Supplementary Tables 1 and 3. * indicates Benjamini-Hochberg adjusted p<0.05, ** indicates p<0.01, and *** indicates p<0.001.

**Supplementary Figure 5:**
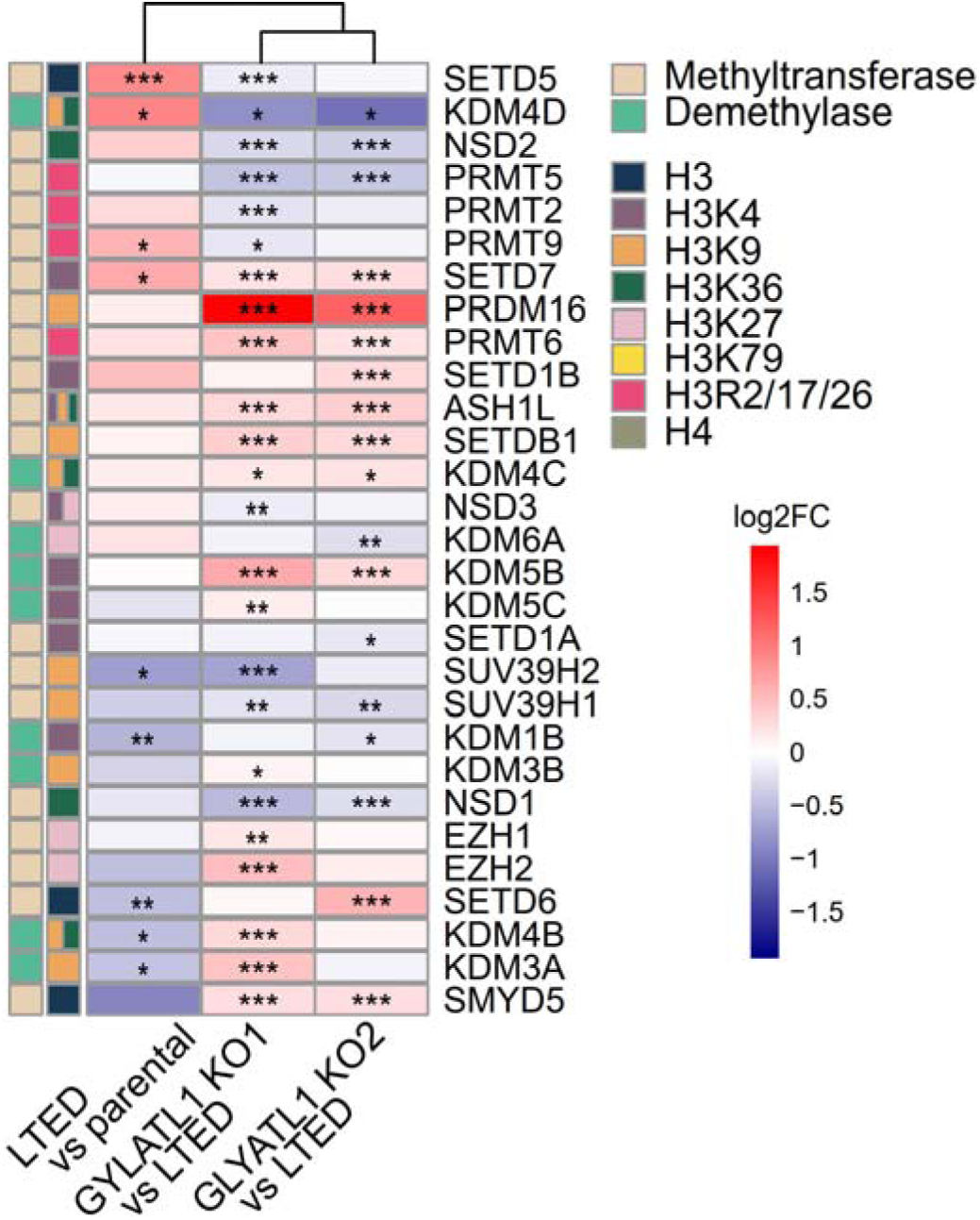
Differentially expressed methyltransferases and demethylases in MCF7 long-term estrogen deprived (LTED) vs. parental cells, and in LTED *GLYATL1* knockout clones KO1 and KO2 vs. LTED cells. Heatmap displays log2 fold-changes (log2FC) in mRNA levels of genes encoding histone modifiers affecting methylation status as measured by RNA sequencing of MCF7 parental and long-term estrogen deprived (LTED) cells, and two LTED *GLYATL1* knockout clones KO1 and KO2, followed by DESeq2 analysis. Respective comparisons are indicated below the heatmaps. Writers of histone marks are indicated in beige and erasers are indicated in green. Affected histone residues are indicated in colors for the respective epigenetic modifiers. * indicates Benjamini Hochberg adjusted p<0.05, ** indicates adjusted p<0.01, and *** indicates adjusted p<0.001.

**Supplementary Figure 6:**
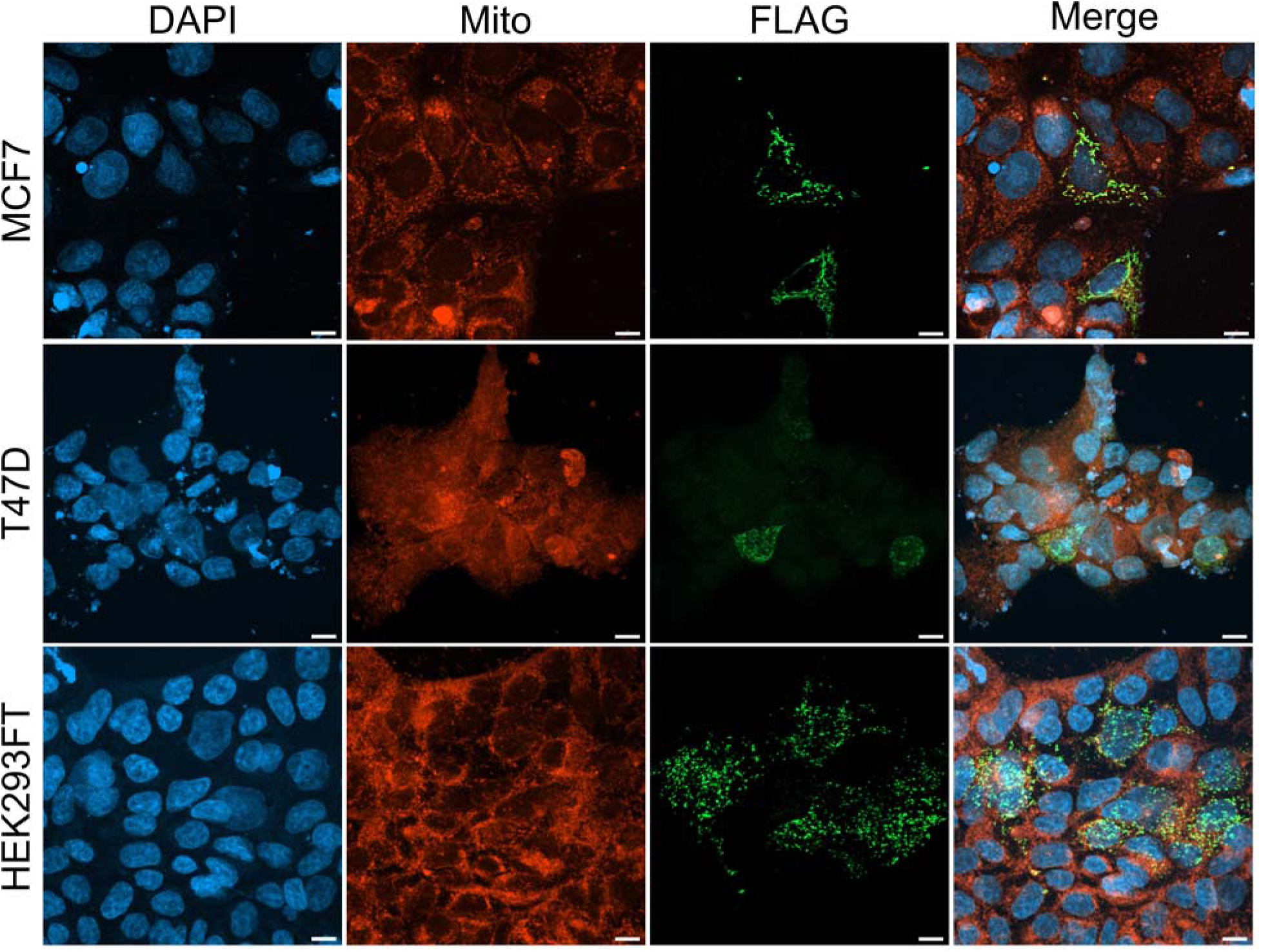
GLYATL1 protein localizes in the mitochondria in MCF7, T47D and HEK293FT cell lines. C-terminally FLAG-tagged GLYATL1 was recombinantly overexpressed in MCF7, T47D and HEK293FT cell lines by transient plasmid transfection. MCF7 and T47D cells were cultivated for 72 hours after transfection and HEK cells for 48 hours, and then incubated with abberior LIVE ORANGE dye to stain mitochondria (Mito). Cells were fixed and incubated with an anti-Flag antibody (FLAG) to detect the GLYATL1-FLAG fusion protein. Finally, nuclei were stained with DAPI and images taken using confocal immunofluorescence microscopy. Image analysis was performed using Zen Blue software and ImageJ (https://imagej.net/ij/). Representative images are shown. Scale bar = 10 µm.

**Supplementary Figure 7:**
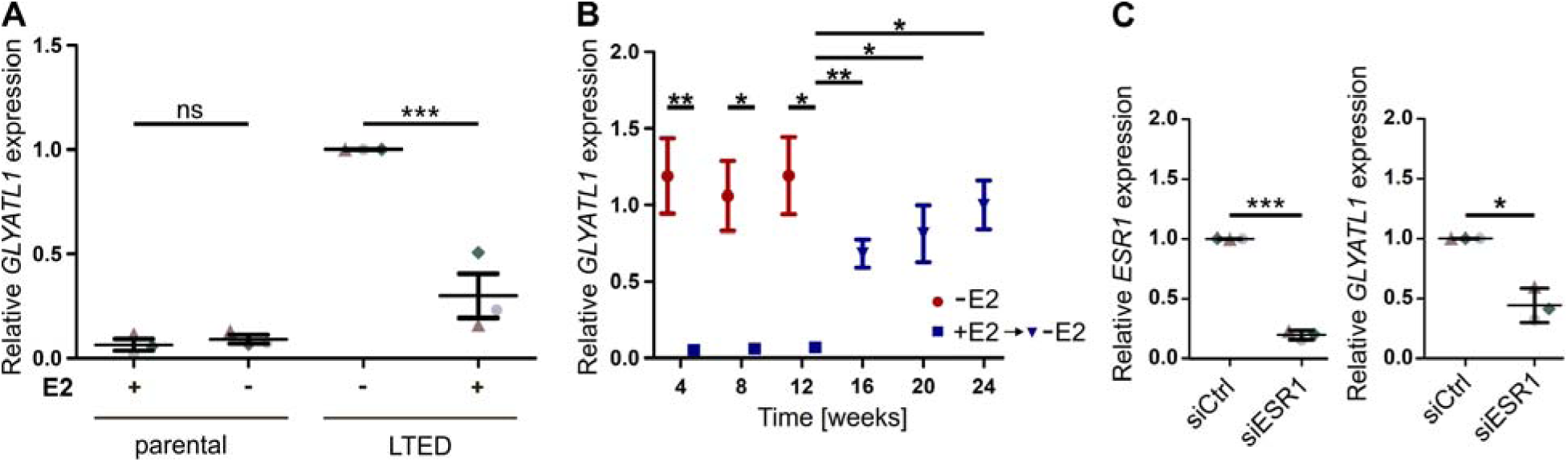
*GLYATL1* expression is regulated by non-canonical estrogen-signaling in long term estrogen deprived T47D cells. (A) *GLYATL1* mRNA levels were assessed via RT-qPCR after 48 hours of estrogen treatment or estrogen deprivation in parental T47D cells (left) and LTED cells (right). Data from all conditions were normalized to LTED cells cultivated in estrogen-depleted media (n=3, with 3 technical replicates each). Statistical significance was assessed using one-way ANOVA with Bonferroni post-test. *** indicates p<0.001, ns: not significant. (B) T47D LTED cells were cultured in the presence (+E2, blue squares) or absence (-E2, red circles) of estrogen for 12 weeks. After these initial 12 weeks, cells were deprived of estrogen again and cultivation was continued for another 12 weeks (+E2 -> -E2, black triangles). mRNA levels were determined by RT-qPCR from cultures harvested at the indicated time points. Relative changes to LTED cultivated in estrogen-deprived media were calculated (n≥4, with 3 technical replicates each). Statistical significance was assessed using unpaired Student’s t-tests. * indicates p<0.05, ** indicates p<0.01. (C) T47D LTED cells were transfected with non-targeting siRNA (siCtrl) or a pool of siRNAs targeting *ESR1* (siESR1) for 72h. Knockdown efficiency (top) and effect on *GLYATL1* expression (bottom) were validated by RT-qPCR and relative levels with respect to control transfected cells were calculated. Values for mRNA expression were normalized to *ACTB* and *PUM1* expression levels. Data are represented by mean ± SEM (n=3). Statistical significance was assessed using paired Student’s t-test, * indicates p<0.05, *** indicates p<0.001.

**Supplementary File:**
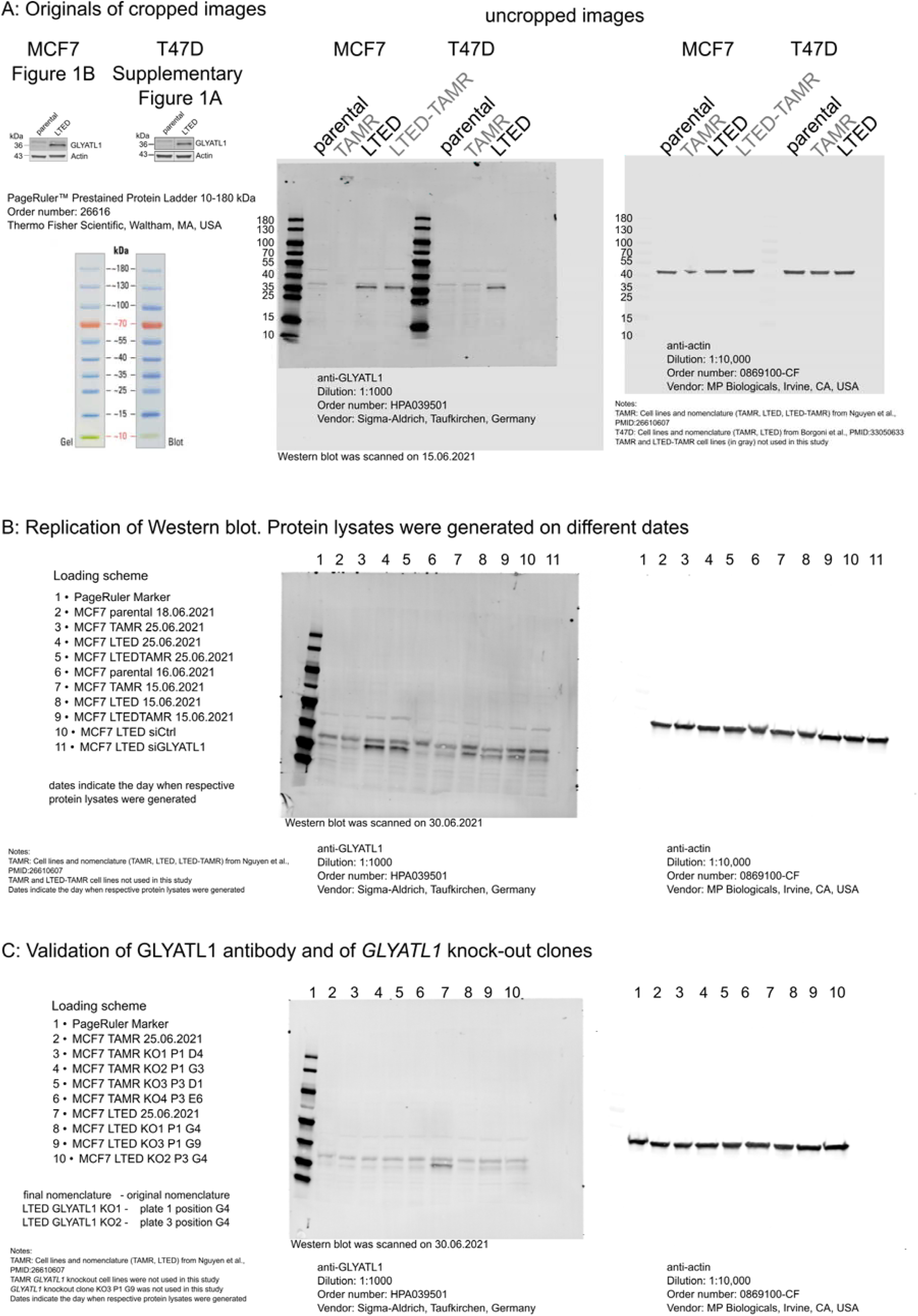
uncropped Western blots, validation of GLYATL1 antibody and initial validation of MCF7 LTED *GLYATL1* knockout clones. Lysates from indicated MCF7 and T47D cell lines were separated by SDS-PAGE and blotted onto a PVDF membranes. The membranes were blocked for unspecific binding of proteins and then incubated with a primary anti GLYATL1 antibodies in blocking buffer. The next day, membranes were washed three times. Then, the blots were incubated with a secondary antibody, which was conjugated with Alexa Fluor™ 680, and washed again. Proteins were visualized with a LI-COR Odyssey scanner using 700 and 800 nm channels. Then, the same blots were reprobed with an anti ß-Actin antibody over night. The next day, the blots were washed and then incubated with a secondary antibody conjugated with DyLight™ 800 4X PEG. Protein bands were again visualized with a LI-COR Odyssey scanner using 700 and 800 nm channels. Original scans and accompanying files are provided in a supplementary ZIP-archive. (A) Cropped images (MCF7 from Figure 1B and T47D from Supplementary Figure 1A) are shown in the top left. The respective uncropped Western blot images of the original gel with indication of molecular weight markers (PageRuler Protein Ladder) are shown on the right. Samples from several MCF7 and T47D cell lines and derivatives were separated on the same gel. Lanes 2 and 4 (MCF7) and 7 and 9 (T47D) were cropped to generate the final images. Other conditions that were tested (i.e., TAMR and LTED-TAMR for MCF7 (19), TAMR for T47D (20)) were not regarded in the current study. (B) Protein lysates of indicated cell lines were generated on the indicated dates and analyzed by SDS-PAGE and Western blot. The same marker was used as in panel A. (C) Protein lysates from *GLYATL1* knockout clones generated from MCF7 TAMR and LTED cells were tested for expression of GLYATL1 protein. Lysates from MCF7 TAMR and LTED cells were used as positive control for GLYATL1 expression. The same marker was used as in panel A.

