## Supplementary Table 1 for "GLYATL1 is associated with metabolic and epigenetic changes and with endocrine resistance in luminal breast cancer"

| ensembl_gene_id | hgnc_symbol | baseMean_LTED_vs_ | log2FoldChange_LTE | lfcSE_LTED_vs_par | stat_LTED_vs_parent | pvalue_LTED_ | padj_LTED_vs_parent |
|---|---|---|---|---|---|---|---|
| ENSG00000072694 | FCGR2B | 484.8762507 | 13.55284072 | 2.768472819 | 10.39808988 | 2.53E-25 | 3.28E-23 |
| ENSG00000234444 | ZNF736 | 626.7989 | 13.45002 | 2.452428 | 10.725 | 7.77E-27 | 1.16E-24 |
| ENSG00000146469 | VIP | 221.3516 | 12.60363 | 4.671253 | 4.112916 | 3.91E-05 | 0.000333 |
| ENSG00000171724 | VAT1L | 2346.3 | 12.38005 | 1.541489 | 8.757199 | NA | NA |
| ENSG00000173093 | CCDC63 | 367.1823 | 12.28203 | 2.445368 | 9.588919 | 8.90E-22 | 8.16E-20 |
| ENSG00000145721 | LIX1 | 211.6637 | 11.87462 | 2.510402 | 9.343228 | 9.34E-21 | 7.52E-19 |
| ENSG00000166840 | GLYATL1 | 271.5448 | 11.78879 | 2.434994 | 9.13442 | 6.58E-20 | 4.79E-18 |
| ENSG00000070882 | OSBPL3 | 565.9964911 | 11.52533514 | 1.34265932 | 9.391422952 | 5.92E-21 | 4.87E-19 |
| ENSG00000268964 | ERVV-2 | 91.95534 | 11.51623 | 2.829341 | 7.168643 | 7.57E-13 | 2.50E-11 |
| ENSG00000182111 | ZNF716 | 78.48767 | 11.42185 | 2.749703 | 8.430582 | 3.44E-17 | 1.92E-15 |
| ENSG00000186448 | ZNF197 | 492.1483 | 11.18076 | 1.281614 | 9.584495 | 9.29E-22 | 8.48E-20 |
| ENSG00000147883 | CDKN2B | 911.6942 | 11.09509 | 1.178178 | 9.322358 | 1.14E-20 | 9.02E-19 |
| ENSG00000197020 | ZNF100 | 95.72002 | 11.05889 | 2.607312 | 8.398348 | 4.53E-17 | 2.49E-15 |
| ENSG00000224244 | | 53.96207 | 10.88691 | 2.635146 | 8.055807 | 7.90E-16 | 3.79E-14 |
| ENSG00000196268 | ZNF493 | 93.08759 | 10.66423 | 2.486491 | 8.300857 | 1.03E-16 | 5.45E-15 |
| ENSG00000196172 | ZNF681 | 140.1867 | 10.62221 | 2.362676 | 8.328718 | 8.17E-17 | 4.36E-15 |
| ENSG00000249568 | | 24.84129 | 10.59873 | 2.919584 | 7.438577 | 1.02E-13 | 3.77E-12 |
| ENSG00000235491 | LINC01889 | 51.6632 | 10.52421 | 2.465372 | 8.00108 | 1.23E-15 | 5.71E-14 |
| ENSG00000151790 | TDO2 | 91.0216 | 10.46172 | 2.444532 | 8.073468 | 6.83E-16 | 3.29E-14 |
| ENSG00000196705 | ZNF431 | 339.7974 | 10.30129 | 1.259017 | 8.872677 | 7.14E-19 | 4.74E-17 |
| ENSG00000123342 | MMP19 | 370.2297398 | 10.24200368 | 1.261678089 | 8.801478996 | 1.35E-18 | 8.67E-17 |
| ENSG00000278318 | ZNF229 | 258.9221 | 10.2394 | 1.273901 | 8.78293 | 1.59E-18 | 1.01E-16 |
| ENSG00000278921 | EPB41L4A-DT | 105.6691 | 10.23464 | 2.330971 | 8.078213 | 6.57E-16 | 3.17E-14 |
| ENSG00000167916 | KRT24 | 41.83543 | 10.19466 | 2.603777 | 7.203577 | 5.87E-13 | 1.97E-11 |
| ENSG00000105750 | ZNF85 | 94.85297445 | 10.14919946 | 2.386407789 | 7.850107092 | 4.16E-15 | 1.79E-13 |
| ENSG00000167232 | ZNF91 | 1357.23 | 10.10188 | 0.8416555 | 11.63809 | 2.64E-31 | 5.70E-29 |
| ENSG00000121413 | ZSCAN18 | 255.4035843 | 10.01303346 | 1.247255156 | 8.683592594 | 3.83E-18 | 2.35E-16 |
| ENSG00000160229 | ZNF66 | 76.52602 | 9.966209 | 2.411113 | 7.64081 | 2.16E-14 | 8.60E-13 |
| ENSG00000211689 | TRGC1 | 1226.522 | 9.884139 | 0.9655504 | 10.61897 | 2.43E-26 | 3.47E-24 |
| ENSG00000267508 | ZNF285 | 39.78754 | 9.679458 | 2.476769 | 7.184448 | 6.75E-13 | 2.25E-11 |
| ENSG00000171476 | HOPX | 225.195 | 9.537842 | 1.527616 | 6.574655 | 4.88E-11 | 1.26E-09 |
| ENSG00000166509 | CLEC3A | 6339.73 | 9.517069 | 0.8643185 | 11.33647 | 8.66E-30 | 1.67E-27 |

| | | | | | | | |
|---|---|---|---|---|---|---|---|
| ENSG00000215146 | | 316.8713 | 9.469539 | 1.260641 | 8.101573 | 5.43E-16 | 2.64E-14 |
| ENSG00000125910 | S1PR4 | 271.3468045 | 9.423044799 | 0.972405703 | 9.28150105 | 1.67E-20 | 1.31E-18 |
| ENSG00000242779 | ZNF702P | 215.5396 | 9.396839 | 2.417926 | 6.933243 | 4.11E-12 | 1.24E-10 |
| ENSG00000215196 | BASP1-AS1 | 288.2615 | 9.293559 | 0.8809052 | 10.42769 | 1.85E-25 | 2.43E-23 |
| ENSG00000213967 | ZNF726 | 103.8441 | 9.291521 | 2.272609 | 7.347787 | 2.02E-13 | 7.20E-12 |
| ENSG00000269994 | | 31.86424 | 9.234941 | 2.774154 | 6.668029 | 2.59E-11 | 7.00E-10 |
| ENSG00000162670 | BRINP3 | 61.98226 | 9.178626 | 1.349494 | 7.510761 | 5.88E-14 | 2.25E-12 |
| ENSG00000124479 | NDP | 31.10284358 | 9.172736532 | 3.055523829 | 4.87316362 | 1.10E-06 | 1.36E-05 |
| ENSG00000188785 | ZNF548 | 286.4139 | 9.148698 | 0.7410429 | 12.11487 | 8.81E-34 | 2.11E-31 |
| ENSG00000278266 | | 32.70973 | 9.131987 | 2.675479 | 6.678376 | 2.42E-11 | 6.55E-10 |
| ENSG00000165092 | ALDH1A1 | 77.93305 | 9.126163 | 2.573215 | 6.427158 | 1.30E-10 | 3.20E-09 |
| ENSG00000237440 | ZNF737 | 265.2971 | 9.091988 | 1.237104 | 7.908714 | 2.60E-15 | 1.16E-13 |
| ENSG00000165029 | ABCA1 | 1217.482 | 9.036087 | 0.7055079 | 12.7782 | 2.17E-37 | 6.48E-35 |
| ENSG00000131845 | ZNF304 | 375.5759062 | 8.947814909 | 0.813471115 | 10.65168944 | 1.71E-26 | 2.50E-24 |
| ENSG00000147889 | CDKN2A | 80.92248 | 8.830092 | 1.322319 | 7.437375 | 1.03E-13 | 3.80E-12 |
| ENSG00000258203 | | 34.97147 | 8.813266 | 2.446927 | 6.626697 | 3.43E-11 | 9.08E-10 |
| ENSG00000105668 | UPK1A | 975.6661373 | 8.786569579 | 1.083351253 | 8.443449284 | 3.08E-17 | 1.73E-15 |
| ENSG00000237174 | | 21.43015 | 8.75489 | 2.401013 | 6.643437 | 3.06E-11 | 8.18E-10 |
| ENSG00000204792 | LINC01291 | 517.7076 | 8.72091 | 0.6697511 | 13.11858 | 2.58E-39 | 8.15E-37 |
| ENSG00000242516 | LINC00960 | 242.1414 | 8.716965 | 0.9557248 | 8.661254 | 4.67E-18 | 2.84E-16 |
| ENSG00000165810 | BTNL9 | 167.8556 | 8.652166 | 0.9673174 | 8.522628 | 1.56E-17 | 8.99E-16 |
| ENSG00000164451 | CALHM4 | 14.24788 | 8.644308 | 2.607737 | 6.030578 | 1.63E-09 | 3.39E-08 |
| ENSG00000275011 | ALOXE3P1 | 80.32729 | 8.600754 | 1.750663 | 5.566417 | 2.60E-08 | 4.39E-07 |
| ENSG00000115594 | IL1R1 | 4175.731873 | 8.494712283 | 0.442964605 | 19.19017166 | 4.47E-82 | 1.70E-78 |
| ENSG00000266602 | | 209.2565 | 8.488385 | 0.9753629 | 8.34285 | 7.25E-17 | 3.88E-15 |
| ENSG00000197128 | ZNF772 | 285.4795 | 8.485003 | 0.7242999 | 11.44444 | 2.51E-30 | 4.97E-28 |
| ENSG00000138653 | NDST4 | 27.25414871 | 8.420232469 | 1.384310892 | 6.807328801 | 9.94E-12 | 2.86E-10 |
| ENSG00000274840 | | 19.79693 | 8.374152 | 2.550269 | 6.217347 | 5.06E-10 | 1.14E-08 |
| ENSG00000230836 | LINC01293 | 1200.828 | 8.314589 | 0.5204146 | 16.10298 | 2.43E-58 | 2.69E-55 |
| ENSG00000211695 | TRGV9 | 292.511 | 8.305188 | 1.082702 | 7.457962 | 8.79E-14 | 3.28E-12 |
| ENSG00000204889 | KRT40 | 14.80727 | 8.258438 | 2.508407 | 5.716811 | 1.09E-08 | 1.98E-07 |
| ENSG00000223523 | | 376.5774 | 8.230725 | 0.6146306 | 13.27313 | 3.31E-40 | 1.09E-37 |
| ENSG00000261776 | | 119.1581 | 8.228726 | 1.702797 | 5.492727 | 3.96E-08 | 6.48E-07 |

| | | | | | | | |
|---|---|---|---|---|---|---|---|
| ENSG00000269586 | CT45A10 | 41.68226 | 8.112472 | 2.497163 | 6.090738 | 1.12E-09 | 2.39E-08 |
| ENSG00000180257 | ZNF816 | 584.9516 | 8.010048 | 0.5383333 | 14.74617 | 3.26E-49 | 1.92E-46 |
| ENSG00000141639 | MAPK4 | 238.4889 | 7.954197 | 1.088319 | 7.741812 | 9.80E-15 | 4.06E-13 |
| ENSG00000185296 | | 42.88617 | 7.944437 | 1.369278 | 6.415927 | 1.40E-10 | 3.43E-09 |
| ENSG00000105388 | CEACAM5 | 1133.643392 | 7.933479399 | 0.951107187 | 8.687715662 | 3.70E-18 | 2.28E-16 |
| ENSG00000225722 | RPL24P9 | 12.71382 | 7.921964 | 2.544575 | 5.795482 | 6.81E-09 | 1.28E-07 |
| ENSG00000183888 | SRARP | 8.019562 | 7.895353 | 2.64277 | 5.157387 | 2.50E-07 | 3.51E-06 |
| ENSG00000287602 | | 77.37948 | 7.869281 | 1.051298 | 7.581298 | 3.42E-14 | 1.34E-12 |
| ENSG00000265296 | FEM1AP2 | 43.30221 | 7.859198 | 1.021908 | 7.541364 | 4.65E-14 | 1.79E-12 |
| ENSG00000179023 | KLHDC7A | 54.8387 | 7.826628 | 2.326301 | 5.80589 | 6.40E-09 | 1.21E-07 |
| ENSG00000224017 | LINC01449 | 20.6754 | 7.781196 | 2.440265 | 5.828658 | 5.59E-09 | 1.07E-07 |
| ENSG00000253369 | | 22.48565 | 7.753447 | 2.540368 | 5.274543 | 1.33E-07 | 1.97E-06 |
| ENSG00000188321 | ZNF559 | 168.3418 | 7.744996 | 0.8186538 | 9.14629 | 5.89E-20 | 4.35E-18 |
| ENSG00000287985 | | 41.93695 | 7.742967 | 1.282585 | 6.63561 | 3.23E-11 | 8.59E-10 |
| ENSG00000253802 | SIRLNT | 81.81028 | 7.687932 | 2.525908 | 5.555599 | 2.77E-08 | 4.64E-07 |
| ENSG00000286206 | | 15.52546 | 7.667371 | 2.332022 | 5.882674 | 4.04E-09 | 7.92E-08 |
| ENSG00000111249 | CUX2 | 131.5669322 | 7.647599906 | 0.813180929 | 9.641429785 | 5.34E-22 | 5.03E-20 |
| ENSG00000221880 | KRTAP1-3 | 9.156212 | 7.613509 | 2.458759 | 5.384424 | 7.27E-08 | 1.13E-06 |
| ENSG00000249199 | | 89.68023 | 7.547934 | 1.008684 | 7.631984 | 2.31E-14 | 9.18E-13 |
| ENSG00000257268 | | 15.31777 | 7.523042 | 2.3842 | 5.677845 | 1.36E-08 | 2.43E-07 |
| ENSG00000139219 | COL2A1 | 207.9511641 | 7.500348616 | 0.724047042 | 10.48332282 | 1.03E-25 | 1.38E-23 |
| ENSG00000227121 | LINC02672 | 129.6446 | 7.498094 | 1.110147 | 7.175202 | 7.22E-13 | 2.39E-11 |
| ENSG00000285205 | | 11.0338 | 7.495504 | 2.529055 | 5.142036 | 2.72E-07 | 3.78E-06 |
| ENSG00000227124 | ZNF717 | 18.1998 | 7.425727 | 1.411832 | 5.868719 | 4.39E-09 | 8.55E-08 |
| ENSG00000187122 | SLIT1 | 5705.075 | 7.408318 | 0.5327752 | 14.08795 | 4.50E-45 | 2.14E-42 |
| ENSG00000242615 | | 7.610404 | 7.381791 | 2.56031 | 5.225779 | 1.73E-07 | 2.51E-06 |
| ENSG00000212901 | KRTAP3-1 | 82.06608 | 7.379487 | 1.065088 | 6.837333 | 8.07E-12 | 2.34E-10 |
| ENSG00000157617 | C2CD2 | 1053.086 | 7.368092 | 0.335921 | 21.90124 | 2.53E-106 | 3.36E-102 |
| ENSG00000250488 | LINC02233 | 44.40315 | 7.360697 | 1.404058 | 5.886569 | 3.94E-09 | 7.74E-08 |
| ENSG00000168542 | COL3A1 | 276.1232 | 7.335692 | 0.9810666 | 7.741581 | 9.82E-15 | 4.06E-13 |
| ENSG00000125245 | GPR18 | 13.73182475 | 7.274159412 | 1.439827499 | 5.667685433 | 1.45E-08 | 2.57E-07 |
| ENSG00000181433 | SAGE1 | 13.83493 | 7.205836 | 2.436441 | 5.246879 | 1.55E-07 | 2.26E-06 |
| ENSG00000088992 | TESC | 305.7693231 | 7.201787623 | 0.595868823 | 12.40581355 | 2.43E-35 | 6.33E-33 |

| | | | | | | | |
|---|---|---|---|---|---|---|---|
| ENSG00000187068 | C3orf70 | 63.33584 | 7.188178 | 0.9956226 | 6.992987 | 2.69E-12 | 8.36E-11 |
| ENSG00000232638 | | 17.33545 | 7.174185 | 2.343244 | 5.440971 | 5.30E-08 | 8.49E-07 |
| ENSG00000135074 | ADAM19 | 38.91880245 | 7.171287198 | 2.244119656 | 5.534046234 | 3.13E-08 | 5.20E-07 |
| ENSG00000262118 | MTCO1P28 | 12.47163 | 7.163394 | 2.590934 | 4.643141 | 3.43E-06 | 3.80E-05 |
| ENSG00000176597 | B3GNT5 | 121.7065 | 7.155636 | 0.9045605 | 7.948776 | 1.88E-15 | 8.50E-14 |
| ENSG00000134569 | LRP4 | 722.9768729 | 6.986765259 | 0.465309245 | 15.26422608 | 1.32E-52 | 8.60E-50 |
| ENSG00000123407 | HOXC12 | 107.6912697 | 6.966463895 | 0.991628672 | 6.78890645 | 1.13E-11 | 3.22E-10 |
| ENSG00000261540 | | 16.82504 | 6.921845 | 2.530697 | 4.521041 | 6.15E-06 | 6.42E-05 |
| ENSG00000168032 | ENTPD3 | 321.7286 | 6.907191 | 0.905663 | 7.878254 | 3.32E-15 | 1.46E-13 |
| ENSG00000267058 | | 7.668322 | 6.906697 | 2.497969 | 4.993481 | 5.93E-07 | 7.75E-06 |
| ENSG00000096006 | CRISP3 | 639.5641255 | 6.905856449 | 0.485245396 | 14.44359705 | 2.75E-47 | 1.43E-44 |
| ENSG00000278862 | | 7.879019 | 6.822114 | 2.602222 | 4.347157 | 1.38E-05 | 0.000131 |
| ENSG00000179477 | ALOX12B | 62.09788 | 6.821162 | 1.377285 | 5.403124 | 6.55E-08 | 1.04E-06 |
| ENSG00000273769 | | 38.80791 | 6.817691 | 2.2994 | 4.962857 | 6.95E-07 | 8.97E-06 |
| ENSG00000279660 | | 11.84547 | 6.806006 | 2.38844 | 4.870007 | 1.12E-06 | 1.38E-05 |
| ENSG00000281091 | | 12.72817 | 6.774929 | 2.298116 | 5.118553 | 3.08E-07 | 4.23E-06 |
| ENSG00000272328 | | 9.245587 | 6.728695 | 2.293577 | 5.097623 | 3.44E-07 | 4.69E-06 |
| ENSG00000156925 | ZIC3 | 297.5869 | 6.71 | 0.6697765 | 10.37077 | 3.37E-25 | 4.26E-23 |
| ENSG00000198054 | DSCR8 | 3.96889 | 6.702104 | 2.889516 | 3.919065 | 8.89E-05 | 0.000689 |
| ENSG00000079841 | RIMS1 | 75.87816622 | 6.672439289 | 0.612985258 | 11.03944016 | 2.47E-28 | 4.17E-26 |
| ENSG00000214999 | | 24.76717 | 6.653191 | 1.541651 | 4.953738 | 7.28E-07 | 9.34E-06 |
| ENSG00000079102 | RUNX1T1 | 133.1968055 | 6.650350003 | 0.743501476 | 9.065968305 | 1.24E-19 | 8.80E-18 |
| ENSG00000257657 | | 10.98514 | 6.615956 | 1.474604 | 5.109333 | 3.23E-07 | 4.43E-06 |
| ENSG00000185483 | ROR1 | 117.5576 | 6.563682 | 1.072294 | 6.107826 | 1.01E-09 | 2.17E-08 |
| ENSG00000286683 | | 5.225983 | 6.560581 | 2.496656 | 4.558619 | 5.15E-06 | 5.48E-05 |
| ENSG00000240522 | RPL7AP10 | 11.56231 | 6.538471 | 2.350973 | 4.883025 | 1.04E-06 | 1.29E-05 |
| ENSG00000146674 | IGFBP3 | 6711.882 | 6.530317 | 0.4590185 | 14.38082 | 6.83E-47 | 3.42E-44 |
| ENSG00000256001 | | 7.575801 | 6.506692 | 2.650591 | 4.401708 | 1.07E-05 | 0.000105 |
| ENSG00000188263 | IL17REL | 154.1584 | 6.476777 | 0.8677451 | 7.850194 | 4.15E-15 | 1.79E-13 |
| ENSG00000086548 | CEACAM6 | 8285.312724 | 6.470712209 | 0.932790688 | 7.398037815 | NA | NA |
| ENSG00000183117 | CSMD1 | 110.5284 | 6.446416 | 0.6311937 | 10.31993 | 5.73E-25 | 7.14E-23 |
| ENSG00000263427 | | 17.96648 | 6.416453 | 1.510548 | 4.958302 | 7.11E-07 | 9.17E-06 |
| ENSG00000064989 | CALCRL | 8.67879219 | 6.395415841 | 1.447909703 | 5.034955407 | 4.78E-07 | 6.36E-06 |

| | | | | | | | |
|---|---|---|---|---|---|---|---|
| ENSG00000269397 | | 17.96905 | 6.391269 | 2.178572 | 5.112528 | 3.18E-07 | 4.36E-06 |
| ENSG00000169347 | GP2 | 17.63449 | 6.372658 | 0.9704262 | 6.657888 | 2.78E-11 | 7.45E-10 |
| ENSG00000282048 | | 42.418 | 6.369048 | 1.421433 | 5.141938 | 2.72E-07 | 3.78E-06 |
| ENSG00000060558 | GNA15 | 5.908412797 | 6.344114548 | 2.404495926 | 4.565986955 | 4.97E-06 | 5.31E-05 |
| ENSG00000129595 | EPB41L4A | 526.9247941 | 6.343099535 | 0.415216092 | 15.4207169 | 1.19E-53 | 8.77E-51 |
| ENSG00000236209 | | 8.961767 | 6.308885 | 2.468497 | 4.439078 | 9.03E-06 | 9.01E-05 |
| ENSG00000106031 | HOXA13 | 24.34614445 | 6.298114881 | 2.393279599 | 4.494175279 | 6.98E-06 | 7.17E-05 |
| ENSG00000109099 | PMP22 | 4938.768138 | 6.281613539 | 0.521964805 | 12.28231131 | 1.13E-34 | 2.75E-32 |
| ENSG00000237187 | NR2F1-AS1 | 127.1024 | 6.269333 | 0.913833 | 6.998639 | 2.58E-12 | 8.05E-11 |
| ENSG00000133116 | KL | 15.5071137 | 6.192036015 | 1.422685311 | 5.014365565 | 5.32E-07 | 7.03E-06 |
| ENSG00000131378 | RFTN1 | 1938.50914 | 6.152905591 | 0.402823008 | 15.48471604 | 4.40E-54 | 3.44E-51 |
| ENSG00000153071 | DAB2 | 862.5747 | 6.144756 | 0.5640777 | 11.15534 | 6.74E-29 | 1.23E-26 |
| ENSG00000178568 | ERBB4 | 202.0668 | 6.139016 | 0.8927362 | 7.261315 | 3.83E-13 | 1.32E-11 |
| ENSG00000171777 | RASGRP4 | 56.04014 | 6.133997 | 0.6382924 | 9.565635 | 1.12E-21 | 9.98E-20 |
| ENSG00000267886 | | 33.04047 | 6.121951 | 1.300268 | 5.265123 | 1.40E-07 | 2.06E-06 |
| ENSG00000176788 | BASP1 | 12702.58 | 6.116147 | 0.3663806 | 16.88099 | 6.21E-64 | 9.70E-61 |
| ENSG00000115252 | PDE1A | 110.5061934 | 6.108658534 | 0.586701329 | 10.55815245 | 4.66E-26 | 6.44E-24 |
| ENSG00000081189 | MEF2C | 445.8271951 | 6.100890847 | 0.713926968 | 8.780183458 | 1.63E-18 | 1.03E-16 |
| ENSG00000281103 | TRG-AS1 | 33.9946 | 6.100439 | 1.260576 | 5.120737 | 3.04E-07 | 4.19E-06 |
| ENSG00000253293 | HOXA10 | 70.53081 | 6.076234 | 0.6221752 | 9.737815 | 2.08E-22 | 2.06E-20 |
| ENSG00000158352 | SHROOM4 | 81.63224 | 6.058029 | 1.029917 | 6.079333 | 1.21E-09 | 2.56E-08 |
| ENSG00000238005 | | 16.52731 | 6.057898 | 2.217061 | 4.720447 | 2.35E-06 | 2.72E-05 |
| ENSG00000140873 | ADAMTS18 | 66.74829 | 6.020514 | 1.225622 | 5.363379 | 8.17E-08 | 1.26E-06 |
| ENSG00000257966 | OLA1P3 | 30.0945 | 6.012025 | 1.294609 | 5.187814 | 2.13E-07 | 3.03E-06 |
| ENSG00000132185 | FCRLA | 9.234425944 | 6.000361321 | 2.394427997 | 4.384223311 | 1.16E-05 | 0.000113 |
| ENSG00000267934 | | 18.19091 | 5.998027 | 2.260022 | 4.518938 | 6.22E-06 | 6.48E-05 |
| ENSG00000272405 | | 14.82239 | 5.951952 | 1.38021 | 4.935189 | 8.01E-07 | 1.02E-05 |
| ENSG00000226320 | LINC01811 | 27.566 | 5.947351 | 1.060443 | 5.599952 | 2.14E-08 | 3.67E-07 |
| ENSG00000135374 | ELF5 | 1486.20996 | 5.943332904 | 0.592087472 | 10.29896376 | 7.12E-25 | 8.68E-23 |
| ENSG00000254101 | LINC02055 | 151.7911 | 5.937862 | 0.4299187 | 14.04886 | 7.83E-45 | 3.59E-42 |
| ENSG00000204128 | C2orf72 | 348.4778 | 5.904038 | 0.6677697 | 9.205709 | 3.39E-20 | 2.59E-18 |
| ENSG00000215262 | KCNU1 | 3.42047 | 5.898328 | 2.812003 | 3.637455 | 0.00027535 | 0.001873 |
| ENSG00000198538 | ZNF28 | 767.8498 | 5.89563 | 0.3267746 | 18.29142 | 9.69E-75 | 1.98E-71 |

| | | | | | | | |
|---|---|---|---|---|---|---|---|
| ENSG00000173267 | SNCG | 779.4688 | 5.878402 | 0.4556413 | 12.96435 | 1.95E-38 | 6.09E-36 |
| ENSG00000137142 | IGFBPL1 | 51.37045442 | 5.873102573 | 0.739840976 | 8.097018296 | 5.63E-16 | 2.74E-14 |
| ENSG00000261083 | LINC02516 | 5.623127 | 5.862018 | 2.332115 | 4.323959 | 1.53E-05 | 0.000144 |
| ENSG00000257591 | ZNF625 | 4.680072 | 5.860873 | 2.502648 | 4.136027 | 3.53E-05 | 0.000304 |
| ENSG00000187689 | AMTN | 74.15408 | 5.857435 | 1.080079 | 5.455656 | 4.88E-08 | 7.88E-07 |
| ENSG00000264080 | | 6.55631 | 5.850949 | 2.26098 | 4.47404 | 7.68E-06 | 7.81E-05 |
| ENSG00000258969 | LINC02307 | 51.83466 | 5.826287 | 1.155671 | 5.549245 | 2.87E-08 | 4.79E-07 |
| ENSG00000180801 | ARSJ | 104.5916 | 5.819263 | 0.5738878 | 10.18634 | 2.28E-24 | 2.68E-22 |
| ENSG00000235786 | ZNRF3-IT1 | 2.439678 | 5.813399 | 2.976523 | 3.46109 | 0.00053799 | 0.003347 |
| ENSG00000231228 | ARPP19P1 | 5.049675 | 5.801245 | 2.520031 | 3.924475 | 8.69E-05 | 0.000676 |
| ENSG00000263177 | MTND1P8 | 7.07363 | 5.798664 | 2.597761 | 3.833924 | 0.00012612 | 0.000942 |
| ENSG00000178229 | ZNF543 | 130.7728 | 5.771467 | 0.4714479 | 12.33765 | 5.68E-35 | 1.42E-32 |
| ENSG00000175745 | NR2F1 | 294.2765 | 5.770995 | 0.6557495 | 9.060897 | 1.29E-19 | 9.19E-18 |
| ENSG00000121361 | KCNJ8 | 421.74947 | 5.76460405 | 0.856785883 | 7.148177117 | 8.79E-13 | 2.88E-11 |
| ENSG00000165617 | DACT1 | 128.6016 | 5.715787 | 0.5190623 | 11.08434 | 1.49E-28 | 2.58E-26 |
| ENSG00000137872 | SEMA6D | 153.0610543 | 5.714877663 | 0.599998687 | 9.581755548 | 9.54E-22 | 8.65E-20 |
| ENSG00000132639 | SNAP25 | 37.10554424 | 5.708676624 | 0.735897277 | 7.866980511 | 3.63E-15 | 1.58E-13 |
| ENSG00000068366 | ACSL4 | 775.5444564 | 5.690595644 | 0.41935899 | 13.74445094 | 5.50E-43 | 2.32E-40 |
| ENSG00000223804 | | 64.88937 | 5.67817 | 0.7194515 | 7.881499 | 3.23E-15 | 1.43E-13 |
| ENSG00000262406 | MMP12 | 9.32591 | 5.660228 | 1.616477 | 4.283666 | 1.84E-05 | 0.000169 |
| ENSG00000183571 | PGPEP1L | 30.35229 | 5.641081 | 0.8835935 | 6.680569 | 2.38E-11 | 6.47E-10 |
| ENSG00000101276 | SLC52A3 | 831.2269215 | 5.632829134 | 0.429092738 | 13.39729393 | 6.27E-41 | 2.14E-38 |
| ENSG00000198681 | MAGEA1 | 52.45407 | 5.612456 | 1.283286 | 4.948138 | 7.49E-07 | 9.57E-06 |
| ENSG00000100368 | CSF2RB | 6.982652006 | 5.606043513 | 2.457358675 | 3.921749052 | 8.79E-05 | 0.000684 |
| ENSG00000287358 | | 7.243639 | 5.601456 | 2.276495 | 4.243921 | 2.20E-05 | 0.000199 |
| ENSG00000168621 | GDNF | 16.51999 | 5.601158 | 1.233451 | 4.885628 | 1.03E-06 | 1.28E-05 |
| ENSG00000198342 | ZNF442 | 65.91105 | 5.593244 | 0.5918748 | 9.474724 | 2.67E-21 | 2.33E-19 |
| ENSG00000078053 | AMPH | 93.62858888 | 5.587434653 | 0.738820242 | 7.864897788 | 3.69E-15 | 1.61E-13 |
| ENSG00000189064 | GAGE2A | 14.24899 | 5.571532 | 2.381388 | 4.227145 | 2.37E-05 | 0.000213 |
| ENSG00000229970 | | 61.9083 | 5.5691 | 0.6497516 | 8.652844 | 5.02E-18 | 3.04E-16 |
| ENSG00000140015 | KCNH5 | 21.00574 | 5.557045 | 2.18866 | 4.38442 | 1.16E-05 | 0.000113 |
| ENSG00000167653 | PSCA | 690.8017 | 5.556358 | 0.5338358 | 10.64355 | 1.87E-26 | 2.70E-24 |
| ENSG00000286791 | | 4.087707 | 5.553051 | 2.642202 | 3.768133 | 0.00016447 | 0.001194 |

| | | | | | | | |
|---|---|---|---|---|---|---|---|
| ENSG00000212719 | LINC02693 | 19.14252 | 5.532425 | 2.230263 | 4.272866 | 1.93E-05 | 0.000177 |
| ENSG00000253541 | SEPTIN10P1 | 14.17416 | 5.528351 | 1.300213 | 4.806534 | 1.54E-06 | 1.85E-05 |
| ENSG00000105810 | CDK6 | 904.4230636 | 5.510520343 | 0.401289768 | 13.8770909 | 8.72E-44 | 3.83E-41 |
| ENSG00000185038 | MROH2A | 22.10503 | 5.505784 | 1.607861 | 4.246977 | 2.17E-05 | 0.000196 |
| ENSG00000148735 | PLEKHS1 | 4919.192 | 5.501883 | 0.5139895 | 11.02519 | 2.89E-28 | 4.86E-26 |
| ENSG00000110375 | UPK2 | 3332.160535 | 5.499984237 | 0.700059474 | 8.240201279 | 1.72E-16 | 8.90E-15 |
| ENSG00000288552 | | 9.314812 | 5.488576 | 2.262671 | 4.186068 | 2.84E-05 | 0.00025 |
| ENSG00000231916 | | 33.15757 | 5.463468 | 1.233564 | 4.947199 | 7.53E-07 | 9.61E-06 |
| ENSG00000106541 | AGR2 | 6750.710945 | 5.436604086 | 0.360606223 | 15.26403373 | 1.33E-52 | 8.60E-50 |
| ENSG00000139329 | LUM | 135.2227 | 5.419474 | 1.366745 | 4.590134 | 4.43E-06 | 4.79E-05 |
| ENSG00000131831 | RAI2 | 413.5901533 | 5.412421108 | 0.37233215 | 14.60904495 | 2.46E-48 | 1.36E-45 |
| ENSG00000231324 | | 5.578388 | 5.383816 | 2.55246 | 3.694703 | 0.00022014 | 0.001543 |
| ENSG00000274265 | | 497.3408 | 5.383032 | 0.4414992 | 12.35578 | 4.53E-35 | 1.15E-32 |
| ENSG00000260880 | HCCAT5 | 3.406232 | 5.365166 | 2.758652 | 3.462686 | 0.00053481 | 0.003331 |
| ENSG00000239572 | | 48.35394 | 5.354028 | 0.9238979 | 5.990451 | 2.09E-09 | 4.27E-08 |
| ENSG00000248485 | PCP4L1 | 37.70257 | 5.353046 | 1.100544 | 4.979162 | 6.39E-07 | 8.29E-06 |
| ENSG00000145113 | MUC4 | 28.95793 | 5.352711 | 0.7067696 | 7.800043 | 6.19E-15 | 2.62E-13 |
| ENSG00000091428 | RAPGEF4 | 398.6799049 | 5.322524175 | 0.405417394 | 13.39311946 | 6.63E-41 | 2.23E-38 |
| ENSG00000288262 | | 2.635025 | 5.297943 | 3.23427 | 3.125059 | 0.0017777 | 0.009379 |
| ENSG00000144063 | MALL | 30.05368 | 5.296286 | 2.255242 | 4.028282 | 5.62E-05 | 0.000459 |
| ENSG00000100362 | PVALB | 39.92971852 | 5.284727295 | 1.285399837 | 4.687454962 | 2.77E-06 | 3.15E-05 |
| ENSG00000249978 | TRGV7 | 7.31939 | 5.280097 | 2.596642 | 3.585664 | 0.00033622 | 0.002235 |
| ENSG00000255367 | | 13.15534 | 5.277602 | 1.401502 | 4.418584 | 9.93E-06 | 9.79E-05 |
| ENSG00000261409 | NA | 88.36255 | 5.271402 | 0.574346 | 9.411239 | 4.90E-21 | 4.11E-19 |
| ENSG00000186417 | GLDN | 870.3851 | 5.27106 | 0.3096153 | 17.25218 | 1.08E-66 | 1.79E-63 |
| ENSG00000087085 | ACHE | 1632.644087 | 5.246875739 | 0.491285551 | 10.93071857 | 8.22E-28 | 1.35E-25 |
| ENSG00000189420 | ZFP92 | 12.35194 | 5.238168 | 1.309714 | 4.591084 | 4.41E-06 | 4.77E-05 |
| ENSG00000157502 | PWWP3B | 116.4691 | 5.22925 | 0.6243269 | 8.659829 | 4.72E-18 | 2.87E-16 |
| ENSG00000251248 | | 6.626676 | 5.213878 | 2.521818 | 3.629086 | 0.00028443 | 0.00193 |
| ENSG00000234373 | SNX18P7 | 10.0626 | 5.211702 | 1.371432 | 4.4351 | 9.20E-06 | 9.15E-05 |
| ENSG00000121966 | CXCR4 | 1342.406363 | 5.197893065 | 0.526607591 | 10.16750068 | 2.77E-24 | 3.23E-22 |
| ENSG00000115590 | IL1R2 | 6.939697115 | 5.190427343 | 2.486448331 | 3.646042127 | 0.00026631 | 0.001823 |
| ENSG00000122787 | AKR1D1 | 64.86957213 | 5.187983225 | 0.905505389 | 6.007081671 | 1.89E-09 | 3.88E-08 |

| | | | | | | | |
|---|---|---|---|---|---|---|---|
| ENSG00000260701 | | 7.16173 | 5.181929 | 2.626599 | 3.528947 | 0.00041722 | 0.002685 |
| ENSG00000128918 | ALDH1A2 | 66.53484529 | 5.178006308 | 0.84870076 | 6.250856761 | 4.08E-10 | 9.36E-09 |
| ENSG00000147606 | SLC26A7 | 76.61348 | 5.174849 | 0.6303345 | 8.425182 | 3.60E-17 | 2.00E-15 |
| ENSG00000248238 | LINC02438 | 2.719529 | 5.174117 | 2.952837 | 3.246373 | 0.00116886 | 0.006544 |
| ENSG00000183566 | BPIFA4P | 256.7067 | 5.173113 | 0.9198708 | 5.891417 | 3.83E-09 | 7.52E-08 |
| ENSG00000248373 | | 65.87521 | 5.172426 | 0.5484776 | 9.588845 | 8.91E-22 | 8.16E-20 |
| ENSG00000275793 | RIMBP3 | 24.75944 | 5.171834 | 0.8162027 | 6.432894 | 1.25E-10 | 3.09E-09 |
| ENSG00000234477 | | 16.29794 | 5.164714 | 1.041468 | 5.310572 | 1.09E-07 | 1.64E-06 |
| ENSG00000215179 | MAPK6P4 | 7.109127 | 5.162336 | 1.375006 | 4.392574 | 1.12E-05 | 0.000109 |
| ENSG00000102243 | VGLL1 | 1612.735221 | 5.161152724 | 1.059551859 | 5.406906685 | 6.41E-08 | 1.01E-06 |
| ENSG00000273674 | | 42.75894 | 5.160089 | 0.6094655 | 8.532026 | 1.44E-17 | 8.40E-16 |
| ENSG00000232196 | MTRNR2L4 | 11.56793 | 5.157776 | 1.589445 | 3.981149 | 6.86E-05 | 0.000547 |
| ENSG00000237457 | LINC01351 | 9.801288 | 5.154622 | 1.45509 | 4.264207 | 2.01E-05 | 0.000183 |
| ENSG00000268678 | | 4.325522 | 5.148195 | 2.306785 | 3.891676 | 9.96E-05 | 0.000762 |
| ENSG00000168675 | LDLRAD4 | 2678.978 | 5.135436 | 0.3110591 | 16.65158 | 2.95E-62 | 3.92E-59 |
| ENSG00000172971 | UNC93B3 | 8.593717 | 5.133413 | 2.342686 | 3.8091 | 0.00013947 | 0.001028 |
| ENSG00000263745 | | 13.57476 | 5.132616 | 1.391621 | 4.342162 | 1.41E-05 | 0.000134 |
| ENSG00000221332 | | 2.476999 | 5.128642 | 2.570771 | 3.561977 | 0.00036807 | 0.002413 |
| ENSG00000269891 | ARHGAP19-SLIT1 | 22.24252 | 5.07668 | 0.9209862 | 5.865994 | 4.46E-09 | 8.68E-08 |
| ENSG00000287824 | | 2.474512 | 5.06653 | 2.588464 | 3.508571 | 0.00045052 | 0.002866 |
| ENSG00000184454 | NCMAP | 30.90165 | 5.057019 | 0.8562402 | 6.25157 | 4.06E-10 | 9.32E-09 |
| ENSG00000133124 | IRS4 | 63.73889128 | 5.056124816 | 0.670948883 | 7.795547045 | 6.41E-15 | 2.71E-13 |
| ENSG00000250753 | | 16.29846 | 5.040886 | 1.055048 | 4.859981 | 1.17E-06 | 1.44E-05 |
| ENSG00000164334 | FAM170A | 3.862075 | 5.037592 | 2.614069 | 3.461383 | 0.00053741 | 0.003344 |
| ENSG00000249937 | LINC02223 | 6.052312 | 5.029362 | 1.485966 | 4.06748 | 4.75E-05 | 0.000396 |
| ENSG00000165061 | ZMAT4 | 137.161 | 5.026774 | 0.4945155 | 10.33271 | 5.01E-25 | 6.31E-23 |
| ENSG00000144278 | GALNT13 | 33.44548 | 5.021929 | 1.129917 | 4.665691 | 3.08E-06 | 3.46E-05 |
| ENSG00000134240 | HMGCS2 | 62.02747316 | 5.018409057 | 1.725846709 | 3.845929223 | 0.0001201 | 0.000901 |
| ENSG00000137875 | BCL2L10 | 5.680403291 | 5.007073228 | 1.585117548 | 3.942574491 | 8.06E-05 | 0.000632 |
| ENSG00000134612 | FOLH1B | 46.22976461 | 5.006616998 | 1.412171941 | 4.181200014 | 2.90E-05 | 0.000255 |
| ENSG00000141469 | SLC14A1 | 77.45098 | 5.00095 | 0.5985296 | 8.579608 | 9.52E-18 | 5.65E-16 |
| ENSG00000245750 | DRAIC | 133.8305 | 4.999705 | 0.5915347 | 8.715295 | 2.90E-18 | 1.81E-16 |
| ENSG00000168874 | ATOH8 | 343.9156 | 4.978323 | 0.9612377 | 5.749243 | 8.96E-09 | 1.66E-07 |

| | | | | | | | |
|---|---|---|---|---|---|---|---|
| ENSG00000233539 | | 2.344575 | 4.956741 | 2.748432 | 3.312948 | 0.00092318 | 0.005329 |
| ENSG00000249345 | LINC02405 | 4.669232 | 4.927829 | 2.773925 | 3.292854 | 0.00099176 | 0.005681 |
| ENSG00000251577 | | 2.415312 | 4.911166 | 2.56849 | 3.45329 | 0.00055379 | 0.003431 |
| ENSG00000265787 | CYP4F35P | 73.80981 | 4.905705 | 0.5906388 | 8.659889 | 4.72E-18 | 2.87E-16 |
| ENSG00000183783 | KCTD8 | 25.34964 | 4.900302 | 0.8617047 | 5.699132 | 1.20E-08 | 2.17E-07 |
| ENSG00000237916 | RPL37P12 | 3.198439 | 4.888801 | 2.498646 | 3.499532 | 0.00046608 | 0.002955 |
| ENSG00000187242 | KRT12 | 4.084208 | 4.876795 | 2.457376 | 3.537426 | 0.00040405 | 0.002613 |
| ENSG00000274253 | | 10.62287 | 4.875559 | 1.132623 | 4.574086 | 4.78E-06 | 5.12E-05 |
| ENSG00000170509 | HSD17B13 | 23.4957 | 4.859602 | 1.058624 | 4.981276 | 6.32E-07 | 8.21E-06 |
| ENSG00000145864 | GABRB2 | 144.9777 | 4.857228 | 0.7563087 | 6.845788 | 7.61E-12 | 2.21E-10 |
| ENSG00000187474 | FPR3 | 4.186622 | 4.848884 | 2.720917 | 3.294915 | 0.00098451 | 0.005645 |
| ENSG00000215018 | COL28A1 | 665.8326 | 4.846561 | 0.5575857 | 9.006246 | 2.13E-19 | 1.50E-17 |
| ENSG00000147408 | CSGALNACT1 | 431.7814 | 4.843207 | 0.4844076 | 10.31541 | 6.00E-25 | 7.45E-23 |
| ENSG00000261154 | LINC02131 | 3.181251 | 4.836831 | 2.60027 | 3.386437 | 0.00070807 | 0.004246 |
| ENSG00000171408 | PDE7B | 45.51463 | 4.823388 | 0.8547058 | 6.05705 | 1.39E-09 | 2.91E-08 |
| ENSG00000274528 | | 65.5897 | 4.813276 | 0.5377766 | 9.139852 | 6.25E-20 | 4.59E-18 |
| ENSG00000259345 | | 42.7633 | 4.809934 | 0.7675351 | 6.742701 | 1.55E-11 | 4.34E-10 |
| ENSG00000007306 | CEACAM7 | 22.04736697 | 4.809082329 | 1.685199772 | 3.734831427 | 0.00018784 | 0.001342 |
| ENSG00000100373 | UPK3A | 794.5814367 | 4.800835301 | 0.938846891 | 5.657901353 | 1.53E-08 | 2.71E-07 |
| ENSG00000237807 | | 632.5548 | 4.79912 | 0.486635 | 10.14121 | 3.63E-24 | 4.15E-22 |
| ENSG00000215861 | | 52.96709 | 4.796736 | 0.6964523 | 7.189475 | 6.50E-13 | 2.17E-11 |
| ENSG00000073792 | IGF2BP2 | 414.9274879 | 4.791693247 | 0.705216249 | 6.839534454 | 7.95E-12 | 2.31E-10 |
| ENSG00000127329 | PTPRB | 79.92174253 | 4.774603802 | 0.918571589 | 5.607129408 | 2.06E-08 | 3.53E-07 |
| ENSG00000269845 | CCNYL6 | 3.596753 | 4.772491 | 2.429335 | 3.520177 | 0.00043126 | 0.002762 |
| ENSG00000079257 | LXN | 3891.919758 | 4.772383417 | 0.496285877 | 9.916272387 | 3.54E-23 | 3.66E-21 |
| ENSG00000180875 | GREM2 | 10.6456 | 4.757803 | 2.364258 | 3.592621 | 0.00032737 | 0.002184 |
| ENSG00000047936 | ROS1 | 10.66667804 | 4.730860566 | 2.223082995 | 3.818049293 | 0.00013451 | 0.000995 |
| ENSG00000136928 | GABBR2 | 1026.634415 | 4.730843545 | 0.779751121 | 6.547306082 | 5.86E-11 | 1.50E-09 |
| ENSG00000101871 | MID1 | 1600.46411 | 4.719947748 | 0.467042373 | 10.39424542 | 2.63E-25 | 3.40E-23 |
| ENSG00000135346 | CGA | 25.49285964 | 4.704846869 | 1.111723507 | 4.609299635 | 4.04E-06 | 4.40E-05 |
| ENSG00000236452 | | 5.434636 | 4.686234 | 2.554064 | 3.361365 | 0.00077558 | 0.004596 |
| ENSG00000230408 | BZW1-AS1 | 2.008596 | 4.676886 | 2.749374 | 3.208447 | 0.00133454 | 0.007358 |
| ENSG00000188828 | GLRA4 | 7.214499 | 4.666342 | 1.386481 | 4.060164 | 4.90E-05 | 0.000407 |

| | | | | | | | |
|---|---|---|---|---|---|---|---|
| ENSG00000285269 | | 2.473477 | 4.654973 | 2.9513 | 3.094902 | 0.00196878 | 0.010215 |
| ENSG00000280225 | | 5.373071 | 4.654386 | 2.330672 | 3.57046 | 0.00035636 | 0.002349 |
| ENSG00000286593 | | 67.42474 | 4.638941 | 0.5094081 | 9.390427 | 5.98E-21 | 4.90E-19 |
| ENSG00000150051 | MKX | 7.115073 | 4.628466 | 1.787604 | 3.481955 | 0.00049777 | 0.003127 |
| ENSG00000254087 | LYN | 181.493 | 4.625439 | 0.647768 | 7.257517 | 3.94E-13 | 1.36E-11 |
| ENSG00000168077 | SCARA3 | 1717.825 | 4.624155 | 0.4863902 | 9.832662 | 8.14E-23 | 8.29E-21 |
| ENSG00000183690 | EFHC2 | 3.515116 | 4.60985 | 2.912781 | 3.116412 | 0.00183066 | 0.009614 |
| ENSG00000183230 | CTNNA3 | 136.0164 | 4.602333 | 0.5819509 | 8.267865 | 1.36E-16 | 7.15E-15 |
| ENSG00000119508 | NR4A3 | 53.82405918 | 4.589136021 | 0.66739818 | 7.162583622 | 7.92E-13 | 2.61E-11 |
| ENSG00000224574 | COL18A1-AS2 | 4.084286 | 4.583549 | 2.326206 | 3.53857 | 0.0004023 | 0.002604 |
| ENSG00000186522 | SEPTIN10 | 850.9536 | 4.573827 | 0.2944468 | 15.73615 | 8.55E-56 | 8.41E-53 |
| ENSG00000248771 | SMIM31 | 14.87124 | 4.557429 | 2.490118 | 3.357654 | 0.00078607 | 0.00465 |
| ENSG00000140798 | ABCC12 | 3.486746 | 4.540439 | 1.86312 | 3.357503 | 0.0007865 | 0.004652 |
| ENSG00000183578 | TNFAIP8L3 | 75.48 | 4.532949 | 0.697053 | 6.852518 | 7.26E-12 | 2.12E-10 |
| ENSG00000226067 | LINC00623 | 175.5906 | 4.523482 | 0.4075735 | 11.32952 | 9.37E-30 | 1.79E-27 |
| ENSG00000259188 | | 8.188214 | 4.5158 | 1.557962 | 3.734477 | 0.00018811 | 0.001342 |
| ENSG00000253596 | | 4.89635 | 4.51546 | 1.39186 | 3.780692 | 0.00015639 | 0.001141 |
| ENSG00000268119 | | 2.585767 | 4.510505 | 2.669071 | 3.199019 | 0.00137896 | 0.00755 |
| ENSG00000274002 | | 25.04707 | 4.508791 | 2.075601 | 3.830975 | 0.00012764 | 0.000951 |
| ENSG00000255189 | GLYATL1P1 | 3.430128 | 4.502521 | 2.329727 | 3.495616 | 0.00047297 | 0.002991 |
| ENSG00000225968 | ELFN1 | 261.8522 | 4.501587 | 0.6286252 | 7.606715 | 2.81E-14 | 1.11E-12 |
| ENSG00000086717 | PPEF1 | 18.90848532 | 4.486345975 | 0.873627885 | 5.384622922 | 7.26E-08 | 1.13E-06 |
| ENSG00000152766 | ANKRD22 | 125.8252 | 4.478052 | 0.7166819 | 6.620241 | 3.59E-11 | 9.46E-10 |
| ENSG00000257588 | | 6.78351 | 4.477418 | 2.755999 | 3.162511 | 0.00156415 | 0.008404 |
| ENSG00000022355 | GABRA1 | 12.43942432 | 4.47062519 | 0.982418232 | 4.907731945 | 9.21E-07 | 1.16E-05 |
| ENSG00000178773 | CPNE7 | 1335.494 | 4.467985 | 0.3325044 | 13.62221 | 2.95E-42 | 1.19E-39 |
| ENSG00000153292 | ADGRF1 | 283.7597 | 4.457847 | 1.69868 | 3.514996 | 0.00043976 | 0.002809 |
| ENSG00000168843 | FSTL5 | 66.78544 | 4.451347 | 0.4884281 | 9.370671 | 7.21E-21 | 5.84E-19 |
| ENSG00000263618 | | 6.527157 | 4.437424 | 1.184649 | 4.097508 | 4.18E-05 | 0.000354 |
| ENSG00000086205 | FOLH1 | 926.913623 | 4.432294604 | 0.890260118 | 5.547073117 | 2.90E-08 | 4.84E-07 |
| ENSG00000165995 | CACNB2 | 142.6525 | 4.431771 | 0.434059 | 10.41683 | 2.08E-25 | 2.71E-23 |
| ENSG00000277301 | | 17.03247 | 4.431479 | 1.134385 | 4.55168 | 5.32E-06 | 5.64E-05 |
| ENSG00000198729 | PPP1R14C | 31.97494 | 4.407947 | 1.342141 | 3.943674 | 8.02E-05 | 0.00063 |

| | | | | | | | |
|---|---|---|---|---|---|---|---|
| ENSG00000257194 | | 24.91001 | 4.385941 | 1.083824 | 4.470034 | 7.82E-06 | 7.94E-05 |
| ENSG00000228714 | | 5.465402 | 4.379358 | 2.430608 | 3.350227 | 0.00080746 | 0.004756 |
| ENSG00000267284 | | 15.90734 | 4.375666 | 2.183111 | 3.630456 | 0.00028292 | 0.001921 |
| ENSG00000160352 | ZNF714 | 288.7789 | 4.370132 | 0.2955235 | 14.98527 | 9.17E-51 | 5.66E-48 |
| ENSG00000226562 | CYP4F26P | 18.34207 | 4.369136 | 0.7896139 | 5.877615 | 4.16E-09 | 8.15E-08 |
| ENSG00000268416 | | 7.510421 | 4.369001 | 1.765625 | 3.366936 | 0.00076008 | 0.004516 |
| ENSG00000163060 | TEKT4 | 12.52686 | 4.367421 | 1.016369 | 4.431771 | 9.35E-06 | 9.28E-05 |
| ENSG00000165078 | CPA6 | 11.21674 | 4.362404 | 0.9534659 | 4.802944 | 1.56E-06 | 1.88E-05 |
| ENSG00000211690 | TRGJ1 | 3.42391 | 4.362274 | 3.006718 | 2.998046 | 0.00271717 | 0.013416 |
| ENSG00000163993 | S100P | 4298.285 | 4.35779 | 0.4770988 | 9.436271 | 3.86E-21 | 3.33E-19 |
| ENSG00000186369 | LINC00643 | 45.26534 | 4.351609 | 2.245294 | 3.485699 | 0.00049085 | 0.003091 |
| ENSG00000103522 | IL21R | 17.41958382 | 4.341428236 | 1.18756113 | 4.293831768 | 1.76E-05 | 0.000163 |
| ENSG00000176049 | JAKMIP2 | 64.31703 | 4.33538 | 0.9063791 | 5.295692 | 1.19E-07 | 1.77E-06 |
| ENSG00000229953 | | 2.216035 | 4.330293 | 2.94569 | 2.988701 | 0.00280166 | 0.013795 |
| ENSG00000130294 | KIF1A | 1458.156254 | 4.307798129 | 0.356760522 | 12.30089547 | 8.96E-35 | 2.20E-32 |
| ENSG00000230506 | | 3.617153 | 4.30678 | 2.470279 | 3.262622 | 0.00110387 | 0.006231 |
| ENSG00000254238 | | 4.094296 | 4.304691 | 1.608419 | 3.48953 | 0.00048387 | 0.003053 |
| ENSG00000258751 | | 9.366391 | 4.301438 | 1.42241 | 3.753473 | 0.0001744 | 0.001256 |
| ENSG00000044459 | CNTLN | 35.02052114 | 4.297337698 | 0.590655052 | 7.44865114 | 9.43E-14 | 3.51E-12 |
| ENSG00000283162 | | 17.67411 | 4.296167 | 1.009715 | 4.386032 | 1.15E-05 | 0.000112 |
| ENSG00000286191 | | 3.466301 | 4.291252 | 2.468957 | 3.258492 | 0.00112006 | 0.00631 |
| ENSG00000286159 | | 121.6978 | 4.284787 | 0.4573869 | 9.591902 | 8.65E-22 | 7.98E-20 |
| ENSG00000113448 | PDE4D | 400.2011567 | 4.282148988 | 0.982669542 | 5.032258841 | 4.85E-07 | 6.44E-06 |
| ENSG00000185477 | GPRIN3 | 219.7199 | 4.277741 | 0.4884011 | 9.033548 | 1.66E-19 | 1.17E-17 |
| ENSG00000232006 | | 3.163108 | 4.27744 | 2.467922 | 3.255763 | 0.00113088 | 0.006362 |
| ENSG00000178033 | CALHM5 | 7.78204 | 4.26946 | 2.687736 | 3.099436 | 0.0019389 | 0.010087 |
| ENSG00000248309 | MEF2C-AS1 | 11.68816 | 4.269401 | 2.314285 | 3.381968 | 0.00071969 | 0.004306 |
| ENSG00000099954 | CECR2 | 366.5183684 | 4.26802484 | 0.238885073 | 17.96900928 | 3.41E-72 | 6.47E-69 |
| ENSG00000121898 | CPXM2 | 36.15758716 | 4.261923401 | 0.754595202 | 6.092659726 | 1.11E-09 | 2.37E-08 |
| ENSG00000272746 | NA | 12.34304 | 4.25959 | 0.9815132 | 4.617828 | 3.88E-06 | 4.24E-05 |
| ENSG00000099260 | PALMD | 36.37499965 | 4.250761954 | 0.744464684 | 5.990332829 | 2.09E-09 | 4.27E-08 |
| ENSG00000111404 | RERGL | 13.18890017 | 4.250047369 | 1.238382177 | 4.182698938 | 2.88E-05 | 0.000254 |
| ENSG00000196074 | SYCP2 | 2716.249 | 4.244027 | 0.2760249 | 15.55533 | 1.46E-54 | 1.25E-51 |

| | | | | | | | |
|---|---|---|---|---|---|---|---|
| ENSG00000068078 | FGFR3 | 3792.57219 | 4.242951761 | 0.757763681 | 6.094575779 | 1.10E-09 | 2.34E-08 |
| ENSG00000129521 | EGLN3 | 5274.794684 | 4.238725523 | 0.487462143 | 8.988101758 | 2.52E-19 | 1.75E-17 |
| ENSG00000277186 | | 14.71532 | 4.230305 | 1.295026 | 3.878665 | 0.00010503 | 0.000799 |
| ENSG00000231131 | LNCAROD | 581.9627 | 4.214379 | 0.4535989 | 9.569627 | 1.07E-21 | 9.63E-20 |
| ENSG00000159640 | ACE | 50.2465 | 4.19623 | 0.552591 | 7.962065 | 1.69E-15 | 7.70E-14 |
| ENSG00000287401 | | 26.70209 | 4.192195 | 0.744836 | 5.86316 | 4.54E-09 | 8.83E-08 |
| ENSG00000135063 | FAM189A2 | 589.5509083 | 4.190099586 | 0.377934777 | 11.31158693 | 1.15E-29 | 2.18E-27 |
| ENSG00000164683 | HEY1 | 46.78087 | 4.186309 | 0.949443 | 5.000757 | 5.71E-07 | 7.50E-06 |
| ENSG00000259438 | MAPK6-DT | 35.76225 | 4.182784 | 0.5311954 | 8.02821 | 9.89E-16 | 4.65E-14 |
| ENSG00000170891 | CYTL1 | 1.721249 | 4.182185 | 2.925002 | 2.962032 | 0.00305616 | 0.014763 |
| ENSG00000231110 | | 2.220659 | 4.181569 | 3.156691 | 2.883099 | 0.00393783 | 0.018209 |
| ENSG00000108551 | RASD1 | 2836.807632 | 4.17911135 | 0.800720003 | 5.744594064 | 9.21E-09 | 1.71E-07 |
| ENSG00000168546 | GFRA2 | 77.29944 | 4.165528 | 1.134504 | 4.426184 | 9.59E-06 | 9.50E-05 |
| ENSG00000266605 | LONRF2P1 | 4.509917 | 4.159835 | 1.332126 | 3.653991 | 0.0002582 | 0.001774 |
| ENSG00000008086 | CDKL5 | 1289.450038 | 4.150542289 | 0.531622926 | 8.146190605 | 3.76E-16 | 1.86E-14 |
| ENSG00000274214 | | 65.551 | 4.142999 | 0.507322 | 8.414853 | 3.93E-17 | 2.18E-15 |
| ENSG00000251473 | | 2.956394 | 4.140368 | 2.697448 | 3.043967 | 0.00233481 | 0.011796 |
| ENSG00000227191 | TRGC2 | 8.977571 | 4.129974 | 1.50549 | 3.422271 | 0.00062101 | 0.003788 |
| ENSG00000264301 | LINC01444 | 3.089957 | 4.127098 | 1.932612 | 3.235048 | 0.00121622 | 0.006777 |
| ENSG00000272068 | | 10.23741 | 4.117778 | 1.094622 | 4.317335 | 1.58E-05 | 0.000148 |
| ENSG00000181449 | SOX2 | 798.6085 | 4.117036 | 0.5124843 | 8.363266 | 6.10E-17 | 3.28E-15 |
| ENSG00000279414 | CDRT15P9 | 13.41612 | 4.109976 | 1.20188 | 3.826729 | 0.00012986 | 0.000965 |
| ENSG00000170927 | PKHD1 | 9.828321 | 4.109498 | 1.132612 | 3.919422 | 8.88E-05 | 0.000688 |
| ENSG00000008311 | AASS | 4306.690677 | 4.098141632 | 0.506204708 | 8.432585621 | 3.38E-17 | 1.89E-15 |
| ENSG00000134258 | VTCN1 | 1246.094116 | 4.096139494 | 0.775782177 | 5.796250597 | 6.78E-09 | 1.28E-07 |
| ENSG00000101842 | VSIG1 | 20.21326851 | 4.095115611 | 0.864239341 | 5.139513981 | 2.75E-07 | 3.82E-06 |
| ENSG00000186777 | ZNF732 | 35.69951 | 4.093326 | 0.5025834 | 8.368231 | 5.85E-17 | 3.16E-15 |
| ENSG00000259377 | | 7.489981 | 4.092576 | 0.9833058 | 4.48187 | 7.40E-06 | 7.56E-05 |
| ENSG00000149289 | ZC3H12C | 106.7296 | 4.086722 | 0.5449147 | 7.711611 | 1.24E-14 | 5.07E-13 |
| ENSG00000285329 | | 3.965585 | 4.086016 | 2.487551 | 3.158205 | 0.00158744 | 0.008503 |
| ENSG00000233441 | CYP2AB1P | 1.847701 | 4.077188 | 3.134442 | 2.878518 | 0.00399548 | 0.018427 |
| ENSG00000138696 | BMPR1B | 156.9449146 | 4.072860666 | 0.506534048 | 8.365161922 | 6.00E-17 | 3.24E-15 |
| ENSG00000229453 | SPINK8 | 5.581297 | 4.057031 | 2.477743 | 3.157822 | 0.00158953 | 0.008511 |

| | | | | | | | |
|---|---|---|---|---|---|---|---|
| ENSG00000271850 | LINC02343 | 24.0252 | 4.055415 | 0.6664406 | 6.31043 | 2.78E-10 | 6.53E-09 |
| ENSG00000283413 | | 20.48499 | 4.038017 | 1.341724 | 3.706115 | 0.00021046 | 0.001484 |
| ENSG00000224565 | LINC01754 | 27.4786 | 4.032303 | 0.6958394 | 6.058783 | 1.37E-09 | 2.89E-08 |
| ENSG00000259306 | | 73.46924 | 4.030047 | 0.406153 | 10.05871 | 8.41E-24 | 9.19E-22 |
| ENSG00000147065 | MSN | 10.37699 | 4.028554 | 2.207513 | 3.365263 | 0.00076471 | 0.00454 |
| ENSG00000180044 | C3orf80 | 28.98936 | 4.025303 | 0.5953187 | 7.08785 | 1.36E-12 | 4.37E-11 |
| ENSG00000253666 | | 5.384273 | 4.019727 | 2.463864 | 3.154965 | 0.00160517 | 0.008583 |
| ENSG00000239801 | DENND6A-AS1 | 9.882243 | 4.005424 | 1.225728 | 3.707521 | 0.0002093 | 0.001477 |
| ENSG00000150667 | FSIP1 | 91.04597 | 3.995282 | 0.5897058 | 7.158505 | 8.16E-13 | 2.68E-11 |
| ENSG00000277883 | NLRP3P1 | 4.240006 | 3.994842 | 1.861882 | 3.135805 | 0.00171383 | 0.00908 |
| ENSG00000172995 | ARPP21 | 3.648724 | 3.994185 | 1.60975 | 3.414495 | 0.000639 | 0.003883 |
| ENSG00000008513 | ST3GAL1 | 3822.609807 | 3.99342095 | 0.264120832 | 15.29360122 | 8.44E-53 | 5.75E-50 |
| ENSG00000235012 | JCADP1 | 303.7181 | 3.993364 | 0.2671074 | 15.01638 | 5.74E-51 | 3.63E-48 |
| ENSG00000237863 | | 29.14283 | 3.991951 | 0.8296348 | 5.31653 | 1.06E-07 | 1.59E-06 |
| ENSG00000117009 | KMO | 33.77972894 | 3.987420001 | 0.87722172 | 4.810081091 | 1.51E-06 | 1.82E-05 |
| ENSG00000281741 | | 6.675357 | 3.983905 | 1.458699 | 3.545028 | 0.00039257 | 0.002553 |
| ENSG00000121101 | TEX14 | 633.9919655 | 3.980701379 | 0.507246544 | 8.185131489 | 2.72E-16 | 1.37E-14 |
| ENSG00000259786 | LINC02109 | 3.033339 | 3.975828 | 2.412166 | 3.16781 | 0.00153592 | 0.008277 |
| ENSG00000178201 | VN1R1 | 16.74497 | 3.969392 | 0.7980231 | 5.312157 | 1.08E-07 | 1.63E-06 |
| ENSG00000183831 | ANKRD45 | 13.48205 | 3.960704 | 1.420701 | 3.440866 | 0.00057986 | 0.00357 |
| ENSG00000285041 | | 2.034538 | 3.95387 | 2.793416 | 2.967003 | 0.00300718 | 0.014587 |
| ENSG00000213075 | RPL31P11 | 6.260332 | 3.944289 | 1.633657 | 3.328477 | 0.00087322 | 0.005088 |
| ENSG00000163898 | LIPH | 423.5181 | 3.936135 | 0.3357001 | 12.00323 | 3.42E-33 | 7.83E-31 |
| ENSG00000087495 | PHACTR3 | 15.35953761 | 3.928477965 | 1.170280679 | 3.925995794 | 8.64E-05 | 0.000672 |
| ENSG00000138134 | STAMBPL1 | 93.13092337 | 3.905248817 | 0.481546896 | 8.387933291 | 4.95E-17 | 2.70E-15 |
| ENSG00000254872 | LINC02688 | 14.2266 | 3.896815 | 1.054698 | 3.945226 | 7.97E-05 | 0.000627 |
| ENSG00000267702 | NA | 22.88549 | 3.887877 | 0.6389903 | 6.377027 | 1.81E-10 | 4.37E-09 |
| ENSG00000259240 | MIR4713HG | 108.4558 | 3.877585 | 0.2904047 | 13.43922 | 3.56E-41 | 1.30E-38 |
| ENSG00000271830 | | 11.14129 | 3.864026 | 1.077601 | 4.186227 | 2.84E-05 | 0.00025 |
| ENSG00000287699 | | 2.238579 | 3.858257 | 2.792303 | 2.925678 | 0.00343706 | 0.016275 |
| ENSG00000139352 | ASCL1 | 57.41778 | 3.858156 | 1.237719 | 3.905091 | 9.42E-05 | 0.000726 |
| ENSG00000156011 | PSD3 | 2035.283 | 3.857838 | 0.3536647 | 11.17025 | 5.70E-29 | 1.04E-26 |
| ENSG00000240138 | EEF1GP4 | 25.40272 | 3.83634 | 0.6616262 | 6.068755 | 1.29E-09 | 2.73E-08 |

| | | | | | | | |
|---|---|---|---|---|---|---|---|
| ENSG00000164684 | ZNF704 | 3113.789 | 3.83329 | 0.276429 | 14.06439 | 6.29E-45 | 2.93E-42 |
| ENSG00000151320 | AKAP6 | 332.7401 | 3.826209 | 0.5237901 | 7.657465 | 1.90E-14 | 7.62E-13 |
| ENSG00000244564 | PRICKLE2-DT | 7.40613 | 3.820351 | 1.357989 | 3.380018 | 0.00072481 | 0.00433 |
| ENSG00000226017 | PRICKLE2-AS3 | 18.37748 | 3.817554 | 0.9399964 | 4.628677 | 3.68E-06 | 4.05E-05 |
| ENSG00000230453 | ANKRD18B | 56.13203 | 3.815481 | 0.5938192 | 6.909923 | 4.85E-12 | 1.45E-10 |
| ENSG00000248515 | | 24.10025 | 3.806999 | 0.5477159 | 7.183644 | 6.79E-13 | 2.26E-11 |
| ENSG00000261553 | | 5.631601 | 3.801924 | 2.276455 | 3.191504 | 0.00141534 | 0.007717 |
| ENSG00000185737 | NRG3 | 150.7221 | 3.799212 | 0.4356275 | 8.997037 | 2.32E-19 | 1.63E-17 |
| ENSG00000185869 | ZNF829 | 54.8585 | 3.798591 | 0.4815729 | 8.127895 | 4.37E-16 | 2.16E-14 |
| ENSG00000104047 | DTWD1 | 4739.166481 | 3.793202384 | 0.204816937 | 18.65210247 | 1.21E-77 | 3.23E-74 |
| ENSG00000130433 | CACNG6 | 20.74677875 | 3.785972487 | 1.271904667 | 3.583560162 | 0.00033894 | 0.00225 |
| ENSG00000198142 | SOWAHC | 914.3585 | 3.785908 | 0.3780812 | 10.30208 | 6.90E-25 | 8.48E-23 |
| ENSG00000287394 | | 18.91325 | 3.784336 | 1.127254 | 3.893074 | 9.90E-05 | 0.000759 |
| ENSG00000258376 | | 90.67508 | 3.783494 | 0.5002531 | 7.870572 | 3.53E-15 | 1.54E-13 |
| ENSG00000268089 | GABRQ | 28.50607 | 3.782644 | 0.578852 | 6.939048 | 3.95E-12 | 1.20E-10 |
| ENSG00000236803 | SDAD1P4 | 7.392585 | 3.778177 | 1.054258 | 4.179493 | 2.92E-05 | 0.000257 |
| ENSG00000175985 | PLEKHD1 | 242.0516 | 3.776521 | 0.4866704 | 8.095062 | 5.72E-16 | 2.78E-14 |
| ENSG00000204334 | ERICH2 | 92.44485 | 3.775202 | 0.4791054 | 8.200104 | 2.40E-16 | 1.22E-14 |
| ENSG00000148677 | ANKRD1 | 1772.851 | 3.769365 | 0.5231332 | 7.571713 | 3.68E-14 | 1.43E-12 |
| ENSG00000218416 | | 154.2685 | 3.76863 | 0.4343742 | 8.918729 | 4.72E-19 | 3.19E-17 |
| ENSG00000269376 | | 23.00745 | 3.752 | 1.014505 | 4.189158 | 2.80E-05 | 0.000247 |
| ENSG00000268628 | | 59.71483 | 3.75154 | 0.8124194 | 5.13767 | 2.78E-07 | 3.86E-06 |
| ENSG00000116774 | OLFML3 | 2618.39881 | 3.748810575 | 0.528971985 | 7.445068601 | 9.69E-14 | 3.61E-12 |
| ENSG00000170396 | ZNF804A | 54.50811 | 3.748121 | 0.702916 | 5.685696 | 1.30E-08 | 2.33E-07 |
| ENSG00000101188 | NTSR1 | 75.61109765 | 3.740979552 | 0.650228936 | 6.041857807 | 1.52E-09 | 3.17E-08 |
| ENSG00000103888 | CEMIP | 454.9390082 | 3.738211623 | 0.412646404 | 9.337118558 | 9.90E-21 | 7.92E-19 |
| ENSG00000175302 | ANKRD30BP1 | 7.808589 | 3.73499 | 1.024679 | 4.065988 | 4.78E-05 | 0.000398 |
| ENSG00000128872 | TMOD2 | 4885.624644 | 3.732117896 | 0.304071745 | 12.48013954 | 9.58E-36 | 2.60E-33 |
| ENSG00000117472 | TSPAN1 | 1155.71779 | 3.732108585 | 0.42047827 | 9.180582904 | 4.29E-20 | 3.23E-18 |
| ENSG00000273693 | CDRT15P10 | 12.09511 | 3.731269 | 1.134802 | 3.665488 | 0.00024687 | 0.001706 |
| ENSG00000184838 | PRR16 | 51.85461 | 3.730553 | 0.7160605 | 5.654468 | 1.56E-08 | 2.75E-07 |
| ENSG00000064787 | BCAS1 | 19937.25764 | 3.728596969 | 0.401316429 | 9.582467243 | 9.48E-22 | 8.62E-20 |
| ENSG00000285102 | | 21.45437 | 3.720105 | 1.4663 | 3.338166 | 0.00084333 | 0.004936 |

| | | | | | | | |
|---|---|---|---|---|---|---|---|
| ENSG00000187848 | P2RX2 | 522.5931 | 3.716097 | 0.48707 | 7.911661 | 2.54E-15 | 1.14E-13 |
| ENSG00000173175 | ADCY5 | 488.2239 | 3.708867 | 0.8960456 | 4.783844 | 1.72E-06 | 2.05E-05 |
| ENSG00000187527 | ATP13A5 | 6.223574 | 3.694418 | 2.913217 | 2.869388 | 0.00411267 | 0.018892 |
| ENSG00000186910 | SERPINA11 | 75.34312 | 3.692786 | 1.064155 | 4.200777 | 2.66E-05 | 0.000236 |
| ENSG00000100078 | PLA2G3 | 535.5797686 | 3.686804845 | 1.160148247 | 3.989759121 | 6.61E-05 | 0.00053 |
| ENSG00000242686 | PDE6B-AS1 | 33.0851 | 3.67983 | 0.4860974 | 7.839196 | 4.53E-15 | 1.94E-13 |
| ENSG00000128849 | CGNL1 | 193.0502967 | 3.676744039 | 0.47026988 | 8.126740391 | 4.41E-16 | 2.17E-14 |
| ENSG00000243838 | PSMC1P7 | 39.03216 | 3.669441 | 0.5263415 | 7.247799 | 4.24E-13 | 1.45E-11 |
| ENSG00000131050 | BPIFA2 | 15.42279593 | 3.666525362 | 2.523969256 | 3.007842296 | 0.0026311 | 0.013062 |
| ENSG00000176406 | RIMS2 | 814.6471 | 3.66277 | 0.3597095 | 10.42875 | 1.83E-25 | 2.41E-23 |
| ENSG00000259194 | | 4.154001 | 3.661891 | 1.560963 | 3.198052 | 0.00138359 | 0.007573 |
| ENSG00000121743 | GJA3 | 19.14665415 | 3.66181679 | 0.790227385 | 5.097358226 | 3.44E-07 | 4.69E-06 |
| ENSG00000048540 | LMO3 | 19.9292577 | 3.658564047 | 0.866106874 | 4.774054886 | 1.81E-06 | 2.14E-05 |
| ENSG00000211691 | TRGJP | 2.332485 | 3.653857 | 3.407156 | 2.763728 | 0.00571452 | 0.024744 |
| ENSG00000213657 | RPL31P44 | 2.954688 | 3.648707 | 2.76775 | 2.852757 | 0.00433418 | 0.019711 |
| ENSG00000137393 | RNF144B | 2465.480751 | 3.63769127 | 0.368496643 | 10.1390355 | 3.71E-24 | 4.23E-22 |
| ENSG00000140287 | HDC | 4.153713 | 3.63541 | 1.624491 | 3.093676 | 0.00197693 | 0.010255 |
| ENSG00000267868 | | 206.1068 | 3.635128 | 0.3510914 | 10.57444 | 3.91E-26 | 5.47E-24 |
| ENSG00000240990 | HOXA11-AS | 52.20936 | 3.62818 | 0.4016581 | 9.159873 | 5.20E-20 | 3.87E-18 |
| ENSG00000259577 | CERNA1 | 119.4368 | 3.625342 | 0.3705147 | 10.08027 | 6.75E-24 | 7.45E-22 |
| ENSG00000180777 | ANKRD30B | 135.3959 | 3.618268 | 0.6710426 | 5.878077 | 4.15E-09 | 8.13E-08 |
| ENSG00000171094 | ALK | 4.121065 | 3.617875 | 2.392046 | 3.033874 | 0.00241436 | 0.012141 |
| ENSG00000138356 | AOX1 | 207.6651014 | 3.616586005 | 0.891664826 | 4.686233194 | 2.78E-06 | 3.16E-05 |
| ENSG00000187672 | ERC2 | 61.85608 | 3.611987 | 0.4109827 | 9.012616 | 2.01E-19 | 1.42E-17 |
| ENSG00000053524 | MCF2L2 | 124.3189987 | 3.60849847 | 0.54319763 | 7.008696714 | 2.41E-12 | 7.51E-11 |
| ENSG00000158055 | GRHL3 | 3673.965 | 3.606318 | 0.7351502 | 5.449383 | 5.05E-08 | 8.12E-07 |
| ENSG00000104490 | NCALD | 84.20217584 | 3.602737302 | 0.597745146 | 6.418018379 | 1.38E-10 | 3.39E-09 |
| ENSG00000126217 | MCF2L | 3989.872598 | 3.602626174 | 0.229482547 | 15.86158449 | 1.17E-56 | 1.19E-53 |
| ENSG00000246985 | SOCS2-AS1 | 163.8557 | 3.598983 | 0.5800228 | 6.60693 | 3.92E-11 | 1.03E-09 |
| ENSG00000196549 | MME | 440.5004 | 3.596983 | 0.5525236 | 6.889664 | 5.59E-12 | 1.66E-10 |
| ENSG00000138378 | STAT4 | 55.14991044 | 3.590627193 | 0.71034506 | 5.503483698 | 3.72E-08 | 6.12E-07 |
| ENSG00000160326 | SLC2A6 | 3626.737 | 3.58946 | 0.4356227 | 8.561686 | 1.11E-17 | 6.57E-16 |
| ENSG00000184564 | SLITRK6 | 860.753 | 3.580267 | 0.4278676 | 8.671799 | 4.25E-18 | 2.60E-16 |

| | | | | | | | |
|---|---|---|---|---|---|---|---|
| ENSG00000108417 | KRT37 | 1.984032213 | 3.574852523 | 2.905529748 | 2.811963266 | 0.00492401 | 0.021877 |
| ENSG00000215057 | PKMP5 | 2.9767 | 3.574127 | 2.429827 | 2.995423 | 0.00274064 | 0.013522 |
| ENSG00000137869 | CYP19A1 | 98.25146901 | 3.569829242 | 0.448192129 | 8.257660504 | 1.49E-16 | 7.75E-15 |
| ENSG00000185352 | HS6ST3 | 647.9231 | 3.569282 | 0.3041916 | 11.93743 | 7.55E-33 | 1.70E-30 |
| ENSG00000265359 | | 3.317338 | 3.569214 | 1.767811 | 3.106064 | 0.00189596 | 0.009889 |
| ENSG00000111961 | SASH1 | 1715.074503 | 3.564984272 | 0.495871398 | 7.545360344 | 4.51E-14 | 1.74E-12 |
| ENSG00000280322 | NA | 6.659677 | 3.56302 | 1.473281 | 3.183628 | 0.00145442 | 0.007899 |
| ENSG00000124721 | DNAH8 | 21.38498412 | 3.562800701 | 0.939107757 | 4.294963098 | 1.75E-05 | 0.000162 |
| ENSG00000198910 | L1CAM | 16266.88 | 3.562747 | 0.5186394 | 7.255801 | 3.99E-13 | 1.38E-11 |
| ENSG00000146013 | GFRA3 | 50.40218 | 3.544991 | 0.8431048 | 4.757507 | 1.96E-06 | 2.31E-05 |
| ENSG00000259678 | | 2.142598 | 3.534535 | 2.950835 | 2.796008 | 0.00517381 | 0.022784 |
| ENSG00000124490 | CRISP2 | 6.179648129 | 3.530401432 | 1.213949412 | 3.548117438 | 0.000388 | 0.002529 |
| ENSG00000166415 | WDR72 | 1743.484 | 3.521904 | 0.2201874 | 16.14365 | 1.26E-58 | 1.45E-55 |
| ENSG00000248455 | LINC02217 | 3.613798 | 3.520138 | 1.531449 | 3.196321 | 0.00139192 | 0.007613 |
| ENSG00000169836 | TACR3 | 26.15286 | 3.517924 | 0.6283715 | 6.050851 | 1.44E-09 | 3.02E-08 |
| ENSG00000239911 | PRKAG2-AS1 | 29.8871 | 3.516705 | 0.7045196 | 5.347582 | 8.91E-08 | 1.36E-06 |
| ENSG00000144218 | AFF3 | 240.2908 | 3.510676 | 0.5034349 | 7.324394 | 2.40E-13 | 8.51E-12 |
| ENSG00000100276 | RASL10A | 80.65316662 | 3.498129981 | 0.395699897 | 9.132520127 | 6.69E-20 | 4.86E-18 |
| ENSG00000280339 | | 81.63872 | 3.494737 | 0.4770419 | 7.654166 | 1.95E-14 | 7.80E-13 |
| ENSG00000276524 | | 10.99747 | 3.49082 | 0.905557 | 4.303723 | 1.68E-05 | 0.000157 |
| ENSG00000189377 | CXCL17 | 380.7613 | 3.488857 | 1.253513 | 3.638659 | 0.00027406 | 0.001867 |
| ENSG00000154646 | TMPRSS15 | 41.92162 | 3.483942 | 0.5124532 | 7.10113 | 1.24E-12 | 3.99E-11 |
| ENSG00000260618 | | 172.7854 | 3.479475 | 0.4052758 | 8.842406 | 9.37E-19 | 6.12E-17 |
| ENSG00000130720 | FIBCD1 | 842.7675056 | 3.476387428 | 0.269834685 | 13.12716519 | 2.30E-39 | 7.37E-37 |
| ENSG00000277840 | | 8.913121 | 3.473674 | 1.021648 | 4.04844 | 5.16E-05 | 0.000426 |
| ENSG00000204186 | ZDBF2 | 294.8596 | 3.468436 | 0.4689396 | 7.736822 | 1.02E-14 | 4.20E-13 |
| ENSG00000126895 | AVPR2 | 27.16189588 | 3.456014031 | 0.682764406 | 5.449753389 | 5.04E-08 | 8.11E-07 |
| ENSG00000227726 | | 14.12984 | 3.454074 | 0.9452739 | 4.015947 | 5.92E-05 | 0.000481 |
| ENSG00000233237 | LINC00472 | 287.4957 | 3.448361 | 0.4991864 | 7.266019 | 3.70E-13 | 1.28E-11 |
| ENSG00000162949 | CAPN13 | 458.8899 | 3.447276 | 0.3675833 | 9.642967 | 5.26E-22 | 4.98E-20 |
| ENSG00000168925 | CTRB1 | 18.49038 | 3.446329 | 1.051402 | 3.807519 | 0.00014037 | 0.001033 |
| ENSG00000280132 | | 14.02065 | 3.445872 | 0.7660853 | 4.903809 | 9.40E-07 | 1.18E-05 |
| ENSG00000057593 | F7 | 74.71006141 | 3.440086168 | 0.707649549 | 5.367935848 | 7.96E-08 | 1.23E-06 |

| | | | | | | | |
|---|---|---|---|---|---|---|---|
| ENSG00000253477 | | 22.28414 | 3.433788 | 0.8385344 | 4.663552 | 3.11E-06 | 3.49E-05 |
| ENSG00000222033 | LINC01124 | 7.642717 | 3.429602 | 2.278066 | 3.018453 | 0.00254069 | 0.012677 |
| ENSG00000146938 | NLGN4X | 64.98913 | 3.423181 | 0.7179696 | 5.006947 | 5.53E-07 | 7.28E-06 |
| ENSG00000125999 | BPIFB1 | 316.7490195 | 3.419189925 | 0.77157044 | 5.018974931 | 5.19E-07 | 6.88E-06 |
| ENSG00000164100 | NDST3 | 108.3819 | 3.417985 | 0.5244747 | 6.808651 | 9.85E-12 | 2.83E-10 |
| ENSG00000271819 | RNU6-94P | 4.514587 | 3.415058 | 1.210752 | 3.358995 | 0.00078226 | 0.004629 |
| ENSG00000172349 | IL16 | 47.14399 | 3.414825 | 0.4342263 | 8.117543 | 4.76E-16 | 2.34E-14 |
| ENSG00000232721 | | 3.27461 | 3.413238 | 1.600997 | 3.022885 | 0.00250378 | 0.012521 |
| ENSG00000187398 | LUZP2 | 398.7087 | 3.409316 | 0.5363515 | 6.744346 | 1.54E-11 | 4.30E-10 |
| ENSG00000265630 | | 2.380076 | 3.403729 | 2.973405 | 2.790176 | 0.00526794 | 0.023127 |
| ENSG00000138379 | MSTN | 26.19153031 | 3.399004125 | 0.865949081 | 4.446147109 | 8.74E-06 | 8.75E-05 |
| ENSG00000003137 | CYP26B1 | 288.4057394 | 3.395319282 | 0.826509737 | 4.733977702 | 2.20E-06 | 2.57E-05 |
| ENSG00000136531 | SCN2A | 18.26133094 | 3.386118948 | 1.0534691 | 3.668412498 | 0.00024406 | 0.00169 |
| ENSG00000166510 | CCDC68 | 410.3791 | 3.38179 | 0.4786541 | 7.432357 | 1.07E-13 | 3.93E-12 |
| ENSG00000114812 | VIPR1 | 599.6839362 | 3.361708127 | 0.435447572 | 8.073138722 | 6.85E-16 | 3.29E-14 |
| ENSG00000106038 | EVX1 | 3.784797895 | 3.355116406 | 2.411698572 | 2.926758924 | 0.00342514 | 0.016227 |
| ENSG00000261713 | SSTR5-AS1 | 4.539007 | 3.348997 | 2.518244 | 2.883837 | 0.00392862 | 0.018175 |
| ENSG00000103647 | CORO2B | 56.95149328 | 3.347254473 | 0.775031419 | 4.899412245 | 9.61E-07 | 1.20E-05 |
| ENSG00000248015 | | 23.23143 | 3.34685 | 0.7753086 | 4.910311 | 9.09E-07 | 1.14E-05 |
| ENSG00000274600 | RIMBP3B | 3.442611 | 3.346633 | 2.554884 | 2.867287 | 0.00414007 | 0.018998 |
| ENSG00000152092 | ASTN1 | 4.317442 | 3.34171 | 1.188729 | 3.537319 | 0.00040421 | 0.002613 |
| ENSG00000156194 | PPEF2 | 2.564238 | 3.336584 | 1.646024 | 2.973524 | 0.00294401 | 0.014368 |
| ENSG00000154269 | ENPP3 | 24.24705 | 3.335786 | 1.179835 | 3.580786 | 0.00034256 | 0.002271 |
| ENSG00000146054 | TRIM7 | 296.1712 | 3.335039 | 0.3754575 | 9.150451 | 5.67E-20 | 4.20E-18 |
| ENSG00000260239 | LINC02533 | 21.69473 | 3.33352 | 0.6609313 | 5.457861 | 4.82E-08 | 7.78E-07 |
| ENSG00000124126 | PREX1 | 105253.9023 | 3.328770248 | 0.314486032 | 10.81729111 | 2.85E-27 | 4.43E-25 |
| ENSG00000211698 | TRGV4 | 2.680469 | 3.326834 | 3.998007 | 2.691815 | 0.00710644 | 0.02952 |
| ENSG00000165091 | TMC1 | 70.07388 | 3.326095 | 0.9247977 | 3.95674 | 7.60E-05 | 0.000601 |
| ENSG00000140285 | FGF7 | 8.843326 | 3.323883 | 0.7299084 | 4.956614 | 7.17E-07 | 9.23E-06 |
| ENSG00000249803 | | 5.012429 | 3.321789 | 2.698033 | 2.778956 | 0.0054534 | 0.023772 |
| ENSG00000270504 | | 50.10897 | 3.319024 | 1.009915 | 3.556312 | 0.0003761 | 0.002462 |
| ENSG00000283692 | | 3.977866 | 3.316342 | 1.689949 | 2.910199 | 0.00361198 | 0.016952 |
| ENSG00000155897 | ADCY8 | 2.27818 | 3.315418 | 2.168836 | 2.831665 | 0.00463064 | 0.020835 |

| | | | | | | | |
|---|---|---|---|---|---|---|---|
| ENSG00000203952 | CCDC160 | 17.44166 | 3.315294 | 1.048151 | 3.846113 | 0.00012001 | 0.000901 |
| ENSG00000064655 | EYA2 | 1235.130647 | 3.312797611 | 0.522458273 | 6.752847449 | 1.45E-11 | 4.07E-10 |
| ENSG00000169031 | COL4A3 | 157.3979 | 3.312399 | 0.4295369 | 8.000355 | 1.24E-15 | 5.73E-14 |
| ENSG00000286033 | | 33.34491 | 3.311981 | 0.6418451 | 5.547434 | 2.90E-08 | 4.84E-07 |
| ENSG00000120833 | SOCS2 | 791.6279937 | 3.300402685 | 0.47130909 | 7.357403619 | 1.88E-13 | 6.71E-12 |
| ENSG00000102683 | SGCG | 2795.336388 | 3.300227357 | 0.384217623 | 8.866552808 | 7.54E-19 | 5.00E-17 |
| ENSG00000268896 | | 4.723093 | 3.297257 | 1.230631 | 3.264332 | 0.00109723 | 0.0062 |
| ENSG00000153956 | CACNA2D1 | 117.3701 | 3.29629 | 0.4428753 | 7.760652 | 8.45E-15 | 3.54E-13 |
| ENSG00000078018 | MAP2 | 1400.924012 | 3.295220186 | 0.456070821 | 7.570775327 | 3.71E-14 | 1.44E-12 |
| ENSG00000122641 | INHBA | 39.82849353 | 3.291133755 | 0.815936422 | 4.643503022 | 3.43E-06 | 3.80E-05 |
| ENSG00000276644 | DACH1 | 12.73577 | 3.290952 | 0.9738133 | 4.041825 | 5.30E-05 | 0.000436 |
| ENSG00000177875 | CCDC184 | 76.53689 | 3.290895 | 0.4929395 | 6.936598 | 4.02E-12 | 1.22E-10 |
| ENSG00000054277 | OPN3 | 306.5913179 | 3.275464521 | 0.35654567 | 9.400815858 | 5.41E-21 | 4.52E-19 |
| ENSG00000233154 | LINC01762 | 11.98183 | 3.274784 | 0.8362569 | 4.488514 | 7.17E-06 | 7.35E-05 |
| ENSG00000143147 | GPR161 | 292.6795 | 3.272473 | 0.4056058 | 8.366588 | 5.93E-17 | 3.20E-15 |
| ENSG00000163053 | SLC16A14 | 734.4679 | 3.267129 | 0.3304511 | 10.14911 | 3.34E-24 | 3.85E-22 |
| ENSG00000154198 | CYP4Z2P | 125.9563 | 3.259702 | 0.7447334 | 4.932392 | 8.12E-07 | 1.03E-05 |
| ENSG00000225793 | TMEM30A-DT | 96.15956 | 3.257828 | 0.5436484 | 6.374741 | 1.83E-10 | 4.43E-09 |
| ENSG00000198478 | SH3BGRL2 | 1357.015 | 3.25233 | 0.2458954 | 13.40194 | 5.89E-41 | 2.03E-38 |
| ENSG00000172602 | RND1 | 1258.49 | 3.250561 | 0.7658407 | 4.84446 | 1.27E-06 | 1.55E-05 |
| ENSG00000276116 | FUT8-AS1 | 122.1232 | 3.249226 | 0.3323796 | 9.990595 | 1.68E-23 | 1.79E-21 |
| ENSG00000173080 | RXFP4 | 28.26824 | 3.245869 | 1.243132 | 3.422689 | 0.00062005 | 0.003784 |
| ENSG00000116032 | GRIN3B | 378.5699193 | 3.244831734 | 0.388017415 | 8.701758751 | 3.27E-18 | 2.03E-16 |
| ENSG00000165028 | NIPSNAP3B | 249.9643 | 3.24357 | 0.3835181 | 8.62767 | 6.26E-18 | 3.75E-16 |
| ENSG00000268041 | ERFL | 33.86551 | 3.242501 | 0.6386824 | 5.427973 | 5.70E-08 | 9.09E-07 |
| ENSG00000176907 | TCIM | 275.3154 | 3.240683 | 0.8640819 | 4.430535 | 9.40E-06 | 9.32E-05 |
| ENSG00000147251 | DOCK11 | 722.8448 | 3.237069 | 0.8615417 | 4.399717 | 1.08E-05 | 0.000106 |
| ENSG00000169856 | ONECUT1 | 179.8302 | 3.236264 | 0.3152392 | 10.52054 | 6.95E-26 | 9.47E-24 |
| ENSG00000178248 | FAM230I | 5.683422 | 3.234177 | 1.578463 | 3.05667 | 0.00223811 | 0.011383 |
| ENSG00000251364 | | 103.6604 | 3.233262 | 0.4158668 | 8.032778 | 9.53E-16 | 4.50E-14 |
| ENSG00000177706 | FAM20C | 4309.799 | 3.228936 | 0.2018592 | 16.16408 | 9.04E-59 | 1.09E-55 |
| ENSG00000261033 | | 7.974493 | 3.225392 | 0.8151128 | 4.365502 | 1.27E-05 | 0.000122 |
| ENSG00000188211 | NCR3LG1 | 979.5883 | 3.225129 | 0.7705807 | 4.766436 | 1.88E-06 | 2.22E-05 |

| | | | | | | | |
|---|---|---|---|---|---|---|---|
| ENSG00000139269 | INHBE | 74.41681 | 3.218053 | 1.179478 | 3.589885 | 0.00033082 | 0.002206 |
| ENSG00000135596 | MICAL1 | 2744.906389 | 3.21769136 | 0.299970396 | 10.9518965 | 6.51E-28 | 1.08E-25 |
| ENSG00000069966 | GNB5 | 2350.86404 | 3.208584586 | 0.209876924 | 15.44423655 | 8.25E-54 | 6.26E-51 |
| ENSG00000233052 | | 172.3791 | 3.206485 | 1.047094 | 3.874458 | 0.00010686 | 0.000811 |
| ENSG00000135362 | PRR5L | 130.5659164 | 3.20564448 | 0.440420488 | 7.536828291 | 4.82E-14 | 1.85E-12 |
| ENSG00000105880 | DLX5 | 17.38909124 | 3.202413324 | 0.906175589 | 4.172896423 | 3.01E-05 | 0.000263 |
| ENSG00000136250 | AOAH | 2.202390131 | 3.197210318 | 1.84072126 | 2.874100036 | 0.00405181 | 0.018653 |
| ENSG00000242575 | TUBAP13 | 13.08402 | 3.192035 | 2.365087 | 2.862867 | 0.00419826 | 0.019209 |
| ENSG00000229558 | SACS-AS1 | 2.953139 | 3.190772 | 3.246094 | 2.677928 | 0.00740792 | 0.03055 |
| ENSG00000275552 | | 14.52543 | 3.181058 | 1.142928 | 3.274498 | 0.0010585 | 0.006009 |
| ENSG00000156049 | GNA14 | 273.8492 | 3.178902 | 0.3470308 | 9.419953 | 4.51E-21 | 3.82E-19 |
| ENSG00000198832 | SELENOM | 1495.946 | 3.178736 | 0.2367819 | 13.60087 | 3.96E-42 | 1.55E-39 |
| ENSG00000206888 | RNU6-48P | 1.474415 | 3.172615 | 2.735377 | 2.756769 | 0.00583755 | NA |
| ENSG00000101144 | BMP7 | 13372.45664 | 3.172322287 | 0.56664216 | 6.040405132 | 1.54E-09 | 3.20E-08 |
| ENSG00000280172 | | 10.16797 | 3.172209 | 0.9371576 | 3.974486 | 7.05E-05 | 0.000561 |
| ENSG00000158786 | PLA2G2F | 48.75459 | 3.168363 | 1.313477 | 3.174691 | 0.00149996 | 0.008112 |
| ENSG00000148926 | ADM | 1242.093 | 3.168035 | 0.4270147 | 7.740992 | 9.86E-15 | 4.07E-13 |
| ENSG00000197619 | ZNF615 | 18.99516 | 3.166163 | 1.099266 | 3.342805 | 0.00082936 | 0.004868 |
| ENSG00000132669 | RIN2 | 679.7722242 | 3.159747486 | 0.39924542 | 8.217496889 | 2.08E-16 | 1.07E-14 |
| ENSG00000250174 | MYLK-AS2 | 2.914348 | 3.158 | 2.688286 | 2.750543 | 0.00594966 | 0.025563 |
| ENSG00000170160 | CCDC144A | 7.748048 | 3.152935 | 1.12179 | 3.454371 | 0.00055158 | 0.00342 |
| ENSG00000150995 | ITPR1 | 1630.051 | 3.152742 | 0.4986726 | 6.706935 | 1.99E-11 | 5.44E-10 |
| ENSG00000253207 | | 2.648783 | 3.152274 | 3.243753 | 2.711978 | 0.0066883 | 0.028055 |
| ENSG00000177398 | UMODL1 | 17.22314 | 3.151191 | 0.7202296 | 4.89023 | 1.01E-06 | 1.25E-05 |
| ENSG00000074803 | SLC12A1 | 24.89814957 | 3.147230909 | 0.724526072 | 4.857014982 | 1.19E-06 | 1.46E-05 |
| ENSG00000254531 | | 727.0755 | 3.135623 | 0.3124985 | 10.28373 | 8.34E-25 | 1.00E-22 |
| ENSG00000226510 | UPK1A-AS1 | 11.81493 | 3.13525 | 2.333688 | 2.884528 | 0.00392001 | 0.018146 |
| ENSG00000266094 | RASSF5 | 210.4717 | 3.134118 | 0.3278268 | 9.836622 | 7.83E-23 | 8.03E-21 |
| ENSG00000214894 | LINC00243 | 87.22626 | 3.132229 | 0.4334366 | 7.551107 | 4.32E-14 | 1.67E-12 |
| ENSG00000154188 | ANGPT1 | 45.61041 | 3.125725 | 0.4608413 | 7.115696 | 1.11E-12 | 3.61E-11 |
| ENSG00000204335 | SP5 | 10.02927 | 3.124908 | 1.089358 | 3.318676 | 0.00090445 | 0.005235 |
| ENSG00000162614 | NEXN | 22.29021 | 3.123236 | 0.7109255 | 4.814891 | 1.47E-06 | 1.78E-05 |
| ENSG00000009765 | IYD | 3.867897513 | 3.123141149 | 2.378316123 | 2.841491006 | 0.00449031 | 0.020286 |

| | | | | | | | |
|---|---|---|---|---|---|---|---|
| ENSG00000124343 | XG | 108.0082697 | 3.118821536 | 0.393779948 | 8.207820864 | 2.25E-16 | 1.15E-14 |
| ENSG00000251539 | SNX18P24 | 2.788566 | 3.117267 | 1.711925 | 2.852655 | 0.00433557 | 0.019711 |
| ENSG00000136205 | TNS3 | 10376.31516 | 3.11284318 | 0.336362282 | 9.521104276 | 1.71E-21 | 1.52E-19 |
| ENSG00000169894 | MUC3A | 720.2583 | 3.103478 | 0.7938982 | 4.561587 | 5.08E-06 | 5.40E-05 |
| ENSG00000248599 | | 54.84231 | 3.102745 | 0.6221984 | 5.372955 | 7.75E-08 | 1.20E-06 |
| ENSG00000255178 | LINC02720 | 7.984227 | 3.092962 | 1.717864 | 2.930999 | 0.00337874 | 0.01605 |
| ENSG00000212766 | EWSAT1 | 49.03332 | 3.09193 | 0.5146205 | 6.368903 | 1.90E-10 | 4.60E-09 |
| ENSG00000151967 | SCHIP1 | 21.12727 | 3.085798 | 1.066592 | 3.337115 | 0.00084653 | 0.004951 |
| ENSG00000183780 | SLC35F3 | 7.352474 | 3.082671 | 1.452045 | 2.994774 | 0.00274648 | 0.013546 |
| ENSG00000182379 | NXPH4 | 406.843 | 3.081019 | 0.3622522 | 8.788597 | 1.51E-18 | 9.67E-17 |
| ENSG00000104381 | GDAP1 | 2435.1132 | 3.078620851 | 0.295591366 | 10.67860523 | 1.28E-26 | 1.89E-24 |
| ENSG00000117020 | AKT3 | 161.0853336 | 3.077201165 | 0.75245035 | 4.465344707 | 7.99E-06 | 8.09E-05 |
| ENSG00000151632 | AKR1C2 | 8854.227 | 3.070167 | 0.7641476 | 4.627976 | 3.69E-06 | 4.06E-05 |
| ENSG00000259556 | | 43.21863 | 3.070078 | 0.4833575 | 6.707401 | 1.98E-11 | 5.43E-10 |
| ENSG00000254966 | | 24.99638 | 3.061748 | 0.7113768 | 4.809894 | 1.51E-06 | 1.82E-05 |
| ENSG00000172061 | LRRC15 | 6.767561 | 3.055775 | 1.274731 | 3.207841 | 0.00133736 | 0.00737 |
| ENSG00000128917 | DLL4 | 46.91370632 | 3.055649774 | 0.60409958 | 5.5220242 | 3.35E-08 | 5.55E-07 |
| ENSG00000267366 | | 4.405202 | 3.054888 | 1.266863 | 3.057942 | 0.00222863 | 0.011341 |
| ENSG00000183844 | FAM3B | 19.29289 | 3.053295 | 0.9521656 | 3.899121 | 9.65E-05 | 0.000742 |
| ENSG00000166262 | FAM227B | 541.7812 | 3.05261 | 0.2700767 | 11.50416 | 1.26E-30 | 2.55E-28 |
| ENSG00000170579 | DLGAP1 | 22.77561 | 3.045289 | 0.8202555 | 4.270828 | 1.95E-05 | 0.000179 |
| ENSG00000250508 | LINC02701 | 1.79098 | 3.045144 | 2.997388 | 2.674725 | 0.00747905 | 0.030798 |
| ENSG00000279289 | | 2.961375 | 3.044468 | 1.493246 | 2.865448 | 0.0041642 | 0.019092 |
| ENSG00000185532 | PRKG1 | 1023.951 | 3.038085 | 0.3871525 | 8.15412 | 3.52E-16 | 1.75E-14 |
| ENSG00000002587 | HS3ST1 | 58.78305483 | 3.035618744 | 0.686625384 | 4.787582387 | 1.69E-06 | 2.01E-05 |
| ENSG00000134594 | RAB33A | 18.14360921 | 3.034566544 | 0.632326525 | 5.143711047 | 2.69E-07 | 3.75E-06 |
| ENSG00000213816 | CNN2P4 | 4.215525 | 3.033464 | 1.763623 | 2.863984 | 0.00418349 | 0.019167 |
| ENSG00000259709 | | 13.3712 | 3.033367 | 0.7299914 | 4.647802 | 3.35E-06 | 3.73E-05 |
| ENSG00000105339 | DENND3 | 895.2704289 | 3.028165637 | 0.280841844 | 11.00945613 | 3.44E-28 | 5.75E-26 |
| ENSG00000169122 | FAM110B | 272.289 | 3.0279 | 0.3777501 | 8.309191 | 9.64E-17 | 5.11E-15 |
| ENSG00000186481 | ANKRD20A5P | 554.6017 | 3.022284 | 0.34218 | 9.097223 | 9.27E-20 | 6.69E-18 |
| ENSG00000126353 | CCR7 | 37.44137021 | 3.019486938 | 0.46832576 | 6.806072743 | 1.00E-11 | 2.88E-10 |
| ENSG00000283526 | PRRT1B | 442.4496 | 3.018509 | 0.3353257 | 9.277065 | 1.74E-20 | 1.36E-18 |

| | | | | | | | |
|---|---|---|---|---|---|---|---|
| ENSG00000198108 | CHSY3 | 137.262 | 3.017885 | 0.3581815 | 8.691001 | 3.59E-18 | 2.22E-16 |
| ENSG00000112964 | GHR | 1807.879367 | 3.016444396 | 0.289625737 | 10.64407798 | 1.86E-26 | 2.70E-24 |
| ENSG00000142319 | SLC6A3 | 36.06764 | 3.013884 | 0.7299662 | 4.606896 | 4.09E-06 | 4.45E-05 |
| ENSG00000172746 | RPL12P13 | 2.250871 | 3.010011 | 2.545059 | 2.759933 | 0.00578132 | 0.025001 |
| ENSG00000260196 | | 247.1087 | 3.004526 | 0.4853634 | 6.565779 | 5.18E-11 | 1.34E-09 |
| ENSG00000132204 | LINC00470 | 9.049463309 | 3.003403927 | 1.236807627 | 3.24292863 | 0.00118308 | 0.006604 |
| ENSG00000179057 | IGSF22 | 55.63754 | 3.000944 | 0.5836205 | 5.564581 | 2.63E-08 | 4.42E-07 |
| ENSG00000184845 | DRD1 | 388.268 | 2.996494 | 0.4210808 | 7.429543 | 1.09E-13 | 4.01E-12 |
| ENSG00000163823 | CCR1 | 26.58797 | 2.991814 | 0.5795739 | 5.641845 | 1.68E-08 | 2.94E-07 |
| ENSG00000104369 | JPH1 | 1237.123336 | 2.985090971 | 0.287053691 | 10.6372942 | 2.00E-26 | 2.87E-24 |
| ENSG00000047346 | FAM214A | 7300.763746 | 2.974514867 | 0.272398083 | 11.14664617 | 7.44E-29 | 1.34E-26 |
| ENSG00000149260 | CAPN5 | 1318.409 | 2.969778 | 0.503325 | 6.314151 | 2.72E-10 | 6.38E-09 |
| ENSG00000245680 | ZNF585B | 308.096 | 2.969053 | 0.2588046 | 11.68013 | 1.61E-31 | 3.51E-29 |
| ENSG00000100290 | BIK | 556.8092125 | 2.966706898 | 0.324456306 | 9.41439279 | 4.76E-21 | 4.00E-19 |
| ENSG00000116016 | EPAS1 | 13619.31508 | 2.965245426 | 0.434969676 | 7.177074839 | 7.12E-13 | 2.37E-11 |
| ENSG00000273328 | | 53.32509 | 2.956151 | 0.719316 | 4.510047 | 6.48E-06 | 6.73E-05 |
| ENSG00000128045 | RASL11B | 7.397924152 | 2.945388854 | 1.591783507 | 2.876022253 | 0.00402721 | 0.018557 |
| ENSG00000259701 | | 6.523383 | 2.94504 | 1.178752 | 3.061414 | 0.00220294 | 0.011234 |
| ENSG00000177707 | NECTIN3 | 575.1793 | 2.943637 | 0.347109 | 8.758833 | 1.97E-18 | 1.25E-16 |
| ENSG00000261330 | PPIAP50 | 5.372258 | 2.942727 | 1.843753 | 2.841242 | 0.00449382 | 0.020296 |
| ENSG00000169783 | LINGO1 | 380.9844 | 2.940893 | 0.4493967 | 6.94188 | 3.87E-12 | 1.19E-10 |
| ENSG00000179097 | HTR1F | 33.96982 | 2.939938 | 0.962061 | 3.773301 | 0.0001611 | 0.001172 |
| ENSG00000139626 | ITGB7 | 55.40867 | 2.938644 | 0.4451725 | 6.804526 | 1.01E-11 | 2.91E-10 |
| ENSG00000286734 | | 4.210974 | 2.937367 | 2.546557 | 2.748512 | 0.00598665 | 0.025684 |
| ENSG00000102287 | GABRE | 510.6764918 | 2.929637517 | 0.883779597 | 3.671533172 | 0.0002411 | 0.001673 |
| ENSG00000118160 | SLC8A2 | 20.76536216 | 2.928097354 | 0.877211424 | 3.813709715 | 0.0001369 | 0.001012 |
| ENSG00000268833 | | 3.071043 | 2.927907 | 2.772389 | 2.703277 | 0.00686595 | 0.028683 |
| ENSG00000285410 | GABPB1-IT1 | 3646.323 | 2.922186 | 0.2207162 | 13.41444 | 4.98E-41 | 1.79E-38 |
| ENSG00000232354 | VIPR1-AS1 | 29.69031 | 2.91972 | 0.6462154 | 5.055473 | 4.29E-07 | 5.77E-06 |
| ENSG00000169059 | VCX3A | 24.92553 | 2.915983 | 2.299692 | 2.791602 | 0.00524479 | 0.02304 |
| ENSG00000226549 | SCDP1 | 664.2595 | 2.915672 | 0.510377 | 6.138348 | 8.34E-10 | 1.81E-08 |
| ENSG00000064195 | DLX3 | 867.2266977 | 2.912765626 | 0.318223195 | 9.428691983 | 4.15E-21 | 3.56E-19 |
| ENSG00000256340 | ABCC6P1 | 5.038562 | 2.912055 | 1.005769 | 3.493717 | 0.00047635 | 0.003008 |

| | | | | | | | |
|---|---|---|---|---|---|---|---|
| ENSG00000177181 | RIMKLA | 360.1091 | 2.90858 | 0.2950872 | 10.09244 | 5.97E-24 | 6.66E-22 |
| ENSG00000176532 | PRR15 | 1426.411 | 2.908491 | 0.2985764 | 9.986626 | 1.74E-23 | 1.85E-21 |
| ENSG00000153064 | BANK1 | 328.5598 | 2.907475 | 0.4535597 | 6.770005 | 1.29E-11 | 3.64E-10 |
| ENSG00000286472 | | 82.83748 | 2.907309 | 0.5549838 | 5.645707 | 1.65E-08 | 2.88E-07 |
| ENSG00000126562 | WNK4 | 60.2745163 | 2.907233371 | 0.456050476 | 6.694490064 | 2.16E-11 | 5.89E-10 |
| ENSG00000257398 | | 12.22186 | 2.906476 | 0.7562322 | 4.299299 | 1.71E-05 | 0.000159 |
| ENSG00000183423 | LRIT3 | 5.933649 | 2.904183 | 1.405581 | 2.910476 | 0.00360879 | 0.01694 |
| ENSG00000236491 | | 7.742966 | 2.903215 | 1.186 | 3.160962 | 0.00157249 | 0.00844 |
| ENSG00000227953 | LINC01341 | 247.8789 | 2.903068 | 0.4308536 | 7.064364 | 1.61E-12 | 5.13E-11 |
| ENSG00000119938 | PPP1R3C | 897.4698807 | 2.902192404 | 0.503232501 | 6.169875594 | 6.83E-10 | 1.51E-08 |
| ENSG00000005073 | HOXA11 | 43.0926937 | 2.896397574 | 0.554425388 | 5.565405973 | 2.62E-08 | 4.41E-07 |
| ENSG00000239467 | | 257.8061 | 2.895703 | 0.3726536 | 8.03045 | 9.71E-16 | 4.58E-14 |
| ENSG00000253552 | HOXA-AS2 | 16.83991 | 2.89493 | 1.070856 | 3.139972 | 0.00168964 | 0.008962 |
| ENSG00000113205 | PCDHB3 | 287.2272653 | 2.89216808 | 0.293245491 | 10.08195667 | 6.64E-24 | 7.35E-22 |
| ENSG00000169026 | SLC49A3 | 40.55115 | 2.889552 | 0.6024388 | 5.301363 | 1.15E-07 | 1.72E-06 |
| ENSG00000258387 | | 2.717501 | 2.885718 | 1.765558 | 2.700122 | 0.00693141 | 0.028892 |
| ENSG00000121753 | ADGRB2 | 756.3772639 | 2.883181296 | 0.357820808 | 8.35436501 | 6.58E-17 | 3.53E-15 |
| ENSG00000251369 | ZNF550 | 418.2978 | 2.881143 | 0.3046621 | 9.697849 | 3.08E-22 | 3.00E-20 |
| ENSG00000264569 | DCXR-DT | 11.77248 | 2.880819 | 0.8365404 | 4.04562 | 5.22E-05 | 0.00043 |
| ENSG00000147509 | RGS20 | 76.04939 | 2.878039 | 0.4958365 | 6.168568 | 6.89E-10 | 1.52E-08 |
| ENSG00000152784 | PRDM8 | 25.91402 | 2.874482 | 0.869194 | 3.795467 | 0.00014737 | 0.00108 |
| ENSG00000142910 | TINAGL1 | 17.71712 | 2.873858 | 1.348618 | 3.076133 | 0.00209705 | 0.010787 |
| ENSG00000255240 | | 25.56628 | 2.863621 | 0.7071616 | 4.605611 | 4.11E-06 | 4.47E-05 |
| ENSG00000212195 | | 17.24416 | 2.862218 | 1.023608 | 3.649156 | 0.0002631 | 0.001806 |
| ENSG00000127831 | VIL1 | 50.15365565 | 2.855807042 | 0.776064461 | 4.287585288 | 1.81E-05 | 0.000167 |
| ENSG00000283154 | IQCJ-SCHIP1 | 92.09654 | 2.853596 | 0.4738838 | 6.405865 | 1.50E-10 | 3.64E-09 |
| ENSG00000137801 | THBS1 | 7263.36434 | 2.85104763 | 0.371774323 | 7.973349575 | 1.54E-15 | 7.06E-14 |
| ENSG00000269825 | | 289.4634 | 2.848922 | 0.4402658 | 6.830474 | 8.46E-12 | 2.45E-10 |
| ENSG00000165495 | PKNOX2 | 4.146617 | 2.848901 | 1.220018 | 2.949576 | 0.0031821 | 0.015258 |
| ENSG00000281692 | PACRG-AS1 | 11.07652 | 2.848158 | 1.125528 | 3.058754 | 0.0022226 | 0.011318 |
| ENSG00000164344 | KLKB1 | 11.18041 | 2.84572 | 0.8815528 | 3.782641 | 0.00015517 | 0.001133 |
| ENSG00000116574 | RHOU | 1549.146468 | 2.842230135 | 0.232187301 | 12.43385716 | 1.71E-35 | 4.50E-33 |
| ENSG00000260228 | | 172.3692 | 2.841749 | 0.5423408 | 5.70234 | 1.18E-08 | 2.14E-07 |

| | | | | | | | |
|---|---|---|---|---|---|---|---|
| ENSG00000235021 | | 14.77361 | 2.840806 | 1.475664 | 2.852374 | 0.00433941 | 0.019718 |
| ENSG00000228252 | COL6A4P2 | 14.89903 | 2.839143 | 0.9078609 | 3.827345 | 0.00012953 | 0.000963 |
| ENSG00000227972 | THAP12P3 | 16.47767 | 2.836741 | 0.7688626 | 4.259154 | 2.05E-05 | 0.000187 |
| ENSG00000278231 | | 139.0384 | 2.834873 | 0.5813741 | 5.375021 | 7.66E-08 | 1.18E-06 |
| ENSG00000263874 | LINC00672 | 153.544 | 2.83442 | 0.3498423 | 8.382785 | 5.17E-17 | 2.82E-15 |
| ENSG00000172201 | ID4 | 152.7478 | 2.833757 | 0.6269229 | 5.052443 | 4.36E-07 | 5.84E-06 |
| ENSG00000077935 | SMC1B | 33.95707173 | 2.83055668 | 0.629309163 | 4.899711238 | 9.60E-07 | 1.20E-05 |
| ENSG00000286705 | | 81.00233 | 2.827845 | 0.3457693 | 8.445403 | 3.03E-17 | 1.70E-15 |
| ENSG00000260118 | | 19.62435 | 2.82193 | 1.017452 | 3.520719 | 0.00043038 | 0.002757 |
| ENSG00000109072 | VTN | 43.7651981 | 2.81504699 | 0.466145535 | 6.401492142 | 1.54E-10 | 3.74E-09 |
| ENSG00000253553 | | 75.8097 | 2.814075 | 0.4791936 | 6.257387 | 3.91E-10 | 9.01E-09 |
| ENSG00000279659 | | 7.94183 | 2.812569 | 1.361065 | 2.954398 | 0.0031328 | 0.015054 |
| ENSG00000144668 | ITGA9 | 10.57354 | 2.811949 | 0.9516651 | 3.632075 | 0.00028115 | 0.001911 |
| ENSG00000238365 | RNU7-57P | 3.603624 | 2.809464 | 1.387768 | 2.804185 | 0.0050444 | 0.022319 |
| ENSG00000189001 | SBSN | 25.44864 | 2.806954 | 0.8481776 | 3.987933 | 6.67E-05 | 0.000533 |
| ENSG00000259868 | | 25.64621 | 2.806227 | 0.6650338 | 4.734438 | 2.20E-06 | 2.56E-05 |
| ENSG00000231119 | | 73.79542 | 2.804877 | 0.6322573 | 4.950227 | 7.41E-07 | 9.50E-06 |
| ENSG00000128989 | ARPP19 | 24377.5269 | 2.800974363 | 0.151373243 | 18.62856955 | 1.88E-77 | 4.55E-74 |
| ENSG00000102048 | ASB9 | 146.083743 | 2.80001681 | 0.50856842 | 5.940366165 | 2.84E-09 | 5.70E-08 |
| ENSG00000272755 | | 8.719559 | 2.797775 | 1.493195 | 2.860208 | 0.00423363 | 0.019334 |
| ENSG00000044524 | EPHA3 | 193.3119417 | 2.797349967 | 0.564712248 | 5.419933651 | 5.96E-08 | 9.47E-07 |
| ENSG00000129467 | ADCY4 | 11.65705516 | 2.793807567 | 0.855991982 | 3.91750961 | 8.95E-05 | 0.000693 |
| ENSG00000236699 | ARHGEF38 | 286.4225 | 2.791808 | 0.4662213 | 6.365066 | 1.95E-10 | 4.71E-09 |
| ENSG00000235280 | MCF2L-AS1 | 227.9474 | 2.787355 | 0.3068393 | 9.297671 | 1.44E-20 | 1.13E-18 |
| ENSG00000115361 | ACADL | 97.00374643 | 2.786166087 | 0.350980902 | 8.18351965 | 2.76E-16 | 1.39E-14 |
| ENSG00000119919 | NKX2-3 | 8.489735223 | 2.784894313 | 1.166096776 | 2.988512369 | 0.00280339 | 0.013801 |
| ENSG00000126262 | FFAR2 | 79.43330334 | 2.779233995 | 0.686730814 | 4.64538954 | 3.39E-06 | 3.77E-05 |
| ENSG00000167105 | TMEM92 | 13.28825 | 2.778833 | 0.8625121 | 3.809579 | 0.0001392 | 0.001027 |
| ENSG00000282511 | CLUHP10 | 2.965009 | 2.778218 | 1.670456 | 2.743885 | 0.00607168 | 0.025969 |
| ENSG00000267751 | | 130.3182 | 2.775625 | 0.3104267 | 9.181073 | 4.27E-20 | 3.22E-18 |
| ENSG00000287751 | | 2.466289 | 2.774355 | 2.863897 | 2.655419 | 0.00792099 | 0.032293 |
| ENSG00000139973 | SYT16 | 728.2894 | 2.774141 | 1.379809 | 3.120379 | 0.00180619 | 0.009512 |
| ENSG00000138592 | USP8 | 9371.886064 | 2.774001235 | 0.180384957 | 15.53600706 | 1.98E-54 | 1.64E-51 |

| | | | | | | | |
|---|---|---|---|---|---|---|---|
| ENSG00000164418 | GRIK2 | 65.96094 | 2.773706 | 0.5099046 | 5.81596 | 6.03E-09 | 1.14E-07 |
| ENSG00000122756 | CNTFR | 31.22423966 | 2.773450491 | 0.541046335 | 5.583273914 | 2.36E-08 | 4.02E-07 |
| ENSG00000280220 | | 2.782726 | 2.772772 | 2.53441 | 2.697762 | 0.00698074 | 0.029048 |
| ENSG00000124140 | SLC12A5 | 745.1002436 | 2.770648961 | 0.203802421 | 13.75955668 | 4.46E-43 | 1.91E-40 |
| ENSG00000278955 | | 2.584469 | 2.768008 | 1.778717 | 2.77611 | 0.00550136 | 0.02395 |
| ENSG00000250334 | LINC00989 | 5.872883 | 2.765333 | 2.313151 | 2.743293 | 0.00608264 | 0.026003 |
| ENSG00000182103 | FAM181B | 55.44895 | 2.764967 | 0.6758082 | 4.572474 | 4.82E-06 | 5.15E-05 |
| ENSG00000135929 | CYP27A1 | 13.8439342 | 2.762167104 | 0.965310092 | 3.433443386 | 0.00059597 | 0.003653 |
| ENSG00000068615 | REEP1 | 371.1368735 | 2.756924461 | 0.283728324 | 9.950110888 | 2.52E-23 | 2.63E-21 |
| ENSG00000251600 | | 21.99216 | 2.75555 | 0.7160218 | 4.374957 | 1.21E-05 | 0.000117 |
| ENSG00000235205 | TATDN2P3 | 697.9817 | 2.754419 | 0.7336684 | 4.339704 | 1.43E-05 | 0.000135 |
| ENSG00000280194 | | 32.57879 | 2.748585 | 0.532058 | 5.57758 | 2.44E-08 | 4.14E-07 |
| ENSG00000088899 | LZTS3 | 188.7383109 | 2.748521234 | 0.346702602 | 8.186567651 | 2.69E-16 | 1.36E-14 |
| ENSG00000107018 | RLN1 | 10.85246487 | 2.739358687 | 0.903027651 | 3.712752168 | 0.00020502 | 0.001449 |
| ENSG00000279042 | | 15.11619 | 2.738632 | 0.7869078 | 4.100986 | 4.11E-05 | 0.000349 |
| ENSG00000259583 | | 98.01338 | 2.736016 | 0.9682437 | 3.363071 | 0.00077081 | 0.00457 |
| ENSG00000198739 | LRRTM3 | 21.65021 | 2.735281 | 1.479725 | 2.794984 | 0.00519024 | 0.022842 |
| ENSG00000070182 | SPTB | 117.3744931 | 2.73425379 | 0.594812457 | 5.089779032 | 3.58E-07 | 4.86E-06 |
| ENSG00000140682 | TGFB1I1 | 347.3392 | 2.734014 | 0.2783302 | 10.06458 | 7.92E-24 | 8.70E-22 |
| ENSG00000251689 | | 2.936361 | 2.733499 | 1.469572 | 2.751859 | 0.00592581 | 0.025484 |
| ENSG00000281655 | | 3.911312 | 2.731153 | 2.100736 | 2.610294 | 0.00904645 | 0.035901 |
| ENSG00000156453 | PCDH1 | 2226.496 | 2.729597 | 0.3493902 | 8.104149 | 5.31E-16 | 2.59E-14 |
| ENSG00000101282 | RSPO4 | 4.44268735 | 2.729099282 | 1.236304591 | 2.870706116 | 0.00409556 | 0.018836 |
| ENSG00000144648 | ACKR2 | 68.54638 | 2.725587 | 0.6104202 | 4.850243 | 1.23E-06 | 1.51E-05 |
| ENSG00000189108 | IL1RAPL2 | 112.7854 | 2.725474 | 0.4643029 | 6.277949 | 3.43E-10 | 7.95E-09 |
| ENSG00000155254 | MARVELD1 | 2808.558 | 2.724554 | 0.3854692 | 7.399041 | 1.37E-13 | 4.99E-12 |
| ENSG00000153823 | PID1 | 216.7556 | 2.721916 | 0.4643713 | 6.263257 | 3.77E-10 | 8.70E-09 |
| ENSG00000279024 | | 3.445673 | 2.721776 | 1.579761 | 2.80287 | 0.00506501 | 0.022391 |
| ENSG00000286836 | | 104.8903 | 2.720391 | 0.5282622 | 5.580313 | 2.40E-08 | 4.08E-07 |
| ENSG00000175264 | CHST1 | 159.9614 | 2.720219 | 0.6504759 | 4.69896 | 2.61E-06 | 2.99E-05 |
| ENSG00000250133 | HOXC-AS2 | 6.882066 | 2.717547 | 1.039812 | 3.351531 | 0.00080366 | 0.004739 |
| ENSG00000287275 | | 46.19597 | 2.714774 | 0.4610496 | 6.22096 | 4.94E-10 | 1.12E-08 |
| ENSG00000229544 | NKX1-2 | 36.52836 | 2.713355 | 0.9974403 | 3.267209 | 0.00108613 | 0.006146 |

| | | | | | | | |
|---|---|---|---|---|---|---|---|
| ENSG00000254024 | | 427.4394 | 2.709849 | 0.3426148 | 8.186884 | 2.68E-16 | 1.36E-14 |
| ENSG00000143365 | RORC | 339.6797 | 2.70854 | 0.5254326 | 5.584111 | 2.35E-08 | 4.00E-07 |
| ENSG00000228412 | LNC-LBCS | 155.7247 | 2.706971 | 1.042969 | 3.483251 | 0.00049536 | 0.003114 |
| ENSG00000259668 | | 174.6157 | 2.705464 | 0.2948273 | 9.42182 | 4.43E-21 | 3.76E-19 |
| ENSG00000170915 | PAQR8 | 172.2443 | 2.705331 | 0.4227578 | 6.790033 | 1.12E-11 | 3.20E-10 |
| ENSG00000118407 | FILIP1 | 18.47898461 | 2.70507063 | 0.759640344 | 4.083038949 | 4.45E-05 | 0.000374 |
| ENSG00000176754 | LINC00303 | 3.657692 | 2.696699 | 2.63157 | 2.661587 | 0.00777732 | 0.031785 |
| ENSG00000255471 | | 73.93502 | 2.696293 | 0.4540935 | 6.34549 | 2.22E-10 | 5.29E-09 |
| ENSG00000232624 | LINC01517 | 17.01797 | 2.695847 | 0.6098051 | 4.911129 | 9.06E-07 | 1.14E-05 |
| ENSG00000156206 | CFAP161 | 12.02136 | 2.695832 | 1.090296 | 3.353586 | 0.00079772 | 0.004708 |
| ENSG00000197535 | MYO5A | 6542.971 | 2.691429 | 0.2006893 | 13.57756 | 5.44E-42 | 2.07E-39 |
| ENSG00000259286 | | 4.091741 | 2.690817 | 1.948969 | 2.738841 | 0.00616561 | 0.026311 |
| ENSG00000258616 | LINC02303 | 68.66875 | 2.690077 | 0.6519198 | 4.660838 | 3.15E-06 | 3.53E-05 |
| ENSG00000133937 | GSC | 96.80665672 | 2.689239965 | 0.510869404 | 5.693389881 | 1.25E-08 | 2.24E-07 |
| ENSG00000215478 | CES5AP1 | 5.471488 | 2.687267 | 1.513251 | 2.779616 | 0.00544232 | 0.023736 |
| ENSG00000123870 | ZNF137P | 95.31447057 | 2.683614196 | 0.532227685 | 5.462142884 | 4.70E-08 | 7.63E-07 |
| ENSG00000108821 | COL1A1 | 2320.092411 | 2.68320447 | 0.407084694 | 6.937904671 | 3.98E-12 | 1.21E-10 |
| ENSG00000100033 | PRODH | 318.8555672 | 2.68224452 | 0.427245618 | 6.658540694 | 2.77E-11 | 7.42E-10 |
| ENSG00000185133 | INPP5J | 1848.204 | 2.680837 | 0.3422657 | 8.126733 | 4.41E-16 | 2.17E-14 |
| ENSG00000162946 | DISC1 | 238.3309 | 2.679326 | 0.2614986 | 10.43597 | 1.70E-25 | 2.25E-23 |
| ENSG00000204131 | NHSL2 | 51.06571 | 2.677789 | 0.5581385 | 5.235508 | 1.65E-07 | 2.39E-06 |
| ENSG00000071282 | LMCD1 | 1827.576173 | 2.676969825 | 0.378622775 | 7.392463142 | 1.44E-13 | 5.22E-12 |
| ENSG00000162458 | FBLIM1 | 739.7659 | 2.67348 | 0.2820403 | 9.692859 | 3.23E-22 | 3.14E-20 |
| ENSG00000170935 | NCBP2L | 32.84071 | 2.672103 | 0.5421372 | 5.383924 | 7.29E-08 | 1.14E-06 |
| ENSG00000118432 | CNR1 | 44.44154657 | 2.671029916 | 0.724867323 | 4.247315823 | 2.16E-05 | 0.000196 |
| ENSG00000198788 | MUC2 | 23.05754 | 2.669256 | 1.511054 | 2.905315 | NA | NA |
| ENSG00000141576 | RNF157 | 409.9479 | 2.667785 | 0.3223253 | 8.595835 | 8.27E-18 | 4.91E-16 |
| ENSG00000171522 | PTGER4 | 1247.442 | 2.662071 | 0.3080208 | 8.911556 | 5.03E-19 | 3.39E-17 |
| ENSG00000167562 | ZNF701 | 280.5978 | 2.65848 | 0.2547703 | 10.61413 | 2.56E-26 | 3.64E-24 |
| ENSG00000005379 | TSPOAP1 | 13.58268644 | 2.65411898 | 1.304818613 | 2.781357983 | 0.0054132 | 0.023632 |
| ENSG00000283294 | | 69.98504 | 2.652527 | 0.6262951 | 4.769428 | 1.85E-06 | 2.19E-05 |
| ENSG00000272668 | | 116.9235 | 2.649497 | 0.5283759 | 5.448107 | 5.09E-08 | 8.17E-07 |
| ENSG00000135338 | LCA5 | 128.930122 | 2.644482187 | 0.441664645 | 6.344610465 | 2.23E-10 | 5.31E-09 |

| | | | | | | | |
|---|---|---|---|---|---|---|---|
| ENSG00000247809 | NR2F2-AS1 | 125.4734 | 2.642935 | 0.3689589 | 7.43827 | 1.02E-13 | 3.77E-12 |
| ENSG00000248473 | LINC01962 | 16.29687 | 2.641433 | 0.7296056 | 4.182024 | 2.89E-05 | 0.000254 |
| ENSG00000126264 | HCST | 3.410562333 | 2.637316878 | 1.547552528 | 2.708626852 | 0.00675623 | 0.028295 |
| ENSG00000286424 | | 14.36458 | 2.63688 | 0.8009897 | 3.947903 | 7.88E-05 | 0.00062 |
| ENSG00000105707 | HPN | 5.03552181 | 2.635946099 | 1.42639819 | 2.728432578 | 0.00636361 | 0.026936 |
| ENSG00000127990 | SGCE | 650.0351449 | 2.635454321 | 0.457359008 | 6.02807692 | 1.66E-09 | 3.43E-08 |
| ENSG00000154262 | ABCA6 | 20.46894 | 2.633526 | 0.6759187 | 4.377287 | 1.20E-05 | 0.000116 |
| ENSG00000219665 | ZNF433-AS1 | 57.11773 | 2.625184 | 0.4117649 | 6.71956 | 1.82E-11 | 5.02E-10 |
| ENSG00000288093 | | 11.6878 | 2.62507 | 0.7268018 | 4.154753 | 3.26E-05 | 0.000283 |
| ENSG00000274499 | | 6.642269 | 2.622716 | 1.443491 | 2.749993 | 0.00595966 | 0.025589 |
| ENSG00000115525 | ST3GAL5 | 2778.100949 | 2.621877321 | 0.410766321 | 6.742185504 | 1.56E-11 | 4.35E-10 |
| ENSG00000115380 | EFEMP1 | 1176.857059 | 2.621681854 | 0.392239875 | 7.030467303 | 2.06E-12 | 6.46E-11 |
| ENSG00000266903 | | 111.5601 | 2.620599 | 0.6524129 | 4.575864 | 4.74E-06 | 5.08E-05 |
| ENSG00000162981 | LRATD1 | 3504.535 | 2.617774 | 0.3803176 | 7.208208 | 5.67E-13 | 1.91E-11 |
| ENSG00000224805 | LINC00853 | 51.27837 | 2.617121 | 0.5032764 | 5.603019 | 2.11E-08 | 3.61E-07 |
| ENSG00000267259 | ERVE-1 | 86.88239 | 2.61084 | 0.5532536 | 5.199007 | 2.00E-07 | 2.87E-06 |
| ENSG00000139970 | RTN1 | 109.0116 | 2.610708 | 0.5359212 | 5.299971 | 1.16E-07 | 1.73E-06 |
| ENSG00000234814 | SVIL2P | 21.49515 | 2.603207 | 0.770899 | 3.916126 | 9.00E-05 | 0.000696 |
| ENSG00000235597 | LINC01102 | 3.625065 | 2.599526 | 1.700084 | 2.637962 | 0.00834058 | 0.033632 |
| ENSG00000112182 | BACH2 | 23.52146224 | 2.597976952 | 0.731418196 | 4.170457924 | 3.04E-05 | 0.000266 |
| ENSG00000157514 | TSC22D3 | 15361.42 | 2.597828 | 0.3015911 | 8.874842 | 7.00E-19 | 4.66E-17 |
| ENSG00000118276 | B4GALT6 | 335.0603876 | 2.596789967 | 0.352518674 | 7.656531176 | 1.91E-14 | 7.67E-13 |
| ENSG00000273123 | | 3.817069 | 2.596239 | 1.705915 | 2.632966 | 0.00846428 | 0.034027 |
| ENSG00000162545 | CAMK2N1 | 2075.251 | 2.591426 | 0.3754831 | 7.232729 | 4.73E-13 | 1.61E-11 |
| ENSG00000164266 | SPINK1 | 22.21718 | 2.591356 | 0.9662781 | 3.431457 | 0.00060035 | 0.003676 |
| ENSG00000182795 | C1orf116 | 376.8257 | 2.589637 | 1.263677 | 3.128771 | 0.00175539 | 0.009278 |
| ENSG00000081479 | LRP2 | 1512.710474 | 2.588402052 | 0.753955469 | 4.104508577 | 4.05E-05 | 0.000344 |
| ENSG00000260337 | | 44.17791 | 2.587829 | 0.7428265 | 4.044894 | 5.23E-05 | 0.000431 |
| ENSG00000138594 | TMOD3 | 7724.565008 | 2.579986019 | 0.155398448 | 16.72493178 | 8.63E-63 | 1.27E-59 |
| ENSG00000149948 | HMGA2 | 24.85982 | 2.578478 | 2.248984 | 2.708824 | 0.00675221 | 0.028283 |
| ENSG00000133135 | RNF128 | 120.3441874 | 2.578329315 | 0.861087996 | 3.649846661 | 0.0002624 | 0.001802 |
| ENSG00000213937 | CLDN9 | 2858.096 | 2.575989 | 0.3044953 | 8.731023 | 2.52E-18 | 1.59E-16 |
| ENSG00000171431 | KRT20 | 108.1889 | 2.57423 | 1.003185 | 3.413511 | 0.00064132 | 0.003895 |

| | | | | | | | |
|---|---|---|---|---|---|---|---|
| ENSG00000259985 | | 35.58738 | 2.573273 | 0.4202737 | 6.454297 | 1.09E-10 | 2.70E-09 |
| ENSG00000270116 | | 13.51384 | 2.572785 | 0.8718564 | 3.651693 | 0.00026052 | 0.00179 |
| ENSG00000287490 | | 2.450064 | 2.571513 | 1.634408 | 2.645248 | 0.00816312 | 0.033047 |
| ENSG00000174469 | CNTNAP2 | 949.8648 | 2.571379 | 0.5536715 | 5.122908 | 3.01E-07 | 4.15E-06 |
| ENSG00000169083 | AR | 5787.367 | 2.568039 | 0.2945892 | 8.98046 | 2.70E-19 | 1.87E-17 |
| ENSG00000107984 | DKK1 | 8517.894175 | 2.566178236 | 0.338608486 | 7.87053736 | 3.53E-15 | 1.54E-13 |
| ENSG00000152818 | UTRN | 1624.745 | 2.565001 | 0.4064715 | 6.664579 | 2.65E-11 | 7.15E-10 |
| ENSG00000144481 | TRPM8 | 32.59946 | 2.564378 | 2.096948 | 2.729374 | 0.00634548 | 0.026891 |
| ENSG00000177432 | NAP1L5 | 73.74855 | 2.559243 | 0.4003804 | 6.65695 | 2.80E-11 | 7.49E-10 |
| ENSG00000256894 | | 7.399831 | 2.55814 | 1.233964 | 2.787706 | 0.00530827 | 0.023254 |
| ENSG00000129596 | CDO1 | 7.548433765 | 2.557371634 | 1.055472023 | 3.154383302 | 0.00160838 | 0.008596 |
| ENSG00000253829 | | 11.24741 | 2.556091 | 0.9927456 | 3.328837 | 0.0008721 | 0.005084 |
| ENSG00000196924 | FLNA | 85439.26 | 2.55601 | 0.2258031 | 11.51545 | 1.10E-30 | 2.27E-28 |
| ENSG00000100336 | APOL4 | 191.9240056 | 2.556009266 | 0.657325045 | 4.446452042 | 8.73E-06 | 8.74E-05 |
| ENSG00000237927 | | 2.916866 | 2.554469 | 1.914046 | 2.586032 | 0.00970879 | 0.037951 |
| ENSG00000234928 | LINC01659 | 20.31054 | 2.553283 | 1.03122 | 3.306399 | 0.00094504 | 0.00544 |
| ENSG00000107014 | RLN2 | 127.8000878 | 2.553257784 | 0.594590049 | 4.82163561 | 1.42E-06 | 1.73E-05 |
| ENSG00000165194 | PCDH19 | 269.6649 | 2.550357 | 0.5128994 | 5.413524 | 6.18E-08 | 9.80E-07 |
| ENSG00000284395 | PERCC1 | 24.25703 | 2.546509 | 0.507833 | 5.375196 | 7.65E-08 | 1.18E-06 |
| ENSG00000107249 | GLIS3 | 41.00662557 | 2.544225926 | 0.609185429 | 4.659939901 | 3.16E-06 | 3.55E-05 |
| ENSG00000176399 | DMRTA1 | 691.3066 | 2.541567 | 0.301038 | 8.7035 | 3.22E-18 | 2.00E-16 |
| ENSG00000011201 | ANOS1 | 254.3605756 | 2.538893106 | 0.357732598 | 7.419578607 | 1.17E-13 | 4.30E-12 |
| ENSG00000092439 | TRPM7 | 17716.98982 | 2.536695239 | 0.239168197 | 10.84225186 | 2.17E-27 | 3.41E-25 |
| ENSG00000019549 | SNAI2 | 70.28604479 | 2.536551158 | 1.153196314 | 3.179321566 | 0.0014762 | 0.007995 |
| ENSG00000165795 | NDRG2 | 1144.176 | 2.52908 | 0.4819204 | 5.68755 | 1.29E-08 | 2.31E-07 |
| ENSG00000213793 | ZNF888 | 587.19 | 2.528945 | 0.3285959 | 8.016324 | 1.09E-15 | 5.10E-14 |
| ENSG00000197847 | SLC22A20P | 91.5412 | 2.527563 | 0.4079245 | 6.554194 | 5.59E-11 | 1.44E-09 |
| ENSG00000069956 | MAPK6 | 9103.016658 | 2.525117431 | 0.226067998 | 11.37755632 | 5.41E-30 | 1.06E-27 |
| ENSG00000214892 | USP8P1 | 256.6481 | 2.524959 | 0.3623535 | 7.272915 | 3.52E-13 | 1.23E-11 |
| ENSG00000188681 | TEKT4P2 | 1170.243 | 2.524031 | 0.2552112 | 10.10695 | 5.15E-24 | 5.79E-22 |
| ENSG00000107282 | APBA1 | 162.4042834 | 2.523824338 | 0.463307687 | 5.831270306 | 5.50E-09 | 1.05E-07 |
| ENSG00000172478 | MAB21L4 | 946.0249 | 2.52348 | 0.9765006 | 3.435696 | NA | NA |
| ENSG00000287500 | | 15.71093 | 2.519887 | 0.8052746 | 3.713889 | 0.0002041 | 0.001444 |

| | | | | | | | |
|---|---|---|---|---|---|---|---|
| ENSG00000258216 | | 20.63148 | 2.518923 | 0.6883436 | 4.160922 | 3.17E-05 | 0.000276 |
| ENSG00000105137 | SYDE1 | 203.9698521 | 2.517959364 | 0.37265036 | 7.077517786 | 1.47E-12 | 4.68E-11 |
| ENSG00000188517 | COL25A1 | 2.752639 | 2.517941 | 1.561989 | 2.633492 | 0.00845118 | 0.033988 |
| ENSG00000235830 | SRGAP3-AS4 | 2.235877 | 2.517725 | 3.472687 | 2.53851 | 0.01113256 | 0.042418 |
| ENSG00000198353 | HOXC4 | 128.6267 | 2.517123 | 0.3156039 | 8.233646 | 1.82E-16 | 9.39E-15 |
| ENSG00000249464 | LINC01091 | 23.64872 | 2.516587 | 0.8679193 | 3.528022 | 0.00041868 | 0.00269 |
| ENSG00000237460 | LINC01661 | 125.6758 | 2.512881 | 0.3861257 | 6.824105 | 8.85E-12 | 2.55E-10 |
| ENSG00000261396 | | 6.77919 | 2.510585 | 1.073589 | 3.09301 | 0.00198138 | 0.010272 |
| ENSG00000233452 | STXBP5-AS1 | 73.2455 | 2.506835 | 0.5025046 | 5.422715 | 5.87E-08 | 9.35E-07 |
| ENSG00000258647 | LINC00930 | 55.69479 | 2.506592 | 0.6768245 | 4.261279 | 2.03E-05 | 0.000185 |
| ENSG00000176928 | GCNT4 | 145.8159 | 2.505976 | 0.3814004 | 6.890612 | 5.56E-12 | 1.65E-10 |
| ENSG00000236658 | | 4.941128 | 2.505365 | 1.244387 | 2.778585 | 0.00545962 | 0.023788 |
| ENSG00000156463 | SH3RF2 | 67.52506 | 2.504471 | 0.5063231 | 5.368469 | 7.94E-08 | 1.22E-06 |
| ENSG00000187479 | C11orf96 | 4.985023 | 2.504212 | 1.562696 | 2.643456 | 0.00820645 | 0.033162 |
| ENSG00000140465 | CYP1A1 | 2073.882 | 2.503179 | 1.603646 | 2.871746 | 0.00408211 | 0.018777 |
| ENSG00000249852 | | 14.43213 | 2.503004 | 0.7129585 | 3.942149 | 8.08E-05 | 0.000633 |
| ENSG00000223715 | LINC01208 | 14.19569 | 2.502619 | 0.7435537 | 3.955866 | 7.63E-05 | 0.000603 |
| ENSG00000168992 | OR7E102P | 19.97312 | 2.502202 | 0.7205478 | 3.97041 | 7.17E-05 | 0.00057 |
| ENSG00000232926 | | 28.71393 | 2.500841 | 0.5693697 | 4.868295 | 1.13E-06 | 1.39E-05 |
| ENSG00000157680 | DGKI | 12.41054 | 2.500388 | 0.9646652 | 3.409432 | 0.00065098 | 0.003945 |
| ENSG00000232504 | ST3GAL5-AS1 | 17.17205 | 2.499538 | 1.245423 | 2.750083 | 0.00595803 | 0.025586 |
| ENSG00000249684 | | 268.7431 | 2.49854 | 0.4129819 | 6.419086 | 1.37E-10 | 3.37E-09 |
| ENSG00000255002 | LINC02324 | 1.899911 | 2.497798 | 2.286005 | 2.574033 | 0.01005208 | 0.039034 |
| ENSG00000169429 | CXCL8 | 31.84114 | 2.49767 | 1.21799 | 3.043313 | 0.00233989 | 0.011817 |
| ENSG00000130528 | HRC | 15.61135126 | 2.497237615 | 0.737752249 | 3.909985379 | 9.23E-05 | 0.000712 |
| ENSG00000257842 | LINC02588 | 113.3556 | 2.488784 | 0.2326555 | 10.8633 | 1.72E-27 | 2.74E-25 |
| ENSG00000277022 | | 53.2833 | 2.484422 | 0.7724689 | 3.862837 | 0.00011208 | 0.000848 |
| ENSG00000123364 | HOXC13 | 2376.645666 | 2.483921288 | 0.244520438 | 10.37581628 | 3.19E-25 | 4.06E-23 |
| ENSG00000203392 | | 131.8636 | 2.480714 | 0.3313192 | 7.75472 | 8.85E-15 | 3.69E-13 |
| ENSG00000166575 | TMEM135 | 3575.265 | 2.480707 | 0.1997709 | 12.59656 | 2.21E-36 | 6.23E-34 |
| ENSG00000225411 | | 21.32377 | 2.480281 | 0.5817752 | 4.721041 | 2.35E-06 | 2.72E-05 |
| ENSG00000182782 | HCAR2 | 306.0363 | 2.480003 | 0.398193 | 6.581147 | 4.67E-11 | 1.21E-09 |
| ENSG00000129682 | FGF13 | 385.8761246 | 2.478354097 | 0.39960387 | 6.54575993 | 5.92E-11 | 1.52E-09 |

| | | | | | | | |
|---|---|---|---|---|---|---|---|
| ENSG00000138600 | SPPL2A | 9476.994157 | 2.477905184 | 0.176619501 | 14.14084011 | 2.13E-45 | 1.03E-42 |
| ENSG00000156113 | KCNMA1 | 93.83838 | 2.476919 | 0.4363425 | 6.060486 | 1.36E-09 | 2.86E-08 |
| ENSG00000140481 | CCDC33 | 20.43853 | 2.475662 | 0.859548 | 3.494622 | 0.00047473 | 0.003001 |
| ENSG00000128641 | MYO1B | 13514.25721 | 2.474967444 | 0.280314164 | 9.077613608 | 1.11E-19 | 7.95E-18 |
| ENSG00000053108 | FSTL4 | 108.7306201 | 2.474539676 | 0.622729796 | 4.506933672 | 6.58E-06 | 6.80E-05 |
| ENSG00000276747 | PADI6 | 9.264557 | 2.473903 | 0.8482724 | 3.611858 | 0.00030401 | 0.002046 |
| ENSG00000121270 | ABCC11 | 51.29008495 | 2.470632043 | 0.551332927 | 4.937029517 | 7.93E-07 | 1.01E-05 |
| ENSG00000261829 | | 4.09733 | 2.465077 | 1.533839 | 2.573572 | 0.01006547 | 0.039075 |
| ENSG00000166477 | LEO1 | 5405.878 | 2.46291 | 0.2166931 | 11.56814 | 5.98E-31 | 1.28E-28 |
| ENSG00000234311 | | 18.23719 | 2.461284 | 0.9498881 | 3.40156 | 0.00067003 | 0.004047 |
| ENSG00000145040 | UCN2 | 10.0567 | 2.457638 | 0.8896021 | 3.437949 | 0.00058614 | 0.003604 |
| ENSG00000215644 | GCGR | 11.66679 | 2.457141 | 0.9937955 | 3.268448 | 0.00108139 | 0.006122 |
| ENSG00000104043 | ATP8B4 | 15.03082985 | 2.454957614 | 0.676821228 | 4.170208355 | 3.04E-05 | 0.000266 |
| ENSG00000128271 | ADORA2A | 20.56258716 | 2.453134148 | 0.877646843 | 3.531906505 | 0.00041258 | 0.00266 |
| ENSG00000232936 | | 23.33761 | 2.452181 | 0.6900313 | 4.174994 | 2.98E-05 | 0.000261 |
| ENSG00000264215 | | 11.33383 | 2.45049 | 1.007812 | 3.208606 | 0.0013338 | 0.007355 |
| ENSG00000181036 | FCRL6 | 1.634623 | 2.448868 | 3.664962 | 2.553349 | 0.01066925 | 0.041032 |
| ENSG00000109472 | CPE | 4859.554853 | 2.44781766 | 0.35224716 | 7.262092608 | 3.81E-13 | 1.32E-11 |
| ENSG00000188868 | ZNF563 | 97.46932 | 2.44725 | 0.2841067 | 8.888154 | 6.21E-19 | 4.17E-17 |
| ENSG00000198183 | BPIFA1 | 51.57541 | 2.443162 | 2.037899 | 2.560876 | 0.01044086 | 0.040285 |
| ENSG00000231504 | NMD3P2 | 10.07715 | 2.442528 | 0.6957817 | 4.013087 | 5.99E-05 | 0.000486 |
| ENSG00000127249 | ATP13A4 | 36.65513424 | 2.44055889 | 0.851556293 | 3.542572847 | 0.00039624 | 0.002572 |
| ENSG00000175497 | DPP10 | 3.834785 | 2.43308 | 3.826084 | 2.552957 | 0.01068126 | 0.041052 |
| ENSG00000250474 | WBP1LP2 | 9.36348 | 2.424009 | 0.8753893 | 3.38948 | 0.00070025 | 0.004205 |
| ENSG00000175906 | ARL4D | 677.3576 | 2.419141 | 0.3663424 | 6.9443 | 3.80E-12 | 1.17E-10 |
| ENSG00000213997 | PGAM1P7 | 74.12436 | 2.413824 | 0.3497405 | 7.207247 | 5.71E-13 | 1.92E-11 |
| ENSG00000148123 | PLPPR1 | 77.11088 | 2.410482 | 0.402619 | 6.341525 | 2.28E-10 | 5.41E-09 |
| ENSG00000259146 | | 277.7852 | 2.405774 | 0.3185468 | 7.835631 | 4.66E-15 | 1.99E-13 |
| ENSG00000257660 | ADCY6-DT | 10.62129 | 2.405023 | 0.8034833 | 3.581207 | 0.00034201 | 0.002269 |
| ENSG00000227755 | | 11.36547 | 2.404613 | 1.267152 | 2.857337 | 0.00427212 | 0.019469 |
| ENSG00000287906 | | 108.6586 | 2.402247 | 0.4097663 | 6.218224 | 5.03E-10 | 1.13E-08 |
| ENSG00000226900 | | 12.83302 | 2.401654 | 0.9194755 | 3.337349 | 0.00084582 | 0.004948 |
| ENSG00000177519 | RPRM | 120.5704 | 2.399334 | 0.5563033 | 4.824769 | 1.40E-06 | 1.70E-05 |

| | | | | | | | |
|---|---|---|---|---|---|---|---|
| ENSG00000124493 | GRM4 | 617.0929092 | 2.398746722 | 0.408253771 | 6.238140314 | 4.43E-10 | 1.01E-08 |
| ENSG00000141527 | CARD14 | 647.8857 | 2.397926 | 0.3618665 | 6.941004 | 3.89E-12 | 1.19E-10 |
| ENSG00000184349 | EFNA5 | 536.8717 | 2.391078 | 0.2558787 | 9.574543 | 1.02E-21 | 9.21E-20 |
| ENSG00000133962 | CATSPERB | 113.1733839 | 2.388859719 | 0.415583341 | 6.088237871 | 1.14E-09 | 2.43E-08 |
| ENSG00000081237 | PTPRC | 5.433697025 | 2.388829059 | 1.589901968 | 2.54444647 | 0.01094511 | 0.041854 |
| ENSG00000189045 | ANKDD1B | 15.11455 | 2.388272 | 0.5833924 | 4.56575 | 4.98E-06 | 5.31E-05 |
| ENSG00000137502 | RAB30 | 1281.178938 | 2.385586421 | 0.43503935 | 5.873142216 | 4.28E-09 | 8.35E-08 |
| ENSG00000272088 | | 7.318413 | 2.382747 | 1.156107 | 2.935812 | 0.00332676 | 0.01584 |
| ENSG00000258101 | | 43.30433 | 2.382667 | 0.5296776 | 4.954434 | 7.25E-07 | 9.32E-06 |
| ENSG00000179111 | HES7 | 17.82891 | 2.380675 | 1.118242 | 3.056014 | 0.00224301 | 0.011399 |
| ENSG00000161642 | ZNF385A | 1948.169 | 2.379921 | 0.284627 | 8.628022 | 6.24E-18 | 3.74E-16 |
| ENSG00000120885 | CLU | 11707.81058 | 2.378570052 | 0.371405506 | 6.738925338 | 1.60E-11 | 4.44E-10 |
| ENSG00000115616 | SLC9A2 | 2595.527243 | 2.378455586 | 0.321748697 | 7.690559419 | 1.46E-14 | 5.96E-13 |
| ENSG00000243004 | | 13.93771 | 2.376945 | 0.9157501 | 3.374147 | 0.00074045 | 0.00441 |
| ENSG00000270604 | HCG17 | 10.48403 | 2.373618 | 0.9165363 | 3.30956 | 0.00093443 | 0.005387 |
| ENSG00000137877 | SPTBN5 | 175.5130493 | 2.372476179 | 0.402236219 | 6.244429062 | 4.25E-10 | 9.73E-09 |
| ENSG00000260011 | | 7.664252 | 2.369899 | 1.063652 | 3.051526 | 0.00227681 | 0.011545 |
| ENSG00000042832 | TG | 12.83163925 | 2.369576468 | 0.716376035 | 3.870621515 | 0.00010856 | 0.000822 |
| ENSG00000228933 | | 20.85451 | 2.368295 | 0.7511611 | 3.703464 | 0.00021268 | 0.001497 |
| ENSG00000259712 | | 10.30657 | 2.368273 | 0.610236 | 4.331846 | 1.48E-05 | 0.00014 |
| ENSG00000170054 | SERPINA9 | 30.91906 | 2.365371 | 0.8203242 | 3.583632 | 0.00033885 | 0.00225 |
| ENSG00000224566 | FAM96AP2 | 11.42197 | 2.36474 | 0.710937 | 3.835139 | 0.00012549 | 0.000937 |
| ENSG00000229483 | LINC00362 | 3.896285 | 2.363023 | 1.455178 | 2.706443 | 0.00680083 | 0.028464 |
| ENSG00000166979 | EVA1C | 54.28671 | 2.360312 | 0.3298931 | 7.425834 | 1.12E-13 | 4.12E-12 |
| ENSG00000020577 | SAMD4A | 4050.772774 | 2.360030018 | 0.345393087 | 7.144835129 | 9.01E-13 | 2.95E-11 |
| ENSG00000149090 | PAMR1 | 16.31627 | 2.358776 | 0.8653579 | 3.451006 | 0.0005585 | 0.003452 |
| ENSG00000168646 | AXIN2 | 274.5701 | 2.358127 | 0.565393 | 4.694158 | 2.68E-06 | 3.06E-05 |
| ENSG00000163393 | SLC22A15 | 310.4001 | 2.357272 | 0.423664 | 5.939534 | 2.86E-09 | 5.73E-08 |
| ENSG00000081014 | AP4E1 | 5318.857867 | 2.353867477 | 0.188976672 | 12.6318105 | 1.41E-36 | 4.12E-34 |
| ENSG00000162461 | SLC25A34 | 37.73112 | 2.352325 | 0.5023331 | 5.107544 | 3.26E-07 | 4.47E-06 |
| ENSG00000137261 | KIAA0319 | 109.8973521 | 2.351523199 | 0.527481757 | 4.946718161 | 7.55E-07 | 9.63E-06 |
| ENSG00000234614 | C2CD4D-AS1 | 85.18482 | 2.350966 | 0.3732589 | 6.610222 | 3.84E-11 | 1.01E-09 |
| ENSG00000151117 | TMEM86A | 520.9282 | 2.34687 | 0.3655449 | 6.743925 | 1.54E-11 | 4.31E-10 |

| | | | | | | | |
|---|---|---|---|---|---|---|---|
| ENSG00000125823 | CSTL1 | 6.78395186 | 2.345898944 | 1.27984208 | 2.635280871 | 0.00840677 | 0.033845 |
| ENSG00000204637 | ARPP19P2 | 8.862275 | 2.342155 | 0.8572735 | 3.41662 | 0.00063404 | 0.003858 |
| ENSG00000101883 | RHOXF1 | 3.364795521 | 2.341629159 | 2.538860454 | 2.569300233 | 0.01019041 | 0.039491 |
| ENSG00000276075 | | 22.98136 | 2.338541 | 0.6587255 | 4.115137 | 3.87E-05 | 0.00033 |
| ENSG00000224251 | | 43.86411 | 2.338479 | 1.042932 | 3.002803 | 0.00267505 | 0.013235 |
| ENSG00000285018 | | 71.29603 | 2.338344 | 0.3075819 | 7.845602 | 4.31E-15 | 1.86E-13 |
| ENSG00000006468 | ETV1 | 10.49234618 | 2.338085154 | 0.829510403 | 3.51401263 | 0.00044139 | 0.002817 |
| ENSG00000118729 | CASQ2 | 5.102997809 | 2.337638997 | 1.662437497 | 2.613600757 | 0.00895937 | 0.035624 |
| ENSG00000135111 | TBX3 | 1716.289437 | 2.337544522 | 0.456493598 | 5.534461049 | 3.12E-08 | 5.19E-07 |
| ENSG00000104953 | TLE6 | 238.3585033 | 2.337346028 | 0.575966768 | 4.620582903 | 3.83E-06 | 4.19E-05 |
| ENSG00000203326 | ZNF525 | 568.7468 | 2.336635 | 0.2395951 | 9.979147 | 1.88E-23 | 1.98E-21 |
| ENSG00000100979 | PLTP | 33.53258375 | 2.335129576 | 0.548795876 | 4.770654578 | 1.84E-06 | 2.18E-05 |
| ENSG00000287377 | | 2.571116 | 2.334861 | 1.71946 | 2.567411 | 0.01024612 | 0.03966 |
| ENSG00000185303 | SFTPA2 | 6.1203 | 2.334852 | 0.9356722 | 3.245529 | 0.00117233 | 0.006557 |
| ENSG00000241622 | RARRES2P1 | 14.34514 | 2.334844 | 0.5951701 | 4.365007 | 1.27E-05 | 0.000122 |
| ENSG00000131711 | MAP1B | 751.1304494 | 2.332927919 | 0.829540114 | 3.554795482 | 0.00037827 | 0.002475 |
| ENSG00000188897 | | 323.4965 | 2.327671 | 0.3508753 | 6.939499 | 3.93E-12 | 1.20E-10 |
| ENSG00000179841 | AKAP5 | 346.9897 | 2.326717 | 0.4298893 | 5.804725 | 6.45E-09 | 1.22E-07 |
| ENSG00000281392 | | 12.81932 | 2.32424 | 0.8657412 | 3.407556 | 0.00065548 | 0.003969 |
| ENSG00000134955 | SLC37A2 | 18.26130766 | 2.323961287 | 0.664569811 | 4.046527044 | 5.20E-05 | 0.000429 |
| ENSG00000102935 | ZNF423 | 129.0732214 | 2.323332315 | 0.585969613 | 4.355211183 | 1.33E-05 | 0.000127 |
| ENSG00000243094 | RPL32P2 | 8.239744 | 2.320201 | 0.8174004 | 3.49654 | 0.00047133 | 0.002985 |
| ENSG00000121552 | CSTA | 134.9061348 | 2.319927013 | 0.579687849 | 4.525532811 | 6.02E-06 | 6.31E-05 |
| ENSG00000148702 | HABP2 | 29.23533 | 2.317708 | 0.6486626 | 4.127726 | 3.66E-05 | 0.000314 |
| ENSG00000207839 | MIR33B | 14.46731 | 2.316126 | 0.7800284 | 3.60096 | 0.00031704 | 0.002124 |
| ENSG00000267239 | | 4.699024 | 2.312977 | 2.717092 | 2.532838 | 0.01131434 | 0.042981 |
| ENSG00000233695 | GAS6-AS1 | 80.43024 | 2.310949 | 0.4902712 | 5.153913 | 2.55E-07 | 3.57E-06 |
| ENSG00000104093 | DMXL2 | 7174.130075 | 2.309015799 | 0.265779436 | 8.923539626 | 4.52E-19 | 3.07E-17 |
| ENSG00000117425 | PTCH2 | 54.45241929 | 2.307753608 | 0.482873784 | 5.197544129 | 2.02E-07 | 2.89E-06 |
| ENSG00000187678 | SPRY4 | 222.4557 | 2.307229 | 0.5753735 | 4.546376 | 5.46E-06 | 5.76E-05 |
| ENSG00000164023 | SGMS2 | 830.5996 | 2.306343 | 0.3375761 | 7.133051 | 9.82E-13 | 3.19E-11 |
| ENSG00000260633 | | 60.49077 | 2.304194 | 0.4960245 | 5.046204 | 4.51E-07 | 6.03E-06 |
| ENSG00000184588 | PDE4B | 137.0912 | 2.301603 | 0.6125754 | 4.306991 | 1.65E-05 | 0.000155 |

| | | | | | | | |
|---|---|---|---|---|---|---|---|
| ENSG00000250835 | LSM3P4 | 1.305252 | 2.301303 | 3.258269 | 2.521178 | 0.01169628 | NA |
| ENSG00000103316 | CRYM | 34.23905413 | 2.300085984 | 0.713453122 | 3.868083032 | 0.00010969 | 0.000831 |
| ENSG00000226102 | SEPTIN7P3 | 4.973441 | 2.29878 | 1.136312 | 2.784553 | 0.00536015 | 0.023446 |
| ENSG00000173727 | | 335.719 | 2.297914 | 0.5896794 | 4.426634 | 9.57E-06 | 9.48E-05 |
| ENSG00000176136 | MC5R | 3.148802 | 2.296919 | 2.138846 | 2.480501 | 0.01311978 | 0.048421 |
| ENSG00000139144 | PIK3C2G | 23.88055569 | 2.296669621 | 0.739548021 | 3.739445618 | 0.00018443 | 0.001319 |
| ENSG00000157613 | CREB3L1 | 260.6303 | 2.295732 | 0.8382271 | 3.48098 | 0.00049958 | 0.003136 |
| ENSG00000175093 | SPSB4 | 63.08059 | 2.293 | 0.5939659 | 4.424068 | 9.69E-06 | 9.57E-05 |
| ENSG00000287051 | | 150.719 | 2.288033 | 0.4932202 | 5.089837 | 3.58E-07 | 4.86E-06 |
| ENSG00000092758 | COL9A3 | 1292.301201 | 2.287163887 | 0.474889858 | 5.246387948 | 1.55E-07 | 2.27E-06 |
| ENSG00000224008 | LINC01441 | 49.63748 | 2.282924 | 0.7015675 | 3.845488 | 0.00012031 | 0.000902 |
| ENSG00000256982 | | 12.1066 | 2.282901 | 0.6625203 | 3.946663 | 7.92E-05 | 0.000623 |
| ENSG00000182118 | FAM89A | 297.2301 | 2.282889 | 0.2637364 | 8.890548 | 6.08E-19 | 4.09E-17 |
| ENSG00000079482 | OPHN1 | 1490.681304 | 2.282171763 | 0.31247366 | 7.591657915 | 3.16E-14 | 1.24E-12 |
| ENSG00000121005 | CRISPLD1 | 296.6118738 | 2.272965884 | 0.343724085 | 6.924416371 | 4.38E-12 | 1.32E-10 |
| ENSG00000205683 | DPF3 | 75.79279 | 2.271307 | 0.3787001 | 6.297563 | 3.02E-10 | 7.05E-09 |
| ENSG00000249326 | | 6.366528 | 2.270965 | 1.392694 | 2.550385 | 0.01076039 | 0.041272 |
| ENSG00000141756 | FKBP10 | 5460.867 | 2.270711 | 0.2992217 | 7.863406 | 3.74E-15 | 1.62E-13 |
| ENSG00000251188 | | 8.702443 | 2.268678 | 0.7908526 | 3.455969 | 0.00054832 | 0.003404 |
| ENSG00000245025 | | 40.04959 | 2.268006 | 0.6155709 | 4.209187 | 2.56E-05 | 0.000229 |
| ENSG00000173253 | DMRT2 | 17.30696 | 2.267234 | 0.6230718 | 4.172539 | 3.01E-05 | 0.000264 |
| ENSG00000248587 | GDNF-AS1 | 15.15012 | 2.265674 | 0.9742695 | 3.153792 | 0.00161164 | 0.008607 |
| ENSG00000184672 | RALYL | 6.222358 | 2.263625 | 1.207997 | 2.84472 | 0.00444505 | 0.020099 |
| ENSG00000170236 | USP50 | 130.7968 | 2.262087 | 0.3981312 | 6.061166 | 1.35E-09 | 2.85E-08 |
| ENSG00000170577 | SIX2 | 646.0662 | 2.261883 | 0.4424455 | 5.527113 | 3.26E-08 | 5.40E-07 |
| ENSG00000234965 | SHISA8 | 20.12038 | 2.259038 | 0.9326462 | 3.206161 | 0.00134519 | 0.007401 |
| ENSG00000155966 | AFF2 | 54.38069 | 2.254711 | 0.6114983 | 4.240077 | 2.23E-05 | 0.000202 |
| ENSG00000189423 | USP32P3 | 256.8087 | 2.254544 | 0.4454609 | 5.463913 | 4.66E-08 | 7.55E-07 |
| ENSG00000169330 | MINAR1 | 471.534 | 2.251756 | 0.4540109 | 5.383245 | 7.32E-08 | 1.14E-06 |
| ENSG00000242732 | RTL5 | 394.3167 | 2.251495 | 0.3183914 | 7.356506 | 1.89E-13 | 6.75E-12 |
| ENSG00000254042 | SLIT3-AS2 | 5.28606 | 2.251081 | 1.20513 | 2.729723 | 0.00633876 | 0.026878 |
| ENSG00000136048 | DRAM1 | 6475.664313 | 2.250162766 | 0.328458987 | 7.151877728 | 8.56E-13 | 2.81E-11 |
| ENSG00000259275 | | 157.9643 | 2.249064 | 0.4510441 | 5.385466 | 7.23E-08 | 1.13E-06 |

| | | | | | | | |
|---|---|---|---|---|---|---|---|
| ENSG00000065609 | SNAP91 | 108.9978183 | 2.248401942 | 0.450690982 | 5.377179581 | 7.57E-08 | 1.17E-06 |
| ENSG00000251141 | MRPS30-DT | 19.27605 | 2.248294 | 0.7390164 | 3.682663 | 0.00023081 | 0.001608 |
| ENSG00000185203 | WASIR1 | 9.782772 | 2.24701 | 0.959075 | 3.120252 | 0.00180697 | 0.009513 |
| ENSG00000223901 | | 168.4854 | 2.244837 | 0.400735 | 5.968856 | 2.39E-09 | 4.85E-08 |
| ENSG00000206538 | VGLL3 | 1227.378 | 2.241065 | 0.983586 | 3.178025 | 0.00148282 | 0.008027 |
| ENSG00000233084 | RPL23AP25 | 3.481687 | 2.240619 | 2.752149 | 2.534909 | 0.01124765 | 0.042799 |
| ENSG00000134256 | CD101 | 61.8771089 | 2.239143236 | 0.427605779 | 5.628897031 | 1.81E-08 | 3.15E-07 |
| ENSG00000061455 | PRDM6 | 440.0254278 | 2.238915671 | 0.263375314 | 8.724864966 | 2.66E-18 | 1.67E-16 |
| ENSG00000033170 | FUT8 | 4751.368741 | 2.23890093 | 0.180164313 | 12.58188945 | 2.66E-36 | 7.43E-34 |
| ENSG00000261707 | | 12.38368 | 2.236487 | 0.7283484 | 3.596883 | 0.00032205 | 0.002153 |
| ENSG00000171016 | PYGO1 | 85.60641 | 2.23373 | 0.6890754 | 3.796041 | 0.00014703 | 0.001077 |
| ENSG00000268100 | ZNF725P | 5.197885 | 2.232727 | 2.933843 | 2.500699 | 0.01239485 | 0.046198 |
| ENSG00000263639 | MSMB | 164.7118 | 2.230026 | 0.9806714 | 3.144008 | 0.00166651 | 0.008852 |
| ENSG00000128610 | FEZF1 | 12.32643058 | 2.228944349 | 0.765791728 | 3.54280249 | 0.0003959 | 0.00257 |
| ENSG00000155760 | FZD7 | 386.0132 | 2.228174 | 0.3318237 | 7.042132 | 1.89E-12 | 5.96E-11 |
| ENSG00000187134 | AKR1C1 | 1521.468 | 2.224226 | 0.4578732 | 5.281647 | 1.28E-07 | 1.90E-06 |
| ENSG00000225062 | CATIP-AS1 | 3.27753 | 2.222913 | 2.7531 | 2.523934 | 0.01160498 | 0.043853 |
| ENSG00000173166 | RAPH1 | 829.7459 | 2.221026 | 0.4337335 | 5.5143 | 3.50E-08 | 5.79E-07 |
| ENSG00000233215 | LINC01687 | 6.066271 | 2.218943 | 1.672836 | 2.46237 | 0.01380223 | 0.050386 |
| ENSG00000196793 | ZNF239 | 1633.889 | 2.218652 | 0.2113033 | 10.70693 | 9.44E-27 | 1.40E-24 |
| ENSG00000253250 | C8orf88 | 4.847133 | 2.218434 | 1.788047 | 2.486201 | 0.0129115 | 0.047811 |
| ENSG00000197308 | GATA3-AS1 | 1733.411 | 2.217894 | 0.312505 | 7.390608 | 1.46E-13 | 5.29E-12 |
| ENSG00000248019 | FAM13A-AS1 | 139.5947 | 2.217658 | 0.4443431 | 5.396149 | 6.81E-08 | 1.07E-06 |
| ENSG00000128739 | SNRPN | 16.81873364 | 2.21599782 | 1.464151315 | 2.520349536 | 0.01172384 | 0.044189 |
| ENSG00000135919 | SERPINE2 | 390.5009306 | 2.21589173 | 0.277761202 | 8.247016043 | 1.62E-16 | 8.44E-15 |
| ENSG00000254427 | LINC02696 | 8.854999 | 2.215312 | 0.8686595 | 3.161952 | 0.00156716 | 0.008418 |
| ENSG00000233232 | NPIPB7 | 1.646332 | 2.212541 | 3.412932 | 2.491822 | 0.01270896 | 0.047199 |
| ENSG00000230118 | | 2.754908 | 2.212526 | 2.474626 | 2.529911 | 0.01140916 | 0.043248 |
| ENSG00000258926 | | 24.12969 | 2.210833 | 0.6324467 | 4.053427 | 5.05E-05 | 0.000418 |
| ENSG00000284883 | | 88.60431 | 2.210738 | 0.5453481 | 4.534714 | 5.77E-06 | 6.06E-05 |
| ENSG00000157654 | PALM2AKAP2 | 301.4209 | 2.208365 | 0.3545195 | 6.542124 | 6.07E-11 | 1.55E-09 |
| ENSG00000167483 | NIBAN3 | 15.46432 | 2.206719 | 0.7513929 | 3.551964 | 0.00038237 | 0.002498 |
| ENSG00000258527 | ASB9P1 | 3.407289 | 2.205656 | 1.133029 | 2.798149 | 0.00513964 | 0.022651 |

| | | | | | | | |
|---|---|---|---|---|---|---|---|
| ENSG00000198569 | SLC34A3 | 219.3279 | 2.205374 | 0.369224 | 6.299329 | 2.99E-10 | 6.99E-09 |
| ENSG00000167037 | SGSM1 | 147.4709 | 2.203284 | 0.2456291 | 9.177705 | 4.40E-20 | 3.31E-18 |
| ENSG00000268970 | | 29.89643 | 2.201624 | 0.6711802 | 3.886643 | 0.00010164 | 0.000776 |
| ENSG00000168928 | CTRB2 | 16.66167 | 2.198155 | 0.8601901 | 3.212715 | 0.00131487 | 0.007263 |
| ENSG00000196542 | SPTSSB | 37055.01 | 2.194808 | 0.3163534 | 7.231117 | 4.79E-13 | 1.63E-11 |
| ENSG00000166289 | PLEKHF1 | 655.3361 | 2.19455 | 0.3542631 | 6.531546 | 6.51E-11 | 1.66E-09 |
| ENSG00000251151 | HOXC-AS3 | 140.8892 | 2.19438 | 0.2819823 | 8.0082 | 1.16E-15 | 5.43E-14 |
| ENSG00000167767 | KRT80 | 33260.77 | 2.194091 | 0.3644231 | 6.360928 | 2.01E-10 | 4.82E-09 |
| ENSG00000198732 | SMOC1 | 9.253587 | 2.193522 | 1.465886 | 2.463753 | 0.01374908 | 0.050213 |
| ENSG00000168993 | CPLX1 | 380.3226 | 2.193226 | 0.2824926 | 8.028952 | 9.83E-16 | 4.63E-14 |
| ENSG00000164430 | CGAS | 339.1831 | 2.191287 | 0.3797252 | 6.120463 | 9.33E-10 | 2.01E-08 |
| ENSG00000154639 | CXADR | 1102.565 | 2.190942 | 0.2645331 | 8.527752 | 1.49E-17 | 8.66E-16 |
| ENSG00000147027 | TMEM47 | 551.8625 | 2.190323 | 0.4080719 | 5.744805 | 9.20E-09 | 1.71E-07 |
| ENSG00000196407 | THEM5 | 79.09659 | 2.187765 | 0.2791368 | 8.091473 | 5.89E-16 | 2.86E-14 |
| ENSG00000143036 | SLC44A3 | 180.7978 | 2.18764 | 0.4363081 | 5.40793 | 6.38E-08 | 1.01E-06 |
| ENSG00000171004 | HS6ST2 | 1246.522 | 2.187345 | 0.3236808 | 7.063141 | 1.63E-12 | 5.17E-11 |
| ENSG00000154118 | JPH3 | 1172.553 | 2.186012 | 0.2922929 | 7.744655 | 9.58E-15 | 3.97E-13 |
| ENSG00000130270 | ATP8B3 | 59.23082686 | 2.185126436 | 0.497937176 | 4.846630338 | 1.26E-06 | 1.53E-05 |
| ENSG00000128833 | MYO5C | 12471.35746 | 2.184539694 | 0.161710114 | 13.65840924 | 1.80E-42 | 7.35E-40 |
| ENSG00000287242 | | 11.76326 | 2.184223 | 0.8669087 | 3.204141 | 0.00135466 | 0.007441 |
| ENSG00000144355 | DLX1 | 1110.955 | 2.183692 | 0.3690217 | 6.264296 | 3.75E-10 | 8.65E-09 |
| ENSG00000228510 | | 8.098478 | 2.183176 | 0.8191039 | 3.350096 | 0.00080783 | 0.004757 |
| ENSG00000142149 | HUNK | 451.2795 | 2.182774 | 0.3694317 | 6.275064 | 3.49E-10 | 8.09E-09 |
| ENSG00000109819 | PPARGC1A | 135.7533627 | 2.180019083 | 0.593383389 | 4.198234404 | 2.69E-05 | 0.000238 |
| ENSG00000243566 | UPK3B | 2101.137 | 2.178282 | 0.5282422 | 4.623833 | 3.77E-06 | 4.13E-05 |
| ENSG00000286536 | | 54.0408 | 2.177786 | 0.4798713 | 4.959352 | 7.07E-07 | 9.12E-06 |
| ENSG00000253302 | STAU2-AS1 | 2.874713 | 2.177413 | 1.522017 | 2.448259 | 0.01435483 | 0.052096 |
| ENSG00000269356 | | 24.71993 | 2.171394 | 0.8963583 | 3.214307 | 0.0013076 | 0.00723 |
| ENSG00000138821 | SLC39A8 | 2505.825938 | 2.170491974 | 0.238170324 | 9.341376804 | 9.51E-21 | 7.63E-19 |
| ENSG00000111799 | COL12A1 | 5661.997919 | 2.170364625 | 0.459924546 | 5.1447382 | 2.68E-07 | 3.74E-06 |
| ENSG00000079337 | RAPGEF3 | 154.9300579 | 2.168318807 | 0.397133589 | 5.823866923 | 5.75E-09 | 1.10E-07 |
| ENSG00000124406 | ATP8A1 | 1551.90741 | 2.167120008 | 0.408206944 | 5.687721057 | 1.29E-08 | 2.31E-07 |
| ENSG00000188130 | MAPK12 | 1591.04 | 2.166894 | 0.2057416 | 10.77765 | 4.39E-27 | 6.74E-25 |

| | | | | | | | |
|---|---|---|---|---|---|---|---|
| ENSG00000099194 | SCD | 177454.8074 | 2.166333666 | 0.470508534 | 5.037084826 | 4.73E-07 | 6.30E-06 |
| ENSG00000100156 | SLC16A8 | 119.3865555 | 2.166032293 | 0.364737917 | 6.254632825 | 3.98E-10 | 9.15E-09 |
| ENSG00000183484 | GPR132 | 48.49457 | 2.16591 | 0.6088919 | 4.097588 | 4.17E-05 | 0.000354 |
| ENSG00000113430 | IRX4 | 407.1771767 | 2.163370009 | 0.305387535 | 7.379791037 | 1.59E-13 | 5.72E-12 |
| ENSG00000287407 | | 40.0868 | 2.162287 | 0.8116045 | 3.400626 | 0.00067232 | 0.004058 |
| ENSG00000100300 | TSPO | 3483.693004 | 2.159493438 | 0.252139317 | 8.816937881 | 1.18E-18 | 7.59E-17 |
| ENSG00000081052 | COL4A4 | 49.47042452 | 2.157323138 | 0.510133355 | 4.673090764 | 2.97E-06 | 3.35E-05 |
| ENSG00000233198 | RNF224 | 241.6521 | 2.154655 | 0.2528414 | 8.756326 | 2.02E-18 | 1.27E-16 |
| ENSG00000134323 | MYCN | 25.93425816 | 2.154377576 | 0.998078705 | 3.070289102 | 0.00213852 | 0.010971 |
| ENSG00000250101 | | 16.47713 | 2.154113 | 0.6456729 | 3.919943 | 8.86E-05 | 0.000687 |
| ENSG00000106789 | CORO2A | 1673.311794 | 2.153157266 | 0.207686435 | 10.55696784 | 4.72E-26 | 6.49E-24 |
| ENSG00000114770 | ABCC5 | 20526.22129 | 2.147901041 | 0.341350824 | 6.607811537 | 3.90E-11 | 1.02E-09 |
| ENSG00000136379 | ABHD17C | 2809.570964 | 2.146990902 | 0.173421151 | 12.56114516 | 3.45E-36 | 9.46E-34 |
| ENSG00000188818 | ZDHHC11 | 374.6478 | 2.146478 | 0.3087335 | 7.237353 | 4.58E-13 | 1.56E-11 |
| ENSG00000131379 | C3orf20 | 16.16325294 | 2.14420276 | 0.546084568 | 4.390898645 | 1.13E-05 | 0.00011 |
| ENSG00000273451 | | 32.26953 | 2.143558 | 0.5337885 | 4.419921 | 9.87E-06 | 9.74E-05 |
| ENSG00000274928 | KRT89P | 77.69516 | 2.142868 | 0.5933423 | 4.148928 | 3.34E-05 | 0.000289 |
| ENSG00000102057 | KCND1 | 211.9295332 | 2.141194636 | 0.535691769 | 4.486792939 | 7.23E-06 | 7.41E-05 |
| ENSG00000143061 | IGSF3 | 14525.13 | 2.140684 | 0.2630674 | 8.390303 | 4.85E-17 | 2.66E-15 |
| ENSG00000128655 | PDE11A | 5.584794449 | 2.14030629 | 1.065806366 | 2.915177926 | 0.00355486 | 0.016737 |
| ENSG00000073605 | GSDMB | 529.7694427 | 2.139923344 | 0.357612432 | 6.317764292 | 2.65E-10 | 6.25E-09 |
| ENSG00000010810 | FYN | 612.1062621 | 2.137643508 | 0.45521088 | 5.120589433 | 3.05E-07 | 4.19E-06 |
| ENSG00000243137 | PSG4 | 52.89408 | 2.136598 | 0.8822236 | 3.207635 | 0.00133831 | 0.007374 |
| ENSG00000082293 | COL19A1 | 2.829400693 | 2.13646388 | 1.598280629 | 2.479762454 | 0.01314699 | 0.048475 |
| ENSG00000252376 | RNA5SP395 | 2.565785 | 2.134297 | 1.913292 | 2.478921 | 0.01317804 | 0.048561 |
| ENSG00000133519 | ZDHHC8P1 | 259.425643 | 2.13285517 | 0.366211772 | 6.160785446 | 7.24E-10 | 1.59E-08 |
| ENSG00000205978 | NYNRIN | 1229.954 | 2.132176 | 0.3621802 | 6.222651 | 4.89E-10 | 1.11E-08 |
| ENSG00000232063 | | 8.803989 | 2.13159 | 0.8692385 | 3.173393 | 0.00150669 | 0.00814 |
| ENSG00000120658 | ENOX1 | 249.9639599 | 2.131297433 | 0.315954967 | 7.037710124 | 1.95E-12 | 6.14E-11 |
| ENSG00000087076 | HSD17B14 | 44.00462122 | 2.129388355 | 0.579859999 | 4.198846442 | 2.68E-05 | 0.000238 |
| ENSG00000273274 | ZBTB8B | 12.04693 | 2.129371 | 1.020707 | 2.914793 | 0.00355925 | 0.016746 |
| ENSG00000244300 | GATA2-AS1 | 32.88228 | 2.128646 | 0.5174369 | 4.524112 | 6.06E-06 | 6.35E-05 |
| ENSG00000149591 | TAGLN | 724.5817 | 2.12853 | 0.2784512 | 7.893099 | 2.95E-15 | 1.31E-13 |

| | | | | | | | |
|---|---|---|---|---|---|---|---|
| ENSG00000286004 | | 100.4244 | 2.127505 | 0.3566163 | 6.286633 | 3.24E-10 | 7.55E-09 |
| ENSG00000122550 | KLHL7 | 1834.865501 | 2.127210125 | 0.174355369 | 12.35952621 | 4.33E-35 | 1.11E-32 |
| ENSG00000225506 | CYP4A22-AS1 | 35.60033 | 2.125831 | 0.4445507 | 5.157542 | 2.50E-07 | 3.51E-06 |
| ENSG00000172399 | MYOZ2 | 8.078191 | 2.125468 | 0.9409616 | 2.895051 | 0.00379097 | 0.01764 |
| ENSG00000253768 | | 19.02454 | 2.125232 | 0.8263871 | 3.295009 | 0.00098419 | 0.005645 |
| ENSG00000286061 | RCAN3AS | 3.887463 | 2.125104 | 1.69483 | 2.519744 | 0.01174403 | 0.04424 |
| ENSG00000116254 | CHD5 | 1341.833809 | 2.120521083 | 0.423285205 | 5.416395391 | 6.08E-08 | 9.65E-07 |
| ENSG00000213777 | | 14.56245 | 2.11839 | 0.7693687 | 3.474421 | 0.00051196 | 0.003201 |
| ENSG00000259531 | | 153.9021 | 2.118096 | 0.2300455 | 9.426028 | 4.26E-21 | 3.64E-19 |
| ENSG00000168994 | PXDC1 | 512.5856 | 2.117418 | 0.4052754 | 5.611912 | 2.00E-08 | 3.44E-07 |
| ENSG00000117318 | ID3 | 3611.27067 | 2.117286915 | 0.580789291 | 4.190324901 | 2.79E-05 | 0.000246 |
| ENSG00000041353 | RAB27B | 3558.901931 | 2.11693592 | 0.353305811 | 6.321447394 | 2.59E-10 | 6.10E-09 |
| ENSG00000174099 | MSRB3 | 111.0274 | 2.116788 | 0.5472286 | 4.372002 | 1.23E-05 | 0.000118 |
| ENSG00000039139 | DNAH5 | 530.7222177 | 2.115976612 | 0.339056405 | 6.557316949 | 5.48E-11 | 1.41E-09 |
| ENSG00000286086 | | 27.75354 | 2.113071 | 0.7186206 | 3.540148 | 0.0003999 | 0.00259 |
| ENSG00000074370 | ATP2A3 | 6888.281826 | 2.112987207 | 0.396330572 | 5.70425247 | 1.17E-08 | 2.11E-07 |
| ENSG00000241527 | CA15P1 | 17.5418 | 2.112438 | 0.5819339 | 4.108731 | 3.98E-05 | 0.000338 |
| ENSG00000260788 | | 3.778357 | 2.109527 | 2.516463 | 2.485684 | 0.01293028 | 0.047861 |
| ENSG00000114279 | FGF12 | 473.1031064 | 2.10884273 | 0.222997435 | 9.660814071 | 4.42E-22 | 4.20E-20 |
| ENSG00000164125 | GASK1B | 359.3687 | 2.108335 | 0.3976911 | 5.671474 | 1.42E-08 | 2.52E-07 |
| ENSG00000286817 | | 3.872317 | 2.106768 | 1.461075 | 2.492183 | 0.01269605 | 0.047179 |
| ENSG00000288545 | | 7.815915 | 2.105425 | 0.9607785 | 2.972856 | 0.00295043 | 0.014394 |
| ENSG00000204248 | COL11A2 | 1000.069 | 2.105302 | 0.3024119 | 7.240283 | 4.48E-13 | 1.54E-11 |
| ENSG00000223547 | ZNF844 | 1.691436 | 2.104808 | 3.427413 | 2.447853 | 0.01437102 | 0.05214 |
| ENSG00000232656 | IDI2-AS1 | 572.8327 | 2.104013 | 0.7501513 | 3.504108 | 0.00045814 | 0.002909 |
| ENSG00000139899 | CBLN3 | 187.9116 | 2.100206 | 0.3803858 | 5.87041 | 4.35E-09 | 8.47E-08 |
| ENSG00000230133 | LINC01721 | 15.17276 | 2.099434 | 1.134102 | 2.550474 | 0.01075766 | 0.041268 |
| ENSG00000153250 | RBMS1 | 2794.434 | 2.097335 | 0.2401241 | 8.965593 | 3.09E-19 | 2.14E-17 |
| ENSG00000223813 | | 114.5596 | 2.096336 | 0.4238739 | 5.339887 | 9.30E-08 | 1.42E-06 |
| ENSG00000205838 | TTC23L | 70.2281 | 2.091237 | 0.4966121 | 4.641844 | 3.45E-06 | 3.82E-05 |
| ENSG00000204044 | SLC12A5-AS1 | 95.98174 | 2.090966 | 0.4280694 | 5.31564 | 1.06E-07 | 1.60E-06 |
| ENSG00000105419 | MEIS3 | 1350.091946 | 2.090530235 | 0.190114245 | 11.17913911 | 5.16E-29 | 9.58E-27 |
| ENSG00000226887 | ERVMER34-1 | 133.934 | 2.090349 | 0.4905485 | 4.742823 | 2.11E-06 | 2.47E-05 |

| | | | | | | | |
|---|---|---|---|---|---|---|---|
| ENSG00000287895 | | 12.50392 | 2.088675 | 0.8654441 | 3.122854 | 0.00179107 | 0.009442 |
| ENSG00000089820 | ARHGAP4 | 1671.342583 | 2.086242133 | 0.248196451 | 8.635223903 | 5.86E-18 | 3.53E-16 |
| ENSG00000164199 | ADGRV1 | 2279.725 | 2.084726 | 0.3146698 | 6.932019 | 4.15E-12 | 1.25E-10 |
| ENSG00000231441 | DST-AS1 | 10.01348 | 2.082231 | 0.6590069 | 3.696644 | 0.00021847 | 0.001534 |
| ENSG00000236283 | | 38.95974 | 2.080462 | 0.5240933 | 4.425536 | 9.62E-06 | 9.52E-05 |
| ENSG00000164938 | TP53INP1 | 8696.884 | 2.080009 | 0.4430003 | 5.112289 | 3.18E-07 | 4.36E-06 |
| ENSG00000204385 | SLC44A4 | 8.396376 | 2.079616 | 1.225976 | 2.584286 | 0.00975808 | 0.03811 |
| ENSG00000187185 | | 20.03374 | 2.079408 | 1.159249 | 2.523072 | 0.01163345 | 0.043926 |
| ENSG00000184343 | SRPK3 | 213.9644 | 2.078722 | 0.3715421 | 5.9488 | 2.70E-09 | 5.45E-08 |
| ENSG00000266145 | RHOT1P1 | 96.64578 | 2.078434 | 0.3625098 | 6.071095 | 1.27E-09 | 2.69E-08 |
| ENSG00000133433 | GSTT2B | 42.04851284 | 2.077631541 | 0.417286865 | 5.356340386 | 8.49E-08 | 1.30E-06 |
| ENSG00000234040 | RPL10P12 | 1.766169 | 2.076181 | 2.871973 | 2.481183 | 0.01309471 | 0.048363 |
| ENSG00000169860 | P2RY1 | 133.7219 | 2.075627 | 0.4307117 | 5.245283 | 1.56E-07 | 2.28E-06 |
| ENSG00000267121 | | 39.02171 | 2.074115 | 0.5143939 | 4.445153 | 8.78E-06 | 8.78E-05 |
| ENSG00000260081 | | 226.5639 | 2.073979 | 0.3122417 | 6.925429 | 4.35E-12 | 1.31E-10 |
| ENSG00000138814 | PPP3CA | 5993.98474 | 2.073186292 | 0.25421091 | 8.401419089 | 4.41E-17 | 2.43E-15 |
| ENSG00000198483 | ANKRD35 | 112.945 | 2.072744 | 0.9538606 | 3.046768 | 0.00231316 | 0.0117 |
| ENSG00000286367 | | 5.645674 | 2.070913 | 1.29287 | 2.44996 | 0.01428723 | 0.051874 |
| ENSG00000232471 | | 1.761057 | 2.070744 | 3.199907 | 2.484095 | 0.01298811 | 0.048028 |
| ENSG00000223861 | | 11.61688 | 2.070053 | 0.7104655 | 3.49577 | 0.0004727 | 0.002991 |
| ENSG00000169213 | RAB3B | 104.1998 | 2.067616 | 0.5729015 | 4.131007 | 3.61E-05 | 0.00031 |
| ENSG00000251273 | LINC02228 | 7.218101 | 2.066815 | 1.271269 | 2.452999 | 0.01416707 | 0.05151 |
| ENSG00000232774 | | 20.17033 | 2.066194 | 0.7183689 | 3.469952 | 0.00052055 | 0.00325 |
| ENSG00000126583 | PRKCG | 39.38627866 | 2.066131655 | 0.622380985 | 3.824041973 | 0.00013128 | 0.000975 |
| ENSG00000205078 | SYCE1L | 220.1916 | 2.063844 | 0.3926938 | 5.627443 | 1.83E-08 | 3.17E-07 |
| ENSG00000171116 | HSFX1 | 9.775545 | 2.06336 | 0.8652767 | 3.103542 | 0.00191219 | 0.009968 |
| ENSG00000260855 | | 270.568 | 2.060986 | 0.4657156 | 4.847436 | 1.25E-06 | 1.53E-05 |
| ENSG00000180353 | HCLS1 | 10.90626 | 2.058394 | 0.6322456 | 3.770435 | 0.00016296 | 0.001185 |
| ENSG00000257830 | | 35.45219 | 2.054359 | 0.8126113 | 3.246528 | 0.00116822 | 0.006544 |
| ENSG00000005243 | COPZ2 | 285.6517853 | 2.052399512 | 0.277366571 | 7.647466297 | 2.05E-14 | 8.18E-13 |
| ENSG00000163704 | PRRT3 | 2032.458 | 2.051907 | 0.2927856 | 7.289025 | 3.12E-13 | 1.09E-11 |
| ENSG00000279579 | LINC01666 | 8.697357 | 2.049809 | 0.7789962 | 3.245073 | 0.0011742 | 0.006563 |
| ENSG00000127954 | STEAP4 | 200.8687774 | 2.048005466 | 0.582388062 | 4.089215311 | 4.33E-05 | 0.000365 |

| | | | | | | | |
|---|---|---|---|---|---|---|---|
| ENSG00000140280 | LYSMD2 | 1593.232 | 2.046474 | 0.1966721 | 10.60077 | 2.96E-26 | 4.15E-24 |
| ENSG00000267767 | LINC01801 | 48.80047 | 2.044825 | 0.742108 | 3.434377 | 0.00059392 | 0.003643 |
| ENSG00000230037 | UBBP1 | 2.695361 | 2.044824 | 1.875781 | 2.484159 | 0.01298579 | 0.048028 |
| ENSG00000287914 | | 181.1791 | 2.043735 | 0.4603772 | 4.864601 | 1.15E-06 | 1.41E-05 |
| ENSG00000228022 | HCG20 | 6.480876 | 2.040636 | 1.092154 | 2.777437 | 0.00547895 | 0.023868 |
| ENSG00000123095 | BHLHE41 | 122.0444774 | 2.040425797 | 0.437351664 | 5.038316126 | 4.70E-07 | 6.26E-06 |
| ENSG00000183111 | ARHGEF37 | 659.7272 | 2.039766 | 0.4079816 | 5.381109 | 7.40E-08 | 1.15E-06 |
| ENSG00000155962 | CLIC2 | 106.4137 | 2.039288 | 0.4565624 | 4.876167 | 1.08E-06 | 1.34E-05 |
| ENSG00000088053 | GP6 | 10.75403802 | 2.035740622 | 1.029237846 | 2.689358278 | 0.00715895 | 0.029701 |
| ENSG00000267882 | | 53.40787 | 2.035066 | 0.4760224 | 4.648256 | 3.35E-06 | 3.73E-05 |
| ENSG00000261754 | | 80.18246 | 2.032424 | 0.4659788 | 4.78269 | 1.73E-06 | 2.06E-05 |
| ENSG00000287629 | | 5.184943 | 2.03239 | 0.9964227 | 2.837478 | 0.00454715 | 0.020508 |
| ENSG00000109758 | HGFAC | 63.95647096 | 2.031142498 | 0.521480636 | 4.374593183 | 1.22E-05 | 0.000117 |
| ENSG00000230882 | | 17.34525 | 2.030944 | 0.6348933 | 3.760411 | 0.00016964 | 0.001227 |
| ENSG00000286311 | | 17.09757 | 2.028896 | 0.7022603 | 3.51109 | 0.00044627 | 0.002843 |
| ENSG00000274414 | | 23.24957 | 2.028235 | 0.6978332 | 3.479464 | 0.00050242 | 0.003151 |
| ENSG00000007062 | PROM1 | 53.49090849 | 2.026436022 | 0.877667828 | 3.113161377 | 0.00185095 | 0.009705 |
| ENSG00000145451 | GLRA3 | 54.10594 | 2.022917 | 0.3240159 | 6.521039 | 6.98E-11 | 1.77E-09 |
| ENSG00000248409 | | 47.06583 | 2.021733 | 0.6033138 | 3.906879 | 9.35E-05 | 0.000721 |
| ENSG00000267920 | SNX6P1 | 9.118156 | 2.020697 | 0.7798362 | 3.194763 | 0.00139946 | 0.007647 |
| ENSG00000249116 | IRX4-AS1 | 54.77466 | 2.018593 | 0.4156228 | 5.24464 | 1.57E-07 | 2.28E-06 |
| ENSG00000170412 | GPRC5C | 1800.12 | 2.016401 | 0.2920613 | 7.180495 | 6.95E-13 | 2.31E-11 |
| ENSG00000204965 | PCDHA5 | 20.91109 | 2.015474 | 0.7049694 | 3.477599 | 0.00050593 | 0.003168 |
| ENSG00000025708 | TYMP | 110.4701409 | 2.015222029 | 0.400060224 | 5.385973605 | 7.21E-08 | 1.13E-06 |
| ENSG00000118922 | KLF12 | 434.5900809 | 2.014820109 | 0.216083739 | 9.524060622 | 1.67E-21 | 1.48E-19 |
| ENSG00000036530 | CYP46A1 | 97.1741518 | 2.013358488 | 0.434396568 | 5.031889455 | 4.86E-07 | 6.45E-06 |
| ENSG00000158560 | DYNC1I1 | 20.45309 | 2.013227 | 0.6778561 | 3.556944 | 0.00037519 | 0.002458 |
| ENSG00000165953 | SERPINA12 | 46.65509 | 2.011487 | 1.495228 | 2.6031 | 0.0092385 | 0.036501 |
| ENSG00000250412 | KLHL2P1 | 30.22124 | 2.010997 | 0.4310254 | 4.988533 | 6.08E-07 | 7.93E-06 |
| ENSG00000069431 | ABCC9 | 7.152979624 | 2.009015291 | 1.545031718 | 2.588214167 | 0.0096475 | 0.037739 |
| ENSG00000148488 | ST8SIA6 | 496.7605 | 2.007909 | 0.571377 | 4.036525 | 5.42E-05 | 0.000445 |
| ENSG00000101210 | EEF1A2 | 36596.2467 | 2.007863254 | 0.204140341 | 10.02979584 | 1.13E-23 | 1.22E-21 |
| ENSG00000287117 | | 1.97239 | 2.006362 | 3.637551 | 2.491337 | 0.01272632 | 0.047224 |

| | | | | | | | |
|---|---|---|---|---|---|---|---|
| ENSG00000250822 | LINC02111 | 3.634872 | 2.001786 | 1.938051 | 2.459558 | 0.01391082 | 0.050713 |
| ENSG00000186197 | EDARADD | 191.5068 | 2.00128 | 0.8680056 | 3.12043 | 0.00180588 | 0.009512 |
| ENSG00000156103 | MMP16 | 1452.82 | 2.000786 | 0.4715556 | 4.686668 | 2.78E-06 | 3.16E-05 |
| ENSG00000163017 | ACTG2 | 16.41924 | 2.000507 | 1.130608 | 2.766409 | 0.00566775 | 0.024565 |
| ENSG00000250331 | LINC01340 | 11.17285 | 1.998676 | 0.7283499 | 3.335756 | 0.00085068 | 0.004972 |
| ENSG00000244879 | GABPB1-AS1 | 8886.842 | 1.997283 | 0.3569104 | 5.936123 | 2.92E-09 | 5.83E-08 |
| ENSG00000227110 | LMCD1-AS1 | 49.67526 | 1.996202 | 0.4538599 | 4.820275 | 1.43E-06 | 1.74E-05 |
| ENSG00000160285 | LSS | 29748.66 | 1.99463 | 0.2607247 | 7.896501 | 2.87E-15 | 1.28E-13 |
| ENSG00000078401 | EDN1 | 1399.705514 | 1.994386168 | 0.47790992 | 4.628469065 | 3.68E-06 | 4.05E-05 |
| ENSG00000112139 | MDGA1 | 120.924836 | 1.994212364 | 1.004832064 | 2.892364755 | 0.00382354 | 0.017765 |
| ENSG00000270035 | | 14.52301 | 1.99373 | 0.7004465 | 3.413386 | 0.00064161 | 0.003896 |
| ENSG00000186212 | SOWAHB | 2501.89 | 1.99205 | 0.3817462 | 5.574356 | 2.48E-08 | 4.20E-07 |
| ENSG00000233029 | | 15.05542 | 1.991838 | 0.8292149 | 3.149816 | 0.00163373 | 0.008699 |
| ENSG00000240567 | LINC02067 | 61.57955 | 1.991063 | 0.4452402 | 4.892475 | 9.96E-07 | 1.24E-05 |
| ENSG00000275880 | | 41.48019 | 1.990881 | 0.6104537 | 3.845111 | 0.0001205 | 0.000903 |
| ENSG00000140470 | ADAMTS17 | 249.7169 | 1.989877 | 0.3154431 | 6.609083 | 3.87E-11 | 1.02E-09 |
| ENSG00000063587 | ZNF275 | 1150.153172 | 1.989471033 | 0.196906996 | 10.29943 | 7.09E-25 | 8.68E-23 |
| ENSG00000123999 | INHA | 202.0754815 | 1.987786202 | 0.332294895 | 6.289767558 | 3.18E-10 | 7.41E-09 |
| ENSG00000256385 | UBE2NP1 | 4.21323 | 1.98715 | 1.442213 | 2.39005 | 0.01684607 | 0.059188 |
| ENSG00000003436 | TFPI | 1391.738906 | 1.986955792 | 0.36110894 | 5.838303309 | 5.27E-09 | 1.01E-07 |
| ENSG00000031081 | ARHGAP31 | 203.0378727 | 1.985731551 | 0.409229404 | 5.232823237 | 1.67E-07 | 2.42E-06 |
| ENSG00000140937 | CDH11 | 80.28189 | 1.98316 | 0.5981885 | 3.884351 | 0.0001026 | 0.000782 |
| ENSG00000003147 | ICA1 | 4697.223775 | 1.982632715 | 0.174569221 | 11.5217547 | 1.03E-30 | 2.13E-28 |
| ENSG00000286017 | | 19.74561 | 1.9818 | 0.5890528 | 3.875701 | 0.00010632 | 0.000808 |
| ENSG00000212993 | POU5F1B | 58.8706 | 1.980993 | 0.3701219 | 5.701544 | 1.19E-08 | 2.14E-07 |
| ENSG00000273464 | | 37.37022 | 1.979146 | 0.6491288 | 3.630888 | 0.00028245 | 0.001918 |
| ENSG00000169851 | PCDH7 | 328.6927 | 1.978252 | 0.2607395 | 7.836754 | 4.62E-15 | 1.98E-13 |
| ENSG00000112530 | PACRG | 58.20960364 | 1.977540454 | 0.437579979 | 4.922056912 | 8.56E-07 | 1.08E-05 |
| ENSG00000007516 | BAIAP3 | 1250.325519 | 1.976435603 | 0.212860206 | 9.486634329 | 2.39E-21 | 2.09E-19 |
| ENSG00000284630 | | 35.77327 | 1.975102 | 0.4975198 | 4.412976 | 1.02E-05 | 0.0001 |
| ENSG00000276107 | | 21.94382 | 1.975086 | 0.9500059 | 2.970358 | 0.00297453 | 0.014472 |
| ENSG00000137819 | PAQR5 | 167.3326256 | 1.974573433 | 0.286234126 | 7.169383551 | 7.53E-13 | 2.49E-11 |
| ENSG00000232551 | MTCO1P14 | 12.7447 | 1.973566 | 0.8166759 | 3.088379 | 0.00201252 | 0.010413 |

| | | | | | | | |
|---|---|---|---|---|---|---|---|
| ENSG00000231703 | | 22.88574 | 1.973467 | 0.6150582 | 3.719015 | 0.0002 | 0.001418 |
| ENSG00000112379 | ARFGEF3 | 6425.058469 | 1.972750765 | 0.326972845 | 6.344583277 | 2.23E-10 | 5.31E-09 |
| ENSG00000152583 | SPARCL1 | 25.3639 | 1.970682 | 0.5276994 | 4.150202 | 3.32E-05 | 0.000288 |
| ENSG00000072182 | ASIC4 | 14.15430171 | 1.968088204 | 0.69359105 | 3.401733052 | 0.0006696 | 0.004045 |
| ENSG00000117643 | MAN1C1 | 236.8276195 | 1.966923952 | 0.352787592 | 5.896628524 | 3.71E-09 | 7.31E-08 |
| ENSG00000102032 | RENBP | 136.7499019 | 1.966898636 | 0.867038965 | 3.061423463 | 0.00220287 | 0.011234 |
| ENSG00000267136 | | 7.777969 | 1.96677 | 0.7741768 | 3.151697 | 0.00162325 | 0.008653 |
| ENSG00000171747 | LGALS4 | 10.32308 | 1.963904 | 0.8475823 | 2.971407 | 0.00296438 | 0.014444 |
| ENSG00000140859 | KIFC3 | 1525.001 | 1.963772 | 0.3892686 | 5.418107 | 6.02E-08 | 9.56E-07 |
| ENSG00000287271 | | 149.6308 | 1.962075 | 0.2860319 | 7.134859 | 9.69E-13 | 3.15E-11 |
| ENSG00000188766 | SPRED3 | 174.8379 | 1.958821 | 0.341387 | 6.042628 | 1.52E-09 | 3.16E-08 |
| ENSG00000262094 | | 9.611647 | 1.958681 | 1.212902 | 2.63318 | 0.00845895 | 0.034011 |
| ENSG00000282935 | DUX4L34 | 2.873621 | 1.9586 | 1.31559 | 2.372919 | 0.01764815 | 0.06143 |
| ENSG00000283433 | RNA5SP532 | 3.90908 | 1.958099 | 1.192265 | 2.594151 | 0.00948248 | 0.03722 |
| ENSG00000180066 | LINC02870 | 32.18492 | 1.955228 | 0.7754615 | 3.246416 | 0.00116868 | 0.006544 |
| ENSG00000100987 | VSX1 | 9.132122391 | 1.955112054 | 1.263638368 | 2.609370463 | 0.0090709 | 0.035971 |
| ENSG00000119535 | CSF3R | 5.049134301 | 1.952047364 | 1.294255091 | 2.539216595 | 0.0111101 | 0.042375 |
| ENSG00000152518 | ZFP36L2 | 8491.33 | 1.952046 | 0.3064154 | 6.671608 | 2.53E-11 | 6.84E-10 |
| ENSG00000261652 | C15orf65 | 56.12991 | 1.95148 | 0.4787812 | 4.520721 | 6.16E-06 | 6.43E-05 |
| ENSG00000114854 | TNNC1 | 58.8297803 | 1.951437092 | 0.562538805 | 3.986074167 | 6.72E-05 | 0.000537 |
| ENSG00000205038 | PKHD1L1 | 5.995381 | 1.950485 | 0.8688138 | 2.950916 | 0.00316833 | 0.015202 |
| ENSG00000130950 | NUTM2F | 7.227254687 | 1.947429623 | 1.075965483 | 2.581560429 | 0.00983548 | 0.038339 |
| ENSG00000260711 | | 143.4499 | 1.94341 | 0.4611804 | 4.646886 | 3.37E-06 | 3.75E-05 |
| ENSG00000271614 | ATP2B1-AS1 | 242.2904 | 1.942321 | 0.3452447 | 5.956204 | 2.58E-09 | 5.22E-08 |
| ENSG00000161405 | IKZF3 | 9.148904 | 1.938864 | 1.100798 | 2.580425 | 0.00986789 | 0.038454 |
| ENSG00000177459 | ERICH5 | 37.56215 | 1.937582 | 0.6892659 | 3.421267 | 0.0006233 | 0.0038 |
| ENSG00000287185 | | 29.29413 | 1.936568 | 0.6963889 | 3.317605 | 0.00090793 | 0.005251 |
| ENSG00000141505 | ASGR1 | 22.42356 | 1.935316 | 0.6209874 | 3.597414 | 0.0003214 | 0.00215 |
| ENSG00000174498 | IGDCC3 | 1027.473 | 1.935058 | 0.2904248 | 6.944081 | 3.81E-12 | 1.17E-10 |
| ENSG00000073146 | MOV10L1 | 13.21814447 | 1.933770272 | 1.183620046 | 2.664230064 | 0.00771648 | 0.031609 |
| ENSG00000104549 | SQLE | 24694.56243 | 1.933306909 | 0.265398166 | 7.548212021 | 4.41E-14 | 1.70E-12 |
| ENSG00000197892 | KIF13B | 1380.913 | 1.933246 | 0.1867177 | 10.55847 | 4.64E-26 | 6.44E-24 |
| ENSG00000162711 | NLRP3 | 4.577581 | 1.932232 | 1.18527 | 2.404927 | 0.01617571 | 0.057219 |

| | | | | | | | |
|---|---|---|---|---|---|---|---|
| ENSG00000275322 | | 70.35362 | 1.931041 | 0.5148658 | 4.249665 | 2.14E-05 | 0.000194 |
| ENSG00000120875 | DUSP4 | 4897.464499 | 1.929879332 | 0.79842395 | 3.170851276 | 0.00151993 | 0.008206 |
| ENSG00000187210 | GCNT1 | 974.4522 | 1.927993 | 0.3672162 | 5.604536 | 2.09E-08 | 3.58E-07 |
| ENSG00000095321 | CRAT | 260.4871348 | 1.92717493 | 0.583628127 | 3.870802924 | 0.00010848 | 0.000822 |
| ENSG00000167772 | ANGPTL4 | 566.0525 | 1.926654 | 0.4231443 | 4.97719 | 6.45E-07 | 8.37E-06 |
| ENSG00000169710 | FASN | 137612.3 | 1.924582 | 0.3546796 | 5.763107 | 8.26E-09 | 1.54E-07 |
| ENSG00000267472 | ARHGAP27P2 | 28.15313 | 1.922044 | 0.8476174 | 3.065501 | 0.00217306 | 0.011109 |
| ENSG00000120278 | PLEKHG1 | 115.5097059 | 1.920655143 | 0.488518905 | 4.295876017 | 1.74E-05 | 0.000162 |
| ENSG00000115474 | KCNJ13 | 12.05448152 | 1.918827547 | 0.654422332 | 3.500982341 | 0.00046355 | 0.00294 |
| ENSG00000114251 | WNT5A | 300.8377103 | 1.917985765 | 0.586901496 | 3.821649511 | 0.00013256 | 0.000982 |
| ENSG00000163508 | EOMES | 380.8822 | 1.91648 | 0.1996965 | 9.790354 | 1.24E-22 | 1.25E-20 |
| ENSG00000140284 | SLC27A2 | 1930.152 | 1.915298 | 0.2477658 | 7.964434 | 1.66E-15 | 7.56E-14 |
| ENSG00000139832 | RAB20 | 1662.343 | 1.914095 | 0.3181336 | 6.328075 | 2.48E-10 | 5.87E-09 |
| ENSG00000241829 | RPL21P54 | 4.133887 | 1.912062 | 1.83126 | 2.372158 | 0.01768454 | 0.061521 |
| ENSG00000156959 | LHFPL4 | 7.038853 | 1.91031 | 0.971302 | 2.792853 | 0.00522455 | 0.022971 |
| ENSG00000135643 | KCNMB4 | 822.8749671 | 1.909937951 | 0.41572596 | 4.998241312 | 5.79E-07 | 7.59E-06 |
| ENSG00000227740 | LINC02803 | 17.41251 | 1.909868 | 0.6117039 | 3.662189 | 0.00025007 | 0.001724 |
| ENSG00000160179 | ABCG1 | 6165.553 | 1.909167 | 0.2582905 | 7.652196 | 1.98E-14 | 7.89E-13 |
| ENSG00000128298 | BAIAP2L2 | 72.40822547 | 1.906405517 | 0.39652681 | 5.164892992 | 2.41E-07 | 3.39E-06 |
| ENSG00000236830 | CBR3-AS1 | 479.4265 | 1.904783 | 0.2530151 | 7.779833 | 7.26E-15 | 3.06E-13 |
| ENSG00000196993 | NPIPB9 | 16.02171 | 1.903413 | 0.8164446 | 3.008346 | 0.00262674 | 0.013045 |
| ENSG00000257261 | | 271.4026 | 1.901336 | 0.1930776 | 10.08715 | 6.30E-24 | 7.00E-22 |
| ENSG00000275897 | | 20.47384 | 1.899659 | 0.5114828 | 4.162375 | 3.15E-05 | 0.000274 |
| ENSG00000111886 | GABRR2 | 93.96743608 | 1.898547402 | 0.685511131 | 3.412271033 | 0.00064424 | 0.00391 |
| ENSG00000176153 | GPX2 | 533.3167 | 1.897949 | 0.5527395 | 3.981232 | 6.86E-05 | 0.000547 |
| ENSG00000170442 | KRT86 | 1506.51 | 1.896851 | 0.4711557 | 4.489275 | 7.15E-06 | 7.33E-05 |
| ENSG00000109814 | UGDH | 5584.303131 | 1.896134747 | 0.168808266 | 11.40030515 | 4.17E-30 | 8.20E-28 |
| ENSG00000184524 | CEND1 | 15.54435 | 1.896116 | 0.8480106 | 2.988825 | 0.00280053 | 0.013792 |
| ENSG00000110934 | BIN2 | 9.006223886 | 1.893386722 | 1.116800359 | 2.596807467 | 0.00940947 | 0.03701 |
| ENSG00000224577 | LINC01117 | 26.82947 | 1.887969 | 0.4380311 | 4.714059 | 2.43E-06 | 2.80E-05 |
| ENSG00000258676 | | 46.82059 | 1.886278 | 0.5613712 | 3.884075 | 0.00010272 | 0.000782 |
| ENSG00000188306 | LRRIQ4 | 3.049366 | 1.882138 | 1.444917 | 2.43254 | 0.01499335 | 0.053898 |
| ENSG00000118156 | ZNF541 | 57.73655285 | 1.882036815 | 0.525789341 | 4.082954362 | 4.45E-05 | 0.000374 |

| | | | | | | | |
|---|---|---|---|---|---|---|---|
| ENSG00000163132 | MSX1 | 116.7679 | 1.88172 | 0.3985967 | 5.112827 | 3.17E-07 | 4.36E-06 |
| ENSG00000280032 | | 13.20996 | 1.880536 | 1.606461 | 2.425566 | 0.01528454 | 0.054722 |
| ENSG00000224081 | SLC44A3-AS1 | 66.5647 | 1.880172 | 0.3854292 | 5.245513 | 1.56E-07 | 2.28E-06 |
| ENSG00000091128 | LAMB4 | 2.907354411 | 1.877939553 | 1.407159479 | 2.390839023 | 0.01680992 | 0.059093 |
| ENSG00000213901 | SLC23A3 | 39.3618 | 1.877768 | 0.5545717 | 3.887122 | 0.00010144 | 0.000775 |
| ENSG00000169116 | PARM1 | 262.6581 | 1.87181 | 0.4652089 | 4.462641 | 8.10E-06 | 8.18E-05 |
| ENSG00000263089 | | 24.73931 | 1.870351 | 0.6397522 | 3.516239 | 0.00043771 | 0.002797 |
| ENSG00000236671 | PRKG1-AS1 | 113.4721 | 1.86757 | 0.3484443 | 5.68463 | 1.31E-08 | 2.34E-07 |
| ENSG00000172497 | ACOT12 | 3.323694 | 1.866219 | 1.851438 | 2.303301 | 0.02126194 | 0.071417 |
| ENSG00000287089 | | 4.783173 | 1.865353 | 0.9594011 | 2.734344 | 0.00625046 | 0.026592 |
| ENSG00000234996 | ACTG1P25 | 195.9669 | 1.864871 | 0.3132483 | 6.264883 | 3.73E-10 | 8.63E-09 |
| ENSG00000004939 | SLC4A1 | 14.56463931 | 1.862829545 | 0.865537514 | 2.919367292 | 0.00350743 | 0.016546 |
| ENSG00000273216 | | 7.312654 | 1.861982 | 0.9072696 | 2.832629 | 0.0046167 | 0.020787 |
| ENSG00000100100 | PIK3IP1 | 1302.798662 | 1.860873883 | 0.457349948 | 4.507705024 | 6.55E-06 | 6.78E-05 |
| ENSG00000166068 | SPRED1 | 579.9189 | 1.860816 | 0.289247 | 6.722838 | 1.78E-11 | 4.93E-10 |
| ENSG00000154479 | CCDC173 | 9.78772 | 1.860656 | 0.9581269 | 2.794689 | 0.00519496 | 0.022859 |
| ENSG00000196220 | SRGAP3 | 493.4742 | 1.858427 | 0.3646142 | 5.436341 | 5.44E-08 | 8.70E-07 |
| ENSG00000172379 | ARNT2 | 1982.945 | 1.85835 | 0.2545164 | 7.550297 | 4.34E-14 | 1.68E-12 |
| ENSG00000235269 | LINC02331 | 37.0171 | 1.85808 | 1.060353 | 2.6814 | 0.00733149 | 0.030294 |
| ENSG00000159263 | SIM2 | 479.5372 | 1.857786 | 0.2034817 | 9.334293 | 1.02E-20 | 8.09E-19 |
| ENSG00000275029 | HMGB1P24 | 2.006441 | 1.856016 | 2.670495 | 2.426192 | 0.01525821 | 0.054647 |
| ENSG00000165996 | HACD1 | 926.7438 | 1.855036 | 0.192324 | 9.834333 | 8.01E-23 | 8.18E-21 |
| ENSG00000172247 | C1QTNF4 | 5.330987 | 1.854904 | 1.814943 | 2.398171 | 0.01647717 | 0.058123 |
| ENSG00000257279 | | 10.44607 | 1.853601 | 0.952406 | 2.843243 | 0.0044657 | 0.020185 |
| ENSG00000184530 | C6orf58 | 9.03906 | 1.853023 | 0.8091315 | 2.964093 | 0.00303577 | 0.014689 |
| ENSG00000143416 | SELENBP1 | 6736.997 | 1.850623 | 0.3869533 | 5.161327 | 2.45E-07 | 3.45E-06 |
| ENSG00000259083 | | 14.0163 | 1.850055 | 0.585506 | 3.682633 | 0.00023084 | 0.001608 |
| ENSG00000099840 | IZUMO4 | 215.5299777 | 1.84987584 | 0.340902157 | 5.754865092 | 8.67E-09 | 1.61E-07 |
| ENSG00000267254 | ZNF790-AS1 | 284.4608 | 1.849333 | 0.3057649 | 6.336271 | 2.35E-10 | 5.58E-09 |
| ENSG00000174640 | SLCO2A1 | 183.5309 | 1.84915 | 0.5260372 | 4.022752 | 5.75E-05 | 0.000469 |
| ENSG00000140279 | DUOX2 | 8.802719 | 1.848605 | 0.9665586 | 2.756169 | 0.00584827 | 0.025216 |
| ENSG00000255129 | | 12.80511 | 1.84855 | 1.067501 | 2.533841 | 0.01128198 | 0.042907 |
| ENSG00000137941 | TTLL7 | 318.1333143 | 1.845443349 | 0.328996564 | 5.933340233 | 2.97E-09 | 5.93E-08 |

| | | | | | | | |
|---|---|---|---|---|---|---|---|
| ENSG00000217026 | RPL10P1 | 7.356472 | 1.842534 | 0.873752 | 2.798908 | 0.00512758 | 0.022609 |
| ENSG00000120211 | INSL4 | 13.56712424 | 1.842360975 | 1.000680998 | 2.72249905 | 0.00647902 | 0.027348 |
| ENSG00000279118 | | 15.83109 | 1.842188 | 1.002115 | 2.675781 | 0.00745554 | 0.030721 |
| ENSG00000237505 | PKN2-AS1 | 33.28762 | 1.842067 | 0.5310964 | 3.999722 | 6.34E-05 | 0.000512 |
| ENSG00000143515 | ATP8B2 | 4791.915 | 1.840758 | 0.3289406 | 5.912399 | 3.37E-09 | 6.70E-08 |
| ENSG00000156886 | ITGAD | 7.606924 | 1.835767 | 0.7693617 | 3.047125 | 0.00231042 | 0.011688 |
| ENSG00000144485 | HES6 | 1448.226 | 1.834988 | 0.3818387 | 5.175904 | 2.27E-07 | 3.22E-06 |
| ENSG00000165272 | AQP3 | 2641.484 | 1.834615 | 0.521796 | 4.018567 | 5.86E-05 | 0.000476 |
| ENSG00000154309 | DISP1 | 243.1908 | 1.832837 | 0.376298 | 5.222231 | 1.77E-07 | 2.55E-06 |
| ENSG00000248858 | | 34.5386 | 1.83213 | 0.4696484 | 4.341731 | 1.41E-05 | 0.000134 |
| ENSG00000166813 | KIF7 | 456.3875 | 1.83207 | 0.2459576 | 7.686256 | 1.52E-14 | 6.14E-13 |
| ENSG00000279127 | | 8.1066 | 1.830306 | 1.064948 | 2.576066 | 0.00999315 | 0.03884 |
| ENSG00000272273 | IER3-AS1 | 7.395564 | 1.829879 | 1.032225 | 2.509565 | 0.01208799 | 0.045272 |
| ENSG00000244405 | ETV5 | 188.6791 | 1.82955 | 0.5506328 | 3.876057 | 0.00010616 | 0.000807 |
| ENSG00000111424 | VDR | 1208.23331 | 1.823806728 | 0.253494683 | 7.443199406 | 9.83E-14 | 3.65E-12 |
| ENSG00000080493 | SLC4A4 | 28.42041763 | 1.823518898 | 1.223957724 | 2.508137912 | 0.01213693 | 0.045405 |
| ENSG00000230074 | | 75.2102 | 1.823109 | 0.3958718 | 4.947711 | 7.51E-07 | 9.59E-06 |
| ENSG00000114270 | COL7A1 | 1228.678689 | 1.821669618 | 0.399807455 | 4.939906746 | 7.82E-07 | 9.95E-06 |
| ENSG00000260731 | KRT8P22 | 13.08763 | 1.821476 | 1.154273 | 2.620697 | 0.00877503 | 0.035049 |
| ENSG00000267475 | | 181.992 | 1.820912 | 0.3768294 | 5.199313 | 2.00E-07 | 2.87E-06 |
| ENSG00000073417 | PDE8A | 3768.206783 | 1.817893683 | 0.230309871 | 8.121210507 | 4.62E-16 | 2.27E-14 |
| ENSG00000167508 | MVD | 15028.68 | 1.817561 | 0.3288114 | 5.854953 | 4.77E-09 | 9.22E-08 |
| ENSG00000246777 | | 4.960477 | 1.817533 | 1.073467 | 2.587669 | 0.00966277 | 0.037788 |
| ENSG00000187244 | BCAM | 8442.715 | 1.814728 | 0.2542432 | 7.388157 | 1.49E-13 | 5.38E-12 |
| ENSG00000137834 | SMAD6 | 1859.022771 | 1.814408855 | 0.452163975 | 4.457169669 | 8.30E-06 | 8.37E-05 |
| ENSG00000267938 | EIF1P6 | 4.544325 | 1.812355 | 0.9638951 | 2.647976 | 0.00809753 | 0.032851 |
| ENSG00000113580 | NR3C1 | 3587.771495 | 1.811297548 | 0.250532431 | 7.488183902 | 6.98E-14 | 2.64E-12 |
| ENSG00000137868 | STRA6 | 122.3957965 | 1.807899961 | 0.837941761 | 2.880108114 | 0.00397539 | 0.018363 |
| ENSG00000105642 | KCNN1 | 8.011852804 | 1.805571207 | 0.789337025 | 2.989023697 | 0.0027987 | 0.013785 |
| ENSG00000166033 | HTRA1 | 677.9376 | 1.805176 | 0.3970098 | 4.939511 | 7.83E-07 | 9.95E-06 |
| ENSG00000067064 | IDI1 | 14399.84935 | 1.804229763 | 0.380661557 | 5.096234767 | 3.46E-07 | 4.71E-06 |
| ENSG00000160282 | FTCD | 74.15685 | 1.804198 | 0.3650975 | 5.269348 | 1.37E-07 | 2.02E-06 |
| ENSG00000133687 | TMTC1 | 328.1741813 | 1.802950826 | 0.53089489 | 3.909327366 | 9.26E-05 | 0.000714 |

| | | | | | | | |
|---|---|---|---|---|---|---|---|
| ENSG00000198753 | PLXNB3 | 1107.687 | 1.802811 | 0.2195174 | 8.423168 | 3.66E-17 | 2.03E-15 |
| ENSG00000078804 | TP53INP2 | 368.5385292 | 1.802105581 | 0.361492546 | 5.337413201 | 9.43E-08 | 1.43E-06 |
| ENSG00000230423 | | 26.33427 | 1.801509 | 0.8138427 | 2.973694 | 0.00294238 | 0.014363 |
| ENSG00000110169 | HPX | 112.2576645 | 1.801215113 | 0.43843991 | 4.518365739 | 6.23E-06 | 6.49E-05 |
| ENSG00000134207 | SYT6 | 6.373692075 | 1.797441648 | 1.536941331 | 2.314485764 | 0.02064109 | 0.069693 |
| ENSG00000270976 | | 6.63769 | 1.796845 | 1.02642 | 2.5393 | 0.01110744 | 0.042371 |
| ENSG00000169064 | ZBBX | 9.364566 | 1.796792 | 0.9964516 | 2.522333 | 0.01165793 | 0.043986 |
| ENSG00000166448 | TMEM130 | 7.605405 | 1.794873 | 1.023915 | 2.552691 | 0.01068942 | 0.041077 |
| ENSG00000274817 | | 21.42949 | 1.793998 | 0.6057385 | 3.492419 | 0.00047867 | 0.003022 |
| ENSG00000132623 | ANKEF1 | 942.8538221 | 1.792257227 | 0.162697648 | 11.18017544 | 5.10E-29 | 9.54E-27 |
| ENSG00000232022 | FAAHP1 | 45.35899 | 1.79213 | 0.5674761 | 3.692903 | 0.00022171 | 0.001553 |
| ENSG00000133943 | DGLUCY | 283.1425314 | 1.791579569 | 0.312028738 | 6.034253047 | 1.60E-09 | 3.31E-08 |
| ENSG00000168071 | CCDC88B | 135.5872 | 1.789016 | 0.3422636 | 5.548917 | 2.87E-08 | 4.80E-07 |
| ENSG00000198189 | HSD17B11 | 387.9787 | 1.788366 | 0.4153714 | 4.707681 | 2.51E-06 | 2.88E-05 |
| ENSG00000109181 | UGT2B10 | 1.722986028 | 1.787664523 | 3.087824481 | 2.359846104 | 0.01828252 | 0.06323 |
| ENSG00000205869 | KRTAP5-1 | 84.30379 | 1.78628 | 0.4182978 | 4.666352 | 3.07E-06 | 3.45E-05 |
| ENSG00000109452 | INPP4B | 9291.103084 | 1.784630429 | 0.197464507 | 9.22889215 | 2.73E-20 | 2.10E-18 |
| ENSG00000148082 | SHC3 | 69.42059 | 1.784066 | 0.4130253 | 4.716879 | 2.39E-06 | 2.77E-05 |
| ENSG00000225422 | RBMS1P1 | 324.3237 | 1.783361 | 0.2890417 | 6.449018 | 1.13E-10 | 2.79E-09 |
| ENSG00000231995 | | 8.320128 | 1.782902 | 0.7499168 | 3.033598 | 0.00241656 | 0.012146 |
| ENSG00000131746 | TNS4 | 66.63900699 | 1.781115547 | 0.923593283 | 2.8032691 | 0.00505874 | 0.022367 |
| ENSG00000133256 | PDE6B | 247.522391 | 1.781009689 | 0.237346814 | 7.721450106 | 1.15E-14 | 4.71E-13 |
| ENSG00000273077 | | 5.506585 | 1.78 | 1.921537 | 2.370225 | 0.01777729 | 0.06179 |
| ENSG00000176716 | OR10AB1P | 2.559071 | 1.779309 | 3.125868 | 2.426803 | 0.01523251 | 0.054588 |
| ENSG00000109705 | NKX3-2 | 116.9865707 | 1.779233685 | 0.440515014 | 4.448417738 | 8.65E-06 | 8.67E-05 |
| ENSG00000178803 | ADORA2A-AS1 | 31.3091 | 1.778985 | 0.9186249 | 2.7824 | 0.00539584 | 0.023575 |
| ENSG00000104899 | AMH | 746.8407347 | 1.778808397 | 0.307100555 | 6.101597815 | 1.05E-09 | 2.24E-08 |
| ENSG00000138798 | EGF | 19.51346736 | 1.776790245 | 0.694019327 | 3.200742404 | 0.00137074 | 0.007514 |
| ENSG00000223652 | | 126.9623 | 1.776162 | 0.2682985 | 6.889486 | 5.60E-12 | 1.66E-10 |
| ENSG00000234405 | | 10.67214 | 1.775653 | 0.8038568 | 2.878415 | 0.0039968 | 0.01843 |
| ENSG00000149269 | PAK1 | 1157.489 | 1.772442 | 0.3045076 | 6.12761 | 8.92E-10 | 1.93E-08 |
| ENSG00000149485 | FADS1 | 7196.969 | 1.772341 | 0.3140008 | 5.947987 | 2.71E-09 | 5.47E-08 |
| ENSG00000196267 | ZNF836 | 283.7078 | 1.770876 | 0.2973876 | 6.242543 | 4.31E-10 | 9.84E-09 |

| | | | | | | | |
|---|---|---|---|---|---|---|---|
| ENSG00000110693 | SOX6 | 160.2592223 | 1.770435264 | 0.453236568 | 4.337888622 | 1.44E-05 | 0.000136 |
| ENSG00000247570 | SDCBPP2 | 4.454675 | 1.770199 | 1.239198 | 2.323712 | 0.02014095 | 0.06836 |
| ENSG00000267128 | RNF157-AS1 | 11.1291 | 1.767363 | 0.7650179 | 2.970794 | 0.00297031 | 0.014464 |
| ENSG00000233073 | GRM3-AS1 | 7.339766 | 1.7662 | 0.8038039 | 2.922743 | 0.00346963 | 0.016392 |
| ENSG00000285872 | | 65.1242 | 1.76545 | 0.2805979 | 6.563524 | 5.26E-11 | 1.36E-09 |
| ENSG00000073150 | PANX2 | 797.3332709 | 1.764195557 | 0.333303467 | 5.625289415 | 1.85E-08 | 3.20E-07 |
| ENSG00000277443 | MARCKS | 8278.574 | 1.763932 | 0.33824 | 5.543779 | 2.96E-08 | 4.92E-07 |
| ENSG00000264201 | MIR4701 | 10.89591 | 1.763655 | 0.7115174 | 3.112381 | 0.00185585 | 0.009724 |
| ENSG00000286379 | | 18.14089 | 1.763609 | 0.6713807 | 3.256689 | 0.0011272 | 0.006346 |
| ENSG00000115457 | IGFBP2 | 2916.552085 | 1.762911009 | 0.362910896 | 5.214687298 | 1.84E-07 | 2.65E-06 |
| ENSG00000231528 | FAM225A | 80.25569 | 1.762447 | 0.4397038 | 4.46031 | 8.18E-06 | 8.26E-05 |
| ENSG00000286337 | | 70.15715 | 1.760686 | 0.4255538 | 4.545437 | 5.48E-06 | 5.78E-05 |
| ENSG00000161835 | TAMALIN | 26.03237 | 1.760486 | 0.6930843 | 3.149857 | 0.00163351 | 0.008699 |
| ENSG00000267778 | | 17.82908 | 1.760344 | 0.5382588 | 3.780519 | 0.0001565 | 0.001141 |
| ENSG00000258689 | LINC01269 | 12.60336 | 1.759947 | 0.9320282 | 2.63866 | 0.00832345 | 0.033583 |
| ENSG00000172382 | PRSS27 | 264.5428 | 1.759896 | 0.3208228 | 5.799083 | 6.67E-09 | 1.26E-07 |
| ENSG00000230798 | FOXD3-AS1 | 183.1084 | 1.759451 | 0.2524926 | 7.196259 | 6.19E-13 | 2.07E-11 |
| ENSG00000121075 | TBX4 | 117.564907 | 1.758643483 | 0.419722177 | 4.587974001 | 4.48E-06 | 4.84E-05 |
| ENSG00000266714 | MYO15B | 4247.557 | 1.758629 | 0.3207701 | 5.793489 | 6.89E-09 | 1.30E-07 |
| ENSG00000185269 | NOTUM | 52.50169 | 1.758242 | 0.8058312 | 2.904068 | 0.00368349 | 0.01721 |
| ENSG00000161243 | FBXO27 | 650.5548 | 1.757514 | 0.2995354 | 6.158555 | 7.34E-10 | 1.61E-08 |
| ENSG00000004776 | HSPB6 | 4.187943457 | 1.757022279 | 1.151078514 | 2.519771275 | 0.01174311 | 0.04424 |
| ENSG00000242021 | | 4.950787 | 1.756209 | 1.413012 | 2.39819 | 0.01647633 | 0.058123 |
| ENSG00000165474 | GJB2 | 93.92771 | 1.7539 | 1.171949 | 2.58515 | 0.00973366 | 0.038031 |
| ENSG00000141449 | GREB1L | 817.6939 | 1.752825 | 0.351237 | 5.33655 | 9.47E-08 | 1.44E-06 |
| ENSG00000271576 | | 30.13549 | 1.752349 | 0.44274 | 4.383716 | 1.17E-05 | 0.000113 |
| ENSG00000132470 | ITGB4 | 11331.45728 | 1.751636966 | 0.269702954 | 6.757446349 | 1.40E-11 | 3.95E-10 |
| ENSG00000166173 | LARP6 | 543.796 | 1.750344 | 0.3236037 | 5.719419 | 1.07E-08 | 1.96E-07 |
| ENSG00000066468 | FGFR2 | 1539.817939 | 1.748925199 | 0.543508131 | 3.743888543 | 0.00018119 | 0.001299 |
| ENSG00000257337 | | 2081.663 | 1.745231 | 0.2744542 | 6.623854 | 3.50E-11 | 9.24E-10 |
| ENSG00000164951 | PDP1 | 1595.904 | 1.745063 | 0.22424 | 8.004613 | 1.20E-15 | 5.57E-14 |
| ENSG00000110975 | SYT10 | 313.6696411 | 1.742023373 | 0.383650967 | 4.902642468 | 9.46E-07 | 1.18E-05 |
| ENSG00000151067 | CACNA1C | 19.17898 | 1.74172 | 0.9351839 | 2.695159 | 0.0070355 | 0.029257 |

| | | | | | | | |
|---|---|---|---|---|---|---|---|
| ENSG00000164056 | SPRY1 | 1500.42 | 1.7416 | 0.2721448 | 6.67524 | 2.47E-11 | 6.68E-10 |
| ENSG00000230212 | | 15.85993 | 1.740922 | 0.6970095 | 3.114347 | 0.00184352 | 0.009674 |
| ENSG00000076641 | PAG1 | 92.09429704 | 1.740599129 | 0.582280091 | 3.553911092 | 0.00037955 | 0.002483 |
| ENSG00000287637 | | 8.219772 | 1.739113 | 0.6371117 | 3.276757 | 0.00105007 | 0.005973 |
| ENSG00000111077 | TNS2 | 1191.481545 | 1.738459782 | 0.294565338 | 6.181997635 | 6.33E-10 | 1.41E-08 |
| ENSG00000225339 | | 65.16253 | 1.737218 | 0.5762975 | 3.555509 | 0.00037725 | 0.002469 |
| ENSG00000166268 | MYRFL | 68.15224 | 1.736546 | 1.315858 | 2.512867 | 0.01197544 | 0.044965 |
| ENSG00000267909 | CCDC177 | 4.018031 | 1.735779 | 1.372866 | 2.442383 | 0.01459065 | 0.052764 |
| ENSG00000145920 | CPLX2 | 4.722039 | 1.735137 | 1.03078 | 2.570043 | 0.0101686 | 0.039429 |
| ENSG00000258871 | | 9.148424 | 1.7338 | 1.039186 | 2.55115 | 0.0107368 | 0.041212 |
| ENSG00000228728 | PPIAP91 | 6.545071 | 1.731643 | 1.179118 | 2.432854 | 0.01498035 | 0.053874 |
| ENSG00000262619 | LINC00621 | 34.06531 | 1.730513 | 0.4860448 | 4.020442 | 5.81E-05 | 0.000473 |
| ENSG00000136243 | NUP42 | 998.0432726 | 1.730478669 | 0.265068669 | 6.779974954 | 1.20E-11 | 3.42E-10 |
| ENSG00000065717 | TLE2 | 5683.952691 | 1.729039826 | 0.169845903 | 10.38796162 | 2.81E-25 | 3.61E-23 |
| ENSG00000151062 | CACNA2D4 | 14.37205 | 1.727446 | 0.5750938 | 3.540381 | 0.00039955 | 0.002589 |
| ENSG00000180818 | HOXC10 | 122.5624 | 1.726775 | 0.242829 | 7.333201 | 2.25E-13 | 7.99E-12 |
| ENSG00000205426 | KRT81 | 7111.851 | 1.726734 | 0.9243388 | 2.751028 | 0.00594087 | 0.025537 |
| ENSG00000258545 | RHOXF1-AS1 | 5.575242 | 1.726585 | 1.052484 | 2.509771 | 0.01208096 | 0.045259 |
| ENSG00000185052 | SLC24A3 | 947.4656 | 1.725047 | 0.3885064 | 4.814025 | 1.48E-06 | 1.79E-05 |
| ENSG00000259388 | | 10.79317 | 1.724116 | 0.6644128 | 3.197804 | 0.00138478 | 0.007578 |
| ENSG00000179846 | NKPD1 | 141.2115 | 1.723736 | 0.4386966 | 4.343901 | 1.40E-05 | 0.000133 |
| ENSG00000222035 | KIAA2012-AS1 | 15.51417 | 1.722409 | 0.5423043 | 3.679027 | 0.00023413 | 0.00163 |
| ENSG00000257829 | | 31.60162 | 1.722383 | 0.4261669 | 4.409947 | 1.03E-05 | 0.000102 |
| ENSG00000174804 | FZD4 | 2655.869 | 1.720392 | 0.269457 | 6.640046 | 3.14E-11 | 8.37E-10 |
| ENSG00000128606 | LRRC17 | 21.56285603 | 1.718996713 | 0.513783238 | 3.802267529 | 0.00014338 | 0.001053 |
| ENSG00000232448 | | 4.156819 | 1.718825 | 1.773471 | 2.2774 | 0.02276236 | 0.075401 |
| ENSG00000235100 | | 30.25506 | 1.71871 | 0.4327576 | 4.360643 | 1.30E-05 | 0.000124 |
| ENSG00000175895 | PLEKHF2 | 5262.976 | 1.717602 | 0.1744733 | 10.02719 | 1.16E-23 | 1.25E-21 |
| ENSG00000136535 | TBR1 | 2.978431428 | 1.717507384 | 1.264290497 | 2.372128281 | 0.01768595 | 0.061521 |
| ENSG00000101400 | SNTA1 | 379.5448001 | 1.717112714 | 0.404658481 | 4.656956891 | 3.21E-06 | 3.59E-05 |
| ENSG00000260727 | SLC7A5P1 | 125.5147 | 1.71686 | 0.2838736 | 6.312519 | 2.75E-10 | 6.44E-09 |
| ENSG00000232762 | | 2.593952 | 1.716703 | 3.097279 | 2.435411 | 0.0148749 | 0.05357 |
| ENSG00000250602 | | 10.05253 | 1.716307 | 0.8250413 | 2.766431 | 0.00566735 | 0.024565 |

| | | | | | | | |
|---|---|---|---|---|---|---|---|
| ENSG00000185585 | OLFML2A | 41.18019 | 1.716039 | 0.634041 | 3.315388 | 0.00091516 | 0.005288 |
| ENSG00000183929 | DUSP5P1 | 8.587951 | 1.715943 | 1.150853 | 2.491413 | 0.0127236 | 0.047223 |
| ENSG00000247982 | LINC00926 | 8.795222 | 1.712229 | 0.752518 | 2.940981 | 0.00327175 | 0.015614 |
| ENSG00000275494 | | 121.2589 | 1.712039 | 0.359797 | 5.102941 | 3.34E-07 | 4.57E-06 |
| ENSG00000226709 | FGF12-AS3 | 7.564278 | 1.711896 | 0.8280948 | 2.804024 | 0.00504691 | 0.022326 |
| ENSG00000261115 | TMEM178B | 1449.235 | 1.710377 | 0.3259602 | 5.56356 | 2.64E-08 | 4.45E-07 |
| ENSG00000095739 | BAMBI | 14937.75703 | 1.70806914 | 0.240148768 | 7.401861849 | 1.34E-13 | 4.89E-12 |
| ENSG00000224063 | | 31.59359 | 1.707849 | 0.7711447 | 2.954486 | 0.0031319 | 0.015054 |
| ENSG00000112972 | HMGCS1 | 9900.56224 | 1.707721047 | 0.342322989 | 5.321130696 | 1.03E-07 | 1.56E-06 |
| ENSG00000197180 | | 351.7454 | 1.70696 | 0.2224601 | 7.952932 | 1.82E-15 | 8.24E-14 |
| ENSG00000135253 | KCP | 25.28669621 | 1.706802977 | 1.004082145 | 2.637191018 | 0.00835957 | 0.033698 |
| ENSG00000144214 | LYG1 | 55.10464 | 1.706685 | 0.7023328 | 3.090571 | 0.00199772 | 0.010342 |
| ENSG00000275212 | LINC02826 | 21.75794 | 1.706381 | 0.515745 | 3.764826 | 0.00016667 | 0.001208 |
| ENSG00000168280 | KIF5C | 3003.967 | 1.705949 | 0.2651692 | 6.697146 | 2.13E-11 | 5.80E-10 |
| ENSG00000175879 | HOXD8 | 28.43081 | 1.705606 | 0.5651288 | 3.541518 | 0.00039783 | 0.00258 |
| ENSG00000172638 | EFEMP2 | 142.9371 | 1.705555 | 0.3016101 | 5.947899 | 2.72E-09 | 5.47E-08 |
| ENSG00000227295 | ELL2P1 | 30.20963 | 1.705276 | 0.4456661 | 4.224612 | 2.39E-05 | 0.000215 |
| ENSG00000139174 | PRICKLE1 | 269.4245346 | 1.705273149 | 0.573280586 | 3.533786871 | 0.00040965 | 0.002645 |
| ENSG00000230069 | LRRC37A15P | 9.753764 | 1.705039 | 0.6468315 | 3.207512 | 0.00133889 | 0.007374 |
| ENSG00000234719 | NPIPB2 | 149.7089 | 1.704487 | 0.4326152 | 4.350546 | 1.36E-05 | 0.000129 |
| ENSG00000265962 | GACAT2 | 20.14323 | 1.702869 | 0.821571 | 2.824446 | 0.00473625 | 0.02121 |
| ENSG00000224769 | MUC20P1 | 120.3672 | 1.702757 | 0.6413078 | 3.255138 | 0.00113337 | 0.006373 |
| ENSG00000108018 | SORCS1 | 28.20706398 | 1.701192889 | 0.619903456 | 3.319860503 | 0.00090062 | 0.005218 |
| ENSG00000260757 | | 17.86993 | 1.700879 | 0.6005175 | 3.367057 | 0.00075975 | 0.004515 |
| ENSG00000146205 | ANO7 | 38.09142 | 1.699724 | 0.358152 | 5.094475 | 3.50E-07 | 4.75E-06 |
| ENSG00000106524 | ANKMY2 | 1132.042069 | 1.699086986 | 0.242034337 | 7.254322667 | 4.04E-13 | 1.39E-11 |
| ENSG00000120896 | SORBS3 | 1784.989575 | 1.698238058 | 0.217816934 | 8.012205087 | 1.13E-15 | 5.26E-14 |
| ENSG00000061918 | GUCY1B1 | 315.8816804 | 1.698108646 | 0.438007529 | 4.309123999 | 1.64E-05 | 0.000153 |
| ENSG00000059377 | TBXAS1 | 52.74506412 | 1.697341537 | 0.528463495 | 3.727278131 | 0.00019356 | 0.001376 |
| ENSG00000279741 | | 22.43076 | 1.697027 | 0.5489031 | 3.597023 | 0.00032188 | 0.002153 |
| ENSG00000256443 | | 112.5272 | 1.696721 | 0.405458 | 4.582699 | 4.59E-06 | 4.94E-05 |
| ENSG00000224216 | | 6.226952 | 1.696512 | 1.019594 | 2.542879 | 0.01099433 | 0.042006 |
| ENSG00000204314 | PRRT1 | 165.5614 | 1.696496 | 0.3141335 | 5.745003 | 9.19E-09 | 1.71E-07 |

| | | | | | | | |
|---|---|---|---|---|---|---|---|
| ENSG00000213468 | FIRRE | 793.9119 | 1.695558 | 0.3154684 | 5.680553 | 1.34E-08 | 2.39E-07 |
| ENSG00000257951 | | 55.24353 | 1.694154 | 0.4157878 | 4.450367 | 8.57E-06 | 8.60E-05 |
| ENSG00000149582 | TMEM25 | 512.1457 | 1.693884 | 0.2186799 | 7.957262 | 1.76E-15 | 7.97E-14 |
| ENSG00000251513 | LIX1-AS1 | 17.27394 | 1.693261 | 0.6301699 | 3.246259 | 0.00116932 | 0.006545 |
| ENSG00000135709 | KIAA0513 | 3929.028347 | 1.690539147 | 0.41022478 | 4.516291869 | 6.29E-06 | 6.55E-05 |
| ENSG00000111834 | RSPH4A | 16.37052274 | 1.685857756 | 0.628173105 | 3.271099894 | 0.0010713 | 0.006069 |
| ENSG00000287527 | | 5.662495 | 1.685262 | 2.336302 | 2.387178 | 0.01697826 | 0.059558 |
| ENSG00000288560 | | 11.28494 | 1.683243 | 0.8105593 | 2.794359 | 0.00520028 | 0.022877 |
| ENSG00000109339 | MAPK10 | 166.1370464 | 1.681535899 | 0.366681655 | 4.930651959 | 8.20E-07 | 1.04E-05 |
| ENSG00000241935 | HOGA1 | 14.62153 | 1.680574 | 0.799051 | 2.863308 | 0.00419243 | 0.019188 |
| ENSG00000282933 | RHOXF1P3 | 44.19293 | 1.67962 | 0.8758657 | 2.620694 | 0.00877509 | 0.035049 |
| ENSG00000169762 | TAPT1 | 1706.499 | 1.675407 | 0.2221224 | 7.758873 | 8.57E-15 | 3.58E-13 |
| ENSG00000162188 | GNG3 | 23.92987 | 1.673874 | 0.5108789 | 3.735633 | 0.00018724 | 0.001338 |
| ENSG00000243422 | RPL23AP49 | 12.97399 | 1.671853 | 0.6476876 | 3.1518 | 0.00162268 | 0.008652 |
| ENSG00000231830 | | 1.906617 | 1.670912 | 3.146044 | 2.420779 | 0.01548727 | 0.055307 |
| ENSG00000004848 | ARX | 114.5078123 | 1.668857068 | 0.305401557 | 5.738966969 | 9.53E-09 | 1.76E-07 |
| ENSG00000254338 | MAFA-AS1 | 57.36145 | 1.668839 | 0.4886445 | 3.891295 | 9.97E-05 | 0.000763 |
| ENSG00000248840 | | 1.541411 | 1.668792 | 2.055939 | 2.252411 | 0.0242963 | NA |
| ENSG00000250237 | | 11.06839 | 1.668771 | 0.8711819 | 2.664074 | 0.00772006 | 0.031619 |
| ENSG00000259153 | | 477.7709 | 1.668055 | 0.2435128 | 7.092181 | 1.32E-12 | 4.24E-11 |
| ENSG00000151715 | TMEM45B | 1375.264 | 1.667928 | 0.4028837 | 4.547448 | 5.43E-06 | 5.73E-05 |
| ENSG00000159753 | CARMIL2 | 154.3407 | 1.666161 | 0.3590454 | 4.996727 | 5.83E-07 | 7.64E-06 |
| ENSG00000147383 | NSDHL | 3612.305 | 1.666126 | 0.2549424 | 6.782459 | 1.18E-11 | 3.36E-10 |
| ENSG00000167766 | ZNF83 | 4190.718 | 1.664548 | 0.2494259 | 6.916552 | 4.63E-12 | 1.38E-10 |
| ENSG00000131171 | SH3BGRL | 8134.067534 | 1.663790388 | 0.348620809 | 5.114782949 | 3.14E-07 | 4.31E-06 |
| ENSG00000164398 | ACSL6 | 3.122251 | 1.663648 | 1.702393 | 2.328475 | 0.01988691 | 0.067652 |
| ENSG00000179104 | TMTC2 | 1468.883 | 1.663345 | 0.2558154 | 6.751553 | 1.46E-11 | 4.10E-10 |
| ENSG00000255693 | LINC02389 | 7.700663 | 1.663029 | 0.8767444 | 2.702795 | 0.00687591 | 0.02872 |
| ENSG00000027075 | PRKCH | 2772.064165 | 1.662776648 | 0.202268818 | 8.404870522 | 4.28E-17 | 2.36E-15 |
| ENSG00000141441 | GAREM1 | 626.4874 | 1.662326 | 0.34111 | 5.201666 | 1.98E-07 | 2.83E-06 |
| ENSG00000168939 | SPRY3 | 207.7925 | 1.661791 | 0.3206746 | 5.480262 | 4.25E-08 | 6.92E-07 |
| ENSG00000253210 | | 169.3371 | 1.661115 | 0.2552674 | 6.741441 | 1.57E-11 | 4.37E-10 |
| ENSG00000112697 | TMEM30A | 14899.46748 | 1.660594817 | 0.204423566 | 8.314469599 | 9.22E-17 | 4.91E-15 |

| | | | | | | | |
|---|---|---|---|---|---|---|---|
| ENSG00000013725 | CD6 | 7.280450996 | 1.659044773 | 1.04144552 | 2.384400157 | 0.01710699 | 0.059892 |
| ENSG00000201134 | | 6.462753 | 1.658906 | 0.9300816 | 2.631688 | 0.00849619 | 0.03412 |
| ENSG00000250942 | ENPP7P11 | 10.45483 | 1.657225 | 0.7132453 | 2.957921 | 0.00309721 | 0.014921 |
| ENSG00000119686 | FLVCR2 | 1105.107981 | 1.655699979 | 0.266038199 | 6.481987715 | 9.05E-11 | 2.26E-09 |
| ENSG00000167964 | RAB26 | 2472.613 | 1.655317 | 0.3033006 | 5.754945 | 8.67E-09 | 1.61E-07 |
| ENSG00000272720 | | 85.71172 | 1.655032 | 0.4213736 | 4.324483 | 1.53E-05 | 0.000144 |
| ENSG00000070371 | CLTCL1 | 587.0166765 | 1.65456608 | 0.36402754 | 4.891571746 | 1.00E-06 | 1.24E-05 |
| ENSG00000288597 | | 20.74021 | 1.654411 | 0.6363309 | 3.174073 | 0.00150316 | 0.008123 |
| ENSG00000279720 | SOWAHCP2 | 52.07877 | 1.651918 | 0.5514649 | 3.514184 | 0.00044111 | 0.002816 |
| ENSG00000067191 | CACNB1 | 597.2460538 | 1.650947703 | 0.251933288 | 6.798265096 | 1.06E-11 | 3.03E-10 |
| ENSG00000256875 | | 2.17715 | 1.650859 | 3.436773 | 2.424962 | 0.01530997 | 0.054806 |
| ENSG00000130477 | UNC13A | 15.7135525 | 1.649829207 | 0.996535418 | 2.529862648 | 0.01141072 | 0.043248 |
| ENSG00000159899 | NPR2 | 261.1467 | 1.648096 | 0.290891 | 5.943265 | 2.79E-09 | 5.61E-08 |
| ENSG00000126561 | STAT5A | 109.9835642 | 1.647312486 | 0.469556797 | 3.95738286 | 7.58E-05 | 0.000599 |
| ENSG00000049089 | COL9A2 | 589.2377615 | 1.646423955 | 0.318410636 | 5.489032961 | 4.04E-08 | 6.60E-07 |
| ENSG00000116701 | NCF2 | 472.2481471 | 1.64571638 | 0.574652255 | 3.419594027 | 0.00062715 | 0.003821 |
| ENSG00000175544 | CABP4 | 4.058541 | 1.643757 | 1.353507 | 2.31749 | 0.02047705 | 0.069253 |
| ENSG00000137312 | FLOT1 | 7050.932764 | 1.642927001 | 0.219671792 | 7.699303289 | 1.37E-14 | 5.58E-13 |
| ENSG00000134716 | CYP2J2 | 748.1736291 | 1.6420187 | 0.204195279 | 8.247236299 | 1.62E-16 | 8.44E-15 |
| ENSG00000250286 | | 32.75955 | 1.639307 | 0.4495259 | 4.071989 | 4.66E-05 | 0.000389 |
| ENSG00000237208 | | 11.43618 | 1.639045 | 0.7316982 | 2.905339 | 0.00366855 | 0.017163 |
| ENSG00000212994 | RPS26P6 | 5.285093 | 1.638602 | 1.18208 | 2.375778 | 0.01751198 | 0.061068 |
| ENSG00000196353 | CPNE4 | 160.098 | 1.637536 | 0.5653665 | 3.446299 | 0.00056832 | 0.003506 |
| ENSG00000030419 | IKZF2 | 1114.737094 | 1.636202927 | 0.333620148 | 5.222165059 | 1.77E-07 | 2.55E-06 |
| ENSG00000187140 | FOXD3 | 78.48057 | 1.635696 | 0.5683579 | 3.419258 | 0.00062792 | 0.003824 |
| ENSG00000185551 | NR2F2 | 7006.204 | 1.635499 | 0.2297098 | 7.317925 | 2.52E-13 | 8.92E-12 |
| ENSG00000197191 | CYSRT1 | 530.1837 | 1.635058 | 0.3261296 | 5.327147 | 9.98E-08 | 1.51E-06 |
| ENSG00000225043 | RPL18AP2 | 7.684664 | 1.634639 | 0.7600379 | 2.803411 | 0.00505652 | 0.022361 |
| ENSG00000116141 | MARK1 | 224.8750043 | 1.633796338 | 0.589024078 | 3.332973716 | 0.00085923 | 0.005012 |
| ENSG00000241359 | SYNPR-AS1 | 82.11228 | 1.633218 | 0.371694 | 4.74057 | 2.13E-06 | 2.49E-05 |
| ENSG00000255050 | | 95.3392 | 1.632476 | 0.4518772 | 4.071346 | 4.67E-05 | 0.00039 |
| ENSG00000170522 | ELOVL6 | 4234.887 | 1.630931 | 0.4156595 | 4.332665 | 1.47E-05 | 0.000139 |
| ENSG00000182568 | SATB1 | 847.1362 | 1.630084 | 0.5463984 | 3.506291 | 0.0004544 | 0.002887 |

| | | | | | | | |
|---|---|---|---|---|---|---|---|
| ENSG00000233429 | HOTAIRM1 | 56.00934 | 1.629809 | 0.5042841 | 3.670264 | 0.0002423 | 0.00168 |
| ENSG00000074590 | NUAK1 | 1893.196218 | 1.62641803 | 0.305422957 | 5.627265579 | 1.83E-08 | 3.17E-07 |
| ENSG00000262223 | | 98.40992 | 1.624727 | 0.4761702 | 3.887295 | 0.00010137 | 0.000775 |
| ENSG00000124006 | OBSL1 | 8187.184004 | 1.624464343 | 0.162039119 | 10.1884286 | 2.23E-24 | 2.64E-22 |
| ENSG00000250995 | TMEM30BP1 | 35.35009 | 1.624381 | 0.4216442 | 4.223908 | 2.40E-05 | 0.000216 |
| ENSG00000272449 | | 251.5979 | 1.624158 | 0.3091897 | 5.551882 | 2.83E-08 | 4.73E-07 |
| ENSG00000179348 | GATA2 | 2120.558 | 1.622589 | 0.1875557 | 8.88715 | 6.27E-19 | 4.20E-17 |
| ENSG00000162746 | FCRLB | 119.4048 | 1.621668 | 0.3048642 | 5.595386 | 2.20E-08 | 3.76E-07 |
| ENSG00000232618 | | 9.730046 | 1.619963 | 0.8296039 | 2.643811 | 0.00819784 | 0.033132 |
| ENSG00000226970 | NEDD8P1 | 12.57462 | 1.619405 | 0.5871807 | 3.265298 | 0.00109349 | 0.006183 |
| ENSG00000115461 | IGFBP5 | 11714.12321 | 1.618648728 | 0.701663536 | 2.982560287 | 0.00285848 | 0.014023 |
| ENSG00000173546 | CSPG4 | 132.2888 | 1.618067 | 0.3674926 | 4.74824 | 2.05E-06 | 2.41E-05 |
| ENSG00000102524 | TNFSF13B | 20.82141604 | 1.617264758 | 0.557033798 | 3.415182858 | 0.00063739 | 0.003877 |
| ENSG00000161298 | ZNF382 | 77.39483 | 1.614951 | 0.5341495 | 3.530397 | 0.00041494 | 0.002673 |
| ENSG00000285331 | | 40.51427 | 1.614842 | 0.4328603 | 4.158397 | 3.20E-05 | 0.000279 |
| ENSG00000154898 | CCDC144CP | 75.4795 | 1.614127 | 0.7102073 | 2.952674 | 0.00315034 | 0.015125 |
| ENSG00000248202 | LINC02234 | 14.12133 | 1.614026 | 0.5753129 | 3.324385 | 0.00088614 | 0.00515 |
| ENSG00000184557 | SOCS3 | 1382.54 | 1.613897 | 0.4207209 | 4.243778 | 2.20E-05 | 0.000199 |
| ENSG00000250820 | | 4.991288 | 1.613216 | 1.223767 | 2.363318 | 0.01811213 | 0.062765 |
| ENSG00000173281 | PPP1R3B | 1110.579 | 1.612516 | 0.2977051 | 5.703151 | 1.18E-08 | 2.13E-07 |
| ENSG00000100985 | MMP9 | 76.88341006 | 1.612348956 | 0.43666156 | 4.131881406 | 3.60E-05 | 0.000309 |
| ENSG00000257137 | LINC02874 | 14.45843 | 1.610263 | 0.7674116 | 2.772149 | 0.00556876 | 0.024196 |
| ENSG00000165966 | PDZRN4 | 2.735644 | 1.609115 | 2.192866 | 2.241626 | 0.02498552 | 0.081075 |
| ENSG00000147526 | TACC1 | 12748.12 | 1.608427 | 0.2785758 | 6.021669 | 1.73E-09 | 3.56E-08 |
| ENSG00000163364 | LINC01116 | 58.07353 | 1.606547 | 0.3818395 | 4.575085 | 4.76E-06 | 5.10E-05 |
| ENSG00000234338 | | 92.79697 | 1.605341 | 0.5483469 | 3.455192 | 0.0005499 | 0.003412 |
| ENSG00000254352 | | 12.68148 | 1.605324 | 0.636062 | 3.108692 | 0.00187917 | 0.009823 |
| ENSG00000128310 | GALR3 | 3.980734171 | 1.604427139 | 1.352433935 | 2.328436921 | 0.01988892 | 0.067652 |
| ENSG00000196684 | HSH2D | 943.7135 | 1.60422 | 0.3287916 | 5.204392 | 1.95E-07 | 2.80E-06 |
| ENSG00000026652 | AGPAT4 | 91.56809499 | 1.603146594 | 0.333382647 | 5.129103398 | 2.91E-07 | 4.02E-06 |
| ENSG00000185070 | FLRT2 | 7.262893 | 1.603117 | 0.9530146 | 2.426629 | 0.01523984 | 0.054607 |
| ENSG00000090534 | THPO | 30.94942161 | 1.602552789 | 0.667044022 | 3.021154309 | 0.00251813 | 0.012583 |
| ENSG00000196139 | AKR1C3 | 2639.252 | 1.601797 | 0.5835675 | 3.303752 | 0.000954 | 0.005486 |

| | | | | | | | |
|---|---|---|---|---|---|---|---|
| ENSG00000211895 | IGHA1 | 11.07097 | 1.597643 | 1.54925 | 2.255636 | 0.02409341 | 0.078796 |
| ENSG00000154237 | LRRK1 | 371.1499 | 1.59733 | 0.2763536 | 6.03038 | 1.64E-09 | 3.39E-08 |
| ENSG00000163297 | ANTXR2 | 558.0813 | 1.596103 | 0.3933272 | 4.438244 | 9.07E-06 | 9.04E-05 |
| ENSG00000167920 | TMEM99 | 673.6121 | 1.595139 | 0.2488951 | 6.643974 | 3.05E-11 | 8.16E-10 |
| ENSG00000092964 | DPYSL2 | 1141.238796 | 1.594546122 | 0.374098884 | 4.633165038 | 3.60E-06 | 3.98E-05 |
| ENSG00000131183 | SLC34A1 | 1.506264548 | 1.593229645 | 3.187725752 | 2.390801886 | 0.01681162 | NA |
| ENSG00000081138 | CDH7 | 90.40238124 | 1.592278791 | 0.554140848 | 3.413558376 | 0.0006412 | 0.003895 |
| ENSG00000081148 | IMPG2 | 21.36709331 | 1.591609848 | 0.649035412 | 3.052712866 | 0.00226783 | 0.011508 |
| ENSG00000232903 | LINC01166 | 59.09508 | 1.590575 | 0.5783225 | 3.274579 | 0.0010582 | 0.006009 |
| ENSG00000010310 | GIPR | 252.4511396 | 1.590379248 | 0.551208385 | 3.422670841 | 0.00062009 | 0.003784 |
| ENSG00000112561 | TFEB | 322.0159979 | 1.590232251 | 0.257845474 | 6.409167862 | 1.46E-10 | 3.57E-09 |
| ENSG00000183625 | CCR3 | 11.46536 | 1.590101 | 0.9333929 | 2.56366 | 0.0103575 | 0.040033 |
| ENSG00000128052 | KDR | 25.03042699 | 1.589878296 | 1.43134683 | 2.181114467 | 0.02917495 | 0.091842 |
| ENSG00000285651 | | 10.21687 | 1.589842 | 0.7951013 | 2.720368 | 0.00652093 | 0.027505 |
| ENSG00000065923 | SLC9A7 | 3848.443691 | 1.589793828 | 0.219960884 | 7.423741327 | 1.14E-13 | 4.18E-12 |
| ENSG00000235770 | LINC00607 | 23.54722 | 1.589729 | 0.6968328 | 2.942793 | 0.00325266 | 0.01554 |
| ENSG00000275454 | | 36.64705 | 1.589699 | 0.5192576 | 3.544509 | 0.00039335 | 0.002557 |
| ENSG00000172461 | FUT9 | 2172.656 | 1.589471 | 0.2776837 | 5.99104 | 2.09E-09 | 4.26E-08 |
| ENSG00000228630 | HOTAIR | 764.9455 | 1.587963 | 0.2712892 | 6.107152 | 1.01E-09 | 2.18E-08 |
| ENSG00000264175 | MIR3189 | 58.24007 | 1.586886 | 0.5483363 | 3.38436 | 0.00071344 | 0.004274 |
| ENSG00000258655 | ARHGAP5-AS1 | 178.2968 | 1.585027 | 0.2418407 | 6.77712 | 1.23E-11 | 3.48E-10 |
| ENSG00000207563 | MIR23B | 6.116801 | 1.584659 | 1.00645 | 2.438767 | 0.01473745 | 0.053165 |
| ENSG00000233449 | MTATP6P14 | 4.654684 | 1.584477 | 1.100617 | 2.286612 | 0.02221846 | 0.074115 |
| ENSG00000217648 | | 77.76683 | 1.58391 | 0.3565185 | 4.760664 | 1.93E-06 | 2.27E-05 |
| ENSG00000187589 | TERF1P2 | 7.547205 | 1.583713 | 0.8882576 | 2.540451 | 0.01107095 | 0.042244 |
| ENSG00000265246 | | 104.4241 | 1.583358 | 0.2796885 | 5.926673 | 3.09E-09 | 6.16E-08 |
| ENSG00000272954 | | 32.3646 | 1.58311 | 0.5021127 | 3.61252 | 0.00030324 | 0.002042 |
| ENSG00000155961 | RAB39B | 1179.549 | 1.582996 | 0.3612827 | 4.730134 | 2.24E-06 | 2.61E-05 |
| ENSG00000070961 | ATP2B1 | 9123.577341 | 1.582110924 | 0.217318897 | 7.491854582 | 6.79E-14 | 2.57E-12 |
| ENSG00000280537 | | 17.20601 | 1.582042 | 0.6328735 | 3.076094 | 0.00209731 | 0.010787 |
| ENSG00000173467 | AGR3 | 19.17383 | 1.580993 | 1.257258 | 2.382797 | 0.01718168 | 0.06009 |
| ENSG00000134765 | DSC1 | 36.96831959 | 1.580984627 | 0.927866528 | 2.4758038 | 0.01329366 | 0.048865 |
| ENSG00000152767 | FARP1 | 2439.211 | 1.580675 | 0.3048343 | 5.49057 | 4.01E-08 | 6.55E-07 |

| | | | | | | | |
|---|---|---|---|---|---|---|---|
| ENSG00000225556 | C2CD4D | 58.30621 | 1.579306 | 0.4891842 | 3.701434 | 0.00021439 | 0.001508 |
| ENSG00000198453 | ZNF568 | 77.35452 | 1.578002 | 0.5531816 | 3.381178 | 0.00072176 | 0.004314 |
| ENSG00000160097 | FNDC5 | 48.83032 | 1.577462 | 0.5667031 | 3.339981 | 0.00083784 | 0.00491 |
| ENSG00000244998 | | 13.39185 | 1.577414 | 0.8629641 | 2.586574 | 0.00969355 | 0.037897 |
| ENSG00000257135 | ODC1-DT | 14.44204 | 1.577043 | 0.9391282 | 2.55316 | 0.01067505 | 0.04104 |
| ENSG00000229914 | | 3.0011 | 1.576488 | 1.540001 | 2.180943 | 0.02918762 | 0.09186 |
| ENSG00000102387 | TAF7L | 71.52160678 | 1.575914966 | 0.407819306 | 4.243557077 | 2.20E-05 | 0.000199 |
| ENSG00000135373 | EHF | 1227.816771 | 1.572385416 | 0.327703241 | 5.13048362 | 2.89E-07 | 4.00E-06 |
| ENSG00000234690 | EPCAM-DT | 13.55069 | 1.571883 | 0.610179 | 3.113108 | 0.00185128 | 0.009705 |
| ENSG00000135926 | TMBIM1 | 603.8085043 | 1.571720769 | 0.523373873 | 3.522289898 | 0.00042784 | 0.002743 |
| ENSG00000134324 | LPIN1 | 6954.757138 | 1.570595796 | 0.325502089 | 5.15708179 | 2.51E-07 | 3.52E-06 |
| ENSG00000071859 | FAM50A | 9498.018103 | 1.56987251 | 0.17795084 | 8.994593551 | 2.37E-19 | 1.65E-17 |
| ENSG00000189409 | MMP23B | 5.093804 | 1.569196 | 1.352799 | 2.213443 | 0.0268671 | 0.085859 |
| ENSG00000258982 | | 23.75628 | 1.567112 | 0.7622058 | 2.782823 | 0.00538883 | 0.023549 |
| ENSG00000184408 | KCND2 | 61.64133 | 1.566487 | 0.8965574 | 2.577709 | 0.00994576 | 0.038689 |
| ENSG00000261504 | LINC01686 | 9.583106 | 1.565822 | 1.032657 | 2.42573 | 0.01527764 | 0.054705 |
| ENSG00000131069 | ACSS2 | 4660.430325 | 1.563069862 | 0.243374599 | 6.64896341 | 2.95E-11 | 7.90E-10 |
| ENSG00000270001 | | 11.01053 | 1.560203 | 0.7789234 | 2.735518 | 0.00622823 | 0.026515 |
| ENSG00000167081 | PBX3 | 1329.726 | 1.559829 | 0.2030528 | 7.881923 | 3.22E-15 | 1.43E-13 |
| ENSG00000138100 | TRIM54 | 27.40784014 | 1.559545499 | 0.643511479 | 3.029002943 | 0.00245362 | 0.012307 |
| ENSG00000280027 | | 40.16305 | 1.559352 | 0.4128585 | 4.156862 | 3.23E-05 | 0.000281 |
| ENSG00000169251 | NMD3 | 16357.25 | 1.558686 | 0.1973224 | 8.090126 | 5.96E-16 | 2.88E-14 |
| ENSG00000272808 | | 1162.302 | 1.558187 | 0.392262 | 4.366828 | 1.26E-05 | 0.000121 |
| ENSG00000067840 | PDZD4 | 1419.158004 | 1.556243423 | 0.234453848 | 6.871728549 | 6.34E-12 | 1.86E-10 |
| ENSG00000138759 | FRAS1 | 2542.406675 | 1.556039013 | 0.434351126 | 3.995397714 | 6.46E-05 | 0.00052 |
| ENSG00000169247 | SH3TC2 | 145.7722 | 1.55598 | 0.4836227 | 3.690797 | 0.00022355 | 0.001564 |
| ENSG00000174473 | GALNTL6 | 103.889 | 1.554569 | 0.3266677 | 5.07535 | 3.87E-07 | 5.22E-06 |
| ENSG00000112902 | SEMA5A | 1828.786668 | 1.554450346 | 0.578615187 | 3.247275473 | 0.00116516 | 0.006534 |
| ENSG00000283235 | | 11.38632 | 1.55395 | 0.8233076 | 2.682758 | 0.00730179 | 0.030185 |
| ENSG00000179532 | DNHD1 | 781.2097 | 1.552964 | 0.3189084 | 5.179694 | 2.22E-07 | 3.15E-06 |
| ENSG00000146216 | TTBK1 | 24.44769 | 1.551567 | 0.5640836 | 3.273313 | 0.00106295 | 0.006032 |
| ENSG00000105974 | CAV1 | 1477.310319 | 1.550873486 | 0.773999142 | 2.747802006 | 0.00599962 | 0.025731 |
| ENSG00000225882 | LINC01456 | 338.6856 | 1.550655 | 0.3268845 | 5.056866 | 4.26E-07 | 5.73E-06 |

| | | | | | | | |
|---|---|---|---|---|---|---|---|
| ENSG00000128923 | MINDY2 | 1066.875301 | 1.55041038 | 0.328963531 | 5.031042333 | 4.88E-07 | 6.47E-06 |
| ENSG00000140993 | TIGD7 | 195.8775 | 1.548173 | 0.3252497 | 5.072341 | 3.93E-07 | 5.29E-06 |
| ENSG00000215386 | MIR99AHG | 285.6404 | 1.547247 | 0.3769632 | 4.463909 | 8.05E-06 | 8.14E-05 |
| ENSG00000185436 | IFNLR1 | 333.8007 | 1.546763 | 0.3304512 | 5.003182 | 5.64E-07 | 7.41E-06 |
| ENSG00000133083 | DCLK1 | 619.2302235 | 1.546454317 | 0.466041481 | 3.745765019 | 0.00017985 | 0.001291 |
| ENSG00000108924 | HLF | 186.6871065 | 1.54542531 | 0.404498951 | 4.199979977 | 2.67E-05 | 0.000237 |
| ENSG00000136695 | IL36RN | 9.08264702 | 1.545342964 | 1.749065954 | 2.202167539 | 0.02765348 | 0.087844 |
| ENSG00000102119 | EMD | 5759.759242 | 1.545127661 | 0.207020823 | 7.687641284 | 1.50E-14 | 6.09E-13 |
| ENSG00000139865 | TTC6 | 550.3018 | 1.544685 | 0.2797607 | 5.793826 | 6.88E-09 | 1.30E-07 |
| ENSG00000278558 | TMEM191B | 164.1599 | 1.544208 | 0.5509638 | 3.344023 | 0.00082573 | 0.004852 |
| ENSG00000186231 | KLHL32 | 32.71378 | 1.543744 | 0.5006005 | 3.54584 | 0.00039137 | 0.002547 |
| ENSG00000009694 | TENM1 | 274.7842533 | 1.543609895 | 0.531769469 | 3.417418301 | 0.00063218 | 0.003848 |
| ENSG00000111860 | CEP85L | 222.9816508 | 1.543098649 | 0.319054869 | 5.144180714 | 2.69E-07 | 3.75E-06 |
| ENSG00000067798 | NAV3 | 42.22122731 | 1.541473774 | 0.958628541 | 2.508743768 | 0.01211613 | 0.045346 |
| ENSG00000130164 | LDLR | 10051.14111 | 1.541473315 | 0.370783977 | 4.514767756 | 6.34E-06 | 6.59E-05 |
| ENSG00000058091 | CDK14 | 372.630249 | 1.540035386 | 0.334005017 | 4.938339389 | 7.88E-07 | 1.00E-05 |
| ENSG00000157064 | NMNAT2 | 831.4824 | 1.5397 | 0.1660292 | 9.429035 | 4.14E-21 | 3.56E-19 |
| ENSG00000167034 | NKX3-1 | 228.3075 | 1.537576 | 0.2333248 | 6.815782 | 9.38E-12 | 2.70E-10 |
| ENSG00000072195 | SPEG | 306.913766 | 1.536721896 | 0.405727122 | 4.180976212 | 2.90E-05 | 0.000255 |
| ENSG00000272732 | | 196.1936 | 1.536254 | 0.3553588 | 4.664913 | 3.09E-06 | 3.47E-05 |
| ENSG00000284606 | | 19.08181 | 1.53597 | 0.6189006 | 3.007324 | 0.00263559 | 0.01308 |
| ENSG00000257002 | | 8.606533 | 1.535384 | 0.8047309 | 2.59506 | 0.00945745 | 0.037154 |
| ENSG00000198892 | SHISA4 | 459.163 | 1.532976 | 0.3206509 | 5.096792 | 3.45E-07 | 4.70E-06 |
| ENSG00000133636 | NTS | 20.8889014 | 1.531324313 | 0.654178042 | 2.918121119 | 0.00352148 | 0.016603 |
| ENSG00000069974 | RAB27A | 899.0737036 | 1.530743902 | 0.161360602 | 9.665117779 | 4.24E-22 | 4.04E-20 |
| ENSG00000186326 | RGS9BP | 87.23955 | 1.529455 | 0.2761941 | 5.792687 | 6.93E-09 | 1.30E-07 |
| ENSG00000065413 | ANKRD44 | 249.4826031 | 1.52912882 | 0.260177582 | 6.152850187 | 7.61E-10 | 1.67E-08 |
| ENSG00000204963 | PCDHA7 | 107.3042 | 1.528301 | 0.2708768 | 5.892645 | 3.80E-09 | 7.48E-08 |
| ENSG00000181649 | PHLDA2 | 1412.06 | 1.528137 | 0.3778191 | 4.416045 | 1.01E-05 | 9.90E-05 |
| ENSG00000225032 | | 62.14148 | 1.528134 | 0.3975916 | 4.222557 | 2.42E-05 | 0.000217 |
| ENSG00000236296 | GUSBP5 | 56.44994 | 1.525967 | 0.5160676 | 3.458611 | 0.00054297 | 0.003375 |
| ENSG00000278540 | ACACA | 11703.33 | 1.525194 | 0.283841 | 5.648216 | 1.62E-08 | 2.84E-07 |
| ENSG00000167554 | ZNF610 | 140.7799 | 1.524982 | 0.3622737 | 4.552761 | 5.29E-06 | 5.61E-05 |

| | | | | | | | |
|---|---|---|---|---|---|---|---|
| ENSG00000279233 | | 233.8028 | 1.524387 | 0.2532183 | 6.278235 | 3.42E-10 | 7.95E-09 |
| ENSG00000176753 | C15orf56 | 16.56521 | 1.524362 | 0.4799515 | 3.609248 | 0.00030709 | 0.002065 |
| ENSG00000232801 | SDCBPP3 | 6.246972 | 1.524355 | 1.036648 | 2.343257 | 0.01911621 | 0.065535 |
| ENSG00000264964 | | 99.83279 | 1.523992 | 0.3806512 | 4.358561 | 1.31E-05 | 0.000125 |
| ENSG00000213903 | LTB4R | 1870.822 | 1.523523 | 0.2744936 | 5.837278 | 5.31E-09 | 1.02E-07 |
| ENSG00000176884 | GRIN1 | 519.6266 | 1.521718 | 0.2839519 | 5.646894 | 1.63E-08 | 2.86E-07 |
| ENSG00000277129 | | 20.98679 | 1.521347 | 0.5953649 | 3.103673 | 0.00191134 | 0.009965 |
| ENSG00000149564 | ESAM | 209.1613 | 1.520819 | 0.4374656 | 3.935972 | 8.29E-05 | 0.000648 |
| ENSG00000286025 | FAM246C | 9.734632 | 1.5207 | 0.9615198 | 2.43256 | 0.01499251 | 0.053898 |
| ENSG00000182732 | RGS6 | 858.1898 | 1.520245 | 0.4200319 | 4.024777 | 5.70E-05 | 0.000465 |
| ENSG00000117407 | ARTN | 386.5766498 | 1.520002624 | 0.383863358 | 4.325342769 | 1.52E-05 | 0.000144 |
| ENSG00000156050 | FAM161B | 728.2302 | 1.519034 | 0.2357805 | 6.663624 | 2.67E-11 | 7.19E-10 |
| ENSG00000100253 | MIOX | 30.47534942 | 1.518483607 | 0.718228646 | 2.789241179 | 0.00528317 | 0.023177 |
| ENSG00000186367 | MINAR2 | 10.81852 | 1.517959 | 0.8235357 | 2.594592 | 0.00947034 | 0.037194 |
| ENSG00000101443 | WFDC2 | 281.2510773 | 1.517939951 | 0.274018616 | 5.797175441 | 6.74E-09 | 1.27E-07 |
| ENSG00000260653 | | 191.3474 | 1.517388 | 0.3414616 | 4.779771 | 1.75E-06 | 2.08E-05 |
| ENSG00000163273 | NPPC | 10.36327 | 1.516797 | 1.198388 | 2.228296 | 0.02586078 | 0.083254 |
| ENSG00000008853 | RHOBTB2 | 823.7605644 | 1.516464725 | 0.352895106 | 4.642348458 | 3.44E-06 | 3.81E-05 |
| ENSG00000182685 | BRICD5 | 810.5231 | 1.51515 | 0.3057884 | 5.25871 | 1.45E-07 | 2.13E-06 |
| ENSG00000249806 | | 3.090548 | 1.514547 | 2.665917 | 2.307536 | 0.02102494 | 0.070791 |
| ENSG00000231672 | DIRC3 | 42.69133 | 1.512039 | 0.6098495 | 3.050864 | 0.00228184 | 0.011565 |
| ENSG00000145569 | OTULINL | 566.9359 | 1.511507 | 0.498662 | 3.536665 | 0.00040521 | 0.002619 |
| ENSG00000183401 | CCDC159 | 625.8009 | 1.511306 | 0.3452454 | 4.704631 | 2.54E-06 | 2.92E-05 |
| ENSG00000259378 | DCAF13P3 | 20.16282 | 1.5111 | 0.5349758 | 3.335011 | 0.00085296 | 0.004981 |
| ENSG00000234409 | CCDC188 | 25.65717 | 1.508748 | 0.5010898 | 3.479394 | 0.00050255 | 0.003151 |
| ENSG00000197183 | NOL4L | 1494.732 | 1.508311 | 0.2614264 | 6.025528 | 1.69E-09 | 3.49E-08 |
| ENSG00000255320 | | 128.3407 | 1.504999 | 0.2812057 | 5.629386 | 1.81E-08 | 3.14E-07 |
| ENSG00000159176 | CSRP1 | 22941.25 | 1.504546 | 0.2776991 | 5.685709 | 1.30E-08 | 2.33E-07 |
| ENSG00000116819 | TFAP2E | 61.87287845 | 1.504325151 | 0.422138634 | 3.9532914 | 7.71E-05 | 0.000608 |
| ENSG00000269978 | | 24.11513 | 1.504268 | 0.568497 | 3.156946 | 0.00159431 | 0.008533 |
| ENSG00000115884 | SDC1 | 5109.406906 | 1.503544485 | 0.287956925 | 5.502404803 | 3.75E-08 | 6.16E-07 |
| ENSG00000143816 | WNT9A | 981.3087 | 1.503538 | 0.2378696 | 6.554054 | 5.60E-11 | 1.44E-09 |
| ENSG00000272512 | | 160.8589 | 1.503157 | 0.6152758 | 3.043957 | 0.00233489 | 0.011796 |

| | | | | | | | |
|---|---|---|---|---|---|---|---|
| ENSG00000080561 | MID2 | 1460.086521 | 1.502762044 | 0.242477202 | 6.428034007 | 1.29E-10 | 3.19E-09 |
| ENSG00000243244 | STON1 | 539.7503 | 1.501926 | 0.3424314 | 4.713441 | 2.44E-06 | 2.81E-05 |
| ENSG00000166592 | RRAD | 4.626313 | 1.501689 | 1.296647 | 2.233149 | 0.02553914 | 0.082458 |
| ENSG00000158747 | NBL1 | 1060.362 | 1.50127 | 0.2767603 | 5.690945 | 1.26E-08 | 2.27E-07 |
| ENSG00000283445 | | 12.30848 | 1.50108 | 0.7800425 | 2.628309 | 0.00858105 | 0.034408 |
| ENSG00000224468 | LAMC1-AS1 | 20.74981 | 1.50105 | 0.7677366 | 2.64467 | 0.00817707 | 0.033083 |
| ENSG00000214708 | | 39.16437 | 1.500285 | 0.4855207 | 3.54382 | 0.00039437 | 0.002562 |
| ENSG00000176933 | TOB2P1 | 64.92788 | 1.500249 | 0.3154505 | 5.034444 | 4.79E-07 | 6.38E-06 |
| ENSG00000130517 | PGPEP1 | 2047.192274 | 1.499877334 | 0.293743444 | 5.385611601 | 7.22E-08 | 1.13E-06 |
| ENSG00000142273 | CBLC | 7.837103 | 1.499225 | 0.9450113 | 2.382994 | 0.01717248 | 0.060065 |
| ENSG00000187808 | SOWAHD | 40.93271 | 1.498467 | 0.5881124 | 3.123794 | 0.00178536 | 0.009415 |
| ENSG00000142197 | DOP1B | 5776.775 | 1.496552 | 0.2181488 | 7.072128 | 1.53E-12 | 4.86E-11 |
| ENSG00000254211 | LINC01485 | 10.53739 | 1.496106 | 0.8630091 | 2.512469 | 0.01198896 | 0.045004 |
| ENSG00000271032 | | 29.38023 | 1.495769 | 0.4866565 | 3.524249 | 0.00042469 | 0.002725 |
| ENSG00000268442 | | 8.121568 | 1.494594 | 1.042566 | 2.230534 | 0.02571201 | 0.082886 |
| ENSG00000278615 | C11orf98 | 26.69662 | 1.493705 | 0.4600628 | 3.689665 | 0.00022455 | 0.001571 |
| ENSG00000226308 | | 15.72174 | 1.492961 | 1.043649 | 2.270858 | 0.02315555 | 0.076374 |
| ENSG00000079819 | EPB41L2 | 1641.33857 | 1.492055448 | 0.269231254 | 5.817260254 | 5.98E-09 | 1.14E-07 |
| ENSG00000150977 | RILPL2 | 1409.751 | 1.491215 | 0.1840966 | 8.280485 | 1.23E-16 | 6.44E-15 |
| ENSG00000265688 | MAFG-DT | 391.8328 | 1.491199 | 0.2626225 | 5.945281 | 2.76E-09 | 5.55E-08 |
| ENSG00000273394 | | 6.538272 | 1.491084 | 1.091286 | 2.282512 | 0.02245916 | 0.074743 |
| ENSG00000121039 | RDH10 | 1141.218239 | 1.490557924 | 0.244746277 | 6.321439712 | 2.59E-10 | 6.10E-09 |
| ENSG00000182263 | FIGN | 412.8522 | 1.490237 | 0.3169289 | 5.012807 | 5.36E-07 | 7.09E-06 |
| ENSG00000187391 | MAGI2 | 151.641 | 1.487797 | 0.2969855 | 5.290137 | 1.22E-07 | 1.82E-06 |
| ENSG00000115414 | FN1 | 2899.992187 | 1.487438066 | 0.398462981 | 4.130872352 | 3.61E-05 | 0.00031 |
| ENSG00000227542 | | 66.54355 | 1.487111 | 0.296968 | 5.287731 | 1.24E-07 | 1.85E-06 |
| ENSG00000226627 | SHANK2-AS1 | 2.966215 | 1.486919 | 1.610235 | 2.167634 | 0.03018658 | 0.094113 |
| ENSG00000204520 | MICA | 2128.225 | 1.485527 | 0.1754025 | 8.632791 | 5.99E-18 | 3.60E-16 |
| ENSG00000175161 | CADM2 | 117.7574 | 1.485201 | 0.5020129 | 3.437171 | 0.00058783 | 0.003612 |
| ENSG00000207751 | | 28.39469 | 1.482633 | 0.4459737 | 3.75581 | 0.00017278 | 0.001246 |
| ENSG00000259426 | | 86.70566 | 1.481738 | 0.4069387 | 4.033321 | 5.50E-05 | 0.00045 |
| ENSG00000279047 | | 4.700222 | 1.481196 | 1.412748 | 2.152108 | 0.03138886 | 0.097177 |
| ENSG00000247287 | | 274.3789 | 1.480482 | 0.3908252 | 4.163048 | 3.14E-05 | 0.000274 |

| | | | | | | | |
|---|---|---|---|---|---|---|---|
| ENSG00000230126 | FGF12-AS2 | 37.81913 | 1.479918 | 0.4617084 | 3.640878 | 0.00027171 | 0.001854 |
| ENSG00000260447 | | 1.457221 | 1.478792 | 2.583394 | 2.35304 | 0.01862065 | NA |
| ENSG00000183696 | UPP1 | 427.8084 | 1.477732 | 0.4450061 | 3.759789 | 0.00017006 | 0.001229 |
| ENSG00000143367 | TUFT1 | 7621.547 | 1.477455 | 0.2372773 | 6.456982 | 1.07E-10 | 2.65E-09 |
| ENSG00000158008 | EXTL1 | 8.678924 | 1.476647 | 0.8648117 | 2.492705 | 0.01267742 | 0.047142 |
| ENSG00000087589 | CASS4 | 11.62451697 | 1.475506176 | 0.824370075 | 2.539535023 | 0.01109999 | 0.042349 |
| ENSG00000183145 | RIPPLY3 | 190.7887 | 1.475283 | 0.4607141 | 3.64904 | 0.00026322 | 0.001806 |
| ENSG00000286333 | | 9.241928 | 1.474823 | 0.9022085 | 2.406351 | 0.01611276 | 0.057042 |
| ENSG00000181781 | ODF3L2 | 5.51226 | 1.474142 | 0.9846817 | 2.360688 | 0.01824108 | 0.06313 |
| ENSG00000154783 | FGD5 | 10.76764 | 1.473966 | 0.8838381 | 2.46116 | 0.01384887 | 0.050529 |
| ENSG00000249797 | LINC02147 | 5.767398 | 1.473399 | 1.207569 | 2.287114 | 0.02218917 | 0.074054 |
| ENSG00000182329 | KIAA2012 | 6.197525 | 1.473334 | 0.8258316 | 2.529386 | 0.01142624 | 0.043295 |
| ENSG00000284128 | | 5.587361 | 1.473002 | 1.107287 | 2.278498 | 0.02269691 | 0.075276 |
| ENSG00000233609 | RPL10P19 | 10.08744 | 1.472931 | 0.8450195 | 2.458221 | 0.01396271 | 0.05086 |
| ENSG00000213906 | LTB4R2 | 369.9204 | 1.472262 | 0.2578896 | 5.960773 | 2.51E-09 | 5.09E-08 |
| ENSG00000249267 | LINC00939 | 115.8942 | 1.471975 | 0.6154786 | 2.977344 | 0.00290758 | 0.014222 |
| ENSG00000215481 | BCRP3 | 6.472911 | 1.4702 | 0.9544742 | 2.390167 | 0.01684072 | 0.059177 |
| ENSG00000272733 | | 14.47116 | 1.46993 | 0.6835027 | 2.759621 | 0.00578685 | 0.02502 |
| ENSG00000184916 | JAG2 | 3824.395 | 1.468913 | 0.2017169 | 7.483267 | 7.25E-14 | 2.73E-12 |
| ENSG00000145349 | CAMK2D | 2645.463 | 1.467318 | 0.2992901 | 5.200083 | 1.99E-07 | 2.86E-06 |
| ENSG00000144152 | FBLN7 | 16.5266 | 1.467 | 0.7015923 | 2.729106 | 0.00635063 | 0.026902 |
| ENSG00000171033 | PKIA | 568.2663 | 1.466505 | 0.5069469 | 3.379762 | 0.00072549 | 0.004333 |
| ENSG00000242866 | STRC | 33.67716 | 1.463539 | 0.6284609 | 2.907314 | 0.00364547 | 0.017084 |
| ENSG00000131386 | GALNT15 | 6.11436866 | 1.463237601 | 0.93532501 | 2.378293691 | 0.01739297 | 0.060733 |
| ENSG00000250786 | SNHG18 | 382.6796 | 1.463032 | 0.4768249 | 3.531846 | 0.00041267 | 0.00266 |
| ENSG00000135604 | STX11 | 24.66163367 | 1.463012636 | 0.815337508 | 2.527165613 | 0.01149873 | 0.043533 |
| ENSG00000187726 | DNAJB13 | 40.61888 | 1.462187 | 0.516453 | 3.318179 | 0.00090607 | 0.005242 |
| ENSG00000186222 | BLOC1S4 | 1116.944 | 1.462097 | 0.2151407 | 7.004372 | 2.48E-12 | 7.74E-11 |
| ENSG00000140876 | NUDT7 | 352.7674 | 1.461811 | 0.2792587 | 5.494873 | 3.91E-08 | 6.41E-07 |
| ENSG00000196337 | CGB7 | 6.5794 | 1.461144 | 0.7703836 | 2.572408 | 0.01009938 | 0.039195 |
| ENSG00000102178 | UBL4A | 5917.680698 | 1.461078361 | 0.269892019 | 5.653556538 | 1.57E-08 | 2.77E-07 |
| ENSG00000267892 | | 51.00683 | 1.459574 | 0.4576193 | 3.59479 | 0.00032465 | 0.002169 |
| ENSG00000182771 | GRID1 | 173.3388 | 1.459168 | 0.4458211 | 3.695692 | 0.00021929 | 0.001538 |

| | | | | | | | |
|---|---|---|---|---|---|---|---|
| ENSG00000235823 | OLMALINC | 2236.275 | 1.458463 | 0.4326478 | 3.794745 | 0.0001478 | 0.001082 |
| ENSG00000089847 | ANKRD24 | 207.8641833 | 1.457494427 | 0.35591133 | 4.428435567 | 9.49E-06 | 9.41E-05 |
| ENSG00000176641 | RNF152 | 1016.155 | 1.456436 | 0.3480604 | 4.518368 | 6.23E-06 | 6.49E-05 |
| ENSG00000261804 | | 99.48784 | 1.455756 | 0.5407425 | 3.209027 | 0.00133185 | 0.007346 |
| ENSG00000227719 | | 138.5687 | 1.455732 | 0.3582808 | 4.394081 | 1.11E-05 | 0.000109 |
| ENSG00000201988 | | 3.03166 | 1.45351 | 1.481703 | 2.15674 | 0.0310259 | 0.0963 |
| ENSG00000272817 | | 18.68634 | 1.452967 | 0.5297078 | 3.227458 | 0.00124895 | 0.006933 |
| ENSG00000137558 | PI15 | 158.5121074 | 1.452907837 | 0.857204365 | 2.503488549 | 0.01229757 | 0.045913 |
| ENSG00000280913 | CTSLP3 | 2.682997 | 1.451876 | 1.468718 | 2.131019 | 0.03308757 | 0.101233 |
| ENSG00000240694 | PNMA2 | 37.02338 | 1.450327 | 0.7484426 | 2.600075 | 0.00932034 | 0.03674 |
| ENSG00000069424 | KCNAB2 | 1158.0433 | 1.450139824 | 0.182827561 | 8.082436307 | 6.35E-16 | 3.07E-14 |
| ENSG00000171346 | KRT15 | 553.1892 | 1.447635 | 0.339982 | 4.586678 | 4.50E-06 | 4.86E-05 |
| ENSG00000272745 | | 3.862233 | 1.446146 | 1.198546 | 2.149916 | 0.03156183 | 0.097587 |
| ENSG00000275540 | | 2.807597 | 1.445916 | 1.47069 | 2.215599 | 0.02671899 | 0.085458 |
| ENSG00000249700 | SRD5A3-AS1 | 37.11256 | 1.445078 | 0.5667594 | 3.066288 | 0.00216734 | 0.011095 |
| ENSG00000146112 | PPP1R18 | 1342.983 | 1.444442 | 0.2501859 | 6.022616 | 1.72E-09 | 3.54E-08 |
| ENSG00000267216 | | 24.20176 | 1.444408 | 0.5464972 | 3.148825 | 0.00163929 | 0.008725 |
| ENSG00000224097 | | 5.040868 | 1.443294 | 0.9773208 | 2.354399 | 0.01855271 | 0.063958 |
| ENSG00000272767 | JMJD1C-AS1 | 76.73464 | 1.443091 | 0.4891545 | 3.424518 | 0.00061589 | 0.00376 |
| ENSG00000135312 | HTR1B | 2.434053866 | 1.442333573 | 2.815559969 | 2.321520626 | 0.02025876 | 0.068702 |
| ENSG00000278212 | | 240.6284 | 1.441589 | 0.2638017 | 5.714815 | 1.10E-08 | 2.00E-07 |
| ENSG00000106829 | TLE4 | 319.9689138 | 1.441463279 | 0.519538672 | 3.269742157 | 0.00107646 | 0.006096 |
| ENSG00000260597 | | 85.82842 | 1.440576 | 0.5163542 | 3.279705 | 0.00103916 | 0.005921 |
| ENSG00000185515 | BRCC3 | 2372.384 | 1.440229 | 0.1720287 | 8.546234 | 1.27E-17 | 7.49E-16 |
| ENSG00000269335 | IKBKG | 646.9981 | 1.439651 | 0.1724029 | 8.527086 | 1.50E-17 | 8.69E-16 |
| ENSG00000196724 | ZNF418 | 228.8434 | 1.439499 | 0.193112 | 7.628878 | 2.37E-14 | 9.39E-13 |
| ENSG00000237487 | VN1R48P | 2.928923 | 1.43942 | 2.403724 | 2.328389 | 0.01989145 | 0.067652 |
| ENSG00000229939 | | 42.81154 | 1.437471 | 0.3071523 | 4.939593 | 7.83E-07 | 9.95E-06 |
| ENSG00000236352 | | 5.70393 | 1.436369 | 1.222051 | 2.138247 | 0.03249666 | 0.099723 |
| ENSG00000168539 | CHRM1 | 372.4759 | 1.435333 | 0.2679984 | 5.615213 | 1.96E-08 | 3.38E-07 |
| ENSG00000175267 | VWA3A | 14.1229 | 1.435286 | 0.5736514 | 3.003069 | 0.00267271 | 0.013229 |
| ENSG00000269893 | SNHG8 | 5471.419 | 1.43478 | 0.2881215 | 5.255939 | 1.47E-07 | 2.16E-06 |
| ENSG00000177788 | | 1405.251 | 1.433425 | 0.3439255 | 4.499066 | 6.83E-06 | 7.03E-05 |

| | | | | | | | |
|---|---|---|---|---|---|---|---|
| ENSG00000170006 | TMEM154 | 728.8778 | 1.432024 | 0.3728669 | 4.197212 | 2.70E-05 | 0.000239 |
| ENSG00000233251 | | 573.0015 | 1.431943 | 0.2932255 | 5.163881 | 2.42E-07 | 3.41E-06 |
| ENSG00000171877 | FRMD5 | 273.7519 | 1.429782 | 0.238591 | 6.219752 | 4.98E-10 | 1.12E-08 |
| ENSG00000251692 | PTX4 | 8.088429 | 1.429722 | 0.7894027 | 2.521029 | 0.01170122 | 0.044117 |
| ENSG00000128218 | VPREB3 | 16.68574002 | 1.429488948 | 0.533214255 | 3.18151548 | 0.00146507 | 0.007942 |
| ENSG00000229869 | | 9.547396 | 1.427541 | 0.9543025 | 2.343027 | 0.01912802 | 0.065567 |
| ENSG00000140519 | RHCG | 70.77328 | 1.427212 | 0.6208231 | 2.89105 | 0.00383957 | 0.017825 |
| ENSG00000213171 | LINGO4 | 2.421838 | 1.426859 | 1.561016 | 2.100879 | 0.03565157 | 0.107116 |
| ENSG00000188042 | ARL4C | 532.2973 | 1.426672 | 0.4979339 | 3.345284 | 0.00082198 | 0.004835 |
| ENSG00000250073 | | 25.87762 | 1.426612 | 0.5602608 | 3.026975 | 0.00247015 | 0.012371 |
| ENSG00000261220 | | 105.0433 | 1.426023 | 0.3384093 | 4.540391 | 5.62E-06 | 5.91E-05 |
| ENSG00000180269 | GPR139 | 41.44998 | 1.425969 | 0.7097639 | 2.662985 | 0.00774509 | 0.031688 |
| ENSG00000234750 | | 8.080389 | 1.425458 | 0.9647586 | 2.316572 | 0.02052707 | 0.069378 |
| ENSG00000163491 | NEK10 | 26.11988 | 1.424719 | 0.4832157 | 3.383319 | 0.00071615 | 0.004288 |
| ENSG00000261471 | | 2.24521 | 1.423977 | 2.710756 | 2.309429 | 0.02091981 | 0.070473 |
| ENSG00000064201 | TSPAN32 | 18.60868518 | 1.423768596 | 0.560866568 | 3.028425929 | 0.00245831 | 0.012327 |
| ENSG00000149654 | CDH22 | 10.5798 | 1.423434 | 1.501332 | 2.172885 | 0.02978896 | 0.09319 |
| ENSG00000141655 | TNFRSF11A | 273.4896 | 1.423177 | 0.2787918 | 5.369455 | 7.90E-08 | 1.22E-06 |
| ENSG00000185220 | PGBD2 | 669.7956 | 1.423144 | 0.2214553 | 6.632759 | 3.29E-11 | 8.74E-10 |
| ENSG00000198535 | C2CD4A | 134.4584 | 1.422871 | 0.9376743 | 2.410652 | 0.01592404 | 0.05651 |
| ENSG00000100504 | PYGL | 3424.443554 | 1.422179717 | 0.43870699 | 3.666656915 | 0.00024574 | 0.001701 |
| ENSG00000269646 | | 21.49875 | 1.421952 | 0.7961734 | 2.476674 | 0.0132613 | 0.048807 |
| ENSG00000174307 | PHLDA3 | 8330.93 | 1.4213 | 0.3284909 | 4.64469 | 3.41E-06 | 3.78E-05 |
| ENSG00000137449 | CPEB2 | 748.083485 | 1.418777444 | 0.337058578 | 4.528590036 | 5.94E-06 | 6.22E-05 |
| ENSG00000114268 | PFKFB4 | 549.2612557 | 1.418545761 | 0.277627328 | 5.379714421 | 7.46E-08 | 1.16E-06 |
| ENSG00000280216 | | 19.70978 | 1.418116 | 0.6093663 | 2.88805 | 0.00387639 | 0.01798 |
| ENSG00000147813 | NAPRT | 946.8545 | 1.418105 | 0.2186936 | 6.700617 | 2.08E-11 | 5.68E-10 |
| ENSG00000140254 | DUOXA1 | 6.146056 | 1.417904 | 1.009594 | 2.275202 | 0.02289384 | 0.07574 |
| ENSG00000258784 | | 1.467081 | 1.417782 | 2.521506 | 2.340996 | 0.01923236 | NA |
| ENSG00000272894 | MIOS-DT | 365.3265 | 1.417557 | 0.3306925 | 4.610799 | 4.01E-06 | 4.37E-05 |
| ENSG00000142556 | ZNF614 | 513.8379 | 1.417539 | 0.2288184 | 6.424655 | 1.32E-10 | 3.25E-09 |
| ENSG00000182718 | ANXA2 | 19081.76 | 1.417383 | 0.2547002 | 5.834939 | 5.38E-09 | 1.03E-07 |
| ENSG00000164742 | ADCY1 | 3593.128 | 1.41727 | 0.3893841 | 4.020862 | 5.80E-05 | 0.000472 |

| | | | | | | | |
|---|---|---|---|---|---|---|---|
| ENSG00000232149 | FERP1 | 7.285637 | 1.41726 | 1.068447 | 2.2991 | 0.02149927 | 0.072114 |
| ENSG00000065809 | FAM107B | 5942.773044 | 1.41709094 | 0.239607201 | 6.152715406 | 7.62E-10 | 1.67E-08 |
| ENSG00000134369 | NAV1 | 647.7942044 | 1.417055034 | 0.308283095 | 4.893678772 | 9.90E-07 | 1.23E-05 |
| ENSG00000278995 | | 22.02345 | 1.416607 | 0.493414 | 3.343359 | 0.00082771 | 0.00486 |
| ENSG00000223502 | | 35.40181 | 1.415678 | 0.6097891 | 2.88868 | 0.00386863 | 0.017957 |
| ENSG00000173511 | VEGFB | 1975.972 | 1.414854 | 0.2302702 | 6.3637 | 1.97E-10 | 4.74E-09 |
| ENSG00000225684 | FAM225B | 1.798067 | 1.414477 | 2.380782 | 2.326362 | 0.01999923 | 0.067923 |
| ENSG00000272079 | | 790.2094 | 1.414285 | 0.259063 | 5.708325 | 1.14E-08 | 2.07E-07 |
| ENSG00000127324 | TSPAN8 | 8.844036373 | 1.414195657 | 1.950972363 | 2.298680886 | 0.02152307 | 0.072166 |
| ENSG00000165959 | CLMN | 3463.276 | 1.413933 | 0.3082985 | 4.885844 | 1.03E-06 | 1.28E-05 |
| ENSG00000169239 | CA5B | 542.3886 | 1.411066 | 0.2587124 | 5.700223 | 1.20E-08 | 2.16E-07 |
| ENSG00000241269 | | 22.14035 | 1.410543 | 0.6017624 | 2.88535 | 0.00390979 | 0.018114 |
| ENSG00000177283 | FZD8 | 1019.045 | 1.409664 | 0.3230732 | 4.685992 | 2.79E-06 | 3.17E-05 |
| ENSG00000143772 | ITPKB | 1015.137 | 1.408974 | 0.1987675 | 7.270373 | 3.58E-13 | 1.25E-11 |
| ENSG00000286193 | | 83.57306 | 1.408554 | 0.4103284 | 3.808331 | 0.00013991 | 0.00103 |
| ENSG00000213445 | SIPA1 | 1118.367 | 1.406798 | 0.2879193 | 5.166719 | 2.38E-07 | 3.37E-06 |
| ENSG00000015520 | NPC1L1 | 20.50007738 | 1.406705188 | 0.636234654 | 2.796303743 | 0.00516908 | 0.022768 |
| ENSG00000223612 | | 130.4443 | 1.406413 | 0.2121671 | 6.847627 | 7.51E-12 | 2.19E-10 |
| ENSG00000273032 | | 95.6535 | 1.406198 | 0.5796045 | 2.980662 | 0.00287626 | 0.014094 |
| ENSG00000231705 | | 6.416657 | 1.405625 | 1.437213 | 2.114224 | 0.03449616 | 0.104555 |
| ENSG00000081842 | PCDHA6 | 174.6808086 | 1.403884299 | 0.267614658 | 5.501636219 | 3.76E-08 | 6.18E-07 |
| ENSG00000285427 | SOD2-OT1 | 23.26361 | 1.403529 | 0.8048771 | 2.510807 | 0.01204555 | 0.045159 |
| ENSG00000205133 | TRIQK | 4183.659 | 1.40312 | 0.2107868 | 6.858468 | 6.96E-12 | 2.04E-10 |
| ENSG00000231651 | DLG3-AS1 | 6.595343 | 1.40301 | 0.7950435 | 2.433943 | 0.01493536 | 0.053755 |
| ENSG00000148120 | AOPEP | 2752.294 | 1.403004 | 0.2192026 | 6.601049 | 4.08E-11 | 1.07E-09 |
| ENSG00000157021 | CIBAR1P1 | 135.047 | 1.402993 | 0.2782381 | 5.316947 | 1.06E-07 | 1.59E-06 |
| ENSG00000115194 | SLC30A3 | 300.7869 | 1.402967147 | 0.403081011 | 3.874651264 | 0.00010678 | 0.00081 |
| ENSG00000249685 | | 10.70832 | 1.402823 | 0.9939005 | 2.220489 | 0.0263856 | 0.084605 |
| ENSG00000185010 | F8 | 332.5159 | 1.401621 | 0.192981 | 7.444364 | 9.74E-14 | 3.62E-12 |
| ENSG00000116991 | SIPA1L2 | 2234.405578 | 1.401524803 | 0.274061393 | 5.3750583 | 7.66E-08 | 1.18E-06 |
| ENSG00000237708 | SPIN2P1 | 2.693927 | 1.400339 | 1.392585 | 2.118214 | 0.03415698 | 0.103716 |
| ENSG00000050767 | COL23A1 | 74.45705001 | 1.399810571 | 0.427876952 | 3.668191906 | 0.00024427 | 0.001691 |
| ENSG00000105464 | GRIN2D | 94.79824576 | 1.399616732 | 0.383682937 | 4.031331581 | 5.55E-05 | 0.000454 |

| | | | | | | | |
|---|---|---|---|---|---|---|---|
| ENSG00000286045 | | 5.456139 | 1.399361 | 0.9378918 | 2.282177 | 0.02247889 | 0.074791 |
| ENSG00000282021 | | 52.95046 | 1.399144 | 0.4177584 | 3.761762 | 0.00016872 | 0.001221 |
| ENSG00000213714 | FAM209B | 8.54371 | 1.398471 | 0.6987089 | 2.621267 | 0.00876037 | 0.035006 |
| ENSG00000179580 | RNF151 | 5.508979 | 1.398056 | 0.8735128 | 2.383205 | 0.01716263 | 0.060047 |
| ENSG00000230490 | | 120.4506 | 1.397878 | 0.6594623 | 2.73616 | 0.00621607 | 0.026476 |
| ENSG00000136114 | THSD1 | 21.23433149 | 1.396739285 | 0.441257501 | 3.574402582 | 0.00035103 | 0.002319 |
| ENSG00000255038 | | 11.50551 | 1.396636 | 0.8824533 | 2.346904 | 0.01893014 | 0.065023 |
| ENSG00000173210 | ABLIM3 | 855.8083 | 1.396546 | 0.2345227 | 6.181977 | 6.33E-10 | 1.41E-08 |
| ENSG00000286543 | | 9.408121 | 1.395969 | 0.9526637 | 2.287225 | 0.02218267 | 0.074042 |
| ENSG00000084731 | KIF3C | 2202.873524 | 1.39406239 | 0.248326681 | 5.860513733 | 4.61E-09 | 8.95E-08 |
| ENSG00000279130 | | 7.624576 | 1.393712 | 1.069124 | 2.22798 | 0.02588183 | 0.083291 |
| ENSG00000145198 | VWA5B2 | 52.86902 | 1.393563 | 0.4790401 | 3.364088 | 0.00076797 | 0.004555 |
| ENSG00000172367 | PDZD3 | 1.575991 | 1.392432 | 2.233689 | 2.287028 | 0.02219418 | 0.074056 |
| ENSG00000114166 | KAT2B | 1507.901063 | 1.392400667 | 0.282616568 | 5.209065867 | 1.90E-07 | 2.73E-06 |
| ENSG00000165164 | CFAP47 | 45.12025 | 1.392318 | 0.4419802 | 3.56325 | 0.00036629 | 0.002404 |
| ENSG00000084710 | EFR3B | 819.7137115 | 1.391820734 | 0.288774824 | 5.107748952 | 3.26E-07 | 4.46E-06 |
| ENSG00000213246 | SUPT4H1 | 3.476351 | 1.391333 | 1.290366 | 2.058611 | 0.03953152 | 0.115839 |
| ENSG00000232710 | | 2.765268 | 1.39023 | 1.215163 | 2.139547 | 0.03239143 | 0.099527 |
| ENSG00000248099 | INSL3 | 19.39413 | 1.389988 | 0.4813393 | 3.319459 | 0.00090192 | 0.005223 |
| ENSG00000151835 | SACS | 645.7013 | 1.389974 | 0.4870198 | 3.325036 | 0.00088407 | 0.005141 |
| ENSG00000119608 | PROX2 | 14.49487773 | 1.388889744 | 0.873716302 | 2.373708201 | 0.01761046 | 0.061339 |
| ENSG00000160781 | PAQR6 | 1325.639 | 1.387851 | 0.2529856 | 5.742751 | 9.32E-09 | 1.72E-07 |
| ENSG00000269001 | | 825.2755 | 1.387518 | 0.2198309 | 6.525628 | 6.77E-11 | 1.72E-09 |
| ENSG00000154358 | OBSCN | 3235.343 | 1.387223 | 0.2787921 | 5.237543 | 1.63E-07 | 2.37E-06 |
| ENSG00000204420 | MPIG6B | 17.41142 | 1.38701 | 0.6762735 | 2.656551 | 0.00789445 | 0.032195 |
| ENSG00000106052 | TAX1BP1 | 5134.533278 | 1.385894221 | 0.152052795 | 9.259245445 | 2.06E-20 | 1.59E-18 |
| ENSG00000072310 | SREBF1 | 43806.67509 | 1.385641562 | 0.296155071 | 4.992098207 | 5.97E-07 | 7.80E-06 |
| ENSG00000100439 | ABHD4 | 2531.005363 | 1.385412196 | 0.191588925 | 7.415357684 | 1.21E-13 | 4.43E-12 |
| ENSG00000100968 | NFATC4 | 646.5946911 | 1.385084135 | 0.297792557 | 4.922887076 | 8.53E-07 | 1.08E-05 |
| ENSG00000102385 | DRP2 | 48.03195075 | 1.384259969 | 0.561076743 | 3.005293601 | 0.00265325 | 0.013145 |
| ENSG00000102181 | CD99L2 | 2293.120407 | 1.383262269 | 0.160200065 | 8.785402559 | 1.56E-18 | 9.93E-17 |
| ENSG00000125337 | KIF25 | 2.854232811 | 1.38264066 | 1.355563773 | 2.127249562 | 0.03339935 | 0.101994 |
| ENSG00000262714 | | 67.63925 | 1.382526 | 0.5286351 | 3.111375 | 0.00186218 | 0.009747 |

| | | | | | | | |
|---|---|---|---|---|---|---|---|
| ENSG00000152910 | CNTNAP4 | 7.550588 | 1.38227 | 1.511495 | 2.105221 | 0.03527207 | 0.106204 |
| ENSG00000180347 | ITPRID1 | 61.64395 | 1.381297 | 0.5669718 | 2.978132 | 0.00290011 | 0.01419 |
| ENSG00000157404 | KIT | 3.465701 | 1.380362 | 1.790432 | 2.184452 | 0.02892902 | 0.091209 |
| ENSG00000129749 | CHRNA10 | 34.85879983 | 1.380227062 | 0.486900097 | 3.263119573 | 0.00110193 | 0.006225 |
| ENSG00000277203 | F8A1 | 1129.215 | 1.379873 | 0.1936953 | 7.312922 | 2.61E-13 | 9.24E-12 |
| ENSG00000162913 | OBSCN-AS1 | 191.4415 | 1.379255 | 0.325528 | 4.53821 | 5.67E-06 | 5.97E-05 |
| ENSG00000148200 | NR6A1 | 491.4427 | 1.378363 | 0.3050155 | 4.814787 | 1.47E-06 | 1.78E-05 |
| ENSG00000232284 | GNG12-AS1 | 35.11281 | 1.375709 | 0.7094771 | 2.605533 | 0.00917315 | 0.036294 |
| ENSG00000107719 | PALD1 | 154.3296872 | 1.375497534 | 0.491418856 | 3.272471052 | 0.00106612 | 0.006046 |
| ENSG00000158186 | MRAS | 325.0042 | 1.375413 | 0.3214654 | 4.592039 | 4.39E-06 | 4.75E-05 |
| ENSG00000152454 | ZNF256 | 257.317 | 1.375209 | 0.2805358 | 5.185543 | 2.15E-07 | 3.06E-06 |
| ENSG00000128578 | STRIP2 | 147.896447 | 1.37476123 | 0.466773073 | 3.394732457 | 0.00068696 | 0.004135 |
| ENSG00000134590 | RTL8C | 4166.49302 | 1.374492599 | 0.149428148 | 9.334492782 | 1.01E-20 | 8.09E-19 |
| ENSG00000183628 | DGCR6 | 313.332 | 1.374378 | 0.2349827 | 6.065728 | 1.31E-09 | 2.77E-08 |
| ENSG00000095383 | TBC1D2 | 1247.636052 | 1.374261352 | 0.249042997 | 5.755122931 | 8.66E-09 | 1.61E-07 |
| ENSG00000230077 | MTAPP2 | 6.221659 | 1.374135 | 0.839732 | 2.357036 | 0.01842147 | 0.063597 |
| ENSG00000251287 | ALG1L2 | 38.17813 | 1.374124 | 0.3104743 | 4.7174 | 2.39E-06 | 2.76E-05 |
| ENSG00000162105 | SHANK2 | 3488.628 | 1.372821 | 0.3302482 | 4.486041 | 7.26E-06 | 7.43E-05 |
| ENSG00000206557 | TRIM71 | 37.76731 | 1.372065 | 0.9825235 | 2.322557 | 0.02020296 | 0.06854 |
| ENSG00000273164 | NA | 12.06936 | 1.371595 | 0.8006231 | 2.445794 | 0.01445334 | 0.05235 |
| ENSG00000141696 | P3H4 | 2315.995 | 1.371582 | 0.1640071 | 8.526753 | 1.51E-17 | 8.69E-16 |
| ENSG00000258673 | LINC01397 | 3.371943 | 1.370865 | 1.2707 | 2.086207 | 0.03695991 | 0.10999 |
| ENSG00000118514 | ALDH8A1 | 8.632547567 | 1.370520833 | 0.782064025 | 2.45831447 | 0.01395909 | 0.050854 |
| ENSG00000136158 | SPRY2 | 410.2607618 | 1.370227086 | 0.290824429 | 4.995154076 | 5.88E-07 | 7.69E-06 |
| ENSG00000231991 | ANXA2P2 | 9724.223 | 1.370113 | 0.2388914 | 5.990571 | 2.09E-09 | 4.27E-08 |
| ENSG00000196391 | ZNF774 | 290.5196 | 1.369558 | 0.2723322 | 5.283029 | 1.27E-07 | 1.89E-06 |
| ENSG00000221923 | ZNF880 | 55.71958 | 1.368691 | 0.5033399 | 3.183222 | 0.00145646 | 0.007907 |
| ENSG00000205885 | C1RL-AS1 | 3.344056 | 1.368681 | 1.609494 | 2.123982 | 0.03367164 | 0.102536 |
| ENSG00000213145 | CRIP1 | 41.22224 | 1.368672 | 0.3751102 | 3.977964 | 6.95E-05 | 0.000554 |
| ENSG00000186496 | ZNF396 | 193.2264 | 1.368391 | 0.2314524 | 6.138815 | 8.31E-10 | 1.81E-08 |
| ENSG00000259937 | | 22.55381 | 1.367798 | 0.4267185 | 3.593048 | 0.00032683 | 0.002181 |
| ENSG00000086159 | AQP6 | 10.05402637 | 1.367180686 | 0.879764702 | 2.354322755 | 0.01855649 | 0.063963 |
| ENSG00000152580 | IGSF10 | 24.09232 | 1.367016 | 0.4845659 | 3.265344 | 0.00109331 | 0.006183 |

| | | | | | | | |
|---|---|---|---|---|---|---|---|
| ENSG00000076685 | NT5C2 | 2591.900404 | 1.366142509 | 0.184248774 | 7.583080002 | 3.37E-14 | 1.32E-12 |
| ENSG00000197558 | SSPOP | 718.4602 | 1.366048 | 0.1985666 | 7.071643 | 1.53E-12 | 4.87E-11 |
| ENSG00000249502 | | 4.650138 | 1.363677 | 1.150782 | 2.18156 | 0.02914202 | 0.091771 |
| ENSG00000270115 | | 35.05143 | 1.362605 | 0.576443 | 2.90145 | 0.0037144 | 0.017338 |
| ENSG00000169057 | MECP2 | 5333.3 | 1.362309 | 0.1951039 | 7.196405 | 6.18E-13 | 2.07E-11 |
| ENSG00000272267 | | 17.59485 | 1.362044 | 0.605592 | 2.809203 | 0.00496644 | 0.022025 |
| ENSG00000013297 | CLDN11 | 19.33540042 | 1.36044702 | 0.897261031 | 2.242969519 | 0.02489878 | 0.080833 |
| ENSG00000227544 | | 3.430218 | 1.359942 | 1.225465 | 2.126312 | 0.03347725 | 0.102159 |
| ENSG00000258477 | PPIAP6 | 3.068363 | 1.35942 | 1.834289 | 2.171965 | 0.02985833 | 0.09333 |
| ENSG00000169126 | ARMC4 | 7.914643 | 1.359029 | 0.6073985 | 2.788391 | 0.00529706 | 0.023216 |
| ENSG00000237433 | RPSAP11 | 6.681962 | 1.35818 | 1.01205 | 2.235667 | 0.02537356 | 0.082003 |
| ENSG00000136842 | TMOD1 | 103.8662032 | 1.356108172 | 0.477848398 | 3.292798488 | 0.00099196 | 0.005681 |
| ENSG00000184956 | MUC6 | 11.24326 | 1.355488 | 0.7830315 | 2.441459 | 0.01462804 | 0.052871 |
| ENSG00000183850 | ZNF730 | 42.06195 | 1.354691 | 0.634716 | 2.753042 | 0.00590444 | 0.025405 |
| ENSG00000077274 | CAPN6 | 5557.358054 | 1.35464798 | 0.646988615 | 2.695778427 | 0.00702244 | 0.029212 |
| ENSG00000258551 | | 7.299104 | 1.354445 | 0.6607427 | 2.653467 | 0.00796696 | 0.032431 |
| ENSG00000110921 | MVK | 3415.331092 | 1.35409947 | 0.393372072 | 3.829814681 | 0.00012824 | 0.000955 |
| ENSG00000278514 | | 2.339471 | 1.354049 | 3.586473 | 2.521331 | 0.01169118 | 0.044091 |
| ENSG00000185662 | SMIM23 | 4.569531 | 1.353941 | 0.9331447 | 2.253237 | 0.02424419 | 0.079207 |
| ENSG00000093100 | | 10.02961364 | 1.353210849 | 0.790486525 | 2.407438615 | 0.01606486 | 0.056918 |
| ENSG00000161249 | DMKN | 5364.903 | 1.353103 | 0.2189429 | 6.399186 | 1.56E-10 | 3.80E-09 |
| ENSG00000286964 | | 133.2176 | 1.352923 | 0.3557098 | 4.134439 | 3.56E-05 | 0.000306 |
| ENSG00000232938 | RPL23AP87 | 117.428 | 1.352613 | 0.4436002 | 3.484056 | 0.00049388 | 0.003106 |
| ENSG00000270882 | H4C14 | 6.388179 | 1.351226 | 0.805147 | 2.385947 | 0.01703521 | 0.059714 |
| ENSG00000101115 | SALL4 | 1355.802348 | 1.350895752 | 0.334745052 | 4.355205146 | 1.33E-05 | 0.000127 |
| ENSG00000011347 | SYT7 | 5010.363444 | 1.349959292 | 0.207602986 | 6.687489494 | 2.27E-11 | 6.17E-10 |
| ENSG00000196421 | C20orf204 | 276.6639 | 1.349907 | 0.3802508 | 3.915108 | 9.04E-05 | 0.000699 |
| ENSG00000183019 | MCEMP1 | 5.367223 | 1.349384 | 1.148639 | 2.199444 | 0.02784639 | 0.088319 |
| ENSG00000241111 | | 177.0694 | 1.349331 | 0.3773453 | 3.917887 | 8.93E-05 | 0.000692 |
| ENSG00000137338 | PGBD1 | 176.4165996 | 1.349320475 | 0.31555371 | 4.57978187 | 4.65E-06 | 5.00E-05 |
| ENSG00000261773 | | 7.985608 | 1.348943 | 0.9549499 | 2.244439 | 0.02480414 | 0.080594 |
| ENSG00000169862 | CTNND2 | 6183.411 | 1.348931 | 0.2861558 | 4.985432 | 6.18E-07 | 8.06E-06 |
| ENSG00000224614 | TNK2-AS1 | 31.0631 | 1.348201 | 0.5139258 | 3.090426 | 0.00199869 | 0.010345 |

| | | | | | | | |
|---|---|---|---|---|---|---|---|
| ENSG00000249921 | | 13.1193 | 1.347177 | 0.7160477 | 2.522011 | 0.0116686 | 0.044012 |
| ENSG00000254275 | LINC00824 | 24.64502 | 1.34579 | 0.4831819 | 3.237077 | 0.00120761 | 0.006731 |
| ENSG00000240914 | RPL15P2 | 9.701505 | 1.345766 | 0.7097979 | 2.546611 | 0.01087747 | 0.041631 |
| ENSG00000171813 | PWWP2B | 3717.938 | 1.345161 | 0.1745327 | 7.856813 | 3.94E-15 | 1.71E-13 |
| ENSG00000287742 | | 8.534494 | 1.344434 | 0.7332554 | 2.48254 | 0.01304495 | 0.048205 |
| ENSG00000075340 | ADD2 | 3.320212589 | 1.343108928 | 1.354861893 | 2.110386126 | 0.03482511 | 0.105228 |
| ENSG00000257038 | | 18.25628 | 1.342965 | 0.6085864 | 2.76242 | 0.00573746 | 0.024823 |
| ENSG00000287806 | | 1.973101 | 1.342653 | 1.660043 | 2.104978 | 0.03529323 | 0.106244 |
| ENSG00000147246 | HTR2C | 559.9481 | 1.342471 | 0.1633361 | 8.37797 | 5.38E-17 | 2.93E-15 |
| ENSG00000185386 | MAPK11 | 1182.365 | 1.341 | 0.2345179 | 5.977471 | 2.27E-09 | 4.61E-08 |
| ENSG00000249767 | ENPP7P10 | 9.760279 | 1.340695 | 1.103823 | 2.197241 | 0.02800322 | 0.088743 |
| ENSG00000257435 | | 3.015328 | 1.340146 | 1.990533 | 2.23607 | 0.0253472 | 0.081948 |
| ENSG00000002834 | LASP1 | 15881.86908 | 1.339477703 | 0.168326135 | 8.048966943 | 8.35E-16 | 3.98E-14 |
| ENSG00000124151 | NCOA3 | 90723.27331 | 1.339022704 | 0.209386092 | 6.592032848 | 4.34E-11 | 1.13E-09 |
| ENSG00000113327 | GABRG2 | 131.9419484 | 1.338354238 | 0.31258256 | 4.57712964 | 4.71E-06 | 5.06E-05 |
| ENSG00000227038 | GTF2IP7 | 5.240038 | 1.337941 | 0.8364218 | 2.340427 | 0.01926169 | 0.065932 |
| ENSG00000274897 | NA | 56.46505 | 1.336083 | 0.424295 | 3.549626 | 0.00038578 | 0.002517 |
| ENSG00000050405 | LIMA1 | 3925.317876 | 1.335924934 | 0.226064743 | 6.122969612 | 9.18E-10 | 1.98E-08 |
| ENSG00000103056 | SMPD3 | 376.658388 | 1.334701628 | 0.395391179 | 3.749554587 | 0.00017715 | 0.001273 |
| ENSG00000245904 | | 55.82489 | 1.334013 | 0.5932222 | 2.80043 | 0.00510345 | 0.022528 |
| ENSG00000163932 | PRKCD | 6524.312 | 1.33331 | 0.1599086 | 8.494138 | 1.99E-17 | 1.14E-15 |
| ENSG00000173890 | GPR160 | 4307.204 | 1.333146 | 0.1587808 | 8.510139 | 1.74E-17 | 9.97E-16 |
| ENSG00000108813 | DLX4 | 314.5930831 | 1.332781225 | 0.197438246 | 6.96045455 | 3.39E-12 | 1.05E-10 |
| ENSG00000133985 | TTC9 | 2507.989909 | 1.332206111 | 0.207728627 | 6.608306143 | 3.89E-11 | 1.02E-09 |
| ENSG00000167371 | PRRT2 | 757.4199 | 1.332073 | 0.3915038 | 3.766107 | 0.00016581 | 0.001202 |
| ENSG00000125257 | ABCC4 | 1506.017861 | 1.330991258 | 0.264620574 | 5.272723646 | 1.34E-07 | 1.99E-06 |
| ENSG00000029993 | HMGB3 | 8514.364714 | 1.330883807 | 0.316366421 | 4.532193695 | 5.84E-06 | 6.12E-05 |
| ENSG00000204282 | NA | 309.2641 | 1.329601 | 0.3513802 | 4.113409 | 3.90E-05 | 0.000332 |
| ENSG00000147394 | ZNF185 | 3302.673 | 1.329484 | 0.296385 | 4.774131 | 1.80E-06 | 2.14E-05 |
| ENSG00000213801 | ZNF321P | 83.83598 | 1.329208 | 0.4420756 | 3.431482 | 0.00060029 | 0.003676 |
| ENSG00000213204 | | 57.57518 | 1.328745 | 0.3837627 | 3.823142 | 0.00013176 | 0.000978 |
| ENSG00000168421 | RHOH | 46.72842 | 1.327329 | 0.5449301 | 2.944738 | 0.00323228 | 0.015457 |
| ENSG00000081377 | CDC14B | 615.0232109 | 1.326866604 | 0.243186859 | 5.679020744 | 1.35E-08 | 2.41E-07 |

| | | | | | | | |
|---|---|---|---|---|---|---|---|
| ENSG00000253197 | | 5.507461 | 1.326174 | 1.015851 | 2.216476 | 0.02665892 | 0.085334 |
| ENSG00000226849 | | 7.979271 | 1.326079 | 0.8283111 | 2.341172 | 0.01922331 | 0.065826 |
| ENSG00000099625 | CBARP | 580.9209424 | 1.325128661 | 0.173269247 | 7.814617195 | 5.51E-15 | 2.35E-13 |
| ENSG00000131389 | SLC6A6 | 8154.8276 | 1.32498465 | 0.2727323 | 5.120743398 | 3.04E-07 | 4.19E-06 |
| ENSG00000180245 | RRH | 6.566236 | 1.324786 | 1.02925 | 2.197139 | 0.02801051 | 0.088744 |
| ENSG00000135740 | SLC9A5 | 531.9415473 | 1.324692674 | 0.233045976 | 5.908452223 | 3.45E-09 | 6.84E-08 |
| ENSG00000262585 | LINC01979 | 24.34647 | 1.324042 | 0.5916872 | 2.778639 | 0.00545872 | 0.023788 |
| ENSG00000236204 | LINC01376 | 252.8184 | 1.323891 | 0.303256 | 4.647058 | 3.37E-06 | 3.74E-05 |
| ENSG00000256029 | | 13.14794 | 1.323362 | 0.6213154 | 2.676693 | 0.00743527 | 0.030651 |
| ENSG00000130997 | POLN | 18.2733366 | 1.322769043 | 0.701973257 | 2.525708506 | 0.01154652 | 0.04367 |
| ENSG00000224141 | MIR548XHG | 394.372 | 1.32201 | 0.2525289 | 5.474332 | 4.39E-08 | 7.15E-07 |
| ENSG00000153707 | PTPRD | 98.88104 | 1.32164 | 0.5947348 | 2.795976 | 0.00517432 | 0.022784 |
| ENSG00000138640 | FAM13A | 456.527004 | 1.321001485 | 0.277594682 | 4.999854083 | 5.74E-07 | 7.54E-06 |
| ENSG00000176834 | VSIG10 | 2676.015 | 1.319855 | 0.1426978 | 9.391611 | 5.91E-21 | 4.87E-19 |
| ENSG00000230316 | FEZF1-AS1 | 116.0942 | 1.319529 | 0.3395968 | 4.202097 | 2.64E-05 | 0.000235 |
| ENSG00000231419 | LINC00689 | 23.93842 | 1.319299 | 0.8096581 | 2.384397 | 0.01710713 | 0.059892 |
| ENSG00000038295 | TLL1 | 11.89211816 | 1.319048015 | 2.146017995 | 2.319511437 | 0.02036732 | 0.068961 |
| ENSG00000172888 | ZNF621 | 2084.988 | 1.318678 | 0.2238685 | 6.102551 | 1.04E-09 | 2.23E-08 |
| ENSG00000234367 | PFN1P3 | 11.14885 | 1.316245 | 0.5807474 | 2.758051 | 0.0058147 | 0.025112 |
| ENSG00000156381 | ANKRD9 | 2515.868 | 1.316132 | 0.2552616 | 5.400877 | 6.63E-08 | 1.04E-06 |
| ENSG00000228873 | | 12.37045 | 1.315854 | 1.069456 | 2.180697 | 0.0292058 | 0.091896 |
| ENSG00000230734 | RPL10P3 | 7.72489 | 1.315756 | 0.9603714 | 2.233475 | 0.02551763 | 0.082429 |
| ENSG00000117245 | KIF17 | 17.64598259 | 1.315748686 | 0.773922621 | 2.411271455 | 0.01589701 | 0.056437 |
| ENSG00000234465 | PINLYP | 28.85276 | 1.315256 | 0.4290961 | 3.465911 | 0.00052844 | 0.003293 |
| ENSG00000095303 | PTGS1 | 341.2083876 | 1.314116726 | 0.864094308 | 2.327563277 | 0.01993531 | 0.067753 |
| ENSG00000259000 | DOCK11P1 | 7.567972 | 1.31403 | 1.79173 | 2.223966 | 0.02615073 | 0.084004 |
| ENSG00000151136 | BTBD11 | 129.4788 | 1.313897 | 0.4328039 | 3.448681 | 0.00056333 | 0.003477 |
| ENSG00000152527 | PLEKHH2 | 648.3498 | 1.313786 | 0.2949839 | 4.71585 | 2.41E-06 | 2.78E-05 |
| ENSG00000162194 | LBHD1 | 1257.425 | 1.31365 | 0.3838376 | 3.791718 | 0.00014961 | 0.001095 |
| ENSG00000128487 | SPECC1 | 3998.851499 | 1.313143698 | 0.200885082 | 6.698851908 | 2.10E-11 | 5.74E-10 |
| ENSG00000276846 | | 90.395 | 1.312913 | 0.3181812 | 4.408455 | 1.04E-05 | 0.000102 |
| ENSG00000159184 | HOXB13 | 21.00826 | 1.312844 | 0.6537129 | 2.615372 | 0.00891303 | 0.035472 |
| ENSG00000131781 | FMO5 | 512.0193164 | 1.312181372 | 0.305433269 | 4.569009443 | 4.90E-06 | 5.24E-05 |

| | | | | | | | |
|---|---|---|---|---|---|---|---|
| ENSG00000257242 | LINC01619 | 12.51778 | 1.311912 | 0.6596619 | 2.600573 | 0.00930682 | 0.036704 |
| ENSG00000116133 | DHCR24 | 35819.65136 | 1.311627887 | 0.337758876 | 4.255928471 | 2.08E-05 | 0.000189 |
| ENSG00000279718 | SNX18P12 | 76.63868 | 1.311188 | 0.4245262 | 3.484776 | 0.00049255 | 0.0031 |
| ENSG00000258183 | LINC02392 | 4.226576 | 1.309525 | 2.019246 | 2.265333 | 0.02349223 | 0.077162 |
| ENSG00000113073 | SLC4A9 | 6.00945249 | 1.30891987 | 0.896006989 | 2.237910181 | 0.02522691 | 0.081669 |
| ENSG00000006327 | TNFRSF12A | 9526.972819 | 1.308669897 | 0.214918458 | 6.314271417 | 2.71E-10 | 6.38E-09 |
| ENSG00000123358 | NR4A1 | 4162.391192 | 1.304244459 | 1.496904852 | 2.213638128 | 0.02685368 | 0.085837 |
| ENSG00000198650 | TAT | 22.22108 | 1.302945 | 0.7968659 | 2.257717 | 0.02396334 | 0.078467 |
| ENSG00000229525 | | 25.19799 | 1.302772 | 0.7102896 | 2.503921 | 0.01228256 | 0.045872 |
| ENSG00000172935 | MRGPRF | 3.814461 | 1.301659 | 1.394864 | 2.138718 | 0.03245855 | 0.099664 |
| ENSG00000260179 | | 17.63279 | 1.301271 | 0.5938682 | 2.703471 | 0.00686195 | 0.028671 |
| ENSG00000152689 | RASGRP3 | 121.7694 | 1.298891 | 0.5489056 | 2.872419 | 0.00407343 | 0.018741 |
| ENSG00000168811 | IL12A | 54.86327 | 1.298315 | 0.6153023 | 2.674156 | 0.00749176 | 0.03084 |
| ENSG00000272146 | ARF4-AS1 | 6.588142 | 1.297025 | 0.8551907 | 2.271724 | 0.02310317 | 0.076268 |
| ENSG00000285486 | | 4.780814 | 1.296837 | 1.075065 | 2.134611 | 0.03279278 | 0.100504 |
| ENSG00000141404 | GNAL | 51.06903 | 1.29677 | 0.3976241 | 3.607413 | 0.00030927 | 0.002078 |
| ENSG00000135973 | GPR45 | 7.365285264 | 1.296266036 | 0.978309429 | 2.190321799 | 0.02850091 | 0.090017 |
| ENSG00000198156 | NPIPB6 | 35.51042 | 1.295561 | 0.600273 | 2.687526 | 0.00719836 | 0.029837 |
| ENSG00000258413 | | 8.338704 | 1.294733 | 0.7885307 | 2.303609 | 0.02124461 | 0.071377 |
| ENSG00000188365 | | 10.8629 | 1.294208 | 0.8345386 | 2.30058 | 0.02141539 | 0.071878 |
| ENSG00000215045 | GRID2IP | 141.749 | 1.293261 | 0.2967106 | 4.648006 | 3.35E-06 | 3.73E-05 |
| ENSG00000205710 | C17orf107 | 12.68406 | 1.293183 | 0.6644113 | 2.514648 | 0.01191514 | 0.044783 |
| ENSG00000108375 | RNF43 | 1661.215538 | 1.293117522 | 0.370625117 | 3.836004655 | 0.00012505 | 0.000935 |
| ENSG00000101049 | SGK2 | 23.56890303 | 1.292232309 | 0.560991963 | 2.810726825 | 0.00494297 | 0.02194 |
| ENSG00000136237 | RAPGEF5 | 475.9004999 | 1.291616325 | 0.319392718 | 4.347227427 | 1.38E-05 | 0.000131 |
| ENSG00000274372 | LINC02804 | 53.90837 | 1.289991 | 0.3186507 | 4.332528 | 1.47E-05 | 0.000139 |
| ENSG00000163462 | TRIM46 | 682.2519 | 1.289783 | 0.2500016 | 5.393902 | 6.89E-08 | 1.08E-06 |
| ENSG00000251056 | ANKRD20A17P | 38.20335 | 1.289585 | 0.4314117 | 3.406997 | 0.00065682 | 0.003976 |
| ENSG00000183615 | FAM167B | 37.88913 | 1.289584 | 0.4345302 | 3.351629 | 0.00080338 | 0.004739 |
| ENSG00000138772 | ANXA3 | 3443.817849 | 1.288300818 | 0.34137945 | 4.095402143 | 4.21E-05 | 0.000356 |
| ENSG00000277879 | | 45.85956 | 1.288062 | 0.4681555 | 3.20966 | 0.00132892 | 0.007333 |
| ENSG00000182963 | GJC1 | 6.57347 | 1.287343 | 1.063832 | 2.15177 | 0.03141546 | 0.097236 |
| ENSG00000273174 | | 33.64012 | 1.286139 | 0.7545314 | 2.410879 | 0.01591412 | 0.056489 |

| | | | | | | | |
|---|---|---|---|---|---|---|---|
| ENSG00000187642 | PERM1 | 56.22144 | 1.285449 | 0.3442004 | 4.036031 | 5.44E-05 | 0.000446 |
| ENSG00000140511 | HAPLN3 | 39.34556 | 1.285311 | 0.7043094 | 2.471207 | 0.01346577 | 0.049396 |
| ENSG00000034677 | RNF19A | 4126.4195 | 1.285052554 | 0.204984716 | 6.466404478 | 1.00E-10 | 2.50E-09 |
| ENSG00000196967 | ZNF585A | 229.8483 | 1.284248 | 0.3649211 | 3.888848 | 0.00010072 | 0.00077 |
| ENSG00000255408 | PCDHA3 | 44.8556 | 1.284013 | 0.4962361 | 3.041303 | 0.00235557 | 0.011887 |
| ENSG00000129951 | PLPPR3 | 94.51220863 | 1.283386293 | 0.577174922 | 2.764670353 | 0.00569804 | 0.024681 |
| ENSG00000154654 | NCAM2 | 3320.754 | 1.28285 | 0.3884126 | 3.661499 | 0.00025074 | 0.001727 |
| ENSG00000176222 | ZNF404 | 85.62285 | 1.282097 | 0.2587086 | 5.194621 | 2.05E-07 | 2.93E-06 |
| ENSG00000141934 | PLPP2 | 3445.246 | 1.280395 | 0.179869 | 7.297694 | 2.93E-13 | 1.03E-11 |
| ENSG00000185275 | CD24P4 | 366.9296 | 1.280047 | 0.3248074 | 4.253707 | 2.10E-05 | 0.000191 |
| ENSG00000246250 | | 12.95432 | 1.279725 | 0.6644215 | 2.531941 | 0.01134332 | 0.043071 |
| ENSG00000136327 | NKX2-8 | 22.9936612 | 1.279672847 | 0.725960326 | 2.368229706 | 0.01787344 | 0.062051 |
| ENSG00000164695 | CHMP4C | 943.3997 | 1.278852 | 0.1920138 | 6.850118 | 7.38E-12 | 2.15E-10 |
| ENSG00000162552 | WNT4 | 247.8223 | 1.278829 | 0.5321371 | 2.912405 | 0.00358657 | 0.016851 |
| ENSG00000279726 | | 9.504009 | 1.278601 | 0.654602 | 2.510233 | 0.01206515 | 0.045213 |
| ENSG00000246526 | LINC02481 | 32.97444 | 1.277852 | 0.4801557 | 3.114712 | 0.00184125 | 0.009666 |
| ENSG00000189196 | LINC00994 | 57.74559 | 1.277497 | 0.5353145 | 2.865984 | 0.00415715 | 0.019066 |
| ENSG00000064666 | CNN2 | 6524.514726 | 1.277264042 | 0.314572197 | 4.355103485 | 1.33E-05 | 0.000127 |
| ENSG00000227288 | ARID3BP1 | 21.7354 | 1.276943 | 0.4313774 | 3.339326 | 0.00083982 | 0.004919 |
| ENSG00000280195 | | 310.2664 | 1.275678 | 0.2510212 | 5.312612 | 1.08E-07 | 1.63E-06 |
| ENSG00000131473 | ACLY | 34018.18021 | 1.27535463 | 0.316383163 | 4.371367901 | 1.23E-05 | 0.000119 |
| ENSG00000171786 | NHLH1 | 6.40982 | 1.274945 | 0.8489395 | 2.215314 | 0.02673851 | 0.0855 |
| ENSG00000188732 | FAM221A | 593.3576 | 1.274762 | 0.246825 | 5.391963 | 6.97E-08 | 1.09E-06 |
| ENSG00000166387 | PPFIBP2 | 1584.118 | 1.274744 | 0.2858001 | 4.734516 | 2.20E-06 | 2.56E-05 |
| ENSG00000100031 | GGT1 | 4.067769314 | 1.274408436 | 1.042502579 | 2.134389391 | 0.03281092 | 0.100537 |
| ENSG00000090339 | ICAM1 | 486.4754749 | 1.274242157 | 0.490050582 | 3.060190004 | 0.00221197 | 0.011276 |
| ENSG00000259935 | | 6.151809 | 1.274097 | 0.9350569 | 2.158045 | 0.03092436 | 0.096086 |
| ENSG00000140839 | CLEC18B | 37.1951 | 1.273613 | 0.6072453 | 2.646938 | 0.00812243 | 0.03293 |
| ENSG00000165929 | TC2N | 5067.756 | 1.273212 | 0.1881862 | 6.922922 | 4.42E-12 | 1.33E-10 |
| ENSG00000230021 | | 22.2149 | 1.272768 | 0.5545253 | 2.804552 | 0.00503865 | 0.022297 |
| ENSG00000180998 | GPR137C | 487.0199 | 1.272644 | 0.4047248 | 3.528732 | 0.00041756 | 0.002686 |
| ENSG00000278385 | | 114.6829 | 1.2722 | 0.3734877 | 3.76934 | 0.00016368 | 0.001189 |
| ENSG00000196440 | ARMCX4 | 25.15491 | 1.271966 | 0.4493736 | 3.246362 | 0.0011689 | 0.006544 |

| | | | | | | | |
|---|---|---|---|---|---|---|---|
| ENSG00000163749 | CCDC158 | 17.54704 | 1.271701 | 0.740164 | 2.386501 | 0.01700954 | 0.059645 |
| ENSG00000272419 | LINC01145 | 368.2675 | 1.271546 | 0.3803711 | 3.693847 | 0.00022089 | 0.001547 |
| ENSG00000108379 | WNT3 | 206.5142592 | 1.271494827 | 0.285343696 | 4.721944769 | 2.34E-06 | 2.71E-05 |
| ENSG00000226508 | LINC01918 | 62.96961 | 1.270572 | 0.5155464 | 2.95181 | 0.00315917 | 0.015162 |
| ENSG00000150281 | CTF1 | 512.4468 | 1.270395 | 0.2271315 | 5.836607 | 5.33E-09 | 1.02E-07 |
| ENSG00000188779 | SKOR1 | 28.28818 | 1.27002 | 0.5842997 | 2.710243 | 0.00672339 | 0.02818 |
| ENSG00000280279 | LINC02887 | 19.63216 | 1.269984 | 0.5276369 | 2.887419 | 0.00388417 | 0.01801 |
| ENSG00000230638 | | 67.0254 | 1.269438 | 0.4216437 | 3.402391 | 0.00066799 | 0.004037 |
| ENSG00000184012 | TMPRSS2 | 2660.103 | 1.269208 | 0.3004579 | 4.507679 | 6.55E-06 | 6.78E-05 |
| ENSG00000272144 | | 50.16766 | 1.269048 | 0.3999626 | 3.528356 | 0.00041815 | 0.002688 |
| ENSG00000255346 | NOX5 | 550.8639 | 1.268721 | 0.2317255 | 5.688888 | 1.28E-08 | 2.29E-07 |
| ENSG00000258072 | | 85.24509 | 1.267912 | 0.4313055 | 3.337482 | 0.00084541 | 0.004946 |
| ENSG00000181358 | CTAGE10P | 12.60378 | 1.267122 | 0.6230651 | 2.596273 | 0.00942413 | 0.037051 |
| ENSG00000122543 | OCM | 11.02656355 | 1.266269983 | 0.700993312 | 2.389353503 | 0.01687805 | 0.059285 |
| ENSG00000269935 | | 10.34152 | 1.265964 | 0.8376347 | 2.251628 | 0.02434579 | 0.079455 |
| ENSG00000124313 | IQSEC2 | 1192.529412 | 1.265938551 | 0.190307696 | 6.822088638 | 8.97E-12 | 2.59E-10 |
| ENSG00000253931 | | 104.7937 | 1.265072 | 0.428316 | 3.357196 | 0.00078737 | 0.004655 |
| ENSG00000089486 | CDIP1 | 61.64662305 | 1.265069772 | 0.361605511 | 3.822309722 | 0.00013221 | 0.00098 |
| ENSG00000134294 | SLC38A2 | 28236.63111 | 1.265064981 | 0.211435373 | 6.178380911 | 6.48E-10 | 1.44E-08 |
| ENSG00000288059 | | 2.707873 | 1.264795 | 1.530699 | 2.055922 | 0.03979006 | 0.116404 |
| ENSG00000204969 | PCDHA2 | 56.22835 | 1.264662 | 0.569663 | 2.745332 | 0.00604498 | 0.025878 |
| ENSG00000287918 | | 33.98922 | 1.264417 | 0.4613458 | 3.146387 | 0.00165301 | 0.008789 |
| ENSG00000235415 | | 26.52817 | 1.264075 | 0.5197429 | 2.888218 | 0.00387432 | 0.017977 |
| ENSG00000185198 | PRSS57 | 18.55146 | 1.264012 | 0.7591579 | 2.35916 | 0.01831636 | 0.063322 |
| ENSG00000228013 | IL6R-AS1 | 12.42156 | 1.263928 | 0.8894347 | 2.205846 | 0.02739476 | 0.08722 |
| ENSG00000187097 | ENTPD5 | 2166.268 | 1.263919 | 0.2326529 | 5.649197 | 1.61E-08 | 2.83E-07 |
| ENSG00000157851 | DPYSL5 | 176.4368 | 1.263779 | 0.9075767 | 2.249577 | 0.02447582 | 0.079762 |
| ENSG00000233912 | | 3.338446 | 1.263511 | 1.562663 | 2.112814 | 0.03461667 | 0.10485 |
| ENSG00000185909 | KLHDC8B | 520.0657 | 1.261899 | 0.3029326 | 4.4485 | 8.65E-06 | 8.67E-05 |
| ENSG00000134247 | PTGFRN | 6052.303929 | 1.261720939 | 0.181026995 | 7.143315383 | 9.11E-13 | 2.98E-11 |
| ENSG00000281100 | | 4.208128 | 1.261497 | 0.9125739 | 2.181998 | 0.02910967 | 0.091691 |
| ENSG00000198125 | MB | 1214.308 | 1.261314 | 0.3818832 | 3.659727 | 0.00025248 | 0.001737 |
| ENSG00000185015 | CA13 | 45.11887 | 1.260819 | 0.4823449 | 3.048261 | 0.0023017 | 0.011646 |

| | | | | | | | |
|---|---|---|---|---|---|---|---|
| ENSG00000256944 | | 21.00093 | 1.260609 | 0.4875674 | 3.017207 | 0.00255116 | 0.012717 |
| ENSG00000213612 | FAM220CP | 6.071341 | 1.259678 | 0.9177502 | 2.178044 | 0.02940279 | 0.09235 |
| ENSG00000233930 | KRTAP5-AS1 | 384.8526 | 1.258608 | 0.3406099 | 4.013167 | 5.99E-05 | 0.000486 |
| ENSG00000218226 | TATDN2P2 | 257.2459 | 1.258197 | 0.4159148 | 3.414934 | 0.00063798 | 0.003878 |
| ENSG00000287234 | | 22.40055 | 1.257879 | 0.5487888 | 2.801262 | 0.00509032 | 0.022485 |
| ENSG00000250295 | RDH10-AS1 | 9.515481 | 1.257671 | 0.8058351 | 2.28315 | 0.02242155 | 0.074661 |
| ENSG00000207157 | RNY3P4 | 5.062409 | 1.256987 | 1.076998 | 2.114897 | 0.03443876 | 0.104417 |
| ENSG00000103269 | RHBDL1 | 460.9340308 | 1.256845025 | 0.27641212 | 4.797533312 | 1.61E-06 | 1.92E-05 |
| ENSG00000180884 | ZNF792 | 379.9015 | 1.256323 | 0.1830549 | 7.044356 | 1.86E-12 | 5.88E-11 |
| ENSG00000011465 | DCN | 4.206038914 | 1.255454228 | 1.886957309 | 2.220438486 | 0.02638902 | 0.084606 |
| ENSG00000249348 | UGDH-AS1 | 115.8803 | 1.255282 | 0.4052904 | 3.467514 | 0.0005253 | 0.003277 |
| ENSG00000143127 | ITGA10 | 125.7408 | 1.255141 | 0.4763263 | 3.075479 | 0.00210165 | 0.010805 |
| ENSG00000235927 | NEXN-AS1 | 2.406438 | 1.25479 | 1.537131 | 2.042858 | 0.04106649 | 0.119454 |
| ENSG00000255741 | | 11.0687 | 1.254339 | 0.53721 | 2.80947 | 0.00496231 | 0.022014 |
| ENSG00000172232 | AZU1 | 10.6746 | 1.253897 | 0.8989713 | 2.208345 | 0.02722026 | 0.086758 |
| ENSG00000272858 | | 33.57773 | 1.253553 | 0.3982053 | 3.510879 | 0.00044663 | 0.002844 |
| ENSG00000197353 | LYPD2 | 9.382931 | 1.252096 | 0.9455257 | 2.102453 | 0.0355136 | 0.106786 |
| ENSG00000213406 | ANXA2P1 | 16.56272 | 1.250986 | 0.6786133 | 2.483615 | 0.01300564 | 0.048073 |
| ENSG00000196159 | FAT4 | 1023.341 | 1.250056 | 0.4229573 | 3.352806 | 0.00079997 | 0.00472 |
| ENSG00000275329 | | 73.84828 | 1.24991 | 0.4970937 | 2.971974 | 0.00295892 | 0.014425 |
| ENSG00000225205 | | 2.912381 | 1.24839 | 1.312527 | 1.99943 | 0.04556188 | 0.12922 |
| ENSG00000179168 | GGN | 23.24547 | 1.247858 | 0.6036026 | 2.611105 | 0.00902502 | 0.035837 |
| ENSG00000055813 | CCDC85A | 122.0869621 | 1.247552822 | 0.454937888 | 3.164655799 | 0.00155267 | 0.008351 |
| ENSG00000287528 | | 12.27258 | 1.246755 | 0.9018287 | 2.116136 | 0.03433322 | 0.104156 |
| ENSG00000138834 | MAPK8IP3 | 7892.351457 | 1.246518703 | 0.248764626 | 5.288072834 | 1.24E-07 | 1.84E-06 |
| ENSG00000140398 | NEIL1 | 1581.117 | 1.246504 | 0.2973802 | 4.459556 | 8.21E-06 | 8.28E-05 |
| ENSG00000080200 | CRYBG3 | 640.8117335 | 1.246259759 | 0.49581471 | 2.984284958 | 0.00284242 | 0.013952 |
| ENSG00000280138 | | 111.548 | 1.246168 | 0.3036679 | 4.384793 | 1.16E-05 | 0.000113 |
| ENSG00000156804 | FBXO32 | 711.247 | 1.24605 | 0.3117207 | 4.289159 | 1.79E-05 | 0.000166 |
| ENSG00000160472 | TMEM190 | 18.02705 | 1.244235 | 0.5624104 | 2.721682 | 0.00649507 | 0.027405 |
| ENSG00000274565 | | 62.23427 | 1.243989 | 0.477044 | 3.055706 | 0.00224531 | 0.011409 |
| ENSG00000240291 | | 41.14271 | 1.24249 | 0.656997 | 2.493819 | 0.0126377 | 0.047014 |
| ENSG00000165511 | ZNF22-AS1 | 19.7052 | 1.242436 | 0.6359732 | 2.515739 | 0.01187831 | 0.04467 |

| | | | | | | | |
|---|---|---|---|---|---|---|---|
| ENSG00000071889 | FAM3A | 3993.010197 | 1.240868471 | 0.129748126 | 9.685306912 | 3.48E-22 | 3.36E-20 |
| ENSG00000137575 | SDCBP | 4009.473223 | 1.240453178 | 0.119257721 | 10.5168282 | 7.23E-26 | 9.80E-24 |
| ENSG00000237772 | LINC02631 | 14.8189 | 1.240062 | 0.5916334 | 2.630399 | 0.00852848 | 0.034219 |
| ENSG00000074966 | TXK | 6.874803077 | 1.239843281 | 0.79511132 | 2.254966352 | 0.02413544 | 0.078914 |
| ENSG00000092607 | TBX15 | 3.396075837 | 1.239709332 | 1.505023523 | 2.059832274 | 0.03941458 | 0.115611 |
| ENSG00000105639 | JAK3 | 211.2270624 | 1.238387595 | 0.362894593 | 3.748805012 | 0.00017768 | 0.001277 |
| ENSG00000268230 | | 85.20438 | 1.238382 | 0.4413471 | 3.209212 | 0.00133099 | 0.007343 |
| ENSG00000189431 | RASSF10 | 10.58704 | 1.236217 | 0.8144854 | 2.20192 | 0.02767099 | 0.08787 |
| ENSG00000152154 | TMEM178A | 38.84825 | 1.236069 | 0.7573054 | 2.322537 | 0.02020406 | 0.06854 |
| ENSG00000254786 | | 3.312954 | 1.235202 | 1.58358 | 2.171776 | 0.02987254 | 0.093353 |
| ENSG00000184305 | CCSER1 | 447.275 | 1.234814 | 0.2651885 | 4.897172 | 9.72E-07 | 1.21E-05 |
| ENSG00000159712 | ANKRD18CP | 13.46259 | 1.234663 | 0.6369678 | 2.523399 | 0.01162266 | 0.043899 |
| ENSG00000173598 | NUDT4 | 3752.799 | 1.23308 | 0.3028362 | 4.358439 | 1.31E-05 | 0.000125 |
| ENSG00000270294 | XIAPP1 | 36.30345 | 1.232954 | 0.5776127 | 2.667722 | 0.00763674 | 0.031336 |
| ENSG00000158220 | ESYT3 | 322.7074 | 1.232496 | 0.2994155 | 4.413424 | 1.02E-05 | 0.0001 |
| ENSG00000257185 | LINC02293 | 54.63807 | 1.232339 | 0.6460634 | 2.498117 | 0.01248548 | 0.046499 |
| ENSG00000196338 | NLGN3 | 77.76896 | 1.23152 | 0.4731894 | 3.034806 | 0.00240691 | 0.012109 |
| ENSG00000086300 | SNX10 | 428.052687 | 1.230831462 | 0.238390284 | 5.379965184 | 7.45E-08 | 1.16E-06 |
| ENSG00000175938 | ORAI3 | 515.0593 | 1.230568 | 0.2520012 | 5.105699 | 3.30E-07 | 4.50E-06 |
| ENSG00000229785 | SLC25A38P1 | 46.67367 | 1.230325 | 0.3609861 | 3.728375 | 0.00019272 | 0.001372 |
| ENSG00000117069 | ST6GALNAC5 | 97.50992725 | 1.229837479 | 0.545577558 | 2.757109871 | 0.00583148 | 0.025168 |
| ENSG00000197442 | MAP3K5 | 1100.275 | 1.229553 | 0.2258732 | 5.655344 | 1.56E-08 | 2.74E-07 |
| ENSG00000245975 | | 27.52372 | 1.229405 | 0.6902805 | 2.430725 | 0.01506863 | 0.05411 |
| ENSG00000127824 | TUBA4A | 681.0083838 | 1.228840053 | 0.467742652 | 3.066480922 | 0.00216595 | 0.01109 |
| ENSG00000286613 | | 3.297857 | 1.228314 | 1.27447 | 2.054184 | 0.03995792 | 0.116754 |
| ENSG00000286431 | | 298.6714 | 1.228246 | 0.3016379 | 4.363235 | 1.28E-05 | 0.000123 |
| ENSG00000272489 | | 3.125709 | 1.227984 | 1.21145 | 1.980102 | 0.04769201 | 0.133659 |
| ENSG00000182040 | USH1G | 39.34469 | 1.227954 | 0.6341089 | 2.513337 | 0.0119595 | 0.044925 |
| ENSG00000204791 | SMPD5 | 25.32975 | 1.227502 | 0.5666777 | 2.686084 | 0.00722949 | 0.029928 |
| ENSG00000286043 | | 8.806544 | 1.227372 | 0.6498752 | 2.466306 | 0.01365147 | 0.049932 |
| ENSG00000272514 | CFAP206 | 7.812834 | 1.226611 | 0.8240393 | 2.202687 | 0.02761679 | 0.087759 |
| ENSG00000125676 | THOC2 | 11591.92951 | 1.226032868 | 0.124836634 | 9.983740133 | 1.80E-23 | 1.90E-21 |
| ENSG00000271086 | NAMA | 4.432929 | 1.225251 | 1.112818 | 2.086622 | 0.03692228 | 0.109919 |

| | | | | | | | |
|---|---|---|---|---|---|---|---|
| ENSG00000179178 | TMEM125 | 1658.656 | 1.225001 | 0.212865 | 5.951145 | 2.66E-09 | 5.38E-08 |
| ENSG00000260398 | | 5.120211 | 1.224196 | 0.8179035 | 2.202228 | 0.02764922 | 0.087841 |
| ENSG00000215319 | EIF5P1 | 56.20086 | 1.223842 | 0.3269417 | 4.058402 | 4.94E-05 | 0.00041 |
| ENSG00000103044 | HAS3 | 64.88739031 | 1.223793353 | 0.403023783 | 3.409315091 | 0.00065126 | 0.003946 |
| ENSG00000184574 | LPAR5 | 10.60187 | 1.222732 | 0.7776794 | 2.232492 | 0.02558246 | 0.082588 |
| ENSG00000224078 | SNHG14 | 49.20764 | 1.222637 | 0.6885365 | 2.387713 | 0.01695356 | 0.059511 |
| ENSG00000284651 | | 18.43856 | 1.222214 | 0.5448834 | 2.728455 | 0.00636317 | 0.026936 |
| ENSG00000281571 | | 62.22299 | 1.221916 | 0.3810714 | 3.564076 | 0.00036514 | 0.002398 |
| ENSG00000267112 | | 3.794198 | 1.221519 | 1.172463 | 2.008468 | 0.04459359 | 0.127214 |
| ENSG00000269807 | | 2.023939 | 1.22139 | 1.58084 | 2.052947 | 0.0400777 | 0.117064 |
| ENSG00000169618 | PROKR1 | 1.746453 | 1.221187 | 2.368134 | 2.358857 | 0.01833134 | 0.06336 |
| ENSG00000184374 | COLEC10 | 47.70058 | 1.22116 | 0.7281733 | 2.35401 | 0.01857213 | 0.064 |
| ENSG00000280739 | EIF1B-AS1 | 58.31123 | 1.220677 | 0.4127012 | 3.33916 | 0.00084032 | 0.004921 |
| ENSG00000176945 | MUC20 | 435.797 | 1.220508 | 0.8025198 | 2.278923 | 0.02267167 | 0.075239 |
| ENSG00000104064 | GABPB1 | 2213.340229 | 1.219179627 | 0.191840497 | 6.536305235 | 6.31E-11 | 1.61E-09 |
| ENSG00000250790 | | 74.06546 | 1.218609 | 0.3800626 | 3.551616 | 0.00038287 | 0.002501 |
| ENSG00000163590 | PPM1L | 1534.932 | 1.218342 | 0.2681923 | 4.793862 | 1.64E-06 | 1.95E-05 |
| ENSG00000206344 | HCG27 | 36.12795 | 1.217698 | 0.4573118 | 3.066895 | 0.00216295 | 0.011079 |
| ENSG00000132031 | MATN3 | 153.7935419 | 1.217023396 | 0.527867988 | 2.795208515 | 0.00518663 | 0.02283 |
| ENSG00000272602 | ZNF595 | 1803.507 | 1.216688 | 0.1690734 | 7.35803 | 1.87E-13 | 6.69E-12 |
| ENSG00000188613 | NANOS1 | 466.6744 | 1.216603 | 0.2698687 | 4.763331 | 1.90E-06 | 2.25E-05 |
| ENSG00000250091 | DNAH10OS | 268.5045 | 1.216569 | 0.3634869 | 3.680866 | 0.00023244 | 0.001619 |
| ENSG00000172380 | GNG12 | 2828.205 | 1.216445 | 0.1723254 | 7.219689 | 5.21E-13 | 1.76E-11 |
| ENSG00000140836 | ZFHX3 | 4457.924 | 1.216047 | 0.3330959 | 3.956604 | 7.60E-05 | 0.000601 |
| ENSG00000269556 | TMEM185A | 1037.134 | 1.215443 | 0.1012775 | 12.08799 | 1.22E-33 | 2.90E-31 |
| ENSG00000136153 | LMO7 | 1969.271615 | 1.215164044 | 0.385773712 | 3.508499106 | 0.00045064 | 0.002866 |
| ENSG00000287024 | | 26.26513 | 1.213805 | 0.5385845 | 2.752516 | 0.00591392 | 0.025442 |
| ENSG00000135333 | EPHA7 | 511.0347222 | 1.213536458 | 0.56219637 | 2.681184002 | 0.00733622 | 0.030304 |
| ENSG00000147400 | CETN2 | 4096.126 | 1.212107 | 0.2155025 | 5.83594 | 5.35E-09 | 1.03E-07 |
| ENSG00000245281 | | 14.5801 | 1.211875 | 0.6745981 | 2.374763 | 0.01756022 | 0.061211 |
| ENSG00000275367 | | 19.99382 | 1.211683 | 0.5842776 | 2.583091 | 0.00979196 | 0.038203 |
| ENSG00000228878 | SEPTIN7-DT | 115.1099 | 1.209832 | 0.3064096 | 4.224589 | 2.39E-05 | 0.000215 |
| ENSG00000139679 | LPAR6 | 5.071807 | 1.209558 | 0.881962 | 2.16054 | 0.03073086 | 0.095585 |

| | | | | | | | |
|---|---|---|---|---|---|---|---|
| ENSG00000254162 | | 10.34645 | 1.209128 | 0.590189 | 2.557782 | 0.01053422 | 0.040592 |
| ENSG00000228634 | | 8.680575 | 1.208786 | 0.7150146 | 2.316198 | 0.02054748 | 0.069417 |
| ENSG00000270372 | | 34.8035 | 1.20865 | 0.3561957 | 3.702844 | 0.0002132 | 0.001501 |
| ENSG00000258245 | RPL10P13 | 3.36361 | 1.207491 | 1.316773 | 1.985602 | 0.04707749 | 0.132343 |
| ENSG00000139540 | SLC39A5 | 8.369149 | 1.207424 | 0.8182693 | 2.198707 | 0.02789877 | 0.088433 |
| ENSG00000236354 | LINC00437 | 2.30444 | 1.20699 | 1.858316 | 2.225747 | 0.02603112 | 0.08365 |
| ENSG00000185973 | TMLHE | 722.2942 | 1.206757 | 0.1692645 | 7.29437 | 3.00E-13 | 1.05E-11 |
| ENSG00000119227 | PIGZ | 642.3883733 | 1.206245271 | 0.190226058 | 6.504407229 | 7.80E-11 | 1.97E-09 |
| ENSG00000104611 | SH2D4A | 1850.37671 | 1.205978419 | 0.322557589 | 4.038146968 | 5.39E-05 | 0.000443 |
| ENSG00000131791 | PRKAB2 | 4861.299611 | 1.205768071 | 0.245822274 | 5.129517173 | 2.90E-07 | 4.01E-06 |
| ENSG00000266554 | LINC01443 | 2.465649 | 1.205069 | 2.71422 | 2.550087 | 0.01076961 | 0.041302 |
| ENSG00000236013 | | 4.488099 | 1.20463 | 1.139579 | 2.069774 | 0.03847353 | 0.113431 |
| ENSG00000277383 | | 68.79991 | 1.204294 | 0.4032686 | 3.350828 | 0.0008057 | 0.004749 |
| ENSG00000257474 | | 4.811437 | 1.203157 | 1.000464 | 2.051481 | 0.04022008 | 0.117404 |
| ENSG00000178988 | MRFAP1L1 | 5236.2 | 1.202965 | 0.1552977 | 7.872358 | 3.48E-15 | 1.53E-13 |
| ENSG00000180549 | FUT7 | 8.590481 | 1.202846 | 0.7894075 | 2.215194 | 0.02674675 | 0.08551 |
| ENSG00000178386 | ZNF223 | 36.26753 | 1.202309 | 0.3992533 | 3.3615 | 0.0007752 | 0.004594 |
| ENSG00000286888 | | 3.493965 | 1.202256 | 1.49237 | 2.023395 | 0.04303246 | 0.123804 |
| ENSG00000224189 | HAGLR | 138.1019 | 1.201742 | 0.374666 | 3.5491 | 0.00038655 | 0.002521 |
| ENSG00000157017 | GHRL | 25.14791 | 1.201604 | 0.5726626 | 2.626923 | 0.00861609 | 0.034528 |
| ENSG00000221818 | EBF2 | 8.820215 | 1.2014 | 0.6314767 | 2.460372 | 0.01387929 | 0.050612 |
| ENSG00000163703 | CRELD1 | 236.7299 | 1.201373 | 0.3080532 | 4.187984 | 2.81E-05 | 0.000248 |
| ENSG00000076864 | RAP1GAP | 1663.509068 | 1.201103786 | 0.289565508 | 4.418403932 | 9.94E-06 | 9.80E-05 |
| ENSG00000184208 | C22orf46 | 1980.813 | 1.20068 | 0.1533061 | 7.982583 | 1.43E-15 | 6.58E-14 |
| ENSG00000270876 | ZNF30-AS1 | 3.963769 | 1.200468 | 1.073505 | 1.986764 | 0.04694853 | 0.132126 |
| ENSG00000249846 | LINC02021 | 131.5121 | 1.200058 | 0.3085864 | 4.179602 | 2.92E-05 | 0.000257 |
| ENSG00000198467 | TPM2 | 781.2458 | 1.199922 | 0.2776659 | 4.576355 | 4.73E-06 | 5.08E-05 |
| ENSG00000196511 | TPK1 | 337.4591 | 1.199444 | 0.2593436 | 4.862762 | 1.16E-06 | 1.42E-05 |
| ENSG00000272369 | | 32.03498 | 1.199434 | 0.499677 | 2.839715 | 0.00451538 | 0.020386 |
| ENSG00000142875 | PRKACB | 1368.366 | 1.198275 | 0.1765086 | 6.940594 | 3.90E-12 | 1.19E-10 |
| ENSG00000110318 | CEP126 | 181.9696782 | 1.198255997 | 0.31380241 | 4.101946957 | 4.10E-05 | 0.000347 |
| ENSG00000279369 | | 28.61235 | 1.197817 | 0.7095241 | 2.291715 | 0.02192211 | 0.073275 |
| ENSG00000154240 | CEP112 | 47.82001 | 1.197198 | 0.3381885 | 3.830838 | 0.00012771 | 0.000951 |

| | | | | | | | |
|---|---|---|---|---|---|---|---|
| ENSG00000288004 | | 43.71075 | 1.196638 | 0.4081009 | 3.308206 | 0.00093896 | 0.00541 |
| ENSG00000185561 | TLCD2 | 746.0385 | 1.196466 | 0.3146745 | 4.089147 | 4.33E-05 | 0.000365 |
| ENSG00000214021 | TTLL3 | 1426.434 | 1.19644 | 0.3348226 | 3.878198 | 0.00010523 | 0.0008 |
| ENSG00000106868 | SUSD1 | 645.3311784 | 1.194308036 | 0.140870917 | 8.610004706 | 7.31E-18 | 4.36E-16 |
| ENSG00000270059 | | 5.15747 | 1.194204 | 0.9780258 | 2.070833 | 0.03837437 | 0.113208 |
| ENSG00000186919 | ZACN | 34.567 | 1.1941 | 0.477049 | 2.926537 | 0.00342758 | 0.016233 |
| ENSG00000272902 | TBC1D8-AS1 | 60.26573 | 1.194095 | 0.3157432 | 4.076516 | 4.57E-05 | 0.000383 |
| ENSG00000272138 | LINC01607 | 8.809207 | 1.193523 | 0.8550945 | 2.157132 | 0.03099541 | 0.09625 |
| ENSG00000068976 | PYGM | 49.12154634 | 1.192986594 | 0.326714801 | 3.930206421 | 8.49E-05 | 0.000662 |
| ENSG00000139631 | CSAD | 1642.994 | 1.192796 | 0.2529918 | 4.943243 | 7.68E-07 | 9.80E-06 |
| ENSG00000188981 | MSANTD1 | 18.5372 | 1.192763 | 0.6546621 | 2.414988 | 0.01573571 | 0.055983 |
| ENSG00000101850 | GPR143 | 280.2683661 | 1.190944915 | 0.318141246 | 4.030582616 | 5.56E-05 | 0.000455 |
| ENSG00000127325 | BEST3 | 2.640100595 | 1.190094574 | 1.610043818 | 2.070965077 | 0.03836206 | 0.113208 |
| ENSG00000109680 | TBC1D19 | 87.35423866 | 1.189948163 | 0.335698283 | 3.845544132 | 0.00012029 | 0.000902 |
| ENSG00000090924 | PLEKHG2 | 2799.489164 | 1.18992088 | 0.305710553 | 4.1719089 | 3.02E-05 | 0.000264 |
| ENSG00000238123 | MID1IP1-AS1 | 41.48719 | 1.189811 | 0.3901579 | 3.392639 | 0.00069223 | 0.004164 |
| ENSG00000251526 | LINC02435 | 2.600714 | 1.189784 | 1.576473 | 2.138986 | 0.03243676 | 0.099631 |
| ENSG00000033327 | GAB2 | 1614.044668 | 1.18952556 | 0.363331884 | 3.60497744 | 0.00031218 | 0.002096 |
| ENSG00000013619 | MAMLD1 | 379.1824778 | 1.188980979 | 0.237220996 | 5.235552511 | 1.64E-07 | 2.39E-06 |
| ENSG00000250997 | | 2.902195 | 1.18812 | 1.639347 | 2.069471 | 0.03850187 | 0.113485 |
| ENSG00000232709 | MARK2P9 | 11.79473 | 1.187889 | 0.6057621 | 2.477764 | 0.01322087 | 0.048699 |
| ENSG00000125266 | EFNB2 | 1176.132946 | 1.187599195 | 0.291692154 | 4.333742874 | 1.47E-05 | 0.000139 |
| ENSG00000117602 | RCAN3 | 678.906933 | 1.187364023 | 0.138658183 | 8.69598989 | 3.44E-18 | 2.13E-16 |
| ENSG00000287979 | | 5.483559 | 1.187084 | 0.9456686 | 2.081615 | 0.03737769 | 0.110935 |
| ENSG00000240338 | | 18.99169 | 1.186585 | 0.8599368 | 2.124905 | 0.03359455 | 0.102383 |
| ENSG00000229853 | | 39.13503 | 1.1864 | 0.3816184 | 3.43856 | 0.00058482 | 0.003598 |
| ENSG00000196369 | SRGAP2B | 486.8278 | 1.185642 | 0.2313182 | 5.347841 | 8.90E-08 | 1.36E-06 |
| ENSG00000257605 | MYG1-AS1 | 25.66691 | 1.185601 | 0.4485649 | 3.038308 | 0.0023791 | 0.011988 |
| ENSG00000177842 | ZNF620 | 360.2865 | 1.18494 | 0.2361427 | 5.236603 | 1.64E-07 | 2.38E-06 |
| ENSG00000229989 | MIR181A1HG | 24.85036 | 1.184731 | 0.45359 | 3.020872 | 0.00252048 | 0.012592 |
| ENSG00000257845 | LINC02294 | 4.440418 | 1.184006 | 1.04073 | 2.042963 | 0.04105607 | 0.119454 |
| ENSG00000176485 | PLAAT3 | 677.7497 | 1.183614 | 0.3654876 | 3.57599 | 0.00034891 | 0.002309 |
| ENSG00000283491 | | 27.84196 | 1.183395 | 0.5179444 | 2.745888 | 0.00603474 | 0.025848 |

| | | | | | | | |
|---|---|---|---|---|---|---|---|
| ENSG00000124333 | VAMP7 | 4213.70941 | 1.182587572 | 0.140372905 | 8.54293445 | 1.31E-17 | 7.69E-16 |
| ENSG00000255020 | | 51.587 | 1.182587 | 0.5989833 | 2.532784 | 0.01131605 | 0.042982 |
| ENSG00000115295 | CLIP4 | 286.9185385 | 1.182555933 | 0.572157161 | 2.592332378 | 0.00953276 | 0.037373 |
| ENSG00000287067 | | 3.069616 | 1.181914 | 1.283362 | 2.012582 | 0.04415862 | 0.126183 |
| ENSG00000180815 | MAP3K15 | 8.685365 | 1.181583 | 0.9258362 | 2.081724 | 0.03736771 | 0.110918 |
| ENSG00000101966 | XIAP | 4550.785045 | 1.181488255 | 0.25841282 | 4.813888347 | 1.48E-06 | 1.79E-05 |
| ENSG00000280310 | | 3.6801 | 1.180972 | 1.355298 | 1.986867 | 0.04693711 | 0.132126 |
| ENSG00000101928 | MOSPD1 | 1924.125877 | 1.18051665 | 0.217728722 | 5.609355485 | 2.03E-08 | 3.49E-07 |
| ENSG00000197857 | ZNF44 | 461.7075 | 1.179925 | 0.169115 | 7.136411 | 9.58E-13 | 3.13E-11 |
| ENSG00000090020 | SLC9A1 | 4437.024379 | 1.179576855 | 0.260031723 | 4.766102374 | 1.88E-06 | 2.22E-05 |
| ENSG00000168026 | TTC21A | 236.6358 | 1.179164 | 0.3608622 | 3.594584 | 0.00032491 | 0.00217 |
| ENSG00000197620 | EOLA1 | 1732.355 | 1.178338 | 0.1905987 | 6.354077 | 2.10E-10 | 5.03E-09 |
| ENSG00000177025 | C19orf18 | 28.14597 | 1.177236 | 0.4450451 | 3.033446 | 0.00241778 | 0.01215 |
| ENSG00000250769 | | 2.748119 | 1.176958 | 1.119702 | 1.975264 | 0.04823823 | 0.134782 |
| ENSG00000063601 | MTMR1 | 3802.350044 | 1.17693143 | 0.158509803 | 7.568667858 | 3.77E-14 | 1.46E-12 |
| ENSG00000090661 | CERS4 | 2539.926873 | 1.176597045 | 0.264925392 | 4.693910989 | 2.68E-06 | 3.06E-05 |
| ENSG00000135862 | LAMC1 | 9275.173355 | 1.176326941 | 0.188627665 | 6.408512524 | 1.47E-10 | 3.58E-09 |
| ENSG00000188167 | TMPPE | 94.78675 | 1.175845 | 0.423689 | 3.127607 | 0.00176236 | 0.009302 |
| ENSG00000205890 | | 651.3152 | 1.175582 | 0.3548303 | 3.649156 | 0.0002631 | 0.001806 |
| ENSG00000239282 | CASTOR1 | 27.66506 | 1.175205 | 0.4235354 | 3.152011 | 0.0016215 | 0.008649 |
| ENSG00000153814 | JAZF1 | 125.0815 | 1.175145 | 0.3650338 | 3.518723 | 0.00043363 | 0.002773 |
| ENSG00000234722 | LINC01287 | 7.74381 | 1.174893 | 0.8433223 | 2.148553 | 0.03166988 | 0.097853 |
| ENSG00000166689 | PLEKHA7 | 3133.906 | 1.174663 | 0.2193548 | 5.54785 | 2.89E-08 | 4.83E-07 |
| ENSG00000270659 | | 91.35395 | 1.174589 | 0.3824759 | 3.425391 | 0.00061392 | 0.003751 |
| ENSG00000213070 | HMGB3P6 | 65.70132 | 1.17398 | 0.4278563 | 3.152557 | 0.00161847 | 0.00864 |
| ENSG00000111412 | SPRING1 | 4777.827502 | 1.173862663 | 0.16580515 | 7.225037489 | 5.01E-13 | 1.70E-11 |
| ENSG00000150630 | VEGFC | 160.9811 | 1.173775 | 0.3743276 | 3.476779 | 0.00050748 | 0.003175 |
| ENSG00000154640 | BTG3 | 842.577 | 1.173525 | 0.2659017 | 4.659331 | 3.17E-06 | 3.55E-05 |
| ENSG00000254274 | TDGF1P5 | 1.872952 | 1.173419 | 1.941858 | 2.258653 | 0.023905 | 0.078295 |
| ENSG00000071205 | ARHGAP10 | 1829.548505 | 1.173245489 | 0.148893473 | 8.016323772 | 1.09E-15 | 5.10E-14 |
| ENSG00000260390 | | 1.524919 | 1.173072 | 2.427177 | 2.450695 | 0.01425808 | NA |
| ENSG00000132801 | ZSWIM3 | 486.7781083 | 1.172231332 | 0.207951718 | 5.819661354 | 5.90E-09 | 1.12E-07 |
| ENSG00000143507 | DUSP10 | 437.1772 | 1.172165 | 0.458995 | 2.976638 | 0.00291428 | 0.014249 |

| | | | | | | | |
|---|---|---|---|---|---|---|---|
| ENSG00000204516 | MICB | 758.9816 | 1.17191 | 0.4322786 | 3.107103 | 0.00188931 | 0.009864 |
| ENSG00000239365 | RPS26P49 | 19.09567 | 1.171137 | 0.5697653 | 2.538628 | 0.0111288 | 0.04241 |
| ENSG00000168899 | VAMP5 | 448.1156 | 1.170611 | 0.2756055 | 4.497465 | 6.88E-06 | 7.08E-05 |
| ENSG00000225828 | FAM229A | 214.7693 | 1.17051 | 0.2342482 | 5.202944 | 1.96E-07 | 2.82E-06 |
| ENSG00000270755 | | 5.119936 | 1.169866 | 0.9526106 | 2.000998 | 0.04539255 | 0.128864 |
| ENSG00000128683 | GAD1 | 284.1512714 | 1.169491638 | 0.437510621 | 3.064132777 | 0.00218302 | 0.011152 |
| ENSG00000258048 | | 5.287686 | 1.169116 | 0.9876046 | 2.020217 | 0.04336084 | 0.124531 |
| ENSG00000126603 | GLIS2 | 1931.48884 | 1.168950554 | 0.1650369 | 7.238119229 | 4.55E-13 | 1.56E-11 |
| ENSG00000167703 | SLC43A2 | 2612.756 | 1.168943 | 0.1935409 | 6.221768 | 4.92E-10 | 1.11E-08 |
| ENSG00000284633 | | 24.21401 | 1.16778 | 0.4869346 | 2.816237 | 0.00485897 | 0.021675 |
| ENSG00000258940 | | 40.68882 | 1.166264 | 0.3742532 | 3.436826 | 0.00058857 | 0.003615 |
| ENSG00000259495 | | 85.46144 | 1.166195 | 0.5388368 | 2.645313 | 0.00816154 | 0.033045 |
| ENSG00000235978 | | 2.447076 | 1.165941 | 1.267271 | 2.021185 | 0.0432606 | 0.124333 |
| ENSG00000101888 | NXT2 | 1015.917171 | 1.165705744 | 0.276557283 | 4.470067067 | 7.82E-06 | 7.94E-05 |
| ENSG00000222612 | RNU2-52P | 89.81186 | 1.163698 | 0.3578304 | 3.572569 | 0.0003535 | 0.002334 |
| ENSG00000138311 | ZNF365 | 97.03760699 | 1.162820137 | 0.290840934 | 4.261301356 | 2.03E-05 | 0.000185 |
| ENSG00000282876 | | 5.02681 | 1.162718 | 0.9715002 | 2.056002 | 0.03978236 | 0.116404 |
| ENSG00000286900 | | 3.526474 | 1.162708 | 1.085525 | 2.038734 | 0.04147656 | 0.120292 |
| ENSG00000171631 | P2RY6 | 347.4489 | 1.162481 | 0.8557582 | 2.146035 | 0.03187022 | 0.098346 |
| ENSG00000271895 | NA | 58.55716 | 1.160919 | 0.3433791 | 3.68675 | 0.00022714 | 0.001587 |
| ENSG00000261150 | EPPK1 | 1815.074 | 1.160807 | 0.529247 | 2.677088 | 0.00742652 | 0.03062 |
| ENSG00000287354 | | 6.701938 | 1.160598 | 0.9622909 | 2.068455 | 0.03859726 | 0.113716 |
| ENSG00000261207 | | 29.38816 | 1.160459 | 0.513341 | 2.713234 | 0.00666301 | 0.02798 |
| ENSG00000170956 | CEACAM3 | 2.13906 | 1.160427 | 1.855554 | 2.196946 | 0.02802427 | 0.088778 |
| ENSG00000230841 | RPL23AP15 | 3.615606 | 1.158557 | 1.19 | 1.919043 | 0.05497893 | 0.149039 |
| ENSG00000158445 | KCNB1 | 27.34677 | 1.158027 | 0.5635152 | 2.56026 | 0.0104594 | 0.040345 |
| ENSG00000198113 | TOR4A | 122.4065 | 1.157629 | 0.3055621 | 4.054088 | 5.03E-05 | 0.000417 |
| ENSG00000284954 | | 13.92708 | 1.157452 | 0.5712927 | 2.500839 | 0.01238996 | 0.046189 |
| ENSG00000117461 | PIK3R3 | 10027.27678 | 1.157323675 | 0.277198429 | 4.423862275 | 9.70E-06 | 9.58E-05 |
| ENSG00000243970 | PPIEL | 108.2052 | 1.156844 | 0.3885494 | 3.323254 | 0.00088974 | 0.005167 |
| ENSG00000139209 | SLC38A4 | 1.121836479 | 1.155654655 | 1.991636404 | 2.276528795 | 0.02281438 | NA |
| ENSG00000143891 | GALM | 908.1434 | 1.155604 | 0.3816035 | 3.374251 | 0.00074017 | 0.004409 |
| ENSG00000169220 | RGS14 | 205.1158 | 1.154234 | 0.2942403 | 4.196382 | 2.71E-05 | 0.00024 |

| | | | | | | | |
|---|---|---|---|---|---|---|---|
| ENSG00000165899 | OTOGL | 8.062017 | 1.154156 | 0.7765911 | 2.194741 | 0.02818215 | 0.089139 |
| ENSG00000225179 | LINC00457 | 2.584218 | 1.153532 | 1.342146 | 1.989877 | 0.04660446 | 0.131406 |
| ENSG00000178458 | H3P16 | 11.38018 | 1.15287 | 0.6844104 | 2.314082 | 0.0206632 | 0.06975 |
| ENSG00000213260 | YWHAZP5 | 33.13878 | 1.15273 | 0.4230646 | 3.10581 | 0.00189759 | 0.009895 |
| ENSG00000279699 | | 6.340917 | 1.152614 | 0.7484987 | 2.178716 | 0.02935277 | 0.092249 |
| ENSG00000169184 | MN1 | 13.73703 | 1.152354 | 0.710015 | 2.268554 | 0.02329544 | 0.076734 |
| ENSG00000114853 | ZBTB47 | 406.7200838 | 1.152281319 | 0.194450414 | 6.108232256 | 1.01E-09 | 2.17E-08 |
| ENSG00000100906 | NFKBIA | 6292.450462 | 1.151264101 | 0.252408723 | 4.812975661 | 1.49E-06 | 1.79E-05 |
| ENSG00000136931 | NR5A1 | 1.983788979 | 1.150984228 | 1.611421612 | 2.111709583 | 0.03471137 | 0.105001 |
| ENSG00000271040 | | 3.242059 | 1.150729 | 1.172525 | 2.020047 | 0.04337853 | 0.124541 |
| ENSG00000236138 | DUX4L26 | 7.644933 | 1.14988 | 0.7308569 | 2.21252 | 0.02693078 | 0.086011 |
| ENSG00000196951 | SCOC-AS1 | 23.81985 | 1.148853 | 0.4346777 | 3.022338 | 0.0025083 | 0.012541 |
| ENSG00000182400 | TRAPPC6B | 812.7308 | 1.148389 | 0.2600324 | 4.650832 | 3.31E-06 | 3.69E-05 |
| ENSG00000183049 | CAMK1D | 2603.804 | 1.147943 | 0.1856574 | 6.351985 | 2.13E-10 | 5.10E-09 |
| ENSG00000148655 | LRMDA | 99.475 | 1.147733 | 0.4262054 | 3.06913 | 0.00214683 | 0.011009 |
| ENSG00000228107 | | 5.419574 | 1.147713 | 1.333921 | 1.997526 | 0.04576805 | 0.129625 |
| ENSG00000179361 | ARID3B | 319.3154 | 1.147099 | 0.1976927 | 5.977264 | 2.27E-09 | 4.62E-08 |
| ENSG00000235522 | | 1.881561 | 1.14694 | 1.445948 | 2.002713 | 0.04520811 | 0.128451 |
| ENSG00000279208 | | 75.00751 | 1.146695 | 0.4190023 | 3.10621 | 0.00189502 | 0.009888 |
| ENSG00000254739 | | 26.71802 | 1.146228 | 0.4753829 | 2.814594 | 0.00488389 | 0.021761 |
| ENSG00000186806 | VSIG10L | 251.9519 | 1.145847 | 0.4576841 | 2.922677 | 0.00347036 | 0.016392 |
| ENSG00000170271 | FAXDC2 | 289.0342 | 1.145773 | 0.3263368 | 3.798831 | 0.00014538 | 0.001067 |
| ENSG00000141506 | PIK3R5 | 25.25936 | 1.145657 | 0.8592965 | 2.119971 | 0.0340085 | 0.10336 |
| ENSG00000255198 | SNHG9 | 360.4613 | 1.145381 | 0.3171994 | 3.900815 | 9.59E-05 | 0.000737 |
| ENSG00000252620 | RNU6ATAC24P | 3.983245 | 1.145191 | 1.08984 | 1.980034 | 0.04769974 | 0.133659 |
| ENSG00000145088 | EAF2 | 64.40182 | 1.144853 | 0.4579999 | 2.924669 | 0.00344823 | 0.01631 |
| ENSG00000163793 | DNAJC5G | 9.784362 | 1.143586 | 0.9496172 | 1.963609 | 0.04957547 | 0.137635 |
| ENSG00000129566 | TEP1 | 2191.540451 | 1.142924442 | 0.261079253 | 4.618878876 | 3.86E-06 | 4.22E-05 |
| ENSG00000160131 | VMA21 | 2901.324 | 1.14266 | 0.1613507 | 7.218962 | 5.24E-13 | 1.77E-11 |
| ENSG00000018236 | CNTN1 | 40.91175816 | 1.142410355 | 0.486740587 | 2.768081136 | 0.00563874 | 0.02446 |
| ENSG00000006071 | ABCC8 | 636.121697 | 1.142368607 | 0.375675594 | 3.379201291 | 0.00072697 | 0.004338 |
| ENSG00000268785 | RPL7P50 | 6.075795 | 1.142034 | 0.8409962 | 2.090874 | 0.0365394 | 0.109007 |
| ENSG00000009724 | MASP2 | 107.6847905 | 1.141951546 | 0.346998373 | 3.599145146 | 0.00031927 | 0.002138 |

| | | | | | | | |
|---|---|---|---|---|---|---|---|
| ENSG00000248375 | | 7.614274 | 1.141892 | 0.8092114 | 2.132932 | 0.03293032 | 0.100814 |
| ENSG00000262823 | | 9.386305 | 1.141464 | 1.351707 | 2.026808 | 0.04268207 | 0.123049 |
| ENSG00000114948 | ADAM23 | 43.12595822 | 1.141228081 | 0.535862731 | 2.608769554 | 0.00908684 | 0.036023 |
| ENSG00000215174 | NLRP2B | 7.543064 | 1.139382 | 0.9662272 | 2.043302 | 0.04102254 | 0.119405 |
| ENSG00000091831 | ESR1 | 13372.65548 | 1.13880655 | 0.310652141 | 3.937985255 | 8.22E-05 | 0.000643 |
| ENSG00000171100 | MTM1 | 868.4325 | 1.138804 | 0.1657172 | 7.028409 | 2.09E-12 | 6.55E-11 |
| ENSG00000168743 | NPNT | 10764.23 | 1.138639 | 0.2999301 | 4.074261 | 4.62E-05 | 0.000386 |
| ENSG00000251667 | BRCC3P1 | 76.85172 | 1.138393 | 0.2530932 | 4.733813 | 2.20E-06 | 2.57E-05 |
| ENSG00000174749 | FAM241A | 1049.464 | 1.138111 | 0.2635229 | 4.55275 | 5.29E-06 | 5.61E-05 |
| ENSG00000129028 | THAP10 | 344.2971571 | 1.137881327 | 0.222177447 | 5.32760057 | 9.95E-08 | 1.51E-06 |
| ENSG00000039523 | RIPOR1 | 2181.084134 | 1.137657911 | 0.194114245 | 6.043891484 | 1.50E-09 | 3.14E-08 |
| ENSG00000104419 | NDRG1 | 1754.729206 | 1.137526263 | 0.349503239 | 3.571088632 | 0.0003555 | 0.002345 |
| ENSG00000258667 | HIF1A-AS3 | 9.499949 | 1.136229 | 0.7144351 | 2.207566 | 0.02727452 | 0.08691 |
| ENSG00000182983 | ZNF662 | 9.371448 | 1.136012 | 1.618888 | 2.146087 | 0.03186601 | 0.098345 |
| ENSG00000213658 | LAT | 22.64936 | 1.135617 | 0.5235319 | 2.626489 | 0.00862709 | 0.034562 |
| ENSG00000029559 | IBSP | 35.25767633 | 1.13538539 | 0.383080215 | 3.289084267 | 0.00100514 | 0.005747 |
| ENSG00000151917 | BEND6 | 393.1231 | 1.134736 | 0.2155008 | 5.460686 | 4.74E-08 | 7.67E-07 |
| ENSG00000268747 | | 7.796108 | 1.134298 | 0.8626499 | 2.104164 | 0.03536418 | 0.106409 |
| ENSG00000232104 | RFX3-AS1 | 36.86122 | 1.134248 | 0.4295458 | 3.016318 | 0.00255865 | 0.012747 |
| ENSG00000248161 | | 14.25842 | 1.133761 | 0.7698061 | 2.150574 | 0.03150985 | 0.09746 |
| ENSG00000198785 | GRIN3A | 10.91226 | 1.133542 | 0.9660407 | 2.050409 | 0.04032455 | 0.117618 |
| ENSG00000227959 | | 18.76108 | 1.133254 | 0.9082958 | 2.052099 | 0.04016009 | 0.117259 |
| ENSG00000280068 | | 12.48486 | 1.13278 | 0.85483 | 2.095639 | 0.03611418 | 0.108224 |
| ENSG00000227051 | C14orf132 | 8654.687 | 1.13275 | 0.5174813 | 2.647674 | 0.00810477 | 0.032876 |
| ENSG00000100852 | ARHGAP5 | 4648.982174 | 1.131361613 | 0.350022132 | 3.563735828 | 0.00036561 | 0.0024 |
| ENSG00000267666 | | 39.03724 | 1.131238 | 0.4685055 | 2.81505 | 0.00487697 | 0.021741 |
| ENSG00000266954 | | 1.630831 | 1.130989 | 2.03753 | 2.302521 | 0.02130581 | 0.071528 |
| ENSG00000197594 | ENPP1 | 1387.262 | 1.130138 | 0.4373245 | 2.969868 | 0.00297928 | 0.014484 |
| ENSG00000234969 | AARSD1P1 | 19.08085 | 1.129696 | 0.4991523 | 2.712096 | 0.00668593 | 0.02805 |
| ENSG00000275936 | | 5.205558 | 1.12968 | 0.9254232 | 2.015547 | 0.04384733 | 0.125591 |
| ENSG00000116833 | NR5A2 | 8.649437151 | 1.129426399 | 1.017756166 | 2.024681349 | 0.0429001 | 0.12353 |
| ENSG00000112246 | SIM1 | 10.2687297 | 1.129270663 | 1.307503119 | 1.962650599 | 0.04968679 | 0.137887 |
| ENSG00000168264 | IRF2BP2 | 4182.482 | 1.127608 | 0.1824358 | 6.33964 | 2.30E-10 | 5.46E-09 |

| | | | | | | | |
|---|---|---|---|---|---|---|---|
| ENSG00000160602 | NEK8 | 1734.287 | 1.126969 | 0.2934008 | 4.11665 | 3.84E-05 | 0.000329 |
| ENSG00000134291 | TMEM106C | 8417.524056 | 1.126895039 | 0.221153018 | 5.291808107 | 1.21E-07 | 1.81E-06 |
| ENSG00000166670 | MMP10 | 7.061288 | 1.12682 | 1.250955 | 2.010103 | 0.04442025 | 0.126835 |
| ENSG00000142871 | CCN1 | 4620.729 | 1.126635 | 0.7441886 | 2.195833 | 0.02810388 | 0.088955 |
| ENSG00000153487 | ING1 | 1117.301 | 1.126365 | 0.2712813 | 4.398951 | 1.09E-05 | 0.000106 |
| ENSG00000279013 | | 13.14121 | 1.125965 | 0.7257325 | 2.172463 | 0.02982074 | 0.093257 |
| ENSG00000213799 | ZNF845 | 640.6049 | 1.125184 | 0.1857716 | 6.224128 | 4.84E-10 | 1.10E-08 |
| ENSG00000005238 | FAM214B | 526.3776412 | 1.124761012 | 0.203151581 | 5.71290118 | 1.11E-08 | 2.02E-07 |
| ENSG00000105679 | GAPDHS | 3.436582754 | 1.12436024 | 1.056712699 | 1.962426398 | 0.04971286 | 0.137945 |
| ENSG00000062524 | LTK | 13.25710392 | 1.124211878 | 0.629482448 | 2.350212777 | 0.01876268 | 0.064557 |
| ENSG00000183323 | CCDC125 | 1541.096 | 1.123841 | 0.2409125 | 4.876886 | 1.08E-06 | 1.33E-05 |
| ENSG00000113070 | HBEGF | 132.1562541 | 1.123654369 | 0.47564311 | 2.78842203 | 0.00529655 | 0.023216 |
| ENSG00000267288 | | 65.73756 | 1.123582 | 0.3578415 | 3.449378 | 0.00056188 | 0.003471 |
| ENSG00000006555 | TTC22 | 30.74662429 | 1.122338873 | 0.570226156 | 2.480958728 | 0.01310295 | 0.048365 |
| ENSG00000131981 | LGALS3 | 9532.365435 | 1.121901173 | 0.349055439 | 3.525043124 | 0.00042341 | 0.002718 |
| ENSG00000159915 | ZNF233 | 292.452 | 1.120807 | 0.2852277 | 4.188786 | 2.80E-05 | 0.000248 |
| ENSG00000153179 | RASSF3 | 1833.667 | 1.119484 | 0.2564687 | 4.599551 | 4.23E-06 | 4.60E-05 |
| ENSG00000235098 | ANKRD65 | 5.386098 | 1.119291 | 0.9440762 | 2.02116 | 0.04326323 | 0.124333 |
| ENSG00000271732 | | 1.685317 | 1.118261 | 1.48254 | 2.004753 | 0.04498944 | 0.127982 |
| ENSG00000278931 | | 121.0099 | 1.117554 | 0.3281445 | 3.696146 | 0.0002189 | 0.001536 |
| ENSG00000114861 | FOXP1 | 746.4526472 | 1.117426227 | 0.268007098 | 4.416077014 | 1.01E-05 | 9.90E-05 |
| ENSG00000272086 | | 226.7147 | 1.117062 | 0.3591769 | 3.437301 | 0.00058754 | 0.003611 |
| ENSG00000172476 | RAB40A | 20.32264 | 1.117056 | 0.4832908 | 2.712958 | 0.00666856 | 0.02799 |
| ENSG00000233718 | MYCNOS | 3.506997 | 1.116774 | 1.326641 | 1.992279 | 0.04634046 | 0.130846 |
| ENSG00000183337 | BCOR | 3382.868 | 1.116734 | 0.2346475 | 4.982129 | 6.29E-07 | 8.18E-06 |
| ENSG00000125378 | BMP4 | 517.5186457 | 1.115360155 | 0.345480369 | 3.533708275 | 0.00040977 | 0.002645 |
| ENSG00000141736 | ERBB2 | 11223.39 | 1.115157 | 0.2003913 | 5.742022 | 9.36E-09 | 1.73E-07 |
| ENSG00000126903 | SLC10A3 | 2867.77947 | 1.114585347 | 0.287659634 | 4.127327954 | 3.67E-05 | 0.000314 |
| ENSG00000154645 | CHODL | 15.95968 | 1.114503 | 0.5655513 | 2.451661 | 0.01421987 | 0.05167 |
| ENSG00000164403 | SHROOM1 | 394.0885 | 1.114491 | 0.3573896 | 3.444257 | 0.00057263 | 0.003529 |
| ENSG00000260244 | | 271.747 | 1.114082 | 0.3982415 | 3.145148 | 0.00166003 | 0.008821 |
| ENSG00000272221 | | 76.94337 | 1.113581 | 0.333621 | 3.627749 | 0.0002859 | 0.001939 |
| ENSG00000287886 | | 587.9895 | 1.112618 | 0.2161914 | 5.337638 | 9.42E-08 | 1.43E-06 |

| | | | | | | | |
|---|---|---|---|---|---|---|---|
| ENSG00000108309 | RUNDC3A | 218.9720565 | 1.111930505 | 0.214328407 | 5.375766729 | 7.63E-08 | 1.18E-06 |
| ENSG00000166311 | SMPD1 | 859.8202 | 1.111926 | 0.2052316 | 5.602345 | 2.11E-08 | 3.62E-07 |
| ENSG00000237515 | SHISA9 | 655.9695 | 1.11171 | 0.1983183 | 5.781507 | 7.40E-09 | 1.39E-07 |
| ENSG00000253395 | | 49.29012 | 1.111529 | 0.5021202 | 2.654254 | 0.0079484 | 0.03237 |
| ENSG00000287595 | | 53.98552 | 1.111109 | 0.6288648 | 2.319462 | 0.02037 | 0.068962 |
| ENSG00000113211 | PCDHB6 | 14.54150073 | 1.110886495 | 0.606919957 | 2.343890894 | 0.01908375 | 0.065466 |
| ENSG00000258175 | LINC02300 | 2.603991 | 1.110766 | 2.239805 | 2.422876 | 0.01539817 | 0.055048 |
| ENSG00000052802 | MSMO1 | 7080.931415 | 1.110115076 | 0.237777946 | 4.878769831 | 1.07E-06 | 1.32E-05 |
| ENSG00000119636 | BBOF1 | 373.8088 | 1.109840248 | 0.25083137 | 4.642689644 | 3.44E-06 | 3.81E-05 |
| ENSG00000269559 | | 7.92127 | 1.108999 | 1.015613 | 1.933564 | 0.05316672 | 0.145482 |
| ENSG00000272055 | RNU6-6P | 6.161821 | 1.108893 | 0.8623692 | 2.050872 | 0.04027946 | 0.117525 |
| ENSG00000184058 | TBX1 | 22.45304 | 1.108755 | 0.6618151 | 2.267851 | 0.0233383 | 0.076828 |
| ENSG00000183742 | MACC1 | 1251.903 | 1.108657 | 0.5190072 | 2.604251 | 0.00920753 | 0.036415 |
| ENSG00000155622 | XAGE2 | 37.42184 | 1.108462 | 1.6922 | 2.196604 | 0.02804871 | 0.088823 |
| ENSG00000229140 | CCDC26 | 26.25482 | 1.107055 | 0.5315902 | 2.533186 | 0.01130308 | 0.042946 |
| ENSG00000229666 | MAST4-AS1 | 35.46388 | 1.107006 | 0.5765062 | 2.428583 | 0.01515796 | 0.054379 |
| ENSG00000150347 | ARID5B | 2295.466 | 1.106768 | 0.312907 | 3.81411 | 0.00013668 | 0.00101 |
| ENSG00000138795 | LEF1 | 36.04163709 | 1.105549843 | 0.520305162 | 2.589243327 | 0.00961871 | 0.037649 |
| ENSG00000224790 | | 97.22984 | 1.105149 | 0.3201095 | 3.728142 | 0.0001929 | 0.001373 |
| ENSG00000132334 | PTPRE | 335.8689868 | 1.104776932 | 0.533954256 | 2.551831835 | 0.01071582 | 0.041161 |
| ENSG00000185950 | IRS2 | 585.7581 | 1.103851 | 0.3882226 | 3.182013 | 0.00146255 | 0.007932 |
| ENSG00000272031 | ANKRD34A | 843.6671 | 1.103786 | 0.2913733 | 4.035883 | 5.44E-05 | 0.000446 |
| ENSG00000237077 | RPL38P2 | 6.299514 | 1.10259 | 0.9349631 | 1.963417 | 0.04959769 | 0.13768 |
| ENSG00000203879 | GDI1 | 12051.56 | 1.10249 | 0.1636909 | 6.873896 | 6.25E-12 | 1.84E-10 |
| ENSG00000164574 | GALNT10 | 6336.022 | 1.102484 | 0.2721635 | 4.294847 | 1.75E-05 | 0.000162 |
| ENSG00000253304 | TMEM200B | 6.117301 | 1.102386 | 1.002953 | 1.955541 | 0.0505193 | 0.139599 |
| ENSG00000242737 | | 5.125605 | 1.101677 | 0.8713316 | 2.044625 | 0.04089182 | 0.119051 |
| ENSG00000122507 | BBS9 | 1034.74246 | 1.100476349 | 0.172787425 | 6.525215625 | 6.79E-11 | 1.72E-09 |
| ENSG00000259763 | LINC02157 | 1.549467 | 1.100395 | 1.873893 | 2.260761 | 0.02377404 | NA |
| ENSG00000102030 | NAA10 | 2666.362589 | 1.100371883 | 0.197505212 | 5.740573919 | 9.44E-09 | 1.74E-07 |
| ENSG00000144935 | TRPC1 | 513.3235 | 1.099756 | 0.2715312 | 4.286741 | 1.81E-05 | 0.000167 |
| ENSG00000274816 | MIR6772 | 3.371724 | 1.099249 | 1.59282 | 2.131706 | 0.033031 | 0.101094 |
| ENSG00000285570 | | 20.94827 | 1.098666 | 0.7725701 | 2.138289 | 0.03249329 | 0.099723 |

| | | | | | | | |
|---|---|---|---|---|---|---|---|
| ENSG00000162843 | WDR64 | 6.23266 | 1.09796 | 0.8078412 | 2.032499 | 0.04210318 | 0.121675 |
| ENSG00000280207 | | 4.703745 | 1.097203 | 1.170872 | 1.97701 | 0.04804052 | 0.134385 |
| ENSG00000166763 | STRCP1 | 12.31118 | 1.097071 | 0.56806 | 2.41065 | 0.0159241 | 0.05651 |
| ENSG00000130827 | PLXNA3 | 12985.81453 | 1.096596026 | 0.285500631 | 4.138229698 | 3.50E-05 | 0.000302 |
| ENSG00000205456 | TP53TG3D | 29.91039 | 1.095249 | 0.5015372 | 2.61577 | 0.00890265 | 0.035449 |
| ENSG00000100558 | PLEK2 | 421.0465773 | 1.094907426 | 0.442734235 | 2.874063401 | 0.00405228 | 0.018653 |
| ENSG00000213160 | KLHL23 | 687.6262 | 1.094592 | 0.2608631 | 4.437276 | 9.11E-06 | 9.08E-05 |
| ENSG00000100422 | CERK | 2409.271991 | 1.094583553 | 0.268051276 | 4.301912927 | 1.69E-05 | 0.000158 |
| ENSG00000109929 | SC5D | 2925.182152 | 1.094552638 | 0.188270282 | 5.976871708 | 2.27E-09 | 4.62E-08 |
| ENSG00000171790 | SLFNL1 | 51.75463 | 1.094337 | 0.3705308 | 3.262802 | 0.00110317 | 0.006228 |
| ENSG00000284654 | | 28.31506 | 1.09332 | 0.4945022 | 2.6447 | 0.00817634 | 0.033083 |
| ENSG00000167107 | ACSF2 | 4405.796 | 1.092931 | 0.2465359 | 4.661158 | 3.14E-06 | 3.53E-05 |
| ENSG00000237892 | KLF7-IT1 | 6.779488 | 1.092347 | 0.8002604 | 2.053153 | 0.04005776 | 0.117033 |
| ENSG00000203778 | FAM229B | 77.84316 | 1.092182 | 0.3419904 | 3.483894 | 0.00049418 | 0.003108 |
| ENSG00000234636 | MED14OS | 12.44205 | 1.092006 | 0.6628715 | 2.234358 | 0.0254595 | 0.082261 |
| ENSG00000164211 | STARD4 | 3449.08 | 1.091849 | 0.376585 | 3.24463 | 0.00117603 | 0.00657 |
| ENSG00000185046 | ANKS1B | 131.3939 | 1.091743 | 0.4210732 | 2.961405 | 0.00306239 | 0.014783 |
| ENSG00000101040 | ZMYND8 | 11260.9908 | 1.091433226 | 0.145895708 | 7.539859483 | 4.70E-14 | 1.81E-12 |
| ENSG00000280351 | | 97.77115 | 1.091283 | 0.379775 | 3.198992 | 0.00137909 | 0.00755 |
| ENSG00000186567 | CEACAM19 | 290.2705 | 1.091061 | 0.4714685 | 2.732293 | 0.00628952 | 0.026707 |
| ENSG00000288538 | | 90.37089 | 1.090896 | 0.4401162 | 2.863385 | 0.00419141 | 0.019187 |
| ENSG00000174137 | FAM53A | 147.9599 | 1.090493 | 0.2088183 | 5.402158 | 6.58E-08 | 1.04E-06 |
| ENSG00000117601 | SERPINC1 | 3.777319269 | 1.090388967 | 0.901987585 | 1.971624631 | 0.04865247 | 0.135668 |
| ENSG00000147459 | DOCK5 | 3136.564 | 1.090016 | 0.2798727 | 4.141339 | 3.45E-05 | 0.000298 |
| ENSG00000184669 | OR7E14P | 275.9998 | 1.089576 | 0.2143485 | 5.266008 | 1.39E-07 | 2.06E-06 |
| ENSG00000227269 | | 2.145025 | 1.089568 | 1.451885 | 1.984752 | 0.04717204 | 0.132514 |
| ENSG00000162571 | TTLL10 | 6.266644 | 1.089548 | 0.8618065 | 1.992266 | 0.04634193 | 0.130846 |
| ENSG00000206120 | EGFEM1P | 15.01126 | 1.089077 | 0.8005013 | 2.071609 | 0.03830192 | 0.113072 |
| ENSG00000213096 | ZNF254 | 604.3234 | 1.088838 | 0.2033053 | 5.525092 | 3.29E-08 | 5.46E-07 |
| ENSG00000250569 | NTAN1P2 | 32.38489 | 1.088783 | 0.3919049 | 3.10847 | 0.00188059 | 0.009828 |
| ENSG00000226944 | RNF207-AS1 | 19.15463 | 1.088524 | 0.6918899 | 2.172077 | 0.02984983 | 0.093315 |
| ENSG00000272398 | CD24 | 43305.88 | 1.08838 | 0.2282846 | 4.997308 | 5.81E-07 | 7.63E-06 |
| ENSG00000029153 | ARNTL2 | 625.2249027 | 1.088041182 | 0.21946842 | 5.159800256 | 2.47E-07 | 3.47E-06 |

| | | | | | | | |
|---|---|---|---|---|---|---|---|
| ENSG00000223776 | LGALS8-AS1 | 25.91721 | 1.086898 | 0.438381 | 2.845308 | 0.00443685 | 0.020076 |
| ENSG00000170846 | | 2937.162 | 1.086865 | 0.2264895 | 5.026278 | 5.00E-07 | 6.63E-06 |
| ENSG00000168502 | MTCL1 | 978.3984 | 1.086411 | 0.2609092 | 4.393066 | 1.12E-05 | 0.000109 |
| ENSG00000231249 | ITPR1-DT | 96.5017 | 1.086288 | 0.4707207 | 2.716709 | 0.00659345 | 0.027745 |
| ENSG00000186973 | FAM183A | 72.18086 | 1.085206 | 0.4928592 | 2.624361 | 0.00868117 | 0.034741 |
| ENSG00000131979 | GCH1 | 1927.521242 | 1.084657407 | 0.159532755 | 6.936926958 | 4.01E-12 | 1.22E-10 |
| ENSG00000164654 | MIOS | 3348.023 | 1.084587 | 0.1705469 | 6.499826 | 8.04E-11 | 2.02E-09 |
| ENSG00000233396 | LINC01719 | 158.1351 | 1.084565 | 0.319771 | 3.669451 | 0.00024307 | 0.001685 |
| ENSG00000279019 | | 14.42237 | 1.084488 | 0.7431769 | 2.111356 | 0.03474169 | 0.105053 |
| ENSG00000272498 | | 17.8382 | 1.084163 | 0.5255835 | 2.516068 | 0.01186723 | 0.044641 |
| ENSG00000272209 | | 27.14316 | 1.082975 | 0.6075278 | 2.312875 | 0.0207295 | 0.069956 |
| ENSG00000135469 | COQ10A | 325.4538633 | 1.082112436 | 0.181271712 | 6.124165799 | 9.12E-10 | 1.97E-08 |
| ENSG00000254614 | | 156.4856 | 1.081873 | 0.2977346 | 3.885268 | 0.00010222 | 0.000779 |
| ENSG00000270231 | NBPF8 | 1314.007 | 1.081813 | 0.2025118 | 5.50682 | 3.65E-08 | 6.01E-07 |
| ENSG00000198157 | HMGN5 | 299.7943 | 1.081793 | 0.3637516 | 3.297141 | 0.00097674 | 0.005607 |
| ENSG00000257869 | | 13.63558 | 1.081236 | 0.5419857 | 2.441767 | 0.01461556 | 0.052833 |
| ENSG00000186479 | RGS7BP | 7.923149 | 1.080975 | 0.6729531 | 2.182315 | 0.02908629 | 0.091628 |
| ENSG00000258290 | BRWD1P2 | 17.50384 | 1.080932 | 0.4901 | 2.619682 | 0.00880119 | 0.035147 |
| ENSG00000281501 | SEPSECS-AS1 | 103.1256 | 1.080681 | 0.299275 | 3.86344 | 0.0001118 | 0.000846 |
| ENSG00000132746 | ALDH3B2 | 15895.36171 | 1.080534621 | 0.307118065 | 3.747369316 | 0.0001787 | 0.001283 |
| ENSG00000143344 | RGL1 | 2613.171 | 1.080399 | 0.4374944 | 2.852482 | 0.00433793 | 0.019715 |
| ENSG00000229487 | ALG13-AS1 | 10.68714 | 1.08008 | 0.8424121 | 2.018101 | 0.04358076 | 0.124962 |
| ENSG00000145431 | PDGFC | 586.7402 | 1.078927 | 0.2601043 | 4.383899 | 1.17E-05 | 0.000113 |
| ENSG00000091409 | ITGA6 | 222.8718292 | 1.078879153 | 0.376068284 | 3.200150953 | 0.00137356 | 0.007526 |
| ENSG00000108387 | SEPTIN4 | 43.93203764 | 1.078733179 | 0.473023988 | 2.710709533 | 0.00671394 | 0.028148 |
| ENSG00000277147 | LINC00869 | 350.2577 | 1.078236 | 0.3225205 | 3.62267 | 0.00029158 | 0.001973 |
| ENSG00000287763 | | 65.96882 | 1.077116 | 0.3663441 | 3.254912 | 0.00113428 | 0.006377 |
| ENSG00000189067 | LITAF | 7454.999 | 1.076597 | 0.1632391 | 6.737343 | 1.61E-11 | 4.48E-10 |
| ENSG00000185813 | PCYT2 | 6878.741 | 1.076253 | 0.2209545 | 5.021098 | 5.14E-07 | 6.80E-06 |
| ENSG00000284707 | | 40.06658 | 1.076236 | 0.4857656 | 2.628433 | 0.00857793 | 0.034401 |
| ENSG00000143320 | CRABP2 | 10824.78 | 1.076065 | 0.472609 | 2.701266 | 0.0069076 | 0.02882 |
| ENSG00000144908 | ALDH1L1 | 19.10792 | 1.075678 | 0.6460458 | 2.186808 | 0.02875656 | 0.090719 |
| ENSG00000227201 | CNN2P1 | 4.509134 | 1.074857 | 0.9319125 | 1.957785 | 0.05025525 | 0.139 |

| | | | | | | | |
|---|---|---|---|---|---|---|---|
| ENSG00000181240 | SLC25A41 | 27.3425 | 1.074318 | 0.5856075 | 2.328811 | 0.0198691 | 0.06761 |
| ENSG00000259165 | DDX18P1 | 25.40094 | 1.074118 | 0.3859883 | 3.113853 | 0.00184662 | 0.009689 |
| ENSG00000214595 | EML6 | 641.5734 | 1.073989 | 0.3182121 | 3.656069 | 0.00025611 | 0.001761 |
| ENSG00000165025 | SYK | 680.5856 | 1.073408 | 0.3948497 | 3.062445 | 0.00219537 | 0.011202 |
| ENSG00000116183 | PAPPA2 | 6.567251363 | 1.072567942 | 0.737110311 | 2.091740401 | 0.03646174 | 0.108849 |
| ENSG00000170899 | GSTA4 | 248.6597 | 1.071901 | 0.3848576 | 3.11893 | 0.00181509 | 0.009548 |
| ENSG00000056998 | GYG2 | 712.2212215 | 1.071017933 | 0.269930483 | 4.203848692 | 2.62E-05 | 0.000233 |
| ENSG00000213277 | MARCKSL1P1 | 7.290149 | 1.069997 | 0.9320661 | 1.964123 | 0.04951583 | 0.137542 |
| ENSG00000267119 | RPL10P15 | 10.79455 | 1.069831 | 0.6381445 | 2.218101 | 0.02654792 | 0.085033 |
| ENSG00000258096 | | 9.082893 | 1.069579 | 0.6164457 | 2.259507 | 0.02385189 | 0.07817 |
| ENSG00000137871 | ZNF280D | 1246.810259 | 1.069149668 | 0.122327075 | 8.849966845 | 8.75E-19 | 5.76E-17 |
| ENSG00000272927 | | 44.21808 | 1.068656 | 0.4336773 | 2.8425 | 0.00447612 | 0.020226 |
| ENSG00000122145 | TBX22 | 1.646474737 | 1.066683454 | 1.695019071 | 2.179977394 | 0.02925914 | 0.092009 |
| ENSG00000207864 | MIR27B | 18.8369 | 1.066442 | 0.5952467 | 2.296835 | 0.02162818 | 0.072464 |
| ENSG00000081181 | ARG2 | 138.5304533 | 1.066388992 | 0.458711125 | 2.731914301 | 0.00629675 | 0.026734 |
| ENSG00000185988 | PLK5 | 45.70507 | 1.066291 | 0.4501174 | 2.74943 | 0.00596991 | 0.025616 |
| ENSG00000270413 | | 3.652658 | 1.065337 | 1.172329 | 1.921222 | 0.05470374 | 0.148551 |
| ENSG00000175115 | PACS1 | 10125.32 | 1.065051 | 0.2686752 | 4.176778 | 2.96E-05 | 0.000259 |
| ENSG00000151552 | QDPR | 929.9852 | 1.064899 | 0.2718943 | 4.153751 | 3.27E-05 | 0.000284 |
| ENSG00000111145 | ELK3 | 739.5326986 | 1.06484794 | 0.225088743 | 4.912501132 | 8.99E-07 | 1.13E-05 |
| ENSG00000099769 | IGFALS | 50.09175385 | 1.064521312 | 0.417282185 | 2.905365827 | 0.00366824 | 0.017163 |
| ENSG00000106852 | LHX6 | 150.0832441 | 1.063820435 | 0.455221218 | 2.742413898 | 0.00609894 | 0.026056 |
| ENSG00000204071 | TCEAL6 | 3.430115 | 1.063042 | 1.273748 | 1.92295 | 0.05448635 | 0.148158 |
| ENSG00000175003 | SLC22A1 | 6.13102 | 1.062638 | 0.8594321 | 1.926023 | 0.05410148 | 0.147428 |
| ENSG00000161653 | NAGS | 87.88235 | 1.060909 | 0.2504848 | 4.451543 | 8.53E-06 | 8.57E-05 |
| ENSG00000233487 | RPSAP69 | 23.56845 | 1.060858 | 0.5830534 | 2.317985 | 0.02045013 | 0.069171 |
| ENSG00000204920 | ZNF155 | 209.598 | 1.05916 | 0.2196016 | 5.016956 | 5.25E-07 | 6.95E-06 |
| ENSG00000167971 | CASKIN1 | 1115.958 | 1.0591 | 0.281977 | 3.992662 | 6.53E-05 | 0.000524 |
| ENSG00000260260 | SNHG19 | 1953.367 | 1.059063 | 0.2294513 | 4.81299 | 1.49E-06 | 1.79E-05 |
| ENSG00000237301 | | 18.56127 | 1.058642 | 0.4315419 | 2.809709 | 0.00495863 | 0.022005 |
| ENSG00000232818 | RPS2P32 | 1.964012 | 1.058388 | 1.418824 | 1.964295 | 0.04949595 | 0.137501 |
| ENSG00000217442 | SYCE3 | 15.75861 | 1.057927 | 0.4995658 | 2.533364 | 0.01129737 | 0.042945 |
| ENSG00000156639 | ZFAND3 | 5388.603 | 1.057412 | 0.131578 | 8.181923 | 2.79E-16 | 1.40E-14 |

| | | | | | | | |
|---|---|---|---|---|---|---|---|
| ENSG00000130513 | GDF15 | 4023.473187 | 1.056671589 | 0.48984559 | 2.588312343 | 0.00964475 | 0.037734 |
| ENSG00000198821 | CD247 | 2.216083 | 1.055853 | 1.459865 | 1.997622 | 0.04575767 | 0.129609 |
| ENSG00000151651 | ADAM8 | 497.2814 | 1.055502 | 0.1920354 | 5.665775 | 1.46E-08 | 2.59E-07 |
| ENSG00000123983 | ACSL3 | 10480.03878 | 1.055336663 | 0.125900918 | 8.490583954 | 2.06E-17 | 1.17E-15 |
| ENSG00000240666 | MME-AS1 | 2.0201 | 1.054838 | 1.471514 | 1.998395 | 0.0456738 | 0.129455 |
| ENSG00000279227 | | 52.22122 | 1.054463 | 0.4121133 | 2.929511 | 0.00339496 | 0.01611 |
| ENSG00000120051 | CFAP58 | 6.732900107 | 1.054248508 | 1.158176588 | 1.85875903 | 0.0630613 | 0.165006 |
| ENSG00000124593 | | 122.5880966 | 1.054041303 | 0.439081422 | 2.777279684 | 0.0054816 | 0.023875 |
| ENSG00000086506 | HBQ1 | 13.26084111 | 1.053825745 | 0.742743237 | 2.067298635 | 0.03870602 | 0.113973 |
| ENSG00000178997 | EXD1 | 3.49148 | 1.053701 | 0.973566 | 1.892179 | 0.05846717 | 0.156138 |
| ENSG00000117834 | SLC5A9 | 3.887968132 | 1.053596597 | 0.972810138 | 1.921467018 | 0.05467286 | 0.148501 |
| ENSG00000187013 | LINC02875 | 442.365 | 1.053585 | 0.2971281 | 3.803101 | 0.0001429 | 0.00105 |
| ENSG00000109686 | SH3D19 | 1599.212314 | 1.053292317 | 0.283264996 | 3.955829429 | 7.63E-05 | 0.000603 |
| ENSG00000241717 | VWFP1 | 2.160852 | 1.053275 | 1.450018 | 1.982302 | 0.04744542 | 0.133158 |
| ENSG00000055955 | ITIH4 | 2.842996953 | 1.052999619 | 1.170625604 | 1.875230853 | 0.06076097 | 0.160505 |
| ENSG00000162069 | BICDL2 | 3935.26 | 1.052466 | 0.2736621 | 4.063002 | 4.84E-05 | 0.000403 |
| ENSG00000205089 | CCNI2 | 50.05805 | 1.052115 | 0.5116292 | 2.523515 | 0.01161881 | 0.043893 |
| ENSG00000132196 | HSD17B7 | 1210.470976 | 1.051825307 | 0.230965726 | 4.754223911 | 1.99E-06 | 2.34E-05 |
| ENSG00000273796 | | 66.60499 | 1.051512 | 0.4369503 | 2.79653 | 0.00516546 | 0.022756 |
| ENSG00000228668 | TRGV5P | 2.045698 | 1.05093 | 1.969135 | 2.376265 | 0.0174889 | 0.061015 |
| ENSG00000280022 | | 8.889372 | 1.050674 | 0.7911306 | 2.009995 | 0.04443168 | 0.126854 |
| ENSG00000077264 | PAK3 | 22.68049594 | 1.050615961 | 0.604387727 | 2.251246587 | 0.02436992 | 0.079504 |
| ENSG00000230665 | THAP12P1 | 2.417862 | 1.050522 | 1.395894 | 1.970449 | 0.04878689 | 0.135943 |
| ENSG00000274024 | | 17.85275 | 1.050416 | 0.6146731 | 2.238869 | 0.02516441 | 0.081506 |
| ENSG00000187098 | MITF | 221.2154 | 1.049945 | 0.2279353 | 4.796393 | 1.62E-06 | 1.93E-05 |
| ENSG00000287578 | | 15.51855 | 1.049309 | 0.6642886 | 2.152338 | 0.03137071 | 0.097139 |
| ENSG00000238145 | TUBAP8 | 3.242248 | 1.049171 | 1.557893 | 2.114161 | 0.03450154 | 0.104559 |
| ENSG00000171388 | APLN | 7.755987 | 1.048967 | 0.6793261 | 2.120285 | 0.03398205 | 0.103303 |
| ENSG00000180089 | TMEM86B | 56.10292 | 1.048849 | 0.3859822 | 3.064785 | 0.00217826 | 0.01113 |
| ENSG00000136830 | NIBAN2 | 11760.45413 | 1.048597767 | 0.220029236 | 4.99228867 | 5.97E-07 | 7.79E-06 |
| ENSG00000234807 | LINC01135 | 391.8705 | 1.048125 | 0.2809535 | 3.976295 | 7.00E-05 | 0.000557 |
| ENSG00000224356 | | 5.023033 | 1.047986 | 0.9484362 | 1.912578 | 0.05580212 | 0.150702 |
| ENSG00000186994 | KANK3 | 32.67978 | 1.047937 | 0.5872247 | 2.277598 | 0.02275055 | 0.075388 |

| | | | | | | | |
|---|---|---|---|---|---|---|---|
| ENSG00000258334 | | 42.95715 | 1.047537 | 0.5787165 | 2.319205 | 0.02038391 | 0.06899 |
| ENSG00000106991 | ENG | 62.66938634 | 1.046958754 | 0.420535842 | 2.850388259 | 0.00436659 | 0.019832 |
| ENSG00000101203 | COL20A1 | 17.47366557 | 1.045149538 | 0.662956956 | 2.12118835 | 0.03390596 | 0.103154 |
| ENSG00000165935 | SMCO2 | 5.591908 | 1.045143 | 0.8545085 | 1.931903 | 0.05337145 | 0.145926 |
| ENSG00000085741 | WNT11 | 14.86944982 | 1.043812513 | 0.489196352 | 2.533651481 | 0.01128809 | 0.042924 |
| ENSG00000164751 | PEX2 | 2293.647 | 1.043298 | 0.1578768 | 6.73984 | 1.59E-11 | 4.42E-10 |
| ENSG00000115221 | ITGB6 | 2335.717076 | 1.043211746 | 0.498626578 | 2.530395739 | 0.01139339 | 0.043201 |
| ENSG00000225981 | | 5.166249 | 1.043114 | 0.9545946 | 1.919942 | 0.05486522 | 0.148853 |
| ENSG00000232412 | | 15.81362 | 1.042914 | 0.5881083 | 2.271306 | 0.02312847 | 0.076316 |
| ENSG00000103528 | SYT17 | 751.2787374 | 1.042282603 | 0.271496048 | 4.068181545 | 4.74E-05 | 0.000395 |
| ENSG00000236772 | | 8.196586 | 1.042203 | 0.8001296 | 2.008263 | 0.0446153 | 0.127255 |
| ENSG00000237939 | IDI1P1 | 4.054894 | 1.042143 | 0.8783648 | 1.921692 | 0.05464458 | 0.148471 |
| ENSG00000185008 | ROBO2 | 42.68016 | 1.040964 | 0.7971702 | 2.006076 | 0.04484811 | 0.127714 |
| ENSG00000227581 | STAU2P1 | 4.986598 | 1.04059 | 0.8244966 | 1.966427 | 0.0492493 | 0.136988 |
| ENSG00000180447 | GAS1 | 7.849583 | 1.040053 | 0.7170654 | 2.045457 | 0.0408098 | 0.118864 |
| ENSG00000237263 | MAPK6P3 | 4.881379 | 1.039739 | 0.8654506 | 1.933337 | 0.0531947 | 0.145531 |
| ENSG00000179528 | LBX2 | 192.9936 | 1.038739 | 0.1814629 | 5.875748 | 4.21E-09 | 8.24E-08 |
| ENSG00000207507 | RNU6-9 | 8.660326 | 1.038447 | 0.7292119 | 2.066687 | 0.03876365 | 0.11413 |
| ENSG00000104427 | ZC2HC1A | 502.2736039 | 1.038288406 | 0.228450505 | 4.732762051 | 2.21E-06 | 2.58E-05 |
| ENSG00000166165 | CKB | 2774.667 | 1.038271 | 0.2412572 | 4.509265 | 6.51E-06 | 6.75E-05 |
| ENSG00000258811 | | 83.0135 | 1.038249 | 0.5652055 | 2.322537 | 0.02020401 | 0.06854 |
| ENSG00000267750 | RUNDC3A-AS1 | 966.1574 | 1.036621 | 0.3145693 | 3.571954 | 0.00035433 | 0.002338 |
| ENSG00000162733 | DDR2 | 30.85808 | 1.036598 | 0.5055403 | 2.483244 | 0.01301919 | 0.048116 |
| ENSG00000260428 | SCX | 300.8093 | 1.035385 | 0.3226329 | 3.484457 | 0.00049314 | 0.003103 |
| ENSG00000081760 | AACS | 5738.393015 | 1.034713757 | 0.172130488 | 6.158097678 | 7.36E-10 | 1.62E-08 |
| ENSG00000001461 | NIPAL3 | 1531.979596 | 1.034050047 | 0.239017026 | 4.522726526 | 6.10E-06 | 6.38E-05 |
| ENSG00000260563 | | 476.9867 | 1.033858 | 0.3034502 | 3.663232 | 0.00024905 | 0.001719 |
| ENSG00000218586 | | 62.43233 | 1.03373 | 0.2931342 | 3.729527 | 0.00019184 | 0.001367 |
| ENSG00000259614 | TUBAP12 | 4.902371 | 1.033577 | 1.252621 | 1.895938 | 0.05796825 | 0.15507 |
| ENSG00000180438 | TPRXL | 17.55499 | 1.032566 | 0.9958025 | 1.772321 | 0.07634125 | 0.190473 |
| ENSG00000185955 | C7orf61 | 8.46891 | 1.032073 | 0.9769309 | 1.915767 | 0.05539472 | 0.149817 |
| ENSG00000285706 | | 4.92632 | 1.032004 | 1.001658 | 1.856739 | 0.06334835 | 0.165529 |
| ENSG00000065361 | ERBB3 | 15260.3147 | 1.031970313 | 0.170820131 | 6.185450022 | 6.19E-10 | 1.38E-08 |

| | | | | | | | |
|---|---|---|---|---|---|---|---|
| ENSG00000115266 | APC2 | 274.7826755 | 1.031768302 | 0.225957884 | 4.759027232 | 1.95E-06 | 2.29E-05 |
| ENSG00000253405 | EVX1-AS | 1.747505 | 1.03158 | 1.738471 | 2.247439 | 0.02461201 | 0.080098 |
| ENSG00000123560 | PLP1 | 6.135370151 | 1.030992695 | 0.736254065 | 2.008446869 | 0.04459583 | 0.127213 |
| ENSG00000186480 | INSIG1 | 12601.22 | 1.030535 | 0.3986549 | 2.969926 | 0.00297872 | 0.014484 |
| ENSG00000196932 | TMEM26 | 1.572216 | 1.029925 | 2.142207 | 2.549566 | 0.01078572 | 0.04134 |
| ENSG00000163141 | BNIPL | 450.5251 | 1.029546 | 0.349313 | 3.246032 | 0.00117026 | 0.006547 |
| ENSG00000149577 | SIDT2 | 1846.045 | 1.027831 | 0.1865534 | 5.664629 | 1.47E-08 | 2.61E-07 |
| ENSG00000163631 | ALB | 25.22663 | 1.027177 | 0.4558019 | 2.636751 | 0.00837042 | 0.033732 |
| ENSG00000170374 | SP7 | 2.366049 | 1.026497 | 1.409815 | 1.967932 | 0.04907591 | 0.136605 |
| ENSG00000106086 | PLEKHA8 | 1336.200059 | 1.026449638 | 0.285179821 | 3.846869179 | 0.00011964 | 0.000899 |
| ENSG00000164972 | C9orf24 | 5.66258 | 1.026263 | 0.753852 | 1.989016 | 0.04669947 | 0.131636 |
| ENSG00000266643 | MIR3677 | 10.09331 | 1.026181 | 0.6557704 | 2.10765 | 0.03506127 | 0.105733 |
| ENSG00000205669 | ACOT6 | 6.187684 | 1.026085 | 1.332815 | 2.030223 | 0.04233387 | 0.122231 |
| ENSG00000258302 | | 40.22879 | 1.026058 | 0.4080214 | 2.846105 | 0.00442575 | 0.020035 |
| ENSG00000185818 | NAT8L | 1287.974 | 1.025539 | 0.2379056 | 4.523496 | 6.08E-06 | 6.36E-05 |
| ENSG00000166750 | SLFN5 | 921.8004 | 1.025422 | 0.5875529 | 2.255768 | 0.02408515 | 0.078779 |
| ENSG00000135678 | CPM | 667.7538866 | 1.025141094 | 0.504814334 | 2.470720886 | 0.0134841 | 0.049442 |
| ENSG00000213920 | MDP1 | 7.684379 | 1.025137 | 0.6625069 | 2.096036 | 0.03607903 | 0.108131 |
| ENSG00000257526 | | 6.751464 | 1.025105 | 0.8908042 | 1.929867 | 0.05362332 | 0.146523 |
| ENSG00000138686 | BBS7 | 981.4433935 | 1.025027693 | 0.204530426 | 5.198583891 | 2.01E-07 | 2.87E-06 |
| ENSG00000070731 | ST6GALNAC2 | 309.1684749 | 1.024853807 | 0.257175338 | 4.203872563 | 2.62E-05 | 0.000233 |
| ENSG00000111266 | DUSP16 | 4080.445758 | 1.024339207 | 0.146667258 | 7.100578886 | 1.24E-12 | 4.00E-11 |
| ENSG00000236409 | NRADDP | 28.36632 | 1.023942 | 0.5217663 | 2.417035 | 0.01564751 | 0.055744 |
| ENSG00000186160 | CYP4Z1 | 47.44582 | 1.023936 | 0.7656535 | 1.934386 | 0.05306573 | 0.145281 |
| ENSG00000129646 | QRICH2 | 324.9369215 | 1.023530397 | 0.372390236 | 3.060105296 | 0.00221259 | 0.011277 |
| ENSG00000165716 | DIPK1B | 282.3098 | 1.023247 | 0.2362771 | 4.536218 | 5.73E-06 | 6.02E-05 |
| ENSG00000181798 | LINC00471 | 54.84694 | 1.023019 | 0.4190793 | 2.794025 | 0.00520564 | 0.022895 |
| ENSG00000139793 | MBNL2 | 1516.785 | 1.02281 | 0.4156166 | 2.817182 | 0.0048447 | 0.021626 |
| ENSG00000112511 | PHF1 | 2661.859802 | 1.022546587 | 0.249043699 | 4.325902961 | 1.52E-05 | 0.000143 |
| ENSG00000287811 | | 10.45818 | 1.021827 | 0.830029 | 1.94794 | 0.05142217 | 0.141522 |
| ENSG00000271119 | | 154.4901 | 1.021792 | 0.2277201 | 4.674256 | 2.95E-06 | 3.34E-05 |
| ENSG00000130962 | PRRG1 | 631.0448147 | 1.021370022 | 0.249102531 | 4.309660116 | 1.64E-05 | 0.000153 |
| ENSG00000154978 | VOPP1 | 1456.961 | 1.021103 | 0.1875738 | 5.607912 | 2.05E-08 | 3.52E-07 |

| | | | | | | | |
|---|---|---|---|---|---|---|---|
| ENSG00000251187 | | 1.734931 | 1.021089 | 1.609529 | 2.153621 | 0.03126991 | 0.096899 |
| ENSG00000241431 | RPL37P6 | 28.57918 | 1.020922 | 0.429173 | 2.73798 | 0.00618178 | 0.026364 |
| ENSG00000169242 | EFNA1 | 9925.38 | 1.019798 | 0.2503942 | 4.304344 | 1.67E-05 | 0.000156 |
| ENSG00000259015 | | 101.5561 | 1.019614 | 0.2982586 | 3.663096 | 0.00024919 | 0.001719 |
| ENSG00000198722 | UNC13B | 2559.148 | 1.019322 | 0.2143226 | 4.934649 | 8.03E-07 | 1.02E-05 |
| ENSG00000234255 | | 46.62639 | 1.018763 | 0.5304672 | 2.392116 | 0.01675153 | 0.058919 |
| ENSG00000167925 | GHDC | 107.5347 | 1.018268 | 0.3588404 | 3.144785 | 0.00166209 | 0.00883 |
| ENSG00000257475 | | 2.235071 | 1.018159 | 1.377638 | 2.054998 | 0.03987923 | 0.116601 |
| ENSG00000264107 | | 15.89987 | 1.017551 | 0.6429446 | 2.157481 | 0.03096821 | 0.096177 |
| ENSG00000196189 | SEMA4A | 1045.503 | 1.017381 | 0.2093146 | 5.033155 | 4.82E-07 | 6.42E-06 |
| ENSG00000249617 | | 2.563062 | 1.017012 | 1.295045 | 1.957728 | 0.05026197 | 0.139004 |
| ENSG00000182372 | CLN8 | 1576.799 | 1.015182 | 0.2523235 | 4.233764 | 2.30E-05 | 0.000207 |
| ENSG00000105877 | DNAH11 | 265.2402293 | 1.01415303 | 0.503337322 | 2.445826865 | 0.01445204 | 0.05235 |
| ENSG00000268756 | | 10.61005 | 1.014115 | 0.729786 | 2.024702 | 0.04289799 | 0.12353 |
| ENSG00000184922 | FMNL1 | 668.3732 | 1.013097 | 0.1671191 | 6.213325 | 5.19E-10 | 1.17E-08 |
| ENSG00000100181 | TPTEP1 | 7.494719225 | 1.013008219 | 0.955519577 | 1.923977299 | 0.05435743 | 0.147913 |
| ENSG00000261442 | | 3.170056 | 1.012939 | 1.34879 | 1.998318 | 0.04568222 | 0.129456 |
| ENSG00000222317 | RNA5SP118 | 2.260364 | 1.012848 | 1.635214 | 2.133742 | 0.03286394 | 0.100657 |
| ENSG00000254013 | MAP2K1P1 | 91.79756 | 1.01282 | 0.2667344 | 4.028553 | 5.61E-05 | 0.000459 |
| ENSG00000156170 | NDUFAF6 | 2069.315 | 1.01269 | 0.1506359 | 6.848102 | 7.48E-12 | 2.18E-10 |
| ENSG00000257433 | | 20.09421 | 1.012611 | 0.4688873 | 2.551022 | 0.01074074 | 0.041221 |
| ENSG00000134531 | EMP1 | 42.54034211 | 1.012318343 | 0.555896048 | 2.294769095 | 0.02174636 | 0.072786 |
| ENSG00000270959 | LPP-AS2 | 100.9126 | 1.012107 | 0.3610545 | 3.128586 | 0.0017565 | 0.00928 |
| ENSG00000147224 | PRPS1 | 3235.267 | 1.01121 | 0.3035832 | 3.589023 | 0.00033192 | 0.002212 |
| ENSG00000228657 | RPL23AP58 | 3.402556 | 1.01051 | 1.217479 | 1.833069 | 0.06679227 | 0.172335 |
| ENSG00000227242 | | 3.321461 | 1.010372 | 1.135412 | 1.856861 | 0.06333089 | 0.1655 |
| ENSG00000272888 | CHASERR | 2522.646 | 1.010367 | 0.2429483 | 4.363619 | 1.28E-05 | 0.000123 |
| ENSG00000278922 | | 211.8718 | 1.009639 | 0.2407091 | 4.392439 | 1.12E-05 | 0.000109 |
| ENSG00000254718 | | 17.9842 | 1.009325 | 0.4340481 | 2.683894 | 0.00727702 | 0.030097 |
| ENSG00000187240 | DYNC2H1 | 1224.529 | 1.009319 | 0.3061983 | 3.552814 | 0.00038113 | 0.002492 |
| ENSG00000235772 | | 63.5264 | 1.009141 | 0.4663037 | 2.569132 | 0.01019536 | 0.039504 |
| ENSG00000230881 | | 1.716799 | 1.008463 | 1.416709 | 2.029865 | 0.04237023 | 0.122292 |
| ENSG00000139651 | ZNF740 | 3633.563 | 1.008032 | 0.181396 | 5.713344 | 1.11E-08 | 2.02E-07 |

| | | | | | | | |
|---|---|---|---|---|---|---|---|
| ENSG00000285863 | | 28.96596 | 1.007856 | 0.7518884 | 1.99325 | 0.04623404 | 0.130624 |
| ENSG00000201998 | SNORA23 | 4.118262 | 1.007838 | 1.323567 | 2.024138 | 0.04295596 | 0.123637 |
| ENSG00000262801 | | 28.65934 | 1.007426 | 0.5210995 | 2.383776 | 0.01713603 | 0.059969 |
| ENSG00000176046 | NUPR1 | 5244.446 | 1.007246 | 1.025399 | 1.906568 | 0.05657651 | 0.152298 |
| ENSG00000102996 | MMP15 | 3118.969341 | 1.006896513 | 0.194710235 | 5.341548457 | 9.22E-08 | 1.41E-06 |
| ENSG00000273355 | | 62.41815 | 1.005987 | 0.3062112 | 3.526183 | 0.0004216 | 0.002707 |
| ENSG00000167981 | ZNF597 | 767.4141 | 1.005865 | 0.1933189 | 5.350371 | 8.78E-08 | 1.34E-06 |
| ENSG00000099812 | MISP | 4315.024067 | 1.00543648 | 0.227268047 | 4.62527991 | 3.74E-06 | 4.11E-05 |
| ENSG00000283342 | NBPF13P | 25.01336 | 1.004987 | 0.4489736 | 2.606583 | 0.00914506 | 0.036222 |
| ENSG00000198208 | RPS6KL1 | 2052.163 | 1.00432 | 0.2572427 | 4.138462 | 3.50E-05 | 0.000302 |
| ENSG00000272361 | | 4.599288 | 1.004282 | 0.9193296 | 1.891162 | 0.05860267 | 0.156377 |
| ENSG00000282572 | | 14.85048 | 1.003898 | 0.7100189 | 2.014869 | 0.04391835 | 0.125726 |
| ENSG00000186594 | MIR22HG | 1100.21 | 1.003489 | 0.1637873 | 6.326218 | 2.51E-10 | 5.93E-09 |
| ENSG00000185920 | PTCH1 | 579.6393 | 1.003484 | 0.287741 | 3.727897 | 0.00019308 | 0.001374 |
| ENSG00000170209 | ANKK1 | 5.852039 | 1.002838 | 0.9219489 | 1.861186 | 0.06271788 | 0.164383 |
| ENSG00000220008 | LINGO3 | 15.11039 | 1.002738 | 0.6693058 | 2.065878 | 0.03884002 | 0.114279 |
| ENSG00000256357 | | 6.584017 | 1.002187 | 1.3718 | 1.981417 | 0.04754455 | 0.133363 |
| ENSG00000140961 | OSGIN1 | 1309.41 | 1.001975 | 0.3445032 | 3.196907 | 0.0013891 | 0.007599 |
| ENSG00000198590 | C3orf35 | 37.07608 | 1.001801 | 0.4172182 | 2.743239 | 0.00608365 | 0.026003 |
| ENSG00000278899 | | 39.87357 | 1.001533 | 0.5220644 | 2.363807 | 0.01808822 | 0.062715 |
| ENSG00000167608 | TMC4 | 4861.232 | 1.001464 | 0.3554745 | 3.122369 | 0.00179402 | 0.009456 |
| ENSG00000257027 | | 248.5922 | 1.001353 | 0.2993212 | 3.608619 | 0.00030783 | 0.00207 |
| ENSG00000240540 | RPL3P9 | 7.211295 | 1.001212 | 0.7610719 | 1.946957 | 0.05153993 | 0.141797 |
| ENSG00000184205 | TSPYL2 | 4974.312 | 1.001202 | 0.2360647 | 4.473246 | 7.70E-06 | 7.84E-05 |
| ENSG00000187531 | SIRT7 | 5156.259 | 1.000933 | 0.1707822 | 5.967217 | 2.41E-09 | 4.89E-08 |
| ENSG00000144031 | ANKRD53 | 8.208703 | 1.000314 | 0.6857687 | 2.041886 | 0.0411628 | 0.119689 |
| ENSG00000283283 | | 11.98879 | 1.000112 | 0.8459913 | 1.934513 | 0.05305012 | 0.145263 |
| ENSG00000180611 | MB21D2 | 987.7776 | 0.9993083 | 0.2912762 | 3.67641 | 0.00023654 | 0.001645 |
| ENSG00000102316 | MAGED2 | 9677.398456 | 0.998691812 | 0.191359248 | 5.330881832 | 9.77E-08 | 1.48E-06 |
| ENSG00000223583 | | 2.280546 | 0.9973036 | 1.15255 | 1.791658 | 0.07318771 | 0.184771 |
| ENSG00000262521 | | 11.04411 | 0.9965041 | 0.7589673 | 1.971047 | 0.04871845 | 0.135809 |
| ENSG00000123388 | HOXC11 | 1188.796014 | 0.996172654 | 0.208788401 | 4.938721234 | 7.86E-07 | 9.99E-06 |
| ENSG00000278952 | | 124.5663 | 0.9959266 | 0.5247585 | 2.346633 | 0.01894392 | 0.065062 |

| | | | | | | | |
|---|---|---|---|---|---|---|---|
| ENSG00000230204 | FTH1P5 | 5.06837 | 0.995847 | 1.274432 | 1.895411 | 0.05803791 | 0.15521 |
| ENSG00000249853 | HS3ST5 | 36.82266 | 0.9958328 | 0.6658082 | 2.069039 | 0.03854243 | 0.113592 |
| ENSG00000184216 | IRAK1 | 15675.02 | 0.9956886 | 0.1638856 | 6.204158 | 5.50E-10 | 1.23E-08 |
| ENSG00000105472 | CLEC11A | 475.1620955 | 0.995325537 | 0.243830777 | 4.288902875 | 1.80E-05 | 0.000166 |
| ENSG00000242578 | | 5.413844 | 0.9953232 | 0.9877776 | 1.843299 | 0.06528533 | 0.169276 |
| ENSG00000230777 | RPS29P5 | 33.05238 | 0.9952695 | 0.4439422 | 2.620957 | 0.00876833 | 0.035032 |
| ENSG00000287315 | | 511.1816 | 0.9949599 | 0.2714213 | 3.896797 | 9.75E-05 | 0.000748 |
| ENSG00000092850 | TEKT2 | 2.037628399 | 0.994573409 | 1.447697833 | 2.063099824 | 0.03910315 | 0.11485 |
| ENSG00000279693 | | 10.67515 | 0.9941404 | 0.877634 | 1.866608 | 0.06195628 | 0.162822 |
| ENSG00000226334 | | 8.454893 | 0.9933927 | 1.394486 | 2.028781 | 0.04248064 | 0.122561 |
| ENSG00000232682 | | 15.73998 | 0.993391 | 0.6018401 | 2.151294 | 0.03145296 | 0.09733 |
| ENSG00000213750 | | 4.050329 | 0.9931166 | 0.9509191 | 1.801223 | 0.0716677 | 0.181866 |
| ENSG00000091262 | ABCC6 | 30.54408152 | 0.993104016 | 0.440146769 | 2.628859035 | 0.00856719 | 0.034363 |
| ENSG00000105509 | HAS1 | 10.46617916 | 0.991353191 | 0.903853652 | 1.829171129 | 0.06737398 | 0.173455 |
| ENSG00000279611 | | 58.37573 | 0.991347 | 0.3661206 | 3.011187 | 0.00260229 | 0.012943 |
| ENSG00000272223 | | 12.12714 | 0.9902628 | 0.5387075 | 2.274742 | 0.02292142 | 0.075777 |
| ENSG00000197816 | CCDC180 | 63.61962 | 0.9900041 | 0.650807 | 2.079013 | 0.03761615 | 0.111431 |
| ENSG00000228172 | | 41.18069 | 0.9897102 | 0.3857633 | 2.878569 | 0.00399483 | 0.018427 |
| ENSG00000125657 | TNFSF9 | 129.3549948 | 0.989140806 | 0.484190669 | 2.44683463 | 0.0144117 | 0.052245 |
| ENSG00000179399 | GPC5 | 145.3175 | 0.9890272 | 0.3713579 | 2.955185 | 0.00312481 | 0.015024 |
| ENSG00000260874 | | 16.25896 | 0.9890204 | 0.6211279 | 2.125929 | 0.03350915 | 0.102199 |
| ENSG00000165424 | ZCCHC24 | 232.2488 | 0.98884 | 0.3406673 | 3.183401 | 0.00145556 | 0.007904 |
| ENSG00000179008 | C14orf39 | 5.681752 | 0.9887448 | 0.8179307 | 1.879771 | 0.0601393 | 0.159306 |
| ENSG00000164506 | STXBP5 | 1821.204 | 0.9884168 | 0.2138635 | 4.796362 | 1.62E-06 | 1.93E-05 |
| ENSG00000134253 | TRIM45 | 1889.875881 | 0.98832799 | 0.180011497 | 5.624725411 | 1.86E-08 | 3.21E-07 |
| ENSG00000183668 | PSG9 | 199.3328 | 0.9876349 | 0.9260067 | 1.90077 | 0.05733212 | 0.153678 |
| ENSG00000269887 | | 28.54425 | 0.9874028 | 0.5422446 | 2.281405 | 0.02252448 | 0.074882 |
| ENSG00000134871 | COL4A2 | 22.74987545 | 0.986837278 | 0.884811875 | 1.793588334 | 0.0728789 | 0.184183 |
| ENSG00000279381 | | 10.41511 | 0.9864517 | 0.6301059 | 2.092544 | 0.03638985 | 0.108732 |
| ENSG00000266970 | | 99.25123 | 0.9863849 | 0.418733 | 2.701908 | 0.00689429 | 0.028787 |
| ENSG00000164823 | OSGIN2 | 4459.009 | 0.9860312 | 0.1939012 | 5.243922 | 1.57E-07 | 2.29E-06 |
| ENSG00000183963 | SMTN | 3071.901 | 0.9858795 | 0.3396561 | 3.191028 | 0.00141768 | 0.007726 |
| ENSG00000268287 | | 2.983597 | 0.9855579 | 1.169014 | 1.838184 | 0.06603529 | 0.17087 |

| | | | | | | | |
|---|---|---|---|---|---|---|---|
| ENSG00000011021 | CLCN6 | 2063.864456 | 0.985436883 | 0.221713963 | 4.624110279 | 3.76E-06 | 4.13E-05 |
| ENSG00000117640 | MTFR1L | 2423.493194 | 0.98508063 | 0.163847368 | 6.151531711 | 7.67E-10 | 1.68E-08 |
| ENSG00000249641 | HOXC13-AS | 493.8533 | 0.9844847 | 0.2640479 | 3.951562 | 7.76E-05 | 0.000612 |
| ENSG00000106689 | LHX2 | 185.4446459 | 0.98442682 | 0.6305436 | 2.107545605 | 0.03507031 | 0.10574 |
| ENSG00000197321 | SVIL | 4465.563 | 0.9841991 | 0.1876041 | 5.401733 | 6.60E-08 | 1.04E-06 |
| ENSG00000261336 | EIF4BP5 | 11.55468 | 0.9839257 | 0.6170969 | 2.101396 | 0.03560621 | 0.106992 |
| ENSG00000260923 | LINC02193 | 8.713474 | 0.9837482 | 0.9025429 | 1.874155 | 0.0609091 | 0.160816 |
| ENSG00000175854 | SWI5 | 1589.431 | 0.9836035 | 0.2856989 | 3.66238 | 0.00024988 | 0.001723 |
| ENSG00000102125 | TAZ | 2651.837418 | 0.983578067 | 0.217671657 | 4.70003598 | 2.60E-06 | 2.98E-05 |
| ENSG00000250317 | SMIM20 | 683.6553 | 0.9831085 | 0.2121365 | 4.805472 | 1.54E-06 | 1.85E-05 |
| ENSG00000225526 | MKRN2OS | 141.2169 | 0.9824557 | 0.3056532 | 3.467131 | 0.00052605 | 0.003281 |
| ENSG00000247363 | | 12.66926 | 0.982405 | 0.7113209 | 1.969061 | 0.04894609 | 0.136301 |
| ENSG00000232536 | | 20.63479 | 0.9822157 | 0.6587304 | 2.036731 | 0.04167697 | 0.120781 |
| ENSG00000106049 | HIBADH | 1756.304644 | 0.982121557 | 0.191679697 | 5.284700355 | 1.26E-07 | 1.87E-06 |
| ENSG00000271253 | | 2.617038 | 0.9816588 | 1.320467 | 1.910244 | 0.05610183 | 0.151327 |
| ENSG00000135407 | AVIL | 328.790601 | 0.981064139 | 0.372669822 | 2.933877516 | 0.00334756 | 0.015919 |
| ENSG00000048471 | SNX29 | 1722.716107 | 0.980903793 | 0.210079527 | 4.849928764 | 1.24E-06 | 1.51E-05 |
| ENSG00000280587 | LINC01348 | 58.56612 | 0.9803978 | 0.55413 | 2.248058 | 0.02457251 | 0.079979 |
| ENSG00000168676 | KCTD19 | 6.75486 | 0.9802277 | 0.7631767 | 1.899918 | 0.05744394 | 0.1539 |
| ENSG00000204934 | ATP6V0E2-AS1 | 217.5357 | 0.9801577 | 0.2387242 | 4.294397 | 1.75E-05 | 0.000163 |
| ENSG00000231359 | | 24.16314 | 0.9801448 | 0.6880101 | 2.003783 | 0.04509332 | 0.128233 |
| ENSG00000258658 | | 4.220195 | 0.9792706 | 1.041748 | 1.834945 | 0.06651382 | 0.171824 |
| ENSG00000283554 | LINC02341 | 3.475661 | 0.9790945 | 1.122407 | 1.779292 | 0.07519187 | 0.188416 |
| ENSG00000180769 | WDFY3-AS2 | 373.4196 | 0.9790094 | 0.206257 | 4.90863 | 9.17E-07 | 1.15E-05 |
| ENSG00000197647 | ZNF433 | 409.9968 | 0.9787227 | 0.2236663 | 4.563855 | 5.02E-06 | 5.35E-05 |
| ENSG00000140939 | NOL3 | 2486.67 | 0.9782164 | 0.1321464 | 7.50074 | 6.35E-14 | 2.42E-12 |
| ENSG00000081386 | ZNF510 | 941.9009164 | 0.978191848 | 0.214956169 | 4.714146631 | 2.43E-06 | 2.80E-05 |
| ENSG00000132906 | CASP9 | 386.965351 | 0.977747864 | 0.247485864 | 4.149617452 | 3.33E-05 | 0.000289 |
| ENSG00000138413 | IDH1 | 8804.648446 | 0.97773879 | 0.241219716 | 4.290127386 | 1.79E-05 | 0.000165 |
| ENSG00000273199 | | 5.367016 | 0.9775803 | 0.9370411 | 1.840857 | 0.06564248 | 0.170036 |
| ENSG00000124224 | PPP4R1L | 1120.116109 | 0.976780388 | 0.195660566 | 5.152320604 | 2.57E-07 | 3.60E-06 |
| ENSG00000140262 | TCF12 | 1729.859 | 0.9764194 | 0.1824069 | 5.494982 | 3.91E-08 | 6.41E-07 |
| ENSG00000273287 | | 3.542403 | 0.9763122 | 1.339474 | 1.939319 | 0.05246255 | 0.143877 |

| | | | | | | | |
|---|---|---|---|---|---|---|---|
| ENSG00000167165 | UGT1A6 | 48.16513 | 0.9760577 | 1.372866 | 2.030918 | 0.04226333 | 0.122072 |
| ENSG00000259884 | NR4A1AS | 125.8734 | 0.9758069 | 1.274113 | 1.984082 | 0.04724675 | 0.132667 |
| ENSG00000129493 | HEATR5A | 1405.650676 | 0.975661481 | 0.254202906 | 4.05634825 | 4.98E-05 | 0.000413 |
| ENSG00000118985 | ELL2 | 739.3114659 | 0.974695586 | 0.207411292 | 4.8688633 | 1.12E-06 | 1.38E-05 |
| ENSG00000083720 | OXCT1 | 4938.376134 | 0.973926826 | 0.289862575 | 3.598382117 | 0.0003202 | 0.002144 |
| ENSG00000127578 | WFIKKN1 | 86.67743712 | 0.973868082 | 0.44182509 | 2.568438468 | 0.01021579 | 0.039572 |
| ENSG00000089335 | ZNF302 | 1702.404247 | 0.973840532 | 0.232166135 | 4.389030204 | 1.14E-05 | 0.000111 |
| ENSG00000105427 | CNFN | 109.6635725 | 0.973633096 | 0.341404425 | 3.139317496 | 0.00169342 | 0.008979 |
| ENSG00000235837 | | 16.08067 | 0.9735842 | 0.5941795 | 2.131237 | 0.0330696 | 0.101189 |
| ENSG00000234390 | USP27X-AS1 | 87.27334 | 0.973536 | 0.3046424 | 3.439468 | 0.00058286 | 0.003587 |
| ENSG00000238054 | | 3.176321 | 0.9732009 | 1.51221 | 2.041031 | 0.04124778 | 0.11984 |
| ENSG00000177173 | NAP1L4P1 | 36.33441 | 0.9723639 | 0.3536739 | 3.050813 | 0.00228223 | 0.011565 |
| ENSG00000101213 | PTK6 | 2270.146793 | 0.972319687 | 0.243009926 | 4.200337625 | 2.67E-05 | 0.000236 |
| ENSG00000259291 | ZNF710-AS1 | 307.7198 | 0.9717275 | 0.2857814 | 3.646284 | 0.00026606 | 0.001822 |
| ENSG00000143867 | OSR1 | 6.201868 | 0.9715006 | 0.8211357 | 1.879622 | 0.06015958 | 0.159344 |
| ENSG00000139537 | CCDC65 | 77.509 | 0.9714356 | 0.3076918 | 3.40536 | 0.00066077 | 0.003997 |
| ENSG00000182168 | UNC5C | 100.1697 | 0.9708053 | 0.5758597 | 2.161709 | 0.03064058 | 0.095338 |
| ENSG00000134245 | WNT2B | 341.7095298 | 0.970654502 | 0.58756392 | 2.141216298 | 0.0322566 | 0.099181 |
| ENSG00000204653 | ASPDH | 24.49391 | 0.9703698 | 0.6578464 | 2.025931 | 0.04277191 | 0.123241 |
| ENSG00000272264 | | 11.38709 | 0.9699134 | 0.6516287 | 2.027074 | 0.04265481 | 0.122984 |
| ENSG00000198589 | LRBA | 9022.721 | 0.9697966 | 0.2211635 | 4.585204 | 4.54E-06 | 4.89E-05 |
| ENSG00000235760 | MSH2-OT1 | 5.907078 | 0.9691169 | 1.005181 | 1.844728 | 0.06507711 | 0.168867 |
| ENSG00000273327 | OR6L2P | 5.372208 | 0.9687598 | 1.014802 | 1.848169 | 0.06457791 | 0.167901 |
| ENSG00000133138 | TBC1D8B | 751.4856614 | 0.968739894 | 0.283889592 | 3.644317167 | 0.0002681 | 0.001833 |
| ENSG00000090539 | CHRD | 859.6580027 | 0.968608179 | 0.526630168 | 2.281123143 | 0.02254116 | 0.074909 |
| ENSG00000280789 | PAGR1 | 1332.898 | 0.9684807 | 0.2385546 | 4.254301 | 2.10E-05 | 0.00019 |
| ENSG00000260630 | SNAI3-AS1 | 447.319 | 0.96808 | 0.2224534 | 4.536572 | 5.72E-06 | 6.01E-05 |
| ENSG00000105327 | BBC3 | 1506.763337 | 0.967931278 | 0.432156299 | 2.590439595 | 0.00958534 | 0.037543 |
| ENSG00000260443 | | 20.23984 | 0.9676006 | 0.4545155 | 2.502885 | 0.01231856 | 0.045968 |
| ENSG00000235939 | | 8.936338 | 0.9673391 | 0.7215208 | 1.943148 | 0.05199824 | 0.142765 |
| ENSG00000213722 | DDAH2 | 4772.306 | 0.9673184 | 0.2124653 | 4.727929 | 2.27E-06 | 2.63E-05 |
| ENSG00000243742 | RPLP0P2 | 44.75415 | 0.9670578 | 0.5166906 | 2.295564 | 0.0217008 | 0.072658 |
| ENSG00000248092 | NNT-AS1 | 857.7968 | 0.966998 | 0.1688469 | 5.857239 | 4.71E-09 | 9.11E-08 |

| | | | | | | | |
|---|---|---|---|---|---|---|---|
| ENSG00000256209 | | 4.453617 | 0.9664148 | 0.9562642 | 1.806968 | 0.07076722 | 0.180132 |
| ENSG00000173991 | TCAP | 27.83072 | 0.965858 | 0.6275469 | 2.063137 | 0.03909963 | 0.11485 |
| ENSG00000250483 | PPM1AP1 | 7.726877 | 0.9656904 | 0.8798698 | 1.864388 | 0.06226728 | 0.16346 |
| ENSG00000162444 | RBP7 | 9.878345 | 0.9655233 | 0.8580077 | 1.854788 | 0.06362656 | 0.166028 |
| ENSG00000172260 | NEGR1 | 5.840894 | 0.9650048 | 0.8606272 | 1.853069 | 0.06387243 | 0.166506 |
| ENSG00000106415 | GLCCI1 | 1458.770797 | 0.964654235 | 0.200313147 | 4.992618819 | 5.96E-07 | 7.78E-06 |
| ENSG00000131094 | C1QL1 | 60.97654871 | 0.963738769 | 0.751812558 | 1.956552582 | 0.0504001 | 0.139342 |
| ENSG00000111371 | SLC38A1 | 44333.09524 | 0.963696828 | 0.241466478 | 4.185092862 | 2.85E-05 | 0.000251 |
| ENSG00000154229 | PRKCA | 1945.874 | 0.9636488 | 0.3160483 | 3.310378 | 0.0009317 | 0.005373 |
| ENSG00000180346 | TIGD2 | 1067.429 | 0.9631013 | 0.2101684 | 4.754132 | 1.99E-06 | 2.34E-05 |
| ENSG00000160111 | CPAMD8 | 417.6822 | 0.9630288 | 0.3521362 | 3.02796 | 0.00246211 | 0.012343 |
| ENSG00000164099 | PRSS12 | 90.55325 | 0.9626877 | 0.6032617 | 2.096174 | 0.0360668 | 0.108107 |
| ENSG00000172893 | DHCR7 | 20426.93 | 0.9622354 | 0.2945768 | 3.550999 | 0.00038377 | 0.002505 |
| ENSG00000262919 | CCNQ | 1632.586 | 0.9619535 | 0.1783506 | 5.550582 | 2.85E-08 | 4.76E-07 |
| ENSG00000261534 | | 201.143 | 0.9618193 | 0.4586614 | 2.473968 | 0.01336215 | 0.049063 |
| ENSG00000136457 | CHAD | 80.2218263 | 0.961595121 | 0.504169029 | 2.318458089 | 0.02042444 | 0.069093 |
| ENSG00000147852 | VLDLR | 553.0774 | 0.9614624 | 0.5126096 | 2.299954 | 0.02145085 | 0.071988 |
| ENSG00000182218 | HHIPL1 | 5.078417 | 0.9614523 | 0.7037334 | 1.928847 | 0.05374986 | 0.146741 |
| ENSG00000119514 | GALNT12 | 794.5162704 | 0.960633477 | 0.260200408 | 3.902552071 | 9.52E-05 | 0.000733 |
| ENSG00000183208 | GDPGP1 | 75.32195 | 0.9593268 | 0.4093588 | 2.677907 | 0.00740838 | 0.03055 |
| ENSG00000185053 | SGCZ | 19.55461 | 0.9591545 | 0.5643887 | 2.154835 | 0.03117475 | 0.096634 |
| ENSG00000231073 | | 25.9355 | 0.9591217 | 0.4656476 | 2.431398 | 0.01504066 | 0.054039 |
| ENSG00000137976 | DNASE2B | 1.967752778 | 0.958806399 | 1.660709702 | 2.257276922 | 0.02399078 | 0.078528 |
| ENSG00000072954 | TMEM38A | 542.9798674 | 0.958616238 | 0.257158439 | 3.934197071 | 8.35E-05 | 0.000653 |
| ENSG00000177854 | TMEM187 | 975.9788 | 0.9584464 | 0.1804146 | 5.46129 | 4.73E-08 | 7.66E-07 |
| ENSG00000270441 | LAMB2P1 | 51.14268 | 0.9581302 | 0.4307921 | 2.571417 | 0.01012834 | 0.039296 |
| ENSG00000163645 | ERICH6 | 15.93533 | 0.9580508 | 0.522258 | 2.251516 | 0.02435285 | 0.079461 |
| ENSG00000251429 | AIDAP2 | 29.99538 | 0.9579903 | 0.4054584 | 2.689955 | 0.00714617 | 0.029662 |
| ENSG00000234695 | TFPI2-DT | 11.3725 | 0.9579855 | 0.6770635 | 1.94894 | 0.05130254 | 0.141323 |
| ENSG00000143797 | MBOAT2 | 3212.819 | 0.9565793 | 0.1757214 | 5.580457 | 2.40E-08 | 4.08E-07 |
| ENSG00000112419 | PHACTR2 | 2660.587817 | 0.956505508 | 0.281592364 | 3.626894213 | 0.00028685 | 0.001945 |
| ENSG00000288075 | | 121.2845 | 0.9557893 | 0.4372426 | 2.541693 | 0.01103169 | 0.042124 |
| ENSG00000109089 | CDR2L | 2242.304789 | 0.955699872 | 0.187180486 | 5.248916524 | 1.53E-07 | 2.24E-06 |

| | | | | | | | |
|---|---|---|---|---|---|---|---|
| ENSG00000165138 | ANKS6 | 3408.173 | 0.9552072 | 0.2158561 | 4.587007 | 4.50E-06 | 4.86E-05 |
| ENSG00000182687 | GALR2 | 25.62169 | 0.9549043 | 0.4057672 | 2.672507 | 0.00752867 | 0.030969 |
| ENSG00000214432 | VPS33B-DT | 22.36386 | 0.9548377 | 0.745541 | 1.917057 | 0.05523071 | 0.149585 |
| ENSG00000153317 | ASAP1 | 8951.494 | 0.9547645 | 0.239483 | 4.208217 | 2.57E-05 | 0.000229 |
| ENSG00000143409 | MINDY1 | 1975.12 | 0.9547034 | 0.2754549 | 3.686022 | 0.00022779 | 0.00159 |
| ENSG00000262587 | | 134.2118 | 0.9541869 | 0.2784328 | 3.647459 | 0.00026485 | 0.001815 |
| ENSG00000198155 | ZNF876P | 55.73992 | 0.9540583 | 0.3779488 | 2.832411 | 0.00461984 | 0.020796 |
| ENSG00000139055 | ERP27 | 138.7460626 | 0.95400947 | 0.379840681 | 2.8114763 | 0.00493147 | 0.021903 |
| ENSG00000066855 | MTFR1 | 2884.157322 | 0.953748477 | 0.171529189 | 5.691706403 | 1.26E-08 | 2.26E-07 |
| ENSG00000253741 | LNCOC1 | 129.5604 | 0.953179 | 0.2895147 | 3.537598 | 0.00040378 | 0.002612 |
| ENSG00000119711 | ALDH6A1 | 2200.432773 | 0.952507827 | 0.316964718 | 3.263040948 | 0.00110224 | 0.006226 |
| ENSG00000285210 | LINC02795 | 7.106895 | 0.9518329 | 1.159047 | 1.868982 | 0.06162526 | 0.162256 |
| ENSG00000263325 | | 51.30221 | 0.9515995 | 0.5911154 | 2.08976 | 0.03663938 | 0.109232 |
| ENSG00000228705 | LINC00659 | 122.2691 | 0.9512332 | 0.5704414 | 2.156954 | 0.03100927 | 0.096278 |
| ENSG00000280106 | | 31.8697 | 0.9504687 | 0.5670425 | 2.146691 | 0.03181792 | 0.098219 |
| ENSG00000198585 | NUDT16 | 3484.802 | 0.9500197 | 0.2202115 | 4.491945 | 7.06E-06 | 7.24E-05 |
| ENSG00000230337 | EXOSC10-AS1 | 20.63102 | 0.9499037 | 0.4525635 | 2.479442 | 0.01315882 | 0.048511 |
| ENSG00000131370 | SH3BP5 | 1707.443996 | 0.949513123 | 0.201295735 | 4.877790788 | 1.07E-06 | 1.33E-05 |
| ENSG00000155099 | PIP4P2 | 269.5994 | 0.9490804 | 0.226581 | 4.372884 | 1.23E-05 | 0.000118 |
| ENSG00000184226 | PCDH9 | 519.0998 | 0.9489516 | 0.4309046 | 2.550663 | 0.01075184 | 0.041251 |
| ENSG00000235584 | | 6.610256 | 0.9488668 | 0.9833274 | 1.821434 | 0.06854099 | 0.175711 |
| ENSG00000287071 | CDRT15P4 | 30.21824 | 0.9486831 | 0.4765437 | 2.368251 | 0.01787242 | 0.062051 |
| ENSG00000104219 | ZDHHC2 | 554.4007295 | 0.948503028 | 0.349415203 | 3.001943557 | 0.00268262 | 0.01327 |
| ENSG00000150471 | ADGRL3 | 87.92102 | 0.9484015 | 0.3754863 | 2.822707 | 0.00476201 | 0.021318 |
| ENSG00000203697 | CAPN8 | 1091.819 | 0.9482475 | 0.814057 | 1.865211 | 0.06215181 | 0.163205 |
| ENSG00000259905 | PWRN1 | 1.788515 | 0.9478565 | 1.536355 | 2.105292 | 0.03526592 | 0.106198 |
| ENSG00000247516 | MIR4458HG | 268.3808 | 0.9475197 | 0.2421685 | 4.112098 | 3.92E-05 | 0.000334 |
| ENSG00000223298 | RNY3P8 | 13.02443 | 0.9470772 | 0.8519115 | 1.866218 | 0.06201082 | 0.162929 |
| ENSG00000262691 | | 145.5606 | 0.9470274 | 0.4021276 | 2.681071 | 0.0073387 | 0.03031 |
| ENSG00000253671 | | 103.2268 | 0.9467951 | 0.4640371 | 2.415333 | 0.01572082 | 0.05594 |
| ENSG00000148219 | ASTN2 | 583.0314 | 0.9463574 | 0.2207466 | 4.468075 | 7.89E-06 | 8.00E-05 |
| ENSG00000254827 | SLC22A18AS | 18.42819 | 0.9459277 | 0.6582085 | 2.004897 | 0.04497403 | 0.127971 |
| ENSG00000161905 | ALOX15 | 546.4574 | 0.9459019 | 0.2521846 | 3.953788 | 7.69E-05 | 0.000607 |

| | | | | | | | |
|---|---|---|---|---|---|---|---|
| ENSG00000137731 | FXYD2 | 15.96521684 | 0.945579256 | 0.711711676 | 1.916866914 | 0.05525484 | 0.149619 |
| ENSG00000183741 | CBX6 | 6177.079 | 0.9452564 | 0.2212787 | 4.448186 | 8.66E-06 | 8.67E-05 |
| ENSG00000282977 | PCBP2-OT1 | 3.521965 | 0.9450253 | 0.9210036 | 1.765431 | 0.07749142 | 0.192598 |
| ENSG00000196810 | CTBP1-DT | 1041.702 | 0.9447902 | 0.1568326 | 6.149777 | 7.76E-10 | 1.70E-08 |
| ENSG00000227765 | MYL12BP2 | 14.53543 | 0.944491 | 0.6934274 | 1.954673 | 0.05062173 | 0.139795 |
| ENSG00000105479 | CCDC114 | 207.8730382 | 0.944181734 | 0.298254517 | 3.422084403 | 0.00062143 | 0.003789 |
| ENSG00000235209 | | 9.171343 | 0.9441024 | 0.7326306 | 1.866815 | 0.06192737 | 0.162813 |
| ENSG00000102878 | HSF4 | 1317.768788 | 0.943670342 | 0.219417797 | 4.49334631 | 7.01E-06 | 7.20E-05 |
| ENSG00000261141 | | 4.064604 | 0.9435829 | 0.9849528 | 1.818511 | 0.06898605 | 0.176579 |
| ENSG00000125409 | TEKT3 | 10.66592452 | 0.943501034 | 0.980193512 | 1.832963821 | 0.06680795 | 0.172349 |
| ENSG00000277128 | | 38.45125 | 0.9431438 | 0.4850119 | 2.338733 | 0.01934924 | 0.06618 |
| ENSG00000170523 | KRT83 | 75.50982 | 0.9427733 | 0.5127446 | 2.270983 | 0.02314801 | 0.076362 |
| ENSG00000171984 | SHLD1 | 82.20129 | 0.9422675 | 0.2808314 | 3.570804 | 0.00035589 | 0.002347 |
| ENSG00000154258 | ABCA9 | 21.7609 | 0.9409316 | 0.640375 | 1.984476 | 0.04720282 | 0.132572 |
| ENSG00000149212 | SESN3 | 1726.471 | 0.9408468 | 0.3482235 | 3.001386 | 0.00268754 | 0.01328 |
| ENSG00000138615 | CILP | 29.92770898 | 0.940829149 | 0.435637823 | 2.505832894 | 0.01221633 | 0.045676 |
| ENSG00000233922 | LINC01694 | 15.64724 | 0.9403541 | 0.9880593 | 1.720106 | 0.08541319 | 0.207398 |
| ENSG00000239344 | | 2.972381 | 0.9399624 | 0.9983719 | 1.751673 | 0.0798301 | 0.1969 |
| ENSG00000279930 | | 18.81836 | 0.9397193 | 0.6976759 | 1.918326 | 0.05506974 | 0.14927 |
| ENSG00000118194 | TNNT2 | 213.6388433 | 0.939628212 | 0.45017886 | 2.444490571 | 0.01450569 | 0.052471 |
| ENSG00000231184 | CCNQP3 | 5.772168 | 0.9394709 | 0.898294 | 1.8084 | 0.07054421 | 0.179667 |
| ENSG00000135334 | AKIRIN2 | 2169.381989 | 0.939156015 | 0.163024072 | 5.915971288 | 3.30E-09 | 6.56E-08 |
| ENSG00000149809 | TM7SF2 | 5613.004 | 0.9389352 | 0.2680275 | 3.71688 | 0.0002017 | 0.001429 |
| ENSG00000254084 | | 2.666105 | 0.9387755 | 1.268536 | 1.974224 | 0.04835633 | 0.135049 |
| ENSG00000261490 | | 68.98929 | 0.9385892 | 0.5874575 | 2.083363 | 0.03721811 | 0.110585 |
| ENSG00000141424 | SLC39A6 | 20289.99 | 0.9381958 | 0.3325429 | 3.084208 | 0.00204095 | 0.010541 |
| ENSG00000159625 | DRC7 | 3.97033 | 0.9381843 | 0.8472751 | 1.805987 | 0.07092039 | 0.180349 |
| ENSG00000120471 | TP53AIP1 | 6.785995693 | 0.93813123 | 0.814902437 | 1.826190591 | 0.06782156 | 0.174337 |
| ENSG00000203288 | TDRKH-AS1 | 391.374 | 0.9378902 | 0.1993287 | 4.862352 | 1.16E-06 | 1.43E-05 |
| ENSG00000286451 | | 720.5449 | 0.9377136 | 0.1632054 | 5.874587 | 4.24E-09 | 8.28E-08 |
| ENSG00000235023 | | 5.028989 | 0.9376312 | 1.34495 | 1.992019 | 0.04636898 | 0.13088 |
| ENSG00000109736 | MFSD10 | 6485.232248 | 0.93718266 | 0.173252743 | 5.545401626 | 2.93E-08 | 4.88E-07 |
| ENSG00000125388 | GRK4 | 211.8739683 | 0.937079711 | 0.224870736 | 4.327412039 | 1.51E-05 | 0.000142 |

| | | | | | | | |
|---|---|---|---|---|---|---|---|
| ENSG00000268262 | NA | 4.069263 | 0.9368719 | 1.14486 | 1.781972 | 0.07475385 | 0.187637 |
| ENSG00000226763 | SRRM5 | 78.54221 | 0.9368557 | 0.3780875 | 2.778823 | 0.00545563 | 0.023778 |
| ENSG00000075461 | CACNG4 | 11607.56057 | 0.936820433 | 0.252639935 | 3.948306868 | 7.87E-05 | 0.000619 |
| ENSG00000155974 | GRIP1 | 908.9024 | 0.9367641 | 0.2734449 | 3.648469 | 0.00026381 | 0.001809 |
| ENSG00000249996 | PPIC-AS1 | 14.25772 | 0.9367281 | 0.5710289 | 2.113048 | 0.03459667 | 0.104824 |
| ENSG00000149084 | HSD17B12 | 841.5377 | 0.9364458 | 0.2830345 | 3.541144 | 0.0003984 | 0.002583 |
| ENSG00000013563 | DNASE1L1 | 1070.20017 | 0.936389696 | 0.134294372 | 7.078982553 | 1.45E-12 | 4.64E-11 |
| ENSG00000130821 | SLC6A8 | 1141.907037 | 0.936290118 | 0.218959956 | 4.450958181 | 8.55E-06 | 8.58E-05 |
| ENSG00000040341 | STAU2 | 1992.277444 | 0.935925246 | 0.115975769 | 8.158240496 | 3.40E-16 | 1.69E-14 |
| ENSG00000243289 | AGAP13P | 9.233763 | 0.935909 | 0.6766563 | 1.945841 | 0.05167381 | 0.14208 |
| ENSG00000248323 | LUCAT1 | 28.94934 | 0.9358179 | 0.5218529 | 2.212161 | 0.02695557 | 0.086067 |
| ENSG00000227398 | KIF9-AS1 | 108.1911 | 0.9353642 | 0.3118056 | 3.246027 | 0.00117028 | 0.006547 |
| ENSG00000272908 | | 4.322295 | 0.9347698 | 1.730972 | 2.450311 | 0.01427328 | 0.051842 |
| ENSG00000100767 | PAPLN | 191.2687961 | 0.934767605 | 0.220810749 | 4.404001059 | 1.06E-05 | 0.000104 |
| ENSG00000084628 | NKAIN1 | 444.1638551 | 0.934534305 | 0.4773361 | 2.344779364 | 0.01903834 | 0.065327 |
| ENSG00000226167 | AP4B1-AS1 | 928.3658 | 0.9345147 | 0.2135021 | 4.542368 | 5.56E-06 | 5.86E-05 |
| ENSG00000132122 | SPATA6 | 204.1561628 | 0.934257599 | 0.265295977 | 3.727734521 | 0.00019321 | 0.001374 |
| ENSG00000253948 | VPS13B-DT | 91.28272 | 0.9331174 | 0.2746216 | 3.615391 | 0.0002999 | 0.002021 |
| ENSG00000287616 | | 5.006027 | 0.9330255 | 0.8939653 | 1.775849 | 0.07575779 | 0.189512 |
| ENSG00000241489 | | 73.01342 | 0.9329162 | 0.3815985 | 2.7453 | 0.00604557 | 0.025878 |
| ENSG00000272841 | MAP3K4-AS1 | 81.47873 | 0.9327749 | 0.4531571 | 2.434154 | 0.01492663 | 0.053737 |
| ENSG00000177873 | ZNF619 | 411.495 | 0.9323709 | 0.1433078 | 6.615084 | 3.71E-11 | 9.79E-10 |
| ENSG00000073910 | FRY | 120.8437287 | 0.932173952 | 0.4726223 | 2.352115659 | 0.01866697 | 0.064261 |
| ENSG00000102053 | ZC3H12B | 33.69152385 | 0.932053558 | 0.590514009 | 2.059064603 | 0.03948805 | 0.115737 |
| ENSG00000179627 | ZBTB42 | 3963.826 | 0.9318086 | 0.2275081 | 4.265438 | 2.00E-05 | 0.000182 |
| ENSG00000164096 | C4orf3 | 5103.249 | 0.9311584 | 0.1328057 | 7.10111 | 1.24E-12 | 3.99E-11 |
| ENSG00000182574 | CSGALNACT2P1 | 2.956498 | 0.9308973 | 1.005842 | 1.763327 | 0.07784536 | 0.193188 |
| ENSG00000234350 | | 10.65754 | 0.9308244 | 0.7043987 | 1.879489 | 0.06017772 | 0.159376 |
| ENSG00000272180 | | 20.1684 | 0.9294353 | 0.578842 | 2.078705 | 0.0376445 | 0.11149 |
| ENSG00000241014 | GPR199P | 12.7412 | 0.9290126 | 0.5028181 | 2.248988 | 0.02451324 | 0.079854 |
| ENSG00000286689 | | 135.3883 | 0.9288674 | 0.3158098 | 3.196991 | 0.00138869 | 0.007598 |
| ENSG00000198689 | SLC9A6 | 1648.062 | 0.9288619 | 0.1336529 | 7.056096 | 1.71E-12 | 5.42E-11 |
| ENSG00000100060 | MFNG | 6.271367864 | 0.928755162 | 0.929072195 | 1.818448674 | 0.06899559 | 0.176586 |

| | | | | | | | |
|---|---|---|---|---|---|---|---|
| ENSG00000161643 | SIGLEC16 | 3.842114 | 0.9287028 | 0.9112012 | 1.785044 | 0.07425422 | 0.186824 |
| ENSG00000114745 | GORASP1 | 3984.052687 | 0.9270974 | 0.127288695 | 7.383207877 | 1.55E-13 | 5.58E-12 |
| ENSG00000261183 | SPINT1-AS1 | 517.3006 | 0.9269442 | 0.1790751 | 5.319778 | 1.04E-07 | 1.57E-06 |
| ENSG00000164040 | PGRMC2 | 5937.852 | 0.9268043 | 0.1323093 | 7.117672 | 1.10E-12 | 3.56E-11 |
| ENSG00000253767 | PCDHGA8 | 22.39995 | 0.9267909 | 0.4916932 | 2.280073 | 0.02260333 | 0.075068 |
| ENSG00000185753 | CXorf38 | 1826.708 | 0.9264685 | 0.2271893 | 4.264697 | 2.00E-05 | 0.000183 |
| ENSG00000217078 | | 4.802288 | 0.9263965 | 1.027275 | 1.770756 | 0.07660127 | 0.190964 |
| ENSG00000253516 | HMGB1P41 | 1.975239 | 0.9258101 | 1.424268 | 2.059587 | 0.03943802 | 0.115629 |
| ENSG00000165175 | MID1IP1 | 5625.015 | 0.9256154 | 0.2578595 | 3.827357 | 0.00012953 | 0.000963 |
| ENSG00000145012 | LPP | 3556.308 | 0.9253414 | 0.2852849 | 3.465755 | 0.00052875 | 0.003295 |
| ENSG00000184451 | CCR10 | 14.34039 | 0.9253154 | 0.6654147 | 1.931398 | 0.05343382 | 0.146049 |
| ENSG00000226950 | DANCR | 2842.964 | 0.9250677 | 0.2146919 | 4.47449 | 7.66E-06 | 7.80E-05 |
| ENSG00000255874 | PRECSIT | 8.059797 | 0.9248629 | 0.6747576 | 1.928349 | 0.05381175 | 0.146849 |
| ENSG00000102886 | GDPD3 | 454.8177986 | 0.924748013 | 0.266583031 | 3.684190676 | 0.00022943 | 0.0016 |
| ENSG00000139344 | AMDHD1 | 41.6131 | 0.9244809 | 0.4291974 | 2.51016 | 0.01206766 | 0.045216 |
| ENSG00000242553 | | 34.62764 | 0.9240079 | 0.4047139 | 2.596877 | 0.00940757 | 0.037008 |
| ENSG00000180336 | MEIOC | 25.20806 | 0.923872 | 0.4436621 | 2.428115 | 0.01517753 | 0.054435 |
| ENSG00000248971 | KRT8P46 | 74.89879 | 0.9237305 | 0.3801113 | 2.728863 | 0.00635532 | 0.026913 |
| ENSG00000273189 | | 4.186533 | 0.9232882 | 0.8956608 | 1.791139 | 0.07327105 | 0.184963 |
| ENSG00000186280 | KDM4D | 40.35525 | 0.9230086 | 0.4205519 | 2.533381 | 0.01129681 | 0.042945 |
| ENSG00000232872 | CTAGE3P | 42.61255 | 0.9229185 | 0.4233859 | 2.508993 | 0.01210759 | 0.045321 |
| ENSG00000184731 | FAM110C | 1439.14 | 0.9227039 | 0.2798224 | 3.514895 | 0.00043993 | 0.00281 |
| ENSG00000112276 | BVES | 28.33814418 | 0.922526813 | 0.50517779 | 2.215352049 | 0.02673591 | 0.085499 |
| ENSG00000203668 | CHML | 2304.938 | 0.9224236 | 0.2418363 | 4.005772 | 6.18E-05 | 0.0005 |
| ENSG00000171843 | MLLT3 | 517.5181 | 0.9221286 | 0.1965496 | 4.848402 | 1.24E-06 | 1.52E-05 |
| ENSG00000172296 | SPTLC3 | 88.34569 | 0.9220409 | 0.5172954 | 2.20353 | 0.02755742 | 0.087633 |
| ENSG00000185278 | ZBTB37 | 1584.566 | 0.921947 | 0.2957916 | 3.351029 | 0.00080512 | 0.004746 |
| ENSG00000119632 | IFI27L2 | 600.4059334 | 0.92164 | 0.331012567 | 3.054506849 | 0.00225431 | 0.011448 |
| ENSG00000218772 | FAM8A6P | 2.229096 | 0.921558 | 1.455631 | 2.107683 | 0.03505844 | 0.105733 |
| ENSG00000278997 | | 45.45964 | 0.9214045 | 0.5265087 | 2.174361 | 0.02967802 | 0.092974 |
| ENSG00000135414 | GDF11 | 1985.793839 | 0.920911687 | 0.228770537 | 4.218713881 | 2.46E-05 | 0.00022 |
| ENSG00000272711 | HK2-DT | 274.9228 | 0.9204819 | 0.2755789 | 3.5688 | 0.00035862 | 0.002363 |
| ENSG00000266446 | | 2.34126 | 0.9193047 | 1.665488 | 2.306622 | 0.02107591 | 0.070927 |

| | | | | | | | |
|---|---|---|---|---|---|---|---|
| ENSG00000258168 | | 3.116794 | 0.9187831 | 1.067252 | 1.725914 | 0.08436296 | 0.205429 |
| ENSG00000205791 | LOH12CR2 | 14.79611 | 0.9184208 | 0.5687687 | 2.068931 | 0.03855259 | 0.11361 |
| ENSG00000197496 | SLC2A10 | 1845.632 | 0.9182703 | 0.1828183 | 5.164969 | 2.40E-07 | 3.39E-06 |
| ENSG00000264491 | | 73.86076 | 0.9180487 | 0.3508943 | 2.897393 | 0.00376278 | 0.017522 |
| ENSG00000273261 | | 6.769105 | 0.9167626 | 0.8782046 | 1.775249 | 0.07585681 | 0.189688 |
| ENSG00000214814 | FER1L6 | 4.577128 | 0.91667 | 1.206352 | 1.804126 | 0.07121159 | 0.180916 |
| ENSG00000152217 | SETBP1 | 285.0546 | 0.9165156 | 0.3128221 | 3.176643 | 0.00148991 | 0.008063 |
| ENSG00000224848 | | 3.38995 | 0.9164428 | 1.146663 | 1.787033 | 0.07393217 | 0.186261 |
| ENSG00000273015 | | 1327.019 | 0.9162993 | 0.190617 | 4.967325 | 6.79E-07 | 8.79E-06 |
| ENSG00000279026 | | 15.56928 | 0.9157296 | 0.4919307 | 2.240308 | 0.02507092 | 0.081312 |
| ENSG00000278948 | | 35.91603 | 0.9155957 | 0.4174963 | 2.512533 | 0.01198679 | 0.045002 |
| ENSG00000112706 | IMPG1 | 37.25455147 | 0.91553692 | 0.425072112 | 2.481587315 | 0.01307987 | 0.04832 |
| ENSG00000105520 | PLPPR2 | 2582.766885 | 0.915199153 | 0.196655023 | 4.804091171 | 1.55E-06 | 1.87E-05 |
| ENSG00000172940 | SLC22A13 | 5.591359 | 0.9151647 | 0.7599529 | 1.812189 | 0.06995695 | 0.178462 |
| ENSG00000266450 | | 2.150067 | 0.9149342 | 1.371981 | 2.012993 | 0.04411532 | 0.126114 |
| ENSG00000155016 | CYP2U1 | 558.3116 | 0.9148313 | 0.3030249 | 3.257449 | 0.00112419 | 0.006331 |
| ENSG00000227473 | TSSK5P | 218.8454 | 0.9147191 | 0.2664197 | 3.640339 | 0.00027228 | 0.001858 |
| ENSG00000159216 | RUNX1 | 2327.973 | 0.9139465 | 0.1898367 | 4.961304 | 7.00E-07 | 9.04E-06 |
| ENSG00000123933 | MXD4 | 5630.100389 | 0.913871157 | 0.265843301 | 3.646126923 | 0.00026622 | 0.001823 |
| ENSG00000249417 | | 3.736005 | 0.9130955 | 1.167634 | 1.830112 | 0.06723316 | 0.17321 |
| ENSG00000105825 | TFPI2 | 1781.879326 | 0.912940913 | 0.393467124 | 2.627078252 | 0.00861215 | 0.034517 |
| ENSG00000272108 | | 35.49651 | 0.9124787 | 0.3865174 | 2.654003 | 0.00795432 | 0.032384 |
| ENSG00000259201 | | 1.671552 | 0.9120446 | 1.377464 | 1.977391 | 0.04799745 | 0.134279 |
| ENSG00000178460 | MCMDC2 | 56.65761 | 0.9120141 | 0.3349412 | 2.980382 | 0.0028789 | 0.014105 |
| ENSG00000164414 | SLC35A1 | 1285.722 | 0.9116309 | 0.1778455 | 5.265479 | 1.40E-07 | 2.06E-06 |
| ENSG00000169314 | C22orf15 | 14.5334 | 0.9114508 | 0.6804196 | 1.906214 | 0.05662242 | 0.15236 |
| ENSG00000144642 | RBMS3 | 41.88746 | 0.9112959 | 0.7454889 | 1.817216 | 0.06918403 | 0.176975 |
| ENSG00000067082 | KLF6 | 5778.26179 | 0.911280684 | 0.528680442 | 2.150974619 | 0.0314782 | 0.097385 |
| ENSG00000231181 | | 8.54528 | 0.9112651 | 0.6650273 | 1.910653 | 0.05604923 | 0.151246 |
| ENSG00000104081 | BMF | 2642.581294 | 0.911233211 | 0.46029774 | 2.361038845 | 0.01822382 | 0.063095 |
| ENSG00000265717 | | 2.976057 | 0.9112042 | 1.469692 | 2.073658 | 0.03811108 | 0.112658 |
| ENSG00000118804 | STBD1 | 248.5839714 | 0.911000822 | 0.220481325 | 4.301734893 | 1.69E-05 | 0.000158 |
| ENSG00000270996 | | 22.23966 | 0.9105098 | 0.6197633 | 1.979587 | 0.0477499 | 0.133755 |

| | | | | | | | |
|---|---|---|---|---|---|---|---|
| ENSG00000125753 | VASP | 6489.1711 | 0.91048028 | 0.164222193 | 5.666742448 | 1.46E-08 | 2.58E-07 |
| ENSG00000196586 | MYO6 | 9335.417 | 0.9102572 | 0.1880654 | 4.985132 | 6.19E-07 | 8.06E-06 |
| ENSG00000182612 | TSPAN10 | 60.81255 | 0.9101522 | 0.4334543 | 2.435352 | 0.0148773 | 0.05357 |
| ENSG00000169507 | SLC38A11 | 8.623771 | 0.9101043 | 0.6815377 | 1.883507 | 0.05963168 | 0.158309 |
| ENSG00000188938 | FAM120AOS | 5113.883 | 0.91008 | 0.1429019 | 6.467849 | 9.94E-11 | 2.48E-09 |
| ENSG00000075643 | MOCOS | 473.809768 | 0.910077078 | 0.306963057 | 3.206657586 | 0.00134287 | 0.007391 |
| ENSG00000262410 | | 52.7677 | 0.9096904 | 0.3628773 | 2.802244 | 0.00507485 | 0.022427 |
| ENSG00000197815 | | 151.7867 | 0.909666 | 0.4182309 | 2.503409 | 0.01230033 | 0.045913 |
| ENSG00000186583 | SPATC1 | 40.69676 | 0.9096292 | 0.5144533 | 2.177446 | 0.02944734 | 0.092426 |
| ENSG00000236212 | | 29.59084 | 0.9095566 | 0.4964383 | 2.226332 | 0.02599197 | 0.083561 |
| ENSG00000170631 | ZNF16 | 898.655 | 0.9092805 | 0.1761865 | 5.301515 | 1.15E-07 | 1.72E-06 |
| ENSG00000271743 | | 4.97687 | 0.908952 | 0.888716 | 1.780479 | 0.07499768 | 0.188053 |
| ENSG00000071553 | ATP6AP1 | 18696.28715 | 0.908925021 | 0.143605257 | 6.418458494 | 1.38E-10 | 3.38E-09 |
| ENSG00000260418 | | 61.45739 | 0.90882 | 0.2998733 | 3.275399 | 0.00105513 | 0.005996 |
| ENSG00000274993 | | 5.256047 | 0.9085274 | 1.319438 | 1.973586 | 0.04842888 | 0.135212 |
| ENSG00000287292 | | 20.24412 | 0.906778 | 0.6596731 | 1.911029 | 0.05600092 | 0.151147 |
| ENSG00000184986 | TMEM121 | 321.5061 | 0.9065942 | 0.2985412 | 3.275094 | 0.00105627 | 0.006 |
| ENSG00000218793 | | 2.529565 | 0.9064636 | 1.164975 | 1.778482 | 0.07532477 | 0.18868 |
| ENSG00000225689 | | 4.112678 | 0.90643 | 1.10026 | 1.806793 | 0.07079463 | 0.180167 |
| ENSG00000267149 | | 1.893662 | 0.906296 | 1.298277 | 1.859211 | 0.06299727 | 0.164888 |
| ENSG00000173442 | EHBP1L1 | 5082.368 | 0.9055968 | 0.2042814 | 4.578432 | 4.68E-06 | 5.04E-05 |
| ENSG00000128346 | C22orf23 | 49.76240081 | 0.90531957 | 0.345070685 | 2.894627954 | 0.00379608 | 0.01766 |
| ENSG00000185432 | METTL7A | 565.5036 | 0.9047456 | 0.6611396 | 1.916725 | 0.05527292 | 0.149623 |
| ENSG00000253991 | | 3.095346 | 0.9042169 | 1.003817 | 1.728552 | 0.08388933 | 0.204546 |
| ENSG00000255236 | | 77.87876 | 0.9041896 | 0.4614068 | 2.328151 | 0.01990411 | 0.067684 |
| ENSG00000243232 | PCDHAC2 | 69.4709 | 0.9039139 | 0.4815055 | 2.254677 | 0.02415363 | 0.078944 |
| ENSG00000163888 | CAMK2N2 | 312.2854 | 0.9037504 | 0.430279 | 2.448198 | 0.01435729 | 0.052098 |
| ENSG00000268105 | | 1.378199 | 0.9033109 | 1.445609 | 2.10651 | 0.03516004 | NA |
| ENSG00000260488 | | 3.137293 | 0.9032763 | 1.32556 | 1.94393 | 0.05190394 | 0.142582 |
| ENSG00000168818 | STX18 | 1166.03 | 0.9030603 | 0.1253175 | 7.310174 | 2.67E-13 | 9.40E-12 |
| ENSG00000138641 | HERC3 | 19.46536359 | 0.90254517 | 0.58474199 | 2.014927437 | 0.04391225 | 0.125722 |
| ENSG00000272323 | TTC23L-AS1 | 124.0795 | 0.9023931 | 0.2494846 | 3.809228 | 0.0001394 | 0.001027 |
| ENSG00000076555 | ACACB | 338.6177998 | 0.902374759 | 0.446449395 | 2.375913385 | 0.01750557 | 0.061053 |

| | | | | | | | |
|---|---|---|---|---|---|---|---|
| ENSG00000270765 | GAS2L2 | 11.76475 | 0.9023158 | 0.6997156 | 1.856639 | 0.06336252 | 0.16555 |
| ENSG00000232229 | LINC00865 | 98.44428 | 0.902223 | 0.248745 | 3.81013 | 0.00013889 | 0.001025 |
| ENSG00000231006 | RPL7P32 | 3.178779 | 0.902092 | 1.039716 | 1.756731 | 0.07896362 | 0.195262 |
| ENSG00000196646 | ZNF136 | 282.1136 | 0.9015202 | 0.2089343 | 4.485219 | 7.28E-06 | 7.45E-05 |
| ENSG00000182472 | CAPN12 | 224.62 | 0.9014365 | 0.375892 | 2.692035 | 0.00710175 | 0.029507 |
| ENSG00000132535 | DLG4 | 591.7913147 | 0.901390802 | 0.338090535 | 2.935326111 | 0.00333197 | 0.015859 |
| ENSG00000100302 | RASD2 | 36.17627101 | 0.901119405 | 0.628915882 | 1.943369315 | 0.05197155 | 0.142722 |
| ENSG00000273335 | | 17.54975 | 0.9007325 | 0.8714094 | 1.779794 | 0.07510968 | 0.188245 |
| ENSG00000231806 | PCAT7 | 829.2416 | 0.9005106 | 0.3549885 | 2.814521 | 0.004885 | 0.021762 |
| ENSG00000249352 | LINC02198 | 8.144026 | 0.8998394 | 0.8247047 | 1.753776 | 0.07946898 | 0.196246 |
| ENSG00000234617 | SNRK-AS1 | 32.3077 | 0.8996855 | 0.5017516 | 2.194308 | 0.0282133 | 0.089217 |
| ENSG00000130066 | SAT1 | 5204.77947 | 0.899374043 | 0.311650221 | 3.117320647 | 0.00182503 | 0.009594 |
| ENSG00000260528 | FAM157C | 11.633 | 0.8992915 | 0.5925747 | 1.978756 | 0.04784348 | 0.133961 |
| ENSG00000164093 | PITX2 | 231.0835 | 0.8992269 | 0.3465988 | 2.868495 | 0.00412429 | 0.018935 |
| ENSG00000186376 | ZNF75D | 3026.854 | 0.8989675 | 0.1871501 | 4.955915 | 7.20E-07 | 9.26E-06 |
| ENSG00000061938 | TNK2 | 4396.768097 | 0.898719528 | 0.221707302 | 4.22313436 | 2.41E-05 | 0.000217 |
| ENSG00000164265 | SCGB3A2 | 1.400238 | 0.8987062 | 1.303007 | 1.941341 | 0.05221698 | NA |
| ENSG00000213830 | CFL1P5 | 21.2312 | 0.8985667 | 0.474494 | 2.269695 | 0.02322611 | 0.076544 |
| ENSG00000100664 | EIF5 | 20158.69727 | 0.898428994 | 0.202221324 | 4.598595494 | 4.25E-06 | 4.62E-05 |
| ENSG00000101307 | SIRPB1 | 2.220206645 | 0.898295408 | 1.357824825 | 2.007335944 | 0.0447139 | 0.127414 |
| ENSG00000234123 | RHBDF1P1 | 2.317954 | 0.8981822 | 1.214155 | 1.853102 | 0.06386769 | 0.166506 |
| ENSG00000159733 | ZFYVE28 | 778.2207 | 0.8981588 | 0.184984 | 4.989661 | 6.05E-07 | 7.89E-06 |
| ENSG00000197021 | EOLA2 | 3121.814 | 0.8965705 | 0.2007195 | 4.621847 | 3.80E-06 | 4.17E-05 |
| ENSG00000280594 | BTG3-AS1 | 9.867104 | 0.8961339 | 0.8017082 | 1.780608 | 0.07497653 | 0.188034 |
| ENSG00000259744 | | 9.011421 | 0.8960133 | 0.7860569 | 1.785587 | 0.07416608 | 0.186655 |
| ENSG00000075223 | SEMA3C | 5371.383475 | 0.895906935 | 0.410168987 | 2.489624546 | 0.01278781 | 0.047413 |
| ENSG00000140931 | CMTM3 | 355.4197 | 0.8958389 | 0.3082507 | 3.150212 | 0.00163152 | 0.00869 |
| ENSG00000286407 | | 61.91614 | 0.8953205 | 0.3862004 | 2.615894 | 0.00889943 | 0.035444 |
| ENSG00000174680 | GRIK1-AS1 | 4.231022 | 0.8950484 | 1.198488 | 1.864985 | 0.06218356 | 0.16327 |
| ENSG00000131788 | PIAS3 | 7079.349209 | 0.894899762 | 0.206136693 | 4.477000792 | 7.57E-06 | 7.71E-05 |
| ENSG00000230333 | | 6.952825 | 0.8937237 | 0.749057 | 1.799941 | 0.07187002 | 0.182206 |
| ENSG00000171462 | DLK2 | 81.75111 | 0.8933528 | 0.3355331 | 2.910738 | 0.00360576 | 0.016929 |
| ENSG00000186395 | KRT10 | 1193.655 | 0.8933021 | 0.2318259 | 4.036058 | 5.44E-05 | 0.000446 |

| | | | | | | | |
|---|---|---|---|---|---|---|---|
| ENSG00000135605 | TEC | 190.4296746 | 0.893289936 | 0.218348094 | 4.248747863 | 2.15E-05 | 0.000195 |
| ENSG00000197406 | DIO3 | 52.44268 | 0.8931715 | 0.6026693 | 1.981679 | 0.04751515 | 0.133294 |
| ENSG00000249896 | LINC02495 | 3.921775 | 0.8931208 | 1.065616 | 1.781762 | 0.07478811 | 0.187705 |
| ENSG00000158815 | FGF17 | 7.065618 | 0.8928615 | 0.9078651 | 1.748899 | 0.08030853 | 0.19786 |
| ENSG00000278376 | | 76.8611 | 0.892839 | 0.3063223 | 3.139165 | 0.0016943 | 0.008982 |
| ENSG00000177335 | C8orf31 | 125.8961 | 0.8925797 | 0.7098886 | 1.844307 | 0.06513846 | 0.168977 |
| ENSG00000130653 | PNPLA7 | 1249.920321 | 0.892314656 | 0.479490372 | 2.242976359 | 0.02489834 | 0.080833 |
| ENSG00000134278 | SPIRE1 | 2400.870068 | 0.892227447 | 0.225728002 | 4.129900837 | 3.63E-05 | 0.000312 |
| ENSG00000226979 | LTA | 2.521121 | 0.8919001 | 0.9794226 | 1.728536 | 0.08389226 | 0.204546 |
| ENSG00000229567 | | 7.740975 | 0.8918148 | 0.6214794 | 1.931877 | 0.05337468 | 0.145926 |
| ENSG00000188343 | CIBAR1 | 1094.87 | 0.8917474 | 0.1689415 | 5.402977 | 6.55E-08 | 1.04E-06 |
| ENSG00000167555 | ZNF528 | 27.78449 | 0.8914948 | 0.6162398 | 1.938918 | 0.05251133 | 0.143966 |
| ENSG00000287913 | | 2.063104 | 0.8905321 | 1.395569 | 2.054238 | 0.03995267 | 0.116754 |
| ENSG00000279821 | | 95.87489 | 0.8905133 | 0.8179751 | 1.783847 | 0.07444857 | 0.187136 |
| ENSG00000138685 | FGF2 | 3.730756515 | 0.890393174 | 0.854261854 | 1.733929685 | 0.08293056 | 0.202834 |
| ENSG00000162078 | ZG16B | 816.7013 | 0.8901345 | 0.2530514 | 3.724036 | 0.00019606 | 0.001393 |
| ENSG00000267026 | | 4.878909 | 0.8896915 | 1.071526 | 1.724562 | 0.08460646 | 0.205871 |
| ENSG00000172348 | RCAN2 | 141.1838 | 0.8895967 | 0.3624902 | 2.732439 | 0.00628673 | 0.0267 |
| ENSG00000139714 | MORN3 | 68.18495 | 0.8895394 | 0.3371892 | 2.891779 | 0.00383067 | 0.01779 |
| ENSG00000259645 | | 2.228962 | 0.889419 | 1.451602 | 2.148819 | 0.03164873 | 0.09781 |
| ENSG00000144655 | CSRNP1 | 1989.755 | 0.8885921 | 0.6355971 | 1.914108 | 0.05560637 | 0.150296 |
| ENSG00000125352 | RNF113A | 1329.957086 | 0.88843625 | 0.151020637 | 5.993081206 | 2.06E-09 | 4.21E-08 |
| ENSG00000179010 | MRFAP1 | 19252.85 | 0.8884107 | 0.1596313 | 5.683755 | 1.32E-08 | 2.35E-07 |
| ENSG00000151617 | EDNRA | 62.17553 | 0.8880043 | 0.3535804 | 2.786811 | 0.00532296 | 0.023303 |
| ENSG00000203837 | PNLIPRP3 | 5.179946 | 0.8879112 | 1.258933 | 1.857922 | 0.06318008 | 0.165187 |
| ENSG00000162009 | SSTR5 | 2.385135 | 0.8878186 | 1.395515 | 2.039786 | 0.0413716 | 0.120055 |
| ENSG00000235314 | LINC00957 | 133.3235 | 0.8877612 | 0.4489874 | 2.330008 | 0.01980574 | 0.067455 |
| ENSG00000275371 | | 8.687832 | 0.8876048 | 0.7919092 | 1.7583 | 0.07869649 | 0.194766 |
| ENSG00000119900 | OGFRL1 | 1341.200871 | 0.88735025 | 0.254990924 | 3.675065659 | 0.00023779 | 0.001653 |
| ENSG00000259685 | IDH2-DT | 4.932993 | 0.8867466 | 0.9612967 | 1.716366 | 0.08609498 | 0.208491 |
| ENSG00000244968 | LIFR-AS1 | 121.5521 | 0.8866979 | 0.2327869 | 3.994687 | 6.48E-05 | 0.00052 |
| ENSG00000123989 | CHPF | 7123.983678 | 0.886466319 | 0.149756452 | 6.04102337 | 1.53E-09 | 3.19E-08 |
| ENSG00000136867 | SLC31A2 | 104.017537 | 0.886418355 | 0.481408644 | 2.216763437 | 0.02663926 | 0.085285 |

| | | | | | | | |
|---|---|---|---|---|---|---|---|
| ENSG00000214274 | ANG | 18.21476 | 0.8863545 | 0.6041785 | 1.94729 | 0.05149999 | 0.141705 |
| ENSG00000117859 | OSBPL9 | 7348.331993 | 0.886059915 | 0.180513886 | 5.032810083 | 4.83E-07 | 6.42E-06 |
| ENSG00000130052 | STARD8 | 46.82203269 | 0.88596877 | 0.475336249 | 2.237169027 | 0.0252753 | 0.081795 |
| ENSG00000101198 | NKAIN4 | 8.151402895 | 0.885823094 | 0.774579855 | 1.749328577 | 0.08023424 | 0.197713 |
| ENSG00000170049 | KCNAB3 | 86.39524 | 0.8857188 | 0.3837746 | 2.613315 | 0.00896686 | 0.035643 |
| ENSG00000250900 | TRIM7-AS1 | 10.26291 | 0.8855599 | 0.7300872 | 1.799098 | 0.07200318 | 0.182439 |
| ENSG00000286024 | | 9.003176 | 0.8854626 | 0.7022083 | 1.828876 | 0.06741825 | 0.173536 |
| ENSG00000260708 | | 321.0979 | 0.8852456 | 0.288217 | 3.295033 | 0.0009841 | 0.005645 |
| ENSG00000162702 | ZNF281 | 2566.182 | 0.8849283 | 0.2072836 | 4.425196 | 9.64E-06 | 9.53E-05 |
| ENSG00000116299 | ELAPOR1 | 4030.114824 | 0.884627496 | 0.431648789 | 2.380835111 | 0.01727344 | 0.060379 |
| ENSG00000215859 | PDZK1P1 | 72.93941 | 0.884384 | 0.3104448 | 3.083114 | 0.00204847 | 0.010573 |
| ENSG00000276809 | | 2.030468 | 0.8842132 | 1.156979 | 1.866718 | 0.06194091 | 0.162813 |
| ENSG00000189127 | ANKRD34B | 265.5623 | 0.8837608 | 0.4129265 | 2.464334 | 0.01372681 | 0.050159 |
| ENSG00000169427 | KCNK9 | 13.02807 | 0.8834775 | 0.6950714 | 1.808543 | 0.070522 | 0.179628 |
| ENSG00000232677 | LINC00665 | 36.47277 | 0.8833913 | 0.4642646 | 2.265415 | 0.02348724 | 0.077155 |
| ENSG00000069943 | PIGB | 555.8559915 | 0.882833418 | 0.183966698 | 4.930945612 | 8.18E-07 | 1.04E-05 |
| ENSG00000174370 | C11orf45 | 162.7448 | 0.882693 | 0.2674514 | 3.506129 | 0.00045468 | 0.002888 |
| ENSG00000185664 | PMEL | 211.2409 | 0.8824454 | 0.3542832 | 2.766099 | 0.00567313 | 0.024585 |
| ENSG00000163531 | NFASC | 32.76379 | 0.8817958 | 0.4341172 | 2.373156 | 0.01763679 | 0.061406 |
| ENSG00000226363 | HAGLROS | 61.01955 | 0.8811841 | 0.329603 | 2.929313 | 0.00339712 | 0.016115 |
| ENSG00000018280 | SLC11A1 | 133.3080595 | 0.881057336 | 0.357634677 | 2.74607158 | 0.00603136 | 0.025846 |
| ENSG00000175567 | UCP2 | 7834.317 | 0.8810325 | 0.2287058 | 4.02668 | 5.66E-05 | 0.000462 |
| ENSG00000010704 | HFE | 293.9587279 | 0.88063962 | 0.287066109 | 3.288042061 | 0.00100887 | 0.005764 |
| ENSG00000235555 | SUMO1P4 | 1.414685 | 0.880425 | 1.42998 | 2.114943 | 0.03443482 | NA |
| ENSG00000225973 | PIGBOS1 | 507.2158 | 0.880298 | 0.2300656 | 3.994893 | 6.47E-05 | 0.00052 |
| ENSG00000102409 | BEX4 | 437.435132 | 0.879944993 | 0.418451193 | 2.430935583 | 0.01505989 | 0.054089 |
| ENSG00000013392 | RWDD2A | 1227.404374 | 0.879287972 | 0.195661625 | 4.65300976 | 3.27E-06 | 3.65E-05 |
| ENSG00000104946 | TBC1D17 | 3574.931316 | 0.879189676 | 0.224395418 | 4.088425572 | 4.34E-05 | 0.000366 |
| ENSG00000104814 | MAP4K1 | 78.50606183 | 0.879163475 | 0.372746638 | 2.644169025 | 0.00818918 | 0.033112 |
| ENSG00000249774 | | 40.01668 | 0.8789754 | 0.3402208 | 2.850084 | 0.00437076 | 0.019847 |
| ENSG00000253540 | FAM86HP | 385.4304 | 0.878675 | 0.1643989 | 5.465796 | 4.61E-08 | 7.48E-07 |
| ENSG00000271930 | | 8.811593 | 0.8785472 | 0.7879856 | 1.752863 | 0.07962558 | 0.196541 |
| ENSG00000188760 | TMEM198 | 361.4373 | 0.878476 | 0.2226446 | 4.109059 | 3.97E-05 | 0.000338 |

| | | | | | | | |
|---|---|---|---|---|---|---|---|
| ENSG00000159763 | PIP | 9.179231 | 0.8784277 | 1.204382 | 1.842162 | 0.06545149 | 0.169657 |
| ENSG00000215068 | | 795.5875 | 0.8783661 | 0.4116678 | 2.455406 | 0.01407254 | 0.051218 |
| ENSG00000078814 | MYH7B | 37.80521613 | 0.878165877 | 0.406073545 | 2.465655653 | 0.01367627 | 0.050009 |
| ENSG00000154370 | TRIM11 | 3552.317 | 0.8781398 | 0.1366348 | 6.528534 | 6.64E-11 | 1.69E-09 |
| ENSG00000261971 | MMP25-AS1 | 878.846 | 0.8779292 | 0.2188014 | 4.186471 | 2.83E-05 | 0.00025 |
| ENSG00000259642 | ST20-AS1 | 176.7712 | 0.8779067 | 0.2089265 | 4.35487 | 1.33E-05 | 0.000127 |
| ENSG00000127947 | PTPN12 | 3786.318473 | 0.877617924 | 0.186941231 | 4.816633624 | 1.46E-06 | 1.77E-05 |
| ENSG00000286550 | | 6.640192 | 0.8774645 | 0.7778167 | 1.742742 | 0.08137864 | 0.199958 |
| ENSG00000214111 | PAFAH1B2P1 | 2.19517 | 0.8772543 | 1.600478 | 2.417383 | 0.01563257 | 0.055699 |
| ENSG00000165195 | PIGA | 897.2887 | 0.8772394 | 0.2457669 | 3.761702 | 0.00016876 | 0.001221 |
| ENSG00000179454 | KLHL28 | 1175.408 | 0.8770002 | 0.2704202 | 3.459429 | 0.00054132 | 0.003366 |
| ENSG00000178177 | LCORL | 652.0296 | 0.8768515 | 0.2112676 | 4.311849 | 1.62E-05 | 0.000152 |
| ENSG00000125648 | SLC25A23 | 3702.438283 | 0.876089816 | 0.272820195 | 3.428443431 | 0.00060705 | 0.003713 |
| ENSG00000257086 | | 47.16328 | 0.8760407 | 0.3222711 | 2.956479 | 0.00311174 | 0.014981 |
| ENSG00000249306 | LINC01411 | 4.307501 | 0.8757989 | 0.8494187 | 1.733405 | 0.0830237 | 0.202968 |
| ENSG00000228501 | RPL15P18 | 13.09867 | 0.8756679 | 0.5862968 | 1.945559 | 0.05170768 | 0.142129 |
| ENSG00000168137 | SETD5 | 15075.66 | 0.8755878 | 0.1535548 | 5.819876 | 5.89E-09 | 1.12E-07 |
| ENSG00000215154 | | 191.3652 | 0.8753713 | 0.3222858 | 2.969968 | 0.00297831 | 0.014484 |
| ENSG00000171943 | SRGAP2C | 425.5811 | 0.8749369 | 0.2253058 | 4.065406 | 4.79E-05 | 0.000399 |
| ENSG00000215375 | MYL5 | 824.1894 | 0.8739648 | 0.221505 | 4.113407 | 3.90E-05 | 0.000332 |
| ENSG00000255014 | ARL6IP1P3 | 2.963082 | 0.8737017 | 1.083254 | 1.764046 | 0.07772426 | 0.192989 |
| ENSG00000116661 | FBXO2 | 1261.148871 | 0.873258363 | 0.406663561 | 2.478257303 | 0.01320259 | 0.048638 |
| ENSG00000076716 | GPC4 | 1231.402648 | 0.873107983 | 0.204748035 | 4.447636326 | 8.68E-06 | 8.69E-05 |
| ENSG00000181666 | ZNF875 | 2102.811 | 0.8729431 | 0.1766133 | 5.071898 | 3.94E-07 | 5.30E-06 |
| ENSG00000184500 | PROS1 | 407.0129 | 0.8727366 | 0.3399098 | 2.82607 | 0.0047123 | 0.021127 |
| ENSG00000206140 | TMEM191C | 43.63418 | 0.8726794 | 0.4239284 | 2.39439 | 0.01664802 | 0.058624 |
| ENSG00000277182 | | 64.32702 | 0.8726469 | 0.3269905 | 2.904783 | 0.00367508 | 0.017182 |
| ENSG00000135298 | ADGRB3 | 9.565566228 | 0.872332006 | 0.616277482 | 1.88445611 | 0.05950331 | 0.158113 |
| ENSG00000223891 | OSER1-DT | 587.2351 | 0.8718067 | 0.2150344 | 4.216678 | 2.48E-05 | 0.000222 |
| ENSG00000125637 | PSD4 | 4965.609992 | 0.871591874 | 0.184525642 | 4.866222735 | 1.14E-06 | 1.40E-05 |
| ENSG00000250910 | | 5.25682 | 0.8715911 | 0.8088411 | 1.741543 | 0.08158849 | 0.200382 |
| ENSG00000235670 | RPL21P40 | 2.395117 | 0.8708945 | 1.074084 | 1.747125 | 0.08061558 | 0.198463 |
| ENSG00000177425 | PAWR | 8492.078 | 0.8708222 | 0.1949159 | 4.57347 | 4.80E-06 | 5.13E-05 |

| | | | | | | | |
|---|---|---|---|---|---|---|---|
| ENSG00000230876 | LINC00486 | 4.305594 | 0.8707717 | 0.9928559 | 1.663811 | 0.09615028 | 0.225972 |
| ENSG00000136099 | PCDH8 | 94.96563925 | 0.870688895 | 0.289845335 | 3.228433804 | 0.0012447 | 0.006917 |
| ENSG00000272574 | | 8.891957 | 0.8703477 | 0.6810198 | 1.830758 | 0.06713674 | 0.173012 |
| ENSG00000270673 | YTHDF3-AS1 | 113.8569 | 0.8700941 | 0.3052752 | 3.085692 | 0.00203079 | 0.010499 |
| ENSG00000169562 | GJB1 | 1.807632 | 0.8692916 | 1.536906 | 2.3062 | 0.02109947 | 0.070979 |
| ENSG00000232140 | | 18.31733 | 0.86905 | 0.6311365 | 1.890389 | 0.05870597 | 0.156555 |
| ENSG00000206625 | RNU6-1 | 5.199978 | 0.8690301 | 0.9093123 | 1.756907 | 0.07893368 | 0.195214 |
| ENSG00000127561 | SYNGR3 | 1191.031456 | 0.868565704 | 0.306032802 | 3.075873091 | 0.00209887 | 0.010793 |
| ENSG00000185261 | KIAA0825 | 77.44209 | 0.8685328 | 0.3493814 | 2.746386 | 0.00602558 | 0.02583 |
| ENSG00000106537 | TSPAN13 | 9000.295858 | 0.868245134 | 0.23575048 | 3.857235897 | 0.00011468 | 0.000865 |
| ENSG00000238099 | LINC01625 | 1.877035 | 0.8681122 | 1.194108 | 1.805578 | 0.07098425 | 0.180442 |
| ENSG00000082684 | SEMA5B | 8.75603281 | 0.868079664 | 1.06132672 | 1.773548517 | 0.07613786 | 0.190087 |
| ENSG00000224014 | | 19.47641 | 0.8676825 | 0.5670116 | 1.975756 | 0.04818242 | 0.134668 |
| ENSG00000196372 | ASB13 | 2456.971 | 0.867677 | 0.1545025 | 5.744116 | 9.24E-09 | 1.71E-07 |
| ENSG00000206965 | RNU6-5P | 2.551647 | 0.8672873 | 0.9573323 | 1.710551 | 0.087164 | 0.210437 |
| ENSG00000152433 | ZNF547 | 116.3007 | 0.8672668 | 0.4112299 | 2.426165 | 0.01525932 | 0.054647 |
| ENSG00000153993 | SEMA3D | 98.1312 | 0.8667249 | 0.529241 | 2.059225 | 0.03947268 | 0.115718 |
| ENSG00000259457 | | 2.301326 | 0.8665998 | 1.509166 | 2.267482 | 0.02336081 | 0.076892 |
| ENSG00000287426 | | 9.299813 | 0.8665205 | 0.8533412 | 1.697705 | 0.08956339 | 0.214902 |
| ENSG00000166188 | ZNF319 | 746.5541 | 0.866446 | 0.212109 | 4.228355 | 2.35E-05 | 0.000212 |
| ENSG00000250256 | | 30.20219 | 0.8663911 | 0.819572 | 1.717747 | 0.08584279 | 0.208155 |
| ENSG00000269318 | | 23.6617 | 0.8658836 | 0.4374183 | 2.317259 | 0.0204896 | 0.069287 |
| ENSG00000244486 | SCARF2 | 96.61622 | 0.8657724 | 0.3459419 | 2.76484 | 0.00569506 | 0.024672 |
| ENSG00000248405 | PRR5-ARHGAP8 | 113.8947 | 0.8657302 | 0.3742839 | 2.600083 | 0.00932011 | 0.03674 |
| ENSG00000162366 | PDZK1IP1 | 29.68315 | 0.8654624 | 1.282882 | 1.925309 | 0.05419076 | 0.147611 |
| ENSG00000267108 | | 6.079706 | 0.8649745 | 0.7843814 | 1.732628 | 0.08316174 | 0.203197 |
| ENSG00000272037 | | 43.38965 | 0.8649556 | 0.4082023 | 2.421613 | 0.0154518 | 0.055202 |
| ENSG00000251136 | | 1560.571 | 0.8649243 | 0.2562758 | 3.56974 | 0.00035734 | 0.002355 |
| ENSG00000113742 | CPEB4 | 1669.974722 | 0.864813342 | 0.325795179 | 2.898576046 | 0.00374861 | 0.017469 |
| ENSG00000090382 | LYZ | 8.843536907 | 0.864803883 | 0.854088214 | 1.730438617 | 0.08355194 | 0.203922 |
| ENSG00000215795 | | 34.24716 | 0.8644917 | 0.345728 | 2.767519 | 0.00564848 | 0.024494 |
| ENSG00000151276 | MAGI1 | 3430.047 | 0.8643088 | 0.2002728 | 4.47605 | 7.60E-06 | 7.75E-05 |
| ENSG00000092096 | SLC22A17 | 829.6467322 | 0.864103586 | 0.327196375 | 2.885139009 | 0.00391241 | 0.018119 |

| | | | | | | | |
|---|---|---|---|---|---|---|---|
| ENSG00000227082 | LINC02798 | 12.74388 | 0.8632987 | 0.503496 | 2.094705 | 0.03619726 | 0.108449 |
| ENSG00000154080 | CHST9 | 8.776301 | 0.8632217 | 0.9358254 | 1.7082 | 0.08759927 | 0.211315 |
| ENSG00000223467 | CALM1P1 | 6.388318 | 0.8631061 | 0.7452414 | 1.772313 | 0.07634268 | 0.190473 |
| ENSG00000267729 | | 21.8303 | 0.8627468 | 0.5312309 | 2.047709 | 0.04058848 | 0.118258 |
| ENSG00000204406 | MBD5 | 1196.869 | 0.8627265 | 0.14677 | 5.996771 | 2.01E-09 | 4.13E-08 |
| ENSG00000204531 | POU5F1 | 19.06594 | 0.8626744 | 0.4984255 | 2.112643 | 0.03463137 | 0.104855 |
| ENSG00000114784 | EIF1B | 1468.00622 | 0.862543629 | 0.18789376 | 4.731690193 | 2.23E-06 | 2.59E-05 |
| ENSG00000166394 | CYB5R2 | 12.13978 | 0.8621865 | 0.6775167 | 1.822316 | 0.06840702 | 0.175469 |
| ENSG00000125746 | EML2 | 1659.427847 | 0.861818293 | 0.310848515 | 2.994805482 | 0.0027462 | 0.013546 |
| ENSG00000241233 | KRTAP5-8 | 11.77565 | 0.8613685 | 0.6653839 | 1.824319 | 0.06810392 | 0.174894 |
| ENSG00000224116 | INHBA-AS1 | 8.092251 | 0.86129 | 0.7413664 | 1.759835 | 0.07843574 | 0.194399 |
| ENSG00000230753 | ZNF341-AS1 | 3.404715 | 0.8612019 | 0.8772868 | 1.69834 | 0.08944353 | 0.214694 |
| ENSG00000104450 | SPAG1 | 1008.355078 | 0.86056103 | 0.162390116 | 5.422224996 | 5.89E-08 | 9.37E-07 |
| ENSG00000182308 | DCAF4L1 | 13.9289 | 0.8605563 | 0.5551412 | 1.969867 | 0.04885358 | 0.13611 |
| ENSG00000157227 | MMP14 | 55.91398 | 0.8604102 | 0.4497615 | 2.256898 | 0.02401442 | 0.078586 |
| ENSG00000110011 | DNAJC4 | 1224.812344 | 0.86031216 | 0.224146144 | 4.007123316 | 6.15E-05 | 0.000498 |
| ENSG00000106443 | PHF14 | 4833.984599 | 0.860277821 | 0.163591333 | 5.380121234 | 7.44E-08 | 1.16E-06 |
| ENSG00000261594 | TPBGL | 22.31512 | 0.8600529 | 0.7462009 | 1.773968 | 0.07606848 | 0.190054 |
| ENSG00000164946 | FREM1 | 9.318581 | 0.859812 | 1.000009 | 1.685274 | 0.09193575 | 0.218875 |
| ENSG00000254548 | | 19.45875 | 0.8597225 | 0.4735685 | 2.169214 | 0.03006647 | 0.093837 |
| ENSG00000287054 | | 4.430241 | 0.859545 | 0.7668981 | 1.733816 | 0.08295069 | 0.202846 |
| ENSG00000105696 | TMEM59L | 70.46961529 | 0.858463301 | 0.524125497 | 2.05128316 | 0.04023938 | 0.117434 |
| ENSG00000282914 | | 3.800654 | 0.8583478 | 0.8751525 | 1.690352 | 0.09096062 | 0.217254 |
| ENSG00000184060 | ADAP2 | 62.341 | 0.8575423 | 0.3042529 | 3.044872 | 0.00232779 | 0.011769 |
| ENSG00000231769 | | 13.90381 | 0.857121 | 0.7342791 | 1.777212 | 0.07553334 | 0.189075 |
| ENSG00000187534 | PRR13P5 | 811.5468 | 0.8570454 | 0.2254127 | 3.97187 | 7.13E-05 | 0.000567 |
| ENSG00000139722 | VPS37B | 1240.96 | 0.8560302 | 0.1447767 | 6.021462 | 1.73E-09 | 3.56E-08 |
| ENSG00000232385 | RPS3AP25 | 5.862124 | 0.8554212 | 1.100992 | 1.823891 | 0.06816849 | 0.175009 |
| ENSG00000163689 | CFAP20DC | 136.4413 | 0.8550318 | 0.3389345 | 2.785718 | 0.00534093 | 0.023366 |
| ENSG00000241697 | TMEFF1 | 2.69167 | 0.854648 | 1.407859 | 2.115086 | 0.03442261 | 0.104384 |
| ENSG00000174004 | NRROS | 2.419829 | 0.8546445 | 1.144253 | 1.862461 | 0.06253813 | 0.164025 |
| ENSG00000186377 | CYP4X1 | 13.18536 | 0.8544592 | 0.6401401 | 1.829479 | 0.06732792 | 0.17337 |
| ENSG00000237115 | | 6.836299 | 0.8542744 | 0.6763561 | 1.792875 | 0.07299291 | 0.184384 |

| | | | | | | | |
|---|---|---|---|---|---|---|---|
| ENSG00000183569 | SERHL2 | 107.3867 | 0.8535279 | 0.3065241 | 3.011214 | 0.00260205 | 0.012943 |
| ENSG00000287539 | | 12.07542 | 0.8532416 | 0.6223195 | 1.848659 | 0.06450709 | 0.167798 |
| ENSG00000213995 | NAXD | 3375.822 | 0.8527389 | 0.1536102 | 5.661369 | 1.50E-08 | 2.65E-07 |
| ENSG00000064651 | SLC12A2 | 4141.103745 | 0.85266113 | 0.256302536 | 3.547773366 | 0.0003885 | 0.002532 |
| ENSG00000258469 | CHMP4BP1 | 9.915258 | 0.8525901 | 0.582598 | 1.906671 | 0.05656326 | 0.152278 |
| ENSG00000230311 | TOMM20P4 | 663.3669 | 0.8523211 | 0.2115111 | 4.212467 | 2.53E-05 | 0.000225 |
| ENSG00000251532 | | 20.9824 | 0.8521441 | 0.46686 | 2.164888 | 0.03039626 | 0.094689 |
| ENSG00000273001 | PFKP-DT | 28.80205 | 0.8520781 | 0.4083707 | 2.408152 | 0.01603351 | 0.056845 |
| ENSG00000283839 | | 5.377752 | 0.8518922 | 1.056784 | 1.704116 | 0.08835946 | 0.212706 |
| ENSG00000197249 | SERPINA1 | 238.5849 | 0.8516942 | 0.5069305 | 2.070796 | 0.03837786 | 0.113208 |
| ENSG00000230084 | | 107.1209 | 0.8512172 | 0.6737152 | 1.809012 | 0.07044915 | 0.179494 |
| ENSG00000182185 | RAD51B | 405.0955 | 0.8504527 | 0.2990658 | 3.073534 | 0.00211539 | 0.010871 |
| ENSG00000039600 | SOX30 | 3.802644375 | 0.850020371 | 0.931666946 | 1.686512566 | 0.09169712 | 0.218505 |
| ENSG00000258884 | LINC02321 | 126.1776 | 0.8497057 | 0.4843971 | 2.140644 | 0.03230275 | 0.0993 |
| ENSG00000147202 | DIAPH2 | 1416.283 | 0.8496641 | 0.1853545 | 4.718493 | 2.38E-06 | 2.75E-05 |
| ENSG00000100890 | PRORP | 396.9450302 | 0.849323199 | 0.197834388 | 4.45009855 | 8.58E-06 | 8.61E-05 |
| ENSG00000163395 | IGFN1 | 8.775848 | 0.8488252 | 0.7553783 | 1.722867 | 0.0849125 | 0.20637 |
| ENSG00000164116 | GUCY1A1 | 803.5122 | 0.8485472 | 0.3405841 | 2.745561 | 0.00604075 | 0.02587 |
| ENSG00000248243 | LINC02014 | 8.862766 | 0.8485117 | 0.6634014 | 1.806651 | 0.07081679 | 0.180172 |
| ENSG00000242861 | | 232.9794 | 0.8483766 | 0.3962807 | 2.440544 | 0.01466517 | 0.052962 |
| ENSG00000167494 | NOS2P2 | 1.438768 | 0.8482604 | 1.5191 | 2.334102 | 0.01959036 | NA |
| ENSG00000267312 | | 155.9328 | 0.8480299 | 0.319378 | 2.902205 | 0.00370546 | 0.017299 |
| ENSG00000279989 | | 16.54183 | 0.8476403 | 0.7465437 | 1.738232 | 0.08217002 | 0.201473 |
| ENSG00000114698 | PLSCR4 | 61.45069122 | 0.847519371 | 0.450983994 | 2.221929425 | 0.02628808 | 0.084333 |
| ENSG00000168118 | RAB4A | 3341.7 | 0.847166 | 0.1346336 | 6.391666 | 1.64E-10 | 3.98E-09 |
| ENSG00000092295 | TGM1 | 305.3550701 | 0.846596773 | 0.368502318 | 2.575671516 | 0.01000456 | 0.038873 |
| ENSG00000270194 | | 138.5699 | 0.846362 | 0.2399354 | 3.70074 | 0.00021497 | 0.001512 |
| ENSG00000287780 | | 15.19441 | 0.8462287 | 0.4959813 | 2.083171 | 0.03723567 | 0.110625 |
| ENSG00000006432 | MAP3K9 | 3075.161301 | 0.845664422 | 0.27332127 | 3.295158261 | 0.00098366 | 0.005645 |
| ENSG00000091181 | IL5RA | 3.498954933 | 0.845663045 | 0.859618694 | 1.67742493 | 0.09345942 | 0.221609 |
| ENSG00000168807 | SNTB2 | 1882.137 | 0.8454573 | 0.2192906 | 4.033978 | 5.48E-05 | 0.000449 |
| ENSG00000269068 | | 6.831338 | 0.8453844 | 0.7762021 | 1.693016 | 0.09045241 | 0.216429 |
| ENSG00000132825 | PPP1R3D | 1591.965348 | 0.845355503 | 0.139313928 | 6.171069554 | 6.78E-10 | 1.50E-08 |

| | | | | | | | |
|---|---|---|---|---|---|---|---|
| ENSG00000230076 | RPL10P6 | 260.806 | 0.8452303 | 0.2200445 | 4.014787 | 5.95E-05 | 0.000483 |
| ENSG00000272183 | | 42.6341 | 0.8445493 | 0.4312058 | 2.275916 | 0.02285102 | 0.075626 |
| ENSG00000152926 | ZNF117 | 3213.258 | 0.8445032 | 0.3200305 | 2.905434 | 0.00366744 | 0.017163 |
| ENSG00000287750 | | 104.1938 | 0.8444733 | 0.3118354 | 2.948671 | 0.00319144 | 0.015292 |
| ENSG00000170525 | PFKFB3 | 6824.158 | 0.8444156 | 0.2049753 | 4.272401 | 1.93E-05 | 0.000177 |
| ENSG00000106004 | HOXA5 | 117.5358138 | 0.844152833 | 0.271438562 | 3.292232388 | 0.00099395 | 0.005691 |
| ENSG00000129518 | EAPP | 2902.859396 | 0.843253422 | 0.159660131 | 5.397766898 | 6.75E-08 | 1.06E-06 |
| ENSG00000287385 | | 21.22029 | 0.8432174 | 0.6921464 | 1.765446 | 0.07748886 | 0.192598 |
| ENSG00000135776 | ABCB10 | 1727.012885 | 0.843107244 | 0.194129285 | 4.47144251 | 7.77E-06 | 7.90E-05 |
| ENSG00000159200 | RCAN1 | 3930.592 | 0.8429429 | 0.4636214 | 2.180892 | 0.02919141 | 0.091862 |
| ENSG00000233588 | CYP51A1P2 | 365.6244 | 0.8424985 | 0.2358591 | 3.753327 | 0.0001745 | 0.001256 |
| ENSG00000148057 | IDNK | 377.449 | 0.8421639 | 0.2671758 | 3.345929 | 0.00082008 | 0.004826 |
| ENSG00000204603 | LINC01257 | 50.42612 | 0.8420937 | 0.6388304 | 1.8357 | 0.06640206 | 0.171568 |
| ENSG00000149300 | C11orf52 | 2.654806 | 0.8420373 | 1.019817 | 1.664308 | 0.09605099 | 0.225839 |
| ENSG00000165895 | ARHGAP42 | 229.6986 | 0.8419417 | 0.3903527 | 2.449876 | 0.01429053 | 0.051877 |
| ENSG00000260361 | | 17.11703 | 0.8418028 | 0.6836356 | 1.776965 | 0.07557403 | 0.189159 |
| ENSG00000214756 | CSKMT | 529.8502 | 0.8413543 | 0.2393082 | 3.69965 | 0.0002159 | 0.001518 |
| ENSG00000287108 | | 1.837347 | 0.8411892 | 1.44447 | 2.222232 | 0.02626765 | 0.084288 |
| ENSG00000091010 | POU4F3 | 8.207135349 | 0.841119917 | 0.685577542 | 1.743595558 | 0.08122964 | 0.199703 |
| ENSG00000140406 | TLNRD1 | 1605.192 | 0.8408015 | 0.1715512 | 5.0398 | 4.66E-07 | 6.22E-06 |
| ENSG00000167460 | TPM4 | 16161.34 | 0.8406977 | 0.2052252 | 4.269692 | 1.96E-05 | 0.000179 |
| ENSG00000277399 | GPR179 | 21.53928 | 0.8406693 | 0.484063 | 2.093079 | 0.03634209 | 0.108713 |
| ENSG00000241769 | LINC00893 | 56.57069 | 0.8404352 | 0.5286153 | 1.993722 | 0.04618241 | 0.130506 |
| ENSG00000069869 | NEDD4 | 1626.202476 | 0.840418233 | 0.20346588 | 4.281568913 | 1.86E-05 | 0.000171 |
| ENSG00000077721 | UBE2A | 5986.02833 | 0.840398301 | 0.10991119 | 7.72360217 | 1.13E-14 | 4.64E-13 |
| ENSG00000207399 | RNU6-1011P | 3.065484 | 0.8402365 | 0.9392966 | 1.681068 | 0.0927498 | 0.220255 |
| ENSG00000077713 | SLC25A43 | 3190.529633 | 0.840159672 | 0.107634557 | 7.885251639 | 3.14E-15 | 1.39E-13 |
| ENSG00000171126 | KCNG3 | 22.75443 | 0.8399424 | 0.589364 | 1.867966 | 0.06176678 | 0.162532 |
| ENSG00000184304 | PRKD1 | 617.6396 | 0.8398537 | 0.2632845 | 3.390868 | 0.00069672 | 0.004188 |
| ENSG00000007384 | RHBDF1 | 4995.286337 | 0.839785811 | 0.186133604 | 4.65036133 | 3.31E-06 | 3.69E-05 |
| ENSG00000207359 | RNU6-925P | 10.18603 | 0.8396329 | 0.59191 | 1.860653 | 0.06279317 | 0.164471 |
| ENSG00000198464 | ZNF480 | 1475.527 | 0.8394737 | 0.1341834 | 6.36472 | 1.96E-10 | 4.71E-09 |
| ENSG00000115758 | ODC1 | 6323.072055 | 0.839383413 | 0.296346651 | 3.041957577 | 0.00235045 | 0.011866 |

| | | | | | | | |
|---|---|---|---|---|---|---|---|
| ENSG00000262362 | | 73.90428 | 0.8393444 | 0.3328886 | 2.75915 | 0.00579519 | 0.025048 |
| ENSG00000242950 | ERVW-1 | 8.868946 | 0.8393322 | 0.7412827 | 1.728792 | 0.08384639 | 0.204509 |
| ENSG00000100647 | SUSD6 | 4082.215305 | 0.838808827 | 0.30700646 | 2.955201284 | 0.00312465 | 0.015024 |
| ENSG00000257270 | | 322.04 | 0.838455 | 0.1982565 | 4.375868 | 1.21E-05 | 0.000117 |
| ENSG00000136261 | BZW2 | 7669.813956 | 0.838232171 | 0.198223546 | 4.369069216 | 1.25E-05 | 0.00012 |
| ENSG00000141314 | RHBDL3 | 47.6296 | 0.8380253 | 0.7284323 | 1.740042 | 0.08185168 | 0.200842 |
| ENSG00000258701 | LINC00638 | 427.4458 | 0.8375872 | 0.3707539 | 2.529164 | 0.01143347 | 0.043316 |
| ENSG00000266993 | | 61.32441 | 0.8375512 | 0.3132413 | 2.905739 | 0.00366387 | 0.01715 |
| ENSG00000164776 | PHKG1 | 30.38266 | 0.8375385 | 0.3964489 | 2.403593 | 0.01623484 | 0.057382 |
| ENSG00000065802 | ASB1 | 1453.533081 | 0.83698159 | 0.137413884 | 6.188124942 | 6.09E-10 | 1.36E-08 |
| ENSG00000200222 | | 5.025043 | 0.8369797 | 1.245605 | 1.919376 | 0.05493674 | 0.149002 |
| ENSG00000173013 | CCDC96 | 67.41267 | 0.8366454 | 0.3565017 | 2.613995 | 0.00894903 | 0.035594 |
| ENSG00000243141 | RPL23AP66 | 1.700017 | 0.8366156 | 1.363579 | 2.110817 | 0.03478801 | 0.10514 |
| ENSG00000167476 | JSRP1 | 9.203355 | 0.8361539 | 0.6533556 | 1.792951 | 0.07298072 | 0.184371 |
| ENSG00000143774 | GUK1 | 8317.02 | 0.8351792 | 0.2038269 | 4.249703 | 2.14E-05 | 0.000194 |
| ENSG00000278685 | IQCA1L | 1.78721 | 0.8341687 | 1.102294 | 1.713685 | 0.0865866 | 0.209405 |
| ENSG00000142188 | TMEM50B | 2831.694 | 0.8341504 | 0.1662828 | 5.154354 | 2.55E-07 | 3.56E-06 |
| ENSG00000262198 | | 31.9617 | 0.8336505 | 0.4334037 | 2.245561 | 0.02473217 | 0.08041 |
| ENSG00000142675 | CNKSR1 | 1445.784 | 0.8332839 | 0.179926 | 4.758605 | 1.95E-06 | 2.30E-05 |
| ENSG00000147231 | RADX | 255.9199 | 0.833189 | 0.4357062 | 2.230207 | 0.02573367 | 0.082925 |
| ENSG00000244682 | FCGR2C | 5.155279 | 0.8327403 | 1.188441 | 1.880417 | 0.06005129 | 0.159184 |
| ENSG00000287078 | | 4.699521 | 0.8325649 | 0.8161955 | 1.664755 | 0.09596174 | 0.225679 |
| ENSG00000170477 | KRT4 | 234.4595 | 0.8325639 | 0.6724547 | 1.794158 | 0.07278798 | 0.184024 |
| ENSG00000118508 | RAB32 | 1778.770956 | 0.83251201 | 0.260902346 | 3.381963794 | 0.0007197 | 0.004306 |
| ENSG00000256249 | | 4.622355 | 0.8323783 | 0.9637474 | 1.641721 | 0.1006477 | 0.233879 |
| ENSG00000266876 | | 62.01165 | 0.8323018 | 0.3253798 | 2.8112 | 0.00493571 | 0.021918 |
| ENSG00000105514 | RAB3D | 2500.988215 | 0.832184796 | 0.145105494 | 5.838237268 | 5.28E-09 | 1.01E-07 |
| ENSG00000143776 | CDC42BPA | 7588.834 | 0.832083 | 0.2258242 | 3.849571 | 0.00011833 | 0.00089 |
| ENSG00000118777 | ABCG2 | 1510.539446 | 0.831767063 | 0.466655971 | 2.139701351 | 0.03237891 | 0.0995 |
| ENSG00000279407 | | 116.3145 | 0.8310148 | 0.3626646 | 2.55904 | 0.01049618 | 0.040475 |
| ENSG00000141179 | PCTP | 1172.534 | 0.8306634 | 0.2114232 | 4.085527 | 4.40E-05 | 0.00037 |
| ENSG00000207524 | RNU6-33P | 7.584999 | 0.8305037 | 0.5667619 | 1.895714 | 0.05799786 | 0.155134 |
| ENSG00000021488 | SLC7A9 | 5.547898684 | 0.830483029 | 0.895887323 | 1.679486645 | 0.09305724 | 0.22083 |

| | | | | | | | |
|---|---|---|---|---|---|---|---|
| ENSG00000279722 | | 37.8961 | 0.8304097 | 0.4054519 | 2.345217 | 0.01901601 | 0.065259 |
| ENSG00000189221 | MAOA | 1552.812 | 0.8303867 | 0.7673318 | 1.706081 | 0.08799308 | 0.212073 |
| ENSG00000135749 | PCNX2 | 1456.966042 | 0.830384747 | 0.298246667 | 3.00114351 | 0.00268968 | 0.013288 |
| ENSG00000107960 | STN1 | 1649.412876 | 0.83031687 | 0.164480664 | 5.119362938 | 3.07E-07 | 4.22E-06 |
| ENSG00000172780 | RAB43 | 214.5097 | 0.8301472 | 0.2313854 | 3.757393 | 0.00017169 | 0.00124 |
| ENSG00000260285 | | 1708.531 | 0.8300838 | 0.3632282 | 2.554276 | 0.01064088 | 0.040937 |
| ENSG00000137968 | SLC44A5 | 222.8523084 | 0.829172869 | 0.596047488 | 1.853270185 | 0.06384364 | 0.16648 |
| ENSG00000225265 | TAF1A-AS1 | 114.6285 | 0.8287754 | 0.2281302 | 3.804042 | 0.00014235 | 0.001046 |
| ENSG00000225449 | RAB6C-AS1 | 69.49762 | 0.8287719 | 0.6473082 | 1.783642 | 0.07448184 | 0.187166 |
| ENSG00000280054 | | 7.609628 | 0.8286826 | 0.7017191 | 1.733625 | 0.08298465 | 0.202891 |
| ENSG00000225154 | TUBAP9 | 2.595379 | 0.8285938 | 1.441772 | 2.278813 | 0.02267819 | 0.075251 |
| ENSG00000233799 | | 14.33872 | 0.8281271 | 0.5098578 | 2.011526 | 0.04426989 | 0.126474 |
| ENSG00000258439 | | 3.042342 | 0.8277741 | 0.9579098 | 1.68425 | 0.09213341 | 0.219188 |
| ENSG00000089692 | LAG3 | 157.063315 | 0.827453641 | 0.503477978 | 2.032284821 | 0.04212483 | 0.12172 |
| ENSG00000140416 | TPM1 | 9948.283 | 0.8274455 | 0.1525566 | 5.533553 | 3.14E-08 | 5.21E-07 |
| ENSG00000244459 | | 26.20472 | 0.8272551 | 0.6455511 | 1.780245 | 0.07503587 | 0.188101 |
| ENSG00000212916 | MAP10 | 737.428 | 0.8272225 | 0.1513984 | 5.560575 | 2.69E-08 | 4.52E-07 |
| ENSG00000237310 | | 97.85547 | 0.8272168 | 0.2841033 | 3.112624 | 0.00185432 | 0.009719 |
| ENSG00000237371 | | 2.977541 | 0.8262283 | 1.110193 | 1.775304 | 0.07584764 | 0.189683 |
| ENSG00000236806 | RPL7AP15 | 4.067267 | 0.8258329 | 0.9236596 | 1.666734 | 0.09556723 | 0.22502 |
| ENSG00000197405 | C5AR1 | 45.43479 | 0.8257662 | 0.481741 | 2.074038 | 0.03807578 | 0.112567 |
| ENSG00000183196 | CHST6 | 4.498324 | 0.8257213 | 0.8128048 | 1.661496 | 0.09661385 | 0.226861 |
| ENSG00000170275 | CRTAP | 6696.089 | 0.8257171 | 0.1200595 | 7.011121 | 2.36E-12 | 7.39E-11 |
| ENSG00000280119 | | 3.683042 | 0.8256824 | 1.091308 | 1.773906 | 0.07607863 | 0.190054 |
| ENSG00000207342 | | 5.622353 | 0.8253758 | 0.7642873 | 1.680185 | 0.09292131 | 0.220549 |
| ENSG00000225787 | LSM3P3 | 8.081692 | 0.8253551 | 0.8796228 | 1.692424 | 0.09056508 | 0.21659 |
| ENSG00000150961 | SEC24D | 3130.277 | 0.8253306 | 0.2195822 | 3.940432 | 8.13E-05 | 0.000637 |
| ENSG00000279568 | | 8.025588 | 0.8252858 | 0.7081333 | 1.713954 | 0.08653715 | 0.209338 |
| ENSG00000063660 | GPC1 | 1937.248167 | 0.825062335 | 0.224501727 | 3.836576495 | 0.00012476 | 0.000933 |
| ENSG00000167588 | GPD1 | 23.42774 | 0.8248411 | 0.5139801 | 1.98173 | 0.04750946 | 0.133292 |
| ENSG00000226853 | | 27.42019 | 0.8248243 | 0.3828902 | 2.424046 | 0.01534866 | 0.054913 |
| ENSG00000156162 | DPY19L4 | 3459.013 | 0.824102 | 0.2007236 | 4.240311 | 2.23E-05 | 0.000202 |
| ENSG00000177989 | ODF3B | 99.7679 | 0.8240635 | 0.4317128 | 2.227694 | 0.02590093 | 0.083343 |

| | | | | | | | |
|---|---|---|---|---|---|---|---|
| ENSG00000173705 | SUSD5 | 9.81709 | 0.8238863 | 0.7122462 | 1.707458 | 0.08773698 | 0.211609 |
| ENSG00000259856 | RAB43P1 | 241.1876 | 0.8237579 | 0.2241549 | 3.850476 | 0.00011789 | 0.000888 |
| ENSG00000140396 | NCOA2 | 3892.304 | 0.8235362 | 0.2563686 | 3.389607 | 0.00069993 | 0.004204 |
| ENSG00000197860 | SGTB | 239.3167 | 0.8234414 | 0.3191977 | 2.812621 | 0.00491395 | 0.021853 |
| ENSG00000139220 | PPFIA2 | 3.149818 | 0.8233156 | 0.8835389 | 1.634929 | 0.102064 | 0.236111 |
| ENSG00000102934 | PLLP | 463.3974529 | 0.823041607 | 0.334375894 | 2.713997229 | 0.00664767 | 0.02793 |
| ENSG00000165030 | NFIL3 | 767.6018 | 0.8227182 | 0.2180358 | 3.934716 | 8.33E-05 | 0.000651 |
| ENSG00000146411 | SLC2A12 | 44.23856 | 0.8224542 | 0.6533535 | 1.745432 | 0.08090974 | 0.199082 |
| ENSG00000198736 | MSRB1 | 2866.932 | 0.8224036 | 0.327275 | 2.77221 | 0.00556771 | 0.024195 |
| ENSG00000113319 | RASGRF2 | 34.12064754 | 0.822385931 | 0.384311392 | 2.412697866 | 0.01583494 | 0.056276 |
| ENSG00000229029 | CDCA4P1 | 2.563492 | 0.8222968 | 1.100653 | 1.740459 | 0.08177845 | 0.200739 |
| ENSG00000234003 | MTATP6P27 | 3.615731 | 0.8221256 | 1.009875 | 1.706176 | 0.08797527 | 0.212049 |
| ENSG00000174227 | PIGG | 3044.069 | 0.8219587 | 0.1692716 | 4.970064 | 6.69E-07 | 8.67E-06 |
| ENSG00000143786 | CNIH3 | 216.1428 | 0.8219313 | 0.2569874 | 3.382408 | 0.00071853 | 0.004301 |
| ENSG00000179455 | MKRN3 | 21.49445 | 0.8214201 | 1.20603 | 1.812427 | 0.06992029 | 0.17842 |
| ENSG00000161594 | KLHL10 | 9.712899 | 0.8213867 | 0.6415964 | 1.781282 | 0.07486636 | 0.187848 |
| ENSG00000185825 | BCAP31 | 19014.1 | 0.8211137 | 0.1632084 | 5.14833 | 2.63E-07 | 3.67E-06 |
| ENSG00000125618 | PAX8 | 151.0773453 | 0.821067063 | 0.194946893 | 4.357175984 | 1.32E-05 | 0.000126 |
| ENSG00000186300 | ZNF555 | 344.5372 | 0.820776 | 0.194691 | 4.354306 | 1.33E-05 | 0.000127 |
| ENSG00000165917 | RAPSN | 19.45141 | 0.8203845 | 0.5418158 | 1.920775 | 0.0547601 | 0.148628 |
| ENSG00000260351 | | 3.650627 | 0.8199463 | 0.8498225 | 1.652799 | 0.09837168 | 0.229831 |
| ENSG00000005471 | ABCB4 | 46.18292391 | 0.819711886 | 0.554629505 | 1.89731893 | 0.05778586 | 0.154676 |
| ENSG00000137331 | IER3 | 4111.253483 | 0.819527177 | 0.338937764 | 2.700762758 | 0.00691807 | 0.028847 |
| ENSG00000163362 | INAVA | 1527.741 | 0.819052 | 0.6584227 | 1.772825 | 0.07625773 | 0.190351 |
| ENSG00000175315 | CST6 | 29.06361 | 0.81853 | 0.9328618 | 1.593794 | 0.1109821 | 0.250888 |
| ENSG00000096395 | MLN | 1.202511198 | 0.818411013 | 1.367483399 | 2.132605996 | 0.03295706 | NA |
| ENSG00000229427 | ANKRD26P4 | 1.188937 | 0.8180415 | 1.323798 | 2.044786 | 0.04087601 | NA |
| ENSG00000167281 | RBFOX3 | 12.04483 | 0.8179421 | 0.6650909 | 1.743748 | 0.08120306 | 0.199656 |
| ENSG00000187446 | CHP1 | 4364.298 | 0.8176869 | 0.1651693 | 5.055041 | 4.30E-07 | 5.78E-06 |
| ENSG00000255153 | TOLLIP-AS1 | 165.6033 | 0.8167976 | 0.2288932 | 3.728062 | 0.00019296 | 0.001373 |
| ENSG00000275223 | | 75.56179 | 0.8167463 | 0.4096717 | 2.29693 | 0.02162275 | 0.072455 |
| ENSG00000261634 | | 3.8929 | 0.8164551 | 0.8099462 | 1.625936 | 0.1039634 | 0.239276 |
| ENSG00000270533 | | 3.913062 | 0.8160878 | 0.8658537 | 1.650125 | 0.09891745 | 0.230781 |

| | | | | | | | |
|---|---|---|---|---|---|---|---|
| ENSG00000271009 | | 76.31664 | 0.8159332 | 0.5238627 | 1.929322 | 0.05369091 | 0.146625 |
| ENSG00000239453 | SIDT1-AS1 | 4.820326 | 0.8158134 | 0.8553941 | 1.657777 | 0.09736255 | 0.228216 |
| ENSG00000236255 | | 78.80905 | 0.8154783 | 0.5756782 | 1.852337 | 0.0639774 | 0.166676 |
| ENSG00000269489 | | 42.59613 | 0.8154035 | 0.5412025 | 1.91576 | 0.05539563 | 0.149817 |
| ENSG00000255883 | FUNDC2P1 | 457.2015 | 0.8153263 | 0.1599486 | 5.230556 | 1.69E-07 | 2.45E-06 |
| ENSG00000110628 | SLC22A18 | 892.6922191 | 0.81511683 | 0.234542852 | 3.64650145 | 0.00026584 | 0.001821 |
| ENSG00000166341 | DCHS1 | 4.151688 | 0.8146618 | 0.9739021 | 1.678162 | 0.09331542 | 0.221307 |
| ENSG00000072858 | SIDT1 | 1952.978642 | 0.814586717 | 0.248642097 | 3.448708457 | 0.00056328 | 0.003477 |
| ENSG00000232533 | | 205.9855 | 0.8145395 | 0.286945 | 3.050605 | 0.0022838 | 0.011569 |
| ENSG00000067829 | IDH3G | 5011.088209 | 0.814219565 | 0.155069018 | 5.344569899 | 9.06E-08 | 1.38E-06 |
| ENSG00000139194 | RBP5 | 43.49187448 | 0.814188459 | 0.575356403 | 1.836216554 | 0.06632564 | 0.171428 |
| ENSG00000158869 | FCER1G | 3.050041 | 0.8140113 | 0.9788469 | 1.67837 | 0.09327499 | 0.22129 |
| ENSG00000237372 | | 3.211829 | 0.8139751 | 1.03192 | 1.670493 | 0.09482189 | 0.22392 |
| ENSG00000175482 | POLD4 | 823.1979 | 0.8139361 | 0.2029311 | 4.163059 | 3.14E-05 | 0.000274 |
| ENSG00000149256 | TENM4 | 145.4086 | 0.8137388 | 0.3438145 | 2.619076 | 0.00881683 | 0.035194 |
| ENSG00000143369 | ECM1 | 95.78686 | 0.8134237 | 0.3777329 | 2.426538 | 0.01524365 | 0.054613 |
| ENSG00000131941 | RHPN2 | 2573.089751 | 0.813363857 | 0.206965791 | 4.075397814 | 4.59E-05 | 0.000385 |
| ENSG00000123124 | WWP1 | 9245.38166 | 0.813297615 | 0.1425279 | 5.786559762 | 7.18E-09 | 1.35E-07 |
| ENSG00000180537 | RNF182 | 7.85784 | 0.8125924 | 0.9752584 | 1.648842 | 0.09917993 | 0.231292 |
| ENSG00000113161 | HMGCR | 10146.92381 | 0.81243846 | 0.276165402 | 3.139481898 | 0.00169247 | 0.008975 |
| ENSG00000231304 | SGO1-AS1 | 6.634571 | 0.812267 | 0.6233916 | 1.758273 | 0.07870101 | 0.194766 |
| ENSG00000235643 | LINC01647 | 2.13301 | 0.8122479 | 1.143265 | 1.866718 | 0.06194105 | 0.162813 |
| ENSG00000079739 | PGM1 | 950.4575639 | 0.812098737 | 0.19712376 | 4.272474482 | 1.93E-05 | 0.000177 |
| ENSG00000235944 | ZNF815P | 344.0617 | 0.8120952 | 0.2113441 | 3.998097 | 6.39E-05 | 0.000515 |
| ENSG00000283268 | TEX54 | 3.572555 | 0.8120641 | 1.005626 | 1.673548 | 0.09421954 | 0.223033 |
| ENSG00000244165 | P2RY11 | 259.3242 | 0.8119824 | 0.2492225 | 3.43085 | 0.00060169 | 0.003683 |
| ENSG00000167861 | HID1 | 12247.78 | 0.811919 | 0.1537097 | 5.380424 | 7.43E-08 | 1.15E-06 |
| ENSG00000151914 | DST | 6631.295 | 0.8114585 | 0.2883392 | 3.021446 | 0.0025157 | 0.012573 |
| ENSG00000163576 | EFHB | 55.10867 | 0.811277 | 0.4096556 | 2.270528 | 0.02317555 | 0.076405 |
| ENSG00000198482 | ZNF808 | 344.9711 | 0.811092 | 0.2423247 | 3.528646 | 0.00041769 | 0.002686 |
| ENSG00000234948 | LINC01524 | 10.49585 | 0.810733 | 0.7057582 | 1.661279 | 0.0966574 | 0.226921 |
| ENSG00000236686 | BZW1P1 | 27.55256 | 0.8105304 | 0.4268841 | 2.221987 | 0.02628416 | 0.084331 |
| ENSG00000254433 | | 12.65093 | 0.8104464 | 0.6347315 | 1.762616 | 0.07796531 | 0.193414 |

| | | | | | | | |
|---|---|---|---|---|---|---|---|
| ENSG00000244274 | DBNDD2 | 97.88912 | 0.8101229 | 0.2350459 | 3.621457 | 0.00029295 | 0.001981 |
| ENSG00000177738 | | 1029.299 | 0.8100345 | 0.1774291 | 4.692458 | 2.70E-06 | 3.08E-05 |
| ENSG00000181830 | SLC35C1 | 3052.868 | 0.8094566 | 0.2170927 | 3.894391 | 9.84E-05 | 0.000755 |
| ENSG00000126822 | PLEKHG3 | 3494.683536 | 0.809254787 | 0.146688649 | 5.594185434 | 2.22E-08 | 3.78E-07 |
| ENSG00000267682 | | 11.70669 | 0.8088572 | 0.6622443 | 1.731756 | 0.08331708 | 0.203479 |
| ENSG00000085276 | MECOM | 618.3378613 | 0.808659789 | 0.391036624 | 2.361061 | 0.01822273 | 0.063095 |
| ENSG00000167657 | DAPK3 | 2436.564 | 0.8085087 | 0.1455226 | 5.653129 | 1.58E-08 | 2.77E-07 |
| ENSG00000286315 | | 15.72953 | 0.8084441 | 0.8838355 | 1.641827 | 0.1006258 | 0.233863 |
| ENSG00000166971 | AKTIP | 2214.901 | 0.8083736 | 0.1608131 | 5.160412 | 2.46E-07 | 3.46E-06 |
| ENSG00000139915 | MDGA2 | 29.79697 | 0.8083627 | 0.5293135 | 1.920029 | 0.05485421 | 0.148838 |
| ENSG00000101197 | BIRC7 | 27.30860775 | 0.808116806 | 0.829412427 | 1.674819716 | 0.09396961 | 0.22259 |
| ENSG00000285215 | | 1.135869 | 0.8077299 | 1.336355 | 2.070175 | 0.03843598 | NA |
| ENSG00000151689 | INPP1 | 245.9512 | 0.807699 | 0.2875824 | 3.01404 | 0.00257794 | 0.012834 |
| ENSG00000136603 | SKIL | 5065.485029 | 0.807430681 | 0.182494994 | 4.558591667 | 5.15E-06 | 5.48E-05 |
| ENSG00000112541 | PDE10A | 256.0477847 | 0.807373031 | 0.364659177 | 2.469407616 | 0.0135337 | 0.049564 |
| ENSG00000262488 | | 2.600648 | 0.8073435 | 1.038241 | 1.66891 | 0.09513524 | 0.224377 |
| ENSG00000007001 | UPP2 | 4.315299815 | 0.807209077 | 0.83330928 | 1.618498292 | 0.10555526 | 0.242059 |
| ENSG00000172915 | NBEA | 1732.417 | 0.8070136 | 0.3130706 | 2.799149 | 0.00512375 | 0.022602 |
| ENSG00000229498 | | 3.16583 | 0.806249 | 1.282822 | 1.986176 | 0.04701379 | 0.132264 |
| ENSG00000196296 | ATP2A1 | 93.31697 | 0.8055026 | 0.3498287 | 2.552191 | 0.01070479 | 0.04113 |
| ENSG00000277287 | | 22.99724 | 0.8053468 | 0.4014979 | 2.280697 | 0.02256636 | 0.074983 |
| ENSG00000243368 | MCCC1-AS1 | 22.70837 | 0.8052364 | 0.5540492 | 1.886655 | 0.05920677 | 0.157524 |
| ENSG00000115290 | GRB14 | 946.0688357 | 0.805078216 | 0.414480074 | 2.246893171 | 0.02464686 | 0.080191 |
| ENSG00000228395 | | 36.17346 | 0.8045114 | 0.374753 | 2.416762 | 0.01565927 | 0.05576 |
| ENSG00000231890 | DARS1-AS1 | 38.30088 | 0.8042357 | 0.341349 | 2.598266 | 0.00936959 | 0.036886 |
| ENSG00000224969 | | 19.1197 | 0.8040611 | 0.5921864 | 1.807031 | 0.07075742 | 0.180124 |
| ENSG00000135119 | RNFT2 | 115.0779161 | 0.804006105 | 0.503097935 | 1.976221694 | 0.04812967 | 0.134578 |
| ENSG00000147234 | FRMPD3 | 100.4458 | 0.8039038 | 0.3410804 | 2.594288 | 0.00947871 | 0.037212 |
| ENSG00000203864 | LINC02868 | 1.693424 | 0.8035282 | 1.41034 | 2.266398 | 0.02342705 | 0.076996 |
| ENSG00000229743 | LINC01159 | 7.642754 | 0.8034205 | 0.863683 | 1.647414 | 0.09947289 | 0.231874 |
| ENSG00000234776 | C11orf94 | 15.39532 | 0.8033969 | 0.6283393 | 1.753873 | 0.0794523 | 0.196241 |
| ENSG00000026508 | CD44 | 1429.626341 | 0.803344553 | 0.433067899 | 2.168558563 | 0.03011621 | 0.093959 |
| ENSG00000118564 | FBXL5 | 1810.876529 | 0.802499057 | 0.098413452 | 8.223041954 | 1.98E-16 | 1.02E-14 |

| | | | | | | | |
|---|---|---|---|---|---|---|---|
| ENSG00000268903 | | 172.6762 | 0.8024013 | 0.3161185 | 2.763324 | 0.00572159 | 0.024767 |
| ENSG00000245205 | EEF1A1P4 | 23.66448 | 0.8023502 | 0.4683007 | 2.052826 | 0.04008942 | 0.117074 |
| ENSG00000013016 | EHD3 | 1139.031946 | 0.802100371 | 0.19986762 | 4.142703832 | 3.43E-05 | 0.000297 |
| ENSG00000197747 | S100A10 | 14842.51 | 0.8017472 | 0.2306183 | 3.705423 | 0.00021104 | 0.001487 |
| ENSG00000279673 | | 18.4109 | 0.8015582 | 0.5910808 | 1.815084 | 0.06951103 | 0.177598 |
| ENSG00000248514 | | 2.099686 | 0.8015568 | 1.235203 | 2.018125 | 0.04357828 | 0.124962 |
| ENSG00000147421 | HMBOX1 | 745.481 | 0.8015084 | 0.2393726 | 3.514233 | 0.00044103 | 0.002816 |
| ENSG00000169635 | HIC2 | 224.7054 | 0.8014145 | 0.2744284 | 3.117749 | 0.00182238 | 0.009582 |
| ENSG00000285053 | TBCE | 392.7427 | 0.8011989 | 0.1908149 | 4.326775 | 1.51E-05 | 0.000143 |
| ENSG00000230510 | PPP5D1 | 17.33525 | 0.8008857 | 0.4783124 | 2.018833 | 0.04350454 | 0.124811 |
| ENSG00000170379 | TCAF2 | 107.4287 | 0.8008442 | 0.459566 | 2.077674 | 0.0377394 | 0.111747 |
| ENSG00000179476 | C14orf28 | 552.5188 | 0.8001856 | 0.189679 | 4.348697 | 1.37E-05 | 0.00013 |
| ENSG00000095752 | IL11 | 144.3391857 | 0.800067337 | 0.504176059 | 1.961951702 | 0.04976811 | 0.13804 |
| ENSG00000272551 | | 2.134441 | 0.799907 | 1.23496 | 1.846867 | 0.06476649 | 0.168242 |
| ENSG00000273230 | | 491.4077 | 0.7998704 | 0.2911342 | 2.968588 | 0.00299171 | 0.014529 |
| ENSG00000287156 | | 4.463477 | 0.7993736 | 0.9839588 | 1.605253 | 0.1084381 | 0.246796 |
| ENSG00000240006 | LINC02004 | 16.01426 | 0.799081 | 0.5497966 | 1.852766 | 0.0639159 | 0.166603 |
| ENSG00000081692 | JMJD4 | 741.7066598 | 0.798684507 | 0.177321239 | 4.629998635 | 3.66E-06 | 4.03E-05 |
| ENSG00000134201 | GSTM5 | 49.69708527 | 0.798658641 | 0.459492065 | 2.064347074 | 0.03898482 | 0.114629 |
| ENSG00000255421 | | 3.156745 | 0.7985068 | 0.9840543 | 1.661241 | 0.09666502 | 0.226921 |
| ENSG00000145819 | ARHGAP26 | 744.7419 | 0.7984049 | 0.2901075 | 2.965017 | 0.00302666 | 0.014661 |
| ENSG00000269421 | | 2.578131 | 0.7982191 | 1.234848 | 1.923494 | 0.05441807 | 0.148039 |
| ENSG00000164649 | CDCA7L | 1529.573 | 0.7982142 | 0.2880383 | 2.96771 | 0.00300027 | 0.014562 |
| ENSG00000232295 | | 6.888794 | 0.7981677 | 0.6718602 | 1.702111 | 0.08873448 | 0.213484 |
| ENSG00000136404 | TM6SF1 | 6.400693511 | 0.7981139 | 0.663043007 | 1.702527249 | 0.08865658 | 0.213323 |
| ENSG00000131724 | IL13RA1 | 5805.245302 | 0.79752529 | 0.228506192 | 3.665951453 | 0.00024642 | 0.001703 |
| ENSG00000146530 | VWDE | 1491.242 | 0.7973721 | 0.3302606 | 2.654798 | 0.00793559 | 0.032333 |
| ENSG00000109606 | DHX15 | 13402.85489 | 0.797309723 | 0.19605387 | 4.21599969 | 2.49E-05 | 0.000223 |
| ENSG00000010295 | IFFO1 | 11.48064544 | 0.797214731 | 0.686783985 | 1.686070892 | 0.09178215 | 0.218587 |
| ENSG00000235897 | TM4SF19-AS1 | 23.18727 | 0.7966976 | 0.5139111 | 1.921162 | 0.05471127 | 0.148556 |
| ENSG00000163702 | IL17RC | 2174.475 | 0.7966046 | 0.1957239 | 4.204426 | 2.62E-05 | 0.000233 |
| ENSG00000226432 | | 420.1018 | 0.7965621 | 0.1712399 | 4.79544 | 1.62E-06 | 1.94E-05 |
| ENSG00000102547 | CAB39L | 948.9306954 | 0.796458178 | 0.393863624 | 2.307486681 | 0.0210277 | 0.070791 |

| | | | | | | | |
|---|---|---|---|---|---|---|---|
| ENSG00000119661 | DNAL1 | 1161.148741 | 0.796388591 | 0.197039153 | 4.201860513 | 2.65E-05 | 0.000235 |
| ENSG00000136286 | MYO1G | 7.169570351 | 0.795474934 | 0.707916917 | 1.682260779 | 0.09251828 | 0.219867 |
| ENSG00000170498 | KISS1 | 74.83412 | 0.7951915 | 0.415588 | 2.212882 | 0.02690581 | 0.085972 |
| ENSG00000129757 | CDKN1C | 35.70269521 | 0.79506999 | 0.542266937 | 1.870800631 | 0.06137272 | 0.161815 |
| ENSG00000204611 | ZNF616 | 813.4061 | 0.7949586 | 0.1689552 | 4.827083 | 1.39E-06 | 1.68E-05 |
| ENSG00000163083 | INHBB | 3060.095 | 0.7948904 | 0.3308043 | 2.644272 | 0.00818668 | 0.033108 |
| ENSG00000287519 | | 8.537552 | 0.7948285 | 0.8693204 | 1.63033 | 0.1030319 | 0.237833 |
| ENSG00000281538 | | 23.7895 | 0.794694 | 0.4714101 | 2.020595 | 0.04332166 | 0.124447 |
| ENSG00000276007 | | 153.7977 | 0.7942404 | 0.4558268 | 2.081828 | 0.03735819 | 0.110914 |
| ENSG00000179954 | SSC5D | 15.06426 | 0.794019 | 0.500506 | 1.952053 | 0.05093196 | 0.140464 |
| ENSG00000235726 | | 10.6227 | 0.7937726 | 0.6166238 | 1.723134 | 0.0848643 | 0.20631 |
| ENSG00000110665 | C11orf21 | 1.544249806 | 0.793438403 | 1.090456555 | 1.679365084 | 0.09308091 | NA |
| ENSG00000216867 | RPL22P12 | 3.534226 | 0.7933689 | 1.069666 | 1.700871 | 0.08896714 | 0.213887 |
| ENSG00000260059 | | 4.152275 | 0.7932231 | 0.8633912 | 1.622891 | 0.1046127 | 0.240375 |
| ENSG00000147174 | GCNA | 177.1991 | 0.7930815 | 0.4562594 | 2.070935 | 0.0383649 | 0.113208 |
| ENSG00000165449 | SLC16A9 | 170.246 | 0.7924116 | 0.4937013 | 1.969657 | 0.04887771 | 0.136154 |
| ENSG00000270761 | | 58.81805 | 0.7923743 | 0.4199516 | 2.190872 | 0.02846107 | 0.089915 |
| ENSG00000249673 | NOP14-AS1 | 852.7143 | 0.7922141 | 0.1896156 | 4.30029 | 1.71E-05 | 0.000159 |
| ENSG00000260916 | CCPG1 | 414.1256 | 0.792196 | 0.2328348 | 3.565443 | 0.00036324 | 0.002387 |
| ENSG00000120149 | MSX2 | 6392.16372 | 0.791542379 | 0.31096939 | 2.783613546 | 0.0053757 | 0.023503 |
| ENSG00000260369 | | 7.114223 | 0.7914965 | 0.7655038 | 1.628195 | 0.1034836 | 0.238502 |
| ENSG00000258648 | UBE2CP1 | 6.960473 | 0.7911672 | 0.8114958 | 1.606111 | 0.1082495 | 0.246493 |
| ENSG00000180953 | ST20 | 168.7792 | 0.7907782 | 0.3619197 | 2.442061 | 0.01460369 | 0.052804 |
| ENSG00000173548 | SNX33 | 3237.565 | 0.7904293 | 0.2281885 | 3.620482 | 0.00029406 | 0.001987 |
| ENSG00000185792 | NLRP9 | 5.14079 | 0.7902822 | 0.7162627 | 1.642886 | 0.1004066 | 0.233477 |
| ENSG00000086544 | ITPKC | 2209.145555 | 0.790087776 | 0.181110792 | 4.480668989 | 7.44E-06 | 7.60E-05 |
| ENSG00000155886 | SLC24A2 | 5.991779 | 0.7897373 | 1.10389 | 1.718971 | 0.08561974 | 0.20771 |
| ENSG00000231414 | RPL39P15 | 18.88061 | 0.7892045 | 0.4447318 | 2.0823 | 0.03731503 | 0.110817 |
| ENSG00000142173 | COL6A2 | 1115.277 | 0.7883082 | 0.3233328 | 2.661362 | 0.00778253 | 0.031802 |
| ENSG00000110721 | CHKA | 3383.131797 | 0.788272912 | 0.183679177 | 4.432130164 | 9.33E-06 | 9.26E-05 |
| ENSG00000152904 | GGPS1 | 1145.121 | 0.788019 | 0.1730547 | 4.670206 | 3.01E-06 | 3.39E-05 |
| ENSG00000250731 | TPM3P6 | 54.51026 | 0.7879707 | 0.3181891 | 2.698481 | 0.00696568 | 0.029004 |
| ENSG00000244153 | WWP1P1 | 223.1755 | 0.7879206 | 0.1714993 | 4.707766 | 2.50E-06 | 2.88E-05 |

| | | | | | | | |
|---|---|---|---|---|---|---|---|
| ENSG00000070081 | NUCB2 | 3672.240574 | 0.787204499 | 0.229106898 | 3.617345427 | 0.00029764 | 0.002008 |
| ENSG00000231369 | | 16.22527 | 0.7869493 | 0.4661484 | 2.030097 | 0.04234663 | 0.122254 |
| ENSG00000197608 | ZNF841 | 915.4861 | 0.7865047 | 0.2189999 | 3.743413 | 0.00018154 | 0.001301 |
| ENSG00000286501 | | 1.736391 | 0.7860964 | 1.22513 | 1.918051 | 0.05510456 | 0.149334 |
| ENSG00000105928 | GSDME | 23.34519696 | 0.785995345 | 0.486631747 | 1.958291631 | 0.0501958 | 0.138864 |
| ENSG00000196110 | ZNF699 | 111.4123 | 0.7856569 | 0.2937798 | 2.883515 | 0.00393263 | 0.01819 |
| ENSG00000197776 | KLHDC1 | 109.6481 | 0.7856204 | 0.4457753 | 2.080824 | 0.03745003 | 0.111098 |
| ENSG00000280219 | | 25.86711 | 0.7855469 | 0.37126 | 2.379189 | 0.01735076 | 0.060609 |
| ENSG00000173726 | TOMM20 | 13233.05 | 0.7854475 | 0.1926504 | 4.209887 | 2.55E-05 | 0.000228 |
| ENSG00000156500 | FAM122C | 253.0576 | 0.7849579 | 0.2242324 | 3.663829 | 0.00024847 | 0.001716 |
| ENSG00000259635 | | 2.942369 | 0.7848719 | 0.9168859 | 1.605832 | 0.1083108 | 0.24659 |
| ENSG00000168056 | LTBP3 | 5449.85 | 0.7847786 | 0.1466276 | 5.451815 | 4.99E-08 | 8.03E-07 |
| ENSG00000157193 | LRP8 | 4709.171 | 0.7846645 | 0.4297872 | 2.156797 | 0.03102147 | 0.096297 |
| ENSG00000256394 | ASIC5 | 2.731026 | 0.784467 | 1.284963 | 2.028268 | 0.04253291 | 0.122686 |
| ENSG00000117758 | STX12 | 3117.728926 | 0.784273208 | 0.152261707 | 5.253676455 | 1.49E-07 | 2.19E-06 |
| ENSG00000254332 | | 652.8713 | 0.7840581 | 0.1984591 | 4.089045 | 4.33E-05 | 0.000365 |
| ENSG00000181626 | ANKRD62 | 1.872528 | 0.7829889 | 1.152416 | 1.772558 | 0.07630205 | 0.190408 |
| ENSG00000232630 | PRPS1P2 | 134.9134 | 0.7829848 | 0.2754888 | 3.03831 | 0.00237909 | 0.011988 |
| ENSG00000162086 | ZNF75A | 2389.466 | 0.7825664 | 0.1450008 | 5.514627 | 3.50E-08 | 5.78E-07 |
| ENSG00000187266 | EPOR | 941.0733 | 0.7825613 | 0.2596076 | 3.145177 | 0.00165987 | 0.008821 |
| ENSG00000110723 | EXPH5 | 368.6435675 | 0.782512031 | 0.26575252 | 3.132226501 | 0.00173486 | 0.009182 |
| ENSG00000236921 | | 16.60035 | 0.7825013 | 0.669479 | 1.668719 | 0.09517305 | 0.22441 |
| ENSG00000109458 | GAB1 | 1743.267885 | 0.782354764 | 0.24699169 | 3.342611722 | 0.00082994 | 0.00487 |
| ENSG00000088854 | C20orf194 | 1318.934199 | 0.782079282 | 0.211636136 | 3.833814425 | 0.00012617 | 0.000942 |
| ENSG00000159399 | HK2 | 5041.39 | 0.7820052 | 0.2464795 | 3.343915 | 0.00082605 | 0.004852 |
| ENSG00000249592 | | 382.6787 | 0.7819805 | 0.2729355 | 3.058741 | 0.00222269 | 0.011318 |
| ENSG00000143882 | ATP6V1C2 | 545.141 | 0.7819783 | 0.4808198 | 1.96769 | 0.04910368 | 0.136668 |
| ENSG00000141738 | GRB7 | 1402.603 | 0.7819542 | 0.2861209 | 2.928066 | 0.00341077 | 0.016168 |
| ENSG00000145476 | CYP4V2 | 1061.445 | 0.7816105 | 0.4294705 | 2.126294 | 0.03347882 | 0.102159 |
| ENSG00000135525 | MAP7 | 4457.083256 | 0.781593599 | 0.109435096 | 7.237979289 | 4.55E-13 | 1.56E-11 |
| ENSG00000237461 | | 10.42222 | 0.7814422 | 0.7479204 | 1.629567 | 0.1031929 | 0.23808 |
| ENSG00000284473 | MIR4720 | 30.19791 | 0.7812077 | 0.3922496 | 2.259414 | 0.02385763 | 0.078179 |
| ENSG00000081818 | PCDHB4 | 57.0024912 | 0.780832293 | 0.368220157 | 2.37305816 | 0.01764149 | 0.061415 |

| | | | | | | | |
|---|---|---|---|---|---|---|---|
| ENSG00000005844 | ITGAL | 6.574893194 | 0.780103892 | 0.849584162 | 1.560030143 | 0.11875276 | 0.263148 |
| ENSG00000166143 | PPP1R14D | 14.62923 | 0.7798574 | 0.5739374 | 1.771916 | 0.07640858 | 0.190602 |
| ENSG00000177125 | ZBTB34 | 842.3259 | 0.7796887 | 0.2561784 | 3.216985 | 0.00129545 | 0.007168 |
| ENSG00000286527 | | 1.34161 | 0.7795271 | 1.183065 | 1.877956 | 0.06038717 | NA |
| ENSG00000197961 | ZNF121 | 2166.922 | 0.7794009 | 0.1728204 | 4.625744 | 3.73E-06 | 4.10E-05 |
| ENSG00000269981 | | 158.8168 | 0.7793629 | 0.3234341 | 2.631511 | 0.00850061 | 0.034132 |
| ENSG00000253686 | LINC01484 | 16.24402 | 0.7792057 | 0.6337909 | 1.695093 | 0.09005775 | 0.215738 |
| ENSG00000166863 | TAC3 | 3.199238 | 0.7788132 | 1.056419 | 1.715638 | 0.08622835 | 0.208747 |
| ENSG00000032219 | ARID4A | 3401.021856 | 0.778715111 | 0.22786353 | 3.57413042 | 0.00035139 | 0.002321 |
| ENSG00000185863 | TMEM210 | 5.227729 | 0.778303 | 0.8556092 | 1.629975 | 0.1031068 | 0.237944 |
| ENSG00000187676 | B3GLCT | 531.8015 | 0.777839 | 0.2977281 | 2.828363 | 0.00467867 | 0.021016 |
| ENSG00000115641 | FHL2 | 1424.493515 | 0.777061989 | 0.451090217 | 2.055276363 | 0.03985232 | 0.116556 |
| ENSG00000213480 | | 11.69823 | 0.7767247 | 0.5832564 | 1.734511 | 0.08282739 | 0.202712 |
| ENSG00000147119 | CHST7 | 187.5931 | 0.7765956 | 0.2499206 | 3.282757 | 0.00102797 | 0.005864 |
| ENSG00000257671 | KRT7-AS | 220.4406 | 0.7765338 | 0.4208363 | 2.154999 | 0.03116189 | 0.096609 |
| ENSG00000265399 | | 58.04686 | 0.7761934 | 0.2802894 | 2.965895 | 0.00301804 | 0.014627 |
| ENSG00000258605 | DYNLL1P2 | 2.030743 | 0.7759002 | 1.021579 | 1.62051 | 0.1051228 | 0.241236 |
| ENSG00000188039 | NWD1 | 7.206384 | 0.7758859 | 0.8815853 | 1.595927 | 0.1105051 | 0.250171 |
| ENSG00000168477 | TNXB | 87.69798 | 0.7755067 | 0.5685364 | 1.782467 | 0.07467308 | 0.187487 |
| ENSG00000178665 | ZNF713 | 82.57603 | 0.7753428 | 0.3820099 | 2.298925 | 0.02150917 | 0.072138 |
| ENSG00000069399 | BCL3 | 1801.531372 | 0.775152703 | 0.255925868 | 3.217151269 | 0.0012947 | 0.007166 |
| ENSG00000212237 | RNA5SP18 | 3.201496 | 0.7747394 | 1.07054 | 1.737057 | 0.08237714 | 0.201852 |
| ENSG00000131725 | WDR44 | 1364.106682 | 0.774556112 | 0.173259941 | 4.595213856 | 4.32E-06 | 4.69E-05 |
| ENSG00000257859 | CASC18 | 2.334777 | 0.7744865 | 1.032103 | 1.64466 | 0.10004 | 0.23291 |
| ENSG00000165972 | CCDC38 | 6.316999 | 0.7744035 | 0.7961813 | 1.585491 | 0.1128549 | 0.253826 |
| ENSG00000186318 | BACE1 | 1519.716 | 0.7742768 | 0.1763482 | 4.49307 | 7.02E-06 | 7.20E-05 |
| ENSG00000225603 | | 5.37431 | 0.7740338 | 1.106152 | 1.835362 | 0.06645208 | 0.171681 |
| ENSG00000287608 | | 71.07514 | 0.773832 | 0.5278329 | 1.879885 | 0.06012382 | 0.159306 |
| ENSG00000071127 | WDR1 | 9690.470591 | 0.773813052 | 0.169081716 | 4.683896627 | 2.81E-06 | 3.20E-05 |
| ENSG00000185710 | SMG1P4 | 170.0284 | 0.7736718 | 0.2851702 | 2.896685 | 0.00377129 | 0.017554 |
| ENSG00000213020 | ZNF611 | 498.7567 | 0.7736081 | 0.2278041 | 3.557196 | 0.00037483 | 0.002456 |
| ENSG00000229582 | | 1.134803 | 0.7732521 | 1.263933 | 2.011699 | 0.04425163 | NA |
| ENSG00000276293 | PIP4K2B | 5126.248 | 0.7731991 | 0.1235917 | 6.340403 | 2.29E-10 | 5.44E-09 |

| | | | | | | | |
|---|---|---|---|---|---|---|---|
| ENSG00000272523 | LINC01023 | 108.5137 | 0.7728938 | 0.35753 | 2.411865 | 0.01587116 | 0.056367 |
| ENSG00000279812 | | 4.570143 | 0.7728397 | 0.9641325 | 1.586631 | 0.1125963 | 0.253447 |
| ENSG00000260955 | | 6.347835 | 0.7725616 | 0.8308967 | 1.580711 | 0.1139442 | 0.255605 |
| ENSG00000264448 | MIR378D2HG | 2.184249 | 0.7725482 | 1.302633 | 2.125874 | 0.03351379 | 0.102199 |
| ENSG00000185437 | SH3BGR | 43.49223 | 0.772487 | 0.3111978 | 2.689004 | 0.00716656 | 0.029719 |
| ENSG00000163347 | CLDN1 | 2424.332 | 0.772377 | 0.4524217 | 2.035652 | 0.04178536 | 0.121002 |
| ENSG00000162783 | IER5 | 4591.585 | 0.7723653 | 0.4265296 | 2.121355 | 0.03389193 | 0.103124 |
| ENSG00000182606 | TRAK1 | 2741.415 | 0.772141 | 0.1862132 | 4.268143 | 1.97E-05 | 0.00018 |
| ENSG00000152595 | MEPE | 2.002607 | 0.7721219 | 1.287171 | 2.057301 | 0.03965724 | 0.116143 |
| ENSG00000162998 | FRZB | 5.3875 | 0.7720863 | 0.837687 | 1.58632 | 0.1126667 | 0.253531 |
| ENSG00000288085 | | 1.500953 | 0.772041 | 1.103342 | 1.778246 | 0.07536348 | NA |
| ENSG00000283674 | | 89.42842 | 0.7713748 | 0.3847826 | 2.275135 | 0.02289783 | 0.075744 |
| ENSG00000240902 | APOOP2 | 6.604358 | 0.7712705 | 1.031124 | 1.688977 | 0.09122377 | 0.217745 |
| ENSG00000069493 | CLEC2D | 483.1649198 | 0.771154432 | 0.275952016 | 2.957197951 | 0.00310449 | 0.014951 |
| ENSG00000242136 | | 15.93768 | 0.770854 | 0.5821722 | 1.752555 | 0.07967839 | 0.196654 |
| ENSG00000185291 | IL3RA | 135.4661 | 0.7707466 | 0.3682079 | 2.346089 | 0.01897155 | 0.065149 |
| ENSG00000144659 | SLC25A38 | 3421.789 | 0.7704726 | 0.1385578 | 5.656856 | 1.54E-08 | 2.72E-07 |
| ENSG00000269289 | | 3.940377 | 0.7701972 | 1.175316 | 1.851124 | 0.06415174 | 0.167037 |
| ENSG00000253582 | | 8.925707 | 0.7700598 | 0.6615416 | 1.645122 | 0.09994469 | 0.232759 |
| ENSG00000139990 | DCAF5 | 6366.688 | 0.7693255 | 0.1902214 | 4.196599 | 2.71E-05 | 0.00024 |
| ENSG00000142512 | SIGLEC10 | 7.830407 | 0.7692478 | 0.8572987 | 1.613714 | 0.1065894 | 0.243778 |
| ENSG00000122644 | ARL4A | 1566.442135 | 0.768994272 | 0.316509888 | 2.647496469 | 0.00810902 | 0.032883 |
| ENSG00000196187 | TMEM63A | 5904.881 | 0.7681647 | 0.290921 | 2.833786 | 0.00460001 | 0.020729 |
| ENSG00000225530 | SP3P | 18.04999 | 0.7681593 | 1.15005 | 1.793981 | 0.07281617 | 0.184077 |
| ENSG00000258524 | NT5CP1 | 5.097324 | 0.7679806 | 0.7937005 | 1.597914 | 0.1100622 | 0.249488 |
| ENSG00000237861 | | 3.077428 | 0.7679336 | 0.9451649 | 1.588752 | 0.1121163 | 0.252696 |
| ENSG00000143418 | CERS2 | 17066.06 | 0.767864 | 0.1764129 | 4.467096 | 7.93E-06 | 8.03E-05 |
| ENSG00000275993 | SIK1B | 853.1324 | 0.7674387 | 0.5143906 | 1.867814 | 0.06178804 | 0.162571 |
| ENSG00000204967 | PCDHA4 | 13.52971 | 0.7670533 | 0.6494067 | 1.656895 | 0.0975407 | 0.228508 |
| ENSG00000186765 | FSCN2 | 212.7317 | 0.7669819 | 0.3125579 | 2.672168 | 0.0075363 | 0.030993 |
| ENSG00000122592 | HOXA7 | 12.35464503 | 0.766829053 | 0.724278759 | 1.638202753 | 0.10137941 | 0.235121 |
| ENSG00000228109 | MELTF-AS1 | 151.9339 | 0.7667949 | 0.2988436 | 2.772977 | 0.00555461 | 0.024154 |
| ENSG00000065833 | ME1 | 3493.670741 | 0.766533862 | 0.285135433 | 2.887281917 | 0.00388586 | 0.018015 |

| | | | | | | | |
|---|---|---|---|---|---|---|---|
| ENSG00000114805 | PLCH1 | 729.7379778 | 0.76651268 | 0.41482693 | 2.141415992 | 0.03224051 | 0.099147 |
| ENSG00000153832 | FBXO36 | 326.4255 | 0.7665057 | 0.1848426 | 4.271923 | 1.94E-05 | 0.000178 |
| ENSG00000087274 | ADD1 | 6368.127076 | 0.766432698 | 0.129623296 | 6.036598925 | 1.57E-09 | 3.27E-08 |
| ENSG00000199906 | RNU5B-2P | 2.718099 | 0.7663747 | 1.018704 | 1.66748 | 0.09541906 | 0.224831 |
| ENSG00000170322 | NFRKB | 3435.862 | 0.7659965 | 0.1942274 | 4.092403 | 4.27E-05 | 0.00036 |
| ENSG00000198075 | SULT1C4 | 1.440716 | 0.7658955 | 1.398261 | 2.38937 | 0.01687729 | NA |
| ENSG00000165775 | FUNDC2 | 3793.017 | 0.7658375 | 0.1413366 | 5.476056 | 4.35E-08 | 7.08E-07 |
| ENSG00000135363 | LMO2 | 193.8447691 | 0.76572719 | 0.316352802 | 2.631905621 | 0.00849075 | 0.034112 |
| ENSG00000119401 | TRIM32 | 1213.759618 | 0.765689117 | 0.171881356 | 4.578251887 | 4.69E-06 | 5.04E-05 |
| ENSG00000196696 | | 563.8015 | 0.765366 | 0.2447144 | 3.305561 | 0.00094787 | 0.005453 |
| ENSG00000285984 | | 3.038448 | 0.7653615 | 0.909242 | 1.572963 | 0.1157275 | 0.258607 |
| ENSG00000153395 | LPCAT1 | 23310.03 | 0.7652991 | 0.1978435 | 4.008446 | 6.11E-05 | 0.000495 |
| ENSG00000224727 | FCF1P7 | 4.66122 | 0.7651443 | 0.7325845 | 1.590888 | 0.1116347 | 0.252063 |
| ENSG00000267023 | LRRC37A16P | 701.4167 | 0.7649645 | 0.221063 | 3.639364 | 0.00027331 | 0.001863 |
| ENSG00000154608 | CEP170P1 | 9.536139 | 0.7647261 | 0.7071909 | 1.610759 | 0.1072323 | 0.244827 |
| ENSG00000162032 | SPSB3 | 42.58935 | 0.7646555 | 0.3368947 | 2.501439 | 0.01236899 | 0.04613 |
| ENSG00000203907 | OOEP | 80.99674 | 0.7645097 | 0.2512391 | 3.2237 | 0.00126546 | 0.007018 |
| ENSG00000232311 | NA | 4.270286 | 0.7644278 | 0.7656713 | 1.578598 | 0.1144284 | 0.256428 |
| ENSG00000177363 | LRRN4CL | 7.339299 | 0.7644173 | 0.6932868 | 1.61675 | 0.1059323 | 0.242645 |
| ENSG00000237118 | CYP2F2P | 42.96599 | 0.7641304 | 0.442165 | 2.034529 | 0.0418983 | 0.12124 |
| ENSG00000164855 | TMEM184A | 5362.77 | 0.7631948 | 0.1756921 | 4.460904 | 8.16E-06 | 8.24E-05 |
| ENSG00000150764 | DIXDC1 | 294.7549 | 0.762978 | 0.3512007 | 2.412055 | 0.01586287 | 0.056345 |
| ENSG00000269937 | | 43.0796 | 0.7629494 | 0.4210672 | 2.109194 | 0.03492784 | 0.105457 |
| ENSG00000143669 | LYST | 682.9406 | 0.7628866 | 0.4030201 | 2.181286 | 0.02916224 | 0.091813 |
| ENSG00000164483 | SAMD3 | 3.300502 | 0.7626974 | 1.385043 | 2.397173 | 0.01652212 | 0.058227 |
| ENSG00000159885 | ZNF222 | 159.7494 | 0.762556 | 0.2266793 | 3.512354 | 0.00044416 | 0.002832 |
| ENSG00000274602 | PI4KAP1 | 602.3574 | 0.7623233 | 0.2002087 | 3.943569 | 8.03E-05 | 0.00063 |
| ENSG00000167807 | | 9.975722 | 0.76232 | 0.6463164 | 1.654812 | 0.0979626 | 0.229238 |
| ENSG00000276570 | | 71.21112 | 0.7622718 | 0.2360295 | 3.394212 | 0.00068826 | 0.004142 |
| ENSG00000275227 | | 17.83662 | 0.762063 | 0.4840732 | 1.921099 | 0.05471929 | 0.148563 |
| ENSG00000112297 | CRYBG1 | 2236.979502 | 0.761723514 | 0.24497957 | 3.288644003 | 0.00100671 | 0.005755 |
| ENSG00000184361 | SPATA32 | 11.59806 | 0.7613184 | 0.5522866 | 1.778218 | 0.07536799 | 0.18875 |
| ENSG00000176894 | PXMP2 | 292.9629 | 0.7609552 | 0.238332 | 3.364578 | 0.00076661 | 0.004548 |

| | | | | | | | |
|---|---|---|---|---|---|---|---|
| ENSG00000160207 | HSF2BP | 100.6429 | 0.760916 | 0.2540195 | 3.159871 | 0.00157839 | 0.00846 |
| ENSG00000287592 | | 1.964893 | 0.760822 | 1.107277 | 1.720928 | 0.08526395 | 0.207073 |
| ENSG00000189269 | DRICH1 | 19.1103 | 0.7606893 | 0.6059285 | 1.700763 | 0.08898742 | 0.213887 |
| ENSG00000230102 | LINC02028 | 22.55732 | 0.760647 | 0.637454 | 1.684459 | 0.09209306 | 0.219134 |
| ENSG00000145860 | RNF145 | 4351.236 | 0.7605504 | 0.1964378 | 4.00324 | 6.25E-05 | 0.000505 |
| ENSG00000123612 | ACVR1C | 38.02070909 | 0.760122452 | 0.458699917 | 1.976244082 | 0.04812714 | 0.134578 |
| ENSG00000118263 | KLF7 | 885.276574 | 0.759755014 | 0.403123391 | 2.168301291 | 0.03013577 | 0.093987 |
| ENSG00000251050 | RBX1P2 | 2.346227 | 0.7595779 | 0.9045688 | 1.570115 | 0.1163883 | 0.259486 |
| ENSG00000181392 | SYNE4 | 3090.232 | 0.7592274 | 0.210391 | 3.766541 | 0.00016553 | 0.001201 |
| ENSG00000117586 | TNFSF4 | 21.11778758 | 0.759122481 | 0.602182922 | 1.693488048 | 0.09036261 | 0.216281 |
| ENSG00000137845 | ADAM10 | 6722.674336 | 0.759081555 | 0.213109026 | 3.69105199 | 0.00022333 | 0.001563 |
| ENSG00000176927 | EFCAB5 | 8.062129 | 0.7590592 | 0.6093092 | 1.678187 | 0.09331058 | 0.221307 |
| ENSG00000158113 | LRRC43 | 1.956492 | 0.7588043 | 1.10844 | 1.692241 | 0.09060006 | 0.216646 |
| ENSG00000168148 | H3-4 | 5.533298 | 0.7584146 | 0.8959709 | 1.574291 | 0.1154203 | 0.25824 |
| ENSG00000107864 | CPEB3 | 330.1569381 | 0.758229757 | 0.278279322 | 2.914945679 | 0.00355751 | 0.016746 |
| ENSG00000111752 | PHC1 | 344.3920333 | 0.758014265 | 0.212325327 | 3.717746317 | 0.00020101 | 0.001425 |
| ENSG00000279286 | | 10.71913 | 0.7579928 | 0.6300557 | 1.672786 | 0.09436937 | 0.223213 |
| ENSG00000250230 | | 11.03543 | 0.7579335 | 0.6367526 | 1.659761 | 0.09696255 | 0.227431 |
| ENSG00000189410 | SH2D5 | 273.3364 | 0.7579312 | 0.3384423 | 2.487459 | 0.01286594 | 0.047687 |
| ENSG00000171606 | ZNF274 | 1639.911 | 0.7578891 | 0.154652 | 5.004685 | 5.60E-07 | 7.36E-06 |
| ENSG00000147614 | ATP6V0D2 | 4.407492 | 0.7576372 | 1.094781 | 1.753775 | 0.07946898 | 0.196246 |
| ENSG00000169193 | CCDC126 | 338.4238 | 0.757517 | 0.2552104 | 3.141189 | 0.00168264 | 0.008932 |
| ENSG00000182747 | SLC35D3 | 1.757661 | 0.7574419 | 1.13513 | 1.779707 | 0.07512399 | 0.188264 |
| ENSG00000258414 | | 113.804 | 0.7573105 | 0.2809875 | 2.88533 | 0.00391003 | 0.018114 |
| ENSG00000278869 | | 34.58433 | 0.7565659 | 0.4300187 | 2.073233 | 0.03815064 | 0.112753 |
| ENSG00000248801 | C8orf34-AS1 | 22.73458 | 0.7564014 | 0.6558521 | 1.636179 | 0.1018022 | 0.235706 |
| ENSG00000215186 | GOLGA6B | 1.923234 | 0.7559583 | 1.296985 | 2.109182 | 0.03492885 | 0.105457 |
| ENSG00000113719 | ERGIC1 | 20341.43985 | 0.755954043 | 0.146954098 | 5.175329804 | 2.28E-07 | 3.22E-06 |
| ENSG00000166927 | MS4A7 | 9.054397 | 0.7557935 | 1.197061 | 1.909511 | 0.0561962 | 0.151505 |
| ENSG00000162383 | SLC1A7 | 2.791187 | 0.7557213 | 1.297426 | 2.135719 | 0.03270236 | 0.100297 |
| ENSG00000109618 | SEPSECS | 1074.72302 | 0.755360264 | 0.203262065 | 3.859492736 | 0.00011362 | 0.000858 |
| ENSG00000261056 | | 4.59333 | 0.7551865 | 0.8543136 | 1.552824 | 0.1204652 | 0.266099 |
| ENSG00000235092 | ID2-AS1 | 289.2154 | 0.7544332 | 0.6746029 | 1.628306 | 0.1034599 | 0.238476 |

| | | | | | | | |
|---|---|---|---|---|---|---|---|
| ENSG00000243069 | ARHGEF26-AS1 | 101.6557 | 0.7543306 | 0.4181695 | 2.092098 | 0.03642974 | 0.108778 |
| ENSG00000169641 | LUZP1 | 1796.515 | 0.7542231 | 0.1846943 | 4.208021 | 2.58E-05 | 0.000229 |
| ENSG00000060140 | STYK1 | 88.07573833 | 0.754154312 | 0.408386654 | 2.143596009 | 0.03206527 | 0.098742 |
| ENSG00000107020 | PLGRKT | 748.9782757 | 0.754117463 | 0.226455032 | 3.481481712 | 0.00049865 | 0.003131 |
| ENSG00000254266 | PKIA-AS1 | 3.295516 | 0.7540949 | 0.9114585 | 1.57759 | 0.1146598 | 0.25682 |
| ENSG00000213640 | EEF1DP4 | 1.757909 | 0.7540234 | 1.054244 | 1.66311 | 0.09629043 | 0.226162 |
| ENSG00000230289 | | 6.289726 | 0.7537667 | 0.648875 | 1.639979 | 0.1010095 | 0.234489 |
| ENSG00000179292 | TMEM151A | 48.16525 | 0.7537177 | 0.5178918 | 1.817195 | 0.0691872 | 0.176975 |
| ENSG00000235005 | | 1.994178 | 0.7536547 | 1.094003 | 1.758968 | 0.07858303 | 0.194555 |
| ENSG00000164175 | SLC45A2 | 4.619058 | 0.7536041 | 1.064353 | 1.710211 | 0.08722689 | 0.210551 |
| ENSG00000187736 | NHEJ1 | 12.57263 | 0.7535224 | 0.4924125 | 1.868584 | 0.06168074 | 0.162386 |
| ENSG00000133935 | ERG28 | 4465.678587 | 0.753325335 | 0.192934013 | 4.020007325 | 5.82E-05 | 0.000473 |
| ENSG00000167046 | | 13.69516 | 0.753292 | 0.5355767 | 1.783774 | 0.07446046 | 0.187148 |
| ENSG00000184584 | STING1 | 196.8995 | 0.7532668 | 0.4342083 | 2.054689 | 0.03990909 | 0.11665 |
| ENSG00000257557 | PPP1R12A-AS1 | 95.61462 | 0.7531796 | 0.253016 | 3.14027 | 0.00168792 | 0.008955 |
| ENSG00000138835 | RGS3 | 1796.890689 | 0.753152197 | 0.306817226 | 2.663520075 | 0.00773278 | 0.031652 |
| ENSG00000274367 | | 30.01191 | 0.7527262 | 0.4770067 | 1.910517 | 0.05606663 | 0.151263 |
| ENSG00000227666 | CYCSP24 | 4.555818 | 0.7525437 | 0.8682912 | 1.588176 | 0.1122465 | 0.252896 |
| ENSG00000093167 | LRRFIP2 | 1406.360544 | 0.752372588 | 0.113605158 | 6.696673042 | 2.13E-11 | 5.81E-10 |
| ENSG00000088340 | FER1L4 | 2006.210778 | 0.752130942 | 0.361525301 | 2.33046889 | 0.01978138 | 0.067398 |
| ENSG00000167123 | CERCAM | 7909.021 | 0.7520526 | 0.1655526 | 4.630534 | 3.65E-06 | 4.02E-05 |
| ENSG00000260136 | | 68.03379 | 0.7517971 | 0.2310287 | 3.400482 | 0.00067267 | 0.004059 |
| ENSG00000171502 | COL24A1 | 53.41126 | 0.7516625 | 0.3955429 | 2.16948 | 0.03004627 | 0.093807 |
| ENSG00000146433 | TMEM181 | 3032.04 | 0.7514219 | 0.2428231 | 3.283364 | 0.00102576 | 0.005853 |
| ENSG00000272129 | | 44.17684 | 0.7509586 | 0.4302158 | 2.049741 | 0.04038968 | 0.117756 |
| ENSG00000231160 | KLF3-AS1 | 102.537 | 0.7509216 | 0.4642969 | 1.942178 | 0.05211559 | 0.143058 |
| ENSG00000173852 | DPY19L1 | 2678.467 | 0.7506738 | 0.2026297 | 3.827488 | 0.00012946 | 0.000963 |
| ENSG00000171931 | FBXW10 | 17.73409 | 0.7505817 | 0.6219963 | 1.645158 | 0.09993725 | 0.232759 |
| ENSG00000155324 | GRAMD2B | 778.1052 | 0.7505103 | 0.247299 | 3.20581 | 0.00134683 | 0.007409 |
| ENSG00000138829 | FBN2 | 492.635962 | 0.750086838 | 0.508536712 | 1.834047523 | 0.06664694 | 0.172084 |
| ENSG00000230005 | SNAP47-AS1 | 2.029861 | 0.7500714 | 1.101081 | 1.788934 | 0.07362538 | 0.185681 |
| ENSG00000261645 | DISC1FP1 | 4.457819 | 0.7498315 | 1.03887 | 1.672659 | 0.0943945 | 0.223228 |
| ENSG00000220305 | HNRNPH1P1 | 9.825442 | 0.7497078 | 0.6609443 | 1.613604 | 0.1066134 | 0.243811 |

| | | | | | | | |
|---|---|---|---|---|---|---|---|
| ENSG00000279443 | | 143.5959 | 0.749707 | 0.3774144 | 2.253102 | 0.02425272 | 0.07921 |
| ENSG00000246582 | | 173.2164 | 0.7494165 | 0.1975938 | 3.929953 | 8.50E-05 | 0.000663 |
| ENSG00000278834 | | 22.1444 | 0.7492269 | 0.4364968 | 2.02025 | 0.0433575 | 0.124531 |
| ENSG00000145014 | TMEM44 | 1063.772 | 0.7491707 | 0.2229251 | 3.509718 | 0.00044858 | 0.002855 |
| ENSG00000232212 | LINC01701 | 11.32165 | 0.7491183 | 0.6047592 | 1.673055 | 0.09431656 | 0.223169 |
| ENSG00000229848 | | 113.0352 | 0.749024 | 0.3847643 | 2.210643 | 0.02706059 | 0.086353 |
| ENSG00000203950 | RTL8A | 4894.228 | 0.7486139 | 0.1643125 | 4.629307 | 3.67E-06 | 4.04E-05 |
| ENSG00000243918 | EIF4BP8 | 2.553805 | 0.7485366 | 1.044587 | 1.628081 | 0.1035076 | 0.238537 |
| ENSG00000231663 | COA6-AS1 | 53.29076 | 0.7485341 | 0.3429421 | 2.412597 | 0.01583934 | 0.056284 |
| ENSG00000277678 | RNU1-153P | 4.52825 | 0.7482994 | 0.8299174 | 1.538741 | 0.1238675 | 0.271538 |
| ENSG00000197576 | HOXA4 | 9.971392 | 0.7479914 | 0.7024788 | 1.58712 | 0.1124854 | 0.253252 |
| ENSG00000100784 | RPS6KA5 | 862.2299843 | 0.747801516 | 0.293536718 | 2.752037024 | 0.00592258 | 0.025475 |
| ENSG00000253736 | | 2.672748 | 0.7477943 | 0.9896647 | 1.617942 | 0.1056752 | 0.242251 |
| ENSG00000161996 | WDR90 | 2564.163 | 0.7476538 | 0.1755432 | 4.330738 | 1.49E-05 | 0.00014 |
| ENSG00000270681 | | 42.5251 | 0.7475286 | 0.3535929 | 2.363941 | 0.01808169 | 0.062701 |
| ENSG00000188158 | NHS | 743.7556 | 0.7472468 | 0.404022 | 2.130971 | 0.03309152 | 0.101233 |
| ENSG00000223802 | CERS1 | 193.6357 | 0.747098 | 0.3350975 | 2.454972 | 0.01408958 | 0.051273 |
| ENSG00000253554 | LINC01414 | 37.08716 | 0.7469453 | 0.3500486 | 2.364919 | 0.01803402 | 0.062544 |
| ENSG00000180914 | OXTR | 86.08822 | 0.7464769 | 0.3875988 | 2.192208 | 0.02836452 | 0.089642 |
| ENSG00000170417 | TMEM182 | 234.5031 | 0.7464683 | 0.2385798 | 3.254522 | 0.00113583 | 0.006384 |
| ENSG00000112981 | NME5 | 14.73078596 | 0.74615977 | 0.596518153 | 1.678298909 | 0.09328876 | 0.221303 |
| ENSG00000247708 | STX18-AS1 | 187.9602 | 0.7460911 | 0.2690234 | 2.948832 | 0.00318977 | 0.015287 |
| ENSG00000144118 | RALB | 3739.88 | 0.7459863 | 0.1794727 | 4.272183 | 1.94E-05 | 0.000178 |
| ENSG00000261744 | | 38.6607 | 0.7456277 | 0.4241988 | 2.04057 | 0.0412936 | 0.119918 |
| ENSG00000196274 | | 9.505204 | 0.7455314 | 0.6302276 | 1.652758 | 0.09838015 | 0.229831 |
| ENSG00000286872 | | 15.68289 | 0.7452318 | 0.5357189 | 1.760787 | 0.07827438 | 0.194036 |
| ENSG00000159239 | | 14.83617 | 0.7451213 | 0.8313236 | 1.559427 | 0.1188954 | 0.26331 |
| ENSG00000271366 | | 17.94539 | 0.7450688 | 0.5059738 | 1.814004 | 0.06967717 | 0.177954 |
| ENSG00000167470 | MIDN | 5082.084 | 0.7450633 | 0.2072727 | 3.714555 | 0.00020356 | 0.001441 |
| ENSG00000229619 | MBNL1-AS1 | 86.07947 | 0.7447342 | 0.3662028 | 2.28928 | 0.02206308 | 0.07367 |
| ENSG00000108786 | HSD17B1 | 234.580688 | 0.744650306 | 0.413884218 | 2.085231479 | 0.03704829 | 0.110176 |
| ENSG00000216740 | ANXA2P3 | 2.711509 | 0.7445551 | 1.051954 | 1.721458 | 0.08516776 | 0.206877 |
| ENSG00000270720 | | 11.69028 | 0.7445054 | 0.5737852 | 1.700317 | 0.08907135 | 0.21405 |

| | | | | | | | |
|---|---|---|---|---|---|---|---|
| ENSG00000119599 | DCAF4 | 4121.05882 | 0.744292581 | 0.182233662 | 4.197854302 | 2.69E-05 | 0.000239 |
| ENSG00000212864 | RNF208 | 1046.05 | 0.7442867 | 0.1742565 | 4.390685 | 1.13E-05 | 0.00011 |
| ENSG00000182827 | ACBD3 | 3821.391 | 0.7442654 | 0.1709084 | 4.46218 | 8.11E-06 | 8.19E-05 |
| ENSG00000155265 | GOLGA7B | 240.0479 | 0.7438664 | 0.3472882 | 2.379869 | 0.01731882 | 0.060513 |
| ENSG00000223797 | ENTPD3-AS1 | 196.8863 | 0.7436429 | 0.2975149 | 2.695549 | 0.00702728 | 0.029228 |
| ENSG00000167378 | IRGQ | 1512.885 | 0.743584 | 0.2081249 | 3.709176 | 0.00020794 | 0.001468 |
| ENSG00000267174 | | 8.29404 | 0.7431534 | 0.6808007 | 1.592715 | 0.1112241 | 0.251306 |
| ENSG00000270276 | H4C15 | 6.777045 | 0.7430857 | 0.6237616 | 1.63547 | 0.1019505 | 0.235931 |
| ENSG00000227087 | RBMX2P5 | 181.1829 | 0.742935 | 0.2083102 | 3.705674 | 0.00021083 | 0.001486 |
| ENSG00000263508 | | 2.960419 | 0.7429134 | 1.041287 | 1.740454 | 0.08177937 | 0.200739 |
| ENSG00000108797 | CNTNAP1 | 140.6561754 | 0.742780699 | 0.399363056 | 2.135247604 | 0.03274079 | 0.100391 |
| ENSG00000140274 | DUOXA2 | 1.292162 | 0.7426744 | 1.229244 | 1.966857 | 0.04919975 | NA |
| ENSG00000248360 | LINC00504 | 20.07212 | 0.7424385 | 0.6603086 | 1.613842 | 0.1065617 | 0.243757 |
| ENSG00000197555 | SIPA1L1 | 4581.526 | 0.742243 | 0.1851943 | 4.129012 | 3.64E-05 | 0.000313 |
| ENSG00000173988 | LRRC63 | 0.9445011 | 0.7422367 | 1.245196 | 2.010992 | 0.04432626 | NA |
| ENSG00000131480 | AOC2 | 59.74441809 | 0.741936238 | 0.43182486 | 2.011375226 | 0.04428584 | 0.126506 |
| ENSG00000218748 | DBIP1 | 3.023707 | 0.7418101 | 1.094238 | 1.840179 | 0.0657419 | 0.170243 |
| ENSG00000279861 | | 77.69224 | 0.7414806 | 0.313021 | 2.576517 | 0.00998014 | 0.0388 |
| ENSG00000261140 | | 3.27803 | 0.7413424 | 0.9677667 | 1.559707 | 0.1188292 | 0.26327 |
| ENSG00000165046 | LETM2 | 76.46346 | 0.7412736 | 0.4074664 | 2.099423 | 0.03577959 | 0.107418 |
| ENSG00000011638 | TMEM159 | 1595.711264 | 0.741055028 | 0.14282502 | 5.296277657 | 1.18E-07 | 1.77E-06 |
| ENSG00000204580 | DDR1 | 23407.82 | 0.7409673 | 0.1932994 | 3.929136 | 8.53E-05 | 0.000665 |
| ENSG00000169604 | ANTXR1 | 2693.906 | 0.7408619 | 0.1972171 | 3.894703 | 9.83E-05 | 0.000754 |
| ENSG00000284621 | | 11.07825 | 0.7408603 | 0.5714481 | 1.699336 | 0.08925593 | 0.214363 |
| ENSG00000162636 | FAM102B | 6043.839 | 0.7404482 | 0.3084817 | 2.592011 | 0.00954168 | 0.037391 |
| ENSG00000186676 | EEF1GP1 | 7.239751 | 0.7404267 | 0.7374152 | 1.538265 | 0.1239838 | 0.271658 |
| ENSG00000266242 | GRAMD4P7 | 1.20085 | 0.7403585 | 1.301069 | 2.219979 | 0.02642022 | NA |
| ENSG00000277051 | | 4.619069 | 0.7402226 | 0.7482124 | 1.539962 | 0.1235697 | 0.271064 |
| ENSG00000219438 | TAFA5 | 364.2222 | 0.7401859 | 0.3029264 | 2.650134 | 0.00804598 | 0.032687 |
| ENSG00000130822 | PNCK | 16.40713626 | 0.740045292 | 0.612474757 | 1.649644441 | 0.09901568 | 0.23099 |
| ENSG00000266306 | | 5.060357 | 0.7399742 | 0.802314 | 1.548518 | 0.1214976 | 0.267756 |
| ENSG00000279948 | | 18.47855 | 0.7394091 | 0.5791974 | 1.699628 | 0.08920091 | 0.214303 |
| ENSG00000231397 | | 2.483285 | 0.739327 | 0.9730891 | 1.560673 | 0.1186009 | 0.262871 |

| | | | | | | | |
|---|---|---|---|---|---|---|---|
| ENSG00000200114 | RNA5SP123 | 3.7967 | 0.7390693 | 0.8407804 | 1.536187 | 0.1244925 | 0.272368 |
| ENSG00000204815 | TTC25 | 115.9037 | 0.7388579 | 0.2380868 | 3.255126 | 0.00113342 | 0.006373 |
| ENSG00000132958 | TPTE2 | 6.028019644 | 0.738764794 | 0.733898245 | 1.547211208 | 0.12181232 | 0.268116 |
| ENSG00000234771 | SLC25A25-AS1 | 964.6995 | 0.7387595 | 0.2675837 | 2.930479 | 0.0033844 | 0.016069 |
| ENSG00000168405 | CMAHP | 9.696303 | 0.7383801 | 0.6177035 | 1.652794 | 0.09837272 | 0.229831 |
| ENSG00000164761 | TNFRSF11B | 1214.301 | 0.7382789 | 0.3874623 | 2.167522 | 0.03019507 | 0.094128 |
| ENSG00000114349 | GNAT1 | 5.305942356 | 0.737820383 | 0.747829887 | 1.552907343 | 0.12044527 | 0.266077 |
| ENSG00000223519 | KIF28P | 77.94116 | 0.7375125 | 0.3698423 | 2.245227 | 0.02475359 | 0.08046 |
| ENSG00000137269 | LRRC1 | 2478.623349 | 0.736993351 | 0.169792034 | 4.450158609 | 8.58E-06 | 8.61E-05 |
| ENSG00000248476 | BACH1-IT1 | 13.83573 | 0.7367673 | 0.5436764 | 1.735467 | 0.0826581 | 0.202447 |
| ENSG00000185215 | TNFAIP2 | 981.032 | 0.7366024 | 0.4916399 | 1.850377 | 0.06425919 | 0.167284 |
| ENSG00000229320 | KRT8P12 | 666.4896 | 0.736426 | 0.2215344 | 3.456818 | 0.00054659 | 0.003395 |
| ENSG00000232065 | LINC01063 | 16.05479 | 0.7362618 | 0.4986532 | 1.819235 | 0.06887562 | 0.176364 |
| ENSG00000131238 | PPT1 | 24912.00987 | 0.735967653 | 0.18279924 | 4.142746747 | 3.43E-05 | 0.000297 |
| ENSG00000251602 | | 114.2951 | 0.7359243 | 0.2235013 | 3.442467 | 0.00057643 | 0.003552 |
| ENSG00000242748 | RPL23AP81 | 8.075767 | 0.7359147 | 0.7440288 | 1.5673 | 0.1170446 | 0.260471 |
| ENSG00000143387 | CTSK | 636.5281 | 0.7359124 | 0.2267744 | 3.379245 | 0.00072685 | 0.004338 |
| ENSG00000171435 | KSR2 | 345.0268 | 0.7357641 | 0.2682849 | 2.91728 | 0.00353099 | 0.01664 |
| ENSG00000198346 | ZNF813 | 546.3653 | 0.7356721 | 0.1957912 | 3.885613 | 0.00010207 | 0.000778 |
| ENSG00000239005 | | 1.536357 | 0.7353578 | 1.129504 | 1.895331 | 0.05804859 | NA |
| ENSG00000230971 | NA | 1.223363 | 0.7349716 | 1.252284 | 2.059513 | 0.03944509 | NA |
| ENSG00000255458 | | 29.45722 | 0.7347123 | 0.3972788 | 2.109357 | 0.03491375 | 0.105436 |
| ENSG00000107771 | CCSER2 | 3346.351229 | 0.734053197 | 0.190299455 | 3.989634353 | 6.62E-05 | 0.00053 |
| ENSG00000237399 | PITRM1-AS1 | 83.6006 | 0.7336921 | 0.2802848 | 2.812837 | 0.00491065 | 0.021843 |
| ENSG00000186056 | MATN1-AS1 | 209.5932 | 0.7334169 | 0.185685 | 4.069843 | 4.70E-05 | 0.000393 |
| ENSG00000130656 | HBZ | 2.92631021 | 0.73317168 | 0.986142785 | 1.592607664 | 0.11124823 | 0.251318 |
| ENSG00000169194 | IL13 | 5.196791 | 0.7326872 | 0.7972886 | 1.543477 | 0.1227151 | 0.269545 |
| ENSG00000261572 | | 189.8 | 0.7324685 | 0.2838508 | 2.77205 | 0.00557045 | 0.024199 |
| ENSG00000158290 | CUL4B | 4999.19 | 0.7318578 | 0.1395913 | 5.331056 | 9.76E-08 | 1.48E-06 |
| ENSG00000100934 | SEC23A | 5796.756883 | 0.731796233 | 0.221223495 | 3.462166659 | 0.00053585 | 0.003336 |
| ENSG00000271869 | | 28.38214 | 0.7317931 | 0.422734 | 2.013334 | 0.04407949 | 0.126052 |
| ENSG00000169689 | CENPX | 8162.637 | 0.7317307 | 0.264275 | 2.955305 | 0.0031236 | 0.015024 |
| ENSG00000182674 | KCNB2 | 1.713356 | 0.7316492 | 1.039871 | 1.733839 | 0.08294657 | 0.202846 |

| | | | | | | | |
|---|---|---|---|---|---|---|---|
| ENSG00000181045 | SLC26A11 | 3149.675 | 0.7315551 | 0.2210154 | 3.452523 | 0.00055537 | 0.003436 |
| ENSG00000022567 | SLC45A4 | 894.4881569 | 0.73153635 | 0.169106939 | 4.43998597 | 9.00E-06 | 8.98E-05 |
| ENSG00000271983 | | 16.46663 | 0.7314402 | 0.6366518 | 1.601862 | 0.109186 | 0.247924 |
| ENSG00000100644 | HIF1A | 9457.852512 | 0.731247172 | 0.135911911 | 5.452944405 | 4.95E-08 | 7.99E-07 |
| ENSG00000279792 | | 16.17752 | 0.7311693 | 0.4660188 | 1.886856 | 0.0591797 | 0.157512 |
| ENSG00000100314 | CABP7 | 6.200450554 | 0.730756653 | 0.698112794 | 1.543188672 | 0.12278501 | 0.269676 |
| ENSG00000181827 | RFX7 | 2581.391 | 0.7305828 | 0.1932136 | 3.895153 | 9.81E-05 | 0.000753 |
| ENSG00000280217 | | 75.60421 | 0.7302419 | 0.5371594 | 1.746977 | 0.08064134 | 0.198495 |
| ENSG00000174233 | ADCY6 | 5938.167 | 0.7300362 | 0.2877169 | 2.73014 | 0.00633073 | 0.026848 |
| ENSG00000271320 | | 1.294027 | 0.7298665 | 1.115831 | 1.755446 | 0.079183 | NA |
| ENSG00000079215 | SLC1A3 | 278.4356268 | 0.729842517 | 0.592677909 | 1.670130124 | 0.09489362 | 0.22403 |
| ENSG00000104140 | RHOV | 1966.271012 | 0.729759847 | 0.340937808 | 2.385524601 | 0.01705478 | 0.059756 |
| ENSG00000204055 | | 19.86675 | 0.7297201 | 0.6276183 | 1.609289 | 0.1075533 | 0.245265 |
| ENSG00000142208 | AKT1 | 18081.19 | 0.7292543 | 0.1859496 | 4.040373 | 5.34E-05 | 0.000439 |
| ENSG00000161791 | FMNL3 | 249.0341 | 0.7291655 | 0.3682212 | 2.230421 | 0.02571953 | 0.08289 |
| ENSG00000069011 | PITX1 | 939.0104671 | 0.729146661 | 0.268028099 | 2.899879425 | 0.00373306 | 0.017413 |
| ENSG00000236483 | MTND2P40 | 4.087299 | 0.7290435 | 0.8698265 | 1.528182 | 0.1264674 | 0.275171 |
| ENSG00000135905 | DOCK10 | 35.00262322 | 0.728986231 | 0.551685159 | 1.710484042 | 0.0871764 | 0.210448 |
| ENSG00000150527 | MIA2 | 2767.829 | 0.7289202 | 0.2202714 | 3.447819 | 0.00056513 | 0.003488 |
| ENSG00000154025 | SLC5A10 | 12.07931 | 0.7288974 | 0.5741574 | 1.666198 | 0.095674 | 0.225251 |
| ENSG00000168314 | MOBP | 2.139942 | 0.7287719 | 1.105316 | 1.732678 | 0.0831529 | 0.203197 |
| ENSG00000172543 | CTSW | 5.619512 | 0.7282573 | 0.6771034 | 1.561965 | 0.1182961 | 0.262432 |
| ENSG00000247675 | LRP4-AS1 | 16.21117 | 0.7281061 | 0.5893305 | 1.650736 | 0.0987925 | 0.230632 |
| ENSG00000153786 | ZDHHC7 | 10094.2 | 0.7280899 | 0.219008 | 3.466471 | 0.00052734 | 0.003287 |
| ENSG00000178187 | ZNF454 | 20.7242 | 0.7277921 | 0.634782 | 1.603594 | 0.1088035 | 0.247281 |
| ENSG00000145287 | PLAC8 | 10.10255 | 0.7276789 | 0.6731364 | 1.570075 | 0.1163978 | 0.259486 |
| ENSG00000232803 | SLCO4A1-AS1 | 24.25686 | 0.727656 | 0.6135768 | 1.63477 | 0.1020973 | 0.236168 |
| ENSG00000254461 | | 117.3561 | 0.7273027 | 0.2746891 | 2.824592 | 0.00473408 | 0.021204 |
| ENSG00000136560 | TANK | 1687.782952 | 0.727206123 | 0.174772272 | 4.266900278 | 1.98E-05 | 0.000181 |
| ENSG00000271858 | | 10.74862 | 0.7267196 | 0.587817 | 1.643001 | 0.1003828 | 0.233442 |
| ENSG00000205634 | LINC00898 | 12.21565 | 0.7266209 | 0.5619155 | 1.691391 | 0.09076209 | 0.216936 |
| ENSG00000149571 | KIRREL3 | 18.67943 | 0.7264625 | 0.543177 | 1.712449 | 0.08681403 | 0.209878 |
| ENSG00000287079 | | 1.039076 | 0.7260745 | 1.165172 | 1.860072 | 0.0628754 | NA |

| | | | | | | | |
|---|---|---|---|---|---|---|---|
| ENSG00000253508 | | 1.274181 | 0.7259574 | 1.185157 | 1.91936 | 0.05493884 | NA |
| ENSG00000141298 | SSH2 | 996.2373 | 0.7258268 | 0.2230184 | 3.401425 | 0.00067036 | 0.004047 |
| ENSG00000170381 | SEMA3E | 900.4907 | 0.7256129 | 0.2997865 | 2.603226 | 0.0092351 | 0.036497 |
| ENSG00000285201 | | 7.676981 | 0.724697 | 1.109573 | 1.75178 | 0.07981168 | 0.196873 |
| ENSG00000123352 | SPATS2 | 2831.607985 | 0.724392543 | 0.227882252 | 3.320334138 | 0.0008991 | 0.00521 |
| ENSG00000104112 | SCG3 | 1.951591081 | 0.72432506 | 1.17256487 | 1.832879955 | 0.06682043 | 0.172364 |
| ENSG00000236420 | CHCHD3P1 | 2.407846 | 0.7239319 | 1.256368 | 2.026261 | 0.04273807 | 0.123157 |
| ENSG00000125968 | ID1 | 5711.593982 | 0.723403569 | 0.7566699 | 1.547351649 | 0.12177847 | 0.268087 |
| ENSG00000256060 | TRAPPC2B | 282.9901 | 0.7233889 | 0.2073797 | 3.626627 | 0.00028715 | 0.001947 |
| ENSG00000151623 | NR3C2 | 481.8616 | 0.7231155 | 0.2693929 | 2.861635 | 0.00421462 | 0.019267 |
| ENSG00000226519 | LINC00390 | 5.136362 | 0.7230951 | 0.8089206 | 1.495223 | 0.1348561 | 0.288138 |
| ENSG00000121577 | POPDC2 | 29.45637229 | 0.723034004 | 0.431877857 | 1.960361827 | 0.04995351 | 0.138367 |
| ENSG00000283656 | MIR4483 | 8.515899 | 0.7227929 | 0.9987415 | 1.653414 | 0.09824668 | 0.229746 |
| ENSG00000125046 | SSUH2 | 8.912206341 | 0.72270932 | 0.791619987 | 1.546230149 | 0.122049 | 0.268469 |
| ENSG00000287032 | | 6.472398 | 0.7224582 | 0.6398791 | 1.568446 | 0.116777 | 0.260028 |
| ENSG00000100197 | CYP2D6 | 49.2327939 | 0.722380023 | 0.325098543 | 2.422700652 | 0.01540562 | 0.055059 |
| ENSG00000163157 | TMOD4 | 91.79574 | 0.7223183 | 0.3645532 | 2.226615 | 0.02597303 | 0.083534 |
| ENSG00000157322 | CLEC18A | 17.21789 | 0.7222589 | 0.5473756 | 1.684133 | 0.0921559 | 0.219222 |
| ENSG00000242880 | | 1.583418 | 0.7221566 | 1.14103 | 1.813592 | 0.06974057 | 0.178086 |
| ENSG00000261528 | CCNYL7 | 1.898066 | 0.7221443 | 1.135341 | 1.806655 | 0.07081609 | 0.180172 |
| ENSG00000127124 | HIVEP3 | 734.8536782 | 0.721555474 | 0.260989347 | 2.934736037 | 0.00333832 | 0.015881 |
| ENSG00000162753 | SLC9C2 | 1.166189 | 0.721395 | 1.245708 | 2.088182 | 0.03678142 | NA |
| ENSG00000269533 | | 1.511836 | 0.7212603 | 1.23517 | 2.062256 | 0.03918333 | NA |
| ENSG00000270571 | | 81.31462 | 0.7212492 | 0.5513065 | 1.691563 | 0.09072934 | 0.216909 |
| ENSG00000261408 | TEN1-CDK3 | 110.5398 | 0.7211707 | 0.2601513 | 2.940458 | 0.00327728 | 0.015635 |
| ENSG00000141968 | VAV1 | 920.7805 | 0.7209437 | 0.2258894 | 3.33527 | 0.00085217 | 0.004978 |
| ENSG00000224682 | SOCS5P2 | 2.855932 | 0.7208948 | 1.025334 | 1.714841 | 0.0863743 | 0.209024 |
| ENSG00000201217 | | 4.623641 | 0.7208152 | 0.8401545 | 1.537775 | 0.1241036 | 0.271876 |
| ENSG00000249167 | SEMA6A-AS2 | 3.270662 | 0.7207109 | 0.8004414 | 1.509684 | 0.131124 | 0.282483 |
| ENSG00000188013 | MEIS3P2 | 63.38647 | 0.7205781 | 0.3499504 | 2.288487 | 0.02210917 | 0.073815 |
| ENSG00000165124 | SVEP1 | 5.92026 | 0.7205367 | 1.107023 | 1.741541 | 0.08158877 | 0.200382 |
| ENSG00000280420 | | 7.852144 | 0.7201701 | 0.7167319 | 1.546288 | 0.1220351 | 0.268469 |
| ENSG00000164082 | GRM2 | 8.518059 | 0.7200045 | 0.7982294 | 1.544907 | 0.1223688 | 0.269074 |

| | | | | | | | |
|---|---|---|---|---|---|---|---|
| ENSG00000268895 | A1BG-AS1 | 197.4687 | 0.7199297 | 0.1995022 | 3.74303 | 0.00018182 | 0.001303 |
| ENSG00000077585 | GPR137B | 1724.25588 | 0.719135507 | 0.292171509 | 2.643157796 | 0.00821368 | 0.033186 |
| ENSG00000230736 | | 8.519036 | 0.7191201 | 0.7036326 | 1.531047 | 0.1257579 | 0.274144 |
| ENSG00000228477 | | 17.06318 | 0.7188903 | 0.5809436 | 1.648923 | 0.0991634 | 0.231274 |
| ENSG00000140548 | ZNF710 | 2696.941 | 0.7188487 | 0.222798 | 3.374393 | 0.00073979 | 0.004408 |
| ENSG00000272529 | | 14.2823 | 0.7186322 | 0.5612784 | 1.663585 | 0.0961953 | 0.226018 |
| ENSG00000125726 | CD70 | 7.347063101 | 0.718475129 | 0.729581103 | 1.535058892 | 0.12476936 | 0.272749 |
| ENSG00000287116 | | 1.706416 | 0.7184315 | 1.199747 | 1.993815 | 0.04617228 | 0.130492 |
| ENSG00000099814 | CEP170B | 8338.427586 | 0.718232483 | 0.147431677 | 4.957873735 | 7.13E-07 | 9.18E-06 |
| ENSG00000204428 | LY6G5C | 221.6928 | 0.7182203 | 0.3254179 | 2.424406 | 0.01533347 | 0.054868 |
| ENSG00000151834 | GABRA2 | 41.49101 | 0.7182133 | 0.8299484 | 1.466938 | 0.1423931 | 0.299805 |
| ENSG00000126821 | SGPP1 | 2258.642292 | 0.717923878 | 0.317458928 | 2.465521351 | 0.0136814 | 0.050021 |
| ENSG00000253368 | TRNP1 | 159.225 | 0.7178483 | 0.3091557 | 2.530317 | 0.01139596 | 0.043205 |
| ENSG00000262712 | | 62.52765 | 0.7176798 | 0.3219888 | 2.417604 | 0.01562305 | 0.055694 |
| ENSG00000207730 | MIR200B | 6.590696 | 0.7176577 | 0.7305398 | 1.527177 | 0.1267169 | 0.275578 |
| ENSG00000153896 | ZNF599 | 307.9632 | 0.7176201 | 0.2082665 | 3.577438 | 0.00034698 | 0.002298 |
| ENSG00000237232 | ZNF295-AS1 | 26.01792 | 0.717572 | 0.4159099 | 2.001861 | 0.04529969 | 0.128641 |
| ENSG00000140564 | FURIN | 5562.436 | 0.7174517 | 0.1701947 | 4.30104 | 1.70E-05 | 0.000158 |
| ENSG00000185043 | CIB1 | 4356.887 | 0.7174427 | 0.2263422 | 3.311063 | 0.00092942 | 0.005362 |
| ENSG00000275532 | | 18.33619 | 0.7172426 | 0.5629399 | 1.663319 | 0.0962485 | 0.226083 |
| ENSG00000259488 | | 60.21484 | 0.7171058 | 0.3047016 | 2.546618 | 0.01087724 | 0.041631 |
| ENSG00000212710 | CTAGE1 | 120.2449 | 0.7169513 | 0.2871938 | 2.679191 | 0.00738003 | 0.030461 |
| ENSG00000273669 | | 1.336146 | 0.7166172 | 1.041726 | 1.629926 | 0.103117 | NA |
| ENSG00000272189 | | 231.9817 | 0.71633 | 0.3329337 | 2.372578 | 0.01766445 | 0.061472 |
| ENSG00000102984 | ZNF821 | 326.0186235 | 0.716233644 | 0.178957153 | 4.114795053 | 3.88E-05 | 0.000331 |
| ENSG00000103479 | RBL2 | 5329.293137 | 0.716131712 | 0.305867754 | 2.538859455 | 0.01112145 | 0.042398 |
| ENSG00000107742 | SPOCK2 | 153.2041866 | 0.716097999 | 0.405725131 | 2.035939753 | 0.04175641 | 0.120945 |
| ENSG00000246082 | NUDT16L2P | 2.560789 | 0.7160129 | 0.9720529 | 1.573861 | 0.1155197 | 0.258354 |
| ENSG00000245146 | MALINC1 | 239.3435 | 0.7158024 | 0.2964012 | 2.60105 | 0.00929389 | 0.036669 |
| ENSG00000237854 | LINC00674 | 588.6768 | 0.7157101 | 0.2588183 | 2.931314 | 0.00337532 | 0.016037 |
| ENSG00000237499 | WAKMAR2 | 9.641618 | 0.7154177 | 0.6605085 | 1.550444 | 0.121035 | 0.266958 |
| ENSG00000234174 | | 2.490352 | 0.7153916 | 1.045014 | 1.665254 | 0.09586201 | 0.225534 |
| ENSG00000267503 | | 2.035097 | 0.715218 | 1.176932 | 1.897622 | 0.05774594 | 0.1546 |

| | | | | | | | |
|---|---|---|---|---|---|---|---|
| ENSG00000205959 | | 9.00906 | 0.7151446 | 0.7611545 | 1.479038 | 0.1391301 | 0.294663 |
| ENSG00000159761 | C16orf86 | 33.42373 | 0.7150638 | 0.4092816 | 2.022763 | 0.04309757 | 0.123964 |
| ENSG00000134897 | BIVM | 783.8758251 | 0.715052195 | 0.244275977 | 3.084789667 | 0.00203696 | 0.010525 |
| ENSG00000225756 | DBH-AS1 | 1.985073 | 0.7143642 | 0.9722087 | 1.585323 | 0.1128929 | 0.253868 |
| ENSG00000166450 | PRTG | 196.9152 | 0.7143625 | 0.471767 | 1.828944 | 0.06740804 | 0.173526 |
| ENSG00000220201 | ZGLP1 | 114.9618 | 0.7140223 | 0.4074783 | 2.02001 | 0.04338238 | 0.124541 |
| ENSG00000260352 | | 11.77832 | 0.7139257 | 0.605665 | 1.60622 | 0.1082256 | 0.24646 |
| ENSG00000235049 | LINC00940 | 5.066341 | 0.7136947 | 0.8501189 | 1.485167 | 0.1374996 | 0.292048 |
| ENSG00000183298 | RPSAP19 | 19.42484 | 0.7134559 | 0.5697261 | 1.659046 | 0.09710662 | 0.227676 |
| ENSG00000145794 | MEGF10 | 14.05066 | 0.7131784 | 0.5322731 | 1.694853 | 0.09010322 | 0.215808 |
| ENSG00000238021 | ARMC4P1 | 8.972435 | 0.7131019 | 0.6638363 | 1.546134 | 0.1220723 | 0.268488 |
| ENSG00000250536 | ABHD17AP3 | 18.4897 | 0.7129472 | 0.5281988 | 1.716154 | 0.08613384 | 0.208556 |
| ENSG00000073598 | FNDC8 | 2.444893767 | 0.712449313 | 0.972603798 | 1.602718045 | 0.10899692 | 0.247601 |
| ENSG00000160211 | G6PD | 22419.46 | 0.7123515 | 0.2715008 | 2.798135 | 0.00513986 | 0.022651 |
| ENSG00000150676 | CCDC83 | 169.3554 | 0.7120538 | 0.3892976 | 2.084668 | 0.03709948 | 0.110282 |
| ENSG00000084636 | COL16A1 | 301.1014174 | 0.712046815 | 0.361817031 | 2.201334508 | 0.02771235 | 0.087968 |
| ENSG00000252192 | | 3.267618 | 0.7120336 | 1.036669 | 1.661071 | 0.09669915 | 0.226961 |
| ENSG00000165914 | TTC7B | 1645.064 | 0.7116937 | 0.182873 | 4.007989 | 6.12E-05 | 0.000496 |
| ENSG00000279306 | | 20.06177 | 0.7116423 | 0.689563 | 1.537501 | 0.1241706 | 0.271947 |
| ENSG00000237575 | PYY2 | 39.08868 | 0.7115793 | 0.3468773 | 2.267875 | 0.02333682 | 0.076828 |
| ENSG00000245552 | | 21.88471 | 0.711538 | 0.4356416 | 1.915997 | 0.05536552 | 0.149797 |
| ENSG00000254582 | PSMA2P1 | 1.560291 | 0.7113227 | 1.10183 | 1.814069 | 0.06966707 | NA |
| ENSG00000094916 | CBX5 | 8511.123231 | 0.710726698 | 0.333040193 | 2.348887017 | 0.01882962 | 0.064745 |
| ENSG00000185917 | SETD4 | 1552 | 0.7107175 | 0.1598703 | 4.554019 | 5.26E-06 | 5.58E-05 |
| ENSG00000273853 | | 17.33148 | 0.7102966 | 0.4792894 | 1.801008 | 0.0717016 | 0.181918 |
| ENSG00000172465 | TCEAL1 | 2111.356 | 0.7102783 | 0.2467316 | 3.062467 | 0.0021952 | 0.011202 |
| ENSG00000285816 | | 5.832952 | 0.7102133 | 0.7723075 | 1.517505 | 0.1291392 | 0.279337 |
| ENSG00000277010 | | 13.56987 | 0.7100188 | 0.8022795 | 1.52788 | 0.1265424 | 0.275244 |
| ENSG00000196427 | NBPF4 | 50.2574 | 0.7099837 | 0.3456216 | 2.252887 | 0.02426631 | 0.079235 |
| ENSG00000115602 | IL1RL1 | 4.066852724 | 0.709875322 | 1.146646895 | 1.866182032 | 0.0620159 | 0.162929 |
| ENSG00000170537 | TMC7 | 277.5816 | 0.7097535 | 0.2545631 | 2.944921 | 0.00323037 | 0.015451 |
| ENSG00000225643 | | 9.596821 | 0.7095643 | 0.7958331 | 1.514926 | 0.1297912 | 0.280405 |
| ENSG00000115085 | ZAP70 | 9.638440406 | 0.709482133 | 0.696188955 | 1.528228175 | 0.1264559 | 0.275168 |

| | | | | | | | |
|---|---|---|---|---|---|---|---|
| ENSG00000143578 | CREB3L4 | 4686.36 | 0.7088552 | 0.2020695 | 3.632486 | 0.0002807 | 0.001908 |
| ENSG00000287778 | | 6.444279 | 0.7088166 | 1.096604 | 1.724582 | 0.08460281 | 0.205871 |
| ENSG00000233021 | | 2.184151 | 0.7087419 | 1.074138 | 1.690668 | 0.09090029 | 0.217168 |
| ENSG00000258870 | EIF4EBP1P1 | 2.974203 | 0.7086187 | 0.9345308 | 1.539492 | 0.1236842 | 0.271225 |
| ENSG00000070540 | WIPI1 | 1936.979339 | 0.708472475 | 0.237367357 | 3.127836222 | 0.00176098 | 0.009298 |
| ENSG00000251081 | | 4.024056 | 0.7083164 | 0.8652287 | 1.487584 | 0.1368606 | 0.291166 |
| ENSG00000254207 | | 12.39654 | 0.7081375 | 0.6651041 | 1.564672 | 0.1176597 | 0.261523 |
| ENSG00000149177 | PTPRJ | 4251.6 | 0.7081065 | 0.2172947 | 3.399383 | 0.00067538 | 0.004073 |
| ENSG00000058335 | RASGRF1 | 145.6165507 | 0.707575049 | 0.540304216 | 1.704104571 | 0.08836155 | 0.212706 |
| ENSG00000152804 | HHEX | 140.2379 | 0.7075235 | 0.3785422 | 2.118537 | 0.03412963 | 0.103671 |
| ENSG00000085719 | CPNE3 | 13556.94181 | 0.707459202 | 0.193378979 | 3.756579884 | 0.00017225 | 0.001243 |
| ENSG00000145390 | USP53 | 2664.653 | 0.7074334 | 0.2498818 | 2.987197 | 0.00281548 | 0.01385 |
| ENSG00000171970 | ZNF57 | 38.87588 | 0.7074105 | 0.366834 | 2.171191 | 0.02991675 | 0.093469 |
| ENSG00000236377 | | 2.245076 | 0.7073819 | 0.9596832 | 1.530924 | 0.1257883 | 0.274187 |
| ENSG00000287861 | | 15.28315 | 0.7072558 | 1.087211 | 1.744833 | 0.08101398 | 0.199262 |
| ENSG00000267344 | | 29.05913 | 0.7066723 | 0.3759382 | 2.127971 | 0.03333946 | 0.101851 |
| ENSG00000115363 | EVA1A | 4.043803192 | 0.706558575 | 0.897908984 | 1.524017354 | 0.12750438 | 0.27686 |
| ENSG00000180834 | MAP6D1 | 37.14649 | 0.7064577 | 0.3681798 | 2.16388 | 0.03047353 | 0.094874 |
| ENSG00000271532 | BBIP1P1 | 2.259719 | 0.7061333 | 0.989822 | 1.577423 | 0.1146981 | 0.256881 |
| ENSG00000228708 | | 2.255763 | 0.7061083 | 1.121756 | 1.786716 | 0.07398344 | 0.186354 |
| ENSG00000251034 | | 18.96697 | 0.7060892 | 0.4791458 | 1.789005 | 0.07361405 | 0.185681 |
| ENSG00000260160 | | 30.02482 | 0.7059772 | 0.5066095 | 1.733149 | 0.08306925 | 0.203061 |
| ENSG00000182768 | NGRN | 2591.817 | 0.7059263 | 0.1562206 | 4.620354 | 3.83E-06 | 4.20E-05 |
| ENSG00000197712 | FAM114A1 | 313.4546 | 0.7057669 | 0.3373535 | 2.30993 | 0.02089201 | 0.070406 |
| ENSG00000134962 | KLB | 54.33588484 | 0.705765278 | 0.423202854 | 1.938200031 | 0.05259882 | 0.144146 |
| ENSG00000279803 | | 1.358121 | 0.7057641 | 1.220476 | 2.095552 | 0.03612195 | NA |
| ENSG00000284648 | | 6.063279 | 0.7056591 | 0.6982729 | 1.521002 | 0.1282592 | 0.278 |
| ENSG00000257817 | | 10.54216 | 0.7056353 | 0.8730017 | 1.516189 | 0.1294717 | 0.279829 |
| ENSG00000181444 | ZNF467 | 2007.165 | 0.7055527 | 0.2515109 | 2.974672 | 0.00293302 | 0.014322 |
| ENSG00000267106 | ZNF561-AS1 | 216.8817 | 0.7055239 | 0.1733956 | 4.169552 | 3.05E-05 | 0.000266 |
| ENSG00000260778 | | 86.11137 | 0.705484 | 0.3279639 | 2.360974 | 0.018227 | 0.063096 |
| ENSG00000155959 | VBP1 | 4249.592 | 0.705257 | 0.226807 | 3.248222 | 0.00116129 | 0.006513 |
| ENSG00000182809 | CRIP2 | 4836.172 | 0.7051801 | 0.277752 | 2.714627 | 0.00663504 | 0.027889 |

| | | | | | | | |
|---|---|---|---|---|---|---|---|
| ENSG00000284966 | | 15.67959 | 0.7050307 | 0.77739 | 1.505056 | 0.1323097 | 0.284505 |
| ENSG00000213918 | DNASE1 | 2469.544 | 0.7049773 | 0.3227288 | 2.393288 | 0.01669812 | 0.058762 |
| ENSG00000234193 | ACSL3-AS1 | 10.22503 | 0.7046958 | 0.6899793 | 1.539558 | 0.123668 | 0.271212 |
| ENSG00000174373 | RALGAPA1 | 1815.646 | 0.7046244 | 0.234408 | 3.14961 | 0.00163489 | 0.008703 |
| ENSG00000266962 | | 36.7957 | 0.7045062 | 0.3349188 | 2.312204 | 0.02076645 | 0.070063 |
| ENSG00000283982 | | 6.890961 | 0.7041332 | 0.7553473 | 1.499776 | 0.1336725 | 0.286408 |
| ENSG00000157540 | DYRK1A | 3115.677 | 0.7040474 | 0.1388375 | 5.140276 | 2.74E-07 | 3.81E-06 |
| ENSG00000204604 | ZNF468 | 1402.78 | 0.7037231 | 0.1516636 | 4.734707 | 2.19E-06 | 2.56E-05 |
| ENSG00000105675 | ATP4A | 3.170124136 | 0.703643893 | 0.904913835 | 1.525833794 | 0.12705126 | 0.276158 |
| ENSG00000106976 | DNM1 | 3005.083019 | 0.703393027 | 0.208953072 | 3.488753439 | 0.00048528 | 0.00306 |
| ENSG00000168702 | LRP1B | 131.8531 | 0.7033747 | 0.5529577 | 1.670308 | 0.09485845 | 0.223966 |
| ENSG00000113532 | ST8SIA4 | 1914.889817 | 0.703212858 | 0.344141824 | 2.261154145 | 0.02374971 | 0.077883 |
| ENSG00000146221 | TCTE1 | 4.779256 | 0.7031375 | 0.6811266 | 1.516466 | 0.1294014 | 0.279722 |
| ENSG00000278959 | | 3.672195 | 0.7029783 | 1.001629 | 1.597161 | 0.1102298 | 0.24974 |
| ENSG00000241322 | CDRT1 | 233.6376 | 0.702802 | 0.4685166 | 1.815128 | 0.0695042 | 0.177597 |
| ENSG00000286828 | | 22.76103 | 0.7027054 | 0.4856445 | 1.765017 | 0.07756093 | 0.192733 |
| ENSG00000118655 | DCLRE1B | 8013.571965 | 0.702702297 | 0.20263296 | 3.594569853 | 0.00032493 | 0.00217 |
| ENSG00000109944 | JHY | 233.2955092 | 0.702688704 | 0.324822367 | 2.362758436 | 0.01813949 | 0.062844 |
| ENSG00000122584 | NXPH1 | 58.29024024 | 0.702597722 | 0.401580985 | 2.014014849 | 0.04400798 | 0.125943 |
| ENSG00000274525 | | 9.747578 | 0.7021173 | 0.7008822 | 1.51789 | 0.1290422 | 0.279241 |
| ENSG00000154359 | LONRF1 | 1441.022 | 0.7020862 | 0.2129357 | 3.432795 | 0.00059739 | 0.003661 |
| ENSG00000234840 | LINC01239 | 199.109 | 0.7020082 | 0.3616006 | 2.174034 | 0.02970261 | 0.093029 |
| ENSG00000159289 | GOLGA6A | 3.142951 | 0.7019782 | 0.8874226 | 1.510918 | 0.1308094 | 0.282074 |
| ENSG00000204311 | PJVK | 76.32116 | 0.701957 | 0.3722882 | 2.126089 | 0.03349582 | 0.102176 |
| ENSG00000170852 | KBTBD2 | 4223.623 | 0.7018998 | 0.1403976 | 5.083369 | 3.71E-07 | 5.02E-06 |
| ENSG00000150672 | DLG2 | 69.16973 | 0.701758 | 0.2960153 | 2.554791 | 0.01062515 | 0.040883 |
| ENSG00000179456 | ZBTB18 | 137.8633 | 0.7008147 | 0.3588159 | 2.175427 | 0.0295981 | 0.092812 |
| ENSG00000175305 | CCNE2 | 734.8254 | 0.700806 | 0.4842855 | 1.761724 | 0.07811586 | 0.193727 |
| ENSG00000186814 | ZSCAN30 | 591.6109 | 0.700778 | 0.3738736 | 2.111336 | 0.03474347 | 0.105053 |
| ENSG00000167580 | AQP2 | 1.905451 | 0.7007482 | 1.176212 | 1.929102 | 0.05371818 | 0.146669 |
| ENSG00000173275 | ZNF449 | 695.5855 | 0.7005666 | 0.3033873 | 2.500129 | 0.01241482 | 0.046256 |
| ENSG00000228612 | HK2P1 | 184.5271 | 0.700401 | 0.3351592 | 2.311047 | 0.02083028 | 0.070225 |
| ENSG00000274833 | | 4.026088 | 0.7002702 | 0.8444833 | 1.487465 | 0.1368921 | 0.291166 |

| | | | | | | | |
|---|---|---|---|---|---|---|---|
| ENSG00000237786 | GFOD1-AS1 | 4.334642 | 0.7001255 | 0.935893 | 1.551407 | 0.1208042 | 0.266626 |
| ENSG00000154143 | PANX3 | 28.56801 | 0.6999394 | 0.5416545 | 1.65955 | 0.09700495 | 0.227498 |
| ENSG00000125731 | SH2D3A | 1477.623573 | 0.69986612 | 0.226797849 | 3.238911897 | 0.00119987 | 0.00669 |
| ENSG00000283288 | SMIM33 | 2.053072 | 0.6997596 | 1.164179 | 1.9003 | 0.05739372 | 0.153797 |
| ENSG00000259540 | | 4.222294 | 0.6994018 | 0.8461358 | 1.499235 | 0.1338126 | 0.286553 |
| ENSG00000260261 | | 765.7718 | 0.6991776 | 0.3269437 | 2.347741 | 0.01888766 | 0.064894 |
| ENSG00000155096 | AZIN1 | 14522.31 | 0.6990643 | 0.1890664 | 3.788431 | 0.0001516 | 0.001108 |
| ENSG00000176024 | ZNF613 | 94.66579 | 0.6987481 | 0.2753582 | 2.71564 | 0.00661477 | 0.027817 |
| ENSG00000099822 | HCN2 | 1178.704198 | 0.698710896 | 0.164727164 | 4.314093803 | 1.60E-05 | 0.00015 |
| ENSG00000273186 | | 59.77214 | 0.6986466 | 0.3178886 | 2.390484 | 0.01682618 | 0.059142 |
| ENSG00000223396 | RPS10P7 | 336.4382 | 0.6980098 | 0.2886537 | 2.602464 | 0.00925564 | 0.036545 |
| ENSG00000246350 | | 3.49532 | 0.6976745 | 0.8756145 | 1.480474 | 0.1387467 | 0.294086 |
| ENSG00000115423 | DNAH6 | 64.33807306 | 0.6976295 | 0.399795617 | 2.000012263 | 0.04549894 | 0.129083 |
| ENSG00000151612 | ZNF827 | 489.8269 | 0.6976216 | 0.3047658 | 2.480309 | 0.01312685 | 0.048426 |
| ENSG00000167550 | RHEBL1 | 125.7343 | 0.6974784 | 0.4547778 | 1.838088 | 0.06604939 | 0.17089 |
| ENSG00000274104 | | 93.11141 | 0.6971864 | 0.3968314 | 2.016273 | 0.04377147 | 0.125401 |
| ENSG00000165140 | FBP1 | 7301.644 | 0.6963622 | 0.2940026 | 2.553329 | 0.01066986 | 0.041032 |
| ENSG00000234521 | | 4.622178 | 0.6963017 | 0.9911693 | 1.60518 | 0.1084542 | 0.246799 |
| ENSG00000138623 | SEMA7A | 518.5705037 | 0.695804536 | 0.366435345 | 2.145626574 | 0.03190278 | 0.098389 |
| ENSG00000104312 | RIPK2 | 2637.603014 | 0.695621974 | 0.16430536 | 4.330959248 | 1.48E-05 | 0.00014 |
| ENSG00000166823 | MESP1 | 29.70484 | 0.6955259 | 0.5380494 | 1.661355 | 0.09664207 | 0.226907 |
| ENSG00000187867 | PALM3 | 1203.318 | 0.6953652 | 0.3989673 | 2.002957 | 0.04518189 | 0.128417 |
| ENSG00000169255 | B3GALNT1 | 2280.809 | 0.6952632 | 0.216303 | 3.343938 | 0.00082598 | 0.004852 |
| ENSG00000035687 | ADSS2 | 3897.967222 | 0.695186758 | 0.175121579 | 4.07791539 | 4.54E-05 | 0.000381 |
| ENSG00000201861 | RNA5SP298 | 1.054931 | 0.6951771 | 1.211954 | 2.073743 | 0.03810317 | NA |
| ENSG00000162522 | KIAA1522 | 18023.52 | 0.6951588 | 0.1422272 | 4.940618 | 7.79E-07 | 9.91E-06 |
| ENSG00000130338 | TULP4 | 2156.488311 | 0.694778006 | 0.27958446 | 2.651373206 | 0.00801652 | 0.032597 |
| ENSG00000276102 | MIR6747 | 2.633631 | 0.6947585 | 0.8676914 | 1.476763 | 0.1397392 | 0.295812 |
| ENSG00000265595 | MIR4756 | 4.971945 | 0.6946484 | 1.082503 | 1.77738 | 0.07550578 | 0.189024 |
| ENSG00000284626 | | 1.713575 | 0.6945198 | 1.133354 | 1.855606 | 0.06350976 | 0.165804 |
| ENSG00000156966 | B3GNT7 | 9.033049 | 0.694003 | 0.5667347 | 1.604186 | 0.108673 | 0.247119 |
| ENSG00000240970 | RPL23AP64 | 111.4605 | 0.6939161 | 0.4330985 | 1.886619 | 0.05921157 | 0.157524 |
| ENSG00000007866 | TEAD3 | 4101.438992 | 0.69389568 | 0.209671398 | 3.436492509 | 0.0005893 | 0.003617 |

| | | | | | | | |
|---|---|---|---|---|---|---|---|
| ENSG00000216906 | | 28.89839 | 0.6938583 | 0.358117 | 2.155526 | 0.03112073 | 0.096515 |
| ENSG00000106333 | PCOLCE | 462.9003765 | 0.693661688 | 0.272257909 | 2.720282332 | 0.00652262 | 0.027508 |
| ENSG00000211683 | | 9.174976 | 0.6936287 | 0.6017974 | 1.562587 | 0.1181497 | 0.262205 |
| ENSG00000204634 | TBC1D8 | 1696.621 | 0.6932453 | 0.1718821 | 4.140303 | 3.47E-05 | 0.000299 |
| ENSG00000274341 | | 1.224451 | 0.6929016 | 1.168563 | 1.923619 | 0.0544024 | NA |
| ENSG00000172006 | ZNF554 | 390.0625 | 0.6928752 | 0.1506973 | 4.699434 | 2.61E-06 | 2.99E-05 |
| ENSG00000164920 | OSR2 | 1629.807 | 0.6925174 | 0.2245318 | 3.222028 | 0.00127287 | 0.007053 |
| ENSG00000272717 | | 25.71145 | 0.6919043 | 0.4467793 | 1.840065 | 0.06575863 | 0.17027 |
| ENSG00000101972 | STAG2 | 7510.568365 | 0.691886974 | 0.171968431 | 4.130873016 | 3.61E-05 | 0.00031 |
| ENSG00000203772 | SPRN | 39.69914 | 0.6917384 | 0.5386989 | 1.648524 | 0.09924523 | 0.231414 |
| ENSG00000170017 | ALCAM | 13531.75 | 0.6916681 | 0.2610191 | 2.811528 | 0.00493068 | 0.021903 |
| ENSG00000225663 | MCRIP1 | 4511.698 | 0.6916568 | 0.1884888 | 3.741102 | 0.00018322 | 0.001312 |
| ENSG00000251013 | GAPDHP62 | 3.525941 | 0.6913651 | 0.862142 | 1.488744 | 0.1365549 | 0.290727 |
| ENSG00000135127 | BICDL1 | 1774.514237 | 0.69130254 | 0.193206496 | 3.702620404 | 0.00021338 | 0.001501 |
| ENSG00000179899 | PHC1P1 | 569.9287 | 0.6911801 | 0.2219109 | 3.252926 | 0.00114223 | 0.006415 |
| ENSG00000278932 | | 222.9028 | 0.6909472 | 0.256669 | 2.847118 | 0.0044117 | 0.019975 |
| ENSG00000277324 | | 13.93808 | 0.6909312 | 0.6499316 | 1.536616 | 0.1243873 | 0.272235 |
| ENSG00000240750 | RN7SL559P | 0.8943534 | 0.6907981 | 1.168712 | 1.951139 | 0.0510405 | NA |
| ENSG00000204173 | LRRC37A5P | 12.24234 | 0.6906942 | 0.6708536 | 1.499623 | 0.1337122 | 0.286408 |
| ENSG00000259993 | | 10.20657 | 0.6904008 | 0.6844836 | 1.511518 | 0.1306565 | 0.281816 |
| ENSG00000116678 | LEPR | 313.6447201 | 0.690113072 | 0.271180025 | 2.717974362 | 0.00656829 | 0.027661 |
| ENSG00000167874 | TMEM88 | 42.6917 | 0.6900448 | 0.402526 | 1.963239 | 0.04961843 | 0.137712 |
| ENSG00000264769 | | 25.86423 | 0.689939 | 0.458589 | 1.80517 | 0.07104814 | 0.180553 |
| ENSG00000227214 | HCG15 | 22.92019 | 0.6892513 | 0.3986302 | 1.979908 | 0.04771382 | 0.133668 |
| ENSG00000149054 | ZNF215 | 1.79514 | 0.6891578 | 1.286419 | 2.45462 | 0.01410336 | 0.051316 |
| ENSG00000126231 | PROZ | 41.40831682 | 0.689125171 | 0.514992813 | 1.669003149 | 0.09511676 | 0.224377 |
| ENSG00000071189 | SNX13 | 3964.386896 | 0.688541437 | 0.195607258 | 3.643660891 | 0.00026879 | 0.001836 |
| ENSG00000237903 | | 3.124674 | 0.6881243 | 0.8726303 | 1.484155 | 0.1377678 | 0.292454 |
| ENSG00000238062 | SPATA3-AS1 | 8.073318 | 0.6876952 | 0.8305092 | 1.512931 | 0.1302973 | 0.28127 |
| ENSG00000275393 | | 12.48695 | 0.687671 | 0.6502114 | 1.515373 | 0.129678 | 0.280229 |
| ENSG00000279406 | | 43.85321 | 0.6871022 | 0.3947905 | 1.977919 | 0.04793784 | 0.134154 |
| ENSG00000236348 | PSMC1P10 | 1.857127 | 0.6870445 | 0.996199 | 1.61983 | 0.1052689 | 0.241507 |
| ENSG00000225614 | ZNF469 | 590.6513 | 0.6862615 | 0.3435454 | 2.223372 | 0.0261907 | 0.084092 |

| | | | | | | | |
|---|---|---|---|---|---|---|---|
| ENSG00000167992 | VWCE | 66.61385 | 0.6859044 | 0.311954 | 2.389587 | 0.01686732 | 0.059255 |
| ENSG00000099875 | MKNK2 | 5725.827369 | 0.685609794 | 0.278748922 | 2.633460777 | 0.00845196 | 0.033988 |
| ENSG00000185105 | MYADML2 | 29.85984 | 0.6855938 | 0.4193944 | 1.89265 | 0.05840444 | 0.156017 |
| ENSG00000072201 | LNX1 | 599.1829908 | 0.685592209 | 0.233103021 | 3.080866361 | 0.00206399 | 0.010644 |
| ENSG00000241015 | TPM3P9 | 943.2653 | 0.6848238 | 0.1969705 | 3.598056 | 0.0003206 | 0.002146 |
| ENSG00000273679 | | 19.99593 | 0.6847291 | 0.4813973 | 1.731282 | 0.08340145 | 0.203611 |
| ENSG00000092820 | EZR | 12706.19763 | 0.684702879 | 0.147662244 | 4.726955859 | 2.28E-06 | 2.64E-05 |
| ENSG00000251143 | | 47.45388 | 0.6846408 | 0.367115 | 2.092765 | 0.0363701 | 0.108713 |
| ENSG00000182957 | SPATA13 | 1353.546 | 0.6845896 | 0.2570861 | 2.848322 | 0.00439504 | 0.019934 |
| ENSG00000011114 | BTBD7 | 4059.829019 | 0.684008943 | 0.225788106 | 3.175795407 | 0.00149426 | 0.008084 |
| ENSG00000051620 | HEBP2 | 4277.081141 | 0.683947128 | 0.109310467 | 6.348140719 | 2.18E-10 | 5.21E-09 |
| ENSG00000279765 | | 14.51171 | 0.6839073 | 0.5116952 | 1.673589 | 0.09421136 | 0.223033 |
| ENSG00000279748 | | 16.03126 | 0.6838187 | 0.5938419 | 1.549808 | 0.1211877 | 0.267184 |
| ENSG00000176723 | ZNF843 | 19.05627 | 0.6835354 | 0.5300205 | 1.644354 | 0.1001032 | 0.232986 |
| ENSG00000132016 | BRME1 | 145.8060135 | 0.683223137 | 0.440156948 | 1.839134309 | 0.06589543 | 0.170591 |
| ENSG00000212747 | RTL8B | 2484.193 | 0.683018 | 0.2186142 | 3.253015 | 0.00114188 | 0.006415 |
| ENSG00000049283 | EPN3 | 5080.585324 | 0.682279122 | 0.263192293 | 2.754767522 | 0.00587339 | 0.025296 |
| ENSG00000169032 | MAP2K1 | 2630.265 | 0.6822225 | 0.1393294 | 4.963927 | 6.91E-07 | 8.93E-06 |
| ENSG00000242628 | | 4.284404 | 0.6821528 | 0.7558551 | 1.462919 | 0.1434896 | 0.301377 |
| ENSG00000163629 | PTPN13 | 4237.421 | 0.6816357 | 0.2436841 | 2.966192 | 0.00301512 | 0.014616 |
| ENSG00000260404 | | 836.2969 | 0.6815086 | 0.1517599 | 4.593376 | 4.36E-06 | 4.73E-05 |
| ENSG00000263126 | | 40.84812 | 0.6814877 | 0.35428 | 2.145048 | 0.03194904 | 0.098492 |
| ENSG00000229349 | ACTG1P9 | 1.86981 | 0.6812029 | 1.098269 | 1.777476 | 0.07548997 | 0.189016 |
| ENSG00000126952 | NXF5 | 3.293085169 | 0.68104289 | 0.860913212 | 1.486178976 | 0.13723179 | 0.291572 |
| ENSG00000147127 | RAB41 | 11.92849 | 0.6806617 | 0.4895217 | 1.7056 | 0.08808254 | 0.212211 |
| ENSG00000267193 | | 2.018196 | 0.6801503 | 1.044628 | 1.623882 | 0.104401 | 0.240075 |
| ENSG00000260259 | LINC02166 | 30.08996 | 0.6801286 | 0.5453433 | 1.613139 | 0.1067143 | 0.243976 |
| ENSG00000267808 | | 1.75841 | 0.680091 | 1.055229 | 1.684578 | 0.09206998 | 0.219134 |
| ENSG00000157985 | AGAP1 | 2969.008 | 0.6800688 | 0.1896035 | 3.678317 | 0.00023478 | 0.001634 |
| ENSG00000272610 | MAGI1-IT1 | 8.948673 | 0.6799335 | 0.6684852 | 1.491931 | 0.1357173 | 0.289387 |
| ENSG00000111725 | PRKAB1 | 2126.487233 | 0.679753786 | 0.146049907 | 4.738349375 | 2.15E-06 | 2.52E-05 |
| ENSG00000244198 | ARHGEF35-AS1 | 65.1545 | 0.6795767 | 0.3472905 | 2.17181 | 0.02986999 | 0.093353 |
| ENSG00000139926 | FRMD6 | 1209.408 | 0.6794844 | 0.2370754 | 3.011789 | 0.00259713 | 0.012922 |

| | | | | | | | |
|---|---|---|---|---|---|---|---|
| ENSG00000168936 | TMEM129 | 2789.776 | 0.679482 | 0.1980484 | 3.549238 | 0.00038635 | 0.00252 |
| ENSG00000222038 | POTEJ | 8.728774 | 0.6792221 | 0.6381335 | 1.510883 | 0.1308182 | 0.282074 |
| ENSG00000204178 | MACO1 | 1989.985 | 0.6792082 | 0.1327027 | 5.195208 | 2.04E-07 | 2.92E-06 |
| ENSG00000130518 | IQCN | 591.2063437 | 0.67913881 | 0.291020772 | 2.522846664 | 0.01164092 | 0.043939 |
| ENSG00000134824 | FADS2 | 798.128837 | 0.67886789 | 0.349169042 | 2.164510112 | 0.03042522 | 0.094768 |
| ENSG00000287165 | | 2.548109 | 0.6785745 | 0.9226839 | 1.511448 | 0.1306742 | 0.281832 |
| ENSG00000100321 | SYNGR1 | 3006.050228 | 0.678041228 | 0.220920689 | 3.205192578 | 0.00134972 | 0.007419 |
| ENSG00000253598 | SLC10A5 | 12.97466 | 0.6777762 | 0.5518973 | 1.58946 | 0.1119566 | 0.252532 |
| ENSG00000135766 | EGLN1 | 311.9780171 | 0.677563155 | 0.29614277 | 2.474946138 | 0.01332563 | 0.048963 |
| ENSG00000225555 | | 2.618987 | 0.6775139 | 0.9283676 | 1.470644 | 0.1413875 | 0.298296 |
| ENSG00000186162 | CIDECP1 | 864.4045 | 0.6774441 | 0.1788288 | 3.895611 | 9.80E-05 | 0.000752 |
| ENSG00000183654 | MARCHF11 | 7.273094 | 0.6773384 | 0.743029 | 1.452232 | 0.146437 | 0.305872 |
| ENSG00000274213 | | 198.4668 | 0.6772514 | 0.3458074 | 2.177959 | 0.02940909 | 0.09235 |
| ENSG00000265445 | | 1.299943 | 0.6770363 | 1.1625 | 1.979997 | 0.04770382 | NA |
| ENSG00000070759 | TESK2 | 1000.798045 | 0.676728154 | 0.389375544 | 1.979005948 | 0.04781534 | 0.133896 |
| ENSG00000125868 | DSTN | 18808.64616 | 0.676515977 | 0.188709942 | 3.657900408 | 0.00025429 | 0.001749 |
| ENSG00000204272 | NBDY | 1964.235 | 0.6762411 | 0.1723143 | 4.052403 | 5.07E-05 | 0.000419 |
| ENSG00000167261 | DPEP2 | 2.097213 | 0.6760051 | 1.053104 | 1.672019 | 0.0945206 | 0.223445 |
| ENSG00000286064 | | 20.87262 | 0.6758658 | 0.3609839 | 2.10014 | 0.03571652 | 0.107287 |
| ENSG00000263531 | | 3.040291 | 0.6757422 | 0.8593294 | 1.460971 | 0.1440235 | 0.302137 |
| ENSG00000132692 | BCAN | 2.364417171 | 0.675246842 | 1.010899649 | 1.622740891 | 0.10464481 | 0.240428 |
| ENSG00000137486 | ARRB1 | 3957.88821 | 0.675136581 | 0.280914418 | 2.577180265 | 0.009961 | 0.038737 |
| ENSG00000270110 | | 24.7165 | 0.6742356 | 0.6282553 | 1.5106 | 0.1308904 | 0.282206 |
| ENSG00000241923 | RPL14P3 | 31.27652 | 0.6742015 | 0.3929541 | 1.964728 | 0.04944578 | 0.137407 |
| ENSG00000172007 | RAB33B | 753.8551 | 0.67402 | 0.2536105 | 2.813113 | 0.00490644 | 0.021833 |
| ENSG00000251580 | LINC02482 | 97.48709 | 0.6740044 | 0.3029497 | 2.404441 | 0.0161972 | 0.057272 |
| ENSG00000228653 | HNRNPCP7 | 22.51594 | 0.6739541 | 0.4677947 | 1.739301 | 0.08198184 | 0.201106 |
| ENSG00000283075 | | 74.00438 | 0.6736857 | 0.352853 | 2.126247 | 0.03348273 | 0.102159 |
| ENSG00000023839 | ABCC2 | 89.53468471 | 0.673536203 | 0.332118466 | 2.226278926 | 0.0259955 | 0.083561 |
| ENSG00000038210 | PI4K2B | 1699.605737 | 0.673455084 | 0.254596614 | 2.803668481 | 0.00505248 | 0.022347 |
| ENSG00000283930 | PLD5P1 | 117.8436 | 0.6733116 | 0.4407718 | 1.809946 | 0.0703042 | 0.179228 |
| ENSG00000272715 | | 7.362624 | 0.6731306 | 0.8034418 | 1.469397 | 0.1417252 | 0.298778 |
| ENSG00000259215 | | 4.316569 | 0.6729899 | 0.7042694 | 1.453489 | 0.1460881 | 0.305359 |

| | | | | | | | |
|---|---|---|---|---|---|---|---|
| ENSG00000198162 | MAN1A2 | 7655.958 | 0.6728458 | 0.2116084 | 3.317378 | 0.00090867 | 0.005254 |
| ENSG00000271590 | | 5.136035 | 0.6728213 | 0.8743782 | 1.462977 | 0.1434738 | 0.301377 |
| ENSG00000224376 | | 29.8186 | 0.6721484 | 0.5365357 | 1.600642 | 0.1094563 | 0.248432 |
| ENSG00000231437 | LINC01750 | 296.3874 | 0.6720327 | 0.3873391 | 1.979261 | 0.04778661 | 0.133844 |
| ENSG00000008516 | MMP25 | 143.4484822 | 0.6719199 | 0.26978128 | 2.662768978 | 0.00775006 | 0.031703 |
| ENSG00000213492 | NT5C3AP1 | 165.728 | 0.6717983 | 0.2227505 | 3.169979 | 0.0015245 | 0.008228 |
| ENSG00000111490 | TBC1D30 | 1095.569393 | 0.671681359 | 0.268126067 | 2.668488897 | 0.00761933 | 0.031284 |
| ENSG00000182986 | ZNF320 | 2.452484 | 0.6715841 | 0.9829695 | 1.619881 | 0.1052578 | 0.241502 |
| ENSG00000215441 | CTAGE12P | 12.43963 | 0.6713977 | 0.5979609 | 1.524451 | 0.127396 | 0.276704 |
| ENSG00000232197 | | 2.407774 | 0.6713262 | 1.187571 | 2.108243 | 0.03500996 | 0.105618 |
| ENSG00000227245 | | 1.595129 | 0.6712874 | 1.010227 | 1.659798 | 0.09695502 | 0.227431 |
| ENSG00000103495 | MAZ | 3572.752379 | 0.671228701 | 0.168469376 | 4.084374659 | 4.42E-05 | 0.000372 |
| ENSG00000132823 | OSER1 | 3149.146634 | 0.671160082 | 0.150548622 | 4.547345839 | 5.43E-06 | 5.73E-05 |
| ENSG00000125386 | FAM193A | 1730.953849 | 0.670961033 | 0.14434359 | 4.708198159 | 2.50E-06 | 2.87E-05 |
| ENSG00000170185 | USP38 | 2752.639 | 0.6709432 | 0.1681009 | 4.07429 | 4.62E-05 | 0.000386 |
| ENSG00000249264 | EEF1A1P9 | 92.27623 | 0.6708461 | 0.3305298 | 2.235393 | 0.02539158 | 0.082052 |
| ENSG00000130711 | PRDM12 | 51.24672339 | 0.670534728 | 0.371435261 | 2.034280983 | 0.04192327 | 0.121283 |
| ENSG00000072042 | RDH11 | 6077.585382 | 0.670534096 | 0.166885996 | 4.152253574 | 3.29E-05 | 0.000286 |
| ENSG00000056558 | TRAF1 | 115.4757661 | 0.670528952 | 0.455581325 | 1.761401279 | 0.07817051 | 0.193814 |
| ENSG00000090316 | MAEA | 3615.89579 | 0.670513134 | 0.183629978 | 3.75966969 | 0.00017014 | 0.001229 |
| ENSG00000088448 | ANKRD10 | 2906.567423 | 0.670449469 | 0.245628674 | 2.877294035 | 0.00401102 | 0.018489 |
| ENSG00000286104 | | 3.096919 | 0.6703727 | 0.8339086 | 1.428682 | 0.1530956 | 0.315758 |
| ENSG00000158286 | RNF207 | 970.6506 | 0.6702588 | 0.2287588 | 3.059516 | 0.00221695 | 0.011295 |
| ENSG00000178567 | EPM2AIP1 | 2728.311 | 0.6700318 | 0.1945523 | 3.532769 | 0.00041123 | 0.002653 |
| ENSG00000253438 | PCAT1 | 257.2975 | 0.6698469 | 0.2372576 | 2.967533 | 0.003002 | 0.014565 |
| ENSG00000114670 | NEK11 | 562.8863276 | 0.669413014 | 0.198712556 | 3.482489062 | 0.00049678 | 0.003122 |
| ENSG00000262686 | GLIS2-AS1 | 8.005628 | 0.6693835 | 0.7165901 | 1.46143 | 0.1438974 | 0.301944 |
| ENSG00000180539 | C9orf139 | 68.5205 | 0.6693654 | 0.4201529 | 1.853657 | 0.06378831 | 0.166368 |
| ENSG00000091106 | NLRC4 | 13.81233373 | 0.669228738 | 0.546833747 | 1.578561949 | 0.11443657 | 0.256428 |
| ENSG00000275160 | | 53.89576 | 0.6691353 | 0.2986592 | 2.417428 | 0.01563061 | 0.055699 |
| ENSG00000233797 | UFL1-AS1 | 18.59222 | 0.6689039 | 0.5617519 | 1.556303 | 0.119636 | 0.264642 |
| ENSG00000197757 | HOXC6 | 339.6192 | 0.6684954 | 0.2499791 | 2.827417 | 0.00469252 | 0.021065 |
| ENSG00000010818 | HIVEP2 | 1699.380735 | 0.668371989 | 0.340199742 | 2.180356369 | 0.02923106 | 0.091937 |

| | | | | | | | |
|---|---|---|---|---|---|---|---|
| ENSG00000091844 | RGS17 | 632.8920081 | 0.668293554 | 0.276831944 | 2.563944287 | 0.01034902 | 0.040012 |
| ENSG00000180673 | EXOC5P1 | 279.6501 | 0.6680048 | 0.2687725 | 2.652842 | 0.00798171 | 0.032468 |
| ENSG00000177465 | ACOT4 | 867.3288 | 0.6676266 | 0.2671293 | 2.653216 | 0.00797289 | 0.03245 |
| ENSG00000263503 | MAPK8IP1P2 | 8.096133 | 0.6675814 | 0.6186895 | 1.493781 | 0.1352329 | 0.288529 |
| ENSG00000123094 | RASSF8 | 1281.53406 | 0.667393749 | 0.219731063 | 3.152339521 | 0.00161968 | 0.008643 |
| ENSG00000158006 | PAFAH2 | 940.6497 | 0.6673913 | 0.2055798 | 3.38132 | 0.00072139 | 0.004314 |
| ENSG00000198455 | ZXDB | 1204.985 | 0.6673475 | 0.2626306 | 2.697917 | 0.00697748 | 0.029048 |
| ENSG00000159423 | ALDH4A1 | 2384.217 | 0.6670503 | 0.349843 | 2.154531 | 0.03119856 | 0.096689 |
| ENSG00000237094 | | 60.567 | 0.6668092 | 0.5085728 | 1.641668 | 0.1006589 | 0.233879 |
| ENSG00000228838 | | 1.199912 | 0.6667651 | 1.106988 | 1.81523 | 0.06948865 | NA |
| ENSG00000258498 | DIO3OS | 67.66109 | 0.6667235 | 0.5956732 | 1.536478 | 0.1244213 | 0.272266 |
| ENSG00000163900 | TMEM41A | 1985.572 | 0.6666804 | 0.1246029 | 5.448073 | 5.09E-08 | 8.17E-07 |
| ENSG00000143257 | NR1I3 | 21.76093 | 0.6665705 | 0.4294026 | 1.815351 | 0.06946994 | 0.177527 |
| ENSG00000144228 | SPOPL | 4557.378 | 0.6665049 | 0.1889718 | 3.638851 | 0.00027386 | 0.001866 |
| ENSG00000274892 | | 3.079442 | 0.6662276 | 0.8749627 | 1.438941 | 0.1501671 | 0.311313 |
| ENSG00000170634 | ACYP2 | 506.5665 | 0.6661615 | 0.1652174 | 4.12303 | 3.74E-05 | 0.00032 |
| ENSG00000237765 | FAM200B | 781.2526 | 0.6658446 | 0.2137519 | 3.245103 | 0.00117408 | 0.006563 |
| ENSG00000229930 | | 5.651465 | 0.6656009 | 0.676036 | 1.447607 | 0.1477269 | 0.30775 |
| ENSG00000213139 | CRYGS | 57.58474 | 0.6655268 | 0.5234597 | 1.612393 | 0.1068765 | 0.244197 |
| ENSG00000175662 | TOM1L2 | 7564.901 | 0.6652748 | 0.2528476 | 2.829257 | 0.00466562 | 0.020964 |
| ENSG00000176320 | | 3.116417 | 0.6651721 | 0.988044 | 1.594395 | 0.1108476 | 0.250712 |
| ENSG00000286977 | | 62.68717 | 0.6651505 | 0.3658844 | 2.039646 | 0.0413856 | 0.12008 |
| ENSG00000185002 | RFX6 | 1.852154 | 0.6649405 | 0.9548432 | 1.493827 | 0.1352209 | 0.288529 |
| ENSG00000128040 | SPINK2 | 7.103216048 | 0.664812013 | 0.80209762 | 1.464521429 | 0.14305153 | 0.300833 |
| ENSG00000275443 | | 1.605521 | 0.6647668 | 1.11874 | 1.861766 | 0.0626361 | 0.164217 |
| ENSG00000077150 | NFKB2 | 2658.346814 | 0.664621321 | 0.18651459 | 3.677341639 | 0.00023568 | 0.00164 |
| ENSG00000105926 | MPP6 | 1161.417173 | 0.664508825 | 0.210846609 | 3.27534069 | 0.00105535 | 0.005996 |
| ENSG00000148218 | ALAD | 4019.965 | 0.6644998 | 0.1880466 | 3.63754 | 0.00027525 | 0.001873 |
| ENSG00000284959 | | 21.8576 | 0.66394 | 0.4903753 | 1.659734 | 0.09696796 | 0.227431 |
| ENSG00000259536 | | 6.342518 | 0.6639092 | 0.6662545 | 1.452372 | 0.1463981 | 0.305814 |
| ENSG00000139914 | FITM1 | 11.47399 | 0.6636875 | 0.5439164 | 1.57151 | 0.1160642 | 0.258961 |
| ENSG00000118596 | SLC16A7 | 964.7360507 | 0.663381254 | 0.321026772 | 2.266634599 | 0.02341256 | 0.076958 |
| ENSG00000267317 | | 223.8814 | 0.6633059 | 0.1889764 | 3.620228 | 0.00029434 | 0.001988 |

| | | | | | | | |
|---|---|---|---|---|---|---|---|
| ENSG00000171488 | LRRC8C | 105.2865 | 0.6631855 | 0.3496904 | 2.111069 | 0.03476636 | 0.105098 |
| ENSG00000198807 | PAX9 | 551.2528 | 0.6630741 | 0.2483547 | 2.817792 | 0.00483552 | 0.021592 |
| ENSG00000105523 | FAM83E | 14.36164103 | 0.663072585 | 0.522371165 | 1.604358396 | 0.10863508 | 0.247084 |
| ENSG00000188916 | INSYN2A | 18.81951 | 0.6629075 | 0.5331333 | 1.598059 | 0.1100299 | 0.249478 |
| ENSG00000121905 | HPCA | 31.67828455 | 0.662719383 | 0.461885886 | 1.733666924 | 0.0829772 | 0.202891 |
| ENSG00000184254 | ALDH1A3 | 1053.94 | 0.6627067 | 0.5387841 | 1.596052 | 0.1104771 | 0.250129 |
| ENSG00000233393 | | 2.957214 | 0.6626761 | 0.9526048 | 1.494022 | 0.13517 | 0.288473 |
| ENSG00000143811 | PYCR2 | 2402.822 | 0.6626213 | 0.1487305 | 4.540562 | 5.61E-06 | 5.90E-05 |
| ENSG00000267394 | | 44.51472 | 0.6625106 | 0.3178813 | 2.265576 | 0.02347734 | 0.077132 |
| ENSG00000078589 | P2RY10 | 1.693499421 | 0.66225171 | 1.209044933 | 2.25310626 | 0.02425245 | 0.07921 |
| ENSG00000164743 | C8orf48 | 16.49079 | 0.6618532 | 0.4663857 | 1.722031 | 0.08506398 | 0.206682 |
| ENSG00000124104 | SNX21 | 1082.976621 | 0.661761132 | 0.202227736 | 3.391953005 | 0.00069396 | 0.004173 |
| ENSG00000117791 | MTARC2 | 967.0954156 | 0.661715387 | 0.184769279 | 3.691915924 | 0.00022257 | 0.001558 |
| ENSG00000258317 | | 33.22294 | 0.6616956 | 0.3716731 | 2.006544 | 0.04479829 | 0.127586 |
| ENSG00000253336 | | 76.90555 | 0.6616321 | 0.3186485 | 2.280202 | 0.02259569 | 0.075052 |
| ENSG00000185022 | MAFF | 353.1939 | 0.6612722 | 0.4277799 | 1.819867 | 0.06877934 | 0.176135 |
| ENSG00000169758 | TMEM266 | 106.7562 | 0.6610049 | 0.2298139 | 3.007981 | 0.0026299 | 0.013058 |
| ENSG00000279342 | | 63.99615 | 0.6609906 | 0.4512971 | 1.756924 | 0.07893083 | 0.195214 |
| ENSG00000129353 | SLC44A2 | 22152.50918 | 0.660957599 | 0.177806559 | 3.822503452 | 0.0001321 | 0.00098 |
| ENSG00000260912 | | 376.4925 | 0.6608899 | 0.2999208 | 2.394494 | 0.01664333 | 0.058616 |
| ENSG00000258630 | LINC02292 | 5.865962 | 0.6606106 | 0.9265901 | 1.469698 | 0.1416435 | 0.298677 |
| ENSG00000141524 | TMC6 | 3504.44 | 0.6605819 | 0.176799 | 3.825185 | 0.00013067 | 0.00097 |
| ENSG00000135245 | HILPDA | 2627.384343 | 0.66051132 | 0.185891083 | 3.6004829 | 0.00031763 | 0.002128 |
| ENSG00000259953 | | 96.62157 | 0.660143 | 0.3455598 | 2.116473 | 0.03430459 | 0.104087 |
| ENSG00000271218 | | 20.90702 | 0.6594047 | 0.4292916 | 1.805334 | 0.07102237 | 0.180522 |
| ENSG00000183153 | GJD3 | 159.7869 | 0.6589197 | 0.2195014 | 3.13712 | 0.00170616 | 0.009041 |
| ENSG00000274798 | | 7.601674 | 0.6588278 | 0.6586118 | 1.465988 | 0.1426516 | 0.300231 |
| ENSG00000165507 | DEPP1 | 203.1032 | 0.6587543 | 0.4703612 | 1.711248 | 0.08703529 | 0.210203 |
| ENSG00000250616 | | 67.24159 | 0.6584543 | 0.2769427 | 2.54161 | 0.01103433 | 0.042128 |
| ENSG00000120162 | MOB3B | 345.1749791 | 0.658185129 | 0.393194676 | 1.921946508 | 0.05461249 | 0.148425 |
| ENSG00000273890 | | 3.642511 | 0.6573789 | 0.8273302 | 1.436671 | 0.1508114 | 0.312335 |
| ENSG00000154263 | ABCA10 | 93.63723 | 0.6573215 | 0.5998137 | 1.4943 | 0.1350972 | 0.288473 |
| ENSG00000225733 | FGD5-AS1 | 11000.67 | 0.6573039 | 0.1592404 | 4.220603 | 2.44E-05 | 0.000219 |

| | | | | | | | |
|---|---|---|---|---|---|---|---|
| ENSG00000143815 | LBR | 3688.43 | 0.6570527 | 0.3260824 | 2.215174 | 0.02674812 | 0.08551 |
| ENSG00000080815 | PSEN1 | 5865.663185 | 0.656901737 | 0.184572005 | 3.662968678 | 0.00024931 | 0.00172 |
| ENSG00000226220 | CICP22 | 4.028712 | 0.656674 | 0.7816239 | 1.405361 | 0.1599139 | 0.326125 |
| ENSG00000120913 | PDLIM2 | 419.3645477 | 0.656516315 | 0.356575362 | 2.066334484 | 0.03879691 | 0.114216 |
| ENSG00000225791 | TRAM2-AS1 | 469.3222 | 0.6564438 | 0.2289969 | 3.006699 | 0.00264101 | 0.013101 |
| ENSG00000204706 | MAMDC2-AS1 | 3.817384 | 0.6563922 | 0.77064 | 1.415457 | 0.1569344 | 0.321777 |
| ENSG00000184292 | TACSTD2 | 25197.07 | 0.6563055 | 0.2612596 | 2.717735 | 0.00657305 | 0.027677 |
| ENSG00000167535 | CACNB3 | 5314.676 | 0.6562838 | 0.156023 | 4.288671 | 1.80E-05 | 0.000166 |
| ENSG00000149926 | TLCD3B | 44.36324 | 0.6560943 | 0.3787501 | 1.963944 | 0.04953663 | 0.137585 |
| ENSG00000223947 | | 7.70449 | 0.6560942 | 0.8065592 | 1.419382 | 0.1557877 | 0.319969 |
| ENSG00000115956 | PLEK | 1.674322158 | 0.656090696 | 1.100544519 | 1.801882254 | 0.07156393 | 0.181638 |
| ENSG00000279511 | | 19.75028 | 0.6560728 | 0.5239842 | 1.569881 | 0.1164427 | 0.259487 |
| ENSG00000226499 | | 3.291009 | 0.6558061 | 0.9598448 | 1.561882 | 0.1183157 | 0.262442 |
| ENSG00000286984 | | 11.99032 | 0.6557653 | 0.578861 | 1.503007 | 0.1328373 | 0.285184 |
| ENSG00000257838 | OTOAP1 | 1.647091 | 0.6553737 | 0.9129519 | 1.469274 | 0.1417584 | 0.2988 |
| ENSG00000262884 | | 4.352261 | 0.655198 | 0.9789204 | 1.599296 | 0.1097549 | 0.249025 |
| ENSG00000260922 | | 2.782016 | 0.6551808 | 1.166673 | 2.094241 | 0.03623854 | 0.10856 |
| ENSG00000263301 | | 39.43407 | 0.655155 | 0.4686323 | 1.698853 | 0.08934691 | 0.214499 |
| ENSG00000286591 | | 1.528931 | 0.6549784 | 1.038395 | 1.729556 | 0.08370957 | NA |
| ENSG00000214544 | GTF2IRD2P1 | 10.57709 | 0.6548676 | 0.6586171 | 1.436937 | 0.1507358 | 0.312224 |
| ENSG00000285545 | | 2.290745 | 0.6547261 | 1.025721 | 1.673603 | 0.09420873 | 0.223033 |
| ENSG00000066248 | NGEF | 168.3351647 | 0.654576282 | 0.268971018 | 2.595169689 | 0.00945442 | 0.037152 |
| ENSG00000250522 | | 10.15727 | 0.6544965 | 0.7516541 | 1.428148 | 0.1532493 | 0.316051 |
| ENSG00000185565 | LSAMP | 41.3844 | 0.6543972 | 0.5285463 | 1.570741 | 0.1162429 | 0.259295 |
| ENSG00000105737 | GRIK5 | 14.76453846 | 0.654262909 | 0.611992958 | 1.484022854 | 0.13780286 | 0.292504 |
| ENSG00000265273 | PGDP1 | 32.10964 | 0.6542407 | 0.4446738 | 1.755638 | 0.07915021 | 0.195641 |
| ENSG00000147862 | NFIB | 1561.828 | 0.6541103 | 0.2183547 | 3.131815 | 0.00173729 | 0.009193 |
| ENSG00000114405 | C3orf14 | 6334.52684 | 0.654051084 | 0.212805208 | 3.160382317 | 0.00157562 | 0.008449 |
| ENSG00000267905 | | 1.965505 | 0.6539183 | 1.074721 | 1.748514 | 0.08037513 | 0.197969 |
| ENSG00000241878 | PISD | 3258.174 | 0.6536783 | 0.1501778 | 4.433376 | 9.28E-06 | 9.21E-05 |
| ENSG00000272501 | | 121.6992 | 0.6535392 | 0.2881809 | 2.439835 | 0.01469397 | 0.053058 |
| ENSG00000163239 | TDRD10 | 1.30748 | 0.653407 | 1.09436 | 1.795875 | 0.07251432 | NA |
| ENSG00000176058 | TPRN | 3250.352 | 0.6533213 | 0.1164647 | 5.684273 | 1.31E-08 | 2.35E-07 |

| | | | | | | | |
|---|---|---|---|---|---|---|---|
| ENSG00000104765 | BNIP3L | 2896.399832 | 0.652655748 | 0.145731456 | 4.580624398 | 4.64E-06 | 4.99E-05 |
| ENSG00000229937 | PRPS1L1 | 1.971781 | 0.6526378 | 1.092983 | 1.8158 | 0.06940101 | 0.177368 |
| ENSG00000148356 | LRSAM1 | 2974.88 | 0.6526046 | 0.1385945 | 4.772378 | 1.82E-06 | 2.16E-05 |
| ENSG00000182814 | FUNDC2P2 | 15.8395 | 0.6525103 | 0.5625692 | 1.535304 | 0.1247092 | 0.272685 |
| ENSG00000104221 | BRF2 | 387.2594736 | 0.652285432 | 0.168817212 | 3.961191526 | 7.46E-05 | 0.000591 |
| ENSG00000232267 | ACTR3P2 | 2.154329 | 0.652259 | 1.00016 | 1.655263 | 0.09787123 | 0.229085 |
| ENSG00000162482 | AKR7A3 | 26.10153 | 0.652136 | 0.4000658 | 1.873975 | 0.06093381 | 0.16085 |
| ENSG00000198780 | FAM169A | 612.7226 | 0.652117 | 0.317566 | 2.244974 | 0.02476978 | 0.080494 |
| ENSG00000133710 | SPINK5 | 54.57354087 | 0.652053245 | 0.422118809 | 1.795953854 | 0.07250186 | 0.18344 |
| ENSG00000229419 | RALGAPA1P1 | 1304.001 | 0.6520288 | 0.2696197 | 2.555654 | 0.01059884 | 0.040816 |
| ENSG00000188997 | KCTD21 | 564.9435 | 0.6514318 | 0.2547618 | 2.7096 | 0.00673644 | 0.028226 |
| ENSG00000171815 | PCDHB1 | 4.025355 | 0.6511819 | 0.8457533 | 1.423924 | 0.1544685 | 0.318022 |
| ENSG00000185482 | STAC3 | 222.9612 | 0.6511548 | 0.3209155 | 2.230991 | 0.02568172 | 0.082817 |
| ENSG00000230317 | LINC01284 | 1.161322 | 0.65112 | 1.156558 | 2.037607 | 0.04158928 | NA |
| ENSG00000168036 | CTNNB1 | 13633.95 | 0.6507336 | 0.1950133 | 3.456695 | 0.00054684 | 0.003395 |
| ENSG00000187550 | SBK2 | 12.20157 | 0.6505684 | 0.7347057 | 1.421649 | 0.1551283 | 0.318939 |
| ENSG00000143450 | OAZ3 | 118.0867 | 0.6505148 | 0.2660842 | 2.603805 | 0.00921951 | 0.036451 |
| ENSG00000114378 | HYAL1 | 17.83628036 | 0.650266313 | 0.520105184 | 1.569098265 | 0.11662504 | 0.25983 |
| ENSG00000278730 | | 2826.417 | 0.6502539 | 0.1414428 | 4.66289 | 3.12E-06 | 3.50E-05 |
| ENSG00000218073 | | 10.86643 | 0.6499964 | 0.6255268 | 1.454479 | 0.1458136 | 0.304881 |
| ENSG00000144724 | PTPRG | 1443.853 | 0.6499523 | 0.2690696 | 2.5677 | 0.01023759 | 0.039633 |
| ENSG00000253327 | RAD21-AS1 | 15.91522 | 0.6498659 | 0.5913732 | 1.499792 | 0.1336684 | 0.286408 |
| ENSG00000233523 | PHBP5 | 2.110244 | 0.64977 | 1.093901 | 1.954896 | 0.05059531 | 0.13975 |
| ENSG00000164010 | ERMAP | 761.0136 | 0.6497399 | 0.2198617 | 3.07712 | 0.00209012 | 0.010756 |
| ENSG00000247157 | LINC01252 | 109.7487 | 0.6493115 | 0.3413856 | 2.107618 | 0.03506407 | 0.105733 |
| ENSG00000168067 | MAP4K2 | 1275.449 | 0.6492675 | 0.194698 | 3.451313 | 0.00055787 | 0.003449 |
| ENSG00000237025 | PARD6BP1 | 1.238979 | 0.6491742 | 1.169379 | 2.120684 | 0.0339484 | NA |
| ENSG00000241563 | CORT | 8.384813 | 0.6484063 | 0.7331321 | 1.413163 | 0.1576079 | 0.32276 |
| ENSG00000115963 | RND3 | 5400.08257 | 0.648405607 | 0.406655767 | 1.848250282 | 0.06456615 | 0.167901 |
| ENSG00000138688 | KIAA1109 | 8706.967967 | 0.648140992 | 0.323058322 | 2.230892332 | 0.02568826 | 0.082819 |
| ENSG00000135838 | NPL | 107.7493115 | 0.648139962 | 0.245743913 | 2.785963969 | 0.00533688 | 0.023352 |
| ENSG00000102174 | PHEX | 308.2502601 | 0.647861137 | 0.280174326 | 2.476152784 | 0.01328067 | 0.048849 |
| ENSG00000130826 | DKC1 | 10494.31044 | 0.647834632 | 0.207947376 | 3.250394151 | 0.00115245 | 0.006468 |

| | | | | | | | |
|---|---|---|---|---|---|---|---|
| ENSG00000088756 | ARHGAP28 | 345.581106 | 0.647652714 | 0.275502193 | 2.512995703 | 0.01197108 | 0.044955 |
| ENSG00000263063 | | 3.905851 | 0.6476189 | 0.9785185 | 1.572326 | 0.115875 | 0.258713 |
| ENSG00000162734 | PEA15 | 5554.765 | 0.6475412 | 0.1720304 | 3.870217 | 0.00010874 | 0.000824 |
| ENSG00000123595 | RAB9A | 1588.801935 | 0.64745704 | 0.203908733 | 3.293310938 | 0.00099015 | 0.005675 |
| ENSG00000249917 | LINC00536 | 6.089293 | 0.6473373 | 0.8044843 | 1.412147 | 0.1579067 | 0.323133 |
| ENSG00000109743 | BST1 | 5.021418752 | 0.647285188 | 0.731665467 | 1.401593098 | 0.16103679 | 0.327735 |
| ENSG00000279074 | | 1.112617 | 0.6472452 | 1.105189 | 1.856896 | 0.06332594 | NA |
| ENSG00000272970 | | 37.20359 | 0.6472442 | 0.4058267 | 1.846759 | 0.06478208 | 0.16825 |
| ENSG00000270624 | | 7.73542 | 0.6467353 | 0.6684852 | 1.437433 | 0.1505949 | 0.311981 |
| ENSG00000159256 | MORC3 | 1619.253 | 0.6466468 | 0.1868654 | 3.566096 | 0.00036234 | 0.002385 |
| ENSG00000213760 | ATP6V1G2 | 64.44829 | 0.6466448 | 0.3936758 | 1.869271 | 0.06158511 | 0.162166 |
| ENSG00000102312 | PORCN | 561.5056864 | 0.64651896 | 0.156751343 | 4.213701582 | 2.51E-05 | 0.000224 |
| ENSG00000168917 | SLC35G2 | 9.331782 | 0.6464798 | 0.6346295 | 1.450393 | 0.146949 | 0.306603 |
| ENSG00000177076 | ACER2 | 503.2569 | 0.646367 | 0.3917366 | 1.89129 | 0.05858564 | 0.15636 |
| ENSG00000206144 | MAPK6P1 | 1.314974 | 0.646355 | 1.064305 | 1.729802 | 0.08366563 | NA |
| ENSG00000105711 | SCN1B | 744.5888157 | 0.646321134 | 0.293288255 | 2.364998913 | 0.01803012 | 0.062538 |
| ENSG00000198914 | POU3F3 | 1.1888 | 0.6459435 | 1.194041 | 2.257888 | 0.02395263 | NA |
| ENSG00000189164 | ZNF527 | 294.0657 | 0.6459349 | 0.1650107 | 4.016451 | 5.91E-05 | 0.00048 |
| ENSG00000160007 | ARHGAP35 | 14330.34 | 0.645827 | 0.1752749 | 3.756302 | 0.00017244 | 0.001244 |
| ENSG00000123091 | RNF11 | 5070.136043 | 0.645715064 | 0.123984319 | 5.259519711 | 1.44E-07 | 2.13E-06 |
| ENSG00000285602 | | 5.89465 | 0.6456288 | 0.5989552 | 1.459838 | 0.1443347 | 0.302627 |
| ENSG00000168907 | PLA2G4F | 250.7335 | 0.6454645 | 0.3422617 | 2.093144 | 0.03633633 | 0.108713 |
| ENSG00000183826 | BTBD9 | 1165.243 | 0.6453593 | 0.2067442 | 3.246432 | 0.00116861 | 0.006544 |
| ENSG00000136848 | DAB2IP | 3329.91614 | 0.644945191 | 0.234471826 | 2.884499693 | 0.00392036 | 0.018146 |
| ENSG00000223198 | RNU2-22P | 4.533827 | 0.6447597 | 0.7208527 | 1.396786 | 0.1624779 | 0.329784 |
| ENSG00000258623 | | 6.073521 | 0.6445969 | 0.7307056 | 1.412471 | 0.1578113 | 0.323077 |
| ENSG00000261104 | | 46.37275 | 0.6444771 | 0.5096593 | 1.600269 | 0.109539 | 0.248598 |
| ENSG00000174482 | LINGO2 | 2.236642 | 0.6443751 | 0.88997 | 1.454682 | 0.1457573 | 0.304847 |
| ENSG00000151553 | FAM160B1 | 1760.887 | 0.64432 | 0.2272883 | 2.964181 | 0.0030349 | 0.014687 |
| ENSG00000263316 | | 38.12506 | 0.6442622 | 0.5146641 | 1.57727 | 0.1147335 | 0.256886 |
| ENSG00000007402 | CACNA2D2 | 192.5080348 | 0.644161053 | 0.240892812 | 2.81287416 | 0.00491009 | 0.021843 |
| ENSG00000151150 | ANK3 | 3225.636 | 0.6439168 | 0.2630494 | 2.600057 | 0.00932084 | 0.03674 |
| ENSG00000283317 | | 4.994125 | 0.6437195 | 0.7498321 | 1.391183 | 0.1641698 | 0.332254 |

| | | | | | | | |
|---|---|---|---|---|---|---|---|
| ENSG00000214199 | EEF1A1P12 | 30.13343 | 0.6436933 | 0.4036762 | 1.843411 | 0.06526909 | 0.169266 |
| ENSG00000119725 | ZNF410 | 49.11109458 | 0.643682594 | 0.359089279 | 2.007239067 | 0.04472421 | 0.127429 |
| ENSG00000240399 | | 20.50742 | 0.6436026 | 0.4973649 | 1.605162 | 0.1084582 | 0.246799 |
| ENSG00000113212 | PCDHB7 | 8.379526058 | 0.643227472 | 0.615588906 | 1.446608999 | 0.14800646 | 0.308229 |
| ENSG00000159314 | ARHGAP27 | 3047.356 | 0.6431884 | 0.1418333 | 4.618772 | 3.86E-06 | 4.22E-05 |
| ENSG00000204947 | ZNF425 | 424.2954 | 0.6430591 | 0.1838802 | 3.601366 | 0.00031655 | 0.002122 |
| ENSG00000204758 | | 31.64434 | 0.6430017 | 0.4988407 | 1.607719 | 0.1078967 | 0.245901 |
| ENSG00000215105 | TTC3P1 | 1946.88 | 0.6426989 | 0.2745214 | 2.501687 | 0.01236031 | 0.046104 |
| ENSG00000236762 | RPL19P16 | 12.62862 | 0.64258 | 0.5167297 | 1.569046 | 0.1166373 | 0.25983 |
| ENSG00000156564 | LRFN2 | 14.87837 | 0.6424959 | 0.7009731 | 1.406355 | 0.1596187 | 0.325723 |
| ENSG00000196183 | RPS2P4 | 30.26689 | 0.6423407 | 0.4113513 | 1.815933 | 0.0693807 | 0.17735 |
| ENSG00000213904 | LIPE-AS1 | 20.45241 | 0.6422434 | 0.468688 | 1.650209 | 0.09890013 | 0.230781 |
| ENSG00000269102 | | 7.581629 | 0.642073 | 0.6906444 | 1.411862 | 0.1579906 | 0.323266 |
| ENSG00000284987 | | 1.415337 | 0.6420325 | 1.065712 | 1.732038 | 0.08326683 | NA |
| ENSG00000174992 | ZG16 | 6.755381 | 0.6418004 | 0.738726 | 1.392843 | 0.1636673 | 0.33159 |
| ENSG00000148814 | LRRC27 | 1329.092 | 0.6417665 | 0.2644903 | 2.578799 | 0.00991444 | 0.038596 |
| ENSG00000215097 | | 7.02933 | 0.6416619 | 0.638258 | 1.432355 | 0.1520424 | 0.314196 |
| ENSG00000157869 | RAB28 | 610.8421 | 0.6404544 | 0.1397877 | 4.657749 | 3.20E-06 | 3.57E-05 |
| ENSG00000236842 | | 1.90236 | 0.640054 | 0.9587036 | 1.496446 | 0.1345375 | 0.287694 |
| ENSG00000227070 | EPS15-AS1 | 78.04818 | 0.6400332 | 0.4180729 | 1.78509 | 0.07424665 | 0.186823 |
| ENSG00000187325 | TAF9B | 2441.881 | 0.6399589 | 0.3293326 | 2.13937 | 0.03240574 | 0.099548 |
| ENSG00000042286 | AIFM2 | 2015.760655 | 0.639925443 | 0.20999817 | 3.169855846 | 0.00152515 | 0.008228 |
| ENSG00000271547 | | 9.898174 | 0.6395331 | 0.6453635 | 1.412535 | 0.1577924 | 0.323077 |
| ENSG00000258056 | | 265.0305 | 0.6392559 | 0.2896251 | 2.375159 | 0.01754137 | 0.061162 |
| ENSG00000261437 | | 341.1026 | 0.6391106 | 0.22769 | 2.942258 | 0.00325828 | 0.015562 |
| ENSG00000241352 | | 22.30458 | 0.6387603 | 0.4459602 | 1.696938 | 0.08970845 | 0.215154 |
| ENSG00000234819 | | 3.657891 | 0.6385994 | 0.7024288 | 1.386082 | 0.165722 | 0.334376 |
| ENSG00000275557 | | 54.7602 | 0.6385775 | 0.7287213 | 1.377805 | 0.1682634 | 0.338041 |
| ENSG00000213236 | YWHAZP2 | 966.6303 | 0.6383007 | 0.1748414 | 3.742707 | 0.00018205 | 0.001304 |
| ENSG00000233913 | RPL10P9 | 5751.738 | 0.63825 | 0.1747822 | 3.73461 | 0.00018801 | 0.001342 |
| ENSG00000248049 | UBA6-AS1 | 349.0565 | 0.6376461 | 0.2113956 | 3.147275 | 0.001648 | 0.008766 |
| ENSG00000224549 | | 7.150439 | 0.637422 | 0.6554711 | 1.400549 | 0.161349 | 0.328245 |
| ENSG00000248746 | ACTN3 | 4.983574 | 0.6373 | 0.8167682 | 1.396737 | 0.1624928 | 0.329789 |

| | | | | | | | |
|---|---|---|---|---|---|---|---|
| ENSG00000215049 | PRDX2P1 | 1.907088 | 0.6367782 | 1.052359 | 1.701414 | 0.08886526 | 0.213709 |
| ENSG00000101882 | NKAP | 1553.961789 | 0.636740047 | 0.158487133 | 4.116096912 | 3.85E-05 | 0.000329 |
| ENSG00000198718 | TOGARAM1 | 2910.768 | 0.6365258 | 0.2421664 | 2.780127 | 0.00543376 | 0.023702 |
| ENSG00000261557 | EEF1A1P38 | 15.04443 | 0.6364326 | 0.5358693 | 1.510294 | 0.1309684 | 0.282306 |
| ENSG00000279205 | | 2.740394 | 0.6357122 | 0.8582992 | 1.408917 | 0.1588598 | 0.324723 |
| ENSG00000274791 | F8A2 | 1.361452 | 0.6356732 | 1.021983 | 1.642417 | 0.1005036 | NA |
| ENSG00000249602 | | 41.52639 | 0.6352895 | 0.3210463 | 2.15562 | 0.03111337 | 0.096515 |
| ENSG00000250332 | | 18.94772 | 0.6352696 | 0.5545683 | 1.497503 | 0.1342625 | 0.287298 |
| ENSG00000136383 | ALPK3 | 163.0787175 | 0.635019567 | 0.351458877 | 2.008107929 | 0.04463182 | 0.127282 |
| ENSG00000240093 | | 2.52721 | 0.6349322 | 0.9888454 | 1.640396 | 0.1009228 | 0.234369 |
| ENSG00000182903 | ZNF721 | 1398.256 | 0.6348927 | 0.1529814 | 4.236536 | 2.27E-05 | 0.000205 |
| ENSG00000100532 | CGRRF1 | 1033.3216 | 0.634801554 | 0.137816518 | 4.681612161 | 2.85E-06 | 3.23E-05 |
| ENSG00000287167 | | 2.039657 | 0.6346823 | 0.8613036 | 1.408234 | 0.1590618 | 0.324986 |
| ENSG00000279953 | | 9.886411 | 0.6345908 | 0.6177744 | 1.444684 | 0.1485467 | 0.30905 |
| ENSG00000146416 | AIG1 | 2128.959 | 0.6345855 | 0.1470304 | 4.394343 | 1.11E-05 | 0.000108 |
| ENSG00000273771 | | 1.567516 | 0.634574 | 1.088425 | 1.825558 | 0.06791693 | NA |
| ENSG00000184774 | MGAT4EP | 28.33263 | 0.6344291 | 0.4813913 | 1.583611 | 0.1132822 | 0.254549 |
| ENSG00000168505 | GBX2 | 26.58946 | 0.6343403 | 0.5518248 | 1.504785 | 0.1323793 | 0.284585 |
| ENSG00000268545 | VN1R107P | 1.704956 | 0.6342313 | 1.020705 | 1.627075 | 0.1037213 | 0.238864 |
| ENSG00000163945 | UVSSA | 1102.473 | 0.6341955 | 0.28561 | 2.384312 | 0.01711111 | 0.059898 |
| ENSG00000232298 | GRHL3-AS1 | 7.774663 | 0.6339782 | 0.8098144 | 1.401976 | 0.1609223 | 0.327534 |
| ENSG00000122557 | HERPUD2 | 1771.315027 | 0.63395287 | 0.17979692 | 3.624354454 | 0.00028968 | 0.001963 |
| ENSG00000157538 | VPS26C | 2583.229 | 0.633695 | 0.1544599 | 4.189723 | 2.79E-05 | 0.000247 |
| ENSG00000244411 | KRTAP5-7 | 9.708244 | 0.6336443 | 0.5309257 | 1.514198 | 0.1299755 | 0.280689 |
| ENSG00000196668 | LINC00173 | 93.76001 | 0.6334339 | 0.7337481 | 1.361265 | 0.17343 | 0.34493 |
| ENSG00000123575 | FAM199X | 6814.296131 | 0.632979604 | 0.156063228 | 4.14259945 | 3.43E-05 | 0.000297 |
| ENSG00000286596 | | 3.709842 | 0.6329633 | 0.7591083 | 1.386182 | 0.1656914 | 0.334344 |
| ENSG00000220695 | | 13.29812 | 0.632856 | 0.487179 | 1.594411 | 0.1108441 | 0.250712 |
| ENSG00000100429 | HDAC10 | 166.4027155 | 0.63280932 | 0.274193008 | 2.461223612 | 0.0138464 | 0.050527 |
| ENSG00000219481 | NBPF1 | 4401.785 | 0.6327378 | 0.2272179 | 2.912767 | 0.00358242 | 0.01684 |
| ENSG00000225193 | RPS12P26 | 9.484958 | 0.6326302 | 0.5427679 | 1.514528 | 0.1298919 | 0.280577 |
| ENSG00000145391 | SETD7 | 9582.58 | 0.6323817 | 0.2227221 | 2.97975 | 0.00288483 | 0.014129 |
| ENSG00000259078 | PTBP1P | 8.617167 | 0.6319594 | 0.6878283 | 1.39711 | 0.1623805 | 0.329662 |

| | | | | | | | |
|---|---|---|---|---|---|---|---|
| ENSG00000167528 | ZNF641 | 1969.688 | 0.6318933 | 0.2725553 | 2.473342 | 0.01338559 | 0.049135 |
| ENSG00000105997 | HOXA3 | 7.402852784 | 0.631820242 | 0.715681324 | 1.377921806 | 0.16822743 | 0.33802 |
| ENSG00000223849 | | 8.679855 | 0.6317299 | 0.7179231 | 1.377444 | 0.1683749 | 0.338163 |
| ENSG00000188765 | TMSB4XP2 | 3.861648 | 0.6314129 | 0.7708393 | 1.389992 | 0.1645313 | 0.332696 |
| ENSG00000223443 | USP17L2 | 2.921489 | 0.6310829 | 0.9012884 | 1.460035 | 0.1442805 | 0.30258 |
| ENSG00000129158 | SERGEF | 607.1184356 | 0.630888148 | 0.204423946 | 3.203288757 | 0.00135868 | 0.007458 |
| ENSG00000287678 | | 32.43489 | 0.63083 | 0.3769482 | 1.893069 | 0.05834874 | 0.155913 |
| ENSG00000167566 | NCKAP5L | 2599.355 | 0.6306988 | 0.1790958 | 3.607801 | 0.0003088 | 0.002075 |
| ENSG00000116903 | EXOC8 | 1175.089162 | 0.630658841 | 0.163721319 | 3.949656279 | 7.83E-05 | 0.000617 |
| ENSG00000245864 | MEF2C-AS2 | 1.714106 | 0.6306197 | 1.090101 | 1.823974 | 0.06815599 | 0.174994 |
| ENSG00000130222 | GADD45G | 631.6299999 | 0.630358824 | 0.233535203 | 2.832617965 | 0.00461685 | 0.020787 |
| ENSG00000284879 | | 6.795512 | 0.6302779 | 0.6811146 | 1.392456 | 0.1637845 | 0.331777 |
| ENSG00000262001 | DLGAP1-AS2 | 194.8234 | 0.6302757 | 0.4947841 | 1.577693 | 0.1146363 | 0.256789 |
| ENSG00000214922 | HLA-F-AS1 | 101.6825 | 0.6300446 | 0.496602 | 1.57295 | 0.1157304 | 0.258607 |
| ENSG00000270095 | | 4.761598 | 0.6299919 | 0.8575504 | 1.424926 | 0.1541785 | 0.317548 |
| ENSG00000171658 | NMRAL2P | 65.43552 | 0.6296988 | 0.8910006 | 1.462085 | 0.1437179 | 0.301705 |
| ENSG00000123427 | EEF1AKMT3 | 571.2440247 | 0.629691755 | 0.220605723 | 2.984662905 | 0.00283891 | 0.013942 |
| ENSG00000137103 | TMEM8B | 397.569993 | 0.629634304 | 0.29362353 | 2.320105588 | 0.02033517 | 0.06887 |
| ENSG00000142583 | SLC2A5 | 3.491812 | 0.6295825 | 0.7797893 | 1.383467 | 0.1665218 | 0.335608 |
| ENSG00000242268 | LINC02082 | 2.703779 | 0.6290001 | 0.883078 | 1.40368 | 0.1604141 | 0.3269 |
| ENSG00000250120 | PCDHA10 | 269.541 | 0.6288319 | 0.3859887 | 1.854841 | 0.06361887 | 0.166024 |
| ENSG00000255389 | | 30.98779 | 0.6286194 | 0.5027826 | 1.554338 | 0.1201038 | 0.2655 |
| ENSG00000241852 | C8orf58 | 415.2304 | 0.6284284 | 0.2414821 | 2.741405 | 0.00611771 | 0.026128 |
| ENSG00000260000 | | 178.1593 | 0.6282322 | 0.2196052 | 2.984503 | 0.0028404 | 0.013947 |
| ENSG00000171385 | KCND3 | 100.835 | 0.6279952 | 0.3419035 | 2.033859 | 0.04196585 | 0.121379 |
| ENSG00000119718 | EIF2B2 | 2999.539838 | 0.627758208 | 0.222488986 | 2.943897958 | 0.00324107 | 0.015493 |
| ENSG00000280287 | | 115.6456 | 0.6277499 | 0.3779989 | 1.875289 | 0.060753 | 0.1605 |
| ENSG00000214559 | | 4.401723 | 0.6275997 | 0.8230966 | 1.39508 | 0.1629918 | 0.33055 |
| ENSG00000183087 | GAS6 | 458.9065 | 0.6274838 | 0.2116381 | 3.087307 | 0.00201979 | 0.010449 |
| ENSG00000141376 | BCAS3 | 37185.07 | 0.6274002 | 0.1795781 | 3.610547 | 0.00030555 | 0.002056 |
| ENSG00000139572 | GPR84 | 2.334697 | 0.6273746 | 1.017791 | 1.625003 | 0.1041618 | 0.23965 |
| ENSG00000139531 | SUOX | 1303.815 | 0.6270537 | 0.1809471 | 3.570005 | 0.00035697 | 0.002353 |
| ENSG00000204116 | CHIC1 | 691.3555 | 0.6269186 | 0.2136154 | 3.051705 | 0.00227546 | 0.011544 |

| | | | | | | | |
|---|---|---|---|---|---|---|---|
| ENSG00000138867 | GUCD1 | 5834.924381 | 0.62686155 | 0.209444375 | 3.110387538 | 0.00186842 | 0.009776 |
| ENSG00000009780 | FAM76A | 609.8146252 | 0.626762355 | 0.14621366 | 4.348222436 | 1.37E-05 | 0.000131 |
| ENSG00000147403 | RPL10 | 25503.9 | 0.6266447 | 0.1586303 | 4.045818 | 5.21E-05 | 0.00043 |
| ENSG00000259079 | | 43.33165 | 0.6266402 | 0.2837557 | 2.372257 | 0.01767977 | 0.061516 |
| ENSG00000285441 | NA | 1.438244 | 0.6266029 | 1.000275 | 1.660838 | 0.09674605 | NA |
| ENSG00000120055 | C10orf95 | 7.602270612 | 0.626550721 | 0.599085915 | 1.425767359 | 0.15393551 | 0.31722 |
| ENSG00000254394 | | 2.874439 | 0.6264272 | 1.050986 | 1.705549 | 0.08809215 | 0.212215 |
| ENSG00000253266 | | 66.8451 | 0.6261147 | 0.5183779 | 1.53038 | 0.1259228 | 0.274391 |
| ENSG00000163482 | STK36 | 3222.113 | 0.626025 | 0.1390579 | 4.532741 | 5.82E-06 | 6.11E-05 |
| ENSG00000131620 | ANO1 | 14.10569048 | 0.625936761 | 0.5212825 | 1.528804041 | 0.12631303 | 0.275029 |
| ENSG00000270179 | | 19.1669 | 0.6258919 | 0.5170113 | 1.525772 | 0.1270668 | 0.276159 |
| ENSG00000001460 | STPG1 | 407.6829618 | 0.625824565 | 0.262907396 | 2.530985181 | 0.01137427 | 0.043147 |
| ENSG00000269907 | PTMAP11 | 4.445363 | 0.6247384 | 0.8465787 | 1.39788 | 0.1621491 | 0.329444 |
| ENSG00000271052 | | 2.653513 | 0.6240572 | 0.9046825 | 1.467073 | 0.1423562 | 0.299751 |
| ENSG00000278987 | | 56.80568 | 0.6236944 | 0.3596342 | 1.944199 | 0.05187146 | 0.142521 |
| ENSG00000130055 | GDPD2 | 4.464398852 | 0.622905405 | 0.772144586 | 1.372891825 | 0.16978598 | 0.340151 |
| ENSG00000171148 | TADA3 | 6750.005 | 0.622809 | 0.1675924 | 3.796477 | 0.00014677 | 0.001076 |
| ENSG00000151876 | FBXO4 | 1230.797 | 0.622487 | 0.2069976 | 3.128569 | 0.0017566 | 0.00928 |
| ENSG00000259797 | | 3.1932 | 0.6224254 | 0.8428086 | 1.394601 | 0.1631361 | 0.330741 |
| ENSG00000171444 | MCC | 976.2909 | 0.62239 | 0.2761942 | 2.400941 | 0.01635298 | 0.057723 |
| ENSG00000163349 | HIPK1 | 34588.35 | 0.6217273 | 0.2473621 | 2.651088 | 0.0080233 | 0.032618 |
| ENSG00000135148 | TRAFD1 | 5360.042549 | 0.621705042 | 0.234621751 | 2.787201702 | 0.00531654 | 0.023279 |
| ENSG00000184697 | CLDN6 | 22.49693 | 0.621524 | 0.5347608 | 1.495697 | 0.1347326 | 0.288009 |
| ENSG00000259673 | IQCH-AS1 | 734.6015 | 0.6215129 | 0.2437278 | 2.689985 | 0.00714553 | 0.029662 |
| ENSG00000147437 | GNRH1 | 49.28438 | 0.6214196 | 0.4877804 | 1.573904 | 0.1155096 | 0.258353 |
| ENSG00000088836 | SLC4A11 | 642.2525173 | 0.621397623 | 0.227905084 | 2.862621563 | 0.00420152 | 0.01922 |
| ENSG00000236078 | LINC01447 | 7.771457 | 0.6212556 | 0.9375423 | 1.550177 | 0.121099 | 0.267033 |
| ENSG00000118096 | IFT46 | 1875.507107 | 0.621142307 | 0.158505129 | 3.998543611 | 6.37E-05 | 0.000514 |
| ENSG00000235903 | CPB2-AS1 | 8.335286 | 0.620968 | 0.6614009 | 1.366982 | 0.171631 | 0.342347 |
| ENSG00000105971 | CAV2 | 799.4306404 | 0.620634548 | 0.488297284 | 1.572157185 | 0.1159141 | 0.258773 |
| ENSG00000063978 | RNF4 | 4064.402263 | 0.62037408 | 0.128313982 | 4.906962329 | 9.25E-07 | 1.16E-05 |
| ENSG00000214796 | TUBA5P | 2137.888 | 0.620315 | 0.3870338 | 1.847654 | 0.06465248 | 0.168012 |
| ENSG00000262312 | | 2.55177 | 0.6202209 | 0.9077015 | 1.482333 | 0.1382516 | 0.2932 |

| | | | | | | | |
|---|---|---|---|---|---|---|---|
| ENSG00000155252 | PI4K2A | 1816.566 | 0.6200028 | 0.1506112 | 4.204132 | 2.62E-05 | 0.000233 |
| ENSG00000233014 | TBC1D3P7 | 1.889639 | 0.6198251 | 1.067947 | 1.763375 | 0.0778373 | 0.193186 |
| ENSG00000161860 | SYCE2 | 30.30145 | 0.6195747 | 0.4178002 | 1.728664 | 0.08386923 | 0.204527 |
| ENSG00000100697 | DICER1 | 7903.029872 | 0.619484967 | 0.223752263 | 2.898091093 | 0.00375442 | 0.017488 |
| ENSG00000258790 | | 5.911052 | 0.619415 | 0.6568211 | 1.370766 | 0.1704479 | 0.341114 |
| ENSG00000106780 | MEGF9 | 1605.960317 | 0.619127233 | 0.218200193 | 2.950795176 | 0.00316957 | 0.015204 |
| ENSG00000184709 | LRRC26 | 37.38173 | 0.6190587 | 0.4676024 | 1.602343 | 0.1090797 | 0.247725 |
| ENSG00000165102 | HGSNAT | 1361.703 | 0.6189024 | 0.1802167 | 3.551994 | 0.00038232 | 0.002498 |
| ENSG00000127081 | ZNF484 | 283.3135675 | 0.618874842 | 0.198323958 | 3.228548768 | 0.0012442 | 0.006916 |
| ENSG00000244125 | | 3.437723 | 0.6186263 | 0.7839132 | 1.370046 | 0.1706725 | 0.341384 |
| ENSG00000225053 | PRPF38AP1 | 1.910501 | 0.6185914 | 0.9895531 | 1.693998 | 0.09026559 | 0.216099 |
| ENSG00000251396 | LINC01301 | 32.17233 | 0.6185411 | 0.3766131 | 1.860209 | 0.06285597 | 0.164599 |
| ENSG00000250312 | ZNF718 | 660.5851 | 0.6184534 | 0.1939307 | 3.288587 | 0.00100692 | 0.005755 |
| ENSG00000258613 | RPL21P7 | 2.097035 | 0.6182709 | 0.909763 | 1.417375 | 0.1563733 | 0.320838 |
| ENSG00000265808 | SEC22B | 6580.603 | 0.6182374 | 0.09293349 | 6.696837 | 2.13E-11 | 5.81E-10 |
| ENSG00000230524 | COL6A4P1 | 44.37534 | 0.6181051 | 0.3672044 | 1.897454 | 0.05776803 | 0.154644 |
| ENSG00000104826 | LHB | 18.51134274 | 0.617982015 | 0.55606933 | 1.470423916 | 0.14144698 | 0.298357 |
| ENSG00000236140 | | 8.577422 | 0.6179286 | 0.6190812 | 1.402827 | 0.1606685 | 0.327262 |
| ENSG00000274070 | CASTOR2 | 864.8723 | 0.6176918 | 0.2459992 | 2.648587 | 0.00808291 | 0.032802 |
| ENSG00000269993 | | 19.59804 | 0.6175865 | 0.6240727 | 1.397675 | 0.1622106 | 0.329478 |
| ENSG00000271803 | | 3.412534 | 0.6174983 | 0.8531097 | 1.412134 | 0.1579104 | 0.323133 |
| ENSG00000099889 | ARVCF | 1369.436853 | 0.617382499 | 0.179835904 | 3.531544598 | 0.00041314 | 0.002662 |
| ENSG00000128564 | VGF | 382.9630224 | 0.617017695 | 0.329369165 | 2.079154253 | 0.03760318 | 0.111431 |
| ENSG00000157036 | EXOG | 513.6386 | 0.6167764 | 0.1823578 | 3.486745 | 0.00048894 | 0.00308 |
| ENSG00000286608 | | 12.35209 | 0.6165385 | 0.7592232 | 1.379755 | 0.1676622 | 0.33719 |
| ENSG00000260082 | | 2.663548 | 0.6161776 | 0.8738639 | 1.409803 | 0.1585977 | 0.324312 |
| ENSG00000243449 | C4orf48 | 382.0142 | 0.6160758 | 0.3057726 | 2.187263 | 0.02872335 | 0.090625 |
| ENSG00000214827 | MTCP1 | 109.9459 | 0.6160387 | 0.2813584 | 2.341937 | 0.01918397 | 0.065708 |
| ENSG00000248138 | | 31.3378 | 0.6157726 | 0.3672628 | 1.89329 | 0.05831928 | 0.1559 |
| ENSG00000253921 | | 2.346111 | 0.6157716 | 0.9571742 | 1.524432 | 0.1274009 | 0.276704 |
| ENSG00000160796 | NBEAL2 | 15577.78 | 0.6157036 | 0.2316471 | 2.862375 | 0.00420479 | 0.019229 |
| ENSG00000137767 | SQOR | 404.2615161 | 0.615690324 | 0.229274639 | 2.816609169 | 0.00485336 | 0.021658 |
| ENSG00000259116 | | 43.7403 | 0.615486 | 0.5504224 | 1.461367 | 0.1439147 | 0.301956 |

| | | | | | | | |
|---|---|---|---|---|---|---|---|
| ENSG00000116748 | AMPD1 | 94.0184976 | 0.61545834 | 0.395638212 | 1.786157026 | 0.07407384 | 0.186529 |
| ENSG00000282508 | LINC01002 | 24.59141 | 0.6150516 | 0.42278 | 1.70436 | 0.0883139 | 0.212653 |
| ENSG00000047056 | WDR37 | 2969.38804 | 0.614974799 | 0.155268597 | 4.031196958 | 5.55E-05 | 0.000454 |
| ENSG00000125775 | SDCBP2 | 55.84708613 | 0.614894967 | 0.36411707 | 1.893192416 | 0.05833229 | 0.155913 |
| ENSG00000125351 | UPF3B | 2070.765091 | 0.614840223 | 0.197779195 | 3.232373896 | 0.00122766 | 0.006831 |
| ENSG00000005108 | THSD7A | 60.28830954 | 0.614545188 | 0.82941939 | 1.400729014 | 0.16129512 | 0.328161 |
| ENSG00000122779 | TRIM24 | 6057.673527 | 0.614532486 | 0.194491498 | 3.287259933 | 0.00101167 | 0.005779 |
| ENSG00000272564 | | 5.359597 | 0.6145065 | 1.015866 | 1.636183 | 0.1018014 | 0.235706 |
| ENSG00000109323 | MANBA | 2211.936619 | 0.614242177 | 0.24777725 | 2.618163332 | 0.00884045 | 0.035257 |
| ENSG00000213762 | ZNF134 | 1327.579 | 0.6140945 | 0.1557833 | 4.034404 | 5.47E-05 | 0.000448 |
| ENSG00000146063 | TRIM41 | 4876.899 | 0.6138465 | 0.1088402 | 5.710756 | 1.12E-08 | 2.04E-07 |
| ENSG00000091317 | CMTM6 | 3352.763192 | 0.613681699 | 0.114031373 | 5.452577929 | 4.96E-08 | 8.00E-07 |
| ENSG00000120437 | ACAT2 | 2290.193791 | 0.613367702 | 0.508709005 | 1.514987726 | 0.12977551 | 0.280394 |
| ENSG00000256433 | | 57.44296 | 0.6130622 | 0.3935206 | 1.789727 | 0.07349775 | 0.18543 |
| ENSG00000180730 | SHISA2 | 2410.748 | 0.6128244 | 0.440079 | 1.653588 | 0.09821121 | 0.229702 |
| ENSG00000254480 | LINC02749 | 14.60251 | 0.6127998 | 0.5647436 | 1.429697 | 0.1528039 | 0.315254 |
| ENSG00000183010 | PYCR1 | 13163.69 | 0.612777 | 0.2112185 | 3.065898 | 0.00217017 | 0.011105 |
| ENSG00000236811 | GAPDHP2 | 2.810148 | 0.6127302 | 0.8581493 | 1.384734 | 0.1661339 | 0.33506 |
| ENSG00000142973 | CYP4B1 | 496.0442 | 0.612694 | 0.321904 | 2.076577 | 0.03784064 | 0.111982 |
| ENSG00000130762 | ARHGEF16 | 2571.432284 | 0.612678974 | 0.165748495 | 3.777219254 | 0.00015859 | 0.001154 |
| ENSG00000279859 | | 4.190352 | 0.6124365 | 0.7644114 | 1.349928 | 0.177039 | 0.35019 |
| ENSG00000073067 | CYP2W1 | 59.97775349 | 0.612397489 | 0.317752448 | 2.102128919 | 0.03554198 | 0.106835 |
| ENSG00000108846 | ABCC3 | 13705.51316 | 0.612209856 | 0.317992226 | 2.109406821 | 0.03490948 | 0.105435 |
| ENSG00000279344 | | 5.251654 | 0.6119754 | 0.7676355 | 1.357455 | 0.1746368 | 0.346523 |
| ENSG00000116729 | WLS | 2190.279279 | 0.611662222 | 0.446734066 | 1.631432285 | 0.10279914 | 0.237417 |
| ENSG00000117036 | ETV3 | 2575.935186 | 0.61148369 | 0.199869769 | 3.168812219 | 0.00153063 | 0.008251 |
| ENSG00000200502 | | 5.569052 | 0.6114364 | 0.7548671 | 1.338019 | 0.1808902 | 0.3554 |
| ENSG00000221890 | NPTXR | 2420.608 | 0.6109674 | 0.1928343 | 3.277576 | 0.00104703 | 0.005959 |
| ENSG00000186205 | MTARC1 | 1064.806 | 0.6105774 | 0.3248121 | 2.060491 | 0.0393516 | 0.115477 |
| ENSG00000253354 | | 1.316155 | 0.6104367 | 1.126168 | 2.081604 | 0.03737863 | NA |
| ENSG00000141437 | SLC25A52 | 2.090516 | 0.6103278 | 0.892757 | 1.406629 | 0.1595374 | 0.325632 |
| ENSG00000188488 | SERPINA5 | 9.862384 | 0.6101163 | 0.7104901 | 1.333457 | 0.182382 | 0.357353 |
| ENSG00000279075 | | 5.931154 | 0.6100313 | 0.7294163 | 1.342218 | 0.1795253 | 0.353607 |

| | | | | | | | |
|---|---|---|---|---|---|---|---|
| ENSG00000168826 | ZBTB49 | 249.8255 | 0.6100246 | 0.1790501 | 3.49819 | 0.00046843 | 0.002967 |
| ENSG00000214243 | | 10.33083 | 0.6097692 | 0.6005375 | 1.40672 | 0.1595106 | 0.325602 |
| ENSG00000149761 | NUDT22 | 1301.495 | 0.6096342 | 0.1860676 | 3.374857 | 0.00073854 | 0.004402 |
| ENSG00000248265 | | 16.44719 | 0.6092663 | 0.5079289 | 1.49801 | 0.1341308 | 0.287072 |
| ENSG00000235806 | | 12.19594 | 0.6091119 | 0.6958175 | 1.348787 | 0.1774053 | 0.350653 |
| ENSG00000147166 | ITGB1BP2 | 50.78811 | 0.6088986 | 0.4227435 | 1.694729 | 0.09012693 | 0.215827 |
| ENSG00000245711 | NADK2-AS1 | 32.18747 | 0.6088303 | 0.4892832 | 1.540465 | 0.1234471 | 0.270817 |
| ENSG00000127526 | SLC35E1 | 3426.84034 | 0.608807569 | 0.16100291 | 3.798076954 | 0.00014582 | 0.00107 |
| ENSG00000118523 | CCN2 | 5007.210015 | 0.608780562 | 0.805154604 | 1.385897411 | 0.16577823 | 0.334439 |
| ENSG00000230023 | LINC02800 | 3.047508 | 0.6087249 | 1.036972 | 1.71193 | 0.0869095 | 0.210033 |
| ENSG00000076662 | ICAM3 | 0.936274287 | 0.608675998 | 1.071996919 | 1.806168217 | 0.07089207 | NA |
| ENSG00000273145 | | 101.0483 | 0.6085272 | 0.2884471 | 2.276339 | 0.02282571 | 0.075571 |
| ENSG00000211688 | TRGJP2 | 1.176104 | 0.6082924 | 1.096361 | 1.938046 | 0.05261766 | NA |
| ENSG00000284708 | | 2.494678 | 0.6079213 | 1.039814 | 1.691296 | 0.09078019 | 0.21696 |
| ENSG00000259793 | | 3.157348 | 0.6078999 | 0.8301029 | 1.377318 | 0.1684139 | 0.338215 |
| ENSG00000279973 | | 13.83217 | 0.6078799 | 0.7170629 | 1.320984 | 0.1865068 | 0.362786 |
| ENSG00000257179 | HMGN2P6 | 1.415501 | 0.6078062 | 1.078497 | 1.884735 | 0.05946568 | NA |
| ENSG00000273381 | | 37.14053 | 0.6076005 | 0.5185112 | 1.488124 | 0.1367181 | 0.290999 |
| ENSG00000144130 | NT5DC4 | 6.546191 | 0.6075084 | 0.7797533 | 1.34923 | 0.1772631 | 0.350477 |
| ENSG00000074621 | SLC24A1 | 1233.847368 | 0.607327094 | 0.209465236 | 3.010816713 | 0.00260546 | 0.012954 |
| ENSG00000167769 | ACER1 | 1.398646 | 0.6072871 | 0.9829948 | 1.569485 | 0.116535 | NA |
| ENSG00000133246 | PRAM1 | 8.902410913 | 0.607183577 | 0.624214589 | 1.364582059 | 0.17238445 | 0.343515 |
| ENSG00000283141 | LINC02666 | 9.902308 | 0.6071585 | 0.5856755 | 1.395147 | 0.1629716 | 0.330534 |
| ENSG00000099864 | PALM | 2145.416294 | 0.607124667 | 0.245147472 | 2.615418603 | 0.00891182 | 0.035472 |
| ENSG00000282826 | FRG1CP | 51.59764 | 0.6069772 | 0.3721085 | 1.838752 | 0.06595168 | 0.170703 |
| ENSG00000198960 | ARMCX6 | 1424.58 | 0.6069266 | 0.1479524 | 4.180335 | 2.91E-05 | 0.000256 |
| ENSG00000158258 | CLSTN2 | 994.9398 | 0.6067925 | 0.4243702 | 1.673591 | 0.09421093 | 0.223033 |
| ENSG00000227173 | MYL6P3 | 4.462888 | 0.6065299 | 0.7602588 | 1.343382 | 0.1791482 | 0.353021 |
| ENSG00000226148 | SLC25A39P1 | 4.223376 | 0.6064796 | 0.7141852 | 1.327021 | 0.1845017 | 0.359808 |
| ENSG00000278374 | | 4.670084 | 0.6064467 | 0.7536892 | 1.357505 | 0.1746208 | 0.346523 |
| ENSG00000166946 | CCNDBP1 | 1914.877 | 0.6063425 | 0.1312284 | 4.683539 | 2.82E-06 | 3.20E-05 |
| ENSG00000087303 | NID2 | 11.27859274 | 0.606326201 | 0.514921953 | 1.484643251 | 0.13763835 | 0.292249 |
| ENSG00000253878 | | 73.76463 | 0.6062777 | 0.3957649 | 1.756708 | 0.07896752 | 0.195262 |

| | | | | | | | |
|---|---|---|---|---|---|---|---|
| ENSG00000120306 | CYSTM1 | 2213.903613 | 0.606190211 | 0.204644199 | 3.085233924 | 0.00203392 | 0.010513 |
| ENSG00000101605 | MYOM1 | 26.79596239 | 0.606055481 | 0.491974584 | 1.518126747 | 0.12898245 | 0.27918 |
| ENSG00000231966 | LINC02818 | 1.445128 | 0.6060327 | 0.9735779 | 1.523727 | 0.1275768 | NA |
| ENSG00000205765 | C5orf51 | 4087.189 | 0.6059613 | 0.2072029 | 3.039198 | 0.00237209 | 0.011961 |
| ENSG00000143702 | CEP170 | 1328.465 | 0.6058366 | 0.2592132 | 2.486142 | 0.01291364 | 0.047813 |
| ENSG00000266963 | | 3.658583 | 0.6057787 | 0.7187326 | 1.335306 | 0.1817762 | 0.356534 |
| ENSG00000239653 | PSMD6-AS2 | 270.5685 | 0.60576 | 0.2787987 | 2.32465 | 0.02009069 | 0.068216 |
| ENSG00000287617 | | 4.771619 | 0.6055649 | 0.7490612 | 1.345368 | 0.1785063 | 0.352201 |
| ENSG00000158856 | DMTN | 857.0413 | 0.6055182 | 0.2086348 | 3.019796 | 0.00252945 | 0.012627 |
| ENSG00000231768 | LINC01354 | 2.062787 | 0.6053986 | 1.037564 | 1.669866 | 0.09494584 | 0.224113 |
| ENSG00000249599 | BMPR1B-DT | 1.244885 | 0.6046002 | 1.068617 | 1.809317 | 0.07040173 | NA |
| ENSG00000090238 | YPEL3 | 3579.326821 | 0.604502252 | 0.267932092 | 2.41565964 | 0.01570674 | 0.055903 |
| ENSG00000058063 | ATP11B | 3055.453674 | 0.604492065 | 0.219848711 | 2.866375594 | 0.00415201 | 0.019046 |
| ENSG00000184347 | SLIT3 | 40.51205 | 0.6044008 | 0.7509465 | 1.349652 | 0.1771278 | 0.350287 |
| ENSG00000277235 | | 4.604328 | 0.604307 | 0.8836027 | 1.420892 | 0.1553481 | 0.319239 |
| ENSG00000276952 | | 36.14089 | 0.604234 | 0.3272893 | 2.034595 | 0.04189167 | 0.12124 |
| ENSG00000267571 | | 41.28408 | 0.6042019 | 0.334441 | 1.991474 | 0.04642879 | 0.131008 |
| ENSG00000143793 | C1orf35 | 2483.039 | 0.6041066 | 0.1894231 | 3.291247 | 0.00099744 | 0.005709 |
| ENSG00000272150 | NBPF25P | 177.288 | 0.6039735 | 0.2297805 | 2.75454 | 0.00587747 | 0.025305 |
| ENSG00000182670 | TTC3 | 14371.03 | 0.6039732 | 0.2014499 | 3.102582 | 0.0019184 | 0.009996 |
| ENSG00000226179 | LINC00685 | 39.55898 | 0.6038272 | 0.5497241 | 1.436509 | 0.1508575 | 0.312379 |
| ENSG00000125879 | OTOR | 1.621855266 | 0.603757482 | 0.988958908 | 1.646405339 | 0.09968034 | 0.232255 |
| ENSG00000165389 | SPTSSA | 4052.841 | 0.6037503 | 0.1544206 | 3.994828 | 6.47E-05 | 0.00052 |
| ENSG00000196859 | KRT39 | 1.230737 | 0.6037155 | 0.9688346 | 1.594147 | 0.1109032 | NA |
| ENSG00000225171 | DUTP6 | 8.993944 | 0.6036938 | 0.6216355 | 1.371162 | 0.1703246 | 0.340919 |
| ENSG00000110427 | KIAA1549L | 24.30023214 | 0.603548044 | 0.614754843 | 1.374176442 | 0.16938691 | 0.33963 |
| ENSG00000230592 | RPSAP8 | 3.07181 | 0.6034131 | 0.8920219 | 1.395197 | 0.1629565 | 0.330528 |
| ENSG00000152439 | ZNF773 | 117.3466 | 0.6028345 | 0.2529788 | 2.517495 | 0.01181926 | 0.044492 |
| ENSG00000285294 | LINC00842 | 51.06884 | 0.6027975 | 0.885118 | 1.436613 | 0.150828 | 0.312342 |
| ENSG00000228917 | NECTIN4-AS1 | 4.491866 | 0.6026275 | 0.8157536 | 1.348554 | 0.1774803 | 0.350751 |
| ENSG00000205106 | LINC02716 | 1.576828 | 0.6025089 | 1.073796 | 1.846381 | 0.06483694 | 0.168343 |
| ENSG00000185669 | SNAI3 | 163.293 | 0.6024126 | 0.3017162 | 2.1615 | 0.03065673 | 0.095366 |
| ENSG00000250506 | CDK3 | 58.8564 | 0.6023419 | 0.2879582 | 2.253211 | 0.02424587 | 0.079207 |

| | | | | | | | |
|---|---|---|---|---|---|---|---|
| ENSG00000280303 | ERICD | 110.19 | 0.602151 | 0.3172575 | 2.072139 | 0.03825253 | 0.112951 |
| ENSG00000197006 | METTL9 | 6073.73 | 0.6020204 | 0.1859146 | 3.336064 | 0.00084974 | 0.004967 |
| ENSG00000166471 | TMEM41B | 1875.347 | 0.6019753 | 0.1488435 | 4.126853 | 3.68E-05 | 0.000315 |
| ENSG00000279878 | | 53.17168 | 0.6019752 | 0.4601109 | 1.583561 | 0.1132937 | 0.254549 |
| ENSG00000235605 | | 3.636403 | 0.6019722 | 0.8232725 | 1.382068 | 0.1669509 | 0.336192 |
| ENSG00000176092 | CRYBG2 | 100.3382 | 0.6018663 | 0.3395889 | 1.960562 | 0.04993012 | 0.138331 |
| ENSG00000156928 | MALSU1 | 1235.25 | 0.6016061 | 0.2388314 | 2.632021 | 0.00848787 | 0.034112 |
| ENSG00000203799 | CCDC162P | 737.8621 | 0.6014209 | 0.3864304 | 1.774868 | 0.07591967 | 0.189721 |
| ENSG00000239218 | RPS20P22 | 8.314112 | 0.6013756 | 0.7101176 | 1.335697 | 0.1816484 | 0.356389 |
| ENSG00000126460 | PRRG2 | 607.9074186 | 0.601346264 | 0.193830075 | 3.20376922 | 0.00135641 | 0.007448 |
| ENSG00000169241 | SLC50A1 | 6610.252 | 0.6011019 | 0.1730635 | 3.588094 | 0.0003331 | 0.002218 |
| ENSG00000198835 | GJC2 | 25.01829 | 0.6009787 | 0.5089436 | 1.457362 | 0.1450164 | 0.303788 |
| ENSG00000241404 | EGFL8 | 111.8058 | 0.6007182 | 0.2920804 | 2.216128 | 0.02668273 | 0.085362 |
| ENSG00000185442 | FAM174B | 4105.652 | 0.6005913 | 0.2448074 | 2.590149 | 0.00959343 | 0.037564 |
| ENSG00000258878 | | 1.614066 | 0.6005802 | 1.046536 | 1.744715 | 0.08103445 | 0.199278 |
| ENSG00000248187 | | 1.560628 | 0.6001384 | 0.9889396 | 1.59172 | 0.1114477 | NA |
| ENSG00000100376 | FAM118A | 2511.472411 | 0.600123153 | 0.210824717 | 2.960944403 | 0.00306697 | 0.0148 |
| ENSG00000125864 | BFSP1 | 201.217478 | 0.600053864 | 0.341911181 | 1.954923292 | 0.05059213 | 0.13975 |
| ENSG00000204147 | ASAH2B | 624.6191 | 0.6000446 | 0.191417 | 3.234737 | 0.00121755 | 0.006782 |
| ENSG00000162415 | ZSWIM5 | 1129.325 | 0.5999679 | 0.2289391 | 2.742797 | 0.00609183 | 0.026034 |
| ENSG00000142530 | FAM71E1 | 118.8518 | 0.5999286 | 0.2246258 | 2.788109 | 0.00530167 | 0.023233 |
| ENSG00000220960 | | 47.71006 | 0.5997435 | 0.4129792 | 1.688505 | 0.09131429 | 0.217869 |
| ENSG00000230847 | OCLNP1 | 16.01329 | 0.599667 | 0.5240051 | 1.451199 | 0.1467245 | 0.306279 |
| ENSG00000101977 | MCF2 | 11.2508879 | 0.599601956 | 0.678719516 | 1.338583998 | 0.18070614 | 0.35517 |
| ENSG00000116035 | VAX2 | 163.9503693 | 0.599492983 | 0.309491666 | 2.109558617 | 0.03489639 | 0.105407 |
| ENSG00000260793 | | 438.2724 | 0.5994067 | 0.2846386 | 2.254729 | 0.02415037 | 0.078943 |
| ENSG00000255500 | | 9.095056 | 0.5992707 | 0.6376089 | 1.357884 | 0.1745005 | 0.346432 |
| ENSG00000240520 | UOX | 0.960653 | 0.5992704 | 1.080785 | 1.914899 | 0.05550532 | NA |
| ENSG00000235084 | CHCHD2P6 | 242.2642 | 0.5992009 | 0.2879944 | 2.240195 | 0.02507826 | 0.081326 |
| ENSG00000119698 | PPP4R4 | 16.2943562 | 0.599154845 | 0.725019348 | 1.318236324 | 0.18742455 | 0.364118 |
| ENSG00000125733 | TRIP10 | 2702.638096 | 0.599064448 | 0.156893549 | 3.901759891 | 9.55E-05 | 0.000735 |
| ENSG00000250722 | SELENOP | 604.8159 | 0.5988249 | 0.6152993 | 1.364461 | 0.1724226 | 0.343541 |
| ENSG00000120910 | PPP3CC | 522.9652777 | 0.598796212 | 0.192158448 | 3.205186258 | 0.00134975 | 0.007419 |

| | | | | | | | |
|---|---|---|---|---|---|---|---|
| ENSG00000164114 | MAP9 | 1605.543 | 0.5987202 | 0.2126855 | 2.941678 | 0.00326439 | 0.015582 |
| ENSG00000279330 | | 14.76173 | 0.5986174 | 0.4737182 | 1.522717 | 0.1278294 | 0.277385 |
| ENSG00000165548 | TMEM63C | 1285.832 | 0.5983257 | 0.3471115 | 1.916762 | 0.05526811 | 0.149623 |
| ENSG00000267681 | | 22.11769 | 0.59829 | 0.4729115 | 1.539411 | 0.1237039 | 0.271246 |
| ENSG00000160190 | SLC37A1 | 2810.628 | 0.5978639 | 0.1826589 | 3.355234 | 0.00079298 | 0.004684 |
| ENSG00000151240 | DIP2C | 3861.287 | 0.5976638 | 0.2207166 | 2.843634 | 0.00446023 | 0.020164 |
| ENSG00000205352 | PRR13 | 3279.366 | 0.5975365 | 0.1952311 | 3.165242 | 0.00154954 | 0.008336 |
| ENSG00000275131 | PDE4DIPP2 | 818.5466 | 0.5975107 | 0.3643665 | 1.859244 | 0.06299262 | 0.164888 |
| ENSG00000175611 | LINC00476 | 307.9186 | 0.5974549 | 0.2108703 | 2.94324 | 0.00324797 | 0.015521 |
| ENSG00000260615 | RPL23AP97 | 4.156658 | 0.5973268 | 0.8427109 | 1.370271 | 0.1706024 | 0.341295 |
| ENSG00000075884 | ARHGAP15 | 4.121971424 | 0.597321517 | 0.786997137 | 1.316481928 | 0.18801234 | 0.364967 |
| ENSG00000283240 | | 1.868167 | 0.5972723 | 0.9407621 | 1.572555 | 0.115822 | 0.258649 |
| ENSG00000116176 | TPSG1 | 29.34207857 | 0.597090794 | 0.564655121 | 1.414587146 | 0.15718958 | 0.322189 |
| ENSG00000273604 | EPOP | 1497.167 | 0.5969157 | 0.2320404 | 2.704335 | 0.00684413 | 0.028609 |
| ENSG00000144711 | IQSEC1 | 6643.118 | 0.596867 | 0.1432038 | 4.258527 | 2.06E-05 | 0.000187 |
| ENSG00000167987 | VPS37C | 3041.862 | 0.5967513 | 0.1082622 | 5.556531 | 2.75E-08 | 4.62E-07 |
| ENSG00000137171 | KLC4 | 1966.690617 | 0.596732844 | 0.225016169 | 2.772353914 | 0.00556525 | 0.024188 |
| ENSG00000187239 | FNBP1 | 3103.664 | 0.5966978 | 0.1335967 | 4.523927 | 6.07E-06 | 6.35E-05 |
| ENSG00000201451 | | 3.034014 | 0.5966843 | 0.9025524 | 1.430634 | 0.1525351 | 0.314871 |
| ENSG00000073803 | MAP3K13 | 1190.700256 | 0.596615342 | 0.13950716 | 4.349521289 | 1.36E-05 | 0.00013 |
| ENSG00000231989 | PPP1R2B | 167.2404 | 0.5966016 | 0.2372987 | 2.637118 | 0.00836137 | 0.0337 |
| ENSG00000249773 | | 2.78942 | 0.5964913 | 0.9163873 | 1.480618 | 0.1387083 | 0.294051 |
| ENSG00000121289 | CEP89 | 2068.97826 | 0.596370882 | 0.144197385 | 4.212225568 | 2.53E-05 | 0.000226 |
| ENSG00000132326 | PER2 | 1008.39368 | 0.596333399 | 0.204490509 | 3.014931187 | 0.00257038 | 0.012801 |
| ENSG00000233621 | LINC01137 | 228.3882 | 0.5960555 | 0.2442695 | 2.566521 | 0.01027244 | 0.039745 |
| ENSG00000150054 | MPP7 | 2080.758 | 0.5959396 | 0.216662 | 2.868603 | 0.00412289 | 0.018932 |
| ENSG00000042813 | ZPBP | 2.069730777 | 0.595688372 | 0.906844331 | 1.402394099 | 0.1607976 | 0.327324 |
| ENSG00000273403 | | 3.831164 | 0.5956584 | 0.8984689 | 1.414236 | 0.1572926 | 0.322362 |
| ENSG00000240991 | RPL23AP67 | 1.935621 | 0.5954637 | 0.9686037 | 1.608532 | 0.1077187 | 0.245579 |
| ENSG00000147697 | GSDMC | 13.80946 | 0.5950733 | 0.7486465 | 1.305053 | 0.1918747 | 0.370153 |
| ENSG00000249425 | LINC02477 | 9.781664 | 0.5950221 | 0.6703765 | 1.327386 | 0.1843809 | 0.359652 |
| ENSG00000213695 | RPS7P14 | 2.731842 | 0.5949564 | 0.8123531 | 1.33537 | 0.1817554 | 0.356534 |
| ENSG00000219451 | RPL23P8 | 302.4391 | 0.5947761 | 0.2448003 | 2.562867 | 0.01038118 | 0.040107 |

| | | | | | | | |
|---|---|---|---|---|---|---|---|
| ENSG00000008517 | IL32 | 28.33088096 | 0.594769965 | 0.679816897 | 1.338403682 | 0.18076488 | 0.355206 |
| ENSG00000065308 | TRAM2 | 2823.314883 | 0.594586218 | 0.21783776 | 2.847920452 | 0.00440059 | 0.019944 |
| ENSG00000235831 | BHLHE40-AS1 | 51.30556 | 0.5942975 | 0.3456031 | 1.905484 | 0.05671717 | 0.152562 |
| ENSG00000197124 | ZNF682 | 250.284 | 0.5942639 | 0.2243236 | 2.770845 | 0.00559111 | 0.024281 |
| ENSG00000197933 | ZNF823 | 376.1496 | 0.5941463 | 0.1653639 | 3.677258 | 0.00023576 | 0.00164 |
| ENSG00000287336 | | 1.439749 | 0.5940713 | 0.9708732 | 1.650692 | 0.09880156 | NA |
| ENSG00000070476 | ZXDC | 3473.882288 | 0.594049161 | 0.143226487 | 4.190765553 | 2.78E-05 | 0.000246 |
| ENSG00000018610 | CXorf56 | 1941.743684 | 0.593879423 | 0.181477172 | 3.378573027 | 0.00072863 | 0.004347 |
| ENSG00000053918 | KCNQ1 | 27.75084973 | 0.593766681 | 0.455512073 | 1.55967084 | 0.11883769 | 0.26327 |
| ENSG00000089916 | GPATCH2L | 4132.568738 | 0.593707647 | 0.28105118 | 2.267051967 | 0.02338705 | 0.076946 |
| ENSG00000276408 | | 12.14248 | 0.5936411 | 0.6391293 | 1.337902 | 0.1809285 | 0.355449 |
| ENSG00000237017 | | 48.17595 | 0.5936405 | 0.3341111 | 1.955666 | 0.05050454 | 0.139573 |
| ENSG00000234911 | TEX21P | 37.48347 | 0.5935144 | 0.344218 | 1.919353 | 0.05493974 | 0.149002 |
| ENSG00000186787 | SPIN2B | 298.5038 | 0.5929913 | 0.2128929 | 2.906413 | 0.00365599 | 0.017126 |
| ENSG00000055208 | TAB2 | 6252.475484 | 0.592933059 | 0.10136498 | 5.898444588 | 3.67E-09 | 7.24E-08 |
| ENSG00000115844 | DLX2 | 317.8745619 | 0.592711008 | 0.406264335 | 1.691424008 | 0.09075585 | 0.216936 |
| ENSG00000160752 | FDPS | 7334.074 | 0.5925269 | 0.2975993 | 2.152424 | 0.03136399 | 0.097134 |
| ENSG00000154305 | MIA3 | 4877.075 | 0.5925179 | 0.2048224 | 3.00433 | 0.00266166 | 0.013181 |
| ENSG00000226986 | PRELID1P5 | 115.9586 | 0.5922395 | 0.3108836 | 2.082278 | 0.03731706 | 0.110817 |
| ENSG00000196436 | NPIPB15 | 284.81 | 0.5919607 | 0.2936818 | 2.174656 | 0.02965593 | 0.092938 |
| ENSG00000157703 | SVOPL | 97.68703 | 0.5917912 | 0.457658 | 1.554295 | 0.120114 | 0.2655 |
| ENSG00000274737 | | 9.062143 | 0.5916245 | 0.6574938 | 1.323027 | 0.1858265 | 0.361749 |
| ENSG00000106006 | HOXA6 | 8.147748154 | 0.591589827 | 0.615512368 | 1.34342651 | 0.17913391 | 0.353021 |
| ENSG00000180211 | | 303.3056 | 0.5914946 | 0.2571081 | 2.437208 | 0.01480118 | 0.053373 |
| ENSG00000255438 | | 106.1047 | 0.59145 | 0.2715419 | 2.319376 | 0.02037464 | 0.068969 |
| ENSG00000274723 | | 2.875953 | 0.5911208 | 0.9488922 | 1.502776 | 0.1328969 | 0.285259 |
| ENSG00000185808 | PIGP | 816.52 | 0.5908819 | 0.1920679 | 3.174707 | 0.00149988 | 0.008112 |
| ENSG00000279838 | | 36.84088 | 0.5908801 | 0.5848797 | 1.371318 | 0.1702759 | 0.340847 |
| ENSG00000166398 | GARRE1 | 2007.751 | 0.5906955 | 0.3200921 | 2.024198 | 0.04294974 | 0.123637 |
| ENSG00000273486 | | 121.5863 | 0.5906441 | 0.3039251 | 2.10988 | 0.03486866 | 0.105336 |
| ENSG00000180481 | GLIPR1L2 | 9.660939 | 0.5903319 | 0.5205974 | 1.443697 | 0.1488242 | 0.309398 |
| ENSG00000234235 | BOK-AS1 | 5.383652 | 0.590087 | 0.8234827 | 1.3495 | 0.1771764 | 0.350357 |
| ENSG00000248919 | ATP5MF-PTCD1 | 3.849726 | 0.5900433 | 0.8659005 | 1.356634 | 0.1748975 | 0.346858 |

| | | | | | | | |
|---|---|---|---|---|---|---|---|
| ENSG00000272588 | | 20.66904 | 0.5899848 | 0.5526094 | 1.406517 | 0.1595707 | 0.325672 |
| ENSG00000159267 | HLCS | 1893.988 | 0.5899767 | 0.2096668 | 2.918611 | 0.00351594 | 0.016581 |
| ENSG00000286915 | | 2.458242 | 0.5898817 | 0.8735474 | 1.3835 | 0.1665117 | 0.335608 |
| ENSG00000185630 | PBX1 | 9354.912 | 0.5897121 | 0.1800239 | 3.360693 | 0.00077747 | 0.004606 |
| ENSG00000233509 | ZNF197-AS1 | 1.074754 | 0.58923 | 1.042443 | 1.706994 | 0.08782316 | NA |
| ENSG00000075151 | EIF4G3 | 7020.763982 | 0.589191184 | 0.147970966 | 4.050146604 | 5.12E-05 | 0.000423 |
| ENSG00000019485 | PRDM11 | 526.2630296 | 0.589159987 | 0.201198802 | 3.036110579 | 0.00239652 | 0.012067 |
| ENSG00000230637 | | 6.067119 | 0.5891591 | 0.6096527 | 1.33733 | 0.1811149 | 0.355657 |
| ENSG00000261512 | | 167.9249 | 0.5891154 | 0.2235543 | 2.753326 | 0.00589931 | 0.025387 |
| ENSG00000258944 | | 3.94698 | 0.5891121 | 0.8322992 | 1.357468 | 0.1746327 | 0.346523 |
| ENSG00000163931 | TKT | 22947 | 0.5890152 | 0.2732638 | 2.288257 | 0.02212254 | 0.07385 |
| ENSG00000226745 | OAZ1P1 | 2.284499 | 0.5889577 | 1.04876 | 1.80784 | 0.07063135 | 0.179839 |
| ENSG00000118495 | PLAGL1 | 272.250043 | 0.588915296 | 0.259646338 | 2.41396098 | 0.01578015 | 0.056111 |
| ENSG00000108785 | HSD17B1P1 | 87.4967395 | 0.588751679 | 0.456840048 | 1.548072883 | 0.12160475 | 0.267859 |
| ENSG00000100139 | MICALL1 | 1881.594551 | 0.588693071 | 0.166660163 | 3.621786745 | 0.00029258 | 0.001979 |
| ENSG00000155275 | TRMT44 | 574.835 | 0.5886753 | 0.1995352 | 3.051637 | 0.00227597 | 0.011545 |
| ENSG00000166128 | RAB8B | 764.2414 | 0.5885332 | 0.2048328 | 2.962707 | 0.00304947 | 0.014736 |
| ENSG00000283050 | GTF2IP12 | 694.9488 | 0.5885102 | 0.250907 | 2.478568 | 0.01319109 | 0.048603 |
| ENSG00000173011 | TADA2B | 1670.066 | 0.5883883 | 0.1448695 | 4.123007 | 3.74E-05 | 0.00032 |
| ENSG00000092445 | TYRO3 | 1967.497723 | 0.588330404 | 0.405239592 | 1.681203573 | 0.09272338 | 0.220255 |
| ENSG00000262580 | | 430.4567 | 0.5882225 | 0.4409892 | 1.589324 | 0.1119873 | 0.25258 |
| ENSG00000146072 | TNFRSF21 | 2582.252 | 0.5880875 | 0.2966189 | 2.166879 | 0.03024407 | 0.094237 |
| ENSG00000259727 | | 3.975718 | 0.5880761 | 0.732752 | 1.302561 | 0.1927246 | 0.371185 |
| ENSG00000002330 | BAD | 1102.684304 | 0.588062996 | 0.248026025 | 2.501996533 | 0.01234951 | 0.046071 |
| ENSG00000254231 | | 318.9127 | 0.587945 | 0.350471 | 1.860938 | 0.062753 | 0.164443 |
| ENSG00000070214 | SLC44A1 | 12736.34819 | 0.58791459 | 0.159581263 | 3.781580135 | 0.00015584 | 0.001137 |
| ENSG00000273523 | | 3.677859 | 0.5878427 | 0.7021934 | 1.301803 | 0.1929838 | 0.371496 |
| ENSG00000224807 | DUX4L9 | 1.238366 | 0.5878382 | 0.990153 | 1.596111 | 0.110464 | NA |
| ENSG00000180488 | MIGA1 | 1157.795 | 0.5876412 | 0.3179529 | 2.020878 | 0.04329238 | 0.124377 |
| ENSG00000236833 | | 3.046221 | 0.5875607 | 0.9015862 | 1.409465 | 0.1586977 | 0.324491 |
| ENSG00000214279 | SCART1 | 280.2523 | 0.5872002 | 0.4720443 | 1.517438 | 0.1291561 | 0.279351 |
| ENSG00000164944 | VIRMA | 7802.303 | 0.5869764 | 0.1337146 | 4.455614 | 8.37E-06 | 8.42E-05 |
| ENSG00000286718 | | 9.129016 | 0.5869071 | 0.5653976 | 1.380534 | 0.1674223 | 0.336846 |

| | | | | | | | |
|---|---|---|---|---|---|---|---|
| ENSG00000164663 | USP49 | 756.422 | 0.58678 | 0.2533316 | 2.453295 | 0.01415542 | 0.051478 |
| ENSG00000214184 | GCC2-AS1 | 23.23768 | 0.5864573 | 0.4169885 | 1.641492 | 0.1006953 | 0.233943 |
| ENSG00000204713 | TRIM27 | 6263.409 | 0.5862416 | 0.1088887 | 5.439533 | 5.34E-08 | 8.56E-07 |
| ENSG00000219702 | | 2.102155 | 0.5859636 | 1.032376 | 1.738298 | 0.08215836 | 0.201466 |
| ENSG00000136854 | STXBP1 | 1704.835179 | 0.585861669 | 0.23034094 | 2.662272642 | 0.0077615 | 0.03173 |
| ENSG00000270513 | | 1.698426 | 0.5858118 | 1.065278 | 1.901792 | 0.05719836 | 0.153397 |
| ENSG00000132359 | RAP1GAP2 | 1757.384626 | 0.585739823 | 0.176305945 | 3.41495675 | 0.00063792 | 0.003878 |
| ENSG00000167720 | SRR | 386.3425 | 0.5856577 | 0.2371569 | 2.603539 | 0.00922666 | 0.036474 |
| ENSG00000127084 | FGD3 | 4749.745721 | 0.585633903 | 0.382725779 | 1.745126686 | 0.08096282 | 0.199194 |
| ENSG00000112406 | HECA | 1219.069576 | 0.585628158 | 0.123000038 | 4.832026324 | 1.35E-06 | 1.64E-05 |
| ENSG00000256294 | ZNF225 | 280.1782 | 0.5850896 | 0.3246219 | 1.979982 | 0.04770553 | 0.133659 |
| ENSG00000198121 | LPAR1 | 46.01343 | 0.5850152 | 0.6443965 | 1.329073 | 0.183824 | 0.359131 |
| ENSG00000135406 | PRPH | 175.7445794 | 0.585008698 | 0.435066493 | 1.586534621 | 0.11261809 | 0.253465 |
| ENSG00000224660 | SH3BP5-AS1 | 1122.143 | 0.584911 | 0.2287637 | 2.666987 | 0.00765346 | 0.031383 |
| ENSG00000267336 | EIF4A2P1 | 2.077004 | 0.5848987 | 0.9148006 | 1.444347 | 0.1486413 | 0.309139 |
| ENSG00000244752 | CRYBB2 | 3.941857 | 0.5847856 | 0.7922989 | 1.307041 | 0.1911988 | 0.36921 |
| ENSG00000279317 | | 2.408552 | 0.5840999 | 0.8968838 | 1.390983 | 0.1642306 | 0.332333 |
| ENSG00000007944 | MYLIP | 2761.56376 | 0.584086843 | 0.172985323 | 3.454192497 | 0.00055194 | 0.003422 |
| ENSG00000270052 | | 1.709285 | 0.5839972 | 0.96993 | 1.612322 | 0.1068919 | 0.244197 |
| ENSG00000129437 | KLK14 | 19.26305875 | 0.583991089 | 0.528229993 | 1.41706561 | 0.15646375 | 0.320986 |
| ENSG00000172939 | OXSR1 | 4241.516 | 0.5839156 | 0.1207966 | 4.892078 | 9.98E-07 | 1.24E-05 |
| ENSG00000254317 | | 21.39618 | 0.5837973 | 0.5731847 | 1.374467 | 0.1692968 | 0.339527 |
| ENSG00000230724 | LINC01001 | 35.05078 | 0.5837831 | 0.4343896 | 1.582465 | 0.1135434 | 0.254844 |
| ENSG00000162241 | SLC25A45 | 301.4709 | 0.5837658 | 0.2409593 | 2.553257 | 0.01067206 | 0.041034 |
| ENSG00000168172 | HOOK3 | 1744.546 | 0.5836631 | 0.2537999 | 2.439767 | 0.01469672 | 0.053061 |
| ENSG00000213401 | MAGEA12 | 36.64664 | 0.5836245 | 0.9866861 | 1.475175 | 0.1401656 | 0.296336 |
| ENSG00000220804 | LINC01881 | 153.4877 | 0.5835286 | 0.3448859 | 1.883671 | 0.05960948 | 0.158281 |
| ENSG00000176165 | FOXG1 | 1.975621 | 0.5834625 | 1.019108 | 1.643953 | 0.1001859 | 0.233127 |
| ENSG00000124641 | MED20 | 1553.363374 | 0.583273166 | 0.127868383 | 4.631946063 | 3.62E-06 | 4.00E-05 |
| ENSG00000105855 | ITGB8 | 195.0541883 | 0.582832045 | 0.405443594 | 1.664733814 | 0.09596587 | 0.225679 |
| ENSG00000285712 | | 5.04426 | 0.5828119 | 0.7793983 | 1.297308 | 0.1945251 | 0.373705 |
| ENSG00000234629 | WDR82P1 | 106.7103 | 0.5827003 | 0.2969992 | 2.120197 | 0.03398944 | 0.103314 |
| ENSG00000177932 | ZNF354C | 717.4521 | 0.582571 | 0.3220763 | 1.984532 | 0.04719662 | 0.132569 |

| | | | | | | | |
|---|---|---|---|---|---|---|---|
| ENSG00000175730 | BAK1P1 | 134.4594 | 0.5824025 | 0.2963689 | 2.127074 | 0.03341393 | 0.10202 |
| ENSG00000259151 | CAP2P1 | 2.875911 | 0.5822778 | 0.7951793 | 1.326175 | 0.1847818 | 0.360222 |
| ENSG00000263644 | | 1.62742 | 0.5822689 | 0.9034301 | 1.411554 | 0.1580813 | 0.323355 |
| ENSG00000270966 | | 6.077168 | 0.582188 | 0.6091191 | 1.330002 | 0.1835177 | 0.358705 |
| ENSG00000166126 | AMN | 18.81357 | 0.5821681 | 0.5331169 | 1.412919 | 0.1576796 | 0.322882 |
| ENSG00000169992 | NLGN2 | 4273.155 | 0.5821313 | 0.2515924 | 2.504626 | 0.0122581 | 0.045793 |
| ENSG00000061273 | HDAC7 | 4981.062917 | 0.582108466 | 0.130394933 | 4.555796896 | 5.22E-06 | 5.55E-05 |
| ENSG00000243779 | | 4.178094 | 0.5820476 | 0.7005805 | 1.287612 | 0.197881 | 0.37796 |
| ENSG00000264148 | | 9.9979 | 0.5819693 | 0.5222713 | 1.41667 | 0.1565793 | 0.321173 |
| ENSG00000198406 | BZW1P2 | 3467.932 | 0.5818302 | 0.2617714 | 2.360936 | 0.0182289 | 0.063096 |
| ENSG00000078399 | HOXA9 | 0.867721537 | 0.581689074 | 1.056626816 | 1.828878318 | 0.06741784 | NA |
| ENSG00000232587 | EEF1A1P3 | 4.445785 | 0.5816346 | 0.73663 | 1.285996 | 0.1984444 | 0.37851 |
| ENSG00000174721 | FGFBP3 | 317.5947 | 0.5814749 | 0.2396969 | 2.55797 | 0.01052853 | 0.040581 |
| ENSG00000250746 | | 6.869686 | 0.581409 | 0.5936628 | 1.339704 | 0.1803417 | 0.354629 |
| ENSG00000162755 | KLHDC9 | 356.909 | 0.5813946 | 0.256963 | 2.398033 | 0.0164834 | 0.058124 |
| ENSG00000229083 | PSMA6P2 | 96.25017 | 0.5813762 | 0.2534008 | 2.432093 | 0.01501185 | 0.053957 |
| ENSG00000113369 | ARRDC3 | 2481.162701 | 0.581200179 | 0.375748408 | 1.75917198 | 0.0785483 | 0.194533 |
| ENSG00000100485 | SOS2 | 2972.087171 | 0.581199672 | 0.248465292 | 2.463942325 | 0.01374182 | 0.0502 |
| ENSG00000279464 | | 10.24977 | 0.5811988 | 0.6329485 | 1.341443 | 0.1797766 | 0.353918 |
| ENSG00000099999 | RNF215 | 1134.129825 | 0.581145029 | 0.15266188 | 3.871098125 | 0.00010835 | 0.000821 |
| ENSG00000271122 | | 1209.362 | 0.5811175 | 0.2056401 | 2.953001 | 0.00314701 | 0.015112 |
| ENSG00000117676 | RPS6KA1 | 2516.960437 | 0.580886847 | 0.152522944 | 3.886418265 | 0.00010173 | 0.000776 |
| ENSG00000182264 | IZUMO1 | 70.98566 | 0.5807769 | 0.3283161 | 1.94343 | 0.05196419 | 0.142716 |
| ENSG00000230013 | CT70 | 6.461074 | 0.5803961 | 0.7781087 | 1.285394 | 0.1986548 | 0.378763 |
| ENSG00000013583 | HEBP1 | 2955.431395 | 0.580190827 | 0.260484368 | 2.332120637 | 0.01969435 | 0.067171 |
| ENSG00000135480 | KRT7 | 659.3383524 | 0.580110974 | 0.386791034 | 1.716634559 | 0.08604596 | 0.208439 |
| ENSG00000139364 | TMEM132B | 136.1872 | 0.5801059 | 0.3967729 | 1.685692 | 0.09185506 | 0.218702 |
| ENSG00000277258 | PCGF2 | 1463.768 | 0.5799587 | 0.163874 | 3.617154 | 0.00029786 | 0.002009 |
| ENSG00000229000 | SEPTIN7P8 | 11.3763 | 0.5798105 | 0.6468763 | 1.296442 | 0.1948231 | 0.37398 |
| ENSG00000115183 | TANC1 | 3487.258445 | 0.57975105 | 0.242853028 | 2.556184071 | 0.01058271 | 0.040761 |
| ENSG00000111450 | STX2 | 456.1232607 | 0.579713262 | 0.170769165 | 3.489411214 | 0.00048409 | 0.003054 |
| ENSG00000178464 | RPL10P16 | 7033.862 | 0.5795327 | 0.1848517 | 3.227911 | 0.00124698 | 0.006928 |
| ENSG00000113657 | DPYSL3 | 40.02435728 | 0.579528001 | 0.596269171 | 1.344335629 | 0.17883988 | 0.352728 |

| | | | | | | | |
|---|---|---|---|---|---|---|---|
| ENSG00000151090 | THRB | 1263.066 | 0.5794911 | 0.352724 | 1.831984 | 0.06695374 | 0.172641 |
| ENSG00000186951 | PPARA | 809.4278 | 0.579344 | 0.2644983 | 2.32108 | 0.0202825 | 0.068753 |
| ENSG00000160469 | BRSK1 | 245.1793 | 0.5790929 | 0.3482056 | 1.852221 | 0.06399409 | 0.166676 |
| ENSG00000172250 | SERHL | 2.617171 | 0.5788297 | 0.9104762 | 1.429753 | 0.1527879 | 0.315246 |
| ENSG00000132911 | NMUR2 | 6.157490152 | 0.578755683 | 0.698525606 | 1.282729021 | 0.19958703 | 0.3798 |
| ENSG00000105219 | CCNP | 117.5519351 | 0.578521314 | 0.366470992 | 1.775860238 | 0.07575596 | 0.189512 |
| ENSG00000178809 | TRIM73 | 11.26514 | 0.5781795 | 0.5985028 | 1.323061 | 0.1858152 | 0.361749 |
| ENSG00000143067 | ZNF697 | 208.2384 | 0.5781379 | 0.2011382 | 2.98289 | 0.0028554 | 0.01401 |
| ENSG00000184047 | DIABLO | 214.3048 | 0.5781282 | 0.199004 | 3.001701 | 0.00268476 | 0.013275 |
| ENSG00000270022 | | 12.36604 | 0.5781261 | 0.5207519 | 1.41213 | 0.1579118 | 0.323133 |
| ENSG00000166313 | APBB1 | 55.94668 | 0.5777739 | 0.359008 | 1.802647 | 0.07144363 | 0.181419 |
| ENSG00000132561 | MATN2 | 2363.564821 | 0.577764279 | 0.314744246 | 2.037816704 | 0.04156827 | 0.120531 |
| ENSG00000136856 | SLC2A8 | 1634.219479 | 0.577576517 | 0.166459913 | 3.587741273 | 0.00033356 | 0.00222 |
| ENSG00000282951 | | 5.354226 | 0.5775472 | 0.7830693 | 1.30273 | 0.192667 | 0.371101 |
| ENSG00000184635 | ZNF93 | 22.33432 | 0.5774675 | 0.4742936 | 1.491992 | 0.1357012 | 0.289387 |
| ENSG00000244124 | ATP1B3-AS1 | 9.778167 | 0.5773176 | 0.6759051 | 1.290902 | 0.1967377 | 0.376675 |
| ENSG00000288533 | | 21.15308 | 0.5772873 | 0.486834 | 1.476395 | 0.1398378 | 0.295859 |
| ENSG00000286996 | | 2.9113 | 0.5772246 | 0.7665966 | 1.282592 | 0.199635 | 0.379841 |
| ENSG00000274272 | | 9.845403 | 0.5771902 | 0.5720439 | 1.357004 | 0.1747799 | 0.346728 |
| ENSG00000157306 | ZFHX2-AS1 | 60.30557 | 0.5771488 | 0.3994366 | 1.660191 | 0.09687606 | 0.227316 |
| ENSG00000104321 | TRPA1 | 1.480103361 | 0.576354165 | 1.058031152 | 1.841312486 | 0.06557578 | NA |
| ENSG00000142657 | PGD | 12668.29 | 0.5762308 | 0.2298882 | 2.62531 | 0.00865702 | 0.034661 |
| ENSG00000280211 | | 29.1616 | 0.5761796 | 0.369772 | 1.75789 | 0.07876624 | 0.194857 |
| ENSG00000174327 | SLC16A13 | 635.8779 | 0.5761164 | 0.1942022 | 3.065587 | 0.00217244 | 0.011108 |
| ENSG00000120899 | PTK2B | 890.2698577 | 0.575857784 | 0.333461468 | 1.89953008 | 0.05749482 | 0.154013 |
| ENSG00000285756 | | 724.7336 | 0.5758285 | 0.1726658 | 3.433705 | 0.00059539 | 0.003651 |
| ENSG00000120457 | KCNJ5 | 1.226582461 | 0.575807515 | 0.913569909 | 1.426162531 | 0.15382143 | NA |
| ENSG00000107104 | KANK1 | 1039.116489 | 0.575793611 | 0.172972987 | 3.440936625 | 0.0005797 | 0.00357 |
| ENSG00000109572 | CLCN3 | 4752.536876 | 0.575623313 | 0.255923913 | 2.405521726 | 0.01614939 | 0.057142 |
| ENSG00000235899 | LINC01564 | 2.05183 | 0.5755915 | 1.016238 | 1.650641 | 0.09881188 | 0.230636 |
| ENSG00000242252 | BGLAP | 10.42145 | 0.575545 | 0.579501 | 1.336618 | 0.1813474 | 0.355903 |
| ENSG00000182544 | MFSD5 | 2281.446 | 0.5754477 | 0.1544845 | 3.793627 | 0.00014846 | 0.001087 |
| ENSG00000175564 | UCP3 | 40.09269 | 0.5754383 | 0.4660237 | 1.493999 | 0.1351759 | 0.288473 |

| | | | | | | | |
|---|---|---|---|---|---|---|---|
| ENSG00000165475 | CRYL1 | 2324.891 | 0.5752486 | 0.1811884 | 3.256154 | 0.00112933 | 0.006356 |
| ENSG00000271525 | ARF4P1 | 15.92508 | 0.575183 | 0.4351764 | 1.5683 | 0.1168112 | 0.260038 |
| ENSG00000207611 | MIR149 | 5.265754 | 0.5747811 | 0.7769015 | 1.290909 | 0.1967354 | 0.376675 |
| ENSG00000281706 | | 167.3057 | 0.5747629 | 0.2579793 | 2.357978 | 0.01837476 | 0.063477 |
| ENSG00000163596 | ICA1L | 340.2939 | 0.5746327 | 0.3043388 | 2.048306 | 0.04053006 | 0.118114 |
| ENSG00000204410 | MSH5 | 138.845 | 0.5746104 | 0.2684512 | 2.279406 | 0.02264296 | 0.075179 |
| ENSG00000237614 | | 11.59849 | 0.5745441 | 0.6015626 | 1.315569 | 0.1883187 | 0.365356 |
| ENSG00000144815 | NXPE3 | 1411.022 | 0.5745349 | 0.221651 | 2.720085 | 0.00652651 | 0.027511 |
| ENSG00000163513 | TGFBR2 | 822.8455 | 0.5744318 | 0.3218358 | 1.957181 | 0.05032621 | 0.139153 |
| ENSG00000149527 | PLCH2 | 70.18114 | 0.5743558 | 0.3691411 | 1.750846 | 0.07997236 | 0.197178 |
| ENSG00000136425 | CIB2 | 17.62114704 | 0.57415198 | 0.733906348 | 1.284669462 | 0.19890781 | 0.379013 |
| ENSG00000228559 | | 3.557365 | 0.5741471 | 1.025977 | 1.718968 | 0.08562015 | 0.20771 |
| ENSG00000167604 | NFKBID | 485.6305 | 0.5740748 | 0.1901674 | 3.115734 | 0.00183488 | 0.009635 |
| ENSG00000213889 | PPM1N | 159.6172 | 0.574019 | 0.2559504 | 2.373533 | 0.01761881 | 0.06136 |
| ENSG00000258561 | | 25.76364 | 0.5739804 | 0.5627902 | 1.352566 | 0.1761944 | 0.348927 |
| ENSG00000145555 | MYO10 | 1782.816 | 0.5736445 | 0.2377192 | 2.53704 | 0.01117943 | 0.042566 |
| ENSG00000198758 | EPS8L3 | 64.43974 | 0.5736306 | 0.3942467 | 1.666824 | 0.09554933 | 0.22502 |
| ENSG00000279203 | | 10.02781 | 0.5736095 | 0.5247921 | 1.398229 | 0.1620443 | 0.329356 |
| ENSG00000151327 | FAM177A1 | 3952.614 | 0.5733509 | 0.1463087 | 3.984358 | 6.77E-05 | 0.000541 |
| ENSG00000152332 | UHMK1 | 9564.186 | 0.572462 | 0.2097372 | 2.883088 | 0.00393798 | 0.018209 |
| ENSG00000164111 | ANXA5 | 12514.5 | 0.5724554 | 0.1995009 | 2.964923 | 0.00302759 | 0.014663 |
| ENSG00000250451 | HOXC-AS1 | 91.17554 | 0.5723063 | 0.4043261 | 1.631394 | 0.1028072 | 0.237417 |
| ENSG00000260078 | | 5.560246 | 0.5722483 | 0.7585192 | 1.304704 | 0.1919938 | 0.370235 |
| ENSG00000227388 | | 73.51026 | 0.5722412 | 0.3665489 | 1.755428 | 0.07918614 | 0.195693 |
| ENSG00000197982 | C1orf122 | 2400.456 | 0.5721139 | 0.1887511 | 3.141571 | 0.00168044 | 0.008922 |
| ENSG00000248329 | APELA | 1.921223 | 0.5720891 | 1.038368 | 1.774888 | 0.07591634 | 0.189721 |
| ENSG00000105483 | CARD8 | 871.0941616 | 0.571913431 | 0.226167118 | 2.646858173 | 0.00812434 | 0.03293 |
| ENSG00000105650 | PDE4C | 3.344184004 | 0.571805127 | 0.890926991 | 1.369451326 | 0.17085824 | 0.341599 |
| ENSG00000180581 | SRP9P1 | 1554.608 | 0.571758 | 0.2012148 | 2.942171 | 0.0032592 | 0.015563 |
| ENSG00000157782 | CABP1 | 5.217517 | 0.5716471 | 0.7338063 | 1.288979 | 0.1974054 | 0.377369 |
| ENSG00000218069 | RSL24D1P1 | 180.0221 | 0.571635 | 0.2022894 | 2.928584 | 0.0034051 | 0.016147 |
| ENSG00000160094 | ZNF362 | 2219.432 | 0.5714253 | 0.1495994 | 3.892662 | 9.92E-05 | 0.00076 |
| ENSG00000144671 | SLC22A14 | 4.076313 | 0.5710917 | 0.7504809 | 1.281632 | 0.1999719 | 0.380346 |

| | | | | | | | |
|---|---|---|---|---|---|---|---|
| ENSG00000185247 | MAGEA11 | 1.577251 | 0.5709335 | 1.027704 | 1.694266 | 0.09021476 | 0.216016 |
| ENSG00000287109 | | 0.9982525 | 0.570792 | 1.057547 | 1.913126 | 0.05573191 | NA |
| ENSG00000225950 | NTF4 | 13.9682 | 0.570705 | 0.5441911 | 1.356656 | 0.1748904 | 0.346858 |
| ENSG00000280193 | | 40.95401 | 0.5706287 | 0.4579194 | 1.495406 | 0.1348084 | 0.288068 |
| ENSG00000229021 | | 19.0388 | 0.5705334 | 0.4113674 | 1.609545 | 0.1074972 | 0.245179 |
| ENSG00000112414 | ADGRG6 | 6029.561741 | 0.570377936 | 0.258352385 | 2.352697902 | 0.01863777 | 0.064202 |
| ENSG00000223697 | | 24.86486 | 0.5703365 | 0.46827 | 1.482666 | 0.1381631 | 0.293059 |
| ENSG00000186577 | SMIM29 | 1729.847 | 0.5702198 | 0.2695855 | 2.238364 | 0.02519734 | 0.081593 |
| ENSG00000134851 | TMEM165 | 4006.67232 | 0.56997971 | 0.159531833 | 3.623115788 | 0.00029108 | 0.00197 |
| ENSG00000273136 | NBPF26 | 997.8141 | 0.5698216 | 0.2379459 | 2.515921 | 0.01187217 | 0.044653 |
| ENSG00000260996 | | 3.870734 | 0.5698112 | 0.8444849 | 1.358032 | 0.1744536 | 0.346391 |
| ENSG00000075043 | KCNQ2 | 16.34750575 | 0.56961332 | 0.505152158 | 1.420456435 | 0.15547484 | 0.319475 |
| ENSG00000230082 | PRRT3-AS1 | 15.79481 | 0.5693771 | 0.6246407 | 1.292375 | 0.1962272 | 0.375969 |
| ENSG00000123685 | BATF3 | 3.37278207 | 0.569028228 | 0.9883364 | 1.588716785 | 0.11212435 | 0.252696 |
| ENSG00000228317 | | 4.379589 | 0.5689575 | 0.8055323 | 1.317379 | 0.1877116 | 0.364543 |
| ENSG00000138613 | APH1B | 775.7650121 | 0.568944897 | 0.326952739 | 1.906296642 | 0.05661173 | 0.15236 |
| ENSG00000258373 | SLC25A3P2 | 2.205672 | 0.5688722 | 0.8389615 | 1.316223 | 0.1880993 | 0.365056 |
| ENSG00000120519 | SLC10A7 | 479.2077019 | 0.568421178 | 0.197943547 | 2.976939015 | 0.00291142 | 0.014238 |
| ENSG00000112245 | PTP4A1 | 29.58355454 | 0.568099752 | 0.545289495 | 1.362384368 | 0.17307661 | 0.344474 |
| ENSG00000272334 | | 1.898047 | 0.5678472 | 0.8890653 | 1.396812 | 0.1624702 | 0.329784 |
| ENSG00000110911 | SLC11A2 | 4688.033598 | 0.567700237 | 0.163222994 | 3.565861767 | 0.00036266 | 0.002386 |
| ENSG00000272137 | | 1.996464 | 0.567658 | 0.930145 | 1.440082 | 0.1498442 | 0.310911 |
| ENSG00000176402 | GJC3 | 18.75853 | 0.5676376 | 0.5502164 | 1.347635 | 0.1777758 | 0.351104 |
| ENSG00000143740 | SNAP47 | 2619.075 | 0.5676002 | 0.1617129 | 3.588404 | 0.00033271 | 0.002215 |
| ENSG00000279700 | | 14.39058 | 0.5675895 | 0.6311158 | 1.28581 | 0.1985095 | 0.378567 |
| ENSG00000131381 | RBSN | 4271.959062 | 0.567542875 | 0.135853959 | 4.242258958 | 2.21E-05 | 0.0002 |
| ENSG00000239213 | NCK1-DT | 294.1322 | 0.5674633 | 0.1723132 | 3.381225 | 0.00072163 | 0.004314 |
| ENSG00000273084 | | 16.61519 | 0.5673678 | 0.6950163 | 1.270669 | 0.2038464 | 0.385453 |
| ENSG00000280173 | | 80.48911 | 0.5672965 | 0.2717941 | 2.226821 | 0.02595925 | 0.0835 |
| ENSG00000174944 | P2RY14 | 2.886534 | 0.5671388 | 0.8237899 | 1.320544 | 0.1866534 | 0.363045 |
| ENSG00000126947 | ARMCX1 | 1108.275428 | 0.567013827 | 0.24149992 | 2.469256774 | 0.0135394 | 0.049573 |
| ENSG00000265282 | | 18.77696 | 0.5669991 | 0.3777105 | 1.698335 | 0.08944453 | 0.214694 |
| ENSG00000277893 | SRD5A2 | 3.855866 | 0.5669219 | 0.7207421 | 1.261614 | 0.2070879 | 0.389453 |

| | | | | | | | |
|---|---|---|---|---|---|---|---|
| ENSG00000256299 | | 1.605274 | 0.5668588 | 1.003802 | 1.644016 | 0.1001728 | 0.233117 |
| ENSG00000277481 | PKD1L3 | 6.517669 | 0.5666643 | 0.6255306 | 1.290047 | 0.1970343 | 0.376972 |
| ENSG00000226342 | NMD3P1 | 9.247175 | 0.5666203 | 0.6928387 | 1.276251 | 0.2018668 | 0.382854 |
| ENSG00000279549 | | 11.30165 | 0.566606 | 0.5212617 | 1.399783 | 0.1615784 | 0.328611 |
| ENSG00000145244 | CORIN | 5.872368 | 0.5662554 | 0.7047632 | 1.273198 | 0.202948 | 0.384356 |
| ENSG00000143878 | RHOB | 14145.06 | 0.5662325 | 0.1472873 | 3.886491 | 0.0001017 | 0.000776 |
| ENSG00000144579 | CTDSP1 | 8178.119 | 0.5661479 | 0.1882499 | 3.11466 | 0.00184157 | 0.009666 |
| ENSG00000182134 | TDRKH | 3396.886 | 0.5660451 | 0.165402 | 3.493814 | 0.00047617 | 0.003008 |
| ENSG00000225031 | EIF4BP7 | 3032.484 | 0.5660401 | 0.2025117 | 2.897307 | 0.00376381 | 0.017522 |
| ENSG00000277103 | | 3.666864 | 0.5660283 | 0.8196844 | 1.305876 | 0.1915946 | 0.369813 |
| ENSG00000139625 | MAP3K12 | 1562.804 | 0.565555 | 0.2414316 | 2.458093 | 0.01396771 | 0.050872 |
| ENSG00000197283 | SYNGAP1 | 2087.432 | 0.5655438 | 0.2521669 | 2.37423 | 0.01758561 | 0.061268 |
| ENSG00000230310 | | 2.269716 | 0.5655009 | 0.8651047 | 1.33872 | 0.1806617 | 0.355109 |
| ENSG00000084070 | SMAP2 | 2392.570983 | 0.565363964 | 0.142142453 | 4.039477875 | 5.36E-05 | 0.00044 |
| ENSG00000118369 | USP35 | 893.995228 | 0.565339867 | 0.283013653 | 2.14897294 | 0.03163655 | 0.097784 |
| ENSG00000117305 | HMGCL | 1571.819324 | 0.565278447 | 0.184502513 | 3.155874924 | 0.00160018 | 0.00856 |
| ENSG00000225347 | SLC25A5P8 | 1.6635 | 0.565089 | 0.9408967 | 1.594875 | 0.1107402 | 0.250618 |
| ENSG00000231188 | | 4.37111 | 0.5648976 | 0.7358547 | 1.259547 | 0.2078327 | 0.39041 |
| ENSG00000147485 | PXDNL | 12.23646 | 0.5648363 | 0.631578 | 1.285906 | 0.1984758 | 0.37853 |
| ENSG00000188827 | SLX4 | 960.5832 | 0.56462 | 0.2639373 | 2.26679 | 0.02340305 | 0.076957 |
| ENSG00000169499 | PLEKHA2 | 879.6162 | 0.5645751 | 0.2180261 | 2.698815 | 0.00695869 | 0.028983 |
| ENSG00000259498 | TPM1-AS | 4.521598 | 0.5645527 | 0.816212 | 1.298745 | 0.1940314 | 0.372863 |
| ENSG00000280407 | | 52.26767 | 0.5644676 | 0.3441968 | 1.82226 | 0.06841553 | 0.175474 |
| ENSG00000263015 | | 1.992093 | 0.5643921 | 0.9537624 | 1.500867 | 0.1333899 | 0.286042 |
| ENSG00000165156 | ZHX1 | 5055.796 | 0.5643414 | 0.201394 | 2.904041 | 0.0036838 | 0.01721 |
| ENSG00000160117 | ANKLE1 | 26.02935 | 0.5641877 | 0.5190128 | 1.38042 | 0.1674575 | 0.336855 |
| ENSG00000268655 | | 21.47649 | 0.5641606 | 0.5640267 | 1.338942 | 0.1805895 | 0.354993 |
| ENSG00000181894 | ZNF329 | 1139.145 | 0.5639447 | 0.1614354 | 3.58032 | 0.00034317 | 0.002274 |
| ENSG00000285581 | | 4.239748 | 0.5637511 | 0.7451599 | 1.284124 | 0.1990985 | 0.379219 |
| ENSG00000144959 | NCEH1 | 793.7143 | 0.5635564 | 0.2377834 | 2.492766 | 0.01267525 | 0.04714 |
| ENSG00000235779 | | 7.213356 | 0.5635372 | 0.7031796 | 1.255906 | 0.20915 | 0.392243 |
| ENSG00000225431 | LINC01671 | 440.6368 | 0.5630656 | 0.6926675 | 1.279057 | 0.2008771 | 0.381549 |
| ENSG00000182324 | KCNJ14 | 64.32668 | 0.5629437 | 0.3027867 | 2.009417 | 0.04449292 | 0.126975 |

| | | | | | | | |
|---|---|---|---|---|---|---|---|
| ENSG00000251571 | DDX3P3 | 1.713336 | 0.5626547 | 0.9328112 | 1.499388 | 0.1337731 | 0.286499 |
| ENSG00000255200 | PGAM1P8 | 68.39054 | 0.5624692 | 0.4335138 | 1.533128 | 0.1252444 | 0.273428 |
| ENSG00000013375 | PGM3 | 3185.449019 | 0.562459276 | 0.183889324 | 3.124608622 | 0.00178042 | 0.009391 |
| ENSG00000243388 | RPL3P3 | 5.432154 | 0.5622229 | 0.6995199 | 1.256439 | 0.208957 | 0.391909 |
| ENSG00000240634 | | 2.95098 | 0.5620424 | 0.8455789 | 1.33324 | 0.182453 | 0.357404 |
| ENSG00000280422 | | 1.859583 | 0.5619585 | 0.8902331 | 1.381164 | 0.1672285 | 0.336675 |
| ENSG00000251313 | | 3.189249 | 0.561817 | 0.7700097 | 1.267379 | 0.2050196 | 0.38701 |
| ENSG00000125633 | CCDC93 | 3357.651472 | 0.561759539 | 0.237400361 | 2.543771683 | 0.01096627 | 0.041916 |
| ENSG00000159692 | CTBP1 | 4006.883 | 0.5617466 | 0.1246368 | 4.631595 | 3.63E-06 | 4.00E-05 |
| ENSG00000245080 | MIR3150BHG | 24.09631 | 0.5615918 | 0.4060516 | 1.603633 | 0.108795 | 0.247281 |
| ENSG00000100528 | CNIH1 | 5860.904055 | 0.561506936 | 0.151433584 | 3.761685416 | 0.00016877 | 0.001221 |
| ENSG00000177202 | SPACA4 | 3.373992 | 0.5614996 | 0.7933396 | 1.288501 | 0.1975715 | 0.377592 |
| ENSG00000233467 | HAX1P1 | 9.455154 | 0.5613433 | 0.6380435 | 1.275542 | 0.2021175 | 0.383247 |
| ENSG00000213309 | RPL9P18 | 3.221967 | 0.5611372 | 0.8091758 | 1.30229 | 0.1928172 | 0.371283 |
| ENSG00000141540 | TTYH2 | 142.2432 | 0.5609757 | 0.2978045 | 2.040133 | 0.04133705 | 0.120005 |
| ENSG00000157168 | NRG1 | 2.595341 | 0.5608547 | 0.936238 | 1.444623 | 0.1485637 | 0.30905 |
| ENSG00000175806 | MSRA | 399.3749 | 0.5604944 | 0.1803783 | 3.202832 | 0.00136083 | 0.007467 |
| ENSG00000158050 | DUSP2 | 1651.705 | 0.5603047 | 0.471833 | 1.450835 | 0.1468259 | 0.306443 |
| ENSG00000253476 | | 10.08341 | 0.5598709 | 0.7338489 | 1.263319 | 0.2064747 | 0.388702 |
| ENSG00000112531 | QKI | 4368.178155 | 0.559770682 | 0.260768978 | 2.284117923 | 0.0223646 | 0.074508 |
| ENSG00000165516 | KLHDC2 | 3704.85 | 0.5596586 | 0.1134329 | 5.021301 | 5.13E-07 | 6.80E-06 |
| ENSG00000276814 | | 8.944791 | 0.5595691 | 0.5829197 | 1.312586 | 0.1893225 | 0.366787 |
| ENSG00000185245 | GP1BA | 81.92503 | 0.5594614 | 0.3768097 | 1.67599 | 0.09374022 | 0.222088 |
| ENSG00000227934 | | 165.5565 | 0.5592963 | 0.1920505 | 2.997606 | 0.00272109 | 0.013433 |
| ENSG00000073584 | SMARCE1 | 750.8832627 | 0.55908324 | 0.190611311 | 3.033084189 | 0.00242068 | 0.012162 |
| ENSG00000150712 | MTMR12 | 3753.55 | 0.5584914 | 0.210715 | 2.753582 | 0.0058947 | 0.025375 |
| ENSG00000241984 | RPL7AP2 | 7.461852 | 0.5584891 | 0.6612447 | 1.264289 | 0.2061263 | 0.388299 |
| ENSG00000196792 | STRN3 | 1945.378 | 0.5584836 | 0.1511341 | 3.759886 | 0.00016999 | 0.001229 |
| ENSG00000271730 | | 5.337316 | 0.558451 | 0.6891693 | 1.245724 | 0.2128657 | 0.39764 |
| ENSG00000188215 | DCUN1D3 | 475.4929 | 0.5584097 | 0.2572318 | 2.289929 | 0.02202543 | 0.073554 |
| ENSG00000240652 | | 12.37853 | 0.5583152 | 0.5288076 | 1.363754 | 0.1726451 | 0.343854 |
| ENSG00000213073 | CHP1P2 | 127.1155 | 0.5580566 | 0.3999809 | 1.603253 | 0.1088787 | 0.247353 |
| ENSG00000167851 | CD300A | 1.828832 | 0.5579457 | 1.015408 | 1.67642 | 0.09365599 | 0.221956 |

| | | | | | | | |
|---|---|---|---|---|---|---|---|
| ENSG00000272989 | LINC02012 | 24.12702 | 0.5578721 | 0.5188122 | 1.376657 | 0.1686183 | 0.338498 |
| ENSG00000123570 | RAB9B | 662.627199 | 0.557631562 | 0.217945991 | 2.680048286 | 0.00736115 | 0.030388 |
| ENSG00000235865 | GSN-AS1 | 6.753781 | 0.5575614 | 0.7038808 | 1.256876 | 0.2087987 | 0.391711 |
| ENSG00000253738 | OTUD6B-AS1 | 1993.368 | 0.5571696 | 0.1484893 | 3.817628 | 0.00013474 | 0.000997 |
| ENSG00000261087 | ZNNT1 | 143.3707 | 0.5569668 | 0.5483099 | 1.328428 | 0.1840368 | 0.359356 |
| ENSG00000260007 | | 161.273 | 0.5569531 | 0.3359451 | 1.831054 | 0.06709251 | 0.172932 |
| ENSG00000235296 | | 6.247209 | 0.5566522 | 0.6275522 | 1.266649 | 0.2052809 | 0.387256 |
| ENSG00000100209 | HSCB | 1164.114126 | 0.556651032 | 0.178638658 | 3.205334243 | 0.00134906 | 0.007418 |
| ENSG00000197258 | EIF4BP6 | 4746.245 | 0.5564411 | 0.1952384 | 2.94094 | 0.00327218 | 0.015614 |
| ENSG00000185168 | LINC00482 | 114.516 | 0.556437 | 0.4350887 | 1.518472 | 0.1288956 | 0.279083 |
| ENSG00000258760 | | 1.529993 | 0.5563124 | 0.9984302 | 1.731068 | 0.0834397 | NA |
| ENSG00000236683 | HMGA1P1 | 50.16665 | 0.5562395 | 0.3246284 | 1.886537 | 0.05922258 | 0.157538 |
| ENSG00000257675 | BTBD10P1 | 12.18703 | 0.5562044 | 0.5127675 | 1.363504 | 0.1727236 | 0.343958 |
| ENSG00000109756 | RAPGEF2 | 3870.862564 | 0.556126012 | 0.3005829 | 2.01093558 | 0.04433226 | 0.126625 |
| ENSG00000213250 | RBMS2P1 | 67.13153 | 0.5558113 | 0.3883647 | 1.636995 | 0.1016315 | 0.235542 |
| ENSG00000197774 | EME2 | 5373.869 | 0.5556492 | 0.2661188 | 2.221024 | 0.02634937 | 0.084509 |
| ENSG00000132356 | PRKAA1 | 5409.052353 | 0.5555548 | 0.207116449 | 2.781827943 | 0.00540537 | 0.023611 |
| ENSG00000279518 | | 95.96668 | 0.5555326 | 0.4474252 | 1.479921 | 0.1388943 | 0.294281 |
| ENSG00000061656 | SPAG4 | 334.5058565 | 0.555460196 | 0.291518201 | 2.041540706 | 0.04119711 | 0.119742 |
| ENSG00000156006 | NAT2 | 4.943105 | 0.5554599 | 0.7682123 | 1.268504 | 0.204618 | 0.386554 |
| ENSG00000218632 | RPL7P28 | 4.518919 | 0.5553979 | 0.9150229 | 1.422563 | 0.154863 | 0.318534 |
| ENSG00000163683 | SMIM14 | 5742.046 | 0.5553919 | 0.1797111 | 3.169901 | 0.00152491 | 0.008228 |
| ENSG00000158828 | PINK1 | 601.0086 | 0.5552044 | 0.2898204 | 2.063272 | 0.03908677 | 0.114835 |
| ENSG00000165084 | C8orf34 | 35.13605 | 0.5549055 | 0.3991072 | 1.599185 | 0.1097796 | 0.249038 |
| ENSG00000138193 | PLCE1 | 231.0885258 | 0.554860348 | 0.415688097 | 1.552371415 | 0.12057338 | 0.266294 |
| ENSG00000087301 | TXNDC16 | 1366.112759 | 0.554746824 | 0.222153605 | 2.610011471 | 0.00905392 | 0.03592 |
| ENSG00000181722 | ZBTB20 | 40.44764 | 0.5545933 | 0.6000577 | 1.30404 | 0.1922199 | 0.370536 |
| ENSG00000271474 | | 12.33415 | 0.5543708 | 0.6059403 | 1.257657 | 0.208516 | 0.391441 |
| ENSG00000223875 | NBEAP3 | 0.8371778 | 0.5543151 | 1.028009 | 1.794453 | 0.07274094 | NA |
| ENSG00000047644 | WWC3 | 8215.996158 | 0.55405126 | 0.247707046 | 2.359809239 | 0.01828434 | 0.06323 |
| ENSG00000088387 | DOCK9 | 2039.333959 | 0.553974819 | 0.173089747 | 3.290350432 | 0.00100063 | 0.005725 |
| ENSG00000265008 | | 1.913472 | 0.5539358 | 0.838209 | 1.311457 | 0.1897033 | 0.367337 |
| ENSG00000224546 | EIF4BP3 | 2022.428 | 0.5537451 | 0.2017285 | 2.847182 | 0.00441081 | 0.019975 |

| | | | | | | | |
|---|---|---|---|---|---|---|---|
| ENSG00000218980 | FTH1P15 | 7.109936 | 0.5537188 | 0.6542553 | 1.257234 | 0.208669 | 0.39159 |
| ENSG00000168334 | XIRP1 | 0.9738504 | 0.5536359 | 1.008901 | 1.695368 | 0.09000554 | NA |
| ENSG00000272325 | NUDT3 | 222.994 | 0.5532293 | 0.2180694 | 2.641903 | 0.00824418 | 0.033289 |
| ENSG00000227885 | | 173.7491 | 0.5531669 | 0.3381562 | 1.81991 | 0.06877272 | 0.176135 |
| ENSG00000178053 | MLF1 | 996.677 | 0.5531386 | 0.2869194 | 2.073042 | 0.03816833 | 0.112777 |
| ENSG00000243156 | MICAL3 | 1744.968 | 0.5529815 | 0.261783 | 2.244151 | 0.0248227 | 0.080622 |
| ENSG00000247735 | | 42.68068 | 0.5529034 | 0.4920643 | 1.398625 | 0.1619253 | 0.32914 |
| ENSG00000273007 | | 31.2459 | 0.5528932 | 0.5543742 | 1.311324 | 0.1897485 | 0.367397 |
| ENSG00000280247 | | 3.120314 | 0.5528725 | 0.8647491 | 1.359901 | 0.1738614 | 0.345577 |
| ENSG00000127419 | TMEM175 | 1369.186278 | 0.552840846 | 0.265550392 | 2.212795468 | 0.02691175 | 0.085978 |
| ENSG00000166428 | PLD4 | 13.76018 | 0.5528257 | 0.6013747 | 1.275461 | 0.2021463 | 0.383274 |
| ENSG00000233822 | H2BC15 | 369.906 | 0.5527896 | 0.2768026 | 2.134893 | 0.03276975 | 0.100457 |
| ENSG00000162825 | NBPF20 | 660.6577 | 0.5527415 | 0.2046766 | 2.802074 | 0.00507753 | 0.022436 |
| ENSG00000205744 | DENND1C | 1026.047 | 0.5526733 | 0.2039434 | 2.80589 | 0.00501777 | 0.022212 |
| ENSG00000232628 | | 0.9762324 | 0.5526222 | 1.028992 | 1.792248 | 0.0730932 | NA |
| ENSG00000213331 | | 2.217482 | 0.5523335 | 0.9155675 | 1.446637 | 0.1479986 | 0.308229 |
| ENSG00000235109 | ZSCAN31 | 253.4745 | 0.5520619 | 0.2163748 | 2.665922 | 0.00767775 | 0.031461 |
| ENSG00000146950 | SHROOM2 | 718.9158 | 0.5519063 | 0.2787297 | 2.108563 | 0.03498232 | 0.105583 |
| ENSG00000067606 | PRKCZ | 1610.165978 | 0.551839258 | 0.163781364 | 3.437880575 | 0.00058629 | 0.003604 |
| ENSG00000271511 | | 1.23745 | 0.5517161 | 0.9459105 | 1.612554 | 0.1068414 | NA |
| ENSG00000266389 | | 19.01708 | 0.5516529 | 0.5369146 | 1.325414 | 0.1850338 | 0.360581 |
| ENSG00000065534 | MYLK | 144.2881019 | 0.551592287 | 0.287892669 | 2.05690795 | 0.03969508 | 0.116229 |
| ENSG00000089723 | OTUB2 | 248.7068585 | 0.551520431 | 0.214097657 | 2.680073626 | 0.0073606 | 0.030388 |
| ENSG00000170356 | OR2A20P | 19.50867 | 0.5514886 | 0.3973142 | 1.595008 | 0.1107105 | 0.250594 |
| ENSG00000101974 | ATP11C | 3455.678276 | 0.551420322 | 0.21404234 | 2.684250892 | 0.00726925 | 0.030079 |
| ENSG00000198019 | FCGR1B | 3.807914 | 0.5514076 | 0.857903 | 1.318383 | 0.1873755 | 0.36405 |
| ENSG00000181291 | TMEM132E | 1.921023 | 0.5513224 | 0.9471977 | 1.542904 | 0.1228541 | 0.269806 |
| ENSG00000215102 | TERF1P4 | 33.78994 | 0.5512814 | 0.3569282 | 1.729982 | 0.08363343 | 0.204064 |
| ENSG00000131943 | C19orf12 | 1354.206877 | 0.551258996 | 0.137058803 | 4.075389549 | 4.59E-05 | 0.000385 |
| ENSG00000126804 | ZBTB1 | 4098.606612 | 0.551211755 | 0.165407731 | 3.447151129 | 0.00056653 | 0.003496 |
| ENSG00000285257 | | 29.93807 | 0.5511061 | 0.4858725 | 1.403074 | 0.1605947 | 0.327187 |
| ENSG00000272092 | | 29.74356 | 0.5509505 | 0.3413058 | 1.78394 | 0.07443342 | 0.187117 |
| ENSG00000228150 | | 21.09814 | 0.5509248 | 0.3827952 | 1.640037 | 0.1009975 | 0.234481 |

| | | | | | | | |
|---|---|---|---|---|---|---|---|
| ENSG00000213131 | YWHAZP4 | 90.45382 | 0.5504874 | 0.2899645 | 2.042201 | 0.04113157 | 0.119617 |
| ENSG00000174226 | SNX31 | 1.690821 | 0.5504491 | 0.9168859 | 1.485748 | 0.1373459 | 0.291745 |
| ENSG00000259711 | | 17.92478 | 0.550383 | 0.448361 | 1.47244 | 0.1409021 | 0.297538 |
| ENSG00000197885 | NKIRAS1 | 820.0868 | 0.5501374 | 0.1598964 | 3.51668 | 0.00043698 | 0.002793 |
| ENSG00000197580 | BCO2 | 20.6771 | 0.5500458 | 0.4925295 | 1.381043 | 0.1672656 | 0.336698 |
| ENSG00000180855 | ZNF443 | 280.2486 | 0.550001 | 0.2300755 | 2.506877 | 0.01218032 | 0.045548 |
| ENSG00000232901 | CYCSP10 | 23.88686 | 0.5499851 | 0.586963 | 1.287021 | 0.1980869 | 0.378124 |
| ENSG00000196683 | TOMM7 | 6132.923 | 0.5499831 | 0.1691533 | 3.299297 | 0.00096927 | 0.005566 |
| ENSG00000239040 | | 4.901427 | 0.5499574 | 0.804421 | 1.28121 | 0.20012 | 0.380492 |
| ENSG00000274512 | TBC1D3L | 263.297 | 0.5498079 | 0.2804881 | 2.101922 | 0.0355601 | 0.106865 |
| ENSG00000225057 | | 3.705224 | 0.5497162 | 0.7744317 | 1.251009 | 0.2109313 | 0.394859 |
| ENSG00000133731 | IMPA1 | 407.9851374 | 0.549574159 | 0.185975002 | 3.051052332 | 0.00228041 | 0.01156 |
| ENSG00000251495 | PPIAP11 | 23.19028 | 0.5494256 | 0.500384 | 1.389862 | 0.1645707 | 0.332736 |
| ENSG00000172795 | DCP2 | 6908.991 | 0.5493127 | 0.2373479 | 2.461987 | 0.01381698 | 0.050426 |
| ENSG00000143761 | ARF1 | 21182.17 | 0.5492147 | 0.1438918 | 3.886329 | 0.00010177 | 0.000776 |
| ENSG00000273253 | | 52.76562 | 0.549077 | 0.3673292 | 1.687194 | 0.0915661 | 0.218288 |
| ENSG00000274514 | | 1.93232 | 0.549043 | 0.9210248 | 1.504055 | 0.1325673 | 0.284874 |
| ENSG00000287737 | | 131.6613 | 0.5489269 | 0.3900489 | 1.626109 | 0.1039264 | 0.239274 |
| ENSG00000135409 | AMHR2 | 1.876121532 | 0.548828883 | 0.919882628 | 1.431850194 | 0.1521867 | 0.314421 |
| ENSG00000244701 | | 19.18007 | 0.5486531 | 0.5888142 | 1.286666 | 0.1982107 | 0.378162 |
| ENSG00000206932 | RNU6-4P | 9.58695 | 0.5485304 | 0.575851 | 1.288312 | 0.1976374 | 0.377663 |
| ENSG00000240401 | | 11.84059 | 0.5481213 | 0.6194211 | 1.249257 | 0.2115711 | 0.395694 |
| ENSG00000249655 | | 5.206953 | 0.5480465 | 0.740373 | 1.259516 | 0.2078441 | 0.39041 |
| ENSG00000197568 | HHLA3 | 231.4423 | 0.5480399 | 0.1985399 | 2.857117 | 0.00427508 | 0.019473 |
| ENSG00000231871 | IPO9-AS1 | 7.749019 | 0.5479402 | 0.6813519 | 1.228255 | 0.2193512 | 0.405478 |
| ENSG00000198598 | MMP17 | 1087.135 | 0.5479357 | 0.2317714 | 2.492256 | 0.01269347 | 0.047179 |
| ENSG00000196923 | PDLIM7 | 3583.531 | 0.5479119 | 0.2549503 | 2.275052 | 0.02290279 | 0.075751 |
| ENSG00000274447 | | 14.51877 | 0.5478918 | 0.6181466 | 1.266356 | 0.2053856 | 0.387358 |
| ENSG00000198221 | AFDN-DT | 30.81925 | 0.5477679 | 0.4746469 | 1.411807 | 0.1580069 | 0.323266 |
| ENSG00000197256 | KANK2 | 2237.923 | 0.5473864 | 0.178715 | 3.163187 | 0.00156052 | 0.008388 |
| ENSG00000233597 | | 10.53084 | 0.5472916 | 0.6002495 | 1.270558 | 0.2038859 | 0.3855 |
| ENSG00000273576 | | 161.9373 | 0.5472578 | 0.3432934 | 1.770484 | 0.07664652 | 0.191034 |
| ENSG00000213981 | | 1.777946 | 0.5472024 | 1.004627 | 1.673699 | 0.09418985 | 0.223033 |

| | | | | | | | |
|---|---|---|---|---|---|---|---|
| ENSG00000127241 | MASP1 | 3.448477734 | 0.547069626 | 0.784142463 | 1.251708655 | 0.21067605 | 0.394604 |
| ENSG00000256061 | DNAAF4 | 15.81505 | 0.5468036 | 0.4405077 | 1.47508 | 0.1401912 | 0.296367 |
| ENSG00000041880 | PARP3 | 376.2155992 | 0.546786309 | 0.194869591 | 2.898868213 | 0.00374512 | 0.017457 |
| ENSG00000188493 | C19orf54 | 1062.629 | 0.5464399 | 0.1921685 | 2.954139 | 0.00313542 | 0.015061 |
| ENSG00000120563 | LYZL1 | 1.891736218 | 0.54613393 | 0.987864911 | 1.616871637 | 0.10590599 | 0.242621 |
| ENSG00000173801 | JUP | 24465.41 | 0.5460961 | 0.1854974 | 3.032827 | 0.00242274 | 0.012171 |
| ENSG00000237976 | RFX5-AS1 | 21.02206 | 0.5459875 | 0.531145 | 1.331468 | 0.1830349 | 0.358032 |
| ENSG00000225527 | | 2.847747 | 0.5459373 | 0.824088 | 1.298698 | 0.1940477 | 0.372863 |
| ENSG00000148584 | A1CF | 1.91881 | 0.545815 | 0.8333469 | 1.280563 | 0.2003471 | 0.380814 |
| ENSG00000116741 | RGS2 | 62.4204401 | 0.545775057 | 0.458562655 | 1.444050369 | 0.14872481 | 0.30924 |
| ENSG00000225405 | RPS15AP17 | 1.287341 | 0.5455643 | 0.943942 | 1.612372 | 0.1068811 | NA |
| ENSG00000228143 | | 4.493258 | 0.5454489 | 0.7131487 | 1.231846 | 0.2180064 | 0.403695 |
| ENSG00000140691 | ARMC5 | 737.274 | 0.5453387 | 0.2485059 | 2.305441 | 0.02114187 | 0.071095 |
| ENSG00000214100 | PLAC9P1 | 1.553819 | 0.5451466 | 0.9847429 | 1.613243 | 0.1066917 | NA |
| ENSG00000250656 | ST3GAL1P1 | 6.128846 | 0.544829 | 0.6410863 | 1.236042 | 0.2164429 | 0.401731 |
| ENSG00000257210 | NACAP8 | 2.888407 | 0.5446228 | 0.8463132 | 1.33153 | 0.1830146 | 0.358032 |
| ENSG00000224155 | | 124.7074 | 0.5445864 | 0.2227287 | 2.5584 | 0.01051552 | 0.040538 |
| ENSG00000151006 | PRSS53 | 140.9937 | 0.5445574 | 0.2528189 | 2.276492 | 0.02281655 | 0.07555 |
| ENSG00000141232 | TOB1 | 9014.296 | 0.5445444 | 0.2197901 | 2.583808 | 0.00977163 | 0.038146 |
| ENSG00000150722 | PPP1R1C | 17.13344 | 0.5444336 | 0.5199296 | 1.335565 | 0.1816916 | 0.356447 |
| ENSG00000143013 | LMO4 | 1152.855 | 0.5443422 | 0.1650243 | 3.372823 | 0.00074402 | 0.004427 |
| ENSG00000091513 | TF | 27.05549362 | 0.544289873 | 0.580609702 | 1.270861106 | 0.20377807 | 0.385378 |
| ENSG00000249741 | | 8.027179 | 0.544226 | 0.6300509 | 1.251509 | 0.210749 | 0.394636 |
| ENSG00000271943 | | 436.5966 | 0.5442188 | 0.1983957 | 2.832676 | 0.00461602 | 0.020787 |
| ENSG00000152495 | CAMK4 | 12.62845 | 0.5437512 | 0.5775129 | 1.272179 | 0.2033095 | 0.384958 |
| ENSG00000164169 | PRMT9 | 980.0901 | 0.5436271 | 0.1636684 | 3.401859 | 0.00066929 | 0.004044 |
| ENSG00000130377 | ACSBG2 | 16.19078119 | 0.543517209 | 0.436063972 | 1.473353092 | 0.14065585 | 0.29716 |
| ENSG00000260519 | | 1.483583 | 0.5433581 | 1.00969 | 1.753747 | 0.07947381 | NA |
| ENSG00000082153 | BZW1 | 7260.939918 | 0.543155336 | 0.247175541 | 2.328681372 | 0.01987595 | 0.067625 |
| ENSG00000284418 | | 7.633257 | 0.5429233 | 0.6516432 | 1.226174 | 0.2201333 | 0.406351 |
| ENSG00000204070 | SYS1 | 1697.63 | 0.542878 | 0.1550127 | 3.521824 | 0.00042859 | 0.002747 |
| ENSG00000110080 | ST3GAL4 | 3573.781892 | 0.542713355 | 0.248674766 | 2.318492095 | 0.02042259 | 0.069093 |
| ENSG00000228323 | MYADM-AS1 | 18.6447 | 0.5426931 | 0.5856396 | 1.270676 | 0.2038438 | 0.385453 |

| | | | | | | | |
|---|---|---|---|---|---|---|---|
| ENSG00000100478 | AP4S1 | 941.838924 | 0.542670977 | 0.164747874 | 3.379879154 | 0.00072518 | 0.004332 |
| ENSG00000273559 | CWC25 | 1281.565 | 0.5424114 | 0.134817 | 4.085487 | 4.40E-05 | 0.00037 |
| ENSG00000165185 | KIAA1958 | 1055.584 | 0.5422561 | 0.242112 | 2.357765 | 0.01838532 | 0.063497 |
| ENSG00000243870 | RN7SL236P | 4.80855 | 0.5418097 | 0.7385397 | 1.231309 | 0.2182072 | 0.403954 |
| ENSG00000168268 | NT5DC2 | 7422.391 | 0.5416469 | 0.1964091 | 2.868812 | 0.00412016 | 0.018923 |
| ENSG00000238287 | | 37.56057 | 0.5416183 | 0.4186022 | 1.508632 | 0.1313928 | 0.282945 |
| ENSG00000187950 | OVCH1 | 1.680057 | 0.5415959 | 0.9823027 | 1.610437 | 0.1073026 | 0.244897 |
| ENSG00000261765 | | 2.034265 | 0.541515 | 0.8607086 | 1.291064 | 0.1966817 | 0.376675 |
| ENSG00000262152 | LINC00514 | 388.5849 | 0.5414242 | 0.3897122 | 1.560642 | 0.1186082 | 0.262871 |
| ENSG00000158109 | TPRG1L | 2434.931 | 0.5413749 | 0.196951 | 2.85799 | 0.00426334 | 0.019439 |
| ENSG00000267733 | | 30.41512 | 0.5413137 | 0.438914 | 1.461776 | 0.1438026 | 0.30184 |
| ENSG00000266934 | | 352.9025 | 0.5408598 | 0.356943 | 1.696491 | 0.08979285 | 0.215258 |
| ENSG00000226310 | | 19.33248 | 0.5408326 | 0.4873083 | 1.367224 | 0.171555 | 0.342255 |
| ENSG00000139725 | RHOF | 272.0833 | 0.5407517 | 0.2027624 | 2.763028 | 0.00572678 | 0.024781 |
| ENSG00000198369 | SPRED2 | 2488.974 | 0.5404453 | 0.2460658 | 2.328869 | 0.01986599 | 0.067609 |
| ENSG00000176623 | RMDN1 | 4584.178 | 0.5403776 | 0.1413616 | 3.89818 | 9.69E-05 | 0.000745 |
| ENSG00000196605 | ZNF846 | 399.0043 | 0.5400685 | 0.2112294 | 2.662619 | 0.00775352 | 0.031712 |
| ENSG00000225871 | | 120.4209 | 0.5397813 | 0.2793384 | 2.06443 | 0.03897693 | 0.114619 |
| ENSG00000213188 | YBX1P4 | 1.713034 | 0.5397232 | 0.929961 | 1.528014 | 0.1265092 | 0.275198 |
| ENSG00000237911 | SRP68P3 | 1.429144 | 0.5397 | 1.033295 | 1.94256 | 0.05206935 | NA |
| ENSG00000254543 | HSPD1P2 | 1.341905 | 0.5395867 | 0.924185 | 1.449719 | 0.147137 | NA |
| ENSG00000273113 | | 35.44609 | 0.5395199 | 0.5634374 | 1.276612 | 0.2017392 | 0.382694 |
| ENSG00000090615 | GOLGA3 | 7635.058788 | 0.539512182 | 0.158004141 | 3.506728178 | 0.00045365 | 0.002883 |
| ENSG00000283709 | FAM238C | 36.68871 | 0.539473 | 0.6487428 | 1.223327 | 0.2212061 | 0.407744 |
| ENSG00000161958 | FGF11 | 43.06067 | 0.5388675 | 0.4976354 | 1.347544 | 0.177805 | 0.351104 |
| ENSG00000272864 | | 7.838604 | 0.5387286 | 0.6112479 | 1.232301 | 0.2178368 | 0.403577 |
| ENSG00000183506 | PI4KAP2 | 443.5789 | 0.5386528 | 0.1588282 | 3.472029 | 0.00051654 | 0.003227 |
| ENSG00000143630 | HCN3 | 1453.101 | 0.5383105 | 0.2517843 | 2.274704 | 0.02292367 | 0.075777 |
| ENSG00000174844 | DNAH12 | 6.181217 | 0.5382535 | 0.6178453 | 1.232904 | 0.2176116 | 0.403301 |
| ENSG00000010270 | STARD3NL | 2060.273223 | 0.538217528 | 0.174872885 | 3.159372047 | 0.0015811 | 0.008471 |
| ENSG00000250740 | | 13.03569 | 0.5382158 | 0.5749626 | 1.260439 | 0.2075112 | 0.390051 |
| ENSG00000199975 | RN7SKP243 | 1.025936 | 0.5381986 | 0.9537286 | 1.527561 | 0.1266216 | NA |
| ENSG00000180264 | ADGRD2 | 28.73709 | 0.5381743 | 0.5675687 | 1.284465 | 0.1989791 | 0.379055 |

| | | | | | | | |
|---|---|---|---|---|---|---|---|
| ENSG00000249328 | | 2.606115 | 0.5379146 | 0.8539537 | 1.300172 | 0.1935419 | 0.372157 |
| ENSG00000153046 | CDYL | 2672.792 | 0.5375781 | 0.1561765 | 3.512668 | 0.00044363 | 0.002829 |
| ENSG00000188002 | | 2269.421 | 0.5374345 | 0.1457763 | 3.757 | 0.00017196 | 0.001242 |
| ENSG00000243055 | GK-AS1 | 6.974299 | 0.5373716 | 0.6257435 | 1.242075 | 0.214209 | 0.399055 |
| ENSG00000125505 | MBOAT7 | 11752.08272 | 0.537281008 | 0.157711957 | 3.478203659 | 0.00050479 | 0.003162 |
| ENSG00000279459 | | 9.898806 | 0.5372646 | 0.6277083 | 1.243935 | 0.2135234 | 0.398308 |
| ENSG00000250081 | | 3.419089 | 0.5372265 | 0.8557848 | 1.333564 | 0.1823469 | 0.357335 |
| ENSG00000113384 | GOLPH3 | 7057.425815 | 0.537196443 | 0.102665487 | 5.283147992 | 1.27E-07 | 1.89E-06 |
| ENSG00000156531 | PHF6 | 3514.173 | 0.537143 | 0.1960919 | 2.810879 | 0.00494064 | 0.021933 |
| ENSG00000133958 | UNC79 | 39.11506012 | 0.537032477 | 0.361341062 | 1.668613763 | 0.09519395 | 0.22442 |
| ENSG00000172057 | ORMDL3 | 2961.847 | 0.5369247 | 0.1377091 | 3.947754 | 7.89E-05 | 0.00062 |
| ENSG00000242337 | INHCAP | 14.9715 | 0.5366313 | 0.5586998 | 1.271109 | 0.2036898 | 0.385321 |
| ENSG00000204257 | HLA-DMA | 186.3408 | 0.5364029 | 0.2706825 | 2.112426 | 0.03464992 | 0.104866 |
| ENSG00000254860 | TMEM9B-AS1 | 128.9407 | 0.5363282 | 0.3602462 | 1.677278 | 0.09348818 | 0.221637 |
| ENSG00000172113 | NME6 | 1380.078 | 0.5362948 | 0.110698 | 4.896791 | 9.74E-07 | 1.21E-05 |
| ENSG00000131067 | GGT7 | 1413.722735 | 0.536050699 | 0.218232874 | 2.544093865 | 0.01095616 | 0.041884 |
| ENSG00000121410 | A1BG | 14.53083577 | 0.535983233 | 0.560949247 | 1.268294039 | 0.20469297 | 0.386626 |
| ENSG00000100441 | KHNYN | 5737.376801 | 0.535980365 | 0.226708375 | 2.518035065 | 0.01180116 | 0.044437 |
| ENSG00000164930 | FZD6 | 4996.744 | 0.5355599 | 0.1822325 | 3.010975 | 0.0026041 | 0.012949 |
| ENSG00000144566 | RAB5A | 5119.002 | 0.5355487 | 0.120194 | 4.544986 | 5.49E-06 | 5.79E-05 |
| ENSG00000259728 | LINC00933 | 35.46019 | 0.5355245 | 0.4807276 | 1.372164 | 0.1700125 | 0.340448 |
| ENSG00000116652 | DLEU2L | 16.98656338 | 0.535291309 | 0.435083111 | 1.456223587 | 0.14533082 | 0.304303 |
| ENSG00000236438 | FAM157A | 76.0131 | 0.5352591 | 0.323048 | 1.818702 | 0.06895688 | 0.176537 |
| ENSG00000233765 | CDRT15P12 | 6.561026 | 0.5351686 | 0.6518008 | 1.209689 | 0.2263984 | 0.413983 |
| ENSG00000137070 | IL11RA | 220.8765375 | 0.5350667 | 0.294957947 | 1.958617174 | 0.05015764 | 0.138788 |
| ENSG00000164442 | CITED2 | 9217.366 | 0.535 | 0.3620655 | 1.667822 | 0.09535109 | 0.22475 |
| ENSG00000137166 | FOXP4 | 2927.967999 | 0.534974395 | 0.204591245 | 2.71255628 | 0.00667664 | 0.028015 |
| ENSG00000112992 | NNT | 5023.164174 | 0.534963157 | 0.167235153 | 3.273692412 | 0.00106152 | 0.006025 |
| ENSG00000241945 | PWP2 | 27.54589 | 0.5345547 | 0.350649 | 1.706508 | 0.0879135 | 0.211958 |
| ENSG00000183856 | IQGAP3 | 3395.79 | 0.5345271 | 0.3788276 | 1.60194 | 0.1091689 | 0.247907 |
| ENSG00000285212 | | 5.337043 | 0.5342674 | 0.6542378 | 1.223865 | 0.2210032 | 0.407398 |
| ENSG00000267064 | UXT-AS1 | 47.34851 | 0.5341296 | 0.4334379 | 1.449713 | 0.1471386 | 0.306927 |
| ENSG00000261143 | ADAMTS7P3 | 2.858255 | 0.5339292 | 0.7682139 | 1.221449 | 0.2219159 | 0.408627 |

| | | | | | | | |
|---|---|---|---|---|---|---|---|
| ENSG00000050555 | LAMC3 | 7.924204447 | 0.533878089 | 0.665353166 | 1.222831935 | 0.22139318 | 0.407976 |
| ENSG00000258441 | LINC00641 | 842.856 | 0.5338444 | 0.3394902 | 1.740954 | 0.0816916 | 0.200579 |
| ENSG00000224536 | | 237.4878 | 0.5338424 | 0.3079378 | 1.886337 | 0.05924956 | 0.157594 |
| ENSG00000124507 | PACSIN1 | 1572.139702 | 0.533769067 | 0.26717832 | 2.124170356 | 0.0336559 | 0.102499 |
| ENSG00000068971 | PPP2R5B | 1128.003023 | 0.53367058 | 0.324900657 | 1.806661825 | 0.07081502 | 0.180172 |
| ENSG00000134597 | RBMX2 | 2533.220896 | 0.533667428 | 0.123231991 | 4.387516594 | 1.15E-05 | 0.000111 |
| ENSG00000106460 | TMEM106B | 3581.266316 | 0.533581 | 0.248476215 | 2.302546382 | 0.02130438 | 0.071528 |
| ENSG00000258365 | | 10.42998 | 0.5334702 | 0.5238694 | 1.300137 | 0.1935541 | 0.372157 |
| ENSG00000126500 | FLRT1 | 46.82380484 | 0.533306283 | 0.474916477 | 1.373873054 | 0.1694811 | 0.339742 |
| ENSG00000171206 | TRIM8 | 3814.03 | 0.5330927 | 0.1749472 | 3.143863 | 0.00166733 | 0.008855 |
| ENSG00000177105 | RHOG | 1127.28 | 0.5330581 | 0.206706 | 2.673251 | 0.007512 | 0.03091 |
| ENSG00000095203 | EPB41L4B | 1951.923862 | 0.533014057 | 0.110012171 | 4.896553312 | 9.75E-07 | 1.21E-05 |
| ENSG00000257809 | | 3.857953 | 0.5329715 | 0.7857673 | 1.237328 | 0.2159653 | 0.401249 |
| ENSG00000100266 | PACSIN2 | 3365.512563 | 0.532836966 | 0.130008737 | 4.159061367 | 3.20E-05 | 0.000278 |
| ENSG00000165105 | RASEF | 3272.413 | 0.5328075 | 0.2595857 | 2.232402 | 0.02558841 | 0.082597 |
| ENSG00000156234 | CXCL13 | 1.091425 | 0.5323669 | 0.9892414 | 1.751499 | 0.07986007 | NA |
| ENSG00000284968 | | 70.93761 | 0.5320843 | 0.3494071 | 1.690717 | 0.09089081 | 0.217165 |
| ENSG00000109654 | TRIM2 | 1693.106624 | 0.531963136 | 0.268876442 | 2.108450487 | 0.03499204 | 0.105588 |
| ENSG00000132406 | TMEM128 | 886.121411 | 0.53189085 | 0.169074436 | 3.223852705 | 0.00126478 | 0.007018 |
| ENSG00000253180 | | 11.65082 | 0.5318428 | 0.5196901 | 1.308853 | 0.1905842 | 0.368613 |
| ENSG00000125772 | GPCPD1 | 567.775493 | 0.53154266 | 0.377842486 | 1.598931843 | 0.10983575 | 0.249123 |
| ENSG00000271664 | | 29.48732 | 0.5313055 | 0.4779197 | 1.372805 | 0.169813 | 0.340177 |
| ENSG00000182158 | CREB3L2 | 4651.159 | 0.5311371 | 0.3486173 | 1.696571 | 0.08977775 | 0.215258 |
| ENSG00000121417 | ZNF211 | 941.2574789 | 0.530642587 | 0.223530891 | 2.479084913 | 0.01317199 | 0.048553 |
| ENSG00000130783 | CCDC62 | 18.29951196 | 0.530619812 | 0.516721043 | 1.301245604 | 0.1931744 | 0.371809 |
| ENSG00000164142 | FAM160A1 | 975.1683 | 0.5300872 | 0.3484181 | 1.705753 | 0.08805412 | 0.212162 |
| ENSG00000277977 | | 1.110965 | 0.5300714 | 0.92909 | 1.522548 | 0.1278719 | NA |
| ENSG00000129480 | DTD2 | 508.1368935 | 0.529449459 | 0.207713543 | 2.651060293 | 0.00802395 | 0.032618 |
| ENSG00000173083 | HPSE | 98.90012 | 0.5294359 | 0.4828081 | 1.362354 | 0.1730863 | 0.344474 |
| ENSG00000217060 | | 58.52953 | 0.5293096 | 0.3033851 | 1.891609 | 0.05854312 | 0.156262 |
| ENSG00000163635 | ATXN7 | 5716.659 | 0.5292447 | 0.21487 | 2.588732 | 0.00963301 | 0.037699 |
| ENSG00000235034 | C19orf81 | 160.7911 | 0.5291108 | 0.1849192 | 2.946862 | 0.00321017 | 0.015366 |
| ENSG00000004799 | PDK4 | 131.7591512 | 0.528960027 | 0.508883842 | 1.319726313 | 0.18692641 | 0.363371 |

| | | | | | | | |
|---|---|---|---|---|---|---|---|
| ENSG00000254909 | | 59.30096 | 0.5289445 | 0.4412349 | 1.431026 | 0.1524227 | 0.314712 |
| ENSG00000287221 | | 11.63209 | 0.5288282 | 0.5874247 | 1.237219 | 0.216006 | 0.401249 |
| ENSG00000136231 | IGF2BP3 | 7.38222474 | 0.528727383 | 0.966474552 | 1.508274009 | 0.13148441 | 0.283119 |
| ENSG00000167619 | TMEM145 | 12.2897 | 0.5286697 | 0.503571 | 1.31634 | 0.1880599 | 0.365006 |
| ENSG00000143742 | SRP9 | 9067.972 | 0.5284488 | 0.1732794 | 3.118976 | 0.00181481 | 0.009548 |
| ENSG00000213994 | | 6.881747 | 0.5283398 | 0.751968 | 1.204465 | 0.22841 | 0.416546 |
| ENSG00000130147 | SH3BP4 | 4174.586674 | 0.528311985 | 0.253733932 | 2.233368525 | 0.02552465 | 0.082438 |
| ENSG00000278616 | BEND3P3 | 55.70969 | 0.5281769 | 0.4641209 | 1.374305 | 0.1693472 | 0.339591 |
| ENSG00000168389 | MFSD2A | 181.574 | 0.52771 | 0.6445305 | 1.216114 | 0.2239417 | 0.41078 |
| ENSG00000224023 | EDRF1-DT | 13.54443 | 0.527503 | 0.4657868 | 1.373479 | 0.1696036 | 0.339937 |
| ENSG00000122085 | MTERF4 | 2345.051253 | 0.52731032 | 0.147971697 | 3.626366535 | 0.00028744 | 0.001948 |
| ENSG00000254682 | | 656.1164 | 0.5267718 | 0.2588944 | 2.155121 | 0.03115237 | 0.096602 |
| ENSG00000214960 | CRPPA | 90.32625 | 0.5266694 | 0.3209865 | 1.798351 | 0.07212138 | 0.182613 |
| ENSG00000268764 | | 3.348736 | 0.5266437 | 0.8036769 | 1.2383 | 0.2156049 | 0.400751 |
| ENSG00000233673 | ANAPC1P1 | 1.876711 | 0.526612 | 0.86761 | 1.334133 | 0.1821604 | 0.357077 |
| ENSG00000225855 | RUSC1-AS1 | 870.331 | 0.5265838 | 0.3517848 | 1.672978 | 0.09433158 | 0.223179 |
| ENSG00000151116 | UEVLD | 2065.21 | 0.5263469 | 0.152932 | 3.501846 | 0.00046205 | 0.002931 |
| ENSG00000104413 | ESRP1 | 10464.0889 | 0.52628078 | 0.14168054 | 3.779722601 | 0.000157 | 0.001144 |
| ENSG00000238055 | | 2.124029 | 0.5260531 | 0.935352 | 1.496288 | 0.1345785 | 0.28773 |
| ENSG00000227192 | | 9.850244 | 0.5258749 | 0.5368416 | 1.284149 | 0.1990898 | 0.379219 |
| ENSG00000162065 | TBC1D24 | 1665.823 | 0.525783 | 0.2014407 | 2.701136 | 0.00691031 | 0.028823 |
| ENSG00000126878 | AIF1L | 4616.422621 | 0.525739561 | 0.289374525 | 1.960880665 | 0.04989295 | 0.138272 |
| ENSG00000221821 | C6orf226 | 396.872 | 0.5257241 | 0.2394992 | 2.316775 | 0.02051601 | 0.06935 |
| ENSG00000124664 | SPDEF | 14031.20375 | 0.52556711 | 0.354002751 | 1.655863488 | 0.09774947 | 0.22886 |
| ENSG00000240137 | ERICH6-AS1 | 142.9098 | 0.5253172 | 0.2600754 | 2.137987 | 0.03251781 | 0.099765 |
| ENSG00000198668 | CALM1 | 22470.66 | 0.5247806 | 0.1937636 | 2.77931 | 0.00544745 | 0.023754 |
| ENSG00000251898 | SCARNA11 | 2.300567 | 0.5247574 | 0.8832013 | 1.356503 | 0.1749393 | 0.346915 |
| ENSG00000214946 | TBC1D26 | 3.68721 | 0.5246361 | 0.8347913 | 1.27403 | 0.2026526 | 0.383878 |
| ENSG00000253520 | | 2.299723 | 0.5246315 | 0.8116572 | 1.241012 | 0.2146014 | 0.399502 |
| ENSG00000253573 | | 10.01971 | 0.5244889 | 0.6198854 | 1.210147 | 0.2262223 | 0.413747 |
| ENSG00000287967 | | 13.27207 | 0.5244143 | 0.606257 | 1.21694 | 0.2236272 | 0.410328 |
| ENSG00000102904 | TSNAXIP1 | 135.5988578 | 0.524338457 | 0.31089003 | 1.837256341 | 0.06617206 | 0.171157 |
| ENSG00000260083 | MIR762HG | 63.15998 | 0.5242887 | 0.3253791 | 1.763398 | 0.07783333 | 0.193186 |

| | | | | | | | |
|---|---|---|---|---|---|---|---|
| ENSG00000234139 | | 1.887634 | 0.5242589 | 0.8739826 | 1.324502 | 0.1853364 | 0.360985 |
| ENSG00000287555 | | 4.448322 | 0.5242151 | 0.7771231 | 1.222041 | 0.2216923 | 0.408329 |
| ENSG00000230154 | AIDAP1 | 52.24504 | 0.5242132 | 0.3911275 | 1.543968 | 0.122596 | 0.269379 |
| ENSG00000253194 | | 13.74977 | 0.5240401 | 0.59346 | 1.227638 | 0.219583 | 0.405649 |
| ENSG00000121895 | TMEM156 | 4.222363259 | 0.523990292 | 1.001923589 | 1.809871843 | 0.07031566 | 0.179236 |
| ENSG00000283047 | FRG1FP | 28.6953 | 0.5239832 | 0.4388368 | 1.416077 | 0.1567529 | 0.321455 |
| ENSG00000163430 | FSTL1 | 129.9294 | 0.5239251 | 0.3798901 | 1.571799 | 0.1159972 | 0.258855 |
| ENSG00000115109 | EPB41L5 | 5925.638525 | 0.523793997 | 0.120685151 | 4.421447953 | 9.80E-06 | 9.68E-05 |
| ENSG00000228486 | C2orf92 | 315.6828 | 0.5236886 | 0.3098946 | 1.841312 | 0.06557583 | 0.169913 |
| ENSG00000130545 | CRB3 | 1291.943536 | 0.523565979 | 0.189733094 | 2.848212359 | 0.00439656 | 0.019937 |
| ENSG00000164953 | TMEM67 | 650.7133 | 0.5235394 | 0.1821217 | 2.956468 | 0.00311185 | 0.014981 |
| ENSG00000225544 | | 39.68873 | 0.5232833 | 0.3569322 | 1.641706 | 0.100651 | 0.233879 |
| ENSG00000054267 | ARID4B | 3977.095979 | 0.523034708 | 0.21696386 | 2.50385504 | 0.01228484 | 0.045874 |
| ENSG00000273154 | | 23.28477 | 0.5230329 | 0.3623443 | 1.617358 | 0.105801 | 0.242434 |
| ENSG00000213463 | SYNJ2BP | 3438.324 | 0.522916 | 0.1583587 | 3.374917 | 0.00073838 | 0.004402 |
| ENSG00000256594 | | 131.4753 | 0.5228506 | 0.3462698 | 1.664942 | 0.09592431 | 0.225621 |
| ENSG00000253384 | | 14.79088 | 0.5227532 | 0.4910354 | 1.324411 | 0.1853665 | 0.360991 |
| ENSG00000198520 | ARMH1 | 244.6308 | 0.5226546 | 0.4930439 | 1.321601 | 0.1863011 | 0.362467 |
| ENSG00000142227 | EMP3 | 90.90345 | 0.5226449 | 0.3112377 | 1.826447 | 0.06778292 | 0.174289 |
| ENSG00000267341 | | 2.734666 | 0.5224896 | 0.7913762 | 1.213128 | 0.2250809 | 0.412113 |
| ENSG00000251787 | RNU7-47P | 0.8879334 | 0.5224856 | 0.9720009 | 1.573113 | 0.1156927 | NA |
| ENSG00000287249 | | 10.5724 | 0.5220356 | 0.6403837 | 1.205368 | 0.2280614 | 0.416108 |
| ENSG00000288398 | | 354.2262 | 0.5215704 | 0.254408 | 2.16918 | 0.03006902 | 0.093837 |
| ENSG00000259366 | | 68.15546 | 0.5214656 | 0.4140389 | 1.465838 | 0.1426926 | 0.300293 |
| ENSG00000254413 | CHKB-CPT1B | 37.34805 | 0.521254 | 0.3062874 | 1.847394 | 0.0646901 | 0.168093 |
| ENSG00000228791 | THRB-AS1 | 20.97069 | 0.5212491 | 0.528463 | 1.271698 | 0.2034805 | 0.385119 |
| ENSG00000102962 | CCL22 | 16.07337649 | 0.521174824 | 0.742911079 | 1.207507464 | 0.22723677 | 0.415088 |
| ENSG00000140025 | EFCAB11 | 553.2254 | 0.5210106 | 0.2578792 | 2.142973 | 0.03211529 | 0.098842 |
| ENSG00000206053 | JPT2 | 13858.67 | 0.5209877 | 0.1845266 | 2.939858 | 0.00328363 | 0.01566 |
| ENSG00000255303 | OR5BA1P | 2.51322 | 0.5209839 | 0.842301 | 1.277664 | 0.2013679 | 0.382144 |
| ENSG00000151846 | PABPC3 | 962.7239 | 0.5209579 | 0.1740522 | 3.065127 | 0.00217578 | 0.011119 |
| ENSG00000238039 | | 12.15093 | 0.5209426 | 0.551689 | 1.238754 | 0.2154365 | 0.400499 |
| ENSG00000250326 | | 32.94465 | 0.5208577 | 0.3832541 | 1.546326 | 0.1220259 | 0.268469 |

| | | | | | | | |
|---|---|---|---|---|---|---|---|
| ENSG00000259113 | | 2.184529 | 0.5207514 | 0.9277442 | 1.484904 | 0.1375691 | 0.292149 |
| ENSG00000277855 | | 6.419047 | 0.5207154 | 0.6012189 | 1.20256 | 0.2291465 | 0.417285 |
| ENSG00000104936 | DMPK | 1570.990878 | 0.52064148 | 0.250420897 | 2.198335123 | 0.02792523 | 0.088506 |
| ENSG00000143674 | MAP3K21 | 1518.337 | 0.5205464 | 0.2123218 | 2.545428 | 0.0109144 | 0.041754 |
| ENSG00000255568 | BRWD1-AS2 | 180.6338 | 0.5203971 | 0.240231 | 2.278537 | 0.0226946 | 0.075276 |
| ENSG00000136888 | ATP6V1G1 | 7360.545117 | 0.520396029 | 0.150904309 | 3.509852955 | 0.00044836 | 0.002854 |
| ENSG00000233108 | GLCCI1-DT | 3.574576 | 0.5202697 | 0.7411633 | 1.191341 | 0.2335198 | 0.422541 |
| ENSG00000152128 | TMEM163 | 13.77751 | 0.5199387 | 0.6559227 | 1.192602 | 0.2330252 | 0.421879 |
| ENSG00000250535 | STK19B | 6.584834 | 0.5199198 | 0.5697628 | 1.220322 | 0.2223427 | 0.409016 |
| ENSG00000235351 | | 23.27569 | 0.5198209 | 0.5099108 | 1.297192 | 0.194565 | 0.373737 |
| ENSG00000249252 | C1QTNF7-AS1 | 3.38763 | 0.5196917 | 0.7468352 | 1.197099 | 0.2312678 | 0.419681 |
| ENSG00000122863 | CHST3 | 1691.937819 | 0.519663612 | 0.211066344 | 2.561428854 | 0.01042426 | 0.040239 |
| ENSG00000114248 | LRRC31 | 6.828995937 | 0.519197645 | 0.862489094 | 1.332931935 | 0.18255414 | 0.357506 |
| ENSG00000223873 | SAP18P2 | 7.169121 | 0.5191662 | 0.5637166 | 1.238226 | 0.2156321 | 0.400751 |
| ENSG00000279541 | | 6.213058 | 0.5190433 | 0.7462874 | 1.193801 | 0.2325557 | 0.421213 |
| ENSG00000198658 | ABHD17AP1 | 134.295 | 0.5190363 | 0.2524506 | 2.176396 | 0.02952563 | 0.092639 |
| ENSG00000177350 | RPL13AP3 | 7.087686 | 0.5190174 | 0.6008233 | 1.203982 | 0.2285966 | 0.416741 |
| ENSG00000259322 | | 13.79167 | 0.5189239 | 0.4513174 | 1.374894 | 0.1691644 | 0.339312 |
| ENSG00000126778 | SIX1 | 775.9877361 | 0.518828283 | 0.21607401 | 2.508141131 | 0.01213682 | 0.045405 |
| ENSG00000272106 | | 323.3894 | 0.5188088 | 0.1972798 | 2.731155 | 0.00631128 | 0.026778 |
| ENSG00000275645 | | 13.54216 | 0.5186854 | 0.5528246 | 1.233601 | 0.2173515 | 0.403012 |
| ENSG00000186106 | ANKRD46 | 1387.47 | 0.5186507 | 0.109586 | 4.781752 | 1.74E-06 | 2.07E-05 |
| ENSG00000166455 | C16orf46 | 1.729907 | 0.5185719 | 0.917087 | 1.440456 | 0.1497385 | 0.31074 |
| ENSG00000270728 | | 45.10103 | 0.5185017 | 0.3304991 | 1.73819 | 0.08217726 | 0.201473 |
| ENSG00000272192 | | 5.470427 | 0.5182024 | 0.8615705 | 1.324857 | 0.1852184 | 0.360835 |
| ENSG00000113555 | PCDH12 | 42.44895613 | 0.518090202 | 0.395927259 | 1.501085583 | 0.13333343 | 0.285965 |
| ENSG00000184545 | DUSP8 | 891.484 | 0.5175869 | 0.2662644 | 2.064147 | 0.03900381 | 0.114672 |
| ENSG00000215712 | TMEM242 | 327.1147 | 0.5174123 | 0.1840201 | 2.894195 | 0.00380132 | 0.01768 |
| ENSG00000167995 | BEST1 | 189.5272 | 0.5173473 | 0.765275 | 1.201614 | 0.229513 | 0.417695 |
| ENSG00000067221 | STOML1 | 573.3850129 | 0.517264777 | 0.205281394 | 2.616942293 | 0.00887213 | 0.035351 |
| ENSG00000109790 | KLHL5 | 1517.416601 | 0.517191636 | 0.30947221 | 1.828569738 | 0.06746409 | 0.17362 |
| ENSG00000184068 | SREBF2-AS1 | 134.9043 | 0.517085 | 0.3213185 | 1.76961 | 0.07679206 | 0.191347 |
| ENSG00000285976 | | 145.7585 | 0.5169973 | 0.3702467 | 1.573447 | 0.1156153 | 0.258459 |

| | | | | | | | |
|---|---|---|---|---|---|---|---|
| ENSG00000067167 | TRAM1 | 10376.13887 | 0.516972278 | 0.159310234 | 3.318583755 | 0.00090475 | 0.005236 |
| ENSG00000168481 | LGI3 | 3.823453 | 0.5169451 | 0.7863705 | 1.20108 | 0.22972 | 0.417929 |
| ENSG00000188738 | FSIP2 | 91.45442 | 0.5168411 | 0.5132486 | 1.27447 | 0.202497 | 0.383693 |
| ENSG00000114480 | GBE1 | 1163.684816 | 0.516822531 | 0.194055695 | 2.757757431 | 0.00581994 | 0.025127 |
| ENSG00000171940 | ZNF217 | 41969.07 | 0.516781 | 0.2480862 | 2.239506 | 0.02512301 | 0.081412 |
| ENSG00000161395 | PGAP3 | 969.8981 | 0.5167673 | 0.195269 | 2.73654 | 0.0062089 | 0.026449 |
| ENSG00000176593 | | 520.3504 | 0.51658 | 0.3217069 | 1.762515 | 0.07798226 | 0.193438 |
| ENSG00000236562 | IFITM3P1 | 2.424185 | 0.5165307 | 0.8033803 | 1.230948 | 0.2183422 | 0.404091 |
| ENSG00000265148 | TSPOAP1-AS1 | 16.96099 | 0.5165143 | 0.5464845 | 1.238242 | 0.2156264 | 0.400751 |
| ENSG00000287641 | | 1.276967 | 0.5164049 | 1.00605 | 1.799083 | 0.07200553 | NA |
| ENSG00000165943 | MOAP1 | 5100.751 | 0.5163304 | 0.1335852 | 3.917674 | 8.94E-05 | 0.000692 |
| ENSG00000174326 | SLC16A11 | 21.43952 | 0.516316 | 0.4693565 | 1.339638 | 0.180363 | 0.354629 |
| ENSG00000240163 | | 6.054799 | 0.5162043 | 0.7232158 | 1.195077 | 0.2320568 | 0.420445 |
| ENSG00000120742 | SERP1 | 14347.78823 | 0.516073155 | 0.143170301 | 3.661710831 | 0.00025054 | 0.001726 |
| ENSG00000205730 | ITPRIPL2 | 1942.279 | 0.5159477 | 0.2506 | 2.181819 | 0.02912288 | 0.091722 |
| ENSG00000107929 | LARP4B | 9375.93157 | 0.515943733 | 0.148401282 | 3.543918491 | 0.00039423 | 0.002562 |
| ENSG00000277452 | RN7SL473P | 8.43364 | 0.5158585 | 0.6981449 | 1.187872 | 0.234884 | 0.424158 |
| ENSG00000198270 | TMEM116 | 849.7325 | 0.5157936 | 0.2008554 | 2.650862 | 0.00802866 | 0.032632 |
| ENSG00000172375 | C2CD2L | 2327.774 | 0.5156969 | 0.17979 | 2.931763 | 0.00337043 | 0.016017 |
| ENSG00000119283 | TRIM67 | 9.030982535 | 0.515500901 | 0.564512221 | 1.225883189 | 0.22024264 | 0.406395 |
| ENSG00000120262 | CCDC170 | 3032.653862 | 0.515461902 | 0.275790203 | 1.994399053 | 0.04610846 | 0.130353 |
| ENSG00000112033 | PPARD | 1777.849734 | 0.515329277 | 0.109726205 | 4.740264534 | 2.13E-06 | 2.49E-05 |
| ENSG00000147100 | SLC16A2 | 892.3739 | 0.5152031 | 0.3868584 | 1.528081 | 0.1264924 | 0.275198 |
| ENSG00000226543 | MYL6P1 | 6.397965 | 0.5150784 | 0.6536443 | 1.177477 | 0.239005 | 0.428947 |
| ENSG00000226919 | | 10.34616 | 0.5149756 | 0.8565116 | 1.361434 | 0.1733766 | 0.344897 |
| ENSG00000049245 | VAMP3 | 3744.929252 | 0.514943836 | 0.144872254 | 3.608500405 | 0.00030797 | 0.00207 |
| ENSG00000226877 | | 1.566809 | 0.5148601 | 0.9158584 | 1.43124 | 0.1523613 | NA |
| ENSG00000182700 | IGIP | 370.6784 | 0.5147121 | 0.3742197 | 1.557504 | 0.1193508 | 0.264209 |
| ENSG00000135932 | CAB39 | 3775.833171 | 0.514670604 | 0.162319334 | 3.223715192 | 0.00126539 | 0.007018 |
| ENSG00000143199 | ADCY10 | 5.669249 | 0.5146009 | 0.6742896 | 1.170196 | 0.2419221 | 0.432487 |
| ENSG00000228283 | KATNBL1P6 | 153.0179 | 0.5145176 | 0.205216 | 2.599606 | 0.0093331 | 0.036771 |
| ENSG00000205579 | DYNLL1P1 | 5.175799 | 0.514308 | 0.6210644 | 1.190825 | 0.2337225 | 0.422779 |
| ENSG00000166530 | HSBP1P2 | 9.956573 | 0.5140013 | 0.6039792 | 1.198809 | 0.2306023 | 0.419046 |

| | | | | | | | |
|---|---|---|---|---|---|---|---|
| ENSG00000243431 | RPL5P30 | 15.82382 | 0.5138283 | 0.5034091 | 1.280771 | 0.2002742 | 0.380703 |
| ENSG00000278757 | | 1.076676 | 0.5138236 | 1.015739 | 1.933076 | 0.05322679 | NA |
| ENSG00000165879 | FRAT1 | 910.5644 | 0.5137648 | 0.2062255 | 2.609775 | 0.00906017 | 0.035934 |
| ENSG00000111540 | RAB5B | 10427.32149 | 0.513145195 | 0.154370588 | 3.400130649 | 0.00067354 | 0.004063 |
| ENSG00000132405 | TBC1D14 | 1650.562987 | 0.512889184 | 0.225991318 | 2.356795944 | 0.01843337 | 0.06363 |
| ENSG00000174776 | WDR49 | 2.716267 | 0.5128374 | 0.775894 | 1.203296 | 0.2288617 | 0.417024 |
| ENSG00000150455 | TIRAP | 218.8483 | 0.5127544 | 0.1879388 | 2.814879 | 0.00487956 | 0.021749 |
| ENSG00000164610 | RP9 | 1684.888 | 0.5126372 | 0.1301358 | 4.000035 | 6.33E-05 | 0.000511 |
| ENSG00000276707 | | 5.527823 | 0.5123957 | 0.7515795 | 1.161826 | 0.245306 | 0.437005 |
| ENSG00000001630 | CYP51A1 | 373.612109 | 0.512220085 | 0.266727674 | 2.052083531 | 0.04016155 | 0.117258 |
| ENSG00000269373 | | 0.9005544 | 0.5121216 | 0.9673245 | 1.559767 | 0.1188148 | NA |
| ENSG00000171552 | BCL2L1 | 4155.469 | 0.5119004 | 0.2190512 | 2.437066 | 0.01480697 | 0.053387 |
| ENSG00000184378 | ACTRT3 | 179.0597 | 0.5118134 | 0.3291208 | 1.711251 | 0.08703486 | 0.210203 |
| ENSG00000259287 | | 25.16348 | 0.5117967 | 0.4094552 | 1.447884 | 0.1476496 | 0.307655 |
| ENSG00000242088 | | 4.822764 | 0.5117055 | 0.7344505 | 1.181134 | 0.2375493 | 0.4272 |
| ENSG00000188451 | SRP72P2 | 14.10261 | 0.5115701 | 0.4395522 | 1.3884 | 0.1650153 | 0.333406 |
| ENSG00000231551 | PDE4DIPP6 | 56.04369 | 0.5114947 | 0.5100993 | 1.266334 | 0.2053936 | 0.387358 |
| ENSG00000105672 | ETV2 | 54.85590878 | 0.51131544 | 0.291020003 | 1.902458441 | 0.05711125 | 0.153179 |
| ENSG00000235175 | RPL26P37 | 1.633834 | 0.5110765 | 0.8909327 | 1.370146 | 0.1706412 | 0.341347 |
| ENSG00000181817 | LSM10 | 1908.473 | 0.5110465 | 0.2444215 | 2.202889 | 0.02760254 | 0.087737 |
| ENSG00000215450 | | 2.534596 | 0.5108289 | 0.9261402 | 1.480256 | 0.138805 | 0.294186 |
| ENSG00000273096 | | 4.8583 | 0.5107806 | 0.7018641 | 1.16947 | 0.2422144 | 0.432835 |
| ENSG00000143355 | LHX9 | 1.91718 | 0.510686 | 0.8388244 | 1.25283 | 0.2102677 | 0.394005 |
| ENSG00000274515 | | 15.36718 | 0.5105489 | 0.5519111 | 1.226856 | 0.2198765 | 0.406021 |
| ENSG00000175206 | NPPA | 13.13085 | 0.5101778 | 0.5999061 | 1.18204 | 0.2371899 | 0.4269 |
| ENSG00000143363 | PRUNE1 | 4412.709 | 0.5099539 | 0.1421008 | 3.639701 | 0.00027296 | 0.001861 |
| ENSG00000256746 | MADD-AS1 | 8.234801 | 0.5098788 | 0.5723989 | 1.204339 | 0.2284587 | 0.416546 |
| ENSG00000280776 | LINC01202 | 66.28739 | 0.5097714 | 0.412702 | 1.448806 | 0.1473918 | 0.307286 |
| ENSG00000233264 | | 6.380305 | 0.5097495 | 0.6317796 | 1.175705 | 0.2397126 | 0.429927 |
| ENSG00000187105 | HEATR4 | 40.87602 | 0.5094858 | 0.4475378 | 1.359485 | 0.1739929 | 0.345786 |
| ENSG00000249786 | EAF1-AS1 | 8.002899 | 0.509299 | 0.6595517 | 1.162549 | 0.2450125 | 0.436719 |
| ENSG00000077943 | ITGA8 | 1.308195708 | 0.50921055 | 0.962818283 | 1.554742333 | 0.12000745 | NA |
| ENSG00000129514 | FOXA1 | 17336.47986 | 0.509109778 | 0.186630269 | 2.741279145 | 0.00612005 | 0.026131 |

| | | | | | | | |
|---|---|---|---|---|---|---|---|
| ENSG00000100503 | NIN | 4361.979164 | 0.509007526 | 0.25020831 | 2.148401267 | 0.03168189 | 0.097879 |
| ENSG00000224389 | C4B | 19.68677 | 0.5089859 | 0.5292579 | 1.234225 | 0.217119 | 0.402669 |
| ENSG00000182952 | HMGN4 | 3710.019 | 0.5089449 | 0.1529568 | 3.39019 | 0.00069844 | 0.004196 |
| ENSG00000168124 | OR1F1 | 7.084552 | 0.5087962 | 0.6924399 | 1.148272 | 0.2508563 | 0.443506 |
| ENSG00000287729 | | 12.04363 | 0.5086779 | 0.6901168 | 1.166002 | 0.2436137 | 0.434839 |
| ENSG00000161267 | BDH1 | 3570.536 | 0.508665 | 0.2875776 | 1.893054 | 0.05835065 | 0.155913 |
| ENSG00000277914 | DDX6P2 | 2.533306 | 0.5085942 | 0.8359492 | 1.263531 | 0.2063985 | 0.388591 |
| ENSG00000157020 | SEC13 | 8750.405 | 0.5085146 | 0.2226074 | 2.367365 | 0.01791524 | 0.062181 |
| ENSG00000255621 | | 3.818153 | 0.5084834 | 0.7384638 | 1.1829 | 0.2368489 | 0.426522 |
| ENSG00000270585 | SNX2P1 | 4.483205 | 0.5084153 | 0.7836119 | 1.220451 | 0.2222939 | 0.408955 |
| ENSG00000263006 | ROCK1P1 | 70.95056 | 0.5083587 | 0.3071007 | 1.797201 | 0.07230369 | 0.182991 |
| ENSG00000103496 | STX4 | 3722.957878 | 0.50832507 | 0.193829269 | 2.699813715 | 0.00693783 | 0.028915 |
| ENSG00000272948 | | 15.84852 | 0.5079599 | 0.4791955 | 1.301951 | 0.1929331 | 0.371425 |
| ENSG00000205085 | FAM71F2 | 39.76144 | 0.5078977 | 0.4732601 | 1.309902 | 0.1902289 | 0.368006 |
| ENSG00000116525 | TRIM62 | 1069.751903 | 0.507828621 | 0.216267748 | 2.446568998 | 0.01442232 | 0.052276 |
| ENSG00000142182 | DNMT3L | 1.781106 | 0.5077747 | 0.9215158 | 1.424841 | 0.1542031 | 0.317574 |
| ENSG00000118557 | PMFBP1 | 35.69517881 | 0.507732543 | 0.456892821 | 1.337493635 | 0.18106156 | 0.355657 |
| ENSG00000228106 | NA | 118.019 | 0.50751 | 0.3810539 | 1.518212 | 0.128961 | 0.279172 |
| ENSG00000240356 | RPL23AP7 | 199.7461 | 0.5073094 | 0.1981878 | 2.655434 | 0.00792064 | 0.032293 |
| ENSG00000174672 | BRSK2 | 1192.073 | 0.5072483 | 0.1954224 | 2.673496 | 0.00750653 | 0.030892 |
| ENSG00000233611 | | 2.456287 | 0.507235 | 0.8522842 | 1.267054 | 0.2051361 | 0.387175 |
| ENSG00000204576 | PRR3 | 1646.92 | 0.5072259 | 0.1882818 | 2.80709 | 0.00499913 | 0.022144 |
| ENSG00000152620 | NADK2 | 1973.874 | 0.5070819 | 0.1754522 | 2.972578 | 0.0029531 | 0.014402 |
| ENSG00000143375 | CGN | 8746.546 | 0.5068033 | 0.1779725 | 2.91483 | 0.00355882 | 0.016746 |
| ENSG00000146267 | FAXC | 290.4023 | 0.5067904 | 0.1919079 | 2.733862 | 0.00625962 | 0.026623 |
| ENSG00000286965 | | 2.588715 | 0.50677 | 0.8757405 | 1.423243 | 0.1546657 | 0.318354 |
| ENSG00000054611 | TBC1D22A | 1223.266996 | 0.506763986 | 0.152506452 | 3.40553844 | 0.00066034 | 0.003996 |
| ENSG00000202337 | RNU6-8 | 68.87145 | 0.5067413 | 0.3415017 | 1.650143 | 0.09891363 | 0.230781 |
| ENSG00000113758 | DBN1 | 9816.719354 | 0.506527857 | 0.219185984 | 2.409646634 | 0.01596798 | 0.056627 |
| ENSG00000104518 | GSDMD | 20.58796874 | 0.506474189 | 0.649907265 | 1.169563947 | 0.24217649 | 0.432797 |
| ENSG00000228028 | | 6.367813 | 0.5064424 | 0.6288126 | 1.172018 | 0.2411899 | 0.431701 |
| ENSG00000181097 | LINC02878 | 63.36258 | 0.5060878 | 0.312051 | 1.764099 | 0.07771534 | 0.192989 |
| ENSG00000255036 | SUGT1P4-STRA6 | 2.504092 | 0.505708 | 0.8346344 | 1.241857 | 0.2142893 | 0.399145 |

| | | | | | | | |
|---|---|---|---|---|---|---|---|
| ENSG00000198865 | CCDC152 | 15.14254 | 0.5055678 | 0.7381598 | 1.175977 | 0.2396042 | 0.429819 |
| ENSG00000138735 | PDE5A | 1953.621295 | 0.505495761 | 0.334773985 | 1.666743941 | 0.09556533 | 0.22502 |
| ENSG00000130957 | FBP2 | 36.60474175 | 0.505118947 | 0.423349016 | 1.398020614 | 0.16210688 | 0.329383 |
| ENSG00000121671 | CRY2 | 2871.152312 | 0.504786793 | 0.221322586 | 2.39810001 | 0.01648037 | 0.058124 |
| ENSG00000099617 | EFNA2 | 57.14724625 | 0.504780448 | 0.445936846 | 1.348351007 | 0.17754551 | 0.350852 |
| ENSG00000227630 | LINC01132 | 24.02205 | 0.5047573 | 0.4598818 | 1.328624 | 0.1839722 | 0.359303 |
| ENSG00000189190 | ZNF600 | 1084.141 | 0.504698 | 0.1729178 | 2.992931 | 0.00276313 | 0.01362 |
| ENSG00000225671 | FCF1P6 | 13.3161 | 0.5042944 | 0.5804216 | 1.18653 | 0.235413 | 0.424853 |
| ENSG00000228701 | TNKS2-AS1 | 65.39151 | 0.5042539 | 0.3635582 | 1.568906 | 0.1166698 | 0.259876 |
| ENSG00000092421 | SEMA6A | 390.6556866 | 0.504151911 | 0.341423603 | 1.638588013 | 0.10129909 | 0.234997 |
| ENSG00000273064 | | 9.118964 | 0.5040731 | 0.7365606 | 1.162803 | 0.2449094 | 0.436565 |
| ENSG00000181450 | ZNF678 | 681.4469 | 0.503989 | 0.2547014 | 2.099079 | 0.03580992 | 0.10749 |
| ENSG00000286406 | | 65.03865 | 0.5039812 | 0.2870902 | 1.883263 | 0.05966468 | 0.158365 |
| ENSG00000160219 | GAB3 | 5.820342 | 0.5039399 | 0.7985687 | 1.224127 | 0.2209042 | 0.407276 |
| ENSG00000079387 | SENP1 | 1260.87167 | 0.503938222 | 0.175250085 | 2.96035591 | 0.00307284 | 0.014825 |
| ENSG00000267980 | | 57.77673 | 0.5038313 | 0.3424862 | 1.639185 | 0.1011748 | 0.234749 |
| ENSG00000269481 | | 21.54661 | 0.5037692 | 0.440582 | 1.356652 | 0.1748918 | 0.346858 |
| ENSG00000180432 | CYP8B1 | 3.490992 | 0.5036113 | 0.9436405 | 1.489595 | 0.1363306 | 0.290447 |
| ENSG00000180878 | C11orf42 | 5.445836 | 0.5035119 | 0.6389601 | 1.152623 | 0.2490652 | 0.441426 |
| ENSG00000124613 | ZNF391 | 381.7865474 | 0.503511093 | 0.232611931 | 2.270139496 | 0.02319912 | 0.076464 |
| ENSG00000169991 | IFFO2 | 861.4309 | 0.5032669 | 0.2384915 | 2.228095 | 0.02587418 | 0.083282 |
| ENSG00000197063 | MAFG | 3345.834 | 0.5031793 | 0.2688767 | 1.993461 | 0.04621095 | 0.130573 |
| ENSG00000160226 | CFAP410 | 865.0344 | 0.5031771 | 0.1655777 | 3.116587 | 0.00182958 | 0.009611 |
| ENSG00000268081 | | 2.685479 | 0.5031599 | 0.9472022 | 1.548109 | 0.1215961 | 0.267859 |
| ENSG00000125457 | MIF4GD | 2659.093135 | 0.502576873 | 0.143366596 | 3.565669953 | 0.00036293 | 0.002386 |
| ENSG00000279140 | | 89.59142 | 0.5025273 | 0.286131 | 1.886767 | 0.05919169 | 0.157518 |
| ENSG00000259251 | | 4.724271 | 0.5020187 | 0.6771551 | 1.143796 | 0.2527081 | 0.445534 |
| ENSG00000160336 | ZNF761 | 2532.176 | 0.501928 | 0.1636916 | 3.127628 | 0.00176223 | 0.009302 |
| ENSG00000265010 | | 4.565025 | 0.5018776 | 0.8100035 | 1.22289 | 0.2213711 | 0.407964 |
| ENSG00000235072 | ARNILA | 11.96206 | 0.5017292 | 0.5483542 | 1.199117 | 0.2304825 | 0.418857 |
| ENSG00000287170 | | 15.68495 | 0.5017189 | 0.4193454 | 1.397783 | 0.1621781 | 0.329478 |
| ENSG00000231999 | LRRC8C-DT | 14.35792 | 0.5016046 | 0.5221805 | 1.250687 | 0.2110489 | 0.394956 |
| ENSG00000183060 | LYSMD4 | 948.7107 | 0.5015083 | 0.2434606 | 2.173501 | 0.02974262 | 0.093115 |

| | | | | | | | |
|---|---|---|---|---|---|---|---|
| ENSG00000178385 | PLEKHM3 | 580.6272 | 0.5013984 | 0.3211629 | 1.712126 | 0.08687352 | 0.210003 |
| ENSG00000144821 | MYH15 | 4.830953 | 0.5012355 | 0.6800729 | 1.139937 | 0.2543124 | 0.447319 |
| ENSG00000259762 | | 2.043231 | 0.5012174 | 0.8282363 | 1.235108 | 0.2167902 | 0.40224 |
| ENSG00000168329 | CX3CR1 | 7.787324 | 0.501171 | 0.6649809 | 1.137572 | 0.2552993 | 0.448674 |
| ENSG00000179766 | ATP8B5P | 4.472275 | 0.5011562 | 0.6454756 | 1.141615 | 0.2536141 | 0.446667 |
| ENSG00000140199 | SLC12A6 | 450.9824 | 0.5011114 | 0.3793728 | 1.510547 | 0.1309038 | 0.282212 |
| ENSG00000177485 | ZBTB33 | 3448.77 | 0.501099 | 0.1925221 | 2.694582 | 0.0070477 | 0.029304 |
| ENSG00000180008 | SOCS4 | 2965.184 | 0.5010062 | 0.1998095 | 2.580022 | 0.0098794 | 0.03848 |
| ENSG00000113638 | TTC33 | 796.1541325 | 0.500942009 | 0.202421722 | 2.572103497 | 0.01010827 | 0.039224 |
| ENSG00000176029 | C11orf16 | 1.067773 | 0.5007941 | 0.9665302 | 1.61683 | 0.105915 | NA |
| ENSG00000260101 | | 17.36372 | 0.5007295 | 0.5244816 | 1.228915 | 0.2191038 | 0.405162 |
| ENSG00000104517 | UBR5 | 18708.2187 | 0.500554562 | 0.206027122 | 2.520172643 | 0.01172973 | 0.044205 |
| ENSG00000241429 | EEF1A1P25 | 3.969917 | 0.5004385 | 0.6623258 | 1.137951 | 0.2551409 | 0.448455 |
| ENSG00000087008 | ACOX3 | 1124.85737 | 0.50026792 | 0.150314609 | 3.389179431 | 0.00070102 | 0.004208 |
| ENSG00000260417 | | 2.354059 | 0.5001434 | 0.8383932 | 1.25659 | 0.2089022 | 0.391861 |
| ENSG00000186026 | ZNF284 | 117.6186 | 0.5000147 | 0.3237747 | 1.69266 | 0.09052017 | 0.216552 |
| ENSG00000157191 | NECAP2 | 2409.426 | 0.4999034 | 0.1492259 | 3.411084 | 0.00064705 | 0.003926 |
| ENSG00000278520 | MIR7851 | 1.478767 | 0.4990939 | 0.8917788 | 1.471145 | 0.141252 | NA |
| ENSG00000232893 | IQCA1-AS1 | 2.066707 | 0.4989867 | 0.9420476 | 1.487235 | 0.1369527 | 0.291212 |
| ENSG00000139718 | SETD1B | 3706.499 | 0.4987166 | 0.2270808 | 2.297211 | 0.02160675 | 0.07241 |
| ENSG00000173480 | ZNF417 | 533.933 | 0.4987124 | 0.2995518 | 1.81243 | 0.0699197 | 0.17842 |
| ENSG00000157734 | SNX22 | 138.3674 | 0.4986727 | 0.214433 | 2.431396 | 0.01504076 | 0.054039 |
| ENSG00000226466 | RPA2P2 | 1.008762 | 0.4985507 | 0.993638 | 1.852898 | 0.06389698 | NA |
| ENSG00000259205 | PRKXP1 | 26.42788 | 0.4985269 | 0.515571 | 1.233484 | 0.2173951 | 0.403012 |
| ENSG00000100271 | TTLL1 | 297.5628794 | 0.498471329 | 0.263256967 | 2.00870619 | 0.0445683 | 0.127162 |
| ENSG00000174996 | KLC2 | 4008.383 | 0.4983218 | 0.1723521 | 2.969548 | 0.00298238 | 0.014492 |
| ENSG00000259422 | | 19.18621 | 0.498152 | 0.4746918 | 1.2809 | 0.2002287 | 0.380667 |
| ENSG00000258081 | | 1.202655 | 0.4980575 | 0.9508493 | 1.618212 | 0.1056168 | NA |
| ENSG00000253320 | MAILR | 392.1087 | 0.4980569 | 0.285229 | 1.870541 | 0.06140871 | 0.161846 |
| ENSG00000254997 | KRTAP5-9 | 43.39107 | 0.4980522 | 0.4303292 | 1.366597 | 0.1717516 | 0.342511 |
| ENSG00000224420 | ADM5 | 177.9224 | 0.49797 | 0.2942858 | 1.829774 | 0.0672837 | 0.173305 |
| ENSG00000180999 | C1orf105 | 1.994768 | 0.4978899 | 0.8606624 | 1.299413 | 0.1938022 | 0.372478 |
| ENSG00000232386 | | 230.3287 | 0.4976264 | 0.3694473 | 1.532934 | 0.1252922 | 0.273487 |

| | | | | | | | |
|---|---|---|---|---|---|---|---|
| ENSG00000179397 | CATSPERE | 54.30371 | 0.4976207 | 0.4457164 | 1.333299 | 0.1824338 | 0.357404 |
| ENSG00000273212 | | 3.709438 | 0.4972434 | 0.7915965 | 1.193106 | 0.2328278 | 0.42162 |
| ENSG00000100815 | TRIP11 | 5002.352742 | 0.49720385 | 0.232968847 | 2.236998649 | 0.02528643 | 0.081821 |
| ENSG00000225518 | LINC01703 | 162.5075 | 0.4970989 | 0.2323469 | 2.2397 | 0.02511041 | 0.081391 |
| ENSG00000131459 | GFPT2 | 252.8449142 | 0.497081616 | 0.279003378 | 1.910035423 | 0.05612865 | 0.151384 |
| ENSG00000154511 | DIPK1A | 306.3065 | 0.4970388 | 0.2518807 | 2.090822 | 0.03654403 | 0.109008 |
| ENSG00000100234 | TIMP3 | 980.5503797 | 0.496957064 | 0.488762242 | 1.261996868 | 0.20694991 | 0.389299 |
| ENSG00000255642 | PABPC1P4 | 233.6103 | 0.4968601 | 0.2517314 | 2.093379 | 0.03631537 | 0.108705 |
| ENSG00000286918 | | 15.41925 | 0.4968005 | 0.5354952 | 1.199663 | 0.2302704 | 0.418557 |
| ENSG00000104964 | TLE5 | 7003.56504 | 0.496771507 | 0.160748377 | 3.160216897 | 0.00157652 | 0.008452 |
| ENSG00000225401 | TGIF2P1 | 2.993833 | 0.4967357 | 0.8258021 | 1.228008 | 0.2194438 | 0.405537 |
| ENSG00000168661 | ZNF30 | 437.9568 | 0.4967002 | 0.1256735 | 4.003933 | 6.23E-05 | 0.000503 |
| ENSG00000240890 | | 8.870076 | 0.4966453 | 0.6336852 | 1.144854 | 0.2522698 | 0.444939 |
| ENSG00000042062 | RIPOR3 | 851.3758018 | 0.496020913 | 0.385550955 | 1.456772413 | 0.14517921 | 0.304033 |
| ENSG00000159674 | SPON2 | 35.41202 | 0.4958703 | 0.3898144 | 1.467355 | 0.1422794 | 0.299661 |
| ENSG00000277013 | | 90.73325 | 0.4958363 | 0.5613303 | 1.18974 | 0.2341485 | 0.423268 |
| ENSG00000175414 | ARL10 | 17.6365 | 0.4957333 | 0.4428468 | 1.336856 | 0.1812698 | 0.355804 |
| ENSG00000116990 | MYCL | 648.3892728 | 0.495725173 | 0.432999985 | 1.358216487 | 0.174395 | 0.346326 |
| ENSG00000152672 | CLEC4F | 9.851706 | 0.4956075 | 0.527438 | 1.202818 | 0.2290469 | 0.417237 |
| ENSG00000185262 | UBALD2 | 2055.574 | 0.4955137 | 0.2721704 | 1.945654 | 0.05169629 | 0.142113 |
| ENSG00000158089 | GALNT14 | 1045.745 | 0.4954458 | 0.3281603 | 1.67275 | 0.09437652 | 0.223213 |
| ENSG00000197355 | UAP1L1 | 3847.269 | 0.495271 | 0.2744868 | 1.932583 | 0.05328755 | 0.145763 |
| ENSG00000169696 | ASPSCR1 | 3322.202 | 0.4949626 | 0.1546701 | 3.27312 | 0.00106368 | 0.006035 |
| ENSG00000005020 | SKAP2 | 884.6778695 | 0.494887065 | 0.20669333 | 2.485250894 | 0.01294602 | 0.047906 |
| ENSG00000271447 | MMP28 | 4.419588 | 0.4945676 | 0.7719602 | 1.165246 | 0.2439196 | 0.435238 |
| ENSG00000285938 | | 33.90732 | 0.4945188 | 0.5033527 | 1.237924 | 0.2157441 | 0.400903 |
| ENSG00000164008 | C1orf50 | 165.0673 | 0.4944816 | 0.2996659 | 1.791867 | 0.07315434 | 0.184756 |
| ENSG00000205560 | CPT1B | 73.85494 | 0.4944378 | 0.3628288 | 1.528006 | 0.1265109 | 0.275198 |
| ENSG00000175265 | GOLGA8A | 3670.044 | 0.4944208 | 0.3571619 | 1.548626 | 0.1214715 | 0.267721 |
| ENSG00000253284 | | 2.978907 | 0.4943906 | 0.7078403 | 1.12795 | 0.2593409 | 0.453676 |
| ENSG00000287683 | | 3.265705 | 0.494179 | 0.768869 | 1.157857 | 0.2469223 | 0.438769 |
| ENSG00000267265 | | 72.46844 | 0.4941612 | 0.3498216 | 1.574157 | 0.1154511 | 0.258266 |
| ENSG00000199036 | MIR219A1 | 3.252136 | 0.49413 | 0.7882424 | 1.175997 | 0.2395961 | 0.429819 |

| | | | | | | | |
|---|---|---|---|---|---|---|---|
| ENSG00000229932 | YWHAZP3 | 2066.478 | 0.4935504 | 0.1762174 | 2.879276 | 0.0039859 | 0.018398 |
| ENSG00000221914 | PPP2R2A | 2735.371 | 0.4934996 | 0.1530323 | 3.277322 | 0.00104797 | 0.005963 |
| ENSG00000249476 | | 24.95194 | 0.493397 | 0.6032276 | 1.150097 | 0.2501039 | 0.442695 |
| ENSG00000158865 | SLC5A11 | 2.946486 | 0.4932499 | 0.8059118 | 1.220907 | 0.2221212 | 0.408875 |
| ENSG00000182541 | LIMK2 | 2889.046 | 0.4931456 | 0.1664784 | 3.032104 | 0.00242856 | 0.012195 |
| ENSG00000224975 | INE1 | 164.8124 | 0.4930674 | 0.2236537 | 2.304148 | 0.02121432 | 0.071284 |
| ENSG00000226823 | SUGT1P1 | 3.309465 | 0.4930359 | 0.8044273 | 1.185617 | 0.2357738 | 0.42515 |
| ENSG00000140848 | CPNE2 | 1061.992 | 0.4928202 | 0.258068 | 2.023235 | 0.04304893 | 0.123838 |
| ENSG00000213965 | NUDT19 | 1634.591 | 0.4927893 | 0.1869557 | 2.741257 | 0.00612046 | 0.026131 |
| ENSG00000268912 | | 72.34606 | 0.4927589 | 0.4178265 | 1.376904 | 0.1685419 | 0.338396 |
| ENSG00000114541 | FRMD4B | 251.0339746 | 0.492740444 | 0.342977279 | 1.602355993 | 0.10907691 | 0.247725 |
| ENSG00000223910 | ZNF32-AS3 | 0.9935834 | 0.4926965 | 0.9509947 | 1.542733 | 0.1228956 | NA |
| ENSG00000108039 | XPNPEP1 | 3360.700956 | 0.49258946 | 0.117401044 | 4.21908592 | 2.45E-05 | 0.00022 |
| ENSG00000163807 | KIAA1143 | 1464.65 | 0.4925714 | 0.1772037 | 2.860566 | 0.00422886 | 0.019319 |
| ENSG00000213213 | CCDC183 | 143.3202 | 0.4922756 | 0.2669101 | 1.961522 | 0.04981811 | 0.13815 |
| ENSG00000036828 | CASR | 1.254146996 | 0.49225739 | 0.917104002 | 1.423642041 | 0.15455012 | NA |
| ENSG00000244585 | RPL12P33 | 1.414736 | 0.4921788 | 0.889712 | 1.3678 | 0.1713747 | NA |
| ENSG00000231519 | | 2.226371 | 0.4921456 | 0.8845858 | 1.35815 | 0.174416 | 0.346342 |
| ENSG00000162510 | MATN1 | 10.86888 | 0.4920707 | 0.4979454 | 1.245025 | 0.2131226 | 0.39784 |
| ENSG00000254428 | | 135.2538 | 0.4919405 | 0.3532818 | 1.558473 | 0.1191212 | 0.263744 |
| ENSG00000103202 | NME4 | 7260.759394 | 0.491821064 | 0.209420185 | 2.439523386 | 0.01470665 | 0.053075 |
| ENSG00000122012 | SV2C | 3.357745261 | 0.49177262 | 0.799810665 | 1.202566167 | 0.22914425 | 0.417285 |
| ENSG00000129422 | MTUS1 | 2004.914165 | 0.491568451 | 0.19396694 | 2.616178203 | 0.00889201 | 0.03542 |
| ENSG00000081320 | STK17B | 602.696633 | 0.491183281 | 0.27186512 | 1.929676268 | 0.05364696 | 0.146565 |
| ENSG00000249375 | CASC11 | 29.32002 | 0.4911025 | 0.4647256 | 1.28775 | 0.197833 | 0.377944 |
| ENSG00000259810 | | 11.08621 | 0.4910203 | 0.7342219 | 1.147689 | 0.2510972 | 0.443843 |
| ENSG00000185658 | BRWD1 | 18939.08 | 0.4909236 | 0.2675801 | 1.952047 | 0.05093265 | 0.140464 |
| ENSG00000142233 | NTN5 | 127.5434 | 0.4909115 | 0.3662001 | 1.508717 | 0.1313712 | 0.282921 |
| ENSG00000116039 | ATP6V1B1 | 33.84976753 | 0.490810023 | 0.488714166 | 1.250968471 | 0.21094598 | 0.394859 |
| ENSG00000164074 | ABHD18 | 1225.483 | 0.4907033 | 0.1835163 | 2.756097 | 0.00584956 | 0.025218 |
| ENSG00000232393 | RPL5P6 | 1.79971 | 0.4906565 | 0.9355685 | 1.463862 | 0.1432315 | 0.301041 |
| ENSG00000228852 | | 18.80376 | 0.4905613 | 0.5867975 | 1.151036 | 0.2497173 | 0.442307 |
| ENSG00000137434 | C6orf52 | 153.7266478 | 0.490373897 | 0.293873906 | 1.802418246 | 0.07147963 | 0.181458 |

| | | | | | | | |
|---|---|---|---|---|---|---|---|
| ENSG00000189144 | ZNF573 | 120.4651 | 0.4902663 | 0.4048651 | 1.402859 | 0.160659 | 0.327262 |
| ENSG00000100614 | PPM1A | 4522.473779 | 0.490210354 | 0.123284506 | 4.011463642 | 6.03E-05 | 0.00049 |
| ENSG00000268333 | | 39.53572 | 0.490155 | 0.3487031 | 1.570338 | 0.1163365 | 0.259416 |
| ENSG00000144635 | DYNC1LI1 | 2102.988 | 0.4900641 | 0.2123852 | 2.414058 | 0.01577593 | 0.05611 |
| ENSG00000237352 | LINC01358 | 2.126348 | 0.4899873 | 0.9379048 | 1.444728 | 0.1485344 | 0.30905 |
| ENSG00000267795 | SMIM22 | 28.85188 | 0.4898557 | 0.5113197 | 1.217582 | 0.223383 | 0.41005 |
| ENSG00000270171 | | 5.953063 | 0.4895462 | 0.6838711 | 1.124287 | 0.2608914 | 0.455578 |
| ENSG00000255152 | MSH5-SAPCD1 | 20.44313 | 0.4895397 | 0.4232151 | 1.357884 | 0.1745004 | 0.346432 |
| ENSG00000139266 | MARCHF9 | 2850.331 | 0.489426 | 0.2214421 | 2.306067 | 0.0211069 | 0.070995 |
| ENSG00000286274 | | 5.275001 | 0.4892181 | 0.6534714 | 1.12142 | 0.2621092 | 0.456894 |
| ENSG00000232027 | | 9.448895 | 0.488854 | 0.5241449 | 1.196782 | 0.2313917 | 0.41982 |
| ENSG00000138161 | CUZD1 | 2.313632775 | 0.488813221 | 0.882010675 | 1.446168929 | 0.14812983 | 0.308413 |
| ENSG00000232871 | SEC1P | 30.52188 | 0.4888116 | 0.5000489 | 1.224122 | 0.2209063 | 0.407276 |
| ENSG00000075399 | VPS9D1 | 1908.271365 | 0.488768808 | 0.174608385 | 2.853633795 | 0.00432223 | 0.019664 |
| ENSG00000272115 | | 20.20616 | 0.4887342 | 0.4500009 | 1.29428 | 0.1955686 | 0.375086 |
| ENSG00000286732 | | 36.22871 | 0.4887169 | 0.4044077 | 1.402451 | 0.1607807 | 0.327324 |
| ENSG00000258875 | | 9.068992 | 0.48857 | 0.5831687 | 1.14241 | 0.2532835 | 0.446223 |
| ENSG00000255213 | NPM1P35 | 2.837399 | 0.4885624 | 0.8032114 | 1.199807 | 0.2302142 | 0.418513 |
| ENSG00000100764 | PSMC1 | 144.1196633 | 0.488410482 | 0.278786629 | 1.878926575 | 0.06025452 | 0.159548 |
| ENSG00000205542 | TMSB4X | 11159.6 | 0.4883922 | 0.2178635 | 2.335785 | 0.01950247 | 0.066619 |
| ENSG00000137745 | MMP13 | 24.80221482 | 0.488245474 | 0.744109949 | 1.151986207 | 0.24932674 | 0.441801 |
| ENSG00000112320 | SOBP | 217.9781177 | 0.48801258 | 0.171278448 | 2.925082138 | 0.00344365 | 0.016298 |
| ENSG00000261431 | | 4.927478 | 0.4877786 | 0.6801716 | 1.119156 | 0.2630737 | 0.458095 |
| ENSG00000187800 | PEAR1 | 4.84753 | 0.4876002 | 0.698127 | 1.133732 | 0.2569069 | 0.450783 |
| ENSG00000273724 | | 10.01407 | 0.4875996 | 0.5950559 | 1.152698 | 0.2490344 | 0.44142 |
| ENSG00000259362 | | 5.116592 | 0.4875064 | 0.8059898 | 1.224532 | 0.2207516 | 0.407076 |
| ENSG00000285802 | | 2.257773 | 0.4874289 | 0.8604487 | 1.287153 | 0.1980409 | 0.378081 |
| ENSG00000155366 | RHOC | 6968.033 | 0.4872811 | 0.195095 | 2.581131 | 0.00984773 | 0.038381 |
| ENSG00000174989 | FBXW8 | 1672.871 | 0.4872337 | 0.2097403 | 2.416743 | 0.01566005 | 0.05576 |
| ENSG00000153560 | UBP1 | 9189.513 | 0.4872316 | 0.2168901 | 2.376244 | 0.01748987 | 0.061015 |
| ENSG00000259408 | | 14.51424 | 0.4870834 | 0.5084142 | 1.207605 | 0.2271994 | 0.415077 |
| ENSG00000204352 | C9orf129 | 2.307268 | 0.4870636 | 0.8017126 | 1.204315 | 0.228468 | 0.416546 |
| ENSG00000170270 | GON7 | 1370.005 | 0.4870503 | 0.1652844 | 3.010071 | 0.00261186 | 0.012981 |

| | | | | | | | |
|---|---|---|---|---|---|---|---|
| ENSG00000176809 | LRRC37A3 | 1000.572 | 0.4869986 | 0.302853 | 1.755646 | 0.07914894 | 0.195641 |
| ENSG00000272479 | | 2.093896 | 0.4869691 | 0.9755774 | 1.706276 | 0.08795663 | 0.212023 |
| ENSG00000267607 | | 1.577987 | 0.4869619 | 0.8973863 | 1.387054 | 0.1654253 | 0.333981 |
| ENSG00000233016 | SNHG7 | 3982.841 | 0.4869446 | 0.2442265 | 2.118337 | 0.03414652 | 0.103696 |
| ENSG00000203644 | | 449.3012 | 0.4866983 | 0.3677256 | 1.497479 | 0.1342685 | 0.287298 |
| ENSG00000250568 | | 156.4655 | 0.4866769 | 0.2877947 | 1.825201 | 0.06797066 | 0.174641 |
| ENSG00000226340 | USP8P2 | 5.096268 | 0.4866662 | 0.730487 | 1.141885 | 0.2535018 | 0.446548 |
| ENSG00000228043 | | 52.91187 | 0.4864407 | 0.3955116 | 1.413752 | 0.1574349 | 0.322555 |
| ENSG00000144426 | NBEAL1 | 1243.029 | 0.4863302 | 0.4324362 | 1.327395 | 0.184378 | 0.359652 |
| ENSG00000238105 | GOLGA2P5 | 2085.678 | 0.4861785 | 0.2228656 | 2.277675 | 0.02274598 | 0.075382 |
| ENSG00000181274 | FRAT2 | 2600.188 | 0.486157 | 0.1467012 | 3.371459 | 0.00074771 | 0.004447 |
| ENSG00000110330 | BIRC2 | 2149.785769 | 0.485917377 | 0.174678225 | 2.829161276 | 0.00466702 | 0.020967 |
| ENSG00000171989 | LDHAL6B | 1.833446 | 0.4858115 | 0.8864408 | 1.338031 | 0.1808863 | 0.3554 |
| ENSG00000286445 | | 28.87971 | 0.4856808 | 0.4875043 | 1.247538 | 0.2122003 | 0.39662 |
| ENSG00000237662 | SOCS2P1 | 1.849557 | 0.4856745 | 0.9483486 | 1.532295 | 0.1254496 | 0.273708 |
| ENSG00000124120 | TTPAL | 2288.116081 | 0.485653635 | 0.119023111 | 4.154979526 | 3.25E-05 | 0.000283 |
| ENSG00000269894 | | 64.35214 | 0.4856208 | 0.3825684 | 1.445603 | 0.1482885 | 0.308671 |
| ENSG00000186340 | THBS2 | 3.897395 | 0.4856142 | 0.821326 | 1.218567 | 0.2230084 | 0.409603 |
| ENSG00000124251 | TP53TG5 | 39.58076587 | 0.485501383 | 0.275971521 | 1.888390319 | 0.05897357 | 0.157127 |
| ENSG00000054282 | SDCCAG8 | 984.3644304 | 0.485422406 | 0.173092565 | 2.873222094 | 0.00406308 | 0.018699 |
| ENSG00000246877 | DNM1P35 | 22.09424 | 0.4852244 | 0.4114822 | 1.373939 | 0.1694607 | 0.339727 |
| ENSG00000272923 | NA | 233.0869 | 0.4851886 | 0.2175048 | 2.32372 | 0.02014053 | 0.06836 |
| ENSG00000259895 | | 15.70154 | 0.4851641 | 0.5515557 | 1.160426 | 0.2458755 | 0.437612 |
| ENSG00000248027 | ARHGAP42-AS1 | 10.68381 | 0.4850064 | 0.7028626 | 1.123536 | 0.2612098 | 0.455805 |
| ENSG00000168158 | OR2C1 | 1.525329 | 0.4846742 | 0.9023056 | 1.385856 | 0.165791 | NA |
| ENSG00000164049 | FBXW12 | 35.18802 | 0.4846291 | 0.4705299 | 1.260848 | 0.2073634 | 0.3898 |
| ENSG00000236257 | EI24P2 | 22.00068 | 0.4845212 | 0.4750568 | 1.248339 | 0.2119069 | 0.396231 |
| ENSG00000162572 | SCNN1D | 293.5032 | 0.4844469 | 0.2354139 | 2.158662 | 0.03087638 | 0.095982 |
| ENSG00000269906 | | 38.1519 | 0.4843592 | 0.3580884 | 1.52134 | 0.1281745 | 0.277839 |
| ENSG00000197530 | MIB2 | 2091.613 | 0.4842548 | 0.2153176 | 2.342929 | 0.019133 | 0.065576 |
| ENSG00000069667 | RORA | 254.1900855 | 0.484226778 | 0.395165608 | 1.411782182 | 0.15801411 | 0.323266 |
| ENSG00000186897 | C1QL4 | 35.26458 | 0.4840443 | 0.5697576 | 1.161398 | 0.2454802 | 0.437172 |
| ENSG00000228089 | PNKDP1 | 5.255009 | 0.4840002 | 0.744585 | 1.160169 | 0.24598 | 0.437722 |

| | | | | | | | |
|---|---|---|---|---|---|---|---|
| ENSG00000186866 | POFUT2 | 2333.551 | 0.4839301 | 0.1549086 | 3.19112 | 0.00141722 | 0.007726 |
| ENSG00000156860 | FBRS | 7021.659 | 0.4838994 | 0.1562468 | 3.161392 | 0.00157017 | 0.008433 |
| ENSG00000259100 | COILP1 | 4.518055 | 0.4836918 | 0.7282439 | 1.136771 | 0.2556338 | 0.449083 |
| ENSG00000287222 | | 495.1192 | 0.4836848 | 0.1809859 | 2.738685 | 0.00616854 | 0.02632 |
| ENSG00000226186 | ELOCP21 | 4.21911 | 0.4836761 | 0.7803895 | 1.153754 | 0.2486008 | 0.440897 |
| ENSG00000037042 | TUBG2 | 1128.70121 | 0.483627219 | 0.185772051 | 2.683997104 | 0.00727477 | 0.030092 |
| ENSG00000278925 | | 1.179315 | 0.4834112 | 0.9547757 | 1.606612 | 0.1081395 | NA |
| ENSG00000250144 | | 32.83029 | 0.4830093 | 0.3573186 | 1.524752 | 0.1273211 | 0.276579 |
| ENSG00000203943 | SAMD13 | 26.51136 | 0.4828785 | 0.4479495 | 1.290742 | 0.1967932 | 0.376755 |
| ENSG00000163874 | ZC3H12A | 1505.802 | 0.4828605 | 0.2481541 | 2.07767 | 0.03773972 | 0.111747 |
| ENSG00000274180 | NATD1 | 918.646 | 0.482858 | 0.3196104 | 1.642762 | 0.1004322 | 0.233516 |
| ENSG00000078900 | TP73 | 429.5917718 | 0.482846995 | 0.339946617 | 1.572653592 | 0.11579905 | 0.258649 |
| ENSG00000198743 | SLC5A3 | 1866.822 | 0.4827644 | 0.316677 | 1.66392 | 0.0961285 | 0.225961 |
| ENSG00000224321 | RPL12P14 | 7.5651 | 0.4827116 | 0.6439713 | 1.112978 | 0.2657177 | 0.461366 |
| ENSG00000187626 | ZKSCAN4 | 550.6711 | 0.4826541 | 0.1332459 | 3.683864 | 0.00022973 | 0.001602 |
| ENSG00000253404 | | 83.42735 | 0.4825886 | 0.3615122 | 1.497969 | 0.1341412 | 0.287072 |
| ENSG00000188659 | SAXO2 | 52.75766 | 0.4825705 | 0.4363298 | 1.307482 | 0.191049 | 0.369082 |
| ENSG00000105443 | CYTH2 | 6002.981602 | 0.482533677 | 0.196506073 | 2.485357569 | 0.01294214 | 0.047898 |
| ENSG00000228966 | HOMER2P1 | 13.19325 | 0.4825115 | 0.7751613 | 1.17797 | 0.2388086 | 0.428738 |
| ENSG00000099866 | MADCAM1 | 60.769062 | 0.482397347 | 0.408974057 | 1.367102803 | 0.17159309 | 0.342297 |
| ENSG00000101187 | SLCO4A1 | 2146.463286 | 0.48237375 | 0.369465645 | 1.480627328 | 0.13870591 | 0.294051 |
| ENSG00000070756 | PABPC1 | 74341.67337 | 0.482178615 | 0.149835584 | 3.281595059 | 0.00103222 | 0.005885 |
| ENSG00000139182 | CLSTN3 | 1824.307671 | 0.481882141 | 0.223157438 | 2.245927225 | 0.02470867 | 0.080363 |
| ENSG00000223263 | RNU6-387P | 2.147246 | 0.4818769 | 0.822549 | 1.222533 | 0.221506 | 0.408127 |
| ENSG00000135094 | SDS | 15.06803696 | 0.481714283 | 0.515132634 | 1.196159703 | 0.23163424 | 0.420038 |
| ENSG00000226125 | LINC01907 | 3.231385 | 0.4814915 | 0.8241651 | 1.226824 | 0.2198887 | 0.406021 |
| ENSG00000135477 | KRT87P | 513.947263 | 0.481470183 | 0.354061992 | 1.523441106 | 0.12764838 | 0.277037 |
| ENSG00000121068 | TBX2 | 13493.80031 | 0.481396216 | 0.165035791 | 2.967545103 | 0.00300188 | 0.014565 |
| ENSG00000126759 | CFP | 17.22996519 | 0.48122016 | 0.449723088 | 1.284610377 | 0.19892847 | 0.379013 |
| ENSG00000239617 | RPL32P27 | 1.190493 | 0.4809213 | 0.9627978 | 1.651152 | 0.09870764 | NA |
| ENSG00000247092 | SNHG10 | 873.0216 | 0.4808657 | 0.1820078 | 2.726662 | 0.00639785 | 0.027055 |
| ENSG00000139644 | TMBIM6 | 43136.27 | 0.4808087 | 0.1041376 | 4.660023 | 3.16E-06 | 3.55E-05 |
| ENSG00000139304 | PTPRQ | 105.8006 | 0.4807415 | 0.442772 | 1.292381 | 0.1962252 | 0.375969 |

| | | | | | | | |
|---|---|---|---|---|---|---|---|
| ENSG00000278464 | | 2.965317 | 0.4802873 | 0.7838712 | 1.167638 | 0.2429529 | 0.433921 |
| ENSG00000071537 | SEL1L | 9112.427723 | 0.480183933 | 0.260056571 | 1.993855728 | 0.04616782 | 0.130492 |
| ENSG00000050393 | MCUR1 | 2625.762249 | 0.480154133 | 0.140184882 | 3.480718044 | 0.00050007 | 0.003138 |
| ENSG00000008441 | NFIX | 936.9567489 | 0.480106533 | 0.338415133 | 1.578322843 | 0.11449146 | 0.256508 |
| ENSG00000116977 | LGALS8 | 5021.847583 | 0.479767479 | 0.127914904 | 3.778291519 | 0.00015791 | 0.00115 |
| ENSG00000250325 | IGBP1P4 | 0.998703 | 0.4797173 | 0.9599626 | 1.65773 | 0.09737202 | NA |
| ENSG00000249961 | TERB1 | 3.57477 | 0.479614 | 0.8336163 | 1.211353 | 0.2257602 | 0.413101 |
| ENSG00000271751 | | 13.56916 | 0.4796122 | 0.4376039 | 1.3007 | 0.1933612 | 0.372007 |
| ENSG00000197062 | ZSCAN26 | 599.2948 | 0.4794031 | 0.1905083 | 2.605774 | 0.0091667 | 0.03628 |
| ENSG00000093183 | SEC22C | 1720.704756 | 0.479357759 | 0.152256998 | 3.205663191 | 0.00134752 | 0.007411 |
| ENSG00000276867 | | 4.067208 | 0.4793186 | 0.7786176 | 1.172444 | 0.2410189 | 0.431512 |
| ENSG00000083642 | PDS5B | 4264.291652 | 0.47910216 | 0.269204211 | 1.894498879 | 0.05815882 | 0.155502 |
| ENSG00000231305 | | 110.4384 | 0.4790205 | 0.2562905 | 1.9742 | 0.04835903 | 0.135049 |
| ENSG00000258590 | NBEAP1 | 11.81977 | 0.478921 | 0.915121 | 1.334589 | 0.1820108 | 0.356915 |
| ENSG00000237172 | B3GNT9 | 577.5179 | 0.4787285 | 0.201821 | 2.450159 | 0.01427933 | 0.051857 |
| ENSG00000206530 | CFAP44 | 1457.633 | 0.4786019 | 0.3352829 | 1.582767 | 0.1134745 | 0.254745 |
| ENSG00000120784 | ZFP30 | 702.2065481 | 0.478553774 | 0.215373031 | 2.311145111 | 0.02082484 | 0.070215 |
| ENSG00000274750 | H3C6 | 1243.114 | 0.4785192 | 0.3045148 | 1.716934 | 0.08599132 | 0.208363 |
| ENSG00000201492 | RNA5SP78 | 2.320591 | 0.4784888 | 0.7844847 | 1.147302 | 0.2512568 | 0.443991 |
| ENSG00000135773 | CAPN9 | 143.2668131 | 0.478385544 | 0.354503356 | 1.509157584 | 0.13125852 | 0.282725 |
| ENSG00000237159 | CNTFR-AS1 | 1.46616 | 0.4783848 | 0.9819992 | 1.863813 | 0.06234801 | NA |
| ENSG00000016082 | ISL1 | 44.76593514 | 0.478223515 | 0.563509837 | 1.145605234 | 0.25195853 | 0.444685 |
| ENSG00000225808 | DNAJC19P5 | 9.401555 | 0.4782043 | 0.7422705 | 1.158942 | 0.2464798 | 0.438306 |
| ENSG00000257611 | | 2.554055 | 0.4779833 | 0.8110247 | 1.202596 | 0.2291328 | 0.417285 |
| ENSG00000229028 | KRT223P | 2.001673 | 0.4779281 | 0.9644057 | 1.725854 | 0.08437371 | 0.205437 |
| ENSG00000161638 | ITGA5 | 423.965 | 0.477916 | 0.3212958 | 1.639615 | 0.1010851 | 0.234582 |
| ENSG00000154914 | USP43 | 1054.274 | 0.477792 | 0.1729789 | 2.834698 | 0.0045869 | 0.020673 |
| ENSG00000232615 | | 113.751 | 0.4777353 | 0.2979304 | 1.725547 | 0.08442895 | 0.205517 |
| ENSG00000244268 | | 2.035439 | 0.477663 | 0.7567011 | 1.125054 | 0.2605661 | 0.45513 |
| ENSG00000233873 | RPL7P44 | 3.166008 | 0.4775036 | 0.7928034 | 1.180833 | 0.2376691 | 0.427295 |
| ENSG00000172986 | GXYLT2 | 128.2331 | 0.4774089 | 0.3658878 | 1.472984 | 0.1407552 | 0.297276 |
| ENSG00000276529 | | 111.7463 | 0.4773698 | 0.2864373 | 1.798398 | 0.07211394 | 0.182613 |
| ENSG00000272141 | | 167.7419 | 0.4772519 | 0.2809555 | 1.824603 | 0.06806096 | 0.174817 |

| | | | | | | | |
|---|---|---|---|---|---|---|---|
| ENSG00000228857 | | 2.464955 | 0.4772022 | 0.8570247 | 1.284615 | 0.198927 | 0.379013 |
| ENSG00000151611 | MMAA | 248.3814 | 0.4771607 | 0.2131873 | 2.331739 | 0.01971443 | 0.067213 |
| ENSG00000226287 | TMEM191A | 740.4442 | 0.4771348 | 0.3697068 | 1.457683 | 0.1449279 | 0.303626 |
| ENSG00000180712 | LINC02363 | 25.17914 | 0.4769124 | 0.4755451 | 1.241821 | 0.2143026 | 0.399145 |
| ENSG00000269392 | | 19.99019 | 0.47681 | 0.5777383 | 1.125195 | 0.2605064 | 0.455057 |
| ENSG00000119574 | ZBTB45 | 1059.662394 | 0.476748731 | 0.161012288 | 3.030956352 | 0.00243781 | 0.012235 |
| ENSG00000167553 | TUBA1C | 8505.237 | 0.4765766 | 0.344033 | 1.587195 | 0.1124684 | 0.253252 |
| ENSG00000149548 | CCDC15 | 106.8317 | 0.4765738 | 0.4539793 | 1.266814 | 0.2052218 | 0.3872 |
| ENSG00000286219 | NOTCH2NLC | 140.2944 | 0.4764771 | 0.3723214 | 1.449629 | 0.1471621 | 0.306952 |
| ENSG00000225345 | SNX18P3 | 152.8487 | 0.4763998 | 0.3503932 | 1.521377 | 0.1281654 | 0.277839 |
| ENSG00000257354 | | 199.708 | 0.4763575 | 0.3584052 | 1.499069 | 0.1338557 | 0.286599 |
| ENSG00000132026 | RTBDN | 2.203882767 | 0.47618389 | 0.83817364 | 1.217621876 | 0.2233677 | 0.41005 |
| ENSG00000230769 | | 1.733749 | 0.4761503 | 0.8685233 | 1.342365 | 0.1794777 | 0.35354 |
| ENSG00000255284 | | 216.1814 | 0.4761322 | 0.221585 | 2.239629 | 0.02511501 | 0.081396 |
| ENSG00000259529 | | 73.69341 | 0.4760833 | 0.3475463 | 1.520651 | 0.1283475 | 0.278146 |
| ENSG00000118873 | RAB3GAP2 | 3167.874557 | 0.476033096 | 0.21971329 | 2.254413275 | 0.02417018 | 0.078979 |
| ENSG00000283627 | | 42.68252 | 0.4760151 | 0.4108773 | 1.343053 | 0.1792549 | 0.353205 |
| ENSG00000149609 | C20orf144 | 23.42394 | 0.4758886 | 0.4004899 | 1.369959 | 0.1706998 | 0.341412 |
| ENSG00000287799 | | 4.355822 | 0.4758707 | 0.7789702 | 1.171596 | 0.2413594 | 0.431917 |
| ENSG00000103111 | MON1B | 5883.554326 | 0.475763925 | 0.139803815 | 3.454704901 | 0.0005509 | 0.003417 |
| ENSG00000283648 | | 1.465052 | 0.475594 | 0.9452985 | 1.509805 | 0.1310932 | NA |
| ENSG00000176531 | PHLDB3 | 1116.438 | 0.4755714 | 0.1425955 | 3.385919 | 0.0007094 | 0.004252 |
| ENSG00000227527 | | 14.77179 | 0.4754995 | 0.4370235 | 1.29012 | 0.197009 | 0.376972 |
| ENSG00000233937 | | 338.4015 | 0.4754632 | 0.3172858 | 1.636112 | 0.1018162 | 0.235706 |
| ENSG00000169410 | PTPN9 | 2365.833 | 0.4752539 | 0.136582 | 3.532154 | 0.00041219 | 0.002658 |
| ENSG00000197386 | HTT | 6952.562 | 0.4752368 | 0.1873231 | 2.608081 | 0.00910515 | 0.03608 |
| ENSG00000048392 | RRM2B | 4615.412659 | 0.47510975 | 0.250866098 | 1.99626096 | 0.04590553 | 0.129931 |
| ENSG00000197070 | ARRDC1 | 5982.612 | 0.4750594 | 0.1482274 | 3.249401 | 0.00115648 | 0.006488 |
| ENSG00000216324 | | 0.8252552 | 0.4750443 | 0.9497272 | 1.586231 | 0.1126869 | NA |
| ENSG00000232977 | LINC00327 | 3.279217 | 0.4748477 | 0.842655 | 1.244115 | 0.2134574 | 0.39828 |
| ENSG00000116604 | MEF2D | 3411.59418 | 0.474800017 | 0.261772956 | 1.925066235 | 0.05422107 | 0.147633 |
| ENSG00000081721 | DUSP12 | 1901.186748 | 0.474732157 | 0.191845238 | 2.549781189 | 0.01077906 | 0.041332 |
| ENSG00000186666 | BCDIN3D | 405.029 | 0.4744986 | 0.2096489 | 2.345595 | 0.01899674 | 0.06521 |

| | | | | | | | |
|---|---|---|---|---|---|---|---|
| ENSG00000171223 | JUNB | 7681.094 | 0.4744512 | 0.3399048 | 1.563494 | 0.1179364 | 0.261863 |
| ENSG00000250986 | LINC02600 | 350.8902 | 0.4744174 | 0.3598569 | 1.491765 | 0.1357608 | 0.289396 |
| ENSG00000175764 | TTLL11 | 322.7108 | 0.4743398 | 0.2520551 | 1.995823 | 0.04595325 | 0.130025 |
| ENSG00000256628 | ZBTB11-AS1 | 251.7285 | 0.4743081 | 0.1785131 | 2.736941 | 0.00620135 | 0.02643 |
| ENSG00000228950 | | 5.173867 | 0.4741339 | 0.6691084 | 1.094324 | 0.2738129 | 0.470565 |
| ENSG00000279607 | | 4.498918 | 0.4739799 | 0.8530643 | 1.277997 | 0.2012504 | 0.381985 |
| ENSG00000138778 | CENPE | 1923.057968 | 0.473912662 | 0.456389485 | 1.25079219 | 0.21101031 | 0.394951 |
| ENSG00000143643 | TTC13 | 1755.176 | 0.4738505 | 0.1648277 | 2.892939 | 0.00381655 | 0.017737 |
| ENSG00000205871 | RPS3AP47 | 27.531 | 0.4737075 | 0.4621242 | 1.25115 | 0.2108796 | 0.39479 |
| ENSG00000100359 | SGSM3 | 138.6543977 | 0.473665882 | 0.335870777 | 1.568099709 | 0.11685787 | 0.26012 |
| ENSG00000105376 | ICAM5 | 580.5214072 | 0.473624063 | 0.253384598 | 1.978680691 | 0.04785197 | 0.13397 |
| ENSG00000259158 | ADAM20P1 | 32.12654 | 0.4735 | 0.5337229 | 1.140179 | 0.2542118 | 0.447236 |
| ENSG00000104343 | UBE2W | 1479.487021 | 0.473430237 | 0.132470922 | 3.618557618 | 0.00029625 | 0.002 |
| ENSG00000196466 | ZNF799 | 227.5401 | 0.4730281 | 0.2690532 | 1.883731 | 0.05960133 | 0.158281 |
| ENSG00000199349 | | 2.048076 | 0.4729028 | 0.7709209 | 1.127055 | 0.2597192 | 0.454069 |
| ENSG00000213085 | CFAP45 | 52.43102 | 0.4728062 | 0.5312619 | 1.155583 | 0.2478518 | 0.440054 |
| ENSG00000237026 | | 1.55256 | 0.472721 | 0.8774024 | 1.318524 | 0.1873282 | NA |
| ENSG00000260001 | TGFBR3L | 115.8465 | 0.4726412 | 0.2170315 | 2.267024 | 0.02338875 | 0.076946 |
| ENSG00000163449 | TMEM169 | 15.40004 | 0.4721878 | 0.4847705 | 1.199279 | 0.2304194 | 0.418771 |
| ENSG00000266921 | | 25.42339 | 0.4721599 | 0.4015912 | 1.349362 | 0.1772208 | 0.350419 |
| ENSG00000116717 | GADD45A | 1125.677064 | 0.472113075 | 0.389001481 | 1.374281526 | 0.1693543 | 0.339591 |
| ENSG00000183154 | | 2.762347 | 0.4719222 | 0.779704 | 1.138559 | 0.2548872 | 0.448068 |
| ENSG00000230415 | LINC01786 | 24.30092 | 0.4716578 | 0.3865438 | 1.396242 | 0.1626417 | 0.330016 |
| ENSG00000226957 | | 2.699095 | 0.4716174 | 0.7097811 | 1.089253 | 0.2760422 | 0.472897 |
| ENSG00000236668 | | 2.273617 | 0.4714693 | 0.8675291 | 1.29875 | 0.1940296 | 0.372863 |
| ENSG00000196655 | TRAPPC4 | 3671.44 | 0.4714345 | 0.160507 | 3.003072 | 0.00267269 | 0.013229 |
| ENSG00000287239 | | 1.122925 | 0.4713474 | 0.9408304 | 1.481566 | 0.1384559 | NA |
| ENSG00000275956 | | 7.48036 | 0.4712221 | 0.6725531 | 1.092412 | 0.2746521 | 0.471519 |
| ENSG00000143595 | AQP10 | 1.644096 | 0.4711647 | 0.8947167 | 1.368103 | 0.1712799 | 0.341929 |
| ENSG00000130787 | HIP1R | 7472.26687 | 0.47111133 | 0.194145297 | 2.504590369 | 0.01225933 | 0.045793 |
| ENSG00000267278 | MAP3K14-AS1 | 548.2376 | 0.470767 | 0.1901837 | 2.568239 | 0.01022167 | 0.039582 |
| ENSG00000165359 | INTS6L | 1326.385 | 0.4707193 | 0.1771961 | 2.733649 | 0.00626368 | 0.026631 |
| ENSG00000254419 | | 59.94568 | 0.4707117 | 0.2570538 | 1.95027 | 0.05114397 | 0.140944 |

| | | | | | | | |
|---|---|---|---|---|---|---|---|
| ENSG00000235954 | TTC28-AS1 | 639.9983 | 0.4706883 | 0.2689947 | 1.86363 | 0.0623737 | 0.163691 |
| ENSG00000230162 | CT45A11P | 1.604905 | 0.4705584 | 0.8862792 | 1.419681 | 0.1557007 | 0.319815 |
| ENSG00000143217 | NECTIN4 | 2568.131 | 0.470514 | 0.2222142 | 2.201277 | 0.02771639 | 0.08797 |
| ENSG00000206562 | METTL6 | 1625.98 | 0.470487 | 0.124047 | 3.847334 | 0.00011941 | 0.000897 |
| ENSG00000234129 | | 33.66351 | 0.4703739 | 0.4615018 | 1.243051 | 0.2138489 | 0.398722 |
| ENSG00000271959 | | 1.27757 | 0.4702311 | 0.8913106 | 1.485748 | 0.1373457 | NA |
| ENSG00000197046 | SIGLEC15 | 114.0906 | 0.4700472 | 0.4666687 | 1.225958 | 0.2202146 | 0.406395 |
| ENSG00000053770 | AP5M1 | 2844.979638 | 0.470010949 | 0.15099523 | 3.174043188 | 0.00150331 | 0.008123 |
| ENSG00000234589 | | 27.55542 | 0.4699784 | 0.4750395 | 1.219335 | 0.2227169 | 0.409386 |
| ENSG00000198924 | DCLRE1A | 1192.334 | 0.4699599 | 0.2902956 | 1.749103 | 0.08027327 | 0.197791 |
| ENSG00000088543 | C3orf18 | 161.4119172 | 0.46995134 | 0.218938616 | 2.238070156 | 0.02521648 | 0.081645 |
| ENSG00000120925 | RNF170 | 1145.971036 | 0.469884684 | 0.244592216 | 2.017807277 | 0.04361134 | 0.125036 |
| ENSG00000211451 | GNRHR2 | 166.221 | 0.4697851 | 0.3050868 | 1.673042 | 0.09431905 | 0.223169 |
| ENSG00000235122 | | 3.336717 | 0.4695789 | 0.7706732 | 1.109865 | 0.2670571 | 0.462817 |
| ENSG00000283020 | DUX4L37 | 28.21959 | 0.4694781 | 0.3749516 | 1.417353 | 0.1563799 | 0.320838 |
| ENSG00000134759 | ELP2 | 6939.824758 | 0.469435189 | 0.232041634 | 2.133792409 | 0.03285978 | 0.100656 |
| ENSG00000256783 | | 1.389161 | 0.4694331 | 0.9611107 | 1.701854 | 0.08878273 | NA |
| ENSG00000168282 | MGAT2 | 60.3927 | 0.4694265 | 0.2807209 | 1.790315 | 0.0734033 | 0.185209 |
| ENSG00000102226 | USP11 | 4763.258013 | 0.469338242 | 0.165604203 | 2.923537642 | 0.00346078 | 0.016358 |
| ENSG00000261003 | | 2.467275 | 0.4693095 | 0.7514408 | 1.120343 | 0.2625678 | 0.457423 |
| ENSG00000258502 | LINC02290 | 0.768915 | 0.469212 | 0.9599658 | 1.703787 | 0.08842091 | NA |
| ENSG00000253899 | | 1.004088 | 0.4691999 | 0.9531084 | 1.65011 | 0.0989205 | NA |
| ENSG00000248103 | | 6.688361 | 0.4690202 | 0.7230643 | 1.109214 | 0.2673378 | 0.463122 |
| ENSG00000272871 | | 10.30018 | 0.4688972 | 0.6191535 | 1.093476 | 0.2741849 | 0.470902 |
| ENSG00000134086 | VHL | 4690.608684 | 0.468747991 | 0.184105413 | 2.619493365 | 0.00880605 | 0.035157 |
| ENSG00000138764 | CCNG2 | 5691.925346 | 0.468697796 | 0.267323839 | 1.858632238 | 0.06307928 | 0.165037 |
| ENSG00000276533 | | 5.673385 | 0.4686536 | 0.6089527 | 1.095731 | 0.2731965 | 0.46984 |
| ENSG00000231852 | CYP21A2 | 13.09452 | 0.4685237 | 0.5226376 | 1.146268 | 0.2516843 | 0.444452 |
| ENSG00000245275 | SAP30L-AS1 | 179.7732 | 0.4684967 | 0.3736653 | 1.418095 | 0.1561631 | 0.320542 |
| ENSG00000161217 | PCYT1A | 5837.813 | 0.4684888 | 0.1018791 | 4.642541 | 3.44E-06 | 3.81E-05 |
| ENSG00000259155 | STK16P1 | 2.60598 | 0.4684755 | 0.7717194 | 1.154544 | 0.2482771 | 0.440587 |
| ENSG00000257376 | | 8.114771 | 0.4683047 | 0.5725984 | 1.112177 | 0.2660621 | 0.461725 |
| ENSG00000259102 | SMARCE1P3 | 1.491307 | 0.4681415 | 0.8755353 | 1.330596 | 0.183322 | NA |

| | | | | | | | |
|---|---|---|---|---|---|---|---|
| ENSG00000206013 | IFITM5 | 1.052548 | 0.4679742 | 0.8807422 | 1.311306 | 0.1897544 | NA |
| ENSG00000276406 | | 1.836734 | 0.4679698 | 0.8871104 | 1.359289 | 0.174055 | 0.345858 |
| ENSG00000164327 | RICTOR | 2677.461 | 0.4675426 | 0.2527587 | 1.946613 | 0.05158112 | 0.141869 |
| ENSG00000153339 | TRAPPC8 | 2383.953 | 0.4675078 | 0.1534782 | 3.126203 | 0.00177079 | 0.009344 |
| ENSG00000133961 | NUMB | 4914.640412 | 0.46748257 | 0.122034274 | 3.918346601 | 8.92E-05 | 0.000691 |
| ENSG00000279108 | | 77.21049 | 0.4674013 | 0.4504618 | 1.245389 | 0.2129888 | 0.397712 |
| ENSG00000158711 | ELK4 | 6817.783 | 0.4673939 | 0.1875782 | 2.563084 | 0.0103747 | 0.040088 |
| ENSG00000286555 | | 34.75809 | 0.4672167 | 0.3891062 | 1.373995 | 0.1694432 | 0.339718 |
| ENSG00000241095 | CYP51A1P1 | 152.3297 | 0.4671235 | 0.2725327 | 1.837065 | 0.0662003 | 0.171197 |
| ENSG00000236939 | BAALC-AS2 | 1.716139 | 0.4670678 | 0.8938254 | 1.376963 | 0.1685238 | 0.338385 |
| ENSG00000101104 | PABPC1L | 7126.325261 | 0.466934785 | 0.327024448 | 1.566670842 | 0.11719165 | 0.260732 |
| ENSG00000185187 | SIGIRR | 4022.741 | 0.4668715 | 0.2194252 | 2.216398 | 0.02666425 | 0.085334 |
| ENSG00000219085 | NPM1P37 | 8.741361 | 0.4667772 | 0.5810213 | 1.114176 | 0.2652039 | 0.460777 |
| ENSG00000218549 | OR4K12P | 4.415882 | 0.4666905 | 0.9023458 | 1.372749 | 0.1698304 | 0.340186 |
| ENSG00000255557 | | 4.464425 | 0.4665294 | 0.8062235 | 1.170061 | 0.2419763 | 0.432555 |
| ENSG00000204929 | | 3.335942 | 0.4663516 | 0.8029607 | 1.147283 | 0.2512645 | 0.443991 |
| ENSG00000100170 | SLC5A1 | 1.059038851 | 0.466117813 | 0.876149169 | 1.316113982 | 0.18813579 | NA |
| ENSG00000146376 | ARHGAP18 | 593.6871 | 0.4660958 | 0.2433858 | 2.01502 | 0.04390258 | 0.125708 |
| ENSG00000104879 | CKM | 2.547668354 | 0.466078919 | 0.71471714 | 1.074975684 | 0.28238562 | 0.480234 |
| ENSG00000239388 | ASB14 | 29.68424 | 0.4660668 | 0.3630118 | 1.44483 | 0.1485056 | 0.30905 |
| ENSG00000265519 | | 176.0521 | 0.4659232 | 0.1993997 | 2.422824 | 0.0154004 | 0.055048 |
| ENSG00000129159 | KCNC1 | 68.85449112 | 0.46588518 | 0.521331336 | 1.15808401 | 0.24682976 | 0.438715 |
| ENSG00000283796 | MIR6510 | 1.040377 | 0.4658823 | 0.9525266 | 1.600022 | 0.1095937 | NA |
| ENSG00000252569 | RNU6-1153P | 1.547662 | 0.4658646 | 0.9021873 | 1.406301 | 0.1596348 | NA |
| ENSG00000113838 | TBCCD1 | 516.2977002 | 0.465569057 | 0.189751368 | 2.531215132 | 0.01136681 | 0.043125 |
| ENSG00000286669 | | 4.600407 | 0.4655545 | 0.6875873 | 1.080973 | 0.2797093 | 0.476996 |
| ENSG00000138363 | ATIC | 11531.47947 | 0.465447026 | 0.130224796 | 3.617217561 | 0.00029779 | 0.002009 |
| ENSG00000229081 | LINC01165 | 20.35562 | 0.46543 | 0.5626422 | 1.11555 | 0.2646148 | 0.460053 |
| ENSG00000167552 | TUBA1A | 7171.298 | 0.465267 | 0.2609585 | 1.891067 | 0.05861543 | 0.156377 |
| ENSG00000172425 | TTC36 | 14.60123 | 0.4652568 | 0.4879395 | 1.177323 | 0.2390665 | 0.429028 |
| ENSG00000233090 | | 1.531557 | 0.4652143 | 0.8583611 | 1.254655 | 0.2096042 | NA |
| ENSG00000163378 | EOGT | 612.6808 | 0.4650509 | 0.1893664 | 2.537684 | 0.01115888 | 0.0425 |
| ENSG00000100344 | PNPLA3 | 99.44830275 | 0.464909086 | 0.413771949 | 1.314928289 | 0.188534 | 0.365678 |

| | | | | | | | |
|---|---|---|---|---|---|---|---|
| ENSG00000146453 | PNLDC1 | 4.935309 | 0.4648978 | 0.7593756 | 1.113659 | 0.2654257 | 0.461102 |
| ENSG00000237418 | | 13.13285 | 0.4648879 | 0.6442745 | 1.076769 | 0.2815833 | 0.479238 |
| ENSG00000118600 | RXYLT1 | 1030.151571 | 0.464844008 | 0.155519217 | 3.054077396 | 0.00225754 | 0.01146 |
| ENSG00000062038 | CDH3 | 6200.804914 | 0.464828217 | 0.13954642 | 3.388680162 | 0.0007023 | 0.004215 |
| ENSG00000084676 | NCOA1 | 2749.516326 | 0.464698482 | 0.226454221 | 2.177508323 | 0.02944266 | 0.092422 |
| ENSG00000155875 | SAXO1 | 3.532226 | 0.4644031 | 0.7321276 | 1.087772 | 0.2766958 | 0.4737 |
| ENSG00000214711 | CAPN14 | 46.03078 | 0.4641771 | 0.7609839 | 1.148594 | 0.2507232 | 0.443359 |
| ENSG00000081853 | PCDHGA2 | 156.4396582 | 0.464171805 | 0.343273734 | 1.498736276 | 0.13394206 | 0.286738 |
| ENSG00000251410 | | 1.824311 | 0.4641252 | 0.9307087 | 1.572357 | 0.1158677 | 0.258713 |
| ENSG00000179886 | TIGD5 | 1123.795 | 0.4637535 | 0.2096957 | 2.296398 | 0.02165316 | 0.072538 |
| ENSG00000259222 | | 1.12811 | 0.4636975 | 0.8964212 | 1.492879 | 0.1354688 | NA |
| ENSG00000272909 | | 6.487896 | 0.4636875 | 0.6124007 | 1.082411 | 0.2790701 | 0.476216 |
| ENSG00000236546 | MYCL-AS1 | 2.33415 | 0.463515 | 0.9005752 | 1.420193 | 0.1555514 | 0.319583 |
| ENSG00000144136 | SLC20A1 | 3904.199 | 0.4635148 | 0.2332219 | 2.085058 | 0.03706406 | 0.110189 |
| ENSG00000100147 | CCDC134 | 175.7217647 | 0.46349101 | 0.194164731 | 2.469469973 | 0.01353134 | 0.049564 |
| ENSG00000108001 | EBF3 | 18.59730661 | 0.463467469 | 0.492774384 | 1.169935363 | 0.24202698 | 0.432617 |
| ENSG00000275585 | PDE4DIPP4 | 1.58067 | 0.4634069 | 0.8722651 | 1.293891 | 0.1957029 | 0.375316 |
| ENSG00000232177 | MTND4P24 | 4.765821 | 0.4632983 | 0.7549289 | 1.125194 | 0.260507 | 0.455057 |
| ENSG00000197045 | GMFB | 7241.542 | 0.4630994 | 0.175034 | 2.698476 | 0.00696577 | 0.029004 |
| ENSG00000157379 | DHRS1 | 1753.302 | 0.4630624 | 0.1862389 | 2.564163 | 0.0103425 | 0.039993 |
| ENSG00000243175 | RPSAP36 | 2.500337 | 0.4628277 | 0.7851928 | 1.144055 | 0.2526006 | 0.445434 |
| ENSG00000159905 | ZNF221 | 18.13459 | 0.4625268 | 0.4731644 | 1.197339 | 0.2311743 | 0.419681 |
| ENSG00000066084 | DIP2B | 4602.946275 | 0.462394338 | 0.209219112 | 2.295049883 | 0.02173027 | 0.072742 |
| ENSG00000243504 | RPS23P1 | 8.260008 | 0.4623746 | 0.7140414 | 1.099272 | 0.2716494 | 0.468058 |
| ENSG00000215604 | ZNF962P | 2.049664 | 0.4622594 | 0.9987195 | 2.390879 | 0.01680808 | 0.059093 |
| ENSG00000249207 | | 10.67434 | 0.4621941 | 0.629068 | 1.087554 | 0.2767921 | 0.473724 |
| ENSG00000236489 | | 2.778689 | 0.4621668 | 0.8380125 | 1.240649 | 0.2147355 | 0.399643 |
| ENSG00000130479 | MAP1S | 812.1791108 | 0.462045845 | 0.210713402 | 2.281127334 | 0.02254091 | 0.074909 |
| ENSG00000171124 | FUT3 | 211.1207 | 0.4620116 | 0.6071155 | 1.085988 | 0.2774842 | 0.474572 |
| ENSG00000234753 | FOXP4-AS1 | 32.2668 | 0.4619052 | 0.4179246 | 1.301245 | 0.1931745 | 0.371809 |
| ENSG00000145337 | PYURF | 3018.101 | 0.4616774 | 0.1535709 | 3.065466 | 0.00217331 | 0.011109 |
| ENSG00000151332 | MBIP | 1590.367 | 0.4616549 | 0.1320116 | 3.547067 | 0.00038955 | 0.002537 |
| ENSG00000278129 | ZNF8 | 663.3089 | 0.4611695 | 0.1785554 | 2.645801 | 0.00814978 | 0.033008 |

| | | | | | | | |
|---|---|---|---|---|---|---|---|
| ENSG00000177337 | DLGAP1-AS1 | 230.876 | 0.4611679 | 0.385545 | 1.36154 | 0.1733432 | 0.344897 |
| ENSG00000251688 | LINC02507 | 2.503346 | 0.4610589 | 0.7761956 | 1.127177 | 0.2596675 | 0.454008 |
| ENSG00000257957 | QRSL1P3 | 1.822468 | 0.4610391 | 0.8342203 | 1.208804 | 0.226738 | 0.414433 |
| ENSG00000246851 | | 1.441748 | 0.4610044 | 0.8378147 | 1.213591 | 0.2249041 | NA |
| ENSG00000266805 | | 1.15485 | 0.4609941 | 0.8719069 | 1.306219 | 0.1914781 | NA |
| ENSG00000170293 | CMTM8 | 509.3072 | 0.4609936 | 0.191397 | 2.493006 | 0.01266668 | 0.047115 |
| ENSG00000168970 | JMJD7-PLA2G4B | 320.673 | 0.4609017 | 0.279021 | 1.766513 | 0.07730988 | 0.19229 |
| ENSG00000285969 | | 2.030036 | 0.4608066 | 0.8695366 | 1.31943 | 0.1870253 | 0.363529 |
| ENSG00000183535 | COL18A1-AS1 | 5.091265 | 0.4603025 | 0.7154546 | 1.085017 | 0.2779141 | 0.475002 |
| ENSG00000280604 | | 30.77499 | 0.4602694 | 0.3747338 | 1.393385 | 0.1635034 | 0.331309 |
| ENSG00000049192 | ADAMTS6 | 78.2204088 | 0.460220705 | 0.437968246 | 1.247229472 | 0.21231337 | 0.396775 |
| ENSG00000151532 | VTI1A | 1664.88 | 0.4601457 | 0.1391255 | 3.365205 | 0.00076487 | 0.00454 |
| ENSG00000240718 | RN7SL851P | 2.837251 | 0.459978 | 0.8717159 | 1.319715 | 0.1869302 | 0.363371 |
| ENSG00000113262 | GRM6 | 4.183826178 | 0.459718938 | 0.663707186 | 1.06237275 | 0.2880665 | 0.486421 |
| ENSG00000137948 | BRDT | 3.510083829 | 0.459616648 | 0.914956773 | 1.537475373 | 0.12417694 | 0.271947 |
| ENSG00000149573 | MPZL2 | 1802.555 | 0.4595649 | 0.4083359 | 1.310167 | 0.1901393 | 0.3679 |
| ENSG00000163637 | PRICKLE2 | 1703.736 | 0.4594713 | 0.2332039 | 2.065092 | 0.03891434 | 0.114447 |
| ENSG00000232765 | | 1.73197 | 0.4593876 | 0.9464965 | 1.583705 | 0.1132608 | 0.254532 |
| ENSG00000184619 | KRBA2 | 21.01206 | 0.4593486 | 0.4872224 | 1.171869 | 0.2412495 | 0.431779 |
| ENSG00000253872 | | 2.156201 | 0.4591231 | 0.9230652 | 1.384152 | 0.1663118 | 0.335337 |
| ENSG00000104325 | DECR1 | 5469.275714 | 0.458893159 | 0.21357075 | 2.233340032 | 0.02552653 | 0.082438 |
| ENSG00000272913 | | 241.7366 | 0.4588363 | 0.287394 | 1.719279 | 0.08556361 | 0.207611 |
| ENSG00000287710 | | 69.92554 | 0.4588053 | 0.3276104 | 1.539286 | 0.1237346 | 0.271291 |
| ENSG00000187049 | TMEM216 | 1037.962 | 0.4587446 | 0.1603417 | 2.924 | 0.00345565 | 0.01634 |
| ENSG00000178449 | COX14 | 2166.428 | 0.4586469 | 0.2011911 | 2.348282 | 0.01886025 | 0.064833 |
| ENSG00000179523 | EIF3J-DT | 654.7159 | 0.4584441 | 0.22443 | 2.134615 | 0.03279251 | 0.100504 |
| ENSG00000184160 | ADRA2C | 1778.669 | 0.4584123 | 0.2697798 | 1.813059 | 0.06982279 | 0.178235 |
| ENSG00000147155 | EBP | 2015.226 | 0.4583834 | 0.3713335 | 1.423197 | 0.154679 | 0.318357 |
| ENSG00000116329 | OPRD1 | 8.15395495 | 0.458340953 | 0.62373833 | 1.079218512 | 0.28049033 | 0.477929 |
| ENSG00000286115 | | 12.79696 | 0.458242 | 0.5176209 | 1.12434 | 0.2608688 | 0.45557 |
| ENSG00000179299 | NSUN7 | 564.1761 | 0.4579088 | 0.1880478 | 2.511592 | 0.01201881 | 0.04509 |
| ENSG00000263142 | LRRC37A17P | 2.996139 | 0.4579045 | 0.7753071 | 1.132527 | 0.2574129 | 0.451389 |
| ENSG00000254634 | SMG1P6 | 17.27055 | 0.4577859 | 0.5474809 | 1.112653 | 0.2658574 | 0.46143 |

| | | | | | | | |
|---|---|---|---|---|---|---|---|
| ENSG00000259502 | | 7.136071 | 0.4576863 | 0.5955944 | 1.07172 | 0.2838457 | 0.4817 |
| ENSG00000204177 | BMS1P1 | 231.1647 | 0.4576218 | 0.2588031 | 1.881634 | 0.05988569 | 0.158824 |
| ENSG00000167972 | ABCA3 | 7758.001 | 0.4575952 | 0.192849 | 2.449944 | 0.01428783 | 0.051874 |
| ENSG00000230650 | | 39.87332 | 0.4575338 | 0.3835436 | 1.366137 | 0.171896 | 0.342747 |
| ENSG00000186174 | BCL9L | 8603.58 | 0.4575276 | 0.266049 | 1.849283 | 0.06441696 | 0.167596 |
| ENSG00000255108 | | 4.211918 | 0.4574359 | 0.6986278 | 1.075994 | 0.2819301 | 0.479644 |
| ENSG00000174599 | TRAM1L1 | 217.4277 | 0.457301 | 0.2375812 | 2.025038 | 0.04286345 | 0.123451 |
| ENSG00000266990 | | 3.326721 | 0.4572035 | 0.8601836 | 1.338434 | 0.1807549 | 0.355206 |
| ENSG00000273481 | | 1.910877 | 0.4571005 | 0.787267 | 1.120415 | 0.2625369 | 0.457412 |
| ENSG00000196233 | LCOR | 3930.917 | 0.4570555 | 0.2104126 | 2.259848 | 0.02383067 | 0.078129 |
| ENSG00000288009 | | 20.36977 | 0.4568654 | 0.5357456 | 1.112696 | 0.2658391 | 0.46143 |
| ENSG00000134070 | IRAK2 | 19.78631697 | 0.456742964 | 0.632498036 | 1.071941328 | 0.28374639 | 0.481562 |
| ENSG00000281731 | METTL14-DT | 25.66242 | 0.4567255 | 0.4418913 | 1.229231 | 0.2189851 | 0.404999 |
| ENSG00000275882 | IKBKGP1 | 1.688476 | 0.4566377 | 0.8827942 | 1.468693 | 0.1419162 | 0.299014 |
| ENSG00000106804 | C5 | 404.3583659 | 0.456554376 | 0.343492668 | 1.47525535 | 0.14014391 | 0.296314 |
| ENSG00000159228 | CBR1 | 2761.379 | 0.4565105 | 0.2732251 | 1.787599 | 0.07384067 | 0.186118 |
| ENSG00000257702 | LBX2-AS1 | 204.5931 | 0.4565026 | 0.2732503 | 1.781001 | 0.07491225 | 0.18791 |
| ENSG00000207009 | | 1.33339 | 0.4563449 | 0.8933397 | 1.508936 | 0.1313151 | NA |
| ENSG00000100075 | SLC25A1 | 13057.72382 | 0.456305514 | 0.237655453 | 2.015973826 | 0.04380271 | 0.125476 |
| ENSG00000270424 | | 1.712425 | 0.4562871 | 0.8454112 | 1.218779 | 0.2229281 | 0.409555 |
| ENSG00000237818 | RPS3AP29 | 9.114097 | 0.4562663 | 0.5444771 | 1.112941 | 0.2657339 | 0.461366 |
| ENSG00000046653 | GPM6B | 138.6176175 | 0.456191436 | 0.354671137 | 1.437453883 | 0.15058907 | 0.311981 |
| ENSG00000228753 | EIF4BP2 | 24.00643 | 0.4561292 | 0.4571466 | 1.200301 | 0.2300226 | 0.418307 |
| ENSG00000226669 | | 2.238712 | 0.4560168 | 0.8098848 | 1.153276 | 0.2487973 | 0.441175 |
| ENSG00000240344 | PPIL3 | 1263.652 | 0.4560113 | 0.1705467 | 2.744794 | 0.00605489 | 0.02591 |
| ENSG00000124549 | BTN2A3P | 114.1032497 | 0.456002375 | 0.285621295 | 1.717316966 | 0.08592127 | 0.20827 |
| ENSG00000265975 | | 22.02169 | 0.4559911 | 0.5440672 | 1.107494 | 0.2680802 | 0.463984 |
| ENSG00000203896 | LIME1 | 438.9517 | 0.4558712 | 0.3074664 | 1.604642 | 0.1085725 | 0.246996 |
| ENSG00000143751 | SDE2 | 1459.855 | 0.455778 | 0.2919743 | 1.687277 | 0.09155002 | 0.218269 |
| ENSG00000144730 | IL17RD | 244.3542 | 0.4556242 | 0.2853172 | 1.713796 | 0.08656622 | 0.209374 |
| ENSG00000164877 | MICALL2 | 3784.931 | 0.4555827 | 0.2489812 | 1.943592 | 0.05194467 | 0.142677 |
| ENSG00000100335 | MIEF1 | 2223.49217 | 0.455404108 | 0.173760667 | 2.6977867 | 0.00698022 | 0.029048 |
| ENSG00000141854 | MISP3 | 1075.768 | 0.4552706 | 0.2478375 | 1.92124 | 0.0547015 | 0.148551 |

| | | | | | | | |
|---|---|---|---|---|---|---|---|
| ENSG00000284773 | | 30.5171 | 0.4549214 | 0.3740873 | 1.375576 | 0.1689529 | 0.339032 |
| ENSG00000151164 | RAD9B | 19.88856 | 0.454891 | 0.4725785 | 1.180857 | 0.2376594 | 0.427295 |
| ENSG00000181856 | SLC2A4 | 71.4784 | 0.4547906 | 0.51161 | 1.141286 | 0.2537511 | 0.446751 |
| ENSG00000228008 | | 1.999017 | 0.4547899 | 0.8945813 | 1.529086 | 0.1262431 | 0.274908 |
| ENSG00000180662 | RPL21P8 | 2.172644 | 0.4547363 | 0.7634181 | 1.098247 | 0.2720965 | 0.468587 |
| ENSG00000164674 | SYTL3 | 56.81782 | 0.4546076 | 0.4085399 | 1.29095 | 0.1967211 | 0.376675 |
| ENSG00000138617 | PARP16 | 993.4320524 | 0.454579759 | 0.155801329 | 2.979079727 | 0.00289116 | 0.014154 |
| ENSG00000227484 | | 2.222114 | 0.4544171 | 0.8505541 | 1.236613 | 0.2162309 | 0.401554 |
| ENSG00000257808 | | 3.315439 | 0.4543402 | 0.8578019 | 1.300459 | 0.1934436 | 0.372085 |
| ENSG00000100221 | JOSD1 | 3494.040483 | 0.45428411 | 0.130955742 | 3.513755212 | 0.00044182 | 0.002819 |
| ENSG00000135750 | KCNK1 | 1643.483682 | 0.454239082 | 0.185307879 | 2.511046319 | 0.01203739 | 0.045145 |
| ENSG00000183808 | RBM12B | 1703.039 | 0.4541857 | 0.2845597 | 1.708367 | 0.08756827 | 0.21126 |
| ENSG00000285108 | | 3.187641 | 0.45417 | 0.7679806 | 1.10831 | 0.2677281 | 0.463617 |
| ENSG00000279456 | | 1.952379 | 0.4539624 | 0.8377237 | 1.220536 | 0.2222617 | 0.408955 |
| ENSG00000182253 | SYNM | 364.1886 | 0.4539573 | 0.3152835 | 1.543876 | 0.1226183 | 0.269397 |
| ENSG00000213512 | GBP7 | 1.18032 | 0.4538832 | 0.9774192 | 2.065645 | 0.03886204 | NA |
| ENSG00000099330 | OCEL1 | 554.0666676 | 0.453814373 | 0.177008456 | 2.639760187 | 0.00829647 | 0.033485 |
| ENSG00000238210 | ETDA | 1.808916 | 0.4536362 | 0.8661453 | 1.29339 | 0.1958761 | 0.375567 |
| ENSG00000170873 | MTSS1 | 629.4688 | 0.453629 | 0.1900113 | 2.446837 | 0.01441159 | 0.052245 |
| ENSG00000215873 | FEN1P1 | 3.44293 | 0.4533508 | 0.7830797 | 1.158414 | 0.2466953 | 0.438554 |
| ENSG00000147649 | MTDH | 12150.07 | 0.4533229 | 0.1599355 | 2.878594 | 0.00399452 | 0.018427 |
| ENSG00000226968 | LINC00423 | 1.375984 | 0.4531486 | 0.9144039 | 1.427663 | 0.1533888 | NA |
| ENSG00000241484 | ARHGAP8 | 534.9834 | 0.4531289 | 0.2702663 | 1.791723 | 0.07317735 | 0.184762 |
| ENSG00000273402 | | 18.32715 | 0.4531015 | 0.4499265 | 1.211704 | 0.2256256 | 0.412911 |
| ENSG00000172346 | CSDC2 | 1.067017 | 0.4529135 | 0.8804788 | 1.450454 | 0.146932 | NA |
| ENSG00000237438 | CECR7 | 12.39033 | 0.4529075 | 0.5489204 | 1.088789 | 0.2762469 | 0.473187 |
| ENSG00000079432 | CIC | 4869.343749 | 0.452766903 | 0.193660567 | 2.414014782 | 0.01577782 | 0.05611 |
| ENSG00000249001 | | 9.159666 | 0.4523028 | 0.6323341 | 1.068112 | 0.2854702 | 0.483469 |
| ENSG00000224509 | MRPS9-AS2 | 5.273699 | 0.4522974 | 0.7265581 | 1.07313 | 0.2832128 | 0.481054 |
| ENSG00000063046 | EIF4B | 30112.17588 | 0.452245661 | 0.150847444 | 3.092230794 | 0.00198658 | 0.010293 |
| ENSG00000269293 | ZSCAN16-AS1 | 109.7632 | 0.4522063 | 0.2998782 | 1.630039 | 0.1030932 | 0.237944 |
| ENSG00000183036 | PCP4 | 714.472 | 0.4519716 | 0.4316997 | 1.236021 | 0.2164507 | 0.401731 |
| ENSG00000182586 | LINC00334 | 3.425111 | 0.4518972 | 0.7117995 | 1.062106 | 0.2881876 | 0.486582 |

| | | | | | | | |
|---|---|---|---|---|---|---|---|
| ENSG00000177000 | MTHFR | 1473.365 | 0.451877 | 0.271312 | 1.761275 | 0.07819186 | 0.193849 |
| ENSG00000221845 | C7orf65 | 1.451904 | 0.4518418 | 0.9260175 | 1.501941 | 0.1331124 | NA |
| ENSG00000244513 | | 48.74631 | 0.4517077 | 0.2925582 | 1.66497 | 0.09591878 | 0.225621 |
| ENSG00000101782 | RIOK3 | 3168.87494 | 0.451627249 | 0.206777723 | 2.268783341 | 0.02328151 | 0.076698 |
| ENSG00000266718 | | 2.201948 | 0.4516144 | 0.8198056 | 1.182896 | 0.2368504 | 0.426522 |
| ENSG00000171357 | LURAP1 | 5.684497 | 0.4515646 | 0.7047307 | 1.053795 | 0.2919769 | 0.490766 |
| ENSG00000106080 | FKBP14 | 475.8605801 | 0.451429163 | 0.188210052 | 2.470725959 | 0.01348391 | 0.049442 |
| ENSG00000277521 | MIR8078 | 1.836982 | 0.4514006 | 0.8557381 | 1.28307 | 0.1994677 | 0.379686 |
| ENSG00000178776 | C5orf46 | 2.775051 | 0.4512913 | 0.8772301 | 1.407414 | 0.1593046 | 0.325307 |
| ENSG00000258633 | | 2.638552 | 0.4511124 | 0.79595 | 1.145219 | 0.2521184 | 0.444819 |
| ENSG00000233967 | | 181.1262 | 0.4509209 | 0.2486342 | 1.913543 | 0.05567854 | 0.150423 |
| ENSG00000059588 | TARBP1 | 4694.959298 | 0.450809186 | 0.223792159 | 2.1286558 | 0.03328275 | 0.101713 |
| ENSG00000140577 | CRTC3 | 2627.767 | 0.4507812 | 0.14698 | 3.117005 | 0.00182699 | 0.009601 |
| ENSG00000184939 | ZFP90 | 3369.817 | 0.4507358 | 0.1642393 | 2.807244 | 0.00499675 | 0.022141 |
| ENSG00000197893 | NRAP | 2.343164 | 0.4503907 | 0.8199758 | 1.163041 | 0.2448131 | 0.436422 |
| ENSG00000288543 | | 1.507472 | 0.4503816 | 0.9270144 | 1.538023 | 0.124043 | NA |
| ENSG00000248213 | CICP16 | 22.82855 | 0.4502009 | 0.6956694 | 1.056067 | 0.2909376 | 0.489522 |
| ENSG00000171757 | LRRC34 | 16.55281 | 0.4499591 | 0.5281399 | 1.100974 | 0.2709082 | 0.467235 |
| ENSG00000249311 | TERF1P3 | 7.448421 | 0.4499556 | 0.6783593 | 1.048668 | 0.2943312 | 0.493381 |
| ENSG00000079101 | CLUL1 | 8.538225479 | 0.449926907 | 0.625982657 | 1.064572474 | 0.28706944 | 0.485373 |
| ENSG00000253513 | | 2.74197 | 0.4498996 | 0.7987466 | 1.142464 | 0.2532614 | 0.446214 |
| ENSG00000251664 | PCDHA12 | 179.696 | 0.4497772 | 0.3231819 | 1.527743 | 0.1265764 | 0.275295 |
| ENSG00000253721 | SUMO2P20 | 1.944506 | 0.449743 | 0.8544137 | 1.354189 | 0.175676 | 0.34809 |
| ENSG00000235513 | L3MBTL2-AS1 | 12.71462 | 0.4494361 | 0.5096514 | 1.115447 | 0.2646589 | 0.460053 |
| ENSG00000013441 | CLK1 | 3815.343319 | 0.449269376 | 0.30157809 | 1.614352911 | 0.1064509 | 0.243592 |
| ENSG00000070367 | EXOC5 | 4897.239216 | 0.449258341 | 0.183417559 | 2.521048892 | 0.01170056 | 0.044117 |
| ENSG00000165023 | DIRAS2 | 15.28984 | 0.4491622 | 0.544875 | 1.082487 | 0.2790363 | 0.476216 |
| ENSG00000197702 | PARVA | 2287.429 | 0.4491192 | 0.213558 | 2.191222 | 0.02843574 | 0.089845 |
| ENSG00000260430 | | 115.7692 | 0.4491159 | 0.2978958 | 1.633739 | 0.1023136 | 0.236524 |
| ENSG00000182931 | WFDC10B | 1.540103 | 0.4490713 | 0.8829985 | 1.301967 | 0.1929277 | NA |
| ENSG00000015592 | STMN4 | 1.182722832 | 0.449068657 | 0.886946709 | 1.344596783 | 0.17875548 | NA |
| ENSG00000128159 | TUBGCP6 | 3381.702287 | 0.448996631 | 0.189152135 | 2.445857253 | 0.01445082 | 0.05235 |
| ENSG00000223529 | EEF1A1P8 | 8.21604 | 0.4489485 | 0.6391884 | 1.058392 | 0.2898766 | 0.488403 |

| | | | | | | | |
|---|---|---|---|---|---|---|---|
| ENSG00000102081 | FMR1 | 4706.941811 | 0.448905231 | 0.118951342 | 3.88723706 | 0.00010139 | 0.000775 |
| ENSG00000271737 | | 122.4766 | 0.4488178 | 0.3298174 | 1.498168 | 0.1340896 | 0.287008 |
| ENSG00000205639 | MFSD2B | 4.959894 | 0.4485977 | 0.6395374 | 1.048581 | 0.2943709 | 0.493416 |
| ENSG00000092148 | HECTD1 | 15409.2844 | 0.44858683 | 0.174613459 | 2.610849618 | 0.00903176 | 0.035858 |
| ENSG00000182310 | SPACA6 | 781.8185 | 0.4483977 | 0.339903 | 1.469306 | 0.1417497 | 0.2988 |
| ENSG00000156273 | BACH1 | 1604.038 | 0.4481929 | 0.1992329 | 2.32742 | 0.01994293 | 0.067753 |
| ENSG00000261216 | | 8.976115 | 0.4480468 | 0.5723231 | 1.072345 | 0.2835651 | 0.481347 |
| ENSG00000265936 | | 1.984598 | 0.4480361 | 0.8007103 | 1.163091 | 0.2447925 | 0.436422 |
| ENSG00000277581 | | 1.633135 | 0.4480214 | 0.8499232 | 1.242276 | 0.2141348 | 0.398973 |
| ENSG00000255398 | HCAR3 | 49.17657 | 0.4479753 | 0.4256542 | 1.240753 | 0.214697 | 0.3996 |
| ENSG00000156030 | MIDEAS | 5869.377 | 0.4479503 | 0.2371574 | 1.982175 | 0.04745965 | 0.133181 |
| ENSG00000125037 | EMC3 | 5744.973533 | 0.447881003 | 0.134317878 | 3.395345537 | 0.00068542 | 0.004127 |
| ENSG00000251161 | | 24.37957 | 0.4478617 | 0.3661744 | 1.373202 | 0.1696896 | 0.340032 |
| ENSG00000286938 | | 4.888651 | 0.4478556 | 0.787397 | 1.142219 | 0.2533631 | 0.446334 |
| ENSG00000132793 | LPIN3 | 3606.624092 | 0.447621195 | 0.239432476 | 1.963789336 | 0.04955453 | 0.137606 |
| ENSG00000134986 | NREP | 143.9506852 | 0.447572081 | 0.244862668 | 1.929423809 | 0.05367827 | 0.146605 |
| ENSG00000259250 | | 36.20961 | 0.4475234 | 0.3940243 | 1.308803 | 0.190601 | 0.368618 |
| ENSG00000198863 | RUNDC1 | 2036.046 | 0.447504 | 0.1776929 | 2.575797 | 0.01000094 | 0.038864 |
| ENSG00000100228 | RAB36 | 243.3872762 | 0.447462137 | 0.377877342 | 1.348551406 | 0.1774811 | 0.350751 |
| ENSG00000100796 | PPP4R3A | 8542.183044 | 0.447329098 | 0.115994794 | 4.001763735 | 6.29E-05 | 0.000508 |
| ENSG00000119866 | BCL11A | 18.91162116 | 0.447042719 | 0.651214229 | 1.044671142 | 0.29617499 | 0.495191 |
| ENSG00000179941 | BBS10 | 941.9864 | 0.446985 | 0.2197983 | 2.088926 | 0.03671438 | 0.109431 |
| ENSG00000236892 | | 0.8590459 | 0.4468795 | 0.9254944 | 1.443652 | 0.148837 | NA |
| ENSG00000271815 | NA | 6.192674 | 0.4468497 | 0.6121534 | 1.051763 | 0.2929082 | 0.491802 |
| ENSG00000142327 | RNPEPL1 | 2702.868 | 0.4468462 | 0.1417289 | 3.200153 | 0.00137355 | 0.007526 |
| ENSG00000250195 | | 0.9419182 | 0.446768 | 0.9420099 | 1.629673 | 0.1031705 | NA |
| ENSG00000257489 | | 137.0338 | 0.4466219 | 0.3213056 | 1.522134 | 0.1279755 | 0.277611 |
| ENSG00000155749 | FLACC1 | 28.41221 | 0.4465904 | 0.5140416 | 1.106061 | 0.2686999 | 0.464722 |
| ENSG00000221269 | MIR1302-8 | 1.070626 | 0.4465396 | 0.8960946 | 1.396702 | 0.1625032 | NA |
| ENSG00000198959 | TGM2 | 232.0557 | 0.4464228 | 0.5998338 | 1.050723 | 0.2933856 | 0.492364 |
| ENSG00000272316 | | 71.27634 | 0.4461734 | 0.4354 | 1.215315 | 0.2242459 | 0.411038 |
| ENSG00000244485 | RPL18P13 | 28.26536 | 0.4461707 | 0.4964483 | 1.123545 | 0.2612062 | 0.455805 |
| ENSG00000207713 | MIR200C | 1.073686 | 0.446153 | 0.9300673 | 1.551441 | 0.1207961 | NA |

| | | | | | | | |
|---|---|---|---|---|---|---|---|
| ENSG00000171574 | ZNF584 | 891.8763 | 0.4461182 | 0.2082912 | 2.216902 | 0.0266298 | 0.085265 |
| ENSG00000094841 | UPRT | 865.8087868 | 0.446024711 | 0.130915772 | 3.452564838 | 0.00055528 | 0.003436 |
| ENSG00000238164 | TNFRSF14-AS1 | 2.124998 | 0.4459194 | 0.8046856 | 1.140512 | 0.2540731 | 0.447081 |
| ENSG00000231274 | SBK3 | 2.59942 | 0.4458478 | 0.856059 | 1.264213 | 0.2061535 | 0.388323 |
| ENSG00000273353 | | 2.520202 | 0.4457928 | 0.8231939 | 1.189163 | 0.2343755 | 0.423527 |
| ENSG00000137843 | PAK6 | 392.3964911 | 0.445753955 | 0.313003856 | 1.570585606 | 0.11627894 | 0.259331 |
| ENSG00000228836 | CT45A5 | 2.86001 | 0.445749 | 0.9293819 | 1.499665 | 0.1337011 | 0.286408 |
| ENSG00000161912 | ADCY10P1 | 102.4858 | 0.4457136 | 0.3998157 | 1.278575 | 0.2010469 | 0.381789 |
| ENSG00000172551 | MUCL1 | 934.9466 | 0.4455134 | 0.4383664 | 1.218547 | 0.2230163 | 0.409603 |
| ENSG00000228817 | BACH1-IT2 | 9.293329 | 0.4455085 | 0.5547742 | 1.064628 | 0.2870441 | 0.485361 |
| ENSG00000196712 | NF1 | 5216.625 | 0.4454782 | 0.2078599 | 2.226256 | 0.02599701 | 0.083561 |
| ENSG00000256124 | LINC01152 | 9.186467 | 0.4454738 | 0.5960516 | 1.043742 | 0.2966046 | 0.495597 |
| ENSG00000236137 | | 29.8115 | 0.4454473 | 0.4966606 | 1.129934 | 0.258504 | 0.452699 |
| ENSG00000215277 | RNF212B | 23.7301 | 0.4454182 | 0.4881086 | 1.147385 | 0.2512227 | 0.443976 |
| ENSG00000251768 | RNA5SP217 | 4.955376 | 0.4453612 | 0.7275329 | 1.064971 | 0.286889 | 0.485253 |
| ENSG00000146232 | NFKBIE | 598.1731 | 0.4453247 | 0.1729413 | 2.604695 | 0.0091956 | 0.036373 |
| ENSG00000223262 | RNA5SP492 | 1.257966 | 0.4451839 | 0.9387986 | 1.616286 | 0.1060324 | NA |
| ENSG00000274423 | SEC22B2P | 2.259667 | 0.4450363 | 0.8546498 | 1.315694 | 0.1882768 | 0.36532 |
| ENSG00000230615 | | 123.0578 | 0.4447814 | 0.308457 | 1.566078 | 0.1173303 | 0.260975 |
| ENSG00000086967 | MYBPC2 | 1.411414849 | 0.444598627 | 0.867701471 | 1.292161843 | 0.1963011 | NA |
| ENSG00000171401 | KRT13 | 136.4452 | 0.4445047 | 0.5836247 | 1.064677 | 0.2870222 | 0.485355 |
| ENSG00000122678 | POLM | 1693.948248 | 0.444266618 | 0.135813902 | 3.321217241 | 0.00089626 | 0.005198 |
| ENSG00000201813 | RNU6-915P | 2.478147 | 0.4442453 | 0.8895222 | 1.390342 | 0.1644252 | 0.332543 |
| ENSG00000167978 | SRRM2 | 46483.16 | 0.4441029 | 0.1814468 | 2.514318 | 0.01192628 | 0.044813 |
| ENSG00000185958 | FAM186A | 9.951007 | 0.4440565 | 0.5947811 | 1.050712 | 0.2933909 | 0.492364 |
| ENSG00000266712 | MIR3149 | 1.325058 | 0.4436843 | 0.9105366 | 1.539113 | 0.1237767 | NA |
| ENSG00000182383 | RPL27AP5 | 6.430583 | 0.4436531 | 0.6907717 | 1.042964 | 0.2969648 | 0.495949 |
| ENSG00000258526 | | 28.41017 | 0.4435957 | 0.3325218 | 1.46283 | 0.143514 | 0.301401 |
| ENSG00000169814 | BTD | 1435.424 | 0.4435881 | 0.247654 | 1.888224 | 0.05899595 | 0.157139 |
| ENSG00000131584 | ACAP3 | 4765.068411 | 0.443466094 | 0.177592476 | 2.561851736 | 0.01041158 | 0.040195 |
| ENSG00000261775 | | 4.979745 | 0.4434655 | 0.7205504 | 1.069238 | 0.2849625 | 0.482948 |
| ENSG00000163512 | AZI2 | 1117.535 | 0.4434015 | 0.11988 | 3.754089 | 0.00017397 | 0.001253 |
| ENSG00000096092 | TMEM14A | 1045.142881 | 0.443322891 | 0.248358716 | 1.886829346 | 0.05918328 | 0.157512 |

| | | | | | | | |
|---|---|---|---|---|---|---|---|
| ENSG00000286795 | | 1.133836 | 0.443314 | 0.9020522 | 1.431291 | 0.1523468 | NA |
| ENSG00000189350 | TOGARAM2 | 30.63666 | 0.4432008 | 0.5550087 | 1.062995 | 0.2877844 | 0.486087 |
| ENSG00000058085 | LAMC2 | 549.5448126 | 0.443093176 | 0.673247018 | 1.045068318 | 0.2959914 | 0.495071 |
| ENSG00000205138 | SDHAF1 | 988.4793 | 0.443018 | 0.1529335 | 2.966225 | 0.0030148 | 0.014616 |
| ENSG00000253958 | CLDN23 | 325.2616 | 0.4429422 | 0.3934694 | 1.294967 | 0.1953315 | 0.374746 |
| ENSG00000286162 | | 36.69899 | 0.4428816 | 0.352496 | 1.403628 | 0.1604295 | 0.3269 |
| ENSG00000205488 | CALML3-AS1 | 42.21529 | 0.442706 | 0.5193616 | 1.097926 | 0.2722367 | 0.468735 |
| ENSG00000176871 | WSB2 | 4171.819 | 0.4426972 | 0.1129521 | 3.971825 | 7.13E-05 | 0.000567 |
| ENSG00000131127 | ZNF141 | 992.4717379 | 0.442694703 | 0.227898758 | 2.034517654 | 0.04189943 | 0.12124 |
| ENSG00000207422 | RNU6-813P | 72.25547 | 0.4426315 | 0.5159848 | 1.083606 | 0.2785396 | 0.475826 |
| ENSG00000244425 | RN7SL268P | 53.92857 | 0.4426214 | 0.3722535 | 1.347238 | 0.1779037 | 0.351273 |
| ENSG00000234277 | LINC01641 | 5.982033 | 0.4424193 | 0.892501 | 1.310168 | 0.1901391 | 0.3679 |
| ENSG00000263527 | MIR4526 | 1.20309 | 0.4422874 | 0.8927813 | 1.377979 | 0.1682097 | NA |
| ENSG00000011405 | PIK3C2A | 5310.481473 | 0.442117371 | 0.265039989 | 1.775662716 | 0.07578853 | 0.189553 |
| ENSG00000243896 | OR2A7 | 10.24825 | 0.4420917 | 0.612499 | 1.044898 | 0.2960701 | 0.495116 |
| ENSG00000269903 | | 8.462975 | 0.4417033 | 0.6202089 | 1.029701 | 0.3031503 | 0.502803 |
| ENSG00000073670 | ADAM11 | 439.327636 | 0.441628689 | 0.3502691 | 1.406480374 | 0.15958153 | 0.325672 |
| ENSG00000168159 | RNF187 | 11802.93 | 0.4414362 | 0.262662 | 1.811413 | 0.07007693 | 0.178734 |
| ENSG00000251194 | | 1.248118 | 0.4413648 | 0.9006427 | 1.482881 | 0.1381062 | NA |
| ENSG00000104497 | SNX16 | 746.5878284 | 0.441358293 | 0.230507241 | 2.008015576 | 0.04464164 | 0.12729 |
| ENSG00000021300 | PLEKHB1 | 700.5014073 | 0.441186063 | 0.307063778 | 1.560109513 | 0.118734 | 0.263128 |
| ENSG00000174652 | ZNF266 | 833.4587 | 0.4410999 | 0.1501285 | 3.001833 | 0.00268359 | 0.013273 |
| ENSG00000081177 | EXD2 | 3636.736823 | 0.441034951 | 0.166031406 | 2.717540748 | 0.00657691 | 0.027688 |
| ENSG00000214561 | RBBP4P4 | 7.739157 | 0.4410027 | 0.6599415 | 1.03027 | 0.3028835 | 0.502455 |
| ENSG00000063322 | MED29 | 3810.500259 | 0.440930471 | 0.116223447 | 3.823048662 | 0.00013181 | 0.000978 |
| ENSG00000164187 | LMBRD2 | 2843.894 | 0.4408937 | 0.2520316 | 1.858453 | 0.0631047 | 0.165058 |
| ENSG00000215199 | YWHAZP6 | 142.6 | 0.4408699 | 0.2117818 | 2.178234 | 0.02938866 | 0.092329 |
| ENSG00000261159 | | 44.15513 | 0.4407331 | 0.3474755 | 1.407765 | 0.1592006 | 0.325169 |
| ENSG00000132541 | RIDA | 2390.189038 | 0.440651752 | 0.270874657 | 1.727532033 | 0.08407216 | 0.204871 |
| ENSG00000261123 | | 25.91972 | 0.44062 | 0.5691025 | 1.054327 | 0.2917333 | 0.490481 |
| ENSG00000103769 | RAB11A | 10492.12824 | 0.440521777 | 0.119560946 | 3.744727056 | 0.00018059 | 0.001296 |
| ENSG00000197989 | SNHG12 | 2074.905 | 0.4404853 | 0.1940774 | 2.331799 | 0.01971129 | 0.067211 |
| ENSG00000141446 | ESCO1 | 2573.799 | 0.4404705 | 0.1482166 | 2.998568 | 0.00271251 | 0.013396 |

| | | | | | | | |
|---|---|---|---|---|---|---|---|
| ENSG00000206573 | THUMPD3-AS1 | 3121.513 | 0.4403849 | 0.1548915 | 2.893633 | 0.00380814 | 0.017704 |
| ENSG00000226571 | | 2.101656 | 0.4401591 | 0.8163817 | 1.146052 | 0.2517736 | 0.444508 |
| ENSG00000162493 | PDPN | 2.806144 | 0.4401115 | 0.8949545 | 1.264914 | 0.2059021 | 0.387987 |
| ENSG00000162923 | WDR26 | 9056.687 | 0.4401004 | 0.1116482 | 3.99781 | 6.39E-05 | 0.000515 |
| ENSG00000237753 | | 149.5571 | 0.4400792 | 0.3207367 | 1.506892 | 0.1318385 | 0.283744 |
| ENSG00000163389 | POGLUT1 | 1644.739 | 0.4400485 | 0.1549958 | 2.899577 | 0.00373667 | 0.017427 |
| ENSG00000168300 | PCMTD1 | 1970.932 | 0.4399665 | 0.2687174 | 1.737392 | 0.08231803 | 0.201744 |
| ENSG00000179058 | C9orf50 | 42.20973 | 0.4398774 | 0.377783 | 1.299827 | 0.1936602 | 0.372274 |
| ENSG00000268001 | CARD8-AS1 | 40.20459 | 0.4398628 | 0.3948363 | 1.283085 | 0.1994624 | 0.379686 |
| ENSG00000254373 | | 164.4706 | 0.4397883 | 0.2193648 | 2.09288 | 0.03635985 | 0.108713 |
| ENSG00000077097 | TOP2B | 16499.64697 | 0.439785427 | 0.130574261 | 3.424678162 | 0.00061553 | 0.003759 |
| ENSG00000147324 | MFHAS1 | 1396.887 | 0.4397214 | 0.1709849 | 2.636247 | 0.00838286 | 0.033772 |
| ENSG00000223458 | LMO7DN-IT1 | 0.9892342 | 0.4396522 | 0.9380843 | 1.664159 | 0.09608062 | NA |
| ENSG00000118257 | NRP2 | 36.28206133 | 0.43954026 | 0.517878033 | 1.099898868 | 0.27137619 | 0.467739 |
| ENSG00000258016 | HIGD1AP1 | 80.06835 | 0.4393711 | 0.34119 | 1.433044 | 0.1518454 | 0.313862 |
| ENSG00000286420 | | 2.808959 | 0.4393635 | 0.8597097 | 1.267758 | 0.2048844 | 0.386865 |
| ENSG00000187653 | TMSB4XP8 | 3231.475 | 0.4393572 | 0.2153333 | 2.127017 | 0.03341867 | 0.102022 |
| ENSG00000156218 | ADAMTSL3 | 49.8299 | 0.4393177 | 0.4788774 | 1.129457 | 0.2587049 | 0.452921 |
| ENSG00000169299 | PGM2 | 2436.253 | 0.4391269 | 0.3040977 | 1.568762 | 0.1167033 | 0.259929 |
| ENSG00000232439 | RPL18AP7 | 2.887833 | 0.4390928 | 0.7657516 | 1.086964 | 0.2770528 | 0.473987 |
| ENSG00000253833 | DSTNP3 | 90.41979 | 0.4390479 | 0.2363558 | 1.953871 | 0.05071642 | 0.140017 |
| ENSG00000166349 | RAG1 | 6.404552 | 0.4388196 | 0.6319897 | 1.030054 | 0.3029845 | 0.502559 |
| ENSG00000273257 | | 3.548372 | 0.438808 | 0.8257644 | 1.200067 | 0.2301132 | 0.418399 |
| ENSG00000267185 | PTP4A2P1 | 117.0683 | 0.4387611 | 0.2443618 | 1.896175 | 0.05793688 | 0.155018 |
| ENSG00000242960 | FTH1P23 | 33.94809 | 0.43867 | 0.4018053 | 1.266014 | 0.2055081 | 0.387492 |
| ENSG00000162402 | USP24 | 5815.143 | 0.4385168 | 0.2174154 | 2.114659 | 0.03445902 | 0.104466 |
| ENSG00000263528 | IKBKE | 665.5906 | 0.4384797 | 0.2677838 | 1.741409 | 0.08161195 | 0.200402 |
| ENSG00000103966 | EHD4 | 2151.283114 | 0.438472767 | 0.193266441 | 2.338036809 | 0.01938534 | 0.06627 |
| ENSG00000175970 | UNC119B | 2942.657 | 0.4384391 | 0.1642608 | 2.713297 | 0.00666174 | 0.027979 |
| ENSG00000106069 | CHN2 | 1.131836256 | 0.438257343 | 0.92408636 | 1.540563625 | 0.12342303 | NA |
| ENSG00000153048 | CARHSP1 | 1975.883 | 0.4382061 | 0.3083218 | 1.534291 | 0.124958 | 0.27298 |
| ENSG00000243830 | | 1.659951 | 0.4381976 | 0.8507997 | 1.343991 | 0.1789514 | 0.35289 |
| ENSG00000137764 | MAP2K5 | 1039.872319 | 0.43819646 | 0.158969887 | 2.808527201 | 0.00497687 | 0.022068 |

| | | | | | | | |
|---|---|---|---|---|---|---|---|
| ENSG00000241007 | SEPTIN7P6 | 62.72009 | 0.4381892 | 0.302366 | 1.57679 | 0.1148438 | 0.257059 |
| ENSG00000228755 | PABPC1P8 | 1.248406 | 0.4379924 | 0.9056593 | 1.375981 | 0.1688274 | NA |
| ENSG00000265982 | | 90.39244 | 0.4379037 | 0.3068625 | 1.546813 | 0.1219083 | 0.268269 |
| ENSG00000162298 | SYVN1 | 3922.457 | 0.4378125 | 0.1873741 | 2.405385 | 0.01615544 | 0.057155 |
| ENSG00000258738 | | 97.63476 | 0.4377535 | 0.347203 | 1.412283 | 0.1578667 | 0.323133 |
| ENSG00000188388 | GOLGA6L3 | 4.737308 | 0.437669 | 0.6684469 | 1.023466 | 0.3060877 | 0.506066 |
| ENSG00000261002 | | 3.706027 | 0.4376239 | 0.6874237 | 1.031389 | 0.3023585 | 0.501834 |
| ENSG00000131374 | TBC1D5 | 7037.423495 | 0.437616518 | 0.209168003 | 2.174954244 | 0.02963355 | 0.092882 |
| ENSG00000146197 | SCUBE3 | 112.9947 | 0.4374613 | 0.4331126 | 1.20466 | 0.2283344 | 0.416492 |
| ENSG00000207607 | MIR200A | 11.63794 | 0.4374611 | 0.5067703 | 1.093034 | 0.2743788 | 0.471141 |
| ENSG00000168904 | LRRC28 | 753.7276 | 0.4374203 | 0.1576966 | 2.845128 | 0.00443936 | 0.020083 |
| ENSG00000270083 | | 20.47683 | 0.4373023 | 0.4796585 | 1.126512 | 0.259949 | 0.454381 |
| ENSG00000114656 | CFAP92 | 276.2123495 | 0.437138453 | 0.267190934 | 1.739589126 | 0.08193119 | 0.201019 |
| ENSG00000130810 | PPAN | 487.1948835 | 0.43672814 | 0.192690573 | 2.342577306 | 0.01915107 | 0.065621 |
| ENSG00000268798 | | 15.83527 | 0.436375 | 0.4564843 | 1.150279 | 0.2500289 | 0.442632 |
| ENSG00000271185 | | 1.343569 | 0.4362893 | 0.8897075 | 1.500203 | 0.1335617 | NA |
| ENSG00000135437 | RDH5 | 28.94891297 | 0.436287712 | 0.520506578 | 1.076045361 | 0.28190697 | 0.479635 |
| ENSG00000228137 | | 54.36822 | 0.4361986 | 0.3794712 | 1.307521 | 0.191036 | 0.369082 |
| ENSG00000175967 | LINC02880 | 2.867803 | 0.436116 | 0.8173047 | 1.171236 | 0.241504 | 0.43206 |
| ENSG00000173542 | MOB1B | 1842.939 | 0.436031 | 0.1863036 | 2.396557 | 0.01654993 | 0.058318 |
| ENSG00000237629 | UQCRHP2 | 1.478381 | 0.4360191 | 0.8289619 | 1.199214 | 0.2304449 | NA |
| ENSG00000218186 | KRT8P43 | 5.472372 | 0.4358256 | 0.6079692 | 1.032755 | 0.3017184 | 0.501179 |
| ENSG00000278570 | NR2E3 | 14.44894 | 0.4357507 | 0.4936151 | 1.106874 | 0.2683483 | 0.464327 |
| ENSG00000113296 | THBS4 | 11.89115761 | 0.435705929 | 0.55279578 | 1.048319114 | 0.29449161 | 0.493525 |
| ENSG00000267281 | ATF7-NPFF | 21.20372 | 0.4356805 | 0.4720377 | 1.133372 | 0.2570583 | 0.450911 |
| ENSG00000257894 | | 14.94281 | 0.4356391 | 0.6265002 | 1.033532 | 0.3013548 | 0.500638 |
| ENSG00000239906 | | 2.260211 | 0.4354926 | 0.822154 | 1.171176 | 0.2415282 | 0.432074 |
| ENSG00000232725 | | 27.78331 | 0.4354266 | 0.4285591 | 1.200873 | 0.2298006 | 0.417993 |
| ENSG00000091536 | MYO15A | 36.64181769 | 0.435395892 | 0.337640643 | 1.425012014 | 0.15415373 | 0.317522 |
| ENSG00000197959 | DNM3 | 56.78531 | 0.4350955 | 0.3667602 | 1.332304 | 0.1827602 | 0.357751 |
| ENSG00000249244 | | 26.51533 | 0.434661 | 0.4722355 | 1.131496 | 0.2578463 | 0.451924 |
| ENSG00000235721 | | 32.86287 | 0.4343541 | 0.4586972 | 1.149267 | 0.250446 | 0.443048 |
| ENSG00000174109 | C16orf91 | 1603.472 | 0.4343455 | 0.2281087 | 1.988956 | 0.04670611 | 0.131636 |

| | | | | | | | |
|---|---|---|---|---|---|---|---|
| ENSG00000090097 | PCBP4 | 1548.762779 | 0.434321761 | 0.203978093 | 2.2029208 | 0.02760033 | 0.087737 |
| ENSG00000064115 | TM7SF3 | 3481.176409 | 0.43424222 | 0.19393687 | 2.256197778 | 0.02405825 | 0.0787 |
| ENSG00000170871 | KIAA0232 | 2556.822 | 0.4341628 | 0.2187776 | 2.063001 | 0.03911253 | 0.114865 |
| ENSG00000166439 | RNF169 | 1856.634 | 0.4341607 | 0.2841195 | 1.636351 | 0.1017662 | 0.235689 |
| ENSG00000173041 | ZNF680 | 897.7462 | 0.4340458 | 0.191983 | 2.337436 | 0.01941655 | 0.066368 |
| ENSG00000229752 | RPL7P10 | 1.378468 | 0.4339569 | 0.8468718 | 1.227675 | 0.219569 | NA |
| ENSG00000111261 | MANSC1 | 1043.648707 | 0.43393776 | 0.181543774 | 2.440835217 | 0.01465334 | 0.052926 |
| ENSG00000234665 | LERFS | 3.019171 | 0.4335815 | 0.8115673 | 1.145759 | 0.2518949 | 0.444632 |
| ENSG00000168016 | TRANK1 | 637.737 | 0.4335406 | 0.4616706 | 1.149945 | 0.2501663 | 0.442695 |
| ENSG00000139620 | KANSL2 | 2287.183 | 0.4334794 | 0.1480459 | 2.987987 | 0.00280822 | 0.013817 |
| ENSG00000105538 | RASIP1 | 95.98315225 | 0.433390437 | 0.276337604 | 1.671736797 | 0.09457624 | 0.223499 |
| ENSG00000182890 | GLUD2 | 1664.117 | 0.4333795 | 0.2149492 | 2.099049 | 0.03581261 | 0.10749 |
| ENSG00000231169 | EEF1B2P1 | 5.855329 | 0.4332153 | 0.6848926 | 1.020828 | 0.3073359 | 0.507559 |
| ENSG00000156427 | FGF18 | 25.11082 | 0.4331994 | 0.5051662 | 1.106246 | 0.2686201 | 0.464645 |
| ENSG00000188739 | RBM34 | 42.12328 | 0.4330762 | 0.2810356 | 1.650143 | 0.09891372 | 0.230781 |
| ENSG00000162004 | CCDC78 | 3472.238 | 0.4328959 | 0.2382972 | 1.914393 | 0.05556996 | 0.150243 |
| ENSG00000256651 | | 2.204052 | 0.4328225 | 0.8647694 | 1.273573 | 0.2028148 | 0.384131 |
| ENSG00000109084 | TMEM97 | 4649.473435 | 0.432700252 | 0.399297557 | 1.271868909 | 0.2034197 | 0.385084 |
| ENSG00000172831 | CES2 | 2480.776 | 0.4326881 | 0.1634591 | 2.691412 | 0.00711503 | 0.029547 |
| ENSG00000254263 | | 7.48366 | 0.432615 | 0.6039282 | 1.024958 | 0.305383 | 0.505213 |
| ENSG00000286485 | | 11.08553 | 0.4324681 | 0.5288514 | 1.060812 | 0.2887755 | 0.487389 |
| ENSG00000287731 | | 3.069707 | 0.4324677 | 0.7750812 | 1.090897 | 0.275318 | 0.472147 |
| ENSG00000236810 | ELOA-AS1 | 35.59164 | 0.4322672 | 0.4776219 | 1.113591 | 0.2654545 | 0.461122 |
| ENSG00000182247 | UBE2E2 | 1170.956 | 0.4322656 | 0.1798303 | 2.468063 | 0.01358464 | 0.049708 |
| ENSG00000275234 | | 289.1898 | 0.4321285 | 0.2637737 | 1.734548 | 0.0828209 | 0.202712 |
| ENSG00000154781 | CCDC174 | 2029.577 | 0.4321116 | 0.1379741 | 3.18244 | 0.0014604 | 0.007923 |
| ENSG00000230026 | | 1.737835 | 0.4320905 | 0.8519784 | 1.241835 | 0.2142973 | 0.399145 |
| ENSG00000260911 | | 19.13257 | 0.4319999 | 0.5158126 | 1.068798 | 0.2851605 | 0.483186 |
| ENSG00000228288 | PCAT6 | 391.1348 | 0.4319891 | 0.2934815 | 1.594565 | 0.1108095 | 0.250712 |
| ENSG00000198939 | ZFP2 | 171.8067 | 0.4318479 | 0.2804477 | 1.646825 | 0.09959408 | 0.232075 |
| ENSG00000179363 | TMEM31 | 30.07593 | 0.4318137 | 0.4078932 | 1.227603 | 0.219596 | 0.405649 |
| ENSG00000229456 | RLIMP1 | 1.573364 | 0.4315911 | 0.8057178 | 1.113718 | 0.2654001 | 0.461088 |
| ENSG00000228525 | | 1.679955 | 0.4315076 | 0.8063698 | 1.122011 | 0.2618578 | 0.456636 |

| | | | | | | | |
|---|---|---|---|---|---|---|---|
| ENSG00000260381 | | 3.046478 | 0.4314205 | 0.740606 | 1.061293 | 0.2885567 | 0.487082 |
| ENSG00000157693 | TMEM268 | 2086.087 | 0.4313756 | 0.2145889 | 2.088862 | 0.03672014 | 0.109436 |
| ENSG00000182796 | TMEM198B | 755.8768 | 0.4312599 | 0.1909062 | 2.325185 | 0.02006206 | 0.068127 |
| ENSG00000112041 | TULP1 | 57.95510481 | 0.430738474 | 0.366087957 | 1.32827295 | 0.18408795 | 0.359356 |
| ENSG00000223599 | | 2.941356 | 0.4307284 | 0.7254921 | 1.03519 | 0.3005802 | 0.499851 |
| ENSG00000183784 | DOCK8-AS1 | 38.3053 | 0.4305768 | 0.3789491 | 1.288946 | 0.197417 | 0.377369 |
| ENSG00000286375 | | 0.9591084 | 0.4305667 | 0.9390189 | 1.711133 | 0.08705665 | NA |
| ENSG00000170122 | FOXD4 | 13.8629 | 0.430469 | 0.5344507 | 1.052612 | 0.2925187 | 0.491366 |
| ENSG00000162591 | MEGF6 | 1270.517 | 0.4303144 | 0.2100446 | 2.12583 | 0.03351742 | 0.102199 |
| ENSG00000266680 | | 112.2779 | 0.4302825 | 0.3295143 | 1.434241 | 0.1515036 | 0.313399 |
| ENSG00000166432 | ZMAT1 | 2193.182 | 0.4299961 | 0.3986406 | 1.24128 | 0.2145024 | 0.399379 |
| ENSG00000242028 | HYPK | 78.2627 | 0.4298005 | 0.4124582 | 1.219584 | 0.2226226 | 0.409344 |
| ENSG00000246922 | UBAP1L | 208.5368 | 0.4297935 | 0.4432547 | 1.157468 | 0.2470812 | 0.438993 |
| ENSG00000183496 | MEX3B | 733.2493 | 0.4296946 | 0.3755133 | 1.288129 | 0.1977011 | 0.377758 |
| ENSG00000180279 | LINC01869 | 16.20344 | 0.4294968 | 0.4937129 | 1.083515 | 0.2785798 | 0.475864 |
| ENSG00000104897 | SF3A2 | 2280.928078 | 0.429465428 | 0.150583155 | 2.907931775 | 0.00363828 | 0.017061 |
| ENSG00000142794 | NBPF3 | 852.6061 | 0.4294091 | 0.1890529 | 2.331809 | 0.01971076 | 0.067211 |
| ENSG00000107105 | ELAVL2 | 76.60101539 | 0.429348674 | 0.558643586 | 1.038226853 | 0.29916445 | 0.498385 |
| ENSG00000110446 | SLC15A3 | 49.60634998 | 0.429114678 | 0.542373285 | 1.045738949 | 0.29568158 | 0.494708 |
| ENSG00000140092 | FBLN5 | 10.47819 | 0.4291102 | 0.7636887 | 1.091142 | 0.2752102 | 0.47205 |
| ENSG00000137876 | RSL24D1 | 7253.490532 | 0.429093628 | 0.206485392 | 2.118528493 | 0.03413033 | 0.103671 |
| ENSG00000131951 | LRRC9 | 1.325064436 | 0.428939423 | 0.866911447 | 1.281648535 | 0.19996597 | NA |
| ENSG00000124812 | CRISP1 | 1.006024714 | 0.42888266 | 0.91544339 | 1.42794548 | 0.15330755 | NA |
| ENSG00000206527 | HACD2 | 9339.101 | 0.4288317 | 0.1447522 | 3.006072 | 0.00264647 | 0.013121 |
| ENSG00000286650 | | 12.1464 | 0.4288202 | 0.6589158 | 1.003532 | 0.3156041 | 0.51659 |
| ENSG00000198246 | SLC29A3 | 1201.765 | 0.4287756 | 0.2684772 | 1.753519 | 0.07951302 | 0.196337 |
| ENSG00000147124 | ZNF41 | 1048.977 | 0.4286533 | 0.1164893 | 3.723201 | 0.00019671 | 0.001397 |
| ENSG00000188322 | SBK1 | 2298.19 | 0.4285756 | 0.2347881 | 1.913278 | 0.05571242 | 0.15049 |
| ENSG00000225496 | RTRAFP1 | 21.78563 | 0.4282315 | 0.4156478 | 1.200724 | 0.2298583 | 0.418037 |
| ENSG00000261609 | GAN | 1080.06 | 0.4281443 | 0.2395061 | 1.866998 | 0.06190192 | 0.162807 |
| ENSG00000188290 | HES4 | 966.0451 | 0.4280772 | 0.2181987 | 2.043155 | 0.04103709 | 0.119434 |
| ENSG00000100311 | PDGFB | 1605.340314 | 0.428015965 | 0.181215272 | 2.428171419 | 0.01517517 | 0.054434 |
| ENSG00000168143 | FAM83B | 1166.104 | 0.4279954 | 0.2915055 | 1.589056 | 0.1120477 | 0.25263 |

| | | | | | | | |
|---|---|---|---|---|---|---|---|
| ENSG00000167994 | RAB3IL1 | 534.017 | 0.4279002 | 0.4315108 | 1.174811 | 0.2400705 | 0.430307 |
| ENSG00000184441 | | 198.0645 | 0.4277982 | 0.2732623 | 1.669496 | 0.09501904 | 0.224226 |
| ENSG00000072864 | NDE1 | 1385.016262 | 0.427783147 | 0.244121255 | 1.831412953 | 0.06703893 | 0.17281 |
| ENSG00000203436 | | 1.404977 | 0.4275337 | 0.8866132 | 1.457208 | 0.145059 | NA |
| ENSG00000259425 | | 1.915383 | 0.4275016 | 0.8854822 | 1.330318 | 0.1834135 | 0.358554 |
| ENSG00000092969 | TGFB2 | 4459.72123 | 0.427461593 | 0.467243932 | 1.121152802 | 0.26222283 | 0.457032 |
| ENSG00000105963 | ADAP1 | 554.4409779 | 0.427352945 | 0.181814874 | 2.419539103 | 0.01554019 | 0.055458 |
| ENSG00000105519 | CAPS | 402.1054562 | 0.427316767 | 0.31225188 | 1.49188106 | 0.13573033 | 0.289387 |
| ENSG00000172318 | B3GALT1 | 4.147077 | 0.4272038 | 0.6483996 | 1.004298 | 0.3152352 | 0.516209 |
| ENSG00000115966 | ATF2 | 1909.82666 | 0.427163144 | 0.235573164 | 1.897867625 | 0.05771352 | 0.154545 |
| ENSG00000162616 | DNAJB4 | 352.8984 | 0.4271462 | 0.2552411 | 1.76951 | 0.07680879 | 0.191367 |
| ENSG00000258769 | RAP1AP | 45.7453 | 0.4269536 | 0.529174 | 1.049795 | 0.2938122 | 0.492822 |
| ENSG00000275479 | | 12.30883 | 0.4268575 | 0.6004399 | 1.007509 | 0.3136903 | 0.514821 |
| ENSG00000008394 | MGST1 | 6098.439899 | 0.426690402 | 0.208908103 | 2.080551971 | 0.03747493 | 0.111112 |
| ENSG00000119986 | AVPI1 | 1564.485468 | 0.426578118 | 0.282268087 | 1.610555708 | 0.10727659 | 0.244865 |
| ENSG00000228139 | LINC01402 | 1.758165 | 0.4265005 | 0.9207762 | 1.645161 | 0.09993666 | 0.232759 |
| ENSG00000228759 | FAF2P1 | 3.778119 | 0.4263554 | 0.6682153 | 0.9948669 | 0.319801 | 0.520733 |
| ENSG00000006118 | TMEM132A | 11181.61497 | 0.426157938 | 0.211667751 | 2.09267368 | 0.03637829 | 0.108713 |
| ENSG00000068903 | SIRT2 | 2908.464777 | 0.426147225 | 0.216697658 | 2.065484793 | 0.03887715 | 0.114363 |
| ENSG00000128656 | CHN1 | 607.930505 | 0.42590134 | 0.223921857 | 1.986756638 | 0.04694937 | 0.132126 |
| ENSG00000134909 | ARHGAP32 | 2052.958368 | 0.425848798 | 0.244199234 | 1.831783658 | 0.06698366 | 0.172701 |
| ENSG00000228158 | TLE1P1 | 154.5546 | 0.4258348 | 0.2556228 | 1.766131 | 0.0773739 | 0.192378 |
| ENSG00000253669 | GASAL1 | 125.6625 | 0.4256877 | 0.3155448 | 1.468825 | 0.1418802 | 0.298962 |
| ENSG00000235706 | DICER1-AS1 | 511.2169 | 0.4256382 | 0.3211767 | 1.454924 | 0.1456903 | 0.304791 |
| ENSG00000104447 | TRPS1 | 7492.959517 | 0.425509119 | 0.243528325 | 1.836523388 | 0.06628029 | 0.17137 |
| ENSG00000264061 | FGF7P1 | 0.8969556 | 0.4254156 | 0.9425205 | 1.802377 | 0.07148608 | NA |
| ENSG00000286952 | | 2.592449 | 0.4253633 | 0.7733638 | 1.068187 | 0.285436 | 0.483442 |
| ENSG00000172466 | ZNF24 | 7063.223 | 0.4253061 | 0.2129577 | 2.072151 | 0.03825134 | 0.112951 |
| ENSG00000243660 | ZNF487 | 249.0289 | 0.4252711 | 0.2148825 | 2.063797 | 0.03903698 | 0.114732 |
| ENSG00000261101 | | 23.06795 | 0.4251962 | 0.4613104 | 1.119789 | 0.2628036 | 0.457774 |
| ENSG00000288109 | | 2.169716 | 0.4251292 | 0.7724864 | 1.070299 | 0.2844845 | 0.482445 |
| ENSG00000178904 | DPY19L3 | 2722.442 | 0.4249736 | 0.2127862 | 2.074272 | 0.03805406 | 0.112515 |
| ENSG00000182107 | TMEM30B | 4908.006 | 0.42491 | 0.1480809 | 2.905094 | 0.00367143 | 0.017171 |

| | | | | | | | |
|---|---|---|---|---|---|---|---|
| ENSG00000233916 | ZDHHC20P1 | 1.033763 | 0.4248495 | 0.8574981 | 1.253023 | 0.2101973 | NA |
| ENSG00000116031 | CD207 | 6.031307001 | 0.42480931 | 0.660890409 | 1.003492401 | 0.31562334 | 0.516589 |
| ENSG00000120500 | ARR3 | 17.58466605 | 0.424792219 | 0.420639508 | 1.185575024 | 0.23579017 | 0.42515 |
| ENSG00000236276 | NDP-AS1 | 1.102577 | 0.4246351 | 0.919477 | 1.486787 | 0.1370711 | NA |
| ENSG00000103671 | TRIP4 | 2052.169508 | 0.424585996 | 0.135952033 | 3.165908546 | 0.00154599 | 0.00832 |
| ENSG00000146535 | GNA12 | 5544.97 | 0.4245214 | 0.146315 | 2.950885 | 0.00316865 | 0.015202 |
| ENSG00000271380 | | 25.25169 | 0.424509 | 0.4086361 | 1.207514 | 0.2272344 | 0.415088 |
| ENSG00000260053 | ABCB10P4 | 6.372019 | 0.4243733 | 0.6714458 | 1.005928 | 0.3144503 | 0.515591 |
| ENSG00000271268 | | 151.0735 | 0.4243411 | 0.2399036 | 1.861605 | 0.06265886 | 0.164254 |
| ENSG00000285103 | | 14.24572 | 0.4243001 | 0.5238637 | 1.040611 | 0.2980561 | 0.497178 |
| ENSG00000162999 | DUSP19 | 29.49446 | 0.4241979 | 0.4452122 | 1.139449 | 0.2545161 | 0.447534 |
| ENSG00000104967 | NOVA2 | 33.99646026 | 0.424061706 | 0.597926165 | 1.008147381 | 0.31338371 | 0.51454 |
| ENSG00000224331 | | 3.240605 | 0.4240302 | 0.6864488 | 0.9958887 | 0.3193042 | 0.520243 |
| ENSG00000281376 | ABALON | 41.81688 | 0.4239805 | 0.4145601 | 1.192288 | 0.2331485 | 0.421999 |
| ENSG00000235529 | AGAP1-IT1 | 7.418851 | 0.4237415 | 0.6721717 | 1.011754 | 0.3116557 | 0.512551 |
| ENSG00000237037 | NDUFA6-DT | 70.01554 | 0.4237175 | 0.2898157 | 1.569364 | 0.1165632 | 0.259726 |
| ENSG00000250072 | SH3TC2-DT | 45.55374 | 0.4236359 | 0.4141715 | 1.194711 | 0.2321999 | 0.420626 |
| ENSG00000152782 | PANK1 | 1151.445 | 0.4236096 | 0.2303162 | 1.924976 | 0.05423233 | 0.147648 |
| ENSG00000220161 | LINC02076 | 2.302028 | 0.4235856 | 0.8031666 | 1.133472 | 0.2570159 | 0.450911 |
| ENSG00000280399 | | 22.07651 | 0.4235742 | 0.4142713 | 1.202511 | 0.2291656 | 0.417287 |
| ENSG00000264290 | | 47.9344 | 0.4235348 | 0.3102037 | 1.480021 | 0.1388677 | 0.294277 |
| ENSG00000255968 | | 4.461486 | 0.4234743 | 0.772215 | 1.085821 | 0.2775582 | 0.474607 |
| ENSG00000256073 | URB1-AS1 | 466.56 | 0.4234412 | 0.2211048 | 1.99787 | 0.04573075 | 0.129553 |
| ENSG00000278998 | | 1.272854 | 0.4232577 | 0.8823503 | 1.364359 | 0.1724545 | NA |
| ENSG00000163666 | HESX1 | 29.13768 | 0.4231911 | 0.4162802 | 1.18381 | 0.2364883 | 0.42607 |
| ENSG00000184307 | ZDHHC23 | 2196.738 | 0.4231425 | 0.2082863 | 2.09668 | 0.03602195 | 0.108033 |
| ENSG00000238090 | | 1.391347 | 0.4231184 | 0.9224999 | 1.616594 | 0.1059659 | NA |
| ENSG00000070031 | SCT | 2.552580375 | 0.423079241 | 0.868226777 | 1.332247919 | 0.18277874 | 0.357761 |
| ENSG00000100345 | MYH9 | 43817.67275 | 0.422987357 | 0.176661772 | 2.45878642 | 0.01394075 | 0.050808 |
| ENSG00000109519 | GRPEL1 | 2933.15064 | 0.422893517 | 0.195919698 | 2.2304382 | 0.02571837 | 0.08289 |
| ENSG00000240057 | | 8.899584 | 0.4228049 | 0.6021165 | 0.9957643 | 0.3193647 | 0.520277 |
| ENSG00000127928 | GNGT1 | 117.871149 | 0.422782554 | 0.252298169 | 1.773774449 | 0.07610047 | 0.190054 |
| ENSG00000229022 | | 4.137191 | 0.4227152 | 0.6359977 | 0.9869535 | 0.3236654 | 0.524581 |

| | | | | | | | |
|---|---|---|---|---|---|---|---|
| ENSG00000260552 | COSMOC | 24.81509 | 0.4227121 | 0.3903447 | 1.243821 | 0.2135654 | 0.398359 |
| ENSG00000227507 | LTB | 60.96746 | 0.4226532 | 0.7634935 | 1.092789 | 0.2744866 | 0.471272 |
| ENSG00000102908 | NFAT5 | 3231.111736 | 0.422551484 | 0.350860597 | 1.358601471 | 0.17427291 | 0.346187 |
| ENSG00000196428 | TSC22D2 | 2723.823 | 0.4224106 | 0.1959858 | 2.226956 | 0.0259502 | 0.083481 |
| ENSG00000183378 | OVCH2 | 2.504705 | 0.4223919 | 0.8838352 | 1.250107 | 0.2112605 | 0.395253 |
| ENSG00000265820 | MIR3177 | 1.632216 | 0.4222614 | 0.8443777 | 1.228118 | 0.2194028 | 0.405518 |
| ENSG00000175600 | SUGCT | 15.98478 | 0.422151 | 0.506185 | 1.056296 | 0.290833 | 0.489462 |
| ENSG00000286675 | | 1.227197 | 0.4221482 | 0.8376367 | 1.195807 | 0.2317718 | NA |
| ENSG00000205642 | VCX3B | 4.682567 | 0.4221441 | 0.9002845 | 1.361855 | 0.1732436 | 0.344736 |
| ENSG00000272368 | | 49.18876 | 0.4221169 | 0.447207 | 1.128272 | 0.2592052 | 0.453528 |
| ENSG00000215270 | TOMM40P2 | 80.5864 | 0.4218574 | 0.2806386 | 1.610673 | 0.107251 | 0.244849 |
| ENSG00000228492 | RAB11FIP1P1 | 135.8497 | 0.4218071 | 0.3401139 | 1.373371 | 0.169637 | 0.339978 |
| ENSG00000135625 | EGR4 | 93.72300721 | 0.421757966 | 0.839246369 | 1.225424828 | 0.2204152 | 0.406653 |
| ENSG00000279331 | RBM12B-AS1 | 51.74763 | 0.4217194 | 0.4629423 | 1.106446 | 0.2685337 | 0.464588 |
| ENSG00000253836 | | 0.9004533 | 0.4216478 | 0.8862095 | 1.494166 | 0.1351322 | NA |
| ENSG00000234942 | GRID1-AS1 | 0.9708414 | 0.421589 | 0.9153385 | 1.522376 | 0.127915 | NA |
| ENSG00000035115 | SH3YL1 | 5167.115329 | 0.4215636 | 0.229001314 | 1.926874105 | 0.05399532 | 0.147184 |
| ENSG00000269858 | EGLN2 | 636.8313 | 0.4215411 | 0.2444572 | 1.820641 | 0.06866146 | 0.175918 |
| ENSG00000164089 | ETNPPL | 7.690895 | 0.421471 | 0.5961533 | 0.9825722 | 0.325818 | 0.527091 |
| ENSG00000185670 | ZBTB3 | 290.9503 | 0.4213985 | 0.1926799 | 2.277389 | 0.02276299 | 0.075401 |
| ENSG00000266104 | MIR4326 | 2.706772 | 0.4211763 | 0.7417141 | 1.038272 | 0.2991436 | 0.498385 |
| ENSG00000145247 | OCIAD2 | 2418.129 | 0.4211043 | 0.1733734 | 2.475988 | 0.0132868 | 0.048857 |
| ENSG00000267834 | | 29.2928 | 0.4209094 | 0.4616456 | 1.103089 | 0.2699885 | 0.466318 |
| ENSG00000153130 | SCOC | 4196.228 | 0.4207199 | 0.1442755 | 2.962911 | 0.00304745 | 0.014729 |
| ENSG00000281327 | LINC01338 | 2.7822 | 0.4206098 | 0.9016763 | 1.331867 | 0.1829039 | 0.357953 |
| ENSG00000013288 | MAN2B2 | 1688.20624 | 0.42054383 | 0.227413832 | 1.927778096 | 0.05388273 | 0.146998 |
| ENSG00000187792 | ZNF70 | 292.8081 | 0.4204826 | 0.2750458 | 1.632595 | 0.1025542 | 0.236957 |
| ENSG00000255142 | | 21.05678 | 0.4203424 | 0.5160893 | 1.0442 | 0.2963929 | 0.49543 |
| ENSG00000260777 | | 8.790239 | 0.4203121 | 0.696474 | 1.007301 | 0.3137902 | 0.514921 |
| ENSG00000276110 | | 2.405703 | 0.4202733 | 0.7650428 | 1.059575 | 0.289338 | 0.487904 |
| ENSG00000249693 | THEGL | 2.33406 | 0.4198629 | 0.8068085 | 1.134021 | 0.2567855 | 0.4506 |
| ENSG00000260075 | NSFP1 | 4.279367 | 0.4195166 | 0.6789995 | 1.004866 | 0.3149613 | 0.516015 |
| ENSG00000079931 | MOXD1 | 1.3468813 | 0.419329193 | 0.859120444 | 1.260946789 | 0.20732802 | NA |

| | | | | | | | |
|---|---|---|---|---|---|---|---|
| ENSG00000159450 | TCHH | 854.0819 | 0.4192964 | 0.2276103 | 1.923028 | 0.05447656 | 0.148146 |
| ENSG00000253171 | | 0.8326467 | 0.4192867 | 0.9217823 | 1.559089 | 0.1189753 | NA |
| ENSG00000163138 | PACRGL | 583.4674 | 0.4192702 | 0.2178813 | 2.00981 | 0.04445135 | 0.126897 |
| ENSG00000239789 | MRPS17 | 247.8704 | 0.4190361 | 0.2343315 | 1.87771 | 0.0604209 | 0.159909 |
| ENSG00000113209 | PCDHB5 | 22.80713657 | 0.418868951 | 0.535460753 | 1.022793058 | 0.3064057 | 0.506432 |
| ENSG00000179362 | HMGN2P46 | 7.125796 | 0.4187919 | 0.5789082 | 1.005094 | 0.3148514 | 0.515898 |
| ENSG00000166546 | BEAN1 | 21.66868 | 0.4187867 | 0.5111987 | 1.040787 | 0.2979743 | 0.497104 |
| ENSG00000282904 | | 0.8863411 | 0.4187285 | 0.8813023 | 1.350553 | 0.1768386 | NA |
| ENSG00000279653 | | 3.741542 | 0.4187204 | 0.6792566 | 0.9912674 | 0.321555 | 0.522329 |
| ENSG00000215769 | ARHGAP27P1-BF | 195.2898 | 0.4186161 | 0.2804094 | 1.597223 | 0.1102161 | 0.24973 |
| ENSG00000266028 | SRGAP2 | 5166.466 | 0.4185492 | 0.2732882 | 1.640784 | 0.1008422 | 0.234244 |
| ENSG00000141258 | SGSM2 | 9247.905 | 0.4182766 | 0.2044299 | 2.116451 | 0.03430646 | 0.104087 |
| ENSG00000180879 | SSR4 | 9685.946 | 0.4180285 | 0.168649 | 2.533201 | 0.01130263 | 0.042946 |
| ENSG00000138785 | INTS12 | 898.0854771 | 0.417999315 | 0.150854792 | 2.812312116 | 0.00491868 | 0.021861 |
| ENSG00000253944 | | 8.190234 | 0.4179635 | 0.6366439 | 0.9907337 | 0.3218156 | 0.52262 |
| ENSG00000197724 | PHF2 | 4546.525 | 0.4178687 | 0.169002 | 2.54122 | 0.01104663 | 0.042169 |
| ENSG00000263327 | TAPT1-AS1 | 129.7534 | 0.4176579 | 0.4954636 | 1.055899 | 0.2910142 | 0.489612 |
| ENSG00000184203 | PPP1R2 | 2674.483 | 0.4176543 | 0.1651756 | 2.615078 | 0.0089207 | 0.035492 |
| ENSG00000258983 | | 4.23054 | 0.4175734 | 0.7371478 | 1.021081 | 0.3072161 | 0.507446 |
| ENSG00000145675 | PIK3R1 | 4360.999 | 0.4174551 | 0.2105595 | 2.068024 | 0.03863781 | 0.113798 |
| ENSG00000211454 | AKR7L | 34.85469 | 0.4173664 | 0.4499344 | 1.119879 | 0.2627652 | 0.457737 |
| ENSG00000250345 | | 1.254365 | 0.4173406 | 0.8813872 | 1.361459 | 0.1733687 | NA |
| ENSG00000083307 | GRHL2 | 8754.289734 | 0.417301167 | 0.136725763 | 3.100429272 | 0.0019324 | 0.010059 |
| ENSG00000101290 | CDS2 | 2227.060363 | 0.417199915 | 0.099289244 | 4.221072683 | 2.43E-05 | 0.000218 |
| ENSG00000234880 | LINC00163 | 2.328516 | 0.4170449 | 0.8033926 | 1.128195 | 0.2592376 | 0.453555 |
| ENSG00000204305 | AGER | 354.8346 | 0.4168156 | 0.3316324 | 1.380475 | 0.1674405 | 0.336846 |
| ENSG00000227815 | | 2.176234 | 0.4167622 | 0.7610408 | 1.050215 | 0.2936195 | 0.492592 |
| ENSG00000174669 | SLC29A2 | 5882.627 | 0.4166905 | 0.1837154 | 2.315251 | 0.02059918 | 0.069569 |
| ENSG00000175105 | ZNF654 | 1320.321 | 0.4163809 | 0.195193 | 2.185367 | 0.02886195 | 0.091008 |
| ENSG00000188848 | BEND4 | 3.465745 | 0.4161809 | 0.7940812 | 1.099755 | 0.2714388 | 0.467786 |
| ENSG00000146021 | KLHL3 | 3.870242 | 0.4159474 | 0.6519486 | 0.9786034 | 0.327776 | 0.529242 |
| ENSG00000115896 | PLCL1 | 24.29099125 | 0.415859464 | 0.562418262 | 0.999263318 | 0.31766715 | 0.518672 |
| ENSG00000141294 | LRRC46 | 170.9991 | 0.4156606 | 0.2935083 | 1.53361 | 0.1251257 | 0.273259 |

| | | | | | | | |
|---|---|---|---|---|---|---|---|
| ENSG00000186517 | ARHGAP30 | 232.6571 | 0.4156599 | 0.3744085 | 1.259522 | 0.207842 | 0.39041 |
| ENSG00000125398 | SOX9 | 435.3446352 | 0.415514246 | 0.420069321 | 1.154043685 | 0.24848226 | 0.440775 |
| ENSG00000183722 | LHFPL6 | 148.8283 | 0.415457 | 0.3613297 | 1.285011 | 0.1987885 | 0.378936 |
| ENSG00000176697 | BDNF | 6.621982 | 0.4150822 | 0.6118274 | 0.9727661 | 0.3306696 | 0.531814 |
| ENSG00000197599 | CCDC154 | 91.7737 | 0.4150458 | 0.4559668 | 1.091362 | 0.2751135 | 0.471945 |
| ENSG00000253973 | | 8.738172 | 0.4149371 | 0.7065185 | 0.993606 | 0.3204147 | 0.521221 |
| ENSG00000213412 | HNRNPA1P33 | 7.227974 | 0.4148287 | 0.7723562 | 1.054244 | 0.2917711 | 0.490513 |
| ENSG00000276085 | CCL3L1 | 1.284489 | 0.4147557 | 0.8470304 | 1.259317 | 0.2079157 | NA |
| ENSG00000210049 | MT-TF | 13.63862 | 0.4146796 | 0.5177366 | 1.034861 | 0.300734 | 0.499981 |
| ENSG00000169084 | DHRSX | 849.4306 | 0.4146506 | 0.2121387 | 2.028525 | 0.04250674 | 0.122623 |
| ENSG00000008277 | ADAM22 | 429.379312 | 0.414555335 | 0.266330007 | 1.654060058 | 0.09811531 | 0.229539 |
| ENSG00000240160 | RN7SL263P | 0.8977104 | 0.4145479 | 0.8805102 | 1.334404 | 0.1820715 | NA |
| ENSG00000173327 | MAP3K11 | 6594.339 | 0.4144845 | 0.153595 | 2.737428 | 0.00619218 | 0.026396 |
| ENSG00000235244 | DANT2 | 18.46012 | 0.4143857 | 0.4960796 | 1.047271 | 0.2949748 | 0.494117 |
| ENSG00000256006 | | 11.38548 | 0.4142929 | 0.5915671 | 0.9844924 | 0.3248735 | 0.525994 |
| ENSG00000255192 | NANOGP8 | 1.005609 | 0.4142266 | 0.8840999 | 1.369551 | 0.1708272 | NA |
| ENSG00000133739 | LRRCC1 | 924.9948163 | 0.414203606 | 0.231036931 | 1.876395296 | 0.06060102 | 0.160178 |
| ENSG00000104881 | PPP1R13L | 2632.489052 | 0.41403688 | 0.177134141 | 2.393631662 | 0.01668249 | 0.058715 |
| ENSG00000119946 | CNNM1 | 1949.522044 | 0.414022338 | 0.280611465 | 1.58106741 | 0.11386263 | 0.255443 |
| ENSG00000138593 | SECISBP2L | 2160.777753 | 0.41389619 | 0.312688392 | 1.442208678 | 0.14924351 | 0.310003 |
| ENSG00000223551 | TMSB4XP4 | 27.48938 | 0.4138877 | 0.4834226 | 1.060086 | 0.2891057 | 0.487729 |
| ENSG00000242100 | RPL9P32 | 9.716996 | 0.4137871 | 0.4843521 | 1.05961 | 0.2893222 | 0.487904 |
| ENSG00000189223 | PAX8-AS1 | 2786.98 | 0.4137752 | 0.1506369 | 2.800524 | 0.00510198 | 0.022525 |
| ENSG00000146094 | DOK3 | 542.2636 | 0.4136548 | 0.4372533 | 1.128148 | 0.2592576 | 0.45356 |
| ENSG00000129515 | SNX6 | 5997.027188 | 0.413559056 | 0.134525454 | 3.118808935 | 0.00181584 | 0.00955 |
| ENSG00000165555 | NOXRED1 | 118.2133 | 0.4133944 | 0.3228122 | 1.404844 | 0.1600676 | 0.326313 |
| ENSG00000245958 | SEPTIN7P14 | 940.1474 | 0.4133574 | 0.1690669 | 2.503415 | 0.01230014 | 0.045913 |
| ENSG00000273203 | | 2.193808 | 0.413233 | 0.8392879 | 1.179693 | 0.2381223 | 0.427883 |
| ENSG00000276550 | HERC2P2 | 3793.396 | 0.4130303 | 0.2922757 | 1.53642 | 0.1244353 | 0.272266 |
| ENSG00000207631 | MIR641 | 9.967778 | 0.412915 | 0.5660435 | 0.9924972 | 0.3209551 | 0.52174 |
| ENSG00000286164 | | 1.942303 | 0.4127555 | 0.8466842 | 1.289792 | 0.1971229 | 0.377112 |
| ENSG00000147799 | ARHGAP39 | 2541.035 | 0.4126808 | 0.1490598 | 2.8224 | 0.00476657 | 0.021331 |
| ENSG00000267077 | | 1.027792 | 0.4126679 | 0.9022046 | 1.421431 | 0.1551914 | NA |

| | | | | | | | |
|---|---|---|---|---|---|---|---|
| ENSG00000147144 | CCDC120 | 2105.976 | 0.4126504 | 0.1822964 | 2.329466 | 0.01983439 | 0.06751 |
| ENSG00000260490 | MYL12BP1 | 60.75689 | 0.412379 | 0.3581387 | 1.292213 | 0.1962834 | 0.376048 |
| ENSG00000105136 | ZNF419 | 703.2580031 | 0.412361603 | 0.209946928 | 2.040999624 | 0.04125086 | 0.11984 |
| ENSG00000228672 | PROB1 | 465.2945 | 0.4123264 | 0.2946762 | 1.49911 | 0.1338452 | 0.286599 |
| ENSG00000130511 | SSBP4 | 2128.937083 | 0.41215355 | 0.197990187 | 2.152696278 | 0.03134256 | 0.09709 |
| ENSG00000099251 | HSD17B7P2 | 353.601137 | 0.412133506 | 0.244036826 | 1.782310073 | 0.07469868 | 0.187534 |
| ENSG00000231993 | EP300-AS1 | 17.26596 | 0.4121248 | 0.4538397 | 1.101644 | 0.2706165 | 0.466884 |
| ENSG00000213478 | CFL1P2 | 6.367112 | 0.412081 | 0.6585715 | 0.9779368 | 0.3281056 | 0.529382 |
| ENSG00000226089 | | 6.596299 | 0.4118199 | 0.7892279 | 1.09449 | 0.2737399 | 0.47048 |
| ENSG00000186051 | TAL2 | 3.675593 | 0.4117382 | 0.7541905 | 1.029096 | 0.3034347 | 0.503086 |
| ENSG00000258311 | | 28.3376 | 0.4116951 | 0.4864708 | 1.050105 | 0.29367 | 0.492645 |
| ENSG00000121310 | ECHDC2 | 2294.049998 | 0.41162684 | 0.258903472 | 1.680835797 | 0.09279481 | 0.220308 |
| ENSG00000198815 | FOXJ3 | 4299.978 | 0.4116211 | 0.1370574 | 3.045732 | 0.00232115 | 0.011738 |
| ENSG00000104356 | POP1 | 2053.000067 | 0.411556882 | 0.29512584 | 1.509570012 | 0.13115318 | 0.282521 |
| ENSG00000114473 | IQCG | 241.5044311 | 0.411446677 | 0.263716447 | 1.655873215 | 0.0977475 | 0.22886 |
| ENSG00000280157 | | 27.55419 | 0.4113898 | 0.4583604 | 1.085471 | 0.2777129 | 0.474841 |
| ENSG00000206195 | DUXAP8 | 1952.496 | 0.4112057 | 0.1871859 | 2.259317 | 0.0238637 | 0.078189 |
| ENSG00000285587 | | 28.14034 | 0.4111297 | 0.6412577 | 0.9485069 | 0.3428715 | 0.544418 |
| ENSG00000251138 | LINC02882 | 88.3394 | 0.4109723 | 0.3559951 | 1.292924 | 0.1960374 | 0.375768 |
| ENSG00000237169 | RPL12P27 | 2.851301 | 0.4109355 | 0.7239576 | 0.9912416 | 0.3215676 | 0.522329 |
| ENSG00000061987 | MON2 | 4963.793809 | 0.410830421 | 0.221744775 | 1.927542378 | 0.05391207 | 0.147048 |
| ENSG00000123268 | ATF1 | 1791.385005 | 0.410802932 | 0.169122562 | 2.470317204 | 0.01349933 | 0.049478 |
| ENSG00000141522 | ARHGDIA | 9336.195 | 0.4107727 | 0.1924729 | 2.182381 | 0.02908145 | 0.091624 |
| ENSG00000238045 | | 233.4848 | 0.4104811 | 0.2987129 | 1.489323 | 0.1364022 | 0.290553 |
| ENSG00000178802 | MPI | 2526.198 | 0.4103928 | 0.1660525 | 2.524755 | 0.01157789 | 0.04377 |
| ENSG00000180758 | GPR157 | 695.0782 | 0.4103641 | 0.1827301 | 2.308289 | 0.02098307 | 0.070659 |
| ENSG00000231616 | | 6.10332 | 0.4103497 | 0.601096 | 0.9741903 | 0.329962 | 0.531029 |
| ENSG00000241336 | LINC01487 | 1.614352 | 0.4102749 | 0.9185844 | 1.612175 | 0.1069238 | 0.244249 |
| ENSG00000248668 | OXCT1-AS1 | 24.93986 | 0.4102701 | 0.3902352 | 1.19344 | 0.2326969 | 0.421411 |
| ENSG00000065613 | SLK | 4843.126543 | 0.410262269 | 0.163041931 | 2.578477359 | 0.00992368 | 0.038626 |
| ENSG00000147647 | DPYS | 1.419428 | 0.4102417 | 0.862318 | 1.382339 | 0.1668675 | NA |
| ENSG00000264655 | PPIAP54 | 25.02904 | 0.410206 | 0.3538287 | 1.29248 | 0.1961911 | 0.375969 |
| ENSG00000136274 | NACAD | 1.317866087 | 0.410190992 | 0.811334863 | 1.135063765 | 0.2563486 | NA |

| | | | | | | | |
|---|---|---|---|---|---|---|---|
| ENSG00000140750 | ARHGAP17 | 1774.873 | 0.4101718 | 0.1360476 | 3.067217 | 0.00216062 | 0.011071 |
| ENSG00000139971 | ARMH4 | 44.60759 | 0.4101388 | 0.4200566 | 1.149183 | 0.2504806 | 0.443048 |
| ENSG00000006282 | SPATA20 | 7681.522913 | 0.410098946 | 0.165927029 | 2.523511979 | 0.01161891 | 0.043893 |
| ENSG00000154316 | TDH | 63.53838 | 0.410049 | 0.4921709 | 1.055588 | 0.2911565 | 0.48979 |
| ENSG00000181135 | ZNF707 | 1836.393 | 0.4099723 | 0.1122484 | 3.688191 | 0.00022585 | 0.00158 |
| ENSG00000245213 | | 105.3658 | 0.4099565 | 0.4088242 | 1.170404 | 0.2418384 | 0.432367 |
| ENSG00000135443 | KRT85 | 1.947447098 | 0.409877147 | 0.900496013 | 1.437342784 | 0.15062062 | 0.31201 |
| ENSG00000144893 | MED12L | 969.7904 | 0.4097751 | 0.2513912 | 1.734253 | 0.08287314 | 0.202757 |
| ENSG00000250049 | | 2.088212 | 0.4096533 | 0.8532579 | 1.329555 | 0.183665 | 0.358888 |
| ENSG00000186960 | LINC01551 | 2.106478 | 0.4096336 | 0.897178 | 1.295242 | 0.1952367 | 0.374639 |
| ENSG00000186825 | CDRT15P3 | 5.837599 | 0.4093905 | 0.7839024 | 1.065276 | 0.286751 | 0.485112 |
| ENSG00000150433 | TMEM218 | 561.375 | 0.409304 | 0.1402545 | 2.966963 | 0.00300757 | 0.014587 |
| ENSG00000181588 | MEX3D | 1275.609 | 0.409248 | 0.2089254 | 2.033116 | 0.04204081 | 0.12153 |
| ENSG00000125851 | PCSK2 | 9.94174567 | 0.409224964 | 0.805086526 | 1.119039278 | 0.26312338 | 0.458121 |
| ENSG00000158092 | NCK1 | 1565.546 | 0.4090775 | 0.1870561 | 2.249784 | 0.02446264 | 0.079729 |
| ENSG00000102078 | SLC25A14 | 933.7280668 | 0.4090715 | 0.131328269 | 3.166577491 | 0.00154244 | 0.008303 |
| ENSG00000279141 | | 1.831499 | 0.409069 | 0.7824324 | 1.035664 | 0.3003589 | 0.499577 |
| ENSG00000204338 | CYP21A1P | 6.27206 | 0.4090369 | 0.6921424 | 0.9732288 | 0.3304396 | 0.53154 |
| ENSG00000248459 | | 1.442979 | 0.4090063 | 0.8356138 | 1.149348 | 0.2504125 | NA |
| ENSG00000166002 | SMCO4 | 286.4993 | 0.4089865 | 0.3535601 | 1.310177 | 0.1901359 | 0.3679 |
| ENSG00000103021 | CCDC113 | 439.9114885 | 0.408969938 | 0.304118317 | 1.448651611 | 0.1474349 | 0.307328 |
| ENSG00000145022 | TCTA | 1549.127 | 0.4089684 | 0.2610198 | 1.663675 | 0.09617745 | 0.226016 |
| ENSG00000226040 | | 3.137047 | 0.4089654 | 0.8027676 | 1.108053 | 0.2678391 | 0.463778 |
| ENSG00000219433 | BTBD10P2 | 2.767195 | 0.4088294 | 0.7727135 | 1.06993 | 0.2846508 | 0.482581 |
| ENSG00000136247 | ZDHHC4 | 2997.364933 | 0.408803586 | 0.153081152 | 2.722467736 | 0.00647964 | 0.027348 |
| ENSG00000285701 | | 0.8987873 | 0.4087762 | 0.8747448 | 1.323378 | 0.1857098 | NA |
| ENSG00000117360 | PRPF3 | 11515.28394 | 0.408712102 | 0.251159573 | 1.71695815 | 0.08598682 | 0.208363 |
| ENSG00000174963 | ZIC4 | 826.5563 | 0.408674 | 0.2969602 | 1.491558 | 0.1358149 | 0.289488 |
| ENSG00000135778 | NTPCR | 2429.403383 | 0.408640536 | 0.144260047 | 2.880672579 | 0.00396828 | 0.018333 |
| ENSG00000100554 | ATP6V1D | 3541.739415 | 0.408575949 | 0.159243943 | 2.618274463 | 0.00883757 | 0.035251 |
| ENSG00000124074 | ENKD1 | 141.6113061 | 0.408519914 | 0.214603963 | 1.985387715 | 0.04710135 | 0.132385 |
| ENSG00000262678 | | 25.4967 | 0.408303 | 0.3763279 | 1.228424 | 0.219288 | 0.405418 |
| ENSG00000060656 | PTPRU | 6501.524466 | 0.408237586 | 0.209267429 | 2.038972778 | 0.04145274 | 0.120236 |

| | | | | | | | |
|---|---|---|---|---|---|---|---|
| ENSG00000259933 | | 23.41187 | 0.4082145 | 0.489008 | 1.044221 | 0.2963834 | 0.49543 |
| ENSG00000177679 | SRRM3 | 1247.61 | 0.4081693 | 0.3124088 | 1.424099 | 0.1544177 | 0.317942 |
| ENSG00000273507 | | 9.47189 | 0.4080714 | 0.5168209 | 1.013279 | 0.3109271 | 0.51174 |
| ENSG00000131375 | CAPN7 | 4633.041691 | 0.407966838 | 0.149652848 | 2.775447014 | 0.00551259 | 0.023991 |
| ENSG00000139977 | NAA30 | 3064.08 | 0.4077869 | 0.1497693 | 2.758633 | 0.00580437 | 0.025072 |
| ENSG00000213082 | PPP1R14BP2 | 1.975277 | 0.4077426 | 0.8235748 | 1.23644 | 0.216295 | 0.401597 |
| ENSG00000050426 | LETMD1 | 4066.426288 | 0.407579134 | 0.207176475 | 2.040596485 | 0.04129095 | 0.119918 |
| ENSG00000259770 | | 0.9027494 | 0.4075198 | 0.9052032 | 1.467592 | 0.142215 | NA |
| ENSG00000242968 | | 6.608512 | 0.4075173 | 0.7055391 | 0.9936681 | 0.3203845 | 0.521203 |
| ENSG00000006756 | ARSD | 2458.543576 | 0.407493155 | 0.262363941 | 1.64313456 | 0.1003551 | 0.233438 |
| ENSG00000243304 | | 3.343599 | 0.4074554 | 0.6866613 | 0.9776334 | 0.3282557 | 0.529422 |
| ENSG00000285737 | LINC02680 | 2.693521 | 0.407299 | 0.7300504 | 0.9975246 | 0.31851 | 0.519681 |
| ENSG00000238035 | | 3.25685 | 0.4072199 | 0.6759054 | 0.9635364 | 0.3352784 | 0.536595 |
| ENSG00000164168 | TMEM184C | 3589.016 | 0.4071636 | 0.1454665 | 2.833654 | 0.00460191 | 0.020734 |
| ENSG00000280388 | | 27.95458 | 0.4071556 | 0.4629968 | 1.068249 | 0.2854083 | 0.483442 |
| ENSG00000230711 | CTAGE13P | 20.68527 | 0.4068959 | 0.4889768 | 1.037111 | 0.2996841 | 0.498891 |
| ENSG00000212329 | RNU6-316P | 1.529835 | 0.4068144 | 0.8611546 | 1.267418 | 0.2050058 | NA |
| ENSG00000224281 | SLC25A5-AS1 | 187.4508 | 0.4067722 | 0.2316215 | 1.837976 | 0.06606601 | 0.170899 |
| ENSG00000181218 | H2AW | 3111.227 | 0.4067702 | 0.2602036 | 1.655389 | 0.09784563 | 0.229065 |
| ENSG00000183475 | ASB7 | 1497.037 | 0.4066963 | 0.1518797 | 2.734003 | 0.00625694 | 0.026616 |
| ENSG00000258466 | | 2.578696 | 0.4065717 | 0.7943782 | 1.100554 | 0.2710908 | 0.467375 |
| ENSG00000124767 | GLO1 | 14534.88203 | 0.40649744 | 0.258277373 | 1.715079125 | 0.08633072 | 0.208938 |
| ENSG00000125249 | RAP2A | 1651.937799 | 0.406424669 | 0.181317478 | 2.295597366 | 0.02169891 | 0.072658 |
| ENSG00000066739 | ATG2B | 5360.648145 | 0.406277789 | 0.242958196 | 1.758648939 | 0.07863716 | 0.194662 |
| ENSG00000255717 | SNHG1 | 8689.266 | 0.4062358 | 0.2302061 | 1.844813 | 0.06506469 | 0.168867 |
| ENSG00000272196 | H2AC19 | 14.75095 | 0.4061486 | 0.5296596 | 0.9996321 | 0.3174886 | 0.518556 |
| ENSG00000137185 | ZSCAN9 | 1022.279389 | 0.406009044 | 0.216418828 | 1.948039472 | 0.05141024 | 0.141517 |
| ENSG00000100105 | PATZ1 | 7227.048028 | 0.406005177 | 0.123324899 | 3.345055819 | 0.00082266 | 0.004836 |
| ENSG00000159579 | RSPRY1 | 1415.337 | 0.405831 | 0.1635852 | 2.48711 | 0.01287854 | 0.047702 |
| ENSG00000213184 | | 2.666034 | 0.4057754 | 0.7125285 | 0.9705484 | 0.3317732 | 0.532911 |
| ENSG00000265293 | ARGFXP2 | 4.918678 | 0.4057687 | 0.6852466 | 0.9606012 | 0.3367527 | 0.537887 |
| ENSG00000123395 | ATG101 | 1962.177732 | 0.405748408 | 0.160110519 | 2.582814681 | 0.0097998 | 0.038222 |
| ENSG00000078618 | NRDC | 13490.7795 | 0.405730169 | 0.092303915 | 4.415235693 | 1.01E-05 | 9.93E-05 |

| | | | | | | | |
|---|---|---|---|---|---|---|---|
| ENSG00000108798 | ABI3 | 1.972056389 | 0.405588424 | 0.734877492 | 0.994718803 | 0.31987305 | 0.520776 |
| ENSG00000186812 | ZNF397 | 1866.636 | 0.4055578 | 0.2357645 | 1.799869 | 0.07188127 | 0.182217 |
| ENSG00000237064 | EIF3IP1 | 0.9754047 | 0.4054852 | 0.8479129 | 1.218723 | 0.2229492 | NA |
| ENSG00000133805 | AMPD3 | 104.1741034 | 0.40531076 | 0.296622803 | 1.47443906 | 0.14036341 | 0.296637 |
| ENSG00000264456 | | 33.23551 | 0.4052987 | 0.3230741 | 1.377575 | 0.1683345 | 0.338107 |
| ENSG00000256637 | LINC01965 | 2.699843 | 0.4051716 | 0.8709716 | 1.396338 | 0.1626128 | 0.329983 |
| ENSG00000254166 | CASC19 | 4.795621 | 0.4051711 | 0.7070885 | 0.9907128 | 0.3218258 | 0.52262 |
| ENSG00000162148 | PPP1R32 | 86.66812 | 0.4051428 | 0.4340924 | 1.103086 | 0.2699899 | 0.466318 |
| ENSG00000226212 | TRGV6 | 1.411236 | 0.405035 | 0.9250205 | 1.743927 | 0.08117175 | NA |
| ENSG00000286413 | | 4.09919 | 0.4050005 | 0.8140425 | 1.115984 | 0.2644289 | 0.459938 |
| ENSG00000277156 | TOMM40P1 | 58.02593 | 0.4047295 | 0.2627105 | 1.639678 | 0.1010722 | 0.234573 |
| ENSG00000244731 | C4A | 63.49445 | 0.4047165 | 0.3649485 | 1.245374 | 0.2129942 | 0.397712 |
| ENSG00000228000 | RPL7AP65 | 1.12503 | 0.4046722 | 0.8240796 | 1.14434 | 0.2524829 | NA |
| ENSG00000254170 | | 7.682866 | 0.4045265 | 0.5229822 | 1.003138 | 0.3157944 | 0.516838 |
| ENSG00000126070 | AGO3 | 1183.045214 | 0.404460261 | 0.204291337 | 2.049890099 | 0.04037516 | 0.11774 |
| ENSG00000130635 | COL5A1 | 5105.532808 | 0.404416687 | 0.338644655 | 1.330632862 | 0.18330984 | 0.358437 |
| ENSG00000226361 | TERF1P5 | 47.75207 | 0.4043517 | 0.3285615 | 1.345894 | 0.1783365 | 0.351892 |
| ENSG00000287513 | | 5.524967 | 0.4042966 | 0.7542301 | 1.050616 | 0.293435 | 0.492376 |
| ENSG00000236234 | | 4.284794 | 0.4042377 | 0.6508963 | 0.9532162 | 0.3404806 | 0.541918 |
| ENSG00000157483 | MYO1E | 1369.245 | 0.4040726 | 0.2104244 | 1.990957 | 0.0464856 | 0.131126 |
| ENSG00000186575 | NF2 | 2526.67 | 0.4037912 | 0.1189143 | 3.466955 | 0.00052639 | 0.003282 |
| ENSG00000236732 | | 31.51514 | 0.4037335 | 0.4674613 | 1.054004 | 0.2918811 | 0.490667 |
| ENSG00000213983 | AP1G2 | 4313.881 | 0.4036656 | 0.199155 | 2.104375 | 0.03534578 | 0.106385 |
| ENSG00000207200 | RNU6-45P | 2.403975 | 0.4036001 | 0.7795478 | 1.037956 | 0.2992903 | 0.498517 |
| ENSG00000185246 | PRPF39 | 4908.906 | 0.4035598 | 0.213726 | 1.929129 | 0.0537148 | 0.146669 |
| ENSG00000179412 | HNRNPCL4 | 3.649229 | 0.4034618 | 0.6864169 | 0.9757061 | 0.3292101 | 0.530313 |
| ENSG00000236456 | | 433.3917 | 0.4034358 | 0.1972597 | 2.115072 | 0.03442383 | 0.104384 |
| ENSG00000251432 | LINC02615 | 395.1787 | 0.4034222 | 0.4730405 | 1.049294 | 0.2940428 | 0.493095 |
| ENSG00000186665 | C17orf58 | 2132.787 | 0.4033885 | 0.1696161 | 2.4247 | 0.01532103 | 0.054838 |
| ENSG00000273711 | | 10.47583 | 0.4033256 | 0.6725689 | 0.9614848 | 0.3363085 | 0.537337 |
| ENSG00000197245 | FAM110D | 4.76462 | 0.4029432 | 0.775021 | 1.05832 | 0.2899095 | 0.488403 |
| ENSG00000282386 | | 93.13612 | 0.4029019 | 0.2557221 | 1.665805 | 0.0957523 | 0.225396 |
| ENSG00000275263 | | 8.021358 | 0.4028839 | 0.5461954 | 0.9756914 | 0.3292174 | 0.530313 |

| | | | | | | | |
|---|---|---|---|---|---|---|---|
| ENSG00000235802 | HCFC1-AS1 | 1.369673 | 0.40278 | 0.8452865 | 1.216871 | 0.2236533 | NA |
| ENSG00000156504 | FAM122B | 4053.227 | 0.4027013 | 0.1864413 | 2.222785 | 0.02623028 | 0.084178 |
| ENSG00000198018 | ENTPD7 | 1100.268 | 0.4025816 | 0.1945113 | 2.152311 | 0.03137288 | 0.097139 |
| ENSG00000254416 | LINC02732 | 8.82562 | 0.402546 | 0.6247069 | 0.9495226 | 0.3423549 | 0.543793 |
| ENSG00000180901 | KCTD2 | 3263.986 | 0.4025169 | 0.1143075 | 3.567802 | 0.00035999 | 0.002371 |
| ENSG00000144649 | GASK1A | 26.533 | 0.4025065 | 0.4789854 | 1.038847 | 0.2988761 | 0.498139 |
| ENSG00000169989 | TIGD4 | 32.28632 | 0.4024633 | 0.5200828 | 0.9970996 | 0.3187162 | 0.519794 |
| ENSG00000287910 | | 9.3777 | 0.4023832 | 0.6589995 | 0.96739 | 0.3333491 | 0.534376 |
| ENSG00000074855 | ANO8 | 2184.264118 | 0.40218215 | 0.176881342 | 2.336640912 | 0.01945786 | 0.066492 |
| ENSG00000132881 | CPLANE2 | 210.5435755 | 0.402149741 | 0.257442926 | 1.666787743 | 0.09555662 | 0.22502 |
| ENSG00000237676 | RPL30P4 | 68.31047 | 0.4021341 | 0.3415923 | 1.307907 | 0.1909048 | 0.368931 |
| ENSG00000262098 | | 2.786598 | 0.4021019 | 0.7760938 | 1.026181 | 0.304806 | 0.504533 |
| ENSG00000236753 | MKLN1-AS | 94.46707 | 0.401958 | 0.3040055 | 1.393819 | 0.1633722 | 0.331144 |
| ENSG00000073712 | FERMT2 | 3611.969894 | 0.401898989 | 0.185989295 | 2.210461677 | 0.02707314 | 0.086381 |
| ENSG00000267141 | | 12.86688 | 0.4017627 | 0.5431033 | 0.9807798 | 0.3267013 | 0.527988 |
| ENSG00000136193 | SCRN1 | 10952.42395 | 0.401734182 | 0.187682058 | 2.203842762 | 0.0275354 | 0.087605 |
| ENSG00000274386 | TMEM269 | 35.76175 | 0.4016208 | 0.333981 | 1.327732 | 0.1842667 | 0.359587 |
| ENSG00000133639 | BTG1 | 3561.456946 | 0.401596582 | 0.218333818 | 1.935060452 | 0.05298288 | 0.145109 |
| ENSG00000232411 | | 1.361404 | 0.4015909 | 0.8966536 | 1.391546 | 0.1640598 | NA |
| ENSG00000238950 | RNU7-173P | 1.281576 | 0.4015882 | 0.8973425 | 1.358831 | 0.1742001 | NA |
| ENSG00000272836 | | 40.28327 | 0.4014771 | 0.4083302 | 1.142596 | 0.2532063 | 0.446146 |
| ENSG00000263540 | MIR5582 | 1.690269 | 0.4014627 | 0.8381638 | 1.249384 | 0.2115247 | 0.395635 |
| ENSG00000268658 | LINC00664 | 0.8656048 | 0.4014422 | 0.8912389 | 1.293594 | 0.1958058 | NA |
| ENSG00000177556 | ATOX1 | 2699.594 | 0.4011133 | 0.1849274 | 2.251189 | 0.02437357 | 0.079507 |
| ENSG00000228327 | | 206.6696 | 0.4010778 | 0.2203554 | 1.907797 | 0.05641742 | 0.151993 |
| ENSG00000116128 | BCL9 | 2883.348314 | 0.401049754 | 0.212306105 | 1.951622446 | 0.05098305 | 0.140585 |
| ENSG00000180535 | BHLHA15 | 4.727413 | 0.4010204 | 0.707174 | 0.9820331 | 0.3260835 | 0.527279 |
| ENSG00000228315 | GUSBP11 | 461.7783 | 0.40096 | 0.346695 | 1.287536 | 0.1979075 | 0.37796 |
| ENSG00000176761 | ZNF285B | 11.25296 | 0.4009584 | 0.5856513 | 0.9526035 | 0.340791 | 0.542132 |
| ENSG00000057252 | SOAT1 | 1830.866794 | 0.400699636 | 0.18051342 | 2.278317911 | 0.02270764 | 0.075302 |
| ENSG00000229601 | | 1.000769 | 0.4006737 | 0.8737994 | 1.317304 | 0.1877369 | NA |
| ENSG00000271499 | | 1.008226 | 0.400569 | 0.9049816 | 1.578464 | 0.114459 | NA |
| ENSG00000175155 | YPEL2 | 2828.036 | 0.400453 | 0.3581436 | 1.251259 | 0.2108399 | 0.394744 |

| | | | | | | | |
|---|---|---|---|---|---|---|---|
| ENSG00000033800 | PIAS1 | 2233.959124 | 0.400451512 | 0.15486988 | 2.642519213 | 0.00822918 | 0.033238 |
| ENSG00000107281 | NPDC1 | 9398.252557 | 0.40038932 | 0.158051032 | 2.596686337 | 0.00941278 | 0.037017 |
| ENSG00000228847 | ATP5MC2P4 | 58.08373 | 0.400056 | 0.2766549 | 1.54735 | 0.1217789 | 0.268087 |
| ENSG00000235330 | RPL12P25 | 7.310999 | 0.3999923 | 0.6303929 | 0.9475559 | 0.3433556 | 0.544861 |
| ENSG00000198911 | SREBF2 | 11748.86 | 0.3999892 | 0.1744573 | 2.358225 | 0.01836257 | 0.063443 |
| ENSG00000113749 | HRH2 | 32.84836922 | 0.399956422 | 0.402685143 | 1.150668916 | 0.24986847 | 0.442437 |
| ENSG00000112282 | MED23 | 2948.584533 | 0.399740648 | 0.261588918 | 1.624006202 | 0.10437449 | 0.240035 |
| ENSG00000170142 | UBE2E1 | 5742.21 | 0.3996568 | 0.1684762 | 2.42062 | 0.01549408 | 0.055316 |
| ENSG00000113924 | HGD | 170.2097038 | 0.399618296 | 0.537406627 | 0.962627112 | 0.33573469 | 0.536872 |
| ENSG00000164188 | RANBP3L | 30.74734 | 0.3995019 | 0.9072407 | 1.680706 | 0.09282004 | 0.220348 |
| ENSG00000241770 | | 2.122452 | 0.3994883 | 0.8030243 | 1.091999 | 0.2748336 | 0.471557 |
| ENSG00000196312 | MFSD14C | 1913.166 | 0.3994699 | 0.2587745 | 1.622653 | 0.1046635 | 0.24045 |
| ENSG00000158882 | TOMM40L | 830.5177 | 0.3994458 | 0.1573657 | 2.593249 | 0.00950739 | 0.037295 |
| ENSG00000228775 | WEE2-AS1 | 147.7312 | 0.3994218 | 0.2999682 | 1.43786 | 0.1504738 | 0.311827 |
| ENSG00000168924 | LETM1 | 3304.68 | 0.3994127 | 0.2200834 | 1.886665 | 0.05920545 | 0.157524 |
| ENSG00000175793 | SFN | 4379.252 | 0.3993849 | 0.2911042 | 1.479969 | 0.1388815 | 0.294277 |
| ENSG00000280620 | SCAANT1 | 126.6715 | 0.399327 | 0.2914981 | 1.47836 | 0.1393114 | 0.295024 |
| ENSG00000112305 | SMAP1 | 575.4335282 | 0.399221431 | 0.163747334 | 2.49162241 | 0.01271611 | 0.047213 |
| ENSG00000132781 | MUTYH | 112.2975744 | 0.399122236 | 0.260944588 | 1.626067761 | 0.10393524 | 0.239273 |
| ENSG00000104432 | IL7 | 41.37494441 | 0.399117527 | 0.393347622 | 1.161255052 | 0.24553819 | 0.437246 |
| ENSG00000198468 | FLVCR1-DT | 71.85252 | 0.3990793 | 0.289544 | 1.485979 | 0.1372847 | 0.291638 |
| ENSG00000162076 | FLYWCH2 | 2387.292 | 0.3989861 | 0.1527255 | 2.630747 | 0.00851973 | 0.034199 |
| ENSG00000260992 | DOCK9-DT | 44.53338 | 0.3989564 | 0.3585535 | 1.251628 | 0.2107056 | 0.394631 |
| ENSG00000272269 | NUP153-AS1 | 19.02158 | 0.3988921 | 0.4757808 | 1.043177 | 0.2968662 | 0.495909 |
| ENSG00000116679 | IVNS1ABP | 10996.63755 | 0.398879289 | 0.223110222 | 1.870893654 | 0.06135982 | 0.161813 |
| ENSG00000042317 | SPATA7 | 829.8629306 | 0.39883439 | 0.220062548 | 1.872818219 | 0.06109349 | 0.161207 |
| ENSG00000254363 | | 75.97511 | 0.3987643 | 0.4195372 | 1.120454 | 0.2625203 | 0.457412 |
| ENSG00000237373 | BRWD1-IT1 | 1.921261 | 0.3985561 | 0.7679952 | 1.032796 | 0.3016992 | 0.501178 |
| ENSG00000134138 | MEIS2 | 445.094873 | 0.398398344 | 0.281169566 | 1.516902261 | 0.12929136 | 0.279592 |
| ENSG00000162620 | LRRIQ3 | 44.55225 | 0.398386 | 0.3579811 | 1.243522 | 0.2136756 | 0.398508 |
| ENSG00000269794 | | 2.707142 | 0.3982651 | 0.7549639 | 1.009637 | 0.3126691 | 0.513716 |
| ENSG00000259924 | | 36.75657 | 0.3981478 | 0.4014857 | 1.146579 | 0.2515555 | 0.444357 |
| ENSG00000264522 | OTUD7B | 4033.375 | 0.3981102 | 0.1357634 | 2.976268 | 0.0029178 | 0.014256 |

| | | | | | | | |
|---|---|---|---|---|---|---|---|
| ENSG00000266338 | NBPF15 | 5334.292 | 0.3980228 | 0.2224861 | 1.866604 | 0.06195696 | 0.162822 |
| ENSG00000135775 | COG2 | 2593.218864 | 0.398005941 | 0.091331079 | 4.393267824 | 1.12E-05 | 0.000109 |
| ENSG00000184319 | RPL23AP82 | 376.0867 | 0.3979952 | 0.2161099 | 1.905031 | 0.05677607 | 0.152603 |
| ENSG00000250403 | | 13.98484 | 0.3979927 | 0.5131819 | 0.9943581 | 0.3200486 | 0.520912 |
| ENSG00000183439 | TRIM61 | 2.87897 | 0.3978717 | 0.720415 | 0.9737029 | 0.3302041 | 0.53129 |
| ENSG00000184730 | APOBR | 138.5332 | 0.3977525 | 0.3173141 | 1.374592 | 0.169258 | 0.339474 |
| ENSG00000271919 | | 1.564837 | 0.3976384 | 0.7995988 | 1.078488 | 0.2808161 | NA |
| ENSG00000162728 | KCNJ9 | 1.736105 | 0.3975907 | 0.7958107 | 1.07477 | 0.2824778 | 0.480302 |
| ENSG00000257202 | | 1.63327 | 0.3974294 | 0.8224883 | 1.189917 | 0.2340789 | 0.423222 |
| ENSG00000226029 | LINC01772 | 48.0243 | 0.3973596 | 0.3401818 | 1.296349 | 0.1948554 | 0.374012 |
| ENSG00000162910 | MRPL55 | 2200.522 | 0.3973252 | 0.2507148 | 1.69607 | 0.08987267 | 0.21543 |
| ENSG00000228314 | CYP4F29P | 132.4303 | 0.3972408 | 0.4997099 | 1.004153 | 0.3153049 | 0.516259 |
| ENSG00000116906 | GNPAT | 3152.024447 | 0.3972206 | 0.137683796 | 2.947410693 | 0.00320447 | 0.015343 |
| ENSG00000287569 | | 23.08927 | 0.3971882 | 0.3838406 | 1.178006 | 0.2387942 | 0.428738 |
| ENSG00000249193 | HSPD1P5 | 13.27064 | 0.3971678 | 0.5019531 | 1.002583 | 0.3160623 | 0.517117 |
| ENSG00000234556 | LINC00701 | 3.094113 | 0.3971411 | 0.889829 | 1.267676 | 0.2049136 | 0.386892 |
| ENSG00000163794 | UCN | 83.7005 | 0.3971326 | 0.4563667 | 1.044902 | 0.2960681 | 0.495116 |
| ENSG00000284719 | | 47.95799 | 0.3970472 | 0.476577 | 1.032137 | 0.302008 | 0.501409 |
| ENSG00000112378 | PERP | 9324.691514 | 0.396910216 | 0.256572654 | 1.673203528 | 0.09428724 | 0.223144 |
| ENSG00000272195 | | 14.17138 | 0.3968082 | 0.5983307 | 0.9460177 | 0.3441396 | 0.545812 |
| ENSG00000256464 | YWHABP2 | 5.704418 | 0.3967423 | 0.5843319 | 0.9462972 | 0.343997 | 0.545651 |
| ENSG00000166707 | ZCCHC18 | 124.3714 | 0.3967005 | 0.4284399 | 1.090843 | 0.2753419 | 0.472154 |
| ENSG00000130167 | TSPAN16 | 2.94049409 | 0.396659034 | 0.742481414 | 1.004522051 | 0.31512705 | 0.516159 |
| ENSG00000272158 | | 2.449404 | 0.3966047 | 0.8131447 | 1.112898 | 0.2657524 | 0.461368 |
| ENSG00000129657 | SEC14L1 | 8810.467831 | 0.396381367 | 0.265136882 | 1.59471588 | 0.11077579 | 0.250677 |
| ENSG00000204366 | ZBTB12 | 779.827 | 0.3963116 | 0.1928934 | 2.123559 | 0.03370704 | 0.102632 |
| ENSG00000235272 | RAMACL | 436.6194 | 0.3962873 | 0.1852072 | 2.195 | 0.02816362 | 0.089102 |
| ENSG00000162817 | C1orf115 | 1045.746 | 0.3962523 | 0.1981395 | 2.068556 | 0.03858774 | 0.113701 |
| ENSG00000215811 | BTNL10 | 5.15014 | 0.3960727 | 0.7040525 | 0.9444168 | 0.3449567 | 0.546537 |
| ENSG00000227486 | | 132.7508 | 0.3960326 | 0.2891788 | 1.473946 | 0.140496 | 0.296893 |
| ENSG00000261670 | | 1.45255 | 0.3959508 | 0.8300512 | 1.145689 | 0.2519239 | NA |
| ENSG00000213462 | ERV3-1 | 1760.775 | 0.3959342 | 0.3746071 | 1.211726 | 0.2256173 | 0.412911 |
| ENSG00000054219 | LY75 | 0.924976879 | 0.395846573 | 0.914626808 | 1.61563792 | 0.10617262 | NA |

| | | | | | | | |
|---|---|---|---|---|---|---|---|
| ENSG00000255282 | WTAPP1 | 10.60217 | 0.3957834 | 0.5604604 | 0.955499 | 0.3393254 | 0.540625 |
| ENSG00000241258 | CRCP | 2428.158 | 0.395752 | 0.1478072 | 2.733306 | 0.0062702 | 0.026642 |
| ENSG00000101489 | CELF4 | 9.482787067 | 0.39570273 | 0.600356404 | 0.922590416 | 0.35622069 | 0.558176 |
| ENSG00000261419 | | 1.129129 | 0.3956798 | 0.9087602 | 1.56058 | 0.1186229 | NA |
| ENSG00000240024 | LINC00888 | 320.4292 | 0.395494 | 0.2221719 | 1.855889 | 0.06346945 | 0.165715 |
| ENSG00000269012 | KRT18P40 | 1.168671 | 0.3954234 | 0.8546535 | 1.297908 | 0.1943191 | NA |
| ENSG00000230979 | | 52.29024 | 0.3953909 | 0.3557163 | 1.239214 | 0.2152662 | 0.400295 |
| ENSG00000197008 | ZNF138 | 1002.017 | 0.3953759 | 0.1553075 | 2.598732 | 0.00935689 | 0.036846 |
| ENSG00000177951 | BET1L | 1989.865 | 0.3953736 | 0.1025372 | 3.899492 | 9.64E-05 | 0.000741 |
| ENSG00000151892 | GFRA1 | 4661.47 | 0.3953712 | 0.5369734 | 0.947325 | 0.3434732 | 0.544991 |
| ENSG00000171824 | EXOSC10 | 3884.612 | 0.3953355 | 0.1565945 | 2.567328 | 0.01024856 | 0.039664 |
| ENSG00000156697 | UTP14A | 2730.362 | 0.3952479 | 0.1958596 | 2.079021 | 0.03761539 | 0.111431 |
| ENSG00000285518 | | 1.777404 | 0.3950226 | 0.7926741 | 1.053893 | 0.2919317 | 0.490721 |
| ENSG00000144035 | NAT8 | 2.55623 | 0.3950129 | 0.728661 | 0.9572746 | 0.3384287 | 0.539775 |
| ENSG00000117000 | RLF | 2093.507747 | 0.394952324 | 0.177493698 | 2.26719561 | 0.02337828 | 0.076931 |
| ENSG00000168615 | ADAM9 | 7368.02 | 0.3948668 | 0.2476104 | 1.723938 | 0.08471898 | 0.20607 |
| ENSG00000244363 | RPL7P23 | 36.67872 | 0.3947269 | 0.4411748 | 1.074513 | 0.282593 | 0.480433 |
| ENSG00000250816 | DCAF13P2 | 0.857008 | 0.3945221 | 0.8838595 | 1.294221 | 0.1955892 | NA |
| ENSG00000169432 | SCN9A | 5.887911 | 0.3945199 | 0.7037025 | 0.9586036 | 0.3377585 | 0.53907 |
| ENSG00000070526 | ST6GALNAC1 | 7.06050735 | 0.394465429 | 0.76608352 | 1.011137624 | 0.31195057 | 0.512916 |
| ENSG00000142627 | EPHA2 | 2653.64 | 0.394439 | 0.3339982 | 1.310388 | 0.1900646 | 0.367869 |
| ENSG00000231544 | RSL24D1P11 | 82.14334 | 0.3944144 | 0.2574041 | 1.622981 | 0.1045934 | 0.240356 |
| ENSG00000169105 | CHST14 | 861.9107 | 0.3943007 | 0.277295 | 1.526664 | 0.1268446 | 0.275788 |
| ENSG00000253930 | TNFRSF10A-AS1 | 35.5941 | 0.3942426 | 0.3773526 | 1.184623 | 0.2361666 | 0.425704 |
| ENSG00000203721 | LINC00862 | 13.97045 | 0.3941651 | 0.5990728 | 0.9417899 | 0.3463002 | 0.547996 |
| ENSG00000112208 | BAG2 | 299.3312485 | 0.393963424 | 0.250812008 | 1.656097888 | 0.097702 | 0.22881 |
| ENSG00000121210 | TMEM131L | 1957.769559 | 0.393871221 | 0.235987214 | 1.753419402 | 0.07953004 | 0.19636 |
| ENSG00000234161 | PABPC5-AS1 | 1.663617 | 0.3938406 | 0.7894431 | 1.045563 | 0.2957627 | 0.494813 |
| ENSG00000081985 | IL12RB2 | 3.272208775 | 0.393819803 | 0.696449204 | 0.950628354 | 0.34179307 | 0.543063 |
| ENSG00000270244 | | 3.908128 | 0.3937098 | 0.7749931 | 1.044871 | 0.2960825 | 0.495116 |
| ENSG00000258424 | | 81.87689 | 0.393455 | 0.3505045 | 1.253858 | 0.2098935 | 0.393332 |
| ENSG00000162813 | BPNT1 | 2319.034 | 0.3933099 | 0.1209164 | 3.313223 | 0.00092228 | 0.005325 |
| ENSG00000011007 | ELOA | 3005.559701 | 0.393260352 | 0.158988322 | 2.515178154 | 0.01189723 | 0.044729 |

| | | | | | | | |
|---|---|---|---|---|---|---|---|
| ENSG00000230943 | LINC02541 | 6.748526 | 0.3932222 | 0.7949654 | 1.135171 | 0.2563036 | 0.450022 |
| ENSG00000089289 | IGBP1 | 4201.689703 | 0.393075519 | 0.177069428 | 2.279309946 | 0.02264865 | 0.075179 |
| ENSG00000117505 | DR1 | 2307.441533 | 0.393005945 | 0.144943198 | 2.763168484 | 0.00572432 | 0.024774 |
| ENSG00000279636 | | 19.71388 | 0.3930047 | 0.4899705 | 1.000422 | 0.3171062 | 0.518187 |
| ENSG00000177374 | HIC1 | 62.70062 | 0.3928608 | 0.492548 | 1.002631 | 0.3160389 | 0.517117 |
| ENSG00000236876 | TMSB4XP1 | 7.421379 | 0.3928224 | 0.5259254 | 0.9680096 | 0.3330396 | 0.534009 |
| ENSG00000288040 | | 1.500179 | 0.3927158 | 0.8897321 | 1.594701 | 0.110779 | NA |
| ENSG00000189171 | S100A13 | 2635.911 | 0.392644 | 0.2537758 | 1.644472 | 0.1000787 | 0.232979 |
| ENSG00000101470 | TNNC2 | 23.07582922 | 0.39259694 | 0.568966758 | 0.945074745 | 0.34462072 | 0.546184 |
| ENSG00000283312 | | 27.85526 | 0.3924934 | 0.5660174 | 0.9416276 | 0.3463833 | 0.548028 |
| ENSG00000170653 | ATF7 | 1017.749 | 0.3924889 | 0.1845385 | 2.207189 | 0.02730088 | 0.086984 |
| ENSG00000131115 | ZNF227 | 626.9127608 | 0.392443739 | 0.183364527 | 2.200524541 | 0.0277697 | 0.088118 |
| ENSG00000196972 | SMIM10L2B | 14.7529 | 0.3923335 | 0.5626427 | 0.9543809 | 0.3398909 | 0.541291 |
| ENSG00000071051 | NCK2 | 1998.269489 | 0.392286018 | 0.165412492 | 2.427773015 | 0.01519185 | 0.054471 |
| ENSG00000148346 | LCN2 | 313.0829 | 0.3922788 | 0.7840148 | 1.080064 | 0.2801137 | 0.47744 |
| ENSG00000113739 | STC2 | 20368.28192 | 0.392278658 | 0.422424147 | 1.087274038 | 0.27691572 | 0.473905 |
| ENSG00000259065 | | 179.0174 | 0.3922427 | 0.2813914 | 1.48727 | 0.1369436 | 0.291212 |
| ENSG00000223203 | RNA5SP221 | 1.04074 | 0.3921962 | 0.8628339 | 1.418614 | 0.1560116 | NA |
| ENSG00000159650 | UROC1 | 1.863674 | 0.3921757 | 0.754315 | 0.999399 | 0.3176015 | 0.518672 |
| ENSG00000252628 | RNU6-453P | 2.074817 | 0.3920341 | 0.8849664 | 1.519032 | 0.1287544 | 0.2788 |
| ENSG00000169446 | MMGT1 | 2988.684 | 0.3919921 | 0.1404739 | 2.813106 | 0.00490655 | 0.021833 |
| ENSG00000251485 | NRBF2P6 | 24.20551 | 0.3918576 | 0.430014 | 1.085108 | 0.277874 | 0.475002 |
| ENSG00000265625 | | 2.376831 | 0.3916977 | 0.7984128 | 1.105075 | 0.269127 | 0.465251 |
| ENSG00000235052 | | 15.17819 | 0.3915194 | 0.5075246 | 0.968747 | 0.3326715 | 0.533703 |
| ENSG00000173875 | ZNF791 | 1188.379 | 0.3912965 | 0.1707272 | 2.330844 | 0.01976159 | 0.067339 |
| ENSG00000270207 | | 1.041369 | 0.3910337 | 0.8904349 | 1.442231 | 0.1492373 | NA |
| ENSG00000134077 | THUMPD3 | 3266.908494 | 0.391031375 | 0.131437016 | 3.010171389 | 0.002611 | 0.012979 |
| ENSG00000178149 | DALRD3 | 1393.656 | 0.391022 | 0.1383113 | 2.874219 | 0.00405029 | 0.01865 |
| ENSG00000169402 | RSPH10B2 | 5.809903 | 0.3910053 | 0.6297043 | 0.9187851 | 0.358208 | 0.5602 |
| ENSG00000279730 | KMT5AP1 | 306.9041 | 0.3908879 | 0.2211115 | 1.845484 | 0.0649671 | 0.168631 |
| ENSG00000187944 | C2orf66 | 5.113859 | 0.3908813 | 0.7508052 | 0.9832798 | 0.3254698 | 0.526767 |
| ENSG00000262165 | | 19.03201 | 0.390879 | 0.5418248 | 0.9582699 | 0.3379267 | 0.539274 |
| ENSG00000287825 | | 31.17575 | 0.3908598 | 0.3343884 | 1.287457 | 0.1979352 | 0.37796 |

| | | | | | | | |
|---|---|---|---|---|---|---|---|
| ENSG00000092108 | SCFD1 | 3834.771917 | 0.390560957 | 0.152022307 | 2.618380867 | 0.00883481 | 0.035245 |
| ENSG00000237351 | ITPK1P1 | 1.63901 | 0.3905601 | 0.7886314 | 1.058733 | 0.2897212 | 0.488241 |
| ENSG00000030110 | BAK1 | 1807.677536 | 0.390399771 | 0.234555355 | 1.751430369 | 0.07987181 | 0.196966 |
| ENSG00000140575 | IQGAP1 | 15986.02 | 0.3902773 | 0.1992691 | 2.013512 | 0.04406079 | 0.126026 |
| ENSG00000186940 | CHCHD2P9 | 724.5597 | 0.390011 | 0.2268944 | 1.798076 | 0.07216496 | 0.182692 |
| ENSG00000103313 | MEFV | 7.350333554 | 0.389905792 | 0.560632142 | 0.940170768 | 0.34712997 | 0.548669 |
| ENSG00000159197 | KCNE2 | 0.830307 | 0.3898858 | 0.8910832 | 1.365386 | 0.1721318 | NA |
| ENSG00000183479 | TREX2 | 85.80562 | 0.3898152 | 0.2278726 | 1.784773 | 0.07429814 | 0.186899 |
| ENSG00000274173 | | 1528.067 | 0.3896588 | 0.3210186 | 1.32263 | 0.1859586 | 0.361932 |
| ENSG00000175505 | CLCF1 | 335.0146 | 0.389657 | 0.3141981 | 1.351947 | 0.1763923 | 0.349184 |
| ENSG00000260442 | ATP2A1-AS1 | 27.65855 | 0.3896288 | 0.4336699 | 1.070972 | 0.2841821 | 0.482086 |
| ENSG00000179981 | TSHZ1 | 900.961 | 0.3896179 | 0.174195 | 2.306246 | 0.02109687 | 0.070979 |
| ENSG00000258845 | | 2.628699 | 0.3894858 | 0.7044344 | 0.9331965 | 0.3507185 | 0.552388 |
| ENSG00000163354 | DCST2 | 229.6202 | 0.3892995 | 0.2620392 | 1.571254 | 0.1161236 | 0.259072 |
| ENSG00000287724 | | 1.456824 | 0.3891818 | 0.8283724 | 1.149933 | 0.2501714 | NA |
| ENSG00000226803 | ZNF451-AS1 | 7.819899 | 0.3890555 | 0.537692 | 0.9467271 | 0.3437778 | 0.545409 |
| ENSG00000164924 | YWHAZ | 49999.85 | 0.389026 | 0.1488704 | 2.661767 | 0.00777317 | 0.031773 |
| ENSG00000108515 | ENO3 | 704.5224443 | 0.388813977 | 0.25738321 | 1.603827111 | 0.10875217 | 0.247246 |
| ENSG00000166278 | C2 | 24.3911 | 0.3887639 | 0.4803618 | 0.9967358 | 0.3188928 | 0.519932 |
| ENSG00000110881 | ASIC1 | 885.6637037 | 0.388666932 | 0.158030073 | 2.531266945 | 0.01136513 | 0.043125 |
| ENSG00000252637 | RNA5SP268 | 2.203274 | 0.3886152 | 0.773831 | 1.039131 | 0.2987437 | 0.497981 |
| ENSG00000257956 | NOP56P3 | 7.512244 | 0.3882994 | 0.6639625 | 0.932139 | 0.3512647 | 0.552921 |
| ENSG00000185518 | SV2B | 2.466471 | 0.3882883 | 0.7946144 | 1.088572 | 0.2763425 | 0.4733 |
| ENSG00000206187 | LINC02346 | 1.122994 | 0.3881714 | 0.9039051 | 1.647314 | 0.0994936 | NA |
| ENSG00000214016 | RPSAP61 | 3.694444 | 0.3877855 | 0.7820017 | 1.051867 | 0.2928607 | 0.491754 |
| ENSG00000113594 | LIFR | 1214.673838 | 0.387684733 | 0.283085724 | 1.467140226 | 0.14233791 | 0.299737 |
| ENSG00000182054 | IDH2 | 13754.78 | 0.3876185 | 0.2880411 | 1.444515 | 0.1485941 | 0.309073 |
| ENSG00000189043 | NDUFA4 | 6636.09 | 0.3875919 | 0.2166567 | 1.862205 | 0.06257426 | 0.164088 |
| ENSG00000257896 | | 14.38095 | 0.3875909 | 0.5974827 | 0.9226152 | 0.3562078 | 0.558176 |
| ENSG00000288593 | | 1.264613 | 0.387574 | 0.8163173 | 1.112155 | 0.2660716 | NA |
| ENSG00000253528 | | 2.276388 | 0.3875095 | 0.7392942 | 0.9653757 | 0.3343567 | 0.535507 |
| ENSG00000142606 | MMEL1 | 79.41841 | 0.3873243 | 0.3220538 | 1.320499 | 0.1866683 | 0.363047 |
| ENSG00000170949 | ZNF160 | 1634.936 | 0.3870401 | 0.1716136 | 2.31237 | 0.02075729 | 0.070041 |

| | | | | | | | |
|---|---|---|---|---|---|---|---|
| ENSG00000256542 | | 3.617268 | 0.3870297 | 0.7388598 | 0.9707293 | 0.3316831 | 0.532838 |
| ENSG00000177410 | ZFAS1 | 3336.714 | 0.3870004 | 0.2534063 | 1.651549 | 0.09862643 | 0.230308 |
| ENSG00000187815 | ZFP69 | 377.5828 | 0.3869674 | 0.187719 | 2.137834 | 0.03253025 | 0.099792 |
| ENSG00000050030 | NEXMIF | 146.5185317 | 0.386900416 | 0.268864437 | 1.530058701 | 0.1260022 | 0.274541 |
| ENSG00000153165 | RGPD3 | 19.70646 | 0.3867576 | 0.4108104 | 1.098257 | 0.2720921 | 0.468587 |
| ENSG00000213272 | RPL7AP9 | 1.157842 | 0.3867223 | 0.8795068 | 1.289313 | 0.1972894 | NA |
| ENSG00000143570 | SLC39A1 | 333.442 | 0.3866997 | 0.187195 | 2.127223 | 0.03340158 | 0.101994 |
| ENSG00000267041 | ZNF850 | 208.5431 | 0.3866271 | 0.2418928 | 1.683699 | 0.09223995 | 0.219402 |
| ENSG00000244722 | RPSAP29 | 1.104856 | 0.3865953 | 0.8441732 | 1.193751 | 0.2325755 | NA |
| ENSG00000280927 | CTBP1-AS | 100.409 | 0.3865692 | 0.3132046 | 1.343552 | 0.1790934 | 0.353004 |
| ENSG00000202382 | | 1.419147 | 0.3865358 | 0.8096322 | 1.102086 | 0.2704241 | NA |
| ENSG00000217555 | CKLF | 8.809607 | 0.3864821 | 0.5362886 | 0.9452202 | 0.3445465 | 0.546099 |
| ENSG00000125503 | PPP1R12C | 5145.223441 | 0.386401986 | 0.157564298 | 2.483790135 | 0.01299924 | 0.048062 |
| ENSG00000283971 | MIR4442 | 7.977195 | 0.386326 | 0.5688596 | 0.9332428 | 0.3506946 | 0.552384 |
| ENSG00000172031 | EPHX4 | 62.4978 | 0.3863144 | 0.3482009 | 1.239272 | 0.2152448 | 0.400292 |
| ENSG00000267370 | | 38.62252 | 0.3863067 | 0.3979833 | 1.115441 | 0.2646616 | 0.460053 |
| ENSG00000196588 | MRTFA | 1705.016 | 0.3862945 | 0.1835032 | 2.138145 | 0.03250501 | 0.099737 |
| ENSG00000172432 | GTPBP2 | 4964.58 | 0.3862749 | 0.2911018 | 1.408601 | 0.1589531 | 0.324889 |
| ENSG00000286357 | | 1.593276 | 0.3862715 | 0.8172032 | 1.101845 | 0.270529 | 0.466782 |
| ENSG00000278549 | MIR6736 | 1.407251 | 0.3862427 | 0.8006026 | 1.087158 | 0.2769668 | NA |
| ENSG00000182473 | EXOC7 | 7241.584 | 0.3862174 | 0.1858839 | 2.142655 | 0.03214079 | 0.098894 |
| ENSG00000269028 | MTRNR2L12 | 75.87988 | 0.3861408 | 0.3847542 | 1.147784 | 0.2510578 | 0.443803 |
| ENSG00000258896 | SCOCP1 | 2.791413 | 0.3860686 | 0.6829755 | 0.9267722 | 0.3540448 | 0.556111 |
| ENSG00000248538 | | 32.19565 | 0.3858403 | 0.4627628 | 1.010567 | 0.3122237 | 0.513254 |
| ENSG00000111644 | ACRBP | 35.95603087 | 0.385758103 | 0.476593877 | 0.994201055 | 0.320125 | 0.520941 |
| ENSG00000065882 | TBC1D1 | 1830.86173 | 0.385658666 | 0.135452449 | 2.892280194 | 0.00382457 | 0.017765 |
| ENSG00000109685 | NSD2 | 7632.479464 | 0.385604131 | 0.24228276 | 1.684844454 | 0.09201859 | 0.219033 |
| ENSG00000265366 | GLUD1P2 | 10.932 | 0.3855984 | 0.493441 | 0.9706496 | 0.3317228 | 0.532862 |
| ENSG00000144597 | EAF1 | 2694.865 | 0.3854989 | 0.1317688 | 2.976458 | 0.00291599 | 0.014251 |
| ENSG00000243020 | RPL7P39 | 6.929611 | 0.3854238 | 0.6432027 | 0.9149183 | 0.3602345 | 0.562244 |
| ENSG00000196417 | ZNF765 | 635.5267 | 0.385383 | 0.1570364 | 2.511762 | 0.01201301 | 0.045075 |
| ENSG00000125629 | INSIG2 | 1959.095038 | 0.385146021 | 0.242972522 | 1.673389665 | 0.09425061 | 0.223086 |
| ENSG00000182873 | PRKCZ-AS1 | 160.8707 | 0.3851106 | 0.3165056 | 1.327488 | 0.1843474 | 0.359644 |

| | | | | | | | |
|---|---|---|---|---|---|---|---|
| ENSG00000139428 | MMAB | 2909.683 | 0.385054 | 0.2184752 | 1.833808 | 0.06668253 | 0.172159 |
| ENSG00000213871 | TAF9BP1 | 2.874647 | 0.3850037 | 0.7104178 | 0.9431958 | 0.3455807 | 0.547085 |
| ENSG00000248714 | | 3.470059 | 0.3850011 | 0.7346875 | 0.9570389 | 0.3385476 | 0.539844 |
| ENSG00000254185 | MAPK6P5 | 0.8621557 | 0.3848455 | 0.9038115 | 1.560833 | 0.1185631 | NA |
| ENSG00000048052 | HDAC9 | 103.6906163 | 0.384830642 | 0.42233663 | 1.063486067 | 0.28756158 | 0.485957 |
| ENSG00000224677 | PDIA3P2 | 7.333534 | 0.3846337 | 0.5967286 | 0.9112761 | 0.3621499 | 0.564172 |
| ENSG00000112146 | FBXO9 | 5116.762796 | 0.38461897 | 0.122289529 | 3.193240935 | 0.00140686 | 0.007682 |
| ENSG00000162763 | LRRC52 | 1.552795 | 0.384597 | 0.8709042 | 1.182919 | 0.2368411 | NA |
| ENSG00000219146 | RPS4XP8 | 8.716645 | 0.3843614 | 0.6062009 | 0.9175125 | 0.3588742 | 0.560812 |
| ENSG00000205918 | PDPK2P | 122.1543 | 0.3843504 | 0.3460335 | 1.237292 | 0.2159786 | 0.401249 |
| ENSG00000230185 | C9orf147 | 25.75735 | 0.3841715 | 0.4137632 | 1.082389 | 0.2790799 | 0.476216 |
| ENSG00000236330 | RPL5P9 | 27.0526 | 0.3841127 | 0.4153604 | 1.083796 | 0.2784554 | 0.475743 |
| ENSG00000223224 | SNORD71 | 1.251382 | 0.3840099 | 0.8387446 | 1.250902 | 0.2109701 | NA |
| ENSG00000229670 | PKP4P1 | 39.8681 | 0.3839618 | 0.3419817 | 1.244752 | 0.2132228 | 0.397971 |
| ENSG00000087266 | SH3BP2 | 2407.329412 | 0.383869394 | 0.189378051 | 2.085058591 | 0.03706398 | 0.110189 |
| ENSG00000273291 | | 1.227527 | 0.3835412 | 0.8797995 | 1.298704 | 0.1940457 | NA |
| ENSG00000085382 | HACE1 | 697.1323437 | 0.383417897 | 0.181778133 | 2.164345304 | 0.03043786 | 0.094796 |
| ENSG00000126246 | IGFLR1 | 13.58824856 | 0.383381999 | 0.523785434 | 0.953565205 | 0.34030378 | 0.541766 |
| ENSG00000075290 | WNT8B | 73.01633809 | 0.383324121 | 0.421343983 | 1.074461484 | 0.2826159 | 0.480442 |
| ENSG00000183248 | PRR36 | 1035.271 | 0.3832783 | 0.1666313 | 2.359933 | 0.01827822 | 0.063225 |
| ENSG00000226969 | | 1.125354 | 0.3831597 | 0.8334868 | 1.220651 | 0.2222183 | NA |
| ENSG00000179277 | MEIS3P1 | 416.4183 | 0.3830127 | 0.199748 | 1.988138 | 0.04679639 | 0.131821 |
| ENSG00000221716 | SNORA11 | 20.85949 | 0.3828629 | 0.4650051 | 1.008346 | 0.3132885 | 0.514416 |
| ENSG00000139637 | MYG1 | 1934.23 | 0.3827872 | 0.1713325 | 2.294103 | 0.02178459 | 0.072859 |
| ENSG00000228358 | LINC02263 | 11.99285 | 0.382755 | 0.9179592 | 1.972216 | 0.0485849 | 0.135537 |
| ENSG00000253593 | | 1.120174 | 0.3826775 | 0.8815299 | 1.25915 | 0.2079763 | NA |
| ENSG00000164073 | MFSD8 | 2298.475 | 0.3826505 | 0.159428 | 2.451437 | 0.01422871 | 0.051695 |
| ENSG00000139289 | PHLDA1 | 1889.829 | 0.3822611 | 0.359045 | 1.195812 | 0.23177 | 0.420162 |
| ENSG00000151468 | CCDC3 | 7.816314 | 0.3822601 | 0.7330388 | 0.9743519 | 0.3298819 | 0.530997 |
| ENSG00000228507 | DAP3P2 | 6.413879 | 0.3821129 | 0.6230246 | 0.9042248 | 0.3658762 | 0.567807 |
| ENSG00000227968 | BUB3P1 | 1.049943 | 0.3821087 | 0.8924633 | 1.461853 | 0.1437815 | NA |
| ENSG00000136940 | PDCL | 1480.161899 | 0.382033476 | 0.179465889 | 2.191302527 | 0.02842991 | 0.089838 |
| ENSG00000168000 | BSCL2 | 108.798 | 0.3820039 | 0.2217343 | 1.79661 | 0.07239751 | 0.183211 |

| | | | | | | | |
|---|---|---|---|---|---|---|---|
| ENSG00000285679 | | 22.91299 | 0.3819748 | 0.4064067 | 1.086884 | 0.277088 | 0.474017 |
| ENSG00000253355 | | 2.055808 | 0.3818667 | 0.8093314 | 1.105146 | 0.2690961 | 0.465251 |
| ENSG00000174282 | ZBTB4 | 5570.875 | 0.3818329 | 0.2004376 | 1.965062 | 0.04940705 | 0.13734 |
| ENSG00000196155 | PLEKHG4 | 2608.518 | 0.3816189 | 0.1789618 | 2.201959 | 0.02766819 | 0.08787 |
| ENSG00000196663 | TECPR2 | 1842.263 | 0.3812469 | 0.227042 | 1.753877 | 0.07945156 | 0.196241 |
| ENSG00000197172 | MAGEA6 | 203.4126 | 0.3811989 | 0.8559043 | 1.042438 | 0.2972085 | 0.496325 |
| ENSG00000217624 | YWHAZP10 | 589.0727 | 0.3811894 | 0.1819818 | 2.17087 | 0.029941 | 0.093534 |
| ENSG00000165912 | PACSIN3 | 3532.309 | 0.3810809 | 0.2680606 | 1.516617 | 0.1293633 | 0.279663 |
| ENSG00000260185 | | 2.106438 | 0.3810758 | 0.7586294 | 0.983059 | 0.3255784 | 0.526911 |
| ENSG00000287492 | | 1.176585 | 0.380879 | 0.8998113 | 1.467573 | 0.1422203 | NA |
| ENSG00000130751 | NPAS1 | 128.1260203 | 0.380871597 | 0.390223693 | 1.117969229 | 0.26358014 | 0.458796 |
| ENSG00000033122 | LRRC7 | 27.36400007 | 0.380837275 | 0.347709284 | 1.222185134 | 0.22163762 | 0.408285 |
| ENSG00000287191 | | 1.254676 | 0.3808349 | 0.8202537 | 1.095987 | 0.2730846 | NA |
| ENSG00000214362 | RPS3AP36 | 1.283988 | 0.3807931 | 0.8370887 | 1.243005 | 0.2138658 | NA |
| ENSG00000245293 | CYP2U1-AS1 | 9.823165 | 0.3806989 | 0.5423633 | 0.9358138 | 0.3493691 | 0.55108 |
| ENSG00000197935 | ZNF311 | 220.8299 | 0.3806325 | 0.2219897 | 1.788961 | 0.07362113 | 0.185681 |
| ENSG00000287104 | | 81.48961 | 0.3804982 | 0.3419201 | 1.231067 | 0.2182976 | 0.404065 |
| ENSG00000215252 | GOLGA8B | 973.108 | 0.3804776 | 0.3524768 | 1.196149 | 0.2316385 | 0.420038 |
| ENSG00000243978 | RTL9 | 2.725924 | 0.3803299 | 0.6809047 | 0.9172032 | 0.3590362 | 0.560966 |
| ENSG00000201239 | | 1.797506 | 0.3803099 | 0.8417401 | 1.207343 | 0.2273002 | 0.415097 |
| ENSG00000265218 | | 17.58005 | 0.3802784 | 0.5213846 | 0.9386871 | 0.3478914 | 0.549663 |
| ENSG00000148297 | MED22 | 4954.575 | 0.380275 | 0.1767732 | 2.223506 | 0.02618172 | 0.084074 |
| ENSG00000125945 | ZNF436 | 871.2705458 | 0.38022157 | 0.231612848 | 1.719703764 | 0.0854863 | 0.207499 |
| ENSG00000227482 | | 0.8641216 | 0.3801197 | 0.8982643 | 1.480017 | 0.1388687 | NA |
| ENSG00000128791 | TWSG1 | 1453.223956 | 0.379967222 | 0.215911682 | 1.82859961 | 0.06745961 | 0.17362 |
| ENSG00000189280 | GJB5 | 26.04931 | 0.3799435 | 0.4208132 | 1.058985 | 0.2896068 | 0.488172 |
| ENSG00000167797 | CDK2AP2 | 3626.171 | 0.3798468 | 0.2239468 | 1.773677 | 0.07611665 | 0.19007 |
| ENSG00000270987 | | 98.94169 | 0.379686 | 0.2738033 | 1.481304 | 0.1385257 | 0.293734 |
| ENSG00000165948 | IFI27L1 | 191.4758 | 0.3795341 | 0.2362137 | 1.688522 | 0.09131108 | 0.217869 |
| ENSG00000261305 | | 11.98436 | 0.3795088 | 0.5865658 | 0.9089746 | 0.3633635 | 0.565433 |
| ENSG00000170456 | DENND5B | 357.4151 | 0.379479 | 0.3359228 | 1.240373 | 0.2148374 | 0.399745 |
| ENSG00000287049 | | 8.492441 | 0.3794357 | 0.5505566 | 0.9166235 | 0.35934 | 0.56121 |
| ENSG00000288044 | | 11.04853 | 0.3794275 | 0.4888035 | 0.9701754 | 0.3319591 | 0.533049 |

| | | | | | | | |
|---|---|---|---|---|---|---|---|
| ENSG00000116095 | PLEKHA3 | 1082.047576 | 0.379329216 | 0.142489861 | 2.703466727 | 0.00686203 | 0.028671 |
| ENSG00000145685 | LHFPL2 | 1127.926 | 0.379184 | 0.3541904 | 1.215454 | 0.2241928 | 0.410997 |
| ENSG00000258700 | LINC00871 | 6.305255 | 0.3791756 | 0.6866137 | 0.9067317 | 0.3645487 | 0.566578 |
| ENSG00000204872 | NAT8B | 2.089491 | 0.3790629 | 0.7960169 | 1.041252 | 0.2977584 | 0.496994 |
| ENSG00000269982 | | 145.6534 | 0.3790248 | 0.2490789 | 1.602493 | 0.1090467 | 0.247693 |
| ENSG00000286974 | | 22.00614 | 0.37874 | 0.524615 | 0.9330423 | 0.3507981 | 0.552408 |
| ENSG00000264273 | | 2.02627 | 0.378581 | 0.7470755 | 0.9656165 | 0.3342361 | 0.535452 |
| ENSG00000213867 | | 17.3121 | 0.378531 | 0.4949601 | 0.9664677 | 0.3338102 | 0.535019 |
| ENSG00000284461 | | 2.832903 | 0.3783887 | 0.7221573 | 0.9428589 | 0.3457531 | 0.547326 |
| ENSG00000196593 | ANKRD20A19P | 3.818423 | 0.3783634 | 0.7284762 | 0.9610472 | 0.3365284 | 0.537624 |
| ENSG00000197111 | PCBP2 | 9648.974 | 0.3782574 | 0.1359793 | 2.820473 | 0.00479529 | 0.021438 |
| ENSG00000248927 | | 2.160822 | 0.3780909 | 0.8537972 | 1.241241 | 0.2145168 | 0.399379 |
| ENSG00000180182 | MED14 | 3265.274 | 0.3779071 | 0.1699906 | 2.281604 | 0.02251274 | 0.074861 |
| ENSG00000287110 | | 64.44657 | 0.3777944 | 0.370168 | 1.154003 | 0.2484988 | 0.440775 |
| ENSG00000235254 | TMEM185AP1 | 1.293017 | 0.377792 | 0.7995858 | 1.074742 | 0.2824901 | NA |
| ENSG00000259270 | | 0.8087547 | 0.3777557 | 0.9071221 | 1.648037 | 0.09934509 | NA |
| ENSG00000225472 | | 1.689705 | 0.3777485 | 0.8319457 | 1.2187 | 0.2229582 | 0.409572 |
| ENSG00000227329 | | 33.04242 | 0.3776866 | 0.3936626 | 1.098826 | 0.2718439 | 0.468303 |
| ENSG00000173889 | PHC3 | 5644.741 | 0.3776167 | 0.2383671 | 1.661589 | 0.09659516 | 0.226837 |
| ENSG00000207650 | MIR570 | 80.76209 | 0.3775897 | 0.256078 | 1.570148 | 0.1163807 | 0.259486 |
| ENSG00000163041 | H3-3A | 452.1143 | 0.377586 | 0.1727582 | 2.246941 | 0.02464383 | 0.080191 |
| ENSG00000053900 | ANAPC4 | 1572.378897 | 0.377539317 | 0.163824595 | 2.358425907 | 0.01835262 | 0.063417 |
| ENSG00000172262 | ZNF131 | 4001.029 | 0.3774237 | 0.1258136 | 3.033611 | 0.00241646 | 0.012146 |
| ENSG00000231443 | | 51.98501 | 0.3773832 | 0.4080199 | 1.079846 | 0.2802109 | 0.477575 |
| ENSG00000069696 | DRD4 | 7.456213338 | 0.377316449 | 0.548732013 | 0.918116215 | 0.35855803 | 0.56045 |
| ENSG00000205268 | PDE7A | 2545.413 | 0.3772773 | 0.1556212 | 2.48116 | 0.01309554 | 0.048363 |
| ENSG00000178623 | GPR35 | 109.2187 | 0.3772114 | 0.4112804 | 1.064997 | 0.2868773 | 0.485253 |
| ENSG00000273828 | | 63.18919 | 0.3771185 | 0.378845 | 1.13218 | 0.2575588 | 0.451569 |
| ENSG00000269915 | | 22.52573 | 0.3770605 | 0.5190077 | 0.9365524 | 0.3489889 | 0.550668 |
| ENSG00000228502 | EEF1A1P11 | 134.2984 | 0.3770312 | 0.2857581 | 1.418812 | 0.1559539 | 0.320237 |
| ENSG00000169228 | RAB24 | 880.58 | 0.3767394 | 0.2878838 | 1.406197 | 0.1596656 | 0.325761 |
| ENSG00000243910 | TUBA4B | 0.9598316 | 0.376712 | 0.8773268 | 1.272185 | 0.2033073 | NA |
| ENSG00000227304 | | 10.29808 | 0.3766814 | 0.5387044 | 0.9199237 | 0.3576127 | 0.559664 |

| | | | | | | | |
|---|---|---|---|---|---|---|---|
| ENSG00000115255 | REEP6 | 2099.479206 | 0.376509432 | 0.260982871 | 1.519358894 | 0.12867218 | 0.27869 |
| ENSG00000080298 | RFX3 | 460.4781732 | 0.376348103 | 0.243300468 | 1.631021953 | 0.10288569 | 0.237576 |
| ENSG00000197142 | ACSL5 | 101.2862 | 0.3763438 | 0.5385126 | 0.9219958 | 0.3565307 | 0.558464 |
| ENSG00000281016 | | 2.188134 | 0.3762287 | 0.7207457 | 0.9340938 | 0.3502555 | 0.551929 |
| ENSG00000173681 | BCLAF3 | 1035.858 | 0.3761881 | 0.2878554 | 1.407824 | 0.1591833 | 0.325159 |
| ENSG00000178974 | FBXO34 | 2404.17 | 0.3758768 | 0.105893 | 3.58404 | 0.00033832 | 0.002247 |
| ENSG00000269834 | ZNF528-AS1 | 3.521306 | 0.3758159 | 0.7681375 | 1.00059 | 0.317025 | 0.518118 |
| ENSG00000234685 | NUS1P2 | 46.08821 | 0.3757968 | 0.3764241 | 1.130212 | 0.258387 | 0.452603 |
| ENSG00000211584 | SLC48A1 | 1136.85 | 0.375769 | 0.2241297 | 1.750604 | 0.08001418 | 0.197247 |
| ENSG00000252414 | RNU6-100P | 1.687256 | 0.3757501 | 0.842386 | 1.240976 | 0.2146146 | 0.399502 |
| ENSG00000203739 | PRDX6-AS1 | 164.241 | 0.375695 | 0.2185504 | 1.792545 | 0.07304577 | 0.1845 |
| ENSG00000245768 | | 3.621225 | 0.3756548 | 0.7038031 | 0.9214574 | 0.3568117 | 0.558805 |
| ENSG00000236682 | MAP3K2-DT | 75.23293 | 0.3753623 | 0.2734327 | 1.461247 | 0.1439477 | 0.302002 |
| ENSG00000149243 | KLHL35 | 599.6136 | 0.37535 | 0.2210238 | 1.769363 | 0.07683332 | 0.19141 |
| ENSG00000254389 | RHPN1-AS1 | 299.2684 | 0.3752182 | 0.1836193 | 2.104301 | 0.03535224 | 0.106385 |
| ENSG00000224172 | | 11.30933 | 0.3750576 | 0.5810064 | 0.9100738 | 0.3627836 | 0.564786 |
| ENSG00000269220 | LINC00528 | 2.53894 | 0.3750457 | 0.7564942 | 0.9761126 | 0.3290087 | 0.530227 |
| ENSG00000067900 | ROCK1 | 6203.20544 | 0.375042067 | 0.197120343 | 1.965175042 | 0.04939398 | 0.137332 |
| ENSG00000010327 | STAB1 | 5.930606201 | 0.375007651 | 0.622020844 | 0.90278772 | 0.36663857 | 0.568565 |
| ENSG00000164123 | C4orf45 | 0.9174488 | 0.374913 | 0.8887681 | 1.447 | 0.1478969 | NA |
| ENSG00000225215 | SMARCE1P1 | 5.616569 | 0.3748892 | 0.6927543 | 0.9119331 | 0.3618039 | 0.563733 |
| ENSG00000173930 | SLCO4C1 | 1447.071 | 0.3748855 | 0.3442133 | 1.196563 | 0.2314768 | 0.419902 |
| ENSG00000175197 | DDIT3 | 1581.789 | 0.3748124 | 0.4471025 | 1.007054 | 0.313909 | 0.515047 |
| ENSG00000276148 | | 11.45133 | 0.3745272 | 0.496332 | 0.9454022 | 0.3444536 | 0.546046 |
| ENSG00000092051 | JPH4 | 2.064132615 | 0.374498557 | 0.813832517 | 1.087182304 | 0.27695625 | 0.473944 |
| ENSG00000100034 | PPM1F | 2220.406023 | 0.374411815 | 0.125262229 | 3.023363609 | 0.00249982 | 0.012503 |
| ENSG00000233834 | | 88.9844 | 0.3743373 | 0.297206 | 1.360337 | 0.1737232 | 0.345379 |
| ENSG00000143171 | RXRG | 1.663319 | 0.3743098 | 0.8358853 | 1.164756 | 0.2441179 | 0.435446 |
| ENSG00000234789 | | 7.374004 | 0.3742809 | 0.6264865 | 0.8902836 | 0.3733136 | 0.575557 |
| ENSG00000158161 | EYA3 | 1563 | 0.3742207 | 0.1811804 | 2.125812 | 0.03351895 | 0.102199 |
| ENSG00000258951 | KRT18P7 | 11.58912 | 0.3741417 | 0.595904 | 0.8995113 | 0.3683804 | 0.570367 |
| ENSG00000121807 | CCR2 | 3.763157485 | 0.374077383 | 0.791904528 | 1.053631094 | 0.29205185 | 0.49083 |
| ENSG00000162627 | SNX7 | 1310.798 | 0.3739987 | 0.1595522 | 2.393884 | 0.01667102 | 0.058698 |

| | | | | | | | |
|---|---|---|---|---|---|---|---|
| ENSG00000286351 | | 1.204719 | 0.3738823 | 0.9165081 | 1.959506 | 0.0500535 | NA |
| ENSG00000188687 | SLC4A5 | 36.00228 | 0.3737574 | 0.4107763 | 1.056131 | 0.2909084 | 0.489522 |
| ENSG00000277873 | | 3.145986 | 0.3737513 | 0.785579 | 1.038789 | 0.2989031 | 0.498153 |
| ENSG00000163606 | CD200R1 | 10.11924 | 0.3737457 | 0.5500864 | 0.9076427 | 0.364067 | 0.566196 |
| ENSG00000224555 | | 16.09854 | 0.3736814 | 0.4252558 | 1.037122 | 0.2996791 | 0.498891 |
| ENSG00000112977 | DAP | 10603.25706 | 0.373643229 | 0.200862622 | 1.929623585 | 0.05365349 | 0.146568 |
| ENSG00000265916 | | 0.930355 | 0.373599 | 0.9012338 | 1.57355 | 0.1155916 | NA |
| ENSG00000165731 | RET | 703.713 | 0.373575 | 0.4724212 | 0.9645778 | 0.3347563 | 0.535953 |
| ENSG00000144362 | PHOSPHO2 | 239.9994 | 0.3735488 | 0.2395034 | 1.643559 | 0.1002673 | 0.233255 |
| ENSG00000260095 | | 18.2038 | 0.3735135 | 0.5066346 | 0.9396441 | 0.3474001 | 0.548984 |
| ENSG00000186272 | ZNF17 | 349.9862 | 0.3734524 | 0.1655954 | 2.310933 | 0.02083656 | 0.070237 |
| ENSG00000146729 | NIPSNAP2 | 2748.469 | 0.3734309 | 0.2085389 | 1.878492 | 0.06031395 | 0.159657 |
| ENSG00000205832 | C16orf96 | 2.905946 | 0.3734132 | 0.7188079 | 0.929034 | 0.3528714 | 0.55476 |
| ENSG00000226747 | FSIP2-AS2 | 2.373274 | 0.3733847 | 0.7755643 | 0.993923 | 0.3202604 | 0.521065 |
| ENSG00000147654 | EBAG9 | 2127.822 | 0.3732552 | 0.1116492 | 3.360565 | 0.00077783 | 0.004606 |
| ENSG00000113391 | FAM172A | 1072.642495 | 0.373181086 | 0.214579964 | 1.787315285 | 0.07388655 | 0.186177 |
| ENSG00000182919 | C11orf54 | 604.2834 | 0.3730815 | 0.2753714 | 1.44621 | 0.1481182 | 0.308413 |
| ENSG00000154001 | PPP2R5E | 4956.157 | 0.373066 | 0.1128296 | 3.326824 | 0.00087842 | 0.005113 |
| ENSG00000241933 | DENND6A-DT | 1.880004 | 0.3730373 | 0.7698894 | 0.992834 | 0.3207909 | 0.521705 |
| ENSG00000066230 | SLC9A3 | 553.6545011 | 0.372845491 | 0.259905134 | 1.51403963 | 0.13001579 | 0.280753 |
| ENSG00000272175 | | 1.198284 | 0.3727671 | 0.8593605 | 1.279764 | 0.200628 | NA |
| ENSG00000114742 | WDR48 | 2098.751446 | 0.372744723 | 0.117027153 | 3.216966308 | 0.00129554 | 0.007168 |
| ENSG00000148429 | USP6NL | 2487.072 | 0.3726539 | 0.210723 | 1.848083 | 0.06459036 | 0.167901 |
| ENSG00000170917 | NUDT6 | 438.8682 | 0.3726169 | 0.1839992 | 2.085931 | 0.03698489 | 0.110039 |
| ENSG00000249471 | ZNF324B | 421.9584 | 0.3725761 | 0.1655418 | 2.308848 | 0.02095203 | 0.070563 |
| ENSG00000187790 | FANCM | 855.8455 | 0.3725553 | 0.2930705 | 1.372265 | 0.1699809 | 0.340422 |
| ENSG00000132872 | SYT4 | 3.339714247 | 0.372507739 | 0.746596837 | 0.943527002 | 0.34541141 | 0.546948 |
| ENSG00000270292 | | 1.155737 | 0.3724494 | 0.8674474 | 1.395316 | 0.1629204 | NA |
| ENSG00000172458 | IL17D | 308.9543 | 0.372392 | 0.1682019 | 2.277845 | 0.0227358 | 0.075358 |
| ENSG00000156127 | BATF | 2109.615 | 0.3722285 | 0.2360153 | 1.651542 | 0.09862786 | 0.230308 |
| ENSG00000139223 | ANP32D | 4.315175 | 0.3721445 | 0.7443972 | 0.9566256 | 0.3387563 | 0.54001 |
| ENSG00000266835 | GAPLINC | 1.970615 | 0.3721181 | 0.8143226 | 1.126319 | 0.2600305 | 0.454463 |
| ENSG00000074706 | IPCEF1 | 5.102345491 | 0.372025044 | 0.735292058 | 0.943698336 | 0.34532383 | 0.546874 |

| | | | | | | | |
|---|---|---|---|---|---|---|---|
| ENSG00000106479 | ZNF862 | 1435.351197 | 0.371913754 | 0.335396048 | 1.228336498 | 0.21932068 | 0.40545 |
| ENSG00000248610 | HSPA8P4 | 3.637148 | 0.371697 | 0.7071756 | 0.91812 | 0.3585561 | 0.56045 |
| ENSG00000159214 | CCDC24 | 734.7358 | 0.3716749 | 0.2118736 | 1.813028 | 0.06982749 | 0.178235 |
| ENSG00000163376 | KBTBD8 | 139.1884 | 0.3716712 | 0.3833059 | 1.107372 | 0.2681332 | 0.464016 |
| ENSG00000243979 | | 2.194456 | 0.3716091 | 0.815328 | 1.123397 | 0.261269 | 0.455874 |
| ENSG00000233170 | | 70.42207 | 0.3715433 | 0.233238 | 1.668931 | 0.09513115 | 0.224377 |
| ENSG00000160838 | LRRC71 | 5.957039 | 0.3714945 | 0.6238152 | 0.8872612 | 0.3749383 | 0.577124 |
| ENSG00000184414 | IRS3P | 1.379337 | 0.3714181 | 0.7938076 | 1.046912 | 0.29514 | NA |
| ENSG00000257191 | | 27.16893 | 0.3712395 | 0.3662043 | 1.137314 | 0.2554071 | 0.448833 |
| ENSG00000252355 | RN7SKP287 | 1.701671 | 0.371155 | 0.7971767 | 1.047761 | 0.2947487 | 0.493863 |
| ENSG00000151014 | NOCT | 350.8816 | 0.3711478 | 0.3082268 | 1.313077 | 0.189157 | 0.366626 |
| ENSG00000116984 | MTR | 4413.061219 | 0.371144121 | 0.225817522 | 1.707276561 | 0.08777066 | 0.211633 |
| ENSG00000261131 | | 2.653979 | 0.3710283 | 0.7439468 | 0.9615419 | 0.3362798 | 0.537337 |
| ENSG00000186472 | PCLO | 514.7973 | 0.3710175 | 0.3412009 | 1.189377 | 0.2342915 | 0.423433 |
| ENSG00000279807 | | 1.743962 | 0.3709441 | 0.7820393 | 1.027755 | 0.304065 | 0.503697 |
| ENSG00000123572 | NRK | 107.8294503 | 0.370936672 | 0.526030206 | 0.917349936 | 0.3589593 | 0.560912 |
| ENSG00000129946 | SHC2 | 587.5458262 | 0.370923221 | 0.468485 | 0.964619493 | 0.33473543 | 0.535952 |
| ENSG00000138744 | NAAA | 1890.525389 | 0.370915263 | 0.157959325 | 2.385859301 | 0.01703927 | 0.059714 |
| ENSG00000250182 | EEF1A1P13 | 219.1661 | 0.3707956 | 0.269773 | 1.473529 | 0.1406085 | 0.297107 |
| ENSG00000198153 | ZNF849P | 7.046105 | 0.3707852 | 0.6921185 | 0.9184731 | 0.3583712 | 0.560323 |
| ENSG00000172867 | KRT2 | 3.476349 | 0.3707473 | 0.7608722 | 1.005096 | 0.3148507 | 0.515898 |
| ENSG00000286303 | | 1.505052 | 0.37053 | 0.8722643 | 1.283478 | 0.1993247 | NA |
| ENSG00000143771 | CNIH4 | 2755.399 | 0.3705008 | 0.1563895 | 2.407115 | 0.01607912 | 0.056961 |
| ENSG00000146278 | PNRC1 | 3031.118 | 0.3704312 | 0.1667361 | 2.271458 | 0.02311927 | 0.076296 |
| ENSG00000275005 | | 2.995332 | 0.3702936 | 0.710866 | 0.9066947 | 0.3645683 | 0.566578 |
| ENSG00000099338 | CATSPERG | 446.43587 | 0.370125513 | 0.304387311 | 1.320105085 | 0.18679994 | 0.363197 |
| ENSG00000185024 | BRF1 | 5009.059 | 0.3700688 | 0.1429239 | 2.630391 | 0.00852867 | 0.034219 |
| ENSG00000253796 | | 1.651114 | 0.3700478 | 0.7669508 | 0.9860451 | 0.324111 | 0.525006 |
| ENSG00000113492 | AGXT2 | 2.669329003 | 0.369946804 | 0.750287487 | 0.929679975 | 0.35253681 | 0.554397 |
| ENSG00000011590 | ZBTB32 | 42.17719113 | 0.369878078 | 0.362022761 | 1.144901699 | 0.25224988 | 0.444939 |
| ENSG00000279415 | | 12.64805 | 0.3698313 | 0.5320096 | 0.9210675 | 0.3570152 | 0.559044 |
| ENSG00000105404 | RABAC1 | 534.6260796 | 0.36980558 | 0.213007038 | 1.804577688 | 0.07114079 | 0.180754 |
| ENSG00000066322 | ELOVL1 | 6834.557485 | 0.369701577 | 0.175141323 | 2.141475345 | 0.03223573 | 0.099147 |

| | | | | | | | |
|---|---|---|---|---|---|---|---|
| ENSG00000200237 | | 1.810736 | 0.369657 | 0.8214779 | 1.115748 | 0.2645302 | 0.459998 |
| ENSG00000253966 | | 6.958442 | 0.3695757 | 0.6566693 | 0.8899651 | 0.3734846 | 0.575609 |
| ENSG00000180229 | HERC2P3 | 16.99396 | 0.3694913 | 0.513568 | 0.9238377 | 0.3555708 | 0.557519 |
| ENSG00000183864 | TOB2 | 6038.75 | 0.3694222 | 0.2642891 | 1.4524 | 0.1463903 | 0.305814 |
| ENSG00000207181 | SNORA14B | 24.44109 | 0.369393 | 0.4606854 | 0.9820418 | 0.3260793 | 0.527279 |
| ENSG00000234702 | VDAC1P3 | 14.92358 | 0.3692746 | 0.4525364 | 0.9809871 | 0.3265991 | 0.52789 |
| ENSG00000188993 | LRRC66 | 3.526823 | 0.3692126 | 0.7394524 | 0.9230036 | 0.3560053 | 0.558084 |
| ENSG00000104863 | LIN7B | 324.2597469 | 0.369126086 | 0.244757695 | 1.597097991 | 0.11024387 | 0.24975 |
| ENSG00000259504 | | 0.9407292 | 0.3691139 | 0.8473811 | 1.196764 | 0.2313987 | NA |
| ENSG00000285568 | | 1.276303 | 0.3690513 | 0.8538412 | 1.292058 | 0.1963369 | NA |
| ENSG00000260816 | | 7.168949 | 0.368817 | 0.6916293 | 0.9056162 | 0.365139 | 0.567001 |
| ENSG00000213706 | RHOG2P | 1.604881 | 0.3688115 | 0.7547308 | 0.9642037 | 0.3349438 | 0.536221 |
| ENSG00000230149 | | 19.4796 | 0.3687475 | 0.4087997 | 1.044148 | 0.296417 | 0.495439 |
| ENSG00000027697 | IFNGR1 | 2174.005099 | 0.368728657 | 0.185736802 | 2.050152452 | 0.04034956 | 0.117678 |
| ENSG00000260176 | | 18.5641 | 0.3685837 | 0.4204558 | 1.034961 | 0.300687 | 0.499966 |
| ENSG00000269904 | MAP2K4P1 | 49.68277 | 0.3685554 | 0.3961654 | 1.064905 | 0.2869189 | 0.485273 |
| ENSG00000210191 | MT-TL2 | 1.487103 | 0.3684462 | 0.8369148 | 1.173452 | 0.2406146 | NA |
| ENSG00000271933 | | 116.2087 | 0.368354 | 0.3629053 | 1.140468 | 0.2540914 | 0.447083 |
| ENSG00000119280 | C1orf198 | 2937.350773 | 0.368275236 | 0.178792128 | 2.116533264 | 0.03429948 | 0.104087 |
| ENSG00000237961 | | 2.702843 | 0.368173 | 0.7254638 | 0.9250553 | 0.3549371 | 0.556901 |
| ENSG00000266473 | | 54.10971 | 0.3681591 | 0.3773197 | 1.10754 | 0.2680605 | 0.46398 |
| ENSG00000235091 | | 2.742958 | 0.3680613 | 0.7645002 | 0.982091 | 0.326055 | 0.527279 |
| ENSG00000232654 | FAM136BP | 3.09404 | 0.367997 | 0.7037284 | 0.9107121 | 0.3624471 | 0.56447 |
| ENSG00000229917 | RPL7P46 | 10.18891 | 0.3679907 | 0.5899357 | 0.8889797 | 0.374014 | 0.576135 |
| ENSG00000038219 | BOD1L1 | 3329.918105 | 0.367922544 | 0.254638456 | 1.572125109 | 0.11592154 | 0.258773 |
| ENSG00000167996 | FTH1 | 5084.625 | 0.3679153 | 0.2432886 | 1.589959 | 0.1118442 | 0.252394 |
| ENSG00000271013 | LRRC37A9P | 28.66507 | 0.3678995 | 0.6439795 | 0.8871544 | 0.3749958 | 0.577146 |
| ENSG00000258733 | LINC02328 | 55.76711 | 0.3677535 | 0.3599408 | 1.141367 | 0.2537174 | 0.446727 |
| ENSG00000213471 | TTLL13P | 3.632392 | 0.3677369 | 0.7097433 | 0.9012801 | 0.3674394 | 0.569506 |
| ENSG00000184007 | PTP4A2 | 12793.72 | 0.3676818 | 0.1258648 | 2.969105 | 0.00298668 | 0.01451 |
| ENSG00000229334 | | 47.51859 | 0.3675943 | 0.3173459 | 1.270138 | 0.2040354 | 0.385728 |
| ENSG00000136895 | GARNL3 | 334.3201453 | 0.367338037 | 0.273778636 | 1.433389114 | 0.15174667 | 0.313755 |
| ENSG00000285873 | | 18.03474 | 0.3673312 | 0.4911498 | 0.937302 | 0.3486032 | 0.550296 |

| | | | | | | | |
|---|---|---|---|---|---|---|---|
| ENSG00000281691 | RBM5-AS1 | 5.875534 | 0.3672039 | 0.5787095 | 0.8811822 | 0.3782192 | 0.580291 |
| ENSG00000242992 | FTH1P4 | 5.966967 | 0.3671604 | 0.656943 | 0.8866442 | 0.3752705 | 0.577435 |
| ENSG00000230953 | | 27.87365 | 0.3670392 | 0.4742867 | 0.9521447 | 0.3410236 | 0.542359 |
| ENSG00000238840 | | 13.4513 | 0.367021 | 0.699854 | 0.9079903 | 0.3638834 | 0.565977 |
| ENSG00000072415 | MPP5 | 3131.879804 | 0.36692095 | 0.157183113 | 2.380357407 | 0.01729585 | 0.060449 |
| ENSG00000200714 | | 5.042564 | 0.3669127 | 0.5960899 | 0.8738232 | 0.3822145 | 0.584531 |
| ENSG00000012822 | CALCOCO1 | 4703.721358 | 0.366791306 | 0.365218806 | 1.132048749 | 0.25761394 | 0.451607 |
| ENSG00000176454 | LPCAT4 | 1620.929 | 0.3667522 | 0.2116977 | 1.801034 | 0.07169757 | 0.181918 |
| ENSG00000134262 | AP4B1 | 4582.449839 | 0.366728907 | 0.153547523 | 2.427268808 | 0.01521298 | 0.054528 |
| ENSG00000269713 | NBPF9 | 4127.873 | 0.3665635 | 0.1803279 | 2.091924 | 0.03644529 | 0.108812 |
| ENSG00000240023 | | 2.297142 | 0.366525 | 0.7518489 | 0.9527824 | 0.3407003 | 0.542072 |
| ENSG00000272969 | | 7.197957 | 0.3662671 | 0.6412948 | 0.8843477 | 0.3765085 | 0.578617 |
| ENSG00000202515 | VTRNA1-3 | 5.83071 | 0.3662657 | 0.6880488 | 0.9098427 | 0.3629055 | 0.564919 |
| ENSG00000271344 | | 8.59316 | 0.3662215 | 0.6763483 | 0.8804908 | 0.3785935 | 0.580732 |
| ENSG00000228802 | | 1.442729 | 0.3662159 | 0.8121178 | 1.177322 | 0.2390668 | NA |
| ENSG00000248544 | | 3.694816 | 0.3662008 | 0.7590191 | 0.9782159 | 0.3279676 | 0.529326 |
| ENSG00000231125 | | 11.6243 | 0.3661731 | 0.592442 | 0.8729024 | 0.3827163 | 0.584963 |
| ENSG00000269246 | | 31.76367 | 0.3661579 | 0.3879318 | 1.079265 | 0.2804695 | 0.477924 |
| ENSG00000147130 | ZMYM3 | 4561.066 | 0.3661553 | 0.1327716 | 2.793067 | 0.00522109 | 0.022959 |
| ENSG00000129003 | VPS13C | 5172.637592 | 0.366100004 | 0.390376156 | 1.069718408 | 0.28474608 | 0.482673 |
| ENSG00000164144 | ARFIP1 | 3173.38 | 0.3660592 | 0.1118355 | 3.30642 | 0.00094496 | 0.00544 |
| ENSG00000224245 | | 1.111005 | 0.366004 | 0.8375052 | 1.171771 | 0.2412889 | NA |
| ENSG00000224893 | LINC02840 | 6.191093 | 0.3656793 | 0.9195637 | 2.655055 | 0.00792954 | 0.032323 |
| ENSG00000188070 | C11orf95 | 1154.041 | 0.3656491 | 0.1797977 | 2.087841 | 0.0368122 | 0.109648 |
| ENSG00000218891 | ZNF579 | 868.9304 | 0.3656335 | 0.1924085 | 1.973552 | 0.04843267 | 0.135212 |
| ENSG00000228570 | NUTM2E | 0.8259875 | 0.3654891 | 0.883985 | 1.431188 | 0.1523763 | NA |
| ENSG00000228169 | PPIAP19 | 2.652956 | 0.3654411 | 0.7206034 | 0.9317226 | 0.3514799 | 0.553161 |
| ENSG00000284624 | | 5.782662 | 0.3653446 | 0.6226981 | 0.8712917 | 0.3835949 | 0.586001 |
| ENSG00000164465 | DCBLD1 | 1088.756 | 0.3653031 | 0.2087986 | 1.817387 | 0.06915794 | 0.176968 |
| ENSG00000170153 | RNF150 | 423.9261 | 0.3652264 | 0.3076296 | 1.283897 | 0.1991781 | 0.379298 |
| ENSG00000285958 | | 2.056902 | 0.3651418 | 0.7590102 | 0.9509024 | 0.3416539 | 0.54292 |
| ENSG00000117016 | RIMS3 | 356.3672466 | 0.365140561 | 0.325070314 | 1.224692983 | 0.22069092 | 0.407046 |
| ENSG00000256747 | | 0.9011069 | 0.3651396 | 0.8236928 | 1.106579 | 0.268476 | NA |

| | | | | | | | |
|---|---|---|---|---|---|---|---|
| ENSG00000161202 | DVL3 | 8406.071 | 0.3650876 | 0.1260945 | 2.929891 | 0.00339081 | 0.016093 |
| ENSG00000105849 | POLR1F | 1534.951455 | 0.365043932 | 0.174712933 | 2.144058135 | 0.03202823 | 0.098662 |
| ENSG00000220785 | MTMR9LP | 196.8353 | 0.3650262 | 0.341298 | 1.180694 | 0.2377243 | 0.427312 |
| ENSG00000224209 | LINC00466 | 1.501957 | 0.3648055 | 0.8056048 | 1.076268 | 0.2818076 | NA |
| ENSG00000179674 | ARL14 | 1.742219 | 0.3647286 | 0.8387124 | 1.180922 | 0.2376336 | 0.427295 |
| ENSG00000082438 | COBLL1 | 1505.722206 | 0.364609882 | 0.279414342 | 1.407069609 | 0.15940675 | 0.32549 |
| ENSG00000160753 | RUSC1 | 8146.4 | 0.3645976 | 0.2710381 | 1.431653 | 0.152243 | 0.314441 |
| ENSG00000186767 | SPIN4 | 738.9106 | 0.3645821 | 0.2954791 | 1.333857 | 0.1822508 | 0.357228 |
| ENSG00000111786 | SRSF9 | 5804.392569 | 0.364534232 | 0.174815799 | 2.167360396 | 0.03020739 | 0.094155 |
| ENSG00000101109 | STK4 | 3154.816019 | 0.364476777 | 0.182890481 | 2.051375862 | 0.04023036 | 0.117421 |
| ENSG00000247081 | BAALC-AS1 | 380.5801 | 0.36439 | 0.1837421 | 2.038204 | 0.04152955 | 0.120432 |
| ENSG00000104228 | TRIM35 | 1574.799066 | 0.364265685 | 0.170230249 | 2.171681266 | 0.02987971 | 0.093364 |
| ENSG00000285473 | | 2.32484 | 0.3640084 | 0.7738759 | 0.9860166 | 0.324125 | 0.525006 |
| ENSG00000043355 | ZIC2 | 234.4186818 | 0.363985752 | 0.301255949 | 1.308298931 | 0.19077194 | 0.368821 |
| ENSG00000172273 | HINFP | 2366.729 | 0.3639399 | 0.1265924 | 2.906518 | 0.00365475 | 0.017123 |
| ENSG00000233823 | | 10.90923 | 0.3636918 | 0.6207062 | 0.8573778 | 0.3912361 | 0.593059 |
| ENSG00000286585 | | 89.95585 | 0.3636432 | 0.2486563 | 1.53973 | 0.1236261 | 0.271143 |
| ENSG00000236432 | MFF-DT | 32.76112 | 0.3633354 | 0.3885609 | 1.064498 | 0.2871033 | 0.485399 |
| ENSG00000135049 | AGTPBP1 | 1271.538915 | 0.363157476 | 0.189242548 | 1.978319526 | 0.04789267 | 0.134042 |
| ENSG00000140463 | BBS4 | 1575.368 | 0.3629672 | 0.1630578 | 2.280421 | 0.02258276 | 0.075019 |
| ENSG00000159596 | TMEM69 | 2528.821 | 0.3628122 | 0.1412244 | 2.612364 | 0.00899186 | 0.035737 |
| ENSG00000099977 | DDT | 7.73816505 | 0.362789775 | 0.608443445 | 0.882431008 | 0.37754377 | 0.579547 |
| ENSG00000277741 | GOLGA6L17P | 7.76392 | 0.362771 | 0.5159193 | 0.9061492 | 0.3648569 | 0.566728 |
| ENSG00000134250 | NOTCH2 | 11329.88396 | 0.362765639 | 0.279325302 | 1.387856513 | 0.16518074 | 0.33369 |
| ENSG00000257175 | | 20.35858 | 0.362601 | 0.4295962 | 1.007539 | 0.3136759 | 0.514821 |
| ENSG00000219928 | | 176.9051 | 0.3625955 | 0.2757634 | 1.405954 | 0.1597377 | 0.32584 |
| ENSG00000239828 | | 0.8340615 | 0.3625465 | 0.8762366 | 1.334118 | 0.1821651 | NA |
| ENSG00000102038 | SMARCA1 | 1480.678809 | 0.3623707 | 0.320744136 | 1.246378878 | 0.21262533 | 0.397274 |
| ENSG00000116918 | TSNAX | 1499.243179 | 0.362363399 | 0.134111249 | 2.735780154 | 0.00622326 | 0.026498 |
| ENSG00000279908 | | 29.70064 | 0.3623365 | 0.4699893 | 0.9479486 | 0.3431556 | 0.544618 |
| ENSG00000262877 | | 363.5356 | 0.3623141 | 0.3100629 | 1.259942 | 0.2076904 | 0.390332 |
| ENSG00000177570 | SAMD12 | 1912.385 | 0.3622844 | 0.2565849 | 1.482707 | 0.1381523 | 0.293059 |
| ENSG00000253313 | C1orf210 | 638.9649 | 0.3622245 | 0.2225656 | 1.6935 | 0.0903604 | 0.216282 |

| | | | | | | | |
|---|---|---|---|---|---|---|---|
| ENSG00000163950 | SLBP | 2446.978 | 0.3618009 | 0.270056 | 1.434084 | 0.1515483 | 0.313435 |
| ENSG00000240793 | UBA52P8 | 7.405845 | 0.3617579 | 0.5983278 | 0.8638294 | 0.3876816 | 0.58987 |
| ENSG00000286089 | | 1.783634 | 0.3617473 | 0.803024 | 1.048488 | 0.294414 | 0.493443 |
| ENSG00000182150 | ERCC6L2 | 1591.66 | 0.3617204 | 0.1489221 | 2.471511 | 0.01345434 | 0.049367 |
| ENSG00000077549 | CAPZB | 6294.852324 | 0.361694342 | 0.242049716 | 1.612444368 | 0.10686528 | 0.244197 |
| ENSG00000182871 | COL18A1 | 8600.439 | 0.3616368 | 0.2095052 | 1.788502 | 0.07369504 | 0.185822 |
| ENSG00000142599 | RERE | 4027.84 | 0.3615732 | 0.1710677 | 2.1669 | 0.03024251 | 0.094237 |
| ENSG00000229380 | PRKAR1B-AS2 | 1.3698 | 0.361504 | 0.8269148 | 1.153692 | 0.2486266 | NA |
| ENSG00000131037 | EPS8L1 | 2554.352616 | 0.361351814 | 0.23491561 | 1.596400415 | 0.11039942 | 0.250081 |
| ENSG00000258045 | | 6.205294 | 0.3612966 | 0.6320109 | 0.8750212 | 0.3815624 | 0.583769 |
| ENSG00000093144 | ECHDC1 | 3376.51469 | 0.361273177 | 0.153705042 | 2.387518593 | 0.01696255 | 0.05952 |
| ENSG00000142444 | TIMM29 | 1513.357 | 0.3612502 | 0.1638757 | 2.259585 | 0.02384705 | 0.078163 |
| ENSG00000256087 | ZNF432 | 397.0431 | 0.3609875 | 0.1825519 | 2.041299 | 0.0412211 | 0.119786 |
| ENSG00000197769 | MAP1LC3C | 1.655493 | 0.3609762 | 0.8588213 | 1.122064 | 0.2618354 | 0.456627 |
| ENSG00000135116 | HRK | 192.2358021 | 0.360897132 | 0.641624142 | 0.857782504 | 0.39101257 | 0.592869 |
| ENSG00000099984 | GSTT2 | 1.330363875 | 0.360871735 | 0.830086073 | 1.139925695 | 0.25431726 | NA |
| ENSG00000128165 | ADM2 | 1255.91234 | 0.360694773 | 0.579794962 | 0.864797429 | 0.38714998 | 0.589456 |
| ENSG00000202290 | RNA5SP37 | 3.2465 | 0.3604575 | 0.7310124 | 0.9265793 | 0.354145 | 0.55617 |
| ENSG00000277621 | | 5.154112 | 0.3604243 | 0.6021651 | 0.8625421 | 0.3883893 | 0.590521 |
| ENSG00000165806 | CASP7 | 3958.077 | 0.3603791 | 0.1593374 | 2.294265 | 0.0217753 | 0.072856 |
| ENSG00000130830 | MPP1 | 5.826585512 | 0.360279581 | 0.763199446 | 0.980368424 | 0.32690429 | 0.528185 |
| ENSG00000100883 | SRP54 | 5991.1764 | 0.360245608 | 0.0891223 | 4.044239864 | 5.25E-05 | 0.000432 |
| ENSG00000280063 | | 183.6944 | 0.3600959 | 0.2570087 | 1.482894 | 0.1381027 | 0.293024 |
| ENSG00000280502 | | 10.04899 | 0.3600352 | 0.566832 | 0.8800595 | 0.3788271 | 0.580989 |
| ENSG00000001617 | SEMA3F | 4718.998221 | 0.359985928 | 0.184324893 | 2.00055979 | 0.04543985 | 0.128961 |
| ENSG00000196597 | ZNF782 | 322.0594 | 0.3598554 | 0.1757182 | 2.10402 | 0.0353767 | 0.106435 |
| ENSG00000168228 | ZCCHC4 | 340.8718 | 0.3598183 | 0.1749556 | 2.115071 | 0.03442388 | 0.104384 |
| ENSG00000109163 | GNRHR | 9.830790344 | 0.359805957 | 0.555742866 | 0.870304739 | 0.38413389 | 0.586492 |
| ENSG00000163961 | RNF168 | 1547.761 | 0.3597872 | 0.224273 | 1.672734 | 0.09437976 | 0.223213 |
| ENSG00000251179 | TMEM92-AS1 | 3.023699 | 0.3596439 | 0.7567161 | 0.9718003 | 0.3311499 | 0.532167 |
| ENSG00000123130 | ACOT9 | 1728.93487 | 0.359605862 | 0.158030803 | 2.320649411 | 0.02030577 | 0.068797 |
| ENSG00000275340 | FGD5P1 | 1.760746 | 0.3594964 | 0.8181836 | 1.093476 | 0.2741847 | 0.470902 |
| ENSG00000162511 | LAPTM5 | 2.851319 | 0.359455 | 0.7895732 | 1.009045 | 0.312953 | 0.514024 |

| | | | | | | | |
|---|---|---|---|---|---|---|---|
| ENSG00000115806 | GORASP2 | 6904.073826 | 0.359419639 | 0.151698447 | 2.333068779 | 0.01964454 | 0.067035 |
| ENSG00000198315 | ZKSCAN8 | 2653.859 | 0.3594086 | 0.2666105 | 1.431664 | 0.1522401 | 0.314441 |
| ENSG00000198933 | TBKBP1 | 1706.349 | 0.3593214 | 0.2507841 | 1.528759 | 0.1263243 | 0.275029 |
| ENSG00000236434 | | 2.429712 | 0.3593155 | 0.7556486 | 0.9555815 | 0.3392837 | 0.540624 |
| ENSG00000241962 | | 15.77286 | 0.3593045 | 0.4743717 | 0.9329105 | 0.3508662 | 0.552425 |
| ENSG00000247572 | CKMT2-AS1 | 418.326 | 0.3592316 | 0.1619349 | 2.279474 | 0.02263892 | 0.075177 |
| ENSG00000248555 | LINC01339 | 0.8616237 | 0.3592102 | 0.8563838 | 1.331917 | 0.1828875 | NA |
| ENSG00000182197 | EXT1 | 1474.517 | 0.3591902 | 0.2090756 | 1.785721 | 0.0741445 | 0.186619 |
| ENSG00000089063 | TMEM230 | 2901.830951 | 0.359185665 | 0.169925289 | 2.172406202 | 0.02982504 | 0.093259 |
| ENSG00000100731 | PCNX1 | 5316.387876 | 0.359082545 | 0.232672625 | 1.616266861 | 0.10603663 | 0.242828 |
| ENSG00000182362 | YBEY | 625.0852 | 0.359072 | 0.2284082 | 1.642052 | 0.1005791 | 0.233777 |
| ENSG00000188846 | RPL14 | 16042.18 | 0.3589585 | 0.170371 | 2.1457 | 0.03189692 | 0.098389 |
| ENSG00000095015 | MAP3K1 | 2576.108776 | 0.358842491 | 0.243208884 | 1.582246631 | 0.11359328 | 0.254925 |
| ENSG00000160305 | DIP2A | 4612.421 | 0.3588339 | 0.201263 | 1.846136 | 0.06487247 | 0.168419 |
| ENSG00000229503 | RPL21P37 | 1.462184 | 0.358833 | 0.801126 | 1.036718 | 0.2998673 | NA |
| ENSG00000173575 | CHD2 | 8584.978 | 0.3587761 | 0.168272 | 2.18256 | 0.02906826 | 0.091593 |
| ENSG00000052723 | SIKE1 | 13188.94149 | 0.358476446 | 0.16050849 | 2.252631889 | 0.02428237 | 0.079277 |
| ENSG00000063169 | BICRA | 1312.893075 | 0.358399171 | 0.191885464 | 1.925242793 | 0.05419899 | 0.147618 |
| ENSG00000137726 | FXYD6 | 14.37354834 | 0.358297117 | 0.602428516 | 0.865704385 | 0.3866523 | 0.589061 |
| ENSG00000246575 | | 195.8989 | 0.3581914 | 0.3617571 | 1.109519 | 0.2672065 | 0.463015 |
| ENSG00000135124 | P2RX4 | 1908.706289 | 0.358174081 | 0.212047305 | 1.756256899 | 0.07904455 | 0.195416 |
| ENSG00000277039 | | 0.909963 | 0.3581346 | 0.8155621 | 1.093226 | 0.2742945 | NA |
| ENSG00000272004 | | 3.173425 | 0.3581263 | 0.7271478 | 0.9095951 | 0.3630361 | 0.565035 |
| ENSG00000245970 | | 350.4504 | 0.3581136 | 0.1954544 | 1.892994 | 0.05835867 | 0.155913 |
| ENSG00000226677 | IGBP1P1 | 42.19713 | 0.3579979 | 0.2658227 | 1.431136 | 0.1523913 | 0.314674 |
| ENSG00000142252 | GEMIN7 | 100.7249 | 0.3579007 | 0.2786745 | 1.372247 | 0.1699867 | 0.340422 |
| ENSG00000267493 | CIRBP-AS1 | 127.9715 | 0.3578723 | 0.2905686 | 1.325955 | 0.1848546 | 0.360311 |
| ENSG00000149600 | COMMD7 | 1363.798 | 0.3578192 | 0.1583798 | 2.306205 | 0.02109918 | 0.070979 |
| ENSG00000250608 | | 35.27018 | 0.3577944 | 0.4046428 | 1.027104 | 0.3043716 | 0.503979 |
| ENSG00000261970 | | 1.356908 | 0.357747 | 0.8065281 | 1.038152 | 0.2991992 | NA |
| ENSG00000155034 | FBXL18 | 2454.155 | 0.3577252 | 0.2275748 | 1.628923 | 0.1033293 | 0.238332 |
| ENSG00000263843 | | 310.8222 | 0.357722 | 0.180455 | 2.040975 | 0.04125332 | 0.11984 |
| ENSG00000257267 | ZNF271P | 1684.648 | 0.3576092 | 0.1391083 | 2.602987 | 0.00924156 | 0.036501 |

| | | | | | | | |
|---|---|---|---|---|---|---|---|
| ENSG00000197540 | GZMM | 6.623501 | 0.3576008 | 0.6236565 | 0.8451619 | 0.3980205 | 0.599117 |
| ENSG00000064393 | HIPK2 | 3356.123383 | 0.357554092 | 0.33480859 | 1.176275178 | 0.23948493 | 0.429692 |
| ENSG00000015171 | ZMYND11 | 11343.96784 | 0.357464478 | 0.152747316 | 2.394945201 | 0.01662285 | 0.058551 |
| ENSG00000273233 | | 1.32471 | 0.3573362 | 0.8218327 | 1.102466 | 0.2702591 | NA |
| ENSG00000126773 | PCNX4 | 2965.507343 | 0.357287642 | 0.162315498 | 2.248231059 | 0.02456146 | 0.079953 |
| ENSG00000287898 | | 1.623507 | 0.3572752 | 0.8327947 | 1.149991 | 0.2501477 | 0.442695 |
| ENSG00000165655 | ZNF503 | 606.2953 | 0.3571476 | 0.2196482 | 1.703143 | 0.08854137 | 0.213065 |
| ENSG00000231903 | | 1.304764 | 0.3570455 | 0.8356169 | 1.295256 | 0.195232 | NA |
| ENSG00000140675 | SLC5A2 | 7.219636 | 0.357043 | 0.6484903 | 0.8505664 | 0.3950103 | 0.596734 |
| ENSG00000158458 | NRG2 | 13.17704 | 0.3568129 | 0.5220785 | 0.8925724 | 0.3720862 | 0.574298 |
| ENSG00000268941 | LINC01711 | 4.149691 | 0.356664 | 0.7087737 | 0.8996715 | 0.3682951 | 0.570302 |
| ENSG00000123349 | PFDN5 | 10441.1018 | 0.356520583 | 0.174909755 | 2.093498219 | 0.0363047 | 0.108705 |
| ENSG00000118292 | C1orf54 | 114.5202525 | 0.356511752 | 0.315999171 | 1.223198052 | 0.2212549 | 0.407778 |
| ENSG00000274286 | ADRA2B | 31.42435 | 0.356425 | 0.4839932 | 0.9459192 | 0.3441898 | 0.54583 |
| ENSG00000275162 | | 2.142221 | 0.3562102 | 0.7919451 | 1.065587 | 0.2866103 | 0.484936 |
| ENSG00000213621 | RPSAP54 | 77.51351 | 0.3561595 | 0.2503454 | 1.495432 | 0.1348016 | 0.288068 |
| ENSG00000130758 | MAP3K10 | 939.1054152 | 0.356017292 | 0.191915124 | 1.913547899 | 0.05567795 | 0.150423 |
| ENSG00000286373 | | 36.89018 | 0.3559668 | 0.4687979 | 0.9335919 | 0.3505145 | 0.55226 |
| ENSG00000072135 | PTPN18 | 2892.6168 | 0.355852448 | 0.155978285 | 2.277465039 | 0.02275847 | 0.075401 |
| ENSG00000214226 | C17orf67 | 398.3095 | 0.3557658 | 0.2992696 | 1.285231 | 0.1987116 | 0.37884 |
| ENSG00000175514 | GPR152 | 0.9663331 | 0.3557499 | 0.823153 | 1.087526 | 0.2768044 | NA |
| ENSG00000256196 | | 20.76721 | 0.3556026 | 0.483816 | 0.9135969 | 0.3609287 | 0.562947 |
| ENSG00000255991 | | 0.7928362 | 0.3553699 | 0.8383229 | 1.150041 | 0.2501268 | NA |
| ENSG00000106799 | TGFBR1 | 2000.092313 | 0.355338093 | 0.238308534 | 1.550989705 | 0.12090415 | 0.266736 |
| ENSG00000166135 | HIF1AN | 4582.288 | 0.3552858 | 0.1265976 | 2.829864 | 0.00465677 | 0.020935 |
| ENSG00000189334 | S100A14 | 13447.19 | 0.3552541 | 0.3601719 | 1.089446 | 0.2759574 | 0.472844 |
| ENSG00000155368 | DBI | 12219.16 | 0.3551269 | 0.3410049 | 1.205909 | 0.2278527 | 0.415813 |
| ENSG00000207151 | | 4.192067 | 0.3550973 | 0.6854537 | 0.8682548 | 0.3852549 | 0.587659 |
| ENSG00000225094 | SETP20 | 7.470241 | 0.355076 | 0.5954443 | 0.8534437 | 0.3934133 | 0.595115 |
| ENSG00000170876 | TMEM43 | 8650.131 | 0.3550344 | 0.189033 | 1.973301 | 0.04846124 | 0.135263 |
| ENSG00000134107 | BHLHE40 | 6793.135961 | 0.354902519 | 0.186675762 | 1.974744097 | 0.0482972 | 0.134904 |
| ENSG00000187118 | CMC1 | 1463.386 | 0.3548854 | 0.1741103 | 2.080608 | 0.03746984 | 0.111112 |
| ENSG00000226824 | | 115.3505 | 0.3548841 | 0.2751182 | 1.383152 | 0.1666181 | 0.335742 |

| | | | | | | | |
|---|---|---|---|---|---|---|---|
| ENSG00000266922 | | 1.298562 | 0.3547066 | 0.847102 | 1.228623 | 0.2192133 | NA |
| ENSG00000236754 | | 2.863301 | 0.3546968 | 0.6598449 | 0.862974 | 0.3881518 | 0.5904 |
| ENSG00000181467 | RAP2B | 2427.153 | 0.3546171 | 0.1759593 | 2.074632 | 0.03802067 | 0.112429 |
| ENSG00000267321 | SNHG30 | 488.6069 | 0.3545897 | 0.2463535 | 1.523711 | 0.1275808 | 0.276981 |
| ENSG00000103489 | XYLT1 | 1212.998189 | 0.354549311 | 0.295660861 | 1.291741982 | 0.19644651 | 0.376281 |
| ENSG00000224597 | SVIL-AS1 | 544.1432 | 0.3545238 | 0.1665549 | 2.175635 | 0.02958258 | 0.092774 |
| ENSG00000258657 | | 1.999748 | 0.3544038 | 0.8035404 | 1.066075 | 0.2863899 | 0.484687 |
| ENSG00000163568 | AIM2 | 16.56231 | 0.3543084 | 0.8437768 | 0.9965963 | 0.3189605 | 0.519945 |
| ENSG00000228521 | | 1.142153 | 0.3542712 | 0.8853917 | 1.55491 | 0.1199675 | NA |
| ENSG00000224183 | SDHDP6 | 4.584873 | 0.3540189 | 0.6741478 | 0.8589733 | 0.3903553 | 0.592276 |
| ENSG00000287200 | | 1.854271 | 0.3539102 | 0.7701803 | 0.951527 | 0.3413369 | 0.542533 |
| ENSG00000154767 | XPC | 3325.548 | 0.3538864 | 0.1979082 | 1.844643 | 0.06508953 | 0.168883 |
| ENSG00000278896 | | 3.085683 | 0.3538864 | 0.7188408 | 0.9030351 | 0.3665073 | 0.568428 |
| ENSG00000144451 | SPAG16 | 0.8617526 | 0.3535913 | 0.8570257 | 1.281236 | 0.2001106 | NA |
| ENSG00000263366 | ABHD17AP5 | 3.7421 | 0.3534879 | 0.650447 | 0.8552267 | 0.3924257 | 0.594061 |
| ENSG00000164941 | INTS8 | 4448.976 | 0.353169 | 0.1815398 | 2.022344 | 0.04314082 | 0.124049 |
| ENSG00000260174 | SYNGR2P1 | 20.14275 | 0.3531071 | 0.4856821 | 0.9121962 | 0.3616655 | 0.563616 |
| ENSG00000130193 | THEM6 | 8879.450208 | 0.352856762 | 0.13325448 | 2.68746769 | 0.00719961 | 0.029837 |
| ENSG00000241635 | UGT1A1 | 30.97752 | 0.3528324 | 0.839533 | 1.018542 | 0.3084206 | 0.508686 |
| ENSG00000115649 | CNPPD1 | 5039.659468 | 0.352749438 | 0.133934163 | 2.673176549 | 0.00751367 | 0.030912 |
| ENSG00000205702 | CYP2D7 | 183.5192 | 0.3526856 | 0.3147514 | 1.215139 | 0.224313 | 0.411132 |
| ENSG00000127220 | ABHD8 | 159.8284194 | 0.352678577 | 0.221479871 | 1.659115468 | 0.09709253 | 0.227663 |
| ENSG00000258682 | | 90.03587 | 0.3526683 | 0.2912076 | 1.300741 | 0.1933472 | 0.372007 |
| ENSG00000143819 | EPHX1 | 5339.084 | 0.3526427 | 0.2166344 | 1.701805 | 0.08879187 | 0.213572 |
| ENSG00000140297 | GCNT3 | 2.692575 | 0.3525409 | 0.7531543 | 0.9382877 | 0.3480966 | 0.549824 |
| ENSG00000265298 | | 632.8246 | 0.352503 | 0.2536277 | 1.479791 | 0.1389291 | 0.294319 |
| ENSG00000165521 | EML5 | 105.8473 | 0.352444 | 0.3585087 | 1.090484 | 0.2755 | 0.472365 |
| ENSG00000100650 | SRSF5 | 15834.61266 | 0.352437993 | 0.133107975 | 2.654029529 | 0.00795369 | 0.032384 |
| ENSG00000214534 | ZNF705E | 14.76996 | 0.3524078 | 0.4998401 | 0.8920825 | 0.3723487 | 0.574503 |
| ENSG00000226791 | LINC02611 | 2.174645 | 0.3523276 | 0.7443615 | 0.930711 | 0.3520031 | 0.553788 |
| ENSG00000224652 | LINC00885 | 142.7883 | 0.3523001 | 0.4098535 | 1.00495 | 0.3149208 | 0.51598 |
| ENSG00000204528 | PSORS1C3 | 1.862051 | 0.3522475 | 0.7602203 | 0.9366055 | 0.3489615 | 0.550666 |
| ENSG00000136816 | TOR1B | 2201.907109 | 0.352236789 | 0.104753037 | 3.39681753 | 0.00068174 | 0.004108 |

| | | | | | | | |
|---|---|---|---|---|---|---|---|
| ENSG00000178462 | TUBAL3 | 26.15566 | 0.3520773 | 0.5948904 | 0.8328233 | 0.4049444 | 0.605695 |
| ENSG00000053254 | FOXN3 | 2945.297147 | 0.351964319 | 0.300172904 | 1.271554904 | 0.20353131 | 0.385142 |
| ENSG00000257513 | NPIPB1P | 2.351693 | 0.351883 | 0.7834354 | 1.005875 | 0.3144759 | 0.515601 |
| ENSG00000225138 | SLC9A3-AS1 | 3794.635 | 0.3518568 | 0.2679583 | 1.3958 | 0.1627749 | 0.330236 |
| ENSG00000232119 | MCTS1 | 3718.423 | 0.3517882 | 0.1603372 | 2.262549 | 0.02366353 | 0.077658 |
| ENSG00000172426 | RSPH9 | 26.98342 | 0.3517008 | 0.4366491 | 0.9569764 | 0.3385791 | 0.539861 |
| ENSG00000234004 | | 10.23649 | 0.3516836 | 0.5071293 | 0.8822899 | 0.3776201 | 0.579564 |
| ENSG00000243746 | EEF1A1P10 | 2.787135 | 0.3516118 | 0.7072053 | 0.8792695 | 0.3792552 | 0.581243 |
| ENSG00000114331 | ACAP2 | 3141.631051 | 0.351530123 | 0.218244789 | 1.66761737 | 0.09539171 | 0.224792 |
| ENSG00000204661 | C5orf60 | 6.570571 | 0.3513737 | 0.6088158 | 0.8424028 | 0.3995625 | 0.600519 |
| ENSG00000227336 | LINC00434 | 0.9744639 | 0.351324 | 0.8382027 | 1.224492 | 0.2207667 | NA |
| ENSG00000213713 | PIGCP1 | 61.45492 | 0.3512436 | 0.3282136 | 1.179304 | 0.2382772 | 0.428074 |
| ENSG00000243696 | | 16.71716 | 0.3512221 | 0.4260906 | 0.9598795 | 0.3371159 | 0.538239 |
| ENSG00000150456 | EEF1AKMT1 | 652.6198 | 0.3510648 | 0.2114992 | 1.73282 | 0.08312759 | 0.203166 |
| ENSG00000238156 | | 1.1569 | 0.3509278 | 0.8543475 | 1.133701 | 0.2569202 | NA |
| ENSG00000229298 | TUBB8P1 | 41.86116 | 0.3508765 | 0.4750905 | 0.9102538 | 0.3626887 | 0.564747 |
| ENSG00000224072 | VEZTP1 | 3.884154 | 0.350415 | 0.6605447 | 0.8551085 | 0.3924911 | 0.59407 |
| ENSG00000157349 | DDX19B | 1080.401 | 0.3502847 | 0.2019589 | 1.795175 | 0.07262583 | 0.183684 |
| ENSG00000162885 | B3GALNT2 | 1451.232 | 0.3501217 | 0.1771821 | 1.985619 | 0.04707567 | 0.132343 |
| ENSG00000107262 | BAG1 | 6679.689979 | 0.350071243 | 0.30142013 | 1.257019284 | 0.20874665 | 0.391652 |
| ENSG00000280231 | | 4.435826 | 0.3500306 | 0.7249916 | 0.8921808 | 0.372296 | 0.574465 |
| ENSG00000186471 | AKAP14 | 1.159214 | 0.3499588 | 0.8703686 | 1.329168 | 0.1837925 | NA |
| ENSG00000235934 | | 6.379801 | 0.3498478 | 0.6530424 | 0.8348118 | 0.4038237 | 0.604652 |
| ENSG00000149743 | TRPT1 | 1561.606 | 0.3498391 | 0.1900789 | 1.870446 | 0.06142189 | 0.161865 |
| ENSG00000279591 | | 1.126861 | 0.3498269 | 0.8702927 | 1.329074 | 0.1838235 | NA |
| ENSG00000286355 | | 2.362275 | 0.3498251 | 0.7973943 | 1.027251 | 0.3043021 | 0.503958 |
| ENSG00000260886 | TAT-AS1 | 7.111426 | 0.3497647 | 0.6002013 | 0.8366844 | 0.40277 | 0.60369 |
| ENSG00000256588 | | 1.779356 | 0.3497644 | 0.7454218 | 0.9229469 | 0.3560349 | 0.558084 |
| ENSG00000280639 | LINC02204 | 1.106721 | 0.3495806 | 0.8274306 | 1.133215 | 0.2571241 | NA |
| ENSG00000237749 | | 15.73976 | 0.3495496 | 0.5231817 | 0.86593 | 0.3865286 | 0.588925 |
| ENSG00000284770 | TBCE | 94.52925 | 0.3495235 | 0.3121962 | 1.220679 | 0.2222076 | 0.408939 |
| ENSG00000261647 | IMPDH1P11 | 1.535541 | 0.3494394 | 0.8381216 | 1.261165 | 0.2072494 | NA |
| ENSG00000168348 | INSM2 | 1.121813 | 0.349435 | 0.8398508 | 1.183456 | 0.2366285 | NA |

| | | | | | | | |
|---|---|---|---|---|---|---|---|
| ENSG00000100522 | GNPNAT1 | 3701.224349 | 0.34933976 | 0.256486978 | 1.4446438 | 0.14855796 | 0.30905 |
| ENSG00000213892 | CEACAM16 | 1.070491 | 0.3492831 | 0.8785147 | 1.445909 | 0.1482028 | NA |
| ENSG00000196866 | H2AC7 | 2.38215 | 0.3492516 | 0.7467947 | 0.9311482 | 0.3517769 | 0.553498 |
| ENSG00000281005 | LINC00921 | 93.51646 | 0.3491934 | 0.3484045 | 1.115949 | 0.2644442 | 0.459938 |
| ENSG00000213701 | SETP22 | 0.8885032 | 0.3491844 | 0.8638399 | 1.238769 | 0.2154311 | NA |
| ENSG00000113905 | HRG | 1.379105523 | 0.349146396 | 0.857149835 | 1.103393322 | 0.2698564 | NA |
| ENSG00000269855 | RNF225 | 1.218545 | 0.349144 | 0.8443702 | 1.269152 | 0.2043868 | NA |
| ENSG00000121851 | POLR3GL | 497.0414733 | 0.349119682 | 0.191553058 | 1.870922249 | 0.06135586 | 0.161813 |
| ENSG00000169957 | ZNF768 | 3934.631 | 0.3489955 | 0.1539846 | 2.3115 | 0.02080528 | 0.070167 |
| ENSG00000277112 | ANKRD20A21P | 35.79449 | 0.3488958 | 0.3920574 | 1.017028 | 0.3091399 | 0.509657 |
| ENSG00000178381 | ZFAND2A | 2085.174 | 0.348792 | 0.1875776 | 1.919522 | 0.05491834 | 0.148982 |
| ENSG00000234353 | | 14.93845 | 0.3487485 | 0.5551247 | 0.8488489 | 0.3959654 | 0.597445 |
| ENSG00000257703 | | 1.209621 | 0.34873 | 0.8135547 | 1.113636 | 0.2654355 | NA |
| ENSG00000203985 | LDLRAD1 | 1.584206 | 0.3486685 | 0.7928278 | 1.039378 | 0.2986288 | 0.497846 |
| ENSG00000157335 | CLEC18C | 3.177725 | 0.3485648 | 0.7466203 | 0.9151992 | 0.360087 | 0.562079 |
| ENSG00000257621 | PSMA3-AS1 | 3975.635 | 0.3483797 | 0.2516995 | 1.476994 | 0.1396774 | 0.295704 |
| ENSG00000100575 | TIMM9 | 2373.499972 | 0.348208062 | 0.193393457 | 1.843358913 | 0.06527663 | 0.169269 |
| ENSG00000257556 | LINC02298 | 6.725527 | 0.3481852 | 0.7861951 | 1.053659 | 0.2920393 | 0.49083 |
| ENSG00000108861 | DUSP3 | 5222.010878 | 0.348137071 | 0.255362156 | 1.440636918 | 0.14968729 | 0.310682 |
| ENSG00000201820 | | 3.847771 | 0.3480983 | 0.6789268 | 0.8512911 | 0.3946077 | 0.596412 |
| ENSG00000259657 | PIGHP1 | 35.53591 | 0.3480669 | 0.3282419 | 1.162003 | 0.2452342 | 0.436968 |
| ENSG00000042088 | TDP1 | 2376.855625 | 0.347948021 | 0.195957515 | 1.838037875 | 0.06605682 | 0.170892 |
| ENSG00000105771 | SMG9 | 2001.632461 | 0.347925234 | 0.204550625 | 1.759641077 | 0.07846868 | 0.19441 |
| ENSG00000153485 | TMEM251 | 1263.266 | 0.347911 | 0.1463999 | 2.413401 | 0.01580442 | 0.056175 |
| ENSG00000267113 | | 4.238106 | 0.347885 | 0.6693714 | 0.8397633 | 0.4010411 | 0.601958 |
| ENSG00000237751 | LINC01143 | 15.03556 | 0.3476105 | 0.4352225 | 0.948422 | 0.3429147 | 0.544454 |
| ENSG00000198131 | ZNF544 | 1702.759 | 0.3476056 | 0.1628901 | 2.184181 | 0.02894894 | 0.091261 |
| ENSG00000228218 | ATF4P3 | 46.32108 | 0.3475361 | 0.4143528 | 0.980517 | 0.326831 | 0.528134 |
| ENSG00000285672 | | 4.708943 | 0.3474373 | 0.6502316 | 0.842636 | 0.3994321 | 0.600486 |
| ENSG00000167363 | FN3K | 808.7461 | 0.3474113 | 0.1642829 | 2.158473 | 0.03089108 | 0.096016 |
| ENSG00000232238 | RPS29P8 | 1.283951 | 0.3473375 | 0.8205637 | 1.097559 | 0.2723969 | NA |
| ENSG00000253706 | | 1.845161 | 0.3473245 | 0.7741632 | 0.9554639 | 0.3393431 | 0.540625 |
| ENSG00000172992 | DCAKD | 2075.065 | 0.347321 | 0.1905585 | 1.867303 | 0.06185931 | 0.162711 |

| | | | | | | | |
|---|---|---|---|---|---|---|---|
| ENSG00000189229 | | 11.73519 | 0.3473155 | 0.8416142 | 0.9632729 | 0.3354106 | 0.536666 |
| ENSG00000262049 | | 428.8298 | 0.3473128 | 0.2449296 | 1.487473 | 0.13689 | 0.291166 |
| ENSG00000229065 | | 4.631169 | 0.3470551 | 0.6745331 | 0.8586598 | 0.3905282 | 0.592403 |
| ENSG00000270614 | | 0.8636202 | 0.3470549 | 0.8736027 | 1.375234 | 0.169059 | NA |
| ENSG00000275070 | | 0.8537194 | 0.3469934 | 0.8638856 | 1.216961 | 0.2236189 | NA |
| ENSG00000229739 | PDC-AS1 | 21.96742 | 0.3469273 | 0.4850946 | 0.8912354 | 0.3728029 | 0.575003 |
| ENSG00000200959 | SNORA74A | 4.37546 | 0.3468879 | 0.6939131 | 0.8573512 | 0.3912508 | 0.593059 |
| ENSG00000112249 | ASCC3 | 4746.804396 | 0.346649189 | 0.191090884 | 1.875836858 | 0.06067769 | 0.160349 |
| ENSG00000214429 | CYCSP6 | 10.80173 | 0.346648 | 0.8404889 | 0.9749172 | 0.3296014 | 0.530716 |
| ENSG00000174527 | MYO1H | 9.86783 | 0.3466062 | 0.601531 | 0.8412253 | 0.4002217 | 0.601197 |
| ENSG00000263011 | | 14.87624 | 0.3466053 | 0.4396098 | 0.938247 | 0.3481175 | 0.549824 |
| ENSG00000101236 | RNF24 | 1494.787195 | 0.346568638 | 0.179175557 | 1.998188071 | 0.04569627 | 0.129477 |
| ENSG00000266865 | | 83.28327 | 0.3465397 | 0.2109838 | 1.705853 | 0.08803543 | 0.212136 |
| ENSG00000163539 | CLASP2 | 2467.307 | 0.3465219 | 0.1931234 | 1.858437 | 0.06310691 | 0.165058 |
| ENSG00000101846 | STS | 1617.039638 | 0.346519546 | 0.306094141 | 1.230811566 | 0.21839335 | 0.40413 |
| ENSG00000214093 | | 2.024373 | 0.3464696 | 0.7640941 | 0.9490965 | 0.3425716 | 0.544072 |
| ENSG00000149932 | TMEM219 | 3312.582 | 0.346411 | 0.2038648 | 1.758037 | 0.07874114 | 0.194832 |
| ENSG00000249288 | SLC25A15P5 | 0.8584717 | 0.3463951 | 0.8517919 | 1.366621 | 0.1717443 | NA |
| ENSG00000154928 | EPHB1 | 33.62706 | 0.3463309 | 0.4948513 | 0.8839225 | 0.376738 | 0.578788 |
| ENSG00000118322 | ATP10B | 43.49681248 | 0.346304883 | 0.524710596 | 0.864144924 | 0.38750828 | 0.589775 |
| ENSG00000170485 | NPAS2 | 540.2637 | 0.3462798 | 0.2600145 | 1.413616 | 0.1574748 | 0.322612 |
| ENSG00000182979 | MTA1 | 8699.899 | 0.3462655 | 0.1249567 | 2.786473 | 0.00532851 | 0.023323 |
| ENSG00000166557 | TMED3 | 11047.77 | 0.3462634 | 0.1500166 | 2.347389 | 0.01890553 | 0.064947 |
| ENSG00000271868 | | 4.087496 | 0.34621 | 0.6462906 | 0.8295997 | 0.4067652 | 0.607427 |
| ENSG00000166199 | ALKBH3 | 679.7063 | 0.3462048 | 0.1818361 | 1.949136 | 0.05127916 | 0.141273 |
| ENSG00000230910 | | 6.732157 | 0.3460953 | 0.5915091 | 0.8281205 | 0.4076022 | 0.608267 |
| ENSG00000169253 | | 23.09762 | 0.3460586 | 0.4569242 | 0.9194745 | 0.3578474 | 0.559815 |
| ENSG00000188710 | QRFP | 14.80075 | 0.3460347 | 0.5054026 | 0.8680886 | 0.3853458 | 0.587757 |
| ENSG00000185033 | SEMA4B | 3688.422 | 0.3460271 | 0.1799003 | 1.964408 | 0.04948275 | 0.137493 |
| ENSG00000163624 | CDS1 | 3556.502 | 0.34597 | 0.1882504 | 1.89768 | 0.0577382 | 0.154595 |
| ENSG00000134668 | SPOCD1 | 380.7554495 | 0.34592987 | 0.428449553 | 0.968412972 | 0.33283816 | 0.533847 |
| ENSG00000010017 | RANBP9 | 3542.033512 | 0.345928602 | 0.172872217 | 2.045146332 | 0.04084043 | 0.11894 |
| ENSG00000271394 | | 1.647963 | 0.345917 | 0.8517734 | 1.124452 | 0.2608211 | 0.455545 |

| | | | | | | | |
|---|---|---|---|---|---|---|---|
| ENSG00000228208 | LINC02869 | 5.224313 | 0.3458977 | 0.7906872 | 1.055534 | 0.2911812 | 0.4898 |
| ENSG00000144229 | THSD7B | 3.680635 | 0.3458409 | 0.7528104 | 0.9322401 | 0.3512124 | 0.552871 |
| ENSG00000237877 | LINC01473 | 27.61251 | 0.3458321 | 0.3664857 | 1.061154 | 0.28862 | 0.487157 |
| ENSG00000100916 | BRMS1L | 1181.756627 | 0.34579201 | 0.185612605 | 1.921456733 | 0.05467415 | 0.148501 |
| ENSG00000257426 | | 2.507807 | 0.3457011 | 0.7666654 | 0.9529607 | 0.34061 | 0.541994 |
| ENSG00000080189 | SLC35C2 | 4776.482901 | 0.34568423 | 0.136263137 | 2.573122716 | 0.01007855 | 0.03912 |
| ENSG00000242852 | ZNF709 | 12.46308 | 0.3454092 | 0.5636283 | 0.8374192 | 0.402357 | 0.603286 |
| ENSG00000250983 | | 10.84906 | 0.3453147 | 0.4525161 | 0.9241284 | 0.3554194 | 0.557414 |
| ENSG00000148848 | ADAM12 | 4.326146 | 0.3452555 | 0.679095 | 0.8505827 | 0.3950012 | 0.596734 |
| ENSG00000198417 | MT1F | 147.1245 | 0.3452103 | 0.4788989 | 0.8895193 | 0.373724 | 0.575822 |
| ENSG00000032742 | IFT88 | 947.6281654 | 0.345166776 | 0.124784878 | 2.792307352 | 0.00523336 | 0.022998 |
| ENSG00000138709 | LARP1B | 2265.755947 | 0.345129674 | 0.167014493 | 2.123046072 | 0.03374999 | 0.10275 |
| ENSG00000122545 | SEPTIN7 | 9502.446064 | 0.34512497 | 0.134723905 | 2.617618049 | 0.00885459 | 0.035297 |
| ENSG00000108602 | ALDH3A1 | 139.2859041 | 0.345061863 | 0.834281254 | 1.012973754 | 0.3110727 | 0.511916 |
| ENSG00000145832 | SLC25A48 | 3.168421 | 0.3450569 | 0.7601092 | 0.9553579 | 0.3393968 | 0.540678 |
| ENSG00000083817 | ZNF416 | 334.8738111 | 0.344977753 | 0.180219599 | 1.96663803 | 0.04922496 | 0.136934 |
| ENSG00000288005 | | 1.433917 | 0.3449621 | 0.8113423 | 1.048246 | 0.2945255 | NA |
| ENSG00000100612 | DHRS7 | 2814.344125 | 0.344931407 | 0.275104435 | 1.337817301 | 0.180956 | 0.355477 |
| ENSG00000232934 | | 2.498028 | 0.3447586 | 0.7683046 | 0.9558095 | 0.3391685 | 0.540484 |
| ENSG00000179119 | SPTY2D1 | 2182.312 | 0.3446369 | 0.2169692 | 1.644413 | 0.100091 | 0.232986 |
| ENSG00000139835 | GRTP1 | 1303.374 | 0.3446067 | 0.1320926 | 2.622209 | 0.00873619 | 0.034925 |
| ENSG00000128951 | DUT | 2370.792546 | 0.344537451 | 0.301230993 | 1.225926138 | 0.22022647 | 0.406395 |
| ENSG00000142867 | BCL10 | 832.7369 | 0.3445137 | 0.2629685 | 1.384172 | 0.1663056 | 0.335337 |
| ENSG00000277782 | | 77.35916 | 0.3444921 | 0.2590139 | 1.413943 | 0.1573787 | 0.322488 |
| ENSG00000271714 | | 8.922288 | 0.3444918 | 0.6174081 | 0.83093 | 0.4060132 | 0.606697 |
| ENSG00000234318 | | 1.33423 | 0.3443798 | 0.8187205 | 1.196064 | 0.2316716 | NA |
| ENSG00000260281 | ITFG1-AS1 | 5.043299 | 0.3443755 | 0.6286971 | 0.8362302 | 0.4030254 | 0.603947 |
| ENSG00000071909 | MYO3B | 4.046428654 | 0.344346173 | 0.722660791 | 0.879057895 | 0.37936989 | 0.581345 |
| ENSG00000250673 | REELD1 | 2.164167 | 0.3443377 | 0.7619296 | 0.931832 | 0.3514234 | 0.553105 |
| ENSG00000244691 | RPL10AP1 | 2.190254 | 0.3443305 | 0.7532477 | 0.9238941 | 0.3555414 | 0.557506 |
| ENSG00000224712 | NPIPA3 | 2.479822 | 0.344281 | 0.7168607 | 0.8738961 | 0.3821748 | 0.584531 |
| ENSG00000280670 | CCDC163 | 194.109 | 0.344209 | 0.2049028 | 1.740247 | 0.08181572 | 0.200772 |
| ENSG00000235192 | | 9.234717 | 0.344202 | 0.567649 | 0.8310662 | 0.4059362 | 0.606687 |

| | | | | | | | |
|---|---|---|---|---|---|---|---|
| ENSG00000167733 | HSD11B1L | 235.1057 | 0.3441809 | 0.3125386 | 1.201323 | 0.2296259 | 0.417815 |
| ENSG00000280319 | | 1.159607 | 0.3440393 | 0.8437436 | 1.273388 | 0.2028805 | NA |
| ENSG00000149557 | FEZ1 | 296.9726 | 0.3439507 | 0.2649525 | 1.378672 | 0.1679959 | 0.337657 |
| ENSG00000104218 | CSPP1 | 1281.016872 | 0.343929819 | 0.160281932 | 2.173492772 | 0.02974325 | 0.093115 |
| ENSG00000185633 | NDUFA4L2 | 165.4876 | 0.3438419 | 0.6894686 | 0.8440559 | 0.3986381 | 0.599605 |
| ENSG00000203321 | CARNMT1-AS1 | 4.445506 | 0.3438345 | 0.6260388 | 0.8283607 | 0.4074662 | 0.608109 |
| ENSG00000279433 | | 1.458035 | 0.3437672 | 0.81402 | 1.06758 | 0.28571 | NA |
| ENSG00000232879 | GLRX5P2 | 9.680353 | 0.3437161 | 0.5040694 | 0.8705951 | 0.3839753 | 0.586322 |
| ENSG00000260461 | | 1.262742 | 0.3436394 | 0.8169646 | 1.07704 | 0.2814624 | NA |
| ENSG00000254615 | NA | 128.1784 | 0.3436106 | 0.2719178 | 1.344168 | 0.1788939 | 0.352808 |
| ENSG00000263558 | RN7SL716P | 4.646215 | 0.3435393 | 0.6288732 | 0.8280268 | 0.4076553 | 0.60828 |
| ENSG00000268307 | LINC02560 | 6.392368 | 0.3433252 | 0.5998945 | 0.8287569 | 0.407242 | 0.607954 |
| ENSG00000160213 | CSTB | 6162.23 | 0.3433025 | 0.1683348 | 2.072509 | 0.03821799 | 0.112874 |
| ENSG00000263847 | | 26.64368 | 0.3432064 | 0.445374 | 0.928333 | 0.3532349 | 0.555192 |
| ENSG00000131669 | NINJ1 | 6369.061941 | 0.34307085 | 0.203462134 | 1.752299683 | 0.07972229 | 0.196744 |
| ENSG00000111875 | ASF1A | 1888.457505 | 0.342996853 | 0.145950131 | 2.395828866 | 0.01658283 | 0.058418 |
| ENSG00000005001 | PRSS22 | 2867.333553 | 0.342981531 | 0.268670872 | 1.359163059 | 0.17409492 | 0.345911 |
| ENSG00000273982 | | 99.7291 | 0.3429686 | 0.310368 | 1.202791 | 0.2290571 | 0.417237 |
| ENSG00000248132 | LINC02101 | 2.162864 | 0.3428935 | 0.8279958 | 1.26373 | 0.206327 | 0.388539 |
| ENSG00000232906 | | 2.278271 | 0.3428872 | 0.7029472 | 0.8611297 | 0.3891666 | 0.591148 |
| ENSG00000105492 | SIGLEC6 | 224.657361 | 0.342883172 | 0.835193114 | 0.996146286 | 0.31917907 | 0.520102 |
| ENSG00000123353 | ORMDL2 | 1007.425069 | 0.342651531 | 0.153780986 | 2.282752455 | 0.02244495 | 0.074711 |
| ENSG00000232531 | KMT5AP2 | 127.4768 | 0.3426504 | 0.1958488 | 1.80921 | 0.07041835 | 0.179433 |
| ENSG00000184163 | C1QTNF12 | 65.30135 | 0.3424653 | 0.291056 | 1.266813 | 0.2052221 | 0.3872 |
| ENSG00000100744 | GSKIP | 2166.0335 | 0.342431778 | 0.124894512 | 2.729569443 | 0.00634171 | 0.026886 |
| ENSG00000187650 | VMAC | 297.4948 | 0.3424242 | 0.2290923 | 1.562609 | 0.1181446 | 0.262205 |
| ENSG00000114859 | CLCN2 | 1111.230699 | 0.342237725 | 0.223603811 | 1.584741435 | 0.11302509 | 0.25408 |
| ENSG00000112078 | KCTD20 | 5041.195558 | 0.342201897 | 0.155799163 | 2.248793597 | 0.02452563 | 0.079875 |
| ENSG00000227370 | | 6.018053 | 0.342111 | 0.6249316 | 0.8312327 | 0.4058422 | 0.606663 |
| ENSG00000253468 | | 0.8117568 | 0.3420721 | 0.8518062 | 1.114237 | 0.2651776 | NA |
| ENSG00000044090 | CUL7 | 3600.370314 | 0.342053092 | 0.165039068 | 2.133622234 | 0.03287372 | 0.100659 |
| ENSG00000177369 | | 1.351798 | 0.3420482 | 0.7871395 | 0.9950294 | 0.319722 | NA |
| ENSG00000265702 | | 68.94656 | 0.3419406 | 0.4575604 | 0.9103993 | 0.362612 | 0.564661 |

| | | | | | | | |
|---|---|---|---|---|---|---|---|
| ENSG00000182831 | C16orf72 | 3337.119 | 0.3418655 | 0.156798 | 2.212125 | 0.026958 | 0.086067 |
| ENSG00000186166 | CENATAC | 1889.007 | 0.3417044 | 0.1913865 | 1.837089 | 0.06619669 | 0.171197 |
| ENSG00000120071 | KANSL1 | 2840.262242 | 0.341691758 | 0.165346555 | 2.106016986 | 0.03520288 | 0.106051 |
| ENSG00000275665 | | 1.649403 | 0.3416576 | 0.8025912 | 1.12984 | 0.2585438 | 0.452699 |
| ENSG00000103343 | ZNF174 | 745.6228725 | 0.341629636 | 0.169318996 | 2.06971483 | 0.03847906 | 0.113431 |
| ENSG00000139624 | CERS5 | 2059.971 | 0.3416186 | 0.1683657 | 2.092778 | 0.03636897 | 0.108713 |
| ENSG00000260368 | | 34.94532 | 0.3416014 | 0.3953243 | 0.9944492 | 0.3200042 | 0.520872 |
| ENSG00000279400 | NA | 11.5357 | 0.3414441 | 0.8388039 | 0.9722093 | 0.3309464 | 0.531937 |
| ENSG00000171759 | PAH | 47.92308 | 0.3413358 | 0.4932935 | 0.866595 | 0.386164 | 0.588471 |
| ENSG00000127585 | FBXL16 | 3180.484695 | 0.34109542 | 0.188151368 | 1.86953283 | 0.06154872 | 0.162086 |
| ENSG00000092529 | CAPN3 | 7.657959701 | 0.34109244 | 0.567279057 | 0.811233772 | 0.41723144 | 0.61716 |
| ENSG00000154734 | ADAMTS1 | 393.7986 | 0.3410378 | 0.4262183 | 0.943935 | 0.3452029 | 0.54678 |
| ENSG00000275022 | MIR6753 | 3.68559 | 0.3409947 | 0.7095549 | 0.8712936 | 0.3835938 | 0.586001 |
| ENSG00000120963 | ZNF706 | 5764.866842 | 0.340970176 | 0.128209352 | 2.728674749 | 0.00635894 | 0.026924 |
| ENSG00000259163 | | 4.020112 | 0.3408714 | 0.7657471 | 0.95301 | 0.340585 | 0.541994 |
| ENSG00000196526 | AFAP1 | 314.3782 | 0.3408452 | 0.3219547 | 1.15866 | 0.2465948 | 0.438422 |
| ENSG00000250917 | | 3.24852 | 0.3407554 | 0.7139146 | 0.8552833 | 0.3923944 | 0.594061 |
| ENSG00000186790 | FOXE3 | 1.621039 | 0.3407467 | 0.7646372 | 0.9480249 | 0.3431168 | 0.544618 |
| ENSG00000160951 | PTGER1 | 3.650182 | 0.3406532 | 0.7080721 | 0.8593391 | 0.3901535 | 0.592145 |
| ENSG00000261468 | | 1.198328 | 0.3406136 | 0.8675256 | 1.302564 | 0.1927237 | NA |
| ENSG00000249456 | | 5.513327 | 0.3405537 | 0.5609509 | 0.8297914 | 0.4066567 | 0.607299 |
| ENSG00000198077 | CYP2A7 | 2.531756 | 0.3405122 | 0.789023 | 0.9971415 | 0.3186958 | 0.519793 |
| ENSG00000180425 | C11orf71 | 347.4165 | 0.3404698 | 0.1772653 | 1.971412 | 0.04867673 | 0.135722 |
| ENSG00000248659 | | 1.335013 | 0.3404339 | 0.816084 | 1.070016 | 0.2846121 | NA |
| ENSG00000172071 | EIF2AK3 | 1580.621 | 0.3404265 | 0.2131963 | 1.664424 | 0.09602781 | 0.225804 |
| ENSG00000133134 | BEX2 | 804.1958259 | 0.340422625 | 0.448332993 | 0.908119246 | 0.36381522 | 0.56593 |
| ENSG00000237101 | | 2.936147 | 0.3404173 | 0.7128593 | 0.869197 | 0.3847394 | 0.587142 |
| ENSG00000180263 | FGD6 | 767.3927 | 0.3403356 | 0.3718441 | 1.036258 | 0.3000817 | 0.499366 |
| ENSG00000131634 | TMEM204 | 12.13507679 | 0.340302867 | 0.507441535 | 0.851341671 | 0.39457958 | 0.596404 |
| ENSG00000228242 | XPC-AS1 | 165.083 | 0.3401962 | 0.2961351 | 1.237484 | 0.2159076 | 0.401178 |
| ENSG00000132549 | VPS13B | 5585.241412 | 0.340156228 | 0.228215131 | 1.55455462 | 0.12005218 | 0.26543 |
| ENSG00000064270 | ATP2C2 | 3838.30948 | 0.340144045 | 0.295498061 | 1.239346124 | 0.21521734 | 0.400288 |
| ENSG00000124459 | ZNF45 | 836.9177894 | 0.340092354 | 0.182695688 | 1.905408773 | 0.05672696 | 0.152562 |

| | | | | | | | |
|---|---|---|---|---|---|---|---|
| ENSG00000166333 | ILK | 41.75421 | 0.3400293 | 0.4289232 | 0.9373563 | 0.3485753 | 0.550285 |
| ENSG00000162931 | TRIM17 | 57.03409 | 0.3398834 | 0.515442 | 0.8511821 | 0.3946682 | 0.596436 |
| ENSG00000105383 | CD33 | 1.041677452 | 0.339861412 | 0.849797823 | 1.095793488 | 0.27316916 | NA |
| ENSG00000165458 | INPPL1 | 19132.76 | 0.339858 | 0.1594104 | 2.167285 | 0.03021317 | 0.094162 |
| ENSG00000183955 | KMT5A | 2497.528 | 0.3397379 | 0.1837207 | 1.905369 | 0.05673208 | 0.152562 |
| ENSG00000236132 | | 1.189169 | 0.3397176 | 0.8243875 | 1.10761 | 0.2680302 | NA |
| ENSG00000244187 | TMEM141 | 1444.759 | 0.3396948 | 0.2449263 | 1.483942 | 0.1378243 | 0.292504 |
| ENSG00000251635 | LINC02278 | 1.121275 | 0.3396569 | 0.8496479 | 1.100634 | 0.2710559 | NA |
| ENSG00000173614 | NMNAT1 | 479.8558 | 0.3395819 | 0.2329607 | 1.526282 | 0.1269396 | 0.275972 |
| ENSG00000270462 | | 1.787388 | 0.3394993 | 0.7758232 | 0.9726185 | 0.3307429 | 0.531835 |
| ENSG00000120137 | PANK3 | 12418.43557 | 0.33943663 | 0.247528716 | 1.442920055 | 0.14904299 | 0.309736 |
| ENSG00000131471 | AOC3 | 26.62808101 | 0.339371404 | 0.473783677 | 0.877636874 | 0.38014081 | 0.582069 |
| ENSG00000161281 | COX7A1 | 3.366175 | 0.3392441 | 0.6508778 | 0.8124213 | 0.4165499 | 0.616495 |
| ENSG00000103429 | BFAR | 4350.266528 | 0.339199922 | 0.14863134 | 2.32742897 | 0.01994245 | 0.067753 |
| ENSG00000237950 | | 132.5892 | 0.3391955 | 0.3525304 | 1.07008 | 0.2845833 | 0.482572 |
| ENSG00000278388 | | 1.031324 | 0.3390977 | 0.850618 | 1.108933 | 0.2674592 | NA |
| ENSG00000258881 | | 4.38215 | 0.3390353 | 0.6261438 | 0.8160722 | 0.4144589 | 0.614598 |
| ENSG00000268516 | | 164.3893 | 0.3389098 | 0.2261454 | 1.562294 | 0.1182188 | 0.262293 |
| ENSG00000125691 | RPL23 | 18087.19652 | 0.3389045 | 0.162714768 | 2.10334078 | 0.03543599 | 0.106601 |
| ENSG00000163602 | RYBP | 3604.087 | 0.3384605 | 0.1357911 | 2.526113 | 0.01153325 | 0.043632 |
| ENSG00000259792 | | 2.94304 | 0.3384307 | 0.753675 | 0.9202816 | 0.3574256 | 0.559503 |
| ENSG00000119640 | ACYP1 | 397.0490731 | 0.338424134 | 0.218924082 | 1.620582693 | 0.10510717 | 0.241236 |
| ENSG00000134152 | KATNBL1 | 1277.074894 | 0.338229696 | 0.1413453 | 2.427292174 | 0.015212 | 0.054528 |
| ENSG00000131400 | NAPSA | 129.4323651 | 0.338191133 | 0.390131587 | 0.991464768 | 0.32145869 | 0.522216 |
| ENSG00000248121 | SMURF2P1 | 8.004807 | 0.338168 | 0.5439545 | 0.8308961 | 0.4060323 | 0.606697 |
| ENSG00000175213 | ZNF408 | 1474.274 | 0.3381034 | 0.1637728 | 2.112822 | 0.03461604 | 0.10485 |
| ENSG00000255650 | FAM222A-AS1 | 2.757132 | 0.3380642 | 0.7502775 | 0.9159403 | 0.3596982 | 0.561604 |
| ENSG00000145016 | RUBCN | 1064.968 | 0.3380178 | 0.2200586 | 1.606698 | 0.1081206 | 0.246263 |
| ENSG00000215424 | MCM3AP-AS1 | 1321.689 | 0.3379734 | 0.2748635 | 1.310709 | 0.1899562 | 0.367719 |
| ENSG00000188611 | ASAH2 | 79.66289 | 0.3379443 | 0.3220313 | 1.147591 | 0.2511375 | 0.443885 |
| ENSG00000261505 | | 355.1299 | 0.337752 | 0.21446 | 1.637351 | 0.1015572 | 0.23541 |
| ENSG00000239377 | ZNF775-AS1 | 7.857725 | 0.3376079 | 0.5983628 | 0.8104678 | 0.4176714 | 0.617501 |
| ENSG00000170786 | SDR16C5 | 78.93345 | 0.3375899 | 0.418318 | 0.9313205 | 0.3516878 | 0.55339 |

| | | | | | | | |
|---|---|---|---|---|---|---|---|
| ENSG00000256943 | | 5.632977 | 0.3374648 | 0.7229358 | 0.8625126 | 0.3884055 | 0.590521 |
| ENSG00000104213 | PDGFRL | 288.1933223 | 0.337411651 | 0.292303945 | 1.242482259 | 0.21405868 | 0.398887 |
| ENSG00000163463 | KRTCAP2 | 418.3789 | 0.3371052 | 0.2649556 | 1.351729 | 0.176462 | 0.349204 |
| ENSG00000283208 | | 9.70836 | 0.337082 | 0.5832981 | 0.8162595 | 0.4143518 | 0.614577 |
| ENSG00000281021 | | 16.17358 | 0.3370615 | 0.5288373 | 0.8357224 | 0.4033111 | 0.604182 |
| ENSG00000224312 | MCCD1P2 | 1.549286 | 0.3370377 | 0.8292752 | 1.234064 | 0.2171789 | NA |
| ENSG00000249898 | MCPH1-AS1 | 58.73868 | 0.337036 | 0.4038287 | 0.9662142 | 0.333937 | 0.535093 |
| ENSG00000254485 | | 5.117793 | 0.3369612 | 0.6285616 | 0.8096408 | 0.4181467 | 0.617934 |
| ENSG00000279162 | | 10.57873 | 0.3368825 | 0.5404799 | 0.8333843 | 0.404628 | 0.605427 |
| ENSG00000108932 | SLC16A6 | 989.8042529 | 0.336867428 | 0.303444739 | 1.195964861 | 0.23171027 | 0.42014 |
| ENSG00000226098 | SEC11B | 42.64985 | 0.3368576 | 0.3580773 | 1.056458 | 0.2907588 | 0.489431 |
| ENSG00000104823 | ECH1 | 3756.335343 | 0.336745018 | 0.229822694 | 1.52151966 | 0.12812948 | 0.277792 |
| ENSG00000136866 | ZFP37 | 0.920881864 | 0.336642517 | 0.873509555 | 1.409653657 | 0.15864198 | NA |
| ENSG00000272795 | GAR1-DT | 3.060249 | 0.3365204 | 0.7344734 | 0.8885186 | 0.3742619 | 0.57635 |
| ENSG00000184470 | TXNRD2 | 1797.637 | 0.3365069 | 0.1697259 | 2.029965 | 0.04236011 | 0.12228 |
| ENSG00000257084 | MIR200CHG | 239.4371 | 0.3364932 | 0.3698195 | 1.026922 | 0.3044571 | 0.504089 |
| ENSG00000105325 | FZR1 | 2710.382477 | 0.336236847 | 0.147288973 | 2.277136532 | 0.02277807 | 0.075441 |
| ENSG00000243649 | CFB | 54.41641 | 0.3362328 | 0.6320148 | 0.8234529 | 0.4102505 | 0.610607 |
| ENSG00000133983 | COX16 | 164.2690188 | 0.335978623 | 0.249202092 | 1.422648588 | 0.15483805 | 0.318534 |
| ENSG00000257221 | | 0.9885959 | 0.3358869 | 0.8615976 | 1.262314 | 0.2068357 | NA |
| ENSG00000243517 | | 36.40281 | 0.3358447 | 0.3969758 | 0.9744862 | 0.3298152 | 0.530954 |
| ENSG00000115107 | STEAP3 | 3194.025338 | 0.335691041 | 0.155152153 | 2.207742282 | 0.02726224 | 0.086881 |
| ENSG00000273209 | | 7.404148 | 0.3356774 | 0.7157383 | 0.8614894 | 0.3889686 | 0.590982 |
| ENSG00000130988 | RGN | 2.174615897 | 0.335483761 | 0.754522539 | 0.909668389 | 0.36299742 | 0.565029 |
| ENSG00000130529 | TRPM4 | 3155.461252 | 0.335402514 | 0.158093294 | 2.1261812 | 0.03348818 | 0.102164 |
| ENSG00000123584 | MAGEA9 | 1.143388998 | 0.335269752 | 0.865193506 | 1.290856307 | 0.19675351 | NA |
| ENSG00000046651 | OFD1 | 2256.105253 | 0.335262857 | 0.159528256 | 2.143063113 | 0.03210803 | 0.098839 |
| ENSG00000084093 | REST | 3359.65536 | 0.335246776 | 0.215059764 | 1.618163864 | 0.10562729 | 0.242162 |
| ENSG00000285818 | | 2.359209 | 0.3352137 | 0.7693876 | 0.927627 | 0.3536011 | 0.555611 |
| ENSG00000288098 | | 31.84805 | 0.3352097 | 0.3802289 | 0.9968621 | 0.3188315 | 0.519919 |
| ENSG00000188895 | MSL1 | 8323.895 | 0.3351129 | 0.1670109 | 2.043111 | 0.04104149 | 0.119434 |
| ENSG00000033867 | SLC4A7 | 2799.805462 | 0.335081579 | 0.22791583 | 1.537302267 | 0.12421931 | 0.27199 |
| ENSG00000274765 | | 1.985809 | 0.3350587 | 0.7518349 | 0.9108183 | 0.3623911 | 0.564416 |

| | | | | | | | |
|---|---|---|---|---|---|---|---|
| ENSG00000251192 | ZNF674 | 263.6386 | 0.334968 | 0.2968914 | 1.218673 | 0.2229683 | 0.409572 |
| ENSG00000175697 | GPR156 | 90.05041 | 0.334836 | 0.3851438 | 0.9989303 | 0.3178284 | 0.518856 |
| ENSG00000112357 | PEX7 | 595.0683989 | 0.33482745 | 0.246826563 | 1.425230039 | 0.15409072 | 0.317416 |
| ENSG00000227256 | MIS18A-AS1 | 12.15267 | 0.3346064 | 0.4805931 | 0.8685033 | 0.3851188 | 0.587542 |
| ENSG00000183889 | | 69.06459 | 0.3345791 | 0.3339335 | 1.109955 | 0.2670184 | 0.46278 |
| ENSG00000088356 | PDRG1 | 2016.152424 | 0.334573935 | 0.187220525 | 1.857002687 | 0.06331077 | 0.16548 |
| ENSG00000117751 | PPP1R8 | 3434.776258 | 0.334553088 | 0.1781612 | 1.936468672 | 0.05281033 | 0.144681 |
| ENSG00000124762 | CDKN1A | 11930.08795 | 0.334468397 | 0.305282867 | 1.215959814 | 0.22400023 | 0.410803 |
| ENSG00000143748 | NVL | 2127.788 | 0.3343007 | 0.1379554 | 2.472098 | 0.01343227 | 0.049293 |
| ENSG00000182557 | SPNS3 | 3.429154 | 0.3342576 | 0.786384 | 1.040853 | 0.2979436 | 0.497084 |
| ENSG00000261427 | | 0.9642798 | 0.3342562 | 0.8747933 | 1.49909 | 0.1338503 | NA |
| ENSG00000179918 | SEPHS2 | 9070.852 | 0.3342388 | 0.1905518 | 1.759611 | 0.07847384 | 0.19441 |
| ENSG00000181061 | HIGD1A | 1386.433 | 0.3341745 | 0.2274352 | 1.519248 | 0.1287 | 0.278718 |
| ENSG00000274021 | | 6.687728 | 0.3341014 | 0.6247015 | 0.8202689 | 0.4120628 | 0.612299 |
| ENSG00000288107 | | 3.267629 | 0.333967 | 0.702907 | 0.8424093 | 0.3995589 | 0.600519 |
| ENSG00000236844 | | 9.840978 | 0.3338731 | 0.5423605 | 0.8178514 | 0.413442 | 0.613478 |
| ENSG00000226701 | RPL15P14 | 3.920253 | 0.3337728 | 0.6955851 | 0.8422845 | 0.3996287 | 0.600585 |
| ENSG00000259940 | | 21.78215 | 0.3337207 | 0.4863588 | 0.8550939 | 0.3924992 | 0.59407 |
| ENSG00000128915 | ICE2 | 2127.557712 | 0.333615217 | 0.139019439 | 2.438805792 | 0.01473589 | 0.053165 |
| ENSG00000156574 | NODAL | 7.384642 | 0.3334852 | 0.6287879 | 0.8102003 | 0.4178251 | 0.617591 |
| ENSG00000203909 | DPPA5 | 1.534638 | 0.3334151 | 0.8185342 | 1.207262 | 0.2273311 | NA |
| ENSG00000244256 | RN7SL130P | 1.403878 | 0.3332223 | 0.8156918 | 1.067416 | 0.285784 | NA |
| ENSG00000260941 | LINC00622 | 1.143816 | 0.3331784 | 0.8293482 | 1.090669 | 0.2754187 | NA |
| ENSG00000182154 | MRPL41 | 5779.283 | 0.3331722 | 0.1772733 | 1.925837 | 0.05412469 | 0.147476 |
| ENSG00000279901 | | 8.564754 | 0.3331152 | 0.6608754 | 0.8123223 | 0.4166067 | 0.616545 |
| ENSG00000128891 | CCDC32 | 466.791947 | 0.333028503 | 0.155954742 | 2.178131911 | 0.02939622 | 0.092342 |
| ENSG00000248429 | FAM198B-AS1 | 193.0634 | 0.3330135 | 0.4904348 | 0.8502249 | 0.3952 | 0.5969 |
| ENSG00000251493 | FOXD1 | 216.4246 | 0.3329976 | 0.2222124 | 1.560723 | 0.1185891 | 0.262871 |
| ENSG00000175832 | ETV4 | 204.211 | 0.3329479 | 0.6443108 | 0.8105021 | 0.4176516 | 0.617501 |
| ENSG00000222046 | DCDC2B | 14.41667 | 0.3328768 | 0.4727611 | 0.8666157 | 0.3861526 | 0.588471 |
| ENSG00000245573 | BDNF-AS | 125.9316 | 0.332816 | 0.3584276 | 1.04192 | 0.2974489 | 0.496571 |
| ENSG00000273599 | | 218.3531 | 0.3327942 | 0.2172123 | 1.593343 | 0.1110833 | 0.251074 |
| ENSG00000228071 | RPL7P47 | 3.456927 | 0.332722 | 0.712534 | 0.8684408 | 0.3851531 | 0.587542 |

| | | | | | | | |
|---|---|---|---|---|---|---|---|
| ENSG00000196781 | TLE1 | 2834.485 | 0.3325845 | 0.2159941 | 1.601089 | 0.1093572 | 0.24825 |
| ENSG00000170558 | CDH2 | 393.0043 | 0.332547 | 0.4559693 | 0.8823617 | 0.3775812 | 0.579547 |
| ENSG00000197714 | ZNF460 | 136.5544 | 0.3324723 | 0.3740079 | 1.001499 | 0.3165856 | 0.517625 |
| ENSG00000110422 | HIPK3 | 3188.001927 | 0.332430234 | 0.247869354 | 1.421777883 | 0.15509074 | 0.318932 |
| ENSG00000164162 | ANAPC10 | 286.2466 | 0.332251 | 0.2295237 | 1.516359 | 0.1294286 | 0.279758 |
| ENSG00000116584 | ARHGEF2 | 3105.50078 | 0.332084522 | 0.303801308 | 1.181285152 | 0.23748946 | 0.427121 |
| ENSG00000224426 | SLC31A1P1 | 1.971235 | 0.3320741 | 0.7827283 | 0.9570473 | 0.3385434 | 0.539844 |
| ENSG00000227379 | PPIAP2 | 1.574711 | 0.3320514 | 0.7506973 | 0.9095808 | 0.3630436 | 0.565035 |
| ENSG00000100417 | PMM1 | 992.2479027 | 0.331918237 | 0.215471189 | 1.583482223 | 0.1133116 | 0.254551 |
| ENSG00000074660 | SCARF1 | 54.66440335 | 0.331888249 | 0.497401124 | 0.840834457 | 0.40044068 | 0.601397 |
| ENSG00000268573 | | 69.35794 | 0.3316769 | 0.3155637 | 1.143808 | 0.2527034 | 0.445534 |
| ENSG00000233200 | LINC02634 | 0.7550807 | 0.3315967 | 0.8695623 | 1.377549 | 0.1683426 | NA |
| ENSG00000229765 | | 2.041975 | 0.3315348 | 0.7233161 | 0.8563817 | 0.3917867 | 0.593601 |
| ENSG00000233015 | SLC25A6P1 | 3.274095 | 0.3315209 | 0.8337635 | 0.9617259 | 0.3361873 | 0.537241 |
| ENSG00000141551 | CSNK1D | 12542.74 | 0.3315081 | 0.1376476 | 2.441411 | 0.01463001 | 0.052871 |
| ENSG00000165915 | SLC39A13 | 3379.094 | 0.3314791 | 0.2101282 | 1.636719 | 0.1016893 | 0.235614 |
| ENSG00000235655 | H3P6 | 4503.115 | 0.3313338 | 0.1666677 | 2.016408 | 0.04375729 | 0.125373 |
| ENSG00000278238 | | 76.61338 | 0.3313254 | 0.3406858 | 1.062781 | 0.2878815 | 0.486199 |
| ENSG00000231107 | LINC01508 | 76.05333 | 0.3312469 | 0.3689281 | 1.010253 | 0.312374 | 0.513454 |
| ENSG00000106355 | LSM5 | 4101.83775 | 0.331207917 | 0.282358223 | 1.255320397 | 0.20936247 | 0.392586 |
| ENSG00000160209 | PDXK | 17998.21 | 0.3310113 | 0.222665 | 1.628291 | 0.1034633 | 0.238476 |
| ENSG00000229965 | RPS3AP7 | 5.975905 | 0.330932 | 0.5587527 | 0.8056526 | 0.4204432 | 0.620287 |
| ENSG00000236345 | SCAT8 | 2.296556 | 0.3309073 | 0.7932427 | 0.9891079 | 0.3226104 | 0.523415 |
| ENSG00000160439 | RDH13 | 1476.577 | 0.3308724 | 0.1438149 | 2.332673 | 0.01966529 | 0.067086 |
| ENSG00000147010 | SH3KBP1 | 1801.552 | 0.3308626 | 0.2479751 | 1.402684 | 0.1607112 | 0.327287 |
| ENSG00000258949 | | 22.43281 | 0.3308624 | 0.4238621 | 0.9186835 | 0.3582611 | 0.560203 |
| ENSG00000263321 | | 4.218605 | 0.3306797 | 0.6828853 | 0.8274835 | 0.4079631 | 0.608532 |
| ENSG00000227401 | RPL37P1 | 4.853436 | 0.3306796 | 0.7330267 | 0.8843124 | 0.3765276 | 0.578617 |
| ENSG00000276855 | | 304.231 | 0.3306695 | 0.2253031 | 1.534736 | 0.1248488 | 0.272856 |
| ENSG00000254990 | | 1.846158 | 0.330579 | 0.7190848 | 0.8446003 | 0.398334 | 0.599392 |
| ENSG00000272263 | | 14.44218 | 0.3303991 | 0.4778735 | 0.8634996 | 0.3878628 | 0.590078 |
| ENSG00000125787 | GNRH2 | 7.586794607 | 0.330394126 | 0.54687719 | 0.806075029 | 0.42019959 | 0.6201 |
| ENSG00000262454 | MIR193BHG | 266.7609 | 0.3303159 | 0.2198806 | 1.564404 | 0.1177227 | 0.261629 |

| | | | | | | | |
|---|---|---|---|---|---|---|---|
| ENSG00000169951 | ZNF764 | 788.885 | 0.3302937 | 0.1755391 | 1.939105 | 0.05248851 | 0.143918 |
| ENSG00000177732 | SOX12 | 2821.475 | 0.3302552 | 0.1710919 | 1.961001 | 0.0498789 | 0.138272 |
| ENSG00000235781 | LINC02569 | 12.91104 | 0.3301595 | 0.5425012 | 0.8178396 | 0.4134488 | 0.613478 |
| ENSG00000123240 | OPTN | 3913.506125 | 0.329896796 | 0.370325555 | 1.020137498 | 0.30766326 | 0.507784 |
| ENSG00000261447 | | 8.270204 | 0.3297333 | 0.7299192 | 0.8680562 | 0.3853636 | 0.587757 |
| ENSG00000214654 | B3GNT10 | 168.0052 | 0.3297153 | 0.2726513 | 1.293254 | 0.1959231 | 0.37563 |
| ENSG00000173409 | ARV1 | 1102.74 | 0.3296639 | 0.1612771 | 2.093036 | 0.03634595 | 0.108713 |
| ENSG00000234432 | | 48.52712 | 0.3295693 | 0.4467362 | 0.8865785 | 0.3753059 | 0.577456 |
| ENSG00000243927 | MRPS6 | 1423.222 | 0.3293538 | 0.2720468 | 1.289753 | 0.1971363 | 0.377112 |
| ENSG00000167779 | IGFBP6 | 40.16901 | 0.3293282 | 0.5620974 | 0.7996781 | 0.4238973 | 0.623903 |
| ENSG00000162300 | ZFPL1 | 336.1327 | 0.3292418 | 0.1753035 | 1.92469 | 0.05426821 | 0.147731 |
| ENSG00000168002 | POLR2G | 6542.867 | 0.3291138 | 0.222896 | 1.538511 | 0.1239237 | 0.271619 |
| ENSG00000273240 | | 3.180417 | 0.3290716 | 0.7044123 | 0.824429 | 0.4096958 | 0.610089 |
| ENSG00000173728 | C1orf100 | 6.493571 | 0.3288127 | 0.6066852 | 0.7948921 | 0.4266763 | 0.626388 |
| ENSG00000103342 | GSPT1 | 15772.01556 | 0.32874737 | 0.1391627 | 2.386409058 | 0.01701382 | 0.059652 |
| ENSG00000172116 | CD8B | 1.12868 | 0.3286852 | 0.8055936 | 1.01282 | 0.3111463 | NA |
| ENSG00000186301 | MST1P2 | 231.1849 | 0.328668 | 0.2727672 | 1.282163 | 0.1997853 | 0.380073 |
| ENSG00000259943 | | 657.7237 | 0.328662 | 0.1674652 | 1.992779 | 0.04628569 | 0.130743 |
| ENSG00000165689 | ENTR1 | 4969.976 | 0.3286587 | 0.1595093 | 2.063876 | 0.03902949 | 0.114722 |
| ENSG00000203523 | TAS2R2P | 5.882022 | 0.3286313 | 0.706291 | 0.8324289 | 0.4051669 | 0.605926 |
| ENSG00000212123 | PRR22 | 143.4087 | 0.3285686 | 0.4222063 | 0.9157575 | 0.3597941 | 0.561721 |
| ENSG00000136371 | MTHFS | 141.2075174 | 0.328452946 | 0.27545969 | 1.270945118 | 0.20374817 | 0.385377 |
| ENSG00000139292 | LGR5 | 3.351793 | 0.328397 | 0.7639474 | 0.9294865 | 0.352637 | 0.55449 |
| ENSG00000214174 | AMZ2P1 | 988.4596 | 0.3283155 | 0.141978 | 2.339574 | 0.01930572 | 0.066057 |
| ENSG00000273416 | | 1.618207 | 0.3282473 | 0.7568905 | 0.8892379 | 0.3738752 | 0.575988 |
| ENSG00000236056 | GAPDHP14 | 1.177232 | 0.3281726 | 0.8615729 | 1.267097 | 0.2051206 | NA |
| ENSG00000267731 | | 13.32943 | 0.3281623 | 0.6146509 | 0.7965423 | 0.4257169 | 0.625567 |
| ENSG00000249196 | TMEM132D-AS1 | 228.2922 | 0.3280447 | 0.8334616 | 0.9371021 | 0.3487061 | 0.550392 |
| ENSG00000251630 | | 3.49071 | 0.3277634 | 0.6667898 | 0.8030962 | 0.4219191 | 0.621809 |
| ENSG00000231684 | EIF1P3 | 286.8935 | 0.3276649 | 0.2358059 | 1.45535 | 0.1455724 | 0.304616 |
| ENSG00000241912 | | 1.270548 | 0.3276518 | 0.855932 | 1.177499 | 0.2389963 | NA |
| ENSG00000275318 | | 17.97192 | 0.3276414 | 0.5313764 | 0.8107711 | 0.4174972 | 0.617347 |
| ENSG00000204231 | RXRB | 3324.778 | 0.3275979 | 0.1509573 | 2.199258 | 0.02785961 | 0.088347 |

| | | | | | | | |
|---|---|---|---|---|---|---|---|
| ENSG00000249669 | CARMN | 0.8272235 | 0.3274964 | 0.8749182 | 1.522711 | 0.127831 | NA |
| ENSG00000258301 | VASH1-AS1 | 10.51171 | 0.3274377 | 0.5272914 | 0.8148893 | 0.4151357 | 0.615074 |
| ENSG00000284618 | | 0.8577502 | 0.3273707 | 0.8293308 | 1.107245 | 0.2681879 | NA |
| ENSG00000178429 | RPS3AP5 | 340.0343 | 0.3273184 | 0.1895971 | 1.78057 | 0.07498275 | 0.188034 |
| ENSG00000233436 | BTBD18 | 2.301053 | 0.3273017 | 0.7286822 | 0.8486729 | 0.3960633 | 0.597471 |
| ENSG00000177807 | KCNJ10 | 7.01365 | 0.3271532 | 0.5842251 | 0.7822884 | 0.4340451 | 0.633034 |
| ENSG00000100902 | PSMA6 | 962.683322 | 0.327120308 | 0.189003703 | 1.789680349 | 0.07350531 | 0.185431 |
| ENSG00000274308 | | 0.7873115 | 0.3270929 | 0.8467669 | 1.061159 | 0.2886178 | NA |
| ENSG00000149201 | CCDC81 | 45.19898 | 0.3270612 | 0.3162891 | 1.130352 | 0.258328 | 0.45256 |
| ENSG00000230274 | PGAM1P3 | 1.053967 | 0.3268565 | 0.8228897 | 1.069418 | 0.2848814 | NA |
| ENSG00000286703 | | 0.9239746 | 0.3268008 | 0.8262547 | 1.209008 | 0.2266599 | NA |
| ENSG00000206417 | H1-10-AS1 | 144.5632 | 0.326774 | 0.3141642 | 1.125223 | 0.2604946 | 0.455057 |
| ENSG00000089351 | GRAMD1A | 4564.485055 | 0.326766864 | 0.204725933 | 1.656862172 | 0.09754734 | 0.228508 |
| ENSG00000235013 | COX20P2 | 2.863451 | 0.3265802 | 0.6679027 | 0.7999893 | 0.423717 | 0.623732 |
| ENSG00000206560 | ANKRD28 | 4919.454 | 0.326474 | 0.1813425 | 1.818967 | 0.06891643 | 0.176452 |
| ENSG00000288025 | | 3.601549 | 0.3264037 | 0.6978366 | 0.8196162 | 0.4124349 | 0.612623 |
| ENSG00000139239 | RPL14P1 | 9350.484 | 0.3263746 | 0.1800187 | 1.899508 | 0.05749778 | 0.154013 |
| ENSG00000202474 | RNA5SP283 | 8.119017 | 0.3262693 | 0.499399 | 0.8260679 | 0.4087656 | 0.609524 |
| ENSG00000187189 | TSPYL4 | 3691.227 | 0.3260823 | 0.125404 | 2.727782 | 0.00637618 | 0.026976 |
| ENSG00000267144 | | 1.101796 | 0.3260786 | 0.8152716 | 1.035127 | 0.3006097 | NA |
| ENSG00000258260 | | 0.9608712 | 0.3260472 | 0.8661229 | 1.394689 | 0.1631096 | NA |
| ENSG00000198723 | TEX45 | 51.6898 | 0.3259963 | 0.384412 | 0.9695497 | 0.332271 | 0.533388 |
| ENSG00000233516 | | 0.8992398 | 0.3259591 | 0.8802701 | 1.690185 | 0.09099248 | NA |
| ENSG00000127452 | FBXL12 | 1245.733653 | 0.325857786 | 0.175573295 | 1.896772543 | 0.05785797 | 0.154853 |
| ENSG00000227183 | HDGFP1 | 0.9517224 | 0.3257793 | 0.8445351 | 1.078056 | 0.2810087 | NA |
| ENSG00000189367 | KIAA0408 | 0.7882919 | 0.3257188 | 0.8464764 | 1.056456 | 0.2907597 | NA |
| ENSG00000274300 | MIR6864 | 1.147759 | 0.3257136 | 0.8211413 | 1.141176 | 0.2537965 | NA |
| ENSG00000279302 | | 2.057669 | 0.3255028 | 0.8450171 | 1.068226 | 0.2854186 | 0.483442 |
| ENSG00000179085 | DPM3 | 1411.817 | 0.3254531 | 0.2209492 | 1.528494 | 0.1263899 | 0.27507 |
| ENSG00000285184 | | 337.1202 | 0.325438 | 0.1998265 | 1.692165 | 0.09061455 | 0.216661 |
| ENSG00000145335 | SNCA | 6.243857 | 0.3252279 | 0.6592223 | 0.7875551 | 0.430957 | 0.630104 |
| ENSG00000249119 | MTND6P4 | 30.35554 | 0.3250912 | 0.4629045 | 0.8624543 | 0.3884376 | 0.590521 |
| ENSG00000135336 | ORC3 | 1728.11995 | 0.325030061 | 0.20497976 | 1.640807907 | 0.1008373 | 0.234243 |

| | | | | | | | |
|---|---|---|---|---|---|---|---|
| ENSG00000068323 | TFE3 | 4305.74851 | 0.324993578 | 0.135038496 | 2.441297468 | 0.0146346 | 0.05288 |
| ENSG00000287813 | | 1.151072 | 0.3248754 | 0.8633194 | 1.542107 | 0.1230475 | NA |
| ENSG00000279555 | | 15.46538 | 0.3248727 | 0.4955017 | 0.8312131 | 0.4058532 | 0.606663 |
| ENSG00000201448 | SNORA63C | 1.415613 | 0.3247052 | 0.8431527 | 1.065114 | 0.2868242 | NA |
| ENSG00000139132 | FGD4 | 569.6376613 | 0.324646833 | 0.228425258 | 1.47665744 | 0.13976749 | 0.295816 |
| ENSG00000258181 | | 4.699722 | 0.3245961 | 0.6932383 | 0.798422 | 0.4246256 | 0.624343 |
| ENSG00000262903 | | 75.37797 | 0.3245609 | 0.3408543 | 1.052938 | 0.2923694 | 0.491239 |
| ENSG00000226174 | TEX22 | 164.4981 | 0.3245316 | 0.282581 | 1.224657 | 0.2207045 | 0.407046 |
| ENSG00000253667 | | 8.218241 | 0.3244946 | 0.520463 | 0.8103194 | 0.4177567 | 0.617591 |
| ENSG00000170473 | PYM1 | 2228.513 | 0.3243395 | 0.2289257 | 1.455415 | 0.1455545 | 0.304603 |
| ENSG00000184560 | SPEM2 | 1.737896 | 0.3242622 | 0.8433109 | 1.076121 | 0.2818731 | 0.479609 |
| ENSG00000258021 | | 1.214525 | 0.3242466 | 0.7844704 | 0.9635176 | 0.3352879 | NA |
| ENSG00000215012 | RTL10 | 2046.088 | 0.3242277 | 0.1478032 | 2.228399 | 0.02585394 | 0.083242 |
| ENSG00000188706 | ZDHHC9 | 5683.323 | 0.3242238 | 0.1742333 | 1.913518 | 0.05568183 | 0.150423 |
| ENSG00000264007 | | 0.7784971 | 0.3241609 | 0.8622697 | 1.323802 | 0.1855687 | NA |
| ENSG00000004961 | HCCS | 2308.267963 | 0.324093165 | 0.15435054 | 2.138855208 | 0.0324474 | 0.099653 |
| ENSG00000203722 | RAET1G | 34.2753 | 0.3240697 | 0.4964363 | 0.8226061 | 0.410732 | 0.61105 |
| ENSG00000120868 | APAF1 | 1177.139243 | 0.324003359 | 0.327540951 | 1.08532197 | 0.27777908 | 0.474924 |
| ENSG00000188026 | RILPL1 | 808.3316 | 0.3239618 | 0.196895 | 1.699111 | 0.08929833 | 0.214421 |
| ENSG00000278147 | | 1.711702 | 0.3239404 | 0.7695832 | 0.9222314 | 0.3564079 | 0.55837 |
| ENSG00000244538 | | 6.291656 | 0.3239118 | 0.6078319 | 0.7897462 | 0.429676 | 0.628915 |
| ENSG00000251073 | NUDT19P5 | 2.239615 | 0.3239084 | 0.7512682 | 0.8826363 | 0.3774328 | 0.57951 |
| ENSG00000085832 | EPS15 | 4316.603336 | 0.323875047 | 0.161162354 | 2.055246463 | 0.03985521 | 0.116556 |
| ENSG00000159409 | CELF3 | 89.83251 | 0.3238641 | 0.3897976 | 0.951915 | 0.3411401 | 0.542447 |
| ENSG00000274653 | | 11.94061 | 0.3237679 | 0.4690898 | 0.8629531 | 0.3881633 | 0.5904 |
| ENSG00000188620 | HMX3 | 3.351035 | 0.3237606 | 0.7012251 | 0.8157295 | 0.4146549 | 0.614649 |
| ENSG00000258044 | | 3.468681 | 0.3234181 | 0.7118309 | 0.8308239 | 0.4060731 | 0.606701 |
| ENSG00000198774 | RASSF9 | 1.23481 | 0.3233898 | 0.8143398 | 1.036388 | 0.3000212 | NA |
| ENSG00000244159 | RPS27AP13 | 0.7535813 | 0.3231567 | 0.8439888 | 1.064607 | 0.2870536 | NA |
| ENSG00000088179 | PTPN4 | 1600.355241 | 0.322982224 | 0.166104366 | 2.003613704 | 0.04511146 | 0.128271 |
| ENSG00000145545 | SRD5A1 | 2998.433 | 0.3229285 | 0.2117566 | 1.578589 | 0.1144303 | 0.256428 |
| ENSG00000177483 | RBM44 | 18.20002 | 0.3227479 | 0.5544235 | 0.7884455 | 0.4304361 | 0.629691 |
| ENSG00000249115 | HAUS5 | 1762.685 | 0.322703 | 0.283898 | 1.222639 | 0.221466 | 0.408082 |

| | | | | | | | |
|---|---|---|---|---|---|---|---|
| ENSG00000237188 | | 5.050076 | 0.3226909 | 0.6437038 | 0.7836133 | 0.4332671 | 0.632237 |
| ENSG00000172037 | LAMB2 | 9492.566 | 0.3226755 | 0.1900774 | 1.74862 | 0.08035675 | 0.197942 |
| ENSG00000215785 | CFL1P6 | 0.98869 | 0.3226564 | 0.8026738 | 1.001786 | 0.316447 | NA |
| ENSG00000253227 | | 9.904762 | 0.3226304 | 0.6124233 | 0.7805778 | 0.4350509 | 0.633814 |
| ENSG00000237273 | RSL24D1P8 | 1.290213 | 0.3225956 | 0.8433017 | 1.072563 | 0.2834674 | NA |
| ENSG00000248275 | TRIM52-AS1 | 724.0936 | 0.3225218 | 0.2638601 | 1.296271 | 0.1948823 | 0.374012 |
| ENSG00000173926 | MARCHF3 | 76.9776 | 0.3225151 | 0.4210426 | 0.9034703 | 0.3662763 | 0.568169 |
| ENSG00000280331 | | 0.89716 | 0.3225027 | 0.8521211 | 1.181012 | 0.237598 | NA |
| ENSG00000135655 | USP15 | 3806.934651 | 0.322457066 | 0.101311871 | 3.208223157 | 0.00133558 | 0.007362 |
| ENSG00000143384 | MCL1 | 38064.1 | 0.3224491 | 0.2343889 | 1.439516 | 0.1500044 | 0.311146 |
| ENSG00000132286 | TIMM10B | 1399.373885 | 0.322428225 | 0.152489258 | 2.129358583 | 0.0332246 | 0.101582 |
| ENSG00000228038 | VN1R51P | 1.621657 | 0.3223874 | 0.7896181 | 0.9777894 | 0.3281785 | 0.529395 |
| ENSG00000237154 | MCFD2P1 | 0.8544098 | 0.3222471 | 0.8516926 | 1.175129 | 0.2399432 | NA |
| ENSG00000164330 | EBF1 | 3.439903 | 0.3221745 | 0.8336558 | 0.9225147 | 0.3562601 | 0.558205 |
| ENSG00000118898 | PPL | 8049.895685 | 0.322163762 | 0.257752016 | 1.307057633 | 0.19119314 | 0.36921 |
| ENSG00000135597 | REPS1 | 2618.475512 | 0.322151536 | 0.105865644 | 3.071693913 | 0.00212848 | 0.010928 |
| ENSG00000223356 | | 6.791205 | 0.3221412 | 0.6335232 | 0.7714293 | 0.4404525 | 0.638603 |
| ENSG00000121406 | ZNF549 | 2.213039554 | 0.322113315 | 0.713790558 | 0.828692718 | 0.40727831 | 0.607954 |
| ENSG00000226348 | VN2R10P | 0.7800036 | 0.3220586 | 0.8621643 | 1.31153 | 0.1896789 | NA |
| ENSG00000154545 | MAGED4 | 4.108349 | 0.321975 | 0.6876227 | 0.7988083 | 0.4244016 | 0.624325 |
| ENSG00000143226 | FCGR2A | 6.911536 | 0.3219059 | 0.716836 | 0.8334774 | 0.4045755 | 0.605416 |
| ENSG00000241278 | ENPP7P4 | 11.20592 | 0.3218839 | 0.5391049 | 0.7892622 | 0.4299588 | 0.629225 |
| ENSG00000214417 | KRT18P13 | 8.498595 | 0.3218405 | 0.5748511 | 0.7875456 | 0.4309626 | 0.630104 |
| ENSG00000076067 | RBMS2 | 642.6940017 | 0.321744568 | 0.23964777 | 1.398863999 | 0.16185377 | 0.329082 |
| ENSG00000286529 | | 11.53778 | 0.3216831 | 0.5345431 | 0.8020135 | 0.4225451 | 0.62255 |
| ENSG00000287322 | | 5.81479 | 0.3216469 | 0.7057883 | 0.8219651 | 0.4110967 | 0.611456 |
| ENSG00000069122 | ADGRF5 | 5.128446667 | 0.321604525 | 0.830654037 | 0.940451252 | 0.34698614 | 0.548568 |
| ENSG00000064687 | ABCA7 | 1330.842961 | 0.321381531 | 0.213661681 | 1.562637962 | 0.11813777 | 0.262205 |
| ENSG00000013588 | GPRC5A | 7171.24748 | 0.321213253 | 0.347160647 | 1.034555036 | 0.30087676 | 0.500062 |
| ENSG00000240253 | FAR2P3 | 1.235363 | 0.3211655 | 0.7807551 | 0.927235 | 0.3538045 | NA |
| ENSG00000259518 | LINC01583 | 0.804455 | 0.3211629 | 0.8543983 | 1.157326 | 0.2471392 | NA |
| ENSG00000280414 | | 0.7633456 | 0.3211388 | 0.8684505 | 1.435665 | 0.1510976 | NA |
| ENSG00000277723 | MIR6827 | 2.117893 | 0.3210979 | 0.7945535 | 0.9862524 | 0.3240092 | 0.524883 |

| | | | | | | | |
|---|---|---|---|---|---|---|---|
| ENSG00000137821 | LRRC49 | 95.05736143 | 0.321093497 | 0.360701608 | 0.999250357 | 0.31767343 | 0.518672 |
| ENSG00000196458 | ZNF605 | 1362.096 | 0.321076 | 0.2007327 | 1.652861 | 0.09835908 | 0.229831 |
| ENSG00000244308 | RN7SL762P | 2.31354 | 0.3209505 | 0.7990518 | 0.9976652 | 0.3184418 | 0.519634 |
| ENSG00000117748 | RPA2 | 3565.393805 | 0.320935265 | 0.248453021 | 1.360267719 | 0.17374522 | 0.345397 |
| ENSG00000279170 | TSTD3 | 309.584 | 0.320921 | 0.2034999 | 1.636675 | 0.1016985 | 0.235614 |
| ENSG00000206069 | TMEM211 | 0.8158869 | 0.3209063 | 0.8439072 | 1.039409 | 0.2986145 | NA |
| ENSG00000279349 | | 1.895191 | 0.3208831 | 0.7928218 | 0.9788988 | 0.32763 | 0.529092 |
| ENSG00000139726 | DENR | 7885.911 | 0.3206746 | 0.1284325 | 2.526996 | 0.01150428 | 0.043547 |
| ENSG00000105552 | BCAT2 | 3309.550262 | 0.320673894 | 0.217360252 | 1.532597863 | 0.12537497 | 0.273578 |
| ENSG00000185013 | NT5C1B | 5.163202 | 0.3206671 | 0.6266014 | 0.7778226 | 0.4366736 | 0.635203 |
| ENSG00000102981 | PARD6A | 423.2989436 | 0.320659038 | 0.274867221 | 1.240506229 | 0.21478821 | 0.399713 |
| ENSG00000164411 | GJB7 | 1.410825 | 0.3205583 | 0.7854593 | 0.9492428 | 0.3424972 | NA |
| ENSG00000279949 | | 7.390268 | 0.3204543 | 0.6559977 | 0.7872385 | 0.4311422 | 0.630262 |
| ENSG00000242134 | RPL5P13 | 0.8145211 | 0.3204128 | 0.8425018 | 1.058802 | 0.2896902 | NA |
| ENSG00000257941 | | 3.377539 | 0.3203993 | 0.6536975 | 0.7835709 | 0.4332919 | 0.632237 |
| ENSG00000282996 | MPP7-DT | 3.515017 | 0.3202744 | 0.6754098 | 0.7839085 | 0.4330938 | 0.632176 |
| ENSG00000274156 | | 1.019185 | 0.3202484 | 0.8426356 | 1.06644 | 0.2862248 | NA |
| ENSG00000075391 | RASAL2 | 1303.057095 | 0.320232551 | 0.200008862 | 1.625100353 | 0.10414118 | 0.239623 |
| ENSG00000261359 | PYCARD-AS1 | 1.162847 | 0.3201388 | 0.7926524 | 0.9684562 | 0.3328166 | NA |
| ENSG00000101608 | MYL12A | 12077.5719 | 0.320133205 | 0.191485809 | 1.691213549 | 0.09079602 | 0.216978 |
| ENSG00000105722 | ERF | 3952.254338 | 0.320072635 | 0.174103341 | 1.870821219 | 0.06136987 | 0.161815 |
| ENSG00000101160 | CTSZ | 1207.116394 | 0.320023439 | 0.209249155 | 1.57997726 | 0.11411207 | 0.255852 |
| ENSG00000231256 | CFAP97D1 | 1.286115 | 0.3197511 | 0.8183521 | 1.142889 | 0.2530846 | NA |
| ENSG00000269843 | | 40.34128 | 0.3196733 | 0.4609412 | 0.842604 | 0.3994499 | 0.600486 |
| ENSG00000250432 | FAM242C | 2.034533 | 0.3195621 | 0.7788003 | 0.9189886 | 0.3581016 | 0.560066 |
| ENSG00000261455 | LINC01003 | 826.487 | 0.319546 | 0.2633599 | 1.284515 | 0.1989618 | 0.379049 |
| ENSG00000137842 | TMEM62 | 836.9055835 | 0.319445425 | 0.147672618 | 2.213521772 | 0.02686169 | 0.085852 |
| ENSG00000189079 | ARID2 | 2389.961 | 0.3193439 | 0.2217014 | 1.503071 | 0.1328208 | 0.285184 |
| ENSG00000104866 | PPP1R37 | 4952.785097 | 0.319341134 | 0.144415371 | 2.238436136 | 0.02519263 | 0.081587 |
| ENSG00000279432 | | 3.4727 | 0.3193192 | 0.6388799 | 0.7805807 | 0.4350491 | 0.633814 |
| ENSG00000105270 | CLIP3 | 334.6766179 | 0.319281636 | 0.365574113 | 0.979530659 | 0.32731785 | 0.528695 |
| ENSG00000248752 | | 1.601166 | 0.3192557 | 0.7822056 | 0.9200796 | 0.3575311 | 0.559578 |
| ENSG00000123066 | MED13L | 10776.5866 | 0.319186732 | 0.324361512 | 1.093312524 | 0.27425661 | 0.470992 |

| | | | | | | | |
|---|---|---|---|---|---|---|---|
| ENSG00000144792 | ZNF660 | 10.63855 | 0.3191058 | 0.4918584 | 0.81611 | 0.4144373 | 0.614598 |
| ENSG00000285693 | | 328.8196 | 0.3190215 | 0.2357891 | 1.422565 | 0.1548624 | 0.318534 |
| ENSG00000267078 | | 4.199491 | 0.3188704 | 0.6088444 | 0.7613029 | 0.4464762 | 0.643407 |
| ENSG00000264395 | MIR3193 | 2.590052 | 0.3188286 | 0.7771748 | 0.9628096 | 0.3356431 | 0.536815 |
| ENSG00000157551 | KCNJ15 | 1.83426 | 0.3187891 | 0.7021383 | 0.8048925 | 0.4208817 | 0.620762 |
| ENSG00000127603 | MACF1 | 7814.167781 | 0.318711228 | 0.232188106 | 1.437538962 | 0.15056491 | 0.311967 |
| ENSG00000211890 | IGHA2 | 2.034451 | 0.3186572 | 0.8431546 | 1.015153 | 0.3100328 | 0.510667 |
| ENSG00000176204 | LRRTM4 | 2.311332 | 0.3186034 | 0.721776 | 0.823113 | 0.4104437 | 0.610826 |
| ENSG00000257225 | | 2.050492 | 0.3185803 | 0.7749184 | 0.9105794 | 0.362517 | 0.564546 |
| ENSG00000259088 | | 19.50968 | 0.3184458 | 0.3917944 | 0.9259052 | 0.3544953 | 0.556358 |
| ENSG00000279382 | | 531.8804 | 0.3183873 | 0.263629 | 1.279197 | 0.2008275 | 0.381509 |
| ENSG00000272861 | | 12.8336 | 0.3182452 | 0.6191998 | 0.7799796 | 0.4354029 | 0.634188 |
| ENSG00000183134 | PTGDR2 | 2.098039 | 0.3182347 | 0.7568111 | 0.9043491 | 0.3658103 | 0.567798 |
| ENSG00000263167 | | 2.126261 | 0.3181362 | 0.8055638 | 1.045787 | 0.2956592 | 0.494708 |
| ENSG00000120322 | PCDHB8 | 99.47318995 | 0.31811656 | 0.261942822 | 1.283967598 | 0.19915329 | 0.379278 |
| ENSG00000246898 | LINC00920 | 0.9339996 | 0.3180925 | 0.8435482 | 1.042769 | 0.2970554 | NA |
| ENSG00000185900 | POMK | 110.5174 | 0.3180742 | 0.3605478 | 0.9865917 | 0.3238429 | 0.524773 |
| ENSG00000215915 | ATAD3C | 91.73762 | 0.3180639 | 0.4898126 | 0.8137507 | 0.4157878 | 0.61561 |
| ENSG00000187955 | COL14A1 | 7.11495 | 0.3180609 | 0.6374257 | 0.7710402 | 0.4406831 | 0.63871 |
| ENSG00000178202 | POGLUT3 | 1558.5 | 0.3179858 | 0.2350238 | 1.420914 | 0.1553418 | 0.319239 |
| ENSG00000231357 | | 15.19005 | 0.3179792 | 0.5137846 | 0.7840842 | 0.4329907 | 0.632095 |
| ENSG00000239455 | | 29.8098 | 0.3179556 | 0.4134136 | 0.8988198 | 0.3687487 | 0.570871 |
| ENSG00000155629 | PIK3AP1 | 40.43018 | 0.3178638 | 0.5301018 | 0.7993839 | 0.4240678 | 0.623976 |
| ENSG00000198816 | ZNF358 | 886.3108 | 0.3178397 | 0.1865672 | 1.759445 | 0.07850199 | 0.194455 |
| ENSG00000256448 | | 1.187669 | 0.317729 | 0.8097903 | 1.027666 | 0.3041071 | NA |
| ENSG00000121644 | DESI2 | 2753.383807 | 0.317718224 | 0.177321073 | 1.847803588 | 0.06463076 | 0.167979 |
| ENSG00000145362 | ANK2 | 203.7312 | 0.3177166 | 0.6228067 | 0.7545179 | 0.4505383 | 0.646801 |
| ENSG00000126247 | CAPNS1 | 14152.28729 | 0.317695441 | 0.190017096 | 1.683126655 | 0.09235058 | 0.219588 |
| ENSG00000152954 | NRSN1 | 1.846284 | 0.3176667 | 0.7790223 | 0.9265817 | 0.3541437 | 0.55617 |
| ENSG00000230666 | CEACAM22P | 2.5371 | 0.3176647 | 0.7350239 | 0.8532517 | 0.3935198 | 0.595147 |
| ENSG00000254999 | BRK1 | 12217.97 | 0.3176551 | 0.1380642 | 2.320496 | 0.02031409 | 0.068816 |
| ENSG00000155428 | TRIM74 | 13.23159 | 0.3175243 | 0.5686555 | 0.766707 | 0.4432557 | 0.640816 |
| ENSG00000130340 | SNX9 | 1712.661012 | 0.317370475 | 0.124424309 | 2.593433488 | 0.00950229 | 0.037281 |

| | | | | | | | |
|---|---|---|---|---|---|---|---|
| ENSG00000287619 | | 28.89776 | 0.3172892 | 0.3999956 | 0.9177055 | 0.3587731 | 0.560753 |
| ENSG00000225246 | RPS2P1 | 0.7827246 | 0.3172633 | 0.8334565 | 1.150621 | 0.2498884 | NA |
| ENSG00000232082 | RPS6KA2-IT1 | 2.54668 | 0.3171936 | 0.7282513 | 0.8327456 | 0.4049882 | 0.605727 |
| ENSG00000197728 | RPS26 | 482.7072 | 0.3171781 | 0.2223851 | 1.488756 | 0.1365516 | 0.290727 |
| ENSG00000224086 | PPM1F-AS1 | 195.5697 | 0.3171227 | 0.1945386 | 1.68964 | 0.09109676 | 0.21754 |
| ENSG00000100592 | DAAM1 | 3950.880655 | 0.317088136 | 0.213350511 | 1.553558767 | 0.1202897 | 0.265822 |
| ENSG00000221883 | ARIH2OS | 85.49047 | 0.3170374 | 0.3227747 | 1.074088 | 0.2827832 | 0.480603 |
| ENSG00000265474 | | 15.51016 | 0.3170105 | 0.4619527 | 0.8358682 | 0.4032291 | 0.604179 |
| ENSG00000161813 | LARP4 | 4638.286 | 0.3169708 | 0.2336428 | 1.419692 | 0.1556975 | 0.319815 |
| ENSG00000076242 | MLH1 | 2110.447063 | 0.316940449 | 0.244608828 | 1.359040883 | 0.17413363 | 0.345963 |
| ENSG00000140474 | ULK3 | 6149.638 | 0.3168385 | 0.1481313 | 2.188712 | 0.02861775 | 0.090324 |
| ENSG00000148672 | GLUD1 | 7114.567 | 0.3168094 | 0.1738066 | 1.881758 | 0.05986893 | 0.158812 |
| ENSG00000168175 | MAPK1IP1L | 4211.761 | 0.316746 | 0.1247939 | 2.570235 | 0.01016296 | 0.039413 |
| ENSG00000087152 | ATXN7L3 | 6738.096442 | 0.31669985 | 0.170764207 | 1.89219187 | 0.05846542 | 0.156138 |
| ENSG00000175262 | C1orf127 | 2.593964 | 0.3166732 | 0.7730284 | 0.9064451 | 0.3647003 | 0.566651 |
| ENSG00000178950 | GAK | 4850.276 | 0.3166513 | 0.1081449 | 2.965362 | 0.00302327 | 0.014647 |
| ENSG00000280053 | | 3.330419 | 0.3165695 | 0.6963828 | 0.7988112 | 0.4243999 | 0.624325 |
| ENSG00000178217 | SH2D4B | 5.642196 | 0.3165618 | 0.5629869 | 0.7727026 | 0.4396984 | 0.638136 |
| ENSG00000277150 | F8A3 | 3.498978 | 0.31654 | 0.7271453 | 0.8591046 | 0.3902828 | 0.592234 |
| ENSG00000286229 | | 2.3262 | 0.3164917 | 0.7366902 | 0.8609347 | 0.389274 | 0.591157 |
| ENSG00000143627 | PKLR | 2.804884 | 0.3164807 | 0.6639146 | 0.7778505 | 0.4366571 | 0.635203 |
| ENSG00000287402 | | 5.442291 | 0.3164699 | 0.6721294 | 0.7905559 | 0.4292032 | 0.628431 |
| ENSG00000142544 | CTU1 | 313.5273 | 0.3163441 | 0.2412124 | 1.379091 | 0.1678666 | 0.337524 |
| ENSG00000286708 | | 1.881495 | 0.3162607 | 0.7365231 | 0.8434863 | 0.3989565 | 0.599914 |
| ENSG00000241954 | | 3.051292 | 0.3162305 | 0.6587163 | 0.7666629 | 0.443282 | 0.640816 |
| ENSG00000105518 | TMEM205 | 2351.911556 | 0.316200513 | 0.196185773 | 1.663877005 | 0.09613701 | 0.225961 |
| ENSG00000249286 | AMD1P3 | 0.932612 | 0.3161759 | 0.8536379 | 1.234788 | 0.2169095 | NA |
| ENSG00000126106 | TMEM53 | 473.8846795 | 0.316097699 | 0.166146651 | 1.954094968 | 0.05068999 | 0.139969 |
| ENSG00000253720 | | 1.937091 | 0.3160715 | 0.8049224 | 1.083683 | 0.2785055 | 0.475798 |
| ENSG00000177380 | PPFIA3 | 2718.568 | 0.316046 | 0.1755774 | 1.85163 | 0.064079 | 0.166867 |
| ENSG00000279569 | | 20.4166 | 0.316032 | 0.3854892 | 0.9353933 | 0.3495857 | 0.551286 |
| ENSG00000207606 | MIR554 | 2.253579 | 0.3160264 | 0.7889513 | 0.9608124 | 0.3366465 | 0.53778 |
| ENSG00000183840 | GPR39 | 72.01218 | 0.3159947 | 0.3377152 | 1.033796 | 0.3012314 | 0.500495 |

| | | | | | | | |
|---|---|---|---|---|---|---|---|
| ENSG00000151414 | NEK7 | 4173.366 | 0.3159756 | 0.1807489 | 1.801964 | 0.07155101 | 0.181622 |
| ENSG00000148225 | WDR31 | 260.8428 | 0.3159543 | 0.2861084 | 1.186596 | 0.2353872 | 0.424836 |
| ENSG00000287855 | | 39.41479 | 0.3159001 | 0.3810043 | 0.9486945 | 0.342776 | 0.544299 |
| ENSG00000285597 | | 2.197301 | 0.3157503 | 0.7534669 | 0.8762293 | 0.3809054 | 0.582932 |
| ENSG00000228499 | TMSB10P1 | 1.41847 | 0.315723 | 0.8395685 | 1.046107 | 0.2955115 | NA |
| ENSG00000249779 | | 4.445395 | 0.3156397 | 0.6914186 | 0.7993796 | 0.4240703 | 0.623976 |
| ENSG00000240392 | RPL9P3 | 2.346883 | 0.3155529 | 0.6866108 | 0.788176 | 0.4305937 | 0.629807 |
| ENSG00000279873 | LINC01126 | 115.8934 | 0.3155138 | 0.2560328 | 1.30083 | 0.1933166 | 0.372002 |
| ENSG00000073849 | ST6GAL1 | 230.249281 | 0.315459331 | 0.328108545 | 1.067351748 | 0.28581303 | 0.483864 |
| ENSG00000138434 | ITPRID2 | 3762.01778 | 0.315447442 | 0.188713006 | 1.721621785 | 0.08513805 | 0.206843 |
| ENSG00000270956 | | 5.460027 | 0.3153816 | 0.6232821 | 0.7602138 | 0.4471268 | 0.643898 |
| ENSG00000254677 | OSBPL9P2 | 2.273761 | 0.3153224 | 0.7392353 | 0.8528372 | 0.3937496 | 0.59542 |
| ENSG00000285676 | | 14.82878 | 0.3152029 | 0.4901722 | 0.8038439 | 0.4214871 | 0.62131 |
| ENSG00000171951 | SCG2 | 5.96086 | 0.3151989 | 0.6342733 | 0.7776962 | 0.4367481 | 0.635252 |
| ENSG00000165650 | PDZD8 | 2164.403 | 0.315146 | 0.2564505 | 1.298666 | 0.1940586 | 0.372863 |
| ENSG00000224550 | NDUFA4P1 | 1382.152 | 0.3151375 | 0.227135 | 1.457041 | 0.1451051 | 0.303926 |
| ENSG00000231227 | | 1.071869 | 0.3149305 | 0.840822 | 1.060531 | 0.2889031 | NA |
| ENSG00000078177 | N4BP2 | 1510.063543 | 0.314919273 | 0.271572353 | 1.227697004 | 0.21956073 | 0.405649 |
| ENSG00000287064 | | 45.11279 | 0.3149002 | 0.7347136 | 0.8459488 | 0.3975813 | 0.598625 |
| ENSG00000151033 | LYZL2 | 1.069952 | 0.3148392 | 0.8400166 | 1.042405 | 0.2972238 | NA |
| ENSG00000084110 | HAL | 21.76663589 | 0.314794044 | 0.413520469 | 0.887194084 | 0.37497442 | 0.577146 |
| ENSG00000251411 | | 8.817534 | 0.3147549 | 0.5251819 | 0.7840121 | 0.433033 | 0.632122 |
| ENSG00000226491 | FTOP1 | 1.81144 | 0.3147492 | 0.7737647 | 0.9089117 | 0.3633967 | 0.565452 |
| ENSG00000223476 | VN1R42P | 5.752692 | 0.3144495 | 0.6458193 | 0.7698423 | 0.4413934 | 0.639234 |
| ENSG00000204121 | ECEL1P1 | 6.383319 | 0.3143378 | 0.8281642 | 0.9267297 | 0.3540669 | 0.556113 |
| ENSG00000137460 | FHDC1 | 1147.439025 | 0.314320781 | 0.201681483 | 1.627468886 | 0.10363755 | 0.238774 |
| ENSG00000286232 | | 0.7910715 | 0.3142096 | 0.8419401 | 1.037083 | 0.2996972 | NA |
| ENSG00000092200 | RPGRIP1 | 84.37759422 | 0.314204554 | 0.321386395 | 1.070633399 | 0.2843343 | 0.482252 |
| ENSG00000267042 | | 1.057047 | 0.3141446 | 0.7863401 | 0.9669158 | 0.3335861 | NA |
| ENSG00000057468 | MSH4 | 6.275926636 | 0.31411096 | 0.651298512 | 0.767772783 | 0.44262219 | 0.640211 |
| ENSG00000266770 | RN7SL619P | 1.076415 | 0.3140112 | 0.8139177 | 1.025366 | 0.3051905 | NA |
| ENSG00000269069 | | 183.1019 | 0.3139664 | 0.3081765 | 1.112685 | 0.2658438 | 0.46143 |
| ENSG00000269892 | | 5.211282 | 0.3139593 | 0.649848 | 0.7646593 | 0.4444744 | 0.641842 |

| | | | | | | | |
|---|---|---|---|---|---|---|---|
| ENSG00000244723 | ASLP1 | 12.37686 | 0.3138721 | 0.5556485 | 0.7700222 | 0.4412867 | 0.639149 |
| ENSG00000236773 | | 370.5655 | 0.3138197 | 0.1494177 | 2.153493 | 0.03127992 | 0.096918 |
| ENSG00000273139 | | 15.56668 | 0.3137889 | 0.5328748 | 0.7718827 | 0.4401839 | 0.638526 |
| ENSG00000238755 | LINC02006 | 1.192642 | 0.3137559 | 0.7753007 | 0.8961003 | 0.3701992 | NA |
| ENSG00000101901 | ALG13 | 3189.57308 | 0.313727619 | 0.157647638 | 2.029835254 | 0.04237329 | 0.122292 |
| ENSG00000110025 | SNX15 | 48.26283466 | 0.313725029 | 0.333652121 | 1.037071713 | 0.29970244 | 0.498891 |
| ENSG00000259876 | | 0.851452 | 0.3136598 | 0.8411029 | 1.06121 | 0.2885944 | NA |
| ENSG00000186230 | ZNF749 | 472.0705 | 0.3135437 | 0.1869079 | 1.716591 | 0.08605391 | 0.208439 |
| ENSG00000240622 | RPL7P15 | 1.190815 | 0.3134612 | 0.768145 | 0.8982661 | 0.3690437 | NA |
| ENSG00000101844 | ATG4A | 1188.439333 | 0.31343279 | 0.139589913 | 2.232181496 | 0.02560297 | 0.082634 |
| ENSG00000244361 | RPL30P7 | 1.223833 | 0.3132272 | 0.7915394 | 0.9565148 | 0.3388122 | NA |
| ENSG00000249635 | | 3.912854 | 0.313206 | 0.6977351 | 0.7915563 | 0.4286195 | 0.62791 |
| ENSG00000124116 | WFDC3 | 6.205080054 | 0.313117801 | 0.622898183 | 0.753200998 | 0.45132914 | 0.647622 |
| ENSG00000088280 | ASAP3 | 1596.11893 | 0.313041504 | 0.192571192 | 1.670926999 | 0.0947361 | 0.223797 |
| ENSG00000152556 | PFKM | 3858.925 | 0.3130402 | 0.1137682 | 2.780732 | 0.00542364 | 0.023666 |
| ENSG00000145425 | RPS3A | 2843.521 | 0.3130339 | 0.1694878 | 1.88824 | 0.05899368 | 0.157139 |
| ENSG00000240005 | STIM2-AS1 | 5.228422 | 0.3129498 | 0.6221809 | 0.7521315 | 0.451972 | 0.648229 |
| ENSG00000253123 | | 0.7905752 | 0.3129381 | 0.8660381 | 1.424958 | 0.1541694 | NA |
| ENSG00000147604 | RPL7 | 2726.884 | 0.3128558 | 0.1775901 | 1.790327 | 0.07340142 | 0.185209 |
| ENSG00000274292 | | 21.32271 | 0.3128318 | 0.3841149 | 0.9259288 | 0.354483 | 0.556358 |
| ENSG00000252026 | RNU6-1262P | 5.405637 | 0.3127905 | 0.6245405 | 0.7570183 | 0.4490389 | 0.645416 |
| ENSG00000090554 | FLT3LG | 13.04720035 | 0.312764352 | 0.528218503 | 0.77247816 | 0.43983128 | 0.638259 |
| ENSG00000173482 | PTPRM | 111.1194 | 0.3126032 | 0.5074906 | 0.792422 | 0.4281147 | 0.627523 |
| ENSG00000248174 | LINC02268 | 0.8123291 | 0.3125243 | 0.812499 | 1.029506 | 0.3032418 | NA |
| ENSG00000261121 | LINC02473 | 1.261649 | 0.3125094 | 0.8488807 | 1.154308 | 0.2483737 | NA |
| ENSG00000257890 | | 0.8036298 | 0.312471 | 0.8483904 | 1.167829 | 0.2428758 | NA |
| ENSG00000119139 | TJP2 | 167.9881923 | 0.312369876 | 0.308382152 | 1.100544717 | 0.27109486 | 0.467375 |
| ENSG00000184154 | LRTOMT | 286.5218 | 0.3122312 | 0.2753874 | 1.209946 | 0.2262995 | 0.41386 |
| ENSG00000281912 | LINC01144 | 74.88626 | 0.3121942 | 0.3375796 | 1.027707 | 0.3040877 | 0.503697 |
| ENSG00000252618 | RNA5SP441 | 1.387353 | 0.3120269 | 0.7549733 | 0.8572153 | 0.3913259 | NA |
| ENSG00000134297 | PLEKHA8P1 | 197.9169017 | 0.312025231 | 0.227418837 | 1.426113785 | 0.1538355 | 0.317038 |
| ENSG00000147571 | CRH | 2.883611 | 0.3119899 | 0.7925871 | 0.9488494 | 0.3426972 | 0.544239 |
| ENSG00000077684 | JADE1 | 1643.025967 | 0.311943391 | 0.180587542 | 1.764976888 | 0.07756765 | 0.192733 |

| | | | | | | | |
|---|---|---|---|---|---|---|---|
| ENSG00000135047 | CTSL | 5248.266417 | 0.311860676 | 0.215164075 | 1.501955948 | 0.13310849 | 0.285621 |
| ENSG00000225563 | | 14.4319 | 0.3117244 | 0.5697209 | 0.7541597 | 0.4507533 | 0.64697 |
| ENSG00000186432 | KPNA4 | 6100.702 | 0.3117071 | 0.1083059 | 2.90229 | 0.00370445 | 0.017298 |
| ENSG00000250222 | | 41.12778 | 0.3114758 | 0.3223038 | 1.055708 | 0.2911017 | 0.489728 |
| ENSG00000100354 | TNRC6B | 1338.868885 | 0.311466498 | 0.210991601 | 1.531776018 | 0.12557772 | 0.273889 |
| ENSG00000251314 | | 1.129462 | 0.311421 | 0.7918406 | 0.954518 | 0.3398215 | NA |
| ENSG00000165861 | ZFYVE1 | 1993.557 | 0.3113273 | 0.2747849 | 1.207378 | 0.2272868 | 0.415097 |
| ENSG00000238405 | RNA5SP311 | 4.143642 | 0.3112526 | 0.6568257 | 0.7437234 | 0.4570438 | 0.652965 |
| ENSG00000274281 | | 1.202727 | 0.3112302 | 0.8735 | 1.734446 | 0.08283898 | NA |
| ENSG00000083828 | ZNF586 | 608.2497743 | 0.311128463 | 0.200276466 | 1.60531966 | 0.10842348 | 0.246783 |
| ENSG00000168792 | ABHD15 | 1740.689 | 0.3110448 | 0.1355952 | 2.349428 | 0.01880228 | 0.064668 |
| ENSG00000233757 | | 40.39847 | 0.3110407 | 0.5711204 | 0.7566452 | 0.4492625 | 0.645517 |
| ENSG00000273132 | | 1.237013 | 0.3109924 | 0.8138003 | 1.028119 | 0.303894 | NA |
| ENSG00000175352 | NRIP3 | 1.126422 | 0.3108906 | 0.8186638 | 1.042643 | 0.2971135 | NA |
| ENSG00000224447 | H3P26 | 0.9098964 | 0.3107152 | 0.8465723 | 1.141489 | 0.2536665 | NA |
| ENSG00000170950 | PGK2 | 4.506775 | 0.3106137 | 0.6540392 | 0.7617969 | 0.4461813 | 0.643188 |
| ENSG00000254664 | | 1.577791 | 0.3105745 | 0.8296357 | 1.161941 | 0.2452596 | 0.436984 |
| ENSG00000181826 | RELL1 | 480.7938 | 0.3105714 | 0.3044259 | 1.104536 | 0.2693608 | 0.465597 |
| ENSG00000267500 | ZNF887P | 17.63826 | 0.310542 | 0.5112548 | 0.7811748 | 0.4346997 | 0.63358 |
| ENSG00000185716 | MOSMO | 691.1235 | 0.3105108 | 0.2645657 | 1.228069 | 0.2194212 | 0.405523 |
| ENSG00000178585 | CTNNBIP1 | 1346.976 | 0.3104607 | 0.1586721 | 1.971211 | 0.04869973 | 0.135772 |
| ENSG00000226051 | ZNF503-AS1 | 5.3878 | 0.3104411 | 0.6257466 | 0.7615868 | 0.4463067 | 0.643333 |
| ENSG00000090612 | ZNF268 | 1261.679982 | 0.31038304 | 0.174997064 | 1.821998082 | 0.06845528 | 0.175542 |
| ENSG00000181191 | PJA1 | 1012.242 | 0.3103314 | 0.1969441 | 1.625941 | 0.1039621 | 0.239276 |
| ENSG00000171450 | CDK5R2 | 12.67901 | 0.310249 | 0.5611837 | 0.758173 | 0.4483474 | 0.64498 |
| ENSG00000261481 | | 2.864127 | 0.3102305 | 0.7605026 | 0.8775499 | 0.380188 | 0.582102 |
| ENSG00000109689 | STIM2 | 1326.204825 | 0.31012437 | 0.146204478 | 2.157632988 | 0.03095638 | 0.096152 |
| ENSG00000225335 | | 77.96654 | 0.3101152 | 0.3577545 | 0.9682161 | 0.3329365 | 0.53394 |
| ENSG00000258727 | | 75.80229 | 0.3100611 | 0.4038854 | 0.8863499 | 0.375429 | 0.577578 |
| ENSG00000232855 | | 12.72185 | 0.3100277 | 0.5297056 | 0.7632355 | 0.445323 | 0.642438 |
| ENSG00000186193 | SAPCD2 | 4587.393 | 0.3099872 | 0.3383952 | 1.010547 | 0.3122331 | 0.513254 |
| ENSG00000155313 | USP25 | 2284.45 | 0.3099826 | 0.168608 | 1.891765 | 0.05852224 | 0.156238 |
| ENSG00000274259 | SYNGAP1-AS1 | 12.13905 | 0.3099417 | 0.5448377 | 0.759069 | 0.4478113 | 0.644453 |

| | | | | | | | |
|---|---|---|---|---|---|---|---|
| ENSG00000252456 | RNA5SP434 | 0.992685 | 0.3098977 | 0.8472821 | 1.164067 | 0.244397 | NA |
| ENSG00000232398 | TMPRSS11CP | 0.8421012 | 0.3097884 | 0.868035 | 1.551469 | 0.1207893 | NA |
| ENSG00000224222 | LINC02636 | 1.267745 | 0.3097664 | 0.7928006 | 0.957455 | 0.3383376 | NA |
| ENSG00000148331 | ASB6 | 2043.688 | 0.3095949 | 0.1232357 | 2.565859 | 0.01029207 | 0.039809 |
| ENSG00000278434 | | 0.8921397 | 0.309482 | 0.848524 | 1.151961 | 0.2493371 | NA |
| ENSG00000170091 | NSG2 | 4.599018 | 0.3094748 | 0.6310782 | 0.7542162 | 0.4507194 | 0.646957 |
| ENSG00000249863 | | 18.29491 | 0.3094598 | 0.4404028 | 0.8349924 | 0.403722 | 0.604548 |
| ENSG00000231313 | CLIC1P1 | 45.08499 | 0.3093905 | 0.4033678 | 0.8960065 | 0.3702493 | 0.572391 |
| ENSG00000184009 | ACTG1 | 90543.35 | 0.3093511 | 0.2008504 | 1.591461 | 0.1115059 | 0.251836 |
| ENSG00000165506 | DNAAF2 | 1899.124 | 0.3091923 | 0.1510432 | 2.077112 | 0.03779121 | 0.111873 |
| ENSG00000187686 | KRT18P59 | 9.50536 | 0.3091279 | 0.5655291 | 0.7595091 | 0.4475481 | 0.644319 |
| ENSG00000226185 | TRIM64FP | 0.8965749 | 0.3090765 | 0.838572 | 1.040899 | 0.2979222 | NA |
| ENSG00000131477 | RAMP2 | 9.366873163 | 0.308818421 | 0.590036245 | 0.751801618 | 0.45217037 | 0.648318 |
| ENSG00000213003 | BTF3P12 | 5.921451 | 0.3087006 | 0.671714 | 0.773447 | 0.4392579 | 0.637775 |
| ENSG00000135205 | CCDC146 | 241.2841699 | 0.308631452 | 0.403577458 | 0.886958023 | 0.3751015 | 0.577241 |
| ENSG00000272788 | | 2.274011 | 0.3086037 | 0.7339556 | 0.8419484 | 0.3998168 | 0.600799 |
| ENSG00000240785 | RPL36AP21 | 0.8650212 | 0.3085777 | 0.8629285 | 1.465577 | 0.1427636 | NA |
| ENSG00000233220 | LINC00167 | 3.480819 | 0.3083451 | 0.7328449 | 0.8354879 | 0.4034431 | 0.604287 |
| ENSG00000272630 | | 16.62066 | 0.3083299 | 0.6704415 | 0.7638469 | 0.4449585 | 0.642221 |
| ENSG00000040731 | CDH10 | 106.1329388 | 0.308038264 | 0.702333079 | 0.794364134 | 0.42698348 | 0.626613 |
| ENSG00000105204 | DYRK1B | 1618.261334 | 0.308027531 | 0.238357922 | 1.351598449 | 0.17650381 | 0.349261 |
| ENSG00000234106 | SRP14P2 | 114.2494 | 0.3079423 | 0.2408969 | 1.347573 | 0.1777957 | 0.351104 |
| ENSG00000197653 | DNAH10 | 101.9524 | 0.3079008 | 0.3529933 | 0.972475 | 0.3308143 | 0.531885 |
| ENSG00000287823 | | 1.665793 | 0.3076905 | 0.8397619 | 1.043112 | 0.2968966 | 0.495923 |
| ENSG00000100483 | VCPKMT | 922.6259591 | 0.307687169 | 0.14138249 | 2.212559509 | 0.02692803 | 0.086011 |
| ENSG00000274276 | CBSL | 71.16611 | 0.3076534 | 0.3370066 | 1.006481 | 0.3141842 | 0.515372 |
| ENSG00000287213 | | 1.338689 | 0.3075992 | 0.7688611 | 0.8933032 | 0.3716948 | NA |
| ENSG00000258282 | BTF3P2 | 13.4367 | 0.3075872 | 0.5415787 | 0.7507629 | 0.4527954 | 0.64892 |
| ENSG00000119699 | TGFB3 | 623.873681 | 0.30751638 | 0.359040955 | 0.969227579 | 0.33243165 | 0.533517 |
| ENSG00000250571 | GLI4 | 1081.813 | 0.3072703 | 0.1628491 | 1.921658 | 0.05464874 | 0.148471 |
| ENSG00000237719 | | 44.90659 | 0.3072199 | 0.4214428 | 0.8612619 | 0.3890938 | 0.591105 |
| ENSG00000159904 | ZNF890P | 16.54403 | 0.3070206 | 0.5440325 | 0.7564604 | 0.4493732 | 0.645547 |
| ENSG00000228451 | SDAD1P1 | 52.90197 | 0.3069934 | 0.3485915 | 0.9765694 | 0.3287824 | 0.529991 |

| | | | | | | | |
|---|---|---|---|---|---|---|---|
| ENSG00000077782 | FGFR1 | 2705.127379 | 0.3069922 | 0.312269006 | 1.070484418 | 0.28440132 | 0.482335 |
| ENSG00000226450 | CYP2D8P | 47.93743 | 0.3069817 | 0.3945156 | 0.8916605 | 0.3725749 | 0.574751 |
| ENSG00000228274 | | 29.60165 | 0.3068549 | 0.4255165 | 0.8524539 | 0.3939622 | 0.595538 |
| ENSG00000128482 | RNF112 | 7.744243058 | 0.30657117 | 0.587017283 | 0.738695007 | 0.46009222 | 0.655523 |
| ENSG00000262663 | | 13.49445 | 0.306485 | 0.4616192 | 0.8167767 | 0.414056 | 0.614241 |
| ENSG00000204110 | LINC02520 | 1.396587 | 0.3064587 | 0.7742304 | 0.9077336 | 0.364019 | NA |
| ENSG00000151135 | TMEM263 | 2027.603 | 0.3064514 | 0.1721659 | 1.798339 | 0.07212328 | 0.182613 |
| ENSG00000229758 | DYNLT3P2 | 1.180854 | 0.3064136 | 0.7640536 | 0.8736539 | 0.3823068 | NA |
| ENSG00000164659 | ELAPOR2 | 1184.065 | 0.3064016 | 0.1162766 | 2.66597 | 0.00767665 | 0.031461 |
| ENSG00000231982 | | 42.30813 | 0.3063636 | 0.3567776 | 0.9625348 | 0.335781 | 0.536902 |
| ENSG00000258556 | LINC02322 | 1.375617 | 0.3063539 | 0.8376373 | 1.045673 | 0.2957122 | NA |
| ENSG00000142046 | TMEM91 | 121.3868 | 0.3063435 | 0.2650751 | 1.225483 | 0.2203932 | 0.406641 |
| ENSG00000253636 | | 17.75247 | 0.3063167 | 0.4420293 | 0.8237882 | 0.4100599 | 0.610397 |
| ENSG00000173715 | C11orf80 | 2244.679 | 0.3062977 | 0.1671129 | 1.879815 | 0.06013323 | 0.159306 |
| ENSG00000261812 | TUBB8P7 | 3.29301 | 0.3062777 | 0.6539451 | 0.7605215 | 0.4469429 | 0.643781 |
| ENSG00000165338 | HECTD2 | 1280.029 | 0.3062719 | 0.2211189 | 1.448172 | 0.1475689 | 0.30754 |
| ENSG00000237872 | POU5F1P4 | 1.460892 | 0.3062146 | 0.7855695 | 0.9833781 | 0.3254214 | NA |
| ENSG00000265182 | SRP72P1 | 15.92575 | 0.3061955 | 0.4503977 | 0.8220697 | 0.4110372 | 0.611401 |
| ENSG00000287472 | | 1.262895 | 0.3060759 | 0.7936042 | 0.9288359 | 0.3529741 | NA |
| ENSG00000005339 | CREBBP | 8071.101106 | 0.306069251 | 0.221353602 | 1.442912345 | 0.14904517 | 0.309736 |
| ENSG00000068383 | INPP5A | 1361.022415 | 0.306012788 | 0.172673064 | 1.817978402 | 0.06906743 | 0.176753 |
| ENSG00000088256 | GNA11 | 3470.136763 | 0.305989402 | 0.176086794 | 1.770066127 | 0.07671613 | 0.191189 |
| ENSG00000248593 | DSTNP2 | 101.007 | 0.305794 | 0.2756726 | 1.186901 | 0.2352666 | 0.424647 |
| ENSG00000143382 | ADAMTSL4 | 227.5141 | 0.305758 | 0.334916 | 1.024814 | 0.3054509 | 0.505294 |
| ENSG00000176988 | FMR1NB | 1.163135 | 0.3056983 | 0.8369697 | 1.199466 | 0.2303467 | NA |
| ENSG00000105613 | MAST1 | 306.4959454 | 0.305617473 | 0.258063315 | 1.24970088 | 0.21140884 | 0.395502 |
| ENSG00000270711 | | 3.091821 | 0.305503 | 0.7080292 | 0.7862434 | 0.4317249 | 0.630906 |
| ENSG00000232987 | LINC01219 | 5.90274 | 0.3054006 | 0.6917632 | 0.7658612 | 0.4437589 | 0.641322 |
| ENSG00000279207 | | 1801.36 | 0.3053247 | 0.2291868 | 1.393769 | 0.1633872 | 0.331149 |
| ENSG00000139874 | SSTR1 | 1.803747 | 0.3052903 | 0.7610814 | 0.8490284 | 0.3958655 | 0.597362 |
| ENSG00000244137 | | 5.160413 | 0.3052226 | 0.611336 | 0.7375912 | 0.4607629 | 0.65574 |
| ENSG00000073008 | PVR | 3661.63925 | 0.305119404 | 0.187149962 | 1.657498959 | 0.09741864 | 0.228287 |
| ENSG00000100577 | GSTZ1 | 2640.73296 | 0.305065583 | 0.259179106 | 1.244866008 | 0.21318101 | 0.397921 |

| | | | | | | | |
|---|---|---|---|---|---|---|---|
| ENSG00000233705 | SLC26A4-AS1 | 2.28018 | 0.3050262 | 0.7928206 | 0.9617912 | 0.3361545 | 0.537224 |
| ENSG00000271133 | | 41.8343 | 0.3049887 | 0.5527711 | 0.7495813 | 0.4535069 | 0.649659 |
| ENSG00000255627 | | 1.090586 | 0.3049185 | 0.8233448 | 1.176109 | 0.2395515 | NA |
| ENSG00000261556 | SMG1P7 | 446.6877 | 0.3048875 | 0.2425353 | 1.320282 | 0.186741 | 0.363109 |
| ENSG00000183067 | IGSF5 | 124.3553 | 0.3048605 | 0.3254711 | 1.020971 | 0.3072683 | 0.507479 |
| ENSG00000225945 | | 8.974989 | 0.3048028 | 0.6012009 | 0.746703 | 0.4552428 | 0.651166 |
| ENSG00000121542 | SEC22A | 849.5928847 | 0.304786353 | 0.121720116 | 2.531234241 | 0.01136619 | 0.043125 |
| ENSG00000100028 | SNRPD3 | 2454.192653 | 0.304776667 | 0.152511643 | 2.036291462 | 0.0417211 | 0.120869 |
| ENSG00000214846 | | 1.148035 | 0.3047477 | 0.8200826 | 1.20148 | 0.229565 | NA |
| ENSG00000213888 | LINC01521 | 479.9058 | 0.3047233 | 0.2705378 | 1.190658 | 0.233788 | 0.422868 |
| ENSG00000214338 | SOGA3 | 5.201081 | 0.3046402 | 0.6051042 | 0.7378399 | 0.4606118 | 0.655647 |
| ENSG00000216588 | IGSF23 | 6.191989 | 0.3046131 | 0.7587 | 0.8825234 | 0.3774938 | 0.579513 |
| ENSG00000113851 | CRBN | 2931.036431 | 0.304355321 | 0.123496868 | 2.491397948 | 0.01272415 | 0.047223 |
| ENSG00000197085 | NPSR1-AS1 | 0.9729707 | 0.3043523 | 0.8215524 | 1.08145 | 0.2794971 | NA |
| ENSG00000248367 | | 115.2442 | 0.3043413 | 0.3388159 | 0.9917767 | 0.3213065 | 0.522096 |
| ENSG00000197782 | ZNF780A | 781.3327 | 0.3042876 | 0.2843514 | 1.146156 | 0.2517304 | 0.444489 |
| ENSG00000267691 | SHC1P2 | 8.837598 | 0.3042822 | 0.5946426 | 0.735657 | 0.4619394 | 0.656639 |
| ENSG00000223575 | RBMX2P3 | 10.67309 | 0.3042796 | 0.5266739 | 0.7605912 | 0.4469013 | 0.643781 |
| ENSG00000200059 | | 1.657505 | 0.3042546 | 0.7835353 | 0.9779739 | 0.3280872 | 0.529382 |
| ENSG00000249042 | | 215.2573 | 0.3042445 | 0.2499933 | 1.282389 | 0.1997061 | 0.379949 |
| ENSG00000267277 | | 19.97843 | 0.3042187 | 0.413825 | 0.8596896 | 0.3899602 | 0.591947 |
| ENSG00000286084 | | 5.482084 | 0.3042151 | 0.6109542 | 0.7391702 | 0.4598037 | 0.655274 |
| ENSG00000188641 | DPYD | 294.1841 | 0.3042018 | 0.473973 | 0.7910314 | 0.4289257 | 0.628128 |
| ENSG00000213574 | LDHAP5 | 12.901 | 0.3040947 | 0.5229632 | 0.7599688 | 0.4472732 | 0.64397 |
| ENSG00000254604 | | 23.51732 | 0.30393 | 0.3325417 | 1.010156 | 0.3124207 | 0.513467 |
| ENSG00000279594 | | 8.336643 | 0.3038935 | 0.5384566 | 0.7465094 | 0.4553598 | 0.651295 |
| ENSG00000186111 | PIP5K1C | 2423.619 | 0.3037548 | 0.1253049 | 2.469236 | 0.0135402 | 0.049573 |
| ENSG00000226937 | CEP164P1 | 10.62253 | 0.3037478 | 0.5931864 | 0.7362416 | 0.4615837 | 0.656292 |
| ENSG00000213440 | H2AZP1 | 2.777461 | 0.303716 | 0.7655742 | 0.8909798 | 0.37294 | 0.575095 |
| ENSG00000111684 | LPCAT3 | 1299.544237 | 0.30361664 | 0.225954972 | 1.417577583 | 0.15631413 | 0.320778 |
| ENSG00000161180 | CCDC116 | 6.948245 | 0.3035765 | 0.5882784 | 0.7430459 | 0.4574539 | 0.653391 |
| ENSG00000127080 | IPPK | 615.1492818 | 0.303513901 | 0.178767389 | 1.749500029 | 0.08020462 | 0.197659 |
| ENSG00000213711 | PHBP7 | 3.827471 | 0.3035077 | 0.6942543 | 0.771098 | 0.4406488 | 0.63871 |

| | | | | | | | |
|---|---|---|---|---|---|---|---|
| ENSG00000287493 | | 0.8543705 | 0.3034147 | 0.8019138 | 1.043192 | 0.2968592 | NA |
| ENSG00000235253 | | 3.894074 | 0.3031628 | 0.6877471 | 0.7744316 | 0.4386756 | 0.637243 |
| ENSG00000137699 | TRIM29 | 64.09242519 | 0.303160235 | 0.576974856 | 0.750506477 | 0.45294972 | 0.649106 |
| ENSG00000122390 | NAA60 | 4249.496759 | 0.303146891 | 0.123883629 | 2.474225188 | 0.01335255 | 0.049048 |
| ENSG00000276696 | | 1.071345 | 0.3031136 | 0.7962761 | 1.071013 | 0.2841636 | NA |
| ENSG00000239719 | | 32.27916 | 0.3030997 | 0.3430355 | 0.9859706 | 0.3241475 | 0.525011 |
| ENSG00000188801 | ZNF322P1 | 5.330932 | 0.303082 | 0.57183 | 0.7346326 | 0.4625633 | 0.656982 |
| ENSG00000123329 | ARHGAP9 | 3.370456478 | 0.302989494 | 0.703130179 | 0.775228291 | 0.43820477 | 0.636782 |
| ENSG00000162836 | ACP6 | 2571.184 | 0.3029804 | 0.1942651 | 1.610957 | 0.1071891 | 0.244771 |
| ENSG00000224048 | LINC02612 | 1.201855 | 0.3028377 | 0.8448415 | 1.085584 | 0.2776631 | NA |
| ENSG00000110925 | CSRNP2 | 2261.352052 | 0.302808981 | 0.22919915 | 1.39192222 | 0.16394595 | 0.331978 |
| ENSG00000212802 | RPL15P3 | 10637.43 | 0.3027402 | 0.1857977 | 1.640154 | 0.1009731 | 0.234466 |
| ENSG00000135093 | USP30 | 1320.735183 | 0.302594333 | 0.155172153 | 1.988991347 | 0.04670216 | 0.131636 |
| ENSG00000259318 | HMGN1P1 | 21.26039 | 0.3025859 | 0.5481653 | 0.7473213 | 0.4548696 | 0.650839 |
| ENSG00000224208 | ZCRB1P1 | 5.660484 | 0.3025822 | 0.6519817 | 0.7457984 | 0.4557892 | 0.651734 |
| ENSG00000280107 | | 1.893001 | 0.3024958 | 0.6981638 | 0.7633112 | 0.4452778 | 0.642438 |
| ENSG00000256091 | MTRF1LP1 | 2.326443 | 0.3024682 | 0.7377293 | 0.823134 | 0.4104318 | 0.610826 |
| ENSG00000181790 | ADGRB1 | 700.9542 | 0.3023409 | 0.2727368 | 1.175582 | 0.2397618 | 0.429979 |
| ENSG00000155269 | GPR78 | 29.56092 | 0.3022796 | 0.3799251 | 0.9046501 | 0.3656508 | 0.567629 |
| ENSG00000255045 | | 5.349022 | 0.302254 | 0.7043431 | 0.7851134 | 0.4323871 | 0.631596 |
| ENSG00000270140 | | 3.252059 | 0.3022283 | 0.7489628 | 0.8532449 | 0.3935235 | 0.595147 |
| ENSG00000154678 | PDE1C | 7.26766 | 0.3022052 | 0.719566 | 0.8079523 | 0.419118 | 0.61902 |
| ENSG00000158122 | PRXL2C | 501.1296 | 0.3021203 | 0.1757026 | 1.764812 | 0.07759536 | 0.192748 |
| ENSG00000207041 | RNU6-3P | 4.898113 | 0.3020202 | 0.6922099 | 0.7697912 | 0.4414238 | 0.639243 |
| ENSG00000130812 | ANGPTL6 | 24.8029941 | 0.301990485 | 0.325339701 | 1.020282699 | 0.30759441 | 0.507721 |
| ENSG00000095585 | BLNK | 717.3060648 | 0.30188672 | 0.546117011 | 0.733860283 | 0.4630339 | 0.657314 |
| ENSG00000148384 | INPP5E | 1858.368 | 0.3018731 | 0.1337857 | 2.282774 | 0.02244368 | 0.074711 |
| ENSG00000068305 | MEF2A | 1932.996545 | 0.301800665 | 0.215331 | 1.458958507 | 0.14457653 | 0.303057 |
| ENSG00000228201 | | 1.222162 | 0.3017733 | 0.7600895 | 0.8528458 | 0.3937448 | NA |
| ENSG00000224049 | | 3.026265 | 0.3017617 | 0.6987354 | 0.7728067 | 0.4396368 | 0.638116 |
| ENSG00000116198 | CEP104 | 2012.981206 | 0.301720167 | 0.204522743 | 1.526770428 | 0.12681811 | 0.275753 |
| ENSG00000270679 | | 3.238245 | 0.3015888 | 0.7334364 | 0.8147836 | 0.4151962 | 0.615074 |
| ENSG00000287687 | | 1.122786 | 0.3015778 | 0.8000854 | 0.9741958 | 0.3299593 | NA |

| | | | | | | | |
|---|---|---|---|---|---|---|---|
| ENSG00000230555 | | 109.7283 | 0.3015293 | 0.3343487 | 0.9970454 | 0.3187425 | 0.519805 |
| ENSG00000150783 | TEX12 | 0.8946683 | 0.3014896 | 0.808721 | 1.004565 | 0.3151065 | NA |
| ENSG00000256229 | ZNF486 | 1.651394 | 0.3014338 | 0.8429206 | 1.09851 | 0.2719819 | 0.468479 |
| ENSG00000280020 | | 8.783262 | 0.3013773 | 0.6092865 | 0.725134 | 0.4683699 | 0.661979 |
| ENSG00000232077 | LINC01031 | 3.78686 | 0.3013393 | 0.653945 | 0.7484453 | 0.4541916 | 0.650324 |
| ENSG00000109787 | KLF3 | 1969.667217 | 0.301322702 | 0.234439746 | 1.322994967 | 0.18583705 | 0.361748 |
| ENSG00000254789 | | 0.7962304 | 0.3013007 | 0.8595853 | 1.384393 | 0.1662381 | NA |
| ENSG00000253971 | CDC42P3 | 3.848832 | 0.3012299 | 0.7190717 | 0.7975437 | 0.4251353 | 0.62485 |
| ENSG00000228766 | RPL7L1P8 | 73.12725 | 0.3011727 | 0.3192401 | 1.035748 | 0.3003199 | 0.499543 |
| ENSG00000171812 | COL8A2 | 1.15241 | 0.3010319 | 0.785097 | 0.9741212 | 0.3299964 | NA |
| ENSG00000103064 | SLC7A6 | 2267.975484 | 0.301017101 | 0.199014048 | 1.583528863 | 0.11330097 | 0.254549 |
| ENSG00000228903 | RASA4CP | 142.8977 | 0.301008 | 0.3643474 | 0.9279277 | 0.3534451 | 0.555432 |
| ENSG00000251155 | SEPTIN14P4 | 3.236361 | 0.3010034 | 0.6830393 | 0.7467575 | 0.4552099 | 0.651166 |
| ENSG00000110218 | PANX1 | 1039.284395 | 0.300858575 | 0.189148736 | 1.636883479 | 0.10165482 | 0.235575 |
| ENSG00000160310 | PRMT2 | 5662.176 | 0.3006773 | 0.1351346 | 2.259001 | 0.02388334 | 0.078234 |
| ENSG00000179088 | C12orf42 | 36.58249 | 0.3004969 | 0.3823721 | 0.8947362 | 0.3709281 | 0.573076 |
| ENSG00000102241 | HTATSF1 | 7956.633757 | 0.300314674 | 0.218403467 | 1.430149976 | 0.15267398 | 0.315084 |
| ENSG00000233837 | EIF3LP2 | 25.49118 | 0.3001606 | 0.4496094 | 0.8076371 | 0.4192995 | 0.619253 |
| ENSG00000255046 | | 10.96279 | 0.3000688 | 0.4878059 | 0.7709622 | 0.4407293 | 0.63871 |
| ENSG00000274105 | | 9.158964 | 0.3000271 | 0.5075221 | 0.7575129 | 0.4487427 | 0.645304 |
| ENSG00000254610 | | 6.062712 | 0.3000141 | 0.7409121 | 0.8258276 | 0.4089019 | 0.609614 |
| ENSG00000157216 | SSBP3 | 2013.155 | 0.2999737 | 0.1421163 | 2.153087 | 0.03131185 | 0.097006 |
| ENSG00000109381 | ELF2 | 2625.882392 | 0.299963351 | 0.10510142 | 2.878647829 | 0.00399384 | 0.018427 |
| ENSG00000010404 | IDS | 741.3014385 | 0.299797485 | 0.206453494 | 1.504613504 | 0.13242347 | 0.284657 |
| ENSG00000111110 | PPM1H | 4226.079594 | 0.29976391 | 0.181579724 | 1.671937061 | 0.09453673 | 0.223445 |
| ENSG00000234921 | | 0.9077817 | 0.2997619 | 0.8355062 | 1.01659 | 0.3093486 | NA |
| ENSG00000139433 | GLTP | 2408.666 | 0.2997472 | 0.1436693 | 2.120444 | 0.03396859 | 0.103291 |
| ENSG00000181523 | SGSH | 3908.459 | 0.2996718 | 0.1299317 | 2.339617 | 0.01930352 | 0.066057 |
| ENSG00000118997 | DNAH7 | 39.84728902 | 0.299540061 | 0.371666617 | 0.912724062 | 0.36138769 | 0.563414 |
| ENSG00000241286 | RPL29P25 | 1.27156 | 0.2995333 | 0.8365345 | 1.011003 | 0.3120152 | NA |
| ENSG00000255974 | CYP2A6 | 17.21945 | 0.2995218 | 0.5129785 | 0.7449429 | 0.4563062 | 0.652157 |
| ENSG00000227001 | NBPF2P | 32.77422 | 0.2994266 | 0.3043349 | 1.065178 | 0.2867952 | 0.485156 |
| ENSG00000135476 | ESPL1 | 2777.927649 | 0.299404688 | 0.461660194 | 0.794469054 | 0.42692242 | 0.626607 |

| | | | | | | | |
|---|---|---|---|---|---|---|---|
| ENSG00000237661 | TNIP2P1 | 0.8382572 | 0.2993251 | 0.8165933 | 1.168445 | 0.2426273 | NA |
| ENSG00000286805 | | 2.547307 | 0.2993137 | 0.8275015 | 0.8490411 | 0.3958584 | 0.597362 |
| ENSG00000160588 | MPZL3 | 921.2578 | 0.2992303 | 0.1446641 | 2.09091 | 0.03653616 | 0.109007 |
| ENSG00000205937 | RNPS1 | 9773.892 | 0.2992035 | 0.1951469 | 1.583428 | 0.1133239 | 0.254554 |
| ENSG00000223703 | | 0.9565043 | 0.2992035 | 0.8419504 | 1.126985 | 0.2597487 | NA |
| ENSG00000198833 | UBE2J1 | 5265.217 | 0.2991996 | 0.133164 | 2.275462 | 0.02287825 | 0.075707 |
| ENSG00000140455 | USP3 | 1695.301 | 0.2991691 | 0.1058421 | 2.853113 | 0.00432933 | 0.019693 |
| ENSG00000272240 | | 55.43655 | 0.2990545 | 0.3957161 | 0.8699746 | 0.3843143 | 0.586695 |
| ENSG00000267651 | | 3.703821 | 0.2989996 | 0.6435326 | 0.731159 | 0.464682 | 0.658636 |
| ENSG00000235277 | | 180.0598 | 0.2988793 | 0.2646875 | 1.198017 | 0.2309103 | 0.419434 |
| ENSG00000204581 | ACOXL-AS1 | 77.39263 | 0.2988661 | 0.4824853 | 0.7722564 | 0.4399626 | 0.638344 |
| ENSG00000286724 | | 70.1477 | 0.29882 | 0.2946013 | 1.090893 | 0.2753201 | 0.472147 |
| ENSG00000175691 | ZNF77 | 378.633 | 0.2987894 | 0.1823753 | 1.690145 | 0.09100013 | 0.217329 |
| ENSG00000163947 | ARHGEF3 | 433.6573 | 0.2987527 | 0.1728564 | 1.773868 | 0.07608496 | 0.190054 |
| ENSG00000228157 | | 1.190455 | 0.2987138 | 0.849943 | 1.203709 | 0.2287021 | NA |
| ENSG00000238186 | | 14.13139 | 0.29871 | 0.4703197 | 0.787671 | 0.4308891 | 0.630066 |
| ENSG00000287997 | | 0.7793423 | 0.2986758 | 0.8532137 | 1.285806 | 0.1985107 | NA |
| ENSG00000161682 | FAM171A2 | 938.0628 | 0.298655 | 0.1614275 | 1.840468 | 0.06569962 | 0.170167 |
| ENSG00000269352 | PTOV1-AS2 | 236.2351 | 0.2986308 | 0.2710136 | 1.16973 | 0.2421098 | 0.432736 |
| ENSG00000231087 | FDPSP7 | 2.475107 | 0.2985411 | 0.7023148 | 0.7672336 | 0.4429426 | 0.64057 |
| ENSG00000240808 | | 2.349414 | 0.2985021 | 0.7105026 | 0.781297 | 0.4346278 | 0.633545 |
| ENSG00000212327 | RNU6-882P | 19.79641 | 0.2984349 | 0.4244592 | 0.8278 | 0.4077838 | 0.608435 |
| ENSG00000241318 | WDR82P2 | 22.02019 | 0.2983285 | 0.4785563 | 0.7743896 | 0.4387004 | 0.637244 |
| ENSG00000270804 | | 67.39558 | 0.2982698 | 0.3582268 | 0.927683 | 0.353572 | 0.555598 |
| ENSG00000165502 | RPL36AL | 8141.734 | 0.2981824 | 0.1967383 | 1.565905 | 0.1173708 | 0.261027 |
| ENSG00000286558 | | 2.147982 | 0.2981704 | 0.7906233 | 0.9703775 | 0.3318583 | 0.532983 |
| ENSG00000207021 | | 1.006906 | 0.2980334 | 0.7967531 | 0.9701596 | 0.331967 | NA |
| ENSG00000287507 | | 32.29457 | 0.2980095 | 0.5162918 | 0.7512297 | 0.4525144 | 0.648657 |
| ENSG00000119688 | ABCD4 | 2817.697973 | 0.297945194 | 0.151689023 | 1.99595075 | 0.04593928 | 0.130011 |
| ENSG00000258819 | LINC02289 | 1.441694 | 0.297893 | 0.8067221 | 1.137914 | 0.2551565 | NA |
| ENSG00000245748 | | 11.97064 | 0.2978793 | 0.5524427 | 0.7302417 | 0.4652424 | 0.659206 |
| ENSG00000238273 | | 7.592721 | 0.2978747 | 0.741363 | 0.8287084 | 0.4072694 | 0.607954 |
| ENSG00000239043 | SNORD127 | 2.914448 | 0.2977437 | 0.7130243 | 0.7804298 | 0.4351379 | 0.633871 |

| | | | | | | | |
|---|---|---|---|---|---|---|---|
| ENSG00000260510 | | 1.922602 | 0.2977316 | 0.7976664 | 1.047699 | 0.2947774 | 0.49388 |
| ENSG00000124615 | MOCS1 | 626.987686 | 0.297667945 | 0.196224267 | 1.567603025 | 0.11697381 | 0.260335 |
| ENSG00000123411 | IKZF4 | 563.8058175 | 0.297540669 | 0.152995333 | 1.983360302 | 0.0473272 | 0.132879 |
| ENSG00000248494 | LNX1-AS2 | 6.73956 | 0.2974618 | 0.6249138 | 0.7191539 | 0.4720461 | 0.66511 |
| ENSG00000259656 | | 6.055178 | 0.2972695 | 0.6619517 | 0.7339985 | 0.4629497 | 0.657314 |
| ENSG00000224945 | | 43.86681 | 0.2972194 | 0.3901302 | 0.8769828 | 0.380496 | 0.582473 |
| ENSG00000277363 | SRCIN1 | 1321.048 | 0.2971547 | 0.3055203 | 1.063301 | 0.2876456 | 0.485978 |
| ENSG00000275232 | | 1.386851 | 0.2969681 | 0.7545507 | 0.8412031 | 0.4002342 | NA |
| ENSG00000197603 | CPLANE1 | 2604.424 | 0.2968767 | 0.335053 | 0.9887318 | 0.3227944 | 0.523534 |
| ENSG00000103599 | IQCH | 188.4214995 | 0.296863564 | 0.22171429 | 1.398845732 | 0.16185925 | 0.329082 |
| ENSG00000171466 | ZNF562 | 1599.497 | 0.2967666 | 0.1718305 | 1.772782 | 0.07626491 | 0.190351 |
| ENSG00000279583 | | 15.93966 | 0.2966414 | 0.4471108 | 0.7923879 | 0.4281345 | 0.627523 |
| ENSG00000201208 | | 1.672378 | 0.2966015 | 0.7595772 | 0.8478196 | 0.3965384 | 0.597631 |
| ENSG00000119650 | IFT43 | 1817.728596 | 0.296561441 | 0.17238419 | 1.783102845 | 0.07456956 | 0.187298 |
| ENSG00000151704 | KCNJ1 | 2.527222 | 0.2965227 | 0.7517921 | 0.8143789 | 0.4154279 | 0.615259 |
| ENSG00000252768 | RNU6-856P | 1.304644 | 0.296401 | 0.7878114 | 0.9206246 | 0.3572465 | NA |
| ENSG00000240038 | AMY2B | 126.2298 | 0.2963892 | 0.6012405 | 0.7139658 | 0.4752483 | 0.667714 |
| ENSG00000260917 | | 283.7234 | 0.2963777 | 0.2500754 | 1.24946 | 0.2114968 | 0.395635 |
| ENSG00000163251 | FZD5 | 933.8468 | 0.29629 | 0.1789906 | 1.700068 | 0.0891182 | 0.214143 |
| ENSG00000088888 | MAVS | 12502.46901 | 0.296225438 | 0.196599037 | 1.554647727 | 0.12002999 | 0.265403 |
| ENSG00000106153 | CHCHD2 | 8380.930346 | 0.296136074 | 0.246913953 | 1.271775421 | 0.20345292 | 0.385119 |
| ENSG00000262313 | | 0.8905198 | 0.2961085 | 0.8335733 | 1.001038 | 0.3168086 | NA |
| ENSG00000124831 | LRRFIP1 | 6316.795052 | 0.296056067 | 0.189488843 | 1.619376465 | 0.10536629 | 0.241674 |
| ENSG00000149179 | C11orf49 | 1617.658 | 0.2960382 | 0.1674728 | 1.809837 | 0.07032104 | 0.179236 |
| ENSG00000267095 | | 1.589121 | 0.295957 | 0.8415993 | 1.111253 | 0.2664594 | 0.462112 |
| ENSG00000138722 | MMRN1 | 1.062036441 | 0.295789613 | 0.803584254 | 1.028904154 | 0.30352472 | NA |
| ENSG00000253223 | | 90.18898 | 0.2957879 | 0.3707063 | 0.8974391 | 0.3694847 | 0.571644 |
| ENSG00000198205 | ZXDA | 304.8257 | 0.2954351 | 0.2764089 | 1.128529 | 0.2590966 | 0.453458 |
| ENSG00000196917 | HCAR1 | 1590.667 | 0.2954326 | 0.3199629 | 1.007532 | 0.3136791 | 0.514821 |
| ENSG00000280206 | | 255.4215 | 0.2954276 | 0.3297526 | 0.98406 | 0.325086 | 0.526274 |
| ENSG00000155744 | FAM126B | 1247.959 | 0.2954153 | 0.283375 | 1.112764 | 0.2658099 | 0.46143 |
| ENSG00000243478 | AOX2P | 1.067216 | 0.2953698 | 0.819379 | 1.093386 | 0.2742243 | NA |
| ENSG00000163958 | ZDHHC19 | 1.214932 | 0.2953422 | 0.7949286 | 1.043114 | 0.2968956 | NA |

| | | | | | | | |
|---|---|---|---|---|---|---|---|
| ENSG00000131323 | TRAF3 | 2529.625231 | 0.295293652 | 0.140058705 | 2.142690951 | 0.03213792 | 0.098894 |
| ENSG00000140553 | UNC45A | 8422.803 | 0.2952575 | 0.1322461 | 2.255108 | 0.02412658 | 0.078895 |
| ENSG00000268043 | NBPF12 | 2417.943 | 0.2952375 | 0.2216038 | 1.345033 | 0.1786145 | 0.352362 |
| ENSG00000164236 | ANKRD33B | 15.24929 | 0.2952101 | 0.6837274 | 0.7563492 | 0.4494399 | 0.645608 |
| ENSG00000231760 | | 11.15459 | 0.2951503 | 0.5296676 | 0.7399151 | 0.4593515 | 0.654889 |
| ENSG00000174405 | LIG4 | 992.9612 | 0.2951493 | 0.294812 | 1.077098 | 0.2814363 | 0.479049 |
| ENSG00000196678 | ERI2 | 1469.228 | 0.2951373 | 0.1982712 | 1.535466 | 0.1246693 | 0.272643 |
| ENSG00000228364 | SAR1AP1 | 1.705228 | 0.2949976 | 0.7914863 | 0.9376611 | 0.3484186 | 0.550162 |
| ENSG00000267060 | PTGES3L | 14.04275 | 0.2949731 | 0.5227231 | 0.7352834 | 0.4621669 | 0.656777 |
| ENSG00000233996 | KDM3AP1 | 1.426963 | 0.2948293 | 0.7646954 | 0.8416337 | 0.399993 | NA |
| ENSG00000273361 | | 1.433853 | 0.2947657 | 0.7598886 | 0.8592825 | 0.3901847 | NA |
| ENSG00000148396 | SEC16A | 13090.24 | 0.2946436 | 0.1811992 | 1.64433 | 0.100108 | 0.232986 |
| ENSG00000272973 | | 4.094781 | 0.2946067 | 0.6782072 | 0.7406421 | 0.4589105 | 0.654576 |
| ENSG00000154813 | DPH3 | 2053.298 | 0.2944464 | 0.167794 | 1.790457 | 0.07338044 | 0.185187 |
| ENSG00000269113 | TRABD2B | 36.21644 | 0.2943934 | 0.5054022 | 0.7523799 | 0.4518227 | 0.648155 |
| ENSG00000236459 | HNRNPA1P22 | 2.151826 | 0.2943432 | 0.7786457 | 0.9834696 | 0.3253764 | 0.52668 |
| ENSG00000112200 | ZNF451 | 1769.527411 | 0.294327664 | 0.139858308 | 2.13848397 | 0.03247748 | 0.09971 |
| ENSG00000266589 | MIR4512 | 4.74644 | 0.2943047 | 0.6253193 | 0.7216348 | 0.4705191 | 0.66397 |
| ENSG00000214548 | MEG3 | 1.248106 | 0.2942318 | 0.8314401 | 1.117712 | 0.2636902 | NA |
| ENSG00000275765 | | 28.96452 | 0.2942279 | 0.4250825 | 0.8133241 | 0.4160323 | 0.615935 |
| ENSG00000286943 | | 1.933937 | 0.2942252 | 0.7366201 | 0.8007118 | 0.4232985 | 0.623285 |
| ENSG00000179965 | ZNF771 | 372.6161 | 0.2942111 | 0.1901305 | 1.600115 | 0.1095731 | 0.248655 |
| ENSG00000212588 | SNORA26 | 35.66343 | 0.2941649 | 0.3644772 | 0.9111977 | 0.3621912 | 0.564204 |
| ENSG00000139910 | NOVA1 | 302.4387 | 0.2941606 | 0.3331395 | 0.9678803 | 0.3331042 | 0.534016 |
| ENSG00000185634 | SHC4 | 14.97962 | 0.2941498 | 0.4708245 | 0.7704619 | 0.4410259 | 0.638841 |
| ENSG00000235106 | BRD3OS | 2246.296 | 0.2938771 | 0.133312 | 2.228405 | 0.0258535 | 0.083242 |
| ENSG00000155849 | ELMO1 | 91.49779 | 0.2937267 | 0.4440676 | 0.8043023 | 0.4212224 | 0.621058 |
| ENSG00000085978 | ATG16L1 | 2159.274677 | 0.293715149 | 0.100396093 | 2.941989816 | 0.00326111 | 0.01557 |
| ENSG00000250107 | CACNA1G-AS1 | 5.618294 | 0.2936801 | 0.6449465 | 0.7266393 | 0.4674469 | 0.661023 |
| ENSG00000102858 | MGRN1 | 3952.917923 | 0.293584513 | 0.138889549 | 2.093599272 | 0.03629569 | 0.108695 |
| ENSG00000115257 | PCSK4 | 291.9635044 | 0.293534699 | 0.241077727 | 1.281380491 | 0.20006005 | 0.380432 |
| ENSG00000269901 | | 13.25893 | 0.2935 | 0.5002538 | 0.7466991 | 0.4552452 | 0.651166 |
| ENSG00000012963 | UBR7 | 3677.643246 | 0.29344082 | 0.258224099 | 1.209372588 | 0.22651974 | 0.414163 |

| | | | | | | | |
|---|---|---|---|---|---|---|---|
| ENSG00000145358 | DDIT4L | 19.01473 | 0.2934111 | 0.5898583 | 0.7027962 | 0.4821828 | 0.673631 |
| ENSG00000223668 | EEF1A1P24 | 4.3228 | 0.2934092 | 0.6578249 | 0.7278125 | 0.4667284 | 0.660515 |
| ENSG00000118200 | CAMSAP2 | 7193.624275 | 0.293379456 | 0.208402419 | 1.457100197 | 0.14508872 | 0.303915 |
| ENSG00000184117 | NIPSNAP1 | 8295.992 | 0.2931417 | 0.1866863 | 1.627916 | 0.1035426 | 0.238597 |
| ENSG00000224383 | PRR29 | 30.32648 | 0.2931323 | 0.5576326 | 0.7327237 | 0.463727 | 0.657836 |
| ENSG00000214318 | ATP5MC1P6 | 1.073772 | 0.2930867 | 0.7960692 | 0.9432201 | 0.3455683 | NA |
| ENSG00000213759 | UGT2B11 | 1.471318 | 0.293084 | 0.8401339 | 1.229643 | 0.2188308 | NA |
| ENSG00000169683 | LRRC45 | 4364.862 | 0.2930719 | 0.2866538 | 1.101163 | 0.2708259 | 0.467142 |
| ENSG00000284391 | | 3.726833 | 0.2929682 | 0.7154765 | 0.7814483 | 0.4345389 | 0.633489 |
| ENSG00000129968 | ABHD17A | 3381.888916 | 0.292936236 | 0.124684931 | 2.360198216 | 0.01826517 | 0.063188 |
| ENSG00000225099 | ATP6V1E1P1 | 63.95094 | 0.2929074 | 0.2825289 | 1.103568 | 0.2697806 | 0.466077 |
| ENSG00000143545 | RAB13 | 924.3862 | 0.2928931 | 0.202023 | 1.50008 | 0.1335936 | 0.286338 |
| ENSG00000119630 | PGF | 14.06695986 | 0.292809114 | 0.598445268 | 0.715930466 | 0.47403429 | 0.666735 |
| ENSG00000287263 | | 58.21906 | 0.2927708 | 0.327995 | 0.9709001 | 0.331598 | 0.532759 |
| ENSG00000102870 | ZNF629 | 4982.105907 | 0.292680249 | 0.100009926 | 2.934957568 | 0.00333593 | 0.015873 |
| ENSG00000165118 | C9orf64 | 1069.648 | 0.2926324 | 0.1661448 | 1.80028 | 0.07181643 | 0.182139 |
| ENSG00000250329 | KDELC1P1 | 1.142621 | 0.2924829 | 0.8334622 | 1.028847 | 0.3035517 | NA |
| ENSG00000233057 | EEF1A1P14 | 2.086745 | 0.2924716 | 0.6942114 | 0.7396025 | 0.4595412 | 0.655019 |
| ENSG00000038382 | TRIO | 11529.4017 | 0.292450937 | 0.239845895 | 1.288975829 | 0.19740649 | 0.377368 |
| ENSG00000152969 | JAKMIP1 | 0.9388823 | 0.2923967 | 0.8349088 | 0.9890554 | 0.3226361 | NA |
| ENSG00000233175 | | 57.67414 | 0.2923962 | 0.3956786 | 0.8482992 | 0.3962714 | 0.597546 |
| ENSG00000246225 | | 35.14518 | 0.2922976 | 0.3860148 | 0.8650567 | 0.3870076 | 0.589349 |
| ENSG00000112367 | FIG4 | 1081.420391 | 0.292294092 | 0.169095609 | 1.776837183 | 0.07559503 | 0.189194 |
| ENSG00000247624 | CPEB2-DT | 3.643205 | 0.2922014 | 0.6693652 | 0.729435 | 0.4657356 | 0.659672 |
| ENSG00000204103 | MAFB | 370.1485 | 0.2921932 | 0.3665532 | 0.8957639 | 0.3703789 | 0.572461 |
| ENSG00000168243 | GNG4 | 232.9397 | 0.2920301 | 0.5143581 | 0.7322929 | 0.4639898 | 0.658076 |
| ENSG00000260685 | | 0.7956557 | 0.2919688 | 0.8455466 | 1.210158 | 0.2262185 | NA |
| ENSG00000253893 | FAM85B | 1.839358 | 0.2919651 | 0.7208277 | 0.7683835 | 0.4422594 | 0.640069 |
| ENSG00000236617 | | 51.33697 | 0.2919447 | 0.3329835 | 0.9701955 | 0.3319491 | 0.533049 |
| ENSG00000103061 | SLC7A6OS | 476.2509474 | 0.291941844 | 0.174755448 | 1.717587098 | 0.08587196 | 0.208188 |
| ENSG00000188822 | CNR2 | 1.029896 | 0.2919407 | 0.8018043 | 0.9862659 | 0.3240026 | NA |
| ENSG00000213190 | MLLT11 | 262.4814 | 0.2919195 | 0.3061689 | 1.035945 | 0.300228 | 0.499484 |
| ENSG00000213600 | | 1.985546 | 0.2919106 | 0.7382143 | 0.8060203 | 0.4202311 | 0.620112 |

| | | | | | | | |
|---|---|---|---|---|---|---|---|
| ENSG00000240895 | | 1.463114 | 0.2918855 | 0.792287 | 0.938035 | 0.3482264 | NA |
| ENSG00000198146 | ZNF770 | 2100.783 | 0.2918172 | 0.3582248 | 0.9169572 | 0.359165 | 0.561104 |
| ENSG00000165288 | BRWD3 | 2343.744 | 0.2917495 | 0.3135964 | 1.011533 | 0.3117614 | 0.512669 |
| ENSG00000211459 | MT-RNR1 | 34760.82 | 0.2916997 | 0.2118192 | 1.499702 | 0.1336916 | 0.286408 |
| ENSG00000180105 | ACER2P1 | 17.60004 | 0.2914711 | 0.5909672 | 0.7136432 | 0.4754478 | 0.667779 |
| ENSG00000114648 | KLHL18 | 1272.839607 | 0.291381256 | 0.157581733 | 1.886832902 | 0.0591828 | 0.157512 |
| ENSG00000233225 | SSBL3P1 | 610.7418 | 0.2913588 | 0.1632989 | 1.834627 | 0.06656103 | 0.171912 |
| ENSG00000149489 | ROM1 | 189.7195 | 0.2912399 | 0.2991285 | 1.040931 | 0.2979077 | 0.497084 |
| ENSG00000162337 | LRP5 | 7273.921 | 0.2912325 | 0.160184 | 1.854162 | 0.06371599 | 0.166228 |
| ENSG00000267282 | | 3.348691 | 0.2912003 | 0.673962 | 0.7314457 | 0.464507 | 0.658522 |
| ENSG00000104388 | RAB2A | 5457.836035 | 0.291155552 | 0.112487888 | 2.602060577 | 0.00926655 | 0.036583 |
| ENSG00000158373 | H2BC5 | 1943.86 | 0.2910214 | 0.2172652 | 1.392092 | 0.1638944 | 0.331924 |
| ENSG00000148841 | ITPRIP | 303.1475 | 0.2909813 | 0.4711217 | 0.7591292 | 0.4477753 | 0.644436 |
| ENSG00000184640 | SEPTIN9 | 14956.85 | 0.2909813 | 0.2319981 | 1.317545 | 0.1876559 | 0.364461 |
| ENSG00000274099 | ABCB10P1 | 3.240383 | 0.290975 | 0.706018 | 0.7568253 | 0.4491546 | 0.645478 |
| ENSG00000206567 | | 701.5368 | 0.2908945 | 0.304838 | 1.03629 | 0.3000667 | 0.499366 |
| ENSG00000256092 | SBNO1-AS1 | 101.0819 | 0.2907892 | 0.3080999 | 1.028888 | 0.3035322 | 0.503157 |
| ENSG00000169692 | AGPAT2 | 3819.655 | 0.2907728 | 0.1807509 | 1.648111 | 0.09932999 | 0.231581 |
| ENSG00000180902 | D2HGDH | 2719.413 | 0.2907571 | 0.1300888 | 2.266802 | 0.02340234 | 0.076957 |
| ENSG00000162378 | ZYG11B | 1974.478 | 0.290664 | 0.1855295 | 1.633245 | 0.1024174 | 0.236702 |
| ENSG00000226851 | | 4.518817 | 0.2906233 | 0.6159217 | 0.7056911 | 0.4803803 | 0.671961 |
| ENSG00000227040 | | 19.40445 | 0.2905449 | 0.4162787 | 0.8205963 | 0.4118762 | 0.612204 |
| ENSG00000224019 | RPL21P32 | 1.328031 | 0.2904903 | 0.7750669 | 0.8753873 | 0.3813632 | NA |
| ENSG00000163719 | MTMR14 | 4383.137 | 0.290424 | 0.1398579 | 2.08746 | 0.0368466 | 0.109726 |
| ENSG00000259557 | HMGN1P26 | 2.957501 | 0.2904082 | 0.7384247 | 0.81492 | 0.4151181 | 0.615074 |
| ENSG00000233055 | | 0.8626059 | 0.2903396 | 0.8044998 | 0.9717961 | 0.331152 | NA |
| ENSG00000175170 | FAM182B | 114.3143 | 0.2903239 | 0.3528889 | 0.9109144 | 0.3623405 | 0.56437 |
| ENSG00000138468 | SENP7 | 1506.140964 | 0.290295654 | 0.241755121 | 1.257087773 | 0.20872185 | 0.391634 |
| ENSG00000258748 | | 25.94561 | 0.2902573 | 0.4541062 | 0.7785825 | 0.4362257 | 0.634829 |
| ENSG00000250966 | H3P14 | 2.94911 | 0.2902224 | 0.6788109 | 0.729263 | 0.4658408 | 0.659777 |
| ENSG00000177455 | CD19 | 1.140081 | 0.2902172 | 0.8463735 | 1.168881 | 0.2424517 | NA |
| ENSG00000237940 | LINC01238 | 53.72961 | 0.290071 | 0.4506923 | 0.7811792 | 0.4346971 | 0.63358 |
| ENSG00000160685 | ZBTB7B | 4977.011 | 0.290056 | 0.1762599 | 1.70524 | 0.08814961 | 0.212334 |

| | | | | | | | |
|---|---|---|---|---|---|---|---|
| ENSG00000116793 | PHTF1 | 2127.31761 | 0.290004582 | 0.259847952 | 1.174448788 | 0.24021531 | 0.430508 |
| ENSG00000143801 | PSEN2 | 1371.651 | 0.2900018 | 0.1959781 | 1.530494 | 0.1258946 | 0.274374 |
| ENSG00000276790 | | 0.9413763 | 0.2899845 | 0.8429526 | 1.275003 | 0.2023081 | NA |
| ENSG00000122643 | NT5C3A | 2578.124236 | 0.28994428 | 0.184096253 | 1.634936552 | 0.10206235 | 0.236111 |
| ENSG00000100393 | EP300 | 4820.816973 | 0.289824824 | 0.258816883 | 1.182388677 | 0.23705151 | 0.426738 |
| ENSG00000254337 | | 252.7551 | 0.289824 | 0.6063476 | 0.6786743 | 0.4973442 | 0.686082 |
| ENSG00000020426 | MNAT1 | 1877.12256 | 0.289744986 | 0.191403723 | 1.559470897 | 0.11888497 | 0.263309 |
| ENSG00000141526 | SLC16A3 | 664.0679 | 0.2897351 | 0.3515297 | 0.9167639 | 0.3592664 | 0.561161 |
| ENSG00000174748 | RPL15 | 37924.21 | 0.2895988 | 0.1758249 | 1.743108 | 0.08131481 | 0.199838 |
| ENSG00000040608 | RTN4R | 376.696329 | 0.289586281 | 0.251728158 | 1.214001132 | 0.22474729 | 0.411786 |
| ENSG00000205643 | CDPF1 | 350.845 | 0.2895593 | 0.1938114 | 1.516659 | 0.1293527 | 0.279662 |
| ENSG00000167257 | RNF214 | 833.5082 | 0.2895236 | 0.159086 | 1.852601 | 0.06393951 | 0.166638 |
| ENSG00000113845 | TIMMDC1 | 4564.662544 | 0.289488927 | 0.147159866 | 1.991600425 | 0.04641492 | 0.130982 |
| ENSG00000261570 | | 3.109297 | 0.2894732 | 0.7326702 | 0.8036263 | 0.4216128 | 0.621426 |
| ENSG00000275945 | EIF3FP1 | 7.802026 | 0.2893651 | 0.5720189 | 0.7092897 | 0.4781447 | 0.670104 |
| ENSG00000260102 | LINC01070 | 0.996723 | 0.2893485 | 0.8320463 | 1.155885 | 0.2477284 | NA |
| ENSG00000267311 | | 4.067724 | 0.289244 | 0.7291183 | 0.7835532 | 0.4333023 | 0.632237 |
| ENSG00000157890 | MEGF11 | 13.01822 | 0.2891831 | 0.4332458 | 0.7929891 | 0.4277842 | 0.627171 |
| ENSG00000187187 | ZNF546 | 234.6113 | 0.2891709 | 0.3367615 | 0.9508859 | 0.3416623 | 0.54292 |
| ENSG00000153246 | PLA2R1 | 479.0591 | 0.289156 | 0.2914642 | 1.074766 | 0.2824797 | 0.480302 |
| ENSG00000273706 | LHX1 | 165.188 | 0.2891136 | 0.3467693 | 0.9125583 | 0.3614749 | 0.563451 |
| ENSG00000278456 | | 6.012168 | 0.2889125 | 0.8219299 | 0.8602941 | 0.3896269 | 0.591577 |
| ENSG00000108984 | MAP2K6 | 175.2822859 | 0.288890266 | 0.524140289 | 0.729637497 | 0.4656118 | 0.659595 |
| ENSG00000069248 | NUP133 | 3650.113887 | 0.288699051 | 0.136978107 | 2.120894356 | 0.03393069 | 0.103206 |
| ENSG00000176700 | SCAND2P | 151.958 | 0.2886828 | 0.1872618 | 1.583688 | 0.1132647 | 0.254532 |
| ENSG00000014914 | MTMR11 | 1752.861377 | 0.288639968 | 0.263374786 | 1.161558231 | 0.24541496 | 0.437085 |
| ENSG00000226581 | LINC02848 | 14.90786 | 0.2886339 | 0.4962127 | 0.7346325 | 0.4625634 | 0.656982 |
| ENSG00000101082 | SLA2 | 2.620695218 | 0.288605965 | 0.676568544 | 0.727950784 | 0.46664371 | 0.66043 |
| ENSG00000136378 | ADAMTS7 | 142.223836 | 0.288604474 | 0.368152136 | 0.898436152 | 0.36895307 | 0.571087 |
| ENSG00000131797 | CLUHP3 | 2216.225001 | 0.288503655 | 0.190516932 | 1.562317823 | 0.11821314 | 0.262293 |
| ENSG00000213110 | RNF11P1 | 1.435188 | 0.2884465 | 0.8383114 | 1.125189 | 0.2605088 | NA |
| ENSG00000285744 | | 10.33166 | 0.288375 | 0.6412342 | 0.7060745 | 0.4801418 | 0.671865 |
| ENSG00000175073 | VCPIP1 | 1261.213 | 0.2883552 | 0.218618 | 1.363044 | 0.1728687 | 0.344222 |

| | | | | | | | |
|---|---|---|---|---|---|---|---|
| ENSG00000180957 | PITPNB | 4243.657 | 0.2882163 | 0.1277645 | 2.285742 | 0.02226938 | 0.074256 |
| ENSG00000099953 | MMP11 | 113.1515576 | 0.288163106 | 0.300006426 | 1.038518772 | 0.2990286 | 0.498268 |
| ENSG00000185730 | ZNF696 | 1221.3 | 0.2881297 | 0.1254826 | 2.324125 | 0.02011877 | 0.068303 |
| ENSG00000126233 | SLURP1 | 13.44331302 | 0.288125264 | 0.51487674 | 0.726782354 | 0.46735929 | 0.661023 |
| ENSG00000249035 | CLMAT3 | 2.507424 | 0.2881132 | 0.6957906 | 0.7338233 | 0.4630564 | 0.657314 |
| ENSG00000236044 | FABP5P2 | 1.771481 | 0.2879651 | 0.7573555 | 0.8585598 | 0.3905834 | 0.592453 |
| ENSG00000122126 | OCRL | 6583.327618 | 0.287888849 | 0.187943737 | 1.571901246 | 0.11597346 | 0.258824 |
| ENSG00000178977 | LINC00324 | 8.582097 | 0.287767 | 0.6081705 | 0.6971077 | 0.4857354 | 0.676779 |
| ENSG00000278281 | MIR7856 | 1.17844 | 0.2877181 | 0.771023 | 0.8698841 | 0.3843637 | NA |
| ENSG00000215812 | ZNF847P | 1.055425 | 0.2877093 | 0.8160983 | 1.171862 | 0.2412524 | NA |
| ENSG00000183562 | | 16.74817 | 0.2876361 | 0.4552899 | 0.7655288 | 0.4439567 | 0.641443 |
| ENSG00000215874 | CAPNS1P1 | 0.9139449 | 0.2876162 | 0.8140613 | 1.169644 | 0.2421445 | NA |
| ENSG00000185019 | UBOX5 | 488.7471 | 0.2874789 | 0.1804493 | 1.636512 | 0.1017325 | 0.235632 |
| ENSG00000240429 | LRRFIP1P1 | 239.5607 | 0.2874735 | 0.2749981 | 1.10996 | 0.2670161 | 0.46278 |
| ENSG00000151726 | ACSL1 | 15051.42 | 0.2874539 | 0.1883 | 1.572534 | 0.1158268 | 0.258649 |
| ENSG00000213328 | | 1.36645 | 0.2873217 | 0.8075682 | 1.013626 | 0.3107614 | NA |
| ENSG00000253972 | MAL2-AS1 | 38.50787 | 0.2873114 | 0.3962192 | 0.8424662 | 0.3995271 | 0.600519 |
| ENSG00000197734 | C14orf178 | 4.325804 | 0.2872138 | 0.6654774 | 0.7131819 | 0.4757332 | 0.668063 |
| ENSG00000241644 | INMT | 0.9555609 | 0.2871603 | 0.8233721 | 1.076453 | 0.2817249 | NA |
| ENSG00000112308 | C6orf62 | 6938.19344 | 0.287088377 | 0.157071507 | 1.903144959 | 0.05702163 | 0.153047 |
| ENSG00000226318 | RPS3AP38 | 7.376336 | 0.2870802 | 0.6440851 | 0.7121528 | 0.4763702 | 0.668675 |
| ENSG00000235582 | | 12.89612 | 0.2870517 | 0.4962363 | 0.734733 | 0.4625021 | 0.656982 |
| ENSG00000067208 | EVI5 | 570.1790248 | 0.28700668 | 0.199198216 | 1.486973029 | 0.13702193 | 0.291313 |
| ENSG00000111879 | FAM184A | 505.5002424 | 0.286996776 | 0.21798089 | 1.3706049 | 0.17049816 | 0.34116 |
| ENSG00000093010 | COMT | 8710.197616 | 0.286993179 | 0.258314272 | 1.183828116 | 0.23648111 | 0.42607 |
| ENSG00000171295 | ZNF440 | 344.7193 | 0.2869438 | 0.2026712 | 1.464208 | 0.1431372 | 0.300943 |
| ENSG00000285095 | | 2.657426 | 0.2868401 | 0.7341614 | 0.7955561 | 0.4262901 | 0.626201 |
| ENSG00000271109 | | 42.03652 | 0.2867817 | 0.4228705 | 0.7991959 | 0.4241769 | 0.624063 |
| ENSG00000248113 | | 48.82265 | 0.2867598 | 0.3924051 | 0.8403491 | 0.4007127 | 0.601703 |
| ENSG00000228544 | CCDC183-AS1 | 237.7162 | 0.2867469 | 0.2248104 | 1.326615 | 0.184636 | 0.360017 |
| ENSG00000227193 | | 1.163894 | 0.2867285 | 0.7679291 | 0.844393 | 0.3984498 | NA |
| ENSG00000224746 | | 6.314021 | 0.2867001 | 0.6689012 | 0.7191176 | 0.4720685 | 0.66511 |
| ENSG00000166170 | BAG5 | 3908.138 | 0.2866599 | 0.1474272 | 2.00657 | 0.04479549 | 0.127586 |

| | | | | | | | |
|---|---|---|---|---|---|---|---|
| ENSG00000259883 | EHD4-AS1 | 2.832843 | 0.2864544 | 0.6787387 | 0.7235774 | 0.4693252 | 0.662848 |
| ENSG00000136950 | ARPC5L | 3958.824126 | 0.286440429 | 0.187792501 | 1.55665113 | 0.11955334 | 0.264525 |
| ENSG00000286696 | | 7.083428 | 0.2861183 | 0.6756123 | 0.7220466 | 0.4702658 | 0.663753 |
| ENSG00000168852 | TPTE2P5 | 37.44102 | 0.2860705 | 0.4224578 | 0.7992795 | 0.4241284 | 0.624026 |
| ENSG00000260267 | | 351.7595 | 0.2860085 | 0.2226213 | 1.340779 | 0.1799922 | 0.354168 |
| ENSG00000157881 | PANK4 | 1144.705 | 0.2859476 | 0.1431033 | 2.027183 | 0.04264374 | 0.122965 |
| ENSG00000167110 | GOLGA2 | 8134.864 | 0.2858944 | 0.1592242 | 1.826193 | 0.06782118 | 0.174337 |
| ENSG00000172738 | TMEM217 | 15.65707 | 0.2858191 | 0.4766926 | 0.7391423 | 0.4598206 | 0.655274 |
| ENSG00000116117 | PARD3B | 924.1855654 | 0.285727967 | 0.314115672 | 0.994092619 | 0.32017778 | 0.520963 |
| ENSG00000257802 | MRS2P2 | 1.314352 | 0.2856682 | 0.7985253 | 1.020371 | 0.3075523 | NA |
| ENSG00000129625 | REEP5 | 15195.93086 | 0.285648923 | 0.138865125 | 1.992910056 | 0.0462713 | 0.130716 |
| ENSG00000255909 | PDCD5P1 | 45.44075 | 0.2855926 | 0.341873 | 0.9300116 | 0.3523651 | 0.554161 |
| ENSG00000168528 | SERINC2 | 984.0358 | 0.2855673 | 0.2400701 | 1.268355 | 0.2046713 | 0.386626 |
| ENSG00000269466 | H3Y1 | 1.375128 | 0.2854848 | 0.8374342 | 1.359232 | 0.1740732 | NA |
| ENSG00000243321 | LINC01998 | 1.808015 | 0.2854319 | 0.7930327 | 0.9565701 | 0.3387843 | 0.54001 |
| ENSG00000276535 | | 7.493566 | 0.2852536 | 0.6057322 | 0.7020661 | 0.4826379 | 0.674019 |
| ENSG00000135631 | RAB11FIP5 | 810.1118558 | 0.285198254 | 0.224739285 | 1.333792932 | 0.18227173 | 0.357242 |
| ENSG00000204241 | LINC02731 | 5.826446 | 0.2851512 | 0.6730541 | 0.7094922 | 0.4780191 | 0.670034 |
| ENSG00000257800 | FNBP1P1 | 86.64481 | 0.2851223 | 0.3670412 | 0.8688491 | 0.3849297 | 0.587331 |
| ENSG00000131849 | ZNF132 | 212.04954 | 0.285080085 | 0.287058335 | 1.064686493 | 0.28701783 | 0.485355 |
| ENSG00000258102 | MAP1LC3B2 | 1032.702 | 0.2849266 | 0.2236439 | 1.324671 | 0.1852804 | 0.360929 |
| ENSG00000175471 | MCTP1 | 98.67106 | 0.2848575 | 0.297893 | 1.034598 | 0.3008565 | 0.50006 |
| ENSG00000216915 | GPR89P | 0.9122018 | 0.2848124 | 0.805331 | 0.9718665 | 0.331117 | NA |
| ENSG00000176018 | LYSMD3 | 2120.288 | 0.2847018 | 0.223222 | 1.324597 | 0.185305 | 0.360951 |
| ENSG00000237350 | CDC42P6 | 1036.157 | 0.2846896 | 0.1907921 | 1.542589 | 0.1229305 | 0.269951 |
| ENSG00000105894 | PTN | 9.377491828 | 0.284675922 | 0.671455047 | 0.731226055 | 0.46464109 | 0.658636 |
| ENSG00000116205 | TCEANC2 | 925.2132902 | 0.284653538 | 0.13612805 | 2.118774606 | 0.03410952 | 0.103631 |
| ENSG00000286112 | | 10.78343 | 0.2846535 | 0.4759495 | 0.7348999 | 0.4624004 | 0.656909 |
| ENSG00000270149 | | 21.57646 | 0.2846146 | 0.4012852 | 0.8215641 | 0.411325 | 0.61159 |
| ENSG00000153029 | MR1 | 1473.192 | 0.2846071 | 0.3086794 | 0.9967616 | 0.3188802 | 0.519932 |
| ENSG00000180015 | | 169.2408 | 0.2845978 | 0.1909029 | 1.536454 | 0.1244272 | 0.272266 |
| ENSG00000270104 | | 2.772969 | 0.2845305 | 0.7237612 | 0.7744487 | 0.4386655 | 0.637243 |
| ENSG00000160746 | ANO10 | 1389.278 | 0.2844761 | 0.1468068 | 1.972167 | 0.04859052 | 0.135538 |

| | | | | | | | |
|---|---|---|---|---|---|---|---|
| ENSG00000247556 | OIP5-AS1 | 4927.961 | 0.2844713 | 0.2038222 | 1.441314 | 0.1494959 | 0.310455 |
| ENSG00000270733 | | 0.826698 | 0.2843937 | 0.7919679 | 0.897902 | 0.3692378 | NA |
| ENSG00000233321 | LINC02669 | 1.930364 | 0.2843477 | 0.7374802 | 0.7937708 | 0.4273289 | 0.626826 |
| ENSG00000164091 | WDR82 | 7798.971 | 0.2843436 | 0.1214929 | 2.368293 | 0.01787037 | 0.062051 |
| ENSG00000165512 | ZNF22 | 1248.209 | 0.2843319 | 0.1894941 | 1.532748 | 0.1253379 | 0.273565 |
| ENSG00000163884 | KLF15 | 44.82027 | 0.2843147 | 0.484063 | 0.7314128 | 0.464527 | 0.658522 |
| ENSG00000271156 | | 5.156296 | 0.2842942 | 0.6091796 | 0.6886299 | 0.4910562 | 0.681494 |
| ENSG00000131748 | STARD3 | 2208.283115 | 0.284042873 | 0.143735886 | 2.009497335 | 0.04448442 | 0.126964 |
| ENSG00000234264 | DEPDC1-AS1 | 18.845 | 0.2839015 | 0.5923445 | 0.6916162 | 0.4891784 | 0.680007 |
| ENSG00000253704 | | 26.29477 | 0.2838854 | 0.4517209 | 0.7571856 | 0.4489387 | 0.645416 |
| ENSG00000226439 | | 1.235931 | 0.2836712 | 0.8101049 | 1.152593 | 0.2490777 | NA |
| ENSG00000112701 | SENP6 | 5902.213457 | 0.283658175 | 0.101123955 | 2.812588735 | 0.00491445 | 0.021853 |
| ENSG00000232545 | | 43.35689 | 0.2835254 | 0.5070781 | 0.7148464 | 0.4747039 | 0.667394 |
| ENSG00000236312 | RPL34P34 | 38.47106 | 0.2835063 | 0.3765786 | 0.8532739 | 0.3935074 | 0.595147 |
| ENSG00000287265 | | 27.12312 | 0.2835022 | 0.3646017 | 0.8716799 | 0.383383 | 0.585745 |
| ENSG00000154814 | OXNAD1 | 1320.051 | 0.2832328 | 0.2081927 | 1.402491 | 0.1607687 | 0.327324 |
| ENSG00000143799 | PARP1 | 10268.37 | 0.2832117 | 0.2781784 | 1.085031 | 0.2779078 | 0.475002 |
| ENSG00000225778 | PROSER2-AS1 | 27.94591 | 0.2831795 | 0.4272427 | 0.7823845 | 0.4339887 | 0.632995 |
| ENSG00000287356 | | 95.95563 | 0.2831703 | 0.3485407 | 0.9001359 | 0.3680479 | 0.570158 |
| ENSG00000230547 | HMGB1P11 | 2.342003 | 0.2830562 | 0.7474411 | 0.8018417 | 0.4226445 | 0.622636 |
| ENSG00000178372 | CALML5 | 592.1596 | 0.2830413 | 0.5797092 | 0.6717282 | 0.5017567 | 0.689342 |
| ENSG00000168214 | RBPJ | 3015.141 | 0.2829307 | 0.1554225 | 1.856389 | 0.06339813 | 0.165627 |
| ENSG00000262188 | LINC01978 | 14.86208 | 0.2827291 | 0.4589473 | 0.753753 | 0.4509975 | 0.64722 |
| ENSG00000259579 | | 1.809987 | 0.2825226 | 0.821573 | 0.8452311 | 0.3979818 | 0.599092 |
| ENSG00000225077 | LINC00337 | 25.09102 | 0.2825086 | 0.529305 | 0.70381 | 0.4815511 | 0.672996 |
| ENSG00000230658 | KLHL7-DT | 18.56368 | 0.2824868 | 0.4160437 | 0.7927476 | 0.4279249 | 0.627319 |
| ENSG00000228794 | LINC01128 | 588.5325 | 0.2824664 | 0.1522001 | 1.900959 | 0.05730739 | 0.153627 |
| ENSG00000262902 | MTCO1P40 | 16.46315 | 0.2824027 | 0.5433676 | 0.7007676 | 0.483448 | 0.67476 |
| ENSG00000219201 | | 527.5217 | 0.2823251 | 0.2546697 | 1.178394 | 0.2386397 | 0.428494 |
| ENSG00000181201 | H2BU2P | 0.9719534 | 0.2821619 | 0.8384473 | 1.064955 | 0.2868963 | NA |
| ENSG00000169967 | MAP3K2 | 3857.156 | 0.2821509 | 0.2046878 | 1.4262 | 0.1538106 | 0.317012 |
| ENSG00000157353 | FCSK | 2273.081 | 0.2820675 | 0.1028618 | 2.769996 | 0.0056057 | 0.024328 |
| ENSG00000229587 | | 2.236056 | 0.2820086 | 0.7040631 | 0.7425817 | 0.457735 | 0.65359 |

| | | | | | | | |
|---|---|---|---|---|---|---|---|
| ENSG00000215414 | PSMA6P1 | 1346.448 | 0.2819951 | 0.2308461 | 1.235037 | 0.2168167 | 0.402249 |
| ENSG00000184207 | PGP | 3403.312 | 0.2819776 | 0.2885134 | 1.056756 | 0.2906232 | 0.489233 |
| ENSG00000166145 | SPINT1 | 11151.49 | 0.2819638 | 0.1256577 | 2.27264 | 0.02304789 | 0.076136 |
| ENSG00000257883 | | 2.356767 | 0.2818799 | 0.7706086 | 0.8316009 | 0.4056342 | 0.606522 |
| ENSG00000250327 | RPSAP70 | 1.830254 | 0.281756 | 0.7594518 | 0.8356302 | 0.403363 | 0.604214 |
| ENSG00000227018 | IL6STP1 | 2.808983 | 0.2817425 | 0.750975 | 0.8226441 | 0.4107104 | 0.61105 |
| ENSG00000276169 | MIR6871 | 2.375396 | 0.2817184 | 0.7062426 | 0.7424673 | 0.4578042 | 0.65359 |
| ENSG00000172840 | PDP2 | 1192.388 | 0.2815723 | 0.2423594 | 1.252714 | 0.2103098 | 0.394029 |
| ENSG00000275812 | | 5.032077 | 0.2814703 | 0.6839171 | 0.718443 | 0.4724842 | 0.665381 |
| ENSG00000170365 | SMAD1 | 1054.624 | 0.281456 | 0.1888541 | 1.534321 | 0.1249507 | 0.27298 |
| ENSG00000163975 | MELTF | 705.3341 | 0.2814489 | 0.2808407 | 1.074224 | 0.2827222 | 0.480585 |
| ENSG00000260386 | LDC1P | 1.280461 | 0.2813726 | 0.7740138 | 0.8739026 | 0.3821713 | NA |
| ENSG00000168412 | MTNR1A | 1.994983 | 0.2812922 | 0.7294393 | 0.7614121 | 0.446411 | 0.643407 |
| ENSG00000262668 | | 11.20922 | 0.2811839 | 0.5945189 | 0.6906741 | 0.4897704 | 0.680368 |
| ENSG00000258199 | | 30.51753 | 0.2811008 | 0.3788274 | 0.8431497 | 0.3991448 | 0.600129 |
| ENSG00000226644 | | 2.727615 | 0.2809164 | 0.7053818 | 0.7409688 | 0.4587124 | 0.654469 |
| ENSG00000269526 | ERVV-1 | 1.445688 | 0.2807687 | 0.8323902 | 0.9576125 | 0.3382582 | NA |
| ENSG00000172239 | PAIP1 | 3616.132 | 0.2806143 | 0.1222087 | 2.357323 | 0.01840721 | 0.063556 |
| ENSG00000172594 | SMPDL3A | 492.6605 | 0.2805927 | 0.2968386 | 1.021697 | 0.3069242 | 0.507068 |
| ENSG00000134489 | HRH4 | 1.299272734 | 0.280512051 | 0.807245646 | 1.068966595 | 0.28508472 | NA |
| ENSG00000177106 | EPS8L2 | 12419.22 | 0.2804374 | 0.2063158 | 1.429896 | 0.1527469 | 0.315186 |
| ENSG00000283209 | | 8.018428 | 0.2804045 | 0.6900625 | 0.7339349 | 0.4629884 | 0.657314 |
| ENSG00000110844 | PRPF40B | 727.4341919 | 0.280379927 | 0.147878612 | 1.928631487 | 0.05377663 | 0.146791 |
| ENSG00000234286 | | 13.77714 | 0.2803594 | 0.4350891 | 0.7648307 | 0.4443723 | 0.641764 |
| ENSG00000272509 | | 28.56883 | 0.2803578 | 0.4424868 | 0.7639967 | 0.4448692 | 0.642138 |
| ENSG00000212829 | RPS26P3 | 4.555542 | 0.2803492 | 0.6766676 | 0.719421 | 0.4718816 | 0.665025 |
| ENSG00000240882 | | 2.048664 | 0.2802058 | 0.72537 | 0.7716359 | 0.4403301 | 0.638603 |
| ENSG00000091039 | OSBPL8 | 4223.752556 | 0.280194446 | 0.234189495 | 1.25149816 | 0.21075279 | 0.394636 |
| ENSG00000103018 | CYB5B | 6029.595679 | 0.280168119 | 0.283360421 | 1.046059151 | 0.29553372 | 0.494669 |
| ENSG00000277463 | | 14.10717 | 0.2800926 | 0.5617317 | 0.6834953 | 0.4942939 | 0.684005 |
| ENSG00000285638 | | 1.404729 | 0.2800729 | 0.8257906 | 1.122367 | 0.2617063 | NA |
| ENSG00000205593 | DENND6B | 555.6657 | 0.2799976 | 0.2552063 | 1.160688 | 0.2457688 | 0.43751 |
| ENSG00000253844 | | 1.243403 | 0.2796801 | 0.7977024 | 1.012042 | 0.3115181 | NA |

| | | | | | | | |
|---|---|---|---|---|---|---|---|
| ENSG00000188549 | CCDC9B | 131.8121 | 0.279677 | 0.5267914 | 0.6891095 | 0.4907544 | 0.681165 |
| ENSG00000260276 | | 75.95951 | 0.2796478 | 0.281992 | 1.057558 | 0.2902569 | 0.488833 |
| ENSG00000236153 | | 1.252258 | 0.2795911 | 0.8019001 | 0.982531 | 0.3258383 | NA |
| ENSG00000078070 | MCCC1 | 3330.177378 | 0.279586143 | 0.110321656 | 2.547128276 | 0.01086135 | 0.041586 |
| ENSG00000114354 | TFG | 8617.366539 | 0.279537538 | 0.10385904 | 2.714861418 | 0.00663035 | 0.027876 |
| ENSG00000203711 | C6orf99 | 105.4112 | 0.2794965 | 0.3070279 | 0.9850309 | 0.3246089 | 0.525598 |
| ENSG00000287887 | | 0.9505785 | 0.2794719 | 0.8302016 | 0.9633992 | 0.3353472 | NA |
| ENSG00000266282 | UBL5P2 | 40.86978 | 0.2793011 | 0.3722229 | 0.8569793 | 0.3914563 | 0.593235 |
| ENSG00000254595 | | 29.74732 | 0.2792875 | 0.4333644 | 0.7671235 | 0.4430081 | 0.640594 |
| ENSG00000106336 | FBXO24 | 23.65367521 | 0.279251607 | 0.384381959 | 0.803160363 | 0.42188205 | 0.621789 |
| ENSG00000284526 | | 31.65171 | 0.2790339 | 0.4058678 | 0.7933789 | 0.4275571 | 0.627044 |
| ENSG00000100652 | SLC10A1 | 4.826961358 | 0.278992164 | 0.59509232 | 0.678699967 | 0.49732799 | 0.686082 |
| ENSG00000272414 | FAM47E-STBD1 | 1.473114 | 0.2789269 | 0.7458038 | 0.7768571 | 0.4372431 | NA |
| ENSG00000228348 | EFCAB14P1 | 36.80786 | 0.2788969 | 0.3463178 | 0.8999637 | 0.3681395 | 0.570215 |
| ENSG00000280213 | UCKL1-AS1 | 188.9708 | 0.2786608 | 0.3690253 | 0.8485092 | 0.3961545 | 0.597526 |
| ENSG00000073737 | DHRS9 | 0.911827281 | 0.278612786 | 0.798128514 | 0.986825872 | 0.32372801 | NA |
| ENSG00000198858 | R3HDM4 | 3524.809 | 0.278468 | 0.1751169 | 1.615135 | 0.1062814 | 0.243304 |
| ENSG00000074319 | TSG101 | 4893.327938 | 0.278287863 | 0.11782078 | 2.388149274 | 0.01693346 | 0.059448 |
| ENSG00000132155 | RAF1 | 11366.84861 | 0.278084721 | 0.130009749 | 2.15600527 | 0.03108325 | 0.096444 |
| ENSG00000266472 | MRPS21 | 8625.918 | 0.2780199 | 0.1740953 | 1.585843 | 0.1127749 | 0.253728 |
| ENSG00000223979 | SMCR2 | 3.420868 | 0.2779808 | 0.6852384 | 0.7109626 | 0.4771074 | 0.669324 |
| ENSG00000188051 | TMEM221 | 31.32683 | 0.2779056 | 0.3798746 | 0.8310019 | 0.4059726 | 0.606687 |
| ENSG00000255224 | | 146.8347 | 0.2775141 | 0.224289 | 1.299762 | 0.1936824 | 0.372274 |
| ENSG00000186086 | NBPF6 | 1.535915 | 0.2774694 | 0.7970782 | 0.914224 | 0.3605992 | NA |
| ENSG00000228767 | KRT8P18 | 2.564202 | 0.2774602 | 0.7442757 | 0.7956103 | 0.4262586 | 0.62619 |
| ENSG00000255960 | | 1.319865 | 0.2774306 | 0.7952545 | 0.9497121 | 0.3422586 | NA |
| ENSG00000270299 | | 2.491574 | 0.2773891 | 0.7160995 | 0.7520479 | 0.4520222 | 0.648231 |
| ENSG00000222112 | RN7SKP16 | 5.85484 | 0.2770552 | 0.6250989 | 0.6784563 | 0.4974824 | 0.686176 |
| ENSG00000241170 | | 58.5719 | 0.2770497 | 0.3602127 | 0.8650643 | 0.3870035 | 0.589349 |
| ENSG00000114857 | NKTR | 5406.380826 | 0.276961573 | 0.224468601 | 1.342400194 | 0.17946627 | 0.35354 |
| ENSG00000259137 | | 38.44569 | 0.2768077 | 0.3626965 | 0.8625697 | 0.3883741 | 0.590521 |
| ENSG00000236824 | BCYRN1 | 3.422455 | 0.276621 | 0.7042731 | 0.7379661 | 0.4605351 | 0.655647 |
| ENSG00000144550 | CPNE9 | 2.233236 | 0.2765748 | 0.699423 | 0.7191185 | 0.4720679 | 0.66511 |

| | | | | | | | |
|---|---|---|---|---|---|---|---|
| ENSG00000224326 | | 6.746449 | 0.2765581 | 0.6518442 | 0.6859131 | 0.4927679 | 0.682788 |
| ENSG00000136710 | CCDC115 | 2169.219467 | 0.276495609 | 0.165765704 | 1.69780813 | 0.08954398 | 0.214875 |
| ENSG00000077044 | DGKD | 2186.868064 | 0.276421801 | 0.142702123 | 1.964719702 | 0.04944669 | 0.137407 |
| ENSG00000258204 | | 43.18402 | 0.2763232 | 0.3131125 | 0.963199 | 0.3354477 | 0.536672 |
| ENSG00000083812 | ZNF324 | 1118.38992 | 0.276319082 | 0.131714097 | 2.12479421 | 0.03360379 | 0.102399 |
| ENSG00000157978 | LDLRAP1 | 1597.536 | 0.2763089 | 0.1962541 | 1.444738 | 0.1485315 | 0.30905 |
| ENSG00000225548 | LINC01980 | 0.8950954 | 0.2762441 | 0.8414262 | 1.15839 | 0.2467049 | NA |
| ENSG00000226137 | BAIAP2-DT | 1703.658 | 0.2761363 | 0.1495341 | 1.873642 | 0.06097981 | 0.160939 |
| ENSG00000125863 | MKKS | 2006.578136 | 0.275993695 | 0.12345049 | 2.259661571 | 0.02384226 | 0.078157 |
| ENSG00000197780 | TAF13 | 1496.979 | 0.2759435 | 0.1387967 | 2.010782 | 0.04434845 | 0.126657 |
| ENSG00000279649 | | 64.24642 | 0.2758914 | 0.422017 | 0.7679048 | 0.4425437 | 0.640164 |
| ENSG00000149435 | GGTLC1 | 3.671577 | 0.2758459 | 0.7883352 | 0.9094577 | 0.3631086 | 0.565103 |
| ENSG00000279259 | | 60.13635 | 0.2758314 | 0.3861991 | 0.8121105 | 0.4167282 | 0.616636 |
| ENSG00000121318 | TAS2R10 | 8.048508048 | 0.275831392 | 0.641966366 | 0.677447633 | 0.49812199 | 0.686635 |
| ENSG00000134899 | ERCC5 | 131.679198 | 0.275702793 | 0.252927025 | 1.154100128 | 0.24845912 | 0.440775 |
| ENSG00000288162 | | 2.165413 | 0.275697 | 0.7669172 | 0.8470624 | 0.3969603 | 0.598029 |
| ENSG00000197332 | | 18.64044 | 0.2755646 | 0.5528313 | 0.6776393 | 0.4980004 | 0.68656 |
| ENSG00000271398 | | 7.156227 | 0.2755352 | 0.6105622 | 0.6687763 | 0.5036382 | 0.690989 |
| ENSG00000213361 | | 1.962001 | 0.2755233 | 0.7584004 | 0.8155693 | 0.4147465 | 0.614726 |
| ENSG00000272030 | | 11.21579 | 0.2755141 | 0.4949983 | 0.7091336 | 0.4782415 | 0.670133 |
| ENSG00000178233 | TMEM151B | 1.329651 | 0.2754959 | 0.7692153 | 0.8508342 | 0.3948615 | NA |
| ENSG00000274902 | | 1.713484 | 0.275444 | 0.7185101 | 0.7446805 | 0.4564649 | 0.652249 |
| ENSG00000225355 | ARL6IP1P2 | 11.51965 | 0.2754224 | 0.5483771 | 0.6836796 | 0.4941775 | 0.683951 |
| ENSG00000135378 | PRRG4 | 953.4622931 | 0.275229659 | 0.148075155 | 1.892987635 | 0.05835951 | 0.155913 |
| ENSG00000162267 | ITIH3 | 1.520078 | 0.2752235 | 0.74526 | 0.7891947 | 0.4299982 | NA |
| ENSG00000046604 | DSG2 | 4098.587758 | 0.275218667 | 0.250475034 | 1.149331342 | 0.25041938 | 0.443048 |
| ENSG00000285664 | | 3.076443 | 0.2752167 | 0.6456927 | 0.6843837 | 0.4937329 | 0.683539 |
| ENSG00000250155 | | 1.545238 | 0.2752082 | 0.7798965 | 0.9465738 | 0.343856 | NA |
| ENSG00000273987 | | 1.472405 | 0.2751988 | 0.7347582 | 0.754806 | 0.4503654 | NA |
| ENSG00000171130 | ATP6V0E2 | 3031.765 | 0.2751131 | 0.2329887 | 1.23699 | 0.2160909 | 0.401379 |
| ENSG00000227811 | INKA2-AS1 | 9.533265 | 0.2750986 | 0.5391895 | 0.678884 | 0.4972114 | 0.68597 |
| ENSG00000230393 | | 1.426841 | 0.2750594 | 0.8076652 | 1.11387 | 0.265335 | NA |
| ENSG00000175606 | TMEM70 | 1437.444 | 0.2749413 | 0.1990186 | 1.425556 | 0.1539965 | 0.317321 |

| | | | | | | | |
|---|---|---|---|---|---|---|---|
| ENSG00000232133 | IMPDH1P10 | 0.9740542 | 0.2748837 | 0.7858651 | 0.9025759 | 0.366751 | NA |
| ENSG00000225972 | MTND1P23 | 10.49113 | 0.2747187 | 0.5685901 | 0.6720039 | 0.5015812 | 0.689314 |
| ENSG00000232883 | | 1.555014 | 0.274669 | 0.7759745 | 0.8879159 | 0.374586 | NA |
| ENSG00000240499 | | 4.727562 | 0.2746563 | 0.6447766 | 0.6812111 | 0.4957379 | 0.684862 |
| ENSG00000213411 | RBM22P2 | 2.466797 | 0.2746554 | 0.6790227 | 0.6939416 | 0.4877188 | 0.678689 |
| ENSG00000137440 | FGFBP1 | 15.96342366 | 0.274572726 | 0.756564197 | 0.830491565 | 0.40626092 | 0.606849 |
| ENSG00000131016 | AKAP12 | 100.4999823 | 0.274527024 | 0.38183087 | 0.806124968 | 0.4201708 | 0.6201 |
| ENSG00000131944 | FAAP24 | 199.2389261 | 0.274418404 | 0.404704184 | 0.783694461 | 0.43321941 | 0.632237 |
| ENSG00000286747 | | 30.95391 | 0.2743948 | 0.4965437 | 0.6953689 | 0.4868241 | 0.677727 |
| ENSG00000091137 | SLC26A4 | 2.406035665 | 0.274384401 | 0.746122248 | 0.781704159 | 0.43438846 | 0.633401 |
| ENSG00000263818 | RDM1P5 | 22.41754 | 0.2743417 | 0.454079 | 0.7365699 | 0.4613839 | 0.656194 |
| ENSG00000126016 | AMOT | 1586.209641 | 0.274337495 | 0.334421821 | 0.906239624 | 0.36480903 | 0.566692 |
| ENSG00000071242 | RPS6KA2 | 674.8283842 | 0.274318024 | 0.582502213 | 0.650336535 | 0.51547486 | 0.700837 |
| ENSG00000228251 | | 7.155104 | 0.2743034 | 0.5781408 | 0.670123 | 0.5027794 | 0.690189 |
| ENSG00000185127 | C6orf120 | 2230.376 | 0.274233 | 0.185826 | 1.513336 | 0.1301943 | 0.281096 |
| ENSG00000217289 | | 95.86113 | 0.2741538 | 0.2299056 | 1.25532 | 0.2093626 | 0.392586 |
| ENSG00000129534 | MIS18BP1 | 3022.625318 | 0.274087409 | 0.271593347 | 1.068723792 | 0.28519415 | 0.483186 |
| ENSG00000150776 | NKAPD1 | 1617.943 | 0.2740159 | 0.1090772 | 2.526745 | 0.01151251 | 0.043572 |
| ENSG00000166908 | PIP4K2C | 4817.683 | 0.2739897 | 0.1125658 | 2.460973 | 0.01385607 | 0.050548 |
| ENSG00000280035 | | 1.414046 | 0.2739367 | 0.7546846 | 0.8176094 | 0.4135802 | NA |
| ENSG00000112599 | GUCA1B | 58.20068822 | 0.273785257 | 0.341664195 | 0.886131649 | 0.37554659 | 0.577691 |
| ENSG00000173212 | MAB21L3 | 2.304976 | 0.273785 | 0.7773295 | 0.9600053 | 0.3370525 | 0.53817 |
| ENSG00000103740 | ACSBG1 | 12.52144285 | 0.273781259 | 0.591258872 | 0.679140944 | 0.49704856 | 0.685888 |
| ENSG00000221676 | RNU6ATAC | 1.913665 | 0.2737208 | 0.7735431 | 0.8842512 | 0.3765606 | 0.578617 |
| ENSG00000228938 | SNRPD2P1 | 5.947236 | 0.2736413 | 0.583772 | 0.6749648 | 0.4996981 | 0.68795 |
| ENSG00000196465 | MYL6B | 2632.096 | 0.2736404 | 0.2093604 | 1.356222 | 0.1750285 | 0.347066 |
| ENSG00000143839 | REN | 10.51137 | 0.2735985 | 0.5062122 | 0.6914595 | 0.4892768 | 0.680076 |
| ENSG00000287155 | | 3.512209 | 0.2735641 | 0.704294 | 0.7195605 | 0.4717956 | 0.665007 |
| ENSG00000245869 | | 1.036531 | 0.2735286 | 0.8172867 | 0.8157735 | 0.4146297 | NA |
| ENSG00000083799 | CYLD | 1128.111637 | 0.273458295 | 0.231430447 | 1.221208696 | 0.22200701 | 0.408738 |
| ENSG00000166800 | LDHAL6A | 7.3992 | 0.2734178 | 0.6361438 | 0.6717209 | 0.5017614 | 0.689342 |
| ENSG00000100601 | ALKBH1 | 1057.507107 | 0.273402323 | 0.153355589 | 1.82072924 | 0.06864803 | 0.175917 |
| ENSG00000065325 | GLP2R | 62.66052179 | 0.273398591 | 0.773837201 | 0.983707123 | 0.32525954 | 0.526523 |

| | | | | | | | |
|---|---|---|---|---|---|---|---|
| ENSG00000273066 | | 165.3036 | 0.27338 | 0.2737927 | 1.060526 | 0.2889055 | 0.487488 |
| ENSG00000121104 | FAM117A | 990.8149839 | 0.273231826 | 0.23312634 | 1.192578807 | 0.23303437 | 0.421879 |
| ENSG00000244332 | | 1.305386 | 0.2732068 | 0.7997187 | 1.082688 | 0.278947 | NA |
| ENSG00000214097 | SMCO1 | 0.974677 | 0.2731999 | 0.8174083 | 0.8181835 | 0.4132524 | NA |
| ENSG00000229237 | HMGN1P37 | 165.2995 | 0.273079 | 0.2596526 | 1.114288 | 0.2651556 | 0.460723 |
| ENSG00000154889 | MPPE1 | 742.1454 | 0.2729854 | 0.150238 | 1.852575 | 0.06394331 | 0.166638 |
| ENSG00000201376 | | 0.8617473 | 0.2729035 | 0.815533 | 1.15129 | 0.2496129 | NA |
| ENSG00000158481 | CD1C | 0.8254204 | 0.2727784 | 0.7894063 | 0.8877985 | 0.3746491 | NA |
| ENSG00000208892 | SNORA49 | 0.8255236 | 0.2727231 | 0.8276473 | 0.9799294 | 0.327121 | NA |
| ENSG00000259330 | INAFM2 | 686.6679 | 0.2726469 | 0.1557291 | 1.778491 | 0.07532332 | 0.18868 |
| ENSG00000276863 | | 1.862994 | 0.2726422 | 0.6931286 | 0.6921236 | 0.4888597 | 0.679677 |
| ENSG00000197272 | IL27 | 3.359914 | 0.2726082 | 0.6470044 | 0.6767432 | 0.4985689 | 0.686951 |
| ENSG00000104154 | SLC30A4 | 168.8596908 | 0.272543108 | 0.325961315 | 0.913747066 | 0.36084977 | 0.562873 |
| ENSG00000141582 | CBX4 | 5902.521 | 0.2722266 | 0.2768821 | 1.04685 | 0.2951689 | 0.49438 |
| ENSG00000287055 | | 0.7565911 | 0.2720684 | 0.8116849 | 0.9853266 | 0.3244637 | NA |
| ENSG00000126458 | RRAS | 768.9344433 | 0.272046254 | 0.269439856 | 1.081111884 | 0.27964735 | 0.47696 |
| ENSG00000109180 | OCIAD1 | 6256.901158 | 0.272016312 | 0.14055331 | 1.958783418 | 0.05013816 | 0.138763 |
| ENSG00000067113 | PLPP1 | 478.0897894 | 0.271829871 | 0.166656806 | 1.65127426 | 0.09868259 | 0.230415 |
| ENSG00000066117 | SMARCD1 | 5086.285251 | 0.271797702 | 0.139974189 | 1.968655381 | 0.04899268 | 0.136402 |
| ENSG00000143633 | C1orf131 | 1329.024 | 0.2717178 | 0.1406651 | 1.966798 | 0.04920646 | 0.136911 |
| ENSG00000236165 | PRADC1P1 | 1.825462 | 0.2716763 | 0.7377132 | 0.7602768 | 0.4470891 | 0.643898 |
| ENSG00000126266 | FFAR1 | 1.172732076 | 0.27167066 | 0.806315335 | 1.015528615 | 0.30985391 | NA |
| ENSG00000224424 | PRKAR2A-AS1 | 49.15943 | 0.2716456 | 0.2733438 | 1.059654 | 0.289302 | 0.487904 |
| ENSG00000166446 | CDYL2 | 6499.321 | 0.2716351 | 0.2729604 | 1.05881 | 0.2896862 | 0.488213 |
| ENSG00000286735 | | 8.076607 | 0.2716021 | 0.7566705 | 0.8217452 | 0.411222 | 0.611539 |
| ENSG00000215035 | FDPSP5 | 20.08987 | 0.2715449 | 0.4584749 | 0.7236781 | 0.4692634 | 0.662831 |
| ENSG00000133740 | E2F5 | 878.5711284 | 0.27154209 | 0.169566294 | 1.639824363 | 0.10104169 | 0.234543 |
| ENSG00000259219 | | 2.311117 | 0.2715308 | 0.7405668 | 0.7760247 | 0.4377344 | 0.636258 |
| ENSG00000285599 | | 1.659713 | 0.2714519 | 0.7183637 | 0.7381659 | 0.4604137 | 0.655647 |
| ENSG00000100625 | SIX4 | 2163.107117 | 0.271392893 | 0.258255897 | 1.109720948 | 0.26711929 | 0.462894 |
| ENSG00000235857 | CTBP2P1 | 1.345094 | 0.2713858 | 0.7844321 | 0.8513039 | 0.3946006 | NA |
| ENSG00000207815 | MIR563 | 2.305588 | 0.2712989 | 0.7599768 | 0.8204814 | 0.4119417 | 0.612233 |
| ENSG00000169733 | RFNG | 3106.855 | 0.2712562 | 0.2068389 | 1.355354 | 0.1753047 | 0.347484 |

| | | | | | | | |
|---|---|---|---|---|---|---|---|
| ENSG00000182752 | PAPPA | 4.541841 | 0.2711836 | 0.5775576 | 0.6596356 | 0.5094877 | 0.69551 |
| ENSG00000228727 | SAPCD1 | 17.78774 | 0.2711419 | 0.5383439 | 0.6725182 | 0.5012539 | 0.689114 |
| ENSG00000262873 | | 10.18296 | 0.2710361 | 0.7023449 | 0.70055 | 0.4835839 | 0.674843 |
| ENSG00000134627 | PIWIL4 | 55.38143668 | 0.270851002 | 0.467581505 | 0.713630633 | 0.47545561 | 0.667779 |
| ENSG00000109956 | B3GAT1 | 1.670953599 | 0.270834321 | 0.735861299 | 0.757042241 | 0.44902456 | 0.645416 |
| ENSG00000224040 | HMGN1P4 | 271.6319 | 0.2707943 | 0.2198335 | 1.287489 | 0.1979238 | 0.37796 |
| ENSG00000163374 | YY1AP1 | 3431.57 | 0.2707244 | 0.1108073 | 2.466786 | 0.01363318 | 0.049879 |
| ENSG00000227220 | | 1.162809 | 0.2706901 | 0.8177574 | 0.8203516 | 0.4120157 | NA |
| ENSG00000102218 | RP2 | 649.3080162 | 0.270674724 | 0.241109102 | 1.181910493 | 0.23724122 | 0.426938 |
| ENSG00000247872 | SPCS2P3 | 1.65136 | 0.2706649 | 0.7825638 | 0.9145648 | 0.3604201 | 0.562434 |
| ENSG00000121766 | ZCCHC17 | 1681.448188 | 0.270620377 | 0.189967543 | 1.464218495 | 0.14313426 | 0.300943 |
| ENSG00000232453 | LINC02777 | 14.17108 | 0.2706075 | 0.5448461 | 0.6633257 | 0.507122 | 0.693592 |
| ENSG00000231093 | TRMT1P1 | 0.8649743 | 0.2705589 | 0.8175186 | 0.8217277 | 0.4112319 | NA |
| ENSG00000239590 | OR1J4 | 0.7902929 | 0.2703538 | 0.8038773 | 0.9557788 | 0.339184 | NA |
| ENSG00000104812 | GYS1 | 3858.54756 | 0.270217366 | 0.152892299 | 1.800157891 | 0.07183571 | 0.182153 |
| ENSG00000286242 | | 6.484369 | 0.2701908 | 0.6565593 | 0.6716145 | 0.5018292 | 0.68937 |
| ENSG00000158773 | USF1 | 3126.665 | 0.2700682 | 0.1730638 | 1.590276 | 0.1117727 | 0.252267 |
| ENSG00000232456 | | 7.205566 | 0.2700239 | 0.6626912 | 0.6953792 | 0.4868177 | 0.677727 |
| ENSG00000224986 | PPP1R8P1 | 0.9284997 | 0.2698654 | 0.8165858 | 0.8106174 | 0.4175854 | NA |
| ENSG00000229586 | TNPO1P3 | 9.587857 | 0.2698519 | 0.4732108 | 0.7113683 | 0.476856 | 0.669145 |
| ENSG00000174628 | IQCK | 322.4855 | 0.2696607 | 0.1795363 | 1.542423 | 0.1229707 | 0.270017 |
| ENSG00000244171 | PBX2P1 | 382.0353 | 0.2696056 | 0.1704849 | 1.611737 | 0.1070192 | 0.244446 |
| ENSG00000269303 | | 1.518548 | 0.269588 | 0.8272412 | 0.9579313 | 0.3380974 | NA |
| ENSG00000196196 | HRCT1 | 4.226327 | 0.2695168 | 0.8151544 | 0.8366086 | 0.4028126 | 0.603696 |
| ENSG00000136878 | USP20 | 2572.31829 | 0.269478095 | 0.156124836 | 1.76603667 | 0.0773897 | 0.192399 |
| ENSG00000121481 | RNF2 | 1067.216159 | 0.269463191 | 0.153393218 | 1.78344127 | 0.0745145 | 0.187231 |
| ENSG00000173811 | CCDC13-AS1 | 3.024015 | 0.2693584 | 0.7167773 | 0.7188674 | 0.4722226 | 0.665243 |
| ENSG00000258620 | | 2.125834 | 0.2693181 | 0.7129043 | 0.7256554 | 0.4680501 | 0.66168 |
| ENSG00000157077 | ZFYVE9 | 1680.436 | 0.2693089 | 0.2135048 | 1.305468 | 0.1917334 | 0.369973 |
| ENSG00000248694 | | 2.43886 | 0.2692652 | 0.720405 | 0.7350913 | 0.4622839 | 0.656814 |
| ENSG00000167543 | TP53I13 | 4060.018 | 0.2692255 | 0.1697724 | 1.620489 | 0.1051272 | 0.241236 |
| ENSG00000273151 | | 806.9112 | 0.269165 | 0.3872412 | 0.788294 | 0.4305248 | 0.629776 |
| ENSG00000287642 | | 38.75285 | 0.2691645 | 0.3694766 | 0.8246356 | 0.4095785 | 0.609996 |

| | | | | | | | |
|---|---|---|---|---|---|---|---|
| ENSG00000075975 | MKRN2 | 3311.605741 | 0.269133949 | 0.142900157 | 1.915914995 | 0.05537592 | 0.149799 |
| ENSG00000224687 | RASAL2-AS1 | 23.83678 | 0.2690404 | 0.3856267 | 0.7942984 | 0.4270217 | 0.626613 |
| ENSG00000241319 | SETP6 | 1.007611 | 0.2690032 | 0.7951499 | 0.9825697 | 0.3258193 | NA |
| ENSG00000257074 | RPL29P33 | 1.557099 | 0.2689608 | 0.7839737 | 0.9004569 | 0.3678771 | NA |
| ENSG00000278962 | | 1.657214 | 0.268929 | 0.7005383 | 0.699181 | 0.4844389 | 0.67556 |
| ENSG00000254852 | NPIPA2 | 3.297244 | 0.2687281 | 0.6697196 | 0.6828968 | 0.494672 | 0.684079 |
| ENSG00000137942 | FNBP1L | 3435.144204 | 0.268683862 | 0.236134599 | 1.188587951 | 0.23460185 | 0.423763 |
| ENSG00000235166 | LINC01440 | 0.9709274 | 0.2686115 | 0.8161682 | 0.8087261 | 0.4186727 | NA |
| ENSG00000155714 | PDZD9 | 1.627144 | 0.2685695 | 0.7391739 | 0.7519373 | 0.4520888 | 0.648257 |
| ENSG00000182853 | VMO1 | 98.47347 | 0.2685674 | 0.2813287 | 1.019978 | 0.3077389 | 0.507877 |
| ENSG00000125844 | RRBP1 | 9132.620075 | 0.268500489 | 0.188406481 | 1.458845814 | 0.14460755 | 0.303099 |
| ENSG00000103319 | EEF2K | 2504.276765 | 0.268430132 | 0.174038139 | 1.540484161 | 0.12344238 | 0.270817 |
| ENSG00000151882 | CCL28 | 201.7543 | 0.2684034 | 0.4405866 | 0.7267421 | 0.4673839 | 0.661023 |
| ENSG00000250751 | | 0.8822718 | 0.268384 | 0.7960815 | 0.9702463 | 0.3319237 | NA |
| ENSG00000076928 | ARHGEF1 | 5695.986815 | 0.268366942 | 0.171544328 | 1.578818889 | 0.11437761 | 0.256361 |
| ENSG00000100578 | KIAA0586 | 1452.260051 | 0.268332343 | 0.179509003 | 1.526856328 | 0.12679674 | 0.275729 |
| ENSG00000231863 | | 4.215505 | 0.2682822 | 0.7325906 | 0.7664631 | 0.4434008 | 0.640918 |
| ENSG00000227992 | | 7.474658 | 0.268185 | 0.6961001 | 0.7105253 | 0.4773785 | 0.669595 |
| ENSG00000186834 | HEXIM1 | 4079.355 | 0.2681601 | 0.1830804 | 1.491936 | 0.1357158 | 0.289387 |
| ENSG00000286153 | | 8.129018 | 0.2681485 | 0.6943652 | 0.7138109 | 0.4753441 | 0.667732 |
| ENSG00000213842 | SUGT1P2 | 3.529074 | 0.2681373 | 0.6302789 | 0.6656442 | 0.5056386 | 0.692419 |
| ENSG00000219545 | UMAD1 | 992.2849 | 0.2680524 | 0.1426512 | 1.939127 | 0.05248584 | 0.143918 |
| ENSG00000283486 | FAM95C | 194.5131 | 0.2680274 | 0.3562036 | 0.8393796 | 0.4012564 | 0.602213 |
| ENSG00000174792 | ODAPH | 12.90057 | 0.2678499 | 0.6234403 | 0.673328 | 0.5007387 | 0.688619 |
| ENSG00000197253 | TPSB2 | 2.015226 | 0.2677861 | 0.7732403 | 0.8494461 | 0.3956331 | 0.597249 |
| ENSG00000240771 | ARHGEF25 | 201.3676 | 0.2677039 | 0.3182306 | 0.916431 | 0.3594409 | 0.561269 |
| ENSG00000119655 | NPC2 | 5832.068455 | 0.267656844 | 0.246579692 | 1.148278781 | 0.2508535 | 0.443506 |
| ENSG00000243402 | | 1.27613 | 0.2675476 | 0.7964234 | 0.9959749 | 0.3192623 | NA |
| ENSG00000166734 | GOLM2 | 3540.415 | 0.2674978 | 0.1200837 | 2.245649 | 0.02472653 | 0.080401 |
| ENSG00000239559 | RPL37P2 | 450.811 | 0.2674848 | 0.2187891 | 1.274681 | 0.2024222 | 0.383606 |
| ENSG00000276404 | MIR6835 | 9.005669 | 0.2674747 | 0.5297075 | 0.664719 | 0.5062302 | 0.692872 |
| ENSG00000268186 | ZNF114-AS1 | 3.246377 | 0.2673515 | 0.7132905 | 0.7146327 | 0.4748361 | 0.667509 |
| ENSG00000248697 | TOX4P1 | 117.3333 | 0.2673059 | 0.2049701 | 1.350317 | 0.1769144 | 0.349969 |

| | | | | | | | |
|---|---|---|---|---|---|---|---|
| ENSG00000156599 | ZDHHC5 | 5002.925 | 0.2672489 | 0.1154415 | 2.339303 | 0.01931974 | 0.066088 |
| ENSG00000263718 | SEPTIN9-DT | 17.35065 | 0.2672119 | 0.5187287 | 0.6722953 | 0.5013957 | 0.689166 |
| ENSG00000154975 | CA10 | 3.051402 | 0.2671129 | 0.6986236 | 0.7047319 | 0.4809771 | 0.672512 |
| ENSG00000122367 | LDB3 | 85.99917113 | 0.267074296 | 0.291168816 | 0.980211107 | 0.32698192 | 0.528217 |
| ENSG00000139946 | PELI2 | 20.38342 | 0.2669983 | 0.5690677 | 0.6610113 | 0.5086051 | 0.694941 |
| ENSG00000178127 | NDUFV2 | 109.6235 | 0.2669848 | 0.3567063 | 0.8370288 | 0.4025764 | 0.603547 |
| ENSG00000102401 | ARMCX3 | 4792.34157 | 0.266961951 | 0.314091646 | 0.922303434 | 0.35637032 | 0.558344 |
| ENSG00000250238 | | 1.285736 | 0.2669353 | 0.7607834 | 0.815829 | 0.414598 | NA |
| ENSG00000148377 | IDI2 | 1.436583 | 0.2669285 | 0.772519 | 0.8656436 | 0.3866857 | NA |
| ENSG00000268938 | | 7.964732 | 0.2669239 | 0.4886434 | 0.6848411 | 0.4934442 | 0.683256 |
| ENSG00000119685 | TTLL5 | 2652.061886 | 0.266920993 | 0.17982487 | 1.51760088 | 0.12911505 | 0.27932 |
| ENSG00000230758 | SNAP23P1 | 2.622666 | 0.266918 | 0.7198977 | 0.7165753 | 0.4736362 | 0.666422 |
| ENSG00000165197 | VEGFD | 2.59658 | 0.2669035 | 0.6997125 | 0.6997241 | 0.4840996 | 0.67535 |
| ENSG00000258501 | EIF3LP1 | 20.29076 | 0.2667375 | 0.4039708 | 0.7651984 | 0.4441534 | 0.641576 |
| ENSG00000184508 | HDDC3 | 858.7925 | 0.2667285 | 0.1800962 | 1.522406 | 0.1279073 | 0.277515 |
| ENSG00000163673 | DCLK3 | 0.9330831 | 0.2667259 | 0.8035433 | 0.9389283 | 0.3477676 | NA |
| ENSG00000152292 | SH2D6 | 4.594874 | 0.2666623 | 0.6481496 | 0.6558329 | 0.5119316 | 0.697889 |
| ENSG00000248493 | | 1.663291 | 0.2666521 | 0.7825797 | 0.8654763 | 0.3867774 | 0.589135 |
| ENSG00000257885 | PHBP18 | 0.9819657 | 0.2666489 | 0.7983313 | 0.9646001 | 0.3347451 | NA |
| ENSG00000170892 | TSEN34 | 5578.023 | 0.266556 | 0.2847084 | 0.997355 | 0.3185922 | 0.519718 |
| ENSG00000001626 | CFTR | 1.2596048 | 0.266511588 | 0.762874118 | 0.809226615 | 0.41838481 | NA |
| ENSG00000272710 | | 2.934865 | 0.2664807 | 0.6752539 | 0.6869447 | 0.4921176 | 0.682311 |
| ENSG00000178082 | TWF1P1 | 865.5174 | 0.2664378 | 0.2274561 | 1.233493 | 0.2173918 | 0.403012 |
| ENSG00000166987 | MBD6 | 4499.087 | 0.2663765 | 0.2277172 | 1.22 | 0.2224649 | 0.409155 |
| ENSG00000256274 | TAS2R64P | 2.39592 | 0.2663745 | 0.7089816 | 0.7060032 | 0.4801861 | 0.671865 |
| ENSG00000115053 | NCL | 38544.24494 | 0.266355945 | 0.235875839 | 1.180708544 | 0.23771853 | 0.427312 |
| ENSG00000283443 | | 1.443841 | 0.2663214 | 0.7742761 | 0.8074155 | 0.4194272 | NA |
| ENSG00000160767 | FAM189B | 3888.249 | 0.2662814 | 0.2821372 | 1.009263 | 0.3128483 | 0.513884 |
| ENSG00000269600 | | 1.575198 | 0.2662754 | 0.7736554 | 0.8737322 | 0.3822641 | 0.58454 |
| ENSG00000137221 | TJAP1 | 1757.804115 | 0.266255973 | 0.161409523 | 1.682573877 | 0.09245761 | 0.219763 |
| ENSG00000259017 | | 1.162615 | 0.2662462 | 0.7997148 | 1.078757 | 0.2806961 | NA |
| ENSG00000160695 | VPS11 | 4933.57 | 0.2662216 | 0.1381575 | 1.987961 | 0.04681598 | 0.131862 |
| ENSG00000163521 | GLB1L | 194.2814 | 0.2662146 | 0.1871309 | 1.463746 | 0.1432634 | 0.301041 |

| | | | | | | | |
|---|---|---|---|---|---|---|---|
| ENSG00000234141 | | 4.948667 | 0.2661996 | 0.6558695 | 0.6669119 | 0.5048284 | 0.691681 |
| ENSG00000100564 | PIGH | 841.8443724 | 0.266140221 | 0.155587775 | 1.744796683 | 0.08102027 | 0.199262 |
| ENSG00000183421 | RIPK4 | 1914.797 | 0.2661043 | 0.2480509 | 1.133298 | 0.2570893 | 0.450911 |
| ENSG00000248740 | LINC02428 | 5.206535 | 0.2661024 | 0.7102856 | 0.6891838 | 0.4907076 | 0.681135 |
| ENSG00000270889 | PPP1R14BP4 | 1.011117 | 0.2660172 | 0.7775187 | 0.9421189 | 0.3461318 | NA |
| ENSG00000279712 | | 10.49711 | 0.2659558 | 0.504226 | 0.6802115 | 0.4963705 | 0.685373 |
| ENSG00000260484 | | 0.9009198 | 0.2658029 | 0.8398034 | 1.18159 | 0.2373682 | NA |
| ENSG00000074800 | ENO1 | 43556.24734 | 0.265651641 | 0.269308933 | 1.046558613 | 0.2953032 | 0.494509 |
| ENSG00000251301 | LINC02384 | 0.9857605 | 0.265575 | 0.7917332 | 0.9289265 | 0.3529272 | NA |
| ENSG00000165410 | CFL2 | 1541.196 | 0.265561 | 0.2298796 | 1.207413 | 0.227273 | 0.415097 |
| ENSG00000213069 | KRT8P40 | 3.786011 | 0.2654961 | 0.6932932 | 0.6892047 | 0.4906944 | 0.681135 |
| ENSG00000225014 | KCTD9P1 | 10.3525 | 0.2654289 | 0.4955323 | 0.686525 | 0.4923821 | 0.682461 |
| ENSG00000250775 | | 3.838107 | 0.2653894 | 0.7450568 | 0.7539 | 0.4509093 | 0.647159 |
| ENSG00000100053 | CRYBB3 | 11.50587197 | 0.265295046 | 0.508452878 | 0.679369371 | 0.49690385 | 0.685724 |
| ENSG00000123297 | TSFM | 39.99925917 | 0.265274673 | 0.273447445 | 1.030955216 | 0.30256182 | 0.502015 |
| ENSG00000187713 | TMEM203 | 2327.706 | 0.2651112 | 0.1928889 | 1.41356 | 0.157491 | 0.32262 |
| ENSG00000126456 | IRF3 | 4911.141641 | 0.265074814 | 0.17018914 | 1.603804532 | 0.10875715 | 0.247246 |
| ENSG00000108639 | SYNGR2 | 15782.41619 | 0.265008934 | 0.18633651 | 1.462377366 | 0.14363783 | 0.301566 |
| ENSG00000215895 | HSPA5P1 | 4.60676 | 0.264911 | 0.6703388 | 0.6773235 | 0.4982007 | 0.686657 |
| ENSG00000172493 | AFF1 | 2395.133 | 0.2648848 | 0.2482049 | 1.128752 | 0.2590025 | 0.45336 |
| ENSG00000288012 | | 16.48158 | 0.2648682 | 0.4026115 | 0.7613414 | 0.4464532 | 0.643407 |
| ENSG00000237211 | SETP4 | 1.851755 | 0.2646884 | 0.7445846 | 0.7790668 | 0.4359403 | 0.634623 |
| ENSG00000213719 | CLIC1 | 13903.72 | 0.2646723 | 0.1761787 | 1.539918 | 0.1235804 | 0.271065 |
| ENSG00000105866 | SP4 | 330.0079112 | 0.26466268 | 0.303365866 | 0.946677576 | 0.3438031 | 0.545409 |
| ENSG00000151093 | OXSM | 1142.584 | 0.2646385 | 0.1367843 | 1.955133 | 0.05056738 | 0.139703 |
| ENSG00000206914 | | 1.984581 | 0.2645789 | 0.7442167 | 0.7707927 | 0.4408298 | 0.638766 |
| ENSG00000255644 | | 0.8971465 | 0.2645658 | 0.8416896 | 1.245055 | 0.2131115 | NA |
| ENSG00000185049 | NELFA | 1826.821 | 0.2645088 | 0.175698 | 1.540577 | 0.1234199 | 0.270803 |
| ENSG00000115977 | AAK1 | 1757.959019 | 0.264485384 | 0.224903056 | 1.224407043 | 0.22079871 | 0.407135 |
| ENSG00000255547 | RPA2P3 | 1.937035 | 0.2644729 | 0.7566078 | 0.818461 | 0.413094 | 0.613152 |
| ENSG00000239219 | | 0.8096822 | 0.264444 | 0.815639 | 0.7952598 | 0.4264624 | NA |
| ENSG00000261728 | | 0.9670658 | 0.2644246 | 0.7884687 | 0.8771004 | 0.3804321 | NA |
| ENSG00000132471 | WBP2 | 7768.094469 | 0.264387702 | 0.164063001 | 1.642305839 | 0.10052664 | 0.233694 |

| | | | | | | | |
|---|---|---|---|---|---|---|---|
| ENSG00000213078 | | 2.557478 | 0.2642871 | 0.7401077 | 0.7714699 | 0.4404285 | 0.638603 |
| ENSG00000228956 | SATB1-AS1 | 32.26355 | 0.2642544 | 0.4225156 | 0.7346126 | 0.4625755 | 0.656982 |
| ENSG00000269473 | | 84.90191 | 0.2641403 | 0.3467986 | 0.8445921 | 0.3983386 | 0.599392 |
| ENSG00000198252 | STYX | 2544.278 | 0.2640555 | 0.1606174 | 1.687357 | 0.09153478 | 0.218253 |
| ENSG00000262298 | | 2.972782 | 0.2639862 | 0.6685503 | 0.6698198 | 0.5029726 | 0.690334 |
| ENSG00000135916 | ITM2C | 1676.462847 | 0.263984069 | 0.308032179 | 0.930477949 | 0.35212368 | 0.553879 |
| ENSG00000223313 | RNU6-516P | 3.118649 | 0.2639698 | 0.678803 | 0.6794976 | 0.4968226 | 0.685678 |
| ENSG00000214359 | RPL18P10 | 1.057319 | 0.2637995 | 0.7863965 | 0.8667681 | 0.3860691 | NA |
| ENSG00000100997 | ABHD12 | 4541.760006 | 0.263777448 | 0.228243267 | 1.214591409 | 0.22452197 | 0.411433 |
| ENSG00000177981 | ASB8 | 1624.674 | 0.2637005 | 0.09650214 | 2.758945 | 0.00579883 | 0.025052 |
| ENSG00000238150 | | 23.34013 | 0.2636948 | 0.3333934 | 0.8706161 | 0.3839638 | 0.586322 |
| ENSG00000219565 | ZNF259P1 | 23.61469 | 0.2636668 | 0.3677648 | 0.8089788 | 0.4185274 | 0.618319 |
| ENSG00000132003 | ZSWIM4 | 190.9744658 | 0.263474401 | 0.259234853 | 1.078647735 | 0.28074479 | 0.478179 |
| ENSG00000258099 | ATXN2-AS | 15.55527 | 0.2634216 | 0.499988 | 0.6723031 | 0.5013908 | 0.689166 |
| ENSG00000160959 | LRRC14 | 2990.48 | 0.2633786 | 0.1852469 | 1.454294 | 0.1458648 | 0.304964 |
| ENSG00000167632 | TRAPPC9 | 2087.679 | 0.2633239 | 0.1403256 | 1.905134 | 0.05676261 | 0.152583 |
| ENSG00000236285 | NPM1P8 | 1.324762 | 0.263179 | 0.7917452 | 1.008145 | 0.3133848 | NA |
| ENSG00000250893 | | 2.406845 | 0.2629732 | 0.7720996 | 0.9227989 | 0.356112 | 0.558137 |
| ENSG00000286366 | | 44.82907 | 0.2629226 | 0.4167566 | 0.7353397 | 0.4621326 | 0.656777 |
| ENSG00000211891 | IGHE | 2.473042 | 0.2627464 | 0.7910636 | 0.9624953 | 0.3358009 | 0.536902 |
| ENSG00000272426 | | 3.29273 | 0.2626826 | 0.6925721 | 0.6828616 | 0.4946943 | 0.684079 |
| ENSG00000113140 | SPARC | 4.498523716 | 0.262675102 | 0.757496624 | 0.799425612 | 0.42404366 | 0.623975 |
| ENSG00000261526 | | 174.9438 | 0.2626558 | 0.2475777 | 1.115338 | 0.2647057 | 0.460092 |
| ENSG00000259982 | CDC37P1 | 1.80663 | 0.2626351 | 0.8070508 | 1.133379 | 0.2570552 | 0.450911 |
| ENSG00000239269 | RPSAP4 | 1.816506 | 0.2625601 | 0.7762905 | 0.9409981 | 0.3467058 | 0.548311 |
| ENSG00000162621 | LRRC53 | 4.300479 | 0.2625251 | 0.8511177 | 1.855843 | NA | NA |
| ENSG00000119411 | BSPRY | 2117.126015 | 0.262494019 | 0.177026884 | 1.515821607 | 0.12956447 | 0.280006 |
| ENSG00000135090 | TAOK3 | 2580.229286 | 0.262386097 | 0.266278471 | 1.043603129 | 0.29666905 | 0.495641 |
| ENSG00000139546 | TARBP2 | 2581.081 | 0.2623315 | 0.2047099 | 1.331577 | 0.1829993 | 0.358032 |
| ENSG00000100038 | TOP3B | 21.57671579 | 0.262105166 | 0.405674602 | 0.747961547 | 0.45448335 | 0.650503 |
| ENSG00000196091 | MYBPC1 | 0.9178248 | 0.2621017 | 0.7965345 | 0.9015644 | 0.3672883 | NA |
| ENSG00000211445 | GPX3 | 1293.82 | 0.262012 | 0.4577858 | 0.6969348 | 0.4858436 | 0.676895 |
| ENSG00000227431 | CSE1L-AS1 | 6.000139 | 0.2618038 | 0.6359868 | 0.6510629 | 0.5150059 | 0.700474 |

| | | | | | | | |
|---|---|---|---|---|---|---|---|
| ENSG00000123171 | CCDC70 | 2.085455192 | 0.261636113 | 0.816486489 | 0.792962198 | 0.42779985 | 0.627171 |
| ENSG00000178761 | FAM219B | 1206.215 | 0.2614238 | 0.2441434 | 1.111857 | 0.2661996 | 0.461852 |
| ENSG00000128383 | APOBEC3A | 2.779826728 | 0.261399058 | 0.69371103 | 0.67428616 | 0.5001294 | 0.688245 |
| ENSG00000258708 | SLC25A21-AS1 | 284.229 | 0.2613748 | 0.3027577 | 0.9393329 | 0.3475599 | 0.549171 |
| ENSG00000252840 | | 1.372684 | 0.2613721 | 0.8140362 | 0.790661 | 0.4291418 | NA |
| ENSG00000070831 | CDC42 | 6739.326059 | 0.261351444 | 0.174545782 | 1.529636998 | 0.12610661 | 0.274678 |
| ENSG00000146085 | MMUT | 2074.548 | 0.2612588 | 0.1534564 | 1.728839 | 0.0838379 | 0.204507 |
| ENSG00000227638 | HNRNPA1P14 | 6.265241 | 0.2612526 | 0.5758395 | 0.6380038 | 0.5234712 | 0.708766 |
| ENSG00000133835 | HSD17B4 | 7898.520466 | 0.2611104 | 0.197617422 | 1.364948448 | 0.17226925 | 0.343337 |
| ENSG00000267038 | | 1.502107 | 0.2609465 | 0.7951352 | 0.9024297 | 0.3668286 | NA |
| ENSG00000286856 | | 1.596143 | 0.2608773 | 0.7493963 | 0.763034 | 0.4454431 | 0.642544 |
| ENSG00000280042 | | 2.217778 | 0.2607551 | 0.7016482 | 0.6920348 | 0.4889155 | 0.679677 |
| ENSG00000267551 | | 0.9845281 | 0.2606076 | 0.831128 | 1.009818 | 0.3125827 | NA |
| ENSG00000259516 | ANP32AP1 | 47.93442 | 0.2604947 | 0.3176033 | 0.8964453 | 0.370015 | 0.572231 |
| ENSG00000286656 | | 11.00071 | 0.2604744 | 0.4674306 | 0.6807706 | 0.4960167 | 0.685033 |
| ENSG00000286331 | | 10.61546 | 0.2604301 | 0.5043681 | 0.6604736 | 0.50895 | 0.695162 |
| ENSG00000256377 | | 2.120265 | 0.2604171 | 0.7148076 | 0.6903767 | 0.4899574 | 0.680533 |
| ENSG00000273509 | CNTNAP3P1 | 2.121795 | 0.2602595 | 0.780358 | 0.9142587 | 0.3605809 | 0.56257 |
| ENSG00000204842 | ATXN2 | 2686.875 | 0.2602462 | 0.2017228 | 1.333148 | 0.1824834 | 0.35742 |
| ENSG00000125508 | SRMS | 112.9255301 | 0.260204862 | 0.312294597 | 0.906986501 | 0.36441394 | 0.566471 |
| ENSG00000253283 | | 0.946436 | 0.260192 | 0.7910687 | 0.9927389 | 0.3208372 | NA |
| ENSG00000213123 | DYNLT2B | 208.6118 | 0.260171 | 0.2759407 | 1.007926 | 0.31349 | 0.514683 |
| ENSG00000259177 | | 0.8118672 | 0.2600047 | 0.8233745 | 0.9441362 | 0.3451001 | NA |
| ENSG00000228606 | | 52.12657 | 0.2599129 | 0.3582838 | 0.8108546 | 0.4174492 | 0.617344 |
| ENSG00000172765 | TMCC1 | 3143.632 | 0.2598443 | 0.1445378 | 1.805665 | 0.07097064 | 0.180425 |
| ENSG00000230221 | | 1.535071 | 0.259766 | 0.7476803 | 0.7727475 | 0.4396718 | NA |
| ENSG00000223945 | | 16.5723 | 0.2596267 | 0.5807145 | 0.6379151 | 0.5235289 | 0.708808 |
| ENSG00000271698 | | 3.288154 | 0.2594492 | 0.6933895 | 0.6858663 | 0.4927974 | 0.682788 |
| ENSG00000147533 | GOLGA7 | 690.4342 | 0.2594362 | 0.1854243 | 1.433442 | 0.1517315 | 0.313749 |
| ENSG00000235088 | | 0.8247787 | 0.259399 | 0.8017559 | 0.9365874 | 0.3489708 | NA |
| ENSG00000257446 | ZNF878 | 0.8185272 | 0.2593855 | 0.8248579 | 0.9345203 | 0.3500356 | NA |
| ENSG00000185847 | LINC01405 | 0.8571367 | 0.2592826 | 0.8463512 | 1.498539 | 0.1339933 | NA |
| ENSG00000107957 | SH3PXD2A | 248.2941004 | 0.259224527 | 0.339637596 | 0.848134051 | 0.39636332 | 0.597607 |

| | | | | | | | |
|---|---|---|---|---|---|---|---|
| ENSG00000248483 | POU5F2 | 1.150102 | 0.2591453 | 0.7782114 | 0.8878829 | 0.3746038 | NA |
| ENSG00000156486 | KCNS2 | 4.877608 | 0.2591439 | 0.6777746 | 0.6735781 | 0.5005796 | 0.688543 |
| ENSG00000243766 | HOTTIP | 1.365775 | 0.2591157 | 0.8363636 | 1.206795 | 0.2275112 | NA |
| ENSG00000242060 | RPS3AP49 | 4.568701 | 0.2590843 | 0.658609 | 0.6534951 | 0.5134371 | 0.69935 |
| ENSG00000246859 | STARD4-AS1 | 183.0066 | 0.2590537 | 0.2948232 | 0.9447375 | 0.3447929 | 0.546359 |
| ENSG00000116731 | PRDM2 | 1624.495662 | 0.259000585 | 0.220305218 | 1.222067573 | 0.22168208 | 0.408329 |
| ENSG00000130956 | HABP4 | 779.1686156 | 0.258980039 | 0.14545726 | 1.80783619 | 0.070632 | 0.179839 |
| ENSG00000143641 | GALNT2 | 4307.051 | 0.2589612 | 0.1614135 | 1.629229 | 0.1032645 | 0.238204 |
| ENSG00000146350 | TBC1D32 | 369.0233 | 0.2588282 | 0.2457381 | 1.102573 | 0.2702128 | 0.466659 |
| ENSG00000267678 | | 0.7668954 | 0.2586587 | 0.8111554 | 1.002359 | 0.3161703 | NA |
| ENSG00000180543 | TSPYL5 | 7316.788 | 0.2585786 | 0.1348892 | 1.940007 | 0.05237883 | 0.143692 |
| ENSG00000223460 | GAPDHP69 | 1.064623 | 0.2584437 | 0.8141661 | 0.7935473 | 0.427459 | NA |
| ENSG00000179364 | PACS2 | 8646.295 | 0.2584033 | 0.1381294 | 1.895422 | 0.05803651 | 0.15521 |
| ENSG00000246366 | LACTB2-AS1 | 17.3473 | 0.2583692 | 0.4669767 | 0.6779835 | 0.4977821 | 0.686461 |
| ENSG00000252690 | | 56.67675 | 0.2583491 | 0.3376935 | 0.8486726 | 0.3960635 | 0.597471 |
| ENSG00000212567 | | 0.9519486 | 0.2583379 | 0.8328818 | 1.108813 | 0.267511 | NA |
| ENSG00000225642 | SNRPEP5 | 3.788319 | 0.2583245 | 0.6957595 | 0.6759527 | 0.4990707 | 0.687464 |
| ENSG00000156467 | UQCRB | 8716.757 | 0.2582196 | 0.1757489 | 1.494072 | 0.1351567 | 0.288473 |
| ENSG00000083750 | RRAGB | 1148.277843 | 0.257969852 | 0.204308351 | 1.306587665 | 0.19135279 | 0.369454 |
| ENSG00000187796 | CARD9 | 208.5659 | 0.2579524 | 0.218826 | 1.229891 | 0.218738 | 0.404598 |
| ENSG00000228665 | | 6.545458 | 0.2578481 | 0.5630609 | 0.6277317 | 0.5301797 | 0.714608 |
| ENSG00000149922 | TBX6 | 276.5814 | 0.2576055 | 0.272302 | 1.006846 | 0.3140089 | 0.515153 |
| ENSG00000227964 | LINC01429 | 1.943366 | 0.2575633 | 0.7784373 | 0.895017 | 0.370778 | 0.572911 |
| ENSG00000203666 | EFCAB2 | 296.1287 | 0.2575542 | 0.193101 | 1.375361 | 0.1690197 | 0.339099 |
| ENSG00000137494 | ANKRD42 | 960.6407629 | 0.257510775 | 0.218055908 | 1.210492716 | 0.22608988 | 0.41359 |
| ENSG00000171425 | ZNF581 | 1675.383 | 0.257487 | 0.1894693 | 1.39221 | 0.1638588 | 0.331877 |
| ENSG00000258581 | | 6.854992 | 0.2574003 | 0.5973805 | 0.6274014 | 0.5303961 | 0.714827 |
| ENSG00000074181 | NOTCH3 | 9148.742905 | 0.257303224 | 0.185102277 | 1.434447167 | 0.1514447 | 0.313302 |
| ENSG00000185619 | PCGF3 | 2977.892 | 0.2572943 | 0.1535942 | 1.709038 | 0.08744388 | 0.210998 |
| ENSG00000255507 | UVRAG-DT | 14.42832 | 0.2572592 | 0.4839343 | 0.6608215 | 0.5087268 | 0.695011 |
| ENSG00000163283 | ALPP | 565.4703 | 0.2572429 | 0.6842204 | 0.7096021 | 0.4779509 | 0.670034 |
| ENSG00000235808 | MYL6P2 | 1.129004 | 0.2571924 | 0.7351164 | 0.7161164 | 0.4739195 | NA |
| ENSG00000233081 | | 5.147956 | 0.2571419 | 0.6745626 | 0.6690141 | 0.5034865 | 0.690861 |

| | | | | | | | |
|---|---|---|---|---|---|---|---|
| ENSG00000153976 | HS3ST3A1 | 19.76395 | 0.25712 | 0.5038704 | 0.6567009 | 0.5113732 | 0.697325 |
| ENSG00000255521 | | 2.407273 | 0.2570299 | 0.6796137 | 0.6488884 | 0.5164105 | 0.701673 |
| ENSG00000288079 | | 4.347386 | 0.2569481 | 0.687949 | 0.6515542 | 0.5146888 | 0.700309 |
| ENSG00000231416 | | 94.97508 | 0.2568835 | 0.2040989 | 1.304742 | 0.1919808 | 0.370235 |
| ENSG00000254760 | | 0.9184996 | 0.2568763 | 0.8235063 | 0.9210735 | 0.357012 | NA |
| ENSG00000154845 | PPP4R1 | 4023.127 | 0.2568389 | 0.1615097 | 1.616487 | 0.1059891 | 0.24274 |
| ENSG00000237149 | ZNF503-AS2 | 188.7083 | 0.256726 | 0.2358493 | 1.139346 | 0.2545589 | 0.44755 |
| ENSG00000157856 | DRC1 | 2.062508 | 0.2567189 | 0.7231945 | 0.7187351 | 0.4723042 | 0.665323 |
| ENSG00000160293 | VAV2 | 5925.041 | 0.2565786 | 0.1388458 | 1.876818 | 0.06054311 | 0.16012 |
| ENSG00000168591 | TMUB2 | 2854.857 | 0.2565633 | 0.1895797 | 1.391621 | 0.1640371 | 0.332061 |
| ENSG00000229212 | | 32.83766 | 0.2564953 | 0.4143045 | 0.7143896 | 0.4749863 | 0.667614 |
| ENSG00000247151 | CSTF3-DT | 2.244555 | 0.2564527 | 0.730612 | 0.7191021 | 0.472078 | 0.66511 |
| ENSG00000172748 | ZNF596 | 535.5054 | 0.2564299 | 0.2335058 | 1.158034 | 0.2468501 | 0.438715 |
| ENSG00000170903 | MSANTD4 | 949.3931 | 0.2563145 | 0.2403187 | 1.122848 | 0.2615021 | 0.456167 |
| ENSG00000275056 | | 11.89369 | 0.2562036 | 0.572659 | 0.6248496 | 0.5320698 | 0.716138 |
| ENSG00000165494 | PCF11 | 7235.607 | 0.2561957 | 0.1952808 | 1.334383 | 0.1820784 | 0.356942 |
| ENSG00000133460 | SLC2A11 | 1662.369827 | 0.256192597 | 0.258595487 | 1.045840392 | 0.29563473 | 0.494708 |
| ENSG00000238165 | | 45.1326 | 0.2561759 | 0.3215762 | 0.8673446 | 0.3857532 | 0.588047 |
| ENSG00000187953 | PMS2CL | 739.0236 | 0.2560593 | 0.1479235 | 1.766353 | 0.07733672 | 0.192316 |
| ENSG00000149311 | ATM | 2536.645 | 0.2560008 | 0.2886177 | 0.9874229 | 0.3234354 | 0.524337 |
| ENSG00000231845 | HMGB3P14 | 2.343191 | 0.2559743 | 0.7398587 | 0.7487626 | 0.4540003 | 0.650094 |
| ENSG00000113504 | SLC12A7 | 5831.107077 | 0.25597327 | 0.176618706 | 1.499375632 | 0.13377621 | 0.286498 |
| ENSG00000157653 | C9orf43 | 80.18314 | 0.2559669 | 0.3037394 | 0.9068701 | 0.3644755 | 0.566533 |
| ENSG00000230623 | | 1.092366 | 0.2559481 | 0.814769 | 0.76102 | 0.4466451 | NA |
| ENSG00000242661 | RPS3AP43 | 1.102612 | 0.255868 | 0.7730623 | 0.8179491 | 0.4133862 | NA |
| ENSG00000064933 | PMS1 | 1131.031027 | 0.25569973 | 0.142728371 | 1.814908629 | 0.06953795 | 0.177649 |
| ENSG00000250244 | | 0.8896294 | 0.2556934 | 0.8114948 | 1.009483 | 0.3127432 | NA |
| ENSG00000184084 | | 1.191608 | 0.2556648 | 0.7865583 | 0.8941526 | 0.3712403 | NA |
| ENSG00000229920 | RPS4XP5 | 1.12852 | 0.2556182 | 0.7957515 | 0.9639371 | 0.3350775 | NA |
| ENSG00000241693 | RN7SL704P | 0.9352165 | 0.2555436 | 0.7581299 | 0.791814 | 0.4284692 | NA |
| ENSG00000225661 | RPL14P5 | 1.3848 | 0.2555398 | 0.7442504 | 0.7496113 | 0.4534888 | NA |
| ENSG00000173039 | RELA | 6655.915 | 0.2555309 | 0.1036734 | 2.488959 | 0.01281176 | 0.047495 |
| ENSG00000142178 | SIK1 | 2.006803 | 0.2554348 | 0.784773 | 0.8605102 | 0.3895078 | 0.591441 |

| | | | | | | | |
|---|---|---|---|---|---|---|---|
| ENSG00000234696 | GPR50-AS1 | 1.588686 | 0.2554202 | 0.7787579 | 0.9074332 | 0.3641778 | 0.566335 |
| ENSG00000145592 | RPL37 | 25449.54 | 0.2554189 | 0.1605334 | 1.590425 | 0.1117391 | 0.252256 |
| ENSG00000253341 | PCBP2P2 | 4489.182 | 0.2553817 | 0.1111404 | 2.303342 | 0.02125959 | 0.071417 |
| ENSG00000168495 | POLR3D | 2243.063 | 0.2553621 | 0.1755908 | 1.494022 | 0.1351699 | 0.288473 |
| ENSG00000157837 | SPPL3 | 3247.567 | 0.2553376 | 0.132994 | 1.949415 | 0.05124591 | 0.141202 |
| ENSG00000133226 | SRRM1 | 3866.233834 | 0.255335806 | 0.124547205 | 2.092930926 | 0.03635532 | 0.108713 |
| ENSG00000197149 | | 5.002746 | 0.2552909 | 0.5967407 | 0.634626 | 0.5256724 | 0.710546 |
| ENSG00000174611 | KY | 3.081745 | 0.2551976 | 0.7235479 | 0.7214125 | 0.4706557 | 0.664057 |
| ENSG00000182484 | WASH6P | 300.6699 | 0.2551145 | 0.2000131 | 1.319269 | 0.1870791 | 0.36357 |
| ENSG00000158423 | RIBC1 | 127.1065 | 0.2551071 | 0.3297379 | 0.8483314 | 0.3962534 | 0.597546 |
| ENSG00000171608 | PIK3CD | 217.6332 | 0.2551023 | 0.2959318 | 0.9296412 | 0.3525569 | 0.554397 |
| ENSG00000260018 | | 293.2818 | 0.2550961 | 0.3099516 | 0.8946101 | 0.3709955 | 0.573147 |
| ENSG00000230551 | | 1320.129 | 0.2549764 | 0.2431749 | 1.100582 | 0.2710785 | 0.467375 |
| ENSG00000137491 | SLCO2B1 | 5.485644156 | 0.254871287 | 0.655654277 | 0.653636343 | 0.51334612 | 0.69927 |
| ENSG00000223727 | | 3.174662 | 0.2548545 | 0.6454447 | 0.6269131 | 0.5307162 | 0.715004 |
| ENSG00000260958 | | 1.121633 | 0.2548047 | 0.8469961 | 1.621522 | 0.1049057 | NA |
| ENSG00000280890 | ELDR | 1.662189 | 0.2547132 | 0.7546951 | 0.76934 | 0.4416915 | 0.639491 |
| ENSG00000145348 | TBCK | 1798.495 | 0.2546384 | 0.1804116 | 1.44417 | 0.1486912 | 0.309195 |
| ENSG00000198964 | SGMS1 | 1538.518 | 0.2546064 | 0.2349833 | 1.142786 | 0.2531273 | 0.446066 |
| ENSG00000214485 | RPL7P1 | 4464.357 | 0.2545832 | 0.1898522 | 1.380297 | 0.1674952 | 0.336905 |
| ENSG00000152270 | PDE3B | 1190.663 | 0.2545241 | 0.2696532 | 0.9972392 | 0.3186484 | 0.519748 |
| ENSG00000107968 | MAP3K8 | 652.8555 | 0.254515964 | 0.289924283 | 0.949607469 | 0.34231174 | 0.543757 |
| ENSG00000256053 | COA8 | 1104.043 | 0.2544539 | 0.1374427 | 1.880297 | 0.06006758 | 0.159211 |
| ENSG00000225533 | PAWRP1 | 2.030791 | 0.2544001 | 0.7374734 | 0.7384074 | 0.4602669 | 0.655631 |
| ENSG00000109670 | FBXW7 | 1701.887147 | 0.254345206 | 0.111835095 | 2.272272883 | 0.02307003 | 0.07618 |
| ENSG00000172350 | ABCG4 | 12.20476 | 0.2543288 | 0.5820944 | 0.6287402 | 0.5295192 | 0.714251 |
| ENSG00000179526 | SHARPIN | 3942.66 | 0.254289 | 0.1388476 | 1.847783 | 0.06463376 | 0.16798 |
| ENSG00000114126 | TFDP2 | 2722.100539 | 0.254271663 | 0.139940333 | 1.858893259 | 0.06304227 | 0.16498 |
| ENSG00000242753 | | 1.459091 | 0.2542345 | 0.8115295 | 0.7657471 | 0.4438268 | NA |
| ENSG00000092203 | TOX4 | 6195.708024 | 0.254192443 | 0.091504481 | 2.789203688 | 0.00528378 | 0.023177 |
| ENSG00000276727 | | 3.816214 | 0.2541704 | 0.6909666 | 0.6675002 | 0.5044527 | 0.691365 |
| ENSG00000256293 | ATP6V1E1P3 | 79.18484 | 0.2541075 | 0.2280166 | 1.161749 | 0.2453372 | 0.437005 |
| ENSG00000114790 | ARHGEF26 | 1977.824573 | 0.254024131 | 0.180472256 | 1.448356708 | 0.14751732 | 0.307475 |

| | | | | | | | |
|---|---|---|---|---|---|---|---|
| ENSG00000134775 | FHOD3 | 42.81214972 | 0.253977593 | 0.501921654 | 0.648366195 | 0.51674813 | 0.70206 |
| ENSG00000177685 | CRACR2B | 1799.493 | 0.2539567 | 0.23762 | 1.115571 | 0.2646059 | 0.460053 |
| ENSG00000279345 | | 12.18613 | 0.2539238 | 0.5509193 | 0.6259976 | 0.5313165 | 0.715504 |
| ENSG00000115310 | RTN4 | 6853.4191 | 0.253897241 | 0.173371605 | 1.486611387 | 0.13711748 | 0.291446 |
| ENSG00000167130 | DOLPP1 | 2011.781 | 0.2538782 | 0.1950498 | 1.336019 | 0.181543 | 0.356235 |
| ENSG00000244371 | PFN1P8 | 17.58734 | 0.2538746 | 0.5191853 | 0.63547 | 0.5251219 | 0.71006 |
| ENSG00000279834 | | 0.8304366 | 0.2538514 | 0.8260054 | 1.010736 | 0.3121428 | NA |
| ENSG00000181333 | HEPHL1 | 63.03954 | 0.2538131 | 0.288828 | 0.9455244 | 0.3443912 | 0.545983 |
| ENSG00000173276 | ZBTB21 | 1887.36 | 0.2537738 | 0.2481657 | 1.056341 | 0.2908123 | 0.489462 |
| ENSG00000101935 | AMMECR1 | 1920.896051 | 0.253705634 | 0.208392795 | 1.263914049 | 0.20626088 | 0.388468 |
| ENSG00000261524 | ABCB10P3 | 1.389287 | 0.2537012 | 0.788797 | 0.8921901 | 0.372291 | NA |
| ENSG00000072786 | STK10 | 3657.677594 | 0.25361903 | 0.155302964 | 1.666791216 | 0.09555593 | 0.22502 |
| ENSG00000151631 | AKR1C6P | 3.814694 | 0.2535953 | 0.6404107 | 0.6328408 | 0.5268376 | 0.711744 |
| ENSG00000238110 | | 55.49812 | 0.2534674 | 0.2885535 | 0.9425991 | 0.345886 | 0.547438 |
| ENSG00000214046 | SMIM7 | 1369.853 | 0.2533525 | 0.1634005 | 1.584739 | 0.1130255 | 0.25408 |
| ENSG00000229887 | HNRNPA1P6 | 6.310243 | 0.253349 | 0.5839412 | 0.6218164 | 0.5340626 | 0.717548 |
| ENSG00000010539 | ZNF200 | 931.3207009 | 0.253346099 | 0.131924114 | 1.926985718 | 0.05398141 | 0.147176 |
| ENSG00000147465 | STAR | 3.148925 | 0.2532195 | 0.6780054 | 0.6431723 | 0.5201123 | 0.705332 |
| ENSG00000102893 | PHKB | 3218.405981 | 0.253156956 | 0.206707328 | 1.272000607 | 0.2033729 | 0.385023 |
| ENSG00000259702 | | 7.129601 | 0.2531217 | 0.6279507 | 0.6352944 | 0.5252364 | 0.710142 |
| ENSG00000140367 | UBE2Q2 | 3532.215 | 0.2529335 | 0.1518838 | 1.667604 | 0.09539437 | 0.224792 |
| ENSG00000154582 | ELOC | 2243.121 | 0.2529285 | 0.2051654 | 1.274943 | 0.2023293 | 0.383512 |
| ENSG00000186281 | GPAT2 | 4.402959 | 0.2528628 | 0.5932445 | 0.6179165 | 0.5366304 | 0.719688 |
| ENSG00000256537 | SMIM10L1 | 1337.866 | 0.2528258 | 0.1168111 | 2.189896 | 0.02853181 | 0.090095 |
| ENSG00000241104 | CEACAMP10 | 3.531219 | 0.2527937 | 0.8112407 | 0.7953674 | 0.4263998 | 0.62628 |
| ENSG00000250371 | | 1.002535 | 0.2527778 | 0.7875618 | 0.8557402 | 0.3921415 | NA |
| ENSG00000287414 | | 1.285992 | 0.2526504 | 0.786755 | 0.9871705 | 0.3235591 | NA |
| ENSG00000071073 | MGAT4A | 4152.383764 | 0.252617614 | 0.199463801 | 1.300549074 | 0.19341285 | 0.372079 |
| ENSG00000237441 | RGL2 | 13098.39 | 0.2525955 | 0.2386293 | 1.107842 | 0.26793 | 0.463846 |
| ENSG00000238098 | ABCA17P | 83.81276 | 0.2525344 | 0.2704459 | 0.9946723 | 0.3198957 | 0.520777 |
| ENSG00000138942 | RNF185 | 2306.819459 | 0.252450223 | 0.184277307 | 1.391300954 | 0.16413419 | 0.332207 |
| ENSG00000274019 | | 3.068817 | 0.2524417 | 0.6774749 | 0.6455445 | 0.5185744 | 0.703857 |
| ENSG00000164047 | CAMP | 4.594373 | 0.2524144 | 0.729001 | 0.7214704 | 0.4706201 | 0.664042 |

| | | | | | | | |
|---|---|---|---|---|---|---|---|
| ENSG00000104231 | ZFAND1 | 3108.854122 | 0.252377814 | 0.234025767 | 1.134963924 | 0.25639043 | 0.450115 |
| ENSG00000166582 | CENPV | 993.1255 | 0.2523584 | 0.2497548 | 1.068577 | 0.2852605 | 0.483268 |
| ENSG00000264029 | | 1.531192 | 0.252355 | 0.7425554 | 0.7379779 | 0.4605279 | NA |
| ENSG00000132718 | SYT11 | 1.804391486 | 0.252292183 | 0.707808967 | 0.683171058 | 0.49449876 | 0.684048 |
| ENSG00000272922 | | 0.9565732 | 0.2522587 | 0.8433 | 1.483697 | 0.1378894 | NA |
| ENSG00000265660 | MIR4664 | 103.8026 | 0.2521808 | 0.3180923 | 0.8638336 | 0.3876793 | 0.58987 |
| ENSG00000187601 | MAGEH1 | 1408.348 | 0.2521221 | 0.1656732 | 1.540575 | 0.1234203 | 0.270803 |
| ENSG00000235191 | NUCB1-AS1 | 0.9669652 | 0.2520905 | 0.8122826 | 0.7663914 | 0.4434435 | NA |
| ENSG00000223393 | | 28.48963 | 0.2520669 | 0.3819105 | 0.7530326 | 0.4514303 | 0.647697 |
| ENSG00000150773 | PIH1D2 | 20.03274 | 0.2517671 | 0.3616712 | 0.7758847 | 0.4378171 | 0.636309 |
| ENSG00000234498 | RPL13AP20 | 28.33778 | 0.2517408 | 0.442538 | 0.6807736 | 0.4960147 | 0.685033 |
| ENSG00000198918 | RPL39 | 1068.452 | 0.2517407 | 0.168097 | 1.53298 | 0.1252807 | 0.273485 |
| ENSG00000242220 | TCP10L | 1.072257 | 0.2517334 | 0.7944755 | 0.9463456 | 0.3439723 | NA |
| ENSG00000105662 | CRTC1 | 819.6490294 | 0.251698197 | 0.199061127 | 1.301131444 | 0.19321347 | 0.371849 |
| ENSG00000119242 | CCDC92 | 2017.132025 | 0.251596229 | 0.15565351 | 1.643049811 | 0.10037263 | 0.233442 |
| ENSG00000286895 | | 2.063688 | 0.251548 | 0.7121629 | 0.681525 | 0.4955394 | 0.684733 |
| ENSG00000158428 | CATIP | 9.452064 | 0.2514721 | 0.6155954 | 0.6285947 | 0.5296144 | 0.714281 |
| ENSG00000097033 | SH3GLB1 | 2509.835915 | 0.25140276 | 0.148657552 | 1.716611535 | 0.08605017 | 0.208439 |
| ENSG00000211692 | TRGJP1 | 1.14035 | 0.2512991 | 0.8222598 | 0.915908 | 0.3597151 | NA |
| ENSG00000205867 | KRTAP5-2 | 0.8509102 | 0.2512957 | 0.7870445 | 0.8453052 | 0.3979405 | NA |
| ENSG00000163481 | RNF25 | 1867.649 | 0.2512728 | 0.2251049 | 1.159541 | 0.2462356 | 0.438038 |
| ENSG00000114120 | SLC25A36 | 10381.67144 | 0.251248688 | 0.219195336 | 1.191063905 | 0.2336285 | 0.422654 |
| ENSG00000225210 | DUXAP9 | 201.1542 | 0.2511789 | 0.2290308 | 1.144938 | 0.2522349 | 0.444939 |
| ENSG00000173153 | ESRRA | 3916.637 | 0.2511722 | 0.1671908 | 1.524775 | 0.1273152 | 0.276579 |
| ENSG00000162888 | C1orf147 | 5.67605 | 0.2511193 | 0.5639446 | 0.6143604 | 0.5389772 | 0.721572 |
| ENSG00000130829 | DUSP9 | 101.1176661 | 0.251020735 | 0.461186489 | 0.668151087 | 0.50403715 | 0.691184 |
| ENSG00000224729 | PCOLCE-AS1 | 5.350305 | 0.250862 | 0.5714152 | 0.615182 | 0.5384345 | 0.721168 |
| ENSG00000005059 | MCUB | 994.132097 | 0.250856552 | 0.312177936 | 0.87902636 | 0.37938698 | 0.581345 |
| ENSG00000197694 | SPTAN1 | 14087.81 | 0.2508328 | 0.1776778 | 1.447334 | 0.1478035 | 0.307879 |
| ENSG00000151413 | NUBPL | 862.0872 | 0.2508118 | 0.1939807 | 1.331793 | 0.1829282 | 0.357975 |
| ENSG00000226981 | ABHD17AP6 | 1.212309 | 0.2506885 | 0.7512433 | 0.7405909 | 0.4589415 | NA |
| ENSG00000149050 | ZNF214 | 3.731166 | 0.2506796 | 0.81484 | 0.7610741 | 0.4466128 | 0.643499 |
| ENSG00000287496 | | 4.19727 | 0.2506245 | 0.6033142 | 0.6106146 | 0.5414548 | 0.723128 |

| | | | | | | | |
|---|---|---|---|---|---|---|---|
| ENSG00000213281 | NRAS | 25028.53 | 0.2505675 | 0.1516555 | 1.726238 | 0.08430463 | 0.205306 |
| ENSG00000151418 | ATP6V1G3 | 0.8816026 | 0.2504318 | 0.8347956 | 1.248023 | 0.2120225 | NA |
| ENSG00000167194 | C16orf92 | 1.549698 | 0.2503617 | 0.7618781 | 0.7710099 | 0.4407011 | NA |
| ENSG00000223044 | RNU6-130P | 1.160968 | 0.2502683 | 0.7734142 | 0.8106897 | 0.4175439 | NA |
| ENSG00000140526 | ABHD2 | 4550.54 | 0.2502503 | 0.2904698 | 0.9241193 | 0.3554242 | 0.557414 |
| ENSG00000228039 | | 1.192469 | 0.2501259 | 0.7979056 | 0.9307893 | 0.3519626 | NA |
| ENSG00000225450 | | 2.543709 | 0.2500881 | 0.7009031 | 0.6736774 | 0.5005165 | 0.688492 |
| ENSG00000119596 | YLPM1 | 6541.84567 | 0.250082729 | 0.155283095 | 1.637464074 | 0.10153354 | 0.235398 |
| ENSG00000214553 | LRRC37A11P | 1.416284 | 0.2499886 | 0.7295975 | 0.7104588 | 0.4774197 | NA |
| ENSG00000153060 | TEKT5 | 18.88064 | 0.2499768 | 0.4536726 | 0.6665224 | 0.5050773 | 0.691829 |
| ENSG00000259732 | | 2.959219 | 0.2498661 | 0.7379402 | 0.731917 | 0.4642192 | 0.658249 |
| ENSG00000126522 | ASL | 1137.911653 | 0.249785083 | 0.190466824 | 1.357512133 | 0.17461854 | 0.346523 |
| ENSG00000118271 | TTR | 1.053798052 | 0.249730222 | 0.800426819 | 0.931578002 | 0.35155466 | NA |
| ENSG00000167815 | PRDX2 | 6702.883 | 0.2497122 | 0.234583 | 1.112999 | 0.2657089 | 0.461366 |
| ENSG00000140521 | POLG | 6545.017 | 0.2496943 | 0.09447384 | 2.652849 | 0.00798155 | 0.032468 |
| ENSG00000187288 | CIDEC | 2.701323 | 0.2496229 | 0.6810206 | 0.6424492 | 0.5205816 | 0.705788 |
| ENSG00000186409 | CCDC30 | 179.1051 | 0.2495898 | 0.3363385 | 0.8180182 | 0.4133468 | 0.613429 |
| ENSG00000242574 | HLA-DMB | 3.073141 | 0.2495386 | 0.7047594 | 0.6745974 | 0.4999316 | 0.688079 |
| ENSG00000248546 | ANP32C | 38.31011 | 0.2494822 | 0.3640186 | 0.7740564 | 0.4388974 | 0.637373 |
| ENSG00000242086 | MUC20-OT1 | 1356.683 | 0.2492269 | 0.2092395 | 1.242883 | 0.2139108 | 0.398736 |
| ENSG00000107959 | PITRM1 | 7869.410264 | 0.249158446 | 0.187109742 | 1.401966064 | 0.16092538 | 0.327534 |
| ENSG00000270025 | BMS1P7 | 0.7824102 | 0.2491522 | 0.8116077 | 0.7598725 | 0.4473308 | NA |
| ENSG00000233614 | DDX11L10 | 1.854361 | 0.2490653 | 0.7470876 | 0.737019 | 0.4611108 | 0.655995 |
| ENSG00000213244 | H3P4 | 0.9627949 | 0.2490549 | 0.8200583 | 1.167569 | 0.2429806 | NA |
| ENSG00000168672 | LRATD2 | 13762.58 | 0.2490203 | 0.1716191 | 1.460483 | 0.1441572 | 0.30237 |
| ENSG00000231028 | LINC00271 | 66.11063 | 0.2488789 | 0.3059246 | 0.8763859 | 0.3808203 | 0.582869 |
| ENSG00000216621 | | 2.456191 | 0.2488755 | 0.7606964 | 0.7761018 | 0.4376889 | 0.636227 |
| ENSG00000109667 | SLC2A9 | 0.903555349 | 0.248826113 | 0.803329753 | 0.957821518 | 0.33815277 | NA |
| ENSG00000283696 | | 5.004293 | 0.2487636 | 0.6911243 | 0.6439606 | 0.519601 | 0.70489 |
| ENSG00000146233 | CYP39A1 | 19.62299 | 0.2486106 | 0.6138987 | 0.6123656 | 0.5402959 | 0.722379 |
| ENSG00000279532 | | 1.122546 | 0.2485884 | 0.7594889 | 0.7811419 | 0.4347191 | NA |
| ENSG00000260211 | | 9.4793 | 0.2485541 | 0.5549938 | 0.6219511 | 0.533974 | 0.717538 |
| ENSG00000110697 | PITPNM1 | 2175.315105 | 0.248484303 | 0.189318391 | 1.353044775 | 0.17604133 | 0.348684 |

| | | | | | | | |
|---|---|---|---|---|---|---|---|
| ENSG00000287839 | | 16.53604 | 0.2483065 | 0.6241722 | 0.6072755 | 0.5436681 | 0.724554 |
| ENSG00000231329 | | 3.071434 | 0.2481896 | 0.6909332 | 0.6606222 | 0.5088546 | 0.695068 |
| ENSG00000130818 | ZNF426 | 21.97364749 | 0.248173115 | 0.432906395 | 0.682668688 | 0.49481622 | 0.684152 |
| ENSG00000226686 | LINC01535 | 0.8451599 | 0.2480668 | 0.8019606 | 0.9121107 | 0.3617104 | NA |
| ENSG00000204524 | ZNF805 | 336.2031 | 0.2480362 | 0.186695 | 1.365771 | 0.1720109 | 0.342899 |
| ENSG00000134115 | CNTN6 | 1.265445678 | 0.247974897 | 0.781656325 | 0.835220817 | 0.40359339 | NA |
| ENSG00000276823 | | 1.833013 | 0.247943 | 0.7271842 | 0.7072772 | 0.4793942 | 0.67143 |
| ENSG00000055163 | CYFIP2 | 2916.80681 | 0.247932944 | 0.21716709 | 1.185923134 | 0.23565266 | 0.42515 |
| ENSG00000284512 | | 17.65412 | 0.2478578 | 0.4969274 | 0.6370135 | 0.524116 | 0.709206 |
| ENSG00000198182 | ZNF607 | 651.7693 | 0.2477935 | 0.1995383 | 1.278441 | 0.2010938 | 0.381824 |
| ENSG00000068024 | HDAC4 | 685.8916931 | 0.247694287 | 0.169257223 | 1.501359111 | 0.1332627 | 0.285929 |
| ENSG00000252782 | RNU6-341P | 0.9502672 | 0.2476387 | 0.763309 | 0.7885355 | 0.4303835 | NA |
| ENSG00000102100 | SLC35A2 | 2877.544052 | 0.247632727 | 0.120757209 | 2.086270732 | 0.03695411 | 0.109985 |
| ENSG00000254550 | OMP | 1.940853 | 0.2476133 | 0.8105108 | 0.7688482 | 0.4419834 | 0.639786 |
| ENSG00000250358 | LINC02200 | 1.003945 | 0.247588 | 0.7651074 | 0.7675558 | 0.4427511 | NA |
| ENSG00000275180 | | 58.65542 | 0.2475871 | 0.3732848 | 0.7475782 | 0.4547146 | 0.650652 |
| ENSG00000271601 | LIX1L | 76.92184 | 0.247533 | 0.3158021 | 0.8559093 | 0.3920479 | 0.593731 |
| ENSG00000131871 | SELENOS | 3630.771745 | 0.247289156 | 0.12919446 | 1.94055605 | 0.05231215 | 0.143553 |
| ENSG00000278630 | LINC02335 | 0.7334737 | 0.2472526 | 0.8368424 | 1.30331 | 0.1924689 | NA |
| ENSG00000118420 | UBE3D | 322.6078485 | 0.247229244 | 0.155446001 | 1.621417014 | 0.10492823 | 0.240933 |
| ENSG00000159882 | ZNF230 | 245.7113 | 0.2472289 | 0.2488884 | 1.035344 | 0.3005084 | 0.499763 |
| ENSG00000227725 | GCOM2 | 99.69957 | 0.2471884 | 0.2561925 | 1.019856 | 0.3077966 | 0.507916 |
| ENSG00000225798 | | 2.507892 | 0.2471177 | 0.6515217 | 0.6253569 | 0.5317369 | 0.715835 |
| ENSG00000183935 | HTR7P1 | 384.4326 | 0.2469886 | 0.2095513 | 1.215361 | 0.2242285 | 0.411034 |
| ENSG00000203441 | LINC00449 | 3.408594 | 0.2469341 | 0.6943519 | 0.6534464 | 0.5134685 | 0.699356 |
| ENSG00000197361 | FBXL22 | 84.14313 | 0.2469248 | 0.254004 | 1.022688 | 0.3064552 | 0.506482 |
| ENSG00000225408 | | 3.254054 | 0.2467739 | 0.739123 | 0.7373575 | 0.460905 | 0.655858 |
| ENSG00000170382 | LRRN2 | 405.408 | 0.2467105 | 0.3898029 | 0.7163928 | 0.4737488 | 0.66651 |
| ENSG00000228541 | | 1.476142 | 0.2466058 | 0.7762332 | 0.9218989 | 0.3565813 | NA |
| ENSG00000256967 | | 16.30548 | 0.2465947 | 0.4811922 | 0.6440011 | 0.5195747 | 0.70489 |
| ENSG00000280160 | | 9.68602 | 0.2465553 | 0.5147926 | 0.6203592 | 0.5350213 | 0.718545 |
| ENSG00000283667 | | 21.48391 | 0.2465529 | 0.4886285 | 0.6317916 | 0.5275231 | 0.712333 |
| ENSG00000235908 | RHOA-IT1 | 3.011669 | 0.2465519 | 0.6499643 | 0.6126474 | 0.5401095 | 0.722316 |

| | | | | | | | |
|---|---|---|---|---|---|---|---|
| ENSG00000232986 | | 3.084181 | 0.2462481 | 0.688849 | 0.6228854 | 0.5333599 | 0.717293 |
| ENSG00000257023 | | 1.326508 | 0.246134 | 0.8113993 | 0.7721374 | 0.440033 | NA |
| ENSG00000180209 | MYLPF | 2.951227 | 0.2460988 | 0.7061041 | 0.6644051 | 0.5064311 | 0.692933 |
| ENSG00000177156 | TALDO1 | 9877.148 | 0.2459237 | 0.220963 | 1.157942 | 0.2468878 | 0.438737 |
| ENSG00000137463 | MGARP | 2.362934954 | 0.245751605 | 0.682423303 | 0.664951705 | 0.50608138 | 0.69274 |
| ENSG00000177565 | TBL1XR1 | 4874.799 | 0.2457495 | 0.1184216 | 2.152509 | 0.03135732 | 0.097124 |
| ENSG00000163938 | GNL3 | 6382.759 | 0.2455697 | 0.1981328 | 1.282695 | 0.1995988 | 0.3798 |
| ENSG00000147894 | C9orf72 | 417.8994 | 0.2455491 | 0.1844175 | 1.369336 | 0.1708944 | 0.341599 |
| ENSG00000284543 | LINC01226 | 3.041383 | 0.2455141 | 0.7075864 | 0.6644654 | 0.5063925 | 0.692933 |
| ENSG00000237501 | | 0.9788758 | 0.2455102 | 0.7589397 | 0.7505867 | 0.4529014 | NA |
| ENSG00000149930 | TAOK2 | 7231.096 | 0.2455045 | 0.1615851 | 1.528636 | 0.1263546 | 0.275049 |
| ENSG00000185433 | LINC00158 | 0.7834554 | 0.2454604 | 0.8077868 | 0.9742995 | 0.3299079 | NA |
| ENSG00000267707 | | 6.47029 | 0.2454368 | 0.5875076 | 0.6023688 | 0.5469287 | 0.726845 |
| ENSG00000261055 | | 2.649691 | 0.2453932 | 0.7242408 | 0.6739089 | 0.5003692 | 0.688436 |
| ENSG00000129680 | MAP7D3 | 1596.933934 | 0.245385749 | 0.195418333 | 1.292174956 | 0.19629656 | 0.376048 |
| ENSG00000278999 | | 1.361591 | 0.2452693 | 0.7643048 | 0.7800896 | 0.4353381 | NA |
| ENSG00000100678 | SLC8A3 | 6.476004714 | 0.24519359 | 0.75430458 | 0.784577857 | 0.43270112 | 0.631846 |
| ENSG00000099968 | BCL2L13 | 3646.253367 | 0.245147129 | 0.120592042 | 2.0561994 | 0.0397633 | 0.116385 |
| ENSG00000163016 | ALMS1P1 | 32.18487 | 0.2451225 | 0.490097 | 0.6218466 | 0.5340427 | 0.717548 |
| ENSG00000286575 | | 1.747224 | 0.245045 | 0.7357525 | 0.7211246 | 0.4708328 | 0.664166 |
| ENSG00000168216 | LMBRD1 | 2078.969 | 0.244975 | 0.1728156 | 1.437677 | 0.1505258 | 0.311911 |
| ENSG00000167136 | ENDOG | 351.9129 | 0.2448948 | 0.2355517 | 1.086578 | 0.2772235 | 0.474187 |
| ENSG00000092330 | TINF2 | 2701.108409 | 0.244880701 | 0.130505493 | 1.903862797 | 0.05692806 | 0.152873 |
| ENSG00000277559 | | 88.47806 | 0.2448591 | 0.3245851 | 0.8247721 | 0.409501 | 0.609936 |
| ENSG00000240731 | | 194.2509 | 0.2447266 | 0.2269249 | 1.127739 | 0.2594303 | 0.453713 |
| ENSG00000176183 | | 2.549299 | 0.2446989 | 0.7350826 | 0.7196775 | 0.4717236 | 0.664963 |
| ENSG00000181873 | IBA57 | 964.8712 | 0.2446752 | 0.2761049 | 0.9502503 | 0.3419851 | 0.543336 |
| ENSG00000276509 | | 1.018088 | 0.2446694 | 0.8098235 | 0.7401118 | 0.4592321 | NA |
| ENSG00000198586 | TLK1 | 3445.515 | 0.2446581 | 0.1252368 | 1.980417 | 0.04765674 | 0.133593 |
| ENSG00000240366 | RPL36AP45 | 1.088359 | 0.244631 | 0.7864575 | 0.8411314 | 0.4002743 | NA |
| ENSG00000259380 | | 0.852927 | 0.2444288 | 0.8300525 | 1.079295 | 0.2804562 | NA |
| ENSG00000159723 | AGRP | 1.113697 | 0.2443809 | 0.8105025 | 0.7554151 | 0.4499999 | NA |
| ENSG00000269836 | | 1.775733 | 0.2442982 | 0.7572712 | 0.7604986 | 0.4469566 | 0.643781 |

| | | | | | | | |
|---|---|---|---|---|---|---|---|
| ENSG00000228643 | | 8.013008 | 0.2441668 | 0.5404619 | 0.6050577 | 0.5451406 | 0.725751 |
| ENSG00000181038 | METTL23 | 1538.599 | 0.2441652 | 0.1678261 | 1.492732 | 0.1355074 | 0.289041 |
| ENSG00000229956 | ZRANB2-AS2 | 33.70868 | 0.244142 | 0.375699 | 0.7307015 | 0.4649615 | 0.658927 |
| ENSG00000160471 | COX6B2 | 3.545849 | 0.2439088 | 0.6621771 | 0.6233988 | 0.5330225 | 0.717057 |
| ENSG00000196182 | STK40 | 3522.87 | 0.2438991 | 0.2100201 | 1.208174 | 0.2269802 | 0.414762 |
| ENSG00000254012 | RPL10P8 | 0.9962794 | 0.2438293 | 0.8052472 | 1.037883 | 0.2993245 | NA |
| ENSG00000213304 | | 8.732971 | 0.2438099 | 0.5804337 | 0.5992719 | 0.5489916 | 0.728397 |
| ENSG00000233013 | FAM157B | 1.01461 | 0.2438002 | 0.8239846 | 1.028475 | 0.3037262 | NA |
| ENSG00000224932 | LINC02262 | 1.553246 | 0.243787 | 0.830929 | 1.170092 | 0.2419639 | NA |
| ENSG00000166801 | FAM111A | 3885.412 | 0.2437397 | 0.269601 | 0.9601729 | 0.3369682 | 0.538165 |
| ENSG00000280968 | | 1.502809 | 0.2437164 | 0.7718726 | 0.8037571 | 0.4215373 | NA |
| ENSG00000111674 | ENO2 | 3188.302055 | 0.24366071 | 0.257567119 | 0.998154688 | 0.31820436 | 0.519342 |
| ENSG00000141425 | RPRD1A | 4794.35 | 0.2435983 | 0.1537763 | 1.583172 | 0.1133823 | 0.254606 |
| ENSG00000164900 | GBX1 | 0.9613741 | 0.243579 | 0.7739495 | 0.8136613 | 0.415839 | NA |
| ENSG00000258384 | RCCD1-AS1 | 14.86438 | 0.2435468 | 0.5466816 | 0.605506 | 0.5448429 | 0.725429 |
| ENSG00000229344 | MTCO2P12 | 106.1725 | 0.2435034 | 0.3956447 | 0.703085 | 0.4820028 | 0.67345 |
| ENSG00000099331 | MYO9B | 3731.016897 | 0.243485577 | 0.198013866 | 1.272109124 | 0.20333435 | 0.384978 |
| ENSG00000286572 | | 23.29783 | 0.2434724 | 0.517101 | 0.6149317 | 0.5385998 | 0.721237 |
| ENSG00000080371 | RAB21 | 3566.31059 | 0.243345529 | 0.111721996 | 2.200307068 | 0.02778512 | 0.088146 |
| ENSG00000140795 | MYLK3 | 37.5986 | 0.2433235 | 0.5581969 | 0.6067049 | 0.5440468 | 0.724914 |
| ENSG00000253661 | ZFHX4-AS1 | 2.243888 | 0.2432923 | 0.8147271 | 0.7120407 | 0.4764396 | 0.668737 |
| ENSG00000272656 | | 18.08936 | 0.2432851 | 0.5117505 | 0.6219625 | 0.5339665 | 0.717538 |
| ENSG00000188581 | KRTAP1-1 | 0.8788645 | 0.2431515 | 0.8110519 | 0.7628983 | 0.445524 | NA |
| ENSG00000276663 | | 5.972082 | 0.2430764 | 0.6319784 | 0.6014345 | 0.5475506 | 0.727321 |
| ENSG00000141458 | NPC1 | 3522.822 | 0.2430734 | 0.1826688 | 1.366044 | 0.1719252 | 0.342766 |
| ENSG00000272008 | | 22.7969 | 0.243035 | 0.4001145 | 0.6990767 | 0.4845041 | 0.67556 |
| ENSG00000139908 | TSSK4 | 248.8963 | 0.2430007 | 0.2046154 | 1.228432 | 0.2192847 | 0.405418 |
| ENSG00000250057 | | 1.478232 | 0.2429821 | 0.7876436 | 0.8964905 | 0.3699909 | NA |
| ENSG00000271283 | | 0.9050682 | 0.2429579 | 0.8108179 | 0.7464671 | 0.4553853 | NA |
| ENSG00000271967 | | 2.240638 | 0.2429359 | 0.6963413 | 0.6417573 | 0.5210308 | 0.706289 |
| ENSG00000120656 | TAF12 | 1815.100869 | 0.242922287 | 0.178934019 | 1.394740844 | 0.163094 | 0.330681 |
| ENSG00000255443 | CD44-AS1 | 3.434783 | 0.2428217 | 0.6918765 | 0.6376131 | 0.5237255 | 0.708884 |
| ENSG00000140945 | CDH13 | 3.515773 | 0.2428059 | 0.7127187 | 0.6673642 | 0.5045395 | 0.691413 |

| | | | | | | | |
|---|---|---|---|---|---|---|---|
| ENSG00000277504 | | 11.3662 | 0.2427115 | 0.495458 | 0.6224109 | 0.5336717 | 0.717377 |
| ENSG00000266155 | | 1.743205 | 0.2425538 | 0.7457517 | 0.7039003 | 0.4814948 | 0.672953 |
| ENSG00000286655 | | 2.821409 | 0.2425279 | 0.727198 | 0.7001901 | 0.4838086 | 0.675015 |
| ENSG00000105619 | TFPT | 1980.345979 | 0.242462047 | 0.218494606 | 1.156841943 | 0.24733694 | 0.439359 |
| ENSG00000124177 | CHD6 | 11174.32953 | 0.242415149 | 0.262076568 | 0.977306478 | 0.32841745 | 0.529602 |
| ENSG00000166483 | WEE1 | 1630.901 | 0.2423342 | 0.3202035 | 0.8226342 | 0.4107161 | 0.61105 |
| ENSG00000035664 | DAPK2 | 465.5186859 | 0.242322621 | 0.310370116 | 0.849424659 | 0.39564504 | 0.597249 |
| ENSG00000014216 | CAPN1 | 16804.66154 | 0.242318065 | 0.143723062 | 1.714532275 | 0.08643102 | 0.209124 |
| ENSG00000225190 | PLEKHM1 | 1553.545 | 0.2422751 | 0.1735295 | 1.435248 | 0.1512165 | 0.312976 |
| ENSG00000179406 | LINC00174 | 717.8735 | 0.2421262 | 0.2891059 | 0.8952276 | 0.3706654 | 0.5728 |
| ENSG00000231177 | LINC00852 | 34.2786 | 0.2419618 | 0.4156669 | 0.6832239 | 0.4944654 | 0.684048 |
| ENSG00000104689 | TNFRSF10A | 709.6210314 | 0.241854224 | 0.220988351 | 1.141557435 | 0.25363803 | 0.446667 |
| ENSG00000172901 | LVRN | 27.09177 | 0.2417241 | 0.4739497 | 0.6316276 | 0.5276303 | 0.7124 |
| ENSG00000141391 | PRELID3A | 432.9294 | 0.2415682 | 0.2304382 | 1.085041 | 0.2779035 | 0.475002 |
| ENSG00000226245 | ZNF32-AS1 | 32.70736 | 0.2415331 | 0.4145366 | 0.6801775 | 0.4963921 | 0.685373 |
| ENSG00000268555 | | 1.029682 | 0.241408 | 0.8095409 | 0.7463654 | 0.4554467 | NA |
| ENSG00000280402 | | 3.355011 | 0.2413754 | 0.7046307 | 0.6478666 | 0.5170713 | 0.702427 |
| ENSG00000140403 | DNAJA4 | 1186.082 | 0.2413506 | 0.2317709 | 1.095553 | 0.2732746 | 0.469944 |
| ENSG00000272447 | | 213.9994 | 0.2412196 | 0.2676765 | 0.9558001 | 0.3391733 | 0.540484 |
| ENSG00000256799 | | 2.878317 | 0.2411659 | 0.6772388 | 0.6283741 | 0.5297589 | 0.714359 |
| ENSG00000226453 | LINC02542 | 114.6877 | 0.2409216 | 0.233555 | 1.091056 | 0.2752481 | 0.472085 |
| ENSG00000140006 | WDR89 | 1222.608 | 0.2408889 | 0.1409981 | 1.734189 | 0.08288455 | 0.202759 |
| ENSG00000037637 | FBXO42 | 1312.485317 | 0.240881076 | 0.125889301 | 1.938433389 | 0.05257037 | 0.144083 |
| ENSG00000249856 | | 0.9897091 | 0.2408758 | 0.8188554 | 0.8690486 | 0.3848205 | NA |
| ENSG00000165312 | OTUD1 | 896.3924 | 0.2408406 | 0.2570897 | 0.9896484 | 0.322346 | 0.523145 |
| ENSG00000092140 | G2E3 | 1845.597342 | 0.240781598 | 0.255887752 | 0.990586807 | 0.32188738 | 0.522633 |
| ENSG00000196236 | XPNPEP3 | 1318.765 | 0.2407284 | 0.1515811 | 1.619554 | 0.1053281 | 0.241622 |
| ENSG00000186868 | MAPT | 3349.926 | 0.2407098 | 0.2370727 | 1.062767 | 0.2878874 | 0.486199 |
| ENSG00000134020 | PEBP4 | 1.835121706 | 0.240696526 | 0.728342188 | 0.659862511 | 0.50934206 | 0.695412 |
| ENSG00000247903 | | 46.39786 | 0.2406141 | 0.3894175 | 0.7065866 | 0.4798234 | 0.671756 |
| ENSG00000159445 | THEM4 | 2419.19 | 0.2404546 | 0.2606853 | 0.9760684 | 0.3290306 | 0.530227 |
| ENSG00000074771 | NOX3 | 1.503607539 | 0.240443555 | 0.831572212 | 1.2235519 | 0.22112131 | NA |
| ENSG00000161533 | ACOX1 | 6824.05 | 0.2404341 | 0.1559033 | 1.579511 | 0.1142188 | 0.256048 |

| | | | | | | | |
|---|---|---|---|---|---|---|---|
| ENSG00000166669 | ATF7IP2 | 757.1922 | 0.2402646 | 0.166991 | 1.45269 | 0.1463098 | 0.305726 |
| ENSG00000171914 | TLN2 | 1608.654 | 0.2401789 | 0.2394183 | 1.07503 | 0.2823613 | 0.480234 |
| ENSG00000260037 | | 6.606882 | 0.2401444 | 0.520531 | 0.6040543 | 0.5458075 | 0.726117 |
| ENSG00000233589 | | 16.79607 | 0.2401417 | 0.5377914 | 0.5970401 | 0.5504806 | 0.729426 |
| ENSG00000170468 | RIOX1 | 1274.534 | 0.2400927 | 0.1756174 | 1.39522 | 0.1629495 | 0.330528 |
| ENSG00000204387 | SNHG32 | 2188.771 | 0.2400779 | 0.3530073 | 0.7634801 | 0.4451772 | 0.642402 |
| ENSG00000152578 | GRIA4 | 0.9581549 | 0.240021 | 0.7674545 | 0.7794255 | 0.4357291 | NA |
| ENSG00000256756 | TIMM17BP1 | 0.9952901 | 0.2398845 | 0.7738066 | 0.8133622 | 0.4160104 | NA |
| ENSG00000011478 | QPCTL | 1197.132383 | 0.239870168 | 0.197747289 | 1.262907935 | 0.20662227 | 0.388847 |
| ENSG00000214176 | PLEKHM1P1 | 3290.939 | 0.2397584 | 0.2439703 | 1.040098 | 0.2982943 | 0.497478 |
| ENSG00000228844 | | 1.288515 | 0.2397473 | 0.7587961 | 0.7628075 | 0.4455782 | NA |
| ENSG00000157933 | SKI | 2188.894 | 0.2397326 | 0.1909563 | 1.289178 | 0.1973364 | 0.377359 |
| ENSG00000231690 | LINC00574 | 2.132142 | 0.2396191 | 0.6588419 | 0.6021804 | 0.5470541 | 0.726916 |
| ENSG00000136052 | SLC41A2 | 1006.659814 | 0.239586603 | 0.226375273 | 1.104234591 | 0.26949139 | 0.465729 |
| ENSG00000267100 | ILF3-DT | 1697.686 | 0.2395805 | 0.1803855 | 1.368376 | 0.1711944 | 0.341836 |
| ENSG00000078596 | ITM2A | 0.777669145 | 0.239550937 | 0.820720673 | 1.16117608 | 0.2455703 | NA |
| ENSG00000241547 | ACTG1P20 | 43.17067 | 0.2395238 | 0.3169243 | 0.8250387 | 0.4093496 | 0.609907 |
| ENSG00000213305 | HNRNPCP6 | 8.655652 | 0.2395124 | 0.5446814 | 0.590289 | 0.5549969 | 0.732691 |
| ENSG00000229214 | LINC00242 | 11.97644 | 0.2394802 | 0.5048906 | 0.6090782 | 0.5424726 | 0.723738 |
| ENSG00000135828 | RNASEL | 617.8922462 | 0.239416737 | 0.265659439 | 0.963385529 | 0.33535409 | 0.536651 |
| ENSG00000094914 | AAAS | 2969.872208 | 0.239385017 | 0.156042947 | 1.569864936 | 0.11644654 | 0.259487 |
| ENSG00000275655 | RN7SL313P | 1.290815 | 0.2393558 | 0.8098975 | 0.7577743 | 0.4485861 | NA |
| ENSG00000117143 | UAP1 | 2612.236877 | 0.239264542 | 0.217578976 | 1.147420229 | 0.25120799 | 0.443976 |
| ENSG00000181982 | CCDC149 | 842.9399 | 0.2392258 | 0.1200746 | 2.010594 | 0.04436831 | 0.126701 |
| ENSG00000251329 | LINC02353 | 1.365522 | 0.2392258 | 0.7738384 | 0.7997158 | 0.4238755 | NA |
| ENSG00000188761 | BCL2L15 | 194.2576 | 0.2391837 | 0.2797953 | 0.9130842 | 0.3611983 | 0.563218 |
| ENSG00000136861 | CDK5RAP2 | 3007.270285 | 0.239165249 | 0.222529742 | 1.097420218 | 0.27245774 | 0.468973 |
| ENSG00000204650 | LINC02210 | 1143.519 | 0.239161 | 0.215996 | 1.129278 | 0.2587805 | 0.453024 |
| ENSG00000237748 | UQCRBP1 | 94.78058 | 0.239154 | 0.3286974 | 0.7986228 | 0.4245091 | 0.624343 |
| ENSG00000178188 | SH2B1 | 4017.704 | 0.2391222 | 0.1848022 | 1.329005 | 0.1838463 | 0.359136 |
| ENSG00000198792 | TMEM184B | 3941.495 | 0.2391178 | 0.1591058 | 1.530674 | 0.12585 | 0.274299 |
| ENSG00000186458 | DEFB132 | 1.299036 | 0.2390397 | 0.7995644 | 0.8750887 | 0.3815256 | NA |
| ENSG00000266980 | | 4.176773 | 0.2389989 | 0.6296339 | 0.5867994 | 0.5573384 | 0.734702 |

| | | | | | | | |
|---|---|---|---|---|---|---|---|
| ENSG00000104327 | CALB1 | 14.58089425 | 0.238973215 | 0.579375643 | 0.606514396 | 0.54417323 | 0.725046 |
| ENSG00000180921 | FAM83H | 12328.53 | 0.2388867 | 0.1670172 | 1.464867 | 0.1429572 | 0.300659 |
| ENSG00000285783 | | 1.076751 | 0.2388254 | 0.7691781 | 0.7758534 | 0.4378355 | NA |
| ENSG00000170779 | CDCA4 | 2137.994 | 0.2388149 | 0.3230186 | 0.8087451 | 0.4186618 | 0.618449 |
| ENSG00000180929 | GPR62 | 1.035157 | 0.238786 | 0.7901675 | 0.8498992 | 0.3953811 | NA |
| ENSG00000218350 | LYPLA1P3 | 295.402 | 0.2387445 | 0.2848024 | 0.8987707 | 0.3687748 | 0.570878 |
| ENSG00000105617 | LENG1 | 619.4682853 | 0.238724289 | 0.157722911 | 1.556736487 | 0.11953306 | 0.264502 |
| ENSG00000268392 | | 1.57592 | 0.2387034 | 0.750591 | 0.7247191 | 0.4686243 | 0.662195 |
| ENSG00000231702 | | 2.284422 | 0.238522 | 0.7160146 | 0.6645091 | 0.5063645 | 0.692933 |
| ENSG00000204628 | RACK1 | 79598.25 | 0.2384 | 0.1454971 | 1.66236 | 0.0964407 | 0.226495 |
| ENSG00000202417 | | 0.8633385 | 0.2383748 | 0.7794775 | 0.8606166 | 0.3894492 | NA |
| ENSG00000166140 | ZFYVE19 | 2977.634 | 0.2382908 | 0.1742671 | 1.39652 | 0.1625578 | 0.329896 |
| ENSG00000242889 | RN7SL449P | 8.033026 | 0.2381694 | 0.5789084 | 0.5829232 | 0.559945 | 0.736239 |
| ENSG00000228065 | LINC01515 | 250.784 | 0.2381281 | 0.2235862 | 1.101874 | 0.2705166 | 0.466782 |
| ENSG00000257604 | | 0.8628665 | 0.2381045 | 0.8172143 | 0.8902386 | 0.3733378 | NA |
| ENSG00000237624 | OXCT2P1 | 4.174614 | 0.2380768 | 0.6584704 | 0.6061156 | 0.544438 | 0.725217 |
| ENSG00000131097 | HIGD1B | 5.252539332 | 0.238026966 | 0.575634721 | 0.58101029 | 0.56123352 | 0.737311 |
| ENSG00000132832 | | 2.909700993 | 0.237988039 | 0.762885288 | 0.749979774 | 0.45326689 | 0.649427 |
| ENSG00000137970 | RPL7P9 | 10649.4648 | 0.237932325 | 0.184249896 | 1.324907074 | 0.18520197 | 0.360829 |
| ENSG00000184083 | FAM120C | 1582.728 | 0.2379309 | 0.188305 | 1.300083 | 0.1935726 | 0.372157 |
| ENSG00000187024 | PTRH1 | 7.66972 | 0.23792 | 0.5487986 | 0.588909 | 0.5559223 | 0.733491 |
| ENSG00000146282 | RARS2 | 2386.293 | 0.2379022 | 0.1640114 | 1.477936 | 0.1394249 | 0.29524 |
| ENSG00000258571 | PTTG4P | 2.18369 | 0.2379002 | 0.7182458 | 0.6738221 | 0.5004244 | 0.688437 |
| ENSG00000225442 | MPRIP-AS1 | 2.171189 | 0.2378778 | 0.6983737 | 0.6389727 | 0.5228407 | 0.708182 |
| ENSG00000189157 | FAM47E | 103.84 | 0.2376971 | 0.3339515 | 0.7800937 | 0.4353357 | 0.634125 |
| ENSG00000227400 | | 2.027407 | 0.237685 | 0.8120904 | 0.7251081 | 0.4683857 | 0.661979 |
| ENSG00000255523 | | 0.8921231 | 0.2376656 | 0.8112211 | 0.7389181 | 0.4599567 | NA |
| ENSG00000129007 | CALML4 | 32.88046638 | 0.237603107 | 0.386374659 | 0.702159296 | 0.48257983 | 0.673973 |
| ENSG00000233741 | RPL27P7 | 1.216011 | 0.2375157 | 0.765683 | 0.7676157 | 0.4427156 | NA |
| ENSG00000199370 | | 1.111169 | 0.2375117 | 0.8296923 | 1.31443 | 0.1887015 | NA |
| ENSG00000225975 | LINC01534 | 201.0388 | 0.2373387 | 0.3152046 | 0.8183849 | 0.4131374 | 0.613152 |
| ENSG00000176054 | RPL23P2 | 21.5206 | 0.237281 | 0.4424526 | 0.643417 | 0.5199536 | 0.705189 |
| ENSG00000091656 | ZFHX4 | 5.75281914 | 0.237255205 | 0.607361343 | 0.581220748 | 0.56109169 | 0.737289 |

| | | | | | | | |
|---|---|---|---|---|---|---|---|
| ENSG00000215182 | MUC5AC | 1.6048 | 0.2371906 | 0.7233876 | 0.6813989 | 0.4956191 | 0.684733 |
| ENSG00000259632 | | 1.175608 | 0.2370309 | 0.7512263 | 0.7150616 | 0.474571 | NA |
| ENSG00000196235 | SUPT5H | 9932.403 | 0.2370032 | 0.1373805 | 1.703924 | 0.08839538 | 0.212753 |
| ENSG00000152953 | STK32B | 16.49155 | 0.2369037 | 0.6215724 | 0.5951353 | 0.5517531 | 0.730566 |
| ENSG00000198563 | DDX39B | 10102.74 | 0.2368888 | 0.1906685 | 1.269531 | 0.2042516 | 0.386054 |
| ENSG00000207752 | MIR199A1 | 1.601922 | 0.2368858 | 0.715608 | 0.6613171 | 0.508409 | 0.694816 |
| ENSG00000214944 | ARHGEF28 | 989.2424 | 0.2367588 | 0.3678117 | 0.7233588 | 0.4694595 | 0.662967 |
| ENSG00000155876 | RRAGA | 2562.248 | 0.2367344 | 0.1673131 | 1.443374 | 0.1489153 | 0.309555 |
| ENSG00000173611 | SCAI | 982.5348 | 0.2366856 | 0.1622428 | 1.495663 | 0.1347416 | 0.288009 |
| ENSG00000173421 | IHO1 | 1.318268 | 0.2365538 | 0.7687615 | 0.7560323 | 0.4496298 | NA |
| ENSG00000250016 | SNX18P23 | 0.8064294 | 0.2364536 | 0.8174855 | 1.065475 | 0.2866611 | NA |
| ENSG00000250116 | RHOQ-AS1 | 27.33089 | 0.2364177 | 0.3969271 | 0.6828866 | 0.4946785 | 0.684079 |
| ENSG00000142459 | EVI5L | 1461.469 | 0.2363872 | 0.1626993 | 1.493774 | 0.1352347 | 0.288529 |
| ENSG00000266124 | MIR5587 | 7.402055 | 0.236197 | 0.551346 | 0.5841673 | 0.5591078 | 0.735831 |
| ENSG00000066629 | EML1 | 942.7027475 | 0.236073277 | 0.312160096 | 0.820349276 | 0.41201702 | 0.612276 |
| ENSG00000263586 | HID1-AS1 | 1.134178 | 0.2359192 | 0.808981 | 0.7463857 | 0.4554345 | NA |
| ENSG00000207047 | SNORD51 | 0.9772133 | 0.2358817 | 0.7489767 | 0.7071306 | 0.4794853 | NA |
| ENSG00000263345 | | 217.7679 | 0.2356682 | 0.3475807 | 0.7573384 | 0.4488471 | 0.645385 |
| ENSG00000263765 | | 1.639089 | 0.2356411 | 0.758127 | 0.7559398 | 0.4496853 | 0.645856 |
| ENSG00000136881 | BAAT | 3.202253509 | 0.235616489 | 0.680138792 | 0.613434179 | 0.53958929 | 0.721961 |
| ENSG00000172500 | FIBP | 4141.67 | 0.2355815 | 0.1539935 | 1.548294 | 0.1215516 | 0.26781 |
| ENSG00000254912 | | 2.248391 | 0.2355258 | 0.7542239 | 0.7069328 | 0.4796082 | 0.671608 |
| ENSG00000182463 | TSHZ2 | 22.40442 | 0.2355092 | 0.4061246 | 0.674752 | 0.4998333 | 0.68798 |
| ENSG00000085563 | ABCB1 | 2.521689676 | 0.235505747 | 0.758514296 | 0.761069477 | 0.44661557 | 0.643499 |
| ENSG00000151693 | ASAP2 | 1984.849 | 0.2353954 | 0.2838307 | 0.8797341 | 0.3790034 | 0.581103 |
| ENSG00000168884 | TNIP2 | 1969.367 | 0.2353713 | 0.1237279 | 1.919286 | 0.05494821 | 0.149002 |
| ENSG00000183283 | DAZAP2 | 12902.64 | 0.2353494 | 0.1271354 | 1.884229 | 0.05953404 | 0.158148 |
| ENSG00000103260 | METRN | 3894.188784 | 0.235348385 | 0.146260281 | 1.636104851 | 0.10181765 | 0.235706 |
| ENSG00000199090 | MIR326 | 0.9915201 | 0.2352371 | 0.8091512 | 0.706239 | 0.4800395 | NA |
| ENSG00000196456 | ZNF775 | 178.6046 | 0.2352183 | 0.2413779 | 1.025865 | 0.3049551 | 0.504662 |
| ENSG00000161132 | | 18.33173 | 0.2351105 | 0.4101431 | 0.6635072 | 0.5070058 | 0.693528 |
| ENSG00000231351 | WBP1P1 | 14.91812 | 0.2350985 | 0.3868385 | 0.6921127 | 0.4888665 | 0.679677 |
| ENSG00000092531 | SNAP23 | 791.0197328 | 0.235097251 | 0.144173995 | 1.655035925 | 0.09791721 | 0.229152 |

| | | | | | | | |
|---|---|---|---|---|---|---|---|
| ENSG00000149295 | DRD2 | 2.47131 | 0.2349707 | 0.7060713 | 0.6375827 | 0.5237454 | 0.708884 |
| ENSG00000225300 | | 0.89401 | 0.2349063 | 0.7864045 | 0.9567372 | 0.3386999 | NA |
| ENSG00000239779 | WBP1 | 987.2081 | 0.2349004 | 0.2545433 | 0.9722967 | 0.330903 | 0.531899 |
| ENSG00000103502 | CDIPT | 5872.510059 | 0.234861585 | 0.144772361 | 1.577252219 | 0.1147375 | 0.256886 |
| ENSG00000183671 | GPR1 | 1.056879 | 0.2348388 | 0.8190117 | 0.8598446 | 0.3898747 | NA |
| ENSG00000237049 | RPL34P1 | 1.319779 | 0.2348247 | 0.7409862 | 0.7077782 | 0.479083 | NA |
| ENSG00000266507 | MIR4479 | 0.8491567 | 0.2347289 | 0.7863194 | 0.8174305 | 0.4136824 | NA |
| ENSG00000172830 | SSH3 | 8016.239 | 0.2346958 | 0.1188308 | 1.980002 | 0.04770334 | 0.133659 |
| ENSG00000258593 | | 25.74718 | 0.2344325 | 0.4224417 | 0.6520944 | 0.5143402 | 0.700097 |
| ENSG00000130202 | NECTIN2 | 9482.924206 | 0.234432048 | 0.159869375 | 1.532273148 | 0.12545504 | 0.273708 |
| ENSG00000176438 | SYNE3 | 345.2383 | 0.2344267 | 0.5500048 | 0.562282 | 0.5739239 | 0.746641 |
| ENSG00000285877 | | 109.8288 | 0.2344145 | 0.3308386 | 0.7858045 | 0.431982 | 0.631212 |
| ENSG00000198496 | NBR2 | 740.2783 | 0.2342804 | 0.3506208 | 0.7442559 | 0.4567217 | 0.65254 |
| ENSG00000122218 | COPA | 12261.63332 | 0.234254514 | 0.141546791 | 1.65530603 | 0.09786244 | 0.229084 |
| ENSG00000179152 | TCAIM | 1182.195 | 0.2342429 | 0.1169696 | 2.026382 | 0.04272563 | 0.123157 |
| ENSG00000205693 | MANSC4 | 2.530964 | 0.2342307 | 0.6987454 | 0.6330179 | 0.5267219 | 0.71168 |
| ENSG00000276103 | | 7.716487 | 0.2341506 | 0.5123833 | 0.5866763 | 0.5574212 | 0.734775 |
| ENSG00000153443 | UBALD1 | 1999.025 | 0.2341456 | 0.1613015 | 1.471014 | 0.1412874 | 0.298163 |
| ENSG00000112715 | VEGFA | 8608.952735 | 0.234072953 | 0.383642128 | 0.691199265 | 0.48944033 | 0.680194 |
| ENSG00000116138 | DNAJC16 | 1182.316907 | 0.23399631 | 0.242573158 | 1.02293766 | 0.30633732 | 0.50635 |
| ENSG00000135535 | CD164 | 16925.33127 | 0.233983518 | 0.165689052 | 1.442742689 | 0.14909297 | 0.309787 |
| ENSG00000100599 | RIN3 | 288.7070565 | 0.233900281 | 0.250521971 | 0.982667233 | 0.32577124 | 0.527062 |
| ENSG00000257264 | | 0.8543515 | 0.2338812 | 0.8086672 | 0.7409371 | 0.4587316 | NA |
| ENSG00000185728 | YTHDF3 | 3524.383 | 0.2338778 | 0.1493621 | 1.598071 | 0.1100272 | 0.249478 |
| ENSG00000270091 | | 29.13918 | 0.2338599 | 0.4096768 | 0.6658323 | 0.5055183 | 0.69229 |
| ENSG00000224993 | RPL29P12 | 7.440713 | 0.2338367 | 0.5707716 | 0.5780221 | 0.5632492 | 0.738828 |
| ENSG00000254479 | SLC25A1P1 | 3.618061 | 0.2338116 | 0.6368739 | 0.5837575 | 0.5593835 | 0.736117 |
| ENSG00000214803 | | 5.779419 | 0.2338015 | 0.5637349 | 0.5772855 | 0.5637466 | 0.739186 |
| ENSG00000248873 | SERBP1P6 | 272.7721 | 0.2337327 | 0.2053075 | 1.183072 | 0.2367808 | 0.426482 |
| ENSG00000180035 | ZNF48 | 1673.231 | 0.2337037 | 0.1084281 | 2.158054 | 0.0309236 | 0.096086 |
| ENSG00000260046 | | 5.83425 | 0.2336793 | 0.5756787 | 0.5719397 | 0.5673628 | 0.741734 |
| ENSG00000188559 | RALGAPA2 | 2741.397 | 0.2335733 | 0.1905492 | 1.261473 | 0.2071386 | 0.389516 |
| ENSG00000108576 | SLC6A4 | 8.820636578 | 0.233562345 | 0.587127314 | 0.574999413 | 0.56529169 | 0.7402 |

| | | | | | | | |
|---|---|---|---|---|---|---|---|
| ENSG00000173486 | FKBP2 | 55.4018 | 0.2334829 | 0.2811359 | 0.8894561 | 0.373758 | 0.575841 |
| ENSG00000199400 | RNY4P19 | 1.042794 | 0.2334451 | 0.7862606 | 0.9574564 | 0.338337 | NA |
| ENSG00000237813 | | 14.55431 | 0.2333921 | 0.6086029 | 0.5832524 | 0.5597234 | 0.73613 |
| ENSG00000226800 | CACTIN-AS1 | 13.87821 | 0.2333512 | 0.4729195 | 0.6144627 | 0.5389096 | 0.721572 |
| ENSG00000281189 | GHET1 | 45.03054 | 0.2332967 | 0.3435287 | 0.7553069 | 0.4500649 | 0.646331 |
| ENSG00000172717 | FAM71D | 9.186282 | 0.232917 | 0.6239858 | 0.5765162 | 0.5642663 | 0.739578 |
| ENSG00000163820 | FYCO1 | 3452.5 | 0.2328508 | 0.2562327 | 0.9588027 | 0.3376581 | 0.538975 |
| ENSG00000183822 | NCF4-AS1 | 0.9900879 | 0.2327758 | 0.8100355 | 0.7296379 | 0.4656116 | NA |
| ENSG00000095596 | CYP26A1 | 102.2571241 | 0.232752439 | 0.508864854 | 0.598107374 | 0.54976829 | 0.728991 |
| ENSG00000115008 | IL1A | 1.245025507 | 0.232654139 | 0.769108551 | 0.759004981 | 0.44784958 | NA |
| ENSG00000170145 | SIK2 | 1511.437 | 0.2326477 | 0.1733395 | 1.369049 | 0.1709839 | 0.341646 |
| ENSG00000180448 | ARHGAP45 | 1637.432 | 0.2326157 | 0.1927185 | 1.220678 | 0.2222081 | 0.408939 |
| ENSG00000283480 | | 1.828214 | 0.2326084 | 0.6851349 | 0.6095029 | 0.5421911 | 0.72353 |
| ENSG00000187741 | FANCA | 3319.194 | 0.2325344 | 0.3118971 | 0.7963052 | 0.4258547 | 0.625711 |
| ENSG00000259542 | | 1.308135 | 0.2325053 | 0.7849035 | 0.9311848 | 0.351758 | NA |
| ENSG00000279434 | | 23.04605 | 0.2324509 | 0.5126785 | 0.5854631 | 0.5582364 | 0.735302 |
| ENSG00000163684 | RPP14 | 725.3188 | 0.2324468 | 0.121119 | 1.943167 | 0.05199601 | 0.142765 |
| ENSG00000087269 | NOP14 | 2732.450786 | 0.232403155 | 0.154736628 | 1.525187162 | 0.12721242 | 0.27643 |
| ENSG00000122299 | ZC3H7A | 4036.972428 | 0.232262545 | 0.098130312 | 2.367358139 | 0.01791559 | 0.062181 |
| ENSG00000131969 | ABHD12B | 5.027271317 | 0.232262532 | 0.75737558 | 0.764227603 | 0.44473162 | 0.642073 |
| ENSG00000272784 | | 5.21726 | 0.2322035 | 0.691057 | 0.6305852 | 0.5283118 | 0.713103 |
| ENSG00000196345 | ZKSCAN7 | 0.7887011 | 0.2321025 | 0.8344127 | 1.350928 | 0.1767185 | NA |
| ENSG00000013364 | MVP | 2079.050356 | 0.232101266 | 0.456789217 | 0.610956534 | 0.54122836 | 0.723008 |
| ENSG00000231064 | THBS3-AS1 | 141.4962 | 0.2320095 | 0.3376941 | 0.7634931 | 0.4451694 | 0.642402 |
| ENSG00000189308 | LIN54 | 1084.415 | 0.2319556 | 0.2694229 | 0.9150862 | 0.3601464 | 0.562139 |
| ENSG00000280067 | | 40.81356 | 0.2318703 | 0.405027 | 0.6646173 | 0.5062953 | 0.692918 |
| ENSG00000258904 | | 8.805985 | 0.2318582 | 0.5350899 | 0.5791954 | 0.5624574 | 0.738154 |
| ENSG00000168724 | DNAJC21 | 4491.073 | 0.2318432 | 0.1129393 | 2.072643 | 0.03820554 | 0.11285 |
| ENSG00000114023 | FAM162A | 2346.874362 | 0.231834176 | 0.258282804 | 0.956550179 | 0.33879435 | 0.54001 |
| ENSG00000269161 | | 3.131852 | 0.2318118 | 0.6649933 | 0.5947203 | 0.5520305 | 0.730751 |
| ENSG00000124374 | PAIP2B | 808.9507542 | 0.231774463 | 0.186861391 | 1.283597299 | 0.1992829 | 0.37947 |
| ENSG00000285907 | | 73.29786 | 0.2317485 | 0.4339831 | 0.630078 | 0.5286436 | 0.713442 |
| ENSG00000135447 | PPP1R1A | 8.046139553 | 0.231743724 | 0.66531475 | 0.602001671 | 0.54717303 | 0.726931 |

| | | | | | | | |
|---|---|---|---|---|---|---|---|
| ENSG00000282870 | FRG1DP | 131.0467 | 0.2317265 | 0.365883 | 0.7137342 | 0.4753916 | 0.66776 |
| ENSG00000174839 | DENND6A | 1335.736 | 0.2317253 | 0.2158772 | 1.121295 | 0.2621621 | 0.456956 |
| ENSG00000255277 | ABCC6P2 | 3.837923 | 0.2317061 | 0.6648515 | 0.5996589 | 0.5487336 | 0.728137 |
| ENSG00000154438 | ASZ1 | 0.9700357 | 0.2316711 | 0.8284222 | 1.140923 | 0.2539018 | NA |
| ENSG00000229384 | HMGB1P16 | 0.899803 | 0.2316604 | 0.7894902 | 0.845866 | 0.3976275 | NA |
| ENSG00000247400 | DNAJC3-DT | 244.7563 | 0.2315593 | 0.279621 | 0.8792297 | 0.3792767 | 0.581243 |
| ENSG00000213363 | RPS3P6 | 22.01129 | 0.2314823 | 0.4672726 | 0.6102337 | 0.541707 | 0.723356 |
| ENSG00000272645 | GTF2IP20 | 809.1496 | 0.2314144 | 0.2999681 | 0.8253004 | 0.409201 | 0.609831 |
| ENSG00000237632 | S100A11P1 | 9.872919 | 0.2313156 | 0.549272 | 0.5761034 | 0.5645453 | 0.739689 |
| ENSG00000171847 | FAM90A1 | 7.492842 | 0.2311888 | 0.5271317 | 0.5720872 | 0.5672629 | 0.741677 |
| ENSG00000229009 | TMPRSS11GP | 0.9324661 | 0.2311632 | 0.7930859 | 0.8306484 | 0.4061723 | NA |
| ENSG00000229089 | ANKRD20A8P | 2.710352 | 0.2311033 | 0.7215444 | 0.6600705 | 0.5092086 | 0.695358 |
| ENSG00000229661 | BUD31P2 | 2.027232 | 0.2310983 | 0.7098382 | 0.6226761 | 0.5334974 | 0.717307 |
| ENSG00000145332 | KLHL8 | 2058.649 | 0.2310624 | 0.207352 | 1.151008 | 0.249729 | 0.442307 |
| ENSG00000148341 | SH3GLB2 | 13025.71 | 0.2309632 | 0.2025044 | 1.151065 | 0.2497055 | 0.442307 |
| ENSG00000130382 | MLLT1 | 4570.132271 | 0.230837531 | 0.162981042 | 1.45199335 | 0.14650345 | 0.305986 |
| ENSG00000162441 | LZIC | 1588.815 | 0.2306485 | 0.1053322 | 2.220569 | 0.02638016 | 0.084598 |
| ENSG00000183090 | FREM3 | 0.8467631 | 0.2306183 | 0.8152212 | 0.8755323 | 0.3812843 | NA |
| ENSG00000116497 | S100PBP | 1740.106395 | 0.230461467 | 0.194082661 | 1.221583595 | 0.22186514 | 0.408619 |
| ENSG00000226648 | PLCG1-AS1 | 188.455 | 0.2304466 | 0.2112078 | 1.1359 | 0.2559984 | 0.449605 |
| ENSG00000224430 | MKRN5P | 2.743114 | 0.2304261 | 0.6899101 | 0.6130203 | 0.5398629 | 0.722091 |
| ENSG00000196498 | NCOR2 | 7952.795 | 0.2303873 | 0.1933079 | 1.220064 | 0.2224407 | 0.409155 |
| ENSG00000136999 | CCN3 | 1.659505081 | 0.23027763 | 0.732513366 | 0.670061283 | 0.50281873 | 0.690194 |
| ENSG00000112031 | MTRF1L | 532.9476608 | 0.23023815 | 0.116923239 | 2.013272508 | 0.04408597 | 0.126057 |
| ENSG00000119402 | FBXW2 | 5078.55213 | 0.230205376 | 0.112838828 | 2.033722572 | 0.04197958 | 0.121406 |
| ENSG00000242247 | ARFGAP3 | 1790.306 | 0.2301892 | 0.180755 | 1.297178 | 0.1945699 | 0.373737 |
| ENSG00000216031 | MIR298 | 4.479601 | 0.2301871 | 0.606751 | 0.5736058 | 0.5662346 | 0.740952 |
| ENSG00000276925 | | 2.077401 | 0.2301781 | 0.7562971 | 0.7568845 | 0.449119 | 0.645478 |
| ENSG00000249222 | ATP5MGL | 22.40598 | 0.2301641 | 0.3882956 | 0.6800505 | 0.4964725 | 0.685403 |
| ENSG00000152380 | FAM151B | 127.3179 | 0.2300677 | 0.2349957 | 1.028887 | 0.3035328 | 0.503157 |
| ENSG00000226476 | LINC01748 | 149.979 | 0.2299414 | 0.2648308 | 0.9200693 | 0.3575366 | 0.559578 |
| ENSG00000205581 | HMGN1 | 23470.19 | 0.2299187 | 0.1578904 | 1.484494 | 0.137678 | 0.29231 |
| ENSG00000278472 | | 5.631673 | 0.2298988 | 0.5885848 | 0.5659976 | 0.5713954 | 0.745129 |

| | | | | | | | |
|---|---|---|---|---|---|---|---|
| ENSG00000103024 | NME3 | 4496.422171 | 0.229815595 | 0.198087856 | 1.195829639 | 0.23176305 | 0.420162 |
| ENSG00000188185 | LINC00265 | 472.7261 | 0.2297988 | 0.2557377 | 0.966244 | 0.3339221 | 0.535093 |
| ENSG00000251775 | | 1.707145 | 0.2297856 | 0.7181271 | 0.6438362 | 0.5196817 | 0.704928 |
| ENSG00000169398 | PTK2 | 10236.21 | 0.2297383 | 0.1305016 | 1.783939 | 0.07443363 | 0.187117 |
| ENSG00000176244 | ACBD7 | 683.1325 | 0.229666 | 0.3908882 | 0.6676684 | 0.5043453 | 0.691254 |
| ENSG00000270479 | BNIP3P37 | 1.012634 | 0.2296164 | 0.7966023 | 0.881989 | 0.3777827 | NA |
| ENSG00000253431 | SRPK2P | 3.416766 | 0.2296069 | 0.6990348 | 0.6254824 | 0.5316545 | 0.71576 |
| ENSG00000265195 | MIR4312 | 2.059458 | 0.2295625 | 0.7196374 | 0.6502183 | 0.5155512 | 0.7009 |
| ENSG00000136279 | DBNL | 8152.760173 | 0.229486512 | 0.142524408 | 1.639705571 | 0.1010664 | 0.234573 |
| ENSG00000072958 | AP1M1 | 1536.669859 | 0.229398473 | 0.231050522 | 1.032474521 | 0.30184988 | 0.501241 |
| ENSG00000250896 | RNPS1P1 | 1282.144 | 0.2293257 | 0.2148819 | 1.080665 | 0.2798461 | 0.477137 |
| ENSG00000123843 | C4BPB | 1.374638778 | 0.229271357 | 0.807214214 | 0.723473205 | 0.46938921 | NA |
| ENSG00000174502 | SLC26A9 | 1.975116 | 0.2291531 | 0.7625164 | 0.7332842 | 0.4633851 | 0.65764 |
| ENSG00000107872 | FBXL15 | 1173.738047 | 0.229053399 | 0.229577506 | 1.043052899 | 0.29692379 | 0.495923 |
| ENSG00000089505 | CMTM1 | 44.19557552 | 0.229042575 | 0.322296415 | 0.776666546 | 0.43735553 | 0.635812 |
| ENSG00000230371 | KARS1P2 | 0.7760316 | 0.2290413 | 0.8072264 | 0.7164839 | 0.4736926 | NA |
| ENSG00000115661 | STK16 | 2021.372946 | 0.228957096 | 0.161050104 | 1.442571275 | 0.14914128 | 0.309839 |
| ENSG00000155729 | KCTD18 | 1410.575 | 0.2289348 | 0.1388734 | 1.671977 | 0.09452893 | 0.223445 |
| ENSG00000253882 | | 4.42483 | 0.2289197 | 0.6544902 | 0.5810287 | 0.5612211 | 0.737311 |
| ENSG00000174106 | LEMD3 | 2809.194 | 0.2288773 | 0.1756251 | 1.341069 | 0.1798981 | 0.354053 |
| ENSG00000147601 | TERF1 | 1196.064 | 0.2288614 | 0.1211888 | 1.925098 | 0.05421712 | 0.147633 |
| ENSG00000261068 | | 77.47494 | 0.2288291 | 0.4801559 | 0.5931821 | 0.5530593 | 0.731456 |
| ENSG00000085185 | BCORL1 | 1003.799594 | 0.228779521 | 0.276851583 | 0.883515284 | 0.37695793 | 0.578992 |
| ENSG00000280333 | | 0.8846553 | 0.2287673 | 0.7762513 | 0.8291069 | 0.4070439 | NA |
| ENSG00000205143 | ARID3C | 7.334717 | 0.228575 | 0.5906611 | 0.5698578 | 0.5687741 | 0.742784 |
| ENSG00000062598 | ELMO2 | 2566.931295 | 0.228526262 | 0.119727008 | 1.927914813 | 0.05386572 | 0.146981 |
| ENSG00000272721 | EIF2B5-DT | 15.72794 | 0.2285119 | 0.4636341 | 0.602365 | 0.5469312 | 0.726845 |
| ENSG00000164190 | NIPBL | 7637.658 | 0.2284613 | 0.2030197 | 1.161591 | 0.2454018 | 0.437085 |
| ENSG00000141985 | SH3GL1 | 4139.265 | 0.2284591 | 0.1935556 | 1.217715 | 0.2233325 | 0.410033 |
| ENSG00000174428 | GTF2IRD2B | 468.3654 | 0.2284437 | 0.3644699 | 0.6985896 | 0.4848086 | 0.675745 |
| ENSG00000248115 | | 2.163259 | 0.2283986 | 0.6998415 | 0.6249964 | 0.5319734 | 0.716045 |
| ENSG00000166828 | SCNN1G | 51.87639 | 0.2283774 | 0.4811875 | 0.5908609 | 0.5546136 | 0.732564 |
| ENSG00000127334 | DYRK2 | 2826.486082 | 0.228300528 | 0.231869375 | 1.007020124 | 0.3139251 | 0.515047 |

| | | | | | | | |
|---|---|---|---|---|---|---|---|
| ENSG00000214717 | ZBED1 | 1260.239 | 0.2281585 | 0.2158166 | 1.097176 | 0.2725645 | 0.469057 |
| ENSG00000275342 | PRAG1 | 1473.679 | 0.2281399 | 0.2411767 | 0.9995676 | 0.3175198 | 0.518575 |
| ENSG00000186416 | NKRF | 1536.62 | 0.2280852 | 0.1762107 | 1.331594 | 0.1829936 | 0.358032 |
| ENSG00000117682 | DHDDS | 1423.06415 | 0.228081648 | 0.13373306 | 1.719455313 | 0.0855315 | 0.207552 |
| ENSG00000214185 | XPOTP1 | 600.9297 | 0.2280401 | 0.3316218 | 0.7570255 | 0.4490346 | 0.645416 |
| ENSG00000271384 | | 5.693172 | 0.2280372 | 0.5831464 | 0.5610622 | 0.5747552 | 0.747097 |
| ENSG00000159658 | EFCAB14 | 6762.718 | 0.228014 | 0.1825294 | 1.264648 | 0.2059977 | 0.388108 |
| ENSG00000156983 | BRPF1 | 3335.68 | 0.2278797 | 0.1543534 | 1.455421 | 0.1455528 | 0.304603 |
| ENSG00000184743 | ATL3 | 2990.656 | 0.2277412 | 0.1403425 | 1.640074 | 0.1009898 | 0.234481 |
| ENSG00000219186 | FTH1P19 | 1.439881 | 0.2277024 | 0.7906336 | 0.8830819 | 0.377192 | NA |
| ENSG00000267416 | | 30.27254 | 0.227639 | 0.3507459 | 0.7234802 | 0.4693849 | 0.662897 |
| ENSG00000160323 | ADAMTS13 | 811.1248 | 0.2275403 | 0.2370547 | 1.0044 | 0.315186 | 0.516196 |
| ENSG00000255529 | POLR2M | 878.6782 | 0.2274472 | 0.1486903 | 1.556796 | 0.1195189 | 0.264493 |
| ENSG00000100280 | AP1B1 | 8446.545595 | 0.227412728 | 0.169129967 | 1.361461928 | 0.17336776 | 0.344897 |
| ENSG00000214784 | RPS3AP21 | 706.5542 | 0.2273892 | 0.1973973 | 1.201727 | 0.2294692 | 0.417644 |
| ENSG00000205609 | EIF3CL | 46.12288 | 0.2273846 | 0.2869271 | 0.8501367 | 0.3952491 | 0.596906 |
| ENSG00000145949 | MYLK4 | 83.57183 | 0.2273547 | 0.4238619 | 0.6309455 | 0.5280761 | 0.712893 |
| ENSG00000173451 | THAP2 | 296.4263 | 0.2271983 | 0.3557702 | 0.714096 | 0.4751678 | 0.667714 |
| ENSG00000274561 | | 215.969 | 0.2270801 | 0.2828326 | 0.8579851 | 0.3909007 | 0.592799 |
| ENSG00000279491 | | 40.06715 | 0.2270108 | 0.4187097 | 0.6360592 | 0.5247378 | 0.709757 |
| ENSG00000100442 | FKBP3 | 4511.72205 | 0.226984898 | 0.17633273 | 1.332715841 | 0.18262507 | 0.357592 |
| ENSG00000204832 | ST8SIA6-AS1 | 55.94231 | 0.2269774 | 0.4212785 | 0.6300932 | 0.5286336 | 0.713442 |
| ENSG00000275389 | | 3.952497 | 0.2267559 | 0.6736468 | 0.5926463 | 0.5534179 | 0.731603 |
| ENSG00000257167 | TMPO-AS1 | 260.7392 | 0.2266949 | 0.3983653 | 0.6549484 | 0.512501 | 0.698397 |
| ENSG00000189136 | UBE2Q2P1 | 266.0758 | 0.2265838 | 0.361125 | 0.70745 | 0.4792869 | 0.671315 |
| ENSG00000238181 | AHCYP2 | 0.7859049 | 0.2263074 | 0.7911058 | 0.8605204 | 0.3895023 | NA |
| ENSG00000112576 | CCND3 | 3819.86296 | 0.226299331 | 0.259560156 | 0.932240209 | 0.3512124 | 0.552871 |
| ENSG00000197841 | ZNF181 | 859.9954 | 0.226274 | 0.1893915 | 1.242768 | 0.2139533 | 0.398746 |
| ENSG00000277720 | H3C5P | 1.099112 | 0.2262486 | 0.7675195 | 0.7732241 | 0.4393898 | NA |
| ENSG00000201302 | SNORA65 | 35.26486 | 0.2262302 | 0.3261536 | 0.7586632 | 0.448054 | 0.644733 |
| ENSG00000232300 | FAM215B | 1.159296 | 0.2262216 | 0.7945682 | 0.865955 | 0.3865148 | NA |
| ENSG00000224121 | ATG12P2 | 0.9096577 | 0.2261294 | 0.8093002 | 0.6844686 | 0.4936793 | NA |
| ENSG00000223559 | | 1124.236 | 0.226106 | 0.286843 | 0.8471034 | 0.3969375 | 0.598029 |

| | | | | | | | |
|---|---|---|---|---|---|---|---|
| ENSG00000257252 | | 5.957691 | 0.226064 | 0.6804421 | 0.5943274 | 0.5522932 | 0.730854 |
| ENSG00000146414 | SHPRH | 884.3882 | 0.2259911 | 0.3621763 | 0.7008019 | 0.4834267 | 0.67476 |
| ENSG00000130311 | DDA1 | 1807.150858 | 0.225945551 | 0.149242766 | 1.54411404 | 0.12256071 | 0.269362 |
| ENSG00000250067 | YJEFN3 | 193.1609 | 0.2258609 | 0.2854712 | 0.8466429 | 0.3971942 | 0.59828 |
| ENSG00000271064 | | 3.82345 | 0.2257406 | 0.6775485 | 0.5842427 | 0.559057 | 0.735831 |
| ENSG00000198198 | SZT2 | 3162.173 | 0.2257383 | 0.2952995 | 0.8188502 | 0.4128719 | 0.612998 |
| ENSG00000179387 | ELMOD2 | 1695.04 | 0.2257033 | 0.1850039 | 1.263286 | 0.2064865 | 0.388702 |
| ENSG00000135473 | PAN2 | 3763.115862 | 0.225681793 | 0.264080172 | 0.904055365 | 0.36596605 | 0.567853 |
| ENSG00000116771 | AGMAT | 284.4279546 | 0.225681174 | 0.322679104 | 0.761286018 | 0.44648625 | 0.643407 |
| ENSG00000162971 | TYW5 | 1149.522 | 0.2256779 | 0.1747808 | 1.328284 | 0.1840842 | 0.359356 |
| ENSG00000181019 | NQO1 | 22264.99 | 0.2256111 | 0.1996224 | 1.166839 | 0.2432756 | 0.434381 |
| ENSG00000239519 | CADM2-AS1 | 1.14462 | 0.2255919 | 0.7539243 | 0.7236115 | 0.4693043 | NA |
| ENSG00000078081 | LAMP3 | 158.174057 | 0.22555872 | 0.476385447 | 0.587169032 | 0.55709019 | 0.734513 |
| ENSG00000261474 | | 85.53955 | 0.2255463 | 0.3016732 | 0.80494 | 0.4208543 | 0.620756 |
| ENSG00000240254 | B4GALT4-AS1 | 2.06998 | 0.2254558 | 0.7557582 | 0.7492135 | 0.4537285 | 0.649906 |
| ENSG00000237709 | EEF1A1P28 | 1.185878 | 0.2254387 | 0.75358 | 0.7250833 | 0.4684009 | NA |
| ENSG00000135924 | DNAJB2 | 3241.18393 | 0.225435923 | 0.23402721 | 1.073321234 | 0.28312701 | 0.480947 |
| ENSG00000223820 | CFL1P1 | 26.76566 | 0.2252545 | 0.4442839 | 0.6088737 | 0.5426082 | 0.723795 |
| ENSG00000129038 | LOXL1 | 80.9716834 | 0.225248868 | 0.38313612 | 0.670537758 | 0.50251505 | 0.690065 |
| ENSG00000213753 | CENPBD1P1 | 1223.263 | 0.225248 | 0.1360936 | 1.667727 | 0.09536998 | 0.224775 |
| ENSG00000213774 | | 1.05706 | 0.2252067 | 0.8156226 | 0.8668798 | 0.3860079 | NA |
| ENSG00000114933 | INO80D | 1018.722489 | 0.225126768 | 0.220439752 | 1.058343598 | 0.28989882 | 0.488403 |
| ENSG00000140986 | RPL3L | 19.98689 | 0.2250347 | 0.4989501 | 0.5766449 | 0.5641794 | 0.739552 |
| ENSG00000240370 | RPL13P5 | 40.45313 | 0.225018 | 0.3344254 | 0.7448857 | 0.4563408 | 0.652171 |
| ENSG00000239402 | CYP4F62P | 1.101191 | 0.2249517 | 0.8065763 | 0.9352989 | 0.3496343 | NA |
| ENSG00000162039 | MEIOB | 23.88371 | 0.2248461 | 0.3648657 | 0.6896622 | 0.4904066 | 0.680949 |
| ENSG00000286161 | | 0.8803952 | 0.2247398 | 0.7569217 | 0.6978363 | 0.4852796 | NA |
| ENSG00000256603 | | 1.378067 | 0.2247344 | 0.7616002 | 0.7142216 | 0.4750902 | NA |
| ENSG00000243886 | | 3.965306 | 0.2247152 | 0.6812704 | 0.6011563 | 0.5477359 | 0.727458 |
| ENSG00000273247 | RAB33B-AS1 | 286.248 | 0.2246605 | 0.1742292 | 1.322721 | 0.1859281 | 0.361899 |
| ENSG00000144747 | TMF1 | 2622.118 | 0.224645 | 0.1661889 | 1.37555 | 0.1689611 | 0.339032 |
| ENSG00000149499 | EML3 | 5055.966 | 0.2245861 | 0.1991003 | 1.182749 | 0.2369085 | 0.426538 |
| ENSG00000245498 | | 58.21088 | 0.2244981 | 0.4292295 | 0.6195265 | 0.5355695 | 0.71889 |

| | | | | | | | |
|---|---|---|---|---|---|---|---|
| ENSG00000231409 | | 27.099 | 0.2244768 | 0.4261054 | 0.62271 | 0.5334751 | 0.717307 |
| ENSG00000152443 | ZNF776 | 1101.274 | 0.2244019 | 0.1886701 | 1.230552 | 0.2184905 | 0.404253 |
| ENSG00000102606 | ARHGEF7 | 2784.180712 | 0.224399818 | 0.139151674 | 1.636570665 | 0.10172021 | 0.235624 |
| ENSG00000141570 | CBX8 | 1898.567 | 0.2243802 | 0.1601164 | 1.42148 | 0.1551773 | 0.318987 |
| ENSG00000198026 | ZNF335 | 2787.807 | 0.2243705 | 0.1442104 | 1.580268 | 0.1140455 | 0.255746 |
| ENSG00000268543 | | 51.32399 | 0.224302 | 0.3399652 | 0.7289248 | 0.4660477 | 0.659868 |
| ENSG00000005022 | SLC25A5 | 24089.74424 | 0.224270227 | 0.242273997 | 0.969854391 | 0.33211908 | 0.533209 |
| ENSG00000163348 | PYGO2 | 5125.421 | 0.2241895 | 0.1577387 | 1.447962 | 0.1476277 | 0.307633 |
| ENSG00000273443 | | 33.26503 | 0.2241884 | 0.3925793 | 0.6582587 | 0.5103719 | 0.696174 |
| ENSG00000255875 | PABPN1P2 | 2.265768 | 0.2241879 | 0.7156537 | 0.6371558 | 0.5240233 | 0.709188 |
| ENSG00000260643 | | 39.59115 | 0.2241318 | 0.324833 | 0.7583217 | 0.4482584 | 0.644887 |
| ENSG00000048707 | VPS13D | 3970.827925 | 0.224129552 | 0.245145826 | 0.959115032 | 0.3375008 | 0.538789 |
| ENSG00000197168 | NEK5 | 95.93578 | 0.2240873 | 0.3340655 | 0.7370769 | 0.4610756 | 0.655995 |
| ENSG00000285601 | | 10.92183 | 0.2239002 | 0.6396758 | 0.5785576 | 0.5628877 | 0.738569 |
| ENSG00000270074 | | 5.623992 | 0.2238458 | 0.6249194 | 0.558489 | 0.5765105 | 0.748537 |
| ENSG00000273080 | | 57.01033 | 0.2237647 | 0.3392509 | 0.7327237 | 0.463727 | 0.657836 |
| ENSG00000260183 | | 2.549747 | 0.2236617 | 0.6596622 | 0.5641567 | 0.5726475 | 0.745897 |
| ENSG00000220412 | | 54.52779 | 0.2236358 | 0.3479985 | 0.7154123 | 0.4743543 | 0.667114 |
| ENSG00000278396 | | 16.81284 | 0.2235921 | 0.4584911 | 0.592877 | 0.5532635 | 0.731539 |
| ENSG00000070770 | CSNK2A2 | 1533.859362 | 0.223579977 | 0.172531945 | 1.326079527 | 0.18481336 | 0.360257 |
| ENSG00000177200 | CHD9 | 3266.6 | 0.2235129 | 0.2845181 | 0.8367983 | 0.4027059 | 0.603678 |
| ENSG00000183281 | PLGLB1 | 49.69773 | 0.2234875 | 0.3822864 | 0.6634391 | 0.5070494 | 0.693529 |
| ENSG00000198711 | SSBP3-AS1 | 35.49224 | 0.2233359 | 0.4692637 | 0.5826938 | 0.5600994 | 0.736369 |
| ENSG00000188786 | MTF1 | 1698.846 | 0.2232528 | 0.2083898 | 1.110432 | 0.2668129 | 0.462635 |
| ENSG00000104164 | BLOC1S6 | 2192.640581 | 0.223248646 | 0.134481862 | 1.681057374 | 0.09275177 | 0.220255 |
| ENSG00000237243 | CREB3L2-AS1 | 0.7865528 | 0.223209 | 0.8038056 | 0.915395 | 0.3599843 | NA |
| ENSG00000182583 | VCX | 45.76983 | 0.2231945 | 0.4323718 | 0.6141137 | 0.5391402 | 0.721597 |
| ENSG00000166676 | TVP23A | 23.65147 | 0.223147 | 0.4374285 | 0.6085096 | 0.5428496 | 0.723863 |
| ENSG00000277895 | | 4.269781 | 0.2231144 | 0.6929267 | 0.6151593 | 0.5384495 | 0.721168 |
| ENSG00000269815 | | 16.68967 | 0.2230774 | 0.4565164 | 0.5921027 | 0.5537819 | 0.731902 |
| ENSG00000255302 | EID1 | 6001.956 | 0.2229556 | 0.1781006 | 1.279577 | 0.2006938 | 0.381303 |
| ENSG00000272172 | | 45.0676 | 0.222765 | 0.3679874 | 0.6857245 | 0.4928868 | 0.682805 |
| ENSG00000144746 | ARL6IP5 | 3970.736 | 0.2227259 | 0.1661746 | 1.393668 | 0.1634179 | 0.331186 |

| | | | | | | | |
|---|---|---|---|---|---|---|---|
| ENSG00000287591 | | 0.8806682 | 0.2227093 | 0.7573164 | 0.7276415 | 0.4668331 | NA |
| ENSG00000235996 | | 0.8191984 | 0.2226495 | 0.8040643 | 1.048768 | 0.2942849 | NA |
| ENSG00000163605 | PPP4R2 | 4127.009 | 0.2226266 | 0.1679873 | 1.353394 | 0.1759297 | 0.348541 |
| ENSG00000116251 | RPL22 | 7391.601725 | 0.222598451 | 0.172762195 | 1.32719514 | 0.18444414 | 0.359749 |
| ENSG00000184612 | RPL7P26 | 12.36104 | 0.2225809 | 0.5216768 | 0.5571771 | 0.5774064 | 0.749261 |
| ENSG00000100941 | PNN | 17574.45323 | 0.222372394 | 0.163897408 | 1.367211097 | 0.17155915 | 0.342255 |
| ENSG00000287698 | | 2.293984 | 0.222317 | 0.6859664 | 0.5905232 | 0.5548399 | 0.732681 |
| ENSG00000186310 | NAP1L3 | 47.12527 | 0.2223101 | 0.6873308 | 0.5855243 | 0.5581952 | 0.735302 |
| ENSG00000166348 | USP54 | 1492.406 | 0.22228 | 0.2444115 | 0.9541306 | 0.3400176 | 0.541405 |
| ENSG00000230321 | MTCO2P27 | 2.115426 | 0.2222798 | 0.7615926 | 0.713906 | 0.4752853 | 0.667717 |
| ENSG00000222043 | | 24.75697 | 0.2222712 | 0.4355035 | 0.6092713 | 0.5423446 | 0.723674 |
| ENSG00000165115 | KIF27 | 536.3975 | 0.2221987 | 0.3140808 | 0.7680449 | 0.4424605 | 0.640164 |
| ENSG00000118855 | MFSD1 | 4780.992547 | 0.222181516 | 0.152200794 | 1.500301186 | 0.1335364 | 0.286239 |
| ENSG00000288547 | C3orf36 | 0.9376676 | 0.2221616 | 0.8150407 | 0.8576244 | 0.3910999 | NA |
| ENSG00000198961 | PJA2 | 5089.014 | 0.2220143 | 0.1973526 | 1.159775 | 0.2461405 | 0.437907 |
| ENSG00000254153 | | 39.35871 | 0.2219606 | 0.4643204 | 0.5865338 | 0.5575168 | 0.734828 |
| ENSG00000165417 | GTF2A1 | 2395.315 | 0.2218846 | 0.2393219 | 0.9758627 | 0.3291325 | 0.530275 |
| ENSG00000171469 | ZNF561 | 2358.44 | 0.221848 | 0.2074133 | 1.116261 | 0.2643102 | 0.459826 |
| ENSG00000114124 | GRK7 | 1.635236325 | 0.221706063 | 0.743564552 | 0.687384517 | 0.49184045 | 0.682066 |
| ENSG00000197882 | OR7E13P | 1.008808 | 0.2216738 | 0.8064827 | 0.7242549 | 0.4689092 | NA |
| ENSG00000230362 | ACTG1P23 | 0.898835 | 0.221645 | 0.7976377 | 0.9442703 | 0.3450315 | NA |
| ENSG00000119723 | COQ6 | 705.1806709 | 0.221614974 | 0.14358799 | 1.570976191 | 0.11618818 | 0.259194 |
| ENSG00000139318 | DUSP6 | 231.0094 | 0.2214113 | 0.487821 | 0.5682283 | 0.5698799 | 0.743744 |
| ENSG00000262434 | | 2.677415 | 0.2213528 | 0.6795524 | 0.5849007 | 0.5586145 | 0.735582 |
| ENSG00000135392 | DNAJC14 | 1262.687948 | 0.22134908 | 0.177275276 | 1.265097958 | 0.20583622 | 0.38791 |
| ENSG00000274020 | LINC01138 | 164.4644 | 0.2213381 | 0.4429086 | 0.6014725 | 0.5475253 | 0.727321 |
| ENSG00000273870 | | 5.123687 | 0.2213125 | 0.6591022 | 0.5672153 | 0.5705679 | 0.744349 |
| ENSG00000275155 | | 5.25963 | 0.2212993 | 0.7599046 | 0.7199331 | 0.4715662 | 0.664789 |
| ENSG00000171291 | ZNF439 | 3.106178 | 0.2212274 | 0.6349469 | 0.5580999 | 0.5767762 | 0.748717 |
| ENSG00000221836 | OR2A5 | 0.8002944 | 0.221148 | 0.8071431 | 0.6909871 | 0.4895737 | NA |
| ENSG00000182512 | GLRX5 | 4528.12 | 0.2211293 | 0.2140235 | 1.072509 | 0.2834916 | 0.481284 |
| ENSG00000072736 | NFATC3 | 1195.891044 | 0.221082577 | 0.230597154 | 0.957698832 | 0.33821465 | 0.539474 |
| ENSG00000226544 | RPL7P22 | 149.2141 | 0.2210823 | 0.2899732 | 0.8192682 | 0.4126334 | 0.612746 |

| | | | | | | | |
|---|---|---|---|---|---|---|---|
| ENSG00000237283 | | 1.111586 | 0.2210765 | 0.810802 | 0.9981408 | 0.3182111 | NA |
| ENSG00000172955 | ADH6 | 8.469374 | 0.221059 | 0.5477747 | 0.549735 | 0.5825011 | 0.753076 |
| ENSG00000258289 | CHURC1 | 4963.159 | 0.2209329 | 0.1716882 | 1.293422 | 0.195865 | 0.375567 |
| ENSG00000134352 | IL6ST | 3040.490829 | 0.220885086 | 0.208211277 | 1.092034755 | 0.27481783 | 0.471557 |
| ENSG00000236611 | LINC02556 | 0.8404545 | 0.2208653 | 0.7964859 | 0.8723761 | 0.3830032 | NA |
| ENSG00000272844 | | 10.51894 | 0.2207629 | 0.5163354 | 0.5567642 | 0.5776886 | 0.749523 |
| ENSG00000271752 | | 1.253112 | 0.2207523 | 0.762473 | 0.7399782 | 0.4593133 | NA |
| ENSG00000147050 | KDM6A | 1987.913 | 0.2206303 | 0.156848 | 1.424331 | 0.1543507 | 0.317829 |
| ENSG00000006534 | ALDH3B1 | 1162.943181 | 0.220535719 | 0.356194924 | 0.700495105 | 0.48361816 | 0.674856 |
| ENSG00000227741 | | 20.51467 | 0.2204024 | 0.4234602 | 0.6079991 | 0.5431881 | 0.724178 |
| ENSG00000255427 | LINC02707 | 1.880701 | 0.2203978 | 0.7099257 | 0.6151326 | 0.5384672 | 0.721168 |
| ENSG00000272347 | | 3.599339 | 0.2202277 | 0.6645082 | 0.5595775 | 0.5757677 | 0.747901 |
| ENSG00000225137 | DYNC1I2P1 | 1187.386 | 0.2202204 | 0.1190187 | 1.872191 | 0.06118024 | 0.161404 |
| ENSG00000267533 | | 465.2636 | 0.2201623 | 0.2398486 | 0.9583754 | 0.3378735 | 0.539222 |
| ENSG00000099904 | ZDHHC8 | 3785.41689 | 0.220134727 | 0.160559372 | 1.389627274 | 0.16464209 | 0.332844 |
| ENSG00000249604 | | 2.595656 | 0.220089 | 0.7061051 | 0.6186603 | 0.5361402 | 0.719256 |
| ENSG00000156831 | NSMCE2 | 2835.881 | 0.2200494 | 0.1949406 | 1.156127 | 0.2476292 | 0.43982 |
| ENSG00000143107 | FNDC7 | 1.006009 | 0.2200448 | 0.76287 | 0.7155918 | 0.4742434 | NA |
| ENSG00000119673 | ACOT2 | 829.0134965 | 0.219891551 | 0.166462544 | 1.382420097 | 0.16684275 | 0.336076 |
| ENSG00000140299 | BNIP2 | 3113.709 | 0.2198829 | 0.1371439 | 1.565895 | 0.1173731 | 0.261027 |
| ENSG00000275207 | MIR6740 | 2.425533 | 0.219858 | 0.7125335 | 0.6142381 | 0.539058 | 0.721572 |
| ENSG00000235028 | HMGN1P30 | 235.6308 | 0.2198304 | 0.2074771 | 1.089565 | 0.2759049 | 0.472815 |
| ENSG00000230590 | FTX | 2383.195 | 0.2198205 | 0.3072967 | 0.7741486 | 0.4388429 | 0.637347 |
| ENSG00000183397 | C19orf71 | 148.7847 | 0.2197656 | 0.2466828 | 0.9359277 | 0.3493104 | 0.55102 |
| ENSG00000164548 | TRA2A | 4353.844 | 0.2196808 | 0.1182099 | 1.914532 | 0.05555224 | 0.15021 |
| ENSG00000023909 | GCLM | 974.1795832 | 0.219650116 | 0.20819092 | 1.095977743 | 0.27308852 | 0.469715 |
| ENSG00000249855 | EEF1A1P19 | 193.5816 | 0.2196499 | 0.1935257 | 1.17126 | 0.2414942 | 0.43206 |
| ENSG00000121022 | COPS5 | 3306.733259 | 0.219634017 | 0.153742337 | 1.434170132 | 0.15152372 | 0.313416 |
| ENSG00000182447 | OTOL1 | 0.9550587 | 0.2195205 | 0.8062024 | 0.7060362 | 0.4801656 | NA |
| ENSG00000131401 | NAPSB | 20.17493357 | 0.21950257 | 0.4905995 | 0.564761425 | 0.57223605 | 0.74572 |
| ENSG00000124587 | PEX6 | 971.9403657 | 0.219480612 | 0.284863042 | 0.836115885 | 0.40308971 | 0.604009 |
| ENSG00000105261 | OVOL3 | 6.463849582 | 0.219405112 | 0.624213773 | 0.545973204 | 0.58508436 | 0.75503 |
| ENSG00000224927 | NDUFA5P10 | 9.34101 | 0.2193685 | 0.5896326 | 0.5375681 | 0.5908753 | 0.759562 |

| | | | | | | | |
|---|---|---|---|---|---|---|---|
| ENSG00000169359 | SLC33A1 | 1046.648 | 0.2193067 | 0.2799391 | 0.8346992 | 0.4038871 | 0.604659 |
| ENSG00000160360 | GPSM1 | 1541.175 | 0.219259 | 0.1490081 | 1.497433 | 0.1342807 | 0.287301 |
| ENSG00000237161 | | 117.4615 | 0.2192377 | 0.3552791 | 0.6913374 | 0.4893535 | 0.680109 |
| ENSG00000166704 | ZNF606 | 954.8891 | 0.2192283 | 0.2010023 | 1.123877 | 0.2610651 | 0.455702 |
| ENSG00000236540 | | 44.22242 | 0.2191676 | 0.3582009 | 0.6823559 | 0.4950139 | 0.684289 |
| ENSG00000286595 | | 3.687437 | 0.2191478 | 0.6419717 | 0.5504608 | 0.5820034 | 0.752579 |
| ENSG00000279103 | | 3.664708 | 0.2191448 | 0.6533707 | 0.5543547 | 0.5793361 | 0.750533 |
| ENSG00000145107 | TM4SF19 | 1.737672 | 0.21914 | 0.7770013 | 0.7939917 | 0.4272003 | 0.626776 |
| ENSG00000188060 | RAB42 | 4.930438 | 0.2191373 | 0.6581133 | 0.5599579 | 0.5755082 | 0.747674 |
| ENSG00000275944 | | 0.8160038 | 0.2191038 | 0.8269491 | 1.171492 | 0.2414012 | NA |
| ENSG00000229178 | | 47.14064 | 0.219056 | 0.3471204 | 0.6994059 | 0.4842984 | 0.67545 |
| ENSG00000271057 | | 0.8153231 | 0.2190026 | 0.7759731 | 0.7398071 | 0.4594171 | NA |
| ENSG00000242262 | | 30.41362 | 0.2189349 | 0.3864909 | 0.6471042 | 0.5175645 | 0.70287 |
| ENSG00000196950 | SLC39A10 | 2574.564 | 0.218931 | 0.2958039 | 0.7983818 | 0.424649 | 0.624343 |
| ENSG00000103197 | TSC2 | 7722.546405 | 0.218924567 | 0.143836387 | 1.535490889 | 0.12466329 | 0.272643 |
| ENSG00000154153 | RETREG1 | 853.6733 | 0.2189213 | 0.2967446 | 0.7962928 | 0.4258618 | 0.625711 |
| ENSG00000090905 | TNRC6A | 4707.832314 | 0.218901212 | 0.199617646 | 1.126557163 | 0.25992976 | 0.454377 |
| ENSG00000244491 | | 1.393636 | 0.2188258 | 0.7145459 | 0.6176557 | 0.5368023 | NA |
| ENSG00000286067 | | 286.6869 | 0.2188223 | 0.2632607 | 0.8818748 | 0.3778445 | 0.57975 |
| ENSG00000142765 | SYTL1 | 2903.814 | 0.2187842 | 0.2845053 | 0.8173369 | 0.4137359 | 0.6138 |
| ENSG00000280248 | | 4.622769 | 0.2187294 | 0.6174305 | 0.5406036 | 0.5887808 | 0.757924 |
| ENSG00000204519 | ZNF551 | 591.6949 | 0.2186432 | 0.2482253 | 0.9346353 | 0.3499763 | 0.551643 |
| ENSG00000156299 | TIAM1 | 2368.076 | 0.2185373 | 0.2663526 | 0.862913 | 0.3881853 | 0.5904 |
| ENSG00000104723 | TUSC3 | 4499.072511 | 0.218501122 | 0.130049352 | 1.695479019 | 0.08998459 | 0.21564 |
| ENSG00000159917 | ZNF235 | 339.2455 | 0.2184679 | 0.2010625 | 1.122654 | 0.2615845 | 0.456249 |
| ENSG00000230595 | RSL24D1P2 | 26.8183 | 0.2184463 | 0.4043078 | 0.6269496 | 0.5306923 | 0.715004 |
| ENSG00000157916 | RER1 | 4452.308 | 0.2183907 | 0.09741886 | 2.22506 | 0.02607719 | 0.083788 |
| ENSG00000103653 | CSK | 5146.096844 | 0.2183813 | 0.170208836 | 1.303936566 | 0.19225521 | 0.370572 |
| ENSG00000270265 | | 9.132684 | 0.2183065 | 0.507394 | 0.5510907 | 0.5815715 | 0.752204 |
| ENSG00000213362 | FTH1P12 | 160.9405 | 0.2182709 | 0.2546959 | 0.9129957 | 0.3612448 | 0.563225 |
| ENSG00000197779 | ZNF81 | 440.3154 | 0.2182245 | 0.3166989 | 0.7645228 | 0.4445558 | 0.641854 |
| ENSG00000161328 | LRRC56 | 577.4116 | 0.2182056 | 0.2325514 | 0.9737073 | 0.3302019 | 0.53129 |
| ENSG00000160691 | SHC1 | 4238.437 | 0.2181643 | 0.1556988 | 1.428019 | 0.1532864 | 0.316103 |

| | | | | | | | |
|---|---|---|---|---|---|---|---|
| ENSG00000174197 | MGA | 4105.606 | 0.2181539 | 0.2978917 | 0.7881935 | 0.4305835 | 0.629807 |
| ENSG00000120937 | NPPB | 1.543970641 | 0.218099034 | 0.807143379 | 0.65344615 | 0.5134687 | NA |
| ENSG00000287284 | | 0.8157332 | 0.2180802 | 0.805848 | 0.7098986 | 0.477767 | NA |
| ENSG00000162438 | CTRC | 2.719401 | 0.2180363 | 0.6755133 | 0.5622943 | 0.5739155 | 0.746641 |
| ENSG00000145687 | SSBP2 | 1795.944 | 0.2179248 | 0.2191323 | 1.037749 | 0.299387 | 0.498616 |
| ENSG00000233560 | KRT8P39 | 25.59218 | 0.2179025 | 0.4287394 | 0.6025597 | 0.5468016 | 0.726798 |
| ENSG00000214121 | PRDX1P1 | 30.20252 | 0.2178184 | 0.3945275 | 0.6400711 | 0.5221263 | 0.707558 |
| ENSG00000249335 | | 0.8585426 | 0.2176489 | 0.7980272 | 0.897958 | 0.369208 | NA |
| ENSG00000237836 | PHKA2-AS1 | 10.62608 | 0.2175975 | 0.5775742 | 0.5337465 | 0.593517 | 0.761353 |
| ENSG00000011105 | TSPAN9 | 1491.992949 | 0.21756388 | 0.189494464 | 1.181883438 | 0.23725195 | 0.426938 |
| ENSG00000255867 | DENND5B-AS1 | 3.858037 | 0.2174853 | 0.6016894 | 0.5392245 | 0.5897319 | 0.758512 |
| ENSG00000230105 | | 3.401899 | 0.2173162 | 0.6611766 | 0.5645857 | 0.5723556 | 0.745771 |
| ENSG00000236723 | | 4.031428 | 0.2172824 | 0.6367119 | 0.5509786 | 0.5816483 | 0.752234 |
| ENSG00000082701 | GSK3B | 7302.610957 | 0.217253689 | 0.179822073 | 1.236525297 | 0.21626337 | 0.401587 |
| ENSG00000214954 | LRRC69 | 55.89846 | 0.2172401 | 0.447359 | 0.5811319 | 0.5611516 | 0.737289 |
| ENSG00000103034 | NDRG4 | 1019.680011 | 0.217185191 | 0.267800837 | 0.857726177 | 0.39104368 | 0.592881 |
| ENSG00000163933 | RFT1 | 1247.407 | 0.2171721 | 0.1567011 | 1.41334 | 0.1575558 | 0.322703 |
| ENSG00000227083 | | 1.170304 | 0.2171058 | 0.7712369 | 0.7331548 | 0.463464 | NA |
| ENSG00000237978 | KCNMB2-AS1 | 284.5337 | 0.2170922 | 0.1788347 | 1.243145 | 0.2138145 | 0.398711 |
| ENSG00000163249 | CCNYL1 | 646.6247 | 0.2170161 | 0.2116816 | 1.066319 | 0.2862795 | 0.484561 |
| ENSG00000285805 | | 2.639637 | 0.2169171 | 0.7518754 | 0.7424819 | 0.4577954 | 0.65359 |
| ENSG00000080573 | COL5A3 | 43.98083942 | 0.216910506 | 0.489563337 | 0.560543966 | 0.57510846 | 0.747374 |
| ENSG00000269242 | | 1.220304 | 0.2168513 | 0.7515105 | 0.6935671 | 0.4879538 | NA |
| ENSG00000077454 | LRCH4 | 888.524284 | 0.216793141 | 0.182493235 | 1.219359831 | 0.22270765 | 0.409386 |
| ENSG00000076770 | MBNL3 | 1267.047428 | 0.216759391 | 0.545173285 | 0.542313222 | 0.58760275 | 0.75702 |
| ENSG00000138660 | AP1AR | 2269.654222 | 0.2167579 | 0.20097524 | 1.100259644 | 0.27121901 | 0.467559 |
| ENSG00000278970 | HEIH | 411.0401 | 0.2166616 | 0.1746267 | 1.262948 | 0.2066079 | 0.388847 |
| ENSG00000213757 | RPS3AP20 | 44.84284 | 0.2165729 | 0.3813455 | 0.6477368 | 0.5171552 | 0.702456 |
| ENSG00000267418 | ELOCP29 | 18.73847 | 0.216478 | 0.3896278 | 0.6407559 | 0.5216813 | 0.707027 |
| ENSG00000286739 | | 1.033542 | 0.2164756 | 0.7525587 | 0.6816232 | 0.4954772 | NA |
| ENSG00000170370 | EMX2 | 0.9113962 | 0.2164655 | 0.8050405 | 0.7053683 | 0.4805811 | NA |
| ENSG00000227825 | SLC9A7P1 | 9.12626 | 0.2164147 | 0.516072 | 0.5415858 | 0.5881038 | 0.757408 |
| ENSG00000248890 | HHIP-AS1 | 10.64044 | 0.2163775 | 0.6531997 | 0.5527495 | 0.580435 | 0.751246 |

| | | | | | | | |
|---|---|---|---|---|---|---|---|
| ENSG00000260088 | DDX59-AS1 | 4.28924 | 0.216358 | 0.589514 | 0.532429 | 0.5944289 | 0.761861 |
| ENSG00000111859 | NEDD9 | 2523.321982 | 0.216310628 | 0.39881738 | 0.628341511 | 0.52978024 | 0.714359 |
| ENSG00000143248 | RGS5 | 22.06694 | 0.2162775 | 0.3758782 | 0.6517899 | 0.5145367 | 0.700237 |
| ENSG00000279022 | | 22.77761 | 0.2162772 | 0.5069385 | 0.5480757 | 0.58364 | 0.753815 |
| ENSG00000103415 | HMOX2 | 4068.46999 | 0.216265324 | 0.205317561 | 1.091726651 | 0.27495327 | 0.471731 |
| ENSG00000247596 | TWF2 | 1080.706 | 0.2160731 | 0.2129355 | 1.036208 | 0.3001052 | 0.499374 |
| ENSG00000234335 | RPS4XP11 | 213.5604 | 0.2160701 | 0.2268255 | 1.010169 | 0.3124141 | 0.513467 |
| ENSG00000226783 | TLK1P1 | 7.908465 | 0.2160529 | 0.5424163 | 0.5392667 | 0.5897029 | 0.758512 |
| ENSG00000232671 | ZNF687-AS1 | 38.12603 | 0.2159923 | 0.3731283 | 0.6591231 | 0.5098167 | 0.695809 |
| ENSG00000134698 | AGO4 | 1543.553205 | 0.215969135 | 0.311708757 | 0.745182051 | 0.45616167 | 0.652028 |
| ENSG00000178028 | DMAP1 | 3582.359 | 0.2158704 | 0.1326727 | 1.643787 | 0.1002202 | 0.233166 |
| ENSG00000112773 | TENT5A | 2435.496121 | 0.215792554 | 0.267657882 | 0.861775632 | 0.388811 | 0.590878 |
| ENSG00000161270 | NPHS1 | 5.076747 | 0.2157224 | 0.72431 | 0.6269527 | 0.5306902 | 0.715004 |
| ENSG00000169981 | ZNF35 | 350.3126 | 0.2156806 | 0.1664075 | 1.328198 | 0.1841128 | 0.359356 |
| ENSG00000188958 | UTS2B | 0.9597467 | 0.2156491 | 0.8045819 | 0.6877907 | 0.4915846 | NA |
| ENSG00000171368 | TPPP | 149.7712 | 0.2156404 | 0.3009619 | 0.7741692 | 0.4388308 | 0.637347 |
| ENSG00000182359 | KBTBD3 | 192.0023 | 0.2156137 | 0.1822257 | 1.220779 | 0.2221697 | 0.408925 |
| ENSG00000222808 | RNU4-47P | 3.493897 | 0.2155558 | 0.6918887 | 0.5857464 | 0.5580459 | 0.735197 |
| ENSG00000171303 | KCNK3 | 2.978697 | 0.2155488 | 0.7139662 | 0.6197429 | 0.5354271 | 0.718872 |
| ENSG00000204291 | COL15A1 | 1.38443 | 0.2154948 | 0.80778 | 0.6800446 | 0.4964763 | NA |
| ENSG00000278486 | YRDCP1 | 2.375011 | 0.2154858 | 0.7049313 | 0.594626 | 0.5520935 | 0.730761 |
| ENSG00000063176 | SPHK2 | 493.1020367 | 0.21548036 | 0.17920299 | 1.232963707 | 0.21758931 | 0.403288 |
| ENSG00000156222 | SLC28A1 | 7.300352 | 0.2154608 | 0.6508972 | 0.5482461 | 0.583523 | 0.753727 |
| ENSG00000119720 | NRDE2 | 1858.051306 | 0.215428581 | 0.128757515 | 1.689248959 | 0.09117173 | 0.217664 |
| ENSG00000228446 | ANKRD49P1 | 3.747487 | 0.2153265 | 0.6862584 | 0.5762227 | 0.5644647 | 0.739657 |
| ENSG00000130844 | ZNF331 | 2418.580583 | 0.21508917 | 0.171345597 | 1.284699831 | 0.1988972 | 0.379013 |
| ENSG00000259093 | | 27.32099 | 0.2148223 | 0.3861776 | 0.6328221 | 0.5268498 | 0.711744 |
| ENSG00000116747 | RO60 | 2405.369568 | 0.214787976 | 0.185224372 | 1.178580416 | 0.23856529 | 0.428418 |
| ENSG00000274252 | GGTLC3 | 2.809355 | 0.2146429 | 0.7749104 | 0.849381 | 0.3956693 | 0.597249 |
| ENSG00000242083 | RPL7AP31 | 64.17112 | 0.2146394 | 0.2958601 | 0.7832301 | 0.433492 | 0.632479 |
| ENSG00000253737 | | 1.374947 | 0.2146022 | 0.7634446 | 0.6921862 | 0.4888204 | NA |
| ENSG00000241101 | PRICKLE2-AS2 | 17.37175 | 0.2144658 | 0.5751436 | 0.5411713 | 0.5883895 | 0.757616 |
| ENSG00000275734 | | 96.33553 | 0.2144251 | 0.3287006 | 0.7178865 | 0.4728273 | 0.665777 |

| | | | | | | | |
|---|---|---|---|---|---|---|---|
| ENSG00000272885 | | 12.57937 | 0.2143443 | 0.5497575 | 0.5293737 | 0.5965462 | 0.763644 |
| ENSG00000124160 | NCOA5 | 4245.776116 | 0.214338172 | 0.163439691 | 1.332855264 | 0.18257931 | 0.357529 |
| ENSG00000040933 | INPP4A | 3354.74664 | 0.214299676 | 0.22282102 | 0.996170576 | 0.31916727 | 0.520102 |
| ENSG00000219790 | OSTCP6 | 5.337525 | 0.21429 | 0.6209839 | 0.5409771 | 0.5885234 | 0.757728 |
| ENSG00000236031 | | 1.010581 | 0.2142766 | 0.7804672 | 0.7999485 | 0.4237406 | NA |
| ENSG00000265574 | WDR45BP1 | 187.8634 | 0.2142533 | 0.2121121 | 1.050938 | 0.2932869 | 0.492283 |
| ENSG00000277764 | | 0.9732894 | 0.2142469 | 0.7841338 | 0.8091148 | 0.4184491 | NA |
| ENSG00000133884 | DPF2 | 3737.986389 | 0.214228577 | 0.144349237 | 1.512407318 | 0.13043027 | 0.281488 |
| ENSG00000214145 | LINC00887 | 2.042886 | 0.2142258 | 0.7515313 | 0.6943748 | 0.4874472 | 0.678417 |
| ENSG00000176148 | TCP11L1 | 682.4862 | 0.2141967 | 0.2627362 | 0.8643457 | 0.387398 | 0.589709 |
| ENSG00000281207 | SLFNL1-AS1 | 68.72248 | 0.2141817 | 0.3029271 | 0.7709392 | 0.4407429 | 0.63871 |
| ENSG00000246203 | | 20.85835 | 0.2141367 | 0.4212552 | 0.5928601 | 0.5532748 | 0.731539 |
| ENSG00000249020 | SNORA58 | 1.765119 | 0.2140563 | 0.721655 | 0.6219086 | 0.5340019 | 0.717539 |
| ENSG00000171189 | GRIK1 | 66.15207 | 0.2139835 | 0.3975857 | 0.6191463 | 0.53582 | 0.719035 |
| ENSG00000248210 | LINC02355 | 0.8668896 | 0.2139786 | 0.8053762 | 0.7042482 | 0.4812782 | NA |
| ENSG00000068885 | IFT80 | 930.9882637 | 0.213822484 | 0.266723633 | 0.835055361 | 0.40368654 | 0.604529 |
| ENSG00000183773 | AIFM3 | 377.8657 | 0.213794 | 0.2809643 | 0.8157314 | 0.4146538 | 0.614649 |
| ENSG00000250848 | | 65.03951 | 0.2137916 | 0.3756809 | 0.6473099 | 0.5174314 | 0.702737 |
| ENSG00000183309 | ZNF623 | 3185.261 | 0.213727 | 0.2253982 | 0.9829012 | 0.3256561 | 0.526946 |
| ENSG00000285588 | | 3.265447 | 0.2137063 | 0.7020343 | 0.5474772 | 0.584051 | 0.754126 |
| ENSG00000174306 | ZHX3 | 4387.96 | 0.2135993 | 0.1728291 | 1.266426 | 0.2053608 | 0.387358 |
| ENSG00000182446 | NPLOC4 | 11164.92 | 0.2135848 | 0.1239434 | 1.746241 | 0.08076901 | 0.198755 |
| ENSG00000205853 | RFPL3S | 6.243221 | 0.2135675 | 0.6344528 | 0.5314512 | 0.5951062 | 0.762472 |
| ENSG00000228868 | MYLKP1 | 0.923869 | 0.2134909 | 0.828867 | 1.412359 | 0.1578443 | NA |
| ENSG00000249784 | SCARNA22 | 11.94583 | 0.213481 | 0.5682064 | 0.5340509 | 0.5933063 | 0.761267 |
| ENSG00000270170 | NCBP2AS2 | 2120.616 | 0.2134366 | 0.22451 | 0.9866587 | 0.32381 | 0.524752 |
| ENSG00000197050 | ZNF420 | 373.2746 | 0.2133104 | 0.2011068 | 1.095143 | 0.2734539 | 0.470191 |
| ENSG00000214389 | RPS3AP26 | 5316.092 | 0.2132988 | 0.19351 | 1.12663 | 0.259899 | 0.454353 |
| ENSG00000163161 | ERCC3 | 6007.763 | 0.2132925 | 0.1306819 | 1.654036 | 0.0981202 | 0.229539 |
| ENSG00000259032 | ENSAP2 | 2253.48 | 0.2132865 | 0.1985268 | 1.107563 | 0.2680507 | 0.46398 |
| ENSG00000132613 | MTSS2 | 2099.777628 | 0.213191918 | 0.178987253 | 1.219108837 | 0.22280289 | 0.409421 |
| ENSG00000204344 | STK19 | 985.9724 | 0.2131342 | 0.1639811 | 1.32543 | 0.1850285 | 0.360581 |
| ENSG00000248196 | | 3.606635 | 0.2131011 | 0.6686936 | 0.5520112 | 0.5809407 | 0.751717 |

| | | | | | | | |
|---|---|---|---|---|---|---|---|
| ENSG00000130544 | ZNF557 | 397.1732567 | 0.213043054 | 0.18824871 | 1.170512766 | 0.24179468 | 0.432347 |
| ENSG00000202111 | VTRNA1-2 | 1.641606 | 0.2130259 | 0.7582213 | 0.7023888 | 0.4824367 | 0.673844 |
| ENSG00000230364 | RPL4P3 | 23.90042 | 0.212931 | 0.4636277 | 0.560165 | 0.5753669 | 0.747542 |
| ENSG00000158792 | SPATA2L | 2102.875 | 0.2128963 | 0.1656579 | 1.311125 | 0.1898157 | 0.367501 |
| ENSG00000137145 | DENND4C | 2341.796503 | 0.212881875 | 0.288448738 | 0.797722519 | 0.42503153 | 0.624767 |
| ENSG00000083223 | TUT7 | 2349.759138 | 0.212857657 | 0.232769451 | 0.953074959 | 0.34055209 | 0.541994 |
| ENSG00000122257 | RBBP6 | 5657.184858 | 0.212806895 | 0.110648182 | 1.927722218 | 0.05388969 | 0.147002 |
| ENSG00000266402 | SNHG25 | 193.9859 | 0.212735 | 0.256496 | 0.8919708 | 0.3724086 | 0.574562 |
| ENSG00000226908 | H2BP6 | 0.8311873 | 0.2125853 | 0.8235902 | 1.190123 | 0.233998 | NA |
| ENSG00000284740 | | 11.27634 | 0.2125224 | 0.6159107 | 0.5258605 | 0.5989851 | 0.765663 |
| ENSG00000207392 | SNORA20 | 6.140983 | 0.2123478 | 0.5815234 | 0.5248727 | 0.5996717 | 0.765998 |
| ENSG00000066777 | ARFGEF1 | 5557.323796 | 0.212274141 | 0.216204225 | 1.010853053 | 0.31208677 | 0.513077 |
| ENSG00000203914 | HSP90B3P | 16.80261 | 0.2121655 | 0.4039749 | 0.610398 | 0.5415982 | 0.723277 |
| ENSG00000130584 | ZBTB46 | 651.1646943 | 0.212121244 | 0.18897822 | 1.15426378 | 0.24839204 | 0.440732 |
| ENSG00000136141 | LRCH1 | 4024.764691 | 0.212094217 | 0.112626295 | 1.884639892 | 0.05947847 | 0.158086 |
| ENSG00000109920 | FNBP4 | 9237.579696 | 0.212054988 | 0.110761622 | 1.931101077 | 0.05347055 | 0.146128 |
| ENSG00000259216 | | 8.499749 | 0.2120518 | 0.5577109 | 0.5298523 | 0.5962143 | 0.763339 |
| ENSG00000198890 | PRMT6 | 16541.57 | 0.2119444 | 0.1793483 | 1.200801 | 0.2298284 | 0.418012 |
| ENSG00000150907 | FOXO1 | 505.6207 | 0.2118887 | 0.202204 | 1.082701 | 0.278941 | 0.476175 |
| ENSG00000219294 | PIP5K1P1 | 13.92807 | 0.2118444 | 0.4798216 | 0.5508841 | 0.5817131 | 0.752277 |
| ENSG00000176746 | MAGEB6 | 1.916883 | 0.2117195 | 0.807412 | 0.674196 | 0.5001867 | 0.688288 |
| ENSG00000248866 | USP46-DT | 139.4933 | 0.2114513 | 0.3087963 | 0.7424323 | 0.4578254 | 0.65359 |
| ENSG00000271659 | | 10.73578 | 0.2114196 | 0.6277 | 0.5264448 | 0.5985792 | 0.76538 |
| ENSG00000145388 | METTL14 | 2051.978 | 0.211335 | 0.1284499 | 1.638306 | 0.1013579 | 0.235092 |
| ENSG00000277283 | | 356.1816 | 0.2111627 | 0.2533967 | 0.8797526 | 0.3789934 | 0.581103 |
| ENSG00000078369 | GNB1 | 14830.75937 | 0.21109522 | 0.135735553 | 1.577206636 | 0.11474799 | 0.256887 |
| ENSG00000254349 | MIR2052HG | 41.65919 | 0.2109025 | 0.4839856 | 0.539066 | 0.5898413 | 0.758616 |
| ENSG00000187775 | DNAH17 | 169.5045 | 0.210869 | 0.2772268 | 0.8148364 | 0.4151659 | 0.615074 |
| ENSG00000079616 | KIF22 | 6031.47657 | 0.210775202 | 0.328677334 | 0.700806828 | 0.48342358 | 0.67476 |
| ENSG00000225329 | LHFPL3-AS2 | 10.17698 | 0.2107747 | 0.5670353 | 0.5229861 | 0.600984 | 0.766786 |
| ENSG00000105255 | FSD1 | 886.9385348 | 0.210647559 | 0.441245685 | 0.577169013 | 0.56382529 | 0.739219 |
| ENSG00000274280 | MIR6069 | 0.8059622 | 0.2106417 | 0.8117754 | 0.8190048 | 0.4127837 | NA |
| ENSG00000133275 | CSNK1G2 | 4498.440943 | 0.210594417 | 0.139204927 | 1.534799851 | 0.124833 | 0.272844 |

| | | | | | | | |
|---|---|---|---|---|---|---|---|
| ENSG00000239830 | RPS4XP22 | 5.015978 | 0.2105392 | 0.5956759 | 0.5210921 | 0.6023026 | 0.767619 |
| ENSG00000283023 | FRG1GP | 117.3962 | 0.2105177 | 0.3396356 | 0.6914558 | 0.4892791 | 0.680076 |
| ENSG00000236166 | | 1.038387 | 0.2104905 | 0.8035798 | 0.6834405 | 0.4943286 | NA |
| ENSG00000214335 | CTAGE16P | 0.9136539 | 0.2104291 | 0.7804574 | 0.7781635 | 0.4364726 | NA |
| ENSG00000259882 | | 1.165189 | 0.2103727 | 0.7995525 | 0.8741183 | 0.3820539 | NA |
| ENSG00000233971 | RPS20P10 | 62.71342 | 0.2103024 | 0.3174271 | 0.7232974 | 0.4694972 | 0.662979 |
| ENSG00000241553 | ARPC4 | 4533.238 | 0.2102783 | 0.2217338 | 0.985136 | 0.3245573 | 0.525578 |
| ENSG00000250645 | | 6.857162 | 0.2101864 | 0.5945422 | 0.5220084 | 0.6016645 | 0.7672 |
| ENSG00000259475 | | 5.296589 | 0.2101079 | 0.6092945 | 0.4992313 | 0.6176164 | 0.77875 |
| ENSG00000250677 | | 1.007638 | 0.2100905 | 0.7745788 | 0.8491594 | 0.3957926 | NA |
| ENSG00000132128 | LRRC41 | 5209.235312 | 0.210049654 | 0.141162161 | 1.472304293 | 0.14093872 | 0.297592 |
| ENSG00000104320 | NBN | 7888.499545 | 0.20998175 | 0.191487839 | 1.133276359 | 0.25709822 | 0.45091 |
| ENSG00000261270 | | 37.87762 | 0.2099642 | 0.411785 | 0.5927565 | 0.5533441 | 0.731542 |
| ENSG00000257831 | | 3.153992 | 0.2099445 | 0.6290259 | 0.523186 | 0.6008448 | 0.766755 |
| ENSG00000075240 | GRAMD4 | 2576.256635 | 0.209684246 | 0.141160035 | 1.507387614 | 0.13171133 | 0.283516 |
| ENSG00000225598 | | 5.849127 | 0.2096606 | 0.6683625 | 0.5581333 | 0.5767534 | 0.748717 |
| ENSG00000242295 | | 0.8974865 | 0.2096543 | 0.8029477 | 0.6613547 | 0.5083849 | NA |
| ENSG00000285564 | | 0.9149841 | 0.2096063 | 0.758802 | 0.6926797 | 0.4885106 | NA |
| ENSG00000249755 | | 1.127256 | 0.2095942 | 0.7598929 | 0.7058702 | 0.4802689 | NA |
| ENSG00000185523 | SPATA45 | 3.33726 | 0.209487 | 0.660866 | 0.5475693 | 0.5839877 | 0.754089 |
| ENSG00000268093 | | 9.260399 | 0.2094755 | 0.4653135 | 0.551101 | 0.5815645 | 0.752204 |
| ENSG00000124098 | FAM210B | 5571.160697 | 0.209393217 | 0.237060366 | 0.96893282 | 0.33257871 | 0.533656 |
| ENSG00000275092 | | 5.04115 | 0.2092365 | 0.7019587 | 0.5928006 | 0.5533146 | 0.731539 |
| ENSG00000228221 | LINC00578 | 44.18489 | 0.2091765 | 0.4826073 | 0.5297213 | 0.5963052 | 0.763419 |
| ENSG00000254670 | RASSF10-DT | 0.9321104 | 0.2091198 | 0.788081 | 0.8354606 | 0.4034584 | NA |
| ENSG00000185065 | | 47.95893 | 0.2090917 | 0.3496063 | 0.6652026 | 0.5059209 | 0.692627 |
| ENSG00000103067 | ESRP2 | 9765.690557 | 0.209076122 | 0.158453135 | 1.337193575 | 0.18115946 | 0.355718 |
| ENSG00000256596 | | 1.418695 | 0.2089162 | 0.7591231 | 0.7234334 | 0.4694136 | NA |
| ENSG00000272211 | | 1.331168 | 0.2088966 | 0.7387617 | 0.655908 | 0.5118833 | NA |
| ENSG00000234797 | RPS3AP6 | 15984.68 | 0.2088568 | 0.1904032 | 1.113303 | 0.2655782 | 0.461263 |
| ENSG00000196459 | TRAPPC2 | 660.7936 | 0.208781 | 0.1600754 | 1.31449 | 0.1886813 | 0.365864 |
| ENSG00000154710 | RABGEF1 | 190.9813 | 0.2087472 | 0.1758102 | 1.217322 | 0.2234816 | 0.410203 |
| ENSG00000168028 | RPSA | 10514.68 | 0.2087153 | 0.2139828 | 1.011204 | 0.3119186 | 0.512896 |

| | | | | | | | |
|---|---|---|---|---|---|---|---|
| ENSG00000186150 | UBL4B | 1.764794 | 0.2086303 | 0.7029356 | 0.5602856 | 0.5752847 | 0.74753 |
| ENSG00000274898 | | 4.915009 | 0.2086039 | 0.6553454 | 0.5356376 | 0.592209 | 0.76052 |
| ENSG00000173744 | AGFG1 | 3067.02 | 0.208561 | 0.164866 | 1.288896 | 0.1974344 | 0.377369 |
| ENSG00000011451 | WIZ | 3288.245054 | 0.208557345 | 0.147576151 | 1.381441965 | 0.16714311 | 0.336528 |
| ENSG00000114573 | ATP6V1A | 6331.333712 | 0.208539798 | 0.118750002 | 1.785746284 | 0.07414035 | 0.186619 |
| ENSG00000128881 | TTBK2 | 510.4323759 | 0.208469198 | 0.371581051 | 0.629638821 | 0.52893092 | 0.713648 |
| ENSG00000113812 | ACTR8 | 1447.922945 | 0.208416329 | 0.121524616 | 1.734308588 | 0.08286335 | 0.202757 |
| ENSG00000267117 | | 1.739962 | 0.2083404 | 0.7252569 | 0.6039473 | 0.5458787 | 0.726117 |
| ENSG00000275445 | | 2.179335 | 0.2082249 | 0.7198027 | 0.6100789 | 0.5418095 | 0.723457 |
| ENSG00000090061 | CCNK | 910.6974891 | 0.208202925 | 0.143430947 | 1.475386904 | 0.14010855 | 0.296287 |
| ENSG00000179342 | | 14.77796 | 0.2081289 | 0.4556035 | 0.5533741 | 0.5800073 | 0.750912 |
| ENSG00000248724 | NPHP3-AS1 | 1.281692 | 0.2081126 | 0.7108511 | 0.568264 | 0.5698557 | NA |
| ENSG00000274294 | | 3.146346 | 0.2080952 | 0.6963885 | 0.5426267 | 0.5873869 | 0.756925 |
| ENSG00000155592 | ZKSCAN2 | 808.7259 | 0.2080813 | 0.1508041 | 1.389968 | 0.1645387 | 0.332696 |
| ENSG00000267522 | LINC01864 | 1.59827 | 0.2080476 | 0.7645261 | 0.6849452 | 0.4933785 | 0.683236 |
| ENSG00000165801 | ARHGEF40 | 432.3235 | 0.2079676 | 0.2555568 | 0.8374261 | 0.4023531 | 0.603286 |
| ENSG00000213654 | GPSM3 | 59.90473 | 0.2079465 | 0.4340558 | 0.5690693 | 0.5693091 | 0.743291 |
| ENSG00000273691 | | 1.624815 | 0.2078914 | 0.7656429 | 0.6835007 | 0.4942905 | 0.684005 |
| ENSG00000105643 | ARRDC2 | 1124.347898 | 0.207888692 | 0.146502477 | 1.45171372 | 0.14658122 | 0.306076 |
| ENSG00000280660 | | 4.834958 | 0.2078575 | 0.7228925 | 0.6046595 | 0.5454053 | 0.725887 |
| ENSG00000232953 | HSPA8P18 | 2.843212 | 0.2078567 | 0.6543658 | 0.5296418 | 0.5963603 | 0.763453 |
| ENSG00000179295 | PTPN11 | 12586.58 | 0.2078303 | 0.1944773 | 1.11008 | 0.2669647 | 0.46278 |
| ENSG00000260669 | | 55.69399 | 0.2076754 | 0.2810908 | 0.7869451 | 0.431314 | 0.630409 |
| ENSG00000175220 | ARHGAP1 | 3924.586 | 0.2076342 | 0.2156719 | 0.9961743 | 0.3191655 | 0.520102 |
| ENSG00000286433 | | 0.8226857 | 0.2076109 | 0.7781363 | 0.7473145 | 0.4548737 | NA |
| ENSG00000188931 | CFAP126 | 195.2921 | 0.2075657 | 0.3282324 | 0.6907972 | 0.489693 | 0.680331 |
| ENSG00000188483 | IER5L | 3342.848 | 0.2075394 | 0.1955207 | 1.112076 | 0.2661054 | 0.46174 |
| ENSG00000140044 | JDP2 | 2926.017 | 0.2075238 | 0.2821165 | 0.788573 | 0.4303616 | 0.629691 |
| ENSG00000234985 | | 1.074656 | 0.2074824 | 0.7715182 | 0.8306421 | 0.4061759 | NA |
| ENSG00000260100 | | 1.279248 | 0.2074822 | 0.7460921 | 0.6539015 | 0.5131753 | NA |
| ENSG00000163625 | WDFY3 | 4611.26 | 0.2074624 | 0.2292703 | 0.9401539 | 0.3471386 | 0.548669 |
| ENSG00000113360 | DROSHA | 5409.630139 | 0.207453034 | 0.197126793 | 1.090096014 | 0.27567085 | 0.472505 |
| ENSG00000255197 | | 5.032508 | 0.2073759 | 0.5654465 | 0.5125683 | 0.6082533 | 0.772056 |

| | | | | | | | |
|---|---|---|---|---|---|---|---|
| ENSG00000105085 | MED26 | 275.2385153 | 0.207347119 | 0.179022873 | 1.202468678 | 0.22918199 | 0.417287 |
| ENSG00000233998 | SETP5 | 2.62242 | 0.2073115 | 0.7291474 | 0.6125563 | 0.5401698 | 0.722353 |
| ENSG00000104660 | LEPROTL1 | 1787.738381 | 0.207266847 | 0.115713455 | 1.800086206 | 0.07184703 | 0.182165 |
| ENSG00000225706 | PTPRD-AS1 | 0.8344356 | 0.207239 | 0.7781077 | 0.7825136 | 0.4339128 | NA |
| ENSG00000232023 | LINC01807 | 67.28135 | 0.2072322 | 0.3807255 | 0.6158874 | 0.5379688 | 0.72101 |
| ENSG00000170946 | DNAJC24 | 516.2905 | 0.2072138 | 0.1940277 | 1.098007 | 0.2722014 | 0.468705 |
| ENSG00000280202 | | 278.5874 | 0.2071558 | 0.3569998 | 0.6815201 | 0.4955425 | 0.684733 |
| ENSG00000128262 | POM121L9P | 4.236507272 | 0.207075981 | 0.575309288 | 0.508157334 | 0.61134301 | 0.774461 |
| ENSG00000257315 | ZBED6 | 118.7143 | 0.2070548 | 0.4484275 | 0.55471 | 0.579093 | 0.750345 |
| ENSG00000216001 | MIR450B | 1.47056 | 0.207021 | 0.7549084 | 0.6795651 | 0.4967799 | NA |
| ENSG00000223791 | UBE2E1-AS1 | 3.472861 | 0.2070183 | 0.6797779 | 0.5536017 | 0.5798515 | 0.750747 |
| ENSG00000137496 | IL18BP | 523.2759975 | 0.206895802 | 0.158451501 | 1.330891347 | 0.18322477 | 0.358344 |
| ENSG00000226241 | | 1.012444 | 0.2068642 | 0.7641535 | 0.7018299 | 0.4827853 | NA |
| ENSG00000169609 | C15orf40 | 1882.323 | 0.2068173 | 0.1942651 | 1.096467 | 0.2728743 | 0.469499 |
| ENSG00000266290 | | 1.01436 | 0.206773 | 0.7267145 | 0.6030448 | 0.5464789 | NA |
| ENSG00000261456 | TUBB8 | 79.89085 | 0.2067578 | 0.378273 | 0.6205451 | 0.534899 | 0.718417 |
| ENSG00000278546 | | 2.353394 | 0.206731 | 0.7308029 | 0.6291136 | 0.5292747 | 0.71404 |
| ENSG00000273891 | | 65.87604 | 0.2067105 | 0.3708194 | 0.6297074 | 0.528886 | 0.713624 |
| ENSG00000176896 | TCEANC | 194.7928 | 0.2066919 | 0.2235695 | 0.9627792 | 0.3356583 | 0.536815 |
| ENSG00000153944 | MSI2 | 14394.42 | 0.2066311 | 0.1607024 | 1.31152 | 0.1896823 | 0.367323 |
| ENSG00000199331 | | 1.769907 | 0.2065876 | 0.7701856 | 0.833681 | 0.4044608 | 0.605347 |
| ENSG00000213772 | EIF1P7 | 12.44762 | 0.2063753 | 0.445869 | 0.5579719 | 0.5768635 | 0.748776 |
| ENSG00000240919 | | 0.8213679 | 0.2063654 | 0.7727561 | 0.8322641 | 0.4052599 | NA |
| ENSG00000254780 | | 6.666213 | 0.2063532 | 0.5425596 | 0.5141178 | 0.6071696 | 0.771197 |
| ENSG00000102572 | STK24 | 2764.250211 | 0.206319963 | 0.166108268 | 1.244026673 | 0.21348975 | 0.39828 |
| ENSG00000170113 | NIPA1 | 1658.458 | 0.206284 | 0.1895545 | 1.090297 | 0.2755822 | 0.472445 |
| ENSG00000122482 | ZNF644 | 1806.834307 | 0.206271682 | 0.186692326 | 1.140908432 | 0.25390803 | 0.446908 |
| ENSG00000169231 | THBS3 | 2628.671 | 0.2062298 | 0.2847595 | 0.7781737 | 0.4364666 | 0.635041 |
| ENSG00000229431 | MED8-AS1 | 41.82549 | 0.2062251 | 0.3396054 | 0.6716872 | 0.5017829 | 0.689342 |
| ENSG00000189143 | CLDN4 | 19114.32 | 0.2062246 | 0.2001165 | 1.057122 | 0.2904557 | 0.489096 |
| ENSG00000188554 | NBR1 | 8880.851 | 0.2061528 | 0.1768262 | 1.195154 | 0.2320268 | 0.420427 |
| ENSG00000286580 | | 3.443843 | 0.2061299 | 0.6971425 | 0.5543517 | 0.5793382 | 0.750533 |
| ENSG00000267137 | | 1.043208 | 0.2061227 | 0.804163 | 0.6600162 | 0.5092434 | NA |

| | | | | | | | |
|---|---|---|---|---|---|---|---|
| ENSG00000260971 | | 5.929814 | 0.2060603 | 0.5832622 | 0.5073564 | 0.6119048 | 0.77473 |
| ENSG00000113916 | BCL6 | 2657.074762 | 0.206011478 | 0.310135997 | 0.721275876 | 0.4707398 | 0.66414 |
| ENSG00000126803 | HSPA2 | 866.5817218 | 0.206011234 | 0.391028197 | 0.606221193 | 0.54436788 | 0.725217 |
| ENSG00000106993 | CDC37L1 | 809.3918348 | 0.205938157 | 0.172017862 | 1.219960422 | 0.22247988 | 0.409155 |
| ENSG00000270177 | | 38.6675 | 0.2058759 | 0.3883178 | 0.6072931 | 0.5436564 | 0.724554 |
| ENSG00000104671 | DCTN6 | 722.4421039 | 0.205575376 | 0.187952894 | 1.119419773 | 0.2629611 | 0.457949 |
| ENSG00000119616 | FCF1 | 2783.831635 | 0.205529115 | 0.137139657 | 1.521662693 | 0.12809362 | 0.277792 |
| ENSG00000286060 | | 2.420352 | 0.2054343 | 0.7664901 | 0.6430625 | 0.5201835 | 0.705393 |
| ENSG00000206936 | RNU4-52P | 4.100801 | 0.2053798 | 0.6486944 | 0.5223111 | 0.6014538 | 0.76709 |
| ENSG00000180370 | PAK2 | 10691.58 | 0.2052852 | 0.2019178 | 1.03842 | 0.2990744 | 0.498314 |
| ENSG00000273950 | | 3.981591 | 0.2052675 | 0.649901 | 0.5260378 | 0.5988619 | 0.765622 |
| ENSG00000152056 | AP1S3 | 235.9052 | 0.2051757 | 0.2068294 | 1.027871 | 0.3040105 | 0.503695 |
| ENSG00000215533 | LINC00189 | 1.289269 | 0.2051226 | 0.8039571 | 0.6581807 | 0.510422 | NA |
| ENSG00000183648 | NDUFB1 | 2348.436 | 0.2049793 | 0.1738699 | 1.224829 | 0.2206398 | 0.407011 |
| ENSG00000159082 | SYNJ1 | 813.4411 | 0.2048948 | 0.2432069 | 0.8840703 | 0.3766582 | 0.578699 |
| ENSG00000268603 | | 2.286547 | 0.204881 | 0.7156942 | 0.591321 | 0.5543054 | 0.732434 |
| ENSG00000197302 | ZNF720 | 1661.992 | 0.2048572 | 0.1199225 | 1.732223 | 0.08323379 | 0.203341 |
| ENSG00000246089 | | 283.2468 | 0.2047632 | 0.1844508 | 1.136475 | 0.2557579 | 0.449242 |
| ENSG00000169885 | CALML6 | 56.19438 | 0.2046397 | 0.2772881 | 0.7922829 | 0.4281957 | 0.627578 |
| ENSG00000048828 | FAM120A | 14604.76063 | 0.204544579 | 0.155525235 | 1.307063656 | 0.19119109 | 0.36921 |
| ENSG00000079691 | CARMIL1 | 1624.767989 | 0.204505114 | 0.174480627 | 1.197887292 | 0.2309609 | 0.419468 |
| ENSG00000134072 | CAMK1 | 57.52456823 | 0.204463093 | 0.259389475 | 0.833632616 | 0.40448804 | 0.605353 |
| ENSG00000196482 | ESRRG | 4.471786 | 0.2044468 | 0.6023909 | 0.5059823 | 0.612869 | 0.775655 |
| ENSG00000164070 | HSPA4L | 2301.174 | 0.204432 | 0.2336048 | 0.904125 | 0.3659291 | 0.567829 |
| ENSG00000187553 | CYP26C1 | 1.040148 | 0.2044126 | 0.7562896 | 0.6667757 | 0.5049154 | NA |
| ENSG00000228399 | | 13.08561 | 0.2043411 | 0.5103747 | 0.5187702 | 0.603921 | 0.768809 |
| ENSG00000211697 | TRGV5 | 1.349966 | 0.2042955 | 0.804159 | 0.6656541 | 0.5056322 | NA |
| ENSG00000241158 | ADAMTS9-AS1 | 24.04206 | 0.2042827 | 0.6215036 | 0.5225489 | 0.6012882 | 0.766989 |
| ENSG00000076604 | TRAF4 | 7323.07866 | 0.204237847 | 0.17959022 | 1.15332236 | 0.2487781 | 0.441175 |
| ENSG00000257307 | | 20.37971 | 0.2041705 | 0.4232787 | 0.5683964 | 0.5697658 | 0.743741 |
| ENSG00000175324 | LSM1 | 847.5196 | 0.2041207 | 0.2107826 | 1.015433 | 0.3098996 | 0.510555 |
| ENSG00000280326 | | 1.125628 | 0.2040834 | 0.7547792 | 0.6692348 | 0.5033457 | NA |
| ENSG00000165355 | FBXO33 | 1346.519 | 0.2040588 | 0.1414032 | 1.455846 | 0.1454351 | 0.304449 |

| | | | | | | | |
|---|---|---|---|---|---|---|---|
| ENSG00000140749 | IGSF6 | 5.232585 | 0.2040176 | 0.5871085 | 0.5077067 | 0.6116591 | 0.774558 |
| ENSG00000267772 | LINC01999 | 1.944906 | 0.2040064 | 0.7352604 | 0.6174971 | 0.5369069 | 0.719986 |
| ENSG00000204271 | SPIN3 | 2153.8 | 0.203797 | 0.2881282 | 0.7667019 | 0.4432587 | 0.640816 |
| ENSG00000199226 | RNU6-50P | 1.492738 | 0.203794 | 0.7530081 | 0.6593856 | 0.5096482 | NA |
| ENSG00000163810 | TGM4 | 1.29023 | 0.2037182 | 0.755134 | 0.6784186 | 0.4975063 | NA |
| ENSG00000185332 | TMEM105 | 231.4421 | 0.2037104 | 0.2967495 | 0.7365392 | 0.4614026 | 0.656194 |
| ENSG00000175376 | EIF1AD | 2397.998 | 0.2036599 | 0.1267936 | 1.623358 | 0.104513 | 0.240229 |
| ENSG00000178750 | STX19 | 5.572621 | 0.203657 | 0.6644391 | 0.5121995 | 0.6085114 | 0.772229 |
| ENSG00000280387 | | 13.89836 | 0.2036219 | 0.5636207 | 0.497367 | 0.6189302 | 0.779563 |
| ENSG00000259916 | | 4.341953 | 0.2036186 | 0.6576313 | 0.522093 | 0.6016056 | 0.7672 |
| ENSG00000258651 | SEC23A-AS1 | 3.337092 | 0.2035967 | 0.6475502 | 0.5177672 | 0.6046207 | 0.76943 |
| ENSG00000260588 | | 91.43896 | 0.2035865 | 0.2911382 | 0.7509832 | 0.4526627 | 0.648791 |
| ENSG00000275609 | | 1.073354 | 0.2035402 | 0.7630807 | 0.6868273 | 0.4921916 | NA |
| ENSG00000135537 | AFG1L | 282.3597106 | 0.203536464 | 0.216174586 | 0.98805051 | 0.32312791 | 0.523902 |
| ENSG00000197102 | DYNC1H1 | 27835.85 | 0.2035343 | 0.249542 | 0.8561723 | 0.3919025 | 0.593641 |
| ENSG00000259235 | | 0.7790951 | 0.2035289 | 0.7973818 | 0.8440192 | 0.3986587 | NA |
| ENSG00000107036 | RIC1 | 1326.287943 | 0.203520452 | 0.216826193 | 0.970720938 | 0.33168726 | 0.532837 |
| ENSG00000226564 | FTH1P20 | 566.1202 | 0.2034791 | 0.2294435 | 0.9272782 | 0.3537821 | 0.555764 |
| ENSG00000258416 | | 1.198859 | 0.2034735 | 0.7682807 | 0.6486053 | 0.5165936 | NA |
| ENSG00000221968 | FADS3 | 1828.382 | 0.2034446 | 0.3280074 | 0.6797142 | 0.4966854 | 0.685565 |
| ENSG00000270540 | | 4.037921 | 0.2032796 | 0.6696461 | 0.5357881 | 0.592105 | 0.760423 |
| ENSG00000231386 | MOB4P1 | 3.502457 | 0.2032783 | 0.6962876 | 0.5653364 | 0.571845 | 0.745437 |
| ENSG00000261096 | | 3.019477 | 0.2032534 | 0.6282427 | 0.506591 | 0.6124418 | 0.775225 |
| ENSG00000165886 | UBTD1 | 1309.677 | 0.2032062 | 0.2152674 | 0.9784313 | 0.3278611 | 0.529299 |
| ENSG00000214867 | SRSF9P1 | 13.04958 | 0.2031814 | 0.5088375 | 0.5138462 | 0.6073595 | 0.771285 |
| ENSG00000264672 | SEPTIN4-AS1 | 25.36981 | 0.2031158 | 0.4645986 | 0.5336754 | 0.5935661 | 0.761364 |
| ENSG00000225792 | | 15.20152 | 0.2030422 | 0.5139108 | 0.5134643 | 0.6076266 | 0.771445 |
| ENSG00000258599 | | 0.8607011 | 0.2029957 | 0.7703607 | 0.8115673 | 0.4170399 | NA |
| ENSG00000160785 | SLC25A44 | 4540.978 | 0.2029307 | 0.1325815 | 1.502378 | 0.1329994 | 0.285427 |
| ENSG00000181991 | MRPS11 | 2004.848 | 0.2028726 | 0.2199564 | 0.9776124 | 0.3282661 | 0.529422 |
| ENSG00000103855 | CD276 | 4897.642891 | 0.202857802 | 0.179287761 | 1.171525834 | 0.24138748 | 0.431939 |
| ENSG00000140612 | SEC11A | 10583.22 | 0.2028516 | 0.1593573 | 1.280249 | 0.2004577 | 0.38097 |
| ENSG00000140265 | ZSCAN29 | 2090.422 | 0.2027885 | 0.2212974 | 0.9563583 | 0.3388912 | 0.540132 |

| | | | | | | | |
|---|---|---|---|---|---|---|---|
| ENSG00000272750 | | 23.18147 | 0.2027265 | 0.4050856 | 0.5811389 | 0.5611468 | 0.737289 |
| ENSG00000168890 | TMEM150A | 1955.187 | 0.2027037 | 0.2489872 | 0.873486 | 0.3823982 | 0.584711 |
| ENSG00000263280 | | 11.90864 | 0.2026448 | 0.4712081 | 0.5279289 | 0.5975487 | 0.764458 |
| ENSG00000278600 | | 488.3325 | 0.2025407 | 0.2825744 | 0.76399 | 0.4448732 | 0.642138 |
| ENSG00000285637 | | 7.985523 | 0.2025336 | 0.6426413 | 0.5033572 | 0.6147132 | 0.776509 |
| ENSG00000182196 | ARL6IP4 | 60.59644 | 0.2025148 | 0.2759097 | 0.7829776 | 0.4336403 | 0.632592 |
| ENSG00000274376 | ADAMTS7P1 | 0.7592223 | 0.2024763 | 0.795676 | 0.8878188 | 0.3746382 | NA |
| ENSG00000216613 | | 1.285472 | 0.2024138 | 0.763984 | 0.6644807 | 0.5063827 | NA |
| ENSG00000149380 | P4HA3 | 9.635707 | 0.2023909 | 0.5785197 | 0.5048152 | 0.6136887 | 0.775861 |
| ENSG00000277135 | | 1.048961 | 0.2022695 | 0.7813305 | 0.7376722 | 0.4607136 | NA |
| ENSG00000255495 | | 7.188146 | 0.2021676 | 0.5489366 | 0.499836 | 0.6171905 | 0.778565 |
| ENSG00000175104 | TRAF6 | 1070.846 | 0.2021028 | 0.1859158 | 1.115695 | 0.2645527 | 0.460007 |
| ENSG00000182973 | CNOT10 | 2038.914 | 0.2021005 | 0.1282862 | 1.592136 | 0.111354 | 0.251536 |
| ENSG00000230581 | ACTG1P14 | 2.468618 | 0.2020654 | 0.7152009 | 0.5883317 | 0.5563097 | 0.73375 |
| ENSG00000103353 | UBFD1 | 7802.715658 | 0.201995132 | 0.121979431 | 1.711800985 | 0.08693335 | 0.21005 |
| ENSG00000163412 | EIF4E3 | 843.6998 | 0.2019296 | 0.19037 | 1.08484 | 0.2779923 | 0.475105 |
| ENSG00000230552 | | 2.06581 | 0.2018174 | 0.7225672 | 0.5567165 | 0.5777212 | 0.749523 |
| ENSG00000091436 | MAP3K20 | 1477.618755 | 0.201805905 | 0.264567007 | 0.810222844 | 0.41781211 | 0.617591 |
| ENSG00000255418 | LINC02718 | 22.63721 | 0.201805 | 0.4675256 | 0.5285344 | 0.5971284 | 0.764085 |
| ENSG00000116171 | SCP2 | 4524.16312 | 0.201744285 | 0.143721412 | 1.41390322 | 0.15739032 | 0.322488 |
| ENSG00000189298 | ZKSCAN3 | 174.0318 | 0.2016703 | 0.2578115 | 0.8240158 | 0.4099306 | 0.610319 |
| ENSG00000259720 | | 0.8708807 | 0.2015975 | 0.7473851 | 0.6491701 | 0.5162285 | NA |
| ENSG00000164830 | OXR1 | 3406.641 | 0.2015299 | 0.1807372 | 1.164832 | 0.2440869 | 0.43542 |
| ENSG00000277476 | | 406.0163 | 0.2015256 | 0.2916824 | 0.7406788 | 0.4588882 | 0.654576 |
| ENSG00000219507 | FTH1P8 | 1771.974 | 0.201514 | 0.2387367 | 0.8804278 | 0.3786276 | 0.58075 |
| ENSG00000240457 | RN7SL472P | 1.600004 | 0.2014833 | 0.7639668 | 0.6674417 | 0.5044901 | 0.691381 |
| ENSG00000258925 | NPM1P5 | 10.24701 | 0.2014784 | 0.4937538 | 0.5210532 | 0.6023297 | 0.767619 |
| ENSG00000189060 | H1-0 | 16973.61 | 0.2014521 | 0.2344709 | 0.8963369 | 0.3700729 | 0.572255 |
| ENSG00000065883 | CDK13 | 4551.425098 | 0.20139153 | 0.148593344 | 1.378063886 | 0.16818356 | 0.337983 |
| ENSG00000213137 | ARF1P2 | 0.7921992 | 0.2013092 | 0.7759139 | 0.7360027 | 0.461729 | NA |
| ENSG00000263393 | | 2.67039 | 0.2013049 | 0.7004847 | 0.5619385 | 0.574158 | 0.746641 |
| ENSG00000185112 | FAM43A | 192.5658 | 0.2012921 | 0.2845219 | 0.7618175 | 0.446169 | 0.643188 |
| ENSG00000260646 | | 8.068144 | 0.2012198 | 0.6343708 | 0.5053907 | 0.6132844 | 0.775778 |

| | | | | | | | |
|---|---|---|---|---|---|---|---|
| ENSG00000141580 | WDR45B | 9426.082 | 0.2011702 | 0.1459673 | 1.390766 | 0.1642965 | 0.332333 |
| ENSG00000276691 | | 1.068117 | 0.2011634 | 0.8016252 | 0.6606006 | 0.5088685 | NA |
| ENSG00000201388 | SNORA68B | 1.795916 | 0.2011457 | 0.7372769 | 0.6344567 | 0.5257828 | 0.710628 |
| ENSG00000224153 | LINC02054 | 0.9534896 | 0.2011109 | 0.768614 | 0.7181905 | 0.4726398 | NA |
| ENSG00000130643 | CALY | 6.627837695 | 0.201080802 | 0.555610758 | 0.49790738 | 0.61854933 | 0.779342 |
| ENSG00000255435 | | 19.98653 | 0.200954 | 0.5160586 | 0.5011094 | 0.6162941 | 0.77773 |
| ENSG00000095066 | HOOK2 | 4654.440287 | 0.200940065 | 0.205658527 | 1.01519542 | 0.31001268 | 0.510667 |
| ENSG00000187486 | KCNJ11 | 734.1511 | 0.2009276 | 0.1520337 | 1.340665 | 0.1800293 | 0.354206 |
| ENSG00000168079 | SCARA5 | 0.8426814 | 0.20092 | 0.7984055 | 1.00796 | 0.3134737 | NA |
| ENSG00000181804 | SLC9A9 | 2.241419 | 0.2008889 | 0.6943966 | 0.5444946 | 0.5861011 | 0.755855 |
| ENSG00000278768 | BACE1-AS | 151.4936 | 0.2007491 | 0.2628482 | 0.8075373 | 0.419357 | 0.619304 |
| ENSG00000266992 | DHX40P1 | 1.73908 | 0.2006921 | 0.7242175 | 0.5915445 | 0.5541556 | 0.73236 |
| ENSG00000237897 | | 6.387168 | 0.2006729 | 0.5209691 | 0.499038 | 0.6177526 | 0.778824 |
| ENSG00000232756 | | 0.9873515 | 0.2006359 | 0.809514 | 0.807521 | 0.4193664 | NA |
| ENSG00000177238 | TRIM72 | 1.748904 | 0.2005816 | 0.6982508 | 0.5564542 | 0.5779004 | 0.749645 |
| ENSG00000131263 | RLIM | 4883.071551 | 0.200445841 | 0.155107384 | 1.318087672 | 0.1874743 | 0.364188 |
| ENSG00000122852 | SFTPA1 | 0.818493415 | 0.20039546 | 0.792786759 | 0.772457716 | 0.43984338 | NA |
| ENSG00000110002 | VWA5A | 28.92330609 | 0.200389041 | 0.470502566 | 0.515233888 | 0.60638959 | 0.770618 |
| ENSG00000123191 | ATP7B | 1828.601451 | 0.20038873 | 0.206112692 | 1.000277321 | 0.31717632 | 0.518204 |
| ENSG00000250869 | | 1.411405 | 0.2003387 | 0.7259177 | 0.601757 | 0.5473359 | NA |
| ENSG00000188092 | GPR89B | 1007.06 | 0.2003167 | 0.1303231 | 1.550286 | 0.1210729 | 0.266997 |
| ENSG00000102034 | ELF4 | 3010.988749 | 0.20030709 | 0.234129299 | 0.893078099 | 0.37181535 | 0.57398 |
| ENSG00000183020 | AP2A2 | 3134.34 | 0.200222 | 0.1223689 | 1.654009 | 0.09812573 | 0.229539 |
| ENSG00000227525 | RPL7P6 | 4.396038 | 0.2002156 | 0.6434618 | 0.5198904 | 0.6031399 | 0.768282 |
| ENSG00000125651 | GTF2F1 | 4880.814365 | 0.200128906 | 0.136285714 | 1.489519607 | 0.13635059 | 0.290467 |
| ENSG00000250012 | | 100.5153 | 0.2000975 | 0.3028502 | 0.722706 | 0.4698605 | 0.663322 |
| ENSG00000140107 | SLC25A47 | 2.31448 | 0.2000822 | 0.6818225 | 0.5255542 | 0.599198 | 0.765684 |
| ENSG00000198976 | MIR429 | 16.69699 | 0.2000308 | 0.4325356 | 0.5498653 | 0.5824118 | 0.752998 |
| ENSG00000101473 | ACOT8 | 1237.698623 | 0.199923044 | 0.157095401 | 1.297037767 | 0.19461819 | 0.373769 |
| ENSG00000126464 | PRR12 | 2753.052415 | 0.199919927 | 0.212756029 | 0.978819036 | 0.3276694 | 0.529102 |
| ENSG00000244607 | CCDC13 | 1.844297 | 0.1998117 | 0.7219846 | 0.568051 | 0.5700004 | 0.743865 |
| ENSG00000235724 | | 19.07008 | 0.1997602 | 0.4138517 | 0.564132 | 0.5726643 | 0.745897 |
| ENSG00000085465 | OVGP1 | 124.7066488 | 0.199747618 | 0.355038855 | 0.620026517 | 0.53524033 | 0.718744 |

| | | | | | | | |
|---|---|---|---|---|---|---|---|
| ENSG00000196507 | TCEAL3 | 3552 | 0.1997389 | 0.2077047 | 0.9816846 | 0.3262552 | 0.52746 |
| ENSG00000165898 | ISCA2 | 1388.37 | 0.1996874 | 0.1662579 | 1.231905 | 0.2179846 | 0.403683 |
| ENSG00000259663 | | 16.85202 | 0.1996645 | 0.4753571 | 0.5168403 | 0.6052676 | 0.769775 |
| ENSG00000159374 | M1AP | 7.108367 | 0.1995779 | 0.5673284 | 0.4963708 | 0.6196329 | 0.780127 |
| ENSG00000269967 | | 5.011765 | 0.1995327 | 0.6004048 | 0.4942189 | 0.6211516 | 0.781145 |
| ENSG00000106771 | TMEM245 | 4954.909304 | 0.199289146 | 0.191620124 | 1.069963582 | 0.2846357 | 0.482581 |
| ENSG00000163399 | ATP1A1 | 23533.44 | 0.1991735 | 0.1309616 | 1.543956 | 0.122599 | 0.269379 |
| ENSG00000184990 | SIVA1 | 6609.157 | 0.1991656 | 0.304649 | 0.6952763 | 0.4868821 | 0.677737 |
| ENSG00000261997 | | 4.324285 | 0.1991272 | 0.6631663 | 0.5203334 | 0.6028312 | 0.767999 |
| ENSG00000151881 | TMEM267 | 1055.474 | 0.1990766 | 0.2750652 | 0.7752116 | 0.4382146 | 0.636782 |
| ENSG00000224713 | | 24.66546 | 0.1990067 | 0.5300241 | 0.4957706 | 0.6200562 | 0.780512 |
| ENSG00000102780 | DGKH | 3849.658421 | 0.198899202 | 0.286708845 | 0.749008749 | 0.45385193 | 0.649978 |
| ENSG00000166928 | MS4A14 | 1.159211 | 0.1988378 | 0.8049641 | 0.6405855 | 0.521792 | NA |
| ENSG00000269608 | OSTCP3 | 23.03038 | 0.1988157 | 0.4001321 | 0.5734792 | 0.5663203 | 0.741028 |
| ENSG00000229191 | | 0.9106107 | 0.1987753 | 0.7777067 | 0.7959119 | 0.4260832 | NA |
| ENSG00000100294 | MCAT | 1147.064822 | 0.198563862 | 0.251946164 | 0.829914793 | 0.40658696 | 0.607229 |
| ENSG00000107679 | PLEKHA1 | 1460.9689 | 0.198533648 | 0.146282921 | 1.382667128 | 0.16676696 | 0.335974 |
| ENSG00000112335 | SNX3 | 7114.857753 | 0.198423159 | 0.129865837 | 1.502914893 | 0.13286099 | 0.285205 |
| ENSG00000186912 | P2RY4 | 1.49793 | 0.198416 | 0.7568368 | 0.6608122 | 0.5087328 | NA |
| ENSG00000259467 | NDUFAF4P1 | 1.037734 | 0.1984079 | 0.7530083 | 0.6543813 | 0.5128662 | NA |
| ENSG00000228626 | | 50.19582 | 0.1983019 | 0.536651 | 0.4874143 | 0.6259648 | 0.784752 |
| ENSG00000167658 | EEF2 | 125864.4 | 0.1982559 | 0.193923 | 1.051602 | 0.2929821 | 0.491833 |
| ENSG00000279785 | | 1.041335 | 0.1981568 | 0.7406646 | 0.6163712 | 0.5376495 | NA |
| ENSG00000274818 | | 20.8869 | 0.1981058 | 0.4611658 | 0.5232832 | 0.6007772 | 0.766755 |
| ENSG00000199719 | RN7SKP74 | 6.743536 | 0.1980954 | 0.6307398 | 0.5019515 | 0.6157017 | 0.777351 |
| ENSG00000271992 | | 1.883291 | 0.1980606 | 0.7346789 | 0.6143177 | 0.5390054 | 0.721572 |
| ENSG00000216921 | FAM240C | 4.099357 | 0.1980175 | 0.639739 | 0.4950172 | 0.620588 | 0.780849 |
| ENSG00000254689 | LINC02235 | 8.847429 | 0.1979499 | 0.561484 | 0.4870559 | 0.6262187 | 0.784923 |
| ENSG00000174738 | NR1D2 | 3565.709 | 0.1979426 | 0.2487362 | 0.8313891 | 0.4057539 | 0.606663 |
| ENSG00000117408 | IPO13 | 3645.815027 | 0.197856936 | 0.162601275 | 1.23958288 | 0.21512972 | 0.400181 |
| ENSG00000085449 | WDFY1 | 1547.077811 | 0.197808726 | 0.130874128 | 1.524306467 | 0.12743217 | 0.276749 |
| ENSG00000253334 | | 1.935856 | 0.1977737 | 0.7373737 | 0.6034716 | 0.546195 | 0.726355 |
| ENSG00000197119 | SLC25A29 | 7536.933 | 0.1977578 | 0.1889047 | 1.068323 | 0.2853749 | 0.483431 |

| | | | | | | | |
|---|---|---|---|---|---|---|---|
| ENSG00000214694 | ARHGEF33 | 3.946788 | 0.1977192 | 0.6277109 | 0.4930026 | 0.6220107 | 0.781677 |
| ENSG00000272695 | GAS6-DT | 9.407751 | 0.197712 | 0.6341528 | 0.5062636 | 0.6126716 | 0.775442 |
| ENSG00000205578 | POM121B | 7.478768 | 0.1976323 | 0.6029292 | 0.4959793 | 0.619909 | 0.78038 |
| ENSG00000160161 | CILP2 | 53.37755 | 0.1975697 | 0.3881105 | 0.5845818 | 0.558829 | 0.735715 |
| ENSG00000229325 | ACAP2-IT1 | 15.91977 | 0.1975645 | 0.4911232 | 0.5083152 | 0.6112323 | 0.774461 |
| ENSG00000196152 | ZNF79 | 375.0792 | 0.1975598 | 0.2200614 | 0.9331498 | 0.3507426 | 0.552394 |
| ENSG00000149346 | SLX4IP | 777.5733 | 0.1975026 | 0.1974174 | 1.032595 | 0.3017934 | 0.50121 |
| ENSG00000223669 | | 3.459123 | 0.1974162 | 0.7588564 | 0.7109593 | 0.4771094 | 0.669324 |
| ENSG00000224834 | BTF3P9 | 12.97547 | 0.1973969 | 0.5122845 | 0.5047526 | 0.6137326 | 0.775861 |
| ENSG00000178226 | PRSS36 | 43.97896 | 0.1973532 | 0.3730213 | 0.5981738 | 0.549724 | 0.728969 |
| ENSG00000134882 | UBAC2 | 3746.26267 | 0.197347116 | 0.184786331 | 1.096178338 | 0.27300074 | 0.469655 |
| ENSG00000256167 | ATF4P4 | 87.76386 | 0.1973027 | 0.2298002 | 0.8962939 | 0.3700958 | 0.572256 |
| ENSG00000244712 | | 1.434042 | 0.1972876 | 0.7386516 | 0.6305897 | 0.5283089 | NA |
| ENSG00000139880 | CDH24 | 1659.973 | 0.1972745 | 0.1931838 | 1.044219 | 0.2963839 | 0.49543 |
| ENSG00000230817 | LINC01362 | 1.143324 | 0.197263 | 0.7721394 | 0.7106413 | 0.4773065 | NA |
| ENSG00000224031 | TCEAL3-AS1 | 2.454105 | 0.1972141 | 0.723899 | 0.5818769 | 0.5606496 | 0.736914 |
| ENSG00000261857 | MIA | 0.7532685 | 0.1971361 | 0.8012332 | 0.8766512 | 0.3806761 | NA |
| ENSG00000257740 | | 31.38101 | 0.1969645 | 0.3438723 | 0.6343136 | 0.5258762 | 0.710682 |
| ENSG00000237506 | RPSAP15 | 603.5133 | 0.1969151 | 0.2279216 | 0.8999437 | 0.3681502 | 0.570215 |
| ENSG00000119004 | CYP20A1 | 723.0823719 | 0.196839205 | 0.218619451 | 0.957732347 | 0.33819774 | 0.539474 |
| ENSG00000105996 | HOXA2 | 3.894897798 | 0.196745968 | 0.717138608 | 0.574897715 | 0.56536048 | 0.740209 |
| ENSG00000231787 | H2BP9 | 3.724437 | 0.1967115 | 0.6429663 | 0.5014105 | 0.6160823 | 0.777647 |
| ENSG00000255145 | STX17-AS1 | 28.31625 | 0.1965523 | 0.3832314 | 0.5809303 | 0.5612874 | 0.737311 |
| ENSG00000274849 | | 22.72383 | 0.1965094 | 0.4507036 | 0.5254604 | 0.5992632 | 0.76573 |
| ENSG00000214578 | HMGN2P15 | 3.911695 | 0.1964835 | 0.6773315 | 0.5247437 | 0.5997614 | 0.765998 |
| ENSG00000107447 | DNTT | 0.842912022 | 0.196461045 | 0.813343299 | 0.892819198 | 0.371954 | NA |
| ENSG00000281491 | DNAJB5-DT | 0.8727449 | 0.1964556 | 0.7910525 | 0.9378345 | 0.3483295 | NA |
| ENSG00000187010 | RHD | 9.612067 | 0.1964478 | 0.6207942 | 0.4887527 | 0.6250168 | 0.783934 |
| ENSG00000141552 | ANAPC11 | 4614.099 | 0.1964315 | 0.2085671 | 0.9825484 | 0.3258297 | 0.527091 |
| ENSG00000121653 | MAPK8IP1 | 1035.92959 | 0.19641026 | 0.172963983 | 1.160601693 | 0.24580392 | 0.437543 |
| ENSG00000248517 | LINC02437 | 0.9886751 | 0.1963926 | 0.8040816 | 0.6342119 | 0.5259425 | NA |
| ENSG00000225724 | ASH2LP3 | 1.062084 | 0.1963625 | 0.7907591 | 0.7869979 | 0.4312831 | NA |
| ENSG00000163812 | ZDHHC3 | 2937.834 | 0.1963613 | 0.1483758 | 1.296738 | 0.1947212 | 0.373892 |

| | | | | | | | |
|---|---|---|---|---|---|---|---|
| ENSG00000163263 | C1orf189 | 2.35632 | 0.1963025 | 0.7106126 | 0.5678207 | 0.5701567 | 0.743996 |
| ENSG00000223500 | RNF14P4 | 0.9257481 | 0.1962825 | 0.7820588 | 0.7128993 | 0.4759081 | NA |
| ENSG00000135390 | ATP5MC2 | 10938.1324 | 0.196211622 | 0.107789257 | 1.85507142 | 0.06358603 | 0.165955 |
| ENSG00000287576 | | 57.28049 | 0.1961738 | 0.3469025 | 0.6287515 | 0.5295118 | 0.714251 |
| ENSG00000147679 | UTP23 | 2570.216 | 0.1961241 | 0.1794695 | 1.106024 | 0.2687163 | 0.464722 |
| ENSG00000162236 | STX5 | 3348.971 | 0.196057 | 0.16578 | 1.213399 | 0.2249772 | 0.412008 |
| ENSG00000248664 | | 5.480643 | 0.1960469 | 0.6459297 | 0.5028546 | 0.6150665 | 0.776882 |
| ENSG00000253238 | | 1.6175 | 0.1960128 | 0.7128937 | 0.5552249 | 0.5787409 | 0.750113 |
| ENSG00000223974 | BNIP3P42 | 1.224423 | 0.1959888 | 0.7317002 | 0.5982315 | 0.5496855 | NA |
| ENSG00000087157 | PGS1 | 2858.448724 | 0.195968199 | 0.198096278 | 1.021053376 | 0.30722915 | 0.507446 |
| ENSG00000259899 | | 0.8808435 | 0.1959617 | 0.8081821 | 0.7866093 | 0.4315106 | NA |
| ENSG00000234616 | JRK | 2928.278 | 0.1959367 | 0.1998206 | 0.9922667 | 0.3210674 | 0.521835 |
| ENSG00000121903 | ZSCAN20 | 312.6579393 | 0.195906127 | 0.215363536 | 0.944118721 | 0.34510898 | 0.546697 |
| ENSG00000149182 | ARFGAP2 | 9265.489 | 0.1957793 | 0.1345541 | 1.475289 | 0.140135 | 0.296314 |
| ENSG00000254387 | MYL12AP1 | 6.779741 | 0.1956795 | 0.5683128 | 0.4840405 | 0.6283571 | 0.786453 |
| ENSG00000262898 | | 3.595852 | 0.1956567 | 0.6203349 | 0.4876914 | 0.6257685 | 0.78456 |
| ENSG00000282911 | DUX4L35 | 1.098258 | 0.195612 | 0.7316188 | 0.57875 | 0.5627579 | NA |
| ENSG00000151789 | ZNF385D | 6.494607 | 0.195558 | 0.717981 | 0.5896781 | 0.5554065 | 0.732922 |
| ENSG00000284031 | MIR4761 | 1.108706 | 0.1955576 | 0.7573995 | 0.658233 | 0.5103884 | NA |
| ENSG00000081277 | PKP1 | 830.6936031 | 0.195538229 | 0.565582627 | 0.468658254 | 0.63931393 | 0.79417 |
| ENSG00000177548 | RABEP2 | 2452.217 | 0.1955249 | 0.161427 | 1.210948 | 0.2259154 | 0.413328 |
| ENSG00000095485 | CWF19L1 | 1652.944301 | 0.195356509 | 0.197386184 | 1.001898692 | 0.31639252 | 0.517466 |
| ENSG00000273455 | | 2.36655 | 0.1953131 | 0.7200745 | 0.565786 | 0.5715393 | 0.745214 |
| ENSG00000092978 | GPATCH2 | 2267.076837 | 0.195272004 | 0.158176529 | 1.25955586 | 0.20782963 | 0.39041 |
| ENSG00000114127 | XRN1 | 3404.850166 | 0.195256762 | 0.285106734 | 0.723901666 | 0.46912611 | 0.662673 |
| ENSG00000123485 | HJURP | 1535.357813 | 0.195149768 | 0.525098169 | 0.484278348 | 0.62818834 | 0.786353 |
| ENSG00000231464 | | 4.590686 | 0.1951201 | 0.6037981 | 0.4859656 | 0.6269916 | 0.785493 |
| ENSG00000142065 | ZFP14 | 482.4526 | 0.1950879 | 0.2714711 | 0.7550669 | 0.4502088 | 0.646466 |
| ENSG00000120334 | CENPL | 987.7054955 | 0.195049283 | 0.346033522 | 0.62353908 | 0.53293033 | 0.716969 |
| ENSG00000152822 | GRM1 | 3.641616 | 0.1949674 | 0.6489132 | 0.5018076 | 0.6158028 | 0.777425 |
| ENSG00000286205 | | 1.963161 | 0.1948505 | 0.7083645 | 0.5425481 | 0.587441 | 0.756958 |
| ENSG00000221843 | C2orf16 | 153.5949 | 0.1947166 | 0.3506845 | 0.6139124 | 0.5392732 | 0.721616 |
| ENSG00000110047 | EHD1 | 4241.028444 | 0.194692716 | 0.224543869 | 0.906461828 | 0.36469146 | 0.56665 |

| | | | | | | | |
|---|---|---|---|---|---|---|---|
| ENSG00000224721 | | 1.910156 | 0.1946792 | 0.7207506 | 0.5729851 | 0.5666548 | 0.741246 |
| ENSG00000115738 | ID2 | 3189.383684 | 0.194638703 | 0.527538282 | 0.480908558 | 0.63058149 | 0.788014 |
| ENSG00000266235 | MIR3176 | 15.14863 | 0.1945397 | 0.5295524 | 0.4889101 | 0.6249053 | 0.783831 |
| ENSG00000198740 | ZNF652 | 2527.068 | 0.1945388 | 0.1369083 | 1.427855 | 0.1533337 | 0.316176 |
| ENSG00000124466 | LYPD3 | 1422.761677 | 0.194489999 | 0.307532441 | 0.676908615 | 0.49846394 | 0.686842 |
| ENSG00000185340 | GAS2L1 | 1859.722 | 0.1944413 | 0.3237333 | 0.6639544 | 0.5067195 | 0.693256 |
| ENSG00000223768 | LINC00205 | 3270.128 | 0.1944316 | 0.2432983 | 0.8367919 | 0.4027095 | 0.603678 |
| ENSG00000237212 | | 1.591441 | 0.1944254 | 0.7773998 | 0.7661421 | 0.4435918 | 0.641124 |
| ENSG00000258256 | | 0.9359296 | 0.1943953 | 0.7640785 | 0.6752379 | 0.4995246 | NA |
| ENSG00000287143 | | 0.8447605 | 0.1943372 | 0.8018648 | 0.6566293 | 0.5114193 | NA |
| ENSG00000286018 | | 64.78676 | 0.1943355 | 0.2801575 | 0.7404384 | 0.459034 | 0.654682 |
| ENSG00000259474 | | 0.9612088 | 0.1943165 | 0.7791458 | 0.7737444 | 0.4390819 | NA |
| ENSG00000139218 | SCAF11 | 7436.251876 | 0.194316238 | 0.174699717 | 1.136618031 | 0.255698 | 0.449166 |
| ENSG00000233602 | ERI3-IT1 | 4.042598 | 0.1942514 | 0.5918907 | 0.4805222 | 0.6308561 | 0.788244 |
| ENSG00000234062 | | 143.6432 | 0.1942228 | 0.2381128 | 0.8532436 | 0.3935242 | 0.595147 |
| ENSG00000248583 | | 14.72245 | 0.1941792 | 0.4788584 | 0.5028096 | 0.6150981 | 0.776885 |
| ENSG00000268723 | | 5.777568 | 0.1940151 | 0.5762741 | 0.4769911 | 0.6333685 | 0.789913 |
| ENSG00000075891 | PAX2 | 1.945270327 | 0.193872003 | 0.719265778 | 0.57478516 | 0.56543661 | 0.740236 |
| ENSG00000229846 | | 3.004552 | 0.1938215 | 0.6996115 | 0.53102 | 0.595405 | 0.762682 |
| ENSG00000229873 | OGFR-AS1 | 18.81893 | 0.1936409 | 0.3796371 | 0.5826205 | 0.5601488 | 0.736398 |
| ENSG00000101474 | APMAP | 8722.59902 | 0.193536715 | 0.115812666 | 1.688196642 | 0.09137349 | 0.217985 |
| ENSG00000083807 | SLC27A5 | 899.0718215 | 0.193462929 | 0.212955748 | 0.941220529 | 0.34659186 | 0.548164 |
| ENSG00000185904 | LINC00839 | 52.48344 | 0.1934158 | 0.7951401 | 0.4449387 | 0.6563641 | 0.806399 |
| ENSG00000235946 | H2BW3P | 9.698822 | 0.1933853 | 0.4879503 | 0.4960419 | 0.6198649 | 0.78038 |
| ENSG00000284686 | | 11.7509 | 0.1933124 | 0.4459769 | 0.5246689 | 0.5998134 | 0.766028 |
| ENSG00000285928 | | 2.352135 | 0.1932259 | 0.6700955 | 0.5068404 | 0.6122668 | 0.775077 |
| ENSG00000242797 | GLYCTK-AS1 | 4.408259 | 0.1932058 | 0.6937828 | 0.5414217 | 0.5882169 | 0.75748 |
| ENSG00000276166 | | 3.83624 | 0.1931582 | 0.6607353 | 0.5152941 | 0.6063475 | 0.770618 |
| ENSG00000221091 | MIR1302-5 | 0.8853349 | 0.1931168 | 0.7604777 | 0.6642864 | 0.506507 | NA |
| ENSG00000286602 | | 5.737203 | 0.1930758 | 0.5675715 | 0.4798493 | 0.6313346 | 0.788593 |
| ENSG00000143889 | HNRNPLL | 3188.594 | 0.192966 | 0.1307929 | 1.495978 | 0.1346593 | 0.287879 |
| ENSG00000217835 | RBMXP5 | 64.47462 | 0.1929503 | 0.3150654 | 0.6681503 | 0.5040377 | 0.691184 |
| ENSG00000171017 | LRRC8E | 99.83624 | 0.1928615 | 0.2481615 | 0.8164549 | 0.41424 | 0.614446 |

| | | | | | | | |
|---|---|---|---|---|---|---|---|
| ENSG00000144283 | PKP4 | 7031.051 | 0.192733 | 0.1381468 | 1.398796 | 0.1618742 | 0.329087 |
| ENSG00000240286 | MEAF6P1 | 3.880674 | 0.1927053 | 0.5885567 | 0.47564 | 0.6343309 | 0.790553 |
| ENSG00000234332 | BCAS2P2 | 521.0729 | 0.192691 | 0.1851249 | 1.071025 | 0.284158 | 0.482076 |
| ENSG00000166012 | TAF1D | 4358.078 | 0.1926801 | 0.1554643 | 1.285458 | 0.1986322 | 0.378763 |
| ENSG00000172534 | HCFC1 | 7167.106 | 0.1925841 | 0.2424 | 0.8320906 | 0.4053578 | 0.606143 |
| ENSG00000115685 | PPP1R7 | 4168.841404 | 0.192582904 | 0.113108096 | 1.712575682 | 0.08679063 | 0.209841 |
| ENSG00000105738 | SIPA1L3 | 2599.644396 | 0.192554335 | 0.194828025 | 1.016916048 | 0.30919337 | 0.509708 |
| ENSG00000162368 | CMPK1 | 6096.919 | 0.192442 | 0.1226705 | 1.586922 | 0.1125304 | 0.253332 |
| ENSG00000244509 | APOBEC3C | 2.082244 | 0.1924385 | 0.7603682 | 0.674481 | 0.5000056 | 0.688145 |
| ENSG00000187954 | CYHR1 | 6152.363 | 0.1924049 | 0.2159665 | 0.9249751 | 0.3549789 | 0.55692 |
| ENSG00000140386 | SCAPER | 824.1247 | 0.1923436 | 0.2433554 | 0.8256032 | 0.4090292 | 0.609694 |
| ENSG00000196505 | GDAP2 | 1415.085 | 0.1921754 | 0.1285369 | 1.488722 | 0.1365606 | 0.290727 |
| ENSG00000086619 | ERO1B | 707.1350346 | 0.192124601 | 0.249392626 | 0.793354082 | 0.42757156 | 0.627044 |
| ENSG00000268670 | | 28.4194 | 0.1921087 | 0.4811076 | 0.4987864 | 0.6179299 | 0.778906 |
| ENSG00000259485 | LINC02253 | 180.6295 | 0.1920156 | 0.2229374 | 0.893091 | 0.3718085 | 0.57398 |
| ENSG00000070061 | ELP1 | 5067.115469 | 0.19200057 | 0.128565975 | 1.519995392 | 0.12851213 | 0.278422 |
| ENSG00000170633 | RNF34 | 1767.501 | 0.1918923 | 0.1476049 | 1.327657 | 0.1842913 | 0.359609 |
| ENSG00000115297 | TLX2 | 2.194175807 | 0.191731718 | 0.701028279 | 0.526090127 | 0.59882559 | 0.765613 |
| ENSG00000250796 | | 3.613606 | 0.191691 | 0.6604454 | 0.4964128 | 0.6196032 | 0.780127 |
| ENSG00000278012 | | 0.841065 | 0.1916724 | 0.8021929 | 0.6362326 | 0.5246248 | NA |
| ENSG00000086666 | ZFAND6 | 3969.918969 | 0.191660671 | 0.138921406 | 1.369365423 | 0.17088508 | 0.341599 |
| ENSG00000080603 | SRCAP | 408.9796327 | 0.191597672 | 0.24798197 | 0.809482129 | 0.41823788 | 0.618029 |
| ENSG00000167487 | KLHL26 | 431.6324 | 0.1915581 | 0.1907378 | 1.034712 | 0.3008032 | 0.500037 |
| ENSG00000244257 | PKD1P1 | 11.26174 | 0.1915103 | 0.5220195 | 0.481233 | 0.6303509 | 0.787909 |
| ENSG00000225578 | NCBP2-AS1 | 34.99161 | 0.1913225 | 0.3989528 | 0.5506598 | 0.5818669 | 0.75244 |
| ENSG00000019995 | ZRANB1 | 1366.372803 | 0.19129358 | 0.153834904 | 1.258685935 | 0.20814379 | 0.390853 |
| ENSG00000277744 | | 21.16298 | 0.19118 | 0.419378 | 0.5327039 | 0.5942386 | 0.761701 |
| ENSG00000130119 | GNL3L | 3690.835824 | 0.191176136 | 0.186916534 | 1.052247169 | 0.29268616 | 0.491554 |
| ENSG00000254762 | | 1.041897 | 0.1911484 | 0.7712746 | 0.8044367 | 0.4211448 | NA |
| ENSG00000104695 | PPP2CB | 2860.918752 | 0.191116825 | 0.094236109 | 2.042337393 | 0.04111808 | 0.119591 |
| ENSG00000103066 | PLA2G15 | 1241.502648 | 0.191092341 | 0.153154626 | 1.270864802 | 0.20377675 | 0.385378 |
| ENSG00000251464 | RPL7L1P13 | 3.01739 | 0.1910855 | 0.6621302 | 0.4886794 | 0.6250687 | 0.783962 |
| ENSG00000232860 | SMG7-AS1 | 95.23683 | 0.1910478 | 0.3019538 | 0.6853772 | 0.4931059 | 0.683001 |

| | | | | | | | |
|---|---|---|---|---|---|---|---|
| ENSG00000126775 | ATG14 | 2149.162344 | 0.191035635 | 0.183332831 | 1.089511053 | 0.27592858 | 0.472825 |
| ENSG00000247746 | USP51 | 276.0844 | 0.191005 | 0.1935433 | 1.016033 | 0.3096138 | 0.510274 |
| ENSG00000079999 | KEAP1 | 5047.017187 | 0.19093823 | 0.223226234 | 0.865564141 | 0.38672923 | 0.589119 |
| ENSG00000204936 | CD177 | 2.228134 | 0.1909073 | 0.7684405 | 0.6500584 | 0.5156545 | 0.700968 |
| ENSG00000206228 | HNRNPA1P4 | 18.05681 | 0.1908801 | 0.441225 | 0.5194242 | 0.603465 | 0.768585 |
| ENSG00000287808 | | 32.51324 | 0.1908487 | 0.368107 | 0.587007 | 0.557199 | 0.734555 |
| ENSG00000285849 | | 3.966016 | 0.1907034 | 0.6097707 | 0.4632754 | 0.643167 | 0.796658 |
| ENSG00000253182 | | 1.463135 | 0.1906094 | 0.8040781 | 0.5969419 | 0.5505462 | NA |
| ENSG00000260317 | | 26.8722 | 0.1906024 | 0.4628055 | 0.5023993 | 0.6153867 | 0.777103 |
| ENSG00000223821 | NA | 1.551989 | 0.1904921 | 0.7602216 | 0.6636692 | 0.506902 | NA |
| ENSG00000252061 | RNU6-415P | 1.904192 | 0.1904788 | 0.7539466 | 0.7101659 | 0.4776013 | 0.669704 |
| ENSG00000249752 | | 2.545503 | 0.1904674 | 0.7404901 | 0.6195176 | 0.5355754 | 0.71889 |
| ENSG00000278917 | | 9.004213 | 0.1904332 | 0.4843372 | 0.4916064 | 0.6229976 | 0.782399 |
| ENSG00000121879 | PIK3CA | 1689.440986 | 0.190424123 | 0.219411539 | 0.904230329 | 0.36587328 | 0.567807 |
| ENSG00000241749 | RPSAP52 | 0.9157484 | 0.1904043 | 0.8032637 | 0.6251658 | 0.5318622 | NA |
| ENSG00000273375 | | 3.59672 | 0.1903961 | 0.697635 | 0.5222701 | 0.6014823 | 0.76709 |
| ENSG00000100219 | XBP1 | 47365.37533 | 0.190361715 | 0.155635214 | 1.244537386 | 0.21330185 | 0.398063 |
| ENSG00000174007 | CEP19 | 106.6487 | 0.1903242 | 0.3001495 | 0.6832654 | 0.4944392 | 0.684048 |
| ENSG00000287125 | | 32.70435 | 0.1902859 | 0.3864383 | 0.560787 | 0.5749427 | 0.747232 |
| ENSG00000101255 | TRIB3 | 6553.441592 | 0.190163343 | 0.293618939 | 0.69892626 | 0.48459812 | 0.675584 |
| ENSG00000196664 | TLR7 | 1.064713 | 0.1900813 | 0.8033521 | 0.6162528 | 0.5377277 | NA |
| ENSG00000144357 | UBR3 | 3632.283 | 0.1900595 | 0.2104173 | 0.940507 | 0.3469575 | 0.548568 |
| ENSG00000234899 | SOX9-AS1 | 6.367704 | 0.1900313 | 0.5683114 | 0.4681079 | 0.6397074 | 0.79435 |
| ENSG00000213859 | KCTD11 | 1025.755 | 0.19003 | 0.1881726 | 1.029592 | 0.3032013 | 0.502856 |
| ENSG00000198551 | ZNF627 | 871.6098 | 0.1899907 | 0.2005437 | 0.9760404 | 0.3290445 | 0.530227 |
| ENSG00000206561 | COLQ | 123.1283 | 0.1899791 | 0.4542719 | 0.5012725 | 0.6161794 | 0.777672 |
| ENSG00000258558 | | 1.511678 | 0.1899707 | 0.7755273 | 0.7704055 | 0.4410594 | NA |
| ENSG00000207034 | | 1.017479 | 0.1899478 | 0.7471177 | 0.6246248 | 0.5322173 | NA |
| ENSG00000239510 | RPL9P5 | 1.577068 | 0.1898793 | 0.7249646 | 0.5698482 | 0.5687807 | 0.742784 |
| ENSG00000119559 | C19orf25 | 1420.288165 | 0.18974715 | 0.166057657 | 1.166169846 | 0.24354578 | 0.434754 |
| ENSG00000186088 | GSAP | 385.3334 | 0.1897235 | 0.3396378 | 0.6130662 | 0.5398326 | 0.722087 |
| ENSG00000270106 | TSNAX-DISC1 | 1.394151 | 0.1896464 | 0.7677141 | 0.6322893 | 0.5271979 | NA |
| ENSG00000155111 | CDK19 | 2503.331 | 0.1896122 | 0.1784601 | 1.087001 | 0.2770365 | 0.473987 |

| | | | | | | | |
|---|---|---|---|---|---|---|---|
| ENSG00000259696 | | 5.841386 | 0.1895814 | 0.5683225 | 0.4662158 | 0.6410611 | 0.795359 |
| ENSG00000231728 | TMSB15B-AS1 | 57.84661 | 0.1895118 | 0.3395993 | 0.613947 | 0.5392504 | 0.721616 |
| ENSG00000136935 | GOLGA1 | 2764.016346 | 0.18949147 | 0.174794963 | 1.107814537 | 0.26794192 | 0.463846 |
| ENSG00000163946 | TASOR | 3959.737 | 0.1894396 | 0.1665876 | 1.160543 | 0.2458278 | 0.437556 |
| ENSG00000263786 | | 28.0736 | 0.1893766 | 0.3468216 | 0.6089156 | 0.5425804 | 0.723795 |
| ENSG00000225235 | INTS6L-AS1 | 2.097953 | 0.1893228 | 0.6840482 | 0.5158495 | 0.6059595 | 0.770322 |
| ENSG00000130779 | CLIP1 | 5451.333677 | 0.189288115 | 0.221909709 | 0.87181182 | 0.38331105 | 0.585736 |
| ENSG00000169379 | ARL13B | 564.2725 | 0.189235 | 0.1866381 | 1.036422 | 0.3000053 | 0.499332 |
| ENSG00000250273 | PSMC1P5 | 0.8325701 | 0.1891926 | 0.812811 | 0.9693806 | 0.3323553 | NA |
| ENSG00000213280 | | 34.40279 | 0.1890157 | 0.3811397 | 0.5639845 | 0.5727647 | 0.745897 |
| ENSG00000271533 | | 318.4127 | 0.1890071 | 0.3075729 | 0.6627825 | 0.5074699 | 0.693818 |
| ENSG00000198838 | RYR3 | 72.73095 | 0.188944 | 0.322098 | 0.6496571 | 0.5159138 | 0.701213 |
| ENSG00000165752 | STK32C | 1158.649 | 0.1888258 | 0.1827821 | 1.05345 | 0.2921348 | 0.490938 |
| ENSG00000127328 | RAB3IP | 2121.614613 | 0.188785781 | 0.135771793 | 1.380890199 | 0.16731272 | 0.336742 |
| ENSG00000261315 | LARP4P | 4.483489 | 0.1887271 | 0.621095 | 0.4715321 | 0.6372608 | 0.792742 |
| ENSG00000240053 | LY6G5B | 255.2348 | 0.1885347 | 0.2216546 | 0.8898171 | 0.3735641 | 0.575609 |
| ENSG00000232380 | ZDHHC20P4 | 195.8569 | 0.1884386 | 0.3771912 | 0.5756304 | 0.564865 | 0.739998 |
| ENSG00000215241 | LINC02449 | 47.8103 | 0.1882431 | 0.3720595 | 0.5695718 | 0.5689681 | 0.742992 |
| ENSG00000279409 | | 1.023508 | 0.1881964 | 0.7802739 | 0.745275 | 0.4561055 | NA |
| ENSG00000197497 | ZNF665 | 0.8560302 | 0.1881806 | 0.7566233 | 0.652043 | 0.5143734 | NA |
| ENSG00000123815 | COQ8B | 1548.158381 | 0.188160689 | 0.168130948 | 1.146978597 | 0.25139047 | 0.444125 |
| ENSG00000165923 | AGBL2 | 142.6001 | 0.1881266 | 0.2924729 | 0.6968041 | 0.4859254 | 0.676902 |
| ENSG00000197670 | | 480.65 | 0.188115 | 0.3465558 | 0.6023086 | 0.5469688 | 0.726845 |
| ENSG00000236494 | | 1.470826 | 0.188094 | 0.7594275 | 0.67309 | 0.50089 | NA |
| ENSG00000157450 | RNF111 | 1491.167 | 0.1880378 | 0.1843161 | 1.025196 | 0.3052708 | 0.505059 |
| ENSG00000233846 | SELENOTP1 | 505.073 | 0.188017 | 0.1467867 | 1.30442 | 0.1920903 | 0.370313 |
| ENSG00000273432 | | 4.667512 | 0.187989 | 0.6227913 | 0.4696668 | 0.6385931 | 0.793879 |
| ENSG00000100330 | MTMR3 | 512.0092777 | 0.187982193 | 0.219802734 | 0.887683957 | 0.37471078 | 0.576907 |
| ENSG00000154144 | TBRG1 | 956.8497 | 0.187979 | 0.1399719 | 1.358364 | 0.1743484 | 0.34626 |
| ENSG00000170516 | COX7B2 | 70.11582 | 0.1879757 | 0.7927537 | 0.4442538 | 0.6568591 | 0.806697 |
| ENSG00000239636 | TFAP2E-AS1 | 5.876164 | 0.1879703 | 0.5978926 | 0.4636252 | 0.6429163 | 0.796464 |
| ENSG00000263326 | | 16.34038 | 0.1878568 | 0.5907266 | 0.4612468 | 0.6446216 | 0.79779 |
| ENSG00000267082 | | 20.14711 | 0.1878491 | 0.4472671 | 0.505484 | 0.6132189 | 0.775778 |

| | | | | | | | |
|---|---|---|---|---|---|---|---|
| ENSG00000260645 | | 41.30285 | 0.1878096 | 0.3765945 | 0.5645805 | 0.5723591 | 0.745771 |
| ENSG00000188647 | PTAR1 | 1936.68 | 0.1877063 | 0.2558821 | 0.7733278 | 0.4393284 | 0.637842 |
| ENSG00000160712 | IL6R | 540.9065 | 0.187585 | 0.5306885 | 0.461065 | 0.644752 | 0.797814 |
| ENSG00000234199 | LINC01191 | 0.823728 | 0.1875647 | 0.7549258 | 0.6329539 | 0.5267638 | NA |
| ENSG00000234801 | MORF4 | 6.098639 | 0.1875596 | 0.5505596 | 0.4669719 | 0.64052 | 0.794898 |
| ENSG00000014164 | ZC3H3 | 1728.053846 | 0.187505961 | 0.152080518 | 1.256524844 | 0.20892574 | 0.391878 |
| ENSG00000044574 | HSPA5 | 26718.06905 | 0.187385591 | 0.220205973 | 0.883193229 | 0.37713188 | 0.579158 |
| ENSG00000186132 | C2orf76 | 431.89 | 0.1873206 | 0.2394451 | 0.8190654 | 0.4127491 | 0.612884 |
| ENSG00000231636 | AGBL5-AS1 | 1.474 | 0.1871728 | 0.7247136 | 0.5584518 | 0.5765359 | NA |
| ENSG00000214212 | C19orf38 | 55.30724 | 0.187172 | 0.3922571 | 0.5396623 | 0.5894299 | 0.758307 |
| ENSG00000198930 | CSAG1 | 19.52996 | 0.1871192 | 0.7931662 | 0.4345656 | 0.6638778 | 0.812014 |
| ENSG00000280401 | | 5.527467 | 0.1870937 | 0.6064918 | 0.4702789 | 0.6381557 | 0.793572 |
| ENSG00000228763 | LIMS1-AS1 | 2.483924 | 0.1870647 | 0.6774054 | 0.5054212 | 0.6132631 | 0.775778 |
| ENSG00000249359 | | 1.021405 | 0.1870399 | 0.7674816 | 0.6390754 | 0.5227739 | NA |
| ENSG00000239254 | | 2.817863 | 0.1870282 | 0.6836503 | 0.5073573 | 0.6119041 | 0.77473 |
| ENSG00000180185 | FAHD1 | 2311.417 | 0.1869702 | 0.1609249 | 1.183362 | 0.2366656 | 0.426332 |
| ENSG00000140941 | MAP1LC3B | 6007.089 | 0.1869098 | 0.2189322 | 0.8859288 | 0.3756559 | 0.577793 |
| ENSG00000278771 | RN7SL3 | 111.4894 | 0.1867115 | 0.4926908 | 0.476664 | 0.6336014 | 0.790039 |
| ENSG00000282950 | | 8.173296 | 0.1866888 | 0.5836399 | 0.4564604 | 0.648059 | 0.800529 |
| ENSG00000170458 | CD14 | 26.29082 | 0.1866173 | 0.5609623 | 0.4676425 | 0.6400402 | 0.794526 |
| ENSG00000239804 | | 1.315033 | 0.1866053 | 0.7556029 | 0.6197393 | 0.5354294 | NA |
| ENSG00000165219 | GAPVD1 | 4676.01 | 0.1865398 | 0.1584681 | 1.200154 | 0.2300796 | 0.418382 |
| ENSG00000243648 | | 1.322666 | 0.1865337 | 0.7648102 | 0.6784971 | 0.4974565 | NA |
| ENSG00000151445 | VIPAS39 | 2498.38 | 0.1865155 | 0.124771 | 1.502668 | 0.1329246 | 0.285296 |
| ENSG00000249825 | THBS4-AS1 | 12.63043 | 0.1864961 | 0.5829181 | 0.4565763 | 0.6479756 | 0.800524 |
| ENSG00000101542 | CDH20 | 1.054744832 | 0.186465243 | 0.766316001 | 0.63457572 | 0.52570517 | NA |
| ENSG00000262766 | | 82.6132 | 0.1864169 | 0.3029438 | 0.6631129 | 0.5072582 | 0.693672 |
| ENSG00000278864 | | 646.606 | 0.1864135 | 0.3246179 | 0.630162 | 0.5285886 | 0.71344 |
| ENSG00000279672 | | 108.0415 | 0.1863938 | 0.2704888 | 0.7318902 | 0.4642356 | 0.658249 |
| ENSG00000239884 | RN7SL608P | 5.823361 | 0.18639 | 0.604615 | 0.4650071 | 0.6419264 | 0.795789 |
| ENSG00000271824 | SMIM32 | 8.591639 | 0.1863664 | 0.6585426 | 0.4950312 | 0.6205781 | 0.780849 |
| ENSG00000233205 | WBP1P2 | 110.9461 | 0.1862836 | 0.2284346 | 0.8481304 | 0.3963653 | 0.597607 |
| ENSG00000099991 | CABIN1 | 5196.132313 | 0.18623543 | 0.123638417 | 1.587140229 | 0.11248089 | 0.253252 |

| | | | | | | | |
|---|---|---|---|---|---|---|---|
| ENSG00000183798 | EMILIN3 | 4.087932 | 0.1861997 | 0.6169201 | 0.4610639 | 0.6447527 | 0.797814 |
| ENSG00000235665 | LINC00298 | 10.49641 | 0.1861324 | 0.7946731 | 0.4135794 | 0.6791822 | 0.821767 |
| ENSG00000258013 | RPL3P13 | 0.8674219 | 0.1859957 | 0.7827672 | 0.7218684 | 0.4703754 | NA |
| ENSG00000204054 | LINC00963 | 2557.947 | 0.1859795 | 0.1976599 | 0.9612605 | 0.3364212 | 0.537485 |
| ENSG00000279852 | | 3.314295 | 0.1859319 | 0.6386613 | 0.4716898 | 0.6371482 | 0.792676 |
| ENSG00000213672 | NCKIPSD | 1646.639 | 0.1859184 | 0.1490644 | 1.268661 | 0.2045621 | 0.386529 |
| ENSG00000242498 | ARPIN | 1318.304 | 0.1858481 | 0.3004615 | 0.6681785 | 0.5040196 | 0.691184 |
| ENSG00000227403 | LINC01806 | 121.8398 | 0.1858278 | 0.3223171 | 0.632292 | 0.5271961 | 0.712031 |
| ENSG00000168297 | PXK | 502.4831 | 0.1858277 | 0.2865607 | 0.7025975 | 0.4823066 | 0.673769 |
| ENSG00000149634 | SPATA25 | 99.50151 | 0.1858228 | 0.3345715 | 0.6160323 | 0.5378732 | 0.720918 |
| ENSG00000276045 | ORAI1 | 1865.461 | 0.1857747 | 0.1844233 | 1.043883 | 0.2965396 | 0.495582 |
| ENSG00000199536 | RNU6-315P | 0.9682874 | 0.185773 | 0.7518944 | 0.6317333 | 0.5275612 | NA |
| ENSG00000168126 | OR2W6P | 1.774911 | 0.1856822 | 0.6993013 | 0.5150998 | 0.6064833 | 0.77062 |
| ENSG00000241030 | RPL29P24 | 3.300246 | 0.1856567 | 0.6416376 | 0.4685967 | 0.639358 | 0.79417 |
| ENSG00000169902 | TPST1 | 266.3884 | 0.1855633 | 0.1845309 | 1.045384 | 0.2958455 | 0.494858 |
| ENSG00000135404 | CD63 | 14895.48024 | 0.18553834 | 0.172642144 | 1.129986576 | 0.25848188 | 0.452699 |
| ENSG00000147535 | PLPP5 | 1326.304 | 0.1854963 | 0.1487819 | 1.265902 | 0.2055481 | 0.387512 |
| ENSG00000077254 | USP33 | 4173.196772 | 0.185300888 | 0.12870397 | 1.429988383 | 0.15272035 | 0.315156 |
| ENSG00000159247 | TUBBP5 | 34.16299 | 0.185271 | 0.4076605 | 0.5328493 | 0.5941379 | 0.761696 |
| ENSG00000100106 | TRIOBP | 2300.031978 | 0.185262207 | 0.137693218 | 1.36524452 | 0.17217621 | 0.343203 |
| ENSG00000225146 | | 4.295938 | 0.1852029 | 0.663046 | 0.4852427 | 0.6275042 | 0.785867 |
| ENSG00000155330 | C16orf87 | 518.7226 | 0.1851984 | 0.1547887 | 1.21816 | 0.2231634 | 0.409817 |
| ENSG00000228007 | | 3.640407 | 0.1850766 | 0.6834889 | 0.5046327 | 0.6138168 | 0.775893 |
| ENSG00000217239 | | 6.340967 | 0.1850561 | 0.6029579 | 0.4639614 | 0.6426754 | 0.796309 |
| ENSG00000231264 | ARAFP1 | 20.37651 | 0.1849783 | 0.7928068 | 0.4286107 | 0.6682065 | 0.815124 |
| ENSG00000104763 | ASAH1 | 6216.433902 | 0.184957741 | 0.241758951 | 0.8014942 | 0.4228456 | 0.622794 |
| ENSG00000267002 | | 1084.531 | 0.1849547 | 0.379728 | 0.5488597 | 0.5831018 | 0.75345 |
| ENSG00000164181 | ELOVL7 | 1878.198 | 0.1849535 | 0.2387866 | 0.8100336 | 0.4179209 | 0.617698 |
| ENSG00000189433 | GJB4 | 4.241379 | 0.1849356 | 0.6649271 | 0.4776837 | 0.6328754 | 0.789647 |
| ENSG00000119862 | LGALSL | 975.2110594 | 0.184901413 | 0.238705626 | 0.833592728 | 0.40451053 | 0.605353 |
| ENSG00000105397 | TYK2 | 6959.096934 | 0.184869834 | 0.122322085 | 1.489103192 | 0.1364602 | 0.290653 |
| ENSG00000186019 | | 8.842209 | 0.1848605 | 0.5481145 | 0.4630779 | 0.6433085 | 0.796684 |
| ENSG00000288579 | | 3.527191 | 0.1847231 | 0.6976004 | 0.5258808 | 0.598971 | 0.765663 |

| | | | | | | | |
|---|---|---|---|---|---|---|---|
| ENSG00000227392 | HPN-AS1 | 0.8693326 | 0.1847084 | 0.8012107 | 0.6195218 | 0.5355727 | NA |
| ENSG00000253394 | LINC00534 | 33.46152 | 0.1847075 | 0.4345195 | 0.5023194 | 0.6154429 | 0.777135 |
| ENSG00000082196 | C1QTNF3 | 177.4456134 | 0.184669037 | 0.35899774 | 0.57008322 | 0.56862126 | 0.742678 |
| ENSG00000277144 | | 1.929013 | 0.1846498 | 0.7558191 | 0.7054123 | 0.4805537 | 0.672168 |
| ENSG00000102699 | PARP4 | 5228.602476 | 0.184640427 | 0.231730562 | 0.830682318 | 0.40615312 | 0.606786 |
| ENSG00000283689 | | 38.34429 | 0.1846059 | 0.3971585 | 0.5353153 | 0.5924319 | 0.760659 |
| ENSG00000144909 | OSBPL11 | 1754.518 | 0.1846056 | 0.1663682 | 1.136282 | 0.2558388 | 0.449354 |
| ENSG00000265355 | MIR3136 | 1.304204 | 0.1844491 | 0.7466018 | 0.6126129 | 0.5401323 | NA |
| ENSG00000217004 | | 0.7817106 | 0.1844181 | 0.7646656 | 0.6544779 | 0.512804 | NA |
| ENSG00000071054 | MAP4K4 | 7069.20016 | 0.184353183 | 0.21619522 | 0.890119054 | 0.37340196 | 0.575572 |
| ENSG00000171496 | OR1L8 | 1.087623 | 0.1842964 | 0.7610206 | 0.6754585 | 0.4993845 | NA |
| ENSG00000118816 | CCNI | 12705.02436 | 0.184272182 | 0.137594069 | 1.348874504 | 0.17737728 | 0.35064 |
| ENSG00000262339 | | 0.919548 | 0.184143 | 0.7720938 | 0.7907939 | 0.4290643 | NA |
| ENSG00000279530 | | 63.97129 | 0.1840686 | 0.509506 | 0.4654857 | 0.6415836 | 0.795667 |
| ENSG00000178951 | ZBTB7A | 2988.721 | 0.1840299 | 0.2041715 | 0.9340842 | 0.3502605 | 0.551929 |
| ENSG00000157214 | STEAP2 | 25.43327 | 0.1839968 | 0.4406991 | 0.4862655 | 0.6267789 | 0.785291 |
| ENSG00000230732 | | 24.22016 | 0.1839723 | 0.4921445 | 0.4756456 | 0.6343269 | 0.790553 |
| ENSG00000278655 | | 2.12474 | 0.1838952 | 0.7185594 | 0.5253907 | 0.5993116 | 0.765755 |
| ENSG00000198843 | SELENOT | 8317.008 | 0.1838501 | 0.1038706 | 1.763712 | 0.0777805 | 0.193081 |
| ENSG00000109083 | IFT20 | 1721.125987 | 0.183820611 | 0.228135694 | 0.819832373 | 0.41231167 | 0.612526 |
| ENSG00000143603 | KCNN3 | 2.506166 | 0.1837731 | 0.668159 | 0.4851849 | 0.6275452 | 0.785881 |
| ENSG00000226276 | | 9.915789 | 0.1837694 | 0.4982518 | 0.4659204 | 0.6412725 | 0.795423 |
| ENSG00000224280 | | 10.13785 | 0.1837629 | 0.5583971 | 0.4569459 | 0.6477099 | 0.800296 |
| ENSG00000276573 | | 1.398052 | 0.1837599 | 0.7487172 | 0.5916357 | 0.5540945 | NA |
| ENSG00000177453 | NIM1K | 30.77803 | 0.1836481 | 0.4040718 | 0.5270139 | 0.5981839 | 0.765087 |
| ENSG00000251595 | ABCA11P | 61.39022 | 0.1836421 | 0.3328336 | 0.6091322 | 0.5424368 | 0.723738 |
| ENSG00000249626 | | 161.2766 | 0.183475 | 0.4402346 | 0.4948292 | 0.6207207 | 0.780967 |
| ENSG00000183018 | SPNS2 | 585.1032 | 0.1834235 | 0.3220684 | 0.6230111 | 0.5332772 | 0.717218 |
| ENSG00000180113 | TDRD6 | 3.555632 | 0.1833532 | 0.6470889 | 0.4660507 | 0.6411792 | 0.795382 |
| ENSG00000259429 | UBE2Q2P2 | 371.7018 | 0.1833157 | 0.3252394 | 0.6205952 | 0.534866 | 0.718409 |
| ENSG00000140678 | ITGAX | 1.756195 | 0.1832762 | 0.7184143 | 0.5412896 | 0.588308 | 0.757561 |
| ENSG00000180938 | ZNF572 | 849.5582 | 0.1832657 | 0.1364874 | 1.36357 | 0.172703 | 0.343943 |
| ENSG00000235225 | RPL31P17 | 24.04669 | 0.183264 | 0.408813 | 0.5201437 | 0.6029634 | 0.768131 |

| | | | | | | | |
|---|---|---|---|---|---|---|---|
| ENSG00000286699 | | 116.1951 | 0.1832023 | 0.3164404 | 0.62982 | 0.5288123 | 0.713591 |
| ENSG00000286128 | | 0.7911407 | 0.183162 | 0.799171 | 0.6173834 | 0.5369818 | NA |
| ENSG00000223546 | LINC00630 | 337.6546 | 0.1831477 | 0.2348244 | 0.8259916 | 0.4088089 | 0.609554 |
| ENSG00000225697 | SLC26A6 | 1611.745 | 0.1831409 | 0.261731 | 0.7430284 | 0.4574645 | 0.653391 |
| ENSG00000144591 | GMPPA | 2709.956 | 0.1831352 | 0.1609576 | 1.145765 | 0.2518926 | 0.444632 |
| ENSG00000177150 | FAM210A | 1384.374 | 0.1831306 | 0.1853713 | 1.013551 | 0.3107972 | 0.511621 |
| ENSG00000259086 | KIAA1143P1 | 94.13606 | 0.1831028 | 0.2893875 | 0.6833335 | 0.4943961 | 0.684048 |
| ENSG00000232237 | ASCL5 | 65.93129 | 0.1831025 | 0.3235425 | 0.6172321 | 0.5370817 | 0.720111 |
| ENSG00000135953 | MFSD9 | 1348.526532 | 0.183072416 | 0.158036154 | 1.169370581 | 0.24225436 | 0.432878 |
| ENSG00000268049 | | 114.7807 | 0.1830112 | 0.2323977 | 0.8217701 | 0.4112077 | 0.611539 |
| ENSG00000004948 | CALCR | 63.39941899 | 0.182988608 | 0.410879363 | 0.51763427 | 0.60471347 | 0.769475 |
| ENSG00000164402 | SEPTIN8 | 8024.259 | 0.1829796 | 0.1648578 | 1.145663 | 0.2519348 | 0.444673 |
| ENSG00000131023 | LATS1 | 2418.152568 | 0.182906626 | 0.249857869 | 0.768385707 | 0.44225807 | 0.640069 |
| ENSG00000267024 | | 2.805157 | 0.1828979 | 0.7175409 | 0.5385175 | 0.5902198 | 0.758944 |
| ENSG00000055211 | GINM1 | 2565.831407 | 0.182866787 | 0.158791008 | 1.170672132 | 0.24173059 | 0.43232 |
| ENSG00000169684 | CHRNA5 | 313.3965 | 0.1828108 | 0.318972 | 0.6282908 | 0.5298134 | 0.714368 |
| ENSG00000274549 | | 2.971146 | 0.1828032 | 0.6879072 | 0.4683635 | 0.6395246 | 0.794189 |
| ENSG00000228986 | PHF10P1 | 16.30471 | 0.1827967 | 0.3850274 | 0.5417516 | 0.5879896 | 0.757298 |
| ENSG00000166548 | TK2 | 250.0734 | 0.1827614 | 0.2223133 | 0.8537155 | 0.3932627 | 0.594921 |
| ENSG00000123444 | KBTBD4 | 602.7379811 | 0.182701595 | 0.187744768 | 1.000510105 | 0.31706371 | 0.518149 |
| ENSG00000104643 | MTMR9 | 652.1487505 | 0.182680652 | 0.200127423 | 0.937607182 | 0.34844632 | 0.550162 |
| ENSG00000213641 | RPL7AP4 | 2.648401 | 0.1826386 | 0.6776908 | 0.4951622 | 0.6204857 | 0.780794 |
| ENSG00000125952 | MAX | 3227.482986 | 0.18261424 | 0.173046488 | 1.082715242 | 0.27893484 | 0.476175 |
| ENSG00000233306 | TRGV2 | 0.790981 | 0.1825855 | 0.8060679 | 0.760612 | 0.4468888 | NA |
| ENSG00000185800 | DMWD | 1573.646 | 0.1825707 | 0.1494123 | 1.250658 | 0.2110594 | 0.394956 |
| ENSG00000287892 | | 1.882567 | 0.1825341 | 0.7381001 | 0.5708365 | 0.5681105 | 0.742456 |
| ENSG00000158793 | NIT1 | 1284.378 | 0.1825319 | 0.1817799 | 1.027765 | 0.3040605 | 0.503697 |
| ENSG00000131015 | ULBP2 | 593.8658923 | 0.182407984 | 0.263454316 | 0.737874013 | 0.46059101 | 0.655646 |
| ENSG00000244693 | CTAGE8 | 31.1424 | 0.1824006 | 0.3929159 | 0.5299577 | 0.5961413 | 0.763339 |
| ENSG00000245112 | SMARCA5-AS1 | 46.50067 | 0.1823478 | 0.3351736 | 0.5973578 | 0.5502685 | 0.729281 |
| ENSG00000189241 | TSPYL1 | 5448.264 | 0.1821127 | 0.1265895 | 1.445956 | 0.1481895 | 0.308513 |
| ENSG00000158022 | TRIM63 | 1.130291 | 0.1820164 | 0.7469667 | 0.5923609 | 0.553609 | NA |
| ENSG00000187021 | PNLIPRP1 | 10.12136 | 0.1819987 | 0.5570819 | 0.4558372 | 0.648507 | 0.800734 |

| | | | | | | | |
|---|---|---|---|---|---|---|---|
| ENSG00000154642 | C21orf91 | 1121.24 | 0.1819165 | 0.1832277 | 1.014865 | 0.3101701 | 0.510797 |
| ENSG00000239265 | CLRN1-AS1 | 9.395954 | 0.1818991 | 0.5531589 | 0.4405817 | 0.6595158 | 0.808465 |
| ENSG00000286741 | | 1.727584 | 0.1818919 | 0.7312319 | 0.5621735 | 0.5739978 | 0.746641 |
| ENSG00000261924 | | 11.35468 | 0.1818586 | 0.5859862 | 0.4418327 | 0.6586103 | 0.807952 |
| ENSG00000269194 | | 4.857952 | 0.1818584 | 0.6107035 | 0.4506337 | 0.6522536 | 0.803679 |
| ENSG00000249092 | PPIAP77 | 1.378464 | 0.1818451 | 0.7743372 | 0.7207733 | 0.471049 | NA |
| ENSG00000120690 | ELF1 | 5116.704835 | 0.181831178 | 0.259216462 | 0.738162033 | 0.46041599 | 0.655646 |
| ENSG00000276997 | | 16.20132 | 0.181783 | 0.4255536 | 0.503058 | 0.6149235 | 0.776738 |
| ENSG00000167800 | TBX10 | 10.427 | 0.1816454 | 0.5574555 | 0.4587395 | 0.6464213 | 0.799272 |
| ENSG00000176473 | WDR25 | 673.8636 | 0.1816179 | 0.1927467 | 0.9770827 | 0.3285282 | 0.529749 |
| ENSG00000277595 | | 6.666894 | 0.1815926 | 0.6742382 | 0.4832351 | 0.6289288 | 0.786761 |
| ENSG00000206815 | RNU6-483P | 0.8424197 | 0.181577 | 0.7573165 | 0.6302075 | 0.5285589 | NA |
| ENSG00000282995 | FRG1EP | 88.4142 | 0.1815123 | 0.3323107 | 0.5983662 | 0.5495956 | 0.728944 |
| ENSG00000279923 | | 3.704767 | 0.1815064 | 0.6035686 | 0.4532882 | 0.6503412 | 0.802434 |
| ENSG00000250892 | | 2.543576 | 0.18145 | 0.6807426 | 0.478899 | 0.6320105 | 0.789069 |
| ENSG00000236905 | | 0.848674 | 0.1813708 | 0.7767378 | 0.749021 | 0.4538445 | NA |
| ENSG00000235045 | RPL7P8 | 4.313131 | 0.1812974 | 0.6524262 | 0.4738476 | 0.6356086 | 0.791501 |
| ENSG00000260761 | | 0.7466917 | 0.1812961 | 0.788516 | 0.7832403 | 0.433486 | NA |
| ENSG00000237852 | | 1.520695 | 0.18127 | 0.7196151 | 0.5315453 | 0.5950409 | NA |
| ENSG00000185345 | PRKN | 30.21652 | 0.1811829 | 0.356648 | 0.5750816 | 0.5652361 | 0.7402 |
| ENSG00000243404 | RPL35AP32 | 1.889111 | 0.1811824 | 0.7346151 | 0.5787916 | 0.5627298 | 0.738475 |
| ENSG00000197948 | FCHSD1 | 1997.26 | 0.1811166 | 0.2388236 | 0.7856631 | 0.4320649 | 0.631216 |
| ENSG00000279179 | | 30.94778 | 0.1810207 | 0.3940829 | 0.5326514 | 0.5942749 | 0.761701 |
| ENSG00000221867 | MAGEA3 | 89.21759 | 0.1809678 | 0.7894986 | 0.4452737 | 0.656122 | 0.806202 |
| ENSG00000180385 | EMC3-AS1 | 382.2248 | 0.1809457 | 0.3326565 | 0.6003431 | 0.5482776 | 0.727909 |
| ENSG00000167548 | KMT2D | 9917.805 | 0.1809304 | 0.3479574 | 0.5736381 | 0.5662127 | 0.740952 |
| ENSG00000267904 | | 34.69322 | 0.1808973 | 0.4248394 | 0.5002098 | 0.6169274 | 0.778339 |
| ENSG00000256940 | PPP1R14B-AS1 | 28.66655 | 0.1807857 | 0.4222717 | 0.5054455 | 0.613246 | 0.775778 |
| ENSG00000179588 | ZFPM1 | 808.8424 | 0.1807553 | 0.1963682 | 0.9691747 | 0.332458 | 0.533527 |
| ENSG00000287503 | | 0.9569011 | 0.1807125 | 0.7597458 | 0.657987 | 0.5105465 | NA |
| ENSG00000237595 | LINC01275 | 1.220089 | 0.1806894 | 0.7624831 | 0.6411413 | 0.5214309 | NA |
| ENSG00000286626 | | 3.961617 | 0.1806745 | 0.6471987 | 0.4717663 | 0.6370936 | 0.792676 |
| ENSG00000244073 | RPS4XP6 | 112.7003 | 0.180671 | 0.2573493 | 0.7411432 | 0.4586066 | 0.654433 |

| | | | | | | | |
|---|---|---|---|---|---|---|---|
| ENSG00000251186 | | 1.220851 | 0.1805558 | 0.7547714 | 0.687099 | 0.4920204 | NA |
| ENSG00000180316 | PNPLA1 | 3.489002 | 0.1805552 | 0.613011 | 0.4475227 | 0.6544977 | 0.805137 |
| ENSG00000128463 | EMC4 | 2399.967039 | 0.180528822 | 0.152428636 | 1.181285698 | 0.23748925 | 0.427121 |
| ENSG00000250266 | LINC01612 | 7.719181 | 0.1805234 | 0.6452857 | 0.4722533 | 0.636746 | 0.79251 |
| ENSG00000218510 | LINC00339 | 110.4029 | 0.180492 | 0.331764 | 0.6050953 | 0.5451157 | 0.725751 |
| ENSG00000100425 | BRD1 | 2573.688686 | 0.180465548 | 0.181864256 | 1.009763842 | 0.31260845 | 0.513648 |
| ENSG00000116539 | ASH1L | 8690.22966 | 0.180443235 | 0.293344092 | 0.658484922 | 0.51022658 | 0.696101 |
| ENSG00000171492 | LRRC8D | 2066.248 | 0.1804303 | 0.1950185 | 0.951916 | 0.3411396 | 0.542447 |
| ENSG00000125734 | GPR108 | 2742.453799 | 0.180408219 | 0.137362438 | 1.310488498 | 0.19003063 | 0.367837 |
| ENSG00000022976 | ZNF839 | 1299.620935 | 0.180398124 | 0.134096702 | 1.372580347 | 0.16988285 | 0.340265 |
| ENSG00000206728 | | 3.088891 | 0.1803942 | 0.6990072 | 0.5054381 | 0.6132512 | 0.775778 |
| ENSG00000284431 | | 3.332108 | 0.1803797 | 0.699017 | 0.5014475 | 0.6160562 | 0.777647 |
| ENSG00000206991 | RNU6-610P | 3.238398 | 0.1803609 | 0.6595408 | 0.4667513 | 0.6406778 | 0.79502 |
| ENSG00000278635 | | 22.58569 | 0.180291 | 0.462699 | 0.4786769 | 0.6321685 | 0.789178 |
| ENSG00000174977 | | 3520.179 | 0.1802638 | 0.1082157 | 1.665692 | 0.09577472 | 0.225429 |
| ENSG00000227742 | CALR4P | 128.8207 | 0.1801082 | 0.3962883 | 0.5192881 | 0.6035598 | 0.768669 |
| ENSG00000258520 | | 7.027364 | 0.1800975 | 0.5616751 | 0.4482613 | 0.6539646 | 0.804895 |
| ENSG00000196166 | C8orf86 | 1.732124 | 0.1800923 | 0.7582355 | 0.5896746 | 0.5554088 | 0.732922 |
| ENSG00000159140 | SON | 16749.45 | 0.1800664 | 0.2005684 | 0.9190704 | 0.3580588 | 0.560032 |
| ENSG00000085662 | AKR1B1 | 1.659281142 | 0.1798324 | 0.755663602 | 0.668385001 | 0.50388787 | 0.691126 |
| ENSG00000234329 | | 18.44789 | 0.1798297 | 0.5454405 | 0.4468357 | 0.6549937 | 0.80534 |
| ENSG00000215093 | EEF1A1P29 | 2.086998 | 0.1798079 | 0.7294482 | 0.5553258 | 0.5786718 | 0.750085 |
| ENSG00000165269 | AQP7 | 5.258836 | 0.1797023 | 0.6482876 | 0.4526392 | 0.6508085 | 0.802697 |
| ENSG00000011198 | ABHD5 | 1085.773184 | 0.17963448 | 0.154221186 | 1.187699684 | 0.23495175 | 0.424204 |
| ENSG00000213370 | RANP6 | 12.26866 | 0.1795896 | 0.5082978 | 0.4570475 | 0.6476369 | 0.800291 |
| ENSG00000232493 | RPL12P11 | 0.8413773 | 0.1795886 | 0.7680125 | 0.6236896 | 0.5328315 | NA |
| ENSG00000131236 | CAP1 | 7032.574697 | 0.179570406 | 0.145610578 | 1.242567197 | 0.21402736 | 0.398856 |
| ENSG00000227942 | FRMD8P1 | 3.85774 | 0.1793 | 0.6815133 | 0.4796182 | 0.6314989 | 0.788593 |
| ENSG00000246523 | FZD4-DT | 28.2146 | 0.1792382 | 0.4063218 | 0.511357 | 0.6091011 | 0.772689 |
| ENSG00000276575 | MIR6895 | 1.232439 | 0.1791951 | 0.7641582 | 0.6103037 | 0.5416606 | NA |
| ENSG00000177103 | DSCAML1 | 134.6818 | 0.1791833 | 0.2311627 | 0.8113427 | 0.4171689 | 0.617102 |
| ENSG00000073756 | PTGS2 | 51.67487459 | 0.179103094 | 0.724091399 | 0.555573758 | 0.57850228 | 0.749955 |
| ENSG00000114735 | HEMK1 | 1715.3183 | 0.179060307 | 0.274182426 | 0.690361989 | 0.48996657 | 0.680533 |

| | | | | | | | |
|---|---|---|---|---|---|---|---|
| ENSG00000129083 | COPB1 | 8795.488689 | 0.178980098 | 0.100259952 | 1.795120882 | 0.07263444 | 0.183688 |
| ENSG00000137310 | TCF19 | 1787.147036 | 0.178913041 | 0.448490353 | 0.468708601 | 0.63927794 | 0.79417 |
| ENSG00000277371 | | 4.136862 | 0.1788825 | 0.6477187 | 0.4697699 | 0.6385195 | 0.793825 |
| ENSG00000274010 | ZFP91P1 | 1.820277 | 0.178865 | 0.7143069 | 0.5144209 | 0.6069577 | 0.77103 |
| ENSG00000197054 | ZNF763 | 11.22016 | 0.1788548 | 0.6064241 | 0.4433518 | 0.6575113 | 0.807092 |
| ENSG00000120509 | PDZD11 | 2680.471047 | 0.178781008 | 0.215058042 | 0.862100551 | 0.38863219 | 0.590708 |
| ENSG00000272979 | | 2.28765 | 0.1787429 | 0.7302078 | 0.5578182 | 0.5769685 | 0.748839 |
| ENSG00000138892 | TTLL8 | 0.830000036 | 0.178725919 | 0.765513732 | 0.653037838 | 0.51373189 | NA |
| ENSG00000118518 | RNF146 | 2028.831343 | 0.178669827 | 0.201911554 | 0.911883262 | 0.36183017 | 0.56374 |
| ENSG00000287878 | | 63.69905 | 0.1786034 | 0.392393 | 0.5204915 | 0.602721 | 0.767932 |
| ENSG00000206754 | SNORD101 | 29.33907 | 0.1786001 | 0.4277887 | 0.4925557 | 0.6223265 | 0.781815 |
| ENSG00000170248 | PDCD6IP | 7853.406 | 0.1785927 | 0.1474802 | 1.227949 | 0.2194662 | 0.40555 |
| ENSG00000136859 | ANGPTL2 | 1.343806321 | 0.178570426 | 0.758980304 | 0.65421155 | 0.51297552 | NA |
| ENSG00000160613 | PCSK7 | 1247.957 | 0.1785526 | 0.1884602 | 0.9542907 | 0.3399365 | 0.541311 |
| ENSG00000155511 | GRIA1 | 1.192124 | 0.1781582 | 0.7607952 | 0.5930878 | 0.5531224 | NA |
| ENSG00000224613 | | 1.156282 | 0.1781243 | 0.7424159 | 0.57125 | 0.5678302 | NA |
| ENSG00000280149 | | 33.83758 | 0.1781001 | 0.3768013 | 0.5327413 | 0.5942127 | 0.761701 |
| ENSG00000242078 | | 9.656094 | 0.1780103 | 0.5006457 | 0.4552709 | 0.6489143 | 0.801162 |
| ENSG00000246308 | | 10.94214 | 0.1780099 | 0.5222883 | 0.447868 | 0.6542485 | 0.80498 |
| ENSG00000169612 | RAMAC | 256.1964 | 0.1779226 | 0.1987346 | 0.9262016 | 0.3543412 | 0.556288 |
| ENSG00000170881 | RNF139 | 6722.863 | 0.1777615 | 0.1376839 | 1.296454 | 0.1948191 | 0.37398 |
| ENSG00000168374 | ARF4 | 6317.568 | 0.1777332 | 0.113715 | 1.643804 | 0.1002166 | 0.233166 |
| ENSG00000108091 | CCDC6 | 8424.871027 | 0.177730882 | 0.10519713 | 1.699580507 | 0.08920986 | 0.214305 |
| ENSG00000151364 | KCTD14 | 3.306272 | 0.177684 | 0.64868 | 0.4559014 | 0.6484609 | 0.800734 |
| ENSG00000172269 | DPAGT1 | 4655.236 | 0.1776688 | 0.1385913 | 1.287726 | 0.1978413 | 0.377944 |
| ENSG00000136149 | RPL13AP25 | 364.4111663 | 0.177653346 | 0.216782069 | 0.84647149 | 0.39728977 | 0.59839 |
| ENSG00000198517 | MAFK | 2736.199 | 0.1775516 | 0.198387 | 0.9220316 | 0.3565121 | 0.558464 |
| ENSG00000232112 | TMA7 | 869.2415 | 0.1775254 | 0.3023781 | 0.6387028 | 0.5230163 | 0.708366 |
| ENSG00000282917 | | 1.517178 | 0.177519 | 0.7254451 | 0.5278177 | 0.5976259 | NA |
| ENSG00000105576 | TNPO2 | 5726.635471 | 0.177494689 | 0.130275649 | 1.368680898 | 0.17109904 | 0.34171 |
| ENSG00000270039 | | 59.47392 | 0.1774203 | 0.31077 | 0.6195266 | 0.5355695 | 0.71889 |
| ENSG00000181852 | RNF41 | 4177.074 | 0.1774067 | 0.2592837 | 0.7206316 | 0.4711362 | 0.664436 |
| ENSG00000228020 | HNRNPA1P46 | 1.325927 | 0.1773652 | 0.7596961 | 0.6162495 | 0.5377299 | NA |

| | | | | | | | |
|---|---|---|---|---|---|---|---|
| ENSG00000143319 | ISG20L2 | 4845.935 | 0.1771627 | 0.1648932 | 1.094598 | 0.2736926 | 0.47045 |
| ENSG00000188991 | SLC15A5 | 0.8406164 | 0.1771541 | 0.7995553 | 0.5895718 | 0.5554778 | NA |
| ENSG00000186073 | CDIN1 | 539.1536 | 0.1770072 | 0.185863 | 0.9784117 | 0.3278707 | 0.529299 |
| ENSG00000130733 | YIPF2 | 3733.239551 | 0.176900189 | 0.16092145 | 1.097456326 | 0.27244196 | 0.468973 |
| ENSG00000271234 | | 1.796314 | 0.1768805 | 0.7344873 | 0.5441322 | 0.5863505 | 0.756114 |
| ENSG00000254837 | | 230.321 | 0.1768706 | 0.1879269 | 0.9680306 | 0.3330291 | 0.534009 |
| ENSG00000227356 | LINC00866 | 2.368463 | 0.176869 | 0.723717 | 0.5219919 | 0.601676 | 0.7672 |
| ENSG00000263893 | | 1.297645 | 0.1768013 | 0.7840729 | 0.6907573 | 0.4897181 | NA |
| ENSG00000252821 | RNU6-388P | 0.8216153 | 0.1767755 | 0.7511307 | 0.5934601 | 0.5528733 | NA |
| ENSG00000125826 | RBCK1 | 6072.569 | 0.176753542 | 0.247608306 | 0.761316551 | 0.44646802 | 0.643407 |
| ENSG00000246982 | | 86.9346 | 0.1767177 | 0.4168996 | 0.4954035 | 0.6203153 | 0.780716 |
| ENSG00000160505 | NLRP4 | 8.310481 | 0.176667 | 0.7904783 | 0.4171929 | 0.6765373 | 0.820122 |
| ENSG00000236018 | | 25.77598 | 0.1766009 | 0.3571658 | 0.5528108 | 0.580393 | 0.751246 |
| ENSG00000166573 | GALR1 | 2.778492 | 0.1765341 | 0.6957752 | 0.4973799 | 0.6189212 | 0.779563 |
| ENSG00000198431 | TXNRD1 | 18585.94 | 0.1765341 | 0.245904 | 0.8013335 | 0.4229386 | 0.622897 |
| ENSG00000261221 | ZNF865 | 1081.585 | 0.176377 | 0.1783086 | 1.00982 | 0.3125817 | 0.513636 |
| ENSG00000231533 | | 0.8488903 | 0.1763559 | 0.7986161 | 0.6238978 | 0.5326947 | NA |
| ENSG00000197324 | LRP10 | 13796.81 | 0.1763493 | 0.1527804 | 1.185954 | 0.2356404 | 0.42515 |
| ENSG00000015475 | BID | 5067.087089 | 0.176319416 | 0.27197568 | 0.687860673 | 0.49154052 | 0.681793 |
| ENSG00000275198 | | 77.52349 | 0.1762905 | 0.3255375 | 0.5929183 | 0.5532358 | 0.731539 |
| ENSG00000225294 | OSTCP2 | 6.500472 | 0.1762704 | 0.5997272 | 0.4392668 | 0.6604683 | 0.809118 |
| ENSG00000255468 | | 254.8362 | 0.176221 | 0.2258755 | 0.8111178 | 0.417298 | 0.617189 |
| ENSG00000286976 | | 5.785707 | 0.1761082 | 0.7936182 | 0.3807657 | 0.7033771 | 0.838592 |
| ENSG00000255150 | EID3 | 24.25871 | 0.1760447 | 0.5379955 | 0.4406276 | 0.6594826 | 0.808462 |
| ENSG00000283691 | | 2.195031 | 0.1758846 | 0.7095751 | 0.5121908 | 0.6085175 | 0.772229 |
| ENSG00000269275 | | 1.444832 | 0.1758182 | 0.7347525 | 0.5443924 | 0.5861715 | NA |
| ENSG00000165695 | AK8 | 85.55621 | 0.1758094 | 0.3266482 | 0.5904292 | 0.5549029 | 0.732685 |
| ENSG00000260565 | ERVK13-1 | 1515.089 | 0.1757521 | 0.323321 | 0.5785394 | 0.5629 | 0.738569 |
| ENSG00000175592 | FOSL1 | 28.90463 | 0.1757453 | 0.4644507 | 0.4646184 | 0.6422047 | 0.795869 |
| ENSG00000267083 | KRT18P61 | 2.082831 | 0.1756873 | 0.6777309 | 0.4703073 | 0.6381355 | 0.793572 |
| ENSG00000099804 | CDC34 | 2996.2635 | 0.175674497 | 0.184035978 | 0.975632223 | 0.32924675 | 0.530328 |
| ENSG00000204065 | TCEAL5 | 1.753688 | 0.17567 | 0.7042453 | 0.5134967 | 0.6076039 | 0.771445 |
| ENSG00000248785 | HIGD1AP14 | 8.996077 | 0.1756662 | 0.5606758 | 0.4414657 | 0.6588759 | 0.808062 |

| | | | | | | | |
|---|---|---|---|---|---|---|---|
| ENSG00000160948 | VPS28 | 6843.121 | 0.175662 | 0.2058909 | 0.8797211 | 0.3790104 | 0.581103 |
| ENSG00000225083 | GRTP1-AS1 | 13.36856 | 0.1755356 | 0.5892077 | 0.4331481 | 0.6649071 | 0.812787 |
| ENSG00000221949 | LINC01465 | 22.72 | 0.1753278 | 0.4667863 | 0.4610493 | 0.6447632 | 0.797814 |
| ENSG00000076554 | TPD52 | 3708.878925 | 0.175276347 | 0.124429128 | 1.432036854 | 0.15213327 | 0.314335 |
| ENSG00000116017 | ARID3A | 3392.088723 | 0.175181919 | 0.131643104 | 1.3345398 | 0.18202701 | 0.356918 |
| ENSG00000226291 | | 1.259473 | 0.1751106 | 0.6952991 | 0.4845854 | 0.6279705 | NA |
| ENSG00000205628 | LINC01446 | 6.329292 | 0.1749721 | 0.7917259 | 0.3979652 | 0.6906558 | 0.829793 |
| ENSG00000225315 | | 0.8839293 | 0.1749387 | 0.7901985 | 0.8285545 | 0.4073565 | NA |
| ENSG00000226661 | | 0.8978239 | 0.1749245 | 0.768402 | 0.6137037 | 0.5394111 | NA |
| ENSG00000207561 | MIR635 | 63.67536 | 0.1748584 | 0.3805664 | 0.5213885 | 0.6020962 | 0.767578 |
| ENSG00000122194 | PLG | 1.07875046 | 0.174808543 | 0.799056728 | 0.586501197 | 0.55753876 | NA |
| ENSG00000265206 | | 0.9923482 | 0.1748024 | 0.7760453 | 0.7173343 | 0.4731679 | NA |
| ENSG00000250365 | | 48.95461 | 0.1746702 | 0.3969935 | 0.5048371 | 0.6136733 | 0.775861 |
| ENSG00000253352 | TUG1 | 16950.96 | 0.1745633 | 0.1880615 | 0.9581261 | 0.3379992 | 0.539325 |
| ENSG00000104679 | R3HCC1 | 1315.325985 | 0.174545374 | 0.204231219 | 0.882658079 | 0.37742104 | 0.57951 |
| ENSG00000043093 | DCUN1D1 | 2982.723832 | 0.1745442 | 0.116654049 | 1.544576384 | 0.12244876 | 0.269183 |
| ENSG00000213212 | NCLP1 | 17.66801 | 0.1745113 | 0.5071917 | 0.4430045 | 0.6577625 | 0.807247 |
| ENSG00000113811 | SELENOK | 1326.841037 | 0.174482712 | 0.193513201 | 0.920228343 | 0.35745345 | 0.559513 |
| ENSG00000237493 | | 718.3107 | 0.1744808 | 0.2096357 | 0.8600078 | 0.3897848 | 0.591749 |
| ENSG00000187609 | EXD3 | 1273.387 | 0.1744577 | 0.2259182 | 0.7788974 | 0.4360401 | 0.634698 |
| ENSG00000272842 | | 9.103255 | 0.1744462 | 0.4756017 | 0.4530422 | 0.6505184 | 0.802546 |
| ENSG00000153558 | FBXL2 | 350.597 | 0.1744401 | 0.3748689 | 0.5172812 | 0.6049599 | 0.769596 |
| ENSG00000285627 | | 1.53852 | 0.1744383 | 0.7435003 | 0.5658475 | 0.5714974 | NA |
| ENSG00000234494 | SP2-AS1 | 349.7333 | 0.174392 | 0.2662909 | 0.6933844 | 0.4880684 | 0.679033 |
| ENSG00000233012 | HDAC1P2 | 1.062281 | 0.1743632 | 0.7457453 | 0.5723944 | 0.5670548 | NA |
| ENSG00000244349 | | 6.034099 | 0.174171 | 0.5813576 | 0.4303964 | 0.6669073 | 0.814371 |
| ENSG00000254224 | | 3.149361 | 0.1741696 | 0.6450155 | 0.4438519 | 0.6571497 | 0.806839 |
| ENSG00000266729 | DSG1-AS1 | 1.158666 | 0.1741637 | 0.8039998 | 0.7340012 | 0.462948 | NA |
| ENSG00000257283 | | 4.193346 | 0.1741043 | 0.6204927 | 0.4373675 | 0.6618449 | 0.810311 |
| ENSG00000231794 | | 2.255259 | 0.1739386 | 0.7070273 | 0.5118777 | 0.6087366 | 0.772377 |
| ENSG00000287844 | | 2.009542 | 0.1738669 | 0.7214342 | 0.513028 | 0.6079317 | 0.771722 |
| ENSG00000212900 | KRTAP3-2 | 0.778264 | 0.1738616 | 0.796198 | 0.9163196 | 0.3594993 | NA |
| ENSG00000111275 | ALDH2 | 47.81013776 | 0.173822298 | 0.502935551 | 0.437146511 | 0.6620051 | 0.810395 |

| | | | | | | | |
|---|---|---|---|---|---|---|---|
| ENSG00000164845 | FAM86FP | 99.6449 | 0.1737009 | 0.3059014 | 0.615319 | 0.5383441 | 0.721137 |
| ENSG00000232678 | SPTLC1P4 | 1.228928 | 0.1736637 | 0.7587773 | 0.673947 | 0.500345 | NA |
| ENSG00000253517 | XRCC6P4 | 2.703298 | 0.1735361 | 0.6659916 | 0.4628903 | 0.643443 | 0.796766 |
| ENSG00000129055 | ANAPC13 | 3325.554229 | 0.173461912 | 0.157857721 | 1.121601737 | 0.26203181 | 0.456791 |
| ENSG00000258798 | | 13.99956 | 0.1733588 | 0.4162313 | 0.4893943 | 0.6245626 | 0.783563 |
| ENSG00000204514 | ZNF814 | 566.5319 | 0.1733251 | 0.3556521 | 0.5432941 | 0.5869273 | 0.756545 |
| ENSG00000092208 | GEMIN2 | 1094.247121 | 0.173309684 | 0.237424912 | 0.762886846 | 0.44553088 | 0.642599 |
| ENSG00000013810 | TACC3 | 3104.006334 | 0.173300598 | 0.517826216 | 0.422696457 | 0.67251674 | 0.817987 |
| ENSG00000188643 | S100A16 | 16650.11 | 0.1731688 | 0.2977329 | 0.6244664 | 0.5323213 | 0.716295 |
| ENSG00000104375 | STK3 | 2419.103542 | 0.173127706 | 0.144505445 | 1.226956643 | 0.21983889 | 0.406021 |
| ENSG00000061936 | SFSWAP | 3946.194094 | 0.173116649 | 0.112248741 | 1.564156018 | 0.11778093 | 0.261692 |
| ENSG00000169020 | ATP5ME | 827.8895 | 0.1730855 | 0.2476283 | 0.7275341 | 0.4668988 | 0.660651 |
| ENSG00000214975 | PPIAP29 | 492.0227 | 0.1730754 | 0.2756894 | 0.6679591 | 0.5041597 | 0.691213 |
| ENSG00000102931 | ARL2BP | 120.6486206 | 0.173054028 | 0.273020488 | 0.673351995 | 0.50072338 | 0.688619 |
| ENSG00000253175 | | 1.211794 | 0.1729825 | 0.7380179 | 0.5526644 | 0.5804932 | NA |
| ENSG00000253328 | SUMO2P19 | 7.154205 | 0.1729387 | 0.5761291 | 0.4239762 | 0.6715832 | 0.81745 |
| ENSG00000152977 | ZIC1 | 1184.283 | 0.1729196 | 0.2036651 | 0.8737358 | 0.3822622 | 0.58454 |
| ENSG00000225173 | | 8.393543 | 0.1728981 | 0.6244745 | 0.436387 | 0.662556 | 0.81092 |
| ENSG00000243187 | CCDC39-AS1 | 1.722708 | 0.172873 | 0.7446732 | 0.5934107 | 0.5529064 | 0.731338 |
| ENSG00000108819 | PPP1R9B | 5915.321217 | 0.172766202 | 0.119376602 | 1.475004343 | 0.14021138 | 0.296386 |
| ENSG00000174579 | MSL2 | 2326.331 | 0.1727602 | 0.1621202 | 1.102394 | 0.2702903 | 0.466715 |
| ENSG00000285830 | | 20.49412 | 0.1727537 | 0.3484982 | 0.5487626 | 0.5831684 | 0.753499 |
| ENSG00000119689 | DLST | 10256.59634 | 0.172690262 | 0.15947558 | 1.083150161 | 0.27874178 | 0.475988 |
| ENSG00000005812 | FBXL3 | 1984.86495 | 0.172648059 | 0.177826495 | 0.995322474 | 0.31957945 | 0.520467 |
| ENSG00000185090 | MANEAL | 1103.089 | 0.1726465 | 0.1850154 | 0.9629132 | 0.3355911 | 0.536804 |
| ENSG00000153208 | MERTK | 136.6672 | 0.17264 | 0.2773437 | 0.6645849 | 0.506316 | 0.692918 |
| ENSG00000278108 | MIR6757 | 0.9876837 | 0.1726339 | 0.7367555 | 0.5126146 | 0.6082209 | NA |
| ENSG00000168487 | BMP1 | 1003.016 | 0.1726279 | 0.2237337 | 0.8148031 | 0.415185 | 0.615074 |
| ENSG00000165322 | ARHGAP12 | 3710.503 | 0.1726278 | 0.189342 | 0.9335043 | 0.3505596 | 0.55226 |
| ENSG00000257674 | | 12.27397 | 0.1726224 | 0.7867703 | 0.43689 | 0.6621911 | 0.810585 |
| ENSG00000231244 | PSMC1P3 | 1.331635 | 0.172577 | 0.7341399 | 0.5418442 | 0.5879258 | NA |
| ENSG00000143457 | GOLPH3L | 3450.098 | 0.1725469 | 0.1758032 | 1.00461 | 0.3150848 | 0.516159 |
| ENSG00000105732 | ZNF574 | 1499.789264 | 0.172432726 | 0.159521189 | 1.101944224 | 0.27048592 | 0.466782 |

| | | | | | | | |
|---|---|---|---|---|---|---|---|
| ENSG00000181101 | ENTR1P2 | 43.9948 | 0.1724134 | 0.3079092 | 0.6099487 | 0.5418958 | 0.723497 |
| ENSG00000112077 | RHAG | 1.217170983 | 0.172377085 | 0.75995223 | 0.633159807 | 0.52662929 | NA |
| ENSG00000246263 | UBR5-AS1 | 142.8163 | 0.1722688 | 0.3020583 | 0.6131864 | 0.5397531 | 0.722087 |
| ENSG00000235787 | RPL35AP8 | 1.886147 | 0.1722574 | 0.7678574 | 0.6226226 | 0.5335325 | 0.717307 |
| ENSG00000235629 | | 1.893371 | 0.1722473 | 0.7121716 | 0.5051593 | 0.613447 | 0.775861 |
| ENSG00000269516 | CYP4F23P | 48.32477 | 0.1722131 | 0.3604552 | 0.5335611 | 0.5936452 | 0.761371 |
| ENSG00000248126 | | 38.24018 | 0.1721191 | 0.3704616 | 0.5233661 | 0.6007195 | 0.766743 |
| ENSG00000143756 | FBXO28 | 3559.638 | 0.1720646 | 0.1621243 | 1.087017 | 0.2770292 | 0.473987 |
| ENSG00000213671 | OLA1P2 | 1.046534 | 0.1720044 | 0.7541516 | 0.6117287 | 0.5407173 | NA |
| ENSG00000237926 | | 0.8642285 | 0.1717514 | 0.7749287 | 0.6553916 | 0.5122156 | NA |
| ENSG00000087253 | LPCAT2 | 392.796407 | 0.171612354 | 0.197743665 | 0.889841708 | 0.37355089 | 0.575609 |
| ENSG00000148358 | GPR107 | 7175.802 | 0.1715902 | 0.1712875 | 1.001496 | 0.3165871 | 0.517625 |
| ENSG00000100292 | HMOX1 | 545.1626453 | 0.171430922 | 0.452117276 | 0.462155651 | 0.6439697 | 0.797169 |
| ENSG00000285650 | | 4.896925 | 0.171396 | 0.5973306 | 0.4268439 | 0.669493 | 0.815897 |
| ENSG00000220105 | | 1.336185 | 0.17137 | 0.7390921 | 0.5400513 | 0.5891617 | NA |
| ENSG00000169371 | SNUPN | 271.1548 | 0.1713686 | 0.2058525 | 0.8718888 | 0.383269 | 0.585705 |
| ENSG00000250526 | CCT6P2 | 23.22085 | 0.1712439 | 0.4489601 | 0.4622094 | 0.6439312 | 0.797158 |
| ENSG00000196576 | PLXNB2 | 18385.39 | 0.1712007 | 0.1473215 | 1.201433 | 0.2295834 | 0.417766 |
| ENSG00000221643 | SNORA77 | 2.707322 | 0.1711759 | 0.6904316 | 0.4747644 | 0.6349549 | 0.791022 |
| ENSG00000134452 | FBH1 | 5897.118595 | 0.171000729 | 0.078556635 | 2.194885382 | 0.02817182 | 0.089117 |
| ENSG00000237338 | FTCD-AS1 | 0.9757833 | 0.1709964 | 0.7664069 | 0.6343253 | 0.5258685 | NA |
| ENSG00000108107 | RPL28 | 17915.17263 | 0.170976159 | 0.155079692 | 1.140818179 | 0.25394559 | 0.446915 |
| ENSG00000212807 | OR2A42 | 4.587798 | 0.170931 | 0.5696146 | 0.4201679 | 0.6743628 | 0.818891 |
| ENSG00000154917 | RAB6B | 1237.083 | 0.1708293 | 0.2541833 | 0.7106571 | 0.4772968 | 0.669516 |
| ENSG00000172575 | RASGRP1 | 330.0174 | 0.1707957 | 0.3947457 | 0.4900826 | 0.6240754 | 0.783271 |
| ENSG00000179021 | C3orf38 | 1463.301 | 0.1707913 | 0.1369344 | 1.252115 | 0.2105281 | 0.39441 |
| ENSG00000120756 | PLS1 | 1563.10522 | 0.170732104 | 0.198504616 | 0.886141403 | 0.37554134 | 0.577691 |
| ENSG00000176681 | LRRC37A | 41.91372 | 0.1707101 | 0.3777404 | 0.5121667 | 0.6085344 | 0.772229 |
| ENSG00000138757 | G3BP2 | 6874.783879 | 0.170647809 | 0.163454534 | 1.063772325 | 0.28743185 | 0.485831 |
| ENSG00000238199 | UBE2V2P3 | 3.515104 | 0.1705896 | 0.6624625 | 0.4506536 | 0.6522393 | 0.803679 |
| ENSG00000269053 | | 2.823299 | 0.1705132 | 0.6863642 | 0.4497396 | 0.6528982 | 0.804137 |
| ENSG00000271676 | | 5.776177 | 0.1704159 | 0.5940379 | 0.4236505 | 0.6718208 | 0.817552 |
| ENSG00000278530 | CHMP1B2P | 0.7255853 | 0.170258 | 0.8105399 | 0.9755369 | 0.329294 | NA |

| | | | | | | | |
|---|---|---|---|---|---|---|---|
| ENSG00000099834 | CDHR5 | 9.782101976 | 0.170170333 | 0.598229794 | 0.425759566 | 0.67028305 | 0.816503 |
| ENSG00000235852 | | 3.733376 | 0.17012 | 0.6240576 | 0.4299255 | 0.6672498 | 0.814638 |
| ENSG00000249850 | KRT18P31 | 6.498121 | 0.1699987 | 0.539435 | 0.4248863 | 0.6709196 | 0.816961 |
| ENSG00000129347 | KRI1 | 3404.983225 | 0.169998688 | 0.185796721 | 0.935040146 | 0.34976761 | 0.551446 |
| ENSG00000279206 | | 1.711526 | 0.1699831 | 0.7526288 | 0.6001946 | 0.5483766 | 0.727909 |
| ENSG00000234107 | TPT1P1 | 2.228178 | 0.1699646 | 0.730379 | 0.5334057 | 0.5937528 | 0.761435 |
| ENSG00000101084 | RAB5IF | 373.1340468 | 0.169923415 | 0.239797236 | 0.742608378 | 0.45771881 | 0.65359 |
| ENSG00000233956 | BTF3P6 | 1.580218 | 0.1696869 | 0.7178857 | 0.5057809 | 0.6130104 | 0.775723 |
| ENSG00000241281 | | 1.985085 | 0.1696457 | 0.7284996 | 0.5302272 | 0.5959544 | 0.763196 |
| ENSG00000231884 | NDUFB1P1 | 124.8337 | 0.1696422 | 0.3221221 | 0.5775446 | 0.5635716 | 0.739049 |
| ENSG00000266918 | | 11.25009 | 0.169621 | 0.5336202 | 0.4212222 | 0.6735929 | 0.818574 |
| ENSG00000241735 | FABP5P3 | 1.144624 | 0.169457 | 0.7314448 | 0.5337398 | 0.5935216 | NA |
| ENSG00000272690 | LINC02018 | 23.82176 | 0.1693559 | 0.38344 | 0.5012007 | 0.6162299 | 0.777685 |
| ENSG00000129315 | CCNT1 | 3450.369237 | 0.169268536 | 0.194566939 | 0.897314423 | 0.36955116 | 0.571647 |
| ENSG00000284994 | | 2.991057 | 0.1692652 | 0.6728788 | 0.4480772 | 0.6540975 | 0.804895 |
| ENSG00000115687 | PASK | 1753.575149 | 0.169230202 | 0.292124701 | 0.632104505 | 0.52731859 | 0.71216 |
| ENSG00000171889 | MIR31HG | 11.25297 | 0.1692005 | 0.7861177 | 0.4310725 | 0.6664156 | 0.813994 |
| ENSG00000268154 | | 0.8063584 | 0.169174 | 0.759575 | 0.6263938 | 0.5310567 | NA |
| ENSG00000228135 | LINC01494 | 1.17541 | 0.1691116 | 0.7445446 | 0.5552237 | 0.5787417 | NA |
| ENSG00000083896 | YTHDC1 | 6717.626141 | 0.169066333 | 0.098974949 | 1.65741461 | 0.09743568 | 0.228287 |
| ENSG00000272674 | PCDHB16 | 93.25156 | 0.1690506 | 0.2643183 | 0.6805051 | 0.4961847 | 0.685194 |
| ENSG00000238009 | | 1.350679 | 0.1689978 | 0.7636186 | 0.5818512 | 0.5606669 | NA |
| ENSG00000180098 | TRNAU1AP | 1158.195 | 0.1689659 | 0.1525259 | 1.1132 | 0.2656226 | 0.461263 |
| ENSG00000254471 | | 2.132621 | 0.1688868 | 0.7536132 | 0.6622536 | 0.5078087 | 0.694174 |
| ENSG00000257761 | | 0.9722274 | 0.1688059 | 0.7559386 | 0.6632804 | 0.507151 | NA |
| ENSG00000085998 | POMGNT1 | 3490.728713 | 0.168795378 | 0.143357151 | 1.178628377 | 0.23854619 | 0.428412 |
| ENSG00000075945 | KIFAP3 | 1394.325816 | 0.168755181 | 0.197348002 | 0.878358955 | 0.37974895 | 0.581698 |
| ENSG00000186566 | GPATCH8 | 3887.905 | 0.1687524 | 0.1933956 | 0.904498 | 0.3657314 | 0.567721 |
| ENSG00000083844 | ZNF264 | 4.036965092 | 0.168734235 | 0.789866416 | 0.388761518 | 0.69745257 | 0.834825 |
| ENSG00000231908 | IDH1-AS1 | 49.25455 | 0.1686471 | 0.3503049 | 0.5360889 | 0.5918971 | 0.760374 |
| ENSG00000161011 | SQSTM1 | 17323.66 | 0.1686094 | 0.2555901 | 0.7280121 | 0.4666062 | 0.660412 |
| ENSG00000225037 | EIF1AX-AS1 | 4.840717 | 0.1685957 | 0.6653018 | 0.4413496 | 0.6589599 | 0.808067 |
| ENSG00000215006 | CHCHD2P2 | 31.59044 | 0.1685813 | 0.3596829 | 0.5256161 | 0.599155 | 0.765665 |

| | | | | | | | |
|---|---|---|---|---|---|---|---|
| ENSG00000259314 | | 3.806607 | 0.1685647 | 0.6298854 | 0.4246219 | 0.6711123 | 0.817101 |
| ENSG00000270460 | | 16.72851 | 0.168542 | 0.4829751 | 0.4332908 | 0.6648035 | 0.812764 |
| ENSG00000134901 | POGLUT2 | 350.6312716 | 0.16850435 | 0.292184739 | 0.619106491 | 0.53584621 | 0.719035 |
| ENSG00000147206 | NXF3 | 1.214739 | 0.168496 | 0.7294688 | 0.5148386 | 0.6066658 | NA |
| ENSG00000186792 | HYAL3 | 216.7614 | 0.168411 | 0.3042689 | 0.5957521 | 0.5513409 | 0.730202 |
| ENSG00000158485 | CD1B | 1.024064 | 0.1682862 | 0.7529222 | 0.5782855 | 0.5630714 | NA |
| ENSG00000107263 | RAPGEF1 | 6812.309399 | 0.16828562 | 0.188243573 | 0.918299377 | 0.35846215 | 0.560387 |
| ENSG00000163577 | EIF5A2 | 473.2971 | 0.1682774 | 0.2063414 | 0.8482837 | 0.39628 | 0.597546 |
| ENSG00000134548 | SPX | 1.463060989 | 0.168259353 | 0.742087395 | 0.532982504 | 0.59404569 | NA |
| ENSG00000182508 | LHFPL1 | 2.585216 | 0.1682481 | 0.6869642 | 0.4485237 | 0.6537753 | 0.804845 |
| ENSG00000100263 | RHBDD3 | 2080.673007 | 0.168184345 | 0.17342895 | 0.992084942 | 0.32115609 | 0.521911 |
| ENSG00000225447 | RPS15AP10 | 5.133977 | 0.168165 | 0.6600058 | 0.4350376 | 0.6635352 | 0.81167 |
| ENSG00000259431 | THTPA | 600.3613 | 0.1681576 | 0.2042902 | 0.8496221 | 0.3955352 | 0.597203 |
| ENSG00000202400 | SNORD82 | 1.120281 | 0.1679485 | 0.7555642 | 0.6595735 | 0.5095276 | NA |
| ENSG00000154124 | OTULIN | 2727.1 | 0.1678972 | 0.2050086 | 0.8475202 | 0.3967052 | 0.59778 |
| ENSG00000172965 | MIR4435-2HG | 991.0285 | 0.1678877 | 0.1962246 | 0.8848994 | 0.3762109 | 0.578446 |
| ENSG00000159871 | LYPD5 | 0.9384642 | 0.1678755 | 0.754508 | 0.6515615 | 0.5146841 | NA |
| ENSG00000256682 | | 1.536785 | 0.1677662 | 0.6925969 | 0.4568387 | 0.647787 | NA |
| ENSG00000274578 | | 0.7829089 | 0.167713 | 0.8047302 | 0.6801407 | 0.4964154 | NA |
| ENSG00000073464 | CLCN4 | 145.5767165 | 0.16762644 | 0.409576691 | 0.468392355 | 0.63950403 | 0.794189 |
| ENSG00000165684 | SNAPC4 | 2951.095 | 0.1675549 | 0.1119908 | 1.506422 | 0.1319589 | 0.283957 |
| ENSG00000173230 | GOLGB1 | 15244.09 | 0.1675381 | 0.2357493 | 0.741133 | 0.4586128 | 0.654433 |
| ENSG00000252577 | SCARNA20 | 11.26257 | 0.1674915 | 0.469964 | 0.4458007 | 0.6557412 | 0.805958 |
| ENSG00000234115 | | 7.152271 | 0.1674845 | 0.6064504 | 0.418123 | 0.6758572 | 0.819835 |
| ENSG00000279584 | | 4.453951 | 0.1674783 | 0.6349188 | 0.4254228 | 0.6705285 | 0.816652 |
| ENSG00000272904 | | 5.724922 | 0.167468 | 0.5511835 | 0.4142431 | 0.678696 | 0.821485 |
| ENSG00000215869 | | 1.906768 | 0.1674499 | 0.7582705 | 0.6311466 | 0.5279447 | 0.712788 |
| ENSG00000132275 | RRP8 | 1026.869051 | 0.167439079 | 0.13162669 | 1.289611382 | 0.19718562 | 0.37718 |
| ENSG00000254088 | SLC2A3P4 | 7.772426 | 0.1673757 | 0.5850797 | 0.4138608 | 0.6789761 | 0.821593 |
| ENSG00000118242 | MREG | 3824.402321 | 0.167334067 | 0.157639559 | 1.080401865 | 0.27996327 | 0.477245 |
| ENSG00000270392 | PFN1P2 | 41.46113 | 0.1670677 | 0.5080219 | 0.4225519 | 0.6726223 | 0.81804 |
| ENSG00000260278 | | 1.585928 | 0.1669972 | 0.71958 | 0.4985109 | 0.618124 | 0.77904 |
| ENSG00000142751 | GPN2 | 1686.156 | 0.1669862 | 0.1248277 | 1.355218 | 0.1753481 | 0.347544 |

| | | | | | | | |
|---|---|---|---|---|---|---|---|
| ENSG00000269313 | MAGIX | 204.3659 | 0.1669211 | 0.2875213 | 0.618523 | 0.5362306 | 0.719265 |
| ENSG00000285244 | DINOL | 7.294552 | 0.1668679 | 0.6102014 | 0.4092233 | 0.6823758 | 0.82413 |
| ENSG00000225462 | FDPSP1 | 41.31831 | 0.1668219 | 0.4633893 | 0.4414567 | 0.6588824 | 0.808062 |
| ENSG00000241529 | RN7SL767P | 4.026502 | 0.1667781 | 0.619433 | 0.4180252 | 0.6759287 | 0.819885 |
| ENSG00000237548 | TTLL11-IT1 | 1.710361 | 0.1667002 | 0.699193 | 0.4784251 | 0.6323476 | 0.789254 |
| ENSG00000034693 | PEX3 | 919.7843135 | 0.166571735 | 0.223730954 | 0.772851331 | 0.43961037 | 0.638112 |
| ENSG00000186063 | AIDA | 866.3656 | 0.1665424 | 0.1993691 | 0.8577802 | 0.3910139 | 0.592869 |
| ENSG00000231113 | | 55.25353 | 0.166542 | 0.307087 | 0.5865999 | 0.5574724 | 0.734806 |
| ENSG00000232852 | CICP4 | 0.9686478 | 0.1665239 | 0.7537622 | 0.5786715 | 0.5628109 | NA |
| ENSG00000219700 | PTCHD3P3 | 6.743013 | 0.1664775 | 0.5170905 | 0.4142678 | 0.678678 | 0.821485 |
| ENSG00000238072 | CDC26P1 | 406.0784 | 0.1664743 | 0.2487385 | 0.7102156 | 0.4775705 | 0.669704 |
| ENSG00000258446 | | 12.36288 | 0.1664432 | 0.5126376 | 0.4185183 | 0.6755682 | 0.819667 |
| ENSG00000134153 | EMC7 | 2807.451212 | 0.166426916 | 0.139611725 | 1.208245064 | 0.22695301 | 0.414741 |
| ENSG00000100994 | PYGB | 8445.368985 | 0.166426385 | 0.135200947 | 1.282726467 | 0.19958792 | 0.3798 |
| ENSG00000105438 | KDELR1 | 9909.721787 | 0.166391611 | 0.138941645 | 1.225877943 | 0.22024461 | 0.406395 |
| ENSG00000109046 | WSB1 | 5935.499715 | 0.166378875 | 0.258644935 | 0.665530406 | 0.5057113 | 0.692483 |
| ENSG00000215210 | RBMXP2 | 191.7459 | 0.1663755 | 0.242379 | 0.7215244 | 0.470587 | 0.66403 |
| ENSG00000178057 | NDUFAF3 | 1205.979 | 0.1663543 | 0.1964531 | 0.8821645 | 0.3776879 | 0.579608 |
| ENSG00000182220 | ATP6AP2 | 7484.828 | 0.1663241 | 0.09686256 | 1.704211 | 0.08834171 | 0.212701 |
| ENSG00000229017 | LINC01277 | 22.83384 | 0.166273 | 0.5092994 | 0.4175018 | 0.6763114 | 0.820059 |
| ENSG00000287721 | | 31.60641 | 0.1662582 | 0.4283777 | 0.457758 | 0.6471263 | 0.799921 |
| ENSG00000130383 | FUT5 | 1.250188243 | 0.166212984 | 0.766040748 | 0.562659844 | 0.57366653 | NA |
| ENSG00000235064 | SLC25A5P2 | 15.4162 | 0.1662059 | 0.500539 | 0.4277004 | 0.6688693 | 0.815566 |
| ENSG00000214102 | WEE2 | 5.117788 | 0.1661297 | 0.6782943 | 0.4466411 | 0.6551342 | 0.805398 |
| ENSG00000114388 | NPRL2 | 1230.199895 | 0.16604098 | 0.188109758 | 0.908087089 | 0.36383221 | 0.56593 |
| ENSG00000224967 | | 1.192607 | 0.1660396 | 0.7639145 | 0.6305494 | 0.5283352 | NA |
| ENSG00000229127 | KANSL1L-AS1 | 22.66283 | 0.1658973 | 0.4757407 | 0.4305753 | 0.6667772 | 0.814361 |
| ENSG00000172081 | MOB3A | 1033.475 | 0.1658936 | 0.1486012 | 1.131433 | 0.2578731 | 0.451941 |
| ENSG00000203546 | | 34.74315 | 0.1658778 | 0.3449755 | 0.5346516 | 0.5928908 | 0.760895 |
| ENSG00000154380 | ENAH | 6160.198 | 0.1658758 | 0.1950636 | 0.8678805 | 0.3854598 | 0.587802 |
| ENSG00000171316 | CHD7 | 3977.894 | 0.1657893 | 0.2191034 | 0.7848997 | 0.4325124 | 0.631709 |
| ENSG00000120458 | MSANTD2 | 588.6613795 | 0.165744279 | 0.28993953 | 0.60109572 | 0.54777623 | 0.727467 |
| ENSG00000259419 | HNRNPCP3 | 2.203272 | 0.1657361 | 0.6676003 | 0.4414676 | 0.6588745 | 0.808062 |

| | | | | | | | |
|---|---|---|---|---|---|---|---|
| ENSG00000067445 | TRO | 11.94195954 | 0.16572408 | 0.51877425 | 0.41534141 | 0.67789199 | 0.820917 |
| ENSG00000227850 | SEPTIN2P1 | 43.89392 | 0.1656859 | 0.3369366 | 0.5418448 | 0.5879255 | 0.757289 |
| ENSG00000175556 | LONRF3 | 29.89759 | 0.1656135 | 0.7845216 | 0.4303419 | 0.666947 | 0.814381 |
| ENSG00000143162 | CREG1 | 5147.965 | 0.1655614 | 0.151947 | 1.108469 | 0.2676595 | 0.463558 |
| ENSG00000220157 | HNRNPA1P12 | 3.513668 | 0.1654549 | 0.6447889 | 0.4212185 | 0.6735955 | 0.818574 |
| ENSG00000186642 | PDE2A | 13.56193 | 0.1654304 | 0.4318077 | 0.4494167 | 0.6531311 | 0.804347 |
| ENSG00000250069 | | 1.186329 | 0.1653117 | 0.7278665 | 0.5170929 | 0.6050913 | NA |
| ENSG00000249129 | SUDS3P1 | 408.4481 | 0.1652647 | 0.2458658 | 0.689781 | 0.4903319 | 0.680949 |
| ENSG00000261801 | LOXL1-AS1 | 384.834 | 0.1651432 | 0.2677082 | 0.6562177 | 0.511684 | 0.697677 |
| ENSG00000129355 | CDKN2D | 397.374239 | 0.165066563 | 0.256938725 | 0.679004189 | 0.49713521 | 0.685915 |
| ENSG00000128591 | FLNC | 2.182688517 | 0.165054611 | 0.686331917 | 0.452236237 | 0.65109881 | 0.802889 |
| ENSG00000103490 | PYCARD | 851.2632289 | 0.165014692 | 0.268783724 | 0.644087464 | 0.51951872 | 0.704851 |
| ENSG00000241506 | PSMC1P1 | 6111.265 | 0.1650103 | 0.2174445 | 0.7880103 | 0.4306907 | 0.629885 |
| ENSG00000142235 | LMTK3 | 1907.05 | 0.1649165 | 0.1723496 | 0.9841637 | 0.325035 | 0.526224 |
| ENSG00000187778 | MCRS1 | 4848.046 | 0.16483 | 0.1699791 | 0.9884435 | 0.3229355 | 0.523686 |
| ENSG00000206603 | SNORA22B | 3.928835 | 0.1648066 | 0.6924621 | 0.462267 | 0.6438899 | 0.797144 |
| ENSG00000250433 | CLSTN2-AS1 | 2.746629 | 0.1647887 | 0.6507981 | 0.4181773 | 0.6758175 | 0.819824 |
| ENSG00000123104 | ITPR2 | 1377.222202 | 0.164774042 | 0.30532706 | 0.584064917 | 0.55917664 | 0.735884 |
| ENSG00000228463 | | 41.12633 | 0.1647424 | 0.3650128 | 0.5049179 | 0.6136165 | 0.775861 |
| ENSG00000171224 | FAM241B | 1397.338 | 0.1647239 | 0.224688 | 0.7593853 | 0.4476221 | 0.644391 |
| ENSG00000146757 | ZNF92 | 605.5776 | 0.1646332 | 0.2173301 | 0.7885513 | 0.4303743 | 0.629691 |
| ENSG00000259654 | | 1.936554 | 0.1646278 | 0.6950437 | 0.4654509 | 0.6416086 | 0.795667 |
| ENSG00000267898 | | 11.00872 | 0.1646142 | 0.4987265 | 0.4192982 | 0.6749982 | 0.819317 |
| ENSG00000259439 | LINC01833 | 7.222589 | 0.1645929 | 0.6587356 | 0.4391319 | 0.6605659 | 0.809118 |
| ENSG00000224271 | | 436.5662 | 0.1645689 | 0.2874195 | 0.6182028 | 0.5364416 | 0.719507 |
| ENSG00000233680 | HNRNPA1P27 | 7.396707 | 0.1644017 | 0.4764933 | 0.428875 | 0.6680142 | 0.815047 |
| ENSG00000234883 | MIR155HG | 1.339253 | 0.1644007 | 0.7539628 | 0.597507 | 0.5501689 | NA |
| ENSG00000135100 | HNF1A | 1.21507953 | 0.164334517 | 0.753680257 | 0.545469424 | 0.5854307 | NA |
| ENSG00000163848 | ZNF148 | 3148.631 | 0.1642858 | 0.1604693 | 1.043771 | 0.2965914 | 0.495597 |
| ENSG00000279801 | | 2.921907 | 0.1642309 | 0.6603844 | 0.4291753 | 0.6677956 | 0.81493 |
| ENSG00000204538 | PSORS1C2 | 12.2831 | 0.1642083 | 0.7036172 | 0.4963739 | 0.6196306 | 0.780127 |
| ENSG00000248592 | STIMATE-MUSTN | 12.50232 | 0.1641479 | 0.4971623 | 0.420813 | 0.6738916 | 0.818684 |
| ENSG00000251330 | | 5.171488 | 0.16412 | 0.5810273 | 0.4067533 | 0.6841892 | 0.825532 |

| | | | | | | | |
|---|---|---|---|---|---|---|---|
| ENSG00000275481 | | 57.68564 | 0.1639864 | 0.3708415 | 0.501319 | 0.6161467 | 0.777672 |
| ENSG00000233578 | EIF4EP1 | 1.815124 | 0.1639164 | 0.7171478 | 0.4813658 | 0.6302566 | 0.787828 |
| ENSG00000271843 | | 142.4652 | 0.1638738 | 0.2521497 | 0.6898121 | 0.4903123 | 0.680949 |
| ENSG00000183837 | PNMA3 | 1.147247 | 0.1638705 | 0.7587713 | 0.6544632 | 0.5128134 | NA |
| ENSG00000185798 | WDR53 | 702.5365 | 0.1638277 | 0.147592 | 1.129637 | 0.2586292 | 0.452818 |
| ENSG00000259648 | | 4.119195 | 0.1637954 | 0.5768508 | 0.402648 | 0.6872072 | 0.82767 |
| ENSG00000228196 | PTPN2P1 | 91.31505 | 0.1636335 | 0.2579849 | 0.6725346 | 0.5012434 | 0.689114 |
| ENSG00000226314 | ZNF192P1 | 7.350079 | 0.1635564 | 0.5742702 | 0.3993589 | 0.6896288 | 0.829057 |
| ENSG00000225850 | | 10.19641 | 0.1635513 | 0.5061861 | 0.4132573 | 0.6794181 | 0.821944 |
| ENSG00000009307 | CSDE1 | 167513.1742 | 0.163494232 | 0.15177284 | 1.095094844 | 0.27347509 | 0.470197 |
| ENSG00000248159 | HSPA8P11 | 2.401599 | 0.1634444 | 0.7023305 | 0.465133 | 0.6418362 | 0.795751 |
| ENSG00000150457 | LATS2 | 1617.344 | 0.1633859 | 0.2116114 | 0.7935593 | 0.427452 | 0.626938 |
| ENSG00000229386 | OR8B9P | 1.023346 | 0.1633351 | 0.7515456 | 0.5640377 | 0.5727285 | NA |
| ENSG00000188257 | PLA2G2A | 11.2568 | 0.1633101 | 0.7858732 | 0.4082393 | 0.683098 | 0.824743 |
| ENSG00000257150 | PGAM1P5 | 8.11051 | 0.1632076 | 0.7859161 | 0.4069114 | 0.684073 | 0.82543 |
| ENSG00000287278 | | 3.861752 | 0.1631521 | 0.7867454 | 0.4010497 | 0.6883836 | 0.828411 |
| ENSG00000070778 | PTPN21 | 1366.824841 | 0.163061214 | 0.35655081 | 0.520969436 | 0.60238806 | 0.767619 |
| ENSG00000185085 | INTS5 | 2322.314 | 0.1630465 | 0.1697626 | 0.9793488 | 0.3274077 | 0.528776 |
| ENSG00000163357 | DCST1 | 97.41336 | 0.1630231 | 0.3054654 | 0.5781878 | 0.5631374 | 0.738718 |
| ENSG00000270433 | H3P37 | 5.030337 | 0.1630022 | 0.6220002 | 0.4046544 | 0.6857315 | 0.82683 |
| ENSG00000234396 | | 0.931081 | 0.1629749 | 0.7833503 | 0.656904 | 0.5112427 | NA |
| ENSG00000137814 | HAUS2 | 1093.450284 | 0.1629462 | 0.24321569 | 0.713282378 | 0.47567104 | 0.668011 |
| ENSG00000130396 | AFDN | 6273.337382 | 0.162882707 | 0.211809139 | 0.791165007 | 0.42884771 | 0.628062 |
| ENSG00000140319 | SRP14 | 10834.94 | 0.1628422 | 0.2017817 | 0.8505443 | 0.3950226 | 0.596734 |
| ENSG00000267592 | | 0.9432069 | 0.1626995 | 0.7571692 | 0.6518975 | 0.5144673 | NA |
| ENSG00000159873 | CCDC117 | 3859.809 | 0.1626857 | 0.2294609 | 0.7481855 | 0.4543483 | 0.65042 |
| ENSG00000113525 | IL5 | 1.043754673 | 0.162660771 | 0.732473825 | 0.518094436 | 0.60439238 | NA |
| ENSG00000218537 | MIF-AS1 | 65.7535 | 0.1626341 | 0.2433712 | 0.7010628 | 0.4832638 | 0.674645 |
| ENSG00000274929 | | 2.5665 | 0.1625633 | 0.6951934 | 0.4486172 | 0.6537078 | 0.804799 |
| ENSG00000159339 | PADI4 | 1.423973 | 0.1625224 | 0.7228018 | 0.4627284 | 0.6435591 | NA |
| ENSG00000236404 | VLDLR-AS1 | 94.20521 | 0.1624721 | 0.6615822 | 0.4115112 | 0.6806977 | 0.823151 |
| ENSG00000168824 | NSG1 | 1.275152 | 0.162407 | 0.7177717 | 0.4930164 | 0.622001 | NA |
| ENSG00000140832 | MARVELD3 | 2315.46 | 0.1623585 | 0.1140475 | 1.437943 | 0.1504503 | 0.311803 |

| | | | | | | | |
|---|---|---|---|---|---|---|---|
| ENSG00000230872 | MFSD13B | 5.601905 | 0.1623069 | 0.5548344 | 0.403732 | 0.6864098 | 0.827298 |
| ENSG00000248996 | | 28.28636 | 0.1622815 | 0.4336388 | 0.4444503 | 0.6567171 | 0.806592 |
| ENSG00000261273 | | 0.9451843 | 0.1622631 | 0.7877756 | 0.7638052 | 0.4449833 | NA |
| ENSG00000288577 | | 1.652899 | 0.1620891 | 0.7006387 | 0.4477101 | 0.6543624 | 0.805045 |
| ENSG00000171735 | CAMTA1 | 1379.509 | 0.1620718 | 0.1825708 | 0.9071241 | 0.3643412 | 0.566424 |
| ENSG00000254851 | | 14.46369 | 0.1620407 | 0.4212705 | 0.4533397 | 0.6503041 | 0.802434 |
| ENSG00000260468 | LINC01290 | 24.50237 | 0.1620105 | 0.4107375 | 0.4558706 | 0.6484831 | 0.800734 |
| ENSG00000078902 | TOLLIP | 2059.627601 | 0.161968754 | 0.22649637 | 0.745837448 | 0.45576562 | 0.651734 |
| ENSG00000288436 | NA | 6.796454 | 0.1618565 | 0.6068867 | 0.3965571 | 0.6916941 | 0.830514 |
| ENSG00000103642 | LACTB | 1100.048242 | 0.161834513 | 0.163946203 | 1.018557471 | 0.3084131 | 0.508686 |
| ENSG00000229897 | SEPTIN7P7 | 13.65155 | 0.1617261 | 0.4410749 | 0.4417477 | 0.6586718 | 0.80799 |
| ENSG00000272692 | | 48.60981 | 0.1616537 | 0.4403782 | 0.4372125 | 0.6619572 | 0.810374 |
| ENSG00000177108 | ZDHHC22 | 75.62531 | 0.1616209 | 0.6438462 | 0.3992622 | 0.6897 | 0.829057 |
| ENSG00000255136 | | 9.409283 | 0.1616177 | 0.5927694 | 0.4059214 | 0.6848003 | 0.826148 |
| ENSG00000082458 | DLG3 | 4418.398728 | 0.161611057 | 0.097295037 | 1.678174138 | 0.09331311 | 0.221307 |
| ENSG00000250366 | TUNAR | 5.319988 | 0.1615974 | 0.7886815 | 0.3747429 | 0.7078517 | 0.841134 |
| ENSG00000235501 | CNN3-DT | 1.196987 | 0.1614006 | 0.7782092 | 0.7097371 | 0.4778672 | NA |
| ENSG00000255670 | | 1.607673 | 0.1613518 | 0.7090115 | 0.4750042 | 0.634784 | 0.790846 |
| ENSG00000104969 | SGTA | 3842.711651 | 0.161330191 | 0.200378833 | 0.81621586 | 0.41437669 | 0.614579 |
| ENSG00000259118 | | 9.358858 | 0.1611905 | 0.5716848 | 0.399667 | 0.6894018 | 0.828952 |
| ENSG00000236976 | | 3.548644 | 0.1610056 | 0.6495269 | 0.4170494 | 0.6766423 | 0.820122 |
| ENSG00000243429 | OR7E29P | 2.281056 | 0.1609796 | 0.6759112 | 0.4284962 | 0.6682899 | 0.815124 |
| ENSG00000286070 | | 5.424801 | 0.1608698 | 0.5383401 | 0.3972342 | 0.6911948 | 0.830171 |
| ENSG00000169155 | ZBTB43 | 1168.333 | 0.1608443 | 0.231463 | 0.686618 | 0.4923235 | 0.682451 |
| ENSG00000148296 | SURF6 | 5447.518 | 0.1608237 | 0.1298431 | 1.254244 | 0.2097535 | 0.393153 |
| ENSG00000175189 | INHBC | 13.38991 | 0.1607729 | 0.5279925 | 0.3962865 | 0.6918937 | 0.830716 |
| ENSG00000271303 | SRXN1 | 36.30304 | 0.1606271 | 0.3336857 | 0.5312548 | 0.5952422 | 0.762609 |
| ENSG00000241860 | | 2.358869 | 0.1605238 | 0.6616329 | 0.4175923 | 0.6762452 | 0.820059 |
| ENSG00000100583 | SAMD15 | 172.6705268 | 0.160511089 | 0.210136245 | 0.790253913 | 0.4293795 | 0.628579 |
| ENSG00000136875 | PRPF4 | 2876.228906 | 0.160508148 | 0.261154469 | 0.709314324 | 0.47812944 | 0.670104 |
| ENSG00000163281 | GNPDA2 | 765.5211 | 0.1605061 | 0.1676107 | 0.9814649 | 0.3263635 | 0.527603 |
| ENSG00000240787 | | 21.05831 | 0.1604414 | 0.3943611 | 0.4691009 | 0.6389975 | 0.794106 |
| ENSG00000164054 | SHISA5 | 7438.76 | 0.1603395 | 0.193604 | 0.8569954 | 0.3914474 | 0.593235 |

| | | | | | | | |
|---|---|---|---|---|---|---|---|
| ENSG00000094963 | FMO2 | 0.803174337 | 0.160249128 | 0.761677157 | 0.614890603 | 0.53862697 | NA |
| ENSG00000039319 | ZFYVE16 | 2791.026758 | 0.160202033 | 0.265907785 | 0.636621968 | 0.52437111 | 0.709369 |
| ENSG00000137841 | PLCB2 | 10.60831344 | 0.160152935 | 0.550417342 | 0.388944926 | 0.69731689 | 0.834703 |
| ENSG00000214812 | | 3.625414 | 0.1601522 | 0.6890273 | 0.4493773 | 0.6531595 | 0.804347 |
| ENSG00000130427 | EPO | 2.419344798 | 0.160130389 | 0.700399213 | 0.463148003 | 0.64325828 | 0.79668 |
| ENSG00000244541 | LINC01213 | 100.8868 | 0.1601237 | 0.3187406 | 0.5175476 | 0.6047739 | 0.769515 |
| ENSG00000228113 | | 2.183398 | 0.1601024 | 0.7868308 | 0.3870381 | 0.698728 | 0.835727 |
| ENSG00000243234 | | 1.306247 | 0.1600961 | 0.7630888 | 0.5875678 | 0.5568224 | NA |
| ENSG00000204256 | BRD2 | 18516.62 | 0.1600583 | 0.1082919 | 1.490655 | 0.136052 | 0.2899 |
| ENSG00000287929 | | 2.428984 | 0.1600262 | 0.7867574 | 0.3872059 | 0.6986038 | 0.835699 |
| ENSG00000285704 | | 0.8818293 | 0.1600167 | 0.734684 | 0.5099123 | 0.6101129 | NA |
| ENSG00000111676 | ATN1 | 9360.537938 | 0.160016436 | 0.185930541 | 0.916765918 | 0.35926532 | 0.561161 |
| ENSG00000286782 | | 1.727388 | 0.1599168 | 0.7397597 | 0.5232166 | 0.6008235 | 0.766755 |
| ENSG00000227105 | PARP1P1 | 34.15961 | 0.1599059 | 0.4010932 | 0.4610099 | 0.6447915 | 0.797814 |
| ENSG00000198400 | NTRK1 | 1.954259 | 0.1598906 | 0.7247949 | 0.5049607 | 0.6135864 | 0.775861 |
| ENSG00000283945 | LINC00032 | 1.314875 | 0.1597893 | 0.7983924 | 0.5551736 | 0.5787759 | NA |
| ENSG00000243663 | RPS4XP14 | 61.70097 | 0.159779 | 0.3357131 | 0.5271648 | 0.5980792 | 0.76499 |
| ENSG00000261762 | | 147.0803 | 0.1596597 | 0.3564235 | 0.4992257 | 0.6176204 | 0.77875 |
| ENSG00000100138 | SNU13 | 5537.716442 | 0.159655393 | 0.197644458 | 0.827488193 | 0.40796041 | 0.608532 |
| ENSG00000133136 | GNG5P2 | 70.29294174 | 0.159645403 | 0.245503333 | 0.686703263 | 0.49226974 | 0.682416 |
| ENSG00000175066 | GK5 | 1526.036 | 0.1595891 | 0.1997278 | 0.8193628 | 0.4125794 | 0.612714 |
| ENSG00000166317 | SYNPO2L | 0.8923906 | 0.1595683 | 0.7352923 | 0.4964913 | 0.6195478 | NA |
| ENSG00000170325 | PRDM10 | 1013.445 | 0.1594769 | 0.1908293 | 0.8579312 | 0.3909305 | 0.59281 |
| ENSG00000259204 | | 1.276749 | 0.1594715 | 0.7630271 | 0.6105556 | 0.5414938 | NA |
| ENSG00000166845 | C18orf54 | 282.7438 | 0.1594296 | 0.3890669 | 0.4660873 | 0.641153 | 0.795382 |
| ENSG00000170545 | SMAGP | 1025.239 | 0.1593487 | 0.2850144 | 0.5974219 | 0.5502257 | 0.729281 |
| ENSG00000117262 | GPR89A | 1060.34093 | 0.159292166 | 0.1033685 | 1.563656446 | 0.11789827 | 0.261822 |
| ENSG00000219992 | | 1.658459 | 0.1592561 | 0.7885774 | 0.3676774 | 0.7131138 | 0.843667 |
| ENSG00000110171 | TRIM3 | 1545.119002 | 0.159212344 | 0.208500364 | 0.779697812 | 0.43556877 | 0.634325 |
| ENSG00000237921 | | 1.262203 | 0.159189 | 0.7553667 | 0.5562317 | 0.5780525 | NA |
| ENSG00000236882 | LINC01554 | 5.997309 | 0.159125 | 0.6197766 | 0.4030705 | 0.6868963 | 0.827596 |
| ENSG00000285314 | | 1.046138 | 0.1591157 | 0.761486 | 0.5856927 | 0.558082 | NA |
| ENSG00000230176 | LINC01433 | 2.151004 | 0.1590372 | 0.7867074 | 0.3868941 | 0.6988346 | 0.835727 |

| | | | | | | | |
|---|---|---|---|---|---|---|---|
| ENSG00000286861 | | 5.221002 | 0.1590335 | 0.5955477 | 0.3887112 | 0.6974898 | 0.834832 |
| ENSG00000130475 | FCHO1 | 749.5163391 | 0.158997428 | 0.249151335 | 0.680177516 | 0.49639207 | 0.685373 |
| ENSG00000243974 | VTI1BP1 | 14.38789 | 0.1589569 | 0.5069189 | 0.4064721 | 0.6843957 | 0.825744 |
| ENSG00000148426 | PROSER2 | 1570.444 | 0.1589434 | 0.2049731 | 0.8061962 | 0.4201297 | 0.6201 |
| ENSG00000230683 | DDTP1 | 0.8721486 | 0.1588889 | 0.7538275 | 0.5549385 | 0.5789367 | NA |
| ENSG00000232176 | | 163.4071 | 0.1588811 | 0.2595636 | 0.6480432 | 0.516957 | 0.702308 |
| ENSG00000287966 | | 76.8952 | 0.1588683 | 0.3111136 | 0.5607175 | 0.5749901 | 0.747257 |
| ENSG00000143437 | ARNT | 6290.257 | 0.1588232 | 0.1321065 | 1.197273 | 0.2312002 | 0.419681 |
| ENSG00000233048 | LINC01722 | 1.379684 | 0.1588101 | 0.7362171 | 0.4986653 | 0.6180152 | NA |
| ENSG00000230366 | DSCR9 | 12.15104 | 0.1587635 | 0.4118441 | 0.4433525 | 0.6575108 | 0.807092 |
| ENSG00000231154 | MORF4L2-AS1 | 142.7854 | 0.1587138 | 0.2662933 | 0.636501 | 0.5244499 | 0.70944 |
| ENSG00000259562 | | 84.43761 | 0.1586812 | 0.2945996 | 0.5776969 | 0.5634688 | 0.739007 |
| ENSG00000181656 | GPR88 | 0.7878987 | 0.1585914 | 0.7985892 | 0.553715 | 0.5797739 | NA |
| ENSG00000176022 | B3GALT6 | 1358.8 | 0.1585322 | 0.1531692 | 1.063916 | 0.2873666 | 0.485752 |
| ENSG00000125744 | RTN2 | 461.6358887 | 0.158496081 | 0.249490393 | 0.665120562 | 0.50597338 | 0.692651 |
| ENSG00000283662 | | 9.138813 | 0.1584569 | 0.5576445 | 0.3957319 | 0.6923028 | 0.83107 |
| ENSG00000275441 | | 13.85103 | 0.1584094 | 0.4449224 | 0.4303955 | 0.666908 | 0.814371 |
| ENSG00000228974 | | 10.84164 | 0.1584077 | 0.5746765 | 0.3941002 | 0.693507 | 0.831714 |
| ENSG00000286019 | NOTCH2NLB | 0.9364291 | 0.1583929 | 0.7697892 | 0.562443 | 0.5738142 | NA |
| ENSG00000198561 | CTNND1 | 463.4338 | 0.1583645 | 0.1959198 | 0.8332077 | 0.4047276 | 0.605497 |
| ENSG00000143622 | RIT1 | 2580.024 | 0.1583444 | 0.1837757 | 0.8927547 | 0.3719886 | 0.574214 |
| ENSG00000183597 | TANGO2 | 874.5228 | 0.1582897 | 0.1394726 | 1.15389 | 0.2485451 | 0.440827 |
| ENSG00000159588 | CCDC17 | 103.031 | 0.1582362 | 0.3633201 | 0.4917296 | 0.6229105 | 0.782326 |
| ENSG00000105131 | EPHX3 | 6.50255695 | 0.158226046 | 0.571097357 | 0.397974622 | 0.69064889 | 0.829793 |
| ENSG00000276298 | | 4.495114 | 0.1582224 | 0.6483659 | 0.4050889 | 0.6854122 | 0.826558 |
| ENSG00000156958 | GALK2 | 1227.861 | 0.1582123 | 0.1635767 | 0.9929596 | 0.3207296 | 0.521669 |
| ENSG00000170745 | KCNS3 | 483.279 | 0.1580834 | 0.2687207 | 0.6142873 | 0.5390255 | 0.721572 |
| ENSG00000129292 | PHF20L1 | 3781.538179 | 0.158075079 | 0.190379575 | 0.842430915 | 0.3995468 | 0.600519 |
| ENSG00000266340 | | 1.724136 | 0.1579734 | 0.7380463 | 0.5110214 | 0.6093361 | 0.772877 |
| ENSG00000226535 | | 7.757156 | 0.1579374 | 0.561238 | 0.3952111 | 0.6926871 | 0.831256 |
| ENSG00000051108 | HERPUD1 | 4068.232491 | 0.157912044 | 0.311308639 | 0.579610734 | 0.56217715 | 0.73793 |
| ENSG00000233087 | RAB6D | 21.83829 | 0.1578099 | 0.4306053 | 0.4351698 | 0.6634392 | 0.81164 |
| ENSG00000269534 | | 5.465215 | 0.1577634 | 0.5632907 | 0.3947438 | 0.693032 | 0.831369 |

| | | | | | | | |
|---|---|---|---|---|---|---|---|
| ENSG00000239474 | KLHL41 | 7.568714 | 0.1576965 | 0.5779863 | 0.3896632 | 0.6967856 | 0.834411 |
| ENSG00000268660 | LETM1P2 | 0.7858115 | 0.1576606 | 0.754449 | 0.5405006 | 0.5888518 | NA |
| ENSG00000236039 | LINC02889 | 20.01822 | 0.1575785 | 0.4472349 | 0.423352 | 0.6720385 | 0.817667 |
| ENSG00000230793 | SMARCE1P5 | 224.4051 | 0.1575627 | 0.1923247 | 0.8389803 | 0.4014804 | 0.602413 |
| ENSG00000248712 | CCDC153 | 116.6243 | 0.1575345 | 0.3276579 | 0.5283457 | 0.5972594 | 0.764199 |
| ENSG00000277453 | | 5.158977 | 0.1575274 | 0.6170763 | 0.4002884 | 0.6889441 | 0.828673 |
| ENSG00000070087 | PFN2 | 12920.96387 | 0.157485645 | 0.152901631 | 0.98998264 | 0.3221826 | 0.522976 |
| ENSG00000168040 | FADD | 539.1679 | 0.1574687 | 0.1755942 | 0.9131407 | 0.3611685 | 0.563205 |
| ENSG00000258090 | | 41.91036 | 0.1573822 | 0.3138808 | 0.5454026 | 0.5854766 | 0.755278 |
| ENSG00000162391 | FAM151A | 1.518127 | 0.1572952 | 0.7871637 | 0.3734518 | 0.7088122 | NA |
| ENSG00000101457 | DNTTIP1 | 2187.575899 | 0.157290355 | 0.184698451 | 0.879622017 | 0.37906411 | 0.581118 |
| ENSG00000268204 | | 44.44751 | 0.1571181 | 0.3938113 | 0.4527671 | 0.6507165 | 0.802697 |
| ENSG00000229809 | ZNF688 | 695.8109 | 0.157074 | 0.2032874 | 0.7902202 | 0.4293992 | 0.628579 |
| ENSG00000166037 | CEP57 | 2457.788 | 0.1570025 | 0.1431588 | 1.111604 | 0.2663084 | 0.461901 |
| ENSG00000156482 | RPL30 | 15572.96 | 0.1568943 | 0.1606227 | 1.002061 | 0.3163142 | 0.517402 |
| ENSG00000134265 | NAPG | 2979.070436 | 0.156882286 | 0.116041432 | 1.387918564 | 0.16516184 | 0.333677 |
| ENSG00000285649 | | 9.080893 | 0.1568787 | 0.5193857 | 0.3924302 | 0.6947404 | 0.832704 |
| ENSG00000156735 | BAG4 | 779.1752 | 0.1568511 | 0.1953658 | 0.8266918 | 0.4084118 | 0.609099 |
| ENSG00000229647 | MYOSLID | 1.004492 | 0.1567572 | 0.7629622 | 0.6168028 | 0.5373648 | NA |
| ENSG00000219703 | RAP1BP3 | 0.8358058 | 0.1566653 | 0.765168 | 0.5877701 | 0.5566866 | NA |
| ENSG00000183161 | FANCF | 1574.208 | 0.1565639 | 0.1108939 | 1.393447 | 0.1634847 | 0.331296 |
| ENSG00000281386 | | 2.750802 | 0.1565572 | 0.6729805 | 0.4108388 | 0.6811907 | 0.823598 |
| ENSG00000204323 | SMIM5 | 379.936 | 0.1565018 | 0.4095211 | 0.4505409 | 0.6523204 | 0.803724 |
| ENSG00000224452 | RSL24D1P6 | 7.088046 | 0.1564823 | 0.5539248 | 0.3894489 | 0.6969441 | 0.834442 |
| ENSG00000141699 | RETREG3 | 2958.557 | 0.1564767 | 0.1465349 | 1.083441 | 0.2786126 | 0.475889 |
| ENSG00000266313 | | 36.48722 | 0.1563729 | 0.3518227 | 0.4945132 | 0.6209438 | 0.781019 |
| ENSG00000188459 | WASF4P | 206.5486 | 0.1562785 | 0.3058683 | 0.5521647 | 0.5808355 | 0.751618 |
| ENSG00000228801 | | 458.4683 | 0.1562781 | 0.1932753 | 0.8354207 | 0.4034809 | 0.604287 |
| ENSG00000161618 | ALDH16A1 | 1449.29 | 0.1562485 | 0.295636 | 0.5659388 | 0.5714354 | 0.745129 |
| ENSG00000240891 | PLCXD2 | 101.9405 | 0.1562331 | 0.3598362 | 0.4835781 | 0.6286853 | 0.786629 |
| ENSG00000287225 | | 2.066133 | 0.1562114 | 0.7886272 | 0.3564529 | 0.7215014 | 0.848466 |
| ENSG00000177721 | ANXA2R | 298.0503 | 0.1560314 | 0.2343119 | 0.6905465 | 0.4898506 | 0.680443 |
| ENSG00000272582 | | 2.079803 | 0.156023 | 0.7019805 | 0.4521534 | 0.6511585 | 0.802925 |

| | | | | | | | |
|---|---|---|---|---|---|---|---|
| ENSG00000164088 | PPM1M | 414.3172 | 0.1560179 | 0.2308352 | 0.7051953 | 0.4806887 | 0.67229 |
| ENSG00000261613 | | 1.321083 | 0.1559846 | 0.7867321 | 0.3738546 | 0.7085125 | NA |
| ENSG00000133193 | FAM104A | 1565.165467 | 0.15589394 | 0.11237405 | 1.389602982 | 0.16464947 | 0.332844 |
| ENSG00000224829 | TOMM20P1 | 7.317774 | 0.1558873 | 0.6301852 | 0.3927504 | 0.6945039 | 0.832535 |
| ENSG00000198752 | CDC42BPB | 10331.62 | 0.1558866 | 0.1761056 | 0.8819237 | 0.3778181 | 0.579743 |
| ENSG00000248279 | LINC02120 | 0.9021718 | 0.155883 | 0.7834899 | 0.7575795 | 0.4487028 | NA |
| ENSG00000070388 | FGF22 | 27.61046716 | 0.155816507 | 0.393797162 | 0.453463876 | 0.65021475 | 0.802395 |
| ENSG00000287723 | | 1.721786 | 0.1558121 | 0.723979 | 0.4775588 | 0.6329643 | 0.789652 |
| ENSG00000181029 | TRAPPC5 | 248.3956 | 0.1558023 | 0.3216518 | 0.5256202 | 0.5991521 | 0.765665 |
| ENSG00000272567 | | 0.8761736 | 0.1557249 | 0.7868933 | 0.8562696 | 0.3918487 | NA |
| ENSG00000251144 | | 1.926867 | 0.155643 | 0.7123216 | 0.4694368 | 0.6387574 | 0.794021 |
| ENSG00000160294 | MCM3AP | 5577.118 | 0.155605 | 0.1461646 | 1.073826 | 0.2829008 | 0.480711 |
| ENSG00000163875 | MEAF6 | 4747.193 | 0.1555453 | 0.2146702 | 0.7402347 | 0.4591576 | 0.654764 |
| ENSG00000260219 | CD2BP2-DT | 134.7146 | 0.1554246 | 0.1860865 | 0.8595147 | 0.3900566 | 0.59206 |
| ENSG00000232294 | | 3.30374 | 0.1554062 | 0.7915738 | 0.3272489 | 0.7434797 | 0.861733 |
| ENSG00000165113 | GKAP1 | 512.2441 | 0.155356 | 0.1879604 | 0.8475535 | 0.3966867 | 0.59778 |
| ENSG00000184575 | XPOT | 24044.85 | 0.155306 | 0.2923046 | 0.566882 | 0.5707943 | 0.744572 |
| ENSG00000199145 | MIR302D | 0.8398861 | 0.1552681 | 0.7646936 | 0.5890032 | 0.5558591 | NA |
| ENSG00000171487 | NLRP5 | 4.065644 | 0.1552431 | 0.7865262 | 0.3734188 | 0.7088368 | 0.841665 |
| ENSG00000273866 | | 2.221485 | 0.1552197 | 0.7260208 | 0.4937961 | 0.6214502 | 0.781231 |
| ENSG00000156471 | PTDSS1 | 6750.26 | 0.1551164 | 0.1551826 | 1.01841 | 0.308483 | 0.508758 |
| ENSG00000255692 | | 1.227389 | 0.155102 | 0.739805 | 0.4957379 | 0.6200794 | NA |
| ENSG00000240497 | | 1.151963 | 0.1550975 | 0.7342075 | 0.4712328 | 0.6374745 | NA |
| ENSG00000204469 | PRRC2A | 20295.63 | 0.1550536 | 0.1505803 | 1.033859 | 0.301202 | 0.500478 |
| ENSG00000124103 | FAM209A | 9.19428059 | 0.154982684 | 0.59099301 | 0.378220405 | 0.70526687 | 0.839876 |
| ENSG00000196247 | ZNF107 | 1599.649 | 0.1549625 | 0.2460906 | 0.6548189 | 0.5125844 | 0.698474 |
| ENSG00000171121 | KCNMB3 | 246.4576 | 0.1549561 | 0.2089732 | 0.7731225 | 0.4394499 | 0.637949 |
| ENSG00000132394 | EEFSEC | 1480.707443 | 0.154945466 | 0.177157432 | 0.895882413 | 0.37031557 | 0.572429 |
| ENSG00000125356 | NDUFA1 | 4583.819898 | 0.154787982 | 0.188618433 | 0.839144076 | 0.40138847 | 0.602377 |
| ENSG00000077092 | RARB | 26.56756573 | 0.154767334 | 0.513994164 | 0.39053505 | 0.69614094 | 0.833781 |
| ENSG00000140332 | TLE3 | 5105.765 | 0.1547213 | 0.172039 | 0.9135761 | 0.3609396 | 0.562947 |
| ENSG00000256827 | | 3.066924 | 0.1546342 | 0.6697429 | 0.4122658 | 0.6801446 | 0.82267 |
| ENSG00000230612 | | 13.1023 | 0.1546006 | 0.5744724 | 0.38274 | 0.7019126 | 0.837622 |

| | | | | | | | |
|---|---|---|---|---|---|---|---|
| ENSG00000100982 | PCIF1 | 2995.902449 | 0.154457533 | 0.201875505 | 0.78843384 | 0.43044298 | 0.629691 |
| ENSG00000010072 | SPRTN | 1164.355022 | 0.154410064 | 0.153980128 | 1.028791364 | 0.30357773 | 0.503198 |
| ENSG00000259707 | | 1.991042 | 0.1543805 | 0.7171906 | 0.4774606 | 0.6330342 | 0.789665 |
| ENSG00000230613 | HM13-AS1 | 6.439185 | 0.1543567 | 0.5788912 | 0.3836144 | 0.7012643 | 0.83727 |
| ENSG00000173273 | TNKS | 2294.077 | 0.1543223 | 0.233766 | 0.6806045 | 0.4961218 | 0.685143 |
| ENSG00000161904 | LEMD2 | 4024.913 | 0.1542655 | 0.1389133 | 1.114716 | 0.2649722 | 0.460465 |
| ENSG00000006625 | GGCT | 2388.444929 | 0.154219307 | 0.231943213 | 0.714440987 | 0.47495454 | 0.667605 |
| ENSG00000139645 | ANKRD52 | 8945.474 | 0.1542192 | 0.200568 | 0.7932226 | 0.4276481 | 0.627062 |
| ENSG00000188033 | ZNF490 | 114.2551 | 0.1541952 | 0.2887062 | 0.5704839 | 0.5683496 | 0.742622 |
| ENSG00000169018 | FEM1B | 4613.213 | 0.1541658 | 0.1664649 | 0.9456548 | 0.3443247 | 0.545915 |
| ENSG00000263293 | | 2.352371 | 0.1541442 | 0.6756697 | 0.4101791 | 0.6816746 | 0.823808 |
| ENSG00000233342 | | 2.740338 | 0.1539998 | 0.7888483 | 0.347853 | 0.7279506 | 0.852261 |
| ENSG00000108950 | FAM20A | 24.31391687 | 0.153999144 | 0.557175461 | 0.377279674 | 0.70596578 | 0.840133 |
| ENSG00000227045 | | 2.768135 | 0.1539562 | 0.667873 | 0.4049516 | 0.6855131 | 0.826642 |
| ENSG00000127511 | SIN3B | 3005.843871 | 0.153942246 | 0.179125755 | 0.884738289 | 0.3762978 | 0.578479 |
| ENSG00000128710 | HOXD10 | 1.025797353 | 0.153917039 | 0.76284467 | 0.590546415 | 0.55482438 | NA |
| ENSG00000197312 | DDI2 | 3032.76 | 0.153867 | 0.2442471 | 0.7243096 | 0.4688757 | 0.662417 |
| ENSG00000100580 | TMED8 | 3465.853858 | 0.153855722 | 0.214377854 | 0.740170758 | 0.45919639 | 0.654773 |
| ENSG00000116649 | SRM | 3908.84269 | 0.153824046 | 0.26078316 | 0.6095853 | 0.54213655 | 0.723521 |
| ENSG00000147804 | SLC39A4 | 1805.323 | 0.1534599 | 0.2557324 | 0.6243396 | 0.5324046 | 0.71637 |
| ENSG00000087206 | UIMC1 | 3201.512513 | 0.153431896 | 0.166875923 | 0.943361653 | 0.34549595 | 0.547016 |
| ENSG00000101557 | USP14 | 5879.519523 | 0.153417028 | 0.209723862 | 0.728540475 | 0.4662828 | 0.660086 |
| ENSG00000237087 | | 3.40452 | 0.1534144 | 0.7860816 | 0.3713507 | 0.7103763 | 0.842335 |
| ENSG00000134900 | TPP2 | 3020.874564 | 0.15338963 | 0.144128505 | 1.077373365 | 0.2813135 | 0.478907 |
| ENSG00000254554 | | 12.07104 | 0.1533699 | 0.53433 | 0.3910147 | 0.6957863 | 0.833469 |
| ENSG00000207697 | MIR573 | 4.762164 | 0.1533685 | 0.6340693 | 0.3890749 | 0.6972207 | 0.834698 |
| ENSG00000224419 | KRT18P27 | 4.676081 | 0.1532697 | 0.6452308 | 0.4040617 | 0.6861673 | 0.827205 |
| ENSG00000099385 | BCL7C | 1664.540576 | 0.153038085 | 0.260333102 | 0.622515096 | 0.53360322 | 0.717366 |
| ENSG00000261606 | | 3.544651 | 0.153005 | 0.6320455 | 0.3787852 | 0.7048474 | 0.839807 |
| ENSG00000287430 | | 83.07952 | 0.1529446 | 0.3858088 | 0.4538608 | 0.649929 | 0.802098 |
| ENSG00000286587 | | 3.08466 | 0.1529441 | 0.7882278 | 0.349427 | 0.7267687 | 0.851739 |
| ENSG00000122203 | KIAA1191 | 7221.274403 | 0.152938331 | 0.117602651 | 1.370659369 | 0.17048117 | 0.341155 |
| ENSG00000203286 | | 9.592685 | 0.1529215 | 0.4686281 | 0.3991673 | 0.68977 | 0.829103 |

| | | | | | | | |
|---|---|---|---|---|---|---|---|
| ENSG00000147548 | NSD3 | 4081.283 | 0.1527595 | 0.17298 | 0.9072593 | 0.3642697 | 0.566379 |
| ENSG00000285020 | | 2.748876 | 0.152685 | 0.6845693 | 0.4223482 | 0.6727709 | 0.818088 |
| ENSG00000237774 | | 0.9306467 | 0.1526772 | 0.7652504 | 0.6011747 | 0.5477236 | NA |
| ENSG00000223361 | FTH1P10 | 354.1245 | 0.1526555 | 0.2105468 | 0.7471076 | 0.4549986 | 0.650989 |
| ENSG00000230282 | RPL7P61 | 0.8203847 | 0.1525858 | 0.7731156 | 0.7317964 | 0.4642929 | NA |
| ENSG00000144674 | GOLGA4 | 7748.476 | 0.1525745 | 0.217352 | 0.727119 | 0.4671531 | 0.660878 |
| ENSG00000203416 | FAM32BP | 1.130662 | 0.1525512 | 0.7708205 | 0.5565873 | 0.5778094 | NA |
| ENSG00000134001 | EIF2S1 | 8746.448395 | 0.152493298 | 0.191540904 | 0.793900229 | 0.42725352 | 0.626798 |
| ENSG00000138780 | GSTCD | 1124.628208 | 0.152466295 | 0.262960552 | 0.614942208 | 0.53859289 | 0.721237 |
| ENSG00000057704 | TMCC3 | 160.0200263 | 0.152457845 | 0.43076054 | 0.443344991 | 0.65751622 | 0.807092 |
| ENSG00000089159 | PXN | 5423.209167 | 0.152435932 | 0.143217204 | 1.083067432 | 0.27877849 | 0.47602 |
| ENSG00000230611 | HMGB1P27 | 0.8854595 | 0.1523812 | 0.801511 | 0.6822463 | 0.4950832 | NA |
| ENSG00000150361 | KLHL1 | 1.382478 | 0.1523655 | 0.7866308 | 0.3629908 | 0.7166118 | NA |
| ENSG00000240063 | | 1.004924 | 0.1523357 | 0.7865713 | 0.3655194 | 0.7147237 | NA |
| ENSG00000255847 | | 1.306986 | 0.1522199 | 0.7302215 | 0.4858771 | 0.6270543 | NA |
| ENSG00000229056 | HECW2-AS1 | 1.058365 | 0.1521898 | 0.7644149 | 0.5904457 | 0.5548919 | NA |
| ENSG00000215063 | RPL21P2 | 1.344236 | 0.1521627 | 0.7266402 | 0.4644049 | 0.6423577 | NA |
| ENSG00000015413 | DPEP1 | 2.31503946 | 0.152152604 | 0.710088133 | 0.439728957 | 0.66013343 | 0.808775 |
| ENSG00000106025 | TSPAN12 | 417.3704931 | 0.152148177 | 0.267656642 | 0.592434466 | 0.55355969 | 0.731731 |
| ENSG00000187051 | RPS19BP1 | 3208.947 | 0.152077 | 0.2053659 | 0.7721273 | 0.440039 | 0.638386 |
| ENSG00000204625 | HCG9 | 1.078999 | 0.1520756 | 0.7586047 | 0.6425256 | 0.520532 | NA |
| ENSG00000268222 | EEF1A1P7 | 5.346789 | 0.1520688 | 0.5928635 | 0.3808024 | 0.7033499 | 0.838592 |
| ENSG00000197467 | COL13A1 | 12.53787 | 0.1520651 | 0.5382509 | 0.3745521 | 0.7079936 | 0.841227 |
| ENSG00000250427 | LINC02148 | 0.7725231 | 0.1520535 | 0.7625555 | 0.541121 | 0.5884242 | NA |
| ENSG00000260815 | PPIAP47 | 1.260576 | 0.1520498 | 0.7195752 | 0.4573854 | 0.6473941 | NA |
| ENSG00000175198 | PCCA | 916.9224 | 0.1520353 | 0.188711 | 0.8294041 | 0.4068758 | 0.607456 |
| ENSG00000151458 | ANKRD50 | 7430.535 | 0.1520137 | 0.2837174 | 0.5701166 | 0.5685986 | 0.742678 |
| ENSG00000225230 | | 2.445767 | 0.1519533 | 0.6606024 | 0.3998861 | 0.6892404 | 0.828804 |
| ENSG00000263821 | | 0.8090856 | 0.1519275 | 0.8091075 | 1.110409 | 0.2668228 | NA |
| ENSG00000107077 | KDM4C | 1528.517513 | 0.151901896 | 0.215924642 | 0.735438627 | 0.46207241 | 0.656759 |
| ENSG00000184481 | FOXO4 | 763.9054 | 0.1518452 | 0.272273 | 0.5962499 | 0.5510083 | 0.729943 |
| ENSG00000273137 | | 67.75351 | 0.1517762 | 0.2324635 | 0.680811 | 0.4959911 | 0.685033 |
| ENSG00000213923 | CSNK1E | 3812.29 | 0.1517553 | 0.1817046 | 0.852714 | 0.3938179 | 0.59549 |

| | | | | | | | |
|---|---|---|---|---|---|---|---|
| ENSG00000076321 | KLHL20 | 1710.978893 | 0.151691314 | 0.211641134 | 0.733644527 | 0.46316542 | 0.657434 |
| ENSG00000169067 | ACTBL2 | 5.655358 | 0.1516824 | 0.7850883 | 0.3754305 | 0.7073403 | 0.840865 |
| ENSG00000234222 | LIX1L-AS1 | 24.47555 | 0.1515215 | 0.3730878 | 0.4626572 | 0.6436101 | 0.796872 |
| ENSG00000166148 | AVPR1A | 1.146527 | 0.1515164 | 0.7973554 | 0.5231155 | 0.6008939 | NA |
| ENSG00000237990 | CNTN4-AS1 | 0.9770455 | 0.1514758 | 0.7699619 | 0.5913115 | 0.5543117 | NA |
| ENSG00000152763 | DNAI4 | 280.3212 | 0.1514676 | 0.4087673 | 0.4274609 | 0.6690437 | 0.81561 |
| ENSG00000278291 | | 188.707 | 0.1514364 | 0.2878832 | 0.562011 | 0.5741085 | 0.746641 |
| ENSG00000059915 | PSD | 474.4662283 | 0.151435641 | 0.179863801 | 0.863920039 | 0.38763182 | 0.589862 |
| ENSG00000107669 | ATE1 | 1568.893089 | 0.15136546 | 0.159687629 | 0.952852035 | 0.34066505 | 0.542049 |
| ENSG00000287364 | | 2.611398 | 0.1513191 | 0.7881362 | 0.3452129 | 0.7299343 | 0.853593 |
| ENSG00000259207 | ITGB3 | 1.32198 | 0.1513064 | 0.7535889 | 0.5555189 | 0.5785398 | NA |
| ENSG00000105058 | FAM32A | 3564.393935 | 0.151186847 | 0.187576086 | 0.825221499 | 0.40924585 | 0.609863 |
| ENSG00000205476 | CCDC85C | 3676.146 | 0.151048 | 0.2125871 | 0.7362228 | 0.4615951 | 0.656292 |
| ENSG00000226251 | LINC02608 | 1.552311 | 0.1510471 | 0.7607264 | 0.6086504 | 0.5427562 | NA |
| ENSG00000287624 | | 1.442382 | 0.1510306 | 0.741763 | 0.4968586 | 0.6192888 | NA |
| ENSG00000274258 | MIR6728 | 1.542212 | 0.1509617 | 0.7257249 | 0.4874496 | 0.6259398 | NA |
| ENSG00000260919 | | 1.921567 | 0.1509378 | 0.690408 | 0.4215271 | 0.6733702 | 0.818528 |
| ENSG00000221995 | TIAF1 | 71.01947 | 0.1508995 | 0.4296917 | 0.4211902 | 0.6736162 | 0.818574 |
| ENSG00000232194 | | 2.315099 | 0.1508224 | 0.788357 | 0.3416044 | 0.7326486 | 0.85508 |
| ENSG00000137831 | UACA | 2218.341381 | 0.150797395 | 0.226839768 | 0.691392531 | 0.4893189 | 0.680096 |
| ENSG00000111897 | SERINC1 | 8869.104808 | 0.150784428 | 0.226805704 | 0.691066123 | 0.48952399 | 0.680239 |
| ENSG00000162231 | NXF1 | 5441.944 | 0.1507244 | 0.1691595 | 0.9088463 | 0.3634313 | 0.565472 |
| ENSG00000277541 | ZNF630-AS1 | 0.8404336 | 0.1507066 | 0.7475463 | 0.4939069 | 0.6213719 | NA |
| ENSG00000261819 | | 34.10405 | 0.1506737 | 0.3273892 | 0.5078799 | 0.6115376 | 0.774478 |
| ENSG00000277999 | | 0.8291976 | 0.1505549 | 0.7685429 | 0.5767305 | 0.5641216 | NA |
| ENSG00000121897 | LIAS | 637.1317763 | 0.150528958 | 0.209584752 | 0.748902455 | 0.453916 | 0.650035 |
| ENSG00000256574 | OR13A1 | 0.9664236 | 0.150403 | 0.7869493 | 0.3531547 | 0.7239725 | NA |
| ENSG00000229409 | | 1.092403 | 0.1503904 | 0.7388794 | 0.5120774 | 0.6085968 | NA |
| ENSG00000112514 | CUTA | 5442.960304 | 0.150371155 | 0.183336856 | 0.8390702 | 0.40142992 | 0.602406 |
| ENSG00000105778 | AVL9 | 10305.55141 | 0.150366121 | 0.184508517 | 0.841412589 | 0.40011684 | 0.601114 |
| ENSG00000073050 | XRCC1 | 2099.256733 | 0.150344567 | 0.198026007 | 0.768583462 | 0.44214063 | 0.639967 |
| ENSG00000246394 | ZNF84-DT | 0.9596332 | 0.1503325 | 0.7506513 | 0.5225687 | 0.6012744 | NA |
| ENSG00000080503 | SMARCA2 | 3686.57227 | 0.150324914 | 0.197030903 | 0.781667392 | 0.43441007 | 0.633401 |

| | | | | | | | |
|---|---|---|---|---|---|---|---|
| ENSG00000132694 | ARHGEF11 | 4204.826371 | 0.150315238 | 0.180993174 | 0.806432893 | 0.41999329 | 0.620032 |
| ENSG00000132702 | HAPLN2 | 6.135482991 | 0.15018145 | 0.571993882 | 0.368299753 | 0.71264974 | 0.843628 |
| ENSG00000082512 | TRAF5 | 2196.10836 | 0.150142672 | 0.201435079 | 0.767141958 | 0.44299712 | 0.640594 |
| ENSG00000236130 | PTCSC2 | 2.507172 | 0.1500869 | 0.6559766 | 0.3998887 | 0.6892385 | 0.828804 |
| ENSG00000204237 | OXLD1 | 2922.378 | 0.1500412 | 0.2186581 | 0.7115171 | 0.4767639 | 0.669086 |
| ENSG00000177383 | MAGEF1 | 4167.967 | 0.1500198 | 0.1681214 | 0.9236195 | 0.3556845 | 0.557664 |
| ENSG00000254473 | | 80.29682 | 0.1500168 | 0.2610432 | 0.6083641 | 0.542946 | 0.723919 |
| ENSG00000106686 | SPATA6L | 47.89227703 | 0.150006659 | 0.313376718 | 0.515159602 | 0.6064415 | 0.770618 |
| ENSG00000144560 | VGLL4 | 2696.584 | 0.1500064 | 0.2199235 | 0.7101222 | 0.4776283 | 0.669704 |
| ENSG00000260064 | | 1.07896 | 0.1499995 | 0.7388932 | 0.4870449 | 0.6262265 | NA |
| ENSG00000279077 | | 1.096395 | 0.1499916 | 0.737495 | 0.4946784 | 0.6208271 | NA |
| ENSG00000185963 | BICD2 | 3192.961 | 0.1499595 | 0.1915172 | 0.8019835 | 0.4225625 | 0.62255 |
| ENSG00000236472 | | 1.392247 | 0.1499086 | 0.7262396 | 0.4811157 | 0.6304343 | NA |
| ENSG00000279246 | | 1.788684 | 0.149755 | 0.7052199 | 0.425432 | 0.6705218 | 0.816652 |
| ENSG00000258359 | PCNPP1 | 23.59777 | 0.149685 | 0.3964875 | 0.4341127 | 0.6642066 | 0.812304 |
| ENSG00000245719 | | 1.469833 | 0.149618 | 0.7303609 | 0.4832189 | 0.6289403 | NA |
| ENSG00000265386 | RN7SL219P | 13.51169 | 0.1495811 | 0.5459068 | 0.3683111 | 0.7126413 | 0.843628 |
| ENSG00000163528 | CHCHD4 | 2242.569 | 0.1495661 | 0.2070297 | 0.7527942 | 0.4515736 | 0.647868 |
| ENSG00000175322 | ZNF519 | 347.2427 | 0.1494651 | 0.2922189 | 0.5432202 | 0.5869782 | 0.756545 |
| ENSG00000180096 | SEPTIN1 | 70.99495 | 0.1494534 | 0.3028497 | 0.5328218 | 0.5941569 | 0.761696 |
| ENSG00000273797 | | 1.100167 | 0.1494448 | 0.7866937 | 0.3525823 | 0.7244016 | NA |
| ENSG00000260698 | | 13.41142 | 0.1493493 | 0.4993243 | 0.3787548 | 0.7048699 | 0.839807 |
| ENSG00000239374 | | 0.8048371 | 0.1492815 | 0.7385323 | 0.4909483 | 0.623463 | NA |
| ENSG00000205808 | PLPP6 | 916.681 | 0.149182 | 0.1671717 | 0.9101718 | 0.3627319 | 0.564781 |
| ENSG00000236064 | | 1.241674 | 0.1491009 | 0.7257105 | 0.4486167 | 0.6537082 | NA |
| ENSG00000213763 | ACTBP2 | 336.2285 | 0.1490672 | 0.2778128 | 0.5733927 | 0.5663788 | 0.741055 |
| ENSG00000170962 | PDGFD | 2.425551 | 0.1490153 | 0.7898152 | 0.3229111 | 0.7467626 | 0.863993 |
| ENSG00000257390 | | 1.661932 | 0.1490068 | 0.697643 | 0.4280661 | 0.668603 | 0.815391 |
| ENSG00000279200 | | 1.136368 | 0.1489894 | 0.7608526 | 0.5878213 | 0.5566522 | NA |
| ENSG00000115165 | CYTIP | 1.206552809 | 0.148918552 | 0.72702559 | 0.477739277 | 0.63283579 | NA |
| ENSG00000143001 | TMEM61 | 10.63246 | 0.1489173 | 0.5943688 | 0.3756991 | 0.7071406 | 0.840791 |
| ENSG00000282206 | | 7.237035 | 0.1487383 | 0.5363112 | 0.3691175 | 0.7120402 | 0.843269 |
| ENSG00000233994 | GDI2P2 | 296.1366 | 0.1487253 | 0.1928526 | 0.795342 | 0.4264146 | 0.62628 |

| | | | | | | | |
|---|---|---|---|---|---|---|---|
| ENSG00000111707 | SUDS3 | 4895.982026 | 0.148621078 | 0.127646267 | 1.155373671 | 0.24793744 | 0.440161 |
| ENSG00000122386 | ZNF205 | 925.8500042 | 0.148608945 | 0.192768499 | 0.790558392 | 0.42920174 | 0.62843 |
| ENSG00000251218 | | 1.202034 | 0.148542 | 0.7863553 | 0.3563098 | 0.7216085 | NA |
| ENSG00000167419 | LPO | 1.988541 | 0.1485364 | 0.6897375 | 0.4157166 | 0.6776174 | 0.820733 |
| ENSG00000274204 | | 3.818433 | 0.148532 | 0.6432915 | 0.3873189 | 0.6985201 | 0.835699 |
| ENSG00000286523 | | 1.064502 | 0.1484705 | 0.7531989 | 0.5264743 | 0.5985587 | NA |
| ENSG00000173110 | HSPA6 | 65.14591 | 0.1484646 | 0.5609357 | 0.3740651 | 0.7083559 | 0.841459 |
| ENSG00000235958 | UBOX5-AS1 | 8.418178 | 0.1483941 | 0.4931686 | 0.3839514 | 0.7010144 | 0.83727 |
| ENSG00000265907 | | 1.035626 | 0.1481724 | 0.7479651 | 0.5124689 | 0.6083229 | NA |
| ENSG00000111832 | RWDD1 | 2581.216919 | 0.148157715 | 0.165713027 | 0.887038223 | 0.37505832 | 0.577208 |
| ENSG00000229951 | | 97.51588 | 0.1480794 | 0.4454487 | 0.4001228 | 0.6890661 | 0.828763 |
| ENSG00000279274 | | 3.254448 | 0.148077 | 0.6597652 | 0.4008715 | 0.6885148 | 0.828457 |
| ENSG00000076108 | BAZ2A | 10083.59684 | 0.148062544 | 0.22719903 | 0.677268935 | 0.49823534 | 0.68667 |
| ENSG00000227479 | | 1.473179 | 0.1480454 | 0.7896924 | 0.3211992 | 0.7480595 | NA |
| ENSG00000269514 | | 1.219765 | 0.1480435 | 0.7883826 | 0.333023 | 0.7391169 | NA |
| ENSG00000163734 | CXCL3 | 4.697674 | 0.1480306 | 0.785002 | 0.36713 | 0.7135221 | 0.843684 |
| ENSG00000225712 | ATP5MC2P1 | 1.980327 | 0.1480152 | 0.7615903 | 0.6020627 | 0.5471324 | 0.726931 |
| ENSG00000213413 | PVRIG | 37.33741 | 0.1479765 | 0.5077505 | 0.3760213 | 0.7069011 | 0.840719 |
| ENSG00000102977 | ACD | 1851.121444 | 0.14795716 | 0.193819021 | 0.806318761 | 0.42005908 | 0.620036 |
| ENSG00000215440 | NPEPL1 | 3037.136 | 0.1479052 | 0.1790581 | 0.8462591 | 0.3974082 | 0.5985 |
| ENSG00000205794 | | 7.260367 | 0.1478708 | 0.568721 | 0.3680388 | 0.7128443 | 0.843639 |
| ENSG00000152700 | SAR1B | 4562.321 | 0.1477544 | 0.1102682 | 1.36247 | 0.1730495 | 0.344474 |
| ENSG00000223534 | HLA-DQB1-AS1 | 1.410009 | 0.1477384 | 0.7872627 | 0.3459519 | 0.7293789 | NA |
| ENSG00000176845 | METRNL | 6601.479 | 0.1477176 | 0.2022495 | 0.7370122 | 0.4611149 | 0.655995 |
| ENSG00000140987 | ZSCAN32 | 1268.212 | 0.1476866 | 0.1566313 | 0.9566457 | 0.3387461 | 0.54001 |
| ENSG00000224758 | LINC01167 | 31.58246 | 0.1476838 | 0.4719191 | 0.3843427 | 0.7007245 | 0.836972 |
| ENSG00000080511 | RDH8 | 1.190153087 | 0.147670111 | 0.801466706 | 0.755869943 | 0.44972718 | NA |
| ENSG00000162804 | SNED1 | 203.8795 | 0.147662 | 0.3092163 | 0.5088187 | 0.6108793 | 0.774202 |
| ENSG00000205231 | TTLL10-AS1 | 1.248222 | 0.1475628 | 0.7303955 | 0.481425 | 0.6302145 | NA |
| ENSG00000175215 | CTDSP2 | 13511.51 | 0.1475238 | 0.1933246 | 0.7846967 | 0.4326314 | 0.631814 |
| ENSG00000010671 | BTK | 3.49437551 | 0.147468331 | 0.6756971 | 0.41191604 | 0.68040096 | 0.82292 |
| ENSG00000232187 | FTH1P7 | 964.4073 | 0.1473608 | 0.2083629 | 0.7195455 | 0.4718049 | 0.665007 |
| ENSG00000233369 | GTF2IP4 | 1081.419 | 0.1473147 | 0.1703951 | 0.8905837 | 0.3731526 | 0.575342 |

| | | | | | | | |
|---|---|---|---|---|---|---|---|
| ENSG00000247011 | | 29.47362 | 0.1472956 | 0.58809 | 0.3682433 | 0.7126918 | 0.843639 |
| ENSG00000228445 | UGT1A2P | 2.933137 | 0.1472241 | 0.7895635 | 0.3200098 | 0.7489609 | 0.865675 |
| ENSG00000287837 | | 178.0816 | 0.1472214 | 0.1955283 | 0.7715751 | 0.4403661 | 0.638603 |
| ENSG00000117481 | NSUN4 | 2339.661415 | 0.147198417 | 0.100583184 | 1.458563115 | 0.14468539 | 0.303214 |
| ENSG00000288544 | NA | 0.8726585 | 0.1470585 | 0.7368599 | 0.4677812 | 0.639941 | NA |
| ENSG00000155636 | RBM45 | 871.7975 | 0.1469336 | 0.1333804 | 1.134605 | 0.256541 | 0.45029 |
| ENSG00000286863 | | 2.140704 | 0.1468851 | 0.7023056 | 0.4172497 | 0.6764957 | 0.820122 |
| ENSG00000254781 | GVINP2 | 0.9485644 | 0.1468586 | 0.8077311 | 1.150039 | 0.2501277 | NA |
| ENSG00000278908 | | 0.8891988 | 0.1467923 | 0.7870652 | 0.3411255 | 0.7330091 | NA |
| ENSG00000078269 | SYNJ2 | 1825.664328 | 0.146769252 | 0.222174448 | 0.681011737 | 0.49586406 | 0.685 |
| ENSG00000155100 | OTUD6B | 2049.77 | 0.1467305 | 0.1774368 | 0.8436173 | 0.3988833 | 0.599838 |
| ENSG00000248079 | DPH6-DT | 10.75321 | 0.1466389 | 0.5118445 | 0.3739978 | 0.708406 | 0.841459 |
| ENSG00000169903 | TM4SF4 | 1.382738 | 0.1466346 | 0.7862418 | 0.3481785 | 0.7277061 | NA |
| ENSG00000149782 | PLCB3 | 4146.185 | 0.1466188 | 0.1539255 | 0.9773689 | 0.3283866 | 0.529584 |
| ENSG00000279421 | | 0.7948018 | 0.1465911 | 0.7566683 | 0.5475219 | 0.5840202 | NA |
| ENSG00000225178 | RPSAP58 | 12794.38 | 0.1465791 | 0.2183092 | 0.7424858 | 0.457793 | 0.65359 |
| ENSG00000202399 | | 2.308298 | 0.1464688 | 0.6928536 | 0.4224985 | 0.6726612 | 0.81805 |
| ENSG00000284677 | | 1.811598 | 0.1464338 | 0.7014168 | 0.4107967 | 0.6812216 | 0.823598 |
| ENSG00000268705 | BNIP3P26 | 4.917341 | 0.1463765 | 0.6417488 | 0.3754403 | 0.707333 | 0.840865 |
| ENSG00000204542 | C6orf15 | 1.081404 | 0.1463425 | 0.7722291 | 0.6046979 | 0.5453797 | NA |
| ENSG00000274034 | MIR6813 | 2.15101 | 0.1463112 | 0.6380697 | 0.3726062 | 0.7094416 | 0.841931 |
| ENSG00000101017 | CD40 | 3.991854701 | 0.146292361 | 0.787053525 | 0.343582697 | 0.73116014 | 0.854161 |
| ENSG00000109205 | ODAM | 34.12602064 | 0.146196567 | 0.679922157 | 0.410047899 | 0.68177081 | 0.823818 |
| ENSG00000275549 | STPG3-AS1 | 42.53782 | 0.1461863 | 0.3635294 | 0.4526924 | 0.6507703 | 0.802697 |
| ENSG00000253915 | MAPRE1P1 | 3.245785 | 0.1460706 | 0.6260335 | 0.3702609 | 0.7111881 | 0.842862 |
| ENSG00000156873 | PHKG2 | 3397.451 | 0.1460508 | 0.1788897 | 0.8439525 | 0.3986959 | 0.599658 |
| ENSG00000268996 | MAN1B1-DT | 323.871 | 0.1460252 | 0.3110857 | 0.5109646 | 0.6093758 | 0.77289 |
| ENSG00000128394 | APOBEC3F | 16.44499372 | 0.145997047 | 0.622124449 | 0.375507443 | 0.70728311 | 0.840865 |
| ENSG00000255181 | CCDC166 | 2.890368 | 0.1459825 | 0.681018 | 0.3766882 | 0.7064054 | 0.840456 |
| ENSG00000121749 | TBC1D15 | 2191.173465 | 0.145941518 | 0.127875541 | 1.152028396 | 0.24930941 | 0.4418 |
| ENSG00000272540 | | 42.30199 | 0.145928 | 0.4199034 | 0.408974 | 0.6825588 | 0.824314 |
| ENSG00000196284 | SUPT3H | 389.2705 | 0.1459197 | 0.2095505 | 0.7226173 | 0.469915 | 0.663331 |
| ENSG00000268199 | | 1.855797 | 0.1458702 | 0.7267098 | 0.4608423 | 0.6449117 | 0.797888 |

| | | | | | | | |
|---|---|---|---|---|---|---|---|
| ENSG00000171243 | SOSTDC1 | 9.384339 | 0.1458306 | 0.5519873 | 0.3586112 | 0.719886 | 0.847625 |
| ENSG00000174938 | SEZ6L2 | 4695.423 | 0.1457647 | 0.202951 | 0.7363785 | 0.4615004 | 0.656252 |
| ENSG00000175582 | RAB6A | 8705.989 | 0.1456992 | 0.1309892 | 1.099296 | 0.2716392 | 0.468058 |
| ENSG00000128039 | SRD5A3 | 1184.026066 | 0.145679182 | 0.199884341 | 0.725136336 | 0.4683684 | 0.661979 |
| ENSG00000227416 | | 0.8613381 | 0.1456664 | 0.7380224 | 0.4770726 | 0.6333104 | NA |
| ENSG00000110108 | TMEM109 | 4613.516786 | 0.145638329 | 0.194107555 | 0.778480032 | 0.43628607 | 0.634848 |
| ENSG00000015285 | WAS | 141.2258737 | 0.145627456 | 0.2781759 | 0.565638555 | 0.57163951 | 0.745294 |
| ENSG00000173418 | NAA20 | 4969.678 | 0.1456171 | 0.1532944 | 0.9572446 | 0.3384438 | 0.539775 |
| ENSG00000160883 | HK3 | 3.452886 | 0.145544 | 0.6401479 | 0.3671543 | 0.7135039 | 0.843684 |
| ENSG00000241472 | PTPRG-AS1 | 434.8487 | 0.1454429 | 0.2963568 | 0.5138482 | 0.6073581 | 0.771285 |
| ENSG00000229926 | IL9RP1 | 1.118036 | 0.1454345 | 0.786592 | 0.3413995 | 0.7328029 | NA |
| ENSG00000256950 | | 2.121363 | 0.1454238 | 0.7243975 | 0.4684931 | 0.639432 | 0.794189 |
| ENSG00000272990 | | 5.49457 | 0.1453988 | 0.6428797 | 0.3715016 | 0.710264 | 0.842335 |
| ENSG00000149925 | ALDOA | 2177.232 | 0.1453671 | 0.212417 | 0.7186431 | 0.4723608 | 0.665367 |
| ENSG00000166263 | STXBP4 | 770.4479 | 0.1452994 | 0.2324013 | 0.6584227 | 0.5102666 | 0.696102 |
| ENSG00000268521 | VN1R83P | 18.16693 | 0.1452813 | 0.5081071 | 0.3696596 | 0.7116361 | 0.843114 |
| ENSG00000282742 | | 1.588881 | 0.1452005 | 0.7248121 | 0.4351547 | 0.6634501 | 0.81164 |
| ENSG00000166272 | WBP1L | 2928.739 | 0.1449088 | 0.134646 | 1.091652 | 0.2749859 | 0.471757 |
| ENSG00000089094 | KDM2B | 2837.630414 | 0.144880736 | 0.191001813 | 0.777870042 | 0.43664563 | 0.635203 |
| ENSG00000279759 | | 232.7065 | 0.1448429 | 0.2848914 | 0.5400241 | 0.5891804 | 0.75806 |
| ENSG00000254542 | NAV2-AS3 | 1.746147 | 0.1447604 | 0.7325478 | 0.4350368 | 0.6635357 | 0.81167 |
| ENSG00000260190 | | 8.647225 | 0.1447179 | 0.5869052 | 0.3615615 | 0.7176798 | 0.846015 |
| ENSG00000279161 | | 6.877548 | 0.1446987 | 0.6259882 | 0.3752207 | 0.7074963 | 0.840975 |
| ENSG00000288534 | | 12.85551 | 0.1446482 | 0.5420291 | 0.3552078 | 0.7224339 | 0.849007 |
| ENSG00000205763 | RP9P | 470.9314 | 0.1445868 | 0.2224671 | 0.6754473 | 0.4993917 | 0.687706 |
| ENSG00000178719 | GRINA | 14778.31 | 0.1445738 | 0.1887598 | 0.7770126 | 0.4371514 | 0.63562 |
| ENSG00000168397 | ATG4B | 5511.862 | 0.1445638 | 0.1556367 | 0.9379129 | 0.3482892 | 0.550029 |
| ENSG00000179403 | VWA1 | 2379.944 | 0.1445569 | 0.2531334 | 0.6099577 | 0.5418898 | 0.723497 |
| ENSG00000235546 | | 4.623952 | 0.1445439 | 0.6947544 | 0.4143489 | 0.6786186 | 0.821485 |
| ENSG00000243243 | | 5.334641 | 0.1445125 | 0.6599202 | 0.3909362 | 0.6958444 | 0.833501 |
| ENSG00000258628 | RHPN2P1 | 2.262722 | 0.1444976 | 0.7323819 | 0.4809059 | 0.6305834 | 0.788014 |
| ENSG00000153187 | HNRNPU | 29074.28 | 0.1444966 | 0.1470759 | 0.9954171 | 0.3195334 | 0.520468 |
| ENSG00000279329 | | 67.69938 | 0.1444758 | 0.3684149 | 0.4410463 | 0.6591795 | 0.808239 |

| | | | | | | | |
|---|---|---|---|---|---|---|---|
| ENSG00000256442 | | 1.018273 | 0.1442615 | 0.7711784 | 0.5510761 | 0.5815815 | NA |
| ENSG00000117616 | RSRP1 | 3207.204451 | 0.144235032 | 0.406239752 | 0.407505848 | 0.6836365 | 0.824997 |
| ENSG00000280384 | | 0.9485372 | 0.1441763 | 0.7868416 | 0.3355041 | 0.7372448 | NA |
| ENSG00000227212 | PFN1P6 | 62.49362 | 0.144138 | 0.432397 | 0.3947915 | 0.6929967 | 0.831365 |
| ENSG00000213015 | ZNF580 | 1411.859 | 0.1441366 | 0.2117282 | 0.6882057 | 0.4913233 | 0.681575 |
| ENSG00000207336 | RNU6-658P | 3.317405 | 0.1441174 | 0.6889824 | 0.4183838 | 0.6756665 | 0.819679 |
| ENSG00000138376 | BARD1 | 1016.25192 | 0.144102978 | 0.361763072 | 0.448329153 | 0.65391566 | 0.804895 |
| ENSG00000156858 | PRR14 | 2959.486 | 0.1440693 | 0.1676353 | 0.8787664 | 0.3795279 | 0.581513 |
| ENSG00000138668 | HNRNPD | 11204.45711 | 0.144032541 | 0.247454836 | 0.609578046 | 0.54214136 | 0.723521 |
| ENSG00000181619 | GPR135 | 227.4601 | 0.1440231 | 0.26237 | 0.5728298 | 0.5667599 | 0.741311 |
| ENSG00000118689 | FOXO3 | 3038.697921 | 0.143964062 | 0.303198301 | 0.517017808 | 0.60514373 | 0.76969 |
| ENSG00000166681 | BEX3 | 8463.717 | 0.143911 | 0.2143201 | 0.7405536 | 0.4589642 | 0.654618 |
| ENSG00000268189 | | 15.09682 | 0.1438722 | 0.4703927 | 0.3782942 | 0.7052121 | 0.839876 |
| ENSG00000261732 | | 18.32871 | 0.1436758 | 0.4655003 | 0.3795248 | 0.7042982 | 0.839351 |
| ENSG00000232377 | | 1.807249 | 0.1436194 | 0.7157727 | 0.439878 | 0.6600255 | 0.808746 |
| ENSG00000140104 | CLBA1 | 2254.49 | 0.1435259 | 0.1340539 | 1.073966 | 0.2828377 | 0.480634 |
| ENSG00000236364 | | 34.08771 | 0.1434902 | 0.3800003 | 0.4272562 | 0.6691927 | 0.815661 |
| ENSG00000231050 | | 15.53082 | 0.1434679 | 0.4817156 | 0.3728209 | 0.7092818 | 0.841817 |
| ENSG00000139263 | LRIG3 | 693.6614 | 0.143417 | 0.2023354 | 0.7335871 | 0.4632004 | 0.657448 |
| ENSG00000182774 | RPS17 | 996.8553 | 0.1433317 | 0.2182159 | 0.6694695 | 0.503196 | 0.690533 |
| ENSG00000124570 | SERPINB6 | 2768.711418 | 0.143302769 | 0.222792028 | 0.661105342 | 0.50854476 | 0.694925 |
| ENSG00000225674 | IPO7P2 | 418.2422 | 0.1431492 | 0.2600929 | 0.5809632 | 0.5612652 | 0.737311 |
| ENSG00000274805 | MIR6768 | 2.358516 | 0.1430794 | 0.6759266 | 0.3914377 | 0.6954737 | 0.83324 |
| ENSG00000007129 | CEACAM21 | 1.325486838 | 0.143005643 | 0.786545075 | 0.334720306 | 0.73783608 | NA |
| ENSG00000227209 | WARS2P1 | 2.54338 | 0.1429625 | 0.6827851 | 0.3513034 | 0.7253607 | 0.850828 |
| ENSG00000207475 | SNORA80E | 4.623537 | 0.142915 | 0.5833981 | 0.3554102 | 0.7222823 | 0.848898 |
| ENSG00000110048 | OSBP | 3483.866931 | 0.142892421 | 0.135375909 | 1.029111903 | 0.30342709 | 0.503086 |
| ENSG00000233231 | HNRNPA1P49 | 33.05328 | 0.1428414 | 0.4321298 | 0.3920262 | 0.6950389 | 0.832987 |
| ENSG00000251459 | | 3.265427 | 0.1428361 | 0.7871814 | 0.3323428 | 0.7396304 | 0.859371 |
| ENSG00000236570 | RAD23BP1 | 45.57622 | 0.1428245 | 0.3486039 | 0.4558719 | 0.6484821 | 0.800734 |
| ENSG00000143420 | ENSA | 23379.7 | 0.1427607 | 0.1426421 | 1.015504 | 0.3098657 | 0.510531 |
| ENSG00000283644 | ETDC | 1.381024 | 0.1427602 | 0.7348033 | 0.4738866 | 0.6355808 | NA |
| ENSG00000077380 | DYNC1I2 | 2873.712432 | 0.142750777 | 0.108601265 | 1.302395226 | 0.19278131 | 0.371241 |

| | | | | | | | |
|---|---|---|---|---|---|---|---|
| ENSG00000239577 | RN7SL388P | 0.9429524 | 0.1427374 | 0.7510729 | 0.489625 | 0.6243992 | NA |
| ENSG00000257635 | | 1.45026 | 0.1427176 | 0.7494647 | 0.4908783 | 0.6235125 | NA |
| ENSG00000265015 | | 1.529883 | 0.1427073 | 0.7476826 | 0.4608276 | 0.6449223 | NA |
| ENSG00000160087 | UBE2J2 | 2381.678 | 0.1426776 | 0.1888635 | 0.7747522 | 0.4384861 | 0.637072 |
| ENSG00000168275 | COA6 | 1086.86 | 0.1426601 | 0.2254666 | 0.660267 | 0.5090825 | 0.695272 |
| ENSG00000158156 | XKR8 | 545.8386 | 0.1426119 | 0.137543 | 1.052862 | 0.2924044 | 0.491267 |
| ENSG00000237494 | | 0.8470152 | 0.1425765 | 0.7529202 | 0.4961236 | 0.6198072 | NA |
| ENSG00000273308 | | 18.7019 | 0.1425498 | 0.4693331 | 0.3759781 | 0.7069332 | 0.840719 |
| ENSG00000162642 | C1orf52 | 1323.315 | 0.1425304 | 0.141474 | 1.025958 | 0.3049115 | 0.504653 |
| ENSG00000127580 | WDR24 | 813.1372563 | 0.142502304 | 0.193552345 | 0.756592821 | 0.44929384 | 0.645516 |
| ENSG00000188820 | CALHM6 | 1.692794 | 0.1424324 | 0.717202 | 0.4166634 | 0.6769246 | 0.820306 |
| ENSG00000286074 | | 3.322244 | 0.1422961 | 0.6348731 | 0.3654195 | 0.7147983 | 0.844633 |
| ENSG00000136059 | VILL | 49.81657308 | 0.142265722 | 0.512161487 | 0.367218683 | 0.7134559 | 0.84368 |
| ENSG00000228623 | ZNF883 | 1.773251 | 0.1422475 | 0.7880078 | 0.3198036 | 0.7491172 | 0.865699 |
| ENSG00000260518 | BMS1P8 | 10.63267 | 0.1422314 | 0.5412764 | 0.3566874 | 0.7213259 | 0.848466 |
| ENSG00000187510 | PLEKHG7 | 1.228932 | 0.1422305 | 0.7858277 | 0.3424316 | 0.7320261 | NA |
| ENSG00000139116 | KIF21A | 2526.819961 | 0.142178156 | 0.215487807 | 0.687138462 | 0.49199548 | 0.68221 |
| ENSG00000176842 | IRX5 | 1737.932 | 0.1421457 | 0.1985804 | 0.7481718 | 0.4543566 | 0.65042 |
| ENSG00000146592 | CREB5 | 6.449751 | 0.1421284 | 0.6277748 | 0.3634835 | 0.7162438 | 0.845492 |
| ENSG00000157152 | SYN2 | 4.215214 | 0.1420368 | 0.6698794 | 0.3913904 | 0.6955087 | 0.83324 |
| ENSG00000135248 | FAM71F1 | 2.609240964 | 0.14200299 | 0.676763411 | 0.38334311 | 0.70146538 | 0.83727 |
| ENSG00000275023 | MLLT6 | 8051.479 | 0.1419571 | 0.1934155 | 0.7624747 | 0.4457767 | 0.642779 |
| ENSG00000283765 | | 4.27505 | 0.1419222 | 0.6189753 | 0.3578571 | 0.7204503 | 0.847936 |
| ENSG00000130182 | ZSCAN10 | 1.173789116 | 0.141818971 | 0.723106767 | 0.408645454 | 0.68279987 | NA |
| ENSG00000287435 | | 2.23605 | 0.1418016 | 0.6848151 | 0.3938489 | 0.6936926 | 0.831861 |
| ENSG00000287035 | | 1.314567 | 0.1417709 | 0.7297577 | 0.4553943 | 0.6488256 | NA |
| ENSG00000254893 | RAP1BL | 1520.598 | 0.1417239 | 0.1417674 | 1.004394 | 0.3151889 | 0.516196 |
| ENSG00000176903 | PNMA1 | 5700.474 | 0.1417054 | 0.1314874 | 1.103593 | 0.2697698 | 0.466077 |
| ENSG00000271131 | PPIAP75 | 1.970379 | 0.1415423 | 0.7218071 | 0.4228826 | 0.6723809 | 0.817934 |
| ENSG00000260132 | | 2.099069 | 0.1415098 | 0.7142439 | 0.4304459 | 0.6668713 | 0.814371 |
| ENSG00000204659 | CBY3 | 16.23356 | 0.1414848 | 0.4399338 | 0.3834447 | 0.7013901 | 0.83727 |
| ENSG00000102225 | CDK16 | 7246.095598 | 0.141464055 | 0.127444667 | 1.109290049 | 0.26730508 | 0.463111 |
| ENSG00000145248 | SLC10A4 | 5.314539 | 0.1414583 | 0.6178063 | 0.3617213 | 0.7175603 | 0.846015 |

| | | | | | | | |
|---|---|---|---|---|---|---|---|
| ENSG00000266777 | SH3GL1P1 | 34.64595 | 0.1414371 | 0.4444088 | 0.3784333 | 0.7051087 | 0.839876 |
| ENSG00000112130 | RNF8 | 1348.16322 | 0.141378097 | 0.145366955 | 0.986316061 | 0.32397804 | 0.524868 |
| ENSG00000266801 | | 1.041922 | 0.1413772 | 0.786544 | 0.3311192 | 0.7405544 | NA |
| ENSG00000085491 | SLC25A24 | 7155.052572 | 0.141344092 | 0.204122083 | 0.70602369 | 0.48017341 | 0.671865 |
| ENSG00000263956 | NBPF11 | 2895.79 | 0.1413394 | 0.2375266 | 0.6192081 | 0.5357793 | 0.719035 |
| ENSG00000120265 | PCMT1 | 5584.49564 | 0.141337445 | 0.241945207 | 0.622405427 | 0.53367531 | 0.717377 |
| ENSG00000286686 | | 1.756233 | 0.1412766 | 0.7363749 | 0.478327 | 0.6324175 | 0.789298 |
| ENSG00000152102 | FAM168B | 10131.81 | 0.1412563 | 0.1554776 | 0.9353583 | 0.3496037 | 0.551286 |
| ENSG00000284669 | | 131.1606 | 0.1412272 | 0.6348058 | 0.3587481 | 0.7197836 | 0.847579 |
| ENSG00000279373 | | 2.714738 | 0.1411973 | 0.6502611 | 0.3675551 | 0.713205 | 0.843667 |
| ENSG00000245156 | | 69.265 | 0.1411043 | 0.3497143 | 0.4468456 | 0.6549865 | 0.80534 |
| ENSG00000273340 | MICE | 43.42143 | 0.1409932 | 0.3967196 | 0.4045411 | 0.6858149 | 0.826893 |
| ENSG00000251503 | CENPS-CORT | 0.8977247 | 0.1409122 | 0.7757307 | 0.6009937 | 0.5478442 | NA |
| ENSG00000203411 | | 6.182628 | 0.1408859 | 0.6151687 | 0.3518282 | 0.7249671 | 0.850636 |
| ENSG00000185522 | LMNTD2 | 500.0409 | 0.140765 | 0.3090083 | 0.4940993 | 0.6212361 | 0.781173 |
| ENSG00000235174 | RPL39P3 | 9164.158 | 0.1407267 | 0.1508759 | 0.9647217 | 0.3346842 | 0.535902 |
| ENSG00000160325 | CACFD1 | 2784.963 | 0.1406978 | 0.265803 | 0.5641302 | 0.5726656 | 0.745897 |
| ENSG00000084092 | NOA1 | 3609.780877 | 0.140665747 | 0.138701769 | 1.028539468 | 0.30369614 | 0.503331 |
| ENSG00000107021 | TBC1D13 | 2721.166726 | 0.140513405 | 0.122531674 | 1.134677116 | 0.25651063 | 0.450266 |
| ENSG00000275457 | | 46.19773 | 0.1405046 | 0.3288433 | 0.4670778 | 0.6404442 | 0.794841 |
| ENSG00000221598 | MIR1249 | 0.816157 | 0.1405038 | 0.7650238 | 0.5839988 | 0.5592211 | NA |
| ENSG00000174132 | FAM174A | 847.2451 | 0.1403809 | 0.1890059 | 0.7627457 | 0.4456151 | 0.642664 |
| ENSG00000120160 | EQTN | 0.938871836 | 0.140290672 | 0.788225304 | 0.312473556 | 0.75468066 | NA |
| ENSG00000133103 | COG6 | 2346.399127 | 0.14028614 | 0.150550135 | 0.954369453 | 0.33989666 | 0.541291 |
| ENSG00000257392 | | 0.8656553 | 0.1402779 | 0.7652029 | 0.5618714 | 0.5742036 | NA |
| ENSG00000197622 | CDC42SE1 | 14271.49 | 0.1401954 | 0.113396 | 1.245454 | 0.2129648 | 0.397712 |
| ENSG00000124444 | ZNF576 | 540.4310642 | 0.140161383 | 0.192120308 | 0.749971594 | 0.45327181 | 0.649427 |
| ENSG00000286121 | | 2.403184 | 0.1401256 | 0.6929511 | 0.4008963 | 0.6884965 | 0.828457 |
| ENSG00000108061 | SHOC2 | 2148.958946 | 0.14006776 | 0.16321015 | 0.877849352 | 0.38002548 | 0.581996 |
| ENSG00000235437 | LINC01278 | 1508.776 | 0.140036 | 0.214196 | 0.6718974 | 0.501649 | 0.689336 |
| ENSG00000175727 | MLXIP | 4460.771 | 0.1399836 | 0.2145575 | 0.6703309 | 0.5026469 | 0.690174 |
| ENSG00000116954 | RRAGC | 1288.541846 | 0.139974208 | 0.201339966 | 0.714267744 | 0.47506163 | 0.667649 |
| ENSG00000115020 | PIKFYVE | 2795.354102 | 0.139973641 | 0.287643688 | 0.525943612 | 0.59892739 | 0.765663 |

| | | | | | | | |
|---|---|---|---|---|---|---|---|
| ENSG00000198944 | SOWAHA | 96.51088 | 0.1399326 | 0.5402088 | 0.3491167 | 0.7270017 | 0.851749 |
| ENSG00000147669 | POLR2K | 3083.599 | 0.1398625 | 0.2200865 | 0.6601036 | 0.5091874 | 0.695358 |
| ENSG00000239198 | RPL5P22 | 4.654912 | 0.1398221 | 0.6662112 | 0.3870063 | 0.6987516 | 0.835727 |
| ENSG00000184313 | MROH7 | 1.33702 | 0.1396828 | 0.7869749 | 0.3253086 | 0.7449476 | NA |
| ENSG00000197471 | SPN | 1.672174 | 0.1396192 | 0.7285045 | 0.4308025 | 0.666612 | 0.814196 |
| ENSG00000267939 | | 13.60506 | 0.1395805 | 0.5276189 | 0.3484616 | 0.7274936 | 0.85203 |
| ENSG00000197181 | PIWIL2 | 2.916508 | 0.1395777 | 0.6503619 | 0.3634605 | 0.7162609 | 0.845492 |
| ENSG00000112039 | FANCE | 609.5574213 | 0.13938349 | 0.196906855 | 0.723607969 | 0.46930645 | 0.662848 |
| ENSG00000165724 | ZMYND19 | 2808.421 | 0.1393572 | 0.2221817 | 0.6193355 | 0.5356954 | 0.719014 |
| ENSG00000280003 | | 0.7957233 | 0.1393484 | 0.7860248 | 0.332744 | 0.7393276 | NA |
| ENSG00000188175 | HEPACAM2 | 2.110553 | 0.1393131 | 0.6996589 | 0.4042211 | 0.6860501 | 0.827102 |
| ENSG00000288526 | | 5.595071 | 0.1392571 | 0.6141597 | 0.351808 | 0.7249823 | 0.850636 |
| ENSG00000215467 | RPL27AP | 1.271286 | 0.1392177 | 0.7358638 | 0.4691625 | 0.6389535 | NA |
| ENSG00000272667 | | 147.5319 | 0.1391766 | 0.2457903 | 0.5979147 | 0.5498969 | 0.729016 |
| ENSG00000163617 | CCDC191 | 510.607 | 0.1391617 | 0.3131331 | 0.4862991 | 0.6267551 | 0.785291 |
| ENSG00000240828 | RPL21P4 | 0.9569823 | 0.1391585 | 0.7337805 | 0.4637619 | 0.6428183 | NA |
| ENSG00000215464 | | 0.8327322 | 0.1391417 | 0.747403 | 0.4821938 | 0.6296683 | NA |
| ENSG00000257735 | | 0.8094806 | 0.139137 | 0.7706882 | 0.5307822 | 0.5955697 | NA |
| ENSG00000159459 | UBR1 | 2107.856 | 0.1390933 | 0.1734558 | 0.8205077 | 0.4119268 | 0.612233 |
| ENSG00000047249 | ATP6V1H | 3012.214377 | 0.139082137 | 0.144975627 | 0.977771639 | 0.32818728 | 0.529395 |
| ENSG00000260898 | ADPGK-AS1 | 4.738713 | 0.138974 | 0.6723884 | 0.3714847 | 0.7102766 | 0.842335 |
| ENSG00000235510 | | 1.000816 | 0.138935 | 0.7868594 | 0.3215096 | 0.7478242 | NA |
| ENSG00000187904 | | 1.075642 | 0.1388373 | 0.7057275 | 0.3965584 | 0.6916931 | NA |
| ENSG00000240463 | RPS19P3 | 24.40353 | 0.1388177 | 0.3944073 | 0.4036999 | 0.6864334 | 0.827298 |
| ENSG00000261267 | | 3.444647 | 0.1388041 | 0.6483014 | 0.3609401 | 0.7181443 | 0.846264 |
| ENSG00000149292 | TTC12 | 581.4852 | 0.1388013 | 0.1740028 | 0.8006287 | 0.4233466 | 0.62329 |
| ENSG00000182108 | DEXI | 284.7565 | 0.1386447 | 0.2597473 | 0.5683105 | 0.5698242 | 0.743744 |
| ENSG00000265806 | MIR4292 | 38.96006 | 0.1385962 | 0.3666187 | 0.4241411 | 0.6714629 | 0.817373 |
| ENSG00000213645 | SLC25A1P3 | 3.374997 | 0.1385648 | 0.7066116 | 0.4030831 | 0.6868871 | 0.827596 |
| ENSG00000279833 | | 83.5985 | 0.1385612 | 0.3020745 | 0.4976711 | 0.6187159 | 0.77949 |
| ENSG00000123810 | B9D2 | 139.2848636 | 0.138559021 | 0.298765374 | 0.499361071 | 0.61752504 | 0.778728 |
| ENSG00000130649 | CYP2E1 | 52.32495544 | 0.138223453 | 0.465589096 | 0.360960678 | 0.71812884 | 0.846264 |
| ENSG00000232202 | CHORDC1P1 | 23.13952 | 0.138207 | 0.4995481 | 0.3735749 | 0.7087206 | 0.841624 |

| | | | | | | | |
|---|---|---|---|---|---|---|---|
| ENSG00000176396 | EID2 | 763.5695 | 0.1381397 | 0.2253447 | 0.6377467 | 0.5236386 | 0.708884 |
| ENSG00000136925 | TSTD2 | 1152.089974 | 0.138103943 | 0.24068139 | 0.593899598 | 0.55257927 | 0.731186 |
| ENSG00000023445 | BIRC3 | 62.60699865 | 0.138081159 | 0.491634269 | 0.356333746 | 0.72159064 | 0.848466 |
| ENSG00000232040 | ZBED9 | 292.6202 | 0.1380802 | 0.2534917 | 0.5710657 | 0.5679551 | 0.742362 |
| ENSG00000112658 | SRF | 4541.686611 | 0.138072095 | 0.251669275 | 0.572668226 | 0.56686936 | 0.741384 |
| ENSG00000140320 | BAHD1 | 3200.547 | 0.1380673 | 0.1492014 | 0.9413817 | 0.3465093 | 0.548075 |
| ENSG00000116791 | CRYZ | 2375.202008 | 0.138042349 | 0.170170728 | 0.830486972 | 0.40626351 | 0.606849 |
| ENSG00000143995 | MEIS1 | 129.245 | 0.1380111 | 0.3992214 | 0.3992786 | 0.689688 | 0.829057 |
| ENSG00000254702 | | 26.02955 | 0.1379215 | 0.3381302 | 0.4468323 | 0.6549961 | 0.80534 |
| ENSG00000214297 | ALDOAP2 | 10.92231 | 0.1378919 | 0.5929792 | 0.3508268 | 0.7257183 | 0.851005 |
| ENSG00000157570 | TSPAN18 | 18.31201 | 0.1378627 | 0.5534366 | 0.3435141 | 0.7312118 | 0.854182 |
| ENSG00000227946 | | 101.1091 | 0.1378567 | 0.2348272 | 0.6223798 | 0.5336922 | 0.717377 |
| ENSG00000275846 | | 1.00543 | 0.1378374 | 0.7437138 | 0.4662065 | 0.6410677 | NA |
| ENSG00000198225 | FKBP1C | 489.002 | 0.137825 | 0.1758925 | 0.7950644 | 0.426576 | 0.626378 |
| ENSG00000258500 | | 1.338696 | 0.1377537 | 0.7249159 | 0.44734 | 0.6546296 | NA |
| ENSG00000227117 | HORMAD2-AS1 | 1.082876 | 0.1376858 | 0.7302369 | 0.4450963 | 0.6562501 | NA |
| ENSG00000219951 | | 4.985537 | 0.1376786 | 0.6067432 | 0.3494824 | 0.7267272 | 0.851739 |
| ENSG00000284636 | | 1.237131 | 0.1376453 | 0.7865293 | 0.3210135 | 0.7482002 | NA |
| ENSG00000153157 | SYCP2L | 10.12271 | 0.1376434 | 0.5281641 | 0.3491499 | 0.7269768 | 0.851749 |
| ENSG00000285777 | | 0.8941454 | 0.1376191 | 0.7983602 | 0.6501372 | 0.5156036 | NA |
| ENSG00000234181 | | 0.9867995 | 0.1375215 | 0.7359567 | 0.457464 | 0.6473376 | NA |
| ENSG00000221970 | OR2A1 | 1.488537 | 0.1374911 | 0.701802 | 0.3953565 | 0.6925798 | NA |
| ENSG00000204859 | ZBTB48 | 954.6297 | 0.1373869 | 0.1080883 | 1.306703 | 0.1913135 | 0.369405 |
| ENSG00000275874 | PICSAR | 0.9770378 | 0.1373219 | 0.7860923 | 0.3241711 | 0.7458085 | NA |
| ENSG00000281883 | | 4.087067 | 0.1372449 | 0.604832 | 0.3415965 | 0.7326546 | 0.85508 |
| ENSG00000235173 | HGH1 | 2780.227 | 0.1372429 | 0.1998272 | 0.7071597 | 0.4794672 | 0.671461 |
| ENSG00000185101 | ANO9 | 3059.278 | 0.137192 | 0.3023195 | 0.5008952 | 0.6164449 | 0.777845 |
| ENSG00000259515 | | 20.90775 | 0.1371301 | 0.4449706 | 0.3675929 | 0.7131768 | 0.843667 |
| ENSG00000080947 | CROCCP3 | 234.3738849 | 0.137098078 | 0.342352683 | 0.438587759 | 0.66096027 | 0.809437 |
| ENSG00000244086 | | 1.816016 | 0.1370925 | 0.7060796 | 0.4092266 | 0.6823734 | 0.82413 |
| ENSG00000271969 | | 6.008577 | 0.1370481 | 0.6277002 | 0.3487599 | 0.7272696 | 0.851826 |
| ENSG00000205041 | | 41.51435 | 0.1370302 | 0.3680365 | 0.4167088 | 0.6768914 | 0.820303 |
| ENSG00000284693 | LINC02606 | 10.07739 | 0.1369756 | 0.544791 | 0.3430023 | 0.7315967 | 0.854371 |

| | | | | | | | |
|---|---|---|---|---|---|---|---|
| ENSG00000231532 | LINC01249 | 9.099541 | 0.1369486 | 0.7397072 | 0.3869485 | 0.6987943 | 0.835727 |
| ENSG00000275066 | SYNRG | 2082.974 | 0.1368703 | 0.1809032 | 0.7592161 | 0.4477233 | 0.644431 |
| ENSG00000115850 | LCT | 3.958664766 | 0.136796602 | 0.67422774 | 0.373797749 | 0.70855479 | 0.841518 |
| ENSG00000156253 | RWDD2B | 1740.621 | 0.1367625 | 0.2155579 | 0.6589885 | 0.5099031 | 0.695852 |
| ENSG00000022840 | RNF10 | 7972.461413 | 0.136641991 | 0.129318598 | 1.039622934 | 0.29851512 | 0.497725 |
| ENSG00000242265 | PEG10 | 917.3313 | 0.1366393 | 0.4136992 | 0.388369 | 0.697743 | 0.835022 |
| ENSG00000238003 | RPL10P4 | 1.224781 | 0.1365978 | 0.7856792 | 0.3283844 | 0.742621 | NA |
| ENSG00000106992 | AK1 | 209.8921542 | 0.136526482 | 0.254898455 | 0.561754573 | 0.57428325 | 0.74674 |
| ENSG00000272491 | | 0.7663045 | 0.1365132 | 0.7997666 | 0.8469046 | 0.3970483 | NA |
| ENSG00000167565 | SERTAD3 | 2331.014 | 0.1364947 | 0.1440555 | 0.9414323 | 0.3464834 | 0.548075 |
| ENSG00000254158 | | 0.928468 | 0.1364416 | 0.7700232 | 0.5556629 | 0.5784413 | NA |
| ENSG00000107593 | PKD2L1 | 1.393559075 | 0.136386551 | 0.704485518 | 0.399039702 | 0.68986395 | NA |
| ENSG00000221826 | PSG3 | 3.126091 | 0.1363678 | 0.7873128 | 0.3110396 | 0.7557705 | 0.869681 |
| ENSG00000187860 | CCDC157 | 150.0881 | 0.1363358 | 0.2623659 | 0.5549408 | 0.5789352 | 0.750255 |
| ENSG00000234936 | | 35.57095 | 0.1362485 | 0.440398 | 0.3724073 | 0.7095896 | 0.842069 |
| ENSG00000183317 | EPHA10 | 709.6644 | 0.1361827 | 0.2075468 | 0.6956469 | 0.48665 | 0.677663 |
| ENSG00000260021 | | 1.221379 | 0.1360796 | 0.7596498 | 0.5883769 | 0.5562793 | NA |
| ENSG00000254676 | | 20.16949 | 0.1360759 | 0.4803368 | 0.3535371 | 0.7236858 | 0.849877 |
| ENSG00000264635 | | 14.98529 | 0.1360468 | 0.5206725 | 0.3430233 | 0.7315809 | 0.854371 |
| ENSG00000223508 | RPL23AP53 | 73.0784 | 0.1360403 | 0.2606262 | 0.5597146 | 0.5756741 | 0.747853 |
| ENSG00000261499 | | 30.19914 | 0.136028 | 0.3902881 | 0.4028046 | 0.687092 | 0.827607 |
| ENSG00000260253 | | 2.158089 | 0.1360092 | 0.7179427 | 0.4387564 | 0.660838 | 0.809377 |
| ENSG00000267422 | | 567.5255 | 0.1359358 | 0.1566761 | 0.8884068 | 0.374322 | 0.576409 |
| ENSG00000104361 | NIPAL2 | 2486.50527 | 0.135917508 | 0.256272529 | 0.556285544 | 0.57801567 | 0.749685 |
| ENSG00000245937 | LINC01184 | 1559.8 | 0.1357087 | 0.1654223 | 0.8405662 | 0.400591 | 0.601554 |
| ENSG00000164690 | SHH | 28.09259 | 0.1356548 | 0.4587269 | 0.3571446 | 0.7209835 | 0.848278 |
| ENSG00000164808 | SPIDR | 4035.571 | 0.1356375 | 0.1453706 | 0.9742479 | 0.3299335 | 0.531028 |
| ENSG00000172900 | ACTE1P | 1.230678 | 0.1356299 | 0.7474144 | 0.4845248 | 0.6280134 | NA |
| ENSG00000188505 | NCCRP1 | 5.355015 | 0.1356129 | 0.5843426 | 0.3411636 | 0.7329804 | 0.855309 |
| ENSG00000154429 | CCSAP | 869.2563 | 0.1354835 | 0.1995957 | 0.6935538 | 0.4879621 | 0.67892 |
| ENSG00000237641 | RPL28P2 | 18.95719 | 0.1354779 | 0.4301778 | 0.3717146 | 0.7101054 | 0.842335 |
| ENSG00000225839 | TRMT2B-AS1 | 1.09325 | 0.1354684 | 0.7801181 | 0.7304499 | 0.4651152 | NA |
| ENSG00000105429 | MEGF8 | 2540.844214 | 0.135446455 | 0.243312228 | 0.58762944 | 0.55678105 | 0.734223 |

| | | | | | | | |
|---|---|---|---|---|---|---|---|
| ENSG00000160055 | TMEM234 | 576.3383 | 0.1353448 | 0.2223079 | 0.6364185 | 0.5245037 | 0.709477 |
| ENSG00000180340 | FZD2 | 1431.699 | 0.1352775 | 0.2475158 | 0.5750468 | 0.5652596 | 0.7402 |
| ENSG00000165323 | FAT3 | 1.387815 | 0.1352648 | 0.7889934 | 0.2931456 | 0.7694108 | NA |
| ENSG00000258471 | | 2.333516 | 0.1352286 | 0.6977937 | 0.3833228 | 0.7014804 | 0.83727 |
| ENSG00000066455 | GOLGA5 | 5436.418965 | 0.135166844 | 0.111627464 | 1.204350277 | 0.22845422 | 0.416546 |
| ENSG00000156172 | C8orf37 | 269.831 | 0.1351424 | 0.2052862 | 0.6839122 | 0.4940306 | 0.683819 |
| ENSG00000165757 | JCAD | 256.2227 | 0.135073 | 0.3871908 | 0.4021214 | 0.6875947 | 0.827949 |
| ENSG00000130005 | GAMT | 810.6608601 | 0.135069861 | 0.236292034 | 0.594320105 | 0.55229803 | 0.730854 |
| ENSG00000255655 | | 1.034295 | 0.135057 | 0.7864244 | 0.3145234 | 0.7531235 | NA |
| ENSG00000257199 | | 19.90485 | 0.1350139 | 0.4631854 | 0.3563666 | 0.721566 | 0.848466 |
| ENSG00000133106 | EPSTI1 | 4.977600917 | 0.134987146 | 0.653338629 | 0.352424259 | 0.72452011 | 0.850449 |
| ENSG00000268070 | | 6.153278 | 0.1349343 | 0.6695795 | 0.3700619 | 0.7113363 | 0.842862 |
| ENSG00000249885 | ARHGEF38-IT1 | 1.163914 | 0.1349328 | 0.7857459 | 0.3234427 | 0.74636 | NA |
| ENSG00000119285 | HEATR1 | 4671.832154 | 0.134890984 | 0.231885459 | 0.617266533 | 0.53705894 | 0.720111 |
| ENSG00000229521 | MYCBP2-AS2 | 1.06611 | 0.1348907 | 0.7597193 | 0.6013004 | 0.5476399 | NA |
| ENSG00000181090 | EHMT1 | 5407.014 | 0.1348712 | 0.1145577 | 1.188319 | 0.2347078 | 0.423897 |
| ENSG00000130881 | LRP3 | 2918.007829 | 0.134858183 | 0.163755792 | 0.844126352 | 0.3985988 | 0.599579 |
| ENSG00000267152 | | 10.80764 | 0.1348447 | 0.5081389 | 0.3418747 | 0.7324452 | 0.854968 |
| ENSG00000140505 | CYP1A2 | 8.329227 | 0.1347923 | 0.692501 | 0.3971143 | 0.6912832 | 0.830171 |
| ENSG00000171490 | RSL1D1 | 13465.95 | 0.134702 | 0.1657012 | 0.8222291 | 0.4109465 | 0.611301 |
| ENSG00000224984 | | 40.41614 | 0.134586 | 0.4702669 | 0.3508037 | 0.7257356 | 0.851005 |
| ENSG00000239407 | | 108.1423 | 0.1345777 | 0.4352853 | 0.3696328 | 0.7116561 | 0.843114 |
| ENSG00000258932 | | 2.014954 | 0.1345776 | 0.7886035 | 0.2970797 | 0.7664057 | 0.876151 |
| ENSG00000175287 | PHYHD1 | 2.057544 | 0.1345689 | 0.7262454 | 0.4479182 | 0.6542122 | 0.804972 |
| ENSG00000251405 | | 0.9094884 | 0.1345057 | 0.7651277 | 0.5487041 | 0.5832086 | NA |
| ENSG00000286873 | | 60.16927 | 0.1344087 | 0.3489269 | 0.4296795 | 0.6674288 | 0.814764 |
| ENSG00000197291 | RAMP2-AS1 | 6.806956 | 0.1343484 | 0.6136893 | 0.3393357 | 0.7343569 | 0.856014 |
| ENSG00000105171 | POP4 | 2437.239184 | 0.134274955 | 0.220764632 | 0.628709927 | 0.52953898 | 0.714251 |
| ENSG00000163785 | RYK | 5629.602 | 0.1342722 | 0.1316123 | 1.009286 | 0.3128376 | 0.513884 |
| ENSG00000254924 | | 1.755889 | 0.1340996 | 0.7281435 | 0.4284814 | 0.6683007 | 0.815124 |
| ENSG00000230965 | SNX18P13 | 18.90678 | 0.1340532 | 0.5019672 | 0.3439776 | 0.7308632 | 0.853987 |
| ENSG00000123728 | RAP2C | 3275.470387 | 0.134050603 | 0.211837193 | 0.685206996 | 0.49321331 | 0.683079 |
| ENSG00000141084 | RANBP10 | 2941.055 | 0.1340094 | 0.1735276 | 0.7942062 | 0.4270754 | 0.626627 |

| | | | | | | | |
|---|---|---|---|---|---|---|---|
| ENSG00000186462 | NAP1L2 | 331.9076 | 0.1338837 | 0.3086629 | 0.4651754 | 0.6418058 | 0.795751 |
| ENSG00000285813 | | 23.41645 | 0.1338347 | 0.4297923 | 0.3678739 | 0.7129672 | 0.843639 |
| ENSG00000104824 | HNRNPL | 6390.60517 | 0.133763674 | 0.192885283 | 0.718300636 | 0.47257194 | 0.665453 |
| ENSG00000272432 | | 5.709987 | 0.1337621 | 0.5632053 | 0.3381345 | 0.7352618 | 0.856543 |
| ENSG00000187091 | PLCD1 | 361.3122 | 0.1336997 | 0.2681573 | 0.5275835 | 0.5977885 | 0.764692 |
| ENSG00000241458 | RPL7P19 | 3.752277 | 0.133656 | 0.6635717 | 0.3617723 | 0.7175222 | 0.846015 |
| ENSG00000184900 | SUMO3 | 11113.76 | 0.1336329 | 0.2030776 | 0.6720878 | 0.5015278 | 0.689312 |
| ENSG00000232434 | AJM1 | 701.9997 | 0.1336295 | 0.1897188 | 0.7291264 | 0.4659243 | 0.659777 |
| ENSG00000198690 | FAN1 | 1955.957 | 0.1335625 | 0.2201512 | 0.6257213 | 0.5314977 | 0.715586 |
| ENSG00000253616 | | 1.999622 | 0.1334856 | 0.7121382 | 0.3971558 | 0.6912526 | 0.830171 |
| ENSG00000234848 | | 1.77254 | 0.1334452 | 0.7265086 | 0.4156518 | 0.6776648 | 0.820733 |
| ENSG00000228677 | TTC3-AS1 | 2.270712 | 0.1334295 | 0.6995982 | 0.3862173 | 0.6993357 | 0.835987 |
| ENSG00000241155 | ARHGAP31-AS1 | 4.166906 | 0.1334046 | 0.6466501 | 0.3497551 | 0.7265225 | 0.851589 |
| ENSG00000139233 | LLPH | 1527.532 | 0.1333173 | 0.1499916 | 0.9134209 | 0.3610212 | 0.563041 |
| ENSG00000231561 | CEACAMP5 | 0.842294 | 0.133317 | 0.7652781 | 0.5458265 | 0.5851852 | NA |
| ENSG00000125458 | NT5C | 1326.281762 | 0.133299462 | 0.154232568 | 0.879359586 | 0.37920634 | 0.581202 |
| ENSG00000224276 | ARHGEF2-AS1 | 1.058692 | 0.1332812 | 0.7269164 | 0.4252867 | 0.6706277 | NA |
| ENSG00000199890 | | 0.9308975 | 0.1332624 | 0.7853991 | 0.3222256 | 0.7472818 | NA |
| ENSG00000254449 | SF3A3P2 | 39.97926 | 0.1332449 | 0.3530714 | 0.4208894 | 0.6738358 | 0.818654 |
| ENSG00000070886 | EPHA8 | 8.39296298 | 0.133244019 | 0.633186415 | 0.348314158 | 0.72760426 | 0.85203 |
| ENSG00000284605 | | 0.8438832 | 0.1331851 | 0.7429984 | 0.448167 | 0.6540327 | NA |
| ENSG00000260555 | | 2.808605 | 0.1331799 | 0.6579705 | 0.3465785 | 0.728908 | 0.853031 |
| ENSG00000286271 | | 1.591103 | 0.133177 | 0.7288082 | 0.3966749 | 0.6916072 | 0.830495 |
| ENSG00000028528 | SNX1 | 8169.500454 | 0.13316004 | 0.128068953 | 1.051605181 | 0.29298073 | 0.491833 |
| ENSG00000270903 | HNRNPA3P9 | 5.460384 | 0.1331061 | 0.6553964 | 0.3481114 | 0.7277565 | 0.852134 |
| ENSG00000255587 | RAB44 | 6.728812 | 0.1330396 | 0.5455698 | 0.3328023 | 0.7392835 | 0.859194 |
| ENSG00000158806 | NPM2 | 2.703218 | 0.1330129 | 0.6668142 | 0.3538667 | 0.7234388 | 0.849774 |
| ENSG00000287965 | | 2.048284 | 0.1329949 | 0.6865719 | 0.3688664 | 0.7122273 | 0.843338 |
| ENSG00000168434 | COG7 | 1615.986 | 0.1329825 | 0.1160748 | 1.153171 | 0.2488402 | 0.441175 |
| ENSG00000235066 | | 1.902997 | 0.1329176 | 0.6771451 | 0.3662585 | 0.7141722 | 0.844077 |
| ENSG00000120337 | TNFSF18 | 0.801939233 | 0.13291277 | 0.797789125 | 0.639747985 | 0.52233645 | NA |
| ENSG00000165240 | ATP7A | 1169.432 | 0.1329071 | 0.3083533 | 0.4641708 | 0.6425254 | 0.796178 |
| ENSG00000124596 | OARD1 | 1602.700927 | 0.132902795 | 0.154175593 | 0.876301794 | 0.38086599 | 0.582905 |

| | | | | | | | |
|---|---|---|---|---|---|---|---|
| ENSG00000116786 | PLEKHM2 | 2654.57967 | 0.132895744 | 0.138000369 | 0.969458326 | 0.33231656 | 0.533405 |
| ENSG00000284711 | | 1.310771 | 0.132894 | 0.7302115 | 0.4305345 | 0.6668069 | NA |
| ENSG00000166949 | SMAD3 | 8691.883 | 0.1328926 | 0.2556762 | 0.5945247 | 0.5521613 | 0.730778 |
| ENSG00000226752 | CUTALP | 3233.085 | 0.1328242 | 0.2780682 | 0.5058417 | 0.6129678 | 0.775706 |
| ENSG00000163597 | SNHG16 | 3204.934 | 0.1327918 | 0.1316017 | 1.006366 | 0.3142394 | 0.515372 |
| ENSG00000118900 | UBN1 | 6127.501657 | 0.132564209 | 0.185069731 | 0.728085739 | 0.4665611 | 0.660384 |
| ENSG00000285692 | | 1.167883 | 0.1325039 | 0.7237643 | 0.4165504 | 0.6770073 | NA |
| ENSG00000062370 | ZNF112 | 259.7774079 | 0.132423582 | 0.287599837 | 0.490572835 | 0.6237286 | 0.782909 |
| ENSG00000236651 | DLX2-DT | 10.44628 | 0.1322073 | 0.6057917 | 0.3356977 | 0.7370988 | 0.85793 |
| ENSG00000257550 | | 1.08414 | 0.132192 | 0.7657919 | 0.5614591 | 0.5744846 | NA |
| ENSG00000258731 | | 1.374643 | 0.132139 | 0.7381508 | 0.4385704 | 0.6609729 | NA |
| ENSG00000091490 | SEL1L3 | 724.5032457 | 0.132136252 | 0.298700147 | 0.473807386 | 0.63563727 | 0.791501 |
| ENSG00000173894 | CBX2 | 4850.88 | 0.132072 | 0.2551645 | 0.5623445 | 0.5738813 | 0.746641 |
| ENSG00000273820 | USP27X | 455.8531 | 0.1320405 | 0.1585812 | 0.8551132 | 0.3924885 | 0.59407 |
| ENSG00000196218 | RYR1 | 7.888371 | 0.1320201 | 0.5750403 | 0.3290879 | 0.7420892 | 0.861061 |
| ENSG00000246316 | | 1.161998 | 0.1318949 | 0.7365879 | 0.4367682 | 0.6622794 | NA |
| ENSG00000188064 | WNT7B | 2580.919 | 0.1317605 | 0.1888894 | 0.7059635 | 0.4802108 | 0.671865 |
| ENSG00000211899 | IGHM | 1.21708 | 0.1317472 | 0.7711416 | 0.4734402 | 0.6358992 | NA |
| ENSG00000170776 | AKAP13 | 3523.302 | 0.1317196 | 0.2511405 | 0.5496548 | 0.5825562 | 0.753111 |
| ENSG00000089091 | DZANK1 | 327.1733524 | 0.131698343 | 0.187347645 | 0.716506312 | 0.47367877 | 0.666446 |
| ENSG00000253030 | MIR2116 | 0.817014 | 0.1315714 | 0.7862632 | 0.3078788 | 0.7581746 | NA |
| ENSG00000102128 | RAB40AL | 1.230944776 | 0.131567617 | 0.711153858 | 0.384012914 | 0.70096888 | NA |
| ENSG00000103549 | RNF40 | 6701.140252 | 0.131497402 | 0.139842865 | 0.952984663 | 0.34059784 | 0.541994 |
| ENSG00000241593 | | 1.816245 | 0.1314422 | 0.7155703 | 0.3889418 | 0.6973192 | 0.834703 |
| ENSG00000235686 | PPIAP20 | 1.189118 | 0.1314132 | 0.738514 | 0.4492897 | 0.6532227 | NA |
| ENSG00000188763 | FZD9 | 21.71379 | 0.1313979 | 0.5909558 | 0.335789 | 0.73703 | 0.857888 |
| ENSG00000179222 | MAGED1 | 9077.365 | 0.1311519 | 0.2128566 | 0.6370119 | 0.5241171 | 0.709206 |
| ENSG00000140057 | AK7 | 1918.357 | 0.1311213 | 0.1454541 | 0.9483469 | 0.3429529 | 0.544483 |
| ENSG00000280080 | TBC1D22A-AS1 | 1.168865 | 0.1310834 | 0.7307403 | 0.4360791 | 0.6627794 | NA |
| ENSG00000258952 | SALRNA1 | 8.237546 | 0.1310396 | 0.5667921 | 0.3280546 | 0.7428704 | 0.861512 |
| ENSG00000243822 | | 1.850906 | 0.1310284 | 0.7865005 | 0.3070673 | 0.7587921 | 0.87145 |
| ENSG00000214612 | RPS19P1 | 87.90568 | 0.1310098 | 0.2821441 | 0.5001315 | 0.6169825 | 0.778339 |
| ENSG00000249565 | SERBP1P5 | 1469.099 | 0.1309861 | 0.2111171 | 0.6445389 | 0.519226 | 0.704562 |

| | | | | | | | |
|---|---|---|---|---|---|---|---|
| ENSG00000240747 | KRBOX1 | 1.025868 | 0.13091 | 0.7872816 | 0.2967631 | 0.7666474 | NA |
| ENSG00000181744 | DIPK2A | 1791.094 | 0.1309082 | 0.22984 | 0.5934302 | 0.5528933 | 0.731338 |
| ENSG00000279138 | | 5.81252 | 0.1309042 | 0.5765061 | 0.3258128 | 0.744566 | 0.862367 |
| ENSG00000189325 | BNIP5 | 0.8064028 | 0.1308283 | 0.7861925 | 0.3059737 | 0.7596246 | NA |
| ENSG00000177406 | NINJ2-AS1 | 227.9345 | 0.1308093 | 0.2866042 | 0.4720836 | 0.6368671 | 0.792549 |
| ENSG00000241890 | RPL13P4 | 0.9538243 | 0.1307026 | 0.7467904 | 0.4473292 | 0.6546374 | NA |
| ENSG00000255363 | LINC02757 | 1.012892 | 0.1306255 | 0.7545618 | 0.4858234 | 0.6270924 | NA |
| ENSG00000088305 | DNMT3B | 980.7144843 | 0.130568332 | 0.451040358 | 0.348391714 | 0.72754603 | 0.85203 |
| ENSG00000179912 | R3HDM2 | 2407.288 | 0.1305213 | 0.2280612 | 0.5924197 | 0.5535695 | 0.731731 |
| ENSG00000260494 | | 3.460111 | 0.130484 | 0.5832605 | 0.3220109 | 0.7474444 | 0.864443 |
| ENSG00000268295 | POLR3GP1 | 6.860549 | 0.130456 | 0.591684 | 0.3277973 | 0.7430649 | 0.861574 |
| ENSG00000109390 | NDUFC1 | 2364.419966 | 0.130387081 | 0.202889185 | 0.660010372 | 0.50924717 | 0.695357 |
| ENSG00000124243 | BCAS4 | 3122.014405 | 0.130361457 | 0.176689705 | 0.753529794 | 0.45113161 | 0.647373 |
| ENSG00000230668 | | 1.578034 | 0.1303432 | 0.7297524 | 0.396636 | 0.6916359 | 0.830495 |
| ENSG00000089050 | RBBP9 | 1142.449135 | 0.130306541 | 0.23808494 | 0.568235002 | 0.56987541 | 0.743744 |
| ENSG00000244144 | | 5.181121 | 0.1302644 | 0.7330514 | 0.4209293 | 0.6738067 | 0.818654 |
| ENSG00000286981 | | 2.674692 | 0.1302532 | 0.7045534 | 0.3828499 | 0.7018311 | 0.837576 |
| ENSG00000235829 | TBCAP2 | 1.267102 | 0.1302428 | 0.7589684 | 0.4903956 | 0.623854 | NA |
| ENSG00000135723 | FHOD1 | 2787.337712 | 0.130227933 | 0.278213174 | 0.509059754 | 0.61071034 | 0.774103 |
| ENSG00000229893 | TAX1BP1-AS1 | 0.8531219 | 0.1302058 | 0.7618655 | 0.5335291 | 0.5936674 | NA |
| ENSG00000204427 | ABHD16A | 182.1694 | 0.1302025 | 0.2360735 | 0.5772476 | 0.5637722 | 0.739186 |
| ENSG00000272677 | HNRNPD-DT | 13.19613 | 0.130199 | 0.4057255 | 0.3717255 | 0.7100972 | 0.842335 |
| ENSG00000213018 | PABPN1P1 | 16.52057 | 0.130127 | 0.387872 | 0.3853615 | 0.6999696 | 0.836549 |
| ENSG00000179958 | DCTPP1 | 4220.947 | 0.1299349 | 0.3284472 | 0.4491021 | 0.653358 | 0.804517 |
| ENSG00000273419 | | 9.051015 | 0.1296665 | 0.57126 | 0.3176563 | 0.7507457 | 0.866601 |
| ENSG00000163866 | SMIM12 | 1906.581 | 0.1296349 | 0.1496861 | 0.8923157 | 0.3722237 | 0.57443 |
| ENSG00000259844 | GNPATP | 6.046884 | 0.1296271 | 0.6997849 | 0.3491871 | 0.7269489 | 0.851749 |
| ENSG00000146731 | CCT6A | 11250.38 | 0.1296029 | 0.2186865 | 0.6149438 | 0.5385918 | 0.721237 |
| ENSG00000240758 | | 7.302071 | 0.1294691 | 0.5444056 | 0.3208177 | 0.7483485 | 0.865221 |
| ENSG00000242267 | SKINT1L | 0.7560088 | 0.1294262 | 0.7987271 | 0.5973681 | 0.5502617 | NA |
| ENSG00000227107 | | 5.051695 | 0.1293881 | 0.6607861 | 0.3590319 | 0.7195712 | 0.847404 |
| ENSG00000129474 | AJUBA | 4824.604524 | 0.129254026 | 0.286457079 | 0.475169552 | 0.63466613 | 0.790773 |
| ENSG00000125818 | PSMF1 | 4374.958201 | 0.129245951 | 0.139748874 | 0.937943113 | 0.34827365 | 0.550029 |

| | | | | | | | |
|---|---|---|---|---|---|---|---|
| ENSG00000224448 | | 1.533251 | 0.1291224 | 0.7135095 | 0.3844137 | 0.7006718 | NA |
| ENSG00000042753 | AP2S1 | 6224.758693 | 0.129099051 | 0.264618726 | 0.514388526 | 0.60698038 | 0.77103 |
| ENSG00000158079 | PTPDC1 | 783.6892 | 0.1290666 | 0.2343703 | 0.5798649 | 0.5620057 | 0.73778 |
| ENSG00000166323 | C11orf65 | 29.31466 | 0.1290569 | 0.3278797 | 0.4311532 | 0.666357 | 0.813994 |
| ENSG00000144036 | EXOC6B | 2521.906 | 0.1289415 | 0.1891576 | 0.6932541 | 0.4881501 | 0.679074 |
| ENSG00000137094 | DNAJB5 | 218.082561 | 0.128733579 | 0.234594513 | 0.576586787 | 0.56421863 | 0.739552 |
| ENSG00000100324 | TAB1 | 1626.359119 | 0.128717162 | 0.159925856 | 0.820908011 | 0.41169867 | 0.612042 |
| ENSG00000278737 | | 0.9715318 | 0.1287014 | 0.7366187 | 0.4100226 | 0.6817893 | NA |
| ENSG00000168575 | SLC20A2 | 1819.499 | 0.1286432 | 0.16617 | 0.7885038 | 0.4304021 | 0.629691 |
| ENSG00000123562 | MORF4L2 | 19194.07332 | 0.128621466 | 0.138296416 | 0.975352955 | 0.32938521 | 0.530455 |
| ENSG00000258634 | | 444.7337 | 0.1285835 | 0.2977238 | 0.4597951 | 0.6456633 | 0.798558 |
| ENSG00000234975 | FTH1P2 | 1341.754 | 0.1285007 | 0.209552 | 0.6386412 | 0.5230563 | 0.708384 |
| ENSG00000175416 | CLTB | 7545.535 | 0.128442 | 0.2282219 | 0.5859133 | 0.5579338 | 0.735159 |
| ENSG00000129472 | RAB2B | 791.3551663 | 0.128433937 | 0.161098069 | 0.812093195 | 0.41673816 | 0.616636 |
| ENSG00000083838 | ZNF446 | 578.7621956 | 0.128265452 | 0.157789612 | 0.843816386 | 0.39877202 | 0.599738 |
| ENSG00000275101 | MIR6766 | 3.334607 | 0.1282269 | 0.6514087 | 0.3440962 | 0.7307739 | 0.853987 |
| ENSG00000266479 | | 2.116767 | 0.1282209 | 0.6940114 | 0.3622032 | 0.7172002 | 0.846015 |
| ENSG00000164929 | BAALC | 9.251342 | 0.1282174 | 0.5739445 | 0.3186289 | 0.750008 | 0.866239 |
| ENSG00000232164 | LINC01873 | 6.485161 | 0.1282079 | 0.6249052 | 0.3342668 | 0.7381782 | 0.858765 |
| ENSG00000233725 | NRAD1 | 17.79045 | 0.1281267 | 0.7297768 | 0.4206444 | 0.6740147 | 0.818723 |
| ENSG00000237292 | LINC01732 | 2.247799 | 0.1280231 | 0.7069463 | 0.3657483 | 0.7145529 | 0.844489 |
| ENSG00000166454 | ATMIN | 8399.839 | 0.1279851 | 0.1214071 | 1.079306 | 0.2804511 | 0.477924 |
| ENSG00000231691 | | 6.051289 | 0.127933 | 0.5853566 | 0.3164607 | 0.7516529 | 0.86703 |
| ENSG00000100418 | DESI1 | 2133.140146 | 0.12787552 | 0.155783504 | 0.815545868 | 0.41475992 | 0.614726 |
| ENSG00000186468 | RPS23 | 7394.766 | 0.1277759 | 0.1797747 | 0.7630316 | 0.4454446 | 0.642544 |
| ENSG00000110042 | DTX4 | 2462.303158 | 0.127757635 | 0.163592353 | 0.800478942 | 0.42343336 | 0.623366 |
| ENSG00000180787 | ZFP3 | 876.0203 | 0.1277519 | 0.2363975 | 0.5766995 | 0.5641425 | 0.739552 |
| ENSG00000204946 | ZNF783 | 2272.155 | 0.1277279 | 0.1775893 | 0.7363493 | 0.4615182 | 0.656252 |
| ENSG00000197153 | H3C12 | 83.60277 | 0.1276866 | 0.2853542 | 0.4764713 | 0.6337387 | 0.790173 |
| ENSG00000143376 | SNX27 | 4968.699 | 0.1276846 | 0.1720445 | 0.7128233 | 0.4759551 | 0.668284 |
| ENSG00000214367 | HAUS3 | 1097.975 | 0.1276384 | 0.1719487 | 0.7565367 | 0.4493275 | 0.645517 |
| ENSG00000031691 | CENPQ | 548.3633645 | 0.127506308 | 0.358245742 | 0.395717607 | 0.69231337 | 0.83107 |
| ENSG00000258750 | | 1.171114 | 0.1275059 | 0.788851 | 0.2746117 | 0.7836145 | NA |

| | | | | | | | |
|---|---|---|---|---|---|---|---|
| ENSG00000271993 | | 8.551468 | 0.1274794 | 0.5427929 | 0.3158845 | 0.7520902 | 0.867249 |
| ENSG00000161542 | PRPSAP1 | 3877.063 | 0.1274781 | 0.1287773 | 1.012158 | 0.3114626 | 0.512399 |
| ENSG00000104960 | PTOV1 | 4494.498034 | 0.127477549 | 0.219668733 | 0.602829766 | 0.54662194 | 0.726642 |
| ENSG00000277639 | | 51.79569 | 0.1273613 | 0.3939671 | 0.3701376 | 0.71128 | 0.842862 |
| ENSG00000143653 | SCCPDH | 11010.94 | 0.1273389 | 0.2068457 | 0.6432606 | 0.520055 | 0.70529 |
| ENSG00000184113 | CLDN5 | 2.627521 | 0.1272784 | 0.6973502 | 0.3772764 | 0.7059682 | 0.840133 |
| ENSG00000274620 | MIR378J | 3.278811 | 0.1272703 | 0.6120969 | 0.3235019 | 0.7463152 | 0.863739 |
| ENSG00000226948 | RPS4XP2 | 253.9832 | 0.1272612 | 0.2335662 | 0.5708535 | 0.568099 | 0.742456 |
| ENSG00000206785 | SNORA15B-2 | 4.249066 | 0.1272285 | 0.6407837 | 0.3289329 | 0.7422064 | 0.861159 |
| ENSG00000276505 | | 9.218865 | 0.1272238 | 0.5341473 | 0.3164698 | 0.751646 | 0.86703 |
| ENSG00000132824 | SERINC3 | 9959.746503 | 0.127160733 | 0.191141965 | 0.652404257 | 0.51414042 | 0.700028 |
| ENSG00000089154 | GCN1 | 15797.33493 | 0.127136775 | 0.133451021 | 0.946401112 | 0.34394404 | 0.5456 |
| ENSG00000217733 | CCT7P1 | 2.750815 | 0.1270917 | 0.6743655 | 0.3537502 | 0.7235261 | 0.849802 |
| ENSG00000248930 | CHP1P1 | 1.17479 | 0.1270597 | 0.7394042 | 0.4348122 | 0.6636987 | NA |
| ENSG00000114503 | NCBP2 | 7096.592011 | 0.127044312 | 0.11087718 | 1.18973032 | 0.2341524 | 0.423268 |
| ENSG00000139636 | LMBR1L | 1216.828 | 0.1269776 | 0.2620917 | 0.5184204 | 0.604165 | 0.769035 |
| ENSG00000206633 | SNORA80B | 3.306149 | 0.1269699 | 0.6810854 | 0.3584175 | 0.7200309 | 0.847677 |
| ENSG00000287012 | | 1.027609 | 0.1269676 | 0.7856001 | 0.3025327 | 0.762246 | NA |
| ENSG00000244097 | RPS4XP17 | 12.29896 | 0.1269144 | 0.4308268 | 0.3488612 | 0.7271935 | 0.851812 |
| ENSG00000280163 | | 6.135667 | 0.126908 | 0.58295 | 0.3193464 | 0.7494638 | 0.865842 |
| ENSG00000128602 | SMO | 995.6146482 | 0.126891928 | 0.18188765 | 0.718577884 | 0.47240104 | 0.665381 |
| ENSG00000217181 | | 2.34323 | 0.1268875 | 0.6609877 | 0.3383075 | 0.7351315 | 0.856543 |
| ENSG00000213752 | | 0.7525721 | 0.1268495 | 0.7863567 | 0.2955371 | 0.7675836 | NA |
| ENSG00000244480 | | 11.3224 | 0.1267549 | 0.5832902 | 0.3074626 | 0.7584913 | 0.87134 |
| ENSG00000167637 | ZNF283 | 479.1157 | 0.126702 | 0.260497 | 0.5150476 | 0.6065198 | 0.770629 |
| ENSG00000239112 | SNORD123 | 5.200473 | 0.1266488 | 0.7354172 | 0.4106324 | 0.6813421 | 0.823668 |
| ENSG00000226478 | UPF3AP1 | 249.6685 | 0.1266297 | 0.1732421 | 0.7513462 | 0.4524443 | 0.648591 |
| ENSG00000242162 | | 1.096331 | 0.1266169 | 0.7859725 | 0.2978244 | 0.7658372 | NA |
| ENSG00000228021 | | 2.734378 | 0.1265885 | 0.625433 | 0.3190377 | 0.7496979 | 0.865959 |
| ENSG00000249709 | ZNF564 | 11.33947 | 0.1265687 | 0.5344052 | 0.3177185 | 0.7506985 | 0.866585 |
| ENSG00000186001 | LRCH3 | 1704.622 | 0.1265615 | 0.1698275 | 0.7575582 | 0.4487155 | 0.645304 |
| ENSG00000278900 | | 1.044634 | 0.126507 | 0.7640171 | 0.5244569 | 0.5999608 | NA |
| ENSG00000279864 | NCOR1P4 | 2.12874 | 0.126409 | 0.8016319 | 0.8787427 | 0.3795408 | 0.581513 |

| | | | | | | | |
|---|---|---|---|---|---|---|---|
| ENSG00000171135 | JAGN1 | 3544.391 | 0.1263738 | 0.168148 | 0.7807307 | 0.4349609 | 0.633766 |
| ENSG00000114956 | DGUOK | 2106.688956 | 0.126351096 | 0.215421783 | 0.604688346 | 0.54538609 | 0.725887 |
| ENSG00000124225 | PMEPA1 | 13505.24681 | 0.126300507 | 0.32217096 | 0.445699222 | 0.65581453 | 0.805976 |
| ENSG00000125384 | PTGER2 | 11.96187639 | 0.126280625 | 0.593186058 | 0.299221638 | 0.76477094 | 0.875128 |
| ENSG00000223138 | RNA5SP450 | 1.721002 | 0.1261714 | 0.7306485 | 0.4325425 | 0.6653472 | 0.813063 |
| ENSG00000204860 | FAM201A | 672.6406 | 0.126171 | 0.1759698 | 0.7327241 | 0.4637267 | 0.657836 |
| ENSG00000164932 | CTHRC1 | 92.09216 | 0.1260994 | 0.3099889 | 0.4413326 | 0.6589723 | 0.808067 |
| ENSG00000235301 | HLA-Z | 0.8682457 | 0.1260651 | 0.7613332 | 0.5831929 | 0.5597635 | NA |
| ENSG00000249572 | | 92.24017 | 0.1260437 | 0.2300533 | 0.5710062 | 0.5679954 | 0.742379 |
| ENSG00000101654 | RNMT | 3545.242927 | 0.126038234 | 0.156881972 | 0.844796346 | 0.39822456 | 0.599322 |
| ENSG00000268047 | | 12.81597 | 0.1259421 | 0.5096857 | 0.31934 | 0.7494687 | 0.865842 |
| ENSG00000288583 | | 1.065362 | 0.1256397 | 0.7872305 | 0.2852354 | 0.7754638 | NA |
| ENSG00000249859 | PVT1 | 3809.23 | 0.12552 | 0.1900514 | 0.6516366 | 0.5146356 | 0.700309 |
| ENSG00000234383 | CTBP2P8 | 78.35328 | 0.1254707 | 0.2374329 | 0.5476176 | 0.5839545 | 0.754089 |
| ENSG00000187855 | ASCL4 | 1.049121 | 0.1254705 | 0.7881502 | 0.2786696 | 0.7804984 | NA |
| ENSG00000254545 | | 2.279943 | 0.1254446 | 0.6728346 | 0.3337129 | 0.7385962 | 0.858824 |
| ENSG00000001561 | ENPP4 | 1348.427966 | 0.125412118 | 0.211193468 | 0.614027881 | 0.5391969 | 0.721616 |
| ENSG00000224415 | | 50.00721 | 0.1252537 | 0.27464 | 0.4852705 | 0.6274845 | 0.785867 |
| ENSG00000158195 | WASF2 | 7408.79 | 0.125075 | 0.1562322 | 0.8142119 | 0.4155236 | 0.615353 |
| ENSG00000213228 | RPL12P38 | 13.3742 | 0.1250571 | 0.4428043 | 0.3411227 | 0.7330112 | 0.855309 |
| ENSG00000266365 | | 4.832827 | 0.1250255 | 0.6395128 | 0.325129 | 0.7450834 | 0.862764 |
| ENSG00000228719 | | 8.047434 | 0.124975 | 0.6117951 | 0.3217037 | 0.7476772 | 0.864637 |
| ENSG00000116752 | BCAS2 | 15348.80854 | 0.124968019 | 0.169081749 | 0.696149522 | 0.4863352 | 0.677366 |
| ENSG00000266869 | | 370.6994 | 0.1249413 | 0.1874617 | 0.6780484 | 0.497741 | 0.686451 |
| ENSG00000266994 | | 3.847979 | 0.1248969 | 0.6260326 | 0.3073119 | 0.758606 | 0.871379 |
| ENSG00000203663 | OR2L2 | 3.370584 | 0.1247604 | 0.6788736 | 0.3414635 | 0.7327547 | 0.855159 |
| ENSG00000244558 | KCNK15-AS1 | 91.20835 | 0.124734 | 0.3276776 | 0.4080245 | 0.6832557 | 0.824763 |
| ENSG00000189419 | SPATA41 | 39.35983 | 0.1247021 | 0.4108651 | 0.3507023 | 0.7258117 | 0.851057 |
| ENSG00000259725 | | 1.630406 | 0.124679 | 0.7223784 | 0.3761654 | 0.7067939 | 0.840667 |
| ENSG00000152592 | DMP1 | 2.20194 | 0.1246728 | 0.6475079 | 0.3080371 | 0.7580541 | 0.871039 |
| ENSG00000286970 | | 1.25355 | 0.1246282 | 0.7293055 | 0.4200652 | 0.6744378 | NA |
| ENSG00000178401 | DNAJC22 | 1396.013 | 0.1246119 | 0.388724 | 0.3622295 | 0.7171806 | 0.846015 |
| ENSG00000260830 | | 7.262491 | 0.1245874 | 0.5727066 | 0.307376 | 0.7585572 | 0.871378 |

| | | | | | | | |
|---|---|---|---|---|---|---|---|
| ENSG00000224820 | BTF3P4 | 27.99716 | 0.1244472 | 0.3823209 | 0.3775819 | 0.7057412 | 0.840057 |
| ENSG00000260269 | | 1.63637 | 0.1244328 | 0.7364137 | 0.4228232 | 0.6724243 | 0.817949 |
| ENSG00000137474 | MYO7A | 164.6534173 | 0.124386722 | 0.41881909 | 0.337244497 | 0.7359326 | 0.85693 |
| ENSG00000117569 | PTBP2 | 853.8813633 | 0.124321779 | 0.181451114 | 0.70117682 | 0.48319267 | 0.674589 |
| ENSG00000166503 | HDGFL3 | 4432.446 | 0.124303 | 0.1639403 | 0.7722164 | 0.4399863 | 0.638344 |
| ENSG00000112309 | B3GAT2 | 21.35952676 | 0.124134768 | 0.379065425 | 0.371493999 | 0.71026963 | 0.842334 |
| ENSG00000260290 | | 1396.658 | 0.1240994 | 0.1135772 | 1.098245 | 0.2720975 | 0.468587 |
| ENSG00000059728 | MXD1 | 1288.982405 | 0.124098273 | 0.221487201 | 0.569194622 | 0.56922407 | 0.743217 |
| ENSG00000255366 | | 2.257265 | 0.1240403 | 0.7162979 | 0.4024477 | 0.6873545 | 0.82781 |
| ENSG00000211818 | TRAV39 | 3.364517 | 0.1240109 | 0.7870348 | 0.2832584 | 0.7769788 | 0.882538 |
| ENSG00000153406 | NMRAL1 | 4157.294 | 0.1239502 | 0.174252 | 0.7266049 | 0.467468 | 0.661023 |
| ENSG00000184274 | LINC00315 | 1.725779 | 0.1238259 | 0.7215238 | 0.3949368 | 0.6928895 | 0.831284 |
| ENSG00000227088 | | 6.668138 | 0.1237934 | 0.732992 | 0.4097428 | 0.6819946 | 0.82392 |
| ENSG00000273716 | | 6.677315 | 0.1237676 | 0.5022971 | 0.3172334 | 0.7510665 | 0.866859 |
| ENSG00000090861 | AARS1 | 25351.29349 | 0.123766658 | 0.254234197 | 0.510103727 | 0.60997879 | 0.773451 |
| ENSG00000223466 | LINC01825 | 4.973468 | 0.1237578 | 0.7414079 | 0.3877156 | 0.6982265 | 0.835488 |
| ENSG00000229393 | | 2.040781 | 0.1236993 | 0.7225042 | 0.3900459 | 0.6965026 | 0.834139 |
| ENSG00000116815 | CD58 | 785.456501 | 0.12369515 | 0.145717614 | 0.862633953 | 0.38833876 | 0.590521 |
| ENSG00000260635 | | 3.726606 | 0.1235849 | 0.6095464 | 0.3125478 | 0.7546242 | 0.86904 |
| ENSG00000259838 | ELOCP2 | 170.9026 | 0.1235049 | 0.2876341 | 0.458834 | 0.6463534 | 0.799226 |
| ENSG00000226578 | | 12.47525 | 0.1235033 | 0.5916637 | 0.3083293 | 0.7578317 | 0.870997 |
| ENSG00000120555 | SEPTIN7P9 | 29.02673525 | 0.123433736 | 0.440480497 | 0.338678456 | 0.73485197 | 0.856328 |
| ENSG00000135314 | KHDC1 | 512.7823494 | 0.123403632 | 0.209068527 | 0.608896627 | 0.54259296 | 0.723795 |
| ENSG00000124486 | USP9X | 9077.153683 | 0.123336405 | 0.142809853 | 0.839781737 | 0.40103078 | 0.601958 |
| ENSG00000240051 | RPL23AP10 | 7.255289 | 0.1233208 | 0.5992436 | 0.3160984 | 0.7519278 | 0.867175 |
| ENSG00000225080 | PFN1P4 | 34.09429 | 0.1232562 | 0.5539868 | 0.3041421 | 0.7610196 | 0.872566 |
| ENSG00000136819 | C9orf78 | 3378.67746 | 0.123238471 | 0.160880795 | 0.787996702 | 0.43069864 | 0.629885 |
| ENSG00000164934 | DCAF13 | 6814.327 | 0.1232216 | 0.1713321 | 0.7184466 | 0.472482 | 0.665381 |
| ENSG00000285867 | | 69.84254 | 0.1231529 | 0.320261 | 0.4260733 | 0.6700544 | 0.8163 |
| ENSG00000271973 | | 4.603715 | 0.1231336 | 0.592131 | 0.3032121 | 0.7617283 | 0.872983 |
| ENSG00000085721 | RRN3 | 4335.980588 | 0.123090639 | 0.178149544 | 0.70408245 | 0.48138143 | 0.672882 |
| ENSG00000176209 | SMIM19 | 1138.662 | 0.123049 | 0.1633867 | 0.7741514 | 0.4388413 | 0.637347 |
| ENSG00000266505 | | 0.910859 | 0.1230293 | 0.787224 | 0.2792401 | 0.7800606 | NA |

| | | | | | | | |
|---|---|---|---|---|---|---|---|
| ENSG00000204120 | GIGYF2 | 4849.863 | 0.1229951 | 0.1820499 | 0.7065482 | 0.4798473 | 0.671756 |
| ENSG00000123636 | BAZ2B | 1739.444893 | 0.122960604 | 0.309128376 | 0.417039364 | 0.67664962 | 0.820122 |
| ENSG00000242477 | | 149.9575 | 0.1229483 | 0.2263905 | 0.5493126 | 0.5827909 | 0.753231 |
| ENSG00000170775 | GPR37 | 531.9096 | 0.1228174 | 0.2467963 | 0.5186463 | 0.6040074 | 0.768871 |
| ENSG00000259726 | CSPG4P11 | 4.305389 | 0.1227463 | 0.5616422 | 0.30212 | 0.7625606 | 0.873673 |
| ENSG00000158106 | RHPN1 | 12087.72 | 0.1226519 | 0.1735441 | 0.7024038 | 0.4824274 | 0.673844 |
| ENSG00000277425 | | 1.106949 | 0.1226418 | 0.7126171 | 0.36884 | 0.712247 | NA |
| ENSG00000206192 | ANKRD20A9P | 0.8507714 | 0.1226381 | 0.7514952 | 0.4478504 | 0.6542612 | NA |
| ENSG00000266036 | | 3.426916 | 0.1226323 | 0.6967229 | 0.3291711 | 0.7420264 | 0.861025 |
| ENSG00000166452 | AKIP1 | 735.0525 | 0.1226275 | 0.1774561 | 0.7111878 | 0.4769679 | 0.669231 |
| ENSG00000164270 | HTR4 | 0.9028094 | 0.1225627 | 0.7868848 | 0.2831429 | 0.7770673 | NA |
| ENSG00000200120 | | 1.00603 | 0.1224642 | 0.7722738 | 0.4989426 | 0.6178198 | NA |
| ENSG00000176095 | IP6K1 | 2379.881 | 0.1224605 | 0.1672307 | 0.7382174 | 0.4603824 | 0.655647 |
| ENSG00000223908 | | 4.343309 | 0.1224327 | 0.592683 | 0.2982496 | 0.7655126 | 0.875583 |
| ENSG00000274038 | | 2.779128 | 0.122364 | 0.710118 | 0.3843402 | 0.7007263 | 0.836972 |
| ENSG00000262979 | | 54.67856 | 0.1223523 | 0.5072539 | 0.3147378 | 0.7529607 | 0.867952 |
| ENSG00000262999 | | 0.8003923 | 0.1223424 | 0.787252 | 0.2774576 | 0.7814288 | NA |
| ENSG00000143079 | CTTNBP2NL | 1436.292 | 0.1223363 | 0.3062572 | 0.4418322 | 0.6586106 | 0.807952 |
| ENSG00000200013 | RNU6-623P | 12.11159 | 0.1223116 | 0.474352 | 0.3187731 | 0.7498986 | 0.86615 |
| ENSG00000232218 | | 3.4352 | 0.12227 | 0.6133834 | 0.3062964 | 0.759379 | 0.871795 |
| ENSG00000273456 | | 90.2271 | 0.1222329 | 0.3409934 | 0.3956019 | 0.6923988 | 0.831098 |
| ENSG00000249068 | | 0.8924323 | 0.1221539 | 0.7603201 | 0.5644939 | 0.572418 | NA |
| ENSG00000021355 | SERPINB1 | 1884.166917 | 0.122124336 | 0.186959499 | 0.690734685 | 0.48973229 | 0.68035 |
| ENSG00000169855 | ROBO1 | 3783.134 | 0.1220456 | 0.2361133 | 0.5452307 | 0.5855948 | 0.755385 |
| ENSG00000198728 | LDB1 | 7143.15 | 0.1220355 | 0.2036302 | 0.6160921 | 0.5378337 | 0.720901 |
| ENSG00000042445 | RETSAT | 2955.552281 | 0.121947331 | 0.142849276 | 0.846119193 | 0.39748626 | 0.59855 |
| ENSG00000186191 | BPIFB4 | 0.8604 | 0.121929 | 0.7871835 | 0.2771698 | 0.7816497 | NA |
| ENSG00000176853 | FAM91A1 | 9523.382 | 0.1218891 | 0.1721082 | 0.7131413 | 0.4757583 | 0.668063 |
| ENSG00000269707 | | 1.719108 | 0.121866 | 0.6968453 | 0.3207791 | 0.7483778 | 0.865221 |
| ENSG00000156990 | RPUSD3 | 3032.7 | 0.1218333 | 0.181854 | 0.6814234 | 0.4956036 | 0.684733 |
| ENSG00000143624 | INTS3 | 8919.17 | 0.1218177 | 0.165869 | 0.7373863 | 0.4608875 | 0.655858 |
| ENSG00000288156 | | 0.7995144 | 0.1217808 | 0.7859259 | 0.2877931 | 0.7735051 | NA |
| ENSG00000236073 | | 0.8811423 | 0.1217624 | 0.7736736 | 0.5477157 | 0.5838872 | NA |

| | | | | | | | |
|---|---|---|---|---|---|---|---|
| ENSG00000278383 | | 48.39051 | 0.1217341 | 0.307876 | 0.4301076 | 0.6671174 | 0.814514 |
| ENSG00000196639 | HRH1 | 3.940679 | 0.1217238 | 0.7457733 | 0.3765988 | 0.7064718 | 0.840456 |
| ENSG00000142687 | KIAA0319L | 5904.084 | 0.1217034 | 0.1654926 | 0.7673818 | 0.4428545 | 0.640477 |
| ENSG00000258386 | | 0.9341217 | 0.121556 | 0.7856043 | 0.2926396 | 0.7697976 | NA |
| ENSG00000155097 | ATP6V1C1 | 7542.48 | 0.1215548 | 0.1264393 | 0.9263249 | 0.3542772 | 0.556279 |
| ENSG00000184708 | EIF4ENIF1 | 1638.784 | 0.1215072 | 0.09852475 | 1.240013 | 0.2149704 | 0.39994 |
| ENSG00000263388 | | 1.615104 | 0.1214236 | 0.7176293 | 0.3731432 | 0.7090419 | 0.841793 |
| ENSG00000160221 | GATD3A | 32.24133 | 0.12141 | 0.4434959 | 0.3275437 | 0.7432567 | 0.861574 |
| ENSG00000198466 | ZNF587 | 1263.838 | 0.1213348 | 0.2800414 | 0.4403609 | 0.6596757 | 0.808573 |
| ENSG00000119953 | SMNDC1 | 1927.791628 | 0.121302592 | 0.13233996 | 0.919360523 | 0.35790703 | 0.559861 |
| ENSG00000090971 | NAT14 | 1670.127236 | 0.121274985 | 0.203603152 | 0.609850852 | 0.54196061 | 0.723513 |
| ENSG00000259750 | NSA2P4 | 6.139447 | 0.1212708 | 0.6049478 | 0.3046048 | 0.7606672 | 0.872369 |
| ENSG00000256128 | LINC00944 | 1.351129 | 0.1212528 | 0.7867354 | 0.2797176 | 0.7796942 | NA |
| ENSG00000238041 | TMEM230P2 | 10.76929 | 0.1211814 | 0.4673659 | 0.3171032 | 0.7511653 | 0.866897 |
| ENSG00000137207 | YIPF3 | 5983.15351 | 0.121167401 | 0.156049814 | 0.791148638 | 0.42885726 | 0.628062 |
| ENSG00000149428 | HYOU1 | 14822.52 | 0.1211416 | 0.1982722 | 0.6279999 | 0.530004 | 0.714449 |
| ENSG00000103494 | RPGRIP1L | 647.3452439 | 0.12113076 | 0.186396526 | 0.672438347 | 0.50130468 | 0.689128 |
| ENSG00000160058 | BSDC1 | 5467.896 | 0.121098 | 0.1405449 | 0.8650559 | 0.3870081 | 0.589349 |
| ENSG00000245910 | SNHG6 | 1995.369 | 0.1210448 | 0.212725 | 0.5887787 | 0.5560098 | 0.733556 |
| ENSG00000096433 | ITPR3 | 7179.543104 | 0.120936899 | 0.229547147 | 0.547627105 | 0.58394797 | 0.754089 |
| ENSG00000149716 | LTO1 | 2181.665 | 0.1208939 | 0.1149237 | 1.060641 | 0.2888532 | 0.487458 |
| ENSG00000233069 | | 0.8159513 | 0.1208491 | 0.7787026 | 0.6057554 | 0.5446772 | NA |
| ENSG00000213708 | SLC16A14P1 | 1.233525 | 0.1207786 | 0.7885469 | 0.2623859 | 0.7930239 | NA |
| ENSG00000228420 | LINC01768 | 66.94512 | 0.1207662 | 0.4455513 | 0.3233389 | 0.7464386 | 0.863806 |
| ENSG00000213152 | RPL7AP60 | 8.180243 | 0.1207638 | 0.584151 | 0.298206 | 0.765546 | 0.875583 |
| ENSG00000148290 | SURF1 | 3721.645 | 0.1207001 | 0.165558 | 0.7228556 | 0.4697686 | 0.663263 |
| ENSG00000231764 | DLX6-AS1 | 1.14916 | 0.1206935 | 0.7867567 | 0.278291 | 0.780789 | NA |
| ENSG00000272247 | | 11.51989 | 0.120684 | 0.6120278 | 0.3108573 | 0.7559091 | 0.869764 |
| ENSG00000222915 | RNU6-564P | 0.8344993 | 0.1206229 | 0.7567827 | 0.456683 | 0.6478989 | NA |
| ENSG00000171885 | AQP4 | 1.145458 | 0.1205741 | 0.7854044 | 0.2897689 | 0.7719931 | NA |
| ENSG00000136936 | XPA | 700.5756916 | 0.120531894 | 0.11846072 | 1.027594174 | 0.30414077 | 0.503754 |
| ENSG00000287029 | | 10.52163 | 0.1203323 | 0.5409712 | 0.2953152 | 0.7677531 | 0.876915 |
| ENSG00000196470 | SIAH1 | 391.7521 | 0.1203213 | 0.1751937 | 0.7049889 | 0.4808171 | 0.67236 |

| | | | | | | | |
|---|---|---|---|---|---|---|---|
| ENSG00000231095 | GYG1P3 | 4.401721 | 0.1203153 | 0.6566265 | 0.3278325 | 0.7430383 | 0.861574 |
| ENSG00000230662 | TNPO1P2 | 7.181502 | 0.120294 | 0.5570221 | 0.2976517 | 0.765969 | 0.875765 |
| ENSG00000104142 | VPS18 | 2415.839357 | 0.120283497 | 0.163778999 | 0.751948982 | 0.45208174 | 0.648257 |
| ENSG00000132254 | ARFIP2 | 5886.883404 | 0.120261862 | 0.151101089 | 0.804593176 | 0.42105448 | 0.620893 |
| ENSG00000165178 | NCF1C | 0.8827736 | 0.1202554 | 0.7513336 | 0.4478723 | 0.6542453 | NA |
| ENSG00000152315 | KCNK13 | 60.39714 | 0.1202512 | 0.5022642 | 0.3101256 | 0.7564654 | 0.869898 |
| ENSG00000270112 | | 1.941123 | 0.120246 | 0.683538 | 0.3276239 | 0.743196 | 0.861574 |
| ENSG00000269956 | MKNK1-AS1 | 11.7248 | 0.1201938 | 0.5238499 | 0.3009266 | 0.7634705 | 0.874228 |
| ENSG00000072609 | CHFR | 3138.385803 | 0.12011073 | 0.125513015 | 0.962970144 | 0.33556251 | 0.536791 |
| ENSG00000170471 | RALGAPB | 4786.17 | 0.1200021 | 0.1773934 | 0.6846777 | 0.4935473 | 0.683363 |
| ENSG00000248240 | | 13.86352 | 0.119957 | 0.6092758 | 0.2943644 | 0.7684794 | 0.877361 |
| ENSG00000226430 | USP17L7 | 1.218182 | 0.1199068 | 0.7249119 | 0.3682554 | 0.7126828 | NA |
| ENSG00000236714 | LINC01844 | 1.033177 | 0.1199055 | 0.7882515 | 0.2631648 | 0.7924236 | NA |
| ENSG00000129654 | FOXJ1 | 44.81769313 | 0.11989663 | 0.669249464 | 0.328104523 | 0.74283263 | 0.861509 |
| ENSG00000125812 | GZF1 | 2145.267862 | 0.119724759 | 0.106605949 | 1.126169753 | 0.26009367 | 0.454544 |
| ENSG00000171564 | FGB | 20.28338 | 0.1196697 | 0.7849598 | 0.2921657 | 0.7701599 | 0.878478 |
| ENSG00000177963 | RIC8A | 5599.894 | 0.1196452 | 0.1425595 | 0.8486549 | 0.3960733 | 0.597471 |
| ENSG00000197617 | VN1R5 | 0.8821992 | 0.1196213 | 0.7436625 | 0.407571 | 0.6835887 | NA |
| ENSG00000283599 | | 0.94005 | 0.1196188 | 0.7621682 | 0.5531019 | 0.5801936 | NA |
| ENSG00000154310 | TNIK | 213.6254 | 0.1195466 | 0.3248665 | 0.4015712 | 0.6879996 | 0.828129 |
| ENSG00000127191 | TRAF2 | 1765.569296 | 0.119508143 | 0.217271401 | 0.570159701 | 0.56856939 | 0.742678 |
| ENSG00000273687 | | 1.64025 | 0.1194379 | 0.7323475 | 0.3669208 | 0.7136781 | 0.843793 |
| ENSG00000285752 | CDC42-AS1 | 37.1859 | 0.1194237 | 0.4689017 | 0.3121148 | 0.7549533 | 0.869057 |
| ENSG00000276261 | | 0.8328359 | 0.119397 | 0.7608382 | 0.4679503 | 0.6398202 | NA |
| ENSG00000284491 | THSD8 | 2.12486 | 0.1192884 | 0.7514522 | 0.4596425 | 0.6457729 | 0.798603 |
| ENSG00000116985 | BMP8B | 1159.321614 | 0.119268283 | 0.231979294 | 0.537294307 | 0.59106434 | 0.759748 |
| ENSG00000100473 | COCH | 214.9683945 | 0.119245533 | 0.292844375 | 0.434896791 | 0.66363734 | 0.811757 |
| ENSG00000270469 | | 3.41565 | 0.1192359 | 0.6578868 | 0.312052 | 0.755001 | 0.869059 |
| ENSG00000228812 | LAMA5-AS1 | 4.181944 | 0.1191617 | 0.6779312 | 0.3395337 | 0.7342077 | 0.855878 |
| ENSG00000169252 | ADRB2 | 3.498759 | 0.1191238 | 0.7415027 | 0.3838197 | 0.7011121 | 0.83727 |
| ENSG00000171873 | ADRA1D | 3.733641 | 0.1191222 | 0.6809068 | 0.3444152 | 0.730534 | 0.853899 |
| ENSG00000081019 | RSBN1 | 3695.828165 | 0.119053553 | 0.198307091 | 0.615381229 | 0.53830299 | 0.721137 |
| ENSG00000258559 | | 55.68729 | 0.119053 | 0.3618379 | 0.3651409 | 0.7150062 | 0.844724 |

| | | | | | | | |
|---|---|---|---|---|---|---|---|
| ENSG00000226386 | PARD3-AS1 | 17.53321 | 0.1190429 | 0.435055 | 0.3269719 | 0.7436891 | 0.861863 |
| ENSG00000155287 | SLC25A28 | 2167.863 | 0.1190171 | 0.1845982 | 0.6513178 | 0.5148414 | 0.700365 |
| ENSG00000269153 | LYPLA2P2 | 88.73599 | 0.1189627 | 0.329725 | 0.40487 | 0.6855731 | 0.826677 |
| ENSG00000277369 | | 29.76494 | 0.1189516 | 0.379102 | 0.3585551 | 0.7199279 | 0.847625 |
| ENSG00000066933 | MYO9A | 3762.804071 | 0.11887412 | 0.300662802 | 0.420890248 | 0.67383523 | 0.818654 |
| ENSG00000064225 | ST3GAL6 | 18.43578844 | 0.118811623 | 0.48270472 | 0.311125739 | 0.75570503 | 0.869643 |
| ENSG00000266420 | RN7SL118P | 1.570375 | 0.1188022 | 0.7282716 | 0.3782995 | 0.7052081 | NA |
| ENSG00000249192 | MTND3P25 | 1.628457 | 0.1187769 | 0.7617632 | 0.5703941 | 0.5684104 | 0.742629 |
| ENSG00000207493 | SNORA46 | 2.345017 | 0.1187569 | 0.663872 | 0.3167956 | 0.7513987 | 0.866978 |
| ENSG00000172785 | CBWD1 | 930.263 | 0.1187104 | 0.2100796 | 0.5832773 | 0.5597067 | 0.73613 |
| ENSG00000205531 | NAP1L4 | 5812.514 | 0.1186721 | 0.1886292 | 0.6614708 | 0.5083104 | 0.694717 |
| ENSG00000260853 | | 508.9607 | 0.118647 | 0.285169 | 0.4496066 | 0.6529941 | 0.804218 |
| ENSG00000131196 | NFATC1 | 564.4571399 | 0.118621714 | 0.163429947 | 0.733965364 | 0.46296985 | 0.657314 |
| ENSG00000217930 | PAM16 | 260.2304 | 0.1185783 | 0.2270329 | 0.5536218 | 0.5798377 | 0.750747 |
| ENSG00000228050 | TOP3BP1 | 11.69185 | 0.1184624 | 0.6140205 | 0.3007871 | 0.7635768 | 0.874249 |
| ENSG00000130522 | JUND | 3337.27937 | 0.118434518 | 0.25450419 | 0.556285897 | 0.57801543 | 0.749685 |
| ENSG00000129295 | LRRC6 | 304.0395916 | 0.11829754 | 0.201469122 | 0.601379549 | 0.54758722 | 0.727333 |
| ENSG00000223774 | | 0.9365514 | 0.1182908 | 0.7358103 | 0.384704 | 0.7004567 | NA |
| ENSG00000280183 | | 3.168144 | 0.1182608 | 0.6658666 | 0.3158954 | 0.7520819 | 0.867249 |
| ENSG00000136932 | TRMO | 673.8902233 | 0.118175935 | 0.199346936 | 0.598475024 | 0.54952302 | 0.72892 |
| ENSG00000222255 | RNU6-101P | 0.8724091 | 0.1180466 | 0.7658305 | 0.5203859 | 0.6027946 | NA |
| ENSG00000100068 | LRP5L | 438.3353365 | 0.117965233 | 0.183051861 | 0.658309988 | 0.51033896 | 0.696164 |
| ENSG00000163535 | SGO2 | 866.3404 | 0.1179429 | 0.4420087 | 0.315235 | 0.7525832 | 0.867632 |
| ENSG00000065000 | AP3D1 | 11049.02496 | 0.117935408 | 0.12382266 | 0.960077695 | 0.33701611 | 0.53817 |
| ENSG00000231770 | TMEM44-AS1 | 553.6481 | 0.1178481 | 0.1627995 | 0.7391047 | 0.4598434 | 0.655274 |
| ENSG00000272911 | | 2.641643 | 0.1178455 | 0.745158 | 0.357591 | 0.7206494 | 0.847998 |
| ENSG00000198932 | GPRASP1 | 335.7964 | 0.1178379 | 0.2854948 | 0.4428485 | 0.6578753 | 0.807348 |
| ENSG00000243680 | RPL37P23 | 505.7364 | 0.1177962 | 0.1679421 | 0.7160926 | 0.4739342 | 0.666661 |
| ENSG00000241631 | RN7SL316P | 0.9878649 | 0.1177914 | 0.7618325 | 0.4836944 | 0.6286028 | NA |
| ENSG00000147118 | ZNF182 | 600.4296 | 0.11768 | 0.2746176 | 0.4526959 | 0.6507678 | 0.802697 |
| ENSG00000106089 | STX1A | 650.0010062 | 0.117601784 | 0.193166512 | 0.620601443 | 0.53486189 | 0.718409 |
| ENSG00000251491 | OR7E28P | 0.7737064 | 0.1175771 | 0.7963835 | 0.5735044 | 0.5663032 | NA |
| ENSG00000271662 | | 64.17305 | 0.1175129 | 0.4140515 | 0.3287581 | 0.7423385 | 0.861237 |

| | | | | | | | |
|---|---|---|---|---|---|---|---|
| ENSG00000204482 | LST1 | 1.476182 | 0.1175036 | 0.7359725 | 0.3995671 | 0.6894754 | NA |
| ENSG00000269374 | | 21.69435 | 0.1173248 | 0.3787488 | 0.3535399 | 0.7236837 | 0.849877 |
| ENSG00000070495 | JMJD6 | 1937.868796 | 0.117287219 | 0.21924953 | 0.556720381 | 0.57771849 | 0.749523 |
| ENSG00000247372 | | 2.194241 | 0.1171468 | 0.7106674 | 0.3738189 | 0.7085391 | 0.841518 |
| ENSG00000233155 | HMGA1P8 | 40.4413 | 0.1171293 | 0.3269255 | 0.3946112 | 0.6931298 | 0.831374 |
| ENSG00000225513 | | 2.266776 | 0.117117 | 0.743837 | 0.3791923 | 0.7045451 | 0.83957 |
| ENSG00000189046 | ALKBH2 | 1840.505 | 0.1170871 | 0.1890241 | 0.6220242 | 0.533926 | 0.717538 |
| ENSG00000166073 | GPR176 | 421.2293 | 0.1170834 | 0.2201389 | 0.5488835 | 0.5830854 | 0.75345 |
| ENSG00000253457 | SMIM18 | 1.395744 | 0.1170703 | 0.7379351 | 0.4165508 | 0.677007 | NA |
| ENSG00000248290 | TNXA | 0.818994 | 0.1170691 | 0.7401245 | 0.3691625 | 0.7120066 | NA |
| ENSG00000280369 | | 3.620448 | 0.1170685 | 0.6762725 | 0.3331263 | 0.739039 | 0.859022 |
| ENSG00000213080 | | 1128.515 | 0.1170634 | 0.2340795 | 0.5156569 | 0.606094 | 0.770457 |
| ENSG00000119509 | INVS | 1035.495238 | 0.1170214 | 0.177833518 | 0.665917673 | 0.50546372 | 0.692251 |
| ENSG00000225194 | LINC00092 | 1.47645 | 0.1169679 | 0.6831013 | 0.3197395 | 0.7491658 | NA |
| ENSG00000258774 | NANOGP7 | 2.548069 | 0.1168906 | 0.7143335 | 0.3713406 | 0.7103838 | 0.842335 |
| ENSG00000253954 | HMGN1P38 | 3777.197 | 0.1168778 | 0.2026532 | 0.5882863 | 0.5563401 | 0.73375 |
| ENSG00000135250 | SRPK2 | 3083.482409 | 0.116815728 | 0.144947775 | 0.800675144 | 0.42331974 | 0.623285 |
| ENSG00000130158 | DOCK6 | 3224.276225 | 0.116811702 | 0.237590475 | 0.523914655 | 0.6003379 | 0.766403 |
| ENSG00000248896 | | 5.091489 | 0.1167903 | 0.5727359 | 0.2937879 | 0.76892 | 0.877553 |
| ENSG00000270165 | | 11.14991 | 0.1167426 | 0.5826531 | 0.2911659 | 0.7709244 | 0.878884 |
| ENSG00000272556 | GTF2IP13 | 467.214 | 0.1166547 | 0.2294529 | 0.5179696 | 0.6044795 | 0.769357 |
| ENSG00000174080 | CTSF | 1436.244 | 0.1166441 | 0.2884184 | 0.4324508 | 0.6654138 | 0.813107 |
| ENSG00000142611 | PRDM16 | 574.4034 | 0.1166108 | 0.261536 | 0.4688148 | 0.639202 | 0.794121 |
| ENSG00000276290 | RBBP4P3 | 3.951943 | 0.1165985 | 0.6162136 | 0.2969326 | 0.766518 | 0.876184 |
| ENSG00000266563 | EIF1P5 | 3.800601 | 0.1165716 | 0.7074024 | 0.351535 | 0.725187 | 0.8507 |
| ENSG00000260144 | | 8.994335 | 0.116563 | 0.4742678 | 0.3036683 | 0.7613806 | 0.872735 |
| ENSG00000158769 | F11R | 5666.791 | 0.1165384 | 0.1167955 | 1.023444 | 0.3060978 | 0.506066 |
| ENSG00000257302 | FAHD2P1 | 1.186564 | 0.1165377 | 0.7879388 | 0.2601359 | 0.7947589 | NA |
| ENSG00000232070 | TMEM253 | 10.42563 | 0.1165003 | 0.4777626 | 0.302061 | 0.7626055 | 0.873687 |
| ENSG00000167011 | NAT16 | 8.952964 | 0.1164955 | 0.6055956 | 0.2979835 | 0.7657157 | 0.875664 |
| ENSG00000145293 | ENOPH1 | 4513.409 | 0.116432 | 0.1966087 | 0.6133131 | 0.5396693 | 0.722014 |
| ENSG00000214765 | SEPTIN7P2 | 1203.475 | 0.1163787 | 0.1686944 | 0.6926912 | 0.4885033 | 0.679425 |
| ENSG00000120992 | LYPLA1 | 4028.037805 | 0.11610149 | 0.196728967 | 0.613886069 | 0.53929062 | 0.721616 |

| | | | | | | | |
|---|---|---|---|---|---|---|---|
| ENSG00000251221 | LINC01337 | 2.754632 | 0.1160701 | 0.6657207 | 0.3158313 | 0.7521306 | 0.867258 |
| ENSG00000276417 | | 1.900898 | 0.116021 | 0.6892804 | 0.3221803 | 0.7473161 | 0.864403 |
| ENSG00000180979 | LRRC57 | 538.9529 | 0.1158825 | 0.1972794 | 0.6069306 | 0.5438969 | 0.724787 |
| ENSG00000268163 | | 1.319964 | 0.115871 | 0.7181029 | 0.3591963 | 0.7194482 | NA |
| ENSG00000224781 | EIF4A2P4 | 81.16876 | 0.1158491 | 0.2853254 | 0.4326541 | 0.6652661 | 0.813001 |
| ENSG00000196976 | LAGE3 | 4275.033 | 0.1158058 | 0.2696969 | 0.4320959 | 0.6656717 | 0.813347 |
| ENSG00000239367 | RN7SL477P | 3.373094 | 0.1157433 | 0.6516421 | 0.3047151 | 0.7605832 | 0.872369 |
| ENSG00000234031 | RPS3AP44 | 4.773535 | 0.1156946 | 0.6089885 | 0.294224 | 0.7685867 | 0.877361 |
| ENSG00000114744 | COMMD2 | 2533.39223 | 0.115676427 | 0.168482856 | 0.712314641 | 0.47626996 | 0.668605 |
| ENSG00000261764 | KRT18P18 | 21.11838 | 0.1155168 | 0.5258553 | 0.2871355 | 0.7740086 | 0.880805 |
| ENSG00000162512 | SDC3 | 2249.724 | 0.1155153 | 0.168337 | 0.6895482 | 0.4904784 | 0.680995 |
| ENSG00000155621 | C9orf85 | 661.2792 | 0.115448 | 0.1951768 | 0.6033498 | 0.546276 | 0.726391 |
| ENSG00000184602 | SNN | 1817.143 | 0.1154364 | 0.1751359 | 0.6828212 | 0.4947198 | 0.684079 |
| ENSG00000267272 | LINC01140 | 5.222216 | 0.1154047 | 0.5927591 | 0.2942697 | 0.7685518 | 0.877361 |
| ENSG00000272812 | | 54.13843 | 0.1153697 | 0.3043369 | 0.4123334 | 0.6800951 | 0.822647 |
| ENSG00000272525 | BTF3-DT | 43.08034 | 0.1153315 | 0.4061633 | 0.3248574 | 0.745289 | 0.86292 |
| ENSG00000233707 | RPL22P11 | 9.504906 | 0.1153007 | 0.4638501 | 0.3042357 | 0.7609483 | 0.872566 |
| ENSG00000270248 | | 7.16208 | 0.1152316 | 0.5695469 | 0.2864189 | 0.7745573 | 0.88111 |
| ENSG00000227910 | | 18.6566 | 0.1152129 | 0.5526275 | 0.2907199 | 0.7712656 | 0.879022 |
| ENSG00000151490 | PTPRO | 1.211277 | 0.1152078 | 0.7393984 | 0.3618386 | 0.7174727 | NA |
| ENSG00000237379 | CBX1P3 | 1.909726 | 0.1151668 | 0.7054394 | 0.345236 | 0.729917 | 0.853593 |
| ENSG00000166924 | NYAP1 | 47.86244 | 0.1151262 | 0.3818345 | 0.3428183 | 0.7317352 | 0.85446 |
| ENSG00000271806 | | 4.07556 | 0.1151063 | 0.6443256 | 0.3008802 | 0.7635059 | 0.874228 |
| ENSG00000183604 | SMG1P5 | 160.1374 | 0.1150287 | 0.239693 | 0.5025941 | 0.6152497 | 0.777039 |
| ENSG00000197930 | ERO1A | 3470.001 | 0.1149695 | 0.180253 | 0.6627857 | 0.5074678 | 0.693818 |
| ENSG00000168092 | PAFAH1B2 | 3271.903 | 0.1146447 | 0.1106332 | 0.9993036 | 0.3176476 | 0.518672 |
| ENSG00000083097 | DOP1A | 1777.858809 | 0.114641345 | 0.247907714 | 0.48001219 | 0.63121873 | 0.788593 |
| ENSG00000214360 | EFCAB9 | 1.9909 | 0.1146239 | 0.6906466 | 0.3294576 | 0.7418099 | 0.860849 |
| ENSG00000114315 | HES1 | 6549.716135 | 0.114579508 | 0.540041715 | 0.276592807 | 0.78209279 | 0.885776 |
| ENSG00000118058 | KMT2A | 7134.232143 | 0.114482143 | 0.323962553 | 0.38243245 | 0.70214063 | 0.837719 |
| ENSG00000237238 | BMS1P10 | 3.173509 | 0.114448 | 0.6105668 | 0.2847383 | 0.7758447 | 0.881951 |
| ENSG00000012983 | MAP4K5 | 3836.666576 | 0.114376728 | 0.146946526 | 0.782677149 | 0.43381672 | 0.632779 |
| ENSG00000232422 | KNOP1P4 | 1.413267 | 0.1143717 | 0.7257119 | 0.3732593 | 0.7089554 | NA |

| | | | | | | | |
|---|---|---|---|---|---|---|---|
| ENSG00000246115 | SUPT4H1P2 | 4.529503 | 0.1142846 | 0.6693952 | 0.3101907 | 0.7564159 | 0.869898 |
| ENSG00000175203 | DCTN2 | 7054.251 | 0.1142527 | 0.1318045 | 0.8308684 | 0.406048 | 0.606697 |
| ENSG00000182580 | EPHB3 | 1035.406 | 0.1142265 | 0.2975135 | 0.4204253 | 0.6741748 | 0.818804 |
| ENSG00000017797 | RALBP1 | 5704.203858 | 0.114214976 | 0.137547533 | 0.841599095 | 0.4000124 | 0.601038 |
| ENSG00000131142 | CCL25 | 1.877665473 | 0.114211731 | 0.745050037 | 0.368921 | 0.71218661 | 0.843338 |
| ENSG00000150459 | SAP18 | 8305.055 | 0.1142051 | 0.1454109 | 0.7724289 | 0.4398604 | 0.638266 |
| ENSG00000114841 | DNAH1 | 641.3206419 | 0.114180699 | 0.418272335 | 0.313522706 | 0.75388357 | 0.868685 |
| ENSG00000271137 | | 1.235761 | 0.1141737 | 0.7601547 | 0.4760565 | 0.6340341 | NA |
| ENSG00000086015 | MAST2 | 3592.827308 | 0.114151363 | 0.172155277 | 0.660685602 | 0.50881396 | 0.695048 |
| ENSG00000261351 | | 90.13409 | 0.1139916 | 0.2092371 | 0.5639425 | 0.5727932 | 0.745897 |
| ENSG00000266897 | | 8.461988 | 0.1139885 | 0.5749155 | 0.2881954 | 0.7731972 | 0.880445 |
| ENSG00000171703 | TCEA2 | 4514.573 | 0.1138994 | 0.1611936 | 0.713807 | 0.4753465 | 0.667732 |
| ENSG00000237880 | LINC01885 | 0.9518127 | 0.1138827 | 0.7884352 | 0.2483409 | 0.8038706 | NA |
| ENSG00000109270 | LAMTOR3 | 2576.039149 | 0.113880016 | 0.164209796 | 0.709674738 | 0.47790586 | 0.670016 |
| ENSG00000198393 | ZNF26 | 1079.002 | 0.1138437 | 0.1405437 | 0.8228187 | 0.4106111 | 0.611041 |
| ENSG00000189159 | JPT1 | 7836.933 | 0.1137518 | 0.2601041 | 0.4652131 | 0.6417789 | 0.795751 |
| ENSG00000134760 | DSG1 | 1.998769179 | 0.113739424 | 0.716949212 | 0.352742841 | 0.72428124 | 0.850275 |
| ENSG00000153132 | CLGN | 686.6466 | 0.1136529 | 0.5724327 | 0.2657874 | 0.790403 | 0.891266 |
| ENSG00000274810 | NPHP3-ACAD11 | 0.9192033 | 0.113635 | 0.7778637 | 0.5435244 | 0.5867688 | NA |
| ENSG00000185158 | LRRC37B | 457.2666 | 0.1135539 | 0.2116823 | 0.5537327 | 0.5797618 | 0.75072 |
| ENSG00000251633 | GYG1P1 | 2.713003 | 0.1135057 | 0.6913735 | 0.3349484 | 0.737664 | 0.858438 |
| ENSG00000189292 | ALKAL2 | 4.112443 | 0.1135007 | 0.6501661 | 0.2944923 | 0.7683818 | 0.877353 |
| ENSG00000226521 | | 2.889258 | 0.1134777 | 0.6755826 | 0.3116027 | 0.7553425 | 0.869377 |
| ENSG00000176973 | FAM89B | 422.9812 | 0.1134764 | 0.2141352 | 0.5518508 | 0.5810506 | 0.751783 |
| ENSG00000198203 | SULT1C2 | 0.8317315 | 0.1134507 | 0.7593731 | 0.4315297 | 0.6660833 | NA |
| ENSG00000114654 | EFCC1 | 82.90359492 | 0.113421579 | 0.32892649 | 0.378893562 | 0.70476691 | 0.839784 |
| ENSG00000103222 | ABCC1 | 6726.149771 | 0.113371961 | 0.177258421 | 0.656729802 | 0.51135467 | 0.697325 |
| ENSG00000273272 | | 2.563488 | 0.1133701 | 0.7136275 | 0.3445361 | 0.7304431 | 0.853899 |
| ENSG00000283399 | | 4.124787 | 0.1133638 | 0.6874612 | 0.3340027 | 0.7383775 | 0.858779 |
| ENSG00000158863 | FAM160B2 | 3557.656 | 0.1132653 | 0.1465421 | 0.7814133 | 0.4345594 | 0.633489 |
| ENSG00000287566 | | 57.30236 | 0.1132216 | 0.6583666 | 0.2892506 | 0.7723896 | 0.879888 |
| ENSG00000214223 | HNRNPA1P10 | 397.0936 | 0.1132096 | 0.161058 | 0.7155782 | 0.4742518 | 0.667005 |
| ENSG00000148935 | GAS2 | 33.47562 | 0.1131851 | 0.4066147 | 0.3196903 | 0.7492031 | 0.865723 |

| | | | | | | | |
|---|---|---|---|---|---|---|---|
| ENSG00000126062 | TMEM115 | 1967.929959 | 0.113155439 | 0.174697906 | 0.662633651 | 0.50756521 | 0.693913 |
| ENSG00000184613 | NELL2 | 56.0978 | 0.1130791 | 0.4621809 | 0.2929809 | 0.7695368 | 0.877917 |
| ENSG00000287264 | | 1.168997 | 0.1130659 | 0.7500453 | 0.413521 | 0.6792249 | NA |
| ENSG00000131096 | PYY | 5.815160913 | 0.113064844 | 0.651616013 | 0.302815689 | 0.76203033 | 0.873216 |
| ENSG00000067225 | PKM | 57145.25694 | 0.112896472 | 0.249136631 | 0.47323257 | 0.63604726 | 0.791825 |
| ENSG00000268940 | CT45A1 | 7.566902 | 0.1128699 | 0.7379194 | 0.3950011 | 0.692842 | 0.831284 |
| ENSG00000226471 | | 110.6702 | 0.1128558 | 0.2979226 | 0.3952791 | 0.6926369 | 0.831233 |
| ENSG00000103404 | USP31 | 1091.292117 | 0.112789961 | 0.214799641 | 0.535109508 | 0.59257414 | 0.760732 |
| ENSG00000286959 | | 1.733632 | 0.1127789 | 0.7498925 | 0.3617273 | 0.7175558 | 0.846015 |
| ENSG00000261673 | | 1.822787 | 0.1127738 | 0.6949129 | 0.3103776 | 0.7562738 | 0.869886 |
| ENSG00000060762 | MPC1 | 997.3673986 | 0.112683698 | 0.19860823 | 0.568930592 | 0.56940324 | 0.743341 |
| ENSG00000238711 | RNY4P25 | 1.93178 | 0.1126787 | 0.746944 | 0.3710512 | 0.7105994 | 0.842553 |
| ENSG00000103266 | STUB1 | 3438.434297 | 0.112655757 | 0.205846763 | 0.560855013 | 0.57489638 | 0.747208 |
| ENSG00000131966 | ACTR10 | 4194.943638 | 0.11264237 | 0.132628077 | 0.846659707 | 0.39718482 | 0.598279 |
| ENSG00000184434 | LRRC19 | 1.887222 | 0.1126383 | 0.6796453 | 0.3049612 | 0.7603957 | 0.872349 |
| ENSG00000153310 | CYRIB | 7203.893 | 0.1125623 | 0.2136008 | 0.5360477 | 0.5919256 | 0.760374 |
| ENSG00000142186 | SCYL1 | 8764.976 | 0.1124552 | 0.1302774 | 0.8702491 | 0.3841643 | 0.5865 |
| ENSG00000185129 | PURA | 2300.698 | 0.1123935 | 0.242396 | 0.4828004 | 0.6292375 | 0.787012 |
| ENSG00000101639 | CEP192 | 2942.208261 | 0.112355029 | 0.199195191 | 0.58582631 | 0.55799224 | 0.735175 |
| ENSG00000233221 | | 2.486308 | 0.1122967 | 0.7512403 | 0.3517239 | 0.7250453 | 0.850636 |
| ENSG00000187952 | HS6ST1P1 | 133.4945 | 0.1122513 | 0.2666936 | 0.4434764 | 0.6574212 | 0.807089 |
| ENSG00000287216 | | 2.666332 | 0.1121958 | 0.6695757 | 0.3134696 | 0.7539239 | 0.868685 |
| ENSG00000185250 | PPIL6 | 108.8499 | 0.1121957 | 0.3331242 | 0.3784972 | 0.7050612 | 0.839876 |
| ENSG00000253228 | NRBF2P4 | 12.77707 | 0.1121873 | 0.477306 | 0.2913848 | 0.770757 | 0.878781 |
| ENSG00000187984 | ANKRD19P | 15.69948 | 0.1121686 | 0.5750166 | 0.2796592 | 0.779739 | 0.884468 |
| ENSG00000145868 | FBXO38 | 2133.984 | 0.1121274 | 0.1575285 | 0.7211945 | 0.4707899 | 0.664156 |
| ENSG00000150403 | TMCO3 | 2390.992 | 0.1121086 | 0.2249545 | 0.5050112 | 0.613551 | 0.775861 |
| ENSG00000089775 | ZBTB25 | 1751.199897 | 0.112058425 | 0.224847662 | 0.517104346 | 0.60508332 | 0.769686 |
| ENSG00000198106 | | 0.9506604 | 0.1120471 | 0.7464674 | 0.4151254 | 0.6780501 | NA |
| ENSG00000180139 | ACTA2-AS1 | 1.203016 | 0.1120178 | 0.7163112 | 0.3497419 | 0.7265324 | NA |
| ENSG00000284830 | | 5.673148 | 0.1119994 | 0.5549887 | 0.2778444 | 0.7811318 | 0.885254 |
| ENSG00000273619 | | 389.6063 | 0.1119504 | 0.2146887 | 0.5422 | 0.5876807 | 0.757083 |
| ENSG00000251288 | | 4.620886 | 0.1117781 | 0.6114392 | 0.2821199 | 0.7778516 | 0.883156 |

| | | | | | | | |
|---|---|---|---|---|---|---|---|
| ENSG00000260895 | | 1.831806 | 0.1117712 | 0.6904755 | 0.3072884 | 0.7586239 | 0.871379 |
| ENSG00000268066 | FMR1-AS1 | 1.79047 | 0.1116502 | 0.6743991 | 0.3015622 | 0.7629858 | 0.873934 |
| ENSG00000214914 | RPL23AP3 | 1.738343 | 0.1116086 | 0.7410869 | 0.4115405 | 0.6806762 | 0.823151 |
| ENSG00000137947 | GTF2B | 1070.603081 | 0.111498597 | 0.155285443 | 0.725572561 | 0.46810085 | 0.661717 |
| ENSG00000227640 | SOX21-AS1 | 1.270712 | 0.1114697 | 0.7368946 | 0.3922558 | 0.6948692 | NA |
| ENSG00000248643 | RBM14-RBM4 | 76.35413 | 0.1114624 | 0.3249053 | 0.3778609 | 0.7055339 | 0.840033 |
| ENSG00000100246 | DNAL4 | 960.7833474 | 0.111436151 | 0.196382136 | 0.56794437 | 0.57007274 | 0.743923 |
| ENSG00000255091 | | 1.038894 | 0.1113377 | 0.787851 | 0.2497211 | 0.8028031 | NA |
| ENSG00000198868 | MTND4LP30 | 2.207949 | 0.111314 | 0.6722493 | 0.3097296 | 0.7567666 | 0.870036 |
| ENSG00000106066 | CPVL | 3.125348992 | 0.111280354 | 0.738880571 | 0.392511013 | 0.69468068 | 0.83267 |
| ENSG00000259407 | | 8.019026 | 0.1112722 | 0.5702653 | 0.2776875 | 0.7812523 | 0.885353 |
| ENSG00000167778 | SPRYD3 | 3613.848 | 0.1111636 | 0.1598919 | 0.7174326 | 0.4731072 | 0.66603 |
| ENSG00000240766 | PLCXD2-AS1 | 0.9588121 | 0.1110218 | 0.724796 | 0.353946 | 0.7233793 | NA |
| ENSG00000156675 | RAB11FIP1 | 2067.528 | 0.1110134 | 0.3031791 | 0.3895274 | 0.6968861 | 0.834442 |
| ENSG00000214249 | CTAGE11P | 9.197012 | 0.1109931 | 0.5326443 | 0.2745834 | 0.7836363 | 0.886724 |
| ENSG00000154222 | CC2D1B | 2588.083 | 0.1109764 | 0.1131861 | 0.9871959 | 0.3235466 | 0.524421 |
| ENSG00000250764 | | 23.21637 | 0.1109147 | 0.4356006 | 0.3058949 | 0.7596847 | 0.871863 |
| ENSG00000137040 | RANBP6 | 1313.915772 | 0.110886666 | 0.153447964 | 0.73785602 | 0.46060195 | 0.655646 |
| ENSG00000273254 | | 5.236685 | 0.1108406 | 0.5741353 | 0.2794124 | 0.7799283 | 0.884569 |
| ENSG00000227407 | | 1.110356 | 0.1108029 | 0.7492138 | 0.3768062 | 0.7063177 | NA |
| ENSG00000131013 | PPIL4 | 2533.630096 | 0.110780661 | 0.121016089 | 0.924619074 | 0.35516409 | 0.557177 |
| ENSG00000141428 | C18orf21 | 845.6864 | 0.1107429 | 0.1853684 | 0.6125183 | 0.5401949 | 0.722353 |
| ENSG00000160200 | CBS | 1622.885 | 0.1106547 | 0.168864 | 0.679078 | 0.4970884 | 0.685907 |
| ENSG00000137962 | ARHGAP29 | 1742.771931 | 0.110642402 | 0.324900474 | 0.369606682 | 0.71167557 | 0.843114 |
| ENSG00000213673 | SLC25A5P3 | 4.810377 | 0.1106277 | 0.6274532 | 0.2850393 | 0.7756141 | 0.881859 |
| ENSG00000271816 | | 38.22719 | 0.1105375 | 0.3996486 | 0.3166116 | 0.7515383 | 0.866998 |
| ENSG00000185379 | RAD51D | 662.1471 | 0.1105343 | 0.1543493 | 0.7364009 | 0.4614868 | 0.656252 |
| ENSG00000200385 | | 1.051557 | 0.1105065 | 0.7386246 | 0.3794687 | 0.7043399 | NA |
| ENSG00000257616 | | 5.269806 | 0.1105035 | 0.5965185 | 0.2824379 | 0.7776078 | 0.882917 |
| ENSG00000134744 | TUT4 | 677.8988881 | 0.110502109 | 0.248290059 | 0.468054711 | 0.63974547 | 0.79435 |
| ENSG00000135801 | TAF5L | 2143.06659 | 0.110440871 | 0.172301917 | 0.652354757 | 0.51417234 | 0.700028 |
| ENSG00000232895 | | 1.786957 | 0.1104056 | 0.7256287 | 0.3513479 | 0.7253273 | 0.850827 |
| ENSG00000105251 | SHD | 0.933836841 | 0.110378179 | 0.784844053 | 0.275005441 | 0.78331206 | NA |

| | | | | | | | |
|---|---|---|---|---|---|---|---|
| ENSG00000159069 | FBXW5 | 7182.799 | 0.1103223 | 0.1648888 | 0.6728874 | 0.5010189 | 0.688898 |
| ENSG00000140471 | LINS1 | 1234.477 | 0.110277 | 0.1295602 | 0.8667762 | 0.3860646 | 0.588387 |
| ENSG00000213730 | POLD2P1 | 1.970197 | 0.1102111 | 0.660027 | 0.2894108 | 0.7722671 | 0.879786 |
| ENSG00000159496 | RGL4 | 24.87431 | 0.1101758 | 0.4500849 | 0.294259 | 0.76856 | 0.877361 |
| ENSG00000224939 | LINC00184 | 5.8957 | 0.1101572 | 0.5476736 | 0.2765973 | 0.7820893 | 0.885776 |
| ENSG00000272431 | ZNF454-DT | 1.004201 | 0.1101329 | 0.757996 | 0.4222745 | 0.6728247 | NA |
| ENSG00000103507 | BCKDK | 3065.896078 | 0.110088776 | 0.214773177 | 0.527398318 | 0.59791701 | 0.764819 |
| ENSG00000023041 | ZDHHC6 | 2831.236734 | 0.110086783 | 0.17626558 | 0.636644522 | 0.52435642 | 0.709369 |
| ENSG00000263924 | | 1.373698 | 0.1100582 | 0.6993556 | 0.3266358 | 0.7439433 | NA |
| ENSG00000270379 | HEATR9 | 1.213171 | 0.1100226 | 0.7378333 | 0.3829624 | 0.7017476 | NA |
| ENSG00000127588 | GNG13 | 25.19557692 | 0.110019958 | 0.47014268 | 0.286870756 | 0.77421128 | 0.880868 |
| ENSG00000173376 | NDNF | 15.9052 | 0.109948 | 0.7379504 | 0.3943665 | 0.6933105 | 0.831553 |
| ENSG00000139974 | SLC38A6 | 582.1827 | 0.1099465 | 0.1478694 | 0.7550284 | 0.4502319 | 0.646466 |
| ENSG00000261257 | | 1.075487 | 0.1098698 | 0.7664243 | 0.4186187 | 0.6754948 | NA |
| ENSG00000271882 | | 21.53886 | 0.1098292 | 0.4719957 | 0.2833252 | 0.7769275 | 0.882522 |
| ENSG00000270503 | YTHDF2P1 | 41.64929 | 0.1096985 | 0.2775446 | 0.4198681 | 0.6745818 | 0.818951 |
| ENSG00000119965 | C10orf88 | 618.1829276 | 0.109684104 | 0.159970908 | 0.699471738 | 0.48425727 | 0.67545 |
| ENSG00000286940 | | 1.26098 | 0.1095694 | 0.7304893 | 0.3626667 | 0.7168539 | NA |
| ENSG00000016864 | GLT8D1 | 1885.322436 | 0.109498305 | 0.103071332 | 1.065995606 | 0.28642563 | 0.484687 |
| ENSG00000222460 | RN7SKP271 | 1.396258 | 0.1094916 | 0.7294778 | 0.3645572 | 0.715442 | NA |
| ENSG00000144744 | UBA3 | 1772.498 | 0.1094746 | 0.1254972 | 0.9004142 | 0.3678998 | 0.570038 |
| ENSG00000118680 | MYL12B | 24292.45672 | 0.109369666 | 0.158122891 | 0.703681825 | 0.48163094 | 0.673037 |
| ENSG00000122877 | EGR2 | 631.4559981 | 0.109325479 | 0.680725604 | 0.290826105 | 0.77118432 | 0.879005 |
| ENSG00000174156 | GSTA3 | 1.626102 | 0.109321 | 0.7992312 | 1.082547 | 0.2790094 | 0.476216 |
| ENSG00000177291 | GJD4 | 0.8089907 | 0.1091889 | 0.7483345 | 0.3591397 | 0.7194906 | NA |
| ENSG00000285531 | | 80.75152 | 0.1091679 | 0.3546105 | 0.3437708 | 0.7310187 | 0.854088 |
| ENSG00000263465 | SRSF8 | 2352.681 | 0.1091653 | 0.1824264 | 0.5922391 | 0.5536905 | 0.731849 |
| ENSG00000224863 | LINC01398 | 3.04734 | 0.1091468 | 0.6690721 | 0.2992018 | 0.7647861 | 0.875128 |
| ENSG00000239415 | | 382.5143 | 0.1091464 | 0.2297775 | 0.4893994 | 0.624559 | 0.783563 |
| ENSG00000163746 | PLSCR2 | 1.452126 | 0.1090509 | 0.7082551 | 0.3250972 | 0.7451075 | NA |
| ENSG00000166189 | HPS6 | 1412.296 | 0.1090463 | 0.1577176 | 0.7039155 | 0.4814854 | 0.672953 |
| ENSG00000287431 | | 461.1774 | 0.1089558 | 0.2823446 | 0.4005824 | 0.6887276 | 0.828482 |
| ENSG00000254369 | HOXA-AS3 | 7.918595 | 0.1089461 | 0.5891886 | 0.2681396 | 0.7885919 | 0.890102 |

| | | | | | | | |
|---|---|---|---|---|---|---|---|
| ENSG00000240809 | CAP1P1 | 83.79686 | 0.1089167 | 0.2501969 | 0.4578537 | 0.6470575 | 0.799892 |
| ENSG00000143952 | VPS54 | 3484.42 | 0.1088321 | 0.1601552 | 0.6740198 | 0.5002987 | 0.688406 |
| ENSG00000287925 | | 1.991211 | 0.1088206 | 0.683016 | 0.3093947 | 0.7570213 | 0.870291 |
| ENSG00000181227 | DLSTP1 | 585.4038 | 0.1088077 | 0.2008612 | 0.5573041 | 0.5773197 | 0.749185 |
| ENSG00000258824 | | 20.40067 | 0.1087985 | 0.4208016 | 0.3013996 | 0.7631098 | 0.874038 |
| ENSG00000249413 | | 17.27681 | 0.1087721 | 0.4634956 | 0.2795208 | 0.7798452 | 0.884534 |
| ENSG00000143258 | USP21 | 1976.183 | 0.1087676 | 0.1853825 | 0.6002801 | 0.5483196 | 0.727909 |
| ENSG00000113318 | MSH3 | 1219.596727 | 0.108714299 | 0.178753214 | 0.628960525 | 0.5293749 | 0.714139 |
| ENSG00000286230 | | 23.48606 | 0.1086663 | 0.5791642 | 0.2656452 | 0.7905125 | 0.891323 |
| ENSG00000283632 | EXOC3L2 | 4.982137 | 0.1086493 | 0.6219153 | 0.2726037 | 0.7851579 | 0.887583 |
| ENSG00000100403 | ZC3H7B | 7039.793833 | 0.1086484 | 0.138366246 | 0.797755473 | 0.42501241 | 0.624767 |
| ENSG00000100596 | SPTLC2 | 9198.854705 | 0.10864635 | 0.222032323 | 0.507189073 | 0.61202216 | 0.774841 |
| ENSG00000286237 | | 6.159327 | 0.1086408 | 0.5179213 | 0.2713251 | 0.786141 | 0.888354 |
| ENSG00000259619 | | 1.563384 | 0.1086207 | 0.731466 | 0.3687919 | 0.7122828 | NA |
| ENSG00000132874 | SLC14A2 | 0.996311643 | 0.108568636 | 0.762527325 | 0.520418579 | 0.60277186 | NA |
| ENSG00000125817 | CENPB | 9409.795106 | 0.108554389 | 0.219358387 | 0.507952285 | 0.61148681 | 0.774478 |
| ENSG00000287588 | | 1.582746 | 0.1085508 | 0.7474068 | 0.3464954 | 0.7289705 | 0.853066 |
| ENSG00000287045 | | 2.690968 | 0.1085372 | 0.710807 | 0.3391244 | 0.734516 | 0.856049 |
| ENSG00000204599 | TRIM39 | 702.3407 | 0.1085205 | 0.1700718 | 0.647711 | 0.5171719 | 0.702456 |
| ENSG00000197601 | FAR1 | 6192.483 | 0.1084534 | 0.1622591 | 0.684019 | 0.4939632 | 0.683761 |
| ENSG00000165591 | FAAH2 | 372.8188 | 0.1084083 | 0.3375052 | 0.3457863 | 0.7295033 | 0.853333 |
| ENSG00000272442 | | 5.276271 | 0.1083916 | 0.6161928 | 0.2753912 | 0.7830157 | 0.886458 |
| ENSG00000135617 | PRADC1 | 1413.214787 | 0.108354837 | 0.307884974 | 0.35268175 | 0.72432704 | 0.850291 |
| ENSG00000223522 | | 8.227977 | 0.1083266 | 0.5223164 | 0.2745726 | 0.7836447 | 0.886724 |
| ENSG00000167183 | PRR15L | 4352.837 | 0.1083246 | 0.2123176 | 0.528538 | 0.597126 | 0.764085 |
| ENSG00000100629 | CEP128 | 412.005425 | 0.108315658 | 0.447969407 | 0.280215381 | 0.77931227 | 0.884135 |
| ENSG00000065491 | TBC1D22B | 1265.559277 | 0.108261769 | 0.110930652 | 0.988796675 | 0.32276263 | 0.523534 |
| ENSG00000173826 | KCNH6 | 2.741161 | 0.1082613 | 0.6990302 | 0.3122899 | 0.7548202 | 0.869057 |
| ENSG00000125848 | FLRT3 | 202.1247923 | 0.108256735 | 0.469176612 | 0.274569996 | 0.78364662 | 0.886724 |
| ENSG00000066923 | STAG3 | 251.4906001 | 0.108217323 | 0.421978828 | 0.296063734 | 0.76718141 | 0.876586 |
| ENSG00000255059 | PPIAP43 | 1.838771 | 0.1081611 | 0.705364 | 0.3336845 | 0.7386177 | 0.858824 |
| ENSG00000100288 | CHKB | 79.4144952 | 0.108084725 | 0.283860062 | 0.408235956 | 0.68310045 | 0.824743 |
| ENSG00000243477 | NAA80 | 404.2941 | 0.1080541 | 0.1994175 | 0.5586168 | 0.5764233 | 0.748533 |

| | | | | | | | |
|---|---|---|---|---|---|---|---|
| ENSG00000254080 | | 1.380872 | 0.1080454 | 0.79916 | 1.024967 | 0.3053788 | NA |
| ENSG00000286071 | | 1.39807 | 0.1080112 | 0.7446298 | 0.3373976 | 0.7358172 | NA |
| ENSG00000075239 | ACAT1 | 2718.879857 | 0.107936104 | 0.18164758 | 0.580015373 | 0.56190425 | 0.737756 |
| ENSG00000182902 | SLC25A18 | 97.86796 | 0.1078975 | 0.4325094 | 0.2879609 | 0.7733767 | 0.880532 |
| ENSG00000141086 | CTRL | 0.7890663 | 0.107843 | 0.7513001 | 0.3675631 | 0.713199 | NA |
| ENSG00000235257 | ITGA9-AS1 | 7.447085 | 0.1077909 | 0.5705991 | 0.2697533 | 0.78735 | 0.889304 |
| ENSG00000167380 | ZNF226 | 1321.55 | 0.1077102 | 0.1750746 | 0.6282369 | 0.5298488 | 0.714379 |
| ENSG00000203811 | H3C14 | 2.141844 | 0.1076547 | 0.6971535 | 0.3098567 | 0.7566699 | 0.870017 |
| ENSG00000168634 | WFDC13 | 0.7835534 | 0.107624 | 0.7390976 | 0.339212 | 0.73445 | NA |
| ENSG00000243824 | RPL12P6 | 6.495639 | 0.1075655 | 0.5912338 | 0.2696036 | 0.7874653 | 0.889371 |
| ENSG00000204498 | NFKBIL1 | 1306.046 | 0.107552 | 0.1967437 | 0.5493949 | 0.5827345 | 0.753231 |
| ENSG00000230479 | | 0.8159529 | 0.1074798 | 0.788498 | 0.234264 | 0.8147801 | NA |
| ENSG00000249138 | | 6.400162 | 0.1074371 | 0.7358824 | 0.4033535 | 0.6866882 | 0.827495 |
| ENSG00000279328 | | 2.447608 | 0.1074241 | 0.6908012 | 0.305909 | 0.759674 | 0.871863 |
| ENSG00000286383 | | 0.9697946 | 0.1073338 | 0.7876512 | 0.2433599 | 0.8077266 | NA |
| ENSG00000196177 | ACADSB | 1370.124 | 0.1072954 | 0.1999793 | 0.5513958 | 0.5813624 | 0.752007 |
| ENSG00000208037 | MIR320A | 1.327665 | 0.1072953 | 0.7282974 | 0.3683783 | 0.7125912 | NA |
| ENSG00000204789 | ZNF204P | 197.0556 | 0.1071882 | 0.3494505 | 0.3378871 | 0.7354482 | 0.856647 |
| ENSG00000224892 | RPS4XP16 | 44.44662 | 0.1071792 | 0.3383205 | 0.3455991 | 0.729644 | 0.853403 |
| ENSG00000277548 | | 5.160742 | 0.1071512 | 0.6116234 | 0.2685708 | 0.78826 | 0.889845 |
| ENSG00000261787 | TCF24 | 80.7004 | 0.1071022 | 0.2824156 | 0.4023783 | 0.6874056 | 0.827834 |
| ENSG00000259354 | | 2.779169 | 0.1069825 | 0.7438758 | 0.3747582 | 0.7078403 | 0.841134 |
| ENSG00000243297 | RPL31P61 | 1.513291 | 0.1069814 | 0.7004463 | 0.3152594 | 0.7525647 | NA |
| ENSG00000235078 | | 9.289781 | 0.1069551 | 0.5329407 | 0.2649091 | 0.7910795 | 0.891628 |
| ENSG00000213235 | EEF1A1P16 | 8.402851 | 0.106911 | 0.5655022 | 0.2649013 | 0.7910855 | 0.891628 |
| ENSG00000227513 | ILRUNP1 | 2.038123 | 0.1068956 | 0.7466959 | 0.3478387 | 0.7279613 | 0.852261 |
| ENSG00000186104 | CYP2R1 | 730.4917 | 0.1068782 | 0.1433134 | 0.7460261 | 0.4556516 | 0.651642 |
| ENSG00000179832 | MROH1 | 8524.293 | 0.106877 | 0.2847261 | 0.3980401 | 0.6906006 | 0.829793 |
| ENSG00000114739 | ACVR2B | 1168.201119 | 0.106852331 | 0.223296799 | 0.48938036 | 0.62457244 | 0.783563 |
| ENSG00000180803 | | 1.512601 | 0.1067685 | 0.7549878 | 0.3401523 | 0.7337418 | NA |
| ENSG00000162384 | CZIB | 1684.939 | 0.1067638 | 0.1368713 | 0.791669 | 0.4285537 | 0.62791 |
| ENSG00000149792 | MRPL49 | 5267.379 | 0.1066133 | 0.1941011 | 0.5622764 | 0.5739277 | 0.746641 |
| ENSG00000134602 | STK26 | 3371.069804 | 0.106569645 | 0.174645254 | 0.630605584 | 0.52829845 | 0.713103 |

| | | | | | | | |
|---|---|---|---|---|---|---|---|
| ENSG00000102239 | BRS3 | 0.78138719 | 0.106508951 | 0.754448022 | 0.336277058 | 0.73666195 | NA |
| ENSG00000163882 | POLR2H | 4711.769 | 0.1063875 | 0.2060582 | 0.53939 | 0.5896178 | 0.758476 |
| ENSG00000151812 | SLC35F4 | 1.510972 | 0.1063472 | 0.7186194 | 0.3404344 | 0.7335294 | NA |
| ENSG00000197213 | ZSCAN5B | 0.8387798 | 0.1062939 | 0.755521 | 0.4037804 | 0.6863742 | NA |
| ENSG00000234546 | LNCTAM34A | 56.62537 | 0.1062542 | 0.3798222 | 0.3168671 | 0.7513445 | 0.866969 |
| ENSG00000108733 | PEX12 | 248.1820499 | 0.106236109 | 0.229940629 | 0.475016884 | 0.63477494 | 0.790846 |
| ENSG00000250202 | | 13.91995 | 0.1062347 | 0.4603606 | 0.281118 | 0.7786199 | 0.883565 |
| ENSG00000205269 | TMEM170B | 800.6128 | 0.1062201 | 0.2477621 | 0.4458697 | 0.6556914 | 0.805934 |
| ENSG00000236887 | | 0.9346724 | 0.1061894 | 0.7481586 | 0.3903166 | 0.6963024 | NA |
| ENSG00000203852 | H3C15 | 2.164895 | 0.1061348 | 0.7260443 | 0.3558692 | 0.7219385 | 0.848725 |
| ENSG00000054523 | KIF1B | 3476.371522 | 0.106090024 | 0.210399596 | 0.524822644 | 0.59970649 | 0.765998 |
| ENSG00000169738 | DCXR | 6598.126 | 0.1060712 | 0.3228254 | 0.3555939 | 0.7221447 | 0.848817 |
| ENSG00000064726 | BTBD1 | 5086.092871 | 0.106036392 | 0.127721478 | 0.839978775 | 0.40092029 | 0.601913 |
| ENSG00000226855 | RPSAP17 | 3.422926 | 0.1059636 | 0.6187134 | 0.2740209 | 0.7840686 | 0.887008 |
| ENSG00000221926 | TRIM16 | 2374.476 | 0.1058782 | 0.2321686 | 0.4796136 | 0.6315022 | 0.788593 |
| ENSG00000103005 | USB1 | 1889.267014 | 0.105840335 | 0.149361635 | 0.700939615 | 0.4833407 | 0.674716 |
| ENSG00000236301 | MRGPRG-AS1 | 2.633156 | 0.1058113 | 0.7104801 | 0.3315185 | 0.7402529 | 0.859716 |
| ENSG00000259045 | MTCO1P2 | 3.071167 | 0.1057693 | 0.63381 | 0.2773159 | 0.7815376 | 0.885492 |
| ENSG00000266733 | TBC1D29P | 0.939321 | 0.1056437 | 0.7481637 | 0.3721351 | 0.7097923 | NA |
| ENSG00000113163 | CERT1 | 3241.637457 | 0.105643313 | 0.181337329 | 0.596827467 | 0.55062258 | 0.729541 |
| ENSG00000125449 | ARMC7 | 1285.493508 | 0.105638452 | 0.208834091 | 0.521939249 | 0.60171263 | 0.767199 |
| ENSG00000259723 | | 128.5112 | 0.1056101 | 0.3116429 | 0.3655215 | 0.7147222 | 0.844614 |
| ENSG00000225361 | PPP1R26-AS1 | 345.7352 | 0.1055577 | 0.2509145 | 0.4388763 | 0.6607512 | 0.809308 |
| ENSG00000014123 | UFL1 | 2874.561051 | 0.105411566 | 0.205114428 | 0.526892089 | 0.59826853 | 0.765158 |
| ENSG00000162869 | PPP1R21 | 1633.226 | 0.1052782 | 0.1142757 | 0.9227088 | 0.356159 | 0.558176 |
| ENSG00000161013 | MGAT4B | 13084.94 | 0.1052675 | 0.1270433 | 0.8379204 | 0.4020754 | 0.603034 |
| ENSG00000257173 | AK6P2 | 1.330074 | 0.1051903 | 0.6993739 | 0.3009506 | 0.7634522 | NA |
| ENSG00000278949 | | 22.43615 | 0.1051562 | 0.4253981 | 0.2899298 | 0.77187 | 0.879447 |
| ENSG00000231881 | | 16.13481 | 0.1051269 | 0.4564755 | 0.2781714 | 0.7808808 | 0.88512 |
| ENSG00000176826 | FKBP9P1 | 2.652623 | 0.105118 | 0.6359547 | 0.2723903 | 0.7853219 | 0.887731 |
| ENSG00000256616 | | 68.11092 | 0.1050772 | 0.2712291 | 0.4095288 | 0.6821517 | 0.823965 |
| ENSG00000160445 | ZER1 | 4745.272 | 0.1050436 | 0.1255265 | 0.8492843 | 0.3957231 | 0.597249 |
| ENSG00000258346 | | 1.381135 | 0.1049701 | 0.7956847 | 0.7368079 | 0.4612392 | NA |

| | | | | | | | |
|---|---|---|---|---|---|---|---|
| ENSG00000246740 | PLA2G4E-AS1 | 4.752334 | 0.104966 | 0.6077372 | 0.264762 | 0.7911928 | 0.891674 |
| ENSG00000218472 | | 0.9677965 | 0.1049658 | 0.7686208 | 0.4968204 | 0.6193157 | NA |
| ENSG00000286346 | | 1.824171 | 0.1049202 | 0.7572535 | 0.3246943 | 0.7454125 | 0.862995 |
| ENSG00000172482 | AGXT | 1.094542 | 0.1049177 | 0.7521868 | 0.3261947 | 0.7442771 | NA |
| ENSG00000135502 | SLC26A10 | 6.468638303 | 0.104855543 | 0.584669064 | 0.259674612 | 0.79511478 | 0.894159 |
| ENSG00000168395 | ING5 | 3364.502 | 0.1047833 | 0.08534934 | 1.220891 | 0.2221274 | 0.408875 |
| ENSG00000262246 | CORO7 | 585.8438 | 0.1047654 | 0.2142077 | 0.5278497 | 0.5976037 | 0.764492 |
| ENSG00000128016 | ZFP36 | 5075.163898 | 0.104752587 | 0.350500726 | 0.359555686 | 0.71917943 | 0.847018 |
| ENSG00000226755 | VPS25P1 | 1.118336 | 0.1047298 | 0.764136 | 0.5395066 | 0.5895374 | NA |
| ENSG00000154079 | SDHAF4 | 449.9844 | 0.1047162 | 0.1819385 | 0.5933982 | 0.5529147 | 0.731338 |
| ENSG00000258171 | LINC02412 | 7.398966 | 0.1047073 | 0.7043375 | 0.3012954 | 0.7631893 | 0.874092 |
| ENSG00000227906 | SNAP25-AS1 | 1.706234 | 0.1046958 | 0.7261406 | 0.3199492 | 0.7490068 | 0.865675 |
| ENSG00000234005 | GAPDHP22 | 6.374811 | 0.1046791 | 0.559607 | 0.2539555 | 0.7995299 | 0.896976 |
| ENSG00000213465 | ARL2 | 1294.502 | 0.104627 | 0.2495492 | 0.4332772 | 0.6648134 | 0.812764 |
| ENSG00000278974 | | 24.40623 | 0.104623 | 0.3948613 | 0.3019736 | 0.7626722 | 0.873725 |
| ENSG00000214413 | BBIP1 | 591.7326 | 0.1046221 | 0.192829 | 0.5528296 | 0.5803801 | 0.751246 |
| ENSG00000266075 | RN7SL574P | 1.356727 | 0.1046207 | 0.7383734 | 0.368395 | 0.7125788 | NA |
| ENSG00000167962 | ZNF598 | 3893.695 | 0.1045584 | 0.1832466 | 0.5704333 | 0.5683839 | 0.742629 |
| ENSG00000223916 | | 2.908939 | 0.1045512 | 0.6487182 | 0.2754376 | 0.7829801 | 0.886458 |
| ENSG00000267073 | | 1.564428 | 0.1045469 | 0.7516462 | 0.3225819 | 0.7470119 | NA |
| ENSG00000170855 | TRIAP1 | 2517.161 | 0.1045349 | 0.2060788 | 0.5217561 | 0.6018402 | 0.767289 |
| ENSG00000261560 | | 69.97043 | 0.104465 | 0.2991414 | 0.3801171 | 0.7038585 | 0.83894 |
| ENSG00000269226 | | 2.714641 | 0.1044111 | 0.6756839 | 0.2813816 | 0.7784178 | 0.883535 |
| ENSG00000274391 | TPTE | 27.4829 | 0.1043168 | 0.4165021 | 0.293052 | 0.7694824 | 0.877917 |
| ENSG00000254122 | PCDHGB7 | 17.76704 | 0.1042708 | 0.4804394 | 0.2731278 | 0.784755 | 0.887316 |
| ENSG00000242808 | SOX2-OT | 9.852299 | 0.1041578 | 0.5165837 | 0.2629242 | 0.792609 | 0.892702 |
| ENSG00000173928 | SWSAP1 | 194.188 | 0.1041464 | 0.2556811 | 0.4293116 | 0.6676965 | 0.814882 |
| ENSG00000115840 | SLC25A12 | 943.8850499 | 0.10413046 | 0.153761741 | 0.693215082 | 0.48817458 | 0.679074 |
| ENSG00000179921 | GPBAR1 | 13.17057 | 0.1040856 | 0.508325 | 0.2740036 | 0.7840819 | 0.887008 |
| ENSG00000158473 | CD1D | 6.186661 | 0.1040529 | 0.6493004 | 0.2885004 | 0.7729637 | 0.880396 |
| ENSG00000224995 | LINC01526 | 1.394168 | 0.1040082 | 0.7517012 | 0.3219888 | 0.7474612 | NA |
| ENSG00000258428 | | 3.549273 | 0.1039989 | 0.6305415 | 0.2665178 | 0.7898405 | 0.890982 |
| ENSG00000167065 | DUSP18 | 111.4283 | 0.1037464 | 0.2703502 | 0.4035932 | 0.6865119 | 0.827321 |

| | | | | | | | |
|---|---|---|---|---|---|---|---|
| ENSG00000251356 | RAB5CP2 | 1.020971 | 0.1037025 | 0.7356321 | 0.3669596 | 0.7136492 | NA |
| ENSG00000143379 | SETDB1 | 6340.264 | 0.1036924 | 0.1468792 | 0.7025471 | 0.482338 | 0.673777 |
| ENSG00000212695 | | 1.579042 | 0.1036533 | 0.7039952 | 0.3184919 | 0.7501119 | 0.866263 |
| ENSG00000104687 | GSR | 5812.386151 | 0.103618941 | 0.234288923 | 0.479893915 | 0.63130283 | 0.788593 |
| ENSG00000166326 | TRIM44 | 8881.745 | 0.1036147 | 0.197106 | 0.5366059 | 0.5915399 | 0.760212 |
| ENSG00000165568 | AKR1E2 | 628.3625 | 0.1035911 | 0.1861371 | 0.5729282 | 0.5666933 | 0.74126 |
| ENSG00000100453 | GZMB | 0.820166615 | 0.103504645 | 0.75497438 | 0.391435354 | 0.69547546 | NA |
| ENSG00000169169 | CPT1C | 50.73691 | 0.1034819 | 0.4021686 | 0.29063 | 0.7713344 | 0.879062 |
| ENSG00000197826 | CFAP299 | 1.273247 | 0.1033822 | 0.7701311 | 0.4816083 | 0.6300843 | NA |
| ENSG00000272691 | | 1.069255 | 0.1033671 | 0.7191353 | 0.3272416 | 0.7434851 | NA |
| ENSG00000183735 | TBK1 | 3566.139 | 0.1032458 | 0.1647129 | 0.6350127 | 0.5254201 | 0.710318 |
| ENSG00000138674 | SEC31A | 9650.234472 | 0.103242755 | 0.11783458 | 0.884413461 | 0.37647306 | 0.578617 |
| ENSG00000213216 | | 4.367765 | 0.1032246 | 0.5957427 | 0.2619763 | 0.7933397 | 0.893115 |
| ENSG00000144559 | TAMM41 | 1428.156 | 0.1031537 | 0.1943476 | 0.5302208 | 0.5959589 | 0.763196 |
| ENSG00000248758 | | 3.151711 | 0.1031496 | 0.6563706 | 0.2732024 | 0.7846976 | 0.887289 |
| ENSG00000125779 | PANK2 | 1148.263032 | 0.102971595 | 0.129713755 | 0.79497115 | 0.42663028 | 0.626378 |
| ENSG00000224470 | ATXN1L | 5810.924 | 0.1029336 | 0.2245311 | 0.4758181 | 0.634204 | 0.790553 |
| ENSG00000280033 | | 21.17256 | 0.1029313 | 0.4248172 | 0.2835403 | 0.7767627 | 0.882518 |
| ENSG00000148300 | REXO4 | 4910.575 | 0.1028635 | 0.1254586 | 0.8104931 | 0.4176568 | 0.617501 |
| ENSG00000164050 | PLXNB1 | 8850.439 | 0.1028275 | 0.181493 | 0.5640646 | 0.5727102 | 0.745897 |
| ENSG00000136536 | MARCHF7 | 5082.59602 | 0.102799837 | 0.138791877 | 0.744827243 | 0.45637616 | 0.652187 |
| ENSG00000198824 | CHAMP1 | 1335.115 | 0.1027839 | 0.219137 | 0.4795863 | 0.6315216 | 0.788593 |
| ENSG00000121900 | TMEM54 | 2713.289676 | 0.102730443 | 0.225078343 | 0.492737732 | 0.62219791 | 0.781764 |
| ENSG00000006831 | ADIPOR2 | 8432.487788 | 0.102650754 | 0.181506365 | 0.585981344 | 0.55788805 | 0.735159 |
| ENSG00000268744 | | 1.180554 | 0.10262 | 0.7551707 | 0.3309751 | 0.7406633 | NA |
| ENSG00000277232 | GTSE1-DT | 60.72228 | 0.1026012 | 0.3376126 | 0.3332988 | 0.7389088 | 0.858927 |
| ENSG00000235142 | LINC02532 | 119.4295 | 0.1025965 | 0.559842 | 0.2722958 | 0.7853946 | 0.887737 |
| ENSG00000160570 | DEDD2 | 2006.195 | 0.1025907 | 0.2016877 | 0.5207842 | 0.6025171 | 0.767746 |
| ENSG00000271426 | | 34.31195 | 0.1024318 | 0.4266901 | 0.2826372 | 0.777455 | 0.882857 |
| ENSG00000275703 | | 3.411862 | 0.1023925 | 0.6609724 | 0.2728506 | 0.7849681 | 0.887468 |
| ENSG00000197818 | SLC9A8 | 2894.633 | 0.1023762 | 0.2114417 | 0.4997941 | 0.61722 | 0.778565 |
| ENSG00000279392 | | 354.738 | 0.1023113 | 0.2672543 | 0.4226011 | 0.6725863 | 0.818034 |
| ENSG00000179151 | EDC3 | 3369.127 | 0.1022349 | 0.1301528 | 0.7828617 | 0.4337083 | 0.632656 |

| | | | | | | | |
|---|---|---|---|---|---|---|---|
| ENSG00000274979 | | 0.7968318 | 0.102186 | 0.7724358 | 0.4518478 | 0.6513786 | NA |
| ENSG00000233476 | EEF1A1P6 | 2597.612 | 0.1021795 | 0.2117277 | 0.4959616 | 0.6199215 | 0.78038 |
| ENSG00000264853 | | 0.7799839 | 0.1021792 | 0.7877796 | 0.2317828 | 0.8167067 | NA |
| ENSG00000239809 | | 4.356065 | 0.1021264 | 0.6310365 | 0.2602573 | 0.7946653 | 0.893984 |
| ENSG00000159348 | CYB5R1 | 6013.955 | 0.1021026 | 0.2050147 | 0.5155018 | 0.6062024 | 0.770549 |
| ENSG00000174365 | SNHG11 | 750.1162 | 0.1020602 | 0.201812 | 0.526776 | 0.5983492 | 0.76521 |
| ENSG00000269300 | | 2.659838 | 0.1020448 | 0.6722351 | 0.2695039 | 0.787542 | 0.889371 |
| ENSG00000110906 | KCTD10 | 3761.367692 | 0.102001912 | 0.123525578 | 0.855459374 | 0.39229689 | 0.594035 |
| ENSG00000242314 | RPL12P32 | 2.286622 | 0.1019491 | 0.6538936 | 0.273578 | 0.7844089 | 0.887189 |
| ENSG00000201659 | RNU12-2P | 1.708171 | 0.1019046 | 0.6972002 | 0.2865903 | 0.7744261 | 0.880999 |
| ENSG00000172689 | MS4A10 | 1.604087 | 0.1017614 | 0.7580145 | 0.3195833 | 0.7492843 | 0.86578 |
| ENSG00000139370 | SLC15A4 | 1452.219 | 0.1017371 | 0.1339823 | 0.7790735 | 0.4359364 | 0.634623 |
| ENSG00000272040 | | 8.215721 | 0.1016748 | 0.5728061 | 0.2541927 | 0.7993467 | 0.896976 |
| ENSG00000197301 | HMGA2-AS1 | 1.011402 | 0.1015503 | 0.789377 | 0.2140779 | 0.8304863 | NA |
| ENSG00000227232 | WASH7P | 83.06532 | 0.1014785 | 0.2968525 | 0.3663401 | 0.7141113 | 0.84406 |
| ENSG00000070159 | PTPN3 | 3369.36459 | 0.101454488 | 0.141006611 | 0.733121234 | 0.46348449 | 0.657711 |
| ENSG00000103852 | TTC23 | 611.0531725 | 0.101436559 | 0.204170378 | 0.509247527 | 0.61057873 | 0.77401 |
| ENSG00000248503 | | 6.97419 | 0.1014336 | 0.5676085 | 0.2553996 | 0.7984145 | 0.896254 |
| ENSG00000255964 | HSPD1P12 | 15.60856 | 0.1013603 | 0.4740339 | 0.2637414 | 0.7919792 | 0.892302 |
| ENSG00000170540 | ARL6IP1 | 9929.33 | 0.1013177 | 0.211243 | 0.4947591 | 0.6207702 | 0.780967 |
| ENSG00000228146 | CASP16P | 91.92554 | 0.1013068 | 0.4325694 | 0.2737824 | 0.7842518 | 0.887087 |
| ENSG00000235308 | | 1.582499 | 0.1012714 | 0.7163387 | 0.3154871 | 0.7523918 | 0.867522 |
| ENSG00000170734 | POLH | 1168.212 | 0.101226 | 0.291209 | 0.3673546 | 0.7133545 | 0.84368 |
| ENSG00000275183 | LENG9 | 750.9224 | 0.1012112 | 0.192526 | 0.5291323 | 0.5967137 | 0.763758 |
| ENSG00000249492 | | 92.47596 | 0.1011031 | 0.2823872 | 0.3852154 | 0.7000778 | 0.836573 |
| ENSG00000286990 | | 7.395962 | 0.1010933 | 0.6403704 | 0.265106 | 0.7909278 | 0.891628 |
| ENSG00000186130 | ZBTB6 | 1053.115 | 0.100971 | 0.1799591 | 0.5721965 | 0.5671888 | 0.741616 |
| ENSG00000225769 | CROCCP1 | 1.863444 | 0.1009569 | 0.7020992 | 0.2858586 | 0.7749864 | 0.881372 |
| ENSG00000122122 | SASH3 | 1.282081028 | 0.100933788 | 0.746428652 | 0.389979467 | 0.69655173 | NA |
| ENSG00000162852 | CNST | 2769.737 | 0.100885 | 0.1955267 | 0.5311701 | 0.5953009 | 0.762648 |
| ENSG00000226982 | CENPCP1 | 39.76517 | 0.1008236 | 0.2985506 | 0.3676757 | 0.713115 | 0.843667 |
| ENSG00000204922 | UQCC3 | 980.3629 | 0.1008048 | 0.1983328 | 0.5234371 | 0.6006701 | 0.766717 |
| ENSG00000176248 | ANAPC2 | 4684.216 | 0.1006963 | 0.09743186 | 1.039358 | 0.2986383 | 0.497846 |

| | | | | | | | |
|---|---|---|---|---|---|---|---|
| ENSG00000250874 | | 34.55798 | 0.1006698 | 0.3676652 | 0.3080906 | 0.7580134 | 0.871039 |
| ENSG00000162365 | CYP4A22 | 1.722997 | 0.1006601 | 0.7415188 | 0.3643698 | 0.7155819 | 0.845179 |
| ENSG00000259096 | SHLD2P2 | 37.00626 | 0.1006329 | 0.4026489 | 0.2881017 | 0.7732689 | 0.880474 |
| ENSG00000237713 | | 0.9885539 | 0.1005328 | 0.7887609 | 0.2189608 | 0.8266806 | NA |
| ENSG00000262959 | RPL23AP86 | 0.7971558 | 0.1004668 | 0.7517924 | 0.3244816 | 0.7455734 | NA |
| ENSG00000188293 | IGFL1 | 91.57049 | 0.1003889 | 0.6681439 | 0.2918517 | 0.7704 | 0.878601 |
| ENSG00000236534 | H3P36 | 5.99782 | 0.1003438 | 0.5851014 | 0.2530046 | 0.8002646 | 0.897349 |
| ENSG00000206622 | SNORA69 | 3.450009 | 0.100253 | 0.6587138 | 0.2711307 | 0.7862905 | 0.88844 |
| ENSG00000251417 | | 35.98659 | 0.1001424 | 0.4647307 | 0.2625402 | 0.792905 | 0.892909 |
| ENSG00000130701 | RBBP8NL | 1273.181778 | 0.100134333 | 0.281972683 | 0.376441991 | 0.70658833 | 0.840497 |
| ENSG00000198258 | UBL5 | 3717.853 | 0.1001308 | 0.2512135 | 0.4173805 | 0.6764001 | 0.820119 |
| ENSG00000287908 | | 3.316684 | 0.1001131 | 0.6437717 | 0.2591062 | 0.7955533 | 0.894387 |
| ENSG00000184227 | ACOT1 | 173.8481 | 0.1001044 | 0.1992178 | 0.5174715 | 0.6048271 | 0.769546 |
| ENSG00000257599 | OVCH1-AS1 | 5.517999 | 0.1000488 | 0.5789741 | 0.2514122 | 0.8014954 | 0.898186 |
| ENSG00000100242 | SUN2 | 2215.419669 | 0.100032833 | 0.121297495 | 0.824801087 | 0.40948453 | 0.609936 |
| ENSG00000253138 | LINC00967 | 0.8292314 | 0.1000275 | 0.7554229 | 0.3282384 | 0.7427314 | NA |
| ENSG00000224906 | SRP14P1 | 5.701873 | 0.1000061 | 0.5504347 | 0.2485129 | 0.8037376 | 0.899554 |
| ENSG00000228882 | CICP9 | 3.493084 | 0.09995936 | 0.6761476 | 0.2669312 | 0.7895221 | 0.890811 |
| ENSG00000005889 | ZFX | 1770.440408 | 0.099957865 | 0.194243765 | 0.525146897 | 0.59948108 | 0.765898 |
| ENSG00000267253 | WHSC1L2P | 4.854805 | 0.0998795 | 0.6240736 | 0.2525594 | 0.8006087 | 0.897578 |
| ENSG00000064961 | HMG20B | 7647.741252 | 0.099844133 | 0.185088945 | 0.552773865 | 0.58041827 | 0.751246 |
| ENSG00000268471 | MIR4453HG | 496.1581 | 0.09982276 | 0.2142037 | 0.4758083 | 0.6342109 | 0.790553 |
| ENSG00000197958 | RPL12 | 11426.43 | 0.09981772 | 0.1623512 | 0.6185176 | 0.5362342 | 0.719265 |
| ENSG00000075413 | MARK3 | 7565.09196 | 0.099785108 | 0.082278561 | 1.237233517 | 0.2160004 | 0.401249 |
| ENSG00000187607 | ZNF286A | 695.8606 | 0.09956613 | 0.1357068 | 0.7450212 | 0.4562589 | 0.652125 |
| ENSG00000178796 | RIIAD1 | 5.911229 | 0.09954047 | 0.5538068 | 0.2445095 | 0.8068363 | 0.900972 |
| ENSG00000163755 | HPS3 | 4600.062 | 0.09952584 | 0.1960297 | 0.52268 | 0.6011969 | 0.766947 |
| ENSG00000198934 | MAGEE1 | 277.2667 | 0.09947636 | 0.2077518 | 0.5008545 | 0.6164735 | 0.777845 |
| ENSG00000100036 | SLC35E4 | 198.0023713 | 0.099473675 | 0.261459938 | 0.400743567 | 0.68860893 | 0.828482 |
| ENSG00000226247 | SUPT4H1P1 | 11.76739 | 0.09943463 | 0.5017153 | 0.2529829 | 0.8002814 | 0.897349 |
| ENSG00000152492 | CCDC50 | 3110.629 | 0.09940787 | 0.158078 | 0.6266482 | 0.5308899 | 0.715129 |
| ENSG00000073331 | ALPK1 | 541.3906862 | 0.099367285 | 0.359351389 | 0.304632548 | 0.76064603 | 0.872369 |
| ENSG00000142409 | ZNF787 | 2070.57 | 0.09925885 | 0.2201306 | 0.4585249 | 0.6465754 | 0.799394 |

| | | | | | | | |
|---|---|---|---|---|---|---|---|
| ENSG00000270499 | MKI67P1 | 0.9525084 | 0.09923927 | 0.7369838 | 0.3397583 | 0.7340386 | NA |
| ENSG00000259590 | LINC02244 | 0.8414374 | 0.09916773 | 0.7471892 | 0.322317 | 0.7472125 | NA |
| ENSG00000278654 | | 1.163173 | 0.09914446 | 0.7562491 | 0.3752199 | 0.7074969 | NA |
| ENSG00000181264 | TLCD5 | 513.8957 | 0.09906801 | 0.2784353 | 0.3784725 | 0.7050796 | 0.839876 |
| ENSG00000137411 | VARS2 | 57.2238135 | 0.09906417 | 0.404457337 | 0.2773107 | 0.78154154 | 0.885492 |
| ENSG00000265462 | MIR3680-1 | 0.972224 | 0.09899884 | 0.7425079 | 0.3636327 | 0.7161323 | NA |
| ENSG00000207357 | RNU6-2 | 14.88451 | 0.09899393 | 0.5373835 | 0.2471856 | 0.8047646 | 0.90021 |
| ENSG00000200742 | | 5.756839 | 0.09898758 | 0.5873935 | 0.2472799 | 0.8046916 | 0.900166 |
| ENSG00000134389 | CFHR5 | 3.760304218 | 0.098834494 | 0.593491227 | 0.243509612 | 0.80761064 | 0.901496 |
| ENSG00000252915 | | 26.19275 | 0.09880605 | 0.3919638 | 0.2883316 | 0.7730929 | 0.880445 |
| ENSG00000240625 | RN7SL403P | 2.844446 | 0.09877465 | 0.6549635 | 0.2578408 | 0.7965298 | 0.89504 |
| ENSG00000188163 | FAM166A | 20.59975 | 0.09873149 | 0.4742094 | 0.2639964 | 0.7917827 | 0.892187 |
| ENSG00000188811 | NHLRC3 | 2547.016 | 0.09865603 | 0.2705322 | 0.4017488 | 0.6878689 | 0.828092 |
| ENSG00000201003 | | 0.900981 | 0.09864279 | 0.7544285 | 0.3073634 | 0.7585668 | NA |
| ENSG00000254198 | | 50.19214 | 0.09859858 | 0.2772759 | 0.3775779 | 0.7057442 | 0.840057 |
| ENSG00000128928 | IVD | 8086.107651 | 0.098471286 | 0.161895702 | 0.609749475 | 0.54202778 | 0.723521 |
| ENSG00000217862 | H4C10P | 24.88926 | 0.09845791 | 0.3775558 | 0.2951696 | 0.7678643 | 0.876989 |
| ENSG00000046647 | GEMIN8 | 808.4785173 | 0.098258406 | 0.166761705 | 0.613895659 | 0.53928428 | 0.721616 |
| ENSG00000265094 | | 1.019435 | 0.09818695 | 0.7377155 | 0.3286006 | 0.7424576 | NA |
| ENSG00000137038 | DMAC1 | 1789.590725 | 0.09813836 | 0.176357708 | 0.565612865 | 0.57165698 | 0.745294 |
| ENSG00000185624 | P4HB | 39814.05 | 0.09806039 | 0.1319311 | 0.7517844 | 0.4521807 | 0.648319 |
| ENSG00000253047 | | 18.93154 | 0.0980443 | 0.4515287 | 0.2600583 | 0.7948188 | 0.894013 |
| ENSG00000165714 | BORCS5 | 279.8168 | 0.0979582 | 0.1929929 | 0.5224483 | 0.6013582 | 0.767005 |
| ENSG00000257696 | | 12.03227 | 0.09789597 | 0.5696771 | 0.2535117 | 0.7998728 | 0.897112 |
| ENSG00000240230 | COX19 | 2305.688 | 0.09785962 | 0.1262855 | 0.8325584 | 0.4050939 | 0.605851 |
| ENSG00000260572 | | 49.17035 | 0.0978545 | 0.2893754 | 0.3587015 | 0.7198184 | 0.847583 |
| ENSG00000287454 | | 1.599156 | 0.09781907 | 0.683881 | 0.278562 | 0.780581 | 0.884969 |
| ENSG00000170222 | ADPRM | 346.5875 | 0.0977694 | 0.2438278 | 0.4223948 | 0.6727368 | 0.818088 |
| ENSG00000180354 | MTURN | 1072.736 | 0.0977618 | 0.290451 | 0.3547231 | 0.7227971 | 0.849321 |
| ENSG00000286376 | | 0.7954906 | 0.09774515 | 0.7645398 | 0.3870453 | 0.6987227 | NA |
| ENSG00000268279 | | 55.09203 | 0.09772041 | 0.2639385 | 0.3904021 | 0.6962392 | 0.833861 |
| ENSG00000204217 | BMPR2 | 5512.048 | 0.09771237 | 0.2127431 | 0.4856433 | 0.6272201 | 0.785585 |
| ENSG00000244482 | LILRA6 | 2.307513 | 0.09768233 | 0.7132162 | 0.3011979 | 0.7632636 | 0.874121 |

| | | | | | | | |
|---|---|---|---|---|---|---|---|
| ENSG00000241057 | | 14.07696 | 0.09766265 | 0.4109122 | 0.2747284 | 0.7835249 | 0.886724 |
| ENSG00000240849 | PEDS1 | 2923.663 | 0.09765981 | 0.1775762 | 0.5364062 | 0.5916778 | 0.760316 |
| ENSG00000258443 | | 1.696484 | 0.09757249 | 0.7188262 | 0.3062522 | 0.7594126 | 0.871795 |
| ENSG00000251102 | CTBP2P4 | 145.6716 | 0.09745806 | 0.225837 | 0.44744 | 0.6545574 | 0.805151 |
| ENSG00000273980 | | 1.993635 | 0.09745399 | 0.7118772 | 0.3080983 | 0.7580075 | 0.871039 |
| ENSG00000122870 | BICC1 | 0.805621603 | 0.097282987 | 0.751073383 | 0.375780754 | 0.7070799 | NA |
| ENSG00000165813 | CCDC186 | 2258.838 | 0.0971182 | 0.2952633 | 0.3608854 | 0.7181852 | 0.846264 |
| ENSG00000278934 | | 0.7875675 | 0.09711583 | 0.7767878 | 0.4206638 | 0.6740006 | NA |
| ENSG00000224635 | | 31.12624 | 0.09711006 | 0.3579785 | 0.2998399 | 0.7642993 | 0.874684 |
| ENSG00000198719 | DLL1 | 717.9746 | 0.09708898 | 0.1471437 | 0.6435953 | 0.5198379 | 0.705104 |
| ENSG00000263515 | MIR548AN | 1.879207 | 0.09708807 | 0.7422793 | 0.3757226 | 0.7071231 | 0.840791 |
| ENSG00000105369 | CD79A | 7.502438422 | 0.097058427 | 0.551117764 | 0.240243103 | 0.8101418 | 0.902923 |
| ENSG00000282757 | DUXB | 0.9974725 | 0.09703214 | 0.7565315 | 0.3280824 | 0.7428493 | NA |
| ENSG00000197562 | RAB40C | 3217.397 | 0.09696557 | 0.139929 | 0.6996046 | 0.4841742 | 0.675419 |
| ENSG00000131089 | ARHGEF9 | 1846.322792 | 0.096882325 | 0.114866954 | 0.838361291 | 0.40182782 | 0.602798 |
| ENSG00000272434 | | 9.412365 | 0.09685124 | 0.4941663 | 0.2417293 | 0.8089899 | 0.902429 |
| ENSG00000198937 | CCDC167 | 576.9459 | 0.09680907 | 0.3324388 | 0.319852 | 0.7490805 | 0.865695 |
| ENSG00000253525 | CPP | 2.36348 | 0.09664981 | 0.748192 | 0.353788 | 0.7234978 | 0.849802 |
| ENSG00000198857 | HSD3BP5 | 1.435793 | 0.09657133 | 0.7530378 | 0.3569085 | 0.7211603 | NA |
| ENSG00000075539 | FRYL | 3636.720615 | 0.09656036 | 0.211446975 | 0.472158534 | 0.63681364 | 0.792549 |
| ENSG00000052795 | FNIP2 | 769.5566912 | 0.09649122 | 0.353827481 | 0.300590535 | 0.76372675 | 0.874293 |
| ENSG00000253730 | | 2.300563 | 0.09645247 | 0.6591458 | 0.2510985 | 0.801738 | 0.898186 |
| ENSG00000234678 | ELF3-AS1 | 587.3451 | 0.09643605 | 0.2588715 | 0.3933029 | 0.6940957 | 0.832232 |
| ENSG00000056097 | ZFR | 8198.976133 | 0.096402311 | 0.168534291 | 0.583054358 | 0.55985671 | 0.736232 |
| ENSG00000022277 | RTF2 | 8867.992291 | 0.09637781 | 0.184672898 | 0.53432941 | 0.59311365 | 0.761093 |
| ENSG00000170242 | USP47 | 2355.353 | 0.09637688 | 0.1439571 | 0.6515444 | 0.5146951 | 0.700309 |
| ENSG00000272769 | | 13.08455 | 0.09637206 | 0.510598 | 0.2411168 | 0.8094646 | 0.902793 |
| ENSG00000161654 | LSM12 | 766.0588 | 0.09629666 | 0.1782156 | 0.5524822 | 0.5806181 | 0.751373 |
| ENSG00000233330 | | 11.92387 | 0.09629566 | 0.5576365 | 0.2389877 | 0.8111151 | 0.903386 |
| ENSG00000050130 | JKAMP | 3071.289917 | 0.096221989 | 0.11087391 | 0.86240685 | 0.38846367 | 0.590521 |
| ENSG00000214719 | | 25.30785 | 0.09613961 | 0.391317 | 0.2788904 | 0.7803289 | 0.884872 |
| ENSG00000207205 | RNVU1-15 | 9.588968 | 0.09605542 | 0.5475948 | 0.237622 | 0.8121743 | 0.904197 |
| ENSG00000264157 | MIR3127 | 1.979204 | 0.09600487 | 0.690021 | 0.2813214 | 0.7784639 | 0.883549 |

| | | | | | | | |
|---|---|---|---|---|---|---|---|
| ENSG00000260751 | | 105.755 | 0.09599647 | 0.2221354 | 0.4460677 | 0.6555483 | 0.805833 |
| ENSG00000126953 | TIMM8A | 1546.147827 | 0.095976116 | 0.191149129 | 0.518890838 | 0.60383687 | 0.768809 |
| ENSG00000269694 | | 4.015948 | 0.09595673 | 0.6330049 | 0.2533127 | 0.8000266 | 0.897214 |
| ENSG00000265417 | | 6.835379 | 0.09593154 | 0.5481326 | 0.2355006 | 0.8138202 | 0.905043 |
| ENSG00000223431 | MTND6P21 | 3.533591 | 0.0959204 | 0.6556265 | 0.2475842 | 0.8044561 | 0.899971 |
| ENSG00000197965 | MPZL1 | 4891.131 | 0.09587287 | 0.1163053 | 0.8491388 | 0.3958041 | 0.597337 |
| ENSG00000100116 | GCAT | 944.9570926 | 0.095831414 | 0.241861389 | 0.419182402 | 0.67508283 | 0.819345 |
| ENSG00000275523 | MIR6508 | 0.7722346 | 0.09583014 | 0.7652514 | 0.5038787 | 0.6143466 | NA |
| ENSG00000226239 | | 5.258148 | 0.09572631 | 0.5845502 | 0.2420187 | 0.8087657 | 0.902255 |
| ENSG00000228236 | TXNP5 | 4.628039 | 0.09572247 | 0.6613377 | 0.257395 | 0.7968739 | 0.895202 |
| ENSG00000102172 | SMS | 3954.583996 | 0.095672863 | 0.231780971 | 0.435704536 | 0.66305111 | 0.811339 |
| ENSG00000198064 | NPIPB13 | 10.47788 | 0.09566816 | 0.4639361 | 0.250269 | 0.8023793 | 0.898677 |
| ENSG00000272288 | ILRUN-AS1 | 133.7409 | 0.09554747 | 0.3168682 | 0.3269779 | 0.7436846 | 0.861863 |
| ENSG00000255524 | NPIPB8 | 3.797101 | 0.09549082 | 0.6280995 | 0.2455056 | 0.806065 | 0.90047 |
| ENSG00000163110 | PDLIM5 | 3023.269 | 0.09541367 | 0.2646605 | 0.3788656 | 0.7047877 | 0.839784 |
| ENSG00000256973 | | 11.29089 | 0.0953954 | 0.5423489 | 0.2366526 | 0.8129263 | 0.90439 |
| ENSG00000114902 | SPCS1 | 3296.269699 | 0.095312078 | 0.161112365 | 0.603599612 | 0.54610987 | 0.726315 |
| ENSG00000272153 | | 40.96625 | 0.09529504 | 0.3205847 | 0.3238946 | 0.7460178 | 0.863507 |
| ENSG00000162601 | MYSM1 | 3760.913 | 0.09526544 | 0.23681 | 0.4160277 | 0.6773897 | 0.820608 |
| ENSG00000240542 | KRTAP9-1 | 1.648016 | 0.09525664 | 0.7055448 | 0.2838823 | 0.7765006 | 0.882376 |
| ENSG00000227676 | LINC01068 | 2.146791 | 0.09520577 | 0.6671012 | 0.2498748 | 0.8026842 | 0.898879 |
| ENSG00000263002 | ZNF234 | 667.73 | 0.09520201 | 0.240257 | 0.4095541 | 0.682133 | 0.823965 |
| ENSG00000242562 | DCAF13P1 | 75.99787 | 0.09508626 | 0.2484035 | 0.4021798 | 0.6875517 | 0.827935 |
| ENSG00000259309 | | 0.7370252 | 0.09506873 | 0.7657762 | 0.3774738 | 0.7058216 | NA |
| ENSG00000286771 | | 0.925868 | 0.0950207 | 0.7542765 | 0.311253 | 0.7556083 | NA |
| ENSG00000253153 | | 1.003575 | 0.09495879 | 0.7560456 | 0.3678119 | 0.7130135 | NA |
| ENSG00000277957 | SENP3-EIF4A1 | 3.282456 | 0.09490656 | 0.6281223 | 0.2374427 | 0.8123134 | 0.9042 |
| ENSG00000198909 | MAP3K3 | 5247.271 | 0.09490511 | 0.148577 | 0.6485451 | 0.5166324 | 0.701939 |
| ENSG00000113108 | APBB3 | 1358.514832 | 0.094809516 | 0.233715847 | 0.41569214 | 0.67763529 | 0.820733 |
| ENSG00000135297 | MTO1 | 2320.256795 | 0.094801764 | 0.14959994 | 0.649703637 | 0.51588367 | 0.701208 |
| ENSG00000184432 | COPB2 | 13892.98 | 0.09472858 | 0.1091919 | 0.8744464 | 0.3818752 | 0.584113 |
| ENSG00000280243 | MTCO1P1 | 4.25655 | 0.09467993 | 0.6536021 | 0.2468632 | 0.8050141 | 0.900308 |
| ENSG00000219682 | | 110.2543 | 0.09461962 | 0.2314477 | 0.4274394 | 0.6690593 | 0.81561 |

| | | | | | | | |
|---|---|---|---|---|---|---|---|
| ENSG00000241217 | RN7SL809P | 9.057429 | 0.09461397 | 0.5352918 | 0.2369503 | 0.8126953 | 0.904263 |
| ENSG00000264279 | MIR4786 | 1.661352 | 0.0945338 | 0.6862578 | 0.2637814 | 0.7919484 | 0.892302 |
| ENSG00000156709 | AIFM1 | 5988.828 | 0.0945183 | 0.220575 | 0.4471491 | 0.6547674 | 0.805245 |
| ENSG00000188856 | RPSAP47 | 297.3255 | 0.09445732 | 0.2077615 | 0.468977 | 0.6390861 | 0.794106 |
| ENSG00000268034 | | 366.8564 | 0.09440488 | 0.2432298 | 0.4077665 | 0.6834451 | 0.824859 |
| ENSG00000225880 | LINC00115 | 42.89031 | 0.09440387 | 0.3457264 | 0.3010153 | 0.7634028 | 0.874223 |
| ENSG00000166401 | SERPINB8 | 151.6993 | 0.09437776 | 0.39427 | 0.2678131 | 0.7888432 | 0.890218 |
| ENSG00000252498 | RNU6-1016P | 11.20919 | 0.0943598 | 0.5230541 | 0.236254 | 0.8132356 | 0.904658 |
| ENSG00000115170 | ACVR1 | 1025.550773 | 0.094307569 | 0.184547839 | 0.516600327 | 0.60543519 | 0.769897 |
| ENSG00000263612 | | 5.691625 | 0.09423459 | 0.5929107 | 0.2324995 | 0.8161501 | 0.906345 |
| ENSG00000213295 | RPL22P10 | 2.574995 | 0.0941522 | 0.688215 | 0.2678098 | 0.7888457 | 0.890218 |
| ENSG00000025800 | KPNA6 | 5322.960976 | 0.094138881 | 0.077102509 | 1.211848711 | 0.2255703 | 0.412895 |
| ENSG00000258791 | LINC00520 | 0.9061904 | 0.09410543 | 0.7467448 | 0.3574655 | 0.7207434 | NA |
| ENSG00000124688 | MAD2L1BP | 1823.026032 | 0.094103858 | 0.163473442 | 0.5961174 | 0.5510968 | 0.730024 |
| ENSG00000196796 | NPIPB10P | 3.42074 | 0.09410112 | 0.6488672 | 0.2526381 | 0.8005479 | 0.897578 |
| ENSG00000238140 | | 1.474463 | 0.09406326 | 0.666602 | 0.2536517 | 0.7997647 | NA |
| ENSG00000253167 | WASHC5-AS1 | 33.46657 | 0.09398563 | 0.3298256 | 0.31026 | 0.7563633 | 0.869898 |
| ENSG00000256250 | | 1.440074 | 0.09391817 | 0.7137627 | 0.2875242 | 0.7737109 | NA |
| ENSG00000163995 | ABLIM2 | 162.1262 | 0.09389059 | 0.280819 | 0.355625 | 0.7221214 | 0.848817 |
| ENSG00000173264 | GPR137 | 1931.789 | 0.09378577 | 0.2363499 | 0.4165763 | 0.6769884 | 0.820346 |
| ENSG00000234356 | | 1.032591 | 0.0937807 | 0.7536429 | 0.3071611 | 0.7587207 | NA |
| ENSG00000204304 | PBX2 | 3594.679 | 0.09375953 | 0.1665744 | 0.588626 | 0.5561122 | 0.733627 |
| ENSG00000226797 | | 7.766189 | 0.09362826 | 0.6209761 | 0.2431634 | 0.8078788 | 0.901644 |
| ENSG00000183977 | PP2D1 | 43.13811 | 0.09360626 | 0.3282503 | 0.3121619 | 0.7549175 | 0.869057 |
| ENSG00000216775 | | 341.8318 | 0.09356989 | 0.2015338 | 0.4833195 | 0.6288689 | 0.786756 |
| ENSG00000235651 | G3BP1P1 | 18.49009 | 0.09354492 | 0.4361321 | 0.2542666 | 0.7992896 | 0.896956 |
| ENSG00000141867 | BRD4 | 2886.198 | 0.09348509 | 0.1303925 | 0.7216673 | 0.4704991 | 0.66397 |
| ENSG00000226650 | KIF4B | 70.38122 | 0.09346779 | 0.5043839 | 0.2401121 | 0.8102434 | 0.902923 |
| ENSG00000167114 | SLC27A4 | 3013.7 | 0.09343443 | 0.1489951 | 0.6157812 | 0.538039 | 0.721067 |
| ENSG00000215717 | TMEM167B | 3326.329 | 0.09340379 | 0.2014777 | 0.475686 | 0.6342981 | 0.790553 |
| ENSG00000116675 | DNAJC6 | 1431.330712 | 0.093399188 | 0.3489813 | 0.293741192 | 0.76895568 | 0.877556 |
| ENSG00000239268 | | 4.853869 | 0.0933812 | 0.6785819 | 0.2456788 | 0.8059309 | 0.90047 |
| ENSG00000220739 | | 0.8453078 | 0.09337435 | 0.7652771 | 0.4801438 | 0.6311251 | NA |

| | | | | | | | |
|---|---|---|---|---|---|---|---|
| ENSG00000277270 | | 0.8242453 | 0.09331209 | 0.7858727 | 0.2327879 | 0.8159261 | NA |
| ENSG00000278370 | | 1.427622 | 0.09328602 | 0.7347693 | 0.3263285 | 0.7441758 | NA |
| ENSG00000226445 | | 65.08172 | 0.09327765 | 0.3259606 | 0.3105242 | 0.7561623 | 0.869868 |
| ENSG00000204118 | NAP1L6P | 116.348 | 0.09327059 | 0.242479 | 0.4014881 | 0.6880608 | 0.828129 |
| ENSG00000258057 | BCDIN3D-AS1 | 65.01712 | 0.09325086 | 0.429839 | 0.2541025 | 0.7994163 | 0.896976 |
| ENSG00000134769 | DTNA | 4525.541281 | 0.093192188 | 0.293426608 | 0.344516511 | 0.73045788 | 0.853899 |
| ENSG00000225978 | HAR1A | 6.38559 | 0.09318169 | 0.6943437 | 0.284718 | 0.7758602 | 0.881951 |
| ENSG00000225051 | HMGB3P22 | 29.08141 | 0.09313245 | 0.4206498 | 0.2584136 | 0.7960877 | 0.894837 |
| ENSG00000253385 | | 1.607684 | 0.09310403 | 0.7364314 | 0.3398548 | 0.7339659 | 0.855711 |
| ENSG00000196943 | NOP9 | 4444.735 | 0.09308908 | 0.1583002 | 0.5996489 | 0.5487402 | 0.728137 |
| ENSG00000229272 | | 7.151341 | 0.09307779 | 0.5895231 | 0.2268266 | 0.8205585 | 0.909023 |
| ENSG00000225727 | | 2.79972 | 0.09305189 | 0.5974303 | 0.2297968 | 0.8182497 | 0.907841 |
| ENSG00000119682 | AREL1 | 5687.693778 | 0.093020131 | 0.109606071 | 0.849797655 | 0.39543759 | 0.597123 |
| ENSG00000237298 | TTN-AS1 | 107.1497 | 0.0929599 | 0.2754605 | 0.3618569 | 0.717459 | 0.846015 |
| ENSG00000183718 | TRIM52 | 1699.493 | 0.09291542 | 0.2900239 | 0.3482505 | 0.7276521 | 0.852049 |
| ENSG00000202358 | RNU6-652P | 9.93355 | 0.09282589 | 0.5518974 | 0.2288754 | 0.8189657 | 0.90837 |
| ENSG00000240303 | ACAD11 | 1.535639 | 0.0928119 | 0.7560967 | 0.3379479 | 0.7354024 | NA |
| ENSG00000239280 | | 2.837149 | 0.09281147 | 0.6695035 | 0.2580363 | 0.7963789 | 0.894975 |
| ENSG00000085224 | ATRX | 8338.302609 | 0.092805353 | 0.22831415 | 0.421517877 | 0.67337696 | 0.818528 |
| ENSG00000243023 | UBA52P3 | 3.826057 | 0.09274027 | 0.6252063 | 0.2356988 | 0.8136664 | 0.904948 |
| ENSG00000279827 | | 0.8979848 | 0.09268509 | 0.7587067 | 0.375142 | 0.7075548 | NA |
| ENSG00000235033 | DAAM2-AS1 | 1.38027 | 0.09267489 | 0.7397251 | 0.3164164 | 0.7516865 | NA |
| ENSG00000123080 | CDKN2C | 810.6149926 | 0.092654638 | 0.340882281 | 0.292387115 | 0.76999066 | 0.87835 |
| ENSG00000198185 | ZNF334 | 1.91054 | 0.09265294 | 0.7553166 | 0.324935 | 0.7452303 | 0.862897 |
| ENSG00000260577 | | 18.26672 | 0.09261238 | 0.5269144 | 0.2355859 | 0.813754 | 0.905008 |
| ENSG00000185641 | | 6867.807 | 0.09258945 | 0.1714553 | 0.5459246 | 0.5851178 | 0.75503 |
| ENSG00000066654 | THUMPD1 | 3399.319755 | 0.092556136 | 0.125898716 | 0.74286551 | 0.45756311 | 0.653496 |
| ENSG00000236054 | SYN3-AS1 | 1.050498 | 0.09247246 | 0.7506078 | 0.3063992 | 0.7593007 | NA |
| ENSG00000143469 | SYT14 | 0.8174332 | 0.09243938 | 0.7710376 | 0.4037937 | 0.6863645 | NA |
| ENSG00000007047 | MARK4 | 1890.649889 | 0.092267695 | 0.148617948 | 0.625737043 | 0.53148743 | 0.715585 |
| ENSG00000131116 | ZNF428 | 989.7743389 | 0.092170412 | 0.222395614 | 0.428975252 | 0.66794124 | 0.814996 |
| ENSG00000167393 | PPP2R3B | 1488.01 | 0.09216973 | 0.1360605 | 0.6866986 | 0.4922727 | 0.682416 |
| ENSG00000227999 | MTND5P1 | 16.36964 | 0.09201469 | 0.5553182 | 0.2286882 | 0.8191113 | 0.908456 |

| | | | | | | | |
|---|---|---|---|---|---|---|---|
| ENSG00000110841 | PPFIBP1 | 1108.307845 | 0.092002724 | 0.225541217 | 0.426868218 | 0.66947532 | 0.815897 |
| ENSG00000122068 | FYTTD1 | 5786.344235 | 0.091787497 | 0.149104625 | 0.628669031 | 0.52956576 | 0.714251 |
| ENSG00000267346 | EIF5AP3 | 0.8755232 | 0.09174817 | 0.746291 | 0.3412958 | 0.7328809 | NA |
| ENSG00000173918 | C1QTNF1 | 5.661384 | 0.09169628 | 0.6677345 | 0.2351742 | 0.8140735 | 0.905136 |
| ENSG00000222881 | | 7.653643 | 0.09165172 | 0.5661734 | 0.2272913 | 0.8201973 | 0.908902 |
| ENSG00000012232 | EXTL3 | 1458.913334 | 0.091615947 | 0.169209273 | 0.541768649 | 0.58797789 | 0.757298 |
| ENSG00000164841 | TMEM74 | 20.99495 | 0.09160793 | 0.5553243 | 0.2252262 | 0.8218033 | 0.909619 |
| ENSG00000122694 | GLIPR2 | 95.7024261 | 0.091565632 | 0.392583713 | 0.268051084 | 0.78866 | 0.890141 |
| ENSG00000074054 | CLASP1 | 3752.956503 | 0.091562191 | 0.172247927 | 0.538691352 | 0.59009984 | 0.758839 |
| ENSG00000130023 | ERMARD | 1884.526388 | 0.091543197 | 0.168210266 | 0.557005812 | 0.57752346 | 0.749376 |
| ENSG00000231133 | HAR1B | 18.7968 | 0.09139579 | 0.6855415 | 0.2874928 | 0.773735 | 0.880665 |
| ENSG00000162222 | TTC9C | 1776.158 | 0.0913799 | 0.1908196 | 0.4891566 | 0.6247308 | 0.783705 |
| ENSG00000212505 | RNA5SP299 | 1.346817 | 0.09124394 | 0.7918119 | 0.6144092 | 0.538945 | NA |
| ENSG00000170689 | HOXB9 | 46.43854 | 0.09108184 | 0.6930233 | 0.2969122 | 0.7665335 | 0.876184 |
| ENSG00000233783 | | 1.012649 | 0.09107532 | 0.7274412 | 0.3000261 | 0.7641572 | NA |
| ENSG00000239345 | HNRNPA1P26 | 63.54519 | 0.09095682 | 0.264412 | 0.3650921 | 0.7150427 | 0.84473 |
| ENSG00000274918 | | 5.211634 | 0.09095681 | 0.6274743 | 0.2322841 | 0.8163174 | 0.906455 |
| ENSG00000136002 | ARHGEF4 | 230.854587 | 0.090940456 | 0.345154533 | 0.298536935 | 0.76529339 | 0.875482 |
| ENSG00000237596 | | 4.60581 | 0.09093098 | 0.6445057 | 0.2273883 | 0.8201218 | 0.908902 |
| ENSG00000139641 | ESYT1 | 11831.25 | 0.09092746 | 0.1570707 | 0.5901578 | 0.5550849 | 0.732713 |
| ENSG00000101417 | PXMP4 | 1747.248892 | 0.090827172 | 0.2286178 | 0.416050577 | 0.67737299 | 0.820608 |
| ENSG00000033030 | ZCCHC8 | 2288.571712 | 0.09076996 | 0.167343971 | 0.560452058 | 0.57517113 | 0.747419 |
| ENSG00000136720 | HS6ST1 | 4310.715725 | 0.090722939 | 0.218299019 | 0.424864004 | 0.67093582 | 0.816961 |
| ENSG00000135845 | PIGC | 2088.447377 | 0.090669684 | 0.145949202 | 0.635510999 | 0.5250952 | 0.71006 |
| ENSG00000179148 | ALOXE3 | 68.28996 | 0.09065802 | 0.4123059 | 0.2504047 | 0.8022744 | 0.898597 |
| ENSG00000118976 | OTUD4P1 | 199.7242051 | 0.090649418 | 0.360587511 | 0.28352903 | 0.77677133 | 0.882518 |
| ENSG00000285646 | | 15.50534 | 0.09064409 | 0.4814822 | 0.2325417 | 0.8161173 | 0.906345 |
| ENSG00000287727 | | 2.053619 | 0.09063982 | 0.7072281 | 0.272823 | 0.7849892 | 0.887468 |
| ENSG00000235989 | MORC2-AS1 | 8.557811 | 0.09062025 | 0.522884 | 0.2270682 | 0.8203707 | 0.908942 |
| ENSG00000230865 | TSEN15P1 | 5.29573 | 0.09058768 | 0.6854367 | 0.2543402 | 0.7992328 | 0.896956 |
| ENSG00000272457 | | 1.903737 | 0.09052457 | 0.7465687 | 0.3504208 | 0.7260229 | 0.851229 |
| ENSG00000223855 | | 481.6311 | 0.09040582 | 0.4158972 | 0.2477567 | 0.8043226 | 0.899897 |
| ENSG00000248827 | | 8.490107 | 0.09038241 | 0.5709244 | 0.2289509 | 0.8189071 | 0.908343 |

| | | | | | | | |
|---|---|---|---|---|---|---|---|
| ENSG00000124733 | MEA1 | 3311.349465 | 0.090370348 | 0.216839735 | 0.427544786 | 0.66898257 | 0.81561 |
| ENSG00000029364 | SLC39A9 | 6055.491335 | 0.090308891 | 0.130309795 | 0.703704551 | 0.48161679 | 0.673037 |
| ENSG00000260583 | | 1.750022 | 0.09023151 | 0.7244094 | 0.3029303 | 0.761943 | 0.873154 |
| ENSG00000270333 | | 1.989939 | 0.09021402 | 0.7105905 | 0.2818743 | 0.7780399 | 0.883295 |
| ENSG00000261738 | MIR3976HG | 1.365972 | 0.09021087 | 0.7452778 | 0.3352852 | 0.7374099 | NA |
| ENSG00000171302 | CANT1 | 6708.748 | 0.09012962 | 0.1531728 | 0.6049684 | 0.5452 | 0.725751 |
| ENSG00000163904 | SENP2 | 3169.815 | 0.09010997 | 0.1169861 | 0.7755701 | 0.4380028 | 0.636544 |
| ENSG00000144233 | AMMECR1L | 3389.734 | 0.09003961 | 0.1625138 | 0.5837196 | 0.559409 | 0.736117 |
| ENSG00000006757 | PNPLA4 | 696.0027261 | 0.089971142 | 0.269951555 | 0.354498854 | 0.72296506 | 0.849406 |
| ENSG00000251389 | YTHDF1P1 | 0.9660051 | 0.08996982 | 0.7297862 | 0.3018723 | 0.7627494 | NA |
| ENSG00000279085 | | 0.8044445 | 0.08996172 | 0.7565261 | 0.2990945 | 0.7648679 | NA |
| ENSG00000169180 | XPO6 | 7542.835 | 0.08992696 | 0.1553198 | 0.5644522 | 0.5724465 | 0.745833 |
| ENSG00000178917 | ZNF852 | 129.3708 | 0.08986083 | 0.2625372 | 0.3533911 | 0.7237952 | 0.84993 |
| ENSG00000227331 | RPL7AP22 | 2.218225 | 0.08985365 | 0.6749223 | 0.2466933 | 0.8051456 | 0.900308 |
| ENSG00000240823 | RN7SL23P | 6.936991 | 0.0898532 | 0.5692626 | 0.2208035 | 0.8252454 | 0.911882 |
| ENSG00000136807 | CDK9 | 2853.509032 | 0.089818429 | 0.149066394 | 0.611817754 | 0.54065834 | 0.722609 |
| ENSG00000273372 | SFTPD-AS1 | 0.8692528 | 0.08968296 | 0.7495268 | 0.3447521 | 0.7302808 | NA |
| ENSG00000118990 | GLRXP3 | 4.427723958 | 0.089619154 | 0.663868366 | 0.250799746 | 0.80196894 | 0.898369 |
| ENSG00000069345 | DNAJA2 | 4305.335087 | 0.089600685 | 0.141704236 | 0.637857438 | 0.52356649 | 0.708822 |
| ENSG00000230756 | RHOQP3 | 126.2511 | 0.089588 | 0.3233449 | 0.3006688 | 0.7636671 | 0.874273 |
| ENSG00000248727 | LINC01948 | 125.1947 | 0.0894607 | 0.4775886 | 0.2117112 | 0.8323323 | 0.915608 |
| ENSG00000002549 | LAP3 | 3625.329195 | 0.089445275 | 0.225582994 | 0.402732605 | 0.68714495 | 0.827633 |
| ENSG00000204160 | ZDHHC18 | 2122.718 | 0.08942919 | 0.1844582 | 0.4981698 | 0.6183643 | 0.779269 |
| ENSG00000260729 | | 11.94767 | 0.08942755 | 0.6143588 | 0.2166943 | 0.8284466 | 0.913464 |
| ENSG00000233613 | DCUN1D2-AS | 5.299617 | 0.08940255 | 0.538929 | 0.2236577 | 0.8230236 | 0.910211 |
| ENSG00000140992 | PDPK1 | 3871.804 | 0.08936091 | 0.2142003 | 0.4197529 | 0.674666 | 0.818951 |
| ENSG00000147140 | NONO | 27990.44 | 0.08935512 | 0.0925432 | 0.9687134 | 0.3326882 | 0.533703 |
| ENSG00000227449 | FGF7P6 | 1.273008 | 0.08930915 | 0.6971274 | 0.2599372 | 0.7949123 | NA |
| ENSG00000260238 | PMF1-BGLAP | 0.7795627 | 0.08927076 | 0.7278697 | 0.2918299 | 0.7704167 | NA |
| ENSG00000267620 | | 5.778735 | 0.08919693 | 0.5925597 | 0.2196815 | 0.8261192 | 0.912152 |
| ENSG00000037757 | MRI1 | 1148.692254 | 0.089178363 | 0.189479493 | 0.488565635 | 0.62514925 | 0.783989 |
| ENSG00000235315 | RPL23AP69 | 1.432438 | 0.08914198 | 0.7233188 | 0.3001507 | 0.7640622 | NA |
| ENSG00000251733 | SCARNA8 | 4.149959 | 0.08910964 | 0.6535081 | 0.2438496 | 0.8073473 | 0.901316 |

| | | | | | | | |
|---|---|---|---|---|---|---|---|
| ENSG00000231826 | LINC01819 | 1.251225 | 0.08909165 | 0.743257 | 0.3678355 | 0.7129959 | NA |
| ENSG00000278993 | | 34.18853 | 0.08905106 | 0.3465952 | 0.2845317 | 0.7760029 | 0.882 |
| ENSG00000079332 | SAR1A | 6155.166924 | 0.089041652 | 0.128111791 | 0.706361223 | 0.47996353 | 0.671801 |
| ENSG00000270574 | | 3.711867 | 0.08903583 | 0.581632 | 0.2097692 | 0.8338479 | 0.91665 |
| ENSG00000186715 | MST1L | 67.54527 | 0.08898576 | 0.3365274 | 0.2881789 | 0.7732098 | 0.880445 |
| ENSG00000169221 | TBC1D10B | 4490.7 | 0.08897906 | 0.1763963 | 0.5190836 | 0.6037025 | 0.768745 |
| ENSG00000267680 | ZNF224 | 840.3268 | 0.08891455 | 0.2515398 | 0.3678667 | 0.7129726 | 0.843639 |
| ENSG00000119414 | PPP6C | 4336.494631 | 0.088855272 | 0.107199876 | 0.829505624 | 0.40681836 | 0.607438 |
| ENSG00000269656 | NOP53-AS1 | 12.59783 | 0.08881047 | 0.5089139 | 0.2222275 | 0.8241368 | 0.911025 |
| ENSG00000235060 | VDAC1P4 | 1.392691 | 0.08879576 | 0.700663 | 0.2469055 | 0.8049814 | NA |
| ENSG00000135454 | B4GALNT1 | 935.325613 | 0.0887414 | 0.390765048 | 0.249329832 | 0.80310566 | 0.899149 |
| ENSG00000213051 | RPL5P5 | 46.99244 | 0.08873064 | 0.3198442 | 0.2957878 | 0.7673921 | 0.876603 |
| ENSG00000197575 | RPS17P2 | 1.188616 | 0.08869079 | 0.7508376 | 0.3339047 | 0.7384515 | NA |
| ENSG00000249148 | | 0.8197506 | 0.08865107 | 0.7673774 | 0.3468924 | 0.7286721 | NA |
| ENSG00000163428 | LRRC58 | 6367.706 | 0.08864597 | 0.2224023 | 0.4175339 | 0.6762879 | 0.820059 |
| ENSG00000185787 | MORF4L1 | 7586.404 | 0.08862879 | 0.2158717 | 0.4237156 | 0.6717733 | 0.817546 |
| ENSG00000264829 | | 1.278082 | 0.08862839 | 0.7290513 | 0.2979107 | 0.7657713 | NA |
| ENSG00000154114 | TBCEL | 730.5774 | 0.08860973 | 0.2113998 | 0.427447 | 0.6690538 | 0.81561 |
| ENSG00000251322 | SHANK3 | 415.9836 | 0.0885351 | 0.2714034 | 0.3337578 | 0.7385623 | 0.858824 |
| ENSG00000143183 | TMCO1 | 5432.968 | 0.08853063 | 0.1424617 | 0.5833434 | 0.5596621 | 0.73613 |
| ENSG00000107186 | MPDZ | 1567.269824 | 0.088491539 | 0.239641056 | 0.382627763 | 0.70199578 | 0.837622 |
| ENSG00000271749 | | 20.31042 | 0.08821956 | 0.5443228 | 0.2193726 | 0.8263598 | 0.912268 |
| ENSG00000196275 | GTF2IRD2 | 199.9216 | 0.0879782 | 0.3840963 | 0.2661138 | 0.7901515 | 0.891256 |
| ENSG00000146122 | DAAM2 | 15.22717 | 0.08781066 | 0.4963717 | 0.2242999 | 0.8225239 | 0.909824 |
| ENSG00000236104 | ZBTB22 | 1918.984 | 0.08778101 | 0.1540208 | 0.5848309 | 0.5586614 | 0.735584 |
| ENSG00000255931 | | 4.694459 | 0.08775052 | 0.6250615 | 0.213367 | 0.8310407 | 0.915116 |
| ENSG00000097007 | ABL1 | 5064.177607 | 0.087713821 | 0.156396146 | 0.565234833 | 0.57191405 | 0.745483 |
| ENSG00000130612 | CYP2G1P | 4.108344063 | 0.087704882 | 0.612745218 | 0.218794467 | 0.82681016 | 0.912571 |
| ENSG00000033178 | UBA6 | 3436.657336 | 0.087699467 | 0.154633291 | 0.565328528 | 0.57185033 | 0.745437 |
| ENSG00000273611 | ZNHIT3 | 1756.651 | 0.08768445 | 0.1870249 | 0.4815406 | 0.6301323 | 0.787755 |
| ENSG00000225093 | RPL3P7 | 415.5728 | 0.08767052 | 0.2158382 | 0.4097247 | 0.6820079 | 0.82392 |
| ENSG00000187583 | PLEKHN1 | 727.1112 | 0.08764942 | 0.2028465 | 0.4364911 | 0.6624804 | 0.810865 |
| ENSG00000235499 | | 147.9062 | 0.08763847 | 0.4684706 | 0.2270209 | 0.8204075 | 0.908942 |

| | | | | | | | |
|---|---|---|---|---|---|---|---|
| ENSG00000177971 | IMP3 | 1896.086 | 0.08758791 | 0.2068947 | 0.4274884 | 0.6690236 | 0.81561 |
| ENSG00000279151 | | 1.901064 | 0.0875592 | 0.6520351 | 0.2330714 | 0.815706 | 0.906139 |
| ENSG00000259644 | | 3.037916 | 0.08753785 | 0.6599048 | 0.2385407 | 0.8114618 | 0.903596 |
| ENSG00000261737 | | 10.65979 | 0.08745492 | 0.5339289 | 0.216171 | 0.8288544 | 0.913655 |
| ENSG00000230448 | LINC00276 | 5.88769 | 0.0874059 | 0.6087792 | 0.2205434 | 0.825448 | 0.911906 |
| ENSG00000107821 | KAZALD1 | 1462.561756 | 0.087392084 | 0.275114426 | 0.337589454 | 0.73567259 | 0.856796 |
| ENSG00000251538 | LINC02201 | 0.7760221 | 0.08733644 | 0.7644878 | 0.2827542 | 0.7773653 | NA |
| ENSG00000112796 | ENPP5 | 755.7715438 | 0.08732456 | 0.245236288 | 0.372265575 | 0.70969513 | 0.842119 |
| ENSG00000259456 | ADNP-AS1 | 117.2089 | 0.08730175 | 0.2687413 | 0.3442813 | 0.7306347 | 0.853923 |
| ENSG00000112787 | FBRSL1 | 4378.223584 | 0.087292888 | 0.156027116 | 0.548677572 | 0.58322674 | 0.753515 |
| ENSG00000215483 | LINC00598 | 11.35039 | 0.08718062 | 0.490175 | 0.2239097 | 0.8228276 | 0.910032 |
| ENSG00000225579 | EDNRB-AS1 | 7.434442 | 0.08712384 | 0.5473123 | 0.2053355 | 0.83731 | 0.918791 |
| ENSG00000119321 | FKBP15 | 3451.840404 | 0.087092488 | 0.098374007 | 0.880903997 | 0.37836978 | 0.580489 |
| ENSG00000021461 | CYP3A43 | 13.55767233 | 0.087081355 | 0.648922181 | 0.216464575 | 0.82862564 | 0.913555 |
| ENSG00000273076 | | 3.585259 | 0.08707244 | 0.6393198 | 0.2291153 | 0.8187793 | 0.908239 |
| ENSG00000265496 | | 4.662026 | 0.08706705 | 0.5648188 | 0.2150903 | 0.8296969 | 0.914205 |
| ENSG00000226647 | | 9.37107 | 0.08706456 | 0.6121389 | 0.2207109 | 0.8253175 | 0.911882 |
| ENSG00000235284 | SNORD62A | 3.28876 | 0.08705371 | 0.6431144 | 0.2303285 | 0.8178365 | 0.907648 |
| ENSG00000092931 | MFSD11 | 1752.896639 | 0.087051749 | 0.103832964 | 0.834591747 | 0.40394761 | 0.604715 |
| ENSG00000228663 | PSMD10P1 | 4.454659 | 0.08696365 | 0.6420429 | 0.2275894 | 0.8199654 | 0.908881 |
| ENSG00000167113 | COQ4 | 3256.649 | 0.08691521 | 0.140456 | 0.6349418 | 0.5254664 | 0.710345 |
| ENSG00000259848 | | 1.853101 | 0.08690093 | 0.7027899 | 0.2618873 | 0.7934084 | 0.89312 |
| ENSG00000229455 | RPS10P18 | 1.270039 | 0.08680887 | 0.7210147 | 0.2743308 | 0.7838304 | NA |
| ENSG00000204104 | TRAF3IP1 | 987.5092 | 0.08680798 | 0.1424871 | 0.6221955 | 0.5338133 | 0.717467 |
| ENSG00000107863 | ARHGAP21 | 4175.157632 | 0.086664531 | 0.214482345 | 0.42316201 | 0.67217706 | 0.817761 |
| ENSG00000268297 | CLEC4GP1 | 1.59052 | 0.08659751 | 0.7469797 | 0.330305 | 0.7411695 | 0.860407 |
| ENSG00000242640 | RPS29P11 | 4.375255 | 0.08647956 | 0.6572549 | 0.233306 | 0.8155238 | 0.90611 |
| ENSG00000124209 | RAB22A | 7513.600129 | 0.086460775 | 0.101352567 | 0.867966676 | 0.38541258 | 0.587798 |
| ENSG00000165702 | GFI1B | 2.983492 | 0.08645426 | 0.6573035 | 0.233919 | 0.8150479 | 0.90584 |
| ENSG00000242242 | NECTIN3-AS1 | 129.7699 | 0.08643751 | 0.2988122 | 0.3092642 | 0.7571205 | 0.870368 |
| ENSG00000232480 | TGFB2-AS1 | 47.62635 | 0.0863848 | 0.5374549 | 0.2211074 | 0.8250088 | 0.9118 |
| ENSG00000276317 | | 15.74571 | 0.08636849 | 0.4710453 | 0.2279055 | 0.8197197 | 0.908827 |
| ENSG00000233924 | RPSAP13 | 2.248044 | 0.08622124 | 0.6716233 | 0.2369539 | 0.8126926 | 0.904263 |

| | | | | | | | |
|---|---|---|---|---|---|---|---|
| ENSG00000278099 | RNVU1-2A | 0.8896435 | 0.08615092 | 0.7357449 | 0.284871 | 0.775743 | NA |
| ENSG00000214725 | CDIPTOSP | 2.50094 | 0.0861057 | 0.7002739 | 0.2665168 | 0.7898412 | 0.890982 |
| ENSG00000278311 | GGNBP2 | 3437.111 | 0.08608156 | 0.08840626 | 0.9519216 | 0.3411368 | 0.542447 |
| ENSG00000275515 | | 3.593257 | 0.08605279 | 0.6024693 | 0.2125356 | 0.8316892 | 0.915261 |
| ENSG00000236111 | | 5.071102 | 0.08601301 | 0.6294119 | 0.2244725 | 0.8223897 | 0.909824 |
| ENSG00000270562 | | 47.59308 | 0.0860093 | 0.4363652 | 0.2327858 | 0.8159277 | 0.906211 |
| ENSG00000200350 | RNU6-1285P | 1.653848 | 0.08595381 | 0.7128795 | 0.2792045 | 0.7800879 | 0.884712 |
| ENSG00000286905 | | 9.533429 | 0.08591535 | 0.5102646 | 0.2114668 | 0.832523 | 0.915659 |
| ENSG00000259612 | EEF1A1P22 | 27.6845 | 0.08589398 | 0.4453939 | 0.2306165 | 0.8176128 | 0.907552 |
| ENSG00000280069 | | 2.956209 | 0.0858452 | 0.6562664 | 0.2300204 | 0.8180759 | 0.907686 |
| ENSG00000240036 | | 175.9471 | 0.08582307 | 0.2108033 | 0.4216025 | 0.6733152 | 0.818528 |
| ENSG00000253633 | | 1.481436 | 0.08579882 | 0.7472543 | 0.3129462 | 0.7543215 | NA |
| ENSG00000272886 | DCP1A | 1332.745 | 0.08574963 | 0.2193379 | 0.4009942 | 0.6884244 | 0.828423 |
| ENSG00000237409 | | 1.326033 | 0.08574701 | 0.7235138 | 0.2724449 | 0.7852799 | NA |
| ENSG00000234283 | LINC01731 | 1.376812 | 0.08574695 | 0.7529304 | 0.3276744 | 0.7431579 | NA |
| ENSG00000105698 | USF2 | 5776.449625 | 0.085745339 | 0.167119308 | 0.523099594 | 0.60090495 | 0.766765 |
| ENSG00000287816 | | 1.649814 | 0.08574077 | 0.6879336 | 0.2448347 | 0.8065844 | 0.90077 |
| ENSG00000271886 | MIR98 | 0.8272171 | 0.08565994 | 0.7488616 | 0.3277817 | 0.7430768 | NA |
| ENSG00000237307 | SRRM1P3 | 4.76479 | 0.08559183 | 0.6228244 | 0.2203734 | 0.8255803 | 0.91192 |
| ENSG00000099246 | RAB18 | 5053.829136 | 0.085570919 | 0.101841855 | 0.834780022 | 0.40384158 | 0.604652 |
| ENSG00000257052 | | 9.167056 | 0.08556214 | 0.5657382 | 0.213072 | 0.8312708 | 0.915137 |
| ENSG00000178502 | KLHL11 | 338.0394 | 0.08554478 | 0.3612431 | 0.2487155 | 0.8035809 | 0.899416 |
| ENSG00000236708 | SDK1-AS1 | 3.924525 | 0.08552929 | 0.6210005 | 0.2145748 | 0.8300988 | 0.914572 |
| ENSG00000219626 | FAM228B | 121.1437 | 0.08551803 | 0.3668321 | 0.2572946 | 0.7969513 | 0.895202 |
| ENSG00000131732 | ZCCHC9 | 1352.084487 | 0.085517704 | 0.147448474 | 0.588158762 | 0.55642573 | 0.733827 |
| ENSG00000272834 | | 2.670243 | 0.08549547 | 0.6813124 | 0.248344 | 0.8038683 | 0.899624 |
| ENSG00000256762 | STH | 1.904403 | 0.08546869 | 0.708989 | 0.263051 | 0.7925113 | 0.892667 |
| ENSG00000237984 | PTENP1 | 760.8162 | 0.08545805 | 0.1606434 | 0.533652 | 0.5935823 | 0.761364 |
| ENSG00000120533 | ENY2 | 5071.693491 | 0.085451903 | 0.190560354 | 0.467310944 | 0.64027743 | 0.794709 |
| ENSG00000127554 | GFER | 811.4001191 | 0.085421835 | 0.158087032 | 0.546801138 | 0.58451536 | 0.754506 |
| ENSG00000170385 | SLC30A1 | 3829.399 | 0.08537028 | 0.3296177 | 0.2869865 | 0.7741227 | 0.880805 |
| ENSG00000109466 | KLHL2 | 779.341483 | 0.085308916 | 0.184612881 | 0.468356018 | 0.63953002 | 0.794189 |
| ENSG00000133055 | MYBPH | 1.638064814 | 0.085239627 | 0.702787068 | 0.244830465 | 0.8065877 | 0.90077 |

| | | | | | | | |
|---|---|---|---|---|---|---|---|
| ENSG00000230912 | | 10.96076 | 0.08523639 | 0.519016 | 0.2137512 | 0.8307411 | 0.914912 |
| ENSG00000205037 | | 2.595657 | 0.08522398 | 0.6398138 | 0.219596 | 0.8261858 | 0.912152 |
| ENSG00000229738 | | 1.337696 | 0.08498579 | 0.7541682 | 0.2812744 | 0.7785 | NA |
| ENSG00000254635 | WAC-AS1 | 2354.222 | 0.08498441 | 0.1483042 | 0.5829394 | 0.5599341 | 0.736239 |
| ENSG00000114786 | ABHD14A-ACY1 | 0.865017378 | 0.084965473 | 0.760502385 | 0.292573561 | 0.76984813 | NA |
| ENSG00000185339 | TCN2 | 15.58993 | 0.08495247 | 0.4111615 | 0.2245244 | 0.8223493 | 0.909824 |
| ENSG00000266521 | | 8.063156 | 0.0848825 | 0.6389675 | 0.2262446 | 0.8210112 | 0.909205 |
| ENSG00000234608 | MAPKAPK5-AS1 | 1060.447 | 0.08480102 | 0.2258229 | 0.3877787 | 0.6981798 | 0.83547 |
| ENSG00000272274 | LINC00551 | 4.642959 | 0.08476706 | 0.6388586 | 0.2234748 | 0.823166 | 0.910331 |
| ENSG00000142669 | SH3BGRL3 | 2029.479 | 0.0847294 | 0.2930115 | 0.3114558 | 0.7554541 | 0.86943 |
| ENSG00000225770 | RPS29P3 | 171.872 | 0.08470388 | 0.2497343 | 0.3519511 | 0.7248749 | 0.850636 |
| ENSG00000237757 | EEF1A1P30 | 2.033378 | 0.08467845 | 0.6975682 | 0.2541349 | 0.7993913 | 0.896976 |
| ENSG00000166192 | SENP8 | 259.3962 | 0.08465634 | 0.1863764 | 0.4530571 | 0.6505076 | 0.802546 |
| ENSG00000102910 | LONP2 | 4106.934911 | 0.084589371 | 0.148586458 | 0.577716477 | 0.56345555 | 0.739007 |
| ENSG00000143614 | GATAD2B | 2933.675 | 0.08449644 | 0.1501134 | 0.5655497 | 0.5716999 | 0.745314 |
| ENSG00000235478 | LINC01664 | 2.243975 | 0.08427012 | 0.6841168 | 0.2402126 | 0.8101654 | 0.902923 |
| ENSG00000259403 | | 2.375419 | 0.08426807 | 0.6559445 | 0.2049735 | 0.8375928 | 0.918934 |
| ENSG00000108510 | MED13 | 19788.09004 | 0.084190542 | 0.250144155 | 0.350642846 | 0.72585631 | 0.851071 |
| ENSG00000099992 | TBC1D10A | 783.4241463 | 0.084172829 | 0.166257621 | 0.511895148 | 0.60872439 | 0.772377 |
| ENSG00000287352 | | 83.35326 | 0.08410079 | 0.2695309 | 0.3285865 | 0.7424683 | 0.861313 |
| ENSG00000272333 | KMT2B | 5515.462 | 0.08407266 | 0.156692 | 0.5328323 | 0.5941497 | 0.761696 |
| ENSG00000123992 | DNPEP | 7405.025097 | 0.083988863 | 0.14525471 | 0.554832076 | 0.57900953 | 0.750278 |
| ENSG00000251535 | | 1.78193 | 0.08396563 | 0.7165332 | 0.2660562 | 0.7901959 | 0.891266 |
| ENSG00000137497 | NUMA1 | 17044.08889 | 0.083944234 | 0.147255948 | 0.583665041 | 0.55944569 | 0.736129 |
| ENSG00000267656 | | 5.637636 | 0.08393908 | 0.6357771 | 0.2176327 | 0.8277153 | 0.913119 |
| ENSG00000275964 | | 423.7105 | 0.08391268 | 0.2711345 | 0.3314786 | 0.740283 | 0.859716 |
| ENSG00000169093 | ASMTL | 1627.906 | 0.08390546 | 0.2400991 | 0.3634872 | 0.716241 | 0.845492 |
| ENSG00000155008 | APOOL | 1770.18 | 0.0838372 | 0.2082707 | 0.4204771 | 0.674137 | 0.818795 |
| ENSG00000243708 | PLA2G4B | 15.10906 | 0.08382319 | 0.5006183 | 0.2091249 | 0.8343508 | 0.916937 |
| ENSG00000206168 | | 17.59939 | 0.08365674 | 0.3763681 | 0.2520143 | 0.80103 | 0.897885 |
| ENSG00000285747 | | 1.181596 | 0.08364034 | 0.7336349 | 0.2867978 | 0.7742672 | NA |
| ENSG00000123454 | DBH | 0.968567992 | 0.083623491 | 0.752682679 | 0.28258805 | 0.77749264 | NA |
| ENSG00000124091 | GCNT7 | 18.75085096 | 0.083557357 | 0.401210032 | 0.240741933 | 0.80975514 | 0.902824 |

| | | | | | | | |
|---|---|---|---|---|---|---|---|
| ENSG00000112079 | STK38 | 5686.183646 | 0.083554536 | 0.172663328 | 0.494452548 | 0.62098661 | 0.781019 |
| ENSG00000105402 | NAPA | 3003.604184 | 0.083498835 | 0.162443015 | 0.533855689 | 0.59344138 | 0.761349 |
| ENSG00000165121 | | 262.7861 | 0.08339449 | 0.3059599 | 0.2914841 | 0.7706811 | 0.878733 |
| ENSG00000232909 | | 25.47467 | 0.08338759 | 0.4468652 | 0.2172631 | 0.8280033 | 0.913361 |
| ENSG00000254270 | ERHP1 | 16.61888 | 0.08332275 | 0.5533508 | 0.2053266 | 0.837317 | 0.918791 |
| ENSG00000260455 | NBAT1 | 1.330572 | 0.08327935 | 0.7597162 | 0.2950076 | 0.7679881 | NA |
| ENSG00000115084 | SLC35F5 | 4740.097064 | 0.083204259 | 0.197353615 | 0.456204011 | 0.64824328 | 0.800631 |
| ENSG00000125355 | TMEM255A | 1.630980884 | 0.083166187 | 0.718892331 | 0.25868176 | 0.79588079 | 0.894642 |
| ENSG00000164031 | DNAJB14 | 2150.516 | 0.0830573 | 0.2125156 | 0.4022057 | 0.6875326 | 0.827935 |
| ENSG00000090060 | PAPOLA | 19882.49203 | 0.083003761 | 0.095027473 | 0.86501815 | 0.38702883 | 0.589349 |
| ENSG00000285850 | | 1.221292 | 0.0829454 | 0.7509682 | 0.3299659 | 0.7414257 | NA |
| ENSG00000204310 | AGPAT1 | 1125.736 | 0.08290993 | 0.1258087 | 0.6667752 | 0.5049157 | 0.69175 |
| ENSG00000260060 | | 5.546724 | 0.08290199 | 0.6038991 | 0.2119421 | 0.8321522 | 0.915581 |
| ENSG00000232139 | LINC00867 | 4.365607 | 0.08290062 | 0.6435936 | 0.2081357 | 0.835123 | 0.917253 |
| ENSG00000065243 | PKN2 | 2796.868324 | 0.082846761 | 0.195325349 | 0.444700969 | 0.65653587 | 0.806416 |
| ENSG00000277972 | CISD3 | 1225.085 | 0.08283469 | 0.1911391 | 0.4448961 | 0.6563948 | 0.806399 |
| ENSG00000230039 | CCNYL5 | 24.59992 | 0.08282309 | 0.4300454 | 0.2331904 | 0.8156135 | 0.90611 |
| ENSG00000230863 | | 11.25583 | 0.08282228 | 0.5720728 | 0.2055071 | 0.837176 | 0.918712 |
| ENSG00000276926 | MIR6797 | 4.84974 | 0.08281323 | 0.5807693 | 0.2057506 | 0.8369857 | 0.918602 |
| ENSG00000139445 | FOXN4 | 5.618621 | 0.08281278 | 0.6988391 | 0.2645272 | 0.7913737 | 0.89184 |
| ENSG00000143753 | DEGS1 | 1941.072 | 0.08278861 | 0.1526789 | 0.5133053 | 0.6077377 | 0.771549 |
| ENSG00000127415 | IDUA | 1477.694573 | 0.082649342 | 0.372034224 | 0.246147528 | 0.80556803 | 0.90031 |
| ENSG00000134575 | ACP2 | 1683.16488 | 0.082648231 | 0.212162024 | 0.402929273 | 0.68700026 | 0.827607 |
| ENSG00000265158 | LRRC37A7P | 6.602068 | 0.08264524 | 0.588811 | 0.1922404 | 0.8475539 | 0.925823 |
| ENSG00000182533 | CAV3 | 1.249626 | 0.08263091 | 0.7156241 | 0.2590875 | 0.7955677 | NA |
| ENSG00000275111 | ZNF2 | 497.7837 | 0.08258427 | 0.1569197 | 0.5344223 | 0.5930494 | 0.761048 |
| ENSG00000186047 | DLEU7 | 2.099836 | 0.08251342 | 0.6867592 | 0.2339693 | 0.8150088 | 0.905834 |
| ENSG00000287456 | | 1.48796 | 0.08245014 | 0.7143231 | 0.269264 | 0.7877266 | NA |
| ENSG00000243144 | | 1.99075 | 0.0824481 | 0.7911828 | 0.380835 | 0.7033257 | 0.838592 |
| ENSG00000223518 | CSNK1A1P1 | 2.409622 | 0.0822119 | 0.6679941 | 0.220138 | 0.8257637 | 0.91192 |
| ENSG00000266445 | NARF-AS1 | 6.20359 | 0.08218831 | 0.5766785 | 0.2030697 | 0.8390805 | 0.920042 |
| ENSG00000143303 | RRNAD1 | 2092.325 | 0.08218183 | 0.1729894 | 0.4815123 | 0.6301525 | 0.787755 |
| ENSG00000127481 | UBR4 | 12010.5236 | 0.082092707 | 0.215266293 | 0.393307578 | 0.69409232 | 0.832232 |

| | | | | | | | |
|---|---|---|---|---|---|---|---|
| ENSG00000286437 | | 5.843891 | 0.08206502 | 0.5640105 | 0.2036348 | 0.8386389 | 0.919709 |
| ENSG00000256351 | | 1.164038 | 0.08199664 | 0.723505 | 0.2602122 | 0.7947001 | NA |
| ENSG00000064999 | ANKS1A | 3401.505525 | 0.08198268 | 0.15403571 | 0.508268958 | 0.61126474 | 0.774461 |
| ENSG00000232442 | MHENCR | 782.5339 | 0.08197433 | 0.2263235 | 0.3672506 | 0.7134321 | 0.84368 |
| ENSG00000092929 | UNC13D | 4408.048008 | 0.081949671 | 0.263635724 | 0.290396007 | 0.7715133 | 0.879191 |
| ENSG00000224232 | KCTD5P1 | 1.571627 | 0.08187327 | 0.700885 | 0.2453503 | 0.8061852 | 0.90051 |
| ENSG00000260658 | | 13.81732 | 0.08184833 | 0.5045802 | 0.2092604 | 0.834245 | 0.916896 |
| ENSG00000100567 | PSMA3 | 8531.392783 | 0.081773782 | 0.232405412 | 0.364866525 | 0.71521104 | 0.844853 |
| ENSG00000241741 | RPL7AP30 | 1470.89 | 0.08177275 | 0.1651807 | 0.4939859 | 0.6213161 | 0.781173 |
| ENSG00000258555 | SPECC1L-ADORA | 2.481554 | 0.08176662 | 0.7211479 | 0.2594155 | 0.7953147 | 0.894202 |
| ENSG00000180284 | CNEP1R1P1 | 1.194714 | 0.08170057 | 0.7064604 | 0.2458902 | 0.8057672 | NA |
| ENSG00000260742 | | 10.59688 | 0.08169085 | 0.5005658 | 0.2078183 | 0.8353708 | 0.917343 |
| ENSG00000283122 | HYMAI | 0.9573814 | 0.08168887 | 0.7573679 | 0.3334047 | 0.7388288 | NA |
| ENSG00000228839 | PIK3IP1-DT | 23.66703 | 0.0815559 | 0.3778553 | 0.2385373 | 0.8114644 | 0.903596 |
| ENSG00000172183 | ISG20 | 320.8338 | 0.08154145 | 0.3553099 | 0.2597688 | 0.7950421 | 0.894115 |
| ENSG00000154473 | BUB3 | 8350.413 | 0.08153228 | 0.138789 | 0.5733242 | 0.5664252 | 0.741055 |
| ENSG00000141380 | SS18 | 5892.845 | 0.08149842 | 0.1087554 | 0.7487528 | 0.4540062 | 0.650094 |
| ENSG00000099364 | FBXL19 | 3336.555283 | 0.081475984 | 0.188544682 | 0.441622183 | 0.65876263 | 0.808062 |
| ENSG00000277619 | | 2.16466 | 0.0814708 | 0.72232 | 0.2618756 | 0.7934173 | 0.89312 |
| ENSG00000258586 | LINC02274 | 1.6229 | 0.08142281 | 0.7404061 | 0.2965221 | 0.7668314 | 0.876374 |
| ENSG00000123473 | STIL | 1988.211251 | 0.081419004 | 0.363854508 | 0.239848265 | 0.81044789 | 0.903034 |
| ENSG00000213522 | RAC1P5 | 1.440788 | 0.08138475 | 0.7247948 | 0.2741879 | 0.7839402 | NA |
| ENSG00000171310 | CHST11 | 981.1197 | 0.08137831 | 0.2014762 | 0.4111665 | 0.6809505 | 0.823382 |
| ENSG00000196600 | SLC22A25 | 0.7849822 | 0.0813604 | 0.7612838 | 0.3256394 | 0.7446972 | NA |
| ENSG00000260032 | NORAD | 13258.97 | 0.08135504 | 0.1856472 | 0.4376528 | 0.661638 | 0.810132 |
| ENSG00000267457 | | 0.7663179 | 0.08134174 | 0.762638 | 0.3264259 | 0.7441021 | NA |
| ENSG00000229704 | EIF2S2P2 | 2.936531 | 0.08132035 | 0.6592976 | 0.2187745 | 0.8268257 | 0.912571 |
| ENSG00000163682 | RPL9 | 1704.029 | 0.08123758 | 0.1691435 | 0.4944511 | 0.6209876 | 0.781019 |
| ENSG00000234618 | RPSAP9 | 39.81038 | 0.08119893 | 0.3716457 | 0.2456641 | 0.8059423 | 0.90047 |
| ENSG00000127054 | INTS11 | 5098.782674 | 0.08112301 | 0.10157829 | 0.798636609 | 0.42450115 | 0.624343 |
| ENSG00000213588 | ZBTB9 | 1405.167 | 0.0810937 | 0.1260208 | 0.6435322 | 0.5198788 | 0.705123 |
| ENSG00000249631 | | 1.242071 | 0.08107447 | 0.7550961 | 0.3002158 | 0.7640125 | NA |
| ENSG00000124164 | VAPB | 11719.21513 | 0.081004605 | 0.138429682 | 0.581872387 | 0.56065264 | 0.736914 |

| | | | | | | | |
|---|---|---|---|---|---|---|---|
| ENSG00000215003 | RPL15P20 | 0.8888088 | 0.0809977 | 0.7526589 | 0.328192 | 0.7427665 | NA |
| ENSG00000279936 | | 0.8694552 | 0.08096365 | 0.735915 | 0.2681763 | 0.7885636 | NA |
| ENSG00000261840 | | 73.09617 | 0.08094115 | 0.2348663 | 0.3566338 | 0.721366 | 0.848466 |
| ENSG00000198001 | IRAK4 | 1259.653 | 0.08093112 | 0.1526072 | 0.5447932 | 0.5858957 | 0.755627 |
| ENSG00000287022 | | 1.149283 | 0.08089128 | 0.7495251 | 0.278239 | 0.7808289 | NA |
| ENSG00000236735 | RPL31P63 | 10.97925 | 0.08087769 | 0.5275056 | 0.2048941 | 0.8376549 | 0.918934 |
| ENSG00000273513 | TBC1D3K | 0.8124491 | 0.08086965 | 0.7540049 | 0.3288172 | 0.7422939 | NA |
| ENSG00000207721 | MIR186 | 27.44136 | 0.08075277 | 0.4000552 | 0.2325076 | 0.8161438 | 0.906345 |
| ENSG00000176915 | ANKLE2 | 6718.433 | 0.08073184 | 0.1294869 | 0.6298936 | 0.5287642 | 0.713569 |
| ENSG00000135720 | DYNC1LI2 | 7172.776757 | 0.080713664 | 0.170911975 | 0.471207658 | 0.63749245 | 0.792993 |
| ENSG00000240376 | | 1209.954 | 0.08060714 | 0.2356286 | 0.3582794 | 0.7201342 | 0.847692 |
| ENSG00000231466 | | 26.07006 | 0.08056347 | 0.383226 | 0.2372308 | 0.8124777 | 0.904236 |
| ENSG00000279135 | | 2.812626 | 0.0804759 | 0.6470219 | 0.2078115 | 0.8353761 | 0.917343 |
| ENSG00000140382 | HMG20A | 1731.287 | 0.08047046 | 0.1458859 | 0.5642099 | 0.5726113 | 0.745897 |
| ENSG00000144476 | ACKR3 | 2524.269 | 0.08040703 | 0.4138087 | 0.2270679 | 0.820371 | 0.908942 |
| ENSG00000151233 | GXYLT1 | 2119.036 | 0.08022764 | 0.187666 | 0.4326903 | 0.6652397 | 0.813001 |
| ENSG00000271259 | | 11.49827 | 0.08020163 | 0.5127517 | 0.2022776 | 0.8396997 | 0.920645 |
| ENSG00000104808 | DHDH | 21.16945755 | 0.080082271 | 0.494277211 | 0.204930617 | 0.83762633 | 0.918934 |
| ENSG00000185591 | SP1 | 9227.434 | 0.08006541 | 0.145507 | 0.557732 | 0.5770274 | 0.748879 |
| ENSG00000286046 | | 46.7959 | 0.08005907 | 0.3642684 | 0.2428863 | 0.8080935 | 0.90177 |
| ENSG00000176182 | MYPOP | 425.7463 | 0.08003079 | 0.2589722 | 0.3246404 | 0.7454532 | 0.863004 |
| ENSG00000112695 | COX7A2 | 5420.052797 | 0.079892905 | 0.190019233 | 0.438561866 | 0.66097903 | 0.809437 |
| ENSG00000259416 | LINC02883 | 2.973386 | 0.07986521 | 0.5935458 | 0.2012628 | 0.8404931 | 0.921114 |
| ENSG00000162520 | SYNC | 10.59736 | 0.07985009 | 0.5013241 | 0.2033812 | 0.8388371 | 0.919851 |
| ENSG00000272047 | GTF2H5 | 877.6094 | 0.07982622 | 0.2218955 | 0.3679569 | 0.7129054 | 0.843639 |
| ENSG00000172009 | THOP1 | 3431.172 | 0.07967518 | 0.2451356 | 0.3340623 | 0.7383325 | 0.858765 |
| ENSG00000287205 | | 13.2383 | 0.07961932 | 0.5146992 | 0.2065109 | 0.8363918 | 0.918117 |
| ENSG00000006638 | TBXA2R | 2.057738336 | 0.079601023 | 0.742419187 | 0.330902841 | 0.74071788 | 0.860071 |
| ENSG00000186908 | ZDHHC17 | 2532.889 | 0.07958559 | 0.2016361 | 0.4038498 | 0.6863232 | 0.827244 |
| ENSG00000213090 | SCYL2P1 | 54.47308 | 0.07957848 | 0.2445057 | 0.3461127 | 0.729258 | 0.853215 |
| ENSG00000104365 | IKBKB | 2919.752579 | 0.079538693 | 0.270900223 | 0.309157907 | 0.75720141 | 0.870423 |
| ENSG00000244230 | RN7SL151P | 1.229876 | 0.07951973 | 0.7926612 | 0.435939 | 0.662881 | NA |
| ENSG00000239857 | GET4 | 194.4136 | 0.07943895 | 0.2351151 | 0.3508657 | 0.7256891 | 0.851005 |

| | | | | | | | |
|---|---|---|---|---|---|---|---|
| ENSG00000257779 | | 0.8693423 | 0.07940809 | 0.741955 | 0.2783008 | 0.7807815 | NA |
| ENSG00000129559 | NEDD8 | 2726.677192 | 0.079389726 | 0.237228559 | 0.369433669 | 0.71180451 | 0.843213 |
| ENSG00000189114 | BLOC1S3 | 1304.841 | 0.07927201 | 0.16885 | 0.4782919 | 0.6324425 | 0.789298 |
| ENSG00000075420 | FNDC3B | 3375.713428 | 0.079216384 | 0.244361196 | 0.37211378 | 0.70980814 | 0.842216 |
| ENSG00000213885 | RPL13AP7 | 323.4212 | 0.07917123 | 0.227949 | 0.3392287 | 0.7344375 | 0.856021 |
| ENSG00000162408 | NOL9 | 1138.844 | 0.07905995 | 0.201221 | 0.4052782 | 0.685273 | 0.826544 |
| ENSG00000279057 | | 8.898651 | 0.0790169 | 0.5445797 | 0.1989947 | 0.8422669 | 0.922165 |
| ENSG00000271318 | MOB4P2 | 7.148134 | 0.07897404 | 0.5635085 | 0.2002433 | 0.8412903 | 0.921607 |
| ENSG00000248383 | PCDHAC1 | 50.49836 | 0.07896457 | 0.4404415 | 0.2099691 | 0.8336918 | 0.916554 |
| ENSG00000106302 | HYAL4 | 1.497577798 | 0.078861833 | 0.735368002 | 0.25567097 | 0.79820491 | NA |
| ENSG00000245662 | LINC02211 | 2.532969 | 0.07881145 | 0.660113 | 0.2129181 | 0.8313908 | 0.915137 |
| ENSG00000156232 | WHAMM | 1202.608 | 0.07880369 | 0.1453916 | 0.5490383 | 0.5829792 | 0.753365 |
| ENSG00000254099 | | 2.015285 | 0.07878563 | 0.681677 | 0.2267985 | 0.8205804 | 0.909023 |
| ENSG00000182117 | NOP10 | 3323.807 | 0.07866498 | 0.2079956 | 0.3919575 | 0.6950896 | 0.832992 |
| ENSG00000133247 | KMT5C | 1332.152581 | 0.078551851 | 0.224000354 | 0.362726539 | 0.71680917 | 0.845914 |
| ENSG00000196205 | EEF1A1P5 | 28248.62 | 0.07845693 | 0.181159 | 0.4420815 | 0.6584302 | 0.807884 |
| ENSG00000163795 | ZNF513 | 135.9917 | 0.07844743 | 0.3107372 | 0.2723245 | 0.7853725 | 0.887737 |
| ENSG00000233893 | EZR-AS1 | 1.325932 | 0.078382 | 0.7509101 | 0.3156117 | 0.7522973 | NA |
| ENSG00000126524 | SBDS | 3755.137017 | 0.078304511 | 0.146386362 | 0.528590731 | 0.59708939 | 0.764085 |
| ENSG00000224217 | PRPF4BP1 | 6.298974 | 0.07829693 | 0.5378588 | 0.1973356 | 0.8435649 | 0.923092 |
| ENSG00000152214 | RIT2 | 1.79091 | 0.07822834 | 0.7301534 | 0.2616253 | 0.7936104 | 0.893218 |
| ENSG00000015479 | MATR3 | 145.2863583 | 0.078158284 | 0.326048854 | 0.264910434 | 0.79107846 | 0.891628 |
| ENSG00000162419 | GMEB1 | 945.7998 | 0.07810759 | 0.1725853 | 0.4636153 | 0.6429234 | 0.796464 |
| ENSG00000232228 | RPL36AP16 | 6.801448 | 0.07803956 | 0.61638 | 0.1981908 | 0.8428958 | 0.92255 |
| ENSG00000258130 | | 5.935681 | 0.07801712 | 0.6819066 | 0.2244931 | 0.8223736 | 0.909824 |
| ENSG00000269800 | PLEKHA3P1 | 16.28083 | 0.07801681 | 0.412175 | 0.2201498 | 0.8257545 | 0.91192 |
| ENSG00000185475 | TMEM179B | 1950.448 | 0.07800146 | 0.1644232 | 0.4940062 | 0.6213018 | 0.781173 |
| ENSG00000246334 | PRR7-AS1 | 75.27474 | 0.07798071 | 0.3045048 | 0.2758726 | 0.7826459 | 0.886221 |
| ENSG00000232858 | RPL34P27 | 5.284186 | 0.07795963 | 0.6084561 | 0.2019959 | 0.83992 | 0.920772 |
| ENSG00000122484 | RPAP2 | 1056.296817 | 0.077941501 | 0.208566588 | 0.383685126 | 0.70121184 | 0.83727 |
| ENSG00000205629 | LCMT1 | 2432.251 | 0.07790078 | 0.1534325 | 0.5088647 | 0.6108471 | 0.774202 |
| ENSG00000271966 | | 72.46289 | 0.07784674 | 0.3283008 | 0.2579626 | 0.7964357 | 0.895001 |
| ENSG00000168005 | SPINDOC | 2157.744 | 0.0778154 | 0.2087413 | 0.3756838 | 0.707152 | 0.840791 |

| | | | | | | | |
|---|---|---|---|---|---|---|---|
| ENSG00000251205 | | 0.8392072 | 0.07778824 | 0.7573251 | 0.2830915 | 0.7771067 | NA |
| ENSG00000226465 | | 112.8668 | 0.07777243 | 0.2839376 | 0.2839125 | 0.7764774 | 0.882376 |
| ENSG00000269139 | | 8.694904 | 0.07775999 | 0.5568188 | 0.1963883 | 0.8443062 | 0.923608 |
| ENSG00000268006 | PTOV1-AS1 | 114.132 | 0.07774542 | 0.3200898 | 0.2626044 | 0.7928555 | 0.892909 |
| ENSG00000036549 | ZZZ3 | 4066.653681 | 0.077712793 | 0.167688515 | 0.470046796 | 0.63832159 | 0.793621 |
| ENSG00000181638 | ZFP41 | 860.8626 | 0.07767665 | 0.1434323 | 0.5377865 | 0.5907245 | 0.759422 |
| ENSG00000137337 | MDC1 | 2524.122726 | 0.07764323 | 0.278411532 | 0.29433209 | 0.76850415 | 0.877361 |
| ENSG00000122432 | SPATA1 | 16.81850782 | 0.077620015 | 0.426230543 | 0.211689577 | 0.83234922 | 0.915608 |
| ENSG00000179743 | | 330.9446 | 0.07759208 | 0.1952596 | 0.4080076 | 0.6832681 | 0.824763 |
| ENSG00000260798 | | 1.218596 | 0.07757097 | 0.7314998 | 0.2590666 | 0.7955839 | NA |
| ENSG00000108239 | TBC1D12 | 1195.551111 | 0.077568469 | 0.135865922 | 0.591060522 | 0.55447987 | 0.732463 |
| ENSG00000115268 | RPS15 | 10008.24658 | 0.077501336 | 0.183965213 | 0.428469589 | 0.66830927 | 0.815124 |
| ENSG00000143028 | SYPL2 | 47.65291 | 0.07738501 | 0.5379065 | 0.1976898 | 0.8432878 | 0.922865 |
| ENSG00000228127 | LINC01649 | 0.761151 | 0.07737406 | 0.7664468 | 0.2480989 | 0.8040579 | NA |
| ENSG00000141338 | ABCA8 | 1.708651 | 0.07736768 | 0.745352 | 0.3384394 | 0.7350321 | 0.8565 |
| ENSG00000163959 | SLC51A | 96.5365 | 0.07718531 | 0.2736568 | 0.2988105 | 0.7650846 | 0.875357 |
| ENSG00000280798 | LINC00294 | 288.4806 | 0.07712799 | 0.2195632 | 0.3700867 | 0.7113179 | 0.842862 |
| ENSG00000263946 | | 3.626507 | 0.07711218 | 0.7044317 | 0.2269798 | 0.8204395 | 0.908942 |
| ENSG00000288103 | | 1.74364 | 0.07697809 | 0.7191278 | 0.248999 | 0.8033616 | 0.899322 |
| ENSG00000225165 | RTRAFP2 | 3.4345 | 0.07690962 | 0.6687733 | 0.2125773 | 0.8316567 | 0.915261 |
| ENSG00000272865 | | 1.220782 | 0.07688376 | 0.7506577 | 0.3410152 | 0.7330922 | NA |
| ENSG00000146425 | DYNLT1 | 3094.931 | 0.07688314 | 0.1933189 | 0.4094948 | 0.6821766 | 0.823965 |
| ENSG00000198925 | ATG9A | 2053.164 | 0.07688219 | 0.1355848 | 0.5753014 | 0.5650875 | 0.7402 |
| ENSG00000182950 | ODF3L1 | 1.033031 | 0.07684696 | 0.7325423 | 0.2644865 | 0.7914051 | NA |
| ENSG00000076650 | GPATCH1 | 1027.296384 | 0.076841915 | 0.127924516 | 0.604429595 | 0.54555807 | 0.725981 |
| ENSG00000244383 | FAM3D-AS1 | 0.9532897 | 0.07679884 | 0.7309493 | 0.2604412 | 0.7945235 | NA |
| ENSG00000254685 | FPGT | 451.6101 | 0.07677365 | 0.2119886 | 0.3714001 | 0.7103396 | 0.842335 |
| ENSG00000253317 | | 8.993983 | 0.07666497 | 0.6110357 | 0.1821578 | 0.8554589 | 0.929979 |
| ENSG00000149646 | CNBD2 | 7.726414 | 0.07664128 | 0.562006 | 0.1879771 | 0.8508946 | 0.927491 |
| ENSG00000225774 | SIRPAP1 | 11.12943 | 0.07655174 | 0.5892012 | 0.177051 | 0.8594683 | 0.932352 |
| ENSG00000157005 | SST | 0.7413412 | 0.07645945 | 0.7707604 | 0.2447205 | 0.8066728 | NA |
| ENSG00000247095 | MIR210HG | 168.1464 | 0.07640903 | 0.2698893 | 0.2936215 | 0.7690471 | 0.877585 |
| ENSG00000242085 | RPS20P33 | 21.39796 | 0.07639473 | 0.4312437 | 0.2122703 | 0.8318962 | 0.915375 |

| | | | | | | | |
|---|---|---|---|---|---|---|---|
| ENSG00000235790 | PEF1-AS1 | 5.775323 | 0.07638788 | 0.5667983 | 0.1824514 | 0.8552285 | 0.929935 |
| ENSG00000233892 | PAIP1P1 | 13.30302 | 0.07636703 | 0.4823813 | 0.1945869 | 0.8457163 | 0.924533 |
| ENSG00000217643 | PTGES3P2 | 1.628165 | 0.07635019 | 0.6858533 | 0.2169629 | 0.8282373 | 0.91345 |
| ENSG00000126705 | AHDC1 | 984.8699837 | 0.076348351 | 0.260887264 | 0.309827083 | 0.75669246 | 0.870017 |
| ENSG00000162819 | BROX | 2909.218 | 0.07621607 | 0.1338909 | 0.5365195 | 0.5915996 | 0.760252 |
| ENSG00000243230 | | 3.264701 | 0.07618774 | 0.6217057 | 0.1969397 | 0.8438748 | 0.923393 |
| ENSG00000169217 | CD2BP2 | 6990.951 | 0.07617905 | 0.1421385 | 0.5455223 | 0.5853943 | 0.755273 |
| ENSG00000287512 | | 1.241712 | 0.07613275 | 0.7368635 | 0.2731953 | 0.7847031 | NA |
| ENSG00000251991 | RNU7-49P | 0.9097052 | 0.07612945 | 0.7525098 | 0.307948 | 0.7581219 | NA |
| ENSG00000220494 | YAP1P1 | 1.437427 | 0.07612404 | 0.741381 | 0.2727176 | 0.7850703 | NA |
| ENSG00000233762 | RPS15P4 | 3441.143 | 0.07606148 | 0.1973206 | 0.3939409 | 0.6936247 | 0.831817 |
| ENSG00000162592 | CCDC27 | 3.424741 | 0.07603898 | 0.616153 | 0.1940396 | 0.8461449 | 0.924846 |
| ENSG00000135637 | CCDC142 | 1473.240947 | 0.076013199 | 0.216334955 | 0.367532418 | 0.71322192 | 0.843666 |
| ENSG00000255062 | | 16.66087 | 0.07597952 | 0.4667501 | 0.1978021 | 0.8431999 | 0.922807 |
| ENSG00000089280 | FUS | 21097.25007 | 0.075976378 | 0.206339569 | 0.378346165 | 0.70517346 | 0.839876 |
| ENSG00000222788 | RNU2-38P | 0.991572 | 0.07594254 | 0.7620754 | 0.2612816 | 0.7938753 | NA |
| ENSG00000233747 | RPL36AP13 | 1.461175 | 0.07589584 | 0.7021874 | 0.2250214 | 0.8219626 | NA |
| ENSG00000232489 | MFAP1P1 | 1.056171 | 0.07582465 | 0.7358687 | 0.2589333 | 0.7956867 | NA |
| ENSG00000142185 | TRPM2 | 5.548508 | 0.07582242 | 0.5819268 | 0.1843168 | 0.853765 | 0.928936 |
| ENSG00000213087 | | 204.8796 | 0.07578717 | 0.2462133 | 0.3266268 | 0.7439502 | 0.862053 |
| ENSG00000286280 | | 1.069555 | 0.07576949 | 0.7619678 | 0.2681294 | 0.7885997 | NA |
| ENSG00000171723 | GPHN | 1905.052 | 0.07574914 | 0.169575 | 0.4635353 | 0.6429807 | 0.796464 |
| ENSG00000205035 | | 9.664215 | 0.07572425 | 0.5348944 | 0.1863349 | 0.8521822 | 0.928062 |
| ENSG00000233276 | GPX1 | 1.241862 | 0.07552696 | 0.7246899 | 0.2572176 | 0.7970108 | NA |
| ENSG00000241288 | LINC02614 | 192.8012 | 0.07551767 | 0.3597066 | 0.2206488 | 0.8253659 | 0.911882 |
| ENSG00000151748 | SAV1 | 1352.946 | 0.07537358 | 0.162932 | 0.4636203 | 0.6429198 | 0.796464 |
| ENSG00000284617 | | 7.627014 | 0.07534064 | 0.6529976 | 0.2140071 | 0.8305415 | 0.914887 |
| ENSG00000133812 | SBF2 | 1390.367207 | 0.075257425 | 0.274431767 | 0.285938415 | 0.77492529 | 0.88134 |
| ENSG00000248508 | SRP14-AS1 | 327.8939 | 0.07516508 | 0.3186152 | 0.2600174 | 0.7948504 | 0.894013 |
| ENSG00000284709 | | 0.9490409 | 0.07513638 | 0.7476471 | 0.2528677 | 0.8003704 | NA |
| ENSG00000074696 | HACD3 | 10291.48871 | 0.075129205 | 0.110570108 | 0.714978548 | 0.4746223 | 0.66735 |
| ENSG00000145990 | GFOD1 | 577.8035 | 0.07511294 | 0.2747322 | 0.2911673 | 0.7709233 | 0.878884 |
| ENSG00000287126 | | 141.1635 | 0.0750617 | 0.2960503 | 0.2712703 | 0.7861831 | 0.888364 |

| | | | | | | | |
|---|---|---|---|---|---|---|---|
| ENSG00000280836 | | 3.618756 | 0.07503144 | 0.6383311 | 0.1916249 | 0.848036 | 0.926147 |
| ENSG00000260948 | | 28.27751 | 0.07492424 | 0.4618035 | 0.192352 | 0.8474665 | 0.925823 |
| ENSG00000257509 | | 5.557086 | 0.07490992 | 0.6218932 | 0.191254 | 0.8483266 | 0.926316 |
| ENSG00000257195 | HNRNPA1P50 | 1.676269 | 0.07450609 | 0.7295773 | 0.2637179 | 0.7919973 | 0.892302 |
| ENSG00000110344 | UBE4A | 3834.516351 | 0.074446188 | 0.25609876 | 0.316032664 | 0.75197772 | 0.867195 |
| ENSG00000120948 | TARDBP | 5594.470787 | 0.074435709 | 0.163263694 | 0.456146888 | 0.64828436 | 0.800645 |
| ENSG00000152348 | ATG10 | 441.3175 | 0.0741614 | 0.1584873 | 0.4826798 | 0.6293231 | 0.787012 |
| ENSG00000171612 | SLC25A33 | 448.8255 | 0.07407223 | 0.2295412 | 0.3353808 | 0.7373378 | 0.858171 |
| ENSG00000186439 | TRDN | 17.13312 | 0.07399115 | 0.5964941 | 0.2088511 | 0.8345645 | 0.917134 |
| ENSG00000259030 | FPGT-TNNI3K | 12.78705 | 0.07396674 | 0.4913241 | 0.1826067 | 0.8551067 | 0.929935 |
| ENSG00000285177 | | 2.006525 | 0.07396321 | 0.6853918 | 0.2138407 | 0.8306713 | 0.914899 |
| ENSG00000047365 | ARAP2 | 1020.207133 | 0.073958144 | 0.272016922 | 0.302680843 | 0.7621331 | 0.873296 |
| ENSG00000228118 | MYL6P5 | 50.05572 | 0.07392097 | 0.3765874 | 0.2206346 | 0.825377 | 0.911882 |
| ENSG00000165509 | MAGEC3 | 32.06864 | 0.07385929 | 0.447946 | 0.1929549 | 0.8469943 | 0.925435 |
| ENSG00000286875 | | 1.676016 | 0.0738203 | 0.702081 | 0.1818291 | 0.8557168 | 0.930183 |
| ENSG00000227502 | MROCKI | 2.857614 | 0.07379161 | 0.7135274 | 0.2487667 | 0.8035413 | 0.89941 |
| ENSG00000228862 | | 3.703837 | 0.07371412 | 0.6119227 | 0.1866208 | 0.851958 | 0.927996 |
| ENSG00000104442 | ARMC1 | 2835.276513 | 0.07360773 | 0.196285684 | 0.399301511 | 0.68967105 | 0.829057 |
| ENSG00000222108 | RNA5SP317 | 10.70487 | 0.07359869 | 0.4768532 | 0.1912375 | 0.8483395 | 0.926316 |
| ENSG00000159788 | RGS12 | 1006.945 | 0.07356898 | 0.1560407 | 0.4876805 | 0.6257762 | 0.78456 |
| ENSG00000241990 | PRR34-AS1 | 89.78643 | 0.07348358 | 0.2580671 | 0.2921218 | 0.7701935 | 0.878478 |
| ENSG00000213205 | STRADBP1 | 1.824978 | 0.0734771 | 0.7228131 | 0.2466799 | 0.8051559 | 0.900308 |
| ENSG00000228928 | KPNA2P2 | 1.812445 | 0.07346134 | 0.7168086 | 0.2511221 | 0.8017197 | 0.898186 |
| ENSG00000068137 | PLEKHH3 | 4334.863921 | 0.073325092 | 0.118826531 | 0.674908795 | 0.49973371 | 0.687949 |
| ENSG00000088766 | CRLS1 | 76.51069342 | 0.073279505 | 0.282269419 | 0.283730136 | 0.7766172 | 0.882471 |
| ENSG00000259565 | KRT8P23 | 1.506061 | 0.07323775 | 0.7256643 | 0.2508473 | 0.8019321 | NA |
| ENSG00000278831 | | 8.636565 | 0.07322839 | 0.5961666 | 0.1836358 | 0.8542992 | 0.92926 |
| ENSG00000236256 | DIAPH2-AS1 | 1.395573 | 0.07322581 | 0.7441753 | 0.3197704 | 0.7491424 | NA |
| ENSG00000278594 | | 30.07135 | 0.07320805 | 0.3652876 | 0.2225967 | 0.8238494 | 0.910821 |
| ENSG00000279225 | | 6.980696 | 0.07319771 | 0.5667232 | 0.1848145 | 0.8533745 | 0.928663 |
| ENSG00000219149 | PPIAP87 | 2.15204 | 0.0731907 | 0.7154023 | 0.2397829 | 0.8104985 | 0.90305 |
| ENSG00000279623 | | 1.146472 | 0.07316244 | 0.7468047 | 0.2874361 | 0.7737784 | NA |
| ENSG00000277901 | | 3.064138 | 0.07315847 | 0.6687544 | 0.1996927 | 0.8417209 | 0.92191 |

| | | | | | | | |
|---|---|---|---|---|---|---|---|
| ENSG00000266743 | | 1.192504 | 0.07311867 | 0.7610895 | 0.2692594 | 0.7877301 | NA |
| ENSG00000165233 | CARD19 | 2860.705 | 0.07310643 | 0.2772647 | 0.2772096 | 0.7816192 | 0.885504 |
| ENSG00000176919 | C8G | 56.05126 | 0.07299354 | 0.3389345 | 0.2284341 | 0.8193088 | 0.908599 |
| ENSG00000115325 | DOK1 | 678.0690872 | 0.072937709 | 0.176673088 | 0.424745951 | 0.67102188 | 0.817029 |
| ENSG00000049759 | NEDD4L | 2726.417691 | 0.072804071 | 0.176305452 | 0.422949477 | 0.67233212 | 0.817912 |
| ENSG00000226045 | | 1.165708 | 0.07279184 | 0.7501456 | 0.3175197 | 0.7508493 | NA |
| ENSG00000116521 | SCAMP3 | 6519.071863 | 0.072765559 | 0.131381395 | 0.536690293 | 0.59148156 | 0.760174 |
| ENSG00000109189 | USP46 | 2617.776649 | 0.072764002 | 0.149318416 | 0.492569958 | 0.62231648 | 0.781814 |
| ENSG00000189129 | PLAC9 | 28.21337 | 0.0727425 | 0.4820999 | 0.1860471 | 0.8524079 | 0.928257 |
| ENSG00000233334 | FAM53B-AS1 | 0.8203727 | 0.07271794 | 0.7246622 | 0.2364782 | 0.8130616 | NA |
| ENSG00000164879 | CA3 | 0.8866205 | 0.0725857 | 0.7545075 | 0.2763699 | 0.782264 | NA |
| ENSG00000227036 | LINC00511 | 46.39612 | 0.07251413 | 0.4444066 | 0.1881093 | 0.850791 | 0.927491 |
| ENSG00000269553 | | 0.8499262 | 0.07248431 | 0.7549507 | 0.2959641 | 0.7672575 | NA |
| ENSG00000231967 | | 1.547867 | 0.0724755 | 0.7464 | 0.2494163 | 0.8030388 | NA |
| ENSG00000236360 | RRAS2P1 | 12.54951 | 0.07246028 | 0.4856536 | 0.1895296 | 0.8496777 | 0.926918 |
| ENSG00000270031 | | 5.417013 | 0.072453 | 0.614507 | 0.181634 | 0.85587 | 0.930273 |
| ENSG00000273188 | | 26.00787 | 0.07243393 | 0.3754442 | 0.2166582 | 0.8284747 | 0.913464 |
| ENSG00000148942 | SLC5A12 | 177.4328 | 0.07242997 | 0.2977665 | 0.2593921 | 0.7953327 | 0.894202 |
| ENSG00000276030 | | 1.676362 | 0.07241272 | 0.7162572 | 0.2333123 | 0.8155189 | 0.90611 |
| ENSG00000226716 | | 1.086966 | 0.07232361 | 0.7277269 | 0.2473974 | 0.8046007 | NA |
| ENSG00000244040 | IL12A-AS1 | 2.201368 | 0.07229605 | 0.6831507 | 0.2023668 | 0.83963 | 0.920606 |
| ENSG00000177688 | SUMO4 | 3.555551 | 0.0722916 | 0.6306695 | 0.1892378 | 0.8499065 | 0.927053 |
| ENSG00000151466 | SCLT1 | 623.5969 | 0.07222084 | 0.2681275 | 0.2842123 | 0.7762477 | 0.882183 |
| ENSG00000206869 | SNORA70F | 1.449646 | 0.07221683 | 0.7209307 | 0.2343808 | 0.8146894 | NA |
| ENSG00000200095 | RNU6-181P | 0.931868 | 0.07216341 | 0.7270562 | 0.234707 | 0.8144361 | NA |
| ENSG00000148335 | NTMT1 | 1557.051 | 0.07209493 | 0.2726669 | 0.2600926 | 0.7947923 | 0.894013 |
| ENSG00000188573 | FBLL1 | 1.341284 | 0.07200064 | 0.7380661 | 0.2318391 | 0.816663 | NA |
| ENSG00000143512 | HHIPL2 | 21.25426 | 0.07195249 | 0.5389012 | 0.1817118 | 0.8558089 | 0.930245 |
| ENSG00000273973 | | 5.746547 | 0.07195065 | 0.6266512 | 0.1873056 | 0.851421 | 0.927829 |
| ENSG00000111321 | LTBR | 6537.892637 | 0.071916269 | 0.169833698 | 0.431091565 | 0.66640179 | 0.813994 |
| ENSG00000269609 | RPARP-AS1 | 1230.693 | 0.07187351 | 0.2226498 | 0.3328943 | 0.739214 | 0.85915 |
| ENSG00000162490 | DRAXIN | 4.430922 | 0.07184979 | 0.578462 | 0.17928 | 0.8577179 | 0.931298 |
| ENSG00000127314 | RAP1B | 2153.436424 | 0.071847548 | 0.116368671 | 0.612423305 | 0.54025773 | 0.722365 |

| | | | | | | | |
|---|---|---|---|---|---|---|---|
| ENSG00000287195 | | 2.58365 | 0.0717639 | 0.7132531 | 0.2450289 | 0.8064341 | 0.900712 |
| ENSG00000165678 | GHITM | 17770.1 | 0.07172864 | 0.1962182 | 0.3418891 | 0.7324344 | 0.854968 |
| ENSG00000273702 | | 492.5247 | 0.07170869 | 0.2084103 | 0.3613953 | 0.717804 | 0.846042 |
| ENSG00000164466 | SFXN1 | 7675.716 | 0.07168901 | 0.236563 | 0.3143481 | 0.7532567 | 0.86819 |
| ENSG00000214541 | RPS4XP3 | 55.571 | 0.07160424 | 0.3702547 | 0.2183765 | 0.8271357 | 0.912821 |
| ENSG00000118454 | ANKRD13C | 1718.425515 | 0.071575549 | 0.133060361 | 0.542436089 | 0.58751812 | 0.75702 |
| ENSG00000213176 | RPL13P6 | 1.46353 | 0.07153799 | 0.7040038 | 0.2211299 | 0.8249913 | NA |
| ENSG00000229613 | LINC01501 | 0.8131562 | 0.07152723 | 0.7663935 | 0.2745507 | 0.7836614 | NA |
| ENSG00000243498 | UBA52P5 | 8.600125 | 0.0715263 | 0.5864246 | 0.1786365 | 0.8582231 | 0.931688 |
| ENSG00000223482 | NUTM2A-AS1 | 1641.541 | 0.07152228 | 0.1800512 | 0.4038493 | 0.6863236 | 0.827244 |
| ENSG00000264188 | | 2.35906 | 0.07149879 | 0.6966918 | 0.2203506 | 0.8255981 | 0.91192 |
| ENSG00000214211 | CTAGE14P | 1.404668 | 0.07147773 | 0.7460323 | 0.2686318 | 0.788213 | NA |
| ENSG00000151338 | MIPOL1 | 365.988 | 0.07145393 | 0.201772 | 0.3427684 | 0.7317727 | 0.85446 |
| ENSG00000247315 | ZCCHC3 | 960.6592 | 0.07142201 | 0.1383334 | 0.5205207 | 0.6027007 | 0.767932 |
| ENSG00000213538 | KRT8P41 | 26.18664 | 0.07138537 | 0.456553 | 0.1891041 | 0.8500112 | 0.927091 |
| ENSG00000237669 | | 8.754051 | 0.07121828 | 0.5607289 | 0.1737795 | 0.8620387 | 0.933808 |
| ENSG00000007171 | NOS2 | 2.32570784 | 0.071209836 | 0.730888645 | 0.256651351 | 0.79744793 | 0.895607 |
| ENSG00000247134 | | 9.805914 | 0.07104426 | 0.5286097 | 0.1697833 | 0.8651806 | 0.935654 |
| ENSG00000227462 | PSME2P6 | 7.938401 | 0.07100676 | 0.6093892 | 0.1814451 | 0.8560182 | 0.930273 |
| ENSG00000058272 | PPP1R12A | 3981.539527 | 0.071006034 | 0.185740092 | 0.384562849 | 0.70056132 | 0.836925 |
| ENSG00000177666 | PNPLA2 | 3374.624 | 0.07098912 | 0.1586391 | 0.466465 | 0.6408827 | 0.7952 |
| ENSG00000001167 | NFYA | 2386.447215 | 0.070988063 | 0.182562205 | 0.397534895 | 0.69097306 | 0.830099 |
| ENSG00000199377 | RNU5F-1 | 1.186653 | 0.07097551 | 0.7153754 | 0.2189462 | 0.8266919 | NA |
| ENSG00000186118 | TEX38 | 6.63626 | 0.070952 | 0.5677705 | 0.1775166 | 0.8591026 | 0.932148 |
| ENSG00000126391 | FRMD8 | 2815.902344 | 0.070917396 | 0.144192699 | 0.511794091 | 0.60879513 | 0.772377 |
| ENSG00000069764 | PLA2G10 | 8.139586003 | 0.07088931 | 0.521228818 | 0.176669489 | 0.85976801 | 0.932527 |
| ENSG00000100319 | ZMAT5 | 584.4845281 | 0.070870097 | 0.208450645 | 0.354347114 | 0.72307876 | 0.849464 |
| ENSG00000254788 | CKLF-CMTM1 | 3.974637 | 0.07070067 | 0.5666037 | 0.1750972 | 0.8610033 | 0.933325 |
| ENSG00000135945 | REV1 | 2870.393639 | 0.070697177 | 0.117891108 | 0.589948107 | 0.55522544 | 0.732862 |
| ENSG00000160271 | RALGDS | 906.9578 | 0.07066895 | 0.1907977 | 0.3714652 | 0.7102911 | 0.842335 |
| ENSG00000135108 | FBXO21 | 3948.206112 | 0.070594812 | 0.152930875 | 0.47196755 | 0.63694995 | 0.792595 |
| ENSG00000233459 | | 4.012655 | 0.07056784 | 0.6882226 | 0.20971 | 0.833894 | 0.916662 |
| ENSG00000207457 | RNU6-476P | 2.800053 | 0.07046462 | 0.6937097 | 0.2213777 | 0.8247984 | 0.911643 |

| | | | | | | | |
|---|---|---|---|---|---|---|---|
| ENSG00000196357 | ZNF565 | 199.4538 | 0.07041511 | 0.2240038 | 0.3270002 | 0.7436677 | 0.861863 |
| ENSG00000234773 | | 6.477827 | 0.07035789 | 0.6135415 | 0.1758113 | 0.8604422 | 0.932877 |
| ENSG00000116922 | C1orf109 | 1741.557591 | 0.070344421 | 0.180849643 | 0.397639312 | 0.69089608 | 0.830044 |
| ENSG00000186141 | POLR3C | 5004.347 | 0.07031276 | 0.1366942 | 0.541145 | 0.5884076 | 0.757616 |
| ENSG00000172757 | CFL1 | 29010.18 | 0.07026867 | 0.2518104 | 0.3299369 | 0.7414476 | 0.860542 |
| ENSG00000272669 | | 63.79248 | 0.0702014 | 0.3598549 | 0.2138615 | 0.8306551 | 0.914899 |
| ENSG00000163156 | SCNM1 | 2992.659 | 0.07018419 | 0.205261 | 0.3517434 | 0.7250307 | 0.850636 |
| ENSG00000100320 | RBFOX2 | 3024.521095 | 0.070174635 | 0.098120476 | 0.708822289 | 0.47843476 | 0.670333 |
| ENSG00000266644 | | 8.034956 | 0.07017062 | 0.5448659 | 0.172071 | 0.8633817 | 0.934425 |
| ENSG00000186204 | CYP4F12 | 64.80872 | 0.07013797 | 0.662709 | 0.2168053 | 0.8283601 | 0.913464 |
| ENSG00000259315 | ACTG1P17 | 9.785881 | 0.07012819 | 0.4654042 | 0.1802271 | 0.8569743 | 0.930826 |
| ENSG00000144589 | STK11IP | 2462.901 | 0.06994102 | 0.1534865 | 0.4278578 | 0.6687546 | 0.815464 |
| ENSG00000254527 | ENPP7P12 | 0.7941846 | 0.06991319 | 0.7510995 | 0.2517185 | 0.8012587 | NA |
| ENSG00000217236 | SP9 | 3.466146 | 0.06991055 | 0.6814108 | 0.1952124 | 0.8452267 | 0.924263 |
| ENSG00000286909 | | 197.9814 | 0.06991012 | 0.2861786 | 0.2603665 | 0.7945811 | 0.893975 |
| ENSG00000188162 | OTOG | 0.8345298 | 0.06988537 | 0.7458077 | 0.2556662 | 0.7982086 | NA |
| ENSG00000122483 | CCDC18 | 394.5553552 | 0.06984995 | 0.312718063 | 0.237367451 | 0.81237174 | 0.904227 |
| ENSG00000145386 | CCNA2 | 2830.642 | 0.06978819 | 0.5315687 | 0.1681319 | 0.8664795 | 0.936211 |
| ENSG00000100711 | ZFYVE21 | 2705.124743 | 0.069724766 | 0.10446384 | 0.670114645 | 0.50278471 | 0.690189 |
| ENSG00000272696 | | 14.13283 | 0.0697154 | 0.4435012 | 0.1870165 | 0.8516477 | 0.927845 |
| ENSG00000184056 | VPS33B | 284.3003 | 0.06971144 | 0.1889152 | 0.3723434 | 0.7096372 | 0.842088 |
| ENSG00000143498 | TAF1A | 316.8329 | 0.06961273 | 0.2387988 | 0.3053996 | 0.7600619 | 0.872089 |
| ENSG00000047578 | KATNIP | 2735.746382 | 0.069606292 | 0.217100399 | 0.31286397 | 0.75438401 | 0.868926 |
| ENSG00000174939 | ASPHD1 | 1128.635 | 0.06959731 | 0.2440206 | 0.3034216 | 0.7615686 | 0.872875 |
| ENSG00000286069 | | 1.436045 | 0.06953954 | 0.7187252 | 0.2337718 | 0.8151622 | NA |
| ENSG00000229980 | TOB1-AS1 | 141.9001 | 0.06949566 | 0.272702 | 0.2748962 | 0.783396 | 0.886723 |
| ENSG00000236878 | MTATP6P26 | 2.687403 | 0.06942803 | 0.6652879 | 0.1811783 | 0.8562277 | 0.930282 |
| ENSG00000240373 | SEC62-AS1 | 9.070041 | 0.06940278 | 0.5559099 | 0.1721638 | 0.8633087 | 0.934398 |
| ENSG00000279917 | | 0.949697 | 0.06938053 | 0.748978 | 0.3202795 | 0.7487565 | NA |
| ENSG00000071655 | MBD3 | 2634.976905 | 0.069332553 | 0.264530568 | 0.274364505 | 0.78380452 | 0.886808 |
| ENSG00000051009 | FAM160A2 | 1605.809931 | 0.069308675 | 0.139512541 | 0.491440146 | 0.62311518 | 0.782509 |
| ENSG00000215878 | MARCKSL1P2 | 2.004831 | 0.0693049 | 0.7033433 | 0.2084864 | 0.8348492 | 0.91716 |
| ENSG00000188522 | FAM83G | 3336.128 | 0.06927812 | 0.1566655 | 0.4081576 | 0.683158 | 0.824763 |

| | | | | | | | |
|---|---|---|---|---|---|---|---|
| ENSG00000152078 | TLCD4 | 506.5732 | 0.06927328 | 0.1968361 | 0.3574888 | 0.7207259 | 0.84805 |
| ENSG00000287206 | | 1.149385 | 0.0692389 | 0.7591167 | 0.2956406 | 0.7675046 | NA |
| ENSG00000166147 | FBN1 | 1063.104 | 0.06919912 | 0.4623761 | 0.1839928 | 0.8540191 | 0.929129 |
| ENSG00000255210 | | 1.080779 | 0.06917421 | 0.7502326 | 0.3329771 | 0.7391516 | NA |
| ENSG00000153294 | ADGRF4 | 153.3134 | 0.0691518 | 0.4999856 | 0.1735943 | 0.8621843 | 0.933895 |
| ENSG00000225762 | LINC01389 | 117.2577 | 0.0691035 | 0.3138743 | 0.2348823 | 0.8143001 | 0.905312 |
| ENSG00000255158 | | 5.356904 | 0.06906022 | 0.5872562 | 0.1685396 | 0.8661588 | 0.936211 |
| ENSG00000287007 | | 0.993259 | 0.06893733 | 0.7507278 | 0.3309681 | 0.7406686 | NA |
| ENSG00000261367 | | 3.78215 | 0.06892641 | 0.6863104 | 0.2070047 | 0.8360062 | 0.917845 |
| ENSG00000156508 | EEF1A1 | 54963.77 | 0.068913 | 0.1740907 | 0.4028867 | 0.6870316 | 0.827607 |
| ENSG00000124743 | KLHL31 | 129.3599498 | 0.068766738 | 0.295495847 | 0.250133497 | 0.80248411 | 0.898756 |
| ENSG00000207956 | MIR579 | 1.607119 | 0.06869156 | 0.7255061 | 0.2407285 | 0.8097655 | 0.902824 |
| ENSG00000198028 | ZNF560 | 0.9796294 | 0.06864994 | 0.7553785 | 0.2922844 | 0.7700692 | NA |
| ENSG00000134853 | PDGFRA | 1.711962926 | 0.068598347 | 0.709518021 | 0.212432909 | 0.83176931 | 0.915311 |
| ENSG00000226810 | | 1.317458 | 0.06858233 | 0.7048239 | 0.2055476 | 0.8371443 | NA |
| ENSG00000226360 | RPL10AP6 | 2577.053 | 0.06855206 | 0.2084893 | 0.3374145 | 0.7358044 | 0.856874 |
| ENSG00000114544 | SLC41A3 | 3802.914649 | 0.068530036 | 0.185625744 | 0.371370465 | 0.71036162 | 0.842334 |
| ENSG00000246214 | | 10.00993 | 0.0685003 | 0.5233333 | 0.1689264 | 0.8658546 | 0.936028 |
| ENSG00000255063 | RBMXP3 | 1.068245 | 0.06843498 | 0.7531325 | 0.2893421 | 0.7723196 | NA |
| ENSG00000236937 | PTGES3P4 | 1.138916 | 0.06837391 | 0.7273973 | 0.2375184 | 0.8122547 | NA |
| ENSG00000270993 | | 5.944128 | 0.06832829 | 0.6110952 | 0.1732652 | 0.862443 | 0.933979 |
| ENSG00000242327 | | 12.41111 | 0.06830507 | 0.4919593 | 0.1713573 | 0.8639429 | 0.934727 |
| ENSG00000276712 | MIR7111 | 22.53442 | 0.06827141 | 0.421885 | 0.1898534 | 0.849424 | 0.92687 |
| ENSG00000222640 | RNU2-51P | 1.938988 | 0.06819218 | 0.6985864 | 0.186383 | 0.8521444 | 0.928062 |
| ENSG00000197024 | ZNF398 | 1752.991 | 0.06818979 | 0.1363783 | 0.4994116 | 0.6174894 | 0.77872 |
| ENSG00000237264 | FTH1P11 | 559.9377 | 0.06817026 | 0.2290851 | 0.3069487 | 0.7588825 | 0.871488 |
| ENSG00000110315 | RNF141 | 1335.368515 | 0.068119878 | 0.208152347 | 0.355374352 | 0.72230916 | 0.848898 |
| ENSG00000287643 | | 2.361166 | 0.06811187 | 0.6548943 | 0.183972 | 0.8540354 | 0.929129 |
| ENSG00000267080 | ASB16-AS1 | 580.6185 | 0.06809853 | 0.2106064 | 0.3318039 | 0.7400373 | 0.859581 |
| ENSG00000227347 | HNRNPKP2 | 477.3709 | 0.0680873 | 0.1665566 | 0.4169642 | 0.6767046 | 0.820152 |
| ENSG00000234418 | | 2.007112 | 0.06803866 | 0.6821256 | 0.196454 | 0.8442548 | 0.923608 |
| ENSG00000066827 | ZFAT | 799.5669812 | 0.067985306 | 0.182196895 | 0.376573802 | 0.70649036 | 0.840456 |
| ENSG00000144231 | POLR2D | 2268.209 | 0.06796187 | 0.1755879 | 0.3946498 | 0.6931013 | 0.831374 |

| | | | | | | | |
|---|---|---|---|---|---|---|---|
| ENSG00000280277 | | 2.379726 | 0.06794961 | 0.6751262 | 0.1880464 | 0.8508403 | 0.927491 |
| ENSG00000185189 | NRBP2 | 2435.918 | 0.06784796 | 0.2770697 | 0.2515861 | 0.801361 | 0.898104 |
| ENSG00000274220 | | 114.7896 | 0.06783817 | 0.2292636 | 0.3107722 | 0.7559738 | 0.869764 |
| ENSG00000285373 | LINC02478 | 1.489862 | 0.06778122 | 0.7205535 | 0.2339288 | 0.8150402 | NA |
| ENSG00000231655 | EMP2P1 | 1.111355 | 0.06774223 | 0.7071586 | 0.2016329 | 0.8402037 | NA |
| ENSG00000120889 | TNFRSF10B | 4651.745532 | 0.067704806 | 0.299491659 | 0.248206483 | 0.80397465 | 0.899636 |
| ENSG00000279669 | | 1.108552 | 0.06769682 | 0.7242851 | 0.1931537 | 0.8468386 | NA |
| ENSG00000143315 | PIGM | 1798.107 | 0.06766375 | 0.1522438 | 0.4391346 | 0.660564 | 0.809118 |
| ENSG00000163630 | SYNPR | 2.080879 | 0.06762592 | 0.677448 | 0.1909192 | 0.8485889 | 0.926453 |
| ENSG00000257218 | GATC | 1470.971 | 0.06760678 | 0.168116 | 0.4080112 | 0.6832655 | 0.824763 |
| ENSG00000234093 | RPS15AP11 | 218.7387 | 0.06758985 | 0.2703583 | 0.2658719 | 0.7903379 | 0.891266 |
| ENSG00000145087 | STXBP5L | 12.06881 | 0.06758608 | 0.4545268 | 0.1783354 | 0.8584596 | 0.93183 |
| ENSG00000173465 | ZNRD2 | 2300.955 | 0.06748994 | 0.2289662 | 0.2866467 | 0.7743828 | 0.880987 |
| ENSG00000169900 | PYDC1 | 1.331716 | 0.06745383 | 0.7518155 | 0.3020821 | 0.7625894 | NA |
| ENSG00000263353 | PPIAL4A | 1.193598 | 0.06743845 | 0.7186146 | 0.2023574 | 0.8396373 | NA |
| ENSG00000214552 | COPS8P2 | 139.8312 | 0.06743025 | 0.2286496 | 0.3126133 | 0.7545744 | 0.86904 |
| ENSG00000111358 | GTF2H3 | 2535.581998 | 0.067407211 | 0.221574313 | 0.312442773 | 0.75470405 | 0.869057 |
| ENSG00000261460 | | 7.301668 | 0.06737111 | 0.5595336 | 0.1659508 | 0.8681957 | 0.936731 |
| ENSG00000169714 | CNBP | 20354.41 | 0.06735302 | 0.1893692 | 0.3982224 | 0.6904663 | 0.829738 |
| ENSG00000028277 | POU2F2 | 25.00906633 | 0.067312065 | 0.503315009 | 0.170650672 | 0.86449845 | 0.935138 |
| ENSG00000253607 | | 0.8822118 | 0.06722303 | 0.7391942 | 0.2466928 | 0.805146 | NA |
| ENSG00000232742 | RHOQP2 | 149.2295 | 0.06721353 | 0.3329284 | 0.2291619 | 0.8187431 | 0.908239 |
| ENSG00000204619 | PPP1R11 | 2888.416 | 0.06718002 | 0.1432614 | 0.469031 | 0.6390475 | 0.794106 |
| ENSG00000089022 | MAPKAPK5 | 2786.717896 | 0.067167263 | 0.098153633 | 0.672420142 | 0.50131626 | 0.689128 |
| ENSG00000243716 | NPIPB5 | 187.638 | 0.06714542 | 0.2374956 | 0.2958755 | 0.7673252 | 0.876599 |
| ENSG00000241370 | RPP21 | 142.2159 | 0.06712818 | 0.2792515 | 0.2550462 | 0.7986874 | 0.896507 |
| ENSG00000133997 | MED6 | 1842.533937 | 0.067125858 | 0.182135918 | 0.376093294 | 0.70684754 | 0.840693 |
| ENSG00000179915 | NRXN1 | 6.522966 | 0.06710459 | 0.6745224 | 0.1805912 | 0.8566884 | 0.930706 |
| ENSG00000125510 | OPRL1 | 27.05010501 | 0.067028102 | 0.3945995 | 0.196103801 | 0.84452892 | 0.923767 |
| ENSG00000287538 | | 0.9689457 | 0.06691534 | 0.7496186 | 0.3108331 | 0.7559275 | NA |
| ENSG00000240972 | MIF | 4.571917 | 0.06691526 | 0.590138 | 0.1626072 | 0.8708277 | 0.938393 |
| ENSG00000185485 | SDHAP1 | 1430.1 | 0.06687398 | 0.1872929 | 0.3751418 | 0.707555 | 0.841007 |
| ENSG00000254815 | LMNTD2-AS1 | 219.7479 | 0.06677984 | 0.3495132 | 0.2115029 | 0.8324949 | 0.915659 |

| | | | | | | | |
|---|---|---|---|---|---|---|---|
| ENSG00000178722 | C5orf64 | 0.9260176 | 0.06676818 | 0.7433303 | 0.2175817 | 0.827755 | NA |
| ENSG00000130749 | ZC3H4 | 3198.663683 | 0.066729063 | 0.192710533 | 0.34395243 | 0.73088206 | 0.853987 |
| ENSG00000206941 | SNORD15A | 0.7952269 | 0.06672244 | 0.7314877 | 0.2301893 | 0.8179447 | NA |
| ENSG00000072518 | MARK2 | 4279.289527 | 0.066680775 | 0.122943373 | 0.526751307 | 0.5983663 | 0.76521 |
| ENSG00000129317 | PUS7L | 1757.409266 | 0.066663746 | 0.180972341 | 0.366319662 | 0.71412656 | 0.84406 |
| ENSG00000177606 | JUN | 15463.93 | 0.06666044 | 0.3964671 | 0.2407698 | 0.8097336 | 0.902824 |
| ENSG00000173848 | NET1 | 7069.2 | 0.0666157 | 0.2799607 | 0.2478554 | 0.8042463 | 0.899858 |
| ENSG00000224680 | PLA2G12AP1 | 75.60083 | 0.06660073 | 0.2802342 | 0.2511419 | 0.8017044 | 0.898186 |
| ENSG00000232859 | LYRM9 | 235.1073 | 0.06658124 | 0.2931839 | 0.2410054 | 0.8095509 | 0.902806 |
| ENSG00000167671 | UBXN6 | 3608.826 | 0.06653472 | 0.1834226 | 0.3731195 | 0.7090595 | 0.841793 |
| ENSG00000240087 | RPSAP12 | 96.34613 | 0.06652205 | 0.2729737 | 0.2573264 | 0.7969268 | 0.895202 |
| ENSG00000267150 | | 5.177773 | 0.06650332 | 0.6542942 | 0.1854325 | 0.8528898 | 0.928429 |
| ENSG00000286092 | | 1.05858 | 0.06643148 | 0.7343517 | 0.2400232 | 0.8103123 | NA |
| ENSG00000200170 | | 0.9625338 | 0.06635808 | 0.7420739 | 0.2260154 | 0.8211894 | NA |
| ENSG00000116580 | GON4L | 4406.980022 | 0.066321409 | 0.151647262 | 0.392994949 | 0.69432321 | 0.832418 |
| ENSG00000181929 | PRKAG1 | 3240.059 | 0.06631715 | 0.1353624 | 0.4843491 | 0.6281382 | 0.786327 |
| ENSG00000172650 | AGAP5 | 21.79236 | 0.06629579 | 0.4862418 | 0.1682281 | 0.8664038 | 0.936211 |
| ENSG00000102805 | CLN5 | 292.8287959 | 0.066245816 | 0.238250208 | 0.295074692 | 0.76793682 | 0.877026 |
| ENSG00000253968 | | 0.997163 | 0.06618115 | 0.7377359 | 0.2388811 | 0.8111978 | NA |
| ENSG00000117450 | PRDX1 | 17654.98807 | 0.06617036 | 0.230062391 | 0.297395606 | 0.7661645 | 0.875951 |
| ENSG00000285231 | OOSP3 | 0.9621908 | 0.06604061 | 0.7466692 | 0.2505794 | 0.8021393 | NA |
| ENSG00000229399 | | 4.226857 | 0.06593705 | 0.6402008 | 0.1764761 | 0.8599199 | 0.932571 |
| ENSG00000139173 | TMEM117 | 343.0469436 | 0.06592719 | 0.175940142 | 0.416266047 | 0.67721533 | 0.820509 |
| ENSG00000160345 | C9orf116 | 434.9427 | 0.06591158 | 0.2398288 | 0.287546 | 0.7736943 | 0.880665 |
| ENSG00000113282 | CLINT1 | 6652.319534 | 0.065904613 | 0.166254609 | 0.442521669 | 0.65811175 | 0.807601 |
| ENSG00000197121 | PGAP1 | 616.0562 | 0.06582955 | 0.3071528 | 0.2400614 | 0.8102827 | 0.902923 |
| ENSG00000132467 | UTP3 | 2693.003795 | 0.06573173 | 0.152333912 | 0.439845538 | 0.66004898 | 0.808746 |
| ENSG00000248245 | | 0.8351517 | 0.06561049 | 0.752041 | 0.3183955 | 0.750185 | NA |
| ENSG00000168066 | SF1 | 12058.34 | 0.06556854 | 0.1485772 | 0.4302656 | 0.6670024 | 0.814411 |
| ENSG00000179335 | CLK3 | 4736.329 | 0.0655663 | 0.1307909 | 0.4878393 | 0.6256637 | 0.78456 |
| ENSG00000226798 | | 1.916261 | 0.06546301 | 0.7002253 | 0.1637973 | 0.8698907 | 0.937688 |
| ENSG00000242899 | RPL7P16 | 4.567194 | 0.06537668 | 0.6264081 | 0.1643138 | 0.8694842 | 0.937402 |
| ENSG00000112852 | PCDHB2 | 326.8342206 | 0.065264289 | 0.204395088 | 0.328440666 | 0.7425785 | 0.861403 |

| | | | | | | | |
|---|---|---|---|---|---|---|---|
| ENSG00000208028 | MIR616 | 62.76989 | 0.0652265 | 0.4860034 | 0.1644942 | 0.8693421 | 0.937389 |
| ENSG00000031003 | FAM13B | 2538.018787 | 0.065209933 | 0.17433956 | 0.385629681 | 0.69977095 | 0.836395 |
| ENSG00000251819 | RNU6-322P | 3.655821 | 0.06518506 | 0.6445198 | 0.1676816 | 0.8668338 | 0.936368 |
| ENSG00000228566 | | 1.314873 | 0.06517319 | 0.7085043 | 0.1865935 | 0.8519794 | NA |
| ENSG00000255437 | | 1.969577 | 0.06513864 | 0.7162084 | 0.1967622 | 0.8440136 | 0.923469 |
| ENSG00000197136 | PCNX3 | 10215.33 | 0.06506691 | 0.1332606 | 0.4934691 | 0.6216811 | 0.78141 |
| ENSG00000284607 | | 1.27019 | 0.06501182 | 0.6994871 | 0.1928321 | 0.8470905 | NA |
| ENSG00000137804 | NUSAP1 | 20.06963331 | 0.064927675 | 0.450771282 | 0.172147215 | 0.8633218 | 0.934398 |
| ENSG00000183172 | SMDT1 | 1944.552 | 0.06492094 | 0.1389063 | 0.4639047 | 0.642716 | 0.796322 |
| ENSG00000142949 | PTPRF | 36901.98 | 0.06486762 | 0.2172689 | 0.3068243 | 0.7589771 | 0.871559 |
| ENSG00000186312 | CA5BP1 | 206.3132 | 0.06481745 | 0.2558192 | 0.2624125 | 0.7930034 | 0.892957 |
| ENSG00000253606 | AFG3L2P1 | 15.14353 | 0.06474318 | 0.4254809 | 0.1793238 | 0.8576835 | 0.931298 |
| ENSG00000106809 | OGN | 0.908829074 | 0.06469092 | 0.750842215 | 0.310488669 | 0.75618938 | NA |
| ENSG00000177479 | ARIH2 | 4206.577 | 0.06468213 | 0.1194588 | 0.5764004 | 0.5643446 | 0.739611 |
| ENSG00000104852 | SNRNP70 | 15664.03605 | 0.064682087 | 0.184858116 | 0.340694859 | 0.73333331 | 0.855398 |
| ENSG00000267088 | | 0.8925077 | 0.06466102 | 0.7435083 | 0.2371377 | 0.81255 | NA |
| ENSG00000197696 | NMB | 583.4048 | 0.06458434 | 0.3833043 | 0.1822984 | 0.8553485 | 0.929935 |
| ENSG00000200674 | RN7SKP160 | 2.540453 | 0.06457505 | 0.6880097 | 0.1903254 | 0.8490542 | 0.926594 |
| ENSG00000009413 | REV3L | 3912.01863 | 0.064554818 | 0.271863964 | 0.248897085 | 0.80344039 | 0.899373 |
| ENSG00000204388 | HSPA1B | 2947.719 | 0.06454636 | 0.353264 | 0.2317637 | 0.8167216 | 0.906752 |
| ENSG00000083099 | LYRM2 | 2930.612568 | 0.064483783 | 0.113155345 | 0.572307483 | 0.56711368 | 0.741591 |
| ENSG00000271874 | | 1.058435 | 0.06440674 | 0.7504721 | 0.3118573 | 0.7551489 | NA |
| ENSG00000253399 | | 1.055604 | 0.06429738 | 0.7449799 | 0.253242 | 0.8000812 | NA |
| ENSG00000167311 | ART5 | 4.890415 | 0.06429218 | 0.6638743 | 0.1781062 | 0.8586396 | 0.93195 |
| ENSG00000011426 | ANLN | 4208.738286 | 0.064281847 | 0.452729952 | 0.169167647 | 0.86566478 | 0.935977 |
| ENSG00000256582 | LINC02390 | 1.41478 | 0.06416925 | 0.7491414 | 0.3114275 | 0.7554757 | NA |
| ENSG00000261024 | | 0.8754998 | 0.06416137 | 0.7404616 | 0.238876 | 0.8112017 | NA |
| ENSG00000151239 | TWF1 | 4693.635 | 0.06414631 | 0.1334049 | 0.485755 | 0.6271409 | 0.78556 |
| ENSG00000163681 | SLMAP | 2633.635 | 0.06408404 | 0.2114325 | 0.3166626 | 0.7514996 | 0.866998 |
| ENSG00000092841 | MYL6 | 21386.59756 | 0.06407862 | 0.179757805 | 0.420302681 | 0.67426435 | 0.818875 |
| ENSG00000223882 | ABCC5-AS1 | 1.811198 | 0.06407045 | 0.6856116 | 0.1824768 | 0.8552086 | 0.929935 |
| ENSG00000125084 | WNT1 | 3.592184931 | 0.064045697 | 0.790182761 | 0.32128279 | 0.7479961 | 0.864893 |
| ENSG00000241532 | AGGF1P3 | 6.670466 | 0.06402421 | 0.5716248 | 0.1566475 | 0.8755227 | 0.941217 |

| | | | | | | | |
|---|---|---|---|---|---|---|---|
| ENSG00000127530 | OR7C1 | 26.58447133 | 0.064022113 | 0.315762666 | 0.223355821 | 0.82325859 | 0.910363 |
| ENSG00000128604 | IRF5 | 916.7927174 | 0.063934052 | 0.208710193 | 0.314323437 | 0.7532754 | 0.86819 |
| ENSG00000249266 | MTHFD2P6 | 1.25058 | 0.06381606 | 0.734395 | 0.2239828 | 0.8227707 | NA |
| ENSG00000279357 | | 1.434198 | 0.06373171 | 0.6924156 | 0.1766676 | 0.8597695 | NA |
| ENSG00000052126 | PLEKHA5 | 1583.828234 | 0.063693591 | 0.168024115 | 0.383661294 | 0.7012295 | 0.83727 |
| ENSG00000213453 | FTH1P3 | 57.08123 | 0.06366293 | 0.2826857 | 0.2392591 | 0.8109047 | 0.903313 |
| ENSG00000197077 | KIAA1671 | 2719.214 | 0.06357715 | 0.2244709 | 0.281414 | 0.7783929 | 0.883535 |
| ENSG00000198169 | ZNF251 | 2910.838 | 0.0635762 | 0.2592111 | 0.258212 | 0.7962433 | 0.894908 |
| ENSG00000235082 | SUMO1P3 | 849.891 | 0.06354167 | 0.198792 | 0.3283901 | 0.7426168 | 0.86141 |
| ENSG00000213290 | PGK1P2 | 45.91138 | 0.06346106 | 0.3901064 | 0.1868905 | 0.8517465 | 0.927845 |
| ENSG00000287070 | | 1.375711 | 0.06343454 | 0.7310264 | 0.2304546 | 0.8177386 | NA |
| ENSG00000274996 | | 0.827674 | 0.06338387 | 0.7579528 | 0.2825322 | 0.7775355 | NA |
| ENSG00000233173 | VN1R34P | 1.45661 | 0.06335243 | 0.7340027 | 0.2153834 | 0.8294684 | NA |
| ENSG00000266640 | MIR4754 | 2.230098 | 0.06333167 | 0.6709474 | 0.1750617 | 0.8610312 | 0.933325 |
| ENSG00000072803 | FBXW11 | 5319.834791 | 0.063312308 | 0.153961604 | 0.418717094 | 0.6754229 | 0.81957 |
| ENSG00000207525 | | 1.200001 | 0.06329078 | 0.7323161 | 0.228158 | 0.8195234 | NA |
| ENSG00000231584 | FAHD2CP | 387.3015 | 0.0632706 | 0.186252 | 0.3478632 | 0.7279429 | 0.852261 |
| ENSG00000198862 | LTN1 | 2719.058 | 0.06321222 | 0.2523746 | 0.2589548 | 0.7956701 | 0.894481 |
| ENSG00000285567 | | 1.116588 | 0.06317196 | 0.7299983 | 0.2137874 | 0.8307128 | NA |
| ENSG00000103051 | COG4 | 5805.924003 | 0.063027473 | 0.144965762 | 0.440091342 | 0.65987095 | 0.808639 |
| ENSG00000135842 | NIBAN1 | 518.017375 | 0.063023495 | 0.678116211 | 0.167424254 | 0.86703624 | 0.936411 |
| ENSG00000122728 | TAF1L | 102.4369257 | 0.063014747 | 0.300828069 | 0.224456741 | 0.82240192 | 0.909824 |
| ENSG00000079435 | LIPE | 7.861396259 | 0.062995854 | 0.571670332 | 0.156327372 | 0.87577498 | 0.941283 |
| ENSG00000266677 | | 3.987885 | 0.06295765 | 0.6491813 | 0.1666245 | 0.8676655 | 0.936695 |
| ENSG00000240616 | RPS6P25 | 124.4124 | 0.06287309 | 0.2975823 | 0.2277767 | 0.8198198 | 0.908881 |
| ENSG00000235734 | HMGN1P36 | 546.6033 | 0.06285777 | 0.2100424 | 0.3062066 | 0.7594473 | 0.871798 |
| ENSG00000213735 | ANAPC10P1 | 19.30226 | 0.06275363 | 0.4571399 | 0.1673173 | 0.8671204 | 0.936413 |
| ENSG00000229057 | RPS3AP54 | 1.814329 | 0.06273105 | 0.7082909 | 0.1851795 | 0.8530882 | 0.928466 |
| ENSG00000134644 | PUM1 | 6754.638598 | 0.062718404 | 0.209377087 | 0.313983212 | 0.75353379 | 0.868386 |
| ENSG00000231802 | DNAJB6P3 | 0.8143219 | 0.06267127 | 0.7394646 | 0.230184 | 0.8179488 | NA |
| ENSG00000252950 | RNA5SP490 | 1.078012 | 0.06265084 | 0.7198051 | 0.2126295 | 0.831616 | NA |
| ENSG00000284402 | MIR7107 | 1.914043 | 0.06263339 | 0.6890914 | 0.1850352 | 0.8532014 | 0.928551 |
| ENSG00000207185 | RNU6-1157P | 1.836043 | 0.06256862 | 0.7296664 | 0.2171274 | 0.8281091 | 0.913402 |

| | | | | | | | |
|---|---|---|---|---|---|---|---|
| ENSG00000081087 | OSTM1 | 1315.9885 | 0.062567184 | 0.240435728 | 0.265754017 | 0.79042866 | 0.891266 |
| ENSG00000105186 | ANKRD27 | 4724.946346 | 0.062551061 | 0.17149215 | 0.354595772 | 0.72289244 | 0.849358 |
| ENSG00000129810 | SGO1 | 503.634264 | 0.062513774 | 0.448323195 | 0.165122796 | 0.86884733 | 0.937133 |
| ENSG00000226890 | | 0.9258754 | 0.06238243 | 0.733854 | 0.2014168 | 0.8403727 | NA |
| ENSG00000284664 | | 7.177916 | 0.0623458 | 0.515243 | 0.158645 | 0.8739485 | 0.940225 |
| ENSG00000261199 | UBE2FP2 | 2.056047 | 0.06231097 | 0.6995273 | 0.1893533 | 0.849816 | 0.926993 |
| ENSG00000286837 | | 2.149923 | 0.06222502 | 0.6639418 | 0.1678467 | 0.8667039 | 0.936304 |
| ENSG00000206897 | SNORA9B | 8.912766 | 0.06221936 | 0.4676994 | 0.1589065 | 0.8737426 | 0.940161 |
| ENSG00000250859 | HNRNPKP1 | 201.0834 | 0.06220373 | 0.1670206 | 0.3811568 | 0.7030869 | 0.838472 |
| ENSG00000233111 | RAB1C | 211.2836 | 0.06218079 | 0.2265177 | 0.2847486 | 0.7758367 | 0.881951 |
| ENSG00000129911 | KLF16 | 1247.644602 | 0.062146541 | 0.194414715 | 0.329967801 | 0.74142429 | 0.860542 |
| ENSG00000175820 | CCDC168 | 5.727189 | 0.06210643 | 0.5921009 | 0.1536565 | 0.8778806 | 0.942366 |
| ENSG00000162576 | MXRA8 | 245.8026 | 0.06209732 | 0.2847898 | 0.2293694 | 0.8185818 | 0.908134 |
| ENSG00000105135 | ILVBL | 3161.601076 | 0.062083777 | 0.184534793 | 0.35838099 | 0.72005822 | 0.847677 |
| ENSG00000234130 | AP2B1P1 | 154.8435 | 0.06204905 | 0.2664217 | 0.2456146 | 0.8059806 | 0.90047 |
| ENSG00000186952 | TMEM232 | 38.82409 | 0.06198824 | 0.4573153 | 0.1636562 | 0.8700018 | 0.937769 |
| ENSG00000215704 | CELA2B | 0.8763977 | 0.06197325 | 0.756009 | 0.2612077 | 0.7939323 | NA |
| ENSG00000259469 | | 1.726756 | 0.06191894 | 0.7286275 | 0.2110445 | 0.8328526 | 0.915897 |
| ENSG00000183723 | CMTM4 | 5601.817 | 0.06182712 | 0.2717079 | 0.2380863 | 0.8118142 | 0.90391 |
| ENSG00000151502 | VPS26B | 3242.581 | 0.06180818 | 0.1067108 | 0.5782408 | 0.5631016 | 0.738708 |
| ENSG00000100865 | CINP | 1747.163786 | 0.061774268 | 0.2073287 | 0.30606207 | 0.75955739 | 0.871849 |
| ENSG00000163877 | SNIP1 | 1316.833 | 0.06168444 | 0.1194276 | 0.5190369 | 0.603735 | 0.768745 |
| ENSG00000171560 | FGA | 4.529135 | 0.06167324 | 0.7879724 | 0.1616448 | 0.8715856 | 0.938753 |
| ENSG00000150637 | CD226 | 3.775179 | 0.06161684 | 0.6386846 | 0.1585091 | 0.8740557 | 0.94027 |
| ENSG00000275519 | MIR6804 | 0.9918742 | 0.06158739 | 0.7347215 | 0.2135183 | 0.8309227 | NA |
| ENSG00000269386 | RAB11B-AS1 | 107.1952 | 0.06158336 | 0.285224 | 0.2221502 | 0.8241969 | 0.911054 |
| ENSG00000255606 | | 4.563501 | 0.06157061 | 0.6186013 | 0.1408004 | 0.8880276 | 0.947572 |
| ENSG00000175216 | CKAP5 | 15248.93 | 0.06156257 | 0.2040468 | 0.3015675 | 0.7629818 | 0.873934 |
| ENSG00000237745 | RPL22P8 | 1.312665 | 0.06148227 | 0.7272038 | 0.2075331 | 0.8355936 | NA |
| ENSG00000104774 | MAN2B1 | 1850.062791 | 0.061358955 | 0.163537626 | 0.361731512 | 0.71755268 | 0.846015 |
| ENSG00000177463 | NR2C2 | 6932.059 | 0.06132244 | 0.2238349 | 0.2801218 | 0.7793841 | 0.884141 |
| ENSG00000248641 | HMGA1P2 | 193.3669 | 0.06131441 | 0.2448269 | 0.2635875 | 0.7920978 | 0.892353 |
| ENSG00000031823 | RANBP3 | 2448.726079 | 0.061310662 | 0.165829705 | 0.387179654 | 0.69862321 | 0.835699 |

| | | | | | | | |
|---|---|---|---|---|---|---|---|
| ENSG00000213293 | | 18.9818 | 0.0612442 | 0.4622491 | 0.1625025 | 0.8709102 | 0.938406 |
| ENSG00000265724 | MIR4284 | 1.44559 | 0.06113701 | 0.7238574 | 0.2141004 | 0.8304688 | NA |
| ENSG00000219932 | RPL12P8 | 7.314449 | 0.06111635 | 0.5406823 | 0.1526224 | 0.8786961 | 0.94286 |
| ENSG00000197934 | CYYR1-AS1 | 1.132108 | 0.06104119 | 0.7164182 | 0.1670566 | 0.8673255 | NA |
| ENSG00000214193 | SH3D21 | 1472.212 | 0.06092718 | 0.2014963 | 0.306697 | 0.7590741 | 0.871595 |
| ENSG00000233586 | | 8.980152 | 0.06092325 | 0.5243741 | 0.1479492 | 0.8823829 | 0.944905 |
| ENSG00000214321 | CBX1P4 | 0.8250196 | 0.06079687 | 0.7281706 | 0.2124625 | 0.8317462 | NA |
| ENSG00000146733 | PSPH | 1641.637 | 0.06078244 | 0.2897498 | 0.2353427 | 0.8139427 | 0.905081 |
| ENSG00000155975 | VPS37A | 1258.797 | 0.06072043 | 0.1781698 | 0.3445387 | 0.7304412 | 0.853899 |
| ENSG00000206149 | HERC2P9 | 788.132 | 0.06071998 | 0.1542229 | 0.3999022 | 0.6892286 | 0.828804 |
| ENSG00000215409 | | 2.07725 | 0.06068476 | 0.6968868 | 0.1834764 | 0.8544242 | 0.929349 |
| ENSG00000273141 | | 10.59453 | 0.06068234 | 0.5182838 | 0.1523556 | 0.8789065 | 0.942968 |
| ENSG00000267526 | | 27.29184 | 0.06064926 | 0.5138337 | 0.1502706 | 0.8805511 | 0.943813 |
| ENSG00000101460 | MAP1LC3A | 0.848876089 | 0.060573194 | 0.754214764 | 0.313230415 | 0.75410561 | NA |
| ENSG00000184209 | SNRNP35 | 1011.797 | 0.06054972 | 0.2063709 | 0.3044934 | 0.760752 | 0.872391 |
| ENSG00000104957 | CCDC130 | 897.7339642 | 0.060492179 | 0.14918976 | 0.406225536 | 0.68457689 | 0.825925 |
| ENSG00000133612 | AGAP3 | 7697.105559 | 0.060361266 | 0.116260969 | 0.556552393 | 0.57783329 | 0.749595 |
| ENSG00000257322 | | 3.189117 | 0.0603102 | 0.692189 | 0.1675537 | 0.8669344 | 0.936384 |
| ENSG00000114626 | ABTB1 | 1319.106283 | 0.060287529 | 0.230795482 | 0.271049804 | 0.78635273 | 0.888442 |
| ENSG00000182318 | ZSCAN22 | 344.5329 | 0.0602493 | 0.2154777 | 0.2876016 | 0.7736517 | 0.880665 |
| ENSG00000084234 | APLP2 | 20700.35853 | 0.060243784 | 0.098304737 | 0.606726843 | 0.54403221 | 0.724914 |
| ENSG00000155906 | RMND1 | 4742.12 | 0.06018388 | 0.1514971 | 0.3971659 | 0.6912451 | 0.830171 |
| ENSG00000182257 | PRR34 | 8.986605 | 0.0601836 | 0.5282652 | 0.1494879 | 0.8811686 | 0.944328 |
| ENSG00000148925 | BTBD10 | 1848.457 | 0.06016905 | 0.1036546 | 0.5744896 | 0.5656365 | 0.740461 |
| ENSG00000076258 | FMO4 | 119.2002353 | 0.060166442 | 0.236998504 | 0.273280561 | 0.78463756 | 0.887289 |
| ENSG00000214460 | TPT1P6 | 17.21005 | 0.06015775 | 0.4184993 | 0.1662105 | 0.8679913 | 0.936731 |
| ENSG00000270419 | CAHM | 42.59028 | 0.06014971 | 0.4256714 | 0.1668749 | 0.8674685 | 0.936559 |
| ENSG00000261407 | | 1.940761 | 0.06012962 | 0.7138631 | 0.1882631 | 0.8506704 | 0.927437 |
| ENSG00000201806 | RNU4-8P | 3.219711 | 0.0601289 | 0.6534586 | 0.1529782 | 0.8784154 | 0.94267 |
| ENSG00000120049 | KCNIP2 | 63.99112122 | 0.060094283 | 0.351451944 | 0.18707399 | 0.85160262 | 0.927845 |
| ENSG00000147676 | MAL2 | 15848.32 | 0.06007034 | 0.155282 | 0.3768457 | 0.7062883 | 0.840366 |
| ENSG00000228294 | BMS1P17 | 4.886332 | 0.0600533 | 0.5472459 | 0.1451166 | 0.8846188 | 0.94582 |
| ENSG00000272476 | | 70.63792 | 0.05997312 | 0.3112181 | 0.2081214 | 0.8351342 | 0.917253 |

| | | | | | | | |
|---|---|---|---|---|---|---|---|
| ENSG00000258279 | LINC00592 | 32.50369 | 0.05995434 | 0.4417911 | 0.1710064 | 0.8642187 | 0.934912 |
| ENSG00000259511 | UBE2Q2L | 0.8668145 | 0.05994719 | 0.762685 | 0.2468072 | 0.8050575 | NA |
| ENSG00000243085 | | 0.9086372 | 0.05993055 | 0.7623022 | 0.2515435 | 0.801394 | NA |
| ENSG00000180806 | HOXC9 | 516.1155 | 0.05990922 | 0.2350332 | 0.2675234 | 0.7890662 | 0.890372 |
| ENSG00000287865 | | 21.87035 | 0.05986785 | 0.4798556 | 0.1488509 | 0.8816713 | 0.944676 |
| ENSG00000131100 | ATP6V1E1 | 5502.027267 | 0.059860288 | 0.160134675 | 0.370079001 | 0.71132363 | 0.842862 |
| ENSG00000130772 | MED18 | 850.6598812 | 0.059840752 | 0.117807575 | 0.511198927 | 0.60921177 | 0.772756 |
| ENSG00000081307 | UBA5 | 2562.254637 | 0.059800931 | 0.172508529 | 0.351692536 | 0.72506886 | 0.850636 |
| ENSG00000116273 | PHF13 | 1545.868075 | 0.059708482 | 0.178506578 | 0.342729812 | 0.73180173 | 0.85446 |
| ENSG00000272483 | | 1.309227 | 0.05961443 | 0.7350304 | 0.2178145 | 0.8275737 | NA |
| ENSG00000225406 | ELF2P4 | 1.400124 | 0.05949109 | 0.7251606 | 0.2007614 | 0.8408851 | NA |
| ENSG00000180423 | HARBI1 | 333.7625 | 0.05945576 | 0.235335 | 0.2462746 | 0.8054696 | 0.90031 |
| ENSG00000239620 | PRR20G | 1.55519 | 0.05934526 | 0.7517804 | 0.2297661 | 0.8182735 | NA |
| ENSG00000219747 | RPL32P16 | 132.16 | 0.05933498 | 0.3298034 | 0.1985659 | 0.8426023 | 0.922267 |
| ENSG00000239605 | STPG4 | 1.647848 | 0.05931022 | 0.7184307 | 0.1958861 | 0.8446993 | 0.923877 |
| ENSG00000225267 | RPL8P2 | 3.313235 | 0.05925655 | 0.6708642 | 0.1652407 | 0.8687545 | 0.937071 |
| ENSG00000227230 | | 5.405449 | 0.05925409 | 0.59569 | 0.1483848 | 0.8820391 | 0.944807 |
| ENSG00000152795 | HNRNPDL | 13908.07 | 0.05924797 | 0.2105628 | 0.2887119 | 0.7728018 | 0.880282 |
| ENSG00000277938 | | 40.81422 | 0.05924442 | 0.4046255 | 0.1688998 | 0.8658755 | 0.936028 |
| ENSG00000229656 | ITGB1-DT | 5.144003 | 0.05914288 | 0.6087463 | 0.1532116 | 0.8782314 | 0.942663 |
| ENSG00000279513 | | 6.140636 | 0.05908804 | 0.5552946 | 0.1433945 | 0.8859786 | 0.946504 |
| ENSG00000232553 | CLK2P1 | 201.2193 | 0.05903696 | 0.2091644 | 0.2883055 | 0.7731129 | 0.880445 |
| ENSG00000001036 | FUCA2 | 3239.190411 | 0.059020516 | 0.182646171 | 0.331284712 | 0.74042944 | 0.859811 |
| ENSG00000162814 | SPATA17 | 319.1718 | 0.0590157 | 0.2295873 | 0.2614473 | 0.7937475 | 0.893302 |
| ENSG00000203804 | ADAMTSL4-AS1 | 66.36817 | 0.0589921 | 0.4183445 | 0.1643554 | 0.8694514 | 0.937402 |
| ENSG00000237977 | EIF4HP2 | 71.46518 | 0.05895281 | 0.413638 | 0.1623272 | 0.8710482 | 0.938517 |
| ENSG00000212497 | RNA5SP465 | 0.8783968 | 0.05878642 | 0.7677355 | 0.2672197 | 0.7893 | NA |
| ENSG00000060303 | RPS17P5 | 1.68367211 | 0.058704066 | 0.68753455 | 0.176150982 | 0.86017533 | 0.932702 |
| ENSG00000237991 | RPL35P1 | 106.4961 | 0.05867526 | 0.2769883 | 0.2283557 | 0.8193698 | 0.908629 |
| ENSG00000228594 | FNDC10 | 1976.746 | 0.05857573 | 0.2699529 | 0.2203425 | 0.8256045 | 0.91192 |
| ENSG00000142396 | ERVK3-1 | 550.6199 | 0.05857531 | 0.1890294 | 0.3233498 | 0.7464303 | 0.863806 |
| ENSG00000119522 | DENND1A | 1695.008603 | 0.058563412 | 0.144138731 | 0.381059747 | 0.70315892 | 0.83852 |
| ENSG00000112624 | BICRAL | 1270.339112 | 0.058551438 | 0.240158983 | 0.256542702 | 0.79753181 | 0.895607 |

| | | | | | | | |
|---|---|---|---|---|---|---|---|
| ENSG00000249751 | ECSCR | 1.002435 | 0.05853789 | 0.7556478 | 0.2307174 | 0.8175344 | NA |
| ENSG00000141026 | MED9 | 1196.099 | 0.05851659 | 0.1556522 | 0.4101816 | 0.6816727 | 0.823808 |
| ENSG00000243926 | TIPARP-AS1 | 166.8693 | 0.05848962 | 0.2661564 | 0.2320971 | 0.8164626 | 0.90654 |
| ENSG00000147570 | DNAJC5B | 0.9898016 | 0.05846949 | 0.7433718 | 0.2064573 | 0.8364337 | NA |
| ENSG00000145888 | GLRA1 | 1.098723 | 0.05843638 | 0.7225761 | 0.1910867 | 0.8484577 | NA |
| ENSG00000223496 | EXOSC6 | 1915.93 | 0.0584356 | 0.1507626 | 0.4099105 | 0.6818716 | 0.823896 |
| ENSG00000203883 | SOX18 | 54.50075 | 0.05843471 | 0.4507395 | 0.1539043 | 0.8776852 | 0.942353 |
| ENSG00000235545 | | 2.531801 | 0.05839892 | 0.671447 | 0.1576527 | 0.8747305 | 0.940788 |
| ENSG00000230435 | | 0.8326724 | 0.05836389 | 0.7646572 | 0.1759621 | 0.8603237 | NA |
| ENSG00000223694 | ADH5P3 | 0.9737129 | 0.05834105 | 0.7294189 | 0.1981766 | 0.8429069 | NA |
| ENSG00000125170 | DOK4 | 835.5306115 | 0.058333437 | 0.211179983 | 0.265964414 | 0.79026662 | 0.891266 |
| ENSG00000240125 | RPL23AP75 | 1.090139 | 0.0583149 | 0.7326639 | 0.1948323 | 0.8455242 | NA |
| ENSG00000221879 | MRPS21P3 | 1.078605 | 0.05826974 | 0.7177626 | 0.1712219 | 0.8640493 | NA |
| ENSG00000137857 | DUOX1 | 84.41256173 | 0.058256934 | 0.450143161 | 0.148612641 | 0.8818593 | 0.944788 |
| ENSG00000262136 | | 39.55637 | 0.05819651 | 0.3529509 | 0.1846045 | 0.8535392 | 0.928728 |
| ENSG00000111269 | CREBL2 | 2030.707214 | 0.058181571 | 0.13756005 | 0.425504471 | 0.67046896 | 0.816652 |
| ENSG00000163466 | ARPC2 | 11824.03 | 0.0581436 | 0.1715215 | 0.3420112 | 0.7323425 | 0.854968 |
| ENSG00000165476 | REEP3 | 3502.87 | 0.05811451 | 0.1509641 | 0.3812705 | 0.7030026 | 0.838447 |
| ENSG00000237514 | PTP4A1P7 | 0.8942101 | 0.05810129 | 0.7609319 | 0.2370548 | 0.8126143 | NA |
| ENSG00000258572 | | 0.7726217 | 0.05803518 | 0.7550155 | 0.1994315 | 0.8419252 | NA |
| ENSG00000280168 | | 4.144744 | 0.05803398 | 0.6715855 | 0.1661389 | 0.8680476 | 0.936731 |
| ENSG00000160678 | S100A1 | 121.224 | 0.05800267 | 0.4312327 | 0.15685 | 0.8753631 | 0.941143 |
| ENSG00000255008 | LINC02739 | 1.252365 | 0.05796703 | 0.7213088 | 0.1887086 | 0.8503212 | NA |
| ENSG00000250742 | LINC02381 | 16.81421 | 0.0579572 | 0.5488025 | 0.1413 | 0.8876329 | 0.94732 |
| ENSG00000277462 | ZNF670 | 357.184 | 0.05792589 | 0.2559151 | 0.2342209 | 0.8148135 | 0.905807 |
| ENSG00000178078 | STAP2 | 3411.935 | 0.05788226 | 0.1841288 | 0.3129993 | 0.7542812 | 0.868871 |
| ENSG00000226457 | RPL22P3 | 21.6961 | 0.05777532 | 0.4447316 | 0.156526 | 0.8756184 | 0.941265 |
| ENSG00000246662 | LINC00535 | 2.666629 | 0.05777279 | 0.6846984 | 0.1618923 | 0.8713907 | 0.938681 |
| ENSG00000278991 | | 25.24112 | 0.05770284 | 0.4992444 | 0.1427235 | 0.8865085 | 0.946696 |
| ENSG00000286733 | | 2.075158 | 0.05766625 | 0.665942 | 0.1542962 | 0.8773762 | 0.94216 |
| ENSG00000184144 | CNTN2 | 2.795065 | 0.05762153 | 0.6894198 | 0.173543 | 0.8622246 | 0.933895 |
| ENSG00000143340 | FAM163A | 3.776852 | 0.05760466 | 0.6358333 | 0.1443124 | 0.8852538 | 0.94607 |
| ENSG00000255298 | OR8G5 | 12.84061 | 0.0575887 | 0.5239699 | 0.143771 | 0.8856813 | 0.946301 |

| | | | | | | | |
|---|---|---|---|---|---|---|---|
| ENSG00000071246 | VASH1 | 1089.113106 | 0.057511682 | 0.225950155 | 0.28419148 | 0.77626364 | 0.882183 |
| ENSG00000135722 | FBXL8 | 946.7174807 | 0.057498989 | 0.166849543 | 0.349942832 | 0.7263816 | 0.851499 |
| ENSG00000135913 | USP37 | 1129.592546 | 0.05728229 | 0.334718429 | 0.181259201 | 0.85616413 | 0.930273 |
| ENSG00000232876 | | 2.127528 | 0.05727119 | 0.6906393 | 0.1731498 | 0.8625337 | 0.934005 |
| ENSG00000218283 | MORF4L1P1 | 7461.49 | 0.05726847 | 0.191089 | 0.3058737 | 0.7597008 | 0.871863 |
| ENSG00000279862 | | 1.020734 | 0.057261 | 0.7322603 | 0.207845 | 0.83535 | NA |
| ENSG00000197768 | STPG3 | 36.13105 | 0.05723856 | 0.3301341 | 0.1903539 | 0.8490318 | 0.926594 |
| ENSG00000010322 | NISCH | 6834.543584 | 0.057229605 | 0.16861056 | 0.343993502 | 0.73085117 | 0.853987 |
| ENSG00000285871 | | 3.27588 | 0.05719703 | 0.6350459 | 0.1514406 | 0.8796282 | 0.943421 |
| ENSG00000133818 | RRAS2 | 1524.427727 | 0.057187717 | 0.147534271 | 0.383066411 | 0.70167052 | 0.837422 |
| ENSG00000172159 | FRMD3 | 257.5584 | 0.05711618 | 0.393959 | 0.1618647 | 0.8714124 | 0.938681 |
| ENSG00000132465 | JCHAIN | 1.501073914 | 0.057027091 | 0.70786486 | 0.174421228 | 0.86153443 | NA |
| ENSG00000112584 | FAM120B | 3449.385574 | 0.056993989 | 0.131017277 | 0.425619759 | 0.67038494 | 0.81659 |
| ENSG00000113615 | SEC24A | 2582.333796 | 0.056967155 | 0.190835194 | 0.307909989 | 0.75815082 | 0.8711 |
| ENSG00000236532 | LINC01695 | 1.86489 | 0.05694762 | 0.6932237 | 0.1431515 | 0.8861705 | 0.946535 |
| ENSG00000125997 | BPIFB9P | 1.162342504 | 0.056867915 | 0.740089427 | 0.212064141 | 0.832057 | NA |
| ENSG00000241671 | | 6.181612 | 0.05680623 | 0.6547346 | 0.1574797 | 0.8748668 | 0.9408 |
| ENSG00000267590 | NDUFA3P1 | 2.642726 | 0.05679572 | 0.6995278 | 0.165494 | 0.8685552 | 0.936894 |
| ENSG00000242169 | | 51.09894 | 0.05670697 | 0.2880666 | 0.2100746 | 0.8336095 | 0.916526 |
| ENSG00000173064 | HECTD4 | 5807.404 | 0.05665923 | 0.2908668 | 0.2059832 | 0.836804 | 0.918493 |
| ENSG00000261646 | | 1.346844 | 0.05656567 | 0.7043009 | 0.162594 | 0.8708381 | NA |
| ENSG00000092020 | PPP2R3C | 879.9816406 | 0.056521891 | 0.174908673 | 0.32879222 | 0.74231274 | 0.861237 |
| ENSG00000171119 | NRTN | 50.98181 | 0.05646896 | 0.3838515 | 0.1685471 | 0.8661529 | 0.936211 |
| ENSG00000111850 | SMIM8 | 350.9748381 | 0.056438468 | 0.188911075 | 0.312560175 | 0.75461484 | 0.86904 |
| ENSG00000221994 | ZNF630 | 349.7744 | 0.05642788 | 0.2419442 | 0.2496424 | 0.8028639 | 0.89903 |
| ENSG00000249363 | | 1.049139 | 0.05639456 | 0.7411005 | 0.2034655 | 0.8387712 | NA |
| ENSG00000144320 | LNPK | 1890.985 | 0.05639168 | 0.1593886 | 0.3679408 | 0.7129173 | 0.843639 |
| ENSG00000175548 | ALG10B | 791.8111 | 0.0563866 | 0.2033348 | 0.2875995 | 0.7736533 | 0.880665 |
| ENSG00000259039 | | 3.342945 | 0.05638584 | 0.6567676 | 0.140544 | 0.8882302 | 0.947729 |
| ENSG00000135040 | NAA35 | 2650.60925 | 0.05635662 | 0.121996522 | 0.474314412 | 0.63527572 | 0.791199 |
| ENSG00000137343 | ATAT1 | 775.5666881 | 0.05622553 | 0.292312571 | 0.208521282 | 0.83482197 | 0.91716 |
| ENSG00000133392 | MYH11 | 56.94417982 | 0.056205357 | 0.357024756 | 0.171491155 | 0.86383759 | 0.934651 |
| ENSG00000197093 | GAL3ST4 | 16.90481 | 0.05612584 | 0.5593355 | 0.1375688 | 0.8905812 | 0.949218 |

| | | | | | | | |
|---|---|---|---|---|---|---|---|
| ENSG00000254535 | PABPC4L | 54.99883 | 0.05605177 | 0.3817034 | 0.1614772 | 0.8717176 | 0.938816 |
| ENSG00000107404 | DVL1 | 4735.371369 | 0.05589823 | 0.154259686 | 0.35641964 | 0.72152632 | 0.848466 |
| ENSG00000176268 | CYCSP34 | 26.68258 | 0.05578938 | 0.4165775 | 0.1518337 | 0.8793181 | 0.943285 |
| ENSG00000264735 | | 1.130359 | 0.0557887 | 0.7317691 | 0.2038475 | 0.8384726 | NA |
| ENSG00000125741 | OPA3 | 1493.344998 | 0.055739167 | 0.152010513 | 0.376454203 | 0.70657926 | 0.840497 |
| ENSG00000118017 | A4GNT | 1.301610512 | 0.05573046 | 0.726420872 | 0.171726085 | 0.86365288 | NA |
| ENSG00000185267 | CDNF | 131.1545 | 0.05564233 | 0.3854987 | 0.1643301 | 0.8694713 | 0.937402 |
| ENSG00000008283 | CYB561 | 23494.47051 | 0.055489157 | 0.142788862 | 0.383613659 | 0.70126481 | 0.83727 |
| ENSG00000237846 | | 1.069665 | 0.05546722 | 0.7318493 | 0.1680577 | 0.8665379 | NA |
| ENSG00000162772 | ATF3 | 1242.129 | 0.05541733 | 0.5695685 | 0.1331622 | 0.8940651 | 0.950882 |
| ENSG00000245648 | KLRK1-AS1 | 5.159965 | 0.05540095 | 0.6297492 | 0.1477804 | 0.8825161 | 0.944952 |
| ENSG00000104885 | DOT1L | 2522.593765 | 0.055369341 | 0.195292935 | 0.289944273 | 0.77185887 | 0.879447 |
| ENSG00000115556 | PLCD4 | 212.8225655 | 0.055333404 | 0.178502915 | 0.334143159 | 0.73827153 | 0.858764 |
| ENSG00000023330 | ALAS1 | 1802.694407 | 0.055316552 | 0.19570315 | 0.277941661 | 0.78105714 | 0.885245 |
| ENSG00000261832 | | 1.677261 | 0.05530338 | 0.7236812 | 0.1870348 | 0.8516333 | 0.927845 |
| ENSG00000273013 | | 1.946635 | 0.05524319 | 0.7162686 | 0.1950419 | 0.8453601 | 0.924333 |
| ENSG00000207729 | MIR556 | 0.719363 | 0.05518262 | 0.7561063 | 0.1845711 | 0.8535654 | NA |
| ENSG00000276805 | | 494.2969 | 0.05515665 | 0.1985514 | 0.2825248 | 0.7775411 | 0.882917 |
| ENSG00000136044 | APPL2 | 3596.907871 | 0.05514427 | 0.157326477 | 0.343366509 | 0.73132275 | 0.854239 |
| ENSG00000197320 | | 8.214159 | 0.05503047 | 0.6377495 | 0.1586073 | 0.8739783 | 0.940225 |
| ENSG00000225285 | LINC01770 | 54.88132 | 0.05498195 | 0.3483783 | 0.1779522 | 0.8587605 | 0.931977 |
| ENSG00000135914 | HTR2B | 1.678496919 | 0.054925459 | 0.716476942 | 0.190370305 | 0.84901897 | 0.926594 |
| ENSG00000225177 | | 274.8642 | 0.05491673 | 0.2552113 | 0.2276103 | 0.8199492 | 0.908881 |
| ENSG00000141434 | MEP1B | 2.166476 | 0.05481788 | 0.681921 | 0.1573563 | 0.8749641 | 0.940866 |
| ENSG00000056586 | RC3H2 | 5061.917516 | 0.054775808 | 0.25809772 | 0.212553393 | 0.83167533 | 0.915261 |
| ENSG00000240776 | | 8.643984 | 0.05466105 | 0.5604969 | 0.1373625 | 0.8907443 | 0.949303 |
| ENSG00000068745 | IP6K2 | 5439.336147 | 0.054646905 | 0.203293463 | 0.276048425 | 0.78251087 | 0.886136 |
| ENSG00000070019 | GUCY2C | 1.687287706 | 0.054625165 | 0.693872353 | 0.154132942 | 0.87750491 | 0.942226 |
| ENSG00000087842 | PIR | 904.7979788 | 0.054615977 | 0.339820213 | 0.174815258 | 0.8612248 | 0.933459 |
| ENSG00000278678 | CYCSP41 | 0.9897518 | 0.05460409 | 0.7528138 | 0.2732797 | 0.7846382 | NA |
| ENSG00000006016 | CRLF1 | 212.3337708 | 0.054546571 | 0.453126585 | 0.148983802 | 0.88156641 | 0.944601 |
| ENSG00000135069 | PSAT1 | 3.728988629 | 0.054524696 | 0.663955196 | 0.128917412 | 0.897423 | 0.952564 |
| ENSG00000158796 | DEDD | 1625.075 | 0.05449061 | 0.1414487 | 0.3491056 | 0.72701 | 0.851749 |

| | | | | | | | |
|---|---|---|---|---|---|---|---|
| ENSG00000265565 | MIR3143 | 1.56267 | 0.05447105 | 0.7091806 | 0.159982 | 0.8728952 | NA |
| ENSG00000186625 | KATNA1 | 918.0947 | 0.05440953 | 0.1432747 | 0.3753616 | 0.7073915 | 0.840888 |
| ENSG00000085644 | ZNF213 | 2111.149753 | 0.054333832 | 0.15408116 | 0.311464095 | 0.75544783 | 0.86943 |
| ENSG00000163331 | DAPL1 | 1.37528 | 0.05431288 | 0.7557183 | 0.3043807 | 0.7608378 | NA |
| ENSG00000225905 | | 1.177859 | 0.05423402 | 0.7178673 | 0.1749163 | 0.8611454 | NA |
| ENSG00000248780 | ARL4AP2 | 12.71795 | 0.05418625 | 0.547523 | 0.132163 | 0.8948554 | 0.951385 |
| ENSG00000136381 | IREB2 | 7833.80918 | 0.054111114 | 0.16392561 | 0.320166516 | 0.74884211 | 0.865607 |
| ENSG00000168803 | ADAL | 510.2961 | 0.05404634 | 0.1633457 | 0.3431613 | 0.7314771 | 0.854344 |
| ENSG00000275374 | | 2.522882 | 0.05397012 | 0.6757581 | 0.1431679 | 0.8861576 | 0.946535 |
| ENSG00000149735 | GPHA2 | 1113.484 | 0.05394502 | 0.6555003 | 0.1733721 | 0.862359 | 0.933926 |
| ENSG00000183354 | KIAA2026 | 1093.555 | 0.05393468 | 0.275204 | 0.2035504 | 0.8387048 | 0.919744 |
| ENSG00000249047 | COX6B1P5 | 1.7119 | 0.05382951 | 0.716054 | 0.1813426 | 0.8560986 | 0.930273 |
| ENSG00000237737 | DCTN1-AS1 | 4.260775 | 0.05374888 | 0.6542678 | 0.1513822 | 0.8796742 | 0.943421 |
| ENSG00000175866 | BAIAP2 | 2714.125 | 0.05373046 | 0.17401 | 0.3254036 | 0.7448757 | 0.862599 |
| ENSG00000285943 | | 56.39483 | 0.05371442 | 0.3294599 | 0.1770465 | 0.8594719 | 0.932352 |
| ENSG00000268751 | SCGB1B2P | 10.77535 | 0.05367962 | 0.5287614 | 0.1314847 | 0.8953919 | 0.951617 |
| ENSG00000106608 | URGCP | 2633.263086 | 0.053567815 | 0.125164992 | 0.443053879 | 0.65772676 | 0.807247 |
| ENSG00000187627 | RGPD1 | 1.642372 | 0.05350737 | 0.707309 | 0.1544254 | 0.8772743 | 0.94213 |
| ENSG00000205771 | CATSPER2P1 | 29.75098 | 0.05346196 | 0.3501085 | 0.1670985 | 0.8672925 | 0.936483 |
| ENSG00000233143 | DIRC3-AS1 | 1.681537 | 0.05345898 | 0.7050048 | 0.1672487 | 0.8671744 | 0.936431 |
| ENSG00000163870 | TPRA1 | 3131.315 | 0.05342617 | 0.1502164 | 0.3629309 | 0.7166565 | 0.845846 |
| ENSG00000162517 | PEF1 | 3971.346 | 0.05340993 | 0.1914515 | 0.2810871 | 0.7786436 | 0.883565 |
| ENSG00000213467 | HMGB1P37 | 2.377749 | 0.0533745 | 0.6701826 | 0.1472812 | 0.8829101 | 0.945233 |
| ENSG00000164086 | DUSP7 | 706.746 | 0.05335531 | 0.1731424 | 0.3082478 | 0.7578938 | 0.87103 |
| ENSG00000254589 | EIF4A2P3 | 42.55678 | 0.05333396 | 0.4035184 | 0.1506507 | 0.8802513 | 0.943649 |
| ENSG00000140157 | NIPA2 | 2033.696 | 0.05329967 | 0.1837003 | 0.2969672 | 0.7664916 | 0.876184 |
| ENSG00000213332 | SLC25A5P6 | 1.046101 | 0.05324795 | 0.7394484 | 0.2067162 | 0.8362315 | NA |
| ENSG00000168096 | ANKS3 | 1533.315 | 0.05324331 | 0.1533343 | 0.3447344 | 0.730294 | 0.853877 |
| ENSG00000126934 | MAP2K2 | 7124.187059 | 0.053233286 | 0.144051178 | 0.378043518 | 0.70539827 | 0.839947 |
| ENSG00000278881 | | 0.8355673 | 0.05309046 | 0.7213893 | 0.1711171 | 0.8641317 | NA |
| ENSG00000145817 | YIPF5 | 2433.781 | 0.05308387 | 0.1248618 | 0.4360971 | 0.6627663 | 0.811103 |
| ENSG00000174243 | DDX23 | 9816.672 | 0.05307394 | 0.1623634 | 0.3351 | 0.7375497 | 0.858342 |
| ENSG00000267692 | TAF9P3 | 4.288477 | 0.05305746 | 0.6477069 | 0.142073 | 0.8870224 | 0.947034 |

| | | | | | | | |
|---|---|---|---|---|---|---|---|
| ENSG00000286672 | | 5.52377 | 0.05303478 | 0.6247297 | 0.1360297 | 0.8917978 | 0.949991 |
| ENSG00000146409 | SLC18B1 | 782.6767 | 0.05299481 | 0.1912372 | 0.2819563 | 0.777977 | 0.883261 |
| ENSG00000173812 | EIF1 | 21820.81 | 0.05297224 | 0.2038719 | 0.2653665 | 0.7907271 | 0.891451 |
| ENSG00000164164 | OTUD4 | 5303.556 | 0.05281317 | 0.1834921 | 0.2977411 | 0.7659008 | 0.875765 |
| ENSG00000287707 | | 15.01659 | 0.05280543 | 0.4927483 | 0.1304477 | 0.8962123 | 0.952078 |
| ENSG00000276710 | CSPG4P10 | 235.1428 | 0.05273266 | 0.2380482 | 0.236751 | 0.81285 | 0.904346 |
| ENSG00000244055 | | 145.4454 | 0.05272027 | 0.5138575 | 0.1556361 | 0.8763199 | 0.941652 |
| ENSG00000112164 | GLP1R | 20.90840862 | 0.052690284 | 0.700631514 | 0.191821611 | 0.84788194 | 0.92613 |
| ENSG00000231752 | EMBP1 | 76.37825 | 0.05266012 | 0.2729482 | 0.2017227 | 0.8401335 | 0.92093 |
| ENSG00000114867 | EIF4G1 | 35284.81701 | 0.052656174 | 0.174475372 | 0.306756962 | 0.75902837 | 0.87158 |
| ENSG00000234648 | | 754.7277 | 0.05260759 | 0.1615609 | 0.3349039 | 0.7376975 | 0.858439 |
| ENSG00000230444 | TFAMP1 | 7.118481 | 0.05258037 | 0.644829 | 0.1292973 | 0.8971224 | 0.952406 |
| ENSG00000254398 | | 1.190086 | 0.05254323 | 0.7019542 | 0.1569772 | 0.8752628 | NA |
| ENSG00000178440 | TIMM23B-AGAP | 29.87074 | 0.05251726 | 0.4011885 | 0.147694 | 0.8825843 | 0.944968 |
| ENSG00000265545 | ELOCP27 | 6.138784 | 0.05251637 | 0.6499383 | 0.1392419 | 0.889259 | 0.948522 |
| ENSG00000139826 | ABHD13 | 852.7463 | 0.05248569 | 0.1480554 | 0.3578678 | 0.7204423 | 0.847936 |
| ENSG00000126790 | L3HYPDH | 1002.246998 | 0.052480606 | 0.167132406 | 0.323046179 | 0.74666027 | 0.863912 |
| ENSG00000155438 | NIFK | 2966.261 | 0.05243057 | 0.1735835 | 0.3000742 | 0.7641206 | 0.874555 |
| ENSG00000173818 | ENDOV | 2282.425 | 0.05238532 | 0.1239561 | 0.4016045 | 0.6879752 | 0.828129 |
| ENSG00000109118 | PHF12 | 4143.722711 | 0.052309071 | 0.10528579 | 0.496524917 | 0.61952411 | 0.780127 |
| ENSG00000169976 | SF3B5 | 4043.113 | 0.05222357 | 0.2376696 | 0.2273795 | 0.8201286 | 0.908902 |
| ENSG00000236591 | UST-AS2 | 4.462908 | 0.05220683 | 0.6637056 | 0.1280905 | 0.8980774 | 0.952878 |
| ENSG00000117281 | CD160 | 36.71881103 | 0.052149219 | 0.411270276 | 0.143170588 | 0.88615546 | 0.946535 |
| ENSG00000152642 | GPD1L | 7875.942 | 0.05204361 | 0.1852934 | 0.2774508 | 0.781434 | 0.885483 |
| ENSG00000269889 | | 32.61818 | 0.05203175 | 0.3609452 | 0.1595842 | 0.8732086 | 0.939968 |
| ENSG00000137218 | FRS3 | 303.9166333 | 0.051942269 | 0.189176375 | 0.280334615 | 0.7792208 | 0.884097 |
| ENSG00000256037 | MRPL40P1 | 6.627714 | 0.05190274 | 0.576347 | 0.1258877 | 0.8998208 | 0.953966 |
| ENSG00000166200 | COPS2 | 4803.488 | 0.05187671 | 0.1861912 | 0.2851203 | 0.775552 | 0.881827 |
| ENSG00000199545 | RNA5SP195 | 0.9208772 | 0.05178261 | 0.7379891 | 0.1826872 | 0.8550435 | NA |
| ENSG00000278276 | | 1.378023 | 0.05177077 | 0.6975468 | 0.1534438 | 0.8780483 | NA |
| ENSG00000224794 | | 3.297309 | 0.05163607 | 0.6611794 | 0.147083 | 0.8830665 | 0.94524 |
| ENSG00000258661 | | 1.38532 | 0.05160585 | 0.7275273 | 0.1552599 | 0.8766164 | NA |
| ENSG00000169925 | BRD3 | 3062.215 | 0.05156466 | 0.2139524 | 0.2363303 | 0.8131764 | 0.90463 |

| | | | | | | | |
|---|---|---|---|---|---|---|---|
| ENSG00000260318 | COX6CP1 | 13.50244 | 0.05155905 | 0.457744 | 0.1349543 | 0.892648 | 0.950211 |
| ENSG00000224628 | | 346.2974 | 0.05155787 | 0.2352618 | 0.2175747 | 0.8277605 | 0.913131 |
| ENSG00000255992 | | 14.97041 | 0.05155103 | 0.4521759 | 0.1336989 | 0.8936407 | 0.950721 |
| ENSG00000237324 | | 0.7184356 | 0.05153248 | 0.76061 | 0.1578457 | 0.8745784 | NA |
| ENSG00000277198 | RN7SL270P | 1.987591 | 0.05153114 | 0.7180623 | 0.1708896 | 0.8643105 | 0.934973 |
| ENSG00000270361 | | 3.692644 | 0.0514943 | 0.6832829 | 0.1504022 | 0.8804473 | 0.943745 |
| ENSG00000267673 | FDX2 | 8.598592 | 0.05148538 | 0.4918926 | 0.1318509 | 0.8951022 | 0.951583 |
| ENSG00000144791 | LIMD1 | 1221.139 | 0.05145399 | 0.2166096 | 0.2440097 | 0.8072233 | 0.901215 |
| ENSG00000254974 | | 2.025958 | 0.05141561 | 0.6896765 | 0.1487465 | 0.8817537 | 0.944726 |
| ENSG00000162426 | SLC45A1 | 8.023462 | 0.05140665 | 0.5334069 | 0.1217783 | 0.9030746 | 0.955125 |
| ENSG00000110321 | EIF4G2 | 38899.42431 | 0.051361676 | 0.173840404 | 0.341945854 | 0.73239164 | 0.854968 |
| ENSG00000261266 | | 0.8528065 | 0.0513511 | 0.7405709 | 0.1643977 | 0.8694181 | NA |
| ENSG00000182095 | TNRC18 | 16369.03 | 0.05127971 | 0.203503 | 0.2521262 | 0.8009435 | 0.897864 |
| ENSG00000070423 | RNF126 | 1854.279787 | 0.051275206 | 0.27323071 | 0.208267049 | 0.83502046 | 0.917253 |
| ENSG00000275674 | | 1.482559 | 0.05122666 | 0.7152084 | 0.1730637 | 0.8626014 | NA |
| ENSG00000256678 | | 2.422913 | 0.05121516 | 0.7172276 | 0.1704191 | 0.8646806 | 0.935259 |
| ENSG00000273444 | | 0.9803106 | 0.05121145 | 0.7384825 | 0.174594 | 0.8613986 | NA |
| ENSG00000125798 | FOXA2 | 1.323790665 | 0.051159994 | 0.78809806 | 0.143288953 | 0.88606198 | NA |
| ENSG00000221953 | C1orf229 | 87.53702 | 0.05112861 | 0.3449715 | 0.1548273 | 0.8769575 | 0.941942 |
| ENSG00000117448 | AKR1A1 | 4499.143909 | 0.051084922 | 0.154404516 | 0.34439424 | 0.73054982 | 0.853899 |
| ENSG00000059122 | FLYWCH1 | 2850.505782 | 0.05104387 | 0.145138898 | 0.356688927 | 0.7213247 | 0.848466 |
| ENSG00000164331 | ANKRA2 | 865.0371 | 0.05099144 | 0.2067132 | 0.2608688 | 0.7941937 | 0.893577 |
| ENSG00000107185 | RGP1 | 1950.092361 | 0.050970999 | 0.170130695 | 0.30117565 | 0.76328056 | 0.874121 |
| ENSG00000158805 | ZNF276 | 2418.277 | 0.05096426 | 0.1341804 | 0.341852 | 0.7324623 | 0.854968 |
| ENSG00000048649 | RSF1 | 4929.19378 | 0.050838503 | 0.175861808 | 0.291704805 | 0.77051234 | 0.878627 |
| ENSG00000167881 | SRP68 | 7693.468 | 0.05082153 | 0.1441196 | 0.362096 | 0.7172803 | 0.846015 |
| ENSG00000169972 | PUSL1 | 970.1927 | 0.05080069 | 0.1618562 | 0.3183625 | 0.75021 | 0.866263 |
| ENSG00000183586 | HMGN3P1 | 8.663291 | 0.05074265 | 0.5469758 | 0.1263848 | 0.8994274 | 0.953788 |
| ENSG00000228224 | NACA4P | 215.2489 | 0.0506943 | 0.1789538 | 0.3101267 | 0.7564646 | 0.869898 |
| ENSG00000160888 | IER2 | 4340.933 | 0.05054641 | 0.3339794 | 0.1859228 | 0.8525053 | 0.928288 |
| ENSG00000288062 | | 3.1958 | 0.05052502 | 0.6538781 | 0.1336908 | 0.893647 | 0.950721 |
| ENSG00000213023 | SYT3 | 82.50903 | 0.05052361 | 0.2869021 | 0.1884713 | 0.8505072 | 0.927366 |
| ENSG00000128731 | HERC2 | 4253.409899 | 0.050491767 | 0.225974915 | 0.229117242 | 0.81877779 | 0.908239 |

| | | | | | | | |
|---|---|---|---|---|---|---|---|
| ENSG00000281756 | C2-AS1 | 2.093061 | 0.05044975 | 0.7091062 | 0.1663393 | 0.8678899 | 0.936731 |
| ENSG00000176102 | CSTF3 | 1906.231 | 0.05043719 | 0.1294402 | 0.3866874 | 0.6989877 | 0.835817 |
| ENSG00000225125 | RANP4 | 16.10586 | 0.05036714 | 0.595253 | 0.1222231 | 0.9027223 | 0.955022 |
| ENSG00000174021 | GNG5 | 654.7617 | 0.0503362 | 0.2000881 | 0.2484499 | 0.8037863 | 0.89957 |
| ENSG00000267212 | | 2.497773 | 0.05032243 | 0.6348774 | 0.1265562 | 0.8992917 | 0.953709 |
| ENSG00000088986 | DYNLL1 | 6604.109457 | 0.050252613 | 0.272421271 | 0.201272765 | 0.8404853 | 0.921114 |
| ENSG00000285219 | HULC | 121.1822 | 0.05015227 | 0.1976734 | 0.2595526 | 0.7952089 | 0.894202 |
| ENSG00000185621 | LMLN | 659.7092 | 0.05010243 | 0.2475958 | 0.2262535 | 0.8210042 | 0.909205 |
| ENSG00000279198 | | 70.6554 | 0.05005098 | 0.3135863 | 0.1765107 | 0.8598928 | 0.932571 |
| ENSG00000178732 | GP5 | 0.8052616 | 0.05003708 | 0.731237 | 0.1704195 | 0.8646802 | NA |
| ENSG00000259736 | CRTC3-AS1 | 20.83914 | 0.05003185 | 0.4112669 | 0.1387378 | 0.8896573 | 0.948692 |
| ENSG00000215030 | RPL13P12 | 14034.97 | 0.05002292 | 0.1929046 | 0.2658529 | 0.7903525 | 0.891266 |
| ENSG00000271401 | | 0.9219151 | 0.04999466 | 0.7430189 | 0.165056 | 0.8688999 | NA |
| ENSG00000221978 | CCNL2 | 13767.72 | 0.04999271 | 0.259234 | 0.1994906 | 0.841879 | 0.922045 |
| ENSG00000147854 | UHRF2 | 2283.329 | 0.04997806 | 0.1745156 | 0.2866896 | 0.77435 | 0.880987 |
| ENSG00000119899 | SLC17A5 | 1444.465125 | 0.049948911 | 0.240027514 | 0.216662722 | 0.8284712 | 0.913464 |
| ENSG00000095139 | ARCN1 | 12125.78775 | 0.049874651 | 0.120439284 | 0.41723358 | 0.67650757 | 0.820122 |
| ENSG00000166105 | GLB1L3 | 0.9943282 | 0.04986727 | 0.7536549 | 0.2132573 | 0.8311263 | NA |
| ENSG00000171603 | CLSTN1 | 8078.614 | 0.04981158 | 0.1637706 | 0.3132454 | 0.7540942 | 0.868768 |
| ENSG00000272646 | | 1.863583 | 0.04976009 | 0.6822871 | 0.1291442 | 0.8972436 | 0.952412 |
| ENSG00000236051 | MYCBP2-AS1 | 1.592806 | 0.04970113 | 0.6655719 | 0.1367454 | 0.8912321 | 0.949585 |
| ENSG00000125304 | TM9SF2 | 10593.34338 | 0.049646953 | 0.134994264 | 0.351619709 | 0.72512348 | 0.850663 |
| ENSG00000264483 | MIR5008 | 1.741505 | 0.04953598 | 0.7329919 | 0.1890561 | 0.8500488 | 0.927094 |
| ENSG00000107611 | CUBN | 1.897128903 | 0.04951434 | 0.709223889 | 0.143012689 | 0.88628016 | 0.946598 |
| ENSG00000134748 | PRPF38A | 2979.010881 | 0.04940361 | 0.163498763 | 0.305292678 | 0.76014325 | 0.872144 |
| ENSG00000282978 | | 187.4565 | 0.04934825 | 0.2768515 | 0.1893794 | 0.8497954 | 0.926993 |
| ENSG00000119231 | SENP5 | 2905.190846 | 0.049220662 | 0.216693943 | 0.227614304 | 0.8199461 | 0.908881 |
| ENSG00000267708 | RDM1P3 | 3.598412 | 0.04920645 | 0.6687975 | 0.1445166 | 0.8850926 | 0.946052 |
| ENSG00000164002 | EXO5 | 291.046 | 0.04918268 | 0.1575401 | 0.3128498 | 0.7543948 | 0.868926 |
| ENSG00000248817 | PMPCAP1 | 2.327878 | 0.04918212 | 0.6747126 | 0.1388997 | 0.8895294 | 0.948689 |
| ENSG00000128228 | SDF2L1 | 1351.131401 | 0.049113923 | 0.287958371 | 0.167885657 | 0.86667323 | 0.936304 |
| ENSG00000111880 | RNGTT | 1974.122562 | 0.049077639 | 0.127779124 | 0.381328492 | 0.70295951 | 0.838433 |
| ENSG00000264314 | MIR548AT | 0.903724 | 0.04906054 | 0.7604127 | 0.2054187 | 0.837245 | NA |

| | | | | | | | |
|---|---|---|---|---|---|---|---|
| ENSG00000279412 | | 19.88513 | 0.0490534 | 0.5041653 | 0.1225274 | 0.9024813 | 0.954999 |
| ENSG00000163636 | PSMD6 | 56745.08 | 0.04904204 | 0.1521279 | 0.3274667 | 0.7433149 | 0.86158 |
| ENSG00000286989 | | 68.39624 | 0.04895196 | 0.3461571 | 0.1567795 | 0.8754186 | 0.941165 |
| ENSG00000105176 | URI1 | 4933.624529 | 0.04892849 | 0.124695848 | 0.413971392 | 0.67889506 | 0.82157 |
| ENSG00000135763 | URB2 | 1769.027226 | 0.048915573 | 0.223271809 | 0.22570237 | 0.82143292 | 0.909357 |
| ENSG00000273557 | | 0.9271151 | 0.04891237 | 0.7171601 | 0.1578586 | 0.8745683 | NA |
| ENSG00000017260 | ATP2C1 | 7360.260501 | 0.048893545 | 0.130460875 | 0.370215301 | 0.71122208 | 0.842862 |
| ENSG00000116871 | MAP7D1 | 3913.690798 | 0.048732078 | 0.175103637 | 0.291648471 | 0.77055541 | 0.878627 |
| ENSG00000258930 | | 0.7658479 | 0.04871461 | 0.7513031 | 0.1773732 | 0.8592152 | NA |
| ENSG00000286403 | | 0.9625854 | 0.04870097 | 0.7394129 | 0.1704059 | 0.8646909 | NA |
| ENSG00000112062 | MAPK14 | 5006.350589 | 0.048587102 | 0.111657062 | 0.428451235 | 0.66832263 | 0.815124 |
| ENSG00000214546 | | 2.204901 | 0.04858134 | 0.6383769 | 0.1205574 | 0.9040417 | 0.95555 |
| ENSG00000175354 | PTPN2 | 2519.626 | 0.048535 | 0.1167997 | 0.4187855 | 0.6753729 | 0.81957 |
| ENSG00000201916 | | 0.8874193 | 0.04844448 | 0.7124378 | 0.1447683 | 0.8848938 | NA |
| ENSG00000166783 | MARF1 | 4228.114 | 0.04836981 | 0.2347827 | 0.2537746 | 0.7996697 | 0.897041 |
| ENSG00000091879 | ANGPT2 | 3.685621283 | 0.04828692 | 0.650243998 | 0.136841279 | 0.89115626 | 0.949554 |
| ENSG00000100722 | ZC3H14 | 6032.096137 | 0.048278342 | 0.134578578 | 0.362045409 | 0.7173181 | 0.846015 |
| ENSG00000206762 | RNU6-418P | 9.012623 | 0.04826417 | 0.5586657 | 0.1215462 | 0.9032584 | 0.955125 |
| ENSG00000082068 | WDR70 | 2068.528171 | 0.04826262 | 0.129697153 | 0.373102643 | 0.70907205 | 0.841793 |
| ENSG00000280239 | | 48.51636 | 0.04817199 | 0.3416075 | 0.1561274 | 0.8759326 | 0.941412 |
| ENSG00000214549 | SDHCP2 | 276.9327 | 0.04814954 | 0.1926221 | 0.2619814 | 0.7933358 | 0.893115 |
| ENSG00000264324 | | 1.226115 | 0.04811969 | 0.7227841 | 0.1716893 | 0.8636818 | NA |
| ENSG00000229348 | HYI-AS1 | 1.918496 | 0.04806769 | 0.6763739 | 0.1279421 | 0.8981948 | 0.952965 |
| ENSG00000286948 | | 11.20687 | 0.04805261 | 0.5327427 | 0.1171099 | 0.906773 | 0.956735 |
| ENSG00000256347 | OR8R1P | 1.911551 | 0.04802371 | 0.7083097 | 0.1537571 | 0.8778012 | 0.942353 |
| ENSG00000084733 | RAB10 | 10886.81444 | 0.047922123 | 0.151088718 | 0.343474121 | 0.73124181 | 0.854182 |
| ENSG00000223188 | | 0.9444012 | 0.04790109 | 0.7392938 | 0.1664108 | 0.8678337 | NA |
| ENSG00000240477 | | 2.114881 | 0.04788811 | 0.7003757 | 0.1423574 | 0.8867977 | 0.946885 |
| ENSG00000214955 | | 3.765419 | 0.04786665 | 0.6382053 | 0.1239664 | 0.9013419 | 0.954626 |
| ENSG00000264589 | MAPT-AS1 | 26.40043 | 0.04773941 | 0.4752074 | 0.1253834 | 0.90022 | 0.954161 |
| ENSG00000173404 | INSM1 | 1.079834 | 0.0477273 | 0.7226115 | 0.1455498 | 0.8842768 | NA |
| ENSG00000106351 | AGFG2 | 954.2258126 | 0.047706954 | 0.218579229 | 0.224951243 | 0.82201721 | 0.909628 |
| ENSG00000278982 | | 2.518548 | 0.0476812 | 0.6986546 | 0.1286421 | 0.8976409 | 0.952681 |

| | | | | | | | |
|---|---|---|---|---|---|---|---|
| ENSG00000047579 | DTNBP1 | 1266.922742 | 0.047669181 | 0.194285069 | 0.248999119 | 0.80336146 | 0.899322 |
| ENSG00000259592 | PRELID1P4 | 3.55739 | 0.04764165 | 0.6344213 | 0.1232425 | 0.9019151 | 0.954853 |
| ENSG00000138448 | ITGAV | 7018.562987 | 0.047633054 | 0.207630328 | 0.232823602 | 0.81589839 | 0.906211 |
| ENSG00000287502 | | 5.375499 | 0.04758395 | 0.551741 | 0.1146627 | 0.9087125 | 0.957395 |
| ENSG00000164615 | CAMLG | 3261.749 | 0.04756794 | 0.159658 | 0.3018678 | 0.7627528 | 0.87378 |
| ENSG00000115339 | GALNT3 | 3455.60274 | 0.047514598 | 0.190851281 | 0.254309295 | 0.79925662 | 0.896956 |
| ENSG00000126856 | PRDM7 | 1.420843704 | 0.047463739 | 0.693193278 | 0.137760077 | 0.89043004 | NA |
| ENSG00000184188 | | 1.554199 | 0.04742946 | 0.7255149 | 0.1739882 | 0.8618747 | NA |
| ENSG00000105701 | FKBP8 | 9081.944807 | 0.047395921 | 0.203511756 | 0.237481291 | 0.81228343 | 0.9042 |
| ENSG00000176318 | FOXN3P1 | 0.8415307 | 0.04737374 | 0.7625365 | 0.2256518 | 0.8214723 | NA |
| ENSG00000146966 | DENND2A | 1.471719 | 0.04736763 | 0.7274458 | 0.1685779 | 0.8661286 | NA |
| ENSG00000286531 | | 1.237514 | 0.04733642 | 0.7435544 | 0.1856713 | 0.8527025 | NA |
| ENSG00000166856 | GPR182 | 1.074781 | 0.04730681 | 0.7265539 | 0.1606138 | 0.8723976 | NA |
| ENSG00000177954 | RPS27 | 2081.772 | 0.04728944 | 0.2616773 | 0.1873619 | 0.8513769 | 0.927819 |
| ENSG00000187172 | BAGE2 | 542.2898 | 0.04726957 | 0.292067 | 0.1590796 | 0.8736062 | 0.940119 |
| ENSG00000115425 | PECR | 416.3149161 | 0.047096762 | 0.273853152 | 0.18113169 | 0.85626421 | 0.930284 |
| ENSG00000148180 | GSN | 2653.435 | 0.04706062 | 0.3733345 | 0.1372341 | 0.8908457 | 0.949337 |
| ENSG00000199805 | RNU1-134P | 0.836115 | 0.0470588 | 0.7370913 | 0.1755159 | 0.8606743 | NA |
| ENSG00000244307 | RN7SL395P | 3.575179 | 0.04702416 | 0.6402534 | 0.1219506 | 0.9029382 | 0.955046 |
| ENSG00000141219 | C17orf80 | 1851.265 | 0.04698593 | 0.1595598 | 0.296093 | 0.767159 | 0.876586 |
| ENSG00000117410 | ATP6V0B | 7701.311135 | 0.046935517 | 0.193716704 | 0.218890013 | 0.82673573 | 0.912569 |
| ENSG00000158301 | GPRASP2 | 904.2531 | 0.04689134 | 0.1635576 | 0.2854073 | 0.7753321 | 0.88169 |
| ENSG00000213316 | LTC4S | 138.4946 | 0.04685438 | 0.2294255 | 0.2118101 | 0.8322552 | 0.915608 |
| ENSG00000101856 | PGRMC1 | 7139.287883 | 0.046831886 | 0.213101772 | 0.226235814 | 0.82101802 | 0.909205 |
| ENSG00000130307 | USHBP1 | 12.71144936 | 0.046804146 | 0.504583085 | 0.119035147 | 0.90524751 | 0.955981 |
| ENSG00000279633 | | 1.972496 | 0.04676988 | 0.7094314 | 0.1333089 | 0.8939491 | 0.950882 |
| ENSG00000048462 | TNFRSF17 | 0.827647326 | 0.046730897 | 0.744347349 | 0.169056574 | 0.86575214 | NA |
| ENSG00000272702 | | 23.39216 | 0.04671788 | 0.4974805 | 0.1229568 | 0.9021413 | 0.95494 |
| ENSG00000207291 | RNU6-30P | 3.29539 | 0.046646 | 0.6434173 | 0.1235918 | 0.9016385 | 0.954802 |
| ENSG00000105549 | THEG | 70.85854313 | 0.046617881 | 0.48035399 | 0.111716583 | 0.91104813 | 0.958677 |
| ENSG00000108960 | MMD | 994.9683776 | 0.046594642 | 0.283066289 | 0.182938786 | 0.85484605 | 0.929731 |
| ENSG00000122778 | KIAA1549 | 1308.84572 | 0.046590107 | 0.212912162 | 0.226098728 | 0.82112463 | 0.909205 |
| ENSG00000279637 | | 1.765009 | 0.04658484 | 0.6555668 | 0.120371 | 0.9041892 | 0.955623 |

| | | | | | | | |
|---|---|---|---|---|---|---|---|
| ENSG00000236901 | MIR600HG | 403.4135 | 0.04654473 | 0.2462788 | 0.1936518 | 0.8464485 | 0.925028 |
| ENSG00000117632 | STMN1 | 7926.157239 | 0.04653797 | 0.376860231 | 0.133726204 | 0.89361908 | 0.950721 |
| ENSG00000280078 | | 0.9279118 | 0.04651355 | 0.741268 | 0.1450464 | 0.8846742 | NA |
| ENSG00000250643 | | 5.113514 | 0.04651226 | 0.5789036 | 0.1000986 | 0.9202661 | 0.962052 |
| ENSG00000135951 | TSGA10 | 503.9805796 | 0.04644816 | 0.216933636 | 0.233001084 | 0.81576056 | 0.906139 |
| ENSG00000164035 | EMCN | 255.7958 | 0.0464411 | 0.2110062 | 0.2331672 | 0.8156316 | 0.90611 |
| ENSG00000096070 | BRPF3 | 3991.307264 | 0.046420739 | 0.223703363 | 0.209140517 | 0.83433855 | 0.916937 |
| ENSG00000262623 | | 1.409948 | 0.04641679 | 0.7004937 | 0.1451434 | 0.8845977 | NA |
| ENSG00000179304 | FAM156B | 1.802359 | 0.04629975 | 0.7215765 | 0.1618693 | 0.8714087 | 0.938681 |
| ENSG00000277602 | | 2.873077 | 0.04629158 | 0.6828619 | 0.1354721 | 0.8922387 | 0.950182 |
| ENSG00000228149 | RPL3P1 | 3.816622 | 0.04621062 | 0.6411506 | 0.1184976 | 0.9056734 | 0.956179 |
| ENSG00000278447 | MIR6781 | 1.18956 | 0.04613283 | 0.708716 | 0.1425251 | 0.8866652 | NA |
| ENSG00000137154 | RPS6 | 31068.8751 | 0.046132034 | 0.181128332 | 0.258796641 | 0.79579215 | 0.89458 |
| ENSG00000163815 | CLEC3B | 1.417403 | 0.04606458 | 0.6781488 | 0.1253214 | 0.9002691 | NA |
| ENSG00000270760 | | 2.664776 | 0.04602507 | 0.6697738 | 0.1300073 | 0.8965606 | 0.952334 |
| ENSG00000204532 | ZSCAN5C | 1.960653 | 0.04602235 | 0.6993367 | 0.1355466 | 0.8921798 | 0.950182 |
| ENSG00000236641 | | 1.794067 | 0.04601226 | 0.7027455 | 0.1465874 | 0.8834577 | 0.945446 |
| ENSG00000230648 | | 2.356137 | 0.04596898 | 0.6922394 | 0.1438389 | 0.8856277 | 0.946282 |
| ENSG00000170502 | NUDT9 | 1826.979 | 0.04591465 | 0.1761208 | 0.2753311 | 0.7830619 | 0.886458 |
| ENSG00000286145 | | 2.117074 | 0.0459049 | 0.6985946 | 0.1473971 | 0.8828186 | 0.945181 |
| ENSG00000172819 | RARG | 6308.382 | 0.0458773 | 0.1498173 | 0.3101028 | 0.7564828 | 0.869898 |
| ENSG00000157426 | AASDH | 967.7038 | 0.04587059 | 0.1382049 | 0.3379097 | 0.7354312 | 0.856647 |
| ENSG00000286158 | | 46.20839 | 0.04585158 | 0.4318323 | 0.1191167 | 0.9051829 | 0.955968 |
| ENSG00000261823 | | 1.034409 | 0.04580881 | 0.7599833 | 0.2758092 | 0.7826946 | NA |
| ENSG00000224287 | MSL3P1 | 32.82747 | 0.04576144 | 0.3476593 | 0.144408 | 0.8851783 | 0.94607 |
| ENSG00000102871 | TRADD | 1738.82367 | 0.045749898 | 0.198544814 | 0.222346652 | 0.82404404 | 0.910998 |
| ENSG00000273056 | | 1.659324 | 0.0456801 | 0.7319903 | 0.1335324 | 0.8937724 | 0.950816 |
| ENSG00000280434 | | 5.574768 | 0.04566776 | 0.5896501 | 0.112465 | 0.9104547 | 0.958167 |
| ENSG00000109171 | SLAIN2 | 2708.861623 | 0.045585657 | 0.151758081 | 0.279496673 | 0.77986369 | 0.884534 |
| ENSG00000198492 | YTHDF2 | 5684.851 | 0.04553707 | 0.09965181 | 0.4106802 | 0.6813071 | 0.823663 |
| ENSG00000274717 | | 1.368037 | 0.0455342 | 0.7205059 | 0.1593752 | 0.8733733 | NA |
| ENSG00000266803 | | 1.655346 | 0.04540587 | 0.6993221 | 0.1375563 | 0.8905911 | 0.949218 |
| ENSG00000251148 | | 1.273065 | 0.04534989 | 0.6931959 | 0.1292901 | 0.8971281 | NA |

| | | | | | | | |
|---|---|---|---|---|---|---|---|
| ENSG00000258984 | UBE2F-SCLY | 1.591546 | 0.0453434 | 0.6779523 | 0.1259325 | 0.8997854 | 0.953966 |
| ENSG00000207708 | MIR141 | 2.888532 | 0.04533443 | 0.671761 | 0.1297516 | 0.896763 | 0.952396 |
| ENSG00000260576 | EIF5A2P1 | 1.219151 | 0.04529843 | 0.7264595 | 0.1557846 | 0.8762028 | NA |
| ENSG00000225282 | | 6.20025 | 0.04529659 | 0.6115416 | 0.1166658 | 0.9071249 | 0.95682 |
| ENSG00000169592 | INO80E | 2449.483 | 0.04523158 | 0.1525986 | 0.2940887 | 0.7686901 | 0.877404 |
| ENSG00000250282 | | 1.764528 | 0.04522432 | 0.6869548 | 0.1277486 | 0.8983479 | 0.953089 |
| ENSG00000126653 | NSRP1 | 1153.03416 | 0.045210361 | 0.154463429 | 0.322826042 | 0.74682699 | 0.86403 |
| ENSG00000212332 | RNU6-780P | 2.281729 | 0.04517628 | 0.6560166 | 0.1191638 | 0.9051456 | 0.955968 |
| ENSG00000197548 | ATG7 | 1812.782 | 0.0451523 | 0.1305746 | 0.3437514 | 0.7310333 | 0.854088 |
| ENSG00000226807 | MROH5 | 1.210821 | 0.04508487 | 0.7315111 | 0.1681146 | 0.8664931 | NA |
| ENSG00000234185 | | 1.864833 | 0.04505565 | 0.7306646 | 0.1665157 | 0.8677512 | 0.936731 |
| ENSG00000147274 | RBMX | 15689.34 | 0.04504862 | 0.1547972 | 0.2468255 | 0.8050432 | 0.900308 |
| ENSG00000244060 | RPS2P41 | 1.315593 | 0.04503016 | 0.7036209 | 0.1412294 | 0.8876887 | NA |
| ENSG00000185834 | RPL12P4 | 3089.695 | 0.04497599 | 0.1615566 | 0.2929388 | 0.7695689 | 0.877917 |
| ENSG00000144554 | FANCD2 | 1997.306 | 0.04496974 | 0.4279075 | 0.1434209 | 0.8859578 | 0.946504 |
| ENSG00000243859 | RPL5P17 | 43.62454 | 0.04494712 | 0.3979639 | 0.1343978 | 0.8930881 | 0.950431 |
| ENSG00000224032 | EPB41L4A-AS1 | 2738.988 | 0.04489124 | 0.2731611 | 0.1768264 | 0.8596448 | 0.932469 |
| ENSG00000252797 | RN7SKP272 | 1.924085 | 0.04484157 | 0.6755617 | 0.1192883 | 0.905047 | 0.955968 |
| ENSG00000276316 | OR8G7P | 1.843552 | 0.04481937 | 0.6788238 | 0.1257863 | 0.8999011 | 0.954013 |
| ENSG00000131910 | NR0B2 | 1.372757695 | 0.044808899 | 0.682723569 | 0.127657587 | 0.89841996 | NA |
| ENSG00000050748 | MAPK9 | 7230.567516 | 0.044785068 | 0.148565411 | 0.287499339 | 0.77373001 | 0.880665 |
| ENSG00000181408 | UTS2R | 1.191072 | 0.04477958 | 0.7320116 | 0.1448897 | 0.884798 | NA |
| ENSG00000234787 | LINC00458 | 8.076645 | 0.04476098 | 0.6756537 | 0.08043678 | 0.9358899 | 0.968364 |
| ENSG00000011243 | AKAP8L | 2652.113328 | 0.04474156 | 0.164606728 | 0.268731069 | 0.78813664 | 0.889844 |
| ENSG00000198604 | BAZ1A | 6115.867 | 0.044729 | 0.1601594 | 0.2845832 | 0.7759635 | 0.881993 |
| ENSG00000278740 | | 9.519103 | 0.0447225 | 0.5646466 | 0.1122558 | 0.9106206 | 0.958265 |
| ENSG00000267927 | | 1.70528 | 0.04465569 | 0.6757352 | 0.1240464 | 0.9012786 | 0.954597 |
| ENSG00000231063 | RPL6P6 | 0.9361909 | 0.04464887 | 0.7401246 | 0.1455526 | 0.8842746 | NA |
| ENSG00000141194 | OR4D1 | 1.158824 | 0.04463615 | 0.7391678 | 0.1439984 | 0.8855017 | NA |
| ENSG00000155657 | TTN | 419.1133 | 0.04459933 | 0.3520423 | 0.1345081 | 0.8930008 | 0.950376 |
| ENSG00000213740 | SERBP1P1 | 1257.928 | 0.04454752 | 0.2366734 | 0.1682022 | 0.8664242 | 0.936211 |
| ENSG00000272537 | | 2.141498 | 0.04453652 | 0.6925925 | 0.108244 | 0.9138022 | 0.959599 |
| ENSG00000100413 | POLR3H | 3873.001928 | 0.0445304 | 0.179044547 | 0.248294086 | 0.80390688 | 0.899629 |

| | | | | | | | |
|---|---|---|---|---|---|---|---|
| ENSG00000205903 | ZNF316 | 5649.928 | 0.04449426 | 0.1244468 | 0.3605343 | 0.7184476 | 0.846494 |
| ENSG00000249159 | | 2.710111 | 0.04435955 | 0.6689862 | 0.1189675 | 0.9053011 | 0.955999 |
| ENSG00000184445 | KNTC1 | 2753.717 | 0.04433372 | 0.3323398 | 0.1418894 | 0.8871673 | 0.947051 |
| ENSG00000116560 | SFPQ | 20052.27726 | 0.044290972 | 0.167274822 | 0.285162636 | 0.77551954 | 0.881827 |
| ENSG00000214919 | | 1.052036 | 0.04423215 | 0.7207555 | 0.124305 | 0.9010738 | NA |
| ENSG00000226091 | LINC00937 | 2.266803 | 0.04422693 | 0.6615133 | 0.1242734 | 0.9010988 | 0.954559 |
| ENSG00000202318 | | 2.44146 | 0.04421467 | 0.7245413 | 0.1616892 | 0.8715506 | 0.938753 |
| ENSG00000234345 | ELF2P1 | 21.87075 | 0.04412521 | 0.4434963 | 0.1174395 | 0.9065118 | 0.95662 |
| ENSG00000257298 | | 5.198123 | 0.04408912 | 0.6105551 | 0.1163982 | 0.907337 | 0.95697 |
| ENSG00000284948 | | 5.620818 | 0.0440747 | 0.6416422 | 0.1126005 | 0.9103473 | 0.958126 |
| ENSG00000168955 | TM4SF20 | 6.574008 | 0.04405549 | 0.5950219 | 0.1039851 | 0.9171812 | 0.960982 |
| ENSG00000111012 | CYP27B1 | 425.2866693 | 0.044028063 | 0.209458858 | 0.208467339 | 0.83486408 | 0.91716 |
| ENSG00000239622 | | 2.136771 | 0.04401844 | 0.7047049 | 0.1301686 | 0.8964331 | 0.952236 |
| ENSG00000276168 | RN7SL1 | 1440.41 | 0.04399647 | 0.4075205 | 0.1127183 | 0.9102539 | 0.958107 |
| ENSG00000273792 | | 0.9603073 | 0.04399628 | 0.7411006 | 0.1716084 | 0.8637454 | NA |
| ENSG00000198648 | STK39 | 2569.746 | 0.04398444 | 0.1611101 | 0.2860867 | 0.7748117 | 0.881324 |
| ENSG00000179141 | MTUS2-AS1 | 1.617443 | 0.04394283 | 0.7012093 | 0.1184345 | 0.9057234 | 0.956179 |
| ENSG00000154102 | C16orf74 | 382.1624 | 0.04387762 | 0.3319652 | 0.1463034 | 0.8836819 | 0.945465 |
| ENSG00000229839 | | 4.131265 | 0.04387143 | 0.5737932 | 0.1063428 | 0.9153104 | 0.960119 |
| ENSG00000176343 | RPL37AP8 | 17.05574 | 0.043859 | 0.4108481 | 0.123244 | 0.9019139 | 0.954853 |
| ENSG00000103657 | HERC1 | 6102.703856 | 0.043847626 | 0.280694297 | 0.122714014 | 0.90233357 | 0.954999 |
| ENSG00000138629 | UBL7 | 1871.589033 | 0.0438388 | 0.218104244 | 0.226079592 | 0.82113952 | 0.909205 |
| ENSG00000107099 | DOCK8 | 414.1762292 | 0.043809986 | 0.226847131 | 0.212999833 | 0.83132709 | 0.915137 |
| ENSG00000274605 | PCCA-DT | 250.7423 | 0.04380773 | 0.2877781 | 0.1618889 | 0.8713934 | 0.938681 |
| ENSG00000267573 | KRT8P5 | 7.414668 | 0.04375475 | 0.550392 | 0.1061514 | 0.9154622 | 0.960119 |
| ENSG00000279806 | | 1.635536 | 0.04374959 | 0.7408669 | 0.161391 | 0.8717854 | 0.938816 |
| ENSG00000113083 | LOX | 73.37950705 | 0.04369281 | 0.409210953 | 0.124686752 | 0.90077155 | 0.954483 |
| ENSG00000265000 | MARCHF10-DT | 1.756905 | 0.04367017 | 0.711131 | 0.1412422 | 0.8876786 | 0.94733 |
| ENSG00000134376 | CRB1 | 2.470296624 | 0.043630245 | 0.644399019 | 0.111567128 | 0.91116664 | 0.958726 |
| ENSG00000182585 | EPGN | 9.690215 | 0.04353139 | 0.652771 | 0.116085 | 0.9075851 | 0.957042 |
| ENSG00000187824 | TMEM220 | 7.532383 | 0.04331266 | 0.5064277 | 0.1103886 | 0.9121012 | 0.958851 |
| ENSG00000260005 | | 67.29381 | 0.04328253 | 0.2901596 | 0.1586602 | 0.8739366 | 0.940225 |
| ENSG00000248415 | GAPDHP61 | 7.078978 | 0.04327512 | 0.5576253 | 0.1134023 | 0.9097116 | 0.957954 |

| | | | | | | | |
|---|---|---|---|---|---|---|---|
| ENSG00000275888 | | 64.27708 | 0.04326588 | 0.3300889 | 0.1460512 | 0.883881 | 0.945465 |
| ENSG00000251455 | | 15.17612 | 0.04325089 | 0.5098584 | 0.1076929 | 0.9142393 | 0.95983 |
| ENSG00000088833 | NSFL1C | 2952.184982 | 0.043094142 | 0.20219234 | 0.21976043 | 0.82605774 | 0.912124 |
| ENSG00000250462 | LRRC37BP1 | 205.7478 | 0.04306393 | 0.2389364 | 0.1986134 | 0.8425652 | 0.922264 |
| ENSG00000261654 | | 40.46481 | 0.0430581 | 0.3766752 | 0.1254857 | 0.900139 | 0.954113 |
| ENSG00000259051 | HNRNPUP1 | 415.5863 | 0.04303855 | 0.1962682 | 0.2244016 | 0.8224448 | 0.909824 |
| ENSG00000270228 | | 7.916613 | 0.04294343 | 0.5887218 | 0.1073297 | 0.9145274 | 0.959909 |
| ENSG00000231086 | TSSK1A | 1.465565 | 0.04287935 | 0.749054 | 0.1727653 | 0.8628359 | NA |
| ENSG00000266947 | | 46.84507 | 0.04284285 | 0.4814361 | 0.11912 | 0.9051803 | 0.955968 |
| ENSG00000198822 | GRM3 | 2.523408 | 0.04281914 | 0.6932012 | 0.127258 | 0.8987362 | 0.953273 |
| ENSG00000235990 | RPL23AP20 | 1.287466 | 0.04273075 | 0.7331159 | 0.1456729 | 0.8841796 | NA |
| ENSG00000250853 | RNF138P1 | 15.77908 | 0.04268724 | 0.4202455 | 0.1167015 | 0.9070966 | 0.95682 |
| ENSG00000184436 | THAP7 | 1900.469 | 0.04267483 | 0.2602617 | 0.1600612 | 0.8728329 | 0.939807 |
| ENSG00000253764 | | 3.354125 | 0.04266629 | 0.6380124 | 0.1058178 | 0.9157269 | 0.960178 |
| ENSG00000167525 | PROCA1 | 459.324 | 0.04262455 | 0.3182375 | 0.1431327 | 0.8861854 | 0.946535 |
| ENSG00000243495 | GMFBP1 | 15.06528 | 0.04261394 | 0.4830368 | 0.1092174 | 0.9130301 | 0.959195 |
| ENSG00000213608 | SLC25A14P1 | 2.841538 | 0.04252694 | 0.6605144 | 0.1133328 | 0.9097667 | 0.957974 |
| ENSG00000105221 | AKT2 | 9705.214903 | 0.042513763 | 0.111696928 | 0.3983342 | 0.69038385 | 0.829729 |
| ENSG00000265137 | MIR3192 | 2.746146 | 0.04248397 | 0.6059803 | 0.103586 | 0.9174979 | 0.961195 |
| ENSG00000105656 | ELL | 899.3319854 | 0.042383401 | 0.145460601 | 0.311695357 | 0.75527206 | 0.869333 |
| ENSG00000143294 | PRCC | 5616.178 | 0.04237678 | 0.1842168 | 0.2081577 | 0.8351058 | 0.917253 |
| ENSG00000130939 | UBE4B | 3187.881719 | 0.042349379 | 0.174007641 | 0.239714872 | 0.81055131 | 0.903071 |
| ENSG00000267169 | | 133.214 | 0.04234044 | 0.3326987 | 0.137112 | 0.8909423 | 0.949364 |
| ENSG00000287515 | | 10.44062 | 0.04227958 | 0.5559833 | 0.1020773 | 0.9186953 | 0.961507 |
| ENSG00000235189 | | 2.257246 | 0.04222767 | 0.6433086 | 0.1099505 | 0.9124486 | 0.95888 |
| ENSG00000236317 | | 1.181925 | 0.0422219 | 0.7357275 | 0.1445196 | 0.8850902 | NA |
| ENSG00000165525 | NEMF | 2908.448 | 0.0422024 | 0.1790386 | 0.2330368 | 0.8157329 | 0.906139 |
| ENSG00000248356 | | 1.374641 | 0.04211803 | 0.7286378 | 0.1551258 | 0.8767222 | NA |
| ENSG00000180422 | LINC00304 | 37.20822 | 0.0420836 | 0.4797589 | 0.09927857 | 0.9209171 | 0.962534 |
| ENSG00000252553 | RNA5SP73 | 1.186983 | 0.0420531 | 0.7362836 | 0.1044629 | 0.916802 | NA |
| ENSG00000114125 | RNF7 | 4148.355599 | 0.042017844 | 0.144319619 | 0.290224672 | 0.77164436 | 0.879303 |
| ENSG00000173231 | TERF1P1 | 1.486982 | 0.04201339 | 0.7206703 | 0.1293403 | 0.8970884 | NA |
| ENSG00000259555 | | 0.8366089 | 0.0419518 | 0.7178333 | 0.1322768 | 0.8947653 | NA |

| | | | | | | | |
|---|---|---|---|---|---|---|---|
| ENSG00000233999 | IGKV3OR2-268 | 0.7790184 | 0.04189606 | 0.7541923 | 0.1602131 | 0.8727132 | NA |
| ENSG00000252464 | RN7SKP70 | 6.971907 | 0.04188391 | 0.5579646 | 0.104406 | 0.9168472 | 0.960822 |
| ENSG00000230626 | | 452.5984 | 0.04186188 | 0.1866179 | 0.2303575 | 0.817814 | 0.907648 |
| ENSG00000108389 | MTMR4 | 9753.448818 | 0.04182637 | 0.201923617 | 0.21286852 | 0.83142952 | 0.915137 |
| ENSG00000184206 | GOLGA6L4 | 7.409271 | 0.04178836 | 0.5505935 | 0.1030427 | 0.9179291 | 0.961208 |
| ENSG00000095261 | PSMD5 | 3017.547638 | 0.041760513 | 0.179643949 | 0.220062246 | 0.82582268 | 0.91194 |
| ENSG00000153391 | INO80C | 491.5777 | 0.04172167 | 0.2970485 | 0.1460814 | 0.8838571 | 0.945465 |
| ENSG00000233183 | | 2.867253 | 0.04170502 | 0.6621648 | 0.1025715 | 0.9183031 | 0.961287 |
| ENSG00000224051 | CPTP | 1328.029 | 0.04163564 | 0.1983806 | 0.2179625 | 0.8274584 | 0.913064 |
| ENSG00000116641 | DOCK7 | 2693.802094 | 0.041591416 | 0.177612026 | 0.23745811 | 0.81230141 | 0.9042 |
| ENSG00000135424 | ITGA7 | 167.7705428 | 0.041531767 | 0.16468462 | 0.254354021 | 0.79922207 | 0.896956 |
| ENSG00000261215 | | 1.173708 | 0.0414613 | 0.7044458 | 0.1274597 | 0.8985766 | NA |
| ENSG00000287743 | | 1.669797 | 0.04145552 | 0.6876315 | 0.1158796 | 0.907748 | 0.957074 |
| ENSG00000232186 | TERF1P7 | 20.65129 | 0.04140526 | 0.4027493 | 0.1162933 | 0.9074201 | 0.95702 |
| ENSG00000170266 | GLB1 | 6306.432 | 0.04138474 | 0.1682387 | 0.2127851 | 0.8314946 | 0.915161 |
| ENSG00000227939 | RPL3P2 | 880.1223 | 0.04137726 | 0.1863446 | 0.2263781 | 0.8209074 | 0.909205 |
| ENSG00000009844 | VTA1 | 3825.982154 | 0.041331125 | 0.163618627 | 0.24815462 | 0.80401478 | 0.899636 |
| ENSG00000137824 | RMDN3 | 3921.07741 | 0.04122226 | 0.114698062 | 0.34941373 | 0.72677873 | 0.851739 |
| ENSG00000258454 | | 5.807528 | 0.04115639 | 0.5584231 | 0.1008239 | 0.9196903 | 0.961781 |
| ENSG00000244586 | WNT5A-AS1 | 13.9841 | 0.04112739 | 0.7020548 | 0.1294862 | 0.896973 | 0.952406 |
| ENSG00000224418 | STK24-AS1 | 11.98418 | 0.04110146 | 0.4820686 | 0.1082822 | 0.9137719 | 0.959599 |
| ENSG00000146049 | KAAG1 | 13.56329 | 0.04106405 | 0.5671517 | 0.09838145 | 0.9216294 | 0.962893 |
| ENSG00000186017 | ZNF566 | 608.5013 | 0.04087537 | 0.1534049 | 0.2690841 | 0.787865 | 0.889621 |
| ENSG00000059145 | UNKL | 1559.579676 | 0.040794507 | 0.211906093 | 0.188638285 | 0.85037632 | 0.927305 |
| ENSG00000065029 | ZNF76 | 2820.688524 | 0.040753441 | 0.151816248 | 0.240153896 | 0.81021095 | 0.902923 |
| ENSG00000178498 | DTX3 | 3457.801 | 0.04074932 | 0.206406 | 0.1886319 | 0.8503813 | 0.927305 |
| ENSG00000153066 | TXNDC11 | 1984.686 | 0.04070959 | 0.1080278 | 0.3773367 | 0.7059234 | 0.840133 |
| ENSG00000266126 | | 1.650473 | 0.04064913 | 0.7244271 | 0.1342252 | 0.8932245 | 0.950538 |
| ENSG00000172014 | ANKRD20A4P | 2.968919 | 0.04050754 | 0.6735927 | 0.1099261 | 0.912468 | 0.95888 |
| ENSG00000201900 | RNY1P13 | 26.52549 | 0.04047697 | 0.4239331 | 0.1090935 | 0.9131284 | 0.959195 |
| ENSG00000241634 | | 43.68687 | 0.04038926 | 0.4386463 | 0.1061745 | 0.9154439 | 0.960119 |
| ENSG00000286095 | SPEGNB | 2.501282 | 0.04035634 | 0.6626735 | 0.1111354 | 0.911509 | 0.95882 |
| ENSG00000287302 | | 0.8650462 | 0.04034944 | 0.7341924 | 0.1454952 | 0.8843199 | NA |

| | | | | | | | |
|---|---|---|---|---|---|---|---|
| ENSG00000274667 | | 21.22232 | 0.04032694 | 0.4194601 | 0.1129155 | 0.9100976 | 0.958056 |
| ENSG00000240371 | RPS4XP13 | 182.6387 | 0.0403078 | 0.2231823 | 0.1868413 | 0.8517851 | 0.927845 |
| ENSG00000276742 | | 1.542895 | 0.04018543 | 0.7148814 | 0.1185802 | 0.9056079 | NA |
| ENSG00000253558 | | 1.250081 | 0.04011108 | 0.6897741 | 0.1118214 | 0.910965 | NA |
| ENSG00000219392 | ZNF602P | 1.101775 | 0.04009507 | 0.7299472 | 0.1466336 | 0.8834213 | NA |
| ENSG00000101421 | CHMP4B | 5450.839548 | 0.04001722 | 0.183525541 | 0.196468524 | 0.84424347 | 0.923608 |
| ENSG00000267006 | | 2.693731 | 0.03997266 | 0.666842 | 0.1100516 | 0.9123685 | 0.958851 |
| ENSG00000282100 | HSP90AB4P | 2.978772 | 0.03990714 | 0.6629939 | 0.1140707 | 0.9091817 | 0.957674 |
| ENSG00000275560 | | 141.1677 | 0.03990702 | 0.2717107 | 0.1537651 | 0.877795 | 0.942353 |
| ENSG00000275901 | | 1.89426 | 0.0398675 | 0.7113664 | 0.1215928 | 0.9032215 | 0.955125 |
| ENSG00000102349 | KLF8 | 372.4612839 | 0.039853628 | 0.358997763 | 0.103997801 | 0.91717109 | 0.960982 |
| ENSG00000196371 | FUT4 | 224.6773 | 0.03984645 | 0.3184952 | 0.135987 | 0.8918315 | 0.949991 |
| ENSG00000257846 | | 21.80734 | 0.03981057 | 0.3631146 | 0.1208929 | 0.9037759 | 0.955376 |
| ENSG00000271828 | | 21.5791 | 0.03974131 | 0.4689826 | 0.09982421 | 0.9204839 | 0.962204 |
| ENSG00000273682 | | 4.435651 | 0.03973256 | 0.585189 | 0.0960532 | 0.9234783 | 0.963616 |
| ENSG00000008710 | PKD1 | 8309.087448 | 0.039619128 | 0.379502612 | 0.112362326 | 0.91053613 | 0.958214 |
| ENSG00000259064 | | 3.444213 | 0.03959712 | 0.6412789 | 0.1092335 | 0.9130173 | 0.959195 |
| ENSG00000087116 | ADAMTS2 | 2.283343984 | 0.039579876 | 0.67069671 | 0.113012447 | 0.91002069 | 0.958028 |
| ENSG00000267160 | | 2.334584 | 0.03954884 | 0.6407568 | 0.1023715 | 0.9184618 | 0.961376 |
| ENSG00000201457 | SNORA55 | 38.60533 | 0.0394893 | 0.3905063 | 0.1152529 | 0.9082447 | 0.957217 |
| ENSG00000168303 | MPLKIP | 1980.307 | 0.03945935 | 0.2188605 | 0.1821664 | 0.8554522 | 0.929979 |
| ENSG00000105127 | AKAP8 | 2114.362966 | 0.039409883 | 0.134745069 | 0.306314335 | 0.75936533 | 0.871794 |
| ENSG00000107937 | GTPBP4 | 9859.044719 | 0.039332789 | 0.224588472 | 0.179583927 | 0.85747922 | 0.931238 |
| ENSG00000267857 | | 1.755446 | 0.03931696 | 0.7093322 | 0.1213687 | 0.903399 | 0.95513 |
| ENSG00000156171 | DRAM2 | 1856.027 | 0.03927462 | 0.2270808 | 0.1738961 | 0.8619472 | 0.9338 |
| ENSG00000255491 | | 7.822976 | 0.03925873 | 0.5762901 | 0.09283678 | 0.9260332 | 0.9644 |
| ENSG00000257103 | LSM14A | 9855.079 | 0.03919275 | 0.1222693 | 0.3472133 | 0.7284311 | 0.852623 |
| ENSG00000285280 | | 6.964607 | 0.03914289 | 0.5991428 | 0.08354302 | 0.9334198 | 0.967249 |
| ENSG00000230373 | GOLGA6L5P | 86.45613 | 0.03909808 | 0.3693281 | 0.1221156 | 0.9028074 | 0.955036 |
| ENSG00000151657 | KIN | 1278.666 | 0.03906372 | 0.1176741 | 0.3257971 | 0.7445778 | 0.862367 |
| ENSG00000269680 | | 35.74059 | 0.03889197 | 0.3556038 | 0.1220051 | 0.902895 | 0.955046 |
| ENSG00000224689 | ZNF812P | 2.332643 | 0.03888231 | 0.6648085 | 0.1018899 | 0.9188441 | 0.961511 |
| ENSG00000272754 | | 1.557562 | 0.03885739 | 0.6790661 | 0.1104206 | 0.9120758 | NA |

| | | | | | | | |
|---|---|---|---|---|---|---|---|
| ENSG00000068438 | FTSJ1 | 2604.071854 | 0.038805395 | 0.174223737 | 0.219219967 | 0.8264787 | 0.912361 |
| ENSG00000287379 | | 3.683503 | 0.03879907 | 0.6433985 | 0.09514625 | 0.9241987 | 0.963623 |
| ENSG00000198755 | RPL10A | 25126.1 | 0.03877047 | 0.1588882 | 0.2372259 | 0.8124815 | 0.904236 |
| ENSG00000259308 | FABP5P8 | 1.471844 | 0.03872495 | 0.6971728 | 0.1210624 | 0.9036416 | NA |
| ENSG00000257624 | | 1.500032 | 0.03870658 | 0.7174498 | 0.1194874 | 0.9048892 | NA |
| ENSG00000204252 | HLA-DOA | 1.962734 | 0.03867289 | 0.6987135 | 0.1248727 | 0.9006244 | 0.954399 |
| ENSG00000217414 | DDX18P3 | 3.901949 | 0.03863008 | 0.6882247 | 0.1223875 | 0.9025921 | 0.954999 |
| ENSG00000015676 | NUDCD3 | 6500.559763 | 0.03861468 | 0.123625728 | 0.327715601 | 0.74312671 | 0.861574 |
| ENSG00000277290 | | 44.11092 | 0.03859191 | 0.336655 | 0.1292172 | 0.8971858 | 0.952412 |
| ENSG00000122733 | PHF24 | 2.993852021 | 0.038550695 | 0.675381108 | 0.093953502 | 0.92514609 | 0.964005 |
| ENSG00000267199 | | 35.85413 | 0.03847609 | 0.4494511 | 0.1017218 | 0.9189775 | 0.961537 |
| ENSG00000234569 | RAD1P1 | 1.041248 | 0.03847276 | 0.7540917 | 0.1744759 | 0.8614915 | NA |
| ENSG00000258521 | | 8.287384 | 0.03837828 | 0.4970072 | 0.09668924 | 0.9229732 | 0.963385 |
| ENSG00000100426 | ZBED4 | 2047.590079 | 0.038334262 | 0.207577813 | 0.186353559 | 0.85216751 | 0.928062 |
| ENSG00000272375 | | 15.01251 | 0.0382976 | 0.49449 | 0.09501099 | 0.9243061 | 0.963692 |
| ENSG00000075624 | ACTB | 78993.78693 | 0.038235914 | 0.268300123 | 0.147056269 | 0.88308761 | 0.94524 |
| ENSG00000202227 | RNU6-282P | 0.9378481 | 0.03820974 | 0.7435024 | 0.1426815 | 0.8865417 | NA |
| ENSG00000111911 | HINT3 | 3020.115403 | 0.038161245 | 0.228442983 | 0.165775911 | 0.8683333 | 0.936731 |
| ENSG00000242791 | | 11.98389 | 0.03816042 | 0.5476782 | 0.09159008 | 0.9270237 | 0.964827 |
| ENSG00000286293 | | 4.593886 | 0.03811095 | 0.6140045 | 0.08333818 | 0.9335826 | 0.967249 |
| ENSG00000135018 | UBQLN1 | 9105.199969 | 0.038099205 | 0.13654109 | 0.25136973 | 0.80152827 | 0.898186 |
| ENSG00000213741 | RPS29 | 9125.902 | 0.03804449 | 0.1564192 | 0.2250502 | 0.8219402 | 0.909628 |
| ENSG00000243725 | TTC4 | 89.36965 | 0.03804084 | 0.1952801 | 0.1991878 | 0.8421158 | 0.922133 |
| ENSG00000274093 | | 4.082204 | 0.03790205 | 0.7298672 | 0.1595099 | 0.8732672 | 0.939993 |
| ENSG00000108406 | DHX40 | 12524.42996 | 0.037892739 | 0.109341925 | 0.377505555 | 0.70579794 | 0.840084 |
| ENSG00000231741 | | 1.109012 | 0.0378845 | 0.7345371 | 0.1443237 | 0.8852448 | NA |
| ENSG00000254035 | | 5.745875 | 0.03784926 | 0.6157905 | 0.0910302 | 0.9274686 | 0.96494 |
| ENSG00000182022 | CHST15 | 4369.496 | 0.03783542 | 0.1831719 | 0.2084994 | 0.834839 | 0.91716 |
| ENSG00000224157 | HCG14 | 2.428922 | 0.03783058 | 0.6604172 | 0.08675226 | 0.9308684 | 0.966437 |
| ENSG00000196557 | CACNA1H | 6111.854 | 0.03778954 | 0.296096 | 0.1337411 | 0.8936073 | 0.950721 |
| ENSG00000122417 | ODF2L | 1113.331923 | 0.037766557 | 0.216437926 | 0.176725966 | 0.85972364 | 0.932517 |
| ENSG00000099917 | MED15 | 3899.072247 | 0.037759036 | 0.138460366 | 0.276856894 | 0.78188999 | 0.88566 |
| ENSG00000273454 | | 3.27213 | 0.03768594 | 0.6457991 | 0.09602159 | 0.9235034 | 0.963616 |

| | | | | | | | |
|---|---|---|---|---|---|---|---|
| ENSG00000138621 | PPCDC | 1104.269359 | 0.037661776 | 0.239393974 | 0.137319853 | 0.89077798 | 0.949303 |
| ENSG00000182199 | SHMT2 | 13256.81 | 0.03762258 | 0.2550957 | 0.1515788 | 0.8795192 | 0.943359 |
| ENSG00000286781 | | 0.816873 | 0.03757923 | 0.7468089 | 0.1383613 | 0.8899549 | NA |
| ENSG00000152749 | GPR180 | 883.4788 | 0.0374589 | 0.2132808 | 0.1813963 | 0.8560565 | 0.930273 |
| ENSG00000073969 | NSF | 3854.81449 | 0.037442082 | 0.122238115 | 0.333618109 | 0.73866775 | 0.858824 |
| ENSG00000178252 | WDR6 | 13509.3 | 0.03741567 | 0.118829 | 0.3163925 | 0.7517046 | 0.86703 |
| ENSG00000100284 | TOM1 | 2019.838302 | 0.037369504 | 0.254460691 | 0.139703816 | 0.88889401 | 0.948284 |
| ENSG00000272650 | | 2.33535 | 0.03700476 | 0.7045929 | 0.1197523 | 0.9046794 | 0.955875 |
| ENSG00000100129 | EIF3L | 7784.296637 | 0.036956897 | 0.182756099 | 0.205260895 | 0.8373683 | 0.918796 |
| ENSG00000107290 | SETX | 5597.258734 | 0.036940661 | 0.187368073 | 0.202065459 | 0.83986555 | 0.920762 |
| ENSG00000102882 | MAPK3 | 6229.82249 | 0.036938725 | 0.201143867 | 0.189770398 | 0.84948905 | 0.926874 |
| ENSG00000261457 | | 4.549795 | 0.03690657 | 0.6103841 | 0.09770656 | 0.9221653 | 0.963092 |
| ENSG00000234981 | | 23.61994 | 0.03683895 | 0.3326696 | 0.1206866 | 0.9039393 | 0.955511 |
| ENSG00000274487 | NPEPPSP1 | 98.27856 | 0.0368047 | 0.2905948 | 0.1334441 | 0.8938422 | 0.950853 |
| ENSG00000225200 | | 1191.103 | 0.03679999 | 0.2008161 | 0.1801837 | 0.8570083 | 0.930826 |
| ENSG00000226309 | SMARCE1P2 | 9.247758 | 0.03678934 | 0.5231717 | 0.09183208 | 0.9268315 | 0.96474 |
| ENSG00000236194 | | 1.6688 | 0.03678695 | 0.6930226 | 0.1069293 | 0.914845 | 0.960049 |
| ENSG00000241981 | | 1.189501 | 0.03677371 | 0.7135961 | 0.1147022 | 0.9086812 | NA |
| ENSG00000134371 | CDC73 | 3768.379714 | 0.036771015 | 0.150717627 | 0.247722322 | 0.80434926 | 0.899897 |
| ENSG00000245322 | H2AZ1-DT | 20.57347 | 0.0367679 | 0.4031766 | 0.1035197 | 0.9175505 | 0.961195 |
| ENSG00000134183 | GNAT2 | 20.29719988 | 0.036697837 | 0.39572471 | 0.105553833 | 0.91593636 | 0.960213 |
| ENSG00000235332 | RPSAP76 | 1.514679 | 0.03666931 | 0.7197337 | 0.1121297 | 0.9107205 | NA |
| ENSG00000172315 | TP53RK | 1830.513 | 0.03666248 | 0.1463774 | 0.2530216 | 0.8002515 | 0.897349 |
| ENSG00000219023 | | 639.5342 | 0.03666158 | 0.207038 | 0.1823274 | 0.8553258 | 0.929935 |
| ENSG00000187144 | SPATA21 | 1.006203 | 0.03664874 | 0.7280296 | 0.1245345 | 0.9008921 | NA |
| ENSG00000288067 | | 2.359325 | 0.03660413 | 0.6822747 | 0.1074206 | 0.9144553 | 0.959909 |
| ENSG00000124659 | TBCC | 1671.054629 | 0.036459629 | 0.153017026 | 0.260226111 | 0.79468936 | 0.893983 |
| ENSG00000272319 | | 0.8452646 | 0.03642615 | 0.7472856 | 0.1196307 | 0.9047757 | NA |
| ENSG00000167977 | KCTD5 | 3126.342 | 0.03635464 | 0.1557922 | 0.2337608 | 0.8151707 | 0.905901 |
| ENSG00000176043 | | 2.059808 | 0.03635008 | 0.6653971 | 0.09527135 | 0.9240993 | 0.963623 |
| ENSG00000101350 | KIF3B | 3443.306204 | 0.036340037 | 0.174660114 | 0.207593388 | 0.83554647 | 0.917492 |
| ENSG00000263154 | | 2.575381 | 0.03633545 | 0.6761314 | 0.09908508 | 0.9210707 | 0.962628 |
| ENSG00000229994 | RPL5P4 | 267.3283 | 0.03632951 | 0.2230857 | 0.1677949 | 0.8667446 | 0.93631 |

| | | | | | | | |
|---|---|---|---|---|---|---|---|
| ENSG00000269019 | HOMER3-AS1 | 9.247724 | 0.03630108 | 0.6260003 | 0.09805024 | 0.9218924 | 0.963032 |
| ENSG00000077463 | SIRT6 | 1406.858926 | 0.036161192 | 0.169237673 | 0.220763299 | 0.82527674 | 0.911882 |
| ENSG00000041802 | LSG1 | 4035.216942 | 0.036161154 | 0.098526871 | 0.365267174 | 0.71491198 | 0.844688 |
| ENSG00000277597 | | 2.322161 | 0.03605644 | 0.7236935 | 0.129638 | 0.8968528 | 0.952406 |
| ENSG00000185986 | SDHAP3 | 1602.811 | 0.03603858 | 0.1896658 | 0.1966739 | 0.8440828 | 0.923507 |
| ENSG00000238283 | TBC1D3P1 | 1.399353 | 0.03597916 | 0.7271133 | 0.09511672 | 0.9242221 | NA |
| ENSG00000142864 | SERBP1 | 17632.63 | 0.03594842 | 0.1957187 | 0.2040407 | 0.8383217 | 0.919437 |
| ENSG00000273248 | | 2.555928 | 0.03588646 | 0.6878824 | 0.1085543 | 0.913556 | 0.95953 |
| ENSG00000269873 | | 2.681979 | 0.03583386 | 0.7120474 | 0.1091517 | 0.9130822 | 0.959195 |
| ENSG00000185651 | UBE2L3 | 4935.783 | 0.03582906 | 0.1725197 | 0.2085107 | 0.8348302 | 0.91716 |
| ENSG00000242445 | RPL7AP11 | 201.4413 | 0.03576353 | 0.1872477 | 0.19532 | 0.8451425 | 0.924242 |
| ENSG00000271555 | | 1.634984 | 0.03574434 | 0.700152 | 0.1017512 | 0.9189542 | 0.961537 |
| ENSG00000250762 | | 4.115527 | 0.0357433 | 0.624181 | 0.0944756 | 0.9247314 | 0.963838 |
| ENSG00000198894 | CIPC | 2585.453 | 0.03562904 | 0.208504 | 0.1750041 | 0.8610764 | 0.933336 |
| ENSG00000239322 | ATP6V1B1-AS1 | 1.83667 | 0.03560765 | 0.6953056 | 0.09515536 | 0.9241914 | 0.963623 |
| ENSG00000197813 | | 21.05888 | 0.03559264 | 0.5468639 | 0.0842086 | 0.9328906 | 0.967215 |
| ENSG00000241886 | | 4.433271 | 0.03557689 | 0.5859566 | 0.08497283 | 0.932283 | 0.966977 |
| ENSG00000118762 | PKD2 | 1598.23811 | 0.035549098 | 0.192270151 | 0.196851544 | 0.84394371 | 0.923431 |
| ENSG00000267198 | | 0.9855571 | 0.03551796 | 0.7406455 | 0.1107731 | 0.9117962 | NA |
| ENSG00000205955 | HSP90AA5P | 1.42423 | 0.03548949 | 0.7024226 | 0.1133689 | 0.9097381 | NA |
| ENSG00000238107 | LINC02806 | 5.312037 | 0.03548823 | 0.6980496 | 0.0962568 | 0.9233166 | 0.963538 |
| ENSG00000287044 | | 0.9257883 | 0.03529418 | 0.7538997 | 0.1456127 | 0.8842271 | NA |
| ENSG00000262484 | CCER2 | 42.15042 | 0.03526489 | 0.3986665 | 0.09770387 | 0.9221674 | 0.963092 |
| ENSG00000149187 | CELF1 | 11993.34 | 0.03525939 | 0.1427102 | 0.228049 | 0.8196081 | 0.908741 |
| ENSG00000004777 | ARHGAP33 | 832.1187637 | 0.035236872 | 0.224306369 | 0.142682685 | 0.88654079 | 0.946696 |
| ENSG00000119446 | RBM18 | 1989.230623 | 0.035176868 | 0.143001166 | 0.263112052 | 0.79246422 | 0.892652 |
| ENSG00000165533 | TTC8 | 1863.877 | 0.03516161 | 0.1125114 | 0.3122309 | 0.754865 | 0.869057 |
| ENSG00000133789 | SWAP70 | 1646.096444 | 0.035134394 | 0.197687244 | 0.180394476 | 0.85684289 | 0.930755 |
| ENSG00000234259 | | 2.636813 | 0.03509611 | 0.6612074 | 0.0955717 | 0.9238607 | 0.963623 |
| ENSG00000201499 | RNU6-312P | 5.642805 | 0.0350886 | 0.5769245 | 0.08525995 | 0.9320547 | 0.966925 |
| ENSG00000200560 | RNU6-288P | 0.8978363 | 0.03504949 | 0.7293422 | 0.1187037 | 0.9055101 | NA |
| ENSG00000153551 | CMTM7 | 16.65084 | 0.03504757 | 0.5063696 | 0.0897221 | 0.9285081 | 0.965308 |
| ENSG00000257449 | | 1.232128 | 0.03497866 | 0.7063258 | 0.09137973 | 0.9271909 | NA |

| | | | | | | | |
|---|---|---|---|---|---|---|---|
| ENSG00000258033 | SNX5P2 | 36.42618 | 0.03495647 | 0.3887211 | 0.1006457 | 0.9198317 | 0.961787 |
| ENSG00000177710 | SLC35G5 | 1.901049 | 0.03487816 | 0.6783692 | 0.09327911 | 0.9256818 | 0.964272 |
| ENSG00000242741 | LINC02005 | 46.87242 | 0.0348675 | 0.3528004 | 0.1107739 | 0.9117956 | 0.958851 |
| ENSG00000266744 | | 0.8779426 | 0.03477601 | 0.7364743 | 0.1091666 | 0.9130704 | NA |
| ENSG00000143858 | SYT2 | 28.36073 | 0.0347215 | 0.5018429 | 0.08598074 | 0.9314817 | 0.966739 |
| ENSG00000148660 | CAMK2G | 4532.862 | 0.03466303 | 0.1612477 | 0.2260591 | 0.8211555 | 0.909205 |
| ENSG00000106113 | CRHR2 | 4.68968069 | 0.034515587 | 0.61453299 | 0.086322686 | 0.93120991 | 0.966609 |
| ENSG00000113712 | CSNK1A1 | 7221.893856 | 0.034473662 | 0.153691307 | 0.222602932 | 0.82384455 | 0.910821 |
| ENSG00000130304 | SLC27A1 | 529.0239996 | 0.03442337 | 0.209923472 | 0.165978257 | 0.86817406 | 0.936731 |
| ENSG00000089737 | DDX24 | 11660.51043 | 0.034314785 | 0.153656214 | 0.227576597 | 0.81997542 | 0.908881 |
| ENSG00000207003 | RNU6-611P | 1.12123 | 0.03424944 | 0.7282201 | 0.1193558 | 0.9049935 | NA |
| ENSG00000236152 | MRPS36P1 | 232.5007 | 0.03423261 | 0.2576263 | 0.1356108 | 0.892129 | 0.950171 |
| ENSG00000214244 | SETP21 | 0.9238486 | 0.03420296 | 0.7558913 | 0.1396257 | 0.8889558 | NA |
| ENSG00000198298 | ZNF485 | 672.1 | 0.0341458 | 0.1351511 | 0.2538784 | 0.7995895 | 0.896989 |
| ENSG00000139405 | RITA1 | 2118.489 | 0.0340765 | 0.2111645 | 0.1701764 | 0.8648714 | 0.935427 |
| ENSG00000147457 | CHMP7 | 2287.146 | 0.0340314 | 0.1409225 | 0.2409867 | 0.8095655 | 0.902806 |
| ENSG00000146476 | ARMT1 | 11369.68 | 0.03400881 | 0.2206166 | 0.1546089 | 0.8771296 | 0.942013 |
| ENSG00000112855 | HARS2 | 3795.435182 | 0.033944039 | 0.164372804 | 0.205786749 | 0.83695749 | 0.918602 |
| ENSG00000269304 | FKBP1AP1 | 1.683784 | 0.03386928 | 0.6957336 | 0.09166156 | 0.9269669 | 0.964805 |
| ENSG00000007545 | CRAMP1 | 1818.404577 | 0.033779193 | 0.179146555 | 0.191182566 | 0.84838256 | 0.926316 |
| ENSG00000179218 | CALR | 18452.85 | 0.03376539 | 0.2555606 | 0.1357453 | 0.8920227 | 0.950096 |
| ENSG00000179750 | APOBEC3B | 302.8049 | 0.0337298 | 0.4166764 | 0.09525487 | 0.9241124 | 0.963623 |
| ENSG00000055483 | USP36 | 5903.198756 | 0.033666519 | 0.158005133 | 0.215701724 | 0.82922026 | 0.913907 |
| ENSG00000230091 | TMEM254-AS1 | 409.6833 | 0.033606 | 0.3167376 | 0.112829 | 0.9101661 | 0.958091 |
| ENSG00000100316 | RPL3 | 47714.08448 | 0.033527199 | 0.160948418 | 0.210045989 | 0.83363178 | 0.916526 |
| ENSG00000232951 | IPO7P1 | 41.54877 | 0.03352401 | 0.2979939 | 0.1235297 | 0.9016876 | 0.954802 |
| ENSG00000232389 | | 231.0789 | 0.03345525 | 0.2585568 | 0.1357739 | 0.892 | 0.950096 |
| ENSG00000248980 | SPCS3-AS1 | 1.134956 | 0.033432 | 0.6987132 | 0.0927479 | 0.9261038 | NA |
| ENSG00000207119 | | 1.937893 | 0.03340351 | 0.7032056 | 0.09666621 | 0.9229915 | 0.963385 |
| ENSG00000077235 | GTF3C1 | 8780.556389 | 0.033382457 | 0.182450609 | 0.168045174 | 0.86654774 | 0.936211 |
| ENSG00000164053 | ATRIP | 912.2441 | 0.03337411 | 0.2376988 | 0.1350292 | 0.8925888 | 0.950211 |
| ENSG00000228629 | | 5.045815 | 0.03336785 | 0.555855 | 0.08009142 | 0.9361645 | 0.968541 |
| ENSG00000258752 | | 6.601597 | 0.03331993 | 0.5595546 | 0.08145041 | 0.9350838 | 0.968074 |

| | | | | | | | |
|---|---|---|---|---|---|---|---|
| ENSG00000172977 | KAT5 | 1196.487 | 0.03323145 | 0.1481941 | 0.2262554 | 0.8210028 | 0.909205 |
| ENSG00000080166 | DCT | 4.188227222 | 0.033226751 | 0.678214199 | 0.090246215 | 0.92809156 | 0.965189 |
| ENSG00000287061 | | 1.440119 | 0.03314071 | 0.7009449 | 0.09144923 | 0.9271356 | NA |
| ENSG00000126091 | ST3GAL3 | 59.63680429 | 0.033077083 | 0.277846353 | 0.123091418 | 0.90203471 | 0.954903 |
| ENSG00000181472 | ZBTB2 | 2582.948 | 0.03306742 | 0.2170978 | 0.1501868 | 0.8806173 | 0.943813 |
| ENSG00000230054 | TEX53 | 1.829906 | 0.03301709 | 0.6604345 | 0.08161349 | 0.9349541 | 0.967999 |
| ENSG00000264553 | MIR4257 | 4.331748 | 0.03301459 | 0.6857255 | 0.1107147 | 0.9118426 | 0.958851 |
| ENSG00000224621 | | 9.623526 | 0.03274901 | 0.6562457 | 0.1051335 | 0.9162699 | 0.96033 |
| ENSG00000263887 | | 3.149923 | 0.03266234 | 0.6720652 | 0.0870163 | 0.9306586 | 0.966414 |
| ENSG00000159840 | ZYX | 3502.356 | 0.03262754 | 0.1929965 | 0.175514 | 0.8606758 | 0.933016 |
| ENSG00000143398 | PIP5K1A | 12106.14 | 0.03258858 | 0.1411949 | 0.2196216 | 0.8261659 | 0.912152 |
| ENSG00000130038 | CRACR2A | 1043.395587 | 0.032567658 | 0.374729849 | 0.10594697 | 0.91562443 | 0.960178 |
| ENSG00000198624 | CCDC69 | 269.5322 | 0.03249229 | 0.3055752 | 0.1154209 | 0.9081115 | 0.957217 |
| ENSG00000038358 | EDC4 | 5465.215674 | 0.03247175 | 0.129978476 | 0.216244515 | 0.82879716 | 0.913655 |
| ENSG00000188419 | CHM | 1542.785 | 0.03245922 | 0.2607974 | 0.128458 | 0.8977865 | 0.95276 |
| ENSG00000205500 | MAPRE3-AS1 | 1.339082 | 0.03245613 | 0.7251264 | 0.1218411 | 0.9030249 | NA |
| ENSG00000163960 | UBXN7 | 3957.378 | 0.03244344 | 0.2312592 | 0.1438839 | 0.8855921 | 0.946282 |
| ENSG00000159921 | GNE | 1159.761 | 0.03241163 | 0.3569051 | 0.1064053 | 0.9152608 | 0.960119 |
| ENSG00000261126 | RBFADN | 145.0723 | 0.03235341 | 0.4283704 | 0.08636935 | 0.9311728 | 0.966608 |
| ENSG00000168779 | SHOX2 | 80.20275 | 0.03231785 | 0.3635492 | 0.09744159 | 0.9223757 | 0.963176 |
| ENSG00000237788 | | 4.157226 | 0.0323134 | 0.5749998 | 0.08143164 | 0.9350987 | 0.968074 |
| ENSG00000230580 | RPL19P5 | 7.913071 | 0.03227686 | 0.5304508 | 0.08091217 | 0.9355118 | 0.968218 |
| ENSG00000281974 | | 1.184971 | 0.0321888 | 0.7050104 | 0.07177184 | 0.9427835 | NA |
| ENSG00000136874 | STX17 | 2889.44835 | 0.032089332 | 0.14649052 | 0.219806634 | 0.82602175 | 0.912122 |
| ENSG00000268649 | | 69.9779 | 0.03197174 | 0.3866399 | 0.09478599 | 0.9244848 | 0.963732 |
| ENSG00000004139 | SARM1 | 337.5331161 | 0.031955317 | 0.297824558 | 0.110373388 | 0.91211326 | 0.958851 |
| ENSG00000242276 | RPL5P3 | 14.91365 | 0.03193497 | 0.4304047 | 0.08681019 | 0.9308224 | 0.966433 |
| ENSG00000242391 | | 8.144 | 0.03192451 | 0.569744 | 0.07681688 | 0.9387692 | 0.969795 |
| ENSG00000227788 | MTCO3P43 | 4.086731 | 0.03190096 | 0.6258587 | 0.07334186 | 0.9415341 | 0.971125 |
| ENSG00000285571 | | 24.08397 | 0.03184543 | 0.3947708 | 0.08936027 | 0.9287956 | 0.965461 |
| ENSG00000215244 | LINC02649 | 70.2984 | 0.03182872 | 0.4503484 | 0.08727223 | 0.9304551 | 0.966278 |
| ENSG00000213307 | RPL18P11 | 1.752547 | 0.03182693 | 0.7183438 | 0.09340478 | 0.925582 | 0.96427 |
| ENSG00000173141 | MRPL57 | 4117.515 | 0.03180978 | 0.2096082 | 0.1648393 | 0.8690705 | 0.937198 |

| | | | | | | | |
|---|---|---|---|---|---|---|---|
| ENSG00000217094 | PPIAP31 | 1202.345 | 0.03174725 | 0.2470328 | 0.1450476 | 0.8846733 | 0.94582 |
| ENSG00000198265 | HELZ | 8777.001 | 0.03169543 | 0.2515102 | 0.1293651 | 0.8970688 | 0.952406 |
| ENSG00000259224 | SLC35G6 | 4.800738 | 0.03169141 | 0.6001692 | 0.07766337 | 0.9380958 | 0.969552 |
| ENSG00000283429 | MIR1244-3 | 2.345554 | 0.03167583 | 0.6421948 | 0.08001282 | 0.9362271 | 0.968541 |
| ENSG00000204136 | GGTA1 | 2.384504 | 0.0315989 | 0.6859324 | 0.08671375 | 0.930899 | 0.966437 |
| ENSG00000006007 | GDE1 | 5035.682788 | 0.031563421 | 0.115411108 | 0.340023122 | 0.73383912 | 0.85569 |
| ENSG00000236148 | RPL23AP37 | 2.516536 | 0.03154648 | 0.6471636 | 0.08063398 | 0.935733 | 0.96828 |
| ENSG00000259867 | | 8.458484 | 0.03149373 | 0.4894025 | 0.08103116 | 0.9354172 | 0.968218 |
| ENSG00000232273 | FTH1P1 | 2.147765 | 0.03147483 | 0.7045345 | 0.1048425 | 0.9165008 | 0.960535 |
| ENSG00000140688 | RUSF1 | 3468.596 | 0.03137459 | 0.1835829 | 0.162538 | 0.8708822 | 0.938406 |
| ENSG00000257341 | | 1.705901 | 0.0312246 | 0.6822491 | 0.090238 | 0.9280981 | 0.965189 |
| ENSG00000111737 | RAB35 | 4682.318144 | 0.031214563 | 0.167552815 | 0.183624569 | 0.85430799 | 0.92926 |
| ENSG00000140374 | ETFA | 13183.35 | 0.03121054 | 0.1857618 | 0.2020518 | 0.8398762 | 0.920762 |
| ENSG00000203635 | | 234.2746 | 0.03117136 | 0.4035953 | 0.0855111 | 0.9318551 | 0.966925 |
| ENSG00000265735 | RN7SL5P | 2.827748 | 0.03115192 | 0.6576657 | 0.07796354 | 0.9378571 | 0.969374 |
| ENSG00000267305 | | 1.30539 | 0.03113139 | 0.7232553 | 0.1075906 | 0.9143204 | NA |
| ENSG00000236713 | | 0.9495986 | 0.03110837 | 0.7314608 | 0.08205223 | 0.9346052 | NA |
| ENSG00000008226 | DLEC1 | 42.46726081 | 0.031072969 | 0.343203182 | 0.09991609 | 0.92041094 | 0.962165 |
| ENSG00000173517 | PEAK1 | 2812.943 | 0.03100747 | 0.2120753 | 0.1469455 | 0.883175 | 0.945296 |
| ENSG00000140153 | WDR20 | 1730.699 | 0.03098601 | 0.1300909 | 0.2442491 | 0.8070379 | 0.901122 |
| ENSG00000110876 | SELPLG | 5.736393803 | 0.030934134 | 0.638339377 | 0.088023724 | 0.92985782 | 0.966006 |
| ENSG00000273821 | | 31.6773 | 0.03090736 | 0.468103 | 0.07948808 | 0.9366444 | 0.968692 |
| ENSG00000236654 | | 3.149695 | 0.03086819 | 0.6886319 | 0.08978217 | 0.9284603 | 0.965301 |
| ENSG00000258653 | | 1.508722 | 0.03086206 | 0.7352709 | 0.1086054 | 0.9135154 | NA |
| ENSG00000196422 | PPP1R26 | 3429.698 | 0.03085589 | 0.1789012 | 0.1790496 | 0.8578987 | 0.931412 |
| ENSG00000142700 | DMRTA2 | 1.145228 | 0.03084824 | 0.7090934 | 0.08981026 | 0.928438 | NA |
| ENSG00000167614 | TTYH1 | 87.3104 | 0.03081983 | 0.3411261 | 0.1041149 | 0.9170782 | 0.960982 |
| ENSG00000254023 | PKMP4 | 4.303459 | 0.03078335 | 0.6468581 | 0.08305038 | 0.9338115 | 0.967249 |
| ENSG00000211645 | IGLV1-50 | 0.9477787 | 0.03076781 | 0.7625363 | 0.1084728 | 0.9136207 | NA |
| ENSG00000108666 | C17orf75 | 895.6438407 | 0.030670224 | 0.151686664 | 0.200993249 | 0.84070385 | 0.921217 |
| ENSG00000140350 | ANP32A | 9279.77 | 0.03066249 | 0.1733545 | 0.1849772 | 0.8532469 | 0.928562 |
| ENSG00000065989 | PDE4A | 543.2065254 | 0.030660037 | 0.445762172 | 0.084122616 | 0.93295894 | 0.967215 |
| ENSG00000276054 | | 1.151043 | 0.03065787 | 0.7398386 | 0.1100317 | 0.9123842 | NA |

| | | | | | | | |
|---|---|---|---|---|---|---|---|
| ENSG00000264943 | SH3GL1P2 | 5.678813 | 0.03063406 | 0.6346678 | 0.07965484 | 0.9365118 | 0.968668 |
| ENSG00000288234 | | 27.26276 | 0.03053498 | 0.4602302 | 0.08315596 | 0.9337275 | 0.967249 |
| ENSG00000077616 | NAALAD2 | 27.09967679 | 0.030477724 | 0.415229128 | 0.083790217 | 0.93322323 | 0.967249 |
| ENSG00000108021 | TASOR2 | 10408.21129 | 0.030440071 | 0.223322092 | 0.138862417 | 0.88955887 | 0.948689 |
| ENSG00000152463 | OLAH | 1.405315 | 0.03040918 | 0.709797 | 0.1007488 | 0.9197498 | NA |
| ENSG00000287879 | | 3.424009 | 0.03032635 | 0.6435046 | 0.0852727 | 0.9320446 | 0.966925 |
| ENSG00000259705 | | 5.892886 | 0.03032524 | 0.6575645 | 0.08335469 | 0.9335695 | 0.967249 |
| ENSG00000285667 | | 36.69337 | 0.03024871 | 0.4169793 | 0.08287873 | 0.933948 | 0.967249 |
| ENSG00000199709 | RNU4-23P | 0.9639842 | 0.0302486 | 0.7228375 | 0.09326749 | 0.9256911 | NA |
| ENSG00000162777 | DENND2D | 3560.693 | 0.03024464 | 0.164663 | 0.2093082 | 0.8342077 | 0.916893 |
| ENSG00000118640 | VAMP8 | 5446.609567 | 0.030238911 | 0.204658 | 0.150552318 | 0.88032888 | 0.943667 |
| ENSG00000263499 | | 1.248524 | 0.03021578 | 0.7279594 | 0.07878881 | 0.9372006 | NA |
| ENSG00000169896 | ITGAM | 14.2677 | 0.03019331 | 0.6293548 | 0.08511543 | 0.9321696 | 0.966963 |
| ENSG00000246339 | EXTL3-AS1 | 49.55121 | 0.03019211 | 0.3683977 | 0.08500855 | 0.9322546 | 0.966977 |
| ENSG00000179101 | H3P47 | 5.948167 | 0.0301808 | 0.6046781 | 0.07509507 | 0.9401391 | 0.97045 |
| ENSG00000103550 | KNOP1 | 1266.466508 | 0.030158042 | 0.188142598 | 0.165846296 | 0.86827791 | 0.936731 |
| ENSG00000260651 | | 2.266535 | 0.0300973 | 0.652815 | 0.08099745 | 0.935444 | 0.968218 |
| ENSG00000102195 | GPR50 | 56.75272442 | 0.030081587 | 0.465706208 | 0.072507932 | 0.94219769 | 0.971637 |
| ENSG00000222248 | RNA5SP201 | 1.414892 | 0.03004887 | 0.7112174 | 0.09858956 | 0.9214642 | NA |
| ENSG00000268288 | | 3.084781 | 0.02983761 | 0.6566467 | 0.0727191 | 0.9420296 | 0.971504 |
| ENSG00000273703 | H2BC14 | 1.952888 | 0.02983282 | 0.6981366 | 0.08416135 | 0.9329281 | 0.967215 |
| ENSG00000178814 | OPLAH | 2367.892 | 0.02979301 | 0.3118286 | 0.1008884 | 0.9196391 | 0.961775 |
| ENSG00000228124 | | 1.507339 | 0.02976945 | 0.7032754 | 0.0829493 | 0.9338919 | NA |
| ENSG00000276436 | | 3.289434 | 0.02975337 | 0.6704748 | 0.07896398 | 0.9370613 | 0.968973 |
| ENSG00000261537 | | 3.011623 | 0.02973085 | 0.6568267 | 0.08386512 | 0.9331637 | 0.967249 |
| ENSG00000288095 | | 59.46013 | 0.02970731 | 0.2932912 | 0.106863 | 0.9148977 | 0.960066 |
| ENSG00000267226 | | 3.539975 | 0.02966934 | 0.6596625 | 0.0823038 | 0.9344051 | 0.967619 |
| ENSG00000267280 | TBX2-AS1 | 3828.134 | 0.02960128 | 0.2584049 | 0.1234303 | 0.9017663 | 0.954833 |
| ENSG00000266707 | PSMD7P1 | 4.380603 | 0.02959585 | 0.6501183 | 0.08525297 | 0.9320603 | 0.966925 |
| ENSG00000141971 | MVB12A | 1672.15 | 0.02952333 | 0.1913402 | 0.1563253 | 0.8757766 | 0.941283 |
| ENSG00000243967 | NBPF5P | 2.852712 | 0.0295096 | 0.6778472 | 0.05556521 | 0.9556882 | 0.978636 |
| ENSG00000288045 | | 16.38965 | 0.02946822 | 0.5970703 | 0.07436309 | 0.9407215 | 0.970675 |
| ENSG00000254409 | | 2.770767 | 0.0293912 | 0.6760634 | 0.08703516 | 0.9306436 | 0.966414 |

| | | | | | | | |
|---|---|---|---|---|---|---|---|
| ENSG00000149547 | EI24 | 5337.104 | 0.02937859 | 0.201776 | 0.1448561 | 0.8848245 | 0.94588 |
| ENSG00000187957 | DNER | 12.86314 | 0.02929022 | 0.6030368 | 0.07917595 | 0.9368927 | 0.968865 |
| ENSG00000239831 | RNF7P1 | 9.490777 | 0.02928793 | 0.4975348 | 0.07349916 | 0.9414089 | 0.971087 |
| ENSG00000227536 | SOCS5P4 | 13.56545 | 0.02927013 | 0.469118 | 0.07029744 | 0.9439569 | 0.972468 |
| ENSG00000198744 | MTCO3P12 | 95.45711 | 0.0292571 | 0.3287452 | 0.1037044 | 0.9174039 | 0.96114 |
| ENSG00000225400 | RAB28P5 | 10.52111 | 0.02923923 | 0.5611327 | 0.06876631 | 0.9451756 | 0.972895 |
| ENSG00000239908 | RN7SL75P | 6.193314 | 0.02923366 | 0.5491935 | 0.07207456 | 0.9425426 | 0.971691 |
| ENSG00000224747 | MTCYBP21 | 10.29016 | 0.02918965 | 0.6292452 | 0.07357373 | 0.9413496 | 0.971064 |
| ENSG00000131368 | MRPS25 | 5971.040091 | 0.029181745 | 0.139012798 | 0.211123497 | 0.83279091 | 0.915867 |
| ENSG00000034533 | ASTE1 | 405.2085014 | 0.029115428 | 0.148404802 | 0.167617575 | 0.86688414 | 0.936384 |
| ENSG00000185359 | HGS | 7548.823 | 0.0290999 | 0.1146286 | 0.2526195 | 0.8005622 | 0.897578 |
| ENSG00000277695 | | 0.7764534 | 0.02908882 | 0.7646174 | 0.104796 | 0.9165377 | NA |
| ENSG00000138823 | MTTP | 2.578630136 | 0.029053814 | 0.650946327 | 0.075451524 | 0.93985547 | 0.970389 |
| ENSG00000233382 | NKAPP1 | 95.96633 | 0.02904081 | 0.2301908 | 0.1350625 | 0.8925625 | 0.950211 |
| ENSG00000119669 | IRF2BPL | 7071.560559 | 0.028945145 | 0.201253278 | 0.154665739 | 0.87708484 | 0.942003 |
| ENSG00000240751 | | 7.376173 | 0.02892875 | 0.5680892 | 0.07368499 | 0.941261 | 0.97101 |
| ENSG00000236474 | GCNT1P1 | 0.9424145 | 0.02891921 | 0.7095449 | 0.08237279 | 0.9343503 | NA |
| ENSG00000235079 | ZRANB2-AS1 | 11.12419 | 0.02890176 | 0.4697037 | 0.07222096 | 0.9424261 | 0.971691 |
| ENSG00000118298 | CA14 | 120.1608098 | 0.028850266 | 0.315902455 | 0.097045631 | 0.92269016 | 0.963243 |
| ENSG00000237972 | TUBG1P | 5.037429 | 0.02881515 | 0.5650454 | 0.06703881 | 0.9465508 | 0.973611 |
| ENSG00000262213 | | 1.883881 | 0.02880948 | 0.7198329 | 0.1072912 | 0.914558 | 0.959909 |
| ENSG00000223571 | DHRSX-IT1 | 1.824358 | 0.02873248 | 0.7027394 | 0.08583079 | 0.9316009 | 0.96675 |
| ENSG00000221909 | FAM200A | 374.864 | 0.02868687 | 0.2093354 | 0.1449852 | 0.8847225 | 0.94582 |
| ENSG00000255326 | | 3.097662 | 0.02866917 | 0.6741512 | 0.0837273 | 0.9332733 | 0.967249 |
| ENSG00000132010 | ZNF20 | 11.43798443 | 0.028451693 | 0.434475636 | 0.076021883 | 0.93940169 | 0.970098 |
| ENSG00000024862 | CCDC28A | 660.7459193 | 0.02843792 | 0.177250069 | 0.155023865 | 0.8768025 | 0.941882 |
| ENSG00000223427 | RBM7P1 | 13.74813 | 0.02837833 | 0.5119187 | 0.07033903 | 0.9439238 | 0.972468 |
| ENSG00000260806 | | 164.2161 | 0.02831379 | 0.2874135 | 0.1032955 | 0.9177285 | 0.961195 |
| ENSG00000269086 | | 15.16683 | 0.02826187 | 0.4066403 | 0.0768348 | 0.938755 | 0.969795 |
| ENSG00000197296 | FITM2 | 1764.922 | 0.02824575 | 0.2290386 | 0.1246554 | 0.9007964 | 0.954483 |
| ENSG00000150628 | SPATA4 | 3.128577 | 0.02823298 | 0.6768583 | 0.08089002 | 0.9355294 | 0.968218 |
| ENSG00000130402 | ACTN4 | 22049.08089 | 0.02813111 | 0.160461924 | 0.179307324 | 0.8576964 | 0.931298 |
| ENSG00000249531 | | 1.535262 | 0.02809632 | 0.7120141 | 0.03331593 | 0.9734226 | NA |

| | | | | | | | |
|---|---|---|---|---|---|---|---|
| ENSG00000215838 | | 1.019677 | 0.02804067 | 0.7171915 | 0.07742972 | 0.9382817 | NA |
| ENSG00000168612 | ZSWIM1 | 882.2651 | 0.02802093 | 0.123157 | 0.2245114 | 0.8223594 | 0.909824 |
| ENSG00000259665 | | 1.772694 | 0.02797115 | 0.6808833 | 0.06671408 | 0.9468093 | 0.973709 |
| ENSG00000231084 | RPL22P24 | 13.57758 | 0.02793102 | 0.4411212 | 0.07449905 | 0.9406133 | 0.970666 |
| ENSG00000181322 | NME9 | 32.27443 | 0.02779472 | 0.4567277 | 0.06974918 | 0.9443933 | 0.972618 |
| ENSG00000178209 | PLEC | 25671.25 | 0.02776127 | 0.2476726 | 0.184639 | 0.8535122 | 0.928728 |
| ENSG00000172936 | MYD88 | 4233.693 | 0.02775472 | 0.1890651 | 0.1457683 | 0.8841043 | 0.945605 |
| ENSG00000102543 | CDADC1 | 304.1513378 | 0.027748601 | 0.203717475 | 0.146842128 | 0.88325663 | 0.945333 |
| ENSG00000262662 | | 1.737046 | 0.02770804 | 0.7018133 | 0.09102133 | 0.9274756 | 0.96494 |
| ENSG00000165805 | C12orf50 | 0.7940437 | 0.02768119 | 0.7307237 | 0.06794226 | 0.9458316 | NA |
| ENSG00000104177 | MYEF2 | 3275.962844 | 0.027605617 | 0.177845667 | 0.150539847 | 0.88033872 | 0.943667 |
| ENSG00000139697 | SBNO1 | 5885.717 | 0.02748864 | 0.1737454 | 0.1664106 | 0.8678338 | 0.936731 |
| ENSG00000272077 | | 22.95421 | 0.02743774 | 0.4443319 | 0.06993538 | 0.9442451 | 0.972607 |
| ENSG00000260613 | | 1.750799 | 0.02741295 | 0.7024918 | 0.06689585 | 0.9466646 | 0.973636 |
| ENSG00000110107 | PRPF19 | 9725.005778 | 0.027302888 | 0.211052882 | 0.121727489 | 0.90311484 | 0.955125 |
| ENSG00000146755 | TRIM50 | 5.645114 | 0.02728712 | 0.5353433 | 0.06494496 | 0.9482178 | 0.974627 |
| ENSG00000217653 | PIMREGP3 | 0.9454327 | 0.02714684 | 0.7241465 | 0.08857065 | 0.9294231 | NA |
| ENSG00000118307 | CFAP94 | 13.26811021 | 0.027085224 | 0.479500133 | 0.067909745 | 0.94585748 | 0.973295 |
| ENSG00000160113 | NR2F6 | 2483.238 | 0.0269967 | 0.2235284 | 0.1243779 | 0.9010161 | 0.954525 |
| ENSG00000223462 | | 1.640544 | 0.02697945 | 0.694285 | 0.07038084 | 0.9438905 | 0.972468 |
| ENSG00000257949 | TEN1 | 2.732386 | 0.02684057 | 0.6456951 | 0.07327791 | 0.941585 | 0.971125 |
| ENSG00000280071 | GATD3B | 490.2991 | 0.02679984 | 0.2055691 | 0.1243589 | 0.9010311 | 0.954525 |
| ENSG00000173531 | MST1 | 655.0622 | 0.026715 | 0.2573692 | 0.09936211 | 0.9208508 | 0.962512 |
| ENSG00000250990 | | 8.286753 | 0.02670276 | 0.6216412 | 0.06711307 | 0.9464917 | 0.973609 |
| ENSG00000177627 | C12orf54 | 1.713953 | 0.02670257 | 0.7000049 | 0.08496121 | 0.9322922 | 0.966977 |
| ENSG00000255970 | LINC02421 | 3.104362 | 0.02661358 | 0.6502676 | 0.07704408 | 0.9385885 | 0.969795 |
| ENSG00000212493 | SNORD19 | 18.37237 | 0.02658219 | 0.4264992 | 0.07372638 | 0.9412281 | 0.97101 |
| ENSG00000163964 | PIGX | 1776.52 | 0.02655575 | 0.09561872 | 0.2769048 | 0.7818532 | 0.885656 |
| ENSG00000171174 | RBKS | 348.9824 | 0.02653887 | 0.1667375 | 0.1600432 | 0.872847 | 0.939807 |
| ENSG00000173805 | HAP1 | 112.9085 | 0.02643099 | 0.2410083 | 0.1101057 | 0.9123255 | 0.958851 |
| ENSG00000263990 | | 19.25442 | 0.02637373 | 0.4933571 | 0.06952215 | 0.944574 | 0.972653 |
| ENSG00000288068 | | 18.60006 | 0.02635218 | 0.5215784 | 0.06248499 | 0.9501766 | 0.975545 |
| ENSG00000228604 | | 2.010238 | 0.02624655 | 0.6992456 | 0.08447598 | 0.932678 | 0.967213 |

| | | | | | | | |
|---|---|---|---|---|---|---|---|
| ENSG00000226234 | KRT18P24 | 2.32531 | 0.02622434 | 0.6702932 | 0.07648181 | 0.9390358 | 0.96992 |
| ENSG00000176542 | USF3 | 1981.856 | 0.02619584 | 0.2296021 | 0.1383009 | 0.8900026 | 0.948936 |
| ENSG00000132463 | GRSF1 | 8910.427992 | 0.026132744 | 0.186034065 | 0.141768227 | 0.88726308 | 0.947081 |
| ENSG00000205423 | CNEP1R1 | 541.6727 | 0.02608023 | 0.1646472 | 0.1800544 | 0.8571099 | 0.930898 |
| ENSG00000120329 | SLC25A2 | 3.189916734 | 0.025984588 | 0.644301073 | 0.067938614 | 0.9458345 | 0.973295 |
| ENSG00000187902 | SHISA7 | 3.901213 | 0.02586806 | 0.5941395 | 0.06320804 | 0.9496008 | 0.975269 |
| ENSG00000121207 | LRAT | 20.60463695 | 0.025833648 | 0.488626151 | 0.063232757 | 0.94958116 | 0.975269 |
| ENSG00000176401 | EID2B | 129.808 | 0.0258118 | 0.2067158 | 0.1293544 | 0.8970772 | 0.952406 |
| ENSG00000133048 | CHI3L1 | 1.064506852 | 0.025741827 | 0.726358833 | 0.070588073 | 0.9437256 | NA |
| ENSG00000127528 | KLF2 | 130.1588096 | 0.025737259 | 0.47643669 | 0.057801669 | 0.95390661 | 0.977602 |
| ENSG00000270839 | | 0.9835396 | 0.02568319 | 0.7401985 | 0.05859066 | 0.9532781 | NA |
| ENSG00000230071 | RPL4P6 | 10.23555 | 0.02565539 | 0.555285 | 0.05907925 | 0.952889 | 0.977059 |
| ENSG00000236088 | COX10-AS1 | 808.4305 | 0.0255709 | 0.2711005 | 0.09859468 | 0.9214601 | 0.962808 |
| ENSG00000144045 | DQX1 | 24.89531 | 0.02552949 | 0.3889698 | 0.07994709 | 0.9362793 | 0.968541 |
| ENSG00000271155 | | 3.227509 | 0.02546219 | 0.6563353 | 0.06926918 | 0.9447754 | 0.972747 |
| ENSG00000280047 | | 19.6641 | 0.02529919 | 0.4971739 | 0.06104683 | 0.9513219 | 0.976178 |
| ENSG00000270822 | | 2.617995 | 0.02515981 | 0.6901835 | 0.08526377 | 0.9320517 | 0.966925 |
| ENSG00000259010 | NUP50P1 | 29.82193 | 0.02514435 | 0.3710587 | 0.0742815 | 0.9407864 | 0.970675 |
| ENSG00000224858 | RPL29P11 | 891.5947 | 0.0251425 | 0.2433104 | 0.1037435 | 0.9173729 | 0.96114 |
| ENSG00000213433 | RPLP1P6 | 5.635758 | 0.02511954 | 0.6060117 | 0.06251284 | 0.9501544 | 0.975545 |
| ENSG00000230155 | CEP250-AS1 | 0.9092837 | 0.02511783 | 0.7328261 | 0.07210289 | 0.94252 | NA |
| ENSG00000144713 | RPL32 | 29034.41 | 0.02511536 | 0.1736931 | 0.1446967 | 0.8849503 | 0.945977 |
| ENSG00000188994 | ZNF292 | 2650.127 | 0.02510686 | 0.2554216 | 0.1140538 | 0.9091951 | 0.957674 |
| ENSG00000258011 | HMGA1P3 | 96.91491 | 0.0250493 | 0.2947354 | 0.09057509 | 0.9278302 | 0.964986 |
| ENSG00000259169 | GNRHR2P1 | 1.431801 | 0.02501904 | 0.7387239 | 0.08110341 | 0.9353597 | NA |
| ENSG00000125912 | NCLN | 4271.255828 | 0.025002793 | 0.21047937 | 0.121484593 | 0.90330722 | 0.955125 |
| ENSG00000230207 | RPL4P5 | 1288.767 | 0.02493594 | 0.1846251 | 0.1282391 | 0.8979597 | 0.952829 |
| ENSG00000232160 | RAP2C-AS1 | 148.5003 | 0.02484279 | 0.3728177 | 0.07123257 | 0.9432127 | 0.972118 |
| ENSG00000140598 | EFL1 | 2300.168 | 0.0247232 | 0.113959 | 0.2114534 | 0.8325335 | 0.915659 |
| ENSG00000136167 | LCP1 | 672.652509 | 0.024692777 | 0.281886349 | 0.085332959 | 0.93199669 | 0.966925 |
| ENSG00000205339 | IPO7 | 11494.96 | 0.02463876 | 0.1896509 | 0.1195147 | 0.9048676 | 0.955922 |
| ENSG00000239523 | MYLK-AS1 | 217.3634 | 0.02463789 | 0.2409607 | 0.1150138 | 0.9084342 | 0.957291 |
| ENSG00000287158 | | 0.7694032 | 0.02453052 | 0.7607005 | 0.08167122 | 0.9349082 | NA |

| | | | | | | | |
|---|---|---|---|---|---|---|---|
| ENSG00000278330 | | 6.16047 | 0.02450944 | 0.5518286 | 0.0585916 | 0.9532774 | 0.977297 |
| ENSG00000166535 | A2ML1 | 5.71146 | 0.02450419 | 0.6451327 | 0.06338079 | 0.9494633 | 0.975269 |
| ENSG00000223663 | NDUFB4P8 | 1.046484 | 0.02434329 | 0.6975502 | 0.06703443 | 0.9465543 | NA |
| ENSG00000178150 | ZNF114 | 329.6569 | 0.02434139 | 0.1968535 | 0.1268745 | 0.8990397 | 0.953529 |
| ENSG00000077458 | FAM76B | 960.3061917 | 0.024324248 | 0.168291404 | 0.144267308 | 0.88528939 | 0.94607 |
| ENSG00000254093 | PINX1 | 394.3041 | 0.0243214 | 0.2957765 | 0.08423919 | 0.9328663 | 0.967215 |
| ENSG00000237036 | ZEB1-AS1 | 252.3845 | 0.02431648 | 0.3077861 | 0.08381699 | 0.9332019 | 0.967249 |
| ENSG00000236914 | LINC01852 | 6.080737 | 0.02411985 | 0.5611609 | 0.05139897 | 0.9590076 | 0.979746 |
| ENSG00000271404 | MZT1P2 | 2.210865 | 0.02406725 | 0.6820454 | 0.0709328 | 0.9434512 | 0.972213 |
| ENSG00000231333 | RPL34P6 | 7.255314 | 0.0240594 | 0.5839102 | 0.06247703 | 0.950183 | 0.975545 |
| ENSG00000265798 | | 5.038422 | 0.02376219 | 0.5254545 | 0.05271521 | 0.9579588 | 0.979347 |
| ENSG00000180104 | EXOC3 | 6508.612 | 0.02371593 | 0.1056424 | 0.2202756 | 0.8256566 | 0.91192 |
| ENSG00000090432 | MUL1 | 1251.787194 | 0.023689033 | 0.163656984 | 0.145625101 | 0.88421735 | 0.945618 |
| ENSG00000105401 | CDC37 | 14249.50714 | 0.023687844 | 0.141343027 | 0.168355775 | 0.8663034 | 0.936211 |
| ENSG00000144369 | FAM171B | 1562.41 | 0.02365399 | 0.2514642 | 0.1014488 | 0.9191942 | 0.96165 |
| ENSG00000171766 | GATM | 15.119 | 0.02359244 | 0.3913361 | 0.06974811 | 0.9443941 | 0.972618 |
| ENSG00000149313 | AASDHPPT | 2240.911 | 0.02358639 | 0.1135752 | 0.2051323 | 0.8374688 | 0.918843 |
| ENSG00000168594 | ADAM29 | 1.298388 | 0.02358311 | 0.7047785 | 0.06982964 | 0.9443293 | NA |
| ENSG00000229190 | | 1.941096 | 0.02351182 | 0.7089226 | 0.07471875 | 0.9404385 | 0.970538 |
| ENSG00000115947 | ORC4 | 2477.029819 | 0.023476209 | 0.115662043 | 0.205231403 | 0.83739134 | 0.918796 |
| ENSG00000204991 | SPIRE2 | 2593.401 | 0.02345027 | 0.142332 | 0.1669838 | 0.8673828 | 0.936531 |
| ENSG00000198160 | MIER1 | 1585.623 | 0.0234374 | 0.1436048 | 0.1692572 | 0.8655943 | 0.935977 |
| ENSG00000234156 | | 38.07668 | 0.02342652 | 0.4387613 | 0.06084381 | 0.9514836 | 0.976211 |
| ENSG00000232702 | | 130.1305 | 0.02339124 | 0.218347 | 0.1152464 | 0.9082498 | 0.957217 |
| ENSG00000100519 | PSMC6 | 4413.605651 | 0.023371917 | 0.177979087 | 0.183916319 | 0.8540791 | 0.929129 |
| ENSG00000139985 | ADAM21 | 6.119777 | 0.02333945 | 0.5619327 | 0.05509143 | 0.9560656 | 0.978774 |
| ENSG00000141337 | ARSG | 1310.117 | 0.02333944 | 0.2157941 | 0.1107562 | 0.9118097 | 0.958851 |
| ENSG00000274444 | | 3.595816 | 0.0232674 | 0.6437296 | 0.05802502 | 0.9537287 | 0.977458 |
| ENSG00000251562 | MALAT1 | 42330.4 | 0.02313959 | 0.3271868 | 0.07208734 | 0.9425324 | 0.971691 |
| ENSG00000240435 | RPS12P27 | 0.8369872 | 0.02311929 | 0.7525245 | 0.07698363 | 0.9386366 | NA |
| ENSG00000286667 | | 25.81286 | 0.02303719 | 0.5403273 | 0.0500305 | 0.9600981 | 0.980097 |
| ENSG00000155755 | TMEM237 | 826.8927 | 0.02300997 | 0.2516098 | 0.09593777 | 0.92357 | 0.963623 |
| ENSG00000225377 | NRSN2-AS1 | 384.1632 | 0.02299281 | 0.3175203 | 0.07215158 | 0.9424813 | 0.971691 |

| | | | | | | | |
|---|---|---|---|---|---|---|---|
| ENSG00000259191 | | 1.901746 | 0.02299129 | 0.7293917 | 0.06213339 | 0.9504566 | 0.975685 |
| ENSG00000173221 | GLRX | 350.7321 | 0.02294404 | 0.3334602 | 0.07183919 | 0.9427299 | 0.971771 |
| ENSG00000123144 | TRIR | 4484.46901 | 0.022941046 | 0.174680415 | 0.130884855 | 0.8958664 | 0.951863 |
| ENSG00000230587 | LINC02580 | 2.029579 | 0.02277956 | 0.6940909 | 0.06402276 | 0.9489521 | 0.975068 |
| ENSG00000258704 | SRP54-AS1 | 36.22952 | 0.0227412 | 0.3642658 | 0.06186548 | 0.95067 | 0.975753 |
| ENSG00000122966 | CIT | 1788.504939 | 0.022737729 | 0.393190356 | 0.061022995 | 0.9513409 | 0.976178 |
| ENSG00000160961 | ZNF333 | 318.2674 | 0.02270921 | 0.3550409 | 0.06699308 | 0.9465872 | 0.973611 |
| ENSG00000249158 | PCDHA11 | 689.612 | 0.02270884 | 0.3183657 | 0.08008471 | 0.9361699 | 0.968541 |
| ENSG00000267637 | | 0.8421071 | 0.02269146 | 0.74924 | 0.08232949 | 0.9343847 | NA |
| ENSG00000236986 | | 5.368699 | 0.02268178 | 0.6759905 | 0.06016713 | 0.9520225 | 0.976538 |
| ENSG00000287868 | | 1.02875 | 0.02267292 | 0.7248568 | 0.04808276 | 0.9616503 | NA |
| ENSG00000078114 | NEBL | 6248.485311 | 0.022645909 | 0.2908622 | 0.076922019 | 0.93868558 | 0.969795 |
| ENSG00000264204 | AGAP7P | 237.4956 | 0.02259364 | 0.2257198 | 0.08921564 | 0.9289105 | 0.965492 |
| ENSG00000068400 | GRIPAP1 | 5210.008369 | 0.022525361 | 0.164928871 | 0.114325744 | 0.90897958 | 0.9576 |
| ENSG00000227155 | | 7.568629 | 0.02247372 | 0.518943 | 0.06113265 | 0.9512536 | 0.976178 |
| ENSG00000148399 | DPH7 | 3239.059 | 0.02243873 | 0.1286958 | 0.1879247 | 0.8509357 | 0.927491 |
| ENSG00000235298 | | 10.83255 | 0.02239733 | 0.4425865 | 0.05805109 | 0.9537079 | 0.977458 |
| ENSG00000136930 | PSMB7 | 7720.797412 | 0.022302436 | 0.231613916 | 0.061535916 | 0.95093241 | 0.975985 |
| ENSG00000229798 | KRT18P26 | 3.292031 | 0.02229212 | 0.6540893 | 0.06319556 | 0.9496108 | 0.975269 |
| ENSG00000102001 | CACNA1F | 0.83760204 | 0.022247915 | 0.74131687 | 0.085227707 | 0.93208036 | NA |
| ENSG00000198680 | TUSC1 | 673.0394 | 0.02223397 | 0.2095825 | 0.1063674 | 0.9152909 | 0.960119 |
| ENSG00000235238 | SUMO2P1 | 845.0967 | 0.02219497 | 0.1578629 | 0.1349703 | 0.8926354 | 0.950211 |
| ENSG00000227962 | | 6.017009 | 0.02218203 | 0.5704988 | 0.05066233 | 0.9595946 | 0.979969 |
| ENSG00000267557 | | 1.094561 | 0.02217829 | 0.7362121 | 0.06473187 | 0.9483875 | NA |
| ENSG00000254340 | | 3.694851 | 0.02207444 | 0.6723047 | 0.0551294 | 0.9560354 | 0.978774 |
| ENSG00000169964 | TMEM42 | 528.3032 | 0.02203704 | 0.2325239 | 0.0944145 | 0.9247799 | 0.963851 |
| ENSG00000236814 | | 2093.424 | 0.02200722 | 0.2204193 | 0.1019952 | 0.9187605 | 0.961511 |
| ENSG00000141569 | TRIM65 | 4304.284 | 0.02193785 | 0.1270701 | 0.172372 | 0.8631451 | 0.934359 |
| ENSG00000102755 | FLT1 | 13.54946112 | 0.021846669 | 0.505546238 | 0.050289268 | 0.95989188 | 0.979994 |
| ENSG00000166246 | C16orf71 | 68.63639 | 0.02182275 | 0.2907907 | 0.07761142 | 0.9381372 | 0.969557 |
| ENSG00000265429 | MIR4782 | 2.815102 | 0.02180828 | 0.6313651 | 0.05610274 | 0.95526 | 0.9785 |
| ENSG00000288031 | | 7.287284 | 0.02179285 | 0.5870314 | 0.04895398 | 0.960956 | 0.980566 |
| ENSG00000198513 | ATL1 | 59.21175 | 0.02178845 | 0.32284 | 0.07746052 | 0.9382572 | 0.969605 |

| | | | | | | | |
|---|---|---|---|---|---|---|---|
| ENSG00000238058 | | 169.087 | 0.0214923 | 0.2111545 | 0.111591 | 0.9111477 | 0.958726 |
| ENSG00000225648 | SBDSP1 | 1534.856 | 0.02146548 | 0.2053175 | 0.1101284 | 0.9123075 | 0.958851 |
| ENSG00000266975 | FARSA-AS1 | 2.270341 | 0.02137734 | 0.6748977 | 0.05904988 | 0.9529124 | 0.977059 |
| ENSG00000279652 | | 4.04264 | 0.0213717 | 0.6466071 | 0.06332582 | 0.9495071 | 0.975269 |
| ENSG00000134982 | APC | 2120.479569 | 0.021263774 | 0.200397304 | 0.10322381 | 0.91778534 | 0.961195 |
| ENSG00000177542 | SLC25A22 | 1709.294 | 0.02119843 | 0.2233022 | 0.09462614 | 0.9246118 | 0.963771 |
| ENSG00000213862 | | 1350.903 | 0.02114366 | 0.1672731 | 0.1310492 | 0.8957364 | 0.951839 |
| ENSG00000269235 | ZNF350-AS1 | 25.14572 | 0.02105786 | 0.5379195 | 0.06320923 | 0.9495999 | 0.975269 |
| ENSG00000137309 | HMGA1 | 9391.958686 | 0.02102187 | 0.24461979 | 0.147084202 | 0.88306556 | 0.94524 |
| ENSG00000198034 | RPS4X | 39611.17 | 0.02096433 | 0.1595805 | 0.1315111 | 0.895371 | 0.951617 |
| ENSG00000132950 | ZMYM5 | 1780.097497 | 0.020923016 | 0.157114953 | 0.123589776 | 0.90164009 | 0.954802 |
| ENSG00000170860 | LSM3 | 3758.385 | 0.0209038 | 0.2631707 | 0.0820464 | 0.9346098 | 0.967756 |
| ENSG00000285736 | | 1.760721 | 0.02087916 | 0.7022148 | 0.05465577 | 0.9564127 | 0.978865 |
| ENSG00000255669 | | 1.020425 | 0.02081491 | 0.7208393 | 0.06646767 | 0.9470055 | NA |
| ENSG00000205534 | SMG1P2 | 204.8398 | 0.02076008 | 0.1968036 | 0.1080404 | 0.9139636 | 0.959693 |
| ENSG00000160953 | PWWP3A | 2216.04 | 0.02073672 | 0.1425443 | 0.1526187 | 0.878699 | 0.94286 |
| ENSG00000135317 | SNX14 | 3819.084396 | 0.02069009 | 0.123345271 | 0.205675486 | 0.83704441 | 0.918605 |
| ENSG00000271153 | RPL23AP88 | 2.317015 | 0.02061271 | 0.6719462 | 0.04914078 | 0.9608071 | 0.980476 |
| ENSG00000274641 | H2BC17 | 53.33064 | 0.02055833 | 0.4427692 | 0.05261558 | 0.9580382 | 0.979347 |
| ENSG00000271788 | | 27.71054 | 0.0205201 | 0.4463311 | 0.05160106 | 0.9588466 | 0.979737 |
| ENSG00000120215 | MLANA | 11.29513277 | 0.020432059 | 0.470716255 | 0.052875196 | 0.95783135 | 0.979347 |
| ENSG00000035928 | RFC1 | 4112.782456 | 0.020333658 | 0.201254595 | 0.098725825 | 0.92135596 | 0.962793 |
| ENSG00000167302 | TEPSIN | 993.3383 | 0.02029083 | 0.2477961 | 0.06847167 | 0.9454102 | 0.973061 |
| ENSG00000263535 | AK4P1 | 265.2858 | 0.02028116 | 0.2995202 | 0.05710047 | 0.9544652 | 0.977986 |
| ENSG00000180357 | ZNF609 | 4882.289 | 0.02028058 | 0.2188332 | 0.09772507 | 0.9221506 | 0.963092 |
| ENSG00000225039 | LINC01058 | 1.695365 | 0.02025268 | 0.7204447 | 0.06054241 | 0.9517236 | 0.976385 |
| ENSG00000230561 | CCDC192 | 20.68097 | 0.02013857 | 0.3884847 | 0.05654925 | 0.9549043 | 0.978285 |
| ENSG00000168101 | NUDT16L1 | 2864.081 | 0.02007134 | 0.2271199 | 0.1008895 | 0.9196382 | 0.961775 |
| ENSG00000162585 | FAAP20 | 1512.623 | 0.02002413 | 0.1763016 | 0.1150558 | 0.9084009 | 0.957291 |
| ENSG00000157106 | SMG1 | 14384.54 | 0.02000684 | 0.3096662 | 0.06530067 | 0.9479346 | 0.974515 |
| ENSG00000283676 | MIR5087 | 3.711714 | 0.01997217 | 0.637774 | 0.05287347 | 0.9578327 | 0.979347 |
| ENSG00000155957 | TMBIM4 | 406.9702 | 0.01988227 | 0.224052 | 0.08996753 | 0.928313 | 0.965223 |
| ENSG00000267852 | | 1.605675 | 0.01986661 | 0.7171583 | 0.06235092 | 0.9502834 | 0.975545 |

| | | | | | | | |
|---|---|---|---|---|---|---|---|
| ENSG00000256262 | USP30-AS1 | 1.658042 | 0.01985739 | 0.6913578 | 0.06007976 | 0.9520921 | 0.976555 |
| ENSG00000237214 | | 185.0453 | 0.01983347 | 0.2493716 | 0.09647864 | 0.9231404 | 0.963465 |
| ENSG00000137822 | TUBGCP4 | 1383.24474 | 0.019814739 | 0.173168102 | 0.114335735 | 0.90897166 | 0.9576 |
| ENSG00000180610 | ZBTB12BP | 3.069865 | 0.01977837 | 0.6556124 | 0.04658286 | 0.9628457 | 0.981577 |
| ENSG00000130811 | EIF3G | 8845.867861 | 0.019769382 | 0.168598556 | 0.102825697 | 0.91810131 | 0.961226 |
| ENSG00000232024 | LSM12P1 | 1130.357 | 0.01975286 | 0.1821936 | 0.1101988 | 0.9122517 | 0.958851 |
| ENSG00000175893 | ZDHHC21 | 1457.223 | 0.01968142 | 0.1812772 | 0.1101377 | 0.9123002 | 0.958851 |
| ENSG00000223723 | TFDP1P2 | 123.1096 | 0.01966218 | 0.2863098 | 0.07129354 | 0.9431641 | 0.972105 |
| ENSG00000213609 | RPL7AP50 | 467.4804 | 0.01965374 | 0.19165 | 0.09536308 | 0.9240264 | 0.963623 |
| ENSG00000180189 | HMGB1P14 | 8.981511 | 0.01963254 | 0.5973019 | 0.04566001 | 0.9635812 | 0.982053 |
| ENSG00000261684 | | 0.908675 | 0.01959308 | 0.7496709 | 0.08138542 | 0.9351354 | NA |
| ENSG00000255329 | H2AZP4 | 2.14705 | 0.01958765 | 0.7045966 | 0.06106247 | 0.9513095 | 0.976178 |
| ENSG00000252334 | RNU6-1337P | 0.7560453 | 0.01958619 | 0.7321401 | 0.04947671 | 0.9605394 | NA |
| ENSG00000251000 | GGCTP1 | 15.19668 | 0.01958443 | 0.5031895 | 0.04864512 | 0.9612021 | 0.980729 |
| ENSG00000117713 | ARID1A | 4917.069199 | 0.019412197 | 0.213142677 | 0.095638176 | 0.92380794 | 0.963623 |
| ENSG00000269296 | | 0.7730335 | 0.01937057 | 0.7311203 | 0.04881305 | 0.9610683 | NA |
| ENSG00000253869 | PIGFP1 | 3.953742 | 0.01933853 | 0.6059976 | 0.04729934 | 0.9622747 | 0.981157 |
| ENSG00000247627 | MTND4P12 | 49.94068 | 0.01933705 | 0.3959376 | 0.06031967 | 0.951901 | 0.976451 |
| ENSG00000254791 | FAR1-IT1 | 1.60286 | 0.01930425 | 0.6913708 | 0.06005464 | 0.9521121 | 0.976555 |
| ENSG00000129084 | PSMA1 | 1441.683913 | 0.019257103 | 0.135975395 | 0.127406764 | 0.89861846 | 0.953206 |
| ENSG00000230325 | | 11.78352 | 0.01925405 | 0.5335219 | 0.04403322 | 0.9648779 | 0.982555 |
| ENSG00000273133 | | 3.219046 | 0.01921203 | 0.6214863 | 0.04443265 | 0.9645595 | 0.982497 |
| ENSG00000188603 | CLN3 | 95.27195 | 0.01917714 | 0.3727318 | 0.05489464 | 0.9562224 | 0.978805 |
| ENSG00000129354 | AP1M2 | 9633.698858 | 0.019149775 | 0.152486094 | 0.069001398 | 0.94498851 | 0.972822 |
| ENSG00000256283 | METTL8P1 | 18.81168 | 0.01911117 | 0.4489855 | 0.04977384 | 0.9603026 | 0.980188 |
| ENSG00000104412 | EMC2 | 3707.383087 | 0.019082576 | 0.142475246 | 0.122989516 | 0.9021154 | 0.95494 |
| ENSG00000144535 | DIS3L2 | 950.6231 | 0.01899112 | 0.1327303 | 0.1354168 | 0.8922824 | 0.950182 |
| ENSG00000143552 | NUP210L | 1650.569 | 0.01894252 | 0.1275822 | 0.1485439 | 0.8819135 | 0.944788 |
| ENSG00000263982 | | 2.733416 | 0.0188874 | 0.6608561 | 0.02207905 | 0.9823849 | 0.990738 |
| ENSG00000184076 | UQCR10 | 2041.214 | 0.01883878 | 0.2439398 | 0.0887839 | 0.9292537 | 0.965722 |
| ENSG00000271127 | | 6.019776 | 0.01879518 | 0.5546322 | 0.0441648 | 0.964773 | 0.982497 |
| ENSG00000104859 | CLASRP | 2871.771366 | 0.018779081 | 0.153914314 | 0.115277589 | 0.9082251 | 0.957217 |
| ENSG00000060971 | ACAA1 | 3142.311014 | 0.018701115 | 0.179660477 | 0.108683827 | 0.91345327 | 0.95946 |

| | | | | | | | |
|---|---|---|---|---|---|---|---|
| ENSG00000275550 | | 1.169957 | 0.01865759 | 0.7456712 | 0.05935816 | 0.9526668 | NA |
| ENSG00000228874 | | 2.088865 | 0.01857849 | 0.6931334 | 0.0413176 | 0.9670427 | 0.983702 |
| ENSG00000182366 | FAM87A | 1.886038 | 0.01855351 | 0.6944944 | 0.05158665 | 0.9588581 | 0.979737 |
| ENSG00000124257 | NEURL2 | 36.59162341 | 0.018540198 | 0.375400303 | 0.053806658 | 0.95708921 | 0.97922 |
| ENSG00000196865 | NHLRC2 | 1811.913 | 0.01853565 | 0.2095818 | 0.08839352 | 0.9295639 | 0.965881 |
| ENSG00000286380 | | 4.611948 | 0.01850121 | 0.6252585 | 0.04693245 | 0.9625671 | 0.981368 |
| ENSG00000130725 | UBE2M | 3007.634421 | 0.018483196 | 0.196475611 | 0.093317021 | 0.92565171 | 0.964272 |
| ENSG00000135451 | TROAP | 2175.347283 | 0.018447481 | 0.440972185 | 0.043886213 | 0.96499511 | 0.982596 |
| ENSG00000125484 | GTF3C4 | 4561.585654 | 0.018437409 | 0.200680254 | 0.09229606 | 0.92646282 | 0.964583 |
| ENSG00000099282 | TSPAN15 | 5251.608852 | 0.018430102 | 0.197086586 | 0.072191021 | 0.94244989 | 0.971691 |
| ENSG00000167578 | RAB4B | 35.8016 | 0.01842468 | 0.3965073 | 0.0525247 | 0.9581106 | 0.979347 |
| ENSG00000205189 | ZBTB10 | 1534.643 | 0.01829171 | 0.2613765 | 0.06897948 | 0.945006 | 0.972822 |
| ENSG00000227653 | ISCA1P6 | 23.8097 | 0.01826283 | 0.4256961 | 0.04937859 | 0.9606176 | 0.980372 |
| ENSG00000185527 | PDE6G | 12.61322 | 0.01820234 | 0.4736143 | 0.04345892 | 0.9653357 | 0.982795 |
| ENSG00000279677 | | 1.021245 | 0.01817062 | 0.749271 | 0.04821162 | 0.9615476 | NA |
| ENSG00000228217 | PNRC2P1 | 82.9904 | 0.01813987 | 0.2377778 | 0.07754615 | 0.9381891 | 0.969573 |
| ENSG00000100888 | CHD8 | 7974.38102 | 0.018138183 | 0.145335184 | 0.121615754 | 0.90320334 | 0.955125 |
| ENSG00000151498 | ACAD8 | 1646.85 | 0.01805613 | 0.1085038 | 0.154992 | 0.8768276 | 0.941882 |
| ENSG00000108312 | UBTF | 10197.09365 | 0.018017337 | 0.109508557 | 0.189687773 | 0.8495538 | 0.926874 |
| ENSG00000152104 | PTPN14 | 1865.071 | 0.0179461 | 0.3266214 | 0.05577296 | 0.9555227 | 0.978579 |
| ENSG00000140379 | BCL2A1 | 1.286748 | 0.01794341 | 0.7023568 | 0.04784944 | 0.9618362 | NA |
| ENSG00000029639 | TFB1M | 391.0159827 | 0.017940436 | 0.159760455 | 0.11623712 | 0.90746462 | 0.957029 |
| ENSG00000229990 | | 3.19975 | 0.01789607 | 0.6699382 | 0.05088199 | 0.9594196 | 0.97996 |
| ENSG00000109625 | CPZ | 9.222417089 | 0.017796422 | 0.55018723 | 0.039341742 | 0.96861793 | 0.983954 |
| ENSG00000137944 | KYAT3 | 1103.02618 | 0.017795032 | 0.205605918 | 0.08851696 | 0.92946581 | 0.965855 |
| ENSG00000198324 | PHETA1 | 565.2515 | 0.01777395 | 0.186039 | 0.09719546 | 0.9225712 | 0.963214 |
| ENSG00000278275 | | 12.4047 | 0.01773212 | 0.5172678 | 0.04210306 | 0.9664165 | 0.983543 |
| ENSG00000228869 | COX4I1P2 | 2.39458 | 0.0176914 | 0.6929538 | 0.04765442 | 0.9619917 | 0.981045 |
| ENSG00000264364 | DYNLL2 | 19923.28 | 0.01768634 | 0.1014124 | 0.1788345 | 0.8580677 | 0.931557 |
| ENSG00000164406 | LEAP2 | 81.39493 | 0.01764848 | 0.3714496 | 0.05282029 | 0.9578751 | 0.979347 |
| ENSG00000236144 | TMEM147-AS1 | 1073.845 | 0.01764653 | 0.2265045 | 0.07914064 | 0.9369208 | 0.968865 |
| ENSG00000102384 | CENPI | 693.1038405 | 0.01762694 | 0.545092676 | 0.027223228 | 0.97828169 | 0.988325 |
| ENSG00000087095 | NLK | 1568.166582 | 0.017604734 | 0.194013415 | 0.086836563 | 0.93080142 | 0.966433 |

| | | | | | | | |
|---|---|---|---|---|---|---|---|
| ENSG00000104848 | KCNA7 | 1.379138723 | 0.017594392 | 0.687551513 | 0.039990392 | 0.96810079 | NA |
| ENSG00000242325 | RPS12P31 | 1.471693 | 0.01753158 | 0.6937894 | 0.0492744 | 0.9607006 | NA |
| ENSG00000279361 | | 1.465958 | 0.01751958 | 0.7074446 | 0.05446773 | 0.9565625 | NA |
| ENSG00000204941 | PSG5 | 11.17385 | 0.01751415 | 0.6358672 | 0.03430059 | 0.9726375 | 0.985921 |
| ENSG00000085872 | CHERP | 1734.725899 | 0.01751122 | 0.171629936 | 0.102530201 | 0.91833584 | 0.961287 |
| ENSG00000223345 | H2BP1 | 0.8674131 | 0.01749604 | 0.7547771 | 0.05463569 | 0.9564287 | NA |
| ENSG00000146677 | RPL32P18 | 3190.032 | 0.01747626 | 0.231947 | 0.07702365 | 0.9386047 | 0.969795 |
| ENSG00000212541 | RNU6-510P | 1.921457 | 0.01746495 | 0.6726729 | 0.0461036 | 0.9632277 | 0.981815 |
| ENSG00000237008 | LAPTM4BP1 | 3.000806 | 0.01731685 | 0.7090957 | 0.05582035 | 0.9554849 | 0.978579 |
| ENSG00000221971 | TTC4P1 | 3.833897 | 0.01731192 | 0.707347 | 0.05849387 | 0.9533552 | 0.977339 |
| ENSG00000280734 | LINC01232 | 490.1035 | 0.01730518 | 0.3394825 | 0.06637968 | 0.9470755 | 0.973832 |
| ENSG00000233406 | | 5.25522 | 0.01730177 | 0.5924116 | 0.0401195 | 0.9679979 | 0.983954 |
| ENSG00000099940 | SNAP29 | 2438.39417 | 0.017295084 | 0.119018813 | 0.159158161 | 0.87354427 | 0.940119 |
| ENSG00000069509 | FUNDC1 | 1093.875463 | 0.017267802 | 0.217431205 | 0.084783371 | 0.93243361 | 0.967048 |
| ENSG00000147364 | FBXO25 | 1497.716 | 0.0170985 | 0.2130161 | 0.05655332 | 0.954901 | 0.978285 |
| ENSG00000231609 | | 25.14092 | 0.01709162 | 0.4522361 | 0.04545384 | 0.9637456 | 0.982108 |
| ENSG00000253605 | HNRNPA1P38 | 2.054488 | 0.01705999 | 0.6686927 | 0.04816779 | 0.9615825 | 0.980809 |
| ENSG00000254505 | CHMP4A | 101.6933 | 0.01693523 | 0.2449833 | 0.07244938 | 0.9422443 | 0.971644 |
| ENSG00000242125 | SNHG3 | 5318.482 | 0.01688325 | 0.1641684 | 0.1032295 | 0.9177808 | 0.961195 |
| ENSG00000185219 | ZNF445 | 1171.644 | 0.01680679 | 0.2210139 | 0.06684458 | 0.9467054 | 0.97364 |
| ENSG00000176444 | CLK2 | 5182.456 | 0.01670332 | 0.2038547 | 0.08361609 | 0.9333617 | 0.967249 |
| ENSG00000261559 | FSCN1P1 | 36.01182 | 0.01662524 | 0.6125595 | 0.05892664 | 0.9530105 | 0.977099 |
| ENSG00000116863 | ADPRS | 2269.834631 | 0.016621019 | 0.212242139 | 0.07557627 | 0.93975622 | 0.970365 |
| ENSG00000205464 | ATP6AP1L | 200.9531 | 0.01658473 | 0.3946489 | 0.04204567 | 0.9664623 | 0.983543 |
| ENSG00000175348 | TMEM9B | 2107.352 | 0.0165359 | 0.1660164 | 0.1010572 | 0.9195051 | 0.961751 |
| ENSG00000166166 | TRMT61A | 3459.596 | 0.01652981 | 0.1516875 | 0.09074137 | 0.9276981 | 0.96494 |
| ENSG00000127540 | UQCR11 | 1142.114154 | 0.016510268 | 0.225433592 | 0.070515361 | 0.94378348 | 0.972468 |
| ENSG00000135837 | CEP350 | 5831.620617 | 0.016366749 | 0.237359547 | 0.067776005 | 0.94596395 | 0.97335 |
| ENSG00000242692 | RPS27AP1 | 48.82896 | 0.01634853 | 0.2831139 | 0.0599072 | 0.9522295 | 0.976637 |
| ENSG00000286586 | | 2.07726 | 0.01633152 | 0.6646955 | 0.03668426 | 0.9707368 | 0.985048 |
| ENSG00000128191 | DGCR8 | 2477.545363 | 0.016330374 | 0.115816701 | 0.147245447 | 0.88293829 | 0.945233 |
| ENSG00000057608 | GDI2 | 16446.15743 | 0.016330206 | 0.180748713 | 0.136713879 | 0.89125696 | 0.949585 |
| ENSG00000147316 | MCPH1 | 782.8142 | 0.01626469 | 0.1609173 | 0.1004638 | 0.9199761 | 0.961824 |

| | | | | | | | |
|---|---|---|---|---|---|---|---|
| ENSG00000278223 | MIR6783 | 1.104791 | 0.01612035 | 0.7472639 | 0.06777539 | 0.9459644 | NA |
| ENSG00000184860 | SDR42E1 | 1049.532 | 0.01605885 | 0.1594961 | 0.100125 | 0.9202451 | 0.962052 |
| ENSG00000047849 | MAP4 | 8466.730093 | 0.016012025 | 0.168897396 | 0.134129998 | 0.89329978 | 0.950572 |
| ENSG00000005801 | ZNF195 | 2123.108249 | 0.016003878 | 0.144759512 | 0.107096485 | 0.91471244 | 0.959948 |
| ENSG00000221988 | PPT2 | 692.5955 | 0.01588592 | 0.2411771 | 0.06486471 | 0.9482817 | 0.974627 |
| ENSG00000100077 | GRK3 | 738.588309 | 0.015863224 | 0.219083958 | 0.075162594 | 0.94008535 | 0.97045 |
| ENSG00000174567 | GOLT1A | 1307.421 | 0.01574122 | 0.2580998 | 0.05560109 | 0.9556596 | 0.978636 |
| ENSG00000232546 | | 31.44633 | 0.01572501 | 0.3607392 | 0.04563415 | 0.9636018 | 0.982053 |
| ENSG00000174953 | DHX36 | 6056.263 | 0.01571041 | 0.1206646 | 0.167315 | 0.8671222 | 0.936413 |
| ENSG00000224839 | RPL12P17 | 1.790694 | 0.01570949 | 0.7043357 | 0.04509487 | 0.9640317 | 0.982258 |
| ENSG00000275756 | | 1.048884 | 0.01565904 | 0.7398124 | 0.03889242 | 0.9689762 | NA |
| ENSG00000112234 | FBXL4 | 1456.680319 | 0.015622393 | 0.223421467 | 0.074928031 | 0.94027197 | 0.97045 |
| ENSG00000105254 | TBCB | 4068.960654 | 0.015526729 | 0.195937486 | 0.078488697 | 0.93743932 | 0.969099 |
| ENSG00000276916 | | 2.012342 | 0.01541874 | 0.6634246 | 0.03598009 | 0.9712982 | 0.985429 |
| ENSG00000285908 | | 1.50506 | 0.01540306 | 0.6624242 | 0.02971138 | 0.9762972 | NA |
| ENSG00000146648 | EGFR | 1849.768 | 0.01536251 | 0.3773374 | 0.04812932 | 0.9616132 | 0.980809 |
| ENSG00000015568 | RGPD5 | 0.834617061 | 0.015345674 | 0.759839663 | 0.054175525 | 0.95679532 | NA |
| ENSG00000160877 | NACC1 | 5115.699 | 0.0152705 | 0.2430583 | 0.06274094 | 0.9499728 | 0.97554 |
| ENSG00000092871 | RFFL | 503.3893412 | 0.015243169 | 0.168034371 | 0.091985578 | 0.9267095 | 0.964715 |
| ENSG00000284376 | | 1.382839 | 0.01522496 | 0.6916982 | 0.04444292 | 0.9645514 | NA |
| ENSG00000229871 | RPSAP20 | 1.076114 | 0.01516162 | 0.7192569 | 0.04050484 | 0.9676907 | NA |
| ENSG00000254004 | ZNF260 | 1409.043 | 0.01510747 | 0.180519 | 0.08740855 | 0.9303468 | 0.966278 |
| ENSG00000253133 | | 1.594024 | 0.01507254 | 0.7091089 | 0.04494181 | 0.9641537 | 0.982345 |
| ENSG00000220744 | RPL5P18 | 24.69015 | 0.01500531 | 0.3282563 | 0.03772428 | 0.9699075 | 0.98462 |
| ENSG00000253562 | | 0.9798549 | 0.0149795 | 0.714336 | 0.03334744 | 0.9733975 | NA |
| ENSG00000224255 | PDCL3P6 | 3.495027 | 0.01490922 | 0.637082 | 0.0342551 | 0.9726737 | 0.985921 |
| ENSG00000253719 | ATXN7L3B | 9577.961 | 0.01478884 | 0.1020766 | 0.1590341 | 0.873642 | 0.940119 |
| ENSG00000145283 | SLC10A6 | 0.7844166 | 0.01478619 | 0.7373284 | 0.04994269 | 0.9601681 | NA |
| ENSG00000163322 | ABRAXAS1 | 539.964 | 0.01477421 | 0.2306484 | 0.06357206 | 0.949311 | 0.975263 |
| ENSG00000180304 | OAZ2 | 4044.099 | 0.01473998 | 0.1781068 | 0.08801314 | 0.9298662 | 0.966007 |
| ENSG00000008294 | SPAG9 | 9477.204832 | 0.014680246 | 0.188649813 | 0.088109136 | 0.92978994 | 0.966003 |
| ENSG00000237793 | RPL18AP16 | 2.647503 | 0.01464919 | 0.6743511 | 0.04728399 | 0.9622869 | 0.981157 |
| ENSG00000287001 | | 11.14302 | 0.01464234 | 0.5541702 | 0.03356456 | 0.9732244 | 0.986254 |

| | | | | | | | |
|---|---|---|---|---|---|---|---|
| ENSG00000236527 | ARF4P2 | 7.711707 | 0.01462711 | 0.5844266 | 0.03489622 | 0.9721625 | 0.985734 |
| ENSG00000054983 | GALC | 2145.589721 | 0.014584766 | 0.243498583 | 0.058148106 | 0.95363066 | 0.977458 |
| ENSG00000157330 | C1orf158 | 0.7914361 | 0.01457384 | 0.7614157 | 0.01988849 | 0.9841323 | NA |
| ENSG00000229122 | AGBL5-IT1 | 9.121941 | 0.01452777 | 0.5244721 | 0.03248808 | 0.9740828 | 0.986555 |
| ENSG00000165119 | HNRNPK | 29233.99 | 0.01446304 | 0.1526667 | 0.02127321 | 0.9830277 | 0.991085 |
| ENSG00000229036 | VDAC1P8 | 205.68 | 0.01442633 | 0.257452 | 0.05087621 | 0.9594242 | 0.97996 |
| ENSG00000122824 | NUDT10 | 5.791255961 | 0.014398901 | 0.585313135 | 0.02986548 | 0.97617434 | 0.987548 |
| ENSG00000213064 | SFT2D2 | 1797.431 | 0.01431381 | 0.1864467 | 0.08074407 | 0.9356455 | 0.968267 |
| ENSG00000163377 | TAFA4 | 3.389148 | 0.01431301 | 0.6691803 | 0.04415587 | 0.9647802 | 0.982497 |
| ENSG00000263603 | | 5.716754 | 0.01429181 | 0.5203234 | 0.03279985 | 0.9738342 | 0.986555 |
| ENSG00000286310 | | 2.930865 | 0.01426368 | 0.6398675 | 0.03231949 | 0.9742173 | 0.986583 |
| ENSG00000063854 | HAGH | 3168.631978 | 0.014209675 | 0.181787629 | 0.082764872 | 0.9340385 | 0.967277 |
| ENSG00000206199 | ANKUB1 | 1.682881 | 0.01420012 | 0.7209755 | 0.04731646 | 0.962261 | 0.981157 |
| ENSG00000236376 | | 2.493462 | 0.01405569 | 0.6878984 | 0.03907396 | 0.9688314 | 0.984054 |
| ENSG00000155666 | KDM8 | 336.2185 | 0.01397646 | 0.2380806 | 0.05303124 | 0.957707 | 0.979347 |
| ENSG00000253773 | C8orf37-AS1 | 9.972184 | 0.01391273 | 0.6045375 | 0.03212237 | 0.9743745 | 0.986622 |
| ENSG00000272070 | | 101.6102 | 0.01384097 | 0.3354576 | 0.02647483 | 0.9788786 | 0.98874 |
| ENSG00000139323 | POC1B | 1553.494 | 0.01383174 | 0.1487182 | 0.08829754 | 0.9296402 | 0.965893 |
| ENSG00000073060 | SCARB1 | 5179.09064 | 0.013822284 | 0.21304151 | 0.073383731 | 0.94150076 | 0.971124 |
| ENSG00000235481 | UBE2R2-AS1 | 7.837876 | 0.0137874 | 0.5577118 | 0.0280739 | 0.9776032 | 0.988186 |
| ENSG00000255993 | PSMC1P9 | 2.357238 | 0.01366446 | 0.7210885 | 0.05138088 | 0.959022 | 0.979746 |
| ENSG00000286365 | | 1.028615 | 0.01364751 | 0.7395871 | 0.01090557 | 0.9912988 | NA |
| ENSG00000238228 | OR7E7P | 9.747187 | 0.01359298 | 0.5523413 | 0.02228176 | 0.9822232 | 0.99065 |
| ENSG00000261798 | | 56.24704 | 0.0135852 | 0.36548 | 0.04324479 | 0.9655064 | 0.982931 |
| ENSG00000198399 | ITSN2 | 3182.443 | 0.01358372 | 0.2225995 | 0.06411943 | 0.9488751 | 0.975041 |
| ENSG00000157470 | FAM81A | 108.5969 | 0.01357537 | 0.3085933 | 0.04432727 | 0.9646435 | 0.982497 |
| ENSG00000279386 | | 7.623672 | 0.01354335 | 0.4783922 | 0.03251723 | 0.9740596 | 0.986555 |
| ENSG00000254923 | | 1.842511 | 0.01353982 | 0.7030331 | 0.02552172 | 0.9796388 | 0.989095 |
| ENSG00000206739 | | 2.779168 | 0.01351065 | 0.6416891 | 0.03714955 | 0.9703658 | 0.984859 |
| ENSG00000230565 | ZNF32-AS2 | 114.645 | 0.01340483 | 0.3417346 | 0.04086187 | 0.967406 | 0.983954 |
| ENSG00000235369 | RPL36AP15 | 2.731209 | 0.01338451 | 0.675915 | 0.03576222 | 0.971472 | 0.985523 |
| ENSG00000261723 | | 3.600266 | 0.01334554 | 0.6520067 | 0.03428652 | 0.9726487 | 0.985921 |
| ENSG00000111058 | ACSS3 | 3185.001439 | 0.013336015 | 0.166745902 | 0.075472439 | 0.93983883 | 0.970389 |

| | | | | | | | |
|---|---|---|---|---|---|---|---|
| ENSG00000187753 | C9orf153 | 1.645186 | 0.01332549 | 0.6866527 | 0.0353057 | 0.971836 | 0.985632 |
| ENSG00000237007 | KRT18P52 | 1.018687 | 0.01327977 | 0.7303669 | 0.03238972 | 0.9741613 | NA |
| ENSG00000196782 | MAML3 | 879.5783 | 0.0132243 | 0.1647519 | 0.08444514 | 0.9327025 | 0.967213 |
| ENSG00000262693 | | 8.324964 | 0.01319555 | 0.5436974 | 0.03160721 | 0.9747853 | 0.986888 |
| ENSG00000003393 | ALS2 | 2385.043797 | 0.013189441 | 0.146058209 | 0.052144789 | 0.95841333 | 0.979576 |
| ENSG00000130714 | POMT1 | 3941.512557 | 0.013123717 | 0.132850733 | 0.098703796 | 0.92137346 | 0.962793 |
| ENSG00000122481 | RWDD3 | 141.6901046 | 0.013106175 | 0.234727482 | 0.053205639 | 0.95756806 | 0.979347 |
| ENSG00000077063 | CTTNBP2 | 2.407684311 | 0.013041094 | 0.680499611 | 0.038562731 | 0.96923902 | 0.984268 |
| ENSG00000285865 | | 6.926932 | 0.01295315 | 0.5486507 | 0.03211877 | 0.9743773 | 0.986622 |
| ENSG00000224236 | MRRFP1 | 8.223604 | 0.01290527 | 0.6039443 | 0.03552036 | 0.9716648 | 0.985632 |
| ENSG00000120738 | EGR1 | 14804.70585 | 0.012861992 | 0.564297014 | -0.026984343 | 0.97847222 | 0.98848 |
| ENSG00000278784 | | 222.128 | 0.01285677 | 0.2707139 | 0.05267917 | 0.9579875 | 0.979347 |
| ENSG00000246477 | | 8.292467 | 0.01269006 | 0.5488077 | 0.03103371 | 0.9752427 | 0.987101 |
| ENSG00000204577 | LILRB3 | 4.805673 | 0.01266838 | 0.634233 | 0.04166118 | 0.9667688 | 0.983576 |
| ENSG00000140650 | PMM2 | 1328.457 | 0.01252641 | 0.1940667 | 0.06909484 | 0.9449141 | 0.972815 |
| ENSG00000184599 | TAFA3 | 3.196088 | 0.01243209 | 0.6297763 | 0.01914756 | 0.9847234 | 0.991967 |
| ENSG00000233927 | RPS28 | 1257.685 | 0.01240986 | 0.1702497 | 0.08665852 | 0.9309429 | 0.966445 |
| ENSG00000238172 | RPS2P35 | 1.122722 | 0.01236439 | 0.6990336 | 0.03225241 | 0.9742708 | NA |
| ENSG00000218976 | IMPDH1P9 | 9.468327 | 0.0123233 | 0.5605777 | 0.03400393 | 0.972874 | 0.986063 |
| ENSG00000196388 | INCA1 | 240.9704 | 0.01230264 | 0.2598989 | 0.04057222 | 0.9676369 | 0.983954 |
| ENSG00000259771 | | 0.8613476 | 0.01226896 | 0.7695891 | -0.06495768 | 0.9482077 | NA |
| ENSG00000230199 | | 2.516649 | 0.01222575 | 0.6742951 | 0.03277379 | 0.973855 | 0.986555 |
| ENSG00000268520 | | 9.559239 | 0.01210732 | 0.518281 | 0.02975671 | 0.9762611 | 0.987598 |
| ENSG00000287342 | | 6.729565 | 0.0120693 | 0.5999058 | 0.0332898 | 0.9734435 | 0.986363 |
| ENSG00000273483 | | 4.670065 | 0.01204918 | 0.6365889 | 0.01829791 | 0.9854012 | 0.992235 |
| ENSG00000250634 | LINC01182 | 2.591897 | 0.01199366 | 0.6596812 | 0.0216705 | 0.9827108 | 0.99092 |
| ENSG00000256338 | RPL41P2 | 50.9172 | 0.01198379 | 0.3129422 | 0.0441871 | 0.9647553 | 0.982497 |
| ENSG00000115694 | STK25 | 8108.575738 | 0.011959811 | 0.123082334 | 0.122750023 | 0.90230505 | 0.954999 |
| ENSG00000206926 | RNU6-1024P | 1.230309 | 0.01193556 | 0.7384664 | 0.0429609 | 0.9657327 | NA |
| ENSG00000114062 | UBE3A | 3351.905014 | 0.011893245 | 0.173745069 | 0.055938766 | 0.95539059 | 0.978579 |
| ENSG00000287202 | | 40.29977 | 0.01180011 | 0.4356977 | 0.02930502 | 0.9766213 | 0.987737 |
| ENSG00000165782 | PIP4P1 | 3666.182 | 0.01173125 | 0.1200608 | 0.1460644 | 0.8838706 | 0.945465 |
| ENSG00000239797 | RPL21P39 | 1.022056 | 0.01168153 | 0.7592141 | 0.03467108 | 0.972342 | NA |

| | | | | | | | |
|---|---|---|---|---|---|---|---|
| ENSG00000266171 | | 14.39232 | 0.01168112 | 0.4676145 | 0.03150586 | 0.9748661 | 0.986901 |
| ENSG00000285897 | | 15.71835 | 0.01166926 | 0.5203751 | 0.02883118 | 0.9769992 | 0.987931 |
| ENSG00000281530 | | 24.53359 | 0.0116012 | 0.3854717 | 0.03145753 | 0.9749047 | 0.986903 |
| ENSG00000104529 | EEF1D | 9909.492337 | 0.011598964 | 0.119822634 | 0.101155297 | 0.91942718 | 0.961751 |
| ENSG00000225885 | | 1.360047 | 0.01151097 | 0.7392094 | 0.0421979 | 0.9663409 | NA |
| ENSG00000214087 | ARL16 | 1565.027 | 0.01149134 | 0.1941683 | 0.05918668 | 0.9528034 | 0.977037 |
| ENSG00000272995 | | 0.7803959 | 0.01147533 | 0.7555732 | 0.02617724 | 0.979116 | NA |
| ENSG00000173706 | HEG1 | 629.2994 | 0.0113888 | 0.5327655 | 0.02637696 | 0.9789567 | 0.988744 |
| ENSG00000171456 | ASXL1 | 4781.367 | 0.01134299 | 0.1731088 | 0.05065964 | 0.9595967 | 0.979969 |
| ENSG00000259020 | | 1.870131 | 0.0113038 | 0.647509 | 0.02610313 | 0.9791751 | 0.988814 |
| ENSG00000122035 | RASL11A | 216.8601387 | 0.011236652 | 0.384787785 | 0.032454125 | 0.9741099 | 0.986555 |
| ENSG00000223549 | MTND5P28 | 4.611735 | 0.01121271 | 0.6100228 | 0.01892224 | 0.9849031 | 0.992063 |
| ENSG00000011009 | LYPLA2 | 2787.559677 | 0.011188525 | 0.202802962 | 0.053738018 | 0.95714389 | 0.97922 |
| ENSG00000132604 | TERF2 | 1800.377593 | 0.011187222 | 0.111932676 | 0.088421757 | 0.92954147 | 0.965881 |
| ENSG00000287822 | | 3.311499 | 0.01118434 | 0.6745799 | 0.03385618 | 0.9729918 | 0.986093 |
| ENSG00000223060 | | 3.570429 | 0.0110863 | 0.5853022 | 0.02426188 | 0.9806437 | 0.989716 |
| ENSG00000117899 | MESD | 4256.567858 | 0.01107168 | 0.126847192 | 0.087884302 | 0.92996863 | 0.966075 |
| ENSG00000238184 | CD81-AS1 | 10.62189 | 0.01103521 | 0.5244892 | 0.03134353 | 0.9749956 | 0.986919 |
| ENSG00000271275 | | 0.8693482 | 0.01093976 | 0.7408497 | 0.03053242 | 0.9756424 | NA |
| ENSG00000234065 | MTND4P26 | 4.662694 | 0.01084076 | 0.6257709 | 0.0184164 | 0.9853067 | 0.992179 |
| ENSG00000168806 | LCMT2 | 955.8262 | 0.01080846 | 0.2552936 | 0.04361039 | 0.965215 | 0.982748 |
| ENSG00000100207 | TCF20 | 3122.084212 | 0.010758911 | 0.174385431 | 0.085170379 | 0.93212594 | 0.966955 |
| ENSG00000258357 | NSA2P2 | 21.01275 | 0.01074384 | 0.4358166 | 0.0278846 | 0.9777542 | 0.988222 |
| ENSG00000228828 | TLK2P2 | 13.79486 | 0.01064134 | 0.4450463 | 0.02825505 | 0.9774587 | 0.988186 |
| ENSG00000134375 | TIMM17A | 5539.936318 | 0.010619867 | 0.240117574 | 0.05198267 | 0.9585425 | 0.979615 |
| ENSG00000160551 | TAOK1 | 7492.944 | 0.01058241 | 0.2755231 | 0.0367075 | 0.9707182 | 0.985048 |
| ENSG00000188283 | ZNF383 | 399.1069 | 0.01057614 | 0.191251 | 0.05280519 | 0.9578871 | 0.979347 |
| ENSG00000167118 | URM1 | 4425.892 | 0.01056788 | 0.1712571 | 0.05294693 | 0.9577742 | 0.979347 |
| ENSG00000095397 | WHRN | 1261.789408 | 0.010482089 | 0.269075776 | 0.051004211 | 0.95932217 | 0.97994 |
| ENSG00000164961 | WASHC5 | 8561.495 | 0.01040695 | 0.1528239 | 0.05539572 | 0.9558232 | 0.978736 |
| ENSG00000213169 | | 27.7346 | 0.01032588 | 0.4368095 | 0.02754106 | 0.9780282 | 0.988291 |
| ENSG00000256223 | ZNF10 | 395.3069 | 0.01031569 | 0.2189375 | 0.04415071 | 0.9647843 | 0.982497 |
| ENSG00000276704 | | 0.943028 | 0.01030803 | 0.7617709 | 0.005670804 | 0.9954754 | NA |

| | | | | | | | |
|---|---|---|---|---|---|---|---|
| ENSG00000248773 | | 16.14945 | 0.01030332 | 0.4728043 | 0.02348019 | 0.9812672 | 0.990118 |
| ENSG00000280114 | | 10.28362 | 0.01027588 | 0.4563075 | 0.02263464 | 0.9819417 | 0.990592 |
| ENSG00000232134 | RPS15AP12 | 59.97951 | 0.01026943 | 0.359713 | 0.03023627 | 0.9758786 | 0.987548 |
| ENSG00000272345 | | 16.9345 | 0.0102542 | 0.534478 | 0.02187239 | 0.9825497 | 0.990866 |
| ENSG00000260602 | HMGN2P40 | 0.9308511 | 0.01016435 | 0.7476 | 0.023516 | 0.9812387 | NA |
| ENSG00000273295 | | 125.2264 | 0.01015844 | 0.322276 | 0.03263779 | 0.9739634 | 0.986555 |
| ENSG00000272154 | | 6.078712 | 0.01011688 | 0.5784784 | 0.0216518 | 0.9827257 | 0.99092 |
| ENSG00000157014 | TATDN2 | 2501.939 | 0.009999545 | 0.1754118 | 0.04955697 | 0.9604754 | 0.980289 |
| ENSG00000119801 | YPEL5 | 4067.650206 | 0.009957823 | 0.23934205 | 0.042164021 | 0.96636794 | 0.983543 |
| ENSG00000178235 | SLITRK1 | 2.034296 | 0.009942043 | 0.6724666 | 0.01398889 | 0.9888388 | 0.994002 |
| ENSG00000268087 | | 33.64254 | 0.009798048 | 0.3413121 | 0.02901634 | 0.9768516 | 0.987895 |
| ENSG00000197971 | MBP | 1305.424 | 0.009794933 | 0.2073524 | 0.04430737 | 0.9646594 | 0.982497 |
| ENSG00000148408 | CACNA1B | 8.593594 | 0.009742102 | 0.5580379 | 0.01948526 | 0.984454 | 0.991846 |
| ENSG00000175309 | PHYKPL | 3893.124 | 0.00970808 | 0.2607822 | 0.03001493 | 0.9760551 | 0.987548 |
| ENSG00000106133 | NSUN5P2 | 33.97832503 | 0.009696792 | 0.370976063 | 0.027623816 | 0.97796219 | 0.988291 |
| ENSG00000188316 | ENO4 | 11.95688 | 0.009694233 | 0.5229662 | 0.01922417 | 0.9846623 | 0.991943 |
| ENSG00000188612 | SUMO2 | 4113.205 | 0.009681153 | 0.1536013 | 0.07194811 | 0.9426432 | 0.971757 |
| ENSG00000278842 | | 2.724195 | 0.009643454 | 0.6445141 | 0.02345374 | 0.9812883 | 0.990118 |
| ENSG00000126003 | PLAGL2 | 1998.862658 | 0.009598109 | 0.200154147 | 0.051005921 | 0.9593208 | 0.97994 |
| ENSG00000261436 | | 0.8093764 | 0.009577908 | 0.7628515 | 0.01221426 | 0.9902547 | NA |
| ENSG00000233690 | EBAG9P1 | 8.833911 | 0.009577753 | 0.5566693 | 0.01718564 | 0.9862885 | 0.992715 |
| ENSG00000244234 | GMCL2 | 44.7455 | 0.0095238 | 0.3399962 | 0.02817832 | 0.9775199 | 0.988186 |
| ENSG00000142632 | ARHGEF19 | 1978.459 | 0.009510944 | 0.2863406 | 0.02674749 | 0.9786611 | 0.988596 |
| ENSG00000226194 | LINC02519 | 1.131092 | 0.009443175 | 0.7210767 | 0.02417035 | 0.9807167 | NA |
| ENSG00000177352 | CCDC71 | 1119.204 | 0.00939249 | 0.2067996 | 0.0442193 | 0.9647296 | 0.982497 |
| ENSG00000235700 | CYCSP52 | 2.828004 | 0.009387534 | 0.6551066 | 0.02109193 | 0.9831723 | 0.991119 |
| ENSG00000158571 | PFKFB1 | 20.54231 | 0.009334085 | 0.3997336 | 0.02370865 | 0.981085 | 0.98999 |
| ENSG00000198618 | PPIAP22 | 17295.7 | 0.009262983 | 0.2282982 | 0.03893411 | 0.9689429 | 0.98413 |
| ENSG00000276957 | | 4.290837 | 0.009250169 | 0.6530698 | 0.02537744 | 0.9797539 | 0.989136 |
| ENSG00000170681 | CAVIN4 | 25.59167 | 0.009197079 | 0.4806075 | 0.0204697 | 0.9836687 | 0.991486 |
| ENSG00000174276 | ZNHIT2 | 771.5324 | 0.00915879 | 0.2272769 | 0.0383776 | 0.9693866 | 0.984317 |
| ENSG00000228366 | | 1.650574 | 0.009144006 | 0.7153951 | 0.00672884 | 0.9946312 | 0.997108 |
| ENSG00000100360 | IFT27 | 1042.071993 | 0.009140367 | 0.163253848 | 0.084467169 | 0.93268501 | 0.967213 |

| | | | | | | | |
|---|---|---|---|---|---|---|---|
| ENSG00000089057 | SLC23A2 | 1657.688407 | 0.009116982 | 0.198877521 | 0.042170352 | 0.9663629 | 0.983543 |
| ENSG00000125821 | DTD1 | 1270.509665 | 0.009095838 | 0.176707366 | 0.049977262 | 0.96014051 | 0.980097 |
| ENSG00000267541 | MTCO2P2 | 19.57759 | 0.008986607 | 0.4872606 | 0.01989864 | 0.9841242 | 0.991697 |
| ENSG00000252641 | RNU6-678P | 1.488001 | 0.008936938 | 0.7178453 | 0.02448032 | 0.9804695 | NA |
| ENSG00000225483 | RPL22P4 | 0.7815012 | 0.008874849 | 0.7445786 | 0.03091471 | 0.9753376 | NA |
| ENSG00000270240 | | 1.417814 | 0.008870348 | 0.7269691 | 0.03244609 | 0.9741163 | NA |
| ENSG00000166337 | TAF10 | 159.3574 | 0.00881765 | 0.2352208 | 0.03935972 | 0.9686036 | 0.983954 |
| ENSG00000159784 | FAM131B | 19.23471 | 0.008785553 | 0.5294441 | 0.02556592 | 0.9796036 | 0.989095 |
| ENSG00000160703 | NLRX1 | 1122.036 | 0.008643594 | 0.1746134 | 0.02529546 | 0.9798193 | 0.989164 |
| ENSG00000206712 | RNU6-26P | 2.865104 | 0.008613051 | 0.6425781 | 0.01695277 | 0.9864743 | 0.992789 |
| ENSG00000236528 | | 1.239629 | 0.008556152 | 0.7332578 | 0.02497899 | 0.9800717 | NA |
| ENSG00000257711 | | 2.076109 | 0.008519774 | 0.6566333 | 0.0210543 | 0.9832023 | 0.991119 |
| ENSG00000168286 | THAP11 | 1889.933 | 0.00847797 | 0.2332655 | 0.01724805 | 0.9862387 | 0.992715 |
| ENSG00000104894 | CD37 | 54.5607362 | 0.008456493 | 0.45624149 | 0.015625883 | 0.98753286 | 0.993181 |
| ENSG00000215301 | DDX3X | 20692.51 | 0.008401612 | 0.1756497 | 0.04623539 | 0.9631226 | 0.981784 |
| ENSG00000224186 | C5orf66 | 78.89588 | 0.008328887 | 0.3183326 | 0.02543382 | 0.9797089 | 0.989128 |
| ENSG00000095380 | NANS | 2542.118498 | 0.008311678 | 0.178894725 | 0.038564384 | 0.9692377 | 0.984268 |
| ENSG00000229896 | | 2.923962 | 0.008275938 | 0.6816215 | 0.02804895 | 0.9776231 | 0.988186 |
| ENSG00000204227 | RING1 | 3222.862 | 0.0082739 | 0.1313901 | 0.05902117 | 0.9529352 | 0.977059 |
| ENSG00000174564 | IL20RB | 85.8869 | 0.008188126 | 0.2463564 | 0.03393391 | 0.9729299 | 0.986068 |
| ENSG00000272944 | | 52.28105 | 0.008093344 | 0.4194016 | 0.01998181 | 0.9840579 | 0.991697 |
| ENSG00000164893 | SLC7A13 | 3.848206 | 0.008062088 | 0.6541728 | -0.00318912 | 0.9974555 | 0.999042 |
| ENSG00000263432 | RN7SL689P | 22.0985 | 0.008017718 | 0.404715 | 0.01985725 | 0.9841573 | 0.991697 |
| ENSG00000219391 | | 7.113262 | 0.008014913 | 0.5921863 | 0.01675234 | 0.9866342 | 0.992852 |
| ENSG00000127870 | RNF6 | 3553.233057 | 0.008007292 | 0.208323205 | 0.033800567 | 0.97303618 | 0.986101 |
| ENSG00000228060 | PABPC4-AS1 | 76.10133 | 0.007963509 | 0.3208595 | 0.03315565 | 0.9735505 | 0.986418 |
| ENSG00000172789 | HOXC5 | 7.854249 | 0.007917756 | 0.5262602 | 0.009951216 | 0.9920602 | 0.995695 |
| ENSG00000215914 | MMP23A | 3.4125 | 0.007848025 | 0.5975377 | 0.01585997 | 0.9873461 | 0.993181 |
| ENSG00000213613 | RPL11P3 | 11.87774 | 0.007763983 | 0.4537 | 0.01808107 | 0.9855742 | 0.992323 |
| ENSG00000234354 | RPS26P47 | 5.262694 | 0.00775555 | 0.637498 | 0.02040095 | 0.9837235 | 0.991486 |
| ENSG00000280273 | | 7.789474 | 0.007719238 | 0.5410732 | 0.01462246 | 0.9883334 | 0.993719 |
| ENSG00000169291 | SHE | 0.8800186 | 0.007716199 | 0.7593412 | -0.000929152 | 0.9992586 | NA |
| ENSG00000108587 | GOSR1 | 1830.348509 | 0.007698108 | 0.1345909 | 0.059562837 | 0.95250382 | 0.976843 |

| | | | | | | | |
|---|---|---|---|---|---|---|---|
| ENSG00000225279 | | 1.262866 | 0.007684522 | 0.7113781 | 0.006164433 | 0.9950815 | NA |
| ENSG00000224765 | | 2.91744 | 0.007679572 | 0.6589198 | 0.01865681 | 0.9851149 | 0.992156 |
| ENSG00000159792 | PSKH1 | 2937.234 | 0.007625412 | 0.1617064 | 0.04391293 | 0.9649738 | 0.982596 |
| ENSG00000161647 | MPP3 | 644.3897 | 0.007527149 | 0.2153105 | 0.03329028 | 0.9734431 | 0.986363 |
| ENSG00000140835 | CHST4 | 1.693661 | 0.007514741 | 0.6749013 | 0.01720033 | 0.9862768 | 0.992715 |
| ENSG00000127666 | TICAM1 | 595.6449052 | 0.007433777 | 0.135307525 | 0.053296956 | 0.95749531 | 0.979347 |
| ENSG00000160223 | ICOSLG | 1.775457 | 0.007414556 | 0.6829642 | 0.01690149 | 0.9865152 | 0.992793 |
| ENSG00000137818 | RPLP1 | 26868.70713 | 0.007411285 | 0.138985917 | 0.030609582 | 0.9755809 | 0.987324 |
| ENSG00000105700 | KXD1 | 1594.744853 | 0.007401325 | 0.144967146 | 0.055026163 | 0.95611762 | 0.978774 |
| ENSG00000090621 | PABPC4 | 12051.84963 | 0.007297674 | 0.194799861 | 0.063310396 | 0.94951934 | 0.975269 |
| ENSG00000235731 | LINC02250 | 1.058033 | 0.007294378 | 0.7190368 | 0.02676633 | 0.9786461 | NA |
| ENSG00000203266 | | 1.098483 | 0.007271952 | 0.7091312 | 0.01570093 | 0.987473 | NA |
| ENSG00000244515 | KRT18P34 | 48.21201 | 0.007215814 | 0.3566898 | 0.02180661 | 0.9826022 | 0.990882 |
| ENSG00000267412 | | 0.9042234 | 0.00720289 | 0.7488085 | 0.01989945 | 0.9841236 | NA |
| ENSG00000264176 | MAGOH2P | 8.356044 | 0.007101248 | 0.5444547 | 0.01485693 | 0.9881463 | 0.993606 |
| ENSG00000186020 | ZNF529 | 533.6159 | 0.007085474 | 0.2330797 | 0.02865521 | 0.9771396 | 0.988036 |
| ENSG00000182149 | IST1 | 11217.41 | 0.007085318 | 0.118272 | 0.08438225 | 0.9327525 | 0.967215 |
| ENSG00000245468 | LINC02447 | 30.76585 | 0.00706343 | 0.3948859 | 0.01876396 | 0.9850294 | 0.992124 |
| ENSG00000134108 | ARL8B | 6511.366366 | 0.007056329 | 0.150573043 | 0.035724327 | 0.97150217 | 0.985523 |
| ENSG00000166233 | ARIH1 | 4635.097 | 0.007029325 | 0.127593 | 0.03857217 | 0.9692315 | 0.984268 |
| ENSG00000119403 | PHF19 | 2134.065883 | 0.007011415 | 0.404382432 | 0.013616296 | 0.9891361 | 0.99405 |
| ENSG00000267096 | | 17.33093 | 0.007000424 | 0.4384074 | 0.01664027 | 0.9867236 | 0.992852 |
| ENSG00000254206 | NPIPB11 | 48.92749 | 0.006986003 | 0.3322139 | 0.0268563 | 0.9785744 | 0.988546 |
| ENSG00000161800 | RACGAP1 | 4232.621 | 0.006967141 | 0.3445784 | 0.0348871 | 0.9721698 | 0.985734 |
| ENSG00000113456 | RAD1 | 2676.426597 | 0.00679806 | 0.206256518 | 0.037542283 | 0.97005263 | 0.984727 |
| ENSG00000117475 | BLZF1 | 2099.495843 | 0.006787306 | 0.190963802 | 0.032648839 | 0.97395462 | 0.986555 |
| ENSG00000108788 | MLX | 4084.244541 | 0.006663469 | 0.158330607 | 0.02419034 | 0.98070078 | 0.989715 |
| ENSG00000068781 | STON1-GTF2A1L | 0.83741469 | 0.006630423 | 0.755772157 | 0.007491962 | 0.99402234 | NA |
| ENSG00000287852 | | 3.288455 | 0.006560365 | 0.6499734 | -0.008406511 | 0.9932927 | 0.996354 |
| ENSG00000228613 | | 170.8386 | 0.006423909 | 0.4477169 | 0.0121159 | 0.9903331 | 0.994751 |
| ENSG00000277588 | MIR6806 | 1.155022 | 0.006390845 | 0.7463953 | 0.01027836 | 0.9917992 | NA |
| ENSG00000254985 | RSF1-IT2 | 2.100521 | 0.006377164 | 0.6750004 | 0.007802277 | 0.9937747 | 0.996626 |
| ENSG00000160973 | FOXH1 | 20.23721 | 0.00635844 | 0.5083311 | 0.0129404 | 0.9896753 | 0.994429 |

| | | | | | | | |
|---|---|---|---|---|---|---|---|
| ENSG00000273838 | | 2.941967 | 0.006335301 | 0.7136037 | 0.01319651 | 0.989471 | 0.994299 |
| ENSG00000135452 | TSPAN31 | 2089.887373 | 0.006328911 | 0.223614121 | 0.022356819 | 0.98216333 | 0.990627 |
| ENSG00000186615 | KTN1-AS1 | 542.3769 | 0.006144897 | 0.2397273 | 0.02848091 | 0.9772786 | 0.988139 |
| ENSG00000270218 | | 1.202006 | 0.006036875 | 0.7186813 | -0.01084729 | 0.9913453 | NA |
| ENSG00000235720 | GABPAP | 141.1838 | 0.006010078 | 0.238724 | 0.02475852 | 0.9802476 | 0.989408 |
| ENSG00000227471 | AKR1B15 | 67.37546 | 0.0059831 | 0.5339129 | 0.02641945 | 0.9789228 | 0.988744 |
| ENSG00000219355 | RPL31P52 | 3.93895 | 0.005873205 | 0.6402478 | 0.01381675 | 0.9889762 | 0.994007 |
| ENSG00000185122 | HSF1 | 6164.247 | 0.005863524 | 0.1169629 | 0.0485326 | 0.9612918 | 0.980745 |
| ENSG00000160075 | SSU72 | 5175.855 | 0.005852815 | 0.187699 | 0.03021731 | 0.9758937 | 0.987548 |
| ENSG00000123684 | LPGAT1 | 2752.098081 | 0.005841327 | 0.191846913 | 0.027943631 | 0.97770711 | 0.988222 |
| ENSG00000110367 | DDX6 | 9334.056387 | 0.005720725 | 0.144789168 | 0.063317284 | 0.94951385 | 0.975269 |
| ENSG00000212242 | RNA5SP219 | 1.528453 | 0.005693058 | 0.6864935 | -0.000367989 | 0.9997064 | NA |
| ENSG00000248564 | | 1.482314 | 0.005692354 | 0.7344383 | -0.002578348 | 0.9979428 | NA |
| ENSG00000127720 | METTL25 | 290.2855376 | 0.005598677 | 0.272164315 | 0.018452451 | 0.98527791 | 0.992179 |
| ENSG00000260689 | HNRNPA3P11 | 2.470316 | 0.005392828 | 0.6966123 | 0.01023442 | 0.9918343 | 0.995543 |
| ENSG00000260672 | | 2.536788 | 0.005379912 | 0.6271599 | 0.01227453 | 0.9902066 | 0.994662 |
| ENSG00000072364 | AFF4 | 10060.6578 | 0.005347782 | 0.274594549 | 0.01628595 | 0.98700627 | 0.993024 |
| ENSG00000053747 | LAMA3 | 300.0774783 | 0.005235657 | 0.290795847 | 0.017218404 | 0.98626238 | 0.992715 |
| ENSG00000153827 | TRIP12 | 9376.942 | 0.005225461 | 0.1397622 | 0.02401424 | 0.9808413 | 0.98982 |
| ENSG00000253868 | FER1L6-AS2 | 1.395188 | 0.005200722 | 0.7312944 | 0.01281558 | 0.9897749 | NA |
| ENSG00000249245 | NCOA4P3 | 12.28202 | 0.005151126 | 0.5023571 | 0.01085601 | 0.9913383 | 0.995309 |
| ENSG00000241511 | RPS15AP24 | 109.5113 | 0.004939367 | 0.3193209 | 0.01670839 | 0.9866693 | 0.992852 |
| ENSG00000197852 | INKA2 | 596.4164 | 0.004925222 | 0.2271068 | 0.02161864 | 0.9827522 | 0.99092 |
| ENSG00000273893 | | 58.64446 | 0.00485885 | 0.2597593 | 0.01704775 | 0.9863985 | 0.992777 |
| ENSG00000265443 | | 2.906874 | 0.004785643 | 0.6019456 | 0.008207185 | 0.9934517 | 0.996394 |
| ENSG00000268062 | SRP14P4 | 1.414262 | 0.004774651 | 0.7042052 | 0.003154317 | 0.9974832 | NA |
| ENSG00000147381 | MAGEA4 | 1.66333 | 0.004711004 | 0.7641961 | -0.07504921 | 0.9401756 | 0.97045 |
| ENSG00000279159 | | 137.1795 | 0.004688521 | 0.274167 | 0.01257776 | 0.9899647 | 0.994603 |
| ENSG00000141753 | IGFBP4 | 7075.744 | 0.004672236 | 0.3607256 | 0.004426984 | 0.9964678 | 0.99846 |
| ENSG00000196611 | MMP1 | 17.05496 | 0.00465518 | 0.6408747 | 0.001799966 | 0.9985638 | 0.99924 |
| ENSG00000233493 | TMEM238 | 332.2332 | 0.004608176 | 0.2393823 | 0.02756419 | 0.9780097 | 0.988291 |
| ENSG00000101126 | ADNP | 10763.12202 | 0.004490958 | 0.127877451 | -0.019610501 | 0.98435409 | 0.99182 |
| ENSG00000251484 | | 6.454789 | 0.004490506 | 0.524304 | 0.006830529 | 0.9945501 | 0.997102 |

| | | | | | | | |
|---|---|---|---|---|---|---|---|
| ENSG00000197363 | ZNF517 | 1516.208 | 0.0044748 | 0.1871261 | 0.01792629 | 0.9856977 | 0.992384 |
| ENSG00000171219 | CDC42BPG | 5605.765 | 0.004434638 | 0.1676174 | 0.02501103 | 0.9800462 | 0.98928 |
| ENSG00000264961 | MIR4730 | 7.371124 | 0.004318775 | 0.544196 | 0.00588698 | 0.9953029 | 0.997518 |
| ENSG00000087302 | RTRAF | 7493.240603 | 0.004302127 | 0.154882746 | 0.044235378 | 0.96471678 | 0.982497 |
| ENSG00000213024 | NUP62 | 3896.075 | 0.004233684 | 0.1861416 | 0.04676489 | 0.9627006 | 0.981466 |
| ENSG00000256222 | MTRNR2L3 | 1.115429 | 0.004216698 | 0.7465032 | 0.01726276 | 0.986227 | NA |
| ENSG00000136827 | TOR1A | 3612.714534 | 0.00419613 | 0.171415654 | 0.039513176 | 0.96848125 | 0.983954 |
| ENSG00000184489 | PTP4A3 | 1020.331 | 0.004180497 | 0.234383 | 0.01149761 | 0.9908264 | 0.994983 |
| ENSG00000261000 | | 1.382745 | 0.004075949 | 0.7074628 | -0.003565599 | 0.9971551 | NA |
| ENSG00000224971 | SUMO2P3 | 2.566491 | 0.004070175 | 0.6511579 | 0.007067837 | 0.9943607 | 0.996987 |
| ENSG00000138375 | SMARCAL1 | 1140.345389 | 0.004007707 | 0.127603655 | 0.030124085 | 0.97596809 | 0.987548 |
| ENSG00000287077 | | 38.37145 | 0.003904121 | 0.3971939 | 0.009164677 | 0.9926877 | 0.996024 |
| ENSG00000085788 | DDHD2 | 2227.720901 | 0.003839247 | 0.145657751 | 0.03102471 | 0.97524983 | 0.987101 |
| ENSG00000100568 | VTI1B | 4146.222049 | 0.003813647 | 0.126757911 | 0.03745213 | 0.97012451 | 0.984727 |
| ENSG00000175063 | UBE2C | 3041.752 | 0.003793537 | 0.4274749 | 0.000195656 | 0.9998439 | 0.999882 |
| ENSG00000120708 | TGFBI | 1049.144203 | 0.003767131 | 0.374026401 | 0.026054939 | 0.97921352 | 0.988815 |
| ENSG00000231851 | MRPS9-AS1 | 14.10328 | 0.003705825 | 0.4991426 | -0.007857235 | 0.9937309 | 0.996619 |
| ENSG00000035862 | TIMP2 | 1643.269505 | 0.003670553 | 0.325105763 | 0.018629791 | 0.98513644 | 0.992156 |
| ENSG00000273218 | | 10.63773 | 0.003669886 | 0.4738657 | 0.003892583 | 0.9968942 | 0.998811 |
| ENSG00000272666 | KLHDC7B-DT | 2.187269 | 0.003648959 | 0.7203575 | -0.01126833 | 0.9910094 | 0.995054 |
| ENSG00000185504 | FAAP100 | 4111.491 | 0.003532451 | 0.1949744 | 0.01170323 | 0.9906624 | 0.994894 |
| ENSG00000117139 | KDM5B | 20640.34502 | 0.003502123 | 0.231057031 | 0.012331281 | 0.99016131 | 0.994654 |
| ENSG00000136709 | WDR33 | 4332.566918 | 0.003496593 | 0.120106503 | 0.041362221 | 0.96700713 | 0.983702 |
| ENSG00000140307 | GTF2A2 | 2381.245 | 0.00348846 | 0.2383389 | 0.02738607 | 0.9781518 | 0.988325 |
| ENSG00000176812 | METTL1P1 | 1.640782 | 0.003480597 | 0.7439575 | 0.02003518 | 0.9840153 | 0.991697 |
| ENSG00000203435 | E2F3P2 | 0.8714759 | 0.003469069 | 0.731814 | -0.02165097 | 0.9827264 | NA |
| ENSG00000103365 | GGA2 | 6263.687142 | 0.003444787 | 0.138206475 | 0.032422414 | 0.97413519 | 0.986555 |
| ENSG00000216624 | GAPDHP72 | 27.37508 | 0.003269706 | 0.4513268 | 0.0137484 | 0.9890307 | 0.994007 |
| ENSG00000118181 | RPS25 | 14707.90402 | 0.003261599 | 0.170794086 | 0.000468839 | 0.99962592 | 0.999814 |
| ENSG00000230914 | KIF19BP | 1.269187 | 0.003256068 | 0.7257642 | 0.008255288 | 0.9934133 | NA |
| ENSG00000099783 | HNRNPM | 13352.7225 | 0.003240492 | 0.241800279 | 0.010329694 | 0.99175824 | 0.995543 |
| ENSG00000130766 | SESN2 | 3706.370107 | 0.003209799 | 0.368615093 | 0.001015041 | 0.99919012 | 0.999453 |
| ENSG00000230409 | TCEA1P2 | 2661.196 | 0.003191422 | 0.2514215 | 0.007531183 | 0.993991 | 0.996683 |

| | | | | | | | |
|---|---|---|---|---|---|---|---|
| ENSG00000203872 | C6orf163 | 33.73932 | 0.003131121 | 0.3908391 | 0.006076544 | 0.9951516 | 0.997404 |
| ENSG00000143158 | MPC2 | 1607.377 | 0.003059297 | 0.2698821 | 0.002863328 | 0.9977154 | 0.999042 |
| ENSG00000218857 | | 0.7621739 | 0.002908479 | 0.7420155 | -0.0283969 | 0.9773456 | NA |
| ENSG00000148985 | PGAP2 | 1257.679 | 0.002890843 | 0.1276627 | 0.01430454 | 0.988587 | 0.993884 |
| ENSG00000226690 | | 2.087749 | 0.002634483 | 0.6864432 | -0.005053831 | 0.9959676 | 0.998034 |
| ENSG00000145147 | SLIT2 | 1067.375 | 0.00254383 | 0.3511107 | 0.007495739 | 0.9940193 | 0.996683 |
| ENSG00000226479 | TMEM185B | 3564.925 | 0.002540658 | 0.147005 | 0.009830271 | 0.9921567 | 0.995717 |
| ENSG00000170476 | MZB1 | 113.0732 | 0.002469412 | 0.307286 | 0.002450704 | 0.9980446 | 0.999173 |
| ENSG00000106404 | CLDN15 | 484.7734656 | 0.002404252 | 0.280422055 | 0.011403539 | 0.99090149 | 0.994983 |
| ENSG00000255337 | | 28.75298 | 0.002379437 | 0.455468 | 0.005083367 | 0.9959441 | 0.998034 |
| ENSG00000206637 | SNORA70H | 1.064028 | 0.00232233 | 0.7691291 | -0.06603017 | 0.9473538 | NA |
| ENSG00000279467 | | 18.83698 | 0.002254816 | 0.4489491 | 0.002259354 | 0.9981973 | 0.999213 |
| ENSG00000143842 | SOX13 | 8174.096 | 0.002227222 | 0.1804804 | 0.02725207 | 0.9782587 | 0.988325 |
| ENSG00000138663 | COPS4 | 2717.230289 | 0.002225461 | 0.250870616 | -0.022510294 | 0.9820409 | 0.990627 |
| ENSG00000168589 | DYNLRB2 | 61.34094 | 0.002222381 | 0.3067414 | 0.006574963 | 0.994754 | 0.997156 |
| ENSG00000276216 | | 18.0028 | 0.002155574 | 0.4383953 | 0.001906638 | 0.9984787 | 0.99924 |
| ENSG00000006453 | BAIAP2L1 | 1187.731561 | 0.002129597 | 0.262505688 | 0.003150328 | 0.99748641 | 0.999042 |
| ENSG00000118046 | STK11 | 4853.180651 | 0.00211918 | 0.136762301 | 0.010188671 | 0.99187076 | 0.995543 |
| ENSG00000225224 | RPS27AP12 | 9.112182 | 0.002091229 | 0.6354868 | -0.001193467 | 0.9990478 | 0.999403 |
| ENSG00000267191 | | 5.261092 | 0.002077831 | 0.596518 | -0.000124433 | 0.9999007 | 0.999901 |
| ENSG00000217716 | RPS10P3 | 425.8899 | 0.00200901 | 0.1855117 | 0.006486164 | 0.9948248 | 0.997167 |
| ENSG00000237773 | | 21.34592 | 0.001988588 | 0.4275391 | 0.00292255 | 0.9976681 | 0.999042 |
| ENSG00000126432 | PRDX5 | 9740.575194 | 0.001935082 | 0.197772256 | 0.018048101 | 0.98560048 | 0.992323 |
| ENSG00000151692 | RNF144A | 707.0358 | 0.001879305 | 0.2754377 | 0.0419897 | 0.9665069 | 0.983543 |
| ENSG00000120253 | NUP43 | 2323.394381 | 0.001834237 | 0.161837419 | -0.002603371 | 0.99792281 | 0.999089 |
| ENSG00000169598 | DFFB | 457.029 | 0.001809604 | 0.1807029 | 0.007506955 | 0.9940104 | 0.996683 |
| ENSG00000172732 | MUS81 | 2844.835 | 0.001803458 | 0.1595589 | -0.000206449 | 0.9998353 | 0.999882 |
| ENSG00000266538 | | 4.818793 | 0.001724108 | 0.632881 | 0.002027848 | 0.998382 | 0.99924 |
| ENSG00000234917 | | 3.458216 | 0.001670573 | 0.590098 | 0.001677159 | 0.9986618 | 0.99924 |
| ENSG00000140043 | PTGR2 | 362.0774 | 0.001562825 | 0.149173 | 0.009764194 | 0.9922094 | 0.99573 |
| ENSG00000158717 | RNF166 | 1876.717 | 0.001511405 | 0.185437 | 0.006467688 | 0.9948396 | 0.997167 |
| ENSG00000273264 | | 6.559503 | 0.001447935 | 0.5795812 | -0.001687189 | 0.9986538 | 0.99924 |
| ENSG00000135185 | TMEM243 | 1262.725067 | 0.001440333 | 0.143256893 | 0.016166529 | 0.98710154 | 0.993082 |

| | | | | | | | |
|---|---|---|---|---|---|---|---|
| ENSG00000182307 | C8orf33 | 5755.424 | 0.001428302 | 0.1971865 | 0.008377127 | 0.9933161 | 0.996354 |
| ENSG00000278811 | LINC00624 | 121.2933 | 0.001320414 | 0.400458 | 0.002373234 | 0.9981064 | 0.999197 |
| ENSG00000188199 | NUTM2B | 58.8633 | 0.001299287 | 0.2530036 | 0.003541127 | 0.9971746 | 0.998979 |
| ENSG00000241582 | RPL23AP8 | 0.8469517 | 0.001262344 | 0.7481411 | -0.02618677 | 0.9791084 | NA |
| ENSG00000236710 | | 1.140551 | 0.001249067 | 0.7296888 | 0.002093653 | 0.9983295 | NA |
| ENSG00000253024 | RNU6-1238P | 2.594217 | 0.001245695 | 0.6846833 | 0.00166585 | 0.9986708 | 0.99924 |
| ENSG00000222179 | RN7SKP26 | 0.859215 | 0.001221496 | 0.7480988 | -0.0246311 | 0.9803492 | NA |
| ENSG00000234515 | PPP1R2P1 | 1.266899 | 0.001145393 | 0.7189396 | 0.007297014 | 0.9941779 | NA |
| ENSG00000160062 | ZBTB8A | 354.5559 | 0.001113835 | 0.1983698 | 0.003050311 | 0.9975662 | 0.999042 |
| ENSG00000077238 | IL4R | 1182.239026 | 0.000863165 | 0.270634093 | 0.004906147 | 0.99608548 | 0.998114 |
| ENSG00000168116 | KIAA1586 | 758.552 | 0.00083392 | 0.2440124 | 0.003619129 | 0.9971124 | 0.998955 |
| ENSG00000104853 | CLPTM1 | 7486.619611 | 0.000809107 | 0.162452503 | 0.003363997 | 0.99731592 | 0.999042 |
| ENSG00000171557 | FGG | 39.49637 | 0.000765828 | 0.6070958 | 0.003672811 | 0.9970695 | 0.998949 |
| ENSG00000243014 | PTMAP8 | 25.39036 | 0.000685525 | 0.4097672 | -0.00156474 | 0.9987515 | 0.999278 |
| ENSG00000152779 | SLC16A12 | 0.8896576 | 0.00064891 | 0.7601645 | -0.08217739 | 0.9345057 | NA |
| ENSG00000232400 | RAD17P1 | 5.132864 | 0.000637949 | 0.5570172 | -0.000212369 | 0.9998306 | 0.999882 |
| ENSG00000117174 | ZNHIT6 | 1048.797765 | 0.000605826 | 0.184770683 | 0.001659317 | 0.99867606 | 0.99924 |
| ENSG00000165501 | LRR1 | 1200.701 | 0.000499548 | 0.3383662 | -0.002097515 | 0.9983264 | 0.99924 |
| ENSG00000105122 | RASAL3 | 1.782514943 | 0.000480064 | 0.728505018 | -0.001881298 | 0.99849894 | 0.99924 |
| ENSG00000242858 | | 0.8473936 | 0.000464609 | 0.755145 | -0.01824563 | 0.9854429 | NA |
| ENSG00000116857 | TMEM9 | 9705.963705 | 0.00046333 | 0.178263361 | -0.022709278 | 0.98188218 | 0.990569 |
| ENSG00000274340 | | 3.955979 | 0.000453414 | 0.6224582 | -0.003220625 | 0.9974303 | 0.999042 |
| ENSG00000279632 | | 2.758763 | 0.000449158 | 0.6481321 | -0.001211227 | 0.9990336 | 0.999403 |
| ENSG00000116754 | SRSF11 | 12050.6215 | 0.000447726 | 0.128049533 | -0.002841275 | 0.99773299 | 0.999042 |
| ENSG00000250979 | | 39.99862 | 0.000434359 | 0.4892522 | -0.008881085 | 0.992914 | 0.996214 |
| ENSG00000145743 | FBXL17 | 1594.315 | 0.000360341 | 0.1837736 | 0.000293389 | 0.9997659 | 0.999882 |
| ENSG00000166352 | IFTAP | 284.9301 | 0.000347292 | 0.2528374 | -0.002802745 | 0.9977637 | 0.999042 |
| ENSG00000166960 | CCDC178 | 1.092988 | 0.00030247 | 0.7407567 | -0.01260524 | 0.9899427 | NA |
| ENSG00000272899 | ATP6V1FNB | 88.53423 | 0.000300465 | 0.3469781 | 0.001372411 | 0.998905 | 0.999356 |
| ENSG00000265262 | TRMT112P3 | 1.33678 | 0.000299231 | 0.7369789 | 0.004051304 | 0.9967675 | NA |
| ENSG00000235677 | NPM1P26 | 20.21679 | 0.000288038 | 0.3859117 | -0.000840054 | 0.9993297 | 0.999556 |
| ENSG00000130313 | PGLS | 2249.259423 | 0.000203874 | 0.157372649 | 0.002273777 | 0.99818579 | 0.999213 |
| ENSG00000286214 | | 111.5805 | 6.72E-05 | 0.3708716 | -0.009720147 | 0.9922446 | 0.99573 |

| | | | | | | | |
|---|---|---|---|---|---|---|---|
| ENSG00000142549 | IGLON5 | 5.50466 | 3.84E-05 | 0.671264 | 0.01052094 | 0.9916057 | 0.99554 |
| ENSG00000232626 | | 7.237746 | 2.88E-05 | 0.4855387 | -0.002860417 | 0.9977177 | 0.999042 |
| ENSG00000233662 | CALM2P4 | 1.130171 | -2.47E-05 | 0.7380695 | -0.006968866 | 0.9944397 | NA |
| ENSG00000280152 | | 127.1778 | -4.17E-05 | 0.39875 | -0.004300722 | 0.9965685 | 0.998523 |
| ENSG00000004534 | RBM6 | 5842.123387 | -0.000100719 | 0.19340532 | 0.001412849 | 0.99887271 | 0.999356 |
| ENSG00000125534 | PPDPF | 11506.38355 | -0.000114923 | 0.194951077 | 0.015636175 | 0.98752465 | 0.993181 |
| ENSG00000126461 | SCAF1 | 3631.637906 | -0.000218677 | 0.139474268 | -0.015268632 | 0.98781787 | 0.993351 |
| ENSG00000261612 | SUB1P3 | 38.8839 | -0.00030454 | 0.4010042 | 0.001045046 | 0.9991662 | 0.999453 |
| ENSG00000232485 | RPL37A-DT | 36.65014 | -0.000385246 | 0.3686762 | -0.002659 | 0.9978784 | 0.999089 |
| ENSG00000240996 | | 31.3235 | -0.000448531 | 0.4184834 | -0.006231954 | 0.9950277 | 0.997317 |
| ENSG00000274691 | | 0.9817322 | -0.000476826 | 0.760855 | -0.08095456 | 0.9354781 | NA |
| ENSG00000254094 | | 2.88997 | -0.000493056 | 0.6663695 | -0.02115836 | 0.9831193 | 0.991119 |
| ENSG00000132698 | RAB25 | 8359.685301 | -0.000562102 | 0.171102379 | -0.008422811 | 0.99327965 | 0.996354 |
| ENSG00000106278 | PTPRZ1 | 1.687939543 | -0.000568161 | 0.681316149 | -0.011446284 | 0.99086739 | 0.994983 |
| ENSG00000103245 | CIAO3 | 2474.324905 | -0.000613226 | 0.169589409 | -0.001868868 | 0.99850886 | 0.99924 |
| ENSG00000282057 | | 4.133948 | -0.000651172 | 0.6272367 | -0.001898921 | 0.9984849 | 0.99924 |
| ENSG00000110046 | ATG2A | 3449.750975 | -0.000660082 | 0.195373409 | -0.002866917 | 0.99771253 | 0.999042 |
| ENSG00000282815 | TEX13C | 2.196839 | -0.000665017 | 0.7551086 | -0.09131434 | 0.9272428 | 0.964903 |
| ENSG00000038427 | VCAN | 2.404094348 | -0.000679178 | 0.680133386 | -0.014728468 | 0.98824881 | 0.993672 |
| ENSG00000229836 | | 45.66924 | -0.000697177 | 0.4357387 | -0.003432934 | 0.9972609 | 0.999028 |
| ENSG00000188800 | TMCO2 | 0.7726061 | -0.000791734 | 0.743978 | -0.01018821 | 0.9918711 | NA |
| ENSG00000088876 | ZNF343 | 453.0136234 | -0.000977362 | 0.18770804 | -0.01119323 | 0.99106928 | 0.995077 |
| ENSG00000236287 | ZBED5 | 1935.197 | -0.001002836 | 0.1533265 | -0.01781248 | 0.9857884 | 0.992437 |
| ENSG00000126777 | KTN1 | 22300.41187 | -0.001011275 | 0.13062525 | -0.021443615 | 0.98289178 | 0.991023 |
| ENSG00000199471 | | 3.552348 | -0.001153161 | 0.7114534 | 0.002644551 | 0.99789 | 0.999089 |
| ENSG00000133398 | MED10 | 2833.421414 | -0.001165278 | 0.202132153 | 0.011760568 | 0.99061664 | 0.994894 |
| ENSG00000032389 | EIPR1 | 2854.657721 | -0.001165387 | 0.20219695 | -0.010355408 | 0.99173773 | 0.995543 |
| ENSG00000138802 | SEC24B | 2667.334564 | -0.001244254 | 0.146127812 | -0.009240002 | 0.99262765 | 0.996002 |
| ENSG00000148943 | LIN7C | 1468.653 | -0.001283857 | 0.1460077 | -0.01042335 | 0.9916835 | 0.995543 |
| ENSG00000137807 | KIF23 | 3654.250077 | -0.001369494 | 0.422418147 | -0.012334335 | 0.99015887 | 0.994654 |
| ENSG00000152284 | TCF7L1 | 509.9149 | -0.001400403 | 0.3144857 | -0.002918003 | 0.9976718 | 0.999042 |
| ENSG00000158813 | EDA | 220.0643 | -0.001447602 | 0.254824 | -0.008553331 | 0.9931755 | 0.996354 |
| ENSG00000259144 | RANBP20P | 5.35068 | -0.001481443 | 0.612657 | -0.005809259 | 0.9953649 | 0.997543 |

| | | | | | | | |
|---|---|---|---|---|---|---|---|
| ENSG00000278861 | | 24.36772 | -0.001496441 | 0.5351623 | -0.007678845 | 0.9938732 | 0.996661 |
| ENSG00000212240 | RNU6-930P | 3.289494 | -0.001505998 | 0.6521822 | -0.00673313 | 0.9946278 | 0.997108 |
| ENSG00000062650 | WAPL | 5897.212108 | -0.001533539 | 0.171715912 | -0.030010008 | 0.97605907 | 0.987548 |
| ENSG00000174607 | UGT8 | 279.4691 | -0.00160112 | 0.304405 | -0.006654078 | 0.9946909 | 0.99713 |
| ENSG00000274767 | | 2.89507 | -0.001628855 | 0.637841 | -0.01644046 | 0.986883 | 0.992964 |
| ENSG00000230447 | CTF2P | 0.8291771 | -0.001744008 | 0.7388435 | -0.0108451 | 0.991347 | NA |
| ENSG00000075826 | SEC31B | 470.6356982 | -0.001762055 | 0.415289094 | -0.018415491 | 0.9853074 | 0.992179 |
| ENSG00000267501 | | 1.127988 | -0.001785988 | 0.7620954 | -0.005134915 | 0.9959029 | NA |
| ENSG00000229867 | STEAP3-AS1 | 17.207 | -0.001878775 | 0.4540706 | -0.005262279 | 0.9958013 | 0.997942 |
| ENSG00000166888 | STAT6 | 3525.561 | -0.001995805 | 0.220284 | -0.04227309 | 0.966281 | 0.983543 |
| ENSG00000149480 | MTA2 | 11825.81 | -0.002043171 | 0.1802012 | -0.03398694 | 0.9728876 | 0.986063 |
| ENSG00000227704 | | 0.8106691 | -0.002093394 | 0.7575908 | -0.08671864 | 0.9308952 | NA |
| ENSG00000175087 | PDIK1L | 802.526 | -0.002117347 | 0.1876823 | -0.01505951 | 0.9879847 | 0.993481 |
| ENSG00000207195 | | 1.600005 | -0.00215619 | 0.726917 | -0.02586714 | 0.9793633 | 0.988892 |
| ENSG00000243417 | RPS3AP18 | 1.010366 | -0.002195692 | 0.7560882 | -0.08932784 | 0.9288214 | NA |
| ENSG00000143393 | PI4KB | 13729.59 | -0.002244815 | 0.1033772 | -0.02422653 | 0.9806719 | 0.989716 |
| ENSG00000254064 | | 14.56373 | -0.002289105 | 0.5147448 | -0.009888537 | 0.9921102 | 0.995708 |
| ENSG00000121053 | EPX | 19.40519767 | -0.002347526 | 0.426287404 | -0.01024883 | 0.99182276 | 0.995543 |
| ENSG00000260403 | | 0.9037556 | -0.002370482 | 0.7592147 | -0.08383419 | 0.9331883 | NA |
| ENSG00000071894 | CPSF1 | 15142.49408 | -0.002393763 | 0.117892417 | -0.022378765 | 0.98214582 | 0.990627 |
| ENSG00000152894 | PTPRK | 5400.829 | -0.002425883 | 0.2221691 | -0.009421472 | 0.9924829 | 0.995932 |
| ENSG00000263627 | PPP4R1-AS1 | 4.074743 | -0.002439642 | 0.6224671 | -0.007007863 | 0.9944086 | 0.996998 |
| ENSG00000215817 | ZC3H11B | 340.351 | -0.002502188 | 0.1926657 | -0.001676035 | 0.9986627 | 0.99924 |
| ENSG00000253102 | | 0.9305994 | -0.002527077 | 0.7434016 | -0.03605944 | 0.971235 | NA |
| ENSG00000275236 | | 7.45672 | -0.00253171 | 0.5762271 | -0.008185388 | 0.9934691 | 0.996394 |
| ENSG00000250220 | | 27.01948 | -0.002599299 | 0.4682444 | -0.008215101 | 0.9934454 | 0.996394 |
| ENSG00000219529 | | 3.620394 | -0.002605891 | 0.6707432 | -0.01640668 | 0.98691 | 0.992964 |
| ENSG00000090659 | CD209 | 0.942492752 | -0.00261673 | 0.756045067 | -0.017630029 | 0.985934 | NA |
| ENSG00000244748 | RN7SL153P | 0.8546391 | -0.002648201 | 0.7624197 | -0.07816061 | 0.9377003 | NA |
| ENSG00000238193 | HADHBP1 | 6.06674 | -0.002676371 | 0.5956645 | -0.01253526 | 0.9899986 | 0.994603 |
| ENSG00000249256 | ATP5MGP3 | 3.867275 | -0.002677435 | 0.6606241 | -0.009343775 | 0.9925449 | 0.995956 |
| ENSG00000100243 | CYB5R3 | 4992.094793 | -0.002684854 | 0.19694548 | -0.0135251 | 0.98920886 | 0.994073 |
| ENSG00000128789 | PSMG2 | 2957.755124 | -0.002688327 | 0.178409347 | -0.025904248 | 0.97933371 | 0.988892 |

| | | | | | | | |
|---|---|---|---|---|---|---|---|
| ENSG00000277058 | HNRNPCL3 | 1.165528 | -0.002713287 | 0.7176128 | -0.03954453 | 0.9684563 | NA |
| ENSG00000125841 | NRSN2 | 2084.45144 | -0.002726443 | 0.186898143 | -0.030874904 | 0.9753693 | 0.987185 |
| ENSG00000264539 | MIR548AR | 3.633838 | -0.002817824 | 0.636743 | -0.01437882 | 0.9885278 | 0.993877 |
| ENSG00000286825 | | 12.46349 | -0.002887518 | 0.4617255 | -0.007664295 | 0.9938848 | 0.996661 |
| ENSG00000164241 | C5orf63 | 385.2015 | -0.002889884 | 0.2885601 | -0.01409542 | 0.9887539 | 0.993954 |
| ENSG00000232260 | BTF3L4P1 | 59.66643 | -0.002919235 | 0.2962597 | -0.01187829 | 0.9905227 | 0.994866 |
| ENSG00000287129 | | 4.554071 | -0.002927213 | 0.5895554 | -0.0104165 | 0.991689 | 0.995543 |
| ENSG00000237781 | ADAMTSL4-AS2 | 22.00504 | -0.00297068 | 0.4489443 | -0.008387118 | 0.9933081 | 0.996354 |
| ENSG00000279080 | | 3.570598 | -0.003046166 | 0.6312532 | -0.01259221 | 0.9899531 | 0.994603 |
| ENSG00000244617 | ASPRV1 | 27.24814 | -0.003076542 | 0.4036791 | -0.01420476 | 0.9886666 | 0.993904 |
| ENSG00000267583 | | 1.016561 | -0.003097438 | 0.7517738 | -0.02696536 | 0.9784874 | NA |
| ENSG00000255165 | | 2.978247 | -0.003179707 | 0.6488317 | -0.01244471 | 0.9900708 | 0.994638 |
| ENSG00000238793 | SNORD124 | 7.313 | -0.003183083 | 0.5538316 | -0.01171437 | 0.9906535 | 0.994894 |
| ENSG00000172775 | PSME3IP1 | 2895.462 | -0.003188268 | 0.1072293 | -0.06818098 | 0.9456416 | 0.973224 |
| ENSG00000278344 | | 1.991617 | -0.003192741 | 0.6563372 | -0.01597628 | 0.9872533 | 0.993181 |
| ENSG00000286588 | | 4.96911 | -0.003210488 | 0.5927979 | -0.008658683 | 0.9930915 | 0.996316 |
| ENSG00000006194 | ZNF263 | 3133.209387 | -0.003220125 | 0.155465058 | -0.022449561 | 0.98208935 | 0.990627 |
| ENSG00000104888 | SLC17A7 | 57.56862487 | -0.003232605 | 0.367360416 | -0.011454865 | 0.99086054 | 0.994983 |
| ENSG00000180138 | CSNK1A1L | 4.989931 | -0.003239054 | 0.6113866 | -0.01271803 | 0.9898528 | 0.994569 |
| ENSG00000172733 | PURG | 1.023815 | -0.003312027 | 0.754044 | -0.09294489 | 0.9259473 | NA |
| ENSG00000076043 | REXO2 | 1100.65332 | -0.003414983 | 0.155730335 | -0.035022273 | 0.97206198 | 0.985715 |
| ENSG00000169490 | TM2D2 | 1948.599 | -0.003419883 | 0.1685835 | -0.03427562 | 0.9726574 | 0.985921 |
| ENSG00000213509 | PPIAP16 | 1.071353 | -0.003448621 | 0.7577319 | -0.08642611 | 0.9311277 | NA |
| ENSG00000161677 | JOSD2 | 1764.314 | -0.003480211 | 0.2282457 | -0.0157675 | 0.9874199 | 0.993181 |
| ENSG00000134318 | ROCK2 | 4046.666658 | -0.003482417 | 0.207895583 | -0.02040835 | 0.98371762 | 0.991486 |
| ENSG00000169857 | AVEN | 644.2544 | -0.003491142 | 0.2389011 | -0.01376522 | 0.9890173 | 0.994007 |
| ENSG00000101161 | PRPF6 | 12346.71098 | -0.003507393 | 0.139297336 | -0.088288031 | 0.92964775 | 0.965893 |
| ENSG00000215251 | FASTKD5 | 888.0672 | -0.003559612 | 0.1571429 | -0.02315459 | 0.981527 | 0.990286 |
| ENSG00000166938 | DIS3L | 1776.508 | -0.003567988 | 0.122337 | -0.0299052 | 0.9761427 | 0.987548 |
| ENSG00000101577 | LPIN2 | 2112.91828 | -0.003607267 | 0.239170715 | -0.019412679 | 0.9845119 | 0.991866 |
| ENSG00000106605 | BLVRA | 3330.301923 | -0.003624363 | 0.206641571 | -0.019875051 | 0.98414305 | 0.991697 |
| ENSG00000140259 | MFAP1 | 1906.787 | -0.003635298 | 0.1343839 | -0.02616476 | 0.9791259 | 0.988814 |
| ENSG00000280145 | | 1.570683 | -0.003698281 | 0.6868326 | -0.01567284 | 0.9874954 | NA |

| | | | | | | | |
|---|---|---|---|---|---|---|---|
| ENSG00000279692 | | 62.01536 | -0.003770358 | 0.2936864 | -0.01818623 | 0.9854903 | 0.992288 |
| ENSG00000172292 | CERS6 | 2902.019 | -0.003936211 | 0.2816962 | 0.001172693 | 0.9990643 | 0.999403 |
| ENSG00000145975 | FAM217A | 14.78825 | -0.003952531 | 0.4476317 | -0.01386743 | 0.9889357 | 0.994007 |
| ENSG00000218490 | FCF1P10 | 0.8136318 | -0.004015372 | 0.7491461 | -0.04838807 | 0.961407 | NA |
| ENSG00000166225 | FRS2 | 2612.439 | -0.00402562 | 0.1985278 | -0.02888197 | 0.9769587 | 0.987931 |
| ENSG00000134905 | CARS2 | 1541.291692 | -0.004105723 | 0.109881982 | -0.041887517 | 0.96658837 | 0.983543 |
| ENSG00000119431 | HDHD3 | 2727.758174 | -0.004191517 | 0.196703384 | -0.003245887 | 0.99741016 | 0.999042 |
| ENSG00000203875 | SNHG5 | 4806.957 | -0.00419155 | 0.2045275 | -0.02727367 | 0.9782415 | 0.988325 |
| ENSG00000270917 | | 9.498897 | -0.004412784 | 0.5450171 | -0.01188766 | 0.9905152 | 0.994866 |
| ENSG00000145216 | FIP1L1 | 1079.929 | -0.004428076 | 0.1227914 | -0.0455794 | 0.9636455 | 0.982053 |
| ENSG00000109610 | SOD3 | 1.91218473 | -0.004463002 | 0.708147475 | -0.037366092 | 0.97019311 | 0.984759 |
| ENSG00000087460 | GNAS | 76852.08716 | -0.00449698 | 0.169877817 | -0.035911063 | 0.97135327 | 0.985448 |
| ENSG00000272986 | | 9.864495 | -0.004506136 | 0.5630207 | -0.008808749 | 0.9929717 | 0.996234 |
| ENSG00000262155 | LINC02175 | 99.6721 | -0.004561706 | 0.4434064 | -0.01888743 | 0.9849309 | 0.992063 |
| ENSG00000254574 | | 0.9194337 | -0.004583122 | 0.7444984 | -0.05218463 | 0.9583816 | NA |
| ENSG00000225528 | | 1.008068 | -0.004590655 | 0.7337967 | -0.02128551 | 0.9830179 | NA |
| ENSG00000285994 | | 0.9926083 | -0.004610756 | 0.7567907 | -0.08795364 | 0.9299135 | NA |
| ENSG00000178403 | NEUROG2 | 4.229649 | -0.004622668 | 0.6836818 | -0.01307407 | 0.9895687 | 0.994359 |
| ENSG00000196890 | H2BU1 | 281.1358 | -0.004631699 | 0.2954569 | -0.03139978 | 0.9749507 | 0.986912 |
| ENSG00000100726 | TELO2 | 5089.095042 | -0.004655998 | 0.240763768 | -0.01383414 | 0.98896231 | 0.994007 |
| ENSG00000278967 | | 0.9472938 | -0.004665782 | 0.7625265 | -0.02313533 | 0.9815423 | NA |
| ENSG00000275120 | | 7.509449 | -0.004692012 | 0.5183333 | -0.01360006 | 0.9891491 | 0.99405 |
| ENSG00000171443 | ZNF524 | 574.8124 | -0.004724801 | 0.2433391 | -0.02458617 | 0.9803851 | 0.98951 |
| ENSG00000227258 | SMIM2-AS1 | 17.26271 | -0.004730985 | 0.483319 | -0.01562117 | 0.9875366 | 0.993181 |
| ENSG00000238290 | | 1.835444 | -0.00477213 | 0.6755183 | -0.02136077 | 0.9829579 | 0.991052 |
| ENSG00000107164 | FUBP3 | 5491.861655 | -0.004773268 | 0.158945915 | -0.03659466 | 0.9708082 | 0.985083 |
| ENSG00000279494 | | 3.774287 | -0.004784469 | 0.6220397 | -0.01573353 | 0.987447 | 0.993181 |
| ENSG00000185347 | TEDC1 | 1681.27 | -0.004805238 | 0.3840408 | -0.01581599 | 0.9873812 | 0.993181 |
| ENSG00000271826 | PLS3-AS1 | 32.1762 | -0.004813035 | 0.4726002 | -0.01667951 | 0.9866923 | 0.992852 |
| ENSG00000276047 | RN7SL711P | 1.40493 | -0.004856796 | 0.7288369 | -0.01145535 | 0.9908602 | NA |
| ENSG00000272010 | | 12.25747 | -0.004902021 | 0.5429836 | -0.01532388 | 0.9877738 | 0.993345 |
| ENSG00000167674 | HDGFL2 | 4365.525 | -0.004907599 | 0.1832661 | -0.02615073 | 0.9791371 | 0.988814 |
| ENSG00000235119 | | 3.087693 | -0.004949641 | 0.5976937 | -0.01701506 | 0.9864246 | 0.992777 |

| | | | | | | | |
|---|---|---|---|---|---|---|---|
| ENSG00000251172 | | 6.788626 | -0.00500391 | 0.5690826 | -0.01577608 | 0.987413 | 0.993181 |
| ENSG00000103254 | ANTKMT | 1515.159175 | -0.005045173 | 0.197919458 | -0.027498871 | 0.97806184 | 0.988291 |
| ENSG00000198738 | SMIM11P1 | 0.7799769 | -0.005061665 | 0.7558403 | -0.08960856 | 0.9285983 | NA |
| ENSG00000244063 | | 1.828219 | -0.005101242 | 0.6871867 | -0.01728874 | 0.9862063 | 0.992715 |
| ENSG00000250290 | NCAPGP1 | 1.144584 | -0.00510414 | 0.7547111 | -0.09172551 | 0.9269161 | NA |
| ENSG00000131778 | CHD1L | 162.8323557 | -0.005113802 | 0.1872436 | -0.02990254 | 0.97614478 | 0.987548 |
| ENSG00000274967 | | 0.7893448 | -0.0052328 | 0.7240639 | -0.03895197 | 0.9689287 | NA |
| ENSG00000156411 | ATP5MPL | 5992.165 | -0.005381368 | 0.2229284 | -0.02661164 | 0.9787695 | 0.988668 |
| ENSG00000172809 | RPL38 | 11484.75 | -0.005419218 | 0.1559874 | -0.06241153 | 0.9502351 | 0.975545 |
| ENSG00000231147 | ARHGAP42P2 | 1.017556 | -0.005441519 | 0.7019275 | -0.04814367 | 0.9616017 | NA |
| ENSG00000156976 | EIF4A2 | 22420.74 | -0.005464687 | 0.1627977 | -0.03675803 | 0.9706779 | 0.985048 |
| ENSG00000147454 | SLC25A37 | 2216.121 | -0.005551756 | 0.1775351 | -0.03460055 | 0.9723983 | 0.985868 |
| ENSG00000146457 | WTAP | 2525.892 | -0.005552374 | 0.1470106 | -0.02992096 | 0.9761301 | 0.987548 |
| ENSG00000082641 | NFE2L1 | 16224.45499 | -0.005561925 | 0.176396499 | -0.056101325 | 0.95526109 | 0.9785 |
| ENSG00000143612 | C1orf43 | 16949.72 | -0.005569698 | 0.1457019 | -0.04122503 | 0.9671165 | 0.983704 |
| ENSG00000180720 | CHRM4 | 4.897863 | -0.005614141 | 0.629536 | -0.01427635 | 0.9886095 | 0.993884 |
| ENSG00000183495 | EP400 | 6199.154 | -0.005650054 | 0.20511 | -0.02883899 | 0.976993 | 0.987931 |
| ENSG00000112473 | SLC39A7 | 10721.26091 | -0.005734955 | 0.123632266 | 0.068661016 | 0.94525945 | 0.972944 |
| ENSG00000271049 | | 1.27368 | -0.005858337 | 0.7197769 | -0.01949368 | 0.9844473 | NA |
| ENSG00000169629 | RGPD8 | 120.3797 | -0.005879009 | 0.1964712 | -0.03342921 | 0.9733323 | 0.986325 |
| ENSG00000260586 | | 1.368382 | -0.005887775 | 0.7070898 | -0.02710655 | 0.9783747 | NA |
| ENSG00000006607 | FARP2 | 2295.810625 | -0.005925258 | 0.20176136 | -0.027817626 | 0.97780761 | 0.988222 |
| ENSG00000189186 | DCAF8L2 | 4.559069 | -0.005977377 | 0.7483097 | -0.1034747 | 0.9175862 | 0.961195 |
| ENSG00000258484 | SPESP1 | 383.1533 | -0.005977637 | 0.2931704 | -0.02240737 | 0.982123 | 0.990627 |
| ENSG00000226636 | | 1.048576 | -0.006001893 | 0.7592541 | -0.08368415 | 0.9333076 | NA |
| ENSG00000210127 | MT-TA | 0.7731088 | -0.006024184 | 0.7549843 | -0.04262946 | 0.9659969 | NA |
| ENSG00000254432 | | 1.135103 | -0.006043502 | 0.7450817 | -0.02753658 | 0.9780318 | NA |
| ENSG00000168802 | CHTF8 | 107.9718 | -0.006154059 | 0.2160602 | -0.02937339 | 0.9765668 | 0.98772 |
| ENSG00000221303 | SNORA79 | 0.9346763 | -0.006158892 | 0.7562449 | -0.0889324 | 0.9291356 | NA |
| ENSG00000126945 | HNRNPH2 | 7452.38035 | -0.0061626 | 0.139070493 | -0.051576923 | 0.95886581 | 0.979737 |
| ENSG00000213028 | | 0.9336431 | -0.006199143 | 0.7465904 | -0.02652656 | 0.9788374 | NA |
| ENSG00000111405 | ENDOU | 2.156027164 | -0.006203603 | 0.659573234 | -0.018998566 | 0.98484225 | 0.992049 |
| ENSG00000197013 | ZNF429 | 567.7244 | -0.006217936 | 0.1540041 | -0.04459457 | 0.9644305 | 0.982497 |

| | | | | | | | |
|---|---|---|---|---|---|---|---|
| ENSG00000233037 | | 1.380213 | -0.006258497 | 0.7538684 | -0.09318046 | 0.9257602 | NA |
| ENSG00000232692 | | 1.906124 | -0.00627357 | 0.6862522 | -0.02731474 | 0.9782087 | 0.988325 |
| ENSG00000267383 | | 0.8398831 | -0.006283512 | 0.7584482 | -0.08527011 | 0.9320467 | NA |
| ENSG00000228955 | | 7.783732 | -0.006342603 | 0.5512898 | -0.0185839 | 0.985173 | 0.992156 |
| ENSG00000172244 | C5orf34 | 316.5968 | -0.006391477 | 0.3419671 | -0.01554084 | 0.9876007 | 0.993208 |
| ENSG00000120159 | CAAP1 | 1170.664894 | -0.006503354 | 0.134161138 | -0.050332231 | 0.95985764 | 0.979994 |
| ENSG00000121892 | PDS5A | 10623.42295 | -0.006507458 | 0.124672136 | -0.090881885 | 0.92758644 | 0.96494 |
| ENSG00000012124 | CD22 | 54.0354473 | -0.006537496 | 0.557292987 | -0.029164405 | 0.97673347 | 0.987813 |
| ENSG00000276662 | | 8.989475 | -0.006551733 | 0.5301532 | -0.02104393 | 0.9832106 | 0.991119 |
| ENSG00000240729 | | 1.484854 | -0.006567823 | 0.6945336 | -0.02216975 | 0.9823125 | NA |
| ENSG00000259523 | | 20.51226 | -0.006585266 | 0.3805067 | -0.02215844 | 0.9823216 | 0.990711 |
| ENSG00000175782 | SLC35E3 | 1059.907 | -0.00668822 | 0.1948468 | -0.03853096 | 0.9692643 | 0.984268 |
| ENSG00000235081 | | 4.704483 | -0.006865552 | 0.5835234 | -0.01990551 | 0.9841188 | 0.991697 |
| ENSG00000280128 | | 1356.84 | -0.006880248 | 0.2931856 | -0.02961584 | 0.9763734 | 0.987603 |
| ENSG00000180628 | PCGF5 | 3286.262 | -0.006883108 | 0.1732217 | -0.04254194 | 0.9660667 | 0.983389 |
| ENSG00000250260 | | 1.017885 | -0.006948961 | 0.7283198 | -0.03996721 | 0.9681193 | NA |
| ENSG00000203801 | LINC00222 | 2.190234 | -0.007012161 | 0.6509904 | -0.01973397 | 0.9842556 | 0.991758 |
| ENSG00000243960 | | 12.28844 | -0.007045398 | 0.452334 | -0.01953971 | 0.9844106 | 0.991839 |
| ENSG00000226400 | | 0.9947519 | -0.007065244 | 0.7662279 | -0.05207743 | 0.958467 | NA |
| ENSG00000254348 | | 0.9502533 | -0.007072864 | 0.7528059 | -0.0950896 | 0.9242437 | NA |
| ENSG00000143554 | SLC27A3 | 2053.898 | -0.007092913 | 0.2169436 | -0.03202242 | 0.9744542 | 0.986622 |
| ENSG00000277067 | | 0.9914599 | -0.007113708 | 0.7183556 | -0.03518033 | 0.9719359 | NA |
| ENSG00000115239 | ASB3 | 876.1661343 | -0.007185164 | 0.14469328 | -0.050473746 | 0.95974487 | 0.979969 |
| ENSG00000265394 | | 1.579464 | -0.007226999 | 0.6780147 | -0.02750282 | 0.9780587 | 0.988291 |
| ENSG00000143373 | ZNF687 | 8628.052 | -0.007248696 | 0.1078433 | -0.06942347 | 0.9446525 | 0.972658 |
| ENSG00000258847 | | 0.8601115 | -0.00729184 | 0.7392481 | -0.0397679 | 0.9682782 | NA |
| ENSG00000153446 | C16orf89 | 4.995484 | -0.007319989 | 0.6108386 | -0.02096168 | 0.9832762 | 0.991148 |
| ENSG00000196387 | ZNF140 | 1020.437 | -0.007376282 | 0.1598692 | -0.04837151 | 0.9614202 | 0.980783 |
| ENSG00000142082 | SIRT3 | 1043.822 | -0.007387765 | 0.1425006 | -0.05339563 | 0.9574167 | 0.979347 |
| ENSG00000197217 | ENTPD4 | 2266.906 | -0.007407192 | 0.2076358 | -0.02941461 | 0.9765339 | 0.98772 |
| ENSG00000166016 | ABTB2 | 1860.916 | -0.007423363 | 0.1560495 | -0.05006846 | 0.9600678 | 0.980097 |
| ENSG00000274373 | | 1.075781 | -0.00743992 | 0.7409356 | -0.04660663 | 0.9628268 | NA |
| ENSG00000256673 | | 31.90952 | -0.007452893 | 0.5276696 | -0.02504843 | 0.9800163 | 0.98928 |

| | | | | | | | |
|---|---|---|---|---|---|---|---|
| ENSG00000220848 | RPS18P9 | 263.4018 | -0.007508295 | 0.1982926 | -0.03976452 | 0.9682809 | 0.983954 |
| ENSG00000241746 | | 0.7687421 | -0.007526003 | 0.7378859 | -0.03305009 | 0.9736346 | NA |
| ENSG00000227500 | SCAMP4 | 3666.362 | -0.007638755 | 0.1481545 | -0.06117497 | 0.9512199 | 0.976178 |
| ENSG00000163788 | SNRK | 558.8616 | -0.00766554 | 0.1392968 | -0.05768657 | 0.9539983 | 0.977659 |
| ENSG00000155858 | LSM11 | 957.6794 | -0.007775423 | 0.18283 | -0.06762754 | 0.9460821 | 0.973414 |
| ENSG00000237668 | RPS15AP38 | 624.9112 | -0.007787032 | 0.1869095 | -0.03981411 | 0.9682413 | 0.983954 |
| ENSG00000112144 | CILK1 | 1897.64508 | -0.00787611 | 0.185709743 | -0.05037556 | 0.95982311 | 0.979994 |
| ENSG00000237456 | ID2B | 0.9783899 | -0.007976247 | 0.7452132 | -0.02816348 | 0.9775318 | NA |
| ENSG00000267116 | PMM2P2 | 0.7809244 | -0.008021237 | 0.7496141 | -0.064847 | 0.9482958 | NA |
| ENSG00000235411 | DRG1P1 | 71.68497 | -0.008035731 | 0.2814594 | -0.02617519 | 0.9791176 | 0.988814 |
| ENSG00000258038 | LINC02327 | 0.8016354 | -0.008064753 | 0.7894715 | 0 | 0.9630965 | NA |
| ENSG00000251490 | | 0.7551998 | -0.008079047 | 0.7894841 | 0 | 0.9639444 | NA |
| ENSG00000211945 | IGHV1-18 | 1.419871 | -0.008081435 | 0.7894831 | 0 | 0.9686595 | NA |
| ENSG00000117385 | P3H1 | 1293.728591 | -0.008099923 | 0.231935346 | -0.041960693 | 0.96653003 | 0.983543 |
| ENSG00000165694 | FRMD7 | 0.9239366 | -0.008141003 | 0.7373333 | -0.05785439 | 0.9538646 | NA |
| ENSG00000132603 | NIP7 | 2955.689108 | -0.008144483 | 0.184050691 | -0.050594231 | 0.95964886 | 0.979969 |
| ENSG00000272627 | | 1.335039 | -0.008153598 | 0.7076993 | -0.02961549 | 0.9763737 | NA |
| ENSG00000287360 | | 0.8142095 | -0.008158389 | 0.7894694 | 0 | 0.9627139 | NA |
| ENSG00000214465 | SMARCE1P6 | 22.03451 | -0.008173727 | 0.3320734 | -0.02832141 | 0.9774058 | 0.988186 |
| ENSG00000287982 | | 1.241452 | -0.008223972 | 0.7570087 | -0.08779992 | 0.9300357 | NA |
| ENSG00000125363 | AMELX | 0.944594072 | -0.008271485 | 0.789449668 | 0 | 0.96082826 | NA |
| ENSG00000163346 | PBXIP1 | 3869.221 | -0.008272477 | 0.3412622 | -0.01932277 | 0.9845836 | 0.991901 |
| ENSG00000277406 | SEC22B4P | 24.35562 | -0.008275786 | 0.4231719 | -0.03063538 | 0.9755603 | 0.987324 |
| ENSG00000244706 | | 1.18386 | -0.008300327 | 0.7894087 | 0 | 0.9570399 | NA |
| ENSG00000229156 | | 0.9871757 | -0.00832558 | 0.7894359 | 0 | 0.9598979 | NA |
| ENSG00000250122 | | 1.022178 | -0.008325651 | 0.7894271 | 0 | 0.9588859 | NA |
| ENSG00000133980 | VRTN | 1.065608806 | -0.008347443 | 0.789430333 | 0 | 0.95948995 | NA |
| ENSG00000260978 | | 1.014072 | -0.008366536 | 0.7894287 | 0 | 0.9590054 | NA |
| ENSG00000259833 | | 1.073468 | -0.008367903 | 0.7894859 | 0 | 0.9666875 | NA |
| ENSG00000248377 | HMGN1P9 | 1.596507 | -0.008368681 | 0.7529341 | -0.09460181 | 0.9246311 | 0.963771 |
| ENSG00000225574 | DRAXINP1 | 1.512956 | -0.008388542 | 0.7894862 | 0 | 0.9686595 | NA |
| ENSG00000260930 | LINC01416 | 1.626435 | -0.008421288 | 0.7894782 | 0 | 0.9686591 | 0.983954 |
| ENSG00000286514 | | 1.09665 | -0.008425902 | 0.7894173 | 0 | 0.9580494 | NA |

| | | | | | | | |
|---|---|---|---|---|---|---|---|
| ENSG00000143156 | NME7 | 607.4787 | -0.008426762 | 0.2087048 | -0.04254208 | 0.9660666 | 0.983389 |
| ENSG00000228203 | GRASLND | 1.17688 | -0.008434905 | 0.7894133 | 0 | 0.9575673 | NA |
| ENSG00000107854 | TNKS2 | 6629.462391 | -0.008442998 | 0.189116654 | -0.050459294 | 0.95975639 | 0.979969 |
| ENSG00000246876 | LINC02466 | 1.673638 | -0.008447763 | 0.7894771 | 0 | 0.968659 | 0.983954 |
| ENSG00000238284 | LINC01448 | 1.000516 | -0.008462533 | 0.789441 | 0 | 0.9596351 | NA |
| ENSG00000239148 | | 2.112134 | -0.008492163 | 0.7894615 | 0 | 0.9686585 | 0.983954 |
| ENSG00000102043 | MTMR8 | 1.766787951 | -0.008522443 | 0.789473678 | 0 | 0.96865885 | 0.983954 |
| ENSG00000205155 | PSENEN | 990.3935 | -0.008532089 | 0.1900996 | -0.0513049 | 0.9590826 | 0.97977 |
| ENSG00000286945 | | 1.855927 | -0.008540735 | 0.7894704 | 0 | 0.9686587 | 0.983954 |
| ENSG00000274835 | | 1.810084 | -0.008554948 | 0.7894724 | 0 | 0.9686588 | 0.983954 |
| ENSG00000067369 | TP53BP1 | 3559.752854 | -0.008569927 | 0.124666532 | -0.083312551 | 0.93360302 | 0.967249 |
| ENSG00000139842 | CUL4A | 5476.744 | -0.008577815 | 0.1138835 | -0.07520559 | 0.9400511 | 0.97045 |
| ENSG00000233693 | PBX1-AS1 | 0.8662658 | -0.008588437 | 0.752564 | -0.09527711 | 0.9240947 | NA |
| ENSG00000213638 | ADAT3 | 162.2141 | -0.008610993 | 0.288606 | -0.02817873 | 0.9775196 | 0.988186 |
| ENSG00000286349 | | 0.8799905 | -0.008617407 | 0.7290244 | -0.04173784 | 0.9667077 | NA |
| ENSG00000211943 | IGHV3-15 | 2.048757 | -0.00862092 | 0.7894645 | 0 | 0.9686584 | 0.983954 |
| ENSG00000276203 | ANKRD20A3P | 1.355428 | -0.008630895 | 0.7893846 | 0 | 0.954931 | NA |
| ENSG00000197323 | TRIM33 | 30997.43 | -0.008665202 | 0.1920873 | -0.06526871 | 0.9479601 | 0.974515 |
| ENSG00000196118 | CCDC189 | 111.5658 | -0.008676371 | 0.2700681 | -0.03696042 | 0.9705166 | 0.984975 |
| ENSG00000204138 | PHACTR4 | 2380.477 | -0.008682375 | 0.1607831 | -0.06256531 | 0.9501127 | 0.975545 |
| ENSG00000167283 | ATP5MG | 2037.512 | -0.008724477 | 0.1948692 | -0.05084035 | 0.9594527 | 0.97996 |
| ENSG00000273365 | | 0.9664875 | -0.008724974 | 0.7365591 | -0.03436668 | 0.9725848 | NA |
| ENSG00000249069 | LINC01033 | 2.98174 | -0.008729815 | 0.6427462 | -0.01761924 | 0.9859426 | 0.992555 |
| ENSG00000228636 | LINC02663 | 1.778518 | -0.008737789 | 0.7893515 | 0 | 0.9502658 | 0.975545 |
| ENSG00000271180 | COX5AP1 | 2.830348 | -0.008743083 | 0.6898099 | -0.02519811 | 0.9798969 | 0.989205 |
| ENSG00000259359 | | 1.053533 | -0.008753375 | 0.751446 | -0.09724695 | 0.9225303 | NA |
| ENSG00000108423 | TUBD1 | 5684.672489 | -0.008766978 | 0.210441107 | -0.03951001 | 0.96848377 | 0.983954 |
| ENSG00000125498 | KIR2DL1 | 2.612959035 | -0.008767678 | 0.789452507 | 0 | 0.96865803 | 0.983954 |
| ENSG00000276724 | | 1.045773 | -0.008775725 | 0.7166878 | -0.04803401 | 0.9616891 | NA |
| ENSG00000253883 | IGHV3-19 | 2.649159 | -0.008779662 | 0.789448 | 0 | 0.9686577 | 0.983954 |
| ENSG00000227578 | RPS3AP53 | 1.806461 | -0.008823268 | 0.7038672 | -0.03527546 | 0.9718601 | 0.985632 |
| ENSG00000250735 | | 2.376964 | -0.008859162 | 0.789456 | 0 | 0.968658 | 0.983954 |
| ENSG00000107295 | SH3GL2 | 2.72622368 | -0.008865048 | 0.789446745 | 0 | 0.96865754 | 0.983954 |

| | | | | | | | |
|---|---|---|---|---|---|---|---|
| ENSG00000286051 | MAGEA6-DT | 2.102724 | -0.008920378 | 0.7894698 | 0 | 0.9686583 | 0.983954 |
| ENSG00000264334 | | 3.414482 | -0.008930024 | 0.7894308 | 0 | 0.9686569 | 0.983954 |
| ENSG00000205659 | LIN52 | 938.2203 | -0.009011164 | 0.1777781 | -0.04887124 | 0.9610219 | 0.980583 |
| ENSG00000261761 | LINC02616 | 4.000963 | -0.009084609 | 0.7894211 | 0 | 0.9686564 | 0.983954 |
| ENSG00000227234 | SPANXB1 | 4.186012 | -0.009099188 | 0.7894222 | 0 | 0.9686566 | 0.983954 |
| ENSG00000205777 | GAGE1 | 2.628776 | -0.009126336 | 0.7892882 | 0 | 0.9427274 | 0.971771 |
| ENSG00000260691 | ANKRD20A1 | 4.238391 | -0.009174716 | 0.7894188 | 0 | 0.9686562 | 0.983954 |
| ENSG00000196961 | AP2A1 | 7174.485 | -0.009182397 | 0.126826 | -0.09788992 | 0.9220197 | 0.963092 |
| ENSG00000264349 | MIR4258 | 21.60269 | -0.009183707 | 0.4260329 | -0.02786385 | 0.9777707 | 0.988222 |
| ENSG00000235236 | | 12.65696 | -0.009208483 | 0.4400269 | -0.02965774 | 0.97634 | 0.987603 |
| ENSG00000132466 | ANKRD17 | 7903.067064 | -0.009212164 | 0.19132113 | -0.057301309 | 0.95430518 | 0.97786 |
| ENSG00000251235 | SNRPCP2 | 23.79823 | -0.009223945 | 0.4261033 | -0.02496392 | 0.9800837 | 0.989281 |
| ENSG00000183304 | FAM9A | 3.170978 | -0.009282849 | 0.7892664 | 0 | 0.9384125 | 0.969728 |
| ENSG00000249642 | | 6.192014 | -0.009366545 | 0.7893943 | 0 | 0.9686551 | 0.983954 |
| ENSG00000277978 | | 16.09755 | -0.00938286 | 0.5056608 | -0.02340947 | 0.9813237 | 0.990118 |
| ENSG00000100325 | ASCC2 | 4190.728837 | -0.009398652 | 0.114919027 | -0.094060773 | 0.92506088 | 0.963954 |
| ENSG00000273058 | | 40.69191 | -0.009459406 | 0.3733419 | -0.0325084 | 0.9740666 | 0.986555 |
| ENSG00000258763 | | 2.841212 | -0.009468158 | 0.7892814 | 0 | 0.9407904 | 0.970675 |
| ENSG00000183570 | PCBP3 | 186.3602 | -0.009489835 | 0.2712782 | -0.04168991 | 0.9667459 | 0.983576 |
| ENSG00000271201 | | 7.762082 | -0.009527533 | 0.7893806 | 0 | 0.9686544 | 0.983954 |
| ENSG00000274502 | ANKRD20A6P | 1.066375 | -0.009564289 | 0.7354782 | -0.05140287 | 0.9590045 | NA |
| ENSG00000144306 | SCRN3 | 1263.597 | -0.009629335 | 0.1131729 | -0.09465065 | 0.9245923 | 0.963771 |
| ENSG00000164520 | RAET1E | 1.828916 | -0.009629775 | 0.7186012 | -0.02376706 | 0.9810384 | 0.989981 |
| ENSG00000188566 | NDOR1 | 1371.766 | -0.009644886 | 0.1560212 | -0.06382326 | 0.949111 | 0.97517 |
| ENSG00000236592 | S100A11P2 | 0.9239183 | -0.009705588 | 0.7483269 | -0.04112444 | 0.9671967 | NA |
| ENSG00000272342 | | 15.93519 | -0.009705611 | 0.502637 | -0.03485953 | 0.9721918 | 0.985734 |
| ENSG00000235488 | JARID2-AS1 | 13.65527 | -0.009721403 | 0.5192648 | -0.02288993 | 0.9817381 | 0.990461 |
| ENSG00000243943 | ZNF512 | 763.1451 | -0.009726797 | 0.2168011 | -0.03795226 | 0.9697257 | 0.984586 |
| ENSG00000183476 | SH2D7 | 0.8021224 | -0.009730091 | 0.7462058 | -0.0526799 | 0.957987 | NA |
| ENSG00000188100 | FAM25A | 2.579928 | -0.009760922 | 0.6927604 | -0.03526399 | 0.9718692 | 0.985632 |
| ENSG00000286548 | | 76.3437 | -0.009863909 | 0.2181782 | -0.04598255 | 0.9633242 | 0.981876 |
| ENSG00000251297 | TUBB7P | 1.061078 | -0.009943306 | 0.7597543 | -0.08292216 | 0.9339134 | NA |
| ENSG00000226372 | DCAF8L1 | 7.68731 | -0.01004075 | 0.7891521 | 0 | 0.9136778 | 0.959592 |

| | | | | | | | |
|---|---|---|---|---|---|---|---|
| ENSG00000178631 | ACTG1P1 | 24.39581 | -0.0100559 | 0.4723358 | -0.03156903 | 0.9748157 | 0.986888 |
| ENSG00000116030 | SUMO1 | 5682.612632 | -0.010131448 | 0.202567839 | -0.047827109 | 0.96185403 | 0.98098 |
| ENSG00000287620 | | 48.98404 | -0.01015887 | 0.4028334 | -0.03245917 | 0.9741059 | 0.986555 |
| ENSG00000227215 | | 1.353245 | -0.01016469 | 0.7074897 | -0.03914116 | 0.9687778 | NA |
| ENSG00000265728 | | 1.747244 | -0.01018426 | 0.7043066 | -0.04772462 | 0.9619357 | 0.981025 |
| ENSG00000172687 | ZNF738 | 11.49155 | -0.01019798 | 0.7891153 | 0 | 0.9085012 | 0.957296 |
| ENSG00000225546 | LINC02476 | 14.40811 | -0.01019867 | 0.7893506 | 0 | 0.9686525 | 0.983954 |
| ENSG00000185774 | KCNIP4 | 4.358811 | -0.01025267 | 0.5556521 | -0.03206569 | 0.9744197 | 0.986622 |
| ENSG00000272758 | WDR5B-DT | 150.2045 | -0.0103481 | 0.2977846 | -0.0281235 | 0.9775637 | 0.988186 |
| ENSG00000152705 | CATSPER3 | 31.65344 | -0.01034821 | 0.354847 | -0.03436743 | 0.9725842 | 0.985921 |
| ENSG00000278965 | | 0.8110434 | -0.01038167 | 0.7601713 | -0.08222105 | 0.9344709 | NA |
| ENSG00000233750 | CICP27 | 9.711103 | -0.01038381 | 0.527921 | -0.0353545 | 0.9717971 | 0.985632 |
| ENSG00000287251 | | 14.18746 | -0.01048688 | 0.7890929 | 0 | 0.9078528 | 0.957096 |
| ENSG00000283103 | | 330.3512 | -0.01049605 | 0.1768338 | -0.06652971 | 0.9469561 | 0.973785 |
| ENSG00000168385 | SEPTIN2 | 19058.72 | -0.01053466 | 0.09393977 | -0.1274279 | 0.8986018 | 0.953206 |
| ENSG00000248767 | FOXL3 | 0.7921253 | -0.01056959 | 0.7599658 | -0.08273637 | 0.9340612 | NA |
| ENSG00000232775 | BMS1P22 | 57.73406 | -0.01062936 | 0.4158942 | -0.03199132 | 0.974479 | 0.986622 |
| ENSG00000130244 | FAM98C | 1897.955768 | -0.010633396 | 0.164593361 | -0.107746271 | 0.91419696 | 0.959824 |
| ENSG00000250654 | | 1.248578 | -0.01069583 | 0.7403911 | -0.05204611 | 0.9584919 | NA |
| ENSG00000176124 | DLEU1 | 394.9062 | -0.01073948 | 0.2079488 | -0.05622223 | 0.9551648 | 0.9785 |
| ENSG00000170571 | EMB | 4645.085 | -0.01076279 | 0.1776561 | -0.05516613 | 0.9560061 | 0.978774 |
| ENSG00000270159 | | 2.873764 | -0.01076808 | 0.6607701 | -0.03312814 | 0.9735724 | 0.986418 |
| ENSG00000267342 | | 40.80702 | -0.01083501 | 0.3018292 | -0.04135055 | 0.9670164 | 0.983702 |
| ENSG00000230392 | | 2.480179 | -0.01083827 | 0.6940671 | -0.02820667 | 0.9774973 | 0.988186 |
| ENSG00000207378 | RNU6-748P | 1.123967 | -0.01088234 | 0.7513627 | -0.09727405 | 0.9225088 | NA |
| ENSG00000254164 | | 4.243737 | -0.01092565 | 0.6613113 | -0.0320356 | 0.9744437 | 0.986622 |
| ENSG00000178015 | GPR150 | 1.13909 | -0.01100498 | 0.7396085 | -0.04079068 | 0.9674628 | NA |
| ENSG00000233229 | CNOT7P1 | 14.27473 | -0.01100909 | 0.4540151 | -0.03433548 | 0.9726096 | 0.985921 |
| ENSG00000196262 | PPIA | 17015.53 | -0.01104614 | 0.2281735 | -0.05399276 | 0.9569409 | 0.979163 |
| ENSG00000259910 | | 1.132446 | -0.01106139 | 0.7141064 | -0.04406137 | 0.9648555 | NA |
| ENSG00000236123 | CEACAMP11 | 1.522191 | -0.01113486 | 0.7512389 | -0.09747254 | 0.9223511 | NA |
| ENSG00000234937 | | 3.416995 | -0.01116302 | 0.641823 | -0.03545776 | 0.9717147 | 0.985632 |
| ENSG00000258230 | | 7.782905 | -0.01118033 | 0.5684885 | -0.02991372 | 0.9761359 | 0.987548 |

| | | | | | | | |
|---|---|---|---|---|---|---|---|
| ENSG00000196914 | ARHGEF12 | 7230.716 | -0.01120334 | 0.2076222 | -0.06981356 | 0.9443421 | 0.972618 |
| ENSG00000165985 | C1QL3 | 24.40774 | -0.0112283 | 0.4625292 | -0.03278907 | 0.9738428 | 0.986555 |
| ENSG00000271643 | | 507.5549 | -0.0112994 | 0.2353619 | -0.05397818 | 0.9569525 | 0.979163 |
| ENSG00000176340 | COX8A | 3620.632 | -0.01131717 | 0.2339856 | -0.04512832 | 0.964005 | 0.982258 |
| ENSG00000233028 | | 1.049461 | -0.01135035 | 0.7265972 | -0.06316352 | 0.9496363 | NA |
| ENSG00000273026 | | 79.3274 | -0.0113594 | 0.264211 | -0.04140283 | 0.9669748 | 0.983702 |
| ENSG00000145780 | FEM1C | 1015.747 | -0.01137405 | 0.1681706 | -0.06207159 | 0.9505058 | 0.975698 |
| ENSG00000278206 | | 30.70065 | -0.01141302 | 0.4267581 | -0.03724973 | 0.9702859 | 0.984816 |
| ENSG00000100196 | KDELR3 | 1035.695708 | -0.011441652 | 0.281199525 | -0.056961678 | 0.95457572 | 0.978062 |
| ENSG00000119487 | MAPKAP1 | 5251.107657 | -0.011453491 | 0.117906411 | -0.092712011 | 0.92613236 | 0.9644 |
| ENSG00000264630 | PRKCA-AS1 | 1.036826 | -0.01146608 | 0.73556 | -0.04044478 | 0.9677385 | NA |
| ENSG00000196562 | SULF2 | 50270.67 | -0.0115196 | 0.1331882 | -0.1090982 | 0.9131246 | 0.959195 |
| ENSG00000286031 | | 0.9503064 | -0.01155169 | 0.7395504 | -0.04087842 | 0.9673928 | NA |
| ENSG00000256312 | | 1.358332 | -0.01159987 | 0.7503665 | -0.09902293 | 0.9211201 | NA |
| ENSG00000126259 | KIRREL2 | 0.816901559 | -0.01164315 | 0.763842386 | -0.054814222 | 0.95628647 | NA |
| ENSG00000136267 | DGKB | 2.247578142 | -0.011643929 | 0.746991033 | -0.105359876 | 0.91609025 | 0.960225 |
| ENSG00000224106 | CYP4F25P | 0.7879847 | -0.01165048 | 0.7891115 | 0 | 0.9639358 | NA |
| ENSG00000103253 | HAGHL | 4494.035316 | -0.011662369 | 0.216678023 | -0.05933495 | 0.95268532 | 0.976954 |
| ENSG00000236559 | BCL2L1-AS1 | 0.8280279 | -0.01168604 | 0.7570673 | -0.08761003 | 0.9301866 | NA |
| ENSG00000196378 | ZNF34 | 549.3784 | -0.01172605 | 0.1603416 | -0.07819059 | 0.9376764 | 0.969232 |
| ENSG00000253591 | KCNIP1-AS1 | 1.954091 | -0.01173051 | 0.7277276 | -0.04857453 | 0.9612584 | 0.980745 |
| ENSG00000099622 | CIRBP | 10140.96369 | -0.011761568 | 0.153765848 | -0.080213641 | 0.93606734 | 0.96851 |
| ENSG00000235552 | RPL6P27 | 10617.54 | -0.01178412 | 0.168544 | -0.09689756 | 0.9228077 | 0.963307 |
| ENSG00000198887 | SMC5 | 2044.738 | -0.01180119 | 0.2077976 | -0.06651943 | 0.9469643 | 0.973785 |
| ENSG00000286910 | | 0.8708673 | -0.01184554 | 0.7890015 | 0 | 0.9546075 | NA |
| ENSG00000254839 | | 0.8482724 | -0.01185708 | 0.7890069 | 0 | 0.9549419 | NA |
| ENSG00000287114 | | 1.713808 | -0.01185928 | 0.7529252 | -0.09493694 | 0.9243649 | 0.963692 |
| ENSG00000233515 | LINC01518 | 4.984949 | -0.01191608 | 0.5898829 | -0.03239979 | 0.9741532 | 0.986555 |
| ENSG00000182070 | OR52A1 | 1.88041 | -0.01193048 | 0.7891447 | 0 | 0.9686567 | 0.983954 |
| ENSG00000213231 | TCL1B | 2.014605 | -0.01193172 | 0.7891508 | 0 | 0.9686565 | 0.983954 |
| ENSG00000230400 | LINC01747 | 0.9389276 | -0.01193652 | 0.7889897 | 0 | 0.9538192 | NA |
| ENSG00000137200 | CMTR1 | 6274.552957 | -0.011982317 | 0.107976696 | -0.133196279 | 0.89403815 | 0.950882 |
| ENSG00000255400 | | 1.138949 | -0.01199993 | 0.7889877 | 0 | 0.954387 | NA |

| | | | | | | | |
|---|---|---|---|---|---|---|---|
| ENSG00000258472 | | 3.026615 | -0.01204852 | 0.6393913 | -0.03912969 | 0.968787 | 0.984046 |
| ENSG00000228527 | | 1.152981 | -0.01205398 | 0.6952345 | -0.04963167 | 0.9604159 | NA |
| ENSG00000280007 | | 28.60428 | -0.01205796 | 0.4082757 | -0.03777324 | 0.9698685 | 0.98462 |
| ENSG00000258710 | LINC01193 | 1.031105 | -0.01209404 | 0.7889734 | 0 | 0.9524631 | NA |
| ENSG00000230424 | EMC1-AS1 | 13.69989 | -0.01209633 | 0.4373019 | -0.03603473 | 0.9712547 | 0.985423 |
| ENSG00000231083 | | 1.071755 | -0.0121106 | 0.7889501 | 0 | 0.9511087 | NA |
| ENSG00000269970 | | 1.351739 | -0.01217615 | 0.7889452 | 0 | 0.9496483 | NA |
| ENSG00000232946 | RPL35AP20 | 2.762638 | -0.01217951 | 0.6991934 | -0.05276089 | 0.9579224 | 0.979347 |
| ENSG00000172020 | GAP43 | 3.014868 | -0.01220976 | 0.7887921 | 0 | 0.9333955 | 0.967249 |
| ENSG00000260093 | | 1.758027 | -0.01221391 | 0.7005354 | -0.03774646 | 0.9698898 | 0.98462 |
| ENSG00000147677 | EIF3H | 14575.38 | -0.01222844 | 0.155525 | -0.1083221 | 0.9137402 | 0.959599 |
| ENSG00000256937 | KRT17P8 | 2.574821 | -0.01222953 | 0.7891305 | 0 | 0.9686558 | 0.983954 |
| ENSG00000146247 | PHIP | 5490.638 | -0.01224433 | 0.2419106 | -0.05474302 | 0.9563432 | 0.978854 |
| ENSG00000137409 | MTCH1 | 11145.02013 | -0.012246323 | 0.141536786 | -0.093024558 | 0.92588405 | 0.964396 |
| ENSG00000278626 | | 1.113683 | -0.01231992 | 0.7889582 | 0 | 0.9511344 | NA |
| ENSG00000249977 | | 1.46425 | -0.01237488 | 0.7134063 | -0.03881557 | 0.9690374 | NA |
| ENSG00000260781 | ARHGAP23P1 | 0.9457196 | -0.01240673 | 0.788956 | 0 | 0.9542641 | NA |
| ENSG00000198796 | ALPK2 | 0.9437299 | -0.01245575 | 0.7889036 | 0 | 0.9463397 | NA |
| ENSG00000227028 | SLC8A1-AS1 | 5.28576 | -0.01245772 | 0.5829285 | -0.03507683 | 0.9720185 | 0.985708 |
| ENSG00000142937 | RPS8 | 33810.82 | -0.01246526 | 0.1745644 | -0.07598951 | 0.9394274 | 0.970098 |
| ENSG00000257636 | G2E3-AS1 | 1.614924 | -0.01248004 | 0.7888881 | 0 | 0.9440929 | 0.972497 |
| ENSG00000121351 | IAPP | 1.685719787 | -0.012548208 | 0.789108186 | 0 | 0.96865469 | 0.983954 |
| ENSG00000249790 | | 1.061531 | -0.01255534 | 0.7888818 | 0 | 0.9440476 | NA |
| ENSG00000248837 | | 0.9670407 | -0.01256798 | 0.7888907 | 0 | 0.9459642 | NA |
| ENSG00000072952 | IRAG1 | 2.155155366 | -0.012580612 | 0.729084258 | -0.054934 | 0.95619104 | 0.978805 |
| ENSG00000170369 | CST2 | 0.9142006 | -0.01261429 | 0.7889398 | 0 | 0.9570207 | NA |
| ENSG00000259256 | LINC01895 | 1.574072 | -0.01263415 | 0.7889062 | 0 | 0.9466042 | 0.973611 |
| ENSG00000063245 | EPN1 | 7226.37323 | -0.012674592 | 0.192755186 | -0.064268835 | 0.94875617 | 0.974994 |
| ENSG00000161992 | PRR35 | 0.8754734 | -0.01270455 | 0.7320094 | -0.04902696 | 0.9608978 | NA |
| ENSG00000258274 | | 1.92191 | -0.01271346 | 0.7510767 | -0.09779433 | 0.9220956 | 0.963092 |
| ENSG00000175294 | CATSPER1 | 1.404469 | -0.0127361 | 0.7314211 | -0.05159267 | 0.9588533 | NA |
| ENSG00000187037 | GPR141 | 2.667519 | -0.01274561 | 0.7888098 | 0 | 0.9351438 | 0.968082 |
| ENSG00000027869 | SH2D2A | 1.994450222 | -0.0127648 | 0.788863839 | 0 | 0.9403534 | 0.970488 |

| | | | | | | | |
|---|---|---|---|---|---|---|---|
| ENSG00000133475 | GGT2 | 2.029775858 | -0.012770993 | 0.788847674 | 0 | 0.93992934 | 0.970428 |
| ENSG00000258903 | | 2.034796 | -0.01279165 | 0.7890905 | 0 | 0.9686535 | 0.983954 |
| ENSG00000207340 | RNVU1-1 | 0.9205984 | -0.01279732 | 0.788871 | 0 | 0.9478019 | NA |
| ENSG00000213976 | | 2.717015 | -0.012883 | 0.7888144 | 0 | 0.9338027 | 0.967249 |
| ENSG00000254272 | BTF3P3 | 15.81135 | -0.01288423 | 0.4152842 | -0.0382171 | 0.9695146 | 0.984409 |
| ENSG00000164576 | SAP30L | 2452.661 | -0.01289366 | 0.200041 | -0.06946415 | 0.9446202 | 0.972658 |
| ENSG00000253676 | TAGLN2P1 | 491.8192 | -0.01291576 | 0.2949793 | -0.04985744 | 0.960236 | 0.980157 |
| ENSG00000249948 | GBA3 | 2.622715 | -0.01295829 | 0.7888116 | 0 | 0.9338588 | 0.967249 |
| ENSG00000183150 | GPR19 | 29.24627 | -0.01304683 | 0.476793 | -0.0420345 | 0.9664712 | 0.983543 |
| ENSG00000168488 | ATXN2L | 11818.28 | -0.01305513 | 0.186713 | -0.07518538 | 0.9400672 | 0.97045 |
| ENSG00000181562 | EDDM3A | 1.246123 | -0.01306405 | 0.7888569 | 0 | 0.9422024 | NA |
| ENSG00000225883 | BMS1P11 | 1.279174 | -0.01316073 | 0.70356 | -0.04638881 | 0.9630004 | NA |
| ENSG00000223564 | CYP4F32P | 3.511125 | -0.01320498 | 0.7887673 | 0 | 0.9274614 | 0.96494 |
| ENSG00000287447 | | 0.9748613 | -0.01321426 | 0.7261522 | -0.04923106 | 0.9607352 | NA |
| ENSG00000250590 | LINC02492 | 3.90692 | -0.01322279 | 0.7887534 | 0 | 0.9260736 | 0.9644 |
| ENSG00000158669 | GPAT4 | 3029.312 | -0.01327651 | 0.09259003 | -0.1572547 | 0.8750441 | 0.940914 |
| ENSG00000145715 | RASA1 | 3128.971 | -0.01330103 | 0.1785935 | -0.06967123 | 0.9444553 | 0.972623 |
| ENSG00000086289 | EPDR1 | 4.592358001 | -0.013388823 | 0.788743815 | 0 | 0.9261339 | 0.9644 |
| ENSG00000173862 | | 0.8905043 | -0.0134633 | 0.7502793 | -0.05371799 | 0.9571598 | NA |
| ENSG00000107187 | LHX3 | 4.352447533 | -0.013463498 | 0.624156376 | -0.037487786 | 0.97009608 | 0.984727 |
| ENSG00000177112 | IRAG1-AS1 | 1.88786 | -0.01346452 | 0.7113677 | -0.05332218 | 0.9574752 | 0.979347 |
| ENSG00000145777 | TSLP | 4.857841 | -0.01347764 | 0.7890433 | 0 | 0.9686506 | 0.983954 |
| ENSG00000225536 | STIP1P3 | 86.45785 | -0.01350653 | 0.2962128 | -0.04816022 | 0.9615886 | 0.980809 |
| ENSG00000106018 | VIPR2 | 24.33848423 | -0.013508225 | 0.53486 | -0.035150874 | 0.97195944 | 0.985686 |
| ENSG00000148291 | SURF2 | 1911.284 | -0.0135318 | 0.1851827 | -0.07865791 | 0.9373047 | 0.96904 |
| ENSG00000201830 | | 0.9873854 | -0.01353289 | 0.7331645 | -0.04922463 | 0.9607403 | NA |
| ENSG00000176160 | HSF5 | 0.8390796 | -0.01355613 | 0.7271225 | -0.06021459 | 0.9519847 | NA |
| ENSG00000236857 | RAP1BP1 | 1.475046 | -0.01358821 | 0.7163816 | -0.04121461 | 0.9671248 | NA |
| ENSG00000231023 | LINC00326 | 1.004408 | -0.01360413 | 0.7889654 | 0 | 0.9545117 | NA |
| ENSG00000123700 | KCNJ2 | 1.222725224 | -0.013619936 | 0.788744057 | 0 | 0.93820593 | NA |
| ENSG00000274552 | MIR6889 | 0.8367684 | -0.01362791 | 0.7362553 | -0.0506004 | 0.9596439 | NA |
| ENSG00000186082 | KRT18P14 | 4.115182 | -0.01365818 | 0.6347913 | -0.03683037 | 0.9706203 | 0.985042 |
| ENSG00000116406 | EDEM3 | 3691.765268 | -0.013659649 | 0.278220843 | -0.044544267 | 0.96447057 | 0.982497 |

| | | | | | | | |
|---|---|---|---|---|---|---|---|
| ENSG00000170889 | RPS9 | 25481.74 | -0.01366189 | 0.1667345 | -0.0842845 | 0.9328302 | 0.967215 |
| ENSG00000266014 | | 0.8140153 | -0.01368068 | 0.7661816 | -0.07162708 | 0.9428987 | NA |
| ENSG00000264717 | NPY4R2 | 3.424127 | -0.01369596 | 0.7887302 | 0 | 0.931259 | 0.966622 |
| ENSG00000180861 | LINC01559 | 2.667606 | -0.01371634 | 0.7890284 | 0 | 0.9686496 | 0.983954 |
| ENSG00000224855 | OPA1-AS1 | 5.828443 | -0.01371689 | 0.5840372 | -0.04173925 | 0.9667066 | 0.983576 |
| ENSG00000112763 | BTN2A1 | 1572.22259 | -0.013737157 | 0.152357464 | -0.109205154 | 0.91303977 | 0.959195 |
| ENSG00000162745 | OLFML2B | 8.204693 | -0.0137431 | 0.535817 | -0.03624708 | 0.9710853 | 0.985289 |
| ENSG00000259081 | | 23.4909 | -0.01375235 | 0.3681477 | -0.04541883 | 0.9637735 | 0.982108 |
| ENSG00000175137 | SH3BP5L | 4010.075 | -0.0137545 | 0.1437151 | -0.1007172 | 0.919775 | 0.961781 |
| ENSG00000213925 | NPM1P33 | 2.663555 | -0.01376636 | 0.6574487 | -0.05191227 | 0.9585986 | 0.979615 |
| ENSG00000197586 | ENTPD6 | 5106.703 | -0.01380433 | 0.1389293 | -0.07380405 | 0.9411663 | 0.97101 |
| ENSG00000102901 | CENPT | 3320.184227 | -0.01383744 | 0.181087738 | -0.084332661 | 0.93279195 | 0.967215 |
| ENSG00000186152 | LILRP1 | 9.291957 | -0.01392417 | 0.7886624 | 0 | 0.9114215 | 0.95882 |
| ENSG00000253363 | | 3.514673 | -0.01392564 | 0.7888752 | 0 | 0.9609683 | 0.980566 |
| ENSG00000258938 | | 42.60254 | -0.0139312 | 0.3607716 | -0.04651113 | 0.9629029 | 0.981597 |
| ENSG00000279181 | | 0.7911524 | -0.01393772 | 0.7523061 | -0.09591239 | 0.9235902 | NA |
| ENSG00000225418 | AKR1C5P | 1.232442 | -0.01405125 | 0.748109 | -0.08330804 | 0.9336066 | NA |
| ENSG00000253455 | | 0.9692886 | -0.0140622 | 0.7625319 | -0.07821373 | 0.937658 | NA |
| ENSG00000272533 | SNORA28 | 9.472162 | -0.01407344 | 0.5310631 | -0.03543813 | 0.9717304 | 0.985632 |
| ENSG00000135486 | HNRNPA1 | 13151.46987 | -0.014162547 | 0.131668362 | -0.115691442 | 0.90789709 | 0.957105 |
| ENSG00000124181 | PLCG1 | 20472.56162 | -0.014177451 | 0.202291322 | -0.029611666 | 0.97637676 | 0.987602 |
| ENSG00000166123 | GPT2 | 5433.815 | -0.01419256 | 0.246669 | -0.04917822 | 0.9607773 | 0.980476 |
| ENSG00000226592 | | 2.724806 | -0.01426731 | 0.7888588 | 0 | 0.9606307 | 0.980372 |
| ENSG00000286512 | | 4.204291 | -0.01427046 | 0.6061523 | -0.04127326 | 0.967078 | 0.983702 |
| ENSG00000276531 | SNX19P4 | 1.593703 | -0.01431743 | 0.7495302 | -0.100472 | 0.9199696 | 0.961824 |
| ENSG00000253982 | | 437.8942 | -0.01434242 | 0.1686873 | -0.08325134 | 0.9336517 | 0.967249 |
| ENSG00000189362 | NEMP2 | 231.5627 | -0.01434539 | 0.3373034 | -0.04796237 | 0.9617462 | 0.980907 |
| ENSG00000226261 | PIMREGP1 | 7.648623 | -0.01435592 | 0.6099236 | -0.03639848 | 0.9709646 | 0.985204 |
| ENSG00000178726 | THBD | 1037.655 | -0.01437165 | 0.2077094 | -0.07044484 | 0.9438396 | 0.972468 |
| ENSG00000236854 | | 8.393827 | -0.01437674 | 0.5854035 | -0.03479745 | 0.9722413 | 0.985746 |
| ENSG00000128242 | GAL3ST1 | 1.177826403 | -0.014441396 | 0.703645217 | -0.057252669 | 0.95434392 | NA |
| ENSG00000111554 | MDM1 | 705.3457773 | -0.014546797 | 0.229203052 | -0.069161036 | 0.94486144 | 0.972798 |
| ENSG00000223834 | | 0.9562296 | -0.01455755 | 0.7360272 | -0.05090377 | 0.9594022 | NA |

| | | | | | | | |
|---|---|---|---|---|---|---|---|
| ENSG00000267388 | | 1.201207 | -0.01461213 | 0.757998 | -0.08949591 | 0.9286878 | NA |
| ENSG00000274813 | | 2.030315 | -0.01461449 | 0.7556909 | -0.0900197 | 0.9282716 | 0.965223 |
| ENSG00000284783 | | 1.153354 | -0.01471815 | 0.7122016 | -0.06058946 | 0.9516862 | NA |
| ENSG00000173559 | NABP1 | 1872.289 | -0.01474969 | 0.3555011 | -0.04182024 | 0.966642 | 0.98356 |
| ENSG00000176381 | PRR18 | 0.958548 | -0.01480912 | 0.7311164 | -0.05200832 | 0.9585221 | NA |
| ENSG00000143252 | SDHC | 3542.487 | -0.01486745 | 0.1537079 | -0.1056134 | 0.9158891 | 0.960213 |
| ENSG00000159479 | MED8 | 1849.366 | -0.01488425 | 0.1515198 | -0.0976225 | 0.9222321 | 0.963117 |
| ENSG00000213194 | CDK7P1 | 31.49648 | -0.0148946 | 0.3272905 | -0.05256697 | 0.9580769 | 0.979347 |
| ENSG00000263276 | | 3.802996 | -0.01491306 | 0.5994048 | -0.04346739 | 0.965329 | 0.982795 |
| ENSG00000230533 | | 1.266733 | -0.01499244 | 0.7889196 | 0 | 0.9686561 | NA |
| ENSG00000273129 | PACERR | 1.02441 | -0.01507986 | 0.7885191 | 0 | 0.9469454 | NA |
| ENSG00000287766 | | 3.3854 | -0.01509498 | 0.6739613 | -0.05277294 | 0.9579128 | 0.979347 |
| ENSG00000088538 | DOCK3 | 353.829438 | -0.015133052 | 0.322282618 | -0.054124613 | 0.95683588 | 0.979162 |
| ENSG00000100401 | RANGAP1 | 9171.895739 | -0.015135198 | 0.189358739 | -0.094870301 | 0.92441785 | 0.9637 |
| ENSG00000058453 | CROCC | 1025.913428 | -0.015150318 | 0.165228056 | -0.097191516 | 0.92257431 | 0.963214 |
| ENSG00000262768 | | 15.64325 | -0.01516237 | 0.3958808 | -0.05402723 | 0.9569135 | 0.979163 |
| ENSG00000062725 | APPBP2 | 39054.07448 | -0.015331761 | 0.126852652 | -0.124545939 | 0.90088303 | 0.954483 |
| ENSG00000267179 | | 1.516921 | -0.01533235 | 0.7086669 | -0.04926377 | 0.9607091 | NA |
| ENSG00000109079 | TNFAIP1 | 4273.55421 | -0.015334775 | 0.143302059 | -0.110746682 | 0.91181723 | 0.958851 |
| ENSG00000100307 | CBX7 | 733.5547245 | -0.015381767 | 0.216364016 | -0.067125052 | 0.94648215 | 0.973609 |
| ENSG00000166046 | TCP11L2 | 340.7737 | -0.01539914 | 0.3183828 | -0.05394794 | 0.9569766 | 0.979163 |
| ENSG00000283538 | | 107.2549 | -0.0154102 | 0.3733543 | -0.04842871 | 0.9613746 | 0.980783 |
| ENSG00000232573 | RPL3P4 | 10971.93 | -0.01541784 | 0.1667167 | -0.1245476 | 0.9008817 | 0.954483 |
| ENSG00000231744 | | 1.541159 | -0.01541982 | 0.6929676 | -0.0535063 | 0.9573285 | NA |
| ENSG00000246422 | DIAPH1-AS1 | 0.9895994 | -0.01551363 | 0.7264854 | -0.06027669 | 0.9519353 | NA |
| ENSG00000175707 | KDF1 | 980.2689 | -0.01552107 | 0.1609286 | -0.1053058 | 0.9161331 | 0.960225 |
| ENSG00000130414 | NDUFA10 | 7321.022453 | -0.015568561 | 0.154933826 | -0.105467123 | 0.91600516 | 0.960213 |
| ENSG00000228414 | LINC01185 | 2.454112 | -0.0156074 | 0.6435556 | -0.05517113 | 0.9560021 | 0.978774 |
| ENSG00000258766 | DIO2-AS1 | 1.055915 | -0.01561444 | 0.78851 | 0 | 0.9440503 | NA |
| ENSG00000270532 | PEBP1P2 | 181.5327 | -0.01561813 | 0.2813393 | -0.06249349 | 0.9501698 | 0.975545 |
| ENSG00000174928 | C3orf33 | 117.8122 | -0.01561916 | 0.2524374 | -0.06775144 | 0.9459835 | 0.97335 |
| ENSG00000249849 | | 1.011367 | -0.01562683 | 0.7421292 | -0.08895944 | 0.9291141 | NA |
| ENSG00000175426 | PCSK1 | 0.7950599 | -0.01568159 | 0.7357765 | -0.06845769 | 0.9454213 | NA |

| | | | | | | | |
|---|---|---|---|---|---|---|---|
| ENSG00000270522 | | 1.275381 | -0.01568258 | 0.7512603 | -0.09769037 | 0.9221782 | NA |
| ENSG00000184995 | IFNE | 0.9201735 | -0.01568648 | 0.7885527 | 0 | 0.9464472 | NA |
| ENSG00000226976 | COX6A1P2 | 3280.203 | -0.01569166 | 0.215589 | -0.08171625 | 0.9348724 | 0.967952 |
| ENSG00000187862 | TTC24 | 1.603245 | -0.01570754 | 0.6862253 | -0.06958717 | 0.9445222 | 0.972637 |
| ENSG00000206791 | RNU1-129P | 0.8729175 | -0.01571811 | 0.7387496 | -0.05971274 | 0.9523844 | NA |
| ENSG00000150656 | CNDP1 | 2.488197 | -0.01574555 | 0.7166171 | -0.03871868 | 0.9691147 | 0.984266 |
| ENSG00000230679 | ENO1-AS1 | 12.35802 | -0.01578366 | 0.5532837 | -0.04188967 | 0.9665866 | 0.983543 |
| ENSG00000246695 | RASSF8-AS1 | 270.3443 | -0.0158092 | 0.3424453 | -0.07168511 | 0.9428525 | 0.971822 |
| ENSG00000262879 | | 1240.075 | -0.01583371 | 0.2405082 | -0.07221923 | 0.9424274 | 0.971691 |
| ENSG00000243547 | HNRNPKP4 | 3168.51 | -0.01583752 | 0.1721347 | -0.09196667 | 0.9267245 | 0.964715 |
| ENSG00000144567 | RETREG2 | 7235.939 | -0.01584791 | 0.1083184 | -0.1566275 | 0.8755384 | 0.941217 |
| ENSG00000265178 | MIR4728 | 2.454963 | -0.01584809 | 0.677978 | -0.04438633 | 0.9645965 | 0.982497 |
| ENSG00000182208 | MOB2 | 1031.107 | -0.01584884 | 0.2163337 | -0.1058363 | 0.9157123 | 0.960178 |
| ENSG00000207750 | MIR553 | 0.8981737 | -0.01585102 | 0.7438764 | -0.07924227 | 0.9368399 | NA |
| ENSG00000234627 | NUS1P3 | 1.165327 | -0.01586317 | 0.7194331 | -0.05666778 | 0.9548098 | NA |
| ENSG00000149503 | INCENP | 2223.755 | -0.01593623 | 0.3795144 | -0.04834704 | 0.9614397 | 0.980783 |
| ENSG00000096093 | EFHC1 | 1686.06391 | -0.015955696 | 0.192590121 | -0.092614154 | 0.9262101 | 0.964433 |
| ENSG00000258638 | | 0.8189278 | -0.01595655 | 0.7885179 | 0 | 0.9448335 | NA |
| ENSG00000122359 | ANXA11 | 6824.627441 | -0.015962765 | 0.151349826 | -0.137584715 | 0.89056864 | 0.949218 |
| ENSG00000197863 | ZNF790 | 252.1164 | -0.0160297 | 0.1790715 | -0.09574137 | 0.923726 | 0.963623 |
| ENSG00000149531 | FRG1BP | 705.712 | -0.01606802 | 0.1889963 | -0.0967592 | 0.9229176 | 0.963383 |
| ENSG00000286839 | | 0.8846249 | -0.01610441 | 0.7886416 | 0 | 0.9590287 | NA |
| ENSG00000186431 | FCAR | 1.475864 | -0.01613438 | 0.7884858 | 0 | 0.9429896 | NA |
| ENSG00000219699 | | 1.172595 | -0.01614066 | 0.7351103 | -0.08300078 | 0.9338509 | NA |
| ENSG00000110696 | C11orf58 | 8849.821737 | -0.016178567 | 0.123247724 | -0.116091253 | 0.90758022 | 0.957042 |
| ENSG00000168569 | TMEM223 | 1472.276 | -0.01619391 | 0.1964528 | -0.08680996 | 0.9308226 | 0.966433 |
| ENSG00000185758 | CLDN24 | 1.047874 | -0.01620283 | 0.7597009 | -0.08331999 | 0.9335971 | NA |
| ENSG00000081154 | PCNP | 5477.144343 | -0.016238154 | 0.131225534 | -0.125528269 | 0.90010535 | 0.954113 |
| ENSG00000235904 | RBMS3-AS3 | 1.638149 | -0.01624563 | 0.7884681 | 0 | 0.9402802 | 0.97045 |
| ENSG00000261807 | LINC02141 | 1.303142 | -0.01626659 | 0.7638463 | -0.07603937 | 0.9393878 | NA |
| ENSG00000251003 | ZFPM2-AS1 | 79.87326 | -0.01627092 | 0.7884009 | 0 | 0.9025292 | 0.954999 |
| ENSG00000169393 | ELSPBP1 | 0.7831822 | -0.01627335 | 0.7884773 | 0 | 0.9441952 | NA |
| ENSG00000284842 | FAM91A2P | 0.8094599 | -0.01628153 | 0.7885457 | 0 | 0.9515975 | NA |

| | | | | | | | |
|---|---|---|---|---|---|---|---|
| ENSG00000280083 | | 0.9921984 | -0.01628855 | 0.7281308 | -0.06762573 | 0.9460836 | NA |
| ENSG00000276980 | | 0.8767816 | -0.01629072 | 0.7886205 | 0 | 0.958571 | NA |
| ENSG00000285766 | | 1.116171 | -0.01635766 | 0.7173622 | -0.07180553 | 0.9427567 | NA |
| ENSG00000207008 | SNORA54 | 1.023104 | -0.01635862 | 0.7237815 | -0.062418 | 0.95023 | NA |
| ENSG00000139133 | ALG10 | 350.022973 | -0.016361335 | 0.276561391 | -0.061991392 | 0.95056969 | 0.975726 |
| ENSG00000172005 | MAL | 1.095055 | -0.01641117 | 0.7884392 | 0 | 0.9391198 | NA |
| ENSG00000285159 | | 0.9764255 | -0.01642898 | 0.7601217 | -0.08257172 | 0.9341921 | NA |
| ENSG00000214857 | SEM1P1 | 2.126627 | -0.01644403 | 0.7068684 | -0.04532884 | 0.9638452 | 0.982143 |
| ENSG00000246379 | | 0.8541928 | -0.0164581 | 0.7885209 | 0 | 0.9497219 | NA |
| ENSG00000224464 | PGAM1P6 | 0.8158087 | -0.01645962 | 0.7423156 | -0.07469956 | 0.9404538 | NA |
| ENSG00000153253 | SCN3A | 1.199579 | -0.01649336 | 0.7884517 | 0 | 0.9427109 | NA |
| ENSG00000272620 | AFAP1-AS1 | 3.890458 | -0.01650653 | 0.598986 | -0.0426914 | 0.9659475 | 0.983343 |
| ENSG00000222020 | HDAC4-AS1 | 22.10457 | -0.0165979 | 0.4415658 | -0.04723117 | 0.962329 | 0.981163 |
| ENSG00000267737 | | 1.158452 | -0.0165999 | 0.7215488 | -0.05363128 | 0.9572289 | NA |
| ENSG00000266708 | | 9.08055 | -0.016645 | 0.5117178 | -0.04384351 | 0.9650291 | 0.982596 |
| ENSG00000129009 | ISLR | 1.043485557 | -0.016689021 | 0.788415534 | 0 | 0.93939843 | NA |
| ENSG00000287437 | | 7.880625 | -0.01669362 | 0.5070146 | -0.05209499 | 0.958453 | 0.979579 |
| ENSG00000212283 | SNORD89 | 8.536485 | -0.01672977 | 0.5515503 | -0.03841526 | 0.9693566 | 0.984317 |
| ENSG00000248705 | | 1.233748 | -0.01677298 | 0.7884304 | 0 | 0.937585 | NA |
| ENSG00000242071 | RPL7AP6 | 12455.56 | -0.01678757 | 0.1557168 | -0.1417632 | 0.8872671 | 0.947081 |
| ENSG00000224057 | EGFR-AS1 | 2.108705 | -0.0168761 | 0.788341 | 0 | 0.9284227 | 0.9653 |
| ENSG00000163935 | SFMBT1 | 1218.097 | -0.01690269 | 0.1327663 | -0.1315361 | 0.8953512 | 0.951617 |
| ENSG00000204174 | NPY4R | 3.632576 | -0.01692573 | 0.7883367 | 0 | 0.9276428 | 0.96494 |
| ENSG00000274618 | H4C6 | 0.9604594 | -0.0169671 | 0.7884147 | 0 | 0.9439838 | NA |
| ENSG00000075415 | SLC25A3 | 21900.73393 | -0.016988521 | 0.163818621 | -0.101245161 | 0.91935585 | 0.961744 |
| ENSG00000225951 | ODF2-AS1 | 3.851097 | -0.017005 | 0.6582057 | -0.05286101 | 0.9578427 | 0.979347 |
| ENSG00000166111 | SVOP | 0.8196984 | -0.01707592 | 0.7883843 | 0 | 0.9382003 | NA |
| ENSG00000137776 | SLTM | 6931.68695 | -0.017112451 | 0.094457526 | -0.189670332 | 0.84956747 | 0.926874 |
| ENSG00000169375 | SIN3A | 5467.09 | -0.01712942 | 0.1576694 | -0.1255653 | 0.9000761 | 0.954113 |
| ENSG00000183666 | GUSBP1 | 745.148 | -0.01714247 | 0.123007 | -0.1381437 | 0.8901268 | 0.948976 |
| ENSG00000136869 | TLR4 | 1.369020551 | -0.017148636 | 0.788372845 | 0 | 0.93822507 | NA |
| ENSG00000265874 | MIR4489 | 2.533229 | -0.01719015 | 0.6642208 | -0.05000364 | 0.9601195 | 0.980097 |
| ENSG00000287561 | | 1.024529 | -0.0172257 | 0.7506191 | -0.07967333 | 0.9364971 | NA |

| | | | | | | | |
|---|---|---|---|---|---|---|---|
| ENSG00000258512 | LINC00239 | 1.218547 | -0.0172276 | 0.7884294 | 0 | 0.9490275 | NA |
| ENSG00000244753 | RPL15P21 | 0.8904365 | -0.01722784 | 0.7882947 | 0 | 0.9311327 | NA |
| ENSG00000273447 | | 24.4452 | -0.0172512 | 0.3648842 | -0.05366775 | 0.9571999 | 0.97924 |
| ENSG00000167749 | KLK4 | 1.274943 | -0.01729617 | 0.7883101 | 0 | 0.9314228 | NA |
| ENSG00000269662 | BNIP3P39 | 0.9810098 | -0.01730128 | 0.788356 | 0 | 0.9394283 | NA |
| ENSG00000280355 | | 30.52705 | -0.01737381 | 0.4586457 | -0.05047899 | 0.9597407 | 0.979969 |
| ENSG00000235586 | | 4.136486 | -0.01738722 | 0.639696 | -0.05049118 | 0.959731 | 0.979969 |
| ENSG00000287467 | | 1.615395 | -0.01739796 | 0.6670829 | -0.0546364 | 0.9564281 | 0.978865 |
| ENSG00000226277 | | 1.044942 | -0.01742846 | 0.7883927 | 0 | 0.9451076 | NA |
| ENSG00000272824 | | 1.157599 | -0.01743831 | 0.788323 | 0 | 0.936373 | NA |
| ENSG00000251439 | KRTAP4-17P | 1.276777 | -0.01744063 | 0.741288 | -0.0628116 | 0.9499165 | NA |
| ENSG00000233926 | | 1.017487 | -0.01744193 | 0.7255399 | -0.05963299 | 0.9524479 | NA |
| ENSG00000261175 | LINC02188 | 1.092419 | -0.01744289 | 0.7883009 | 0 | 0.9355933 | NA |
| ENSG00000236582 | PRPF38AP2 | 9.136046 | -0.01744627 | 0.5486664 | -0.04826356 | 0.9615062 | 0.980809 |
| ENSG00000224815 | | 0.9543837 | -0.01747042 | 0.7884601 | 0 | 0.9534761 | NA |
| ENSG00000273176 | | 1.42833 | -0.01748564 | 0.7178679 | -0.05717344 | 0.954407 | NA |
| ENSG00000226079 | | 0.7801987 | -0.01749206 | 0.7883898 | 0 | 0.9410465 | NA |
| ENSG00000274297 | | 0.8407603 | -0.01749224 | 0.7358902 | -0.06290828 | 0.9498395 | NA |
| ENSG00000229782 | | 0.8472281 | -0.01752083 | 0.7883635 | 0 | 0.9405595 | NA |
| ENSG00000121797 | CCRL2 | 1.030139818 | -0.017554714 | 0.78831526 | 0 | 0.92897518 | NA |
| ENSG00000280029 | | 0.9193497 | -0.01758495 | 0.7883414 | 0 | 0.9405708 | NA |
| ENSG00000203667 | COX20 | 105.1979 | -0.01760061 | 0.3081375 | -0.0649865 | 0.9481847 | 0.974627 |
| ENSG00000198846 | TOX | 1.126432 | -0.01761799 | 0.7883062 | 0 | 0.9354465 | NA |
| ENSG00000100121 | GGTLC2 | 4.941288699 | -0.017646804 | 0.788223756 | 0 | 0.91791922 | 0.961208 |
| ENSG00000267218 | | 2.917831 | -0.01766986 | 0.6861486 | -0.05480627 | 0.9562928 | 0.97884 |
| ENSG00000120068 | HOXB8 | 2.028961054 | -0.017673417 | 0.788268381 | 0 | 0.9236892 | 0.963623 |
| ENSG00000089220 | PEBP1 | 16814.07571 | -0.017691876 | 0.20922078 | -0.09074817 | 0.9276927 | 0.96494 |
| ENSG00000150048 | CLEC1A | 1.005955 | -0.01769838 | 0.7883527 | 0 | 0.9385381 | NA |
| ENSG00000253952 | HIGD1AP18 | 1.292026 | -0.01771125 | 0.7181707 | -0.05695391 | 0.9545819 | NA |
| ENSG00000100938 | GMPR2 | 6122.527744 | -0.017715468 | 0.144902466 | -0.13144894 | 0.89542018 | 0.951617 |
| ENSG00000251537 | | 13.81337 | -0.01777748 | 0.5021975 | -0.04479045 | 0.9642743 | 0.98243 |
| ENSG00000241120 | HMGN1P8 | 6.828633 | -0.01777947 | 0.5450651 | -0.04730871 | 0.9622672 | 0.981157 |
| ENSG00000259179 | UBE2CP4 | 4.312988 | -0.01782705 | 0.6814563 | -0.05436007 | 0.9566483 | 0.979015 |

| | | | | | | | |
|---|---|---|---|---|---|---|---|
| ENSG00000112210 | RAB23 | 411.283701 | -0.017835217 | 0.294899619 | -0.070279883 | 0.94397089 | 0.972468 |
| ENSG00000142541 | RPL13A | 30060.93 | -0.01783591 | 0.1873172 | -0.09521157 | 0.9241468 | 0.963623 |
| ENSG00000136273 | HUS1 | 981.1669468 | -0.017857565 | 0.132291419 | -0.145458508 | 0.88434887 | 0.945712 |
| ENSG00000211815 | TRAV36DV7 | 1.022858 | -0.01789782 | 0.7882643 | 0 | 0.9312199 | NA |
| ENSG00000121390 | PSPC1 | 4779.411891 | -0.017957689 | 0.156598977 | -0.105629879 | 0.91587602 | 0.960213 |
| ENSG00000251520 | | 7.000992 | -0.01798747 | 0.5433089 | -0.05062786 | 0.9596221 | 0.979969 |
| ENSG00000167625 | ZNF526 | 961.8217 | -0.01811448 | 0.1534451 | -0.110656 | 0.9118891 | 0.958851 |
| ENSG00000125686 | MED1 | 3315.074192 | -0.018130105 | 0.22589741 | -0.082961204 | 0.93388239 | 0.967249 |
| ENSG00000156052 | GNAQ | 5537.972 | -0.01813646 | 0.1499216 | -0.1388867 | 0.8895397 | 0.948689 |
| ENSG00000116691 | MIIP | 1182.25387 | -0.018175145 | 0.23503922 | -0.085620359 | 0.93176821 | 0.966886 |
| ENSG00000223969 | | 1.60433 | -0.01822704 | 0.7882345 | 0 | 0.9267374 | 0.964715 |
| ENSG00000231358 | | 2.817838 | -0.01827289 | 0.7880897 | 0 | 0.9195437 | 0.961751 |
| ENSG00000243959 | RN7SL684P | 1.150895 | -0.0182781 | 0.7259981 | -0.05946541 | 0.9525814 | NA |
| ENSG00000163145 | C1QTNF7 | 7.515155 | -0.01828107 | 0.743296 | -0.1130745 | 0.9099715 | 0.958028 |
| ENSG00000141349 | G6PC3 | 3724.486 | -0.01831209 | 0.1822284 | -0.1063518 | 0.9153033 | 0.960119 |
| ENSG00000270184 | | 48.33698 | -0.01834916 | 0.3765183 | -0.05872488 | 0.9531712 | 0.977226 |
| ENSG00000187721 | GTF2IP3 | 1.93634 | -0.01835396 | 0.7882465 | 0 | 0.936476 | 0.968668 |
| ENSG00000240445 | FOXO3B | 480.347 | -0.01835786 | 0.2878586 | -0.0702534 | 0.943992 | 0.972468 |
| ENSG00000216639 | | 1.304448 | -0.01837413 | 0.7140331 | -0.06175816 | 0.9507554 | NA |
| ENSG00000199716 | | 0.9628154 | -0.01837589 | 0.7237282 | -0.07407706 | 0.9409491 | NA |
| ENSG00000288037 | | 1.135006 | -0.01838039 | 0.7369918 | -0.06171711 | 0.9507881 | NA |
| ENSG00000274211 | SOCS7 | 2158.972 | -0.01839245 | 0.2556146 | -0.07997257 | 0.9362591 | 0.968541 |
| ENSG00000234377 | OBI1-AS1 | 0.947435 | -0.01841442 | 0.7079984 | -0.06520595 | 0.94801 | NA |
| ENSG00000178695 | KCTD12 | 853.3398 | -0.0184218 | 0.251239 | -0.08287081 | 0.9339543 | 0.967249 |
| ENSG00000258986 | TMEM179 | 6.313998 | -0.01842674 | 0.7425511 | -0.1131413 | 0.9099185 | 0.95802 |
| ENSG00000228335 | | 56.72557 | -0.01843254 | 0.2934214 | -0.07019134 | 0.9440414 | 0.972481 |
| ENSG00000201118 | | 1.717201 | -0.01843957 | 0.6641362 | -0.06568365 | 0.9476297 | 0.974289 |
| ENSG00000287306 | | 47.28358 | -0.01850808 | 0.3688036 | -0.05661199 | 0.9548543 | 0.978285 |
| ENSG00000089876 | DHX32 | 2767.990885 | -0.018511926 | 0.134547932 | -0.143651993 | 0.88577528 | 0.946363 |
| ENSG00000147687 | TATDN1 | 4054.095 | -0.01851853 | 0.1766221 | -0.1079147 | 0.9140634 | 0.959721 |
| ENSG00000275769 | | 0.8376074 | -0.01852511 | 0.755496 | -0.09078799 | 0.9276611 | NA |
| ENSG00000121741 | ZMYM2 | 9948.406222 | -0.018541283 | 0.160997962 | -0.124197364 | 0.90115901 | 0.954585 |
| ENSG00000279066 | HEXD-IT1 | 19.82551 | -0.01857791 | 0.4354138 | -0.05508665 | 0.9560694 | 0.978774 |

| | | | | | | | |
|---|---|---|---|---|---|---|---|
| ENSG00000282381 | | 6.625698 | -0.01858638 | 0.5813853 | -0.05580584 | 0.9554965 | 0.978579 |
| ENSG00000287133 | | 7.451537 | -0.01859869 | 0.5851594 | -0.05193736 | 0.9585786 | 0.979615 |
| ENSG00000108671 | PSMD11 | 5014.836379 | -0.018622779 | 0.241016226 | -0.051494088 | 0.95893181 | 0.979741 |
| ENSG00000106105 | GARS1 | 9199.45415 | -0.018665175 | 0.273146543 | -0.076854729 | 0.93873911 | 0.969795 |
| ENSG00000090776 | EFNB1 | 555.7227586 | -0.018665315 | 0.216552491 | -0.092248298 | 0.92650077 | 0.964585 |
| ENSG00000274718 | | 1.745229 | -0.01870763 | 0.788144 | 0 | 0.9276663 | 0.96494 |
| ENSG00000225217 | HSPA7 | 2.672837 | -0.01870789 | 0.7881526 | 0 | 0.9209448 | 0.962534 |
| ENSG00000287475 | | 3.486837 | -0.01871705 | 0.6322679 | -0.05247208 | 0.9581525 | 0.979347 |
| ENSG00000212939 | | 2.167228 | -0.01876317 | 0.7881306 | 0 | 0.9169919 | 0.960936 |
| ENSG00000255341 | OR8Q1P | 5.602678 | -0.01884437 | 0.5513647 | -0.0495683 | 0.9604664 | 0.980289 |
| ENSG00000218175 | | 2797.02 | -0.01887093 | 0.1859122 | -0.1139279 | 0.909295 | 0.957705 |
| ENSG00000251669 | FAM86EP | 314.2582 | -0.01887856 | 0.1928443 | -0.1027629 | 0.9181511 | 0.96124 |
| ENSG00000166887 | VPS39 | 4568.845 | -0.01889243 | 0.14484 | -0.1432984 | 0.8860545 | 0.946535 |
| ENSG00000229604 | MTATP8P2 | 2.24725 | -0.01894145 | 0.7171276 | -0.06087366 | 0.9514598 | 0.976211 |
| ENSG00000241413 | RN7SL441P | 0.9464298 | -0.01894432 | 0.7595436 | -0.08359485 | 0.9333786 | NA |
| ENSG00000129696 | TTI2 | 506.9420552 | -0.018958521 | 0.194719183 | -0.103321924 | 0.91770748 | 0.961195 |
| ENSG00000205632 | LINC01310 | 2.011224 | -0.0189688 | 0.7881401 | 0 | 0.9227107 | 0.963243 |
| ENSG00000132570 | PCBD2 | 1119.796555 | -0.018980667 | 0.161697591 | -0.119095376 | 0.90519979 | 0.955968 |
| ENSG00000174371 | EXO1 | 993.9003 | -0.01905538 | 0.5468493 | -0.05502673 | 0.9561172 | 0.978774 |
| ENSG00000261054 | | 1.960779 | -0.01913762 | 0.7106393 | -0.0531353 | 0.9576241 | 0.979347 |
| ENSG00000212135 | SNORD67 | 10.55711 | -0.01915045 | 0.5115883 | -0.05164393 | 0.9588124 | 0.979737 |
| ENSG00000124523 | SIRT5 | 1190.832059 | -0.019157173 | 0.142342384 | -0.153119284 | 0.87830421 | 0.94267 |
| ENSG00000265107 | GJA5 | 7.147091 | -0.01922879 | 0.7882285 | 0 | NA | NA |
| ENSG00000184887 | BTBD6 | 4090.983 | -0.01924134 | 0.1733432 | -0.1329979 | 0.894195 | 0.950906 |
| ENSG00000272931 | LRRC8D-DT | 79.12452 | -0.01927547 | 0.3273816 | -0.06743215 | 0.9462377 | 0.973536 |
| ENSG00000197976 | AKAP17A | 2873.205 | -0.01927782 | 0.1298109 | -0.1658051 | 0.8683103 | 0.936731 |
| ENSG00000249509 | NEUROG2-AS1 | 2.315045 | -0.01931702 | 0.7880782 | 0 | 0.9189 | 0.961532 |
| ENSG00000168427 | KLHL30 | 1.255975 | -0.01931721 | 0.72996 | -0.07475161 | 0.9404123 | NA |
| ENSG00000165409 | TSHR | 3.559981 | -0.01933492 | 0.6829518 | -0.04615246 | 0.9631887 | 0.981813 |
| ENSG00000282906 | | 0.8440558 | -0.01938947 | 0.7692925 | -0.0663342 | 0.9471118 | NA |
| ENSG00000155918 | RAET1L | 3.067749 | -0.01938969 | 0.7140854 | -0.04557969 | 0.9636453 | 0.982053 |
| ENSG00000119408 | NEK6 | 2626.507835 | -0.019459252 | 0.136276743 | -0.142671547 | 0.88654959 | 0.946696 |
| ENSG00000149806 | FAU | 7848.811 | -0.0195242 | 0.1796212 | -0.09322054 | 0.9257284 | 0.964272 |

| | | | | | | | |
|---|---|---|---|---|---|---|---|
| ENSG00000107223 | EDF1 | 10364.12703 | -0.019526348 | 0.184265779 | -0.187559025 | 0.85122235 | 0.927727 |
| ENSG00000130312 | MRPL34 | 2107.516568 | -0.019538838 | 0.234591591 | -0.093551942 | 0.92546509 | 0.964262 |
| ENSG00000156413 | FUT6 | 5.292951 | -0.01954122 | 0.7174003 | -0.06841697 | 0.9454537 | 0.973068 |
| ENSG00000157884 | CIB4 | 1.470084 | -0.01954509 | 0.7557857 | -0.09028031 | 0.9280645 | NA |
| ENSG00000184492 | FOXD4L1 | 22.62344 | -0.01955953 | 0.5011725 | -0.05617859 | 0.9551995 | 0.9785 |
| ENSG00000126107 | HECTD3 | 2851.130115 | -0.019597732 | 0.111989273 | -0.184631132 | 0.85351837 | 0.928728 |
| ENSG00000249986 | YWHAQP6 | 2.794197 | -0.01960016 | 0.6969756 | -0.06882874 | 0.9451259 | 0.972882 |
| ENSG00000277493 | | 2.0109 | -0.01960663 | 0.7880844 | 0 | 0.9255609 | 0.96427 |
| ENSG00000288473 | | 1.118327 | -0.01963061 | 0.703676 | -0.07029495 | 0.9439589 | NA |
| ENSG00000285774 | | 2.225319 | -0.01965693 | 0.7274057 | -0.05249909 | 0.958131 | 0.979347 |
| ENSG00000261420 | | 21.70185 | -0.01969219 | 0.4464701 | -0.05323174 | 0.9575473 | 0.979347 |
| ENSG00000110713 | NUP98 | 5464.028603 | -0.019721368 | 0.14618059 | -0.159101724 | 0.87358873 | 0.940119 |
| ENSG00000171551 | ECEL1 | 1.503396 | -0.01973714 | 0.7560881 | -0.08974248 | 0.9284919 | NA |
| ENSG00000214652 | ZNF727 | 30.82062 | -0.01974691 | 0.7880186 | 0 | 0.9050002 | 0.955968 |
| ENSG00000261461 | UBE2MP1 | 778.6727 | -0.01981744 | 0.2069797 | -0.1029062 | 0.9180374 | 0.961226 |
| ENSG00000258837 | | 2.272402 | -0.01982073 | 0.788066 | 0 | 0.926877 | 0.96475 |
| ENSG00000261898 | | 1.460266 | -0.01982802 | 0.7189182 | -0.06097112 | 0.9513822 | NA |
| ENSG00000133142 | TCEAL4 | 11533.19103 | -0.019839202 | 0.200849418 | -0.105331354 | 0.91611288 | 0.960225 |
| ENSG00000102189 | EEA1 | 2502.076887 | -0.019843892 | 0.268099831 | -0.078657525 | 0.93730503 | 0.96904 |
| ENSG00000252503 | RNU6-531P | 4.745929 | -0.0199183 | 0.5860031 | -0.05376101 | 0.9571256 | 0.979221 |
| ENSG00000215009 | ACSM4 | 3.366079 | -0.01992784 | 0.7436047 | -0.1108751 | 0.9117154 | 0.958851 |
| ENSG00000252516 | RNA5SP82 | 18.43299 | -0.01997132 | 0.3677941 | -0.06367671 | 0.9492277 | 0.975253 |
| ENSG00000205929 | C21orf62 | 2.314418 | -0.02000782 | 0.6498952 | -0.06362909 | 0.9492656 | 0.975254 |
| ENSG00000244094 | SPRR2F | 0.890343 | -0.02002586 | 0.761265 | -0.08068382 | 0.9356934 | NA |
| ENSG00000140497 | SCAMP2 | 5834.96 | -0.02005009 | 0.1338503 | -0.153798 | 0.877769 | 0.942353 |
| ENSG00000109445 | ZNF330 | 2457.492944 | -0.020088752 | 0.116012242 | -0.176216552 | 0.86012382 | 0.932684 |
| ENSG00000236478 | | 1.380763 | -0.02009143 | 0.7293381 | -0.06679328 | 0.9467463 | NA |
| ENSG00000115486 | GGCX | 3280.834997 | -0.020097599 | 0.15274559 | -0.18724759 | 0.85146651 | 0.927841 |
| ENSG00000176533 | GNG7 | 243.148 | -0.02019793 | 0.2678768 | -0.08188133 | 0.9347411 | 0.967854 |
| ENSG00000236992 | RPL12P12 | 55.01301 | -0.02027303 | 0.3137362 | -0.06283891 | 0.9498948 | 0.975523 |
| ENSG00000176358 | TAC4 | 4.079725 | -0.02029291 | 0.6133493 | -0.05446239 | 0.9565668 | 0.978969 |
| ENSG00000254397 | | 2.916027 | -0.02037075 | 0.6330288 | -0.05805851 | 0.953702 | 0.977458 |
| ENSG00000085274 | MYNN | 1645.156924 | -0.02040891 | 0.115841758 | -0.190391713 | 0.84900219 | 0.926594 |

| | | | | | | | |
|---|---|---|---|---|---|---|---|
| ENSG00000169509 | CRCT1 | 1.040963 | -0.02041444 | 0.788037 | 0 | 0.9336601 | NA |
| ENSG00000206848 | RNU6-890P | 1.454791 | -0.02046215 | 0.7385232 | -0.08887722 | 0.9291795 | NA |
| ENSG00000287009 | | 18.81578 | -0.02047485 | 0.3714693 | -0.06572253 | 0.9475987 | 0.974289 |
| ENSG00000116132 | PRRX1 | 1.057233415 | -0.020487977 | 0.78798631 | 0 | 0.93407176 | NA |
| ENSG00000223390 | | 1.249424 | -0.02050616 | 0.7111708 | -0.07488446 | 0.9403066 | NA |
| ENSG00000265491 | RNF115 | 5086.946 | -0.02052551 | 0.1130934 | -0.1871182 | 0.851568 | 0.927845 |
| ENSG00000214203 | RPS4XP1 | 95.64847 | -0.02056365 | 0.2745196 | -0.08346787 | 0.9334795 | 0.967249 |
| ENSG00000187605 | TET3 | 6475.587 | -0.02063104 | 0.2388408 | -0.09409375 | 0.9250347 | 0.963954 |
| ENSG00000267497 | NFE2L3P1 | 0.9865471 | -0.02064474 | 0.788014 | 0 | 0.9370051 | NA |
| ENSG00000250318 | | 13.09688 | -0.02064924 | 0.5150415 | -0.05751413 | 0.9541356 | 0.977724 |
| ENSG00000266578 | | 2.216848 | -0.0206587 | 0.7242078 | -0.05761703 | 0.9540537 | 0.977678 |
| ENSG00000272002 | | 1.280889 | -0.02071426 | 0.7880414 | 0 | 0.9406838 | NA |
| ENSG00000118482 | PHF3 | 4795.131781 | -0.02073117 | 0.171565365 | -0.149216217 | 0.88138302 | 0.944481 |
| ENSG00000152034 | MCHR2 | 1.930856 | -0.02074502 | 0.7879604 | 0 | 0.9301401 | 0.966178 |
| ENSG00000229375 | USP24P1 | 3.648254 | -0.02077426 | 0.7880516 | 0 | 0.9179391 | 0.961208 |
| ENSG00000013293 | SLC7A14 | 0.946288366 | -0.020780718 | 0.75579178 | -0.090141782 | 0.92817455 | NA |
| ENSG00000105246 | EBI3 | 3.086117414 | -0.020828523 | 0.753568821 | -0.09408541 | 0.92504131 | 0.963954 |
| ENSG00000083168 | KAT6A | 2998.839356 | -0.020838834 | 0.272633038 | -0.106600031 | 0.91510629 | 0.960119 |
| ENSG00000109193 | SULT1E1 | 1.983087497 | -0.020913182 | 0.787947774 | 0 | 0.92820164 | 0.965223 |
| ENSG00000230537 | | 4.954487 | -0.02094064 | 0.6186933 | -0.05308055 | 0.9576677 | 0.979347 |
| ENSG00000265984 | | 1.063675 | -0.02098421 | 0.787963 | 0 | 0.9298456 | NA |
| ENSG00000286769 | | 0.9521691 | -0.02102657 | 0.7880581 | 0 | 0.9460576 | NA |
| ENSG00000253896 | | 0.8855169 | -0.02103836 | 0.7880907 | 0 | 0.9481376 | NA |
| ENSG00000111775 | COX6A1 | 2032.974228 | -0.0210625 | 0.223518792 | -0.123570413 | 0.90165542 | 0.954802 |
| ENSG00000238258 | | 0.9535235 | -0.02106355 | 0.7880117 | 0 | 0.9408404 | NA |
| ENSG00000275663 | H4C7 | 1.181423 | -0.02108823 | 0.7297074 | -0.0714417 | 0.9430462 | NA |
| ENSG00000258615 | | 0.904518 | -0.02109217 | 0.7879561 | 0 | 0.934487 | NA |
| ENSG00000213707 | HMGB1P10 | 14.53299 | -0.02113436 | 0.5423283 | -0.05828757 | 0.9535196 | 0.977458 |
| ENSG00000224728 | IMPDH1P8 | 56.66953 | -0.02113968 | 0.3354944 | -0.07271588 | 0.9420322 | 0.971504 |
| ENSG00000230584 | CCT5P2 | 56.50253 | -0.02116377 | 0.3208362 | -0.07093776 | 0.9434473 | 0.972213 |
| ENSG00000255856 | | 0.9409844 | -0.0211645 | 0.7324222 | -0.08547729 | 0.9318819 | NA |
| ENSG00000169891 | REPS2 | 691.6704 | -0.02120373 | 0.2221393 | -0.1028592 | 0.9180748 | 0.961226 |
| ENSG00000117620 | SLC35A3 | 1365.188482 | -0.021217408 | 0.18940236 | -0.130506572 | 0.89616566 | 0.952066 |

| | | | | | | | |
|---|---|---|---|---|---|---|---|
| ENSG00000243885 | | 3.018133 | -0.0212234 | 0.7445559 | -0.1091205 | 0.9131069 | 0.959195 |
| ENSG00000136918 | WDR38 | 2.763103768 | -0.02123103 | 0.685129001 | -0.059375141 | 0.95265331 | 0.976954 |
| ENSG00000100908 | EMC9 | 1283.944874 | -0.021360886 | 0.196557719 | -0.118601844 | 0.9055908 | 0.956153 |
| ENSG00000275379 | H3C11 | 3.050881 | -0.02140677 | 0.6832018 | -0.06053886 | 0.9517265 | 0.976385 |
| ENSG00000214563 | GAPDHP15 | 0.8819658 | -0.02143974 | 0.7879673 | 0 | 0.9388328 | NA |
| ENSG00000201643 | SNORA14A | 0.9681001 | -0.02145478 | 0.7879796 | 0 | 0.9411892 | NA |
| ENSG00000287834 | | 1.338605 | -0.02145766 | 0.7879454 | 0 | 0.934366 | NA |
| ENSG00000148343 | MIGA2 | 3310.727 | -0.02145827 | 0.1178185 | -0.1876168 | 0.8511771 | 0.927716 |
| ENSG00000139579 | NABP2 | 2878.257 | -0.02154319 | 0.2465824 | -0.0378826 | 0.9697813 | 0.984605 |
| ENSG00000265489 | | 1.215617 | -0.02155012 | 0.7408305 | -0.07612183 | 0.9393222 | NA |
| ENSG00000174444 | RPL4 | 58142.4 | -0.02157142 | 0.1806242 | -0.1259315 | 0.8997862 | 0.953966 |
| ENSG00000136492 | BRIP1 | 10663.19772 | -0.021647638 | 0.426192861 | -0.068105508 | 0.94570165 | 0.973226 |
| ENSG00000258745 | | 1.495766 | -0.02168138 | 0.7500046 | -0.1005113 | 0.9199384 | NA |
| ENSG00000242435 | UPK3BP1 | 1.21837 | -0.02169375 | 0.7325008 | -0.07203254 | 0.942576 | NA |
| ENSG00000285016 | | 0.8341418 | -0.02174924 | 0.7878881 | 0 | 0.9237917 | NA |
| ENSG00000168522 | FNTA | 2421.56 | -0.02175844 | 0.1889866 | -0.1179708 | 0.9060908 | 0.956276 |
| ENSG00000177182 | CLVS1 | 0.8810301 | -0.02177206 | 0.7485187 | -0.1002914 | 0.920113 | NA |
| ENSG00000244176 | | 1.298233 | -0.02179086 | 0.78788 | 0 | 0.9288543 | NA |
| ENSG00000237417 | XRCC6P1 | 4.537824 | -0.02180341 | 0.6090549 | -0.0582256 | 0.9535689 | 0.977458 |
| ENSG00000283072 | | 3.687163 | -0.02192692 | 0.7878319 | 0 | 0.9253109 | 0.964139 |
| ENSG00000132002 | DNAJB1 | 7260.255566 | -0.021936724 | 0.181705583 | -0.140774507 | 0.88804808 | 0.947572 |
| ENSG00000227404 | KRT8P20 | 3.543864 | -0.0219712 | 0.6803504 | -0.06272555 | 0.9499851 | 0.97554 |
| ENSG00000256538 | | 2.493661 | -0.02202384 | 0.6460373 | -0.06399447 | 0.9489746 | 0.975068 |
| ENSG00000105755 | ETHE1 | 501.6767459 | -0.022034839 | 0.180443323 | -0.129759358 | 0.89675682 | 0.952396 |
| ENSG00000244151 | | 45.34275 | -0.0220432 | 0.3227139 | -0.06717549 | 0.946442 | 0.973609 |
| ENSG00000214584 | PGGT1BP2 | 33.70191 | -0.02204441 | 0.3331814 | -0.07557233 | 0.9397593 | 0.970365 |
| ENSG00000269054 | | 53.30768 | -0.02206931 | 0.3645856 | -0.06894781 | 0.9450312 | 0.972822 |
| ENSG00000267452 | LINC02073 | 9.136543 | -0.02215417 | 0.6966477 | -0.04758065 | 0.9620505 | 0.981067 |
| ENSG00000231426 | FILNC1 | 0.9030843 | -0.02219754 | 0.7577345 | -0.0871435 | 0.9305575 | NA |
| ENSG00000211817 | TRAV38-2DV8 | 1.303753 | -0.02224164 | 0.7879045 | 0 | 0.9374202 | NA |
| ENSG00000149308 | NPAT | 841.6934 | -0.0222608 | 0.1850391 | -0.1263717 | 0.8994377 | 0.953788 |
| ENSG00000044446 | PHKA2 | 2469.973768 | -0.022269161 | 0.179664362 | -0.129818317 | 0.89671017 | 0.952396 |
| ENSG00000254143 | | 1.538594 | -0.0222864 | 0.787849 | 0 | 0.9323904 | NA |

| | | | | | | | |
|---|---|---|---|---|---|---|---|
| ENSG00000214114 | MYCBP | 1499.676 | -0.02228821 | 0.3196355 | -0.07495712 | 0.9402488 | 0.97045 |
| ENSG00000206838 | SNORA5A | 23.72096 | -0.02232425 | 0.386053 | -0.06728561 | 0.9463543 | 0.973609 |
| ENSG00000043143 | JADE2 | 4676.612354 | -0.022334741 | 0.144902427 | -0.171698211 | 0.86367479 | 0.934551 |
| ENSG00000267325 | LINC01415 | 1.472583 | -0.02237178 | 0.7877806 | 0 | 0.9196726 | NA |
| ENSG00000227802 | DNAJB3 | 3.856157 | -0.02237236 | 0.7878023 | 0 | 0.9183407 | 0.961287 |
| ENSG00000235032 | BMP7-AS1 | 1.227347 | -0.02238312 | 0.7540187 | -0.09323939 | 0.9257134 | NA |
| ENSG00000100348 | TXN2 | 3839.175152 | -0.022384019 | 0.189474895 | -0.095241964 | 0.92412264 | 0.963623 |
| ENSG00000242352 | FAM91A3P | 196.9675 | -0.02241595 | 0.2648565 | -0.08876946 | 0.9292651 | 0.965722 |
| ENSG00000236480 | PKMP1 | 39.31623 | -0.022421 | 0.3803794 | -0.06990037 | 0.944273 | 0.972607 |
| ENSG00000253886 | | 1.542901 | -0.02242119 | 0.6892063 | -0.07005119 | 0.9441529 | NA |
| ENSG00000147133 | TAF1 | 5288.261 | -0.0224221 | 0.2517214 | -0.09740417 | 0.9224054 | 0.963176 |
| ENSG00000225705 | | 1.338109 | -0.02247742 | 0.7877839 | 0 | 0.9252156 | NA |
| ENSG00000136244 | IL6 | 2.511401348 | -0.022482021 | 0.724920612 | -0.061129283 | 0.95125625 | 0.976178 |
| ENSG00000006606 | CCL26 | 1.100443452 | -0.022485649 | 0.78782288 | 0 | 0.92909068 | NA |
| ENSG00000258001 | | 1.228376 | -0.02252123 | 0.7549358 | -0.09164249 | 0.9269821 | NA |
| ENSG00000124440 | HIF3A | 1.038425495 | -0.022525641 | 0.787798154 | 0 | 0.92682554 | NA |
| ENSG00000204439 | C6orf47 | 1203.193 | -0.0225371 | 0.1349322 | -0.1717246 | 0.863654 | 0.934551 |
| ENSG00000269191 | | 6.44945 | -0.0225408 | 0.6065027 | -0.06043975 | 0.9518054 | 0.976391 |
| ENSG00000134398 | ERN2 | 1.486237115 | -0.022543482 | 0.787767678 | 0 | 0.92317296 | NA |
| ENSG00000282542 | | 3.562583 | -0.02255909 | 0.6950119 | -0.06584172 | 0.9475038 | 0.974235 |
| ENSG00000278524 | MIR6810 | 1.206559 | -0.02256852 | 0.7878237 | 0 | 0.931397 | NA |
| ENSG00000196188 | CTSE | 2.169329 | -0.02257961 | 0.7877483 | 0 | 0.9260221 | 0.9644 |
| ENSG00000203857 | HSD3B1 | 13.39831 | -0.02258488 | 0.7877681 | 0 | 0.9112216 | 0.958746 |
| ENSG00000175164 | ABO | 8.043422 | -0.02258732 | 0.7877686 | 0 | 0.9119409 | 0.958851 |
| ENSG00000171649 | ZIK1 | 0.8594082 | -0.02259468 | 0.732668 | -0.08831357 | 0.9296275 | NA |
| ENSG00000286215 | | 2.161133 | -0.02260758 | 0.6643606 | -0.05817717 | 0.9536075 | 0.977458 |
| ENSG00000132432 | SEC61G | 3224.272029 | -0.022628819 | 0.19513933 | -0.139798378 | 0.8888193 | 0.948243 |
| ENSG00000258464 | | 1.374154 | -0.02264026 | 0.7031131 | -0.07219776 | 0.9424445 | NA |
| ENSG00000113520 | IL4 | 1.152975552 | -0.02266866 | 0.741443272 | -0.081428888 | 0.93510088 | NA |
| ENSG00000174446 | SNAPC5 | 1617.018 | -0.0227143 | 0.1471702 | -0.1594468 | 0.8733169 | 0.940008 |
| ENSG00000267709 | | 2.466648 | -0.0227157 | 0.7515093 | -0.09779897 | 0.9220919 | 0.963092 |
| ENSG00000259420 | | 2.213786 | -0.02272914 | 0.7467353 | -0.1062527 | 0.9153818 | 0.960119 |
| ENSG00000130234 | ACE2 | 2.159480712 | -0.022776572 | 0.787736399 | 0 | 0.91668708 | 0.960692 |

| | | | | | | | |
|---|---|---|---|---|---|---|---|
| ENSG00000233153 | UBE2E2-AS1 | 1.068088 | -0.02278101 | 0.7569653 | -0.08821598 | 0.929705 | NA |
| ENSG00000130717 | UCK1 | 3424.228193 | -0.02280129 | 0.205836802 | -0.12378311 | 0.90148701 | 0.954742 |
| ENSG00000212402 | SNORA74B | 2.827113 | -0.02281139 | 0.6594402 | -0.06046521 | 0.9517851 | 0.976391 |
| ENSG00000277710 | NBEAP5 | 1.010128 | -0.02281497 | 0.7437414 | -0.1264251 | 0.8993954 | NA |
| ENSG00000203588 | IGBP1-AS1 | 7.354128 | -0.02283923 | 0.5605361 | -0.05968403 | 0.9524073 | 0.976782 |
| ENSG00000240270 | RPL12P37 | 1.250242 | -0.02287775 | 0.722555 | -0.1071183 | 0.9146951 | NA |
| ENSG00000237523 | LINC00857 | 11.45016 | -0.02290993 | 0.7877254 | 0 | 0.9060207 | 0.956276 |
| ENSG00000151883 | PARP8 | 1203.127 | -0.02292836 | 0.2366728 | -0.1477577 | 0.882534 | 0.944952 |
| ENSG00000047457 | CP | 112.6363933 | -0.022950299 | 0.591395509 | -0.055855381 | 0.95545702 | 0.978579 |
| ENSG00000229256 | ST13P13 | 9.852535 | -0.02303223 | 0.5333066 | -0.06188046 | 0.950658 | 0.975753 |
| ENSG00000005700 | IBTK | 4913.624218 | -0.023055878 | 0.155298803 | -0.164757279 | 0.86913503 | 0.937215 |
| ENSG00000227118 | BTF3P13 | 3.01676 | -0.02307066 | 0.6588133 | -0.06648288 | 0.9469934 | 0.973785 |
| ENSG00000198040 | ZNF84 | 3183.434 | -0.02307401 | 0.2633944 | -0.106391 | 0.9152722 | 0.960119 |
| ENSG00000170162 | VGLL2 | 0.8581849 | -0.0230771 | 0.7492847 | -0.1252154 | 0.9003531 | NA |
| ENSG00000224814 | | 2.22379 | -0.02309039 | 0.7877442 | 0 | 0.9271365 | 0.96485 |
| ENSG00000244476 | ERVFRD-1 | 0.874846 | -0.02310492 | 0.7538871 | -0.09356144 | 0.9254575 | NA |
| ENSG00000185596 | WASH3P | 614.8418 | -0.02310732 | 0.1663173 | -0.1440746 | 0.8854415 | 0.946159 |
| ENSG00000085514 | PILRA | 16.78416602 | -0.023119689 | 0.508716858 | -0.064918027 | 0.94823927 | 0.974627 |
| ENSG00000005194 | CIAPIN1 | 3051.8549 | -0.023122463 | 0.212050843 | -0.11523978 | 0.90825507 | 0.957217 |
| ENSG00000167384 | ZNF180 | 841.1916 | -0.02312584 | 0.1910966 | -0.1197642 | 0.90467 | 0.955875 |
| ENSG00000159363 | ATP13A2 | 2477.45 | -0.02313324 | 0.1549122 | -0.155871 | 0.8761347 | 0.941553 |
| ENSG00000238324 | RN7SKP198 | 1.000828 | -0.0232426 | 0.7432404 | -0.09825041 | 0.9217335 | NA |
| ENSG00000231848 | KRTCAP2P1 | 294.7541 | -0.02330233 | 0.1848558 | -0.1328917 | 0.894279 | 0.950906 |
| ENSG00000284194 | SCO2 | 36.91259 | -0.02336116 | 0.3083776 | -0.07876151 | 0.9372223 | 0.96904 |
| ENSG00000231964 | | 13.24389 | -0.02341863 | 0.7876846 | 0 | 0.9078088 | 0.957088 |
| ENSG00000070729 | CNGB1 | 3.146560713 | -0.023422842 | 0.746878637 | -0.106070972 | 0.91552604 | 0.960119 |
| ENSG00000226954 | | 3.521784 | -0.02342532 | 0.7449103 | -0.1087269 | 0.9134191 | 0.95946 |
| ENSG00000143207 | COP1 | 3615.333 | -0.02343254 | 0.1534652 | -0.173962 | 0.8618953 | 0.9338 |
| ENSG00000234945 | GTF3C2-AS1 | 10.00916 | -0.02343307 | 0.4788894 | -0.06498496 | 0.948186 | 0.974627 |
| ENSG00000105290 | APLP1 | 367.1248744 | -0.023437602 | 0.370092447 | -0.073693572 | 0.94125421 | 0.97101 |
| ENSG00000164746 | C7orf57 | 1.129634 | -0.02345073 | 0.7199137 | -0.08332898 | 0.93359 | NA |
| ENSG00000259381 | | 7.259852 | -0.02358067 | 0.5698543 | -0.06413828 | 0.9488601 | 0.975041 |
| ENSG00000109065 | NAT9 | 4339.716358 | -0.02365182 | 0.193056774 | -0.134991887 | 0.89261829 | 0.95021 |

| | | | | | | | |
|---|---|---|---|---|---|---|---|
| ENSG00000170367 | CST5 | 1.521095 | -0.02366603 | 0.7877825 | 0 | 0.9374492 | NA |
| ENSG00000175334 | BANF1 | 7508.94 | -0.02367646 | 0.2240844 | -0.1240883 | 0.9012454 | 0.954597 |
| ENSG00000178723 | GLULP4 | 1.633114 | -0.02368581 | 0.6707462 | -0.06808621 | 0.945717 | 0.973226 |
| ENSG00000074603 | DPP8 | 3854.245365 | -0.023855704 | 0.153817938 | -0.189141344 | 0.84998204 | 0.927091 |
| ENSG00000090470 | PDCD7 | 2004.718917 | -0.02396288 | 0.155051232 | -0.188084108 | 0.85081072 | 0.927491 |
| ENSG00000160216 | AGPAT3 | 8678.729 | -0.02401596 | 0.09768185 | -0.2620331 | 0.7932959 | 0.893115 |
| ENSG00000260796 | | 6.694132 | -0.02412354 | 0.560035 | -0.06091359 | 0.951428 | 0.976211 |
| ENSG00000207186 | RNA5SP122 | 1.248416 | -0.02416687 | 0.7108347 | -0.08045467 | 0.9358757 | NA |
| ENSG00000268836 | | 4.76415 | -0.02417173 | 0.6752092 | -0.07327005 | 0.9415912 | 0.971125 |
| ENSG00000204584 | | 8.860326 | -0.02417658 | 0.5648997 | -0.0646828 | 0.9484266 | 0.974693 |
| ENSG00000267504 | | 1.276296 | -0.02418176 | 0.7262771 | -0.07881441 | 0.9371802 | NA |
| ENSG00000214562 | NUTM2D | 69.53718 | -0.02431053 | 0.3604157 | -0.08106033 | 0.935394 | 0.968218 |
| ENSG00000166342 | NETO1 | 4.583279 | -0.02434947 | 0.7875938 | 0 | 0.9187812 | 0.961511 |
| ENSG00000164463 | CREBRF | 1358.783 | -0.02440531 | 0.3921843 | -0.07852972 | 0.9374067 | 0.9691 |
| ENSG00000187690 | EZHIP | 197.4989 | -0.02447407 | 0.7379951 | -0.1227103 | 0.9023365 | 0.954999 |
| ENSG00000174951 | FUT1 | 745.5422 | -0.02450935 | 0.204172 | -0.1375621 | 0.8905865 | 0.949218 |
| ENSG00000253307 | | 1.243985 | -0.02457627 | 0.7495553 | -0.1010074 | 0.9195446 | NA |
| ENSG00000242419 | PCDHGC4 | 9.542151 | -0.02459648 | 0.5445168 | -0.06480946 | 0.9483257 | 0.974627 |
| ENSG00000164825 | DEFB1 | 29.47697 | -0.02461096 | 0.7875699 | 0 | 0.904791 | 0.955917 |
| ENSG00000111670 | GNPTAB | 5142.475848 | -0.024653506 | 0.175802036 | -0.171632448 | 0.8637265 | 0.934569 |
| ENSG00000234206 | | 2.492401 | -0.02468438 | 0.7224925 | -0.07872579 | 0.9372507 | 0.96904 |
| ENSG00000249302 | FTH1P24 | 1.211422 | -0.02469958 | 0.7348765 | -0.08646176 | 0.9310994 | NA |
| ENSG00000110887 | DAO | 3.109736821 | -0.024727811 | 0.73022419 | -0.095261772 | 0.92410691 | 0.963623 |
| ENSG00000166035 | LIPC | 8.20664 | -0.02473521 | 0.7409057 | -0.1158688 | 0.9077566 | 0.957074 |
| ENSG00000255040 | MORF4L1P3 | 1.810788 | -0.02474661 | 0.6744248 | -0.07104048 | 0.9433655 | 0.972202 |
| ENSG00000261242 | | 8.602323 | -0.02487146 | 0.5132871 | -0.06482312 | 0.9483148 | 0.974627 |
| ENSG00000242876 | RN7SL812P | 1.199615 | -0.02492099 | 0.7019693 | -0.07750354 | 0.938223 | NA |
| ENSG00000286904 | | 1.580887 | -0.02495658 | 0.7270966 | -0.07792599 | 0.9378869 | 0.969374 |
| ENSG00000274995 | | 0.9142993 | -0.02498683 | 0.7605932 | -0.08221898 | 0.9344726 | NA |
| ENSG00000239544 | | 1.875144 | -0.02499503 | 0.6915095 | -0.07659538 | 0.9389454 | 0.969864 |
| ENSG00000213178 | RPL22P1 | 3415.731 | -0.02499668 | 0.1670385 | -0.1332192 | 0.8940201 | 0.950882 |
| ENSG00000145214 | DGKQ | 1725.899 | -0.0250293 | 0.1554472 | -0.1770082 | 0.859502 | 0.932352 |
| ENSG00000159377 | PSMB4 | 24009.89 | -0.02505431 | 0.1806154 | -0.1590013 | 0.8736678 | 0.940119 |

| | | | | | | | |
|---|---|---|---|---|---|---|---|
| ENSG00000259326 | | 2.480912 | -0.02511839 | 0.6882852 | -0.08058518 | 0.9357719 | 0.96828 |
| ENSG00000263072 | ZNF213-AS1 | 1310.755 | -0.02517983 | 0.2178929 | -0.1200533 | 0.9044409 | 0.95578 |
| ENSG00000261469 | | 26.72357 | -0.02520416 | 0.3627031 | -0.08073855 | 0.9356499 | 0.968267 |
| ENSG00000111300 | NAA25 | 4157.08298 | -0.025275339 | 0.141066268 | -0.222060665 | 0.82426666 | 0.911093 |
| ENSG00000157399 | ARSL | 0.9352042 | -0.02533184 | 0.7308104 | -0.1035339 | 0.9175392 | NA |
| ENSG00000204923 | FBXO48 | 57.87736 | -0.02535531 | 0.3542779 | -0.08219917 | 0.9344883 | 0.967668 |
| ENSG00000230189 | | 2.594201 | -0.02536058 | 0.6613753 | -0.06724349 | 0.9463879 | 0.973609 |
| ENSG00000167461 | RAB8A | 1019.89 | -0.0254351 | 0.2232214 | -0.1221287 | 0.9027971 | 0.955036 |
| ENSG00000169302 | STK32A | 9.89846 | -0.02545309 | 0.5849006 | -0.06530031 | 0.9479349 | 0.974515 |
| ENSG00000178125 | PPP1R42 | 3.052423 | -0.02546776 | 0.6202171 | -0.07500718 | 0.940209 | 0.97045 |
| ENSG00000141295 | SCRN2 | 3276.985 | -0.02554717 | 0.1777217 | -0.144282 | 0.8852778 | 0.94607 |
| ENSG00000269404 | SPIB | 1.319021 | -0.0256135 | 0.7238834 | -0.09492654 | 0.9243732 | NA |
| ENSG00000141052 | MYOCD | 1.568235 | -0.02561949 | 0.7489133 | -0.1023186 | 0.9185038 | NA |
| ENSG00000169551 | CT55 | 1.227664 | -0.02562177 | 0.7378372 | -0.08901873 | 0.929067 | NA |
| ENSG00000213516 | RBMXL1 | 1215.538 | -0.02564382 | 0.1793273 | -0.1675283 | 0.8669544 | 0.936384 |
| ENSG00000116703 | PDC | 1.345453922 | -0.025657301 | 0.691032683 | -0.087786211 | 0.9300466 | NA |
| ENSG00000091592 | NLRP1 | 43.18992074 | -0.02568795 | 0.72545943 | -0.05563084 | 0.9556359 | 0.978636 |
| ENSG00000182722 | SEPHS1P1 | 1.014644 | -0.02573208 | 0.7236786 | -0.08665151 | 0.9309485 | NA |
| ENSG00000200816 | SNORA38 | 1.098684 | -0.02574087 | 0.7172364 | -0.0898477 | 0.9284082 | NA |
| ENSG00000178556 | CKS1BP6 | 19.68454 | -0.02575664 | 0.418329 | -0.07447191 | 0.9406349 | 0.970666 |
| ENSG00000235313 | HM13-IT1 | 20.68086 | -0.02576878 | 0.3989513 | -0.076875 | 0.938723 | 0.969795 |
| ENSG00000164244 | PRRC1 | 5657.656 | -0.02581969 | 0.163618 | -0.1722595 | 0.8632336 | 0.934398 |
| ENSG00000187999 | HNRNPA1P61 | 0.9568621 | -0.02588491 | 0.7344524 | -0.1038469 | 0.9172909 | NA |
| ENSG00000225521 | SNED1-AS1 | 4.229433 | -0.02591338 | 0.7429252 | -0.1129948 | 0.9100347 | 0.958028 |
| ENSG00000232403 | RPL27P5 | 1.080196 | -0.02600923 | 0.7418067 | -0.11905 | 0.9052358 | NA |
| ENSG00000065054 | SLC9A3R2 | 5221.975304 | -0.026027879 | 0.225504614 | -0.123401944 | 0.90178882 | 0.954833 |
| ENSG00000161509 | GRIN2C | 146.6642 | -0.02605792 | 0.2553091 | -0.1210245 | 0.9036716 | 0.955342 |
| ENSG00000175886 | RPL7AP66 | 605.312 | -0.02608342 | 0.1922835 | -0.1414512 | 0.8875135 | 0.947268 |
| ENSG00000137098 | SPAG8 | 30.11564792 | -0.026120803 | 0.428371303 | -0.074344062 | 0.94073662 | 0.970675 |
| ENSG00000241868 | RN7SL434P | 1.654933 | -0.0261883 | 0.7517811 | -0.09721238 | 0.9225577 | 0.963214 |
| ENSG00000254370 | | 1.366063 | -0.02623277 | 0.715838 | -0.1005333 | 0.9199209 | NA |
| ENSG00000226912 | ISCA2P1 | 1.367475 | -0.02630657 | 0.7321044 | -0.1264826 | 0.8993499 | NA |
| ENSG00000273252 | OR7E39P | 2.40121 | -0.02631057 | 0.7551048 | -0.09244443 | 0.9263449 | 0.964498 |

| | | | | | | | |
|---|---|---|---|---|---|---|---|
| ENSG00000139613 | SMARCC2 | 9276.559 | -0.02632295 | 0.1654665 | -0.183867 | 0.8541178 | 0.929129 |
| ENSG00000279588 | | 4.112936 | -0.02641114 | 0.7874222 | 0 | 0.9138679 | 0.95963 |
| ENSG00000223963 | THAP12P8 | 5.441063 | -0.02641297 | 0.7874559 | 0 | 0.9192903 | 0.961713 |
| ENSG00000177058 | SLC38A9 | 1050.887 | -0.02642529 | 0.1751364 | -0.1546825 | 0.8770717 | 0.942003 |
| ENSG00000263740 | RN7SL4P | 11.37293 | -0.02643277 | 0.5993249 | -0.06965079 | 0.9444716 | 0.972623 |
| ENSG00000213060 | | 1.250521 | -0.02646328 | 0.7343341 | -0.136022 | 0.8918039 | NA |
| ENSG00000198643 | FAM3D | 2.524803 | -0.02654594 | 0.7220748 | -0.08365922 | 0.9333274 | 0.967249 |
| ENSG00000008130 | NADK | 3957.611819 | -0.02657333 | 0.130873974 | -0.215762301 | 0.82917304 | 0.913893 |
| ENSG00000203356 | LINC01562 | 1.472917 | -0.02658417 | 0.7313858 | -0.09534567 | 0.9240403 | NA |
| ENSG00000247626 | MARS2 | 1058.62 | -0.02660991 | 0.2038491 | -0.1772196 | 0.8593359 | 0.932325 |
| ENSG00000198837 | DENND4B | 8051.049 | -0.0266901 | 0.1787736 | -0.1650391 | 0.8689132 | 0.937166 |
| ENSG00000246560 | UBE2D3-AS1 | 35.36204 | -0.02671643 | 0.4128435 | -0.07665474 | 0.9388982 | 0.969853 |
| ENSG00000226367 | ST7-AS2 | 0.9287469 | -0.02673144 | 0.72641 | -0.09623288 | 0.9233356 | NA |
| ENSG00000279794 | | 72.20842 | -0.02673672 | 0.3889161 | -0.08104718 | 0.9354044 | 0.968218 |
| ENSG00000180764 | PIPSL | 1564.279 | -0.02678016 | 0.1622612 | -0.1691466 | 0.8656814 | 0.935977 |
| ENSG00000230499 | | 2.066908 | -0.02678016 | 0.7496873 | -0.1007356 | 0.9197603 | 0.961781 |
| ENSG00000227440 | ATP5MC1P4 | 28.96853 | -0.02682215 | 0.519957 | -0.07240674 | 0.9422782 | 0.971644 |
| ENSG00000244280 | ECEL1P2 | 3.500238 | -0.02682818 | 0.7875562 | 0 | 0.9171284 | 0.960982 |
| ENSG00000266676 | MIR4297 | 1.226301 | -0.02683089 | 0.7208781 | -0.1011287 | 0.9194483 | NA |
| ENSG00000186732 | MPPED1 | 166.4665 | -0.02683534 | 0.7405879 | -0.1166235 | 0.9071584 | 0.95682 |
| ENSG00000267523 | | 109.8689 | -0.02687645 | 0.3113015 | -0.09628586 | 0.9232935 | 0.963538 |
| ENSG00000244122 | UGT1A7 | 10.22714 | -0.0268909 | 0.78745 | 0 | 0.9094996 | 0.957844 |
| ENSG00000214062 | RPL7P17 | 1.421008 | -0.0269551 | 0.7059486 | -0.08911645 | 0.9289894 | NA |
| ENSG00000174715 | PPIAP72 | 2.157495 | -0.02696886 | 0.7011677 | -0.08857919 | 0.9294163 | 0.965841 |
| ENSG00000185803 | SLC52A2 | 4113.387 | -0.02697112 | 0.1735879 | -0.1743889 | 0.8615598 | 0.933594 |
| ENSG00000170854 | RIOX2 | 2074.832 | -0.02697338 | 0.1805143 | -0.1597218 | 0.8731002 | 0.939929 |
| ENSG00000224533 | TMLHE-AS1 | 10.93815 | -0.02698988 | 0.5102148 | -0.07030473 | 0.9439511 | 0.972468 |
| ENSG00000102007 | PLP2 | 2593.891259 | -0.027069725 | 0.188691644 | -0.159011822 | 0.87365956 | 0.940119 |
| ENSG00000268533 | | 0.7919592 | -0.02709876 | 0.7466622 | -0.1172134 | 0.9066909 | NA |
| ENSG00000253683 | | 1054.466 | -0.02713423 | 0.2633578 | -0.1183521 | 0.9057887 | 0.956179 |
| ENSG00000287408 | | 8.651175 | -0.02723814 | 0.5372659 | -0.07174111 | 0.9428079 | 0.971814 |
| ENSG00000252906 | SCARNA3 | 0.7986245 | -0.02724038 | 0.7546296 | -0.09262897 | 0.9261983 | NA |
| ENSG00000271755 | | 4.821035 | -0.02726494 | 0.580915 | -0.07103774 | 0.9433677 | 0.972202 |

| | | | | | | | |
|---|---|---|---|---|---|---|---|
| ENSG00000226984 | | 1.817373 | -0.02726512 | 0.7261117 | -0.1019192 | 0.9188208 | 0.961511 |
| ENSG00000240395 | RPL5P23 | 142.1714 | -0.02726703 | 0.2507024 | -0.1062107 | 0.9154152 | 0.960119 |
| ENSG00000211816 | TRAV38-1 | 5.304688 | -0.02727981 | 0.7873763 | 0 | 0.9185779 | 0.961449 |
| ENSG00000250397 | | 101.5918 | -0.02728112 | 0.3389936 | -0.0831034 | 0.9337693 | 0.967249 |
| ENSG00000204595 | DPRX | 1.763255 | -0.02740454 | 0.7596828 | -0.08387553 | 0.9331554 | 0.967249 |
| ENSG00000253559 | OSGEPL1-AS1 | 9.556104 | -0.02741004 | 0.4917876 | -0.07686373 | 0.938732 | 0.969795 |
| ENSG00000277245 | | 1.511975 | -0.02746974 | 0.7252647 | -0.1192108 | 0.9051084 | NA |
| ENSG00000164897 | TMUB1 | 3128.813 | -0.0275186 | 0.1967928 | -0.1407902 | 0.8880357 | 0.947572 |
| ENSG00000286939 | | 1.67066 | -0.02762828 | 0.721645 | -0.08407402 | 0.9329976 | 0.967217 |
| ENSG00000189166 | TNRC18P3 | 1.019426 | -0.02768552 | 0.7215386 | -0.1088521 | 0.9133198 | NA |
| ENSG00000108958 | | 343.6210416 | -0.027693776 | 0.192489209 | -0.155710736 | 0.87626103 | 0.941651 |
| ENSG00000239382 | ALKBH6 | 332.9731 | -0.02778682 | 0.2582798 | -0.1172793 | 0.9066387 | 0.956652 |
| ENSG00000245466 | | 2.383014 | -0.02782365 | 0.7470486 | -0.1054563 | 0.9160137 | 0.960213 |
| ENSG00000145375 | SPATA5 | 548.496 | -0.02792169 | 0.232921 | -0.1373943 | 0.8907192 | 0.949303 |
| ENSG00000257759 | | 1.322104 | -0.02797275 | 0.7137803 | -0.09390893 | 0.9251815 | NA |
| ENSG00000218418 | | 381.4155 | -0.02800918 | 0.2581929 | -0.1098567 | 0.912523 | 0.958888 |
| ENSG00000229862 | | 1.810977 | -0.02801503 | 0.6944889 | -0.0955899 | 0.9238463 | 0.963623 |
| ENSG00000248932 | | 353.6653 | -0.02804445 | 0.2897834 | -0.1071863 | 0.9146412 | 0.959911 |
| ENSG00000162068 | NTN3 | 16.21847 | -0.02808395 | 0.4762263 | -0.07673007 | 0.9388383 | 0.969829 |
| ENSG00000157625 | TAB3 | 3689.212 | -0.02809396 | 0.2186286 | -0.1427926 | 0.886454 | 0.946696 |
| ENSG00000270402 | | 1.421471 | -0.02811181 | 0.7267059 | -0.1091058 | 0.9131186 | NA |
| ENSG00000158578 | ALAS2 | 3.289306 | -0.02811773 | 0.6037585 | -0.07832476 | 0.9375697 | 0.969167 |
| ENSG00000178607 | ERN1 | 2645.88 | -0.0281265 | 0.3662389 | -0.08938342 | 0.9287772 | 0.965461 |
| ENSG00000224080 | UBE2FP1 | 20.9805 | -0.02812993 | 0.3885674 | -0.08731209 | 0.9304234 | 0.966278 |
| ENSG00000275449 | MIR6859-3 | 1.056062 | -0.02813561 | 0.7211344 | -0.1238531 | 0.9014316 | NA |
| ENSG00000105501 | SIGLEC5 | 12.55909466 | -0.028163976 | 0.787355817 | 0 | 0.91023011 | 0.958107 |
| ENSG00000229852 | | 142.5771 | -0.02816598 | 0.2333502 | -0.1349929 | 0.8926175 | 0.950211 |
| ENSG00000287580 | | 22.47909 | -0.02818805 | 0.3790568 | -0.08751789 | 0.9302599 | 0.966227 |
| ENSG00000241081 | RPL22P2 | 1166.417 | -0.0281954 | 0.17993 | -0.1674068 | 0.86705 | 0.936411 |
| ENSG00000090530 | P3H2 | 393.4338439 | -0.028196926 | 0.537234801 | -0.051479931 | 0.95894309 | 0.979741 |
| ENSG00000230383 | | 339.9914 | -0.02831237 | 0.1946697 | -0.1520902 | 0.8791158 | 0.943155 |
| ENSG00000229638 | RPL4P4 | 10967.06 | -0.02840135 | 0.1969479 | -0.1214646 | 0.903323 | 0.955125 |
| ENSG00000118785 | SPP1 | 56.3317581 | -0.02840244 | 0.739724907 | -0.119136877 | 0.90516692 | 0.955968 |

| | | | | | | | |
|---|---|---|---|---|---|---|---|
| ENSG00000278914 | | 1.308048 | -0.02842238 | 0.7318936 | -0.116456 | 0.9072911 | NA |
| ENSG00000225180 | PVALEF | 9.853534 | -0.02843247 | 0.5172267 | -0.07607043 | 0.9393631 | 0.970098 |
| ENSG00000006611 | USH1C | 4.191097889 | -0.028439628 | 0.66685308 | -0.083143263 | 0.93373763 | 0.967249 |
| ENSG00000151575 | TEX9 | 147.2685 | -0.02845944 | 0.2194821 | -0.1285684 | 0.8976992 | 0.952705 |
| ENSG00000133874 | RNF122 | 152.7224235 | -0.028467767 | 0.236547032 | -0.128173055 | 0.89801202 | 0.952847 |
| ENSG00000270672 | MTRNR2L6 | 1.065476 | -0.02848688 | 0.7527168 | -0.09595842 | 0.9235536 | NA |
| ENSG00000144891 | AGTR1 | 0.8784827 | -0.02852075 | 0.7475092 | -0.1553042 | 0.8765815 | NA |
| ENSG00000287769 | | 329.5782 | -0.02852373 | 0.3510243 | -0.09078686 | 0.927662 | 0.96494 |
| ENSG00000234841 | | 0.9638194 | -0.02856551 | 0.7342097 | -0.1323726 | 0.8946896 | NA |
| ENSG00000023287 | RB1CC1 | 4150.170362 | -0.028642222 | 0.202796285 | -0.144972213 | 0.88473281 | 0.94582 |
| ENSG00000243568 | | 0.8228487 | -0.02864515 | 0.7405096 | -0.1059322 | 0.9156362 | NA |
| ENSG00000166819 | PLIN1 | 46.817 | -0.02867069 | 0.5262837 | -0.07949225 | 0.9366411 | 0.968692 |
| ENSG00000218996 | ARL4AP5 | 6.910622 | -0.02867279 | 0.5311245 | -0.07616658 | 0.9392866 | 0.970066 |
| ENSG00000162607 | USP1 | 4444.006 | -0.0287343 | 0.3090473 | -0.1055845 | 0.915912 | 0.960213 |
| ENSG00000224950 | | 61.25791 | -0.02896706 | 0.533786 | -0.076317 | 0.9391669 | 0.96998 |
| ENSG00000153201 | RANBP2 | 8765.991 | -0.02896962 | 0.1952227 | -0.1491009 | 0.881474 | 0.944541 |
| ENSG00000101146 | RAE1 | 8998.14411 | -0.029065585 | 0.206265239 | -0.166147475 | 0.86804089 | 0.936731 |
| ENSG00000182010 | RTKN2 | 902.2221 | -0.0290868 | 0.4154458 | -0.08593619 | 0.9315171 | 0.966739 |
| ENSG00000218233 | NEPNP | 0.9055998 | -0.0291096 | 0.7347727 | -0.1037145 | 0.9173959 | NA |
| ENSG00000184867 | ARMCX2 | 1144.823 | -0.02912634 | 0.2182704 | -0.1462258 | 0.8837432 | 0.945465 |
| ENSG00000225814 | GRPEL2P2 | 8.13821 | -0.02914358 | 0.5108923 | -0.07831481 | 0.9375776 | 0.969167 |
| ENSG00000170743 | SYT9 | 1.5476 | -0.02915567 | 0.7287768 | -0.09138567 | 0.9271861 | NA |
| ENSG00000111885 | MAN1A1 | 2271.096589 | -0.029156154 | 0.25787404 | -0.124070437 | 0.90125951 | 0.954597 |
| ENSG00000162704 | ARPC5 | 7947.758 | -0.029185 | 0.1282168 | -0.2332593 | 0.8155601 | 0.90611 |
| ENSG00000275651 | MIR6851 | 1.345065 | -0.02921105 | 0.7300489 | -0.09679522 | 0.922889 | NA |
| ENSG00000173213 | TUBB8B | 1.688528 | -0.02925166 | 0.7218754 | -0.09894791 | 0.9211796 | 0.962704 |
| ENSG00000148303 | RPL7A | 38342.65 | -0.02930213 | 0.1491003 | -0.2400622 | 0.810282 | 0.902923 |
| ENSG00000111011 | RSRC2 | 4744.170443 | -0.029303907 | 0.185005062 | -0.165871934 | 0.86825774 | 0.936731 |
| ENSG00000116670 | MAD2L2 | 1398.770977 | -0.029338446 | 0.231883453 | -0.132578304 | 0.89452689 | 0.951124 |
| ENSG00000162413 | KLHL21 | 2145.573 | -0.0293424 | 0.1829129 | -0.1683093 | 0.86634 | 0.936211 |
| ENSG00000279311 | | 1.061662 | -0.02946756 | 0.6940905 | -0.09723619 | 0.9225388 | NA |
| ENSG00000275166 | MIR6814 | 2.747778 | -0.02950296 | 0.7198764 | -0.08611107 | 0.9313781 | 0.966708 |
| ENSG00000068489 | PRR11 | 4774.001064 | -0.029515661 | 0.391732125 | -0.106119598 | 0.91548746 | 0.960119 |

| | | | | | | | |
|---|---|---|---|---|---|---|---|
| ENSG00000258725 | PRC1-AS1 | 28.69919 | -0.02952503 | 0.383731 | -0.09119646 | 0.9273365 | 0.96494 |
| ENSG00000281904 | | 3.107127 | -0.02954208 | 0.6279038 | -0.07977921 | 0.9364129 | 0.968641 |
| ENSG00000151092 | NGLY1 | 2061.665 | -0.02959614 | 0.1683671 | -0.1838775 | 0.8541095 | 0.929129 |
| ENSG00000224714 | LINC02671 | 3.19477 | -0.02963619 | 0.744917 | -0.1060779 | 0.9155206 | 0.960119 |
| ENSG00000235652 | FBXO30-DT | 97.4573 | -0.02968155 | 0.3853139 | -0.09108958 | 0.9274214 | 0.96494 |
| ENSG00000244026 | FAM86DP | 678.8844 | -0.02974851 | 0.183901 | -0.1688877 | 0.865885 | 0.936028 |
| ENSG00000235018 | | 0.8248828 | -0.02975451 | 0.7663199 | -0.07283503 | 0.9419374 | NA |
| ENSG00000175356 | SCUBE2 | 2225.576 | -0.02976321 | 0.4495147 | -0.09142447 | 0.9271553 | 0.96485 |
| ENSG00000228318 | | 1.26791 | -0.02978901 | 0.744689 | -0.1300767 | 0.8965057 | NA |
| ENSG00000198363 | ASPH | 6617.822 | -0.02979268 | 0.2492058 | -0.119318 | 0.9050234 | 0.955968 |
| ENSG00000251185 | | 1.132765 | -0.02990332 | 0.7610984 | -0.08218301 | 0.9345012 | NA |
| ENSG00000119729 | RHOQ | 3680.54095 | -0.029985311 | 0.240353486 | -0.134531765 | 0.89298209 | 0.950376 |
| ENSG00000135387 | CAPRIN1 | 16616.91915 | -0.030020253 | 0.12604948 | -0.243187518 | 0.80786014 | 0.901644 |
| ENSG00000275965 | | 2.629702 | -0.03005503 | 0.6835482 | -0.08968241 | 0.9285396 | 0.965308 |
| ENSG00000261465 | | 1.128144 | -0.03017264 | 0.7527326 | -0.09661026 | 0.9230359 | NA |
| ENSG00000183828 | NUDT14 | 1573.248 | -0.03017382 | 0.2418524 | -0.1388142 | 0.889597 | 0.948692 |
| ENSG00000134419 | RPS15A | 6750.796331 | -0.030192061 | 0.169181213 | -0.185280858 | 0.85300875 | 0.928455 |
| ENSG00000163286 | ALPG | 26.08898 | -0.03020627 | 0.7871068 | 0 | 0.9098334 | 0.958006 |
| ENSG00000271784 | | 10.46758 | -0.03022462 | 0.5240512 | -0.08091585 | 0.9355089 | 0.968218 |
| ENSG00000130921 | C12orf65 | 1270.918471 | -0.030255676 | 0.161070275 | -0.190216086 | 0.84913981 | 0.926636 |
| ENSG00000105852 | PON3 | 1.508719802 | -0.030270088 | 0.756652075 | -0.090131701 | 0.92818256 | NA |
| ENSG00000128645 | HOXD1 | 1.128383682 | -0.030309334 | 0.726520432 | -0.120616697 | 0.90399464 | NA |
| ENSG00000140395 | WDR61 | 4205.516 | -0.03033602 | 0.1842817 | -0.1771042 | 0.8594266 | 0.932352 |
| ENSG00000007908 | SELE | 18.47317721 | -0.030376253 | 0.742611305 | -0.112598375 | 0.91034897 | 0.958125 |
| ENSG00000235636 | NUS1P1 | 711.4336 | -0.03041905 | 0.2164741 | -0.1511922 | 0.8798241 | 0.943451 |
| ENSG00000160199 | PKNOX1 | 1169.7 | -0.03043855 | 0.1395858 | -0.2133974 | 0.831017 | 0.915116 |
| ENSG00000286721 | | 2.465405 | -0.03045406 | 0.641365 | -0.08771843 | 0.9301005 | 0.966174 |
| ENSG00000239122 | RNU2-11P | 5.036086 | -0.03045869 | 0.6134948 | -0.07635561 | 0.9391362 | 0.96998 |
| ENSG00000235485 | | 5.738928 | -0.03049197 | 0.5308874 | -0.07917558 | 0.936893 | 0.968865 |
| ENSG00000180071 | ANKRD18A | 735.2405 | -0.03050195 | 0.2288959 | -0.1471977 | 0.882976 | 0.945235 |
| ENSG00000214003 | ATP5PBP3 | 2.693561 | -0.03061632 | 0.6245422 | -0.08754698 | 0.9302367 | 0.966227 |
| ENSG00000178694 | NSUN3 | 862.8771 | -0.03065218 | 0.1348604 | -0.2262085 | 0.8210393 | 0.909205 |
| ENSG00000279727 | LINC02033 | 0.814638 | -0.03065767 | 0.7611539 | -0.1659305 | 0.8682117 | NA |

| | | | | | | | |
|---|---|---|---|---|---|---|---|
| ENSG00000279041 | | 78.65396 | -0.0307837 | 0.3730144 | -0.09575579 | 0.9237145 | 0.963623 |
| ENSG00000280543 | ASAP1-IT2 | 2.558594 | -0.03079387 | 0.7045297 | -0.08541805 | 0.931929 | 0.966925 |
| ENSG00000225071 | RPS26P58 | 2.072354 | -0.03081423 | 0.6710112 | -0.09957924 | 0.9206784 | 0.962369 |
| ENSG00000166851 | PLK1 | 2422.839 | -0.03088804 | 0.5282045 | -0.08328617 | 0.933624 | 0.967249 |
| ENSG00000128699 | ORMDL1 | 2989.086802 | -0.030894583 | 0.194294144 | -0.166629182 | 0.86766183 | 0.936695 |
| ENSG00000213432 | RPL17P34 | 4.61608 | -0.03090036 | 0.599212 | -0.08588817 | 0.9315553 | 0.966741 |
| ENSG00000061676 | NCKAP1 | 12356.14899 | -0.030911025 | 0.1650933 | -0.213397247 | 0.83101713 | 0.915116 |
| ENSG00000075673 | ATP12A | 15.2644958 | -0.030920814 | 0.600693927 | -0.080606793 | 0.93575466 | 0.96828 |
| ENSG00000235924 | ZNRF2P3 | 0.8221752 | -0.03096645 | 0.7219598 | -0.1196171 | 0.9047865 | NA |
| ENSG00000265646 | TUFMP1 | 9.002571 | -0.03097546 | 0.5940459 | -0.08330619 | 0.9336081 | 0.967249 |
| ENSG00000060718 | COL11A1 | 3.86970596 | -0.030984391 | 0.718839518 | -0.087320894 | 0.93041645 | 0.966278 |
| ENSG00000181704 | YIPF6 | 3934.274 | -0.03100277 | 0.1610256 | -0.198617 | 0.8425624 | 0.922264 |
| ENSG00000280649 | | 50.19388 | -0.03104196 | 0.787022 | 0 | 0.9041749 | 0.955623 |
| ENSG00000243199 | | 3810.248 | -0.03104756 | 0.1812922 | -0.1814545 | 0.8560109 | 0.930273 |
| ENSG00000265407 | MIR4324 | 1.120959 | -0.03106488 | 0.7380281 | -0.1249125 | 0.9005928 | NA |
| ENSG00000054793 | ATP9A | 13547.2798 | -0.031064951 | 0.262511054 | -0.129322759 | 0.89710226 | 0.952406 |
| ENSG00000215841 | | 8.665061 | -0.03107925 | 0.5804027 | -0.07865252 | 0.937309 | 0.96904 |
| ENSG00000108448 | TRIM16L | 1984.683528 | -0.03116125 | 0.247486252 | -0.117955847 | 0.90610264 | 0.956276 |
| ENSG00000204351 | SKIV2L | 4087.981 | -0.03116886 | 0.1339384 | -0.2340758 | 0.8149262 | 0.905821 |
| ENSG00000274986 | MIR6750 | 1.975074 | -0.03118871 | 0.6754596 | -0.09425707 | 0.924905 | 0.963943 |
| ENSG00000157107 | FCHO2 | 1486.746 | -0.03123769 | 0.2029217 | -0.1738335 | 0.8619963 | 0.9338 |
| ENSG00000139168 | ZCRB1 | 2342.325192 | -0.031270637 | 0.136617164 | -0.24352955 | 0.8075952 | 0.901496 |
| ENSG00000270528 | FAM133EP | 1.02412 | -0.03137174 | 0.7284735 | -0.143452 | 0.8859332 | NA |
| ENSG00000186451 | SPATA12 | 2.655866 | -0.03139738 | 0.6605773 | -0.07956154 | 0.936586 | 0.968692 |
| ENSG00000178537 | SLC25A20 | 652.0062 | -0.03141787 | 0.1900387 | -0.1779511 | 0.8587614 | 0.931977 |
| ENSG00000237784 | RPS20P12 | 0.9579634 | -0.03142171 | 0.7591657 | -0.08534668 | 0.9319858 | NA |
| ENSG00000265542 | | 0.9851465 | -0.03142374 | 0.7299565 | -0.1063325 | 0.9153185 | NA |
| ENSG00000240720 | LRRD1 | 2.421379 | -0.03144892 | 0.6774655 | -0.091442 | 0.9271414 | 0.96485 |
| ENSG00000235413 | KRT18P63 | 7.149512 | -0.0315326 | 0.521511 | -0.08918498 | 0.9289349 | 0.965492 |
| ENSG00000271228 | | 3.18562 | -0.03153394 | 0.6706656 | -0.09523778 | 0.924126 | 0.963623 |
| ENSG00000269867 | | 64.39325 | -0.03154827 | 0.2889211 | -0.1159557 | 0.9076876 | 0.957074 |
| ENSG00000172725 | CORO1B | 14068.14 | -0.0315846 | 0.1425162 | -0.2391375 | 0.8109989 | 0.903381 |
| ENSG00000141564 | RPTOR | 2865.18 | -0.03159253 | 0.169145 | -0.1980073 | 0.8430394 | 0.922669 |

| | | | | | | | |
|---|---|---|---|---|---|---|---|
| ENSG00000141448 | GATA6 | 163.4997 | -0.03163879 | 0.2207343 | -0.1510215 | 0.8799587 | 0.943516 |
| ENSG00000244289 | RPS3AP35 | 2.998137 | -0.0316933 | 0.6372942 | -0.08606139 | 0.9314176 | 0.966711 |
| ENSG00000227908 | | 5.587696 | -0.03172221 | 0.593992 | -0.08685812 | 0.9307843 | 0.966433 |
| ENSG00000227376 | FTH1P16 | 127.8275 | -0.03173472 | 0.2670873 | -0.1286539 | 0.8976315 | 0.952681 |
| ENSG00000262655 | SPON1 | 17.81675 | -0.03179679 | 0.3832784 | -0.09758294 | 0.9222635 | 0.963117 |
| ENSG00000226971 | | 8.118713 | -0.03185493 | 0.5497845 | -0.08644939 | 0.9311092 | 0.96658 |
| ENSG00000201725 | RNU6-304P | 1.148532 | -0.03190622 | 0.73226 | -0.1051485 | 0.9162579 | NA |
| ENSG00000277268 | LHX1-DT | 94.76139 | -0.03192731 | 0.3488493 | -0.1147645 | 0.9086318 | 0.957348 |
| ENSG00000113648 | MACROH2A1 | 12538.19779 | -0.031946925 | 0.119159784 | -0.249138557 | 0.80325361 | 0.899277 |
| ENSG00000218014 | KRT19P1 | 381.6477 | -0.03195977 | 0.1884564 | -0.1627437 | 0.8707202 | 0.938353 |
| ENSG00000145103 | ILDR1 | 690.7145 | -0.03196699 | 0.2065465 | -0.1744162 | 0.8615384 | 0.933594 |
| ENSG00000277498 | | 1.209766 | -0.03198913 | 0.7373133 | -0.1177696 | 0.9062502 | NA |
| ENSG00000253877 | LINC01608 | 26.96922 | -0.03204243 | 0.7016372 | -0.1079517 | 0.914034 | 0.959721 |
| ENSG00000168878 | SFTPB | 1.620816 | -0.0321056 | 0.7149785 | -0.09190776 | 0.9267713 | 0.964715 |
| ENSG00000269125 | | 4.461637 | -0.03211704 | 0.6238711 | -0.09249511 | 0.9263047 | 0.964494 |
| ENSG00000239881 | RPS27P25 | 1.127413 | -0.03216609 | 0.7440992 | -0.1220475 | 0.9028614 | NA |
| ENSG00000280113 | | 1.042225 | -0.03222049 | 0.7576857 | -0.08794067 | 0.9299238 | NA |
| ENSG00000084652 | TXLNA | 5532.039081 | -0.032230343 | 0.163380227 | -0.200477303 | 0.84110731 | 0.921503 |
| ENSG00000253499 | | 1.524524 | -0.03234579 | 0.6899951 | -0.09680547 | 0.9228809 | NA |
| ENSG00000142192 | APP | 32502.9 | -0.03235015 | 0.1387642 | -0.2511356 | 0.8017093 | 0.898186 |
| ENSG00000204569 | PPP1R10 | 4975.623 | -0.032367 | 0.1899284 | -0.1782314 | 0.8585413 | 0.931881 |
| ENSG00000165300 | SLITRK5 | 261.5668 | -0.03239046 | 0.3144026 | -0.1058238 | 0.9157221 | 0.960178 |
| ENSG00000248554 | C5orf34-AS1 | 27.98191 | -0.03242039 | 0.4208159 | -0.09528908 | 0.9240852 | 0.963623 |
| ENSG00000162994 | CLHC1 | 227.6802 | -0.03246788 | 0.2885479 | -0.1324512 | 0.8946274 | 0.951193 |
| ENSG00000254454 | RCC2P6 | 21.74917 | -0.03257875 | 0.4130359 | -0.09662015 | 0.9230281 | 0.963385 |
| ENSG00000143337 | TOR1AIP1 | 3666.538 | -0.03258488 | 0.1415492 | -0.2783692 | 0.7807289 | 0.885062 |
| ENSG00000087884 | AAMDC | 277.3129875 | -0.032589078 | 0.174819394 | -0.200619749 | 0.84099591 | 0.921419 |
| ENSG00000253258 | | 1.82355 | -0.03264122 | 0.724283 | -0.1134373 | 0.9096839 | 0.957954 |
| ENSG00000175029 | CTBP2 | 5025.441 | -0.03267611 | 0.1242631 | -0.2706098 | 0.7866912 | 0.888711 |
| ENSG00000149929 | HIRIP3 | 899.0259 | -0.03268058 | 0.2043798 | -0.1692476 | 0.8656019 | 0.935977 |
| ENSG00000173868 | PHOSPHO1 | 4.275979 | -0.03270585 | 0.689429 | -0.1010688 | 0.9194958 | 0.961751 |
| ENSG00000141560 | FN3KRP | 2750.024 | -0.03271752 | 0.2735953 | -0.1231531 | 0.9019859 | 0.95489 |
| ENSG00000215284 | CHMP5P1 | 39.82842 | -0.03273417 | 0.3060464 | -0.1195629 | 0.9048294 | 0.955919 |

| | | | | | | | |
|---|---|---|---|---|---|---|---|
| ENSG00000198610 | AKR1C4 | 5.568038 | -0.03274186 | 0.6871855 | -0.08481932 | 0.932405 | 0.967048 |
| ENSG00000279773 | | 1.537796 | -0.03276554 | 0.7811891 | -0.3246878 | 0.7454174 | NA |
| ENSG00000229605 | RPL21P93 | 4.258866 | -0.03288294 | 0.5951169 | -0.08948464 | 0.9286968 | 0.965433 |
| ENSG00000234925 | ATP5PDP4 | 75.80083 | -0.03289374 | 0.3173793 | -0.1166279 | 0.9071549 | 0.95682 |
| ENSG00000234232 | PDE4DIPP7 | 0.9602338 | -0.03291527 | 0.7602196 | -0.08392053 | 0.9331196 | NA |
| ENSG00000223375 | | 1.102397 | -0.03292101 | 0.75077 | -0.09983019 | 0.9204791 | NA |
| ENSG00000251018 | HMMR-AS1 | 1.309733 | -0.03292285 | 0.7042274 | -0.1132356 | 0.9098438 | NA |
| ENSG00000183813 | CCR4 | 1.173769 | -0.03294402 | 0.7546212 | -0.09332743 | 0.9256434 | NA |
| ENSG00000214293 | APTR | 583.9213 | -0.03299274 | 0.1947554 | -0.1682123 | 0.8664163 | 0.936211 |
| ENSG00000251791 | SCARNA6 | 2.220559 | -0.03305769 | 0.6549541 | -0.08912776 | 0.9289804 | 0.965502 |
| ENSG00000130684 | ZNF337 | 348.8627709 | -0.033228032 | 0.272619758 | -0.129342295 | 0.89708681 | 0.952406 |
| ENSG00000286276 | | 4.177631 | -0.03322978 | 0.6576395 | -0.09001376 | 0.9282763 | 0.965223 |
| ENSG00000271092 | TLCD4-RWDD3 | 15.31143 | -0.03322994 | 0.4337445 | -0.09326103 | 0.9256962 | 0.964272 |
| ENSG00000100593 | ISM2 | 6.411700315 | -0.033239075 | 0.663520732 | -0.090676044 | 0.92775001 | 0.96494 |
| ENSG00000224358 | TMCO1-AS1 | 23.9136 | -0.03325331 | 0.4477817 | -0.09270013 | 0.9261418 | 0.9644 |
| ENSG00000244791 | | 1.977155 | -0.03326617 | 0.7040456 | -0.09492589 | 0.9243737 | 0.963692 |
| ENSG00000270236 | | 3.030348 | -0.03330181 | 0.6380577 | -0.0981471 | 0.9218155 | 0.96299 |
| ENSG00000225022 | UBE2D3P1 | 904.1681 | -0.03333052 | 0.138163 | -0.2456163 | 0.8059793 | 0.90047 |
| ENSG00000258758 | | 1.090827 | -0.03342745 | 0.7556945 | -0.09149254 | 0.9271012 | NA |
| ENSG00000141098 | GFOD2 | 1769.939 | -0.03347798 | 0.1667789 | -0.2400794 | 0.8102687 | 0.902923 |
| ENSG00000236608 | EIF4A1P6 | 1.50105 | -0.03349698 | 0.7062799 | -0.1006182 | 0.9198535 | NA |
| ENSG00000095932 | SMIM24 | 1.063904884 | -0.033536278 | 0.718707434 | -0.116571359 | 0.90719974 | NA |
| ENSG00000287579 | | 0.8517045 | -0.03355923 | 0.7600579 | -0.0842446 | 0.932862 | NA |
| ENSG00000275017 | | 11.24281 | -0.03359609 | 0.5960356 | -0.09069184 | 0.9277375 | 0.96494 |
| ENSG00000232599 | | 1.262768 | -0.03360327 | 0.7135233 | -0.1080538 | 0.913953 | NA |
| ENSG00000167967 | E4F1 | 3626.951 | -0.03367402 | 0.1097402 | -0.3037086 | 0.7613499 | 0.872735 |
| ENSG00000080546 | SESN1 | 1116.756169 | -0.033732745 | 0.22824246 | -0.15322612 | 0.87821996 | 0.942663 |
| ENSG00000179219 | LINC00311 | 1.225739 | -0.03381432 | 0.7488387 | -0.1032357 | 0.9177759 | NA |
| ENSG00000149506 | ZP1 | 1.556401 | -0.03393087 | 0.6982514 | -0.1040861 | 0.9171011 | NA |
| ENSG00000198901 | PRC1 | 130.157 | -0.03393661 | 0.3414639 | -0.1149178 | 0.9085103 | 0.957296 |
| ENSG00000198081 | ZBTB14 | 960.916 | -0.033946 | 0.1270999 | -0.27715 | 0.7816649 | 0.885518 |
| ENSG00000232739 | | 0.8408062 | -0.03399437 | 0.7575474 | -0.08865843 | 0.9293534 | NA |
| ENSG00000207784 | MIR542 | 1.913883 | -0.03408365 | 0.6979168 | -0.1072335 | 0.9146037 | 0.959909 |

| | | | | | | | |
|---|---|---|---|---|---|---|---|
| ENSG00000087128 | TMPRSS11E | 3.6611515 | -0.034137354 | 0.691501082 | -0.110422168 | 0.91207457 | 0.958851 |
| ENSG00000215005 | HSPD1P7 | 0.8072207 | -0.03415 | 0.731409 | -0.1279628 | 0.8981784 | NA |
| ENSG00000236262 | SHANK2-AS2 | 0.9697397 | -0.03418525 | 0.7314139 | -0.13882 | 0.8895924 | NA |
| ENSG00000148483 | TMEM236 | 2.98437 | -0.03418668 | 0.625216 | -0.09588212 | 0.9236142 | 0.963623 |
| ENSG00000165821 | SALL2 | 460.2008 | -0.0342134 | 0.3211092 | -0.1131425 | 0.9099176 | 0.95802 |
| ENSG00000232004 | CAP1P2 | 32.53026 | -0.03422446 | 0.3671299 | -0.1181979 | 0.9059108 | 0.956236 |
| ENSG00000231552 | IGBP1P3 | 1.674524 | -0.03426041 | 0.7217517 | -0.134916 | 0.8926783 | 0.950211 |
| ENSG00000278677 | H2AC17 | 19.22079 | -0.03428141 | 0.3959122 | -0.102936 | 0.9180138 | 0.961226 |
| ENSG00000167536 | DHRS13 | 492.7721 | -0.03433167 | 0.2356983 | -0.164822 | 0.8690841 | 0.937198 |
| ENSG00000130150 | MOSPD2 | 1222.414115 | -0.03433331 | 0.27665296 | -0.134885838 | 0.89270214 | 0.95021 |
| ENSG00000259751 | TUBAP11 | 3.164139 | -0.03434444 | 0.5894935 | -0.09280528 | 0.9260583 | 0.9644 |
| ENSG00000235438 | ESRRAP2 | 2.242998 | -0.03438859 | 0.6989391 | -0.1104536 | 0.9120496 | 0.958851 |
| ENSG00000185838 | GNB1L | 779.1808 | -0.03443641 | 0.1934451 | -0.1864812 | 0.8520674 | 0.928062 |
| ENSG00000262319 | | 1.26944 | -0.03448901 | 0.7306859 | -0.1270837 | 0.8988742 | NA |
| ENSG00000232406 | | 4.536906 | -0.03460921 | 0.6411356 | -0.08918875 | 0.9289319 | 0.965492 |
| ENSG00000277775 | H3C7 | 19.58785 | -0.03464213 | 0.4376521 | -0.09845419 | 0.9215716 | 0.962886 |
| ENSG00000099974 | DDTL | 163.6383738 | -0.034698018 | 0.271929434 | -0.139287507 | 0.88922296 | 0.948521 |
| ENSG00000249506 | ZEB2P1 | 24.2161 | -0.03473158 | 0.3473493 | -0.1154737 | 0.9080697 | 0.957217 |
| ENSG00000101439 | CST3 | 6121.778121 | -0.034741265 | 0.270306531 | -0.140297301 | 0.8884251 | 0.947898 |
| ENSG00000268743 | | 6.036541 | -0.0347569 | 0.604073 | -0.09001173 | 0.9282779 | 0.965223 |
| ENSG00000277367 | NEK2P1 | 0.9441245 | -0.03479127 | 0.7618184 | -0.1888202 | 0.8502337 | NA |
| ENSG00000232083 | RPL31P7 | 2.582525 | -0.03482274 | 0.643232 | -0.09707805 | 0.9226644 | 0.963243 |
| ENSG00000260452 | TPRKBP2 | 1.328015 | -0.03482736 | 0.7204364 | -0.1195508 | 0.904839 | NA |
| ENSG00000136631 | VPS45 | 5625.186561 | -0.03483186 | 0.11230866 | -0.346026728 | 0.72932263 | 0.853215 |
| ENSG00000272354 | | 1.285058 | -0.03488516 | 0.7166374 | -0.1353069 | 0.8923692 | NA |
| ENSG00000286611 | | 1.83076 | -0.03490033 | 0.7145309 | -0.1262653 | 0.8995219 | 0.953839 |
| ENSG00000089009 | RPL6 | 12770.71883 | -0.034976361 | 0.161379215 | -0.242202179 | 0.80862351 | 0.902134 |
| ENSG00000077232 | DNAJC10 | 6936.259455 | -0.034986997 | 0.240615284 | -0.156440696 | 0.87568566 | 0.941283 |
| ENSG00000279762 | | 1.349431 | -0.03506998 | 0.7254916 | -0.1172618 | 0.9066526 | NA |
| ENSG00000181513 | ACBD4 | 841.1242 | -0.03508756 | 0.2388207 | -0.1818708 | 0.8556841 | 0.930183 |
| ENSG00000215515 | IFIT1P1 | 2.670439 | -0.03509734 | 0.684366 | -0.1055304 | 0.9159549 | 0.960213 |
| ENSG00000197705 | KLHL14 | 2.839089 | -0.0352224 | 0.6716996 | -0.1111579 | 0.9114912 | 0.95882 |
| ENSG00000274678 | | 1.099478 | -0.03523507 | 0.7412986 | -0.1529539 | 0.8784346 | NA |

| | | | | | | | |
|---|---|---|---|---|---|---|---|
| ENSG00000188917 | TRMT2B | 2396.348 | -0.03534563 | 0.1468271 | -0.2520437 | 0.8010073 | 0.897885 |
| ENSG00000001629 | ANKIB1 | 4963.699635 | -0.035369686 | 0.185384484 | -0.21293585 | 0.831377 | 0.915137 |
| ENSG00000138760 | SCARB2 | 10053.96104 | -0.035437056 | 0.153394011 | -0.237626543 | 0.81217076 | 0.904197 |
| ENSG00000179044 | EXOC3L1 | 94.64744 | -0.03544707 | 0.3213317 | -0.1199129 | 0.9045521 | 0.955816 |
| ENSG00000179776 | CDH5 | 59.01822 | -0.03553439 | 0.7422401 | -0.1134876 | 0.909644 | 0.957954 |
| ENSG00000139117 | CPNE8 | 12.50226882 | -0.035590125 | 0.743973182 | -0.111190621 | 0.91146519 | 0.95882 |
| ENSG00000252614 | RNU6-807P | 4.923242 | -0.03559105 | 0.621196 | -0.09737504 | 0.9224286 | 0.963176 |
| ENSG00000234337 | | 361.7089 | -0.03564625 | 0.2445909 | -0.1553739 | 0.8765266 | 0.941708 |
| ENSG00000240288 | GHRLOS | 35.45801 | -0.03568858 | 0.376153 | -0.1102399 | 0.9122191 | 0.958851 |
| ENSG00000122971 | ACADS | 513.3840844 | -0.035706905 | 0.254744168 | -0.146037683 | 0.88389164 | 0.945465 |
| ENSG00000204560 | DHX16 | 4022.695 | -0.03578949 | 0.1101974 | -0.3318674 | 0.7399894 | 0.859581 |
| ENSG00000260647 | | 6.589388 | -0.03581376 | 0.5752303 | -0.09343374 | 0.925559 | 0.96427 |
| ENSG00000270690 | | 3.18827 | -0.03582829 | 0.6675647 | -0.1064356 | 0.9152367 | 0.960119 |
| ENSG00000134283 | PPHLN1 | 3341.412602 | -0.035842086 | 0.129777358 | -0.28886064 | 0.77268803 | 0.88019 |
| ENSG00000130165 | ELOF1 | 2345.437912 | -0.035929196 | 0.198741407 | -0.192233643 | 0.84755919 | 0.925823 |
| ENSG00000084072 | PPIE | 4165.173485 | -0.035946255 | 0.161388215 | -0.234087588 | 0.81491698 | 0.905821 |
| ENSG00000132879 | FBXO44 | 1734.066493 | -0.035954491 | 0.256114464 | -0.151370733 | 0.87968328 | 0.943421 |
| ENSG00000279590 | | 2.020147 | -0.0359579 | 0.7231669 | -0.1095632 | 0.9127558 | 0.959069 |
| ENSG00000267890 | | 2.586001 | -0.03597183 | 0.6706856 | -0.09818937 | 0.9217819 | 0.96299 |
| ENSG00000184992 | BRI3BP | 3572.666 | -0.03597327 | 0.3444807 | -0.1213728 | 0.9033957 | 0.95513 |
| ENSG00000232333 | RPS27AP2 | 7.02081 | -0.03609532 | 0.5572901 | -0.09625295 | 0.9233197 | 0.963538 |
| ENSG00000228879 | RPL35AP6 | 3.572123 | -0.03613863 | 0.6664008 | -0.1010515 | 0.9195095 | 0.961751 |
| ENSG00000226057 | PHF2P2 | 36.7913 | -0.03614088 | 0.7412564 | -0.1151672 | 0.9083126 | 0.957239 |
| ENSG00000207062 | SNORA15B-1 | 3.524433 | -0.03615321 | 0.5812587 | -0.09420563 | 0.9249458 | 0.963948 |
| ENSG00000162669 | HFM1 | 8.789833 | -0.03617414 | 0.5018928 | -0.09875227 | 0.921335 | 0.962793 |
| ENSG00000169752 | NRG4 | 32.12214 | -0.03618076 | 0.4537864 | -0.1006985 | 0.9197898 | 0.961781 |
| ENSG00000271554 | | 1.428563 | -0.03620889 | 0.7393254 | -0.1562165 | 0.8758624 | NA |
| ENSG00000277892 | MIR6746 | 2.497365 | -0.03625159 | 0.6131217 | -0.09862807 | 0.9214336 | 0.962808 |
| ENSG00000219487 | | 51.75357 | -0.03626802 | 0.389608 | -0.1063923 | 0.9152711 | 0.960119 |
| ENSG00000213891 | RPL3P6 | 1.534722 | -0.03633795 | 0.6907367 | -0.1073115 | 0.9145418 | NA |
| ENSG00000146859 | TMEM140 | 542.823 | -0.03636012 | 0.2421083 | -0.1587171 | 0.8738918 | 0.940225 |
| ENSG00000258548 | LINC00645 | 2.667146 | -0.03637252 | 0.7347023 | -0.1944494 | 0.845824 | 0.924612 |
| ENSG00000178852 | EFCAB13 | 201.4919 | -0.03641805 | 0.4819581 | -0.08284556 | 0.9339743 | 0.967249 |

| | | | | | | | |
|---|---|---|---|---|---|---|---|
| ENSG00000163930 | BAP1 | 6565.659 | -0.03647274 | 0.130025 | -0.2871336 | 0.77401 | 0.880805 |
| ENSG00000179943 | FIZ1 | 879.1766 | -0.03653952 | 0.1312402 | -0.284987 | 0.7756541 | 0.881867 |
| ENSG00000127589 | TUBBP1 | 2009.341799 | -0.036601503 | 0.367358122 | -0.125696676 | 0.89997203 | 0.95405 |
| ENSG00000147059 | SPIN2A | 4.526457 | -0.0366343 | 0.6141404 | -0.09835464 | 0.9216507 | 0.962893 |
| ENSG00000165685 | TMEM52B | 84.68386 | -0.03674027 | 0.7866618 | 0 | 0.9071457 | 0.95682 |
| ENSG00000237758 | BANF1P3 | 814.2203 | -0.03674227 | 0.2048638 | -0.1956641 | 0.8448731 | 0.923991 |
| ENSG00000286511 | | 6.941933 | -0.03682164 | 0.6063882 | -0.1015962 | 0.9190772 | 0.961604 |
| ENSG00000233380 | RPS2P48 | 1.885183 | -0.03687391 | 0.7217739 | -0.1101654 | 0.9122782 | 0.958851 |
| ENSG00000164749 | HNF4G | 155.2731 | -0.03687467 | 0.4003999 | -0.1148446 | 0.9085683 | 0.957319 |
| ENSG00000232352 | SEMA3B-AS1 | 47.46675 | -0.03689874 | 0.3152048 | -0.1332761 | 0.893975 | 0.950882 |
| ENSG00000196757 | ZNF700 | 460.3815 | -0.03692378 | 0.2386835 | -0.1664588 | 0.8677959 | 0.936731 |
| ENSG00000267560 | | 8.205489 | -0.03695621 | 0.5475933 | -0.09605623 | 0.9234759 | 0.963616 |
| ENSG00000235267 | | 1.253155 | -0.03702536 | 0.7117053 | -0.1265181 | 0.8993219 | NA |
| ENSG00000224059 | HSPA8P16 | 0.8488194 | -0.03711551 | 0.7567219 | -0.09021718 | 0.9281146 | NA |
| ENSG00000235912 | | 339.4001 | -0.03712507 | 0.2431298 | -0.1619972 | 0.871308 | 0.938681 |
| ENSG00000267203 | SNRPGP4 | 20.42457 | -0.03716962 | 0.4598159 | -0.1032325 | 0.9177784 | 0.961195 |
| ENSG00000243104 | MTND4LP14 | 0.9659827 | -0.03717882 | 0.7648883 | -0.2364668 | 0.8130705 | NA |
| ENSG00000226243 | RPL37AP1 | 167.1307 | -0.03725921 | 0.2896101 | -0.1359606 | 0.8918524 | 0.949991 |
| ENSG00000153975 | ZUP1 | 937.8413 | -0.03726385 | 0.1355246 | -0.2814383 | 0.7783742 | 0.883535 |
| ENSG00000232952 | | 357.642 | -0.03727547 | 0.1596226 | -0.2455854 | 0.8060032 | 0.90047 |
| ENSG00000255384 | | 0.8599895 | -0.03728274 | 0.7376799 | -0.1401465 | 0.8885443 | NA |
| ENSG00000238741 | SCARNA7 | 2.090617 | -0.03729024 | 0.7212437 | -0.1273822 | 0.8986379 | 0.953206 |
| ENSG00000245017 | LINC02453 | 1.999537 | -0.03731128 | 0.6754635 | -0.1125597 | 0.9103797 | 0.958126 |
| ENSG00000084073 | ZMPSTE24 | 4949.856493 | -0.037316048 | 0.144132674 | -0.24142114 | 0.80922873 | 0.90262 |
| ENSG00000249234 | | 1.322919 | -0.03740194 | 0.7081355 | -0.1230473 | 0.9020697 | NA |
| ENSG00000122378 | PRXL2A | 8188.651392 | -0.037468828 | 0.156298946 | -0.274670983 | 0.78356903 | 0.886724 |
| ENSG00000267786 | | 1.353577 | -0.03754702 | 0.7005913 | -0.1254673 | 0.9001536 | NA |
| ENSG00000198912 | C1orf174 | 961.0343 | -0.03754868 | 0.1677741 | -0.2303567 | 0.8178146 | 0.907648 |
| ENSG00000122477 | LRRC39 | 34.46633281 | -0.037554174 | 0.369329239 | -0.118853867 | 0.90539113 | 0.956018 |
| ENSG00000152932 | RAB3C | 0.8345981 | -0.03758257 | 0.7540446 | -0.2067378 | 0.8362146 | NA |
| ENSG00000198797 | BRINP2 | 91.93988 | -0.03760668 | 0.5312845 | -0.1102344 | 0.9122235 | 0.958851 |
| ENSG00000203497 | PDCD4-AS1 | 299.0914 | -0.03760788 | 0.2995108 | -0.1463653 | 0.883633 | 0.945465 |
| ENSG00000239521 | CASTOR3 | 630.282 | -0.03761404 | 0.3447044 | -0.1224077 | 0.9025761 | 0.954999 |

| | | | | | | | |
|---|---|---|---|---|---|---|---|
| ENSG00000011304 | PTBP1 | 17881.04641 | -0.03768455 | 0.237809302 | -0.169832325 | 0.865142 | 0.935654 |
| ENSG00000155307 | SAMSN1 | 1.59079 | -0.03772325 | 0.6735404 | -0.1169478 | 0.9069014 | 0.956815 |
| ENSG00000178026 | LRRC75B | 550.7238 | -0.03773692 | 0.2648696 | -0.15542 | 0.8764902 | 0.941707 |
| ENSG00000163050 | COQ8A | 3015.545 | -0.03774343 | 0.1242509 | -0.3057222 | 0.7598161 | 0.87192 |
| ENSG00000169607 | CKAP2L | 587.7982 | -0.03786397 | 0.3926971 | -0.1170883 | 0.9067901 | 0.956735 |
| ENSG00000255447 | | 2.47241 | -0.03786904 | 0.7006068 | -0.1025931 | 0.9182859 | 0.961287 |
| ENSG00000162227 | TAF6L | 1107.701 | -0.03787802 | 0.2129496 | -0.185932 | 0.8524981 | 0.928288 |
| ENSG00000244004 | RPS3AP19 | 2.607826 | -0.03788383 | 0.653915 | -0.1005245 | 0.9199279 | 0.961824 |
| ENSG00000143632 | ACTA1 | 5.135034 | -0.03790945 | 0.5930621 | -0.09565496 | 0.9237946 | 0.963623 |
| ENSG00000223730 | | 1.143921 | -0.03806159 | 0.7148198 | -0.1599655 | 0.8729082 | NA |
| ENSG00000243829 | | 145.2554 | -0.03811986 | 0.2220412 | -0.1734857 | 0.8622697 | 0.933906 |
| ENSG00000266498 | | 3.202276 | -0.03814689 | 0.6065089 | -0.1113816 | 0.9113138 | 0.958767 |
| ENSG00000267348 | GEMIN7-AS1 | 50.808 | -0.03815475 | 0.3280421 | -0.1276011 | 0.8984646 | 0.953175 |
| ENSG00000279144 | | 2.290445 | -0.03821994 | 0.7194171 | -0.1244397 | 0.9009672 | 0.954525 |
| ENSG00000282393 | | 9.776716 | -0.03823654 | 0.5381901 | -0.1034535 | 0.9176031 | 0.961195 |
| ENSG00000140990 | NDUFB10 | 4811.791 | -0.03824273 | 0.2344597 | -0.1640641 | 0.8696807 | 0.93751 |
| ENSG00000273342 | | 1.01486 | -0.0384171 | 0.7294251 | -0.1407942 | 0.8880325 | NA |
| ENSG00000134056 | MRPS36 | 1157.268253 | -0.03842335 | 0.215859128 | -0.198759094 | 0.8424512 | 0.922253 |
| ENSG00000229562 | ZFYVE9P1 | 7.094166 | -0.0385774 | 0.5961539 | -0.1021923 | 0.918604 | 0.961449 |
| ENSG00000284862 | CCDC39 | 17.40942 | -0.03861445 | 0.4413551 | -0.1098257 | 0.9125476 | 0.958888 |
| ENSG00000105011 | ASF1B | 1864.581278 | -0.03861617 | 0.550412471 | -0.107274743 | 0.91457102 | 0.959909 |
| ENSG00000224409 | | 1.193556 | -0.03863068 | 0.7405236 | -0.1566868 | 0.8754917 | NA |
| ENSG00000116350 | SRSF4 | 5882.96537 | -0.038636572 | 0.153220016 | -0.291889073 | 0.77037144 | 0.878601 |
| ENSG00000286629 | | 3.948731 | -0.03869989 | 0.6819593 | -0.1189117 | 0.9053453 | 0.956008 |
| ENSG00000128000 | ZNF780B | 850.4778876 | -0.038726173 | 0.305281089 | -0.137847019 | 0.89036132 | 0.949126 |
| ENSG00000074317 | SNCB | 2.445487238 | -0.038767213 | 0.698212197 | -0.139160623 | 0.88932322 | 0.948552 |
| ENSG00000279314 | | 0.9751434 | -0.03880906 | 0.7376124 | -0.1402087 | 0.8884951 | NA |
| ENSG00000123472 | ATPAF1 | 4124.798029 | -0.038844869 | 0.209472562 | -0.193873071 | 0.84627527 | 0.924915 |
| ENSG00000163320 | CGGBP1 | 8119.296 | -0.03886715 | 0.1702057 | -0.1915883 | 0.8480647 | 0.926147 |
| ENSG00000163806 | SPDYA | 23.41069 | -0.03889755 | 0.3935085 | -0.116637 | 0.9071478 | 0.95682 |
| ENSG00000198176 | TFDP1 | 5745.745 | -0.03891057 | 0.2280953 | -0.1764387 | 0.8599493 | 0.932571 |
| ENSG00000253141 | | 1.610837 | -0.03898379 | 0.7133965 | -0.1386825 | 0.8897011 | 0.948692 |
| ENSG00000099949 | LZTR1 | 1142.941915 | -0.03898818 | 0.208140103 | -0.199342177 | 0.84199509 | 0.922096 |

| | | | | | | | |
|---|---|---|---|---|---|---|---|
| ENSG00000166347 | CYB5A | 2896.85 | -0.03905109 | 0.2369187 | -0.1931545 | 0.8468379 | 0.92531 |
| ENSG00000160014 | CALM3 | 10084.91 | -0.03913826 | 0.2840301 | -0.1542551 | 0.8774086 | 0.94216 |
| ENSG00000198663 | C6orf89 | 6238.71 | -0.03914763 | 0.1791533 | -0.2320132 | 0.8165278 | 0.906574 |
| ENSG00000212396 | RNA5SP323 | 2.649588 | -0.03922717 | 0.7210992 | -0.138298 | 0.8900049 | 0.948936 |
| ENSG00000105355 | PLIN3 | 2964.586641 | -0.039231461 | 0.176791592 | -0.249385024 | 0.80306297 | 0.899149 |
| ENSG00000232346 | | 1834.037 | -0.03927426 | 0.2044822 | -0.2103824 | 0.8333693 | 0.916351 |
| ENSG00000179902 | C1orf194 | 16.57786 | -0.03928332 | 0.4180218 | -0.1136991 | 0.9094763 | 0.957844 |
| ENSG00000081803 | CADPS2 | 1599.173015 | -0.039289488 | 0.210261582 | -0.180530918 | 0.85673578 | 0.93072 |
| ENSG00000152601 | MBNL1 | 4993.222 | -0.03929126 | 0.1560481 | -0.2410888 | 0.8094863 | 0.902793 |
| ENSG00000160050 | CCDC28B | 198.0003 | -0.03940062 | 0.3623504 | -0.1246356 | 0.900812 | 0.954483 |
| ENSG00000167986 | DDB1 | 22146.54 | -0.03945365 | 0.1733214 | -0.2359275 | 0.8134889 | 0.904826 |
| ENSG00000083845 | RPS5 | 28659.94289 | -0.039477237 | 0.194966797 | -0.243299947 | 0.80777305 | 0.901601 |
| ENSG00000157593 | SLC35B2 | 6231.596 | -0.03948455 | 0.1115144 | -0.3469953 | 0.7285949 | 0.852739 |
| ENSG00000100814 | CCNB1IP1 | 3182.276037 | -0.039490031 | 0.14675143 | -0.323088941 | 0.74662789 | 0.863912 |
| ENSG00000173391 | OLR1 | 120.1587 | -0.03953596 | 0.786675 | 0 | 0.9056954 | 0.956179 |
| ENSG00000076053 | RBM7 | 451.0261859 | -0.03959621 | 0.147424427 | -0.282918012 | 0.77723969 | 0.88265 |
| ENSG00000224387 | | 1.894289 | -0.03961352 | 0.72479 | -0.1250185 | 0.9005089 | 0.954353 |
| ENSG00000159079 | CFAP298 | 477.3548 | -0.03969886 | 0.1997087 | -0.2152783 | 0.8295503 | 0.914195 |
| ENSG00000108231 | LGI1 | 8.551584212 | -0.039733706 | 0.66629547 | -0.120144903 | 0.90436836 | 0.955774 |
| ENSG00000162927 | PUS10 | 491.357 | -0.03984736 | 0.263965 | -0.1597194 | 0.8731021 | 0.939929 |
| ENSG00000259904 | ACTG1P15 | 3.945834 | -0.03987292 | 0.6584847 | -0.1142055 | 0.9090749 | 0.957663 |
| ENSG00000109220 | CHIC2 | 465.6786441 | -0.039884712 | 0.161319154 | -0.262145101 | 0.79320958 | 0.893113 |
| ENSG00000124942 | AHNAK | 42146.02928 | -0.039895018 | 0.311384238 | -0.146387376 | 0.88361559 | 0.945465 |
| ENSG00000261595 | | 2.236266 | -0.04009193 | 0.6750519 | -0.1222241 | 0.9027216 | 0.955022 |
| ENSG00000267924 | | 2.681131 | -0.04013325 | 0.6871258 | -0.1073919 | 0.9144781 | 0.959909 |
| ENSG00000272054 | | 80.61876 | -0.04030067 | 0.2927109 | -0.1517562 | 0.8793792 | 0.943285 |
| ENSG00000284138 | ATP6V0CP4 | 1.043096 | -0.04032231 | 0.7345607 | -0.1462372 | 0.8837341 | NA |
| ENSG00000168913 | ENHO | 35.7075 | -0.04034202 | 0.5195147 | -0.1104384 | 0.9120617 | 0.958851 |
| ENSG00000221823 | PPP3R1 | 1923.811 | -0.04041081 | 0.1517527 | -0.2755854 | 0.7828665 | 0.886426 |
| ENSG00000261239 | ANKRD26P1 | 1.097983 | -0.04045055 | 0.7187938 | -0.1565733 | 0.8755812 | NA |
| ENSG00000177191 | B3GNT8 | 59.68399 | -0.04048112 | 0.4024258 | -0.1209305 | 0.9037461 | 0.955376 |
| ENSG00000066136 | NFYC | 2269.161212 | -0.040499978 | 0.162584538 | -0.264412953 | 0.79146173 | 0.891863 |
| ENSG00000169744 | LDB2 | 12.60252 | -0.04050583 | 0.5512304 | -0.1064709 | 0.9152087 | 0.960119 |

| | | | | | | | |
|---|---|---|---|---|---|---|---|
| ENSG00000136014 | USP44 | 1.145765705 | -0.040523047 | 0.728881504 | -0.167258174 | 0.86716691 | NA |
| ENSG00000152061 | RABGAP1L | 1535.168 | -0.04055785 | 0.1712094 | -0.2367471 | 0.812853 | 0.904346 |
| ENSG00000259340 | | 0.7988366 | -0.04056559 | 0.7382011 | -0.1827835 | 0.8549679 | NA |
| ENSG00000223635 | | 2.956707 | -0.04061738 | 0.7098564 | -0.1442253 | 0.8853226 | 0.94607 |
| ENSG00000115419 | GLS | 3641.549146 | -0.04063795 | 0.231991831 | -0.172537313 | 0.86301513 | 0.934294 |
| ENSG00000237586 | | 2.246156 | -0.0406758 | 0.7280748 | -0.1420451 | 0.8870444 | 0.947034 |
| ENSG00000254477 | | 7.471717 | -0.04068084 | 0.6149924 | -0.101473 | 0.919175 | 0.96165 |
| ENSG00000261303 | | 2.662936 | -0.04069494 | 0.6508123 | -0.1158644 | 0.90776 | 0.957074 |
| ENSG00000171161 | ZNF672 | 2362.568 | -0.040705 | 0.1676085 | -0.2466329 | 0.8051923 | 0.900308 |
| ENSG00000259818 | | 2.217297 | -0.04072841 | 0.702513 | -0.1313991 | 0.8954596 | 0.951621 |
| ENSG00000173432 | SAA1 | 5.355635 | -0.04074085 | 0.732047 | -0.1628984 | 0.8705984 | 0.93826 |
| ENSG00000196632 | WNK3 | 7.132617 | -0.04075491 | 0.5465511 | -0.1083892 | 0.913687 | 0.959592 |
| ENSG00000110074 | FOXRED1 | 1439.939983 | -0.040754991 | 0.309259447 | -0.13514778 | 0.89249503 | 0.95021 |
| ENSG00000224806 | ARL5AP4 | 2.122805 | -0.04083145 | 0.6927 | -0.1154081 | 0.9081217 | 0.957217 |
| ENSG00000197585 | | 1.536451 | -0.0408596 | 0.7181061 | -0.1351157 | 0.8925204 | NA |
| ENSG00000114646 | CSPG5 | 127.938397 | -0.040921959 | 0.401747235 | -0.120046639 | 0.9044462 | 0.95578 |
| ENSG00000156931 | VPS8 | 3321.608 | -0.04100335 | 0.2174001 | -0.1988976 | 0.8423429 | 0.922211 |
| ENSG00000150991 | UBC | 34436.22 | -0.04113321 | 0.1955112 | -0.2295551 | 0.8184375 | 0.908012 |
| ENSG00000239791 | | 3.667954 | -0.04118267 | 0.6384601 | -0.1074538 | 0.914429 | 0.959909 |
| ENSG00000152582 | SPEF2 | 703.7226 | -0.0412326 | 0.3451625 | -0.1425165 | 0.886672 | 0.946788 |
| ENSG00000279168 | | 10.45085 | -0.04126609 | 0.5555087 | -0.1113912 | 0.9113061 | 0.958767 |
| ENSG00000115507 | OTX1 | 298.4865439 | -0.041291592 | 0.176039362 | -0.244073509 | 0.80717389 | 0.901198 |
| ENSG00000269842 | | 2.8393 | -0.04129552 | 0.7491175 | -0.103182 | 0.9178186 | 0.961195 |
| ENSG00000270120 | | 2.21606 | -0.04134299 | 0.6761488 | -0.1363695 | 0.8915291 | 0.949761 |
| ENSG00000255343 | | 2.293529 | -0.04140169 | 0.689084 | -0.1354355 | 0.8922675 | 0.950182 |
| ENSG00000258895 | | 232.9189 | -0.04144963 | 0.2820288 | -0.1554218 | 0.8764888 | 0.941707 |
| ENSG00000230721 | | 3.388953 | -0.04145305 | 0.6368465 | -0.1100813 | 0.9123449 | 0.958851 |
| ENSG00000285923 | | 21.89843 | -0.04149834 | 0.4172616 | -0.1219559 | 0.902934 | 0.955046 |
| ENSG00000285696 | | 2.111022 | -0.04151665 | 0.7246947 | -0.1250677 | 0.90047 | 0.95435 |
| ENSG00000167394 | ZNF668 | 334.737 | -0.04153837 | 0.1705154 | -0.2444099 | 0.8069134 | 0.901021 |
| ENSG00000247240 | UBL7-AS1 | 227.9047 | -0.04158897 | 0.3011339 | -0.148139 | 0.8822331 | 0.944859 |
| ENSG00000090006 | LTBP4 | 2974.513955 | -0.041665162 | 0.172132548 | -0.239671079 | 0.81058526 | 0.903071 |
| ENSG00000232750 | | 1.663848 | -0.04167147 | 0.7168596 | -0.1317073 | 0.8952158 | 0.951596 |

| | | | | | | | |
|---|---|---|---|---|---|---|---|
| ENSG00000100109 | TFIP11 | 1666.76726 | -0.041720314 | 0.12959944 | -0.312801674 | 0.75443134 | 0.86893 |
| ENSG00000227477 | STK4-AS1 | 28.72509 | -0.04172828 | 0.4283955 | -0.1223951 | 0.9025861 | 0.954999 |
| ENSG00000171862 | PTEN | 6012.381 | -0.04178508 | 0.09939413 | -0.4267798 | 0.6695397 | 0.815897 |
| ENSG00000112592 | TBP | 1406.92096 | -0.041785886 | 0.116614492 | -0.374997746 | 0.70766214 | 0.841059 |
| ENSG00000054179 | ENTPD2 | 766.6133227 | -0.041825137 | 0.267541836 | -0.170511531 | 0.86460787 | 0.935218 |
| ENSG00000111596 | CNOT2 | 4576.663174 | -0.041906451 | 0.092740315 | -0.467113625 | 0.64041859 | 0.794841 |
| ENSG00000278610 | NA | 0.9176 | -0.04196723 | 0.7392448 | -0.157166 | 0.8751141 | NA |
| ENSG00000265888 | DSCAS | 3.325546 | -0.041973 | 0.6466493 | -0.1173526 | 0.9065806 | 0.956628 |
| ENSG00000107651 | SEC23IP | 3857.689659 | -0.0419758 | 0.128673024 | -0.370827179 | 0.71076626 | 0.842606 |
| ENSG00000128272 | ATF4 | 22001.83298 | -0.042012204 | 0.165518543 | -0.268599817 | 0.78823765 | 0.889844 |
| ENSG00000188152 | NUTM2G | 29.36975 | -0.04207965 | 0.3633886 | -0.1329983 | 0.8941947 | 0.950906 |
| ENSG00000149781 | FERMT3 | 75.99218 | -0.04208469 | 0.2723283 | -0.1655907 | 0.8684791 | 0.93685 |
| ENSG00000259760 | | 0.7628182 | -0.04214872 | 0.761755 | -0.2230947 | 0.8234618 | NA |
| ENSG00000130487 | KLHDC7B | 252.3902848 | -0.042192128 | 0.64487388 | -0.131077681 | 0.89571386 | 0.951839 |
| ENSG00000187959 | CPSF4L | 9.265025 | -0.04225585 | 0.5326622 | -0.10737 | 0.9144954 | 0.959909 |
| ENSG00000254470 | AP5B1 | 3049.766 | -0.04230085 | 0.2086517 | -0.212948 | 0.8313675 | 0.915137 |
| ENSG00000183580 | FBXL7 | 61.11792 | -0.04230604 | 0.5077033 | -0.1161659 | 0.9075211 | 0.957042 |
| ENSG00000274560 | | 3.104458 | -0.0424022 | 0.6875251 | -0.1349097 | 0.8926833 | 0.950211 |
| ENSG00000183682 | BMP8A | 167.9314 | -0.04244646 | 0.1844847 | -0.2433484 | 0.8077355 | 0.901598 |
| ENSG00000219642 | BMPR1AP1 | 244.7052 | -0.04246561 | 0.2186406 | -0.2088038 | 0.8346014 | 0.917136 |
| ENSG00000110660 | SLC35F2 | 1217.589406 | -0.042496094 | 0.149088145 | -0.304688651 | 0.76060329 | 0.872369 |
| ENSG00000275476 | | 5.228022 | -0.04257579 | 0.6520797 | -0.1174083 | 0.9065365 | 0.95662 |
| ENSG00000215217 | C5orf49 | 67.01818 | -0.04260207 | 0.3960797 | -0.1296573 | 0.8968376 | 0.952406 |
| ENSG00000226015 | CCT8P1 | 421.4553 | -0.04266827 | 0.2228488 | -0.2039707 | 0.8383764 | 0.919459 |
| ENSG00000225330 | | 3.498598 | -0.04269458 | 0.6392081 | -0.1316997 | 0.8952218 | 0.951596 |
| ENSG00000250770 | | 25.09176 | -0.04283201 | 0.4327614 | -0.1386788 | 0.889704 | 0.948692 |
| ENSG00000279789 | | 172.4196 | -0.04293294 | 0.3400432 | -0.1525696 | 0.8787377 | 0.942863 |
| ENSG00000201183 | RNVU1-3 | 4.677116 | -0.04294608 | 0.686859 | -0.1249617 | 0.9005538 | 0.954362 |
| ENSG00000236780 | LINC01829 | 19.49498 | -0.04295029 | 0.392432 | -0.1283758 | 0.8978516 | 0.95278 |
| ENSG00000269980 | | 2.76001 | -0.04296728 | 0.6679647 | -0.1181849 | 0.9059211 | 0.956236 |
| ENSG00000173545 | ZNF622 | 3419.74 | -0.04319257 | 0.1304678 | -0.3490328 | 0.7270647 | 0.851749 |
| ENSG00000261318 | | 0.9374111 | -0.04321053 | 0.7607907 | -0.2245359 | 0.8223403 | NA |
| ENSG00000276968 | | 1.477071 | -0.04325645 | 0.7255646 | -0.1389946 | 0.8894545 | NA |

| | | | | | | | |
|---|---|---|---|---|---|---|---|
| ENSG00000152936 | LMNTD1 | 2.372318 | -0.04327221 | 0.7124268 | -0.1895524 | 0.8496599 | 0.926918 |
| ENSG00000167701 | GPT | 265.0617 | -0.04327288 | 0.2920302 | -0.158623 | 0.8739659 | 0.940225 |
| ENSG00000204618 | RNF39 | 192.9916 | -0.04327327 | 0.2351705 | -0.1887267 | 0.850307 | 0.9273 |
| ENSG00000158710 | TAGLN2 | 12762.8 | -0.04328049 | 0.2861841 | -0.1661504 | 0.8680386 | 0.936731 |
| ENSG00000237206 | IMPDH1P4 | 1.041122 | -0.04330875 | 0.7117408 | -0.1444024 | 0.8851827 | NA |
| ENSG00000085433 | WDR47 | 1184.353418 | -0.043316218 | 0.224945438 | -0.2083627 | 0.83494578 | 0.917211 |
| ENSG00000261527 | | 1.223159 | -0.04334176 | 0.7243245 | -0.1521253 | 0.8790881 | NA |
| ENSG00000213950 | RPS10P2 | 1.800857 | -0.04339355 | 0.7009265 | -0.1309043 | 0.895851 | 0.951863 |
| ENSG00000242087 | RPL36AP41 | 0.8381581 | -0.0434475 | 0.7443407 | -0.1871663 | 0.8515303 | NA |
| ENSG00000108592 | FTSJ3 | 14924.33617 | -0.04349951 | 0.10977582 | -0.400577922 | 0.6887309 | 0.828482 |
| ENSG00000105953 | OGDH | 8625.04104 | -0.043507978 | 0.205280965 | -0.216938796 | 0.82825604 | 0.91345 |
| ENSG00000267595 | BRCA1P1 | 22.34657 | -0.04352833 | 0.4612937 | -0.1199944 | 0.9044876 | 0.955786 |
| ENSG00000268967 | | 1.378172 | -0.04353802 | 0.7013183 | -0.1492274 | 0.8813742 | NA |
| ENSG00000274184 | | 6.804742 | -0.043547 | 0.6754929 | -0.1395753 | 0.8889955 | 0.948317 |
| ENSG00000126254 | RBM42 | 5086.746415 | -0.043548808 | 0.228597182 | -0.158038441 | 0.87442651 | 0.940555 |
| ENSG00000078140 | UBE2K | 5097.669708 | -0.043585349 | 0.165713916 | -0.331359326 | 0.74037309 | 0.859783 |
| ENSG00000239932 | RN7SL606P | 1.620982 | -0.04358715 | 0.7294143 | -0.1509424 | 0.8800211 | 0.943516 |
| ENSG00000260277 | | 9.261068 | -0.04365923 | 0.4985245 | -0.110831 | 0.9117504 | 0.958851 |
| ENSG00000104408 | EIF3E | 26539.35299 | -0.043681167 | 0.150193567 | -0.309794699 | 0.75671708 | 0.870017 |
| ENSG00000143412 | ANXA9 | 2842.88 | -0.04370591 | 0.2231962 | -0.2123356 | 0.8318452 | 0.915357 |
| ENSG00000213942 | | 26.51475 | -0.04375929 | 0.3422668 | -0.1454159 | 0.8843825 | 0.945712 |
| ENSG00000268423 | NA | 3.779901 | -0.04382597 | 0.5915611 | -0.1168534 | 0.9069763 | 0.95682 |
| ENSG00000196123 | KIAA0895L | 1614.138 | -0.04383852 | 0.2011354 | -0.2251821 | 0.8218376 | 0.909619 |
| ENSG00000180776 | ZDHHC20 | 6710.036 | -0.04392268 | 0.182843 | -0.249358 | 0.8030839 | 0.899149 |
| ENSG00000286787 | | 20.11083 | -0.0439387 | 0.4506339 | -0.1205487 | 0.9040485 | 0.95555 |
| ENSG00000163705 | FANCD2OS | 4.341214 | -0.04394322 | 0.6253721 | -0.118697 | 0.9055154 | 0.956112 |
| ENSG00000244535 | | 2.536557 | -0.04399093 | 0.6933968 | -0.1251152 | 0.9004324 | 0.954348 |
| ENSG00000099785 | MARCHF2 | 508.7088106 | -0.044010844 | 0.145157744 | -0.304193464 | 0.7609805 | 0.872566 |
| ENSG00000227454 | MTND4P30 | 0.7727088 | -0.04417025 | 0.7222897 | -0.1681932 | 0.8664313 | NA |
| ENSG00000277998 | | 2463.171 | -0.04422006 | 0.186822 | -0.2407026 | 0.8097856 | 0.902824 |
| ENSG00000286113 | | 1.47858 | -0.04431806 | 0.7139629 | -0.1554952 | 0.876431 | NA |
| ENSG00000174842 | GLMN | 374.8189 | -0.04440516 | 0.2891064 | -0.1530235 | 0.8783798 | 0.94267 |
| ENSG00000181220 | ZNF746 | 1732.511 | -0.04444742 | 0.1298479 | -0.3708207 | 0.7107711 | 0.842606 |

| | | | | | | | |
|---|---|---|---|---|---|---|---|
| ENSG00000274922 | | 67.92582 | -0.04444876 | 0.3311638 | -0.1512246 | 0.8797986 | 0.943451 |
| ENSG00000125755 | SYMPK | 5686.781602 | -0.044614094 | 0.115625078 | -0.372928574 | 0.7092016 | 0.841795 |
| ENSG00000179447 | SLC24A3-AS1 | 1.069237 | -0.04461637 | 0.7427657 | -0.1889943 | 0.8500973 | NA |
| ENSG00000131653 | TRAF7 | 9337.534565 | -0.044659099 | 0.131800765 | -0.362825629 | 0.71673515 | 0.845902 |
| ENSG00000232693 | | 1.327494 | -0.04466455 | 0.7103281 | -0.1411045 | 0.8877874 | NA |
| ENSG00000249014 | HMGN2P4 | 108.1677 | -0.04469959 | 0.3256023 | -0.1552206 | 0.8766474 | 0.9418 |
| ENSG00000199568 | RNU5A-1 | 5.12132 | -0.04479483 | 0.6297834 | -0.1219226 | 0.9029603 | 0.955046 |
| ENSG00000178935 | ZNF552 | 927.0856 | -0.04494508 | 0.1242062 | -0.3732407 | 0.7089693 | 0.841784 |
| ENSG00000136877 | FPGS | 3548.302629 | -0.04496487 | 0.19095482 | -0.240485103 | 0.80995421 | 0.902923 |
| ENSG00000257331 | RACGAP1P1 | 2.483506 | -0.04497152 | 0.730853 | -0.1419355 | 0.8871309 | 0.947051 |
| ENSG00000142408 | CACNG8 | 47.87737 | -0.04510611 | 0.4210532 | -0.1223382 | 0.9026312 | 0.955002 |
| ENSG00000260860 | | 1.16655 | -0.04511135 | 0.7494893 | -0.1821622 | 0.8554555 | NA |
| ENSG00000272518 | | 23.76605 | -0.0451398 | 0.4974967 | -0.1140972 | 0.9091607 | 0.957674 |
| ENSG00000279786 | | 1.761857 | -0.04516327 | 0.7003157 | -0.1329187 | 0.8942577 | 0.950906 |
| ENSG00000261633 | | 2.286139 | -0.04517441 | 0.6625847 | -0.1309109 | 0.8958458 | 0.951863 |
| ENSG00000196381 | ZNF781 | 16.84741 | -0.0452144 | 0.5378962 | -0.1196222 | 0.9047824 | 0.955917 |
| ENSG00000104915 | STX10 | 1801.716457 | -0.045231844 | 0.191925867 | -0.241915296 | 0.8088458 | 0.902306 |
| ENSG00000123908 | AGO2 | 8112.099956 | -0.045264994 | 0.250907644 | -0.194057958 | 0.8461305 | 0.924846 |
| ENSG00000232043 | | 19.09925 | -0.04531029 | 0.4828769 | -0.1180297 | 0.9060441 | 0.956276 |
| ENSG00000270714 | MICOS10P2 | 37.45648 | -0.04532191 | 0.4045082 | -0.1345274 | 0.8929856 | 0.950376 |
| ENSG00000224401 | RPL7P57 | 29.52973 | -0.04533069 | 0.3291505 | -0.1531059 | 0.8783147 | 0.94267 |
| ENSG00000224431 | | 12.56323 | -0.04539115 | 0.545112 | -0.1183438 | 0.9057952 | 0.956179 |
| ENSG00000159259 | CHAF1B | 823.2813 | -0.04540191 | 0.4650995 | -0.1233043 | 0.9018661 | 0.954853 |
| ENSG00000170191 | NANP | 1106.919 | -0.04541932 | 0.164029 | -0.2878002 | 0.7734997 | 0.880624 |
| ENSG00000231507 | LINC01353 | 1.985497 | -0.04544068 | 0.7039306 | -0.1350724 | 0.8925546 | 0.950211 |
| ENSG00000196547 | MAN2A2 | 6741.761 | -0.0454533 | 0.2037339 | -0.2142309 | 0.830367 | 0.914791 |
| ENSG00000204673 | AKT1S1 | 5859.598 | -0.04566612 | 0.1915677 | -0.230095 | 0.818018 | 0.907686 |
| ENSG00000148337 | CIZ1 | 9290.665 | -0.04567015 | 0.1193792 | -0.3873001 | 0.698534 | 0.835699 |
| ENSG00000197870 | PRB3 | 4.900404 | -0.04568145 | 0.6141074 | -0.1216906 | 0.903144 | 0.955125 |
| ENSG00000174429 | ABRA | 0.7754938 | -0.04569494 | 0.7691083 | -0.2721313 | 0.7855211 | NA |
| ENSG00000253618 | GRPEL2-AS1 | 8.070902 | -0.04581461 | 0.4966738 | -0.1226187 | 0.9024091 | 0.954999 |
| ENSG00000160799 | CCDC12 | 1512.574 | -0.0458195 | 0.2159677 | -0.2230656 | 0.8234844 | 0.910531 |
| ENSG00000140718 | FTO | 1943.654 | -0.04582824 | 0.1797427 | -0.2702554 | 0.7869638 | 0.888944 |

| | | | | | | | |
|---|---|---|---|---|---|---|---|
| ENSG00000264012 | | 0.9986971 | -0.04583092 | 0.7304086 | -0.1576019 | 0.8747705 | NA |
| ENSG00000244226 | ILF2P1 | 1.076728 | -0.04584629 | 0.7477527 | -0.1833668 | 0.8545102 | NA |
| ENSG00000232341 | RPL4P2 | 20.73103 | -0.04589796 | 0.4170269 | -0.1321327 | 0.8948794 | 0.951385 |
| ENSG00000277969 | | 167.5852 | -0.0459337 | 0.2571267 | -0.1906425 | 0.8488057 | 0.926586 |
| ENSG00000099899 | TRMT2A | 2919.526257 | -0.045965893 | 0.159012529 | -0.305772376 | 0.75977796 | 0.871913 |
| ENSG00000143224 | PPOX | 1034.726 | -0.04597031 | 0.2478824 | -0.2032319 | 0.8389538 | 0.919941 |
| ENSG00000204386 | NEU1 | 7019.093 | -0.04604273 | 0.2451711 | -0.1795134 | 0.8575346 | 0.931245 |
| ENSG00000263466 | | 0.9544512 | -0.04608312 | 0.7200675 | -0.1596411 | 0.8731638 | NA |
| ENSG00000255000 | | 1.164154 | -0.04620323 | 0.735696 | -0.1631843 | 0.8703733 | NA |
| ENSG00000163191 | S100A11 | 21306.83 | -0.04625322 | 0.1929267 | -0.2631468 | 0.7924375 | 0.892652 |
| ENSG00000270050 | | 2.158694 | -0.04629108 | 0.6819582 | -0.1381582 | 0.8901154 | 0.948976 |
| ENSG00000165983 | PTER | 816.693 | -0.04629869 | 0.2012607 | -0.2576957 | 0.7966418 | 0.895081 |
| ENSG00000070269 | TMEM260 | 1560.214141 | -0.046331938 | 0.266045072 | -0.194924807 | 0.84545181 | 0.924357 |
| ENSG00000224273 | | 0.8494465 | -0.04633523 | 0.7271656 | -0.1810032 | 0.8563651 | NA |
| ENSG00000267632 | | 40.0377 | -0.04641158 | 0.4388488 | -0.1295936 | 0.8968879 | 0.952406 |
| ENSG00000259802 | | 200.2247 | -0.0464132 | 0.3010779 | -0.1691273 | 0.8656965 | 0.935977 |
| ENSG00000115365 | LANCL1 | 3178.361184 | -0.046432787 | 0.179353451 | -0.268643609 | 0.78820395 | 0.889844 |
| ENSG00000164022 | AIMP1 | 5215.719 | -0.04643886 | 0.1382482 | -0.3405264 | 0.7334602 | 0.855457 |
| ENSG00000260578 | | 5.977283 | -0.04646978 | 0.6096784 | -0.1174304 | 0.906519 | 0.95662 |
| ENSG00000144736 | SHQ1 | 1251.409 | -0.04648803 | 0.2382376 | -0.2068054 | 0.8361618 | 0.91794 |
| ENSG00000177669 | MBOAT4 | 1.287669 | -0.04650247 | 0.7158118 | -0.1604016 | 0.8725647 | NA |
| ENSG00000272825 | | 16.9604 | -0.0465043 | 0.4565856 | -0.1268512 | 0.8990582 | 0.953529 |
| ENSG00000257443 | | 1.301645 | -0.04652299 | 0.7091924 | -0.1458656 | 0.8840275 | NA |
| ENSG00000232969 | | 2.60822 | -0.04656767 | 0.6729262 | -0.1316469 | 0.8952636 | 0.951603 |
| ENSG00000115274 | INO80B | 91.42514273 | -0.046573531 | 0.250805889 | -0.197458277 | 0.84346893 | 0.923025 |
| ENSG00000120727 | PAIP2 | 3713.964565 | -0.046695899 | 0.129507457 | -0.383653649 | 0.70123517 | 0.83727 |
| ENSG00000261094 | | 150.0495 | -0.04670216 | 0.3323367 | -0.1497659 | 0.8809493 | 0.944131 |
| ENSG00000267011 | | 8.421598 | -0.04671298 | 0.5043819 | -0.1212369 | 0.9035034 | 0.955202 |
| ENSG00000232690 | HSPE1P9 | 3.966388 | -0.04677052 | 0.6882439 | -0.1395995 | 0.8889765 | 0.948317 |
| ENSG00000287310 | | 0.76241 | -0.04680842 | 0.732203 | -0.1702482 | 0.8648149 | NA |
| ENSG00000164185 | ZNF474 | 3.005173 | -0.04685298 | 0.6506728 | -0.1261127 | 0.8996427 | 0.953929 |
| ENSG00000280311 | | 4.612909 | -0.04686657 | 0.6473999 | -0.1317928 | 0.8951482 | 0.951594 |
| ENSG00000025770 | NCAPH2 | 2259.793706 | -0.046895745 | 0.313965036 | -0.160441311 | 0.87253345 | 0.939577 |

| | | | | | | | |
|---|---|---|---|---|---|---|---|
| ENSG00000182347 | PDSS1P1 | 0.9495991 | -0.04691138 | 0.7174209 | -0.1605923 | 0.8724145 | NA |
| ENSG00000130254 | SAFB2 | 3871.212089 | -0.046926161 | 0.149989335 | -0.310013859 | 0.75655042 | 0.869938 |
| ENSG00000185567 | AHNAK2 | 929.5605 | -0.04694925 | 0.2323122 | -0.2100602 | 0.8336207 | 0.916526 |
| ENSG00000143543 | JTB | 1214.54 | -0.04700035 | 0.1774072 | -0.2749011 | 0.7833922 | 0.886723 |
| ENSG00000174343 | CHRNA9 | 1.268484 | -0.04701351 | 0.722675 | -0.18396 | 0.8540449 | NA |
| ENSG00000182551 | ADI1 | 3996.748 | -0.04702168 | 0.2650898 | -0.1681034 | 0.8665019 | 0.936211 |
| ENSG00000224988 | | 1.672528 | -0.04703321 | 0.7219258 | -0.1735687 | 0.8622044 | 0.933895 |
| ENSG00000272682 | | 1.234224 | -0.04709882 | 0.7150904 | -0.1622761 | 0.8710884 | NA |
| ENSG00000006042 | TMEM98 | 24.07697695 | -0.047149542 | 0.469159981 | -0.130553344 | 0.89612865 | 0.952065 |
| ENSG00000140830 | TXNL4B | 1410.551 | -0.04719143 | 0.1641904 | -0.2957842 | 0.7673949 | 0.876603 |
| ENSG00000214269 | LGMNP1 | 29.25451 | -0.04723862 | 0.3593745 | -0.1485375 | 0.8819186 | 0.944788 |
| ENSG00000206949 | RNU6-1324P | 1.516911 | -0.04728604 | 0.7869624 | -0.5600362 | 0.5754548 | NA |
| ENSG00000272226 | | 6.595853 | -0.04732283 | 0.6398889 | -0.1226132 | 0.9024134 | 0.954999 |
| ENSG00000173898 | SPTBN2 | 9467.209 | -0.04743532 | 0.2240923 | -0.2060439 | 0.8367566 | 0.918479 |
| ENSG00000214558 | | 186.6597 | -0.04747559 | 0.2531691 | -0.1899954 | 0.8493127 | 0.926786 |
| ENSG00000176731 | RBIS | 1124.771 | -0.04752366 | 0.1706223 | -0.2884947 | 0.7729681 | 0.880396 |
| ENSG00000166997 | CNPY4 | 427.3944 | -0.04755125 | 0.2341207 | -0.2151579 | 0.8296442 | 0.914205 |
| ENSG00000100227 | POLDIP3 | 4809.29092 | -0.047557355 | 0.094576184 | -0.524018976 | 0.60026534 | 0.766374 |
| ENSG00000213226 | | 1.91896 | -0.04764963 | 0.6786388 | -0.1456178 | 0.8842231 | 0.945618 |
| ENSG00000273487 | | 1.475425 | -0.04765875 | 0.7201813 | -0.1746422 | 0.8613608 | NA |
| ENSG00000253854 | | 92.50251 | -0.04770774 | 0.3749302 | -0.1509851 | 0.8799875 | 0.943516 |
| ENSG00000136280 | CCM2 | 2321.193721 | -0.047826329 | 0.22371633 | -0.233716147 | 0.81520535 | 0.905901 |
| ENSG00000175595 | ERCC4 | 710.6548 | -0.0478643 | 0.3077133 | -0.1829551 | 0.8548332 | 0.929731 |
| ENSG00000285000 | | 2.126231 | -0.04786754 | 0.6690888 | -0.1480796 | 0.88228 | 0.944871 |
| ENSG00000077984 | CST7 | 3.067813984 | -0.047943687 | 0.656112388 | -0.122604846 | 0.90242002 | 0.954999 |
| ENSG00000279048 | | 73.47655 | -0.04806851 | 0.4313741 | -0.1371579 | 0.890906 | 0.949364 |
| ENSG00000200788 | | 3.486139 | -0.04807187 | 0.6667571 | -0.1413013 | 0.8876319 | 0.94732 |
| ENSG00000068028 | RASSF1 | 771.9104983 | -0.048076679 | 0.17405388 | -0.283444989 | 0.77683575 | 0.882518 |
| ENSG00000020256 | ZFP64 | 1419.870921 | -0.048081506 | 0.117891935 | -0.421596308 | 0.6733197 | 0.818528 |
| ENSG00000162929 | KIAA1841 | 721.7022 | -0.04814257 | 0.2268315 | -0.2239183 | 0.8228209 | 0.910032 |
| ENSG00000233836 | | 174.2467 | -0.04817169 | 0.6953783 | -0.1140099 | 0.9092299 | 0.957674 |
| ENSG00000226496 | LINC00323 | 13.4944 | -0.04822663 | 0.611547 | -0.1102716 | 0.912194 | 0.958851 |
| ENSG00000171222 | SCAND1 | 2466.511 | -0.04823385 | 0.2306501 | -0.2193995 | 0.8263389 | 0.912268 |

| | | | | | | | |
|---|---|---|---|---|---|---|---|
| ENSG00000237550 | | 8965.218 | -0.04835848 | 0.1929747 | -0.2744736 | 0.7837207 | 0.886751 |
| ENSG00000230079 | STK24P1 | 1.694082 | -0.04841252 | 0.708879 | -0.1460236 | 0.8839028 | 0.945465 |
| ENSG00000138769 | CDKL2 | 31.23187979 | -0.048426431 | 0.461514363 | -0.138115779 | 0.89014892 | 0.948975 |
| ENSG00000083814 | ZNF671 | 7.415733221 | -0.048445605 | 0.552742736 | -0.125992426 | 0.89973792 | 0.953966 |
| ENSG00000137509 | PRCP | 2395.547132 | -0.048455512 | 0.182000995 | -0.283499831 | 0.77679371 | 0.882518 |
| ENSG00000119714 | GPR68 | 305.0940426 | -0.048596663 | 0.362197271 | -0.162659811 | 0.87078629 | 0.938387 |
| ENSG00000159231 | CBR3 | 100.8619 | -0.04860957 | 0.3988341 | -0.1443674 | 0.8852103 | 0.94607 |
| ENSG00000167186 | COQ7 | 935.0432 | -0.04861069 | 0.1410605 | -0.3564746 | 0.7214852 | 0.848466 |
| ENSG00000225591 | | 3.99565 | -0.0487735 | 0.6426116 | -0.1340939 | 0.8933283 | 0.950572 |
| ENSG00000099381 | SETD1A | 3247.46142 | -0.04883149 | 0.192405951 | -0.266583005 | 0.78979024 | 0.890982 |
| ENSG00000226049 | TLK2P1 | 2701.567 | -0.04892502 | 0.1416143 | -0.3844183 | 0.7006684 | 0.836972 |
| ENSG00000160392 | C19orf47 | 741.9247 | -0.04893852 | 0.2304169 | -0.2464246 | 0.8053535 | 0.90031 |
| ENSG00000278576 | | 14.54102 | -0.04900936 | 0.5071516 | -0.1292868 | 0.8971307 | 0.952406 |
| ENSG00000005302 | MSL3 | 2274.203473 | -0.049016154 | 0.15981237 | -0.333897836 | 0.73845665 | 0.858824 |
| ENSG00000143178 | TBX19 | 108.8699 | -0.04910682 | 0.2932729 | -0.18235 | 0.8553081 | 0.929935 |
| ENSG00000196214 | ZNF766 | 1321.883 | -0.04912402 | 0.1441583 | -0.343054 | 0.7315578 | 0.854371 |
| ENSG00000138758 | SEPTIN11 | 11000.41316 | -0.049129808 | 0.248827977 | -0.211618537 | 0.83240465 | 0.915631 |
| ENSG00000254858 | MPV17L2 | 847.0796 | -0.04915216 | 0.2294645 | -0.2313024 | 0.8170799 | 0.907036 |
| ENSG00000151292 | CSNK1G3 | 2532.256 | -0.04926397 | 0.1307261 | -0.4101136 | 0.6817226 | 0.823818 |
| ENSG00000221869 | CEBPD | 866.4233 | -0.04927378 | 0.2853934 | -0.1855884 | 0.8527676 | 0.928421 |
| ENSG00000115520 | COQ10B | 1790.698076 | -0.049274408 | 0.176872404 | -0.299057906 | 0.76489586 | 0.875216 |
| ENSG00000161265 | U2AF1L4 | 202.6638 | -0.04927465 | 0.2441369 | -0.1998184 | 0.8416226 | 0.92184 |
| ENSG00000073350 | LLGL2 | 23152.82501 | -0.04927515 | 0.197231946 | -0.29618978 | 0.76708515 | 0.876551 |
| ENSG00000028839 | TBPL1 | 702.76816 | -0.049288548 | 0.19047602 | -0.268565905 | 0.78826375 | 0.889844 |
| ENSG00000262248 | | 8.452995 | -0.04929189 | 0.5256551 | -0.1302786 | 0.896346 | 0.952182 |
| ENSG00000111328 | CDK2AP1 | 6695.10562 | -0.049384295 | 0.262209087 | -0.20727203 | 0.83579742 | 0.91773 |
| ENSG00000231612 | | 1.791054 | -0.04947677 | 0.6830993 | -0.1644629 | 0.8693667 | 0.937389 |
| ENSG00000147121 | KRBOX4 | 840.9932 | -0.04948174 | 0.1787769 | -0.2873905 | 0.7738133 | 0.880717 |
| ENSG00000204262 | COL5A2 | 445.6472 | -0.04949917 | 0.3269481 | -0.1907548 | 0.8487177 | 0.926556 |
| ENSG00000155903 | RASA2 | 1198.257 | -0.04952941 | 0.213218 | -0.2573531 | 0.7969062 | 0.895202 |
| ENSG00000197756 | RPL37A | 19720.78 | -0.04953939 | 0.1571354 | -0.3444888 | 0.7304787 | 0.853899 |
| ENSG00000066185 | ZMYND12 | 25.27909339 | -0.049572814 | 0.425642213 | -0.140892341 | 0.88795499 | 0.947572 |
| ENSG00000200913 | SNORD46 | 35.93767 | -0.04962293 | 0.3040865 | -0.177895 | 0.8588055 | 0.931977 |

| | | | | | | | |
|---|---|---|---|---|---|---|---|
| ENSG00000143549 | TPM3 | 16009.96 | -0.04966125 | 0.2286812 | -0.2302786 | 0.8178753 | 0.907653 |
| ENSG00000260790 | | 2.318776 | -0.04972001 | 0.6726989 | -0.1331381 | 0.8940842 | 0.950882 |
| ENSG00000131507 | NDFIP1 | 5353.473333 | -0.049782016 | 0.174063389 | -0.305049267 | 0.76032863 | 0.872319 |
| ENSG00000285091 | | 17.18406 | -0.04978951 | 0.518439 | -0.1283426 | 0.8978779 | 0.95278 |
| ENSG00000258565 | BLZF2P | 1.456275 | -0.0497936 | 0.6993827 | -0.1529722 | 0.8784202 | NA |
| ENSG00000263335 | | 21.91834 | -0.04987817 | 0.3979917 | -0.1483347 | 0.8820786 | 0.944807 |
| ENSG00000213959 | | 1.717716 | -0.04988231 | 0.7390327 | -0.1912814 | 0.8483051 | 0.926316 |
| ENSG00000166925 | TSC22D4 | 3783.964 | -0.04993009 | 0.1796325 | -0.2830471 | 0.7771407 | 0.882585 |
| ENSG00000166743 | ACSM1 | 4.431007 | -0.04996767 | 0.6011575 | -0.1291679 | 0.8972248 | 0.952412 |
| ENSG00000210082 | MT-RNR2 | 161836.8 | -0.04997382 | 0.1808115 | -0.287017 | 0.7740993 | 0.880805 |
| ENSG00000223505 | RPL35AP5 | 6.537359 | -0.04999157 | 0.6233992 | -0.1328823 | 0.8942865 | 0.950906 |
| ENSG00000109586 | GALNT7 | 2987.03576 | -0.050116094 | 0.23878019 | -0.21782989 | 0.82756166 | 0.913064 |
| ENSG00000100347 | SAMM50 | 2202.52774 | -0.050206661 | 0.20475591 | -0.256200021 | 0.79779639 | 0.895734 |
| ENSG00000232363 | | 2.243301 | -0.05021526 | 0.7140968 | -0.1549883 | 0.8768306 | 0.941882 |
| ENSG00000236015 | | 57.35951 | -0.0502223 | 0.3711607 | -0.1563343 | 0.8757695 | 0.941283 |
| ENSG00000185883 | ATP6V0C | 20.26271 | -0.05028701 | 0.4207493 | -0.141542 | 0.8874417 | 0.94723 |
| ENSG00000240509 | RPL34P18 | 859.1137 | -0.05031743 | 0.1895146 | -0.2966534 | 0.7667311 | 0.876327 |
| ENSG00000170837 | GPR27 | 789.2723 | -0.05032012 | 0.1484262 | -0.3379111 | 0.7354302 | 0.856647 |
| ENSG00000160741 | CRTC2 | 3995.303 | -0.05034687 | 0.1409919 | -0.332484 | 0.7395238 | 0.859371 |
| ENSG00000182405 | PGBD4 | 284.5506 | -0.05046095 | 0.1913763 | -0.2754075 | 0.7830031 | 0.886458 |
| ENSG00000235740 | PHACTR2-AS1 | 0.9139797 | -0.05048342 | 0.7355995 | -0.2099145 | 0.8337344 | NA |
| ENSG00000125447 | GGA3 | 4998.283443 | -0.050493025 | 0.131434778 | -0.47759504 | 0.63293846 | 0.789652 |
| ENSG00000154217 | PITPNC1 | 3310.326 | -0.05069406 | 0.1120595 | -0.4615626 | 0.644395 | 0.797658 |
| ENSG00000212663 | | 0.7523421 | -0.05078027 | 0.7467072 | -0.2123695 | 0.8318188 | NA |
| ENSG00000261423 | TMEM202-AS1 | 58.13757 | -0.05080786 | 0.3405756 | -0.1683778 | 0.8662861 | 0.936211 |
| ENSG00000214222 | TUBBP2 | 2.556312 | -0.05083192 | 0.6875516 | -0.1570787 | 0.8751828 | 0.941025 |
| ENSG00000286503 | | 8.337955 | -0.0509219 | 0.5273227 | -0.130597 | 0.8960941 | 0.952065 |
| ENSG00000262481 | TMEM256-PLSCI | 1.919338 | -0.05096187 | 0.686154 | -0.1559179 | 0.8760978 | 0.941552 |
| ENSG00000172667 | ZMAT3 | 1871.67 | -0.05105645 | 0.3596686 | -0.1379658 | 0.8902675 | 0.949064 |
| ENSG00000165934 | CPSF2 | 5601.288 | -0.05106989 | 0.1604349 | -0.3285942 | 0.7424624 | 0.861313 |
| ENSG00000134461 | ANKRD16 | 822.4354304 | -0.051078574 | 0.163175852 | -0.329602034 | 0.74170069 | 0.860798 |
| ENSG00000189057 | FAM111B | 1223.634 | -0.05114736 | 0.5820313 | -0.1422664 | 0.8868696 | 0.946923 |
| ENSG00000170619 | COMMD5 | 2914.861 | -0.05114819 | 0.1832502 | -0.2985545 | 0.76528 | 0.875483 |

| | | | | | | | |
|---|---|---|---|---|---|---|---|
| ENSG00000259759 | | 0.9451875 | -0.05125258 | 0.7487137 | -0.2648892 | 0.7910948 | NA |
| ENSG00000112851 | ERBIN | 5503.253464 | -0.05130074 | 0.150956902 | -0.351157482 | 0.72547021 | 0.850907 |
| ENSG00000184661 | CDCA2 | 748.3147 | -0.05132527 | 0.4583462 | -0.1451136 | 0.8846212 | 0.94582 |
| ENSG00000132676 | DAP3 | 9331.422653 | -0.051342446 | 0.119909471 | -0.395120898 | 0.69275367 | 0.831284 |
| ENSG00000185418 | TARS3 | 728.9888 | -0.05137703 | 0.1697173 | -0.3197431 | 0.7491631 | 0.865715 |
| ENSG00000236451 | | 7.496683 | -0.05139753 | 0.5630223 | -0.1366036 | 0.8913442 | 0.949602 |
| ENSG00000286874 | | 30.41973 | -0.0514385 | 0.4171868 | -0.1466427 | 0.883414 | 0.945437 |
| ENSG00000150477 | KIAA1328 | 328.5382 | -0.05145657 | 0.1686324 | -0.3222634 | 0.7472532 | 0.864403 |
| ENSG00000172086 | KRCC1 | 2287.061 | -0.05147344 | 0.1939724 | -0.2718745 | 0.7857185 | 0.88799 |
| ENSG00000272248 | | 5.17454 | -0.05149821 | 0.5798409 | -0.129787 | 0.896735 | 0.952396 |
| ENSG00000057757 | PITHD1 | 1766.746614 | -0.051521676 | 0.198339702 | -0.270919592 | 0.78645288 | 0.888518 |
| ENSG00000119396 | RAB14 | 6895.939674 | -0.05158036 | 0.102992428 | -0.507100833 | 0.61208407 | 0.774883 |
| ENSG00000023516 | AKAP11 | 5322.331493 | -0.051593884 | 0.280458778 | -0.201203575 | 0.8405394 | 0.921114 |
| ENSG00000049239 | H6PD | 2233.997404 | -0.051609709 | 0.249165184 | -0.217661112 | 0.82769317 | 0.913119 |
| ENSG00000134852 | CLOCK | 3112.237236 | -0.051717879 | 0.190056126 | -0.284632845 | 0.77592545 | 0.881987 |
| ENSG00000074071 | MRPS34 | 5611.558112 | -0.051723704 | 0.197585218 | -0.297244146 | 0.76628012 | 0.876045 |
| ENSG00000250299 | MRPS31P4 | 48.75247 | -0.05178525 | 0.2849075 | -0.1937069 | 0.8464054 | 0.925019 |
| ENSG00000231731 | | 63.55311 | -0.05180619 | 0.2661811 | -0.2079493 | 0.8352686 | 0.917301 |
| ENSG00000284575 | MIR4793 | 0.9042618 | -0.05188194 | 0.7371364 | -0.2039699 | 0.838377 | NA |
| ENSG00000165490 | DDIAS | 886.7835 | -0.0519372 | 0.5141763 | -0.1456417 | 0.8842042 | 0.945618 |
| ENSG00000142655 | PEX14 | 802.8697 | -0.05203158 | 0.1978889 | -0.2738428 | 0.7842054 | 0.887074 |
| ENSG00000213253 | RPL12P42 | 11.76509 | -0.05213996 | 0.5266556 | -0.1288712 | 0.8974595 | 0.952565 |
| ENSG00000223650 | UHRF2P1 | 2.76462 | -0.05222758 | 0.6852879 | -0.1508162 | 0.8801207 | 0.943585 |
| ENSG00000103423 | DNAJA3 | 7317.048764 | -0.05226307 | 0.160432413 | -0.325956024 | 0.74445763 | 0.862302 |
| ENSG00000287884 | | 1.08192 | -0.0524271 | 0.7413764 | -0.2129089 | 0.831398 | NA |
| ENSG00000216802 | | 0.7848991 | -0.05245809 | 0.7394004 | -0.2114024 | 0.8325732 | NA |
| ENSG00000102898 | NUTF2 | 5906.680328 | -0.052540439 | 0.190715388 | -0.280686879 | 0.77895057 | 0.883841 |
| ENSG00000253848 | | 1.235385 | -0.05266374 | 0.7206452 | -0.1741877 | 0.861718 | NA |
| ENSG00000037749 | MFAP3 | 1443.581998 | -0.052670281 | 0.305198053 | -0.204232437 | 0.83817186 | 0.919311 |
| ENSG00000200169 | RNU5D-1 | 0.8612932 | -0.05275511 | 0.7339584 | -0.1985122 | 0.8426444 | NA |
| ENSG00000241146 | RPL7P41 | 1.242476 | -0.05278218 | 0.7587812 | -0.2794645 | 0.7798884 | NA |
| ENSG00000135677 | GNS | 12498.75889 | -0.052809317 | 0.281487237 | -0.201870275 | 0.84001814 | 0.920842 |
| ENSG00000272459 | FAM193B-DT | 5.381269 | -0.05284662 | 0.5704284 | -0.1348357 | 0.8927418 | 0.950215 |

| | | | | | | | |
|---|---|---|---|---|---|---|---|
| ENSG00000171467 | ZNF318 | 2771.956 | -0.05289147 | 0.2412056 | -0.2337933 | 0.8151455 | 0.905901 |
| ENSG00000273033 | LINC02035 | 337.5508 | -0.0528973 | 0.3211813 | -0.1824107 | 0.8552605 | 0.929935 |
| ENSG00000099810 | MTAP | 26.22715134 | -0.052922541 | 0.502834421 | -0.148010723 | 0.88233431 | 0.944891 |
| ENSG00000278493 | | 3.135532 | -0.05296315 | 0.6723526 | -0.1543027 | 0.8773711 | 0.94216 |
| ENSG00000129103 | SUMF2 | 5300.123331 | -0.052990745 | 0.129979447 | -0.418968029 | 0.6752395 | 0.81946 |
| ENSG00000231563 | | 22.92767 | -0.05301285 | 0.4537236 | -0.1428765 | 0.8863877 | 0.946675 |
| ENSG00000236859 | NIFK-AS1 | 274.6106 | -0.05303236 | 0.1784835 | -0.3037448 | 0.7613224 | 0.872735 |
| ENSG00000137770 | CTDSPL2 | 2018.02899 | -0.053039955 | 0.213667926 | -0.263393953 | 0.79224695 | 0.892483 |
| ENSG00000213540 | TXN2P1 | 1.652795 | -0.05305808 | 0.73842 | -0.2159119 | 0.8290564 | 0.913802 |
| ENSG00000274124 | | 3.458828 | -0.05308038 | 0.6724113 | -0.1460935 | 0.8838476 | 0.945465 |
| ENSG00000259070 | LINC00639 | 7.015458 | -0.05308711 | 0.5961264 | -0.1450832 | 0.8846452 | 0.94582 |
| ENSG00000071967 | CYBRD1 | 169.4035425 | -0.053115385 | 0.489459009 | -0.127467911 | 0.89857007 | 0.953206 |
| ENSG00000005448 | WDR54 | 1967.52817 | -0.053128271 | 0.264589257 | -0.215339562 | 0.82950259 | 0.91418 |
| ENSG00000275675 | | 0.7920897 | -0.05320849 | 0.7529667 | -0.2530902 | 0.8001985 | NA |
| ENSG00000034152 | MAP2K3 | 3196.762811 | -0.053258616 | 0.209376874 | -0.26500703 | 0.79100405 | 0.891628 |
| ENSG00000270482 | | 8.932529 | -0.05329261 | 0.5569768 | -0.1366606 | 0.8912991 | 0.949592 |
| ENSG00000112319 | EYA4 | 74.91085356 | -0.05336247 | 0.44526781 | -0.150207453 | 0.88060095 | 0.943813 |
| ENSG00000277152 | | 3.990255 | -0.05339962 | 0.6426452 | -0.1402492 | 0.8884631 | 0.947901 |
| ENSG00000287030 | | 2.068432 | -0.05341744 | 0.6564851 | -0.1483491 | 0.8820673 | 0.944807 |
| ENSG00000283828 | | 44.32755 | -0.05345428 | 0.3218015 | -0.1740608 | 0.8618177 | 0.933759 |
| ENSG00000101911 | PRPS2 | 2278.212256 | -0.053521009 | 0.279340096 | -0.209785643 | 0.83383498 | 0.91665 |
| ENSG00000112053 | SLC26A8 | 7.486575465 | -0.053548979 | 0.655973944 | -0.133856724 | 0.89351587 | 0.950721 |
| ENSG00000270190 | | 2.63243 | -0.05356481 | 0.7226458 | -0.1710417 | 0.864191 | 0.934912 |
| ENSG00000146223 | RPL7L1 | 10379.6 | -0.05357331 | 0.1614477 | -0.347573 | 0.7281609 | 0.852439 |
| ENSG00000109332 | UBE2D3 | 11323.99669 | -0.053575131 | 0.099470793 | -0.56494014 | 0.57211448 | 0.745598 |
| ENSG00000231345 | BEND3P1 | 270.1748 | -0.05358828 | 0.3353189 | -0.2079912 | 0.8352358 | 0.917301 |
| ENSG00000069849 | ATP1B3 | 5579.109677 | -0.053590509 | 0.184465277 | -0.304853708 | 0.76047757 | 0.872349 |
| ENSG00000255355 | | 2.169152 | -0.05359593 | 0.6630075 | -0.1604044 | 0.8725625 | 0.939577 |
| ENSG00000226706 | | 25.50432 | -0.05367835 | 0.3516752 | -0.1728833 | 0.8627431 | 0.9341 |
| ENSG00000227857 | | 26.31765 | -0.05376086 | 0.4360724 | -0.1511545 | 0.8798538 | 0.943451 |
| ENSG00000275506 | H3P25 | 0.762804 | -0.05387323 | 0.7480036 | -0.227034 | 0.8203973 | NA |
| ENSG00000173335 | CST9 | 14.85451 | -0.05387996 | 0.7830034 | -0.5436801 | 0.5866616 | 0.756284 |
| ENSG00000269044 | | 113.261 | -0.05401999 | 0.3144 | -0.1939981 | 0.8461774 | 0.924846 |

| | | | | | | | |
|---|---|---|---|---|---|---|---|
| ENSG00000258744 | | 233.9928 | -0.05403232 | 0.2581797 | -0.2187555 | 0.8268405 | 0.912571 |
| ENSG00000213047 | DENND1B | 2309.6 | -0.05403931 | 0.2498978 | -0.2314661 | 0.8169527 | 0.906933 |
| ENSG00000259594 | | 14.13323 | -0.05405393 | 0.4695812 | -0.1468115 | 0.8832808 | 0.945333 |
| ENSG00000256843 | | 9.644904 | -0.05406133 | 0.5564242 | -0.1418924 | 0.887165 | 0.947051 |
| ENSG00000239223 | RPL34P31 | 28.27675 | -0.0541297 | 0.3806608 | -0.158453 | 0.8740999 | 0.940279 |
| ENSG00000155816 | FMN2 | 3.986092 | -0.0542129 | 0.6532122 | -0.1569484 | 0.8752855 | 0.941098 |
| ENSG00000229835 | KHSRPP1 | 244.59 | -0.05436324 | 0.2645427 | -0.2206145 | 0.8253926 | 0.911882 |
| ENSG00000228084 | | 4.665922 | -0.05438269 | 0.6392008 | -0.1445601 | 0.8850582 | 0.946052 |
| ENSG00000183978 | COA3 | 2093.842 | -0.05441677 | 0.2207672 | -0.2629537 | 0.7925862 | 0.892702 |
| ENSG00000105258 | POLR2I | 361.7127488 | -0.05445402 | 0.257545645 | -0.228162097 | 0.81952023 | 0.908698 |
| ENSG00000235843 | | 1.239198 | -0.05449089 | 0.7438302 | -0.2091629 | 0.8343211 | NA |
| ENSG00000272186 | | 24.17622 | -0.0545786 | 0.3644098 | -0.1723822 | 0.8631371 | 0.934359 |
| ENSG00000078795 | PKD2L2 | 2.491429783 | -0.054588575 | 0.638213021 | -0.147778093 | 0.8825179 | 0.944952 |
| ENSG00000176108 | CHMP6 | 1374.183 | -0.05463062 | 0.2368966 | -0.2370119 | 0.8126476 | 0.904263 |
| ENSG00000146556 | WASH2P | 415.0056 | -0.05468254 | 0.1666276 | -0.3303718 | 0.7411191 | 0.860407 |
| ENSG00000153093 | ACOXL | 269.3745 | -0.05488486 | 0.4860198 | -0.1268166 | 0.8990856 | 0.953529 |
| ENSG00000160563 | MED27 | 650.6737 | -0.05491966 | 0.2334591 | -0.2426285 | 0.8082932 | 0.901868 |
| ENSG00000277952 | | 50.5443 | -0.05498528 | 0.3529349 | -0.1758379 | 0.8604213 | 0.932877 |
| ENSG00000108255 | CRYBA1 | 1.493432893 | -0.055036656 | 0.703630055 | -0.17556126 | 0.86063864 | NA |
| ENSG00000254481 | PTP4A2P2 | 12.80184 | -0.05505273 | 0.5015992 | -0.1461271 | 0.8838211 | 0.945465 |
| ENSG00000196656 | | 1292.273 | -0.05507375 | 0.2572022 | -0.2139872 | 0.830557 | 0.914887 |
| ENSG00000173540 | GMPPB | 1190.18 | -0.0551057 | 0.1858608 | -0.3088516 | 0.7574344 | 0.870653 |
| ENSG00000144596 | GRIP2 | 2.447008 | -0.05511267 | 0.6438325 | -0.1511879 | 0.8798275 | 0.943451 |
| ENSG00000157637 | SLC38A10 | 8197.045 | -0.05511698 | 0.1446127 | -0.4147759 | 0.678306 | 0.821268 |
| ENSG00000120526 | NUDCD1 | 4172.320375 | -0.055172708 | 0.235588732 | -0.248192239 | 0.80398567 | 0.899636 |
| ENSG00000272573 | MUSTN1 | 0.9287147 | -0.05517403 | 0.7423403 | -0.2336094 | 0.8152882 | NA |
| ENSG00000167941 | SOST | 1.800573 | -0.05523818 | 0.789402 | -0.6891984 | 0.4906984 | 0.681135 |
| ENSG00000266826 | IGBP1P2 | 4.449077 | -0.0553111 | 0.6716575 | -0.1450082 | 0.8847044 | 0.94582 |
| ENSG00000103326 | CAPN15 | 4144.579091 | -0.055311235 | 0.152860636 | -0.374886762 | 0.70774469 | 0.841108 |
| ENSG00000110514 | MADD | 7641.953111 | -0.055312039 | 0.124673211 | -0.420155811 | 0.67437163 | 0.818891 |
| ENSG00000179195 | ZNF664 | 15365.09 | -0.05532129 | 0.1558075 | -0.3754516 | 0.7073247 | 0.840865 |
| ENSG00000163312 | HELQ | 915.6641 | -0.05533685 | 0.09994643 | -0.5841777 | 0.5591007 | 0.735831 |
| ENSG00000247317 | LY6E-DT | 189.491 | -0.05535161 | 0.410822 | -0.1583247 | 0.8742009 | 0.94035 |

| | | | | | | | |
|---|---|---|---|---|---|---|---|
| ENSG00000223825 | DAZAP2P1 | 20.48363 | -0.05537401 | 0.3947327 | -0.1640511 | 0.8696909 | 0.93751 |
| ENSG00000213606 | AKR1B10P1 | 1.278505 | -0.05539302 | 0.790484 | -0.221859 | 0.8244237 | NA |
| ENSG00000131873 | CHSY1 | 2762.503284 | -0.055413292 | 0.154249839 | -0.370711198 | 0.71085265 | 0.842627 |
| ENSG00000169946 | ZFPM2 | 22.95438 | -0.05544999 | 0.67018 | -0.1549335 | 0.8768737 | 0.94189 |
| ENSG00000187634 | SAMD11 | 2377.66 | -0.05547935 | 0.3406907 | -0.1773648 | 0.8592218 | 0.932239 |
| ENSG00000143442 | POGZ | 13764.49 | -0.05550513 | 0.2266033 | -0.2594084 | 0.7953202 | 0.894202 |
| ENSG00000211513 | MIR320E | 3.08524 | -0.05552097 | 0.6341416 | -0.1506614 | 0.8802428 | 0.943649 |
| ENSG00000166913 | YWHAB | 17893.81 | -0.055531 | 0.1530077 | -0.3953457 | 0.6925877 | 0.831229 |
| ENSG00000228532 | SUMO2P21 | 1467.199 | -0.05558573 | 0.1738683 | -0.3299412 | 0.7414444 | 0.860542 |
| ENSG00000152133 | GPATCH11 | 518.8581 | -0.0556723 | 0.2442916 | -0.2430136 | 0.8079948 | 0.901698 |
| ENSG00000154485 | MMP21 | 26.99693 | -0.05568949 | 0.5001238 | -0.1463615 | 0.8836361 | 0.945465 |
| ENSG00000130702 | LAMA5 | 26163.84022 | -0.055692877 | 0.175772226 | -0.349345303 | 0.72683009 | 0.851749 |
| ENSG00000241549 | GUSBP2 | 100.6685 | -0.05571542 | 0.371214 | -0.1741457 | 0.861751 | 0.933725 |
| ENSG00000241556 | | 28.80827 | -0.05572646 | 0.3689533 | -0.1728553 | 0.8627652 | 0.9341 |
| ENSG00000235245 | | 121.3978 | -0.05582273 | 0.2369265 | -0.2369317 | 0.8127098 | 0.904263 |
| ENSG00000224647 | | 4.280619 | -0.05585908 | 0.633714 | -0.15386 | 0.8777201 | 0.942353 |
| ENSG00000112659 | CUL9 | 3334.974195 | -0.055982934 | 0.175818941 | -0.334627729 | 0.73790593 | 0.858607 |
| ENSG00000229589 | ACVR2B-AS1 | 131.2879 | -0.05600774 | 0.2637084 | -0.2269802 | 0.8204391 | 0.908942 |
| ENSG00000236953 | ZDHHC20-IT1 | 2.850751 | -0.05614582 | 0.6245079 | -0.1520137 | 0.8791761 | 0.943181 |
| ENSG00000249936 | RAC1P2 | 3847.174 | -0.05617544 | 0.1257006 | -0.4564582 | 0.6480605 | 0.800529 |
| ENSG00000213386 | | 2.087352 | -0.05621964 | 0.6751061 | -0.1599811 | 0.8728959 | 0.939821 |
| ENSG00000233045 | RPL15P5 | 9.251392 | -0.05625639 | 0.5023375 | -0.1458546 | 0.8840362 | 0.94557 |
| ENSG00000244527 | | 3.000818 | -0.05627564 | 0.6422557 | -0.1575379 | 0.8748209 | 0.940788 |
| ENSG00000207923 | MIR559 | 20.3573 | -0.05628884 | 0.377026 | -0.1729898 | 0.8626594 | 0.934093 |
| ENSG00000229950 | TFAP2A-AS1 | 71.56866 | -0.0563351 | 0.3070062 | -0.2006617 | 0.8409631 | 0.921419 |
| ENSG00000267309 | | 60.85425 | -0.05644214 | 0.2392802 | -0.2512888 | 0.8015908 | 0.898186 |
| ENSG00000203740 | METTL11B | 2.161713 | -0.0564726 | 0.6982612 | -0.180387 | 0.8568488 | 0.930755 |
| ENSG00000175048 | ZDHHC14 | 324.164 | -0.05657092 | 0.2057641 | -0.2870286 | 0.7740904 | 0.880805 |
| ENSG00000268108 | | 1.831197 | -0.05664863 | 0.7206276 | -0.1911653 | 0.8483961 | 0.926316 |
| ENSG00000263846 | CIAPIN1P | 68.6734 | -0.05667181 | 0.291089 | -0.2105289 | 0.8332549 | 0.91629 |
| ENSG00000182195 | LDOC1 | 6881.842 | -0.05668685 | 0.20011 | -0.290742 | 0.7712487 | 0.879022 |
| ENSG00000105298 | CACTIN | 1024.433277 | -0.0567951 | 0.149504271 | -0.377647632 | 0.70569238 | 0.840057 |
| ENSG00000234009 | RPL5P34 | 1471.97 | -0.05680562 | 0.1886247 | -0.3084435 | 0.7577449 | 0.870972 |

| | | | | | | | |
|---|---|---|---|---|---|---|---|
| ENSG00000261329 | | 8.634111 | -0.05695583 | 0.5348758 | -0.1483932 | 0.8820324 | 0.944807 |
| ENSG00000236552 | RPL13AP5 | 25188 | -0.05700954 | 0.1891082 | -0.3054205 | 0.7600459 | 0.872089 |
| ENSG00000231024 | | 1.313932 | -0.05709519 | 0.7409173 | -0.2300139 | 0.818081 | NA |
| ENSG00000269986 | | 1.244329 | -0.05710882 | 0.7309923 | -0.1964942 | 0.8442234 | NA |
| ENSG00000182141 | ZNF708 | 319.2082 | -0.05716377 | 0.249291 | -0.2380059 | 0.8118765 | 0.903941 |
| ENSG00000157500 | APPL1 | 2409.608 | -0.05717116 | 0.1688865 | -0.3500859 | 0.7262742 | 0.851411 |
| ENSG00000206822 | | 1.131405 | -0.05726035 | 0.6980512 | -0.1874971 | 0.8512709 | NA |
| ENSG00000234741 | GAS5 | 11245.27 | -0.0572664 | 0.2178713 | -0.2764819 | 0.7821779 | 0.885835 |
| ENSG00000228808 | HMGB3P4 | 1.650547 | -0.05732533 | 0.6880525 | -0.1760715 | 0.8602377 | 0.932732 |
| ENSG00000103832 | GOLGA8UP | 3.15552441 | -0.057343384 | 0.680499088 | -0.17761922 | 0.85902203 | 0.932098 |
| ENSG00000090686 | USP48 | 3880.18925 | -0.057378743 | 0.143522765 | -0.416121837 | 0.67732085 | 0.820599 |
| ENSG00000196922 | ZNF252P | 2403.593 | -0.05737911 | 0.1925566 | -0.3107811 | 0.7559671 | 0.869764 |
| ENSG00000177576 | C18orf32 | 137.7894 | -0.05743558 | 0.3161261 | -0.2012599 | 0.8404954 | 0.921114 |
| ENSG00000158526 | TSR2 | 3382.731 | -0.05744374 | 0.1563126 | -0.3772371 | 0.7059974 | 0.840133 |
| ENSG00000235888 | | 2.89481 | -0.05746595 | 0.6351462 | -0.1597545 | 0.8730745 | 0.939929 |
| ENSG00000261433 | | 1.27564 | -0.05751492 | 0.7005477 | -0.1968927 | 0.8439115 | NA |
| ENSG00000115145 | STAM2 | 1932.920478 | -0.057568016 | 0.168173985 | -0.341033041 | 0.73307871 | 0.855314 |
| ENSG00000213079 | SCAF8 | 3637.385 | -0.0576026 | 0.1663937 | -0.3635745 | 0.7161758 | 0.845492 |
| ENSG00000272663 | PPP1R21-DT | 12.52763 | -0.05762407 | 0.5320677 | -0.1482269 | 0.8821637 | 0.944859 |
| ENSG00000036054 | TBC1D23 | 2492.941954 | -0.05762444 | 0.12313391 | -0.470041205 | 0.63832558 | 0.793621 |
| ENSG00000120129 | DUSP1 | 6199.98033 | -0.057664236 | 0.458971863 | -0.161813778 | 0.8714525 | 0.938686 |
| ENSG00000179051 | RCC2 | 10448.84 | -0.05770123 | 0.1253135 | -0.5212604 | 0.6021854 | 0.767619 |
| ENSG00000147471 | PLPBP | 1706.846 | -0.057719 | 0.1292221 | -0.4597404 | 0.6457025 | 0.79857 |
| ENSG00000127743 | IL17B | 1.424859405 | -0.05778276 | 0.724797095 | -0.206428039 | 0.83645657 | NA |
| ENSG00000259845 | HERC2P10 | 8.021125 | -0.05783738 | 0.5539454 | -0.1536526 | 0.8778837 | 0.942366 |
| ENSG00000116885 | OSCP1 | 355.0812589 | -0.057843027 | 0.21242831 | -0.27870484 | 0.78047135 | 0.884964 |
| ENSG00000151690 | MFSD6 | 2207.352 | -0.05788039 | 0.2096248 | -0.2949327 | 0.7680453 | 0.877045 |
| ENSG00000253816 | GUSBP14 | 104.6043 | -0.05792123 | 0.354393 | -0.1858368 | 0.8525728 | 0.928323 |
| ENSG00000110442 | COMMD9 | 2254.561464 | -0.058076348 | 0.179933889 | -0.333073268 | 0.73907898 | 0.859031 |
| ENSG00000261441 | | 9.520022 | -0.05813066 | 0.5554597 | -0.1530183 | 0.8783838 | 0.94267 |
| ENSG00000286321 | | 1.410922 | -0.05822161 | 0.7904061 | -0.7461065 | 0.4556031 | NA |
| ENSG00000229780 | UBE2Q1-AS1 | 19.16064 | -0.05825845 | 0.5136324 | -0.1517886 | 0.8793537 | 0.943285 |
| ENSG00000129214 | SHBG | 27.92921388 | -0.058353487 | 0.394158723 | -0.172687645 | 0.86289696 | 0.934204 |

| | | | | | | | |
|---|---|---|---|---|---|---|---|
| ENSG00000260325 | HSPB9 | 5.631144 | -0.05842194 | 0.6016679 | -0.1516835 | 0.8794366 | 0.943308 |
| ENSG00000237970 | TMEM161BP1 | 9.337548 | -0.05845236 | 0.5271297 | -0.1524762 | 0.8788114 | 0.942904 |
| ENSG00000226312 | CFLAR-AS1 | 6.241268 | -0.0585158 | 0.6050424 | -0.1526771 | 0.8786529 | 0.94286 |
| ENSG00000156345 | CDK20 | 1055.121 | -0.0585401 | 0.16517 | -0.3581524 | 0.7202293 | 0.847766 |
| ENSG00000228484 | | 1.504944 | -0.05855349 | 0.7200559 | -0.2375896 | 0.8121995 | NA |
| ENSG00000115524 | SF3B1 | 24704.8506 | -0.058579412 | 0.131739416 | -0.452976961 | 0.65056533 | 0.802566 |
| ENSG00000276077 | | 0.9700604 | -0.05858413 | 0.7275791 | -0.2113067 | 0.8326479 | NA |
| ENSG00000205572 | SERF1B | 12.7311 | -0.05859617 | 0.430934 | -0.1633938 | 0.8702084 | 0.937954 |
| ENSG00000067704 | IARS2 | 9579.97697 | -0.058624778 | 0.204391555 | -0.322125708 | 0.74735746 | 0.864403 |
| ENSG00000112306 | RPS12 | 17878.51372 | -0.0586482 | 0.167195145 | -0.363239549 | 0.71642595 | 0.845649 |
| ENSG00000206885 | SNORA75 | 18.00729 | -0.05868294 | 0.4726856 | -0.1590846 | 0.8736022 | 0.940119 |
| ENSG00000233771 | CICP5 | 0.8531817 | -0.05869701 | 0.7551077 | -0.2715789 | 0.7859458 | NA |
| ENSG00000100416 | TRMU | 2290.022138 | -0.058699498 | 0.180542311 | -0.332407923 | 0.73958126 | 0.859371 |
| ENSG00000120709 | FAM53C | 3235.637286 | -0.05879141 | 0.175962208 | -0.337352794 | 0.73585097 | 0.856891 |
| ENSG00000105329 | TGFB1 | 1777.571467 | -0.058832717 | 0.289963184 | -0.216067107 | 0.82893544 | 0.913707 |
| ENSG00000116830 | TTF2 | 4932.452518 | -0.058864318 | 0.270861356 | -0.246567146 | 0.80524323 | 0.900308 |
| ENSG00000150401 | DCUN1D2 | 953.179 | -0.05886442 | 0.1287955 | -0.4679652 | 0.6398095 | 0.79435 |
| ENSG00000134453 | RBM17 | 9300.464029 | -0.058942063 | 0.142146231 | -0.422726934 | 0.6724945 | 0.817987 |
| ENSG00000114547 | ROPN1B | 7.164742457 | -0.058995687 | 0.567966644 | -0.149320409 | 0.88130081 | 0.944431 |
| ENSG00000066422 | ZBTB11 | 3481.348517 | -0.059071025 | 0.22380913 | -0.278379718 | 0.78072089 | 0.885062 |
| ENSG00000227237 | | 5.427762 | -0.05907358 | 0.6303298 | -0.1630817 | 0.8704541 | 0.938143 |
| ENSG00000158517 | NCF1 | 0.8089801 | -0.05908373 | 0.7538169 | -0.2683335 | 0.7884426 | NA |
| ENSG00000259087 | | 8.397667 | -0.05914847 | 0.5528175 | -0.1481593 | 0.882217 | 0.944859 |
| ENSG00000102804 | TSC22D1 | 8413.10202 | -0.059258331 | 0.139513755 | -0.462525176 | 0.64370475 | 0.796952 |
| ENSG00000181555 | SETD2 | 4673.211 | -0.05928458 | 0.1479399 | -0.3873338 | 0.6985091 | 0.835699 |
| ENSG00000287214 | | 1.195431 | -0.05929251 | 0.7281036 | -0.1990854 | 0.842196 | NA |
| ENSG00000276853 | | 32.83529 | -0.0593209 | 0.4090909 | -0.1717638 | 0.8636232 | 0.934551 |
| ENSG00000126883 | NUP214 | 3507.8965 | -0.059351596 | 0.158271516 | -0.384803941 | 0.70038268 | 0.836787 |
| ENSG00000186314 | PRELID2 | 94.09834 | -0.05938002 | 0.2577671 | -0.2583174 | 0.7961619 | 0.894882 |
| ENSG00000124920 | MYRF | 79.35381473 | -0.059435199 | 0.389469087 | -0.180356201 | 0.85687294 | 0.930755 |
| ENSG00000238207 | | 2.725285 | -0.05945631 | 0.6747272 | -0.1738506 | 0.8619829 | 0.9338 |
| ENSG00000110060 | PUS3 | 443.2509641 | -0.059485119 | 0.184587363 | -0.340635508 | 0.733378 | 0.855398 |
| ENSG00000269846 | | 1.77771 | -0.0595139 | 0.7277831 | -0.1986584 | 0.84253 | 0.922264 |

| | | | | | | | |
|---|---|---|---|---|---|---|---|
| ENSG00000085871 | MGST2 | 1017.278349 | -0.05952123 | 0.224147293 | -0.279086614 | 0.78017836 | 0.884739 |
| ENSG00000274695 | | 10.83584 | -0.05953455 | 0.5835392 | -0.157975 | 0.8744765 | 0.94057 |
| ENSG00000207425 | | 95.68161 | -0.05958346 | 0.2581089 | -0.2407697 | 0.8097337 | 0.902824 |
| ENSG00000111802 | TDP2 | 2608.105064 | -0.059583979 | 0.124700553 | -0.489135324 | 0.62474589 | 0.783705 |
| ENSG00000284976 | | 1355.724 | -0.05960322 | 0.206133 | -0.2988168 | 0.7650798 | 0.875357 |
| ENSG00000222345 | SNORD19C | 23.21228 | -0.05965229 | 0.4214218 | -0.1669522 | 0.8674077 | 0.936531 |
| ENSG00000224405 | LINC00572 | 1.586561 | -0.05981889 | 0.7139867 | -0.2016377 | 0.8402 | 0.920965 |
| ENSG00000179922 | ZNF784 | 388.7505 | -0.05982679 | 0.2692612 | -0.2374848 | 0.8122807 | 0.9042 |
| ENSG00000126226 | PCID2 | 3104.269769 | -0.059833346 | 0.146682201 | -0.445415175 | 0.65601975 | 0.806151 |
| ENSG00000279520 | | 106.46 | -0.05983584 | 0.2512114 | -0.2498614 | 0.8026946 | 0.898879 |
| ENSG00000244050 | DEFB109F | 2.308948 | -0.05987304 | 0.6845998 | -0.1691447 | 0.8656829 | 0.935977 |
| ENSG00000232642 | | 0.9160009 | -0.05991788 | 0.7334301 | -0.2164997 | 0.8285982 | NA |
| ENSG00000126787 | DLGAP5 | 1916.798001 | -0.059926827 | 0.517840777 | -0.164975141 | 0.86896355 | 0.937183 |
| ENSG00000186442 | KRT3 | 3.942975 | -0.05993984 | 0.7250442 | -0.1947602 | 0.8455807 | 0.924422 |
| ENSG00000224334 | | 6.657726 | -0.06003301 | 0.5483753 | -0.1575721 | 0.874794 | 0.940788 |
| ENSG00000120053 | GOT1 | 3949.254177 | -0.060119579 | 0.174172886 | -0.364127394 | 0.71576288 | 0.845317 |
| ENSG00000279711 | | 1.274676 | -0.06017922 | 0.737409 | -0.2643991 | 0.7914724 | NA |
| ENSG00000166847 | DCTN5 | 8082.772 | -0.06024843 | 0.1137689 | -0.534636 | 0.5929016 | 0.760895 |
| ENSG00000227495 | KIF1C-AS1 | 6.67909 | -0.06025808 | 0.5448477 | -0.1556201 | 0.8763325 | 0.941652 |
| ENSG00000246889 | | 86.45359 | -0.0604686 | 0.2359651 | -0.2714208 | 0.7860674 | 0.888314 |
| ENSG00000241282 | RPL34P33 | 89.17613 | -0.06051154 | 0.2536854 | -0.2469668 | 0.804934 | 0.900308 |
| ENSG00000228238 | | 1.38602 | -0.06060255 | 0.7058431 | -0.2084934 | 0.8348437 | NA |
| ENSG00000093072 | ADA2 | 15.99323788 | -0.060685805 | 0.464413308 | -0.165856676 | 0.86826974 | 0.936731 |
| ENSG00000278554 | | 2.312841 | -0.06073464 | 0.7208176 | -0.2244686 | 0.8223927 | 0.909824 |
| ENSG00000145428 | RNF175 | 0.8121941 | -0.06073619 | 0.7574487 | -0.2828325 | 0.7773053 | NA |
| ENSG00000259358 | DEPDC1P1 | 27.81388 | -0.06075739 | 0.5061894 | -0.1576002 | 0.8747719 | 0.940788 |
| ENSG00000135930 | EIF4E2 | 2650.014933 | -0.060762551 | 0.137249859 | -0.44846043 | 0.65382094 | 0.804863 |
| ENSG00000267340 | | 22.97883 | -0.06081415 | 0.4650423 | -0.1619041 | 0.8713814 | 0.938681 |
| ENSG00000228696 | ARL17B | 13.43616 | -0.06082703 | 0.4619183 | -0.1641626 | 0.8696031 | 0.937492 |
| ENSG00000281133 | | 2.477457 | -0.06083796 | 0.6412266 | -0.1729531 | 0.8626883 | 0.934093 |
| ENSG00000287074 | | 7.253323 | -0.06090851 | 0.5958171 | -0.1615324 | 0.8716741 | 0.93881 |
| ENSG00000144895 | EIF2A | 7981.606 | -0.06098993 | 0.1603172 | -0.4007676 | 0.6885912 | 0.828482 |
| ENSG00000186635 | ARAP1 | 3418.522 | -0.06099815 | 0.173276 | -0.3728884 | 0.7092315 | 0.841795 |

| | | | | | | | |
|---|---|---|---|---|---|---|---|
| ENSG00000282855 | | 17.6111 | -0.06106125 | 0.4722113 | -0.1648897 | 0.8690308 | 0.937198 |
| ENSG00000248449 | PCDHGB8P | 1.472198 | -0.06107487 | 0.7002525 | -0.1864736 | 0.8520733 | NA |
| ENSG00000285979 | | 153.9681 | -0.061163 | 0.3386393 | -0.2009889 | 0.8407072 | 0.921217 |
| ENSG00000236268 | LINC01361 | 2.006986 | -0.06116904 | 0.6499813 | -0.1743982 | 0.8615526 | 0.933594 |
| ENSG00000205746 | PKD1P4 | 445.4246 | -0.06118209 | 0.3931467 | -0.1854938 | 0.8528418 | 0.928426 |
| ENSG00000269949 | | 7.496085 | -0.06122694 | 0.4912769 | -0.159618 | 0.873182 | 0.939968 |
| ENSG00000204396 | VWA7 | 46.63214 | -0.06134388 | 0.3216766 | -0.2137374 | 0.8307519 | 0.914912 |
| ENSG00000265845 | | 2.847353 | -0.06141859 | 0.6832575 | -0.1745262 | 0.861452 | 0.933591 |
| ENSG00000196642 | RABL6 | 8854.909 | -0.06142826 | 0.1496812 | -0.4222873 | 0.6728154 | 0.818088 |
| ENSG00000108557 | RAI1 | 7960.109046 | -0.061451748 | 0.180102258 | -0.351127981 | 0.72549234 | 0.850907 |
| ENSG00000284308 | C2orf81 | 104.9826 | -0.06147183 | 0.3060876 | -0.2212049 | 0.8249329 | 0.911754 |
| ENSG00000252174 | RNU7-18P | 3.92835 | -0.06155254 | 0.6481994 | -0.1689799 | 0.8658125 | 0.936028 |
| ENSG00000118007 | STAG1 | 3477.749048 | -0.061583808 | 0.208828112 | -0.308022241 | 0.75806541 | 0.871039 |
| ENSG00000108256 | NUFIP2 | 11268.40876 | -0.061587355 | 0.245920504 | -0.266945015 | 0.7895115 | 0.890811 |
| ENSG00000115446 | UNC50 | 1782.114239 | -0.06170021 | 0.193389275 | -0.32627847 | 0.74421368 | 0.862245 |
| ENSG00000273384 | | 1.715118 | -0.06179172 | 0.7159458 | -0.2167635 | 0.8283927 | 0.913464 |
| ENSG00000113732 | ATP6V0E1 | 9889.935548 | -0.061797813 | 0.171297286 | -0.377618778 | 0.70571381 | 0.840057 |
| ENSG00000159352 | PSMD4 | 15287.65 | -0.06180711 | 0.202345 | -0.3212935 | 0.747988 | 0.864893 |
| ENSG00000079950 | STX7 | 2377.519931 | -0.061850668 | 0.171235596 | -0.379593023 | 0.70424754 | 0.839341 |
| ENSG00000235408 | SNORA71B | 29.55227 | -0.06186911 | 0.3759073 | -0.1906302 | 0.8488153 | 0.926586 |
| ENSG00000091073 | DTX2 | 619.2033596 | -0.061894255 | 0.219192953 | -0.300439689 | 0.76384179 | 0.874321 |
| ENSG00000235297 | FAUP1 | 2077.599 | -0.06195485 | 0.1774162 | -0.3597728 | 0.719017 | 0.846902 |
| ENSG00000259548 | | 2.503911 | -0.06195861 | 0.6617887 | -0.1711803 | 0.864082 | 0.93484 |
| ENSG00000171421 | MRPL36 | 2803.542 | -0.0619718 | 0.2494313 | -0.2257439 | 0.8214006 | 0.909357 |
| ENSG00000240106 | RN7SL146P | 1.820157 | -0.06203807 | 0.6964695 | -0.1917858 | 0.84791 | 0.92613 |
| ENSG00000066379 | POLR1H | 621.7717561 | -0.062130517 | 0.203472488 | -0.318450905 | 0.75014293 | 0.866263 |
| ENSG00000254245 | PCDHGA3 | 43.34095 | -0.06226159 | 0.4603422 | -0.1554536 | 0.8764637 | 0.941707 |
| ENSG00000142330 | CAPN10 | 1368.65 | -0.0623014 | 0.1855384 | -0.3457311 | 0.7295448 | 0.853333 |
| ENSG00000130755 | GMFG | 3.792625341 | -0.062378455 | 0.589291886 | -0.167099152 | 0.86729203 | 0.936483 |
| ENSG00000259605 | | 41.70566 | -0.06242339 | 0.3800411 | -0.1874562 | 0.8513029 | 0.927777 |
| ENSG00000083093 | PALB2 | 1627.585423 | -0.062459352 | 0.167162753 | -0.394226672 | 0.6934137 | 0.831639 |
| ENSG00000268069 | | 4.061206 | -0.06247876 | 0.6238841 | -0.1631429 | 0.8704059 | 0.938129 |
| ENSG00000188315 | C3orf62 | 323.1418 | -0.06253072 | 0.2674098 | -0.2522133 | 0.8008762 | 0.897826 |

| | | | | | | | |
|---|---|---|---|---|---|---|---|
| ENSG00000020129 | NCDN | 4598.360447 | -0.062613746 | 0.120503019 | -0.489828082 | 0.62425556 | 0.783439 |
| ENSG00000125843 | AP5S1 | 1121.246852 | -0.062617078 | 0.202614919 | -0.31685838 | 0.75135106 | 0.866969 |
| ENSG00000260671 | | 1.344666 | -0.06267255 | 0.7209852 | -0.2287045 | 0.8190986 | NA |
| ENSG00000263731 | | 46.11141 | -0.062726 | 0.3996138 | -0.1852152 | 0.8530603 | 0.928466 |
| ENSG00000278484 | | 4.575265 | -0.06276339 | 0.6343777 | -0.1777291 | 0.8589357 | 0.932062 |
| ENSG00000104714 | ERICH1 | 1057.333368 | -0.062814175 | 0.136283142 | -0.466202423 | 0.64107062 | 0.795359 |
| ENSG00000229473 | RGS17P1 | 4.641916 | -0.06284551 | 0.6610613 | -0.1824536 | 0.8552268 | 0.929935 |
| ENSG00000268945 | | 9.366607 | -0.06285488 | 0.5201296 | -0.1614212 | 0.8717617 | 0.938816 |
| ENSG00000268947 | | 73.01163 | -0.06291624 | 0.3651872 | -0.1963557 | 0.8443318 | 0.923608 |
| ENSG00000258427 | RBM8B | 1519.074 | -0.06298539 | 0.1977765 | -0.3340994 | 0.7383046 | 0.858765 |
| ENSG00000101134 | DOK5 | 1.006579528 | -0.062992903 | 0.763532897 | -0.314485198 | 0.75315256 | NA |
| ENSG00000176289 | IDSP1 | 6.185838 | -0.06302061 | 0.5454482 | -0.1660852 | 0.8680899 | 0.936731 |
| ENSG00000063244 | U2AF2 | 14715.41663 | -0.063046234 | 0.182471752 | -0.357435373 | 0.7207659 | 0.848059 |
| ENSG00000108878 | CACNG1 | 71.24597632 | -0.063049125 | 0.431960243 | -0.173421467 | 0.86232015 | 0.933922 |
| ENSG00000114302 | PRKAR2A | 3023.221208 | -0.063052462 | 0.22341603 | -0.301776804 | 0.76282222 | 0.873822 |
| ENSG00000215492 | HNRNPA1P7 | 5270.12 | -0.06306801 | 0.1504289 | -0.4441189 | 0.6569566 | 0.80678 |
| ENSG00000180900 | SCRIB | 10997.67 | -0.0631556 | 0.1200105 | -0.5218284 | 0.6017898 | 0.767261 |
| ENSG00000148153 | INIP | 2035.692 | -0.06321116 | 0.1697746 | -0.3777788 | 0.705595 | 0.840057 |
| ENSG00000162063 | CCNF | 1562.602 | -0.06325685 | 0.4214894 | -0.1777065 | 0.8589535 | 0.932062 |
| ENSG00000089234 | BRAP | 1645.797925 | -0.063339545 | 0.09766296 | -0.67807553 | 0.49772381 | 0.686451 |
| ENSG00000274659 | LINC02371 | 1.614752 | -0.06335555 | 0.7020008 | -0.19041 | 0.8489879 | 0.926594 |
| ENSG00000162341 | TPCN2 | 856.473 | -0.06339666 | 0.1349101 | -0.4798181 | 0.6313568 | 0.788593 |
| ENSG00000280039 | RN7SKP23 | 11.19381 | -0.06344731 | 0.530077 | -0.1659225 | 0.8682179 | 0.936731 |
| ENSG00000128596 | CCDC136 | 76.38609618 | -0.063524238 | 0.345675022 | -0.204826572 | 0.83770763 | 0.918954 |
| ENSG00000280300 | | 15.34372 | -0.06355377 | 0.387566 | -0.1855249 | 0.8528173 | 0.928426 |
| ENSG00000259295 | CSPG4P12 | 118.2068 | -0.0636614 | 0.266429 | -0.2553816 | 0.7984284 | 0.896254 |
| ENSG00000184616 | SPDYE12P | 35.77073 | -0.06367788 | 0.3577833 | -0.2043545 | 0.8380764 | 0.919244 |
| ENSG00000239246 | | 412.6809 | -0.06375881 | 0.2329102 | -0.3026087 | 0.7621881 | 0.873321 |
| ENSG00000264644 | KRT18P8 | 17.76561 | -0.06379113 | 0.5362396 | -0.1682473 | 0.8663887 | 0.936211 |
| ENSG00000199132 | MIR450A1 | 1.362689 | -0.06386687 | 0.7403552 | -0.2430713 | 0.8079502 | NA |
| ENSG00000244119 | PDCL3P4 | 180.6531 | -0.06387772 | 0.1742348 | -0.3619306 | 0.7174039 | 0.846015 |
| ENSG00000070193 | FGF10 | 4.017463119 | -0.06387899 | 0.792635083 | -0.464809755 | 0.6420677 | 0.795852 |
| ENSG00000136542 | GALNT5 | 4.487241195 | -0.063905903 | 0.701206066 | -0.188254529 | 0.85067713 | 0.927437 |

| | | | | | | | |
|---|---|---|---|---|---|---|---|
| ENSG00000230454 | | 146.2075 | -0.06407191 | 0.3447517 | -0.2093584 | 0.8341685 | 0.916888 |
| ENSG00000137177 | KIF13A | 3072.192112 | -0.064157929 | 0.222519508 | -0.310452844 | 0.75621661 | 0.869886 |
| ENSG00000188305 | PEAK3 | 3.326743 | -0.06416838 | 0.6138653 | -0.1721865 | 0.8632909 | 0.934398 |
| ENSG00000265136 | | 17.315 | -0.06420779 | 0.383498 | -0.1963331 | 0.8443495 | 0.923608 |
| ENSG00000225066 | MDM4P1 | 4.94559 | -0.06428261 | 0.6602345 | -0.1869433 | 0.8517051 | 0.927845 |
| ENSG00000256263 | DDX11L8 | 3.234069 | -0.06434731 | 0.6431127 | -0.1731455 | 0.8625371 | 0.934005 |
| ENSG00000187988 | KCTD9P3 | 13.8568 | -0.06437918 | 0.544951 | -0.1660703 | 0.8681017 | 0.936731 |
| ENSG00000134109 | EDEM1 | 3192.778459 | -0.064430977 | 0.222870292 | -0.317240573 | 0.75106107 | 0.866859 |
| ENSG00000182923 | CEP63 | 1913.204 | -0.06444026 | 0.1047432 | -0.6595095 | 0.5095686 | 0.695578 |
| ENSG00000116191 | RALGPS2 | 2771.731258 | -0.064492461 | 0.125427734 | -0.524865102 | 0.59967697 | 0.765998 |
| ENSG00000257815 | PRANCR | 86.19017 | -0.06449581 | 0.330497 | -0.2179589 | 0.8274611 | 0.913064 |
| ENSG00000240342 | RPS2P5 | 16503.6 | -0.06449784 | 0.1939326 | -0.3459182 | 0.7294042 | 0.853273 |
| ENSG00000231889 | TRAF3IP2-AS1 | 80.59765 | -0.06451563 | 0.2314812 | -0.2931154 | 0.7694339 | 0.877913 |
| ENSG00000166396 | SERPINB7 | 1.415278 | -0.06451635 | 0.7928787 | -0.4648149 | 0.642064 | NA |
| ENSG00000136811 | ODF2 | 3890.618558 | -0.064532846 | 0.239591954 | -0.288244934 | 0.77315926 | 0.880445 |
| ENSG00000176407 | KCMF1 | 4253.396 | -0.06457767 | 0.1321632 | -0.5078841 | 0.6115346 | 0.774478 |
| ENSG00000234817 | ECI2-DT | 4.651326 | -0.06457799 | 0.5919394 | -0.1697759 | 0.8651864 | 0.935654 |
| ENSG00000248648 | | 11.089 | -0.06472523 | 0.4712502 | -0.1745334 | 0.8614462 | 0.933591 |
| ENSG00000122406 | RPL5 | 27154.7731 | -0.064768223 | 0.166926562 | -0.396621 | 0.69164696 | 0.830495 |
| ENSG00000205309 | NT5M | 283.6789 | -0.06478752 | 0.2793708 | -0.2536125 | 0.7997949 | 0.897068 |
| ENSG00000272693 | | 52.99816 | -0.06478781 | 0.2729999 | -0.2554459 | 0.7983787 | 0.896254 |
| ENSG00000228709 | LINC02575 | 3.630156 | -0.06489644 | 0.7079725 | -0.1926985 | 0.8471951 | 0.925578 |
| ENSG00000228063 | LYPLAL1-DT | 1.00694 | -0.06492137 | 0.7323776 | -0.2432801 | 0.8077884 | NA |
| ENSG00000244627 | TPTEP2 | 93.77869 | -0.06493056 | 0.4276605 | -0.1795655 | 0.8574937 | 0.931239 |
| ENSG00000101190 | TCFL5 | 3285.451666 | -0.06495604 | 0.145601534 | -0.523460286 | 0.60065398 | 0.766716 |
| ENSG00000230897 | RPS18P12 | 952.4377 | -0.06505326 | 0.2154173 | -0.3142931 | 0.7532984 | 0.86819 |
| ENSG00000113387 | SUB1 | 8867.992501 | -0.065082038 | 0.196604913 | -0.337681318 | 0.73560336 | 0.856753 |
| ENSG00000127483 | HP1BP3 | 9110.059162 | -0.065135195 | 0.18902311 | -0.368691286 | 0.71235785 | 0.843395 |
| ENSG00000185864 | NPIPB4 | 263.1674 | -0.06514338 | 0.2433254 | -0.280149 | 0.7793632 | 0.884141 |
| ENSG00000123374 | CDK2 | 2616.220508 | -0.065186772 | 0.349794251 | -0.225005272 | 0.82197517 | 0.909628 |
| ENSG00000146858 | ZC3HAV1L | 264.4992 | -0.06521722 | 0.3228464 | -0.2239702 | 0.8227805 | 0.910032 |
| ENSG00000257964 | | 5.943057 | -0.06522451 | 0.5977812 | -0.1681964 | 0.8664288 | 0.936211 |
| ENSG00000137692 | DCUN1D5 | 1955.841697 | -0.065251291 | 0.182838673 | -0.372969712 | 0.70917099 | 0.841795 |

| | | | | | | | |
|---|---|---|---|---|---|---|---|
| ENSG00000229152 | ANKRD10-IT1 | 342.7272 | -0.0653958 | 0.318331 | -0.2261442 | 0.8210893 | 0.909205 |
| ENSG00000177731 | FLII | 16168.01 | -0.06545932 | 0.1147025 | -0.5458352 | 0.5851792 | 0.755032 |
| ENSG00000124541 | RRP36 | 3413.080461 | -0.065529835 | 0.221829011 | -0.306509401 | 0.75921682 | 0.871684 |
| ENSG00000166716 | ZNF592 | 5407.199 | -0.06557987 | 0.1658418 | -0.4839856 | 0.6283961 | 0.786465 |
| ENSG00000095906 | NUBP2 | 4643.745337 | -0.065649685 | 0.194275714 | -0.352082843 | 0.72477614 | 0.850631 |
| ENSG00000130985 | UBA1 | 23009.01469 | -0.065688794 | 0.152026203 | -0.444414886 | 0.65674265 | 0.806592 |
| ENSG00000134242 | PTPN22 | 77.51138059 | -0.065692194 | 0.303624531 | -0.236940007 | 0.81270333 | 0.904263 |
| ENSG00000178397 | FAM220A | 1.006468 | -0.06570473 | 0.7449111 | -0.2499715 | 0.8026094 | NA |
| ENSG00000150316 | CWC15 | 1890.885 | -0.06572722 | 0.1782818 | -0.3854711 | 0.6998884 | 0.836497 |
| ENSG00000264548 | | 97.56955 | -0.06576549 | 0.3563688 | -0.217194 | 0.8280572 | 0.913383 |
| ENSG00000084112 | SSH1 | 3714.883683 | -0.065820596 | 0.233538881 | -0.298412251 | 0.76538854 | 0.875498 |
| ENSG00000198168 | SVIP | 2581.525 | -0.06588298 | 0.190499 | -0.3675359 | 0.7132193 | 0.843667 |
| ENSG00000168256 | NKIRAS2 | 2615.992 | -0.06602961 | 0.128775 | -0.5212176 | 0.6022152 | 0.767619 |
| ENSG00000273179 | | 2.817053 | -0.06605554 | 0.6683139 | -0.1853154 | 0.8529817 | 0.928455 |
| ENSG00000171345 | KRT19 | 51878.58 | -0.06610347 | 0.1829577 | -0.3750937 | 0.7075908 | 0.841012 |
| ENSG00000231841 | FAM192BP | 305.1093 | -0.06633571 | 0.1562755 | -0.436039 | 0.6628085 | 0.811117 |
| ENSG00000196812 | ZSCAN16 | 458.1136 | -0.06634124 | 0.1846835 | -0.3768758 | 0.7062659 | 0.840366 |
| ENSG00000173660 | UQCRH | 5623.418 | -0.06634204 | 0.229047 | -0.2967628 | 0.7666476 | 0.876277 |
| ENSG00000114209 | PDCD10 | 3742.050775 | -0.066358948 | 0.149614896 | -0.452029748 | 0.65124755 | 0.802998 |
| ENSG00000239483 | RPS15AP16 | 3.25438 | -0.06641181 | 0.6702361 | -0.1911241 | 0.8484284 | 0.926316 |
| ENSG00000074047 | GLI2 | 9.726531071 | -0.066414956 | 0.628803988 | -0.176490659 | 0.85990849 | 0.932571 |
| ENSG00000092847 | AGO1 | 3213.686386 | -0.0664993 | 0.140010715 | -0.484925143 | 0.62772945 | 0.786038 |
| ENSG00000101346 | POFUT1 | 3560.425298 | -0.06666922 | 0.168520927 | -0.39843082 | 0.69031264 | 0.829715 |
| ENSG00000023191 | RNH1 | 7570.816214 | -0.06675679 | 0.131294724 | -0.524352396 | 0.60003346 | 0.766235 |
| ENSG00000099910 | KLHL22 | 1600.547087 | -0.066784541 | 0.146949499 | -0.465395641 | 0.64164816 | 0.795666 |
| ENSG00000146676 | PURB | 6782.233 | -0.06683017 | 0.1719689 | -0.4291021 | 0.6678489 | 0.814958 |
| ENSG00000177663 | IL17RA | 2028.382 | -0.06687702 | 0.1388389 | -0.4940072 | 0.6213011 | 0.781173 |
| ENSG00000239305 | RNF103 | 1457.615 | -0.06698016 | 0.1366333 | -0.5137686 | 0.6074138 | 0.771285 |
| ENSG00000287960 | | 0.7876481 | -0.06700048 | 0.7650583 | -0.3163282 | 0.7517534 | NA |
| ENSG00000166860 | ZBTB39 | 1018.565 | -0.06705805 | 0.2098338 | -0.3332775 | 0.7389248 | 0.858927 |
| ENSG00000163872 | YEATS2 | 2839.679 | -0.06707852 | 0.2024917 | -0.3372224 | 0.7359492 | 0.85693 |
| ENSG00000102271 | KLHL4 | 0.955070377 | -0.067103078 | 0.779565117 | -0.500263606 | 0.61688948 | NA |
| ENSG00000229944 | EIF4EP2 | 1450.482 | -0.06714266 | 0.1893221 | -0.3709459 | 0.7106778 | 0.842599 |

| | | | | | | | |
|---|---|---|---|---|---|---|---|
| ENSG00000066427 | ATXN3 | 1833.048751 | -0.067219816 | 0.139266935 | -0.496374556 | 0.61963017 | 0.780127 |
| ENSG00000213625 | LEPROT | 4235.437 | -0.0672549 | 0.1740994 | -0.3973893 | 0.6910804 | 0.830171 |
| ENSG00000265791 | | 6.849196 | -0.06725554 | 0.5239855 | -0.1746274 | 0.8613724 | 0.933581 |
| ENSG00000147164 | SNX12 | 4470.101 | -0.06726367 | 0.1053999 | -0.6396097 | 0.5224264 | 0.707748 |
| ENSG00000258509 | | 7.213834 | -0.06737842 | 0.6005836 | -0.1718507 | 0.8635549 | 0.934536 |
| ENSG00000224775 | BRAFP1 | 2.953474 | -0.0674538 | 0.6123013 | -0.1779216 | 0.8587845 | 0.931977 |
| ENSG00000273156 | | 14.98974 | -0.06747441 | 0.5446512 | -0.1755443 | 0.860652 | 0.933016 |
| ENSG00000181126 | HLA-V | 5.512009 | -0.06748068 | 0.5254283 | -0.178363 | 0.8584379 | 0.93183 |
| ENSG00000258815 | LINC02820 | 99.5009 | -0.06749824 | 0.3053869 | -0.2398454 | 0.8104501 | 0.903034 |
| ENSG00000213149 | CNN2P9 | 4.18701 | -0.06759192 | 0.6031508 | -0.1755282 | 0.8606646 | 0.933016 |
| ENSG00000231185 | SPRY4-AS1 | 9.498895 | -0.06760314 | 0.6244308 | -0.1720087 | 0.8634307 | 0.93444 |
| ENSG00000261632 | | 0.8254911 | -0.06773272 | 0.771497 | -0.3693826 | 0.7118426 | NA |
| ENSG00000178691 | SUZ12 | 2658.984 | -0.06785096 | 0.1963444 | -0.3617831 | 0.7175141 | 0.846015 |
| ENSG00000006652 | IFRD1 | 3163.562968 | -0.067913271 | 0.195318253 | -0.366603558 | 0.71391475 | 0.843961 |
| ENSG00000125611 | CHCHD5 | 1083.593371 | -0.067980654 | 0.2165215 | -0.330069229 | 0.74134765 | 0.860542 |
| ENSG00000100462 | PRMT5 | 4174.19409 | -0.068012594 | 0.208719202 | -0.347023024 | 0.72857402 | 0.852739 |
| ENSG00000253092 | | 2.99316 | -0.0680266 | 0.66757 | -0.1868566 | 0.8517731 | 0.927845 |
| ENSG00000206652 | RNU1-1 | 1.790769 | -0.0680468 | 0.7060092 | -0.2302039 | 0.8179333 | 0.90768 |
| ENSG00000261522 | | 1.12782 | -0.068157 | 0.74093 | -0.2578041 | 0.7965581 | NA |
| ENSG00000235207 | TUBBP6 | 1.059015 | -0.06816248 | 0.758865 | -0.3022978 | 0.7624251 | NA |
| ENSG00000080618 | CPB2 | 4.147195988 | -0.068215211 | 0.670616428 | -0.185785797 | 0.85261274 | 0.928328 |
| ENSG00000171988 | JMJD1C | 4040.271 | -0.06828801 | 0.2368085 | -0.306013 | 0.7595948 | 0.871854 |
| ENSG00000271482 | | 1.581139 | -0.06829738 | 0.7266108 | -0.227705 | 0.8198756 | 0.908881 |
| ENSG00000267796 | LIN37 | 231.4582 | -0.06831371 | 0.2508844 | -0.2898524 | 0.7719292 | 0.879476 |
| ENSG00000134058 | CDK7 | 2187.024763 | -0.068363876 | 0.146675765 | -0.476973564 | 0.63338095 | 0.789913 |
| ENSG00000174903 | RAB1B | 8868.323 | -0.06838826 | 0.1821433 | -0.392091 | 0.694991 | 0.832967 |
| ENSG00000227881 | ASS1P5 | 2.470222 | -0.06840381 | 0.6829619 | -0.2178414 | 0.8275527 | 0.913064 |
| ENSG00000173218 | VANGL1 | 4404.946 | -0.06847222 | 0.1775436 | -0.3688416 | 0.7122458 | 0.843338 |
| ENSG00000212743 | LINC02656 | 36.41775 | -0.06849964 | 0.5320153 | -0.168061 | 0.8665353 | 0.936211 |
| ENSG00000128694 | OSGEPL1 | 359.9598681 | -0.068552538 | 0.195691483 | -0.364623134 | 0.71539274 | 0.84503 |
| ENSG00000161203 | AP2M1 | 23582.96 | -0.06860794 | 0.1708238 | -0.4473629 | 0.6546131 | 0.805151 |
| ENSG00000125703 | ATG4C | 578.5043259 | -0.068625133 | 0.22215841 | -0.324378336 | 0.74565162 | 0.863159 |
| ENSG00000275582 | | 26.43693 | -0.06865782 | 0.4265686 | -0.1957377 | 0.8448155 | 0.923966 |

| | | | | | | | |
|---|---|---|---|---|---|---|---|
| ENSG00000221983 | UBA52 | 12409.71 | -0.06866522 | 0.1815931 | -0.3460902 | 0.7292749 | 0.853215 |
| ENSG00000236896 | | 7.64181 | -0.06873694 | 0.5835319 | -0.1743028 | 0.8616275 | 0.933629 |
| ENSG00000237328 | RAI1-AS1 | 0.9609824 | -0.06881709 | 0.7486956 | -0.2851969 | 0.7754933 | NA |
| ENSG00000088812 | ATRN | 3751.005775 | -0.068827293 | 0.20109136 | -0.356265022 | 0.7216421 | 0.848489 |
| ENSG00000187688 | TRPV2 | 8.367385 | -0.06889964 | 0.5461751 | -0.1783456 | 0.8584516 | 0.93183 |
| ENSG00000263400 | TMEM220-AS1 | 2.445734 | -0.06890159 | 0.7239909 | -0.2387577 | 0.8112935 | 0.903519 |
| ENSG00000153237 | CCDC148 | 54.34764 | -0.06892113 | 0.3021956 | -0.2449808 | 0.8064713 | 0.900716 |
| ENSG00000147099 | HDAC8 | 463.4794 | -0.06897038 | 0.1947588 | -0.3678622 | 0.712976 | 0.843639 |
| ENSG00000278725 | | 1.453773 | -0.06898938 | 0.6891346 | -0.2100795 | 0.8336057 | NA |
| ENSG00000258591 | PPIAP4 | 6.700168 | -0.069023 | 0.5761148 | -0.1762873 | 0.8600683 | 0.932662 |
| ENSG00000177197 | PCNPP5 | 219.5793 | -0.06908557 | 0.2037908 | -0.3535919 | 0.7236447 | 0.849877 |
| ENSG00000234284 | ZNF879 | 330.2227 | -0.06915021 | 0.2222507 | -0.3261164 | 0.7443363 | 0.862303 |
| ENSG00000146426 | TIAM2 | 222.6107 | -0.06927054 | 0.2012431 | -0.3653864 | 0.714823 | 0.844633 |
| ENSG00000254065 | | 2.89359 | -0.06929058 | 0.6494536 | -0.1903087 | 0.8490672 | 0.926594 |
| ENSG00000259245 | LINC02853 | 1.203113 | -0.06931323 | 0.7440779 | -0.2548571 | 0.7988334 | NA |
| ENSG00000257727 | CNPY2 | 495.1862 | -0.06937591 | 0.138316 | -0.5091066 | 0.6106775 | 0.774098 |
| ENSG00000100124 | ANKRD54 | 1791.613757 | -0.069384198 | 0.177277269 | -0.402820556 | 0.68708024 | 0.827607 |
| ENSG00000287274 | | 0.9259244 | -0.06940976 | 0.735202 | -0.2775646 | 0.7813466 | NA |
| ENSG00000183921 | SDR42E2 | 2.231029 | -0.06946122 | 0.6756528 | -0.1990766 | 0.8422028 | 0.922133 |
| ENSG00000233426 | EIF3FP3 | 1466.268 | -0.06947801 | 0.1584833 | -0.4500712 | 0.6526591 | 0.803955 |
| ENSG00000108946 | PRKAR1A | 20634.74872 | -0.069496776 | 0.092438999 | -0.804645064 | 0.42102453 | 0.620893 |
| ENSG00000135968 | GCC2 | 2626.800883 | -0.069579422 | 0.228121608 | -0.313038352 | 0.75425153 | 0.86887 |
| ENSG00000267507 | | 2.306407 | -0.06960731 | 0.6483625 | -0.193239 | 0.8467718 | 0.925306 |
| ENSG00000246451 | | 173.7267 | -0.06963744 | 0.3540474 | -0.2243148 | 0.8225124 | 0.909824 |
| ENSG00000169418 | NPR1 | 1.858487 | -0.06964028 | 0.6858176 | -0.2065655 | 0.8363492 | 0.918108 |
| ENSG00000100216 | TOMM22 | 2453.296473 | -0.069648758 | 0.222733875 | -0.328346574 | 0.74264963 | 0.86141 |
| ENSG00000273749 | CYFIP1 | 6582.773 | -0.06972468 | 0.1637163 | -0.4648135 | 0.642065 | 0.795852 |
| ENSG00000154146 | NRGN | 169.8821 | -0.06974017 | 0.5724464 | -0.1812995 | 0.8561325 | 0.930273 |
| ENSG00000162373 | BEND5 | 20.85548 | -0.06978428 | 0.5143908 | -0.1863202 | 0.8521937 | 0.928062 |
| ENSG00000237489 | C10orf143 | 228.7352 | -0.06982233 | 0.2671962 | -0.2849274 | 0.7756998 | 0.881881 |
| ENSG00000149639 | SOGA1 | 4415.323 | -0.06983929 | 0.2766968 | -0.281395 | 0.7784074 | 0.883535 |
| ENSG00000102974 | CTCF | 6093.789318 | -0.069840163 | 0.15739524 | -0.464642653 | 0.64218738 | 0.795869 |
| ENSG00000182888 | RPS27AP20 | 1.459798 | -0.06984519 | 0.717863 | -0.2430774 | 0.8079455 | NA |

| | | | | | | | |
|---|---|---|---|---|---|---|---|
| ENSG00000263823 | | 26.36651 | -0.06990004 | 0.4174422 | -0.1991082 | 0.8421781 | 0.922133 |
| ENSG00000149823 | VPS51 | 4358.879 | -0.06992262 | 0.1825819 | -0.3918694 | 0.6951547 | 0.833013 |
| ENSG00000115540 | MOB4 | 1164.051653 | -0.069956152 | 0.220248504 | -0.333341837 | 0.73887626 | 0.858927 |
| ENSG00000063177 | RPL18 | 25346.54162 | -0.069958481 | 0.178438745 | -0.405188583 | 0.6853389 | 0.826544 |
| ENSG00000178715 | | 144.1832 | -0.06998893 | 0.285195 | -0.2694758 | 0.7875636 | 0.889371 |
| ENSG00000287663 | | 1.47374 | -0.07000795 | 0.7121083 | -0.2241515 | 0.8226394 | NA |
| ENSG00000107862 | GBF1 | 5830.419727 | -0.07007301 | 0.151308938 | -0.480854709 | 0.63061976 | 0.788023 |
| ENSG00000080845 | DLGAP4 | 2266.201573 | -0.070129435 | 0.144209389 | -0.504836312 | 0.61367381 | 0.775861 |
| ENSG00000202314 | SNORD6 | 19.13073 | -0.07021671 | 0.4321192 | -0.1958986 | 0.8446896 | 0.923877 |
| ENSG00000156973 | PDE6D | 799.0982 | -0.0702202 | 0.2562584 | -0.2932795 | 0.7693085 | 0.877808 |
| ENSG00000253716 | MINCR | 580.2088 | -0.07022049 | 0.255265 | -0.2917837 | 0.770452 | 0.878622 |
| ENSG00000283098 | | 6.734334 | -0.07023341 | 0.5905913 | -0.1812603 | 0.8561633 | 0.930273 |
| ENSG00000100058 | CRYBB2P1 | 824.6281191 | -0.07029407 | 0.152873869 | -0.472115533 | 0.63684433 | 0.792549 |
| ENSG00000088808 | PPP1R13B | 2632.323241 | -0.070308815 | 0.134838461 | -0.562784569 | 0.57358159 | 0.746595 |
| ENSG00000110987 | BCL7A | 1968.333715 | -0.070314679 | 0.162722722 | -0.450078631 | 0.65265375 | 0.803955 |
| ENSG00000270605 | | 227.5973 | -0.07032818 | 0.4161877 | -0.2096319 | 0.833955 | 0.916692 |
| ENSG00000184678 | H2BC21 | 10606.9 | -0.07036866 | 0.2951097 | -0.2611289 | 0.7939931 | 0.893416 |
| ENSG00000285934 | | 0.7466155 | -0.07037256 | 0.794873 | -0.9230905 | 0.3559601 | NA |
| ENSG00000280378 | | 9.273848 | -0.07049765 | 0.55604 | -0.1814726 | 0.8559966 | 0.930273 |
| ENSG00000261546 | | 6.083671 | -0.0705193 | 0.5648192 | -0.1792344 | 0.8577536 | 0.931298 |
| ENSG00000214783 | POLR2J4 | 49.23255 | -0.07054453 | 0.3906089 | -0.2128564 | 0.831439 | 0.915137 |
| ENSG00000223973 | PAFAH1B1P2 | 2.54458 | -0.07055832 | 0.694083 | -0.2162141 | 0.8288209 | 0.913655 |
| ENSG00000257698 | GIHCG | 132.6503 | -0.07057338 | 0.3201258 | -0.2372272 | 0.8124806 | 0.904236 |
| ENSG00000233527 | ZNF529-AS1 | 178.6344 | -0.0706029 | 0.3530782 | -0.2208735 | 0.8251909 | 0.911882 |
| ENSG00000082213 | C5orf22 | 2247.957823 | -0.070646518 | 0.177711048 | -0.410405505 | 0.68150851 | 0.823757 |
| ENSG00000165660 | ABRAXAS2 | 1431.175 | -0.0706829 | 0.1065524 | -0.6557616 | 0.5119775 | 0.697897 |
| ENSG00000250642 | | 1.090371 | -0.07075757 | 0.7512667 | -0.3624636 | 0.7170056 | NA |
| ENSG00000262227 | | 29.87925 | -0.07092225 | 0.3408383 | -0.2401261 | 0.8102325 | 0.902923 |
| ENSG00000143322 | ABL2 | 2649.553 | -0.07098732 | 0.2313118 | -0.3265809 | 0.7439849 | 0.862056 |
| ENSG00000131165 | CHMP1A | 10534.67328 | -0.071035256 | 0.200324067 | -0.371910029 | 0.70995984 | 0.842283 |
| ENSG00000213197 | NUDCP1 | 13.06728 | -0.07105184 | 0.482312 | -0.1903116 | 0.849065 | 0.926594 |
| ENSG00000224973 | LARGE-AS1 | 0.9723684 | -0.07110172 | 0.7111091 | -0.2310098 | 0.8173072 | NA |
| ENSG00000244005 | NFS1 | 1881.919 | -0.07111058 | 0.2442085 | -0.3121365 | 0.7549368 | 0.869057 |

| | | | | | | | |
|---|---|---|---|---|---|---|---|
| ENSG00000213174 | | 1.933862 | -0.07111695 | 0.689223 | -0.2117709 | 0.8322858 | 0.915608 |
| ENSG00000070915 | SLC12A3 | 1.206360273 | -0.071151797 | 0.795177582 | -1.058037672 | 0.29003827 | NA |
| ENSG00000115009 | CCL20 | 3.553687524 | -0.071210999 | 0.723729277 | -0.246709202 | 0.80513328 | 0.900308 |
| ENSG00000169442 | CD52 | 1.377694 | -0.07123358 | 0.7952182 | -1.071143 | 0.284105 | NA |
| ENSG00000227115 | LINC01630 | 2.204227 | -0.07124676 | 0.7953501 | -1.21608 | 0.2239543 | 0.41078 |
| ENSG00000201078 | RN7SKP214 | 1.38581 | -0.07124812 | 0.7952323 | -1.083875 | 0.27842 | NA |
| ENSG00000264247 | LINC00909 | 652.7288 | -0.07128239 | 0.1866893 | -0.3896545 | 0.696792 | 0.834411 |
| ENSG00000163207 | IVL | 39.85135 | -0.07130621 | 0.7119806 | -0.1928797 | 0.8470531 | 0.925461 |
| ENSG00000239944 | TRBV8-2 | 1.375583 | -0.07131383 | 0.7952424 | -1.095288 | 0.2733903 | NA |
| ENSG00000228686 | | 1.628766 | -0.07131448 | 0.6934728 | -0.2321119 | 0.8164511 | 0.90654 |
| ENSG00000279359 | | 1.363544 | -0.07132239 | 0.7952414 | -1.093053 | 0.2743704 | NA |
| ENSG00000141198 | TOM1L1 | 4700.452 | -0.07132388 | 0.1246936 | -0.5292255 | 0.596649 | 0.763712 |
| ENSG00000287382 | | 1.642471 | -0.07133819 | 0.795296 | -1.145551 | 0.2519808 | 0.444695 |
| ENSG00000231290 | APCDD1L-DT | 13.49952 | -0.07134583 | 0.5063733 | -0.1854001 | 0.8529152 | 0.928429 |
| ENSG00000254213 | PRXL2AP2 | 2.475205 | -0.07135865 | 0.6706689 | -0.1999881 | 0.8414899 | 0.921735 |
| ENSG00000248795 | | 49.86014 | -0.07137168 | 0.3047049 | -0.2562688 | 0.7977433 | 0.895734 |
| ENSG00000268529 | CYP2T3P | 1.031465 | -0.07141471 | 0.7296151 | -0.2562234 | 0.7977784 | NA |
| ENSG00000121989 | ACVR2A | 553.7145195 | -0.071442139 | 0.159142625 | -0.451501733 | 0.65162798 | 0.803243 |
| ENSG00000134504 | KCTD1 | 577.9763596 | -0.071445497 | 0.136510627 | -0.533043791 | 0.59400327 | 0.76161 |
| ENSG00000230585 | PHBP12 | 4.648237 | -0.07144668 | 0.6281536 | -0.1916319 | 0.8480305 | 0.926147 |
| ENSG00000176654 | NANOGP1 | 1.555409 | -0.07146398 | 0.7952976 | -1.129728 | 0.2585908 | NA |
| ENSG00000108381 | ASPA | 1.590115897 | -0.071488384 | 0.795306801 | -1.124338442 | 0.26086948 | 0.45557 |
| ENSG00000111647 | UHRF1BP1L | 1438.463268 | -0.071488458 | 0.255987217 | -0.300769911 | 0.76358995 | 0.874249 |
| ENSG00000285724 | | 1.631439 | -0.07149953 | 0.7953166 | -1.143815 | 0.2527005 | 0.445534 |
| ENSG00000214289 | RPL39P5 | 7.79172 | -0.07154063 | 0.6089513 | -0.1792274 | 0.8577591 | 0.931298 |
| ENSG00000214535 | RPS15AP1 | 360.614 | -0.07156378 | 0.2590132 | -0.3032766 | 0.7616791 | 0.872964 |
| ENSG00000237922 | | 0.9039708 | -0.07158606 | 0.7697669 | -0.4116455 | 0.6805993 | NA |
| ENSG00000286062 | | 2.013518 | -0.07158688 | 0.7953778 | -1.202917 | 0.2290084 | 0.417234 |
| ENSG00000164764 | SBSPON | 24.06737 | -0.07159395 | 0.4082916 | -0.2070195 | 0.8359946 | 0.917845 |
| ENSG00000227418 | PCGEM1 | 1.909857 | -0.07162928 | 0.7953721 | -1.18802 | 0.2348254 | 0.424081 |
| ENSG00000286889 | | 1.791673 | -0.07164347 | 0.7953555 | -1.154156 | 0.2484364 | 0.440775 |
| ENSG00000284825 | | 2.108706 | -0.07169031 | 0.7954018 | -1.2159 | 0.224023 | 0.410803 |
| ENSG00000253660 | | 2.060104 | -0.07170666 | 0.7953992 | -1.209352 | 0.2265278 | 0.414163 |

| | | | | | | | |
|---|---|---|---|---|---|---|---|
| ENSG00000279897 | BIRC6-AS2 | 1.107573 | -0.07175282 | 0.7217755 | -0.2376928 | 0.8121194 | NA |
| ENSG00000283400 | | 3.828779 | -0.07179106 | 0.7955107 | -1.383967 | 0.1663687 | 0.335376 |
| ENSG00000272727 | LINC02266 | 2.264965 | -0.07182688 | 0.7954347 | -1.23599 | 0.2164623 | 0.401731 |
| ENSG00000136897 | MRPL50 | 1749.318644 | -0.071855182 | 0.213410036 | -0.351936323 | 0.72488602 | 0.850636 |
| ENSG00000237574 | LINC01856 | 1.153843 | -0.07186922 | 0.7261307 | -0.2734684 | 0.7844932 | NA |
| ENSG00000137392 | CLPS | 4.93937997 | -0.071962115 | 0.795562852 | -1.450111187 | 0.14702752 | 0.306719 |
| ENSG00000234537 | | 10.19995 | -0.0719634 | 0.495001 | -0.1896712 | 0.8495668 | 0.926874 |
| ENSG00000241975 | ELOCP19 | 20.74702 | -0.07197136 | 0.4465423 | -0.1990771 | 0.8422024 | 0.922133 |
| ENSG00000203995 | ZYG11A | 900.3173 | -0.07198255 | 0.2264435 | -0.3408514 | 0.7332155 | 0.855322 |
| ENSG00000111817 | DSE | 676.7881002 | -0.07201083 | 0.197958678 | -0.378077308 | 0.70537317 | 0.839947 |
| ENSG00000187581 | COX8C | 2.480509 | -0.07201827 | 0.6544414 | -0.1991273 | 0.8421632 | 0.922133 |
| ENSG00000088038 | CNOT3 | 4153.800155 | -0.072062189 | 0.142166702 | -0.514405214 | 0.60696872 | 0.77103 |
| ENSG00000264010 | MIR4429 | 3.936521 | -0.07210093 | 0.7955562 | -1.3917 | 0.1640133 | 0.332049 |
| ENSG00000272779 | | 14.74772 | -0.07210667 | 0.4164372 | -0.2057234 | 0.837007 | 0.918602 |
| ENSG00000255121 | CENATAC-DT | 127.6915 | -0.0721076 | 0.3637268 | -0.2251302 | 0.821878 | 0.909626 |
| ENSG00000196126 | HLA-DRB1 | 165.0167 | -0.07211629 | 0.4250204 | -0.2012734 | 0.8404848 | 0.921114 |
| ENSG00000211716 | TRBV9 | 3.981539 | -0.07214364 | 0.7955635 | -1.394933 | 0.1630362 | 0.330614 |
| ENSG00000280444 | | 4.728684 | -0.07215961 | 0.795584 | -1.443347 | 0.1489228 | 0.309555 |
| ENSG00000277411 | | 0.7717872 | -0.07216442 | 0.7548385 | -0.305222 | 0.7601971 | NA |
| ENSG00000160808 | MYL3 | 4.993864 | -0.07220186 | 0.7955963 | -1.450676 | 0.14687 | 0.306487 |
| ENSG00000005884 | ITGA3 | 7506.999445 | -0.072271604 | 0.231655983 | -0.290902227 | 0.7711261 | 0.878976 |
| ENSG00000138463 | SLC49A4 | 1561.363371 | -0.072435752 | 0.113209924 | -0.658735366 | 0.51006572 | 0.695971 |
| ENSG00000124614 | RPS10 | 167.7401037 | -0.072457419 | 0.240034357 | -0.319418555 | 0.74940914 | 0.865842 |
| ENSG00000253309 | SERPINE3 | 5.968998 | -0.07249357 | 0.5535668 | -0.1856286 | 0.8527361 | 0.928421 |
| ENSG00000255394 | C8orf49 | 2.475209 | -0.07265007 | 0.7955647 | -1.261166 | 0.2072491 | 0.389641 |
| ENSG00000223501 | VPS52 | 3388.952 | -0.07271177 | 0.1220549 | -0.6097012 | 0.5420598 | 0.723521 |
| ENSG00000228436 | | 64.95857 | -0.07272163 | 0.4977905 | -0.1923053 | 0.8475031 | 0.925823 |
| ENSG00000254855 | | 10.0004 | -0.07277866 | 0.5301405 | -0.1885487 | 0.8504465 | 0.927338 |
| ENSG00000156466 | GDF6 | 6.616444 | -0.07280321 | 0.6875318 | -0.236179 | 0.8132938 | 0.904685 |
| ENSG00000276250 | | 3.252714 | -0.07284115 | 0.6336211 | -0.1915955 | 0.8480591 | 0.926147 |
| ENSG00000277759 | MIR6719 | 2.069948 | -0.07284492 | 0.6738486 | -0.2203012 | 0.8256366 | 0.91192 |
| ENSG00000223729 | LINC02247 | 8.784553 | -0.07287118 | 0.7957284 | -1.61764 | 0.1057401 | 0.242358 |
| ENSG00000223884 | | 3.731549 | -0.07290109 | 0.6670528 | -0.2084827 | 0.8348521 | 0.91716 |

| | | | | | | | |
|---|---|---|---|---|---|---|---|
| ENSG00000052841 | TTC17 | 4431.479053 | -0.072956929 | 0.228995684 | -0.320353531 | 0.74870035 | 0.865481 |
| ENSG00000111906 | HDDC2 | 3504.159829 | -0.072977103 | 0.187497034 | -0.408794001 | 0.68269084 | 0.824398 |
| ENSG00000070047 | PHRF1 | 3165.861788 | -0.073148677 | 0.137039739 | -0.486518586 | 0.62659953 | 0.785215 |
| ENSG00000082781 | ITGB5 | 4525.643753 | -0.073200895 | 0.231225662 | -0.29206025 | 0.77024056 | 0.878494 |
| ENSG00000172171 | TEFM | 315.3805 | -0.0732156 | 0.1864336 | -0.4074838 | 0.6836527 | 0.824997 |
| ENSG00000256591 | | 18.90242 | -0.07323345 | 0.4055682 | -0.2151575 | 0.8296446 | 0.914205 |
| ENSG00000181773 | GPR3 | 42.76693 | -0.07329085 | 0.5794808 | -0.1883095 | 0.850634 | 0.927437 |
| ENSG00000117523 | PRRC2C | 19116.75558 | -0.073341358 | 0.199896749 | -0.426400504 | 0.66981603 | 0.816084 |
| ENSG00000130731 | METTL26 | 4114.700842 | -0.073378293 | 0.203405487 | -0.377885021 | 0.70551601 | 0.840033 |
| ENSG00000021776 | AQR | 3584.278197 | -0.073378797 | 0.152377882 | -0.49448561 | 0.62096327 | 0.781019 |
| ENSG00000114779 | ABHD14B | 2415.937211 | -0.073395911 | 0.169555237 | -0.439136438 | 0.66056268 | 0.809118 |
| ENSG00000269427 | | 18.57174 | -0.07347438 | 0.4512876 | -0.201198 | 0.8405437 | 0.921114 |
| ENSG00000188873 | RPL10AP2 | 78.83131 | -0.07355244 | 0.2944271 | -0.2677438 | 0.7888965 | 0.890218 |
| ENSG00000144524 | COPS7B | 2135.992 | -0.07355258 | 0.1526096 | -0.4919116 | 0.6227818 | 0.782218 |
| ENSG00000126733 | DACH2 | 15.66872657 | -0.073583627 | 0.451691982 | -0.204575259 | 0.83790399 | 0.919131 |
| ENSG00000198574 | SH2D1B | 1.887912 | -0.07363052 | 0.7268575 | -0.2830361 | 0.7771491 | 0.882585 |
| ENSG00000184232 | OAF | 582.9284 | -0.07369331 | 0.2365345 | -0.3252567 | 0.7449868 | 0.86269 |
| ENSG00000112640 | PPP2R5D | 1866.746463 | -0.073697665 | 0.195758052 | -0.397287619 | 0.69115538 | 0.830171 |
| ENSG00000116544 | DLGAP3 | 66.7260802 | -0.073810485 | 0.456600049 | -0.18272565 | 0.85501329 | 0.929875 |
| ENSG00000196911 | KPNA5 | 842.2278 | -0.0738148 | 0.1548692 | -0.4877601 | 0.6257198 | 0.78456 |
| ENSG00000285589 | | 97.89303 | -0.07384887 | 0.2634927 | -0.3000809 | 0.7641154 | 0.874555 |
| ENSG00000224599 | BMS1P12 | 1.250719 | -0.07387832 | 0.7128984 | -0.2417663 | 0.8089612 | NA |
| ENSG00000258092 | | 1.056892 | -0.07392386 | 0.7079116 | -0.2433352 | 0.8077457 | NA |
| ENSG00000156603 | MED19 | 739.4049 | -0.07395524 | 0.1338907 | -0.5611806 | 0.5746744 | 0.747029 |
| ENSG00000105610 | KLF1 | 9.744884359 | -0.073956164 | 0.495345762 | -0.194788107 | 0.84555883 | 0.924422 |
| ENSG00000125966 | MMP24 | 72.87017911 | -0.07400822 | 0.403684161 | -0.21386803 | 0.83064997 | 0.914899 |
| ENSG00000278023 | RDM1 | 65.29197 | -0.07409901 | 0.4677006 | -0.2003428 | 0.8412125 | 0.921581 |
| ENSG00000261349 | | 14.08184 | -0.07411881 | 0.5971883 | -0.1942903 | 0.8459486 | 0.92471 |
| ENSG00000145632 | PLK2 | 10856.79 | -0.0741205 | 0.3638596 | -0.2112794 | 0.8326693 | 0.915771 |
| ENSG00000266412 | NCOA4 | 10252.15 | -0.07424541 | 0.1314543 | -0.5751524 | 0.5651882 | 0.7402 |
| ENSG00000283849 | | 4.207501 | -0.0742455 | 0.6121105 | -0.1994287 | 0.8419274 | 0.92206 |
| ENSG00000260500 | | 7.000348 | -0.07424978 | 0.5389862 | -0.1812331 | 0.8561846 | 0.930273 |
| ENSG00000229688 | CRPPA-AS1 | 1.826618 | -0.07427608 | 0.7094491 | -0.2414444 | 0.8092107 | 0.90262 |

| | | | | | | | |
|---|---|---|---|---|---|---|---|
| ENSG00000100632 | ERH | 4868.10015 | -0.074284745 | 0.201671258 | -0.39619676 | 0.69195988 | 0.830758 |
| ENSG00000128254 | C22orf24 | 9.482174302 | -0.074348853 | 0.524405942 | -0.187926456 | 0.8509343 | 0.927491 |
| ENSG00000253908 | | 16.08309 | -0.07445279 | 0.4655852 | -0.1988218 | 0.8424022 | 0.922238 |
| ENSG00000123838 | C4BPA | 5.974179694 | -0.074520163 | 0.705813411 | -0.225968212 | 0.82122614 | 0.909245 |
| ENSG00000287218 | | 4.796754 | -0.07454866 | 0.6192018 | -0.1932454 | 0.8467668 | 0.925306 |
| ENSG00000199804 | RNA5SP383 | 16.28704 | -0.07455648 | 0.4081996 | -0.2147723 | 0.8299449 | 0.91444 |
| ENSG00000054118 | THRAP3 | 10751.42982 | -0.074610024 | 0.146967073 | -0.482668739 | 0.62933096 | 0.787012 |
| ENSG00000185467 | KPNA7 | 14.10004 | -0.07463303 | 0.5816814 | -0.1868826 | 0.8517527 | 0.927845 |
| ENSG00000107485 | GATA3 | 33949.88941 | -0.07473551 | 0.155352529 | -0.598262223 | 0.54966498 | 0.728963 |
| ENSG00000131408 | NR1H2 | 3857.003092 | -0.074743882 | 0.157218566 | -0.487080512 | 0.62620128 | 0.784923 |
| ENSG00000267908 | ZSCAN5DP | 1.034485 | -0.07481735 | 0.7275454 | -0.2581534 | 0.7962886 | NA |
| ENSG00000265052 | RN7SL622P | 0.9254433 | -0.07495385 | 0.7428616 | -0.3030707 | 0.761836 | NA |
| ENSG00000185947 | ZNF267 | 915.2736 | -0.07497111 | 0.224228 | -0.3565101 | 0.7214586 | 0.848466 |
| ENSG00000175061 | SNHG29 | 15484.64 | -0.07500277 | 0.2061537 | -0.3884354 | 0.6976938 | 0.835001 |
| ENSG00000181524 | RPL24P4 | 3332.087 | -0.07505375 | 0.2081592 | -0.3784217 | 0.7051174 | 0.839876 |
| ENSG00000066044 | ELAVL1 | 5054.685888 | -0.07505691 | 0.169939271 | -0.463565751 | 0.6429589 | 0.796464 |
| ENSG00000232388 | SMIM26 | 861.1402 | -0.07511324 | 0.2051821 | -0.38336 | 0.7014528 | 0.83727 |
| ENSG00000150756 | ATPSCKMT | 1935.33 | -0.07512454 | 0.1496864 | -0.4832826 | 0.6288951 | 0.786756 |
| ENSG00000131242 | RAB11FIP4 | 2478.548381 | -0.075132927 | 0.116556583 | -0.626015515 | 0.53130477 | 0.715504 |
| ENSG00000279500 | | 59.29769 | -0.07519073 | 0.4173018 | -0.2151099 | 0.8296816 | 0.914205 |
| ENSG00000082258 | CCNT2 | 3109.20518 | -0.075209407 | 0.17947077 | -0.433773308 | 0.66445307 | 0.812493 |
| ENSG00000241907 | RPS20P4 | 4.194806 | -0.07523725 | 0.5735468 | -0.1952816 | 0.8451725 | 0.924242 |
| ENSG00000183513 | COA5 | 2649.841 | -0.07525914 | 0.1229728 | -0.6400247 | 0.5221565 | 0.707562 |
| ENSG00000226539 | MLXP1 | 1.721981 | -0.07527891 | 0.6778626 | -0.2177637 | 0.8276132 | 0.913083 |
| ENSG00000234127 | TRIM26 | 4343.52 | -0.0754228 | 0.1342576 | -0.5660733 | 0.5713439 | 0.745106 |
| ENSG00000259649 | | 0.9739242 | -0.07542602 | 0.7431814 | -0.2918207 | 0.7704237 | NA |
| ENSG00000215271 | HOMEZ | 1531.972 | -0.07547176 | 0.2149527 | -0.3687797 | 0.712292 | 0.843355 |
| ENSG00000006125 | AP2B1 | 8275.524046 | -0.075537695 | 0.171372679 | -0.457381799 | 0.64739664 | 0.800098 |
| ENSG00000238249 | HMGN2P17 | 0.8024948 | -0.07558383 | 0.7409618 | -0.2910087 | 0.7710447 | NA |
| ENSG00000198870 | STKLD1 | 155.1282 | -0.07561086 | 0.323807 | -0.273207 | 0.7846941 | 0.887289 |
| ENSG00000234213 | FHP1 | 6.144635 | -0.07563304 | 0.5827508 | -0.193145 | 0.8468454 | 0.92531 |
| ENSG00000260339 | HEXA-AS1 | 55.73401 | -0.07564507 | 0.3846983 | -0.2262147 | 0.8210345 | 0.909205 |
| ENSG00000260382 | | 3.901409 | -0.07566466 | 0.6528464 | -0.2080928 | 0.8351565 | 0.917253 |

| | | | | | | | |
|---|---|---|---|---|---|---|---|
| ENSG00000228998 | | 34.2633 | -0.07574191 | 0.3965898 | -0.2270222 | 0.8204064 | 0.908942 |
| ENSG00000227868 | TEX46 | 1.07473 | -0.07576225 | 0.7444446 | -0.293561 | 0.7690934 | NA |
| ENSG00000185104 | FAF1 | 3308.708 | -0.07581916 | 0.129996 | -0.5929875 | 0.5531895 | 0.731539 |
| ENSG00000124702 | KLHDC3 | 7292.693313 | -0.076016392 | 0.20144797 | -0.3857948 | 0.69964865 | 0.836286 |
| ENSG00000266236 | NARF-IT1 | 16.8398 | -0.07606283 | 0.4362746 | -0.2121877 | 0.8319606 | 0.915408 |
| ENSG00000125531 | FNDC11 | 6.803914707 | -0.076077422 | 0.541128956 | -0.194940026 | 0.8454399 | 0.924357 |
| ENSG00000251022 | THAP9-AS1 | 2125.988 | -0.07613477 | 0.2015358 | -0.3895211 | 0.6968907 | 0.834442 |
| ENSG00000107789 | MINPP1 | 1772.624656 | -0.076204533 | 0.205268262 | -0.384866145 | 0.70033659 | 0.836769 |
| ENSG00000124222 | STX16 | 13395.31834 | -0.076230558 | 0.189718525 | -0.414177637 | 0.67874402 | 0.821499 |
| ENSG00000230667 | SETSIP | 68.42503 | -0.07624562 | 0.2782412 | -0.2976705 | 0.7659547 | 0.875765 |
| ENSG00000103199 | ZNF500 | 1032.779103 | -0.076250025 | 0.127255835 | -0.607654566 | 0.5434166 | 0.724367 |
| ENSG00000011332 | DPF1 | 270.4738462 | -0.076313634 | 0.387609197 | -0.223349556 | 0.82326346 | 0.910363 |
| ENSG00000243989 | ACY1 | 56.73388 | -0.07631512 | 0.3438085 | -0.2465473 | 0.8052586 | 0.900308 |
| ENSG00000271763 | | 9.058688 | -0.07633009 | 0.5611959 | -0.1951607 | 0.8452671 | 0.92427 |
| ENSG00000221500 | SNORD100 | 33.76027 | -0.07635108 | 0.3521513 | -0.2425531 | 0.8083516 | 0.901868 |
| ENSG00000279375 | | 6.701858 | -0.07636473 | 0.6049847 | -0.2002236 | 0.8413057 | 0.921607 |
| ENSG00000231278 | OR9S24P | 0.9674461 | -0.0764565 | 0.7957739 | -1.033323 | 0.3014526 | NA |
| ENSG00000269425 | | 10.26323 | -0.07653932 | 0.4852797 | -0.2008154 | 0.8408429 | 0.921328 |
| ENSG00000240674 | | 2.700532 | -0.07655462 | 0.7143618 | -0.2411387 | 0.8094476 | 0.902793 |
| ENSG00000136367 | ZFHX2 | 344.9883573 | -0.076604687 | 0.359867866 | -0.240424578 | 0.81000113 | 0.902923 |
| ENSG00000260475 | | 22.67322 | -0.07661572 | 0.4202417 | -0.2167777 | 0.8283816 | 0.913464 |
| ENSG00000165171 | METTL27 | 76.22321 | -0.07666793 | 0.301701 | -0.2773132 | 0.7815396 | 0.885492 |
| ENSG00000217455 | | 0.938143 | -0.07667743 | 0.7489284 | -0.3325906 | 0.7394434 | NA |
| ENSG00000162396 | PARS2 | 578.0842 | -0.0766918 | 0.1226681 | -0.6341875 | 0.5259585 | 0.710757 |
| ENSG00000236801 | RPL24P8 | 2528.375 | -0.07677231 | 0.2030032 | -0.3916647 | 0.695306 | 0.833119 |
| ENSG00000154822 | PLCL2 | 12.23995 | -0.07687266 | 0.5748133 | -0.1888895 | 0.8501794 | 0.927199 |
| ENSG00000107798 | LIPA | 6271.616141 | -0.076954118 | 0.178504733 | -0.481490214 | 0.63016813 | 0.787755 |
| ENSG00000278963 | | 3.164414 | -0.07703482 | 0.6459021 | -0.2104976 | 0.8332793 | 0.91629 |
| ENSG00000257913 | DDN-AS1 | 12.32364 | -0.07707221 | 0.5117744 | -0.1999854 | 0.841492 | 0.921735 |
| ENSG00000100030 | MAPK1 | 6985.797369 | -0.077147581 | 0.130819512 | -0.59822169 | 0.54969202 | 0.728963 |
| ENSG00000249465 | RBMXP4 | 114.4071 | -0.07720359 | 0.2793669 | -0.2964522 | 0.7668847 | 0.876397 |
| ENSG00000130349 | MTRES1 | 815.654079 | -0.077359927 | 0.193520629 | -0.416528147 | 0.67702358 | 0.820351 |
| ENSG00000234315 | OSTCP5 | 1.357006 | -0.07736539 | 0.6884806 | -0.2281725 | 0.8195121 | NA |

| | | | | | | | |
|---|---|---|---|---|---|---|---|
| ENSG00000254973 | | 226.6508 | -0.07740394 | 0.27716 | -0.2999843 | 0.7641891 | 0.874596 |
| ENSG00000221518 | RNU6ATAC16P | 1.214543 | -0.07744063 | 0.7457623 | -0.3130828 | 0.7542177 | NA |
| ENSG00000236813 | BTF3P8 | 3.505572 | -0.07751622 | 0.6366352 | -0.2044851 | 0.8379745 | 0.91917 |
| ENSG00000162437 | RAVER2 | 1966.966 | -0.07765183 | 0.1287723 | -0.6131204 | 0.5397967 | 0.722087 |
| ENSG00000237992 | LINC01808 | 1.009148 | -0.077682 | 0.7362033 | -0.3141641 | 0.7533964 | NA |
| ENSG00000143514 | TP53BP2 | 2563.212 | -0.0778383 | 0.1438696 | -0.5560695 | 0.5781633 | 0.749803 |
| ENSG00000169727 | GPS1 | 9649.398 | -0.07792057 | 0.1729704 | -0.4693391 | 0.6388273 | 0.794021 |
| ENSG00000123505 | AMD1 | 5277.990318 | -0.077921254 | 0.216610854 | -0.377305561 | 0.70594654 | 0.840133 |
| ENSG00000235660 | LINC00345 | 8.297307 | -0.07800248 | 0.5114309 | -0.2086059 | 0.8347559 | 0.91716 |
| ENSG00000260163 | | 1.836805 | -0.07801845 | 0.7222732 | -0.2625307 | 0.7929123 | 0.892909 |
| ENSG00000232450 | | 69.96738 | -0.07802828 | 0.4015695 | -0.2256624 | 0.821464 | 0.909357 |
| ENSG00000182899 | RPL35A | 15845.66 | -0.07810847 | 0.1602194 | -0.5080517 | 0.6114171 | 0.774478 |
| ENSG00000106399 | RPA3 | 2244.283245 | -0.078121044 | 0.362008943 | -0.250839044 | 0.80193855 | 0.898369 |
| ENSG00000169249 | ZRSR2 | 712.8385 | -0.07818448 | 0.1911595 | -0.4177365 | 0.6761398 | 0.820028 |
| ENSG00000114904 | NEK4 | 1495.587122 | -0.078200693 | 0.174226455 | -0.470197751 | 0.63821374 | 0.793593 |
| ENSG00000163909 | HEYL | 302.621 | -0.07831893 | 0.2548699 | -0.3309068 | 0.7407149 | 0.860071 |
| ENSG00000227218 | | 130.8844 | -0.07832912 | 0.3840227 | -0.2396041 | 0.8106372 | 0.903091 |
| ENSG00000198218 | QRICH1 | 4144.634 | -0.0783345 | 0.1189573 | -0.6677557 | 0.5042896 | 0.691213 |
| ENSG00000156711 | MAPK13 | 2726.66 | -0.07835482 | 0.157515 | -0.5082047 | 0.6113098 | 0.774461 |
| ENSG00000261618 | LINC02605 | 4.866568 | -0.07837864 | 0.6447394 | -0.2132595 | 0.8311246 | 0.915137 |
| ENSG00000131061 | ZNF341 | 421.9706116 | -0.078394942 | 0.222587302 | -0.361550527 | 0.71768794 | 0.846015 |
| ENSG00000279148 | | 28.8737 | -0.07840731 | 0.4123854 | -0.2232774 | 0.8233196 | 0.910387 |
| ENSG00000197162 | ZNF785 | 702.9656 | -0.07841747 | 0.279063 | -0.3037157 | 0.7613445 | 0.872735 |
| ENSG00000217512 | | 1.104512 | -0.07842698 | 0.7539551 | -0.3338424 | 0.7384985 | NA |
| ENSG00000239704 | CDRT4 | 17.08495 | -0.07850332 | 0.4288856 | -0.2202316 | 0.8256908 | 0.91192 |
| ENSG00000257303 | | 58.06384 | -0.07850899 | 0.269277 | -0.3121543 | 0.7549232 | 0.869057 |
| ENSG00000196670 | ZFP62 | 3597.595 | -0.07853089 | 0.2172628 | -0.3815628 | 0.7027856 | 0.838263 |
| ENSG00000067057 | PFKP | 12277.29288 | -0.078717388 | 0.217119077 | -0.364066119 | 0.71580864 | 0.845328 |
| ENSG00000184261 | KCNK12 | 52.80325 | -0.07885655 | 0.4282789 | -0.2201305 | 0.8257695 | 0.91192 |
| ENSG00000125551 | PLGLB2 | 32.62252289 | -0.078898754 | 0.438566088 | -0.220950379 | 0.82513107 | 0.911882 |
| ENSG00000068354 | TBC1D25 | 902.6602627 | -0.0789254 | 0.150989925 | -0.535124686 | 0.59256364 | 0.760732 |
| ENSG00000174915 | PTDSS2 | 2276.532 | -0.07892772 | 0.1686208 | -0.4773017 | 0.6331473 | 0.789733 |
| ENSG00000110944 | IL23A | 26.54691824 | -0.078936674 | 0.405740196 | -0.228750209 | 0.81906307 | 0.90844 |

| | | | | | | | |
|---|---|---|---|---|---|---|---|
| ENSG00000166343 | MSS51 | 120.1128 | -0.07894374 | 0.2343455 | -0.3553858 | 0.7223006 | 0.848898 |
| ENSG00000166169 | POLL | 1766.369 | -0.07911698 | 0.1042598 | -0.769974 | 0.4413153 | 0.639156 |
| ENSG00000242307 | RPS26P52 | 2.167608 | -0.07913684 | 0.7024459 | -0.239037 | 0.8110769 | 0.903386 |
| ENSG00000188338 | SLC38A3 | 4.552963 | -0.07915848 | 0.6443236 | -0.2178741 | 0.8275272 | 0.913064 |
| ENSG00000110801 | PSMD9 | 540.5215966 | -0.079165758 | 0.200561862 | -0.414281008 | 0.67866832 | 0.821485 |
| ENSG00000170011 | MYRIP | 57.84322 | -0.07921847 | 0.4736572 | -0.218082 | 0.8273652 | 0.913037 |
| ENSG00000260470 | | 3.799595 | -0.07927813 | 0.6415041 | -0.2129723 | 0.8313486 | 0.915137 |
| ENSG00000099942 | CRKL | 5202.083836 | -0.079333137 | 0.158819709 | -0.531682842 | 0.59494568 | 0.762303 |
| ENSG00000206775 | SNORD37 | 2.126829 | -0.07933753 | 0.6801332 | -0.225441 | 0.8216362 | 0.90951 |
| ENSG00000271181 | | 0.8347698 | -0.07933855 | 0.7483644 | -0.3130328 | 0.7542557 | NA |
| ENSG00000139291 | TMEM19 | 707.3399 | -0.07935321 | 0.2672007 | -0.319956 | 0.7490017 | 0.865675 |
| ENSG00000278462 | | 1.653783 | -0.07941589 | 0.7252913 | -0.2601314 | 0.7947624 | 0.894013 |
| ENSG00000166025 | AMOTL1 | 4040.099 | -0.0794251 | 0.3392482 | -0.2566028 | 0.7974854 | 0.895607 |
| ENSG00000182093 | GET1 | 1915.539 | -0.079434 | 0.1608578 | -0.5056909 | 0.6130736 | 0.775766 |
| ENSG00000167595 | PROSER3 | 749.4357 | -0.07949497 | 0.1706393 | -0.4774936 | 0.6330107 | 0.789665 |
| ENSG00000250948 | | 0.8694394 | -0.07952134 | 0.7636265 | -0.4000886 | 0.6890912 | NA |
| ENSG00000204463 | BAG6 | 12786.6 | -0.07953468 | 0.1431867 | -0.5999986 | 0.5485072 | 0.727973 |
| ENSG00000178934 | LGALS7B | 1.556074 | -0.07956463 | 0.7962906 | -1.089672 | 0.2758576 | NA |
| ENSG00000225675 | LINC01771 | 24.31428 | -0.07958067 | 0.3685664 | -0.2475498 | 0.8044828 | 0.899971 |
| ENSG00000071082 | RPL31 | 19223.04208 | -0.079646953 | 0.169487903 | -0.476730825 | 0.63355381 | 0.790017 |
| ENSG00000105556 | MIER2 | 941.4148994 | -0.079684887 | 0.180538261 | -0.46541814 | 0.64163205 | 0.795666 |
| ENSG00000239650 | GUSBP4 | 1.025061 | -0.07974478 | 0.7428476 | -0.3288514 | 0.742268 | NA |
| ENSG00000167549 | CORO6 | 21.69379 | -0.07978186 | 0.4055885 | -0.2328134 | 0.8159063 | 0.906211 |
| ENSG00000185238 | PRMT3 | 1803.612 | -0.07982774 | 0.1980137 | -0.4237348 | 0.6717593 | 0.817546 |
| ENSG00000121764 | HCRTR1 | 3.130531724 | -0.079833309 | 0.635219594 | -0.213241574 | 0.83113854 | 0.915137 |
| ENSG00000177943 | MAMDC4 | 1242.507 | -0.07985279 | 0.2192518 | -0.3915401 | 0.695398 | 0.833192 |
| ENSG00000124574 | ABCC10 | 2485.655724 | -0.079869748 | 0.180735683 | -0.450166931 | 0.65259008 | 0.803944 |
| ENSG00000287180 | | 1.300404 | -0.079905 | 0.6911929 | -0.2467695 | 0.8050866 | NA |
| ENSG00000281183 | NPTN-IT1 | 183.8164 | -0.07991634 | 0.3629878 | -0.2511428 | 0.8017037 | 0.898186 |
| ENSG00000157999 | ANKRD61 | 100.3377 | -0.07992572 | 0.2972266 | -0.292941 | 0.7695673 | 0.877917 |
| ENSG00000171155 | C1GALT1C1 | 1890.851 | -0.07995726 | 0.1586393 | -0.5438462 | 0.5865473 | 0.756247 |
| ENSG00000279187 | | 14.10241 | -0.08005118 | 0.5118206 | -0.2070181 | 0.8359957 | 0.917845 |
| ENSG00000126970 | ZC4H2 | 300.4162516 | -0.080122685 | 0.293900114 | -0.295040633 | 0.76796284 | 0.877026 |

| | | | | | | | |
|---|---|---|---|---|---|---|---|
| ENSG00000221947 | XKR9 | 74.99667 | -0.08012809 | 0.3687558 | -0.2461067 | 0.8055997 | 0.90031 |
| ENSG00000233060 | | 0.8195447 | -0.08017768 | 0.7392349 | -0.3021146 | 0.7625647 | NA |
| ENSG00000241293 | PPATP1 | 66.31276 | -0.08027001 | 0.3294873 | -0.2747099 | 0.7835392 | 0.886724 |
| ENSG00000284738 | | 27.96986 | -0.08043543 | 0.4033593 | -0.233182 | 0.8156201 | 0.90611 |
| ENSG00000198856 | OSTC | 4510.133 | -0.08043576 | 0.1632664 | -0.5161103 | 0.6057774 | 0.770165 |
| ENSG00000088682 | COQ9 | 2533.884933 | -0.080462785 | 0.178411412 | -0.49287662 | 0.62209977 | 0.781714 |
| ENSG00000167702 | KIFC2 | 8171.741 | -0.08064382 | 0.3031404 | -0.2934853 | 0.7691512 | 0.877666 |
| ENSG00000104133 | SPG11 | 4306.04466 | -0.080648888 | 0.225496699 | -0.380640406 | 0.7034701 | 0.838628 |
| ENSG00000271382 | | 0.8344506 | -0.08073714 | 0.731481 | -0.2844251 | 0.7760846 | NA |
| ENSG00000266490 | | 28.45553 | -0.08074069 | 0.3745732 | -0.2455916 | 0.8059984 | 0.90047 |
| ENSG00000227615 | | 6074.957 | -0.08075953 | 0.1763159 | -0.4692339 | 0.6389025 | 0.794041 |
| ENSG00000128185 | DGCR6L | 3176.778966 | -0.080808509 | 0.21748419 | -0.391358605 | 0.69553219 | 0.83324 |
| ENSG00000254732 | | 4.593523 | -0.08085223 | 0.624569 | -0.2068454 | 0.8361306 | 0.91794 |
| ENSG00000122687 | MRM2 | 71.02351652 | -0.080888946 | 0.238071811 | -0.360049734 | 0.71880994 | 0.846874 |
| ENSG00000177426 | TGIF1 | 5382.826 | -0.08089325 | 0.2038447 | -0.4073911 | 0.6837207 | 0.825042 |
| ENSG00000279608 | | 1.289581 | -0.08095383 | 0.7069519 | -0.2612188 | 0.7939238 | NA |
| ENSG00000146555 | SDK1 | 1561.248 | -0.08099583 | 0.3561697 | -0.2694021 | 0.7876203 | 0.889383 |
| ENSG00000215218 | UBE2QL1 | 397.3373 | -0.08111005 | 0.3737314 | -0.246418 | 0.8053587 | 0.90031 |
| ENSG00000284946 | | 301.6887 | -0.0811221 | 0.3794072 | -0.2536219 | 0.7997877 | 0.897068 |
| ENSG00000286035 | | 1.840724 | -0.0811875 | 0.6649161 | -0.2301527 | 0.8179731 | 0.907686 |
| ENSG00000116688 | MFN2 | 5685.314759 | -0.081304454 | 0.133983163 | -0.627619801 | 0.53025303 | 0.71467 |
| ENSG00000270062 | | 22.57487 | -0.08132563 | 0.4632858 | -0.218969 | 0.8266742 | 0.912539 |
| ENSG00000226427 | HMGN2P7 | 1.243283 | -0.08141104 | 0.7433748 | -0.3238592 | 0.7460446 | NA |
| ENSG00000286614 | | 21.32602 | -0.08141849 | 0.398858 | -0.2392967 | 0.8108755 | 0.903313 |
| ENSG00000011454 | RABGAP1 | 6255.70326 | -0.081505069 | 0.180500027 | -0.469237207 | 0.63890009 | 0.794041 |
| ENSG00000165699 | TSC1 | 4036.27 | -0.08151574 | 0.2577239 | -0.338187 | 0.7352223 | 0.856543 |
| ENSG00000239557 | PPP2R5CP | 245.9933 | -0.08151966 | 0.1948338 | -0.4296366 | 0.66746 | 0.814764 |
| ENSG00000163754 | GYG1 | 5743.989 | -0.08152498 | 0.2127928 | -0.4158532 | 0.6775175 | 0.820725 |
| ENSG00000227543 | SPAG5-AS1 | 96.59185 | -0.08158878 | 0.2638962 | -0.3276135 | 0.7432039 | 0.861574 |
| ENSG00000253945 | | 4.594161 | -0.08160466 | 0.6628582 | -0.232362 | 0.8162568 | 0.906425 |
| ENSG00000137055 | PLAA | 1687.174011 | -0.081658873 | 0.137266088 | -0.601012426 | 0.54783171 | 0.727476 |
| ENSG00000263004 | | 320.8332 | -0.08169239 | 0.2897445 | -0.3045566 | 0.7607039 | 0.872373 |
| ENSG00000286846 | | 5.481041 | -0.08173925 | 0.6032436 | -0.2141863 | 0.8304018 | 0.914792 |

| | | | | | | | |
|---|---|---|---|---|---|---|---|
| ENSG00000234241 | | 514.2869 | -0.08174748 | 0.2377607 | -0.3673292 | 0.7133735 | 0.84368 |
| ENSG00000260737 | LINC01227 | 3.987199 | -0.0817902 | 0.6265441 | -0.2163973 | 0.8286781 | 0.913575 |
| ENSG00000239264 | TXNDC5 | 112.679 | -0.08183425 | 0.3444319 | -0.2644288 | 0.7914496 | 0.891863 |
| ENSG00000114316 | USP4 | 1975.389555 | -0.081870347 | 0.135273043 | -0.616976931 | 0.53724994 | 0.7203 |
| ENSG00000236444 | UBE2L5 | 1.363549 | -0.08188566 | 0.7196162 | -0.265102 | 0.7909309 | NA |
| ENSG00000133773 | CCDC59 | 2263.219416 | -0.081960315 | 0.162387755 | -0.561180164 | 0.57467472 | 0.747029 |
| ENSG00000183077 | AFMID | 1079.5 | -0.08197277 | 0.1459072 | -0.5650318 | 0.5720521 | 0.745554 |
| ENSG00000143569 | UBAP2L | 13811.25 | -0.08200338 | 0.153456 | -0.5483625 | 0.583443 | 0.753696 |
| ENSG00000198573 | SPANXC | 1.902915 | -0.08206204 | 0.7953353 | -0.6955122 | 0.4867343 | 0.677727 |
| ENSG00000270108 | | 13.66148 | -0.08209891 | 0.5121457 | -0.2118236 | 0.8322446 | 0.915608 |
| ENSG00000236848 | RPL23AP95 | 1.510572 | -0.08212003 | 0.7226616 | -0.2753506 | 0.7830469 | NA |
| ENSG00000067334 | DNTTIP2 | 2600.054708 | -0.082143966 | 0.21905671 | -0.420642853 | 0.6740159 | 0.818723 |
| ENSG00000029534 | ANK1 | 74.55625138 | -0.082177081 | 0.387402091 | -0.243740641 | 0.8074317 | 0.901372 |
| ENSG00000123607 | TTC21B | 920.1818539 | -0.082193867 | 0.202675443 | -0.436194889 | 0.66269534 | 0.811053 |
| ENSG00000177733 | HNRNPA0 | 11382.93 | -0.08219528 | 0.1956718 | -0.4190622 | 0.6751707 | 0.819414 |
| ENSG00000246465 | | 116.0836 | -0.08220846 | 0.3677234 | -0.2706628 | 0.7866504 | 0.888703 |
| ENSG00000117592 | PRDX6 | 17558.57436 | -0.082214846 | 0.194356277 | -0.448155932 | 0.65404066 | 0.804895 |
| ENSG00000204444 | APOM | 99.93184 | -0.08233563 | 0.3057384 | -0.2960012 | 0.7672292 | 0.876588 |
| ENSG00000144040 | SFXN5 | 1622.464 | -0.0823738 | 0.2892025 | -0.3106029 | 0.7561025 | 0.869868 |
| ENSG00000223711 | | 1.422778 | -0.08249509 | 0.702665 | -0.2521431 | 0.8009304 | NA |
| ENSG00000221539 | SNORD99 | 50.47951 | -0.08255341 | 0.3459491 | -0.2568698 | 0.7972793 | 0.895456 |
| ENSG00000100225 | FBXO7 | 4783.552085 | -0.082768526 | 0.102778942 | -0.815915064 | 0.41454872 | 0.614649 |
| ENSG00000234740 | | 2.310923 | -0.0827797 | 0.6940057 | -0.2466507 | 0.8051786 | 0.900308 |
| ENSG00000281106 | TMEM272 | 3.663591 | -0.08282853 | 0.6568337 | -0.2316068 | 0.8168435 | 0.906849 |
| ENSG00000137815 | RTF1 | 4703.070665 | -0.082840196 | 0.164662771 | -0.540339926 | 0.58896263 | 0.757928 |
| ENSG00000156384 | SFR1 | 162.4223 | -0.08290854 | 0.3544629 | -0.2654536 | 0.7906601 | 0.891421 |
| ENSG00000169246 | NPIPB3 | 221.9822 | -0.08291057 | 0.2618795 | -0.3416957 | 0.7325799 | 0.855068 |
| ENSG00000227922 | SPTLC1P5 | 2.672205 | -0.082953 | 0.7131766 | -0.268972 | 0.7879512 | 0.889681 |
| ENSG00000177133 | PRDM16-DT | 42.11305 | -0.08297751 | 0.4545113 | -0.2222392 | 0.8241277 | 0.911025 |
| ENSG00000280486 | | 23.55907 | -0.08298901 | 0.3930286 | -0.2461845 | 0.8055394 | 0.90031 |
| ENSG00000279800 | BCLAF1P2 | 1001.29 | -0.08301744 | 0.2219435 | -0.3881241 | 0.6979242 | 0.835202 |
| ENSG00000010379 | SLC6A13 | 1.528947982 | -0.083022158 | 0.678188706 | -0.240628977 | 0.80984269 | NA |
| ENSG00000269938 | | 12.33932 | -0.08304376 | 0.5831492 | -0.2131746 | 0.8311908 | 0.915137 |

| | | | | | | | |
|---|---|---|---|---|---|---|---|
| ENSG00000176912 | TYMSOS | 74.09667 | -0.08306738 | 0.3590473 | -0.2613806 | 0.793799 | 0.893322 |
| ENSG00000230629 | RPS23P8 | 3796.792 | -0.08306957 | 0.1759157 | -0.4785302 | 0.6322729 | 0.789227 |
| ENSG00000286493 | | 0.9451392 | -0.08308272 | 0.7387664 | -0.3191555 | 0.7496086 | NA |
| ENSG00000257954 | | 5.384819 | -0.08315795 | 0.624536 | -0.216467 | 0.8286238 | 0.913555 |
| ENSG00000235920 | THRAP3P3 | 2.41605 | -0.08317957 | 0.6628633 | -0.2425933 | 0.8083205 | 0.901868 |
| ENSG00000228929 | RPS13P2 | 2045.279 | -0.08320768 | 0.1671697 | -0.5331241 | 0.5939477 | 0.761575 |
| ENSG00000100603 | SNW1 | 5618.596004 | -0.083225092 | 0.093965182 | -0.898014756 | 0.36917768 | 0.571335 |
| ENSG00000228793 | | 6.219147 | -0.08327085 | 0.6725467 | -0.2273172 | 0.8201771 | 0.908902 |
| ENSG00000205111 | CDKL4 | 2.858473 | -0.08327225 | 0.6566576 | -0.2352658 | 0.8140025 | 0.905094 |
| ENSG00000249007 | | 269.7491 | -0.08339043 | 0.4489602 | -0.228189 | 0.8194993 | 0.908698 |
| ENSG00000231942 | HNRNPA1P36 | 3.371574 | -0.08344261 | 0.6071391 | -0.2143016 | 0.8303119 | 0.914768 |
| ENSG00000108799 | EZH1 | 2858.17409 | -0.083546705 | 0.226836892 | -0.388983938 | 0.69728803 | 0.834703 |
| ENSG00000179988 | PSTK | 255.35 | -0.08358144 | 0.2245951 | -0.4028601 | 0.6870511 | 0.827607 |
| ENSG00000214820 | MPRIPP1 | 220.8485 | -0.08360622 | 0.2601121 | -0.3431748 | 0.731467 | 0.854344 |
| ENSG00000272831 | | 2.345077 | -0.08368443 | 0.6794721 | -0.2460127 | 0.8056724 | 0.900354 |
| ENSG00000250043 | | 2.247199 | -0.08371613 | 0.6747993 | -0.2575086 | 0.7967862 | 0.895202 |
| ENSG00000227954 | TARID | 1.213492 | -0.08374241 | 0.7269521 | -0.3001143 | 0.76409 | NA |
| ENSG00000166797 | CIAO2A | 3183.901 | -0.08376919 | 0.2165176 | -0.4052169 | 0.6853181 | 0.826544 |
| ENSG00000130770 | ATP5IF1 | 5390.685326 | -0.084021544 | 0.175281868 | -0.490880012 | 0.62351132 | 0.782785 |
| ENSG00000236662 | | 4.438965 | -0.08402914 | 0.6116432 | -0.2208364 | 0.8252198 | 0.911882 |
| ENSG00000186716 | BCR | 4823.998 | -0.08409197 | 0.1435431 | -0.5934075 | 0.5529085 | 0.731338 |
| ENSG00000125124 | BBS2 | 1819.413187 | -0.084206141 | 0.159566416 | -0.581438595 | 0.56094489 | 0.737225 |
| ENSG00000266909 | SLC25A6P4 | 1.051651 | -0.08424863 | 0.7258546 | -0.28596 | 0.7749088 | NA |
| ENSG00000114391 | RPL24 | 15451.37432 | -0.084277142 | 0.176663873 | -0.430461515 | 0.66685996 | 0.814371 |
| ENSG00000140525 | FANCI | 6870.933 | -0.08436058 | 0.4389255 | -0.2340285 | 0.8149628 | 0.905821 |
| ENSG00000160447 | PKN3 | 1385.816 | -0.08437204 | 0.3245821 | -0.2695115 | 0.7875361 | 0.889371 |
| ENSG00000205940 | HSP90AB2P | 35.16885 | -0.08450743 | 0.3422818 | -0.2770923 | 0.7817092 | 0.885531 |
| ENSG00000282886 | PTMAP12 | 19.51782 | -0.08451011 | 0.4204076 | -0.238595 | 0.8114196 | 0.903596 |
| ENSG00000197372 | ZNF675 | 683.5346 | -0.08451567 | 0.1564827 | -0.5532876 | 0.5800665 | 0.750952 |
| ENSG00000159556 | ISL2 | 57.21054 | -0.08453845 | 0.3704628 | -0.2593636 | 0.7953547 | 0.894202 |
| ENSG00000103363 | ELOB | 6435.573411 | -0.084554521 | 0.210829333 | -0.445533518 | 0.65593424 | 0.806083 |
| ENSG00000188375 | H3-5 | 219.4373 | -0.08457832 | 0.2909218 | -0.3182148 | 0.750322 | 0.866301 |
| ENSG00000183291 | SELENOF | 5934.934 | -0.08461017 | 0.1050138 | -0.8149072 | 0.4151254 | 0.615074 |

| | | | | | | | |
|---|---|---|---|---|---|---|---|
| ENSG00000272862 | | 0.9412383 | -0.0846238 | 0.7204166 | -0.2752132 | 0.7831525 | NA |
| ENSG00000271075 | | 9.57588 | -0.08465156 | 0.5220422 | -0.218566 | 0.8269881 | 0.912696 |
| ENSG00000123737 | EXOSC9 | 1808.010428 | -0.084714194 | 0.300919531 | -0.307619482 | 0.75837189 | 0.871241 |
| ENSG00000104522 | GFUS | 5860.847793 | -0.084744425 | 0.185333471 | -0.476825559 | 0.63348634 | 0.789998 |
| ENSG00000229618 | | 1.269511 | -0.08480139 | 0.7332144 | -0.3309619 | 0.7406733 | NA |
| ENSG00000100241 | SBF1 | 5041.417648 | -0.084833919 | 0.15658952 | -0.559533898 | 0.5757974 | 0.747903 |
| ENSG00000107175 | CREB3 | 2222.718372 | -0.084862325 | 0.145657526 | -0.593332819 | 0.55295844 | 0.73136 |
| ENSG00000179889 | PDXDC1 | 11975.59 | -0.0849158 | 0.1137173 | -0.73289 | 0.4636255 | 0.657806 |
| ENSG00000163291 | PAQR3 | 728.0964 | -0.08503313 | 0.226862 | -0.3930381 | 0.6942913 | 0.832418 |
| ENSG00000226268 | | 274.3498 | -0.08505674 | 0.2343627 | -0.3785123 | 0.7050501 | 0.839876 |
| ENSG00000261451 | | 1.538316 | -0.08515768 | 0.7308191 | -0.2941145 | 0.7686704 | NA |
| ENSG00000206585 | RNVU1-7 | 1.705372 | -0.08517109 | 0.7444152 | -0.3365649 | 0.7364449 | 0.85732 |
| ENSG00000231646 | FSIP2-AS1 | 3.483962 | -0.08518801 | 0.6816031 | -0.2466805 | 0.8051555 | 0.900308 |
| ENSG00000185404 | SP140L | 645.964 | -0.08522171 | 0.2078536 | -0.4264649 | 0.6697692 | 0.816064 |
| ENSG00000186364 | NUDT17 | 623.6245 | -0.08524365 | 0.2178917 | -0.4098152 | 0.6819415 | 0.82392 |
| ENSG00000270723 | RPL23AP92 | 0.9094911 | -0.0852792 | 0.7481787 | -0.3307091 | 0.7408642 | NA |
| ENSG00000286645 | | 1.233626 | -0.0853001 | 0.7145377 | -0.2733645 | 0.784573 | NA |
| ENSG00000123159 | GIPC1 | 9487.304384 | -0.085336042 | 0.183402333 | -0.488318977 | 0.62532392 | 0.784171 |
| ENSG00000266373 | | 26.15346 | -0.08546921 | 0.3864779 | -0.2562051 | 0.7977925 | 0.895734 |
| ENSG00000229721 | | 1.018601 | -0.08547306 | 0.7500021 | -0.3346057 | 0.7379225 | NA |
| ENSG00000256646 | | 0.771107 | -0.08556824 | 0.7552199 | -0.3759844 | 0.7069285 | NA |
| ENSG00000272308 | NA | 4.026493 | -0.08562271 | 0.6637616 | -0.238118 | 0.8117896 | 0.90391 |
| ENSG00000119383 | PTPA | 15433.38348 | -0.085796355 | 0.164181629 | -0.592822233 | 0.55330013 | 0.731539 |
| ENSG00000251400 | ALDH7A1P1 | 664.4631 | -0.0857998 | 0.3038879 | -0.319034 | 0.7497007 | 0.865959 |
| ENSG00000206924 | RNU6-689P | 1.683363 | -0.08581333 | 0.6769106 | -0.2452557 | 0.8062584 | 0.900554 |
| ENSG00000128654 | MTX2 | 2088.676191 | -0.085959685 | 0.170977246 | -0.510084232 | 0.60999245 | 0.773451 |
| ENSG00000114200 | BCHE | 3.922195033 | -0.086002239 | 0.623914106 | -0.231125596 | 0.81721723 | 0.90715 |
| ENSG00000163697 | APBB2 | 6496.656 | -0.08605686 | 0.2208186 | -0.4104569 | 0.6814708 | 0.823757 |
| ENSG00000113790 | EHHADH | 263.2166832 | -0.086087645 | 0.267996799 | -0.339943206 | 0.7338993 | 0.855706 |
| ENSG00000272402 | | 10.57753 | -0.08611145 | 0.4944853 | -0.2273241 | 0.8201718 | 0.908902 |
| ENSG00000121089 | NACA3P | 1012.599368 | -0.086173158 | 0.152918813 | -0.580693031 | 0.56144736 | 0.73741 |
| ENSG00000255464 | SEPTIN14P8 | 1.735629 | -0.08618525 | 0.701553 | -0.2647805 | 0.7911786 | 0.891674 |
| ENSG00000100097 | LGALS1 | 2633.139022 | -0.086246811 | 0.53310871 | -0.233440684 | 0.81541922 | 0.906101 |

| | | | | | | | |
|---|---|---|---|---|---|---|---|
| ENSG00000279713 | | 7.375302 | -0.08626416 | 0.5931661 | -0.2229833 | 0.8235485 | 0.910564 |
| ENSG00000225656 | | 1.178765 | -0.08628066 | 0.7972792 | -1.152389 | 0.2491613 | NA |
| ENSG00000221990 | EXOC3-AS1 | 907.3421 | -0.08628701 | 0.1745235 | -0.5349776 | 0.5926653 | 0.760812 |
| ENSG00000127837 | AAMP | 10613.64354 | -0.086385572 | 0.166943527 | -0.561925862 | 0.57416653 | 0.746641 |
| ENSG00000274051 | | 3.504606 | -0.08639725 | 0.6450368 | -0.2285003 | 0.8192573 | 0.90858 |
| ENSG00000228053 | | 1.950354 | -0.08647443 | 0.6969182 | -0.2547325 | 0.7989297 | 0.896704 |
| ENSG00000255737 | AGAP2-AS1 | 164.0006 | -0.08648273 | 0.2137101 | -0.4213645 | 0.673489 | 0.818569 |
| ENSG00000279982 | | 9.591606 | -0.08655409 | 0.5149894 | -0.2249919 | 0.8219855 | 0.909628 |
| ENSG00000273489 | | 1.204106 | -0.08657138 | 0.7600396 | -0.3661573 | 0.7142477 | NA |
| ENSG00000189091 | SF3B3 | 16104.45 | -0.08660641 | 0.2381542 | -0.3848849 | 0.7003227 | 0.83677 |
| ENSG00000142347 | MYO1F | 124.0177 | -0.08667002 | 0.2373137 | -0.38215 | 0.7023501 | 0.837894 |
| ENSG00000170044 | ZPLD1 | 1.075119 | -0.08671065 | 0.7404361 | -0.3436047 | 0.7311436 | NA |
| ENSG00000186648 | CARMIL3 | 319.2039 | -0.0868213 | 0.2623154 | -0.3530183 | 0.7240747 | 0.850074 |
| ENSG00000268205 | | 658.465 | -0.08685505 | 0.3082906 | -0.3121205 | 0.7549489 | 0.869057 |
| ENSG00000182858 | ALG12 | 771.97 | -0.08689904 | 0.1563501 | -0.5717723 | 0.5674762 | 0.741846 |
| ENSG00000231887 | PRH1 | 19.34831 | -0.08692065 | 0.4251893 | -0.2468051 | 0.805059 | 0.900308 |
| ENSG00000112339 | HBS1L | 2859.40227 | -0.086926232 | 0.163289228 | -0.553713994 | 0.5797746 | 0.75072 |
| ENSG00000268015 | | 0.9430447 | -0.08696628 | 0.7472763 | -0.3453827 | 0.7298066 | NA |
| ENSG00000138767 | CNOT6L | 1973.024868 | -0.087090817 | 0.161376134 | -0.563649072 | 0.57299298 | 0.746011 |
| ENSG00000261195 | | 0.7853225 | -0.08710668 | 0.7431952 | -0.3347813 | 0.7377901 | NA |
| ENSG00000180596 | H2BC4 | 300.1959 | -0.0872954 | 0.2835394 | -0.3341405 | 0.7382735 | 0.858765 |
| ENSG00000263826 | | 213.7927 | -0.08729656 | 0.303194 | -0.3211606 | 0.7480887 | 0.864962 |
| ENSG00000286786 | | 13.86395 | -0.08730869 | 0.5125668 | -0.225691 | 0.8214418 | 0.909357 |
| ENSG00000250015 | | 4.526957 | -0.08735493 | 0.6456173 | -0.2350198 | 0.8141934 | 0.905231 |
| ENSG00000247950 | SEC24B-AS1 | 56.27206 | -0.08737104 | 0.3212232 | -0.2992665 | 0.7647367 | 0.875128 |
| ENSG00000286622 | | 0.9108295 | -0.08738321 | 0.7472917 | -0.3455644 | 0.7296701 | NA |
| ENSG00000282221 | | 120.5798 | -0.08750763 | 0.2808313 | -0.341362 | 0.7328311 | 0.855211 |
| ENSG00000244313 | | 13604.95 | -0.08751166 | 0.187855 | -0.482211 | 0.629656 | 0.7873 |
| ENSG00000271817 | | 4.791619 | -0.087516 | 0.6216001 | -0.2242834 | 0.8225368 | 0.909824 |
| ENSG00000233030 | | 231.1069 | -0.08752323 | 0.4637713 | -0.2353256 | 0.813956 | 0.905081 |
| ENSG00000287733 | | 11.74545 | -0.08754924 | 0.5458085 | -0.2266371 | 0.8207059 | 0.909124 |
| ENSG00000100154 | TTC28 | 520.8847117 | -0.087715852 | 0.274268991 | -0.335324655 | 0.7373802 | 0.858183 |
| ENSG00000217241 | CBX3P9 | 862.3209 | -0.08775342 | 0.2667493 | -0.3475506 | 0.7281777 | 0.852439 |

| | | | | | | | |
|---|---|---|---|---|---|---|---|
| ENSG00000196295 | GARS1-DT | 974.5906 | -0.08781782 | 0.3056375 | -0.3147518 | 0.7529501 | 0.867952 |
| ENSG00000198087 | CD2AP | 3364.795 | -0.08784082 | 0.177865 | -0.4954133 | 0.6203084 | 0.780716 |
| ENSG00000064886 | CHI3L2 | 10.18061214 | -0.087920578 | 0.588559957 | -0.228143458 | 0.81953472 | 0.908698 |
| ENSG00000198502 | HLA-DRB5 | 25.93609 | -0.08793664 | 0.5975795 | -0.2169934 | 0.8282135 | 0.91345 |
| ENSG00000224745 | | 6.101681 | -0.0879427 | 0.5408185 | -0.2275353 | 0.8200075 | 0.908881 |
| ENSG00000119927 | GPAM | 1007.253935 | -0.087948073 | 0.236305565 | -0.392965619 | 0.69434488 | 0.832418 |
| ENSG00000286335 | | 1.04161 | -0.08795639 | 0.7632455 | -0.4160394 | 0.6773812 | NA |
| ENSG00000248916 | NUP210P3 | 0.9590982 | -0.08802696 | 0.7342908 | -0.3019443 | 0.7626946 | NA |
| ENSG00000173638 | SLC19A1 | 3007.748 | -0.08810231 | 0.2522382 | -0.3714895 | 0.710273 | 0.842335 |
| ENSG00000224578 | HNRNPA1P48 | 6342.689 | -0.08811149 | 0.1638015 | -0.5516502 | 0.5811881 | 0.751891 |
| ENSG00000171160 | MORN4 | 872.9893 | -0.08813355 | 0.2446077 | -0.3816804 | 0.7026984 | 0.838197 |
| ENSG00000162244 | RPL29 | 16969.68 | -0.0881425 | 0.1864745 | -0.4923654 | 0.622461 | 0.781947 |
| ENSG00000274937 | | 1.07465 | -0.08814374 | 0.7415767 | -0.3206462 | 0.7484785 | NA |
| ENSG00000264187 | | 13.18205 | -0.08823491 | 0.4345717 | -0.2455037 | 0.8060664 | 0.90047 |
| ENSG00000207927 | MIR302A | 0.858752 | -0.0883545 | 0.7600321 | -0.3678109 | 0.7130143 | NA |
| ENSG00000122042 | UBL3 | 10426.52017 | -0.088430696 | 0.252422616 | -0.374600155 | 0.70795786 | 0.841222 |
| ENSG00000196998 | WDR45 | 1769.38 | -0.08846608 | 0.2301583 | -0.4150895 | 0.6780764 | 0.821028 |
| ENSG00000086848 | ALG9 | 197.7084996 | -0.088512535 | 0.196113232 | -0.466057525 | 0.64117433 | 0.795382 |
| ENSG00000248966 | BCLAF1P1 | 18.08237 | -0.08862308 | 0.4354433 | -0.2454401 | 0.8061157 | 0.90047 |
| ENSG00000213790 | OLA1P1 | 729.3054 | -0.08877522 | 0.1654682 | -0.555779 | 0.578362 | 0.749951 |
| ENSG00000155026 | RSPH10B | 5.501441 | -0.08881971 | 0.5617976 | -0.2253869 | 0.8216783 | 0.909518 |
| ENSG00000268864 | | 117.5147 | -0.08886692 | 0.4336281 | -0.2506086 | 0.8021168 | 0.898497 |
| ENSG00000265855 | MIR4514 | 1.525356 | -0.08889123 | 0.7262688 | -0.3060734 | 0.7595487 | NA |
| ENSG00000246640 | PICART1 | 16.20725 | -0.08889919 | 0.4544258 | -0.2389627 | 0.8111345 | 0.903386 |
| ENSG00000267919 | | 1.54278 | -0.08898855 | 0.6994973 | -0.2869243 | 0.7741703 | NA |
| ENSG00000200087 | SNORA73B | 243.6044 | -0.08909673 | 0.2391248 | -0.3917798 | 0.6952209 | 0.833055 |
| ENSG00000147853 | AK3 | 3435.259 | -0.08915248 | 0.1326066 | -0.7184239 | 0.4724959 | 0.665381 |
| ENSG00000252621 | | 0.8508079 | -0.08930187 | 0.796794 | -0.8465385 | 0.3972524 | NA |
| ENSG00000271557 | YPEL5P2 | 2.353805 | -0.08949615 | 0.6765964 | -0.2536792 | 0.7997434 | 0.897068 |
| ENSG00000233247 | UBE2E4P | 184.5757 | -0.0894967 | 0.2674758 | -0.3546588 | 0.7228452 | 0.84934 |
| ENSG00000157181 | ODR4 | 2416.162 | -0.08955378 | 0.1308991 | -0.6924458 | 0.4886574 | 0.679603 |
| ENSG00000259238 | | 1.482581 | -0.08960326 | 0.7214744 | -0.3000852 | 0.7641122 | NA |
| ENSG00000247228 | | 1.071084 | -0.08967474 | 0.7264402 | -0.3012847 | 0.7631974 | NA |

| | | | | | | | |
|---|---|---|---|---|---|---|---|
| ENSG00000146006 | LRRTM2 | 2.492353 | -0.08972101 | 0.6390113 | -0.2403966 | 0.8100228 | 0.902923 |
| ENSG00000124391 | IL17C | 32.34689335 | -0.089789917 | 0.474617979 | -0.246367447 | 0.8053978 | 0.90031 |
| ENSG00000177051 | FBXO46 | 932.6621 | -0.08982027 | 0.1893317 | -0.4910131 | 0.6234172 | 0.782759 |
| ENSG00000111452 | ADGRD1 | 22.44777877 | -0.089881563 | 0.534807452 | -0.230027797 | 0.81807017 | 0.907686 |
| ENSG00000136870 | ZNF189 | 958.4671726 | -0.08995188 | 0.212770821 | -0.441238626 | 0.65904025 | 0.808106 |
| ENSG00000267072 | NAGPA-AS1 | 2.527928 | -0.08999804 | 0.6995403 | -0.2684189 | 0.7883769 | 0.889897 |
| ENSG00000263235 | | 54.22736 | -0.09000106 | 0.3641043 | -0.2812036 | 0.7785543 | 0.883564 |
| ENSG00000229324 | NUTF2P7 | 1.033688 | -0.09004714 | 0.7304628 | -0.3109922 | 0.7558065 | NA |
| ENSG00000176953 | NFATC2IP | 4551.678 | -0.09005743 | 0.1643434 | -0.5565601 | 0.577828 | 0.749595 |
| ENSG00000050628 | PTGER3 | 272.6425625 | -0.090065561 | 0.432502265 | -0.256312845 | 0.79770928 | 0.895734 |
| ENSG00000227321 | MTND4P15 | 4.721908 | -0.09008669 | 0.5941071 | -0.234361 | 0.8147048 | 0.905724 |
| ENSG00000137198 | GMPR | 85.8761313 | -0.090155376 | 0.325627296 | -0.307826439 | 0.7582144 | 0.871135 |
| ENSG00000279196 | | 56.89676 | -0.09022528 | 0.3628955 | -0.2733324 | 0.7845977 | 0.887289 |
| ENSG00000197385 | ZNF860 | 452.1201 | -0.0902947 | 0.2725456 | -0.3615466 | 0.7176909 | 0.846015 |
| ENSG00000137364 | TPMT | 944.4971887 | -0.090369445 | 0.20248628 | -0.473304693 | 0.63599582 | 0.791799 |
| ENSG00000164076 | CAMKV | 3.324272 | -0.09039107 | 0.6936879 | -0.2730464 | 0.7848175 | 0.887349 |
| ENSG00000103978 | TMEM87A | 2503.279695 | -0.090470051 | 0.118113423 | -0.810772311 | 0.41749644 | 0.617347 |
| ENSG00000259804 | | 12.09708 | -0.09052191 | 0.5477861 | -0.2305698 | 0.817649 | 0.907554 |
| ENSG00000145002 | FAM86B2 | 11.64577 | -0.09055907 | 0.4722915 | -0.2441177 | 0.8071397 | 0.901197 |
| ENSG00000238103 | RPL9P7 | 10.43975 | -0.09057882 | 0.5056076 | -0.2353704 | 0.8139213 | 0.905081 |
| ENSG00000224604 | | 6.926544 | -0.0906655 | 0.6621586 | -0.2796725 | 0.7797288 | 0.884468 |
| ENSG00000269091 | | 33.35624 | -0.09084194 | 0.3990911 | -0.267829 | 0.788831 | 0.890218 |
| ENSG00000244560 | | 359.4829 | -0.09087415 | 0.300092 | -0.3315142 | 0.7402561 | 0.859716 |
| ENSG00000207568 | MIR203A | 0.8099173 | -0.090915 | 0.7336296 | -0.3234775 | 0.7463336 | NA |
| ENSG00000284820 | | 15.4928 | -0.09091912 | 0.4358176 | -0.2539527 | 0.7995321 | 0.896976 |
| ENSG00000211452 | DIO1 | 127.8503 | -0.09099483 | 0.4202309 | -0.2617213 | 0.7935363 | 0.893178 |
| ENSG00000232905 | | 2.890798 | -0.09108647 | 0.6832019 | -0.2625107 | 0.7929278 | 0.892909 |
| ENSG00000227097 | RPS28P7 | 5050.535 | -0.0911124 | 0.1710876 | -0.5638158 | 0.5728795 | 0.745937 |
| ENSG00000128626 | MRPS12 | 3281.54719 | -0.091116244 | 0.267755059 | -0.348975386 | 0.72710779 | 0.851749 |
| ENSG00000055950 | MRPL43 | 2612.637992 | -0.091210726 | 0.196752518 | -0.475375034 | 0.63451969 | 0.790628 |
| ENSG00000221044 | | 5.360968 | -0.09123595 | 0.6045539 | -0.2370785 | 0.8125959 | 0.904263 |
| ENSG00000102580 | DNAJC3 | 2653.551703 | -0.091311033 | 0.158488289 | -0.580395402 | 0.56164801 | 0.737529 |
| ENSG00000111843 | TMEM14C | 4133.153515 | -0.091410782 | 0.166058124 | -0.541903264 | 0.58788515 | 0.757273 |

| | | | | | | | |
|---|---|---|---|---|---|---|---|
| ENSG00000119335 | SET | 24281.21534 | -0.091445157 | 0.175372652 | -0.576594486 | 0.56421343 | 0.739552 |
| ENSG00000144401 | METTL21A | 854.1602 | -0.09148215 | 0.164045 | -0.5726647 | 0.5668718 | 0.741384 |
| ENSG00000106330 | MOSPD3 | 749.5025303 | -0.091485345 | 0.185497496 | -0.503442771 | 0.61465301 | 0.776509 |
| ENSG00000089053 | ANAPC5 | 9692.359078 | -0.091486099 | 0.137032319 | -0.680021701 | 0.49649072 | 0.685403 |
| ENSG00000186185 | KIF18B | 1550.289 | -0.09153417 | 0.5277281 | -0.2389562 | 0.8111395 | 0.903386 |
| ENSG00000258354 | | 8.104464 | -0.09154178 | 0.5759666 | -0.2340494 | 0.8149466 | 0.905821 |
| ENSG00000101745 | ANKRD12 | 2967.767829 | -0.091586762 | 0.265459185 | -0.36537248 | 0.71483338 | 0.844633 |
| ENSG00000275332 | | 1.387391 | -0.09159946 | 0.7290825 | -0.3358239 | 0.7370036 | NA |
| ENSG00000271327 | | 45.03818 | -0.0916275 | 0.4945808 | -0.2404203 | 0.8100045 | 0.902923 |
| ENSG00000242485 | MRPL20 | 2982.679 | -0.09168999 | 0.1991717 | -0.4743759 | 0.6352319 | 0.791181 |
| ENSG00000258828 | KRT8P2 | 9.209487 | -0.0919583 | 0.5734265 | -0.235857 | 0.8135436 | 0.904849 |
| ENSG00000003402 | CFLAR | 2342.625851 | -0.091972523 | 0.217940886 | -0.445142648 | 0.65621667 | 0.806281 |
| ENSG00000250346 | EEF1GP2 | 1.357002 | -0.0919875 | 0.7221263 | -0.2981788 | 0.7655667 | NA |
| ENSG00000263179 | HNRNPCP4 | 32.96052 | -0.09202024 | 0.3211449 | -0.3164345 | 0.7516727 | 0.86703 |
| ENSG00000146872 | TLK2 | 7087.312 | -0.09204393 | 0.08920987 | -1.079652 | 0.2802973 | 0.477692 |
| ENSG00000172053 | QARS1 | 8153.624 | -0.09206562 | 0.1119213 | -0.8079897 | 0.4190965 | 0.61902 |
| ENSG00000071626 | DAZAP1 | 8177.553937 | -0.092186975 | 0.172087235 | -0.551886388 | 0.5810262 | 0.751783 |
| ENSG00000269961 | | 26.04596 | -0.09230881 | 0.3638324 | -0.2860024 | 0.7748763 | 0.881331 |
| ENSG00000222898 | RN7SKP97 | 4.160802 | -0.09231066 | 0.6705062 | -0.2539369 | 0.7995443 | 0.896976 |
| ENSG00000266097 | MIR5192 | 0.987066 | -0.09233705 | 0.7737339 | -0.4816316 | 0.6300677 | NA |
| ENSG00000170265 | ZNF282 | 3475.843 | -0.09238048 | 0.1140326 | -0.8555114 | 0.3922681 | 0.594025 |
| ENSG00000287262 | | 58.89585 | -0.09239767 | 0.3791095 | -0.282475 | 0.7775793 | 0.882917 |
| ENSG00000104883 | PEX11G | 222.4253338 | -0.092417702 | 0.271642907 | -0.363987694 | 0.7158672 | 0.845328 |
| ENSG00000197483 | ZNF628 | 403.3489 | -0.09249276 | 0.2382908 | -0.4112954 | 0.6808559 | 0.823305 |
| ENSG00000180846 | CSNK1G2-AS1 | 4.180201 | -0.09257372 | 0.6027888 | -0.2408351 | 0.8096829 | 0.902824 |
| ENSG00000230188 | | 1.732316 | -0.09266368 | 0.7201957 | -0.2978724 | 0.7658006 | 0.875723 |
| ENSG00000198791 | CNOT7 | 4020.692 | -0.09278168 | 0.1590404 | -0.6007534 | 0.5480042 | 0.727632 |
| ENSG00000266896 | | 54.20482 | -0.09288322 | 0.3786923 | -0.28322 | 0.7770082 | 0.882538 |
| ENSG00000133243 | BTBD2 | 4090.995102 | -0.092950595 | 0.149611014 | -0.637539542 | 0.52377346 | 0.708886 |
| ENSG00000197978 | GOLGA6L9 | 663.3077 | -0.09300633 | 0.2637266 | -0.3793817 | 0.7044044 | 0.83944 |
| ENSG00000188107 | EYS | 5.720425 | -0.09303931 | 0.6253072 | -0.2426425 | 0.8082823 | 0.901868 |
| ENSG00000234175 | | 1.836451 | -0.09304632 | 0.7125353 | -0.3005051 | 0.7637919 | 0.874321 |
| ENSG00000207313 | SNORA2B | 4.348991 | -0.09308786 | 0.6412669 | -0.2499433 | 0.8026312 | 0.898879 |

| | | | | | | | |
|---|---|---|---|---|---|---|---|
| ENSG00000087299 | L2HGDH | 2054.069745 | -0.093108157 | 0.244987158 | -0.395076049 | 0.69278676 | 0.831284 |
| ENSG00000158435 | CNOT11 | 4636.982 | -0.09314013 | 0.1131191 | -0.8486153 | 0.3960954 | 0.597471 |
| ENSG00000261357 | | 2.355601 | -0.09314307 | 0.7141915 | -0.2895248 | 0.7721798 | 0.879724 |
| ENSG00000214700 | C12orf71 | 1.275835 | -0.09320802 | 0.7212964 | -0.328804 | 0.7423039 | NA |
| ENSG00000261513 | | 11.41681 | -0.09324599 | 0.5497377 | -0.2360871 | 0.8133651 | 0.904727 |
| ENSG00000067560 | RHOA | 13656.58119 | -0.093334481 | 0.128895576 | -0.735848949 | 0.46182263 | 0.656545 |
| ENSG00000162782 | TDRD5 | 1031.786 | -0.09334546 | 0.2180736 | -0.4471695 | 0.6547527 | 0.805245 |
| ENSG00000121067 | SPOP | 4351.80224 | -0.093414576 | 0.14635071 | -0.663025113 | 0.50731447 | 0.693713 |
| ENSG00000279570 | | 0.9577854 | -0.09348482 | 0.7531706 | -0.4148121 | 0.6782794 | NA |
| ENSG00000020219 | CCT8L1P | 0.881283389 | -0.093514557 | 0.742071067 | -0.339127035 | 0.73451403 | NA |
| ENSG00000285955 | | 2.797718 | -0.09359605 | 0.6702554 | -0.2618068 | 0.7934704 | 0.893142 |
| ENSG00000230542 | LINC00102 | 0.8945322 | -0.09362366 | 0.7584146 | -0.4060952 | 0.6846726 | NA |
| ENSG00000176142 | TMEM39A | 2820.124 | -0.09366233 | 0.2298775 | -0.4339711 | 0.6643094 | 0.812355 |
| ENSG00000250548 | LINC01303 | 1.183931 | -0.09366462 | 0.7349249 | -0.3210688 | 0.7481582 | NA |
| ENSG00000179630 | LACC1 | 584.2339 | -0.09373587 | 0.2043379 | -0.4784975 | 0.6322962 | 0.789227 |
| ENSG00000138777 | PPA2 | 4472.713243 | -0.093810318 | 0.206628098 | -0.474522181 | 0.63512759 | 0.791166 |
| ENSG00000233980 | FDPSP2 | 1.794199 | -0.09385952 | 0.6947678 | -0.2750331 | 0.7832908 | 0.886679 |
| ENSG00000163453 | IGFBP7 | 9.621299 | -0.09396895 | 0.6412623 | -0.2426474 | 0.8082786 | 0.901868 |
| ENSG00000204389 | HSPA1A | 6392.066 | -0.09398934 | 0.3080243 | -0.3348479 | 0.7377398 | 0.858451 |
| ENSG00000156970 | BUB1B | 1705.859 | -0.09399978 | 0.4231504 | -0.2746507 | 0.7835846 | 0.886724 |
| ENSG00000163798 | SLC4A1AP | 2727.82 | -0.09402199 | 0.1046934 | -0.9168519 | 0.3592203 | 0.561155 |
| ENSG00000245060 | LINC00847 | 460.5817 | -0.09402928 | 0.2341473 | -0.4292721 | 0.6677252 | 0.814882 |
| ENSG00000273450 | | 1.202748 | -0.0940807 | 0.7354666 | -0.3442406 | 0.7306654 | NA |
| ENSG00000080822 | CLDND1 | 2438.81966 | -0.094174064 | 0.182779548 | -0.553019902 | 0.58024978 | 0.751153 |
| ENSG00000228175 | GEMIN8P4 | 26.87093 | -0.09421053 | 0.4144974 | -0.2704353 | 0.7868254 | 0.888825 |
| ENSG00000230202 | | 6960.858 | -0.09421924 | 0.1730416 | -0.5643157 | 0.5725393 | 0.745896 |
| ENSG00000135482 | ZC3H10 | 613.0442763 | -0.094283141 | 0.187563089 | -0.52286841 | 0.60106583 | 0.766828 |
| ENSG00000215022 | | 19.28251 | -0.09431017 | 0.444045 | -0.2582008 | 0.796252 | 0.894908 |
| ENSG00000255122 | | 1.209215 | -0.09436142 | 0.717179 | -0.3044262 | 0.7608032 | NA |
| ENSG00000239494 | RN7SL333P | 12.78561 | -0.09442927 | 0.5290111 | -0.2454673 | 0.8060946 | 0.90047 |
| ENSG00000162542 | TMCO4 | 722.6701 | -0.09448083 | 0.1455423 | -0.6635149 | 0.5070009 | 0.693528 |
| ENSG00000231829 | | 21.08751 | -0.09449696 | 0.5138937 | -0.2488394 | 0.803485 | 0.899385 |
| ENSG00000256742 | KDM2B-DT | 8.817608 | -0.09451341 | 0.5207626 | -0.243111 | 0.8079194 | 0.901651 |

| | | | | | | | |
|---|---|---|---|---|---|---|---|
| ENSG00000188385 | JAKMIP3 | 199.7246 | -0.09456096 | 0.2522727 | -0.3825092 | 0.7020837 | 0.837689 |
| ENSG00000259461 | ANP32BP3 | 0.985783 | -0.09460628 | 0.7256953 | -0.3304939 | 0.7410268 | NA |
| ENSG00000162144 | CYB561A3 | 1666.156 | -0.09463525 | 0.1402401 | -0.6794627 | 0.4968447 | 0.685678 |
| ENSG00000007520 | TSR3 | 2172.831548 | -0.094667092 | 0.168121466 | -0.580235286 | 0.56175596 | 0.737634 |
| ENSG00000214135 | | 1162.793 | -0.09467312 | 0.2749844 | -0.3652141 | 0.7149516 | 0.844697 |
| ENSG00000250886 | | 4.403188 | -0.09467317 | 0.6121346 | -0.2558367 | 0.7980769 | 0.896011 |
| ENSG00000105426 | PTPRS | 4994.029751 | -0.094729156 | 0.179962238 | -0.553853358 | 0.57967921 | 0.75067 |
| ENSG00000223822 | EEF1A1P1 | 11.57663 | -0.09472923 | 0.4566344 | -0.2598256 | 0.7949983 | 0.894104 |
| ENSG00000266594 | MIR4766 | 1.053129 | -0.0948007 | 0.7552705 | -0.4073094 | 0.6837807 | NA |
| ENSG00000126267 | COX6B1 | 6761.755718 | -0.094805997 | 0.214796971 | -0.461476484 | 0.64445679 | 0.79766 |
| ENSG00000179284 | DAND5 | 2.032292 | -0.0948165 | 0.6819443 | -0.2782898 | 0.7807899 | 0.885093 |
| ENSG00000087338 | GMCL1 | 2481.148266 | -0.094820928 | 0.164658297 | -0.604935888 | 0.5452216 | 0.725751 |
| ENSG00000140391 | TSPAN3 | 6536.016 | -0.09482461 | 0.1649039 | -0.6087255 | 0.5427064 | 0.723848 |
| ENSG00000230280 | HNRNPA1P59 | 116.3207 | -0.09482848 | 0.3623927 | -0.2980951 | 0.7656305 | 0.875613 |
| ENSG00000100029 | PES1 | 6014.613858 | -0.094838922 | 0.186026743 | -0.534216973 | 0.59319143 | 0.761156 |
| ENSG00000110324 | IL10RA | 3.856989762 | -0.094920221 | 0.593375153 | -0.246284684 | 0.80546187 | 0.90031 |
| ENSG00000137168 | PPIL1 | 2720.847903 | -0.094925464 | 0.235944738 | -0.415633132 | 0.67767848 | 0.820733 |
| ENSG00000010278 | CD9 | 11885.75726 | -0.094976894 | 0.176859122 | -0.55852135 | 0.57648843 | 0.748537 |
| ENSG00000234558 | API5P1 | 421.6291 | -0.09502085 | 0.2194841 | -0.450637 | 0.6522512 | 0.803679 |
| ENSG00000272505 | | 22.17494 | -0.09503481 | 0.4385279 | -0.2622203 | 0.7931516 | 0.893085 |
| ENSG00000074582 | BCS1L | 1813.077771 | -0.095167258 | 0.140163192 | -0.677162905 | 0.4983026 | 0.686727 |
| ENSG00000272366 | | 0.9796507 | -0.09522628 | 0.7577658 | -0.3894785 | 0.6969222 | NA |
| ENSG00000136240 | KDELR2 | 15837.95718 | -0.095259362 | 0.08188744 | -1.202384532 | 0.22921458 | 0.417287 |
| ENSG00000240003 | RPL7P24 | 3.456856 | -0.09527416 | 0.6620505 | -0.2663564 | 0.7899647 | 0.891083 |
| ENSG00000271964 | | 22.73489 | -0.09527813 | 0.5369225 | -0.2593867 | 0.7953369 | 0.894202 |
| ENSG00000227006 | | 16.40534 | -0.0952861 | 0.4154739 | -0.2740635 | 0.7840358 | 0.887008 |
| ENSG00000166816 | LDHD | 3476.645 | -0.09531171 | 0.3513375 | -0.2984094 | 0.7653907 | 0.875498 |
| ENSG00000285287 | | 12.9275 | -0.09531228 | 0.5029843 | -0.246236 | 0.8054996 | 0.90031 |
| ENSG00000070444 | MNT | 1321.247832 | -0.095313153 | 0.185490859 | -0.521953063 | 0.60170301 | 0.767199 |
| ENSG00000155719 | OTOA | 0.913721 | -0.09532742 | 0.7397717 | -0.3707288 | 0.7108395 | NA |
| ENSG00000245571 | FAM111A-DT | 340.9587 | -0.09534168 | 0.3137988 | -0.3322226 | 0.7397212 | 0.859439 |
| ENSG00000099308 | MAST3 | 665.3888875 | -0.095358791 | 0.211315588 | -0.468909696 | 0.63913419 | 0.794106 |
| ENSG00000101843 | PSMD10 | 4255.8989 | -0.095426487 | 0.119431003 | -0.814362597 | 0.41543726 | 0.615259 |

| | | | | | | | |
|---|---|---|---|---|---|---|---|
| ENSG00000110448 | CD5 | 25.25569659 | -0.095432449 | 0.457570643 | -0.252543002 | 0.80062138 | 0.897578 |
| ENSG00000161204 | ABCF3 | 5867.067 | -0.09544609 | 0.1150145 | -0.8193544 | 0.4125843 | 0.612714 |
| ENSG00000153147 | SMARCA5 | 9160.47 | -0.09550256 | 0.1148882 | -0.7931782 | 0.427674 | 0.627062 |
| ENSG00000264570 | SNX19P3 | 10.50654 | -0.09551985 | 0.4899396 | -0.2519686 | 0.8010653 | 0.897887 |
| ENSG00000287351 | | 1.296925 | -0.09567571 | 0.7208341 | -0.3086411 | 0.7575946 | NA |
| ENSG00000105229 | PIAS4 | 1819.198153 | -0.095868812 | 0.161664762 | -0.604382361 | 0.54558946 | 0.725986 |
| ENSG00000258012 | | 6.272999 | -0.09593603 | 0.5775021 | -0.2447314 | 0.8066644 | 0.900818 |
| ENSG00000277215 | SPANXA2-OT1 | 37.06948 | -0.09598546 | 0.4018878 | -0.2833752 | 0.7768892 | 0.882518 |
| ENSG00000168701 | TMEM208 | 2105.402 | -0.09604905 | 0.2028193 | -0.4907197 | 0.6236247 | 0.782851 |
| ENSG00000126214 | KLC1 | 1868.074536 | -0.096072527 | 0.122830516 | -0.795026912 | 0.42659785 | 0.626378 |
| ENSG00000237617 | | 0.9975997 | -0.09620418 | 0.7645046 | -0.4432523 | 0.6575832 | NA |
| ENSG00000241743 | XACT | 1.152685 | -0.0962525 | 0.7200756 | -0.3108928 | 0.7558821 | NA |
| ENSG00000127125 | PPCS | 3236.900798 | -0.096306725 | 0.115538893 | -0.845497316 | 0.39783323 | 0.59897 |
| ENSG00000160072 | ATAD3B | 1941.332 | -0.09633217 | 0.1833958 | -0.5440714 | 0.5863924 | 0.75612 |
| ENSG00000196535 | MYO18A | 4304.662 | -0.0963948 | 0.1782771 | -0.5834477 | 0.559592 | 0.73613 |
| ENSG00000179115 | FARSA | 3970.589 | -0.09641997 | 0.2259576 | -0.4709413 | 0.6376826 | 0.793155 |
| ENSG00000072121 | ZFYVE26 | 4289.370844 | -0.096434026 | 0.2620543 | -0.397338853 | 0.6911176 | 0.830171 |
| ENSG00000162399 | BSND | 1.028067 | -0.09653094 | 0.7119628 | -0.3172693 | 0.7510393 | NA |
| ENSG00000237222 | LINC01968 | 1.388818 | -0.09655096 | 0.7141947 | -0.3022352 | 0.7624728 | NA |
| ENSG00000100084 | HIRA | 960.6796354 | -0.096661774 | 0.153513327 | -0.613105016 | 0.5398069 | 0.722087 |
| ENSG00000178104 | PDE4DIP | 6344.442 | -0.09671876 | 0.2360198 | -0.4127763 | 0.6797705 | 0.822292 |
| ENSG00000119778 | ATAD2B | 2026.162819 | -0.096746306 | 0.21203641 | -0.476265975 | 0.6338849 | 0.790244 |
| ENSG00000157911 | PEX10 | 1156.284 | -0.09677012 | 0.164075 | -0.6493648 | 0.5161026 | 0.701398 |
| ENSG00000181789 | COPG1 | 12562.86 | -0.0967883 | 0.1127218 | -0.7080524 | 0.4789127 | 0.670968 |
| ENSG00000262265 | | 1.877225 | -0.09683516 | 0.668866 | -0.2721787 | 0.7854847 | 0.887801 |
| ENSG00000242931 | RPL7P49 | 1.232588 | -0.09685582 | 0.718942 | -0.3111608 | 0.7556784 | NA |
| ENSG00000163479 | SSR2 | 9515.49 | -0.09686247 | 0.1117586 | -0.9023902 | 0.3668496 | 0.568859 |
| ENSG00000142748 | FCN3 | 4.203301 | -0.09688313 | 0.6319964 | -0.2571136 | 0.7970911 | 0.895283 |
| ENSG00000286244 | | 11.59342 | -0.09691824 | 0.5486493 | -0.2464069 | 0.8053673 | 0.90031 |
| ENSG00000229007 | EXOSC3P1 | 1.27752 | -0.09695462 | 0.7259297 | -0.3438736 | 0.7309414 | NA |
| ENSG00000176101 | SSNA1 | 3560.406 | -0.09695597 | 0.2456709 | -0.419864 | 0.6745848 | 0.818951 |
| ENSG00000259305 | ZHX1-C8orf76 | 30.48625 | -0.09698359 | 0.3074143 | -0.3420598 | 0.7323059 | 0.854968 |
| ENSG00000070413 | DGCR2 | 7663.060309 | -0.097028418 | 0.138547578 | -0.708944917 | 0.47835866 | 0.670262 |

| | | | | | | | |
|---|---|---|---|---|---|---|---|
| ENSG00000198542 | ITGBL1 | 0.8756331 | -0.09716887 | 0.7899407 | -0.3277277 | 0.7431176 | NA |
| ENSG00000237742 | | 0.7976125 | -0.09717833 | 0.7877531 | -0.7491636 | 0.4537586 | NA |
| ENSG00000143149 | ALDH9A1 | 2804.877 | -0.09720712 | 0.211536 | -0.484076 | 0.6283319 | 0.786453 |
| ENSG00000143368 | SF3B4 | 10247.49 | -0.09724899 | 0.1938186 | -0.5204246 | 0.6027677 | 0.767955 |
| ENSG00000106631 | MYL7 | 0.874067061 | -0.097267896 | 0.762366766 | -0.412022471 | 0.68032295 | NA |
| ENSG00000231858 | | 0.7792964 | -0.09729457 | 0.7579528 | -0.3903176 | 0.6963017 | NA |
| ENSG00000263264 | NA | 2.964991 | -0.09732446 | 0.665201 | -0.2667333 | 0.7896745 | 0.890945 |
| ENSG00000171241 | SHCBP1 | 957.1887 | -0.09735383 | 0.5400254 | -0.2548381 | 0.7988482 | 0.89665 |
| ENSG00000239705 | | 1.931223 | -0.09735517 | 0.6957367 | -0.2949804 | 0.7680088 | 0.877041 |
| ENSG00000251615 | | 136.2995 | -0.09735922 | 0.4125574 | -0.2811779 | 0.7785739 | 0.883564 |
| ENSG00000235328 | | 3.276077 | -0.09736307 | 0.697153 | -0.2938989 | 0.7688352 | 0.877494 |
| ENSG00000214832 | UPF3AP2 | 153.7043 | -0.09739941 | 0.1954429 | -0.5143421 | 0.6070128 | 0.771034 |
| ENSG00000260296 | | 32.27115 | -0.09743882 | 0.4357762 | -0.2733135 | 0.7846122 | 0.887289 |
| ENSG00000236526 | | 20.84605 | -0.09750081 | 0.4975935 | -0.2505124 | 0.8021911 | 0.898542 |
| ENSG00000123901 | GPR83 | 3.141956277 | -0.097504268 | 0.613654003 | -0.253980762 | 0.79951042 | 0.896976 |
| ENSG00000187555 | USP7 | 9789.795 | -0.09753907 | 0.1502302 | -0.6886038 | 0.4910726 | 0.681494 |
| ENSG00000260526 | | 58.05393 | -0.0976116 | 0.3169494 | -0.3378288 | 0.7354922 | 0.856661 |
| ENSG00000140983 | RHOT2 | 9249.554 | -0.09761473 | 0.1457917 | -0.6832805 | 0.4944296 | 0.684048 |
| ENSG00000222032 | | 0.9722447 | -0.09761935 | 0.7949998 | -0.990963 | 0.3217036 | NA |
| ENSG00000260265 | LINC02562 | 0.8455581 | -0.09769119 | 0.7904837 | -0.321535 | 0.747805 | NA |
| ENSG00000231527 | FAM27C | 1.267489 | -0.0977332 | 0.772005 | -0.4830388 | 0.6290682 | NA |
| ENSG00000258479 | LINC00640 | 1.965443 | -0.09776099 | 0.67284 | -0.2782263 | 0.7808387 | 0.88511 |
| ENSG00000225695 | HNRNPA1P35 | 34.66901 | -0.09779839 | 0.3264695 | -0.3282234 | 0.7427427 | 0.861443 |
| ENSG00000130227 | XPO7 | 5557.615288 | -0.097810552 | 0.14093944 | -0.704063001 | 0.48139354 | 0.672882 |
| ENSG00000225419 | RPL21P3 | 2.550206 | -0.09781442 | 0.7035208 | -0.2959318 | 0.7672822 | 0.876588 |
| ENSG00000017483 | SLC38A5 | 8.946917158 | -0.097843962 | 0.58732877 | -0.25116985 | 0.80168279 | 0.898186 |
| ENSG00000221184 | | 59.43619 | -0.09784976 | 0.3103307 | -0.3451523 | 0.7299799 | 0.853608 |
| ENSG00000232706 | NUTM2HP | 1.164805 | -0.09785029 | 0.7535392 | -0.4121021 | 0.6802646 | NA |
| ENSG00000166484 | MAPK7 | 973.1074 | -0.09785196 | 0.1846099 | -0.542368 | 0.587565 | 0.75702 |
| ENSG00000110871 | COQ5 | 2271.151443 | -0.097859035 | 0.173037234 | -0.594650013 | 0.55207744 | 0.730761 |
| ENSG00000161914 | ZNF653 | 218.8016 | -0.09785956 | 0.2306458 | -0.4401809 | 0.6598061 | 0.808635 |
| ENSG00000174485 | DENND4A | 1896.419 | -0.09787698 | 0.214301 | -0.4786862 | 0.6321619 | 0.789178 |
| ENSG00000268412 | TRMT112P6 | 562.7627 | -0.09789404 | 0.2119008 | -0.4824503 | 0.6294861 | 0.787127 |

| | | | | | | | |
|---|---|---|---|---|---|---|---|
| ENSG00000248830 | ZNF807P | 1.783518 | -0.09792507 | 0.7184004 | -0.3183634 | 0.7502093 | 0.866263 |
| ENSG00000267544 | HIKESHIP2 | 122.9573 | -0.09795181 | 0.2807732 | -0.3771236 | 0.7060817 | 0.840196 |
| ENSG00000274569 | | 1.192745 | -0.09796069 | 0.7289496 | -0.3343818 | 0.7380914 | NA |
| ENSG00000206899 | RNU6-36P | 3.683806 | -0.09799158 | 0.5863846 | -0.2530099 | 0.8002606 | 0.897349 |
| ENSG00000144455 | SUMF1 | 2936.508 | -0.09805055 | 0.1601611 | -0.611406 | 0.5409308 | 0.722792 |
| ENSG00000172728 | FUT10 | 326.0473 | -0.09805983 | 0.2378461 | -0.4332576 | 0.6648276 | 0.812764 |
| ENSG00000232590 | | 0.7002479 | -0.09808545 | 0.7926621 | -0.8772118 | 0.3803716 | NA |
| ENSG00000136868 | SLC31A1 | 4212.998594 | -0.098098967 | 0.234143431 | -0.441324549 | 0.65897806 | 0.808067 |
| ENSG00000170667 | RASA4B | 50.98348 | -0.0981398 | 0.3917232 | -0.2923557 | 0.7700147 | 0.87835 |
| ENSG00000254929 | | 5.499193 | -0.09820351 | 0.5850167 | -0.2494473 | 0.8030148 | 0.899149 |
| ENSG00000275774 | HNRNPCL2 | 3.477147 | -0.09821012 | 0.6024155 | -0.2565209 | 0.7975486 | 0.895607 |
| ENSG00000213729 | RPS16P2 | 5.830315 | -0.09821256 | 0.6022634 | -0.25734 | 0.7969163 | 0.895202 |
| ENSG00000183340 | JRKL | 526.0418 | -0.09825704 | 0.2609498 | -0.3972749 | 0.6911648 | 0.830171 |
| ENSG00000152990 | ADGRA3 | 2158.279 | -0.09835197 | 0.1284376 | -0.775083 | 0.4382906 | 0.636836 |
| ENSG00000154227 | CERS3 | 0.8800904 | -0.09845386 | 0.7481509 | -0.3898534 | 0.696645 | NA |
| ENSG00000111361 | EIF2B1 | 3970.728807 | -0.098505311 | 0.144157525 | -0.696888544 | 0.48587255 | 0.676899 |
| ENSG00000179041 | RRS1 | 2412.747 | -0.0985276 | 0.2054831 | -0.5025453 | 0.615284 | 0.777045 |
| ENSG00000279928 | DDX11L17 | 0.8795637 | -0.0985343 | 0.7348496 | -0.3631835 | 0.7164678 | NA |
| ENSG00000103723 | AP3B2 | 339.8185271 | -0.098610838 | 0.248596382 | -0.422292376 | 0.67281162 | 0.818088 |
| ENSG00000235281 | | 0.9500654 | -0.09865528 | 0.790512 | -0.3236645 | 0.7461921 | NA |
| ENSG00000259589 | | 0.8340678 | -0.09886119 | 0.7690984 | -0.4229814 | 0.6723088 | NA |
| ENSG00000261777 | | 46.75477 | -0.09888384 | 0.3297208 | -0.3294125 | 0.7418439 | 0.860851 |
| ENSG00000270019 | | 5.713001 | -0.0989415 | 0.6375295 | -0.2612065 | 0.7939333 | 0.893398 |
| ENSG00000100823 | APEX1 | 12699.03225 | -0.099033855 | 0.109524635 | -0.912661133 | 0.36142079 | 0.563416 |
| ENSG00000129991 | TNNI3 | 1.737657583 | -0.09909885 | 0.737963438 | -0.355598775 | 0.72214106 | 0.848817 |
| ENSG00000036257 | CUL3 | 3116.905112 | -0.099137981 | 0.138465958 | -0.752131287 | 0.45197211 | 0.648229 |
| ENSG00000068308 | OTUD5 | 4487.863259 | -0.099154617 | 0.112160153 | -0.914722822 | 0.36033716 | 0.562338 |
| ENSG00000177239 | MAN1B1 | 6537.113 | -0.09916453 | 0.1012514 | -1.010899 | 0.3120646 | 0.513072 |
| ENSG00000153291 | SLC25A27 | 149.7008 | -0.09922727 | 0.3892279 | -0.2942492 | 0.7685675 | 0.877361 |
| ENSG00000233033 | CASK-AS1 | 0.9577325 | -0.0992459 | 0.7459263 | -0.3979839 | 0.690642 | NA |
| ENSG00000158163 | DZIP1L | 44.83879 | -0.09927509 | 0.4078877 | -0.2901651 | 0.7716899 | 0.879317 |
| ENSG00000233503 | HNRNPLP1 | 15.66496 | -0.09930923 | 0.5101215 | -0.2577867 | 0.7965715 | 0.89504 |
| ENSG00000223714 | LINC02601 | 5.532211 | -0.09931589 | 0.6114833 | -0.2519021 | 0.8011168 | 0.897906 |

| | | | | | | | |
|---|---|---|---|---|---|---|---|
| ENSG00000262294 | | 2.633744 | -0.09932002 | 0.6849131 | -0.3004281 | 0.7638507 | 0.874321 |
| ENSG00000284735 | | 4.559253 | -0.09932952 | 0.5979485 | -0.2554093 | 0.798407 | 0.896254 |
| ENSG00000255769 | GOLGA2P10 | 395.2385 | -0.0993809 | 0.1956883 | -0.5261939 | 0.5987535 | 0.765557 |
| ENSG00000136463 | TACO1 | 4786.66493 | -0.099430497 | 0.2846384 | -0.36667215 | 0.71386358 | 0.843961 |
| ENSG00000186076 | | 737.6535 | -0.09943543 | 0.3003841 | -0.3612011 | 0.7179491 | 0.846132 |
| ENSG00000147475 | ERLIN2 | 2858.309 | -0.09946652 | 0.20219 | -0.5321418 | 0.5946278 | 0.762079 |
| ENSG00000258839 | MC1R | 492.5918 | -0.09948398 | 0.2204809 | -0.498962 | 0.6178061 | 0.778824 |
| ENSG00000173200 | PARP15 | 1.287036 | -0.0994928 | 0.7215563 | -0.3246272 | 0.7454632 | NA |
| ENSG00000277020 | | 5.387775 | -0.09958341 | 0.5435052 | -0.2516175 | 0.8013367 | 0.898104 |
| ENSG00000182628 | SKA2 | 9513.434 | -0.09964032 | 0.2820948 | -0.3959648 | 0.692131 | 0.830926 |
| ENSG00000075218 | GTSE1 | 1110.642281 | -0.099659363 | 0.530217188 | -0.259961105 | 0.79489378 | 0.894024 |
| ENSG00000272359 | RNU4-89P | 0.8235357 | -0.09967866 | 0.7355528 | -0.3567943 | 0.7212458 | NA |
| ENSG00000024422 | EHD2 | 622.7405953 | -0.099757229 | 0.269500208 | -0.401411143 | 0.68811745 | 0.828128 |
| ENSG00000227518 | MIR1302-9HG | 3.562661 | -0.09984481 | 0.6256576 | -0.2649922 | 0.7910155 | 0.891628 |
| ENSG00000223877 | RPS8P10 | 8.535138 | -0.09985693 | 0.4956317 | -0.2578177 | 0.7965476 | 0.89504 |
| ENSG00000287439 | | 0.7868909 | -0.09987232 | 0.7595514 | -0.4341084 | 0.6642097 | NA |
| ENSG00000256075 | | 3.738508 | -0.099874 | 0.6726948 | -0.2778593 | 0.7811204 | 0.885254 |
| ENSG00000200579 | | 1.66617 | -0.09990118 | 0.7341724 | -0.3518516 | 0.7249496 | 0.850636 |
| ENSG00000183814 | LIN9 | 700.9968 | -0.0999048 | 0.3057994 | -0.3598093 | 0.7189897 | 0.846902 |
| ENSG00000168209 | DDIT4 | 6108.271 | -0.09990643 | 0.3230664 | -0.3439722 | 0.7308672 | 0.853987 |
| ENSG00000179632 | MAF1 | 9197.654 | -0.09991554 | 0.1618629 | -0.610708 | 0.5413929 | 0.723118 |
| ENSG00000169189 | NSMCE1 | 1912.415 | -0.1000504 | 0.1840942 | -0.5620276 | 0.5740972 | 0.746641 |
| ENSG00000267365 | KCNJ2-AS1 | 0.8973709 | -0.1000508 | 0.7905256 | -0.3271363 | 0.7435648 | NA |
| ENSG00000115226 | FNDC4 | 11.86226617 | -0.10009358 | 0.504943343 | -0.269458685 | 0.78757673 | 0.889371 |
| ENSG00000116478 | HDAC1 | 11287.68687 | -0.100094666 | 0.15131033 | -0.681730994 | 0.49540907 | 0.684728 |
| ENSG00000242353 | RPL12P30 | 0.8849341 | -0.1001411 | 0.7905542 | -0.3269647 | 0.7436946 | NA |
| ENSG00000139890 | REM2 | 88.61489 | -0.1001667 | 0.3745257 | -0.3070349 | 0.7588168 | 0.87145 |
| ENSG00000234835 | PHBP13 | 2.303994 | -0.1001889 | 0.709315 | -0.3171511 | 0.7511289 | 0.866893 |
| ENSG00000271266 | LAPTM4BP2 | 97.58912 | -0.100224 | 0.2165828 | -0.4845281 | 0.6280112 | 0.786205 |
| ENSG00000239900 | ADSL | 320.2068 | -0.1002713 | 0.1731749 | -0.5833377 | 0.559666 | 0.73613 |
| ENSG00000132424 | PNISR | 9602.480983 | -0.100283006 | 0.252123086 | -0.42377417 | 0.67173052 | 0.817546 |
| ENSG00000237672 | KRR1P1 | 9.305157 | -0.1003272 | 0.5324191 | -0.2534751 | 0.7999011 | 0.897112 |
| ENSG00000235497 | | 0.797251 | -0.1003316 | 0.7427652 | -0.3809617 | 0.7032316 | NA |

| | | | | | | | |
|---|---|---|---|---|---|---|---|
| ENSG00000273747 | | 9.3298 | -0.1003352 | 0.5058378 | -0.2613152 | 0.7938494 | 0.893341 |
| ENSG00000090889 | KIF4A | 2086.235332 | -0.100385263 | 0.46315258 | -0.263689584 | 0.79201912 | 0.892302 |
| ENSG00000226756 | | 3.056338 | -0.1004309 | 0.6494261 | -0.2720157 | 0.7856099 | 0.887905 |
| ENSG00000236691 | NDUFA4P2 | 1.519408 | -0.1004601 | 0.7184257 | -0.317022 | 0.7512269 | NA |
| ENSG00000213935 | RPL22P16 | 33.31373 | -0.1004711 | 0.3404571 | -0.3295458 | 0.7417432 | 0.86081 |
| ENSG00000254873 | | 9.301163 | -0.1004766 | 0.4965445 | -0.2619681 | 0.7933461 | 0.893115 |
| ENSG00000119844 | AFTPH | 3887.17693 | -0.100544741 | 0.18959764 | -0.567590548 | 0.57031303 | 0.744126 |
| ENSG00000151948 | GLT1D1 | 1.606389 | -0.100671 | 0.7336398 | -0.353995 | 0.7233427 | 0.849699 |
| ENSG00000257043 | SRSF3P1 | 63.22216 | -0.1007553 | 0.3314995 | -0.3402577 | 0.7336625 | 0.85558 |
| ENSG00000120860 | WASHC3 | 1053.538327 | -0.100793539 | 0.16968497 | -0.606040869 | 0.54448762 | 0.725223 |
| ENSG00000271424 | GLRX5P3 | 2.895172 | -0.1008025 | 0.6526536 | -0.2684693 | 0.7883381 | 0.889891 |
| ENSG00000049323 | LTBP1 | 5972.841895 | -0.100806205 | 0.23514711 | -0.420115608 | 0.674401 | 0.818891 |
| ENSG00000179431 | FJX1 | 1106.169 | -0.1008151 | 0.2192242 | -0.4828152 | 0.6292269 | 0.787012 |
| ENSG00000278041 | | 0.9238218 | -0.1009016 | 0.7727948 | -0.4998198 | 0.6172019 | NA |
| ENSG00000201340 | | 0.8965906 | -0.1009293 | 0.7605585 | -0.4406248 | 0.6594846 | NA |
| ENSG00000237868 | | 53.29324 | -0.1010095 | 0.3040523 | -0.357729 | 0.7205461 | 0.847936 |
| ENSG00000133703 | KRAS | 1826.64163 | -0.101034426 | 0.151086753 | -0.685306727 | 0.49315039 | 0.683027 |
| ENSG00000108298 | RPL19 | 26886.12281 | -0.101059723 | 0.167192102 | -0.611707657 | 0.5407312 | 0.722634 |
| ENSG00000167700 | MFSD3 | 2914.102 | -0.1011285 | 0.2408683 | -0.4184817 | 0.675595 | 0.819667 |
| ENSG00000171307 | ZDHHC16 | 4159.895 | -0.1012329 | 0.1380992 | -0.7705328 | 0.4409839 | 0.638839 |
| ENSG00000251402 | FAM90A25P | 2.062104 | -0.101276 | 0.7121577 | -0.3303093 | 0.7411663 | 0.860407 |
| ENSG00000185519 | FAM131C | 101.9106 | -0.1013756 | 0.3973927 | -0.2959701 | 0.7672529 | 0.876588 |
| ENSG00000267498 | | 4.149758 | -0.1013764 | 0.6452288 | -0.2665953 | 0.7897808 | 0.890982 |
| ENSG00000278505 | C17orf78 | 1.420476 | -0.1013775 | 0.7157672 | -0.3241757 | 0.745805 | NA |
| ENSG00000185837 | HDHD5-AS1 | 17.95761 | -0.1014119 | 0.4822783 | -0.271415 | 0.7860719 | 0.888314 |
| ENSG00000276272 | | 2.227012 | -0.1014193 | 0.6858725 | -0.2910874 | 0.7709845 | 0.878884 |
| ENSG00000285614 | | 0.9551004 | -0.1014382 | 0.7542215 | -0.4218294 | 0.6731495 | NA |
| ENSG00000243016 | | 3.844814 | -0.1015421 | 0.7098982 | -0.3083912 | 0.7577847 | 0.87098 |
| ENSG00000167615 | LENG8 | 10136.69 | -0.1017144 | 0.2531438 | -0.4282228 | 0.6684889 | 0.81529 |
| ENSG00000236991 | EDRF1-AS1 | 38.23267 | -0.1017258 | 0.3474209 | -0.3280159 | 0.7428996 | 0.861512 |
| ENSG00000111678 | C12orf57 | 1252.201525 | -0.101726067 | 0.214208739 | -0.48417561 | 0.62826125 | 0.786407 |
| ENSG00000230092 | | 7.537719 | -0.1018116 | 0.5251146 | -0.2602702 | 0.7946554 | 0.893984 |
| ENSG00000267414 | SETBP1-DT | 5.782834 | -0.1018966 | 0.6238951 | -0.26586 | 0.790347 | 0.891266 |

| | | | | | | | |
|---|---|---|---|---|---|---|---|
| ENSG00000111796 | KLRB1 | 1.264556113 | -0.101941696 | 0.731984114 | -0.342738628 | 0.7317951 | NA |
| ENSG00000105173 | CCNE1 | 910.1361981 | -0.101949778 | 0.257447606 | -0.420008332 | 0.67447937 | 0.818912 |
| ENSG00000143164 | DCAF6 | 3481.76 | -0.1020086 | 0.20702 | -0.513546 | 0.6075694 | 0.771445 |
| ENSG00000137404 | NRM | 524.0264843 | -0.102146643 | 0.453325311 | -0.277489063 | 0.7814046 | 0.885483 |
| ENSG00000203327 | | 3.768004 | -0.1021851 | 0.6237507 | -0.2654447 | 0.7906669 | 0.891421 |
| ENSG00000100897 | DCAF11 | 7482.728983 | -0.10221048 | 0.090037193 | -1.158670266 | 0.24659062 | 0.438422 |
| ENSG00000259774 | | 1.981777 | -0.102213 | 0.6785409 | -0.2940075 | 0.7687522 | 0.877437 |
| ENSG00000285857 | | 8.608351 | -0.1022744 | 0.5991419 | -0.261588 | 0.7936391 | 0.893218 |
| ENSG00000272305 | | 7.718485 | -0.1023707 | 0.5608293 | -0.2610981 | 0.7940169 | 0.893416 |
| ENSG00000283078 | | 78.96732 | -0.1023872 | 0.2699817 | -0.4039865 | 0.6862227 | 0.827235 |
| ENSG00000260634 | | 1.083166 | -0.1023944 | 0.7486304 | -0.4211194 | 0.6736679 | NA |
| ENSG00000100811 | YY1 | 8519.791948 | -0.102435493 | 0.114644616 | -0.904802904 | 0.36556981 | 0.567562 |
| ENSG00000185414 | MRPL30 | 2906.476 | -0.1024755 | 0.1593223 | -0.6497598 | 0.5158474 | 0.701195 |
| ENSG00000224905 | | 2.9867 | -0.1024952 | 0.6265423 | -0.2758645 | 0.7826522 | 0.886221 |
| ENSG00000270871 | | 48.19485 | -0.1025061 | 0.3641978 | -0.3194816 | 0.7493613 | 0.865831 |
| ENSG00000286507 | | 0.8586472 | -0.1025355 | 0.7603413 | -0.4304851 | 0.6668428 | NA |
| ENSG00000280767 | | 2.655904 | -0.1026355 | 0.6703533 | -0.287567 | 0.7736782 | 0.880665 |
| ENSG00000255753 | | 2.251428 | -0.1027621 | 0.650039 | -0.2833858 | 0.7768811 | 0.882518 |
| ENSG00000241438 | TDGF1P6 | 1.003602 | -0.1027883 | 0.7184384 | -0.3477777 | 0.7280072 | NA |
| ENSG00000151963 | ZNF37CP | 1.693143 | -0.1027915 | 0.7279396 | -0.3741197 | 0.7083152 | 0.841459 |
| ENSG00000185338 | SOCS1 | 126.1473 | -0.1028548 | 0.3316372 | -0.3461228 | 0.7292504 | 0.853215 |
| ENSG00000244556 | ODCP | 8.81718 | -0.1029815 | 0.565715 | -0.2580388 | 0.796377 | 0.894975 |
| ENSG00000263220 | | 1.957287 | -0.1030052 | 0.6952362 | -0.3023715 | 0.7623689 | 0.873491 |
| ENSG00000178605 | GTPBP6 | 3575.069 | -0.1030215 | 0.1977397 | -0.5525021 | 0.5806044 | 0.751373 |
| ENSG00000141858 | SAMD1 | 2368.716 | -0.103033 | 0.2377759 | -0.4578324 | 0.6470729 | 0.799892 |
| ENSG00000245849 | RAD51-AS1 | 710.4761 | -0.1030625 | 0.353963 | -0.3267488 | 0.7438579 | 0.862021 |
| ENSG00000232316 | LINC02518 | 42.93991 | -0.1031867 | 0.5444631 | -0.2572489 | 0.7969867 | 0.895203 |
| ENSG00000078687 | TNRC6C | 1223.944283 | -0.103213228 | 0.280775381 | -0.398014976 | 0.69061915 | 0.829793 |
| ENSG00000284484 | | 1.161179 | -0.103309 | 0.7187651 | -0.3324484 | 0.7395507 | NA |
| ENSG00000261739 | GOLGA8S | 1.188003 | -0.1033183 | 0.730834 | -0.3669429 | 0.7136616 | NA |
| ENSG00000110195 | FOLR1 | 2.422216102 | -0.103360009 | 0.696553324 | -0.311281016 | 0.755587 | 0.869545 |
| ENSG00000185130 | H2BC13 | 31.59751 | -0.1034043 | 0.4019026 | -0.2980851 | 0.7656382 | 0.875613 |
| ENSG00000024526 | DEPDC1 | 1275.400958 | -0.103407279 | 0.436863773 | -0.290448059 | 0.77147348 | 0.879183 |

| | | | | | | | |
|---|---|---|---|---|---|---|---|
| ENSG00000136715 | SAP130 | 2375.35684 | -0.10348352 | 0.189738119 | -0.567250991 | 0.57054367 | 0.744349 |
| ENSG00000224690 | UBE2D3P3 | 2.147876 | -0.1035031 | 0.7029921 | -0.3108531 | 0.7559123 | 0.869764 |
| ENSG00000233242 | | 4.658654 | -0.1035113 | 0.6461266 | -0.2734148 | 0.7845344 | 0.887289 |
| ENSG00000203499 | IQANK1 | 5260.59 | -0.1035485 | 0.1936043 | -0.5552978 | 0.578691 | 0.750085 |
| ENSG00000231105 | ECE1-AS1 | 1.345942 | -0.103575 | 0.721851 | -0.3343804 | 0.7380925 | NA |
| ENSG00000198673 | TAFA2 | 21.05722 | -0.1036434 | 0.3945356 | -0.3061551 | 0.7594866 | 0.871805 |
| ENSG00000134516 | DOCK2 | 9.274699045 | -0.103713105 | 0.636322489 | -0.270054552 | 0.78711829 | 0.88908 |
| ENSG00000199691 | RN7SKP173 | 19.6158 | -0.1038091 | 0.4038012 | -0.3041206 | 0.761036 | 0.872566 |
| ENSG00000146109 | ABT1 | 2235.288 | -0.1038622 | 0.2156376 | -0.5033801 | 0.614697 | 0.776509 |
| ENSG00000179978 | NAIPP2 | 45.38945 | -0.103907 | 0.3497202 | -0.3318361 | 0.740013 | 0.859581 |
| ENSG00000267565 | | 25.29179 | -0.1039453 | 0.3462216 | -0.3336092 | 0.7386745 | 0.858824 |
| ENSG00000196476 | C20orf96 | 417.5965 | -0.1039684 | 0.2118174 | -0.5248253 | 0.5997047 | 0.765998 |
| ENSG00000233121 | VN1R20P | 3.355406 | -0.1039808 | 0.6831 | -0.2911046 | 0.7709713 | 0.878884 |
| ENSG00000223537 | | 1.517239 | -0.104033 | 0.7317368 | -0.3721357 | 0.7097918 | NA |
| ENSG00000110851 | PRDM4 | 3086.536931 | -0.104122083 | 0.180208293 | -0.597469687 | 0.55019384 | 0.729281 |
| ENSG00000110700 | RPS13 | 14312.20437 | -0.104128166 | 0.164517574 | -0.654756644 | 0.51262445 | 0.698493 |
| ENSG00000222920 | RNA5SP29 | 2.772922 | -0.1041326 | 0.6818722 | -0.2956536 | 0.7674946 | 0.87668 |
| ENSG00000266877 | | 1.469142 | -0.104139 | 0.6970454 | -0.3196126 | 0.749262 | NA |
| ENSG00000184897 | H1-10 | 4093.224 | -0.1041806 | 0.2685322 | -0.4204881 | 0.6741289 | 0.818795 |
| ENSG00000103275 | UBE2I | 6363.193434 | -0.104198209 | 0.135135558 | -0.789128006 | 0.4300372 | 0.629305 |
| ENSG00000259124 | | 0.9344634 | -0.1043776 | 0.7896802 | -0.7916528 | 0.4285632 | NA |
| ENSG00000013306 | SLC25A39 | 21649.68519 | -0.104391421 | 0.173904514 | -0.618049643 | 0.53654262 | 0.719606 |
| ENSG00000103274 | NUBP1 | 722.1046604 | -0.104401341 | 0.214904215 | -0.507674108 | 0.61168191 | 0.774558 |
| ENSG00000238141 | BRWD1-AS1 | 31.78094 | -0.1044097 | 0.3726975 | -0.3191254 | 0.7496314 | 0.865959 |
| ENSG00000164209 | SLC25A46 | 3935.525 | -0.1044206 | 0.1306568 | -0.7841883 | 0.4329296 | 0.632041 |
| ENSG00000228661 | STIM1-AS1 | 4.049893 | -0.1044212 | 0.6214692 | -0.2760705 | 0.7824939 | 0.886136 |
| ENSG00000105550 | FGF21 | 6.644954152 | -0.104431838 | 0.671545023 | -0.315232381 | 0.75258522 | 0.867632 |
| ENSG00000223984 | HNRNPRP1 | 44.87658 | -0.1044577 | 0.3479087 | -0.3358918 | 0.7369525 | 0.857835 |
| ENSG00000114019 | AMOTL2 | 14015.65033 | -0.104512774 | 0.325242708 | -0.339600414 | 0.73415747 | 0.855857 |
| ENSG00000257803 | PIGAP1 | 2.751572 | -0.1046143 | 0.631996 | -0.2710964 | 0.7863169 | 0.88844 |
| ENSG00000197019 | SERTAD1 | 858.9211 | -0.1046518 | 0.2035647 | -0.5359261 | 0.5920096 | 0.760374 |
| ENSG00000248098 | BCKDHA | 72.17441 | -0.1046904 | 0.244937 | -0.4523859 | 0.650991 | 0.802793 |
| ENSG00000162687 | KCNT2 | 11.31897 | -0.104697 | 0.5221491 | -0.273841 | 0.7842068 | 0.887074 |

| | | | | | | | |
|---|---|---|---|---|---|---|---|
| ENSG00000184381 | PLA2G6 | 640.9962 | -0.1047462 | 0.2424603 | -0.4512025 | 0.6518436 | 0.80336 |
| ENSG00000271680 | | 9.568039 | -0.104761 | 0.7994828 | -1.367662 | 0.1714178 | 0.34215 |
| ENSG00000239827 | SUGT1P3 | 38.62059 | -0.1049201 | 0.3853764 | -0.3153665 | 0.7524834 | 0.86759 |
| ENSG00000130177 | CDC16 | 3258.013073 | -0.105006576 | 0.11261473 | -0.94611683 | 0.344089 | 0.545764 |
| ENSG00000170419 | VSTM2A | 11.56679 | -0.105074 | 0.5652457 | -0.2786116 | 0.7805429 | 0.884969 |
| ENSG00000231784 | DBIL5P | 29.98474 | -0.1050913 | 0.3852933 | -0.3170002 | 0.7512435 | 0.86695 |
| ENSG00000228075 | BOD1L2 | 0.8352213 | -0.1051314 | 0.7305502 | -0.3927167 | 0.6945287 | NA |
| ENSG00000105321 | CCDC9 | 1319.459912 | -0.10519967 | 0.141908662 | -0.757540662 | 0.44872601 | 0.645304 |
| ENSG00000233985 | LINC01681 | 1.019665 | -0.1052919 | 0.7906787 | -0.3390765 | 0.7345521 | NA |
| ENSG00000277738 | | 7.192665 | -0.1053321 | 0.8011759 | -1.850178 | 0.06428785 | 0.167337 |
| ENSG00000288032 | | 3.433546 | -0.105372 | 0.639539 | -0.2811741 | 0.7785769 | 0.883564 |
| ENSG00000165392 | WRN | 1129.639 | -0.1054083 | 0.1905676 | -0.5731549 | 0.5665398 | 0.741132 |
| ENSG00000213560 | | 15.18842 | -0.1054173 | 0.4863115 | -0.2791617 | 0.7801207 | 0.884712 |
| ENSG00000196700 | ZNF512B | 11868.58 | -0.1054333 | 0.141566 | -0.7581273 | 0.4483748 | 0.644985 |
| ENSG00000227077 | | 6.345229 | -0.1054614 | 0.5457804 | -0.265812 | 0.790384 | 0.891266 |
| ENSG00000278662 | GOLGA6L10 | 240.8668 | -0.1054887 | 0.292637 | -0.3906749 | 0.6960376 | 0.833695 |
| ENSG00000231341 | VDAC1P6 | 118.9593 | -0.1054906 | 0.2478608 | -0.4517619 | 0.6514406 | 0.803115 |
| ENSG00000272140 | | 85.75486 | -0.1056213 | 0.2627022 | -0.428768 | 0.6680921 | 0.815105 |
| ENSG00000266667 | | 1.515495 | -0.1056498 | 0.715714 | -0.3338881 | 0.738464 | NA |
| ENSG00000226237 | GAS1RR | 0.9640669 | -0.1057742 | 0.7340569 | -0.3820767 | 0.7024044 | NA |
| ENSG00000280062 | | 2.994336 | -0.10588 | 0.6143583 | -0.2772625 | 0.7815786 | 0.885496 |
| ENSG00000261587 | TMEM249 | 11.29689 | -0.1059027 | 0.5313085 | -0.2736643 | 0.7843426 | 0.887152 |
| ENSG00000140988 | RPS2 | 31849.7 | -0.105976 | 0.1943178 | -0.5979666 | 0.5498622 | 0.729016 |
| ENSG00000263069 | RNF213-AS1 | 274.0901 | -0.1060752 | 0.3240065 | -0.3619744 | 0.7173711 | 0.846015 |
| ENSG00000232757 | ETF1P1 | 1.802435 | -0.1060861 | 0.7273972 | -0.351865 | 0.7249395 | 0.850636 |
| ENSG00000213563 | C8orf82 | 1977.7 | -0.106122 | 0.1816535 | -0.6121577 | 0.5404334 | 0.722434 |
| ENSG00000254502 | FNTAP1 | 6.764212 | -0.1061334 | 0.596925 | -0.2817416 | 0.7781417 | 0.883372 |
| ENSG00000188032 | C19orf67 | 4.009763 | -0.1061436 | 0.6430954 | -0.2786056 | 0.7805475 | 0.884969 |
| ENSG00000175602 | CCDC85B | 2530.986 | -0.1061961 | 0.2309017 | -0.5012576 | 0.6161898 | 0.777672 |
| ENSG00000236017 | ASMTL-AS1 | 146.9289 | -0.1062135 | 0.3173739 | -0.3689439 | 0.7121695 | 0.843338 |
| ENSG00000260084 | | 0.8340355 | -0.1062421 | 0.7347832 | -0.3670706 | 0.7135663 | NA |
| ENSG00000127362 | TAS2R3 | 2.338301324 | -0.106274074 | 0.722836768 | -0.344753952 | 0.73027935 | 0.853877 |
| ENSG00000270048 | | 3.770466 | -0.106326 | 0.6421953 | -0.2844262 | 0.7760838 | 0.882054 |

| | | | | | | | |
|---|---|---|---|---|---|---|---|
| ENSG00000235508 | RPS2P7 | 27.58615 | -0.1063281 | 0.39126 | -0.3134881 | 0.7539098 | 0.868685 |
| ENSG00000156469 | MTERF3 | 1388.559 | -0.106478 | 0.1633927 | -0.6684676 | 0.5038352 | 0.691125 |
| ENSG00000181625 | SLX1B | 2.713601 | -0.1065029 | 0.6892947 | -0.3168492 | 0.751358 | 0.866969 |
| ENSG00000162433 | AK4 | 2783.468 | -0.1065272 | 0.2694188 | -0.4272745 | 0.6691794 | 0.815661 |
| ENSG00000229700 | | 2.824209 | -0.1065905 | 0.6454708 | -0.2833736 | 0.7768904 | 0.882518 |
| ENSG00000210151 | MT-TS1 | 1.264486 | -0.1066102 | 0.7247377 | -0.3531964 | 0.7239412 | NA |
| ENSG00000200090 | | 34.02548 | -0.1066122 | 0.3780881 | -0.3245099 | 0.745552 | 0.863081 |
| ENSG00000130561 | SAG | 1.07822877 | -0.106686553 | 0.738126682 | -0.377837953 | 0.70555098 | NA |
| ENSG00000167325 | RRM1 | 4400.844 | -0.1067974 | 0.3945098 | -0.3336353 | 0.7386548 | 0.858824 |
| ENSG00000251298 | | 2.9822 | -0.1068088 | 0.6205071 | -0.2767904 | 0.781941 | 0.88568 |
| ENSG00000153363 | LINC00467 | 470.6148 | -0.1068177 | 0.1813998 | -0.6085832 | 0.5428008 | 0.723848 |
| ENSG00000260625 | | 15.86014 | -0.1068218 | 0.5207598 | -0.274548 | 0.7836636 | 0.886724 |
| ENSG00000089041 | P2RX7 | 2.124584438 | -0.10683715 | 0.672569154 | -0.296226388 | 0.7670572 | 0.876551 |
| ENSG00000162073 | PAQR4 | 1388.751 | -0.1068789 | 0.2459556 | -0.4896777 | 0.624362 | 0.783483 |
| ENSG00000231010 | | 6.970441 | -0.1069267 | 0.5607987 | -0.2726032 | 0.7851582 | 0.887583 |
| ENSG00000132005 | RFX1 | 918.1029571 | -0.106929227 | 0.180787737 | -0.611505999 | 0.54086465 | 0.72274 |
| ENSG00000189343 | RPS2P46 | 5578.48 | -0.1069693 | 0.2044054 | -0.5405912 | 0.5887894 | 0.757924 |
| ENSG00000253439 | CDC42P5 | 2.241211 | -0.1070594 | 0.7052906 | -0.3220995 | 0.7473773 | 0.864403 |
| ENSG00000115760 | BIRC6 | 9302.890209 | -0.107103603 | 0.285012834 | -0.407501711 | 0.68363954 | 0.824997 |
| ENSG00000279924 | | 0.929769 | -0.107127 | 0.7459014 | -0.4247826 | 0.6709952 | NA |
| ENSG00000091947 | TMEM101 | 2420.750802 | -0.107127726 | 0.170493725 | -0.646085822 | 0.51822378 | 0.703529 |
| ENSG00000181788 | SIAH2 | 4920.4 | -0.1072143 | 0.2472675 | -0.4683586 | 0.6395282 | 0.794189 |
| ENSG00000127838 | PNKD | 6792.552671 | -0.107251781 | 0.292832462 | -0.362470094 | 0.71700077 | 0.846015 |
| ENSG00000171763 | SPATA5L1 | 717.8491 | -0.1072935 | 0.1800325 | -0.6109642 | 0.5412233 | 0.723008 |
| ENSG00000170581 | STAT2 | 3754.863 | -0.1073109 | 0.2361925 | -0.467398 | 0.6402151 | 0.794706 |
| ENSG00000136197 | C7orf25 | 29.43940887 | -0.10733377 | 0.327021839 | -0.358335778 | 0.72009205 | 0.847679 |
| ENSG00000228981 | | 22.76424 | -0.1073487 | 0.3886665 | -0.3205524 | 0.7485496 | 0.865382 |
| ENSG00000285644 | | 98.84679 | -0.1074162 | 0.3058609 | -0.3845648 | 0.7005599 | 0.836925 |
| ENSG00000227758 | | 1.342002 | -0.107426 | 0.7274406 | -0.3810091 | 0.7031965 | NA |
| ENSG00000137860 | SLC28A2 | 1.440706887 | -0.10746455 | 0.672352938 | -0.307497409 | 0.7584648 | NA |
| ENSG00000269896 | | 104.7287 | -0.1074881 | 0.2055035 | -0.5366957 | 0.5914779 | 0.760174 |
| ENSG00000104979 | C19orf53 | 3090.732058 | -0.107519112 | 0.212199514 | -0.531868098 | 0.59481735 | 0.762212 |
| ENSG00000170684 | ZNF296 | 272.6124 | -0.1075689 | 0.1894377 | -0.5947485 | 0.5520116 | 0.730751 |

| | | | | | | | |
|---|---|---|---|---|---|---|---|
| ENSG00000183255 | PTTG1IP | 13951.27 | -0.1075859 | 0.1378382 | -0.8284927 | 0.4073915 | 0.608055 |
| ENSG00000197446 | CYP2F1 | 1.180075 | -0.107621 | 0.7300791 | -0.375805 | 0.7070619 | NA |
| ENSG00000273226 | | 14.48639 | -0.1077102 | 0.5255974 | -0.2753545 | 0.7830439 | 0.886458 |
| ENSG00000213373 | LINC00671 | 0.9447828 | -0.1077793 | 0.7794139 | -0.6429543 | 0.5202538 | NA |
| ENSG00000228044 | | 1.152215 | -0.1077923 | 0.7656065 | -0.5253257 | 0.5993568 | NA |
| ENSG00000173567 | ADGRF3 | 25.92124 | -0.1078309 | 0.3988963 | -0.3166014 | 0.7515461 | 0.866998 |
| ENSG00000196230 | TUBB | 27077.37 | -0.1078946 | 0.3742397 | -0.2677741 | 0.7888732 | 0.890218 |
| ENSG00000226124 | FTCDNL1 | 242.0458 | -0.1078951 | 0.2312899 | -0.48482 | 0.627804 | 0.786057 |
| ENSG00000183458 | PKD1P3 | 907.7769 | -0.1079319 | 0.3770876 | -0.323147 | 0.7465839 | 0.863912 |
| ENSG00000023171 | GRAMD1B | 2.802476524 | -0.107955653 | 0.702798031 | -0.313747194 | 0.75371305 | 0.868555 |
| ENSG00000111530 | CAND1 | 11978.87464 | -0.108053276 | 0.172691846 | -0.603007971 | 0.54650339 | 0.726642 |
| ENSG00000255823 | MTRNR2L8 | 17.03945 | -0.1080668 | 0.5026118 | -0.2806829 | 0.7789536 | 0.883841 |
| ENSG00000068097 | HEATR6 | 17383.82236 | -0.1080727 | 0.130380511 | -0.866951711 | 0.38596847 | 0.588308 |
| ENSG00000232699 | BDH2P1 | 51.23866 | -0.1080887 | 0.3811432 | -0.3261184 | 0.7443348 | 0.862303 |
| ENSG00000102144 | PGK1 | 26974.47518 | -0.108142021 | 0.212857249 | -0.53097546 | 0.59543579 | 0.762682 |
| ENSG00000094755 | GABRP | 1843.976417 | -0.108153194 | 0.646072764 | -0.278698556 | 0.78047617 | 0.884964 |
| ENSG00000217557 | | 1.307241 | -0.1081879 | 0.7345896 | -0.3688943 | 0.7122065 | NA |
| ENSG00000196071 | OR2L13 | 5.275571 | -0.1083713 | 0.6624639 | -0.3162718 | 0.7517962 | 0.867075 |
| ENSG00000166965 | RCCD1 | 1799.166 | -0.1084239 | 0.1596338 | -0.7006907 | 0.483496 | 0.674792 |
| ENSG00000261838 | | 1.467125 | -0.1085183 | 0.7444505 | -0.396747 | 0.6915541 | NA |
| ENSG00000153989 | NUS1 | 3138.47 | -0.108594 | 0.1629195 | -0.700193 | 0.4838068 | 0.675015 |
| ENSG00000278499 | NPAP1L | 0.9126923 | -0.1086263 | 0.7459784 | -0.4267243 | 0.6695801 | NA |
| ENSG00000189042 | ZNF567 | 587.7304 | -0.1086553 | 0.19489 | -0.5811193 | 0.5611601 | 0.737289 |
| ENSG00000180198 | RCC1 | 10599.69 | -0.1087163 | 0.2487735 | -0.4414616 | 0.6588788 | 0.808062 |
| ENSG00000254495 | | 0.8296397 | -0.1087333 | 0.7602666 | -0.4376906 | 0.6616106 | NA |
| ENSG00000213071 | LPAL2 | 42.52081 | -0.1087698 | 0.4332573 | -0.3034299 | 0.7615623 | 0.872875 |
| ENSG00000070614 | NDST1 | 9307.869827 | -0.108782769 | 0.152199013 | -0.762490452 | 0.44576734 | 0.642779 |
| ENSG00000041515 | MYO16 | 83.61821614 | -0.108828916 | 0.490861602 | -0.285992475 | 0.77488389 | 0.881331 |
| ENSG00000217416 | ISCA1P1 | 284.8442 | -0.1088324 | 0.1889625 | -0.588575 | 0.5561464 | 0.733627 |
| ENSG00000279811 | | 0.7729246 | -0.1088594 | 0.7625483 | -0.4830393 | 0.6290679 | NA |
| ENSG00000240489 | SETP14 | 2130.476 | -0.1089072 | 0.1949135 | -0.5792038 | 0.5624517 | 0.738154 |
| ENSG00000269688 | | 83.97378 | -0.1089498 | 0.2330297 | -0.4908206 | 0.6235533 | 0.7828 |
| ENSG00000165182 | CXorf58 | 1.699704 | -0.1089571 | 0.6739769 | -0.306598 | 0.7591494 | 0.871644 |

| | | | | | | | |
|---|---|---|---|---|---|---|---|
| ENSG00000129473 | BCL2L2 | 3198.335518 | -0.10896518 | 0.239193156 | -0.479685777 | 0.63145084 | 0.788593 |
| ENSG00000180458 | | 1.544924 | -0.1089921 | 0.7030922 | -0.3255082 | 0.7447965 | NA |
| ENSG00000202522 | | 7.474087 | -0.1090715 | 0.5208577 | -0.2936942 | 0.7689916 | 0.877559 |
| ENSG00000151572 | ANO4 | 4.370339 | -0.1090848 | 0.8018493 | -1.709558 | 0.08734759 | 0.210806 |
| ENSG00000103145 | HCFC1R1 | 3049.77552 | -0.109289016 | 0.219443916 | -0.524201912 | 0.60013811 | 0.766332 |
| ENSG00000238260 | | 20.4382 | -0.1092942 | 0.5744165 | -0.2780866 | 0.7809459 | 0.885156 |
| ENSG00000136206 | SPDYE1 | 42.04227539 | -0.109319666 | 0.393303825 | -0.321879547 | 0.74754395 | 0.864521 |
| ENSG00000196900 | TEX43 | 0.8128654 | -0.1093502 | 0.7176026 | -0.351969 | 0.7248615 | NA |
| ENSG00000248898 | | 1.960705 | -0.1093538 | 0.669 | -0.3031131 | 0.7618036 | 0.873031 |
| ENSG00000269506 | | 0.844284 | -0.1093656 | 0.7832533 | -0.6410979 | 0.5214591 | NA |
| ENSG00000124214 | STAU1 | 12103.17576 | -0.109368776 | 0.111469199 | -0.971928188 | 0.33108628 | 0.532097 |
| ENSG00000263033 | | 14.78378 | -0.1093719 | 0.4744224 | -0.2870053 | 0.7741082 | 0.880805 |
| ENSG00000213876 | RPL7AP64 | 0.9884509 | -0.1094143 | 0.7768796 | -0.5887226 | 0.5560473 | NA |
| ENSG00000215014 | | 18.63361 | -0.1094146 | 0.4777825 | -0.2916847 | 0.7705277 | 0.878627 |
| ENSG00000153879 | CEBPG | 4397.742 | -0.1094487 | 0.3278221 | -0.3664162 | 0.7140545 | 0.84405 |
| ENSG00000174038 | C9orf131 | 1.441545 | -0.1094874 | 0.7298385 | -0.3651887 | 0.7149705 | NA |
| ENSG00000126581 | BECN1 | 3692.063124 | -0.109516908 | 0.134717236 | -0.824560789 | 0.40962099 | 0.610012 |
| ENSG00000198039 | ZNF273 | 360.0724 | -0.1096209 | 0.2711891 | -0.4322854 | 0.665534 | 0.813216 |
| ENSG00000258136 | | 31.95919 | -0.1096506 | 0.3707743 | -0.3401584 | 0.7337372 | 0.85563 |
| ENSG00000183250 | LINC01547 | 683.7288 | -0.1097463 | 0.2084267 | -0.5525205 | 0.5805918 | 0.751373 |
| ENSG00000158042 | MRPL17 | 2779.424 | -0.1097574 | 0.2558874 | -0.4339955 | 0.6642917 | 0.812355 |
| ENSG00000115942 | ORC2 | 2884.083512 | -0.109770966 | 0.168328058 | -0.666669091 | 0.50498353 | 0.691783 |
| ENSG00000224877 | NDUFAF8 | 1530.506 | -0.109825 | 0.1937115 | -0.5931422 | 0.553086 | 0.731456 |
| ENSG00000166441 | RPL27A | 10575.17 | -0.1098314 | 0.1648785 | -0.7051886 | 0.4806929 | 0.67229 |
| ENSG00000246763 | RGMB-AS1 | 132.8181 | -0.109838 | 0.3024687 | -0.3953273 | 0.6926013 | 0.831229 |
| ENSG00000254440 | PBOV1 | 0.9584232 | -0.1098887 | 0.7310952 | -0.3666314 | 0.713894 | NA |
| ENSG00000048991 | R3HDM1 | 2876.025886 | -0.10993361 | 0.199244559 | -0.574811529 | 0.56541877 | 0.740236 |
| ENSG00000144161 | ZC3H8 | 1524.668 | -0.1099542 | 0.1446541 | -0.7654659 | 0.4439942 | 0.641462 |
| ENSG00000164983 | TMEM65 | 2600.434 | -0.1099635 | 0.1838225 | -0.6207715 | 0.5347499 | 0.718326 |
| ENSG00000242208 | RPL5P29 | 42.82128 | -0.1099661 | 0.378928 | -0.3337883 | 0.7385393 | 0.858824 |
| ENSG00000079805 | DNM2 | 12476.74017 | -0.109975449 | 0.155068327 | -0.726676093 | 0.4674244 | 0.661023 |
| ENSG00000250903 | GMDS-DT | 249.4863 | -0.1099807 | 0.3503535 | -0.3530137 | 0.7240782 | 0.850074 |
| ENSG00000168772 | CXXC4 | 78.06227 | -0.1099856 | 0.3637244 | -0.3392157 | 0.7344473 | 0.856021 |

| | | | | | | | |
|---|---|---|---|---|---|---|---|
| ENSG00000114353 | GNAI2 | 6991.967392 | -0.109993418 | 0.204444161 | -0.551813446 | 0.58107617 | 0.751783 |
| ENSG00000213529 | RPL32P22 | 2.177664 | -0.1100137 | 0.6993896 | -0.3436804 | 0.7310867 | 0.854113 |
| ENSG00000091483 | FH | 3865.241372 | -0.110057761 | 0.227661815 | -0.500720542 | 0.61656781 | 0.777927 |
| ENSG00000274615 | | 1.929971 | -0.1100969 | 0.729198 | -0.3622619 | 0.7171563 | 0.846015 |
| ENSG00000266957 | | 0.9015026 | -0.1101451 | 0.7666029 | -0.5171455 | 0.6050546 | NA |
| ENSG00000255423 | EBLN2 | 111.0918 | -0.1101875 | 0.328371 | -0.3744075 | 0.7081011 | 0.841263 |
| ENSG00000154162 | CDH12 | 6.5793 | -0.1102173 | 0.7013272 | -0.3275158 | 0.7432778 | 0.861574 |
| ENSG00000241061 | RPL5P1 | 857.8108 | -0.1102591 | 0.1875821 | -0.6074845 | 0.5435294 | 0.724479 |
| ENSG00000063761 | ADCK1 | 933.0405097 | -0.110276357 | 0.149547933 | -0.750313485 | 0.45306592 | 0.649237 |
| ENSG00000187735 | TCEA1 | 3720.308 | -0.1102996 | 0.1934974 | -0.5792594 | 0.5624142 | 0.738154 |
| ENSG00000257576 | HSPD1P4 | 314.7175 | -0.1103022 | 0.250258 | -0.4654232 | 0.6416284 | 0.795667 |
| ENSG00000270977 | | 55.92609 | -0.1103698 | 0.5116255 | -0.2854469 | 0.7753018 | 0.88169 |
| ENSG00000167642 | SPINT2 | 12740.22 | -0.110385 | 0.1589788 | -0.7057124 | 0.480367 | 0.671961 |
| ENSG00000166086 | JAM3 | 1.333067 | -0.1103946 | 0.7008812 | -0.3320319 | 0.7398652 | NA |
| ENSG00000131779 | PEX11B | 3617.436769 | -0.110414487 | 0.135022967 | -0.841340448 | 0.40015724 | 0.601141 |
| ENSG00000286634 | | 10.07408 | -0.1104248 | 0.5785602 | -0.2830406 | 0.7771457 | 0.882585 |
| ENSG00000279110 | | 4.813838 | -0.1104352 | 0.5913738 | -0.2803023 | 0.7792456 | 0.884097 |
| ENSG00000256393 | RPL41P5 | 376.8974 | -0.1104452 | 0.1787327 | -0.6323073 | 0.5271861 | 0.712031 |
| ENSG00000241343 | RPL36A | 817.5216 | -0.1104848 | 0.2960858 | -0.4080017 | 0.6832725 | 0.824763 |
| ENSG00000119979 | DENND10 | 1259.078211 | -0.110545001 | 0.152189604 | -0.745248468 | 0.45612153 | 0.652028 |
| ENSG00000278739 | MIR6859-4 | 1.415956 | -0.1105776 | 0.7147194 | -0.3404942 | 0.7334844 | NA |
| ENSG00000200344 | | 0.9999001 | -0.110597 | 0.7195103 | -0.3643859 | 0.7155699 | NA |
| ENSG00000235776 | | 453.9341 | -0.1105971 | 0.1821122 | -0.6264127 | 0.5310443 | 0.715301 |
| ENSG00000125485 | DDX31 | 1638.495565 | -0.110650973 | 0.10623363 | -1.053373975 | 0.29216963 | 0.490965 |
| ENSG00000130958 | SLC35D2 | 261.7486902 | -0.110687644 | 0.264846952 | -0.443957351 | 0.65707342 | 0.806839 |
| ENSG00000160284 | SPATC1L | 574.9951 | -0.1107777 | 0.220351 | -0.5191034 | 0.6036886 | 0.768745 |
| ENSG00000133872 | SARAF | 7095.621669 | -0.110777881 | 0.099198664 | -1.139788031 | 0.25437462 | 0.447374 |
| ENSG00000278986 | | 22.48242 | -0.1108467 | 0.4180304 | -0.314786 | 0.7529241 | 0.867952 |
| ENSG00000263257 | | 0.832686 | -0.1108585 | 0.7597418 | -0.443339 | 0.6575206 | NA |
| ENSG00000262140 | | 0.8472698 | -0.1108914 | 0.7414315 | -0.4035216 | 0.6865646 | NA |
| ENSG00000257512 | | 104.9604 | -0.110915 | 0.3504521 | -0.3569015 | 0.7211655 | 0.848435 |
| ENSG00000183751 | TBL3 | 3696.826 | -0.110932 | 0.1811067 | -0.6257554 | 0.5314754 | 0.715586 |
| ENSG00000125462 | MIR9-1HG | 51.92467982 | -0.110965145 | 0.424082493 | -0.313365049 | 0.75400334 | 0.868735 |

| | | | | | | | |
|---|---|---|---|---|---|---|---|
| ENSG00000217130 | | 445.2542 | -0.1109991 | 0.241314 | -0.4824476 | 0.629488 | 0.787127 |
| ENSG00000228540 | | 0.8084331 | -0.1110016 | 0.790198 | -0.3619521 | 0.7173878 | NA |
| ENSG00000264931 | MIR3138 | 1.010803 | -0.1110091 | 0.7637402 | -0.4642622 | 0.6424599 | NA |
| ENSG00000136527 | TRA2B | 8841.279893 | -0.111073744 | 0.213120499 | -0.546330143 | 0.58483902 | 0.75485 |
| ENSG00000151247 | EIF4E | 1264.593 | -0.1113489 | 0.1948834 | -0.5913122 | 0.5543112 | 0.732434 |
| ENSG00000267219 | | 19.10656 | -0.1113648 | 0.474652 | -0.2986191 | 0.7652307 | 0.875483 |
| ENSG00000179820 | MYADM | 1380.328 | -0.1113978 | 0.4139368 | -0.3263384 | 0.7441684 | 0.862231 |
| ENSG00000166685 | COG1 | 2636.388 | -0.1114205 | 0.139411 | -0.8137476 | 0.4157896 | 0.61561 |
| ENSG00000229625 | | 1.692532 | -0.1114646 | 0.7210512 | -0.3568814 | 0.7211806 | 0.848435 |
| ENSG00000125149 | C16orf70 | 1418.195394 | -0.111474985 | 0.116480175 | -0.967910473 | 0.33308908 | 0.534016 |
| ENSG00000215493 | | 104.8853 | -0.1115027 | 0.2347929 | -0.4911952 | 0.6232884 | 0.78269 |
| ENSG00000131435 | PDLIM4 | 0.834766305 | -0.111502905 | 0.762063959 | -0.494031082 | 0.62128423 | NA |
| ENSG00000102158 | MAGT1 | 5128.539148 | -0.111510886 | 0.156659063 | -0.740690262 | 0.45888127 | 0.654576 |
| ENSG00000233672 | RNASEH2B-AS1 | 19.28831 | -0.1115302 | 0.4765581 | -0.2952977 | 0.7677665 | 0.876915 |
| ENSG00000286234 | | 2.452144 | -0.1115842 | 0.702213 | -0.3408555 | 0.7332124 | 0.855322 |
| ENSG00000117242 | PINK1-AS | 137.7833026 | -0.111613544 | 0.237977782 | -0.496457749 | 0.61957149 | 0.780127 |
| ENSG00000257594 | GALNT4 | 9.441107 | -0.1116406 | 0.476166 | -0.3009019 | 0.7634893 | 0.874228 |
| ENSG00000125652 | ALKBH7 | 1402.272316 | -0.111646928 | 0.200129953 | -0.595536727 | 0.55148476 | 0.730319 |
| ENSG00000173757 | STAT5B | 3038.164 | -0.1116771 | 0.1312474 | -0.8646062 | 0.387255 | 0.589525 |
| ENSG00000224016 | | 4.988537 | -0.111692 | 0.587192 | -0.2861122 | 0.7747922 | 0.881324 |
| ENSG00000227733 | | 2.877537 | -0.1117462 | 0.6890246 | -0.3306283 | 0.7409252 | 0.860274 |
| ENSG00000224437 | PIGUP1 | 44.37212 | -0.1117486 | 0.3150778 | -0.3805801 | 0.7035148 | 0.838644 |
| ENSG00000197744 | PTMAP2 | 2036.677 | -0.1117718 | 0.2508857 | -0.4541146 | 0.6497463 | 0.802003 |
| ENSG00000164124 | TMEM144 | 489.3183 | -0.1117956 | 0.3161252 | -0.3866017 | 0.6990511 | 0.835817 |
| ENSG00000268583 | | 6.550209 | -0.1118175 | 0.5888978 | -0.2851516 | 0.775528 | 0.881827 |
| ENSG00000136319 | TTC5 | 1361.761478 | -0.111833376 | 0.163187536 | -0.710736334 | 0.47724764 | 0.669482 |
| ENSG00000248124 | RRN3P1 | 1.097727 | -0.1119346 | 0.7418076 | -0.4207783 | 0.673917 | NA |
| ENSG00000248374 | | 1.908973 | -0.111974 | 0.7594274 | -0.4910831 | 0.6233677 | 0.782752 |
| ENSG00000173113 | TRMT112 | 5565.485 | -0.111975 | 0.2047436 | -0.5673066 | 0.5705059 | 0.744342 |
| ENSG00000228247 | UBBP2 | 1.021137 | -0.1119799 | 0.7441871 | -0.4303779 | 0.6669207 | NA |
| ENSG00000146722 | | 0.8112261 | -0.1120301 | 0.7415174 | -0.414948 | 0.67818 | NA |
| ENSG00000169756 | LIMS1 | 1860.311 | -0.1120499 | 0.2051639 | -0.5701372 | 0.5685846 | 0.742678 |
| ENSG00000204956 | PCDHGA1 | 86.23811 | -0.1121047 | 0.351384 | -0.3532993 | 0.7238641 | 0.849973 |

| | | | | | | | |
|---|---|---|---|---|---|---|---|
| ENSG00000162909 | CAPN2 | 2775.278 | -0.1121308 | 0.3706362 | -0.3472348 | 0.7284149 | 0.852623 |
| ENSG00000136813 | ECPAS | 8558.34325 | -0.112153794 | 0.131668252 | -0.882608361 | 0.37744791 | 0.57951 |
| ENSG00000276302 | | 0.8552265 | -0.1121967 | 0.7593539 | -0.4472802 | 0.6546728 | NA |
| ENSG00000253649 | PRSS51 | 6.588053 | -0.1122816 | 0.5647326 | -0.2882605 | 0.7731474 | 0.880445 |
| ENSG00000179262 | RAD23A | 3350.512 | -0.1123088 | 0.1506059 | -0.7602715 | 0.4470923 | 0.643898 |
| ENSG00000270890 | | 53.41638 | -0.1123674 | 0.3702591 | -0.3463132 | 0.7291073 | 0.853151 |
| ENSG00000287405 | | 0.8582391 | -0.112373 | 0.7632857 | -0.468442 | 0.6394685 | NA |
| ENSG00000108306 | FBXL20 | 1147.358727 | -0.112391902 | 0.183339856 | -0.626714759 | 0.53084623 | 0.715107 |
| ENSG00000106723 | SPIN1 | 6660.976349 | -0.112535632 | 0.173503158 | -0.667829126 | 0.50424267 | 0.691213 |
| ENSG00000105640 | RPL18A | 3121.866083 | -0.112538945 | 0.21168419 | -0.556409854 | 0.5779307 | 0.749648 |
| ENSG00000226889 | | 0.8199876 | -0.1127534 | 0.769968 | -0.5362896 | 0.5917584 | NA |
| ENSG00000259917 | HNRNPLP2 | 1655.638 | -0.1128218 | 0.2724834 | -0.4459315 | 0.6556468 | 0.805916 |
| ENSG00000114923 | SLC4A3 | 527.6363825 | -0.11283298 | 0.226786179 | -0.525724095 | 0.59907992 | 0.765665 |
| ENSG00000216285 | | 1232.54 | -0.1128499 | 0.2796553 | -0.4367811 | 0.6622701 | 0.810645 |
| ENSG00000199879 | RNVU1-22 | 6.313155 | -0.1128692 | 0.5856977 | -0.2879487 | 0.773386 | 0.880532 |
| ENSG00000236871 | LINC00106 | 106.6363 | -0.1129528 | 0.3451859 | -0.367723 | 0.7130798 | 0.843667 |
| ENSG00000143458 | GABPB2 | 1928.402 | -0.1129561 | 0.2015292 | -0.5834878 | 0.5595649 | 0.73613 |
| ENSG00000256282 | | 14.39955 | -0.1129817 | 0.5611716 | -0.2870619 | 0.7740649 | 0.880805 |
| ENSG00000258609 | LINC-ROR | 0.8312426 | -0.11309 | 0.7336428 | -0.3878177 | 0.698151 | NA |
| ENSG00000286408 | | 87.55501 | -0.1131318 | 0.3423708 | -0.36807 | 0.712821 | 0.843639 |
| ENSG00000237073 | | 2.819662 | -0.1131347 | 0.6528675 | -0.3182362 | 0.7503058 | 0.866301 |
| ENSG00000215845 | TSTD1 | 2131.054 | -0.113289 | 0.1760145 | -0.684881 | 0.493419 | 0.683256 |
| ENSG00000163159 | VPS72 | 6782.833 | -0.1132954 | 0.1349342 | -0.8675593 | 0.3856357 | 0.587969 |
| ENSG00000205250 | E2F4 | 5815.53 | -0.1133526 | 0.1273608 | -0.9062468 | 0.3648053 | 0.566692 |
| ENSG00000231340 | ACTG1P10 | 7.150095 | -0.1133857 | 0.6548871 | -0.3121489 | 0.7549273 | 0.869057 |
| ENSG00000286363 | | 1.25504 | -0.113398 | 0.7471844 | -0.4475828 | 0.6544543 | NA |
| ENSG00000223382 | LINC01778 | 8.135404 | -0.1134026 | 0.6144398 | -0.2945286 | 0.768354 | 0.877353 |
| ENSG00000234324 | RPL9P2 | 1.66421 | -0.1134307 | 0.7269356 | -0.3758597 | 0.7070212 | 0.840749 |
| ENSG00000275854 | | 18.96618 | -0.1135275 | 0.4568677 | -0.3080963 | 0.758009 | 0.871039 |
| ENSG00000162613 | FUBP1 | 8492.324 | -0.1135471 | 0.1550112 | -0.7561603 | 0.4495531 | 0.645736 |
| ENSG00000174775 | HRAS | 2479.49 | -0.113548 | 0.1853405 | -0.6273039 | 0.53046 | 0.714877 |
| ENSG00000161526 | SAP30BP | 6661.521 | -0.1135541 | 0.1391141 | -0.8398111 | 0.4010143 | 0.601958 |
| ENSG00000234369 | TATDN1P1 | 651.466 | -0.1135822 | 0.2205859 | -0.5303348 | 0.5958798 | 0.763169 |

| | | | | | | | |
|---|---|---|---|---|---|---|---|
| ENSG00000188157 | AGRN | 19203.08 | -0.1136439 | 0.1616917 | -0.7269902 | 0.467232 | 0.660946 |
| ENSG00000214182 | PTMAP5 | 1469.599 | -0.1136456 | 0.2675149 | -0.4517325 | 0.6514617 | 0.803115 |
| ENSG00000173867 | | 0.8058874 | -0.1138783 | 0.7535069 | -0.4532328 | 0.6503811 | NA |
| ENSG00000132321 | IQCA1 | 6.158006882 | -0.113928216 | 0.612857967 | -0.294169654 | 0.76862827 | 0.877371 |
| ENSG00000220793 | RPL21P119 | 7.420533 | -0.1139437 | 0.5365973 | -0.291051 | 0.7710123 | 0.878884 |
| ENSG00000175573 | C11orf68 | 2082.817 | -0.1140671 | 0.1561745 | -0.7514971 | 0.4523536 | 0.648496 |
| ENSG00000284652 | | 0.9464308 | -0.1141151 | 0.7269079 | -0.3860926 | 0.6994281 | NA |
| ENSG00000205584 | | 0.7630924 | -0.1142236 | 0.7712763 | -0.5189709 | 0.603781 | NA |
| ENSG00000189167 | ZAR1L | 3.764511 | -0.1142299 | 0.64855 | -0.3131981 | 0.7541302 | 0.868772 |
| ENSG00000187514 | PTMA | 21931.18 | -0.114246 | 0.2266067 | -0.5298531 | 0.5962138 | 0.763339 |
| ENSG00000159720 | ATP6V0D1 | 6521.605 | -0.1142591 | 0.1578658 | -0.7584737 | 0.4481675 | 0.644826 |
| ENSG00000221986 | MYBPHL | 2.969875 | -0.1142635 | 0.6233472 | -0.3077047 | 0.7583071 | 0.871204 |
| ENSG00000274712 | | 689.5068 | -0.1142683 | 0.2534235 | -0.4847603 | 0.6278464 | 0.786065 |
| ENSG00000231977 | | 0.8495401 | -0.1144687 | 0.7914 | -0.8332261 | 0.4047172 | NA |
| ENSG00000228115 | | 1.364068 | -0.1144872 | 0.7183344 | -0.3706363 | 0.7109085 | NA |
| ENSG00000100024 | UPB1 | 2.418721262 | -0.114564443 | 0.671470627 | -0.332015362 | 0.73987766 | 0.859546 |
| ENSG00000105323 | HNRNPUL1 | 14746.47232 | -0.114684998 | 0.155722122 | -0.766687073 | 0.44326759 | 0.640816 |
| ENSG00000184507 | NUTM1 | 0.8373124 | -0.1147828 | 0.7618513 | -0.4795783 | 0.6315272 | NA |
| ENSG00000278084 | | 1.222597 | -0.1147828 | 0.7218092 | -0.3791756 | 0.7045575 | NA |
| ENSG00000214770 | | 43.5447 | -0.1148036 | 0.3639833 | -0.3566112 | 0.7213829 | 0.848466 |
| ENSG00000132680 | KHDC4 | 5267.174378 | -0.114829784 | 0.361619475 | -0.347367595 | 0.72831518 | 0.852562 |
| ENSG00000236086 | HMGN2P28 | 2.964367 | -0.1148611 | 0.6829504 | -0.3277379 | 0.7431099 | 0.861574 |
| ENSG00000267053 | | 1.078506 | -0.1149233 | 0.7534102 | -0.4403334 | 0.6596956 | NA |
| ENSG00000229043 | ZFAND2A-DT | 1129.092 | -0.1149513 | 0.1201455 | -0.9753525 | 0.3293855 | 0.530455 |
| ENSG00000077147 | TM9SF3 | 16056.97099 | -0.114966519 | 0.133255838 | -0.953338689 | 0.3404185 | 0.541851 |
| ENSG00000228285 | LYPLA2P1 | 352.657 | -0.1150267 | 0.2188368 | -0.5495229 | 0.5826466 | 0.753191 |
| ENSG00000179933 | C14orf119 | 3756.637 | -0.11509 | 0.1659696 | -0.7152165 | 0.4744753 | 0.667178 |
| ENSG00000205913 | SRRM2-AS1 | 267.102 | -0.1151031 | 0.2807036 | -0.4407497 | 0.6593942 | 0.808407 |
| ENSG00000154175 | ABI3BP | 5.461061 | -0.1151972 | 0.5476346 | -0.2983899 | 0.7654056 | 0.875498 |
| ENSG00000231074 | HCG18 | 2534.58 | -0.1152131 | 0.2551904 | -0.4600014 | 0.6455152 | 0.798412 |
| ENSG00000283769 | | 0.8833382 | -0.1153163 | 0.8021817 | -1.221761 | 0.2217979 | NA |
| ENSG00000237938 | | 16.15201 | -0.1154135 | 0.4536584 | -0.3103355 | 0.7563059 | 0.869886 |
| ENSG00000025772 | TOMM34 | 4490.558836 | -0.115458892 | 0.21226877 | -0.571437908 | 0.56770284 | 0.742106 |

| | | | | | | | |
|---|---|---|---|---|---|---|---|
| ENSG00000134709 | HOOK1 | 3540.906169 | -0.115481821 | 0.200536213 | -0.595361559 | 0.55160182 | 0.730402 |
| ENSG00000278492 | | 1.222432 | -0.1155296 | 0.7141109 | -0.3722898 | 0.7096771 | NA |
| ENSG00000260274 | | 135.3806 | -0.1155559 | 0.237775 | -0.5038669 | 0.6143549 | 0.776315 |
| ENSG00000158985 | CDC42SE2 | 2782.983 | -0.1155596 | 0.1208011 | -0.9776608 | 0.3282421 | 0.529422 |
| ENSG00000148229 | POLE3 | 6680.489 | -0.1155927 | 0.2787871 | -0.4668961 | 0.6405742 | 0.794928 |
| ENSG00000235974 | VN2R19P | 1.070487 | -0.115692 | 0.7288983 | -0.3784945 | 0.7050633 | NA |
| ENSG00000222017 | | 30.49228 | -0.1157178 | 0.3615755 | -0.3636283 | 0.7161356 | 0.845492 |
| ENSG00000135999 | EPC2 | 1861.184793 | -0.115724822 | 0.1496567 | -0.780850654 | 0.43489034 | 0.633754 |
| ENSG00000235065 | RPL24P2 | 529.9079 | -0.1157571 | 0.2405258 | -0.5139309 | 0.6073003 | 0.771285 |
| ENSG00000170677 | SOCS6 | 1098.369 | -0.1159437 | 0.1612502 | -0.7352157 | 0.4622081 | 0.656777 |
| ENSG00000272335 | | 451.9765 | -0.1159909 | 0.273107 | -0.4465505 | 0.6551996 | 0.805441 |
| ENSG00000186474 | KLK12 | 0.8067198 | -0.1160802 | 0.7913171 | -0.3606488 | 0.718362 | NA |
| ENSG00000000457 | SCYL3 | 1168.86545 | -0.116096344 | 0.143615053 | -0.824951807 | 0.40939895 | 0.609907 |
| ENSG00000273568 | | 93.40451 | -0.1161912 | 0.2144999 | -0.562077 | 0.5740635 | 0.746641 |
| ENSG00000168958 | MFF | 2417.599 | -0.1161928 | 0.1161371 | -1.012015 | 0.3115309 | 0.512448 |
| ENSG00000158552 | ZFAND2B | 1837.31 | -0.116332 | 0.1817754 | -0.6603653 | 0.5090195 | 0.695221 |
| ENSG00000240096 | | 5.093736 | -0.116337 | 0.6563255 | -0.3179656 | 0.750511 | 0.866443 |
| ENSG00000223478 | | 163.2023 | -0.1164129 | 0.4083707 | -0.334086 | 0.7383147 | 0.858765 |
| ENSG00000158636 | EMSY | 2715.65 | -0.1164205 | 0.2191663 | -0.5609043 | 0.5748628 | 0.747201 |
| ENSG00000125730 | C3 | 668.6533149 | -0.116451865 | 0.542347934 | -0.297696042 | 0.76593516 | 0.875765 |
| ENSG00000259364 | | 2.924087 | -0.116465 | 0.6507632 | -0.318344 | 0.750224 | 0.866263 |
| ENSG00000216937 | CCDC7 | 204.7066 | -0.116491 | 0.2742076 | -0.4512804 | 0.6517875 | 0.80336 |
| ENSG00000237200 | ZBTB40-IT1 | 10.05283 | -0.1165376 | 0.5345363 | -0.2966194 | 0.7667571 | 0.876327 |
| ENSG00000250076 | MAPRE1P2 | 1.951463 | -0.116565 | 0.732107 | -0.4005745 | 0.6887335 | 0.828482 |
| ENSG00000229999 | SGSM3-AS1 | 1.173141 | -0.1165756 | 0.6894048 | -0.3406237 | 0.7333869 | NA |
| ENSG00000232912 | RERE-AS1 | 1.465864 | -0.1166175 | 0.7473084 | -0.4319095 | 0.6658072 | NA |
| ENSG00000148842 | CNNM2 | 750.1106 | -0.116629 | 0.2424764 | -0.5085342 | 0.6110788 | 0.774324 |
| ENSG00000177990 | DPY19L2 | 2.109344 | -0.1166401 | 0.6786055 | -0.3400044 | 0.7338532 | 0.85569 |
| ENSG00000259768 | | 294.2301 | -0.1167286 | 0.2921468 | -0.4401214 | 0.6598492 | 0.808639 |
| ENSG00000015153 | YAF2 | 658.0031552 | -0.11677345 | 0.13752879 | -0.864161407 | 0.38749923 | 0.589775 |
| ENSG00000285561 | | 1.30618 | -0.1168631 | 0.7061583 | -0.3591324 | 0.7194961 | NA |
| ENSG00000218426 | | 6743.311 | -0.1169815 | 0.1644658 | -0.7352428 | 0.4621916 | 0.656777 |
| ENSG00000249485 | RBBP4P1 | 748.2178 | -0.116997 | 0.1413629 | -0.8468493 | 0.3970791 | 0.598174 |

| | | | | | | | |
|---|---|---|---|---|---|---|---|
| ENSG00000187498 | COL4A1 | 2.089932 | -0.1170281 | 0.72304 | -0.3614116 | 0.7177918 | 0.846042 |
| ENSG00000168283 | BMI1 | 2618.173 | -0.117047 | 0.1229711 | -0.961787 | 0.3361566 | 0.537224 |
| ENSG00000252892 | RNU6-548P | 0.9165652 | -0.1171087 | 0.7452103 | -0.4410059 | 0.6592087 | NA |
| ENSG00000109475 | RPL34 | 12693.03963 | -0.117173949 | 0.169569071 | -0.712259927 | 0.47630383 | 0.668617 |
| ENSG00000175390 | EIF3F | 4637.432 | -0.1171936 | 0.155821 | -0.7417851 | 0.4582175 | 0.654009 |
| ENSG00000140534 | TICRR | 1573.939 | -0.1172573 | 0.5270211 | -0.3039145 | 0.761193 | 0.87267 |
| ENSG00000198915 | RASGEF1A | 107.3472 | -0.117279 | 0.4418917 | -0.3203938 | 0.7486698 | 0.865481 |
| ENSG00000223658 | C1GALT1C1L | 73.85626 | -0.1172865 | 0.3671484 | -0.3600178 | 0.7188338 | 0.846874 |
| ENSG00000261371 | PECAM1 | 5.9404 | -0.1173246 | 0.6377037 | -0.3048433 | 0.7604855 | 0.872349 |
| ENSG00000205740 | | 67.12 | -0.1173701 | 0.3664046 | -0.3606938 | 0.7183283 | 0.846391 |
| ENSG00000160679 | CHTOP | 7549.001 | -0.1173862 | 0.1187536 | -1.00735 | 0.3137668 | 0.514915 |
| ENSG00000250988 | SNHG21 | 110.4773 | -0.1174982 | 0.2597712 | -0.4796633 | 0.6314669 | 0.788593 |
| ENSG00000169515 | CCDC8 | 0.9669916 | -0.1175413 | 0.7264958 | -0.3961848 | 0.6919687 | NA |
| ENSG00000010244 | ZNF207 | 7294.509489 | -0.117674193 | 0.098447783 | -1.219483717 | 0.22266065 | 0.409374 |
| ENSG00000234176 | HSPA8P1 | 52.93616 | -0.1177334 | 0.3223053 | -0.4039285 | 0.6862653 | 0.827244 |
| ENSG00000105379 | ETFB | 1129.951611 | -0.11773437 | 0.28774617 | -0.425116199 | 0.67075197 | 0.816887 |
| ENSG00000276584 | MIR6737 | 1.055607 | -0.1177937 | 0.7493902 | -0.443151 | 0.6576565 | NA |
| ENSG00000171159 | C9orf16 | 2660.354 | -0.1178105 | 0.1847367 | -0.6558128 | 0.5119446 | 0.697889 |
| ENSG00000261519 | | 0.7810659 | -0.1178191 | 0.7904189 | -0.3790413 | 0.7046572 | NA |
| ENSG00000276449 | | 22.55734 | -0.1178253 | 0.4395045 | -0.3259672 | 0.7444492 | 0.862303 |
| ENSG00000287207 | | 1.75427 | -0.1178989 | 0.7567324 | -0.4212976 | 0.6735378 | 0.818574 |
| ENSG00000182327 | GLTPD2 | 172.6499 | -0.1179634 | 0.2295467 | -0.5402969 | 0.5889923 | 0.757928 |
| ENSG00000120088 | CRHR1 | 0.947224159 | -0.117990427 | 0.763507282 | -0.473603535 | 0.63578265 | NA |
| ENSG00000142676 | RPL11 | 22354.68 | -0.1180094 | 0.1632012 | -0.7679867 | 0.4424951 | 0.640164 |
| ENSG00000232335 | | 1.685717 | -0.1180637 | 0.7711168 | -0.5805557 | 0.5615399 | 0.737424 |
| ENSG00000270127 | | 75.12618 | -0.118078 | 0.4724973 | -0.3166287 | 0.7515253 | 0.866998 |
| ENSG00000033627 | ATP6V0A1 | 3646.343232 | -0.11810378 | 0.204284691 | -0.597246753 | 0.55034265 | 0.729316 |
| ENSG00000227109 | CRIP1P3 | 2.002951 | -0.1181271 | 0.7286801 | -0.3804315 | 0.7036251 | 0.838737 |
| ENSG00000101132 | PFDN4 | 12074.03591 | -0.118131581 | 0.199257389 | -0.614331378 | 0.53899637 | 0.721572 |
| ENSG00000259820 | | 581.3672 | -0.1183174 | 0.2174682 | -0.5651722 | 0.5719566 | 0.745502 |
| ENSG00000120370 | GORAB | 830.1410036 | -0.118319821 | 0.101536197 | -1.169658699 | 0.24213835 | 0.432758 |
| ENSG00000112029 | FBXO5 | 952.6749436 | -0.118343348 | 0.391247459 | -0.361727001 | 0.71755605 | 0.846015 |
| ENSG00000141542 | RAB40B | 1542.384 | -0.1183636 | 0.146923 | -0.821693 | 0.4112517 | 0.611549 |

| | | | | | | | |
|---|---|---|---|---|---|---|---|
| ENSG00000128245 | YWHAH | 3400.200919 | -0.118389944 | 0.199803456 | -0.620262366 | 0.53508507 | 0.718595 |
| ENSG00000228352 | | 7.77766 | -0.1183991 | 0.5752622 | -0.3006557 | 0.7636771 | 0.874273 |
| ENSG00000261067 | | 55.26539 | -0.1184344 | 0.4153565 | -0.3375304 | 0.7357171 | 0.85681 |
| ENSG00000232328 | | 1.717037 | -0.1184386 | 0.6900648 | -0.3463509 | 0.729079 | 0.853151 |
| ENSG00000259062 | ACTN1-AS1 | 4.696355 | -0.1184597 | 0.6331684 | -0.3070451 | 0.7588091 | 0.87145 |
| ENSG00000124357 | NAGK | 3867.597152 | -0.118496773 | 0.192093672 | -0.644354922 | 0.51934531 | 0.704687 |
| ENSG00000101294 | HM13 | 10299.61261 | -0.118595335 | 0.118197107 | -1.015598979 | 0.30982039 | 0.510488 |
| ENSG00000255425 | OR8G3P | 5.38732 | -0.1186233 | 0.5986657 | -0.3071667 | 0.7587165 | 0.871448 |
| ENSG00000167880 | EVPL | 6901.865 | -0.1186258 | 0.1527362 | -0.8380804 | 0.4019856 | 0.602933 |
| ENSG00000234145 | NAP1L4P3 | 304.8882 | -0.1186523 | 0.1661033 | -0.7315949 | 0.4644158 | 0.658469 |
| ENSG00000166004 | CEP295 | 983.353 | -0.1187044 | 0.2645066 | -0.4836004 | 0.6286695 | 0.786629 |
| ENSG00000221406 | MIR320B2 | 1.028816 | -0.1187307 | 0.7364758 | -0.42682 | 0.6695104 | NA |
| ENSG00000198855 | FICD | 447.8351 | -0.1187443 | 0.271029 | -0.4688093 | 0.6392059 | 0.794121 |
| ENSG00000239291 | | 1.375482 | -0.1187659 | 0.6889914 | -0.356147 | 0.7217305 | NA |
| ENSG00000196544 | BORCS6 | 368.0034 | -0.118773 | 0.2024772 | -0.6111038 | 0.5411309 | 0.72297 |
| ENSG00000186493 | C5orf38 | 1657.893 | -0.1187775 | 0.2077685 | -0.5914661 | 0.5542082 | 0.732393 |
| ENSG00000265566 | RN7SL605P | 4.35512 | -0.1187856 | 0.5876354 | -0.3039182 | 0.7611902 | 0.87267 |
| ENSG00000225942 | LINC01875 | 0.9829857 | -0.1188131 | 0.8030299 | -1.273369 | 0.202887 | NA |
| ENSG00000130669 | PAK4 | 4851.03708 | -0.118824691 | 0.211332739 | -0.608664759 | 0.54274667 | 0.723848 |
| ENSG00000125834 | STK35 | 1699.007338 | -0.118832597 | 0.165033338 | -0.749613045 | 0.45348779 | 0.649659 |
| ENSG00000170858 | LILRP2 | 1.369058 | -0.1188716 | 0.7911944 | -0.3710125 | 0.7106283 | NA |
| ENSG00000182518 | FAM104B | 854.8612 | -0.1190208 | 0.1889843 | -0.6526468 | 0.513984 | 0.699915 |
| ENSG00000165629 | ATP5F1C | 13219.44 | -0.1190989 | 0.2100431 | -0.5935637 | 0.5528039 | 0.731338 |
| ENSG00000032444 | PNPLA6 | 2539.699016 | -0.119110465 | 0.241971811 | -0.52103092 | 0.60234523 | 0.767619 |
| ENSG00000158691 | ZSCAN12 | 248.6361 | -0.119113 | 0.2733627 | -0.4702629 | 0.6381672 | 0.793572 |
| ENSG00000232608 | TIMM9P2 | 10.573 | -0.1191475 | 0.5582539 | -0.3048819 | 0.7604561 | 0.872349 |
| ENSG00000185888 | PRSS38 | 0.788146 | -0.1191804 | 0.7901691 | -0.3877454 | 0.6982045 | NA |
| ENSG00000224004 | ATP5F1CP1 | 144.2585 | -0.1192305 | 0.2542923 | -0.4859007 | 0.6270376 | 0.785493 |
| ENSG00000154957 | ZNF18 | 569.1434 | -0.1193168 | 0.2263019 | -0.5689802 | 0.5693696 | 0.743334 |
| ENSG00000157259 | GATAD1 | 2826.047 | -0.1193287 | 0.1929793 | -0.6375953 | 0.5237371 | 0.708884 |
| ENSG00000252743 | RNU6-850P | 7.650073 | -0.1193329 | 0.54374 | -0.3046202 | 0.7606554 | 0.872369 |
| ENSG00000136738 | STAM | 1479.665552 | -0.119353486 | 0.147392096 | -0.824114404 | 0.40987456 | 0.610287 |
| ENSG00000112893 | MAN2A1 | 4418.047258 | -0.11942064 | 0.217871438 | -0.570254989 | 0.56850477 | 0.742678 |

| | | | | | | | |
|---|---|---|---|---|---|---|---|
| ENSG00000178343 | SHISA3 | 2.318207 | -0.1194207 | 0.7122872 | -0.3692732 | 0.7119241 | 0.843269 |
| ENSG00000065135 | GNAI3 | 8000.065057 | -0.119458781 | 0.21326083 | -0.585914269 | 0.55793313 | 0.735159 |
| ENSG00000160714 | UBE2Q1 | 7427.769 | -0.1195773 | 0.109701 | -1.102145 | 0.2703986 | 0.466781 |
| ENSG00000167685 | ZNF444 | 3471.403 | -0.1196115 | 0.1752188 | -0.706328 | 0.4799842 | 0.671801 |
| ENSG00000259351 | | 2.117323 | -0.119612 | 0.7566368 | -0.4729891 | 0.6362209 | 0.792005 |
| ENSG00000107672 | NSMCE4A | 2243.379741 | -0.119617035 | 0.175804778 | -0.701475747 | 0.48300617 | 0.674391 |
| ENSG00000237481 | | 1.969196 | -0.1196233 | 0.7289327 | -0.3852761 | 0.7000329 | 0.836557 |
| ENSG00000250075 | | 2.226335 | -0.1196609 | 0.6601177 | -0.3282513 | 0.7427216 | 0.861443 |
| ENSG00000178503 | NECAP1P1 | 0.8059244 | -0.1197996 | 0.7769625 | -0.5721527 | 0.5672185 | NA |
| ENSG00000110429 | FBXO3 | 2316.386382 | -0.119877601 | 0.154517334 | -0.78619785 | 0.43175159 | 0.63091 |
| ENSG00000225416 | | 4.145046 | -0.1198947 | 0.5909778 | -0.3055559 | 0.7599428 | 0.872027 |
| ENSG00000280721 | LINC01943 | 2.552223 | -0.1199047 | 0.6539746 | -0.3318465 | 0.7400052 | 0.859581 |
| ENSG00000140545 | MFGE8 | 789.3198 | -0.1199066 | 0.3422999 | -0.3885627 | 0.6975997 | 0.834926 |
| ENSG00000124508 | BTN2A2 | 526.0925562 | -0.119949021 | 0.255605728 | -0.497879211 | 0.61856918 | 0.779342 |
| ENSG00000230498 | | 1.130655 | -0.1201358 | 0.7306411 | -0.4198177 | 0.6746186 | NA |
| ENSG00000254165 | | 18.72317 | -0.1202356 | 0.4003867 | -0.3521575 | 0.7247202 | 0.850603 |
| ENSG00000116288 | PARK7 | 8161.090238 | -0.120269989 | 0.198765547 | -0.651029576 | 0.5150274 | 0.700474 |
| ENSG00000178896 | EXOSC4 | 1540.833 | -0.1202873 | 0.2487878 | -0.5168395 | 0.6052682 | 0.769775 |
| ENSG00000039987 | BEST2 | 2.731735912 | -0.120336122 | 0.635282224 | -0.317408723 | 0.75093349 | 0.86678 |
| ENSG00000211892 | IGHG4 | 0.9405874 | -0.120349 | 0.7387315 | -0.4288554 | 0.6680284 | NA |
| ENSG00000182450 | KCNK4 | 0.939605 | -0.1203519 | 0.7560967 | -0.4774117 | 0.633069 | NA |
| ENSG00000231130 | HLA-T | 0.8680589 | -0.1203673 | 0.7556022 | -0.432034 | 0.6657167 | NA |
| ENSG00000285399 | NA | 347.5354 | -0.1203777 | 0.2249691 | -0.558236 | 0.5766832 | 0.748688 |
| ENSG00000262172 | | 20.41719 | -0.1203848 | 0.4896212 | -0.317856 | 0.7505942 | 0.866502 |
| ENSG00000181016 | LSMEM1 | 91.31456 | -0.1203963 | 0.3415368 | -0.3927058 | 0.6945367 | 0.832535 |
| ENSG00000196704 | AMZ2 | 8121.012 | -0.1204398 | 0.1474369 | -0.8453141 | 0.3979355 | 0.599057 |
| ENSG00000146833 | TRIM4 | 2613.489 | -0.1205059 | 0.2277289 | -0.5300443 | 0.5960812 | 0.763316 |
| ENSG00000120029 | ARMH3 | 2191.796707 | -0.120539372 | 0.109326695 | -1.123604139 | 0.261181 | 0.455805 |
| ENSG00000286899 | | 0.9444291 | -0.1205395 | 0.7620353 | -0.4931869 | 0.6218806 | NA |
| ENSG00000137414 | FAM8A1 | 2411.740308 | -0.120666136 | 0.161175026 | -0.771448515 | 0.44044113 | 0.638603 |
| ENSG00000176920 | FUT2 | 54.95632 | -0.1206815 | 0.4081593 | -0.3483541 | 0.7275742 | 0.85203 |
| ENSG00000102359 | SRPX2 | 4.911191251 | -0.12068664 | 0.700259625 | -0.355972067 | 0.72186148 | 0.848672 |
| ENSG00000245928 | SDAD1-AS1 | 1.011444 | -0.120715 | 0.7331112 | -0.4238929 | 0.6716439 | NA |

| | | | | | | | |
|---|---|---|---|---|---|---|---|
| ENSG00000226149 | | 1.63678 | -0.1207771 | 0.7392445 | -0.4103609 | 0.6815413 | 0.823759 |
| ENSG00000056277 | ZNF280C | 617.1200933 | -0.120796544 | 0.202396903 | -0.620011006 | 0.53525054 | 0.718744 |
| ENSG00000166024 | R3HCC1L | 1399.851 | -0.1208044 | 0.1350241 | -0.9413697 | 0.3465154 | 0.548075 |
| ENSG00000213312 | RPL7AP34 | 9.588194 | -0.1208358 | 0.5186775 | -0.3103317 | 0.7563087 | 0.869886 |
| ENSG00000203616 | RHOT1P2 | 17.02335 | -0.1208451 | 0.4308208 | -0.3371652 | 0.7359923 | 0.856943 |
| ENSG00000170909 | OSCAR | 53.06387 | -0.120881 | 0.3180166 | -0.4184378 | 0.6756271 | 0.819668 |
| ENSG00000114030 | KPNA1 | 3668.769525 | -0.120890871 | 0.135468245 | -0.911593408 | 0.36198279 | 0.563945 |
| ENSG00000225507 | | 100.9401 | -0.1209922 | 0.2436233 | -0.506669 | 0.6123871 | 0.775192 |
| ENSG00000226913 | BSN-DT | 21.89774 | -0.1210022 | 0.4903249 | -0.3199173 | 0.749031 | 0.865675 |
| ENSG00000197808 | ZNF461 | 430.4299 | -0.1210432 | 0.2191407 | -0.5711591 | 0.5678918 | 0.742316 |
| ENSG00000167524 | RSKR | 467.1182 | -0.1210541 | 0.2646901 | -0.4897685 | 0.6242977 | 0.783439 |
| ENSG00000133107 | TRPC4 | 9.234593985 | -0.121145392 | 0.518175074 | -0.323924562 | 0.74599515 | 0.863507 |
| ENSG00000115705 | TPO | 249.3228758 | -0.121176299 | 0.401063979 | -0.334088518 | 0.73831276 | 0.858764 |
| ENSG00000270948 | MTDHP1 | 5.877953 | -0.1211773 | 0.5923022 | -0.314167 | 0.7533942 | 0.868263 |
| ENSG00000207468 | SNORA19 | 2.406694 | -0.1211916 | 0.6909834 | -0.3510443 | 0.7255551 | 0.850944 |
| ENSG00000126746 | ZNF384 | 1860.60506 | -0.121274644 | 0.148952762 | -0.862920019 | 0.38818144 | 0.5904 |
| ENSG00000251634 | NAIPP4 | 36.07109 | -0.1213327 | 0.3803821 | -0.3673182 | 0.7133817 | 0.84368 |
| ENSG00000275910 | | 17.38487 | -0.1213901 | 0.5221504 | -0.3105515 | 0.7561416 | 0.869868 |
| ENSG00000169188 | APEX2 | 2463.352 | -0.1214019 | 0.2292237 | -0.5348581 | 0.592748 | 0.760881 |
| ENSG00000041988 | THAP3 | 565.2473676 | -0.121408933 | 0.171510498 | -0.72710959 | 0.46715882 | 0.660878 |
| ENSG00000284657 | | 4.699454 | -0.121412 | 0.611575 | -0.3162553 | 0.7518088 | 0.867075 |
| ENSG00000236065 | | 1.394299 | -0.1214617 | 0.7169308 | -0.376678 | 0.7064129 | NA |
| ENSG00000117477 | CCDC181 | 13.81469957 | -0.121488999 | 0.433580933 | -0.337013889 | 0.73610643 | 0.857038 |
| ENSG00000235454 | HAUS6P3 | 5.159598 | -0.1215028 | 0.5873761 | -0.314308 | 0.7532871 | 0.86819 |
| ENSG00000223878 | PPIAP53 | 11.90319 | -0.1215032 | 0.548459 | -0.3120976 | 0.7549664 | 0.869057 |
| ENSG00000117533 | VAMP4 | 688.2141975 | -0.121506865 | 0.185186185 | -0.677016648 | 0.49839539 | 0.686783 |
| ENSG00000252206 | RNU7-40P | 1.309318 | -0.1215603 | 0.7282226 | -0.3891514 | 0.6971641 | NA |
| ENSG00000213344 | PCNPP3 | 1.449259 | -0.1215746 | 0.6889913 | -0.3628372 | 0.7167265 | NA |
| ENSG00000213269 | RPS6P26 | 2.743343 | -0.1215818 | 0.648965 | -0.3230464 | 0.7466601 | 0.863912 |
| ENSG00000233594 | BTF3P5 | 28.03316 | -0.1216477 | 0.371703 | -0.3729396 | 0.7091934 | 0.841795 |
| ENSG00000237412 | PRSS56 | 1.019054 | -0.1216593 | 0.7352602 | -0.4247554 | 0.671015 | NA |
| ENSG00000107882 | SUFU | 1257.959115 | -0.121676253 | 0.144399579 | -0.860496183 | 0.38951559 | 0.591441 |
| ENSG00000213630 | BOLA3P3 | 1.276342 | -0.1216837 | 0.7622961 | -0.492986 | 0.6220225 | NA |

| | | | | | | | |
|---|---|---|---|---|---|---|---|
| ENSG00000001084 | GCLC | 3849.37336 | -0.121752046 | 0.161891253 | -0.769644948 | 0.44151053 | 0.639299 |
| ENSG00000261060 | | 1.544491 | -0.1217528 | 0.7092293 | -0.376069 | 0.7068656 | NA |
| ENSG00000128908 | INO80 | 3358.632662 | -0.121768705 | 0.14481676 | -0.868716908 | 0.38500199 | 0.587408 |
| ENSG00000164904 | ALDH7A1 | 4229.551 | -0.1218474 | 0.2954325 | -0.4482397 | 0.6539802 | 0.804895 |
| ENSG00000277479 | | 1.456186 | -0.1218492 | 0.7413885 | -0.4382869 | 0.6611783 | NA |
| ENSG00000279589 | | 0.9581481 | -0.1219139 | 0.7357465 | -0.4181865 | 0.6758108 | NA |
| ENSG00000237931 | CLIC4P3 | 34.85066 | -0.1219313 | 0.3642977 | -0.3802956 | 0.703726 | 0.83882 |
| ENSG00000161847 | RAVER1 | 1952.809 | -0.1221759 | 0.2608718 | -0.4689475 | 0.6391072 | 0.794106 |
| ENSG00000216412 | RPL12P2 | 0.8396281 | -0.1222663 | 0.7568355 | -0.4746232 | 0.6350556 | NA |
| ENSG00000273466 | NA | 3.704748 | -0.1222951 | 0.6351687 | -0.3215278 | 0.7478104 | 0.864754 |
| ENSG00000179909 | ZNF154 | 11.81036 | -0.1222961 | 0.4710539 | -0.3235563 | 0.746274 | 0.863729 |
| ENSG00000138175 | ARL3 | 1773.948947 | -0.12234451 | 0.218283174 | -0.591229523 | 0.55436664 | 0.732434 |
| ENSG00000162695 | SLC30A7 | 1189.157 | -0.1223491 | 0.180271 | -0.6893489 | 0.4906037 | 0.681098 |
| ENSG00000269486 | ERVK9-11 | 37.20202 | -0.1223966 | 0.3967963 | -0.3620414 | 0.7173211 | 0.846015 |
| ENSG00000168288 | MMADHC | 5451.138 | -0.1224575 | 0.1852142 | -0.6739047 | 0.5003719 | 0.688436 |
| ENSG00000259500 | KRT8P24 | 2.496338 | -0.1225384 | 0.6615975 | -0.3325606 | 0.739466 | 0.859368 |
| ENSG00000277692 | | 1.249664 | -0.1225438 | 0.7186661 | -0.3883276 | 0.6977736 | NA |
| ENSG00000166595 | CIAO2B | 2369.804 | -0.1225472 | 0.2242205 | -0.591214 | 0.554377 | 0.732434 |
| ENSG00000231482 | | 20.63844 | -0.1226161 | 0.5033547 | -0.3191106 | 0.7496427 | 0.865959 |
| ENSG00000078124 | ACER3 | 1122.687287 | -0.122703252 | 0.199341019 | -0.639857452 | 0.52226528 | 0.707599 |
| ENSG00000283683 | MYOCOS | 1.826623 | -0.122793 | 0.6948508 | -0.3634951 | 0.716235 | 0.845492 |
| ENSG00000006451 | RALA | 4479.860186 | -0.122823763 | 0.149427849 | -0.868878045 | 0.38491384 | 0.587331 |
| ENSG00000248696 | | 0.8441829 | -0.1229149 | 0.746825 | -0.4549686 | 0.6491318 | NA |
| ENSG00000221930 | DENND10P1 | 817.8897 | -0.1229325 | 0.1671329 | -0.7516131 | 0.4522838 | 0.648431 |
| ENSG00000259915 | | 1.23213 | -0.122957 | 0.7274031 | -0.3931068 | 0.6942406 | NA |
| ENSG00000142686 | C1orf216 | 1691.145 | -0.1229898 | 0.1598674 | -0.786898 | 0.4313416 | 0.630415 |
| ENSG00000265241 | RBM8A | 7215.59 | -0.1230271 | 0.1245839 | -1.001931 | 0.3163769 | 0.517466 |
| ENSG00000213303 | | 36.32481 | -0.1231276 | 0.3358007 | -0.4083089 | 0.6830469 | 0.824743 |
| ENSG00000257183 | | 0.9090442 | -0.1231517 | 0.7209155 | -0.4004523 | 0.6888234 | NA |
| ENSG00000109320 | NFKB1 | 2228.649365 | -0.123254655 | 0.163071169 | -0.779271606 | 0.43581974 | 0.634517 |
| ENSG00000276744 | | 21.67335 | -0.1232577 | 0.4165489 | -0.3454408 | 0.729763 | 0.853505 |
| ENSG00000202461 | | 1.404683 | -0.1232812 | 0.7208541 | -0.3897814 | 0.6966982 | NA |
| ENSG00000214425 | LRRC37A4P | 13.53965 | -0.1233233 | 0.4191584 | -0.350323 | 0.7260963 | 0.85124 |

| | | | | | | | |
|---|---|---|---|---|---|---|---|
| ENSG00000227375 | DLG1-AS1 | 6.837379 | -0.1233327 | 0.5983031 | -0.3181712 | 0.7503551 | 0.866301 |
| ENSG00000016602 | CLCA4 | 2.659511936 | -0.123465291 | 0.646912264 | -0.332078226 | 0.73983019 | 0.859528 |
| ENSG00000271500 | | 1.620662 | -0.1235003 | 0.700598 | -0.3670074 | 0.7136135 | 0.843754 |
| ENSG00000094975 | SUCO | 5002.288883 | -0.123554222 | 0.173020905 | -0.747913387 | 0.4545124 | 0.650503 |
| ENSG00000230799 | RPS2P16 | 3.680455 | -0.1235624 | 0.642383 | -0.324822 | 0.7453158 | 0.86292 |
| ENSG00000197279 | ZNF165 | 535.1473 | -0.1235657 | 0.2093509 | -0.6134145 | 0.5396023 | 0.721961 |
| ENSG00000237735 | | 0.9307005 | -0.1236038 | 0.7938873 | -0.9170952 | 0.3590927 | NA |
| ENSG00000235613 | NSRP1P1 | 2.094759 | -0.123643 | 0.7347895 | -0.4164043 | 0.6771142 | 0.820424 |
| ENSG00000220583 | RPL35P2 | 4.795014 | -0.1236874 | 0.5678281 | -0.3133263 | 0.7540328 | 0.868735 |
| ENSG00000167112 | TRUB2 | 3796.531 | -0.1237226 | 0.1887317 | -0.6869332 | 0.4921248 | 0.682311 |
| ENSG00000224870 | MRPL20-AS1 | 1442.357 | -0.1237916 | 0.1434135 | -0.8749027 | 0.3816269 | 0.583834 |
| ENSG00000255182 | | 326.7835 | -0.1238021 | 0.2281997 | -0.5705583 | 0.5682991 | 0.742593 |
| ENSG00000188529 | SRSF10 | 8185.625 | -0.1238289 | 0.1415674 | -0.8821269 | 0.3777082 | 0.579608 |
| ENSG00000135316 | SYNCRIP | 14920.22313 | -0.12386855 | 0.186369529 | -0.682634731 | 0.49483768 | 0.684152 |
| ENSG00000259595 | | 1.289989 | -0.1238787 | 0.7455282 | -0.4490497 | 0.6533958 | NA |
| ENSG00000119684 | MLH3 | 1888.965506 | -0.123942749 | 0.150136355 | -0.859921275 | 0.38983244 | 0.591787 |
| ENSG00000186998 | EMID1 | 602.0477 | -0.1240499 | 0.2859058 | -0.468034 | 0.6397603 | 0.79435 |
| ENSG00000231035 | RPL7L1P9 | 5.47865 | -0.1240733 | 0.6238367 | -0.318401 | 0.7501808 | 0.866263 |
| ENSG00000287817 | | 56.05376 | -0.1241055 | 0.25746 | -0.5131352 | 0.6078568 | 0.771664 |
| ENSG00000185839 | | 1365.211 | -0.1241287 | 0.1948176 | -0.6610699 | 0.5085675 | 0.694925 |
| ENSG00000232093 | DCST1-AS1 | 60.8523 | -0.1241328 | 0.3530093 | -0.3953263 | 0.6926021 | 0.831229 |
| ENSG00000166558 | SLC38A8 | 1.015893 | -0.124186 | 0.7600044 | -0.5007225 | 0.6165665 | NA |
| ENSG00000183186 | C2CD4C | 126.3763 | -0.124191 | 0.5052837 | -0.3225866 | 0.7470083 | 0.864202 |
| ENSG00000261235 | | 104.6061 | -0.1242115 | 0.2707819 | -0.4929421 | 0.6220535 | 0.781693 |
| ENSG00000117010 | ZNF684 | 120.7902738 | -0.124324871 | 0.324291516 | -0.424125549 | 0.67147425 | 0.817373 |
| ENSG00000266495 | | 1.34424 | -0.1243371 | 0.7297756 | -0.4224052 | 0.6727293 | NA |
| ENSG00000204392 | LSM2 | 1704.312 | -0.1243798 | 0.2726803 | -0.4909927 | 0.6234316 | 0.782759 |
| ENSG00000007314 | SCN4A | 1.285940138 | -0.124405282 | 0.732095106 | -0.436455419 | 0.66250634 | NA |
| ENSG00000234080 | | 1.23991 | -0.1244392 | 0.7313018 | -0.4102845 | 0.6815973 | NA |
| ENSG00000196531 | NACA | 15725.65 | -0.1244542 | 0.1383521 | -0.9194771 | 0.3578461 | 0.559815 |
| ENSG00000126243 | LRFN3 | 800.3937334 | -0.124463921 | 0.170492149 | -0.76327513 | 0.44529933 | 0.642438 |
| ENSG00000129351 | ILF3 | 27518.6444 | -0.124523915 | 0.135070349 | -0.965008381 | 0.33454061 | 0.535769 |
| ENSG00000264112 | | 1271.958 | -0.124524 | 0.321416 | -0.4263147 | 0.6698786 | 0.816123 |

| | | | | | | | |
|---|---|---|---|---|---|---|---|
| ENSG00000102383 | ZDHHC15 | 0.802226802 | -0.124578967 | 0.756433894 | -0.479887022 | 0.63130773 | NA |
| ENSG00000202523 | | 0.9652661 | -0.1246328 | 0.7394979 | -0.4512641 | 0.6517992 | NA |
| ENSG00000265480 | KRT18P55 | 14.83624 | -0.1247027 | 0.4630982 | -0.3363555 | 0.7366028 | 0.857466 |
| ENSG00000129932 | DOHH | 913.5121505 | -0.124719712 | 0.218077806 | -0.598239165 | 0.54968036 | 0.728963 |
| ENSG00000260660 | | 1.715888 | -0.12481 | 0.7069424 | -0.3807918 | 0.7033578 | 0.838592 |
| ENSG00000279452 | | 51.61152 | -0.1248672 | 0.3915933 | -0.3690903 | 0.7120604 | 0.843269 |
| ENSG00000265800 | | 21.79275 | -0.1248894 | 0.405323 | -0.35988 | 0.7189369 | 0.846902 |
| ENSG00000138698 | RAP1GDS1 | 2262.755211 | -0.124924315 | 0.135812098 | -0.936481652 | 0.34902526 | 0.550668 |
| ENSG00000142168 | SOD1 | 12694.38 | -0.1249611 | 0.2385223 | -0.5427144 | 0.5873264 | 0.756921 |
| ENSG00000125656 | CLPP | 2278.513172 | -0.124963449 | 0.229940818 | -0.572348937 | 0.5670856 | 0.741591 |
| ENSG00000287862 | | 1.77393 | -0.125161 | 0.7900952 | -0.4100409 | 0.681776 | 0.823818 |
| ENSG00000244681 | MTHFD2P1 | 0.9879903 | -0.1251707 | 0.8043788 | -1.309843 | 0.190249 | NA |
| ENSG00000213530 | MTHFD2P5 | 1.448143 | -0.1251879 | 0.6973713 | -0.3742548 | 0.7082147 | NA |
| ENSG00000143919 | CAMKMT | 390.2375 | -0.1252203 | 0.1619027 | -0.7794069 | 0.4357401 | 0.634435 |
| ENSG00000138107 | ACTR1A | 5771.443504 | -0.125265599 | 0.149322723 | -0.910051567 | 0.36279532 | 0.564786 |
| ENSG00000204952 | FBXO47 | 2.01736 | -0.1252692 | 0.751506 | -0.4606206 | 0.6450708 | 0.798011 |
| ENSG00000172530 | BANP | 1168.73 | -0.1252837 | 0.2386771 | -0.554081 | 0.5795234 | 0.750615 |
| ENSG00000183576 | SETD3 | 5940.405 | -0.1252956 | 0.1183028 | -1.073741 | 0.2829386 | 0.480732 |
| ENSG00000132323 | ILKAP | 1164.053071 | -0.12537929 | 0.188880444 | -0.68654755 | 0.49236789 | 0.682461 |
| ENSG00000270036 | | 1.068862 | -0.125403 | 0.7564269 | -0.5006449 | 0.616621 | NA |
| ENSG00000238423 | SNORD42B | 5.306866 | -0.1257178 | 0.5854244 | -0.3221623 | 0.7473297 | 0.864403 |
| ENSG00000134970 | TMED7 | 3709.984085 | -0.125719999 | 0.196564275 | -0.670756275 | 0.50237581 | 0.690014 |
| ENSG00000008988 | RPS20 | 16470.9819 | -0.125724099 | 0.160573982 | -0.809335985 | 0.41832191 | 0.618103 |
| ENSG00000264235 | | 46.62432 | -0.1257278 | 0.4721591 | -0.3369252 | 0.7361733 | 0.857078 |
| ENSG00000107779 | BMPR1A | 2653.172761 | -0.125887677 | 0.122570126 | -1.041256469 | 0.29775653 | 0.496994 |
| ENSG00000223701 | RAET1E-AS1 | 1.23197 | -0.1258954 | 0.7293721 | -0.4019911 | 0.6876905 | NA |
| ENSG00000224079 | | 2.506427 | -0.1260234 | 0.6759092 | -0.3597892 | 0.7190048 | 0.846902 |
| ENSG00000253408 | | 0.8628843 | -0.1260446 | 0.7679365 | -0.515383 | 0.6062854 | NA |
| ENSG00000150337 | FCGR1A | 89.83163 | -0.1260455 | 0.5174352 | -0.3275926 | 0.7432197 | 0.861574 |
| ENSG00000232259 | | 1.960683 | -0.1261053 | 0.7009831 | -0.3637938 | 0.716012 | 0.845424 |
| ENSG00000123415 | SMUG1 | 2093.387342 | -0.126108985 | 0.171865203 | -0.758497259 | 0.44815335 | 0.644826 |
| ENSG00000198597 | ZNF536 | 0.8241604 | -0.1261614 | 0.7740699 | -0.6092262 | 0.5423745 | NA |
| ENSG00000170074 | FAM153A | 1.34009 | -0.1262285 | 0.693527 | -0.3694127 | 0.7118201 | NA |

| | | | | | | | |
|---|---|---|---|---|---|---|---|
| ENSG00000227339 | THRAP3P1 | 9.611937 | -0.1262555 | 0.5345927 | -0.3260516 | 0.7443853 | 0.862303 |
| ENSG00000115464 | USP34 | 9088.662634 | -0.126323173 | 0.241068548 | -0.553924486 | 0.57963053 | 0.75067 |
| ENSG00000263513 | FAM72C | 9.911068 | -0.1263393 | 0.6344681 | -0.3365738 | 0.7364382 | 0.85732 |
| ENSG00000274501 | | 2.989931 | -0.1263551 | 0.6652023 | -0.3410313 | 0.73308 | 0.855314 |
| ENSG00000131409 | LRRC4B | 18.94024197 | -0.12644149 | 0.497957202 | -0.33239506 | 0.73959097 | 0.859371 |
| ENSG00000168393 | DTYMK | 2702.643 | -0.1264571 | 0.3738328 | -0.3894855 | 0.696917 | 0.834442 |
| ENSG00000259299 | | 8.485208 | -0.1264671 | 0.4758422 | -0.3345179 | 0.7379888 | 0.858665 |
| ENSG00000287952 | | 10.14362 | -0.1265041 | 0.6233158 | -0.3642863 | 0.7156443 | 0.845215 |
| ENSG00000241597 | | 2.321612 | -0.1265056 | 0.6940717 | -0.3705806 | 0.7109499 | 0.842705 |
| ENSG00000073921 | PICALM | 9998.677165 | -0.126562508 | 0.106481383 | -1.162155653 | 0.24517224 | 0.436916 |
| ENSG00000055118 | KCNH2 | 411.6506688 | -0.126594647 | 0.347568646 | -0.408284923 | 0.68306451 | 0.824743 |
| ENSG00000213214 | ARHGEF35 | 345.1813 | -0.1267039 | 0.1659894 | -0.7815484 | 0.43448 | 0.633469 |
| ENSG00000174236 | REP15 | 0.8426353 | -0.1267082 | 0.7600433 | -0.5031485 | 0.6148598 | NA |
| ENSG00000286369 | | 29.76426 | -0.126723 | 0.428436 | -0.3541008 | 0.7232634 | 0.849643 |
| ENSG00000174720 | LARP7 | 2105.26 | -0.1267356 | 0.1696243 | -0.7681045 | 0.4424251 | 0.640164 |
| ENSG00000287248 | | 1.552979 | -0.1268179 | 0.7915891 | -0.390474 | 0.6961861 | NA |
| ENSG00000160766 | GBAP1 | 1516.39 | -0.1268431 | 0.2231853 | -0.5904449 | 0.5548924 | 0.732685 |
| ENSG00000109854 | HTATIP2 | 3710.03819 | -0.126977373 | 0.139082934 | -0.928938316 | 0.35292105 | 0.554805 |
| ENSG00000285859 | | 1.197608 | -0.1269909 | 0.791881 | -0.3861118 | 0.6994139 | NA |
| ENSG00000283051 | LINC01668 | 1.306314 | -0.1269957 | 0.7564977 | -0.4822982 | 0.6295941 | NA |
| ENSG00000242583 | KMT5AP3 | 0.9364419 | -0.1270583 | 0.7610372 | -0.5069142 | 0.612215 | NA |
| ENSG00000255585 | | 47.84199 | -0.1271103 | 0.3824525 | -0.3838374 | 0.701099 | 0.83727 |
| ENSG00000270178 | NA | 2.199627 | -0.1271205 | 0.7061982 | -0.4117071 | 0.6805541 | 0.823053 |
| ENSG00000215190 | LINC00680 | 703.6425 | -0.1271503 | 0.1755777 | -0.7573648 | 0.4488313 | 0.645385 |
| ENSG00000214132 | | 6.237687 | -0.1271514 | 0.5463292 | -0.3254979 | 0.7448043 | 0.862554 |
| ENSG00000268201 | | 21.60532 | -0.1271996 | 0.4677546 | -0.3388088 | 0.7347538 | 0.856251 |
| ENSG00000154721 | JAM2 | 21.15337 | -0.1272243 | 0.5143998 | -0.325748 | 0.7446151 | 0.862372 |
| ENSG00000270598 | | 3.399024 | -0.1272336 | 0.6782407 | -0.3523877 | 0.7245475 | 0.850449 |
| ENSG00000285533 | RELA-DT | 472.3249 | -0.1272723 | 0.23465 | -0.5667281 | 0.5708989 | 0.744672 |
| ENSG00000100258 | LMF2 | 2538.091002 | -0.127274994 | 0.189149469 | -0.674993227 | 0.49968007 | 0.687949 |
| ENSG00000259417 | CTXND1 | 4.528556 | -0.1273201 | 0.5963059 | -0.3223575 | 0.7471819 | 0.864365 |
| ENSG00000169621 | APLF | 11.39109 | -0.1273686 | 0.5387847 | -0.3259769 | 0.7444418 | 0.862303 |
| ENSG00000114439 | BBX | 3188.41565 | -0.127387449 | 0.30563222 | -0.456984957 | 0.64768186 | 0.800296 |

| | | | | | | | |
|---|---|---|---|---|---|---|---|
| ENSG00000254662 | | 1.722472 | -0.1273984 | 0.6801763 | -0.3626157 | 0.716892 | 0.845974 |
| ENSG00000238231 | | 1.17529 | -0.1274011 | 0.7167582 | -0.4003163 | 0.6889236 | NA |
| ENSG00000155827 | RNF20 | 3221.01 | -0.1274021 | 0.09647372 | -1.343533 | 0.1790994 | 0.353004 |
| ENSG00000223309 | RNU6-722P | 4.479453 | -0.1275782 | 0.666363 | -0.3490041 | 0.7270862 | 0.851749 |
| ENSG00000254812 | | 29.26578 | -0.1275871 | 0.3474164 | -0.4140733 | 0.6788205 | 0.821518 |
| ENSG00000286562 | | 1.707706 | -0.1275925 | 0.7283735 | -0.4118991 | 0.6804134 | 0.82292 |
| ENSG00000134061 | CD180 | 1.82603445 | -0.127631683 | 0.728947803 | -0.413911242 | 0.67893911 | 0.821586 |
| ENSG00000070404 | FSTL3 | 3113.427856 | -0.127828245 | 0.258837772 | -0.52408589 | 0.6002188 | 0.766361 |
| ENSG00000230870 | FBXW11P1 | 25.20578 | -0.1279208 | 0.4613281 | -0.3467722 | 0.7287624 | 0.852898 |
| ENSG00000076944 | STXBP2 | 1986.615818 | -0.128012301 | 0.189507921 | -0.705778318 | 0.48032601 | 0.671956 |
| ENSG00000265753 | RN7SL444P | 4.195553 | -0.1280718 | 0.6552749 | -0.3382626 | 0.7351653 | 0.856543 |
| ENSG00000002919 | SNX11 | 2090.076355 | -0.128077925 | 0.11592869 | -1.117764733 | 0.26366749 | 0.458918 |
| ENSG00000231104 | | 3.389151 | -0.1281051 | 0.6358239 | -0.3391995 | 0.7344595 | 0.856021 |
| ENSG00000280186 | | 24.08125 | -0.1281248 | 0.5188896 | -0.3332838 | 0.73892 | 0.858927 |
| ENSG00000123154 | WDR83 | 262.0210559 | -0.128144253 | 0.196434846 | -0.675580705 | 0.4993069 | 0.687683 |
| ENSG00000224479 | | 324.2409 | -0.1281472 | 0.2196469 | -0.6096241 | 0.5421109 | 0.723521 |
| ENSG00000183530 | PRR14L | 3654.826 | -0.128249 | 0.2301275 | -0.5864072 | 0.5576019 | 0.734904 |
| ENSG00000279951 | | 2.056221 | -0.1283102 | 0.6713028 | -0.3558187 | 0.7219763 | 0.848732 |
| ENSG00000173137 | ADCK5 | 1186.873 | -0.1283672 | 0.1986152 | -0.6680734 | 0.5040867 | 0.691184 |
| ENSG00000164631 | ZNF12 | 3897.78 | -0.1283835 | 0.2039611 | -0.6653788 | 0.5058082 | 0.692569 |
| ENSG00000253534 | TRBV6-8 | 0.8306319 | -0.128435 | 0.8034599 | -1.017616 | 0.3088606 | NA |
| ENSG00000276397 | LINC01879 | 2.41158 | -0.1284363 | 0.7200609 | -0.4524449 | 0.6509485 | 0.802778 |
| ENSG00000244342 | LINC00698 | 3.527885 | -0.1284524 | 0.605234 | -0.327563 | 0.7432421 | 0.861574 |
| ENSG00000135956 | TMEM127 | 6086.463683 | -0.128488568 | 0.155362227 | -0.855239272 | 0.3924187 | 0.594061 |
| ENSG00000280336 | | 1.651927 | -0.1285693 | 0.7373292 | -0.4430275 | 0.6577458 | 0.807247 |
| ENSG00000104856 | RELB | 230.0869128 | -0.128594251 | 0.422239404 | -0.340652239 | 0.7333654 | 0.855398 |
| ENSG00000278719 | MCM8-AS1 | 0.8771882 | -0.128691 | 0.7596541 | -0.5075782 | 0.6117492 | NA |
| ENSG00000229807 | XIST | 1.272836 | -0.1287492 | 0.7120129 | -0.3910027 | 0.6957952 | NA |
| ENSG00000251175 | GSTCD-AS1 | 2.870701 | -0.1287543 | 0.6462277 | -0.3423353 | 0.7320986 | 0.854769 |
| ENSG00000284697 | | 1.936049 | -0.1288406 | 0.7049102 | -0.3870607 | 0.6987113 | 0.835727 |
| ENSG00000276445 | | 7.238755 | -0.1288748 | 0.624978 | -0.3335256 | 0.7387376 | 0.85886 |
| ENSG00000270975 | MAGOH3P | 0.8598795 | -0.1289093 | 0.7783223 | -0.6362753 | 0.524597 | NA |
| ENSG00000071991 | CDH19 | 0.879339526 | -0.128910157 | 0.793197307 | -0.374141682 | 0.70829891 | NA |

| | | | | | | | |
|---|---|---|---|---|---|---|---|
| ENSG00000172046 | USP19 | 3710.632 | -0.1289365 | 0.1060028 | -1.245426 | 0.2129753 | 0.397712 |
| ENSG00000213058 | | 5.688033 | -0.1289652 | 0.5910088 | -0.3315564 | 0.7402242 | 0.859716 |
| ENSG00000244203 | FOXP1-AS1 | 1.229883 | -0.1289766 | 0.7383024 | -0.4402582 | 0.6597501 | NA |
| ENSG00000112312 | GMNN | 1496.012254 | -0.12900818 | 0.513070862 | -0.330552139 | 0.7409828 | 0.860303 |
| ENSG00000235400 | | 12.75925 | -0.1290683 | 0.4747897 | -0.3409448 | 0.7331452 | 0.855321 |
| ENSG00000270863 | DDX55P1 | 1.95979 | -0.1290854 | 0.6921502 | -0.3743839 | 0.7081187 | 0.841263 |
| ENSG00000213061 | PFN1P11 | 3.591774 | -0.1291938 | 0.6490077 | -0.3383016 | 0.7351359 | 0.856543 |
| ENSG00000216866 | RPS2P55 | 748.4171 | -0.1292416 | 0.2377962 | -0.5849963 | 0.5585502 | 0.73557 |
| ENSG00000162877 | PM20D1 | 5.591614 | -0.1292692 | 0.5928228 | -0.3266336 | 0.743945 | 0.862053 |
| ENSG00000169139 | UBE2V2 | 4144.88 | -0.1292835 | 0.2054568 | -0.6572476 | 0.5110217 | 0.696953 |
| ENSG00000264044 | | 8.030641 | -0.1292975 | 0.585183 | -0.3237701 | 0.7461121 | 0.863579 |
| ENSG00000169435 | RASSF6 | 3.61864 | -0.1294637 | 0.6945005 | -0.3729614 | 0.7091772 | 0.841795 |
| ENSG00000232369 | PPIAP35 | 6.492899 | -0.1295415 | 0.5845315 | -0.3299826 | 0.7414131 | 0.860542 |
| ENSG00000230641 | USP12-AS2 | 0.8872299 | -0.1295663 | 0.7622812 | -0.5316075 | 0.5949979 | NA |
| ENSG00000105053 | VRK3 | 1849.248946 | -0.129594072 | 0.171305171 | -0.768291254 | 0.44231417 | 0.640081 |
| ENSG00000255381 | | 1.130712 | -0.1296728 | 0.7162381 | -0.4039622 | 0.6862405 | NA |
| ENSG00000277806 | | 4.477844 | -0.1297788 | 0.6532581 | -0.3508638 | 0.7256905 | 0.851005 |
| ENSG00000253919 | THAP12P7 | 347.594 | -0.129901 | 0.2117186 | -0.6371096 | 0.5240534 | 0.709193 |
| ENSG00000261872 | | 0.9020987 | -0.129903 | 0.7594511 | -0.5101104 | 0.6099741 | NA |
| ENSG00000271321 | CTAGE6 | 1.173657 | -0.1299796 | 0.7386236 | -0.4517615 | 0.6514408 | NA |
| ENSG00000287081 | | 2.300284 | -0.1300517 | 0.6850289 | -0.3835409 | 0.7013187 | 0.83727 |
| ENSG00000066032 | CTNNA2 | 3.801551819 | -0.130107043 | 0.624005724 | -0.342736817 | 0.73179646 | 0.85446 |
| ENSG00000102103 | PQBP1 | 4393.165943 | -0.130134561 | 0.161666914 | -0.82661663 | 0.40845439 | 0.609128 |
| ENSG00000176155 | CCDC57 | 3240.896 | -0.1301656 | 0.1539773 | -0.8697064 | 0.3844608 | 0.586839 |
| ENSG00000130559 | CAMSAP1 | 4530.297443 | -0.130205389 | 0.18373917 | -0.73847498 | 0.46022587 | 0.655608 |
| ENSG00000233775 | | 1.759165 | -0.1302077 | 0.7440603 | -0.4551478 | 0.6490029 | 0.801234 |
| ENSG00000285904 | | 20.22326 | -0.1302242 | 0.4970086 | -0.3398086 | 0.7340006 | 0.855711 |
| ENSG00000101440 | ASIP | 1.861065661 | -0.130287801 | 0.685451193 | -0.373429004 | 0.70882918 | 0.841664 |
| ENSG00000251247 | ZNF345 | 278.0978 | -0.1303198 | 0.24863 | -0.5399579 | 0.5892261 | 0.758082 |
| ENSG00000273180 | | 0.9262303 | -0.1304379 | 0.790653 | -0.4181753 | 0.6758189 | NA |
| ENSG00000136628 | EPRS1 | 16148.46044 | -0.130451512 | 0.222083344 | -0.615695198 | 0.53809571 | 0.721092 |
| ENSG00000163714 | U2SURP | 11355.21 | -0.1304649 | 0.159496 | -0.8376789 | 0.4022111 | 0.603203 |
| ENSG00000236636 | | 1.231868 | -0.1305193 | 0.7492961 | -0.5072605 | 0.6119721 | NA |

| | | | | | | | |
|---|---|---|---|---|---|---|---|
| ENSG00000024048 | UBR2 | 3431.66824 | -0.130550772 | 0.193728809 | -0.694326785 | 0.4874773 | 0.678424 |
| ENSG00000279255 | | 35.28276 | -0.1305853 | 0.4062471 | -0.3759166 | 0.7069789 | 0.840736 |
| ENSG00000123739 | PLA2G12A | 1886.198134 | -0.130601064 | 0.206853042 | -0.673720195 | 0.50048922 | 0.68849 |
| ENSG00000141568 | FOXK2 | 7090.475 | -0.1306645 | 0.1332837 | -1.018176 | 0.3085943 | 0.50891 |
| ENSG00000135441 | BLOC1S1 | 309.796107 | -0.130669981 | 0.22051905 | -0.619550462 | 0.53555379 | 0.71889 |
| ENSG00000241058 | NSUN6 | 512.2637 | -0.1307164 | 0.22517 | -0.6211243 | 0.5345178 | 0.718051 |
| ENSG00000238251 | NSA2P7 | 72.70899 | -0.1307539 | 0.313118 | -0.4575621 | 0.6472671 | 0.800058 |
| ENSG00000257818 | C1GALT1P1 | 17.84485 | -0.1307956 | 0.5680897 | -0.3323699 | 0.73961 | 0.859371 |
| ENSG00000142453 | CARM1 | 5490.79 | -0.1308565 | 0.1557884 | -0.9426704 | 0.3458495 | 0.547413 |
| ENSG00000196666 | FAM180B | 1.300816 | -0.1308982 | 0.7614147 | -0.5501826 | 0.5821942 | NA |
| ENSG00000251381 | LINC00958 | 1.394724 | -0.1309343 | 0.7664429 | -0.5301304 | 0.5960215 | NA |
| ENSG00000095059 | DHPS | 250.1758779 | -0.131010653 | 0.280546517 | -0.493834555 | 0.62142303 | 0.781231 |
| ENSG00000198355 | PIM3 | 5211.741 | -0.1310396 | 0.2250426 | -0.6189293 | 0.5359629 | 0.719083 |
| ENSG00000224609 | | 6.073083 | -0.1310507 | 0.6516394 | -0.3490992 | 0.7270148 | 0.851749 |
| ENSG00000008128 | CDK11A | 210.4047247 | -0.131091193 | 0.283482743 | -0.498602906 | 0.61805916 | 0.778995 |
| ENSG00000117280 | RAB29 | 5041.421201 | -0.131138942 | 0.297521811 | -0.464694174 | 0.64215048 | 0.795869 |
| ENSG00000132600 | PRMT7 | 2148.418077 | -0.131180297 | 0.195217197 | -0.696830909 | 0.48590862 | 0.676902 |
| ENSG00000243370 | RN7SL775P | 0.9904572 | -0.1312813 | 0.7685517 | -0.5732011 | 0.5665085 | NA |
| ENSG00000228887 | EEF1DP1 | 283.3944 | -0.1312897 | 0.2629013 | -0.5332893 | 0.5938333 | 0.761502 |
| ENSG00000091986 | CCDC80 | 61.38015858 | -0.131315626 | 0.399334858 | -0.383326682 | 0.70147756 | 0.83727 |
| ENSG00000131188 | PRR7 | 1608.310981 | -0.131376402 | 0.232227274 | -0.597326841 | 0.55028919 | 0.729281 |
| ENSG00000150636 | CCDC102B | 29.58121 | -0.1313876 | 0.4391555 | -0.3673025 | 0.7133934 | 0.84368 |
| ENSG00000196302 | GUSBP15 | 2.11926 | -0.1314167 | 0.6571613 | -0.353153 | 0.7239737 | 0.850065 |
| ENSG00000213347 | MXD3 | 1560.58 | -0.1315043 | 0.3337253 | -0.4312663 | 0.6662747 | 0.813934 |
| ENSG00000215559 | ANKRD20A11P | 73.19883 | -0.1315251 | 0.467606 | -0.3534318 | 0.7237648 | 0.84993 |
| ENSG00000104825 | NFKBIB | 1252.664122 | -0.131540786 | 0.220307006 | -0.597663291 | 0.55006462 | 0.729166 |
| ENSG00000084764 | MAPRE3 | 1053.380338 | -0.131580974 | 0.214441545 | -0.641015909 | 0.52151235 | 0.70687 |
| ENSG00000227200 | | 1.290286 | -0.13164 | 0.7584086 | -0.476267 | 0.6338842 | NA |
| ENSG00000234061 | | 1.155427 | -0.1317384 | 0.7165002 | -0.4338874 | 0.6643702 | NA |
| ENSG00000156787 | TBC1D31 | 828.9246 | -0.1317649 | 0.2776379 | -0.5033854 | 0.6146933 | 0.776509 |
| ENSG00000285710 | | 1.306913 | -0.1317919 | 0.7314959 | -0.4267985 | 0.6695261 | NA |
| ENSG00000256751 | PLBD1-AS1 | 67.20653 | -0.1317964 | 0.3467576 | -0.4269763 | 0.6693966 | 0.815872 |
| ENSG00000250056 | LINC01018 | 7.637439 | -0.1318117 | 0.5264467 | -0.3452641 | 0.7298959 | 0.853593 |

| | | | | | | | |
|---|---|---|---|---|---|---|---|
| ENSG00000237254 | TRBV30 | 1.225291 | -0.1318352 | 0.7587316 | -0.5165791 | 0.60545 | NA |
| ENSG00000150760 | DOCK1 | 2649.327 | -0.1319775 | 0.1831969 | -0.7426684 | 0.4576824 | 0.65359 |
| ENSG00000230712 | GGTLC4P | 1.983111 | -0.1320129 | 0.7455906 | -0.4776527 | 0.6328974 | 0.789647 |
| ENSG00000163435 | ELF3 | 16133.28 | -0.132024 | 0.3046926 | -0.4563442 | 0.6481425 | 0.800544 |
| ENSG00000149483 | TMEM138 | 1638.094 | -0.1320609 | 0.1713125 | -0.7999254 | 0.423754 | 0.623752 |
| ENSG00000231296 | | 1.410816 | -0.132091 | 0.7432545 | -0.4747111 | 0.6349929 | NA |
| ENSG00000007080 | CCDC124 | 1895.419842 | -0.132124656 | 0.246541457 | -0.562416308 | 0.57383241 | 0.746641 |
| ENSG00000247077 | PGAM5 | 3884.563 | -0.132151 | 0.263751 | -0.5384893 | 0.5902393 | 0.758944 |
| ENSG00000251361 | | 1.220018 | -0.1321629 | 0.7921844 | -0.398782 | 0.6900538 | NA |
| ENSG00000278700 | | 1.154686 | -0.1322083 | 0.7441553 | -0.4696703 | 0.6385906 | NA |
| ENSG00000287121 | | 0.8308462 | -0.1322567 | 0.7910899 | -0.4162927 | 0.6771959 | NA |
| ENSG00000254003 | SRI-AS1 | 2.091455 | -0.132298 | 0.6968904 | -0.3826339 | 0.7019912 | 0.837622 |
| ENSG00000073536 | NLE1 | 1379.506332 | -0.132375556 | 0.220924145 | -0.627206901 | 0.53052362 | 0.714924 |
| ENSG00000101493 | ZNF516 | 1095.991784 | -0.132424883 | 0.364678819 | -0.417947393 | 0.67598558 | 0.819916 |
| ENSG00000177736 | FXNP2 | 1.150081 | -0.1324469 | 0.7367581 | -0.446919 | 0.6549336 | NA |
| ENSG00000265222 | | 1.644249 | -0.1324694 | 0.7103781 | -0.4031384 | 0.6868464 | 0.827596 |
| ENSG00000277368 | | 23.49882 | -0.1325059 | 0.4933392 | -0.3460683 | 0.7292914 | 0.853215 |
| ENSG00000240869 | RN7SL128P | 1.932457 | -0.1325175 | 0.6998987 | -0.3872785 | 0.69855 | 0.835699 |
| ENSG00000204380 | PKP4-AS1 | 27.0895 | -0.1325371 | 0.4006459 | -0.3859058 | 0.6995665 | 0.836225 |
| ENSG00000245479 | LINC01585 | 4.952273 | -0.1325457 | 0.6087309 | -0.339839 | 0.7339778 | 0.855711 |
| ENSG00000164039 | BDH2 | 244.3762 | -0.132554 | 0.2951106 | -0.4856588 | 0.6272091 | 0.785585 |
| ENSG00000203737 | GPR52 | 1.151913 | -0.1326016 | 0.7340708 | -0.4503593 | 0.6524514 | NA |
| ENSG00000136834 | OR1J1 | 2.93565704 | -0.132611391 | 0.669042191 | -0.364504695 | 0.71548117 | 0.845097 |
| ENSG00000183726 | TMEM50A | 3447.913 | -0.1326174 | 0.111874 | -1.19615 | 0.231638 | 0.420038 |
| ENSG00000138801 | PAPSS1 | 4434.113795 | -0.132654071 | 0.127498153 | -1.07143727 | 0.28397286 | 0.481824 |
| ENSG00000286623 | | 4.420251 | -0.1326956 | 0.6286841 | -0.3503601 | 0.7260685 | 0.85124 |
| ENSG00000225465 | RFPL1S | 2.169994 | -0.1327346 | 0.665577 | -0.3692437 | 0.7119461 | 0.843269 |
| ENSG00000286081 | | 97.8414 | -0.1327694 | 0.4376458 | -0.3620842 | 0.7172891 | 0.846015 |
| ENSG00000158164 | TMSB15A | 41.11229 | -0.1328077 | 0.6088192 | -0.3411918 | 0.7329592 | 0.855309 |
| ENSG00000055147 | FAM114A2 | 979.853388 | -0.132816672 | 0.189244646 | -0.729111212 | 0.46593364 | 0.659777 |
| ENSG00000198695 | MT-ND6 | 30505.2 | -0.132926 | 0.204059 | -0.6774073 | 0.4981476 | 0.686635 |
| ENSG00000076984 | MAP2K7 | 2146.209344 | -0.132936442 | 0.116910817 | -1.152330433 | 0.24918532 | 0.441609 |
| ENSG00000179172 | HNRNPCL1 | 9.997061 | -0.132954 | 0.5071972 | -0.3404789 | 0.7334959 | 0.855461 |

| | | | | | | | |
|---|---|---|---|---|---|---|---|
| ENSG00000139675 | HNRNPA1L2 | 818.5153 | -0.1329826 | 0.1872535 | -0.7357368 | 0.4618909 | 0.656607 |
| ENSG00000225573 | RPL35P5 | 663.8797 | -0.1330015 | 0.2425994 | -0.5907877 | 0.5546626 | 0.732593 |
| ENSG00000200737 | | 2.787582 | -0.1330089 | 0.6733124 | -0.3627725 | 0.7167748 | 0.845911 |
| ENSG00000256116 | | 3.565551 | -0.133009 | 0.6017433 | -0.338173 | 0.7352328 | 0.856543 |
| ENSG00000088247 | KHSRP | 9053.624422 | -0.133030256 | 0.183290968 | -0.768288929 | 0.44231555 | 0.640081 |
| ENSG00000213598 | | 714.0612 | -0.1331079 | 0.2235691 | -0.626738 | 0.530831 | 0.715107 |
| ENSG00000006747 | SCIN | 10274.19919 | -0.133147089 | 0.428808716 | -0.376584994 | 0.70648204 | 0.840456 |
| ENSG00000250479 | CHCHD10 | 1036.311 | -0.1331646 | 0.2732404 | -0.5035085 | 0.6146068 | 0.776509 |
| ENSG00000271172 | | 0.8286881 | -0.1331739 | 0.7718341 | -0.5975422 | 0.5501455 | NA |
| ENSG00000212443 | SNORA53 | 9.526449 | -0.1332214 | 0.4928962 | -0.3564602 | 0.7214959 | 0.848466 |
| ENSG00000136238 | RAC1 | 16162.58269 | -0.133223111 | 0.138606145 | -1.039658323 | 0.29849867 | 0.497725 |
| ENSG00000285688 | | 1.959839 | -0.1332867 | 0.6539337 | -0.3598764 | 0.7189396 | 0.846902 |
| ENSG00000114098 | ARMC8 | 3241.715778 | -0.133328132 | 0.125481605 | -1.020465421 | 0.30750778 | 0.507653 |
| ENSG00000106636 | YKT6 | 8479.111443 | -0.13337361 | 0.116793283 | -1.093884619 | 0.27400559 | 0.470756 |
| ENSG00000207168 | SNORA15 | 1.34497 | -0.1334206 | 0.7252766 | -0.4494053 | 0.6531393 | NA |
| ENSG00000028310 | BRD9 | 4722.075684 | -0.133438377 | 0.191141677 | -0.710218463 | 0.47756867 | 0.669704 |
| ENSG00000167077 | MEI1 | 8.696381 | -0.1334545 | 0.5328273 | -0.3341397 | 0.7382741 | 0.858765 |
| ENSG00000141429 | GALNT1 | 5227.133 | -0.1335646 | 0.1500399 | -0.9043244 | 0.3658234 | 0.567798 |
| ENSG00000173821 | RNF213 | 33657.22 | -0.1335795 | 0.283775 | -0.4596186 | 0.64579 | 0.798603 |
| ENSG00000267855 | NDUFA7 | 21.25769 | -0.1336678 | 0.4283788 | -0.3736907 | 0.7086344 | 0.841575 |
| ENSG00000115561 | CHMP3 | 4141.960509 | -0.133678544 | 0.16086004 | -0.846073492 | 0.39751175 | 0.598554 |
| ENSG00000168872 | DDX19A | 880.5603 | -0.1337128 | 0.1682725 | -0.831009 | 0.4059685 | 0.606687 |
| ENSG00000271924 | RNA5SP108 | 2.159425 | -0.1337759 | 0.681645 | -0.3811626 | 0.7030826 | 0.838472 |
| ENSG00000280116 | | 1.237562 | -0.1338246 | 0.7384443 | -0.4709755 | 0.6376582 | NA |
| ENSG00000078747 | ITCH | 3540.56929 | -0.133868564 | 0.170774977 | -0.798635975 | 0.42450152 | 0.624343 |
| ENSG00000138750 | NUP54 | 2388.465797 | -0.133869928 | 0.214567391 | -0.650461918 | 0.51539389 | 0.700793 |
| ENSG00000253414 | | 4.283124 | -0.1338801 | 0.6345841 | -0.3656214 | 0.7146476 | 0.844563 |
| ENSG00000169760 | NLGN1 | 1.109752 | -0.1338846 | 0.7161174 | -0.4185227 | 0.675565 | NA |
| ENSG00000122696 | SLC25A51 | 427.5011302 | -0.133887589 | 0.142514632 | -0.949636337 | 0.34229707 | 0.543757 |
| ENSG00000132612 | VPS4A | 3566.835024 | -0.133986862 | 0.106276887 | -1.287007204 | 0.19809178 | 0.378123 |
| ENSG00000188610 | FAM72B | 256.1693 | -0.1340009 | 0.4601153 | -0.3491954 | 0.7269426 | 0.851749 |
| ENSG00000256278 | | 6.208238 | -0.1340146 | 0.5782277 | -0.3390169 | 0.734597 | 0.856106 |
| ENSG00000133318 | RTN3 | 7903.091283 | -0.134053145 | 0.146572336 | -0.937779426 | 0.34835778 | 0.550105 |

| | | | | | | | |
|---|---|---|---|---|---|---|---|
| ENSG00000078304 | PPP2R5C | 6606.719001 | -0.13407592 | 0.099643752 | -1.364729307 | 0.17233815 | 0.343448 |
| ENSG00000260619 | | 16.72943 | -0.1343256 | 0.4496464 | -0.3672603 | 0.7134249 | 0.84368 |
| ENSG00000256045 | MTRNR2L10 | 4.288456 | -0.1344067 | 0.6603989 | -0.3550066 | 0.7225847 | 0.849109 |
| ENSG00000286410 | | 0.8570488 | -0.1344075 | 0.7584561 | -0.5207506 | 0.6025405 | NA |
| ENSG00000133056 | PIK3C2B | 6714.423805 | -0.134429607 | 0.192595051 | -0.714343011 | 0.4750151 | 0.667619 |
| ENSG00000243007 | RPL12P35 | 8.330879 | -0.1345136 | 0.5858545 | -0.3457223 | 0.7295515 | 0.853333 |
| ENSG00000270813 | NANOGNBP3 | 12.41658 | -0.1345435 | 0.5146947 | -0.3444785 | 0.7304865 | 0.853899 |
| ENSG00000133401 | PDZD2 | 172.5535639 | -0.13455381 | 0.348504529 | -0.433577872 | 0.66459501 | 0.812629 |
| ENSG00000149257 | SERPINH1 | 10892.05 | -0.1345585 | 0.1438994 | -0.9523515 | 0.3409187 | 0.542258 |
| ENSG00000161082 | CELF5 | 166.1404 | -0.1345603 | 0.3222005 | -0.460853 | 0.6449041 | 0.797888 |
| ENSG00000188690 | UROS | 3175.467 | -0.1345901 | 0.1389491 | -0.9886197 | 0.3228492 | 0.523578 |
| ENSG00000277170 | | 9.867942 | -0.1346279 | 0.5965713 | -0.3409377 | 0.7331505 | 0.855321 |
| ENSG00000280381 | | 0.8434229 | -0.1347541 | 0.7770598 | -0.6042874 | 0.5456526 | NA |
| ENSG00000238018 | | 35.88537 | -0.1348118 | 0.3763168 | -0.4052243 | 0.6853126 | 0.826544 |
| ENSG00000236114 | | 3.991755 | -0.134861 | 0.635561 | -0.3495324 | 0.7266897 | 0.851739 |
| ENSG00000145495 | MARCHF6 | 17084.87 | -0.1348824 | 0.1931624 | -0.7272324 | 0.4670836 | 0.660877 |
| ENSG00000039560 | RAI14 | 2428.372175 | -0.135067068 | 0.216433393 | -0.651961766 | 0.51442584 | 0.700152 |
| ENSG00000282885 | | 8.721948 | -0.1350985 | 0.5187021 | -0.3483394 | 0.7275853 | 0.85203 |
| ENSG00000087365 | SF3B2 | 18953.60962 | -0.135121085 | 0.18337323 | -0.771659688 | 0.44031601 | 0.638603 |
| ENSG00000008300 | CELSR3 | 5619.82055 | -0.135314576 | 0.230673059 | -0.668394503 | 0.5038818 | 0.691126 |
| ENSG00000164151 | ICE1 | 6390.21 | -0.1353788 | 0.2024843 | -0.6976496 | 0.4853964 | 0.676413 |
| ENSG00000271550 | BNIP3P11 | 54.63716 | -0.1353974 | 0.3315014 | -0.4517302 | 0.6514633 | 0.803115 |
| ENSG00000287917 | | 15.06659 | -0.1354493 | 0.4772559 | -0.3564898 | 0.7214738 | 0.848466 |
| ENSG00000213365 | | 0.9786481 | -0.1354777 | 0.7574138 | -0.5340863 | 0.5932818 | NA |
| ENSG00000143155 | TIPRL | 2741.757 | -0.1354925 | 0.09325189 | -1.467932 | 0.1421228 | 0.299402 |
| ENSG00000253333 | | 2.058045 | -0.1355332 | 0.6522285 | -0.3615847 | 0.7176624 | 0.846015 |
| ENSG00000280143 | | 1.312825 | -0.1355488 | 0.7207992 | -0.4285282 | 0.6682666 | NA |
| ENSG00000234933 | CDC42P1 | 0.891859 | -0.1355702 | 0.7916121 | -0.4185827 | 0.6755211 | NA |
| ENSG00000149599 | DUSP15 | 27.86829 | -0.135612 | 0.4466627 | -0.3720668 | 0.7098431 | 0.84222 |
| ENSG00000153214 | TMEM87B | 3273.606 | -0.1356136 | 0.1725949 | -0.7967509 | 0.4255957 | 0.625423 |
| ENSG00000273729 | | 90.12368 | -0.135725 | 0.2375897 | -0.6015264 | 0.5474894 | 0.727312 |
| ENSG00000198783 | ZNF830 | 563.8553 | -0.1357251 | 0.1592861 | -0.8707801 | 0.3838743 | 0.586259 |
| ENSG00000117362 | APH1A | 24524.9088 | -0.135758407 | 0.153821696 | -0.905713788 | 0.36508736 | 0.566956 |

| | | | | | | | |
|---|---|---|---|---|---|---|---|
| ENSG00000183684 | ALYREF | 5502.38 | -0.1358349 | 0.3000909 | -0.4938075 | 0.6214421 | 0.781231 |
| ENSG00000138768 | USO1 | 4610.805018 | -0.13587894 | 0.153994797 | -0.902265629 | 0.36691578 | 0.568915 |
| ENSG00000165527 | ARF6 | 8007.106 | -0.1359041 | 0.1339308 | -1.054999 | 0.2914259 | 0.49015 |
| ENSG00000214655 | ZSWIM8 | 4280.839 | -0.1359501 | 0.2006892 | -0.704322 | 0.4812322 | 0.672798 |
| ENSG00000226833 | | 40.53045 | -0.1359813 | 0.465105 | -0.365029 | 0.7150897 | 0.844748 |
| ENSG00000080709 | KCNN2 | 137.4737676 | -0.135988078 | 0.2556748 | -0.562328049 | 0.57389253 | 0.746641 |
| ENSG00000052344 | PRSS8 | 6588.784946 | -0.136040866 | 0.2991543 | -0.495236454 | 0.62043322 | 0.780792 |
| ENSG00000140368 | PSTPIP1 | 1.347972 | -0.1362478 | 0.7241228 | -0.4403077 | 0.6597142 | NA |
| ENSG00000151806 | GUF1 | 2168.825 | -0.1362766 | 0.1570423 | -0.8873762 | 0.3748764 | 0.577124 |
| ENSG00000198786 | MT-ND5 | 98310.67 | -0.136307 | 0.1902819 | -0.7413681 | 0.4584703 | 0.654335 |
| ENSG00000163125 | RPRD2 | 6733.226 | -0.1363869 | 0.1578862 | -0.8889346 | 0.3740382 | 0.576139 |
| ENSG00000008438 | PGLYRP1 | 2.443641585 | -0.136390238 | 0.654505824 | -0.369207499 | 0.71197307 | 0.843269 |
| ENSG00000259630 | FABP5P9 | 0.8285862 | -0.1364756 | 0.7339798 | -0.4579843 | 0.6469637 | NA |
| ENSG00000139921 | TMX1 | 4031.718 | -0.1365167 | 0.1613624 | -0.8616119 | 0.3889011 | 0.590947 |
| ENSG00000198682 | PAPSS2 | 1691.05 | -0.1365886 | 0.3803329 | -0.3912262 | 0.69563 | 0.83332 |
| ENSG00000257058 | | 5.749364 | -0.1365918 | 0.5686586 | -0.3403507 | 0.7335924 | 0.855536 |
| ENSG00000214455 | RCN1P2 | 5.975767 | -0.1366128 | 0.5896851 | -0.3447183 | 0.7303061 | 0.853877 |
| ENSG00000169062 | UPF3A | 2232.282 | -0.1366227 | 0.1019548 | -1.368265 | 0.1712292 | 0.341879 |
| ENSG00000129535 | NRL | 130.8554856 | -0.136685624 | 0.258403697 | -0.561955541 | 0.57414631 | 0.746641 |
| ENSG00000231875 | | 1.289247 | -0.1368163 | 0.7604077 | -0.522373 | 0.6014106 | NA |
| ENSG00000276282 | | 0.8071177 | -0.1368188 | 0.7932512 | -0.3975203 | 0.6909838 | NA |
| ENSG00000215237 | | 1.107297 | -0.1368788 | 0.7258429 | -0.4368778 | 0.6621999 | NA |
| ENSG00000278035 | | 2.100741 | -0.1369149 | 0.654318 | -0.3709132 | 0.7107022 | 0.842599 |
| ENSG00000196689 | TRPV1 | 291.4427 | -0.1369707 | 0.3105482 | -0.4934998 | 0.6216595 | 0.78141 |
| ENSG00000014824 | SLC30A9 | 4716.40217 | -0.137036409 | 0.123112024 | -1.13431969 | 0.25666047 | 0.45047 |
| ENSG00000212128 | TAS2R13 | 1.332381 | -0.1370494 | 0.7164688 | -0.4276468 | 0.6689083 | NA |
| ENSG00000204572 | KRTAP5-10 | 23.04035 | -0.1371062 | 0.4865923 | -0.3595603 | 0.719176 | 0.847018 |
| ENSG00000277795 | | 2.044079 | -0.137125 | 0.7044365 | -0.39955 | 0.689488 | 0.828952 |
| ENSG00000008323 | PLEKHG6 | 627.2175298 | -0.137144825 | 0.26760613 | -0.544826285 | 0.585873 | 0.755627 |
| ENSG00000107938 | EDRF1 | 1796.522623 | -0.13733227 | 0.202440899 | -0.704180265 | 0.48132052 | 0.672851 |
| ENSG00000260252 | | 9.985126 | -0.1373608 | 0.4938362 | -0.3560343 | 0.7218148 | 0.848655 |
| ENSG00000242068 | | 24.42407 | -0.1374062 | 0.369787 | -0.4231792 | 0.6721645 | 0.817761 |
| ENSG00000285991 | | 10.5175 | -0.1374293 | 0.573697 | -0.3443507 | 0.7305825 | 0.8539 |

| | | | | | | | |
|---|---|---|---|---|---|---|---|
| ENSG00000006712 | PAF1 | 4643.683395 | -0.137508779 | 0.252820465 | -0.563983223 | 0.57276555 | 0.745897 |
| ENSG00000239602 | RPL35AP16 | 0.920363 | -0.1375921 | 0.7928202 | -0.4072463 | 0.6838271 | NA |
| ENSG00000151366 | NDUFC2 | 1222.591 | -0.1376928 | 0.1748176 | -0.8063985 | 0.4200131 | 0.620032 |
| ENSG00000165434 | PGM2L1 | 415.6697 | -0.1377498 | 0.4656448 | -0.3613881 | 0.7178093 | 0.846042 |
| ENSG00000205879 | FAM90A2P | 2.361097 | -0.1377593 | 0.7097459 | -0.4197789 | 0.6746469 | 0.818951 |
| ENSG00000279453 | | 119.8407 | -0.1377768 | 0.3290949 | -0.472637 | 0.6364722 | 0.792243 |
| ENSG00000230122 | ECEL1P3 | 2.39974 | -0.1378033 | 0.7053306 | -0.4156452 | 0.6776696 | 0.820733 |
| ENSG00000138430 | OLA1 | 6221.721442 | -0.137804996 | 0.14820561 | -0.972362199 | 0.33087039 | 0.531899 |
| ENSG00000225964 | NRIR | 3.795811 | -0.1378179 | 0.740644 | -0.4798383 | 0.6313424 | 0.788593 |
| ENSG00000234793 | | 5.510312 | -0.1378817 | 0.5588412 | -0.3487647 | 0.727266 | 0.851826 |
| ENSG00000138286 | FAM149B1 | 1555.740633 | -0.137890829 | 0.132225763 | -1.062463308 | 0.28802541 | 0.486401 |
| ENSG00000177034 | MTX3 | 2271.853 | -0.1379378 | 0.208467 | -0.6859705 | 0.4927317 | 0.682788 |
| ENSG00000136111 | TBC1D4 | 1801.747433 | -0.138003058 | 0.200437427 | -0.716953526 | 0.47340277 | 0.666199 |
| ENSG00000156853 | ZNF689 | 944.718 | -0.1380169 | 0.1770525 | -0.7948266 | 0.4267144 | 0.626409 |
| ENSG00000144028 | SNRNP200 | 27396.44 | -0.1381231 | 0.1334725 | -1.050868 | 0.2933194 | 0.492306 |
| ENSG00000257222 | | 8.58564 | -0.1381625 | 0.5615096 | -0.3543578 | 0.7230707 | 0.849464 |
| ENSG00000275055 | | 4.74716 | -0.1381808 | 0.6241207 | -0.3530791 | 0.7240292 | 0.850074 |
| ENSG00000233217 | MROH3P | 11.49631 | -0.138238 | 0.6217039 | -0.3441299 | 0.7307486 | 0.853987 |
| ENSG00000112365 | ZBTB24 | 1053.539373 | -0.138259466 | 0.150997269 | -0.935589769 | 0.34948444 | 0.551196 |
| ENSG00000185627 | PSMD13 | 6087.164 | -0.1382754 | 0.2244942 | -0.6424627 | 0.5205728 | 0.705788 |
| ENSG00000111671 | SPSB2 | 568.6791179 | -0.138292303 | 0.1844849 | -0.773173173 | 0.4394199 | 0.63794 |
| ENSG00000163885 | CFAP100 | 2.37443 | -0.1383771 | 0.6523847 | -0.3694087 | 0.7118231 | 0.843213 |
| ENSG00000242163 | | 1.552912 | -0.1384266 | 0.7346491 | -0.4572978 | 0.647457 | NA |
| ENSG00000108828 | VAT1 | 10420.61637 | -0.13843129 | 0.241069282 | -0.607073152 | 0.54380237 | 0.724697 |
| ENSG00000269888 | | 2.461434 | -0.1384501 | 0.660982 | -0.368164 | 0.712751 | 0.843639 |
| ENSG00000213414 | | 1.656347 | -0.1384553 | 0.7199677 | -0.4403345 | 0.6596949 | 0.808573 |
| ENSG00000272660 | | 2.87295 | -0.1384808 | 0.6710263 | -0.3810134 | 0.7031933 | 0.838524 |
| ENSG00000115808 | STRN | 2706.241042 | -0.138510556 | 0.209646197 | -0.686900778 | 0.49214526 | 0.682311 |
| ENSG00000113269 | RNF130 | 4493.314905 | -0.138516486 | 0.166378829 | -0.847314779 | 0.39681969 | 0.597919 |
| ENSG00000233295 | FAM90A20P | 0.7910841 | -0.138532 | 0.7766988 | -0.6131115 | 0.5398026 | NA |
| ENSG00000279145 | | 6.867223 | -0.1386037 | 0.5899645 | -0.3608805 | 0.7181888 | 0.846264 |
| ENSG00000231401 | | 0.9471492 | -0.1386585 | 0.7564208 | -0.4975071 | 0.6188315 | NA |
| ENSG00000148248 | SURF4 | 18387.37 | -0.1387289 | 0.120286 | -1.137009 | 0.2555346 | 0.448968 |

| | | | | | | | |
|---|---|---|---|---|---|---|---|
| ENSG00000047315 | POLR2B | 8881.45316 | -0.138735718 | 0.136128025 | -1.076607678 | 0.28165557 | 0.4793 |
| ENSG00000146576 | C7orf26 | 2776.487 | -0.1387954 | 0.1483437 | -0.9439435 | 0.3451985 | 0.54678 |
| ENSG00000286440 | | 1.049898 | -0.1388047 | 0.7533997 | -0.520074 | 0.603012 | NA |
| ENSG00000235563 | | 1.630085 | -0.1389268 | 0.7912052 | -0.4409277 | 0.6592653 | 0.808307 |
| ENSG00000178409 | BEND3 | 714.3904 | -0.1389601 | 0.2163737 | -0.6719234 | 0.5016325 | 0.689336 |
| ENSG00000116690 | PRG4 | 5.611852063 | -0.138987148 | 0.632084908 | -0.363903913 | 0.71592976 | 0.845364 |
| ENSG00000286834 | | 12.27079 | -0.1390311 | 0.511651 | -0.3620031 | 0.7173497 | 0.846015 |
| ENSG00000239726 | RN7SL688P | 6.487888 | -0.1390902 | 0.5679846 | -0.3497629 | 0.7265166 | 0.851589 |
| ENSG00000178966 | RMI1 | 967.0561 | -0.1391008 | 0.3370266 | -0.4603166 | 0.645289 | 0.798207 |
| ENSG00000202260 | RN7SKP69 | 0.8832845 | -0.1391039 | 0.7567125 | -0.4962947 | 0.6196865 | NA |
| ENSG00000124588 | NQO2 | 1992.00395 | -0.139173714 | 0.174181577 | -0.82042385 | 0.41197452 | 0.612247 |
| ENSG00000175130 | MARCKSL1 | 11231.53 | -0.1392196 | 0.1876194 | -0.7688351 | 0.4419912 | 0.639786 |
| ENSG00000274799 | | 1.188499 | -0.13923 | 0.7338718 | -0.4592692 | 0.6460409 | NA |
| ENSG00000163075 | CFAP221 | 6.956165 | -0.1393018 | 0.529417 | -0.3577276 | 0.7205472 | 0.847936 |
| ENSG00000249609 | | 0.8456901 | -0.1393219 | 0.7510399 | -0.5307361 | 0.5956016 | NA |
| ENSG00000241185 | | 2.267329 | -0.1393261 | 0.7098234 | -0.4207051 | 0.6739705 | 0.818723 |
| ENSG00000225298 | LINC00113 | 4.183023 | -0.1393266 | 0.742067 | -0.4781502 | 0.6325433 | 0.789387 |
| ENSG00000203908 | KHDC3L | 1.234516 | -0.1393533 | 0.757885 | -0.540276 | 0.5890067 | NA |
| ENSG00000170638 | TRABD | 2581.652 | -0.1394626 | 0.1953545 | -0.7329822 | 0.4635693 | 0.657796 |
| ENSG00000132821 | VSTM2L | 3.041313943 | -0.139502888 | 0.682483143 | -0.382635819 | 0.70198981 | 0.837622 |
| ENSG00000153495 | TEX29 | 4.62075 | -0.1395904 | 0.5607601 | -0.3550254 | 0.7225705 | 0.849109 |
| ENSG00000125962 | ARMCX5 | 1072.761643 | -0.13960143 | 0.205224162 | -0.701597879 | 0.48292998 | 0.674356 |
| ENSG00000184792 | OSBP2 | 825.5097 | -0.1396342 | 0.1684651 | -0.8524702 | 0.3939532 | 0.595538 |
| ENSG00000232578 | | 1.114084 | -0.1396425 | 0.7262362 | -0.4577836 | 0.6471079 | NA |
| ENSG00000270926 | | 0.9769505 | -0.1396764 | 0.7698476 | -0.6417025 | 0.5210664 | NA |
| ENSG00000272205 | | 9.208224 | -0.1397057 | 0.5466118 | -0.352375 | 0.7245571 | 0.850449 |
| ENSG00000277149 | TYW1B | 94.82383 | -0.1397451 | 0.2529653 | -0.5829648 | 0.559917 | 0.736239 |
| ENSG00000162769 | FLVCR1 | 1013.909 | -0.1398134 | 0.2160957 | -0.6705569 | 0.5025029 | 0.690065 |
| ENSG00000203724 | C1orf53 | 179.6927 | -0.13982 | 0.3010673 | -0.5047855 | 0.6137095 | 0.775861 |
| ENSG00000224668 | IPO8P1 | 7.01606 | -0.1399177 | 0.5586182 | -0.3588546 | 0.7197038 | 0.847523 |
| ENSG00000284678 | LINC02811 | 1.476261 | -0.1399419 | 0.7546921 | -0.509375 | 0.6104894 | NA |
| ENSG00000090989 | EXOC1 | 2058.802201 | -0.139967335 | 0.190863453 | -0.765657626 | 0.44388005 | 0.641412 |
| ENSG00000258876 | TGFB3-AS1 | 1.420039 | -0.1400153 | 0.7101946 | -0.4294786 | 0.6675749 | NA |

| | | | | | | | |
|---|---|---|---|---|---|---|---|
| ENSG00000246863 | GPR176-DT | 3.720194 | -0.1401036 | 0.6006237 | -0.3577517 | 0.7205291 | 0.847936 |
| ENSG00000271871 | | 7.761014 | -0.1401089 | 0.633557 | -0.3795789 | 0.704258 | 0.839341 |
| ENSG00000271937 | | 1.147578 | -0.1401672 | 0.7549785 | -0.5080467 | 0.6114206 | NA |
| ENSG00000088726 | TMEM40 | 121.6670251 | -0.140167783 | 0.498293386 | -0.370259524 | 0.71118913 | 0.842862 |
| ENSG00000154059 | IMPACT | 2342.709 | -0.1402207 | 0.2036231 | -0.7175435 | 0.4730388 | 0.665969 |
| ENSG00000179271 | GADD45GIP1 | 1971.297 | -0.1402518 | 0.2904562 | -0.5288799 | 0.5968887 | 0.763945 |
| ENSG00000288586 | | 17.00529 | -0.140292 | 0.4281535 | -0.3865363 | 0.6990995 | 0.835818 |
| ENSG00000213650 | | 3.936578 | -0.1403172 | 0.6481516 | -0.372654 | 0.709406 | 0.841926 |
| ENSG00000114446 | IFT57 | 2607.295653 | -0.140339524 | 0.208775887 | -0.722198867 | 0.47017222 | 0.663656 |
| ENSG00000224888 | | 73.8323 | -0.1404808 | 0.3651292 | -0.4353111 | 0.6633366 | 0.811584 |
| ENSG00000166492 | FAM86GP | 8.890888 | -0.1405702 | 0.5303611 | -0.3585255 | 0.7199501 | 0.847625 |
| ENSG00000134825 | TMEM258 | 2257.120548 | -0.140617647 | 0.195547115 | -0.743377442 | 0.4572532 | 0.65317 |
| ENSG00000272455 | MRPL20-DT | 252.9451 | -0.1406339 | 0.2592098 | -0.573358 | 0.5664023 | 0.741055 |
| ENSG00000264296 | | 1.919775 | -0.1406591 | 0.7193454 | -0.4440101 | 0.6570353 | 0.806839 |
| ENSG00000167280 | ENGASE | 2195.944 | -0.1406654 | 0.2090341 | -0.7034642 | 0.4817665 | 0.673156 |
| ENSG00000251216 | | 1.143384 | -0.1406675 | 0.7413047 | -0.519262 | 0.6035781 | NA |
| ENSG00000276984 | | 2.069942 | -0.1406921 | 0.6770812 | -0.3863964 | 0.6992031 | 0.835904 |
| ENSG00000114346 | ECT2 | 5076.120974 | -0.140707395 | 0.361439279 | -0.443305451 | 0.65754481 | 0.807092 |
| ENSG00000114416 | FXR1 | 6951.197112 | -0.14073316 | 0.11797807 | -1.155121539 | 0.24804066 | 0.440285 |
| ENSG00000116194 | ANGPTL1 | 1.509672372 | -0.140760866 | 0.693866945 | -0.41440957 | 0.67857418 | NA |
| ENSG00000186660 | ZFP91 | 3639.282 | -0.1407899 | 0.1221578 | -1.185506 | 0.2358176 | 0.42515 |
| ENSG00000230772 | VN1R108P | 0.9172382 | -0.1408902 | 0.7603425 | -0.5180925 | 0.6043937 | NA |
| ENSG00000276476 | | 1.625261 | -0.1409774 | 0.791331 | -0.4468476 | 0.6549851 | 0.80534 |
| ENSG00000250377 | | 0.8017239 | -0.1409871 | 0.7400089 | -0.4996894 | 0.6172938 | NA |
| ENSG00000255052 | FAM66D | 74.51322 | -0.1409956 | 0.260781 | -0.5749356 | 0.5653348 | 0.740209 |
| ENSG00000231625 | SLC47A1P2 | 2.975626 | -0.1411174 | 0.6248943 | -0.369969 | 0.7114056 | 0.842907 |
| ENSG00000188486 | H2AX | 6637.685 | -0.1411312 | 0.3930558 | -0.4202166 | 0.6743272 | 0.818891 |
| ENSG00000229676 | ZNF492 | 37.38716 | -0.1411793 | 0.5508926 | -0.3616295 | 0.7176289 | 0.846015 |
| ENSG00000118579 | MED28 | 2730.82405 | -0.141291291 | 0.136956999 | -1.045760874 | 0.29567145 | 0.494708 |
| ENSG00000175701 | MTLN | 984.0679 | -0.1412927 | 0.2297062 | -0.645575 | 0.5185546 | 0.703857 |
| ENSG00000257718 | CPNE8-AS1 | 2.128395 | -0.1412968 | 0.7937832 | -0.4078449 | 0.6833876 | 0.824859 |
| ENSG00000131697 | NPHP4 | 1205.010674 | -0.141334769 | 0.171686451 | -0.877962925 | 0.37996384 | 0.581993 |
| ENSG00000268869 | ESPNP | 27.50501 | -0.1413662 | 0.433617 | -0.3956664 | 0.6923511 | 0.831078 |

| | | | | | | | |
|---|---|---|---|---|---|---|---|
| ENSG00000006740 | ARHGAP44 | 1231.639197 | -0.14146126 | 0.126918031 | -1.1683721 | 0.2426567 | 0.433538 |
| ENSG00000227066 | | 0.9852876 | -0.141522 | 0.7544585 | -0.5132328 | 0.6077885 | NA |
| ENSG00000132849 | PATJ | 5360.716525 | -0.141584312 | 0.257840698 | -0.590265269 | 0.55501282 | 0.732691 |
| ENSG00000223839 | FAM95B1 | 1.928136 | -0.1415851 | 0.7914216 | -0.4477813 | 0.654311 | 0.805019 |
| ENSG00000177595 | PIDD1 | 1559.393 | -0.1415937 | 0.2119714 | -0.7068692 | 0.4796478 | 0.671608 |
| ENSG00000204962 | PCDHA8 | 30.82578 | -0.1416782 | 0.4661648 | -0.3739914 | 0.7084107 | 0.841459 |
| ENSG00000267783 | | 1.943611 | -0.1417007 | 0.701827 | -0.4200523 | 0.6744473 | 0.81891 |
| ENSG00000285398 | | 1.980024 | -0.1417094 | 0.7507922 | -0.5172489 | 0.6049824 | 0.769596 |
| ENSG00000224914 | LINC00863 | 441.3174 | -0.1417679 | 0.1822252 | -0.8044109 | 0.4211597 | 0.621 |
| ENSG00000232593 | KANTR | 837.9159 | -0.1418977 | 0.253061 | -0.5983978 | 0.5495745 | 0.728944 |
| ENSG00000279483 | | 559.0573 | -0.1419632 | 0.2411059 | -0.6162707 | 0.5377159 | 0.720816 |
| ENSG00000163660 | CCNL1 | 11463.23 | -0.1419801 | 0.1688298 | -0.8778676 | 0.3800156 | 0.581996 |
| ENSG00000286521 | | 0.7968818 | -0.1419946 | 0.7973251 | -0.5550851 | 0.5788365 | NA |
| ENSG00000178913 | TAF7 | 9675.68 | -0.142013 | 0.1078062 | -1.331458 | 0.1830385 | 0.358032 |
| ENSG00000124201 | ZNFX1 | 7427.001351 | -0.142031109 | 0.264524395 | -0.590720465 | 0.55470773 | 0.732616 |
| ENSG00000197584 | KCNMB2 | 5.295955 | -0.1420336 | 0.5584874 | -0.3577031 | 0.7205655 | 0.847936 |
| ENSG00000236675 | MTX1P1 | 283.4033 | -0.1420668 | 0.2045902 | -0.7170633 | 0.473335 | 0.666174 |
| ENSG00000005206 | SPPL2B | 2703.692825 | -0.142141636 | 0.153811485 | -0.950903153 | 0.34165354 | 0.54292 |
| ENSG00000110711 | AIP | 3385.703135 | -0.142173956 | 0.216433823 | -0.685902026 | 0.49277489 | 0.682788 |
| ENSG00000260192 | LINC02240 | 0.8154267 | -0.142184 | 0.7576994 | -0.5739001 | 0.5660354 | NA |
| ENSG00000079134 | THOC1 | 2046.828676 | -0.142196076 | 0.198683751 | -0.747704189 | 0.4546386 | 0.650614 |
| ENSG00000234949 | | 134.3134 | -0.1422522 | 0.2501014 | -0.6056065 | 0.5447761 | 0.725409 |
| ENSG00000112237 | CCNC | 3506.650491 | -0.1423054 | 0.208298512 | -0.733232627 | 0.46341656 | 0.65765 |
| ENSG00000248208 | WDR45P1 | 10.2388 | -0.1423416 | 0.5407337 | -0.3613271 | 0.717855 | 0.846058 |
| ENSG00000202078 | | 1.496938 | -0.1423654 | 0.7568946 | -0.4991282 | 0.6176891 | NA |
| ENSG00000277496 | | 57.19564 | -0.1424411 | 0.3372721 | -0.4707869 | 0.6377929 | 0.793254 |
| ENSG00000276071 | | 26.44352 | -0.1425244 | 0.4141971 | -0.4058894 | 0.6848239 | 0.826148 |
| ENSG00000106624 | AEBP1 | 206.4641983 | -0.142532033 | 0.284696393 | -0.542148611 | 0.58771613 | 0.757092 |
| ENSG00000213199 | ASIC3 | 345.5608 | -0.1426234 | 0.2954374 | -0.5252424 | 0.5994147 | 0.76585 |
| ENSG00000101216 | GMEB2 | 2290.30088 | -0.142669524 | 0.166806264 | -0.927321201 | 0.35375979 | 0.555762 |
| ENSG00000280028 | | 1.267903 | -0.1426883 | 0.7620404 | -0.5678629 | 0.570128 | NA |
| ENSG00000226210 | WASH8P | 1235.667 | -0.1426921 | 0.19957 | -0.7443818 | 0.4566455 | 0.652467 |
| ENSG00000271105 | SCML2P2 | 10.14252 | -0.1426955 | 0.5517871 | -0.3617993 | 0.717502 | 0.846015 |

| | | | | | | | |
|---|---|---|---|---|---|---|---|
| ENSG00000213526 | SETP8 | 18.5093 | -0.1427115 | 0.4002743 | -0.4187325 | 0.6754117 | 0.81957 |
| ENSG00000265437 | | 1.657241 | -0.1427906 | 0.7257297 | -0.4435072 | 0.6573989 | 0.807089 |
| ENSG00000198830 | HMGN2 | 5798.395 | -0.1427993 | 0.2534335 | -0.5967509 | 0.5506737 | 0.729573 |
| ENSG00000227879 | PSPC1P1 | 2.503577 | -0.1428444 | 0.6988683 | -0.4235642 | 0.6718837 | 0.817591 |
| ENSG00000255397 | NASPP1 | 103.9997 | -0.1428519 | 0.422705 | -0.4036606 | 0.6864623 | 0.827298 |
| ENSG00000183911 | RPL21P132 | 6.479884 | -0.1429995 | 0.5697784 | -0.3591105 | 0.7195124 | 0.847373 |
| ENSG00000240322 | RN7SL481P | 6.483891 | -0.1431086 | 0.580252 | -0.3631688 | 0.7164788 | 0.845674 |
| ENSG00000079313 | REXO1 | 1355.849581 | -0.143124201 | 0.187867528 | -0.776911177 | 0.43721118 | 0.635672 |
| ENSG00000204623 | ZNRD1ASP | 480.9105 | -0.1431645 | 0.4131233 | -0.4104417 | 0.681482 | 0.823757 |
| ENSG00000287511 | | 3.491332 | -0.1431921 | 0.6574929 | -0.3865798 | 0.6990673 | 0.835817 |
| ENSG00000010803 | SCMH1 | 2216.898271 | -0.143202571 | 0.192333445 | -0.779820796 | 0.43549636 | 0.634254 |
| ENSG00000213553 | RPLP0P6 | 19120.1 | -0.14323 | 0.1903558 | -0.7783699 | 0.436351 | 0.634907 |
| ENSG00000260979 | | 5.583144 | -0.1432682 | 0.5469618 | -0.3666022 | 0.7139157 | 0.843961 |
| ENSG00000181004 | BBS12 | 156.9466 | -0.1433704 | 0.3224347 | -0.4906803 | 0.6236526 | 0.782851 |
| ENSG00000232470 | | 29.65006 | -0.1433728 | 0.4333394 | -0.4031866 | 0.6868109 | 0.827596 |
| ENSG00000059573 | ALDH18A1 | 9966.351143 | -0.143404542 | 0.138470858 | -1.116851585 | 0.26405779 | 0.459447 |
| ENSG00000196072 | BLOC1S2 | 2702.079 | -0.1434153 | 0.1791021 | -0.8311777 | 0.4058733 | 0.606663 |
| ENSG00000213352 | GAPDHP44 | 2.244369 | -0.1434319 | 0.7103907 | -0.4273979 | 0.6690895 | 0.81561 |
| ENSG00000093000 | NUP50 | 3081.009487 | -0.143473363 | 0.1934965 | -0.771797142 | 0.44023459 | 0.638565 |
| ENSG00000248254 | | 0.9046836 | -0.1434926 | 0.7317525 | -0.4677203 | 0.6399846 | NA |
| ENSG00000273004 | | 15.29374 | -0.1436236 | 0.4930298 | -0.3743963 | 0.7081095 | 0.841263 |
| ENSG00000124159 | MATN4 | 10.54495081 | -0.143714288 | 0.576140763 | -0.364017129 | 0.71584522 | 0.845328 |
| ENSG00000229635 | | 3.204143 | -0.1437196 | 0.5948443 | -0.3665258 | 0.7139728 | 0.843991 |
| ENSG00000271580 | | 29.24719 | -0.1437237 | 0.3881725 | -0.4279413 | 0.6686939 | 0.815427 |
| ENSG00000138380 | CARF | 636.4107925 | -0.143827854 | 0.289515802 | -0.553871964 | 0.57966647 | 0.75067 |
| ENSG00000196455 | PIK3R4 | 2424.826 | -0.1438313 | 0.1457579 | -1.025707 | 0.3050297 | 0.504729 |
| ENSG00000279466 | | 0.8358357 | -0.1439265 | 0.7654426 | -0.6047083 | 0.5453729 | NA |
| ENSG00000278234 | | 1.130529 | -0.143947 | 0.761148 | -0.5738186 | 0.5660906 | NA |
| ENSG00000176563 | CNTD1 | 72.60113 | -0.1439984 | 0.2672388 | -0.5757219 | 0.5648032 | 0.739975 |
| ENSG00000105193 | RPS16 | 20926.05904 | -0.144009001 | 0.172344454 | -0.859121419 | 0.39027353 | 0.592234 |
| ENSG00000260774 | | 79.47032 | -0.1440258 | 0.2431949 | -0.6243 | 0.5324306 | 0.71637 |
| ENSG00000099290 | WASHC2A | 5554.644529 | -0.14403414 | 0.219753933 | -0.690960689 | 0.48959025 | 0.680283 |
| ENSG00000065526 | SPEN | 4859.251915 | -0.144058464 | 0.229842669 | -0.651927851 | 0.51444772 | 0.700152 |

| | | | | | | | |
|---|---|---|---|---|---|---|---|
| ENSG00000235042 | SMARCAL1-AS1 | 1.266525 | -0.1440897 | 0.7597424 | -0.5741442 | 0.5658702 | NA |
| ENSG00000170848 | PSG6 | 1.043824 | -0.1441388 | 0.79951 | -0.6270789 | 0.5306075 | NA |
| ENSG00000203709 | MIR29B2CHG | 305.3159 | -0.1441928 | 0.3636413 | -0.4481691 | 0.6540312 | 0.804895 |
| ENSG00000286042 | LCAL1 | 1.519612 | -0.1441968 | 0.7586364 | -0.5413628 | 0.5882576 | NA |
| ENSG00000270858 | VDAC1P5 | 4.480467 | -0.1442388 | 0.6373953 | -0.3782084 | 0.7052758 | 0.839876 |
| ENSG00000239883 | PARGP1 | 552.9598 | -0.1442437 | 0.2582961 | -0.5943152 | 0.5523013 | 0.730854 |
| ENSG00000278267 | MIR6859-1 | 19.87986 | -0.1442749 | 0.4407581 | -0.3949891 | 0.6928509 | 0.831284 |
| ENSG00000146281 | PM20D2 | 1774.677 | -0.1443487 | 0.2524505 | -0.6064626 | 0.5442076 | 0.725055 |
| ENSG00000206755 | SNORA30 | 1.060618 | -0.1443909 | 0.7565831 | -0.5428864 | 0.587208 | NA |
| ENSG00000163104 | SMARCAD1 | 2653.456 | -0.1444649 | 0.1628199 | -0.9330112 | 0.3508142 | 0.552408 |
| ENSG00000274162 | SNX18P1Y | 2.718785 | -0.1444903 | 0.6843654 | -0.4015493 | 0.6880158 | 0.828129 |
| ENSG00000008118 | CAMK1G | 2.278076588 | -0.144505243 | 0.75722506 | -0.540297416 | 0.58899194 | 0.757928 |
| ENSG00000213183 | RPS7P8 | 0.8488184 | -0.1445397 | 0.7597798 | -0.5254835 | 0.5992471 | NA |
| ENSG00000170390 | DCLK2 | 137.6408 | -0.1445516 | 0.4430171 | -0.3995793 | 0.6894664 | 0.828952 |
| ENSG00000108443 | RPS6KB1 | 41049.45099 | -0.144637883 | 0.115682582 | -1.083156869 | 0.2787388 | 0.475988 |
| ENSG00000121274 | TENT4B | 1250.49133 | -0.14464649 | 0.184801575 | -0.794944335 | 0.42664588 | 0.626378 |
| ENSG00000229901 | FOXO6-AS1 | 2.690088 | -0.1446655 | 0.6534953 | -0.3853273 | 0.699995 | 0.836549 |
| ENSG00000234424 | | 0.8362403 | -0.1446995 | 0.8032309 | -0.7720775 | 0.4400685 | NA |
| ENSG00000174574 | AKIRIN1 | 3460.116 | -0.1447277 | 0.1573425 | -0.9175463 | 0.3588565 | 0.560812 |
| ENSG00000082074 | FYB1 | 0.854214634 | -0.144748554 | 0.792288793 | -0.441651227 | 0.65874161 | NA |
| ENSG00000166848 | TERF2IP | 4884.142 | -0.1447784 | 0.1265228 | -1.15158 | 0.2494938 | 0.442038 |
| ENSG00000287090 | | 5.690625 | -0.1448915 | 0.6068072 | -0.3703591 | 0.711115 | 0.842862 |
| ENSG00000180660 | MAB21L1 | 0.8222236 | -0.1449341 | 0.7920534 | -0.4480546 | 0.6541138 | NA |
| ENSG00000246228 | CASC8 | 14.89181 | -0.144949 | 0.478585 | -0.382012 | 0.7024524 | 0.837979 |
| ENSG00000253161 | LINC01605 | 2.255326 | -0.1449744 | 0.6882179 | -0.423699 | 0.6717854 | 0.817546 |
| ENSG00000072756 | TRNT1 | 2997.274038 | -0.144987884 | 0.140946591 | -1.052539667 | 0.29255202 | 0.49139 |
| ENSG00000230495 | | 2.36062 | -0.1450996 | 0.696216 | -0.421496 | 0.6733929 | 0.818528 |
| ENSG00000110031 | LPXN | 238.3329535 | -0.145103909 | 0.289058496 | -0.542773952 | 0.58728545 | 0.756905 |
| ENSG00000257083 | | 0.9230686 | -0.1451042 | 0.7485457 | -0.5147929 | 0.6066977 | NA |
| ENSG00000164933 | SLC25A32 | 11.71161 | -0.1451971 | 0.5302851 | -0.3719623 | 0.7099209 | 0.842274 |
| ENSG00000163655 | GMPS | 7177.32 | -0.1454898 | 0.2167833 | -0.7063224 | 0.4799876 | 0.671801 |
| ENSG00000272465 | | 4.496441 | -0.1455447 | 0.655686 | -0.3890813 | 0.697216 | 0.834698 |
| ENSG00000233668 | | 8.165206 | -0.1455609 | 0.5138199 | -0.3691072 | 0.7120478 | 0.843269 |

| | | | | | | | |
|---|---|---|---|---|---|---|---|
| ENSG00000115286 | NDUFS7 | 732.0715781 | -0.145627669 | 0.20971382 | -0.721176503 | 0.47080093 | 0.664156 |
| ENSG00000280214 | | 12.92796 | -0.1456497 | 0.5413527 | -0.3680659 | 0.7128241 | 0.843639 |
| ENSG00000173678 | SPDYE2B | 4.073223 | -0.1457689 | 0.6518942 | -0.3869189 | 0.6988162 | 0.835727 |
| ENSG00000285303 | | 1.049816 | -0.1457961 | 0.7936615 | -0.4239001 | 0.6716387 | NA |
| ENSG00000183432 | ZBTB8OSP1 | 9.272683 | -0.145832 | 0.5034876 | -0.378257 | 0.7052397 | 0.839876 |
| ENSG00000265321 | MIR4263 | 11.06804 | -0.1458325 | 0.4691259 | -0.3866376 | 0.6990245 | 0.835817 |
| ENSG00000268324 | LRRC2-AS1 | 0.8386329 | -0.1459445 | 0.7757628 | -0.6358265 | 0.5248895 | NA |
| ENSG00000274353 | | 1.191828 | -0.1459464 | 0.8096371 | -1.470355 | 0.1414657 | NA |
| ENSG00000221520 | MIR1285-1 | 2.242538 | -0.1459747 | 0.6633229 | -0.399889 | 0.6892382 | 0.828804 |
| ENSG00000234006 | DDX39B-AS1 | 2.708952 | -0.1460287 | 0.6997899 | -0.4142508 | 0.6786904 | 0.821485 |
| ENSG00000201746 | RNU6-828P | 0.7951367 | -0.1460638 | 0.7757776 | -0.6359817 | 0.5247884 | NA |
| ENSG00000104626 | ERI1 | 986.100523 | -0.14611334 | 0.268618245 | -0.580768187 | 0.5613967 | 0.73741 |
| ENSG00000172954 | LCLAT1 | 1291.408 | -0.1461145 | 0.1614247 | -0.9361216 | 0.3492106 | 0.550928 |
| ENSG00000132740 | IGHMBP2 | 1719.028189 | -0.146136179 | 0.188266711 | -0.81114429 | 0.41728281 | 0.617189 |
| ENSG00000221944 | TIGD1 | 774.1446 | -0.1462007 | 0.2670571 | -0.5887525 | 0.5560273 | 0.733556 |
| ENSG00000242366 | UGT1A8 | 2.038745 | -0.1462389 | 0.7941393 | -0.4209475 | 0.6737934 | 0.818654 |
| ENSG00000224100 | | 1.749233 | -0.1462478 | 0.7382 | -0.4920823 | 0.6226612 | 0.782124 |
| ENSG00000139410 | SDSL | 599.6212 | -0.1462538 | 0.2835856 | -0.5555685 | 0.5785059 | 0.749955 |
| ENSG00000134255 | CEPT1 | 1822.204942 | -0.146416063 | 0.181067203 | -0.831151338 | 0.40588814 | 0.606663 |
| ENSG00000140105 | WARS1 | 9919.033 | -0.1464491 | 0.3159911 | -0.5112035 | 0.6092086 | 0.772756 |
| ENSG00000166803 | PCLAF | 631.3498 | -0.1464546 | 0.5277218 | -0.3833488 | 0.7014612 | 0.83727 |
| ENSG00000108175 | ZMIZ1 | 6763.377081 | -0.146483317 | 0.214775584 | -0.714660563 | 0.47481881 | 0.667509 |
| ENSG00000131747 | TOP2A | 12183.43108 | -0.146524342 | 0.39207519 | -0.437480758 | 0.66176273 | 0.810248 |
| ENSG00000078061 | ARAF | 3950.739746 | -0.146573455 | 0.148771736 | -0.995013892 | 0.31972951 | 0.52068 |
| ENSG00000112578 | BYSL | 2161.930565 | -0.146617222 | 0.194489157 | -0.791543792 | 0.42862673 | 0.62791 |
| ENSG00000272383 | | 1.508997 | -0.1466403 | 0.730827 | -0.4677004 | 0.6399989 | NA |
| ENSG00000256043 | CTSO | 580.4819 | -0.1466419 | 0.3206025 | -0.5040801 | 0.6142051 | 0.776236 |
| ENSG00000245532 | NEAT1 | 28154.21 | -0.1466855 | 0.2835968 | -0.5594412 | 0.5758607 | 0.747949 |
| ENSG00000000971 | CFH | 8.491109685 | -0.146689226 | 0.700197442 | -0.420899474 | 0.67382849 | 0.818654 |
| ENSG00000118520 | ARG1 | 5.323108797 | -0.146776642 | 0.564644723 | -0.371723886 | 0.71009844 | 0.842334 |
| ENSG00000251665 | | 1.348853 | -0.146803 | 0.7547749 | -0.5682735 | 0.5698493 | NA |
| ENSG00000170515 | PA2G4 | 6478.171 | -0.1468124 | 0.2373433 | -0.6503308 | 0.5154786 | 0.700837 |
| ENSG00000131469 | RPL27 | 17090.31802 | -0.146817617 | 0.179817713 | -0.866485973 | 0.38622372 | 0.588528 |

| | | | | | | | |
|---|---|---|---|---|---|---|---|
| ENSG00000146670 | CDCA5 | 2894.74 | -0.1469344 | 0.5370215 | -0.3735503 | 0.7087389 | 0.841624 |
| ENSG00000167178 | ISLR2 | 1.648576 | -0.1469488 | 0.7517266 | -0.5353178 | 0.5924301 | 0.760659 |
| ENSG00000102921 | N4BP1 | 1987.481 | -0.14695529 | 0.158093069 | -0.944373455 | 0.34497884 | 0.546537 |
| ENSG00000256826 | ATP5MFP4 | 1.079681 | -0.146959 | 0.7934663 | -0.8833797 | 0.3770312 | NA |
| ENSG00000108556 | CHRNE | 4.927372573 | -0.147000702 | 0.596832641 | -0.378456581 | 0.70509144 | 0.839876 |
| ENSG00000260735 | | 19.50461 | -0.1470643 | 0.4056898 | -0.4250729 | 0.6707835 | 0.816888 |
| ENSG00000225008 | TWF1P2 | 11.70596 | -0.1471177 | 0.4741597 | -0.3872609 | 0.6985631 | 0.835699 |
| ENSG00000265996 | MIR3671 | 2.10035 | -0.1471411 | 0.6886184 | -0.4341799 | 0.6641578 | 0.812282 |
| ENSG00000134717 | BTF3L4 | 4102.436601 | -0.147217893 | 0.115706796 | -1.284862681 | 0.19884027 | 0.379008 |
| ENSG00000238288 | MTND3P23 | 2.364424 | -0.1472561 | 0.7259 | -0.4693458 | 0.6388225 | 0.794021 |
| ENSG00000214872 | SMTNL1 | 10.64547 | -0.1472694 | 0.6021461 | -0.3684744 | 0.7125195 | 0.843549 |
| ENSG00000089157 | RPLP0 | 44583.20932 | -0.147275449 | 0.18092651 | -0.839401205 | 0.40124421 | 0.602213 |
| ENSG00000130803 | ZNF317 | 1823.896829 | -0.147360295 | 0.09657374 | -1.523749967 | 0.12757118 | 0.276981 |
| ENSG00000079150 | FKBP7 | 49.08785385 | -0.147365142 | 0.348243767 | -0.476793733 | 0.63350901 | 0.789998 |
| ENSG00000287042 | | 19.17077 | -0.1474035 | 0.5271941 | -0.3776302 | 0.7057053 | 0.840057 |
| ENSG00000125971 | DYNLRB1 | 409.0734007 | -0.147458364 | 0.206194946 | -0.749252151 | 0.45370524 | 0.649906 |
| ENSG00000136021 | SCYL2 | 4881.039467 | -0.147464193 | 0.17017492 | -0.890130553 | 0.37339579 | 0.575572 |
| ENSG00000049883 | PTCD2 | 1177.354225 | -0.147476468 | 0.179515937 | -0.851137145 | 0.39469317 | 0.59644 |
| ENSG00000131019 | ULBP3 | 210.2329724 | -0.147489148 | 0.313340576 | -0.516536704 | 0.60547961 | 0.769897 |
| ENSG00000167258 | CDK12 | 4164.482 | -0.1475264 | 0.1631608 | -0.9544531 | 0.3398543 | 0.541291 |
| ENSG00000251718 | RNU2-13P | 0.7683948 | -0.147593 | 0.8091964 | -1.290153 | 0.1969974 | NA |
| ENSG00000214530 | STARD10 | 16963.02 | -0.147624 | 0.2222633 | -0.6967544 | 0.4859565 | 0.67691 |
| ENSG00000268364 | SMC5-AS1 | 25.53819 | -0.1476437 | 0.3965142 | -0.4341876 | 0.6641522 | 0.812282 |
| ENSG00000108582 | CPD | 2747.448682 | -0.147660765 | 0.20245599 | -0.76089494 | 0.44671982 | 0.643615 |
| ENSG00000257769 | | 1.135239 | -0.1477066 | 0.7477119 | -0.5248575 | 0.5996823 | NA |
| ENSG00000129455 | KLK8 | 0.935823001 | -0.147808806 | 0.793710114 | -0.431032701 | 0.66644459 | NA |
| ENSG00000235339 | GEMIN2P2 | 2.898934 | -0.1478208 | 0.6785688 | -0.4130763 | 0.6795507 | 0.822064 |
| ENSG00000127952 | STYXL1 | 1621.197111 | -0.147864466 | 0.15802397 | -0.941611924 | 0.34639136 | 0.548028 |
| ENSG00000285728 | | 235.526 | -0.1478782 | 0.2942287 | -0.5578181 | 0.5769686 | 0.748839 |
| ENSG00000007541 | PIGQ | 2899.358399 | -0.147887826 | 0.172121548 | -0.862801327 | 0.38824671 | 0.590459 |
| ENSG00000279231 | | 2.436357 | -0.1479027 | 0.6710029 | -0.4043229 | 0.6859753 | 0.827049 |
| ENSG00000127616 | SMARCA4 | 14797.22416 | -0.147938587 | 0.166061877 | -0.975726483 | 0.32920003 | 0.530313 |
| ENSG00000205116 | TMEM88B | 1.202055 | -0.1479659 | 0.7507924 | -0.5456177 | 0.5853287 | NA |

| | | | | | | | |
|---|---|---|---|---|---|---|---|
| ENSG00000236861 | BET1-AS1 | 7.321007 | -0.1479916 | 0.5957693 | -0.3707339 | 0.7108357 | 0.842627 |
| ENSG00000285458 | | 5.7613 | -0.1480432 | 0.5696104 | -0.3748574 | 0.7077666 | 0.841108 |
| ENSG00000196277 | GRM7 | 1.921267 | -0.1480554 | 0.7096727 | -0.4438425 | 0.6571565 | 0.806839 |
| ENSG00000222287 | RNU6-1043P | 1.303971 | -0.1482298 | 0.7275584 | -0.4687889 | 0.6392206 | NA |
| ENSG00000119185 | ITGB1BP1 | 3758.163837 | -0.148313768 | 0.222488724 | -0.697267983 | 0.4856351 | 0.676675 |
| ENSG00000070950 | RAD18 | 2076.591538 | -0.148367056 | 0.349341349 | -0.486598032 | 0.62654322 | 0.785181 |
| ENSG00000241127 | YAE1 | 833.2024 | -0.1483831 | 0.1834438 | -0.8357127 | 0.4033166 | 0.604182 |
| ENSG00000147789 | ZNF7 | 2363.332 | -0.1484024 | 0.1004945 | -1.497231 | 0.1343333 | 0.287354 |
| ENSG00000207789 | MIR26A2 | 1.399501 | -0.1484422 | 0.7203557 | -0.4631533 | 0.6432545 | NA |
| ENSG00000272312 | | 0.8162994 | -0.1484816 | 0.7925874 | -0.4518015 | 0.651412 | NA |
| ENSG00000277052 | MIR6763 | 4.355029 | -0.1484832 | 0.5905262 | -0.3738209 | 0.7085376 | 0.841518 |
| ENSG00000168610 | STAT3 | 10526.84 | -0.1485873 | 0.2545757 | -0.5851332 | 0.5584581 | 0.735521 |
| ENSG00000252212 | RNU2-58P | 0.9713773 | -0.1485873 | 0.7644647 | -0.5623795 | 0.5738575 | NA |
| ENSG00000273045 | C2orf15 | 496.9163 | -0.1486163 | 0.1450416 | -1.046618 | 0.2952758 | 0.494509 |
| ENSG00000144026 | ZNF514 | 2191.896 | -0.1487739 | 0.2965376 | -0.545431 | 0.5854572 | 0.755278 |
| ENSG00000095574 | IKZF5 | 1184.857521 | -0.148801211 | 0.156288071 | -0.972998695 | 0.33055395 | 0.53166 |
| ENSG00000167173 | C15orf39 | 2510.63 | -0.1488511 | 0.2172646 | -0.7310586 | 0.4647434 | 0.658653 |
| ENSG00000261173 | | 5.564315 | -0.1489209 | 0.6000515 | -0.3806464 | 0.7034657 | 0.838628 |
| ENSG00000158941 | CCAR2 | 4647.438 | -0.1489391 | 0.1556725 | -0.9768873 | 0.3286249 | 0.52984 |
| ENSG00000246130 | | 1.982957 | -0.1489444 | 0.6891625 | -0.4245661 | 0.671153 | 0.817113 |
| ENSG00000106100 | NOD1 | 331.8461878 | -0.148947723 | 0.140577542 | -1.07285194 | 0.28333755 | 0.481141 |
| ENSG00000268030 | | 74.71785 | -0.1489694 | 0.2772878 | -0.5763408 | 0.5643849 | 0.739624 |
| ENSG00000131043 | AAR2 | 2602.502248 | -0.149049849 | 0.146968653 | -1.036657738 | 0.2998954 | 0.499181 |
| ENSG00000135046 | ANXA1 | 208.2609782 | -0.14905792 | 0.593833927 | -0.381845647 | 0.70257586 | 0.838088 |
| ENSG00000186687 | LYRM7 | 929.8742 | -0.1490932 | 0.2119541 | -0.7378377 | 0.4606131 | 0.655647 |
| ENSG00000140264 | SERF2 | 8447.631 | -0.1491439 | 0.1615124 | -0.9700831 | 0.3320051 | 0.53309 |
| ENSG00000263098 | | 1.249446 | -0.1491559 | 0.7302512 | -0.4904162 | 0.6238394 | NA |
| ENSG00000164796 | CSMD3 | 1.379205 | -0.1491943 | 0.7311288 | -0.4948605 | 0.6206986 | NA |
| ENSG00000178927 | CYBC1 | 5817.238 | -0.149236 | 0.1694722 | -0.9057438 | 0.3650715 | 0.566956 |
| ENSG00000136450 | SRSF1 | 6009.549833 | -0.149260382 | 0.163298503 | -0.926071448 | 0.35440885 | 0.556354 |
| ENSG00000136169 | SETDB2 | 645.9270344 | -0.149290021 | 0.203997531 | -0.778488209 | 0.43628126 | 0.634848 |
| ENSG00000232927 | USP12PY | 3.622165 | -0.1492929 | 0.6520746 | -0.4001025 | 0.689081 | 0.828763 |
| ENSG00000231079 | KIF5C-AS1 | 0.8347824 | -0.1493159 | 0.7608114 | -0.5813082 | 0.5610327 | NA |

| | | | | | | | |
|---|---|---|---|---|---|---|---|
| ENSG00000101435 | CST9L | 0.896822633 | -0.149333404 | 0.80318562 | -1.172872382 | 0.24084699 | NA |
| ENSG00000134287 | ARF3 | 11625.37179 | -0.149355559 | 0.160703429 | -0.91930019 | 0.35793858 | 0.559877 |
| ENSG00000262470 | TVP23CP2 | 7.498524 | -0.1494351 | 0.610519 | -0.3823148 | 0.7022279 | 0.837786 |
| ENSG00000221491 | SNORA2C | 2.459287 | -0.1494597 | 0.6638362 | -0.400546 | 0.6887544 | 0.828482 |
| ENSG00000271707 | ATP1B3P1 | 22.99606 | -0.1494696 | 0.3612057 | -0.4648302 | 0.6420531 | 0.795852 |
| ENSG00000134809 | TIMM10 | 1688.722409 | -0.149477026 | 0.292717425 | -0.554140323 | 0.57948282 | 0.750599 |
| ENSG00000254016 | ALG1L10P | 5.361567 | -0.1495457 | 0.5776862 | -0.376285 | 0.706705 | 0.840598 |
| ENSG00000012211 | PRICKLE3 | 960.0259838 | -0.149553903 | 0.174473993 | -0.870808043 | 0.38385897 | 0.586259 |
| ENSG00000170921 | TANC2 | 14534.41 | -0.1495643 | 0.2821957 | -0.605569 | 0.544801 | 0.725409 |
| ENSG00000217801 | | 736.7789 | -0.1495869 | 0.2754572 | -0.5862283 | 0.5577221 | 0.735026 |
| ENSG00000101951 | PAGE4 | 2.274339614 | -0.149614186 | 0.794443285 | -0.429432783 | 0.6676083 | 0.814814 |
| ENSG00000253301 | LINC01606 | 1.163782 | -0.1496261 | 0.7935576 | -0.4422929 | 0.6582773 | NA |
| ENSG00000197081 | IGF2R | 10649.26 | -0.1497036 | 0.2991018 | -0.5453927 | 0.5854835 | 0.755278 |
| ENSG00000198633 | ZNF534 | 8.50871 | -0.1497901 | 0.5277467 | -0.3862959 | 0.6992775 | 0.835955 |
| ENSG00000139517 | LNX2 | 1982.35 | -0.1498007 | 0.2115151 | -0.7383033 | 0.4603302 | 0.655647 |
| ENSG00000274494 | MIR6832 | 3.66025 | -0.1498498 | 0.6455114 | -0.4006747 | 0.6886597 | 0.828482 |
| ENSG00000232034 | | 0.8520884 | -0.149866 | 0.7924425 | -0.4617749 | 0.6442428 | NA |
| ENSG00000188747 | NOXA1 | 1765.886 | -0.1499603 | 0.277843 | -0.5739273 | 0.566017 | 0.74074 |
| ENSG00000167202 | TBC1D2B | 2762.708 | -0.1499696 | 0.1728468 | -0.8942959 | 0.3711636 | 0.573273 |
| ENSG00000229894 | GK3P | 11.99366 | -0.1499879 | 0.6121484 | -0.3868265 | 0.6988847 | 0.835749 |
| ENSG00000158234 | FAIM | 799.7612 | -0.1501767 | 0.229207 | -0.6946645 | 0.4872656 | 0.6782 |
| ENSG00000260545 | | 1.982523 | -0.1501924 | 0.7143826 | -0.4517596 | 0.6514422 | 0.803115 |
| ENSG00000273733 | | 5.867793 | -0.150193 | 0.5310873 | -0.383659 | 0.7012312 | 0.83727 |
| ENSG00000180869 | LINC01555 | 1.246637 | -0.1502701 | 0.7361381 | -0.4951664 | 0.6204826 | NA |
| ENSG00000231747 | | 19.33046 | -0.1503015 | 0.4300369 | -0.4176559 | 0.6761987 | 0.820059 |
| ENSG00000245059 | | 3.3631 | -0.1505048 | 0.6608626 | -0.4014425 | 0.6880944 | 0.828129 |
| ENSG00000137441 | FGFBP2 | 26.33364512 | -0.150552509 | 0.564147337 | -0.392714844 | 0.69453011 | 0.832535 |
| ENSG00000251609 | SETP12 | 2.47115 | -0.1505969 | 0.7061451 | -0.4418357 | 0.6586081 | 0.807952 |
| ENSG00000104872 | PIH1D1 | 3879.053317 | -0.150609722 | 0.21951094 | -0.71948461 | 0.47184238 | 0.665025 |
| ENSG00000140632 | GLYR1 | 7920.744 | -0.1506326 | 0.1130331 | -1.348107 | 0.1776238 | 0.350937 |
| ENSG00000251191 | LINC00589 | 6.025692 | -0.1506756 | 0.691136 | -0.4234829 | 0.671943 | 0.817626 |
| ENSG00000258788 | CKS1BP1 | 9.891307 | -0.150795 | 0.5062104 | -0.391936 | 0.6951055 | 0.832992 |
| ENSG00000236530 | KPNA2P1 | 6.276275 | -0.1509015 | 0.6475196 | -0.3983922 | 0.6903411 | 0.829715 |

| | | | | | | | |
|---|---|---|---|---|---|---|---|
| ENSG00000213442 | RPL18AP3 | 12417.6 | -0.1509139 | 0.2059944 | -0.7714154 | 0.4404608 | 0.638603 |
| ENSG00000126785 | RHOJ | 2.03112439 | -0.150916151 | 0.792586169 | -0.463133596 | 0.64326861 | 0.79668 |
| ENSG00000167664 | TMIGD2 | 0.9608705 | -0.1510063 | 0.7948679 | -0.4271085 | 0.6693003 | NA |
| ENSG00000286682 | | 18.12247 | -0.1510285 | 0.3839706 | -0.4498297 | 0.6528332 | 0.804114 |
| ENSG00000147419 | CCDC25 | 2099.06 | -0.1510357 | 0.1802285 | -0.8655362 | 0.3867446 | 0.589119 |
| ENSG00000206418 | RAB12 | 2017.294 | -0.1510374 | 0.1214837 | -1.250622 | 0.2110724 | 0.394956 |
| ENSG00000152475 | ZNF837 | 305.9446 | -0.1510471 | 0.280691 | -0.5807192 | 0.5614297 | 0.73741 |
| ENSG00000066651 | TRMT11 | 799.5284291 | -0.151130471 | 0.169252119 | -0.918659913 | 0.35827348 | 0.560203 |
| ENSG00000257875 | | 1.096861 | -0.1511704 | 0.7688394 | -0.6506365 | 0.5152812 | NA |
| ENSG00000267546 | | 24.4453 | -0.1512119 | 0.5839961 | -0.385164 | 0.7001159 | 0.836581 |
| ENSG00000104904 | OAZ1 | 13105.64893 | -0.151508396 | 0.194963617 | -0.81981192 | 0.41232333 | 0.612526 |
| ENSG00000006025 | OSBPL7 | 3522.881144 | -0.151525866 | 0.211998549 | -0.756861355 | 0.44913293 | 0.645477 |
| ENSG00000228526 | MIR34AHG | 664.1409 | -0.1515681 | 0.4128417 | -0.4456963 | 0.6558166 | 0.805976 |
| ENSG00000151229 | SLC2A13 | 408.981 | -0.1516401 | 0.2606881 | -0.6227838 | 0.5334266 | 0.717307 |
| ENSG00000259211 | | 5.170333 | -0.1516481 | 0.5890279 | -0.3844639 | 0.7006346 | 0.836972 |
| ENSG00000214380 | NFYBP1 | 8.347247 | -0.1516768 | 0.5294451 | -0.3831507 | 0.701608 | 0.837385 |
| ENSG00000273035 | | 0.9514567 | -0.1517725 | 0.7443896 | -0.5542306 | 0.579421 | NA |
| ENSG00000140854 | KATNB1 | 1685.528 | -0.151816 | 0.2088177 | -0.7627302 | 0.4456243 | 0.642664 |
| ENSG00000250727 | | 0.8052331 | -0.151831 | 0.771871 | -0.6361072 | 0.5247066 | NA |
| ENSG00000187838 | PLSCR3 | 39.85097 | -0.1518372 | 0.3264338 | -0.5085192 | 0.6110893 | 0.774324 |
| ENSG00000029363 | BCLAF1 | 8911.809544 | -0.151839872 | 0.145380761 | -1.080591359 | 0.27987893 | 0.477163 |
| ENSG00000264741 | MIR4505 | 1.038325 | -0.1518732 | 0.7698431 | -0.6271195 | 0.5305809 | NA |
| ENSG00000169764 | UGP2 | 5564.578 | -0.1518927 | 0.1527927 | -1.008405 | 0.3132601 | 0.514401 |
| ENSG00000260206 | | 0.8552846 | -0.1518989 | 0.7460142 | -0.5347226 | 0.5928417 | NA |
| ENSG00000286141 | | 0.9558358 | -0.1519512 | 0.8051383 | -1.197787 | 0.231 | NA |
| ENSG00000157823 | AP3S2 | 1351.369 | -0.1521542 | 0.1662822 | -0.9655232 | 0.3342828 | 0.535452 |
| ENSG00000188487 | INSC | 0.8127743 | -0.1522457 | 0.7784445 | -0.6752633 | 0.4995085 | NA |
| ENSG00000247131 | | 4.905358 | -0.1523622 | 0.6585493 | -0.4029789 | 0.6869637 | 0.827607 |
| ENSG00000251229 | | 2.005737 | -0.1523689 | 0.7079062 | -0.4542733 | 0.6496321 | 0.801899 |
| ENSG00000249955 | | 1.152772 | -0.1524089 | 0.741683 | -0.5165215 | 0.6054902 | NA |
| ENSG00000272275 | | 13.38449 | -0.1524273 | 0.4828605 | -0.3981968 | 0.6904852 | 0.829738 |
| ENSG00000078808 | SDF4 | 5712.874277 | -0.152644226 | 0.127752855 | -1.231699134 | 0.21806149 | 0.403718 |
| ENSG00000283959 | | 52.29233 | -0.1527218 | 0.3444447 | -0.4946331 | 0.6208591 | 0.781011 |

| | | | | | | | |
|---|---|---|---|---|---|---|---|
| ENSG00000285967 | NIPBL-DT | 1330.879 | -0.1527631 | 0.2421303 | -0.6671975 | 0.504646 | 0.691523 |
| ENSG00000253433 | | 5.630766 | -0.1528021 | 0.6508287 | -0.4019951 | 0.6876876 | 0.828024 |
| ENSG00000114520 | SNX4 | 3404.114472 | -0.152846895 | 0.118939416 | -1.310840676 | 0.1899116 | 0.36766 |
| ENSG00000067141 | NEO1 | 5047.222332 | -0.152867509 | 0.199740166 | -0.800910237 | 0.42318361 | 0.623154 |
| ENSG00000173914 | RBM4B | 2283.257 | -0.1528851 | 0.184372 | -0.8443015 | 0.3985009 | 0.599534 |
| ENSG00000111325 | OGFOD2 | 160.1196151 | -0.152898151 | 0.201693605 | -0.784229869 | 0.43290524 | 0.63204 |
| ENSG00000101333 | PLCB4 | 1961.539043 | -0.152937575 | 0.216111102 | -0.740222356 | 0.45916508 | 0.654764 |
| ENSG00000007392 | LUC7L | 5139.11946 | -0.153005625 | 0.201114968 | -0.790289878 | 0.4293585 | 0.628579 |
| ENSG00000157368 | IL34 | 1.401696 | -0.1532349 | 0.7328868 | -0.502843 | 0.6150746 | NA |
| ENSG00000234371 | RPSAP31 | 1.57952 | -0.1533417 | 0.709956 | -0.4532522 | 0.6503672 | 0.802434 |
| ENSG00000262500 | MAPK8IP1P1 | 1.209613 | -0.1533449 | 0.7714597 | -0.6472035 | 0.5175002 | NA |
| ENSG00000237804 | HNRNPA1P39 | 1.411858 | -0.1533785 | 0.7354572 | -0.5123044 | 0.608438 | NA |
| ENSG00000243701 | DUBR | 345.2937 | -0.1533977 | 0.2978575 | -0.5628693 | 0.5735239 | 0.746593 |
| ENSG00000154451 | GBP5 | 0.9478511 | -0.1534153 | 0.7604859 | -0.5873328 | 0.5569802 | NA |
| ENSG00000198380 | GFPT1 | 5542.305 | -0.1534245 | 0.1877658 | -0.8022752 | 0.4223938 | 0.622405 |
| ENSG00000144802 | NFKBIZ | 1267.611 | -0.1535404 | 0.3504866 | -0.4800031 | 0.6312252 | 0.788593 |
| ENSG00000286878 | | 1.712979 | -0.1535651 | 0.7094164 | -0.4585188 | 0.6465798 | 0.799394 |
| ENSG00000214141 | ACTN4P2 | 1.226553 | -0.1535775 | 0.7317744 | -0.512159 | 0.6085398 | NA |
| ENSG00000121057 | AKAP1 | 10117.19145 | -0.153698722 | 0.177644331 | -0.898301209 | 0.36902499 | 0.571164 |
| ENSG00000260025 | CRIM1-DT | 120.9066 | -0.1537291 | 0.3394233 | -0.4994859 | 0.6174371 | 0.77871 |
| ENSG00000145741 | BTF3 | 22370.53 | -0.1537683 | 0.143662 | -1.100007 | 0.2713293 | 0.46771 |
| ENSG00000223803 | RPS20P14 | 1536.986 | -0.1537796 | 0.1681631 | -0.9302272 | 0.3522534 | 0.554018 |
| ENSG00000158553 | POM121L2 | 1.315105 | -0.1538996 | 0.7544943 | -0.5651395 | 0.5719789 | NA |
| ENSG00000079277 | MKNK1 | 1584.697543 | -0.153984779 | 0.145163933 | -1.06878188 | 0.28516797 | 0.483186 |
| ENSG00000230844 | ZNF674-AS1 | 332.6906 | -0.153989 | 0.1974502 | -0.8063119 | 0.420063 | 0.620036 |
| ENSG00000137478 | FCHSD2 | 1585.109165 | -0.153999574 | 0.122611979 | -1.265677632 | 0.20562852 | 0.387609 |
| ENSG00000072832 | CRMP1 | 0.840595118 | -0.154013913 | 0.794658449 | -0.918327711 | 0.35844732 | NA |
| ENSG00000169718 | DUS1L | 11269.32 | -0.1541106 | 0.1601919 | -0.9915032 | 0.3214399 | 0.522216 |
| ENSG00000178636 | | 3.607201 | -0.1541114 | 0.6180095 | -0.4019023 | 0.6877559 | 0.828047 |
| ENSG00000185495 | SEPTIN7P13 | 272.3511 | -0.154221 | 0.3291383 | -0.5241149 | 0.6001986 | 0.766361 |
| ENSG00000148737 | TCF7L2 | 547.5085 | -0.1542621 | 0.3015692 | -0.5593167 | 0.5759456 | 0.748023 |
| ENSG00000163093 | BBS5 | 367.5369 | -0.154431 | 0.1971644 | -0.8138197 | 0.4157482 | 0.61561 |
| ENSG00000254431 | | 1.136713 | -0.1544369 | 0.7341677 | -0.5020245 | 0.6156503 | NA |

| | | | | | | | |
|---|---|---|---|---|---|---|---|
| ENSG00000200677 | | 1.621716 | -0.1545072 | 0.7138218 | -0.4809725 | 0.6305361 | 0.788014 |
| ENSG00000244036 | | 0.8272319 | -0.1545141 | 0.7556895 | -0.5915104 | 0.5541785 | NA |
| ENSG00000178038 | ALS2CL | 1063.338 | -0.1545174 | 0.3568178 | -0.5023977 | 0.6153878 | 0.777103 |
| ENSG00000272473 | | 38.71057 | -0.1546107 | 0.4227739 | -0.433161 | 0.6648978 | 0.812787 |
| ENSG00000238832 | | 4.59355 | -0.1546248 | 0.6968755 | -0.4332885 | 0.6648052 | 0.812764 |
| ENSG00000231707 | PABPC1P1 | 7.069453 | -0.1546465 | 0.5390831 | -0.3949251 | 0.6928981 | 0.831284 |
| ENSG00000113971 | NPHP3 | 511.043715 | -0.154656939 | 0.253018406 | -0.658951949 | 0.50992663 | 0.695852 |
| ENSG00000280070 | | 2.936054 | -0.1546786 | 0.6649414 | -0.4185089 | 0.6755751 | 0.819667 |
| ENSG00000259138 | | 1.556354 | -0.1547315 | 0.7311726 | -0.4889551 | 0.6248735 | NA |
| ENSG00000255185 | PDXDC2P | 2.674958 | -0.1548546 | 0.6273673 | -0.4018845 | 0.687769 | 0.828047 |
| ENSG00000153936 | HS2ST1 | 2670.048 | -0.1548836 | 0.1572027 | -0.9797818 | 0.3271939 | 0.528527 |
| ENSG00000145050 | MANF | 2296.086 | -0.1549136 | 0.2371014 | -0.6885698 | 0.4910941 | 0.681494 |
| ENSG00000161940 | BCL6B | 1.448168 | -0.1549387 | 0.7102867 | -0.4612324 | 0.6446318 | NA |
| ENSG00000234664 | HMGN2P5 | 2140.308 | -0.1549389 | 0.2892115 | -0.5751636 | 0.5651806 | 0.7402 |
| ENSG00000105699 | LSR | 6711.783512 | -0.154949649 | 0.179771821 | -0.880542643 | 0.37856542 | 0.580722 |
| ENSG00000196933 | RPS26P11 | 0.9007722 | -0.1549538 | 0.7826088 | -0.7599561 | 0.4472808 | NA |
| ENSG00000184014 | DENND5A | 2502.123 | -0.1549602 | 0.1769656 | -0.8990279 | 0.3686378 | 0.570732 |
| ENSG00000272940 | | 6.620795 | -0.1549957 | 0.5566069 | -0.3934791 | 0.6939657 | 0.832151 |
| ENSG00000224616 | RTCA-AS1 | 77.53997 | -0.1550286 | 0.456951 | -0.4214483 | 0.6734278 | 0.818533 |
| ENSG00000066135 | KDM4A | 2731.111941 | -0.155065386 | 0.186918905 | -0.849315709 | 0.39570564 | 0.597249 |
| ENSG00000204188 | GGNBP1 | 1.684722 | -0.1552596 | 0.6716505 | -0.4382181 | 0.6612282 | 0.809705 |
| ENSG00000259209 | | 13.77724 | -0.1552644 | 0.5280369 | -0.3996166 | 0.6894389 | 0.828952 |
| ENSG00000267308 | LINC01764 | 1.089073 | -0.1552981 | 0.7558879 | -0.5587995 | 0.5762986 | NA |
| ENSG00000184795 | UNC93B5 | 0.8795937 | -0.1553059 | 0.7932972 | -0.4680115 | 0.6397764 | NA |
| ENSG00000259956 | RBM15B | 3805.99 | -0.1553966 | 0.1665763 | -0.9566944 | 0.3387215 | 0.54001 |
| ENSG00000198354 | DCAF12L2 | 88.70753 | -0.1553992 | 0.3748722 | -0.4736064 | 0.6357806 | 0.791605 |
| ENSG00000234493 | RHOXF1P1 | 5.410706 | -0.1554585 | 0.7554638 | -0.5732083 | 0.5665037 | 0.741122 |
| ENSG00000225918 | RPL7P59 | 1.307066 | -0.1554942 | 0.7302605 | -0.4916507 | 0.6229663 | NA |
| ENSG00000224944 | CASC6 | 0.937446 | -0.1555002 | 0.796048 | -0.4246658 | 0.6710803 | NA |
| ENSG00000153714 | LURAP1L | 935.4185 | -0.1555239 | 0.3831212 | -0.4745064 | 0.6351388 | 0.791166 |
| ENSG00000167315 | ACAA2 | 3731.414 | -0.1556244 | 0.3146858 | -0.5437576 | 0.5866083 | 0.756284 |
| ENSG00000228474 | OST4 | 3653.017 | -0.1556795 | 0.1729511 | -0.9352258 | 0.349672 | 0.551328 |
| ENSG00000243176 | | 142.6795 | -0.1557183 | 0.1989434 | -0.8102038 | 0.417823 | 0.617591 |

| | | | | | | | |
|---|---|---|---|---|---|---|---|
| ENSG00000261236 | BOP1 | 7343.96 | -0.1557609 | 0.1499051 | -1.074151 | 0.2827549 | 0.480586 |
| ENSG00000267127 | | 41.4063 | -0.1557893 | 0.4845443 | -0.4088896 | 0.6826207 | 0.824351 |
| ENSG00000231355 | | 0.9226621 | -0.1557899 | 0.7353958 | -0.5171856 | 0.6050266 | NA |
| ENSG00000267769 | | 13.03589 | -0.1558106 | 0.4859776 | -0.4055743 | 0.6850554 | 0.82639 |
| ENSG00000180389 | ATP5F1EP2 | 352.0522 | -0.1558246 | 0.2678068 | -0.6280648 | 0.5299615 | 0.714449 |
| ENSG00000175147 | TMEM51-AS1 | 79.96703 | -0.1559278 | 0.3855426 | -0.4717121 | 0.6371323 | 0.792676 |
| ENSG00000154478 | GPR26 | 1.488717 | -0.155991 | 0.7435187 | -0.5726564 | 0.5668774 | NA |
| ENSG00000174780 | SRP72 | 9397.177 | -0.156047 | 0.1736806 | -0.9341696 | 0.3502164 | 0.551925 |
| ENSG00000175591 | P2RY2 | 1740.996 | -0.1560543 | 0.1895086 | -0.8564093 | 0.3917714 | 0.593601 |
| ENSG00000266296 | ARIH2P1 | 90.4246 | -0.156068 | 0.2260681 | -0.7226146 | 0.4699167 | 0.663331 |
| ENSG00000232811 | | 63.55009 | -0.1560862 | 0.5119061 | -0.4051029 | 0.6854019 | 0.826558 |
| ENSG00000167863 | ATP5PD | 5054.968 | -0.1561408 | 0.2059039 | -0.785686 | 0.4320515 | 0.631216 |
| ENSG00000249915 | PDCD6 | 1614.552 | -0.1562504 | 0.194121 | -0.8334261 | 0.4046044 | 0.605426 |
| ENSG00000171722 | SPATA46 | 26.64754 | -0.1562955 | 0.4817663 | -0.4140717 | 0.6788216 | 0.821518 |
| ENSG00000198646 | NCOA6 | 3704.528 | -0.1563093 | 0.1876436 | -0.8576457 | 0.3910881 | 0.592914 |
| ENSG00000125875 | TBC1D20 | 2685.728994 | -0.156310621 | 0.227361032 | -0.722734451 | 0.46984305 | 0.663322 |
| ENSG00000198712 | MT-CO2 | 95518.55 | -0.1563827 | 0.1561176 | -1.163075 | 0.244799 | 0.436422 |
| ENSG00000163939 | PBRM1 | 2600.802 | -0.1564161 | 0.2244023 | -0.7469236 | 0.4551097 | 0.651112 |
| ENSG00000246067 | RAB30-DT | 433.207 | -0.1564524 | 0.2496295 | -0.6668944 | 0.5048396 | 0.691681 |
| ENSG00000169100 | SLC25A6 | 23786.74 | -0.1565723 | 0.1995428 | -0.814585 | 0.4153099 | 0.615174 |
| ENSG00000243445 | | 14.92388 | -0.1566594 | 0.4094964 | -0.4482594 | 0.653966 | 0.804895 |
| ENSG00000224413 | | 1.072674 | -0.156684 | 0.7755203 | -0.7113587 | 0.476862 | NA |
| ENSG00000285825 | CCDC15-DT | 28.57731 | -0.1567176 | 0.485583 | -0.4108078 | 0.6812135 | 0.823598 |
| ENSG00000220924 | OSTCP4 | 71.56286 | -0.1568545 | 0.2386191 | -0.6937422 | 0.4878439 | 0.678827 |
| ENSG00000163472 | TMEM79 | 1047.713 | -0.1568688 | 0.1553205 | -1.029495 | 0.3032471 | 0.502869 |
| ENSG00000136449 | MYCBPAP | 5.562747726 | -0.156935878 | 0.567855239 | -0.394645823 | 0.69310429 | 0.831374 |
| ENSG00000279716 | | 89.16639 | -0.1569584 | 0.3709157 | -0.4838305 | 0.6285062 | 0.786528 |
| ENSG00000198130 | HIBCH | 1474.981 | -0.1569687 | 0.2194257 | -0.7542577 | 0.4506945 | 0.646956 |
| ENSG00000163611 | SPICE1 | 219.2099 | -0.1569761 | 0.2177865 | -0.7531066 | 0.4513859 | 0.647668 |
| ENSG00000224897 | POT1-AS1 | 311.142 | -0.1569924 | 0.3553864 | -0.4976039 | 0.6187633 | 0.779513 |
| ENSG00000156009 | MAGEA8 | 4.706088 | -0.1570077 | 0.6723133 | -0.432906 | 0.665083 | 0.81289 |
| ENSG00000280832 | GSEC | 171.1054 | -0.1570901 | 0.2745845 | -0.6173084 | 0.5370313 | 0.720111 |
| ENSG00000122565 | CBX3 | 13338.46685 | -0.157094682 | 0.22567866 | -0.730216559 | 0.46525782 | 0.659206 |

| | | | | | | | |
|---|---|---|---|---|---|---|---|
| ENSG00000229539 | PANK2-AS1 | 22.26516 | -0.1571529 | 0.3693884 | -0.4820792 | 0.6297497 | 0.78738 |
| ENSG00000155761 | SPAG17 | 38.65604 | -0.157228 | 0.4780329 | -0.417119 | 0.6765913 | 0.820122 |
| ENSG00000074416 | MGLL | 1228.091274 | -0.157228425 | 0.328105836 | -0.524880663 | 0.59966615 | 0.765998 |
| ENSG00000141933 | TPGS1 | 267.4689 | -0.1573182 | 0.2241199 | -0.7391775 | 0.4597993 | 0.655274 |
| ENSG00000196502 | SULT1A1 | 748.1256 | -0.1573394 | 0.2436035 | -0.6829919 | 0.494612 | 0.684079 |
| ENSG00000160868 | CYP3A4 | 1.815347 | -0.1573954 | 0.7275202 | -0.5020474 | 0.6156342 | 0.77734 |
| ENSG00000138459 | SLC35A5 | 1067.891622 | -0.157461373 | 0.165457852 | -0.975859224 | 0.32913423 | 0.530275 |
| ENSG00000153234 | NR4A2 | 612.8082 | -0.1574732 | 0.4643117 | -0.4358411 | 0.662952 | 0.811255 |
| ENSG00000214881 | TMEM14DP | 379.7928 | -0.1575048 | 0.2173407 | -0.7591501 | 0.4477628 | 0.644436 |
| ENSG00000142230 | SAE1 | 10023.53 | -0.1575446 | 0.2551908 | -0.656476 | 0.5115179 | 0.697486 |
| ENSG00000254759 | NAP1L1P1 | 134.2308 | -0.1575534 | 0.1947352 | -0.8375705 | 0.402272 | 0.60326 |
| ENSG00000231424 | | 4.207048 | -0.1575548 | 0.5684557 | -0.3982669 | 0.6904335 | 0.829738 |
| ENSG00000142039 | CCDC97 | 2186.178 | -0.1575774 | 0.1154887 | -1.382716 | 0.1667521 | 0.33597 |
| ENSG00000262953 | EIF4A1P9 | 1.205881 | -0.1575948 | 0.7840576 | -0.7585053 | 0.4481485 | NA |
| ENSG00000250906 | | 5.444728 | -0.1576166 | 0.6550756 | -0.4267845 | 0.6695363 | 0.815897 |
| ENSG00000119314 | PTBP3 | 7319.102617 | -0.157654013 | 0.173427562 | -0.958945032 | 0.33758644 | 0.538893 |
| ENSG00000164038 | SLC9B2 | 640.497 | -0.1577616 | 0.2440387 | -0.679832 | 0.4966108 | 0.685505 |
| ENSG00000102595 | UGGT2 | 1072.804741 | -0.157763878 | 0.225713285 | -0.71991896 | 0.47157489 | 0.664789 |
| ENSG00000248498 | ASNSP1 | 259.3771 | -0.1578097 | 0.6220645 | -0.4151757 | 0.6780133 | 0.820998 |
| ENSG00000284934 | | 1.013448 | -0.1578691 | 0.7935804 | -0.4739863 | 0.6355097 | NA |
| ENSG00000258920 | FOXN3-AS1 | 172.4898 | -0.1579018 | 0.2195451 | -0.7509528 | 0.452681 | 0.648791 |
| ENSG00000257497 | | 72.12511 | -0.1579224 | 0.2785343 | -0.6076747 | 0.5434032 | 0.724367 |
| ENSG00000183647 | ZNF530 | 301.1472 | -0.1579588 | 0.3078341 | -0.5580789 | 0.5767905 | 0.748717 |
| ENSG00000156475 | PPP2R2B | 303.3992 | -0.1579924 | 0.5680525 | -0.4077749 | 0.683439 | 0.824859 |
| ENSG00000234268 | | 18.2985 | -0.1580114 | 0.3890418 | -0.4651514 | 0.6418231 | 0.795751 |
| ENSG00000226548 | | 1.217469 | -0.1580375 | 0.7464436 | -0.5622864 | 0.5739209 | NA |
| ENSG00000239774 | | 2.961035 | -0.158153 | 0.6911105 | -0.4568746 | 0.6477612 | 0.800296 |
| ENSG00000144580 | CNOT9 | 3674.863 | -0.1581725 | 0.1532199 | -1.051998 | 0.2928006 | 0.491715 |
| ENSG00000186628 | FSD2 | 4.495328 | -0.1582362 | 0.6246479 | -0.4054887 | 0.6851183 | 0.826428 |
| ENSG00000280169 | | 1.382476 | -0.1583926 | 0.7150534 | -0.4864967 | 0.626615 | NA |
| ENSG00000138231 | DBR1 | 1224.147353 | -0.158408291 | 0.208068985 | -0.789316439 | 0.42992708 | 0.629213 |
| ENSG00000242457 | RBBP4P2 | 251.4852 | -0.1584903 | 0.1795036 | -0.9124886 | 0.3615115 | 0.563475 |
| ENSG00000138182 | KIF20B | 2247.740491 | -0.158504949 | 0.391464917 | -0.444697207 | 0.65653859 | 0.806416 |

| | | | | | | | |
|---|---|---|---|---|---|---|---|
| ENSG00000258429 | PDF | 243.1764 | -0.1585064 | 0.215045 | -0.7768564 | 0.4372435 | 0.635684 |
| ENSG00000273437 | | 16.23508 | -0.158518 | 0.5313565 | -0.4018346 | 0.6878057 | 0.828053 |
| ENSG00000235010 | | 1.230728 | -0.1585184 | 0.7243532 | -0.5069129 | 0.6122159 | NA |
| ENSG00000104331 | BPNT2 | 5735.907214 | -0.158528601 | 0.145028034 | -1.108490962 | 0.26764984 | 0.463558 |
| ENSG00000121064 | SCPEP1 | 3755.926151 | -0.158565887 | 0.242448443 | -0.701528695 | 0.48297313 | 0.674381 |
| ENSG00000264647 | | 5.20656 | -0.1585818 | 0.6398877 | -0.4102047 | 0.6816558 | 0.823808 |
| ENSG00000174225 | ARL13A | 10.57299 | -0.1586334 | 0.4725555 | -0.4177371 | 0.6761394 | 0.820028 |
| ENSG00000180479 | ZNF571 | 243.3995 | -0.1586621 | 0.2393794 | -0.7006308 | 0.4835335 | 0.674808 |
| ENSG00000235601 | BARX1-DT | 4.27014 | -0.1586881 | 0.6676177 | -0.4285714 | 0.6682351 | 0.815124 |
| ENSG00000286804 | | 1.678303 | -0.158784 | 0.694351 | -0.4487054 | 0.6536442 | 0.804795 |
| ENSG00000225292 | | 1.608217 | -0.1587886 | 0.7156555 | -0.485062 | 0.6276324 | 0.785953 |
| ENSG00000197912 | SPG7 | 10758.07 | -0.1589183 | 0.1389487 | -1.16616 | 0.2435499 | 0.434754 |
| ENSG00000278200 | LINC01971 | 1.404444 | -0.1589549 | 0.7379951 | -0.5363444 | 0.5917205 | NA |
| ENSG00000286312 | | 1.818331 | -0.1589738 | 0.7055534 | -0.4673494 | 0.6402499 | 0.794709 |
| ENSG00000135835 | KIAA1614 | 190.7979256 | -0.159005075 | 0.237871038 | -0.707962485 | 0.47896855 | 0.671011 |
| ENSG00000148700 | ADD3 | 1579.617 | -0.1590188 | 0.2603888 | -0.64898 | 0.5163513 | 0.701628 |
| ENSG00000006837 | CDKL3 | 50.88174399 | -0.159021694 | 0.367864469 | -0.489799098 | 0.62427607 | 0.783439 |
| ENSG00000165698 | SPACA9 | 246.8658 | -0.1590284 | 0.2061259 | -0.798541 | 0.4245566 | 0.624343 |
| ENSG00000235568 | NFAM1 | 3.113494 | -0.1590444 | 0.663062 | -0.4233567 | 0.672035 | 0.817667 |
| ENSG00000187837 | H1-2 | 3113.538 | -0.1590602 | 0.2272439 | -0.7204792 | 0.47123 | 0.664514 |
| ENSG00000128059 | PPAT | 2377.441689 | -0.159069514 | 0.184624285 | -0.894536298 | 0.37103502 | 0.573175 |
| ENSG00000233885 | YEATS2-AS1 | 70.76628 | -0.159161 | 0.3026646 | -0.5686429 | 0.5695985 | 0.743559 |
| ENSG00000220267 | ACTBP8 | 1.154497 | -0.1592372 | 0.7952272 | -0.4512383 | 0.6518178 | NA |
| ENSG00000196570 | PFN3 | 0.9719671 | -0.1593806 | 0.7523548 | -0.5879121 | 0.5565913 | NA |
| ENSG00000168619 | ADAM18 | 0.7407731 | -0.1594444 | 0.8027272 | -1.109292 | 0.2673042 | NA |
| ENSG00000273387 | | 13.76156 | -0.1594538 | 0.4403629 | -0.4420363 | 0.6584629 | 0.807884 |
| ENSG00000252147 | RNY3P16 | 3.46068 | -0.1594557 | 0.666192 | -0.4285404 | 0.6682577 | 0.815124 |
| ENSG00000266265 | KLF14 | 1.815967 | -0.1594558 | 0.7956753 | -0.4493048 | 0.6532118 | 0.804375 |
| ENSG00000044115 | CTNNA1 | 24914.77538 | -0.159684804 | 0.099020328 | -1.590335627 | 0.11175917 | 0.252267 |
| ENSG00000167447 | SMG8 | 4654.929 | -0.1597177 | 0.1151787 | -1.418583 | 0.1560206 | 0.320344 |
| ENSG00000197889 | MEIG1 | 47.23445 | -0.1597277 | 0.3615774 | -0.4994642 | 0.6174524 | 0.77871 |
| ENSG00000055070 | SZRD1 | 4436.508091 | -0.159770206 | 0.212238775 | -0.787125456 | 0.43120843 | 0.630292 |
| ENSG00000175193 | PARL | 482.1713 | -0.159796 | 0.2843917 | -0.6231946 | 0.5331566 | 0.717129 |

| | | | | | | | |
|---|---|---|---|---|---|---|---|
| ENSG00000272918 | | 24.06446 | -0.1598151 | 0.421173 | -0.4510639 | 0.6519435 | 0.803446 |
| ENSG00000230457 | PA2G4P4 | 641.7264 | -0.1599265 | 0.2587791 | -0.6542739 | 0.5129353 | 0.698845 |
| ENSG00000199787 | SNORA80C | 1.715451 | -0.1599456 | 0.7536361 | -0.5435796 | 0.5867308 | 0.756337 |
| ENSG00000257613 | NA | 1.95221 | -0.1599502 | 0.6743371 | -0.4379046 | 0.6614554 | 0.809946 |
| ENSG00000198060 | MARCHF5 | 4683.04 | -0.1599547 | 0.0940041 | -1.762924 | 0.07791335 | 0.193321 |
| ENSG00000135587 | SMPD2 | 1003.264203 | -0.159979781 | 0.16149574 | -1.014859252 | 0.31017292 | 0.510797 |
| ENSG00000218208 | RPS27AP11 | 46.23802 | -0.1599807 | 0.3641882 | -0.4989738 | 0.6177979 | 0.778824 |
| ENSG00000106459 | NRF1 | 1195.405485 | -0.160033953 | 0.146905236 | -1.098156393 | 0.2721362 | 0.468623 |
| ENSG00000228223 | HCG11 | 594.4508 | -0.1601012 | 0.2385604 | -0.698954 | 0.4845807 | 0.675584 |
| ENSG00000008018 | PSMB1 | 11256.12685 | -0.160132729 | 0.188316031 | -0.878522825 | 0.37966006 | 0.581595 |
| ENSG00000158987 | RAPGEF6 | 545.5142 | -0.160159 | 0.1558389 | -1.059275 | 0.2894744 | 0.488042 |
| ENSG00000101306 | MYLK2 | 11.41420926 | -0.160201004 | 0.605436106 | -0.409688291 | 0.68203462 | 0.82392 |
| ENSG00000163352 | LENEP | 7.627175 | -0.1602241 | 0.5961205 | -0.4014567 | 0.6880839 | 0.828129 |
| ENSG00000253506 | NACA2 | 481.2616 | -0.1602273 | 0.1653063 | -0.9925447 | 0.3209319 | 0.52174 |
| ENSG00000158955 | WNT9B | 0.7785167 | -0.1602392 | 0.8076488 | -0.9207552 | 0.3571782 | NA |
| ENSG00000233954 | UQCRHL | 3057.795 | -0.1602394 | 0.1933991 | -0.8582903 | 0.3907322 | 0.592628 |
| ENSG00000228133 | | 1.411667 | -0.1603644 | 0.6975293 | -0.4731992 | 0.636071 | NA |
| ENSG00000287508 | | 1.365126 | -0.1605339 | 0.7222585 | -0.5160754 | 0.6058018 | NA |
| ENSG00000206384 | COL6A6 | 1.739744 | -0.1605615 | 0.703785 | -0.4744348 | 0.6351898 | 0.791166 |
| ENSG00000138115 | CYP2C8 | 67.43269672 | -0.160763706 | 0.341343499 | -0.523092544 | 0.60090985 | 0.766765 |
| ENSG00000260743 | | 3.130199 | -0.1607726 | 0.6381385 | -0.4197751 | 0.6746497 | 0.818951 |
| ENSG00000111481 | COPZ1 | 7814.603229 | -0.160890591 | 0.134137285 | -1.244069923 | 0.21347383 | 0.39828 |
| ENSG00000244471 | | 8.96373 | -0.1609208 | 0.6039484 | -0.4192255 | 0.6750513 | 0.819344 |
| ENSG00000227619 | | 2.493763 | -0.161095 | 0.7489904 | -0.5372079 | 0.591124 | 0.759788 |
| ENSG00000265345 | MIR5188 | 1.142731 | -0.1610965 | 0.7510034 | -0.562969 | 0.573456 | NA |
| ENSG00000280353 | | 1.497418 | -0.1611729 | 0.735485 | -0.53875 | 0.5900594 | NA |
| ENSG00000287186 | | 0.8648349 | -0.1612096 | 0.776517 | -0.7083941 | 0.4787005 | NA |
| ENSG00000101057 | MYBL2 | 6339.953548 | -0.161251526 | 0.60813395 | -0.424105231 | 0.67148907 | 0.817373 |
| ENSG00000200376 | RNU5E-10P | 3.381401 | -0.1613243 | 0.6735334 | -0.4385805 | 0.6609655 | 0.809437 |
| ENSG00000138303 | ASCC1 | 2092.362328 | -0.161332061 | 0.183970731 | -0.902019298 | 0.36704662 | 0.569065 |
| ENSG00000280296 | | 0.7968921 | -0.1613617 | 0.7585907 | -0.6054105 | 0.5449063 | NA |
| ENSG00000251586 | TET2-AS1 | 1.028126 | -0.1613976 | 0.774622 | -0.6903229 | 0.4899912 | NA |
| ENSG00000142945 | KIF2C | 2318.249 | -0.1614483 | 0.4539597 | -0.4392195 | 0.6605025 | 0.809118 |

| | | | | | | | |
|---|---|---|---|---|---|---|---|
| ENSG00000237588 | | 8.958574 | -0.1615319 | 0.6166867 | -0.4154323 | 0.6778255 | 0.820874 |
| ENSG00000288028 | | 1.152737 | -0.1615458 | 0.752895 | -0.6016432 | 0.5474117 | NA |
| ENSG00000236397 | DDX11L2 | 77.71613 | -0.1615697 | 0.3484147 | -0.5190638 | 0.6037163 | 0.768745 |
| ENSG00000164117 | FBXO8 | 825.5682 | -0.1616265 | 0.1574677 | -1.046239 | 0.2954506 | 0.494633 |
| ENSG00000203279 | | 11.0845 | -0.1617085 | 0.4734432 | -0.4315524 | 0.6660668 | 0.813792 |
| ENSG00000100056 | ESS2 | 1505.345351 | -0.161991078 | 0.20033241 | -0.837317456 | 0.40241415 | 0.603337 |
| ENSG00000279281 | | 1.057575 | -0.1620367 | 0.7597797 | -0.6000319 | 0.548485 | NA |
| ENSG00000178467 | P4HTM | 4061.291 | -0.1620569 | 0.1813028 | -0.922815 | 0.3561036 | 0.558137 |
| ENSG00000139351 | SYCP3 | 12.54632 | -0.1620799 | 0.5313831 | -0.4137516 | 0.679056 | 0.821652 |
| ENSG00000256898 | ARPC3P4 | 4.031915 | -0.1622171 | 0.6125043 | -0.4172446 | 0.6764995 | 0.820122 |
| ENSG00000282840 | | 2.415478 | -0.1624054 | 0.7356726 | -0.516394 | 0.6055792 | 0.769986 |
| ENSG00000136942 | RPL35 | 16547.16658 | -0.162413917 | 0.203534094 | -0.844937512 | 0.39814573 | 0.599237 |
| ENSG00000183665 | TRMT12 | 2665.968 | -0.1625278 | 0.09507281 | -1.735062 | 0.08272976 | 0.202548 |
| ENSG00000261204 | | 10.99314 | -0.1625391 | 0.472785 | -0.431341 | 0.6662205 | 0.813905 |
| ENSG00000188242 | | 14.10505 | -0.1626027 | 0.5476601 | -0.4096722 | 0.6820465 | 0.82392 |
| ENSG00000179967 | PPP1R14BP3 | 2945.857 | -0.1626435 | 0.2341965 | -0.7481897 | 0.4543457 | 0.65042 |
| ENSG00000273012 | | 2.504468 | -0.162807 | 0.7560313 | -0.5784947 | 0.5629302 | 0.738569 |
| ENSG00000180172 | RPS12P23 | 11.89438 | -0.1628826 | 0.5002343 | -0.4168768 | 0.6767685 | 0.820192 |
| ENSG00000225921 | NOL7 | 3476.15 | -0.1629015 | 0.1401948 | -1.188602 | 0.2345963 | 0.423764 |
| ENSG00000159592 | GPBP1L1 | 6958.848 | -0.1629101 | 0.1433169 | -1.16007 | 0.2460203 | 0.437723 |
| ENSG00000235795 | | 7.345931 | -0.1629461 | 0.5937267 | -0.4144657 | 0.6785331 | 0.821485 |
| ENSG00000286094 | NA | 1.509021 | -0.1631195 | 0.7189388 | -0.5153093 | 0.6063369 | NA |
| ENSG00000131475 | VPS25 | 3792.211193 | -0.16312062 | 0.227656687 | -0.75570859 | 0.44982393 | 0.64602 |
| ENSG00000258644 | SYNJ2BP-COX16 | 0.8814364 | -0.1631296 | 0.7696765 | -0.6560363 | 0.5118008 | NA |
| ENSG00000142609 | CFAP74 | 15.49356 | -0.163138 | 0.4695906 | -0.4397462 | 0.660121 | 0.808775 |
| ENSG00000172404 | DNAJB7 | 2.659935 | -0.1631741 | 0.686719 | -0.4554584 | 0.6487795 | 0.801033 |
| ENSG00000286579 | | 11.10006 | -0.1631793 | 0.5083289 | -0.4142362 | 0.6787011 | 0.821485 |
| ENSG00000235681 | | 21.16068 | -0.1632354 | 0.4634249 | -0.4372139 | 0.6619562 | 0.810374 |
| ENSG00000273872 | SLC25A15P4 | 22.16007 | -0.1632416 | 0.4294854 | -0.4564029 | 0.6481003 | 0.800529 |
| ENSG00000073734 | ABCB11 | 0.930282678 | -0.163274494 | 0.795172274 | -0.472807452 | 0.63635056 | NA |
| ENSG00000255007 | LINC02489 | 7.786238 | -0.1633452 | 0.6496092 | -0.4297071 | 0.6674087 | 0.814764 |
| ENSG00000269343 | ZNF587B | 877.6316 | -0.1634224 | 0.2675627 | -0.6525958 | 0.5140169 | 0.699924 |
| ENSG00000254701 | | 48.16364 | -0.163424 | 0.332297 | -0.5351715 | 0.5925313 | 0.760732 |

| | | | | | | | |
|---|---|---|---|---|---|---|---|
| ENSG00000172456 | FGGY | 353.0069 | -0.1634458 | 0.2553194 | -0.7050297 | 0.4807917 | 0.672359 |
| ENSG00000239556 | | 17.26453 | -0.1634509 | 0.4253063 | -0.4467001 | 0.6550916 | 0.805383 |
| ENSG00000267662 | | 1.215479 | -0.1635621 | 0.7951464 | -0.4731141 | 0.6361318 | NA |
| ENSG00000285636 | | 0.8635143 | -0.1635847 | 0.7717634 | -0.6891274 | 0.4907431 | NA |
| ENSG00000255933 | | 6.764954 | -0.1636353 | 0.5653426 | -0.4151648 | 0.6780213 | 0.820998 |
| ENSG00000271199 | | 2.049167 | -0.1636861 | 0.7204232 | -0.5064722 | 0.6125252 | 0.775293 |
| ENSG00000285768 | LINC02849 | 1.023757 | -0.1637365 | 0.755526 | -0.6012793 | 0.547654 | NA |
| ENSG00000244229 | RPL26P35 | 1.456197 | -0.1637846 | 0.7421827 | -0.5617269 | 0.5743021 | NA |
| ENSG00000275294 | LINC02340 | 4.951754 | -0.1638906 | 0.6195059 | -0.4170753 | 0.6766233 | 0.820122 |
| ENSG00000138138 | ATAD1 | 3766.269344 | -0.163948138 | 0.186256693 | -0.914819879 | 0.3602862 | 0.562291 |
| ENSG00000234902 | | 1.594976 | -0.1639638 | 0.7425445 | -0.5659231 | 0.571446 | 0.745129 |
| ENSG00000182944 | EWSR1 | 17706.36 | -0.1639699 | 0.1505827 | -1.17079 | 0.2416833 | 0.432264 |
| ENSG00000137500 | CCDC90B | 3130.558201 | -0.164026882 | 0.112360255 | -1.475639558 | 0.14004068 | 0.29619 |
| ENSG00000240471 | PHBP8 | 14.65119 | -0.1640649 | 0.4877004 | -0.429444 | 0.6676001 | 0.814814 |
| ENSG00000123384 | LRP1 | 1503.660221 | -0.164127296 | 0.2442956 | -0.725460327 | 0.46816968 | 0.661779 |
| ENSG00000131931 | THAP1 | 487.1306462 | -0.164146034 | 0.160836322 | -1.035904426 | 0.30024673 | 0.499484 |
| ENSG00000135902 | CHRND | 7.204975732 | -0.164179377 | 0.579676858 | -0.413399176 | 0.67931418 | 0.82189 |
| ENSG00000266261 | | 4.382639 | -0.1641841 | 0.5891731 | -0.4132537 | 0.6794208 | 0.821944 |
| ENSG00000111339 | ART4 | 0.988240693 | -0.164208183 | 0.801018656 | -0.645367775 | 0.51868888 | NA |
| ENSG00000173226 | IQCB1 | 2555.348 | -0.1642296 | 0.1744413 | -0.9932656 | 0.3205805 | 0.521459 |
| ENSG00000124172 | ATP5F1E | 12352.66319 | -0.164244764 | 0.190063969 | -0.893339801 | 0.37167523 | 0.573863 |
| ENSG00000228415 | PTMAP1 | 2.45582 | -0.1642554 | 0.7041628 | -0.4715727 | 0.6372318 | 0.792742 |
| ENSG00000198089 | SFI1 | 2186.588 | -0.1642746 | 0.1996759 | -0.8582702 | 0.3907433 | 0.592628 |
| ENSG00000117614 | SYF2 | 2064.56552 | -0.164276707 | 0.187074408 | -0.926502998 | 0.35418464 | 0.556199 |
| ENSG00000198873 | GRK5 | 180.4467 | -0.1642934 | 0.287881 | -0.6197445 | 0.535426 | 0.718872 |
| ENSG00000144285 | SCN1A | 1.981245 | -0.1645766 | 0.7936866 | -0.5123614 | 0.6083981 | 0.772203 |
| ENSG00000267062 | | 1.02505 | -0.1645877 | 0.7563102 | -0.6002503 | 0.5483394 | NA |
| ENSG00000163694 | RBM47 | 4506.655 | -0.1646318 | 0.1546595 | -1.081474 | 0.2794861 | 0.476799 |
| ENSG00000049130 | KITLG | 2364.61798 | -0.164698395 | 0.231538869 | -0.752280189 | 0.45188258 | 0.648206 |
| ENSG00000167216 | KATNAL2 | 287.3465 | -0.1647344 | 0.1975658 | -0.861894 | 0.3887458 | 0.590813 |
| ENSG00000161652 | IZUMO2 | 0.844093 | -0.1648058 | 0.7942113 | -0.4958842 | 0.6199761 | NA |
| ENSG00000204084 | INPP5B | 1616.268 | -0.1648445 | 0.232378 | -0.7458526 | 0.4557565 | 0.651734 |
| ENSG00000064835 | POU1F1 | 0.86080388 | -0.164870572 | 0.747342647 | -0.582369104 | 0.56031809 | NA |

| | | | | | | | |
|---|---|---|---|---|---|---|---|
| ENSG00000213493 | ACTN4P1 | 84.15345 | -0.1648805 | 0.3250992 | -0.5583876 | 0.5765798 | 0.74859 |
| ENSG00000151422 | FER | 838.394 | -0.1649275 | 0.1998499 | -0.8566849 | 0.391619 | 0.593414 |
| ENSG00000114026 | OGG1 | 1570.680855 | -0.164937072 | 0.139383855 | -1.143041597 | 0.25302132 | 0.445909 |
| ENSG00000235674 | LDHAP2 | 3.368649 | -0.16496 | 0.6697594 | -0.4480467 | 0.6541195 | 0.804895 |
| ENSG00000237436 | CAMTA1-DT | 27.15653 | -0.1649675 | 0.4698576 | -0.4295638 | 0.667513 | 0.814773 |
| ENSG00000073282 | TP63 | 204.403605 | -0.16496954 | 0.589323657 | -0.433073058 | 0.6649617 | 0.812816 |
| ENSG00000156136 | DCK | 993.3092 | -0.1649768 | 0.2727922 | -0.6492423 | 0.5161817 | 0.701434 |
| ENSG00000241464 | RPL39P38 | 3.708636 | -0.1649816 | 0.6347646 | -0.4296158 | 0.6674752 | 0.814764 |
| ENSG00000100813 | ACIN1 | 17566.65739 | -0.165000507 | 0.138870462 | -1.224544463 | 0.2207469 | 0.407076 |
| ENSG00000175455 | CCDC14 | 4327.304 | -0.1650071 | 0.1810311 | -0.9399484 | 0.3472441 | 0.54877 |
| ENSG00000279837 | | 9.297375 | -0.1650319 | 0.5864933 | -0.4174904 | 0.6763197 | 0.820059 |
| ENSG00000251574 | | 6.023676 | -0.1650693 | 0.5713714 | -0.4264713 | 0.6697645 | 0.816064 |
| ENSG00000225126 | CAMTA1-AS1 | 1.113334 | -0.1650847 | 0.7774571 | -0.7434331 | 0.4572195 | NA |
| ENSG00000275409 | | 2.695525 | -0.1651108 | 0.6481018 | -0.4468591 | 0.6549768 | 0.80534 |
| ENSG00000130723 | NA | 17516.7163 | -0.165137115 | 0.228327387 | -0.75931764 | 0.44766257 | 0.644412 |
| ENSG00000254719 | | 281.9084 | -0.1651671 | 0.21659 | -0.8065615 | 0.4199192 | 0.619962 |
| ENSG00000270706 | PRMT1P1 | 4.484481 | -0.1651769 | 0.6317494 | -0.4279469 | 0.6686898 | 0.815427 |
| ENSG00000273369 | | 39.70793 | -0.1653727 | 0.3406228 | -0.5432334 | 0.5869691 | 0.756545 |
| ENSG00000267858 | MZF1-AS1 | 235.1423 | -0.1654105 | 0.3155443 | -0.5751982 | 0.5651573 | 0.7402 |
| ENSG00000157510 | AFAP1L1 | 1.992055 | -0.1654512 | 0.6790897 | -0.4473389 | 0.6546304 | 0.805151 |
| ENSG00000258568 | RHOQP1 | 53.08792 | -0.1654557 | 0.27386 | -0.6474705 | 0.5173275 | 0.702631 |
| ENSG00000278845 | MRPL45 | 2496.268 | -0.1654982 | 0.1327455 | -1.271695 | 0.2034816 | 0.385119 |
| ENSG00000239912 | RPL39P36 | 9.173475 | -0.165526 | 0.4701773 | -0.4353024 | 0.663343 | 0.811584 |
| ENSG00000234118 | RPL13AP6 | 12.22919 | -0.1655399 | 0.4518441 | -0.4476301 | 0.6544202 | 0.805079 |
| ENSG00000244462 | RBM12 | 4864.03 | -0.1655808 | 0.1802403 | -0.9538276 | 0.3401709 | 0.541587 |
| ENSG00000278467 | | 20.5188 | -0.1656226 | 0.5265039 | -0.4218893 | 0.6731058 | 0.818366 |
| ENSG00000113761 | ZNF346 | 2076.842583 | -0.165661085 | 0.109540151 | -1.534449171 | 0.12491919 | 0.272969 |
| ENSG00000107537 | PHYH | 1438.497906 | -0.165687942 | 0.204158263 | -0.84028357 | 0.40074941 | 0.601724 |
| ENSG00000286978 | | 1.139156 | -0.1657183 | 0.7671066 | -0.671805 | 0.5017078 | NA |
| ENSG00000284015 | MIR1281 | 3.422311 | -0.1657817 | 0.6752503 | -0.4573493 | 0.64742 | 0.800098 |
| ENSG00000285879 | | 1.339442 | -0.1658139 | 0.7492315 | -0.5816626 | 0.560794 | NA |
| ENSG00000259098 | | 2.901986 | -0.1658973 | 0.7938835 | -0.5152664 | 0.6063668 | 0.770618 |
| ENSG00000029725 | RABEP1 | 3809.043925 | -0.165945851 | 0.160542181 | -1.044202706 | 0.29639162 | 0.49543 |

| | | | | | | | |
|---|---|---|---|---|---|---|---|
| ENSG00000227172 | | 1.869579 | -0.1660705 | 0.7274744 | -0.5094824 | 0.6104142 | 0.773875 |
| ENSG00000233540 | DNM3-IT1 | 0.7995249 | -0.1661726 | 0.7630663 | -0.6475254 | 0.5172919 | NA |
| ENSG00000264968 | | 0.9174369 | -0.1662257 | 0.7374557 | -0.5593908 | 0.575895 | NA |
| ENSG00000246430 | LINC00968 | 0.8220474 | -0.1662459 | 0.795731 | -0.4750521 | 0.6347498 | NA |
| ENSG00000214188 | ST7-OT4 | 1.141582 | -0.1663005 | 0.7586481 | -0.6214577 | 0.5342985 | NA |
| ENSG00000259363 | | 1.057777 | -0.1663861 | 0.7964363 | -0.4679835 | 0.6397964 | NA |
| ENSG00000168263 | KCNV2 | 3.057882 | -0.1664101 | 0.6723114 | -0.4580303 | 0.6469307 | 0.799791 |
| ENSG00000130764 | LRRC47 | 2833.938121 | -0.166481205 | 0.146853429 | -1.155632684 | 0.24783143 | 0.440054 |
| ENSG00000072071 | ADGRL1 | 4185.506703 | -0.166534365 | 0.145211547 | -1.206485355 | 0.2276304 | 0.41555 |
| ENSG00000104805 | NUCB1 | 6680.905325 | -0.166537989 | 0.130905311 | -1.300063181 | 0.19357932 | 0.372157 |
| ENSG00000197785 | ATAD3A | 1822.213 | -0.1665756 | 0.2481719 | -0.709147 | 0.4782333 | 0.670133 |
| ENSG00000197879 | MYO1C | 14529.16 | -0.1665828 | 0.1566299 | -1.073649 | 0.28298 | 0.480753 |
| ENSG00000171298 | GAA | 12814.09 | -0.1666033 | 0.1422176 | -1.197148 | 0.231249 | 0.419681 |
| ENSG00000231170 | | 2.271588 | -0.1666066 | 0.7518041 | -0.5637744 | 0.5729077 | 0.745937 |
| ENSG00000214144 | | 10.34066 | -0.1666572 | 0.465755 | -0.4438721 | 0.6571351 | 0.806839 |
| ENSG00000137193 | PIM1 | 1181.593354 | -0.166813627 | 0.416536349 | -0.471948638 | 0.63696345 | 0.792595 |
| ENSG00000187164 | SHTN1 | 5491.533 | -0.1668482 | 0.2166449 | -0.8249271 | 0.409413 | 0.609907 |
| ENSG00000250166 | | 1.700489 | -0.1668494 | 0.720946 | -0.4980228 | 0.6184679 | 0.779308 |
| ENSG00000109906 | ZBTB16 | 1.342537509 | -0.166908675 | 0.815747123 | -1.607885209 | 0.10786032 | NA |
| ENSG00000110931 | CAMKK2 | 4463.514947 | -0.167008793 | 0.118175774 | -1.427487325 | 0.15343948 | 0.31632 |
| ENSG00000260121 | | 327.2642 | -0.167009 | 0.205835 | -0.8450081 | 0.3981063 | 0.599212 |
| ENSG00000213394 | RPSAP46 | 1.119896 | -0.1670285 | 0.7839785 | -0.7834514 | 0.4333621 | NA |
| ENSG00000223442 | TH2LCRR | 13.12945 | -0.1670591 | 0.4844147 | -0.4403871 | 0.6596568 | 0.808573 |
| ENSG00000167397 | VKORC1 | 640.1707 | -0.167085 | 0.2223343 | -0.7775749 | 0.4368197 | 0.635253 |
| ENSG00000117399 | CDC20 | 3402.042195 | -0.167149575 | 0.519759247 | -0.429043252 | 0.66789175 | 0.814973 |
| ENSG00000162961 | DPY30 | 2729.548 | -0.1673161 | 0.242863 | -0.7256621 | 0.4680459 | 0.66168 |
| ENSG00000165632 | TAF3 | 1447.058 | -0.1673395 | 0.193594 | -0.914355 | 0.3605304 | 0.56254 |
| ENSG00000110104 | CCDC86 | 1847.285501 | -0.167405385 | 0.23099937 | -0.7725336 | 0.43979845 | 0.638246 |
| ENSG00000115233 | PSMD14 | 5490.306436 | -0.167406938 | 0.195978377 | -0.877507849 | 0.38021086 | 0.582103 |
| ENSG00000228212 | OFD1P17 | 16.52567 | -0.1674542 | 0.4740109 | -0.4447548 | 0.656497 | 0.806416 |
| ENSG00000198251 | CYP2A7P2 | 0.8517406 | -0.1674698 | 0.7732652 | -0.7055693 | 0.480456 | NA |
| ENSG00000102096 | PIM2 | 897.9834573 | -0.167481838 | 0.245018242 | -0.717245068 | 0.4732229 | 0.666087 |
| ENSG00000236358 | | 6.335779 | -0.1675118 | 0.57383 | -0.4222274 | 0.672859 | 0.818103 |

| | | | | | | | |
|---|---|---|---|---|---|---|---|
| ENSG00000241357 | | 0.9514126 | -0.1675248 | 0.7598277 | -0.6273668 | 0.5304188 | NA |
| ENSG00000121380 | BCL2L14 | 4.371109 | -0.167584148 | 0.667010467 | -0.440198415 | 0.65979341 | 0.808635 |
| ENSG00000238132 | CASC4P1 | 11.31409 | -0.1676516 | 0.506818 | -0.4330244 | 0.664997 | 0.812822 |
| ENSG00000240682 | ISY1 | 1035.61 | -0.1676889 | 0.1562854 | -1.087625 | 0.2767606 | 0.4737 |
| ENSG00000148308 | GTF3C5 | 4782.81 | -0.1677191 | 0.2275541 | -0.76765 | 0.4426952 | 0.640281 |
| ENSG00000117419 | ERI3 | 3293.831585 | -0.167733597 | 0.187659868 | -0.92623331 | 0.35432475 | 0.556288 |
| ENSG00000125122 | LRRC29 | 80.92479966 | -0.167739722 | 0.302495073 | -0.606142273 | 0.54442028 | 0.725217 |
| ENSG00000184515 | BEX5 | 123.608 | -0.1677448 | 0.2877783 | -0.633503 | 0.5264052 | 0.711324 |
| ENSG00000171451 | DSEL | 268.8088 | -0.1678714 | 0.3918043 | -0.4991719 | 0.6176583 | 0.77875 |
| ENSG00000226262 | PHKBP1 | 1.431246 | -0.1678802 | 0.7015636 | -0.4836711 | 0.6286193 | NA |
| ENSG00000167676 | PLIN4 | 272.893 | -0.1678916 | 0.3950595 | -0.4986104 | 0.6180539 | 0.778995 |
| ENSG00000165794 | SLC39A2 | 1.07003 | -0.1678996 | 0.7686713 | -0.6476318 | 0.5172231 | NA |
| ENSG00000174840 | PDE12 | 1566.481 | -0.1680338 | 0.1374068 | -1.244561 | 0.2132932 | 0.398063 |
| ENSG00000265749 | | 65.19886 | -0.1680395 | 0.3400828 | -0.5484842 | 0.5833595 | 0.753636 |
| ENSG00000019169 | MARCO | 1.162407935 | -0.16804741 | 0.795449113 | -0.493998258 | 0.62130741 | NA |
| ENSG00000173436 | MICOS10 | 353.265 | -0.1680856 | 0.1686011 | -1.02796 | 0.3039688 | 0.503658 |
| ENSG00000241973 | PI4KA | 8972.776 | -0.1681689 | 0.1653595 | -1.048151 | 0.2945691 | 0.493624 |
| ENSG00000172243 | CLEC7A | 104.706 | -0.1681734 | 0.7079428 | -0.4626922 | 0.643585 | 0.796872 |
| ENSG00000242372 | EIF6 | 3928.013 | -0.168202 | 0.2188205 | -0.8496306 | 0.3955305 | 0.597203 |
| ENSG00000207127 | | 0.853821 | -0.1682407 | 0.7970134 | -0.4645148 | 0.642279 | NA |
| ENSG00000227711 | | 1.067258 | -0.1682423 | 0.7528705 | -0.596735 | 0.5506843 | NA |
| ENSG00000239419 | RN7SL535P | 9.20801 | -0.1682624 | 0.5740504 | -0.4265751 | 0.6696889 | 0.816041 |
| ENSG00000116237 | ICMT | 4732.629517 | -0.168263754 | 0.143501258 | -1.201525164 | 0.22954755 | 0.417729 |
| ENSG00000248774 | | 4.911416 | -0.1684304 | 0.57866 | -0.4249517 | 0.6708719 | 0.816958 |
| ENSG00000173599 | PC | 2349.546 | -0.1684534 | 0.3645402 | -0.5332321 | 0.593873 | 0.761516 |
| ENSG00000203730 | TEDDM1 | 0.9224669 | -0.1684535 | 0.7853007 | -0.7557476 | 0.4498006 | NA |
| ENSG00000148110 | MFSD14B | 3184.527 | -0.1684855 | 0.1049408 | -1.622078 | 0.1047866 | 0.240671 |
| ENSG00000250959 | GLUD1P3 | 124.8264 | -0.1685025 | 0.3158756 | -0.5761028 | 0.5645457 | 0.739689 |
| ENSG00000188277 | C15orf62 | 96.89697 | -0.168524 | 0.3033575 | -0.6028189 | 0.5466292 | 0.726642 |
| ENSG00000288596 | C8orf44 | 172.814 | -0.1685889 | 0.3379121 | -0.5543384 | 0.5793472 | 0.750533 |
| ENSG00000181215 | C4orf50 | 1.004321 | -0.1687062 | 0.7641217 | -0.6545385 | 0.5127649 | NA |
| ENSG00000186010 | NDUFA13 | 79.48802 | -0.168714 | 0.2915557 | -0.6230623 | 0.5332436 | 0.717209 |
| ENSG00000227063 | RPL41P1 | 550.4868 | -0.1687317 | 0.195929 | -0.8959693 | 0.3702691 | 0.572391 |

| | | | | | | | |
|---|---|---|---|---|---|---|---|
| ENSG00000258640 | RPL21P5 | 1.202024 | -0.1687978 | 0.7342584 | -0.544336 | 0.5862103 | NA |
| ENSG00000232387 | SKA2P1 | 489.9664 | -0.1688157 | 0.2955812 | -0.6127129 | 0.5400662 | 0.722316 |
| ENSG00000144712 | CAND2 | 29.08297 | -0.1688609 | 0.3421044 | -0.5492675 | 0.5828219 | 0.753235 |
| ENSG00000225715 | PSMA1P1 | 1.606769 | -0.1688698 | 0.6982377 | -0.4788832 | 0.6320217 | 0.789069 |
| ENSG00000175336 | APOF | 1.86066 | -0.16892 | 0.7334992 | -0.5459655 | 0.5850896 | 0.75503 |
| ENSG00000228110 | ST13P19 | 247.9243 | -0.1689208 | 0.1485099 | -1.159457 | 0.2462701 | 0.438038 |
| ENSG00000075711 | DLG1 | 5066.609701 | -0.168921937 | 0.14725527 | -1.172094744 | 0.24115902 | 0.431684 |
| ENSG00000167985 | SDHAF2 | 544.3808 | -0.1689699 | 0.2722397 | -0.6608715 | 0.5086947 | 0.695011 |
| ENSG00000255154 | HTD2 | 4.059013 | -0.1689865 | 0.6510603 | -0.4473493 | 0.6546229 | 0.805151 |
| ENSG00000236867 | | 2.280017 | -0.1689886 | 0.6920844 | -0.4778734 | 0.6327403 | 0.789558 |
| ENSG00000169087 | HSPBAP1 | 433.2758 | -0.1690824 | 0.1981741 | -0.8802091 | 0.3787461 | 0.580899 |
| ENSG00000203993 | ARRDC1-AS1 | 830.1859 | -0.1691659 | 0.1718557 | -1.019847 | 0.3078008 | 0.507916 |
| ENSG00000101152 | DNAJC5 | 12358.84416 | -0.169232925 | 0.112649197 | -1.39477969 | 0.16308229 | 0.330681 |
| ENSG00000233929 | MT1XP1 | 9.939936 | -0.1692824 | 0.6555595 | -0.4564237 | 0.6480853 | 0.800529 |
| ENSG00000262408 | | 1.322197 | -0.1692868 | 0.739513 | -0.5498022 | 0.5824551 | NA |
| ENSG00000259462 | CPEB1-AS1 | 2.841312 | -0.1693159 | 0.7102267 | -0.5095335 | 0.6103783 | 0.773866 |
| ENSG00000119328 | ABITRAM | 1702.024421 | -0.169361829 | 0.13215135 | -1.3079028 | 0.19090628 | 0.368931 |
| ENSG00000158125 | XDH | 1.280449 | -0.1693771 | 0.778868 | -0.7122423 | 0.4763148 | NA |
| ENSG00000287857 | | 1.173329 | -0.1694194 | 0.7445943 | -0.5841794 | 0.5590996 | NA |
| ENSG00000133393 | CEP20 | 3845.801573 | -0.169467379 | 0.149101278 | -1.169258355 | 0.24229956 | 0.432929 |
| ENSG00000163923 | RPL39L | 748.0237 | -0.1695062 | 0.3157399 | -0.5909946 | 0.554524 | 0.732482 |
| ENSG00000271425 | NBPF10 | 699.3114 | -0.1695436 | 0.2223721 | -0.811491 | 0.4170838 | 0.617035 |
| ENSG00000270629 | NBPF14 | 1118.819 | -0.1695689 | 0.1968572 | -0.8913394 | 0.3727471 | 0.57495 |
| ENSG00000121774 | KHDRBS1 | 10132.49147 | -0.169607625 | 0.163485785 | -1.090384864 | 0.27554365 | 0.472409 |
| ENSG00000157895 | C12orf43 | 1044.522 | -0.1696198 | 0.1967277 | -0.8877639 | 0.3746678 | 0.576875 |
| ENSG00000280109 | PLAC4 | 6.859412 | -0.1696307 | 0.5874188 | -0.4242669 | 0.6713712 | 0.817342 |
| ENSG00000235076 | GAPDHP52 | 7.676092 | -0.1696409 | 0.5137129 | -0.4355518 | 0.6631619 | 0.811437 |
| ENSG00000214049 | UCA1 | 289.5069 | -0.1696769 | 0.4477634 | -0.4467595 | 0.6550487 | 0.805367 |
| ENSG00000272023 | | 4.071164 | -0.1697783 | 0.6593283 | -0.4525055 | 0.6509049 | 0.802762 |
| ENSG00000144134 | RABL2A | 556.0796 | -0.1698178 | 0.3040531 | -0.602301 | 0.5469738 | 0.726845 |
| ENSG00000170448 | NFXL1 | 1436.918 | -0.1698644 | 0.1595398 | -1.08583 | 0.2775542 | 0.474607 |
| ENSG00000271455 | | 1.148 | -0.1699145 | 0.7512759 | -0.6150745 | 0.5385056 | NA |
| ENSG00000112739 | PRPF4B | 6892.407226 | -0.169918287 | 0.162608152 | -1.083298609 | 0.2786759 | 0.475967 |

| | | | | | | | |
|---|---|---|---|---|---|---|---|
| ENSG00000204348 | DXO | 1462.081 | -0.1699232 | 0.1812686 | -0.965522 | 0.3342834 | 0.535452 |
| ENSG00000220541 | PPIHP2 | 20.44479 | -0.1699687 | 0.4413714 | -0.4679485 | 0.6398214 | 0.79435 |
| ENSG00000107140 | TESK1 | 1748.787736 | -0.170010757 | 0.173860439 | -1.005717653 | 0.31455141 | 0.515665 |
| ENSG00000088832 | FKBP1A | 4780.773435 | -0.170030067 | 0.155965048 | -1.14960391 | 0.25030705 | 0.442888 |
| ENSG00000213185 | FAM24B | 194.58 | -0.170033 | 0.2003027 | -0.870916 | 0.3838 | 0.586214 |
| ENSG00000149089 | APIP | 1127.726 | -0.1700453 | 0.1931264 | -0.9138365 | 0.3608028 | 0.562833 |
| ENSG00000169045 | HNRNPH1 | 36931.11 | -0.1700513 | 0.1715945 | -1.076964 | 0.2814962 | 0.479121 |
| ENSG00000167900 | TK1 | 5167.566 | -0.1700733 | 0.5789752 | -0.4489545 | 0.6534645 | 0.804611 |
| ENSG00000120498 | TEX11 | 1.048222887 | -0.17009199 | 0.724434502 | -0.544199701 | 0.58630406 | NA |
| ENSG00000083290 | ULK2 | 2140.220501 | -0.170110442 | 0.15213592 | -1.172562434 | 0.24097132 | 0.431455 |
| ENSG00000129538 | RNASE1 | 1.591384114 | -0.170145477 | 0.795927666 | -0.493007701 | 0.62200715 | 0.781676 |
| ENSG00000237133 | RASGRP3-AS1 | 1.150111 | -0.1702517 | 0.7563617 | -0.6215021 | 0.5342693 | NA |
| ENSG00000230330 | HMGN2P3 | 955.1407 | -0.1703324 | 0.2932752 | -0.6316589 | 0.5276098 | 0.7124 |
| ENSG00000264304 | | 18.62447 | -0.1703475 | 0.4593729 | -0.4551034 | 0.6490348 | 0.801236 |
| ENSG00000271018 | | 2.348395 | -0.1703813 | 0.7227886 | -0.520977 | 0.6023828 | 0.767619 |
| ENSG00000275562 | SEPTIN7P12 | 2.275389 | -0.1704641 | 0.6566701 | -0.4546904 | 0.649332 | 0.801566 |
| ENSG00000187559 | FOXD4L3 | 1.680982 | -0.170511 | 0.7420683 | -0.5911607 | 0.5544127 | 0.732445 |
| ENSG00000004779 | NDUFAB1 | 4368.666997 | -0.170649543 | 0.262133815 | -0.689641071 | 0.49041993 | 0.680949 |
| ENSG00000279943 | | 14.98725 | -0.1706737 | 0.5101101 | -0.4568875 | 0.6477519 | 0.800296 |
| ENSG00000241983 | RN7SL566P | 2.289003 | -0.1707298 | 0.6895854 | -0.4869932 | 0.6262631 | 0.784941 |
| ENSG00000145354 | CISD2 | 1342.823 | -0.170778 | 0.155764 | -1.121804 | 0.2619458 | 0.456718 |
| ENSG00000089685 | BIRC5 | 3414.494506 | -0.170788276 | 0.533363515 | -0.440731251 | 0.65940757 | 0.808407 |
| ENSG00000057663 | ATG5 | 2353.952062 | -0.170867186 | 0.123051533 | -1.421647718 | 0.15512855 | 0.318939 |
| ENSG00000184465 | WDR27 | 2354.073 | -0.1708675 | 0.2251273 | -0.7996668 | 0.4239038 | 0.623903 |
| ENSG00000214019 | RBM39P1 | 348.0531 | -0.1708691 | 0.188315 | -0.9352636 | 0.3496525 | 0.551328 |
| ENSG00000256625 | | 4.488262 | -0.1708747 | 0.659274 | -0.4516642 | 0.6515109 | 0.803136 |
| ENSG00000263675 | MIR5581 | 17.29188 | -0.170879 | 0.5287516 | -0.4259025 | 0.6701789 | 0.816414 |
| ENSG00000216657 | GLRX3P2 | 371.5792 | -0.1708791 | 0.2395877 | -0.7605384 | 0.4469328 | 0.643781 |
| ENSG00000130363 | RSPH3 | 516.4872887 | -0.170889678 | 0.209604943 | -0.848753808 | 0.3960183 | 0.597471 |
| ENSG00000115514 | TXNDC9 | 1868.932912 | -0.17091133 | 0.184897783 | -0.941875089 | 0.3462566 | 0.547959 |
| ENSG00000108840 | HDAC5 | 2119.614594 | -0.17094058 | 0.294105736 | -0.631786026 | 0.52752671 | 0.712333 |
| ENSG00000214941 | ZSWIM7 | 1410.814 | -0.1709494 | 0.1600697 | -1.105107 | 0.2691132 | 0.465251 |
| ENSG00000272128 | | 2.307272 | -0.1709637 | 0.6856394 | -0.4707464 | 0.6378218 | 0.793254 |

| | | | | | | | |
|---|---|---|---|---|---|---|---|
| ENSG00000064601 | CTSA | 8103.422828 | -0.170971332 | 0.228989193 | -0.783613121 | 0.43326715 | 0.632237 |
| ENSG00000260871 | | 3.724619 | -0.1709901 | 0.6815636 | -0.4735238 | 0.6358395 | 0.791641 |
| ENSG00000257452 | | 0.8805595 | -0.1710245 | 0.8012064 | -0.6233483 | 0.5330556 | NA |
| ENSG00000103035 | PSMD7 | 5035.912168 | -0.17110052 | 0.17269159 | -1.027158761 | 0.30434571 | 0.503968 |
| ENSG00000255893 | | 8.342832 | -0.1711314 | 0.5626411 | -0.4313494 | 0.6662143 | 0.813905 |
| ENSG00000271427 | | 55.62179 | -0.1711628 | 0.4022192 | -0.4956289 | 0.6201563 | 0.780601 |
| ENSG00000141076 | UTP4 | 4457.166 | -0.1711923 | 0.1900094 | -0.9073227 | 0.3642362 | 0.566364 |
| ENSG00000258355 | | 1.853433 | -0.1712138 | 0.7574351 | -0.6099101 | 0.5419214 | 0.723497 |
| ENSG00000108262 | GIT1 | 6607.469724 | -0.171229218 | 0.13063766 | -1.317919185 | 0.1875307 | 0.364271 |
| ENSG00000260884 | | 49.79483 | -0.1712691 | 0.4834256 | -0.4502658 | 0.6525188 | 0.803931 |
| ENSG00000234146 | AKAP8P1 | 1.14762 | -0.1713174 | 0.7472341 | -0.5960755 | 0.5511248 | NA |
| ENSG00000105063 | PPP6R1 | 7661.731085 | -0.171319286 | 0.157467719 | -1.119526465 | 0.26291561 | 0.457909 |
| ENSG00000186260 | MRTFB | 3893.01 | -0.1713225 | 0.2826806 | -0.6593481 | 0.5096723 | 0.695684 |
| ENSG00000166478 | ZNF143 | 831.0812 | -0.1713422 | 0.1392726 | -1.25733 | 0.2086341 | 0.391565 |
| ENSG00000203593 | | 0.9705783 | -0.1713699 | 0.7495195 | -0.5997207 | 0.5486924 | NA |
| ENSG00000141385 | AFG3L2 | 4558.3 | -0.171386 | 0.1809678 | -0.9828935 | 0.3256599 | 0.526946 |
| ENSG00000219361 | RPSAP72 | 2.393403 | -0.1714777 | 0.6756843 | -0.4603372 | 0.6452742 | 0.798207 |
| ENSG00000104901 | DKKL1 | 29.17527891 | -0.171591987 | 0.436982558 | -0.461031553 | 0.64477597 | 0.797814 |
| ENSG00000261334 | | 5.265151 | -0.1715966 | 0.6366409 | -0.442414 | 0.6581896 | 0.807659 |
| ENSG00000212371 | | 8.013227 | -0.1716196 | 0.5837998 | -0.432798 | 0.6651615 | 0.812948 |
| ENSG00000015133 | CCDC88C | 4944.433946 | -0.171649043 | 0.232221244 | -0.755076016 | 0.45020337 | 0.646466 |
| ENSG00000143153 | ATP1B1 | 6447.591 | -0.1716933 | 0.353946 | -0.5482166 | 0.5835432 | 0.753727 |
| ENSG00000075618 | FSCN1 | 2875.593496 | -0.171700958 | 0.385625758 | -0.510016502 | 0.6100399 | 0.773474 |
| ENSG00000220925 | IGBP1-AS2 | 0.9534107 | -0.1717426 | 0.7609163 | -0.6537902 | 0.513247 | NA |
| ENSG00000172716 | SLFN11 | 1.234203 | -0.1718395 | 0.7753081 | -0.7413779 | 0.4584643 | NA |
| ENSG00000028203 | VEZT | 4555.816822 | -0.171868068 | 0.124662397 | -1.366730949 | 0.17170965 | 0.342464 |
| ENSG00000257511 | | 998.9863 | -0.1718691 | 0.1494633 | -1.165204 | 0.2439365 | 0.435239 |
| ENSG00000275383 | | 3.41008 | -0.1718742 | 0.6659368 | -0.4645819 | 0.6422309 | 0.795869 |
| ENSG00000261644 | | 2.576032 | -0.1719084 | 0.6630744 | -0.45262 | 0.6508224 | 0.802697 |
| ENSG00000177868 | SVBP | 515.9916 | -0.1719451 | 0.1762887 | -1.005236 | 0.3147832 | 0.515872 |
| ENSG00000278828 | H3C10 | 1635.609 | -0.1719466 | 0.254394 | -0.7172682 | 0.4732086 | 0.666087 |
| ENSG00000253819 | LINC01151 | 1.396211 | -0.1719777 | 0.7497487 | -0.6087247 | 0.542707 | NA |
| ENSG00000267800 | | 3.747611 | -0.1720325 | 0.6970419 | -0.4862662 | 0.6267785 | 0.785291 |

| | | | | | | | |
|---|---|---|---|---|---|---|---|
| ENSG00000217539 | IQCB2P | 15.65713 | -0.1720624 | 0.4481031 | -0.4679241 | 0.6398389 | 0.79435 |
| ENSG00000088367 | EPB41L1 | 3119.455963 | -0.17209666 | 0.166478117 | -1.072702979 | 0.2834044 | 0.481197 |
| ENSG00000257758 | KRT18P20 | 20.61329 | -0.1721116 | 0.4304553 | -0.4790311 | 0.6319165 | 0.789012 |
| ENSG00000207033 | RNU6-154P | 2.550246 | -0.1721279 | 0.7186295 | -0.5361038 | 0.5918868 | 0.760374 |
| ENSG00000184281 | TSSC4 | 1793.619 | -0.1721771 | 0.2033433 | -0.8755561 | 0.3812714 | 0.583425 |
| ENSG00000264743 | DPRXP4 | 4.247519 | -0.1722649 | 0.6329149 | -0.4486348 | 0.6536951 | 0.804799 |
| ENSG00000139656 | SMIM2 | 1.227887 | -0.1722673 | 0.7575823 | -0.6333525 | 0.5265035 | NA |
| ENSG00000222022 | | 1.069946 | -0.1723237 | 0.8168229 | -1.498902 | 0.1338992 | NA |
| ENSG00000261839 | | 2.095898 | -0.1724327 | 0.7046245 | -0.4980004 | 0.6184838 | 0.779308 |
| ENSG00000231548 | OR55B1P | 0.8751828 | -0.172437 | 0.7712139 | -0.6984298 | 0.4849085 | NA |
| ENSG00000274031 | | 1.24702 | -0.1726406 | 0.751897 | -0.616007 | 0.5378899 | NA |
| ENSG00000088035 | ALG6 | 938.3141685 | -0.172650242 | 0.222950694 | -0.824995737 | 0.40937401 | 0.609907 |
| ENSG00000272810 | | 1.245393 | -0.1726563 | 0.7709697 | -0.6948042 | 0.487178 | NA |
| ENSG00000286000 | | 1.661025 | -0.1727103 | 0.6992285 | -0.4995208 | 0.6174125 | 0.77871 |
| ENSG00000130348 | QRSL1 | 1790.985244 | -0.172816263 | 0.149601327 | -1.178399117 | 0.23863753 | 0.428494 |
| ENSG00000164970 | FAM219A | 1474.189 | -0.1728281 | 0.1291046 | -1.359514 | 0.1739838 | 0.345786 |
| ENSG00000237039 | RPS28P4 | 8.231037 | -0.1728536 | 0.6147278 | -0.4420348 | 0.658464 | 0.807884 |
| ENSG00000134313 | KIDINS220 | 5283.370133 | -0.172859322 | 0.259433595 | -0.693277851 | 0.4881352 | 0.679074 |
| ENSG00000278153 | | 4.286227 | -0.1728773 | 0.6545014 | -0.4498082 | 0.6528487 | 0.804114 |
| ENSG00000270806 | C17orf50 | 4.267116 | -0.1729827 | 0.6343675 | -0.4453146 | 0.6560924 | 0.806202 |
| ENSG00000021826 | CPS1 | 1173.861282 | -0.172994191 | 0.521706786 | -0.45712264 | 0.6475829 | 0.800262 |
| ENSG00000137504 | CREBZF | 9735.941231 | -0.173089219 | 0.171114085 | -1.054403649 | 0.29169815 | 0.490456 |
| ENSG00000236816 | ANKRD20A7P | 2.23808 | -0.1731437 | 0.7971267 | -0.4896168 | 0.6244051 | 0.7835 |
| ENSG00000251370 | SEMA5A-AS1 | 0.953857 | -0.1731853 | 0.796678 | -0.4912535 | 0.6232472 | NA |
| ENSG00000234043 | NUDT9P1 | 7.85167 | -0.1733157 | 0.6087797 | -0.4399578 | 0.6599676 | 0.808721 |
| ENSG00000183617 | MRPL54 | 1149.972 | -0.173521 | 0.2453725 | -0.738908 | 0.4599628 | 0.655374 |
| ENSG00000188636 | RTL6 | 3195.417 | -0.1735708 | 0.1584232 | -1.115902 | 0.2644643 | 0.459943 |
| ENSG00000157870 | PRXL2B | 1878.365 | -0.1736672 | 0.1942602 | -0.925695 | 0.3546045 | 0.556497 |
| ENSG00000288573 | | 46.47932 | -0.1736786 | 0.3396961 | -0.5702318 | 0.5685205 | 0.742678 |
| ENSG00000203965 | EFCAB7 | 205.7528 | -0.1737043 | 0.2151714 | -0.8441681 | 0.3985755 | 0.599578 |
| ENSG00000261873 | SMIM36 | 2.051822 | -0.1737618 | 0.6893864 | -0.4885975 | 0.6251267 | 0.783989 |
| ENSG00000157554 | ERG | 18.70514 | -0.173767 | 0.6875702 | -0.4612506 | 0.6446188 | 0.79779 |
| ENSG00000273027 | | 51.86069 | -0.1737755 | 0.3638359 | -0.5436975 | 0.5866497 | 0.756284 |

| | | | | | | | |
|---|---|---|---|---|---|---|---|
| ENSG00000284490 | | 0.9965692 | -0.1737803 | 0.7745597 | -0.7101188 | 0.4776305 | NA |
| ENSG00000232401 | LINC00112 | 2.841443 | -0.1738026 | 0.7502462 | -0.5847019 | 0.5587482 | 0.735648 |
| ENSG00000110092 | CCND1 | 16698.68822 | -0.173820141 | 0.176534249 | -1.025698965 | 0.3050335 | 0.504729 |
| ENSG00000234393 | | 6.003881 | -0.1739632 | 0.65882 | -0.463124 | 0.6432755 | 0.796681 |
| ENSG00000256146 | | 1.033518 | -0.1739656 | 0.7417487 | -0.5696358 | 0.5689247 | NA |
| ENSG00000236542 | MED28P7 | 0.8141772 | -0.1740315 | 0.8091427 | -0.8761448 | 0.3809513 | NA |
| ENSG00000269925 | | 24.95861 | -0.1740649 | 0.4506888 | -0.4710802 | 0.6375835 | 0.793069 |
| ENSG00000226107 | RPL36AP54 | 9.283453 | -0.1741003 | 0.5704699 | -0.4448239 | 0.656447 | 0.806415 |
| ENSG00000111913 | RIPOR2 | 1.495561329 | -0.174325729 | 0.730094183 | -0.543098415 | 0.58706204 | NA |
| ENSG00000247765 | | 0.8635899 | -0.1743894 | 0.771024 | -0.6914386 | 0.48929 | NA |
| ENSG00000101310 | SEC23B | 5740.225464 | -0.174429817 | 0.125666404 | -1.400760253 | 0.16128578 | 0.328161 |
| ENSG00000198917 | SPOUT1 | 22.91731 | -0.1744325 | 0.3920149 | -0.518757 | 0.6039302 | 0.768809 |
| ENSG00000148950 | IMMP1L | 437.5035 | -0.1745107 | 0.214759 | -0.8492986 | 0.3957152 | 0.597249 |
| ENSG00000279935 | | 0.9514256 | -0.1745133 | 0.8172234 | -1.476605 | 0.1397816 | NA |
| ENSG00000196453 | ZNF777 | 1488.981 | -0.1745186 | 0.1553877 | -1.148716 | 0.2506733 | 0.4433 |
| ENSG00000115827 | DCAF17 | 1519.88373 | -0.174597518 | 0.200831037 | -0.902874475 | 0.36659252 | 0.568527 |
| ENSG00000163608 | NEPRO | 3771.51 | -0.174648 | 0.123085 | -1.435118 | 0.1512534 | 0.313028 |
| ENSG00000286129 | | 36.90661 | -0.1746649 | 0.3795984 | -0.5228555 | 0.6010748 | 0.766828 |
| ENSG00000287972 | | 0.993942 | -0.1747063 | 0.777732 | -0.7705672 | 0.4409635 | NA |
| ENSG00000229023 | RAB1AP1 | 5.574721 | -0.1747207 | 0.6297586 | -0.4480658 | 0.6541057 | 0.804895 |
| ENSG00000132341 | RAN | 20214.71028 | -0.174743841 | 0.225335122 | -0.813089859 | 0.41616654 | 0.616065 |
| ENSG00000118217 | ATF6 | 3452.708826 | -0.174758793 | 0.225004023 | -0.815764519 | 0.41463483 | 0.614649 |
| ENSG00000110583 | NAA40 | 2767.45691 | -0.174778549 | 0.192652652 | -0.947937379 | 0.34316133 | 0.544618 |
| ENSG00000236431 | | 53.52639 | -0.1748234 | 0.4516565 | -0.4685862 | 0.6393655 | 0.79417 |
| ENSG00000126216 | TUBGCP3 | 1433.373953 | -0.174835638 | 0.18121398 | -0.998010414 | 0.31827431 | 0.519393 |
| ENSG00000270157 | | 211.0524 | -0.1749044 | 0.3915866 | -0.5145781 | 0.6068479 | 0.771009 |
| ENSG00000105372 | RPS19 | 28242.84909 | -0.175004433 | 0.169585832 | -1.059639339 | 0.28930871 | 0.487904 |
| ENSG00000272576 | | 13.45026 | -0.1751891 | 0.4372813 | -0.4835506 | 0.6287048 | 0.786629 |
| ENSG00000095787 | WAC | 13246.26124 | -0.175192574 | 0.135030843 | -1.392993241 | 0.16362184 | 0.331524 |
| ENSG00000230927 | TMBIM7P | 2.186412 | -0.1752302 | 0.7238727 | -0.5338294 | 0.5934596 | 0.761349 |
| ENSG00000270427 | NRBF2P5 | 7.347281 | -0.1753859 | 0.5359325 | -0.4448843 | 0.6564034 | 0.806399 |
| ENSG00000095464 | PDE6C | 17.23872029 | -0.175474683 | 0.526158266 | -0.45075236 | 0.65216804 | 0.803679 |
| ENSG00000166912 | MTMR10 | 1030.285 | -0.1755143 | 0.201645 | -0.9036667 | 0.3661722 | 0.568097 |

| | | | | | | | |
|---|---|---|---|---|---|---|---|
| ENSG00000279696 | | 56.85944 | -0.1756404 | 0.3333187 | -0.5870759 | 0.5571527 | 0.73453 |
| ENSG00000188735 | TMEM120B | 1193.089 | -0.1756448 | 0.1647075 | -1.092095 | 0.2747915 | 0.471557 |
| ENSG00000278829 | | 1.191269 | -0.175646 | 0.7786439 | -0.7617654 | 0.4462001 | NA |
| ENSG00000163743 | RCHY1 | 1297.853 | -0.1756571 | 0.1371187 | -1.303343 | 0.1924578 | 0.370846 |
| ENSG00000189283 | FHIT | 337.8224 | -0.1756839 | 0.2932363 | -0.6438431 | 0.5196772 | 0.704928 |
| ENSG00000144224 | UBXN4 | 11126.42 | -0.1756932 | 0.1569722 | -1.140869 | 0.2539244 | 0.446908 |
| ENSG00000110274 | CEP164 | 2354.617612 | -0.175704481 | 0.109659761 | -1.592959736 | 0.11116922 | 0.251225 |
| ENSG00000151773 | CCDC122 | 514.5139 | -0.1757214 | 0.1961419 | -0.9283027 | 0.3532506 | 0.555192 |
| ENSG00000215198 | | 2.210746 | -0.1757556 | 0.7268979 | -0.5556193 | 0.5784711 | 0.749955 |
| ENSG00000259212 | | 29.55204 | -0.1758614 | 0.4617894 | -0.4678584 | 0.6398859 | 0.794371 |
| ENSG00000273314 | | 161.762 | -0.1758856 | 0.3180352 | -0.6056988 | 0.5447148 | 0.725367 |
| ENSG00000177054 | ZDHHC13 | 1634.528 | -0.1758994 | 0.1049217 | -1.696507 | 0.08978997 | 0.215258 |
| ENSG00000236618 | PITPNA-AS1 | 89.58565 | -0.1759412 | 0.2336804 | -0.7894692 | 0.4298378 | 0.629117 |
| ENSG00000051825 | MPHOSPH9 | 1635.767238 | -0.175945096 | 0.257384402 | -0.719030739 | 0.47212198 | 0.665136 |
| ENSG00000263338 | | 4.390609 | -0.1759473 | 0.6496735 | -0.4659906 | 0.6412222 | 0.795398 |
| ENSG00000165837 | ERICH6B | 11.73234 | -0.1762239 | 0.4748777 | -0.4650088 | 0.6419252 | 0.795789 |
| ENSG00000147168 | IL2RG | 0.828631 | -0.176233 | 0.8125652 | -1.029892 | 0.3030605 | NA |
| ENSG00000226085 | UQCRFS1P1 | 295.4599 | -0.1762777 | 0.2778852 | -0.6985797 | 0.4848147 | 0.675745 |
| ENSG00000169131 | ZNF354A | 2141.255 | -0.1763383 | 0.1728185 | -1.04106 | 0.2978477 | 0.497049 |
| ENSG00000124226 | RNF114 | 13901.5897 | -0.176357234 | 0.147102796 | -1.222302438 | 0.22159328 | 0.408231 |
| ENSG00000232150 | ST13P4 | 487.1182 | -0.1763931 | 0.150311 | -1.195282 | 0.2319768 | 0.420365 |
| ENSG00000250891 | LINC02208 | 12.69475 | -0.1765441 | 0.5275733 | -0.4533267 | 0.6503135 | 0.802434 |
| ENSG00000271420 | | 0.9298435 | -0.1765451 | 0.7629419 | -0.6668233 | 0.504885 | NA |
| ENSG00000235123 | DSCAM-AS1 | 15269.76 | -0.1766211 | 0.3404747 | -0.5843291 | 0.5589989 | 0.735831 |
| ENSG00000231856 | | 95.63009 | -0.176732 | 0.2775506 | -0.677956 | 0.4977996 | 0.686461 |
| ENSG00000172731 | LRRC20 | 1047.773 | -0.1767441 | 0.2184347 | -0.8480528 | 0.3964086 | 0.597631 |
| ENSG00000202379 | | 10.47522 | -0.176783 | 0.5357725 | -0.4502112 | 0.6525582 | 0.803942 |
| ENSG00000110172 | CHORDC1 | 2949.831152 | -0.176803702 | 0.265026037 | -0.711219776 | 0.47694806 | 0.669231 |
| ENSG00000275395 | FCGBP | 85.20155 | -0.176817 | 0.4173607 | -0.5039127 | 0.6143227 | 0.776311 |
| ENSG00000275339 | | 8.014418 | -0.176877 | 0.5166374 | -0.4538393 | 0.6499445 | 0.802098 |
| ENSG00000185305 | ARL15 | 1206.838 | -0.1769431 | 0.1742047 | -1.045484 | 0.2957995 | 0.494816 |
| ENSG00000135823 | STX6 | 5377.548381 | -0.176955783 | 0.083024544 | -2.163987044 | 0.03046534 | 0.094868 |
| ENSG00000170035 | UBE2E3 | 2027.121 | -0.1769826 | 0.2496748 | -0.732919 | 0.4636078 | 0.657806 |

| | | | | | | | |
|---|---|---|---|---|---|---|---|
| ENSG00000124496 | TRERF1 | 2021.410746 | -0.177003506 | 0.256134899 | -0.735618314 | 0.46196301 | 0.656639 |
| ENSG00000174460 | ZCCHC12 | 5.896057 | -0.1770807 | 0.7173315 | -0.5108342 | 0.6094672 | 0.772932 |
| ENSG00000249249 | | 27.76046 | -0.1770946 | 0.4155657 | -0.5009022 | 0.61644 | 0.777845 |
| ENSG00000256981 | | 0.7912426 | -0.1771242 | 0.7665451 | -0.6723989 | 0.5013298 | NA |
| ENSG00000229534 | HNRNPA1P53 | 2.299022 | -0.1772039 | 0.7517995 | -0.5801092 | 0.561841 | 0.73771 |
| ENSG00000131051 | RBM39 | 19834.60568 | -0.17720541 | 0.165922299 | -1.119541346 | 0.26290926 | 0.457909 |
| ENSG00000196437 | ZNF569 | 255.1145 | -0.1772295 | 0.2853213 | -0.6710905 | 0.5021629 | 0.689793 |
| ENSG00000235763 | SNRPGP5 | 3.669945 | -0.1773295 | 0.6573602 | -0.4725446 | 0.6365381 | 0.792288 |
| ENSG00000103227 | LMF1 | 1257.743949 | -0.177367631 | 0.190417863 | -0.964931623 | 0.33457906 | 0.535798 |
| ENSG00000266501 | ATG12P1 | 76.49672 | -0.1774723 | 0.2442943 | -0.7678705 | 0.4425641 | 0.640164 |
| ENSG00000136986 | DERL1 | 8411.747435 | -0.177528158 | 0.081010426 | -2.206778435 | 0.02732954 | 0.087065 |
| ENSG00000101220 | C20orf27 | 5816.477869 | -0.177578903 | 0.253384465 | -0.73969508 | 0.45948503 | 0.655009 |
| ENSG00000270342 | | 7.220339 | -0.1775802 | 0.6229388 | -0.4512312 | 0.6518229 | 0.80336 |
| ENSG00000242667 | | 3.75743 | -0.1776086 | 0.6704436 | -0.4796493 | 0.6314768 | 0.788593 |
| ENSG00000272829 | | 26.02165 | -0.1777192 | 0.4437442 | -0.4862903 | 0.6267614 | 0.785291 |
| ENSG00000143436 | MRPL9 | 4702.837 | -0.1777234 | 0.1882615 | -0.976523 | 0.3288053 | 0.529991 |
| ENSG00000237422 | | 1.140437 | -0.1777944 | 0.7836142 | -0.7597444 | 0.4474074 | NA |
| ENSG00000226088 | HHLA3-AS1 | 2.959215 | -0.1778088 | 0.6671564 | -0.4791355 | 0.6318422 | 0.788957 |
| ENSG00000151491 | EPS8 | 2446 | -0.1778524 | 0.1820486 | -1.00621 | 0.3143147 | 0.515464 |
| ENSG00000145241 | CENPC | 1224.73 | -0.1778725 | 0.1694364 | -1.072259 | 0.2836037 | 0.481382 |
| ENSG00000226147 | TUBBP10 | 0.9289754 | -0.1778883 | 0.7486498 | -0.5984832 | 0.5495176 | NA |
| ENSG00000082126 | MPP4 | 1.780501037 | -0.177981873 | 0.70423445 | -0.513789373 | 0.60739926 | 0.771285 |
| ENSG00000259664 | LINC02254 | 7.946257 | -0.1780082 | 0.5943438 | -0.4513021 | 0.6517719 | 0.80336 |
| ENSG00000141027 | NCOR1 | 9465.914 | -0.1781029 | 0.1688897 | -1.087628 | 0.2767594 | 0.4737 |
| ENSG00000136451 | VEZF1 | 6607.382462 | -0.178219043 | 0.227142741 | -0.823716117 | 0.41010088 | 0.610419 |
| ENSG00000086232 | EIF2AK1 | 16918.34372 | -0.178279407 | 0.169044451 | -1.088559543 | 0.27634817 | 0.473299 |
| ENSG00000263219 | RYKP1 | 8.442588 | -0.1782939 | 0.4959928 | -0.4615003 | 0.6444397 | 0.79766 |
| ENSG00000132664 | POLR3F | 1185.438352 | -0.178330028 | 0.168622271 | -1.087937442 | 0.27662273 | 0.473648 |
| ENSG00000257138 | TAS2R38 | 2.794795 | -0.1783872 | 0.7956902 | -0.5486623 | 0.5832372 | 0.753515 |
| ENSG00000182326 | C1S | 46.8792 | -0.1784671 | 0.3575878 | -0.5596731 | 0.5757024 | 0.747853 |
| ENSG00000167074 | TEF | 2325.288 | -0.1785235 | 0.1693254 | -1.087628 | 0.2767595 | 0.4737 |
| ENSG00000287760 | | 58.40126 | -0.1785317 | 0.3465799 | -0.5749899 | 0.5652981 | 0.7402 |
| ENSG00000271971 | | 128.7662 | -0.1785959 | 0.3187636 | -0.6124289 | 0.540254 | 0.722365 |

| | | | | | | | |
|---|---|---|---|---|---|---|---|
| ENSG00000257743 | MGAM2 | 4.557449 | -0.1787022 | 0.6740281 | -0.4941836 | 0.6211765 | 0.781145 |
| ENSG00000228384 | | 1.648957 | -0.1787253 | 0.7277202 | -0.5557168 | 0.5784044 | 0.749955 |
| ENSG00000143033 | MTF2 | 1278.169 | -0.1787291 | 0.1511075 | -1.20851 | 0.2268511 | 0.414583 |
| ENSG00000148843 | PDCD11 | 5263.454 | -0.178753 | 0.159269 | -1.141358 | 0.253721 | 0.446727 |
| ENSG00000157343 | ARMC12 | 51.91416 | -0.1787541 | 0.4583986 | -0.4858716 | 0.6270582 | 0.785493 |
| ENSG00000240231 | RPS27P29 | 2.423009 | -0.1787602 | 0.7054115 | -0.5154694 | 0.606225 | 0.770549 |
| ENSG00000179938 | GOLGA8J | 0.777913 | -0.1788086 | 0.8023802 | -1.055925 | 0.2910026 | NA |
| ENSG00000135638 | EMX1 | 5.961766937 | -0.178838096 | 0.716724208 | -0.524757458 | 0.59975181 | 0.765998 |
| ENSG00000198408 | OGA | 10230.93 | -0.1788932 | 0.2136937 | -0.8744916 | 0.3818506 | 0.584109 |
| ENSG00000167526 | RPL13 | 46053.72 | -0.1789516 | 0.1718881 | -1.031118 | 0.3024853 | 0.501951 |
| ENSG00000129244 | ATP1B2 | 1.952960142 | -0.179224321 | 0.76438312 | -0.642128493 | 0.52078976 | 0.706034 |
| ENSG00000230896 | | 35.53509 | -0.1792729 | 0.3277664 | -0.6039534 | 0.5458746 | 0.726117 |
| ENSG00000287920 | | 1.530211 | -0.1792878 | 0.7115381 | -0.5268962 | 0.5982657 | NA |
| ENSG00000182378 | PLCXD1 | 954.7837 | -0.1793033 | 0.2151934 | -0.871696 | 0.3833743 | 0.585745 |
| ENSG00000286563 | | 6.087487 | -0.1793517 | 0.5782403 | -0.4539628 | 0.6498556 | 0.802063 |
| ENSG00000228253 | MT-ATP8 | 7803.357 | -0.1793592 | 0.1762134 | -1.063645 | 0.2874894 | 0.485875 |
| ENSG00000229630 | LINC01264 | 0.9585382 | -0.1793731 | 0.7857289 | -0.8322422 | 0.4052722 | NA |
| ENSG00000156515 | HK1 | 12334.04 | -0.1793991 | 0.1502703 | -1.265818 | 0.2055783 | 0.387542 |
| ENSG00000151012 | SLC7A11 | 2847.792 | -0.1794886 | 0.6178925 | -0.4693655 | 0.6388084 | 0.794021 |
| ENSG00000151500 | THYN1 | 902.8994 | -0.1795832 | 0.1950767 | -0.9581468 | 0.3379888 | 0.539325 |
| ENSG00000254877 | | 2.867869 | -0.1796003 | 0.6943443 | -0.5104905 | 0.6097079 | 0.773201 |
| ENSG00000248936 | | 6.501694 | -0.1796308 | 0.5349095 | -0.4591852 | 0.6461012 | 0.798951 |
| ENSG00000167799 | NUDT8 | 1217.042 | -0.1796597 | 0.3687856 | -0.5587781 | 0.5763132 | 0.748427 |
| ENSG00000286999 | | 1.432026 | -0.1797321 | 0.7312879 | -0.5741511 | 0.5658655 | NA |
| ENSG00000130940 | CASZ1 | 1449.599505 | -0.179813927 | 0.193088723 | -0.963247108 | 0.33542353 | 0.536666 |
| ENSG00000138604 | GLCE | 2135.485007 | -0.179889834 | 0.20647235 | -0.907775883 | 0.36399664 | 0.56612 |
| ENSG00000268650 | | 1.47394 | -0.1799148 | 0.7431467 | -0.5927994 | 0.5533154 | NA |
| ENSG00000141101 | NOB1 | 3757.346 | -0.1799258 | 0.170467 | -1.08936 | 0.2759954 | 0.472878 |
| ENSG00000145020 | AMT | 4.407495 | -0.1800001 | 0.6295857 | -0.460801 | 0.6449413 | 0.797888 |
| ENSG00000181315 | ZNF322 | 583.7781 | -0.1800081 | 0.2390336 | -0.7938844 | 0.4272627 | 0.626798 |
| ENSG00000214917 | | 2.709554 | -0.1800549 | 0.6811201 | -0.4918931 | 0.6227949 | 0.782218 |
| ENSG00000163380 | LMOD3 | 4.117493 | -0.1802849 | 0.67741 | -0.4774111 | 0.6330694 | 0.789672 |
| ENSG00000081059 | TCF7 | 20.54533666 | -0.180295602 | 0.427658314 | -0.504344131 | 0.61401957 | 0.776113 |

| | | | | | | | |
|---|---|---|---|---|---|---|---|
| ENSG00000182511 | FES | 263.3344 | -0.1803412 | 0.2471095 | -0.7765799 | 0.4374066 | 0.635852 |
| ENSG00000231556 | RSL24D1P3 | 0.8385133 | -0.180352 | 0.7742987 | -0.722287 | 0.470118 | NA |
| ENSG00000261008 | LINC01572 | 131.773 | -0.1803643 | 0.2542416 | -0.7542925 | 0.4506736 | 0.646956 |
| ENSG00000230295 | GTF2IP23 | 37.96895 | -0.1804223 | 0.3698214 | -0.5562038 | 0.5780715 | 0.749721 |
| ENSG00000113194 | FAF2 | 8934.431223 | -0.18042504 | 0.091697461 | -1.988386941 | 0.04676891 | 0.131757 |
| ENSG00000271963 | | 1.484316 | -0.1805503 | 0.7384156 | -0.5856311 | 0.5581234 | NA |
| ENSG00000201573 | | 1.88934 | -0.1805631 | 0.7210767 | -0.5358373 | 0.592071 | 0.760417 |
| ENSG00000101150 | TPD52L2 | 16482.16033 | -0.180661231 | 0.13425385 | -1.367772877 | 0.17138318 | 0.34211 |
| ENSG00000152931 | PART1 | 5.058333 | -0.180674 | 0.6505564 | -0.4757748 | 0.6342349 | 0.790553 |
| ENSG00000238043 | | 1.888652 | -0.1807018 | 0.6989489 | -0.5195529 | 0.6033752 | 0.768545 |
| ENSG00000113312 | TTC1 | 3072.099026 | -0.180718967 | 0.141506336 | -1.300260914 | 0.19351156 | 0.372157 |
| ENSG00000065618 | COL17A1 | 6.357902474 | -0.180815641 | 0.667093083 | -0.478503936 | 0.63229157 | 0.789227 |
| ENSG00000260927 | | 8.889682 | -0.1809575 | 0.6602596 | -0.4753923 | 0.6345074 | 0.790628 |
| ENSG00000185684 | EP400P1 | 1475.571 | -0.1809821 | 0.1296812 | -1.416109 | 0.1567437 | 0.321455 |
| ENSG00000261837 | | 2.277443 | -0.1810092 | 0.7046828 | -0.5308566 | 0.5955182 | 0.762742 |
| ENSG00000152670 | DDX4 | 1.75351 | -0.1810331 | 0.7485864 | -0.5979274 | 0.5498884 | 0.729016 |
| ENSG00000270589 | | 29.29556 | -0.1810383 | 0.4286295 | -0.5053331 | 0.6133249 | 0.775788 |
| ENSG00000248527 | MTATP6P1 | 6871.118 | -0.1810439 | 0.1938032 | -0.9688101 | 0.33264 | 0.53369 |
| ENSG00000018189 | RUFY3 | 1146.357832 | -0.18108709 | 0.165544881 | -1.129867197 | 0.25853219 | 0.452699 |
| ENSG00000159173 | TNNI1 | 301.8944 | -0.181162 | 0.2816135 | -0.688392 | 0.491206 | 0.681575 |
| ENSG00000273245 | | 2.41805 | -0.1811648 | 0.6868133 | -0.5041044 | 0.614188 | 0.776236 |
| ENSG00000279845 | | 4.954179 | -0.1811948 | 0.5934897 | -0.4540455 | 0.6497961 | 0.802027 |
| ENSG00000276248 | | 30.58186 | -0.1812203 | 0.4202393 | -0.5121814 | 0.608524 | 0.772229 |
| ENSG00000176476 | SGF29 | 807.7626 | -0.1812662 | 0.1766805 | -1.047941 | 0.2946657 | 0.493755 |
| ENSG00000237053 | FUCA1P1 | 1.48239 | -0.1812728 | 0.7450298 | -0.6252665 | 0.5317962 | NA |
| ENSG00000270257 | | 6.705974 | -0.1813326 | 0.5392825 | -0.4629493 | 0.6434007 | 0.796761 |
| ENSG00000223745 | CCDC18-AS1 | 607.3978 | -0.1813756 | 0.4380635 | -0.5054566 | 0.6132382 | 0.775778 |
| ENSG00000263307 | | 14.29893 | -0.1813915 | 0.5231512 | -0.4641501 | 0.6425402 | 0.796178 |
| ENSG00000197056 | ZMYM1 | 1015.689 | -0.1814737 | 0.173279 | -1.077645 | 0.2811923 | 0.478787 |
| ENSG00000105991 | HOXA1 | 30.37578666 | -0.18153524 | 0.531506464 | -0.456093317 | 0.64832288 | 0.800655 |
| ENSG00000197858 | GPAA1 | 7864.432 | -0.1815365 | 0.1603918 | -1.172573 | 0.240967 | 0.431455 |
| ENSG00000204316 | MRPL38 | 211.4684 | -0.1815644 | 0.2178529 | -0.8694671 | 0.3845917 | 0.586984 |
| ENSG00000102897 | LYRM1 | 1174.588881 | -0.181598814 | 0.172674299 | -1.08466402 | 0.27807049 | 0.475208 |

| | | | | | | | |
|---|---|---|---|---|---|---|---|
| ENSG00000239920 | | 1.42399 | -0.1817536 | 0.7963363 | -0.5511631 | 0.5815218 | NA |
| ENSG00000287931 | | 1.061081 | -0.181758 | 0.7467987 | -0.6017357 | 0.5473501 | NA |
| ENSG00000145506 | NKD2 | 359.4355 | -0.1817584 | 0.3978455 | -0.5401012 | 0.5891273 | 0.758028 |
| ENSG00000131650 | KREMEN2 | 753.5745695 | -0.181817761 | 0.392576032 | -0.537956554 | 0.59060704 | 0.759344 |
| ENSG00000240661 | | 18.62434 | -0.1818888 | 0.4343141 | -0.4966615 | 0.6194278 | 0.780127 |
| ENSG00000233040 | FAM204BP | 7.44236 | -0.1819023 | 0.553187 | -0.4574253 | 0.6473654 | 0.800098 |
| ENSG00000169436 | COL22A1 | 0.800206 | -0.1819354 | 0.7605935 | -0.6536736 | 0.5133221 | NA |
| ENSG00000238083 | LRRC37A2 | 220.2463 | -0.1819387 | 0.2673693 | -0.7310815 | 0.4647293 | 0.658653 |
| ENSG00000132199 | ENOSF1 | 4634.825382 | -0.18194782 | 0.20390686 | -0.933683682 | 0.35046709 | 0.552222 |
| ENSG00000171150 | SOCS5 | 1108.42 | -0.1819844 | 0.1867957 | -1.005514 | 0.3146494 | 0.515758 |
| ENSG00000153233 | PTPRR | 38.53526 | -0.1820437 | 0.6763995 | -0.4755354 | 0.6344054 | 0.790579 |
| ENSG00000250064 | | 0.8214309 | -0.1821891 | 0.809486 | -1.222548 | 0.2215004 | NA |
| ENSG00000287041 | | 1.832351 | -0.1822371 | 0.7119108 | -0.5439571 | 0.586471 | 0.756185 |
| ENSG00000253106 | | 234.3064 | -0.1822534 | 0.2867108 | -0.6861924 | 0.4925918 | 0.682645 |
| ENSG00000267395 | DM1-AS | 45.26233 | -0.182294 | 0.3739627 | -0.5542525 | 0.579406 | 0.750572 |
| ENSG00000264343 | NOTCH2NLA | 23.56597 | -0.1823066 | 0.4652794 | -0.4847272 | 0.6278699 | 0.786065 |
| ENSG00000137561 | TTPA | 51.61268912 | -0.182356052 | 0.439043381 | -0.504778222 | 0.61371461 | 0.775861 |
| ENSG00000189319 | FAM53B | 2135.24 | -0.1824331 | 0.1154295 | -1.612328 | 0.1068907 | 0.244197 |
| ENSG00000178035 | IMPDH2 | 11611.72 | -0.1825012 | 0.1407769 | -1.331181 | 0.1831296 | 0.358184 |
| ENSG00000174013 | FBXO45 | 2614.991 | -0.1825441 | 0.1724486 | -1.091284 | 0.2751478 | 0.471974 |
| ENSG00000197273 | GUCA2A | 1.421695 | -0.1825572 | 0.8092621 | -0.839954 | 0.4009342 | NA |
| ENSG00000134884 | ARGLU1 | 8787.21709 | -0.182564261 | 0.292917717 | -0.676132111 | 0.49895677 | 0.687392 |
| ENSG00000219329 | | 6.108699 | -0.1825779 | 0.6383884 | -0.4763614 | 0.633817 | 0.790234 |
| ENSG00000056972 | TRAF3IP2 | 1432.15154 | -0.182583428 | 0.198042301 | -0.962314121 | 0.33589184 | 0.53693 |
| ENSG00000262904 | TMPOP2 | 29.43168 | -0.1825998 | 0.6168319 | -0.4690265 | 0.6390507 | 0.794106 |
| ENSG00000230993 | RPL12P15 | 0.8794237 | -0.1826687 | 0.7733377 | -0.7198983 | 0.4715876 | NA |
| ENSG00000277299 | | 1.199274 | -0.1826963 | 0.7598894 | -0.6653129 | 0.5058504 | NA |
| ENSG00000070501 | POLB | 791.2004441 | -0.182713437 | 0.162829632 | -1.14327078 | 0.25292618 | 0.445771 |
| ENSG00000262420 | | 7.248951 | -0.1827555 | 0.5445049 | -0.4573533 | 0.6474171 | 0.800098 |
| ENSG00000204253 | HNRNPCP2 | 4406.56 | -0.1827846 | 0.156944 | -1.189627 | 0.2341931 | 0.423313 |
| ENSG00000256030 | CBX3P4 | 1.971982 | -0.182793 | 0.7483299 | -0.6011783 | 0.5477213 | 0.727458 |
| ENSG00000141837 | CACNA1A | 2.386155 | -0.1827993 | 0.6786827 | -0.4928017 | 0.6221527 | 0.781744 |
| ENSG00000286618 | | 1.234777 | -0.1829178 | 0.7412035 | -0.6093516 | 0.5422914 | NA |

| | | | | | | | |
|---|---|---|---|---|---|---|---|
| ENSG00000132376 | INPP5K | 2088.156878 | -0.182941631 | 0.19647564 | -0.969249352 | 0.33242079 | 0.533517 |
| ENSG00000213693 | SEC14L1P1 | 52.91234 | -0.182954 | 0.2895401 | -0.6789875 | 0.4971458 | 0.685915 |
| ENSG00000115216 | NRBP1 | 10957.89949 | -0.182965508 | 0.146034831 | -1.277501075 | 0.2014254 | 0.382208 |
| ENSG00000229915 | | 2.940044 | -0.1830816 | 0.7098729 | -0.5317693 | 0.5948858 | 0.762263 |
| ENSG00000189266 | PNRC2 | 4025.687 | -0.1831467 | 0.1848583 | -1.026197 | 0.3047985 | 0.504533 |
| ENSG00000272502 | | 19.37577 | -0.1832224 | 0.488166 | -0.4755138 | 0.6344208 | 0.790579 |
| ENSG00000170959 | DCDC1 | 4.401104 | -0.1832348 | 0.6423556 | -0.47365 | 0.6357495 | 0.791603 |
| ENSG00000251303 | CAB39P1 | 1.351552 | -0.183249 | 0.7642186 | -0.692926 | 0.488356 | NA |
| ENSG00000168065 | SLC22A11 | 0.8798536 | -0.1832765 | 0.758493 | -0.6640423 | 0.5066632 | NA |
| ENSG00000186153 | WWOX | 3472.257 | -0.1832907 | 0.2819001 | -0.7003667 | 0.4836984 | 0.674932 |
| ENSG00000272277 | | 142.4629 | -0.1833043 | 0.2875467 | -0.6824694 | 0.4949422 | 0.684225 |
| ENSG00000260036 | | 9.072086 | -0.1833284 | 0.6377531 | -0.4742515 | 0.6353206 | 0.791217 |
| ENSG00000286118 | | 5.797731 | -0.1833924 | 0.5977049 | -0.4647193 | 0.6421325 | 0.795869 |
| ENSG00000159388 | BTG2 | 7411.241 | -0.1833991 | 0.2894706 | -0.6845888 | 0.4936034 | 0.683405 |
| ENSG00000205795 | CYS1 | 30.3654 | -0.1834159 | 0.5598494 | -0.4602611 | 0.6453289 | 0.798219 |
| ENSG00000174586 | ZNF497 | 238.6169 | -0.1834222 | 0.1995276 | -0.9524993 | 0.3408438 | 0.542171 |
| ENSG00000064489 | BORCS8-MEF2B | 1.268983094 | -0.183439344 | 0.75063894 | -0.620007799 | 0.53525265 | NA |
| ENSG00000037965 | HOXC8 | 146.5965569 | -0.183531872 | 0.27010459 | -0.728510464 | 0.46630117 | 0.660086 |
| ENSG00000157110 | RBPMS | 1234.271 | -0.183555 | 0.2619227 | -0.7367013 | 0.461304 | 0.656181 |
| ENSG00000232060 | SLC4A1APP1 | 305.3279 | -0.1836528 | 0.2133487 | -0.8910085 | 0.3729246 | 0.575095 |
| ENSG00000124232 | RBPJL | 1.24623631 | -0.183658319 | 0.735962401 | -0.584392053 | 0.55895657 | NA |
| ENSG00000144741 | SLC25A26 | 1180.905 | -0.1837165 | 0.1819985 | -1.02772 | 0.3040815 | 0.503697 |
| ENSG00000223416 | RPS26P15 | 1.806597 | -0.1837444 | 0.7772691 | -0.741057 | 0.4586589 | 0.654461 |
| ENSG00000204700 | OR2J2 | 1.19354 | -0.1837512 | 0.7992245 | -0.5025402 | 0.6152876 | NA |
| ENSG00000164080 | RAD54L2 | 1424.87 | -0.1837697 | 0.2148596 | -0.8923686 | 0.3721954 | 0.57443 |
| ENSG00000273628 | | 1.79443 | -0.183882 | 0.7680929 | -0.699439 | 0.4842778 | 0.67545 |
| ENSG00000164385 | LINC01600 | 2.370192 | -0.1839344 | 0.66048 | -0.4838449 | 0.6284959 | 0.786528 |
| ENSG00000277681 | MIR6739 | 1.124526 | -0.1839818 | 0.764606 | -0.7010593 | 0.483266 | NA |
| ENSG00000165675 | ENOX2 | 1111.205 | -0.1840848 | 0.1537383 | -1.232043 | 0.2179329 | 0.403647 |
| ENSG00000135436 | FAM186B | 43.55535376 | -0.184162748 | 0.422590757 | -0.517891565 | 0.60453393 | 0.769357 |
| ENSG00000140323 | DISP2 | 161.4048 | -0.1841741 | 0.3342151 | -0.6096454 | 0.5420967 | 0.723521 |
| ENSG00000227582 | ADGRF5P1 | 19.30538 | -0.1842316 | 0.5244281 | -0.4744762 | 0.6351604 | 0.791166 |
| ENSG00000089682 | RBM41 | 1204.628105 | -0.184249328 | 0.218257121 | -0.884249044 | 0.37656179 | 0.578617 |

| | | | | | | | |
|---|---|---|---|---|---|---|---|
| ENSG00000236603 | RANP1 | 1825.713 | -0.184257 | 0.2523647 | -0.7917007 | 0.4285352 | 0.62791 |
| ENSG00000287975 | | 1.616432 | -0.1843169 | 0.7967109 | -0.5601818 | 0.5753554 | 0.747542 |
| ENSG00000204539 | CDSN | 1.065249 | -0.1843239 | 0.820666 | -1.558379 | 0.1191434 | NA |
| ENSG00000100023 | PPIL2 | 3817.09936 | -0.184366462 | 0.116035079 | -1.620464343 | 0.10513257 | 0.241236 |
| ENSG00000114796 | KLHL24 | 2165.764679 | -0.184384218 | 0.389178105 | -0.56219841 | 0.57398085 | 0.746641 |
| ENSG00000228369 | TXNDC12-AS1 | 2.212917 | -0.1843913 | 0.69085 | -0.5194927 | 0.6034172 | 0.768562 |
| ENSG00000105568 | PPP2R1A | 14462.81748 | -0.184448437 | 0.193817729 | -0.977028855 | 0.32855487 | 0.529759 |
| ENSG00000158019 | BABAM2 | 1167.584 | -0.1844974 | 0.1402333 | -1.328875 | 0.1838893 | 0.359194 |
| ENSG00000139436 | GIT2 | 930.5868 | -0.1845298 | 0.1939736 | -0.9808359 | 0.3266737 | 0.527976 |
| ENSG00000169564 | PCBP1 | 1130.503 | -0.1845414 | 0.1653105 | -1.165853 | 0.2436738 | 0.434917 |
| ENSG00000100014 | SPECC1L | 2033.192489 | -0.184541875 | 0.140653013 | -1.339770077 | 0.18032011 | 0.354629 |
| ENSG00000065150 | IPO5 | 7167.411912 | -0.18456589 | 0.223424568 | -0.864769802 | 0.38716515 | 0.589456 |
| ENSG00000162947 | LINC01931 | 1.83316 | -0.1846925 | 0.7992169 | -0.508781 | 0.6109057 | 0.774202 |
| ENSG00000259022 | DNAJC8P1 | 36.37803 | -0.1846949 | 0.3019698 | -0.6642688 | 0.5065183 | 0.693016 |
| ENSG00000204710 | SPDYC | 11.83985 | -0.1847564 | 0.5026413 | -0.4810421 | 0.6304865 | 0.788004 |
| ENSG00000149476 | TKFC | 3087.704 | -0.1847659 | 0.1803435 | -1.056057 | 0.2909421 | 0.489522 |
| ENSG00000235165 | CDK5P1 | 0.9385731 | -0.1848221 | 0.806591 | -0.7169693 | 0.473393 | NA |
| ENSG00000223305 | RN7SKP30 | 1.853497 | -0.1848242 | 0.7383279 | -0.5954646 | 0.5515329 | 0.730347 |
| ENSG00000233379 | | 5.619073 | -0.1848275 | 0.7971685 | -0.5449713 | 0.5857732 | 0.755542 |
| ENSG00000228126 | FALEC | 45.9488 | -0.1848738 | 0.4068048 | -0.5326629 | 0.594267 | 0.761701 |
| ENSG00000188986 | NELFB | 5569.099 | -0.184877 | 0.1845881 | -1.034497 | 0.3009041 | 0.500076 |
| ENSG00000182584 | ACTL10 | 193.2241 | -0.1848891 | 0.2920474 | -0.6777544 | 0.4979275 | 0.68653 |
| ENSG00000219027 | RPS3AP2 | 3.908044 | -0.1849125 | 0.6412555 | -0.4780474 | 0.6326165 | 0.789441 |
| ENSG00000058668 | ATP2B4 | 2737.206402 | -0.184934439 | 0.280631183 | -0.733369729 | 0.46333296 | 0.657601 |
| ENSG00000168938 | PPIC | 2687.65 | -0.1849781 | 0.1600486 | -1.168302 | 0.2426849 | 0.433559 |
| ENSG00000272034 | SNORD14A | 58.39552 | -0.1849817 | 0.3467104 | -0.5997915 | 0.5486452 | 0.72812 |
| ENSG00000244398 | | 13829.82 | -0.1850291 | 0.1681347 | -1.137575 | 0.255298 | 0.448674 |
| ENSG00000185236 | RAB11B | 1946.581 | -0.1850381 | 0.2219954 | -0.9067049 | 0.3645629 | 0.566578 |
| ENSG00000134365 | CFHR4 | 3.476201151 | -0.185054767 | 0.674927187 | -0.503654747 | 0.61450401 | 0.776466 |
| ENSG00000287859 | | 2.113549 | -0.1850578 | 0.6779821 | -0.5020045 | 0.6156644 | 0.777341 |
| ENSG00000166359 | WDR88 | 94.98444 | -0.1852221 | 0.2545997 | -0.7629569 | 0.4454891 | 0.642573 |
| ENSG00000136152 | COG3 | 3726.237654 | -0.185226848 | 0.157475301 | -1.186945782 | 0.23524901 | 0.424647 |
| ENSG00000182934 | SRPRA | 5480.133 | -0.1852578 | 0.1150427 | -1.622972 | 0.1045953 | 0.240356 |

| | | | | | | | |
|---|---|---|---|---|---|---|---|
| ENSG00000272033 | | 2.044439 | -0.1853335 | 0.7327469 | -0.5777366 | 0.5634419 | 0.739007 |
| ENSG00000172590 | MRPL52 | 2590.331 | -0.185343 | 0.195977 | -0.9806518 | 0.3267645 | 0.528058 |
| ENSG00000244357 | RN7SL145P | 1.197792 | -0.1853921 | 0.7708179 | -0.7127393 | 0.4760071 | NA |
| ENSG00000233297 | RASA4DP | 8.635376 | -0.1854042 | 0.5067207 | -0.4771332 | 0.6332672 | 0.789845 |
| ENSG00000273565 | | 8.157353 | -0.185584 | 0.5411351 | -0.4657684 | 0.6413813 | 0.795521 |
| ENSG00000273851 | | 1.755683 | -0.1855944 | 0.7045169 | -0.5339666 | 0.5933646 | 0.761305 |
| ENSG00000184634 | MED12 | 4877.749 | -0.1856248 | 0.1700336 | -1.120543 | 0.2624826 | 0.457412 |
| ENSG00000108395 | TRIM37 | 57971.37963 | -0.185634533 | 0.112309329 | -1.672268631 | 0.09447136 | 0.22337 |
| ENSG00000214772 | | 1.287896 | -0.1857277 | 0.732217 | -0.58435 | 0.5589848 | NA |
| ENSG00000242952 | RPSAP3 | 2.542206 | -0.1857482 | 0.7241942 | -0.5663678 | 0.5711438 | 0.744898 |
| ENSG00000035403 | VCL | 4868.930003 | -0.185760466 | 0.180856231 | -1.046363894 | 0.29539306 | 0.494568 |
| ENSG00000272699 | | 0.9491528 | -0.1857897 | 0.7800137 | -0.7888987 | 0.4301712 | NA |
| ENSG00000276334 | | 44.82433 | -0.185819 | 0.4227563 | -0.5230232 | 0.6009581 | 0.766786 |
| ENSG00000145536 | ADAMTS16 | 15.02812 | -0.1858683 | 0.676906 | -0.4983185 | 0.6182596 | 0.779174 |
| ENSG00000238118 | SLC25A24P2 | 1.405004 | -0.1859351 | 0.7206012 | -0.5775375 | 0.5635764 | NA |
| ENSG00000276545 | PCDHGB9P | 2.863214 | -0.1859443 | 0.6593696 | -0.4946259 | 0.6208642 | 0.781011 |
| ENSG00000124772 | CPNE5 | 1.408141855 | -0.185972284 | 0.757068441 | -0.655459752 | 0.51217179 | NA |
| ENSG00000105287 | PRKD2 | 3137.6017 | -0.186078226 | 0.167407182 | -1.138699479 | 0.25482852 | 0.447994 |
| ENSG00000129167 | TPH1 | 7.598190044 | -0.186100569 | 0.550225319 | -0.47280483 | 0.63635243 | 0.792131 |
| ENSG00000034510 | TMSB10 | 14569.63482 | -0.186111172 | 0.23880397 | -0.821424328 | 0.41140461 | 0.611674 |
| ENSG00000136754 | ABI1 | 3396.086365 | -0.186144446 | 0.095351447 | -1.971873023 | 0.0486241 | 0.135618 |
| ENSG00000233337 | UBE2FP3 | 20.28489 | -0.1861783 | 0.4903614 | -0.4848339 | 0.6277942 | 0.786057 |
| ENSG00000236439 | | 1124.958 | -0.1861807 | 0.1730085 | -1.117458 | 0.2637984 | 0.459086 |
| ENSG00000111863 | ADTRP | 5.940567875 | -0.186223872 | 0.67484654 | -0.518807737 | 0.60389482 | 0.768809 |
| ENSG00000261404 | | 7.240864 | -0.1862682 | 0.5761106 | -0.4628606 | 0.6434643 | 0.796766 |
| ENSG00000228737 | | 9.797595 | -0.1863386 | 0.5001599 | -0.4826597 | 0.6293374 | 0.787012 |
| ENSG00000167264 | DUS2 | 1186.033 | -0.18637 | 0.2315003 | -0.8609668 | 0.3892564 | 0.591157 |
| ENSG00000249791 | | 1.248101 | -0.1863896 | 0.7334065 | -0.6207693 | 0.5347514 | NA |
| ENSG00000213315 | | 136.4349 | -0.1863898 | 0.2941472 | -0.687157 | 0.4919838 | 0.68221 |
| ENSG00000162302 | RPS6KA4 | 2456.173 | -0.186407 | 0.1927676 | -1.003743 | 0.3155025 | 0.516487 |
| ENSG00000250030 | | 15.87691 | -0.1864155 | 0.4437573 | -0.5086679 | 0.6109851 | 0.774266 |
| ENSG00000163116 | STPG2 | 4.94289 | -0.1864797 | 0.570898 | -0.4700838 | 0.6382952 | 0.793621 |
| ENSG00000254867 | | 1.763027 | -0.1864841 | 0.7531432 | -0.6283738 | 0.5297591 | 0.714359 |

| | | | | | | | |
|---|---|---|---|---|---|---|---|
| ENSG00000150967 | ABCB9 | 1062.54 | -0.1865206 | 0.1995286 | -0.9733333 | 0.3303877 | 0.531517 |
| ENSG00000204172 | AGAP9 | 370.8095 | -0.1866276 | 0.192989 | -1.0006 | 0.3170204 | 0.518118 |
| ENSG00000125247 | TMTC4 | 1219.679477 | -0.186766045 | 0.169571883 | -1.134274684 | 0.25667935 | 0.450473 |
| ENSG00000100387 | RBX1 | 2132.370596 | -0.186782847 | 0.241125767 | -0.814980919 | 0.41508323 | 0.615074 |
| ENSG00000243414 | TICAM2 | 1.393136 | -0.1867975 | 0.7416628 | -0.6042921 | 0.5456495 | NA |
| ENSG00000288568 | | 29.51752 | -0.1868095 | 0.4280622 | -0.5174271 | 0.6048581 | 0.769548 |
| ENSG00000132423 | COQ3 | 558.8306142 | -0.186825053 | 0.355855952 | -0.589712348 | 0.55538352 | 0.732922 |
| ENSG00000233309 | RPS6P12 | 13.12173 | -0.1868363 | 0.4208054 | -0.5245021 | 0.5999294 | 0.766139 |
| ENSG00000176720 | BOK | 3064.569 | -0.1869095 | 0.1741161 | -1.111737 | 0.2662514 | 0.461872 |
| ENSG00000274944 | | 0.8642815 | -0.1869488 | 0.7848705 | -0.8160661 | 0.4144623 | NA |
| ENSG00000105723 | GSK3A | 2978.7203 | -0.186949278 | 0.165107474 | -1.155562003 | 0.24786036 | 0.440054 |
| ENSG00000198836 | OPA1 | 6136.909 | -0.1869496 | 0.1915065 | -0.9953546 | 0.3195638 | 0.520468 |
| ENSG00000267159 | | 10.69274 | -0.1869783 | 0.5902926 | -0.4739584 | 0.6355296 | 0.791441 |
| ENSG00000237649 | KIFC1 | 2834.371 | -0.1870067 | 0.4393728 | -0.5179007 | 0.6045275 | 0.769357 |
| ENSG00000249379 | | 11.66253 | -0.1870799 | 0.4627494 | -0.4947848 | 0.6207521 | 0.780967 |
| ENSG00000237276 | ANO7L1 | 59.10743 | -0.1871047 | 0.3376532 | -0.6113154 | 0.5409908 | 0.722836 |
| ENSG00000162735 | PEX19 | 2023.711 | -0.1871697 | 0.1323844 | -1.435033 | 0.1512776 | 0.313035 |
| ENSG00000158062 | UBXN11 | 1155.899 | -0.1872041 | 0.2100482 | -0.9304201 | 0.3521536 | 0.553894 |
| ENSG00000075785 | RAB7A | 16434.03568 | -0.1872496 | 0.129827686 | -1.464481903 | 0.14306232 | 0.300833 |
| ENSG00000277396 | | 2.754263 | -0.1872696 | 0.7072993 | -0.5375448 | 0.5908913 | 0.759562 |
| ENSG00000106304 | SPAM1 | 32.21102162 | -0.187322191 | 0.485523597 | -0.466609959 | 0.64077896 | 0.795108 |
| ENSG00000121579 | NAA50 | 12903.13976 | -0.187401506 | 0.198177892 | -0.980332094 | 0.32692222 | 0.528185 |
| ENSG00000232234 | | 2.177588 | -0.1874384 | 0.7987352 | -0.5293433 | 0.5965673 | 0.763644 |
| ENSG00000147872 | PLIN2 | 497.9943 | -0.1874576 | 0.308947 | -0.663141 | 0.5072403 | 0.693672 |
| ENSG00000168685 | IL7R | 0.9568048 | -0.1875498 | 0.7983225 | -0.5454445 | 0.5854478 | NA |
| ENSG00000109184 | DCUN1D4 | 2895.679879 | -0.18755879 | 0.134515818 | -1.451795227 | 0.14655855 | 0.306076 |
| ENSG00000260240 | APOOP5 | 5.372173 | -0.187759 | 0.6646263 | -0.5040083 | 0.6142555 | 0.776263 |
| ENSG00000277159 | | 25.23938 | -0.1877972 | 0.5251125 | -0.4756262 | 0.6343407 | 0.790553 |
| ENSG00000272654 | | 127.8836 | -0.187802 | 0.2453778 | -0.8058638 | 0.4203214 | 0.620177 |
| ENSG00000134339 | SAA2 | 2.93924389 | -0.187812044 | 0.797706786 | -0.553945601 | 0.57961608 | 0.75067 |
| ENSG00000226286 | | 1.995382 | -0.1878492 | 0.7987564 | -0.5320414 | 0.5946973 | 0.762132 |
| ENSG00000248256 | OCIAD1-AS1 | 6.609259 | -0.1879268 | 0.5167181 | -0.4811626 | 0.630401 | 0.787935 |
| ENSG00000259397 | | 29.4085 | -0.1879436 | 0.3543886 | -0.5935105 | 0.5528395 | 0.731338 |

| | | | | | | | |
|---|---|---|---|---|---|---|---|
| ENSG00000230068 | CDC42-IT1 | 1.044608 | -0.1880696 | 0.7413323 | -0.6099979 | 0.5418632 | NA |
| ENSG00000233868 | LLPHP3 | 380.9627 | -0.1882153 | 0.2082831 | -0.9436526 | 0.3453472 | 0.546879 |
| ENSG00000213514 | | 67.42654 | -0.1884085 | 0.432768 | -0.522563 | 0.6012784 | 0.766989 |
| ENSG00000126067 | PSMB2 | 5981.135272 | -0.188454955 | 0.232318809 | -0.848018534 | 0.39642765 | 0.597631 |
| ENSG00000163634 | THOC7 | 17528.21 | -0.1884677 | 0.1651013 | -1.170557 | 0.2417768 | 0.432344 |
| ENSG00000175224 | ATG13 | 5720.878 | -0.1885222 | 0.113937 | -1.663597 | 0.09619304 | 0.226018 |
| ENSG00000152359 | POC5 | 996.6137 | -0.1885271 | 0.1887775 | -1.025881 | 0.3049477 | 0.504662 |
| ENSG00000162604 | TM2D1 | 1151.774 | -0.188669 | 0.1589518 | -1.236236 | 0.216371 | 0.401702 |
| ENSG00000255520 | | 1.560782 | -0.1888734 | 0.7548856 | -0.6458481 | 0.5183777 | NA |
| ENSG00000256994 | LINC02378 | 1.000502 | -0.1888976 | 0.821876 | -1.531828 | 0.125565 | NA |
| ENSG00000231665 | OGFOD1P1 | 1.03244 | -0.1889169 | 0.7560712 | -0.670372 | 0.5026206 | NA |
| ENSG00000007952 | NOX1 | 36.09628408 | -0.188919085 | 0.293433405 | -0.69596541 | 0.48645049 | 0.677456 |
| ENSG00000072062 | PRKACA | 3566.506359 | -0.18892978 | 0.220016369 | -0.901417335 | 0.36736647 | 0.569461 |
| ENSG00000214045 | IMPDH1P2 | 4.775393 | -0.1890041 | 0.6302341 | -0.4828277 | 0.6292181 | 0.787012 |
| ENSG00000135636 | DYSF | 12.9167876 | -0.189016863 | 0.528558963 | -0.482716315 | 0.62929718 | 0.787012 |
| ENSG00000202079 | | 1.659418 | -0.1890579 | 0.735196 | -0.606184 | 0.5443926 | 0.725217 |
| ENSG00000271936 | | 50.34889 | -0.189065 | 0.482643 | -0.4991701 | 0.6176596 | 0.77875 |
| ENSG00000149115 | TNKS1BP1 | 7395.886 | -0.1891345 | 0.1482243 | -1.334447 | 0.1820574 | 0.356928 |
| ENSG00000269930 | | 1.055765 | -0.1892132 | 0.7602294 | -0.6729467 | 0.5009812 | NA |
| ENSG00000040199 | PHLPP2 | 2874.393705 | -0.189219429 | 0.188630643 | -1.054684333 | 0.29156972 | 0.490317 |
| ENSG00000233124 | LINC00456 | 7.771996 | -0.1892443 | 0.5465395 | -0.4718201 | 0.6370552 | 0.792672 |
| ENSG00000172845 | SP3 | 5610.439 | -0.1892907 | 0.1635881 | -1.196559 | 0.2314783 | 0.419902 |
| ENSG00000162521 | RBBP4 | 5878.317 | -0.1893033 | 0.1215515 | -1.555908 | 0.1197299 | 0.264805 |
| ENSG00000253632 | | 1.475458 | -0.1893604 | 0.7830952 | -0.8139825 | 0.415655 | NA |
| ENSG00000165874 | SHLD2P1 | 2.840458 | -0.1893842 | 0.6514284 | -0.4936601 | 0.6215463 | 0.781315 |
| ENSG00000148411 | NACC2 | 3555.041 | -0.1894898 | 0.2014449 | -0.9647689 | 0.3346606 | 0.535897 |
| ENSG00000102317 | RBM3 | 11808.45439 | -0.189492966 | 0.207792548 | -0.95530547 | 0.33942324 | 0.540688 |
| ENSG00000162398 | LEXM | 1.799187 | -0.1896286 | 0.7288113 | -0.578405 | 0.5629907 | 0.738599 |
| ENSG00000177776 | RPS2P2 | 0.965058 | -0.1896858 | 0.7824372 | -0.8149455 | 0.4151035 | NA |
| ENSG00000260809 | VPS35P1 | 0.8438482 | -0.1897735 | 0.7768877 | -0.729552 | 0.4656641 | NA |
| ENSG00000287700 | | 1.362282 | -0.1897864 | 0.8072972 | -1.116674 | 0.2641336 | NA |
| ENSG00000137693 | YAP1 | 3492.817018 | -0.189915283 | 0.155435373 | -1.26614222 | 0.20546217 | 0.387433 |
| ENSG00000104290 | FZD3 | 1201.590041 | -0.19002823 | 0.137935255 | -1.399826586 | 0.16156525 | 0.32861 |

| | | | | | | | |
|---|---|---|---|---|---|---|---|
| ENSG00000107829 | FBXW4 | 2783.224237 | -0.190045344 | 0.161365558 | -1.202093767 | 0.2293272 | 0.417414 |
| ENSG00000167523 | SPATA33 | 783.9902 | -0.1900563 | 0.2032891 | -0.975456 | 0.3293341 | 0.530436 |
| ENSG00000253653 | | 1.495672 | -0.1900769 | 0.7548955 | -0.64064 | 0.5217566 | NA |
| ENSG00000226043 | | 3.88653 | -0.190102 | 0.6819761 | -0.5212472 | 0.6021946 | 0.767619 |
| ENSG00000259442 | | 1.266651 | -0.1901127 | 0.7484266 | -0.625471 | 0.531662 | NA |
| ENSG00000263164 | | 4.21314 | -0.1901571 | 0.6196142 | -0.4833288 | 0.6288623 | 0.786756 |
| ENSG00000186184 | POLR1D | 7830.164 | -0.1902379 | 0.1607456 | -1.220451 | 0.2222938 | 0.408955 |
| ENSG00000224908 | TIMM8BP2 | 5.271819 | -0.1902406 | 0.5649852 | -0.4776484 | 0.6329005 | 0.789647 |
| ENSG00000169203 | NPIPB12 | 26.47272 | -0.1902847 | 0.36529 | -0.5893255 | 0.5556429 | 0.733158 |
| ENSG00000165671 | NSD1 | 12253.75 | -0.1903397 | 0.2193885 | -0.9073178 | 0.3642388 | 0.566364 |
| ENSG00000162430 | SELENON | 2212.849 | -0.1903614 | 0.1750916 | -1.129862 | 0.2585343 | 0.452699 |
| ENSG00000219410 | | 11.63673 | -0.1903681 | 0.5577414 | -0.4800222 | 0.6312116 | 0.788593 |
| ENSG00000261302 | | 1.232267 | -0.1903969 | 0.750773 | -0.6341849 | 0.5259602 | NA |
| ENSG00000157456 | CCNB2 | 2256.795 | -0.1904408 | 0.5388509 | -0.487204 | 0.6261138 | 0.784865 |
| ENSG00000258988 | SF3B4P1 | 4.333861 | -0.1905269 | 0.6219664 | -0.4893787 | 0.6245736 | 0.783563 |
| ENSG00000177889 | UBE2N | 6552.955 | -0.1905787 | 0.2269214 | -0.8571295 | 0.3913733 | 0.5932 |
| ENSG00000163257 | DCAF16 | 1418.75 | -0.1908379 | 0.2005916 | -0.9908151 | 0.3217759 | 0.522603 |
| ENSG00000252835 | SCARNA21 | 9.796095 | -0.1909025 | 0.57571 | -0.4763012 | 0.6338598 | 0.790244 |
| ENSG00000173825 | TIGD3 | 110.6716 | -0.1910394 | 0.3019382 | -0.6864108 | 0.4924541 | 0.68249 |
| ENSG00000080819 | CPOX | 1205.951501 | -0.191041625 | 0.209151236 | -0.951596366 | 0.34130173 | 0.542509 |
| ENSG00000188021 | UBQLN2 | 2646.519 | -0.1910594 | 0.171195 | -1.145885 | 0.2518427 | 0.444599 |
| ENSG00000255035 | SDHCP4 | 185.9988 | -0.1910627 | 0.2074354 | -0.9580681 | 0.3380284 | 0.539339 |
| ENSG00000230929 | | 7.627217 | -0.1911006 | 0.5388651 | -0.4805275 | 0.6308524 | 0.788244 |
| ENSG00000277948 | | 1.436874 | -0.1912989 | 0.7437725 | -0.6143208 | 0.5390033 | NA |
| ENSG00000273765 | | 1.350717 | -0.1913631 | 0.7180997 | -0.5523295 | 0.5807226 | NA |
| ENSG00000240549 | THOC7-AS1 | 14.22497 | -0.1913728 | 0.4561416 | -0.5108666 | 0.6094444 | 0.772932 |
| ENSG00000234219 | CDCA4P4 | 1.727479 | -0.1915059 | 0.7100832 | -0.5556556 | 0.5784463 | 0.749955 |
| ENSG00000225082 | DAP3P1 | 3.478704 | -0.1915183 | 0.6050463 | -0.4837589 | 0.628557 | 0.786555 |
| ENSG00000240964 | RN7SL751P | 1.181802 | -0.1915224 | 0.7659263 | -0.7128785 | 0.475921 | NA |
| ENSG00000100218 | RSPH14 | 5.116550403 | -0.191563228 | 0.657563636 | -0.499526185 | 0.61740875 | 0.77871 |
| ENSG00000137501 | SYTL2 | 15385.96142 | -0.19160517 | 0.270047854 | -0.760142098 | 0.44716965 | 0.643914 |
| ENSG00000231500 | RPS18 | 25847.77 | -0.1916299 | 0.1770129 | -1.127536 | 0.2595159 | 0.453833 |
| ENSG00000233469 | ST6GALNAC4P1 | 2.648373 | -0.1916567 | 0.7016842 | -0.5441193 | 0.5863594 | 0.756114 |

| | | | | | | | |
|---|---|---|---|---|---|---|---|
| ENSG00000197849 | OR8G1 | 3.084933 | -0.191739 | 0.6816805 | -0.5163064 | 0.6056404 | 0.770027 |
| ENSG00000160318 | CLDND2 | 56.87608 | -0.1917552 | 0.5136427 | -0.4908962 | 0.6234999 | 0.782785 |
| ENSG00000279285 | | 1.526832 | -0.1917579 | 0.7461249 | -0.640379 | 0.5219263 | NA |
| ENSG00000181924 | COA4 | 4857.196 | -0.1917971 | 0.1865764 | -1.078768 | 0.2806914 | 0.478149 |
| ENSG00000005483 | KMT2E | 2978.395119 | -0.191835518 | 0.294107808 | -0.70233454 | 0.48247056 | 0.673856 |
| ENSG00000120256 | LRP11 | 168.014361 | -0.191857007 | 0.237888815 | -0.847874966 | 0.39650761 | 0.597631 |
| ENSG00000163612 | FAM86KP | 4.417408 | -0.1919679 | 0.6222839 | -0.4932861 | 0.6218104 | 0.781536 |
| ENSG00000166104 | | 1.0056 | -0.192042 | 0.807414 | -1.118642 | 0.2632927 | NA |
| ENSG00000270277 | | 7.322728 | -0.1920645 | 0.6325639 | -0.4953747 | 0.6203356 | 0.780716 |
| ENSG00000232486 | DEPDC1P2 | 23.95521 | -0.1920714 | 0.5520174 | -0.4859188 | 0.6270247 | 0.785493 |
| ENSG00000236872 | PPIAP36 | 3.164139 | -0.1921042 | 0.6727182 | -0.5237251 | 0.6004698 | 0.766534 |
| ENSG00000119760 | SUPT7L | 4074.797447 | -0.192177973 | 0.16959719 | -1.162489726 | 0.2450366 | 0.436733 |
| ENSG00000232940 | HCG25 | 53.55347 | -0.192267 | 0.447133 | -0.5210578 | 0.6023265 | 0.767619 |
| ENSG00000114115 | RBP1 | 1.032865191 | -0.192319361 | 0.758992659 | -0.673878762 | 0.5003884 | NA |
| ENSG00000185924 | RTN4RL1 | 1217.51 | -0.1923777 | 0.300546 | -0.6977735 | 0.4853189 | 0.676341 |
| ENSG00000214915 | | 1.532341 | -0.1924024 | 0.7319195 | -0.6035228 | 0.546161 | NA |
| ENSG00000135315 | CEP162 | 434.5797151 | -0.192451306 | 0.214868874 | -0.932955321 | 0.35084304 | 0.552421 |
| ENSG00000134532 | SOX5 | 16.12500247 | -0.192550818 | 0.564771423 | -0.505926459 | 0.61290828 | 0.775667 |
| ENSG00000225313 | | 60.27903 | -0.1926239 | 0.3092139 | -0.6775998 | 0.4980255 | 0.68656 |
| ENSG00000184863 | RBM33 | 3511.758 | -0.1926602 | 0.1858169 | -1.078973 | 0.2805998 | 0.478085 |
| ENSG00000254339 | | 1.101614 | -0.1926742 | 0.7556831 | -0.6597028 | 0.5094446 | NA |
| ENSG00000138495 | COX17 | 1784.154464 | -0.192704655 | 0.233163936 | -0.875034825 | 0.38155496 | 0.583769 |
| ENSG00000213055 | EEF1B2P7 | 0.8996342 | -0.1927178 | 0.7873869 | -0.8085775 | 0.4187582 | NA |
| ENSG00000163344 | PMVK | 2273.007 | -0.1927791 | 0.14882 | -1.321771 | 0.1862445 | 0.362435 |
| ENSG00000174456 | C12orf76 | 573.796 | -0.1927811 | 0.2455876 | -0.830623 | 0.4061866 | 0.606802 |
| ENSG00000267677 | | 2.990649 | -0.1927898 | 0.6219502 | -0.4925974 | 0.6222971 | 0.781815 |
| ENSG00000169914 | OTUD3 | 1117.277 | -0.1927931 | 0.2128076 | -0.9694488 | 0.3323213 | 0.533405 |
| ENSG00000109911 | ELP4 | 950.7430858 | -0.19279937 | 0.230053575 | -0.879942149 | 0.37889065 | 0.58102 |
| ENSG00000085552 | IGSF9 | 2199.304189 | -0.192804231 | 0.246494288 | -0.809314301 | 0.41833438 | 0.618103 |
| ENSG00000163626 | COX18 | 704.5357 | -0.1928064 | 0.1721716 | -1.153184 | 0.248835 | 0.441175 |
| ENSG00000261208 | | 1.252222 | -0.1928423 | 0.7546994 | -0.6719785 | 0.5015974 | NA |
| ENSG00000227694 | RPL23AP74 | 1.167571 | -0.1928627 | 0.7766344 | -0.7798929 | 0.4354539 | NA |
| ENSG00000147257 | GPC3 | 31.20374 | -0.1928644 | 0.5177619 | -0.4923017 | 0.6225061 | 0.781966 |

| | | | | | | | |
|---|---|---|---|---|---|---|---|
| ENSG00000225558 | UBE2D3P2 | 246.4661 | -0.1929171 | 0.1778341 | -1.11602 | 0.2644136 | 0.459938 |
| ENSG00000185344 | ATP6V0A2 | 1778.423 | -0.1930121 | 0.127199 | -1.547671 | 0.1217014 | 0.267985 |
| ENSG00000229117 | RPL41 | 5490.037 | -0.1930314 | 0.1884992 | -0.9956645 | 0.3194132 | 0.520324 |
| ENSG00000138435 | CHRNA1 | 5.213934833 | -0.193083835 | 0.631125057 | -0.493070304 | 0.62196291 | 0.781676 |
| ENSG00000224883 | | 1.002115 | -0.1931469 | 0.76659 | -0.7117058 | 0.476647 | NA |
| ENSG00000102978 | POLR2C | 3523.20872 | -0.193174392 | 0.203631419 | -0.981834096 | 0.32618159 | 0.527405 |
| ENSG00000205209 | SCGB2B2 | 9.950024 | -0.1933023 | 0.4547194 | -0.523989 | 0.6002862 | 0.766374 |
| ENSG00000168386 | FILIP1L | 74.95114 | -0.1933319 | 0.5580941 | -0.4869258 | 0.6263109 | 0.784964 |
| ENSG00000231713 | | 19.29407 | -0.1933631 | 0.5200301 | -0.507988 | 0.6114617 | 0.774478 |
| ENSG00000249858 | SNX5P1 | 72.50463 | -0.1934235 | 0.3142555 | -0.6749887 | 0.499683 | 0.68795 |
| ENSG00000267298 | | 1.390261 | -0.1934586 | 0.7548443 | -0.6592558 | 0.5097315 | NA |
| ENSG00000167889 | MGAT5B | 281.1596 | -0.1935038 | 0.3033193 | -0.693143 | 0.4882198 | 0.679101 |
| ENSG00000276523 | | 50.30217 | -0.1935309 | 0.432971 | -0.5360741 | 0.5919074 | 0.760374 |
| ENSG00000212789 | ST13P5 | 255.8766 | -0.1935314 | 0.1739017 | -1.144857 | 0.2522684 | 0.444939 |
| ENSG00000227081 | | 5257.776 | -0.1935577 | 0.1604675 | -1.239258 | 0.2152499 | 0.400292 |
| ENSG00000177803 | YWHAQP4 | 3.022354 | -0.1936022 | 0.6286546 | -0.4988686 | 0.6178719 | 0.77887 |
| ENSG00000188186 | LAMTOR4 | 3513.133 | -0.1936346 | 0.2049762 | -0.9827407 | 0.3257351 | 0.527036 |
| ENSG00000110057 | UNC93B1 | 4669.595239 | -0.193668114 | 0.212402743 | -0.948167429 | 0.34304423 | 0.544595 |
| ENSG00000148634 | HERC4 | 3924.847 | -0.1936945 | 0.1394129 | -1.425338 | 0.1540595 | 0.317377 |
| ENSG00000207696 | MIR659 | 5.088031 | -0.1937007 | 0.5845331 | -0.4876721 | 0.6257821 | 0.78456 |
| ENSG00000131467 | PSME3 | 8808.285638 | -0.193760337 | 0.17011761 | -1.175089469 | 0.23995892 | 0.430165 |
| ENSG00000106803 | SEC61B | 4586.830004 | -0.193834506 | 0.184976975 | -1.102187549 | 0.27038014 | 0.466781 |
| ENSG00000217165 | ANKRD18EP | 465.1702 | -0.1938591 | 0.2158402 | -0.9406053 | 0.3469072 | 0.548564 |
| ENSG00000116266 | STXBP3 | 3126.334811 | -0.193888658 | 0.126407836 | -1.567646487 | 0.11696366 | 0.260334 |
| ENSG00000256254 | | 1.096673 | -0.1939157 | 0.7997325 | -0.9452628 | 0.3445247 | NA |
| ENSG00000271161 | BOLA2P2 | 5.541217 | -0.1939459 | 0.6297428 | -0.4903096 | 0.6239148 | 0.783106 |
| ENSG00000145725 | PPIP5K2 | 2666.137 | -0.1939966 | 0.2023424 | -0.9999155 | 0.3173514 | 0.518364 |
| ENSG00000237571 | LINC02862 | 1.948617 | -0.1940259 | 0.7580176 | -0.714892 | 0.4746758 | 0.66739 |
| ENSG00000173846 | PLK3 | 319.4496 | -0.1941064 | 0.3278952 | -0.6585928 | 0.5101573 | 0.69606 |
| ENSG00000224165 | DNAJC27-AS1 | 134.43 | -0.194147 | 0.3627143 | -0.6060266 | 0.5444971 | 0.725223 |
| ENSG00000146666 | LINC00525 | 2.630207 | -0.1941702 | 0.7998954 | -0.5458794 | 0.5851488 | 0.75503 |
| ENSG00000148357 | HMCN2 | 4.794754 | -0.1942561 | 0.5769407 | -0.4890548 | 0.6248029 | 0.78374 |
| ENSG00000197429 | IPP | 485.8936 | -0.1942569 | 0.2130233 | -0.9518271 | 0.3411847 | 0.542486 |

| | | | | | | | |
|---|---|---|---|---|---|---|---|
| ENSG00000158480 | SPATA2 | 2453.24 | -0.1943304 | 0.1627708 | -1.217166 | 0.2235411 | 0.41023 |
| ENSG00000287691 | | 2.924338 | -0.1943454 | 0.7054969 | -0.5514042 | 0.5813567 | 0.752007 |
| ENSG00000147650 | LRP12 | 1194.531 | -0.1943534 | 0.2235064 | -0.9082828 | 0.3637288 | 0.565869 |
| ENSG00000232203 | SLC25A6P2 | 6.384899 | -0.1944901 | 0.5910899 | -0.4866604 | 0.626499 | 0.785163 |
| ENSG00000173578 | XCR1 | 1.255669 | -0.1944938 | 0.7204514 | -0.576111 | 0.5645402 | NA |
| ENSG00000135144 | DTX1 | 33.84413601 | -0.194502126 | 0.4020308 | -0.563530843 | 0.57307346 | 0.746043 |
| ENSG00000123201 | GUCY1B2 | 625.7805655 | -0.194735321 | 0.36409801 | -0.61036374 | 0.54162088 | 0.723277 |
| ENSG00000068120 | COASY | 3931.799359 | -0.194735994 | 0.184709205 | -1.089865969 | 0.27577219 | 0.472648 |
| ENSG00000110536 | PTPMT1 | 822.3693902 | -0.194776046 | 0.183238102 | -1.08422896 | 0.27826329 | 0.475445 |
| ENSG00000196504 | PRPF40A | 8815.429 | -0.1948943 | 0.1440404 | -1.378725 | 0.1679794 | 0.337649 |
| ENSG00000213498 | TPT1P13 | 1.318495 | -0.1950754 | 0.7805154 | -0.8118821 | 0.4168593 | NA |
| ENSG00000100714 | MTHFD1 | 8314.385423 | -0.195084262 | 0.290215933 | -0.727143212 | 0.46713823 | 0.660878 |
| ENSG00000228034 | HMGN2P21 | 10.64879 | -0.1951793 | 0.5332426 | -0.4973393 | 0.6189498 | 0.779563 |
| ENSG00000101751 | POLI | 2330.151009 | -0.195310866 | 0.240045042 | -0.839994423 | 0.40091151 | 0.601913 |
| ENSG00000167904 | TMEM68 | 1119.028 | -0.1953211 | 0.1815441 | -1.132405 | 0.2574642 | 0.451433 |
| ENSG00000160410 | SHKBP1 | 2625.541 | -0.195371 | 0.1631751 | -1.230166 | 0.2186349 | 0.404436 |
| ENSG00000282936 | | 173.1988 | -0.1954211 | 0.3307582 | -0.6538473 | 0.5132102 | 0.699148 |
| ENSG00000171533 | MAP6 | 2.94877 | -0.195433 | 0.6704347 | -0.5170639 | 0.6051116 | 0.769686 |
| ENSG00000120008 | WDR11 | 3670.643506 | -0.19553213 | 0.124486947 | -1.577386897 | 0.11470653 | 0.256881 |
| ENSG00000236241 | | 0.9870728 | -0.1955456 | 0.8053075 | -1.074303 | 0.282687 | NA |
| ENSG00000118507 | AKAP7 | 98.57008835 | -0.195569931 | 0.283557438 | -0.748221736 | 0.45432642 | 0.65042 |
| ENSG00000151287 | TEX30 | 373.7696 | -0.1955922 | 0.3000884 | -0.7115853 | 0.4767216 | 0.669062 |
| ENSG00000101222 | SPEF1 | 29.43963046 | -0.195610129 | 0.404264023 | -0.56442822 | 0.57246274 | 0.745833 |
| ENSG00000206612 | SNORA2A | 3.082047 | -0.1956179 | 0.6487673 | -0.5041157 | 0.6141801 | 0.776236 |
| ENSG00000266598 | | 1.062158 | -0.1956305 | 0.8171 | -1.036514 | 0.2999624 | NA |
| ENSG00000255151 | GLYATL1B | 10.03377 | -0.1957793 | 0.5208998 | -0.4964757 | 0.6195588 | 0.780127 |
| ENSG00000086758 | HUWE1 | 30904.49274 | -0.195876794 | 0.232133306 | -0.887857716 | 0.37461729 | 0.57683 |
| ENSG00000225492 | GBP1P1 | 2.048957 | -0.195882 | 0.7545725 | -0.6520954 | 0.5143397 | 0.700097 |
| ENSG00000259075 | POC1B-GALNT4 | 5.021924 | -0.195968 | 0.627028 | -0.4980939 | 0.6184178 | 0.779299 |
| ENSG00000255128 | HSPD1P3 | 2.338598 | -0.1960173 | 0.7346495 | -0.6141892 | 0.5390903 | 0.721572 |
| ENSG00000168439 | STIP1 | 15300.32 | -0.1960295 | 0.1624249 | -1.315805 | 0.1882394 | 0.365274 |
| ENSG00000118193 | KIF14 | 1891.634817 | -0.196088131 | 0.443010135 | -0.540476351 | 0.58886857 | 0.757928 |
| ENSG00000271200 | | 5.286639 | -0.1961141 | 0.6183105 | -0.4974251 | 0.6188893 | 0.779563 |

| | | | | | | | |
|---|---|---|---|---|---|---|---|
| ENSG00000198691 | ABCA4 | 591.7739 | -0.1961604 | 0.5693945 | -0.4873728 | 0.6259942 | 0.784752 |
| ENSG00000275381 | | 3.128748 | -0.1962276 | 0.7040721 | -0.5548393 | 0.5790046 | 0.750278 |
| ENSG00000203325 | | 1.417119 | -0.1962632 | 0.7177584 | -0.574463 | 0.5656545 | NA |
| ENSG00000233538 | | 1.638692 | -0.1962915 | 0.7336497 | -0.6168618 | 0.5373259 | 0.720366 |
| ENSG00000279738 | | 40.65739 | -0.1963051 | 0.37707 | -0.6019986 | 0.5471751 | 0.726931 |
| ENSG00000182173 | TSEN54 | 9183.878 | -0.1963074 | 0.172954 | -1.17417 | 0.2403271 | 0.430592 |
| ENSG00000145861 | C1QTNF2 | 1.629674 | -0.1963375 | 0.7272805 | -0.6040669 | 0.5457992 | 0.726117 |
| ENSG00000248333 | CDK11B | 2103.106 | -0.1963695 | 0.1730718 | -1.154719 | 0.2482056 | 0.44049 |
| ENSG00000287593 | | 0.8690881 | -0.196386 | 0.7675424 | -0.7321264 | 0.4640914 | NA |
| ENSG00000180210 | F2 | 1.527451 | -0.1964554 | 0.7055424 | -0.5598214 | 0.5756013 | NA |
| ENSG00000154945 | ANKRD40 | 3370.686 | -0.1964698 | 0.1893767 | -1.077366 | 0.2813166 | 0.478907 |
| ENSG00000023228 | NDUFS1 | 7209.462529 | -0.196504944 | 0.180745809 | -1.102061224 | 0.27043506 | 0.466782 |
| ENSG00000269959 | SPACA6P-AS | 156.1755 | -0.1965538 | 0.607045 | -0.5078472 | 0.6115605 | 0.774478 |
| ENSG00000178741 | COX5A | 5177.294 | -0.1968061 | 0.193795 | -1.063298 | 0.2876468 | 0.485978 |
| ENSG00000138018 | SELENOI | 5043.160402 | -0.196846526 | 0.2123161 | -0.965484785 | 0.33430205 | 0.535452 |
| ENSG00000166164 | BRD7 | 2589.227 | -0.1968706 | 0.1043311 | -1.907599 | 0.05644302 | 0.152016 |
| ENSG00000162641 | AKNAD1 | 6.505918 | -0.1969418 | 0.630165 | -0.5078818 | 0.6115362 | 0.774478 |
| ENSG00000260593 | | 14.85049 | -0.1970109 | 0.579588 | -0.4815592 | 0.6301191 | 0.787755 |
| ENSG00000234076 | TPRG1-AS1 | 0.8317873 | -0.1970297 | 0.8226729 | -1.36592 | 0.1719641 | NA |
| ENSG00000234782 | TPT1P9 | 737.6298 | -0.1970691 | 0.2266289 | -0.9129959 | 0.3612447 | 0.563225 |
| ENSG00000278705 | H4C2 | 6.889028 | -0.1970831 | 0.6180792 | -0.4943086 | 0.6210882 | 0.781108 |
| ENSG00000226472 | | 4.18229 | -0.1972294 | 0.6356466 | -0.5050325 | 0.613536 | 0.775861 |
| ENSG00000080644 | CHRNA3 | 54.04959689 | -0.197283738 | 0.367927178 | -0.608573025 | 0.54280749 | 0.723848 |
| ENSG00000227413 | | 3.032041 | -0.1972867 | 0.7028953 | -0.565107 | 0.572001 | 0.745523 |
| ENSG00000162878 | PKDCC | 43.25954 | -0.1973718 | 0.4795664 | -0.5151431 | 0.6064531 | 0.770618 |
| ENSG00000207870 | MIR221 | 2.222923 | -0.1974006 | 0.8003497 | -0.5587952 | 0.5763015 | 0.748427 |
| ENSG00000120802 | TMPO | 9772.509958 | -0.197517658 | 0.330975842 | -0.663204623 | 0.50719951 | 0.693663 |
| ENSG00000259782 | | 1.42686 | -0.1975763 | 0.7717442 | -0.7386697 | 0.4601076 | NA |
| ENSG00000226540 | GAPDHP73 | 2.897954 | -0.1976917 | 0.7069931 | -0.5601476 | 0.5753788 | 0.747542 |
| ENSG00000150779 | TIMM8B | 1326.139 | -0.1977866 | 0.2143814 | -0.9630983 | 0.3354982 | 0.53672 |
| ENSG00000234287 | | 3931.114 | -0.1978324 | 0.1423554 | -1.412198 | 0.1578915 | 0.323133 |
| ENSG00000238278 | ALG1L6P | 134.2308 | -0.1978505 | 0.3313903 | -0.66159 | 0.508234 | 0.694648 |
| ENSG00000240710 | | 13.61878 | -0.1978974 | 0.5913368 | -0.5016468 | 0.615916 | 0.777511 |

| | | | | | | | |
|---|---|---|---|---|---|---|---|
| ENSG00000242697 | RPL5P12 | 12.34901 | -0.1979569 | 0.5230804 | -0.5001409 | 0.6169759 | 0.778339 |
| ENSG00000213371 | NAP1L1P3 | 323.8977 | -0.1979821 | 0.1701877 | -1.190257 | 0.2339453 | 0.423066 |
| ENSG00000157578 | LCA5L | 392.948 | -0.1980337 | 0.1695682 | -1.197141 | 0.2312516 | 0.419681 |
| ENSG00000285155 | | 1.654529 | -0.1981649 | 0.7994109 | -0.5872372 | 0.5570444 | 0.734509 |
| ENSG00000231831 | MTHFD1P1 | 117.9762 | -0.1982074 | 0.3724841 | -0.6043173 | 0.5456327 | 0.726007 |
| ENSG00000277050 | | 16.91984 | -0.1982272 | 0.5133598 | -0.5094301 | 0.6104508 | 0.773884 |
| ENSG00000113966 | ARL6 | 188.3168703 | -0.198260724 | 0.28864703 | -0.742621946 | 0.4577106 | 0.65359 |
| ENSG00000232665 | PHBP10 | 4.419512 | -0.1982785 | 0.6784822 | -0.5359572 | 0.5919882 | 0.760374 |
| ENSG00000196834 | POTEI | 27.99584 | -0.1982875 | 0.4595125 | -0.5310925 | 0.5953547 | 0.76268 |
| ENSG00000101052 | IFT52 | 1473.002434 | -0.19832342 | 0.158439451 | -1.289406967 | 0.19725664 | 0.377234 |
| ENSG00000287387 | | 17.08632 | -0.1983297 | 0.5502591 | -0.4952063 | 0.6204545 | 0.780792 |
| ENSG00000013374 | NUB1 | 2981.055157 | -0.198537913 | 0.156464092 | -1.321249486 | 0.18641819 | 0.362667 |
| ENSG00000235246 | | 1.180594 | -0.1985695 | 0.7536505 | -0.6764733 | 0.4987402 | NA |
| ENSG00000282728 | PGR-AS1 | 8.519759 | -0.1985708 | 0.6291832 | -0.5360011 | 0.5919578 | 0.760374 |
| ENSG00000232600 | TONSL-AS1 | 37.64346 | -0.1986344 | 0.3434374 | -0.6470761 | 0.5175827 | 0.70287 |
| ENSG00000257763 | OR5BK1P | 2.811567 | -0.1987502 | 0.6851831 | -0.5408369 | 0.58862 | 0.757816 |
| ENSG00000218180 | SLC25A5P7 | 1.11435 | -0.1988009 | 0.7778337 | -0.7897585 | 0.4296688 | NA |
| ENSG00000233524 | RANP8 | 2.336991 | -0.1988528 | 0.706995 | -0.5677374 | 0.5702133 | 0.744033 |
| ENSG00000279422 | | 5.804267 | -0.1988967 | 0.6206319 | -0.5053859 | 0.6132878 | 0.775778 |
| ENSG00000184100 | BRD7P2 | 693.591 | -0.198907 | 0.113023 | -1.787535 | 0.07385109 | 0.186127 |
| ENSG00000287574 | | 79.89754 | -0.1989756 | 0.3134498 | -0.6921139 | 0.4888658 | 0.679677 |
| ENSG00000230699 | | 174.3282 | -0.1990071 | 0.2689077 | -0.7915026 | 0.4286508 | 0.62791 |
| ENSG00000078668 | VDAC3 | 4157.942636 | -0.199010336 | 0.277322885 | -0.784297359 | 0.43286565 | 0.632017 |
| ENSG00000143157 | POGK | 7001.364 | -0.1990408 | 0.1557834 | -1.324155 | 0.1854517 | 0.361131 |
| ENSG00000198612 | COPS8 | 1713.89 | -0.1990865 | 0.1909601 | -1.089577 | 0.2758995 | 0.472815 |
| ENSG00000274015 | | 39.3604 | -0.1991162 | 0.294278 | -0.7282612 | 0.4664537 | 0.660267 |
| ENSG00000172073 | TEX37 | 1.504495 | -0.1991441 | 0.8005694 | -0.5680234 | 0.5700191 | NA |
| ENSG00000287097 | | 11.75638 | -0.1991453 | 0.7488895 | -0.6330831 | 0.5266794 | 0.711659 |
| ENSG00000279413 | | 38.47938 | -0.1991716 | 0.3008369 | -0.7202068 | 0.4713977 | 0.664715 |
| ENSG00000261114 | | 28.17092 | -0.1992012 | 0.4993813 | -0.515909 | 0.6059179 | 0.770306 |
| ENSG00000228413 | | 3.133479 | -0.199203 | 0.6976828 | -0.5574774 | 0.5772013 | 0.749068 |
| ENSG00000267567 | | 8.250683 | -0.1993228 | 0.5503656 | -0.5017853 | 0.6158186 | 0.777425 |
| ENSG00000179997 | RPA3P1 | 0.9470706 | -0.1993236 | 0.7721453 | -0.7605271 | 0.4469396 | NA |

| | | | | | | | |
|---|---|---|---|---|---|---|---|
| ENSG00000276757 | RN7SL192P | 8.262377 | -0.1993359 | 0.5149007 | -0.503559 | 0.6145713 | 0.776509 |
| ENSG00000206559 | ZCWPW2 | 32.59781 | -0.1994705 | 0.3938513 | -0.5872234 | 0.5570537 | 0.734509 |
| ENSG00000119737 | GPR75 | 51.64087513 | -0.199486399 | 0.461508474 | -0.53537945 | 0.5923875 | 0.760659 |
| ENSG00000229204 | PTGES3P3 | 295.7501 | -0.1995327 | 0.2067369 | -0.9949537 | 0.3197588 | 0.520696 |
| ENSG00000115137 | DNAJC27 | 556.9632344 | -0.199562743 | 0.198329526 | -1.046096684 | 0.2955164 | 0.494669 |
| ENSG00000164106 | SCRG1 | 4.042519 | -0.1995689 | 0.6503731 | -0.5179037 | 0.6045255 | 0.769357 |
| ENSG00000108469 | RECQL5 | 4480.983739 | -0.199678921 | 0.149226651 | -1.369249797 | 0.17092121 | 0.341599 |
| ENSG00000248690 | HAS2-AS1 | 0.7960596 | -0.1996915 | 0.7771335 | -0.7920355 | 0.42834 | NA |
| ENSG00000091542 | ALKBH5 | 5583.344709 | -0.199696666 | 0.139093783 | -1.458220614 | 0.14477974 | 0.303364 |
| ENSG00000173402 | DAG1 | 10052.08 | -0.1997043 | 0.1689915 | -1.216191 | 0.2239123 | 0.410766 |
| ENSG00000256618 | MTRNR2L1 | 1.732914 | -0.1997055 | 0.7523095 | -0.6625684 | 0.507607 | 0.693934 |
| ENSG00000119326 | CTNNAL1 | 1417.003501 | -0.199727757 | 0.400855844 | -0.578479427 | 0.56294048 | 0.738569 |
| ENSG00000230387 | | 1.086466 | -0.1997574 | 0.7809972 | -0.8064654 | 0.4199745 | NA |
| ENSG00000187762 | HSPE1P11 | 1.313685 | -0.1997624 | 0.7690852 | -0.7481526 | 0.4543681 | NA |
| ENSG00000120685 | PROSER1 | 5130.322551 | -0.199765462 | 0.145956782 | -1.397026804 | 0.16240551 | 0.329688 |
| ENSG00000260459 | FTLP14 | 35.10039 | -0.1999032 | 0.322694 | -0.6784776 | 0.4974689 | 0.686176 |
| ENSG00000206127 | GOLGA8O | 1.134484 | -0.1999214 | 0.7700897 | -0.7480297 | 0.4544422 | NA |
| ENSG00000168484 | SFTPC | 0.9021749 | -0.199982 | 0.7767244 | -0.7956442 | 0.4262389 | NA |
| ENSG00000279964 | | 0.875698 | -0.2000193 | 0.7750725 | -0.7679522 | 0.4425156 | NA |
| ENSG00000100983 | GSS | 3471.659755 | -0.2000291 | 0.19349151 | -1.078253681 | 0.28092056 | 0.478414 |
| ENSG00000237415 | NRBF2P3 | 1.46964 | -0.2000438 | 0.7385499 | -0.6159815 | 0.5379068 | NA |
| ENSG00000260316 | | 4.531701 | -0.2000571 | 0.606556 | -0.5047073 | 0.6137644 | 0.775864 |
| ENSG00000234203 | | 2.016668 | -0.200105 | 0.6715758 | -0.5361316 | 0.5918676 | 0.760374 |
| ENSG00000223284 | RNU6-195P | 2.568052 | -0.2001378 | 0.6731407 | -0.5335746 | 0.5936359 | 0.761371 |
| ENSG00000100364 | KIAA0930 | 2846.743541 | -0.200168211 | 0.2096223 | -0.998411442 | 0.31807989 | 0.519203 |
| ENSG00000244578 | LINC01391 | 1.028566 | -0.2001776 | 0.7288712 | -0.6182866 | 0.5363864 | NA |
| ENSG00000112304 | ACOT13 | 1142.134139 | -0.200178322 | 0.211130443 | -0.987751634 | 0.3232743 | 0.524108 |
| ENSG00000125871 | MGME1 | 2296.288608 | -0.200186412 | 0.14554955 | -1.391682538 | 0.16401855 | 0.332049 |
| ENSG00000006015 | REX1BD | 1624.954463 | -0.20022312 | 0.147430528 | -1.383284394 | 0.16657768 | 0.335695 |
| ENSG00000196876 | SCN8A | 650.3876 | -0.2002791 | 0.2804634 | -0.771142 | 0.4406228 | 0.63871 |
| ENSG00000187699 | C2orf88 | 22.79851 | -0.200284 | 0.4969734 | -0.5188643 | 0.6038554 | 0.768809 |
| ENSG00000277246 | | 2.086462 | -0.2003079 | 0.7247715 | -0.6222838 | 0.5337553 | 0.717425 |
| ENSG00000226332 | | 243.8708 | -0.2005146 | 0.292084 | -0.7411386 | 0.4586094 | 0.654433 |

| | | | | | | | |
|---|---|---|---|---|---|---|---|
| ENSG00000100949 | RABGGTA | 2028.274141 | -0.200540307 | 0.170942017 | -1.204893679 | 0.22824435 | 0.416385 |
| ENSG00000272343 | | 3.077825 | -0.2005478 | 0.694211 | -0.5546743 | 0.5791174 | 0.750345 |
| ENSG00000123545 | NDUFAF4 | 1191.995995 | -0.200560995 | 0.199193616 | -1.047504473 | 0.29486697 | 0.493998 |
| ENSG00000138036 | DYNC2LI1 | 753.0405647 | -0.200565161 | 0.167051866 | -1.230462014 | 0.21852414 | 0.404287 |
| ENSG00000266258 | LINC01909 | 5.136808 | -0.2006039 | 0.6490645 | -0.5138116 | 0.6073837 | 0.771285 |
| ENSG00000131470 | PSMC3IP | 446.4613885 | -0.200606652 | 0.319284693 | -0.691086189 | 0.48951138 | 0.680239 |
| ENSG00000102796 | DHRS12 | 203.554776 | -0.200642281 | 0.239648561 | -0.888873434 | 0.37407111 | 0.576156 |
| ENSG00000235358 | | 4.273331 | -0.2006561 | 0.6422203 | -0.5173321 | 0.6049243 | 0.769596 |
| ENSG00000227963 | RBM15-AS1 | 14.94197 | -0.200691 | 0.4503891 | -0.5458874 | 0.5851433 | 0.75503 |
| ENSG00000213189 | BTF3L4P2 | 1193.178 | -0.2007259 | 0.1232724 | -1.655148 | 0.09789445 | 0.229119 |
| ENSG00000261341 | | 2.7828 | -0.2007421 | 0.6573902 | -0.5258189 | 0.599014 | 0.765663 |
| ENSG00000171853 | TRAPPC12 | 1987.924 | -0.2007445 | 0.09071542 | -2.229378 | 0.02578879 | 0.083083 |
| ENSG00000287276 | | 1.178025 | -0.2008123 | 0.7661366 | -0.7408987 | 0.4587549 | NA |
| ENSG00000256269 | HMBS | 3640.133 | -0.2008249 | 0.2469375 | -0.8642478 | 0.3874518 | 0.589757 |
| ENSG00000259244 | | 8.944538 | -0.2008834 | 0.4790296 | -0.5264352 | 0.5985859 | 0.76538 |
| ENSG00000233502 | | 1.306523 | -0.2009135 | 0.7525536 | -0.6742983 | 0.5001217 | NA |
| ENSG00000243775 | OSTCP1 | 4.612268 | -0.2009183 | 0.6116028 | -0.5081797 | 0.6113273 | 0.774461 |
| ENSG00000247728 | | 5.789339 | -0.2009422 | 0.6008312 | -0.4973944 | 0.618911 | 0.779563 |
| ENSG00000251988 | RNU4ATAC18P | 3.551664 | -0.2009479 | 0.6306981 | -0.5117921 | 0.6087965 | 0.772377 |
| ENSG00000255533 | | 0.7483205 | -0.2009703 | 0.8118259 | -1.160196 | 0.2459692 | NA |
| ENSG00000112110 | MRPL18 | 2366.565949 | -0.201046352 | 0.227521511 | -0.925954345 | 0.35446971 | 0.556358 |
| ENSG00000229996 | | 2.299028 | -0.2010501 | 0.7678672 | -0.7301571 | 0.4652941 | 0.659223 |
| ENSG00000259493 | TFDP1P3 | 4.943224 | -0.2010994 | 0.6204185 | -0.5101417 | 0.6099522 | 0.773451 |
| ENSG00000263727 | | 2.539878 | -0.2011611 | 0.7029141 | -0.5619069 | 0.5741794 | 0.746641 |
| ENSG00000133422 | MORC2 | 6293.943322 | -0.201178228 | 0.137136297 | -1.494019651 | 0.13517048 | 0.288473 |
| ENSG00000233635 | | 1.069166 | -0.2011968 | 0.7456676 | -0.6412094 | 0.5213866 | NA |
| ENSG00000269420 | PLA2G4C-AS1 | 1.620348 | -0.2013734 | 0.7340724 | -0.6155755 | 0.5381747 | 0.721105 |
| ENSG00000150768 | DLAT | 2560.276 | -0.2014213 | 0.2598966 | -0.8280248 | 0.4076565 | 0.60828 |
| ENSG00000145782 | ATG12 | 2946.628 | -0.2014406 | 0.1554583 | -1.333213 | 0.1824618 | 0.357404 |
| ENSG00000172869 | DMXL1 | 3136.099 | -0.2015221 | 0.2743737 | -0.80976 | 0.4180781 | 0.617896 |
| ENSG00000226862 | | 6.266478 | -0.2015299 | 0.594781 | -0.5049029 | 0.613627 | 0.775861 |
| ENSG00000272049 | | 5.384933 | -0.2015357 | 0.6373022 | -0.5165557 | 0.6054663 | 0.769897 |
| ENSG00000062716 | VMP1 | 53810.18311 | -0.20156371 | 0.309416969 | -0.713565464 | 0.47549592 | 0.667801 |

| | | | | | | | |
|---|---|---|---|---|---|---|---|
| ENSG00000170832 | USP32 | 39471 | -0.2015643 | 0.1451311 | -1.438817 | 0.1502023 | 0.311362 |
| ENSG00000108344 | PSMD3 | 6984.089266 | -0.201651764 | 0.208352242 | -1.002414746 | 0.31614332 | 0.517186 |
| ENSG00000273606 | MIR6776 | 1.326089 | -0.2016564 | 0.7710179 | -0.7068301 | 0.4796721 | NA |
| ENSG00000120942 | UBIAD1 | 1298.033315 | -0.201741961 | 0.183338552 | -1.13404128 | 0.25677723 | 0.4506 |
| ENSG00000137747 | TMPRSS13 | 1858.16001 | -0.20177276 | 0.287602381 | -0.771382894 | 0.44048001 | 0.638603 |
| ENSG00000130332 | LSM7 | 2446.011774 | -0.201797743 | 0.208190478 | -1.014512662 | 0.31033819 | 0.510993 |
| ENSG00000138614 | INTS14 | 3785.080572 | -0.202051696 | 0.127213702 | -1.676803544 | 0.09358091 | 0.221836 |
| ENSG00000168152 | THAP9 | 303.1799 | -0.2020786 | 0.1483381 | -1.386267 | 0.1656655 | 0.334344 |
| ENSG00000227588 | CNTN4-AS2 | 2.106339 | -0.2021105 | 0.747148 | -0.6707932 | 0.5023523 | 0.690014 |
| ENSG00000166747 | AP1G1 | 11974.44 | -0.2021585 | 0.1296002 | -1.599216 | 0.1097726 | 0.249038 |
| ENSG00000236797 | SPA17P1 | 2.194154 | -0.2022531 | 0.7082893 | -0.585813 | 0.5580012 | 0.735175 |
| ENSG00000182872 | RBM10 | 5084.751 | -0.2022703 | 0.1693199 | -1.222484 | 0.2215247 | 0.408133 |
| ENSG00000229931 | ATXN1-AS1 | 55.93139 | -0.2023431 | 0.4666774 | -0.5403565 | 0.5889512 | 0.757928 |
| ENSG00000240859 | | 174.9581 | -0.2024039 | 0.2614413 | -0.8226064 | 0.4107319 | 0.61105 |
| ENSG00000232519 | | 6.319082 | -0.2024051 | 0.5654119 | -0.5114276 | 0.6090517 | 0.772664 |
| ENSG00000120798 | NR2C1 | 1877.450615 | -0.202420809 | 0.188887673 | -1.102034671 | 0.2704466 | 0.466782 |
| ENSG00000176890 | TYMS | 943.3392 | -0.2024247 | 0.4127481 | -0.5795716 | 0.5622035 | 0.73793 |
| ENSG00000173692 | PSMD1 | 5242.417 | -0.2024633 | 0.2423711 | -0.8778393 | 0.3800309 | 0.581996 |
| ENSG00000002586 | CD99 | 2977.288505 | -0.202472612 | 0.144080209 | -1.435554084 | 0.15112927 | 0.312829 |
| ENSG00000275468 | | 6.893883 | -0.202498 | 0.6445333 | -0.523193 | 0.6008399 | 0.766755 |
| ENSG00000285978 | | 1.48643 | -0.2024981 | 0.7431699 | -0.6569333 | 0.5112238 | NA |
| ENSG00000277423 | | 4.783064 | -0.2025861 | 0.6124777 | -0.5167345 | 0.6053415 | 0.769832 |
| ENSG00000252643 | RNU6-1136P | 1.045702 | -0.2026279 | 0.7832821 | -0.8014267 | 0.4228847 | NA |
| ENSG00000164134 | NAA15 | 5122.314 | -0.2027362 | 0.228664 | -0.9244008 | 0.3552777 | 0.557303 |
| ENSG00000204130 | RUFY2 | 1377.123 | -0.2027532 | 0.1418599 | -1.453107 | 0.1461941 | 0.305532 |
| ENSG00000196136 | SERPINA3 | 6.224506 | -0.2027766 | 0.6632353 | -0.5666197 | 0.5709726 | 0.744731 |
| ENSG00000215267 | AKR1C7P | 1.514247 | -0.2027986 | 0.7615218 | -0.7261968 | 0.4677182 | NA |
| ENSG00000182836 | PLCXD3 | 17.64107 | -0.2028184 | 0.6102114 | -0.50757 | 0.611755 | 0.774614 |
| ENSG00000272764 | | 31.03902 | -0.2028996 | 0.3755995 | -0.615533 | 0.5382028 | 0.721105 |
| ENSG00000184905 | TCEAL2 | 0.8900291 | -0.2029001 | 0.794671 | -0.9496972 | 0.3422661 | NA |
| ENSG00000112218 | GPR63 | 124.951734 | -0.202956082 | 0.290838688 | -0.745173435 | 0.45616688 | 0.652028 |
| ENSG00000257564 | | 50.80581 | -0.2030422 | 0.3622057 | -0.6348747 | 0.5255101 | 0.710368 |
| ENSG00000131626 | PPFIA1 | 5226.080159 | -0.20304971 | 0.159439094 | -1.302168065 | 0.19285894 | 0.371309 |

| | | | | | | | |
|---|---|---|---|---|---|---|---|
| ENSG00000255966 | | 13.10255 | -0.2030609 | 0.5108764 | -0.5177113 | 0.6046597 | 0.769443 |
| ENSG00000133030 | MPRIP | 12170.88015 | -0.20311642 | 0.215753554 | -0.982429914 | 0.3258881 | 0.527091 |
| ENSG00000274963 | RN7SL600P | 7.992536 | -0.203127 | 0.5933084 | -0.5089951 | 0.6107556 | 0.774123 |
| ENSG00000162600 | OMA1 | 852.6033 | -0.2031991 | 0.1702629 | -1.228984 | 0.2190777 | 0.405142 |
| ENSG00000166866 | MYO1A | 1.271367 | -0.2032573 | 0.7420181 | -0.6494291 | 0.516061 | NA |
| ENSG00000174206 | C12orf66 | 1011.88 | -0.2033317 | 0.1690249 | -1.23169 | 0.2180648 | 0.403718 |
| ENSG00000275895 | U2AF1L5 | 1.346365 | -0.2034918 | 0.7661446 | -0.7414928 | 0.4583947 | NA |
| ENSG00000229598 | PRDX3P1 | 9.432546 | -0.2035097 | 0.5583372 | -0.512964 | 0.6079765 | 0.771742 |
| ENSG00000213590 | | 3.166659 | -0.2035131 | 0.6642146 | -0.5395991 | 0.5894736 | 0.758327 |
| ENSG00000267283 | | 2.40666 | -0.2035317 | 0.6633516 | -0.5360355 | 0.591934 | 0.760374 |
| ENSG00000264006 | AKR1C8P | 1.38484 | -0.2035825 | 0.7493782 | -0.679477 | 0.4968357 | NA |
| ENSG00000184517 | ZFP1 | 1991.448 | -0.2036945 | 0.1505661 | -1.389009 | 0.1648301 | 0.333133 |
| ENSG00000226086 | EIF3LP3 | 9.909284 | -0.2037037 | 0.4997667 | -0.5257848 | 0.5990378 | 0.765663 |
| ENSG00000248327 | NOL8P1 | 1.220279 | -0.2037182 | 0.7600997 | -0.6928359 | 0.4884126 | NA |
| ENSG00000158315 | RHBDL2 | 19.43371 | -0.203734 | 0.6434418 | -0.5285148 | 0.597142 | 0.764085 |
| ENSG00000101413 | RPRD1B | 3429.541279 | -0.203739107 | 0.107696303 | -1.90306653 | 0.05703187 | 0.153059 |
| ENSG00000163867 | ZMYM6 | 724.2868 | -0.2038015 | 0.1758063 | -1.188692 | 0.2345609 | 0.423747 |
| ENSG00000187243 | MAGED4B | 3.831368 | -0.203831 | 0.5880999 | -0.5151732 | 0.606432 | 0.770618 |
| ENSG00000228470 | | 5.155811 | -0.2038713 | 0.6157747 | -0.5118664 | 0.6087445 | 0.772377 |
| ENSG00000271147 | ARMCX5-GPRAS | 430.0298 | -0.203906 | 0.1879525 | -1.12369 | 0.2611447 | 0.455781 |
| ENSG00000223823 | LINC01342 | 4.857686 | -0.203936 | 0.7215293 | -0.6044343 | 0.5455549 | 0.725981 |
| ENSG00000170890 | PLA2G1B | 0.8743192 | -0.203976 | 0.7630055 | -0.7102451 | 0.4775522 | NA |
| ENSG00000260027 | HOXB7 | 260.1419 | -0.2039787 | 0.2854115 | -0.7746214 | 0.4385634 | 0.63715 |
| ENSG00000116151 | MORN1 | 374.1379027 | -0.204171719 | 0.168560069 | -1.24642638 | 0.2126079 | 0.39727 |
| ENSG00000197980 | LEKR1 | 155.0871 | -0.2041944 | 0.2594058 | -0.8366615 | 0.4027829 | 0.60369 |
| ENSG00000100239 | PPP6R2 | 3916.924353 | -0.204196684 | 0.140245215 | -1.474833335 | 0.14025736 | 0.296436 |
| ENSG00000146243 | IRAK1BP1 | 202.76 | -0.2042645 | 0.2755391 | -0.7970061 | 0.4254475 | 0.62524 |
| ENSG00000161021 | MAML1 | 6262.907 | -0.2042668 | 0.1778361 | -1.184068 | 0.2363861 | 0.425944 |
| ENSG00000266651 | | 64.36878 | -0.2043661 | 0.3361707 | -0.6686168 | 0.50374 | 0.691066 |
| ENSG00000123143 | PKN1 | 6471.875864 | -0.204369455 | 0.182570413 | -1.159470039 | 0.24626464 | 0.438038 |
| ENSG00000285884 | | 16.88904 | -0.2044131 | 0.4526705 | -0.5494291 | 0.582711 | 0.753231 |
| ENSG00000174796 | THAP6 | 947.1173 | -0.2044365 | 0.2120371 | -1.000367 | 0.3171331 | 0.518191 |
| ENSG00000285727 | | 1.605979 | -0.204438 | 0.7422001 | -0.65144 | 0.5147625 | 0.700365 |

| | | | | | | | |
|---|---|---|---|---|---|---|---|
| ENSG00000230823 | CBX1P1 | 38.13873 | -0.2044531 | 0.3174107 | -0.7072226 | 0.4794282 | 0.671442 |
| ENSG00000228554 | | 3.166188 | -0.204541 | 0.6791793 | -0.5514461 | 0.5813279 | 0.752007 |
| ENSG00000225092 | | 2.710897 | -0.2045969 | 0.6694139 | -0.5414404 | 0.5882041 | 0.75748 |
| ENSG00000127445 | PIN1 | 2099.349474 | -0.204664973 | 0.228044911 | -0.947315159 | 0.34347821 | 0.544991 |
| ENSG00000100461 | RBM23 | 4898.898695 | -0.204705598 | 0.153374529 | -1.36976366 | 0.17076069 | 0.341457 |
| ENSG00000250330 | | 1.744276 | -0.20479 | 0.7814186 | -0.8249368 | 0.4094075 | 0.609907 |
| ENSG00000177469 | CAVIN1 | 1460.47 | -0.2047963 | 0.3072858 | -0.7242781 | 0.468895 | 0.662417 |
| ENSG00000177138 | FAM9B | 1.101628 | -0.2048139 | 0.8279272 | -1.648817 | 0.09918506 | NA |
| ENSG00000228561 | | 11.41882 | -0.2048361 | 0.5405677 | -0.5143994 | 0.6069728 | 0.77103 |
| ENSG00000236554 | ASNSP3 | 2.514244 | -0.2048612 | 0.6475412 | -0.5407555 | 0.5886761 | 0.757851 |
| ENSG00000228280 | | 1.183147 | -0.2049527 | 0.7843561 | -0.8394763 | 0.4012021 | NA |
| ENSG00000095794 | CREM | 913.5795597 | -0.204958964 | 0.131464833 | -1.584911917 | 0.11298635 | 0.254035 |
| ENSG00000135974 | C2orf49 | 1887.05669 | -0.205016683 | 0.212537247 | -1.004071083 | 0.31534435 | 0.516292 |
| ENSG00000177084 | POLE | 6987.115 | -0.205018 | 0.3733476 | -0.6042169 | 0.5456994 | 0.72606 |
| ENSG00000224791 | KRT18P39 | 34.25075 | -0.2051812 | 0.4018814 | -0.5912847 | 0.5543297 | 0.732434 |
| ENSG00000262468 | LINC01569 | 472.6334 | -0.2052578 | 0.1925294 | -1.109444 | 0.2672387 | 0.463041 |
| ENSG00000100201 | DDX17 | 24257.75163 | -0.205282432 | 0.245363707 | -0.885375847 | 0.37595396 | 0.578152 |
| ENSG00000285820 | | 1.366792 | -0.2052859 | 0.78247 | -0.8094561 | 0.4182528 | NA |
| ENSG00000124217 | MOCS3 | 1785.262601 | -0.20529978 | 0.138249281 | -1.507819232 | 0.1316008 | 0.283324 |
| ENSG00000132436 | FIGNL1 | 1654.023185 | -0.205560038 | 0.375600086 | -0.631055646 | 0.52800414 | 0.712832 |
| ENSG00000179157 | RPS2P28 | 3.629921 | -0.205569 | 0.6341688 | -0.5199677 | 0.6030861 | 0.76825 |
| ENSG00000266648 | SETP3 | 32.09589 | -0.2055944 | 0.3892982 | -0.6106358 | 0.5414407 | 0.723128 |
| ENSG00000109534 | GAR1 | 1892.756683 | -0.205621465 | 0.193880528 | -1.102304476 | 0.27032932 | 0.466752 |
| ENSG00000273017 | | 4.564815 | -0.2056685 | 0.637409 | -0.5256931 | 0.5991015 | 0.765665 |
| ENSG00000198840 | MT-ND3 | 17401.34 | -0.2056963 | 0.1676881 | -1.276413 | 0.2018097 | 0.3828 |
| ENSG00000242299 | | 9754.093 | -0.2057975 | 0.1852396 | -1.151357 | 0.2495852 | 0.44217 |
| ENSG00000186704 | DTX2P1 | 1.967761 | -0.2058443 | 0.7150367 | -0.5828417 | 0.5599998 | 0.736275 |
| ENSG00000105248 | YJU2 | 1230.325532 | -0.205919148 | 0.225754241 | -0.954356145 | 0.3399034 | 0.541291 |
| ENSG00000097046 | CDC7 | 866.845908 | -0.205967854 | 0.401417929 | -0.598993019 | 0.54917754 | 0.728535 |
| ENSG00000229677 | | 1.879029 | -0.2059777 | 0.7355734 | -0.636955 | 0.5241542 | 0.70922 |
| ENSG00000064995 | TAF11 | 3569.49725 | -0.206015423 | 0.18010118 | -1.148458883 | 0.25077918 | 0.443428 |
| ENSG00000100281 | HMGXB4 | 1821.787535 | -0.206050548 | 0.190673213 | -1.119094956 | 0.26309963 | 0.45811 |
| ENSG00000257496 | | 1.444651 | -0.2060955 | 0.7057151 | -0.5847935 | 0.5586866 | NA |

| | | | | | | | |
|---|---|---|---|---|---|---|---|
| ENSG00000232460 | BMPR1AP2 | 169.9474 | -0.2061325 | 0.1959624 | -1.102464 | 0.27026 | 0.466693 |
| ENSG00000119707 | RBM25 | 12370.61159 | -0.206136275 | 0.159474374 | -1.326705282 | 0.18460619 | 0.359985 |
| ENSG00000182600 | SNORC | 150.2134 | -0.2061854 | 0.2786766 | -0.7921788 | 0.4282564 | 0.627598 |
| ENSG00000107338 | SHB | 573.7919446 | -0.206205146 | 0.190188173 | -1.120398845 | 0.26254384 | 0.457412 |
| ENSG00000224764 | | 1.346654 | -0.2062089 | 0.7262148 | -0.6346142 | 0.52568 | NA |
| ENSG00000258659 | TRIM34 | 3.315295 | -0.2062241 | 0.800937 | -0.6118741 | 0.5406211 | 0.722596 |
| ENSG00000112159 | MDN1 | 8522.780952 | -0.20622974 | 0.247869113 | -0.863635739 | 0.38778803 | 0.589998 |
| ENSG00000266074 | BAHCC1 | 2176.308 | -0.2063921 | 0.2429041 | -0.8975431 | 0.3694292 | 0.571644 |
| ENSG00000153933 | DGKE | 2770.894 | -0.2064195 | 0.2579733 | -0.8488941 | 0.3959402 | 0.597441 |
| ENSG00000286859 | NA | 16.59773 | -0.2064265 | 0.4362046 | -0.5724533 | 0.5670149 | 0.741535 |
| ENSG00000286388 | | 611.7677 | -0.2064506 | 0.1966184 | -1.092189 | 0.2747499 | 0.471557 |
| ENSG00000283525 | | 0.8176392 | -0.2065297 | 0.8093873 | -0.730389 | 0.4651524 | NA |
| ENSG00000141034 | GID4 | 1215.856 | -0.206539 | 0.2081592 | -1.016333 | 0.3094707 | 0.510102 |
| ENSG00000231120 | BTF3P10 | 206.2229 | -0.2065519 | 0.2479462 | -0.8731668 | 0.3825721 | 0.584842 |
| ENSG00000186714 | CCDC73 | 16.60499 | -0.2066622 | 0.41605 | -0.5854831 | 0.5582229 | 0.735302 |
| ENSG00000255094 | | 2.023608 | -0.2067766 | 0.7006056 | -0.5898476 | 0.5552928 | 0.732896 |
| ENSG00000238198 | LRIG2-DT | 3.236983 | -0.206869 | 0.7096811 | -0.5973292 | 0.5502876 | 0.729281 |
| ENSG00000126088 | UROD | 2421.326629 | -0.206998307 | 0.186736286 | -1.152760951 | 0.24900852 | 0.441414 |
| ENSG00000144061 | NPHP1 | 192.4157 | -0.2070155 | 0.2168342 | -1.006366 | 0.3142395 | 0.515372 |
| ENSG00000286087 | | 2.723941 | -0.2070834 | 0.7182535 | -0.604614 | 0.5454355 | 0.72589 |
| ENSG00000279332 | | 10.56626 | -0.2071344 | 0.4881035 | -0.5401968 | 0.5890613 | 0.75798 |
| ENSG00000169826 | CSGALNACT2 | 978.9428 | -0.2072429 | 0.1817716 | -1.174564 | 0.2401691 | 0.430454 |
| ENSG00000236337 | FMR1-IT1 | 27.02183 | -0.2072517 | 0.388717 | -0.616104 | 0.5378258 | 0.720901 |
| ENSG00000233144 | | 1.916272 | -0.207265 | 0.6889672 | -0.5670667 | 0.5706688 | 0.744445 |
| ENSG00000143390 | RFX5 | 7293.618 | -0.2072986 | 0.169001 | -1.263943 | 0.2062503 | 0.388468 |
| ENSG00000260022 | | 13.65034 | -0.2073078 | 0.4993934 | -0.5353356 | 0.5924178 | 0.760659 |
| ENSG00000073169 | SELENOO | 1695.851918 | -0.20732365 | 0.192263955 | -1.121094749 | 0.26224753 | 0.457045 |
| ENSG00000140968 | IRF8 | 7.070117 | -0.2073768 | 0.8011222 | -0.6247475 | 0.5321368 | 0.716192 |
| ENSG00000137727 | ARHGAP20 | 1.085058375 | -0.207391391 | 0.745550769 | -0.663390761 | 0.50708032 | NA |
| ENSG00000169905 | TOR1AIP2 | 2860.944 | -0.207452 | 0.1668171 | -1.280171 | 0.2004849 | 0.380994 |
| ENSG00000206647 | | 1.002671 | -0.2074669 | 0.8170034 | -1.281978 | 0.1998503 | NA |
| ENSG00000285509 | TBCEL-TECTA | 7.653426 | -0.2075537 | 0.5928653 | -0.5334235 | 0.5937405 | 0.761435 |
| ENSG00000229211 | KCTD9P6 | 14.64408 | -0.2075605 | 0.4911377 | -0.5392527 | 0.5897125 | 0.758512 |

| | | | | | | | |
|---|---|---|---|---|---|---|---|
| ENSG00000273320 | | 2.328919 | -0.207617 | 0.8012778 | -0.6121395 | 0.5404455 | 0.722434 |
| ENSG00000189195 | BTBD8 | 130.9352 | -0.207719 | 0.274226 | -0.8060817 | 0.4201957 | 0.6201 |
| ENSG00000267940 | | 14.26923 | -0.2079194 | 0.5030412 | -0.5309654 | 0.5954428 | 0.762682 |
| ENSG00000201563 | | 2.305697 | -0.2079323 | 0.6967249 | -0.5762218 | 0.5644653 | 0.739657 |
| ENSG00000111252 | SH2B3 | 416.9252181 | -0.207987781 | 0.233963862 | -0.922152102 | 0.35644924 | 0.558402 |
| ENSG00000236146 | | 4.345134 | -0.2080036 | 0.7031048 | -0.5977043 | 0.5500372 | 0.729166 |
| ENSG00000226377 | | 25.32836 | -0.2080507 | 0.4121612 | -0.5956791 | 0.5513896 | 0.73023 |
| ENSG00000223692 | DIP2A-IT1 | 5.619556 | -0.2080535 | 0.6334373 | -0.5347677 | 0.5928105 | 0.760888 |
| ENSG00000253166 | | 1.190211 | -0.2080539 | 0.7632614 | -0.7160512 | 0.4739598 | NA |
| ENSG00000151849 | CENPJ | 1361.861 | -0.2080975 | 0.3523908 | -0.6644247 | 0.5064185 | 0.692933 |
| ENSG00000230356 | NCAPD2P1 | 1.085604 | -0.2081116 | 0.7762686 | -0.7713785 | 0.4404826 | NA |
| ENSG00000185163 | DDX51 | 2798.603 | -0.2081159 | 0.1345692 | -1.57256 | 0.1158207 | 0.258649 |
| ENSG00000023572 | GLRX2 | 1503.300016 | -0.208150225 | 0.151690183 | -1.415833941 | 0.15682413 | 0.321576 |
| ENSG00000104131 | EIF3J | 5579.837146 | -0.208218449 | 0.129767072 | -1.631893232 | 0.10270198 | 0.237243 |
| ENSG00000137210 | TMEM14B | 1806.470057 | -0.2082917 | 0.221806202 | -0.9809844 | 0.32660044 | 0.527889 |
| ENSG00000237945 | LINC00649 | 374.1897 | -0.2083313 | 0.339382 | -0.6857298 | 0.4928835 | 0.682805 |
| ENSG00000212296 | SNORD72 | 9.102605 | -0.2083547 | 0.6028201 | -0.5264517 | 0.5985744 | 0.76538 |
| ENSG00000253392 | | 12.51525 | -0.2083755 | 0.4553583 | -0.5551423 | 0.5787973 | 0.75015 |
| ENSG00000115073 | ACTR1B | 5829.371104 | -0.208381779 | 0.142515091 | -1.53515969 | 0.12474461 | 0.272718 |
| ENSG00000173662 | TAS1R1 | 1.142151 | -0.2083858 | 0.7725933 | -0.7616586 | 0.4462638 | NA |
| ENSG00000104147 | OIP5 | 145.636875 | -0.208657028 | 0.447519825 | -0.570049746 | 0.56864396 | 0.742678 |
| ENSG00000204574 | ABCF1 | 7468.106 | -0.2086664 | 0.1580094 | -1.370569 | 0.1705095 | 0.34116 |
| ENSG00000260773 | | 12.54958 | -0.2086819 | 0.411796 | -0.6046682 | 0.5453995 | 0.725887 |
| ENSG00000213857 | ACTBP15 | 1.575301 | -0.2087472 | 0.7478264 | -0.6756517 | 0.4992618 | 0.687656 |
| ENSG00000216316 | | 0.9546199 | -0.2088363 | 0.8006833 | -0.9583265 | 0.3378982 | NA |
| ENSG00000241399 | CD302 | 33.1907 | -0.2089625 | 0.4289619 | -0.5834643 | 0.5595808 | 0.73613 |
| ENSG00000210144 | MT-TY | 14.96046 | -0.2089773 | 0.4315977 | -0.5799668 | 0.561937 | 0.737763 |
| ENSG00000264270 | | 1.595073 | -0.2090027 | 0.7164109 | -0.6073013 | 0.543651 | 0.724554 |
| ENSG00000255308 | CSRP3-AS1 | 4.466092 | -0.2090857 | 0.6395242 | -0.5426474 | 0.5873726 | 0.756925 |
| ENSG00000038945 | MSR1 | 1.020760866 | -0.209122197 | 0.76907426 | -0.780157629 | 0.4352981 | NA |
| ENSG00000260805 | | 133.3587 | -0.2091613 | 0.2442286 | -0.9019646 | 0.3670757 | 0.569077 |
| ENSG00000173992 | CCS | 2103.87 | -0.20918 | 0.1968465 | -1.10129 | 0.2707704 | 0.467089 |
| ENSG00000251414 | | 47.79884 | -0.2092233 | 0.3266525 | -0.7036183 | 0.4816705 | 0.673057 |

| | | | | | | | |
|---|---|---|---|---|---|---|---|
| ENSG00000134910 | STT3A | 6009.825133 | -0.209264705 | 0.108175815 | -1.955158788 | 0.05056433 | 0.139703 |
| ENSG00000239835 | | 1.082847 | -0.2094143 | 0.7557036 | -0.6847792 | 0.4934833 | NA |
| ENSG00000163632 | C3orf49 | 79.71424 | -0.2094169 | 0.2394174 | -0.9206732 | 0.3572211 | 0.559314 |
| ENSG00000086504 | MRPL28 | 5316.40666 | -0.209430564 | 0.267844469 | -0.835382761 | 0.40350223 | 0.604287 |
| ENSG00000143179 | UCK2 | 2552.975 | -0.2094624 | 0.1702495 | -1.263876 | 0.2062744 | 0.388468 |
| ENSG00000278627 | | 14.44526 | -0.2094654 | 0.5104334 | -0.5382353 | 0.5904146 | 0.759133 |
| ENSG00000252367 | | 0.8264256 | -0.2095159 | 0.7750895 | -0.7827501 | 0.4337739 | NA |
| ENSG00000261635 | | 2.20657 | -0.2095719 | 0.6938077 | -0.5806533 | 0.5614741 | 0.73741 |
| ENSG00000180251 | SLC9A4 | 283.7852 | -0.2095897 | 0.5404532 | -0.5225086 | 0.6013162 | 0.766989 |
| ENSG00000096080 | MRPS18A | 890.0315714 | -0.209600157 | 0.205811436 | -1.072461409 | 0.28351284 | 0.481289 |
| ENSG00000265843 | LINC01029 | 24.39572 | -0.2096265 | 0.8020922 | -0.6501496 | 0.5155956 | 0.700924 |
| ENSG00000118503 | TNFAIP3 | 126.6955943 | -0.209671437 | 0.469230308 | -0.54833364 | 0.58346283 | 0.753696 |
| ENSG00000267178 | PHF5CP | 26.24494 | -0.2097531 | 0.4854305 | -0.5493322 | 0.5827775 | 0.753231 |
| ENSG00000261575 | | 11.99408 | -0.2097618 | 0.5024959 | -0.5403758 | 0.5889379 | 0.757928 |
| ENSG00000131462 | TUBG1 | 3586.387792 | -0.209825612 | 0.383490388 | -0.628462829 | 0.52970079 | 0.714359 |
| ENSG00000234832 | NXT1-AS1 | 11.61293 | -0.2098321 | 0.5465211 | -0.5298735 | 0.5961996 | 0.763339 |
| ENSG00000134690 | CDCA8 | 2053.630776 | -0.209843038 | 0.439160706 | -0.584816041 | 0.55867142 | 0.735584 |
| ENSG00000105649 | RAB3A | 241.2583797 | -0.20985273 | 0.240891459 | -0.921639137 | 0.35671684 | 0.558722 |
| ENSG00000228195 | RPL5P27 | 0.785037 | -0.2099318 | 0.7824268 | -0.8199941 | 0.4122195 | NA |
| ENSG00000139998 | RAB15 | 9454.616 | -0.2099498 | 0.1834743 | -1.178794 | 0.2384804 | 0.428323 |
| ENSG00000270781 | | 2.843438 | -0.2099656 | 0.6827681 | -0.5624029 | 0.5738415 | 0.746641 |
| ENSG00000259974 | LINC00261 | 23.3717 | -0.2100367 | 0.8021193 | -0.6389872 | 0.5228312 | 0.708182 |
| ENSG00000205981 | DNAJC19 | 1200.435 | -0.210103 | 0.1575852 | -1.353171 | 0.1760009 | 0.34863 |
| ENSG00000272420 | | 13.15149 | -0.2101452 | 0.5322138 | -0.5337926 | 0.593485 | 0.761349 |
| ENSG00000247877 | | 5.485523 | -0.2101631 | 0.660692 | -0.5558465 | 0.5783158 | 0.749928 |
| ENSG00000253875 | | 2.028589 | -0.2103153 | 0.7165371 | -0.6116186 | 0.5407901 | 0.722677 |
| ENSG00000251333 | RTN3P1 | 597.5039 | -0.2103184 | 0.1832779 | -1.18052 | 0.2377934 | 0.427378 |
| ENSG00000178074 | C2orf69 | 951.8765 | -0.2104356 | 0.1897994 | -1.14605 | 0.2517744 | 0.444508 |
| ENSG00000229336 | SREK1IP1P1 | 2.422975 | -0.2104575 | 0.7297202 | -0.6459609 | 0.5183047 | 0.703563 |
| ENSG00000198700 | IPO9 | 4912.784 | -0.210464 | 0.2236043 | -0.9802891 | 0.3269434 | 0.528187 |
| ENSG00000259347 | | 22.49934 | -0.2104854 | 0.4622201 | -0.5620031 | 0.5741139 | 0.746641 |
| ENSG00000236790 | LINC00299 | 14.92655 | -0.2106087 | 0.5777981 | -0.5279289 | 0.5975487 | 0.764458 |
| ENSG00000177600 | RPLP2 | 15239.81 | -0.2106209 | 0.1715055 | -1.274347 | 0.2025405 | 0.38373 |

| | | | | | | | |
|---|---|---|---|---|---|---|---|
| ENSG00000287193 | | 12.60523 | -0.2106382 | 0.5371911 | -0.5318908 | 0.5948017 | 0.762212 |
| ENSG00000161999 | JMJD8 | 1079.343 | -0.2106785 | 0.1406049 | -1.528737 | 0.1263296 | 0.275029 |
| ENSG00000165202 | OR1Q1 | 1.012619 | -0.2106803 | 0.7649493 | -0.7408663 | 0.4587745 | NA |
| ENSG00000234896 | OR7E62P | 2.863272 | -0.2107975 | 0.8031113 | -0.6029398 | 0.5465488 | 0.726642 |
| ENSG00000256235 | SMIM3 | 411.2694 | -0.2108476 | 0.2527075 | -0.8776298 | 0.3801446 | 0.582069 |
| ENSG00000254598 | CSNK2A3 | 482.8017 | -0.2108676 | 0.1592252 | -1.354447 | 0.1755937 | 0.347979 |
| ENSG00000059769 | DNAJC25 | 464.9502319 | -0.210905264 | 0.145846582 | -1.470884096 | 0.14132246 | 0.298213 |
| ENSG00000110917 | MLEC | 10992.01867 | -0.210967983 | 0.141894672 | -1.537227296 | 0.12423766 | 0.27199 |
| ENSG00000135070 | ISCA1 | 2192.975576 | -0.210971106 | 0.142424448 | -1.497336641 | 0.13430569 | 0.287331 |
| ENSG00000047230 | CTPS2 | 2283.496357 | -0.211016648 | 0.103294629 | -2.062637966 | 0.03914704 | 0.114953 |
| ENSG00000163069 | SGCB | 884.2503 | -0.2110243 | 0.1536473 | -1.40407 | 0.1602981 | 0.326733 |
| ENSG00000171566 | PLRG1 | 3775.308 | -0.2110503 | 0.09632528 | -2.218172 | 0.0265431 | 0.085028 |
| ENSG00000141367 | CLTC | 75235.81 | -0.2111038 | 0.1243737 | -1.707335 | 0.08775976 | 0.211625 |
| ENSG00000011132 | APBA3 | 625.5731218 | -0.211112003 | 0.172175112 | -1.262153166 | 0.20689368 | 0.389248 |
| ENSG00000110435 | PDHX | 2350.791053 | -0.21112928 | 0.157265122 | -1.39022664 | 0.16446007 | 0.332588 |
| ENSG00000105393 | BABAM1 | 1108.214162 | -0.211191801 | 0.228409292 | -0.974363619 | 0.32987603 | 0.530997 |
| ENSG00000187109 | NAP1L1 | 23049.21 | -0.2111951 | 0.1223495 | -1.721506 | 0.08515905 | 0.206875 |
| ENSG00000168813 | ZNF507 | 2364.803 | -0.2112301 | 0.2047032 | -1.072816 | 0.2833535 | 0.481141 |
| ENSG00000137955 | RABGGTB | 4898.885385 | -0.211260342 | 0.202288621 | -1.073716699 | 0.28294968 | 0.480732 |
| ENSG00000087053 | MTMR2 | 1628.653117 | -0.211279678 | 0.180117668 | -1.207331823 | 0.22730438 | 0.415097 |
| ENSG00000185298 | CCDC137 | 3624.397 | -0.2113231 | 0.2121657 | -1.030087 | 0.3029692 | 0.502559 |
| ENSG00000250471 | GMPSP1 | 354.1901 | -0.2113696 | 0.2847925 | -0.8017486 | 0.4226984 | 0.622681 |
| ENSG00000119878 | CRIPT | 804.3941115 | -0.211421379 | 0.179046487 | -1.217032832 | 0.22359172 | 0.410291 |
| ENSG00000160716 | CHRNB2 | 13.43347 | -0.2114275 | 0.4634625 | -0.56146 | 0.574484 | 0.746928 |
| ENSG00000242531 | | 4.445545 | -0.2115041 | 0.6271376 | -0.5346895 | 0.5928646 | 0.760895 |
| ENSG00000235366 | LINC01055 | 2.131382 | -0.2115107 | 0.7233227 | -0.62365 | 0.5328575 | 0.716908 |
| ENSG00000207975 | MIR181B1 | 1.49602 | -0.2115121 | 0.8035265 | -0.5954387 | 0.5515502 | NA |
| ENSG00000258172 | | 0.9054843 | -0.2115811 | 0.7711631 | -0.7680499 | 0.4424576 | NA |
| ENSG00000115091 | ACTR3 | 8645.271221 | -0.211692264 | 0.116783993 | -1.823122396 | 0.06828486 | 0.175243 |
| ENSG00000164068 | RNF123 | 2402.814 | -0.2117495 | 0.1653135 | -1.319117 | 0.1871299 | 0.363626 |
| ENSG00000224316 | GTF2IP5 | 7.868759 | -0.2118314 | 0.560266 | -0.5281821 | 0.597373 | 0.764307 |
| ENSG00000243759 | ST13P15 | 272.6184 | -0.2119484 | 0.1912724 | -1.139926 | 0.2543169 | 0.447319 |
| ENSG00000264924 | | 1.574288 | -0.21196 | 0.712972 | -0.6035432 | 0.5461474 | 0.726329 |

| | | | | | | | |
|---|---|---|---|---|---|---|---|
| ENSG00000206535 | LNP1 | 125.49 | -0.2120861 | 0.3036889 | -0.756549 | 0.4493201 | 0.645517 |
| ENSG00000236117 | LINC01639 | 152.2219 | -0.2120959 | 0.803724 | -0.5948493 | 0.5519442 | 0.730746 |
| ENSG00000280997 | CCAT2 | 3.435052 | -0.2121545 | 0.6495097 | -0.5491156 | 0.5829261 | 0.753333 |
| ENSG00000277661 | | 1.228587 | -0.2121879 | 0.8091171 | -0.7100218 | 0.4776906 | NA |
| ENSG00000286577 | | 1.040707 | -0.2121879 | 0.7669852 | -0.7479927 | 0.4544645 | NA |
| ENSG00000168038 | ULK4 | 154.276 | -0.2123013 | 0.2482408 | -0.9062333 | 0.3648124 | 0.566692 |
| ENSG00000203972 | GLYATL3 | 22.65845 | -0.2123398 | 0.8021752 | -0.6523694 | 0.5141629 | 0.700028 |
| ENSG00000283761 | | 4.306892 | -0.2123798 | 0.6373495 | -0.54505 | 0.5857191 | 0.755509 |
| ENSG00000237729 | | 7.29527 | -0.2125558 | 0.5476757 | -0.5286558 | 0.5970442 | 0.764085 |
| ENSG00000197223 | C1D | 502.426 | -0.2125587 | 0.1691424 | -1.284655 | 0.1989128 | 0.379013 |
| ENSG00000238280 | | 2.583307 | -0.2126775 | 0.7312573 | -0.6466402 | 0.5178648 | 0.703146 |
| ENSG00000269958 | | 407.8896 | -0.212678 | 0.2050275 | -1.076145 | 0.2818625 | 0.479609 |
| ENSG00000173457 | PPP1R14B | 2412.601 | -0.2126906 | 0.2411975 | -0.8999379 | 0.3681533 | 0.570215 |
| ENSG00000171448 | ZBTB26 | 1184.187 | -0.212737 | 0.2513374 | -0.8973573 | 0.3695283 | 0.571647 |
| ENSG00000286847 | | 3.259994 | -0.2127742 | 0.6032239 | -0.5377853 | 0.5907253 | 0.759422 |
| ENSG00000196352 | CD55 | 5430.455 | -0.2127909 | 0.2650152 | -0.8366563 | 0.4027858 | 0.60369 |
| ENSG00000177692 | DNAJC28 | 62.87813 | -0.2128079 | 0.3657982 | -0.6619171 | 0.5080243 | 0.694398 |
| ENSG00000234816 | H2AC5P | 2.093601 | -0.2128135 | 0.6936529 | -0.5906492 | 0.5547555 | 0.732642 |
| ENSG00000205758 | CRYZL1 | 793.0813 | -0.2128443 | 0.1323777 | -1.635745 | 0.1018931 | 0.235819 |
| ENSG00000286285 | | 1.729002 | -0.2128859 | 0.7154356 | -0.6085801 | 0.5428028 | 0.723848 |
| ENSG00000164902 | PHAX | 3058.223 | -0.212893 | 0.1607254 | -1.353418 | 0.1759221 | 0.348541 |
| ENSG00000241680 | RPL31P49 | 67.36234 | -0.2129411 | 0.3789728 | -0.6399006 | 0.5222372 | 0.707599 |
| ENSG00000253174 | | 1.981364 | -0.2129597 | 0.688024 | -0.5871365 | 0.557112 | 0.734513 |
| ENSG00000005189 | REXO5 | 427.7188676 | -0.212979718 | 0.380264292 | -0.644548597 | 0.51921976 | 0.704562 |
| ENSG00000238247 | RDXP3 | 5.468409 | -0.2131156 | 0.6169579 | -0.5362095 | 0.5918138 | 0.760374 |
| ENSG00000136783 | NIPSNAP3A | 1570.608488 | -0.213379653 | 0.158651839 | -1.383126385 | 0.16662612 | 0.335742 |
| ENSG00000279968 | CCDC28A-AS1 | 8.306444 | -0.213418 | 0.5613141 | -0.534804 | 0.5927854 | 0.760888 |
| ENSG00000143553 | SNAPIN | 2115.868 | -0.2134938 | 0.1705312 | -1.287248 | 0.198008 | 0.378072 |
| ENSG00000177508 | IRX3 | 2009.874 | -0.2135269 | 0.2650335 | -0.8705839 | 0.3839814 | 0.586322 |
| ENSG00000116209 | TMEM59 | 11388.91631 | -0.21352875 | 0.152495829 | -1.44111677 | 0.1495517 | 0.310546 |
| ENSG00000236739 | CLIC4P1 | 159.9389 | -0.2136036 | 0.3871481 | -0.6398192 | 0.5222901 | 0.707599 |
| ENSG00000106261 | ZKSCAN1 | 8937.614574 | -0.21361551 | 0.352357425 | -0.682527111 | 0.49490571 | 0.68421 |
| ENSG00000188368 | PRR19 | 120.2221 | -0.2136428 | 0.3486037 | -0.6784452 | 0.4974895 | 0.686176 |

| | | | | | | | |
|---|---|---|---|---|---|---|---|
| ENSG00000151687 | ANKAR | 110.2529 | -0.2136608 | 0.2695726 | -0.8510373 | 0.3947486 | 0.596456 |
| ENSG00000286011 | | 25.8363 | -0.2137442 | 0.3861677 | -0.6383722 | 0.5232314 | 0.708521 |
| ENSG00000278009 | | 18.36435 | -0.2137506 | 0.4217483 | -0.6002257 | 0.5483558 | 0.727909 |
| ENSG00000201121 | RNY1P12 | 0.9662798 | -0.2137786 | 0.7815286 | -0.8375489 | 0.4022841 | NA |
| ENSG00000177640 | CASC2 | 95.66114 | -0.2138838 | 0.2832907 | -0.814542 | 0.4153345 | 0.615176 |
| ENSG00000264281 | | 30.1479 | -0.213936 | 0.3406746 | -0.6991057 | 0.484486 | 0.67556 |
| ENSG00000186526 | CYP4F8 | 10.70659 | -0.2139509 | 0.8028374 | -0.6259963 | 0.5313174 | 0.715504 |
| ENSG00000274378 | | 4.059388 | -0.2139569 | 0.6220732 | -0.5499176 | 0.5823759 | 0.752988 |
| ENSG00000236229 | VEZF1P1 | 17.90747 | -0.213961 | 0.5168088 | -0.5461273 | 0.5849784 | 0.754993 |
| ENSG00000147586 | MRPS28 | 331.5389 | -0.2141915 | 0.2762434 | -0.8237821 | 0.4100634 | 0.610397 |
| ENSG00000188542 | DUSP28 | 374.6101 | -0.2142015 | 0.1665117 | -1.330624 | 0.1833129 | 0.358437 |
| ENSG00000253203 | GUSBP3 | 5.894461 | -0.2142174 | 0.5526155 | -0.538899 | 0.5899566 | 0.758691 |
| ENSG00000232640 | | 71.90804 | -0.2142342 | 0.2418716 | -0.928056 | 0.3533785 | 0.55536 |
| ENSG00000215311 | NPM1P12 | 10.81789 | -0.2142782 | 0.5470729 | -0.5423507 | 0.5875769 | 0.75702 |
| ENSG00000229833 | PET100 | 217.5105 | -0.214305 | 0.2234654 | -1.009512 | 0.312729 | 0.513751 |
| ENSG00000180697 | C3orf22 | 1.099715 | -0.2143144 | 0.7722954 | -0.774276 | 0.4387676 | NA |
| ENSG00000117133 | RPF1 | 1980.720952 | -0.214331224 | 0.148561939 | -1.472031291 | 0.14101243 | 0.297677 |
| ENSG00000075073 | TACR2 | 18.71506748 | -0.214357904 | 0.456218997 | -0.575470526 | 0.56497312 | 0.740103 |
| ENSG00000214081 | CYP4F30P | 1.056436 | -0.2144064 | 0.7936833 | -0.9191733 | 0.3580049 | NA |
| ENSG00000269243 | | 14.83884 | -0.2144427 | 0.4174617 | -0.6026876 | 0.5467165 | 0.726722 |
| ENSG00000115652 | UXS1 | 2827.137707 | -0.214468706 | 0.136520826 | -1.607654944 | 0.10791077 | 0.245908 |
| ENSG00000182389 | CACNB4 | 26.0999 | -0.2145552 | 0.5328776 | -0.5472577 | 0.5842017 | 0.754247 |
| ENSG00000237452 | MEIOSIN | 2.12825 | -0.2145903 | 0.7297351 | -0.6414946 | 0.5212014 | 0.706484 |
| ENSG00000262074 | SNORD3B-2 | 7.693076 | -0.2146561 | 0.6553711 | -0.5621671 | 0.5740022 | 0.746641 |
| ENSG00000103994 | ZNF106 | 4351.213291 | -0.214725223 | 0.243936051 | -0.926808395 | 0.35402603 | 0.556111 |
| ENSG00000113946 | CLDN16 | 4.554570912 | -0.214730836 | 0.802868715 | -0.627837022 | 0.53011071 | 0.714551 |
| ENSG00000133619 | KRBA1 | 925.8185103 | -0.214763402 | 0.166894232 | -1.315331035 | 0.18839867 | 0.365476 |
| ENSG00000214401 | KANSL1-AS1 | 44.59453 | -0.2147889 | 0.3755003 | -0.6522852 | 0.5142172 | 0.700036 |
| ENSG00000183199 | HSP90AB3P | 1781.283 | -0.2147908 | 0.2343791 | -0.9605988 | 0.3367539 | 0.537887 |
| ENSG00000179165 | PXT1 | 1.916304 | -0.2148176 | 0.7163664 | -0.608391 | 0.5429282 | 0.723919 |
| ENSG00000197329 | PELI1 | 444.7422 | -0.2148603 | 0.2021375 | -1.102005 | 0.2704596 | 0.466782 |
| ENSG00000125319 | HROB | 548.6424469 | -0.21495381 | 0.439803192 | -0.563564939 | 0.57305025 | 0.746043 |
| ENSG00000065060 | UHRF1BP1 | 4602.363299 | -0.214976885 | 0.253570206 | -0.899724986 | 0.36826662 | 0.570291 |

| | | | | | | | |
|---|---|---|---|---|---|---|---|
| ENSG00000264177 | | 1.073668 | -0.2149938 | 0.7474519 | -0.6980631 | 0.4851377 | NA |
| ENSG00000274322 | | 1.318294 | -0.2150165 | 0.769507 | -0.7613678 | 0.4464374 | NA |
| ENSG00000239183 | SNORA84 | 0.8449358 | -0.2150766 | 0.7907714 | -0.9029009 | 0.3665785 | NA |
| ENSG00000203581 | OR1F2P | 2.010537 | -0.215099 | 0.7048127 | -0.6164518 | 0.5375964 | 0.720692 |
| ENSG00000136699 | SMPD4 | 6065.491762 | -0.215119865 | 0.271542823 | -0.849404825 | 0.39565607 | 0.597249 |
| ENSG00000179950 | PUF60 | 12619.99 | -0.2151775 | 0.2373153 | -0.9549926 | 0.3395814 | 0.540908 |
| ENSG00000205267 | DGAT2L7P | 0.9087294 | -0.215259 | 0.8114772 | -1.157576 | 0.2470371 | NA |
| ENSG00000125630 | POLR1B | 3423.525714 | -0.215279619 | 0.146463284 | -1.51223509 | 0.13047407 | 0.28156 |
| ENSG00000138658 | ZGRF1 | 1126.089224 | -0.215561908 | 0.37801949 | -0.641794562 | 0.52100658 | 0.706289 |
| ENSG00000122376 | SHLD2 | 3614.250725 | -0.215578243 | 0.142475768 | -1.535347836 | 0.12469841 | 0.272684 |
| ENSG00000265033 | RN7SL262P | 2.060554 | -0.215593 | 0.6835464 | -0.5812614 | 0.5610643 | 0.737289 |
| ENSG00000258741 | H2AZ2P1 | 475.7986 | -0.2156049 | 0.2813068 | -0.8294443 | 0.406853 | 0.607456 |
| ENSG00000173557 | FAM166C | 1.28411 | -0.2156213 | 0.7816023 | -0.8441229 | 0.3986007 | NA |
| ENSG00000227175 | | 0.9048054 | -0.215672 | 0.8072069 | -1.040568 | 0.298076 | NA |
| ENSG00000187513 | GJA4 | 1.677995 | -0.2157279 | 0.7047833 | -0.6029536 | 0.5465395 | 0.726642 |
| ENSG00000180398 | MCFD2 | 6350.999 | -0.2157426 | 0.1402625 | -1.565253 | 0.1175237 | 0.261274 |
| ENSG00000231549 | ATP5MDP1 | 22.90893 | -0.2157869 | 0.4255744 | -0.6076519 | 0.5434183 | 0.724367 |
| ENSG00000107372 | ZFAND5 | 5258.85745 | -0.215822121 | 0.156053636 | -1.406999764 | 0.15942746 | 0.325507 |
| ENSG00000214875 | MED28P1 | 1.857962 | -0.2158343 | 0.7170209 | -0.6220816 | 0.5338882 | 0.717531 |
| ENSG00000286223 | | 19.0678 | -0.2158489 | 0.4227156 | -0.6039784 | 0.545858 | 0.726117 |
| ENSG00000113300 | CNOT6 | 7754.63531 | -0.215869479 | 0.152840651 | -1.463756999 | 0.14326036 | 0.301041 |
| ENSG00000266010 | GATA6-AS1 | 5.675904 | -0.2158841 | 0.6043122 | -0.5389636 | 0.589912 | 0.75867 |
| ENSG00000177885 | GRB2 | 13504.84 | -0.2159266 | 0.1393668 | -1.566298 | 0.1172788 | 0.260904 |
| ENSG00000133019 | CHRM3 | 365.1556231 | -0.216075652 | 0.236421542 | -0.970452879 | 0.3318208 | 0.532955 |
| ENSG00000114491 | UMPS | 4433.533151 | -0.216128689 | 0.223609277 | -1.011762234 | 0.31165175 | 0.512551 |
| ENSG00000264775 | PPIAP14 | 1.291208 | -0.2162682 | 0.8142296 | -1.189677 | 0.2341732 | NA |
| ENSG00000118197 | DDX59 | 1590.579097 | -0.216296847 | 0.137764551 | -1.597913688 | 0.11006219 | 0.249488 |
| ENSG00000100526 | CDKN3 | 1321.214166 | -0.216357459 | 0.527567323 | -0.547061322 | 0.5843366 | 0.754311 |
| ENSG00000179913 | B3GNT3 | 379.9738 | -0.2165877 | 0.3618778 | -0.675815 | 0.4991581 | 0.687549 |
| ENSG00000080007 | DDX43 | 576.6282726 | -0.216609512 | 0.189556588 | -1.175317165 | 0.23986785 | 0.430085 |
| ENSG00000099849 | RASSF7 | 3843.592434 | -0.216618301 | 0.187750083 | -1.189788746 | 0.23412943 | 0.423268 |
| ENSG00000108947 | EFNB3 | 1324.141933 | -0.216799355 | 0.229294491 | -0.990119657 | 0.32211564 | 0.522899 |
| ENSG00000082996 | RNF13 | 3379.674171 | -0.216949898 | 0.157365135 | -1.377759341 | 0.1682776 | 0.338044 |

| | | | | | | | |
|---|---|---|---|---|---|---|---|
| ENSG00000181481 | RNF135 | 902.0182 | -0.2169831 | 0.1964526 | -1.143593 | 0.2527926 | 0.445654 |
| ENSG00000067992 | PDK3 | 2576.06249 | -0.217045578 | 0.241001651 | -0.941638763 | 0.34637762 | 0.548028 |
| ENSG00000163501 | IHH | 1.149557 | -0.2172234 | 0.7727638 | -0.7806796 | 0.434991 | NA |
| ENSG00000237899 | | 20.98258 | -0.2173867 | 0.4458508 | -0.5903956 | 0.5549255 | 0.732685 |
| ENSG00000225616 | | 107.0976 | -0.2174139 | 0.3021579 | -0.7789214 | 0.436026 | 0.634698 |
| ENSG00000269921 | | 17.19424 | -0.217418 | 0.426099 | -0.6059477 | 0.5445495 | 0.725256 |
| ENSG00000170315 | UBB | 10577.21 | -0.2174685 | 0.1604401 | -1.417678 | 0.1562849 | 0.320743 |
| ENSG00000104313 | EYA1 | 4.827493715 | -0.21747615 | 0.613762311 | -0.54036252 | 0.58894705 | 0.757928 |
| ENSG00000224152 | | 112.452 | -0.2174814 | 0.2607284 | -0.8851164 | 0.3760938 | 0.5783 |
| ENSG00000232713 | RPS12P3 | 26.52445 | -0.2174977 | 0.3746462 | -0.6570589 | 0.5111431 | 0.697082 |
| ENSG00000187682 | ERAS | 2.179087 | -0.2175242 | 0.6951799 | -0.5947661 | 0.5519999 | 0.730751 |
| ENSG00000165480 | SKA3 | 1298.425 | -0.2175555 | 0.5674194 | -0.547766 | 0.5838526 | 0.754053 |
| ENSG00000281849 | | 6.129436 | -0.2176628 | 0.5636535 | -0.547148 | 0.584277 | 0.754308 |
| ENSG00000110075 | PPP6R3 | 9981.018106 | -0.217693933 | 0.136941471 | -1.617678218 | 0.10573196 | 0.242358 |
| ENSG00000233287 | | 9.670067 | -0.2176962 | 0.5201226 | -0.5472773 | 0.5841882 | 0.754247 |
| ENSG00000213380 | COG8 | 893.1835 | -0.2177627 | 0.1440653 | -1.528575 | 0.1263699 | 0.275049 |
| ENSG00000189306 | RRP7A | 2944.239 | -0.2177776 | 0.2505716 | -0.9082377 | 0.3637526 | 0.565873 |
| ENSG00000163738 | MTHFD2L | 603.009 | -0.2177859 | 0.185464 | -1.210193 | 0.2262047 | 0.413743 |
| ENSG00000255503 | | 5.851014 | -0.2177971 | 0.6754699 | -0.5833223 | 0.5596763 | 0.73613 |
| ENSG00000272541 | | 0.8991507 | -0.2178147 | 0.7951914 | -0.9120344 | 0.3617506 | NA |
| ENSG00000164620 | RELL2 | 970.9073 | -0.2178203 | 0.1900058 | -1.190603 | 0.2338095 | 0.422879 |
| ENSG00000171208 | NETO2 | 1886.768 | -0.217844 | 0.1270273 | -1.719989 | 0.08543451 | 0.207429 |
| ENSG00000141384 | TAF4B | 654.5053 | -0.2178499 | 0.2297715 | -0.9946503 | 0.3199064 | 0.520777 |
| ENSG00000235162 | C12orf75 | 1203.644 | -0.2178514 | 0.3298089 | -0.7262533 | 0.4676835 | 0.661268 |
| ENSG00000141744 | PNMT | 3.047876 | -0.217904 | 0.7313246 | -0.6701078 | 0.5027891 | 0.690189 |
| ENSG00000261790 | | 108.1187 | -0.2179829 | 0.2131988 | -1.067903 | 0.2855641 | 0.483597 |
| ENSG00000287757 | | 7.242365 | -0.2179986 | 0.5582079 | -0.5475607 | 0.5839936 | 0.754089 |
| ENSG00000171475 | WIPF2 | 2218.206 | -0.218051 | 0.1622191 | -1.374957 | 0.1691446 | 0.339298 |
| ENSG00000237864 | LINC00322 | 10.66684 | -0.2180675 | 0.6379918 | -0.5541984 | 0.579443 | 0.750584 |
| ENSG00000256103 | ATP5MFP5 | 0.8128036 | -0.2180904 | 0.7766918 | -0.8094686 | 0.4182457 | NA |
| ENSG00000124608 | AARS2 | 1546.321391 | -0.218218837 | 0.166011946 | -1.344710898 | 0.17871861 | 0.352541 |
| ENSG00000184702 | SEPTIN5 | 22.68865 | -0.2182481 | 0.4571377 | -0.5849231 | 0.5585994 | 0.735582 |
| ENSG00000261061 | | 329.4926 | -0.2182629 | 0.4322642 | -0.6092572 | 0.542354 | 0.723674 |

| | | | | | | | |
|---|---|---|---|---|---|---|---|
| ENSG00000096384 | HSP90AB1 | 76786.49963 | -0.218295256 | 0.208012118 | -1.092144361 | 0.27476965 | 0.471557 |
| ENSG00000196814 | MVB12B | 900.2127 | -0.2183268 | 0.2144521 | -1.037125 | 0.2996776 | 0.498891 |
| ENSG00000197147 | LRRC8B | 1164.472 | -0.2184531 | 0.1900743 | -1.194172 | 0.2324107 | 0.420979 |
| ENSG00000145919 | BOD1 | 8100.372 | -0.2184545 | 0.1695054 | -1.333255 | 0.1824481 | 0.357404 |
| ENSG00000153113 | CAST | 8592.378 | -0.218469 | 0.1737815 | -1.283323 | 0.1993788 | 0.379598 |
| ENSG00000100296 | THOC5 | 3101.95857 | -0.218513892 | 0.165288155 | -1.35198514 | 0.17638007 | 0.349184 |
| ENSG00000160633 | SAFB | 5705.295 | -0.2185251 | 0.1764776 | -1.268611 | 0.2045797 | 0.386529 |
| ENSG00000169740 | ZNF32 | 1878.188 | -0.2186063 | 0.1827452 | -1.238014 | 0.2157108 | 0.400869 |
| ENSG00000127311 | HELB | 141.1376709 | -0.218628374 | 0.232370951 | -0.988377532 | 0.32296779 | 0.523707 |
| ENSG00000141741 | MIEN1 | 972.1446 | -0.218706 | 0.2109258 | -1.078216 | 0.2809372 | 0.478414 |
| ENSG00000138180 | CEP55 | 2293.107417 | -0.218836311 | 0.546766485 | -0.546742951 | 0.58455534 | 0.754521 |
| ENSG00000221916 | C19orf73 | 77.53005 | -0.218854 | 0.2542611 | -0.9195259 | 0.3578206 | 0.559815 |
| ENSG00000230065 | | 0.7789948 | -0.2188541 | 0.7915216 | -0.8909698 | 0.3729454 | NA |
| ENSG00000182676 | PPP1R27 | 7.575503 | -0.2188773 | 0.5416134 | -0.5500645 | 0.5822752 | 0.752894 |
| ENSG00000102763 | VWA8 | 1945.881618 | -0.218938612 | 0.185255556 | -1.217689707 | 0.22334191 | 0.410033 |
| ENSG00000112081 | SRSF3 | 15119.55233 | -0.218961454 | 0.223302494 | -1.026461126 | 0.30467428 | 0.504417 |
| ENSG00000174912 | METTL15P1 | 120.8158 | -0.2189726 | 0.2325103 | -0.9987205 | 0.3179301 | 0.51899 |
| ENSG00000265802 | RN7SL49P | 4.142794 | -0.2189914 | 0.6693122 | -0.5739882 | 0.5659758 | 0.740723 |
| ENSG00000232838 | PET117 | 86.76453 | -0.2190418 | 0.213949 | -1.066238 | 0.2863163 | 0.484593 |
| ENSG00000168876 | ANKRD49 | 773.1759 | -0.2190764 | 0.2291419 | -0.9942549 | 0.3200988 | 0.52093 |
| ENSG00000278224 | PRICKLE4 | 8.183441 | -0.2190874 | 0.5408227 | -0.5509743 | 0.5816513 | 0.752234 |
| ENSG00000167323 | STIM1 | 1289.383 | -0.2191297 | 0.1327438 | -1.691536 | 0.09073448 | 0.216909 |
| ENSG00000185721 | DRG1 | 3445.889 | -0.2192089 | 0.1739799 | -1.304626 | 0.1920202 | 0.370235 |
| ENSG00000227492 | | 7.021598 | -0.2192098 | 0.5573538 | -0.5470624 | 0.5843358 | 0.754312 |
| ENSG00000170791 | CHCHD7 | 1229.714 | -0.2192295 | 0.1928477 | -1.17598 | 0.239603 | 0.429819 |
| ENSG00000187231 | SESTD1 | 1705.3 | -0.2192454 | 0.2076472 | -1.10895 | 0.2674519 | 0.463289 |
| ENSG00000201176 | RNU6-853P | 1.092038 | -0.2192761 | 0.7402326 | -0.6746623 | 0.4998903 | NA |
| ENSG00000086102 | NFX1 | 3021.522869 | -0.21928062 | 0.120278886 | -1.85611745 | 0.06343682 | 0.165679 |
| ENSG00000254038 | | 1.187693 | -0.2193311 | 0.8165312 | -1.222519 | 0.2215116 | NA |
| ENSG00000140443 | IGF1R | 5412.271 | -0.2195251 | 0.3066002 | -0.7862917 | 0.4316966 | 0.630899 |
| ENSG00000196756 | SNHG17 | 1640.52 | -0.2195299 | 0.1755693 | -1.28542 | 0.1986455 | 0.378763 |
| ENSG00000013561 | RNF14 | 2935.737466 | -0.219547773 | 0.138316067 | -1.646987078 | 0.0995607 | 0.232017 |
| ENSG00000130724 | CHMP2A | 2794.891646 | -0.219577043 | 0.172955923 | -1.310356419 | 0.19007529 | 0.367869 |

| | | | | | | | |
|---|---|---|---|---|---|---|---|
| ENSG00000272910 | | 5.297268 | -0.2196224 | 0.6091002 | -0.5550801 | 0.5788399 | 0.750168 |
| ENSG00000287141 | | 1.36814 | -0.2196275 | 0.7294558 | -0.650883 | 0.515122 | NA |
| ENSG00000129675 | ARHGEF6 | 273.3891769 | -0.219663489 | 0.378631991 | -0.672033771 | 0.5015622 | 0.689314 |
| ENSG00000213962 | API5P2 | 5.493683 | -0.2196834 | 0.6921897 | -0.5986957 | 0.5493758 | 0.728762 |
| ENSG00000118260 | CREB1 | 2714.977546 | -0.219704606 | 0.177996336 | -1.285196553 | 0.19872361 | 0.37884 |
| ENSG00000188312 | CENPP | 302.333 | -0.2199224 | 0.3278739 | -0.7452337 | 0.4561305 | 0.652028 |
| ENSG00000108669 | CYTH1 | 3041.726173 | -0.220008766 | 0.149283697 | -1.517578288 | 0.12912074 | 0.27932 |
| ENSG00000163209 | SPRR3 | 6.104091 | -0.2200168 | 0.8048323 | -0.6252801 | 0.5317872 | 0.715866 |
| ENSG00000279236 | | 4.13958 | -0.2200259 | 0.6674435 | -0.5806054 | 0.5615064 | 0.737416 |
| ENSG00000285791 | | 1.575555 | -0.2200286 | 0.72103 | -0.636789 | 0.5242623 | 0.70931 |
| ENSG00000132024 | CC2D1A | 3573.164414 | -0.220067523 | 0.144975521 | -1.572073065 | 0.11593361 | 0.258778 |
| ENSG00000109805 | NCAPG | 1050.918545 | -0.220135805 | 0.543505689 | -0.561266722 | 0.57461572 | 0.747026 |
| ENSG00000245148 | ARAP1-AS2 | 8.015007 | -0.2202288 | 0.5828548 | -0.5554899 | 0.5785596 | 0.749988 |
| ENSG00000130173 | ANGPTL8 | 170.2560013 | -0.220251048 | 0.293091698 | -0.812214548 | 0.41666853 | 0.616602 |
| ENSG00000182504 | CEP97 | 709.6955 | -0.2203256 | 0.2754765 | -0.8559032 | 0.3920513 | 0.593731 |
| ENSG00000146038 | DCDC2 | 3788.208 | -0.2204429 | 0.3268829 | -0.7433729 | 0.457256 | 0.65317 |
| ENSG00000271392 | | 3.385557 | -0.2204473 | 0.6724533 | -0.589551 | 0.5554917 | 0.732995 |
| ENSG00000163527 | STT3B | 11840.48 | -0.2204487 | 0.1319725 | -1.696936 | 0.08970874 | 0.215154 |
| ENSG00000260063 | | 23.96037 | -0.220451 | 0.4527355 | -0.5960517 | 0.5511407 | 0.730046 |
| ENSG00000184949 | FAM227A | 265.7036 | -0.2204997 | 0.3398748 | -0.7249436 | 0.4684866 | 0.662086 |
| ENSG00000259520 | | 2.943453 | -0.2205578 | 0.6939599 | -0.6025028 | 0.5468395 | 0.726812 |
| ENSG00000270068 | | 4.244257 | -0.2205627 | 0.6401017 | -0.5620544 | 0.5740789 | 0.746641 |
| ENSG00000218739 | CEBPZOS | 1833.127 | -0.2206501 | 0.1664531 | -1.358763 | 0.1742217 | 0.346112 |
| ENSG00000245848 | CEBPA | 615.7562 | -0.2208285 | 0.2410322 | -0.9709341 | 0.3315811 | 0.532759 |
| ENSG00000179873 | NLRP11 | 1.200397 | -0.2208677 | 0.8116125 | -0.7730933 | 0.4394672 | NA |
| ENSG00000215908 | CROCCP2 | 1412.979 | -0.2208975 | 0.1584635 | -1.426587 | 0.1536991 | 0.316806 |
| ENSG00000258325 | ITFG2-AS1 | 1.682247 | -0.2209437 | 0.7286241 | -0.6596399 | 0.5094849 | 0.69551 |
| ENSG00000251821 | RNU6-583P | 4.568948 | -0.2209571 | 0.6924929 | -0.6079888 | 0.5431949 | 0.724178 |
| ENSG00000076248 | UNG | 5930.0408 | -0.221006676 | 0.370892588 | -0.66348016 | 0.50702308 | 0.693528 |
| ENSG00000023892 | DEF6 | 290.3299118 | -0.221007986 | 0.311998132 | -0.782258915 | 0.43406243 | 0.633034 |
| ENSG00000225357 | RPF2P1 | 50.37382 | -0.2210946 | 0.2715692 | -0.8689405 | 0.3848797 | 0.587322 |
| ENSG00000138639 | ARHGAP24 | 68.69287568 | -0.22111925 | 0.445517286 | -0.604988523 | 0.54518662 | 0.725751 |
| ENSG00000111615 | KRR1 | 3068.868955 | -0.221130194 | 0.142180372 | -1.588969277 | 0.11206733 | 0.252653 |

| | | | | | | | |
|---|---|---|---|---|---|---|---|
| ENSG00000210194 | MT-TE | 2.999005 | -0.2211981 | 0.6894435 | -0.5937504 | 0.552679 | 0.731245 |
| ENSG00000246548 | LINC02288 | 7.217856 | -0.2212221 | 0.6231053 | -0.5560098 | 0.5782041 | 0.74982 |
| ENSG00000164219 | PGGT1B | 1549.43 | -0.221232 | 0.121662 | -1.841167 | 0.06559711 | 0.169951 |
| ENSG00000278903 | | 2.923574 | -0.2212788 | 0.6510902 | -0.5742996 | 0.5657651 | 0.740563 |
| ENSG00000105808 | RASA4 | 31.59852255 | -0.221282026 | 0.3786415 | -0.670328249 | 0.50264856 | 0.690174 |
| ENSG00000184371 | CSF1 | 327.2884 | -0.221299 | 0.3782681 | -0.6666146 | 0.5050184 | 0.691784 |
| ENSG00000229119 | | 2082.523 | -0.221491 | 0.1910989 | -1.207179 | 0.227363 | 0.415147 |
| ENSG00000109066 | TMEM104 | 1435.820469 | -0.221503042 | 0.208228626 | -1.104490952 | 0.26938023 | 0.465597 |
| ENSG00000183775 | KCTD16 | 3.360122 | -0.2215236 | 0.6582449 | -0.5613117 | 0.5745851 | 0.747023 |
| ENSG00000171236 | LRG1 | 168.3744 | -0.2215791 | 0.4099018 | -0.6577214 | 0.5107172 | 0.696609 |
| ENSG00000161981 | SNRNP25 | 1559.473 | -0.2216138 | 0.4575791 | -0.5979917 | 0.5498455 | 0.729016 |
| ENSG00000254248 | | 3.492745 | -0.2216826 | 0.6703361 | -0.5756979 | 0.5648194 | 0.739975 |
| ENSG00000162694 | EXTL2 | 277.8516 | -0.2217093 | 0.2918666 | -0.8227542 | 0.4106478 | 0.61105 |
| ENSG00000254814 | | 1.931914 | -0.2218336 | 0.7215441 | -0.6527436 | 0.5139216 | 0.699866 |
| ENSG00000248884 | | 0.7488181 | -0.2219417 | 0.8179123 | -0.8961094 | 0.3701943 | NA |
| ENSG00000236394 | | 3.385429 | -0.2219428 | 0.6773038 | -0.5880664 | 0.5564877 | 0.733872 |
| ENSG00000233017 | | 27.89479 | -0.2220383 | 0.4872093 | -0.5765906 | 0.564216 | 0.739552 |
| ENSG00000273010 | | 16.71613 | -0.2221656 | 0.4120516 | -0.6345903 | 0.5256956 | 0.710546 |
| ENSG00000167785 | ZNF558 | 1067.081 | -0.222173 | 0.2799345 | -0.8549391 | 0.3925849 | 0.594132 |
| ENSG00000090520 | DNAJB11 | 2395.319633 | -0.222185741 | 0.145377369 | -1.55062417 | 0.12099178 | 0.266885 |
| ENSG00000124134 | KCNS1 | 3.029768436 | -0.222351121 | 0.668641063 | -0.584544643 | 0.55885394 | 0.735715 |
| ENSG00000167635 | ZNF146 | 7432.28 | -0.2223677 | 0.1356385 | -1.681437 | 0.09267807 | 0.220188 |
| ENSG00000225151 | GOLGA2P7 | 52.28861 | -0.2223791 | 0.2600853 | -0.9052895 | 0.365312 | 0.567203 |
| ENSG00000206989 | SNORD63 | 2.784273 | -0.2223941 | 0.6254055 | -0.5626468 | 0.5736754 | 0.746641 |
| ENSG00000143190 | POU2F1 | 1739.509 | -0.2224577 | 0.1995273 | -1.165098 | 0.2439795 | 0.435272 |
| ENSG00000117335 | CD46 | 16728.60344 | -0.22246608 | 0.194178362 | -1.197226108 | 0.23121844 | 0.419681 |
| ENSG00000117862 | TXNDC12 | 399.6173331 | -0.222481073 | 0.191496292 | -1.202799046 | 0.22905409 | 0.417237 |
| ENSG00000240135 | PSMD12P1 | 42.43524 | -0.2225803 | 0.3718577 | -0.6829214 | 0.4946565 | 0.684079 |
| ENSG00000065154 | OAT | 5281.797557 | -0.22261594 | 0.227025065 | -1.028469294 | 0.30372913 | 0.503354 |
| ENSG00000080293 | SCTR | 5.439018144 | -0.222648648 | 0.682123186 | -0.581210109 | 0.56109886 | 0.737289 |
| ENSG00000287378 | | 1.031891 | -0.2226751 | 0.7748544 | -0.7761206 | 0.4376778 | NA |
| ENSG00000104067 | TJP1 | 4166.165971 | -0.222698439 | 0.1879623 | -1.247889666 | 0.21207147 | 0.396407 |
| ENSG00000147536 | GINS4 | 334.5131 | -0.2227116 | 0.4735833 | -0.5844413 | 0.5589234 | 0.73577 |

| | | | | | | | |
|---|---|---|---|---|---|---|---|
| ENSG00000122176 | FMOD | 14.17861307 | -0.222717428 | 0.80517733 | -0.646010606 | 0.51827249 | 0.703555 |
| ENSG00000202058 | RN7SKP80 | 4.407613 | -0.2227268 | 0.6053451 | -0.5584904 | 0.5765096 | 0.748537 |
| ENSG00000205485 | | 172.5303 | -0.2227821 | 0.1925469 | -1.192416 | 0.233098 | 0.421965 |
| ENSG00000025423 | HSD17B6 | 121.0085698 | -0.222790122 | 0.412129223 | -0.632433693 | 0.52710352 | 0.711981 |
| ENSG00000245888 | NSMCE1-DT | 77.20886 | -0.2229109 | 0.4310774 | -0.6186518 | 0.5361458 | 0.719256 |
| ENSG00000234073 | | 1.601135 | -0.2229136 | 0.7442201 | -0.693036 | 0.4882869 | 0.679159 |
| ENSG00000287953 | | 1.047056 | -0.2229986 | 0.8344046 | -1.65583 | 0.09775623 | NA |
| ENSG00000168298 | H1-4 | 41.41906 | -0.2230295 | 0.436142 | -0.6119503 | 0.5405706 | 0.722565 |
| ENSG00000248216 | KCTD9P5 | 4.324183 | -0.2230719 | 0.6347605 | -0.571942 | 0.5673612 | 0.741734 |
| ENSG00000267197 | | 15.13502 | -0.2230772 | 0.5503096 | -0.5627986 | 0.5735721 | 0.746595 |
| ENSG00000132716 | DCAF8 | 2252.476635 | -0.223086324 | 0.167481684 | -1.386659212 | 0.1655457 | 0.334198 |
| ENSG00000058262 | SEC61A1 | 23466.33233 | -0.223109468 | 0.108127293 | -2.08519522 | 0.03705158 | 0.110176 |
| ENSG00000236711 | SMAD9-IT1 | 1.277457 | -0.2231805 | 0.7846633 | -0.8662259 | 0.3863663 | NA |
| ENSG00000269535 | | 4.120691 | -0.2232348 | 0.6431183 | -0.5682698 | 0.5698518 | 0.743744 |
| ENSG00000204556 | | 14.93292 | -0.2232355 | 0.4263051 | -0.6195985 | 0.5355222 | 0.71889 |
| ENSG00000120063 | GNA13 | 10287.06229 | -0.223265389 | 0.146252402 | -1.580297465 | 0.11403876 | 0.255746 |
| ENSG00000237082 | COX5BP6 | 7.231594 | -0.2233525 | 0.6089381 | -0.5616173 | 0.5743768 | 0.746825 |
| ENSG00000275438 | | 8.296558 | -0.223388 | 0.5542774 | -0.5640952 | 0.5726894 | 0.745897 |
| ENSG00000205105 | COX17P1 | 341.755 | -0.2234431 | 0.252557 | -0.9384718 | 0.348002 | 0.549739 |
| ENSG00000272202 | | 2.765566 | -0.2234463 | 0.7444127 | -0.6754317 | 0.4994016 | 0.687706 |
| ENSG00000225580 | PA2G4P5 | 630.7071 | -0.2236466 | 0.2470153 | -0.9615139 | 0.3362939 | 0.537337 |
| ENSG00000179031 | NSA2P3 | 13.40988 | -0.2236523 | 0.5393646 | -0.5637955 | 0.5728933 | 0.745937 |
| ENSG00000234263 | MAP3K5-AS1 | 0.97609 | -0.2236561 | 0.7597978 | -0.7545672 | 0.4505087 | NA |
| ENSG00000207234 | RNU6-125P | 1.072916 | -0.2236977 | 0.7859462 | -0.8726938 | 0.38283 | NA |
| ENSG00000204397 | CARD16 | 16.4394 | -0.2237066 | 0.8050899 | -0.658735 | 0.510066 | 0.695971 |
| ENSG00000008324 | SS18L2 | 769.8131327 | -0.223718965 | 0.232073131 | -1.016078708 | 0.30959191 | 0.51027 |
| ENSG00000142731 | PLK4 | 1608.621 | -0.2237278 | 0.4528144 | -0.5922038 | 0.5537142 | 0.731849 |
| ENSG00000237248 | LINC00987 | 4.707247 | -0.2237385 | 0.7164062 | -0.6509848 | 0.5150563 | 0.700478 |
| ENSG00000166473 | PKD1L2 | 157.5139 | -0.2237427 | 0.3375568 | -0.726718 | 0.4673987 | 0.661023 |
| ENSG00000198435 | NRARP | 971.3132 | -0.2238562 | 0.2829412 | -0.8568416 | 0.3915324 | 0.593317 |
| ENSG00000228041 | EIF3JP1 | 0.8482295 | -0.2238696 | 0.8083987 | -1.058575 | 0.2897934 | NA |
| ENSG00000146143 | PRIM2 | 862.8365 | -0.2239363 | 0.3724142 | -0.6814654 | 0.495577 | 0.684733 |
| ENSG00000273061 | CDC37L1-DT | 39.01897 | -0.2239886 | 0.3374057 | -0.7381999 | 0.460393 | 0.655647 |

| | | | | | | | |
|---|---|---|---|---|---|---|---|
| ENSG00000251017 | | 79.5714 | -0.2240701 | 0.2243503 | -1.045883 | 0.2956148 | 0.494708 |
| ENSG00000171817 | ZNF540 | 27.92622 | -0.2240746 | 0.5879348 | -0.5645884 | 0.5723538 | 0.745771 |
| ENSG00000092199 | HNRNPC | 25992.16664 | -0.224097219 | 0.159897497 | -1.434601951 | 0.15140056 | 0.313235 |
| ENSG00000203791 | EEF1AKMT2 | 351.6258 | -0.2242003 | 0.1652988 | -1.387345 | 0.1653366 | 0.333928 |
| ENSG00000125835 | SNRPB | 6439.61693 | -0.224220299 | 0.296068028 | -0.820240923 | 0.41207878 | 0.612299 |
| ENSG00000286722 | | 2.060925 | -0.2242323 | 0.7357842 | -0.6702032 | 0.5027283 | 0.690189 |
| ENSG00000230783 | RPS3AP13 | 1.0548 | -0.2242942 | 0.7900863 | -0.918162 | 0.358534 | NA |
| ENSG00000158201 | ABHD3 | 841.82 | -0.224324 | 0.2282193 | -1.02425 | 0.3057172 | 0.50564 |
| ENSG00000224976 | PARP4P2 | 0.8449452 | -0.2243368 | 0.7777631 | -0.8261024 | 0.408746 | NA |
| ENSG00000280254 | | 411.7835 | -0.2243417 | 0.2089879 | -1.113256 | 0.2655983 | 0.461263 |
| ENSG00000167088 | SNRPD1 | 3768.21 | -0.2243556 | 0.2389115 | -0.9887157 | 0.3228022 | 0.523534 |
| ENSG00000261578 | | 319.5667 | -0.2243958 | 0.4026586 | -0.6558469 | 0.5119226 | 0.697889 |
| ENSG00000262558 | | 1.582306 | -0.2244744 | 0.7171476 | -0.6551593 | 0.5123652 | 0.698247 |
| ENSG00000215784 | FAM72D | 210.0455 | -0.2244868 | 0.4286636 | -0.6281273 | 0.5299206 | 0.71444 |
| ENSG00000119523 | ALG2 | 2574.419503 | -0.224529525 | 0.163558327 | -1.408505822 | 0.15898134 | 0.324896 |
| ENSG00000237390 | | 2.139272 | -0.2246973 | 0.7220874 | -0.6553129 | 0.5122663 | 0.69822 |
| ENSG00000164535 | DAGLB | 2089.945 | -0.2247217 | 0.1202293 | -1.866822 | 0.06192645 | 0.162813 |
| ENSG00000231300 | EZH2P1 | 9.831879 | -0.2247347 | 0.6126681 | -0.5694858 | 0.5690265 | 0.743032 |
| ENSG00000130300 | PLVAP | 3.818503417 | -0.224782878 | 0.741032618 | -0.667811876 | 0.50425369 | 0.691213 |
| ENSG00000159884 | CCDC107 | 56.2134 | -0.2248289 | 0.3341139 | -0.7410191 | 0.4586819 | 0.654461 |
| ENSG00000284070 | | 23.2905 | -0.2248428 | 0.4978786 | -0.5823056 | 0.5603608 | 0.736636 |
| ENSG00000174442 | ZWILCH | 1883.155 | -0.2249 | 0.4319632 | -0.6141828 | 0.5390946 | 0.721572 |
| ENSG00000242199 | | 5.313645 | -0.2250468 | 0.6551445 | -0.5775682 | 0.5635557 | 0.739049 |
| ENSG00000231650 | RFESDP1 | 3.625972 | -0.2250472 | 0.6934322 | -0.6152914 | 0.5383623 | 0.721137 |
| ENSG00000100926 | TM9SF1 | 396.1764762 | -0.225068143 | 0.18344876 | -1.269482832 | 0.20426891 | 0.386059 |
| ENSG00000204175 | GPRIN2 | 1.230337 | -0.2250684 | 0.7778278 | -0.8466385 | 0.3971967 | NA |
| ENSG00000253880 | | 1.507973 | -0.2251013 | 0.817157 | -0.8736285 | 0.3823206 | NA |
| ENSG00000169504 | CLIC4 | 7511.186 | -0.2251058 | 0.3101274 | -0.7952909 | 0.4264443 | 0.626283 |
| ENSG00000230956 | CAPZA1P1 | 91.45162 | -0.2252032 | 0.2375729 | -1.005346 | 0.31473 | 0.515827 |
| ENSG00000215866 | LINC01356 | 1.444532 | -0.2252196 | 0.7895691 | -0.9041436 | 0.3659193 | NA |
| ENSG00000230312 | SLC25A53P1 | 2.586047 | -0.2252428 | 0.7297945 | -0.668332 | 0.5039217 | 0.691137 |
| ENSG00000101365 | IDH3B | 3515.542649 | -0.225301585 | 0.198291171 | -1.179395294 | 0.23824081 | 0.428067 |
| ENSG00000087995 | METTL2A | 4390.944134 | -0.225396074 | 0.094933003 | -2.423864944 | 0.01535632 | 0.054928 |

| | | | | | | | |
|---|---|---|---|---|---|---|---|
| ENSG00000161980 | POLR3K | 1413.589 | -0.2254482 | 0.2859303 | -0.8445394 | 0.3983681 | 0.599402 |
| ENSG00000162928 | PEX13 | 2687.372 | -0.2254823 | 0.1497498 | -1.567224 | 0.1170625 | 0.260489 |
| ENSG00000011258 | MBTD1 | 5133.626346 | -0.2254892 | 0.173600439 | -1.342535867 | 0.17942231 | 0.353509 |
| ENSG00000142528 | ZNF473 | 1205 | -0.2255706 | 0.1609449 | -1.434057 | 0.1515561 | 0.313435 |
| ENSG00000197275 | RAD54B | 214.481 | -0.225593 | 0.4826784 | -0.5910574 | 0.5544819 | 0.732463 |
| ENSG00000163154 | TNFAIP8L2 | 2.428072 | -0.2256499 | 0.6677522 | -0.6000585 | 0.5484672 | 0.727956 |
| ENSG00000160188 | RSPH1 | 225.1448 | -0.2257226 | 0.2838275 | -0.8554029 | 0.3923281 | 0.594048 |
| ENSG00000126814 | TRMT5 | 1984.55604 | -0.225737073 | 0.15302844 | -1.514805213 | 0.12982174 | 0.280448 |
| ENSG00000070018 | LRP6 | 2197.837931 | -0.225763005 | 0.226571156 | -1.041126589 | 0.2978168 | 0.497029 |
| ENSG00000285870 | | 0.759107 | -0.2257838 | 0.8127287 | -1.14047 | 0.2540904 | NA |
| ENSG00000203812 | H2AC18 | 39.45631 | -0.2257851 | 0.4258309 | -0.6328962 | 0.5268014 | 0.711744 |
| ENSG00000235768 | BRD7P5 | 66.41605 | -0.2257987 | 0.3217803 | -0.7786056 | 0.4362121 | 0.634829 |
| ENSG00000166889 | PATL1 | 2306.326 | -0.225851 | 0.1435936 | -1.60434 | 0.1086392 | 0.247084 |
| ENSG00000198768 | APCDD1L | 0.9258734 | -0.2258985 | 0.7857021 | -0.8818535 | 0.377856 | NA |
| ENSG00000116514 | RNF19B | 889.7575351 | -0.225908668 | 0.183133949 | -1.266971814 | 0.20516537 | 0.387175 |
| ENSG00000287151 | C2orf27A | 559.9778 | -0.22593 | 0.1718494 | -1.357665 | 0.1745699 | 0.346523 |
| ENSG00000092853 | CLSPN | 940.1373064 | -0.225967211 | 0.584229822 | -0.574130222 | 0.56587968 | 0.740633 |
| ENSG00000178718 | RPP25 | 2115.33 | -0.2259681 | 0.2109698 | -1.153169 | 0.2488408 | 0.441175 |
| ENSG00000139668 | WDFY2 | 606.5963 | -0.2261322 | 0.219688 | -1.081366 | 0.2795342 | 0.47685 |
| ENSG00000259031 | | 6.165712 | -0.2263437 | 0.5931264 | -0.5692353 | 0.5691965 | 0.743217 |
| ENSG00000047188 | YTHDC2 | 2874.417645 | -0.22638854 | 0.165043657 | -1.392364123 | 0.16381216 | 0.331808 |
| ENSG00000204564 | C6orf136 | 1126.247 | -0.2264102 | 0.196035 | -1.195511 | 0.2318873 | 0.420289 |
| ENSG00000137802 | MAPKBP1 | 766.075186 | -0.226421072 | 0.206121756 | -1.141575615 | 0.25363047 | 0.446667 |
| ENSG00000003987 | MTMR7 | 135.2982808 | -0.226431399 | 0.238899429 | -0.997317398 | 0.31861047 | 0.519718 |
| ENSG00000279557 | | 0.8537593 | -0.2266164 | 0.7603555 | -0.745189 | 0.4561575 | NA |
| ENSG00000243279 | PRAF2 | 393.9548 | -0.2266516 | 0.1947381 | -1.202277 | 0.2292561 | 0.417313 |
| ENSG00000276043 | UHRF1 | 1176.694 | -0.2266543 | 0.5792659 | -0.5742923 | 0.56577 | 0.740563 |
| ENSG00000230508 | RPL19P21 | 10.44204 | -0.2266832 | 0.5141154 | -0.5763965 | 0.5643472 | 0.739611 |
| ENSG00000100027 | YPEL1 | 85.97955536 | -0.226734961 | 0.397105726 | -0.66363331 | 0.50692503 | 0.693501 |
| ENSG00000231007 | CDC20P1 | 77.3094 | -0.2268137 | 0.5006237 | -0.5841662 | 0.5591085 | 0.735831 |
| ENSG00000223486 | | 0.9136896 | -0.2268529 | 0.7883836 | -0.900405 | 0.3679047 | NA |
| ENSG00000136938 | ANP32B | 9605.235451 | -0.226935048 | 0.17871589 | -1.318552263 | 0.18731884 | 0.363966 |
| ENSG00000276234 | TADA2A | 839.223 | -0.2269757 | 0.1683343 | -1.38411 | 0.1663248 | 0.335338 |

| | | | | | | | |
|---|---|---|---|---|---|---|---|
| ENSG00000255112 | CHMP1B | 2805.335 | -0.227001 | 0.1942148 | -1.207838 | 0.2271098 | 0.414942 |
| ENSG00000279348 | | 91.13741 | -0.2270212 | 0.3493066 | -0.7546647 | 0.4504502 | 0.64671 |
| ENSG00000209645 | SNORD105 | 1.192309 | -0.2270331 | 0.7711079 | -0.8083638 | 0.4188812 | NA |
| ENSG00000259121 | | 1.148754 | -0.2270545 | 0.7751614 | -0.7847289 | 0.4326126 | NA |
| ENSG00000138756 | BMP2K | 354.398129 | -0.227181183 | 0.241485311 | -0.995956799 | 0.31927114 | 0.52022 |
| ENSG00000280212 | | 2.085522 | -0.2271835 | 0.7394676 | -0.6992603 | 0.4843894 | 0.675542 |
| ENSG00000179603 | GRM8 | 1.530911 | -0.2271954 | 0.7518225 | -0.7142904 | 0.4750476 | NA |
| ENSG00000111445 | RFC5 | 1832.240043 | -0.227203505 | 0.376239211 | -0.70637378 | 0.47995572 | 0.671801 |
| ENSG00000155093 | PTPRN2 | 69.38365 | -0.2272474 | 0.4607408 | -0.6020058 | 0.5471703 | 0.726931 |
| ENSG00000129562 | DAD1 | 5534.36306 | -0.227263787 | 0.152805663 | -1.532672293 | 0.12535662 | 0.273578 |
| ENSG00000143862 | ARL8A | 5975.075 | -0.2273323 | 0.1771228 | -1.326495 | 0.1846758 | 0.360042 |
| ENSG00000228339 | AMD1P1 | 2.190385 | -0.2273891 | 0.7636719 | -0.7837371 | 0.4331944 | 0.632237 |
| ENSG00000253173 | H2AZP2 | 20.5807 | -0.2274108 | 0.4189315 | -0.6376811 | 0.5236813 | 0.708884 |
| ENSG00000163001 | CFAP36 | 2300.52 | -0.2274988 | 0.176803 | -1.337359 | 0.1811054 | 0.355657 |
| ENSG00000226632 | UBE2V1P1 | 2.227186 | -0.2275549 | 0.7516201 | -0.7123615 | 0.476241 | 0.6686 |
| ENSG00000253695 | | 2.45577 | -0.227655 | 0.7253629 | -0.6662572 | 0.5052467 | 0.691989 |
| ENSG00000244009 | B3GAT3P1 | 44.13829 | -0.2277046 | 0.344417 | -0.7351133 | 0.4622705 | 0.656814 |
| ENSG00000169660 | HEXD | 2380.693 | -0.2277335 | 0.190932 | -1.232246 | 0.2178572 | 0.403577 |
| ENSG00000177030 | DEAF1 | 2088.904 | -0.227756 | 0.1840736 | -1.279128 | 0.2008521 | 0.381529 |
| ENSG00000214961 | NT5DC1P2 | 11.90239 | -0.2277748 | 0.4848599 | -0.5903503 | 0.5549558 | 0.732688 |
| ENSG00000140932 | CMTM2 | 3.018291 | -0.2278338 | 0.7113482 | -0.6320469 | 0.5273562 | 0.712175 |
| ENSG00000152256 | PDK1 | 518.125 | -0.227872 | 0.2736059 | -0.884132 | 0.3766249 | 0.578681 |
| ENSG00000286261 | NA | 56.04408 | -0.2278874 | 0.411 | -0.6513558 | 0.5148169 | 0.700365 |
| ENSG00000256771 | ZNF253 | 331.0877 | -0.2278883 | 0.2759669 | -0.8844828 | 0.3764356 | 0.578617 |
| ENSG00000108474 | PIGL | 804.5007048 | -0.227928877 | 0.163757111 | -1.42534517 | 0.15405745 | 0.317377 |
| ENSG00000254864 | | 8.466005 | -0.2279839 | 0.6051085 | -0.5742284 | 0.5658133 | 0.740583 |
| ENSG00000131446 | MGAT1 | 5682.12346 | -0.227989308 | 0.123104127 | -1.86022634 | 0.06285351 | 0.164599 |
| ENSG00000218027 | | 1.106115 | -0.228021 | 0.7563107 | -0.736672 | 0.4613218 | NA |
| ENSG00000213033 | AURKAP1 | 2.731233 | -0.2280243 | 0.7106417 | -0.6463037 | 0.5180827 | 0.703405 |
| ENSG00000275410 | HNF1B | 1.013407 | -0.2280253 | 0.7788491 | -0.8465577 | 0.3972417 | NA |
| ENSG00000134551 | PRH2 | 3.568704605 | -0.228039059 | 0.621359142 | -0.577525553 | 0.56358448 | 0.739049 |
| ENSG00000215193 | PEX26 | 3776.152 | -0.2281936 | 0.1591091 | -1.468886 | 0.1418638 | 0.298951 |
| ENSG00000199990 | VTRNA1-1 | 0.8809328 | -0.2282866 | 0.7743658 | -0.8106725 | 0.4175538 | NA |

| | | | | | | | |
|---|---|---|---|---|---|---|---|
| ENSG00000121864 | ZNF639 | 2463.196193 | -0.22830611 | 0.160851686 | -1.462912254 | 0.14349139 | 0.301377 |
| ENSG00000204536 | CCHCR1 | 1288.155 | -0.2283209 | 0.3053131 | -0.8461558 | 0.3974658 | 0.59855 |
| ENSG00000115568 | ZNF142 | 2181.162156 | -0.228330559 | 0.194774472 | -1.207080167 | 0.22740127 | 0.415189 |
| ENSG00000230989 | HSBP1 | 12388.55 | -0.2283725 | 0.2016096 | -1.202618 | 0.2291242 | 0.417285 |
| ENSG00000126012 | KDM5C | 9417.193981 | -0.228380668 | 0.111806645 | -2.065828239 | 0.0388447 | 0.11428 |
| ENSG00000145191 | EIF2B5 | 5514.794 | -0.2284431 | 0.1699773 | -1.384913 | 0.1660792 | 0.334995 |
| ENSG00000166974 | MAPRE2 | 132.6836 | -0.2285412 | 0.3742798 | -0.6985379 | 0.4848409 | 0.675746 |
| ENSG00000167005 | NUDT21 | 7428.714 | -0.2287667 | 0.1628762 | -1.441063 | 0.1495668 | 0.310553 |
| ENSG00000221955 | SLC12A8 | 517.0801 | -0.2287754 | 0.1904949 | -1.231731 | 0.2180497 | 0.403718 |
| ENSG00000165792 | METTL17 | 2471.349 | -0.2287794 | 0.1640758 | -1.435593 | 0.1511183 | 0.312829 |
| ENSG00000279766 | | 5.375612 | -0.2288143 | 0.6019735 | -0.5663486 | 0.5711568 | 0.744898 |
| ENSG00000260335 | | 0.893802 | -0.2288349 | 0.7976322 | -0.9620276 | 0.3360357 | NA |
| ENSG00000132855 | ANGPTL3 | 1.352629629 | -0.228878948 | 0.768540737 | -0.797628976 | 0.42508583 | NA |
| ENSG00000262497 | FAM187B2P | 2.259372 | -0.2289252 | 0.7370456 | -0.6857383 | 0.4928782 | 0.682805 |
| ENSG00000255571 | MIR9-3HG | 469.3027 | -0.2289362 | 0.3513171 | -0.7350383 | 0.4623162 | 0.656825 |
| ENSG00000262583 | TMEM231P1 | 23.94902 | -0.2289431 | 0.4565282 | -0.6126428 | 0.5401126 | 0.722316 |
| ENSG00000113597 | TRAPPC13 | 1525.48286 | -0.228979548 | 0.119656521 | -1.938581263 | 0.05255235 | 0.144054 |
| ENSG00000174417 | TRHR | 1.787315 | -0.229046 | 0.7287693 | -0.6837322 | 0.4941443 | 0.68394 |
| ENSG00000253361 | | 186.3535 | -0.2290531 | 0.3865183 | -0.6820559 | 0.4952036 | 0.684515 |
| ENSG00000106952 | TNFSF8 | 6.13134132 | -0.229073379 | 0.840937636 | -2.490978982 | 0.01273917 | 0.047252 |
| ENSG00000259066 | | 6.023233 | -0.229156 | 0.6034655 | -0.5706775 | 0.5682183 | 0.742524 |
| ENSG00000273336 | OR7M1P | 1.003361 | -0.2292433 | 0.818137 | -1.228008 | 0.2194439 | NA |
| ENSG00000131508 | UBE2D2 | 5451.700513 | -0.229251225 | 0.17507924 | -1.341702335 | 0.17969253 | 0.353832 |
| ENSG00000185088 | RPS27L | 3696.052 | -0.2292819 | 0.2639793 | -0.9215407 | 0.3567682 | 0.55877 |
| ENSG00000207325 | RNY1P4 | 0.9314408 | -0.2293018 | 0.7771976 | -0.8388675 | 0.4015437 | NA |
| ENSG00000173915 | ATP5MD | 2796.142 | -0.2293104 | 0.2416095 | -1.001106 | 0.3167757 | 0.517901 |
| ENSG00000151893 | CACUL1 | 4190.599 | -0.2293118 | 0.1139591 | -2.033496 | 0.04200246 | 0.121459 |
| ENSG00000285966 | | 3.214154 | -0.2293661 | 0.6658372 | -0.6037223 | 0.5460283 | 0.726243 |
| ENSG00000233266 | HMGB1P31 | 4.295935 | -0.2293889 | 0.6680614 | -0.5994217 | 0.5488917 | 0.728301 |
| ENSG00000235635 | | 1.843012 | -0.2294519 | 0.7251831 | -0.6684803 | 0.5038271 | 0.691125 |
| ENSG00000094804 | CDC6 | 1954.253281 | -0.229500225 | 0.591764031 | -0.582268524 | 0.56038582 | 0.736636 |
| ENSG00000279425 | | 26.36913 | -0.2295331 | 0.424505 | -0.6232131 | 0.5331445 | 0.717129 |
| ENSG00000119906 | SLF2 | 2664.504161 | -0.229542711 | 0.20340945 | -1.192363041 | 0.23311893 | 0.421974 |

| | | | | | | | |
|---|---|---|---|---|---|---|---|
| ENSG00000115677 | HDLBP | 34933.45017 | -0.229724686 | 0.103358822 | -2.24420273 | 0.02481936 | 0.080622 |
| ENSG00000127774 | EMC6 | 124.1026224 | -0.229786952 | 0.305001225 | -0.818882722 | 0.41285333 | 0.612998 |
| ENSG00000196420 | S100A5 | 1.982876 | -0.2298021 | 0.7239995 | -0.6652071 | 0.505918 | 0.692627 |
| ENSG00000203876 | ADD3-AS1 | 1.543385 | -0.2299381 | 0.7743568 | -0.8029641 | 0.4219955 | NA |
| ENSG00000233098 | CCDC144NL-AS1 | 1.944769 | -0.2299642 | 0.7201427 | -0.6647875 | 0.5061864 | 0.692848 |
| ENSG00000255455 | | 156.5027 | -0.2300227 | 0.3356784 | -0.7626753 | 0.4456571 | 0.642676 |
| ENSG00000237126 | | 41.52823 | -0.2300684 | 0.2784477 | -0.8909223 | 0.3729708 | 0.575095 |
| ENSG00000183066 | WBP2NL | 34.91538 | -0.2300742 | 0.5072192 | -0.5905868 | 0.5547973 | 0.732661 |
| ENSG00000115718 | PROC | 8.664808568 | -0.230152693 | 0.623552647 | -0.577262776 | 0.56376196 | 0.739186 |
| ENSG00000085982 | USP40 | 2297.73784 | -0.230212468 | 0.188364935 | -1.2750208 | 0.20230188 | 0.383487 |
| ENSG00000138606 | SHF | 107.3344674 | -0.230263983 | 0.317629591 | -0.793170517 | 0.42767848 | 0.627062 |
| ENSG00000207171 | | 1.121274 | -0.2303512 | 0.8127627 | -0.7557174 | 0.4498187 | NA |
| ENSG00000187566 | NHLRC1 | 362.2722 | -0.2305213 | 0.2145725 | -1.106747 | 0.2684033 | 0.464392 |
| ENSG00000280432 | | 0.8337087 | -0.230564 | 0.8022275 | -1.007331 | 0.3137757 | NA |
| ENSG00000258890 | CEP95 | 6115.429 | -0.2306231 | 0.1456648 | -1.638533 | 0.1013105 | 0.235002 |
| ENSG00000198053 | SIRPA | 172.5975 | -0.2306581 | 0.3312871 | -0.7694627 | 0.4416187 | 0.639421 |
| ENSG00000199237 | RNU6-834P | 1.864319 | -0.2306658 | 0.7367488 | -0.6879955 | 0.4914556 | 0.681711 |
| ENSG00000231940 | RPS7P3 | 12.94723 | -0.2307158 | 0.4932289 | -0.5992095 | 0.5490332 | 0.728408 |
| ENSG00000211460 | TSN | 6870.944 | -0.2308476 | 0.1762927 | -1.357374 | 0.1746625 | 0.346547 |
| ENSG00000280800 | | 1.678023 | -0.2309878 | 0.7475181 | -0.7462901 | 0.4554922 | 0.651449 |
| ENSG00000232864 | NUCKS1P1 | 4.476821 | -0.2309974 | 0.7003452 | -0.6398731 | 0.5222551 | 0.707599 |
| ENSG00000233286 | MTND3P10 | 1.848095 | -0.2310185 | 0.741024 | -0.7093293 | 0.4781202 | 0.670104 |
| ENSG00000250980 | NSA2P6 | 1.238272 | -0.2310318 | 0.7702688 | -0.8042797 | 0.4212355 | NA |
| ENSG00000229497 | | 1.577667 | -0.2311244 | 0.7429683 | -0.7140694 | 0.4751843 | 0.667714 |
| ENSG00000276651 | | 5.376823 | -0.2311621 | 0.5568537 | -0.5751152 | 0.5652134 | 0.7402 |
| ENSG00000272977 | | 66.10846 | -0.2312585 | 0.4269535 | -0.645133 | 0.518841 | 0.704147 |
| ENSG00000101447 | FAM83D | 1254.854107 | -0.231277892 | 0.451457077 | -0.622796455 | 0.53341829 | 0.717307 |
| ENSG00000166831 | RBPMS2 | 83.011 | -0.2313332 | 0.5489968 | -0.5786583 | 0.5628198 | 0.738557 |
| ENSG00000213326 | RPS7P11 | 686.353 | -0.2313631 | 0.1893218 | -1.261194 | 0.2072389 | 0.389641 |
| ENSG00000174521 | TTC9B | 17.51229 | -0.2313707 | 0.504232 | -0.5965157 | 0.5508308 | 0.729744 |
| ENSG00000259790 | ANP32BP1 | 272.87 | -0.231384 | 0.2193277 | -1.103798 | 0.2696808 | 0.465976 |
| ENSG00000249409 | | 2.402247 | -0.2314115 | 0.7085705 | -0.6468036 | 0.5177591 | 0.703038 |
| ENSG00000213533 | STIMATE | 40.04835 | -0.2314302 | 0.3331676 | -0.7707181 | 0.440874 | 0.638795 |

| | | | | | | | |
|---|---|---|---|---|---|---|---|
| ENSG00000221241 | SNORD88A | 6.247408 | -0.2314842 | 0.6633352 | -0.602828 | 0.5466231 | 0.726642 |
| ENSG00000229519 | | 5.577503 | -0.2315226 | 0.6405953 | -0.5958258 | 0.5512916 | 0.730173 |
| ENSG00000286339 | | 0.7763951 | -0.2316561 | 0.8303267 | -1.232973 | 0.217586 | NA |
| ENSG00000108774 | RAB5C | 2875.844138 | -0.231683784 | 0.185889211 | -1.28775097 | 0.19783267 | 0.377944 |
| ENSG00000100600 | LGMN | 1769.204028 | -0.231690996 | 0.262697753 | -0.940082265 | 0.34717537 | 0.548694 |
| ENSG00000275091 | | 3.976468 | -0.2317656 | 0.6650936 | -0.6038426 | 0.5459483 | 0.726173 |
| ENSG00000261916 | | 1.664028 | -0.2317725 | 0.7451969 | -0.7029618 | 0.4820795 | 0.673522 |
| ENSG00000237015 | | 50.58611 | -0.2317966 | 0.4027901 | -0.6717715 | 0.5017292 | 0.689342 |
| ENSG00000242061 | | 0.8835388 | -0.2318114 | 0.7955952 | -0.9211592 | 0.3569673 | NA |
| ENSG00000125454 | SLC25A19 | 880.2531018 | -0.231816096 | 0.249411819 | -0.979478917 | 0.3273434 | 0.528704 |
| ENSG00000100626 | GALNT16 | 240.6190519 | -0.231822095 | 0.460831854 | -0.627168201 | 0.53054899 | 0.714924 |
| ENSG00000130255 | RPL36 | 13182.7651 | -0.23183604 | 0.174392151 | -1.387755612 | 0.16521148 | 0.333726 |
| ENSG00000236325 | RPL32P5 | 1.074588 | -0.2318491 | 0.7660166 | -0.7734342 | 0.4392654 | NA |
| ENSG00000172139 | SLC9C1 | 3.015565 | -0.2318618 | 0.7350899 | -0.6836227 | 0.4942135 | 0.683965 |
| ENSG00000111679 | PTPN6 | 2414.534733 | -0.231877613 | 0.199376777 | -1.219290089 | 0.22273411 | 0.409386 |
| ENSG00000253159 | PCDHGA12 | 5.058362 | -0.231913 | 0.7005292 | -0.6325629 | 0.5270191 | 0.711937 |
| ENSG00000047932 | GOPC | 2754.608213 | -0.231928925 | 0.170720484 | -1.397630968 | 0.16222391 | 0.329478 |
| ENSG00000048544 | MRPS10 | 3562.869763 | -0.231948192 | 0.14013793 | -1.696983167 | 0.08969984 | 0.215154 |
| ENSG00000227617 | CERS6-AS1 | 1.621626 | -0.2319781 | 0.7440291 | -0.7209504 | 0.47094 | 0.664246 |
| ENSG00000179134 | SAMD4B | 5060.462 | -0.2320023 | 0.161893 | -1.482734 | 0.138145 | 0.293059 |
| ENSG00000141371 | C17orf64 | 8.743686 | -0.2320209 | 0.6154972 | -0.5799183 | 0.5619697 | 0.737769 |
| ENSG00000198899 | MT-ATP6 | 65976.41 | -0.2320231 | 0.1784209 | -1.408506 | 0.1589813 | 0.324896 |
| ENSG00000232809 | VDAC1P10 | 0.8252375 | -0.2320506 | 0.8269386 | -1.106418 | 0.2685458 | NA |
| ENSG00000204620 | | 32.08632 | -0.2320603 | 0.3585271 | -0.7311872 | 0.4646648 | 0.658636 |
| ENSG00000225423 | TNPO1P1 | 8.43591 | -0.2320635 | 0.5295892 | -0.5835246 | 0.5595402 | 0.73613 |
| ENSG00000285560 | | 6.157347 | -0.2321345 | 0.5448111 | -0.5834051 | 0.5596206 | 0.73613 |
| ENSG00000074935 | TUBE1 | 936.2149436 | -0.232165319 | 0.419035621 | -0.646081784 | 0.51822639 | 0.703529 |
| ENSG00000261655 | | 2.75552 | -0.2321749 | 0.7480498 | -0.7243173 | 0.4688709 | 0.662417 |
| ENSG00000264937 | | 7.788106 | -0.2321945 | 0.5933877 | -0.5796989 | 0.5621177 | 0.737891 |
| ENSG00000252947 | SCARNA1 | 2.989591 | -0.2322373 | 0.6827153 | -0.615346 | 0.5383263 | 0.721137 |
| ENSG00000164287 | CDC20B | 48.96076 | -0.2322829 | 0.7165549 | -0.6468623 | 0.517721 | 0.703022 |
| ENSG00000079459 | FDFT1 | 14760.04746 | -0.232365996 | 0.255295541 | -0.967161834 | 0.33346314 | 0.534527 |
| ENSG00000251680 | | 5.447407 | -0.2323763 | 0.5592338 | -0.5831999 | 0.5597587 | 0.73614 |

| | | | | | | | |
|---|---|---|---|---|---|---|---|
| ENSG00000177464 | GPR4 | 1.468255 | -0.2323781 | 0.7558258 | -0.7445407 | 0.4565494 | NA |
| ENSG00000275025 | | 1.406965 | -0.2324731 | 0.7724035 | -0.7740057 | 0.4389274 | NA |
| ENSG00000268499 | | 2.222884 | -0.2324872 | 0.7315454 | -0.6738221 | 0.5004244 | 0.688437 |
| ENSG00000136937 | NCBP1 | 4427.382007 | -0.232508169 | 0.133361136 | -1.774908612 | 0.07591298 | 0.189721 |
| ENSG00000219891 | ZSCAN12P1 | 149.3393 | -0.2325358 | 0.3954563 | -0.6875396 | 0.4917428 | 0.681967 |
| ENSG00000261669 | | 6.862038 | -0.2325735 | 0.5682667 | -0.5815198 | 0.5608902 | 0.73719 |
| ENSG00000171055 | FEZ2 | 2714.722 | -0.2326513 | 0.1739074 | -1.393903 | 0.1633468 | 0.331118 |
| ENSG00000258823 | | 1.658616 | -0.2326937 | 0.7459359 | -0.7208922 | 0.4709758 | 0.664261 |
| ENSG00000267138 | | 1.408648 | -0.2327347 | 0.7480258 | -0.7220741 | 0.4702489 | NA |
| ENSG00000117595 | IRF6 | 1919.916854 | -0.232805307 | 0.179503623 | -1.368659321 | 0.17110579 | 0.34171 |
| ENSG00000104341 | LAPTM4B | 19746.78114 | -0.232805394 | 0.194261308 | -1.240425915 | 0.2148179 | 0.399741 |
| ENSG00000257285 | | 89.47003 | -0.2329014 | 0.3920356 | -0.6856491 | 0.4929344 | 0.68282 |
| ENSG00000138794 | CASP6 | 1154.07066 | -0.232952991 | 0.147815822 | -1.613780242 | 0.1065751 | 0.243767 |
| ENSG00000122566 | HNRNPA2B1 | 46074.73724 | -0.23301121 | 0.240699753 | -1.021749549 | 0.30689945 | 0.507059 |
| ENSG00000203387 | SNF8P1 | 1.952129 | -0.2330964 | 0.7039689 | -0.6512686 | 0.5148731 | 0.700372 |
| ENSG00000151348 | EXT2 | 3587.406 | -0.2331797 | 0.07456975 | -3.122954 | 0.00179045 | 0.00944 |
| ENSG00000220937 | HNRNPA1P41 | 0.8015754 | -0.2332359 | 0.7898497 | -0.9080203 | 0.3638675 | NA |
| ENSG00000223540 | RPS15AP5 | 7.479501 | -0.2333559 | 0.5885514 | -0.570709 | 0.5681969 | 0.742524 |
| ENSG00000286885 | | 8.099222 | -0.2334321 | 0.5366179 | -0.5852328 | 0.5583912 | 0.73547 |
| ENSG00000132330 | SCLY | 103.8536112 | -0.233438471 | 0.231649206 | -1.056256634 | 0.29085098 | 0.489462 |
| ENSG00000140365 | COMMD4 | 4107.859 | -0.2334402 | 0.2326829 | -1.06333 | 0.2876325 | 0.485978 |
| ENSG00000168769 | TET2 | 2261.165 | -0.2335611 | 0.2602815 | -0.9578847 | 0.3381209 | 0.539454 |
| ENSG00000216676 | RPL8P1 | 1.051425 | -0.2335695 | 0.787108 | -0.8526252 | 0.3938672 | NA |
| ENSG00000147082 | CCNB3 | 13.78848 | -0.233582 | 0.4387567 | -0.6429478 | 0.520258 | 0.705458 |
| ENSG00000008952 | SEC62 | 7587.820886 | -0.233757598 | 0.13649568 | -1.76482282 | 0.07759355 | 0.192748 |
| ENSG00000198818 | SFT2D1 | 1817.853 | -0.2337788 | 0.1575311 | -1.523531 | 0.1276259 | 0.277037 |
| ENSG00000103510 | KAT8 | 2204.521065 | -0.233786232 | 0.130449292 | -1.820661223 | 0.06865837 | 0.175918 |
| ENSG00000006576 | PHTF2 | 2723.700308 | -0.233818878 | 0.191544959 | -1.254533104 | 0.2096483 | 0.393011 |
| ENSG00000238227 | TMEM250 | 3291.88 | -0.2338489 | 0.1315671 | -1.806084 | 0.07090521 | 0.180327 |
| ENSG00000253894 | | 1.999616 | -0.2338828 | 0.7433148 | -0.7102738 | 0.4775344 | 0.669704 |
| ENSG00000242110 | AMACR | 160.7145 | -0.2340119 | 0.2043454 | -1.187801 | 0.2349119 | 0.424179 |
| ENSG00000225739 | NPM1P18 | 32.20624 | -0.2340293 | 0.4260588 | -0.6552643 | 0.5122976 | 0.69822 |
| ENSG00000251015 | SLC25A30-AS1 | 2.404981 | -0.2340297 | 0.7420244 | -0.7342238 | 0.4628124 | 0.657178 |

| | | | | | | | |
|---|---|---|---|---|---|---|---|
| ENSG00000201524 | RNU6-450P | 2.170688 | -0.2340612 | 0.6880036 | -0.6252042 | 0.531837 | 0.715897 |
| ENSG00000133858 | ZFC3H1 | 4593.856018 | -0.234115678 | 0.342888151 | -0.764052055 | 0.44483622 | 0.642138 |
| ENSG00000230701 | FBXW4P1 | 19.45257 | -0.234177 | 0.3934871 | -0.6893551 | 0.4905998 | 0.681098 |
| ENSG00000135966 | TGFBRAP1 | 2839.467301 | -0.234230416 | 0.127427995 | -1.875340706 | 0.06074587 | 0.160497 |
| ENSG00000101638 | ST8SIA5 | 3.884951817 | -0.234233595 | 0.654540442 | -0.599178017 | 0.54905418 | 0.728408 |
| ENSG00000228816 | AK3P5 | 3.176379 | -0.2343326 | 0.6978272 | -0.6355718 | 0.5250556 | 0.710054 |
| ENSG00000243819 | RN7SL832P | 40.51529 | -0.2344216 | 0.4178163 | -0.6650938 | 0.5059905 | 0.692651 |
| ENSG00000282961 | PRNCR1 | 1.894633 | -0.2344384 | 0.7422655 | -0.695329 | 0.4868491 | 0.677727 |
| ENSG00000287113 | | 2.397533 | -0.2344686 | 0.6844166 | -0.6244699 | 0.532319 | 0.716295 |
| ENSG00000251675 | | 4.15211 | -0.2345182 | 0.6585836 | -0.5971172 | 0.5504291 | 0.729394 |
| ENSG00000215840 | | 4.722243 | -0.2346668 | 0.6298343 | -0.5938138 | 0.5526367 | 0.731225 |
| ENSG00000260604 | | 250.8064 | -0.2347094 | 0.5791864 | -0.572257 | 0.5671479 | 0.741599 |
| ENSG00000184844 | CYCSP45 | 11.39651 | -0.2347105 | 0.5385462 | -0.5945574 | 0.5521393 | 0.730778 |
| ENSG00000253923 | | 1.393471 | -0.2347247 | 0.7404103 | -0.7081821 | 0.4788322 | NA |
| ENSG00000100146 | SOX10 | 1.933327741 | -0.234734626 | 0.729095318 | -0.677892602 | 0.49783979 | 0.68648 |
| ENSG00000187840 | EIF4EBP1 | 3524.152 | -0.2347827 | 0.2612866 | -0.9624114 | 0.335843 | 0.536916 |
| ENSG00000273338 | | 7.917387 | -0.2348401 | 0.5912102 | -0.5885487 | 0.556164 | 0.733627 |
| ENSG00000158615 | PPP1R15B | 12301.66 | -0.2349887 | 0.2000314 | -1.205378 | 0.2280574 | 0.416108 |
| ENSG00000139163 | ETNK1 | 1904.898 | -0.234991919 | 0.188541941 | -1.292558151 | 0.19616392 | 0.375956 |
| ENSG00000232928 | DDX3P1 | 1.46671 | -0.2351625 | 0.7148356 | -0.6749728 | 0.499693 | NA |
| ENSG00000242756 | RHOT1P3 | 58.77505 | -0.2351667 | 0.2820944 | -0.8958206 | 0.3703486 | 0.572447 |
| ENSG00000260536 | | 1.224417 | -0.235278 | 0.7507263 | -0.7558661 | 0.4497295 | NA |
| ENSG00000247473 | CARS1-AS1 | 6.631927 | -0.2353088 | 0.5939387 | -0.5883119 | 0.5563229 | 0.73375 |
| ENSG00000153012 | LGI2 | 37.30465 | -0.2353371 | 0.550273 | -0.5856831 | 0.5580885 | 0.735217 |
| ENSG00000063015 | SEZ6 | 4.904791881 | -0.235369393 | 0.633795105 | -0.589827427 | 0.55530635 | 0.732896 |
| ENSG00000142513 | ACP4 | 9.799968 | -0.2354275 | 0.5091135 | -0.6008911 | 0.5479125 | 0.727547 |
| ENSG00000234500 | | 20.01234 | -0.2354604 | 0.4511916 | -0.6354083 | 0.5251622 | 0.710078 |
| ENSG00000267871 | ZNF460-AS1 | 46.83101 | -0.2354746 | 0.4340576 | -0.6511121 | 0.5149741 | 0.700473 |
| ENSG00000232732 | | 2.992469 | -0.2355959 | 0.627173 | -0.6002306 | 0.5483526 | 0.727909 |
| ENSG00000260903 | XKR7 | 12.4719 | -0.2356154 | 0.5545721 | -0.5850068 | 0.5585431 | 0.73557 |
| ENSG00000183048 | SLC25A10 | 1184.901 | -0.2357739 | 0.3496968 | -0.7719933 | 0.4401184 | 0.638466 |
| ENSG00000042429 | MED17 | 231.9209921 | -0.235784286 | 0.154205267 | -1.559544988 | 0.11886745 | 0.263307 |
| ENSG00000166266 | CUL5 | 1908.162 | -0.2358997 | 0.1637927 | -1.458798 | 0.1446208 | 0.303103 |

| | | | | | | | |
|---|---|---|---|---|---|---|---|
| ENSG00000285554 | | 1.075156 | -0.2360028 | 0.7765432 | -0.8351985 | 0.403606 | NA |
| ENSG00000272854 | | 1.790824 | -0.2360153 | 0.7289486 | -0.6849815 | 0.4933556 | 0.683236 |
| ENSG00000253837 | | 1.44777 | -0.236122 | 0.7474365 | -0.7306965 | 0.4649646 | NA |
| ENSG00000183444 | OR7E38P | 16.7421 | -0.2362192 | 0.4788707 | -0.619185 | 0.5357945 | 0.719035 |
| ENSG00000183396 | TMEM89 | 2.172309 | -0.2362323 | 0.7585964 | -0.7570728 | 0.4490062 | 0.645416 |
| ENSG00000143198 | MGST3 | 1455.797 | -0.2362593 | 0.1908339 | -1.279598 | 0.2006867 | 0.381303 |
| ENSG00000162736 | NCSTN | 4459.464 | -0.2362867 | 0.1684191 | -1.456831 | 0.1451629 | 0.304023 |
| ENSG00000162385 | MAGOH | 1815.199 | -0.2363237 | 0.2111095 | -1.166445 | 0.2434347 | 0.434607 |
| ENSG00000049769 | PPP1R3F | 467.6958221 | -0.2363392 | 0.228055599 | -1.088244826 | 0.27648704 | 0.473476 |
| ENSG00000282458 | WASH5P | 4.406143 | -0.2363552 | 0.6179501 | -0.5950844 | 0.5517871 | 0.730574 |
| ENSG00000142784 | WDTC1 | 2742.219 | -0.2363556 | 0.1365187 | -1.768351 | 0.07700219 | 0.191723 |
| ENSG00000237882 | PPIAP13 | 0.8793714 | -0.2364461 | 0.8160284 | -0.7967737 | 0.4255825 | NA |
| ENSG00000226676 | LINC02679 | 1.353018 | -0.2364768 | 0.8144982 | -0.765298 | 0.4440941 | NA |
| ENSG00000286886 | | 2.701559 | -0.2364973 | 0.6803522 | -0.6211907 | 0.5344742 | 0.718051 |
| ENSG00000130024 | PHF10 | 5894.969402 | -0.236510147 | 0.122838805 | -1.906935996 | 0.05652888 | 0.152216 |
| ENSG00000257113 | | 1.710896 | -0.236519 | 0.7743884 | -0.813923 | 0.4156891 | 0.615564 |
| ENSG00000182625 | | 7.806497 | -0.2365327 | 0.5861879 | -0.5859539 | 0.5579065 | 0.735159 |
| ENSG00000126882 | FAM78A | 349.5127546 | -0.236536935 | 0.28358242 | -0.903598124 | 0.36620854 | 0.568097 |
| ENSG00000261596 | | 12.39518 | -0.2365751 | 0.5735775 | -0.5902234 | 0.5550409 | 0.732691 |
| ENSG00000080986 | NDC80 | 1106.624703 | -0.236625072 | 0.45819957 | -0.647747265 | 0.51714843 | 0.702456 |
| ENSG00000284024 | | 1866.517 | -0.2366377 | 0.1222699 | -1.934138 | 0.05309621 | 0.145315 |
| ENSG00000111605 | CPSF6 | 7295.85156 | -0.236648784 | 0.134163544 | -1.79077707 | 0.07332908 | 0.185092 |
| ENSG00000112761 | CCN6 | 1.784887579 | -0.236680159 | 0.746306927 | -0.737902405 | 0.46057376 | 0.655646 |
| ENSG00000101266 | CSNK2A1 | 4408.471377 | -0.236681421 | 0.108041044 | -2.241129474 | 0.02501769 | 0.081159 |
| ENSG00000286401 | | 3.916478 | -0.2368847 | 0.6492528 | -0.6085678 | 0.5428109 | 0.723848 |
| ENSG00000134574 | DDB2 | 3236.101112 | -0.236886103 | 0.178927415 | -1.368185653 | 0.17125397 | 0.341903 |
| ENSG00000271029 | | 2.090987 | -0.2369494 | 0.7173724 | -0.6705344 | 0.5025172 | 0.690065 |
| ENSG00000142166 | IFNAR1 | 3531.403 | -0.2369642 | 0.1485541 | -1.6236 | 0.1044612 | 0.240172 |
| ENSG00000287562 | | 131.7134 | -0.2370374 | 0.3877936 | -0.7051501 | 0.4807168 | 0.67229 |
| ENSG00000230183 | CNOT6LP1 | 52.19958 | -0.237049 | 0.3755678 | -0.7201343 | 0.4714423 | 0.664722 |
| ENSG00000100312 | ACR | 3.24437905 | -0.237081541 | 0.795043227 | -0.95976978 | 0.3371711 | 0.538295 |
| ENSG00000232362 | ATP5MGP2 | 45.10042 | -0.2371342 | 0.3470002 | -0.7647117 | 0.4444432 | 0.641831 |
| ENSG00000235183 | SRP14P3 | 1.20142 | -0.2371344 | 0.7504761 | -0.7333162 | 0.4633656 | NA |

| | | | | | | | |
|---|---|---|---|---|---|---|---|
| ENSG00000136908 | DPM2 | 2515.126721 | -0.237236262 | 0.204228602 | -1.207032909 | 0.22741947 | 0.415193 |
| ENSG00000112941 | TENT4A | 4095.457329 | -0.237271267 | 0.166008542 | -1.467863505 | 0.1421413 | 0.299417 |
| ENSG00000232888 | RPS11P5 | 2174.035 | -0.23752 | 0.1635921 | -1.496798 | 0.1344459 | 0.287539 |
| ENSG00000219863 | RPL31P28 | 1.210031 | -0.2375531 | 0.7635487 | -0.7751659 | 0.4382416 | NA |
| ENSG00000000938 | FGR | 1.210551068 | -0.237565676 | 0.818664493 | -0.87830984 | 0.3797756 | NA |
| ENSG00000085377 | PREP | 2773.754425 | -0.237647811 | 0.186950549 | -1.307695405 | 0.19097664 | 0.368996 |
| ENSG00000265112 | MIR3153 | 34.06319 | -0.2376564 | 0.4785781 | -0.6245405 | 0.5322727 | 0.716295 |
| ENSG00000090863 | GLG1 | 10996.31745 | -0.237732147 | 0.167123795 | -1.473194489 | 0.1406986 | 0.297203 |
| ENSG00000081870 | HSPB11 | 1712.680501 | -0.237747908 | 0.309136853 | -0.842917044 | 0.39927484 | 0.60029 |
| ENSG00000107130 | NCS1 | 2653.958234 | -0.237754804 | 0.215242574 | -1.162057641 | 0.24521205 | 0.436958 |
| ENSG00000118137 | APOA1 | 14.54840296 | -0.237777648 | 0.845460022 | -2.952462591 | 0.0031525 | 0.015133 |
| ENSG00000163319 | MRPS18C | 711.4723 | -0.2378981 | 0.2281268 | -1.094477 | 0.2737459 | 0.47048 |
| ENSG00000080854 | IGSF9B | 577.4287771 | -0.237921001 | 0.287193119 | -0.895473645 | 0.37053395 | 0.572634 |
| ENSG00000198754 | OXCT2 | 27.82982 | -0.2380209 | 0.3886471 | -0.7062015 | 0.4800629 | 0.671835 |
| ENSG00000149532 | CPSF7 | 6029.377 | -0.2380454 | 0.1558271 | -1.587421 | 0.1124172 | 0.253184 |
| ENSG00000211643 | IGLV5-52 | 0.9046957 | -0.2380506 | 0.8310264 | -1.163137 | 0.2447741 | NA |
| ENSG00000182087 | TMEM259 | 10501.94 | -0.2380749 | 0.149556 | -1.624213 | 0.1043304 | 0.239966 |
| ENSG00000283041 | | 32264.42 | -0.238163 | 0.1606776 | -1.568574 | 0.1167471 | 0.259986 |
| ENSG00000205090 | TMEM240 | 17.7326 | -0.2382029 | 0.5276451 | -0.5996657 | 0.5487291 | 0.728137 |
| ENSG00000064313 | TAF2 | 2840.270832 | -0.238306217 | 0.190806827 | -1.290964077 | 0.19671613 | 0.376675 |
| ENSG00000198799 | LRIG2 | 1430.813 | -0.2385779 | 0.1606444 | -1.519977 | 0.1285168 | 0.278422 |
| ENSG00000160789 | LMNA | 12869.85 | -0.2388358 | 0.1305347 | -1.858084 | 0.06315712 | 0.165147 |
| ENSG00000224763 | FDPSP8 | 1.585593 | -0.2389794 | 0.7656603 | -0.7982827 | 0.4247064 | 0.624393 |
| ENSG00000186283 | TOR3A | 5427.705 | -0.2390231 | 0.1831831 | -1.346367 | 0.1781843 | 0.3517 |
| ENSG00000269821 | KCNQ1OT1 | 537.4033 | -0.2390637 | 0.3939553 | -0.7066445 | 0.4797874 | 0.671756 |
| ENSG00000125743 | SNRPD2 | 8066.145551 | -0.239276783 | 0.211455999 | -1.187453266 | 0.23504888 | 0.42434 |
| ENSG00000067715 | SYT1 | 1041.947067 | -0.239325305 | 0.2742146 | -0.937218031 | 0.34864642 | 0.550332 |
| ENSG00000230583 | GTF2IRD1P1 | 16.72901 | -0.2393809 | 0.418229 | -0.6705475 | 0.5025088 | 0.690065 |
| ENSG00000084623 | EIF3I | 9915.631402 | -0.239412041 | 0.199908131 | -1.22631132 | 0.22008154 | 0.406291 |
| ENSG00000198134 | PTMAP9 | 91.22794 | -0.239444 | 0.3134416 | -0.8340975 | 0.4042261 | 0.605064 |
| ENSG00000088451 | TGDS | 635.5806028 | -0.239466662 | 0.142498966 | -1.707407366 | 0.08774636 | 0.211612 |
| ENSG00000270553 | | 6.095677 | -0.2394803 | 0.6019828 | -0.6000911 | 0.5484455 | 0.727956 |
| ENSG00000168566 | SNRNP48 | 1288.943 | -0.2396375 | 0.1849624 | -1.340377 | 0.1801229 | 0.354338 |

| | | | | | | | |
|---|---|---|---|---|---|---|---|
| ENSG00000167272 | POP5 | 1600.064 | -0.2396667 | 0.1564248 | -1.549479 | 0.1212665 | 0.267314 |
| ENSG00000178146 | GK4P | 153.291 | -0.2396854 | 0.3945004 | -0.7048377 | 0.4809112 | 0.672456 |
| ENSG00000115657 | ABCB6 | 262.9587073 | -0.23972254 | 0.287930666 | -0.890030451 | 0.37344954 | 0.575599 |
| ENSG00000198727 | MT-CYB | 84903.46 | -0.2397545 | 0.1573075 | -1.612792 | 0.1067897 | 0.24411 |
| ENSG00000178307 | TMEM11 | 2193.433 | -0.2397575 | 0.211494 | -1.18162 | 0.2373565 | 0.427013 |
| ENSG00000187556 | NANOS3 | 6.45282 | -0.2397975 | 0.550223 | -0.6034006 | 0.5462422 | 0.726382 |
| ENSG00000259921 | | 2.837931 | -0.2400125 | 0.7324915 | -0.6990978 | 0.4844909 | 0.67556 |
| ENSG00000167395 | ZNF646 | 2018.212 | -0.2401177 | 0.1956812 | -1.270997 | 0.2037296 | 0.385369 |
| ENSG00000233862 | | 3.026235 | -0.2402129 | 0.6515896 | -0.6156712 | 0.5381115 | 0.721092 |
| ENSG00000159166 | LAD1 | 8214.728 | -0.2402804 | 0.2590087 | -1.002587 | 0.3160601 | 0.517117 |
| ENSG00000268350 | FAM156A | 11.41481 | -0.2403168 | 0.4562792 | -0.6492864 | 0.5161533 | 0.701431 |
| ENSG00000243710 | CFAP57 | 14.61871 | -0.2403726 | 0.4624773 | -0.6401997 | 0.5220428 | 0.70748 |
| ENSG00000236319 | | 9.999907 | -0.2404047 | 0.6298884 | -0.6002883 | 0.5483141 | 0.727909 |
| ENSG00000199121 | MIR26B | 2.125893 | -0.2404289 | 0.7015242 | -0.6589875 | 0.5099038 | 0.695852 |
| ENSG00000229585 | RPL21P44 | 1.407008 | -0.2404472 | 0.7655244 | -0.7836989 | 0.4332168 | NA |
| ENSG00000086730 | LAT2 | 30.19524391 | -0.240472887 | 0.381036136 | -0.724696355 | 0.46863833 | 0.662195 |
| ENSG00000259274 | | 0.7954433 | -0.2405099 | 0.7899011 | -0.9322394 | 0.3512128 | NA |
| ENSG00000209042 | SNORD12C | 32.67095 | -0.2405534 | 0.5252555 | -0.6057824 | 0.5446592 | 0.72533 |
| ENSG00000135870 | RC3H1 | 2639.636571 | -0.240624699 | 0.194275079 | -1.290310457 | 0.19694289 | 0.376932 |
| ENSG00000175467 | SART1 | 5752.403 | -0.2406873 | 0.141538 | -1.723811 | 0.08474204 | 0.206102 |
| ENSG00000287860 | | 2.549568 | -0.2407368 | 0.7660308 | -0.8010307 | 0.4231139 | 0.623086 |
| ENSG00000205323 | SARNP | 5.751986 | -0.2407993 | 0.5994486 | -0.5969498 | 0.5505409 | 0.729469 |
| ENSG00000228409 | CCT6P1 | 476.0356 | -0.2408123 | 0.1538456 | -1.583318 | 0.1133491 | 0.254571 |
| ENSG00000234518 | PTGES3P1 | 1953.2 | -0.2408172 | 0.2115519 | -1.191292 | 0.2335391 | 0.422541 |
| ENSG00000164081 | TEX264 | 2250.974 | -0.2408259 | 0.1754982 | -1.413464 | 0.1575194 | 0.322653 |
| ENSG00000112242 | E2F3 | 2315.334369 | -0.240924314 | 0.1879625 | -1.315884338 | 0.18821287 | 0.365249 |
| ENSG00000231700 | | 13.13035 | -0.2409904 | 0.4733941 | -0.6355586 | 0.5250642 | 0.710054 |
| ENSG00000232794 | HNRNPDP1 | 7.602548 | -0.2410325 | 0.6161211 | -0.609062 | 0.5424833 | 0.723738 |
| ENSG00000159128 | IFNGR2 | 887.3581 | -0.2410872 | 0.1755192 | -1.422793 | 0.1547962 | 0.318524 |
| ENSG00000181222 | POLR2A | 9138.872 | -0.241119 | 0.2116214 | -1.200868 | 0.2298024 | 0.417993 |
| ENSG00000164011 | ZNF691 | 681.0016 | -0.2411544 | 0.1601387 | -1.544576 | 0.1224488 | 0.269183 |
| ENSG00000268575 | | 141.8361 | -0.2412265 | 0.2673463 | -0.9602893 | 0.3369096 | 0.538104 |
| ENSG00000132840 | BHMT2 | 1.155277295 | -0.241326371 | 0.81458615 | -1.127322182 | 0.25960629 | NA |

| | | | | | | | |
|---|---|---|---|---|---|---|---|
| ENSG00000163803 | PLB1 | 3.201116 | -0.2413325 | 0.6720829 | -0.6259342 | 0.5313581 | 0.715506 |
| ENSG00000129636 | ITFG1 | 2167.413434 | -0.241401533 | 0.146631901 | -1.696509791 | 0.08978938 | 0.215258 |
| ENSG00000176055 | MBLAC2 | 804.1277 | -0.2414755 | 0.2163584 | -1.180335 | 0.2378668 | 0.427481 |
| ENSG00000248347 | | 6.022401 | -0.2415816 | 0.6450004 | -0.6178237 | 0.5366916 | 0.719733 |
| ENSG00000156239 | N6AMT1 | 880.9862 | -0.2416253 | 0.2053472 | -1.223276 | 0.2212256 | 0.407752 |
| ENSG00000279954 | | 8.889369 | -0.2417239 | 0.5807437 | -0.595945 | 0.551212 | 0.730104 |
| ENSG00000111364 | DDX55 | 2602.440997 | -0.241771537 | 0.118499553 | -2.067717706 | 0.03866658 | 0.11387 |
| ENSG00000130299 | GTPBP3 | 1735.39262 | -0.241787099 | 0.15459627 | -1.594079879 | 0.11091815 | 0.250786 |
| ENSG00000117411 | B4GALT2 | 4329.79456 | -0.241802957 | 0.212949689 | -1.184230938 | 0.23632166 | 0.425886 |
| ENSG00000248846 | LINC02065 | 1.544296 | -0.2418462 | 0.7485673 | -0.7395132 | 0.4595954 | NA |
| ENSG00000261079 | | 19.90384 | -0.2418747 | 0.4047468 | -0.7000561 | 0.4838923 | 0.675096 |
| ENSG00000125820 | NKX2-2 | 1.59450515 | -0.241935486 | 0.782196116 | -0.897089264 | 0.36967129 | 0.571799 |
| ENSG00000228373 | | 9.467514 | -0.2420191 | 0.509678 | -0.6185692 | 0.5362002 | 0.719265 |
| ENSG00000252542 | SNORD36C | 9.406185 | -0.2420682 | 0.4898068 | -0.6246627 | 0.5321925 | 0.71623 |
| ENSG00000197713 | RPE | 4091.32 | -0.2421326 | 0.1467336 | -1.693355 | 0.09038786 | 0.216314 |
| ENSG00000261884 | | 112.9232 | -0.2421419 | 0.3062731 | -0.8609165 | 0.3892841 | 0.591157 |
| ENSG00000203620 | | 0.7868711 | -0.2421702 | 0.8194536 | -1.177068 | 0.2391684 | NA |
| ENSG00000276462 | | 1.033512 | -0.2422663 | 0.81181 | -1.100833 | 0.2709694 | NA |
| ENSG00000196365 | LONP1 | 7469.099 | -0.2422776 | 0.2083481 | -1.195712 | 0.2318089 | 0.420204 |
| ENSG00000255689 | | 0.9242538 | -0.2422859 | 0.8352315 | -1.293744 | 0.195754 | NA |
| ENSG00000276975 | HYDIN2 | 1.586206 | -0.2423206 | 0.7351814 | -0.7074766 | 0.4792704 | 0.671315 |
| ENSG00000236060 | HSPB1P1 | 53.66433 | -0.2423474 | 0.3724844 | -0.7343412 | 0.4627408 | 0.657147 |
| ENSG00000161179 | YDJC | 3057.689 | -0.2424309 | 0.2451082 | -1.035503 | 0.3004343 | 0.499671 |
| ENSG00000229719 | MIR194-2HG | 5.889636 | -0.2424752 | 0.6491346 | -0.614347 | 0.538986 | 0.721572 |
| ENSG00000272341 | | 156.2912 | -0.2425218 | 0.4399489 | -0.6800821 | 0.4964525 | 0.685403 |
| ENSG00000179564 | LSMEM2 | 3.767748 | -0.2425366 | 0.6574911 | -0.6297866 | 0.5288342 | 0.713591 |
| ENSG00000276593 | | 1.455538 | -0.2426353 | 0.7694544 | -0.8136052 | 0.4158712 | NA |
| ENSG00000175756 | AURKAIP1 | 4339.723 | -0.2426848 | 0.2672507 | -0.9698911 | 0.3321008 | 0.533209 |
| ENSG00000127423 | AUNIP | 446.3469361 | -0.24270017 | 0.347876075 | -0.782070503 | 0.43417314 | 0.63316 |
| ENSG00000174891 | RSRC1 | 1351.761 | -0.2427187 | 0.2629941 | -0.9850608 | 0.3245942 | 0.525598 |
| ENSG00000227361 | RPS24P7 | 6.910238 | -0.2428828 | 0.5479456 | -0.6232525 | 0.5331186 | 0.717129 |
| ENSG00000260123 | CARMAL | 1.56697 | -0.2429304 | 0.8191071 | -0.8404421 | 0.4006605 | NA |
| ENSG00000023608 | SNAPC1 | 454.8154762 | -0.243165379 | 0.228274189 | -1.124015449 | 0.26100647 | 0.455629 |

| | | | | | | | |
|---|---|---|---|---|---|---|---|
| ENSG00000248576 | | 2.947109 | -0.2432528 | 0.6958705 | -0.6532571 | 0.5135905 | 0.699451 |
| ENSG00000252311 | RNU1-103P | 10.12096 | -0.2433513 | 0.524833 | -0.6155704 | 0.5381781 | 0.721105 |
| ENSG00000141378 | PTRH2 | 5257.439 | -0.2433526 | 0.1620123 | -1.568381 | 0.1167923 | 0.26003 |
| ENSG00000185164 | NOMO2 | 1653.964 | -0.2433861 | 0.2022882 | -1.261411 | 0.2071607 | 0.38953 |
| ENSG00000096654 | ZNF184 | 454.2202856 | -0.243396981 | 0.165995582 | -1.503436995 | 0.13272639 | 0.285069 |
| ENSG00000103591 | AAGAB | 5675.070552 | -0.243424217 | 0.127224188 | -1.940977347 | 0.05226103 | 0.143428 |
| ENSG00000257365 | FNTB | 394.2452 | -0.2434625 | 0.1990321 | -1.269309 | 0.204331 | 0.386149 |
| ENSG00000162191 | UBXN1 | 7128.456 | -0.2435752 | 0.2043142 | -1.259533 | 0.2078377 | 0.39041 |
| ENSG00000105647 | PIK3R2 | 7.66034915 | -0.243741086 | 0.506740913 | -0.619002835 | 0.53591449 | 0.719054 |
| ENSG00000104524 | PYCR3 | 2307.612218 | -0.243748775 | 0.256946412 | -1.02444799 | 0.30562373 | 0.505517 |
| ENSG00000201592 | | 1.437262 | -0.2437733 | 0.7756882 | -0.8454581 | 0.3978551 | NA |
| ENSG00000173258 | ZNF483 | 454.5973 | -0.2437737 | 0.2486116 | -1.032546 | 0.3018163 | 0.501216 |
| ENSG00000272078 | | 35.0121 | -0.2438228 | 0.3889439 | -0.7240871 | 0.4690122 | 0.662547 |
| ENSG00000101166 | PRELID3B | 14272.64485 | -0.243903863 | 0.152987647 | -1.66704443 | 0.09550557 | 0.224994 |
| ENSG00000280878 | | 0.906513 | -0.2439069 | 0.8423 | -1.643799 | 0.1002177 | NA |
| ENSG00000269653 | | 1.301656 | -0.2439589 | 0.7747026 | -0.8308127 | 0.4060794 | NA |
| ENSG00000100479 | POLE2 | 483.9896654 | -0.244060005 | 0.476369222 | -0.651622909 | 0.51464447 | 0.700309 |
| ENSG00000252916 | RNU6-762P | 1.87617 | -0.2440773 | 0.7057073 | -0.6827992 | 0.4947337 | 0.684079 |
| ENSG00000141556 | TBCD | 7428.162 | -0.2440913 | 0.2548385 | -1.075675 | 0.2820728 | 0.479821 |
| ENSG00000253953 | PCDHGB4 | 5.79528 | -0.2441317 | 0.5401637 | -0.6108824 | 0.5412774 | 0.72302 |
| ENSG00000117228 | GBP1 | 147.0669782 | -0.244156009 | 0.516187893 | -0.60106414 | 0.54779727 | 0.727467 |
| ENSG00000278683 | | 0.9023484 | -0.2442338 | 0.7825612 | -0.8611822 | 0.3891377 | NA |
| ENSG00000095564 | BTAF1 | 4823.31966 | -0.244236903 | 0.312877166 | -0.856172547 | 0.39190235 | 0.593641 |
| ENSG00000177225 | GATD1 | 3382.602 | -0.244255 | 0.1918914 | -1.32372 | 0.1855959 | 0.361359 |
| ENSG00000172366 | MCRIP2 | 3468.276 | -0.2442595 | 0.1877834 | -1.346686 | 0.1780815 | 0.351545 |
| ENSG00000252874 | | 4.214472 | -0.2443297 | 0.6518387 | -0.6226586 | 0.5335089 | 0.717307 |
| ENSG00000272006 | | 3.436933 | -0.24435 | 0.654135 | -0.6279938 | 0.530008 | 0.714449 |
| ENSG00000180988 | OR52N2 | 1.663547 | -0.2444921 | 0.7854937 | -0.8974724 | 0.3694669 | 0.571644 |
| ENSG00000099956 | SMARCB1 | 6315.863941 | -0.244532293 | 0.205423205 | -1.239523595 | 0.21515166 | 0.400194 |
| ENSG00000232724 | TRIM80P | 1.631474 | -0.2446855 | 0.7608653 | -0.7829781 | 0.43364 | 0.632592 |
| ENSG00000276124 | MIR6855 | 0.82545 | -0.2446956 | 0.8180522 | -0.815258 | 0.4149246 | NA |
| ENSG00000156642 | NPTN | 5210.812 | -0.2447334 | 0.1176849 | -2.081915 | 0.0373502 | 0.110903 |
| ENSG00000227775 | | 39.99129 | -0.2447899 | 0.4698583 | -0.6453122 | 0.5187249 | 0.704025 |

| | | | | | | | |
|---|---|---|---|---|---|---|---|
| ENSG00000188383 | GPAT2P2 | 3.941544 | -0.2449163 | 0.624769 | -0.6122069 | 0.5404009 | 0.722434 |
| ENSG00000230067 | HSPD1P6 | 35.63716 | -0.2450039 | 0.3876378 | -0.7275979 | 0.4668598 | 0.660651 |
| ENSG00000254648 | | 1.262581 | -0.2450602 | 0.7489413 | -0.748849 | 0.4539482 | NA |
| ENSG00000092036 | HAUS4 | 1119.211738 | -0.245071979 | 0.179755785 | -1.406224315 | 0.15965753 | 0.325761 |
| ENSG00000101997 | CCDC22 | 1885.430537 | -0.245084455 | 0.123194027 | -2.022546881 | 0.04311989 | 0.124002 |
| ENSG00000130726 | TRIM28 | 22336.51161 | -0.245165574 | 0.223987503 | -1.146417154 | 0.25162259 | 0.444446 |
| ENSG00000231776 | LINC01611 | 19.36546 | -0.2451991 | 0.3432644 | -0.7980517 | 0.4248405 | 0.624555 |
| ENSG00000242660 | | 5.013726 | -0.2452107 | 0.613429 | -0.6110812 | 0.5411458 | 0.72297 |
| ENSG00000198804 | MT-CO1 | 192493 | -0.2452129 | 0.1593213 | -1.575476 | 0.1151466 | 0.257693 |
| ENSG00000138190 | EXOC6 | 1424.147614 | -0.245226659 | 0.164570787 | -1.529106823 | 0.12623796 | 0.274908 |
| ENSG00000234742 | | 38.30468 | -0.2452411 | 0.3379154 | -0.8059385 | 0.4202783 | 0.620147 |
| ENSG00000212385 | RNU6-817P | 10.48991 | -0.2452499 | 0.5036413 | -0.6260728 | 0.5312672 | 0.715504 |
| ENSG00000273551 | | 3.12212 | -0.2453824 | 0.700042 | -0.6496163 | 0.5159401 | 0.701213 |
| ENSG00000285911 | | 2.485271 | -0.2453999 | 0.7422006 | -0.7139491 | 0.4752587 | 0.667714 |
| ENSG00000259056 | DUXAP1 | 0.7809728 | -0.2454246 | 0.7932127 | -0.9227415 | 0.3561419 | NA |
| ENSG00000239272 | RPL21P10 | 8.313215 | -0.2454937 | 0.5819573 | -0.6108599 | 0.5412923 | 0.72302 |
| ENSG00000116663 | FBXO6 | 745.5323461 | -0.245545202 | 0.217178513 | -1.181865701 | 0.23725899 | 0.426938 |
| ENSG00000229018 | PMS2P7 | 5.374475 | -0.2457061 | 0.5308281 | -0.621138 | 0.5345088 | 0.718051 |
| ENSG00000072506 | HSD17B10 | 5786.771654 | -0.245751701 | 0.225213043 | -1.12521273 | 0.2604989 | 0.455057 |
| ENSG00000101558 | VAPA | 6069.373828 | -0.245912716 | 0.13922718 | -1.795409679 | 0.07258845 | 0.183606 |
| ENSG00000274756 | | 12.38355 | -0.245981 | 0.5524539 | -0.6117761 | 0.5406859 | 0.72261 |
| ENSG00000178199 | ZC3H12D | 13.39793 | -0.2459829 | 0.5447728 | -0.6154902 | 0.538231 | 0.721106 |
| ENSG00000256211 | SKP1P2 | 96.44949 | -0.2459892 | 0.274532 | -0.957784 | 0.3381717 | 0.539471 |
| ENSG00000254028 | | 10.80006 | -0.2460279 | 0.5564869 | -0.609554 | 0.5421573 | 0.723521 |
| ENSG00000249637 | | 57.84362 | -0.2460856 | 0.3032718 | -0.8796368 | 0.3790561 | 0.581118 |
| ENSG00000233256 | | 1.621722 | -0.2461238 | 0.8176576 | -0.8295119 | 0.4068148 | 0.607438 |
| ENSG00000135341 | MAP3K7 | 3798.585794 | -0.246184022 | 0.123209421 | -2.023906431 | 0.04297978 | 0.123679 |
| ENSG00000074527 | NTN4 | 1950.485243 | -0.246277004 | 0.59473725 | -0.605892062 | 0.54458643 | 0.725269 |
| ENSG00000269976 | | 0.9060973 | -0.2463347 | 0.8139301 | -1.115823 | 0.2644978 | NA |
| ENSG00000286670 | | 0.933575 | -0.2465168 | 0.7990084 | -0.9604253 | 0.3368412 | NA |
| ENSG00000051180 | RAD51 | 921.2800566 | -0.246686462 | 0.464990067 | -0.65572159 | 0.51200328 | 0.697897 |
| ENSG00000060339 | CCAR1 | 6868.147269 | -0.246962677 | 0.155962074 | -1.623795197 | 0.10441953 | 0.240097 |
| ENSG00000108424 | KPNB1 | 22101.47312 | -0.24702752 | 0.224196508 | -1.154078181 | 0.24846812 | 0.440775 |

| | | | | | | | |
|---|---|---|---|---|---|---|---|
| ENSG00000261794 | GOLGA8H | 10.24511 | -0.2470468 | 0.4717617 | -0.6545866 | 0.512734 | 0.698607 |
| ENSG00000273305 | | 56.11559 | -0.2471192 | 0.3702221 | -0.7578651 | 0.4485318 | 0.645138 |
| ENSG00000086827 | ZW10 | 1208.438305 | -0.247129377 | 0.198102012 | -1.303249 | 0.19248977 | 0.370867 |
| ENSG00000108187 | PBLD | 655.7731594 | -0.247129431 | 0.2015065 | -1.273636844 | 0.20279215 | 0.384115 |
| ENSG00000164366 | CCDC127 | 1636.249 | -0.2473269 | 0.187085 | -1.342415 | 0.1794614 | 0.35354 |
| ENSG00000105641 | SLC5A5 | 58.80345998 | -0.247354465 | 0.700892565 | -0.668781434 | 0.50363491 | 0.690989 |
| ENSG00000267474 | | 3.526312 | -0.2473842 | 0.649726 | -0.6259774 | 0.5313297 | 0.715504 |
| ENSG00000276851 | | 1.969283 | -0.2474014 | 0.7212433 | -0.6987904 | 0.484683 | 0.675667 |
| ENSG00000226784 | PGAM4 | 232.7281 | -0.2474027 | 0.266649 | -0.9890024 | 0.322662 | 0.523427 |
| ENSG00000253799 | LINC01030 | 0.9325 | -0.2474076 | 0.839623 | -1.399692 | 0.1616056 | NA |
| ENSG00000136802 | LRRC8A | 2333.17112 | -0.247424433 | 0.189068763 | -1.346360316 | 0.17818634 | 0.3517 |
| ENSG00000203734 | ECT2L | 6.315977 | -0.2474263 | 0.5984958 | -0.6122818 | 0.5403514 | 0.722417 |
| ENSG00000261187 | | 10.8215 | -0.2474336 | 0.5438733 | -0.6192033 | 0.5357824 | 0.719035 |
| ENSG00000067596 | DHX8 | 4892.621279 | -0.247455031 | 0.130564091 | -1.923384842 | 0.05443174 | 0.148039 |
| ENSG00000149273 | RPS3 | 59830.13 | -0.2475214 | 0.1649347 | -1.538424 | 0.123945 | 0.271619 |
| ENSG00000138398 | PPIG | 4597.114766 | -0.247552705 | 0.144559686 | -1.766950691 | 0.07723649 | 0.19216 |
| ENSG00000147684 | NDUFB9 | 7060.052 | -0.2475561 | 0.1958766 | -1.286432 | 0.1982922 | 0.378289 |
| ENSG00000167965 | MLST8 | 2797.475 | -0.2476523 | 0.2267635 | -1.139903 | 0.2543265 | 0.447319 |
| ENSG00000067365 | METTL22 | 1242.113866 | -0.247688471 | 0.131426235 | -1.914162407 | 0.05559941 | 0.150296 |
| ENSG00000258643 | BCL2L2-PABPN1 | 30.60609 | -0.2478517 | 0.3884184 | -0.7366749 | 0.4613201 | 0.656181 |
| ENSG00000276292 | | 2.821171 | -0.2479238 | 0.6886361 | -0.6607081 | 0.5087995 | 0.695048 |
| ENSG00000125107 | CNOT1 | 12221.63141 | -0.247985502 | 0.21434572 | -1.206403825 | 0.22766182 | 0.415579 |
| ENSG00000207370 | | 1.849004 | -0.2480869 | 0.7278858 | -0.7153602 | 0.4743865 | 0.667117 |
| ENSG00000165804 | ZNF219 | 3678.319 | -0.2481163 | 0.2452299 | -1.059842 | 0.2892165 | 0.487854 |
| ENSG00000228205 | | 1266.687 | -0.2481292 | 0.2122069 | -1.218871 | 0.2228932 | 0.409547 |
| ENSG00000271980 | | 2.121302 | -0.2481307 | 0.7128704 | -0.6908089 | 0.4896856 | 0.680331 |
| ENSG00000119705 | SLIRP | 2541.744318 | -0.248182223 | 0.262706378 | -1.006097777 | 0.31436854 | 0.51552 |
| ENSG00000070785 | EIF2B3 | 1278.792374 | -0.248241157 | 0.276784503 | -0.963699359 | 0.33519668 | 0.536561 |
| ENSG00000267184 | | 2.442619 | -0.2483913 | 0.746417 | -0.749091 | 0.4538024 | 0.649942 |
| ENSG00000272787 | | 57.11016 | -0.2484006 | 0.3516713 | -0.8024335 | 0.4223022 | 0.622339 |
| ENSG00000086598 | TMED2 | 11328.82581 | -0.248434099 | 0.142958608 | -1.754211752 | 0.07939423 | 0.19614 |
| ENSG00000239930 | | 1.488147 | -0.2484364 | 0.7510265 | -0.761436 | 0.4463967 | NA |
| ENSG00000177082 | WDR73 | 935.6474 | -0.2485567 | 0.1223003 | -2.057554 | 0.03963297 | 0.116085 |

| | | | | | | | |
|---|---|---|---|---|---|---|---|
| ENSG00000276255 | LINC02809 | 9.169829 | -0.2485749 | 0.6549852 | -0.6343322 | 0.525864 | 0.710682 |
| ENSG00000158528 | PPP1R9A | 1420.767 | -0.2486056 | 0.3222819 | -0.8733445 | 0.3824753 | 0.584762 |
| ENSG00000101940 | WDR13 | 3735.780378 | -0.248635738 | 0.191019524 | -1.344359942 | 0.17883202 | 0.352728 |
| ENSG00000259232 | | 1.107074 | -0.2487275 | 0.7886569 | -0.926625 | 0.3541213 | NA |
| ENSG00000264859 | DSG2-AS1 | 6.194571 | -0.2487355 | 0.5744394 | -0.6190324 | 0.535895 | 0.719054 |
| ENSG00000105185 | PDCD5 | 3082.167276 | -0.248759289 | 0.19171949 | -1.34823756 | 0.17758199 | 0.350898 |
| ENSG00000227487 | NCAM1-AS1 | 1.304063 | -0.248802 | 0.8428895 | -1.550615 | 0.120994 | NA |
| ENSG00000224481 | | 20.46652 | -0.2488105 | 0.5217682 | -0.6324308 | 0.5271054 | 0.711981 |
| ENSG00000276023 | DUSP14 | 1166.858 | -0.2488219 | 0.1261757 | -2.006911 | 0.04475917 | 0.127515 |
| ENSG00000104951 | IL4I1 | 10.89340481 | -0.24895926 | 0.531479932 | -0.630652672 | 0.52826765 | 0.713103 |
| ENSG00000239392 | | 19.47849 | -0.2492076 | 0.3835201 | -0.7421957 | 0.4579688 | 0.653696 |
| ENSG00000147123 | NDUFB11 | 5428.758 | -0.2492841 | 0.1961349 | -1.308256 | 0.1907866 | 0.368821 |
| ENSG00000248569 | | 246.9788 | -0.2493237 | 0.2621529 | -1.012852 | 0.311131 | 0.511981 |
| ENSG00000211714 | TRBV7-3 | 1.319468 | -0.2493809 | 0.8471003 | -1.877278 | 0.06048003 | NA |
| ENSG00000108433 | GOSR2 | 3593.261544 | -0.249400764 | 0.14831199 | -1.711703392 | 0.08695134 | 0.210051 |
| ENSG00000182325 | FBXL6 | 1994.871 | -0.2494314 | 0.1742226 | -1.466692 | 0.1424599 | 0.299922 |
| ENSG00000260009 | LINC02130 | 1.023823 | -0.2495189 | 0.8357962 | -1.485507 | 0.1374096 | NA |
| ENSG00000167182 | SP2 | 1903.965 | -0.2495932 | 0.1921887 | -1.347544 | 0.1778052 | 0.351104 |
| ENSG00000198793 | MTOR | 4939.587 | -0.2496783 | 0.1630102 | -1.5636 | 0.1179114 | 0.261829 |
| ENSG00000240579 | | 1.600949 | -0.2497104 | 0.7816921 | -0.8794879 | 0.3791368 | 0.581158 |
| ENSG00000175202 | HIGD2B | 2.677435 | -0.2498081 | 0.6786568 | -0.6522639 | 0.514231 | 0.700036 |
| ENSG00000133059 | DSTYK | 5054.219237 | -0.249846489 | 0.215023803 | -1.234605011 | 0.21697754 | 0.402463 |
| ENSG00000266844 | | 0.8342272 | -0.2498673 | 0.7993216 | -0.9990242 | 0.317783 | NA |
| ENSG00000076201 | PTPN23 | 2786.067788 | -0.249885124 | 0.142286331 | -1.808643554 | 0.0705064 | 0.179606 |
| ENSG00000278665 | | 1.03132 | -0.2499685 | 0.7807898 | -0.8703354 | 0.3841172 | NA |
| ENSG00000071994 | PDCD2 | 5913.518935 | -0.249994391 | 0.111386147 | -2.286099339 | 0.02224845 | 0.074196 |
| ENSG00000184182 | UBE2F | 431.9932 | -0.2500903 | 0.2049951 | -1.267203 | 0.2050826 | 0.387101 |
| ENSG00000235500 | SNX19P2 | 10.51507 | -0.2501481 | 0.5131178 | -0.6357241 | 0.5249562 | 0.70998 |
| ENSG00000145220 | LYAR | 1055.568 | -0.2502107 | 0.2839761 | -0.9467098 | 0.3437867 | 0.545409 |
| ENSG00000159202 | UBE2Z | 13872.69 | -0.2502273 | 0.1097388 | -2.305013 | 0.02116586 | 0.071157 |
| ENSG00000229927 | RHEBP1 | 4.031592 | -0.2502455 | 0.6208349 | -0.6198577 | 0.5353515 | 0.718843 |
| ENSG00000187147 | RNF220 | 3290.574 | -0.2503433 | 0.1347915 | -1.907676 | 0.05643305 | 0.152013 |
| ENSG00000007255 | TRAPPC6A | 2290.852216 | -0.250347057 | 0.206906721 | -1.257485287 | 0.20857796 | 0.391529 |

| | | | | | | | |
|---|---|---|---|---|---|---|---|
| ENSG00000254528 | | 1.449063 | -0.2503842 | 0.7544275 | -0.7855324 | 0.4321415 | NA |
| ENSG00000153885 | KCTD15 | 4611.629 | -0.2504642 | 0.1858256 | -1.390772 | 0.1642946 | 0.332333 |
| ENSG00000126453 | BCL2L12 | 1434.425286 | -0.250517158 | 0.369786495 | -0.756092116 | 0.44959397 | 0.64576 |
| ENSG00000152457 | DCLRE1C | 1193.739 | -0.2505491 | 0.1697877 | -1.526023 | 0.127004 | 0.27609 |
| ENSG00000196747 | H2AC13 | 315.0892 | -0.2505877 | 0.2859912 | -0.9434537 | 0.3454489 | 0.546974 |
| ENSG00000068784 | SRBD1 | 1150.228663 | -0.250590639 | 0.265918877 | -1.004319544 | 0.31522462 | 0.516209 |
| ENSG00000181819 | KCTD9P2 | 49.02115 | -0.2506654 | 0.3420487 | -0.8184298 | 0.4131118 | 0.613152 |
| ENSG00000239470 | | 8.530564 | -0.2507758 | 0.5288505 | -0.6269329 | 0.5307032 | 0.715004 |
| ENSG00000118412 | CASP8AP2 | 1965.672975 | -0.25085214 | 0.239955236 | -1.10630103 | 0.26859625 | 0.464635 |
| ENSG00000241782 | | 1.717933 | -0.2509215 | 0.7164857 | -0.6994434 | 0.484275 | 0.67545 |
| ENSG00000152082 | MZT2B | 2253.116 | -0.2509765 | 0.253242 | -1.043627 | 0.2966579 | 0.495641 |
| ENSG00000108468 | CBX1 | 13846.87844 | -0.25101929 | 0.203049416 | -1.283107407 | 0.19945445 | 0.379686 |
| ENSG00000149798 | CDC42EP2 | 1.303282 | -0.2510404 | 0.7611906 | -0.7859352 | 0.4319054 | NA |
| ENSG00000262652 | | 21.5198 | -0.2510649 | 0.4155945 | -0.7092383 | 0.4781766 | 0.670113 |
| ENSG00000265871 | MIR3174 | 1.556757 | -0.2510871 | 0.769368 | -0.8372817 | 0.4024342 | NA |
| ENSG00000282988 | | 39.51371 | -0.2511433 | 0.2888766 | -0.9355327 | 0.3495139 | 0.55121 |
| ENSG00000151746 | BICD1 | 403.2061 | -0.2512948 | 0.2714751 | -0.9897699 | 0.3222866 | 0.523081 |
| ENSG00000201616 | RNU1-91P | 1.61806 | -0.2512995 | 0.7626363 | -0.8004589 | 0.4234449 | 0.623366 |
| ENSG00000279149 | | 1.300239 | -0.2512995 | 0.739188 | -0.749955 | 0.4532818 | NA |
| ENSG00000272716 | SLC30A6-DT | 34.06296 | -0.2513472 | 0.3314557 | -0.8374452 | 0.4023424 | 0.603286 |
| ENSG00000136051 | WASHC4 | 4341.349536 | -0.251425554 | 0.235750602 | -1.10704629 | 0.26827392 | 0.464229 |
| ENSG00000134758 | RNF138 | 1434.942791 | -0.251528489 | 0.195101277 | -1.334799188 | 0.18194208 | 0.356807 |
| ENSG00000010438 | PRSS3 | 0.940172663 | -0.251572102 | 0.789637333 | -0.928067314 | 0.35337265 | NA |
| ENSG00000260676 | LINC01541 | 60.84575 | -0.2516011 | 0.7097325 | -0.692246 | 0.4887828 | 0.679677 |
| ENSG00000220506 | | 1.036876 | -0.2516349 | 0.7860976 | -0.9045711 | 0.3656927 | NA |
| ENSG00000252652 | | 1.595676 | -0.2517981 | 0.7585607 | -0.7796285 | 0.4356096 | 0.63435 |
| ENSG00000213779 | | 0.8025463 | -0.2518601 | 0.8011573 | -0.9805077 | 0.3268356 | NA |
| ENSG00000246323 | | 4.885155 | -0.2518618 | 0.5983086 | -0.629547 | 0.528991 | 0.713693 |
| ENSG00000136161 | RCBTB2 | 46.23512955 | -0.251977175 | 0.428668474 | -0.709721573 | 0.47787681 | 0.670011 |
| ENSG00000287226 | | 2.183486 | -0.2520018 | 0.779612 | -0.8616644 | 0.3888722 | 0.590937 |
| ENSG00000183873 | SCN5A | 35.04156 | -0.2521069 | 0.5198494 | -0.6380036 | 0.5234713 | 0.708766 |
| ENSG00000122025 | FLT3 | 2.052217396 | -0.252118935 | 0.737844734 | -0.726923748 | 0.46727267 | 0.660968 |
| ENSG00000230191 | IFITM3P4 | 3.285399 | -0.2521325 | 0.7378945 | -0.7402658 | 0.4591387 | 0.654764 |

| | | | | | | | |
|---|---|---|---|---|---|---|---|
| ENSG00000066557 | LRRC40 | 814.0092588 | -0.252194646 | 0.249834953 | -1.071328586 | 0.28402171 | 0.481876 |
| ENSG00000142546 | NOSIP | 2724.093 | -0.2522469 | 0.2033286 | -1.308574 | 0.1906785 | 0.368741 |
| ENSG00000162377 | COA7 | 3081.001 | -0.2523373 | 0.1818399 | -1.430436 | 0.1525918 | 0.314957 |
| ENSG00000140400 | MAN2C1 | 5315.799 | -0.252357 | 0.252192 | -1.057835 | 0.2901307 | 0.488652 |
| ENSG00000128708 | HAT1 | 4002.379411 | -0.252387629 | 0.301560024 | -0.93739015 | 0.34855791 | 0.550285 |
| ENSG00000269951 | | 20.92218 | -0.2524109 | 0.407493 | -0.730495 | 0.4650877 | 0.659071 |
| ENSG00000250254 | PTTG2 | 1.915153 | -0.2525046 | 0.7364752 | -0.7423124 | 0.4578981 | 0.653659 |
| ENSG00000271918 | | 111.3472 | -0.2525171 | 0.339027 | -0.8273363 | 0.4080465 | 0.608622 |
| ENSG00000260657 | | 1.674193 | -0.2525801 | 0.7459427 | -0.7750674 | 0.4382999 | 0.636836 |
| ENSG00000118894 | EEF2KMT | 770.8335715 | -0.252626633 | 0.206898367 | -1.276684788 | 0.20171355 | 0.38268 |
| ENSG00000143222 | UFC1 | 3083.607 | -0.2527526 | 0.1489564 | -1.731231 | 0.08341054 | 0.203614 |
| ENSG00000229659 | RPL26P6 | 19.9231 | -0.2527681 | 0.4388527 | -0.6909334 | 0.4896074 | 0.680283 |
| ENSG00000224810 | | 1.114654 | -0.2529286 | 0.7750853 | -0.8528307 | 0.3937532 | NA |
| ENSG00000131233 | GJA9 | 2.063775501 | -0.25299395 | 0.733124728 | -0.728832386 | 0.4661042 | 0.659913 |
| ENSG00000229251 | HNRNPA1P8 | 7.935595 | -0.2530731 | 0.5513315 | -0.6359209 | 0.524828 | 0.709843 |
| ENSG00000226964 | RHEBP2 | 38.27847 | -0.2531216 | 0.3574479 | -0.7944329 | 0.4269435 | 0.626607 |
| ENSG00000277161 | PIGW | 811.8706 | -0.2531222 | 0.2971422 | -0.925939 | 0.3544777 | 0.556358 |
| ENSG00000287559 | | 0.8340429 | -0.2532446 | 0.8321284 | -1.36589 | 0.1719735 | NA |
| ENSG00000154723 | ATP5PF | 2878.497 | -0.2532791 | 0.2363866 | -1.127246 | 0.2596385 | 0.453994 |
| ENSG00000224003 | YES1P1 | 5.082877 | -0.2533409 | 0.7004381 | -0.6814348 | 0.4955964 | 0.684733 |
| ENSG00000271926 | | 12.54549 | -0.2533687 | 0.5098735 | -0.644698 | 0.5191229 | 0.704494 |
| ENSG00000267676 | THA1P | 18.09099 | -0.2534331 | 0.455054 | -0.677816 | 0.4978883 | 0.686512 |
| ENSG00000234743 | EIF5AP4 | 40.16561 | -0.2534928 | 0.3795609 | -0.7665902 | 0.4433252 | 0.640844 |
| ENSG00000271626 | H3P42 | 0.8721168 | -0.2535564 | 0.8141539 | -1.13637 | 0.2558019 | NA |
| ENSG00000122729 | ACO1 | 1765.520599 | -0.253561412 | 0.151345606 | -1.711762504 | 0.08694045 | 0.21005 |
| ENSG00000211581 | MIR765 | 0.8361917 | -0.2535725 | 0.8215496 | -0.8750688 | 0.3815365 | NA |
| ENSG00000049656 | CLPTM1L | 10710.66953 | -0.25359521 | 0.180494847 | -1.463753312 | 0.14326136 | 0.301041 |
| ENSG00000264610 | MIR4685 | 3.725437 | -0.2536164 | 0.6253578 | -0.6389532 | 0.5228534 | 0.708182 |
| ENSG00000198171 | DDRGK1 | 2682.323 | -0.2538154 | 0.1681249 | -1.559776 | 0.1188127 | 0.263259 |
| ENSG00000242659 | | 3.31932 | -0.2538192 | 0.7140733 | -0.7044137 | 0.4811752 | 0.672754 |
| ENSG00000287934 | | 33.75577 | -0.2538256 | 0.3807419 | -0.7651713 | 0.4441695 | 0.641576 |
| ENSG00000160218 | TRAPPC10 | 3041.379 | -0.2538262 | 0.1944888 | -1.366713 | 0.1717154 | 0.342464 |
| ENSG00000158023 | CFAP251 | 93.55707 | -0.253846 | 0.2759873 | -0.9779665 | 0.3280909 | 0.529382 |

| | | | | | | | |
|---|---|---|---|---|---|---|---|
| ENSG00000166181 | API5 | 6451.298 | -0.2538551 | 0.1530548 | -1.698204 | 0.08946921 | 0.214715 |
| ENSG00000087191 | PSMC5 | 15815.08025 | -0.253913883 | 0.167843105 | -1.557020917 | 0.11946552 | 0.264396 |
| ENSG00000149100 | EIF3M | 7991.986 | -0.2539617 | 0.1630637 | -1.595148 | 0.1106791 | 0.250544 |
| ENSG00000166405 | RIC3 | 0.9123179 | -0.2540074 | 0.7942559 | -0.9379437 | 0.3482733 | NA |
| ENSG00000230778 | ANKRD63 | 9.06638 | -0.2540301 | 0.6054653 | -0.6351826 | 0.5253093 | 0.710205 |
| ENSG00000229262 | | 0.911554 | -0.2540459 | 0.813784 | -1.140773 | 0.2539645 | NA |
| ENSG00000213376 | GAPDHP71 | 13.70302 | -0.254072 | 0.4993238 | -0.6508725 | 0.5151288 | 0.70054 |
| ENSG00000125870 | SNRPB2 | 3927.497246 | -0.254100916 | 0.214241607 | -1.258499309 | 0.20821124 | 0.390952 |
| ENSG00000233806 | LINC01237 | 52.86006 | -0.2541221 | 0.3958733 | -0.740663 | 0.4588978 | 0.654576 |
| ENSG00000116698 | SMG7 | 10154.97393 | -0.254134023 | 0.140653482 | -1.884611208 | 0.05948235 | 0.158086 |
| ENSG00000182759 | MAFA | 92.19335 | -0.2541757 | 0.2794821 | -0.9745627 | 0.3297772 | 0.530954 |
| ENSG00000069275 | NUCKS1 | 55927.67045 | -0.254212985 | 0.24072148 | -1.114532809 | 0.26505068 | 0.460571 |
| ENSG00000198625 | MDM4 | 7071.906 | -0.2542597 | 0.1771635 | -1.483976 | 0.1378154 | 0.292504 |
| ENSG00000135503 | ACVR1B | 2803.932385 | -0.254279532 | 0.188154513 | -1.397196854 | 0.16235438 | 0.329634 |
| ENSG00000069329 | VPS35 | 7582.692406 | -0.254348394 | 0.127007084 | -2.036444714 | 0.04170572 | 0.120851 |
| ENSG00000255139 | | 17.07907 | -0.254433 | 0.3635982 | -0.7914889 | 0.4286587 | 0.62791 |
| ENSG00000243646 | IL10RB | 46.82873 | -0.2544724 | 0.2607983 | -1.036037 | 0.3001849 | 0.499444 |
| ENSG00000054116 | TRAPPC3 | 3940.151569 | -0.254496868 | 0.162036791 | -1.63741711 | 0.10154335 | 0.235399 |
| ENSG00000112619 | PRPH2 | 1.56465514 | -0.254497156 | 0.711100162 | -0.69770154 | 0.48536387 | NA |
| ENSG00000159593 | NAE1 | 3218.213 | -0.2545558 | 0.2208592 | -1.231187 | 0.218253 | 0.404011 |
| ENSG00000280233 | | 8.390714 | -0.2546027 | 0.5416584 | -0.6386003 | 0.523083 | 0.708384 |
| ENSG00000181904 | C5orf24 | 3222.629 | -0.2546179 | 0.2310172 | -1.185566 | 0.2357938 | 0.42515 |
| ENSG00000202272 | | 1.777941 | -0.2546843 | 0.8276653 | -1.007771 | 0.3135642 | 0.514709 |
| ENSG00000132383 | RPA1 | 5632.176845 | -0.254868004 | 0.270513648 | -1.00784937 | 0.31352678 | 0.514709 |
| ENSG00000136810 | TXN | 8798.264234 | -0.254917148 | 0.225652779 | -1.174384566 | 0.24024102 | 0.430525 |
| ENSG00000184608 | FAM167A-AS1 | 1.111308 | -0.2550279 | 0.7868432 | -0.9442979 | 0.3450174 | NA |
| ENSG00000083123 | BCKDHB | 940.4142774 | -0.25504898 | 0.170693402 | -1.53145557 | 0.12565684 | 0.274028 |
| ENSG00000229800 | ATP8A2P2 | 1.276726 | -0.2550654 | 0.7895455 | -0.9240766 | 0.3554464 | NA |
| ENSG00000198373 | WWP2 | 4079.674 | -0.2550847 | 0.1523163 | -1.692408 | 0.0905683 | 0.21659 |
| ENSG00000153283 | CD96 | 8.645491 | -0.2550981 | 0.5418591 | -0.636771 | 0.524274 | 0.70931 |
| ENSG00000089902 | RCOR1 | 4087.085464 | -0.255120903 | 0.164536139 | -1.593022718 | 0.1111551 | 0.251215 |
| ENSG00000075914 | EXOSC7 | 1262.994252 | -0.255257666 | 0.221629417 | -1.211258125 | 0.22579649 | 0.413139 |
| ENSG00000127948 | POR | 5939.547215 | -0.25531623 | 0.184368932 | -1.459730356 | 0.1443642 | 0.30266 |

| | | | | | | | |
|---|---|---|---|---|---|---|---|
| ENSG00000102743 | SLC25A15 | 2387.377678 | -0.255357178 | 0.249690083 | -1.085860244 | 0.27754083 | 0.474607 |
| ENSG00000235672 | | 2.422602 | -0.255362 | 0.6915947 | -0.6761985 | 0.4989147 | 0.687392 |
| ENSG00000279031 | | 11.43131 | -0.2555517 | 0.5330786 | -0.6392632 | 0.5226517 | 0.708017 |
| ENSG00000268366 | | 51.27772 | -0.2555534 | 0.3253742 | -0.8677112 | 0.3855524 | 0.58791 |
| ENSG00000261118 | | 59.91981 | -0.2555594 | 0.4224355 | -0.7169725 | 0.4733911 | 0.666199 |
| ENSG00000250734 | | 1.360821 | -0.2555743 | 0.7588472 | -0.7943049 | 0.427018 | NA |
| ENSG00000287107 | | 1.984658 | -0.2555838 | 0.788014 | -0.8998866 | 0.3681806 | 0.570224 |
| ENSG00000286659 | | 8.286953 | -0.2558195 | 0.5430918 | -0.6441364 | 0.519487 | 0.704844 |
| ENSG00000224822 | THRB-IT1 | 1.282335 | -0.2559768 | 0.8194883 | -0.8085212 | 0.4187906 | NA |
| ENSG00000278875 | | 4.635022 | -0.2560015 | 0.7408288 | -0.7342417 | 0.4628015 | 0.657178 |
| ENSG00000065357 | DGKA | 131.4212426 | -0.25603743 | 0.288044637 | -0.953348656 | 0.34041345 | 0.541851 |
| ENSG00000114650 | SCAP | 4058.646044 | -0.256046956 | 0.117592015 | -2.20915541 | 0.02716383 | 0.086609 |
| ENSG00000252933 | RNU6-1223P | 2.59641 | -0.2561181 | 0.7246541 | -0.7145336 | 0.4748973 | 0.66756 |
| ENSG00000176428 | VPS37D | 275.0356 | -0.2561247 | 0.239256 | -1.131181 | 0.2579788 | 0.452067 |
| ENSG00000122642 | FKBP9 | 4549.800938 | -0.256134367 | 0.154432087 | -1.705054052 | 0.08818434 | 0.212399 |
| ENSG00000108439 | PNPO | 2252.450546 | -0.256180703 | 0.126311945 | -2.059634178 | 0.03943353 | 0.115629 |
| ENSG00000272417 | | 1.731396 | -0.2562002 | 0.7489922 | -0.7772319 | 0.4370219 | 0.635466 |
| ENSG00000133895 | MEN1 | 3929.827205 | -0.256246738 | 0.138496729 | -1.893640638 | 0.05827273 | 0.155791 |
| ENSG00000188010 | MORN2 | 613.5219 | -0.2562554 | 0.2080987 | -1.286672 | 0.1982085 | 0.378162 |
| ENSG00000071564 | TCF3 | 9143.45337 | -0.256353583 | 0.206016878 | -1.290049217 | 0.19703357 | 0.376972 |
| ENSG00000088205 | DDX18 | 6831.657866 | -0.256535609 | 0.213190702 | -1.262493152 | 0.20677139 | 0.3891 |
| ENSG00000265981 | MIR544B | 2.077835 | -0.2565718 | 0.74902 | -0.7640816 | 0.4448186 | 0.642138 |
| ENSG00000198554 | WDHD1 | 1518.343 | -0.2565985 | 0.4221363 | -0.7385411 | 0.4601857 | 0.655608 |
| ENSG00000008196 | TFAP2B | 6.856002281 | -0.25660226 | 0.689976185 | -0.669522142 | 0.50316246 | 0.690525 |
| ENSG00000258713 | C20orf141 | 1.445839 | -0.2568284 | 0.8509769 | -1.938246 | 0.05259321 | NA |
| ENSG00000164051 | CCDC51 | 1098.373 | -0.2568988 | 0.1876276 | -1.407439 | 0.1592973 | 0.325307 |
| ENSG00000161664 | ASB16 | 149.0519 | -0.2569836 | 0.3240137 | -0.8786976 | 0.3795653 | 0.581517 |
| ENSG00000136643 | RPS6KC1 | 1210.361769 | -0.256996808 | 0.137330868 | -1.905178207 | 0.05675691 | 0.152583 |
| ENSG00000148824 | MTG1 | 565.6695 | -0.2570089 | 0.2551311 | -1.074909 | 0.2824156 | 0.480255 |
| ENSG00000286729 | | 29.15095 | -0.2570169 | 0.5191827 | -0.6536263 | 0.5133526 | 0.69927 |
| ENSG00000177757 | FAM87B | 7.748654 | -0.2570211 | 0.5188876 | -0.6533848 | 0.5135082 | 0.699375 |
| ENSG00000101138 | CSTF1 | 5133.117961 | -0.257030759 | 0.131657908 | -2.004741982 | 0.04499064 | 0.127982 |
| ENSG00000224292 | | 4.307513 | -0.2570746 | 0.6137646 | -0.6409597 | 0.5215489 | 0.706883 |

| | | | | | | | |
|---|---|---|---|---|---|---|---|
| ENSG00000196396 | PTPN1 | 7161.372 | -0.2571067 | 0.1570199 | -1.68772 | 0.09146508 | 0.218165 |
| ENSG00000113494 | PRLR | 7705.800674 | -0.257117333 | 0.384908287 | -0.780881581 | 0.43487215 | 0.633754 |
| ENSG00000229447 | | 10.58609 | -0.2571624 | 0.5418487 | -0.6505449 | 0.5153403 | 0.700756 |
| ENSG00000078487 | ZCWPW1 | 412.6073843 | -0.257169241 | 0.329689934 | -0.864095511 | 0.38753543 | 0.589783 |
| ENSG00000132938 | MTUS2 | 41.84816788 | -0.257202187 | 0.353416502 | -0.815798462 | 0.41461541 | 0.614649 |
| ENSG00000258851 | | 5.318356 | -0.257415 | 0.6131919 | -0.6383634 | 0.5232371 | 0.708521 |
| ENSG00000273142 | LINC02604 | 487.0365 | -0.2575167 | 0.2475399 | -1.105969 | 0.2687398 | 0.464732 |
| ENSG00000116106 | EPHA4 | 2797.311346 | -0.257647151 | 0.423039964 | -0.733872473 | 0.46302647 | 0.657314 |
| ENSG00000234459 | | 4.323875 | -0.2576624 | 0.689483 | -0.6718485 | 0.5016801 | 0.689342 |
| ENSG00000270612 | | 1.287857 | -0.2576861 | 0.7881182 | -0.9428733 | 0.3457457 | NA |
| ENSG00000157827 | FMNL2 | 2417.318 | -0.2577151 | 0.2125814 | -1.245067 | 0.2131071 | 0.397839 |
| ENSG00000202279 | | 0.9248273 | -0.2577303 | 0.8353245 | -1.157551 | 0.2470475 | NA |
| ENSG00000120693 | SMAD9 | 151.7510958 | -0.25777984 | 0.355618794 | -0.819996079 | 0.41221834 | 0.612438 |
| ENSG00000019102 | VSIG2 | 5.672294154 | -0.257824631 | 0.649923883 | -0.652072448 | 0.51435444 | 0.700096 |
| ENSG00000278709 | NKILA | 711.9868 | -0.2578479 | 0.3435863 | -0.8363951 | 0.4029327 | 0.603842 |
| ENSG00000088827 | SIGLEC1 | 2.012423421 | -0.257967455 | 0.779906373 | -0.866798414 | 0.38605247 | 0.588387 |
| ENSG00000278727 | | 5.063517 | -0.2580806 | 0.6321188 | -0.6426199 | 0.5204708 | 0.70571 |
| ENSG00000276058 | STMN1P1 | 4.198791 | -0.2581454 | 0.6836054 | -0.6936063 | 0.4879292 | 0.67891 |
| ENSG00000272071 | | 4.797362 | -0.2581465 | 0.6359039 | -0.6505719 | 0.5153229 | 0.700756 |
| ENSG00000011028 | MRC2 | 1892.303007 | -0.258172729 | 0.456513693 | -0.688191661 | 0.4913321 | 0.681575 |
| ENSG00000181741 | FDX1P1 | 2.082251 | -0.2581831 | 0.7406187 | -0.7633064 | 0.4452807 | 0.642438 |
| ENSG00000287726 | | 2.958398 | -0.2582111 | 0.6660224 | -0.6584284 | 0.5102629 | 0.696102 |
| ENSG00000280161 | | 72.16306 | -0.2582519 | 0.3747824 | -0.7877005 | 0.4308719 | 0.630066 |
| ENSG00000121058 | COIL | 3102.963946 | -0.258258644 | 0.108635646 | -2.402460544 | 0.01628519 | 0.057522 |
| ENSG00000265539 | MIR3164 | 2.994102 | -0.2582935 | 0.6666282 | -0.659965 | 0.5092763 | 0.695358 |
| ENSG00000102452 | NALCN | 1.632684369 | -0.258293768 | 0.831516471 | -1.052770965 | 0.29244598 | 0.491306 |
| ENSG00000101752 | MIB1 | 4396.108202 | -0.258441043 | 0.218227575 | -1.234170174 | 0.2171395 | 0.402679 |
| ENSG00000109016 | DHRS7B | 1372.758734 | -0.258485254 | 0.1943003 | -1.381115385 | 0.16724349 | 0.336679 |
| ENSG00000076924 | XAB2 | 2738.889656 | -0.258486593 | 0.156797369 | -1.714299534 | 0.08647373 | 0.209208 |
| ENSG00000256069 | A2MP1 | 34.18233 | -0.2585504 | 0.7363673 | -0.7658318 | 0.4437764 | 0.641322 |
| ENSG00000124766 | SOX4 | 7365.774197 | -0.258590485 | 0.295766984 | -0.945331832 | 0.34448949 | 0.546046 |
| ENSG00000103351 | CLUAP1 | 1053.157881 | -0.258636814 | 0.187828374 | -1.415172598 | 0.1570179 | 0.321874 |
| ENSG00000234585 | CCT6P3 | 260.4129 | -0.2587081 | 0.2311166 | -1.173876 | 0.2404447 | 0.430745 |

| | | | | | | | |
|---|---|---|---|---|---|---|---|
| ENSG00000196981 | WDR5B | 1062.485 | -0.258734 | 0.2250896 | -1.216753 | 0.2236981 | 0.41043 |
| ENSG00000108773 | KAT2A | 5822.239392 | -0.258774807 | 0.171520249 | -1.589459505 | 0.11195669 | 0.252532 |
| ENSG00000164867 | NOS3 | 37.27993 | -0.2588127 | 0.6248323 | -0.6552317 | 0.5123186 | 0.69822 |
| ENSG00000276488 | | 40.73185 | -0.2588158 | 0.2976734 | -0.9405003 | 0.346961 | 0.548568 |
| ENSG00000259659 | | 11.8638 | -0.2588532 | 0.4485798 | -0.6989377 | 0.484591 | 0.675584 |
| ENSG00000244134 | RPS12P20 | 2.329148 | -0.2589618 | 0.7851867 | -0.886093 | 0.3755674 | 0.577691 |
| ENSG00000138279 | ANXA7 | 6697.480523 | -0.258992963 | 0.154905971 | -1.718267316 | 0.08574787 | 0.207977 |
| ENSG00000234322 | ST13P18 | 193.7877 | -0.2590767 | 0.1887072 | -1.410803 | 0.1583027 | 0.323758 |
| ENSG00000162688 | AGL | 1332.069 | -0.2591797 | 0.2195022 | -1.236759 | 0.2161767 | 0.401482 |
| ENSG00000175634 | RPS6KB2 | 4387.506 | -0.2593148 | 0.178978 | -1.491863 | 0.1357351 | 0.289387 |
| ENSG00000124422 | USP22 | 11158.35923 | -0.259384398 | 0.184958398 | -1.454024209 | 0.14593959 | 0.305084 |
| ENSG00000171314 | PGAM1 | 3862.982 | -0.2593891 | 0.3034609 | -0.9286548 | 0.353068 | 0.55497 |
| ENSG00000162775 | RBM15 | 1063.174 | -0.2594631 | 0.2213251 | -1.233359 | 0.2174419 | 0.403071 |
| ENSG00000285043 | | 80.93677 | -0.2595408 | 0.3725907 | -0.7936527 | 0.4273977 | 0.626892 |
| ENSG00000188428 | BLOC1S5 | 670.3605 | -0.2595905 | 0.2070113 | -1.308207 | 0.1908031 | 0.368821 |
| ENSG00000256690 | | 21.48435 | -0.2596697 | 0.4171051 | -0.7326928 | 0.4637458 | 0.657836 |
| ENSG00000174453 | VWC2L | 1.783186 | -0.259689 | 0.7318977 | -0.7650927 | 0.4442164 | 0.641608 |
| ENSG00000242020 | RN7SL68P | 0.8508713 | -0.2597145 | 0.8281101 | -1.292874 | 0.1960545 | NA |
| ENSG00000152049 | KCNE4 | 31.73515 | -0.2598069 | 0.7200804 | -0.7348199 | 0.4624492 | 0.656943 |
| ENSG00000104756 | KCTD9 | 1164.220349 | -0.2598991 | 0.185356324 | -1.455520149 | 0.14552532 | 0.304603 |
| ENSG00000115211 | EIF2B4 | 3203.20665 | -0.259927454 | 0.133759229 | -1.985765351 | 0.04705939 | 0.132337 |
| ENSG00000167693 | NXN | 300.4672 | -0.2599956 | 0.3966427 | -0.7600741 | 0.4472103 | 0.643937 |
| ENSG00000261864 | | 4.52437 | -0.2600105 | 0.6896403 | -0.6702447 | 0.5027018 | 0.690189 |
| ENSG00000287136 | | 1.412988 | -0.2600209 | 0.7723395 | -0.8453369 | 0.3979228 | NA |
| ENSG00000213839 | TMX2P1 | 194.3295 | -0.2600981 | 0.2386423 | -1.156998 | 0.247273 | 0.439275 |
| ENSG00000164256 | PRDM9 | 4.520724 | -0.260113 | 0.639509 | -0.6540017 | 0.5131107 | 0.699048 |
| ENSG00000176390 | CRLF3 | 359.7828 | -0.2601429 | 0.1989913 | -1.35477 | 0.1754907 | 0.3478 |
| ENSG00000204590 | GNL1 | 4888.322 | -0.2601817 | 0.1789157 | -1.519786 | 0.1285648 | 0.278503 |
| ENSG00000132964 | CDK8 | 1478.001211 | -0.260274677 | 0.15537502 | -1.711865139 | 0.08692152 | 0.210043 |
| ENSG00000168461 | RAB31 | 1007.299 | -0.2603143 | 0.2140448 | -1.242957 | 0.2138836 | 0.398728 |
| ENSG00000132837 | DMGDH | 76.26669131 | -0.260328981 | 0.691781886 | -0.707547628 | 0.47922623 | 0.671301 |
| ENSG00000143575 | HAX1 | 6844.187 | -0.2603799 | 0.155666 | -1.722547 | 0.08497056 | 0.206474 |
| ENSG00000203684 | IBA57-DT | 10.15511 | -0.2604162 | 0.503681 | -0.659628 | 0.5094926 | 0.69551 |

| | | | | | | | |
|---|---|---|---|---|---|---|---|
| ENSG00000187764 | SEMA4D | 1349.028 | -0.2604923 | 0.1441049 | -1.833356 | 0.06674961 | 0.172282 |
| ENSG00000229611 | LINC02851 | 2.288068 | -0.2605456 | 0.7718191 | -0.8337454 | 0.4044245 | 0.605327 |
| ENSG00000229891 | LINC01315 | 34.38646 | -0.2605762 | 0.3920201 | -0.7408559 | 0.4587808 | 0.654532 |
| ENSG00000206913 | | 2.277593 | -0.260596 | 0.6829028 | -0.6831609 | 0.4945052 | 0.684048 |
| ENSG00000240441 | | 1.724978 | -0.2606411 | 0.7443341 | -0.7705763 | 0.4409581 | 0.638839 |
| ENSG00000168070 | MAJIN | 2.64161 | -0.2607037 | 0.7847336 | -0.89436 | 0.3711293 | 0.573254 |
| ENSG00000197771 | MCMBP | 4644.427 | -0.2607158 | 0.2476284 | -1.109959 | 0.2670167 | 0.46278 |
| ENSG00000138078 | PREPL | 3053.40365 | -0.260727229 | 0.249579421 | -1.049103052 | 0.29413069 | 0.4932 |
| ENSG00000112699 | GMDS | 1291.841101 | -0.26075012 | 0.221410511 | -1.228168909 | 0.21938357 | 0.40551 |
| ENSG00000255857 | PXN-AS1 | 274.4636 | -0.2607634 | 0.1789101 | -1.499652 | 0.1337047 | 0.286408 |
| ENSG00000215835 | | 1851.183 | -0.2607661 | 0.200922 | -1.34524 | 0.1785477 | 0.352256 |
| ENSG00000285931 | | 1.51332 | -0.2608089 | 0.7622723 | -0.8218607 | 0.4111562 | NA |
| ENSG00000137492 | THAP12 | 5522.149135 | -0.260810124 | 0.173629445 | -1.553904502 | 0.1202072 | 0.265662 |
| ENSG00000283498 | MIR1244-2 | 69.81883 | -0.2609483 | 0.3829041 | -0.7830181 | 0.4336165 | 0.632592 |
| ENSG00000236035 | NSA2P1 | 16.28823 | -0.2609616 | 0.4345295 | -0.7177674 | 0.4729007 | 0.665845 |
| ENSG00000235698 | PA2G4P2 | 249.72 | -0.2611842 | 0.2900946 | -0.9760883 | 0.3290207 | 0.530227 |
| ENSG00000235316 | DUSP8P5 | 150.045 | -0.2612207 | 0.3037336 | -0.9310478 | 0.3518288 | 0.553547 |
| ENSG00000185252 | ZNF74 | 1575.849 | -0.2612632 | 0.1582805 | -1.681963 | 0.09257599 | 0.219979 |
| ENSG00000020181 | ADGRA2 | 292.358426 | -0.261319614 | 0.287340733 | -1.017021087 | 0.3091434 | 0.509657 |
| ENSG00000073905 | VDAC1P1 | 1072.284574 | -0.261338948 | 0.17514908 | -1.522395371 | 0.12791004 | 0.277514 |
| ENSG00000125450 | NUP85 | 5284.380264 | -0.261345984 | 0.258288141 | -1.06736533 | 0.2858069 | 0.483864 |
| ENSG00000231302 | RPL36P2 | 1.632617 | -0.2614991 | 0.7484731 | -0.7846361 | 0.432667 | 0.631831 |
| ENSG00000122912 | SLC25A16 | 1087.125829 | -0.261505126 | 0.230583632 | -1.190070202 | 0.2340188 | 0.423171 |
| ENSG00000287016 | | 38.45871 | -0.2616623 | 0.4871253 | -0.6794771 | 0.4968356 | 0.685678 |
| ENSG00000111728 | ST8SIA1 | 75.89561666 | -0.261675876 | 0.559394093 | -0.660005642 | 0.50925021 | 0.695357 |
| ENSG00000288007 | | 3.446586 | -0.2616857 | 0.7007526 | -0.6954503 | 0.4867731 | 0.677727 |
| ENSG00000236890 | | 1.716277 | -0.2617797 | 0.7487266 | -0.7705042 | 0.4410009 | 0.638839 |
| ENSG00000179131 | | 3.711561 | -0.2617951 | 0.6850012 | -0.680834 | 0.4959765 | 0.685033 |
| ENSG00000138385 | SSB | 6493.894368 | -0.261833255 | 0.17360418 | -1.573963251 | 0.11549595 | 0.258344 |
| ENSG00000204435 | CSNK2B | 1046.146 | -0.2618506 | 0.1886944 | -1.432161 | 0.1520977 | 0.314286 |
| ENSG00000261783 | | 28.21088 | -0.2619645 | 0.3937707 | -0.7713472 | 0.4405012 | 0.638603 |
| ENSG00000269439 | | 58.56929 | -0.2619657 | 0.3735432 | -0.7955032 | 0.4263209 | 0.626212 |
| ENSG00000214192 | UBE2V1P2 | 287.8773 | -0.2619821 | 0.2595708 | -1.07163 | 0.2838864 | 0.481731 |

| | | | | | | | |
|---|---|---|---|---|---|---|---|
| ENSG00000152683 | SLC30A6 | 2301.987 | -0.2620431 | 0.1293972 | -2.072801 | 0.03819078 | 0.112831 |
| ENSG00000138002 | IFT172 | 1475.59721 | -0.262061441 | 0.207108518 | -1.314601105 | 0.188644 | 0.365845 |
| ENSG00000136682 | CBWD2 | 1537.678722 | -0.262100351 | 0.16360546 | -1.648042301 | 0.09934399 | 0.231593 |
| ENSG00000085840 | ORC1 | 817.2854021 | -0.262155579 | 0.620443729 | -0.6687447 | 0.50365835 | 0.690989 |
| ENSG00000264580 | MIR5692B | 1.070663 | -0.2621859 | 0.8227842 | -0.8525406 | 0.3939141 | NA |
| ENSG00000070601 | FRMPD1 | 1.828590246 | -0.262236124 | 0.70557002 | -0.71027656 | 0.47753265 | 0.669704 |
| ENSG00000108883 | EFTUD2 | 7561.007872 | -0.262285171 | 0.204719427 | -1.333012567 | 0.18252768 | 0.35748 |
| ENSG00000266824 | | 1.433524 | -0.2623133 | 0.7511987 | -0.7819222 | 0.4342603 | NA |
| ENSG00000141956 | PRDM15 | 1405.839 | -0.2623523 | 0.165543 | -1.617392 | 0.1057937 | 0.242434 |
| ENSG00000161547 | SRSF2 | 10063.16 | -0.2624122 | 0.1968779 | -1.390957 | 0.1642384 | 0.332333 |
| ENSG00000204866 | IGFL2 | 1.133318 | -0.2624203 | 0.8259254 | -0.9233571 | 0.3558211 | NA |
| ENSG00000259954 | IL21R-AS1 | 1.921236 | -0.2624495 | 0.6984662 | -0.7011671 | 0.4831987 | 0.674589 |
| ENSG00000235631 | RNF148 | 5.489452 | -0.2624698 | 0.6580264 | -0.6678274 | 0.5042438 | 0.691213 |
| ENSG00000213399 | RPS2P17 | 14.59976 | -0.2624875 | 0.4861144 | -0.681507 | 0.4955508 | 0.684733 |
| ENSG00000214391 | TUBAP2 | 332.0018 | -0.2624984 | 0.444499 | -0.7184311 | 0.4724915 | 0.665381 |
| ENSG00000287087 | | 2.70155 | -0.2625222 | 0.70695 | -0.7217628 | 0.4704403 | 0.663929 |
| ENSG00000047410 | TPR | 13546.05052 | -0.262587541 | 0.168210881 | -1.60727449 | 0.10799417 | 0.246017 |
| ENSG00000116874 | WARS2 | 974.7667702 | -0.262610748 | 0.202291038 | -1.347193732 | 0.17791785 | 0.351274 |
| ENSG00000173905 | GOLIM4 | 2262.464 | -0.2626433 | 0.2109427 | -1.292406 | 0.1962166 | 0.375969 |
| ENSG00000144848 | ATG3 | 3491.449 | -0.2627082 | 0.1379291 | -1.949396 | 0.05124816 | 0.141202 |
| ENSG00000110958 | PTGES3 | 15250.50976 | -0.262752528 | 0.202630342 | -1.346132955 | 0.17825964 | 0.351782 |
| ENSG00000263368 | | 0.8395606 | -0.2628257 | 0.806823 | -1.073129 | 0.2832131 | NA |
| ENSG00000233250 | | 18.19845 | -0.2628633 | 0.4149619 | -0.7499983 | 0.4532557 | 0.649427 |
| ENSG00000227512 | | 4.266868 | -0.2629508 | 0.7053505 | -0.6950198 | 0.4870429 | 0.677926 |
| ENSG00000253729 | PRKDC | 19242.84 | -0.2630476 | 0.260813 | -1.037091 | 0.2996935 | 0.498891 |
| ENSG00000272173 | | 40.44482 | -0.2631024 | 0.3701078 | -0.8051386 | 0.4207397 | 0.620621 |
| ENSG00000113068 | PFDN1 | 2439.459266 | -0.263115478 | 0.247711961 | -1.13204743 | 0.25761449 | 0.451607 |
| ENSG00000188693 | CYP51A1-AS1 | 6.997414 | -0.2631529 | 0.6087407 | -0.6513562 | 0.5148166 | 0.700365 |
| ENSG00000215131 | C16orf90 | 0.8611893 | -0.2632097 | 0.8038328 | -1.046902 | 0.2951447 | NA |
| ENSG00000260103 | | 3.110835 | -0.2632476 | 0.7028942 | -0.7119886 | 0.4764718 | 0.668747 |
| ENSG00000100380 | ST13 | 8262.123466 | -0.263324869 | 0.110079085 | -2.422297085 | 0.01542274 | 0.055113 |
| ENSG00000275278 | | 0.7885023 | -0.2633843 | 0.8484381 | -1.558358 | 0.1191483 | NA |
| ENSG00000132205 | EMILIN2 | 26.60193067 | -0.263393074 | 0.504178686 | -0.673052938 | 0.50091361 | 0.688789 |

| | | | | | | | |
|---|---|---|---|---|---|---|---|
| ENSG00000224679 | MED15P4 | 1.718103 | -0.2634066 | 0.7791222 | -0.8901945 | 0.3733614 | 0.575572 |
| ENSG00000226699 | | 0.9438575 | -0.26341 | 0.85132 | -1.721619 | 0.0851385 | NA |
| ENSG00000227888 | FAM66A | 5.883847 | -0.2634331 | 0.6251539 | -0.6608093 | 0.5087346 | 0.695011 |
| ENSG00000272320 | | 29.26133 | -0.2634404 | 0.6134664 | -0.656015 | 0.5118145 | 0.697819 |
| ENSG00000129460 | NGDN | 2837.966016 | -0.263509984 | 0.200767606 | -1.360221565 | 0.17375982 | 0.345401 |
| ENSG00000287735 | | 20.69738 | -0.2635122 | 0.4770525 | -0.6960759 | 0.4863813 | 0.677395 |
| ENSG00000260441 | | 2.45817 | -0.2635166 | 0.7137361 | -0.7109745 | 0.4771 | 0.669324 |
| ENSG00000223300 | | 3.621344 | -0.2635361 | 0.6582075 | -0.6733382 | 0.5007321 | 0.688619 |
| ENSG00000170703 | TTLL6 | 3.59418 | -0.2635904 | 0.6383096 | -0.6653504 | 0.5058264 | 0.692569 |
| ENSG00000279879 | | 6.621611 | -0.2636771 | 0.4941467 | -0.6809105 | 0.4959281 | 0.685033 |
| ENSG00000267345 | | 2.951337 | -0.2636879 | 0.7159079 | -0.7193749 | 0.47191 | 0.665025 |
| ENSG00000223343 | | 18.48462 | -0.2637057 | 0.3598774 | -0.8251552 | 0.4092835 | 0.609885 |
| ENSG00000260410 | | 3.589579 | -0.2637062 | 0.6406737 | -0.6591722 | 0.5097852 | 0.695802 |
| ENSG00000177613 | CSTF2T | 2147.44 | -0.2637448 | 0.1392815 | -1.92688 | 0.05399458 | 0.147184 |
| ENSG00000242190 | | 1.484866 | -0.2637687 | 0.7750985 | -0.8624124 | 0.3884606 | NA |
| ENSG00000230397 | SPTLC1P1 | 99.2735 | -0.2638473 | 0.2405807 | -1.150449 | 0.2499589 | 0.442537 |
| ENSG00000213761 | MT1P1 | 0.9195629 | -0.2638549 | 0.7806463 | -0.8833573 | 0.3770433 | NA |
| ENSG00000227436 | FCF1P1 | 1.854047 | -0.2638741 | 0.7371222 | -0.7700802 | 0.4412523 | 0.639134 |
| ENSG00000183091 | NEB | 817.1705 | -0.263894 | 0.3273581 | -0.8939225 | 0.3713634 | 0.573448 |
| ENSG00000224819 | | 3.862064 | -0.2639513 | 0.6781831 | -0.6920608 | 0.4888992 | 0.679677 |
| ENSG00000188037 | CLCN1 | 0.7747202 | -0.2639689 | 0.8227441 | -0.864626 | 0.3872441 | NA |
| ENSG00000007923 | DNAJC11 | 2487.28063 | -0.264003696 | 0.141353602 | -1.904431676 | 0.05685399 | 0.152766 |
| ENSG00000267439 | | 2.322883 | -0.2640188 | 0.643767 | -0.6677934 | 0.5042655 | 0.691213 |
| ENSG00000167695 | TLCD3A | 1112.954 | -0.2640919 | 0.1638392 | -1.64205 | 0.1005797 | 0.233777 |
| ENSG00000177311 | ZBTB38 | 1158.635 | -0.264105 | 0.2839968 | -1.000929 | 0.316861 | 0.517945 |
| ENSG00000225259 | ST13P6 | 988.9056 | -0.264107 | 0.1435412 | -1.87072 | 0.06138395 | 0.161822 |
| ENSG00000137992 | DBT | 1074.072971 | -0.264133777 | 0.199655191 | -1.367610912 | 0.1714339 | 0.34215 |
| ENSG00000243071 | | 6.043626 | -0.2644247 | 0.5746706 | -0.6576003 | 0.510795 | 0.696679 |
| ENSG00000025796 | SEC63 | 5825.219105 | -0.264432792 | 0.1546891 | -1.762757626 | 0.07794138 | 0.193372 |
| ENSG00000233328 | PFN1P1 | 923.9326 | -0.2644942 | 0.2744074 | -1.032641 | 0.3017718 | 0.501206 |
| ENSG00000103540 | CCP110 | 1826.1305 | -0.264542128 | 0.264461974 | -1.070053067 | 0.28459542 | 0.482572 |
| ENSG00000241709 | RN7SL265P | 1.850469 | -0.2645857 | 0.8002354 | -0.9962529 | 0.3191273 | 0.520102 |
| ENSG00000223374 | | 35.72782 | -0.2645926 | 0.3763386 | -0.7987182 | 0.4244538 | 0.624343 |

| | | | | | | | |
|---|---|---|---|---|---|---|---|
| ENSG00000230022 | FNTAP2 | 2.725047 | -0.2646033 | 0.6641802 | -0.677598 | 0.4980266 | 0.68656 |
| ENSG00000257884 | | 3.145797 | -0.2646493 | 0.6727279 | -0.6856254 | 0.4929493 | 0.68282 |
| ENSG00000180525 | DIP2C-AS1 | 2.379246 | -0.264766 | 0.7192625 | -0.7292082 | 0.4658743 | 0.659777 |
| ENSG00000270343 | UNGP3 | 0.9807342 | -0.2648315 | 0.8478875 | -1.498107 | 0.1341054 | NA |
| ENSG00000180626 | ZNF594 | 954.5987 | -0.2648916 | 0.2586352 | -1.089262 | 0.2760383 | 0.472897 |
| ENSG00000271798 | SNORA51 | 2.356225 | -0.2649723 | 0.6708972 | -0.6815478 | 0.4955249 | 0.684733 |
| ENSG00000158545 | ZC3H18 | 5335.74 | -0.2650494 | 0.128838 | -2.097934 | 0.03591095 | 0.10773 |
| ENSG00000118961 | LDAH | 1470.776516 | -0.265170391 | 0.157528388 | -1.727186859 | 0.08413411 | 0.204985 |
| ENSG00000152219 | ARL14EP | 1351.95 | -0.2652058 | 0.1611086 | -1.681086 | 0.09274613 | 0.220255 |
| ENSG00000108465 | CDK5RAP3 | 3845.783158 | -0.265235225 | 0.258625685 | -1.090563863 | 0.27546484 | 0.472335 |
| ENSG00000170836 | PPM1D | 14563.67 | -0.2652366 | 0.1668088 | -1.643008 | 0.1003814 | 0.233442 |
| ENSG00000138449 | SLC40A1 | 1.131858741 | -0.265351592 | 0.760742105 | -0.840485574 | 0.40063619 | NA |
| ENSG00000100299 | ARSA | 1039.963053 | -0.265366197 | 0.29962241 | -0.966303215 | 0.33389248 | 0.535093 |
| ENSG00000163633 | C4orf36 | 15.79094 | -0.2653764 | 0.4270986 | -0.7378211 | 0.4606232 | 0.655647 |
| ENSG00000164284 | GRPEL2 | 2246.683 | -0.2654032 | 0.1826734 | -1.505958 | 0.1320779 | 0.284098 |
| ENSG00000136160 | EDNRB | 39.71714177 | -0.26541866 | 0.822793026 | -0.817838758 | 0.41344926 | 0.613478 |
| ENSG00000234171 | RNASEH1-AS1 | 454.2081 | -0.2654751 | 0.2227885 | -1.240916 | 0.2146366 | 0.399515 |
| ENSG00000103942 | HOMER2 | 3181.942516 | -0.265494767 | 0.190807043 | -1.4399609 | 0.14987846 | 0.310933 |
| ENSG00000288087 | | 0.9449677 | -0.2655003 | 0.7838513 | -0.9006292 | 0.3677855 | NA |
| ENSG00000237730 | RPS5P8 | 12.12238 | -0.2656233 | 0.5446415 | -0.6620634 | 0.5079306 | 0.694305 |
| ENSG00000138075 | ABCG5 | 1.032436821 | -0.265713386 | 0.787619527 | -0.917420237 | 0.35892247 | NA |
| ENSG00000079462 | PAFAH1B3 | 2007.782973 | -0.265737458 | 0.220930496 | -1.275088508 | 0.20227792 | 0.383469 |
| ENSG00000157184 | CPT2 | 860.9373 | -0.2657481 | 0.2533304 | -1.107411 | 0.2681164 | 0.464016 |
| ENSG00000287564 | | 0.819808 | -0.2657835 | 0.8315455 | -1.024361 | 0.3056649 | NA |
| ENSG00000261338 | | 34.84902 | -0.2659829 | 0.4463138 | -0.7230294 | 0.4696618 | 0.663147 |
| ENSG00000142279 | WTIP | 594.0429 | -0.2660253 | 0.1905316 | -1.439972 | 0.1498752 | 0.310933 |
| ENSG00000183624 | HMCES | 5529.672 | -0.2660574 | 0.2991279 | -0.9600825 | 0.3370137 | 0.53817 |
| ENSG00000234036 | TXNP6 | 362.97 | -0.2660799 | 0.2612059 | -1.084562 | 0.2781157 | 0.475224 |
| ENSG00000254561 | | 0.851521 | -0.266126 | 0.8263811 | -0.9384783 | 0.3479987 | NA |
| ENSG00000233859 | ADH5P4 | 269.3206 | -0.2661305 | 0.2023776 | -1.364191 | 0.1725076 | 0.343683 |
| ENSG00000253754 | | 3.967639 | -0.2661645 | 0.6828906 | -0.6941439 | 0.487592 | 0.678548 |
| ENSG00000231944 | PHKA1-AS1 | 0.8294836 | -0.2661916 | 0.8083226 | -1.044046 | 0.2964641 | NA |
| ENSG00000238221 | | 1.11372 | -0.2661946 | 0.7792477 | -0.8799595 | 0.3788813 | NA |

| | | | | | | | |
|---|---|---|---|---|---|---|---|
| ENSG00000230131 | LINC02641 | 1.456399 | -0.2662843 | 0.7596196 | -0.8062046 | 0.4201249 | NA |
| ENSG00000149541 | B3GAT3 | 3449.465 | -0.2663839 | 0.1997655 | -1.389352 | 0.1647256 | 0.332973 |
| ENSG00000277566 | | 5.752023 | -0.2664681 | 0.6694723 | -0.6954465 | 0.4867755 | 0.677727 |
| ENSG00000278053 | DDX52 | 1228.869 | -0.2666332 | 0.1082157 | -2.475918 | 0.01328942 | 0.048857 |
| ENSG00000251409 | | 1.475092 | -0.266659 | 0.7894297 | -0.9561032 | 0.3390201 | NA |
| ENSG00000139112 | GABARAPL1 | 654.7486166 | -0.266689852 | 0.305858346 | -0.95173362 | 0.34123209 | 0.542509 |
| ENSG00000050820 | BCAR1 | 7497.010948 | -0.266703711 | 0.196356046 | -1.423604014 | 0.15456113 | 0.318163 |
| ENSG00000254278 | | 0.9174654 | -0.2667042 | 0.8240264 | -0.8628877 | 0.3881992 | NA |
| ENSG00000107831 | FGF8 | 1.72335758 | -0.266721066 | 0.785809455 | -0.932090512 | 0.35128975 | 0.552928 |
| ENSG00000283010 | | 0.8243347 | -0.26676 | 0.8014397 | -0.9990619 | 0.3177647 | NA |
| ENSG00000238024 | DDX39BP2 | 0.8122105 | -0.2668901 | 0.8020196 | -0.9901342 | 0.3221085 | NA |
| ENSG00000272024 | | 10.44188 | -0.2668989 | 0.5625679 | -0.6628594 | 0.5074206 | 0.693818 |
| ENSG00000148090 | AUH | 577.5552 | -0.2669337 | 0.1388671 | -1.959489 | 0.05005556 | 0.138606 |
| ENSG00000125375 | DMAC2L | 910.5687039 | -0.266961436 | 0.209144473 | -1.328642726 | 0.18396587 | 0.359303 |
| ENSG00000165244 | ZNF367 | 1083.834 | -0.2669773 | 0.4501967 | -0.7248174 | 0.4685641 | 0.662161 |
| ENSG00000090565 | RAB11FIP3 | 5373.455775 | -0.266983894 | 0.145241414 | -1.928566322 | 0.05378472 | 0.146791 |
| ENSG00000160446 | ZDHHC12 | 2173.753 | -0.2670018 | 0.2977669 | -0.9966836 | 0.3189181 | 0.519932 |
| ENSG00000115523 | GNLY | 0.963711112 | -0.267036942 | 0.802414855 | -0.998519342 | 0.31802759 | NA |
| ENSG00000196290 | NIF3L1 | 1883.019 | -0.2670579 | 0.2150102 | -1.294891 | 0.1953579 | 0.374763 |
| ENSG00000227692 | MED28P3 | 2.078347 | -0.2671244 | 0.7197116 | -0.7400498 | 0.4592698 | 0.654843 |
| ENSG00000267065 | LINC02080 | 0.9539889 | -0.2671368 | 0.7776259 | -0.8771839 | 0.3803867 | NA |
| ENSG00000278017 | | 6.295573 | -0.2671441 | 0.551633 | -0.6612235 | 0.508469 | 0.694862 |
| ENSG00000075303 | SLC25A40 | 1197.195137 | -0.267216519 | 0.168739747 | -1.630009071 | 0.10309958 | 0.237944 |
| ENSG00000272953 | | 65.25713 | -0.2673838 | 0.4013024 | -0.7728527 | 0.4396096 | 0.638112 |
| ENSG00000173933 | RBM4 | 2309.836 | -0.2674727 | 0.1647462 | -1.672808 | 0.0943651 | 0.223213 |
| ENSG00000252218 | | 1.037912 | -0.2674866 | 0.8228052 | -1.174691 | 0.2401182 | NA |
| ENSG00000221340 | RNU6ATAC18P | 1.356429 | -0.267528 | 0.8243956 | -0.8627708 | 0.3882635 | NA |
| ENSG00000113658 | SMAD5 | 3716.395968 | -0.267867219 | 0.187979208 | -1.475656822 | 0.14003604 | 0.29619 |
| ENSG00000155189 | AGPAT5 | 2296.624 | -0.2678982 | 0.1698288 | -1.625415 | 0.1040741 | 0.239489 |
| ENSG00000100991 | TRPC4AP | 5795.358292 | -0.267909656 | 0.127852441 | -2.121547143 | 0.03387579 | 0.103086 |
| ENSG00000266308 | RN7SL510P | 0.9851418 | -0.2679195 | 0.7739328 | -0.8664387 | 0.3862496 | NA |
| ENSG00000259553 | | 0.8377051 | -0.2679599 | 0.8386094 | -1.181863 | 0.2372602 | NA |
| ENSG00000206601 | RNU6-431P | 2.986027 | -0.2680186 | 0.6881851 | -0.7077451 | 0.4791036 | 0.671164 |

| | | | | | | | |
|---|---|---|---|---|---|---|---|
| ENSG00000198874 | TYW1 | 1303.95 | -0.2680554 | 0.1457204 | -1.874764 | 0.0608252 | 0.160611 |
| ENSG00000198242 | RPL23A | 2994.751 | -0.2680835 | 0.170195 | -1.627109 | 0.103714 | 0.238864 |
| ENSG00000241478 | HSPA8P9 | 17.73688 | -0.2681241 | 0.4750413 | -0.7060969 | 0.4801279 | 0.671865 |
| ENSG00000101000 | PROCR | 119.0123122 | -0.268145442 | 0.366730111 | -0.831214397 | 0.40585252 | 0.606663 |
| ENSG00000248122 | APOOP4 | 0.9455353 | -0.2681711 | 0.8089572 | -1.085144 | 0.2778578 | NA |
| ENSG00000163739 | CXCL1 | 3.816768 | -0.2682218 | 0.8288534 | -0.9739523 | 0.3300802 | 0.531187 |
| ENSG00000202441 | RNY4P10 | 21.81143 | -0.2682662 | 0.4854199 | -0.6957721 | 0.4865716 | 0.677589 |
| ENSG00000261542 | | 17.88178 | -0.2682888 | 0.5331969 | -0.6744104 | 0.5000504 | 0.688172 |
| ENSG00000213661 | | 1.894844 | -0.268324 | 0.758137 | -0.8055538 | 0.4205001 | 0.620303 |
| ENSG00000144381 | HSPD1 | 31727.06 | -0.2683999 | 0.220783 | -1.271197 | 0.2036585 | 0.385317 |
| ENSG00000183655 | KLHL25 | 803.3417 | -0.2684131 | 0.2361221 | -1.203615 | 0.2287385 | 0.416914 |
| ENSG00000212907 | MT-ND4L | 17060.33 | -0.2684502 | 0.1794105 | -1.540853 | 0.1233526 | 0.270699 |
| ENSG00000203705 | TATDN3 | 370.9585 | -0.2684543 | 0.2031727 | -1.369844 | 0.1707357 | 0.341433 |
| ENSG00000180747 | SMG1P3 | 258.8409 | -0.2685035 | 0.3059404 | -0.9561369 | 0.3390031 | 0.540278 |
| ENSG00000287853 | | 4.956133 | -0.2686045 | 0.7160761 | -0.7296049 | 0.4656317 | 0.659595 |
| ENSG00000125482 | TTF1 | 1155.572702 | -0.268622144 | 0.162645894 | -1.694087523 | 0.09024865 | 0.216078 |
| ENSG00000155465 | SLC7A7 | 35.2055 | -0.2686925 | 0.4446889 | -0.7303821 | 0.4651567 | 0.659133 |
| ENSG00000183287 | CCBE1 | 1.316067 | -0.2687107 | 0.7882037 | -0.9271404 | 0.3538537 | NA |
| ENSG00000054965 | FAM168A | 5421.269009 | -0.268768994 | 0.130391934 | -2.086946462 | 0.03689297 | 0.109852 |
| ENSG00000239454 | LINC02047 | 1.518541 | -0.2687709 | 0.7436993 | -0.7866774 | 0.4314707 | NA |
| ENSG00000109832 | DDX25 | 1.15098294 | -0.268856084 | 0.799494348 | -0.992335695 | 0.3210338 | NA |
| ENSG00000286535 | | 15.00514 | -0.2688881 | 0.4115738 | -0.7655573 | 0.4439398 | 0.641443 |
| ENSG00000249026 | CTNNA1P1 | 11.31275 | -0.2689348 | 0.4650674 | -0.712793 | 0.4759738 | 0.668284 |
| ENSG00000273784 | | 3.060804 | -0.2690771 | 0.643004 | -0.6828684 | 0.49469 | 0.684079 |
| ENSG00000170340 | B3GNT2 | 1559.31 | -0.2690792 | 0.1845241 | -1.503404 | 0.132735 | 0.285069 |
| ENSG00000225402 | | 6.40127 | -0.2690979 | 0.5974149 | -0.6725497 | 0.5012338 | 0.689114 |
| ENSG00000228051 | WBP11P3 | 0.8642356 | -0.2691426 | 0.8401526 | -1.206237 | 0.227726 | NA |
| ENSG00000281593 | | 4.635128 | -0.2691517 | 0.6281321 | -0.6670721 | 0.5047261 | 0.691597 |
| ENSG00000232623 | MRAP-AS1 | 1.414411 | -0.2691519 | 0.7529618 | -0.80157 | 0.4228017 | NA |
| ENSG00000279267 | | 21.69074 | -0.269196 | 0.3683969 | -0.8259499 | 0.4088325 | 0.609555 |
| ENSG00000213742 | ZNF337-AS1 | 364.4202 | -0.2693621 | 0.2854315 | -1.017796 | 0.308775 | 0.509176 |
| ENSG00000228407 | | 27.98353 | -0.269367 | 0.370774 | -0.8255564 | 0.4090558 | 0.609694 |
| ENSG00000168061 | SAC3D1 | 1118.274 | -0.2693949 | 0.3026982 | -0.9720054 | 0.3310478 | 0.532068 |

| | | | | | | | |
|---|---|---|---|---|---|---|---|
| ENSG00000205790 | DPP9-AS1 | 15.26872 | -0.2693986 | 0.4593786 | -0.7199802 | 0.4715372 | 0.664789 |
| ENSG00000270182 | | 1.191533 | -0.2694231 | 0.7997068 | -1.024166 | 0.3057569 | NA |
| ENSG00000109113 | RAB34 | 25.01511592 | -0.26947351 | 0.611988568 | -0.669285255 | 0.50331353 | 0.690659 |
| ENSG00000100304 | TTLL12 | 4837.798236 | -0.269541254 | 0.218623872 | -1.287181701 | 0.19803096 | 0.378081 |
| ENSG00000219806 | ATP5PBP6 | 6.025659 | -0.269547 | 0.5980042 | -0.6666309 | 0.5050079 | 0.691784 |
| ENSG00000183688 | RFLNB | 123.8279 | -0.2696406 | 0.4137607 | -0.767906 | 0.4425431 | 0.640164 |
| ENSG00000153037 | SRP19 | 1035.42 | -0.2697639 | 0.1104279 | -2.469815 | 0.01351829 | 0.049534 |
| ENSG00000236852 | SSBL2P | 34.90898 | -0.2697958 | 0.3867156 | -0.8175243 | 0.4136289 | 0.61371 |
| ENSG00000100505 | TRIM9 | 24.64862535 | -0.269796803 | 0.600397591 | -0.669865807 | 0.50294334 | 0.690329 |
| ENSG00000273192 | | 3.52992 | -0.2698029 | 0.6391789 | -0.6760867 | 0.4989856 | 0.687392 |
| ENSG00000158482 | SNX29P1 | 7.76339 | -0.2698817 | 0.5305656 | -0.6760765 | 0.4989921 | 0.687392 |
| ENSG00000176749 | CDK5R1 | 377.1224 | -0.2700183 | 0.2210547 | -1.274333 | 0.2025455 | 0.38373 |
| ENSG00000138315 | OIT3 | 0.912063216 | -0.270060724 | 0.797681052 | -0.966489297 | 0.3337994 | NA |
| ENSG00000278873 | | 2.065368 | -0.2700742 | 0.7405094 | -0.7804606 | 0.4351198 | 0.633871 |
| ENSG00000189339 | SLC35E2B | 3363.894 | -0.2701084 | 0.2247544 | -1.266264 | 0.2054185 | 0.387378 |
| ENSG00000257545 | | 3.669647 | -0.2701296 | 0.6364842 | -0.6695202 | 0.5031637 | 0.690525 |
| ENSG00000199933 | RNY1P16 | 3.262568 | -0.2702266 | 0.6390392 | -0.6840928 | 0.4939166 | 0.683732 |
| ENSG00000008735 | MAPK8IP2 | 267.0991797 | -0.270259823 | 0.413352092 | -0.777673546 | 0.43676149 | 0.635252 |
| ENSG00000009830 | POMT2 | 2817.550051 | -0.270310887 | 0.14959117 | -1.832512048 | 0.06687517 | 0.172472 |
| ENSG00000250909 | | 16.33797 | -0.270325 | 0.4164634 | -0.7661587 | 0.4435819 | 0.641124 |
| ENSG00000110900 | TSPAN11 | 6.819502568 | -0.270353624 | 0.644790674 | -0.679823274 | 0.49661637 | 0.685505 |
| ENSG00000232118 | BACH1-AS1 | 107.9591 | -0.2703739 | 0.4002248 | -0.7919062 | 0.4284153 | 0.627796 |
| ENSG00000170430 | MGMT | 1963.039 | -0.2705355 | 0.2179063 | -1.294957 | 0.1953351 | 0.374746 |
| ENSG00000281404 | LINC01176 | 93.44309 | -0.2706523 | 0.3212995 | -0.9255305 | 0.35469 | 0.556598 |
| ENSG00000153789 | CIBAR2 | 1.005631 | -0.2706927 | 0.8261075 | -0.9069436 | 0.3644366 | NA |
| ENSG00000165259 | HDX | 193.6214 | -0.2706997 | 0.2349873 | -1.192893 | 0.2329111 | 0.421713 |
| ENSG00000063127 | SLC6A16 | 34.84325645 | -0.270732998 | 0.434455798 | -0.747988357 | 0.45446718 | 0.650503 |
| ENSG00000151503 | NCAPD3 | 2716.236 | -0.2707469 | 0.3346859 | -0.8948283 | 0.3708789 | 0.573033 |
| ENSG00000215883 | CYB5RL | 422.4632 | -0.2708167 | 0.216399 | -1.305676 | 0.1916629 | 0.369891 |
| ENSG00000149196 | HIKESHI | 1946.784 | -0.270842 | 0.1679651 | -1.673181 | 0.09429177 | 0.223144 |
| ENSG00000173120 | KDM2A | 10829.44 | -0.2708505 | 0.1342468 | -2.050561 | 0.04030974 | 0.117591 |
| ENSG00000148468 | FAM171A1 | 296.8399 | -0.2708997 | 0.3349804 | -0.8961458 | 0.3701749 | 0.572345 |
| ENSG00000158296 | SLC13A3 | 282.7048 | -0.2709238 | 0.272 | -1.063101 | 0.2877359 | 0.486067 |

| | | | | | | | |
|---|---|---|---|---|---|---|---|
| ENSG00000269570 | | 13.53279 | -0.2709484 | 0.5609009 | -0.6749113 | 0.4997321 | 0.68795 |
| ENSG00000262039 | | 1.317249 | -0.2709527 | 0.7483467 | -0.800734 | 0.4232856 | NA |
| ENSG00000272832 | | 4.377089 | -0.2710076 | 0.625765 | -0.6833797 | 0.494367 | 0.684048 |
| ENSG00000116809 | ZBTB17 | 1436.492541 | -0.271119408 | 0.159587803 | -1.75200287 | 0.07977331 | 0.196801 |
| ENSG00000164494 | PDSS2 | 1311.368 | -0.2711797 | 0.2096081 | -1.348849 | 0.1773855 | 0.35064 |
| ENSG00000250131 | | 19.87837 | -0.2712445 | 0.423576 | -0.7578278 | 0.4485541 | 0.645138 |
| ENSG00000172932 | ANKRD13D | 4717.858 | -0.2713183 | 0.139122 | -2.000544 | 0.04544156 | 0.128961 |
| ENSG00000238713 | | 2.813701 | -0.2713518 | 0.6821697 | -0.7018196 | 0.4827917 | 0.674198 |
| ENSG00000232917 | HSPE1P6 | 3.762006 | -0.2713588 | 0.6936311 | -0.7062056 | 0.4800603 | 0.671835 |
| ENSG00000106211 | HSPB1 | 35968.79772 | -0.271381886 | 0.211854384 | -1.334503693 | 0.18203884 | 0.356918 |
| ENSG00000215790 | SLC35E2A | 979.074 | -0.2713961 | 0.2709492 | -1.078694 | 0.2807241 | 0.478174 |
| ENSG00000272084 | | 10.72684 | -0.2714881 | 0.5442342 | -0.6749231 | 0.4997246 | 0.68795 |
| ENSG00000242375 | | 4.064193 | -0.2715137 | 0.5863625 | -0.6679067 | 0.5041932 | 0.691213 |
| ENSG00000253347 | | 5.124022 | -0.271527 | 0.6239031 | -0.6663452 | 0.5051905 | 0.691948 |
| ENSG00000159023 | EPB41 | 4698.7 | -0.271599 | 0.1959459 | -1.445367 | 0.148355 | 0.308785 |
| ENSG00000189376 | C8orf76 | 404.4085 | -0.2716526 | 0.1931764 | -1.458265 | 0.1447675 | 0.303362 |
| ENSG00000254996 | ANKHD1-EIF4EBI | 101.1801 | -0.2717788 | 0.3367572 | -0.890939 | 0.3729619 | 0.575095 |
| ENSG00000179796 | LRRC3B | 0.9920951 | -0.271801 | 0.809097 | -1.088854 | 0.2762184 | NA |
| ENSG00000250974 | LINC02196 | 1.274798 | -0.2718775 | 0.7841542 | -0.9254147 | 0.3547502 | NA |
| ENSG00000174695 | TMEM167A | 5511.446 | -0.2719243 | 0.1230874 | -2.189018 | 0.0285955 | 0.090275 |
| ENSG00000120254 | MTHFD1L | 4042.564281 | -0.271947574 | 0.201198841 | -1.438317102 | 0.1503441 | 0.311607 |
| ENSG00000174684 | B4GAT1 | 1794.264 | -0.2719628 | 0.2035523 | -1.391712 | 0.1640096 | 0.332049 |
| ENSG00000250846 | EPHA5-AS1 | 0.8371268 | -0.271965 | 0.8189118 | -1.158056 | 0.246841 | NA |
| ENSG00000109436 | TBC1D9 | 9436.491244 | -0.272031539 | 0.292184457 | -1.009558406 | 0.31270691 | 0.513747 |
| ENSG00000108509 | CAMTA2 | 2259.456466 | -0.272086918 | 0.187092963 | -1.514430094 | 0.12991679 | 0.280608 |
| ENSG00000257384 | | 1.373406 | -0.2720954 | 0.7745299 | -0.8837339 | 0.3768399 | NA |
| ENSG00000220920 | | 3.227758 | -0.2721208 | 0.63281 | -0.6747799 | 0.4998156 | 0.68798 |
| ENSG00000165688 | PMPCA | 6661.074 | -0.2722348 | 0.183934 | -1.547279 | 0.1217959 | 0.268102 |
| ENSG00000139567 | ACVRL1 | 1.050807 | -0.2724335 | 0.799807 | -1.029621 | 0.3031878 | NA |
| ENSG00000240857 | RDH14 | 1183.833 | -0.2724503 | 0.1321514 | -2.097869 | 0.03591674 | 0.10773 |
| ENSG00000051596 | THOC3 | 2425.793164 | -0.272483148 | 0.276916754 | -1.059193168 | 0.28951182 | 0.488058 |
| ENSG00000214753 | HNRNPUL2 | 2854.123 | -0.2725301 | 0.1699335 | -1.647358 | 0.09948454 | 0.23188 |
| ENSG00000180190 | TDRP | 4.078884 | -0.2725392 | 0.65299 | -0.6771022 | 0.4983411 | 0.686744 |

| | | | | | | | |
|---|---|---|---|---|---|---|---|
| ENSG00000125995 | ROMO1 | 2063.413889 | -0.272628739 | 0.247842314 | -1.158841396 | 0.24652084 | 0.438349 |
| ENSG00000184787 | UBE2G2 | 10339.48 | -0.2726439 | 0.2188916 | -1.308237 | 0.190793 | 0.368821 |
| ENSG00000279006 | | 1.157298 | -0.2726802 | 0.7893328 | -0.9330885 | 0.3507743 | NA |
| ENSG00000198805 | PNP | 3551.108 | -0.2726846 | 0.2803479 | -1.060408 | 0.2889591 | 0.487544 |
| ENSG00000238082 | NUDCP2 | 50.59364 | -0.2727112 | 0.3004065 | -0.982897 | 0.3256581 | 0.526946 |
| ENSG00000258412 | | 0.8088878 | -0.272718 | 0.7867185 | -0.9429993 | 0.3456813 | NA |
| ENSG00000249142 | | 199.7128 | -0.2727317 | 0.2248246 | -1.274775 | 0.2023889 | 0.383598 |
| ENSG00000271396 | | 8.036464 | -0.2727334 | 0.5786745 | -0.6754511 | 0.4993892 | 0.687706 |
| ENSG00000089123 | TASP1 | 1009.607943 | -0.272790335 | 0.146780762 | -1.893138963 | 0.05833939 | 0.155913 |
| ENSG00000187323 | DCC | 11.06932 | -0.2727937 | 0.5931266 | -0.6680961 | 0.5040722 | 0.691184 |
| ENSG00000105497 | ZNF175 | 48.42941989 | -0.272842496 | 0.52912254 | -0.674853228 | 0.49976902 | 0.687962 |
| ENSG00000227034 | | 15.28442 | -0.2728528 | 0.5298821 | -0.6864189 | 0.492449 | 0.68249 |
| ENSG00000236384 | LINC00479 | 11.09461 | -0.2728925 | 0.7113349 | -0.7243662 | 0.468841 | 0.662417 |
| ENSG00000244502 | HDAC11-AS1 | 2.881485 | -0.2729319 | 0.6382502 | -0.6843549 | 0.4937511 | 0.683539 |
| ENSG00000280276 | | 1.25073 | -0.273013 | 0.8238333 | -0.8547278 | 0.3927018 | NA |
| ENSG00000286850 | | 3.38903 | -0.2730632 | 0.6192665 | -0.67739 | 0.4981585 | 0.686635 |
| ENSG00000254507 | DEFB131E | 0.9690293 | -0.2730715 | 0.8487639 | -1.415956 | 0.1567885 | NA |
| ENSG00000225726 | | 0.9381887 | -0.2731309 | 0.7880235 | -0.9309403 | 0.3518845 | NA |
| ENSG00000213559 | HNRNPA1P64 | 2.85198 | -0.2733147 | 0.7439868 | -0.7705802 | 0.4409558 | 0.638839 |
| ENSG00000176386 | CDC26 | 281.7926 | -0.2733284 | 0.2146742 | -1.322336 | 0.1860563 | 0.362096 |
| ENSG00000166762 | CATSPER2 | 291.4264 | -0.2733362 | 0.3018271 | -0.986527 | 0.3238746 | 0.524793 |
| ENSG00000204475 | NCR3 | 2.098591 | -0.2734575 | 0.7213304 | -0.7678678 | 0.4425657 | 0.640164 |
| ENSG00000273302 | | 9.170018 | -0.2734652 | 0.5827838 | -0.6735366 | 0.500606 | 0.688544 |
| ENSG00000011275 | RNF216 | 6163.381525 | -0.273706313 | 0.126031284 | -2.202879695 | 0.02760323 | 0.087737 |
| ENSG00000214810 | CYCSP55 | 22.69746 | -0.2737543 | 0.389496 | -0.8114618 | 0.4171005 | 0.617035 |
| ENSG00000264529 | DNAJB6P8 | 0.9096938 | -0.2737718 | 0.8260458 | -0.8789895 | 0.379407 | NA |
| ENSG00000102710 | SUPT20H | 4249.862038 | -0.273776731 | 0.163887416 | -1.715464729 | 0.08626006 | 0.208805 |
| ENSG00000148572 | NRBF2 | 1422.552 | -0.2737818 | 0.1250724 | -2.21215 | 0.02695632 | 0.086067 |
| ENSG00000241537 | | 2.486717 | -0.2738275 | 0.7513869 | -0.8187472 | 0.4129307 | 0.61302 |
| ENSG00000232098 | | 644.8844 | -0.2738728 | 0.222952 | -1.303322 | 0.1924648 | 0.370846 |
| ENSG00000222014 | RAB6C | 84.68978 | -0.273873 | 0.3601914 | -0.8619319 | 0.388725 | 0.590813 |
| ENSG00000141543 | EIF4A3 | 8635.681 | -0.2740215 | 0.2197876 | -1.302857 | 0.1926236 | 0.371045 |
| ENSG00000134186 | PRPF38B | 7895.832172 | -0.274040962 | 0.132145245 | -2.107052973 | 0.03511299 | 0.105845 |

| | | | | | | | |
|---|---|---|---|---|---|---|---|
| ENSG00000286048 | | 0.7808321 | -0.2740577 | 0.8323528 | -1.312286 | 0.1894236 | NA |
| ENSG00000225434 | LINC01504 | 35.99735 | -0.2741001 | 0.5214869 | -0.6964203 | 0.4861657 | 0.677166 |
| ENSG00000226352 | PSPC1-AS2 | 54.88403 | -0.2741089 | 0.3869685 | -0.8131813 | 0.4161141 | 0.616021 |
| ENSG00000100099 | HPS4 | 2440.961741 | -0.274166464 | 0.093036111 | -2.963625867 | 0.00304038 | 0.014706 |
| ENSG00000178982 | EIF3K | 6967.364 | -0.2742304 | 0.1694972 | -1.686387 | 0.09172121 | 0.218521 |
| ENSG00000231799 | PA2G4P6 | 67.32279 | -0.2742727 | 0.2987174 | -0.9958219 | 0.3193367 | 0.520264 |
| ENSG00000108010 | GLRX3 | 3477.535377 | -0.274303465 | 0.21380598 | -1.35745285 | 0.17463737 | 0.346523 |
| ENSG00000258034 | | 0.9462972 | -0.2744565 | 0.8035795 | -1.010944 | 0.3120432 | NA |
| ENSG00000171320 | ESCO2 | 493.6025 | -0.2744706 | 0.5816668 | -0.688244 | 0.4912991 | 0.681575 |
| ENSG00000275632 | | 7.710419 | -0.2744731 | 0.5329912 | -0.6886262 | 0.4910586 | 0.681494 |
| ENSG00000286818 | | 0.7636793 | -0.2744857 | 0.8084932 | -1.038958 | 0.2988241 | NA |
| ENSG00000154342 | WNT3A | 56.81518 | -0.2745126 | 0.5023742 | -0.6920648 | 0.4888967 | 0.679677 |
| ENSG00000253263 | | 6.795887 | -0.2746541 | 0.5789792 | -0.6818927 | 0.4953068 | 0.684622 |
| ENSG00000269910 | | 15.59581 | -0.2746889 | 0.4201159 | -0.7709701 | 0.4407246 | 0.63871 |
| ENSG00000180019 | | 0.8999441 | -0.2746998 | 0.8195164 | -1.168229 | 0.2427143 | NA |
| ENSG00000166482 | MFAP4 | 9.954732 | -0.2747735 | 0.4942168 | -0.7095128 | 0.4780063 | 0.670034 |
| ENSG00000243802 | | 53.387 | -0.2747912 | 0.3574239 | -0.8615387 | 0.3889414 | 0.590975 |
| ENSG00000004455 | AK2 | 7644.982395 | -0.274805088 | 0.148284211 | -1.891663199 | 0.05853587 | 0.156258 |
| ENSG00000214653 | HNRNPA3P3 | 118.5215 | -0.274907 | 0.24053 | -1.203847 | 0.2286488 | 0.416808 |
| ENSG00000169570 | DTWD2 | 935.5761 | -0.2750424 | 0.236979 | -1.226925 | 0.2198507 | 0.406021 |
| ENSG00000270344 | POC1B-AS1 | 266.7362 | -0.2751629 | 0.3466372 | -0.8808399 | 0.3784045 | 0.580509 |
| ENSG00000106633 | GCK | 1.248166483 | -0.27519003 | 0.767556923 | -0.85175109 | 0.39435226 | NA |
| ENSG00000181544 | FANCB | 199.6848 | -0.2753157 | 0.4406239 | -0.7580387 | 0.4484278 | 0.645026 |
| ENSG00000270480 | | 2.627154 | -0.2753705 | 0.6956998 | -0.7210527 | 0.4708771 | 0.664193 |
| ENSG00000170178 | HOXD12 | 9.410331 | -0.2753744 | 0.5654847 | -0.6875637 | 0.4917276 | 0.681967 |
| ENSG00000198938 | MT-CO3 | 101765.5 | -0.2754125 | 0.1752575 | -1.62864 | 0.1033893 | 0.238368 |
| ENSG00000235371 | | 6.083658 | -0.2754199 | 0.5403261 | -0.687656 | 0.4916695 | 0.681936 |
| ENSG00000283516 | LITAFD | 1.055662 | -0.2755708 | 0.7741068 | -0.8790588 | 0.3793694 | NA |
| ENSG00000188672 | RHCE | 13.83341 | -0.2756576 | 0.486226 | -0.716177 | 0.4738821 | 0.666661 |
| ENSG00000153531 | ADPRHL1 | 863.9998 | -0.2756671 | 0.2087985 | -1.373204 | 0.1696888 | 0.340032 |
| ENSG00000215477 | RCN1P1 | 7.240984 | -0.2757399 | 0.5835748 | -0.6730786 | 0.5008973 | 0.688789 |
| ENSG00000054654 | SYNE2 | 9409.812468 | -0.275747537 | 0.266982553 | -1.103463998 | 0.26982572 | 0.466125 |
| ENSG00000177697 | CD151 | 7399.075 | -0.2757507 | 0.13049 | -2.145668 | 0.03189947 | 0.098389 |

| | | | | | | | |
|---|---|---|---|---|---|---|---|
| ENSG00000126838 | PZP | 2.628765887 | -0.27585706 | 0.826007823 | -0.894157798 | 0.37123747 | 0.573296 |
| ENSG00000255434 | | 1.352719 | -0.2759229 | 0.7685902 | -0.8489624 | 0.3959022 | NA |
| ENSG00000186827 | TNFRSF4 | 2.212622 | -0.2760015 | 0.7268382 | -0.7638128 | 0.4449788 | 0.642221 |
| ENSG00000115750 | TAF1B | 1206.655744 | -0.276024348 | 0.169101949 | -1.671110905 | 0.09469977 | 0.223731 |
| ENSG00000125571 | IL37 | 2.432045225 | -0.276030735 | 0.763502989 | -0.852499845 | 0.39393673 | 0.595538 |
| ENSG00000258603 | | 0.9525547 | -0.2760869 | 0.8349633 | -1.045243 | 0.2959105 | NA |
| ENSG00000154174 | TOMM70 | 7558.036 | -0.2761116 | 0.1254778 | -2.201557 | 0.0276966 | 0.087928 |
| ENSG00000238390 | | 15.06585 | -0.2762128 | 0.4064131 | -0.7958298 | 0.426131 | 0.626037 |
| ENSG00000234912 | SNHG20 | 1030.116 | -0.276254 | 0.2643283 | -1.118101 | 0.2635238 | 0.458728 |
| ENSG00000278590 | RN7SL113P | 3.025841 | -0.2762646 | 0.6763236 | -0.7058414 | 0.4802868 | 0.671936 |
| ENSG00000104613 | INTS10 | 2844.923162 | -0.27637885 | 0.11293264 | -2.479756268 | 0.01314722 | 0.048475 |
| ENSG00000026025 | VIM | 47.05972208 | -0.276379125 | 0.530651434 | -0.688275154 | 0.49127953 | 0.681575 |
| ENSG00000281087 | | 6.506497 | -0.2763832 | 0.5523018 | -0.6896985 | 0.4903838 | 0.680949 |
| ENSG00000259336 | | 1.714389 | -0.2764068 | 0.7565201 | -0.8217828 | 0.4112005 | 0.611539 |
| ENSG00000286352 | | 1.533135 | -0.2764583 | 0.7880859 | -0.9233696 | 0.3558146 | NA |
| ENSG00000234769 | WASH4P | 28.32944 | -0.276462 | 0.4325174 | -0.7636958 | 0.4450486 | 0.642287 |
| ENSG00000141622 | RNF165 | 212.8053 | -0.2764762 | 0.2865196 | -1.040531 | 0.2980932 | 0.497209 |
| ENSG00000285677 | | 17.39004 | -0.2765383 | 0.3908256 | -0.8174627 | 0.413664 | 0.613728 |
| ENSG00000164405 | UQCRQ | 4378.863 | -0.2767939 | 0.2562767 | -1.156006 | 0.2476787 | 0.439849 |
| ENSG00000276641 | MIR6769A | 2.047658 | -0.276795 | 0.7244077 | -0.7605651 | 0.4469169 | 0.643781 |
| ENSG00000100804 | PSMB5 | 11513.44921 | -0.276807311 | 0.228157141 | -1.272338179 | 0.20325299 | 0.384878 |
| ENSG00000282944 | | 2.026673 | -0.2769222 | 0.6891054 | -0.7265765 | 0.4674854 | 0.661023 |
| ENSG00000258450 | | 14.3145 | -0.2769237 | 0.461017 | -0.7377341 | 0.4606761 | 0.655687 |
| ENSG00000158104 | HPD | 24.65789 | -0.2769468 | 0.3573552 | -0.8678961 | 0.3854512 | 0.587802 |
| ENSG00000253200 | | 96.57624 | -0.2769603 | 0.230993 | -1.258757 | 0.2081181 | 0.390832 |
| ENSG00000013523 | ANGEL1 | 4224.232557 | -0.276997539 | 0.152204127 | -1.87838471 | 0.06032856 | 0.15968 |
| ENSG00000231468 | PRDX3P2 | 9.673277 | -0.2770283 | 0.5411235 | -0.6882769 | 0.4912784 | 0.681575 |
| ENSG00000219986 | BTF3P7 | 20.96077 | -0.2770644 | 0.3775948 | -0.8389983 | 0.4014703 | 0.602413 |
| ENSG00000217027 | TPT1P4 | 152.4112 | -0.277082 | 0.3005233 | -1.00365 | 0.3155472 | 0.516529 |
| ENSG00000257488 | LINC02354 | 0.8353622 | -0.2772258 | 0.8047608 | -1.027945 | 0.3039756 | NA |
| ENSG00000244128 | LINC01322 | 103.8924 | -0.2773002 | 0.3689336 | -0.8498328 | 0.395418 | 0.597123 |
| ENSG00000110497 | AMBRA1 | 3205.738544 | -0.277453377 | 0.120333641 | -2.336330556 | 0.01947402 | 0.06653 |
| ENSG00000276380 | UBE2NL | 4.79196 | -0.2774564 | 0.6224137 | -0.692349 | 0.4887182 | 0.679652 |

| | | | | | | | |
|---|---|---|---|---|---|---|---|
| ENSG00000162623 | TYW3 | 1895.505 | -0.2775177 | 0.1816309 | -1.572024 | 0.115945 | 0.258782 |
| ENSG00000099817 | POLR2E | 6367.103228 | -0.277545742 | 0.228683448 | -1.219113276 | 0.2228012 | 0.409421 |
| ENSG00000013503 | POLR3B | 920.907317 | -0.277617859 | 0.223737103 | -1.303903013 | 0.19226665 | 0.370572 |
| ENSG00000215796 | | 3.164374 | -0.2776874 | 0.7411751 | -0.8047232 | 0.4209794 | 0.620872 |
| ENSG00000094631 | HDAC6 | 3005.990511 | -0.277711114 | 0.182402901 | -1.572731692 | 0.11578096 | 0.258649 |
| ENSG00000242622 | | 313.0386 | -0.2777378 | 0.2302179 | -1.284097 | 0.199108 | 0.379219 |
| ENSG00000260828 | HMGB3P32 | 1.386073 | -0.2777851 | 0.8000967 | -1.018576 | 0.3084042 | NA |
| ENSG00000261678 | SCRT1 | 9.051262 | -0.2779018 | 0.5039293 | -0.7101156 | 0.4776324 | 0.669704 |
| ENSG00000230741 | | 1.561663 | -0.2779683 | 0.7659582 | -0.8464533 | 0.3972999 | NA |
| ENSG00000198841 | KTI12 | 283.7806 | -0.2779728 | 0.2544331 | -1.163553 | 0.2446051 | 0.436203 |
| ENSG00000131437 | KIF3A | 1550.757415 | -0.278075716 | 0.19202117 | -1.501112405 | 0.13332649 | 0.285965 |
| ENSG00000175550 | DRAP1 | 4012.654 | -0.278223 | 0.1802313 | -1.58879 | 0.1121079 | 0.252696 |
| ENSG00000021762 | OSBPL5 | 1017.398619 | -0.278292181 | 0.159541926 | -1.796262042 | 0.07245285 | 0.183333 |
| ENSG00000161558 | TMEM143 | 399.2705 | -0.2783586 | 0.2201813 | -1.319245 | 0.1870873 | 0.36357 |
| ENSG00000206778 | RNU6-213P | 1.88909 | -0.2783623 | 0.7354907 | -0.7794242 | 0.4357299 | 0.634435 |
| ENSG00000273024 | INTS4P2 | 30.40112 | -0.2784586 | 0.394948 | -0.8153853 | 0.4148518 | 0.614804 |
| ENSG00000090857 | PDPR | 4131.444372 | -0.278485419 | 0.180120525 | -1.590776511 | 0.11165988 | 0.252098 |
| ENSG00000265451 | | 4.851374 | -0.2785084 | 0.6955069 | -0.7381098 | 0.4604477 | 0.655647 |
| ENSG00000155329 | ZCCHC10 | 1174.379 | -0.2785615 | 0.1361234 | -2.082959 | 0.03725496 | 0.110669 |
| ENSG00000116337 | AMPD2 | 4621.506373 | -0.278678586 | 0.169936908 | -1.67440874 | 0.0940503 | 0.222731 |
| ENSG00000232807 | | 42.965 | -0.2786949 | 0.3536669 | -0.8835223 | 0.3769542 | 0.578992 |
| ENSG00000114107 | CEP70 | 1250.965891 | -0.278703541 | 0.190542907 | -1.509985822 | 0.13104704 | 0.282422 |
| ENSG00000276945 | | 4.336316 | -0.2787666 | 0.6982452 | -0.7446737 | 0.456469 | 0.652249 |
| ENSG00000287002 | | 0.9115968 | -0.278784 | 0.8522266 | -1.442355 | 0.1492023 | NA |
| ENSG00000272372 | | 1.672341 | -0.2788689 | 0.8166054 | -1.140416 | 0.254113 | 0.447092 |
| ENSG00000106701 | FSD1L | 261.1839288 | -0.278877553 | 0.216894733 | -1.331727458 | 0.18294976 | 0.35799 |
| ENSG00000232472 | EEF1B2P3 | 7242.511 | -0.2788911 | 0.1731265 | -1.664024 | 0.09610754 | 0.225952 |
| ENSG00000224490 | TTC21B-AS1 | 1.391855 | -0.2790402 | 0.7051328 | -0.7413408 | 0.4584868 | NA |
| ENSG00000135679 | MDM2 | 5144.689044 | -0.27905063 | 0.221893829 | -1.317673811 | 0.18761287 | 0.364404 |
| ENSG00000133250 | ZNF414 | 392.7078566 | -0.279094267 | 0.266872135 | -1.111803568 | 0.26622262 | 0.461852 |
| ENSG00000239503 | MARK2P8 | 1.543049 | -0.2791592 | 0.764356 | -0.8354905 | 0.4034416 | NA |
| ENSG00000173406 | DAB1 | 5.084603 | -0.2792022 | 0.7415996 | -0.784359 | 0.4328295 | 0.631999 |
| ENSG00000113810 | SMC4 | 6644.938705 | -0.279386614 | 0.359208852 | -0.830341216 | 0.40634589 | 0.606903 |

| | | | | | | | |
|---|---|---|---|---|---|---|---|
| ENSG00000254092 | | 7.384232 | -0.2794266 | 0.7821698 | -0.9060192 | 0.3649257 | 0.566802 |
| ENSG00000256238 | SUPT16HP1 | 428.8497 | -0.2794784 | 0.2609869 | -1.141203 | 0.2537855 | 0.446782 |
| ENSG00000172270 | BSG | 21578.5 | -0.2795638 | 0.1771151 | -1.649035 | 0.09914042 | 0.231241 |
| ENSG00000105373 | NOP53 | 14031.26525 | -0.279582583 | 0.247020391 | -1.19725267 | 0.23120809 | 0.419681 |
| ENSG00000119048 | UBE2B | 2984.132721 | -0.279583123 | 0.119446556 | -2.371637945 | 0.01770944 | 0.061587 |
| ENSG00000139354 | GAS2L3 | 935.9295 | -0.2796574 | 0.3325672 | -0.9402205 | 0.3471045 | 0.548669 |
| ENSG00000114638 | UPK1B | 2.096491114 | -0.279668108 | 0.826367733 | -0.860041052 | 0.38976641 | 0.591749 |
| ENSG00000226790 | HNRNPA3P1 | 13.86102 | -0.2797705 | 0.4212465 | -0.7849382 | 0.4324898 | 0.631709 |
| ENSG00000246705 | H2AJ | 5133.63 | -0.2799277 | 0.2596442 | -1.134983 | 0.2563823 | 0.450115 |
| ENSG00000236609 | ZNF853 | 25.16556 | -0.2799811 | 0.4837511 | -0.7290607 | 0.4659645 | 0.659785 |
| ENSG00000183579 | ZNRF3 | 1182.865 | -0.2799827 | 0.218196 | -1.34799 | 0.1776617 | 0.350977 |
| ENSG00000119782 | FKBP1B | 5.935574395 | -0.280031887 | 0.663799954 | -0.714005698 | 0.47522366 | 0.667714 |
| ENSG00000067601 | PMS2P4 | 100.6590509 | -0.280094482 | 0.226374349 | -1.286772947 | 0.19817344 | 0.378162 |
| ENSG00000199038 | MIR210 | 5.087757 | -0.2802111 | 0.6005516 | -0.6884492 | 0.4911699 | 0.681563 |
| ENSG00000125877 | ITPA | 2499.451093 | -0.280303011 | 0.159455029 | -1.803912043 | 0.0712451 | 0.180967 |
| ENSG00000252010 | SCARNA5 | 2.311406 | -0.2803163 | 0.7734915 | -0.8794529 | 0.3791558 | 0.581158 |
| ENSG00000186684 | CYP27C1 | 31.90367 | -0.2803745 | 0.8270377 | -0.8538768 | 0.3931732 | 0.59482 |
| ENSG00000276410 | H2BC3 | 1.557911 | -0.2804009 | 0.8015157 | -1.00881 | 0.3130659 | NA |
| ENSG00000226383 | LINC01876 | 1.342551 | -0.2804348 | 0.816444 | -1.110957 | 0.2665869 | NA |
| ENSG00000234718 | | 0.8919307 | -0.2805399 | 0.8053186 | -1.024119 | 0.3057793 | NA |
| ENSG00000251204 | | 13.92743 | -0.2805659 | 0.4706856 | -0.7396119 | 0.4595355 | 0.655019 |
| ENSG00000090581 | GNPTG | 2581.632547 | -0.28057621 | 0.249558017 | -1.184566231 | 0.236189 | 0.425704 |
| ENSG00000166275 | BORCS7 | 709.7084 | -0.2806331 | 0.1843717 | -1.573685 | 0.1155603 | 0.258423 |
| ENSG00000256663 | | 556.0625 | -0.2806392 | 0.6063076 | -0.704189 | 0.4813151 | 0.672851 |
| ENSG00000228886 | | 1.014142 | -0.2807329 | 0.7882179 | -0.9309542 | 0.3518773 | NA |
| ENSG00000254536 | | 84.52963 | -0.2807519 | 0.3022485 | -1.01216 | 0.3114614 | 0.512399 |
| ENSG00000142534 | RPS11 | 26336.79 | -0.2807679 | 0.1608098 | -1.787281 | 0.07389208 | 0.186177 |
| ENSG00000140030 | GPR65 | 8.990137 | -0.2807837 | 0.5459496 | -0.7176226 | 0.47299 | 0.665936 |
| ENSG00000286189 | | 1.592087 | -0.2807983 | 0.7883196 | -0.9620376 | 0.3360307 | 0.537087 |
| ENSG00000255538 | OR10V2P | 1.054146 | -0.2808484 | 0.7781154 | -0.9149189 | 0.3602342 | NA |
| ENSG00000108518 | PFN1 | 14630.90376 | -0.280866299 | 0.254559413 | -1.17090413 | 0.24163732 | 0.432211 |
| ENSG00000070366 | SMG6 | 1568.774319 | -0.28089027 | 0.159486069 | -1.801654803 | 0.07159973 | 0.181711 |
| ENSG00000116044 | NFE2L2 | 5517.681238 | -0.280939809 | 0.14130615 | -2.014013443 | 0.04400812 | 0.125943 |

| | | | | | | | |
|---|---|---|---|---|---|---|---|
| ENSG00000105677 | TMEM147 | 4266.515697 | -0.28096539 | 0.184063582 | -1.584493781 | 0.1130814 | 0.254184 |
| ENSG00000115944 | COX7A2L | 7631.351695 | -0.28099916 | 0.164158351 | -1.793786462 | 0.07284726 | 0.184121 |
| ENSG00000236471 | | 1.783294 | -0.2810545 | 0.7274328 | -0.7794548 | 0.4357118 | 0.634435 |
| ENSG00000281195 | | 46.9467 | -0.2810723 | 0.4214609 | -0.7902471 | 0.4293835 | 0.628579 |
| ENSG00000277047 | | 18.9686 | -0.2810763 | 0.3650478 | -0.8729012 | 0.3827169 | 0.584963 |
| ENSG00000141698 | NT5C3B | 2900.174 | -0.2811356 | 0.1902658 | -1.505774 | 0.1321252 | 0.284177 |
| ENSG00000222489 | SNORA79B | 7.413209 | -0.2813428 | 0.5745605 | -0.6981097 | 0.4851086 | 0.676083 |
| ENSG00000272148 | | 13.96297 | -0.2813698 | 0.4681816 | -0.745384 | 0.4560396 | 0.652022 |
| ENSG00000164109 | MAD2L1 | 2841.306 | -0.2814733 | 0.4325679 | -0.7836813 | 0.4332271 | 0.632237 |
| ENSG00000230438 | SERPINB9P1 | 2.378107 | -0.2815473 | 0.7012509 | -0.7453423 | 0.4560648 | 0.652023 |
| ENSG00000220585 | DDX18P6 | 5.020417 | -0.2816666 | 0.6618461 | -0.7160857 | 0.4739384 | 0.666661 |
| ENSG00000256525 | POLG2 | 3252.071 | -0.281771 | 0.247266 | -1.205204 | 0.2281245 | 0.416194 |
| ENSG00000233975 | LINC02574 | 2.231925 | -0.2818882 | 0.8435017 | -1.449125 | 0.1473027 | 0.307148 |
| ENSG00000228421 | | 0.9651996 | -0.2819705 | 0.8301557 | -0.9306186 | 0.3520509 | NA |
| ENSG00000286168 | | 6.804033 | -0.2820063 | 0.589359 | -0.6896599 | 0.4904081 | 0.680949 |
| ENSG00000235449 | | 72.58417 | -0.2820537 | 0.2732835 | -1.100614 | 0.2710648 | 0.467375 |
| ENSG00000230445 | LRRC37A6P | 111.9989 | -0.2820627 | 0.2516966 | -1.184094 | 0.2363759 | 0.425944 |
| ENSG00000096746 | HNRNPH3 | 9977.835248 | -0.282199286 | 0.126453228 | -2.28281297 | 0.02244139 | 0.074711 |
| ENSG00000155545 | MIER3 | 1767.309 | -0.2822092 | 0.2171666 | -1.362739 | 0.1729647 | 0.344387 |
| ENSG00000178952 | TUFM | 14814.25 | -0.282221 | 0.1985608 | -1.48879 | 0.1365426 | 0.290727 |
| ENSG00000286607 | | 5.141037 | -0.2823115 | 0.6387127 | -0.7297192 | 0.4655619 | 0.659567 |
| ENSG00000177340 | | 2.949236 | -0.28235 | 0.7392215 | -0.8068602 | 0.419747 | 0.619777 |
| ENSG00000181751 | MACIR | 1661.989 | -0.2824184 | 0.1476938 | -1.94841 | 0.05136591 | 0.141424 |
| ENSG00000125846 | ZNF133 | 998.4201262 | -0.282530457 | 0.131370919 | -2.175682079 | 0.02957904 | 0.092774 |
| ENSG00000231550 | PTCHD3P2 | 3.004995 | -0.2826002 | 0.7606216 | -0.8442566 | 0.398526 | 0.599538 |
| ENSG00000259424 | IRAIN | 30.78301 | -0.282615 | 0.4465247 | -0.7780788 | 0.4365226 | 0.635088 |
| ENSG00000224800 | RPS27AP6 | 1.289637 | -0.2827155 | 0.7694408 | -0.8690298 | 0.3848309 | NA |
| ENSG00000241316 | SUCLG2-AS1 | 62.7833 | -0.2827193 | 0.4284317 | -0.7898324 | 0.4296257 | 0.628876 |
| ENSG00000230849 | GOT2P2 | 46.87361 | -0.2827385 | 0.3631329 | -0.8825356 | 0.3774872 | 0.579513 |
| ENSG00000120907 | ADRA1A | 1.546079918 | -0.282742678 | 0.838420893 | -1.092771967 | 0.27449393 | NA |
| ENSG00000279356 | | 20.25197 | -0.2827712 | 0.4131401 | -0.8039765 | 0.4214106 | 0.621232 |
| ENSG00000131187 | F12 | 1060.978111 | -0.282929696 | 0.226809081 | -1.303380167 | 0.192445 | 0.370846 |
| ENSG00000166855 | CLPX | 4200.708 | -0.2830013 | 0.1481104 | -1.948638 | 0.05133862 | 0.141364 |

| | | | | | | | |
|---|---|---|---|---|---|---|---|
| ENSG00000160049 | DFFA | 2929.952 | -0.2830303 | 0.178529 | -1.63746 | 0.1015344 | 0.235399 |
| ENSG00000176978 | DPP7 | 8178.512 | -0.2830712 | 0.2021419 | -1.45597 | 0.145401 | 0.304402 |
| ENSG00000287033 | | 2.530361 | -0.2831538 | 0.6910965 | -0.736566 | 0.4613863 | 0.656194 |
| ENSG00000153767 | GTF2E1 | 1519.179 | -0.2831984 | 0.1584077 | -1.827597 | 0.06761 | 0.173945 |
| ENSG00000239462 | | 2.80112 | -0.2833302 | 0.7360037 | -0.7994044 | 0.424056 | 0.623976 |
| ENSG00000184719 | RNLS | 545.1535 | -0.2833814 | 0.1892665 | -1.551449 | 0.120794 | 0.266626 |
| ENSG00000274471 | | 68.05454 | -0.2833884 | 0.3129111 | -0.994124 | 0.3201625 | 0.520963 |
| ENSG00000000419 | DPM1 | 5260.486783 | -0.283400579 | 0.153870111 | -1.883768624 | 0.05959628 | 0.158281 |
| ENSG00000228624 | HDAC2-AS2 | 25.69303 | -0.2834323 | 0.4366981 | -0.7773476 | 0.4369537 | 0.635402 |
| ENSG00000133316 | WDR74 | 3098.770713 | -0.283441294 | 0.200026156 | -1.46833342 | 0.14201368 | 0.299196 |
| ENSG00000120314 | WDR55 | 3108.395462 | -0.283465772 | 0.143187214 | -2.01853365 | 0.04353571 | 0.124887 |
| ENSG00000260448 | LCMT1-AS1 | 12.39081 | -0.2835593 | 0.5133604 | -0.7166915 | 0.4735645 | 0.666374 |
| ENSG00000165891 | E2F7 | 736.2931 | -0.2835701 | 0.5600217 | -0.7095612 | 0.4779763 | 0.670034 |
| ENSG00000198356 | GET3 | 4044.094 | -0.2836242 | 0.1942608 | -1.517801 | 0.1290646 | 0.279267 |
| ENSG00000267091 | CTBP2P7 | 2.411076 | -0.2836818 | 0.6892261 | -0.739928 | 0.4593437 | 0.654889 |
| ENSG00000139505 | MTMR6 | 2577.604 | -0.2836902 | 0.1779853 | -1.653031 | 0.09832457 | 0.229822 |
| ENSG00000261124 | | 3.533314 | -0.2836941 | 0.6587233 | -0.7127651 | 0.4759911 | 0.668284 |
| ENSG00000287100 | | 5.926244 | -0.2837605 | 0.6399072 | -0.6986533 | 0.4847687 | 0.675745 |
| ENSG00000287192 | | 2.165728 | -0.2838114 | 0.7628141 | -0.8511846 | 0.3946668 | 0.596436 |
| ENSG00000154727 | GABPA | 1387.649 | -0.2838723 | 0.1607962 | -1.802437 | 0.07147675 | 0.181458 |
| ENSG00000250302 | LINC01618 | 2.603969 | -0.2840014 | 0.7565418 | -0.8382133 | 0.4019109 | 0.602855 |
| ENSG00000100523 | DDHD1 | 2932.245844 | -0.284051771 | 0.15029692 | -1.960832121 | 0.04989861 | 0.138272 |
| ENSG00000167106 | FAM102A | 6804.817 | -0.284099 | 0.2850629 | -1.082463 | 0.2790469 | 0.476216 |
| ENSG00000171262 | FAM98B | 1847.222 | -0.2842125 | 0.2603132 | -1.135386 | 0.2562137 | 0.449924 |
| ENSG00000110619 | CARS1 | 6760.001012 | -0.284405449 | 0.284562776 | -1.080763495 | 0.27980233 | 0.477093 |
| ENSG00000060688 | SNRNP40 | 2015.97779 | -0.284456156 | 0.24013544 | -1.252048489 | 0.2105522 | 0.394427 |
| ENSG00000122873 | CISD1 | 1370.368718 | -0.284506262 | 0.206915156 | -1.440910367 | 0.14961001 | 0.310594 |
| ENSG00000228328 | PGK1P1 | 60.72172 | -0.2845477 | 0.3610413 | -0.886418 | 0.3753923 | 0.577555 |
| ENSG00000234694 | | 45.98137 | -0.2845729 | 0.3645423 | -0.8848332 | 0.3762466 | 0.578467 |
| ENSG00000255565 | | 0.8822353 | -0.2846132 | 0.8372642 | -1.35706 | 0.174762 | NA |
| ENSG00000198826 | ARHGAP11A | 2115.005 | -0.2846221 | 0.4308237 | -0.7856376 | 0.4320798 | 0.631216 |
| ENSG00000214147 | | 1.014363 | -0.284715 | 0.8227698 | -1.191059 | 0.2336305 | NA |
| ENSG00000156521 | TYSND1 | 4028.806 | -0.284759 | 0.1601353 | -1.826269 | 0.06780971 | 0.174337 |

| | | | | | | | |
|---|---|---|---|---|---|---|---|
| ENSG00000253406 | | 0.8735755 | -0.2847931 | 0.8081602 | -1.086584 | 0.2772208 | NA |
| ENSG00000243335 | KCTD7 | 418.2151 | -0.284815 | 0.3442406 | -0.924384 | 0.3552864 | 0.557303 |
| ENSG00000251323 | LINC02728 | 20.36028 | -0.2848511 | 0.5877675 | -0.706536 | 0.4798549 | 0.671756 |
| ENSG00000256280 | ATP5MGP1 | 1.280897 | -0.2848972 | 0.7850384 | -0.9460635 | 0.3441162 | NA |
| ENSG00000177946 | CENPBD1 | 1358.999 | -0.2849869 | 0.1454917 | -1.995051 | 0.04603733 | 0.130193 |
| ENSG00000232645 | LINC01431 | 36.17209 | -0.2849915 | 0.3550876 | -0.8997737 | 0.3682407 | 0.570284 |
| ENSG00000107949 | BCCIP | 3370.586574 | -0.285012428 | 0.215094119 | -1.397617589 | 0.16222793 | 0.329478 |
| ENSG00000218305 | CDC14C | 11.8583 | -0.2851696 | 0.6061108 | -0.6974534 | 0.4855191 | 0.676549 |
| ENSG00000271639 | PPFIA1P1 | 1.451284 | -0.2854073 | 0.7861237 | -0.9384234 | 0.3480268 | NA |
| ENSG00000106615 | RHEB | 2692.481172 | -0.285425824 | 0.149649755 | -1.948637595 | 0.05133872 | 0.141364 |
| ENSG00000112996 | MRPS30 | 3640.649916 | -0.285444462 | 0.095148219 | -3.066985218 | 0.0021623 | 0.011078 |
| ENSG00000201785 | SNORD117 | 7.17763 | -0.2854473 | 0.6105252 | -0.7068908 | 0.4796344 | 0.671608 |
| ENSG00000187630 | DHRS4L2 | 945.0434 | -0.2855122 | 0.2336242 | -1.290106 | 0.197014 | 0.376972 |
| ENSG00000214821 | HMGB1P4 | 1.046789 | -0.2855732 | 0.7714419 | -0.8809824 | 0.3783273 | NA |
| ENSG00000243224 | | 26.35973 | -0.2856381 | 0.3650571 | -0.883063 | 0.3772022 | 0.579233 |
| ENSG00000237973 | MTCO1P12 | 2541.998 | -0.2856387 | 0.1961458 | -1.509839 | 0.1310844 | 0.282455 |
| ENSG00000165916 | PSMC3 | 9097.146 | -0.2856754 | 0.2912971 | -1.058577 | 0.2897925 | 0.48833 |
| ENSG00000235750 | KIAA0040 | 1346.598 | -0.2856935 | 0.2539271 | -1.174265 | 0.2402889 | 0.430553 |
| ENSG00000186710 | CFAP73 | 11.82015 | -0.2857485 | 0.5059627 | -0.7344786 | 0.4626571 | 0.657063 |
| ENSG00000137161 | CNPY3 | 3194.220639 | -0.2858398 | 0.166831831 | -1.758946732 | 0.07858656 | 0.194555 |
| ENSG00000251774 | RNU6-377P | 1.931136 | -0.2858703 | 0.7514619 | -0.839815 | 0.4010121 | 0.601958 |
| ENSG00000091129 | NRCAM | 2855.288568 | -0.286040562 | 0.395017284 | -0.85030809 | 0.39515382 | 0.596864 |
| ENSG00000227632 | | 12.28442 | -0.286177 | 0.4891481 | -0.7373327 | 0.4609201 | 0.655858 |
| ENSG00000286106 | NOTCH2NLR | 3.458923 | -0.2864607 | 0.6967487 | -0.7522185 | 0.4519196 | 0.648224 |
| ENSG00000186645 | SPDYE17 | 10.54727 | -0.2864712 | 0.5303682 | -0.7193625 | 0.4719176 | 0.665025 |
| ENSG00000277654 | | 114.8443 | -0.2865194 | 0.3048457 | -1.022196 | 0.306688 | 0.506836 |
| ENSG00000064932 | SBNO2 | 4435.119269 | -0.286536676 | 0.153150426 | -1.911520429 | 0.05593773 | 0.150992 |
| ENSG00000089060 | SLC8B1 | 641.8413485 | -0.286597848 | 0.218967229 | -1.367491928 | 0.17147117 | 0.342182 |
| ENSG00000230216 | HSPB1P2 | 117.27 | -0.2866752 | 0.2872358 | -1.069919 | 0.2846556 | 0.482581 |
| ENSG00000260891 | | 3.149426 | -0.2866787 | 0.7022751 | -0.759955 | 0.4472815 | 0.64397 |
| ENSG00000203307 | FAM78B-AS1 | 0.8933362 | -0.2867247 | 0.7817603 | -0.9173354 | 0.3589669 | NA |
| ENSG00000089356 | FXYD3 | 169.2807712 | -0.286742116 | 0.30918187 | -1.015164282 | 0.31002752 | 0.510667 |
| ENSG00000132434 | LANCL2 | 1590.903143 | -0.286770916 | 0.252780205 | -1.191280144 | 0.23354363 | 0.422541 |

| | | | | | | | |
|---|---|---|---|---|---|---|---|
| ENSG00000100591 | AHSA1 | 11542.57977 | -0.286787416 | 0.189375639 | -1.564341157 | 0.11773747 | 0.261629 |
| ENSG00000178162 | FAR2P2 | 7.48618 | -0.2868556 | 0.6793411 | -0.7390294 | 0.4598891 | 0.655304 |
| ENSG00000112651 | MRPL2 | 271.4100766 | -0.286968906 | 0.208437452 | -1.422631209 | 0.15484309 | 0.318534 |
| ENSG00000137806 | NDUFAF1 | 558.097463 | -0.287014614 | 0.22585535 | -1.34107349 | 0.17989659 | 0.354053 |
| ENSG00000260953 | | 1.539155 | -0.2870244 | 0.7588615 | -0.8500473 | 0.3952988 | NA |
| ENSG00000113558 | SKP1 | 8596.240077 | -0.287107709 | 0.168127438 | -1.793376357 | 0.07291277 | 0.184251 |
| ENSG00000286931 | | 94.49568 | -0.2872335 | 0.2307149 | -1.296291 | 0.1948751 | 0.374012 |
| ENSG00000116898 | MRPS15 | 4430.836864 | -0.287328581 | 0.234359013 | -1.288913054 | 0.19742832 | 0.377368 |
| ENSG00000286835 | | 53.79441 | -0.2873835 | 0.3497958 | -0.9210444 | 0.3570273 | 0.559044 |
| ENSG00000228804 | | 3.275588 | -0.2873968 | 0.6696142 | -0.7320366 | 0.4641462 | 0.658228 |
| ENSG00000100749 | VRK1 | 1957.138016 | -0.287477303 | 0.3782189 | -0.892168839 | 0.37230244 | 0.574464 |
| ENSG00000226608 | FTLP3 | 1816.544 | -0.2874978 | 0.2156109 | -1.390864 | 0.1642666 | 0.332333 |
| ENSG00000270210 | | 2.217793 | -0.2875063 | 0.8124589 | -1.056965 | 0.2905276 | 0.489115 |
| ENSG00000137709 | POU2F3 | 242.2686467 | -0.287517656 | 0.375970089 | -0.8773143 | 0.38031595 | 0.582231 |
| ENSG00000197358 | BNIP3P1 | 385.6269 | -0.2875235 | 0.1690667 | -1.747532 | 0.08054511 | 0.198314 |
| ENSG00000141873 | SLC39A3 | 763.2783 | -0.287584 | 0.1884688 | -1.576832 | 0.1148341 | 0.257059 |
| ENSG00000144867 | SRPRB | 5093.286 | -0.2877012 | 0.1382405 | -2.125489 | 0.03354583 | 0.10227 |
| ENSG00000171169 | NAIF1 | 536.2089 | -0.2877496 | 0.1477848 | -1.985973 | 0.04703627 | 0.1323 |
| ENSG00000283766 | | 35.67933 | -0.2878158 | 0.5056918 | -0.7352828 | 0.4621673 | 0.656777 |
| ENSG00000129219 | PLD2 | 2203.644904 | -0.287880504 | 0.113565399 | -2.583232748 | 0.00978793 | 0.038199 |
| ENSG00000188266 | HYKK | 233.9656 | -0.2880575 | 0.2913046 | -1.069764 | 0.2847255 | 0.482669 |
| ENSG00000129173 | E2F8 | 837.1776143 | -0.288070341 | 0.55661122 | -0.716055836 | 0.47395687 | 0.666661 |
| ENSG00000271390 | | 0.8090126 | -0.2880885 | 0.8612594 | -1.65896 | 0.09712395 | NA |
| ENSG00000163832 | ELP6 | 1280.391 | -0.2882463 | 0.2767011 | -1.116927 | 0.2640256 | 0.459421 |
| ENSG00000279372 | | 4.870963 | -0.2882489 | 0.6742228 | -0.7380621 | 0.4604767 | 0.655647 |
| ENSG00000155367 | PPM1J | 150.8279 | -0.2882796 | 0.3509415 | -0.9204156 | 0.3573556 | 0.55943 |
| ENSG00000092470 | WDR76 | 736.3206703 | -0.288385393 | 0.453111056 | -0.781403143 | 0.43456542 | 0.633489 |
| ENSG00000079974 | RABL2B | 1001.642357 | -0.288478998 | 0.180942812 | -1.639349075 | 0.10114058 | 0.23469 |
| ENSG00000183474 | GTF2H2C | 514.2721 | -0.2884894 | 0.1980428 | -1.508077 | 0.1315349 | 0.283205 |
| ENSG00000258515 | | 101.5208 | -0.2887344 | 0.3043758 | -1.048699 | 0.2943167 | 0.493381 |
| ENSG00000124795 | DEK | 8619.130574 | -0.288737627 | 0.311683607 | -1.04119786 | 0.29778373 | 0.497005 |
| ENSG00000237934 | | 11.69271 | -0.2887595 | 0.5272929 | -0.7232638 | 0.4695178 | 0.662979 |
| ENSG00000274227 | | 11.25715 | -0.288765 | 0.4921543 | -0.7454286 | 0.4560126 | 0.652018 |

| | | | | | | | |
|---|---|---|---|---|---|---|---|
| ENSG00000131828 | PDHA1 | 6417.705023 | -0.288789242 | 0.197045648 | -1.523347314 | 0.12767184 | 0.277066 |
| ENSG00000159346 | ADIPOR1 | 14090.97 | -0.2888083 | 0.1169893 | -2.418668 | 0.01557744 | 0.055561 |
| ENSG00000274356 | | 29.60657 | -0.2888146 | 0.4484218 | -0.7759406 | 0.437784 | 0.636296 |
| ENSG00000101442 | ACTR5 | 793.2690482 | -0.288818454 | 0.213988792 | -1.407939235 | 0.15914906 | 0.325114 |
| ENSG00000139278 | GLIPR1 | 29.93366 | -0.2888863 | 0.3932511 | -0.8478378 | 0.3965283 | 0.597631 |
| ENSG00000238317 | SNORD11 | 8.456941 | -0.2890292 | 0.5511112 | -0.7170865 | 0.4733207 | 0.666174 |
| ENSG00000172613 | RAD9A | 2444.485 | -0.2890417 | 0.224664 | -1.354345 | 0.1756265 | 0.348018 |
| ENSG00000249889 | ALG1L11P | 2.340915 | -0.2890423 | 0.7115567 | -0.7722567 | 0.4399624 | 0.638344 |
| ENSG00000184277 | TM2D3 | 2297.159 | -0.2890453 | 0.1549489 | -1.932265 | 0.05332683 | 0.145852 |
| ENSG00000231607 | DLEU2 | 133.6915 | -0.2891114 | 0.2765569 | -1.121599 | 0.2620328 | 0.456791 |
| ENSG00000125901 | MRPS26 | 4125.953975 | -0.289253167 | 0.195355559 | -1.542111937 | 0.1230464 | 0.270161 |
| ENSG00000258569 | | 0.9686636 | -0.2892612 | 0.8573972 | -1.482393 | 0.1382357 | NA |
| ENSG00000224965 | LINC02586 | 0.9919985 | -0.2894181 | 0.845774 | -1.423058 | 0.1547194 | NA |
| ENSG00000162645 | GBP2 | 402.3632 | -0.2894418 | 0.3974629 | -0.8505797 | 0.3950029 | 0.596734 |
| ENSG00000284731 | | 4.310813 | -0.2894728 | 0.6998088 | -0.7571639 | 0.4489517 | 0.645416 |
| ENSG00000109501 | WFS1 | 2061.172448 | -0.289511759 | 0.261059408 | -1.17780067 | 0.23887608 | 0.428802 |
| ENSG00000220205 | VAMP2 | 1472.116 | -0.2897001 | 0.2254566 | -1.347653 | 0.1777699 | 0.351104 |
| ENSG00000279370 | | 0.8826012 | -0.2897407 | 0.8395523 | -1.0741 | 0.2827778 | NA |
| ENSG00000060642 | PIGV | 924.0565195 | -0.289758838 | 0.136616929 | -2.15480372 | 0.0311772 | 0.096634 |
| ENSG00000164161 | HHIP | 5.249855 | -0.2897712 | 0.6352428 | -0.7153272 | 0.4744069 | 0.667117 |
| ENSG00000105447 | GRWD1 | 3529.057749 | -0.289863353 | 0.238324638 | -1.280866126 | 0.20024069 | 0.380666 |
| ENSG00000040487 | SLC66A1 | 987.7375629 | -0.289894444 | 0.162489084 | -1.826950965 | 0.06770714 | 0.174171 |
| ENSG00000125459 | MSTO1 | 1246.068091 | -0.289961994 | 0.172500837 | -1.722668677 | 0.08494846 | 0.206439 |
| ENSG00000127399 | LRRC61 | 782.4468489 | -0.290024265 | 0.222116456 | -1.360508653 | 0.17366902 | 0.345298 |
| ENSG00000087086 | FTL | 19055.91304 | -0.290031163 | 0.22274518 | -1.362453907 | 0.17305468 | 0.344474 |
| ENSG00000267470 | ZNF571-AS1 | 53.52174 | -0.2900363 | 0.3043761 | -1.038207 | 0.2991737 | 0.498385 |
| ENSG00000252759 | | 4.124611 | -0.2900444 | 0.6280363 | -0.7114547 | 0.4768025 | 0.669105 |
| ENSG00000217702 | | 9.593941 | -0.2900636 | 0.5325643 | -0.729436 | 0.465735 | 0.659672 |
| ENSG00000103876 | FAH | 1335.282994 | -0.290088323 | 0.269637507 | -1.170448112 | 0.24182068 | 0.432364 |
| ENSG00000241416 | KRT8P13 | 156.9573 | -0.2900899 | 0.2066649 | -1.459848 | 0.1443317 | 0.302627 |
| ENSG00000178297 | TMPRSS9 | 47.07262 | -0.2900917 | 0.3788909 | -0.874562 | 0.3818123 | 0.584084 |
| ENSG00000116213 | WRAP73 | 1892.210954 | -0.290167474 | 0.144641177 | -2.081764347 | 0.037364 | 0.110918 |
| ENSG00000224061 | SRSF3P4 | 2.17234 | -0.2901982 | 0.7278762 | -0.7942638 | 0.4270419 | 0.626613 |

| | | | | | | | |
|---|---|---|---|---|---|---|---|
| ENSG00000248787 | | 15.17353 | -0.2902108 | 0.4237042 | -0.8120131 | 0.4167841 | 0.61667 |
| ENSG00000155256 | ZFYVE27 | 3405.127 | -0.2902577 | 0.1423034 | -2.105995 | 0.03520477 | 0.106052 |
| ENSG00000087586 | AURKA | 5460.3138 | -0.290296973 | 0.397126448 | -0.864034361 | 0.38756902 | 0.5898 |
| ENSG00000160688 | FLAD1 | 3935.774 | -0.2905953 | 0.154613 | -1.90425 | 0.05687765 | 0.152799 |
| ENSG00000020922 | MRE11 | 1424.639472 | -0.29069699 | 0.227114695 | -1.339410197 | 0.18043717 | 0.354746 |
| ENSG00000255815 | KRT8P11 | 68.17887 | -0.2908196 | 0.2934446 | -1.072906 | 0.2833135 | 0.481141 |
| ENSG00000249965 | CDC42P4 | 2.56265 | -0.2910577 | 0.7461573 | -0.8265616 | 0.4084856 | 0.60914 |
| ENSG00000078674 | PCM1 | 5870.547856 | -0.291103357 | 0.136973646 | -2.156923962 | 0.03101159 | 0.096278 |
| ENSG00000244052 | RPL5P24 | 5.191556 | -0.2911321 | 0.6082231 | -0.7166704 | 0.4735775 | 0.666374 |
| ENSG00000131148 | EMC8 | 2848.071636 | -0.291189297 | 0.240742181 | -1.26594378 | 0.20553321 | 0.387511 |
| ENSG00000173960 | UBXN2A | 1566.081 | -0.2912704 | 0.124384 | -2.354251 | 0.01856009 | 0.063967 |
| ENSG00000120451 | SNX19 | 3297.646428 | -0.291274843 | 0.213878253 | -1.424383419 | 0.15433551 | 0.317822 |
| ENSG00000115129 | TP53I3 | 1352.800519 | -0.291275825 | 0.158341089 | -1.848180428 | 0.06457625 | 0.167901 |
| ENSG00000165819 | METTL3 | 3711.276 | -0.2912783 | 0.18282 | -1.631878 | 0.1027051 | 0.237243 |
| ENSG00000220875 | H3C9P | 1.03059 | -0.2912839 | 0.783812 | -0.9276735 | 0.353577 | NA |
| ENSG00000113643 | RARS1 | 5796.858105 | -0.291305685 | 0.168659698 | -1.778274298 | 0.07535882 | 0.188745 |
| ENSG00000228809 | | 5.504415 | -0.2913542 | 0.6149226 | -0.717303 | 0.4731871 | 0.666087 |
| ENSG00000283341 | | 56.87419 | -0.2913838 | 0.5892286 | -0.7201175 | 0.4714527 | 0.664722 |
| ENSG00000264573 | RN7SL15P | 4.080308 | -0.2914598 | 0.6915504 | -0.7421883 | 0.4579732 | 0.653696 |
| ENSG00000251720 | RNU7-123P | 1.640178 | -0.2914617 | 0.7380597 | -0.8206579 | 0.4118412 | 0.612186 |
| ENSG00000137413 | TAF8 | 1143.077333 | -0.291462373 | 0.142460762 | -2.083854991 | 0.03717336 | 0.110476 |
| ENSG00000259692 | | 74.24896 | -0.2914701 | 0.3354573 | -0.9600125 | 0.3370489 | 0.53817 |
| ENSG00000283929 | MIR5010 | 6.238169 | -0.2915951 | 0.5496897 | -0.7206097 | 0.4711497 | 0.664436 |
| ENSG00000237361 | TUSC8 | 3.878966 | -0.2916394 | 0.6419715 | -0.7322972 | 0.4639872 | 0.658076 |
| ENSG00000236577 | SNRPGP14 | 2.126253 | -0.2917204 | 0.737476 | -0.8042594 | 0.4212472 | 0.62106 |
| ENSG00000198382 | UVRAG | 1782.8 | -0.2917397 | 0.1164332 | -2.539095 | 0.01111397 | 0.042378 |
| ENSG00000129988 | LBP | 6.02463947 | -0.291747951 | 0.588160079 | -0.726176149 | 0.46773079 | 0.661299 |
| ENSG00000076344 | RGS11 | 203.2763184 | -0.29177776 | 0.413083668 | -0.835833945 | 0.40324833 | 0.604179 |
| ENSG00000166986 | MARS1 | 14323.63 | -0.2917909 | 0.2949457 | -1.073092 | 0.2832298 | 0.481054 |
| ENSG00000124839 | RAB17 | 4365.112623 | -0.291904112 | 0.321389433 | -0.977767166 | 0.32818949 | 0.529395 |
| ENSG00000213261 | EEF1B2P6 | 1496.352 | -0.2919592 | 0.1705217 | -1.76856 | 0.0769673 | 0.191659 |
| ENSG00000171863 | RPS7 | 11387.3 | -0.2919937 | 0.1694705 | -1.770739 | 0.07660411 | 0.190964 |
| ENSG00000226711 | FAM66C | 121.8611 | -0.2920009 | 0.2751915 | -1.139488 | 0.2544998 | 0.447534 |

| | | | | | | | |
|---|---|---|---|---|---|---|---|
| ENSG00000230837 | RPL31P2 | 15.50957 | -0.2921248 | 0.4747609 | -0.7537479 | 0.4510006 | 0.64722 |
| ENSG00000278599 | TBC1D3E | 5.641893 | -0.292278 | 0.6138158 | -0.7139617 | 0.4752509 | 0.667714 |
| ENSG00000241313 | WWTR1-AS1 | 35.24629 | -0.2923229 | 0.4495566 | -0.7909871 | 0.4289515 | 0.628131 |
| ENSG00000132670 | PTPRA | 3699.997697 | -0.292339816 | 0.099233725 | -2.97378184 | 0.00294154 | 0.014361 |
| ENSG00000267922 | | 9.795764 | -0.2925161 | 0.7774997 | -0.906405 | 0.3647215 | 0.566651 |
| ENSG00000225531 | | 49.75526 | -0.2926218 | 0.3118116 | -1.023336 | 0.3061488 | 0.50607 |
| ENSG00000107874 | CUEDC2 | 2661.130259 | -0.292645877 | 0.202793229 | -1.504188175 | 0.13253292 | 0.284846 |
| ENSG00000068912 | ERLEC1 | 3332.918432 | -0.2926478 | 0.124875678 | -2.37473367 | 0.01756162 | 0.061211 |
| ENSG00000259581 | TYRO3P | 10.03982 | -0.2927354 | 0.5460063 | -0.7217846 | 0.4704269 | 0.663929 |
| ENSG00000219693 | FGF7P8 | 1.97992 | -0.2928963 | 0.7348963 | -0.8157756 | 0.4146285 | 0.614649 |
| ENSG00000170802 | FOXN2 | 1046.449 | -0.2929041 | 0.2514071 | -1.236043 | 0.2164425 | 0.401731 |
| ENSG00000244703 | CD46P1 | 1.928803 | -0.2929542 | 0.7800025 | -0.925399 | 0.3547584 | 0.556672 |
| ENSG00000188295 | ZNF669 | 833.6855 | -0.2929685 | 0.14786 | -2.020157 | 0.04336712 | 0.124531 |
| ENSG00000167193 | CRK | 4688.156 | -0.292971 | 0.1130389 | -2.625953 | 0.00864068 | 0.034611 |
| ENSG00000131373 | HACL1 | 1371.74019 | -0.293098472 | 0.218069795 | -1.405219908 | 0.1599559 | 0.32616 |
| ENSG00000139496 | NUP58 | 5247.772 | -0.2932677 | 0.180469 | -1.69897 | 0.08932482 | 0.214465 |
| ENSG00000152684 | PELO | 1414.939 | -0.2934049 | 0.1206668 | -2.463265 | 0.0137678 | 0.050268 |
| ENSG00000075131 | TIPIN | 673.4734918 | -0.293643521 | 0.364421225 | -0.916953908 | 0.35916679 | 0.561104 |
| ENSG00000159958 | TNFRSF13C | 28.71085 | -0.2936897 | 0.5625697 | -0.7275698 | 0.466877 | 0.660651 |
| ENSG00000264538 | SUZ12P1 | 663.4697 | -0.2937469 | 0.1979781 | -1.535888 | 0.1245659 | 0.272484 |
| ENSG00000232059 | | 9.460162 | -0.2938221 | 0.7126413 | -0.7727591 | 0.439665 | 0.638122 |
| ENSG00000285852 | | 13.56928 | -0.2938604 | 0.5660361 | -0.7291548 | 0.465907 | 0.659777 |
| ENSG00000263232 | ATP5F1AP3 | 12.71829 | -0.2938709 | 0.5604818 | -0.7315555 | 0.4644399 | 0.658469 |
| ENSG00000132912 | DCTN4 | 5017.27995 | -0.293879563 | 0.110318822 | -2.706217694 | 0.00680544 | 0.028464 |
| ENSG00000108349 | CASC3 | 5878.126834 | -0.29390874 | 0.125472996 | -2.383668611 | 0.01714103 | 0.059979 |
| ENSG00000263266 | RPS7P1 | 9504.413 | -0.2939512 | 0.1788742 | -1.689155 | 0.09118977 | 0.217684 |
| ENSG00000255220 | DDX18P5 | 37.46339 | -0.293983 | 0.3352781 | -0.9732976 | 0.3304054 | 0.531517 |
| ENSG00000132622 | HSPA12B | 48.38316762 | -0.294008517 | 0.430779801 | -0.816116329 | 0.41443361 | 0.614598 |
| ENSG00000213771 | KRT8P37 | 34.47656 | -0.2940163 | 0.3998805 | -0.8563236 | 0.3918188 | 0.593615 |
| ENSG00000268866 | | 1.150042 | -0.2940326 | 0.8061257 | -1.039275 | 0.2986768 | NA |
| ENSG00000099326 | MZF1 | 1911.460743 | -0.294143596 | 0.242553655 | -1.287525337 | 0.19791124 | 0.37796 |
| ENSG00000101278 | RPS10P5 | 18.43446193 | -0.294194623 | 0.396110557 | -0.858877242 | 0.39040825 | 0.592289 |
| ENSG00000231969 | MMADHC-DT | 10.43185 | -0.2942403 | 0.4759722 | -0.7645712 | 0.4445269 | 0.641854 |

| | | | | | | | |
|---|---|---|---|---|---|---|---|
| ENSG00000117500 | TMED5 | 2217.348085 | -0.29435733 | 0.149936683 | -2.002188346 | 0.04526448 | 0.128582 |
| ENSG00000261476 | | 6.956341 | -0.2943597 | 0.5402088 | -0.7360417 | 0.4617053 | 0.656413 |
| ENSG00000279817 | | 2.260502 | -0.2943685 | 0.666414 | -0.752413 | 0.4518027 | 0.648155 |
| ENSG00000243257 | MTCO2P29 | 1.332794 | -0.2944438 | 0.8074489 | -1.041687 | 0.2975567 | NA |
| ENSG00000258608 | DNAJC19P9 | 552.8605 | -0.2946429 | 0.1857499 | -1.634536 | 0.1021465 | 0.23624 |
| ENSG00000213026 | CFL1P4 | 1.940425 | -0.2947096 | 0.7837942 | -0.9401672 | 0.3471318 | 0.548669 |
| ENSG00000164062 | APEH | 7204.693 | -0.2947114 | 0.2053315 | -1.491356 | 0.135868 | 0.289578 |
| ENSG00000100083 | GGA1 | 4372.872078 | -0.29472351 | 0.133573356 | -2.239933169 | 0.02509526 | 0.081351 |
| ENSG00000225385 | | 9.898322 | -0.2947374 | 0.5471024 | -0.7324586 | 0.4638887 | 0.658003 |
| ENSG00000176087 | SLC35A4 | 5877.933 | -0.2947537 | 0.1748188 | -1.735389 | 0.0826719 | 0.202462 |
| ENSG00000097021 | ACOT7 | 1947.517076 | -0.294771096 | 0.372159283 | -0.8937424 | 0.37145973 | 0.573564 |
| ENSG00000108370 | RGS9 | 86.789634 | -0.294802063 | 0.455541404 | -0.787330625 | 0.43108835 | 0.630241 |
| ENSG00000138678 | GPAT3 | 212.5919486 | -0.294848536 | 0.448842228 | -0.802342935 | 0.42235461 | 0.622382 |
| ENSG00000243678 | NME2 | 354.6643 | -0.2948695 | 0.172964 | -1.754747 | 0.07930262 | 0.195963 |
| ENSG00000234297 | PSIP1P1 | 96.91614 | -0.2948833 | 0.2890583 | -1.096068 | 0.2730491 | 0.469678 |
| ENSG00000282608 | ADORA3 | 1.768521 | -0.2948955 | 0.8638847 | -1.860894 | 0.06275916 | 0.164443 |
| ENSG00000286681 | | 198.3565 | -0.2949283 | 0.2234546 | -1.387191 | 0.1653836 | 0.333944 |
| ENSG00000111785 | RIC8B | 664.4830541 | -0.294951436 | 0.209814306 | -1.46664124 | 0.14247368 | 0.299927 |
| ENSG00000130520 | LSM4 | 4258.529134 | -0.294952181 | 0.299702528 | -1.070690041 | 0.28430882 | 0.48224 |
| ENSG00000230433 | | 1.114269 | -0.2949543 | 0.8326272 | -0.9250773 | 0.3549257 | NA |
| ENSG00000138381 | ASNSD1 | 1280.673423 | -0.294978172 | 0.193774722 | -1.582495937 | 0.1135364 | 0.254844 |
| ENSG00000113575 | PPP2CA | 8501.113076 | -0.295101474 | 0.167813309 | -1.81226463 | 0.06994532 | 0.178451 |
| ENSG00000101247 | NDUFAF5 | 835.8120663 | -0.295126471 | 0.189422537 | -1.610407895 | 0.10730884 | 0.244897 |
| ENSG00000212232 | SNORD17 | 31.71452 | -0.2952576 | 0.4881549 | -0.7692414 | 0.44175 | 0.639541 |
| ENSG00000173653 | RCE1 | 2390.586 | -0.2952743 | 0.1705443 | -1.781046 | 0.07490493 | 0.18791 |
| ENSG00000237357 | | 2.420015 | -0.295281 | 0.718668 | -0.7947492 | 0.4267594 | 0.62644 |
| ENSG00000134864 | GGACT | 210.3987896 | -0.295451188 | 0.244083895 | -1.277865837 | 0.20129674 | 0.382045 |
| ENSG00000269190 | FBXO17 | 676.804 | -0.295531 | 0.2608096 | -1.203572 | 0.2287551 | 0.416916 |
| ENSG00000214650 | | 4.998664 | -0.2955904 | 0.6747865 | -0.7478728 | 0.4545369 | 0.650503 |
| ENSG00000183431 | SF3A3 | 6084.973 | -0.2955988 | 0.1805668 | -1.686507 | 0.09169816 | 0.218505 |
| ENSG00000250321 | | 84.67435 | -0.2957886 | 0.4191613 | -0.8354454 | 0.403467 | 0.604287 |
| ENSG00000236463 | LINC00427 | 0.9602487 | -0.2957948 | 0.8404796 | -1.369918 | 0.1707125 | NA |
| ENSG00000165669 | FAM204A | 1898.312 | -0.2958867 | 0.1523098 | -1.995667 | 0.04597014 | 0.130059 |

| | | | | | | | |
|---|---|---|---|---|---|---|---|
| ENSG00000279089 | | 213.1354 | -0.2958933 | 0.2308566 | -1.343939 | 0.1789682 | 0.35289 |
| ENSG00000248206 | | 5.590853 | -0.2959632 | 0.6473351 | -0.7384804 | 0.4602226 | 0.655608 |
| ENSG00000287963 | | 1.93621 | -0.2960079 | 0.7373877 | -0.8216269 | 0.4112893 | 0.611571 |
| ENSG00000177675 | CD163L1 | 50.20174 | -0.2961207 | 0.4772359 | -0.7652184 | 0.4441415 | 0.641576 |
| ENSG00000275401 | | 6.42199 | -0.2962782 | 0.5743162 | -0.732107 | 0.4641033 | 0.658202 |
| ENSG00000179163 | FUCA1 | 1156.139 | -0.2963814 | 0.1706818 | -1.783935 | 0.07443425 | 0.187117 |
| ENSG00000278133 | | 357.5884 | -0.2964632 | 0.187006 | -1.631514 | 0.1027819 | 0.2374 |
| ENSG00000130703 | OSBPL2 | 6361.149842 | -0.296470062 | 0.166605229 | -1.83033169 | 0.06720036 | 0.173143 |
| ENSG00000236819 | LINC01563 | 1.028151 | -0.2965858 | 0.805464 | -1.066198 | 0.286334 | NA |
| ENSG00000103194 | USP10 | 9000.589054 | -0.296699143 | 0.135010968 | -2.262987585 | 0.02363646 | 0.077598 |
| ENSG00000273382 | TMEM167B-DT | 335.0788 | -0.2967134 | 0.1937711 | -1.589231 | 0.1120083 | 0.252606 |
| ENSG00000249532 | MIR302CHG | 3.676122 | -0.2967267 | 0.627011 | -0.7319837 | 0.4641785 | 0.658239 |
| ENSG00000175931 | UBE2O | 5168.358 | -0.2967404 | 0.1418218 | -2.140427 | 0.03232031 | 0.099343 |
| ENSG00000258378 | | 2.080421 | -0.2967451 | 0.7297413 | -0.7994495 | 0.4240298 | 0.623976 |
| ENSG00000285803 | | 32.73183 | -0.2967502 | 0.373771 | -0.9047724 | 0.365586 | 0.567562 |
| ENSG00000118690 | ARMC2 | 100.1784965 | -0.296837003 | 0.24384845 | -1.278217717 | 0.20117267 | 0.381866 |
| ENSG00000124198 | ARFGEF2 | 8944.433789 | -0.29688966 | 0.243704027 | -1.286663552 | 0.19821158 | 0.378162 |
| ENSG00000170604 | IRF2BP1 | 2004.395 | -0.2969445 | 0.1728606 | -1.7731 | 0.0762122 | 0.190255 |
| ENSG00000226696 | LENG8-AS1 | 192.7956 | -0.296966 | 0.3371971 | -0.9810965 | 0.3265451 | 0.527864 |
| ENSG00000236670 | KRT18P5 | 10.47002 | -0.2970251 | 0.6009573 | -0.7286935 | 0.4661892 | 0.659998 |
| ENSG00000203780 | FANK1 | 229.9325 | -0.2971168 | 0.2615725 | -1.206346 | 0.227684 | 0.415591 |
| ENSG00000072110 | ACTN1 | 12495.87793 | -0.29715319 | 0.153196396 | -1.972547048 | 0.04854719 | 0.13546 |
| ENSG00000277459 | | 19.24666 | -0.297201 | 0.5002718 | -0.7612583 | 0.4465028 | 0.643407 |
| ENSG00000213339 | QTRT1 | 3158.038 | -0.2972321 | 0.1722852 | -1.775765 | 0.07577168 | 0.189529 |
| ENSG00000133678 | TMEM254 | 1778.585221 | -0.297270574 | 0.179147232 | -1.710561606 | 0.08716207 | 0.210437 |
| ENSG00000229814 | RPL35AP21 | 12.55167 | -0.2972865 | 0.5125783 | -0.7491111 | 0.4537903 | 0.649942 |
| ENSG00000167747 | C19orf48 | 8595.23 | -0.2972991 | 0.2259548 | -1.377709 | 0.1682932 | 0.33805 |
| ENSG00000168778 | TCTN2 | 303.7194 | -0.297319 | 0.2415343 | -1.295165 | 0.1952632 | 0.374662 |
| ENSG00000266232 | MIR3178 | 2.748589 | -0.2973743 | 0.6895412 | -0.7618142 | 0.4461709 | 0.643188 |
| ENSG00000182866 | LCK | 4.274897 | -0.2973946 | 0.6067961 | -0.7371694 | 0.4610193 | 0.655964 |
| ENSG00000114942 | EEF1B2 | 10952.70508 | -0.297530646 | 0.16884487 | -1.806830695 | 0.07078868 | 0.180167 |
| ENSG00000136718 | IMP4 | 5511.715352 | -0.297556606 | 0.210022102 | -1.476631247 | 0.13977451 | 0.295816 |
| ENSG00000185880 | TRIM69 | 597.0424 | -0.2975618 | 0.2957113 | -1.086843 | 0.2771061 | 0.474017 |

| | | | | | | | |
|---|---|---|---|---|---|---|---|
| ENSG00000143106 | PSMA5 | 10674.71 | -0.2976282 | 0.197544 | -1.557945 | 0.1192462 | 0.263999 |
| ENSG00000171657 | GPR82 | 1.050265 | -0.2976474 | 0.7708909 | -0.8942894 | 0.371167 | NA |
| ENSG00000138443 | ABI2 | 4099.233956 | -0.297653235 | 0.12420379 | -2.405518717 | 0.01614953 | 0.057142 |
| ENSG00000155463 | OXA1L | 9190.528 | -0.2976685 | 0.1064221 | -2.829583 | 0.00466087 | 0.020946 |
| ENSG00000072210 | ALDH3A2 | 5532.364326 | -0.297699413 | 0.162437318 | -1.905431324 | 0.05672403 | 0.152562 |
| ENSG00000162434 | JAK1 | 9269.367 | -0.2977832 | 0.190576 | -1.598568 | 0.1099166 | 0.249285 |
| ENSG00000163646 | CLRN1 | 1.169214 | -0.2978015 | 0.8062431 | -1.064596 | 0.2870588 | NA |
| ENSG00000227782 | | 33.91366 | -0.2978662 | 0.3847455 | -0.8914332 | 0.3726968 | 0.574906 |
| ENSG00000262681 | | 0.8774907 | -0.2979277 | 0.8520937 | -1.291759 | 0.1964406 | NA |
| ENSG00000160256 | FAM207A | 1471.178 | -0.2980899 | 0.1690113 | -1.798316 | 0.07212697 | 0.182613 |
| ENSG00000137714 | FDX1 | 922.5311652 | -0.298266379 | 0.199286863 | -1.551291683 | 0.1208318 | 0.26663 |
| ENSG00000196323 | ZBTB44 | 1898.61 | -0.2982803 | 0.1546455 | -1.969416 | 0.04890534 | 0.136216 |
| ENSG00000068650 | ATP11A | 3089.396235 | -0.298307633 | 0.265501039 | -1.202354589 | 0.22922618 | 0.417287 |
| ENSG00000175110 | MRPS22 | 2250.347 | -0.2983658 | 0.1844715 | -1.66925 | 0.09506783 | 0.224321 |
| ENSG00000162139 | NEU3 | 1392.423 | -0.2984185 | 0.2011694 | -1.538404 | 0.1239499 | 0.271619 |
| ENSG00000143537 | ADAM15 | 11820.76 | -0.2985582 | 0.1768627 | -1.745046 | 0.08097688 | 0.199206 |
| ENSG00000158683 | PKD1L1 | 14.92595 | -0.2986644 | 0.5066693 | -0.7588431 | 0.4479464 | 0.644613 |
| ENSG00000167984 | NLRC3 | 78.00479 | -0.2986898 | 0.3612669 | -0.9364886 | 0.3490217 | 0.550668 |
| ENSG00000148187 | MRRF | 3313.747 | -0.2987099 | 0.1418265 | -2.170637 | 0.0299586 | 0.093577 |
| ENSG00000215630 | GUSBP9 | 25.46184 | -0.2987434 | 0.4335063 | -0.8283474 | 0.4074738 | 0.608109 |
| ENSG00000213585 | VDAC1 | 12959.28 | -0.298776 | 0.1780378 | -1.727058 | 0.08415725 | 0.205022 |
| ENSG00000258891 | | 3.618103 | -0.2988469 | 0.5928662 | -0.7375951 | 0.4607605 | 0.65574 |
| ENSG00000113583 | C5orf15 | 3932.90506 | -0.298922331 | 0.171435757 | -1.810883038 | 0.07015896 | 0.178909 |
| ENSG00000268942 | CKS1BP3 | 271.282 | -0.2989299 | 0.3810392 | -0.9006728 | 0.3677623 | 0.569908 |
| ENSG00000259585 | RBM17P4 | 11.2443 | -0.2989547 | 0.5152234 | -0.7571155 | 0.4489807 | 0.645416 |
| ENSG00000231128 | | 10.90105 | -0.2989992 | 0.5412364 | -0.7476611 | 0.4546646 | 0.650616 |
| ENSG00000228331 | RPL17P43 | 0.8024273 | -0.2993797 | 0.8433807 | -1.089568 | 0.2759037 | NA |
| ENSG00000287543 | | 9.501826 | -0.2994367 | 0.6379624 | -0.7367364 | 0.4612826 | 0.656181 |
| ENSG00000242571 | RPL21P11 | 6.487696 | -0.2995292 | 0.5957776 | -0.7397653 | 0.4594424 | 0.654983 |
| ENSG00000276188 | | 2.753333 | -0.2995377 | 0.7092951 | -0.7922309 | 0.4282261 | 0.627588 |
| ENSG00000266950 | | 1.424664 | -0.2995608 | 0.7686427 | -0.882921 | 0.3772789 | NA |
| ENSG00000196305 | IARS1 | 15408.88 | -0.2995798 | 0.2955819 | -1.099792 | 0.2714229 | 0.467786 |
| ENSG00000151923 | TIAL1 | 5716.751 | -0.2996046 | 0.130293 | -2.373189 | 0.01763525 | 0.061406 |

| | | | | | | | |
|---|---|---|---|---|---|---|---|
| ENSG00000155506 | LARP1 | 21660.21 | -0.2996128 | 0.1830412 | -1.692517 | 0.09054742 | 0.216579 |
| ENSG00000100065 | CARD10 | 462.84606 | -0.299662633 | 0.365315034 | -0.931388083 | 0.35165285 | 0.553368 |
| ENSG00000215187 | FAM166B | 9.460203 | -0.2997425 | 0.6563948 | -0.7602013 | 0.4471343 | 0.643898 |
| ENSG00000225872 | LINC01529 | 37.59836 | -0.2997444 | 0.4554797 | -0.8068868 | 0.4197317 | 0.619777 |
| ENSG00000104886 | PLEKHJ1 | 2550.637784 | -0.299763685 | 0.182965486 | -1.688493367 | 0.09131656 | 0.217869 |
| ENSG00000237353 | PATE4 | 0.809526 | -0.2997726 | 0.8213642 | -1.135533 | 0.2561521 | NA |
| ENSG00000223023 | | 0.9221521 | -0.2998184 | 0.8172092 | -1.125281 | 0.26047 | NA |
| ENSG00000143970 | ASXL2 | 3028.02 | -0.29986 | 0.3113741 | -1.054398 | 0.2917006 | 0.490456 |
| ENSG00000181938 | GINS3 | 435.8973 | -0.2999446 | 0.425854 | -0.844409 | 0.3984409 | 0.599478 |
| ENSG00000169118 | CSNK1G1 | 2203.086 | -0.300019 | 0.1539159 | -1.995554 | 0.04598252 | 0.13008 |
| ENSG00000276174 | | 0.8384099 | -0.3000961 | 0.8432288 | -1.095321 | 0.2733761 | NA |
| ENSG00000261386 | | 17.72281 | -0.3001031 | 0.3856975 | -0.8922907 | 0.3722371 | 0.57443 |
| ENSG00000174917 | MICOS13 | 1715.15 | -0.3001184 | 0.1984119 | -1.56624 | 0.1172924 | 0.260913 |
| ENSG00000072501 | SMC1A | 8365.58711 | -0.30012791 | 0.268606814 | -1.197362445 | 0.23116532 | 0.419681 |
| ENSG00000204529 | GUCY2EP | 3.563489 | -0.3001506 | 0.7892692 | -0.9872588 | 0.3235158 | 0.524403 |
| ENSG00000270647 | TAF15 | 7705.265 | -0.3001698 | 0.1676512 | -1.850148 | 0.06429224 | 0.167337 |
| ENSG00000101199 | ARFGAP1 | 9409.375219 | -0.300199839 | 0.119380014 | -2.543661244 | 0.01096974 | 0.041918 |
| ENSG00000252755 | RNU6-703P | 2.974613 | -0.3002563 | 0.7158854 | -0.8010424 | 0.4231071 | 0.623086 |
| ENSG00000096872 | IFT74 | 531.5009508 | -0.300331442 | 0.154801697 | -1.985608747 | 0.04707678 | 0.132343 |
| ENSG00000267838 | | 17.76732 | -0.3003653 | 0.5797957 | -0.739125 | 0.4598311 | 0.655274 |
| ENSG00000125967 | NECAB3 | 2553.094411 | -0.300418632 | 0.152804374 | -2.016957304 | 0.04369997 | 0.12525 |
| ENSG00000125970 | RALY | 6796.110724 | -0.300490994 | 0.193322927 | -1.607396627 | 0.10796739 | 0.245998 |
| ENSG00000123552 | USP45 | 914.1317683 | -0.300493696 | 0.168954329 | -1.823230816 | 0.06826844 | 0.175232 |
| ENSG00000238197 | PAXBP1-AS1 | 240.1192 | -0.3005019 | 0.2510793 | -1.265446 | 0.2057114 | 0.387737 |
| ENSG00000178301 | AQP11 | 230.4668 | -0.3005792 | 0.1962269 | -1.586484 | 0.1126296 | 0.253469 |
| ENSG00000077312 | SNRPA | 3276.512857 | -0.300610051 | 0.28661502 | -1.138250876 | 0.25501574 | 0.448264 |
| ENSG00000270878 | GCATP1 | 27.46988 | -0.3007092 | 0.5014871 | -0.7708584 | 0.4407908 | 0.638744 |
| ENSG00000279253 | | 42.91869 | -0.3007303 | 0.4717565 | -0.7912797 | 0.4287808 | 0.628054 |
| ENSG00000266401 | | 1.566886 | -0.3007693 | 0.8343244 | -0.9426833 | 0.3458429 | NA |
| ENSG00000225721 | | 6.328881 | -0.3007932 | 0.6002217 | -0.7433646 | 0.457261 | 0.65317 |
| ENSG00000214027 | ARPC3P5 | 28.33229 | -0.3008113 | 0.4549487 | -0.8088256 | 0.4186155 | 0.618415 |
| ENSG00000256050 | NA | 1.19564 | -0.3008839 | 0.7769211 | -0.9325051 | 0.3510756 | NA |
| ENSG00000105223 | PLD3 | 5106.790711 | -0.300982034 | 0.199371994 | -1.564988963 | 0.1175855 | 0.26139 |

| | | | | | | | |
|---|---|---|---|---|---|---|---|
| ENSG00000196116 | TDRD7 | 1493.991 | -0.3009964 | 0.1643453 | -1.887914 | 0.0590375 | 0.157218 |
| ENSG00000119392 | GLE1 | 3998.243673 | -0.301066567 | 0.184171968 | -1.685953345 | 0.09180479 | 0.218622 |
| ENSG00000272983 | | 20.95671 | -0.3012167 | 0.4680468 | -0.7942935 | 0.4270246 | 0.626613 |
| ENSG00000100395 | L3MBTL2 | 1466.990275 | -0.301224102 | 0.181007998 | -1.712017061 | 0.08689352 | 0.210013 |
| ENSG00000096401 | CDC5L | 3837.714758 | -0.301343994 | 0.160474631 | -1.921633157 | 0.05465193 | 0.148471 |
| ENSG00000065518 | NDUFB4 | 3572.162303 | -0.301357007 | 0.142554599 | -2.160102269 | 0.03076475 | 0.095657 |
| ENSG00000162639 | HENMT1 | 2525.097 | -0.3013607 | 0.212767 | -1.4627 | 0.1435494 | 0.301451 |
| ENSG00000167333 | TRIM68 | 218.1533 | -0.3014045 | 0.2126884 | -1.474863 | 0.1402493 | 0.296436 |
| ENSG00000279419 | | 3.462185 | -0.3014624 | 0.7000331 | -0.7697335 | 0.441458 | 0.639258 |
| ENSG00000110768 | GTF2H1 | 3877.015088 | -0.301488 | 0.141935307 | -2.219958402 | 0.02642159 | 0.0847 |
| ENSG00000277728 | | 23.13982 | -0.3015836 | 0.4284976 | -0.841543 | 0.4000438 | 0.601039 |
| ENSG00000258017 | | 10.05358 | -0.3016622 | 0.4920096 | -0.7786624 | 0.4361786 | 0.634829 |
| ENSG00000229689 | | 426.7938 | -0.3016717 | 0.3585374 | -0.9375888 | 0.3484558 | 0.550162 |
| ENSG00000236565 | HNRNPA3P5 | 1344.173 | -0.30201 | 0.1972821 | -1.586611 | 0.1126008 | 0.253447 |
| ENSG00000278002 | | 4.112153 | -0.3020617 | 0.6966169 | -0.7871234 | 0.4312096 | 0.630292 |
| ENSG00000249489 | GAPDHP70 | 13.36797 | -0.3021048 | 0.5560075 | -0.7520573 | 0.4520166 | 0.648231 |
| ENSG00000158859 | ADAMTS4 | 2.500927 | -0.3023993 | 0.6668722 | -0.7712962 | 0.4405314 | 0.638612 |
| ENSG00000039650 | PNKP | 2395.117259 | -0.302443762 | 0.161554153 | -1.914115143 | 0.05560545 | 0.150296 |
| ENSG00000223843 | EFCAB6-AS1 | 1.585133 | -0.3025107 | 0.8365511 | -0.9780243 | 0.3280623 | 0.529382 |
| ENSG00000159335 | PTMS | 12191.74 | -0.3025239 | 0.2200342 | -1.399259 | 0.1617352 | 0.328895 |
| ENSG00000110888 | CAPRIN2 | 535.4599321 | -0.302551996 | 0.264450581 | -1.22130722 | 0.22196972 | 0.408698 |
| ENSG00000214457 | RPL7P29 | 0.9748702 | -0.3025972 | 0.8518777 | -1.49567 | 0.1347397 | NA |
| ENSG00000125885 | MCM8 | 361.3951104 | -0.302745179 | 0.32110481 | -1.037765959 | 0.29937903 | 0.498616 |
| ENSG00000215349 | MRPL3P1 | 140.8022 | -0.3027856 | 0.3271881 | -1.01453 | 0.31033 | 0.510993 |
| ENSG00000120217 | CD274 | 13.60390355 | -0.302868245 | 0.559836138 | -0.747885841 | 0.45452902 | 0.650503 |
| ENSG00000107560 | RAB11FIP2 | 679.7294249 | -0.302894056 | 0.184771532 | -1.690470121 | 0.09093805 | 0.21722 |
| ENSG00000254964 | | 10.2313 | -0.3029189 | 0.52971 | -0.7619702 | 0.4460778 | 0.643143 |
| ENSG00000203362 | POLH-AS1 | 13.57945 | -0.3031416 | 0.5589134 | -0.7549036 | 0.4503068 | 0.646539 |
| ENSG00000179094 | PER1 | 1397.277 | -0.3031568 | 0.260601 | -1.242224 | 0.214154 | 0.39898 |
| ENSG00000260360 | | 9.43631 | -0.3031861 | 0.491809 | -0.7808028 | 0.4349185 | 0.63376 |
| ENSG00000280023 | | 3.893191 | -0.3032105 | 0.7234462 | -0.8209982 | 0.4116473 | 0.612 |
| ENSG00000116704 | SLC35D1 | 1050.467114 | -0.303294302 | 0.222822274 | -1.422698418 | 0.1548236 | 0.318534 |
| ENSG00000268686 | | 5.022758 | -0.3033675 | 0.6800452 | -0.766889 | 0.4431475 | 0.640761 |

| | | | | | | | |
|---|---|---|---|---|---|---|---|
| ENSG00000230513 | THAP7-AS1 | 176.6358 | -0.303385 | 0.2483121 | -1.290423 | 0.1969037 | 0.376885 |
| ENSG00000220472 | | 1190.961 | -0.3034121 | 0.2258415 | -1.406869 | 0.1594661 | 0.325561 |
| ENSG00000187951 | | 560.2573 | -0.3035706 | 0.3614178 | -0.9555464 | 0.3393014 | 0.540624 |
| ENSG00000269933 | | 1.409943 | -0.3036289 | 0.7679503 | -0.9039119 | 0.3660421 | NA |
| ENSG00000257097 | CLIP1-AS1 | 1.362615 | -0.3037716 | 0.8159003 | -1.121488 | 0.2620803 | NA |
| ENSG00000196862 | RGPD4 | 5.33362 | -0.303804 | 0.585005 | -0.7495887 | 0.4535025 | 0.649659 |
| ENSG00000103150 | MLYCD | 1698.115371 | -0.303862227 | 0.145460778 | -2.129192246 | 0.03323836 | 0.101607 |
| ENSG00000183943 | PRKX | 1718.651 | -0.3039148 | 0.1962134 | -1.60362 | 0.1087978 | 0.247281 |
| ENSG00000101004 | NINL | 2426.12077 | -0.303934727 | 0.158299076 | -1.962059981 | 0.0497555 | 0.138021 |
| ENSG00000106609 | TMEM248 | 4597.910825 | -0.303953207 | 0.107560252 | -2.857194097 | 0.00427404 | 0.019473 |
| ENSG00000277288 | LINC02881 | 0.9433861 | -0.3040461 | 0.8382327 | -1.288932 | 0.1974218 | NA |
| ENSG00000272849 | | 98.22328 | -0.3040888 | 0.2960818 | -1.11225 | 0.2660308 | 0.461701 |
| ENSG00000118162 | KPTN | 547.7118369 | -0.304133276 | 0.241555753 | -1.328527811 | 0.1840038 | 0.359338 |
| ENSG00000218265 | RPS4XP7 | 12.54055 | -0.3041707 | 0.5050574 | -0.7739401 | 0.4389662 | 0.637421 |
| ENSG00000232882 | PHKA1P1 | 113.7281 | -0.3041983 | 0.3393924 | -0.990216 | 0.3220685 | 0.522855 |
| ENSG00000213860 | RPL21P75 | 19.95434 | -0.3042424 | 0.4813163 | -0.7932286 | 0.4276446 | 0.627062 |
| ENSG00000110200 | ANAPC15 | 532.4880647 | -0.304281815 | 0.293916372 | -1.127848759 | 0.2593838 | 0.453691 |
| ENSG00000011485 | PPP5C | 4415.497917 | -0.304296539 | 0.231986961 | -1.373006987 | 0.16975018 | 0.340128 |
| ENSG00000171634 | BPTF | 15998.22 | -0.3043259 | 0.203572 | -1.574174 | 0.1154472 | 0.258266 |
| ENSG00000245694 | CRNDE | 4072.525 | -0.3043377 | 0.2788223 | -1.174313 | 0.2402696 | 0.430547 |
| ENSG00000251670 | | 0.8025052 | -0.3043603 | 0.8177404 | -1.143913 | 0.2526597 | NA |
| ENSG00000099624 | ATP5F1D | 4432.913592 | -0.304420347 | 0.185196396 | -1.700910661 | 0.08895976 | 0.213887 |
| ENSG00000256028 | | 8.877641 | -0.304425 | 0.5187792 | -0.7691482 | 0.4418054 | 0.639586 |
| ENSG00000228330 | | 2.742867 | -0.3044958 | 0.6938838 | -0.7817563 | 0.4343578 | 0.633395 |
| ENSG00000173401 | GLIPR1L1 | 1.935267 | -0.3046266 | 0.8108934 | -1.090202 | 0.2756241 | 0.472486 |
| ENSG00000143951 | WDPCP | 317.51 | -0.3046301 | 0.2061191 | -1.528241 | 0.1264528 | 0.275168 |
| ENSG00000211450 | SELENOH | 299.512 | -0.3047038 | 0.1801608 | -1.743528 | 0.08124136 | 0.199713 |
| ENSG00000132481 | TRIM47 | 452.2579402 | -0.304789369 | 0.328945693 | -1.021589186 | 0.30697538 | 0.507121 |
| ENSG00000182272 | B4GALNT4 | 5067.578 | -0.3048022 | 0.189288 | -1.670329 | 0.09485425 | 0.223966 |
| ENSG00000179818 | PCBP1-AS1 | 933.0672 | -0.304855 | 0.2468851 | -1.305032 | 0.191882 | 0.370153 |
| ENSG00000269973 | | 178.1106 | -0.3049168 | 0.2393482 | -1.350458 | 0.176869 | 0.349906 |
| ENSG00000218896 | TUBB8P2 | 1.537564 | -0.3050046 | 0.7793134 | -0.9349493 | 0.3498144 | NA |
| ENSG00000248559 | | 6.496766 | -0.3050309 | 0.5805006 | -0.746745 | 0.4552175 | 0.651166 |

| | | | | | | | |
|---|---|---|---|---|---|---|---|
| ENSG00000183207 | RUVBL2 | 7440.19 | -0.3051572 | 0.2276941 | -1.405279 | 0.1599383 | 0.326149 |
| ENSG00000133112 | TPT1 | 43051.81385 | -0.305170464 | 0.149191191 | -2.09482683 | 0.03618639 | 0.108429 |
| ENSG00000146410 | MTFR2 | 310.8836 | -0.3052014 | 0.5707788 | -0.7566891 | 0.4492361 | 0.645517 |
| ENSG00000201772 | SNORA5C | 49.21882 | -0.3052593 | 0.3056927 | -1.083002 | 0.2788074 | 0.476038 |
| ENSG00000279689 | | 40.86019 | -0.3053997 | 0.3707226 | -0.9404396 | 0.3469921 | 0.548568 |
| ENSG00000267274 | | 3.914024 | -0.3054331 | 0.6774195 | -0.7656458 | 0.4438871 | 0.641412 |
| ENSG00000255980 | | 3.607864 | -0.3054687 | 0.6840666 | -0.7885392 | 0.4303814 | 0.629691 |
| ENSG00000226031 | FGF13-AS1 | 12.98279 | -0.3054759 | 0.535257 | -0.7681815 | 0.4423793 | 0.640138 |
| ENSG00000273654 | | 557.3802 | -0.305502 | 0.1755318 | -1.787907 | 0.07379097 | 0.186011 |
| ENSG00000109107 | ALDOC | 1014.989809 | -0.305559122 | 0.325850021 | -1.013326154 | 0.3109044 | 0.51174 |
| ENSG00000111224 | PARP11 | 358.3645777 | -0.305603985 | 0.218923688 | -1.450621055 | 0.14688541 | 0.306495 |
| ENSG00000146067 | FAM193B | 4832.939 | -0.3056648 | 0.1958516 | -1.61672 | 0.1059388 | 0.242645 |
| ENSG00000279640 | | 3.962008 | -0.3056735 | 0.6994431 | -0.7924394 | 0.4281045 | 0.627523 |
| ENSG00000163607 | GTPBP8 | 490.1011 | -0.3059053 | 0.2221388 | -1.439229 | 0.1500857 | 0.311193 |
| ENSG00000264056 | MIR5685 | 1.365505 | -0.3059128 | 0.7582491 | -0.8708339 | 0.3838449 | NA |
| ENSG00000266117 | FBXO36P1 | 4.42506 | -0.3059379 | 0.6514135 | -0.7652293 | 0.444135 | 0.641576 |
| ENSG00000231920 | NEBL-AS1 | 44.80695 | -0.3059895 | 0.322534 | -1.044699 | 0.2961622 | 0.495191 |
| ENSG00000010165 | EEF1AKNMT | 5341.078223 | -0.306048718 | 0.139647812 | -2.23198827 | 0.02561574 | 0.082655 |
| ENSG00000229133 | RPS7P4 | 124.1266 | -0.3060497 | 0.2342716 | -1.373718 | 0.1695294 | 0.339813 |
| ENSG00000233432 | MTIF2P1 | 6.56657 | -0.3060544 | 0.6387789 | -0.7511409 | 0.4525679 | 0.648699 |
| ENSG00000269540 | | 1.68894 | -0.3060649 | 0.8121342 | -1.072632 | 0.2834364 | 0.481221 |
| ENSG00000124562 | SNRPC | 5493.112309 | -0.306080509 | 0.238877445 | -1.349679283 | 0.17711888 | 0.350287 |
| ENSG00000115762 | PLEKHB2 | 8506.87637 | -0.306086134 | 0.141185563 | -2.206416819 | 0.02735483 | 0.087135 |
| ENSG00000204194 | RPL12P1 | 1.230075 | -0.3060931 | 0.7831221 | -0.964835 | 0.3346275 | NA |
| ENSG00000125977 | EIF2S2 | 4911.248724 | -0.306173219 | 0.186783616 | -1.697100814 | 0.0896776 | 0.215154 |
| ENSG00000213601 | KRT18P19 | 1.025095 | -0.3061782 | 0.8104427 | -1.102136 | 0.2704027 | NA |
| ENSG00000229180 | | 979.8669 | -0.3061841 | 0.1707114 | -1.829736 | 0.06728949 | 0.173305 |
| ENSG00000113441 | LNPEP | 4919.375353 | -0.306352004 | 0.208601986 | -1.551809185 | 0.12070789 | 0.266547 |
| ENSG00000225857 | LINC02816 | 2.331719 | -0.3064017 | 0.6938878 | -0.7855061 | 0.4321569 | 0.631294 |
| ENSG00000286677 | | 5.217218 | -0.3064747 | 0.6510144 | -0.7633267 | 0.4452686 | 0.642438 |
| ENSG00000254631 | | 0.8161615 | -0.3065343 | 0.8229213 | -1.164418 | 0.2442546 | NA |
| ENSG00000150551 | LYPD1 | 350.713 | -0.3067021 | 0.3521673 | -0.9716098 | 0.3312447 | 0.532288 |
| ENSG00000152402 | GUCY1A2 | 691.4114 | -0.3067599 | 0.4509291 | -0.8357087 | 0.4033188 | 0.604182 |

| | | | | | | | |
|---|---|---|---|---|---|---|---|
| ENSG00000204904 | LINC01545 | 3.011702 | -0.3067986 | 0.6958554 | -0.7911617 | 0.4288496 | 0.628062 |
| ENSG00000260612 | | 1.131798 | -0.3068775 | 0.7936103 | -0.9827616 | 0.3257248 | NA |
| ENSG00000152558 | TMEM123 | 8413.095 | -0.3069872 | 0.1345336 | -2.320908 | 0.02029181 | 0.068758 |
| ENSG00000287570 | | 3.620932 | -0.3072062 | 0.7883046 | -1.008624 | 0.3131551 | 0.514324 |
| ENSG00000110955 | ATP5F1B | 34296.18342 | -0.307245642 | 0.210863053 | -1.517637832 | 0.12910572 | 0.27932 |
| ENSG00000167862 | MRPL58 | 2353.067 | -0.3073475 | 0.2489484 | -1.307197 | 0.1911458 | 0.36921 |
| ENSG00000232995 | | 38.67674 | -0.3073707 | 0.4039203 | -0.8847928 | 0.3762684 | 0.578467 |
| ENSG00000287088 | | 2.039439 | -0.307447 | 0.7577842 | -0.8834016 | 0.3770193 | 0.579052 |
| ENSG00000085511 | MAP3K4 | 1481.834827 | -0.307450174 | 0.177511825 | -1.786040917 | 0.07409263 | 0.186559 |
| ENSG00000204434 | POTEKP | 638.1339 | -0.3074715 | 0.4108342 | -0.8873028 | 0.3749159 | 0.577124 |
| ENSG00000151148 | UBE3B | 4058.692 | -0.3075213 | 0.164855 | -1.942596 | 0.05206504 | 0.142934 |
| ENSG00000105618 | PRPF31 | 2755.217163 | -0.30756903 | 0.211207096 | -1.525615899 | 0.12710555 | 0.27622 |
| ENSG00000229912 | | 1.025282 | -0.3076489 | 0.8434407 | -1.329922 | 0.1835441 | NA |
| ENSG00000278891 | | 0.8699824 | -0.3076513 | 0.8724766 | -1.75481 | 0.07929189 | NA |
| ENSG00000226878 | GSTM3P2 | 1.02937 | -0.3076662 | 0.8147853 | -1.108164 | 0.267791 | NA |
| ENSG00000197182 | MIRLET7BHG | 92.36376 | -0.3076674 | 0.3384734 | -1.010059 | 0.3124671 | 0.513511 |
| ENSG00000119681 | LTBP2 | 537.924079 | -0.307714916 | 0.342870972 | -1.00220954 | 0.3162424 | 0.517316 |
| ENSG00000107581 | EIF3A | 18831.20865 | -0.307771217 | 0.158597471 | -1.988560485 | 0.04674974 | 0.131717 |
| ENSG00000282022 | | 1.031762 | -0.3078226 | 0.8304733 | -1.20341 | 0.2288176 | NA |
| ENSG00000268949 | MRPS17P1 | 8.306293 | -0.3079286 | 0.6106191 | -0.7566019 | 0.4492884 | 0.645517 |
| ENSG00000185842 | DNAH14 | 923.1685 | -0.3079395 | 0.2350362 | -1.38093 | 0.1673006 | 0.336742 |
| ENSG00000234676 | IFT74-AS1 | 1.174683 | -0.3079418 | 0.8287258 | -1.188574 | 0.2346072 | NA |
| ENSG00000160783 | PMF1 | 901.0635 | -0.3079441 | 0.323511 | -1.049501 | 0.2939474 | 0.493018 |
| ENSG00000259563 | | 3.910017 | -0.3079612 | 0.6651579 | -0.7740372 | 0.4389088 | 0.637373 |
| ENSG00000266111 | | 1.793952 | -0.3079805 | 0.7316677 | -0.8331792 | 0.4047436 | 0.605497 |
| ENSG00000273514 | FOXD4L6 | 1.131901 | -0.3081001 | 0.850789 | -1.450902 | 0.1468072 | NA |
| ENSG00000225830 | ERCC6 | 1451.527 | -0.3082034 | 0.2155644 | -1.488943 | 0.1365025 | 0.290697 |
| ENSG00000265666 | RARA-AS1 | 4.433349 | -0.3082079 | 0.5963302 | -0.7583713 | 0.4482287 | 0.644879 |
| ENSG00000235527 | HIPK1-AS1 | 129.7451 | -0.308222 | 0.2863626 | -1.159169 | 0.2463875 | 0.4382 |
| ENSG00000249140 | PRDX2P3 | 4.423253 | -0.3083735 | 0.6087229 | -0.7592797 | 0.4476853 | 0.644412 |
| ENSG00000253425 | HSPA8P13 | 11.77279 | -0.30856 | 0.6870355 | -0.7946418 | 0.4268219 | 0.626498 |
| ENSG00000236105 | PRELID3BP10 | 23.6064 | -0.3085871 | 0.4056787 | -0.8901043 | 0.3734099 | 0.575572 |
| ENSG00000249176 | | 120.0169 | -0.308614 | 0.3161926 | -1.071598 | 0.2839004 | 0.481731 |

| | | | | | | | |
|---|---|---|---|---|---|---|---|
| ENSG00000212694 | LINC01089 | 551.6899 | -0.3088515 | 0.35467 | -0.9758624 | 0.3291327 | 0.530275 |
| ENSG00000213063 | RPL29P7 | 1.861091 | -0.3090086 | 0.8657926 | -1.509679 | 0.1311252 | 0.282483 |
| ENSG00000142856 | ITGB3BP | 725.6912 | -0.309032 | 0.3136263 | -1.081856 | 0.2793167 | 0.476541 |
| ENSG00000129194 | SOX15 | 47.98905246 | -0.30905993 | 0.371592055 | -0.945915334 | 0.34419177 | 0.54583 |
| ENSG00000287164 | | 1.722767 | -0.3092279 | 0.8089231 | -1.052625 | 0.2925129 | 0.491366 |
| ENSG00000241157 | RPL32P32 | 19.1627 | -0.3092767 | 0.451695 | -0.8303984 | 0.4063136 | 0.606889 |
| ENSG00000030066 | NUP160 | 6027.980869 | -0.309299157 | 0.236169569 | -1.380720127 | 0.16736503 | 0.336771 |
| ENSG00000280758 | | 9.964827 | -0.3093068 | 0.5092459 | -0.7879633 | 0.4307182 | 0.629885 |
| ENSG00000130054 | FAM155B | 1277.086703 | -0.309454639 | 0.130814803 | -2.403139118 | 0.016255 | 0.057446 |
| ENSG00000260678 | | 2.461605 | -0.3094816 | 0.7331474 | -0.8463302 | 0.3973686 | 0.598474 |
| ENSG00000123975 | CKS2 | 3149.21439 | -0.309504342 | 0.314985819 | -1.137024132 | 0.2555282 | 0.448968 |
| ENSG00000237821 | ST13P7 | 251.4909 | -0.3095414 | 0.1776132 | -1.777443 | 0.07549534 | 0.189016 |
| ENSG00000160949 | TONSL | 4021.625 | -0.3095987 | 0.3693465 | -0.9345969 | 0.3499961 | 0.551643 |
| ENSG00000188000 | OR7D2 | 1.045548 | -0.3096797 | 0.815969 | -1.117456 | 0.2637993 | NA |
| ENSG00000177674 | AGTRAP | 1215.206 | -0.3096927 | 0.1911825 | -1.669933 | 0.09493266 | 0.224102 |
| ENSG00000124356 | STAMBP | 3199.126598 | -0.309696575 | 0.117064601 | -2.686700225 | 0.00721617 | 0.029892 |
| ENSG00000012048 | BRCA1 | 1946.4381 | -0.309730512 | 0.424713381 | -0.891789255 | 0.3725059 | 0.574678 |
| ENSG00000143324 | XPR1 | 2196.678 | -0.3097336 | 0.1309255 | -2.40264 | 0.01627722 | 0.057509 |
| ENSG00000143590 | EFNA3 | 774.783 | -0.3098238 | 0.3067951 | -1.105572 | 0.2689117 | 0.464999 |
| ENSG00000215016 | RPL24P7 | 53.84566 | -0.3098976 | 0.355117 | -0.9783607 | 0.327896 | 0.529301 |
| ENSG00000087088 | BAX | 4736.695426 | -0.309913049 | 0.215683674 | -1.50112117 | 0.13332422 | 0.285965 |
| ENSG00000060749 | QSER1 | 3417.96822 | -0.309920372 | 0.20809795 | -1.520309773 | 0.12843314 | 0.278286 |
| ENSG00000126858 | RHOT1 | 2014.134157 | -0.309942865 | 0.115780308 | -2.706362479 | 0.00680248 | 0.028464 |
| ENSG00000150593 | PDCD4 | 6519.023 | -0.3100361 | 0.2049891 | -1.591835 | 0.1114218 | 0.251668 |
| ENSG00000126261 | UBA2 | 8434.075548 | -0.310128844 | 0.160643745 | -1.972491679 | 0.04855351 | 0.135463 |
| ENSG00000225693 | LAGE3P1 | 4.291322 | -0.3101467 | 0.6945459 | -0.7985051 | 0.4245774 | 0.624343 |
| ENSG00000057935 | MTA3 | 2690.730033 | -0.310161534 | 0.134111543 | -2.365583527 | 0.01800168 | 0.062456 |
| ENSG00000160191 | PDE9A | 871.1939 | -0.3101697 | 0.2916506 | -1.147068 | 0.2513537 | 0.444096 |
| ENSG00000137817 | PARP6 | 6510.061501 | -0.310218696 | 0.212941978 | -1.521590553 | 0.12811171 | 0.277792 |
| ENSG00000118496 | FBXO30 | 708.0967938 | -0.310249203 | 0.265562964 | -1.245420518 | 0.21297722 | 0.397712 |
| ENSG00000226469 | ADAM1B | 6.952441 | -0.3102592 | 0.5827217 | -0.7624044 | 0.4458187 | 0.642804 |
| ENSG00000108784 | NAGLU | 2310.488667 | -0.310296051 | 0.234684824 | -1.377825582 | 0.16825715 | 0.338041 |
| ENSG00000132952 | USPL1 | 2264.049763 | -0.310314715 | 0.215564779 | -1.504833633 | 0.13236686 | 0.284581 |

| | | | | | | | |
|---|---|---|---|---|---|---|---|
| ENSG00000176371 | ZSCAN2 | 513.2094 | -0.31038 | 0.1927185 | -1.680444 | 0.09287105 | 0.22045 |
| ENSG00000251107 | PDCD5P2 | 8.884375 | -0.3103804 | 0.5006436 | -0.7983827 | 0.4246485 | 0.624343 |
| ENSG00000179698 | WDR97 | 2120.001 | -0.3105497 | 0.1407407 | -2.249896 | 0.02445557 | 0.079725 |
| ENSG00000206077 | ZDHHC11B | 109.2603 | -0.3105556 | 0.3245297 | -1.046549 | 0.2953076 | 0.494509 |
| ENSG00000286176 | | 36.66566 | -0.3105974 | 0.3743707 | -0.9478691 | 0.3431961 | 0.544641 |
| ENSG00000237281 | CATIP-AS2 | 1.510668 | -0.310598 | 0.8388964 | -0.990784 | 0.3217911 | NA |
| ENSG00000250031 | | 12.79443 | -0.3106241 | 0.5757532 | -0.7645509 | 0.444539 | 0.641854 |
| ENSG00000164167 | LSM6 | 848.5193 | -0.3106794 | 0.2179299 | -1.496351 | 0.1345622 | 0.287718 |
| ENSG00000172508 | CARNS1 | 116.869 | -0.3106812 | 0.3557199 | -0.9833561 | 0.3254322 | 0.526738 |
| ENSG00000132661 | NXT1 | 1291.985988 | -0.310897611 | 0.257837595 | -1.278910932 | 0.20092842 | 0.381614 |
| ENSG00000116685 | KIAA2013 | 2780.896599 | -0.310909682 | 0.166793769 | -1.907333088 | 0.05647747 | 0.152093 |
| ENSG00000286457 | | 1.23097 | -0.3109596 | 0.7845536 | -0.9623641 | 0.3358667 | NA |
| ENSG00000228889 | UBAC2-AS1 | 129.2407 | -0.3109682 | 0.2268163 | -1.433156 | 0.1518132 | 0.31382 |
| ENSG00000196366 | C9orf163 | 154.8261 | -0.3109947 | 0.2723659 | -1.212113 | 0.2254691 | 0.412739 |
| ENSG00000171700 | RGS19 | 1666.966 | -0.3109998 | 0.2634258 | -1.242295 | 0.2141278 | 0.398973 |
| ENSG00000111144 | LTA4H | 6737.832075 | -0.311051112 | 0.16579744 | -1.931639706 | 0.05340399 | 0.145991 |
| ENSG00000101158 | NELFCD | 19920.51896 | -0.311231584 | 0.176613346 | -1.836102947 | 0.06634243 | 0.171431 |
| ENSG00000280269 | | 3.07183 | -0.3113345 | 0.721371 | -0.8200165 | 0.4122067 | 0.612438 |
| ENSG00000279529 | | 67.35746 | -0.3113501 | 0.35874 | -0.9764966 | 0.3288184 | 0.529991 |
| ENSG00000178654 | PPIAP33 | 9.503811 | -0.3114444 | 0.5477355 | -0.780715 | 0.4349702 | 0.633766 |
| ENSG00000228057 | SEC63P1 | 284.8479 | -0.3114986 | 0.2007318 | -1.609373 | 0.1075349 | 0.245244 |
| ENSG00000153561 | RMND5A | 5870.979 | -0.311517 | 0.2048444 | -1.581145 | 0.113845 | 0.255425 |
| ENSG00000251867 | | 80.59204 | -0.3115324 | 0.3790334 | -0.9305302 | 0.3520966 | 0.55387 |
| ENSG00000189275 | LINC01164 | 1.574148 | -0.3116064 | 0.7884489 | -0.9872606 | 0.3235149 | 0.524403 |
| ENSG00000152147 | GEMIN6 | 1248.52 | -0.3118078 | 0.2527877 | -1.311788 | 0.1895915 | 0.367174 |
| ENSG00000117906 | RCN2 | 5872.685994 | -0.311831138 | 0.149753018 | -2.146258469 | 0.03185236 | 0.098314 |
| ENSG00000232549 | SRD5A1P1 | 13.23609 | -0.311837 | 0.5329004 | -0.7798421 | 0.4354838 | 0.634254 |
| ENSG00000139687 | RB1 | 2442.495 | -0.3119093 | 0.1819131 | -1.768094 | 0.07704523 | 0.191812 |
| ENSG00000148773 | MKI67 | 8794.309 | -0.311915 | 0.5593689 | -0.7775619 | 0.4368273 | 0.635253 |
| ENSG00000181359 | HSP90AA6P | 21.76271 | -0.3119162 | 0.48035 | -0.8117683 | 0.4169246 | 0.616843 |
| ENSG00000165704 | HPRT1 | 3056.17 | -0.3119659 | 0.2466174 | -1.312188 | 0.1894569 | 0.366967 |
| ENSG00000250138 | | 28.51009 | -0.3119742 | 0.4537177 | -0.8386481 | 0.4016668 | 0.602625 |
| ENSG00000225806 | | 51.98698 | -0.3120205 | 0.4278772 | -0.8676583 | 0.3855814 | 0.58792 |

| | | | | | | | |
|---|---|---|---|---|---|---|---|
| ENSG00000286362 | | 4.624979 | -0.3121765 | 0.606818 | -0.7649356 | 0.4443099 | 0.641709 |
| ENSG00000166578 | IQCD | 111.4719 | -0.3121956 | 0.285842 | -1.175844 | 0.2396572 | 0.429885 |
| ENSG00000273951 | | 12.73194 | -0.3122149 | 0.5253163 | -0.7810813 | 0.4347547 | 0.633626 |
| ENSG00000252391 | RNU6-638P | 0.9883542 | -0.312251 | 0.8303524 | -1.220855 | 0.2221411 | NA |
| ENSG00000273175 | | 1.812873 | -0.3122788 | 0.7855735 | -0.9620533 | 0.3360228 | 0.537087 |
| ENSG00000166436 | TRIM66 | 1649.75 | -0.312337 | 0.2624689 | -1.266925 | 0.2051822 | 0.387179 |
| ENSG00000223734 | LAPTM4A-DT | 1.120747 | -0.3123633 | 0.8249849 | -1.155457 | 0.2479034 | NA |
| ENSG00000243273 | | 1.958371 | -0.3124029 | 0.7102194 | -0.8239958 | 0.4099419 | 0.610319 |
| ENSG00000117480 | FAAH | 2752.277151 | -0.312411736 | 0.179784606 | -1.787980778 | 0.07377911 | 0.185998 |
| ENSG00000144749 | LRIG1 | 1813.407 | -0.3124873 | 0.2996696 | -1.140773 | 0.2539646 | 0.446919 |
| ENSG00000112293 | GPLD1 | 135.9209911 | -0.31254234 | 0.280800727 | -1.193727843 | 0.23258445 | 0.421236 |
| ENSG00000288071 | | 1.669691 | -0.3127371 | 0.8134406 | -1.102549 | 0.2702229 | 0.466659 |
| ENSG00000104880 | ARHGEF18 | 835.4118021 | -0.312752266 | 0.166516204 | -1.928597719 | 0.05378082 | 0.146791 |
| ENSG00000104635 | SLC39A14 | 3155.226378 | -0.312828845 | 0.23856959 | -1.382317717 | 0.16687417 | 0.336089 |
| ENSG00000131080 | EDA2R | 991.837819 | -0.312902916 | 0.421893138 | -0.891129969 | 0.37285945 | 0.575057 |
| ENSG00000227470 | RPL39P16 | 3.524636 | -0.312905 | 0.7137158 | -0.8246201 | 0.4095873 | 0.609996 |
| ENSG00000105829 | BET1 | 888.6284859 | -0.313093667 | 0.154460196 | -2.089906811 | 0.03662617 | 0.109204 |
| ENSG00000101337 | TM9SF4 | 5357.920347 | -0.313144688 | 0.136846779 | -2.370744872 | 0.01775228 | 0.061719 |
| ENSG00000259343 | TMC3-AS1 | 27.88677 | -0.3132064 | 0.5288092 | -0.7915405 | 0.4286287 | 0.62791 |
| ENSG00000198886 | MT-ND4 | 150507.7 | -0.313213 | 0.1774157 | -1.817075 | 0.06920568 | 0.177005 |
| ENSG00000235169 | SMIM1 | 60.70275 | -0.3132616 | 0.3905743 | -0.926342 | 0.3542683 | 0.556279 |
| ENSG00000233559 | LINC00513 | 2.216455 | -0.313356 | 0.7933174 | -0.9965446 | 0.3189856 | 0.519945 |
| ENSG00000167895 | TMC8 | 67.19523 | -0.3133906 | 0.4102855 | -0.8973433 | 0.3695357 | 0.571647 |
| ENSG00000241956 | | 2.038156 | -0.3134022 | 0.7529783 | -0.9030371 | 0.3665062 | 0.568428 |
| ENSG00000274698 | | 40.72705 | -0.3134394 | 0.402451 | -0.8963358 | 0.3700734 | 0.572255 |
| ENSG00000136770 | DNAJC1 | 2641.704326 | -0.313474341 | 0.152953858 | -2.088049282 | 0.03679339 | 0.109605 |
| ENSG00000185056 | C5orf47 | 2.855583 | -0.3136071 | 0.6969023 | -0.80818 | 0.418987 | 0.618895 |
| ENSG00000147036 | LANCL3 | 1.542562 | -0.3136315 | 0.7896088 | -0.9909393 | 0.3217152 | NA |
| ENSG00000124657 | OR2B6 | 40.61988231 | -0.313733774 | 0.503790907 | -0.80457694 | 0.42106385 | 0.620893 |
| ENSG00000137288 | UQCC2 | 2048.479834 | -0.313739817 | 0.193897815 | -1.672076757 | 0.09450919 | 0.22344 |
| ENSG00000267255 | | 2.131173 | -0.3138526 | 0.761996 | -0.9121204 | 0.3617054 | 0.563645 |
| ENSG00000223725 | | 2.969743 | -0.3139366 | 0.7849223 | -0.9516072 | 0.3412962 | 0.54251 |
| ENSG00000185115 | NSMCE3 | 508.5412 | -0.3139497 | 0.1960385 | -1.659763 | 0.09696216 | 0.227431 |

| | | | | | | | |
|---|---|---|---|---|---|---|---|
| ENSG00000279059 | | 479.4346 | -0.3139984 | 0.2115206 | -1.54818 | 0.121579 | 0.267847 |
| ENSG00000279536 | | 3.757228 | -0.3140351 | 0.6888377 | -0.8092559 | 0.418368 | 0.618118 |
| ENSG00000140740 | UQCRC2 | 10986.82 | -0.314098 | 0.1469448 | -2.111672 | 0.03471462 | 0.105002 |
| ENSG00000125850 | OVOL2 | 390.2785872 | -0.314208947 | 0.281971419 | -1.198264524 | 0.23081405 | 0.419287 |
| ENSG00000131174 | COX7B | 4465.002426 | -0.314214431 | 0.245313779 | -1.339700708 | 0.18034267 | 0.354629 |
| ENSG00000247271 | ZBED5-AS1 | 142.3362 | -0.3142936 | 0.2398988 | -1.382318 | 0.1668741 | 0.336089 |
| ENSG00000198556 | ZNF789 | 994.3318 | -0.3143109 | 0.2156053 | -1.528581 | 0.1263682 | 0.275049 |
| ENSG00000124102 | PI3 | 6.780383619 | -0.314345147 | 0.875710709 | -1.906243912 | 0.05661857 | 0.15236 |
| ENSG00000155115 | GTF3C6 | 1311.232 | -0.3143628 | 0.2866537 | -1.186086 | 0.2355884 | 0.425125 |
| ENSG00000173473 | SMARCC1 | 7671.47 | -0.3143962 | 0.1703353 | -1.902896 | 0.05705412 | 0.153103 |
| ENSG00000168653 | NDUFS5 | 5281.492 | -0.3144895 | 0.2427661 | -1.355741 | 0.1751816 | 0.347291 |
| ENSG00000240759 | RPS27P21 | 1.016305 | -0.3145941 | 0.8255425 | -1.175877 | 0.2396438 | NA |
| ENSG00000163956 | LRPAP1 | 4754.365 | -0.314733 | 0.1591417 | -2.012925 | 0.04412256 | 0.126121 |
| ENSG00000124789 | NUP153 | 4578.105451 | -0.314740872 | 0.212522107 | -1.54621218 | 0.12205333 | 0.268469 |
| ENSG00000196150 | ZNF250 | 1331.756 | -0.314868 | 0.2156073 | -1.522329 | 0.1279268 | 0.277528 |
| ENSG00000235397 | EPN2-AS1 | 6.144663 | -0.3148778 | 0.5987194 | -0.7736141 | 0.439159 | 0.637666 |
| ENSG00000163026 | WDCP | 462.0131 | -0.3149089 | 0.223226 | -1.47659 | 0.1397857 | 0.295816 |
| ENSG00000261588 | | 13.18429 | -0.3150098 | 0.4566468 | -0.8454063 | 0.397884 | 0.599013 |
| ENSG00000101246 | ARFRP1 | 3481.7193 | -0.315041698 | 0.174998083 | -1.852222665 | 0.06399386 | 0.166676 |
| ENSG00000173065 | FAM222B | 1131.623 | -0.3151313 | 0.1771739 | -1.836173 | 0.06633215 | 0.171428 |
| ENSG00000135365 | PHF21A | 2028.124302 | -0.315161394 | 0.28444286 | -1.210654689 | 0.22602778 | 0.413505 |
| ENSG00000134686 | PHC2 | 3680.961746 | -0.315210363 | 0.112037008 | -2.821693287 | 0.00477708 | 0.021367 |
| ENSG00000171806 | METTL18 | 563.0262 | -0.315298 | 0.1850384 | -1.75803 | 0.07874249 | 0.194832 |
| ENSG00000225976 | CKS1BP2 | 29.51697 | -0.315345 | 0.4613433 | -0.8422139 | 0.3996682 | 0.60061 |
| ENSG00000261040 | WFDC21P | 33.72933 | -0.3153725 | 0.4180198 | -0.8853745 | 0.3759547 | 0.578153 |
| ENSG00000167770 | OTUB1 | 5807.824 | -0.3153828 | 0.1754062 | -1.8538 | 0.06376778 | 0.166347 |
| ENSG00000256381 | | 0.8837472 | -0.3153962 | 0.8676984 | -1.47178 | 0.1410802 | NA |
| ENSG00000254532 | | 3.736177 | -0.3153988 | 0.6481158 | -0.7714532 | 0.4404384 | 0.638603 |
| ENSG00000137812 | KNL1 | 1174.860081 | -0.315609192 | 0.466149557 | -0.850172278 | 0.39522931 | 0.596906 |
| ENSG00000117298 | ECE1 | 4268.3456 | -0.315790781 | 0.219821551 | -1.494985196 | 0.13491831 | 0.288247 |
| ENSG00000058673 | ZC3H11A | 58.67757442 | -0.315793464 | 0.31303811 | -1.101349118 | 0.27074474 | 0.467075 |
| ENSG00000105467 | SYNGR4 | 5.410072968 | -0.315816562 | 0.600072688 | -0.778772878 | 0.43611352 | 0.63477 |
| ENSG00000284691 | | 165.1204 | -0.3158481 | 0.2645311 | -1.271538 | 0.2035374 | 0.385142 |

| | | | | | | | |
|---|---|---|---|---|---|---|---|
| ENSG00000168438 | CDC40 | 1788.147 | -0.3158577 | 0.1760426 | -1.8369 | 0.06622461 | 0.171243 |
| ENSG00000101193 | GID8 | 11038.16677 | -0.315861536 | 0.161852582 | -1.995438236 | 0.0459951 | 0.130102 |
| ENSG00000175772 | LINC01106 | 76.1477 | -0.3158891 | 0.3332592 | -1.043751 | 0.2966008 | 0.495597 |
| ENSG00000196553 | CCDC196 | 3.000433 | -0.3160292 | 0.7108066 | -0.8257995 | 0.4089178 | 0.609614 |
| ENSG00000226617 | RPL21P110 | 2.400698 | -0.3160554 | 0.7006528 | -0.8183959 | 0.4131312 | 0.613152 |
| ENSG00000228300 | FAM174C | 1591.032 | -0.3161498 | 0.2169574 | -1.521553 | 0.1281211 | 0.277792 |
| ENSG00000108848 | LUC7L3 | 17706.71643 | -0.316221863 | 0.220432987 | -1.499943175 | 0.13362912 | 0.286387 |
| ENSG00000278399 | | 4.445226 | -0.3162907 | 0.6608031 | -0.7857505 | 0.4320137 | 0.631216 |
| ENSG00000171109 | MFN1 | 3736.828 | -0.316479 | 0.1144647 | -2.817891 | 0.00483402 | 0.021589 |
| ENSG00000168496 | FEN1 | 3671.465 | -0.3165104 | 0.4402648 | -0.8693613 | 0.3846496 | 0.587039 |
| ENSG00000179091 | CYC1 | 12052.08 | -0.3165851 | 0.2179715 | -1.51683 | 0.1293096 | 0.279592 |
| ENSG00000086589 | RBM22 | 3272.461834 | -0.31676644 | 0.148329206 | -2.199227068 | 0.02786178 | 0.088347 |
| ENSG00000124228 | DDX27 | 7339.571978 | -0.31679655 | 0.121926622 | -2.605492239 | 0.00917424 | 0.036294 |
| ENSG00000174898 | CATSPERD | 2.514149 | -0.3168139 | 0.7659714 | -0.9182738 | 0.3584755 | 0.560387 |
| ENSG00000213430 | HSPD1P1 | 1674.573 | -0.3168366 | 0.2399563 | -1.400201 | 0.1614531 | 0.328432 |
| ENSG00000175166 | PSMD2 | 16230.53 | -0.3168395 | 0.2435954 | -1.276673 | 0.2017176 | 0.38268 |
| ENSG00000272043 | | 21.0227 | -0.3169531 | 0.4798442 | -0.826808 | 0.4083459 | 0.609035 |
| ENSG00000151576 | QTRT2 | 2724.733 | -0.3170081 | 0.1557539 | -2.107389 | 0.03508383 | 0.105769 |
| ENSG00000136270 | TBRG4 | 7162.061608 | -0.317098166 | 0.223927306 | -1.465390054 | 0.14281453 | 0.300454 |
| ENSG00000243988 | RPS24P17 | 2.536001 | -0.3171048 | 0.7166254 | -0.8432587 | 0.3990838 | 0.600071 |
| ENSG00000108599 | AKAP10 | 2408.068132 | -0.317243118 | 0.144808676 | -2.240850343 | 0.02503577 | 0.081208 |
| ENSG00000184983 | NDUFA6 | 2180.86 | -0.3172582 | 0.2791613 | -1.226022 | 0.2201906 | 0.406395 |
| ENSG00000162664 | ZNF326 | 2134.761 | -0.3172991 | 0.154767 | -2.096216 | 0.03606304 | 0.108107 |
| ENSG00000197114 | ZGPAT | 751.7045 | -0.3173352 | 0.1666089 | -1.947934 | 0.05142288 | 0.141522 |
| ENSG00000065548 | ZC3H15 | 5808.364016 | -0.317382021 | 0.18214707 | -1.77168467 | 0.07644692 | 0.19068 |
| ENSG00000100410 | PHF5A | 2386.931388 | -0.317428927 | 0.256847267 | -1.312791457 | 0.18925323 | 0.366706 |
| ENSG00000205100 | HSP90AA4P | 13.85924 | -0.3174432 | 0.4540747 | -0.8543245 | 0.3929252 | 0.594546 |
| ENSG00000124216 | SNAI1 | 69.38041938 | -0.31751814 | 0.410423364 | -0.909305246 | 0.36318902 | 0.565195 |
| ENSG00000065183 | WDR3 | 4067.164149 | -0.3175399 | 0.183463325 | -1.790570916 | 0.07336218 | 0.185158 |
| ENSG00000096063 | SRPK1 | 3323.388589 | -0.317550043 | 0.187009645 | -1.773769493 | 0.07610129 | 0.190054 |
| ENSG00000253204 | | 177.5153 | -0.3175827 | 0.2861217 | -1.201002 | 0.2297506 | 0.417956 |
| ENSG00000005249 | PRKAR2B | 305.9274397 | -0.317605529 | 0.264627406 | -1.282776524 | 0.19957038 | 0.3798 |
| ENSG00000261592 | | 9.568183 | -0.3176739 | 0.6400943 | -0.7873045 | 0.4311036 | 0.630241 |

| | | | | | | | |
|---|---|---|---|---|---|---|---|
| ENSG00000165169 | DYNLT3 | 1564.665 | -0.3177322 | 0.1447098 | -2.23095 | 0.02568446 | 0.082817 |
| ENSG00000163584 | RPL22L1 | 4054.972 | -0.3180313 | 0.232589 | -1.402957 | 0.1606297 | 0.327233 |
| ENSG00000243339 | RN7SL738P | 6.260429 | -0.3183269 | 0.6282141 | -0.7776441 | 0.4367789 | 0.635252 |
| ENSG00000162972 | MAIP1 | 1368.375 | -0.3183273 | 0.1976052 | -1.672602 | 0.09440558 | 0.223234 |
| ENSG00000265055 | | 25.68935 | -0.3183889 | 0.3353925 | -1.049428 | 0.2939812 | 0.493043 |
| ENSG00000112294 | ALDH5A1 | 4082.869708 | -0.318421036 | 0.101954585 | -3.156646279 | 0.00159595 | 0.00854 |
| ENSG00000205726 | ITSN1 | 1332.493 | -0.3184265 | 0.1984754 | -1.668995 | 0.09511846 | 0.224377 |
| ENSG00000123416 | TUBA1B | 9019.811327 | -0.318509205 | 0.433918245 | -0.865307136 | 0.38687024 | 0.589243 |
| ENSG00000116221 | MRPL37 | 5593.233908 | -0.318574672 | 0.254580109 | -1.328105699 | 0.18414319 | 0.359373 |
| ENSG00000206531 | CD200R1L | 1.376934 | -0.3186192 | 0.7934049 | -1.004319 | 0.315225 | NA |
| ENSG00000254986 | DPP3 | 5896.161 | -0.3187786 | 0.1782864 | -1.860642 | 0.06279479 | 0.164471 |
| ENSG00000174136 | RGMB | 749.4303 | -0.3188093 | 0.2093763 | -1.587533 | 0.1123921 | 0.25317 |
| ENSG00000276791 | | 192.0283 | -0.3189099 | 0.2784743 | -1.227197 | 0.2197485 | 0.405902 |
| ENSG00000229727 | | 1.004784 | -0.3189434 | 0.8311452 | -1.229288 | 0.2189637 | NA |
| ENSG00000273294 | C1QTNF3-AMAC | 5.261819 | -0.3191635 | 0.6894284 | -0.8110447 | 0.41734 | 0.617217 |
| ENSG00000064300 | NGFR | 19.12650192 | -0.319163738 | 0.526097489 | -0.801590868 | 0.42278966 | 0.622781 |
| ENSG00000077157 | PPP1R12B | 4395.669032 | -0.319213824 | 0.262254048 | -1.308212793 | 0.19080114 | 0.368821 |
| ENSG00000146587 | RBAK | 3180.51 | -0.3192171 | 0.1358445 | -2.397234 | 0.01651936 | 0.058225 |
| ENSG00000231715 | COX6CP2 | 1.859157 | -0.319271 | 0.7403814 | -0.8838456 | 0.3767796 | 0.578818 |
| ENSG00000083312 | TNPO1 | 9063.48123 | -0.319273262 | 0.184251381 | -1.803787378 | 0.07126465 | 0.180999 |
| ENSG00000273473 | | 25.88908 | -0.3192991 | 0.4423114 | -0.8710137 | 0.3837467 | 0.586166 |
| ENSG00000286502 | | 2.281521 | -0.3193598 | 0.7534612 | -0.8942476 | 0.3711894 | 0.57328 |
| ENSG00000222588 | | 1.284458 | -0.3193653 | 0.7886658 | -0.9706406 | 0.3317273 | NA |
| ENSG00000206832 | RNU6V | 0.9131347 | -0.3193765 | 0.8539893 | -1.189007 | 0.2344369 | NA |
| ENSG00000285948 | | 3.867519 | -0.3194041 | 0.645689 | -0.7972047 | 0.4253322 | 0.625105 |
| ENSG00000108651 | UTP6 | 2612.902941 | -0.319527538 | 0.15587972 | -2.102623492 | 0.0354987 | 0.106753 |
| ENSG00000173020 | GRK2 | 10705.3 | -0.3195561 | 0.1370878 | -2.406179 | 0.01612038 | 0.057054 |
| ENSG00000115866 | DARS1 | 4333.335398 | -0.319636297 | 0.154201536 | -2.124424456 | 0.03363467 | 0.102463 |
| ENSG00000197299 | BLM | 979.391 | -0.3196397 | 0.539422 | -0.8036403 | 0.4216047 | 0.621426 |
| ENSG00000285596 | | 25.43464 | -0.3196644 | 0.3814599 | -0.9569303 | 0.3386024 | 0.539866 |
| ENSG00000197798 | FAM118B | 514.3773 | -0.3198158 | 0.1436451 | -2.2618 | 0.02370978 | 0.077761 |
| ENSG00000260585 | | 2.515571 | -0.3198328 | 0.7816851 | -0.9514842 | 0.3413586 | 0.542535 |
| ENSG00000267201 | LINC01775 | 8.29772 | -0.3199714 | 0.6550726 | -0.7950254 | 0.4265987 | 0.626378 |

| | | | | | | | |
|---|---|---|---|---|---|---|---|
| ENSG00000146386 | ABRACL | 2203.234 | -0.3201133 | 0.1513158 | -2.118436 | 0.03413816 | 0.103683 |
| ENSG00000165644 | COMTD1 | 1477.126 | -0.3201616 | 0.2146141 | -1.564056 | 0.1178043 | 0.2617 |
| ENSG00000156671 | SAMD8 | 1614.499 | -0.3203225 | 0.2152594 | -1.551315 | 0.1208263 | 0.26663 |
| ENSG00000140694 | PARN | 2984.574 | -0.3203686 | 0.08974498 | -3.58943 | 0.0003314 | 0.002209 |
| ENSG00000217824 | SNRPEP6 | 2.711533 | -0.3204062 | 0.7319804 | -0.8407515 | 0.4004871 | 0.601432 |
| ENSG00000273243 | | 13.49637 | -0.3204337 | 0.496161 | -0.8248345 | 0.4094656 | 0.609936 |
| ENSG00000249096 | LINC02362 | 2.831186 | -0.3204611 | 0.6883352 | -0.8067601 | 0.4198047 | 0.619827 |
| ENSG00000230928 | | 1.029778 | -0.3205392 | 0.8608408 | -1.53754 | 0.1241612 | NA |
| ENSG00000176595 | KBTBD11 | 342.6043 | -0.3205406 | 0.1870249 | -1.759627 | 0.07847101 | 0.19441 |
| ENSG00000227627 | | 22.33086 | -0.3205785 | 0.4561802 | -0.8525305 | 0.3939197 | 0.595538 |
| ENSG00000091164 | TXNL1 | 3713.757101 | -0.320848148 | 0.160194386 | -2.051532677 | 0.0402151 | 0.117402 |
| ENSG00000224631 | RPS27AP16 | 7920.044 | -0.3209127 | 0.1749547 | -1.877551 | 0.06044267 | 0.15995 |
| ENSG00000231892 | | 1.898587 | -0.320915 | 0.7988661 | -1.044103 | 0.2964379 | 0.495443 |
| ENSG00000065970 | FOXJ2 | 1315.200671 | -0.320938424 | 0.202529976 | -1.644690468 | 0.10003366 | 0.23291 |
| ENSG00000180329 | CCDC43 | 1596.115 | -0.320982 | 0.1481077 | -2.206146 | 0.02737381 | 0.087174 |
| ENSG00000104884 | ERCC2 | 2259.920991 | -0.321019768 | 0.112896676 | -2.89736131 | 0.00376316 | 0.017522 |
| ENSG00000183022 | TPM3P8 | 42.7952 | -0.3210319 | 0.364295 | -0.9953444 | 0.3195688 | 0.520468 |
| ENSG00000235077 | | 1.68796 | -0.3210619 | 0.8248826 | -1.160753 | 0.2457423 | 0.437492 |
| ENSG00000125631 | HTR5BP | 3.253172701 | -0.32107115 | 0.735448545 | -0.84752966 | 0.39669996 | 0.59778 |
| ENSG00000113569 | NUP155 | 4436.013607 | -0.321155912 | 0.282346357 | -1.230959092 | 0.21833817 | 0.404091 |
| ENSG00000101596 | SMCHD1 | 3736.272102 | -0.321280916 | 0.225457324 | -1.494018835 | 0.1351707 | 0.288473 |
| ENSG00000212432 | | 1.423047 | -0.3213624 | 0.8297758 | -1.189682 | 0.2341715 | NA |
| ENSG00000125434 | SLC25A35 | 197.1665302 | -0.321548882 | 0.222629665 | -1.50981346 | 0.13109103 | 0.282455 |
| ENSG00000173207 | CKS1B | 1672.26 | -0.3215502 | 0.3470289 | -1.023383 | 0.3061268 | 0.506066 |
| ENSG00000127184 | COX7C | 10318.84963 | -0.321628614 | 0.191666105 | -1.753010003 | 0.07960029 | 0.196497 |
| ENSG00000131914 | LIN28A | 1.194105683 | -0.321791744 | 0.762797552 | -0.908555802 | 0.36358465 | NA |
| ENSG00000120438 | TCP1 | 9343.550997 | -0.321802142 | 0.227848837 | -1.480914417 | 0.13862938 | 0.293931 |
| ENSG00000160208 | RRP1B | 5037.377 | -0.3218687 | 0.2283453 | -1.460299 | 0.1442079 | 0.302452 |
| ENSG00000224425 | SSBP3P2 | 4.873305 | -0.3218961 | 0.7201438 | -0.8479621 | 0.3964591 | 0.597631 |
| ENSG00000249493 | ANKRD20A18P | 15.97554 | -0.3219024 | 0.426053 | -0.8974948 | 0.3694549 | 0.571644 |
| ENSG00000286208 | | 509.8442 | -0.321951 | 0.4839501 | -0.8384957 | 0.4017524 | 0.602719 |
| ENSG00000254681 | PKD1P5 | 152.2271 | -0.3219754 | 0.358569 | -1.011723 | 0.3116704 | 0.512551 |
| ENSG00000226874 | | 7.221674 | -0.3221863 | 0.6045877 | -0.7847522 | 0.4325989 | 0.631801 |

| | | | | | | | |
|---|---|---|---|---|---|---|---|
| ENSG00000132780 | NASP | 7391.717033 | -0.3224081 | 0.346529104 | -1.02521774 | 0.30526046 | 0.505059 |
| ENSG00000108452 | ZNF29P | 5.26412855 | -0.322421364 | 0.687098358 | -0.816454409 | 0.41424029 | 0.614446 |
| ENSG00000230267 | HERC2P4 | 3.054016 | -0.3224323 | 0.7826885 | -0.9516359 | 0.3412817 | 0.54251 |
| ENSG00000257093 | DENND11 | 4712.062 | -0.3224359 | 0.2680339 | -1.281931 | 0.1998668 | 0.380201 |
| ENSG00000248008 | NRAV | 629.744 | -0.3225335 | 0.1265936 | -2.58868 | 0.00963444 | 0.037699 |
| ENSG00000081923 | ATP8B1 | 4150.190573 | -0.322537357 | 0.194593826 | -1.721313822 | 0.08519389 | 0.206922 |
| ENSG00000196517 | SLC6A9 | 3295.104 | -0.3226437 | 0.3939213 | -0.9360798 | 0.3492321 | 0.550929 |
| ENSG00000229468 | RPL39P18 | 3.739347 | -0.3227415 | 0.6964943 | -0.8238911 | 0.4100014 | 0.610373 |
| ENSG00000170089 | | 1709.498 | -0.3228368 | 0.2982278 | -1.177559 | 0.2389726 | 0.428917 |
| ENSG00000267515 | | 1.060728 | -0.3228614 | 0.8061201 | -1.069165 | 0.2849952 | NA |
| ENSG00000170234 | PWWP2A | 1380.146 | -0.3229166 | 0.1418197 | -2.328115 | 0.01990598 | 0.067684 |
| ENSG00000141252 | VPS53 | 2071.85 | -0.3229185 | 0.101383 | -3.224012 | 0.00126408 | 0.007016 |
| ENSG00000224295 | OLFM5P | 1.156989 | -0.3229355 | 0.8321067 | -1.233971 | 0.2172138 | NA |
| ENSG00000218676 | BRD7P4 | 21.91476 | -0.3229575 | 0.365636 | -0.9983557 | 0.3181069 | 0.519215 |
| ENSG00000185420 | SMYD3 | 1201.075 | -0.3229786 | 0.2830419 | -1.232834 | 0.2176378 | 0.403322 |
| ENSG00000050327 | ARHGEF5 | 788.8941305 | -0.323031957 | 0.176322619 | -1.885591106 | 0.05935008 | 0.157845 |
| ENSG00000225595 | LINC01623 | 1.777729 | -0.3232435 | 0.8005458 | -1.038999 | 0.2988051 | 0.498052 |
| ENSG00000088970 | KIZ | 602.5614037 | -0.323247136 | 0.177062514 | -1.876553931 | 0.06057926 | 0.160158 |
| ENSG00000227896 | | 117.0155 | -0.3233014 | 0.3704906 | -0.9928757 | 0.3207706 | 0.521704 |
| ENSG00000114978 | MOB1A | 9526.682357 | -0.323354528 | 0.100415436 | -3.246987526 | 0.00116634 | 0.006539 |
| ENSG00000237125 | HAND2-AS1 | 49.92705 | -0.3234684 | 0.3851605 | -0.9658198 | 0.3341344 | 0.535377 |
| ENSG00000059758 | CDK17 | 1605.659232 | -0.323529979 | 0.177525195 | -1.87510335 | 0.06077851 | 0.160519 |
| ENSG00000284634 | | 33.59236 | -0.3236233 | 0.4878127 | -0.8411932 | 0.4002397 | 0.601197 |
| ENSG00000166133 | RPUSD2 | 718.2681 | -0.3236336 | 0.2121236 | -1.590278 | 0.1117721 | 0.252267 |
| ENSG00000155229 | MMS19 | 7562.775 | -0.3237174 | 0.09250149 | -3.529474 | 0.00041639 | 0.002681 |
| ENSG00000070761 | CFAP20 | 1434.548709 | -0.324031622 | 0.175658188 | -1.878520077 | 0.06031005 | 0.159657 |
| ENSG00000259972 | | 419.1484 | -0.3240409 | 0.3359113 | -1.070912 | 0.284209 | 0.482101 |
| ENSG00000103154 | NECAB2 | 125.834883 | -0.324054436 | 0.346842678 | -1.042013923 | 0.29740522 | 0.49656 |
| ENSG00000180573 | H2AC6 | 5030.161 | -0.3240879 | 0.2356867 | -1.423092 | 0.1547095 | 0.318386 |
| ENSG00000231579 | RPL7P21 | 125.039 | -0.3244007 | 0.2717942 | -1.279547 | 0.2007046 | 0.381303 |
| ENSG00000092201 | SUPT16H | 14594.24464 | -0.324464785 | 0.211471364 | -1.598518645 | 0.1099276 | 0.249289 |
| ENSG00000115648 | MLPH | 13879.75319 | -0.32449857 | 0.359070761 | -1.023999014 | 0.30583574 | 0.505742 |
| ENSG00000239149 | SNORA59A | 6.745451 | -0.3245049 | 0.5987995 | -0.7952574 | 0.4264638 | 0.626283 |

| | | | | | | | |
|---|---|---|---|---|---|---|---|
| ENSG00000130638 | ATXN10 | 6206.154909 | -0.324568988 | 0.209161226 | -1.618926315 | 0.10546312 | 0.241869 |
| ENSG00000067842 | ATP2B3 | 1.753740847 | -0.32458519 | 0.739531138 | -0.894433718 | 0.37108988 | 0.573226 |
| ENSG00000164291 | ARSK | 271.3611 | -0.3246384 | 0.2582157 | -1.335332 | 0.1817676 | 0.356534 |
| ENSG00000267049 | | 14.57991 | -0.3246436 | 0.4226106 | -0.9120146 | 0.361761 | 0.563699 |
| ENSG00000232344 | | 10.93188 | -0.3246634 | 0.4967745 | -0.8328344 | 0.4049382 | 0.605695 |
| ENSG00000235602 | POU5F1P3 | 6.209235 | -0.3247087 | 0.652211 | -0.7960056 | 0.4260288 | 0.625921 |
| ENSG00000250708 | LINC02269 | 21.40239 | -0.3247445 | 0.3756074 | -0.9856868 | 0.3242868 | 0.525204 |
| ENSG00000231167 | YBX1P2 | 723.3994 | -0.3248418 | 0.2634635 | -1.314144 | 0.1887977 | 0.366063 |
| ENSG00000180867 | PDIA3P1 | 3920.917 | -0.3248446 | 0.1308085 | -2.543728 | 0.01096766 | 0.041916 |
| ENSG00000277687 | | 44.73097 | -0.3248583 | 0.435865 | -0.8956027 | 0.370465 | 0.57256 |
| ENSG00000257501 | PAFAH1B2P2 | 1.642144 | -0.3248978 | 0.8649876 | -1.549258 | 0.1213197 | 0.267409 |
| ENSG00000100889 | PCK2 | 9521.124625 | -0.32489951 | 0.374630954 | -0.992495691 | 0.32095578 | 0.52174 |
| ENSG00000053371 | AKR7A2 | 2258.957711 | -0.324966007 | 0.219547909 | -1.547626635 | 0.12171222 | 0.267985 |
| ENSG00000196289 | BECN2 | 0.8387237 | -0.3250008 | 0.8580359 | -1.215926 | 0.2240131 | NA |
| ENSG00000144647 | POMGNT2 | 514.7979 | -0.3250188 | 0.2839183 | -1.232969 | 0.2175873 | 0.403288 |
| ENSG00000287603 | | 6.58282 | -0.3251124 | 0.5764803 | -0.7937921 | 0.4273165 | 0.626826 |
| ENSG00000138031 | ADCY3 | 2462.003158 | -0.325156414 | 0.18549516 | -1.820104507 | 0.06874309 | 0.176076 |
| ENSG00000234509 | | 8.486026 | -0.3253289 | 0.5520699 | -0.805701 | 0.4204153 | 0.620281 |
| ENSG00000214533 | KRT18P33 | 14.70275 | -0.3253595 | 0.4858801 | -0.8415604 | 0.4000341 | 0.601039 |
| ENSG00000270016 | | 1.424331 | -0.3253821 | 0.7730977 | -0.9614231 | 0.3363395 | NA |
| ENSG00000144191 | CNGA3 | 1.282928 | -0.3254023 | 0.7735062 | -0.9543957 | 0.3398834 | NA |
| ENSG00000160972 | PPP1R16A | 2493.436 | -0.3254883 | 0.184263 | -1.82519 | 0.06797243 | 0.174641 |
| ENSG00000139597 | N4BP2L1 | 452.6982 | -0.325555 | 0.3815802 | -0.9767264 | 0.3287046 | 0.529904 |
| ENSG00000225420 | | 1.658439 | -0.3255624 | 0.8443727 | -1.020258 | 0.3076062 | 0.507721 |
| ENSG00000001497 | LAS1L | 4671.170246 | -0.325647641 | 0.201190116 | -1.686162588 | 0.09176449 | 0.218565 |
| ENSG00000136045 | PWP1 | 4741.198518 | -0.325659133 | 0.237137763 | -1.439318547 | 0.15006029 | 0.311165 |
| ENSG00000254044 | | 1.033118 | -0.3257419 | 0.8683483 | -1.595846 | 0.1105233 | NA |
| ENSG00000115839 | RAB3GAP1 | 4343.836569 | -0.325764469 | 0.145567894 | -2.286433032 | 0.02222894 | 0.07414 |
| ENSG00000138738 | PRDM5 | 446.9285304 | -0.325837333 | 0.21462509 | -1.589068933 | 0.11204483 | 0.25263 |
| ENSG00000223685 | LINC00571 | 23.9372 | -0.3259492 | 0.4623075 | -0.8669957 | 0.3859444 | 0.588305 |
| ENSG00000055609 | KMT2C | 7636.67042 | -0.326029247 | 0.308754759 | -1.154425673 | 0.2483257 | 0.440644 |
| ENSG00000225913 | | 6.392564 | -0.3260519 | 0.6972863 | -0.8253636 | 0.4091652 | 0.609812 |
| ENSG00000134317 | GRHL1 | 2457.354087 | -0.326133988 | 0.260772479 | -1.337376172 | 0.18109988 | 0.355657 |

| | | | | | | | |
|---|---|---|---|---|---|---|---|
| ENSG00000154803 | FLCN | 2410.774 | -0.3261744 | 0.2908631 | -1.213673 | 0.2248726 | 0.411928 |
| ENSG00000233974 | | 5.395024 | -0.3262354 | 0.5910704 | -0.7985026 | 0.4245789 | 0.624343 |
| ENSG00000204390 | HSPA1L | 73.99402 | -0.3262625 | 0.3331678 | -1.084598 | 0.2780999 | 0.475224 |
| ENSG00000062194 | GPBP1 | 6455.623455 | -0.326282983 | 0.10239166 | -3.192300846 | 0.00141144 | 0.007702 |
| ENSG00000213013 | RPS15AP36 | 5.499779 | -0.3262977 | 0.6800045 | -0.8242003 | 0.4098258 | 0.610248 |
| ENSG00000037474 | NSUN2 | 9683.71956 | -0.326316027 | 0.18654004 | -1.805197756 | 0.07104374 | 0.180553 |
| ENSG00000262482 | | 11.40269 | -0.3263884 | 0.4600572 | -0.8697414 | 0.3844418 | 0.586839 |
| ENSG00000234488 | RPL27P6 | 8.598486 | -0.3263887 | 0.5725101 | -0.8021493 | 0.4224666 | 0.622478 |
| ENSG00000164404 | GDF9 | 73.31277 | -0.3264007 | 0.2850973 | -1.234377 | 0.2170623 | 0.402592 |
| ENSG00000267248 | | 225.6855 | -0.3264415 | 0.3515842 | -1.041949 | 0.2974354 | 0.496571 |
| ENSG00000148840 | PPRC1 | 4187.925 | -0.3264885 | 0.1518812 | -2.168692 | 0.03010606 | 0.093939 |
| ENSG00000230797 | YY2 | 96.33052 | -0.3265627 | 0.2665418 | -1.304655 | 0.1920103 | 0.370235 |
| ENSG00000166228 | PCBD1 | 3210.016 | -0.3266458 | 0.1380066 | -2.433232 | 0.01496469 | 0.053835 |
| ENSG00000281420 | | 16.79608 | -0.3266666 | 0.4243395 | -0.9083813 | 0.3636768 | 0.565821 |
| ENSG00000099797 | TECR | 2050.324279 | -0.326784341 | 0.218590068 | -1.561013335 | 0.11852061 | 0.262765 |
| ENSG00000156795 | NTAQ1 | 1405.16 | -0.3268113 | 0.2206634 | -1.551943 | 0.1206758 | 0.266498 |
| ENSG00000255967 | HADHAP2 | 87.77105 | -0.3268347 | 0.2329792 | -1.472526 | 0.140879 | 0.297513 |
| ENSG00000287168 | | 122.4356 | -0.3268576 | 0.3416427 | -1.072008 | 0.2837165 | 0.481542 |
| ENSG00000256268 | LINC02454 | 2.299709 | -0.326935 | 0.8193723 | -1.146248 | 0.2516927 | 0.444452 |
| ENSG00000184967 | NOC4L | 1265.447 | -0.3269406 | 0.2409769 | -1.430733 | 0.1525068 | 0.314837 |
| ENSG00000140829 | DHX38 | 6955.008 | -0.3271145 | 0.1336268 | -2.500269 | 0.01240989 | 0.046244 |
| ENSG00000128268 | MGAT3 | 10.24478893 | -0.327131848 | 0.649415851 | -0.797618291 | 0.42509203 | 0.624821 |
| ENSG00000158427 | TMSB15B | 23.76133 | -0.3272244 | 0.3317483 | -1.092249 | 0.2747238 | 0.471557 |
| ENSG00000259772 | | 3.754576 | -0.3273082 | 0.7019413 | -0.8478249 | 0.3965355 | 0.597631 |
| ENSG00000266174 | MIR4666A | 5.962992 | -0.3273416 | 0.5719141 | -0.8041307 | 0.4213215 | 0.621135 |
| ENSG00000093217 | XYLB | 502.8474262 | -0.327396581 | 0.18348578 | -1.841324466 | 0.06557403 | 0.169913 |
| ENSG00000198677 | TTC37 | 4503.811 | -0.3274441 | 0.2136519 | -1.599075 | 0.1098038 | 0.249072 |
| ENSG00000249275 | | 0.9329302 | -0.3274546 | 0.8403012 | -1.299308 | 0.1938382 | NA |
| ENSG00000168259 | DNAJC7 | 5109.46 | -0.3274805 | 0.1577388 | -2.126968 | 0.0334227 | 0.102023 |
| ENSG00000185974 | GRK1 | 1.09183 | -0.327545 | 0.8529285 | -1.114197 | 0.2651945 | NA |
| ENSG00000270607 | | 1.11206 | -0.3275497 | 0.8473396 | -1.054749 | 0.2915401 | NA |
| ENSG00000188933 | USP32P1 | 3.826828 | -0.3275691 | 0.661079 | -0.809635 | 0.41815 | 0.617934 |
| ENSG00000028116 | VRK2 | 701.3391033 | -0.327630911 | 0.178683425 | -1.887219481 | 0.05913081 | 0.157435 |

| | | | | | | | |
|---|---|---|---|---|---|---|---|
| ENSG00000180509 | KCNE1 | 1.288365 | -0.3276397 | 0.8577062 | -1.204831 | 0.2282687 | NA |
| ENSG00000126218 | F10 | 15.08811573 | -0.327853206 | 0.457208704 | -0.876578092 | 0.38071584 | 0.582742 |
| ENSG00000109511 | ANXA10 | 0.969358018 | -0.327864427 | 0.869454592 | -1.396621407 | 0.16252745 | NA |
| ENSG00000254186 | | 1.887259 | -0.3278942 | 0.764559 | -0.9453262 | 0.3444924 | 0.546046 |
| ENSG00000148053 | NTRK2 | 1.911052 | -0.3278996 | 0.8461116 | -1.023582 | 0.3060327 | 0.506036 |
| ENSG00000225568 | | 26.91074 | -0.3279112 | 0.3745318 | -0.9991535 | 0.3177203 | 0.518712 |
| ENSG00000236311 | TLX1NB | 1.986418 | -0.3279165 | 0.7390863 | -0.9066107 | 0.3646127 | 0.566614 |
| ENSG00000132646 | PCNA | 4466.060396 | -0.327959235 | 0.428069626 | -0.915427866 | 0.35996706 | 0.561958 |
| ENSG00000260949 | | 9.418633 | -0.3280027 | 0.6612087 | -0.8124759 | 0.4165186 | 0.616483 |
| ENSG00000241612 | | 5.164736 | -0.3281266 | 0.6703225 | -0.8155176 | 0.4147761 | 0.614726 |
| ENSG00000111653 | ING4 | 1642.542893 | -0.328141049 | 0.260067426 | -1.347777317 | 0.17773002 | 0.351086 |
| ENSG00000262700 | | 1.362802 | -0.3283373 | 0.8221336 | -1.168144 | 0.2427487 | NA |
| ENSG00000176946 | THAP4 | 3283.537 | -0.3285399 | 0.1799772 | -1.889242 | 0.05885942 | 0.156917 |
| ENSG00000236200 | KDM4A-AS1 | 143.8877 | -0.3285462 | 0.2583639 | -1.353321 | 0.1759532 | 0.348561 |
| ENSG00000119318 | RAD23B | 13741.61148 | -0.328597128 | 0.170019691 | -1.938564997 | 0.05255433 | 0.144054 |
| ENSG00000258345 | | 0.8523452 | -0.3286021 | 0.8261294 | -1.168888 | 0.2424489 | NA |
| ENSG00000135829 | DHX9 | 17532.5459 | -0.328630225 | 0.222322079 | -1.546787472 | 0.1219145 | 0.268269 |
| ENSG00000165238 | WNK2 | 1045.388 | -0.3286364 | 0.2209476 | -1.541917 | 0.1230937 | 0.27022 |
| ENSG00000158169 | FANCC | 1221.693 | -0.3287062 | 0.2079067 | -1.659419 | 0.0970313 | 0.22754 |
| ENSG00000105281 | SLC1A5 | 18062.38983 | -0.328709598 | 0.192339385 | -1.773565752 | 0.07613501 | 0.190087 |
| ENSG00000090054 | SPTLC1 | 4333.376201 | -0.328835833 | 0.13215273 | -2.525839316 | 0.01154223 | 0.04366 |
| ENSG00000138771 | SHROOM3 | 4368.427759 | -0.328867685 | 0.25399106 | -1.338473786 | 0.18074204 | 0.355206 |
| ENSG00000197894 | ADH5 | 5979.854 | -0.3290313 | 0.1816382 | -1.86798 | 0.06176482 | 0.162532 |
| ENSG00000267480 | | 11.94916 | -0.3290573 | 0.4781286 | -0.8589304 | 0.3903789 | 0.592278 |
| ENSG00000232499 | | 621.8736 | -0.3290953 | 0.1609675 | -2.090621 | 0.03656204 | 0.109038 |
| ENSG00000252448 | | 1.251452 | -0.3291729 | 0.8433736 | -1.294453 | 0.1955089 | NA |
| ENSG00000188177 | ZC3H6 | 625.9484 | -0.3292787 | 0.2512065 | -1.39748 | 0.1622694 | 0.329502 |
| ENSG00000171163 | ZNF692 | 4391.133 | -0.3293502 | 0.2636009 | -1.332117 | 0.1828217 | 0.357819 |
| ENSG00000254831 | KRT18P58 | 3.07162 | -0.3293638 | 0.691095 | -0.8312311 | 0.4058431 | 0.606663 |
| ENSG00000101003 | GINS1 | 1700.236715 | -0.329388711 | 0.47666359 | -0.879570651 | 0.37909194 | 0.581127 |
| ENSG00000119929 | CUTC | 1237.570953 | -0.329489916 | 0.249423247 | -1.399234624 | 0.16174264 | 0.328895 |
| ENSG00000112167 | SAYSD1 | 1375.430488 | -0.329576474 | 0.182495082 | -1.852540991 | 0.06394818 | 0.166638 |
| ENSG00000005810 | MYCBP2 | 3492.610375 | -0.329626592 | 0.244176718 | -1.427195659 | 0.15352351 | 0.316469 |

| | | | | | | | |
|---|---|---|---|---|---|---|---|
| ENSG00000122986 | HVCN1 | 6.03444135 | -0.329654748 | 0.594976464 | -0.801495271 | 0.42284498 | 0.622794 |
| ENSG00000283672 | MIR4678 | 2.969291 | -0.3297138 | 0.7129744 | -0.8609819 | 0.389248 | 0.591157 |
| ENSG00000233381 | AK4P3 | 11.9202 | -0.3297172 | 0.5565673 | -0.818703 | 0.4129559 | 0.61302 |
| ENSG00000196741 | LINC01560 | 149.1932 | -0.3297268 | 0.2222294 | -1.54969 | 0.1212158 | 0.267224 |
| ENSG00000286220 | | 1.599811 | -0.3297411 | 0.8186742 | -1.139685 | 0.2544177 | 0.44742 |
| ENSG00000145740 | SLC30A5 | 3245.836 | -0.3297559 | 0.1024567 | -3.259088 | 0.00111771 | 0.006301 |
| ENSG00000236495 | | 1.664914 | -0.329773 | 0.7827522 | -0.9895278 | 0.322405 | 0.523209 |
| ENSG00000145979 | TBC1D7 | 1216.866 | -0.3298373 | 0.2307809 | -1.512833 | 0.1303221 | 0.281281 |
| ENSG00000265100 | | 41.36049 | -0.3298787 | 0.3364469 | -1.082821 | 0.2788879 | 0.476145 |
| ENSG00000262814 | MRPL12 | 962.6783 | -0.3299365 | 0.2988099 | -1.198727 | 0.2306342 | 0.419075 |
| ENSG00000062485 | CS | 11148.89317 | -0.330028463 | 0.162118836 | -2.091189436 | 0.03651109 | 0.108971 |
| ENSG00000132485 | ZRANB2 | 5161.402385 | -0.330165839 | 0.199324025 | -1.719603955 | 0.08550446 | 0.207524 |
| ENSG00000183763 | TRAIP | 502.6176 | -0.3301966 | 0.4457363 | -0.8898919 | 0.373524 | 0.575609 |
| ENSG00000142207 | URB1 | 3583.82 | -0.3302478 | 0.219605 | -1.568568 | 0.1167486 | 0.259986 |
| ENSG00000237054 | PRMT5-AS1 | 16.03896 | -0.3303619 | 0.4137573 | -0.9382584 | 0.3481116 | 0.549824 |
| ENSG00000105486 | LIG1 | 3243.83189 | -0.33041278 | 0.421849179 | -0.93059216 | 0.35206457 | 0.553852 |
| ENSG00000230530 | LIMD1-AS1 | 27.90001 | -0.3304555 | 0.3621666 | -1.027209 | 0.3043219 | 0.50396 |
| ENSG00000185324 | CDK10 | 7219.874 | -0.3304557 | 0.1647338 | -2.065433 | 0.03888207 | 0.114365 |
| ENSG00000260571 | BNIP3P5 | 2.094041 | -0.3304748 | 0.7333081 | -0.8868258 | 0.3751727 | 0.577318 |
| ENSG00000271047 | | 2.511477 | -0.3305018 | 0.7582467 | -0.9347738 | 0.3499049 | 0.551565 |
| ENSG00000278879 | | 7.869561 | -0.3306065 | 0.6624718 | -0.8193467 | 0.4125886 | 0.612714 |
| ENSG00000258553 | | 1.30705 | -0.3306252 | 0.8110126 | -1.081136 | 0.2796368 | NA |
| ENSG00000172943 | PHF8 | 3252.19 | -0.3306732 | 0.1825442 | -1.866416 | 0.06198323 | 0.162875 |
| ENSG00000240409 | MTATP8P1 | 1.225636 | -0.3307008 | 0.8574875 | -1.178895 | 0.2384399 | NA |
| ENSG00000233791 | LINC01136 | 13.83204 | -0.33076 | 0.5566224 | -0.80746 | 0.4194015 | 0.619335 |
| ENSG00000108591 | DRG2 | 2360.558625 | -0.330771758 | 0.111395001 | -3.012138 | 0.00259415 | 0.01291 |
| ENSG00000213300 | HNRNPA3P6 | 5313.842 | -0.3307756 | 0.1792395 | -1.906012 | 0.05664861 | 0.152399 |
| ENSG00000280238 | | 1.083073 | -0.3307798 | 0.7971433 | -1.018283 | 0.3085436 | NA |
| ENSG00000150787 | PTS | 599.3286 | -0.3307857 | 0.2183481 | -1.580329 | 0.1140315 | 0.255746 |
| ENSG00000245685 | FRG1-DT | 83.11315 | -0.3308446 | 0.3058184 | -1.173172 | 0.240727 | 0.431105 |
| ENSG00000188229 | TUBB4B | 17296.43 | -0.3308516 | 0.3628385 | -1.001707 | 0.316485 | 0.517522 |
| ENSG00000101624 | CEP76 | 589.3861209 | -0.330972442 | 0.231522089 | -1.517933132 | 0.12903126 | 0.27924 |
| ENSG00000176393 | RNPEP | 4454.918 | -0.3310268 | 0.1915804 | -1.798792 | 0.07205151 | 0.182544 |

| | | | | | | | |
|---|---|---|---|---|---|---|---|
| ENSG00000035681 | NSMAF | 3104.354914 | -0.331087824 | 0.141062792 | -2.401712749 | 0.01631852 | 0.057623 |
| ENSG00000173950 | XXYLT1 | 1646.482 | -0.3311674 | 0.187046 | -1.804077 | 0.07121919 | 0.180918 |
| ENSG00000121486 | TRMT1L | 1440.395418 | -0.331254972 | 0.176779598 | -1.932231105 | 0.05333099 | 0.145852 |
| ENSG00000260156 | | 2.012526 | -0.3312781 | 0.7157301 | -0.8587622 | 0.3904717 | 0.592352 |
| ENSG00000176105 | YES1 | 4823.951 | -0.3313845 | 0.1410343 | -2.39363 | 0.01668256 | 0.058715 |
| ENSG00000081041 | CXCL2 | 7.882910098 | -0.33138585 | 0.843585026 | -0.992049994 | 0.32117314 | 0.521911 |
| ENSG00000138382 | METTL5 | 1553.93176 | -0.331435565 | 0.172723562 | -1.960756336 | 0.04990745 | 0.138283 |
| ENSG00000125445 | MRPS7 | 6048.343654 | -0.331466346 | 0.246037889 | -1.417761134 | 0.15626052 | 0.320717 |
| ENSG00000137880 | GCHFR | 50.1713867 | -0.331647981 | 0.333797463 | -1.101819879 | 0.27053999 | 0.466782 |
| ENSG00000204954 | C12orf73 | 880.4 | -0.3316551 | 0.1814597 | -1.882565 | 0.05975933 | 0.158553 |
| ENSG00000198945 | L3MBTL3 | 283.488 | -0.3316854 | 0.2809463 | -1.268581 | 0.2045904 | 0.386529 |
| ENSG00000085733 | CTTN | 22632.86107 | -0.331749714 | 0.150850281 | -2.263990191 | 0.02357472 | 0.077424 |
| ENSG00000188523 | CFAP77 | 2.005475 | -0.3318012 | 0.7531008 | -0.9165059 | 0.3594016 | 0.56124 |
| ENSG00000196199 | MPHOSPH8 | 4248.775 | -0.3318082 | 0.1630743 | -2.094198 | 0.03624234 | 0.10856 |
| ENSG00000133195 | SLC39A11 | 2056.9252 | -0.331844283 | 0.128175951 | -2.640441664 | 0.00827981 | 0.033427 |
| ENSG00000243160 | | 1.844374 | -0.3318725 | 0.7943081 | -1.007802 | 0.3135497 | 0.514709 |
| ENSG00000164078 | MST1R | 124.143 | -0.3319393 | 0.4529062 | -0.8932581 | 0.371719 | 0.573898 |
| ENSG00000196968 | FUT11 | 437.5808 | -0.3319835 | 0.2032483 | -1.695263 | 0.09002548 | 0.215699 |
| ENSG00000213275 | IFITM9P | 1.100289 | -0.3319836 | 0.8578271 | -1.183214 | 0.2367244 | NA |
| ENSG00000120733 | KDM3B | 8071.733561 | -0.331989295 | 0.156035774 | -2.189224555 | 0.02858052 | 0.090238 |
| ENSG00000269463 | | 6.918521 | -0.3320102 | 0.6896436 | -0.8382992 | 0.4018627 | 0.602817 |
| ENSG00000145833 | DDX46 | 8170.904 | -0.3320781 | 0.1922697 | -1.814798 | 0.06955503 | 0.177676 |
| ENSG00000124399 | NDUFB4P12 | 276.2685439 | -0.332089392 | 0.221045164 | -1.565515394 | 0.11746211 | 0.261159 |
| ENSG00000219102 | HNRNPA3P12 | 142.3868 | -0.332102 | 0.22487 | -1.551433 | 0.120798 | 0.266626 |
| ENSG00000071794 | HLTF | 4250.058824 | -0.332249052 | 0.234350196 | -1.452647364 | 0.14632169 | 0.305727 |
| ENSG00000273356 | LINC02019 | 28.75354 | -0.3322543 | 0.6033364 | -0.8197214 | 0.412375 | 0.612568 |
| ENSG00000141905 | NFIC | 3096.854 | -0.3324568 | 0.1939387 | -1.784696 | 0.07431068 | 0.186913 |
| ENSG00000146083 | RNF44 | 6851.675 | -0.3325915 | 0.1603561 | -2.158085 | 0.03092123 | 0.096086 |
| ENSG00000221164 | SNORA11F | 28.11087 | -0.3326324 | 0.358565 | -1.040883 | 0.2979301 | 0.497084 |
| ENSG00000287601 | | 3.200561 | -0.3326528 | 0.7290218 | -0.8721088 | 0.383149 | 0.585556 |
| ENSG00000198276 | UCKL1 | 4281.437 | -0.3327087 | 0.1406991 | -2.401951 | 0.01630787 | 0.057595 |
| ENSG00000223621 | AK4P4 | 2.15345 | -0.3327876 | 0.7554674 | -0.9230289 | 0.3559922 | 0.558084 |
| ENSG00000184857 | TMEM186 | 871.8291 | -0.3328477 | 0.1261517 | -2.705942 | 0.0068111 | 0.02848 |

| | | | | | | | |
|---|---|---|---|---|---|---|---|
| ENSG00000227799 | CDPF1P1 | 91.84791 | -0.3330287 | 0.2245256 | -1.54473 | 0.1224115 | 0.269145 |
| ENSG00000275576 | | 8.620622 | -0.3330645 | 0.5703886 | -0.8146383 | 0.4152794 | 0.615163 |
| ENSG00000070669 | ASNS | 153.2362265 | -0.333092225 | 0.615769348 | -0.812608282 | 0.41644268 | 0.616405 |
| ENSG00000227852 | RPS29P17 | 5.470077 | -0.3330992 | 0.6318978 | -0.805593 | 0.4204776 | 0.620303 |
| ENSG00000240311 | | 3.48301 | -0.3331796 | 0.6838329 | -0.8286206 | 0.4073191 | 0.607981 |
| ENSG00000107518 | ATRNL1 | 38.18384405 | -0.333254755 | 0.533024473 | -0.840873224 | 0.40041896 | 0.601397 |
| ENSG00000177888 | ZBTB41 | 3103.689 | -0.3334058 | 0.2879937 | -1.248505 | 0.2118463 | 0.396181 |
| ENSG00000267323 | SLC25A1P5 | 1.116157 | -0.3334281 | 0.8142549 | -1.085223 | 0.2778227 | NA |
| ENSG00000091651 | ORC6 | 1078.285775 | -0.333523785 | 0.473327066 | -0.88527425 | 0.37600874 | 0.578202 |
| ENSG00000164669 | INTS4P1 | 21.11322 | -0.3335466 | 0.3941918 | -0.9739095 | 0.3301015 | 0.531189 |
| ENSG00000235376 | RPEL1 | 217.5571 | -0.3336202 | 0.2396436 | -1.465465 | 0.1427941 | 0.300435 |
| ENSG00000170759 | KIF5B | 13039.76 | -0.3336489 | 0.1916948 | -1.799753 | 0.07189972 | 0.182246 |
| ENSG00000260077 | | 77.81806 | -0.3336998 | 0.3779596 | -1.013079 | 0.3110225 | 0.511866 |
| ENSG00000037280 | FLT4 | 101.7271732 | -0.333878011 | 0.529602349 | -0.834747945 | 0.40385964 | 0.604652 |
| ENSG00000235251 | | 2.767358 | -0.3338839 | 0.7590809 | -0.9139615 | 0.3607371 | 0.562763 |
| ENSG00000156265 | MAP3K7CL | 1.582289 | -0.3340113 | 0.7543668 | -0.9153115 | 0.3600281 | 0.56202 |
| ENSG00000112096 | SOD2 | 2019.950517 | -0.334025651 | 0.178155748 | -1.937470857 | 0.05268781 | 0.144375 |
| ENSG00000233997 | LINC01425 | 7.767439 | -0.3341303 | 0.6637266 | -0.8331167 | 0.4047789 | 0.605516 |
| ENSG00000135211 | TMEM60 | 805.0461492 | -0.33437399 | 0.174234698 | -1.969836449 | 0.04885712 | 0.13611 |
| ENSG00000234884 | | 0.99133 | -0.3344581 | 0.8670002 | -1.568304 | 0.1168103 | NA |
| ENSG00000169813 | HNRNPF | 22338.63 | -0.3345059 | 0.1500483 | -2.275416 | 0.02288097 | 0.075707 |
| ENSG00000105655 | ISYNA1 | 4849.653081 | -0.334552731 | 0.26860535 | -1.336641735 | 0.18133961 | 0.355903 |
| ENSG00000247345 | | 1.792013 | -0.3346323 | 0.8449461 | -0.9967132 | 0.3189038 | 0.519932 |
| ENSG00000148516 | ZEB1 | 8.654761 | -0.3347002 | 0.5462794 | -0.8332299 | 0.4047151 | 0.605497 |
| ENSG00000006715 | VPS41 | 4595.153572 | -0.334705658 | 0.215484415 | -1.612559118 | 0.10684033 | 0.244163 |
| ENSG00000121578 | B4GALT4 | 2300.97656 | -0.334714406 | 0.183301943 | -1.899029243 | 0.05756063 | 0.154151 |
| ENSG00000171962 | DRC3 | 188.0931 | -0.3348055 | 0.4042713 | -0.9785615 | 0.3277967 | 0.529243 |
| ENSG00000267147 | LINC01842 | 4.986869 | -0.334861 | 0.6360744 | -0.8255655 | 0.4090506 | 0.609694 |
| ENSG00000286584 | | 30.29503 | -0.3349147 | 0.4197882 | -0.9432229 | 0.3455669 | 0.547085 |
| ENSG00000107438 | PDLIM1 | 3483.495827 | -0.334999131 | 0.257427465 | -1.361989895 | 0.17320107 | 0.344677 |
| ENSG00000104738 | MCM4 | 8377.630337 | -0.335007354 | 0.387613062 | -0.998110474 | 0.31822579 | 0.519345 |
| ENSG00000278259 | MYO19 | 3663.067 | -0.335047 | 0.2267185 | -1.554112 | 0.1201577 | 0.265575 |
| ENSG00000058600 | POLR3E | 3722.724211 | -0.335142011 | 0.143752973 | -2.374722288 | 0.01756216 | 0.061211 |

| | | | | | | | |
|---|---|---|---|---|---|---|---|
| ENSG00000269997 | | 15.00391 | -0.3351549 | 0.5653445 | -0.8206825 | 0.4118271 | 0.612186 |
| ENSG00000136997 | MYC | 14302.48286 | -0.335222123 | 0.286086541 | -1.264612762 | 0.20601018 | 0.388108 |
| ENSG00000235695 | HIGD2AP2 | 2.199554 | -0.3352674 | 0.7463532 | -0.9001419 | 0.3680447 | 0.570158 |
| ENSG00000287425 | | 4.136852 | -0.3353155 | 0.6957008 | -0.8571022 | 0.3913884 | 0.5932 |
| ENSG00000257900 | | 5.601577 | -0.3353452 | 0.5411999 | -0.8276416 | 0.4078735 | 0.608532 |
| ENSG00000132017 | DCAF15 | 1261.870219 | -0.335416848 | 0.249584384 | -1.429514071 | 0.15285653 | 0.315339 |
| ENSG00000183960 | KCNH8 | 61.95984 | -0.3354235 | 0.4814878 | -0.8777237 | 0.3800937 | 0.582058 |
| ENSG00000283973 | | 1.076147 | -0.3354433 | 0.8145925 | -1.106781 | 0.2683885 | NA |
| ENSG00000170364 | SETMAR | 945.8807 | -0.3354588 | 0.1514862 | -2.261982 | 0.02369854 | 0.077734 |
| ENSG00000157927 | RADIL | 156.219 | -0.335504 | 0.2842325 | -1.274621 | 0.2024434 | 0.383619 |
| ENSG00000135686 | KLHL36 | 6797.205031 | -0.33553497 | 0.205294536 | -1.708887002 | 0.08747188 | 0.211046 |
| ENSG00000182810 | DDX28 | 1356.277 | -0.3355541 | 0.279854 | -1.288338 | 0.1976282 | 0.377663 |
| ENSG00000030582 | GRN | 28264.12398 | -0.335618459 | 0.241103378 | -1.367254671 | 0.1715455 | 0.342255 |
| ENSG00000021645 | NRXN3 | 450.8597791 | -0.335623166 | 0.358841192 | -1.035068623 | 0.30063687 | 0.499914 |
| ENSG00000196141 | SPATS2L | 2368.181 | -0.3356592 | 0.281142 | -1.27712 | 0.2015597 | 0.382435 |
| ENSG00000103160 | HSDL1 | 3858.726827 | -0.335701095 | 0.159406649 | -2.174516622 | 0.02966636 | 0.09296 |
| ENSG00000103042 | SLC38A7 | 1316.87043 | -0.335728409 | 0.253461116 | -1.403371678 | 0.16050603 | 0.327031 |
| ENSG00000264932 | | 4.569493 | -0.335763 | 0.7009693 | -0.8589991 | 0.390341 | 0.592276 |
| ENSG00000100836 | PABPN1 | 4609.86115 | -0.335854449 | 0.173249811 | -1.995912529 | 0.04594345 | 0.130011 |
| ENSG00000170348 | TMED10 | 19322.79 | -0.3359299 | 0.1135857 | -2.978061 | 0.00290079 | 0.014191 |
| ENSG00000255508 | | 340.6609 | -0.3359482 | 0.2274001 | -1.541052 | 0.1233042 | 0.270615 |
| ENSG00000188219 | POTEE | 126.5108 | -0.3359583 | 0.4027761 | -0.9745047 | 0.329806 | 0.530954 |
| ENSG00000263317 | LINC01982 | 0.8821661 | -0.3359613 | 0.8669211 | -1.313151 | 0.1891322 | NA |
| ENSG00000229759 | MRPS18AP1 | 1.967175 | -0.3360518 | 0.7559849 | -0.9330546 | 0.3507918 | 0.552408 |
| ENSG00000120645 | IQSEC3 | 0.957683887 | -0.336144602 | 0.800298031 | -1.051775912 | 0.29290237 | NA |
| ENSG00000175749 | EIF3KP1 | 1.814433 | -0.3361454 | 0.7901848 | -1.030414 | 0.3028158 | 0.502374 |
| ENSG00000158470 | B4GALT5 | 5516.54 | -0.3361949 | 0.178359 | -1.940347 | 0.05233755 | 0.1436 |
| ENSG00000248863 | | 3.064263 | -0.3362309 | 0.6308718 | -0.819469 | 0.4125189 | 0.612713 |
| ENSG00000260034 | LCMT1-AS2 | 15.32802 | -0.3363255 | 0.517761 | -0.8484602 | 0.3961817 | 0.597534 |
| ENSG00000288061 | | 2.293537 | -0.3364485 | 0.7000966 | -0.8543423 | 0.3929154 | 0.594546 |
| ENSG00000233901 | LINC01503 | 248.5322 | -0.3365542 | 0.2888504 | -1.257311 | 0.2086411 | 0.391565 |
| ENSG00000169495 | HTRA4 | 43.07226 | -0.3365974 | 0.4783397 | -0.8836078 | 0.3769079 | 0.578982 |
| ENSG00000232884 | | 3.170478 | -0.3367057 | 0.7519702 | -0.9323469 | 0.3511573 | 0.55285 |

| | | | | | | | |
|---|---|---|---|---|---|---|---|
| ENSG00000105856 | HBP1 | 2520.472495 | -0.336725172 | 0.201898379 | -1.732865595 | 0.08311956 | 0.203165 |
| ENSG00000261186 | LINC01238 | 26.6327 | -0.3367968 | 0.3968218 | -0.9779046 | 0.3281215 | 0.529382 |
| ENSG00000106348 | IMPDH1 | 6657.531377 | -0.336807341 | 0.166732365 | -2.070213397 | 0.03843237 | 0.113356 |
| ENSG00000063241 | ISOC2 | 3819.962552 | -0.336848033 | 0.285557933 | -1.288877714 | 0.1974406 | 0.377368 |
| ENSG00000171067 | C11orf24 | 1839.352 | -0.336911 | 0.1969167 | -1.780233 | 0.07503787 | 0.188101 |
| ENSG00000269737 | | 51.27209 | -0.336977 | 0.3666951 | -1.039346 | 0.2986439 | 0.497846 |
| ENSG00000200534 | SNORA33 | 88.59735 | -0.337047 | 0.3009016 | -1.217996 | 0.2232253 | 0.409902 |
| ENSG00000244218 | RN7SL81P | 5.503133 | -0.3371082 | 0.6668798 | -0.818724 | 0.4129439 | 0.61302 |
| ENSG00000231407 | GORAB-AS1 | 21.16535 | -0.3371457 | 0.4440013 | -0.9165306 | 0.3593887 | 0.56124 |
| ENSG00000259470 | KRT8P9 | 111.3688 | -0.3371875 | 0.2434121 | -1.461434 | 0.1438964 | 0.301944 |
| ENSG00000133624 | ZNF767P | 1376.828391 | -0.337189254 | 0.377803185 | -1.023382162 | 0.30612719 | 0.506066 |
| ENSG00000280327 | | 19.20682 | -0.3372096 | 0.5142686 | -0.8530475 | 0.393633 | 0.595278 |
| ENSG00000268902 | CSAG2 | 1.78769 | -0.3372419 | 0.8725996 | -1.368519 | 0.1711495 | 0.341772 |
| ENSG00000237406 | NDUFA9P1 | 1.147575 | -0.3372618 | 0.7953195 | -1.037467 | 0.2995184 | NA |
| ENSG00000181404 | WASHC1 | 155.295 | -0.3373606 | 0.2361214 | -1.503672 | 0.1326659 | 0.284971 |
| ENSG00000034713 | GABARAPL2 | 6040.877108 | -0.337418581 | 0.158708449 | -2.174941302 | 0.02963452 | 0.092882 |
| ENSG00000273343 | | 49.99604 | -0.337528 | 0.3765781 | -1.02194 | 0.3068092 | 0.507004 |
| ENSG00000268061 | NAPA-AS1 | 38.83168 | -0.3376096 | 0.3599893 | -1.05844 | 0.2898548 | 0.488403 |
| ENSG00000111276 | CDKN1B | 4786.809737 | -0.337621132 | 0.184968736 | -1.833044301 | 0.06679598 | 0.172335 |
| ENSG00000235325 | SRSF3P5 | 1.854498 | -0.3377194 | 0.7987287 | -1.030997 | 0.3025422 | 0.502014 |
| ENSG00000176974 | SHMT1 | 3045.873 | -0.3377392 | 0.3221918 | -1.149257 | 0.2504502 | 0.443048 |
| ENSG00000135821 | GLUL | 11862.4899 | -0.33775751 | 0.151325169 | -2.29941229 | 0.02148154 | 0.072073 |
| ENSG00000170373 | CST1 | 3.712654 | -0.3377611 | 0.8486221 | -1.051636 | 0.2929666 | 0.491833 |
| ENSG00000136010 | ALDH1L2 | 2270.704415 | -0.337870906 | 0.698814424 | -0.872491897 | 0.38294009 | 0.58527 |
| ENSG00000258734 | | 2.266242 | -0.3378762 | 0.7529351 | -0.9370637 | 0.3487258 | 0.550392 |
| ENSG00000249459 | ZNF286B | 197.6093 | -0.3378818 | 0.2156002 | -1.621144 | 0.1049867 | 0.240984 |
| ENSG00000258388 | PPT2-EGFL8 | 35.04266 | -0.3378915 | 0.38478 | -1.00601 | 0.3144107 | 0.515558 |
| ENSG00000236281 | NDUFB9P2 | 39.10225 | -0.3379104 | 0.3798624 | -1.020556 | 0.3074649 | 0.507632 |
| ENSG00000153914 | SREK1 | 6836.674 | -0.3379463 | 0.1640929 | -2.11777 | 0.03419458 | 0.103795 |
| ENSG00000286444 | | 1.122361 | -0.3380277 | 0.8498443 | -1.046771 | 0.2952053 | NA |
| ENSG00000204764 | RANBP17 | 1362.899 | -0.3380446 | 0.1736878 | -2.003446 | 0.04512941 | 0.128309 |
| ENSG00000131269 | ABCB7 | 1322.880933 | -0.338074514 | 0.145639468 | -2.369267077 | 0.01782338 | 0.061934 |
| ENSG00000226328 | NUP50-DT | 547.9635 | -0.3380749 | 0.2309898 | -1.538125 | 0.124018 | 0.271711 |

| | | | | | | | |
|---|---|---|---|---|---|---|---|
| ENSG00000183426 | NPIPA1 | 524.336 | -0.3381073 | 0.2792294 | -1.301103 | 0.193223 | 0.371849 |
| ENSG00000257953 | | 2.508413 | -0.3382034 | 0.7112128 | -0.8734429 | 0.3824217 | 0.584713 |
| ENSG00000124571 | XPO5 | 6670.173025 | -0.338205019 | 0.120222036 | -2.831305168 | 0.00463585 | 0.020844 |
| ENSG00000257851 | HNRNPA3P10 | 516.3954 | -0.338373 | 0.2418193 | -1.513071 | 0.1302616 | 0.281216 |
| ENSG00000171056 | SOX7 | 6.968504 | -0.3385268 | 0.5780002 | -0.8255432 | 0.4090633 | 0.609694 |
| ENSG00000255031 | | 35.10781 | -0.3385882 | 0.4000624 | -0.9894373 | 0.3224492 | 0.523217 |
| ENSG00000128294 | TPST2 | 674.3649827 | -0.338619623 | 0.199328181 | -1.759242652 | 0.0785363 | 0.194521 |
| ENSG00000260400 | | 34.58589 | -0.3386244 | 0.4628453 | -0.8960766 | 0.3702119 | 0.572369 |
| ENSG00000130204 | TOMM40 | 5916.574016 | -0.338745554 | 0.314875983 | -1.179987177 | 0.23800532 | 0.427701 |
| ENSG00000169245 | CXCL10 | 1.710934 | -0.3387485 | 0.8669816 | -1.29197 | 0.1963675 | 0.376156 |
| ENSG00000144468 | RHBDD1 | 971.7827 | -0.3387801 | 0.2912173 | -1.259162 | 0.207972 | 0.390585 |
| ENSG00000255445 | | 3.394701 | -0.3388108 | 0.6976068 | -0.8610046 | 0.3892355 | 0.591157 |
| ENSG00000119013 | NDUFB3 | 1862.92527 | -0.338867729 | 0.224074576 | -1.597471427 | 0.11016066 | 0.249647 |
| ENSG00000101441 | CST4 | 3.761170994 | -0.338935829 | 0.849313893 | -1.05105646 | 0.29323266 | 0.492223 |
| ENSG00000141503 | MINK1 | 7253.199 | -0.3390543 | 0.1217666 | -2.813594 | 0.00489911 | 0.02181 |
| ENSG00000234707 | SEC61G-DT | 2.461488 | -0.3390776 | 0.7678935 | -0.9443561 | 0.3449877 | 0.546537 |
| ENSG00000146090 | RASGEF1C | 25.7138 | -0.3390827 | 0.5529648 | -0.841116 | 0.4002829 | 0.601228 |
| ENSG00000223940 | KRT8P8 | 6.677713 | -0.3391674 | 0.590989 | -0.8225625 | 0.4107568 | 0.611053 |
| ENSG00000131495 | NDUFA2 | 1765.488307 | -0.339192546 | 0.246409641 | -1.454809213 | 0.14572209 | 0.304809 |
| ENSG00000205864 | KRTAP5-6 | 0.8919149 | -0.3391992 | 0.8041806 | -1.071425 | 0.2839785 | NA |
| ENSG00000214029 | ZNF891 | 474.4086 | -0.3392404 | 0.361786 | -1.05466 | 0.2915806 | 0.490317 |
| ENSG00000234420 | ZNF37BP | 2249.859 | -0.3392607 | 0.1867249 | -1.876494 | 0.0605875 | 0.160158 |
| ENSG00000164675 | IQUB | 3.728455 | -0.3392742 | 0.6695002 | -0.8275258 | 0.4079391 | 0.608532 |
| ENSG00000168509 | HJV | 1.119452 | -0.3393967 | 0.8608732 | -1.418381 | 0.1560794 | NA |
| ENSG00000256452 | | 1.388183 | -0.3394219 | 0.7879006 | -0.9997393 | 0.3174367 | NA |
| ENSG00000206113 | CFAP99 | 28.01552 | -0.3395976 | 0.471502 | -0.8982624 | 0.3690457 | 0.571164 |
| ENSG00000219395 | HSPA8P15 | 78.51074 | -0.3396676 | 0.3014106 | -1.227651 | 0.2195779 | 0.405649 |
| ENSG00000215692 | | 2.887979 | -0.339747 | 0.7145929 | -0.8662818 | 0.3863357 | 0.588665 |
| ENSG00000166825 | ANPEP | 3.31953 | -0.3398053 | 0.6511895 | -0.8321745 | 0.4053105 | 0.606107 |
| ENSG00000214013 | GANC | 363.9391 | -0.3398307 | 0.1880279 | -1.86338 | 0.06240886 | 0.163751 |
| ENSG00000244245 | | 105.1684 | -0.339899 | 0.2487837 | -1.446265 | 0.148103 | 0.308406 |
| ENSG00000183023 | SLC8A1 | 362.0809 | -0.340018 | 0.2587888 | -1.397455 | 0.1622768 | 0.329502 |
| ENSG00000127863 | TNFRSF19 | 40.6850031 | -0.340018846 | 0.640922609 | -0.812964899 | 0.41623818 | 0.616136 |

| | | | | | | | |
|---|---|---|---|---|---|---|---|
| ENSG00000083937 | CHMP2B | 4134.914019 | -0.340104577 | 0.072952687 | -4.699392206 | 2.61E-06 | 2.99E-05 |
| ENSG00000166579 | NDEL1 | 1994.963 | -0.3401727 | 0.1501804 | -2.318479 | 0.0204233 | 0.069093 |
| ENSG00000232712 | KIZ-AS1 | 4.868203 | -0.3401941 | 0.6148784 | -0.8158792 | 0.4145692 | 0.614649 |
| ENSG00000267156 | TPMTP1 | 54.78293 | -0.3402014 | 0.2750139 | -1.32654 | 0.184661 | 0.36004 |
| ENSG00000160299 | PCNT | 7163.063 | -0.3402087 | 0.2071359 | -1.76218 | 0.07803883 | 0.19356 |
| ENSG00000199664 | RNU6-1266P | 1.301979 | -0.3403079 | 0.8189682 | -1.122161 | 0.2617942 | NA |
| ENSG00000119912 | IDE | 5044.435709 | -0.340311045 | 0.162663636 | -2.159236396 | 0.03083183 | 0.095854 |
| ENSG00000132510 | KDM6B | 3009.724249 | -0.340323183 | 0.295946929 | -1.254896773 | 0.20951624 | 0.392819 |
| ENSG00000104783 | KCNN4 | 200.5990557 | -0.340348792 | 0.623514131 | -0.827580611 | 0.40790805 | 0.608532 |
| ENSG00000147439 | BIN3 | 527.2005 | -0.340401 | 0.1647394 | -2.109927 | 0.03486463 | 0.105335 |
| ENSG00000183273 | CCDC60 | 2.143735 | -0.3404964 | 0.7908099 | -1.005552 | 0.3146312 | 0.515758 |
| ENSG00000205544 | TMEM256 | 273.8346 | -0.3405151 | 0.3154923 | -1.180798 | 0.2376828 | 0.427295 |
| ENSG00000087448 | KLHL42 | 1808.726221 | -0.340587752 | 0.166645828 | -2.08550439 | 0.03702354 | 0.110117 |
| ENSG00000185009 | AP3M1 | 3336.505 | -0.3408882 | 0.1340881 | -2.580178 | 0.00987495 | 0.03847 |
| ENSG00000141452 | RMC1 | 1052.798 | -0.3408945 | 0.105383 | -3.276808 | 0.00104988 | 0.005973 |
| ENSG00000166526 | ZNF3 | 2425.742 | -0.3410507 | 0.1385274 | -2.504584 | 0.01225957 | 0.045793 |
| ENSG00000250486 | FAM218A | 197.5693 | -0.3411289 | 0.3003318 | -1.230322 | 0.2185767 | 0.404356 |
| ENSG00000114021 | NIT2 | 4717.889087 | -0.341231118 | 0.138575742 | -2.505454152 | 0.01222943 | 0.045719 |
| ENSG00000025293 | PHF20 | 2254.226808 | -0.341254603 | 0.121549571 | -2.85968965 | 0.00424056 | 0.019362 |
| ENSG00000105227 | PRX | 424.1057955 | -0.341276146 | 0.205727514 | -1.723985207 | 0.08471051 | 0.206068 |
| ENSG00000181754 | AMIGO1 | 1244.612 | -0.3413568 | 0.244245 | -1.49103 | 0.1359535 | 0.289737 |
| ENSG00000100297 | MCM5 | 3201.259996 | -0.341458276 | 0.555466912 | -0.848363193 | 0.39623574 | 0.597546 |
| ENSG00000196652 | ZKSCAN5 | 1058.035 | -0.3414692 | 0.1533441 | -2.272535 | 0.02305419 | 0.076147 |
| ENSG00000244306 | | 1315.466 | -0.3416088 | 0.2571563 | -1.379394 | 0.1677734 | 0.337388 |
| ENSG00000285711 | LINC02835 | 1.071804 | -0.3416764 | 0.8101682 | -1.082647 | 0.278965 | NA |
| ENSG00000172771 | EFCAB12 | 11.39443 | -0.3416899 | 0.483389 | -0.8872965 | 0.3749193 | 0.577124 |
| ENSG00000070010 | UFD1 | 3211.477857 | -0.341697784 | 0.238286283 | -1.505603168 | 0.13216908 | 0.284248 |
| ENSG00000203865 | ATP1A1-AS1 | 308.9784 | -0.341756 | 0.2187015 | -1.630821 | 0.1029281 | 0.237635 |
| ENSG00000268104 | SLC6A14 | 192.2595 | -0.3418131 | 0.364263 | -1.063252 | 0.2876676 | 0.485982 |
| ENSG00000101986 | ABCD1 | 1882.776831 | -0.34186311 | 0.19985388 | -1.768642594 | 0.07695354 | 0.191655 |
| ENSG00000181690 | PLAG1 | 137.1533 | -0.3419893 | 0.4969452 | -0.8630729 | 0.3880974 | 0.5904 |
| ENSG00000166900 | STX3 | 4236.499 | -0.3420301 | 0.2672151 | -1.368685 | 0.1710979 | 0.34171 |
| ENSG00000242259 | C22orf39 | 1540.787 | -0.3422343 | 0.1408806 | -2.477074 | 0.01324646 | 0.048766 |

| | | | | | | | |
|---|---|---|---|---|---|---|---|
| ENSG00000269984 | | 69.87621 | -0.3422345 | 0.3897509 | -1.013616 | 0.3107658 | 0.511602 |
| ENSG00000136758 | YME1L1 | 13214.68073 | -0.342300362 | 0.127811644 | -2.709627229 | 0.00673589 | 0.028226 |
| ENSG00000130734 | ATG4D | 1457.61049 | -0.342466519 | 0.198703153 | -1.783692768 | 0.0744736 | 0.187163 |
| ENSG00000204843 | DCTN1 | 9417.584 | -0.3425015 | 0.1363724 | -2.579681 | 0.00988915 | 0.038509 |
| ENSG00000278075 | | 6.904101 | -0.3426247 | 0.6255971 | -0.8341241 | 0.404211 | 0.605064 |
| ENSG00000059691 | GATB | 1494.521295 | -0.34263552 | 0.235621615 | -1.529820372 | 0.1260612 | 0.274647 |
| ENSG00000161692 | DBF4B | 896.928 | -0.3428209 | 0.365529 | -1.068725 | 0.2851937 | 0.483186 |
| ENSG00000273521 | | 13.52551 | -0.3428282 | 0.5880929 | -0.8388527 | 0.401552 | 0.602487 |
| ENSG00000250240 | | 29.42074 | -0.3428473 | 0.4194072 | -0.9577858 | 0.3381708 | 0.539471 |
| ENSG00000272076 | | 1.760673 | -0.3429009 | 0.7959864 | -1.031923 | 0.3021082 | 0.501482 |
| ENSG00000272799 | | 4.219591 | -0.3429458 | 0.6919361 | -0.8581064 | 0.3908337 | 0.592731 |
| ENSG00000135698 | MPHOSPH6 | 1769.6327 | -0.342982321 | 0.246234232 | -1.471265652 | 0.14121929 | 0.298066 |
| ENSG00000249550 | LINC01234 | 5.672223 | -0.3430375 | 0.7979302 | -1.037188 | 0.2996482 | 0.498891 |
| ENSG00000123219 | CENPK | 579.293349 | -0.3431047 | 0.424139507 | -0.965562993 | 0.3342629 | 0.535452 |
| ENSG00000249395 | CASC9 | 286.6746 | -0.3431228 | 0.1738629 | -2.016572 | 0.04374023 | 0.125352 |
| ENSG00000241520 | | 4.333933 | -0.3432919 | 0.6787156 | -0.8524828 | 0.3939462 | 0.595538 |
| ENSG00000163468 | CCT3 | 30859.36 | -0.3433685 | 0.1407651 | -2.538245 | 0.01114098 | 0.042444 |
| ENSG00000232042 | | 1.596216 | -0.3434455 | 0.8204891 | -1.127756 | 0.2594231 | 0.453713 |
| ENSG00000285669 | | 78.77294 | -0.3435608 | 0.4519354 | -0.9250386 | 0.3549459 | 0.556901 |
| ENSG00000281649 | EBLN3P | 3802.661 | -0.3435683 | 0.2438665 | -1.473013 | 0.1407475 | 0.297276 |
| ENSG00000148444 | COMMD3 | 833.1282 | -0.3436182 | 0.2270357 | -1.594224 | 0.1108858 | 0.250777 |
| ENSG00000204301 | NOTCH4 | 22.72028 | -0.343647 | 0.450909 | -0.928725 | 0.3530316 | 0.554946 |
| ENSG00000272384 | | 92.90473 | -0.3437159 | 0.3201702 | -1.178859 | 0.2384544 | 0.428305 |
| ENSG00000196975 | ANXA4 | 4352.992 | -0.3437298 | 0.1621725 | -2.173153 | 0.02976879 | 0.093149 |
| ENSG00000236090 | LDHAP3 | 22.27487 | -0.3437383 | 0.4580151 | -0.9204114 | 0.3573578 | 0.55943 |
| ENSG00000233230 | | 22.43152 | -0.3438572 | 0.477053 | -0.8970395 | 0.3696978 | 0.571807 |
| ENSG00000265917 | MIR3685 | 15.56066 | -0.3439133 | 0.4730744 | -0.9038062 | 0.3660982 | 0.568025 |
| ENSG00000225159 | NPM1P39 | 1544.569 | -0.3440766 | 0.2175934 | -1.649566 | 0.09903177 | 0.231008 |
| ENSG00000222383 | RNA5SP203 | 3.582498 | -0.3441681 | 0.6894463 | -0.855332 | 0.3923674 | 0.594061 |
| ENSG00000145901 | TNIP1 | 4029.049 | -0.3442865 | 0.1180838 | -2.95576 | 0.003119 | 0.015004 |
| ENSG00000173641 | HSPB7 | 4.179283 | -0.3443083 | 0.6879145 | -0.8575916 | 0.391118 | 0.592926 |
| ENSG00000148339 | SLC25A25 | 1674.349 | -0.344371 | 0.2477867 | -1.465264 | 0.142849 | 0.300498 |
| ENSG00000267741 | UBE2L4 | 11.79728 | -0.3444347 | 0.5205579 | -0.8673464 | 0.3857522 | 0.588047 |

| | | | | | | | |
|---|---|---|---|---|---|---|---|
| ENSG00000101367 | MAPRE1 | 6338.960382 | -0.344490927 | 0.22220995 | -1.621651459 | 0.104878 | 0.240839 |
| ENSG00000164197 | RNF180 | 124.997 | -0.3445884 | 0.3589395 | -1.082463 | 0.279047 | 0.476216 |
| ENSG00000214160 | ALG3 | 2852.921 | -0.3446147 | 0.1377909 | -2.579989 | 0.00988036 | 0.03848 |
| ENSG00000173214 | MFSD4B | 819.7712 | -0.3447322 | 0.2342873 | -1.548355 | 0.1215369 | 0.26781 |
| ENSG00000137266 | SLC22A23 | 1605.52074 | -0.344774807 | 0.236850085 | -1.511088787 | 0.13076583 | 0.282006 |
| ENSG00000233327 | USP32P2 | 8.693646 | -0.3448242 | 0.573727 | -0.8447193 | 0.3982676 | 0.599353 |
| ENSG00000244952 | | 5.512739 | -0.3448735 | 0.6749736 | -0.8510855 | 0.3947219 | 0.596449 |
| ENSG00000220323 | H2BC19P | 1638.338 | -0.3448747 | 0.2342706 | -1.551614 | 0.1207547 | 0.2666 |
| ENSG00000229376 | CICP3 | 3.727869 | -0.3449445 | 0.7333117 | -0.9031624 | 0.3664397 | 0.568389 |
| ENSG00000263834 | MIR4635 | 4.45254 | -0.3450267 | 0.7257119 | -0.8951921 | 0.3706844 | 0.5728 |
| ENSG00000234170 | LINC02645 | 1.419882 | -0.3450714 | 0.9006594 | -2.11091 | 0.03478006 | NA |
| ENSG00000224559 | LINC01087 | 344.4143 | -0.3450777 | 0.3881061 | -1.028706 | 0.3036179 | 0.503233 |
| ENSG00000229425 | | 1.599807 | -0.3450932 | 0.887484 | -1.786889 | 0.07395536 | 0.186301 |
| ENSG00000006283 | CACNA1G | 11.14772001 | -0.345122071 | 0.664834289 | -0.854192674 | 0.39299824 | 0.594622 |
| ENSG00000141959 | PFKL | 14173.37 | -0.3451674 | 0.1520664 | -2.307217 | 0.02104272 | 0.070833 |
| ENSG00000269924 | | 1.776823 | -0.3452346 | 0.7405393 | -0.9124162 | 0.3615497 | 0.563502 |
| ENSG00000132635 | PCED1A | 2081.53162 | -0.345294887 | 0.18466197 | -1.946922687 | 0.051544 | 0.141797 |
| ENSG00000164024 | METAP1 | 4192.083 | -0.3453174 | 0.196218 | -1.838689 | 0.06596092 | 0.170711 |
| ENSG00000232727 | YWHAEP1 | 15.3851 | -0.3453379 | 0.4406411 | -0.9458275 | 0.3442366 | 0.545836 |
| ENSG00000267296 | CEBPA-DT | 272.1343 | -0.3453625 | 0.2426628 | -1.502347 | 0.1330075 | 0.285427 |
| ENSG00000226666 | HSPA9P1 | 210.3084 | -0.3454603 | 0.254171 | -1.440494 | 0.1497278 | 0.31074 |
| ENSG00000113645 | WWC1 | 6894.272427 | -0.34558911 | 0.143480146 | -2.447170568 | 0.01439827 | 0.052222 |
| ENSG00000225392 | MRPL36P1 | 2.15281 | -0.3456003 | 0.7984318 | -1.03857 | 0.2990046 | 0.49826 |
| ENSG00000113240 | CLK4 | 1850.260229 | -0.3456956 | 0.27474619 | -1.375306323 | 0.1690365 | 0.339107 |
| ENSG00000287686 | | 48.91569 | -0.3457992 | 0.3184215 | -1.187685 | 0.2349575 | 0.424204 |
| ENSG00000108176 | DNAJC12 | 212.0590824 | -0.345857172 | 0.430421783 | -0.962469556 | 0.33581379 | 0.536902 |
| ENSG00000256612 | CYP2B7P | 22.37554 | -0.3458589 | 0.5973496 | -0.8471122 | 0.3969326 | 0.598029 |
| ENSG00000174238 | PITPNA | 4992.919 | -0.3459018 | 0.1304591 | -2.689027 | 0.00716606 | 0.029719 |
| ENSG00000048162 | NOP16 | 2483.50938 | -0.345936984 | 0.234494503 | -1.552967761 | 0.12043084 | 0.266067 |
| ENSG00000079335 | CDC14A | 272.6740458 | -0.345950288 | 0.254559724 | -1.456117374 | 0.14536018 | 0.30434 |
| ENSG00000246731 | | 6.741414 | -0.3460323 | 0.5553512 | -0.8560424 | 0.3919743 | 0.593682 |
| ENSG00000287127 | | 1.653944 | -0.3460693 | 0.8404808 | -1.232233 | 0.2178621 | 0.403577 |
| ENSG00000286156 | BMS1P23 | 297.8444 | -0.3460893 | 0.2368638 | -1.544194 | 0.1225414 | 0.269342 |

| | | | | | | | |
|---|---|---|---|---|---|---|---|
| ENSG00000229268 | PES1P2 | 7.072161 | -0.3461255 | 0.6798154 | -0.8550209 | 0.3925396 | 0.594098 |
| ENSG00000213005 | PTTG3P | 4.850701 | -0.3461411 | 0.6857543 | -0.8702996 | 0.3841367 | 0.586492 |
| ENSG00000145734 | BDP1 | 3711.742 | -0.3461651 | 0.2726368 | -1.390355 | 0.1644211 | 0.332543 |
| ENSG00000227311 | | 4.371797 | -0.346189 | 0.6881073 | -0.8649558 | 0.3870631 | 0.589368 |
| ENSG00000166197 | NOLC1 | 12338.62 | -0.3461972 | 0.2462688 | -1.486508 | 0.1371447 | 0.291481 |
| ENSG00000264520 | | 3.930957 | -0.3462644 | 0.7078 | -0.8831946 | 0.3771311 | 0.579158 |
| ENSG00000286813 | | 20.49981 | -0.3462753 | 0.5335161 | -0.8623665 | 0.3884859 | 0.590521 |
| ENSG00000040633 | PHF23 | 3058.613264 | -0.346316631 | 0.190494189 | -1.877094179 | 0.06050519 | 0.160068 |
| ENSG00000196081 | ZNF724 | 52.47262 | -0.3463384 | 0.4740505 | -0.9055186 | 0.3651907 | 0.567048 |
| ENSG00000235262 | KDM5C-IT1 | 1.407492 | -0.3463404 | 0.7833102 | -0.9843433 | 0.3249467 | NA |
| ENSG00000134184 | GSTM1 | 1.243032505 | -0.346408401 | 0.826584479 | -1.201486788 | 0.22956243 | NA |
| ENSG00000176783 | RUFY1 | 4342.025 | -0.3464512 | 0.1302635 | -2.658985 | 0.00783764 | 0.031992 |
| ENSG00000159111 | MRPL10 | 5679.309 | -0.3464651 | 0.1658735 | -2.144069 | 0.03202738 | 0.098662 |
| ENSG00000248150 | LINC02150 | 1.414308 | -0.3464739 | 0.8382065 | -1.22674 | 0.2199204 | NA |
| ENSG00000254900 | | 4.390059 | -0.346693 | 0.6617947 | -0.8562359 | 0.3918673 | 0.593641 |
| ENSG00000108854 | SMURF2 | 4109.296525 | -0.346731629 | 0.166194098 | -2.142958818 | 0.03211641 | 0.098842 |
| ENSG00000273542 | H4C12 | 58.44831 | -0.3468219 | 0.3684082 | -1.065994 | 0.2864265 | 0.484687 |
| ENSG00000174943 | KCTD13 | 1665.508 | -0.3468965 | 0.1748717 | -2.028954 | 0.04246298 | 0.122524 |
| ENSG00000149474 | KAT14 | 1456.287 | -0.3469052 | 0.1594181 | -2.262106 | 0.02369085 | 0.077727 |
| ENSG00000005175 | RPAP3 | 2133.616244 | -0.347043113 | 0.194968997 | -1.852296177 | 0.0639833 | 0.166676 |
| ENSG00000169174 | PCSK9 | 15.87551 | -0.3471172 | 0.8940523 | -1.782043 | 0.07474221 | 0.187625 |
| ENSG00000235016 | SEMA3F-AS1 | 68.06589 | -0.3471588 | 0.262665 | -1.409118 | 0.1588004 | 0.324631 |
| ENSG00000234752 | LINC02676 | 8.726771 | -0.3471999 | 0.5462504 | -0.8593327 | 0.390157 | 0.592145 |
| ENSG00000173456 | RNF26 | 2195.006 | -0.3473154 | 0.278857 | -1.342093 | 0.179566 | 0.353635 |
| ENSG00000132763 | MMACHC | 766.9698309 | -0.347484683 | 0.157136539 | -2.262856035 | 0.02364457 | 0.077605 |
| ENSG00000149357 | LAMTOR1 | 7472.335 | -0.3475243 | 0.1509901 | -2.365693 | 0.01799638 | 0.062446 |
| ENSG00000175221 | MED16 | 2065.812 | -0.3475697 | 0.1932052 | -1.856056 | 0.06344564 | 0.165686 |
| ENSG00000101746 | NOL4 | 1.820013877 | -0.34760362 | 0.753997949 | -0.945016465 | 0.34465047 | 0.546199 |
| ENSG00000261416 | | 1.421052 | -0.3476118 | 0.8208097 | -1.1668 | 0.2432913 | NA |
| ENSG00000226355 | | 0.8072098 | -0.3476282 | 0.8459679 | -1.26873 | 0.2045373 | NA |
| ENSG00000178602 | OTOS | 1.245199 | -0.3477012 | 0.8358994 | -1.239857 | 0.2150284 | NA |
| ENSG00000113595 | TRIM23 | 997.4535994 | -0.347744488 | 0.235825329 | -1.551091514 | 0.12087976 | 0.266704 |
| ENSG00000220666 | RCC2P7 | 1.064569 | -0.3478024 | 0.8186502 | -1.146483 | 0.2515955 | NA |

| | | | | | | | |
|---|---|---|---|---|---|---|---|
| ENSG00000278619 | MRM1 | 466.7705 | -0.3478524 | 0.1823796 | -1.965092 | 0.04940358 | 0.13734 |
| ENSG00000288060 | | 5.435719 | -0.3479572 | 0.6689912 | -0.8436705 | 0.3988536 | 0.599827 |
| ENSG00000160932 | LY6E | 17382.53 | -0.3481686 | 0.3369265 | -1.124027 | 0.2610016 | 0.45563 |
| ENSG00000117598 | PLPPR5 | 163.9825516 | -0.348248372 | 0.25799491 | -1.434990952 | 0.15128968 | 0.313035 |
| ENSG00000255005 | | 6.334002 | -0.3483287 | 0.5781677 | -0.8543282 | 0.3929232 | 0.594546 |
| ENSG00000285532 | LINC02788 | 0.9519799 | -0.3483802 | 0.8487807 | -1.346708 | 0.1780743 | NA |
| ENSG00000274191 | | 5.267306 | -0.3484369 | 0.6542427 | -0.8541319 | 0.3930319 | 0.59464 |
| ENSG00000276096 | | 5.395728 | -0.3484472 | 0.6792009 | -0.8646439 | 0.3872343 | 0.589525 |
| ENSG00000160870 | CYP3A7 | 2.139562 | -0.3484823 | 0.77706 | -0.9685135 | 0.332788 | 0.533799 |
| ENSG00000279853 | | 2.189096 | -0.3485003 | 0.7132495 | -0.8885514 | 0.3742442 | 0.57635 |
| ENSG00000100918 | REC8 | 679.5777791 | -0.348594982 | 0.342809275 | -1.132864058 | 0.25727135 | 0.451184 |
| ENSG00000135446 | CDK4 | 9068.06014 | -0.3486422 | 0.170542344 | -2.092701572 | 0.0363758 | 0.108713 |
| ENSG00000164587 | RPS14 | 26577.98 | -0.3486927 | 0.1822591 | -1.99785 | 0.04573297 | 0.129553 |
| ENSG00000160584 | SIK3 | 1035.92 | -0.3487203 | 0.1874062 | -1.916116 | 0.05535038 | 0.149777 |
| ENSG00000226777 | FAM30A | 50.0206 | -0.3487283 | 0.9109354 | -3.099484 | 0.00193858 | 0.010087 |
| ENSG00000135439 | AGAP2 | 617.3858173 | -0.348797101 | 0.259250261 | -1.429399006 | 0.15288958 | 0.315382 |
| ENSG00000231312 | MAP4K3-DT | 367.2329 | -0.3488057 | 0.2808074 | -1.332474 | 0.1827046 | 0.357685 |
| ENSG00000121680 | PEX16 | 2332.087838 | -0.348852741 | 0.202567967 | -1.816232939 | 0.06933465 | 0.177249 |
| ENSG00000265095 | FTLP12 | 8.494936 | -0.3488591 | 0.5902113 | -0.850462 | 0.3950683 | 0.596769 |
| ENSG00000149636 | DSN1 | 1315.399 | -0.3488663 | 0.4688373 | -0.9183377 | 0.3584421 | 0.560387 |
| ENSG00000103047 | TANGO6 | 408.8382301 | -0.348886364 | 0.254481359 | -1.467537257 | 0.14222996 | 0.29958 |
| ENSG00000153140 | CETN3 | 1253.43 | -0.3489094 | 0.1921804 | -1.881475 | 0.05990732 | 0.158866 |
| ENSG00000186105 | LRRC70 | 2.011015 | -0.3490996 | 0.789863 | -1.015129 | 0.3100443 | 0.510667 |
| ENSG00000126215 | XRCC3 | 1309.430876 | -0.34911237 | 0.317678716 | -1.214728985 | 0.22446947 | 0.411391 |
| ENSG00000111181 | SLC6A12 | 27.31546372 | -0.349202391 | 0.795958205 | -1.058858628 | 0.28966417 | 0.488207 |
| ENSG00000264229 | RNU4ATAC | 8.177706 | -0.3492047 | 0.6239693 | -0.8514314 | 0.3945297 | 0.596362 |
| ENSG00000013275 | PSMC4 | 6710.667999 | -0.34924436 | 0.197565049 | -1.832718825 | 0.0668444 | 0.172409 |
| ENSG00000218690 | H2AC10P | 31.91164 | -0.3492766 | 0.5017242 | -0.8891826 | 0.3739049 | 0.576001 |
| ENSG00000233254 | RPL21P134 | 2.005852 | -0.3493675 | 0.7508775 | -0.9332933 | 0.3506686 | 0.552375 |
| ENSG00000198952 | SMG5 | 13382.4 | -0.3494052 | 0.1031937 | -3.403661 | 0.00066489 | 0.00402 |
| ENSG00000135736 | CCDC102A | 371.1126053 | -0.34941378 | 0.204945886 | -1.766877968 | 0.07724867 | 0.19216 |
| ENSG00000233483 | EFCAB15P | 50.48944 | -0.3495153 | 0.4235067 | -0.9742326 | 0.3299411 | 0.531028 |
| ENSG00000226553 | IMPDH1P7 | 25.31646 | -0.34975 | 0.3799636 | -1.05245 | 0.2925931 | 0.491428 |

| | | | | | | | |
|---|---|---|---|---|---|---|---|
| ENSG00000260331 | | 4.579886 | -0.3498047 | 0.6923173 | -0.8738619 | 0.3821935 | 0.584531 |
| ENSG00000214433 | GOLGA2P8 | 12.13852 | -0.349863 | 0.4854749 | -0.9011187 | 0.3675252 | 0.569574 |
| ENSG00000102738 | MRPS31 | 1347.310776 | -0.349903692 | 0.182025759 | -1.982800985 | 0.04738966 | 0.133041 |
| ENSG00000281469 | | 2.165413 | -0.3499332 | 0.786573 | -1.013284 | 0.3109244 | 0.51174 |
| ENSG00000227008 | RPS24P19 | 2068.585 | -0.3502328 | 0.175605 | -2.054784 | 0.03989983 | 0.116635 |
| ENSG00000135972 | MRPS9 | 1521.038713 | -0.350245653 | 0.202052673 | -1.806192309 | 0.07088831 | 0.180302 |
| ENSG00000067177 | PHKA1 | 1762.159598 | -0.350264815 | 0.179974559 | -2.005477128 | 0.04491206 | 0.127841 |
| ENSG00000245317 | | 164.8888 | -0.350267 | 0.2581571 | -1.43933 | 0.150057 | 0.311165 |
| ENSG00000253549 | CA3-AS1 | 23.29878 | -0.3504967 | 0.4688736 | -0.9240118 | 0.3554801 | 0.557443 |
| ENSG00000164916 | FOXK1 | 3229.128 | -0.350534 | 0.1825036 | -1.975712 | 0.04818744 | 0.134668 |
| ENSG00000185100 | ADSS1 | 551.052 | -0.3505952 | 0.1568607 | -2.281817 | 0.02250014 | 0.074829 |
| ENSG00000269883 | | 0.8924805 | -0.3506255 | 0.8561579 | -1.392507 | 0.163769 | NA |
| ENSG00000128692 | EIF2S2P4 | 2391.709472 | -0.35063813 | 0.203723415 | -1.813408771 | 0.0697688 | 0.178126 |
| ENSG00000138796 | HADH | 4418.264242 | -0.350739972 | 0.230715646 | -1.56102175 | 0.11851862 | 0.262765 |
| ENSG00000235848 | RMDN2-AS1 | 2.656302 | -0.350882 | 0.7030239 | -0.8886898 | 0.3741698 | 0.576275 |
| ENSG00000227579 | | 6.152791 | -0.3509861 | 0.6495189 | -0.875275 | 0.3814243 | 0.583625 |
| ENSG00000151065 | DCP1B | 618.0886 | -0.3510076 | 0.2456201 | -1.535619 | 0.1246319 | 0.272606 |
| ENSG00000121060 | TRIM25 | 10736.93837 | -0.351105078 | 0.158195032 | -2.30984563 | 0.0208967 | 0.070413 |
| ENSG00000196449 | YRDC | 1460.258 | -0.3512864 | 0.2394227 | -1.546042 | 0.1220944 | 0.268515 |
| ENSG00000281641 | SAMD12-AS1 | 31.90528 | -0.3514073 | 0.4180683 | -0.9863107 | 0.3239807 | 0.524869 |
| ENSG00000261616 | | 17.31479 | -0.3514979 | 0.469757 | -0.9206065 | 0.3572559 | 0.559336 |
| ENSG00000103248 | MTHFSD | 1552.296908 | -0.351507038 | 0.157637193 | -2.305994171 | 0.02111096 | 0.071 |
| ENSG00000236915 | CLCA4-AS1 | 4.672612 | -0.351576 | 0.6682764 | -0.8656871 | 0.3866618 | 0.589061 |
| ENSG00000163818 | LZTFL1 | 784.5349 | -0.3517743 | 0.1872757 | -1.934165 | 0.05309287 | 0.145315 |
| ENSG00000254205 | | 1.498617 | -0.3518089 | 0.8034401 | -1.05813 | 0.289996 | NA |
| ENSG00000217889 | KRT18P48 | 205.4498 | -0.3518976 | 0.2931245 | -1.296965 | 0.1946431 | 0.373769 |
| ENSG00000125895 | TMEM74B | 2.774513673 | -0.352213284 | 0.854209862 | -1.049280011 | 0.29404926 | 0.493095 |
| ENSG00000104983 | CCDC61 | 385.5754631 | -0.352216025 | 0.209472724 | -1.750597787 | 0.08001522 | 0.197247 |
| ENSG00000242017 | ALG1L15P | 96.76252 | -0.3522419 | 0.2368928 | -1.563984 | 0.1178212 | 0.2617 |
| ENSG00000266959 | | 1.256251 | -0.3522863 | 0.8646864 | -1.466733 | 0.1424487 | NA |
| ENSG00000273234 | OR2A13P | 1.961058 | -0.3523126 | 0.7861731 | -1.005208 | 0.3147966 | 0.515872 |
| ENSG00000058799 | YIPF1 | 2205.485216 | -0.352502558 | 0.10924257 | -3.291715792 | 0.00099578 | 0.005701 |
| ENSG00000235194 | PPP1R3E | 1044.503 | -0.3525929 | 0.3577809 | -1.111626 | 0.2662991 | 0.461901 |

| | | | | | | | |
|---|---|---|---|---|---|---|---|
| ENSG00000172661 | WASHC2C | 4465.602 | -0.3527205 | 0.2241787 | -1.652102 | 0.09851373 | 0.230102 |
| ENSG00000249541 | | 2.357885 | -0.3527491 | 0.7936668 | -1.02067 | 0.3074109 | 0.507632 |
| ENSG00000230899 | MAGEA8-AS1 | 1.405319 | -0.3527518 | 0.8588053 | -1.36482 | 0.1723097 | NA |
| ENSG00000186501 | TMEM222 | 1608.4 | -0.3529364 | 0.1479838 | -2.42725 | 0.01521378 | 0.054528 |
| ENSG00000180011 | ZADH2 | 2718.201 | -0.3530672 | 0.1164997 | -3.065608 | 0.00217228 | 0.011108 |
| ENSG00000183155 | RABIF | 3435.406 | -0.3531357 | 0.1181838 | -3.033853 | 0.00241452 | 0.012141 |
| ENSG00000273428 | | 6.520047 | -0.353213 | 0.6026898 | -0.8626358 | 0.3883377 | 0.590521 |
| ENSG00000102781 | KATNAL1 | 431.4682953 | -0.353318545 | 0.321964191 | -1.208721046 | 0.22677003 | 0.414463 |
| ENSG00000274973 | BSNDP2 | 2.350325 | -0.3534335 | 0.7484148 | -0.9445359 | 0.3448959 | 0.54649 |
| ENSG00000245526 | LINC00461 | 162.0323 | -0.3535016 | 0.2358441 | -1.589941 | 0.111848 | 0.252394 |
| ENSG00000185031 | SLC2A3P2 | 1.640606 | -0.3535953 | 0.8051379 | -1.073314 | 0.2831302 | 0.480947 |
| ENSG00000106635 | BCL7B | 2364.041265 | -0.353683531 | 0.19244391 | -1.904125861 | 0.05689379 | 0.152827 |
| ENSG00000164849 | GPR146 | 25.75736 | -0.3537595 | 0.3696168 | -1.088031 | 0.2765812 | 0.473607 |
| ENSG00000225163 | NA | 23.9538 | -0.3537996 | 0.4024794 | -1.024115 | 0.305781 | 0.505714 |
| ENSG00000162062 | TEDC2 | 1098.375 | -0.3538145 | 0.3791531 | -1.083216 | 0.2787126 | 0.475988 |
| ENSG00000186350 | RXRA | 8833.393 | -0.3538185 | 0.1782931 | -2.040224 | 0.04132804 | 0.119997 |
| ENSG00000177839 | PCDHB9 | 72.85012 | -0.3538328 | 0.3495685 | -1.128484 | 0.2591155 | 0.453461 |
| ENSG00000173209 | AHSA2P | 4871.359 | -0.3538334 | 0.314741 | -1.232744 | 0.2176715 | 0.403356 |
| ENSG00000100220 | RTCB | 5194.276316 | -0.353836515 | 0.187113866 | -1.953170303 | 0.05079942 | 0.140169 |
| ENSG00000186094 | AGBL4 | 27.35601 | -0.3538741 | 0.4097236 | -1.014101 | 0.3105347 | 0.511253 |
| ENSG00000257403 | | 3.345047 | -0.3538823 | 0.6800334 | -0.8717499 | 0.3833448 | 0.585745 |
| ENSG00000234017 | | 1.062288 | -0.3539235 | 0.837398 | -1.255408 | 0.2093307 | NA |
| ENSG00000261064 | LINC02256 | 8.156485 | -0.3539459 | 0.5330329 | -0.8823907 | 0.3775656 | 0.579547 |
| ENSG00000261766 | | 24.80455 | -0.3539961 | 0.3865769 | -1.050623 | 0.2934315 | 0.492376 |
| ENSG00000175470 | PPP2R2D | 2167.385 | -0.353997 | 0.1255512 | -2.861823 | 0.00421212 | 0.019259 |
| ENSG00000227176 | NSA2P5 | 9.162714 | -0.35403 | 0.5669652 | -0.8612008 | 0.3891274 | 0.591122 |
| ENSG00000174292 | TNK1 | 1108.412 | -0.3540525 | 0.161179 | -2.254499 | 0.02416481 | 0.078971 |
| ENSG00000225511 | LINC00475 | 30.33567 | -0.3541893 | 0.4188659 | -0.9973297 | 0.3186045 | 0.519718 |
| ENSG00000250441 | | 7.362928 | -0.3542419 | 0.5857387 | -0.8674147 | 0.3857148 | 0.588047 |
| ENSG00000249215 | NCOA4P4 | 1.14404 | -0.354245 | 0.8622 | -1.137356 | 0.2553897 | NA |
| ENSG00000146463 | ZMYM4 | 4879.92 | -0.3544088 | 0.135016 | -2.669698 | 0.00759196 | 0.031196 |
| ENSG00000263624 | | 6.015744 | -0.3544314 | 0.6608651 | -0.8613029 | 0.3890712 | 0.591104 |
| ENSG00000265203 | RBP3 | 1.13394 | -0.3544417 | 0.8421444 | -1.27705 | 0.2015847 | NA |

| | | | | | | | |
|---|---|---|---|---|---|---|---|
| ENSG00000236698 | EIF1AXP1 | 1241.076 | -0.3544606 | 0.235019 | -1.587175 | 0.1124729 | 0.253252 |
| ENSG00000166167 | BTRC | 1558.779 | -0.3544613 | 0.1525519 | -2.368737 | 0.01784892 | 0.062007 |
| ENSG00000167645 | YIF1B | 2790.643 | -0.3545121 | 0.2078019 | -1.781401 | 0.07484689 | 0.187817 |
| ENSG00000139154 | AEBP2 | 1314.147997 | -0.354611033 | 0.13860359 | -2.599587249 | 0.0093336 | 0.036771 |
| ENSG00000082269 | FAM135A | 168.4146299 | -0.354699534 | 0.297725494 | -1.303157178 | 0.19252111 | 0.370901 |
| ENSG00000228002 | DHX9P1 | 26.88297 | -0.3547199 | 0.3820538 | -1.0654 | 0.2866952 | 0.485049 |
| ENSG00000164180 | TMEM161B | 2159.223 | -0.3547802 | 0.2429936 | -1.544506 | 0.1224657 | 0.269198 |
| ENSG00000270076 | | 10.11441 | -0.3548145 | 0.7423429 | -0.901242 | 0.3674597 | 0.569506 |
| ENSG00000145494 | NDUFS6 | 3970.465 | -0.3549915 | 0.1927908 | -1.90917 | 0.05624015 | 0.151592 |
| ENSG00000255647 | | 10.22569 | -0.3551152 | 0.5693126 | -0.8560723 | 0.3919578 | 0.593682 |
| ENSG00000265142 | MIR133A1HG | 1.235734 | -0.3552887 | 0.8133977 | -1.094035 | 0.2739397 | NA |
| ENSG00000226416 | MRPL23-AS1 | 38.10991 | -0.355302 | 0.4017849 | -1.028325 | 0.3037968 | 0.503404 |
| ENSG00000285077 | ARHGAP11B | 117.0409 | -0.3553433 | 0.3929505 | -1.044848 | 0.2960933 | 0.495116 |
| ENSG00000271849 | MAN2A1-DT | 5.447845 | -0.3554256 | 0.6340088 | -0.8592279 | 0.3902148 | 0.592198 |
| ENSG00000149923 | PPP4C | 8614.134 | -0.3554503 | 0.2159981 | -1.725542 | 0.0844299 | 0.205517 |
| ENSG00000249753 | | 2.066799 | -0.3554976 | 0.8039533 | -1.059691 | 0.2892851 | 0.487904 |
| ENSG00000270137 | | 2.146014 | -0.355511 | 0.7819083 | -1.004543 | 0.3151167 | 0.516159 |
| ENSG00000275484 | | 24.17134 | -0.3555158 | 0.4569625 | -0.9427611 | 0.3458031 | 0.547372 |
| ENSG00000005893 | LAMP2 | 10682.46463 | -0.355577871 | 0.158634247 | -2.296057635 | 0.02167259 | 0.072585 |
| ENSG00000157764 | BRAF | 1342.824 | -0.3556756 | 0.2216634 | -1.677378 | 0.09346855 | 0.22161 |
| ENSG00000241411 | | 2.623399 | -0.3556941 | 0.8581253 | -1.392033 | 0.1639124 | 0.331935 |
| ENSG00000286100 | | 1.42491 | -0.3558108 | 0.7788831 | -0.9891169 | 0.3226059 | NA |
| ENSG00000241361 | SLC25A24P1 | 54.36586 | -0.3559584 | 0.3837377 | -1.067756 | 0.2856305 | 0.483679 |
| ENSG00000247679 | | 408.761 | -0.3559637 | 0.3338059 | -1.182853 | 0.2368673 | 0.426522 |
| ENSG00000186815 | TPCN1 | 2444.363 | -0.3560005 | 0.2661342 | -1.45271 | 0.1463042 | 0.305726 |
| ENSG00000248923 | MTND5P11 | 43.8806 | -0.356031 | 0.4133485 | -1.012073 | 0.3115034 | 0.512435 |
| ENSG00000267612 | | 3.488075 | -0.3561311 | 0.7397684 | -0.9261927 | 0.3543458 | 0.556288 |
| ENSG00000089006 | SNX5 | 8432.565485 | -0.356336128 | 0.238451701 | -1.57437906 | 0.11539984 | 0.258216 |
| ENSG00000227834 | | 1.155437 | -0.3563706 | 0.8857016 | -1.509703 | 0.1311193 | NA |
| ENSG00000275620 | | 0.9894951 | -0.3564352 | 0.8355853 | -1.214034 | 0.2247347 | NA |
| ENSG00000068796 | KIF2A | 3118.597963 | -0.356498169 | 0.177560735 | -2.071806784 | 0.03828346 | 0.11303 |
| ENSG00000280828 | | 27.91384 | -0.3565574 | 0.4860872 | -0.9194562 | 0.357857 | 0.559815 |
| ENSG00000066279 | ASPM | 5120.22273 | -0.356576873 | 0.415948531 | -1.012750623 | 0.31117929 | 0.512028 |

| | | | | | | | |
|---|---|---|---|---|---|---|---|
| ENSG00000144840 | RABL3 | 2107.717 | -0.3566302 | 0.1799159 | -2.052913 | 0.04008097 | 0.117064 |
| ENSG00000149136 | SSRP1 | 8821.745 | -0.3567562 | 0.2649203 | -1.442658 | 0.1491167 | 0.309812 |
| ENSG00000165275 | TRMT10B | 462.7948 | -0.3567734 | 0.1818742 | -2.020919 | 0.04328816 | 0.124377 |
| ENSG00000176302 | FOXR1 | 3.566412 | -0.3568694 | 0.6912032 | -0.8729554 | 0.3826874 | 0.584963 |
| ENSG00000130021 | PUDP | 1027.938996 | -0.356897617 | 0.23457136 | -1.609837846 | 0.10743326 | 0.245075 |
| ENSG00000272807 | | 29.36778 | -0.3570624 | 0.4059956 | -1.026225 | 0.3047857 | 0.504533 |
| ENSG00000248677 | LINC02102 | 9.509587 | -0.3572295 | 0.5806785 | -0.8623639 | 0.3884873 | 0.590521 |
| ENSG00000186743 | TPI1P3 | 1.295903 | -0.3573169 | 0.8242814 | -1.176543 | 0.2393781 | NA |
| ENSG00000130175 | PRKCSH | 10890.26792 | -0.357487572 | 0.118945602 | -3.044073154 | 0.00233398 | 0.011796 |
| ENSG00000220842 | RPL21P16 | 2030.708 | -0.3575049 | 0.1776555 | -2.069753 | 0.03847548 | 0.113431 |
| ENSG00000007341 | ST7L | 706.8484146 | -0.357556153 | 0.213897218 | -1.74769984 | 0.08051602 | 0.19826 |
| ENSG00000198746 | GPATCH3 | 604.4437 | -0.3577272 | 0.1773928 | -2.080784 | 0.03745366 | 0.111098 |
| ENSG00000103485 | QPRT | 5522.986898 | -0.357807988 | 0.436414692 | -1.004198617 | 0.31528289 | 0.516255 |
| ENSG00000212978 | | 298.4285 | -0.3578268 | 0.1818911 | -2.025641 | 0.04280156 | 0.1233 |
| ENSG00000176040 | TMPRSS7 | 6.611419 | -0.3578706 | 0.6655477 | -0.8696814 | 0.3844745 | 0.586839 |
| ENSG00000248318 | | 1.043973 | -0.3579127 | 0.8164549 | -1.15239 | 0.2491609 | NA |
| ENSG00000267302 | RNFT1-DT | 24.44851 | -0.3579555 | 0.3619676 | -1.114717 | 0.2649719 | 0.460465 |
| ENSG00000153107 | ANAPC1 | 3524.59 | -0.3580052 | 0.1982438 | -1.89179 | 0.05851899 | 0.156238 |
| ENSG00000170482 | SLC23A1 | 48.19396 | -0.3581213 | 0.3367702 | -1.181598 | 0.2373651 | 0.427013 |
| ENSG00000236047 | RPL13AP12 | 4.755778 | -0.358127 | 0.6913208 | -0.8842493 | 0.3765617 | 0.578617 |
| ENSG00000232874 | | 16.3479 | -0.3581329 | 0.4966994 | -0.9195248 | 0.3578211 | 0.559815 |
| ENSG00000273424 | | 5.759933 | -0.3581503 | 0.7061992 | -0.8975062 | 0.3694489 | 0.571644 |
| ENSG00000259319 | | 238.8302 | -0.3581551 | 0.2694838 | -1.422528 | 0.1548729 | 0.318534 |
| ENSG00000172216 | CEBPB | 2828.906 | -0.3582374 | 0.3302294 | -1.199294 | 0.2304136 | 0.418771 |
| ENSG00000108588 | CCDC47 | 20765.83341 | -0.358281733 | 0.156558721 | -2.348550016 | 0.01884667 | 0.064795 |
| ENSG00000268157 | | 5.906825 | -0.3583166 | 0.6798731 | -0.8757443 | 0.381169 | 0.583302 |
| ENSG00000113328 | CCNG1 | 6183.904326 | -0.358365394 | 0.187379244 | -1.979162553 | 0.04779771 | 0.133861 |
| ENSG00000107331 | ABCA2 | 13969.01268 | -0.358378797 | 0.201305308 | -1.860920751 | 0.06275537 | 0.164443 |
| ENSG00000231201 | | 1.29254 | -0.3583879 | 0.8180822 | -1.115874 | 0.2644761 | NA |
| ENSG00000118418 | HMGN3 | 1981.287353 | -0.35870595 | 0.259378481 | -1.463012943 | 0.14346384 | 0.301377 |
| ENSG00000260105 | AOC4P | 1.024532 | -0.3587736 | 0.8279543 | -1.169467 | 0.2422155 | NA |
| ENSG00000230314 | ELOVL2-AS1 | 23.20281 | -0.3588679 | 0.8247045 | -1.19732 | 0.231182 | 0.419681 |
| ENSG00000214041 | PGAM1P4 | 1.348634 | -0.3589252 | 0.8449802 | -1.252978 | 0.2102137 | NA |

| | | | | | | | |
|---|---|---|---|---|---|---|---|
| ENSG00000267811 | | 16.13852 | -0.3589335 | 0.4249731 | -0.9973961 | 0.3185723 | 0.519718 |
| ENSG00000055732 | MCOLN3 | 830.7470444 | -0.358939181 | 0.372618467 | -1.092331661 | 0.27468735 | 0.471549 |
| ENSG00000065057 | NTHL1 | 1696.417575 | -0.358993936 | 0.237725517 | -1.59608458 | 0.11046991 | 0.250129 |
| ENSG00000116001 | TIA1 | 6253.193995 | -0.359101095 | 0.219971983 | -1.706293209 | 0.0879535 | 0.212023 |
| ENSG00000106692 | FKTN | 2206.900683 | -0.359120369 | 0.198511105 | -1.876709781 | 0.06055788 | 0.160144 |
| ENSG00000266405 | CBX3P2 | 37.10933 | -0.3591226 | 0.4515921 | -0.9688716 | 0.3326093 | 0.533673 |
| ENSG00000124380 | SNRNP27 | 989.3900412 | -0.359128905 | 0.179118962 | -2.072713187 | 0.03819898 | 0.112843 |
| ENSG00000170542 | SERPINB9 | 37.91144 | -0.3591704 | 0.3585689 | -1.132505 | 0.2574222 | 0.451389 |
| ENSG00000158411 | MITD1 | 1225.707 | -0.3592018 | 0.2347517 | -1.611249 | 0.1071254 | 0.244667 |
| ENSG00000231226 | TRIM31-AS1 | 2.711528 | -0.3593684 | 0.7912641 | -1.028339 | 0.3037905 | 0.503404 |
| ENSG00000156802 | ATAD2 | 9155.545 | -0.3594608 | 0.4242998 | -1.00849 | 0.3132195 | 0.514366 |
| ENSG00000202343 | | 8.390097 | -0.3594709 | 0.6603698 | -0.8684349 | 0.3851563 | 0.587542 |
| ENSG00000168795 | ZBTB5 | 1810.109 | -0.3595162 | 0.1724127 | -2.145914 | 0.03187984 | 0.098364 |
| ENSG00000137710 | RDX | 2917.65037 | -0.359676737 | 0.179112458 | -2.063956764 | 0.03902182 | 0.114712 |
| ENSG00000260733 | | 1.530565 | -0.3597649 | 0.8666572 | -1.175595 | 0.2397568 | NA |
| ENSG00000230305 | | 1.029323 | -0.3599966 | 0.8329006 | -1.193215 | 0.2327852 | NA |
| ENSG00000134815 | DHX34 | 2603.22901 | -0.360002624 | 0.140589531 | -2.594273202 | 0.00947911 | 0.037212 |
| ENSG00000121940 | CLCC1 | 2049.867705 | -0.360039997 | 0.132833803 | -2.754712826 | 0.00587437 | 0.025296 |
| ENSG00000104299 | INTS9 | 869.9571359 | -0.360116593 | 0.128636627 | -2.862852595 | 0.00419846 | 0.019209 |
| ENSG00000179604 | CDC42EP4 | 4002.535 | -0.360151 | 0.2224208 | -1.695562 | 0.08996879 | 0.215622 |
| ENSG00000135541 | AHI1 | 1165.962884 | -0.360239685 | 0.224897554 | -1.676719875 | 0.09359727 | 0.221836 |
| ENSG00000253295 | | 2.296377 | -0.3603847 | 0.7405724 | -0.941589 | 0.3464031 | 0.548028 |
| ENSG00000074276 | CDHR2 | 141.6064059 | -0.36042761 | 0.287245678 | -1.352274214 | 0.17628761 | 0.349067 |
| ENSG00000121964 | GTDC1 | 938.3347055 | -0.360510123 | 0.154035005 | -2.392961017 | 0.01671302 | 0.058799 |
| ENSG00000112290 | WASF1 | 1158.55992 | -0.360718139 | 0.171861858 | -2.160223162 | 0.0307554 | 0.095646 |
| ENSG00000131759 | RARA | 3030.99836 | -0.360838924 | 0.189991846 | -1.965764215 | 0.04932585 | 0.137157 |
| ENSG00000167291 | TBC1D16 | 7900.697 | -0.360903 | 0.2537794 | -1.50997 | 0.1310512 | 0.282422 |
| ENSG00000213225 | NOC2LP1 | 1024.332 | -0.3609314 | 0.1694795 | -2.188239 | 0.02865222 | 0.090422 |
| ENSG00000273181 | | 10.09072 | -0.3610289 | 0.5921534 | -0.8711886 | 0.3836512 | 0.586054 |
| ENSG00000196363 | WDR5 | 5382.022 | -0.3610359 | 0.1638836 | -2.279301 | 0.02264917 | 0.075179 |
| ENSG00000065978 | YBX1 | 17172.24563 | -0.361237193 | 0.235408707 | -1.614758549 | 0.10636299 | 0.24347 |
| ENSG00000137216 | TMEM63B | 4281.315281 | -0.361474215 | 0.14534247 | -2.568200127 | 0.01022281 | 0.039582 |
| ENSG00000272081 | | 20.61536 | -0.3616146 | 0.4465431 | -0.972309 | 0.3308969 | 0.531899 |

| | | | | | | | |
|---|---|---|---|---|---|---|---|
| ENSG00000103043 | VAC14 | 3177.813268 | -0.361637579 | 0.116516456 | -3.154505092 | 0.00160771 | 0.008595 |
| ENSG00000138069 | RAB1A | 10585.88688 | -0.361755943 | 0.113727842 | -3.194747644 | 0.00139953 | 0.007647 |
| ENSG00000232490 | OSBPL10-AS1 | 0.8340922 | -0.3618965 | 0.8905716 | -1.538111 | 0.1240216 | NA |
| ENSG00000197170 | PSMD12 | 10894.34 | -0.3619877 | 0.2395795 | -1.592637 | 0.1112417 | 0.251318 |
| ENSG00000267293 | | 16.46869 | -0.3620339 | 0.5070515 | -0.9240878 | 0.3554406 | 0.557414 |
| ENSG00000184984 | CHRM5 | 8.296781 | -0.3620917 | 0.5397619 | -0.8985603 | 0.3688869 | 0.571018 |
| ENSG00000130640 | TUBGCP2 | 5115.458412 | -0.36218269 | 0.142224117 | -2.616359074 | 0.0088873 | 0.035407 |
| ENSG00000121931 | LRIF1 | 1408.571751 | -0.362245847 | 0.274556261 | -1.443901009 | 0.14876682 | 0.309303 |
| ENSG00000274215 | | 1.329285 | -0.3622466 | 0.8161247 | -1.116882 | 0.2640446 | NA |
| ENSG00000185000 | DGAT1 | 2090.375 | -0.3623251 | 0.209305 | -1.856907 | 0.06332435 | 0.165499 |
| ENSG00000197498 | RPF2 | 1999.854 | -0.3623695 | 0.1747869 | -2.138426 | 0.03248222 | 0.099713 |
| ENSG00000164880 | INTS1 | 12895.56 | -0.3623962 | 0.1879158 | -1.985796 | 0.04705595 | 0.132337 |
| ENSG00000233514 | | 140.0002 | -0.3624639 | 0.3153037 | -1.258098 | 0.2083561 | 0.391168 |
| ENSG00000166226 | CCT2 | 17426.78 | -0.3624765 | 0.2277027 | -1.669652 | 0.09498816 | 0.224173 |
| ENSG00000126698 | DNAJC8 | 7179.899916 | -0.36258748 | 0.18143585 | -2.060213131 | 0.03937817 | 0.11553 |
| ENSG00000274559 | | 0.8459095 | -0.3625876 | 0.8450787 | -1.261527 | 0.2071189 | NA |
| ENSG00000254907 | | 8.182761 | -0.3626548 | 0.5884887 | -0.8767195 | 0.380639 | 0.582658 |
| ENSG00000256576 | LINC02361 | 36.57605 | -0.3627108 | 0.4289663 | -1.005347 | 0.3147299 | 0.515827 |
| ENSG00000228536 | LYPLAL1-AS1 | 4.40973 | -0.3627604 | 0.6360661 | -0.8786148 | 0.3796102 | 0.581552 |
| ENSG00000164045 | CDC25A | 692.5106 | -0.3627646 | 0.5883288 | -0.8927078 | 0.3720137 | 0.574219 |
| ENSG00000151470 | C4orf33 | 269.8215 | -0.3627975 | 0.2680233 | -1.444175 | 0.1486898 | 0.309195 |
| ENSG00000237176 | | 19.43483 | -0.3628318 | 0.5324161 | -0.9003945 | 0.3679104 | 0.570038 |
| ENSG00000267009 | | 23.23734 | -0.3628475 | 0.4485865 | -0.9779073 | 0.3281202 | 0.529382 |
| ENSG00000122952 | ZWINT | 5124.277806 | -0.362850549 | 0.563000233 | -0.902240825 | 0.36692895 | 0.568915 |
| ENSG00000211456 | SACM1L | 1614.065 | -0.3629245 | 0.1659716 | -2.242158 | 0.02495119 | 0.080983 |
| ENSG00000100104 | SRRD | 830.1370049 | -0.362996362 | 0.156171495 | -2.370447683 | 0.01776656 | 0.061761 |
| ENSG00000132589 | FLOT2 | 6817.706056 | -0.3630606 | 0.163317968 | -2.290909242 | 0.02196866 | 0.073401 |
| ENSG00000188372 | ZP3 | 982.7011 | -0.363063 | 0.2476069 | -1.550114 | 0.1211142 | 0.267044 |
| ENSG00000144445 | KANSL1L | 746.4997 | -0.3630743 | 0.3101652 | -1.314042 | 0.1888322 | 0.366077 |
| ENSG00000109062 | SLC9A3R1 | 15627.41922 | -0.363211962 | 0.213589425 | -1.773757314 | 0.0761033 | 0.190054 |
| ENSG00000236529 | | 18.72031 | -0.3632414 | 0.5124398 | -0.9172822 | 0.3589948 | 0.560935 |
| ENSG00000250492 | INTS6P1 | 67.40608 | -0.3634102 | 0.3000611 | -1.312865 | 0.1892283 | 0.366684 |
| ENSG00000254862 | LGR4-AS1 | 2.695947 | -0.3634223 | 0.6765955 | -0.9004913 | 0.3678588 | 0.570025 |

| | | | | | | | |
|---|---|---|---|---|---|---|---|
| ENSG00000243155 | | 74.56932 | -0.363442 | 0.2974697 | -1.3216 | 0.1863014 | 0.362467 |
| ENSG00000239917 | RPS10P16 | 1.090789 | -0.3634653 | 0.836955 | -1.233123 | 0.2175297 | NA |
| ENSG00000261308 | FIGNL2 | 2.26575 | -0.3635256 | 0.7350666 | -0.9415439 | 0.3464262 | 0.548032 |
| ENSG00000172878 | METAP1D | 687.7222 | -0.3635369 | 0.2463057 | -1.563975 | 0.1178235 | 0.2617 |
| ENSG00000237828 | PA2G4P1 | 129.2017 | -0.3635782 | 0.3153967 | -1.265068 | 0.205847 | 0.387911 |
| ENSG00000214026 | MRPL23 | 1717.108 | -0.363602 | 0.2346625 | -1.622401 | 0.1047176 | 0.240554 |
| ENSG00000230319 | TTLL1-AS1 | 1.497221 | -0.3636243 | 0.8339334 | -1.184231 | 0.2363218 | NA |
| ENSG00000048140 | TSPAN17 | 5566.980865 | -0.363679297 | 0.128727462 | -2.878006687 | 0.00400197 | 0.01845 |
| ENSG00000277701 | | 2.507124 | -0.3637519 | 0.7730538 | -1.019428 | 0.3079997 | 0.508181 |
| ENSG00000231811 | | 5.606144 | -0.363769 | 0.738995 | -0.9636435 | 0.3352247 | 0.536574 |
| ENSG00000287124 | | 1.591972 | -0.3639393 | 0.7793568 | -1.020533 | 0.3074759 | 0.507632 |
| ENSG00000207523 | SNORA66 | 14.7129 | -0.3640121 | 0.5173414 | -0.9175725 | 0.3588427 | 0.560812 |
| ENSG00000132294 | EFR3A | 4956.820428 | -0.364048558 | 0.176511186 | -2.112527943 | 0.03464119 | 0.104855 |
| ENSG00000120805 | ARL1 | 4470.093065 | -0.364095018 | 0.114538807 | -3.233599372 | 0.00122241 | 0.006806 |
| ENSG00000104907 | TRMT1 | 2795.737153 | -0.36411639 | 0.137777479 | -2.694254748 | 0.00705462 | 0.029328 |
| ENSG00000259561 | | 0.9529866 | -0.3641649 | 0.855822 | -1.327551 | 0.1843266 | NA |
| ENSG00000132153 | DHX30 | 4933.249642 | -0.364241914 | 0.181622302 | -2.083493551 | 0.03720626 | 0.110562 |
| ENSG00000229326 | RPL17P15 | 1.663256 | -0.3643425 | 0.7969211 | -1.062357 | 0.2880736 | 0.486421 |
| ENSG00000275426 | | 7.799762 | -0.3643871 | 0.5262552 | -0.914238 | 0.3605918 | 0.56257 |
| ENSG00000136826 | KLF4 | 1491.250422 | -0.364455689 | 0.313614702 | -1.267895848 | 0.20483515 | 0.386799 |
| ENSG00000124783 | SSR1 | 7596.816028 | -0.364551417 | 0.080116017 | -4.578223353 | 4.69E-06 | 5.04E-05 |
| ENSG00000222019 | URAHP | 126.7805 | -0.3645597 | 0.2210091 | -1.724125 | 0.08468522 | 0.206044 |
| ENSG00000186532 | SMYD4 | 863.479 | -0.3646165 | 0.1636008 | -2.291037 | 0.02196126 | 0.073386 |
| ENSG00000248954 | | 25.96654 | -0.3647369 | 0.4892484 | -0.9395825 | 0.3474318 | 0.549002 |
| ENSG00000213999 | MEF2B | 7.442518 | -0.3649256 | 0.6073314 | -0.8732561 | 0.3825235 | 0.584802 |
| ENSG00000276556 | | 1.967742 | -0.3651452 | 0.7869736 | -1.03987 | 0.2984001 | 0.497596 |
| ENSG00000181284 | TMEM102 | 509.0224 | -0.3651619 | 0.1750214 | -2.145271 | 0.03193121 | 0.098454 |
| ENSG00000155393 | HEATR3 | 1690.868 | -0.3651862 | 0.1176017 | -3.195459 | 0.00139609 | 0.007634 |
| ENSG00000111790 | FGFR1OP2 | 1489.819961 | -0.365204521 | 0.162057631 | -2.319744257 | 0.02035472 | 0.068927 |
| ENSG00000196204 | RNF216P1 | 885.9912 | -0.3652129 | 0.13943 | -2.669513 | 0.00759614 | 0.031208 |
| ENSG00000229153 | EPHA1-AS1 | 1.86832 | -0.3653219 | 0.8332712 | -1.213494 | 0.224941 | 0.411971 |
| ENSG00000114554 | PLXNA1 | 13021.16596 | -0.365346038 | 0.195270709 | -1.937065965 | 0.05273728 | 0.144496 |
| ENSG00000173237 | C11orf86 | 0.9863311 | -0.365392 | 0.9040602 | -1.784587 | 0.07432841 | NA |

| | | | | | | | |
|---|---|---|---|---|---|---|---|
| ENSG00000143621 | ILF2 | 15407.6 | -0.3654477 | 0.2360274 | -1.60863 | 0.1076972 | 0.245551 |
| ENSG00000231948 | HS1BP3-IT1 | 1.534126 | -0.3654776 | 0.8181627 | -1.118881 | 0.2631909 | NA |
| ENSG00000225725 | FAM66E | 1.794929 | -0.3655657 | 0.8198079 | -1.123818 | 0.2610903 | 0.455716 |
| ENSG00000277437 | | 2.234613 | -0.3656785 | 0.8338598 | -1.203705 | 0.2287035 | 0.416879 |
| ENSG00000125166 | GOT2 | 7572.506506 | -0.365685313 | 0.216846504 | -1.738367263 | 0.08214612 | 0.201466 |
| ENSG00000254233 | LINC02365 | 13.07216 | -0.3657841 | 0.6750174 | -0.8966888 | 0.369885 | 0.572063 |
| ENSG00000105197 | TIMM50 | 3908.573626 | -0.365799105 | 0.208597001 | -1.824861288 | 0.06802196 | 0.174734 |
| ENSG00000171827 | ZNF570 | 194.0902 | -0.3658062 | 0.2279996 | -1.682968 | 0.09238128 | 0.219625 |
| ENSG00000197381 | ADARB1 | 2774.705 | -0.3658506 | 0.2035139 | -1.870694 | 0.06138753 | 0.161822 |
| ENSG00000129187 | DCTD | 3873.293305 | -0.365880809 | 0.139434244 | -2.681486541 | 0.00732959 | 0.030291 |
| ENSG00000276966 | H4C5 | 294.4362 | -0.3660078 | 0.4667683 | -0.9662327 | 0.3339278 | 0.535093 |
| ENSG00000166295 | ANAPC16 | 5792.961 | -0.366125 | 0.1529824 | -2.444607 | 0.01450102 | 0.052462 |
| ENSG00000105364 | MRPL4 | 3728.833966 | -0.366133612 | 0.250145697 | -1.571680027 | 0.11602478 | 0.258895 |
| ENSG00000230916 | MTCO1P53 | 4.414966 | -0.3662044 | 0.6375427 | -0.8799525 | 0.378885 | 0.58102 |
| ENSG00000269148 | | 17.02849 | -0.3663941 | 0.4256144 | -1.015977 | 0.3096403 | 0.510286 |
| ENSG00000144410 | CPO | 5.533121 | -0.3664636 | 0.7418275 | -0.9686667 | 0.3327115 | 0.533708 |
| ENSG00000101945 | SUV39H1 | 734.3236073 | -0.366487429 | 0.428965247 | -1.024601342 | 0.30555133 | 0.505429 |
| ENSG00000271870 | | 210.8524 | -0.3666446 | 0.2977581 | -1.3282 | 0.184112 | 0.359356 |
| ENSG00000145907 | G3BP1 | 11816.3 | -0.3666558 | 0.1523483 | -2.47413 | 0.01335609 | 0.04905 |
| ENSG00000173124 | ACSM6 | 7.341198 | -0.3666945 | 0.7031065 | -0.9017135 | 0.3672091 | 0.56925 |
| ENSG00000189366 | ALG1L | 372.0317 | -0.3667558 | 0.3026489 | -1.325345 | 0.1850566 | 0.360599 |
| ENSG00000188725 | SMIM15 | 3400.257 | -0.366765 | 0.1491729 | -2.51688 | 0.01183992 | 0.044545 |
| ENSG00000127463 | EMC1 | 3199.503701 | -0.366903983 | 0.12686182 | -2.930643819 | 0.0033826 | 0.016063 |
| ENSG00000132475 | H3-3B | 48419.29556 | -0.366915803 | 0.205801155 | -1.853732228 | 0.06377748 | 0.166356 |
| ENSG00000222005 | LINC01118 | 8.033293 | -0.3669164 | 0.6263089 | -0.8822604 | 0.377636 | 0.579564 |
| ENSG00000101280 | ANGPT4 | 3.806629141 | -0.366927492 | 0.655572807 | -0.894147672 | 0.37124289 | 0.573296 |
| ENSG00000262833 | | 0.9480734 | -0.3669487 | 0.8714798 | -1.221994 | 0.2217099 | NA |
| ENSG00000177045 | SIX5 | 518.079 | -0.3670369 | 0.1796757 | -2.102636 | 0.03549761 | 0.106753 |
| ENSG00000099995 | SF3A1 | 5549.438052 | -0.367044662 | 0.185210467 | -2.05515395 | 0.03986414 | 0.116569 |
| ENSG00000004700 | RECQL | 866.7027504 | -0.367110405 | 0.281671537 | -1.402662 | 0.16071765 | 0.327287 |
| ENSG00000203760 | CENPW | 532.4119 | -0.3672055 | 0.5559424 | -0.9100445 | 0.3627991 | 0.564786 |
| ENSG00000232021 | LEF1-AS1 | 1.169202 | -0.3672597 | 0.836791 | -1.241613 | 0.2143795 | NA |
| ENSG00000142449 | FBN3 | 2.847575 | -0.3673658 | 0.7152936 | -0.9290612 | 0.3528574 | 0.55476 |

| | | | | | | | |
|---|---|---|---|---|---|---|---|
| ENSG00000100802 | C14orf93 | 680.9557927 | -0.367403621 | 0.205512219 | -1.858074111 | 0.06315849 | 0.165147 |
| ENSG00000047634 | SCML1 | 1676.980036 | -0.367416556 | 0.182671745 | -2.073199313 | 0.03815373 | 0.112753 |
| ENSG00000179256 | SMCO3 | 1.0896 | -0.367575 | 0.8780595 | -1.2904 | 0.1969117 | NA |
| ENSG00000149658 | YTHDF1 | 6899.32 | -0.3676148 | 0.1137177 | -3.213476 | 0.00131139 | 0.007247 |
| ENSG00000217128 | FNIP1 | 2224.932 | -0.3677796 | 0.3209342 | -1.261826 | 0.2070116 | 0.389387 |
| ENSG00000106078 | COBL | 1671.998986 | -0.368086961 | 0.237730606 | -1.628527308 | 0.10341312 | 0.238402 |
| ENSG00000228919 | | 0.990597 | -0.3681629 | 0.8746662 | -1.479431 | 0.1390253 | NA |
| ENSG00000090674 | MCOLN1 | 723.9206078 | -0.368240244 | 0.2275962 | -1.692549405 | 0.09054127 | 0.216579 |
| ENSG00000171858 | RPS21 | 16118.11 | -0.3682958 | 0.1817609 | -2.10221 | 0.03553488 | 0.106826 |
| ENSG00000273448 | | 25.39689 | -0.3684641 | 0.576682 | -0.9001273 | 0.3680525 | 0.570158 |
| ENSG00000152404 | CWF19L2 | 495.686 | -0.3685673 | 0.2075095 | -1.84808 | 0.06459077 | 0.167901 |
| ENSG00000113578 | FGF1 | 1.384722818 | -0.368579528 | 0.842774927 | -1.277467353 | 0.2014373 | NA |
| ENSG00000129071 | MBD4 | 3593.07915 | -0.368682644 | 0.139844283 | -2.690500329 | 0.0071345 | 0.029623 |
| ENSG00000160194 | NDUFV3 | 2952.704 | -0.3687438 | 0.125968 | -2.97037 | 0.00297442 | 0.014472 |
| ENSG00000109917 | ZPR1 | 2110.06002 | -0.368779821 | 0.17479492 | -2.175650408 | 0.02958141 | 0.092774 |
| ENSG00000134827 | TCN1 | 280.9260616 | -0.368795615 | 0.763997643 | -1.048676884 | 0.29432686 | 0.493381 |
| ENSG00000152503 | TRIM36 | 396.4947 | -0.3688419 | 0.2678132 | -1.472235 | 0.1409573 | 0.297608 |
| ENSG00000260409 | | 1.01746 | -0.368849 | 0.8529213 | -1.297734 | 0.1943788 | NA |
| ENSG00000083444 | PLOD1 | 3176.855465 | -0.368868132 | 0.186881004 | -2.05798241 | 0.03959182 | 0.115977 |
| ENSG00000160908 | ZNF394 | 1257.731 | -0.3689637 | 0.1421893 | -2.658047 | 0.0078595 | 0.032077 |
| ENSG00000134007 | ADAM20 | 2.620067968 | -0.368981042 | 0.7109257 | -0.929211751 | 0.35277936 | 0.554681 |
| ENSG00000241360 | PDXP | 39.88035 | -0.3689996 | 0.3629586 | -1.150211 | 0.2500569 | 0.442652 |
| ENSG00000206634 | SNORA22 | 11.55451 | -0.3690756 | 0.4890824 | -0.9508341 | 0.3416886 | 0.54293 |
| ENSG00000261549 | | 47.60163 | -0.3691288 | 0.3287611 | -1.241667 | 0.2143594 | 0.399223 |
| ENSG00000035499 | DEPDC1B | 803.7264435 | -0.369189292 | 0.558558004 | -0.907138641 | 0.36433349 | 0.566424 |
| ENSG00000258457 | | 93.4962 | -0.3693292 | 0.4436579 | -1.000954 | 0.3168488 | 0.517945 |
| ENSG00000167930 | FAM234A | 5649.224 | -0.3694622 | 0.1676499 | -2.270745 | 0.02316243 | 0.076374 |
| ENSG00000271734 | | 5.09278 | -0.3695056 | 0.6295827 | -0.8842582 | 0.3765569 | 0.578617 |
| ENSG00000100211 | CBY1 | 674.4875654 | -0.369527605 | 0.183233023 | -2.076535532 | 0.03784445 | 0.111982 |
| ENSG00000230224 | PHBP9 | 79.10964 | -0.3697975 | 0.3878952 | -1.099605 | 0.2715042 | 0.467868 |
| ENSG00000186907 | RTN4RL2 | 59.85073 | -0.369813 | 0.4763801 | -0.972484 | 0.3308098 | 0.531885 |
| ENSG00000244038 | DDOST | 10880.52 | -0.3698213 | 0.1438031 | -2.629306 | 0.00855593 | 0.034323 |
| ENSG00000204961 | PCDHA9 | 8.497532 | -0.3698308 | 0.6235784 | -0.8856029 | 0.3758316 | 0.57803 |

| | | | | | | | |
|---|---|---|---|---|---|---|---|
| ENSG00000186871 | ERCC6L | 524.7393 | -0.3700036 | 0.5897652 | -0.9131642 | 0.3611562 | 0.563205 |
| ENSG00000263606 | CHORDC1P4 | 201.6207 | -0.3702459 | 0.317618 | -1.274222 | 0.2025848 | 0.383777 |
| ENSG00000115282 | TTC31 | 2915.595264 | -0.370373867 | 0.136178379 | -2.773564 | 0.00554459 | 0.024126 |
| ENSG00000196151 | WDSUB1 | 711.1032 | -0.3704074 | 0.2395957 | -1.653402 | 0.09824905 | 0.229746 |
| ENSG00000185608 | MRPL40 | 1438.555 | -0.3704797 | 0.2406006 | -1.610856 | 0.1072112 | 0.2448 |
| ENSG00000260038 | | 5.575353 | -0.3705163 | 0.6251431 | -0.8898544 | 0.3735441 | 0.575609 |
| ENSG00000000003 | TSPAN6 | 3533.305759 | -0.370593339 | 0.19971534 | -1.922391307 | 0.05455654 | 0.148318 |
| ENSG00000108511 | HOXB6 | 13.78311239 | -0.370595224 | 0.799162702 | -1.095293884 | 0.27338791 | 0.470108 |
| ENSG00000155304 | HSPA13 | 2164.008 | -0.3706191 | 0.2670392 | -1.482016 | 0.1383359 | 0.293355 |
| ENSG00000272686 | WASL-DT | 455.4541 | -0.3706337 | 0.2320864 | -1.689241 | 0.09117329 | 0.217664 |
| ENSG00000256399 | | 0.9358542 | -0.3708006 | 0.8744061 | -1.2608 | 0.2073809 | NA |
| ENSG00000153774 | CFDP1 | 3062.042 | -0.370904 | 0.1882873 | -2.040205 | 0.04132992 | 0.119997 |
| ENSG00000229048 | DUTP1 | 1.329682 | -0.3709311 | 0.8943201 | -1.529994 | 0.1260182 | NA |
| ENSG00000130713 | EXOSC2 | 2757.496706 | -0.370975318 | 0.239849125 | -1.627419395 | 0.10364806 | 0.238778 |
| ENSG00000254027 | | 12.48914 | -0.3709914 | 0.4997942 | -0.9438832 | 0.3452294 | 0.54679 |
| ENSG00000283031 | | 10.8592 | -0.3709966 | 0.55 | -0.9198333 | 0.3576599 | 0.559705 |
| ENSG00000197345 | MRPL21 | 2953.74 | -0.3710412 | 0.239764 | -1.616208 | 0.1060493 | 0.242836 |
| ENSG00000170100 | ZNF778 | 2906.287 | -0.3711325 | 0.2559667 | -1.532619 | 0.1253698 | 0.273578 |
| ENSG00000261582 | | 6.512876 | -0.3711397 | 0.554257 | -0.9036144 | 0.3661999 | 0.568097 |
| ENSG00000226525 | RPS7P10 | 540.9456 | -0.3711471 | 0.2339859 | -1.671758 | 0.09457204 | 0.223499 |
| ENSG00000256021 | ELOCP31 | 0.8310103 | -0.3711921 | 0.8515359 | -1.315533 | 0.188331 | NA |
| ENSG00000167748 | KLK1 | 4.043446 | -0.3713362 | 0.8629012 | -1.116964 | 0.2640099 | 0.459421 |
| ENSG00000230262 | LINC02603 | 9.645516 | -0.3714047 | 0.5562529 | -0.9070279 | 0.364392 | 0.56647 |
| ENSG00000139160 | ETFBKMT | 48.00768978 | -0.371505032 | 0.29331214 | -1.370389434 | 0.17056537 | 0.341246 |
| ENSG00000198825 | INPP5F | 923.9059 | -0.3715132 | 0.1769648 | -2.16917 | 0.03006978 | 0.093837 |
| ENSG00000069812 | HES2 | 271.824571 | -0.371520225 | 0.413253732 | -1.045849639 | 0.29563046 | 0.494708 |
| ENSG00000235618 | FAM21EP | 77.907 | -0.3715741 | 0.3246917 | -1.259811 | 0.2077374 | 0.390365 |
| ENSG00000223503 | | 6.40818 | -0.3717027 | 0.6808307 | -0.901333 | 0.3674113 | 0.569497 |
| ENSG00000240086 | | 347.5693 | -0.3717831 | 0.2404324 | -1.628716 | 0.1033732 | 0.238351 |
| ENSG00000178821 | TMEM52 | 148.7697 | -0.3718209 | 0.4255018 | -1.034896 | 0.3007174 | 0.499981 |
| ENSG00000279368 | | 57.92522 | -0.371821 | 0.4197496 | -1.046182 | 0.2954772 | 0.494646 |
| ENSG00000120837 | NFYB | 3925.258912 | -0.371822397 | 0.17614962 | -2.183751888 | 0.02898049 | 0.091338 |
| ENSG00000235374 | SSR4P1 | 202.7043 | -0.3719144 | 0.2088564 | -1.846549 | 0.06481253 | 0.168312 |

| | | | | | | | |
|---|---|---|---|---|---|---|---|
| ENSG00000230417 | LINC00595 | 1.283984 | -0.3719284 | 0.796127 | -1.069611 | 0.2847944 | NA |
| ENSG00000115504 | EHBP1 | 1828.145916 | -0.372100657 | 0.17061837 | -2.277910089 | 0.02273194 | 0.075355 |
| ENSG00000228037 | | 1.510136 | -0.3721973 | 0.784968 | -1.027812 | 0.3040383 | NA |
| ENSG00000236533 | RPS12P4 | 9.619958 | -0.3722054 | 0.5814921 | -0.8996297 | 0.3683174 | 0.570303 |
| ENSG00000154265 | ABCA5 | 3141.617 | -0.3722391 | 0.2567167 | -1.532224 | 0.1254672 | 0.273712 |
| ENSG00000134780 | DAGLA | 451.0167757 | -0.372245873 | 0.182978602 | -2.11067494 | 0.03480026 | 0.105165 |
| ENSG00000272768 | | 16.61217 | -0.3723343 | 0.5137171 | -0.9315698 | 0.3515589 | 0.553253 |
| ENSG00000126756 | UXT | 1858.171385 | -0.372384935 | 0.129429034 | -2.936717233 | 0.00331706 | 0.015803 |
| ENSG00000214794 | KRT18P42 | 1.962315 | -0.3724344 | 0.819047 | -1.134835 | 0.2564444 | 0.45018 |
| ENSG00000236493 | EIF2S2P3 | 118.0861 | -0.372455 | 0.2384944 | -1.648502 | 0.09924975 | 0.231414 |
| ENSG00000260509 | | 3.929602 | -0.3724579 | 0.6866062 | -0.9229451 | 0.3560358 | 0.558084 |
| ENSG00000137642 | SORL1 | 3706.620308 | -0.372500728 | 0.257033364 | -1.539240957 | 0.12374548 | 0.271292 |
| ENSG00000122548 | KIAA0087 | 0.943853292 | -0.372579498 | 0.882188662 | -1.56207254 | 0.1182709 | NA |
| ENSG00000235124 | KRT8P17 | 15.84626 | -0.372586 | 0.4103935 | -1.060748 | 0.2888043 | 0.487407 |
| ENSG00000107815 | TWNK | 1966.407436 | -0.372612142 | 0.142995297 | -2.654905962 | 0.00793305 | 0.032332 |
| ENSG00000112118 | MCM3 | 8198.854501 | -0.372723582 | 0.423007394 | -1.046521025 | 0.29532054 | 0.494509 |
| ENSG00000275464 | | 688.8843 | -0.3727603 | 0.2067014 | -1.875791 | 0.06068401 | 0.160349 |
| ENSG00000109919 | MTCH2 | 7104.269392 | -0.372865403 | 0.184624798 | -2.089262445 | 0.03668411 | 0.109353 |
| ENSG00000184923 | NUTM2A | 24.57146 | -0.3729594 | 0.4052566 | -1.072819 | 0.2833522 | 0.481141 |
| ENSG00000106266 | SNX8 | 3207.57716 | -0.373006356 | 0.1714768 | -2.237503179 | 0.02525347 | 0.081735 |
| ENSG00000236498 | | 15.81879 | -0.3730752 | 0.572552 | -0.9057112 | 0.3650888 | 0.566956 |
| ENSG00000227911 | LINC02344 | 1.199719 | -0.3731057 | 0.8422509 | -1.252739 | 0.2103007 | NA |
| ENSG00000207280 | SNORD20 | 13.50839 | -0.3731097 | 0.4897303 | -0.9664864 | 0.3338008 | 0.535019 |
| ENSG00000112425 | EPM2A | 442.8661915 | -0.373176573 | 0.145115702 | -2.62145738 | 0.00875547 | 0.034992 |
| ENSG00000109846 | CRYAB | 27.0761447 | -0.373330475 | 0.797742407 | -1.059173238 | 0.28952089 | 0.488058 |
| ENSG00000233384 | CNIH3-AS2 | 4.339585 | -0.3734874 | 0.7109355 | -0.940817 | 0.3467987 | 0.548425 |
| ENSG00000033100 | CHPF2 | 4485.067906 | -0.373882064 | 0.208731618 | -1.863866802 | 0.06234041 | 0.16362 |
| ENSG00000133114 | GPALPP1 | 2096.814759 | -0.374016291 | 0.137319096 | -2.770674696 | 0.00559403 | 0.024289 |
| ENSG00000260298 | ACTG1P16 | 0.9191315 | -0.3740582 | 0.9105319 | -1.809812 | 0.07032497 | NA |
| ENSG00000262692 | | 29.32668 | -0.3740663 | 0.4950149 | -0.9623719 | 0.3358628 | 0.536916 |
| ENSG00000176601 | MAP3K19 | 1.171511 | -0.3741776 | 0.8242109 | -1.157671 | 0.2469982 | NA |
| ENSG00000283463 | HSFX4 | 1.948192 | -0.3742747 | 0.746046 | -0.9749661 | 0.3295771 | 0.530716 |
| ENSG00000143947 | RPS27A | 11815.39 | -0.374283 | 0.188708 | -2.033354 | 0.04201676 | 0.121474 |

| | | | | | | | |
|---|---|---|---|---|---|---|---|
| ENSG00000251169 | LINC01843 | 13.59546 | -0.374411 | 0.5710801 | -0.9110124 | 0.3622888 | 0.564323 |
| ENSG00000234473 | ZNF101P2 | 1.670401 | -0.3745126 | 0.7564849 | -0.9926857 | 0.3208632 | 0.52174 |
| ENSG00000180708 | OR10K2 | 15.39902 | -0.3748547 | 0.5488126 | -0.9186868 | 0.3582594 | 0.560203 |
| ENSG00000066697 | MSANTD3 | 743.1312088 | -0.374860087 | 0.194273813 | -1.99628364 | 0.04590306 | 0.129931 |
| ENSG00000257773 | ST13P3 | 115.3791 | -0.3750554 | 0.208697 | -1.867421 | 0.06184277 | 0.162699 |
| ENSG00000113621 | TXNDC15 | 1396.169533 | -0.37513747 | 0.149940114 | -2.577788286 | 0.00994349 | 0.038686 |
| ENSG00000237296 | SMG1P1 | 79.67288 | -0.375149 | 0.3241194 | -1.278215 | 0.2011735 | 0.381866 |
| ENSG00000080345 | RIF1 | 5081.926745 | -0.375191784 | 0.213633127 | -1.834328632 | 0.06660523 | 0.171993 |
| ENSG00000178764 | ZHX2 | 2947.993 | -0.3752118 | 0.1895836 | -2.036343 | 0.04171595 | 0.120867 |
| ENSG00000213100 | KRT18P68 | 14.61419 | -0.3753782 | 0.5122594 | -0.9447428 | 0.3447902 | 0.546359 |
| ENSG00000234439 | KRT18P2 | 2.492084 | -0.3753978 | 0.8112601 | -1.097641 | 0.2723614 | 0.468889 |
| ENSG00000184384 | MAML2 | 370.9698 | -0.3754181 | 0.4727659 | -0.9904843 | 0.3219375 | 0.522674 |
| ENSG00000092068 | SLC7A8 | 226.5066959 | -0.375437474 | 0.487815272 | -0.973093116 | 0.33050703 | 0.531616 |
| ENSG00000135517 | MIP | 3.19659622 | -0.375573133 | 0.691061169 | -0.914494078 | 0.36045729 | 0.562459 |
| ENSG00000137522 | RNF121 | 2245.661049 | -0.375722229 | 0.11369872 | -3.365851193 | 0.00076308 | 0.004532 |
| ENSG00000072818 | ACAP1 | 192.0355063 | -0.3757524 | 0.380617783 | -1.13335414 | 0.25706556 | 0.45091 |
| ENSG00000277151 | | 7.156512 | -0.3758046 | 0.6334578 | -0.9041925 | 0.3658934 | 0.567807 |
| ENSG00000235847 | LDHAP7 | 48.69856 | -0.3758079 | 0.3578217 | -1.18429 | 0.2362983 | 0.425872 |
| ENSG00000244041 | LINC01011 | 112.3856 | -0.3758135 | 0.2365438 | -1.680841 | 0.09279386 | 0.220308 |
| ENSG00000128944 | KNSTRN | 1637.192184 | -0.375865599 | 0.390599454 | -1.106207123 | 0.26863688 | 0.464645 |
| ENSG00000149554 | CHEK1 | 1005.27 | -0.3759098 | 0.4591354 | -0.9992437 | 0.3176767 | 0.518672 |
| ENSG00000132688 | NES | 224.9548044 | -0.375968263 | 0.699952936 | -0.943740224 | 0.34530241 | 0.546873 |
| ENSG00000271021 | | 2.800691 | -0.3759933 | 0.8038159 | -1.058273 | 0.2899309 | 0.488408 |
| ENSG00000136485 | DCAF7 | 21749.56093 | -0.376135938 | 0.134689936 | -2.839090534 | 0.00452423 | 0.020419 |
| ENSG00000286010 | | 15.58009 | -0.3761383 | 0.4906443 | -0.9690407 | 0.3325249 | 0.533602 |
| ENSG00000100445 | SDR39U1 | 697.9420309 | -0.37623137 | 0.240524924 | -1.653272094 | 0.0982755 | 0.229768 |
| ENSG00000119812 | FAM98A | 3520.586532 | -0.376255522 | 0.147310887 | -2.606242177 | 0.00915417 | 0.036252 |
| ENSG00000135900 | MRPL44 | 1269.556519 | -0.376287987 | 0.214643637 | -1.833259997 | 0.06676392 | 0.172302 |
| ENSG00000279903 | SLC7A2-IT1 | 0.9118965 | -0.3763209 | 0.8891717 | -1.624249 | 0.1043227 | NA |
| ENSG00000280057 | | 14.7227 | -0.3763516 | 0.5964604 | -0.9123329 | 0.3615935 | 0.563537 |
| ENSG00000188227 | ZNF793 | 621.4047 | -0.3764662 | 0.2038778 | -1.917206 | 0.05521171 | 0.149548 |
| ENSG00000198908 | BHLHB9 | 2.158399 | -0.3765203 | 0.7484434 | -0.9727047 | 0.3307001 | 0.53183 |
| ENSG00000253898 | LINC01419 | 6.193657 | -0.3767207 | 0.8641913 | -1.111822 | 0.2662148 | 0.461852 |

| | | | | | | | |
|---|---|---|---|---|---|---|---|
| ENSG00000012174 | MBTPS2 | 1744.36104 | -0.376836067 | 0.229187824 | -1.719456608 | 0.08553126 | 0.207552 |
| ENSG00000272572 | | 34.4106 | -0.3768363 | 0.3787442 | -1.14013 | 0.2542322 | 0.447242 |
| ENSG00000180667 | YOD1 | 57.46163 | -0.3769167 | 0.2991038 | -1.369175 | 0.1709446 | 0.341599 |
| ENSG00000169567 | HINT1 | 8901.534 | -0.3770158 | 0.1844026 | -2.111141 | 0.03476019 | 0.105091 |
| ENSG00000287021 | | 1.049478 | -0.3770188 | 0.875423 | -1.242512 | 0.2140477 | NA |
| ENSG00000260439 | LMF1-AS1 | 5.98983 | -0.3770297 | 0.7122402 | -0.9342161 | 0.3501924 | 0.55192 |
| ENSG00000251587 | LDHAP1 | 1.533938 | -0.3771753 | 0.8394485 | -1.21975 | 0.2225598 | NA |
| ENSG00000155542 | SETD9 | 245.7772 | -0.3771823 | 0.3029325 | -1.356688 | 0.1748803 | 0.346858 |
| ENSG00000255725 | TDGP1 | 241.3469 | -0.3771926 | 0.3377093 | -1.245993 | 0.2127669 | 0.397483 |
| ENSG00000285524 | | 0.8261321 | -0.3773473 | 0.9043466 | -1.635121 | 0.1020236 | NA |
| ENSG00000130816 | DNMT1 | 8860.342291 | -0.377368098 | 0.314374938 | -1.316353238 | 0.18805551 | 0.365006 |
| ENSG00000081791 | DELE1 | 4488.976036 | -0.3774196 | 0.114368211 | -3.346543386 | 0.00081826 | 0.004816 |
| ENSG00000214300 | SPDYE3 | 230.1419 | -0.377435 | 0.2091089 | -1.88321 | 0.05967191 | 0.158368 |
| ENSG00000122335 | SERAC1 | 486.2289727 | -0.377616778 | 0.155514859 | -2.480370703 | 0.01312459 | 0.048425 |
| ENSG00000285591 | | 1.229937 | -0.3777497 | 0.8144672 | -1.126399 | 0.2599966 | NA |
| ENSG00000177098 | SCN4B | 2.615992 | -0.3778312 | 0.7178987 | -0.9628323 | 0.3356317 | 0.536815 |
| ENSG00000100150 | DEPDC5 | 752.1068561 | -0.377835203 | 0.164323533 | -2.357868996 | 0.01838018 | 0.063487 |
| ENSG00000116983 | HPCAL4 | 2.932010811 | -0.378191973 | 0.702433293 | -0.927573749 | 0.35362872 | 0.555622 |
| ENSG00000177700 | POLR2L | 4311.226 | -0.3782177 | 0.2252061 | -1.767573 | 0.07713232 | 0.191953 |
| ENSG00000275835 | TUBGCP5 | 1253.395 | -0.3782228 | 0.1908423 | -2.04733 | 0.04062568 | 0.118354 |
| ENSG00000183765 | CHEK2 | 1703.704 | -0.3782654 | 0.2981444 | -1.379178 | 0.1678399 | 0.337496 |
| ENSG00000244192 | | 16.17474 | -0.3783166 | 0.4474427 | -1.015751 | 0.3097481 | 0.510401 |
| ENSG00000259570 | | 6.5542 | -0.3783959 | 0.568108 | -0.9226705 | 0.3561789 | 0.558176 |
| ENSG00000182749 | PAQR7 | 567.4999 | -0.3784028 | 0.2326783 | -1.710612 | 0.08715269 | 0.210437 |
| ENSG00000261098 | | 19.36951 | -0.3787467 | 0.4536313 | -1.016634 | 0.3093278 | 0.509898 |
| ENSG00000287655 | | 7.243571 | -0.3787773 | 0.6162096 | -0.9064048 | 0.3647217 | 0.566651 |
| ENSG00000224429 | LINC00539 | 63.61803 | -0.3788665 | 0.2760644 | -1.469114 | 0.1418018 | 0.298868 |
| ENSG00000200209 | | 3.549272 | -0.3788688 | 0.6911127 | -0.9334749 | 0.3505748 | 0.55226 |
| ENSG00000153162 | BMP6 | 10.65221 | -0.379043 | 0.6706396 | -0.92131 | 0.3568886 | 0.558893 |
| ENSG00000163554 | SPTA1 | 30.19469 | -0.3790536 | 0.8643252 | -1.089166 | 0.2760806 | 0.472933 |
| ENSG00000163516 | ANKZF1 | 5386.08 | -0.3791251 | 0.2744476 | -1.498406 | 0.1340278 | 0.286898 |
| ENSG00000263563 | UBBP4 | 98.26636 | -0.3791455 | 0.3321808 | -1.259851 | 0.2077232 | 0.390365 |
| ENSG00000003756 | RBM5 | 5184.22858 | -0.379208641 | 0.201329249 | -1.948830142 | 0.05131571 | 0.141333 |

| | | | | | | | |
|---|---|---|---|---|---|---|---|
| ENSG00000088826 | SMOX | 328.2642689 | -0.379248423 | 0.262121364 | -1.537348731 | 0.12420793 | 0.27199 |
| ENSG00000271989 | | 1.548793 | -0.3793603 | 0.8500094 | -1.30608 | 0.1915254 | NA |
| ENSG00000269444 | | 2.864496 | -0.3794516 | 0.7750832 | -1.028884 | 0.3035342 | 0.503157 |
| ENSG00000166444 | DENND2B | 275.4524 | -0.3795263 | 0.2214977 | -1.798588 | 0.07208381 | 0.182591 |
| ENSG00000228506 | | 205.1243 | -0.3797358 | 0.2292718 | -1.732619 | 0.08316333 | 0.203197 |
| ENSG00000267532 | MIR497HG | 7.265779 | -0.379775 | 0.568725 | -0.9241474 | 0.3554096 | 0.557414 |
| ENSG00000164048 | ZNF589 | 626.7671 | -0.3797903 | 0.214053 | -1.850058 | 0.06430519 | 0.167355 |
| ENSG00000242607 | RPS3AP34 | 4.949369 | -0.3798612 | 0.64995 | -0.9166768 | 0.359312 | 0.561199 |
| ENSG00000230097 | ME2P1 | 0.9909066 | -0.3799358 | 0.8812613 | -1.532074 | 0.1255041 | NA |
| ENSG00000100379 | KCTD17 | 830.6296712 | -0.380024223 | 0.225217818 | -1.776701414 | 0.07561738 | 0.189232 |
| ENSG00000283418 | | 9.229152 | -0.3801141 | 0.5524828 | -0.9335047 | 0.3505594 | 0.55226 |
| ENSG00000186854 | TRABD2A | 194.567 | -0.3801493 | 0.4098436 | -1.092002 | 0.2748321 | 0.471557 |
| ENSG00000132361 | CLUH | 10380.20976 | -0.380179863 | 0.191259407 | -2.075794294 | 0.03791298 | 0.112148 |
| ENSG00000271916 | | 15.593 | -0.3802602 | 0.4693811 | -1.002488 | 0.3161081 | 0.51716 |
| ENSG00000158417 | EIF5B | 11177.95 | -0.380273 | 0.1807835 | -2.16608 | 0.03030505 | 0.094415 |
| ENSG00000187994 | RINL | 65.09102 | -0.3803633 | 0.4177466 | -1.074192 | 0.2827367 | 0.480585 |
| ENSG00000267244 | | 93.10977 | -0.3803714 | 0.3730853 | -1.163909 | 0.2444609 | 0.435999 |
| ENSG00000124788 | ATXN1 | 1193.209006 | -0.380406478 | 0.403634558 | -1.104062139 | 0.26956618 | 0.465828 |
| ENSG00000213626 | LBH | 39.81723 | -0.38041 | 0.5987115 | -0.9161915 | 0.3595665 | 0.561432 |
| ENSG00000103226 | NOMO3 | 792.2989767 | -0.380415996 | 0.231462295 | -1.723736136 | 0.08475549 | 0.206102 |
| ENSG00000091622 | PITPNM3 | 436.4677808 | -0.380441262 | 0.27625032 | -1.480477125 | 0.13874596 | 0.294085 |
| ENSG00000280433 | | 201.2218 | -0.3804569 | 0.2826671 | -1.448163 | 0.1475714 | 0.30754 |
| ENSG00000134987 | WDR36 | 3084.024904 | -0.380951386 | 0.186534739 | -2.108494995 | 0.0349882 | 0.105588 |
| ENSG00000161381 | PLXDC1 | 17.16974 | -0.3810496 | 0.3948599 | -1.115787 | 0.2645134 | 0.459998 |
| ENSG00000204371 | EHMT2 | 5013.25 | -0.3810938 | 0.1358345 | -2.917097 | 0.00353306 | 0.016647 |
| ENSG00000242888 | | 1.114102 | -0.3811048 | 0.8668244 | -1.411722 | 0.1580319 | NA |
| ENSG00000176261 | ZBTB8OS | 1178.604 | -0.3811474 | 0.1581511 | -2.47612 | 0.01328189 | 0.048849 |
| ENSG00000279900 | | 14.30729 | -0.3812815 | 0.6029496 | -0.9195492 | 0.3578084 | 0.559815 |
| ENSG00000130748 | TMEM160 | 516.9033735 | -0.381301786 | 0.272270958 | -1.497213056 | 0.13433783 | 0.287354 |
| ENSG00000172172 | MRPL13 | 2605.029 | -0.381321 | 0.2718592 | -1.501283 | 0.1332824 | 0.285948 |
| ENSG00000237988 | OR2I1P | 6.47612 | -0.3813824 | 0.870938 | -1.18309 | 0.2367733 | 0.426482 |
| ENSG00000149016 | TUT1 | 937.7433 | -0.3814382 | 0.1628075 | -2.399319 | 0.01642561 | 0.057964 |
| ENSG00000233585 | | 148.604 | -0.3815351 | 0.338502 | -1.252753 | 0.2102956 | 0.394029 |

| | | | | | | | |
|---|---|---|---|---|---|---|---|
| ENSG00000241985 | WWTR1-IT1 | 3.319681 | -0.3815658 | 0.7322922 | -0.9726226 | 0.3307409 | 0.531835 |
| ENSG00000249216 | | 3.507325 | -0.3815895 | 0.8669607 | -1.107801 | 0.2679478 | 0.463846 |
| ENSG00000287390 | | 23.02982 | -0.3816046 | 0.4588868 | -1.017493 | 0.308919 | 0.50935 |
| ENSG00000181026 | AEN | 3386.048 | -0.3818082 | 0.1972669 | -2.004871 | 0.04497679 | 0.127971 |
| ENSG00000011260 | UTP18 | 5193.747368 | -0.381828421 | 0.209041237 | -1.90405488 | 0.05690304 | 0.152836 |
| ENSG00000271270 | TMCC1-AS1 | 266.3513 | -0.3819065 | 0.4094772 | -1.093954 | 0.2739753 | 0.470753 |
| ENSG00000271888 | | 53.59488 | -0.3820133 | 0.2908431 | -1.420005 | 0.1556061 | 0.319671 |
| ENSG00000230982 | DSTNP1 | 80.2582 | -0.3820749 | 0.3797909 | -1.144381 | 0.2524657 | 0.445225 |
| ENSG00000166411 | IDH3A | 2052.495 | -0.3822372 | 0.1897642 | -2.086055 | 0.03697366 | 0.110018 |
| ENSG00000100412 | ACO2 | 5564.031579 | -0.3823961 | 0.136494097 | -2.851422973 | 0.0043524 | 0.019771 |
| ENSG00000075234 | TTC38 | 1059.685234 | -0.382474797 | 0.136785021 | -2.85442932 | 0.00431142 | 0.019621 |
| ENSG00000213866 | YBX1P10 | 2422.465 | -0.382587 | 0.2468286 | -1.633159 | 0.1024356 | 0.236724 |
| ENSG00000224861 | YBX1P1 | 3084.567 | -0.3828494 | 0.2413 | -1.665265 | 0.0958599 | 0.225534 |
| ENSG00000224114 | | 184.2327 | -0.3828713 | 0.2542945 | -1.589714 | 0.1118992 | 0.252467 |
| ENSG00000153982 | GDPD1 | 365.627 | -0.383029 | 0.4394207 | -1.054676 | 0.2915735 | 0.490317 |
| ENSG00000110066 | KMT5B | 4403.886975 | -0.383129905 | 0.134000977 | -2.908435474 | 0.00363242 | 0.017039 |
| ENSG00000169519 | METTL15 | 536.7954 | -0.3831997 | 0.1971252 | -2.045702 | 0.04078571 | 0.118807 |
| ENSG00000187866 | FAM122A | 660.778 | -0.3832663 | 0.2071541 | -1.921984 | 0.05460779 | 0.148425 |
| ENSG00000112149 | CD83 | 89.66686364 | -0.383430181 | 0.324242418 | -1.302448916 | 0.19276297 | 0.371232 |
| ENSG00000140022 | STON2 | 224.5961 | -0.3834686 | 0.286434 | -1.443224 | 0.1489574 | 0.309603 |
| ENSG00000134755 | DSC2 | 691.5440476 | -0.38362076 | 0.317364085 | -1.330385412 | 0.18339132 | 0.358537 |
| ENSG00000087237 | CETP | 1.737680454 | -0.383654317 | 0.840904026 | -1.22495572 | 0.22059191 | 0.406951 |
| ENSG00000114850 | SSR3 | 8353.003436 | -0.383894026 | 0.150799383 | -2.598952146 | 0.00935088 | 0.036828 |
| ENSG00000109111 | SUPT6H | 13051.30595 | -0.384059049 | 0.101680988 | -3.833794916 | 0.00012618 | 0.000942 |
| ENSG00000101343 | CRNKL1 | 3022.440898 | -0.384064219 | 0.186097601 | -2.138753073 | 0.03245567 | 0.099664 |
| ENSG00000114423 | CBLB | 1210.627192 | -0.384092348 | 0.17655137 | -2.236426242 | 0.02532386 | 0.081894 |
| ENSG00000187961 | KLHL17 | 2095.501 | -0.3840933 | 0.1541271 | -2.547091 | 0.0108625 | 0.041586 |
| ENSG00000127980 | PEX1 | 1259.7864 | -0.384141706 | 0.152195566 | -2.576416889 | 0.00998302 | 0.038806 |
| ENSG00000166780 | BMERB1 | 420.4187 | -0.3842539 | 0.3504606 | -1.233517 | 0.2173831 | 0.403012 |
| ENSG00000235701 | PCBP2P1 | 6.102417 | -0.3844907 | 0.6740972 | -0.9349546 | 0.3498117 | 0.551483 |
| ENSG00000254244 | PAICSP4 | 310.5776 | -0.3845117 | 0.2331276 | -1.744003 | 0.08115857 | 0.199565 |
| ENSG00000259404 | EFL1P1 | 6.541568 | -0.3846454 | 0.6605613 | -0.9480786 | 0.3430894 | 0.544618 |
| ENSG00000241932 | RPL35AP34 | 4.810329 | -0.3846872 | 0.6816831 | -0.9126419 | 0.3614309 | 0.563416 |

| | | | | | | | |
|---|---|---|---|---|---|---|---|
| ENSG00000174173 | TRMT10C | 2517.12 | -0.3847527 | 0.268616 | -1.531848 | 0.12556 | 0.273889 |
| ENSG00000171848 | RRM2 | 6142.315 | -0.3847723 | 0.6422322 | -0.9634745 | 0.3353094 | 0.536612 |
| ENSG00000053501 | USE1 | 682.4456302 | -0.384774052 | 0.18480042 | -2.156551733 | 0.03104061 | 0.096334 |
| ENSG00000113721 | PDGFRB | 2.30029018 | -0.384971742 | 0.757177167 | -1.014841886 | 0.3101812 | 0.510797 |
| ENSG00000254109 | RBPMS-AS1 | 44.86945 | -0.3850273 | 0.2819789 | -1.467282 | 0.1422993 | 0.299679 |
| ENSG00000175581 | MRPL48 | 2068.129 | -0.3850328 | 0.2116465 | -1.912353 | 0.05583093 | 0.150765 |
| ENSG00000227337 | RPL23AP43 | 2.575797 | -0.385034 | 0.7708064 | -1.014134 | 0.3105189 | 0.511253 |
| ENSG00000208772 | SNORD94 | 26.36885 | -0.3851158 | 0.4347347 | -1.063378 | 0.2876103 | 0.485978 |
| ENSG00000237682 | PRKCIP1 | 264.6964 | -0.3851588 | 0.3048479 | -1.37826 | 0.1681231 | 0.337887 |
| ENSG00000173171 | MTX1 | 1522.68 | -0.3851943 | 0.148843 | -2.644255 | 0.00818709 | 0.033108 |
| ENSG00000236779 | | 2.757559 | -0.3852383 | 0.7727282 | -1.031319 | 0.3023912 | 0.501837 |
| ENSG00000172828 | CES3 | 75.81729 | -0.3852486 | 0.3775801 | -1.170577 | 0.2417688 | 0.432344 |
| ENSG00000230124 | ACBD6 | 4014.077 | -0.3852886 | 0.1306615 | -3.001616 | 0.00268551 | 0.013275 |
| ENSG00000253712 | LINC02886 | 0.8820613 | -0.3853459 | 0.8800879 | -1.257604 | 0.2085352 | NA |
| ENSG00000161016 | RPL8 | 61815.56 | -0.3853959 | 0.1797631 | -2.251094 | 0.0243796 | 0.079517 |
| ENSG00000249087 | ZNF436-AS1 | 73.35493 | -0.3854108 | 0.496181 | -0.989825 | 0.3222597 | 0.523069 |
| ENSG00000226054 | MEMO1P1 | 862.0748 | -0.3854367 | 0.146904 | -2.684011 | 0.00727448 | 0.030092 |
| ENSG00000164346 | NSA2 | 6266.091 | -0.3855684 | 0.1684307 | -2.349094 | 0.01881915 | 0.064717 |
| ENSG00000230358 | SPDYE21 | 6.952356 | -0.385802 | 0.5841481 | -0.9356557 | 0.3494505 | 0.551175 |
| ENSG00000116586 | LAMTOR2 | 2816.396458 | -0.385900014 | 0.175073849 | -2.277905795 | 0.02273219 | 0.075355 |
| ENSG00000170312 | CDK1 | 3461.262 | -0.3860006 | 0.4594513 | -1.018562 | 0.308411 | 0.508686 |
| ENSG00000180917 | CMTR2 | 2994.698 | -0.3860063 | 0.1870323 | -2.13728 | 0.03257521 | 0.099918 |
| ENSG00000065457 | ADAT1 | 2993.005067 | -0.386155118 | 0.181992208 | -2.19718332 | 0.02800736 | 0.088744 |
| ENSG00000227141 | | 17.76737 | -0.3862091 | 0.4853299 | -1.002749 | 0.3159821 | 0.517081 |
| ENSG00000279675 | | 2.08699 | -0.3863424 | 0.789934 | -1.067637 | 0.2856841 | 0.483708 |
| ENSG00000144021 | CIAO1 | 8999.854 | -0.3864572 | 0.09011426 | -4.319704 | 1.56E-05 | 0.000147 |
| ENSG00000286273 | | 2.097912 | -0.3865828 | 0.8032369 | -1.094909 | 0.2735565 | 0.470307 |
| ENSG00000268707 | | 3.866186 | -0.3866048 | 0.6961912 | -0.9500156 | 0.3421043 | 0.543493 |
| ENSG00000258730 | ITPK1-AS1 | 1.880247 | -0.3866119 | 0.7224145 | -0.9680618 | 0.3330135 | 0.534009 |
| ENSG00000131044 | TTLL9 | 32.43135418 | -0.386646273 | 0.405639925 | -1.115430596 | 0.26466595 | 0.460053 |
| ENSG00000224195 | | 36.59087 | -0.3868256 | 0.386367 | -1.143502 | 0.2528304 | 0.445691 |
| ENSG00000250039 | | 1.28664 | -0.3868624 | 0.8162569 | -1.11657 | 0.2641783 | NA |
| ENSG00000229356 | LRRC3-DT | 1.325847 | -0.3868784 | 0.8038114 | -1.081094 | 0.2796553 | NA |

| | | | | | | | |
|---|---|---|---|---|---|---|---|
| ENSG00000158402 | CDC25C | 569.7267 | -0.386996 | 0.4709603 | -1.030834 | 0.3026185 | 0.502078 |
| ENSG00000127337 | YEATS4 | 1525.929293 | -0.387019836 | 0.220320653 | -1.829669845 | 0.06729932 | 0.173314 |
| ENSG00000267534 | S1PR2 | 155.8292 | -0.3873114 | 0.2874433 | -1.451495 | 0.1466421 | 0.306155 |
| ENSG00000166529 | ZSCAN21 | 323.7221 | -0.3873183 | 0.2106871 | -1.911852 | 0.05589514 | 0.150907 |
| ENSG00000115275 | MOGS | 6513.228332 | -0.387335458 | 0.151651339 | -2.610028522 | 0.00905347 | 0.03592 |
| ENSG00000119041 | GTF3C3 | 3072.753136 | -0.38743327 | 0.146531485 | -2.701729952 | 0.00689798 | 0.028798 |
| ENSG00000213130 | EEF1DP5 | 1.017959 | -0.3874432 | 0.8678464 | -1.389383 | 0.1647162 | NA |
| ENSG00000144048 | DUSP11 | 1678.876 | -0.3874679 | 0.122926 | -3.182155 | 0.00146184 | 0.00793 |
| ENSG00000271002 | | 1.098394 | -0.3876276 | 0.8339859 | -1.189872 | 0.2340967 | NA |
| ENSG00000137574 | TGS1 | 1218.807401 | -0.387750943 | 0.176259904 | -2.231080884 | 0.02567577 | 0.082809 |
| ENSG00000251046 | ZNF969P | 0.8121904 | -0.3877922 | 0.9003673 | -1.481934 | 0.1383579 | NA |
| ENSG00000240132 | ETF1P2 | 123.4696 | -0.3878035 | 0.2516679 | -1.633563 | 0.1023506 | 0.236589 |
| ENSG00000262209 | PCDHGB3 | 33.56526 | -0.3880995 | 0.4733905 | -1.019588 | 0.3079237 | 0.508088 |
| ENSG00000255374 | TAS2R43 | 1.654197 | -0.3881097 | 0.7514054 | -1.001775 | 0.3164523 | 0.5175 |
| ENSG00000179038 | | 1.600967 | -0.3882641 | 0.7772313 | -1.037287 | 0.2996022 | 0.498891 |
| ENSG00000249437 | NAIP | 52.29614 | -0.3882887 | 0.3177857 | -1.348094 | 0.1776282 | 0.350937 |
| ENSG00000258344 | | 8.609822 | -0.3884838 | 0.5974659 | -0.9348111 | 0.3498856 | 0.551565 |
| ENSG00000237801 | AMD1P2 | 3.686352 | -0.3884856 | 0.7346007 | -0.9824298 | 0.3258882 | 0.527091 |
| ENSG00000277662 | | 12.01741 | -0.3885104 | 0.4997149 | -0.9924671 | 0.3209697 | 0.52174 |
| ENSG00000138246 | DNAJC13 | 6909.205138 | -0.38873475 | 0.16797715 | -2.383023082 | 0.01717112 | 0.060065 |
| ENSG00000115761 | NOL10 | 2405.197026 | -0.388794272 | 0.18722911 | -2.14710044 | 0.03178529 | 0.09813 |
| ENSG00000134046 | MBD2 | 2923.788115 | -0.388844276 | 0.180290673 | -2.216146898 | 0.02668144 | 0.085362 |
| ENSG00000224738 | | 1378.257 | -0.388926 | 0.3604979 | -1.213806 | 0.2248217 | 0.411866 |
| ENSG00000263120 | | 3.615001 | -0.3890257 | 0.6990256 | -0.9487472 | 0.3427492 | 0.544289 |
| ENSG00000200924 | RNU6-1048P | 6.139653 | -0.3890319 | 0.5682988 | -0.9424686 | 0.3459528 | 0.547511 |
| ENSG00000159618 | ADGRG5 | 155.0267 | -0.3891254 | 0.390522 | -1.149921 | 0.2501763 | 0.442695 |
| ENSG00000257271 | KIRREL3-AS1 | 1.188313 | -0.3891705 | 0.9265848 | -2.003617 | 0.04511107 | NA |
| ENSG00000121621 | KIF18A | 569.4183317 | -0.389180486 | 0.469105839 | -1.036039672 | 0.30018364 | 0.499444 |
| ENSG00000227113 | | 7.748425 | -0.3891917 | 0.6450151 | -0.936615 | 0.3489566 | 0.550666 |
| ENSG00000144852 | NR1I2 | 1.877003 | -0.3891979 | 0.7498085 | -1.006782 | 0.3140394 | 0.515171 |
| ENSG00000163902 | RPN1 | 17376.43 | -0.3892828 | 0.1624763 | -2.454303 | 0.0141158 | 0.051355 |
| ENSG00000213222 | TOMM40P4 | 537.9767 | -0.3893639 | 0.237303 | -1.728251 | 0.08394325 | 0.204614 |
| ENSG00000104472 | CHRAC1 | 1805.614775 | -0.389370544 | 0.203836634 | -1.997100235 | 0.0458143 | 0.129714 |

| | | | | | | | |
|---|---|---|---|---|---|---|---|
| ENSG00000100151 | PICK1 | 1621.740045 | -0.389394128 | 0.155567389 | -2.555189035 | 0.01061301 | 0.040848 |
| ENSG00000261335 | | 8.59412 | -0.3895809 | 0.5996276 | -0.9274794 | 0.3536777 | 0.555666 |
| ENSG00000284642 | | 14.1253 | -0.389675 | 0.4416243 | -1.064316 | 0.2871856 | 0.485492 |
| ENSG00000107551 | RASSF4 | 52.25938526 | -0.389694546 | 0.431979081 | -1.088502554 | 0.27637331 | 0.473312 |
| ENSG00000237517 | DGCR5 | 64.63326 | -0.3897353 | 0.4209006 | -1.096277 | 0.2729576 | 0.469611 |
| ENSG00000251611 | FAM160A1-DT | 24.345 | -0.3899426 | 0.4300745 | -1.078079 | 0.2809986 | 0.478488 |
| ENSG00000261202 | | 2.180468 | -0.3900832 | 0.7383646 | -0.9853991 | 0.3244281 | 0.525401 |
| ENSG00000008382 | MPND | 926.0355242 | -0.390089462 | 0.208517838 | -1.950468684 | 0.05112028 | 0.140923 |
| ENSG00000249353 | NPM1P27 | 13604.78 | -0.3903561 | 0.1901337 | -2.102759 | 0.03548685 | 0.106742 |
| ENSG00000233547 | | 2.851956 | -0.3903968 | 0.7267566 | -0.9916026 | 0.3213914 | 0.522202 |
| ENSG00000064703 | DDX20 | 1998.459398 | -0.390532981 | 0.160563328 | -2.476807662 | 0.01325633 | 0.048796 |
| ENSG00000260608 | | 8.86063 | -0.3905755 | 0.6645489 | -0.9456493 | 0.3443275 | 0.545915 |
| ENSG00000067955 | CBFB | 2368.063631 | -0.390578009 | 0.229962902 | -1.778476589 | 0.07532561 | 0.18868 |
| ENSG00000198794 | SCAMP5 | 2961.62 | -0.3905794 | 0.163807 | -2.439553 | 0.01470543 | 0.053075 |
| ENSG00000213853 | EMP2 | 11963.73 | -0.3906326 | 0.1289389 | -3.081011 | 0.00206299 | 0.010641 |
| ENSG00000136271 | DDX56 | 8416.511797 | -0.390636813 | 0.175837493 | -2.286672529 | 0.02221494 | 0.074112 |
| ENSG00000175395 | ZNF25 | 652.4952 | -0.3906503 | 0.1951384 | -2.071415 | 0.03832001 | 0.113112 |
| ENSG00000153563 | CD8A | 240.8641 | -0.3906594 | 0.2902639 | -1.465119 | 0.1428885 | 0.300539 |
| ENSG00000271198 | VDAC3P1 | 20.78324 | -0.390681 | 0.4478637 | -1.060222 | 0.2890437 | 0.487656 |
| ENSG00000273321 | | 6.343046 | -0.3907691 | 0.7052013 | -0.9635438 | 0.3352747 | 0.536595 |
| ENSG00000177646 | ACAD9 | 4140.958 | -0.3908927 | 0.15962 | -2.496396 | 0.01254624 | 0.046713 |
| ENSG00000253219 | KRT18P41 | 8.2423 | -0.3909661 | 0.616246 | -0.9385246 | 0.3479749 | 0.549729 |
| ENSG00000279662 | | 400.5693 | -0.390967 | 0.3611399 | -1.213509 | 0.2249354 | 0.411971 |
| ENSG00000199172 | MIR331 | 2.427664 | -0.391014 | 0.7648735 | -1.007075 | 0.3138986 | 0.515047 |
| ENSG00000276763 | | 29.54018 | -0.3911058 | 0.4941651 | -1.006375 | 0.3142353 | 0.515372 |
| ENSG00000188112 | C6orf132 | 2099.396 | -0.3913187 | 0.261376 | -1.603813 | 0.1087554 | 0.247246 |
| ENSG00000117222 | RBBP5 | 3733.222149 | -0.391327132 | 0.155250562 | -2.578404921 | 0.00992576 | 0.038629 |
| ENSG00000140905 | GCSH | 159.134 | -0.3913317 | 0.3241977 | -1.332449 | 0.1827128 | 0.357685 |
| ENSG00000283537 | | 14.4319 | -0.3913533 | 0.5901949 | -0.9368563 | 0.3488325 | 0.550528 |
| ENSG00000247595 | SPTY2D1OS | 78.73963 | -0.3914011 | 0.3361868 | -1.289555 | 0.1972052 | 0.377186 |
| ENSG00000115970 | THADA | 2792.8301 | -0.391509652 | 0.155534481 | -2.582271841 | 0.00981522 | 0.038274 |
| ENSG00000113407 | TARS1 | 13542.57207 | -0.391617647 | 0.222716201 | -1.840258261 | 0.06573033 | 0.17023 |
| ENSG00000120832 | MTERF2 | 915.911692 | -0.39177799 | 0.221560634 | -1.863006334 | 0.06246138 | 0.163872 |

| | | | | | | | |
|---|---|---|---|---|---|---|---|
| ENSG00000266456 | | 8.046819 | -0.3918373 | 0.5274046 | -0.9678895 | 0.3330995 | 0.534016 |
| ENSG00000107897 | ACBD5 | 3116.618751 | -0.391921828 | 0.166576628 | -2.406617912 | 0.01610101 | 0.057008 |
| ENSG00000112679 | DUSP22 | 1209.412246 | -0.392003916 | 0.223796812 | -1.836155116 | 0.06633472 | 0.171428 |
| ENSG00000161091 | MFSD12 | 2572.571 | -0.3921633 | 0.1600972 | -2.520002 | 0.0117354 | 0.04422 |
| ENSG00000278637 | H4C1 | 3.773465 | -0.3922012 | 0.7565637 | -0.9938514 | 0.3202952 | 0.52109 |
| ENSG00000127884 | ECHS1 | 9577.546486 | -0.39223497 | 0.274908242 | -1.534440948 | 0.12492121 | 0.272969 |
| ENSG00000228818 | | 386.3588 | -0.3922905 | 0.2474022 | -1.676735 | 0.09359436 | 0.221836 |
| ENSG00000124193 | SRSF6 | 9069.795033 | -0.392297897 | 0.153952845 | -2.59607711 | 0.00942949 | 0.037066 |
| ENSG00000153006 | SREK1IP1 | 1105.401 | -0.392299 | 0.1253098 | -3.167167 | 0.00153932 | 0.008292 |
| ENSG00000180992 | MRPL14 | 3343.394 | -0.392389 | 0.2608772 | -1.582447 | 0.1135476 | 0.254844 |
| ENSG00000271265 | | 3.753793 | -0.392395 | 0.7663315 | -1.026152 | 0.30482 | 0.504533 |
| ENSG00000260293 | | 54.94577 | -0.3924594 | 0.357936 | -1.232474 | 0.2177719 | 0.403514 |
| ENSG00000129925 | PGAP6 | 5396.996949 | -0.392601356 | 0.192733971 | -2.108382411 | 0.03499793 | 0.105594 |
| ENSG00000174483 | BBS1 | 75.89626 | -0.3927436 | 0.3759759 | -1.189508 | 0.2342398 | 0.423369 |
| ENSG00000288063 | | 2.557443 | -0.3929174 | 0.7114593 | -0.9768145 | 0.328661 | 0.529866 |
| ENSG00000169021 | UQCRFS1 | 5078.802 | -0.3929555 | 0.2417094 | -1.715244 | 0.08630043 | 0.208884 |
| ENSG00000256667 | KLRA1P | 103.8904 | -0.3929582 | 0.3272613 | -1.327143 | 0.1844613 | 0.359756 |
| ENSG00000265656 | | 1.988405 | -0.392979 | 0.8300091 | -1.182874 | 0.236859 | 0.426522 |
| ENSG00000270638 | | 8.005743 | -0.3930907 | 0.6907729 | -0.963315 | 0.3353895 | 0.536666 |
| ENSG00000278090 | LUNAR1 | 5.141364 | -0.3931122 | 0.6516296 | -0.9522787 | 0.3409556 | 0.542284 |
| ENSG00000248530 | BCL2L12P1 | 14.88187 | -0.3931427 | 0.5548129 | -0.9640398 | 0.335026 | 0.53632 |
| ENSG00000258785 | LINC01580 | 2.676419 | -0.3931717 | 0.8210079 | -1.171475 | 0.2414077 | 0.431946 |
| ENSG00000187773 | DIPK1C | 6.028759 | -0.3932197 | 0.7381672 | -1.001789 | 0.3164457 | 0.5175 |
| ENSG00000166833 | NAV2 | 261.2185 | -0.3932383 | 0.338807 | -1.296583 | 0.1947749 | 0.373968 |
| ENSG00000278876 | | 42.36497 | -0.3933309 | 0.3914627 | -1.155763 | 0.247778 | 0.439996 |
| ENSG00000270049 | | 73.3422 | -0.3933391 | 0.4284183 | -1.085154 | 0.2778535 | 0.475002 |
| ENSG00000121073 | SLC35B1 | 5428.2515 | -0.393565908 | 0.16493461 | -2.445526799 | 0.01446407 | 0.052364 |
| ENSG00000272100 | | 4.521369 | -0.393621 | 0.6788131 | -0.9525883 | 0.3407987 | 0.542132 |
| ENSG00000118849 | RARRES1 | 1.755663275 | -0.393660246 | 0.76821254 | -1.038164338 | 0.29919355 | 0.498387 |
| ENSG00000244402 | RN7SL314P | 1.002514 | -0.3936855 | 0.9142829 | -1.681648 | 0.09263713 | NA |
| ENSG00000154134 | ROBO3 | 254.2353 | -0.3937404 | 0.2924192 | -1.450348 | 0.1469614 | 0.306605 |
| ENSG00000250410 | | 3.655837 | -0.3937718 | 0.7676738 | -1.019164 | 0.3081253 | 0.508357 |
| ENSG00000007968 | E2F2 | 780.6112772 | -0.39392764 | 0.477620206 | -1.024045699 | 0.30581369 | 0.505737 |

| | | | | | | | |
|---|---|---|---|---|---|---|---|
| ENSG00000082898 | XPO1 | 16555.8485 | -0.393936564 | 0.157052502 | -2.59662083 | 0.00941458 | 0.037019 |
| ENSG00000272563 | | 0.8479446 | -0.394069 | 0.9192468 | -1.740739 | 0.08172941 | NA |
| ENSG00000164318 | EGFLAM | 1.534354 | -0.3941271 | 0.8978797 | -1.576493 | 0.1149122 | NA |
| ENSG00000115042 | FAHD2A | 1572.039475 | -0.394199105 | 0.162718732 | -2.48412152 | 0.01298715 | 0.048028 |
| ENSG00000226493 | RPL26P27 | 2.001074 | -0.3942225 | 0.7713062 | -1.040673 | 0.2980275 | 0.497162 |
| ENSG00000114988 | LMAN2L | 2739.346788 | -0.394274944 | 0.162208225 | -2.494153456 | 0.0126258 | 0.046976 |
| ENSG00000120697 | ALG5 | 2373.055025 | -0.394292659 | 0.152925076 | -2.638221506 | 0.00833421 | 0.033611 |
| ENSG00000099377 | HSD3B7 | 623.7986656 | -0.39440531 | 0.21567276 | -1.906676915 | 0.05656244 | 0.152278 |
| ENSG00000224903 | RNF32-AS1 | 24.95942 | -0.3944256 | 0.4917725 | -1.012246 | 0.3114204 | 0.512393 |
| ENSG00000247121 | | 95.29915 | -0.3944808 | 0.2635032 | -1.58884 | 0.1120964 | 0.252696 |
| ENSG00000226009 | KCNIP2-AS1 | 5.072916 | -0.3945732 | 0.6579221 | -0.9458562 | 0.3442219 | 0.545836 |
| ENSG00000273472 | | 345.8353 | -0.3946622 | 0.3794403 | -1.19177 | 0.2333514 | 0.422309 |
| ENSG00000167670 | CHAF1A | 1694.657 | -0.394672 | 0.4165887 | -1.110197 | 0.2669143 | 0.46278 |
| ENSG00000204370 | SDHD | 1947.363 | -0.3947013 | 0.2184737 | -1.888887 | 0.05890701 | 0.156981 |
| ENSG00000223553 | SMPD4P1 | 1.925078 | -0.3949266 | 0.8603657 | -1.323792 | 0.185572 | 0.361338 |
| ENSG00000274363 | | 6.644443 | -0.3949782 | 0.7122383 | -0.9782914 | 0.3279302 | 0.529301 |
| ENSG00000072274 | TFRC | 19315.50187 | -0.394988739 | 0.213477768 | -1.961472133 | 0.04982397 | 0.138152 |
| ENSG00000108506 | INTS2 | 5174.03454 | -0.395031365 | 0.204355538 | -2.026271104 | 0.04273701 | 0.123157 |
| ENSG00000249668 | KRT18P56 | 3.535183 | -0.3952185 | 0.7819234 | -1.064295 | 0.287195 | 0.485492 |
| ENSG00000100142 | POLR2F | 1703.345563 | -0.395469263 | 0.250040784 | -1.674535889 | 0.09402533 | 0.222692 |
| ENSG00000126005 | MMP24OS | 825.4477781 | -0.395571335 | 0.193516673 | -2.143175825 | 0.03209898 | 0.098822 |
| ENSG00000273549 | | 3.163977 | -0.3956672 | 0.8018111 | -1.101138 | 0.2708365 | 0.467142 |
| ENSG00000100490 | CDKL1 | 132.4018251 | -0.395747144 | 0.241157756 | -1.741479628 | 0.08159954 | 0.20039 |
| ENSG00000247970 | | 4.393807 | -0.3958089 | 0.617259 | -0.9479484 | 0.3431557 | 0.544618 |
| ENSG00000023902 | PLEKHO1 | 900.5325903 | -0.396008358 | 0.443047841 | -1.093828626 | 0.27403015 | 0.470756 |
| ENSG00000186306 | OR10T2 | 12.41877 | -0.3960924 | 0.6114182 | -0.9516492 | 0.3412749 | 0.54251 |
| ENSG00000264425 | MIR4653 | 13.08988 | -0.3961261 | 0.4836858 | -1.021259 | 0.3071317 | 0.507348 |
| ENSG00000144227 | NXPH2 | 6.285441 | -0.3961345 | 0.873848 | -1.145135 | 0.2521531 | 0.444851 |
| ENSG00000112983 | BRD8 | 5501.204575 | -0.396153847 | 0.196382731 | -2.093078494 | 0.03634215 | 0.108713 |
| ENSG00000205018 | | 186.567 | -0.3961704 | 0.2405522 | -1.736878 | 0.08240879 | 0.201911 |
| ENSG00000090372 | STRN4 | 7646.285282 | -0.396214961 | 0.147774768 | -2.732489789 | 0.00628576 | 0.0267 |
| ENSG00000100162 | CENPM | 726.287935 | -0.396391566 | 0.483611005 | -1.021798523 | 0.30687627 | 0.507052 |
| ENSG00000141028 | CDRT15P1 | 50.48231 | -0.3964541 | 0.314326 | -1.380489 | 0.167436 | 0.336846 |

| | | | | | | | |
|---|---|---|---|---|---|---|---|
| ENSG00000204315 | FKBPL | 583.8643 | -0.3964647 | 0.2155108 | -1.919313 | 0.05494475 | 0.149002 |
| ENSG00000131351 | HAUS8 | 640.9356027 | -0.39655408 | 0.435978074 | -1.069400436 | 0.28488927 | 0.482885 |
| ENSG00000233554 | B4GALT1-AS1 | 13.11721 | -0.3967398 | 0.5935879 | -0.9586059 | 0.3377573 | 0.53907 |
| ENSG00000145882 | PCYOX1L | 1263.527 | -0.396757 | 0.2116503 | -1.954855 | 0.05060023 | 0.13975 |
| ENSG00000241255 | | 224.2882 | -0.3968602 | 0.2255367 | -1.844739 | 0.06507554 | 0.168867 |
| ENSG00000187535 | IFT140 | 3049.522 | -0.3969961 | 0.1881803 | -2.186022 | 0.02881397 | 0.090878 |
| ENSG00000223117 | RN7SKP296 | 0.9665621 | -0.3971403 | 0.8633808 | -1.371142 | 0.1703308 | NA |
| ENSG00000188396 | TCTEX1D4 | 3.069952 | -0.3971639 | 0.8854395 | -1.473251 | 0.1406833 | 0.297195 |
| ENSG00000230825 | | 7.415785 | -0.3974878 | 0.6298506 | -0.9534381 | 0.3403681 | 0.541836 |
| ENSG00000055917 | PUM2 | 9125.461486 | -0.397691204 | 0.180039301 | -2.281462008 | 0.02252112 | 0.07488 |
| ENSG00000104129 | DNAJC17 | 800.8253478 | -0.397783309 | 0.241493392 | -1.759594537 | 0.07847658 | 0.19441 |
| ENSG00000002933 | TMEM176A | 1.025770139 | -0.397850639 | 0.821080799 | -1.16602672 | 0.24360364 | NA |
| ENSG00000256148 | | 10.21747 | -0.3978667 | 0.6112282 | -0.9540929 | 0.3400366 | 0.541405 |
| ENSG00000234289 | H2BS1 | 1.29211 | -0.3979004 | 0.8389354 | -1.251329 | 0.2108143 | NA |
| ENSG00000275202 | | 38.01553 | -0.3979798 | 0.2881774 | -1.487705 | 0.1368286 | 0.291158 |
| ENSG00000156298 | TSPAN7 | 5.734183 | -0.3980342 | 0.6836512 | -0.9627243 | 0.3356859 | 0.536827 |
| ENSG00000213856 | VDAC1P2 | 631.6474 | -0.398222 | 0.1650503 | -2.468125 | 0.01358227 | 0.049708 |
| ENSG00000243147 | MRPL33 | 2246.85 | -0.39823 | 0.1580794 | -2.585363 | 0.00972765 | 0.038019 |
| ENSG00000281560 | LSINCT5 | 1.091333 | -0.3982611 | 0.8862451 | -1.282757 | 0.1995772 | NA |
| ENSG00000108100 | CCNY | 5063.725291 | -0.398315903 | 0.114658992 | -3.52857869 | 0.0004178 | 0.002686 |
| ENSG00000217769 | | 1.749563 | -0.3983678 | 0.798863 | -1.099978 | 0.2713418 | 0.46771 |
| ENSG00000170027 | YWHAG | 10730.98 | -0.3984122 | 0.1495482 | -2.66707 | 0.00765158 | 0.031382 |
| ENSG00000287335 | | 57.95233 | -0.3984399 | 0.5113671 | -1.000337 | 0.3171476 | 0.518191 |
| ENSG00000256737 | RBBP4P5 | 41.9289 | -0.3984488 | 0.3148955 | -1.386176 | 0.1656933 | 0.334344 |
| ENSG00000200403 | RNU6-1099P | 1.443446 | -0.3985008 | 0.8151761 | -1.131925 | 0.257666 | NA |
| ENSG00000171517 | LPAR3 | 10.10681 | -0.3986081 | 0.6174961 | -0.9492783 | 0.3424791 | 0.543958 |
| ENSG00000089195 | TRMT6 | 1084.744106 | -0.398650614 | 0.232267308 | -1.808641153 | 0.07050677 | 0.179606 |
| ENSG00000170043 | TRAPPC1 | 2070.02 | -0.3986749 | 0.2095234 | -1.980296 | 0.04767025 | 0.133617 |
| ENSG00000227057 | WDR46 | 4904.037 | -0.3986812 | 0.1724562 | -2.380842 | 0.0172731 | 0.060379 |
| ENSG00000178338 | ZNF354B | 1246.979 | -0.3987421 | 0.1890932 | -2.182664 | 0.02906053 | 0.09158 |
| ENSG00000277721 | | 1.387696 | -0.3989425 | 0.8103101 | -1.130646 | 0.2582041 | NA |
| ENSG00000064102 | INTS13 | 1409.263139 | -0.398976351 | 0.207258412 | -2.00208372 | 0.04527573 | 0.128587 |
| ENSG00000287495 | | 1.634715 | -0.3992502 | 0.8197266 | -1.131636 | 0.2577875 | 0.451851 |

| | | | | | | | |
|---|---|---|---|---|---|---|---|
| ENSG00000169372 | CRADD | 527.5974 | -0.3993116 | 0.1655659 | -2.472804 | 0.01340578 | 0.049203 |
| ENSG00000116750 | UCHL5 | 3338.403873 | -0.39932033 | 0.212424376 | -1.963859837 | 0.04954635 | 0.137598 |
| ENSG00000139719 | VPS33A | 851.1614 | -0.3993387 | 0.1644743 | -2.487426 | 0.01286712 | 0.047687 |
| ENSG00000254331 | CKS1BP7 | 112.9525 | -0.3994112 | 0.410892 | -1.13693 | 0.2555675 | 0.448996 |
| ENSG00000135547 | HEY2 | 29.56523858 | -0.399431834 | 0.410108055 | -1.148782531 | 0.25064566 | 0.443281 |
| ENSG00000039068 | CDH1 | 33379.34293 | -0.399441146 | 0.126348205 | -3.207595538 | 0.0013385 | 0.007374 |
| ENSG00000171246 | NPTX1 | 23.35781 | -0.3996098 | 0.5342254 | -0.9922805 | 0.3210607 | 0.521835 |
| ENSG00000182348 | ZNF804B | 0.8621394 | -0.3996588 | 0.9210323 | -1.738281 | 0.08216132 | NA |
| ENSG00000251712 | RNU7-20P | 3.035054 | -0.399705 | 0.6798978 | -0.9696672 | 0.3322124 | 0.533326 |
| ENSG00000230918 | DPP4-DT | 1.203008 | -0.3997326 | 0.848068 | -1.258157 | 0.208335 | NA |
| ENSG00000178234 | GALNT11 | 1869.613 | -0.3997445 | 0.1659272 | -2.452971 | 0.01416816 | 0.05151 |
| ENSG00000178980 | SELENOW | 5807.544 | -0.3998064 | 0.1522335 | -2.711201 | 0.00670399 | 0.028112 |
| ENSG00000136521 | NDUFB5 | 4143.495843 | -0.399973822 | 0.175999015 | -2.353809323 | 0.01858214 | 0.064027 |
| ENSG00000110455 | ACCS | 1055.189312 | -0.400025063 | 0.268303894 | -1.597305727 | 0.11019758 | 0.249709 |
| ENSG00000136003 | ISCU | 4030.975329 | -0.400029172 | 0.154954041 | -2.646302322 | 0.00813771 | 0.032977 |
| ENSG00000057019 | DCBLD2 | 3459.671116 | -0.400051362 | 0.176905652 | -2.336911905 | 0.01944377 | 0.066452 |
| ENSG00000109133 | TMEM33 | 3116.301567 | -0.400103341 | 0.116079475 | -3.502196752 | 0.00046144 | 0.002928 |
| ENSG00000206828 | RNVU1-30 | 29.73633 | -0.4002012 | 0.5144704 | -1.011964 | 0.3115553 | 0.512457 |
| ENSG00000274997 | H2AC12 | 6.795671 | -0.4003077 | 0.5978724 | -0.9656296 | 0.3342296 | 0.535452 |
| ENSG00000143971 | ETAA1 | 1006.599 | -0.4003099 | 0.1614157 | -2.531686 | 0.01135157 | 0.043092 |
| ENSG00000231587 | SNORD62B | 25.73961 | -0.4003948 | 0.3406477 | -1.305328 | 0.191781 | 0.370012 |
| ENSG00000133069 | TMCC2 | 112.8374275 | -0.40040254 | 0.46351591 | -1.061440635 | 0.2884897 | 0.486999 |
| ENSG00000257534 | | 6.429142 | -0.4006102 | 0.6699322 | -0.9526105 | 0.3407875 | 0.542132 |
| ENSG00000151151 | IPMK | 884.8692 | -0.4006326 | 0.2353099 | -1.795019 | 0.07265075 | 0.183712 |
| ENSG00000275437 | | 249.9947 | -0.400649 | 0.2544255 | -1.674797 | 0.09397407 | 0.22259 |
| ENSG00000149231 | CCDC82 | 490.7655 | -0.4006493 | 0.2785756 | -1.541064 | 0.1233013 | 0.270615 |
| ENSG00000008869 | HEATR5B | 2841.825389 | -0.400753466 | 0.218093447 | -1.9208142 | 0.05475514 | 0.148628 |
| ENSG00000266717 | | 3.081544 | -0.4007678 | 0.7972604 | -1.111576 | 0.2663205 | 0.461901 |
| ENSG00000258646 | | 16.12968 | -0.4007924 | 0.4010085 | -1.158014 | 0.2468585 | 0.438715 |
| ENSG00000253846 | PCDHGA10 | 15.62701 | -0.4008153 | 0.5274766 | -0.9889712 | 0.3226772 | 0.523427 |
| ENSG00000156026 | MCU | 1588.054 | -0.4008522 | 0.2449361 | -1.728843 | 0.08383728 | 0.204507 |
| ENSG00000238178 | | 1.546177 | -0.4009616 | 0.8560858 | -1.320903 | 0.1865336 | NA |
| ENSG00000235957 | COX7CP1 | 275.0505 | -0.4010551 | 0.2184649 | -1.923385 | 0.05443167 | 0.148039 |

| | | | | | | | |
|---|---|---|---|---|---|---|---|
| ENSG00000198231 | DDX42 | 25134.88 | -0.4010731 | 0.1556076 | -2.635098 | 0.00841131 | 0.033855 |
| ENSG00000205084 | TMEM231 | 938.075 | -0.4012185 | 0.1529549 | -2.689513 | 0.00715563 | 0.029692 |
| ENSG00000196227 | FAM217B | 2172.999 | -0.4013553 | 0.1969417 | -2.111989 | 0.03468743 | 0.104943 |
| ENSG00000276571 | | 7.361931 | -0.4013673 | 0.5540803 | -0.974905 | 0.3296074 | 0.530716 |
| ENSG00000207554 | MIR647 | 51.76003 | -0.4013773 | 0.3038744 | -1.435829 | 0.1510511 | 0.312731 |
| ENSG00000134333 | LDHA | 20512.74208 | -0.401392691 | 0.266625174 | -1.6076189 | 0.10791867 | 0.245908 |
| ENSG00000272662 | TIMMDC1-DT | 5.169628 | -0.4014401 | 0.725381 | -0.9965256 | 0.3189948 | 0.519945 |
| ENSG00000286319 | | 3.748888 | -0.4014446 | 0.6706357 | -0.9571389 | 0.3384972 | 0.539828 |
| ENSG00000132388 | UBE2G1 | 2472.172341 | -0.401497039 | 0.106529423 | -3.820485719 | 0.00013319 | 0.000987 |
| ENSG00000234072 | | 948.7283 | -0.4015369 | 0.1872827 | -2.216999 | 0.02662316 | 0.085254 |
| ENSG00000105245 | NUMBL | 1173.593662 | -0.401601741 | 0.197583593 | -2.119410562 | 0.03405579 | 0.10348 |
| ENSG00000213186 | TRIM59 | 841.3993 | -0.4016101 | 0.2245788 | -1.861938 | 0.06261184 | 0.16417 |
| ENSG00000158555 | GDPD5 | 750.117 | -0.4016435 | 0.3091202 | -1.414195 | 0.1573047 | 0.322362 |
| ENSG00000239608 | RUVBL1-AS1 | 2.266304 | -0.401645 | 0.8006219 | -1.114725 | 0.2649684 | 0.460465 |
| ENSG00000130590 | SAMD10 | 1315.404666 | -0.401700371 | 0.19557276 | -2.12880988 | 0.03326999 | 0.101685 |
| ENSG00000126351 | THRA | 2497.901673 | -0.401746276 | 0.205332331 | -2.033946558 | 0.04195698 | 0.121367 |
| ENSG00000108219 | TSPAN14 | 6311.839869 | -0.401796351 | 0.112102263 | -3.629808824 | 0.00028363 | 0.001925 |
| ENSG00000237842 | SMU1P1 | 131.4099 | -0.4018826 | 0.2183339 | -1.92724 | 0.05394967 | 0.147106 |
| ENSG00000169908 | TM4SF1 | 1308.114 | -0.4018881 | 0.5735768 | -0.9386005 | 0.3479359 | 0.5497 |
| ENSG00000254694 | | 20.1388 | -0.4018919 | 0.5388371 | -0.9926345 | 0.3208881 | 0.52174 |
| ENSG00000274925 | ZKSCAN2-DT | 257.583 | -0.4019091 | 0.3620458 | -1.250684 | 0.2110498 | 0.394956 |
| ENSG00000235419 | | 1.275543 | -0.4019376 | 0.9267856 | -1.809768 | 0.07033173 | NA |
| ENSG00000147905 | ZCCHC7 | 1146.144 | -0.4020107 | 0.1501603 | -2.732488 | 0.00628581 | 0.0267 |
| ENSG00000140948 | ZCCHC14 | 5091.165 | -0.4021004 | 0.2277563 | -1.842929 | 0.06533944 | 0.169399 |
| ENSG00000267658 | | 2.588285 | -0.4021893 | 0.7616375 | -1.043565 | 0.2966868 | 0.495641 |
| ENSG00000125246 | CLYBL | 73.36945981 | -0.40221476 | 0.399757713 | -1.159432759 | 0.24627983 | 0.438038 |
| ENSG00000196787 | H2AC11 | 966.6724 | -0.4023546 | 0.2046712 | -2.044687 | 0.04088573 | 0.119046 |
| ENSG00000121691 | CAT | 3899.788946 | -0.402425103 | 0.138926951 | -2.958778681 | 0.00308861 | 0.014887 |
| ENSG00000160051 | IQCC | 438.9294 | -0.4024257 | 0.2930929 | -1.487583 | 0.136861 | 0.291166 |
| ENSG00000159842 | ABR | 5732.487 | -0.4025526 | 0.1252166 | -3.327621 | 0.00087591 | 0.005101 |
| ENSG00000163510 | CWC22 | 1664.855 | -0.402643 | 0.1240687 | -3.275931 | 0.00105314 | 0.005987 |
| ENSG00000223784 | LINP1 | 11.64836 | -0.4027598 | 0.8758401 | -1.145247 | 0.2521068 | 0.444819 |
| ENSG00000120158 | RCL1 | 1509.180181 | -0.402803202 | 0.194217537 | -2.155045407 | 0.03115828 | 0.096609 |

| | | | | | | | |
|---|---|---|---|---|---|---|---|
| ENSG00000122741 | DCAF10 | 2200.892149 | -0.402840738 | 0.16957536 | -2.445569491 | 0.01446236 | 0.052364 |
| ENSG00000174514 | MFSD4A | 63.50953 | -0.4028545 | 0.3730385 | -1.23222 | 0.2178669 | 0.403577 |
| ENSG00000228264 | PSMD8P1 | 14.78013 | -0.4028893 | 0.4946789 | -1.03186 | 0.3021377 | 0.501499 |
| ENSG00000263050 | | 26.43224 | -0.4029267 | 0.4494234 | -1.081101 | 0.2796523 | 0.47696 |
| ENSG00000265452 | MIR3682 | 6.440586 | -0.4029732 | 0.6346131 | -0.9715158 | 0.3312915 | 0.532331 |
| ENSG00000166532 | RIMKLB | 1693.705 | -0.4031346 | 0.2453334 | -1.737894 | 0.08222945 | 0.201583 |
| ENSG00000234851 | RPL23AP42 | 653.0461 | -0.4031575 | 0.1832379 | -2.27416 | 0.02295637 | 0.075871 |
| ENSG00000113716 | HMGXB3 | 5201.565184 | -0.403206557 | 0.137766699 | -3.03887159 | 0.00237466 | 0.011972 |
| ENSG00000103152 | MPG | 2182.161692 | -0.403238707 | 0.170593194 | -2.440984561 | 0.01464728 | 0.052919 |
| ENSG00000143977 | SNRPG | 1463.906 | -0.4033757 | 0.2724383 | -1.587926 | 0.112303 | 0.252991 |
| ENSG00000274427 | | 3.656661 | -0.4034981 | 0.7250819 | -0.99207 | 0.3211634 | 0.521911 |
| ENSG00000214078 | CPNE1 | 4597.797 | -0.4035159 | 0.2037128 | -2.034876 | 0.04186332 | 0.121175 |
| ENSG00000106245 | BUD31 | 2037.091892 | -0.403663836 | 0.125740859 | -3.262830612 | 0.00110305 | 0.006228 |
| ENSG00000240583 | AQP1 | 13.08527 | -0.4037479 | 0.5550978 | -0.9823414 | 0.3259317 | 0.527129 |
| ENSG00000164896 | FASTK | 6174.595 | -0.4038203 | 0.1616787 | -2.562484 | 0.01039263 | 0.04014 |
| ENSG00000232956 | SNHG15 | 3316.325 | -0.4038353 | 0.2837664 | -1.503694 | 0.1326601 | 0.284971 |
| ENSG00000164542 | KIAA0895 | 608.1139 | -0.403911 | 0.1971256 | -2.125105 | 0.03357784 | 0.102344 |
| ENSG00000138160 | KIF11 | 3569.563739 | -0.403918066 | 0.550914128 | -0.996506373 | 0.31900417 | 0.519945 |
| ENSG00000129691 | ASH2L | 2139.112491 | -0.403935558 | 0.195244661 | -2.144985593 | 0.031954 | 0.098492 |
| ENSG00000207755 | MIR450A2 | 1.721942 | -0.4039722 | 0.8838235 | -1.469623 | 0.141664 | 0.298696 |
| ENSG00000266173 | STRADA | 759.9703 | -0.4039912 | 0.1955069 | -2.14109 | 0.03226679 | 0.099201 |
| ENSG00000272848 | | 47.68838 | -0.4040173 | 0.4721255 | -1.058867 | 0.2896604 | 0.488207 |
| ENSG00000268496 | | 3.179605 | -0.4041037 | 0.7761973 | -1.05988 | 0.2891993 | 0.487854 |
| ENSG00000160539 | PLPP7 | 6.669823 | -0.4042846 | 0.5808098 | -0.9824573 | 0.3258746 | 0.527091 |
| ENSG00000124786 | SLC35B3 | 1327.155991 | -0.404285426 | 0.217964345 | -1.941889664 | 0.05215046 | 0.143139 |
| ENSG00000068394 | GPKOW | 2075.696004 | -0.404303521 | 0.174498875 | -2.412154643 | 0.01585855 | 0.056337 |
| ENSG00000075568 | TMEM131 | 4567.414852 | -0.40439422 | 0.145180433 | -2.870466279 | 0.00409867 | 0.018843 |
| ENSG00000287181 | | 3.573238 | -0.4045362 | 0.7738244 | -1.077242 | 0.2813721 | 0.478971 |
| ENSG00000099901 | RANBP1 | 4380.341892 | -0.404546582 | 0.319271243 | -1.378966804 | 0.16790499 | 0.337576 |
| ENSG00000215571 | GRK6P1 | 12.39071 | -0.404618 | 0.5010224 | -1.03197 | 0.3020861 | 0.501476 |
| ENSG00000165802 | NSMF | 5151.11 | -0.4046923 | 0.1651003 | -2.526604 | 0.01151712 | 0.043583 |
| ENSG00000214050 | FBXO16 | 112.0677 | -0.4047548 | 0.2188586 | -1.932104 | 0.05334668 | 0.14588 |
| ENSG00000115306 | SPTBN1 | 7209.970474 | -0.404887256 | 0.276399488 | -1.572537034 | 0.11582606 | 0.258649 |

| | | | | | | | |
|---|---|---|---|---|---|---|---|
| ENSG00000251553 | DDX18P4 | 3.101477 | -0.4049772 | 0.7293893 | -0.9975807 | 0.3184827 | 0.519669 |
| ENSG00000213569 | GTF3C6P2 | 159.2218 | -0.4050209 | 0.3293817 | -1.362484 | 0.1730452 | 0.344474 |
| ENSG00000280852 | | 95.98703 | -0.4050868 | 0.3017745 | -1.460555 | 0.1441376 | 0.302352 |
| ENSG00000088727 | KIF9 | 278.5526561 | -0.405100931 | 0.238409246 | -1.785491525 | 0.07418163 | 0.186677 |
| ENSG00000074356 | NCBP3 | 3825.534426 | -0.405134561 | 0.152713275 | -2.71483291 | 0.00663092 | 0.027876 |
| ENSG00000075292 | ZNF638 | 8992.370237 | -0.405274772 | 0.150664266 | -2.746113345 | 0.00603059 | 0.025846 |
| ENSG00000227076 | | 3.21634 | -0.4053255 | 0.8037949 | -1.112086 | 0.266101 | 0.46174 |
| ENSG00000186523 | FAM86B1 | 148.0186 | -0.4053273 | 0.2816886 | -1.54829 | 0.1215524 | 0.26781 |
| ENSG00000182481 | KPNA2 | 18435.79 | -0.405379 | 0.3677568 | -1.300065 | 0.1935789 | 0.372157 |
| ENSG00000162650 | ATXN7L2 | 938.4261 | -0.4055095 | 0.2911687 | -1.506269 | 0.1319981 | 0.284018 |
| ENSG00000126768 | TIMM17B | 2927.384768 | -0.405868552 | 0.2066438 | -2.03987172 | 0.0413631 | 0.120054 |
| ENSG00000091157 | WDR7 | 894.7304047 | -0.40596221 | 0.171097087 | -2.438560247 | 0.0147459 | 0.053188 |
| ENSG00000163558 | PRKCI | 5194.439 | -0.4060543 | 0.1603419 | -2.593983 | 0.00948713 | 0.037232 |
| ENSG00000272980 | | 15.05922 | -0.406075 | 0.5567018 | -0.9914927 | 0.3214451 | 0.522216 |
| ENSG00000279457 | WASH9P | 368.0517 | -0.4062098 | 0.2048585 | -2.070795 | 0.03837792 | 0.113208 |
| ENSG00000258836 | | 1.52799 | -0.4062235 | 0.8931314 | -1.549374 | 0.1212919 | NA |
| ENSG00000275720 | | 6.102575 | -0.4062277 | 0.6765388 | -0.9701811 | 0.3319562 | 0.533049 |
| ENSG00000164077 | MON1A | 457.9727 | -0.4063276 | 0.1684622 | -2.48045 | 0.01312168 | 0.048421 |
| ENSG00000234323 | LINC01505 | 2.696257 | -0.4063346 | 0.7730972 | -1.05943 | 0.2894041 | 0.487954 |
| ENSG00000242841 | KRT8P35 | 52.4209 | -0.4065407 | 0.3163514 | -1.410433 | 0.1584117 | 0.323956 |
| ENSG00000176171 | BNIP3 | 2351.786 | -0.4066104 | 0.1410912 | -2.959912 | 0.00307727 | 0.014844 |
| ENSG00000165832 | TRUB1 | 1883.376 | -0.4066568 | 0.1997052 | -2.112807 | 0.03461731 | 0.10485 |
| ENSG00000182621 | PLCB1 | 1938.718 | -0.406722 | 0.2545311 | -1.675971 | 0.09374388 | 0.222088 |
| ENSG00000266370 | MIR3657 | 2.299433 | -0.4067372 | 0.7582058 | -1.03264 | 0.3017722 | 0.501206 |
| ENSG00000160214 | RRP1 | 4514.731 | -0.406815 | 0.2263542 | -1.884857 | 0.05944912 | 0.158029 |
| ENSG00000213409 | C1QBPP2 | 3.560125 | -0.4069139 | 0.7619725 | -1.050433 | 0.2935189 | 0.492485 |
| ENSG00000175182 | FAM131A | 825.9418 | -0.4069862 | 0.2579867 | -1.663417 | 0.09622895 | 0.226057 |
| ENSG00000132953 | XPO4 | 3422.213128 | -0.407068031 | 0.220473905 | -1.917607853 | 0.05516075 | 0.149441 |
| ENSG00000230042 | AK3P3 | 13.94201 | -0.4071478 | 0.5643886 | -0.9894556 | 0.3224403 | 0.523217 |
| ENSG00000224831 | TMEM183B | 4045.604 | -0.4073376 | 0.1980091 | -2.147681 | 0.03173914 | 0.098015 |
| ENSG00000136731 | UGGT1 | 5948.549511 | -0.407369772 | 0.225532563 | -1.892266382 | 0.0584555 | 0.156138 |
| ENSG00000238728 | MIR1972-1 | 1.684215 | -0.4077743 | 0.8020846 | -1.100857 | 0.2709591 | 0.467262 |
| ENSG00000224837 | GCSHP5 | 1557.379 | -0.4078507 | 0.2501351 | -1.729799 | 0.08366617 | 0.204126 |

| | | | | | | | |
|---|---|---|---|---|---|---|---|
| ENSG00000075407 | ZNF37A | 2246.520519 | -0.407867961 | 0.221958841 | -1.927313854 | 0.05394053 | 0.147106 |
| ENSG00000142677 | IL22RA1 | 12.25777 | -0.4079812 | 0.6825782 | -0.9782876 | 0.3279321 | 0.529301 |
| ENSG00000143924 | EML4 | 3802.591 | -0.4079861 | 0.1786247 | -2.350381 | 0.01875422 | 0.064544 |
| ENSG00000264017 | RN7SL336P | 2.654033 | -0.4080904 | 0.8035955 | -1.09278 | 0.2744906 | 0.471272 |
| ENSG00000164611 | PTTG1 | 1528.452 | -0.4081131 | 0.5985449 | -0.9869145 | 0.3236846 | 0.524581 |
| ENSG00000273249 | WDR5-DT | 14.38238 | -0.4083328 | 0.4620497 | -1.073999 | 0.2828233 | 0.480634 |
| ENSG00000266184 | | 5.061056 | -0.4085783 | 0.7387206 | -1.020583 | 0.307452 | 0.507632 |
| ENSG00000175106 | TVP23C | 125.8686 | -0.408649 | 0.2421051 | -1.78151 | 0.0748291 | 0.18779 |
| ENSG00000053372 | MRTO4 | 2659.914857 | -0.408657329 | 0.295038547 | -1.536602735 | 0.12439062 | 0.272235 |
| ENSG00000278918 | | 84.85176 | -0.4087509 | 0.2588635 | -1.680008 | 0.09295567 | 0.220611 |
| ENSG00000103381 | CPPED1 | 598.6400001 | -0.408809291 | 0.282733149 | -1.55277913 | 0.12047591 | 0.266101 |
| ENSG00000176986 | SEC24C | 4581.193 | -0.4089507 | 0.1979966 | -2.144977 | 0.03195469 | 0.098492 |
| ENSG00000264350 | SNRPGP2 | 726.0665 | -0.4089531 | 0.3068676 | -1.470442 | 0.141442 | 0.298357 |
| ENSG00000229118 | | 0.9038153 | -0.409026 | 0.8934283 | -1.327331 | 0.1843992 | NA |
| ENSG00000286864 | | 3.066615 | -0.4091511 | 0.7753545 | -1.045479 | 0.2958017 | 0.494816 |
| ENSG00000235962 | RPL7AP53 | 5.908842 | -0.4091978 | 0.5784231 | -0.9883175 | 0.3229972 | 0.523722 |
| ENSG00000105499 | PLA2G4C | 221.3009762 | -0.409397704 | 0.5075205 | -1.032052758 | 0.3020474 | 0.501443 |
| ENSG00000108479 | GALK1 | 3586.97159 | -0.409443265 | 0.276670827 | -1.609988938 | 0.10740027 | 0.245021 |
| ENSG00000112186 | CAP2 | 1794.081915 | -0.409539859 | 0.251689239 | -1.690789656 | 0.09087698 | 0.217152 |
| ENSG00000172345 | STARD5 | 72.09841 | -0.4095847 | 0.3539338 | -1.309135 | 0.1904886 | 0.368455 |
| ENSG00000254783 | | 18.45586 | -0.4096458 | 0.436635 | -1.12574 | 0.2602754 | 0.454772 |
| ENSG00000154518 | ATP5MC3 | 8495.973 | -0.4096708 | 0.2244407 | -1.911547 | 0.05593436 | 0.150992 |
| ENSG00000108829 | LRRC59 | 12908.09945 | -0.409703926 | 0.210158558 | -2.035387722 | 0.04181188 | 0.121062 |
| ENSG00000170004 | CHD3 | 17554.26 | -0.4097066 | 0.182921 | -2.31209 | 0.02077275 | 0.070075 |
| ENSG00000107147 | KCNT1 | 12.58031334 | -0.409839068 | 0.624498612 | -0.980437153 | 0.32687038 | 0.528165 |
| ENSG00000166261 | ZNF202 | 1009.692 | -0.4098863 | 0.1684927 | -2.49177 | 0.01271084 | 0.0472 |
| ENSG00000215114 | UBXN2B | 1759.969 | -0.4100333 | 0.1286881 | -3.232279 | 0.00122807 | 0.006831 |
| ENSG00000131771 | PPP1R1B | 2.763413118 | -0.41006291 | 0.70890458 | -1.00015849 | 0.31723381 | 0.518204 |
| ENSG00000147044 | CASK | 2754.941 | -0.4101134 | 0.1513207 | -2.768889 | 0.00562478 | 0.024403 |
| ENSG00000198010 | DLGAP2 | 5.288223 | -0.4102929 | 0.6550511 | -0.9683285 | 0.3328803 | 0.533883 |
| ENSG00000102900 | NUP93 | 2646.530867 | -0.4103507 | 0.268192954 | -1.645100023 | 0.09994919 | 0.232759 |
| ENSG00000197061 | H4C3 | 15.96094 | -0.4103972 | 0.4963595 | -1.038626 | 0.2989786 | 0.498248 |
| ENSG00000260493 | | 4.768816 | -0.4105068 | 0.617952 | -0.9723678 | 0.3308676 | 0.531899 |

| | | | | | | | |
|---|---|---|---|---|---|---|---|
| ENSG00000168894 | RNF181 | 3987.834 | -0.410594 | 0.2068186 | -2.068367 | 0.03860555 | 0.113728 |
| ENSG00000228476 | CSTP1 | 2.315442 | -0.4106206 | 0.9445222 | -2.228507 | 0.02584676 | 0.083239 |
| ENSG00000204568 | MRPS18B | 3927.368 | -0.4106869 | 0.162012 | -2.608488 | 0.00909432 | 0.036042 |
| ENSG00000258315 | C17orf49 | 36.09127 | -0.4106893 | 0.3916939 | -1.205503 | 0.2280093 | 0.41607 |
| ENSG00000184402 | SS18L1 | 3904.895 | -0.4107069 | 0.1764672 | -2.399864 | 0.01640116 | 0.057886 |
| ENSG00000163013 | FBXO41 | 2913.768 | -0.4107126 | 0.1317954 | -3.185417 | 0.00144546 | 0.007863 |
| ENSG00000280870 | | 3.904287 | -0.4107458 | 0.6300588 | -0.9889891 | 0.3226685 | 0.523427 |
| ENSG00000065559 | MAP2K4 | 2593.313698 | -0.410843448 | 0.087409025 | -4.738873695 | 2.15E-06 | 2.51E-05 |
| ENSG00000181163 | NPM1 | 27744.78 | -0.4109565 | 0.1983419 | -2.148228 | 0.03169567 | 0.09791 |
| ENSG00000112742 | TTK | 1312.927318 | -0.410984367 | 0.463342902 | -1.103783754 | 0.26968695 | 0.465976 |
| ENSG00000240219 | | 60.12864 | -0.4110242 | 0.5044852 | -1.042162 | 0.2973364 | 0.496476 |
| ENSG00000175792 | RUVBL1 | 7261.201 | -0.411051 | 0.2142865 | -2.005789 | 0.0448788 | 0.127774 |
| ENSG00000201544 | SNORA16B | 2.214187 | -0.4110753 | 0.8679764 | -1.360674 | 0.1736169 | 0.34522 |
| ENSG00000264772 | | 8.315736 | -0.4111241 | 0.4894984 | -1.057023 | 0.290501 | 0.489115 |
| ENSG00000121281 | ADCY7 | 284.1427124 | -0.411146825 | 0.279481289 | -1.578340505 | 0.11448741 | 0.256508 |
| ENSG00000277200 | | 25.2686 | -0.4111676 | 0.469889 | -1.081261 | 0.2795808 | 0.476899 |
| ENSG00000268266 | | 2.655839 | -0.4112103 | 0.7479795 | -1.046983 | 0.2951073 | 0.494308 |
| ENSG00000254995 | STX16-NPEPL1 | 174.7153 | -0.4112333 | 0.3150633 | -1.432381 | 0.1520348 | 0.314196 |
| ENSG00000108604 | SMARCD2 | 26709.78524 | -0.411276397 | 0.129981401 | -3.213251822 | 0.00131241 | 0.007251 |
| ENSG00000236908 | LINC02827 | 9.34317 | -0.4114081 | 0.7067515 | -0.9942906 | 0.3200814 | 0.52093 |
| ENSG00000154548 | SRSF12 | 5.165885 | -0.4114697 | 0.7430281 | -1.032184 | 0.301986 | 0.501404 |
| ENSG00000229806 | RPS15P5 | 6.280415 | -0.411581 | 0.6181207 | -0.9788728 | 0.3276428 | 0.529092 |
| ENSG00000287091 | | 6.108787 | -0.4116705 | 0.5792029 | -1.000858 | 0.3168953 | 0.517969 |
| ENSG00000175575 | PAAF1 | 2215.918 | -0.4117639 | 0.127183 | -3.28289 | 0.00102749 | 0.005862 |
| ENSG00000283378 | CNTNAP3C | 1.524692 | -0.4117681 | 0.9513019 | -2.29491 | 0.02173826 | NA |
| ENSG00000265908 | | 3.414758 | -0.4117802 | 0.7723561 | -1.057894 | 0.2901036 | 0.488637 |
| ENSG00000167094 | TTC16 | 4.067238 | -0.4117956 | 0.6126751 | -0.9864557 | 0.3239095 | 0.524817 |
| ENSG00000256436 | TAS2R31 | 7.121089 | -0.4120001 | 0.623943 | -0.9817805 | 0.326208 | 0.527416 |
| ENSG00000284610 | | 10.0778 | -0.4120233 | 0.6768378 | -0.994671 | 0.3198963 | 0.520777 |
| ENSG00000269934 | | 2.592538 | -0.4120246 | 0.7243165 | -1.017685 | 0.3088277 | 0.509231 |
| ENSG00000236975 | LINC02814 | 29.81816 | -0.4121439 | 0.4911964 | -1.063055 | 0.287757 | 0.486071 |
| ENSG00000196860 | TOMM20L | 15.39844 | -0.4121885 | 0.5062936 | -1.040063 | 0.2983106 | 0.497478 |
| ENSG00000139767 | SRRM4 | 1.175408 | -0.4123388 | 0.9468293 | -2.125802 | 0.03351971 | NA |

| | | | | | | | |
|---|---|---|---|---|---|---|---|
| ENSG00000232233 | LINC02043 | 10.59194 | -0.4124746 | 0.594338 | -0.9905763 | 0.3218925 | 0.522633 |
| ENSG00000277534 | | 198.4176 | -0.4124787 | 0.2031258 | -2.124378 | 0.03363856 | 0.102463 |
| ENSG00000280129 | | 34.84896 | -0.412777 | 0.410819 | -1.175705 | 0.2397129 | 0.429927 |
| ENSG00000102445 | RUBCNL | 19.34870877 | -0.412799601 | 0.571147979 | -0.992751448 | 0.3208311 | 0.521738 |
| ENSG00000124145 | SDC4 | 9892.303058 | -0.412826853 | 0.190033116 | -2.262062574 | 0.02369354 | 0.077727 |
| ENSG00000163808 | KIF15 | 860.995 | -0.4128783 | 0.47496 | -1.077548 | 0.2812354 | 0.47883 |
| ENSG00000264546 | | 1.476521 | -0.41314 | 0.7926188 | -1.084336 | 0.2782159 | NA |
| ENSG00000158850 | B4GALT3 | 2830.313 | -0.4131617 | 0.156229 | -2.749611 | 0.00596661 | 0.025609 |
| ENSG00000235821 | IFITM4P | 1.702544 | -0.413216 | 0.8910256 | -1.464069 | 0.1431751 | 0.300975 |
| ENSG00000199325 | RNU4-39P | 2.933802 | -0.4132753 | 0.7411961 | -1.040309 | 0.2981964 | 0.49735 |
| ENSG00000203485 | INF2 | 7380.056 | -0.4133809 | 0.2279545 | -1.884447 | 0.05950453 | 0.158113 |
| ENSG00000204882 | GPR20 | 1.219408 | -0.413403 | 0.827739 | -1.197589 | 0.231077 | NA |
| ENSG00000215909 | BRWD1P1 | 3.791438 | -0.413409 | 0.7316986 | -1.018941 | 0.3082309 | 0.5085 |
| ENSG00000079263 | SP140 | 5.428101251 | -0.413424392 | 0.643320013 | -0.973543495 | 0.33028325 | 0.531385 |
| ENSG00000137275 | RIPK1 | 1740.920202 | -0.413454169 | 0.154690333 | -2.727034678 | 0.00639063 | 0.027028 |
| ENSG00000189007 | ADAT2 | 630.8029 | -0.4134749 | 0.1969913 | -2.172484 | 0.02981916 | 0.093257 |
| ENSG00000267131 | | 635.1773 | -0.4134905 | 0.2951113 | -1.521884 | 0.1280381 | 0.277724 |
| ENSG00000104973 | MED25 | 2462.409788 | -0.41352802 | 0.133237535 | -3.169939552 | 0.00152471 | 0.008228 |
| ENSG00000270802 | | 0.8980663 | -0.4135451 | 0.9066679 | -1.446818 | 0.147948 | NA |
| ENSG00000133063 | CHIT1 | 1.793362007 | -0.413558745 | 0.787990717 | -1.104370469 | 0.26943247 | 0.465657 |
| ENSG00000233008 | LINC01725 | 23.67451 | -0.4135959 | 0.4036577 | -1.185591 | 0.235784 | 0.42515 |
| ENSG00000049246 | PER3 | 1816.23903 | -0.413632175 | 0.233492048 | -1.856133167 | 0.06343458 | 0.165679 |
| ENSG00000274317 | LINC02334 | 15.80793 | -0.413678 | 0.6461513 | -0.9911358 | 0.3216193 | 0.522381 |
| ENSG00000132004 | FBXW9 | 786.7423509 | -0.413768266 | 0.218034665 | -2.000919188 | 0.0454011 | 0.128874 |
| ENSG00000238084 | | 166.4919 | -0.4137974 | 0.2471881 | -1.769601 | 0.07679371 | 0.191347 |
| ENSG00000198900 | TOP1 | 12160.47 | -0.4139564 | 0.1263522 | -3.32437 | 0.00088619 | 0.00515 |
| ENSG00000149657 | LSM14B | 4881.004 | -0.4139581 | 0.1202482 | -3.412399 | 0.00064394 | 0.003909 |
| ENSG00000239983 | | 4.505035 | -0.4140757 | 0.6970422 | -1.000168 | 0.317229 | 0.518204 |
| ENSG00000224531 | SMIM13 | 950.8551 | -0.4141359 | 0.1933145 | -2.218348 | 0.02653112 | 0.085 |
| ENSG00000279155 | | 17.45433 | -0.4141813 | 0.4922501 | -1.063635 | 0.2874943 | 0.485875 |
| ENSG00000131503 | ANKHD1 | 193.7982784 | -0.414353644 | 0.220344524 | -1.964923901 | 0.04942305 | 0.13737 |
| ENSG00000146826 | MAP11 | 1449.566 | -0.4147271 | 0.1848587 | -2.313406 | 0.02070031 | 0.069866 |
| ENSG00000136933 | RABEPK | 1320.638943 | -0.414817316 | 0.149480287 | -2.835139363 | 0.00458057 | 0.020648 |

| | | | | | | | |
|---|---|---|---|---|---|---|---|
| ENSG00000228666 | KRT18P1 | 52.04935 | -0.4148575 | 0.3696407 | -1.275765 | 0.2020387 | 0.383152 |
| ENSG00000168887 | C2orf68 | 3098.516 | -0.4148615 | 0.1258488 | -3.330727 | 0.0008662 | 0.005051 |
| ENSG00000102098 | SCML2 | 447.2436089 | -0.41487939 | 0.254892131 | -1.726855932 | 0.08419354 | 0.205054 |
| ENSG00000231395 | ARL4AP4 | 3.197326 | -0.4149868 | 0.7418009 | -1.034311 | 0.3009908 | 0.500189 |
| ENSG00000139350 | NEDD1 | 1948.861 | -0.4151746 | 0.2333907 | -1.869857 | 0.06150363 | 0.162016 |
| ENSG00000122783 | CYREN | 1047.685902 | -0.415259742 | 0.191940053 | -2.231542119 | 0.02564524 | 0.082731 |
| ENSG00000187066 | TMEM262 | 9.156214 | -0.4153238 | 0.5110297 | -1.043039 | 0.2969305 | 0.495923 |
| ENSG00000198326 | TMEM239 | 1.919967 | -0.4153602 | 0.9576882 | -2.446881 | 0.01440985 | 0.052245 |
| ENSG00000142634 | EFHD2 | 1666.382 | -0.4153796 | 0.2287883 | -1.909827 | 0.05615545 | 0.151425 |
| ENSG00000263829 | SINHCAFP1 | 9.469273 | -0.4154877 | 0.6565313 | -0.9891378 | 0.3225957 | 0.523415 |
| ENSG00000158716 | DUSP23 | 1185.006 | -0.4154972 | 0.2793707 | -1.614661 | 0.1063842 | 0.243498 |
| ENSG00000125520 | SLC2A4RG | 7008.423828 | -0.415564368 | 0.205562803 | -2.101947632 | 0.03555786 | 0.106865 |
| ENSG00000183171 | | 388.8507 | -0.4158026 | 0.2276194 | -1.918209 | 0.05508455 | 0.149295 |
| ENSG00000125650 | PSPN | 95.18086921 | -0.415824491 | 0.276253158 | -1.608406697 | 0.10774614 | 0.245598 |
| ENSG00000100372 | SLC25A17 | 1339.622921 | -0.416033134 | 0.206437439 | -2.092334836 | 0.03640857 | 0.108763 |
| ENSG00000182083 | OR6B2 | 0.8974614 | -0.4161046 | 0.9342112 | -1.778075 | 0.07539154 | NA |
| ENSG00000114767 | RRP9 | 1804.206915 | -0.416116258 | 0.207013432 | -2.098653218 | 0.03584748 | 0.107571 |
| ENSG00000258735 | LINC00637 | 3.898327 | -0.4161509 | 0.6697538 | -0.9945144 | 0.3199725 | 0.520852 |
| ENSG00000185666 | SYN3 | 7.87357 | -0.4161597 | 0.637382 | -0.9906369 | 0.3218629 | 0.522633 |
| ENSG00000004864 | SLC25A13 | 1902.417018 | -0.416166241 | 0.216589316 | -2.005313692 | 0.04492952 | 0.127877 |
| ENSG00000267115 | | 1.070983 | -0.4161949 | 0.9033151 | -1.555767 | 0.1197635 | NA |
| ENSG00000161010 | MRNIP | 3772.225 | -0.4162323 | 0.2494706 | -1.723261 | 0.08484135 | 0.206292 |
| ENSG00000165125 | TRPV6 | 163.2802 | -0.4162966 | 0.4784484 | -1.080866 | 0.2797566 | 0.477046 |
| ENSG00000120333 | MRPS14 | 2107.918041 | -0.416326959 | 0.137183054 | -3.059792194 | 0.00221491 | 0.011287 |
| ENSG00000235916 | MRPS18CP7 | 85.573 | -0.4163495 | 0.3090077 | -1.472157 | 0.1409784 | 0.297629 |
| ENSG00000047648 | ARHGAP6 | 238.0761062 | -0.416376136 | 0.5121227 | -1.032248399 | 0.30195577 | 0.501385 |
| ENSG00000244945 | RUFY1-AS1 | 52.13819 | -0.4166243 | 0.2990078 | -1.515161 | 0.1297316 | 0.280322 |
| ENSG00000073614 | KDM5A | 3273.605713 | -0.416762392 | 0.188646157 | -2.282017399 | 0.02248831 | 0.074799 |
| ENSG00000105607 | GCDH | 1041.566687 | -0.416799739 | 0.247152268 | -1.795707204 | 0.0725411 | 0.183522 |
| ENSG00000262732 | | 4.153758 | -0.4169207 | 0.7322652 | -1.03726 | 0.2996148 | 0.498891 |
| ENSG00000163507 | CIP2A | 1433.688 | -0.4171049 | 0.4259996 | -1.143299 | 0.2529144 | 0.445771 |
| ENSG00000111877 | MCM9 | 1139.725304 | -0.417117477 | 0.120247294 | -3.522340768 | 0.00042775 | 0.002743 |
| ENSG00000287083 | | 2.211262 | -0.4171681 | 0.7665325 | -1.082364 | 0.2790909 | 0.476216 |

| | | | | | | | |
|---|---|---|---|---|---|---|---|
| ENSG00000163686 | ABHD6 | 182.0322 | -0.4173838 | 0.2159232 | -2.018324 | 0.04355753 | 0.124936 |
| ENSG00000254208 | | 3.942113 | -0.4174054 | 0.6878779 | -1.004527 | 0.3151247 | 0.516159 |
| ENSG00000120696 | KBTBD7 | 1133.871595 | -0.417516599 | 0.16916238 | -2.534879897 | 0.01124859 | 0.042799 |
| ENSG00000114686 | MRPL3 | 5791.371828 | -0.417719787 | 0.216867319 | -1.998401846 | 0.04567311 | 0.129455 |
| ENSG00000235910 | APOA1-AS | 1.105829 | -0.4178647 | 0.8744081 | -1.354802 | 0.1754807 | NA |
| ENSG00000134248 | LAMTOR5 | 3481.113977 | -0.417922379 | 0.161134301 | -2.657101578 | 0.00788157 | 0.032147 |
| ENSG00000272356 | | 41.03705 | -0.4180039 | 0.3709969 | -1.275372 | 0.2021776 | 0.383307 |
| ENSG00000074211 | PPP2R2C | 3294.326614 | -0.418007499 | 0.137182412 | -3.102398264 | 0.0019196 | 0.01 |
| ENSG00000122692 | SMU1 | 3083.115802 | -0.418029568 | 0.143937576 | -2.945785789 | 0.00322135 | 0.015413 |
| ENSG00000272168 | CASC15 | 20.62673 | -0.4180488 | 0.4721261 | -1.096603 | 0.272815 | 0.469427 |
| ENSG00000104755 | ADAM2 | 3.986170138 | -0.418105267 | 0.821478185 | -1.195422171 | 0.23192213 | 0.420323 |
| ENSG00000259479 | SORD2P | 166.7268 | -0.4181812 | 0.2627537 | -1.71392 | 0.08654347 | 0.209338 |
| ENSG00000162676 | GFI1 | 128.1571 | -0.4182051 | 0.2866492 | -1.580426 | 0.1140094 | 0.255729 |
| ENSG00000261058 | | 1.101081 | -0.4182187 | 0.9057734 | -1.40601 | 0.1597212 | NA |
| ENSG00000225912 | RPL26P36 | 249.7617 | -0.4182512 | 0.2230852 | -1.960049 | 0.04999008 | 0.138454 |
| ENSG00000110756 | HPS5 | 1655.396795 | -0.418270824 | 0.169224994 | -2.535120956 | 0.01124085 | 0.042788 |
| ENSG00000102109 | PCSK1N | 1913.004272 | -0.418346185 | 0.282680317 | -1.605111872 | 0.10846919 | 0.246803 |
| ENSG00000213970 | | 4.503498 | -0.4183487 | 0.7302179 | -1.009982 | 0.312504 | 0.51354 |
| ENSG00000171928 | TVP23B | 2320.377 | -0.4184098 | 0.1922163 | -2.273006 | 0.02302581 | 0.076082 |
| ENSG00000269940 | | 36.91197 | -0.418436 | 0.3040133 | -1.495389 | 0.1348128 | 0.288068 |
| ENSG00000169282 | KCNAB1 | 20.79675 | -0.4185145 | 0.4245203 | -1.161803 | 0.2453156 | 0.437005 |
| ENSG00000260932 | | 5.985822 | -0.4185976 | 0.659606 | -0.9924881 | 0.3209595 | 0.52174 |
| ENSG00000275572 | GRIFIN | 2.88484 | -0.4186154 | 0.7880478 | -1.092148 | 0.2747682 | 0.471557 |
| ENSG00000146828 | SLC12A9 | 4523.032 | -0.4186984 | 0.1507346 | -2.831508 | 0.00463291 | 0.020835 |
| ENSG00000151304 | SRFBP1 | 717.1706 | -0.4187244 | 0.2073534 | -2.099429 | 0.03577913 | 0.107418 |
| ENSG00000132963 | POMP | 5646.733092 | -0.418789359 | 0.202995998 | -2.143910834 | 0.03204003 | 0.098675 |
| ENSG00000183778 | B3GALT5 | 905.6355 | -0.4188794 | 0.5680078 | -1.033657 | 0.3012967 | 0.500572 |
| ENSG00000141497 | ZMYND15 | 297.5746 | -0.4189178 | 0.2942513 | -1.534146 | 0.1249938 | 0.273016 |
| ENSG00000143727 | ACP1 | 6484.04 | -0.4189288 | 0.1297481 | -3.284442 | 0.00102185 | 0.005832 |
| ENSG00000122547 | EEPD1 | 153.1198039 | -0.418984693 | 0.265950235 | -1.681933587 | 0.09258172 | 0.219979 |
| ENSG00000224324 | THAP5P1 | 6.591452 | -0.4191638 | 0.5600797 | -1.021811 | 0.3068702 | 0.507052 |
| ENSG00000248180 | GAPDHP60 | 65.79235 | -0.4192285 | 0.4116771 | -1.195007 | 0.2320844 | 0.420445 |
| ENSG00000104324 | CPQ | 539.4090982 | -0.419311512 | 0.308991965 | -1.482905058 | 0.13809964 | 0.293024 |

| | | | | | | | |
|---|---|---|---|---|---|---|---|
| ENSG00000171045 | TSNARE1 | 1509.789 | -0.4193476 | 0.1954937 | -2.22579 | 0.02602823 | 0.08365 |
| ENSG00000269399 | | 66.73021 | -0.4195264 | 0.5382025 | -1.034196 | 0.3010444 | 0.500247 |
| ENSG00000066027 | PPP2R5A | 3082.055552 | -0.419591313 | 0.110430125 | -3.835634872 | 0.00012524 | 0.000936 |
| ENSG00000184110 | EIF3C | 1452.458 | -0.4197886 | 0.1736244 | -2.495764 | 0.01256863 | 0.046786 |
| ENSG00000205707 | ETFRF1 | 437.7181 | -0.4198001 | 0.1391569 | -3.063646 | 0.00218658 | 0.011164 |
| ENSG00000261326 | LINC01355 | 314.5162 | -0.419954 | 0.4812715 | -1.078814 | 0.2806708 | 0.478145 |
| ENSG00000274001 | | 7.139459 | -0.4200006 | 0.6154185 | -1.000949 | 0.3168513 | 0.517945 |
| ENSG00000259038 | | 48.46492 | -0.4201364 | 0.5967242 | -1.00022 | 0.3172042 | 0.518204 |
| ENSG00000288598 | | 15.8833 | -0.4201385 | 0.5115185 | -1.056296 | 0.2908332 | 0.489462 |
| ENSG00000122705 | CLTA | 6426.319038 | -0.420331194 | 0.146505102 | -2.924986918 | 0.00344471 | 0.0163 |
| ENSG00000106638 | TBL2 | 2354.71607 | -0.420380035 | 0.116208298 | -3.663197599 | 0.00024909 | 0.001719 |
| ENSG00000171729 | TMEM51 | 549.677 | -0.4203866 | 0.2308438 | -1.916788 | 0.05526491 | 0.149623 |
| ENSG00000261188 | | 409.5632 | -0.4204469 | 0.1729641 | -2.47516 | 0.01331764 | 0.04894 |
| ENSG00000132773 | TOE1 | 757.970069 | -0.420741267 | 0.368380538 | -1.277636426 | 0.20137765 | 0.382144 |
| ENSG00000253213 | USP12P1 | 15.30851 | -0.4208882 | 0.5363032 | -1.042305 | 0.2972702 | 0.496397 |
| ENSG00000240541 | TM4SF1-AS1 | 38.06355 | -0.4209867 | 0.5252578 | -1.046031 | 0.2955468 | 0.494669 |
| ENSG00000088930 | XRN2 | 8410.745144 | -0.421037095 | 0.137313611 | -3.1356881 | 0.00171451 | 0.009082 |
| ENSG00000226121 | AHCTF1P1 | 1784.868 | -0.4210607 | 0.3081075 | -1.494006 | 0.1351742 | 0.288473 |
| ENSG00000247774 | PCED1B-AS1 | 3.541067 | -0.4211735 | 0.6274312 | -1.008499 | 0.3132148 | 0.514366 |
| ENSG00000139083 | ETV6 | 626.1301595 | -0.421371513 | 0.230967804 | -1.915901464 | 0.05537764 | 0.149799 |
| ENSG00000067182 | TNFRSF1A | 6294.214581 | -0.421436131 | 0.135345745 | -3.178015278 | 0.00148287 | 0.008027 |
| ENSG00000260394 | LINC02867 | 19.87953 | -0.4214757 | 0.4395859 | -1.14634 | 0.2516546 | 0.444452 |
| ENSG00000234806 | RPL26P29 | 4.790703 | -0.4214771 | 0.7817984 | -1.107831 | 0.267935 | 0.463846 |
| ENSG00000239468 | RN7SL569P | 1.677311 | -0.4215379 | 0.8982107 | -1.543834 | 0.1226286 | 0.269397 |
| ENSG00000143321 | HDGF | 28759.08 | -0.4216336 | 0.1945142 | -2.316378 | 0.02053766 | 0.0694 |
| ENSG00000168734 | PKIG | 801.5673 | -0.4216652 | 0.2513194 | -1.78041 | 0.07500891 | 0.188064 |
| ENSG00000235214 | FAM83C-AS1 | 2.489295 | -0.4218702 | 0.8090682 | -1.156205 | 0.2475974 | 0.439793 |
| ENSG00000189227 | C15orf61 | 416.1726 | -0.4218948 | 0.1860871 | -2.347957 | 0.01887668 | 0.064865 |
| ENSG00000181915 | ADO | 2289.576 | -0.4219131 | 0.117345 | -3.640296 | 0.00027233 | 0.001858 |
| ENSG00000284726 | | 8.025689 | -0.4221262 | 0.7127227 | -1.027543 | 0.3041648 | 0.503762 |
| ENSG00000177169 | ULK1 | 7522.967 | -0.4221346 | 0.2608601 | -1.73563 | 0.08262923 | 0.202395 |
| ENSG00000249173 | LINC01093 | 1.15831 | -0.4224467 | 0.9457719 | -1.929469 | 0.05367263 | NA |
| ENSG00000174886 | NDUFA11 | 730.731 | -0.4224965 | 0.2586116 | -1.734042 | 0.08291065 | 0.202804 |

| | | | | | | | |
|---|---|---|---|---|---|---|---|
| ENSG00000178531 | CTXN1 | 1139.542 | -0.422501 | 0.3049208 | -1.512027 | 0.130527 | 0.281628 |
| ENSG00000228187 | | 63.16147 | -0.4227182 | 0.2901465 | -1.573521 | 0.1155981 | 0.258454 |
| ENSG00000164976 | MYORG | 2113.938 | -0.4227203 | 0.2804795 | -1.652847 | 0.09836209 | 0.229831 |
| ENSG00000213667 | PGGT1BP1 | 18.67758 | -0.4228384 | 0.3978542 | -1.230905 | 0.2183586 | 0.404093 |
| ENSG00000111665 | CDCA3 | 1469.394918 | -0.422882236 | 0.59492697 | -1.020328599 | 0.30757264 | 0.507721 |
| ENSG00000125347 | IRF1 | 1467.95752 | -0.422895225 | 0.283364855 | -1.599785319 | 0.10964622 | 0.248799 |
| ENSG00000099203 | TMED1 | 691.9020278 | -0.422931393 | 0.198334904 | -2.211598611 | 0.02699441 | 0.086162 |
| ENSG00000237522 | NONOP2 | 49.60543 | -0.4230056 | 0.3670862 | -1.304523 | 0.1920554 | 0.370272 |
| ENSG00000198003 | CCDC151 | 47.87815 | -0.4230194 | 0.4543624 | -1.133599 | 0.2569629 | 0.450852 |
| ENSG00000170088 | TMEM192 | 3464.023 | -0.4232278 | 0.212295 | -2.080084 | 0.03751786 | 0.111214 |
| ENSG00000273117 | INSIG1-DT | 298.6345 | -0.423385 | 0.3412455 | -1.385702 | 0.1658379 | 0.334534 |
| ENSG00000155158 | TTC39B | 1054.681 | -0.4234146 | 0.4415774 | -1.164077 | 0.244393 | 0.435907 |
| ENSG00000140451 | PIF1 | 620.4159 | -0.4234895 | 0.4975317 | -1.067675 | 0.285667 | 0.483708 |
| ENSG00000276728 | | 118.9683 | -0.4235286 | 0.3218627 | -1.445771 | 0.1482414 | 0.308597 |
| ENSG00000229184 | ATP5PDP2 | 5.817629 | -0.4236014 | 0.6663753 | -1.003039 | 0.3158422 | 0.516884 |
| ENSG00000234800 | PCMTD1P3 | 12.28278 | -0.4237486 | 0.4845786 | -1.094004 | 0.273953 | 0.470745 |
| ENSG00000204152 | TIMM23B | 211.2666 | -0.4238666 | 0.291937 | -1.574483 | 0.1153757 | 0.258184 |
| ENSG00000088325 | TPX2 | 5701.960781 | -0.423933831 | 0.427726817 | -1.180586632 | 0.23776698 | 0.42736 |
| ENSG00000235138 | | 26.71736 | -0.4239799 | 0.4337381 | -1.161045 | 0.2456236 | 0.437339 |
| ENSG00000114383 | TUSC2 | 1778.325863 | -0.424033755 | 0.216260763 | -2.041862465 | 0.04116518 | 0.119689 |
| ENSG00000169258 | GPRIN1 | 1103.909 | -0.4240522 | 0.3402935 | -1.387286 | 0.1653547 | 0.33394 |
| ENSG00000269743 | SLC25A53 | 189.4569 | -0.4240767 | 0.2397327 | -1.869575 | 0.06154285 | 0.162086 |
| ENSG00000271855 | | 26.96116 | -0.4240768 | 0.4921355 | -1.087799 | 0.2766841 | 0.4737 |
| ENSG00000240452 | MTCO1P29 | 1.226596 | -0.4242053 | 0.8160332 | -1.170676 | 0.2417292 | NA |
| ENSG00000111602 | TIMELESS | 5641.666217 | -0.424460048 | 0.420931143 | -1.175501534 | 0.23979412 | 0.429985 |
| ENSG00000107902 | LHPP | 1048.231919 | -0.424502823 | 0.240902994 | -1.858571784 | 0.06308786 | 0.165043 |
| ENSG00000205302 | SNX2 | 5808.023 | -0.4245074 | 0.150289 | -2.919609 | 0.00350471 | 0.016536 |
| ENSG00000258539 | | 7.573856 | -0.4246142 | 0.5734014 | -1.020554 | 0.3074659 | 0.507632 |
| ENSG00000152382 | TADA1 | 1504.359 | -0.4246185 | 0.182762 | -2.397685 | 0.01649903 | 0.058161 |
| ENSG00000263105 | | 23.19241 | -0.4247844 | 0.4428228 | -1.149909 | 0.2501814 | 0.442695 |
| ENSG00000099795 | NDUFB7 | 2484.80585 | -0.424869446 | 0.246763717 | -1.814745969 | 0.06956296 | 0.177679 |
| ENSG00000107362 | ABHD17B | 726.0569949 | -0.424884722 | 0.181328278 | -2.415316623 | 0.01572154 | 0.05594 |
| ENSG00000012061 | ERCC1 | 2204.934728 | -0.424900273 | 0.186163996 | -2.35477756 | 0.0185338 | 0.06391 |

| | | | | | | | |
|---|---|---|---|---|---|---|---|
| ENSG00000252534 | | 4.308608 | -0.4249157 | 0.6833203 | -1.005712 | 0.314554 | 0.515665 |
| ENSG00000117625 | RCOR3 | 2185.631022 | -0.425023861 | 0.167803945 | -2.60301326 | 0.00924084 | 0.036501 |
| ENSG00000167840 | ZNF232 | 602.7438 | -0.4250882 | 0.1809874 | -2.436362 | 0.01483582 | 0.053474 |
| ENSG00000130508 | PXDN | 9236.744539 | -0.425096405 | 0.171335007 | -2.547706884 | 0.01084335 | 0.04153 |
| ENSG00000287362 | | 4.613586 | -0.4251064 | 0.6892393 | -1.004 | 0.3153784 | 0.516316 |
| ENSG00000244754 | N4BP2L2 | 7299.241 | -0.4255929 | 0.2901231 | -1.585724 | 0.1128018 | 0.253728 |
| ENSG00000133111 | RFXAP | 596.5528583 | -0.425623829 | 0.20822011 | -2.125131491 | 0.03357564 | 0.102344 |
| ENSG00000187186 | | 6.644354 | -0.4257158 | 0.6436254 | -1.015929 | 0.3096633 | 0.510292 |
| ENSG00000160606 | TLCD1 | 1045.125 | -0.4257959 | 0.2536444 | -1.774938 | 0.0759081 | 0.189721 |
| ENSG00000213104 | NPM1P46 | 72.8997 | -0.4258338 | 0.240903 | -1.865601 | 0.06209726 | 0.163078 |
| ENSG00000125898 | FAM110A | 983.712931 | -0.425969687 | 0.196452846 | -2.248835598 | 0.02452296 | 0.079875 |
| ENSG00000152242 | C18orf25 | 2069.382 | -0.4261498 | 0.1560589 | -2.804821 | 0.00503445 | 0.022282 |
| ENSG00000126767 | ELK1 | 2960.047468 | -0.426208641 | 0.156036445 | -2.788491465 | 0.00529541 | 0.023216 |
| ENSG00000274370 | | 143.9701 | -0.4263907 | 0.4320029 | -1.181483 | 0.2374107 | 0.427037 |
| ENSG00000206992 | RNU6-574P | 2.053989 | -0.4264034 | 0.7701173 | -1.093473 | 0.2741864 | 0.470902 |
| ENSG00000143228 | NUF2 | 1297.358 | -0.4264171 | 0.449906 | -1.150668 | 0.2498689 | 0.442437 |
| ENSG00000286537 | | 2.412466 | -0.4267877 | 0.7747987 | -1.086967 | 0.2770515 | 0.473987 |
| ENSG00000066583 | ISOC1 | 6187.319384 | -0.42694221 | 0.241042795 | -1.869985213 | 0.06148587 | 0.161985 |
| ENSG00000142102 | PGGHG | 2478.066 | -0.4271082 | 0.2746115 | -1.670521 | 0.09481626 | 0.22392 |
| ENSG00000105982 | RNF32 | 126.618404 | -0.427458382 | 0.213280947 | -2.085824494 | 0.03699452 | 0.110056 |
| ENSG00000245384 | CXXC4-AS1 | 4.536189 | -0.4278956 | 0.6747333 | -1.018763 | 0.3083154 | 0.508608 |
| ENSG00000087087 | SRRT | 8607.004816 | -0.42801639 | 0.214085633 | -2.096544322 | 0.03603393 | 0.108045 |
| ENSG00000147162 | OGT | 29498.23 | -0.4282037 | 0.3954553 | -1.257152 | 0.2086985 | 0.391617 |
| ENSG00000139618 | BRCA2 | 1218.735 | -0.4283775 | 0.5649029 | -1.055224 | 0.291323 | 0.490008 |
| ENSG00000258858 | | 1.382351 | -0.4284442 | 0.9485773 | -1.887177 | 0.05913649 | NA |
| ENSG00000152484 | USP12 | 3222.977 | -0.4285364 | 0.1707093 | -2.575262 | 0.01001641 | 0.038907 |
| ENSG00000114999 | TTL | 2450.382048 | -0.428641632 | 0.178749858 | -2.470330912 | 0.01349881 | 0.049478 |
| ENSG00000165280 | VCP | 21466.84 | -0.4287155 | 0.1808224 | -2.448908 | 0.01432901 | 0.052009 |
| ENSG00000258153 | HSPE1P4 | 177.3605 | -0.4287263 | 0.2917292 | -1.593903 | 0.1109578 | 0.250854 |
| ENSG00000260989 | | 5.701386 | -0.4287875 | 0.7287052 | -1.049953 | 0.2937398 | 0.492731 |
| ENSG00000101452 | DHX35 | 797.0216655 | -0.428866875 | 0.166591478 | -2.642395156 | 0.0082322 | 0.033245 |
| ENSG00000230702 | RPL31P30 | 1.01644 | -0.4289117 | 0.8902582 | -1.445005 | 0.1484565 | NA |
| ENSG00000238061 | | 2.232384 | -0.4289585 | 0.7868491 | -1.11005 | 0.2669777 | 0.46278 |

| | | | | | | | |
|---|---|---|---|---|---|---|---|
| ENSG00000279744 | | 51.51015 | -0.429023 | 0.3519815 | -1.368906 | 0.1710285 | 0.341692 |
| ENSG00000146540 | C7orf50 | 9126.266 | -0.4290874 | 0.1949892 | -2.278137 | 0.0227184 | 0.075328 |
| ENSG00000151665 | PIGF | 625.2597 | -0.4291728 | 0.1192533 | -3.654294 | 0.00025789 | 0.001773 |
| ENSG00000159086 | PAXBP1 | 3479.249 | -0.4292434 | 0.1477398 | -2.963182 | 0.00304477 | 0.014722 |
| ENSG00000251131 | | 41.54386 | -0.4292818 | 0.3206945 | -1.470683 | 0.141377 | 0.298296 |
| ENSG00000223773 | CD99P1 | 162.3838 | -0.4293373 | 0.2606078 | -1.751992 | 0.07977518 | 0.196801 |
| ENSG00000119820 | YIPF4 | 3135.279393 | -0.429341916 | 0.202013719 | -2.210421223 | 0.02707594 | 0.086381 |
| ENSG00000250221 | KRT8P32 | 140.4775 | -0.4293554 | 0.2837207 | -1.634966 | 0.1020561 | 0.236111 |
| ENSG00000054148 | PHPT1 | 4472.033035 | -0.429365117 | 0.161114995 | -2.718382873 | 0.00656019 | 0.027631 |
| ENSG00000230992 | FAM201B | 12.76915 | -0.4294382 | 0.4595867 | -1.144426 | 0.252447 | 0.445222 |
| ENSG00000184164 | CRELD2 | 2072.394 | -0.4294631 | 0.2657552 | -1.724705 | 0.08458073 | 0.205846 |
| ENSG00000182287 | AP1S2 | 562.9679 | -0.429486 | 0.1435498 | -3.050216 | 0.00228677 | 0.011579 |
| ENSG00000163071 | SPATA18 | 732.8248 | -0.4295661 | 0.40634 | -1.228475 | 0.2192688 | 0.405418 |
| ENSG00000261320 | | 2.043812 | -0.4299399 | 0.9297347 | -1.612699 | 0.1068099 | 0.244136 |
| ENSG00000232334 | | 3.144662 | -0.4300355 | 0.7833826 | -1.108424 | 0.2676789 | 0.463562 |
| ENSG00000099341 | PSMD8 | 10056.60288 | -0.430183444 | 0.186190436 | -2.407974301 | 0.01604131 | 0.056857 |
| ENSG00000261167 | | 111.1652 | -0.4302508 | 0.3188012 | -1.482423 | 0.1382277 | 0.293173 |
| ENSG00000272009 | | 12.30753 | -0.430319 | 0.5169285 | -1.061631 | 0.2884035 | 0.486902 |
| ENSG00000179242 | CDH4 | 91.90285 | -0.4303279 | 0.3750793 | -1.306319 | 0.191444 | 0.369603 |
| ENSG00000103512 | NOMO1 | 4436.455596 | -0.430369034 | 0.148339651 | -2.958734166 | 0.00308905 | 0.014887 |
| ENSG00000229593 | SUCLA2P3 | 4.806158 | -0.4304432 | 0.7002066 | -1.03632 | 0.3000531 | 0.499366 |
| ENSG00000213943 | KRT18P17 | 572.6967 | -0.4305446 | 0.2094358 | -2.143192 | 0.03209769 | 0.098822 |
| ENSG00000272145 | NFYC-AS1 | 175.9566 | -0.430596 | 0.2321293 | -1.945018 | 0.05177286 | 0.142279 |
| ENSG00000206147 | RPL23AP57 | 0.8700355 | -0.4306505 | 0.9306875 | -1.609944 | 0.1074102 | NA |
| ENSG00000136891 | TEX10 | 2507.824365 | -0.430917799 | 0.126907198 | -3.440666586 | 0.00058028 | 0.003572 |
| ENSG00000238116 | RAD21P1 | 6.723328 | -0.4311048 | 0.6638193 | -1.029215 | 0.3033786 | 0.503056 |
| ENSG00000148688 | RPP30 | 1881.582 | -0.4312314 | 0.2883988 | -1.614159 | 0.1064928 | 0.243662 |
| ENSG00000173698 | ADGRG2 | 52.19011 | -0.4312424 | 0.4241585 | -1.199915 | 0.2301722 | 0.418465 |
| ENSG00000275963 | | 1.874287 | -0.4315039 | 0.7762818 | -1.094723 | 0.2736378 | 0.470415 |
| ENSG00000173295 | FAM86B3P | 344.6387 | -0.4315236 | 0.2654121 | -1.73682 | 0.08241889 | 0.201917 |
| ENSG00000145681 | HAPLN1 | 56.86359 | -0.431745 | 0.8570828 | -1.300157 | 0.1935471 | 0.372157 |
| ENSG00000138326 | RPS24 | 29293.5777 | -0.431831527 | 0.156931091 | -2.821335079 | 0.00478242 | 0.021388 |
| ENSG00000251728 | | 2.281838 | -0.4318335 | 0.7989686 | -1.126349 | 0.260018 | 0.454463 |

| | | | | | | | |
|---|---|---|---|---|---|---|---|
| ENSG00000238363 | SNORA13 | 32.92282 | -0.432216 | 0.414372 | -1.214587 | 0.2245236 | 0.411433 |
| ENSG00000000460 | C1orf112 | 612.0831085 | -0.432324422 | 0.389653805 | -1.275619425 | 0.20209008 | 0.383223 |
| ENSG00000060237 | WNK1 | 8221.860534 | -0.432512802 | 0.202763918 | -2.21842781 | 0.02652568 | 0.084993 |
| ENSG00000127022 | CANX | 52082.32819 | -0.432549818 | 0.151839 | -2.986059114 | 0.00282598 | 0.013889 |
| ENSG00000251271 | ALG1L7P | 5.601547 | -0.4325718 | 0.5615287 | -1.041674 | 0.2975629 | 0.49673 |
| ENSG00000149927 | DOC2A | 1404.363 | -0.4326317 | 0.2164805 | -2.118123 | 0.03416464 | 0.103728 |
| ENSG00000130695 | CEP85 | 1421.549765 | -0.432767607 | 0.233235957 | -1.970838843 | 0.04874231 | 0.135848 |
| ENSG00000150753 | CCT5 | 25996.01 | -0.4328372 | 0.223943 | -2.024152 | 0.04295448 | 0.123637 |
| ENSG00000052749 | RRP12 | 5182.687451 | -0.433143316 | 0.130409321 | -3.405499625 | 0.00066043 | 0.003996 |
| ENSG00000124279 | FASTKD3 | 1295.854439 | -0.433161467 | 0.183819952 | -2.43320734 | 0.01496573 | 0.053835 |
| ENSG00000288018 | | 1.650741 | -0.4331886 | 0.865607 | -1.314536 | 0.1886658 | 0.365861 |
| ENSG00000078142 | PIK3C3 | 1415.339311 | -0.43341586 | 0.19988472 | -2.249894083 | 0.02445567 | 0.079725 |
| ENSG00000118733 | OLFM3 | 1.232313424 | -0.433429477 | 0.947438514 | -1.816957408 | 0.06922364 | NA |
| ENSG00000234825 | XRCC6P2 | 804.6365 | -0.4335113 | 0.2820263 | -1.653582 | 0.09821255 | 0.229702 |
| ENSG00000287600 | | 6.134559 | -0.4336947 | 0.6928913 | -1.048466 | 0.2944238 | 0.493443 |
| ENSG00000225768 | LINC02620 | 2.617438 | -0.4337444 | 0.8425167 | -1.246014 | 0.2127592 | 0.397483 |
| ENSG00000235558 | GUSBP17 | 10.63953 | -0.4337701 | 0.4747534 | -1.130213 | 0.2583866 | 0.452603 |
| ENSG00000262402 | MCUR1P1 | 1.938759 | -0.4337955 | 0.8329538 | -1.213636 | 0.2248867 | 0.411928 |
| ENSG00000174516 | PELI3 | 940.0611 | -0.4338283 | 0.1735886 | -2.563717 | 0.01035579 | 0.040032 |
| ENSG00000232894 | MRPS31P2 | 2.967654 | -0.433848 | 0.8013543 | -1.12179 | 0.2619517 | 0.456718 |
| ENSG00000038532 | CLEC16A | 2696.785295 | -0.433870861 | 0.164429029 | -2.681213828 | 0.00733556 | 0.030304 |
| ENSG00000204019 | CT83 | 873.0191 | -0.4339555 | 0.2100599 | -2.156404 | 0.03105212 | 0.096359 |
| ENSG00000137656 | BUD13 | 813.3815598 | -0.434063917 | 0.231365164 | -1.975067679 | 0.04826047 | 0.13483 |
| ENSG00000168356 | SCN11A | 2.83789 | -0.4340925 | 0.7120035 | -1.051873 | 0.2928579 | 0.491754 |
| ENSG00000273082 | LARGE1-AS1 | 1.374511 | -0.4341594 | 0.8837492 | -1.419487 | 0.1557571 | NA |
| ENSG00000271851 | | 44.50464 | -0.4341762 | 0.4053902 | -1.242824 | 0.2139326 | 0.398736 |
| ENSG00000173714 | WFIKKN2 | 1.037271 | -0.4342094 | 0.8732192 | -1.358534 | 0.1742944 | NA |
| ENSG00000198954 | KIFBP | 2567.1 | -0.4342316 | 0.1863659 | -2.406905 | 0.01608837 | 0.056982 |
| ENSG00000235919 | ASH1L-AS1 | 306.3857 | -0.4342858 | 0.2108623 | -2.144559 | 0.03198812 | 0.098584 |
| ENSG00000142920 | AZIN2 | 116.8881 | -0.4344884 | 0.3669033 | -1.339912 | 0.180274 | 0.354583 |
| ENSG00000205622 | | 9.996267 | -0.4345212 | 0.5556125 | -1.057096 | 0.2904679 | 0.489096 |
| ENSG00000143333 | RGS16 | 995.2267 | -0.4345707 | 0.4518943 | -1.173737 | 0.2405004 | 0.430803 |
| ENSG00000198919 | DZIP3 | 2261.864 | -0.4346302 | 0.2139901 | -2.120421 | 0.03397052 | 0.103291 |

| | | | | | | | |
|---|---|---|---|---|---|---|---|
| ENSG00000168237 | GLYCTK | 184.6186 | -0.4349159 | 0.4490602 | -1.173546 | 0.240577 | 0.430895 |
| ENSG00000178115 | GOLGA8Q | 2.798145 | -0.4349538 | 0.730614 | -1.060521 | 0.2889077 | 0.487488 |
| ENSG00000261799 | | 175.9829 | -0.4352229 | 0.3021194 | -1.565848 | 0.1173842 | 0.261029 |
| ENSG00000217325 | PRELID1P1 | 85.95007 | -0.4352272 | 0.3141247 | -1.519672 | 0.1285936 | 0.278543 |
| ENSG00000162066 | AMDHD2 | 1754.212 | -0.4352275 | 0.2147231 | -2.112644 | 0.03463123 | 0.104855 |
| ENSG00000258813 | | 26.33337 | -0.4352958 | 0.3806652 | -1.307875 | 0.1909156 | 0.368931 |
| ENSG00000213619 | NDUFS3 | 5051.333 | -0.4353865 | 0.2674777 | -1.737756 | 0.08225376 | 0.201624 |
| ENSG00000114982 | KANSL3 | 3794.007472 | -0.435480194 | 0.140803299 | -3.13546675 | 0.00171581 | 0.009087 |
| ENSG00000133216 | EPHB2 | 651.696826 | -0.435583455 | 0.284381951 | -1.640608825 | 0.10087864 | 0.234308 |
| ENSG00000128536 | CDHR3 | 96.46431811 | -0.435590266 | 0.268452825 | -1.727415472 | 0.08409307 | 0.204904 |
| ENSG00000170037 | CNTROB | 2670.332 | -0.4356217 | 0.1566163 | -2.847597 | 0.00440506 | 0.019959 |
| ENSG00000156256 | USP16 | 2457.044 | -0.4356281 | 0.1484324 | -2.998584 | 0.00271238 | 0.013396 |
| ENSG00000273474 | | 2.488543 | -0.4356424 | 0.8313742 | -1.193019 | 0.2328618 | 0.421652 |
| ENSG00000197165 | SULT1A2 | 24.1888 | -0.435845 | 0.520774 | -1.093859 | 0.2740166 | 0.470756 |
| ENSG00000168781 | PPIP5K1 | 1341.583 | -0.4358859 | 0.2041917 | -2.216349 | 0.0266676 | 0.085334 |
| ENSG00000197106 | SLC6A17 | 2.869843 | -0.4359049 | 0.8245074 | -1.179341 | 0.2382622 | 0.428074 |
| ENSG00000274080 | | 7.956727 | -0.4359072 | 0.6246413 | -1.02955 | 0.3032211 | 0.502858 |
| ENSG00000275700 | AATF | 2990.867 | -0.4360069 | 0.1318333 | -3.35964 | 0.00078044 | 0.004619 |
| ENSG00000240184 | PCDHGC3 | 1101.865 | -0.4360846 | 0.2259787 | -2.022634 | 0.04311087 | 0.123989 |
| ENSG00000130560 | UBAC1 | 3518.202924 | -0.436162205 | 0.188832157 | -2.393072506 | 0.01670794 | 0.058789 |
| ENSG00000286281 | | 1.46844 | -0.4361674 | 0.8405373 | -1.238708 | 0.2154536 | NA |
| ENSG00000103091 | WDR59 | 7229.497952 | -0.436241959 | 0.108661306 | -4.057085355 | 4.97E-05 | 0.000412 |
| ENSG00000164975 | SNAPC3 | 1253.303 | -0.436341 | 0.2886372 | -1.630652 | 0.1029638 | 0.237696 |
| ENSG00000128886 | ELL3 | 87.61607889 | -0.436448738 | 0.313649471 | -1.523444525 | 0.12764753 | 0.277037 |
| ENSG00000110076 | NRXN2 | 2.36873184 | -0.436515315 | 0.802821204 | -1.154846917 | 0.24815312 | 0.440426 |
| ENSG00000128590 | DNAJB9 | 1565.814449 | -0.436860691 | 0.265382201 | -1.752052764 | 0.07976474 | 0.196801 |
| ENSG00000057657 | PRDM1 | 4.34217517 | -0.436887793 | 0.84582678 | -1.267005321 | 0.20515339 | 0.387175 |
| ENSG00000100353 | EIF3D | 9687.618679 | -0.436933852 | 0.159684443 | -2.812167766 | 0.00492088 | 0.021867 |
| ENSG00000117519 | CNN3 | 2.666724852 | -0.436988666 | 0.762648963 | -1.097310706 | 0.27250559 | 0.468986 |
| ENSG00000254951 | | 2.2655 | -0.4371654 | 0.8342344 | -1.201194 | 0.2296759 | 0.417877 |
| ENSG00000128513 | POT1 | 1077.06964 | -0.437190238 | 0.155877943 | -2.875859912 | 0.00402929 | 0.018563 |
| ENSG00000103356 | EARS2 | 3129.441123 | -0.437376632 | 0.170116525 | -2.644867394 | 0.00817229 | 0.033074 |
| ENSG00000229820 | | 16.79521 | -0.4374922 | 0.4450966 | -1.185716 | 0.2357343 | 0.42515 |

| | | | | | | | |
|---|---|---|---|---|---|---|---|
| ENSG00000196628 | TCF4 | 21.3331 | -0.437539 | 0.7266997 | -1.031304 | 0.3023981 | 0.501837 |
| ENSG00000170291 | ELP5 | 1674.678 | -0.4376218 | 0.1870679 | -2.419015 | 0.01556261 | 0.055516 |
| ENSG00000242272 | AK2P2 | 24.76511 | -0.4377243 | 0.35615 | -1.381813 | 0.1670292 | 0.336324 |
| ENSG00000212452 | SNORD69 | 15.55997 | -0.4379562 | 0.4502063 | -1.173503 | 0.2405941 | 0.430896 |
| ENSG00000160124 | CCDC58 | 827.6881 | -0.437958 | 0.1941935 | -2.334725 | 0.0195578 | 0.066782 |
| ENSG00000172409 | CLP1 | 639.2491 | -0.4380178 | 0.1609069 | -2.789269 | 0.00528271 | 0.023177 |
| ENSG00000219133 | | 4.883303 | -0.4380247 | 0.6784091 | -1.034666 | 0.300825 | 0.500039 |
| ENSG00000262703 | | 33.06911 | -0.4380426 | 0.4944358 | -1.122512 | 0.2616448 | 0.456324 |
| ENSG00000198105 | ZNF248 | 1609.124 | -0.4380974 | 0.2545632 | -1.798348 | 0.07212181 | 0.182613 |
| ENSG00000103264 | FBXO31 | 2769.2263 | -0.438114105 | 0.129485113 | -3.436829149 | 0.00058857 | 0.003615 |
| ENSG00000286150 | | 76.24937 | -0.4381463 | 0.2766209 | -1.695788 | 0.08992599 | 0.215538 |
| ENSG00000255624 | C10orf88B | 137.3316 | -0.4381588 | 0.2324075 | -1.986656 | 0.04696052 | 0.132136 |
| ENSG00000131504 | DIAPH1 | 12575.02738 | -0.4381853 | 0.220079808 | -2.068259017 | 0.03861567 | 0.113745 |
| ENSG00000101079 | NDRG3 | 2016.239972 | -0.43828408 | 0.154843026 | -2.895696403 | 0.00378318 | 0.017606 |
| ENSG00000242951 | | 404.371 | -0.4382985 | 0.2589208 | -1.809231 | 0.0704151 | 0.179433 |
| ENSG00000224856 | RPL35AP36 | 11.78116 | -0.4384159 | 0.5677048 | -1.047413 | 0.2949091 | 0.494038 |
| ENSG00000188352 | FOCAD | 2541.022 | -0.4384292 | 0.1540173 | -2.911205 | 0.00360038 | 0.01691 |
| ENSG00000171403 | KRT9 | 3.636173 | -0.438447 | 0.9329942 | -1.756413 | 0.07901794 | 0.195368 |
| ENSG00000127993 | RBM48 | 818.0542998 | -0.438693881 | 0.197602028 | -2.297863694 | 0.02156955 | 0.072304 |
| ENSG00000108654 | DDX5 | 86329.18364 | -0.438846042 | 0.131241577 | -3.397231237 | 0.00068071 | 0.004104 |
| ENSG00000105968 | H2AZ2 | 9512.671227 | -0.438858648 | 0.198743808 | -2.292013844 | 0.02190484 | 0.073234 |
| ENSG00000272626 | | 1.154354 | -0.4389028 | 0.9546038 | -1.865988 | 0.06204303 | NA |
| ENSG00000241112 | RPL29P14 | 3.67126 | -0.4389937 | 0.8141378 | -1.185278 | 0.2359074 | 0.425283 |
| ENSG00000161277 | THAP8 | 587.6334 | -0.4392291 | 0.2881739 | -1.645343 | 0.09989901 | 0.232724 |
| ENSG00000186918 | ZNF395 | 1303.227 | -0.4392767 | 0.1203982 | -3.704781 | 0.00021157 | 0.001491 |
| ENSG00000213755 | | 1.135164 | -0.4393282 | 0.9642231 | -2.046714 | 0.0406862 | NA |
| ENSG00000233122 | CTAGE7P | 88.29675 | -0.4393537 | 0.4007058 | -1.271127 | 0.2036836 | 0.385321 |
| ENSG00000113272 | THG1L | 1276.625913 | -0.439355225 | 0.177536541 | -2.545168518 | 0.0109225 | 0.041779 |
| ENSG00000254111 | | 1.741178 | -0.4393629 | 0.8662519 | -1.351813 | 0.176435 | 0.349192 |
| ENSG00000067836 | ROGDI | 1231.98091 | -0.43945539 | 0.189014088 | -2.403878708 | 0.01622216 | 0.057351 |
| ENSG00000185513 | L3MBTL1 | 227.6975 | -0.4396224 | 0.3547846 | -1.397633 | 0.1622233 | 0.329478 |
| ENSG00000143479 | DYRK3 | 278.4208 | -0.4396835 | 0.3708188 | -1.350564 | 0.1768351 | 0.349865 |
| ENSG00000226359 | ACTG1P24 | 6.497758 | -0.4397531 | 0.6736822 | -1.04095 | 0.2978987 | 0.497084 |

| | | | | | | | |
|---|---|---|---|---|---|---|---|
| ENSG00000109762 | SNX25 | 1371.300173 | -0.439760415 | 0.197325073 | -2.311699223 | 0.02079426 | 0.070139 |
| ENSG00000175387 | SMAD2 | 3910.306 | -0.4400099 | 0.103713 | -4.274683 | 1.91E-05 | 0.000176 |
| ENSG00000154743 | TSEN2 | 1774.179 | -0.4402321 | 0.1673051 | -2.697821 | 0.0069795 | 0.029048 |
| ENSG00000286854 | | 1.187651 | -0.4402721 | 0.8333415 | -1.232477 | 0.2177708 | NA |
| ENSG00000183520 | UTP11 | 2323.45 | -0.4403539 | 0.1456944 | -3.096416 | 0.00195876 | 0.010174 |
| ENSG00000266967 | AARSD1 | 117.1812 | -0.4403668 | 0.2191699 | -2.104328 | 0.03534988 | 0.106385 |
| ENSG00000221817 | PPP3CB-AS1 | 175.2275 | -0.4404616 | 0.2955328 | -1.607295 | 0.1079897 | 0.246017 |
| ENSG00000137700 | SLC37A4 | 3254.378831 | -0.440670997 | 0.172168823 | -2.631917248 | 0.00849045 | 0.034112 |
| ENSG00000230426 | LINC01036 | 1.988052 | -0.4407089 | 0.8740659 | -1.388916 | 0.1648584 | 0.333156 |
| ENSG00000162407 | PLPP3 | 894.1382 | -0.440736 | 0.2322918 | -1.990393 | 0.0465477 | 0.131287 |
| ENSG00000132305 | IMMT | 7211.608274 | -0.440737621 | 0.213871343 | -2.149159962 | 0.03162172 | 0.09775 |
| ENSG00000277142 | LINC00235 | 279.4314 | -0.4408286 | 0.2458268 | -1.900619 | 0.0573519 | 0.153704 |
| ENSG00000285632 | | 10.25423 | -0.4408542 | 0.5638789 | -1.064773 | 0.2869785 | 0.485343 |
| ENSG00000167131 | CCDC103 | 2.087203 | -0.4408675 | 0.7612121 | -1.105214 | 0.2690668 | 0.465237 |
| ENSG00000258768 | | 45.41113 | -0.440941 | 0.4631136 | -1.166903 | 0.2432495 | 0.434364 |
| ENSG00000131143 | COX4I1 | 15826.15376 | -0.441021746 | 0.175442901 | -2.588019613 | 0.00965295 | 0.037755 |
| ENSG00000253784 | | 2.31735 | -0.4410304 | 0.9333971 | -1.582134 | 0.1136191 | 0.254962 |
| ENSG00000162980 | ARL5A | 1012.742 | -0.4411047 | 0.2246636 | -2.076464 | 0.03785102 | 0.111989 |
| ENSG00000255517 | | 224.4946 | -0.4411136 | 0.2218117 | -2.079985 | 0.03752686 | 0.111228 |
| ENSG00000239472 | RN7SL221P | 2.276884 | -0.441275 | 0.7509437 | -1.096072 | 0.2730474 | 0.469678 |
| ENSG00000250651 | PABPC1P7 | 2.048012 | -0.4413057 | 0.7749307 | -1.123118 | 0.2613873 | 0.456025 |
| ENSG00000135372 | NAT10 | 5847.458549 | -0.441371926 | 0.133324784 | -3.383217923 | 0.00071642 | 0.004289 |
| ENSG00000272578 | | 71.5258 | -0.4413828 | 0.29982 | -1.597572 | 0.1101383 | 0.249621 |
| ENSG00000124713 | GNMT | 35.85968562 | -0.441390633 | 0.351008654 | -1.410908895 | 0.15827148 | 0.323719 |
| ENSG00000143374 | TARS2 | 4938.407 | -0.4414118 | 0.1962641 | -2.329687 | 0.01982271 | 0.067478 |
| ENSG00000255222 | SETP17 | 1.450679 | -0.4414559 | 0.8980277 | -1.465878 | 0.1426816 | NA |
| ENSG00000165209 | STRBP | 4020.041 | -0.4415922 | 0.1459626 | -3.088635 | 0.00201079 | 0.010406 |
| ENSG00000186723 | OR10H1 | 1.381105 | -0.4416135 | 0.8890731 | -1.446847 | 0.1479398 | NA |
| ENSG00000188501 | LCTL | 18.36723 | -0.4416341 | 0.4353881 | -1.211753 | 0.225607 | 0.412911 |
| ENSG00000183760 | ACP7 | 59.98315 | -0.4416715 | 0.3944821 | -1.297087 | 0.1946012 | 0.373769 |
| ENSG00000270580 | | 54.76232 | -0.4418517 | 0.3269848 | -1.487512 | 0.1368796 | 0.291166 |
| ENSG00000215347 | SLC25A5P1 | 9.21223 | -0.4420426 | 0.6459678 | -1.035824 | 0.3002841 | 0.499515 |
| ENSG00000248854 | HNRNPH1P3 | 11.32759 | -0.4422046 | 0.5995657 | -1.050228 | 0.2936132 | 0.492592 |

| | | | | | | | |
|---|---|---|---|---|---|---|---|
| ENSG00000111581 | NUP107 | 4288.210256 | -0.442204693 | 0.273112106 | -1.731972609 | 0.08327844 | 0.203404 |
| ENSG00000267251 | | 7.403724 | -0.4422473 | 0.5637668 | -1.069012 | 0.2850642 | 0.483089 |
| ENSG00000251023 | | 4.36074 | -0.4425134 | 0.6746726 | -1.034708 | 0.3008051 | 0.500037 |
| ENSG00000233622 | CYP2T1P | 471.6635 | -0.4425535 | 0.4849598 | -1.147877 | 0.2510194 | 0.443764 |
| ENSG00000250659 | | 3.306634 | -0.4427101 | 0.6444243 | -1.046446 | 0.2953553 | 0.494536 |
| ENSG00000142961 | MOB3C | 882.2226 | -0.4427534 | 0.1771445 | -2.566426 | 0.01027526 | 0.03975 |
| ENSG00000147003 | CLTRN | 16.29029 | -0.4430326 | 0.4822483 | -1.141523 | 0.2536525 | 0.446667 |
| ENSG00000196368 | NUDT11 | 275.309 | -0.4432082 | 0.2768435 | -1.718488 | 0.08570766 | 0.207904 |
| ENSG00000134146 | DPH6 | 358.9725689 | -0.443295455 | 0.218513763 | -2.120566674 | 0.03395829 | 0.103278 |
| ENSG00000198715 | GLMP | 1663.318 | -0.4434068 | 0.2017928 | -2.278703 | 0.02268474 | 0.075264 |
| ENSG00000204308 | RNF5 | 2185.268 | -0.4436959 | 0.1945505 | -2.362066 | 0.01817339 | 0.062953 |
| ENSG00000101230 | ISM1 | 34.17843263 | -0.443714616 | 0.452280611 | -1.188992441 | 0.23444264 | 0.423591 |
| ENSG00000275052 | PPP4R3B | 6806.935 | -0.4438374 | 0.1507977 | -3.00164 | 0.00268529 | 0.013275 |
| ENSG00000175054 | ATR | 4075.704 | -0.4440587 | 0.2011472 | -2.296191 | 0.02166498 | 0.072569 |
| ENSG00000176208 | ATAD5 | 530.0923 | -0.4441886 | 0.4276152 | -1.243048 | 0.21385 | 0.398722 |
| ENSG00000100226 | GTPBP1 | 2112.686672 | -0.444362412 | 0.160019057 | -2.865857349 | 0.00415882 | 0.019071 |
| ENSG00000177447 | CBX3P1 | 24.39601 | -0.4444572 | 0.3864519 | -1.324462 | 0.1853496 | 0.360985 |
| ENSG00000277954 | | 4.309959 | -0.444531 | 0.7539992 | -1.082045 | 0.2792324 | 0.476427 |
| ENSG00000101574 | METTL4 | 688.2781853 | -0.444546587 | 0.182584737 | -2.507933346 | 0.01214396 | 0.045425 |
| ENSG00000233830 | EIF4HP1 | 1857.721 | -0.444658 | 0.1569523 | -2.908076 | 0.0036366 | 0.017056 |
| ENSG00000115386 | REG1A | 2.450375251 | -0.44480511 | 0.87352949 | -1.349733843 | 0.17710137 | 0.350287 |
| ENSG00000172572 | PDE3A | 5.487345 | -0.4448196 | 0.8458821 | -1.263617 | 0.2063675 | 0.388588 |
| ENSG00000213938 | SEPHS1P6 | 34.00036 | -0.4451447 | 0.331581 | -1.484914 | 0.1375666 | 0.292149 |
| ENSG00000104728 | ARHGEF10 | 77.04876807 | -0.445264896 | 0.445291068 | -1.204312759 | 0.22846872 | 0.416546 |
| ENSG00000271778 | | 6.290596 | -0.44528 | 0.6989503 | -1.057504 | 0.2902814 | 0.488844 |
| ENSG00000141627 | DYM | 2889.492 | -0.4452926 | 0.1530823 | -2.972095 | 0.00295775 | 0.014422 |
| ENSG00000122515 | ZMIZ2 | 9590.758885 | -0.445365596 | 0.164728348 | -2.799511007 | 0.00511801 | 0.022581 |
| ENSG00000250284 | | 10.41151 | -0.4455719 | 0.7076457 | -1.048805 | 0.2942679 | 0.493368 |
| ENSG00000097096 | SYDE2 | 1261.204151 | -0.445608618 | 0.288818228 | -1.665465622 | 0.0958199 | 0.225495 |
| ENSG00000156374 | PCGF6 | 663.6968 | -0.4456971 | 0.2637736 | -1.815818 | 0.06939822 | 0.177368 |
| ENSG00000223695 | | 1.839494 | -0.4457162 | 0.8482682 | -1.243338 | 0.2137432 | 0.398606 |
| ENSG00000224093 | BCAR3-AS1 | 4.964232 | -0.4458684 | 0.7001139 | -1.059517 | 0.2893642 | 0.487918 |
| ENSG00000198301 | SDAD1 | 3513.079 | -0.4458734 | 0.1427834 | -3.181445 | 0.00146542 | 0.007942 |

| | | | | | | | |
|---|---|---|---|---|---|---|---|
| ENSG00000223760 | MED15P9 | 1.896207 | -0.4459735 | 0.8563568 | -1.29295 | 0.1960283 | 0.375768 |
| ENSG00000122008 | POLK | 683.9933988 | -0.44602512 | 0.265680587 | -1.793330181 | 0.07292014 | 0.184253 |
| ENSG00000177728 | TMEM94 | 10176.26 | -0.4460502 | 0.1713837 | -2.706186 | 0.0068061 | 0.028464 |
| ENSG00000152939 | MARVELD2 | 2211.966 | -0.4460578 | 0.1781108 | -2.57759 | 0.00994921 | 0.038697 |
| ENSG00000269176 | | 44.98732 | -0.4461316 | 0.3678296 | -1.36917 | 0.1709461 | 0.341599 |
| ENSG00000100206 | DMC1 | 84.40097173 | -0.446278063 | 0.465743411 | -1.192237337 | 0.2331682 | 0.422006 |
| ENSG00000163541 | SUCLG1 | 6549.516 | -0.4463752 | 0.1647535 | -2.800547 | 0.0051016 | 0.022525 |
| ENSG00000169019 | COMMD8 | 716.7437 | -0.4463805 | 0.2078848 | -2.233169 | 0.0255378 | 0.082458 |
| ENSG00000122033 | MTIF3 | 2004.003586 | -0.446470852 | 0.152585807 | -2.984447297 | 0.00284091 | 0.013947 |
| ENSG00000145365 | TIFA | 366.8942 | -0.446484 | 0.2749915 | -1.747899 | 0.08048145 | 0.198194 |
| ENSG00000165646 | SLC18A2 | 11.31677 | -0.446546 | 0.6037819 | -1.05695 | 0.2905344 | 0.489115 |
| ENSG00000215421 | ZNF407 | 779.2352 | -0.4467414 | 0.1632157 | -2.807115 | 0.00499874 | 0.022144 |
| ENSG00000188234 | AGAP4 | 426.2728 | -0.4467993 | 0.1679322 | -2.727843 | 0.006375 | 0.026975 |
| ENSG00000184270 | H2AC21 | 86.81666 | -0.4468438 | 0.4061091 | -1.281632 | 0.1999717 | 0.380346 |
| ENSG00000260086 | | 8.658661 | -0.4468959 | 0.6079542 | -1.061608 | 0.2884138 | 0.486902 |
| ENSG00000239498 | | 6.824538 | -0.4470212 | 0.6861113 | -1.066951 | 0.285994 | 0.48414 |
| ENSG00000132478 | UNK | 3168.389055 | -0.44707973 | 0.125435815 | -3.614807581 | 0.00030057 | 0.002025 |
| ENSG00000172336 | POP7 | 2094.487 | -0.4471754 | 0.2861254 | -1.695168 | 0.09004358 | 0.215723 |
| ENSG00000101019 | UQCC1 | 2444.896141 | -0.447338548 | 0.148610469 | -3.074944932 | 0.00210541 | 0.010822 |
| ENSG00000214439 | FAM185BP | 53.25707 | -0.4474705 | 0.3236597 | -1.521575 | 0.1281157 | 0.277792 |
| ENSG00000049167 | ERCC8 | 566.8650296 | -0.447570174 | 0.228070606 | -2.063713035 | 0.03904493 | 0.114742 |
| ENSG00000236058 | RPL17P36 | 140.0696 | -0.4475844 | 0.2348539 | -2.002762 | 0.0452028 | 0.128451 |
| ENSG00000023734 | STRAP | 5763.895538 | -0.447679946 | 0.190872077 | -2.42464777 | 0.01532325 | 0.054839 |
| ENSG00000261963 | CCDC92B | 4.40061 | -0.4476844 | 0.8023951 | -1.171093 | 0.2415616 | 0.432105 |
| ENSG00000220771 | BOLA2P3 | 4.792269 | -0.4477152 | 0.7542541 | -1.097408 | 0.2724632 | 0.468973 |
| ENSG00000236794 | BCRP8 | 3.337835 | -0.4477432 | 0.7319243 | -1.081019 | 0.2796886 | 0.476991 |
| ENSG00000124575 | H1-3 | 111.1466895 | -0.447797928 | 0.344640071 | -1.452387273 | 0.14639395 | 0.305814 |
| ENSG00000273555 | MIR6812 | 1.867612 | -0.4479917 | 0.7882585 | -1.131788 | 0.2577237 | 0.451769 |
| ENSG00000242114 | MTFP1 | 150.711 | -0.4481118 | 0.306863 | -1.594158 | 0.1109006 | 0.250786 |
| ENSG00000212158 | SNORD66 | 2.1888 | -0.4484266 | 0.7744841 | -1.122845 | 0.2615032 | 0.456167 |
| ENSG00000106588 | PSMA2 | 289.9955249 | -0.448431811 | 0.241313947 | -1.958680062 | 0.05015027 | 0.138782 |
| ENSG00000136828 | RALGPS1 | 1512.252981 | -0.448547954 | 0.274257238 | -1.752008809 | 0.07977229 | 0.196801 |
| ENSG00000235554 | SRSF6P2 | 366.0297 | -0.4485502 | 0.1684154 | -2.7338 | 0.00626081 | 0.026623 |

| | | | | | | | |
|---|---|---|---|---|---|---|---|
| ENSG00000204394 | VARS1 | 6688.611 | -0.4485511 | 0.1884861 | -2.478978 | 0.01317595 | 0.04856 |
| ENSG00000162889 | MAPKAPK2 | 13778.35 | -0.4486804 | 0.1386607 | -3.302675 | 0.00095767 | 0.005505 |
| ENSG00000121775 | TMEM39B | 808.3902375 | -0.448809479 | 0.154554878 | -2.968901822 | 0.00298866 | 0.014517 |
| ENSG00000248568 | KRT8P48 | 44.98654 | -0.4488523 | 0.3373343 | -1.477819 | 0.1394563 | 0.295283 |
| ENSG00000099860 | GADD45B | 3660.756715 | -0.448909527 | 0.254771113 | -1.850585481 | 0.06422921 | 0.167223 |
| ENSG00000086696 | HSD17B2 | 3.017531805 | -0.449101389 | 0.795538736 | -1.139391216 | 0.25454002 | 0.447546 |
| ENSG00000178971 | CTC1 | 1622.105 | -0.4491387 | 0.2075513 | -2.251962 | 0.02432466 | 0.079406 |
| ENSG00000278916 | CEP83-DT | 48.74008 | -0.4492046 | 0.2906721 | -1.669191 | 0.09507958 | 0.224329 |
| ENSG00000286606 | | 13.32093 | -0.4493686 | 0.4451311 | -1.211505 | 0.2257021 | 0.413023 |
| ENSG00000054598 | FOXC1 | 305.4904554 | -0.449395778 | 0.227041506 | -2.075854783 | 0.03790738 | 0.112144 |
| ENSG00000235363 | SNRPGP10 | 902.1368 | -0.4495236 | 0.2799955 | -1.725553 | 0.08442794 | 0.205517 |
| ENSG00000151838 | CCDC175 | 0.922582 | -0.4495357 | 0.9675166 | -1.969056 | 0.04894662 | NA |
| ENSG00000239808 | RN7SL255P | 1.474293 | -0.4495693 | 0.9432463 | -1.756757 | 0.0789592 | NA |
| ENSG00000231057 | | 1.094331 | -0.4496591 | 0.9304249 | -1.526146 | 0.1269736 | NA |
| ENSG00000249212 | ATP1B1P1 | 3.573232 | -0.4497682 | 0.6803539 | -1.057913 | 0.2900949 | 0.488637 |
| ENSG00000286360 | | 48.13305 | -0.4498253 | 0.3605311 | -1.405862 | 0.1597651 | 0.325871 |
| ENSG00000238286 | SLC35E1P1 | 20.81137 | -0.4499173 | 0.469411 | -1.179251 | 0.2382984 | 0.428083 |
| ENSG00000250132 | | 137.5705 | -0.4499825 | 0.3566188 | -1.418207 | 0.1561305 | 0.3205 |
| ENSG00000158604 | TMED4 | 6240.242 | -0.4500004 | 0.1548245 | -2.964317 | 0.00303355 | 0.014686 |
| ENSG00000266920 | ACTBP9 | 2.528968 | -0.4500008 | 0.8094851 | -1.166676 | 0.2433411 | 0.434469 |
| ENSG00000198553 | KCNRG | 7.654529 | -0.4501005 | 0.629186 | -1.06575 | 0.2865367 | 0.484842 |
| ENSG00000196090 | PTPRT | 115.749 | -0.450162 | 0.6638131 | -1.066395 | 0.2862453 | 0.484534 |
| ENSG00000214870 | | 23.21157 | -0.4503317 | 0.4566035 | -1.199726 | 0.2302457 | 0.418541 |
| ENSG00000014919 | COX15 | 1954.075611 | -0.450410421 | 0.133089838 | -3.459865385 | 0.00054045 | 0.003361 |
| ENSG00000237080 | EHMT2-AS1 | 17.92 | -0.4504305 | 0.4164744 | -1.268167 | 0.2047382 | 0.386644 |
| ENSG00000258445 | MAD2L1P1 | 72.46603 | -0.4504709 | 0.4643187 | -1.190401 | 0.2338887 | 0.422993 |
| ENSG00000177302 | TOP3A | 3567.268 | -0.4509337 | 0.1812887 | -2.554859 | 0.01062307 | 0.040881 |
| ENSG00000143119 | CD53 | 1.586642 | -0.4509389 | 0.8526311 | -1.303032 | 0.1925638 | 0.370956 |
| ENSG00000273983 | H3C8 | 310.9653 | -0.4509529 | 0.4051584 | -1.293012 | 0.1960068 | 0.375763 |
| ENSG00000217261 | POM121L4P | 2.463906 | -0.4511537 | 0.7791749 | -1.141062 | 0.2538439 | 0.446825 |
| ENSG00000260803 | | 2.057309 | -0.451163 | 0.8342338 | -1.213299 | 0.2250155 | 0.412022 |
| ENSG00000268736 | MTCO3P39 | 39.65816 | -0.4511753 | 0.5650495 | -1.094685 | 0.2736548 | 0.470415 |
| ENSG00000261373 | VPS9D1-AS1 | 1094.358 | -0.4512702 | 0.2048257 | -2.291657 | 0.02192545 | 0.073275 |

| | | | | | | | |
|---|---|---|---|---|---|---|---|
| ENSG00000198888 | MT-ND1 | 46497.07 | -0.451382 | 0.1510512 | -3.085045 | 0.00203521 | 0.010518 |
| ENSG00000113013 | HSPA9 | 29348.06419 | -0.451599331 | 0.179294464 | -2.594435407 | 0.00947464 | 0.037205 |
| ENSG00000243592 | RPL17P22 | 2.317326 | -0.4516142 | 0.7921718 | -1.145226 | 0.2521157 | 0.444819 |
| ENSG00000095951 | HIVEP1 | 1272.571692 | -0.451650429 | 0.273965781 | -1.76866115 | 0.07695044 | 0.191655 |
| ENSG00000279283 | | 29.64302 | -0.4516722 | 0.4930515 | -1.154858 | 0.2481487 | 0.440426 |
| ENSG00000266984 | POLR3GP2 | 3.326089 | -0.4518376 | 0.8237678 | -1.217688 | 0.2233427 | 0.410033 |
| ENSG00000008405 | CRY1 | 1524.058587 | -0.451967669 | 0.17337538 | -2.685639246 | 0.00723912 | 0.029964 |
| ENSG00000265817 | FSBP | 135.3038 | -0.4520109 | 0.5173947 | -1.150032 | 0.2501307 | 0.442695 |
| ENSG00000172586 | CHCHD1 | 2926.598 | -0.4520628 | 0.2060523 | -2.283731 | 0.02238735 | 0.074566 |
| ENSG00000224520 | KRT8P45 | 2170.693 | -0.4521483 | 0.2172963 | -2.186352 | 0.02878988 | 0.090813 |
| ENSG00000171960 | PPIH | 986.3733 | -0.4523335 | 0.3185916 | -1.560915 | 0.1185437 | 0.262794 |
| ENSG00000143443 | C1orf56 | 1011.685 | -0.4524054 | 0.1741506 | -2.67005 | 0.007584 | 0.031168 |
| ENSG00000180228 | PRKRA | 1106.538 | -0.4524273 | 0.1447781 | -3.185963 | 0.00144273 | 0.007852 |
| ENSG00000280087 | | 2.901132 | -0.4524277 | 0.7842571 | -1.141087 | 0.2538337 | 0.446825 |
| ENSG00000237135 | DDX10P1 | 0.9792691 | -0.4525048 | 0.8881405 | -1.445462 | 0.1483281 | NA |
| ENSG00000178412 | | 1.274935 | -0.4525234 | 0.9094475 | -1.544678 | 0.1224242 | NA |
| ENSG00000278766 | | 1.310665 | -0.452618 | 0.8596784 | -1.272007 | 0.2033707 | NA |
| ENSG00000121858 | TNFSF10 | 63.90268845 | -0.452632382 | 0.670443573 | -1.053112422 | 0.29228947 | 0.491136 |
| ENSG00000138439 | FAM117B | 815.4338862 | -0.452643127 | 0.214288018 | -2.202534535 | 0.02762757 | 0.087782 |
| ENSG00000196119 | OR8A1 | 30.6951 | -0.4528242 | 0.462124 | -1.198289 | 0.2308046 | 0.419288 |
| ENSG00000101361 | NOP56 | 7859.236688 | -0.452912784 | 0.204564469 | -2.301727385 | 0.02135055 | 0.071669 |
| ENSG00000104889 | RNASEH2A | 15.84233677 | -0.452961493 | 0.618486648 | -1.074698544 | 0.28250972 | 0.480323 |
| ENSG00000172167 | MTBP | 574.2232 | -0.4529847 | 0.3084579 | -1.606231 | 0.1082232 | 0.24646 |
| ENSG00000089101 | CFAP61 | 4.685358649 | -0.453080014 | 0.688321674 | -1.078863436 | 0.28064861 | 0.478137 |
| ENSG00000279032 | | 0.7995805 | -0.453213 | 0.9604065 | -1.819562 | 0.06882571 | NA |
| ENSG00000226468 | | 21.70748 | -0.4532798 | 0.4954026 | -1.143356 | 0.2528909 | 0.445768 |
| ENSG00000273002 | ARHGEF2-AS2 | 34.13093 | -0.4535467 | 0.5570579 | -1.097696 | 0.2723373 | 0.468878 |
| ENSG00000081913 | PHLPP1 | 1010.653597 | -0.453555092 | 0.139189721 | -3.321740753 | 0.00089458 | 0.00519 |
| ENSG00000164543 | STK17A | 1194.989 | -0.4535556 | 0.1476799 | -3.128462 | 0.00175724 | 0.009282 |
| ENSG00000175463 | TBC1D10C | 100.2754 | -0.4535804 | 0.3554146 | -1.433181 | 0.1518061 | 0.31382 |
| ENSG00000149418 | ST14 | 7961.727 | -0.4536662 | 0.2030862 | -2.335088 | 0.01953882 | 0.066734 |
| ENSG00000133065 | SLC41A1 | 8123.356798 | -0.453681354 | 0.202807339 | -2.334773524 | 0.01955526 | 0.066782 |
| ENSG00000136504 | KAT7 | 5123.955331 | -0.453744541 | 0.128843129 | -3.588691957 | 0.00033234 | 0.002214 |

| | | | | | | | |
|---|---|---|---|---|---|---|---|
| ENSG00000286616 | | 2.134085 | -0.4537949 | 0.7757888 | -1.135242 | 0.256274 | 0.45 |
| ENSG00000107816 | LZTS2 | 4333.538903 | -0.453823265 | 0.159659318 | -2.92664754 | 0.00342637 | 0.01623 |
| ENSG00000277693 | | 50.93616 | -0.4538944 | 0.3891004 | -1.340766 | 0.1799965 | 0.354168 |
| ENSG00000083520 | DIS3 | 3026.373788 | -0.453902432 | 0.140855804 | -3.278850364 | 0.00104231 | 0.005935 |
| ENSG00000283167 | | 5.337308 | -0.4539867 | 0.7450785 | -1.097333 | 0.2724961 | 0.468986 |
| ENSG00000133302 | SLF1 | 618.8394817 | -0.454000659 | 0.386658478 | -1.341458915 | 0.1797715 | 0.353918 |
| ENSG00000187475 | H1-6 | 2.61717 | -0.4541418 | 0.8000838 | -1.153129 | 0.2488574 | 0.441175 |
| ENSG00000261716 | H2BC20P | 7537.537 | -0.4541716 | 0.294322 | -1.671125 | 0.09469691 | 0.223731 |
| ENSG00000106603 | COA1 | 3919.234857 | -0.454263548 | 0.122609768 | -3.754077062 | 0.00017398 | 0.001253 |
| ENSG00000153815 | CMIP | 2203.17 | -0.4544472 | 0.164287 | -2.852682 | 0.00433519 | 0.019711 |
| ENSG00000278177 | | 6.113055 | -0.4544703 | 0.6308513 | -1.069279 | 0.284944 | 0.482947 |
| ENSG00000280055 | TMEM75 | 12.88932 | -0.4544729 | 0.6303866 | -1.07564 | 0.2820881 | 0.479821 |
| ENSG00000237605 | DYRK3-AS1 | 9.19404 | -0.4545443 | 0.5913361 | -1.086089 | 0.2774398 | 0.474527 |
| ENSG00000231925 | TAPBP | 7993.287 | -0.4546356 | 0.2263313 | -2.105012 | 0.0352903 | 0.106244 |
| ENSG00000273899 | NOL12 | 164.7074 | -0.4546461 | 0.3361846 | -1.503677 | 0.1326646 | 0.284971 |
| ENSG00000111727 | HCFC2 | 747.7229222 | -0.454684469 | 0.187328283 | -2.511759706 | 0.01201309 | 0.045075 |
| ENSG00000111215 | PRR4 | 17.40271163 | -0.454699121 | 0.422814125 | -1.271560901 | 0.20352918 | 0.385142 |
| ENSG00000242474 | | 782.0692 | -0.4548209 | 0.2276651 | -2.093437 | 0.03631013 | 0.108705 |
| ENSG00000062582 | MRPS24 | 40.33483047 | -0.455026772 | 0.370535406 | -1.392584575 | 0.16374545 | 0.331723 |
| ENSG00000137203 | TFAP2A | 1952.505764 | -0.455067512 | 0.175589013 | -2.671775967 | 0.0075451 | 0.031016 |
| ENSG00000229644 | NAMPTP1 | 1289.512 | -0.4551409 | 0.201142 | -2.350896 | 0.01872825 | 0.064463 |
| ENSG00000164305 | CASP3 | 1431.008 | -0.455212 | 0.2346438 | -2.049089 | 0.04045341 | 0.117929 |
| ENSG00000148498 | PARD3 | 4961.44 | -0.4552201 | 0.1251181 | -3.638219 | 0.00027453 | 0.00187 |
| ENSG00000259376 | | 6.919523 | -0.455502 | 0.568746 | -1.093157 | 0.2743248 | 0.471079 |
| ENSG00000164066 | INTU | 394.2282 | -0.4555078 | 0.2518789 | -1.921846 | 0.05462518 | 0.148444 |
| ENSG00000115548 | KDM3A | 3296.887362 | -0.455518585 | 0.169797094 | -2.760685902 | 0.00576801 | 0.024947 |
| ENSG00000287190 | | 10.30137 | -0.4556247 | 0.6504294 | -1.078572 | 0.2807785 | 0.478205 |
| ENSG00000224843 | LINC00240 | 40.20285 | -0.4556301 | 0.3417456 | -1.484261 | 0.1377398 | 0.292418 |
| ENSG00000262119 | | 2.898177 | -0.4556576 | 0.7837527 | -1.150574 | 0.2499076 | 0.442476 |
| ENSG00000159147 | DONSON | 850.9985 | -0.4557112 | 0.3720089 | -1.364458 | 0.1724234 | 0.343541 |
| ENSG00000121152 | NCAPH | 1957.113102 | -0.455814594 | 0.56639145 | -1.110567569 | 0.26675453 | 0.462564 |
| ENSG00000261502 | | 12.82062 | -0.455968 | 0.4917748 | -1.167769 | 0.2429001 | 0.433882 |
| ENSG00000272430 | LINC02637 | 3.977636 | -0.4559945 | 0.7023723 | -1.076678 | 0.2816241 | 0.479277 |

| | | | | | | | |
|---|---|---|---|---|---|---|---|
| ENSG00000271880 | AGAP11 | 6.860208 | -0.4560445 | 0.7231341 | -1.080362 | 0.2799812 | 0.477245 |
| ENSG00000273306 | | 72.08227 | -0.4561142 | 0.2853928 | -1.723738 | 0.0847552 | 0.206102 |
| ENSG00000215867 | KRT18P57 | 1.913986 | -0.4562942 | 0.9344456 | -1.49475 | 0.1349796 | 0.288342 |
| ENSG00000249307 | LINC01088 | 2.402802 | -0.456301 | 0.728816 | -1.087747 | 0.2767066 | 0.4737 |
| ENSG00000175643 | RMI2 | 1183.476 | -0.4563458 | 0.4284353 | -1.254597 | 0.2096251 | 0.392995 |
| ENSG00000119977 | TCTN3 | 3857.196882 | -0.45646905 | 0.112931356 | -4.075172039 | 4.60E-05 | 0.000385 |
| ENSG00000204622 | HLA-J | 17.20575 | -0.456582 | 0.5315346 | -1.123362 | 0.2612838 | 0.455874 |
| ENSG00000273760 | | 90.89722 | -0.4567651 | 0.6854033 | -1.049062 | 0.2941497 | 0.493201 |
| ENSG00000229160 | | 3.075048 | -0.456825 | 0.8525597 | -1.292771 | 0.1960904 | 0.375842 |
| ENSG00000184786 | DYNLT2 | 283.2187 | -0.4568375 | 0.2672748 | -1.823539 | 0.06822176 | 0.175129 |
| ENSG00000164794 | KCNV1 | 127.2453 | -0.4568756 | 0.3741359 | -1.395373 | 0.1629034 | 0.330471 |
| ENSG00000269929 | MIRLET7A1HG | 410.5264 | -0.4569513 | 0.2205759 | -2.178932 | 0.02933674 | 0.09221 |
| ENSG00000261019 | | 10.61984 | -0.4571028 | 0.5032851 | -1.151736 | 0.2494297 | 0.441954 |
| ENSG00000167969 | ECI1 | 3624.445 | -0.4571432 | 0.1689013 | -2.783769 | 0.00537313 | 0.023495 |
| ENSG00000130706 | ADRM1 | 6538.262728 | -0.457170049 | 0.219709173 | -2.1783218 | 0.02938209 | 0.092323 |
| ENSG00000286576 | | 1.091212 | -0.457182 | 0.8637315 | -1.306126 | 0.1915097 | NA |
| ENSG00000233056 | ERVH48-1 | 7.95753 | -0.4572489 | 0.634624 | -1.075813 | 0.282011 | 0.479751 |
| ENSG00000213780 | GTF2H4 | 30.66537 | -0.4573663 | 0.4101837 | -1.302211 | 0.1928443 | 0.371308 |
| ENSG00000227827 | PKD1P2 | 18.52473 | -0.4575819 | 0.4786487 | -1.185635 | 0.2357665 | 0.42515 |
| ENSG00000157326 | DHRS4 | 508.2147 | -0.458033 | 0.2010302 | -2.365537 | 0.01800396 | 0.062456 |
| ENSG00000157224 | CLDN12 | 1090.192 | -0.4580358 | 0.2653225 | -1.812596 | 0.06989415 | 0.178388 |
| ENSG00000269837 | IPO5P1 | 319.4173 | -0.4581175 | 0.2214923 | -2.160196 | 0.03075752 | 0.095646 |
| ENSG00000279384 | | 7.161635 | -0.4581303 | 0.7292836 | -1.10216 | 0.2703919 | 0.466781 |
| ENSG00000286615 | | 21.25494 | -0.4581607 | 0.4033973 | -1.321661 | 0.1862812 | 0.362467 |
| ENSG00000078319 | PMS2P1 | 574.555042 | -0.458327562 | 0.146413722 | -3.193323474 | 0.00140645 | 0.007681 |
| ENSG00000159210 | SNF8 | 3287.116 | -0.4584922 | 0.1413131 | -3.325862 | 0.00088146 | 0.005127 |
| ENSG00000262096 | PCDHB19P | 8.940732 | -0.4585172 | 0.5398529 | -1.12404 | 0.2609959 | 0.45563 |
| ENSG00000101811 | CSTF2 | 2267.257005 | -0.45855842 | 0.241693182 | -2.002205554 | 0.04526263 | 0.128582 |
| ENSG00000164332 | UBLCP1 | 1490.064 | -0.4586098 | 0.1296993 | -3.575514 | 0.00034954 | 0.002311 |
| ENSG00000233416 | RPS25P3 | 1.637257 | -0.4587342 | 0.9130961 | -1.487239 | 0.1369516 | 0.291212 |
| ENSG00000163214 | DHX57 | 1476.117 | -0.4588479 | 0.2018313 | -2.369016 | 0.01783549 | 0.061968 |
| ENSG00000235961 | PNMA6A | 55.13518 | -0.4589391 | 0.4222672 | -1.283324 | 0.1993786 | 0.379598 |
| ENSG00000214860 | EVPLL | 2.480511 | -0.4589795 | 0.8388403 | -1.248071 | 0.2120052 | 0.396339 |

| | | | | | | | |
|---|---|---|---|---|---|---|---|
| ENSG00000259248 | USP3-AS1 | 68.04296 | -0.4590036 | 0.2602318 | -1.876521 | 0.06058372 | 0.160158 |
| ENSG00000034053 | APBA2 | 197.4305832 | -0.45904087 | 0.285698281 | -1.734733573 | 0.08278801 | 0.202653 |
| ENSG00000128973 | CLN6 | 2116.858785 | -0.459068892 | 0.201337919 | -2.363204802 | 0.01811765 | 0.062776 |
| ENSG00000115307 | AUP1 | 9383.777772 | -0.459097708 | 0.164395744 | -2.882610923 | 0.00394394 | 0.01823 |
| ENSG00000278341 | | 91.59772 | -0.4591894 | 0.3575507 | -1.440795 | 0.1496426 | 0.310614 |
| ENSG00000138032 | PPM1B | 863.5217099 | -0.459222624 | 0.161519234 | -2.914359069 | 0.0035642 | 0.016761 |
| ENSG00000286760 | | 124.779 | -0.45925 | 0.2867296 | -1.724051 | 0.08469859 | 0.206058 |
| ENSG00000251548 | | 30.01764 | -0.4593751 | 0.3741008 | -1.401203 | 0.1611533 | 0.327936 |
| ENSG00000251079 | BMS1P2 | 21.69754 | -0.4594245 | 0.3442981 | -1.487822 | 0.1367979 | 0.291116 |
| ENSG00000162723 | SLAMF9 | 1.488673 | -0.4594801 | 0.8987848 | -1.437388 | 0.1506077 | NA |
| ENSG00000206964 | | 1.637223 | -0.4596281 | 0.8194133 | -1.195899 | 0.231736 | 0.420158 |
| ENSG00000257507 | LINC02373 | 2.321269 | -0.4598383 | 0.8736669 | -1.329859 | 0.1835647 | 0.358771 |
| ENSG00000166922 | SCG5 | 10.73828 | -0.4598893 | 0.6341411 | -1.090162 | 0.2756419 | 0.472486 |
| ENSG00000170421 | KRT8 | 163694.2 | -0.4600892 | 0.1969592 | -2.421392 | 0.0154612 | 0.055228 |
| ENSG00000213523 | SRA1 | 1648.181 | -0.460112 | 0.1661279 | -2.840823 | 0.00449973 | 0.020318 |
| ENSG00000196811 | CHRNG | 20.41209 | -0.4602617 | 0.4977388 | -1.160947 | 0.2456635 | 0.437381 |
| ENSG00000279347 | | 15.17897 | -0.4604645 | 0.673681 | -1.094202 | 0.2738662 | 0.470626 |
| ENSG00000164978 | NUDT2 | 1347.25 | -0.4607942 | 0.173767 | -2.720114 | 0.00652595 | 0.027511 |
| ENSG00000239096 | | 2.519388 | -0.4608025 | 0.8058377 | -1.159058 | 0.2464326 | 0.438251 |
| ENSG00000253873 | PCDHGA11 | 6.481989 | -0.4608646 | 0.6364042 | -1.075017 | 0.2823673 | 0.480234 |
| ENSG00000132932 | ATP8A2 | 1527.456824 | -0.460893234 | 0.354232566 | -1.457884929 | 0.14487226 | 0.303534 |
| ENSG00000105289 | TJP3 | 2723.056694 | -0.460933981 | 0.215436004 | -2.285619866 | 0.02227651 | 0.074271 |
| ENSG00000189180 | ZNF33A | 4006.879 | -0.4610016 | 0.2112523 | -2.272205 | 0.02307414 | 0.076185 |
| ENSG00000089597 | GANAB | 29207.85279 | -0.461042797 | 0.129321713 | -3.620224584 | 0.00029435 | 0.001988 |
| ENSG00000079246 | XRCC5 | 17110.65456 | -0.461112321 | 0.211602539 | -2.274680742 | 0.02292509 | 0.075777 |
| ENSG00000271727 | | 3.416785 | -0.4611167 | 0.7199384 | -1.0805 | 0.2799197 | 0.477202 |
| ENSG00000251613 | | 3.142106 | -0.4611374 | 0.7288629 | -1.098676 | 0.2719093 | 0.468384 |
| ENSG00000233654 | | 75.20788 | -0.4611736 | 0.2807338 | -1.761498 | 0.0781541 | 0.193792 |
| ENSG00000157954 | WIPI2 | 6615.327 | -0.4614607 | 0.1254247 | -3.734707 | 0.00018793 | 0.001342 |
| ENSG00000286654 | | 2.256127 | -0.4616119 | 1.002843 | -2.631744 | 0.00849479 | 0.034119 |
| ENSG00000121957 | GPSM2 | 1931.535598 | -0.461651939 | 0.360322854 | -1.473408518 | 0.14064092 | 0.297152 |
| ENSG00000157778 | PSMG3 | 2658.502 | -0.4616742 | 0.2741086 | -1.830399 | 0.06719032 | 0.173134 |
| ENSG00000156875 | MFSD14A | 110.0435 | -0.461733 | 0.2446915 | -1.991417 | 0.04643512 | 0.131011 |

| | | | | | | | |
|---|---|---|---|---|---|---|---|
| ENSG00000104980 | TIMM44 | 2711.568757 | -0.461768851 | 0.159282962 | -2.964188515 | 0.00303482 | 0.014687 |
| ENSG00000137571 | SLCO5A1 | 14.69260761 | -0.461952845 | 0.692357679 | -1.101888005 | 0.27051036 | 0.466782 |
| ENSG00000260425 | | 19.14903 | -0.4620147 | 0.5342506 | -1.130706 | 0.2581788 | 0.452358 |
| ENSG00000123836 | PFKFB2 | 3426.90679 | -0.462031515 | 0.309184182 | -1.616853976 | 0.10590981 | 0.242621 |
| ENSG00000243849 | CFAP44-AS1 | 1.213455 | -0.4620874 | 0.9237694 | -1.408596 | 0.1589546 | NA |
| ENSG00000261868 | MFSD1P1 | 2.435718 | -0.4621476 | 0.7522809 | -1.117677 | 0.263705 | 0.458953 |
| ENSG00000011143 | MKS1 | 2660.734775 | -0.46233049 | 0.14529279 | -3.245035798 | 0.00117436 | 0.006563 |
| ENSG00000261584 | | 1.448375 | -0.4624714 | 0.8438955 | -1.253999 | 0.2098424 | NA |
| ENSG00000272994 | C2orf49-DT | 446.3588 | -0.4625228 | 0.1632436 | -2.904983 | 0.00367273 | 0.017174 |
| ENSG00000141040 | ZNF287 | 731.6591 | -0.4626352 | 0.1017193 | -4.591765 | 4.40E-06 | 4.76E-05 |
| ENSG00000185453 | ZSWIM9 | 739.7147 | -0.4628471 | 0.1676856 | -2.831564 | 0.00463209 | 0.020835 |
| ENSG00000162959 | MEMO1 | 147.3188 | -0.4629607 | 0.3202985 | -1.591345 | 0.1115319 | 0.251874 |
| ENSG00000110651 | CD81 | 7851.52368 | -0.46296322 | 0.149030328 | -3.19150989 | 0.00141531 | 0.007717 |
| ENSG00000197226 | TBC1D9B | 22179.49 | -0.4630172 | 0.09901957 | -4.729454 | 2.25E-06 | 2.62E-05 |
| ENSG00000181652 | ATG9B | 40.85259 | -0.4630246 | 0.5665995 | -1.113234 | 0.2656079 | 0.461263 |
| ENSG00000270270 | BNIP3P10 | 1.318234 | -0.4632261 | 0.9717342 | -1.873887 | 0.06094602 | NA |
| ENSG00000264386 | MIR4513 | 1.430156 | -0.4632682 | 0.8174728 | -1.197593 | 0.2310753 | NA |
| ENSG00000106125 | MINDY4 | 13.68244824 | -0.463391161 | 0.555943051 | -1.108554009 | 0.26762263 | 0.463555 |
| ENSG00000213639 | PPP1CB | 7402.191 | -0.4635935 | 0.1817445 | -2.634001 | 0.00843854 | 0.03395 |
| ENSG00000228716 | DHFR | 1749.539 | -0.4639244 | 0.353422 | -1.49547 | 0.1347918 | 0.288068 |
| ENSG00000213872 | RPL32P11 | 5.41947 | -0.4639683 | 0.6233125 | -1.093475 | 0.2741852 | 0.470902 |
| ENSG00000105835 | NAMPT | 3294.416887 | -0.464052744 | 0.171411532 | -2.776970538 | 0.00548682 | 0.02389 |
| ENSG00000184224 | C11orf72 | 4.719413 | -0.4640945 | 0.8223164 | -1.185529 | 0.2358083 | 0.42515 |
| ENSG00000163728 | TTC14 | 2992.896 | -0.4649904 | 0.2862028 | -1.753179 | 0.07957122 | 0.196444 |
| ENSG00000120210 | INSL6 | 1.28820094 | -0.465033897 | 0.835660501 | -1.236557415 | 0.21625144 | NA |
| ENSG00000006744 | ELAC2 | 7291.493062 | -0.465049695 | 0.215440093 | -2.246389929 | 0.02467904 | 0.080276 |
| ENSG00000260969 | WWOX-AS1 | 1.209402 | -0.465071 | 0.9386511 | -1.521556 | 0.1281203 | NA |
| ENSG00000033011 | ALG1 | 1300.083654 | -0.465166249 | 0.158608419 | -3.004997883 | 0.00265583 | 0.013155 |
| ENSG00000260279 | | 340.013 | -0.4652666 | 0.2608478 | -1.917953 | 0.05511695 | 0.149353 |
| ENSG00000274976 | | 3.228758 | -0.4653686 | 0.8647762 | -1.315249 | 0.1884261 | 0.365503 |
| ENSG00000231531 | HINT1P1 | 4.785501 | -0.4656513 | 0.6658525 | -1.095769 | 0.27318 | 0.46984 |
| ENSG00000241479 | MECOM-AS1 | 1.351488 | -0.4657114 | 0.9258085 | -1.415618 | 0.1568873 | NA |
| ENSG00000254620 | | 4.773084 | -0.4660411 | 0.7860693 | -1.128626 | 0.2590558 | 0.453416 |

| | | | | | | | |
|---|---|---|---|---|---|---|---|
| ENSG00000156510 | HKDC1 | 4.560689 | -0.4661278 | 0.7633632 | -1.141521 | 0.2536532 | 0.446667 |
| ENSG00000207741 | MIR590 | 19.2587 | -0.4663072 | 0.422492 | -1.308287 | 0.1907761 | 0.368821 |
| ENSG00000224543 | SNRPGP15 | 179.8462 | -0.4664527 | 0.3088072 | -1.653193 | 0.09829163 | 0.229785 |
| ENSG00000103037 | SETD6 | 1495.289306 | -0.466583528 | 0.144198698 | -3.300220735 | 0.00096609 | 0.00555 |
| ENSG00000280010 | | 9.954235 | -0.4665908 | 0.6281409 | -1.098924 | 0.271801 | 0.468289 |
| ENSG00000131153 | GINS2 | 2635.436627 | -0.466632799 | 0.6425515 | -1.118195918 | 0.26348333 | 0.458687 |
| ENSG00000231365 | WARS2-AS1 | 763.4266 | -0.466695 | 0.2401518 | -2.041427 | 0.04120838 | 0.119762 |
| ENSG00000199753 | SNORD104 | 683.2365 | -0.4667206 | 0.2510157 | -1.97528 | 0.04823642 | 0.134782 |
| ENSG00000213881 | NPM1P6 | 1355.144 | -0.4667627 | 0.2093155 | -2.338077 | 0.01938324 | 0.06627 |
| ENSG00000217159 | LARP1P1 | 5.973572 | -0.4668978 | 0.7648022 | -1.149184 | 0.2504803 | 0.443048 |
| ENSG00000244694 | PTCHD4 | 56.83438 | -0.4670106 | 0.4209934 | -1.304966 | 0.1919046 | 0.370169 |
| ENSG00000180574 | EIF2S3B | 2547.928 | -0.4670449 | 0.1995818 | -2.429879 | 0.01510385 | 0.054222 |
| ENSG00000072133 | RPS6KA6 | 1150.969904 | -0.467087092 | 0.185444566 | -2.600033012 | 0.00932148 | 0.03674 |
| ENSG00000238222 | MKRN4P | 2.118023 | -0.4672458 | 0.8911236 | -1.3884 | 0.1650153 | 0.333406 |
| ENSG00000251211 | | 2.942957 | -0.4673503 | 0.8072367 | -1.167733 | 0.2429143 | 0.433882 |
| ENSG00000238275 | HOMER2P2 | 1.467445 | -0.4673563 | 0.8549076 | -1.293741 | 0.1957548 | NA |
| ENSG00000171311 | EXOSC1 | 2533.346 | -0.4674053 | 0.1679999 | -2.870418 | 0.00409929 | 0.018843 |
| ENSG00000259746 | HSPE1P3 | 290.3673 | -0.4674126 | 0.2740628 | -1.827633 | 0.0676046 | 0.173945 |
| ENSG00000145736 | GTF2H2 | 424.1162 | -0.4674441 | 0.1992626 | -2.432548 | 0.01499302 | 0.053898 |
| ENSG00000143195 | ILDR2 | 10.92316 | -0.4674691 | 0.6079981 | -1.106334 | 0.2685818 | 0.464635 |
| ENSG00000139684 | ESD | 11710.31 | -0.4674847 | 0.1660879 | -2.89811 | 0.00375419 | 0.017488 |
| ENSG00000184752 | NDUFA12 | 2045.248 | -0.4675 | 0.2128791 | -2.320989 | 0.02028741 | 0.068753 |
| ENSG00000230593 | PPIAP40 | 5.308028 | -0.4675111 | 0.6135009 | -1.098824 | 0.2718448 | 0.468303 |
| ENSG00000179029 | TMEM107 | 262.8836 | -0.4676643 | 0.330692 | -1.570686 | 0.1162557 | 0.259301 |
| ENSG00000102390 | PBDC1 | 2047.637022 | -0.467719143 | 0.254577525 | -1.950307723 | 0.05113945 | 0.140944 |
| ENSG00000138172 | CALHM2 | 772.5485392 | -0.467794879 | 0.266661377 | -1.864119954 | 0.06230486 | 0.163543 |
| ENSG00000120686 | UFM1 | 4571.083684 | -0.467817065 | 0.169213667 | -2.849860948 | 0.00437384 | 0.019853 |
| ENSG00000163444 | TMEM183A | 6413.114 | -0.4678172 | 0.1496174 | -3.227491 | 0.00124881 | 0.006933 |
| ENSG00000201151 | | 2.072506 | -0.4679407 | 0.7776276 | -1.151116 | 0.2496844 | 0.442307 |
| ENSG00000118946 | PCDH17 | 66.34718809 | -0.467972659 | 0.508100005 | -1.183653321 | 0.23655032 | 0.426153 |
| ENSG00000157315 | TMED6 | 25.67004 | -0.4679846 | 0.3592953 | -1.4656 | 0.1427573 | 0.300382 |
| ENSG00000163002 | NUP35 | 890.488 | -0.4680229 | 0.2736345 | -1.83316 | 0.06677875 | 0.172324 |
| ENSG00000250033 | SLC7A11-AS1 | 17.75207 | -0.4680846 | 0.6984788 | -1.113502 | 0.2654931 | 0.461159 |

| | | | | | | | |
|---|---|---|---|---|---|---|---|
| ENSG00000167792 | NDUFV1 | 12625.25 | -0.4681133 | 0.1570727 | -3.068278 | 0.00215296 | 0.011035 |
| ENSG00000248925 | | 3.519844 | -0.468132 | 0.7455261 | -1.128742 | 0.2590066 | 0.45336 |
| ENSG00000111642 | CHD4 | 1309.100779 | -0.468445382 | 0.199359993 | -2.445143388 | 0.01447946 | 0.052405 |
| ENSG00000173674 | EIF1AX | 2887.852 | -0.4684579 | 0.1839777 | -2.630567 | 0.00852426 | 0.034212 |
| ENSG00000279685 | MAPT-IT1 | 2.926094 | -0.4685791 | 0.8205296 | -1.181511 | 0.2373997 | 0.427037 |
| ENSG00000279044 | | 2.804408 | -0.4686003 | 0.8494302 | -1.255201 | 0.2094058 | 0.39264 |
| ENSG00000287957 | | 8.787661 | -0.4686121 | 0.6437965 | -1.097125 | 0.2725868 | 0.469065 |
| ENSG00000265559 | RN7SL652P | 1.497434 | -0.4687249 | 0.8972512 | -1.451247 | 0.1467111 | NA |
| ENSG00000255734 | HNRNPABP1 | 7.574834 | -0.4687276 | 0.6082675 | -1.110055 | 0.2669755 | 0.46278 |
| ENSG00000166839 | ANKDD1A | 332.0706 | -0.4688751 | 0.3062286 | -1.668847 | 0.09514762 | 0.224377 |
| ENSG00000105516 | DBP | 1119.729038 | -0.468987497 | 0.240154053 | -2.059623371 | 0.03943456 | 0.115629 |
| ENSG00000196715 | VKORC1L1 | 3082.045 | -0.4690206 | 0.1436359 | -3.333184 | 0.00085858 | 0.005009 |
| ENSG00000134697 | GNL2 | 4123.568676 | -0.469068265 | 0.17788207 | -2.714093834 | 0.00664573 | 0.02793 |
| ENSG00000168301 | KCTD6 | 251.5494 | -0.4692536 | 0.284834 | -1.777469 | 0.07549114 | 0.189016 |
| ENSG00000141030 | COPS3 | 5107.845 | -0.4694446 | 0.2504706 | -1.985568 | 0.0470813 | 0.132343 |
| ENSG00000188807 | TMEM201 | 1153.299 | -0.4694598 | 0.2532347 | -1.966717 | 0.04921581 | 0.136923 |
| ENSG00000054967 | RELT | 544.1261048 | -0.469511317 | 0.286136126 | -1.770836192 | 0.07658795 | 0.19096 |
| ENSG00000235863 | B3GALT4 | 117.2857 | -0.469689 | 0.3221574 | -1.606767 | 0.1081054 | 0.246249 |
| ENSG00000123064 | DDX54 | 5916.888623 | -0.469747951 | 0.200362422 | -2.403838996 | 0.01622392 | 0.057351 |
| ENSG00000232075 | MRPL35P2 | 7.337231 | -0.4698283 | 0.5716427 | -1.118688 | 0.2632733 | 0.458352 |
| ENSG00000240204 | SMKR1 | 361.7174 | -0.4698965 | 0.2686035 | -1.871448 | 0.06128302 | 0.161643 |
| ENSG00000176998 | HCG4 | 6.56077 | -0.47022 | 0.631205 | -1.100912 | 0.2709348 | 0.467251 |
| ENSG00000240498 | CDKN2B-AS1 | 1182.341 | -0.4703366 | 0.2851853 | -1.781002 | 0.07491205 | 0.18791 |
| ENSG00000102054 | RBBP7 | 9279.697232 | -0.470357466 | 0.223474196 | -2.204276174 | 0.02750493 | 0.087529 |
| ENSG00000180219 | FAM71C | 1.174104 | -0.47048 | 0.9824976 | -1.948357 | 0.05137222 | NA |
| ENSG00000202031 | SNORD38A | 8.938324 | -0.4705222 | 0.4949691 | -1.197736 | 0.2310198 | 0.419518 |
| ENSG00000011566 | MAP4K3 | 2588.751797 | -0.470606452 | 0.159655838 | -3.019760817 | 0.00252974 | 0.012627 |
| ENSG00000161103 | | 0.8663261 | -0.4707984 | 0.9659768 | -1.743476 | 0.08125055 | NA |
| ENSG00000222222 | RNU2-17P | 2.745222 | -0.4708021 | 0.8355167 | -1.217204 | 0.2235266 | 0.41023 |
| ENSG00000267586 | LINC00907 | 5.017805 | -0.4708372 | 0.6501314 | -1.103162 | 0.2699569 | 0.466318 |
| ENSG00000099821 | POLRMT | 2633.238121 | -0.470939873 | 0.12319925 | -3.888767638 | 0.00010076 | 0.00077 |
| ENSG00000111237 | VPS29 | 2867.722541 | -0.470942749 | 0.13675774 | -3.506755449 | 0.00045361 | 0.002883 |
| ENSG00000184677 | ZBTB40 | 2942.941 | -0.4709536 | 0.1855208 | -2.624722 | 0.00867199 | 0.03471 |

| | | | | | | | |
|---|---|---|---|---|---|---|---|
| ENSG00000272872 | | 145.5984 | -0.4709741 | 0.319617 | -1.621224 | 0.1049697 | 0.240979 |
| ENSG00000023318 | ERP44 | 2840.208077 | -0.470985758 | 0.146097508 | -3.289627595 | 0.0010032 | 0.005737 |
| ENSG00000245330 | | 5.794932 | -0.4709928 | 0.6801453 | -1.104516 | 0.2693693 | 0.465597 |
| ENSG00000172469 | MANEA | 670.9006 | -0.4710508 | 0.2480944 | -2.025413 | 0.04282493 | 0.123354 |
| ENSG00000250584 | LINC01511 | 1.607776 | -0.4714214 | 0.8318894 | -1.220542 | 0.2222596 | 0.408955 |
| ENSG00000115541 | HSPE1 | 1699.583699 | -0.471504798 | 0.225318994 | -2.196831477 | 0.02803248 | 0.088782 |
| ENSG00000236756 | DNAJC9-AS1 | 67.91082 | -0.4717059 | 0.298453 | -1.711675 | 0.08695653 | 0.210051 |
| ENSG00000237476 | LINC01637 | 132.9443 | -0.471739 | 0.2625417 | -1.910593 | 0.05605687 | 0.151252 |
| ENSG00000218018 | RBM38-AS1 | 94.14006 | -0.4717658 | 0.2335304 | -2.122469 | 0.03379838 | 0.102874 |
| ENSG00000267114 | | 8.585926 | -0.4718785 | 0.5113642 | -1.18175 | 0.2373049 | 0.426991 |
| ENSG00000270060 | | 62.85288 | -0.4719222 | 0.3073267 | -1.682899 | 0.09239458 | 0.219632 |
| ENSG00000215807 | KRT18P65 | 73.61911 | -0.4720277 | 0.3648491 | -1.46579 | 0.1427055 | 0.300296 |
| ENSG00000137878 | GCOM1 | 48.61237707 | -0.47214673 | 0.511738213 | -1.182809402 | 0.23688469 | 0.426524 |
| ENSG00000254180 | | 20.77202 | -0.4721587 | 0.4485677 | -1.267404 | 0.205011 | 0.38701 |
| ENSG00000131652 | THOC6 | 2300.006378 | -0.472166235 | 0.165621911 | -2.924828822 | 0.00344646 | 0.016305 |
| ENSG00000273901 | | 19.38165 | -0.4722885 | 0.5778815 | -1.127863 | 0.2593777 | 0.453691 |
| ENSG00000284154 | MIR3605 | 1.243765 | -0.472395 | 0.9299843 | -1.425054 | 0.1541416 | NA |
| ENSG00000055130 | CUL1 | 5432.540237 | -0.472461169 | 0.155061283 | -3.204315715 | 0.00135384 | 0.007438 |
| ENSG00000197037 | ZSCAN25 | 1202.616 | -0.472539 | 0.1969077 | -2.480969 | 0.01310259 | 0.048365 |
| ENSG00000261424 | ATP5PBP7 | 132.4103 | -0.4727868 | 0.2601382 | -1.934529 | 0.05304807 | 0.145263 |
| ENSG00000185829 | ARL17A | 140.3658 | -0.4728141 | 0.2799189 | -1.812262 | 0.0699458 | 0.178451 |
| ENSG00000135045 | C9orf40 | 699.7109262 | -0.472856058 | 0.325055569 | -1.596237523 | 0.11043577 | 0.2501 |
| ENSG00000261824 | LINC00662 | 1265.69 | -0.4730724 | 0.3571185 | -1.492942 | 0.1354524 | 0.288949 |
| ENSG00000148334 | PTGES2 | 4580.242 | -0.4730851 | 0.1974512 | -2.48726 | 0.01287314 | 0.047692 |
| ENSG00000099365 | STX1B | 114.9466504 | -0.473183563 | 0.393187782 | -1.386439827 | 0.16561264 | 0.334308 |
| ENSG00000180773 | SLC36A4 | 531.9544 | -0.4732902 | 0.3085838 | -1.687537 | 0.09150013 | 0.218229 |
| ENSG00000242802 | AP5Z1 | 6947.398 | -0.4732939 | 0.1838659 | -2.691541 | 0.00711228 | 0.02954 |
| ENSG00000145982 | FARS2 | 554.2716 | -0.4733025 | 0.2302027 | -2.167867 | 0.03016881 | 0.094079 |
| ENSG00000129250 | KIF1C | 13708.7157 | -0.473303706 | 0.137122636 | -3.431429567 | 0.00060041 | 0.003676 |
| ENSG00000181396 | OGFOD3 | 2457.421 | -0.4735134 | 0.172316 | -2.827402 | 0.00469274 | 0.021065 |
| ENSG00000143850 | PLEKHA6 | 9438.321 | -0.4735247 | 0.1845876 | -2.674999 | 0.00747295 | 0.030778 |
| ENSG00000145692 | BHMT | 3.451449 | -0.4736514 | 0.862515 | -1.325421 | 0.1850315 | 0.360581 |
| ENSG00000126457 | PRMT1 | 11930.5005 | -0.47373987 | 0.236106014 | -2.112368244 | 0.03465488 | 0.104868 |

| | | | | | | | |
|---|---|---|---|---|---|---|---|
| ENSG00000235043 | TECRP1 | 1109.594 | -0.4739415 | 0.2762877 | -1.838913 | 0.06592798 | 0.170659 |
| ENSG00000225493 | LINC01107 | 1.320202 | -0.473962 | 0.8776612 | -1.359001 | 0.1741463 | NA |
| ENSG00000139734 | DIAPH3 | 538.3775 | -0.4739946 | 0.5045622 | -1.206089 | 0.2277833 | 0.415715 |
| ENSG00000103148 | NPRL3 | 2963.046114 | -0.474008132 | 0.153345369 | -3.164416771 | 0.00155394 | 0.008356 |
| ENSG00000076351 | SLC46A1 | 474.6276458 | -0.474207977 | 0.247235936 | -2.025656714 | 0.04279998 | 0.1233 |
| ENSG00000235151 | | 6.022924 | -0.4742841 | 0.6749976 | -1.109268 | 0.2673145 | 0.463112 |
| ENSG00000175536 | LIPT2 | 275.8246 | -0.4742849 | 0.4058508 | -1.363792 | 0.1726331 | 0.343854 |
| ENSG00000134014 | ELP3 | 1141.765233 | -0.474355666 | 0.178562718 | -2.736716669 | 0.00620557 | 0.026444 |
| ENSG00000124535 | WRNIP1 | 4087.403692 | -0.474404732 | 0.137245438 | -3.521539566 | 0.00042905 | 0.002749 |
| ENSG00000236349 | SUCLG2P2 | 347.3072 | -0.4744436 | 0.2144437 | -2.314818 | 0.02062291 | 0.06964 |
| ENSG00000272853 | SUV39H2-DT | 96.14571 | -0.474508 | 0.2460316 | -2.0334 | 0.04201216 | 0.121474 |
| ENSG00000188626 | GOLGA8M | 1.339438 | -0.4746677 | 0.9480808 | -1.54804 | 0.1216128 | NA |
| ENSG00000273183 | | 27.02103 | -0.4747556 | 0.5307329 | -1.165565 | 0.2437905 | 0.435067 |
| ENSG00000157557 | ETS2 | 2651.383 | -0.4748224 | 0.2224391 | -2.231168 | 0.02566997 | 0.0828 |
| ENSG00000248313 | CYP4F27P | 2.518756 | -0.4748639 | 0.8591962 | -1.313723 | 0.1889396 | 0.366232 |
| ENSG00000151461 | UPF2 | 3992.045 | -0.4749389 | 0.1449981 | -3.345112 | 0.00082249 | 0.004836 |
| ENSG00000253710 | ALG11 | 596.4949 | -0.475039 | 0.2576943 | -1.957885 | 0.05024348 | 0.138982 |
| ENSG00000167196 | FBXO22 | 3551.56 | -0.4751718 | 0.1342062 | -3.597418 | 0.00032139 | 0.00215 |
| ENSG00000133313 | CNDP2 | 5550.814342 | -0.475212745 | 0.169134148 | -2.894114232 | 0.0038023 | 0.01768 |
| ENSG00000051382 | PIK3CB | 3245.338045 | -0.475283937 | 0.135468098 | -3.589965254 | 0.00033072 | 0.002206 |
| ENSG00000286163 | | 3.91996 | -0.4753061 | 0.7124286 | -1.121778 | 0.2619568 | 0.456718 |
| ENSG00000241651 | | 1.32981 | -0.4753565 | 0.8671218 | -1.314243 | 0.1887645 | NA |
| ENSG00000267221 | C17orf113 | 254.543 | -0.4753961 | 0.210433 | -2.352637 | 0.01864081 | 0.064204 |
| ENSG00000183137 | CEP57L1 | 426.3664 | -0.4754637 | 0.2451488 | -2.055987 | 0.03978381 | 0.116404 |
| ENSG00000278601 | | 5.619481 | -0.4758738 | 0.6723681 | -1.116367 | 0.2642649 | 0.459777 |
| ENSG00000136982 | DSCC1 | 922.3055405 | -0.475944565 | 0.495553382 | -1.218223386 | 0.2231391 | 0.4098 |
| ENSG00000139324 | TMTC3 | 1978.686 | -0.4759485 | 0.2044008 | -2.426387 | 0.01525001 | 0.054628 |
| ENSG00000213985 | | 351.0468 | -0.4759927 | 0.2563938 | -1.984423 | 0.04720873 | 0.132575 |
| ENSG00000197927 | NBEAP2 | 32.47872 | -0.4760041 | 0.3943426 | -1.394232 | 0.1632475 | 0.330942 |
| ENSG00000167711 | SERPINF2 | 11.32259 | -0.476078 | 0.5165565 | -1.188537 | 0.2346219 | 0.423771 |
| ENSG00000091009 | RBM27 | 1219.611501 | -0.476115267 | 0.173226039 | -2.826410126 | 0.0047073 | 0.021112 |
| ENSG00000252244 | RNU7-3P | 3.758492 | -0.4761289 | 0.7894534 | -1.184585 | 0.2361815 | 0.425704 |
| ENSG00000287935 | | 12.81999 | -0.476153 | 0.7020868 | -1.13 | 0.2584763 | 0.452699 |

| | | | | | | | |
|---|---|---|---|---|---|---|---|
| ENSG00000130699 | TAF4 | 2322.534734 | -0.47616585 | 0.161405712 | -3.029446299 | 0.00245003 | 0.012291 |
| ENSG00000218902 | PTMAP3 | 4.792709 | -0.4762055 | 0.6438601 | -1.120445 | 0.262524 | 0.457412 |
| ENSG00000214424 | FAM149B1P1 | 18.08549 | -0.4762827 | 0.3979131 | -1.387154 | 0.1653948 | 0.333944 |
| ENSG00000277053 | GTF2IP1 | 280.0386 | -0.4766983 | 0.1580753 | -3.078127 | 0.00208306 | 0.010728 |
| ENSG00000096717 | SIRT1 | 2030.620317 | -0.476720665 | 0.17107579 | -2.875220012 | 0.00403746 | 0.018594 |
| ENSG00000124467 | PSG8 | 5.26939363 | -0.47674599 | 0.85441039 | -1.263182908 | 0.20652345 | 0.388744 |
| ENSG00000078246 | TULP3 | 1463.669256 | -0.476775963 | 0.157310856 | -3.098841446 | 0.00194279 | 0.010099 |
| ENSG00000184985 | SORCS2 | 81.70574 | -0.4770614 | 0.5584612 | -1.146282 | 0.2516786 | 0.444452 |
| ENSG00000170469 | SPATA24 | 194.6333 | -0.4772811 | 0.2506747 | -1.998683 | 0.0456427 | 0.129436 |
| ENSG00000184178 | SCFD2 | 896.1332 | -0.4772845 | 0.2044452 | -2.460624 | 0.01386956 | 0.050583 |
| ENSG00000234785 | EEF1GP5 | 10.25751 | -0.4773305 | 0.6284574 | -1.119392 | 0.2629731 | 0.457949 |
| ENSG00000253641 | | 6.690712 | -0.4776604 | 0.5576266 | -1.150731 | 0.249843 | 0.442437 |
| ENSG00000124155 | PIGT | 6121.158763 | -0.477737174 | 0.151094614 | -3.22213364 | 0.0012724 | 0.007052 |
| ENSG00000117525 | F3 | 139.6139376 | -0.477776421 | 0.340073599 | -1.559108569 | 0.11897069 | 0.263455 |
| ENSG00000216360 | | 1.419402 | -0.4778835 | 0.8757223 | -1.375979 | 0.1688282 | NA |
| ENSG00000279058 | AGAP14P | 30.23713 | -0.4779765 | 0.3371946 | -1.570413 | 0.116319 | 0.259399 |
| ENSG00000115350 | POLE4 | 1939.078277 | -0.478055715 | 0.246917211 | -2.050900179 | 0.04027667 | 0.117525 |
| ENSG00000101849 | TBL1X | 5971.249151 | -0.478087652 | 0.17152382 | -2.865287524 | 0.00416631 | 0.019098 |
| ENSG00000240950 | | 275.9837 | -0.4782171 | 0.2293714 | -2.183827 | 0.02897495 | 0.091332 |
| ENSG00000250251 | PKD1P6 | 892.3347 | -0.4782938 | 0.3510491 | -1.520479 | 0.1283906 | 0.278216 |
| ENSG00000258645 | HSPE1P2 | 597.3814 | -0.4783367 | 0.2597805 | -1.967633 | 0.04911028 | 0.136672 |
| ENSG00000248671 | | 151.7258 | -0.4783709 | 0.2057931 | -2.42338 | 0.01537683 | 0.054979 |
| ENSG00000187942 | LDLRAD2 | 2.22515 | -0.4783968 | 0.9404632 | -1.506816 | 0.1318578 | 0.283762 |
| ENSG00000213700 | RPL17P50 | 5.086946 | -0.4784724 | 0.6213903 | -1.116193 | 0.2643395 | 0.459847 |
| ENSG00000124171 | PARD6B | 10186.50443 | -0.478529879 | 0.146027962 | -3.355508436 | 0.00079219 | 0.00468 |
| ENSG00000140995 | DEF8 | 5551.346 | -0.4786668 | 0.1370288 | -3.580865 | 0.00034246 | 0.002271 |
| ENSG00000230935 | RPS3P1 | 2.153909 | -0.4792076 | 0.873482 | -1.328172 | 0.1841211 | 0.359356 |
| ENSG00000103241 | FOXF1 | 2.157060755 | -0.479319095 | 0.931700871 | -1.414565912 | 0.15719581 | 0.322189 |
| ENSG00000171659 | GPR34 | 1.531277 | -0.4793353 | 0.8941192 | -1.40122 | 0.1611483 | NA |
| ENSG00000198720 | ANKRD13B | 1618.37 | -0.4794008 | 0.1607046 | -3.056411 | 0.00224004 | 0.011388 |
| ENSG00000175279 | CENPS | 74.14026 | -0.4794275 | 0.3952544 | -1.402464 | 0.1607766 | 0.327324 |
| ENSG00000231767 | RPS27AP5 | 653.6623 | -0.4795002 | 0.1806116 | -2.73757 | 0.00618949 | 0.026392 |
| ENSG00000176994 | SMCR8 | 4816.917 | -0.4795224 | 0.2599347 | -1.963732 | 0.04956117 | 0.13761 |

| | | | | | | | |
|---|---|---|---|---|---|---|---|
| ENSG00000266969 | | 6.458021 | -0.4795478 | 0.6998686 | -1.12585 | 0.260229 | 0.45475 |
| ENSG00000235204 | | 7.224889 | -0.4795769 | 0.6223672 | -1.111202 | 0.2664813 | 0.46212 |
| ENSG00000284629 | SCYGR1 | 3.639618 | -0.4796151 | 0.7670243 | -1.160135 | 0.2459938 | 0.437722 |
| ENSG00000262412 | | 34.90737 | -0.479665 | 0.3337392 | -1.587437 | 0.1124137 | 0.253184 |
| ENSG00000285836 | | 3.98635 | -0.4797118 | 0.8162715 | -1.207245 | 0.2273377 | 0.41513 |
| ENSG00000202415 | RN7SKP269 | 2.079984 | -0.4797277 | 0.7987058 | -1.195364 | 0.2319448 | 0.420336 |
| ENSG00000027847 | B4GALT7 | 1973.186995 | -0.479733477 | 0.121492723 | -3.99497351 | 6.47E-05 | 0.00052 |
| ENSG00000198848 | CES1 | 2.018513 | -0.4798249 | 1.008209 | -2.265625 | 0.02347437 | 0.077132 |
| ENSG00000177096 | PHETA2 | 13.18734 | -0.4798387 | 0.5306609 | -1.182246 | 0.2371081 | 0.426811 |
| ENSG00000165487 | MICU2 | 3691.95 | -0.4798818 | 0.1171987 | -4.140586 | 3.46E-05 | 0.000299 |
| ENSG00000163515 | RETNLB | 3.528526 | -0.479893 | 0.8841943 | -1.404491 | 0.1601728 | 0.326502 |
| ENSG00000230568 | SF3A3P1 | 50.14612 | -0.479965 | 0.2892006 | -1.797296 | 0.07228869 | 0.18297 |
| ENSG00000165973 | NELL1 | 2.262326 | -0.4801027 | 0.8558575 | -1.320371 | 0.186711 | 0.363077 |
| ENSG00000266933 | | 12.12717 | -0.4801169 | 0.6070318 | -1.124207 | 0.2609253 | 0.455608 |
| ENSG00000134684 | YARS1 | 11714.75719 | -0.48020835 | 0.272015301 | -1.89071208 | 0.05866279 | 0.156457 |
| ENSG00000169679 | BUB1 | 2146.856 | -0.4802339 | 0.5636168 | -1.17165 | 0.2413375 | 0.431907 |
| ENSG00000134215 | VAV3 | 1307.944025 | -0.480264106 | 0.411618975 | -1.375842245 | 0.16887048 | 0.338902 |
| ENSG00000085415 | SEH1L | 3074.490285 | -0.480348438 | 0.207696272 | -2.407572865 | 0.01605896 | 0.056905 |
| ENSG00000236484 | RRM2P2 | 1.342081 | -0.4803692 | 0.9913166 | -1.977841 | 0.04794659 | NA |
| ENSG00000141510 | TP53 | 5710.223 | -0.4804875 | 0.1760734 | -2.82796 | 0.00468456 | 0.021035 |
| ENSG00000089248 | ERP29 | 9472.182711 | -0.480519702 | 0.143449684 | -3.424844848 | 0.00061515 | 0.003757 |
| ENSG00000256166 | LINC02571 | 49.03828 | -0.4806371 | 0.5106474 | -1.200049 | 0.2301202 | 0.418399 |
| ENSG00000202533 | | 2.428956 | -0.4806423 | 0.8231552 | -1.198561 | 0.2306986 | 0.419164 |
| ENSG00000274425 | | 10.27998 | -0.4806573 | 0.5813283 | -1.146797 | 0.2514657 | 0.444228 |
| ENSG00000141401 | IMPA2 | 1573.096 | -0.4807031 | 0.2768756 | -1.870576 | 0.06140384 | 0.161846 |
| ENSG00000168078 | PBK | 1290.009 | -0.4807999 | 0.5866226 | -1.158398 | 0.2467018 | 0.438554 |
| ENSG00000263815 | RN7SL426P | 1.105383 | -0.4809844 | 0.9865278 | -1.884991 | 0.059431 | NA |
| ENSG00000182165 | TP53TG1 | 309.1479 | -0.4810863 | 0.2302865 | -2.195986 | 0.02809292 | 0.088931 |
| ENSG00000244687 | UBE2V1 | 287.6454 | -0.4811366 | 0.1733974 | -2.85655 | 0.00428273 | 0.019498 |
| ENSG00000133067 | LGR6 | 117.3870709 | -0.481223271 | 0.595347375 | -1.125805608 | 0.26024781 | 0.454753 |
| ENSG00000276434 | | 17.64711 | -0.4812416 | 0.56073 | -1.157772 | 0.2469572 | 0.438802 |
| ENSG00000164032 | H2AZ1 | 13807.52 | -0.4812523 | 0.2729293 | -1.889639 | 0.05880632 | 0.156791 |
| ENSG00000277373 | | 10.80082 | -0.4813417 | 0.678613 | -1.130868 | 0.2581105 | 0.452268 |

| | | | | | | | |
|---|---|---|---|---|---|---|---|
| ENSG00000224221 | RPS6P8 | 6.618086 | -0.4813523 | 0.6873982 | -1.130606 | 0.2582209 | 0.452402 |
| ENSG00000274964 | | 19.3007 | -0.4813692 | 0.4964278 | -1.226144 | 0.2201445 | 0.406351 |
| ENSG00000083457 | ITGAE | 977.3721811 | -0.481389325 | 0.301964823 | -1.742885247 | 0.08135366 | 0.199915 |
| ENSG00000228519 | RANBP1P1 | 819.5558 | -0.4814151 | 0.3450272 | -1.559516 | 0.1188744 | 0.263308 |
| ENSG00000156398 | SFXN2 | 1020.741 | -0.4815009 | 0.1975153 | -2.530548 | 0.01138846 | 0.043189 |
| ENSG00000137076 | TLN1 | 7640.994628 | -0.481568529 | 0.188142218 | -2.638375877 | 0.00833042 | 0.033601 |
| ENSG00000254333 | NDST1-AS1 | 2.623775 | -0.4816568 | 0.9044241 | -1.476487 | 0.1398133 | 0.295851 |
| ENSG00000078699 | CBFA2T2 | 2031.682302 | -0.481905055 | 0.226902025 | -2.229454028 | 0.02578371 | 0.083076 |
| ENSG00000267809 | NDUFV2P1 | 1078.65 | -0.4819211 | 0.1958187 | -2.55198 | 0.01071127 | 0.041149 |
| ENSG00000270021 | | 41.54478 | -0.4820068 | 0.464411 | -1.266772 | 0.205237 | 0.3872 |
| ENSG00000159131 | GART | 5392.923 | -0.4820184 | 0.1353029 | -3.60997 | 0.00030623 | 0.00206 |
| ENSG00000141577 | CEP131 | 2445.312 | -0.4820494 | 0.1863906 | -2.669322 | 0.00760045 | 0.031221 |
| ENSG00000277282 | IGHV1OR21-1 | 2.323189 | -0.4821476 | 0.9569963 | -1.604263 | 0.1086561 | 0.247101 |
| ENSG00000230964 | | 1.866455 | -0.4822155 | 0.8448339 | -1.278877 | 0.2009404 | 0.381614 |
| ENSG00000249494 | | 4.292236 | -0.4824241 | 0.6969279 | -1.122744 | 0.261546 | 0.456212 |
| ENSG00000250539 | KRT8P33 | 1677.937 | -0.4824902 | 0.216455 | -2.339487 | 0.01931023 | 0.066064 |
| ENSG00000111057 | KRT18 | 81607.78882 | -0.482510412 | 0.20806806 | -2.413601167 | 0.01579574 | 0.056152 |
| ENSG00000007376 | RPUSD1 | 4386.268547 | -0.482536241 | 0.234916703 | -2.150007167 | 0.03155465 | 0.097576 |
| ENSG00000111640 | GAPDH | 52772.79431 | -0.482650984 | 0.21490777 | -2.343404867 | 0.01910864 | 0.065519 |
| ENSG00000251880 | RNU7-75P | 7.252797 | -0.4828358 | 0.6369952 | -1.131277 | 0.2579387 | 0.452027 |
| ENSG00000209482 | SNORD83A | 41.64913 | -0.4830026 | 0.3547764 | -1.524972 | 0.127266 | 0.276524 |
| ENSG00000271779 | | 1.86618 | -0.4830878 | 0.905741 | -1.436666 | 0.1508129 | 0.312335 |
| ENSG00000111344 | RASAL1 | 384.4750996 | -0.483142545 | 0.23952091 | -2.126528901 | 0.03345925 | 0.102123 |
| ENSG00000048740 | CELF2 | 1.088295238 | -0.483189757 | 0.903837666 | -1.481858883 | 0.13837785 | NA |
| ENSG00000100350 | FOXRED2 | 3151.217318 | -0.483256216 | 0.205776179 | -2.446093421 | 0.01444136 | 0.052331 |
| ENSG00000136478 | TEX2 | 6618.078324 | -0.483394034 | 0.132743041 | -3.708916977 | 0.00020815 | 0.001469 |
| ENSG00000214264 | KCTD9P4 | 53.77593 | -0.4833973 | 0.4463168 | -1.307048 | 0.1911964 | 0.36921 |
| ENSG00000116668 | SWT1 | 280.3336303 | -0.483478281 | 0.283048607 | -1.841525588 | 0.06554457 | 0.169865 |
| ENSG00000224928 | KRT8P30 | 11.08931 | -0.4835796 | 0.5463965 | -1.163536 | 0.244612 | 0.436203 |
| ENSG00000130741 | EIF2S3 | 12396.60618 | -0.483582106 | 0.179185906 | -2.781807158 | 0.00540572 | 0.023611 |
| ENSG00000142733 | MAP3K6 | 1202.52 | -0.4836841 | 0.1835751 | -2.72931 | 0.0063467 | 0.026891 |
| ENSG00000146834 | MEPCE | 3796.24 | -0.483696 | 0.2235239 | -2.269409 | 0.02324346 | 0.076591 |
| ENSG00000127564 | PKMYT1 | 2581.290921 | -0.483738331 | 0.467339293 | -1.296456675 | 0.1948182 | 0.37398 |

| | | | | | | | |
|---|---|---|---|---|---|---|---|
| ENSG00000010361 | FUZ | 860.8333641 | -0.48379462 | 0.208849703 | -2.411277002 | 0.01589677 | 0.056437 |
| ENSG00000231381 | RNF2P1 | 6.855371 | -0.4838948 | 0.7189247 | -1.124062 | 0.2609867 | 0.45563 |
| ENSG00000226393 | IFNA20P | 4.389828 | -0.4839313 | 0.6511946 | -1.128387 | 0.2591567 | 0.453503 |
| ENSG00000212163 | SNORD91A | 2.461031 | -0.4839463 | 0.8075305 | -1.209837 | 0.2263413 | 0.413908 |
| ENSG00000188878 | FBF1 | 431.188 | -0.4839571 | 0.2471963 | -2.073179 | 0.03815566 | 0.112753 |
| ENSG00000229870 | RPL21P89 | 4.922797 | -0.4839882 | 0.7712719 | -1.17523 | 0.2399027 | 0.430093 |
| ENSG00000166822 | TMEM170A | 1777.287 | -0.4840151 | 0.1673325 | -2.966517 | 0.00301194 | 0.014606 |
| ENSG00000214140 | PRCD | 21.97208 | -0.4840201 | 0.5448905 | -1.176739 | 0.2392997 | 0.429389 |
| ENSG00000257122 | RRN3P3 | 352.4998 | -0.4842654 | 0.1823932 | -2.744241 | 0.00606511 | 0.025949 |
| ENSG00000280184 | | 13.59155 | -0.4842876 | 0.5982735 | -1.145277 | 0.2520945 | 0.444819 |
| ENSG00000223718 | | 1.480881 | -0.4843063 | 0.8546726 | -1.284641 | 0.1989176 | NA |
| ENSG00000167914 | GSDMA | 1.821624 | -0.484511 | 0.8834536 | -1.384725 | 0.1661367 | 0.33506 |
| ENSG00000268858 | | 849.9187 | -0.484512 | 0.1951611 | -2.569717 | 0.01017815 | 0.039461 |
| ENSG00000253981 | ALG1L13P | 39.4014 | -0.4845366 | 0.3766276 | -1.464164 | 0.1431491 | 0.300944 |
| ENSG00000108106 | UBE2S | 2251.516151 | -0.48471072 | 0.385868843 | -1.447777943 | 0.14767916 | 0.307692 |
| ENSG00000275807 | | 81.6657 | -0.4847603 | 0.2908767 | -1.788229 | 0.07373905 | 0.185915 |
| ENSG00000213683 | | 343.7193 | -0.4848127 | 0.5070857 | -1.216619 | 0.2237492 | 0.410495 |
| ENSG00000218359 | SUMO2P13 | 1.838245 | -0.4849666 | 0.8392998 | -1.258382 | 0.2082535 | 0.391003 |
| ENSG00000172803 | SNX32 | 75.12099 | -0.4849757 | 0.3280689 | -1.635774 | 0.101887 | 0.235819 |
| ENSG00000270933 | | 19.24728 | -0.4849833 | 0.5919811 | -1.142686 | 0.2531691 | 0.44611 |
| ENSG00000107625 | DDX50 | 2885.130055 | -0.485003484 | 0.186511757 | -2.687312711 | 0.00720295 | 0.029846 |
| ENSG00000260213 | CENPN-AS1 | 6.389454 | -0.4850043 | 0.6737327 | -1.135687 | 0.2560874 | 0.449731 |
| ENSG00000263862 | LINC01543 | 1.538503 | -0.4850257 | 0.9142616 | -1.494441 | 0.1350605 | NA |
| ENSG00000255135 | | 438.4548 | -0.4850291 | 0.2072523 | -2.436977 | 0.01481063 | 0.053393 |
| ENSG00000164754 | RAD21 | 18912.93 | -0.4850624 | 0.2177339 | -2.327394 | 0.01994428 | 0.067753 |
| ENSG00000162714 | ZNF496 | 6418.477 | -0.485207 | 0.1527322 | -3.243498 | 0.00118072 | 0.006593 |
| ENSG00000116489 | CAPZA1 | 9461.251611 | -0.485236575 | 0.127135226 | -3.873898112 | 0.00010711 | 0.000812 |
| ENSG00000134444 | RELCH | 2275.791411 | -0.485247942 | 0.152395277 | -3.254309585 | 0.00113668 | 0.006388 |
| ENSG00000101353 | MROH8 | 79.55938181 | -0.485304821 | 0.326172715 | -1.639029722 | 0.10120707 | 0.234804 |
| ENSG00000099219 | ERMP1 | 8161.785226 | -0.485430291 | 0.19329602 | -2.608564143 | 0.0090923 | 0.03604 |
| ENSG00000174231 | PRPF8 | 21391.66 | -0.4856696 | 0.14551 | -3.387124 | 0.0007063 | 0.004236 |
| ENSG00000185028 | LRRC14B | 1.292134 | -0.4858112 | 0.9275636 | -1.579745 | 0.1141653 | NA |
| ENSG00000060491 | OGFR | 7896.103055 | -0.485844711 | 0.215288443 | -2.356250124 | 0.01846048 | 0.06369 |

| | | | | | | | |
|---|---|---|---|---|---|---|---|
| ENSG00000273073 | | 36.20961 | -0.4858767 | 0.4185317 | -1.359367 | 0.1740304 | 0.345835 |
| ENSG00000243795 | LINC02044 | 2.074148 | -0.4861949 | 0.7776992 | -1.1624 | 0.2450729 | 0.436768 |
| ENSG00000267213 | | 51.13721 | -0.4862389 | 0.3386624 | -1.589137 | 0.1120294 | 0.25263 |
| ENSG00000174177 | CTU2 | 1863.456 | -0.4862552 | 0.153386 | -3.249522 | 0.00115599 | 0.006486 |
| ENSG00000164626 | KCNK5 | 376.5147 | -0.486262 | 0.3379725 | -1.594396 | 0.1108473 | 0.250712 |
| ENSG00000113460 | BRIX1 | 2999.843479 | -0.486371093 | 0.235625751 | -2.17512788 | 0.02962054 | 0.092871 |
| ENSG00000160201 | U2AF1 | 352.8488 | -0.4863753 | 0.2186452 | -2.327648 | 0.01993082 | 0.067753 |
| ENSG00000186226 | LCE1E | 1.680461 | -0.4865593 | 0.89182 | -1.387494 | 0.1652911 | 0.333862 |
| ENSG00000250130 | GAPDHP43 | 1.439934 | -0.4866302 | 0.945445 | -1.657176 | 0.09748396 | NA |
| ENSG00000286443 | | 12.98133 | -0.486777 | 0.4492812 | -1.310016 | 0.1901904 | 0.367958 |
| ENSG00000251550 | | 2.109681 | -0.486841 | 0.8819056 | -1.361339 | 0.1734067 | 0.34493 |
| ENSG00000214998 | CCNB2P1 | 3.002747 | -0.4870194 | 0.7574896 | -1.16733 | 0.2430771 | 0.4341 |
| ENSG00000163072 | NOSTRIN | 75.77593 | -0.487069 | 0.4835017 | -1.256852 | 0.2088071 | 0.391711 |
| ENSG00000244926 | ALKBH3-AS1 | 9.263046 | -0.4871078 | 0.6349556 | -1.127236 | 0.2596425 | 0.453994 |
| ENSG00000284747 | | 33.9724 | -0.4871729 | 0.3920405 | -1.425475 | 0.1540199 | 0.317344 |
| ENSG00000264563 | MIR4633 | 2.104308 | -0.4872221 | 0.8348875 | -1.262924 | 0.2066163 | 0.388847 |
| ENSG00000233184 | | 425.9758 | -0.4872953 | 0.2083489 | -2.435337 | 0.01487793 | 0.05357 |
| ENSG00000277440 | | 5.395572 | -0.487475 | 0.7586849 | -1.175551 | 0.2397743 | 0.429979 |
| ENSG00000287548 | | 5.821006 | -0.4875856 | 0.6276401 | -1.13726 | 0.2554295 | 0.448843 |
| ENSG00000166130 | IKBIP | 627.0201 | -0.4876042 | 0.1284568 | -3.854102 | 0.00011616 | 0.000875 |
| ENSG00000197265 | GTF2E2 | 1577.278 | -0.487662 | 0.171895 | -2.920669 | 0.0034928 | 0.016486 |
| ENSG00000225234 | TRAPPC12-AS1 | 30.07882 | -0.4876621 | 0.5448155 | -1.185771 | 0.2357127 | 0.42515 |
| ENSG00000127586 | CHTF18 | 3425.686855 | -0.487703633 | 0.326180911 | -1.650189339 | 0.09890422 | 0.230781 |
| ENSG00000198730 | CTR9 | 2572.301 | -0.4877334 | 0.1345619 | -3.687834 | 0.00022617 | 0.001581 |
| ENSG00000275216 | | 1.502227 | -0.4878563 | 0.9578557 | -1.562613 | 0.1181437 | NA |
| ENSG00000121211 | MND1 | 389.6221373 | -0.487870079 | 0.489551808 | -1.245278497 | 0.2130294 | 0.397722 |
| ENSG00000285230 | RALY-AS1 | 359.867 | -0.4879135 | 0.2355818 | -2.188763 | 0.02861405 | 0.090323 |
| ENSG00000225484 | NUTM2B-AS1 | 1004.363 | -0.4879141 | 0.1313249 | -3.772732 | 0.00016147 | 0.001174 |
| ENSG00000256745 | ARPC3P3 | 1808.978 | -0.487993 | 0.2068787 | -2.454145 | 0.01412199 | 0.05137 |
| ENSG00000229522 | LINC01523 | 0.9958342 | -0.4882688 | 0.8940947 | -1.395092 | 0.1629882 | NA |
| ENSG00000270195 | SLBP-DT | 35.78793 | -0.4882695 | 0.4641893 | -1.285977 | 0.198451 | 0.37851 |
| ENSG00000226864 | ATE1-AS1 | 4.2321 | -0.4884818 | 0.7881549 | -1.189964 | 0.2340607 | 0.423218 |
| ENSG00000131398 | KCNC3 | 3121.848489 | -0.488599509 | 0.248647142 | -2.086348459 | 0.03694707 | 0.109976 |

| | | | | | | | |
|---|---|---|---|---|---|---|---|
| ENSG00000010256 | UQCRC1 | 6417.726486 | -0.488713123 | 0.258471465 | -2.008084535 | 0.04463431 | 0.127282 |
| ENSG00000205277 | MUC12 | 9.665019 | -0.4888192 | 0.8723983 | -1.379839 | 0.1676363 | 0.337163 |
| ENSG00000229816 | DDX50P1 | 56.19798 | -0.4890276 | 0.3191654 | -1.695335 | 0.09001193 | 0.215686 |
| ENSG00000025434 | NR1H3 | 741.6958112 | -0.489106566 | 0.251893593 | -2.063631256 | 0.03905269 | 0.114752 |
| ENSG00000164362 | TERT | 67.82141 | -0.4891079 | 0.7356925 | -1.172177 | 0.2411261 | 0.431674 |
| ENSG00000219284 | | 0.9978591 | -0.4891826 | 0.9615493 | -1.614686 | 0.1063787 | NA |
| ENSG00000197763 | TXNRD3 | 835.8052 | -0.4892189 | 0.2361459 | -2.227048 | 0.02594407 | 0.083472 |
| ENSG00000251201 | TMED7-TICAM2 | 2.292062 | -0.4892394 | 0.8267958 | -1.23884 | 0.2154047 | 0.400483 |
| ENSG00000139438 | FAM222A | 399.6137 | -0.4892506 | 0.451446 | -1.317226 | 0.187763 | 0.364616 |
| ENSG00000106628 | POLD2 | 7557.886958 | -0.489358969 | 0.227448527 | -2.256696677 | 0.02402704 | 0.078618 |
| ENSG00000213676 | ATF6B | 5132.321 | -0.4894916 | 0.142594 | -3.502273 | 0.00046131 | 0.002928 |
| ENSG00000235587 | GAPDHP65 | 923.1302 | -0.4895405 | 0.2394892 | -2.162628 | 0.0305698 | 0.095134 |
| ENSG00000110395 | CBL | 2549.312968 | -0.489623786 | 0.292040696 | -1.817026882 | 0.069213 | 0.177006 |
| ENSG00000283654 | LMLN2 | 2.531697 | -0.4896415 | 0.7771301 | -1.189257 | 0.2343385 | 0.423489 |
| ENSG00000228305 | PRELID1P6 | 1082.461 | -0.4899093 | 0.2769438 | -1.902542 | 0.0571004 | 0.153165 |
| ENSG00000273344 | PAXIP1-DT | 1038.109 | -0.4900385 | 0.1906653 | -2.654728 | 0.00793724 | 0.032334 |
| ENSG00000270930 | GRAMD4P8 | 21.96019 | -0.490077 | 0.6217149 | -1.150885 | 0.2497795 | 0.442367 |
| ENSG00000136352 | NKX2-1 | 4.068454361 | -0.490100335 | 0.943593911 | -1.469514619 | 0.14169326 | 0.298734 |
| ENSG00000279880 | | 15.79865 | -0.4901191 | 0.4972135 | -1.239141 | 0.2152932 | 0.400317 |
| ENSG00000115364 | MRPL19 | 4101.210362 | -0.490307034 | 0.233083603 | -2.212054585 | 0.02696289 | 0.086072 |
| ENSG00000166920 | C15orf48 | 27.57552 | -0.4903258 | 0.5838256 | -1.160112 | 0.2460034 | 0.437722 |
| ENSG00000134480 | CCNH | 1447.332117 | -0.490369955 | 0.168866971 | -2.988200093 | 0.00280626 | 0.01381 |
| ENSG00000239763 | | 36.83366 | -0.4904089 | 0.4674455 | -1.286793 | 0.1981664 | 0.378162 |
| ENSG00000109265 | CRACD | 581.999161 | -0.490684841 | 0.428701392 | -1.361257957 | 0.17343218 | 0.34493 |
| ENSG00000261360 | | 71.84706 | -0.4906976 | 0.4344508 | -1.343674 | 0.1790538 | 0.352993 |
| ENSG00000261286 | ATP2C2-AS1 | 70.07914 | -0.490779 | 0.2583488 | -2.020134 | 0.04336952 | 0.124531 |
| ENSG00000004487 | KDM1A | 5668.883107 | -0.490782368 | 0.200201731 | -2.5516304 | 0.01072202 | 0.041173 |
| ENSG00000095319 | NUP188 | 6420.501158 | -0.49078749 | 0.225395691 | -2.274838994 | 0.02291559 | 0.075774 |
| ENSG00000101407 | TTI1 | 2028.000188 | -0.490963408 | 0.212558202 | -2.439678298 | 0.01470035 | 0.053067 |
| ENSG00000213698 | CAPZA1P3 | 23.38619 | -0.4910683 | 0.4194182 | -1.382095 | 0.1669425 | 0.336192 |
| ENSG00000246273 | SBF2-AS1 | 227.0714 | -0.4910873 | 0.2933496 | -1.817205 | 0.06918572 | 0.176975 |
| ENSG00000164985 | PSIP1 | 2765.87 | -0.4910939 | 0.1979057 | -2.570658 | 0.01015055 | 0.039371 |
| ENSG00000136448 | NMT1 | 6397.171859 | -0.491097056 | 0.147255698 | -3.380789562 | 0.00072278 | 0.004319 |

| | | | | | | | |
|---|---|---|---|---|---|---|---|
| ENSG00000251139 | | 4.285549 | -0.491128 | 0.7452687 | -1.147059 | 0.2513573 | 0.444096 |
| ENSG00000269487 | | 3.729688 | -0.4912419 | 0.728644 | -1.152672 | 0.2490451 | 0.44142 |
| ENSG00000258674 | | 20.13704 | -0.4913016 | 0.5341547 | -1.204299 | 0.2284741 | 0.416546 |
| ENSG00000106462 | EZH2 | 3880.761197 | -0.491372624 | 0.265436375 | -1.977572758 | 0.04797693 | 0.134249 |
| ENSG00000240216 | CPHL1P | 331.1586 | -0.4914446 | 0.4691118 | -1.280279 | 0.2004471 | 0.38097 |
| ENSG00000223509 | | 78.92368 | -0.4915067 | 0.2529846 | -2.066099 | 0.03881911 | 0.114248 |
| ENSG00000048342 | CC2D2A | 491.8787092 | -0.491546439 | 0.194561026 | -2.614770409 | 0.00892875 | 0.035518 |
| ENSG00000151694 | ADAM17 | 2088.908 | -0.4916838 | 0.1879915 | -2.707316 | 0.00678298 | 0.028399 |
| ENSG00000107731 | UNC5B | 345.6590824 | -0.49172596 | 0.391302757 | -1.456328187 | 0.14530192 | 0.304266 |
| ENSG00000254294 | IMPDH1P6 | 4.121266 | -0.4919615 | 0.7782311 | -1.19567 | 0.2318252 | 0.420205 |
| ENSG00000244733 | | 31.06849 | -0.4921412 | 0.3349017 | -1.623088 | 0.1045706 | 0.240341 |
| ENSG00000164306 | PRIMPOL | 405.5855 | -0.4921815 | 0.1403227 | -3.580444 | 0.00034301 | 0.002273 |
| ENSG00000240048 | DDX50P2 | 21.71193 | -0.4921977 | 0.4436776 | -1.332517 | 0.1826904 | 0.357685 |
| ENSG00000170175 | CHRNB1 | 321.1006 | -0.4924339 | 0.2629427 | -2.000393 | 0.04545786 | 0.128983 |
| ENSG00000231827 | | 5.598038 | -0.4926071 | 0.6675959 | -1.128275 | 0.259204 | 0.453528 |
| ENSG00000132677 | RHBG | 431.6438619 | -0.49263358 | 0.242346476 | -2.145366134 | 0.03192359 | 0.098442 |
| ENSG00000113734 | BNIP1 | 1032.077373 | -0.492679506 | 0.251173351 | -2.092256153 | 0.03641561 | 0.108769 |
| ENSG00000154874 | CCDC144B | 188.8908 | -0.4926935 | 0.5238226 | -1.221534 | 0.221884 | 0.408619 |
| ENSG00000260288 | | 10.79293 | -0.4927552 | 0.529189 | -1.223942 | 0.220974 | 0.407373 |
| ENSG00000246790 | | 3.750679 | -0.4927555 | 0.7513433 | -1.163506 | 0.2446245 | 0.436203 |
| ENSG00000161970 | RPL26 | 3176.157 | -0.492883 | 0.1731343 | -2.920395 | 0.00349589 | 0.016498 |
| ENSG00000277791 | PSMB3 | 3997.188 | -0.4930611 | 0.2541836 | -2.03978 | 0.04137223 | 0.120055 |
| ENSG00000232457 | SLC16A6P1 | 18.17648 | -0.4931013 | 0.4918382 | -1.259218 | 0.2079516 | 0.390575 |
| ENSG00000230497 | EEF1B2P8 | 1.801374 | -0.4931763 | 1.006062 | -2.142438 | 0.03215826 | 0.098936 |
| ENSG00000085760 | MTIF2 | 2657.503564 | -0.493255526 | 0.153737672 | -3.279232907 | 0.0010409 | 0.00593 |
| ENSG00000133328 | PLAAT2 | 21.93009566 | -0.49330239 | 0.553179827 | -1.196544596 | 0.23148411 | 0.419902 |
| ENSG00000252361 | RNU6-118P | 5.930402 | -0.4933665 | 0.6814095 | -1.149101 | 0.2505142 | 0.443078 |
| ENSG00000123200 | ZC3H13 | 5946.673282 | -0.493399306 | 0.177333342 | -2.863797588 | 0.00418595 | 0.019172 |
| ENSG00000228797 | FAM207BP | 497.3797 | -0.4935551 | 0.1987898 | -2.595429 | 0.00944731 | 0.037131 |
| ENSG00000164715 | LMTK2 | 2170.481 | -0.4935695 | 0.2447439 | -2.090637 | 0.03656058 | 0.109038 |
| ENSG00000235448 | LURAP1L-AS1 | 5.81911 | -0.4935858 | 0.7382213 | -1.173714 | 0.2405097 | 0.430803 |
| ENSG00000082805 | ERC1 | 1781.966343 | -0.493592195 | 0.20085416 | -2.551514599 | 0.01072558 | 0.04118 |
| ENSG00000227769 | | 2.47046 | -0.4936526 | 0.7874705 | -1.203116 | 0.2289316 | 0.417123 |

| | | | | | | | |
|---|---|---|---|---|---|---|---|
| ENSG00000258284 | POLR2KP1 | 2.869792 | -0.4936573 | 0.7942943 | -1.185555 | 0.2357981 | 0.42515 |
| ENSG00000086065 | CHMP5 | 3821.373893 | -0.493785733 | 0.158683967 | -3.210385364 | 0.00132557 | 0.007317 |
| ENSG00000015532 | XYLT2 | 3702.723191 | -0.493900458 | 0.110626162 | -4.510421489 | 6.47E-06 | 6.72E-05 |
| ENSG00000286370 | | 2.215613 | -0.4939994 | 0.8246778 | -1.244434 | 0.2133398 | 0.398106 |
| ENSG00000084754 | HADHA | 13375.74441 | -0.494108679 | 0.122244157 | -4.096095534 | 4.20E-05 | 0.000355 |
| ENSG00000235159 | | 21.03673 | -0.494252 | 0.441464 | -1.335096 | 0.1818449 | 0.356643 |
| ENSG00000242140 | | 5.994947 | -0.4945682 | 0.6167405 | -1.165288 | 0.2439024 | 0.435237 |
| ENSG00000123500 | COL10A1 | 3.745368667 | -0.494659202 | 0.762188275 | -1.165610637 | 0.24377191 | 0.435063 |
| ENSG00000106546 | AHR | 875.3389556 | -0.494889194 | 0.295879606 | -1.813580914 | 0.06974227 | 0.178086 |
| ENSG00000167858 | TEKT1 | 1.846529 | -0.4949932 | 0.9668966 | -1.730199 | 0.08359471 | 0.203989 |
| ENSG00000187416 | LHFPL3 | 1.116037 | -0.4950208 | 0.9333633 | -1.565942 | 0.1173623 | NA |
| ENSG00000173085 | COQ2 | 1215.335 | -0.4950873 | 0.3657831 | -1.535211 | 0.1247321 | 0.272713 |
| ENSG00000226284 | ARPC3P1 | 324.9201 | -0.4951889 | 0.2103401 | -2.444847 | 0.01449136 | 0.052441 |
| ENSG00000114395 | CYB561D2 | 673.2205668 | -0.495227418 | 0.227185875 | -2.285487665 | 0.02228425 | 0.074287 |
| ENSG00000231210 | COMETT | 0.9330049 | -0.4954715 | 0.9853537 | -1.776598 | 0.07563448 | NA |
| ENSG00000188817 | SNTN | 84.90669 | -0.4955431 | 0.2941028 | -1.820814 | 0.06863512 | 0.175901 |
| ENSG00000182057 | OGFRP1 | 298.6342 | -0.4956856 | 0.2182369 | -2.371435 | 0.01771918 | 0.061612 |
| ENSG00000207344 | SNORA22C | 2.113759 | -0.4957555 | 0.8405683 | -1.274682 | 0.2024219 | 0.383606 |
| ENSG00000170445 | HARS1 | 5211.779 | -0.4957908 | 0.2103456 | -2.452809 | 0.01417457 | 0.051526 |
| ENSG00000222009 | BTBD19 | 12.96056 | -0.4958471 | 0.7062492 | -1.161754 | 0.2453353 | 0.437005 |
| ENSG00000264745 | TTC39C-AS1 | 5.311912 | -0.4959537 | 0.5725197 | -1.181701 | 0.2373244 | 0.426997 |
| ENSG00000240877 | RN7SL521P | 13.81242 | -0.4959838 | 0.5114388 | -1.235541 | 0.2166293 | 0.402013 |
| ENSG00000270300 | PHACTR2P1 | 2.88501 | -0.4961044 | 0.7390266 | -1.165077 | 0.2439877 | 0.435272 |
| ENSG00000267092 | | 14.0806 | -0.4962104 | 0.4739529 | -1.289012 | 0.1973938 | 0.377369 |
| ENSG00000267287 | | 2.415896 | -0.4962793 | 0.751329 | -1.163096 | 0.2447908 | 0.436422 |
| ENSG00000228192 | | 48.85753 | -0.4962958 | 0.4621507 | -1.312517 | 0.1893457 | 0.366805 |
| ENSG00000167004 | PDIA3 | 12243.91 | -0.496356 | 0.1678228 | -3.065688 | 0.0021717 | 0.011108 |
| ENSG00000173334 | TRIB1 | 4002.259 | -0.4966119 | 0.4625791 | -1.298227 | 0.1942094 | 0.373125 |
| ENSG00000280417 | | 38.23022 | -0.4966447 | 0.337648 | -1.629764 | 0.1031514 | 0.238005 |
| ENSG00000278867 | | 13.0916 | -0.4969217 | 0.5310468 | -1.219264 | 0.2227439 | 0.409386 |
| ENSG00000112874 | NUDT12 | 1402.701077 | -0.497002692 | 0.219503703 | -2.384158627 | 0.01711822 | 0.059915 |
| ENSG00000273763 | | 183.6614 | -0.4970581 | 0.1695959 | -3.007297 | 0.00263582 | 0.01308 |
| ENSG00000269983 | | 7.608219 | -0.4974678 | 0.5321194 | -1.217161 | 0.2235431 | 0.41023 |

| | | | | | | | |
|---|---|---|---|---|---|---|---|
| ENSG00000106346 | USP42 | 1980.947677 | -0.497538442 | 0.171967817 | -2.97119741 | 0.00296641 | 0.014451 |
| ENSG00000201264 | SNORD73B | 1.615466 | -0.497569 | 0.8913326 | -1.427672 | 0.1533862 | 0.316259 |
| ENSG00000040531 | CTNS | 1315.643745 | -0.497628647 | 0.193432757 | -2.64996787 | 0.00804994 | 0.032695 |
| ENSG00000236846 | | 1.042365 | -0.4976302 | 1.015888 | -2.120661 | 0.03395038 | NA |
| ENSG00000243364 | EFNA4 | 1038.386 | -0.4976366 | 0.2023381 | -2.552998 | 0.01068002 | 0.041052 |
| ENSG00000172340 | SUCLG2 | 2357.524 | -0.4978227 | 0.169344 | -3.023437 | 0.00249921 | 0.012503 |
| ENSG00000106948 | AKNA | 725.383432 | -0.498124705 | 0.33110086 | -1.665025306 | 0.09590771 | 0.225621 |
| ENSG00000160172 | FAM86C2P | 671.0311 | -0.4981387 | 0.1597707 | -3.19474 | 0.00139957 | 0.007647 |
| ENSG00000227890 | PSMA2P3 | 27.94452 | -0.4981589 | 0.4655147 | -1.312927 | 0.1892074 | 0.36667 |
| ENSG00000226055 | PAICSP1 | 8.786682 | -0.498269 | 0.5776549 | -1.182642 | 0.2369511 | 0.426586 |
| ENSG00000272217 | | 16.23367 | -0.4983012 | 0.6437279 | -1.157097 | 0.2472328 | 0.439233 |
| ENSG00000175877 | TMEM270 | 13.28834 | -0.4983091 | 0.6234265 | -1.161167 | 0.245574 | 0.43728 |
| ENSG00000164252 | AGGF1 | 1391.389 | -0.4983157 | 0.180012 | -2.848127 | 0.00439774 | 0.019939 |
| ENSG00000253955 | | 29.43204 | -0.4985044 | 0.4135514 | -1.408452 | 0.1589973 | 0.324904 |
| ENSG00000116353 | MECR | 1429.923244 | -0.49856971 | 0.202008521 | -2.562174005 | 0.01040192 | 0.04017 |
| ENSG00000148481 | MINDY3 | 1549.008 | -0.4986075 | 0.1540984 | -3.307039 | 0.00094288 | 0.005431 |
| ENSG00000243389 | | 4.402652 | -0.4986125 | 0.7151073 | -1.167802 | 0.2428865 | 0.433882 |
| ENSG00000235872 | | 2.002125 | -0.498667 | 0.8573661 | -1.286908 | 0.1981263 | 0.378162 |
| ENSG00000229020 | AKR7A2P1 | 1.724617 | -0.4986744 | 0.9009728 | -1.43347 | 0.1517237 | 0.313749 |
| ENSG00000285130 | | 15.17462 | -0.4986861 | 0.6080173 | -1.167309 | 0.2430857 | 0.4341 |
| ENSG00000082146 | STRADB | 828.5965531 | -0.498719789 | 0.181883808 | -2.828281579 | 0.00467986 | 0.021017 |
| ENSG00000258777 | HIF1A-AS1 | 2.910204 | -0.4987945 | 0.8767434 | -1.346194 | 0.1782401 | 0.35178 |
| ENSG00000271976 | | 165.0107 | -0.4988323 | 0.3128414 | -1.746378 | 0.08074531 | 0.198715 |
| ENSG00000285856 | | 2.102948 | -0.4989059 | 0.9138161 | -1.494313 | 0.1350939 | 0.288473 |
| ENSG00000011376 | LARS2 | 1455.387705 | -0.498910077 | 0.251299722 | -2.108905788 | 0.03495271 | 0.105517 |
| ENSG00000164304 | CAGE1 | 5.898405 | -0.4989653 | 0.6534415 | -1.155267 | 0.2479813 | 0.440209 |
| ENSG00000256325 | | 4.64677 | -0.4989783 | 0.7491578 | -1.173918 | 0.2404277 | 0.430743 |
| ENSG00000160233 | LRRC3 | 416.4973 | -0.4991391 | 0.3111322 | -1.755514 | 0.07917142 | 0.195675 |
| ENSG00000284428 | | 99.92686 | -0.4991666 | 0.2950025 | -1.825731 | 0.06789079 | 0.174482 |
| ENSG00000074201 | CLNS1A | 7595.394418 | -0.499183358 | 0.121556959 | -4.157092268 | 3.22E-05 | 0.00028 |
| ENSG00000102145 | GATA1 | 2.642243672 | -0.499237632 | 0.823973619 | -1.262104426 | 0.20691121 | 0.389253 |
| ENSG00000214012 | KRT18P38 | 499.8272 | -0.4992438 | 0.2598315 | -2.06136 | 0.03926868 | 0.11526 |
| ENSG00000196591 | HDAC2 | 7292.47 | -0.4992599 | 0.1616741 | -3.160607 | 0.00157441 | 0.008447 |

| | | | | | | | |
|---|---|---|---|---|---|---|---|
| ENSG00000143256 | PFDN2 | 2329.063 | -0.4993035 | 0.2223647 | -2.355888 | 0.01847849 | 0.063727 |
| ENSG00000235609 | | 159.9204 | -0.4993517 | 0.4529557 | -1.336264 | 0.1814631 | 0.356104 |
| ENSG00000171097 | KYAT1 | 7.942167 | -0.4993769 | 0.5872168 | -1.179044 | 0.2383808 | 0.428202 |
| ENSG00000149262 | INTS4 | 1568.595 | -0.4994104 | 0.1296291 | -3.910549 | 9.21E-05 | 0.000711 |
| ENSG00000073111 | MCM2 | 4547.020171 | -0.499556712 | 0.463617519 | -1.314898143 | 0.18854414 | 0.365678 |
| ENSG00000182841 | RRP7BP | 388.3042 | -0.4996567 | 0.1631396 | -3.131604 | 0.00173854 | 0.009198 |
| ENSG00000179761 | PIPOX | 29.63694 | -0.4996788 | 0.4397922 | -1.361492 | 0.1733583 | 0.344897 |
| ENSG00000168014 | C2CD3 | 2016.175 | -0.4997998 | 0.1766865 | -2.922702 | 0.00347008 | 0.016392 |
| ENSG00000215086 | NPM1P24 | 729.3131 | -0.4998262 | 0.1943118 | -2.662461 | 0.00775716 | 0.031717 |
| ENSG00000110675 | ELMOD1 | 3.066610664 | -0.499857146 | 0.915887921 | -1.477623697 | 0.13950853 | 0.29537 |
| ENSG00000106305 | AIMP2 | 2163.241669 | -0.499938391 | 0.214810526 | -2.429438059 | 0.01512225 | 0.054273 |
| ENSG00000143486 | EIF2D | 7792.616 | -0.500028 | 0.1760367 | -2.925349 | 0.00344071 | 0.016287 |
| ENSG00000287338 | | 2.854133 | -0.5000823 | 0.7929965 | -1.219296 | 0.222732 | 0.409386 |
| ENSG00000248268 | | 10.44996 | -0.5001742 | 0.5403392 | -1.219092 | 0.2228094 | 0.409421 |
| ENSG00000128965 | CHAC1 | 1162.731954 | -0.500315441 | 0.631281448 | -1.180826024 | 0.23767184 | 0.427295 |
| ENSG00000133477 | FAM83F | 12.79691377 | -0.500493747 | 0.641225444 | -1.156058815 | 0.2476571 | 0.43984 |
| ENSG00000151005 | TKTL2 | 1.726063 | -0.5004947 | 0.9952035 | -1.902726 | 0.05707629 | 0.153147 |
| ENSG00000176624 | MEX3C | 2539.197 | -0.5005981 | 0.1734368 | -2.969668 | 0.00298122 | 0.014491 |
| ENSG00000165006 | UBAP1 | 2673.873 | -0.500713 | 0.1093752 | -4.64271 | 3.44E-06 | 3.81E-05 |
| ENSG00000138092 | CENPO | 1006.006724 | -0.500719958 | 0.434544165 | -1.371428157 | 0.17024153 | 0.340827 |
| ENSG00000165304 | MELK | 1380.321 | -0.5007356 | 0.6310835 | -1.177931 | 0.2388242 | 0.428738 |
| ENSG00000204220 | PFDN6 | 2230.549 | -0.5007375 | 0.1998593 | -2.601926 | 0.0092702 | 0.036592 |
| ENSG00000259349 | | 369.8566 | -0.500756 | 0.2668147 | -2.012665 | 0.04414986 | 0.126183 |
| ENSG00000111229 | ARPC3 | 6161.020227 | -0.501017389 | 0.186780384 | -2.78330255 | 0.00538086 | 0.023518 |
| ENSG00000101444 | AHCY | 13146.90128 | -0.501158635 | 0.195513193 | -2.649161073 | 0.00806919 | 0.032756 |
| ENSG00000171084 | FAM86JP | 423.8259 | -0.5012998 | 0.2835516 | -1.904363 | 0.05686295 | 0.152775 |
| ENSG00000090487 | SPG21 | 7571.693993 | -0.501362697 | 0.111612604 | -4.523846968 | 6.07E-06 | 6.35E-05 |
| ENSG00000233966 | UBE2SP1 | 361.1256 | -0.5014595 | 0.3770883 | -1.518003 | 0.1290138 | 0.279225 |
| ENSG00000255449 | | 3.249797 | -0.5017164 | 0.7837698 | -1.216049 | 0.2239664 | 0.41078 |
| ENSG00000270964 | | 103.9884 | -0.5017178 | 0.2887777 | -1.875107 | 0.06077796 | 0.160519 |
| ENSG00000110063 | DCPS | 693.8214327 | -0.501861332 | 0.21104627 | -2.476159009 | 0.01328044 | 0.048849 |
| ENSG00000205356 | TECPR1 | 837.7575 | -0.5023269 | 0.1947195 | -2.667413 | 0.00764377 | 0.03136 |
| ENSG00000174010 | KLHL15 | 455.6725 | -0.5023449 | 0.290375 | -1.86562 | 0.06209462 | 0.163078 |

| | | | | | | | |
|---|---|---|---|---|---|---|---|
| ENSG00000119421 | NDUFA8 | 2609.764601 | -0.502390301 | 0.20754815 | -2.530616615 | 0.01138622 | 0.043186 |
| ENSG00000128829 | EIF2AK4 | 3076.152826 | -0.502434278 | 0.143611797 | -3.565641858 | 0.00036297 | 0.002386 |
| ENSG00000234705 | HMGA1P4 | 35.52019 | -0.5025073 | 0.452422 | -1.340105 | 0.1802111 | 0.354485 |
| ENSG00000168140 | VASN | 817.1898 | -0.5025598 | 0.2388626 | -2.223634 | 0.02617311 | 0.084066 |
| ENSG00000223657 | KRT8P51 | 6.776339 | -0.5025892 | 0.6315109 | -1.158033 | 0.2468504 | 0.438715 |
| ENSG00000123009 | NME2P1 | 714.1802225 | -0.502664059 | 0.221269621 | -2.376865007 | 0.01746048 | 0.060936 |
| ENSG00000111206 | FOXM1 | 3038.151716 | -0.502691999 | 0.577600577 | -1.213318411 | 0.22500811 | 0.412022 |
| ENSG00000006062 | MAP3K14 | 914.8155896 | -0.502700988 | 0.259799677 | -2.058077717 | 0.03958267 | 0.115963 |
| ENSG00000232368 | FTLP2 | 27.11135 | -0.5027678 | 0.4027488 | -1.441406 | 0.14947 | 0.310425 |
| ENSG00000286632 | | 2.368736 | -0.5027923 | 0.9427553 | -1.40934 | 0.1587347 | 0.324542 |
| ENSG00000236778 | INTS6-AS1 | 78.08458 | -0.502797 | 0.5124031 | -1.262186 | 0.206882 | 0.389248 |
| ENSG00000169230 | PRELID1 | 3001.534 | -0.5028059 | 0.2553387 | -2.085572 | 0.03701745 | 0.110112 |
| ENSG00000127663 | KDM4B | 2853.473367 | -0.502902606 | 0.212683479 | -2.468071519 | 0.01358432 | 0.049708 |
| ENSG00000064309 | CDON | 557.0334271 | -0.502942739 | 0.430837088 | -1.390832003 | 0.16427638 | 0.332333 |
| ENSG00000041357 | PSMA4 | 11391.2056 | -0.503029068 | 0.244611037 | -2.17348401 | 0.02974391 | 0.093115 |
| ENSG00000241627 | UBQLN4P1 | 838.0136 | -0.5031586 | 0.1899228 | -2.739522 | 0.00615285 | 0.026261 |
| ENSG00000285467 | | 14.39244 | -0.5031766 | 0.5380513 | -1.234835 | 0.216892 | 0.40236 |
| ENSG00000228906 | | 18.34061 | -0.5032685 | 0.5577969 | -1.210336 | 0.22615 | 0.413672 |
| ENSG00000188985 | DHFRP1 | 966.7167 | -0.5034341 | 0.4086812 | -1.434982 | 0.1512923 | 0.313035 |
| ENSG00000263155 | MYZAP | 39.18735 | -0.5034792 | 0.6512825 | -1.172082 | 0.2411642 | 0.431684 |
| ENSG00000243403 | | 1.737406 | -0.5036549 | 0.8891274 | -1.378745 | 0.1679735 | 0.337649 |
| ENSG00000163166 | IWS1 | 3518.505 | -0.5036689 | 0.09835013 | -5.185803 | 2.15E-07 | 3.06E-06 |
| ENSG00000079385 | CEACAM1 | 108.886241 | -0.503743081 | 0.548260862 | -1.202394883 | 0.22921057 | 0.417287 |
| ENSG00000213221 | DNLZ | 11.2766 | -0.5038423 | 0.6648153 | -1.177627 | 0.2389453 | 0.428897 |
| ENSG00000239528 | RPS14P8 | 2.620191 | -0.5040872 | 0.7974148 | -1.229846 | 0.2187548 | 0.404601 |
| ENSG00000265458 | | 123.7244 | -0.5041112 | 0.3704724 | -1.546756 | 0.1219221 | 0.268269 |
| ENSG00000241837 | ATP5PO | 1443.546 | -0.5041497 | 0.2061905 | -2.538131 | 0.01114463 | 0.042452 |
| ENSG00000258806 | OR11H7 | 2.082414 | -0.5041839 | 0.8180114 | -1.247076 | 0.2123698 | 0.396853 |
| ENSG00000243181 | | 11.89881 | -0.5042638 | 0.5784823 | -1.198517 | 0.2307158 | 0.419166 |
| ENSG00000102221 | JADE3 | 2287.921576 | -0.504264159 | 0.155582795 | -3.319501503 | 0.00090178 | 0.005223 |
| ENSG00000170144 | HNRNPA3 | 13244.22 | -0.5045218 | 0.1225757 | -4.169717 | 3.05E-05 | 0.000266 |
| ENSG00000227345 | PARG | 2434.635 | -0.5046023 | 0.125765 | -4.073105 | 4.64E-05 | 0.000388 |
| ENSG00000236430 | KRT8P29 | 1.799272 | -0.5046795 | 0.8599743 | -1.294695 | 0.1954254 | 0.374838 |

| | | | | | | | |
|---|---|---|---|---|---|---|---|
| ENSG00000261395 | HSPE1P5 | 14.20893 | -0.5046842 | 0.5891934 | -1.196961 | 0.2313217 | 0.41975 |
| ENSG00000235286 | | 0.9959575 | -0.5047803 | 0.9929856 | -1.773865 | 0.07608547 | NA |
| ENSG00000196419 | XRCC6 | 15028.73 | -0.5048425 | 0.2891049 | -1.88701 | 0.05915895 | 0.157494 |
| ENSG00000272072 | | 20.32859 | -0.5048579 | 0.5700541 | -1.206111 | 0.2277747 | 0.415715 |
| ENSG00000137100 | DCTN3 | 2258.066118 | -0.504874179 | 0.181708311 | -2.870323593 | 0.00410052 | 0.018846 |
| ENSG00000171453 | POLR1C | 2053.359 | -0.504928 | 0.1402477 | -3.666329 | 0.00024606 | 0.001702 |
| ENSG00000219200 | RNASEK | 108.8646 | -0.504975 | 0.3018683 | -1.818662 | 0.06896301 | 0.176537 |
| ENSG00000143921 | ABCG8 | 2.879182 | -0.5050032 | 0.7270217 | -1.172732 | 0.2409034 | 0.431392 |
| ENSG00000090273 | NUDC | 4452.488912 | -0.505017618 | 0.20516004 | -2.603826486 | 0.00921894 | 0.036451 |
| ENSG00000011295 | TTC19 | 4348.110147 | -0.505033177 | 0.133658402 | -3.849212421 | 0.0001185 | 0.000891 |
| ENSG00000126368 | NR1D1 | 644.4470853 | -0.505183987 | 0.248389561 | -2.145690887 | 0.03189765 | 0.098389 |
| ENSG00000158158 | CNNM4 | 5982.314 | -0.5053715 | 0.1688294 | -3.084158 | 0.00204129 | 0.010541 |
| ENSG00000197766 | CFD | 303.9063 | -0.505399 | 0.2648273 | -2.037753 | 0.04157469 | 0.120537 |
| ENSG00000231307 | RPS3P2 | 4.147575 | -0.5054972 | 0.7461142 | -1.188796 | 0.23452 | 0.423702 |
| ENSG00000109576 | AADAT | 963.6028961 | -0.505616922 | 0.175470044 | -2.962262905 | 0.00305387 | 0.014755 |
| ENSG00000222365 | SNORD12B | 3.062499 | -0.5056182 | 0.8132154 | -1.218831 | 0.2229084 | 0.409547 |
| ENSG00000279086 | | 2.006096 | -0.5057897 | 0.801802 | -1.203303 | 0.2288589 | 0.417024 |
| ENSG00000167680 | SEMA6B | 320.9067 | -0.5063106 | 0.4343739 | -1.38406 | 0.16634 | 0.335343 |
| ENSG00000122026 | RPL21 | 841.6348736 | -0.506359107 | 0.213105839 | -2.476260781 | 0.01327666 | 0.048849 |
| ENSG00000152942 | RAD17 | 2478.343 | -0.506367 | 0.1241189 | -4.136691 | 3.52E-05 | 0.000303 |
| ENSG00000187595 | ZNF385C | 91.76233 | -0.5067054 | 0.4216494 | -1.421643 | 0.15513 | 0.318939 |
| ENSG00000188869 | TMC3 | 5.732763 | -0.506729 | 0.6715091 | -1.17706 | 0.2391715 | 0.429187 |
| ENSG00000274364 | | 1.807286 | -0.5068085 | 0.9215772 | -1.476691 | 0.1397584 | 0.295816 |
| ENSG00000287688 | | 20.33902 | -0.5068789 | 0.4751202 | -1.320451 | 0.1866845 | 0.363052 |
| ENSG00000213366 | GSTM2 | 197.8828 | -0.5069441 | 0.4347024 | -1.38433 | 0.1662574 | 0.335278 |
| ENSG00000100055 | CYTH4 | 1.567428852 | -0.507049306 | 0.966690169 | -1.587856665 | 0.11231876 | NA |
| ENSG00000261211 | | 109.7033 | -0.5072546 | 0.365504 | -1.562808 | 0.1180977 | 0.262193 |
| ENSG00000278236 | | 0.8718528 | -0.5073461 | 0.9692425 | -1.608475 | 0.1077312 | NA |
| ENSG00000008838 | MED24 | 4699.8013 | -0.50736938 | 0.15399847 | -3.367350324 | 0.00075894 | 0.004511 |
| ENSG00000213820 | RPL13P2 | 1.457282 | -0.507461 | 0.9682942 | -1.583726 | 0.1132562 | NA |
| ENSG00000287457 | | 7.252346 | -0.5075929 | 0.6691484 | -1.175281 | 0.2398823 | 0.430085 |
| ENSG00000159322 | ADPGK | 4644.698 | -0.5076421 | 0.08693382 | -5.871665 | 4.31E-09 | 8.42E-08 |
| ENSG00000268615 | | 17.34804 | -0.5077255 | 0.5043614 | -1.265214 | 0.2057946 | 0.387867 |

| | | | | | | | |
|---|---|---|---|---|---|---|---|
| ENSG00000242094 | FOXP1-IT1 | 1.843331 | -0.5078118 | 0.9501185 | -1.454608 | 0.1457779 | 0.304854 |
| ENSG00000215572 | ESRRAP1 | 4.838479 | -0.5081085 | 0.6665645 | -1.160257 | 0.2459444 | 0.437705 |
| ENSG00000168060 | NAALADL1 | 3.611209 | -0.5081114 | 0.7895806 | -1.213841 | 0.2248084 | 0.411866 |
| ENSG00000117013 | KCNQ4 | 401.2690952 | -0.508118402 | 0.294990462 | -1.877471868 | 0.06045345 | 0.159963 |
| ENSG00000036448 | MYOM2 | 3.565151938 | -0.508129888 | 0.782043703 | -1.215520972 | 0.22416746 | 0.410997 |
| ENSG00000102468 | HTR2A | 6.590359359 | -0.508131169 | 0.862994716 | -1.341172585 | 0.17986443 | 0.354039 |
| ENSG00000182180 | MRPS16 | 5236.602 | -0.5082706 | 0.1727969 | -3.02839 | 0.00245861 | 0.012327 |
| ENSG00000287644 | | 3.02296 | -0.5084941 | 0.7124886 | -1.19104 | 0.2336378 | 0.422654 |
| ENSG00000259891 | | 223.4216 | -0.508532 | 0.3069129 | -1.805823 | 0.07094606 | 0.180397 |
| ENSG00000172663 | TMEM134 | 2921.753 | -0.5086271 | 0.1708356 | -3.057257 | 0.00223373 | 0.011365 |
| ENSG00000147724 | FAM135B | 2.806104 | -0.5089747 | 0.9097846 | -1.449574 | 0.1471773 | 0.306959 |
| ENSG00000146263 | MMS22L | 1413.539 | -0.5090233 | 0.3419955 | -1.657413 | 0.09743596 | 0.228287 |
| ENSG00000159167 | STC1 | 742.6949 | -0.5090629 | 0.5106618 | -1.278324 | 0.2011353 | 0.381866 |
| ENSG00000152944 | MED21 | 1245.627 | -0.5091088 | 0.1938193 | -2.731334 | 0.00630786 | 0.026776 |
| ENSG00000260924 | LINC01311 | 168.4348 | -0.5091641 | 0.2881811 | -1.909708 | 0.05617083 | 0.151452 |
| ENSG00000259120 | SMIM6 | 7.740607 | -0.5092481 | 0.6082513 | -1.182169 | 0.2371385 | 0.426837 |
| ENSG00000198042 | MAK16 | 1170.381 | -0.5092996 | 0.1882139 | -2.798895 | 0.00512779 | 0.022609 |
| ENSG00000253372 | | 5.151443 | -0.5093326 | 0.5931599 | -1.197842 | 0.2309784 | 0.419471 |
| ENSG00000242999 | RN7SL239P | 3.507432 | -0.509408 | 0.7642926 | -1.20909 | 0.2266281 | 0.414261 |
| ENSG00000140326 | CDAN1 | 1251.851 | -0.5094583 | 0.1492694 | -3.485273 | 0.00049164 | 0.003095 |
| ENSG00000150510 | FAM124A | 30.21201 | -0.5094888 | 0.5341291 | -1.249396 | 0.2115204 | 0.395635 |
| ENSG00000241494 | | 140.0726 | -0.5095375 | 0.2760698 | -1.982708 | 0.04740009 | 0.133056 |
| ENSG00000146007 | ZMAT2 | 6502.108 | -0.5095669 | 0.1808005 | -2.936781 | 0.00331638 | 0.015802 |
| ENSG00000251002 | | 6.393106 | -0.5097056 | 0.9526032 | -1.454004 | 0.1459452 | 0.305084 |
| ENSG00000111652 | COPS7A | 3348.154433 | -0.50976487 | 0.131633531 | -3.941188556 | 8.11E-05 | 0.000635 |
| ENSG00000213593 | TMX2 | 3202.856 | -0.5097677 | 0.1274081 | -4.022057 | 5.77E-05 | 0.00047 |
| ENSG00000219470 | | 27.36973 | -0.5097928 | 0.3397779 | -1.665944 | 0.09572456 | 0.225351 |
| ENSG00000259826 | | 57.26009 | -0.5099025 | 0.3711588 | -1.554646 | 0.1200303 | 0.265403 |
| ENSG00000261888 | | 838.4188 | -0.5099548 | 0.3265007 | -1.686431 | 0.09171278 | 0.21852 |
| ENSG00000225630 | MTND2P28 | 1523.589 | -0.5099696 | 0.1808764 | -2.90729 | 0.00364575 | 0.017084 |
| ENSG00000108523 | RNF167 | 4963.230242 | -0.510371555 | 0.145186646 | -3.576947692 | 0.00034763 | 0.002301 |
| ENSG00000133026 | MYH10 | 4637.708476 | -0.510435996 | 0.184162698 | -2.859628537 | 0.00424138 | 0.019362 |
| ENSG00000253417 | LINC02159 | 3.098439 | -0.5104631 | 0.864333 | -1.330848 | 0.1832392 | 0.358345 |

| | | | | | | | |
|---|---|---|---|---|---|---|---|
| ENSG00000156304 | SCAF4 | 3382.993 | -0.5105375 | 0.1379525 | -3.765901 | 0.00016595 | 0.001203 |
| ENSG00000141279 | NPEPPS | 9318.83 | -0.5106423 | 0.1141443 | -4.501973 | 6.73E-06 | 6.94E-05 |
| ENSG00000200550 | RNU6-137P | 1.054854 | -0.5106647 | 0.9425036 | -1.608551 | 0.1077145 | NA |
| ENSG00000227373 | RABGAP1L-DT | 9.362235 | -0.51075 | 0.5830626 | -1.215895 | 0.224025 | 0.410803 |
| ENSG00000265261 | | 0.8567394 | -0.5108212 | 0.9500139 | -1.6126 | 0.1068315 | NA |
| ENSG00000267724 | | 7.042978 | -0.5110859 | 0.7042969 | -1.186068 | 0.2355956 | 0.425125 |
| ENSG00000279088 | | 830.8777 | -0.5111298 | 0.2383419 | -2.262493 | 0.02366696 | 0.077659 |
| ENSG00000148834 | GSTO1 | 2293.747 | -0.5114725 | 0.293214 | -1.889033 | 0.05888741 | 0.156961 |
| ENSG00000285665 | TOP1P1 | 440.8457 | -0.511507 | 0.1704512 | -3.086813 | 0.00202315 | 0.010462 |
| ENSG00000171617 | ENC1 | 6242.56 | -0.5116022 | 0.3060988 | -1.785851 | 0.07412339 | 0.186601 |
| ENSG00000287575 | | 2.897248 | -0.5116888 | 0.7143827 | -1.174981 | 0.2400024 | 0.430213 |
| ENSG00000200879 | SNORD14E | 30.26493 | -0.5117535 | 0.4857247 | -1.309154 | 0.1904821 | 0.368455 |
| ENSG00000253330 | | 13.32658 | -0.5118681 | 0.6505582 | -1.186937 | 0.2352523 | 0.424647 |
| ENSG00000178096 | BOLA1 | 2986.485 | -0.5121042 | 0.184353 | -2.870486 | 0.00409841 | 0.018843 |
| ENSG00000163918 | RFC4 | 1285.175 | -0.5123226 | 0.4194965 | -1.439139 | 0.1501113 | 0.311222 |
| ENSG00000260193 | LINC02846 | 45.94385 | -0.5124129 | 0.4108116 | -1.454835 | 0.1457149 | 0.30481 |
| ENSG00000261342 | | 27.7823 | -0.5124163 | 0.3762536 | -1.551583 | 0.1207621 | 0.2666 |
| ENSG00000141562 | NARF | 3293.563 | -0.5124935 | 0.2204813 | -2.458906 | 0.01393613 | 0.050798 |
| ENSG00000188460 | ACTBP11 | 4.401463 | -0.5126019 | 0.7424032 | -1.189083 | 0.2344069 | 0.423555 |
| ENSG00000073578 | SDHA | 1605.156366 | -0.51263796 | 0.229019506 | -2.348014951 | 0.01887376 | 0.064863 |
| ENSG00000174437 | ATP2A2 | 13178.91 | -0.5126929 | 0.1645379 | -3.19474 | 0.00139957 | 0.007647 |
| ENSG00000273783 | | 127.5419 | -0.5127031 | 0.3240936 | -1.740544 | 0.08176348 | 0.200737 |
| ENSG00000227624 | SNRPEP3 | 8.15212 | -0.5127689 | 0.6122359 | -1.197123 | 0.2312586 | 0.419681 |
| ENSG00000236680 | | 7.138055 | -0.512843 | 0.6118923 | -1.196473 | 0.2315121 | 0.419924 |
| ENSG00000132749 | TESMIN | 2054.088479 | -0.512930045 | 0.19287175 | -2.783331237 | 0.00538039 | 0.023518 |
| ENSG00000286207 | | 53.23152 | -0.5130577 | 0.4463189 | -1.37649 | 0.1686701 | 0.33856 |
| ENSG00000230750 | SDAD1P2 | 21.47261 | -0.513083 | 0.4139463 | -1.451741 | 0.1465736 | 0.306076 |
| ENSG00000204616 | TRIM31 | 40.2774 | -0.5132253 | 0.6128504 | -1.197938 | 0.2309411 | 0.419461 |
| ENSG00000234183 | LINC01952 | 2.838895 | -0.5132868 | 0.7874749 | -1.226652 | 0.2199534 | 0.406094 |
| ENSG00000277218 | | 13.79806 | -0.5133312 | 0.6112382 | -1.196219 | 0.2316111 | 0.420038 |
| ENSG00000124334 | IL9R | 2.923143903 | -0.513530081 | 0.841395534 | -1.281421021 | 0.20004582 | 0.380432 |
| ENSG00000128408 | RIBC2 | 118.9403706 | -0.513614738 | 0.477074252 | -1.329053885 | 0.18383019 | 0.359131 |
| ENSG00000254459 | | 1.827529 | -0.5136326 | 0.9178846 | -1.449491 | 0.1472005 | 0.306984 |

| | | | | | | | |
|---|---|---|---|---|---|---|---|
| ENSG00000236829 | | 203.6574 | -0.5136844 | 0.4870014 | -1.314079 | 0.1888197 | 0.366077 |
| ENSG00000179799 | OR7E22P | 4.21968 | -0.513737 | 0.7620005 | -1.226637 | 0.2199591 | 0.406094 |
| ENSG00000164591 | MYOZ3 | 1.585413 | -0.5138268 | 0.8551087 | -1.308146 | 0.1908237 | 0.368834 |
| ENSG00000172215 | CXCR6 | 14.50201 | -0.5138362 | 0.4872119 | -1.312416 | 0.1893799 | 0.366844 |
| ENSG00000287896 | | 8.15134 | -0.5138432 | 0.5922278 | -1.21233 | 0.2253862 | 0.412615 |
| ENSG00000223404 | LINC00397 | 1.32993 | -0.5139117 | 0.9823163 | -1.74789 | 0.08048311 | NA |
| ENSG00000267319 | SELENOKP1 | 16.42163 | -0.5139374 | 0.5023274 | -1.296052 | 0.1949576 | 0.37413 |
| ENSG00000189120 | SP6 | 1003.637 | -0.513941 | 0.4390535 | -1.397544 | 0.1622499 | 0.329498 |
| ENSG00000235368 | SAPCD2P2 | 3.68649 | -0.5139586 | 0.8416332 | -1.306183 | 0.1914905 | 0.369666 |
| ENSG00000244021 | | 3.172631 | -0.5140615 | 0.7962037 | -1.238818 | 0.2154129 | 0.400483 |
| ENSG00000158966 | CACHD1 | 326.0615 | -0.5140898 | 0.3022351 | -1.847299 | 0.06470378 | 0.168096 |
| ENSG00000234788 | HSPA8P3 | 2.997902 | -0.5141746 | 0.8180912 | -1.261753 | 0.2070376 | 0.389408 |
| ENSG00000259494 | MRPL46 | 441.5964 | -0.5143933 | 0.180703 | -2.935052 | 0.00333492 | 0.015871 |
| ENSG00000107443 | CCNJ | 1520.097878 | -0.514448473 | 0.178303924 | -2.965720544 | 0.00301975 | 0.014633 |
| ENSG00000253570 | RNF5P1 | 551.5222 | -0.514503 | 0.173157 | -3.050508 | 0.00228455 | 0.01157 |
| ENSG00000202347 | RNU1-16P | 1.663567 | -0.5145497 | 0.8984463 | -1.403664 | 0.1604188 | 0.3269 |
| ENSG00000233786 | CDC27P1 | 26.84326 | -0.5145903 | 0.3588756 | -1.611049 | 0.1071691 | 0.244746 |
| ENSG00000176422 | SPRYD4 | 1145.257 | -0.5146448 | 0.1510351 | -3.474647 | 0.00051153 | 0.003199 |
| ENSG00000274776 | | 2.605451 | -0.5149262 | 0.7941041 | -1.234732 | 0.2169304 | 0.402403 |
| ENSG00000168010 | ATG16L2 | 2555.052 | -0.5149761 | 0.2100321 | -2.561064 | 0.0104352 | 0.040275 |
| ENSG00000224046 | | 25.38449 | -0.5150278 | 0.4684534 | -1.343389 | 0.1791462 | 0.353021 |
| ENSG00000212607 | SNORA3B | 23.59131 | -0.5150369 | 0.466031 | -1.352111 | 0.1763399 | 0.349145 |
| ENSG00000093009 | CDC45 | 1723.407029 | -0.515427102 | 0.625524245 | -1.209237656 | 0.22657156 | 0.414197 |
| ENSG00000213434 | VTI1BP2 | 5.235161 | -0.5155734 | 0.589767 | -1.21441 | 0.2245912 | 0.411529 |
| ENSG00000131876 | SNRPA1 | 2436.76677 | -0.515637874 | 0.316536479 | -1.788728106 | 0.07365861 | 0.185748 |
| ENSG00000266066 | POLRMTP1 | 292.7433 | -0.5156436 | 0.1648664 | -3.214112 | 0.00130848 | 0.007234 |
| ENSG00000130347 | RTN4IP1 | 470.8888147 | -0.515644771 | 0.355887416 | -1.630982945 | 0.10289392 | 0.237576 |
| ENSG00000145041 | DCAF1 | 2132.779 | -0.5157419 | 0.1548317 | -3.409544 | 0.00065072 | 0.003945 |
| ENSG00000164850 | GPER1 | 1271.443 | -0.5157912 | 0.5730404 | -1.247972 | 0.2120414 | 0.396379 |
| ENSG00000122140 | MRPS2 | 3076.634444 | -0.515973064 | 0.260484124 | -2.096286701 | 0.03605677 | 0.108101 |
| ENSG00000188976 | NOC2L | 7743.159 | -0.5159956 | 0.2197905 | -2.463994 | 0.01373982 | 0.0502 |
| ENSG00000179542 | SLITRK4 | 263.332 | -0.5162037 | 0.3196279 | -1.816978 | 0.06922042 | 0.177008 |
| ENSG00000278983 | | 1.923052 | -0.5162485 | 0.8923849 | -1.420946 | 0.1553326 | 0.319239 |

| | | | | | | | |
|---|---|---|---|---|---|---|---|
| ENSG00000224786 | CETN4P | 1.297503 | -0.5162862 | 0.9804477 | -1.790237 | 0.0734158 | NA |
| ENSG00000178425 | NT5DC1 | 1925.846 | -0.5164241 | 0.1947499 | -2.76093 | 0.0057637 | 0.024933 |
| ENSG00000224985 | | 35.78036 | -0.5165061 | 0.3446828 | -1.668832 | 0.09515065 | 0.224377 |
| ENSG00000009790 | TRAF3IP3 | 4.030598925 | -0.516533492 | 0.641102933 | -1.185633977 | 0.23576688 | 0.42515 |
| ENSG00000285658 | | 2.253483 | -0.5165443 | 0.9124759 | -1.438328 | 0.150341 | 0.311607 |
| ENSG00000249031 | SUMO2P6 | 4.494227 | -0.5167197 | 0.7837843 | -1.221122 | 0.22204 | 0.408771 |
| ENSG00000087470 | DNM1L | 4029.779074 | -0.516813449 | 0.111928382 | -4.67240421 | 2.98E-06 | 3.36E-05 |
| ENSG00000286757 | | 10.50376 | -0.5168725 | 0.6189508 | -1.191496 | 0.2334588 | 0.422474 |
| ENSG00000137462 | TLR2 | 139.4430302 | -0.517024851 | 0.410528821 | -1.470127471 | 0.14152723 | 0.298479 |
| ENSG00000196497 | IPO4 | 105.5764 | -0.517231 | 0.2130636 | -2.533176 | 0.01130341 | 0.042946 |
| ENSG00000117153 | KLHL12 | 6188.861919 | -0.5174053 | 0.153963458 | -3.422501814 | 0.00062048 | 0.003785 |
| ENSG00000124532 | MRS2 | 3499.477855 | -0.517446997 | 0.177092469 | -3.036100459 | 0.0023966 | 0.012067 |
| ENSG00000251054 | | 1.273769 | -0.517447 | 0.998993 | -1.766076 | 0.07738312 | NA |
| ENSG00000275719 | | 27.40359 | -0.5178393 | 0.4275529 | -1.435963 | 0.1510128 | 0.312676 |
| ENSG00000274210 | RNVU1-27 | 9.705251 | -0.5178512 | 0.577322 | -1.224785 | 0.2206562 | 0.407013 |
| ENSG00000010932 | FMO1 | 7.173996386 | -0.517881165 | 0.569339488 | -1.235089814 | 0.21679708 | 0.40224 |
| ENSG00000208036 | MIR106B | 9.42876 | -0.5179194 | 0.530155 | -1.266399 | 0.2053701 | 0.387358 |
| ENSG00000145808 | ADAMTS19 | 1544.974 | -0.5180423 | 0.290725 | -1.933551 | 0.05316834 | 0.145482 |
| ENSG00000014138 | POLA2 | 439.2558381 | -0.51805304 | 0.477772535 | -1.337397328 | 0.18109298 | 0.355657 |
| ENSG00000198000 | NOL8 | 3117.768 | -0.5181094 | 0.1195017 | -4.383433 | 1.17E-05 | 0.000113 |
| ENSG00000144034 | TPRKB | 1310.021 | -0.5181559 | 0.2177547 | -2.491952 | 0.01270433 | 0.047193 |
| ENSG00000272161 | | 1.337024 | -0.5181641 | 0.9515839 | -1.577306 | 0.1147252 | NA |
| ENSG00000265683 | SYPL1P2 | 34.04018 | -0.5183797 | 0.4298315 | -1.430406 | 0.1526007 | 0.314957 |
| ENSG00000115204 | MPV17 | 2612.746423 | -0.518457975 | 0.15658473 | -3.395644792 | 0.00068467 | 0.004124 |
| ENSG00000007168 | PAFAH1B1 | 5250.447199 | -0.518850329 | 0.124628004 | -4.172389423 | 3.01E-05 | 0.000264 |
| ENSG00000167100 | SAMD14 | 22.60057 | -0.5189715 | 0.6146625 | -1.220016 | 0.2224588 | 0.409155 |
| ENSG00000161681 | SHANK1 | 11.69279 | -0.5190167 | 0.517969 | -1.281331 | 0.2000776 | 0.380438 |
| ENSG00000198951 | NAGA | 1458.453 | -0.51902 | 0.2225818 | -2.444022 | 0.01452456 | 0.052532 |
| ENSG00000007372 | PAX6 | 3.26107224 | -0.519349473 | 0.668396853 | -1.196826936 | 0.23137402 | 0.419817 |
| ENSG00000226221 | RPL26P19 | 4645.539 | -0.5194412 | 0.2045489 | -2.642823 | 0.0082218 | 0.033213 |
| ENSG00000161955 | TNFSF13 | 90.86282 | -0.5195965 | 0.2204548 | -2.463788 | 0.01374774 | 0.050213 |
| ENSG00000269388 | | 53.10737 | -0.5198535 | 0.3376995 | -1.68248 | 0.09247576 | 0.219786 |
| ENSG00000198642 | KLHL9 | 2056.978 | -0.5198612 | 0.2175993 | -2.498327 | 0.0124781 | 0.046478 |

| | | | | | | | |
|---|---|---|---|---|---|---|---|
| ENSG00000166451 | CENPN | 1586.87 | -0.5198687 | 0.4577389 | -1.407443 | 0.159296 | 0.325307 |
| ENSG00000138050 | THUMPD2 | 870.0552295 | -0.519888581 | 0.146691395 | -3.601130774 | 0.00031684 | 0.002124 |
| ENSG00000279605 | | 74.14953 | -0.5199669 | 0.2650044 | -2.092168 | 0.03642349 | 0.108771 |
| ENSG00000231203 | KRT8P10 | 810.8436 | -0.5199699 | 0.2270772 | -2.445692 | 0.01445746 | 0.052354 |
| ENSG00000197953 | AADACL2 | 2.661227 | -0.5199776 | 1.069624 | -2.815377 | 0.00487201 | 0.021723 |
| ENSG00000005007 | UPF1 | 5193.759321 | -0.520065471 | 0.154025992 | -3.47482664 | 0.00051118 | 0.003197 |
| ENSG00000235027 | | 125.1529 | -0.5201682 | 0.3443305 | -1.690506 | 0.0909312 | 0.21722 |
| ENSG00000251584 | | 1.512272 | -0.5202048 | 1.05419 | -2.398329 | 0.01647007 | NA |
| ENSG00000167080 | B4GALNT2 | 23.23043 | -0.5205146 | 0.4853981 | -1.330542 | 0.1833397 | 0.358463 |
| ENSG00000070610 | GBA2 | 3547.625462 | -0.52083686 | 0.168727069 | -3.185110738 | 0.00144699 | 0.00787 |
| ENSG00000203761 | MSTO2P | 474.8318 | -0.5209166 | 0.2222185 | -2.45264 | 0.01418121 | 0.051543 |
| ENSG00000254221 | PCDHGB1 | 134.6032 | -0.5209437 | 0.3916774 | -1.533908 | 0.1250524 | 0.273121 |
| ENSG00000138101 | DTNB | 897.6688924 | -0.520960329 | 0.160876121 | -3.318097517 | 0.00090633 | 0.005243 |
| ENSG00000213117 | MEMO1P2 | 13.81631 | -0.521056 | 0.5235417 | -1.284702 | 0.1988964 | 0.379013 |
| ENSG00000218502 | H2AZP3 | 33.67356 | -0.5211511 | 0.4322809 | -1.435539 | 0.1511337 | 0.312829 |
| ENSG00000272438 | | 34.30361 | -0.5212118 | 0.3525096 | -1.656703 | 0.09757957 | 0.228543 |
| ENSG00000131848 | ZSCAN5A | 216.9637309 | -0.521239095 | 0.268125675 | -2.079026712 | 0.0376149 | 0.111431 |
| ENSG00000117122 | MFAP2 | 304.1284047 | -0.521332859 | 0.508278807 | -1.312962088 | 0.18919572 | 0.36667 |
| ENSG00000197822 | OCLN | 2582.312 | -0.5215108 | 0.1701076 | -3.184551 | 0.00144979 | 0.00788 |
| ENSG00000110237 | ARHGEF17 | 4013.994588 | -0.521807279 | 0.133804449 | -3.969222433 | 7.21E-05 | 0.000572 |
| ENSG00000005882 | PDK2 | 1579.841724 | -0.521827608 | 0.197157269 | -2.742069636 | 0.00610534 | 0.026079 |
| ENSG00000144290 | SLC4A10 | 128.2663 | -0.521904 | 0.4957624 | -1.342142 | 0.1795499 | 0.353629 |
| ENSG00000122490 | SLC66A2 | 2172.69027 | -0.52196973 | 0.18255494 | -2.94817719 | 0.00319654 | 0.015311 |
| ENSG00000280181 | | 1.206615 | -0.5220365 | 1.015137 | -1.891844 | 0.05851181 | NA |
| ENSG00000117054 | ACADM | 2382.288272 | -0.522052268 | 0.134007942 | -3.970929689 | 7.16E-05 | 0.000569 |
| ENSG00000287584 | | 19.16161 | -0.522078 | 0.6426073 | -1.2034 | 0.2288215 | 0.417008 |
| ENSG00000037241 | RPL26L1 | 1893.137155 | -0.522137778 | 0.264696158 | -2.105603699 | 0.03523879 | 0.10614 |
| ENSG00000167775 | CD320 | 2098.184 | -0.5221704 | 0.2570428 | -2.155857 | 0.03109481 | 0.096469 |
| ENSG00000249839 | | 28.34755 | -0.5221726 | 0.443673 | -1.411744 | 0.1580254 | 0.323266 |
| ENSG00000185946 | RNPC3 | 783.8912 | -0.5223066 | 0.3050697 | -1.864948 | 0.06218874 | 0.16327 |
| ENSG00000069998 | HDHD5 | 4284.22084 | -0.522449585 | 0.200107026 | -2.713144451 | 0.00666481 | 0.02798 |
| ENSG00000122884 | P4HA1 | 2503.50858 | -0.5225728 | 0.220912507 | -2.479790937 | 0.01314594 | 0.048475 |
| ENSG00000197563 | PIGN | 1311.804 | -0.5226794 | 0.2203721 | -2.485764 | 0.01292737 | 0.047857 |

| | | | | | | | |
|---|---|---|---|---|---|---|---|
| ENSG00000200795 | RNU4-1 | 5.221631 | -0.5228617 | 0.760186 | -1.204562 | 0.2283725 | 0.416533 |
| ENSG00000071539 | TRIP13 | 2313.534299 | -0.522885972 | 0.581676956 | -1.251332794 | 0.21081309 | 0.394721 |
| ENSG00000174744 | BRMS1 | 3977.949 | -0.5230828 | 0.2162353 | -2.527463 | 0.01148899 | 0.043508 |
| ENSG00000236184 | TCEA1P4 | 5.889364 | -0.5231632 | 0.7381985 | -1.207878 | 0.227094 | 0.414941 |
| ENSG00000104731 | KLHDC4 | 4856.136921 | -0.523251001 | 0.133432203 | -3.994921778 | 6.47E-05 | 0.00052 |
| ENSG00000225356 | | 678.7224 | -0.5232577 | 0.1633459 | -3.302846 | 0.00095709 | 0.005503 |
| ENSG00000008256 | CYTH3 | 2001.286064 | -0.523401656 | 0.279233365 | -2.013414863 | 0.044071 | 0.126041 |
| ENSG00000250982 | GAPDHP35 | 10.18412 | -0.5234291 | 0.5930785 | -1.232038 | 0.2179351 | 0.403647 |
| ENSG00000235370 | DNM1P51 | 11.81719 | -0.5234772 | 0.7023411 | -1.215463 | 0.2241895 | 0.410997 |
| ENSG00000113368 | LMNB1 | 6858.494755 | -0.523606893 | 0.442719562 | -1.422048035 | 0.15501231 | 0.318796 |
| ENSG00000111962 | UST | 561.7141478 | -0.523638582 | 0.313045636 | -1.8308535 | 0.06712241 | 0.172992 |
| ENSG00000174788 | PCP2 | 47.46212 | -0.5237973 | 0.4208956 | -1.462563 | 0.1435869 | 0.301507 |
| ENSG00000204949 | FAM83A-AS1 | 10.1773 | -0.5242453 | 0.7741708 | -1.195039 | 0.2320718 | 0.420445 |
| ENSG00000203280 | KIAA1671-AS1 | 2.187031 | -0.5244433 | 0.8110876 | -1.269925 | 0.2041114 | 0.385844 |
| ENSG00000240764 | PCDHGC5 | 9.376176 | -0.5244632 | 0.5588534 | -1.25415 | 0.2097876 | 0.393161 |
| ENSG00000135521 | LTV1 | 1515.800346 | -0.524525018 | 0.181738656 | -2.976420954 | 0.00291634 | 0.014251 |
| ENSG00000124207 | CSE1L | 13883.80226 | -0.524804364 | 0.313024053 | -1.836327565 | 0.06630923 | 0.171428 |
| ENSG00000236539 | HNRNPA1P54 | 2.368499 | -0.5250535 | 0.9909443 | -1.686941 | 0.09161469 | 0.218384 |
| ENSG00000257524 | | 5.581654 | -0.5252353 | 0.6140659 | -1.220507 | 0.2222728 | 0.408955 |
| ENSG00000148719 | DNAJB12 | 3510.879 | -0.5253728 | 0.1188163 | -4.472306 | 7.74E-06 | 7.87E-05 |
| ENSG00000108826 | MRPL27 | 3776.855727 | -0.525472218 | 0.249031936 | -2.228619279 | 0.02583925 | 0.083225 |
| ENSG00000187045 | TMPRSS6 | 34.17081 | -0.5254768 | 0.4364645 | -1.439851 | 0.1499095 | 0.310973 |
| ENSG00000258244 | OSTF1P1 | 28.27317 | -0.5255546 | 0.3361734 | -1.734347 | 0.0828565 | 0.202757 |
| ENSG00000176349 | | 1.809964 | -0.5256764 | 0.9263481 | -1.470613 | 0.1413957 | 0.298296 |
| ENSG00000127995 | CASD1 | 1382.032282 | -0.525732604 | 0.217524305 | -2.531930553 | 0.01134365 | 0.043071 |
| ENSG00000148908 | RGS10 | 1302.547 | -0.5257453 | 0.2214893 | -2.490509 | 0.01275602 | 0.047301 |
| ENSG00000148158 | SNX30 | 2685.047 | -0.5258105 | 0.2504897 | -2.223502 | 0.02618198 | 0.084074 |
| ENSG00000137601 | NEK1 | 910.2332272 | -0.52601028 | 0.176209657 | -3.069796262 | 0.00214205 | 0.010987 |
| ENSG00000176714 | CCDC121 | 224.1917 | -0.5260314 | 0.2198513 | -2.501346 | 0.01237222 | 0.046136 |
| ENSG00000177427 | MIEF2 | 1340.354 | -0.5261445 | 0.1565503 | -3.44564 | 0.00056971 | 0.003514 |
| ENSG00000261886 | | 3.24077 | -0.5261698 | 0.7513856 | -1.233891 | 0.2172435 | 0.402844 |
| ENSG00000183032 | SLC25A21 | 87.8284 | -0.526173 | 0.3435897 | -1.709553 | 0.08734856 | 0.210806 |
| ENSG00000261310 | | 5.206726 | -0.5262957 | 0.6477496 | -1.209221 | 0.2265778 | 0.414197 |

| | | | | | | | |
|---|---|---|---|---|---|---|---|
| ENSG00000160408 | ST6GALNAC6 | 6.511016 | -0.5264503 | 0.5676832 | -1.251998 | 0.2105706 | 0.394434 |
| ENSG00000092094 | OSGEP | 2388.302144 | -0.526759159 | 0.170753842 | -3.166879167 | 0.00154084 | 0.008299 |
| ENSG00000213209 | | 2.167016 | -0.5269005 | 0.9783187 | -1.683248 | 0.09232703 | 0.21957 |
| ENSG00000221025 | MIR1250 | 1.167916 | -0.527056 | 1.034716 | -2.035674 | 0.04178307 | NA |
| ENSG00000170260 | ZNF212 | 1207.826 | -0.5273192 | 0.1537481 | -3.510556 | 0.00044717 | 0.002847 |
| ENSG00000235621 | LINC00494 | 58.41709 | -0.5273244 | 0.540394 | -1.290424 | 0.1969035 | 0.376885 |
| ENSG00000135052 | GOLM1 | 6912.739036 | -0.527393242 | 0.214736813 | -2.562589747 | 0.01038947 | 0.040133 |
| ENSG00000179562 | GCC1 | 1755.389 | -0.5274405 | 0.1879745 | -2.907775 | 0.0036401 | 0.017063 |
| ENSG00000038274 | MAT2B | 2963.547229 | -0.527451353 | 0.104981908 | -5.077599903 | 3.82E-07 | 5.16E-06 |
| ENSG00000217950 | NOC2LP2 | 122.7639 | -0.5274739 | 0.2842425 | -2.000385 | 0.0454587 | 0.128983 |
| ENSG00000152253 | SPC25 | 580.0422 | -0.5275229 | 0.6332606 | -1.240339 | 0.2148501 | 0.399745 |
| ENSG00000132768 | DPH2 | 2885.922301 | -0.527544241 | 0.1552756 | -3.458408681 | 0.00054338 | 0.003377 |
| ENSG00000279758 | | 2.812096 | -0.5277287 | 0.7871065 | -1.251534 | 0.2107399 | 0.394636 |
| ENSG00000136144 | RCBTB1 | 1216.84357 | -0.527900711 | 0.340618896 | -1.728700869 | 0.08386265 | 0.204527 |
| ENSG00000025039 | RRAGD | 1578.494123 | -0.527947214 | 0.490722135 | -1.341244043 | 0.17984123 | 0.35402 |
| ENSG00000214322 | CBX1P2 | 22.85875 | -0.5280039 | 0.3988466 | -1.536041 | 0.1245283 | 0.272424 |
| ENSG00000272707 | | 3.558062 | -0.5281291 | 0.8130089 | -1.260923 | 0.2073366 | 0.389778 |
| ENSG00000254285 | KRT8P3 | 10966.07 | -0.5281296 | 0.2299547 | -2.393803 | 0.01667469 | 0.058703 |
| ENSG00000196376 | SLC35F1 | 63.82096 | -0.5282759 | 0.3011355 | -1.90923 | 0.05623241 | 0.151587 |
| ENSG00000082397 | EPB41L3 | 14.76400578 | -0.528293186 | 0.695749391 | -1.204845098 | 0.22826311 | 0.41639 |
| ENSG00000127364 | TAS2R4 | 40.0893331 | -0.528308146 | 0.581136571 | -1.248268658 | 0.21193269 | 0.396231 |
| ENSG00000127914 | AKAP9 | 4322.336953 | -0.528434099 | 0.283548428 | -2.017320611 | 0.04366207 | 0.125155 |
| ENSG00000232454 | | 66.99841 | -0.528646 | 0.3016699 | -1.903961 | 0.05691523 | 0.152854 |
| ENSG00000255237 | | 11.18267 | -0.5286802 | 0.5756105 | -1.241468 | 0.214433 | 0.399332 |
| ENSG00000272655 | POLR2J4 | 255.4003 | -0.5287089 | 0.2549097 | -2.199164 | 0.02786628 | 0.088351 |
| ENSG00000183128 | CALHM3 | 3.517343 | -0.5287596 | 0.7386827 | -1.219613 | 0.2226115 | 0.409344 |
| ENSG00000235590 | GNAS-AS1 | 88.00148 | -0.5288745 | 0.3043912 | -1.889729 | 0.05879427 | 0.156775 |
| ENSG00000237190 | CDKN2AIPNL | 1309.444 | -0.5289443 | 0.1897795 | -2.888537 | 0.00387039 | 0.017962 |
| ENSG00000092098 | RNF31 | 130.228898 | -0.528954559 | 0.266840587 | -2.115457116 | 0.03439101 | 0.10432 |
| ENSG00000260942 | CAPN10-DT | 247.9148 | -0.5291306 | 0.2683299 | -2.112155 | 0.03467319 | 0.104912 |
| ENSG00000101349 | PAK5 | 7.019866478 | -0.529332918 | 0.704670961 | -1.226822778 | 0.21988921 | 0.406021 |
| ENSG00000005436 | GCFC2 | 895.1269308 | -0.529353586 | 0.196778465 | -2.788395556 | 0.00529698 | 0.023216 |
| ENSG00000238364 | RNU7-140P | 2.321781 | -0.5295321 | 0.9158052 | -1.442851 | 0.1490625 | 0.309748 |

| | | | | | | | |
|---|---|---|---|---|---|---|---|
| ENSG00000253785 | | 1385.099 | -0.5296688 | 0.1808387 | -3.020174 | 0.0025263 | 0.012617 |
| ENSG00000133265 | HSPBP1 | 2638.982141 | -0.529749432 | 0.260311643 | -2.177680064 | 0.02942986 | 0.092393 |
| ENSG00000176855 | KRT18P28 | 224.4885 | -0.5299481 | 0.2413379 | -2.321457 | 0.02026217 | 0.068702 |
| ENSG00000137996 | RTCA | 1109.379204 | -0.530255232 | 0.167190898 | -3.252905509 | 0.00114232 | 0.006415 |
| ENSG00000082014 | SMARCD3 | 1004.84447 | -0.530279576 | 0.243520871 | -2.297807532 | 0.02157275 | 0.072306 |
| ENSG00000124782 | RREB1 | 2977.256309 | -0.530279759 | 0.216772962 | -2.560149061 | 0.01046273 | 0.040352 |
| ENSG00000288066 | | 286.1427 | -0.530322 | 0.3269221 | -1.799489 | 0.07194139 | 0.182311 |
| ENSG00000099800 | TIMM13 | 1608.360742 | -0.530375311 | 0.200483513 | -2.751406096 | 0.005934 | 0.025511 |
| ENSG00000261740 | BOLA2-SMG1P6 | 13.50129 | -0.5304036 | 0.4879513 | -1.35179 | 0.1764424 | 0.349192 |
| ENSG00000087903 | RFX2 | 390.289661 | -0.530471056 | 0.217429961 | -2.558804195 | 0.01050329 | 0.040497 |
| ENSG00000105767 | CADM4 | 638.6904655 | -0.530486281 | 0.285757854 | -1.973779926 | 0.04840677 | 0.135168 |
| ENSG00000129195 | PIMREG | 803.6123025 | -0.530602597 | 0.664848686 | -1.238415747 | 0.21556195 | 0.400704 |
| ENSG00000271754 | | 8.803287 | -0.5307415 | 0.5381675 | -1.29017 | 0.1969916 | 0.376972 |
| ENSG00000135622 | SEMA4F | 629.6735426 | -0.530852113 | 0.162258983 | -3.354367588 | 0.00079547 | 0.004697 |
| ENSG00000214331 | | 416.4457 | -0.5309062 | 0.2816659 | -2.041588 | 0.0411924 | 0.119742 |
| ENSG00000167306 | MYO5B | 4539.608 | -0.5310139 | 0.2615568 | -2.162604 | 0.03057164 | 0.095134 |
| ENSG00000180376 | CCDC66 | 686.9299 | -0.5310935 | 0.1692734 | -3.221845 | 0.00127368 | 0.007056 |
| ENSG00000288559 | | 196.0328 | -0.5312228 | 0.2874964 | -1.995072 | 0.04603504 | 0.130193 |
| ENSG00000065615 | CYB5R4 | 1220.658815 | -0.53153425 | 0.20291342 | -2.73077303 | 0.0063186 | 0.026801 |
| ENSG00000172062 | SMN1 | 574.0414 | -0.5315483 | 0.2140712 | -2.590414 | 0.00958605 | 0.037543 |
| ENSG00000112685 | EXOC2 | 1988.954123 | -0.531579276 | 0.095021845 | -5.647814462 | 1.63E-08 | 2.85E-07 |
| ENSG00000281097 | LINC01395 | 6.281726 | -0.5315854 | 0.6831782 | -1.212847 | 0.2251884 | 0.412282 |
| ENSG00000256849 | TCP1P3 | 15.41888 | -0.531602 | 0.4913559 | -1.357302 | 0.1746852 | 0.346566 |
| ENSG00000273618 | | 1.916953 | -0.5317835 | 0.88335 | -1.343917 | 0.1789753 | 0.35289 |
| ENSG00000108384 | RAD51C | 7465.213015 | -0.531837174 | 0.308498897 | -1.882300033 | 0.05979529 | 0.158632 |
| ENSG00000181610 | MRPS23 | 8216.109 | -0.5322173 | 0.2106709 | -2.630783 | 0.00851884 | 0.034199 |
| ENSG00000230291 | | 98.47868 | -0.53235 | 0.3345838 | -1.764502 | 0.07764754 | 0.192859 |
| ENSG00000235095 | | 3.253814 | -0.5324385 | 0.7917941 | -1.257337 | 0.2086317 | 0.391565 |
| ENSG00000072849 | DERL2 | 1769.827208 | -0.532472553 | 0.145712778 | -3.728641326 | 0.00019252 | 0.001371 |
| ENSG00000278126 | | 7.764718 | -0.5324732 | 0.6644354 | -1.221502 | 0.221896 | 0.408619 |
| ENSG00000170606 | HSPA4 | 12516.83 | -0.5327915 | 0.2043511 | -2.756705 | 0.00583869 | 0.025195 |
| ENSG00000234028 | EIF2AK3-DT | 157.694 | -0.5327929 | 0.209116 | -2.6548 | 0.00793555 | 0.032333 |
| ENSG00000166794 | PPIB | 5827.948 | -0.5328796 | 0.1773963 | -3.093352 | 0.00197909 | 0.010262 |

| | | | | | | | |
|---|---|---|---|---|---|---|---|
| ENSG00000155530 | LRGUK | 21.67366 | -0.5329788 | 0.5959769 | -1.250567 | 0.2110924 | 0.394966 |
| ENSG00000231574 | LINC02015 | 38.44464 | -0.53324 | 0.6750949 | -1.215772 | 0.2240718 | 0.41086 |
| ENSG00000187522 | HSPA14 | 1779.679 | -0.5332535 | 0.3146378 | -1.857758 | 0.0632034 | 0.165216 |
| ENSG00000244716 | | 2899.245 | -0.5333002 | 0.1780437 | -3.091091 | 0.00199422 | 0.010326 |
| ENSG00000128050 | PAICS | 12591.66134 | -0.533327192 | 0.226544841 | -2.438500908 | 0.01474832 | 0.05319 |
| ENSG00000250197 | HMGN1P15 | 1.715929 | -0.5334214 | 0.8413853 | -1.290626 | 0.1968333 | 0.376804 |
| ENSG00000174501 | ANKRD36C | 411.1995 | -0.5335982 | 0.4553005 | -1.41634 | 0.1566761 | 0.321347 |
| ENSG00000213786 | NHP2P2 | 3.311366 | -0.5336654 | 0.818226 | -1.293824 | 0.1957261 | 0.375334 |
| ENSG00000235914 | MACROD2-AS1 | 0.9312952 | -0.5339656 | 0.9893684 | -1.640096 | 0.1009851 | NA |
| ENSG00000012223 | LTF | 2.037453744 | -0.533969446 | 0.997821233 | -1.824484118 | 0.06807892 | 0.174847 |
| ENSG00000224094 | RPS24P8 | 610.8588 | -0.5342035 | 0.2033077 | -2.734879 | 0.00624032 | 0.026558 |
| ENSG00000270951 | | 8.925199 | -0.5342376 | 0.7370789 | -1.236428 | 0.2162994 | 0.401597 |
| ENSG00000135164 | DMTF1 | 3655.61178 | -0.534787031 | 0.186722057 | -2.978325874 | 0.00289828 | 0.014184 |
| ENSG00000214089 | RPL9P21 | 2.325099 | -0.5350744 | 0.8482536 | -1.337091 | 0.181193 | 0.355758 |
| ENSG00000141977 | CIB3 | 15.39941 | -0.5351727 | 0.6330114 | -1.242996 | 0.2138695 | 0.398728 |
| ENSG00000161671 | EMC10 | 4810.369 | -0.5352034 | 0.1874376 | -2.95001 | 0.00317763 | 0.015239 |
| ENSG00000115112 | TFCP2L1 | 1.145320925 | -0.535482565 | 1.030590654 | -1.921835173 | 0.0546265 | NA |
| ENSG00000171793 | CTPS1 | 6068.963 | -0.5355439 | 0.2356958 | -2.413694 | 0.01579171 | 0.056145 |
| ENSG00000163644 | PPM1K | 1114.312 | -0.5355754 | 0.2838635 | -2.032917 | 0.04206087 | 0.121575 |
| ENSG00000115875 | SRSF7 | 10995.76541 | -0.535772271 | 0.143528996 | -3.786465905 | 0.00015281 | 0.001116 |
| ENSG00000228363 | | 52.3848 | -0.535943 | 0.4026939 | -1.544927 | 0.1223638 | 0.269074 |
| ENSG00000270061 | | 11.75464 | -0.5360182 | 0.648202 | -1.231917 | 0.2179803 | 0.403683 |
| ENSG00000119915 | ELOVL3 | 17.27635748 | -0.536023904 | 0.648145777 | -1.236917123 | 0.21611785 | 0.401401 |
| ENSG00000106009 | BRAT1 | 6996.8305 | -0.536211127 | 0.17071017 | -3.227763803 | 0.00124762 | 0.006929 |
| ENSG00000106571 | GLI3 | 1056.234247 | -0.536272977 | 0.243239103 | -2.327364457 | 0.01994588 | 0.067753 |
| ENSG00000226774 | HIKESHIP1 | 1.032591 | -0.5362906 | 1.007447 | -1.748661 | 0.08034963 | NA |
| ENSG00000198522 | GPN1 | 2295.741 | -0.5367286 | 0.1799372 | -3.106081 | 0.00189585 | 0.009889 |
| ENSG00000278535 | DHRS11 | 549.1141 | -0.5367356 | 0.2892324 | -2.001719 | 0.04531494 | 0.128671 |
| ENSG00000178075 | GRAMD1C | 131.3533 | -0.5368582 | 0.3729061 | -1.640502 | 0.1009008 | 0.234339 |
| ENSG00000207445 | SNORD15B | 48.12366 | -0.5368984 | 0.320331 | -1.844408 | 0.06512374 | 0.168955 |
| ENSG00000236264 | RPL26P30 | 520.164 | -0.5370297 | 0.2731007 | -2.120943 | 0.03392659 | 0.103205 |
| ENSG00000116161 | CACYBP | 6141.41572 | -0.537064782 | 0.260984994 | -2.17796164 | 0.02940889 | 0.09235 |
| ENSG00000172531 | PPP1CA | 12635.55 | -0.5371965 | 0.2398699 | -2.361833 | 0.01818485 | 0.062984 |

| | | | | | | | |
|---|---|---|---|---|---|---|---|
| ENSG00000226081 | USP12PX | 34.83651 | -0.5373077 | 0.4721814 | -1.399935 | 0.1615327 | 0.328569 |
| ENSG00000165215 | CLDN3 | 8591.517 | -0.5373453 | 0.2216184 | -2.505005 | 0.01224495 | 0.045758 |
| ENSG00000256361 | | 4.207245 | -0.5373644 | 0.7883696 | -1.260046 | 0.2076529 | 0.390289 |
| ENSG00000266289 | | 4.364642 | -0.5373849 | 0.8395922 | -1.268217 | 0.2047205 | 0.386638 |
| ENSG00000185480 | PARPBP | 900.1018 | -0.5373943 | 0.4170823 | -1.511735 | 0.1306013 | 0.281743 |
| ENSG00000099284 | MACROH2A2 | 2047.647541 | -0.537426302 | 0.155309279 | -3.529466341 | 0.0004164 | 0.002681 |
| ENSG00000198763 | MT-ND2 | 67562.11 | -0.5374494 | 0.164373 | -3.341266 | 0.00083397 | 0.004889 |
| ENSG00000133641 | C12orf29 | 968.875941 | -0.537453429 | 0.174636298 | -3.189774393 | 0.00142384 | 0.007758 |
| ENSG00000105875 | WDR91 | 958.9599863 | -0.537582712 | 0.147091081 | -3.728015211 | 0.00019299 | 0.001373 |
| ENSG00000270591 | | 4.595705 | -0.5376504 | 0.7144468 | -1.227859 | 0.2194999 | 0.405584 |
| ENSG00000136444 | RSAD1 | 5813.487983 | -0.537762326 | 0.171851589 | -3.219127686 | 0.00128581 | 0.007122 |
| ENSG00000279081 | | 1.414638 | -0.5379435 | 0.9562155 | -1.616888 | 0.1059025 | NA |
| ENSG00000244582 | RPL21P120 | 4.672427 | -0.5382657 | 0.6813843 | -1.219569 | 0.2226285 | 0.409344 |
| ENSG00000233355 | CHRM3-AS2 | 1.221897 | -0.5383488 | 0.9533363 | -1.604948 | 0.1085052 | NA |
| ENSG00000160180 | TFF3 | 1189.255 | -0.5385958 | 0.7305978 | -1.261606 | 0.2070906 | 0.389453 |
| ENSG00000227344 | HAUS6P1 | 96.16363 | -0.5387599 | 0.288959 | -2.01819 | 0.04357146 | 0.124962 |
| ENSG00000139372 | TDG | 4236.413 | -0.5387935 | 0.2041584 | -2.745372 | 0.00604424 | 0.025878 |
| ENSG00000106526 | ACTR3C | 231.8123266 | -0.538810884 | 0.216360923 | -2.601581642 | 0.0092795 | 0.036623 |
| ENSG00000287654 | | 3.047423 | -0.5390564 | 0.7959742 | -1.276353 | 0.2018308 | 0.382813 |
| ENSG00000228010 | | 19.9242 | -0.5390628 | 0.5854884 | -1.272591 | 0.2031632 | 0.384736 |
| ENSG00000283156 | | 3.495444 | -0.5391204 | 0.749322 | -1.230676 | 0.2184441 | 0.404195 |
| ENSG00000103335 | PIEZO1 | 34115.49032 | -0.539192965 | 0.145403261 | -3.780969371 | 0.00015622 | 0.00114 |
| ENSG00000070718 | AP3M2 | 1261.212123 | -0.539375432 | 0.133965591 | -4.097323398 | 4.18E-05 | 0.000354 |
| ENSG00000233360 | PDXP-DT | 5.258319 | -0.5395687 | 0.6660236 | -1.239715 | 0.2150808 | 0.400118 |
| ENSG00000169136 | ATF5 | 2381.178 | -0.5397045 | 0.29548 | -1.980822 | 0.04761123 | 0.133479 |
| ENSG00000225606 | | 18.53349 | -0.5399692 | 0.6846535 | -1.235153 | 0.2167735 | 0.40224 |
| ENSG00000079785 | DDX1 | 8228.148691 | -0.539972604 | 0.201388606 | -2.792651948 | 0.00522779 | 0.022977 |
| ENSG00000275894 | | 33.33605 | -0.5400516 | 0.4918501 | -1.371602 | 0.1701875 | 0.340747 |
| ENSG00000247498 | GPRC5D-AS1 | 242.6962 | -0.5405461 | 0.2264103 | -2.513022 | 0.0119702 | 0.044955 |
| ENSG00000119950 | MXI1 | 1593.085483 | -0.540649039 | 0.263007514 | -2.169191648 | 0.03006814 | 0.093837 |
| ENSG00000244381 | SDHDP3 | 6.452945 | -0.5407657 | 0.6943155 | -1.244017 | 0.2134932 | 0.39828 |
| ENSG00000277075 | H2AC8 | 482.7365 | -0.5407934 | 0.332146 | -1.806437 | 0.07085015 | 0.180239 |
| ENSG00000265706 | SNORD53B | 2.015803 | -0.5408541 | 0.9252874 | -1.46642 | 0.142534 | 0.300007 |

| | | | | | | | |
|---|---|---|---|---|---|---|---|
| ENSG00000278112 | | 37.44717 | -0.5408746 | 0.352175 | -1.717408 | 0.08590466 | 0.208248 |
| ENSG00000232586 | KIAA1614-AS1 | 6.567585 | -0.5409179 | 0.7254243 | -1.241353 | 0.2144754 | 0.399379 |
| ENSG00000085063 | CD59 | 3326.031777 | -0.540963673 | 0.20754847 | -2.713124721 | 0.0066652 | 0.02798 |
| ENSG00000103121 | CMC2 | 2011.487174 | -0.541031252 | 0.309038922 | -1.916098473 | 0.05535256 | 0.149777 |
| ENSG00000223653 | | 5.257654 | -0.5414123 | 0.6413774 | -1.245309 | 0.2130182 | 0.397722 |
| ENSG00000196301 | HLA-DRB9 | 5.314231 | -0.5414343 | 0.8801944 | -1.346109 | 0.1782674 | 0.351782 |
| ENSG00000118363 | SPCS2 | 1825.492572 | -0.541438496 | 0.091660072 | -5.952117805 | 2.65E-09 | 5.35E-08 |
| ENSG00000257243 | PARK7P1 | 9.017533 | -0.5414654 | 0.676636 | -1.236151 | 0.2164024 | 0.401731 |
| ENSG00000169439 | SDC2 | 2848.669 | -0.5416165 | 0.2372226 | -2.408721 | 0.01600854 | 0.056764 |
| ENSG00000265678 | | 48.55261 | -0.5417501 | 0.2631276 | -2.195138 | 0.02815372 | 0.089092 |
| ENSG00000106328 | FSCN3 | 2.037645838 | -0.54225993 | 0.809506525 | -1.287555717 | 0.19790066 | 0.37796 |
| ENSG00000031698 | SARS1 | 18029.74111 | -0.542270673 | 0.322246768 | -1.853562519 | 0.06380177 | 0.166387 |
| ENSG00000147416 | ATP6V1B2 | 3094.568 | -0.5425565 | 0.1285984 | -4.288772 | 1.80E-05 | 0.000166 |
| ENSG00000120688 | WBP4 | 1324.445061 | -0.542679813 | 0.135037263 | -4.085793475 | 4.39E-05 | 0.00037 |
| ENSG00000148362 | PAXX | 4920.472 | -0.5427492 | 0.1913852 | -2.932943 | 0.00335765 | 0.015962 |
| ENSG00000158486 | DNAH3 | 222.0991 | -0.5427681 | 0.2384355 | -2.396146 | 0.01656848 | 0.058375 |
| ENSG00000181123 | | 1.011602 | -0.5427786 | 1.063561 | -2.196316 | 0.02806936 | NA |
| ENSG00000246145 | RRS1-AS1 | 3.481647 | -0.5429586 | 0.7176632 | -1.241236 | 0.2145185 | 0.399379 |
| ENSG00000268140 | PCGF7P | 91.88714 | -0.5429607 | 0.3067348 | -1.933316 | 0.05319731 | 0.145531 |
| ENSG00000101384 | JAG1 | 694.4601937 | -0.543003749 | 0.316991732 | -1.880072503 | 0.0600982 | 0.159261 |
| ENSG00000117971 | CHRNB4 | 49.8257758 | -0.543136898 | 0.384356345 | -1.617412209 | 0.10578933 | 0.242434 |
| ENSG00000143353 | LYPLAL1 | 1115.934 | -0.543153 | 0.2069004 | -2.733412 | 0.00626819 | 0.026639 |
| ENSG00000233723 | LINC01122 | 13.51438 | -0.5432478 | 0.5278055 | -1.329603 | 0.1836492 | 0.358883 |
| ENSG00000137267 | TUBB2A | 301.1647458 | -0.543272958 | 0.381910485 | -1.629297092 | 0.10325014 | 0.238192 |
| ENSG00000235354 | RPS29P16 | 27.67228 | -0.5433124 | 0.4281344 | -1.50532 | 0.1322419 | 0.284382 |
| ENSG00000125991 | ERGIC3 | 10535.39944 | -0.543482131 | 0.156717253 | -3.565509933 | 0.00036315 | 0.002387 |
| ENSG00000187624 | C17orf97 | 93.05291 | -0.5435545 | 0.2933854 | -2.007413 | 0.04470569 | 0.127414 |
| ENSG00000172354 | GNB2 | 12265.97 | -0.5439257 | 0.2042253 | -2.768958 | 0.00562359 | 0.024402 |
| ENSG00000257176 | | 65.92934 | -0.543988 | 0.4532178 | -1.450713 | 0.1468597 | 0.306487 |
| ENSG00000172818 | OVOL1 | 1061.151 | -0.5440657 | 0.2134317 | -2.663768 | 0.00772708 | 0.031638 |
| ENSG00000198331 | HYLS1 | 573.3071 | -0.5442751 | 0.2741029 | -2.127837 | 0.03335063 | 0.101873 |
| ENSG00000168827 | GFM1 | 4688.424 | -0.5443135 | 0.1528796 | -3.641578 | 0.00027097 | 0.00185 |
| ENSG00000232931 | LINC00342 | 1284.662 | -0.5445235 | 0.4399617 | -1.483776 | 0.1378684 | 0.292574 |

| | | | | | | | |
|---|---|---|---|---|---|---|---|
| ENSG00000278703 | | 2.854039 | -0.5445795 | 0.8020162 | -1.279944 | 0.2005649 | 0.381119 |
| ENSG00000123600 | METTL8 | 1037.438557 | -0.54464571 | 0.207860667 | -2.727717973 | 0.00637741 | 0.026977 |
| ENSG00000167220 | HDHD2 | 78.63582 | -0.5447279 | 0.3401271 | -1.783227 | 0.07454943 | 0.187283 |
| ENSG00000087074 | PPP1R15A | 2501.066689 | -0.54474319 | 0.341594431 | -1.772616772 | 0.07629223 | 0.190401 |
| ENSG00000175274 | TP53I11 | 7689.992 | -0.5447934 | 0.2158114 | -2.636293 | 0.00838174 | 0.033772 |
| ENSG00000248399 | | 2.455877 | -0.5448834 | 0.8188467 | -1.294718 | 0.1954174 | 0.374838 |
| ENSG00000035141 | FAM136A | 6128.490574 | -0.545003411 | 0.182494554 | -3.079491257 | 0.00207355 | 0.010687 |
| ENSG00000162526 | TSSK3 | 18.13822 | -0.5450695 | 0.6824952 | -1.248291 | 0.2119246 | 0.396231 |
| ENSG00000229404 | LINC00858 | 436.1013 | -0.5451998 | 0.2775879 | -2.106805 | 0.03513446 | 0.105898 |
| ENSG00000107951 | MTPAP | 2063.322612 | -0.545232975 | 0.186163513 | -3.024272655 | 0.00249232 | 0.012471 |
| ENSG00000286177 | | 61.45239 | -0.5452612 | 0.407358 | -1.557258 | 0.1194093 | 0.264316 |
| ENSG00000174165 | ZDHHC24 | 1996.837 | -0.5454078 | 0.1776143 | -3.157291 | 0.00159243 | 0.008525 |
| ENSG00000268564 | | 8.928317 | -0.5454911 | 0.5565544 | -1.310146 | 0.1901465 | 0.3679 |
| ENSG00000141127 | PRPSAP2 | 2486.146 | -0.5455748 | 0.1467301 | -3.800451 | 0.00014443 | 0.00106 |
| ENSG00000227689 | SRP68P2 | 1.70496 | -0.5458607 | 0.9327646 | -1.498791 | 0.1339279 | 0.286731 |
| ENSG00000279070 | | 2.180992 | -0.5459396 | 0.7978987 | -1.264759 | 0.2059576 | 0.388064 |
| ENSG00000117400 | MPL | 19.65034408 | -0.545992102 | 0.47773022 | -1.408158971 | 0.159084 | 0.325006 |
| ENSG00000160957 | RECQL4 | 5173.775 | -0.5459951 | 0.3981791 | -1.588604 | 0.1121498 | 0.25271 |
| ENSG00000225470 | JPX | 187.7929 | -0.5461382 | 0.228134 | -2.509385 | 0.01209416 | 0.045283 |
| ENSG00000223254 | | 1.994661 | -0.5461454 | 0.8911263 | -1.404929 | 0.1600423 | 0.3263 |
| ENSG00000204685 | STARD7-AS1 | 300.6768 | -0.5461651 | 0.2011351 | -2.818124 | 0.00483052 | 0.021577 |
| ENSG00000175768 | TOMM5 | 200.495 | -0.5462654 | 0.3239348 | -1.859478 | 0.06295939 | 0.164821 |
| ENSG00000221886 | ZBED8 | 315.7491 | -0.5463386 | 0.2857596 | -2.063245 | 0.03908934 | 0.114835 |
| ENSG00000170615 | SLC26A5 | 10.58305 | -0.5463514 | 0.7053633 | -1.245518 | 0.2129414 | 0.397712 |
| ENSG00000163536 | SERPINI1 | 63.52458 | -0.5467024 | 0.3361195 | -1.811167 | 0.07011507 | 0.178814 |
| ENSG00000164087 | POC1A | 785.3373 | -0.5472806 | 0.4239707 | -1.516861 | 0.1293019 | 0.279592 |
| ENSG00000279278 | | 126.2846 | -0.5473088 | 0.4119808 | -1.550883 | 0.1209298 | 0.266771 |
| ENSG00000254222 | | 2.279113 | -0.5473495 | 0.8541123 | -1.351926 | 0.1763991 | 0.349184 |
| ENSG00000165572 | KBTBD6 | 1454.248 | -0.5474586 | 0.181977 | -3.099008 | 0.0019417 | 0.010096 |
| ENSG00000141002 | TCF25 | 9080.42 | -0.5475018 | 0.1434503 | -3.860474 | 0.00011317 | 0.000854 |
| ENSG00000101544 | ADNP2 | 2917.848436 | -0.547523095 | 0.208245316 | -2.756360026 | 0.00584486 | 0.025206 |
| ENSG00000075213 | SEMA3A | 32.42754979 | -0.547607374 | 0.349228922 | -1.743442277 | 0.08125639 | 0.199732 |
| ENSG00000117543 | DPH5 | 1427.786966 | -0.547855465 | 0.160288088 | -3.499472423 | 0.00046618 | 0.002955 |

| | | | | | | | |
|---|---|---|---|---|---|---|---|
| ENSG00000113649 | TCERG1 | 6458.946689 | -0.548044254 | 0.196830142 | -2.883851615 | 0.00392844 | 0.018175 |
| ENSG00000006695 | COX10 | 1315.911992 | -0.548062618 | 0.189400222 | -2.99707158 | 0.00272587 | 0.013452 |
| ENSG00000106012 | IQCE | 6123.570595 | -0.548299759 | 0.154888222 | -3.666395867 | 0.00024599 | 0.001702 |
| ENSG00000229512 | | 42.11394 | -0.5483187 | 0.4353713 | -1.496436 | 0.1345402 | 0.287694 |
| ENSG00000137054 | POLR1E | 1431.86246 | -0.548611407 | 0.2205838 | -2.59691395 | 0.00940655 | 0.037008 |
| ENSG00000160813 | PPP1R35 | 784.9258 | -0.5486626 | 0.2049422 | -2.79036 | 0.00526494 | 0.023118 |
| ENSG00000106591 | MRPL32 | 3203.375932 | -0.548952914 | 0.190151858 | -2.985315616 | 0.00283286 | 0.013918 |
| ENSG00000113396 | SLC27A6 | 490.9930067 | -0.548977874 | 0.314049029 | -1.914055422 | 0.05561308 | 0.150298 |
| ENSG00000101191 | DIDO1 | 9459.804417 | -0.549051986 | 0.169415483 | -3.345140797 | 0.00082241 | 0.004836 |
| ENSG00000165637 | VDAC2 | 2815.295 | -0.5490683 | 0.1878501 | -3.042123 | 0.00234916 | 0.011861 |
| ENSG00000163382 | NAXE | 4693.845 | -0.5491663 | 0.2147533 | -2.719623 | 0.00653564 | 0.027541 |
| ENSG00000125944 | HNRNPR | 9450.002821 | -0.549189623 | 0.169066445 | -3.339501334 | 0.00083929 | 0.004917 |
| ENSG00000254034 | INTS9-AS1 | 3.182053 | -0.5493322 | 0.7303792 | -1.242828 | 0.2139314 | 0.398736 |
| ENSG00000235703 | LINC00894 | 213.485 | -0.5493886 | 0.4772732 | -1.415194 | 0.1570116 | 0.321874 |
| ENSG00000167468 | GPX4 | 9501.039 | -0.5496057 | 0.1697248 | -3.323633 | 0.00088853 | 0.005162 |
| ENSG00000173786 | CNP | 4890.221 | -0.549686 | 0.2112065 | -2.755515 | 0.00585997 | 0.025257 |
| ENSG00000115241 | PPM1G | 11460.44322 | -0.549818623 | 0.23965776 | -2.41902882 | 0.01556201 | 0.055516 |
| ENSG00000124701 | APOBEC2 | 3.163864154 | -0.550078813 | 0.872334836 | -1.367330501 | 0.17152174 | 0.342255 |
| ENSG00000251279 | SMIM15-AS1 | 3.569779 | -0.5502179 | 0.724466 | -1.254159 | 0.2097844 | 0.393161 |
| ENSG00000250215 | CIR1P2 | 56.41623 | -0.5504876 | 0.2982751 | -1.998532 | 0.04565897 | 0.129454 |
| ENSG00000134996 | OSTF1 | 1372.620353 | -0.550510486 | 0.127474539 | -4.379755698 | 1.19E-05 | 0.000115 |
| ENSG00000185739 | SRL | 8.230144 | -0.5505668 | 0.6199663 | -1.268275 | 0.2046999 | 0.386626 |
| ENSG00000146830 | GIGYF1 | 9133.85 | -0.5506312 | 0.2085831 | -2.743919 | 0.00607105 | 0.025969 |
| ENSG00000227268 | KLLN | 193.6019 | -0.550648 | 0.2491495 | -2.349991 | 0.01877389 | 0.064587 |
| ENSG00000285796 | | 589.1188 | -0.5506703 | 0.3218434 | -1.888244 | 0.05899315 | 0.157139 |
| ENSG00000250697 | | 3.992913 | -0.550764 | 0.7171477 | -1.243556 | 0.213663 | 0.398508 |
| ENSG00000113141 | IK | 4915.544739 | -0.550767292 | 0.129855311 | -4.286086151 | 1.82E-05 | 0.000168 |
| ENSG00000255566 | | 47.31426 | -0.5508329 | 0.4860197 | -1.409109 | 0.1588028 | 0.324631 |
| ENSG00000172339 | ALG14 | 286.6599 | -0.5509201 | 0.2769927 | -2.134268 | 0.03282081 | 0.100555 |
| ENSG00000155893 | PXYLP1 | 938.6485 | -0.5509992 | 0.2194174 | -2.649902 | 0.0080515 | 0.032695 |
| ENSG00000238213 | TARDBPP2 | 9.965423 | -0.5511336 | 0.652485 | -1.262304 | 0.2068394 | 0.389201 |
| ENSG00000186487 | MYT1L | 3.71706 | -0.551239 | 0.9804438 | -1.525772 | 0.1270666 | 0.276159 |
| ENSG00000131730 | CKMT2 | 28.36177685 | -0.551445912 | 0.390333467 | -1.619363148 | 0.10536915 | 0.241674 |

| | | | | | | | |
|---|---|---|---|---|---|---|---|
| ENSG00000101181 | MTG2 | 2618.879271 | -0.551472412 | 0.1264068 | -4.425668705 | 9.61E-06 | 9.52E-05 |
| ENSG00000152464 | RPP38 | 783.7642 | -0.551508 | 0.1597898 | -3.533283 | 0.00041043 | 0.002649 |
| ENSG00000149091 | DGKZ | 4996.484 | -0.551578 | 0.1810318 | -3.151601 | 0.00162378 | 0.008654 |
| ENSG00000176658 | MYO1D | 5209.43 | -0.5515945 | 0.1785913 | -3.184855 | 0.00144826 | 0.007875 |
| ENSG00000123136 | DDX39A | 4578.932701 | -0.55175128 | 0.262937689 | -2.236791076 | 0.0253 | 0.081855 |
| ENSG00000157741 | UBN2 | 2011.315 | -0.5518037 | 0.3157384 | -1.919158 | 0.05496429 | 0.149015 |
| ENSG00000228589 | SPCS2P4 | 1767.546 | -0.551854 | 0.09533254 | -5.850272 | 4.91E-09 | 9.47E-08 |
| ENSG00000278985 | | 9.414021 | -0.5519398 | 0.5811563 | -1.300464 | 0.1934419 | 0.372085 |
| ENSG00000262777 | | 5.155169 | -0.5520465 | 0.6170604 | -1.26986 | 0.2041343 | 0.38586 |
| ENSG00000130309 | COLGALT1 | 3668.656177 | -0.552403033 | 0.206081328 | -2.786412873 | 0.00532949 | 0.023324 |
| ENSG00000111271 | ACAD10 | 1638.439576 | -0.552424193 | 0.190280185 | -2.997444377 | 0.00272254 | 0.013438 |
| ENSG00000132581 | SDF2 | 3204.27653 | -0.552428334 | 0.147972632 | -3.813514276 | 0.00013701 | 0.001012 |
| ENSG00000127989 | MTERF1 | 605.2623327 | -0.552442335 | 0.156743386 | -3.605074767 | 0.00031206 | 0.002095 |
| ENSG00000160818 | GPATCH4 | 4291.912 | -0.5525922 | 0.1317153 | -4.267831 | 1.97E-05 | 0.000181 |
| ENSG00000231245 | C1DP1 | 50.47115 | -0.5526157 | 0.2646372 | -2.226342 | 0.0259913 | 0.083561 |
| ENSG00000213699 | SLC35F6 | 3056.729 | -0.5527037 | 0.2300746 | -2.523007 | 0.0116356 | 0.043926 |
| ENSG00000260273 | | 17.84807 | -0.5527708 | 0.5645581 | -1.321065 | 0.1864796 | 0.36276 |
| ENSG00000287809 | | 15.24259 | -0.5528439 | 0.5447914 | -1.333526 | 0.1823591 | 0.357335 |
| ENSG00000286830 | | 1.694572 | -0.5529111 | 0.9877303 | -1.703546 | 0.08846602 | 0.212903 |
| ENSG00000119888 | EPCAM | 12045.5821 | -0.553105688 | 0.108244864 | -5.165382539 | 2.40E-07 | 3.38E-06 |
| ENSG00000254469 | | 109.848 | -0.5531476 | 0.3590542 | -1.734744 | 0.08278618 | 0.202653 |
| ENSG00000280120 | | 124.9229 | -0.5532792 | 0.3688309 | -1.704384 | 0.08830938 | 0.212653 |
| ENSG00000224725 | CEP57L1P1 | 28.66098 | -0.5533012 | 0.4516705 | -1.47937 | 0.1390414 | 0.294499 |
| ENSG00000135002 | RFK | 1455.610025 | -0.553498378 | 0.170543336 | -3.336146394 | 0.00084948 | 0.004967 |
| ENSG00000266469 | | 9.149819 | -0.5535092 | 0.6081041 | -1.27829 | 0.2011473 | 0.381866 |
| ENSG00000223864 | NPM1P19 | 24.19528 | -0.5535569 | 0.3991795 | -1.605333 | 0.1084205 | 0.246784 |
| ENSG00000257950 | P2RX5-TAX1BP3 | 60.12574 | -0.5535792 | 0.2631093 | -2.239963 | 0.02509329 | 0.081351 |
| ENSG00000130449 | ZSWIM6 | 1012.961639 | -0.553923906 | 0.180252041 | -3.159406613 | 0.00158091 | 0.008471 |
| ENSG00000237686 | | 21.09561 | -0.5540046 | 0.6728568 | -1.271233 | 0.2036458 | 0.385317 |
| ENSG00000129197 | RPAIN | 1975.298529 | -0.554346476 | 0.11613552 | -4.833805126 | 1.34E-06 | 1.63E-05 |
| ENSG00000119703 | ZC2HC1C | 137.2889924 | -0.554420984 | 0.243579364 | -2.397941873 | 0.01648748 | 0.058128 |
| ENSG00000226002 | GTF2IP14 | 16.66808 | -0.5544312 | 0.5278791 | -1.355991 | 0.1751022 | 0.34716 |
| ENSG00000259321 | | 71.68575 | -0.5544733 | 0.3163828 | -1.90523 | 0.05675012 | 0.15258 |

| | | | | | | | |
|---|---|---|---|---|---|---|---|
| ENSG00000111726 | CMAS | 2211.828221 | -0.554543483 | 0.213930047 | -2.708284834 | 0.0067632 | 0.02832 |
| ENSG00000177370 | TIMM22 | 1257.144 | -0.5545484 | 0.2045082 | -2.825255 | 0.0047243 | 0.021171 |
| ENSG00000230911 | PPIHP1 | 146.9488 | -0.5546102 | 0.3069855 | -1.977402 | 0.04799622 | 0.134279 |
| ENSG00000120705 | ETF1 | 12361.97792 | -0.554621695 | 0.163474676 | -3.455300165 | 0.00054968 | 0.003411 |
| ENSG00000186564 | FOXD2 | 59.71393 | -0.5547168 | 0.3351118 | -1.82605 | 0.06784271 | 0.174375 |
| ENSG00000232810 | TNF | 9.474245 | -0.5548175 | 0.734973 | -1.26355 | 0.2063917 | 0.388591 |
| ENSG00000119787 | ATL2 | 3625.098614 | -0.554839937 | 0.187218345 | -3.068759135 | 0.0021495 | 0.011021 |
| ENSG00000134802 | SLC43A3 | 6.723745293 | -0.554995414 | 0.612218099 | -1.28666989 | 0.19820937 | 0.378162 |
| ENSG00000083535 | PIBF1 | 609.0251965 | -0.555032281 | 0.189736978 | -3.027105455 | 0.00246908 | 0.012371 |
| ENSG00000102753 | KPNA3 | 2570.060834 | -0.555111389 | 0.166233929 | -3.427115704 | 0.00061003 | 0.003729 |
| ENSG00000256955 | | 26.3286 | -0.555131 | 0.448548 | -1.492081 | 0.1356779 | 0.289382 |
| ENSG00000213341 | CHUK | 1988.812 | -0.5551989 | 0.1614468 | -3.512235 | 0.00044436 | 0.002833 |
| ENSG00000135506 | OS9 | 10754.75115 | -0.555407723 | 0.16374547 | -3.51884711 | 0.00043343 | 0.002773 |
| ENSG00000131668 | BARX1 | 8.403577017 | -0.555440715 | 0.656776721 | -1.247239953 | 0.21230953 | 0.396775 |
| ENSG00000080802 | CNOT4 | 1208.419068 | -0.555450806 | 0.192599506 | -2.986725554 | 0.00281983 | 0.013866 |
| ENSG00000124299 | PEPD | 2477.065113 | -0.555485195 | 0.215111281 | -2.704155789 | 0.00684782 | 0.02862 |
| ENSG00000165097 | KDM1B | 1965.276 | -0.5555139 | 0.1811143 | -3.17413 | 0.00150286 | 0.008123 |
| ENSG00000271781 | | 169.0629 | -0.5555198 | 0.3034528 | -1.991793 | 0.04639379 | 0.130937 |
| ENSG00000227308 | LINC02832 | 4.30826 | -0.5557127 | 0.7509397 | -1.254348 | 0.2097156 | 0.39311 |
| ENSG00000271072 | PHBP20 | 4.108749 | -0.5558026 | 0.7150729 | -1.259501 | 0.2078493 | 0.39041 |
| ENSG00000279254 | | 2.492897 | -0.5558122 | 0.8553687 | -1.355468 | 0.1752684 | 0.347438 |
| ENSG00000256356 | HSPA8P5 | 18.62291 | -0.5558779 | 0.4095577 | -1.582949 | 0.1134332 | 0.254674 |
| ENSG00000287106 | | 1.533755 | -0.5560345 | 1.001345 | -1.623417 | 0.1045003 | NA |
| ENSG00000137074 | APTX | 1820.018817 | -0.55605825 | 0.132766344 | -4.255829541 | 2.08E-05 | 0.000189 |
| ENSG00000120616 | EPC1 | 1882.966901 | -0.556099228 | 0.140900491 | -4.000890594 | 6.31E-05 | 0.000509 |
| ENSG00000286340 | | 28.87035 | -0.5561657 | 0.4161909 | -1.564115 | 0.1177905 | 0.261692 |
| ENSG00000234737 | KRT18P15 | 874.1729 | -0.5563868 | 0.2422666 | -2.456009 | 0.01404895 | 0.051146 |
| ENSG00000144645 | OSBPL10 | 2263.652 | -0.5566056 | 0.1867402 | -3.084314 | 0.00204022 | 0.01054 |
| ENSG00000253121 | | 1.403405 | -0.5566165 | 0.9805489 | -1.682329 | 0.09250516 | NA |
| ENSG00000050344 | NFE2L3 | 640.054261 | -0.556620497 | 0.184033556 | -3.134880981 | 0.00171924 | 0.009103 |
| ENSG00000182224 | CYB5D1 | 30.18075 | -0.5566631 | 0.3435486 | -1.802465 | 0.07147226 | 0.181458 |
| ENSG00000272472 | | 1.872012 | -0.5566885 | 1.107778 | -2.644408 | 0.0081834 | 0.033104 |
| ENSG00000257108 | NHLRC4 | 2.05192 | -0.5567351 | 0.8503744 | -1.325259 | 0.1850852 | 0.360628 |

| | | | | | | | |
|---|---|---|---|---|---|---|---|
| ENSG00000276390 | | 27.33975 | -0.5568347 | 0.4538748 | -1.486234 | 0.1372172 | 0.291565 |
| ENSG00000089818 | NECAP1 | 1738.702355 | -0.556884685 | 0.096606605 | -5.810941568 | 6.21E-09 | 1.18E-07 |
| ENSG00000132182 | NUP210 | 16914.92494 | -0.557103808 | 0.183093476 | -3.129911538 | 0.00174859 | 0.009249 |
| ENSG00000276291 | FRG1HP | 318.6955 | -0.5571323 | 0.3020469 | -1.997103 | 0.04581405 | 0.129714 |
| ENSG00000237289 | CKMT1B | 837.462 | -0.5571869 | 0.3260631 | -1.888988 | 0.05889347 | 0.156961 |
| ENSG00000219553 | | 23.6484 | -0.5576541 | 0.4170504 | -1.573184 | 0.1156763 | 0.25853 |
| ENSG00000144043 | TEX261 | 7908.271 | -0.5579797 | 0.1118405 | -5.046736 | 4.49E-07 | 6.02E-06 |
| ENSG00000272870 | SAP30-DT | 54.1368 | -0.5582588 | 0.417201 | -1.572876 | 0.1157476 | 0.258624 |
| ENSG00000114631 | PODXL2 | 5241.247516 | -0.558347435 | 0.20894087 | -2.781524184 | 0.00541043 | 0.023627 |
| ENSG00000137752 | CASP1 | 16.01042836 | -0.558359216 | 0.683394046 | -1.24837461 | 0.2118939 | 0.396231 |
| ENSG00000187621 | TCL6 | 1.367614 | -0.5584227 | 1.018305 | -1.722438 | 0.08499025 | NA |
| ENSG00000230832 | SSBL4P | 37.70971 | -0.5584534 | 0.4003438 | -1.621197 | 0.1049753 | 0.240979 |
| ENSG00000253771 | TPTE2P1 | 14.64294 | -0.5584597 | 0.6951669 | -1.267391 | 0.2050153 | 0.38701 |
| ENSG00000273888 | FRMD6-AS1 | 90.64887 | -0.5584802 | 0.2990252 | -2.029221 | 0.04243578 | 0.122459 |
| ENSG00000168273 | SMIM4 | 615.649 | -0.5585524 | 0.1760927 | -3.260204 | 0.00111332 | 0.006278 |
| ENSG00000089693 | MLF2 | 11370.76069 | -0.558743673 | 0.192822709 | -3.040506311 | 0.00236181 | 0.011914 |
| ENSG00000155085 | AK9 | 563.9311 | -0.5591772 | 0.3082663 | -1.992202 | 0.04634891 | 0.130852 |
| ENSG00000072163 | LIMS2 | 5.941755246 | -0.559285089 | 0.823047862 | -1.291030825 | 0.19669299 | 0.376675 |
| ENSG00000178996 | SNX18 | 1715.472 | -0.5597478 | 0.1900897 | -3.048698 | 0.00229835 | 0.011631 |
| ENSG00000176383 | B3GNT4 | 117.6652 | -0.5597978 | 0.2513589 | -2.360595 | 0.01824564 | 0.063137 |
| ENSG00000085999 | RAD54L | 859.0888667 | -0.560242754 | 0.478382499 | -1.451653412 | 0.146598 | 0.306087 |
| ENSG00000004399 | PLXND1 | 3178.535161 | -0.560349049 | 0.291191935 | -2.080557908 | 0.03747439 | 0.111112 |
| ENSG00000165807 | PPP1R36 | 44.29262 | -0.5605481 | 0.4223629 | -1.563821 | 0.1178596 | 0.261758 |
| ENSG00000248968 | | 4.543287 | -0.5605597 | 0.7792864 | -1.282849 | 0.1995451 | 0.3798 |
| ENSG00000235290 | HLA-W | 11.69687 | -0.5605632 | 0.6430932 | -1.269088 | 0.2044097 | 0.38627 |
| ENSG00000280162 | | 1.454785 | -0.5607562 | 0.946535 | -1.562344 | 0.118207 | NA |
| ENSG00000264500 | MIR3124 | 2.706608 | -0.5608909 | 0.7477526 | -1.270476 | 0.2039153 | 0.385528 |
| ENSG00000226138 | | 8.354665 | -0.5613982 | 0.6373102 | -1.281426 | 0.2000442 | 0.380432 |
| ENSG00000196734 | LCE1B | 1.221811 | -0.561699 | 1.060785 | -1.9836 | 0.04730044 | NA |
| ENSG00000248487 | ABHD14A | 100.4348 | -0.5617379 | 0.1970347 | -2.953245 | 0.00314452 | 0.015102 |
| ENSG00000235275 | KRT18P16 | 87.86426 | -0.561927 | 0.3135814 | -1.959369 | 0.0500696 | 0.138631 |
| ENSG00000111639 | MRPL51 | 5001.507033 | -0.562018266 | 0.24606922 | -2.433791787 | 0.01494159 | 0.05377 |
| ENSG00000255409 | RSF1-IT1 | 2.490784 | -0.5620968 | 0.8377124 | -1.340618 | 0.1800447 | 0.35421 |

| | | | | | | | |
|---|---|---|---|---|---|---|---|
| ENSG00000108094 | CUL2 | 3383.644158 | -0.56227846 | 0.138045174 | -4.153416196 | 3.28E-05 | 0.000284 |
| ENSG00000227590 | ATP5MC1P5 | 0.9948051 | -0.5623855 | 1.043558 | -1.87186 | 0.06122603 | NA |
| ENSG00000228925 | | 26.10861 | -0.5625606 | 0.4573073 | -1.486395 | 0.1371747 | 0.291498 |
| ENSG00000260401 | | 1187.344 | -0.5625691 | 0.2089874 | -2.807956 | 0.00498571 | 0.0221 |
| ENSG00000160803 | UBQLN4 | 6282.512 | -0.5626333 | 0.1330467 | -4.293092 | 1.76E-05 | 0.000163 |
| ENSG00000128011 | LRFN1 | 194.4056863 | -0.562637944 | 0.307561855 | -1.992150535 | 0.04635454 | 0.130854 |
| ENSG00000265354 | TIMM23 | 4373.36 | -0.5630456 | 0.2408953 | -2.46543 | 0.01368489 | 0.050027 |
| ENSG00000197905 | TEAD4 | 1546.605 | -0.5631449 | 0.2783733 | -2.170028 | 0.03000473 | 0.0937 |
| ENSG00000079156 | OSBPL6 | 678.070695 | -0.563209455 | 0.325228908 | -1.914749804 | 0.05552442 | 0.15015 |
| ENSG00000224936 | SUCLA2P1 | 12.84644 | -0.5632912 | 0.4862897 | -1.440403 | 0.1497535 | 0.310747 |
| ENSG00000250461 | | 149.6459 | -0.5633225 | 0.2286202 | -2.590123 | 0.00959418 | 0.037564 |
| ENSG00000141642 | ELAC1 | 238.6188 | -0.563416 | 0.2955707 | -2.06608 | 0.03882096 | 0.114248 |
| ENSG00000274403 | | 12.81687 | -0.5638122 | 0.5335669 | -1.371799 | 0.170126 | 0.340649 |
| ENSG00000272265 | | 1.529446 | -0.5640424 | 1.008201 | -1.718775 | 0.08565535 | NA |
| ENSG00000260495 | | 48.35326 | -0.564368 | 0.6036272 | -1.312741 | 0.1892702 | 0.366712 |
| ENSG00000125814 | NAPB | 1215.708234 | -0.564484567 | 0.183586572 | -3.17411383 | 0.00150295 | 0.008123 |
| ENSG00000184831 | APOO | 751.906 | -0.5644943 | 0.2630525 | -2.284653 | 0.0223332 | 0.074432 |
| ENSG00000152520 | PAN3 | 3539.489 | -0.5645316 | 0.177942 | -3.275041 | 0.00105647 | 0.006 |
| ENSG00000228487 | | 46.59959 | -0.5646203 | 0.3601668 | -1.750867 | 0.07996882 | 0.197178 |
| ENSG00000229312 | | 27.84489 | -0.5647286 | 0.455353 | -1.503695 | 0.1326599 | 0.284971 |
| ENSG00000163661 | PTX3 | 4.43006 | -0.5648888 | 0.7354518 | -1.278477 | 0.2010814 | 0.381824 |
| ENSG00000170906 | NDUFA3 | 2209.738 | -0.5649092 | 0.2123115 | -2.802461 | 0.00507144 | 0.022416 |
| ENSG00000176619 | LMNB2 | 9147.149 | -0.5650444 | 0.2784949 | -2.179425 | 0.02930008 | 0.092105 |
| ENSG00000115526 | CHST10 | 786.4986217 | -0.565096882 | 0.177509277 | -3.278847672 | 0.00104232 | 0.005935 |
| ENSG00000229413 | | 141.7747 | -0.5651287 | 0.2638262 | -2.282494 | 0.0224602 | 0.074743 |
| ENSG00000135931 | ARMC9 | 887.8177116 | -0.565271244 | 0.149611247 | -3.853049844 | 0.00011666 | 0.000879 |
| ENSG00000234450 | | 13.98199 | -0.5653637 | 0.7130764 | -1.286361 | 0.1983172 | 0.378309 |
| ENSG00000204429 | | 3.597468 | -0.5654188 | 0.8281822 | -1.317177 | 0.1877794 | 0.364621 |
| ENSG00000068654 | POLR1A | 6218.472064 | -0.565453344 | 0.203339876 | -2.904299951 | 0.00368075 | 0.017202 |
| ENSG00000106733 | NMRK1 | 940.6136656 | -0.565630087 | 0.153698469 | -3.756930815 | 0.00017201 | 0.001242 |
| ENSG00000175826 | CTDNEP1 | 2139.541 | -0.5657434 | 0.1612485 | -3.585778 | 0.00033607 | 0.002235 |
| ENSG00000204421 | LY6G6C | 28.37054 | -0.5657891 | 0.4772017 | -1.448817 | 0.1473887 | 0.307286 |
| ENSG00000144029 | MRPS5 | 5084.198 | -0.5659964 | 0.1530487 | -3.791479 | 0.00014975 | 0.001095 |

| | | | | | | | |
|---|---|---|---|---|---|---|---|
| ENSG00000170893 | TRH | 1.969655 | -0.5660562 | 0.9249569 | -1.487441 | 0.1368983 | 0.291166 |
| ENSG00000174851 | YIF1A | 5020.286 | -0.5660709 | 0.1714368 | -3.391437 | 0.00069527 | 0.00418 |
| ENSG00000235319 | | 9.08876 | -0.5661201 | 0.5602852 | -1.339215 | 0.1805008 | 0.354845 |
| ENSG00000271367 | | 3.131531 | -0.5664356 | 0.7904068 | -1.305798 | 0.1916213 | 0.369838 |
| ENSG00000077279 | DCX | 8.217050303 | -0.566613454 | 0.809391919 | -1.36090966 | 0.17354224 | 0.345097 |
| ENSG00000134243 | SORT1 | 9047.532413 | -0.56718192 | 0.152006207 | -3.77787898 | 0.00015817 | 0.001152 |
| ENSG00000113522 | RAD50 | 363.5759142 | -0.567235185 | 0.276392673 | -2.200028073 | 0.0278049 | 0.088198 |
| ENSG00000105220 | GPI | 20579.90698 | -0.567266764 | 0.22708649 | -2.619639254 | 0.00880228 | 0.035147 |
| ENSG00000155508 | CNOT8 | 3771.691 | -0.5674401 | 0.1578563 | -3.686194 | 0.00022763 | 0.00159 |
| ENSG00000188191 | PRKAR1B | 4348.681 | -0.5674516 | 0.2542989 | -2.368418 | 0.01786436 | 0.062044 |
| ENSG00000105784 | RUNDC3B | 165.8733175 | -0.567562821 | 0.378112656 | -1.700847693 | 0.08897159 | 0.213887 |
| ENSG00000105048 | TNNT1 | 1896.156536 | -0.567624301 | 0.343654779 | -1.888500262 | 0.05895882 | 0.157103 |
| ENSG00000186510 | CLCNKA | 8.723798 | -0.567722 | 0.603287 | -1.313729 | 0.1889374 | 0.366232 |
| ENSG00000238961 | SNORA47 | 6.180077 | -0.567735 | 0.7440351 | -1.296985 | 0.1946363 | 0.373769 |
| ENSG00000140955 | ADAD2 | 1.125816 | -0.5677414 | 0.9000546 | -1.395012 | 0.1630122 | NA |
| ENSG00000144158 | SNRPA1P1 | 674.1027 | -0.5677814 | 0.5064486 | -1.416768 | 0.1565509 | 0.32114 |
| ENSG00000231458 | RNASEH1P2 | 46.66348 | -0.5677853 | 0.3138131 | -1.982294 | 0.04744638 | 0.133158 |
| ENSG00000003509 | NDUFAF7 | 1116.977704 | -0.56787906 | 0.150159163 | -3.860502265 | 0.00011315 | 0.000854 |
| ENSG00000106682 | EIF4H | 13447.64225 | -0.568003974 | 0.157636399 | -3.686126449 | 0.00022769 | 0.00159 |
| ENSG00000263766 | | 21.22355 | -0.5680439 | 0.4263553 | -1.56989 | 0.1164407 | 0.259487 |
| ENSG00000167842 | MIS12 | 1843.43 | -0.5680744 | 0.2064717 | -2.856918 | 0.00427776 | 0.019482 |
| ENSG00000251441 | RTEL1P1 | 2.591488 | -0.5681235 | 0.7912152 | -1.300711 | 0.1933575 | 0.372007 |
| ENSG00000175787 | ZNF169 | 347.2028 | -0.5681827 | 0.3614355 | -1.769062 | 0.07688353 | 0.191517 |
| ENSG00000170584 | NUDCD2 | 2181.355 | -0.5682056 | 0.1916566 | -3.076361 | 0.00209544 | 0.010781 |
| ENSG00000187742 | SECISBP2 | 5115.474 | -0.568206 | 0.221865 | -2.686237 | 0.00722619 | 0.029924 |
| ENSG00000131844 | MCCC2 | 10780.01357 | -0.568255713 | 0.170309569 | -3.435283947 | 0.00059193 | 0.003632 |
| ENSG00000133794 | ARNTL | 959.3578006 | -0.568380018 | 0.18884203 | -3.11326151 | 0.00185032 | 0.009704 |
| ENSG00000273204 | | 8.402535 | -0.568488 | 0.666654 | -1.28952 | 0.1972173 | 0.377186 |
| ENSG00000226744 | | 4.449639 | -0.568567 | 0.8268106 | -1.335881 | 0.1815883 | 0.356297 |
| ENSG00000085231 | AK6 | 1297.786351 | -0.568635059 | 0.151723745 | -3.831721732 | 0.00012725 | 0.000948 |
| ENSG00000163781 | TOPBP1 | 6071.717 | -0.5688117 | 0.2188273 | -2.727906 | 0.00637378 | 0.026974 |
| ENSG00000016391 | CHDH | 1938.981283 | -0.569036714 | 0.15728041 | -3.698456134 | 0.00021692 | 0.001523 |
| ENSG00000113048 | MRPS27 | 5000.774215 | -0.569072599 | 0.210348392 | -2.818766208 | 0.00482086 | 0.021538 |

| | | | | | | | |
|---|---|---|---|---|---|---|---|
| ENSG00000259840 | | 18.05619 | -0.5692317 | 0.496653 | -1.431133 | 0.1523922 | 0.314674 |
| ENSG00000146574 | CCZ1B | 1989.121 | -0.5692747 | 0.1230596 | -4.691699 | 2.71E-06 | 3.09E-05 |
| ENSG00000122882 | ECD | 2632.384749 | -0.569276341 | 0.114787962 | -5.046341689 | 4.50E-07 | 6.03E-06 |
| ENSG00000228204 | | 2.895962 | -0.5692868 | 0.828533 | -1.305608 | 0.1916859 | 0.369909 |
| ENSG00000260464 | | 17.05914 | -0.5697551 | 0.5256679 | -1.398034 | 0.1621027 | 0.329383 |
| ENSG00000226688 | ENTPD1-AS1 | 140.0207 | -0.5697637 | 0.2370886 | -2.528307 | 0.01146141 | 0.04341 |
| ENSG00000259514 | | 5.747931 | -0.5698167 | 0.7580285 | -1.299788 | 0.1936736 | 0.372274 |
| ENSG00000132141 | CCT6B | 140.1972579 | -0.56983274 | 0.261122938 | -2.322826949 | 0.02018845 | 0.068513 |
| ENSG00000110799 | VWF | 11.00546242 | -0.569833199 | 0.630410696 | -1.304802005 | 0.19196028 | 0.370235 |
| ENSG00000182791 | CCDC87 | 54.11882 | -0.5699889 | 0.3350652 | -1.883452 | 0.0596392 | 0.158313 |
| ENSG00000272259 | LINC01749 | 1.108483 | -0.5699924 | 1.024403 | -1.839909 | 0.06578161 | NA |
| ENSG00000231878 | SNRPFP1 | 81.27035 | -0.5700543 | 0.4134831 | -1.615875 | 0.1061214 | 0.24298 |
| ENSG00000183734 | ASCL2 | 398.7345 | -0.5701194 | 0.2437037 | -2.477376 | 0.01323524 | 0.048732 |
| ENSG00000273363 | | 2.466041 | -0.5701759 | 0.9278209 | -1.454661 | 0.145763 | 0.304847 |
| ENSG00000026103 | FAS | 434.0928488 | -0.570269192 | 0.327124995 | -1.921011787 | 0.05473022 | 0.148577 |
| ENSG00000228779 | | 3.90093 | -0.5703064 | 0.7935024 | -1.306047 | 0.1915364 | 0.369728 |
| ENSG00000259076 | | 2.822471 | -0.5704809 | 0.8597832 | -1.36408 | 0.1725423 | 0.343726 |
| ENSG00000224411 | HSP90AA2P | 2655.965 | -0.5705395 | 0.2431535 | -2.464452 | 0.01372229 | 0.05015 |
| ENSG00000197451 | HNRNPAB | 14269.95 | -0.5705439 | 0.2496077 | -2.331323 | 0.01973632 | 0.067279 |
| ENSG00000151779 | NBAS | 2086.165 | -0.5706649 | 0.1708598 | -3.461444 | 0.00053729 | 0.003344 |
| ENSG00000219607 | PPP1R3G | 2.266053 | -0.570682 | 0.8252804 | -1.329776 | 0.183592 | 0.358798 |
| ENSG00000136872 | ALDOB | 1.263713144 | -0.570795775 | 0.998799037 | -1.687068059 | 0.09159027 | NA |
| ENSG00000112984 | KIF20A | 2858.109663 | -0.571047706 | 0.603867208 | -1.343706716 | 0.17904325 | 0.352992 |
| ENSG00000112212 | TSPO2 | 1.962229624 | -0.571100698 | 0.901012157 | -1.387198648 | 0.1653812 | 0.333944 |
| ENSG00000157796 | WDR19 | 1249.018 | -0.5712406 | 0.2678562 | -2.275973 | 0.0228476 | 0.075624 |
| ENSG00000154447 | SH3RF1 | 1823.505 | -0.5713292 | 0.1553972 | -3.753563 | 0.00017434 | 0.001256 |
| ENSG00000260423 | LINC02367 | 21.17949 | -0.57139 | 0.6704006 | -1.307709 | 0.1909719 | 0.368996 |
| ENSG00000261189 | | 162.7991 | -0.5715248 | 0.278971 | -2.199005 | 0.02787753 | 0.088376 |
| ENSG00000204592 | HLA-E | 9069.879 | -0.5715366 | 0.2108376 | -2.824166 | 0.00474038 | 0.021225 |
| ENSG00000250504 | KRT18P51 | 2.237277 | -0.5715616 | 0.9672349 | -1.600949 | 0.1093881 | 0.248298 |
| ENSG00000239351 | NPM1P29 | 3.312777 | -0.5715862 | 0.7955936 | -1.311932 | 0.1895431 | 0.367107 |
| ENSG00000250115 | AK3P2 | 2.38928 | -0.5717406 | 1.006775 | -1.757885 | 0.078767 | 0.194857 |
| ENSG00000280367 | | 14.83039 | -0.571848 | 0.5837531 | -1.329232 | 0.1837714 | 0.359069 |

| | | | | | | | |
|---|---|---|---|---|---|---|---|
| ENSG00000146066 | HIGD2A | 2858.627 | -0.571932 | 0.1829885 | -3.234725 | 0.0012176 | 0.006782 |
| ENSG00000106290 | TAF6 | 1709.592526 | -0.572013043 | 0.183092711 | -3.231202643 | 0.00123271 | 0.006855 |
| ENSG00000278000 | | 52.38373 | -0.572018 | 0.4480838 | -1.531189 | 0.1257226 | 0.274119 |
| ENSG00000143499 | SMYD2 | 3816.564 | -0.5720756 | 0.1999401 | -2.969534 | 0.00298252 | 0.014492 |
| ENSG00000172766 | NAA16 | 1608.594 | -0.5723621 | 0.1668344 | -3.511702 | 0.00044525 | 0.002837 |
| ENSG00000080824 | HSP90AA1 | 96236.12147 | -0.572388269 | 0.211630073 | -2.819074967 | 0.00481623 | 0.021521 |
| ENSG00000286688 | | 5.099576 | -0.5723958 | 0.6426523 | -1.30462 | 0.1920221 | 0.370235 |
| ENSG00000214353 | VAC14-AS1 | 10.74566 | -0.5725556 | 0.5566155 | -1.367585 | 0.171442 | 0.34215 |
| ENSG00000042980 | ADAM28 | 3.161882169 | -0.572593373 | 0.962585974 | -1.594104535 | 0.11091262 | 0.250786 |
| ENSG00000238133 | MAP3K20-AS1 | 7.619501 | -0.5726564 | 0.736063 | -1.305376 | 0.1917648 | 0.370007 |
| ENSG00000115421 | PAPOLG | 1075.562885 | -0.572661575 | 0.228253186 | -2.633529314 | 0.00845025 | 0.033988 |
| ENSG00000260367 | | 50.64093 | -0.5727873 | 0.3670578 | -1.763733 | 0.077777 | 0.193081 |
| ENSG00000188243 | COMMD6 | 1945.755 | -0.5728081 | 0.1835577 | -3.147425 | 0.00164716 | 0.008763 |
| ENSG00000253130 | | 1.3785 | -0.5728289 | 1.038047 | -1.902128 | 0.05715437 | NA |
| ENSG00000122435 | TRMT13 | 701.9291862 | -0.572976241 | 0.244564153 | -2.477621071 | 0.01322615 | 0.048712 |
| ENSG00000155868 | MED7 | 894.1453 | -0.5730153 | 0.1522996 | -3.841972 | 0.00012205 | 0.000914 |
| ENSG00000279495 | | 138.9261 | -0.5730543 | 0.3123503 | -2.009654 | 0.04446785 | 0.12693 |
| ENSG00000102786 | INTS6 | 2189.891076 | -0.573238187 | 0.197175966 | -3.015648073 | 0.00256431 | 0.012773 |
| ENSG00000272382 | | 32.39715 | -0.5733522 | 0.4154232 | -1.614421 | 0.1064361 | 0.243592 |
| ENSG00000229222 | KRT18P4 | 37.07789 | -0.5733758 | 0.3504164 | -1.824207 | 0.06812085 | 0.174921 |
| ENSG00000177855 | CACYBPP2 | 1726.711 | -0.5734508 | 0.2842304 | -2.173346 | 0.02975431 | 0.093126 |
| ENSG00000250938 | | 9.464221 | -0.5735927 | 0.5140914 | -1.412512 | 0.1577992 | 0.323077 |
| ENSG00000119969 | HELLS | 2400.950126 | -0.573593596 | 0.482769721 | -1.479771444 | 0.13893425 | 0.294319 |
| ENSG00000249064 | KRT18P25 | 24.6829 | -0.5736499 | 0.459626 | -1.511889 | 0.1305621 | 0.281681 |
| ENSG00000102471 | NDFIP2 | 1949.491323 | -0.573883545 | 0.155148827 | -3.789165274 | 0.00015115 | 0.001105 |
| ENSG00000037897 | METTL1 | 1266.529905 | -0.573885654 | 0.234609417 | -2.582247983 | 0.0098159 | 0.038274 |
| ENSG00000233178 | | 2.847788 | -0.5739891 | 0.951959 | -1.556184 | 0.1196643 | 0.264682 |
| ENSG00000155846 | PPARGC1B | 877.2441 | -0.5740688 | 0.3107371 | -2.019554 | 0.04342966 | 0.124636 |
| ENSG00000230170 | HNRNPA1P28 | 1.205608 | -0.5740811 | 0.9938598 | -1.676199 | 0.09369931 | NA |
| ENSG00000161960 | EIF4A1 | 649.8294 | -0.5745448 | 0.2096671 | -2.85832 | 0.00425891 | 0.019422 |
| ENSG00000278733 | | 4.197529 | -0.5745834 | 0.7996298 | -1.328388 | 0.1840499 | 0.359356 |
| ENSG00000171202 | TMEM126A | 1000.845 | -0.57459 | 0.2315311 | -2.612119 | 0.0089983 | 0.035752 |
| ENSG00000270497 | | 2.492699 | -0.5749442 | 0.8180744 | -1.356021 | 0.1750926 | 0.34716 |

| | | | | | | | |
|---|---|---|---|---|---|---|---|
| ENSG00000117395 | EBNA1BP2 | 5074.634135 | -0.574994563 | 0.248241766 | -2.402845196 | 0.01626807 | 0.057485 |
| ENSG00000286257 | | 2.677155 | -0.5752601 | 0.8794363 | -1.40674 | 0.1595045 | 0.325602 |
| ENSG00000243667 | WDR92 | 313.2094 | -0.5753499 | 0.1739102 | -3.415138 | 0.0006375 | 0.003877 |
| ENSG00000196313 | POM121 | 2793.773 | -0.5757102 | 0.1923487 | -3.101165 | 0.00192761 | 0.010038 |
| ENSG00000159055 | MIS18A | 887.9497 | -0.5757873 | 0.388102 | -1.694885 | 0.09009722 | 0.215808 |
| ENSG00000119333 | DYNC2I2 | 8343.108975 | -0.575841346 | 0.349963759 | -1.842367177 | 0.06542146 | 0.169595 |
| ENSG00000270808 | | 5.228273 | -0.5758786 | 0.6383464 | -1.316953 | 0.1878544 | 0.36474 |
| ENSG00000111142 | METAP2 | 6867.310333 | -0.575960218 | 0.229395755 | -2.617794171 | 0.00885002 | 0.035285 |
| ENSG00000257704 | INAFM1 | 408.3094 | -0.5760058 | 0.27819 | -2.228073 | 0.02587568 | 0.083282 |
| ENSG00000259767 | | 8.140797 | -0.576053 | 0.5127834 | -1.433169 | 0.1518095 | 0.31382 |
| ENSG00000229097 | CALM2P2 | 494.6259 | -0.5761095 | 0.23101 | -2.624133 | 0.00868699 | 0.034744 |
| ENSG00000278588 | H2BC10 | 7.921608 | -0.5763161 | 0.6246426 | -1.316616 | 0.1879673 | 0.364933 |
| ENSG00000204856 | FAM216A | 559.3973 | -0.5763338 | 0.3443194 | -1.86815 | 0.06174118 | 0.162497 |
| ENSG00000145996 | CDKAL1 | 474.0545 | -0.5766015 | 0.1646363 | -3.587126 | 0.00033434 | 0.002225 |
| ENSG00000280441 | | 2.489563 | -0.5766321 | 0.9430227 | -1.494396 | 0.1350721 | 0.288473 |
| ENSG00000234134 | | 23.42706 | -0.5767179 | 0.4419815 | -1.554338 | 0.1201037 | 0.2655 |
| ENSG00000156110 | ADK | 4567.369 | -0.5767499 | 0.1985559 | -3.034421 | 0.00240998 | 0.012122 |
| ENSG00000103534 | TMC5 | 197.7662847 | -0.576785384 | 0.326175854 | -1.94573855 | 0.05168615 | 0.142099 |
| ENSG00000286136 | | 10.3702 | -0.5768631 | 0.5729709 | -1.352537 | 0.1762037 | 0.348927 |
| ENSG00000263001 | GTF2I | 3580.219 | -0.5769649 | 0.1579433 | -3.73945 | 0.00018442 | 0.001319 |
| ENSG00000166788 | SAAL1 | 975.5189 | -0.5772819 | 0.3300176 | -1.934303 | 0.05307594 | 0.145289 |
| ENSG00000226779 | NAALADL2-AS2 | 2.400574 | -0.5777829 | 1.14297 | -2.815796 | 0.00486566 | 0.021698 |
| ENSG00000136436 | CALCOCO2 | 6935.988204 | -0.577791739 | 0.193436453 | -3.109470869 | 0.00187423 | 0.009801 |
| ENSG00000197903 | H2BC12 | 3631.299 | -0.5777979 | 0.1741127 | -3.425306 | 0.00061411 | 0.003752 |
| ENSG00000221829 | FANCG | 1958.527 | -0.578068 | 0.4330129 | -1.581814 | 0.113692 | 0.255104 |
| ENSG00000286327 | | 4.893101 | -0.5781333 | 0.8414671 | -1.363872 | 0.1726077 | 0.343831 |
| ENSG00000239332 | LINC01119 | 13.81094 | -0.578172 | 0.5531523 | -1.375474 | 0.1689846 | 0.339054 |
| ENSG00000164949 | GEM | 173.8813 | -0.5784237 | 0.5125012 | -1.448665 | 0.1474311 | 0.307328 |
| ENSG00000230697 | SAP18P3 | 1.513915 | -0.5784764 | 0.9779085 | -1.627329 | 0.1036673 | NA |
| ENSG00000166260 | COX11 | 4165.482 | -0.5785118 | 0.1407866 | -4.186294 | 2.84E-05 | 0.00025 |
| ENSG00000177042 | TMEM80 | 1850.86 | -0.5787672 | 0.290005 | -2.157935 | 0.0309329 | 0.09609 |
| ENSG00000188959 | C9orf152 | 606.9534 | -0.5787806 | 0.293606 | -2.135563 | 0.03271501 | 0.100324 |
| ENSG00000018699 | TTC27 | 1312.236596 | -0.578850175 | 0.212299447 | -2.884711626 | 0.00391772 | 0.01814 |

| | | | | | | | |
|---|---|---|---|---|---|---|---|
| ENSG00000177683 | THAP5 | 1189.687 | -0.5789292 | 0.1715883 | -3.471409 | 0.00051774 | 0.003234 |
| ENSG00000266936 | | 9.076171 | -0.5790932 | 0.5705898 | -1.365161 | 0.1722026 | 0.34323 |
| ENSG00000189337 | KAZN | 619.4924 | -0.5791073 | 0.2014089 | -2.984927 | 0.00283646 | 0.013933 |
| ENSG00000218582 | GAPDHP63 | 72.03713 | -0.5791521 | 0.3999672 | -1.679245 | 0.09310424 | 0.220904 |
| ENSG00000131899 | LLGL1 | 3130.463681 | -0.579190568 | 0.220559247 | -2.745914113 | 0.00603426 | 0.025848 |
| ENSG00000132967 | HMGB1P5 | 4310.235156 | -0.579257082 | 0.342440219 | -1.885130988 | 0.05941216 | 0.157947 |
| ENSG00000281333 | | 24.53281 | -0.5793331 | 0.5650649 | -1.37139 | 0.1702534 | 0.340828 |
| ENSG00000218281 | H2AC9P | 27.60501 | -0.5795231 | 0.3978964 | -1.683474 | 0.09228328 | 0.219486 |
| ENSG00000269038 | | 20.12615 | -0.5795421 | 0.4169028 | -1.625864 | 0.1039786 | 0.23929 |
| ENSG00000273345 | | 11.29803 | -0.5796417 | 0.5441819 | -1.396179 | 0.1626607 | 0.330029 |
| ENSG00000110436 | SLC1A2 | 261.5305898 | -0.579791091 | 0.639118918 | -1.330249223 | 0.18343617 | 0.358572 |
| ENSG00000140943 | MBTPS1 | 15549.5 | -0.5799099 | 0.1894318 | -3.167191 | 0.00153919 | 0.008292 |
| ENSG00000182898 | TCHHL1 | 63.82207 | -0.5799311 | 0.6051935 | -1.336905 | 0.1812536 | 0.355804 |
| ENSG00000239335 | LLPH-DT | 48.37785 | -0.5801147 | 0.3199403 | -1.985063 | 0.04713752 | 0.132459 |
| ENSG00000109971 | HSPA8 | 54957.84018 | -0.580138554 | 0.128660035 | -4.597882985 | 4.27E-06 | 4.63E-05 |
| ENSG00000213383 | | 384.3231 | -0.5805548 | 0.2186295 | -2.781481 | 0.00541116 | 0.023627 |
| ENSG00000229267 | SNHG31 | 38.33375 | -0.5805787 | 0.4238804 | -1.614343 | 0.1064531 | 0.243592 |
| ENSG00000172175 | MALT1 | 1094.816 | -0.5807083 | 0.1800438 | -3.32093 | 0.00089718 | 0.005202 |
| ENSG00000163170 | BOLA3 | 768.2628 | -0.5808229 | 0.3101523 | -2.049795 | 0.04038446 | 0.117754 |
| ENSG00000187079 | TEAD1 | 5941.96 | -0.5808975 | 0.2706413 | -2.295534 | 0.02170252 | 0.072658 |
| ENSG00000105072 | C19orf44 | 160.966373 | -0.580922808 | 0.227042695 | -2.675268362 | 0.00746695 | 0.030758 |
| ENSG00000272468 | | 51.91733 | -0.5811811 | 0.5299726 | -1.421291 | 0.1552322 | 0.319075 |
| ENSG00000161888 | SPC24 | 861.0363 | -0.5811996 | 0.6131933 | -1.357641 | 0.1745775 | 0.346523 |
| ENSG00000215475 | SIAH3 | 2.086053 | -0.5815039 | 0.8861444 | -1.402435 | 0.1607855 | 0.327324 |
| ENSG00000278743 | | 7.895883 | -0.5815197 | 0.712818 | -1.319887 | 0.1868727 | 0.363312 |
| ENSG00000248445 | SEMA6A-AS1 | 41.84921 | -0.581725 | 0.4277118 | -1.601781 | 0.1092041 | 0.247944 |
| ENSG00000105202 | FBL | 9344.011904 | -0.581780271 | 0.218993517 | -2.787533616 | 0.00531109 | 0.023263 |
| ENSG00000260896 | ARLNC1 | 1.404519 | -0.5818102 | 1.021749 | -1.656252 | 0.09767089 | NA |
| ENSG00000267742 | SINHCAFP2 | 126.6839 | -0.5820267 | 0.3478477 | -1.874063 | 0.0609217 | 0.160834 |
| ENSG00000142892 | PIGK | 1041.278 | -0.582061 | 0.2048536 | -2.954346 | 0.00313332 | 0.015054 |
| ENSG00000065665 | SEC61A2 | 636.8280124 | -0.582163833 | 0.168666895 | -3.541476615 | 0.00039789 | 0.00258 |
| ENSG00000075702 | WDR62 | 1847.09229 | -0.582167119 | 0.488938289 | -1.494058205 | 0.13516041 | 0.288473 |
| ENSG00000154832 | CXXC1 | 3153.053 | -0.5822262 | 0.1571967 | -3.803494 | 0.00014267 | 0.001048 |

| | | | | | | | |
|---|---|---|---|---|---|---|---|
| ENSG00000255652 | | 5.409621 | -0.5822527 | 0.6550401 | -1.32825 | 0.1840956 | 0.359356 |
| ENSG00000213025 | COX20P1 | 348.4144 | -0.5822545 | 0.2119031 | -2.867327 | 0.00413955 | 0.018998 |
| ENSG00000287978 | | 115.2173 | -0.5822877 | 0.3798893 | -1.74874 | 0.08033598 | 0.197909 |
| ENSG00000230606 | | 15.56541 | -0.5824024 | 0.5722691 | -1.376843 | 0.1685609 | 0.338408 |
| ENSG00000162757 | C1orf74 | 280.7831 | -0.5824408 | 0.2292948 | -2.666348 | 0.00766804 | 0.03143 |
| ENSG00000171804 | WDR87 | 10.13188 | -0.5824945 | 0.6482743 | -1.319331 | 0.1870586 | 0.363567 |
| ENSG00000168517 | HEXIM2 | 157.4156 | -0.5826571 | 0.2534753 | -2.420164 | 0.01551349 | 0.05537 |
| ENSG00000197461 | PDGFA | 1181.324 | -0.5826995 | 0.2508947 | -2.458728 | 0.01394301 | 0.050809 |
| ENSG00000100911 | PSME2 | 6438.447985 | -0.583044727 | 0.208604594 | -2.91185902 | 0.00359285 | 0.016877 |
| ENSG00000221887 | HMSD | 3.153299 | -0.5831445 | 0.8049438 | -1.343584 | 0.179083 | 0.353004 |
| ENSG00000270124 | | 3.767224 | -0.5832495 | 0.8462075 | -1.38076 | 0.1673527 | 0.336771 |
| ENSG00000167513 | CDT1 | 3913.622 | -0.5837176 | 0.3851234 | -1.745013 | 0.08098265 | 0.199206 |
| ENSG00000147874 | HAUS6 | 1323.404 | -0.58372 | 0.2201967 | -2.772911 | 0.00555573 | 0.024155 |
| ENSG00000141646 | SMAD4 | 1894.319 | -0.5837891 | 0.09468277 | -6.223504 | 4.86E-10 | 1.10E-08 |
| ENSG00000127922 | SEM1 | 3538.004302 | -0.58387048 | 0.200756261 | -3.032260183 | 0.0024273 | 0.012191 |
| ENSG00000175283 | DOLK | 1533.935 | -0.5840217 | 0.1421848 | -4.180316 | 2.91E-05 | 0.000256 |
| ENSG00000287349 | | 1.580297 | -0.5840796 | 1.056732 | -1.956347 | 0.05042426 | 0.139395 |
| ENSG00000159307 | SCUBE1 | 4.587541 | -0.5841352 | 0.8380704 | -1.341025 | 0.1799125 | 0.354055 |
| ENSG00000232748 | | 25.86199 | -0.5841742 | 0.5113877 | -1.453301 | 0.1461401 | 0.305443 |
| ENSG00000197566 | ZNF624 | 476.8038 | -0.5842258 | 0.1957431 | -3.095912 | 0.00196209 | 0.010186 |
| ENSG00000106263 | EIF3B | 26991.87492 | -0.584274801 | 0.191650999 | -3.153789954 | 0.00161165 | 0.008607 |
| ENSG00000243964 | RPL23AP65 | 73.5808 | -0.5843092 | 0.3103559 | -2.05617 | 0.03976615 | 0.116385 |
| ENSG00000189212 | DPY19L2P1 | 77.51456 | -0.5843206 | 0.3427346 | -1.890708 | 0.05866332 | 0.156457 |
| ENSG00000174015 | CBY2 | 38.84376 | -0.5844788 | 0.3520403 | -1.849745 | 0.06435034 | 0.167456 |
| ENSG00000114993 | RTKN | 3000.701442 | -0.58453675 | 0.152509786 | -3.916341819 | 8.99E-05 | 0.000695 |
| ENSG00000132507 | EIF5A | 10709.56 | -0.584562251 | 0.285634981 | -2.21021318 | 0.02709037 | 0.086417 |
| ENSG00000200769 | | 3.527965 | -0.584568 | 0.7579286 | -1.303677 | 0.1923436 | 0.370693 |
| ENSG00000106853 | PTGR1 | 1414.512511 | -0.584570179 | 0.227475539 | -2.701483073 | 0.0069031 | 0.028815 |
| ENSG00000253910 | PCDHGB2 | 61.01595 | -0.5847268 | 0.2893516 | -2.181355 | 0.0291572 | 0.091808 |
| ENSG00000078967 | UBE2D4 | 341.4539897 | -0.584820701 | 0.183822848 | -3.28591552 | 0.00101652 | 0.005804 |
| ENSG00000154930 | ACSS1 | 1808.575 | -0.5848376 | 0.3165167 | -2.035257 | 0.04182504 | 0.121078 |
| ENSG00000280073 | | 19.52271 | -0.5849274 | 0.6408559 | -1.341548 | 0.1797425 | 0.353904 |
| ENSG00000080608 | PUM3 | 2050.684538 | -0.585088483 | 0.215267498 | -2.842996356 | 0.00446916 | 0.020197 |

| | | | | | | | |
|---|---|---|---|---|---|---|---|
| ENSG00000250258 | | 8.786175 | -0.5852244 | 0.5675455 | -1.38628 | 0.1656615 | 0.334344 |
| ENSG00000276672 | | 132.2495 | -0.585393 | 0.5451769 | -1.41536 | 0.1569629 | 0.321811 |
| ENSG00000239969 | | 1.539774 | -0.5856299 | 0.9589209 | -1.561315 | 0.1184493 | NA |
| ENSG00000262772 | LINC01977 | 298.7652 | -0.5856869 | 0.2752491 | -2.271694 | 0.02310502 | 0.076268 |
| ENSG00000280278 | | 10.84393 | -0.5857893 | 0.6637509 | -1.316571 | 0.1879826 | 0.364936 |
| ENSG00000213930 | GALT | 637.2966 | -0.5858825 | 0.1980044 | -3.079243 | 0.00207527 | 0.010694 |
| ENSG00000256968 | SNRPEP2 | 1872.805 | -0.5859578 | 0.2229585 | -2.750646 | 0.00594778 | 0.025562 |
| ENSG00000267543 | | 6.651007 | -0.5860725 | 0.8128641 | -1.347044 | 0.1779661 | 0.351344 |
| ENSG00000286522 | H3C2 | 99.19601 | -0.586247 | 0.4152354 | -1.653304 | 0.09826903 | 0.229768 |
| ENSG00000271383 | NBPF19 | 18.08174 | -0.5862962 | 0.5773672 | -1.377941 | 0.1682215 | 0.33802 |
| ENSG00000159685 | CHCHD6 | 938.7362 | -0.5864122 | 0.2326188 | -2.648962 | 0.00807394 | 0.032771 |
| ENSG00000244921 | MTCYBP18 | 12.94165 | -0.5865839 | 0.618101 | -1.348959 | 0.1773501 | 0.350622 |
| ENSG00000166136 | NDUFB8 | 509.3378 | -0.5865875 | 0.2359548 | -2.619018 | 0.00881833 | 0.035195 |
| ENSG00000074755 | ZZEF1 | 5330.366234 | -0.586687308 | 0.241975049 | -2.575400986 | 0.01001239 | 0.038897 |
| ENSG00000286001 | | 6.977981 | -0.5867595 | 0.7626562 | -1.30718 | 0.1911515 | 0.36921 |
| ENSG00000118939 | UCHL3 | 114.2152202 | -0.586782982 | 0.324764799 | -1.992297438 | 0.04633843 | 0.130846 |
| ENSG00000234684 | SDCBP2-AS1 | 171.5405 | -0.5868807 | 0.2617042 | -2.378891 | 0.01736483 | 0.060642 |
| ENSG00000161921 | CXCL16 | 3780.924 | -0.5868944 | 0.163974 | -3.668532 | 0.00024395 | 0.00169 |
| ENSG00000174808 | BTC | 513.2604 | -0.5869885 | 0.2079263 | -2.938665 | 0.00329629 | 0.015718 |
| ENSG00000275132 | RN7SL663P | 2.557299 | -0.5873928 | 0.8979251 | -1.377087 | 0.1684854 | 0.338333 |
| ENSG00000255690 | TRIL | 27.63315 | -0.5875124 | 0.5946344 | -1.362414 | 0.1730674 | 0.344474 |
| ENSG00000270722 | RNVU1-31 | 4.800575 | -0.5875476 | 0.7977607 | -1.339634 | 0.1803645 | 0.354629 |
| ENSG00000158864 | NDUFS2 | 5900.58 | -0.5875488 | 0.1588404 | -3.809497 | 0.00013925 | 0.001027 |
| ENSG00000147689 | FAM83A | 294.3575 | -0.5875754 | 0.4181339 | -1.646026 | 0.09975844 | 0.232417 |
| ENSG00000277589 | | 3.082861 | -0.5876787 | 0.8719327 | -1.391309 | 0.1641317 | 0.332207 |
| ENSG00000213539 | YBX1P6 | 3.174809 | -0.5878817 | 0.7677377 | -1.315558 | 0.1883226 | 0.365356 |
| ENSG00000265395 | MIR3944 | 1.673766 | -0.588199 | 0.967912 | -1.594893 | 0.1107361 | 0.250618 |
| ENSG00000111845 | PAK1IP1 | 1398.276014 | -0.588422332 | 0.223825018 | -2.756651993 | 0.00583965 | 0.025195 |
| ENSG00000259920 | | 5.320324 | -0.5884239 | 0.8406252 | -1.369167 | 0.1709472 | 0.341599 |
| ENSG00000237883 | DGUOK-AS1 | 133.0417 | -0.5885237 | 0.2555446 | -2.451141 | 0.0142404 | 0.05173 |
| ENSG00000106610 | STAG3L4 | 1016.413223 | -0.588566179 | 0.162452228 | -3.713214723 | 0.00020464 | 0.001447 |
| ENSG00000176909 | MAMSTR | 61.23753 | -0.5886286 | 0.2983936 | -2.13498 | 0.03276263 | 0.100447 |
| ENSG00000164609 | SLU7 | 3793.125 | -0.5887699 | 0.1278269 | -4.674219 | 2.95E-06 | 3.34E-05 |

| | | | | | | | |
|---|---|---|---|---|---|---|---|
| ENSG00000277117 | | 155.4599 | -0.5894322 | 0.2886978 | -2.196875 | 0.02802935 | 0.088782 |
| ENSG00000189056 | RELN | 1.092386 | -0.5895706 | 1.115183 | -2.214762 | 0.02677638 | NA |
| ENSG00000185674 | LYG2 | 8.267924 | -0.5898343 | 0.584378 | -1.369888 | 0.1707219 | 0.341431 |
| ENSG00000227671 | ZNF731P | 3574.017 | -0.5899905 | 0.2705904 | -2.338567 | 0.01935788 | 0.066193 |
| ENSG00000133028 | SCO1 | 3475.839113 | -0.590287883 | 0.185232452 | -3.290006585 | 0.00100185 | 0.00573 |
| ENSG00000014641 | MDH1 | 9748.972259 | -0.590602585 | 0.243976198 | -2.557041656 | 0.01055666 | 0.040673 |
| ENSG00000283064 | | 25.27586 | -0.590707 | 0.4335771 | -1.610333 | 0.1073251 | 0.244913 |
| ENSG00000276180 | H4C9 | 1302.156 | -0.5907287 | 0.165385 | -3.672477 | 0.00024021 | 0.001668 |
| ENSG00000112667 | DNPH1 | 2720.764832 | -0.590792978 | 0.229364764 | -2.701227597 | 0.00690841 | 0.02882 |
| ENSG00000049618 | ARID1B | 2845.026387 | -0.59081615 | 0.16267366 | -3.737042385 | 0.0001862 | 0.001331 |
| ENSG00000231162 | COX11P1 | 50.91605 | -0.5909969 | 0.3418145 | -1.929845 | 0.05362601 | 0.146523 |
| ENSG00000214263 | RPSAP53 | 1.174262 | -0.5910407 | 1.037928 | -1.728079 | 0.08397399 | NA |
| ENSG00000115993 | TRAK2 | 5100.190156 | -0.591110507 | 0.295331543 | -2.172337938 | 0.02983018 | 0.093264 |
| ENSG00000101391 | CDK5RAP1 | 1987.127028 | -0.591223283 | 0.179367277 | -3.409284084 | 0.00065134 | 0.003946 |
| ENSG00000175518 | UBQLNL | 3.191382 | -0.5913581 | 0.9617311 | -1.547642 | 0.1217085 | 0.267985 |
| ENSG00000165271 | NOL6 | 3985.097 | -0.591538 | 0.1693925 | -3.578584 | 0.00034546 | 0.002288 |
| ENSG00000223572 | CKMT1A | 675.3592 | -0.5916824 | 0.3311731 | -1.981277 | 0.04756026 | 0.133393 |
| ENSG00000226259 | GTF2H2B | 44.2574 | -0.5917109 | 0.2792529 | -2.269133 | 0.02326023 | 0.076637 |
| ENSG00000201555 | | 4.483672 | -0.5919091 | 0.7493648 | -1.327603 | 0.1843093 | 0.359618 |
| ENSG00000160404 | TOR2A | 1040.067 | -0.5919475 | 0.1966236 | -3.121414 | 0.00179985 | 0.009482 |
| ENSG00000063180 | CA11 | 744.2613783 | -0.59217986 | 0.355530146 | -1.878600536 | 0.06029906 | 0.15965 |
| ENSG00000166377 | ATP9B | 724.128 | -0.5922008 | 0.2673002 | -2.363392 | 0.0181085 | 0.062761 |
| ENSG00000265778 | | 73.59884 | -0.5923307 | 0.3629572 | -1.846429 | 0.06482997 | 0.168341 |
| ENSG00000261758 | | 5.598038 | -0.5925361 | 0.7964725 | -1.336888 | 0.1812592 | 0.355804 |
| ENSG00000101363 | MANBAL | 2171.405506 | -0.59255989 | 0.16421062 | -3.699517638 | 0.00021601 | 0.001518 |
| ENSG00000226330 | | 110.9599 | -0.5925694 | 0.3592905 | -1.854298 | 0.06369654 | 0.166194 |
| ENSG00000158467 | AHCYL2 | 1591.208 | -0.5926267 | 0.1708842 | -3.564258 | 0.00036489 | 0.002397 |
| ENSG00000169895 | SYAP1 | 7119.627 | -0.5926791 | 0.1046473 | -5.716736 | 1.09E-08 | 1.98E-07 |
| ENSG00000228350 | LINC02585 | 15.4081 | -0.5927274 | 0.4691409 | -1.53838 | 0.1239557 | 0.271619 |
| ENSG00000120539 | MASTL | 2040.633967 | -0.592867364 | 0.402488947 | -1.717066523 | 0.08596702 | 0.208361 |
| ENSG00000144182 | LIPT1 | 184.7079 | -0.5929602 | 0.2034967 | -3.026347 | 0.00247528 | 0.012395 |
| ENSG00000210135 | MT-TN | 106.5084 | -0.5930168 | 0.2250533 | -2.756467 | 0.00584295 | 0.025205 |
| ENSG00000138442 | WDR12 | 2171.215058 | -0.593174916 | 0.152523129 | -3.974512647 | 7.05E-05 | 0.000561 |

| | | | | | | | |
|---|---|---|---|---|---|---|---|
| ENSG00000240668 | KRT8P36 | 34.34608 | -0.5932174 | 0.4114748 | -1.684424 | 0.09209973 | 0.219134 |
| ENSG00000237798 | | 1.56279 | -0.5936886 | 1.030485 | -1.725822 | 0.0843795 | NA |
| ENSG00000214207 | KRT18P10 | 452.2081 | -0.5937167 | 0.3075997 | -2.108006 | 0.03503045 | 0.105668 |
| ENSG00000123146 | ADGRE5 | 715.2198962 | -0.593720595 | 0.186078224 | -3.300101312 | 0.0009665 | 0.005551 |
| ENSG00000204356 | NELFE | 3951.996 | -0.5937294 | 0.2180233 | -2.852293 | 0.00434051 | 0.01972 |
| ENSG00000242689 | CNTF | 31.10274 | -0.5939123 | 0.4151839 | -1.676055 | 0.09372741 | 0.222088 |
| ENSG00000240974 | RPS27AP10 | 0.9745094 | -0.5941141 | 1.05235 | -1.77921 | 0.07520539 | NA |
| ENSG00000101189 | MRGBP | 4524.161587 | -0.594145541 | 0.219246988 | -2.837923314 | 0.00454081 | 0.020486 |
| ENSG00000214282 | KRT8P14 | 66.43979 | -0.5943592 | 0.2961468 | -2.172976 | 0.0297821 | 0.09318 |
| ENSG00000212664 | | 433.3245 | -0.5945874 | 0.2252891 | -2.765092 | 0.00569067 | 0.024657 |
| ENSG00000237667 | LINC01115 | 11.75402 | -0.5946297 | 0.4730734 | -1.537548 | 0.1241591 | 0.271947 |
| ENSG00000147592 | LACTB2 | 884.8373 | -0.5951432 | 0.1636513 | -3.723731 | 0.0001963 | 0.001394 |
| ENSG00000166963 | MAP1A | 47.2617 | -0.595145 | 0.4300209 | -1.647052 | 0.09954727 | 0.232006 |
| ENSG00000279133 | | 75.30482 | -0.595271 | 0.4533428 | -1.584037 | 0.1131852 | 0.254396 |
| ENSG00000255120 | OVOL1-AS1 | 34.36632 | -0.5954405 | 0.5181917 | -1.462417 | 0.1436271 | 0.301566 |
| ENSG00000178105 | DDX10 | 1263.109 | -0.595463 | 0.1279482 | -4.715638 | 2.41E-06 | 2.78E-05 |
| ENSG00000154719 | MRPL39 | 1320.385 | -0.5955548 | 0.2982613 | -2.168347 | 0.03013228 | 0.093987 |
| ENSG00000167600 | CYP2S1 | 21.65676 | -0.5955625 | 0.5231867 | -1.459829 | 0.144337 | 0.302627 |
| ENSG00000144504 | ANKMY1 | 1194.892 | -0.595619 | 0.1579017 | -3.857742 | 0.00011444 | 0.000864 |
| ENSG00000137101 | CD72 | 35.23454893 | -0.595800323 | 0.561071583 | -1.412256639 | 0.15787441 | 0.323133 |
| ENSG00000004766 | VPS50 | 1335.946412 | -0.595815764 | 0.177782794 | -3.452642875 | 0.00055512 | 0.003436 |
| ENSG00000163162 | RNF149 | 2947.937 | -0.5961491 | 0.1410263 | -4.312878 | 1.61E-05 | 0.000151 |
| ENSG00000271267 | | 14.16264 | -0.5962088 | 0.6337663 | -1.352968 | 0.1760659 | 0.348706 |
| ENSG00000165076 | PRSS37 | 1.384552 | -0.5964364 | 1.042479 | -1.849048 | 0.06445095 | NA |
| ENSG00000232842 | KAT7P1 | 82.51199 | -0.5965314 | 0.2603393 | -2.440854 | 0.01465256 | 0.052926 |
| ENSG00000188599 | NPIPP1 | 156.7448 | -0.5966387 | 0.2486049 | -2.542074 | 0.01101967 | 0.042097 |
| ENSG00000224126 | UBE2SP2 | 267.2784 | -0.5966603 | 0.3847615 | -1.783378 | 0.07452479 | 0.187239 |
| ENSG00000226926 | PDZPH1P | 1.609039 | -0.5968796 | 0.9793801 | -1.633768 | 0.1023077 | 0.236524 |
| ENSG00000184925 | LCN12 | 117.6008 | -0.5969808 | 0.3097601 | -2.106229 | 0.03518447 | 0.106036 |
| ENSG00000230360 | DDX10P2 | 3.979263 | -0.5971181 | 0.7456382 | -1.336976 | 0.1812305 | 0.355804 |
| ENSG00000112655 | PTK7 | 2742.490397 | -0.597126706 | 0.202201256 | -3.06825685 | 0.00215312 | 0.011035 |
| ENSG00000074219 | TEAD2 | 2696.501739 | -0.597190101 | 0.254662991 | -2.498484984 | 0.01247254 | 0.046464 |
| ENSG00000116285 | ERRFI1 | 2316.105988 | -0.597200684 | 0.359270143 | -1.869572628 | 0.06154319 | 0.162086 |

| | | | | | | | |
|---|---|---|---|---|---|---|---|
| ENSG00000182376 | | 7.901801 | -0.5973742 | 0.6850086 | -1.32748 | 0.1843499 | 0.359644 |
| ENSG00000286545 | | 97.96872 | -0.5976093 | 0.3539605 | -1.903384 | 0.05699043 | 0.152994 |
| ENSG00000156261 | CCT8 | 10296.86 | -0.5976223 | 0.2354448 | -2.700177 | 0.00693027 | 0.028892 |
| ENSG00000185650 | ZFP36L1 | 13223.21 | -0.5977252 | 0.2727343 | -2.345651 | 0.0189939 | 0.065209 |
| ENSG00000237491 | LINC01409 | 120.1707 | -0.5980492 | 0.4249097 | -1.656784 | 0.09756311 | 0.228525 |
| ENSG00000127804 | METTL16 | 1801.348335 | -0.598591136 | 0.124584698 | -4.881666276 | 1.05E-06 | 1.30E-05 |
| ENSG00000144115 | THNSL2 | 1384.272 | -0.5987991 | 0.2103679 | -2.959104 | 0.00308535 | 0.014877 |
| ENSG00000260804 | LINC01963 | 272.1825 | -0.598974 | 0.2881004 | -2.2455 | 0.02473603 | 0.080413 |
| ENSG00000138081 | FBXO11 | 2037.739845 | -0.598996516 | 0.165527989 | -3.712653788 | 0.0002051 | 0.00145 |
| ENSG00000199961 | SNORD1B | 5.624958 | -0.5991697 | 0.6167976 | -1.361041 | 0.1735009 | 0.345041 |
| ENSG00000103184 | SEC14L5 | 19.8588715 | -0.59922701 | 0.489834961 | -1.518194071 | 0.12896548 | 0.279172 |
| ENSG00000242338 | BMS1P4 | 49.98554 | -0.5992877 | 0.4762305 | -1.551274 | 0.1208361 | 0.26663 |
| ENSG00000228513 | | 1.380676 | -0.5994037 | 1.105714 | -2.074364 | 0.03804552 | NA |
| ENSG00000236496 | GPS2P1 | 150.9239 | -0.5994714 | 0.287142 | -2.250823 | 0.02439673 | 0.079553 |
| ENSG00000165997 | ARL5B | 2705.334 | -0.5994867 | 0.2365249 | -2.668571 | 0.00761746 | 0.031281 |
| ENSG00000138095 | LRPPRC | 19129.49467 | -0.599499865 | 0.161025214 | -3.815887752 | 0.00013569 | 0.001003 |
| ENSG00000248578 | NPM1P21 | 30.59024 | -0.5995819 | 0.4422487 | -1.624678 | 0.1042311 | 0.239788 |
| ENSG00000134490 | TMEM241 | 954.8261464 | -0.599594891 | 0.161498765 | -3.808831239 | 0.00013963 | 0.001028 |
| ENSG00000268916 | CSAG3 | 2.345702 | -0.5997558 | 1.036323 | -1.685035 | 0.09198174 | 0.218965 |
| ENSG00000143933 | CALM2 | 13929.44 | -0.6000258 | 0.2219185 | -2.829725 | 0.00465881 | 0.020941 |
| ENSG00000258303 | | 1.709933 | -0.6000275 | 1.010715 | -1.715918 | 0.08617706 | 0.208642 |
| ENSG00000058056 | USP13 | 2453.042385 | -0.600120612 | 0.1618501 | -3.800965447 | 0.00014413 | 0.001058 |
| ENSG00000134940 | ACRV1 | 13.0795979 | -0.600301111 | 0.720898898 | -1.351820917 | 0.17643261 | 0.349192 |
| ENSG00000111981 | ULBP1 | 809.5870155 | -0.60033861 | 0.493196494 | -1.524185465 | 0.12746239 | 0.276792 |
| ENSG00000165055 | METTL2B | 1275.708 | -0.6006069 | 0.09821523 | -6.165992 | 7.00E-10 | 1.55E-08 |
| ENSG00000280383 | | 23.85323 | -0.6011103 | 0.4414271 | -1.614109 | 0.1065037 | 0.243666 |
| ENSG00000065485 | PDIA5 | 1827.182144 | -0.601213091 | 0.156072349 | -3.920454593 | 8.84E-05 | 0.000686 |
| ENSG00000153207 | AHCTF1 | 6840.005 | -0.6012155 | 0.1908008 | -3.262156 | 0.00110568 | 0.00624 |
| ENSG00000198920 | KIAA0753 | 1329.643 | -0.601344 | 0.1178744 | -5.166097 | 2.39E-07 | 3.37E-06 |
| ENSG00000136108 | CKAP2 | 3171.039762 | -0.601473231 | 0.243538415 | -2.592143929 | 0.00953799 | 0.037388 |
| ENSG00000213036 | | 53.70451 | -0.6015132 | 0.3395986 | -1.975743 | 0.04818385 | 0.134668 |
| ENSG00000259984 | | 3.294182 | -0.6017264 | 0.8389802 | -1.368692 | 0.1710957 | 0.34171 |
| ENSG00000172974 | VDAC2P5 | 3417.372 | -0.6020513 | 0.191926 | -3.243527 | 0.0011806 | 0.006593 |

| | | | | | | | |
|---|---|---|---|---|---|---|---|
| ENSG00000215912 | TTC34 | 61.59977 | -0.6020989 | 0.4982635 | -1.512826 | 0.1303238 | 0.281281 |
| ENSG00000101665 | SMAD7 | 1024.378447 | -0.602099605 | 0.271127437 | -2.379019272 | 0.01735877 | 0.060629 |
| ENSG00000123643 | SLC36A1 | 1490.616639 | -0.602103149 | 0.246326546 | -2.584567043 | 0.00975014 | 0.038085 |
| ENSG00000198814 | GK | 141.6625 | -0.6021592 | 0.3881558 | -1.782966 | 0.07459185 | 0.187336 |
| ENSG00000140263 | SORD | 3723.424 | -0.6024687 | 0.2038206 | -3.07281 | 0.00212054 | 0.010893 |
| ENSG00000116062 | MSH6 | 4719.892312 | -0.602486637 | 0.284108817 | -2.284280344 | 0.02235506 | 0.074486 |
| ENSG00000239553 | RN7SL12P | 3.980099 | -0.6025017 | 0.7317457 | -1.333558 | 0.1823489 | 0.357335 |
| ENSG00000205571 | SMN2 | 425.1963 | -0.6026191 | 0.1577054 | -3.921068 | 8.82E-05 | 0.000685 |
| ENSG00000158748 | HTR6 | 23.22499 | -0.6026438 | 0.6557007 | -1.369272 | 0.1709141 | 0.341599 |
| ENSG00000276600 | RAB7B | 61.22801 | -0.6029104 | 0.3484373 | -1.935455 | 0.05293452 | 0.144992 |
| ENSG00000141519 | CCDC40 | 432.0041 | -0.6032283 | 0.3906296 | -1.766462 | 0.07731836 | 0.192294 |
| ENSG00000231822 | SMC3P1 | 169.677 | -0.6033465 | 0.2354367 | -2.697417 | 0.00698798 | 0.029074 |
| ENSG00000210196 | MT-TP | 1874.36 | -0.6033491 | 0.201399 | -3.110814 | 0.00186573 | 0.009764 |
| ENSG00000160185 | UBASH3A | 2.162676 | -0.6033492 | 1.062127 | -1.872864 | 0.06108723 | 0.161207 |
| ENSG00000204149 | AGAP6 | 121.8277 | -0.6033987 | 0.2402549 | -2.652053 | 0.00800039 | 0.032537 |
| ENSG00000123965 | PMS2P5 | 39.51115885 | -0.603698477 | 0.324288556 | -2.044747172 | 0.04087979 | 0.119042 |
| ENSG00000259781 | HMGB1P6 | 2246.739 | -0.6037724 | 0.3533707 | -1.907898 | 0.0564044 | 0.151973 |
| ENSG00000242588 | | 258.9787 | -0.604047 | 0.4480599 | -1.626257 | 0.103895 | 0.239222 |
| ENSG00000224224 | HAUS1P2 | 1.227981 | -0.6042224 | 1.08125 | -1.962808 | 0.04966851 | NA |
| ENSG00000143093 | STRIP1 | 3436.387 | -0.6043289 | 0.1374867 | -4.479589 | 7.48E-06 | 7.63E-05 |
| ENSG00000215089 | KRT18P11 | 946.9965 | -0.6043853 | 0.2868932 | -2.282151 | 0.02248043 | 0.074791 |
| ENSG00000137073 | UBAP2 | 2821.015107 | -0.604452111 | 0.13412036 | -4.591569639 | 4.40E-06 | 4.76E-05 |
| ENSG00000236008 | LINC01814 | 123.5224 | -0.6045239 | 0.3262682 | -2.041715 | 0.04117982 | 0.119718 |
| ENSG00000129245 | FXR2 | 2984.925194 | -0.604687533 | 0.147793232 | -4.163039125 | 3.14E-05 | 0.000274 |
| ENSG00000233347 | ERP29P1 | 68.04144 | -0.6047328 | 0.352 | -1.923408 | 0.0544288 | 0.148039 |
| ENSG00000226180 | | 97.46167 | -0.6051068 | 0.316548 | -2.09113 | 0.03651642 | 0.108975 |
| ENSG00000084693 | AGBL5 | 1793.115257 | -0.605258662 | 0.139959243 | -4.40765318 | 1.04E-05 | 0.000102 |
| ENSG00000232729 | | 9.49606 | -0.6053191 | 0.6850621 | -1.351553 | 0.1765185 | 0.349264 |
| ENSG00000006530 | AGK | 1823.713406 | -0.605357757 | 0.147939805 | -4.179180338 | 2.93E-05 | 0.000257 |
| ENSG00000271797 | | 6.515866 | -0.6054524 | 0.6475358 | -1.369404 | 0.1708731 | 0.341599 |
| ENSG00000108963 | DPH1 | 1563.809135 | -0.605643653 | 0.149482423 | -4.129759352 | 3.63E-05 | 0.000312 |
| ENSG00000009954 | BAZ1B | 7586.984784 | -0.605733676 | 0.169715727 | -3.669201824 | 0.00024331 | 0.001687 |
| ENSG00000198929 | NOS1AP | 446.1357 | -0.6059808 | 0.3557127 | -1.913756 | 0.05565134 | 0.150387 |

| | | | | | | | |
|---|---|---|---|---|---|---|---|
| ENSG00000115816 | CEBPZ | 2229.119952 | -0.606117514 | 0.186753992 | -3.350408455 | 0.00080693 | 0.004754 |
| ENSG00000164296 | TIGD6 | 391.9084 | -0.6061357 | 0.1709432 | -3.641495 | 0.00027106 | 0.00185 |
| ENSG00000168476 | REEP4 | 2140.186 | -0.6062774 | 0.2583183 | -2.491237 | 0.01272991 | 0.047231 |
| ENSG00000250363 | KRT18P21 | 12.86365 | -0.6064504 | 0.5930055 | -1.408327 | 0.1590344 | 0.324955 |
| ENSG00000236782 | | 10.53177 | -0.606702 | 0.5420151 | -1.463828 | 0.1432409 | 0.301041 |
| ENSG00000136146 | MED4 | 1263.830227 | -0.606731363 | 0.171832801 | -3.630969617 | 0.00028236 | 0.001918 |
| ENSG00000184428 | TOP1MT | 2972.774 | -0.6067678 | 0.159885 | -3.920204 | 8.85E-05 | 0.000687 |
| ENSG00000286430 | | 21.86571 | -0.606975 | 0.3918315 | -1.780664 | 0.07496731 | 0.18803 |
| ENSG00000255362 | LINC02761 | 4.198667 | -0.6071673 | 0.7393675 | -1.358475 | 0.1743129 | 0.346241 |
| ENSG00000124383 | MPHOSPH10 | 3807.895091 | -0.607213438 | 0.1358198 | -4.501564863 | 6.75E-06 | 6.95E-05 |
| ENSG00000251206 | | 2.83736 | -0.607306 | 1.060936 | -1.860714 | 0.06278453 | 0.164471 |
| ENSG00000211455 | STK38L | 1398.839 | -0.6073656 | 0.2264617 | -2.822417 | 0.00476632 | 0.021331 |
| ENSG00000248489 | LINC02062 | 49.2452 | -0.6075027 | 0.3313475 | -2.020915 | 0.04328856 | 0.124377 |
| ENSG00000240211 | PPP1R35-AS1 | 7.742761 | -0.6076498 | 0.593815 | -1.406018 | 0.1597188 | 0.325827 |
| ENSG00000230010 | | 12.28586 | -0.6077071 | 0.6593454 | -1.366025 | 0.1719311 | 0.342766 |
| ENSG00000148730 | EIF4EBP2 | 8413.107 | -0.6077137 | 0.2344771 | -2.726363 | 0.00640366 | 0.027075 |
| ENSG00000231689 | LINC01090 | 1.361096 | -0.6077199 | 1.095003 | -2.047051 | 0.04065309 | NA |
| ENSG00000255265 | TMA16P1 | 2.38609 | -0.6077446 | 0.9287108 | -1.512178 | 0.1304885 | 0.281568 |
| ENSG00000247137 | | 173.8457 | -0.6078116 | 0.3704466 | -1.858875 | 0.06304486 | 0.16498 |
| ENSG00000255622 | PCDHB17P | 10.25982 | -0.6078397 | 0.5444873 | -1.447595 | 0.1477302 | 0.30775 |
| ENSG00000183258 | DDX41 | 8128.159 | -0.6080397 | 0.1720403 | -3.637219 | 0.0002756 | 0.001875 |
| ENSG00000260899 | | 2.418211 | -0.6084294 | 1.009114 | -1.668265 | 0.09526315 | 0.224563 |
| ENSG00000113593 | PPWD1 | 1765.708246 | -0.608458939 | 0.132848451 | -4.65663354 | 3.21E-06 | 3.59E-05 |
| ENSG00000115318 | LOXL3 | 586.4763033 | -0.608555941 | 0.356880457 | -1.91652563 | 0.05529822 | 0.149661 |
| ENSG00000166435 | XRRA1 | 392.799 | -0.6089966 | 0.2118302 | -2.987132 | 0.00281608 | 0.013851 |
| ENSG00000062822 | POLD1 | 2814.938801 | -0.609355594 | 0.377903151 | -1.838492545 | 0.06598986 | 0.170769 |
| ENSG00000197362 | ZNF786 | 1025.072 | -0.609595 | 0.1089784 | -5.639546 | 1.70E-08 | 2.97E-07 |
| ENSG00000122512 | PMS2 | 2268.306797 | -0.609723498 | 0.153775865 | -4.05879752 | 4.93E-05 | 0.00041 |
| ENSG00000141447 | OSBPL1A | 1363.065 | -0.6097551 | 0.2363682 | -2.717442 | 0.00657887 | 0.027692 |
| ENSG00000173269 | MMRN2 | 64.78673 | -0.609775 | 0.3907481 | -1.77141 | 0.07649251 | 0.190775 |
| ENSG00000183793 | NPIPA5 | 7.061653 | -0.6098316 | 0.6003956 | -1.398025 | 0.1621055 | 0.329383 |
| ENSG00000260810 | | 23.08219 | -0.609867 | 0.4121184 | -1.737189 | 0.08235374 | 0.201813 |
| ENSG00000152465 | NMT2 | 1058.555 | -0.6103595 | 0.1421934 | -4.351992 | 1.35E-05 | 0.000129 |

| | | | | | | | |
|---|---|---|---|---|---|---|---|
| ENSG00000242759 | LINC00882 | 8.869128 | -0.6108424 | 0.5171757 | -1.504132 | 0.1325475 | 0.284854 |
| ENSG00000230913 | NPM1P51 | 4.156266 | -0.6110488 | 0.8022071 | -1.376459 | 0.1686797 | 0.33856 |
| ENSG00000177853 | ZNF518A | 1910.518 | -0.6114052 | 0.2191179 | -2.916343 | 0.00354161 | 0.016681 |
| ENSG00000004142 | POLDIP2 | 8667.214882 | -0.611443462 | 0.206680192 | -3.079193721 | 0.00207562 | 0.010694 |
| ENSG00000234630 | | 25.72956 | -0.6114464 | 0.6118433 | -1.406167 | 0.1596744 | 0.325761 |
| ENSG00000131591 | C1orf159 | 1162.705054 | -0.611661241 | 0.149364686 | -4.170441903 | 3.04E-05 | 0.000266 |
| ENSG00000252050 | | 2.276102 | -0.6118646 | 0.8893237 | -1.442145 | 0.1492614 | 0.310016 |
| ENSG00000228140 | | 1.733554 | -0.611873 | 0.9476951 | -1.53724 | 0.1242345 | 0.27199 |
| ENSG00000228782 | MRPL45P2 | 426.034 | -0.6120023 | 0.2114499 | -3.017318 | 0.00255022 | 0.012715 |
| ENSG00000231066 | NPM1P9 | 9.875162 | -0.6121725 | 0.5844146 | -1.431516 | 0.1522823 | 0.314496 |
| ENSG00000115128 | SF3B6 | 4644.556307 | -0.6123305 | 0.19167706 | -3.306009207 | 0.00094635 | 0.005446 |
| ENSG00000136104 | RNASEH2B | 1186.558734 | -0.612344513 | 0.179077653 | -3.528228157 | 0.00041835 | 0.002689 |
| ENSG00000089048 | ESF1 | 1811.909652 | -0.612765521 | 0.178771565 | -3.547120497 | 0.00038947 | 0.002537 |
| ENSG00000133640 | LRRIQ1 | 9.813368451 | -0.612829868 | 0.665528979 | -1.368853382 | 0.17104511 | 0.341692 |
| ENSG00000251365 | LINC02236 | 57.80672 | -0.61287 | 0.4449585 | -1.650657 | 0.09880864 | 0.230636 |
| ENSG00000102967 | DHODH | 1211.025189 | -0.613036349 | 0.187225721 | -3.388382163 | 0.00070306 | 0.004219 |
| ENSG00000230873 | STMND1 | 3.368573 | -0.6130929 | 0.780673 | -1.36625 | 0.1718604 | 0.342702 |
| ENSG00000164889 | SLC4A2 | 11962.37 | -0.6131247 | 0.1150968 | -5.390391 | 7.03E-08 | 1.10E-06 |
| ENSG00000072401 | UBE2D1 | 1369.871838 | -0.613251977 | 0.171733451 | -3.670448764 | 0.00024213 | 0.00168 |
| ENSG00000273165 | | 1.331208 | -0.613507 | 1.055101 | -1.734926 | 0.08275397 | NA |
| ENSG00000283390 | | 63.67554 | -0.6139016 | 0.3174118 | -2.122624 | 0.03378534 | 0.102846 |
| ENSG00000171540 | OTP | 6.592303 | -0.6141106 | 0.7313778 | -1.370779 | 0.1704438 | 0.341114 |
| ENSG00000163121 | NEURL3 | 29.64365 | -0.6141253 | 0.7818562 | -1.34201 | 0.1795927 | 0.353661 |
| ENSG00000205592 | MUC19 | 25.37708 | -0.6142235 | 0.5549684 | -1.465231 | 0.1428579 | 0.300498 |
| ENSG00000160396 | HIPK4 | 8.431939 | -0.6142924 | 0.5914109 | -1.418319 | 0.1560977 | 0.320458 |
| ENSG00000171823 | FBXL14 | 736.1642 | -0.6143368 | 0.2145618 | -2.992489 | 0.00276713 | 0.013638 |
| ENSG00000226084 | | 1735.103 | -0.6144641 | 0.1711655 | -3.688061 | 0.00022597 | 0.00158 |
| ENSG00000231240 | KLF2P1 | 163.5866 | -0.6145034 | 0.4218765 | -1.719932 | 0.08544472 | 0.207429 |
| ENSG00000236764 | COX7A2P2 | 14.6717 | -0.6145104 | 0.5138603 | -1.505988 | 0.1320703 | 0.284098 |
| ENSG00000231439 | WASIR2 | 46.63636 | -0.6145211 | 0.4272358 | -1.692781 | 0.0904972 | 0.216517 |
| ENSG00000232850 | | 29.62625 | -0.6145855 | 0.5642893 | -1.449197 | 0.1472826 | 0.307131 |
| ENSG00000103249 | CLCN7 | 5186.024429 | -0.614903806 | 0.32342685 | -2.092737851 | 0.03637256 | 0.108713 |
| ENSG00000204438 | GPANK1 | 1016.119 | -0.6149432 | 0.1812524 | -3.495733 | 0.00047276 | 0.002991 |

| | | | | | | | |
|---|---|---|---|---|---|---|---|
| ENSG00000173163 | COMMD1 | 866.2234 | -0.6149714 | 0.2258875 | -2.858749 | 0.00425315 | 0.019399 |
| ENSG00000233325 | MIPEPP3 | 66.50888 | -0.6150495 | 0.2703185 | -2.423665 | 0.01536475 | 0.054943 |
| ENSG00000204950 | LRRC10B | 81.04032 | -0.6151707 | 0.4656277 | -1.612564 | 0.1068392 | 0.244163 |
| ENSG00000268584 | | 70.07884 | -0.6152246 | 0.3134463 | -2.143941 | 0.03203761 | 0.098675 |
| ENSG00000103196 | CRISPLD2 | 766.9241901 | -0.615296901 | 0.373391054 | -1.872596731 | 0.06112409 | 0.161272 |
| ENSG00000235105 | | 231.3629 | -0.6153589 | 0.2373486 | -2.730872 | 0.0063167 | 0.026797 |
| ENSG00000168310 | IRF2 | 1324.538 | -0.6154063 | 0.2076761 | -3.082227 | 0.00205458 | 0.010602 |
| ENSG00000265490 | | 9.723765 | -0.6157919 | 0.5577208 | -1.451329 | 0.1466881 | 0.306228 |
| ENSG00000196550 | FAM72A | 328.7505 | -0.6161122 | 0.4606605 | -1.616906 | 0.1058987 | 0.242621 |
| ENSG00000243753 | HLA-L | 121.9198 | -0.6162376 | 0.4033394 | -1.767539 | 0.07713803 | 0.191953 |
| ENSG00000277758 | | 95.08035 | -0.6164397 | 0.4798051 | -1.578176 | 0.1145251 | 0.256561 |
| ENSG00000222071 | MIR1915 | 3.098716 | -0.6165683 | 0.9636586 | -1.556506 | 0.1195877 | 0.264558 |
| ENSG00000116127 | ALMS1 | 2367.693407 | -0.616593909 | 0.225820599 | -2.863688203 | 0.0041874 | 0.019175 |
| ENSG00000087263 | OGFOD1 | 2081.792684 | -0.616660171 | 0.151903519 | -4.151671359 | 3.30E-05 | 0.000286 |
| ENSG00000260326 | PHBP21 | 21.42045 | -0.6167906 | 0.4598228 | -1.626536 | 0.1038356 | 0.239106 |
| ENSG00000257350 | | 4.871731 | -0.6168255 | 0.7663528 | -1.380841 | 0.1673278 | 0.336747 |
| ENSG00000138670 | RASGEF1B | 86.85916293 | -0.616867913 | 0.43228199 | -1.693468949 | 0.09036624 | 0.216281 |
| ENSG00000284505 | LYNX1-SLURP2 | 2.188696 | -0.6169833 | 0.9389036 | -1.491776 | 0.1357578 | 0.289396 |
| ENSG00000261659 | | 62.08391 | -0.6172385 | 0.4284636 | -1.702086 | 0.08873931 | 0.213484 |
| ENSG00000080839 | RBL1 | 815.7935723 | -0.617752155 | 0.396520118 | -1.785967392 | 0.07410454 | 0.186571 |
| ENSG00000172361 | CFAP53 | 86.96506 | -0.6177568 | 0.3347093 | -2.035705 | 0.04178003 | 0.121 |
| ENSG00000248973 | | 3.804938 | -0.6178021 | 0.8695613 | -1.423622 | 0.154556 | 0.318163 |
| ENSG00000128510 | CPA4 | 3.333259567 | -0.617813411 | 1.078585817 | -1.810119665 | 0.07027723 | 0.179176 |
| ENSG00000102309 | PIN4 | 1226.05538 | -0.617836482 | 0.207473041 | -3.096051533 | 0.00196116 | 0.010184 |
| ENSG00000198546 | ZNF511 | 479.4938 | -0.617914 | 0.313279 | -2.151451 | 0.03144063 | 0.097303 |
| ENSG00000083857 | FAT1 | 8166.294572 | -0.618130719 | 0.305833393 | -2.201761297 | 0.02768217 | 0.087893 |
| ENSG00000171103 | TRMT61B | 820.2545 | -0.6181827 | 0.1386361 | -4.544715 | 5.50E-06 | 5.80E-05 |
| ENSG00000053438 | NNAT | 47.29458048 | -0.61819883 | 0.418579774 | -1.728256059 | 0.08394233 | 0.204614 |
| ENSG00000203880 | PCMTD2 | 6664.955 | -0.6183754 | 0.223735 | -2.891452 | 0.00383466 | 0.017806 |
| ENSG00000110090 | CPT1A | 10799.85735 | -0.619157451 | 0.169948772 | -3.787806613 | 0.00015198 | 0.001111 |
| ENSG00000094880 | CDC23 | 3584.739948 | -0.619157478 | 0.179244455 | -3.574994327 | 0.00035024 | 0.002314 |
| ENSG00000116199 | FAM20B | 4226.051503 | -0.619264058 | 0.14437099 | -4.394379085 | 1.11E-05 | 0.000108 |
| ENSG00000214517 | PPME1 | 2523.549 | -0.6193462 | 0.1826913 | -3.496178 | 0.00047197 | 0.002988 |

| | | | | | | | |
|---|---|---|---|---|---|---|---|
| ENSG00000261663 | | 108.1765 | -0.6194086 | 0.2864319 | -2.332643 | 0.01966689 | 0.067086 |
| ENSG00000227598 | | 4.088399 | -0.6195471 | 0.7727628 | -1.376433 | 0.1686876 | 0.33856 |
| ENSG00000137133 | HINT2 | 37.07073635 | -0.619576927 | 0.363730252 | -1.922172949 | 0.054584 | 0.148378 |
| ENSG00000231043 | | 1222.734 | -0.6195899 | 0.1363284 | -4.658447 | 3.19E-06 | 3.57E-05 |
| ENSG00000225678 | | 5.097559 | -0.6196059 | 0.7421111 | -1.358434 | 0.174326 | 0.346241 |
| ENSG00000203650 | LINC01285 | 7.223361 | -0.6197298 | 0.8908933 | -1.398697 | 0.1619039 | 0.329122 |
| ENSG00000119541 | VPS4B | 2665.961027 | -0.619806097 | 0.167673485 | -3.770067721 | 0.0001632 | 0.001186 |
| ENSG00000265681 | RPL17 | 693.2371 | -0.6198125 | 0.2518159 | -2.617509 | 0.00885742 | 0.035303 |
| ENSG00000229019 | | 1.173203 | -0.620054 | 1.01606 | -1.682608 | 0.09245091 | NA |
| ENSG00000112759 | SLC29A1 | 2330.630301 | -0.620092532 | 0.359230973 | -1.939396341 | 0.0524531 | 0.143866 |
| ENSG00000259881 | | 62.46085 | -0.6202165 | 0.4033719 | -1.783167 | 0.07455907 | 0.187289 |
| ENSG00000283635 | | 3.068676 | -0.6203047 | 0.8142685 | -1.40118 | 0.1611602 | 0.327936 |
| ENSG00000067248 | DHX29 | 2989.11147 | -0.620681739 | 0.142210514 | -4.451310206 | 8.53E-06 | 8.57E-05 |
| ENSG00000115289 | PCGF1 | 1374.648056 | -0.621122313 | 0.189531474 | -3.394083644 | 0.00068859 | 0.004143 |
| ENSG00000247373 | TMED2-DT | 190.6997 | -0.6213298 | 0.3187439 | -2.129164 | 0.03324068 | 0.101607 |
| ENSG00000231840 | TMEM139-AS1 | 4.174549 | -0.6217246 | 0.9452763 | -1.547663 | 0.1217034 | 0.267985 |
| ENSG00000122970 | IFT81 | 675.4586144 | -0.621877077 | 0.13442854 | -4.710881279 | 2.47E-06 | 2.84E-05 |
| ENSG00000204010 | IFIT1B | 1.054329 | -0.6219442 | 1.077741 | -1.807567 | 0.07067399 | NA |
| ENSG00000162779 | AXDND1 | 8.430825 | -0.621951 | 0.6780595 | -1.386202 | 0.1656851 | 0.334344 |
| ENSG00000111711 | GOLT1B | 1702.2819 | -0.62207418 | 0.139102197 | -4.550439163 | 5.35E-06 | 5.67E-05 |
| ENSG00000233270 | SNRPEP4 | 2629.672 | -0.6222987 | 0.2303239 | -2.837069 | 0.00455298 | 0.020531 |
| ENSG00000213939 | | 52.26237 | -0.622308 | 0.3836171 | -1.855352 | 0.06354597 | 0.165883 |
| ENSG00000136068 | FLNB | 11485.43478 | -0.622352557 | 0.298689607 | -2.263327333 | 0.02361552 | 0.077538 |
| ENSG00000126602 | TRAP1 | 6589.683827 | -0.622618971 | 0.246723861 | -2.668422504 | 0.00762084 | 0.031285 |
| ENSG00000273768 | RNVU1-29 | 3.76192 | -0.622704 | 0.7217687 | -1.372886 | 0.1697877 | 0.340152 |
| ENSG00000222726 | RNU2-7P | 5.828514 | -0.6227937 | 0.6382375 | -1.403901 | 0.1603485 | 0.32681 |
| ENSG00000092010 | PSME1 | 13492.04806 | -0.622919221 | 0.176721375 | -3.634982896 | 0.000278 | 0.00189 |
| ENSG00000139746 | RBM26 | 3050.44 | -0.6231237 | 0.1492519 | -4.263335 | 2.01E-05 | 0.000184 |
| ENSG00000176715 | ACSF3 | 2785.641 | -0.6232227 | 0.1201972 | -5.251946 | 1.51E-07 | 2.21E-06 |
| ENSG00000152117 | SMPD4BP | 2897.078 | -0.6232279 | 0.1891278 | -3.404189 | 0.00066361 | 0.004014 |
| ENSG00000002079 | MYH16 | 14.24391396 | -0.623343319 | 0.45691519 | -1.635973204 | 0.1018452 | 0.235749 |
| ENSG00000176225 | RTTN | 492.3253 | -0.6235816 | 0.3402482 | -2.042925 | 0.04105986 | 0.119454 |
| ENSG00000182870 | GALNT9 | 2.776355 | -0.6235971 | 0.9261688 | -1.468977 | 0.1418391 | 0.298923 |

| | | | | | | | |
|---|---|---|---|---|---|---|---|
| ENSG00000157429 | ZNF19 | 223.1662 | -0.6237309 | 0.1852365 | -3.47716 | 0.00050676 | 0.003172 |
| ENSG00000118960 | HS1BP3 | 2140.068594 | -0.623862073 | 0.263849425 | -2.520477378 | 0.01171958 | 0.04418 |
| ENSG00000177192 | PUS1 | 2610.87 | -0.6240024 | 0.1597154 | -3.990463 | 6.59E-05 | 0.000528 |
| ENSG00000115207 | GTF3C2 | 4096.856638 | -0.624016437 | 0.138837478 | -4.584986756 | 4.54E-06 | 4.89E-05 |
| ENSG00000127366 | TAS2R5 | 22.14796744 | -0.624066777 | 0.553383879 | -1.487154255 | 0.13697408 | 0.291234 |
| ENSG00000119777 | TMEM214 | 5067.014772 | -0.624144564 | 0.132264097 | -4.808523547 | 1.52E-06 | 1.83E-05 |
| ENSG00000271862 | | 34.86824 | -0.6245643 | 0.3410034 | -2.039212 | 0.04142884 | 0.120179 |
| ENSG00000100665 | SERPINA4 | 6.087407007 | -0.624722728 | 0.824901328 | -1.451082951 | 0.14675676 | 0.306323 |
| ENSG00000231402 | WASF5P | 16.32048 | -0.62478 | 0.5649846 | -1.462932 | 0.1434859 | 0.301377 |
| ENSG00000279320 | | 8.469515 | -0.6249628 | 0.7487948 | -1.388889 | 0.1648665 | 0.333156 |
| ENSG00000180233 | ZNRF2 | 534.2248 | -0.6253014 | 0.2120814 | -3.063979 | 0.00218414 | 0.011153 |
| ENSG00000271789 | | 7.849018 | -0.6254428 | 0.7968439 | -1.382462 | 0.16683 | 0.336076 |
| ENSG00000150893 | FREM2 | 8564.587 | -0.6255981 | 0.3866974 | -1.855069 | 0.06358635 | 0.165955 |
| ENSG00000179611 | DGKZP1 | 3142.766 | -0.6258795 | 0.186958 | -3.453133 | 0.00055412 | 0.003433 |
| ENSG00000286811 | | 3.414494 | -0.6262326 | 0.8807049 | -1.461586 | 0.1438546 | 0.301925 |
| ENSG00000169288 | MRPL1 | 1319.956 | -0.6262617 | 0.2906625 | -2.330319 | 0.01978932 | 0.067412 |
| ENSG00000170624 | SGCD | 2.533963 | -0.6263714 | 1.076455 | -1.86275 | 0.0624975 | 0.163951 |
| ENSG00000008282 | SYPL1 | 5713.134619 | -0.626642096 | 0.133769298 | -4.799734521 | 1.59E-06 | 1.90E-05 |
| ENSG00000103932 | RPAP1 | 1852.719477 | -0.626840374 | 0.134985577 | -4.722667409 | 2.33E-06 | 2.70E-05 |
| ENSG00000080031 | PTPRH | 21.2687389 | -0.626849623 | 0.755116098 | -1.383516792 | 0.16650646 | 0.335608 |
| ENSG00000095002 | MSH2 | 4442.016411 | -0.627020875 | 0.307855626 | -2.23658833 | 0.02531326 | 0.081888 |
| ENSG00000249421 | ADAMTS19-AS1 | 206.6848 | -0.6270519 | 0.2692743 | -2.492121 | 0.01269827 | 0.047179 |
| ENSG00000254503 | | 6.975592 | -0.6272492 | 0.8059211 | -1.368881 | 0.1710365 | 0.341692 |
| ENSG00000171365 | CLCN5 | 908.9956 | -0.6274671 | 0.2852734 | -2.369348 | 0.01781948 | 0.061929 |
| ENSG00000180817 | PPA1 | 8728.938 | -0.6279828 | 0.1668029 | -3.862475 | 0.00011224 | 0.000848 |
| ENSG00000197579 | TOPORS | 1289.454 | -0.6281375 | 0.1380148 | -4.631857 | 3.62E-06 | 4.00E-05 |
| ENSG00000150938 | CRIM1 | 4542.572 | -0.6282211 | 0.316169 | -2.187905 | 0.02867656 | 0.090488 |
| ENSG00000102393 | GLA | 2178.312245 | -0.628281365 | 0.30139213 | -2.266777517 | 0.02340382 | 0.076957 |
| ENSG00000279296 | PRAL | 3.799268 | -0.6283235 | 1.171595 | -2.311286 | 0.02081704 | 0.070198 |
| ENSG00000274383 | | 11.97143 | -0.6284087 | 0.5071662 | -1.555189 | 0.119901 | 0.265162 |
| ENSG00000275713 | H2BC9 | 517.8604 | -0.6284281 | 0.3964655 | -1.826898 | 0.06771504 | 0.174171 |
| ENSG00000272181 | | 38.58824 | -0.6285206 | 0.3909847 | -1.845572 | 0.06495439 | 0.168615 |
| ENSG00000116761 | CTH | 118.0322921 | -0.628683063 | 0.429581541 | -1.732142565 | 0.08324818 | 0.203348 |

| | | | | | | | |
|---|---|---|---|---|---|---|---|
| ENSG00000266189 | MIR3186 | 9.891757 | -0.628685 | 0.6525305 | -1.420224 | 0.1555425 | 0.319583 |
| ENSG00000143869 | GDF7 | 16.4757 | -0.6294091 | 0.6165319 | -1.439341 | 0.1500538 | 0.311165 |
| ENSG00000188389 | PDCD1 | 1.25711 | -0.6297886 | 1.022892 | -1.66303 | 0.09630651 | NA |
| ENSG00000205863 | C1QTNF9B | 1.560006 | -0.6298225 | 1.060024 | -1.771124 | 0.07654016 | NA |
| ENSG00000274066 | MIR6514 | 3.126493 | -0.6300153 | 0.8291413 | -1.405713 | 0.1598093 | 0.325936 |
| ENSG00000175766 | EIF4E1B | 2.100305 | -0.6301594 | 1.182656 | -2.40618 | 0.01612033 | 0.057054 |
| ENSG00000144120 | TMEM177 | 781.3305 | -0.6301742 | 0.2109415 | -3.1119 | 0.00185888 | 0.009736 |
| ENSG00000172890 | NADSYN1 | 9394.198 | -0.6303554 | 0.2382377 | -2.792776 | 0.0052258 | 0.022972 |
| ENSG00000134779 | TPGS2 | 4331.379931 | -0.630573197 | 0.210544465 | -3.111582888 | 0.00186087 | 0.009744 |
| ENSG00000267397 | | 4.381832 | -0.6306032 | 0.737709 | -1.378768 | 0.1679663 | 0.337649 |
| ENSG00000269952 | | 6.379714 | -0.6307655 | 0.752442 | -1.390768 | 0.1642958 | 0.332333 |
| ENSG00000213089 | PDCL3P5 | 179.4682 | -0.6310187 | 0.2117592 | -3.107764 | 0.00188508 | 0.00985 |
| ENSG00000124802 | EEF1E1 | 446.332556 | -0.631138871 | 0.216551259 | -3.061979681 | 0.00219878 | 0.011217 |
| ENSG00000175711 | B3GNTL1 | 358.5785 | -0.6313807 | 0.1617403 | -3.995315 | 6.46E-05 | 0.00052 |
| ENSG00000236643 | | 8.990973 | -0.6314994 | 0.6240537 | -1.417431 | 0.1563569 | 0.320838 |
| ENSG00000251259 | | 14.07414 | -0.6316153 | 0.5783619 | -1.470074 | 0.1415417 | 0.298486 |
| ENSG00000235560 | | 76.9778 | -0.6316643 | 0.2562613 | -2.610769 | 0.00903389 | 0.035862 |
| ENSG00000215388 | ACTG1P3 | 8.573879 | -0.6318304 | 0.6747235 | -1.398189 | 0.1620563 | 0.329356 |
| ENSG00000187193 | MT1X | 164.6592 | -0.631854 | 0.7679471 | -1.419004 | 0.1558979 | 0.320146 |
| ENSG00000244045 | TMEM199 | 723.9896 | -0.6320448 | 0.1385874 | -4.645224 | 3.40E-06 | 3.77E-05 |
| ENSG00000172799 | ZBTB8OSP2 | 141.8582 | -0.6320955 | 0.2670838 | -2.522763 | 0.01164369 | 0.043943 |
| ENSG00000262576 | PCDHGA4 | 63.6117 | -0.6321516 | 0.4387763 | -1.716474 | 0.08607535 | 0.208472 |
| ENSG00000285761 | | 4.397177 | -0.632358 | 0.8422981 | -1.423066 | 0.154717 | 0.318386 |
| ENSG00000132522 | GPS2 | 71.40092879 | -0.63239148 | 0.293717166 | -2.32759588 | 0.01993357 | 0.067753 |
| ENSG00000121988 | ZRANB3 | 680.1280286 | -0.632466323 | 0.234089577 | -2.844997127 | 0.00444118 | 0.020085 |
| ENSG00000198056 | PRIM1 | 650.0321 | -0.6325261 | 0.4057346 | -1.82568 | 0.06789847 | 0.174484 |
| ENSG00000171865 | RNASEH1 | 1892.915 | -0.6327069 | 0.2122594 | -3.107486 | 0.00188686 | 0.009857 |
| ENSG00000223764 | LINC02593 | 1905.942 | -0.6327554 | 0.3242339 | -2.139385 | 0.03240447 | 0.099548 |
| ENSG00000270823 | | 3.651544 | -0.6328052 | 0.9444427 | -1.488113 | 0.1367212 | 0.290999 |
| ENSG00000137513 | NARS2 | 1789.934072 | -0.63289728 | 0.196801744 | -3.334289553 | 0.00085518 | 0.00499 |
| ENSG00000277959 | | 24.28618 | -0.6333129 | 0.4989764 | -1.591107 | 0.1115855 | 0.251973 |
| ENSG00000239672 | NME1 | 3840.716 | -0.6339817 | 0.3150947 | -2.206068 | 0.02737925 | 0.087181 |
| ENSG00000218803 | GSTM2P1 | 90.22293 | -0.6340826 | 0.4795801 | -1.625942 | 0.103962 | 0.239276 |

| | | | | | | | |
|---|---|---|---|---|---|---|---|
| ENSG00000276092 | | 1.155913 | -0.6345541 | 1.048035 | -1.746836 | 0.08066574 | NA |
| ENSG00000239247 | RN7SL589P | 2.277699 | -0.6347043 | 0.8587438 | -1.404907 | 0.1600488 | 0.3263 |
| ENSG00000163006 | CCDC138 | 379.6311 | -0.6348691 | 0.4220811 | -1.766868 | 0.07725033 | 0.19216 |
| ENSG00000228702 | | 5.134054 | -0.6349555 | 0.7157663 | -1.385927 | 0.1657693 | 0.334439 |
| ENSG00000164258 | NDUFS4 | 1839.569 | -0.6350479 | 0.2173059 | -3.0581 | 0.00222745 | 0.011337 |
| ENSG00000266900 | | 1.414409 | -0.6350846 | 1.071819 | -1.833885 | 0.06667103 | NA |
| ENSG00000160145 | KALRN | 246.383 | -0.6350923 | 0.2602947 | -2.591209 | 0.00956393 | 0.037473 |
| ENSG00000259959 | | 230.0827 | -0.6352601 | 0.1889325 | -3.47695 | 0.00050715 | 0.003174 |
| ENSG00000184924 | PTRHD1 | 656.1307 | -0.6355594 | 0.2186803 | -3.041331 | 0.00235535 | 0.011887 |
| ENSG00000137507 | LRRC32 | 2.525573474 | -0.635610615 | 0.921986967 | -1.500354865 | 0.1335225 | 0.286239 |
| ENSG00000283897 | | 55.39658 | -0.6358666 | 0.3044681 | -2.270732 | 0.02316319 | 0.076374 |
| ENSG00000204052 | LRRC73 | 174.2232 | -0.6363927 | 0.3684053 | -1.958815 | 0.05013449 | 0.138763 |
| ENSG00000178878 | APOLD1 | 256.4353 | -0.6364162 | 0.2294055 | -2.899923 | 0.00373254 | 0.017413 |
| ENSG00000182459 | TEX19 | 54.93809 | -0.6364403 | 0.5826227 | -1.489049 | 0.1364744 | 0.290661 |
| ENSG00000272689 | | 2.209243 | -0.6364488 | 0.885582 | -1.461538 | 0.1438678 | 0.301929 |
| ENSG00000225956 | | 5.610995 | -0.6365577 | 0.7101586 | -1.402733 | 0.1606966 | 0.327287 |
| ENSG00000248590 | GLDCP1 | 23.89669 | -0.6366271 | 0.7743707 | -1.389152 | 0.1647866 | 0.333071 |
| ENSG00000160991 | ORAI2 | 2454.901 | -0.6366958 | 0.1502825 | -4.31929 | 1.57E-05 | 0.000147 |
| ENSG00000283355 | | 4.643071 | -0.6366959 | 0.8617114 | -1.440841 | 0.1496297 | 0.310611 |
| ENSG00000228232 | GAPDHP1 | 2891.551 | -0.6369239 | 0.2447651 | -2.733558 | 0.0062654 | 0.026634 |
| ENSG00000164919 | COX6C | 9502.299 | -0.6371013 | 0.252707 | -2.674117 | 0.00749264 | 0.03084 |
| ENSG00000227199 | ST7-AS1 | 54.95286 | -0.6372629 | 0.4153849 | -1.795643 | 0.07255139 | 0.18353 |
| ENSG00000169248 | CXCL11 | 1.402339 | -0.6373712 | 1.082635 | -1.798834 | 0.07204501 | NA |
| ENSG00000058729 | RIOK2 | 2186.354902 | -0.637740216 | 0.117169651 | -5.512503289 | 3.54E-08 | 5.83E-07 |
| ENSG00000173621 | LRFN4 | 3810.27 | -0.6381945 | 0.185188 | -3.565784 | 0.00036277 | 0.002386 |
| ENSG00000133704 | IPO8 | 2697.672699 | -0.638214727 | 0.202481944 | -3.271479204 | 0.00106987 | 0.006063 |
| ENSG00000085365 | SCAMP1 | 4495.900148 | -0.638468466 | 0.174305122 | -3.769595309 | 0.00016351 | 0.001188 |
| ENSG00000115368 | WDR75 | 3462.784954 | -0.638469068 | 0.176021788 | -3.735803205 | 0.00018712 | 0.001337 |
| ENSG00000153015 | CWC27 | 1798.02 | -0.6385114 | 0.1778534 | -3.703604 | 0.00021256 | 0.001497 |
| ENSG00000223576 | | 2.787828 | -0.6387159 | 0.8073412 | -1.413272 | 0.1575758 | 0.322719 |
| ENSG00000110717 | NDUFS8 | 5456.68273 | -0.638773846 | 0.199823993 | -3.315137295 | 0.00091598 | 0.005292 |
| ENSG00000175984 | DENND2C | 945.9536 | -0.6389105 | 0.3166374 | -2.212764 | 0.02691394 | 0.085978 |
| ENSG00000178295 | GEN1 | 1954.296 | -0.6389426 | 0.4246565 | -1.763127 | 0.07787907 | 0.193254 |

| | | | | | | | |
|---|---|---|---|---|---|---|---|
| ENSG00000116260 | QSOX1 | 9295.53879 | -0.63902424 | 0.213455371 | -3.111947724 | 0.00185857 | 0.009736 |
| ENSG00000272800 | | 41.41676 | -0.6390504 | 0.3693422 | -1.956098 | 0.05045361 | 0.139447 |
| ENSG00000068878 | PSME4 | 5311.366089 | -0.639342804 | 0.194502395 | -3.409788847 | 0.00065013 | 0.003942 |
| ENSG00000198707 | CEP290 | 1868.389 | -0.6393707 | 0.1963518 | -3.387434 | 0.0007055 | 0.004232 |
| ENSG00000270130 | | 11.54978 | -0.6396037 | 0.5438734 | -1.521509 | 0.1281322 | 0.277792 |
| ENSG00000266053 | NDUFV2-AS1 | 62.71279 | -0.6396071 | 0.2967114 | -2.333336 | 0.01963052 | 0.066996 |
| ENSG00000164300 | SERINC5 | 4120.096 | -0.6398734 | 0.2653802 | -2.592921 | 0.00951647 | 0.03732 |
| ENSG00000235062 | BCRP5 | 2.615937 | -0.6403851 | 0.9089651 | -1.502996 | 0.1328402 | 0.285184 |
| ENSG00000155066 | PROM2 | 4716.389 | -0.6406964 | 0.165174 | -3.995945 | 6.44E-05 | 0.000519 |
| ENSG00000227014 | FKBP14-AS1 | 23.33289 | -0.6409659 | 0.557731 | -1.513332 | 0.1301954 | 0.281096 |
| ENSG00000173145 | NOC3L | 1999.061 | -0.6412133 | 0.2049773 | -3.251782 | 0.00114684 | 0.006438 |
| ENSG00000156509 | FBXO43 | 33.80384 | -0.6412459 | 0.4698572 | -1.666935 | 0.09552725 | 0.22502 |
| ENSG00000165730 | STOX1 | 261.0128 | -0.6413023 | 0.2956813 | -2.342362 | 0.01916212 | 0.065642 |
| ENSG00000275710 | | 6.844337 | -0.6414293 | 0.5994498 | -1.466572 | 0.1424926 | 0.299943 |
| ENSG00000181035 | SLC25A42 | 454.4706 | -0.6414462 | 0.1930385 | -3.429246 | 0.00060526 | 0.003703 |
| ENSG00000129048 | ACKR4 | 6.366354151 | -0.641565741 | 0.670008258 | -1.430738525 | 0.15250517 | 0.314837 |
| ENSG00000153922 | CHD1 | 3589.321 | -0.6419554 | 0.1900437 | -3.519657 | 0.00043211 | 0.002767 |
| ENSG00000229880 | IMMTP1 | 55.37244 | -0.6420154 | 0.2659609 | -2.577073 | 0.0099641 | 0.038744 |
| ENSG00000075188 | NUP37 | 1549.898938 | -0.642096364 | 0.269931282 | -2.531392951 | 0.01136105 | 0.043122 |
| ENSG00000177770 | CDKN2AIPNLP1 | 15.56174 | -0.6421206 | 0.5178477 | -1.580209 | 0.1140591 | 0.255755 |
| ENSG00000172301 | COPRS | 1542.311 | -0.6421413 | 0.1983956 | -3.360618 | 0.00077768 | 0.004606 |
| ENSG00000145242 | EPHA5 | 4.757658 | -0.6423151 | 0.8401161 | -1.455491 | 0.1455333 | 0.304603 |
| ENSG00000236792 | | 134.0764 | -0.6424552 | 0.2695016 | -2.542005 | 0.01102187 | 0.042099 |
| ENSG00000004975 | DVL2 | 4687.596265 | -0.642484584 | 0.164295815 | -4.023012455 | 5.75E-05 | 0.000468 |
| ENSG00000139508 | SLC46A3 | 1325.102 | -0.6427426 | 0.2346285 | -2.882554 | 0.00394465 | 0.01823 |
| ENSG00000187801 | ZFP69B | 214.5559 | -0.6428424 | 0.3111854 | -2.260846 | 0.02376879 | 0.077936 |
| ENSG00000170075 | GPR37L1 | 2175.472 | -0.6431081 | 0.2941172 | -2.361255 | 0.01821322 | 0.063074 |
| ENSG00000177303 | CASKIN2 | 5808.966 | -0.6432534 | 0.1691351 | -3.91994 | 8.86E-05 | 0.000687 |
| ENSG00000108578 | BLMH | 1689.958264 | -0.643323307 | 0.340565609 | -2.097874234 | 0.03591626 | 0.10773 |
| ENSG00000271711 | | 8.470427 | -0.6433302 | 0.6881181 | -1.427586 | 0.1534112 | 0.316286 |
| ENSG00000021574 | SPAST | 1456.718626 | -0.643438545 | 0.161087097 | -4.091393625 | 4.29E-05 | 0.000362 |
| ENSG00000262147 | | 21.43272 | -0.6434818 | 0.4105219 | -1.822433 | 0.0683893 | 0.17544 |
| ENSG00000242616 | GNG10 | 29.8948 | -0.6438547 | 0.3271372 | -2.163635 | 0.03049241 | 0.094921 |

| | | | | | | | |
|---|---|---|---|---|---|---|---|
| ENSG00000234478 | ACBD3-AS1 | 7.461634 | -0.6440774 | 0.6439872 | -1.442495 | 0.1491628 | 0.30986 |
| ENSG00000120699 | EXOSC8 | 2422.742089 | -0.644253871 | 0.189759325 | -3.512973892 | 0.00044312 | 0.002827 |
| ENSG00000279123 | | 4.511103 | -0.644321 | 1.144947 | -2.05908 | 0.03948654 | 0.115737 |
| ENSG00000275580 | | 14.67294 | -0.6443329 | 0.6157591 | -1.471058 | 0.1412753 | 0.298161 |
| ENSG00000235347 | | 1.031202 | -0.6443716 | 1.019366 | -1.675872 | 0.09376319 | NA |
| ENSG00000270264 | NDUFB8P2 | 56.17342 | -0.6445335 | 0.3692635 | -1.970661 | 0.04876266 | 0.13589 |
| ENSG00000140853 | NLRC5 | 434.3377 | -0.6447169 | 0.725506 | -1.432933 | 0.1518769 | 0.313903 |
| ENSG00000184840 | TMED9 | 14488.72 | -0.6448754 | 0.1866078 | -3.550875 | 0.00038395 | 0.002506 |
| ENSG00000171217 | CLDN20 | 13.41182 | -0.6452483 | 0.5363028 | -1.553448 | 0.1203161 | 0.265858 |
| ENSG00000133661 | SFTPD | 121.4168311 | -0.645269739 | 0.299932022 | -2.329731965 | 0.01982032 | 0.067478 |
| ENSG00000155011 | DKK2 | 1.420114 | -0.6453143 | 0.9435399 | -1.534671 | 0.1248646 | NA |
| ENSG00000138433 | CIR1 | 1638.4318 | -0.645331214 | 0.145348936 | -4.550153493 | 5.36E-06 | 5.67E-05 |
| ENSG00000137075 | RNF38 | 1369.359336 | -0.645400491 | 0.172849067 | -3.840425715 | 0.00012282 | 0.000919 |
| ENSG00000106299 | WASL | 3732.462102 | -0.645555792 | 0.164163801 | -4.031733621 | 5.54E-05 | 0.000453 |
| ENSG00000263272 | | 215.475 | -0.6456215 | 0.3228328 | -2.203604 | 0.02755222 | 0.087627 |
| ENSG00000254271 | ANK3-DT | 13.737 | -0.6457799 | 0.5703934 | -1.504568 | 0.1324353 | 0.284659 |
| ENSG00000141644 | MBD1 | 4108.173 | -0.6461311 | 0.1140288 | -5.735536 | 9.72E-09 | 1.79E-07 |
| ENSG00000215283 | HMGB3P24 | 18.2001 | -0.6466818 | 0.5294028 | -1.565806 | 0.117394 | 0.261029 |
| ENSG00000214988 | RPL7AP26 | 6.633374 | -0.64688 | 0.7437593 | -1.431713 | 0.152226 | 0.314441 |
| ENSG00000259877 | | 415.5829 | -0.6468929 | 0.2087427 | -3.223709 | 0.00126542 | 0.007018 |
| ENSG00000107562 | CXCL12 | 633.0454668 | -0.646898747 | 0.646131729 | -1.470327119 | 0.14147318 | 0.298389 |
| ENSG00000164163 | ABCE1 | 5917.901 | -0.646934 | 0.1841727 | -3.624033 | 0.00029005 | 0.001965 |
| ENSG00000070190 | DAPP1 | 134.2856165 | -0.647023979 | 0.765063971 | -1.418549179 | 0.15603049 | 0.320344 |
| ENSG00000196367 | TRRAP | 7901.44 | -0.6472442 | 0.1988531 | -3.383535 | 0.00071559 | 0.004286 |
| ENSG00000166153 | DEPDC4 | 100.3503 | -0.6474651 | 0.2429016 | -2.80959 | 0.00496047 | 0.02201 |
| ENSG00000129484 | PARP2 | 1889.816513 | -0.647562121 | 0.216578079 | -3.117087198 | 0.00182648 | 0.0096 |
| ENSG00000264057 | | 37.72964 | -0.6475656 | 0.5228065 | -1.583393 | 0.1133318 | 0.254554 |
| ENSG00000137720 | C11orf1 | 361.1471254 | -0.647685954 | 0.261137448 | -2.643996705 | 0.00819335 | 0.033119 |
| ENSG00000288104 | | 1.583664 | -0.6477988 | 0.9675469 | -1.573617 | 0.1155759 | 0.258436 |
| ENSG00000215529 | EFCAB8 | 4.592642 | -0.6478747 | 0.7015246 | -1.421533 | 0.1551617 | 0.31898 |
| ENSG00000255959 | | 1.831228 | -0.6479976 | 0.9908656 | -1.608478 | 0.1077305 | 0.245585 |
| ENSG00000220749 | RPL21P28 | 87.85985 | -0.6481626 | 0.2834321 | -2.459678 | 0.01390619 | 0.050703 |
| ENSG00000236869 | ZKSCAN7-AS1 | 4.135846 | -0.6482241 | 0.6946365 | -1.43165 | 0.152244 | 0.314441 |

| | | | | | | | |
|---|---|---|---|---|---|---|---|
| ENSG00000198876 | DCAF12 | 4311.721 | -0.6482855 | 0.1066579 | -6.145019 | 8.00E-10 | 1.74E-08 |
| ENSG00000136156 | ITM2B | 6708.667734 | -0.648341389 | 0.167976245 | -3.997146848 | 6.41E-05 | 0.000516 |
| ENSG00000170919 | TPT1-AS1 | 483.3271 | -0.6484307 | 0.4179988 | -1.810179 | 0.07026802 | 0.17917 |
| ENSG00000179071 | CCDC89 | 25.84561 | -0.6484789 | 0.587583 | -1.500319 | 0.1335319 | 0.286239 |
| ENSG00000287625 | | 71.01703 | -0.6486313 | 0.3138296 | -2.257463 | 0.02397918 | 0.078509 |
| ENSG00000095539 | SEMA4G | 241.7679658 | -0.648658425 | 0.237485299 | -2.869980112 | 0.00410498 | 0.018863 |
| ENSG00000139197 | PEX5 | 2127.168212 | -0.649206627 | 0.110800406 | -5.94085629 | 2.84E-09 | 5.69E-08 |
| ENSG00000165526 | RPUSD4 | 1299.092 | -0.6492835 | 0.1320474 | -4.997221 | 5.82E-07 | 7.63E-06 |
| ENSG00000111052 | LIN7A | 2628.140581 | -0.649298494 | 0.300499417 | -2.352517253 | 0.01864683 | 0.064214 |
| ENSG00000179935 | LINC00652 | 29.56463 | -0.6494877 | 0.3867004 | -1.916876 | 0.05525375 | 0.149619 |
| ENSG00000132842 | AP3B1 | 3452.06324 | -0.649608235 | 0.139162031 | -4.754404074 | 1.99E-06 | 2.34E-05 |
| ENSG00000160298 | C21orf58 | 1594.731 | -0.6498618 | 0.2836886 | -2.462207 | 0.0138085 | 0.050402 |
| ENSG00000275450 | | 8.108065 | -0.6499568 | 0.6146526 | -1.475859 | 0.1399817 | 0.296113 |
| ENSG00000253485 | PCDHGA5 | 41.66794 | -0.6499601 | 0.4505926 | -1.734235 | 0.08287639 | 0.202757 |
| ENSG00000086475 | SEPHS1 | 6575.89583 | -0.649970939 | 0.162380815 | -4.095526077 | 4.21E-05 | 0.000356 |
| ENSG00000259316 | | 1.416659 | -0.6504233 | 1.087208 | -1.77408 | 0.07604998 | NA |
| ENSG00000135940 | COX5B | 6722.197868 | -0.650536836 | 0.197739509 | -3.453637337 | 0.00055308 | 0.003428 |
| ENSG00000145331 | TRMT10A | 633.9038 | -0.6505393 | 0.1735391 | -3.85139 | 0.00011745 | 0.000884 |
| ENSG00000236021 | | 16.97903 | -0.6507432 | 0.5532679 | -1.541958 | 0.1230839 | 0.27022 |
| ENSG00000259706 | HSP90B2P | 99.029 | -0.6512038 | 0.2612137 | -2.650707 | 0.00803234 | 0.032642 |
| ENSG00000253626 | EIF5AL1 | 3007.222 | -0.6512754 | 0.2841638 | -2.460624 | 0.01386956 | 0.050583 |
| ENSG00000251072 | LMNB1-DT | 4.238879 | -0.6514673 | 0.8433527 | -1.440995 | 0.1495862 | 0.310569 |
| ENSG00000154122 | ANKH | 1508.697 | -0.651519 | 0.2403065 | -2.860902 | 0.00422438 | 0.019305 |
| ENSG00000252213 | SNORA74D | 2.113498 | -0.6516342 | 1.086466 | -1.860162 | 0.06286264 | 0.1646 |
| ENSG00000243323 | PTPRVP | 200.713 | -0.6517173 | 0.4443024 | -1.731687 | 0.08332932 | 0.203491 |
| ENSG00000103174 | NAGPA | 1006.740759 | -0.651790959 | 0.167400516 | -3.996698367 | 6.42E-05 | 0.000517 |
| ENSG00000087502 | ERGIC2 | 2024.159899 | -0.651840103 | 0.128839046 | -5.136004168 | 2.81E-07 | 3.89E-06 |
| ENSG00000199477 | SNORA31 | 16.936 | -0.651878 | 0.4721675 | -1.684409 | 0.0921026 | 0.219134 |
| ENSG00000272360 | | 2.277643 | -0.6519303 | 1.105492 | -1.906334 | 0.05660693 | 0.15236 |
| ENSG00000224259 | LINC01133 | 24.73683 | -0.6520455 | 1.058749 | -1.660347 | 0.09684466 | 0.227263 |
| ENSG00000164118 | CEP44 | 636.3946 | -0.6520785 | 0.1743806 | -3.846983 | 0.00011958 | 0.000899 |
| ENSG00000279799 | | 61.0978 | -0.6521455 | 0.3501226 | -2.078903 | 0.0376263 | 0.111449 |
| ENSG00000169583 | CLIC3 | 2677.08 | -0.6523468 | 0.4109284 | -1.863497 | 0.06239232 | 0.163724 |

| | | | | | | | |
|---|---|---|---|---|---|---|---|
| ENSG00000232332 | ARL5AP5 | 3.481604 | -0.6523852 | 0.8546688 | -1.455043 | 0.1456575 | 0.304746 |
| ENSG00000103995 | CEP152 | 744.5475547 | -0.65241169 | 0.397566669 | -1.910323694 | 0.05609155 | 0.151314 |
| ENSG00000077348 | EXOSC5 | 1877.032702 | -0.652533061 | 0.212464941 | -3.2027854 | 0.00136105 | 0.007467 |
| ENSG00000229692 | SOS1-IT1 | 79.71168 | -0.6525465 | 0.3794136 | -1.952966 | 0.05082365 | 0.140207 |
| ENSG00000242882 | RPL5P11 | 7.164291 | -0.6528609 | 0.796044 | -1.419166 | 0.1558505 | 0.320074 |
| ENSG00000227709 | | 3.086617 | -0.652942 | 0.9037377 | -1.4901 | 0.1361979 | 0.290188 |
| ENSG00000224885 | EIPR1-IT1 | 11.85699 | -0.6530369 | 0.5952929 | -1.500865 | 0.1333906 | 0.286042 |
| ENSG00000075856 | SART3 | 4462.422058 | -0.6532159 | 0.123258215 | -5.361053198 | 8.27E-08 | 1.27E-06 |
| ENSG00000271602 | KRT8P49 | 27.49057 | -0.6533403 | 0.5351749 | -1.57734 | 0.1147173 | 0.256884 |
| ENSG00000083635 | NUFIP1 | 880.8104935 | -0.653490594 | 0.211881024 | -3.21491234 | 0.00130484 | 0.007218 |
| ENSG00000119772 | DNMT3A | 4815.131682 | -0.653514047 | 0.124318036 | -5.3000702 | 1.16E-07 | 1.73E-06 |
| ENSG00000165072 | MAMDC2 | 5.237201 | -0.6535263 | 0.8172521 | -1.454518 | 0.1458027 | 0.304881 |
| ENSG00000170264 | FAM161A | 257.1318 | -0.6535955 | 0.4226062 | -1.821918 | 0.06846744 | 0.175556 |
| ENSG00000196821 | ILRUN | 9482.503 | -0.6536629 | 0.1506377 | -4.435802 | 9.17E-06 | 9.13E-05 |
| ENSG00000170500 | LONRF2 | 4221.763 | -0.6538881 | 0.3690614 | -2.008707 | 0.04456818 | 0.127162 |
| ENSG00000174032 | SLC25A30 | 1635.369 | -0.6539892 | 0.2264756 | -3.040335 | 0.00236315 | 0.011918 |
| ENSG00000231344 | | 19.0201 | -0.6540598 | 0.4389354 | -1.775184 | 0.07586745 | 0.189697 |
| ENSG00000183638 | RP1L1 | 5.794554 | -0.6541321 | 0.7349277 | -1.429054 | 0.1529888 | 0.315563 |
| ENSG00000040275 | SPDL1 | 1898.415493 | -0.654318891 | 0.386850639 | -1.924471243 | 0.05429554 | 0.147775 |
| ENSG00000135912 | TTLL4 | 1517.259808 | -0.654341375 | 0.183255081 | -3.680566064 | 0.00023272 | 0.001621 |
| ENSG00000123131 | PRDX4 | 4098.322379 | -0.654376111 | 0.213052419 | -3.204336265 | 0.00135374 | 0.007438 |
| ENSG00000124370 | MCEE | 505.6297499 | -0.654502762 | 0.185112859 | -3.64560399 | 0.00026677 | 0.001825 |
| ENSG00000168135 | KCNJ4 | 128.6461 | -0.6549903 | 0.4415295 | -1.764062 | 0.07772158 | 0.192989 |
| ENSG00000115539 | PDCL3 | 1492.557477 | -0.655060703 | 0.222545881 | -3.092470217 | 0.00198498 | 0.010287 |
| ENSG00000111731 | C2CD5 | 2058.490718 | -0.655192666 | 0.134124341 | -4.964296655 | 6.90E-07 | 8.91E-06 |
| ENSG00000182004 | SNRPE | 2979.036 | -0.6555837 | 0.2423722 | -2.845378 | 0.00443587 | 0.020076 |
| ENSG00000231628 | | 34.92381 | -0.6555893 | 0.4300624 | -1.800204 | 0.07182838 | 0.182152 |
| ENSG00000145723 | GIN1 | 517.9341 | -0.6558184 | 0.1435399 | -4.650168 | 3.32E-06 | 3.70E-05 |
| ENSG00000165733 | BMS1 | 6859.763 | -0.6559758 | 0.2094455 | -3.268013 | 0.00108305 | 0.00613 |
| ENSG00000102890 | ELMO3 | 3880.822929 | -0.656418746 | 0.153976807 | -4.356466051 | 1.32E-05 | 0.000126 |
| ENSG00000286409 | | 2.203922 | -0.656514 | 0.9462666 | -1.534996 | 0.1247849 | 0.272761 |
| ENSG00000105612 | DNASE2 | 3357.753449 | -0.656563944 | 0.169011965 | -3.992140568 | 6.55E-05 | 0.000525 |
| ENSG00000227359 | | 20.77276 | -0.6567234 | 0.4053596 | -1.870117 | 0.06146764 | 0.161953 |

| | | | | | | | |
|---|---|---|---|---|---|---|---|
| ENSG00000228434 | | 16.00155 | -0.6568468 | 0.5306349 | -1.584968 | 0.1129737 | 0.254028 |
| ENSG00000256463 | SALL3 | 2.110017 | -0.6569627 | 1.147649 | -2.027837 | 0.04257689 | 0.122786 |
| ENSG00000263968 | RN7SL381P | 60.73096 | -0.6569958 | 0.443565 | -1.770534 | 0.0766383 | 0.191031 |
| ENSG00000136295 | TTYH3 | 5263.434013 | -0.65713481 | 0.15149609 | -4.438542244 | 9.06E-06 | 9.03E-05 |
| ENSG00000126870 | DYNC2I1 | 1540.721022 | -0.657247345 | 0.147019768 | -4.550395595 | 5.35E-06 | 5.67E-05 |
| ENSG00000116459 | ATP5PB | 9434.953231 | -0.657404198 | 0.216886506 | -3.16968874 | 0.00152602 | 0.008231 |
| ENSG00000278958 | | 1.809304 | -0.6576502 | 1.036056 | -1.711168 | 0.08705004 | 0.210219 |
| ENSG00000168883 | USP39 | 5673.431 | -0.6581863 | 0.1545712 | -4.373584 | 1.22E-05 | 0.000118 |
| ENSG00000234709 | UPF3AP3 | 11.48776 | -0.6583606 | 0.6700209 | -1.444502 | 0.1485979 | 0.309073 |
| ENSG00000248794 | | 11.37099 | -0.658542 | 0.5239295 | -1.597564 | 0.11014 | 0.249621 |
| ENSG00000143314 | MRPL24 | 4406.959 | -0.6586644 | 0.1885067 | -3.637522 | 0.00027527 | 0.001873 |
| ENSG00000197360 | ZNF98 | 3.991656 | -0.6587581 | 0.9575328 | -1.553288 | 0.1203544 | 0.265921 |
| ENSG00000144452 | ABCA12 | 3382.762 | -0.6588882 | 0.6586628 | -1.497017 | 0.1343888 | 0.28744 |
| ENSG00000135241 | PNPLA8 | 1999.713703 | -0.659118579 | 0.172920209 | -3.916939005 | 8.97E-05 | 0.000694 |
| ENSG00000184260 | H2AC20 | 359.2473 | -0.6591506 | 0.2918956 | -2.444631 | 0.01450006 | 0.052462 |
| ENSG00000183801 | OLFML1 | 2.14426 | -0.6592164 | 1.114839 | -1.841721 | 0.06551604 | 0.169807 |
| ENSG00000134508 | CABLES1 | 2351.82113 | -0.659262611 | 0.167241542 | -4.049362395 | 5.14E-05 | 0.000424 |
| ENSG00000270006 | | 52.30857 | -0.6595766 | 0.372764 | -2.00658 | 0.04479444 | 0.127586 |
| ENSG00000043514 | TRIT1 | 1568.796105 | -0.659640214 | 0.139655554 | -4.810880888 | 1.50E-06 | 1.81E-05 |
| ENSG00000215333 | KRT18P23 | 27.36838 | -0.6597925 | 0.463957 | -1.720457 | 0.08534938 | 0.207262 |
| ENSG00000171503 | ETFDH | 1163.78 | -0.6598584 | 0.1563031 | -4.317085 | 1.58E-05 | 0.000148 |
| ENSG00000167721 | TSR1 | 3897.465 | -0.6601181 | 0.1706377 | -3.977566 | 6.96E-05 | 0.000555 |
| ENSG00000150990 | DHX37 | 2820.985 | -0.6603776 | 0.2050566 | -3.351552 | 0.0008036 | 0.004739 |
| ENSG00000148606 | POLR3A | 3142.411 | -0.6604274 | 0.1432804 | -4.727127 | 2.28E-06 | 2.64E-05 |
| ENSG00000273802 | H2BC8 | 1607.259 | -0.6605296 | 0.2737556 | -2.583003 | 0.00979445 | 0.038207 |
| ENSG00000139737 | SLAIN1 | 721.8333 | -0.6609476 | 0.2073703 | -3.315679 | 0.00091421 | 0.005285 |
| ENSG00000262519 | TXNP4 | 99.04831 | -0.6611241 | 0.3058801 | -2.343396 | 0.01910908 | 0.065519 |
| ENSG00000118705 | RPN2 | 16009.83264 | -0.661408355 | 0.167797946 | -4.046984335 | 5.19E-05 | 0.000428 |
| ENSG00000272945 | | 9.56534 | -0.6614614 | 0.6590326 | -1.479597 | 0.1389809 | 0.294394 |
| ENSG00000075089 | ACTR6 | 2332.224075 | -0.661545471 | 0.209345694 | -3.271617062 | 0.00106934 | 0.006062 |
| ENSG00000227540 | | 98.19476 | -0.6616146 | 0.3014609 | -2.386735 | 0.01699876 | 0.059615 |
| ENSG00000260747 | | 635.1611 | -0.6616965 | 0.2196909 | -3.154177 | 0.00160951 | 0.008601 |
| ENSG00000273271 | | 75.19104 | -0.6618465 | 0.3312968 | -2.205342 | 0.0274301 | 0.087322 |

| | | | | | | | |
|---|---|---|---|---|---|---|---|
| ENSG00000165732 | DDX21 | 14390.12 | -0.6618623 | 0.2444679 | -2.849861 | 0.00437383 | 0.019853 |
| ENSG00000226659 | | 15.49514 | -0.6621173 | 0.5342754 | -1.596148 | 0.1104557 | 0.250123 |
| ENSG00000226396 | | 102.4059 | -0.6621952 | 0.2911022 | -2.457139 | 0.01400484 | 0.050993 |
| ENSG00000214309 | MBLAC1 | 91.02905 | -0.6625272 | 0.3453271 | -2.13368 | 0.03286897 | 0.100657 |
| ENSG00000164828 | SUN1 | 15250.77 | -0.6625633 | 0.1687425 | -4.053797 | 5.04E-05 | 0.000417 |
| ENSG00000197748 | CFAP43 | 22.66298 | -0.662711 | 0.4782033 | -1.684527 | 0.09207991 | 0.219134 |
| ENSG00000245522 | LINC02709 | 18.66617 | -0.6627148 | 0.5796993 | -1.544061 | 0.1225735 | 0.269368 |
| ENSG00000146700 | SSC4D | 53.63715 | -0.6627857 | 0.3030514 | -2.374471 | 0.01757412 | 0.061244 |
| ENSG00000203414 | BTBD7P1 | 67.31734 | -0.6628438 | 0.398944 | -1.920216 | 0.05483061 | 0.148804 |
| ENSG00000137106 | GRHPR | 4525.242479 | -0.663023372 | 0.155730474 | -4.353456656 | 1.34E-05 | 0.000128 |
| ENSG00000179583 | CIITA | 208.4461 | -0.6632517 | 0.3574667 | -2.088537 | 0.03674939 | 0.109489 |
| ENSG00000108953 | YWHAE | 18089.02315 | -0.663307283 | 0.199792097 | -3.445038953 | 0.00057098 | 0.003521 |
| ENSG00000232296 | | 4.039555 | -0.6639296 | 0.9142588 | -1.459289 | 0.1444855 | 0.302891 |
| ENSG00000240875 | LINC00886 | 1177.731 | -0.6639904 | 0.397019 | -1.920144 | 0.05483968 | 0.148814 |
| ENSG00000141994 | DUS3L | 1407.516 | -0.6641322 | 0.2181506 | -3.186746 | 0.00143883 | 0.007834 |
| ENSG00000278638 | | 1.40033 | -0.6643487 | 0.9755756 | -1.544014 | 0.122585 | NA |
| ENSG00000145863 | GABRA6 | 3.999597 | -0.6645426 | 0.792651 | -1.444994 | 0.1484595 | 0.308979 |
| ENSG00000143994 | ABHD1 | 44.12376 | -0.6646402 | 0.3275859 | -2.241548 | 0.02499062 | 0.081081 |
| ENSG00000273841 | TAF9 | 3334.198 | -0.6646571 | 0.2115685 | -3.273036 | 0.00106399 | 0.006035 |
| ENSG00000107643 | MAPK8 | 2368.056969 | -0.664752274 | 0.110740911 | -6.047391755 | 1.47E-09 | 3.07E-08 |
| ENSG00000068697 | LAPTM4A | 10698.16929 | -0.664952694 | 0.151072957 | -4.494367141 | 6.98E-06 | 7.17E-05 |
| ENSG00000215041 | NEURL4 | 2410.762 | -0.6649989 | 0.1070068 | -6.27869 | 3.41E-10 | 7.93E-09 |
| ENSG00000144406 | UNC80 | 1.981642 | -0.6652965 | 0.8899153 | -1.503228 | 0.1327802 | 0.285124 |
| ENSG00000198948 | MFAP3L | 796.5263 | -0.6653844 | 0.2978742 | -2.420699 | 0.01549069 | 0.055311 |
| ENSG00000236993 | GAPDHP21 | 7.507657 | -0.6656569 | 0.5981964 | -1.51705 | 0.1292541 | 0.27954 |
| ENSG00000119042 | SATB2 | 1541.854737 | -0.66566149 | 0.224873906 | -3.092685297 | 0.00198354 | 0.010281 |
| ENSG00000276698 | | 49.02531 | -0.6656777 | 0.3243466 | -2.251745 | 0.02433838 | 0.079441 |
| ENSG00000178999 | AURKB | 1700.444 | -0.6663474 | 0.6398678 | -1.531134 | 0.1257363 | 0.274119 |
| ENSG00000253305 | PCDHGB6 | 20.51362 | -0.6663537 | 0.3785963 | -1.994507 | 0.04609663 | 0.130333 |
| ENSG00000267374 | MIR924HG | 168.8217 | -0.666893 | 0.5084327 | -1.656031 | 0.09771553 | 0.228821 |
| ENSG00000132591 | ERAL1 | 4018.790619 | -0.666946932 | 0.214205572 | -3.246737273 | 0.00116736 | 0.006542 |
| ENSG00000226200 | SGMS1-AS1 | 368.9938 | -0.6669913 | 0.4192901 | -1.867316 | 0.06185746 | 0.162711 |
| ENSG00000280385 | | 115.1244 | -0.667153 | 0.2821811 | -2.534719 | 0.01125377 | 0.042812 |

| | | | | | | | |
|---|---|---|---|---|---|---|---|
| ENSG00000236305 | SLC12A9-AS1 | 46.1237 | -0.6672401 | 0.3474302 | -2.1414 | 0.0322418 | 0.099147 |
| ENSG00000196912 | ANKRD36B | 241.3462 | -0.6672733 | 0.3273734 | -2.25084 | 0.02439569 | 0.079553 |
| ENSG00000173588 | CEP83 | 936.658 | -0.6675981 | 0.2310733 | -3.03206 | 0.00242891 | 0.012195 |
| ENSG00000138792 | ENPEP | 505.0531733 | -0.667710173 | 0.212490562 | -3.280712426 | 0.00103545 | 0.005901 |
| ENSG00000240889 | NDUFB2-AS1 | 23.64157 | -0.6678092 | 0.3642006 | -2.062492 | 0.03916096 | 0.114975 |
| ENSG00000234492 | RPL34-DT | 2.619913 | -0.6679912 | 0.9452184 | -1.556504 | 0.1195883 | 0.264558 |
| ENSG00000164182 | NDUFAF2 | 1103.605 | -0.6682855 | 0.2192609 | -3.18915 | 0.00142692 | 0.007774 |
| ENSG00000163817 | SLC6A20 | 3.678353 | -0.6683554 | 1.126928 | -1.927787 | 0.05388159 | 0.146998 |
| ENSG00000026036 | RTEL1-TNFRSF6E | 534.2153124 | -0.668444766 | 0.291780238 | -2.47627827 | 0.01327601 | 0.048849 |
| ENSG00000152291 | TGOLN2 | 14450.87 | -0.668506 | 0.2154952 | -3.165731 | 0.00154694 | 0.008323 |
| ENSG00000107796 | ACTA2 | 181.6208113 | -0.668506743 | 0.387309261 | -1.972912882 | 0.04850549 | 0.135372 |
| ENSG00000167377 | ZNF23 | 121.0858 | -0.668527 | 0.2493614 | -2.814179 | 0.0048902 | 0.021778 |
| ENSG00000221639 | | 1.92437 | -0.668844 | 0.9044849 | -1.484651 | 0.1376362 | 0.292249 |
| ENSG00000132640 | BTBD3 | 2327.364142 | -0.668907626 | 0.151613028 | -4.50833528 | 6.53E-06 | 6.77E-05 |
| ENSG00000252473 | | 2.54025 | -0.6691651 | 0.8885647 | -1.471313 | 0.1412064 | 0.298063 |
| ENSG00000105374 | NKG7 | 7.497807453 | -0.66937662 | 0.786228007 | -1.463210411 | 0.14340981 | 0.301325 |
| ENSG00000177551 | NHLH2 | 4.780347 | -0.6695283 | 0.8091945 | -1.446901 | 0.1479247 | 0.308107 |
| ENSG00000204228 | HSD17B8 | 629.4951 | -0.6696825 | 0.2041429 | -3.401419 | 0.00067037 | 0.004047 |
| ENSG00000276141 | WHAMMP3 | 20.87002 | -0.6698193 | 0.4117794 | -1.891123 | 0.05860793 | 0.156377 |
| ENSG00000279469 | | 2.776439 | -0.6699626 | 0.8777611 | -1.488347 | 0.1366595 | 0.290915 |
| ENSG00000264895 | | 1.56217 | -0.6700024 | 1.028618 | -1.694151 | 0.09023668 | NA |
| ENSG00000136155 | SCEL | 4.572285965 | -0.670065144 | 0.935830726 | -1.486413525 | 0.13716977 | 0.291498 |
| ENSG00000248946 | MTND3P22 | 22.57478 | -0.6701529 | 0.58016 | -1.558868 | 0.1190276 | 0.263559 |
| ENSG00000188649 | CC2D2B | 10.51418 | -0.6703735 | 0.6396074 | -1.500348 | 0.1335243 | 0.286239 |
| ENSG00000230715 | | 412.817 | -0.6704337 | 0.2719871 | -2.638489 | 0.00832764 | 0.033595 |
| ENSG00000138587 | MNS1 | 113.3353595 | -0.670478953 | 0.493071735 | -1.689398948 | 0.091143 | 0.217631 |
| ENSG00000204588 | LINC01123 | 105.0218 | -0.6705876 | 0.3681059 | -2.058102 | 0.0395803 | 0.115963 |
| ENSG00000258525 | | 10.09298 | -0.6707903 | 0.6764655 | -1.490931 | 0.1359796 | 0.289769 |
| ENSG00000161956 | SENP3 | 1067.946 | -0.6708965 | 0.1820278 | -3.796392 | 0.00014682 | 0.001076 |
| ENSG00000152127 | MGAT5 | 3769.114 | -0.6709623 | 0.1523625 | -4.504464 | 6.65E-06 | 6.87E-05 |
| ENSG00000128567 | PODXL | 1592.630203 | -0.670998041 | 0.252404591 | -2.77924065 | 0.00544861 | 0.023755 |
| ENSG00000284602 | | 29.78095 | -0.6710118 | 0.5260455 | -1.627814 | 0.1035643 | 0.238626 |
| ENSG00000225067 | RPL23AP2 | 43.94844 | -0.6710302 | 0.2982821 | -2.439041 | 0.01472629 | 0.053139 |

| | | | | | | | |
|---|---|---|---|---|---|---|---|
| ENSG00000105341 | DMAC2 | 2615.429786 | -0.671420588 | 0.210086193 | -3.322759711 | 0.00089132 | 0.005175 |
| ENSG00000260233 | ZNRD2-AS1 | 212.3274 | -0.6714427 | 0.3856923 | -1.989972 | 0.04659398 | 0.13139 |
| ENSG00000122861 | PLAU | 372.9017151 | -0.671451085 | 0.472109281 | -1.738293578 | 0.0821591 | 0.201466 |
| ENSG00000224773 | HSPA8P7 | 72.18384 | -0.6716541 | 0.2762283 | -2.6137 | 0.00895676 | 0.035619 |
| ENSG00000253537 | PCDHGA7 | 22.17663 | -0.6716757 | 0.3665514 | -2.056787 | 0.0397067 | 0.116241 |
| ENSG00000103168 | TAF1C | 6611.258199 | -0.671784635 | 0.134465931 | -5.0794238 | 3.79E-07 | 5.12E-06 |
| ENSG00000225342 | LRRK2-DT | 4.195602 | -0.6718575 | 0.846439 | -1.486633 | 0.1371119 | 0.291446 |
| ENSG00000224899 | LINC02830 | 92.84258 | -0.6719381 | 0.4694341 | -1.740266 | 0.08181231 | 0.200772 |
| ENSG00000279029 | | 2.431524 | -0.672055 | 1.002356 | -1.603304 | 0.1088676 | 0.247349 |
| ENSG00000171132 | PRKCE | 1160.888 | -0.6723378 | 0.2095514 | -3.328637 | 0.00087272 | 0.005086 |
| ENSG00000198580 | | 21.3552 | -0.672437 | 0.6495892 | -1.500753 | 0.1334195 | 0.28608 |
| ENSG00000115392 | FANCL | 775.0096595 | -0.672571005 | 0.176139473 | -3.927276051 | 8.59E-05 | 0.00067 |
| ENSG00000288065 | | 26.69535 | -0.6726578 | 0.5320461 | -1.621226 | 0.1049693 | 0.240979 |
| ENSG00000115163 | CENPA | 824.8197895 | -0.672778116 | 0.480273738 | -1.72302634 | 0.08488376 | 0.206319 |
| ENSG00000177879 | AP3S1 | 3111.634 | -0.6729126 | 0.1483317 | -4.650565 | 3.31E-06 | 3.69E-05 |
| ENSG00000196850 | PPTC7 | 1403.013 | -0.6729401 | 0.2057604 | -3.395298 | 0.00068554 | 0.004127 |
| ENSG00000165264 | NDUFB6 | 1838.764 | -0.6730035 | 0.2519943 | -2.832352 | 0.0046207 | 0.020796 |
| ENSG00000099250 | NRP1 | 6927.908828 | -0.673058731 | 0.394507707 | -1.961348735 | 0.04983836 | 0.138178 |
| ENSG00000196581 | AJAP1 | 11.63026 | -0.6734035 | 0.9448606 | -1.507465 | 0.1316914 | 0.283496 |
| ENSG00000276931 | | 13.96724 | -0.6734506 | 0.6033941 | -1.534254 | 0.1249671 | 0.27298 |
| ENSG00000135702 | CHST5 | 4.34722251 | -0.673481124 | 0.808042581 | -1.462052566 | 0.14372681 | 0.301705 |
| ENSG00000233080 | LINC01399 | 5.498391 | -0.6735966 | 0.8887462 | -1.501137 | 0.13332 | 0.285965 |
| ENSG00000196730 | DAPK1 | 10.22508 | -0.6737318 | 0.6702691 | -1.476388 | 0.1398398 | 0.295859 |
| ENSG00000258917 | ZMYND19P1 | 2.323478 | -0.6737876 | 0.9695931 | -1.53177 | 0.1255793 | 0.273889 |
| ENSG00000108759 | KRT32 | 4.210034547 | -0.673823609 | 0.937746745 | -1.569994112 | 0.11641648 | 0.259486 |
| ENSG00000088002 | SULT2B1 | 1195.826676 | -0.673840531 | 0.29199436 | -2.500667763 | 0.01239594 | 0.046198 |
| ENSG00000224785 | | 30.78515 | -0.6738926 | 0.383802 | -2.004698 | 0.04499531 | 0.127982 |
| ENSG00000207757 | MIR93 | 16.00831 | -0.6740862 | 0.5092922 | -1.665356 | 0.09584183 | 0.225527 |
| ENSG00000288585 | | 129.4094 | -0.6741954 | 0.4377222 | -1.829374 | 0.06734358 | 0.173394 |
| ENSG00000224067 | | 3.182917 | -0.6743783 | 1.017873 | -1.610267 | 0.1073395 | 0.244921 |
| ENSG00000103187 | COTL1 | 5018.628246 | -0.674616306 | 0.421738458 | -1.858402189 | 0.06311192 | 0.165058 |
| ENSG00000189403 | HMGB1 | 14270.95 | -0.6746547 | 0.3151305 | -2.344289 | 0.01906339 | 0.065405 |
| ENSG00000280198 | | 16.17243 | -0.6746581 | 0.4788341 | -1.719906 | 0.0854495 | 0.207429 |

| | | | | | | | |
|---|---|---|---|---|---|---|---|
| ENSG00000284753 | EEF1AKMT4 | 555.2282 | -0.674673 | 0.2704321 | -2.666954 | 0.00765422 | 0.031383 |
| ENSG00000166340 | TPP1 | 2431.579 | -0.674791 | 0.3485027 | -2.161534 | 0.03065408 | 0.095366 |
| ENSG00000257159 | | 6.171893 | -0.6748827 | 0.6346519 | -1.503735 | 0.1326495 | 0.284971 |
| ENSG00000137124 | ALDH1B1 | 1384.255902 | -0.675012006 | 0.347505384 | -2.157993932 | 0.0309283 | 0.096087 |
| ENSG00000204011 | COL5A1-AS1 | 1.826273 | -0.6751031 | 0.983678 | -1.570036 | 0.1164068 | 0.259486 |
| ENSG00000162881 | OXER1 | 5.194903 | -0.675868 | 0.9223882 | -1.56154 | 0.1183964 | 0.262577 |
| ENSG00000273367 | | 5.996686 | -0.6760158 | 0.9589424 | -1.58578 | 0.1127893 | 0.253728 |
| ENSG00000183779 | ZNF703 | 3201.171 | -0.6760708 | 0.237889 | -2.986612 | 0.00282087 | 0.013869 |
| ENSG00000188833 | ENTPD8 | 113.9231 | -0.6761685 | 0.5473332 | -1.61836 | 0.105585 | 0.242106 |
| ENSG00000145604 | SKP2 | 1585.689 | -0.6762912 | 0.3371538 | -2.226352 | 0.0259906 | 0.083561 |
| ENSG00000106367 | AP1S1 | 6024.51622 | -0.676688648 | 0.183338045 | -3.830332563 | 0.00012797 | 0.000953 |
| ENSG00000173838 | MARCHF10 | 78.49262 | -0.6767442 | 0.3998418 | -1.945227 | 0.05174764 | 0.142224 |
| ENSG00000236972 | FABP5P1 | 44.61356 | -0.6767936 | 0.4383161 | -1.833449 | 0.0667358 | 0.172263 |
| ENSG00000176771 | NCKAP5 | 215.6629 | -0.6768988 | 0.5000071 | -1.699298 | 0.0892631 | 0.214363 |
| ENSG00000276547 | PCDHGB5 | 667.6538 | -0.6770517 | 0.2614124 | -2.757999 | 0.00581564 | 0.025112 |
| ENSG00000197993 | KEL | 2.921122 | -0.677238 | 0.9289154 | -1.537151 | 0.1242564 | 0.272009 |
| ENSG00000119771 | KLHL29 | 466.3876322 | -0.677255999 | 0.242388622 | -2.947552263 | 0.00320301 | 0.015339 |
| ENSG00000136522 | MRPL47 | 2208.58904 | -0.677296672 | 0.278306695 | -2.624026458 | 0.0086897 | 0.03475 |
| ENSG00000198515 | CNGA1 | 234.57 | -0.6773927 | 0.351933 | -2.147658 | 0.03174091 | 0.098015 |
| ENSG00000279129 | | 2.299429 | -0.6774487 | 1.16138 | -1.953263 | 0.05078839 | 0.140167 |
| ENSG00000232653 | GOLGA8N | 44.28367 | -0.6775347 | 0.3136945 | -2.35211 | 0.01866723 | 0.064261 |
| ENSG00000166896 | ATP23 | 488.1148 | -0.6776026 | 0.2870283 | -2.546296 | 0.01088727 | 0.041663 |
| ENSG00000137936 | BCAR3 | 977.3417061 | -0.677610846 | 0.210571735 | -3.362941307 | 0.00077117 | 0.004571 |
| ENSG00000184675 | AMER1 | 1073.138 | -0.6778683 | 0.2648681 | -2.738392 | 0.00617405 | 0.026339 |
| ENSG00000215256 | DHRS4-AS1 | 2226.171 | -0.6780101 | 0.1895882 | -3.69419 | 0.00022059 | 0.001546 |
| ENSG00000273162 | | 100.411 | -0.6780381 | 0.3590515 | -2.129443 | 0.03321763 | 0.101572 |
| ENSG00000106244 | PDAP1 | 6490.587182 | -0.678201698 | 0.191078653 | -3.672035085 | 0.00024063 | 0.001671 |
| ENSG00000236266 | | 6.153413 | -0.6782169 | 0.6153828 | -1.523514 | 0.1276302 | 0.277037 |
| ENSG00000274225 | | 26.72141 | -0.6782684 | 0.5829893 | -1.569037 | 0.1166393 | 0.25983 |
| ENSG00000140464 | PML | 2275.519 | -0.6783082 | 0.174751 | -3.99264 | 6.53E-05 | 0.000524 |
| ENSG00000179240 | GVQW3 | 948.0835 | -0.6784089 | 0.339604 | -2.216359 | 0.02666695 | 0.085334 |
| ENSG00000120820 | GLT8D2 | 24.87056397 | -0.678461292 | 0.408642706 | -1.926607862 | 0.05402852 | 0.147244 |
| ENSG00000101412 | E2F1 | 2229.382017 | -0.6784763 | 0.499845613 | -1.707916319 | 0.08765187 | 0.211423 |

| | | | | | | | |
|---|---|---|---|---|---|---|---|
| ENSG00000177917 | ARL6IP6 | 1057.874 | -0.678517 | 0.1885381 | -3.716894 | 0.00020169 | 0.001429 |
| ENSG00000213390 | ARHGAP19 | 1556.552 | -0.678584 | 0.2729401 | -2.659565 | 0.00782416 | 0.031942 |
| ENSG00000166562 | SEC11C | 1748.583 | -0.6786325 | 0.1220852 | -5.641277 | 1.69E-08 | 2.94E-07 |
| ENSG00000268355 | | 3.060607 | -0.6786483 | 0.9859617 | -1.579789 | 0.1141551 | 0.255927 |
| ENSG00000236942 | | 8.124459 | -0.6787359 | 0.5993834 | -1.543796 | 0.1226377 | 0.269397 |
| ENSG00000124602 | UNC5CL | 37.84935919 | -0.678897057 | 0.363997427 | -2.093390336 | 0.03631432 | 0.108705 |
| ENSG00000178700 | DHFR2 | 502.9528 | -0.6790336 | 0.2740685 | -2.652981 | 0.00797845 | 0.032467 |
| ENSG00000151376 | ME3 | 8.309387 | -0.6791151 | 0.7767354 | -1.439396 | 0.1500384 | 0.311165 |
| ENSG00000119715 | ESRRB | 10.28213312 | -0.679298492 | 0.675475042 | -1.503368997 | 0.13274392 | 0.285069 |
| ENSG00000263080 | | 6.779849 | -0.6794866 | 0.7605072 | -1.476333 | 0.1398546 | 0.295867 |
| ENSG00000158270 | COLEC12 | 46.39813 | -0.6798178 | 1.082881 | -1.723077 | 0.08487461 | 0.206316 |
| ENSG00000260310 | | 4.409949 | -0.6799579 | 0.8705546 | -1.509957 | 0.1310543 | 0.282422 |
| ENSG00000120725 | SIL1 | 2119.765892 | -0.680229462 | 0.195865272 | -3.586146328 | 0.0003356 | 0.002232 |
| ENSG00000112877 | CEP72 | 685.1393311 | -0.680319677 | 0.271608881 | -2.680074803 | 0.00736057 | 0.030388 |
| ENSG00000273311 | DGCR11 | 110.3624 | -0.6804266 | 0.4244928 | -1.88532 | 0.0593867 | 0.157895 |
| ENSG00000088298 | EDEM2 | 1108.817025 | -0.680434302 | 0.165245573 | -4.21435841 | 2.50E-05 | 0.000224 |
| ENSG00000171204 | TMEM126B | 2842.229 | -0.680746 | 0.1308474 | -5.294532 | 1.19E-07 | 1.78E-06 |
| ENSG00000197375 | SLC22A5 | 3370.677 | -0.6812313 | 0.2004056 | -3.545152 | 0.00039239 | 0.002552 |
| ENSG00000171792 | RHNO1 | 1826.989 | -0.6823615 | 0.2660022 | -2.731223 | 0.00630998 | 0.026777 |
| ENSG00000152223 | EPG5 | 2402.957 | -0.6824394 | 0.2572623 | -2.814647 | 0.00488309 | 0.021761 |
| ENSG00000005486 | RHBDD2 | 2799.206366 | -0.682494512 | 0.192995102 | -3.659757275 | 0.00025245 | 0.001737 |
| ENSG00000271538 | LINC02427 | 10.76454 | -0.6825166 | 0.7271038 | -1.480004 | 0.1388722 | 0.294277 |
| ENSG00000116473 | RAP1A | 2940.33054 | -0.682625901 | 0.12715796 | -5.427545333 | 5.71E-08 | 9.11E-07 |
| ENSG00000273160 | | 2.083361 | -0.6828416 | 0.8697219 | -1.494732 | 0.1349844 | 0.288342 |
| ENSG00000245149 | RNF139-AS1 | 513.5642 | -0.6830468 | 0.2424268 | -2.971054 | 0.0029678 | 0.014455 |
| ENSG00000221819 | GAS8-AS1 | 139.0605 | -0.6831298 | 0.3293385 | -2.287009 | 0.02219532 | 0.074056 |
| ENSG00000251474 | RPL32P3 | 1542.168 | -0.683251 | 0.3684178 | -2.092661 | 0.03637944 | 0.108713 |
| ENSG00000047621 | C12orf4 | 590.1063332 | -0.683263492 | 0.208572629 | -3.408718239 | 0.00065269 | 0.003953 |
| ENSG00000171720 | HDAC3 | 3864.244 | -0.6833009 | 0.1667474 | -4.214321 | 2.51E-05 | 0.000224 |
| ENSG00000086189 | DIMT1 | 1136.96119 | -0.683426167 | 0.171748784 | -4.093449323 | 4.25E-05 | 0.000359 |
| ENSG00000114738 | MAPKAPK3 | 2560.906211 | -0.683547242 | 0.158494295 | -4.413401892 | 1.02E-05 | 0.0001 |
| ENSG00000241764 | | 91.08272 | -0.6839582 | 0.289648 | -2.54452 | 0.01094279 | 0.041851 |
| ENSG00000122674 | CCZ1 | 769.4223267 | -0.683982464 | 0.125980427 | -5.520124371 | 3.39E-08 | 5.61E-07 |

| | | | | | | | |
|---|---|---|---|---|---|---|---|
| ENSG00000122034 | GTF3A | 7834.319754 | -0.684062617 | 0.163366961 | -4.293685461 | 1.76E-05 | 0.000163 |
| ENSG00000163762 | TM4SF18 | 155.0561 | -0.6841979 | 0.5562252 | -1.610555 | 0.1072767 | 0.244865 |
| ENSG00000151465 | CDC123 | 3776.175 | -0.6842989 | 0.2017424 | -3.51947 | 0.00043241 | 0.002768 |
| ENSG00000197969 | VPS13A | 3878.138 | -0.684448 | 0.2674644 | -2.731246 | 0.00630954 | 0.026777 |
| ENSG00000277767 | | 7.076177 | -0.6845038 | 0.7984394 | -1.486067 | 0.1372613 | 0.291612 |
| ENSG00000158483 | FAM86C1P | 585.2772 | -0.6846195 | 0.1631475 | -4.306408 | 1.66E-05 | 0.000155 |
| ENSG00000145491 | ROPN1L | 12.63442 | -0.6847837 | 0.6572583 | -1.519224 | 0.1287062 | 0.278718 |
| ENSG00000143401 | ANP32E | 5583.848 | -0.6849587 | 0.3957921 | -1.985076 | 0.04713601 | 0.132459 |
| ENSG00000265692 | LINC01970 | 27.27913 | -0.6851106 | 0.4002168 | -1.966274 | 0.04926696 | 0.137008 |
| ENSG00000266983 | | 17.74091 | -0.6855227 | 0.4181079 | -1.905247 | 0.05674792 | 0.15258 |
| ENSG00000001631 | KRIT1 | 379.9492167 | -0.685698738 | 0.177919802 | -3.964680024 | 7.35E-05 | 0.000583 |
| ENSG00000256897 | | 10.40221 | -0.685726 | 0.6612238 | -1.506198 | 0.1320164 | 0.284035 |
| ENSG00000272416 | | 7.305039 | -0.6860165 | 0.8111808 | -1.482766 | 0.1381366 | 0.293059 |
| ENSG00000140009 | ESR2 | 27.04899 | -0.6860871 | 0.5299719 | -1.657447 | 0.09742913 | 0.228287 |
| ENSG00000268883 | PNMA6B | 10.8213 | -0.6862571 | 0.7294142 | -1.487922 | 0.1367713 | 0.291083 |
| ENSG00000004897 | CDC27 | 5255.671435 | -0.68635861 | 0.169095275 | -4.178801112 | 2.93E-05 | 0.000257 |
| ENSG00000168906 | MAT2A | 20532.7 | -0.6865283 | 0.1734263 | -4.097165 | 4.18E-05 | 0.000354 |
| ENSG00000163811 | WDR43 | 5363.848 | -0.6867601 | 0.212657 | -3.368638 | 0.00075541 | 0.004492 |
| ENSG00000242798 | | 36.87396 | -0.6868014 | 0.4395995 | -1.857781 | 0.06320013 | 0.165216 |
| ENSG00000223799 | IL10RB-DT | 11.26485 | -0.6869454 | 0.5562587 | -1.610233 | 0.1073469 | 0.244921 |
| ENSG00000137760 | ALKBH8 | 431.5401783 | -0.687283752 | 0.16596037 | -4.246066587 | 2.18E-05 | 0.000197 |
| ENSG00000089163 | SIRT4 | 46.0117073 | -0.687297031 | 0.365842205 | -2.111560749 | 0.03472414 | 0.105018 |
| ENSG00000064012 | CASP8 | 1509.040686 | -0.687317895 | 0.178613086 | -3.961601251 | 7.44E-05 | 0.00059 |
| ENSG00000163827 | LRRC2 | 3.260784 | -0.6873295 | 0.9608252 | -1.60592 | 0.1082915 | 0.246568 |
| ENSG00000096696 | DSP | 20381.93068 | -0.68740166 | 0.220370194 | -3.261420941 | 0.00110855 | 0.006254 |
| ENSG00000115234 | SNX17 | 7404.531815 | -0.687431541 | 0.120198226 | -5.779547477 | 7.49E-09 | 1.40E-07 |
| ENSG00000225076 | WWC3-AS1 | 1.882607 | -0.6874407 | 1.057346 | -1.704071 | 0.08836781 | 0.212706 |
| ENSG00000128585 | MKLN1 | 2096.173565 | -0.687787049 | 0.220941094 | -3.25607379 | 0.00112964 | 0.006356 |
| ENSG00000166598 | HSP90B1 | 25040.58 | -0.6878504 | 0.1634811 | -4.312013 | 1.62E-05 | 0.000152 |
| ENSG00000005469 | CROT | 845.0852933 | -0.688356537 | 0.317427171 | -2.384479254 | 0.01710331 | 0.059892 |
| ENSG00000176732 | PFN4 | 46.23291 | -0.688386 | 0.3273396 | -2.309367 | 0.02092323 | 0.070475 |
| ENSG00000090013 | BLVRB | 6724.400574 | -0.688403729 | 0.248271254 | -2.932628727 | 0.00336106 | 0.015975 |
| ENSG00000286138 | | 62.69088 | -0.6884268 | 0.3088622 | -2.416733 | 0.01566051 | 0.05576 |

| | | | | | | | |
|---|---|---|---|---|---|---|---|
| ENSG00000114529 | C3orf52 | 768.4086255 | -0.68856627 | 0.328205049 | -2.320985555 | 0.02028762 | 0.068753 |
| ENSG00000277310 | | 8.064172 | -0.6886525 | 0.7321902 | -1.486675 | 0.1371007 | 0.291446 |
| ENSG00000204681 | GABBR1 | 1895.725 | -0.688694 | 0.2959306 | -2.523371 | 0.01162356 | 0.043899 |
| ENSG00000287672 | | 15.39037 | -0.6887375 | 0.5359402 | -1.654421 | 0.09804208 | 0.229404 |
| ENSG00000215158 | | 486.7379 | -0.6889304 | 0.3155089 | -2.391774 | 0.01676717 | 0.058966 |
| ENSG00000224881 | | 4.096534 | -0.6889767 | 0.753706 | -1.492938 | 0.1354534 | 0.288949 |
| ENSG00000177830 | CHID1 | 2911.7 | -0.68914 | 0.1741507 | -4.078154 | 4.54E-05 | 0.000381 |
| ENSG00000223705 | NSUN5P1 | 1892.697 | -0.689143 | 0.2570204 | -2.813338 | 0.00490301 | 0.021824 |
| ENSG00000136147 | PHF11 | 1275.335441 | -0.689250029 | 0.224810696 | -3.206716924 | 0.00134259 | 0.007391 |
| ENSG00000084090 | STARD7 | 11227.02091 | -0.689335865 | 0.186669919 | -3.811179722 | 0.00013831 | 0.001021 |
| ENSG00000183044 | ABAT | 5734.752 | -0.6894821 | 0.2696474 | -2.733392 | 0.00626858 | 0.026639 |
| ENSG00000132313 | MRPL35 | 2694.119276 | -0.689574159 | 0.211596018 | -3.396889967 | 0.00068156 | 0.004108 |
| ENSG00000259370 | | 17.48844 | -0.6896897 | 0.5883959 | -1.588668 | 0.1121353 | 0.252699 |
| ENSG00000241839 | PLEKHO2 | 256.3342 | -0.6897008 | 0.2123473 | -3.385123 | 0.00071146 | 0.004263 |
| ENSG00000116455 | WDR77 | 2209.123668 | -0.689781904 | 0.226953006 | -3.19429866 | 0.00140171 | 0.007657 |
| ENSG00000202508 | | 2.371179 | -0.6899508 | 0.9644553 | -1.562457 | 0.1181804 | 0.262251 |
| ENSG00000242396 | | 1.069781 | -0.6899795 | 1.045321 | -1.728169 | 0.08395798 | NA |
| ENSG00000178538 | CA8 | 670.5117 | -0.6902846 | 0.4074076 | -1.967361 | 0.04914164 | 0.136745 |
| ENSG00000286997 | | 9.369784 | -0.690341 | 0.6355747 | -1.533192 | 0.1252285 | 0.273416 |
| ENSG00000283005 | CDC27P3 | 4.245968 | -0.690366 | 0.8487758 | -1.509004 | 0.1312977 | 0.282786 |
| ENSG00000259865 | | 415.8666 | -0.6909514 | 0.2933478 | -2.548593 | 0.01081585 | 0.041437 |
| ENSG00000274460 | | 84.5876 | -0.6909657 | 0.5631418 | -1.621943 | 0.1048156 | 0.240716 |
| ENSG00000167157 | PRRX2 | 15.8824 | -0.6909728 | 0.8147835 | -1.520739 | 0.1283254 | 0.27812 |
| ENSG00000138085 | ATRAID | 4513.256957 | -0.690991516 | 0.157108326 | -4.498883917 | 6.83E-06 | 7.03E-05 |
| ENSG00000157423 | HYDIN | 2.455604 | -0.6910547 | 0.9596564 | -1.529651 | 0.1261032 | 0.274678 |
| ENSG00000102003 | SYP | 100.3615557 | -0.691418269 | 0.314610219 | -2.401464357 | 0.0163296 | 0.057648 |
| ENSG00000270012 | | 195.9901 | -0.6915268 | 0.2049217 | -3.507348 | 0.0004526 | 0.002877 |
| ENSG00000215374 | FAM66B | 42.81362 | -0.6915798 | 0.3365739 | -2.266701 | 0.02340848 | 0.076957 |
| ENSG00000160401 | CFAP157 | 184.8052 | -0.69185 | 0.2718211 | -2.722071 | 0.00648742 | 0.027377 |
| ENSG00000248626 | GAPDHP40 | 90.83446 | -0.6919021 | 0.437627 | -1.876948 | 0.06052517 | 0.160105 |
| ENSG00000055044 | NOP58 | 5217.673623 | -0.691984501 | 0.182064933 | -3.919393996 | 8.88E-05 | 0.000688 |
| ENSG00000260920 | | 222.4805 | -0.6922046 | 0.2931284 | -2.551422 | 0.01072843 | 0.041185 |
| ENSG00000272892 | | 19.52918 | -0.6925246 | 0.5392277 | -1.651588 | 0.09861852 | 0.230308 |

| | | | | | | | |
|---|---|---|---|---|---|---|---|
| ENSG00000135624 | CCT7 | 21712.23328 | -0.692635009 | 0.217840959 | -3.320399656 | 0.00089889 | 0.00521 |
| ENSG00000130943 | PKDREJ | 21.71547933 | -0.692747956 | 0.525855367 | -1.675963956 | 0.09374525 | 0.222088 |
| ENSG00000236048 | SSBL5P | 32.34902 | -0.6928323 | 0.3341701 | -2.294433 | 0.02176564 | 0.072833 |
| ENSG00000227682 | ATP5F1AP2 | 9.064168 | -0.6929378 | 0.6425585 | -1.547685 | 0.1216982 | 0.267985 |
| ENSG00000249593 | | 11.87222 | -0.6929782 | 0.5312282 | -1.668677 | 0.09518135 | 0.22441 |
| ENSG00000213239 | NPM1P32 | 10.56646 | -0.6930857 | 0.6586094 | -1.537489 | 0.1241735 | 0.271947 |
| ENSG00000132256 | TRIM5 | 369.5369773 | -0.693271068 | 0.317734858 | -2.381743193 | 0.01723091 | 0.060254 |
| ENSG00000128512 | DOCK4 | 111.0687228 | -0.693403412 | 0.396392565 | -2.013616267 | 0.04404984 | 0.126008 |
| ENSG00000136122 | BORA | 359.0470018 | -0.693527504 | 0.365523651 | -2.144287996 | 0.03200982 | 0.098639 |
| ENSG00000109927 | TECTA | 20.86447918 | -0.69357223 | 0.465961776 | -1.800288346 | 0.07181512 | 0.182139 |
| ENSG00000136026 | CKAP4 | 2710.771121 | -0.693687018 | 0.145145741 | -4.871939062 | 1.11E-06 | 1.36E-05 |
| ENSG00000205930 | C21orf62-AS1 | 45.1836 | -0.6937823 | 0.6331487 | -1.561437 | 0.1184206 | 0.262609 |
| ENSG00000148450 | MSRB2 | 1361.715 | -0.6939589 | 0.1920579 | -3.742616 | 0.00018211 | 0.001304 |
| ENSG00000137225 | CAPN11 | 1.349528629 | -0.6939673 | 1.100371059 | -1.780427069 | 0.0750061 | NA |
| ENSG00000163913 | IFT122 | 1884.505 | -0.6940078 | 0.1362838 | -5.18596 | 2.15E-07 | 3.06E-06 |
| ENSG00000122958 | VPS26A | 4834.379494 | -0.69403783 | 0.133976384 | -5.263642291 | 1.41E-07 | 2.08E-06 |
| ENSG00000282772 | | 1.797272 | -0.6940549 | 0.9548285 | -1.518184 | 0.128968 | 0.279172 |
| ENSG00000077420 | APBB1IP | 6.168762817 | -0.694066825 | 0.852966175 | -1.455172387 | 0.14562154 | 0.304695 |
| ENSG00000103707 | MTFMT | 992.0575196 | -0.694067539 | 0.166072338 | -4.286403441 | 1.82E-05 | 0.000168 |
| ENSG00000167699 | GLOD4 | 3662.678 | -0.6942923 | 0.1630493 | -4.354384 | 1.33E-05 | 0.000127 |
| ENSG00000076513 | ANKRD13A | 3378.056356 | -0.69434668 | 0.277552841 | -2.682673561 | 0.00730362 | 0.030188 |
| ENSG00000285799 | | 41.61064 | -0.6945098 | 0.554632 | -1.636305 | 0.1017759 | 0.235691 |
| ENSG00000287804 | | 3.414358 | -0.6948954 | 0.8412568 | -1.511594 | 0.1306371 | 0.281797 |
| ENSG00000250148 | KRT8P31 | 4.690631 | -0.6949838 | 0.8176811 | -1.475513 | 0.1400746 | 0.296238 |
| ENSG00000270084 | GAS5-AS1 | 38.58597 | -0.6950947 | 0.2994123 | -2.51741 | 0.01182211 | 0.044497 |
| ENSG00000182489 | XKRX | 25.03218 | -0.6950962 | 0.4045947 | -1.986135 | 0.04701829 | 0.132264 |
| ENSG00000136490 | LIMD2 | 592.5813897 | -0.695166496 | 0.393086342 | -2.032485533 | 0.04210453 | 0.121674 |
| ENSG00000165060 | FXN | 299.2275 | -0.6954818 | 0.2813198 | -2.659718 | 0.00782061 | 0.031933 |
| ENSG00000286302 | | 2.218826 | -0.6957008 | 1.005176 | -1.585325 | 0.1128926 | 0.253868 |
| ENSG00000224011 | | 2.633433 | -0.6958196 | 0.9327595 | -1.499908 | 0.1336381 | 0.286387 |
| ENSG00000279544 | | 2.204814 | -0.6958237 | 1.0313 | -1.682958 | 0.09238315 | 0.219625 |
| ENSG00000261291 | | 1.958082 | -0.6960269 | 1.035335 | -1.686753 | 0.09165095 | 0.218451 |
| ENSG00000168246 | UBTD2 | 2663.634 | -0.6971009 | 0.1269284 | -5.572138 | 2.52E-08 | 4.26E-07 |

| | | | | | | | |
|---|---|---|---|---|---|---|---|
| ENSG00000076003 | MCM6 | 6219.263971 | -0.69721259 | 0.438903012 | -1.883693402 | 0.05960646 | 0.158281 |
| ENSG00000101182 | PSMA7 | 14459.72326 | -0.69727528 | 0.240498095 | -3.056191088 | 0.00224168 | 0.011395 |
| ENSG00000188921 | HACD4 | 10.23213 | -0.6972963 | 0.6177894 | -1.562684 | 0.1181269 | 0.262205 |
| ENSG00000167085 | PHB | 9667.661 | -0.6980974 | 0.2523377 | -2.914429 | 0.0035634 | 0.01676 |
| ENSG00000167716 | WDR81 | 2303.701 | -0.6986536 | 0.1561992 | -4.572773 | 4.81E-06 | 5.15E-05 |
| ENSG00000130305 | NSUN5 | 2246.425763 | -0.698670414 | 0.17243054 | -4.157076924 | 3.22E-05 | 0.00028 |
| ENSG00000207217 | SNORA80D | 4.006158 | -0.698818 | 0.8914668 | -1.529656 | 0.1261019 | 0.274678 |
| ENSG00000268218 | | 118.3602 | -0.6992735 | 0.2285143 | -3.206784 | 0.00134228 | 0.007391 |
| ENSG00000272142 | | 270.4113 | -0.6993025 | 0.3180176 | -2.416643 | 0.01566437 | 0.055767 |
| ENSG00000104691 | UBXN8 | 586.3122303 | -0.699339896 | 0.213565502 | -3.415967926 | 0.00063556 | 0.003867 |
| ENSG00000253251 | SHLD3 | 178.4448 | -0.6993467 | 0.2305954 | -3.186042 | 0.00144234 | 0.007851 |
| ENSG00000141750 | STAC2 | 1.903842 | -0.6998438 | 1.011188 | -1.605586 | 0.1083649 | 0.246693 |
| ENSG00000163406 | SLC15A2 | 258.2046 | -0.699874 | 0.227305 | -3.228942 | 0.00124249 | 0.006908 |
| ENSG00000226856 | THORLNC | 16.04535 | -0.7000138 | 0.508359 | -1.716804 | 0.08601501 | 0.208402 |
| ENSG00000227474 | RPL6P24 | 1.233568 | -0.7000148 | 1.19437 | -1.993722 | 0.04618243 | NA |
| ENSG00000108055 | SMC3 | 4072.697208 | -0.700096824 | 0.216669018 | -3.37271476 | 0.00074431 | 0.004428 |
| ENSG00000214146 | LINC02026 | 6.990008 | -0.7002432 | 0.6949648 | -1.514282 | 0.1299544 | 0.280666 |
| ENSG00000168944 | CEP120 | 1365.483 | -0.7002647 | 0.1593898 | -4.496067 | 6.92E-06 | 7.12E-05 |
| ENSG00000077514 | POLD3 | 1859.619463 | -0.700363917 | 0.308119918 | -2.48100969 | 0.01310108 | 0.048365 |
| ENSG00000286049 | | 3.159764 | -0.7003938 | 0.8654028 | -1.510366 | 0.1309502 | 0.282289 |
| ENSG00000241889 | | 115.0366 | -0.7004521 | 0.3083561 | -2.490597 | 0.01275285 | 0.047296 |
| ENSG00000196735 | HLA-DQA1 | 3.705535 | -0.7005228 | 0.9692578 | -1.538414 | 0.1239474 | 0.271619 |
| ENSG00000235472 | EIF4A1P7 | 121.7021 | -0.7005752 | 0.2381508 | -3.095515 | 0.00196471 | 0.010197 |
| ENSG00000117877 | POLR1G | 430.3970071 | -0.70057691 | 0.291804534 | -2.593622482 | 0.00949707 | 0.037266 |
| ENSG00000280332 | | 53.46177 | -0.7006104 | 0.45337 | -1.853158 | 0.06385967 | 0.166505 |
| ENSG00000249884 | RNF103-CHMP3 | 7.678206 | -0.700637 | 0.640738 | -1.54698 | 0.1218681 | 0.268217 |
| ENSG00000270820 | | 98.58029 | -0.7007667 | 0.3325215 | -2.320297 | 0.02032484 | 0.068844 |
| ENSG00000139514 | SLC7A1 | 13799.69 | -0.7008067 | 0.3170748 | -2.422078 | 0.01543202 | 0.055139 |
| ENSG00000196935 | SRGAP1 | 1293.146 | -0.7012242 | 0.2940718 | -2.579006 | 0.0099085 | 0.038578 |
| ENSG00000136143 | SUCLA2 | 2148.826635 | -0.701269061 | 0.204814375 | -3.561403703 | 0.00036888 | 0.002418 |
| ENSG00000091640 | SPAG7 | 2572.881371 | -0.701553881 | 0.184564782 | -3.92085296 | 8.82E-05 | 0.000685 |
| ENSG00000267655 | | 17.19895 | -0.7016159 | 0.4440199 | -1.876665 | 0.06056405 | 0.160144 |
| ENSG00000115904 | SOS1 | 4091.547572 | -0.701752 | 0.329169703 | -2.331239604 | 0.01974073 | 0.067286 |

| | | | | | | | |
|---|---|---|---|---|---|---|---|
| ENSG00000083544 | TDRD3 | 420.5263598 | -0.701764572 | 0.201699543 | -3.610806064 | 0.00030525 | 0.002054 |
| ENSG00000283016 | ATP5MC1P2 | 12.63487 | -0.7024152 | 0.5091462 | -1.727532 | 0.08407218 | 0.204872 |
| ENSG00000267033 | | 4.691678 | -0.7027465 | 0.7349181 | -1.494006 | 0.135174 | 0.288473 |
| ENSG00000128253 | RFPL2 | 3.577226326 | -0.702805384 | 0.897423908 | -1.533514797 | 0.12514906 | 0.273287 |
| ENSG00000229509 | | 6.809445 | -0.7028375 | 0.5794062 | -1.623389 | 0.1045062 | 0.240229 |
| ENSG00000134049 | IER3IP1 | 1136.201191 | -0.70316576 | 0.171233491 | -4.21248361 | 2.53E-05 | 0.000225 |
| ENSG00000239959 | ENPP7P2 | 4.273817 | -0.7034442 | 0.805277 | -1.505992 | 0.1320694 | 0.284098 |
| ENSG00000176840 | MIR7-3HG | 1.315931 | -0.7034757 | 1.132284 | -1.852555 | 0.0639461 | NA |
| ENSG00000274828 | | 403.8275 | -0.7038385 | 0.25655 | -2.905768 | 0.00366353 | 0.01715 |
| ENSG00000286881 | | 3.353039 | -0.7038691 | 0.9914005 | -1.628782 | 0.1033592 | 0.23834 |
| ENSG00000240695 | | 84.45253 | -0.7039907 | 0.2683068 | -2.801048 | 0.00509369 | 0.022496 |
| ENSG00000116212 | LRRC42 | 2599.111656 | -0.704032321 | 0.243472015 | -3.049926645 | 0.00228897 | 0.011588 |
| ENSG00000169223 | LMAN2 | 9409.258 | -0.7041417 | 0.1660107 | -4.3446 | 1.40E-05 | 0.000133 |
| ENSG00000244468 | | 1071.043 | -0.7042849 | 0.6137579 | -1.586027 | 0.1127331 | 0.253659 |
| ENSG00000189430 | NCR1 | 3.149215 | -0.70429 | 1.194324 | -1.998301 | 0.04568401 | 0.129456 |
| ENSG00000081026 | MAGI3 | 2941.765884 | -0.70429129 | 0.222069511 | -3.314469522 | 0.00091817 | 0.005302 |
| ENSG00000051341 | POLQ | 1429.089856 | -0.705071948 | 0.47291527 | -1.820886436 | 0.06862412 | 0.17589 |
| ENSG00000135269 | TES | 3266.499606 | -0.705147458 | 0.259942805 | -2.886762419 | 0.00389228 | 0.018042 |
| ENSG00000234084 | | 26.04832 | -0.705194 | 0.357796 | -2.210093 | 0.02709871 | 0.086433 |
| ENSG00000124813 | RUNX2 | 505.7921803 | -0.705295578 | 0.303282568 | -2.511377547 | 0.0120261 | 0.045111 |
| ENSG00000273008 | CSGALNACT2-DT | 62.79271 | -0.7053021 | 0.3776357 | -2.11254 | 0.03464012 | 0.104855 |
| ENSG00000148019 | CEP78 | 2120.905 | -0.7054396 | 0.336622 | -2.321372 | 0.0202668 | 0.068709 |
| ENSG00000115998 | C2orf42 | 718.3256823 | -0.705544207 | 0.258378541 | -2.89854538 | 0.00374898 | 0.017469 |
| ENSG00000183598 | H3C13 | 147.401 | -0.7056729 | 0.4217367 | -1.959283 | 0.05007968 | 0.138644 |
| ENSG00000224934 | | 234.4231 | -0.7057759 | 0.4669267 | -1.846809 | 0.06477484 | 0.168247 |
| ENSG00000230807 | | 2.057257 | -0.70614 | 1.035499 | -1.671335 | 0.09465552 | 0.223666 |
| ENSG00000065621 | GSTO2 | 1565.037 | -0.706406465 | 0.165482784 | -4.376156631 | 1.21E-05 | 0.000117 |
| ENSG00000103710 | RASL12 | 2.554540813 | -0.706590282 | 1.019929849 | -1.683123385 | 0.09235121 | 0.219588 |
| ENSG00000231789 | PIK3CD-AS2 | 9.530534 | -0.7068825 | 0.9631243 | -1.589855 | 0.1118674 | 0.252417 |
| ENSG00000263412 | NFE2L1-DT | 124.8238 | -0.7071539 | 0.452404 | -1.873884 | 0.06094641 | 0.160867 |
| ENSG00000167434 | CA4 | 843.1323 | -0.7073666 | 0.3136467 | -2.465235 | 0.01369236 | 0.050047 |
| ENSG00000215302 | | 34.76054 | -0.7074194 | 0.3423679 | -2.299289 | 0.02148855 | 0.072087 |
| ENSG00000108852 | MPP2 | 458.342191 | -0.707455556 | 0.395123356 | -2.061023682 | 0.03930078 | 0.115341 |

| | | | | | | | |
|---|---|---|---|---|---|---|---|
| ENSG00000134057 | CCNB1 | 6046.48814 | -0.70781582 | 0.439359971 | -1.912159801 | 0.0558557 | 0.150816 |
| ENSG00000226754 | | 63.95181 | -0.7078796 | 0.3248606 | -2.390379 | 0.016831 | 0.059151 |
| ENSG00000258352 | IFITM3P6 | 1.435582 | -0.7079296 | 1.051447 | -1.700637 | 0.08901115 | NA |
| ENSG00000137713 | PPP2R1B | 1207.80366 | -0.708404858 | 0.238123023 | -3.137566725 | 0.00170357 | 0.009029 |
| ENSG00000171105 | INSR | 1727.291 | -0.7085061 | 0.2230787 | -3.328244 | 0.00087395 | 0.005091 |
| ENSG00000227354 | RBM26-AS1 | 80.2534 | -0.7088693 | 0.3042058 | -2.527162 | 0.01149884 | 0.043533 |
| ENSG00000286892 | | 1.741708 | -0.709424 | 1.074931 | -1.750364 | 0.08005559 | 0.197328 |
| ENSG00000253196 | | 6.224529 | -0.7095623 | 0.7124428 | -1.533394 | 0.1251789 | 0.27333 |
| ENSG00000176410 | DNAJC30 | 597.0244 | -0.7098254 | 0.1524624 | -4.758564 | 1.95E-06 | 2.30E-05 |
| ENSG00000248557 | | 16.68565 | -0.7099592 | 0.5125513 | -1.742445 | 0.08143068 | 0.20004 |
| ENSG00000111837 | MAK | 18.32484441 | -0.710102365 | 0.646694762 | -1.566942706 | 0.11712808 | 0.260613 |
| ENSG00000257114 | LINC02450 | 11.26043 | -0.7103918 | 0.7974331 | -1.504993 | 0.1323258 | 0.284516 |
| ENSG00000146373 | RNF217 | 3.261001 | -0.7110065 | 0.8473623 | -1.534319 | 0.1249512 | 0.27298 |
| ENSG00000215021 | PHB2 | 13034.02 | -0.7112668 | 0.186491 | -3.926948 | 8.60E-05 | 0.00067 |
| ENSG00000128563 | PRKRIP1 | 2685.790013 | -0.711390504 | 0.128327035 | -5.632482423 | 1.78E-08 | 3.09E-07 |
| ENSG00000271034 | | 11.77434 | -0.7117409 | 0.5809783 | -1.636619 | 0.1017102 | 0.235621 |
| ENSG00000259699 | HMGB1P8 | 9.81303 | -0.711784 | 0.748123 | -1.541778 | 0.1231276 | 0.270273 |
| ENSG00000148484 | RSU1 | 4250.775 | -0.7119788 | 0.1301837 | -5.552936 | 2.81E-08 | 4.71E-07 |
| ENSG00000101162 | TUBB1 | 54.52834557 | -0.711989279 | 0.519066942 | -1.726905016 | 0.08418472 | 0.205052 |
| ENSG00000164329 | TENT2 | 2908.13 | -0.7123359 | 0.1891634 | -3.891034 | 9.98E-05 | 0.000764 |
| ENSG00000258169 | LINC00485 | 0.993249 | -0.7123643 | 1.225422 | -2.079971 | 0.03752819 | NA |
| ENSG00000074842 | MYDGF | 3277.744888 | -0.71239806 | 0.147239783 | -4.912759295 | 8.98E-07 | 1.13E-05 |
| ENSG00000124260 | MAGEA10 | 41.90576223 | -0.712583591 | 1.113753762 | -1.774996973 | 0.07589839 | 0.189721 |
| ENSG00000095637 | SORBS1 | 163.3236983 | -0.713281965 | 0.456099021 | -1.868147784 | 0.06174148 | 0.162497 |
| ENSG00000231749 | ABCA9-AS1 | 1.414835 | -0.7133601 | 1.272926 | -2.24547 | 0.02473798 | NA |
| ENSG00000123106 | CCDC91 | 922.9861942 | -0.713540618 | 0.172943249 | -4.24495041 | 2.19E-05 | 0.000198 |
| ENSG00000121716 | PILRB | 343.88787 | -0.713540623 | 0.279259772 | -2.740792496 | 0.00612912 | 0.026164 |
| ENSG00000148835 | TAF5 | 463.432 | -0.7136565 | 0.3886586 | -2.097997 | 0.03590539 | 0.10773 |
| ENSG00000138071 | ACTR2 | 17065.95769 | -0.713855853 | 0.15201538 | -4.842402548 | 1.28E-06 | 1.56E-05 |
| ENSG00000205559 | CHKB-DT | 171.7859 | -0.7140373 | 0.2622536 | -2.888049 | 0.0038764 | 0.01798 |
| ENSG00000106683 | LIMK1 | 2086.071728 | -0.714122688 | 0.158429237 | -4.616246376 | 3.91E-06 | 4.27E-05 |
| ENSG00000279865 | | 20.04465 | -0.7143394 | 0.5094967 | -1.764018 | 0.07772892 | 0.192989 |
| ENSG00000224953 | SRIP3 | 2.176259 | -0.7143396 | 1.180699 | -1.962271 | 0.04973092 | 0.137981 |

| | | | | | | | |
|---|---|---|---|---|---|---|---|
| ENSG00000152413 | HOMER1 | 1333.76 | -0.7147764 | 0.2005265 | -3.698566 | 0.00021682 | 0.001523 |
| ENSG00000254708 | MMADHCP2 | 4.257116 | -0.7149161 | 0.821882 | -1.507321 | 0.1317284 | 0.28353 |
| ENSG00000236516 | KLF2P4 | 4.222534 | -0.7149416 | 0.9374666 | -1.575672 | 0.1151013 | 0.257613 |
| ENSG00000163467 | TSACC | 81.27374 | -0.7150429 | 0.2693282 | -2.826797 | 0.00470161 | 0.021094 |
| ENSG00000125354 | SEPTIN6 | 232.0079044 | -0.715091099 | 0.436886305 | -1.934371288 | 0.05306749 | 0.145281 |
| ENSG00000185896 | LAMP1 | 10821.1 | -0.7151082 | 0.1654382 | -4.434673 | 9.22E-06 | 9.16E-05 |
| ENSG00000117984 | CTSD | 7653.978016 | -0.715468402 | 0.229357901 | -3.272247693 | 0.00106696 | 0.00605 |
| ENSG00000135048 | CEMIP2 | 2305.481164 | -0.715531507 | 0.254004669 | -2.957581121 | 0.00310063 | 0.014935 |
| ENSG00000113119 | TMCO6 | 1089.834628 | -0.715907282 | 0.14844155 | -4.921233956 | 8.60E-07 | 1.08E-05 |
| ENSG00000240854 | PSMC2P1 | 5.932182 | -0.7159585 | 0.7800584 | -1.531428 | 0.1256636 | 0.274028 |
| ENSG00000136011 | STAB2 | 3.15384862 | -0.716025821 | 0.743597785 | -1.529222732 | 0.12620924 | 0.274879 |
| ENSG00000054690 | PLEKHH1 | 2393.782252 | -0.716245781 | 0.222845249 | -3.357399041 | 0.0007868 | 0.004653 |
| ENSG00000166507 | NDST2 | 43.02597 | -0.7163636 | 0.3810098 | -2.132915 | 0.03293167 | 0.100814 |
| ENSG00000168564 | CDKN2AIP | 1332.849 | -0.7164201 | 0.1452695 | -5.035185 | 4.77E-07 | 6.36E-06 |
| ENSG00000159199 | ATP5MC1 | 4069.127 | -0.7170983 | 0.1721705 | -4.28413 | 1.83E-05 | 0.000169 |
| ENSG00000254577 | | 6.520015 | -0.7172705 | 0.7575121 | -1.536634 | 0.1243829 | 0.272235 |
| ENSG00000112394 | SLC16A10 | 63.56138823 | -0.717305914 | 0.510721339 | -1.767617242 | 0.07712491 | 0.191953 |
| ENSG00000213958 | KRT18P29 | 234.9058 | -0.7173545 | 0.2366459 | -3.184097 | 0.00145206 | 0.007888 |
| ENSG00000090263 | MRPS33 | 2284.581232 | -0.717363921 | 0.155741996 | -4.710160565 | 2.48E-06 | 2.85E-05 |
| ENSG00000257543 | | 9.164367 | -0.7176652 | 0.5947226 | -1.624511 | 0.1042667 | 0.23985 |
| ENSG00000227073 | SDHDP2 | 25.25075 | -0.7176719 | 0.5095961 | -1.771922 | 0.07640753 | 0.190602 |
| ENSG00000184363 | PKP3 | 6970.464 | -0.71768 | 0.1708887 | -4.328011 | 1.50E-05 | 0.000142 |
| ENSG00000155754 | C2CD6 | 4.598925 | -0.7178233 | 0.7224675 | -1.531158 | 0.1257304 | 0.274119 |
| ENSG00000165609 | NUDT5 | 4378.574 | -0.7178306 | 0.1571114 | -4.672737 | 2.97E-06 | 3.36E-05 |
| ENSG00000285600 | | 9.059471 | -0.7178374 | 0.9019369 | -1.585733 | 0.1128 | 0.253728 |
| ENSG00000236924 | | 15.54155 | -0.7179188 | 0.6304126 | -1.597965 | 0.1100509 | 0.249488 |
| ENSG00000215482 | CALM2P3 | 132.6038 | -0.7179311 | 0.3062372 | -2.549021 | 0.01080259 | 0.041398 |
| ENSG00000261889 | | 89.32933 | -0.7180794 | 0.3608957 | -2.234044 | 0.02548017 | 0.082318 |
| ENSG00000247828 | TMEM161B-AS1 | 1213.5 | -0.7181264 | 0.441833 | -1.926749 | 0.05401096 | 0.147211 |
| ENSG00000138111 | MFSD13A | 313.0168874 | -0.71862734 | 0.264003581 | -2.902911961 | 0.00369711 | 0.017266 |
| ENSG00000101464 | PIGU | 1351.886314 | -0.718681954 | 0.19085862 | -3.892421854 | 9.92E-05 | 0.00076 |
| ENSG00000226266 | | 26.06179 | -0.7187135 | 0.3566267 | -2.259036 | 0.02388116 | 0.078234 |
| ENSG00000262333 | HNRNPA1P16 | 103.5678 | -0.7187593 | 0.3023604 | -2.578156 | 0.00993292 | 0.038651 |

| | | | | | | | |
|---|---|---|---|---|---|---|---|
| ENSG00000128309 | MPST | 3439.756092 | -0.718792696 | 0.179193206 | -4.13243295 | 3.59E-05 | 0.000309 |
| ENSG00000116005 | PCYOX1 | 4986.20502 | -0.718821449 | 0.188947063 | -3.931571145 | 8.44E-05 | 0.000659 |
| ENSG00000115902 | SLC1A4 | 2338.955863 | -0.718851106 | 0.353946427 | -2.271585707 | 0.02311155 | 0.07628 |
| ENSG00000197472 | ZNF695 | 362.7201 | -0.7188986 | 0.4608104 | -1.88422 | 0.05953524 | 0.158148 |
| ENSG00000115484 | CCT4 | 11253.68245 | -0.719352822 | 0.207363248 | -3.615152439 | 0.00030017 | 0.002023 |
| ENSG00000273107 | | 1.384352 | -0.71963 | 1.27298 | -2.217445 | 0.02659272 | NA |
| ENSG00000005100 | DHX33 | 3295.953091 | -0.719650436 | 0.203731844 | -3.675318699 | 0.00023755 | 0.001651 |
| ENSG00000255559 | ZNF252P-AS1 | 55.43234 | -0.7196792 | 0.4672276 | -1.86241 | 0.06254526 | 0.164028 |
| ENSG00000227698 | | 22.62716 | -0.7198498 | 0.5425439 | -1.709225 | 0.08740935 | 0.210934 |
| ENSG00000169750 | RAC3 | 3794.589 | -0.7200563 | 0.3129819 | -2.515418 | 0.01188914 | 0.044705 |
| ENSG00000226161 | EIF4A1P5 | 21.26333 | -0.7201547 | 0.3829915 | -2.140191 | 0.03233932 | 0.09939 |
| ENSG00000261168 | | 6.474082 | -0.7203164 | 0.81359 | -1.530445 | 0.1259065 | 0.274377 |
| ENSG00000149679 | CABLES2 | 2679.738 | -0.7204158 | 0.1664826 | -4.43555 | 9.18E-06 | 9.14E-05 |
| ENSG00000134308 | YWHAQ | 14985.24998 | -0.720578755 | 0.262805727 | -2.919214104 | 0.00350915 | 0.016552 |
| ENSG00000225075 | | 13.23366 | -0.7208508 | 0.6115922 | -1.624189 | 0.1043354 | 0.239966 |
| ENSG00000279114 | | 11.58647 | -0.7209814 | 0.6362227 | -1.592777 | 0.1112101 | 0.251296 |
| ENSG00000172197 | MBOAT1 | 969.4427 | -0.7210568 | 0.2828879 | -2.745009 | 0.00605093 | 0.025897 |
| ENSG00000131149 | GSE1 | 19630.1697 | -0.721149452 | 0.18855511 | -3.933910897 | 8.36E-05 | 0.000653 |
| ENSG00000264812 | | 6.782025 | -0.7212656 | 0.795481 | -1.551664 | 0.1207426 | 0.2666 |
| ENSG00000127527 | EPS15L1 | 2363.047185 | -0.721334522 | 0.119267906 | -6.134575887 | 8.54E-10 | 1.85E-08 |
| ENSG00000214706 | IFRD2 | 3929.898 | -0.7220842 | 0.1988668 | -3.767268 | 0.00016504 | 0.001197 |
| ENSG00000213551 | DNAJC9 | 1361.655 | -0.7224617 | 0.3609887 | -2.239346 | 0.02513344 | 0.081427 |
| ENSG00000228960 | OR2A9P | 6.188259 | -0.7226055 | 0.6338115 | -1.596288 | 0.1104245 | 0.250095 |
| ENSG00000249526 | | 1.675092 | -0.7226813 | 1.141367 | -1.833456 | 0.06673485 | 0.172263 |
| ENSG00000267040 | | 43.04311 | -0.7229763 | 0.3764574 | -2.177214 | 0.02946461 | 0.092469 |
| ENSG00000116120 | FARSB | 3463.257009 | -0.723086771 | 0.220138821 | -3.435208092 | 0.0005921 | 0.003632 |
| ENSG00000117697 | NSL1 | 948.2939823 | -0.723149798 | 0.200008829 | -3.752868117 | 0.00017482 | 0.001258 |
| ENSG00000186654 | PRR5 | 632.8684 | -0.7234959 | 0.1945652 | -3.84678 | 0.00011968 | 0.000899 |
| ENSG00000257790 | EIF4A1P4 | 205.8941 | -0.7242667 | 0.2251758 | -3.36824 | 0.0007565 | 0.004498 |
| ENSG00000188050 | RNF133 | 4.121344 | -0.7243964 | 0.9348852 | -1.568358 | 0.1167977 | 0.26003 |
| ENSG00000100285 | NEFH | 3.629200859 | -0.72463425 | 0.872488497 | -1.524745342 | 0.12732263 | 0.276579 |
| ENSG00000123178 | SPRYD7 | 646.7647462 | -0.724766156 | 0.163385177 | -4.549723074 | 5.37E-06 | 5.68E-05 |
| ENSG00000258458 | | 41.00166 | -0.7247741 | 0.3691854 | -2.208668 | 0.02719771 | 0.086707 |

| | | | | | | | |
|---|---|---|---|---|---|---|---|
| ENSG00000256686 | | 2.82372 | -0.7248521 | 0.8996766 | -1.603323 | 0.1088634 | 0.247349 |
| ENSG00000215305 | VPS16 | 1876.221 | -0.7250659 | 0.1155922 | -6.34041 | 2.29E-10 | 5.44E-09 |
| ENSG00000243302 | | 759.7061 | -0.7252966 | 0.2777842 | -2.809331 | 0.00496446 | 0.02202 |
| ENSG00000232611 | | 51.63226 | -0.7254601 | 0.4164475 | -2.03218 | 0.04213548 | 0.121738 |
| ENSG00000099337 | KCNK6 | 4548.14395 | -0.725493516 | 0.202228606 | -3.726444566 | 0.0001942 | 0.00138 |
| ENSG00000172115 | CYCS | 6453.478 | -0.7255701 | 0.2168119 | -3.495354 | 0.00047343 | 0.002993 |
| ENSG00000286194 | | 1.88251 | -0.7255865 | 1.220867 | -2.02342 | 0.04302988 | 0.123804 |
| ENSG00000151718 | WWC2 | 3477.698 | -0.7259649 | 0.2708805 | -2.861434 | 0.00421729 | 0.019276 |
| ENSG00000002016 | RAD52 | 654.2418248 | -0.726136513 | 0.32794156 | -2.43627182 | 0.01483953 | 0.053474 |
| ENSG00000280187 | | 294.518 | -0.7261797 | 0.179087 | -4.171522 | 3.03E-05 | 0.000265 |
| ENSG00000196693 | ZNF33B | 2988.196 | -0.7266465 | 0.279969 | -2.798619 | 0.00513216 | 0.022624 |
| ENSG00000220685 | | 7.09222 | -0.7272617 | 0.7043833 | -1.573239 | 0.1156635 | 0.258523 |
| ENSG00000141480 | ARRB2 | 1815.745 | -0.727858 | 0.2028097 | -3.722077 | 0.00019759 | 0.001402 |
| ENSG00000214992 | AKAP17BP | 1.77992 | -0.7279196 | 1.056399 | -1.732202 | 0.08323764 | 0.203341 |
| ENSG00000058404 | CAMK2B | 1753.941198 | -0.728187881 | 0.26994653 | -2.876508888 | 0.00402101 | 0.018532 |
| ENSG00000141499 | WRAP53 | 1013.731 | -0.7282321 | 0.2727454 | -2.859158 | 0.00424767 | 0.019384 |
| ENSG00000177628 | GBA | 4368.101 | -0.7288425 | 0.1770498 | -4.218559 | 2.46E-05 | 0.00022 |
| ENSG00000250899 | | 435.8815 | -0.7289398 | 0.2882643 | -2.729299 | 0.00634692 | 0.026891 |
| ENSG00000198015 | MRPL42 | 3127.489 | -0.7289533 | 0.1925025 | -3.913728 | 9.09E-05 | 0.000702 |
| ENSG00000187257 | RSBN1L | 1501.173 | -0.7290508 | 0.1692257 | -4.411779 | 1.03E-05 | 0.000101 |
| ENSG00000185304 | RGPD2 | 40.55358 | -0.7291926 | 0.3310224 | -2.429048 | 0.01513854 | 0.054317 |
| ENSG00000019505 | SYT13 | 1.744545092 | -0.729405785 | 1.174772192 | -1.903224804 | 0.05701122 | 0.153034 |
| ENSG00000224505 | | 16.63631 | -0.7296951 | 0.5625044 | -1.698256 | 0.08945952 | 0.214711 |
| ENSG00000010318 | PHF7 | 262.4019525 | -0.729741941 | 0.217968563 | -3.494061456 | 0.00047573 | 0.003006 |
| ENSG00000217897 | HSPE1P8 | 3.099033 | -0.7299953 | 1.087067 | -1.701271 | 0.08889209 | 0.213755 |
| ENSG00000259677 | | 7.761789 | -0.730129 | 0.6729118 | -1.594419 | 0.1108421 | 0.250712 |
| ENSG00000174151 | CYB561D1 | 2144.679 | -0.7301493 | 0.1993078 | -3.79679 | 0.00014658 | 0.001075 |
| ENSG00000107890 | ANKRD26 | 1262.288478 | -0.730255006 | 0.201300933 | -3.747562628 | 0.00017856 | 0.001283 |
| ENSG00000168234 | TTC39C | 832.8543 | -0.7303881 | 0.1568896 | -4.766216 | 1.88E-06 | 2.22E-05 |
| ENSG00000102554 | KLF5 | 2832.598717 | -0.730409689 | 0.290176638 | -2.749653467 | 0.00596583 | 0.025609 |
| ENSG00000272386 | | 4.572736 | -0.730623 | 0.9050158 | -1.573498 | 0.1156036 | 0.258454 |
| ENSG00000261934 | PCDHGA9 | 18.31649 | -0.7307155 | 0.5533927 | -1.712972 | 0.08671766 | 0.209703 |
| ENSG00000188707 | ZBED6CL | 1101.744 | -0.7311696 | 0.1866643 | -4.043574 | 5.26E-05 | 0.000433 |

| | | | | | | | |
|---|---|---|---|---|---|---|---|
| ENSG00000121454 | LHX4 | 570.4034689 | -0.731572152 | 0.296655936 | -2.671809417 | 0.00754435 | 0.031016 |
| ENSG00000288021 | | 5.39726 | -0.7317252 | 0.7605966 | -1.557172 | 0.1194297 | 0.264339 |
| ENSG00000120662 | MTRF1 | 484.1101001 | -0.732028739 | 0.190409676 | -3.953903444 | 7.69E-05 | 0.000607 |
| ENSG00000168710 | AHCYL1 | 8369.098 | -0.7320547 | 0.1523152 | -4.914505 | 8.90E-07 | 1.12E-05 |
| ENSG00000225963 | | 15.82908 | -0.7321605 | 0.7926833 | -1.562779 | 0.1181046 | 0.262193 |
| ENSG00000150093 | ITGB1 | 13633.73 | -0.7321632 | 0.2045671 | -3.717125 | 0.0002015 | 0.001428 |
| ENSG00000204977 | TRIM13 | 785.2994 | -0.7322865 | 0.143959 | -5.183913 | 2.17E-07 | 3.09E-06 |
| ENSG00000068724 | TTC7A | 1783.244562 | -0.732341495 | 0.19903245 | -3.818491886 | 0.00013427 | 0.000994 |
| ENSG00000207031 | SNORD59A | 28.64381 | -0.7326786 | 0.3739007 | -2.222911 | 0.02622177 | 0.084165 |
| ENSG00000179862 | CITED4 | 622.4239 | -0.7331276 | 0.3358135 | -2.418404 | 0.01558876 | 0.055594 |
| ENSG00000166171 | DPCD | 814.6639 | -0.7332078 | 0.2213108 | -3.469321 | 0.00052178 | 0.003257 |
| ENSG00000168291 | PDHB | 2240.891 | -0.7333794 | 0.2632617 | -2.970347 | 0.00297464 | 0.014472 |
| ENSG00000180822 | PSMG4 | 1239.398 | -0.7336893 | 0.1904819 | -3.988287 | 6.66E-05 | 0.000533 |
| ENSG00000268282 | | 15.69943 | -0.7337798 | 0.4950081 | -1.822914 | 0.06831644 | 0.175287 |
| ENSG00000273284 | | 32.5957 | -0.7346008 | 0.4659616 | -1.906166 | 0.05662869 | 0.152361 |
| ENSG00000198742 | SMURF1 | 2269.575 | -0.7349598 | 0.1421617 | -5.266922 | 1.39E-07 | 2.05E-06 |
| ENSG00000273018 | FAM106A | 151.6627 | -0.7350185 | 0.4952366 | -1.840921 | 0.06563318 | 0.170028 |
| ENSG00000260329 | | 16.02465 | -0.7353736 | 0.5351901 | -1.747097 | 0.08062054 | 0.198463 |
| ENSG00000144827 | ABHD10 | 1345.85 | -0.7354705 | 0.1652418 | -4.554822 | 5.24E-06 | 5.57E-05 |
| ENSG00000278840 | | 1.901558 | -0.7355454 | 1.113112 | -1.822943 | 0.06831198 | 0.175287 |
| ENSG00000061794 | MRPS35 | 3296.734898 | -0.735829227 | 0.154439793 | -4.868526554 | 1.12E-06 | 1.39E-05 |
| ENSG00000265590 | CFAP298-TCP10l | 20.9109 | -0.7359033 | 0.4248306 | -2.030385 | 0.04231737 | 0.122197 |
| ENSG00000116096 | SPR | 3878.369864 | -0.73610888 | 0.171726511 | -4.420125439 | 9.86E-06 | 9.74E-05 |
| ENSG00000128266 | GNAZ | 300.8446818 | -0.736184002 | 0.284440474 | -2.784207704 | 0.00536587 | 0.023468 |
| ENSG00000152455 | SUV39H2 | 1221.819 | -0.736272 | 0.3019565 | -2.648286 | 0.00809011 | 0.032826 |
| ENSG00000187905 | LRRC74B | 4.336046 | -0.7362849 | 0.864146 | -1.583165 | 0.113384 | 0.254606 |
| ENSG00000170324 | FRMPD2 | 5.632466 | -0.7368266 | 0.6953375 | -1.588156 | 0.1122512 | 0.252896 |
| ENSG00000140543 | DET1 | 116.6543 | -0.7370309 | 0.3827678 | -2.193308 | 0.0282852 | 0.089423 |
| ENSG00000272391 | POM121C | 3040.07 | -0.737076 | 0.1697035 | -4.480727 | 7.44E-06 | 7.60E-05 |
| ENSG00000225969 | ABHD11-AS1 | 70.99826 | -0.73733 | 0.4925593 | -1.843566 | 0.06524642 | 0.169224 |
| ENSG00000261668 | | 3.006425 | -0.7374464 | 0.8652225 | -1.564649 | 0.1176653 | 0.261523 |
| ENSG00000139343 | SNRPF | 3106.015 | -0.7374821 | 0.2663593 | -2.930871 | 0.00338014 | 0.016054 |
| ENSG00000064547 | LPAR2 | 1037.54587 | -0.737527295 | 0.23391469 | -3.324382752 | 0.00088615 | 0.00515 |

| | | | | | | | |
|---|---|---|---|---|---|---|---|
| ENSG00000261105 | LMO7-AS1 | 27.96328 | -0.7376485 | 0.4700607 | -1.895988 | 0.05796163 | 0.155068 |
| ENSG00000274270 | | 37.49524 | -0.7381325 | 0.5511383 | -1.739484 | 0.08194962 | 0.201045 |
| ENSG00000123179 | EBPL | 1472.257447 | -0.738211131 | 0.206651321 | -3.713347629 | 0.00020454 | 0.001447 |
| ENSG00000276934 | | 3.352608 | -0.7382121 | 0.8629628 | -1.561088 | 0.1185029 | 0.262765 |
| ENSG00000136824 | SMC2 | 2216.102711 | -0.738450354 | 0.453383615 | -1.960839773 | 0.04989772 | 0.138272 |
| ENSG00000122965 | RBM19 | 2677.414349 | -0.738562856 | 0.089296339 | -8.348168344 | 6.93E-17 | 3.71E-15 |
| ENSG00000253677 | UBE2HP1 | 91.0861 | -0.7386443 | 0.2227846 | -3.462462 | 0.00053526 | 0.003333 |
| ENSG00000066926 | FECH | 2591.125717 | -0.738894866 | 0.261874973 | -2.991407628 | 0.00277695 | 0.013683 |
| ENSG00000171953 | ATPAF2 | 1307.88 | -0.7393445 | 0.2023806 | -3.795179 | 0.00014754 | 0.00108 |
| ENSG00000168090 | COPS6 | 5058.992 | -0.739457 | 0.1966615 | -3.899221 | 9.65E-05 | 0.000742 |
| ENSG00000200394 | SNORA38B | 3.886645 | -0.7395235 | 0.8223138 | -1.550372 | 0.1210523 | 0.266974 |
| ENSG00000159208 | CIART | 1918.047 | -0.7395862 | 0.2603966 | -3.020307 | 0.00252519 | 0.012614 |
| ENSG00000130429 | ARPC1B | 127.8862286 | -0.73978914 | 0.223090068 | -3.470966294 | 0.00051859 | 0.003238 |
| ENSG00000228819 | AGKP2 | 8.463688 | -0.7401383 | 0.6467699 | -1.628883 | 0.1033378 | 0.238332 |
| ENSG00000117118 | SDHB | 3157.6086 | -0.740298648 | 0.205637293 | -3.744565697 | 0.00018071 | 0.001296 |
| ENSG00000212512 | | 2.287413 | -0.7405243 | 1.191875 | -1.883056 | 0.05969278 | 0.158408 |
| ENSG00000239887 | C1orf226 | 1595.996 | -0.7406046 | 0.361789 | -2.298703 | 0.02152179 | 0.072166 |
| ENSG00000105357 | MYH14 | 13139.29384 | -0.740624111 | 0.230117978 | -3.375937237 | 0.00073565 | 0.004386 |
| ENSG00000270558 | | 46.56051 | -0.7406999 | 0.4365691 | -2.00737 | 0.0447103 | 0.127414 |
| ENSG00000179859 | RNF227 | 162.688 | -0.740739 | 0.312425 | -2.585282 | 0.00972993 | 0.038022 |
| ENSG00000281344 | HELLPAR | 37.51199 | -0.7408546 | 0.3804932 | -2.217295 | 0.02660295 | 0.085199 |
| ENSG00000135423 | GLS2 | 21.05235721 | -0.741245928 | 0.454113089 | -1.951587742 | 0.05098717 | 0.140585 |
| ENSG00000166224 | SGPL1 | 9097.741 | -0.7412898 | 0.1618108 | -4.689268 | 2.74E-06 | 3.12E-05 |
| ENSG00000161920 | MED11 | 704.9374 | -0.7416821 | 0.2369567 | -3.301372 | 0.00096213 | 0.005528 |
| ENSG00000005156 | LIG3 | 1693.796509 | -0.741965906 | 0.141922211 | -5.340604589 | 9.26E-08 | 1.41E-06 |
| ENSG00000135077 | HAVCR2 | 2.982837527 | -0.742009505 | 0.866314094 | -1.564320995 | 0.1177422 | 0.261629 |
| ENSG00000183891 | TTC32 | 434.7275 | -0.7422016 | 0.1690839 | -4.506862 | 6.58E-06 | 6.80E-05 |
| ENSG00000259783 | LINC02259 | 1.350844 | -0.7423333 | 1.200793 | -1.954432 | 0.05065016 | NA |
| ENSG00000286819 | | 4.27128 | -0.742378 | 0.8077497 | -1.561944 | 0.1183011 | 0.262432 |
| ENSG00000230922 | | 5.95397 | -0.7427175 | 0.7005213 | -1.604359 | 0.1086348 | 0.247084 |
| ENSG00000197238 | H4C11 | 13.69024 | -0.7429003 | 0.5320717 | -1.770986 | 0.07656303 | 0.190915 |
| ENSG00000246090 | | 77.23567 | -0.7430995 | 0.3559158 | -2.336414 | 0.01946967 | 0.066524 |
| ENSG00000172209 | GPR22 | 1.161263 | -0.7431483 | 1.229877 | -1.990161 | 0.0465732 | NA |

| | | | | | | | |
|---|---|---|---|---|---|---|---|
| ENSG00000249899 | | 1.210526 | -0.7432633 | 1.209889 | -2.018381 | 0.04355158 | NA |
| ENSG00000166902 | MRPL16 | 2526.015 | -0.7434444 | 0.2790501 | -2.857528 | 0.00426955 | 0.019461 |
| ENSG00000233647 | NENFP1 | 2.57079 | -0.7435131 | 1.003719 | -1.633482 | 0.1023676 | 0.236608 |
| ENSG00000279894 | | 11.24457 | -0.743631 | 0.6798114 | -1.613565 | 0.1066219 | 0.243811 |
| ENSG00000132819 | RBM38 | 3911.171986 | -0.743939477 | 0.245133274 | -3.179371717 | 0.00147595 | 0.007995 |
| ENSG00000179598 | PLD6 | 765.8432 | -0.7442443 | 0.211571 | -3.663265 | 0.00024902 | 0.001719 |
| ENSG00000271991 | | 5.369573 | -0.7443687 | 0.9070123 | -1.601159 | 0.1093417 | 0.248235 |
| ENSG00000126861 | OMG | 1.44684199 | -0.74444465 | 1.156354325 | -1.894170598 | 0.05820237 | NA |
| ENSG00000266910 | | 25.84737 | -0.7446253 | 0.5924944 | -1.696677 | 0.08975767 | 0.215252 |
| ENSG00000218890 | NUFIP1P1 | 68.35234 | -0.7446476 | 0.2671661 | -2.975891 | 0.00292139 | 0.014271 |
| ENSG00000164347 | GFM2 | 2728.23 | -0.7447688 | 0.1285993 | -5.861257 | 4.59E-09 | 8.92E-08 |
| ENSG00000164061 | BSN | 122.9061 | -0.7447885 | 0.2323501 | -3.364204 | 0.00076765 | 0.004554 |
| ENSG00000243317 | STMP1 | 3425.498 | -0.7448374 | 0.1736913 | -4.411096 | 1.03E-05 | 0.000101 |
| ENSG00000260051 | | 94.46203 | -0.7453008 | 0.3809599 | -2.223202 | 0.02620215 | 0.084119 |
| ENSG00000253658 | LINC01592 | 2.810046 | -0.7453822 | 1.1486 | -1.889019 | 0.05888924 | 0.156961 |
| ENSG00000010219 | DYRK4 | 236.6613046 | -0.745619096 | 0.262987373 | -3.002880606 | 0.00267437 | 0.013234 |
| ENSG00000172889 | EGFL7 | 1973.264 | -0.7456256 | 0.3015794 | -2.678917 | 0.00738607 | 0.030475 |
| ENSG00000007237 | GAS7 | 1.429180449 | -0.745767328 | 1.12565824 | -1.842792611 | 0.0653593 | NA |
| ENSG00000256084 | | 1.589604 | -0.745857 | 1.278009 | -2.131879 | 0.03301682 | 0.101063 |
| ENSG00000164338 | UTP15 | 1393.519 | -0.7461291 | 0.224505 | -3.47892 | 0.00050344 | 0.003155 |
| ENSG00000071462 | BUD23 | 4840.537578 | -0.74621013 | 0.157571368 | -4.858010165 | 1.19E-06 | 1.46E-05 |
| ENSG00000279791 | | 6.384008 | -0.7471885 | 0.7589981 | -1.573376 | 0.1156317 | 0.258474 |
| ENSG00000286488 | | 4.911905 | -0.7475416 | 0.8326229 | -1.5896 | 0.1119251 | 0.252504 |
| ENSG00000278195 | SSTR3 | 2.993011 | -0.7476639 | 1.41901 | -2.728986 | 0.00635294 | 0.026908 |
| ENSG00000255234 | | 3.890509 | -0.7477082 | 0.8542978 | -1.572695 | 0.1157895 | 0.258649 |
| ENSG00000095587 | TLL2 | 20.25358226 | -0.747738089 | 0.516437784 | -1.822714182 | 0.06834669 | 0.175348 |
| ENSG00000227848 | SUCLA2-AS1 | 9.064466 | -0.7478566 | 0.6646281 | -1.627246 | 0.1036848 | 0.238821 |
| ENSG00000181027 | FKRP | 1084.176 | -0.748066 | 0.1327074 | -5.717349 | 1.08E-08 | 1.98E-07 |
| ENSG00000285416 | | 62.34175 | -0.7482876 | 0.2904934 | -2.773348 | 0.00554827 | 0.024134 |
| ENSG00000140527 | WDR93 | 20.18813 | -0.7484209 | 0.4346821 | -2.027946 | 0.04256574 | 0.122767 |
| ENSG00000186976 | EFCAB6 | 48.1443 | -0.7484676 | 0.4340752 | -2.026335 | 0.04273044 | 0.123157 |
| ENSG00000170409 | | 8.004553 | -0.7487266 | 0.6943807 | -1.603486 | 0.1088275 | 0.247301 |
| ENSG00000076826 | CAMSAP3 | 2810.063668 | -0.748805402 | 0.194451346 | -3.983277262 | 6.80E-05 | 0.000543 |

| | | | | | | | |
|---|---|---|---|---|---|---|---|
| ENSG00000274251 | | 6.058502 | -0.7490619 | 0.7103471 | -1.603567 | 0.1088096 | 0.247281 |
| ENSG00000260954 | | 9.824664 | -0.7491685 | 0.6258104 | -1.669654 | 0.09498775 | 0.224173 |
| ENSG00000198626 | RYR2 | 3.499605 | -0.7492552 | 0.8353223 | -1.561754 | 0.1183461 | 0.262487 |
| ENSG00000225131 | PSME2P2 | 794.6476 | -0.7497259 | 0.2612621 | -3.049513 | 0.00229213 | 0.011602 |
| ENSG00000272843 | | 5.210116 | -0.7498889 | 0.9266588 | -1.605044 | 0.108484 | 0.246816 |
| ENSG00000138399 | FASTKD1 | 1350.352851 | -0.750086705 | 0.147109896 | -5.19421466 | 2.06E-07 | 2.94E-06 |
| ENSG00000131095 | GFAP | 4.939913809 | -0.750246578 | 0.666011443 | -1.634227716 | 0.10221105 | 0.236328 |
| ENSG00000261600 | | 40.65415 | -0.7503655 | 0.3741273 | -2.256916 | 0.02401331 | 0.078586 |
| ENSG00000261278 | | 3.798932 | -0.7504745 | 0.8059691 | -1.578985 | 0.1143395 | 0.256297 |
| ENSG00000130935 | NOL11 | 10245.84604 | -0.751074629 | 0.189706885 | -4.121322103 | 3.77E-05 | 0.000322 |
| ENSG00000092621 | PHGDH | 11091.59315 | -0.751110003 | 0.448233014 | -2.00210811 | 0.04527311 | 0.128587 |
| ENSG00000115756 | HPCAL1 | 2193.800205 | -0.751237001 | 0.250151453 | -3.181394642 | 0.00146568 | 0.007942 |
| ENSG00000264920 | | 223.2184 | -0.7513242 | 0.2308602 | -3.418068 | 0.00063068 | 0.00384 |
| ENSG00000178395 | CCDC185 | 1.986654 | -0.7514081 | 0.980851 | -1.653039 | 0.09832291 | 0.229822 |
| ENSG00000109738 | GLRB | 361.8833569 | -0.75181721 | 0.29709711 | -2.725490077 | 0.00642061 | 0.027138 |
| ENSG00000231473 | RB1-DT | 1.757967 | -0.7519084 | 1.118523 | -1.759895 | 0.07842556 | 0.194392 |
| ENSG00000154920 | EME1 | 644.8256 | -0.7519732 | 0.4472278 | -2.015057 | 0.04389864 | 0.125708 |
| ENSG00000117724 | CENPF | 5238.777875 | -0.752045949 | 0.494530875 | -1.896441328 | 0.05790171 | 0.15495 |
| ENSG00000277511 | | 10.47366 | -0.752338 | 0.7385068 | -1.596355 | 0.1104096 | 0.250083 |
| ENSG00000239300 | | 10.44199 | -0.752351 | 0.6488084 | -1.652597 | 0.09841299 | 0.229887 |
| ENSG00000113552 | GNPDA1 | 3870.205419 | -0.752712743 | 0.224505397 | -3.508621519 | 0.00045044 | 0.002866 |
| ENSG00000235001 | EIF4A1P2 | 835.848 | -0.7531479 | 0.2087634 | -3.750235 | 0.00017667 | 0.00127 |
| ENSG00000172059 | KLF11 | 1285.995 | -0.753304 | 0.2724536 | -2.9617 | 0.00305946 | 0.014774 |
| ENSG00000277449 | CEBPB-AS1 | 178.5271 | -0.75419 | 0.2990166 | -2.750533 | 0.00594984 | 0.025563 |
| ENSG00000287005 | | 11.20049 | -0.7543693 | 0.5977043 | -1.700666 | 0.08900574 | 0.213912 |
| ENSG00000128607 | KLHDC10 | 3114.048266 | -0.754696467 | 0.203859284 | -3.853579338 | 0.0001164 | 0.000877 |
| ENSG00000266601 | | 19.36507 | -0.7547992 | 0.4540404 | -1.987746 | 0.04683977 | 0.131895 |
| ENSG00000140743 | CDR2 | 1712.749 | -0.7548974 | 0.1941422 | -4.023616 | 5.73E-05 | 0.000468 |
| ENSG00000128335 | APOL2 | 1534.820703 | -0.754943216 | 0.299148605 | -2.754710132 | 0.00587442 | 0.025296 |
| ENSG00000214575 | CPEB1 | 28.28259 | -0.7549619 | 0.621902 | -1.688893 | 0.09123985 | 0.217764 |
| ENSG00000108528 | SLC25A11 | 3413.429872 | -0.755009441 | 0.222450247 | -3.545922107 | 0.00039124 | 0.002547 |
| ENSG00000104892 | KLC3 | 326.0482861 | -0.755288043 | 0.220250598 | -3.586081852 | 0.00033568 | 0.002232 |
| ENSG00000160710 | ADAR | 32019.1 | -0.7553989 | 0.1880764 | -4.148598 | 3.35E-05 | 0.00029 |

| | | | | | | | |
|---|---|---|---|---|---|---|---|
| ENSG00000262943 | ALOX12P2 | 742.0069 | -0.7554703 | 0.2710499 | -2.972959 | 0.00294943 | 0.014392 |
| ENSG00000280099 | | 202.1024 | -0.7560399 | 0.3762818 | -2.278594 | 0.02269122 | 0.075276 |
| ENSG00000214198 | TTC41P | 22.2576 | -0.756201 | 0.4328192 | -2.06221 | 0.03918774 | 0.115035 |
| ENSG00000207231 | | 1.149223 | -0.7565688 | 1.228468 | -1.965017 | 0.04941222 | NA |
| ENSG00000226082 | | 3.015804 | -0.756762 | 0.9579294 | -1.622321 | 0.1047346 | 0.240572 |
| ENSG00000144488 | ESPNL | 1.470548 | -0.7568229 | 1.393944 | -2.496874 | 0.01252934 | NA |
| ENSG00000241468 | ATP5MF | 2217.831 | -0.757106 | 0.2383916 | -3.338231 | 0.00084314 | 0.004936 |
| ENSG00000272103 | | 11.2706 | -0.7574209 | 0.7003852 | -1.623388 | 0.1045065 | 0.240229 |
| ENSG00000111912 | NCOA7 | 1184.149709 | -0.757460793 | 0.437111977 | -2.048523335 | 0.04050875 | 0.118078 |
| ENSG00000196415 | PRTN3 | 6.794878 | -0.7574674 | 0.8026489 | -1.594398 | 0.110847 | 0.250712 |
| ENSG00000279418 | LINC00244 | 2.495089 | -0.7577792 | 1.160104 | -1.880703 | 0.06001238 | 0.159113 |
| ENSG00000108839 | ALOX12 | 130.8450132 | -0.757876106 | 0.343211557 | -2.447389574 | 0.01438952 | 0.0522 |
| ENSG00000280088 | | 194.732 | -0.758355 | 0.2554505 | -3.146537 | 0.00165216 | 0.008786 |
| ENSG00000127946 | HIP1 | 1357.426402 | -0.758387577 | 0.309479536 | -2.671744002 | 0.00754582 | 0.031016 |
| ENSG00000286282 | | 1.453431 | -0.7585604 | 1.198931 | -1.962926 | 0.04965482 | NA |
| ENSG00000248468 | | 3.259835 | -0.7586414 | 1.021058 | -1.693153 | 0.09042639 | 0.216386 |
| ENSG00000107341 | UBE2R2 | 3120.806576 | -0.758651475 | 0.083858855 | -9.151282313 | 5.63E-20 | 4.18E-18 |
| ENSG00000186591 | UBE2H | 6561.686 | -0.7586716 | 0.18578 | -4.221549 | 2.43E-05 | 0.000218 |
| ENSG00000144395 | CCDC150 | 282.2319 | -0.7587106 | 0.4927627 | -1.901456 | 0.05724233 | 0.153468 |
| ENSG00000138073 | PREB | 5160.698416 | -0.758871218 | 0.148134266 | -5.235140349 | 1.65E-07 | 2.40E-06 |
| ENSG00000259448 | LINC02352 | 15.4451 | -0.7591044 | 0.6563269 | -1.658893 | 0.09713728 | 0.227728 |
| ENSG00000267446 | | 4.042266 | -0.759229 | 1.072816 | -1.742656 | 0.08139377 | 0.199977 |
| ENSG00000138400 | MDH1B | 104.2759855 | -0.759523244 | 0.30007747 | -2.743411641 | 0.00608044 | 0.025998 |
| ENSG00000143631 | FLG | 1.471407 | -0.7595434 | 1.230948 | -2.02554 | 0.04281195 | NA |
| ENSG00000204839 | MROH6 | 1445.653 | -0.7597472 | 0.313024 | -2.635726 | 0.00839575 | 0.033808 |
| ENSG00000257918 | | 2.366764 | -0.7599314 | 1.0214 | -1.67945 | 0.09306443 | 0.22083 |
| ENSG00000262115 | | 10.17974 | -0.7600383 | 0.5772334 | -1.742422 | 0.08143463 | 0.20004 |
| ENSG00000177119 | ANO6 | 2364.563 | -0.7600407 | 0.2718865 | -2.982407 | 0.00285991 | 0.014027 |
| ENSG00000014257 | ACP3 | 61.14691324 | -0.760356698 | 0.439419311 | -2.044462566 | 0.04090787 | 0.119084 |
| ENSG00000271452 | | 30.44013 | -0.7605119 | 0.3292865 | -2.536196 | 0.01120641 | 0.042663 |
| ENSG00000286808 | | 81.74884 | -0.7605376 | 0.3188025 | -2.607845 | 0.00911143 | 0.036099 |
| ENSG00000227621 | PHBP11 | 21.84877 | -0.761175 | 0.5781099 | -1.754199 | 0.07939636 | 0.19614 |
| ENSG00000115041 | KCNIP3 | 321.446512 | -0.761185311 | 0.321219083 | -2.598095618 | 0.00937424 | 0.036898 |

| | | | | | | | |
|---|---|---|---|---|---|---|---|
| ENSG00000196620 | UGT2B15 | 50.11099 | -0.7616437 | 1.008198 | -1.716949 | 0.08598848 | 0.208363 |
| ENSG00000283409 | MIR3662 | 2.075172 | -0.7623645 | 0.9496554 | -1.618201 | 0.1056192 | 0.242162 |
| ENSG00000273920 | | 3.163695 | -0.7624368 | 0.8683057 | -1.614056 | 0.1065153 | 0.243672 |
| ENSG00000269895 | | 2.447636 | -0.762649 | 1.055225 | -1.660516 | 0.09681066 | 0.227203 |
| ENSG00000269720 | CCDC194 | 13.20271 | -0.7627066 | 1.364314 | -2.283787 | 0.02238408 | 0.074564 |
| ENSG00000148948 | LRRC4C | 70.2984 | -0.7627546 | 0.6390348 | -1.686591 | 0.09168194 | 0.218505 |
| ENSG00000129204 | USP6 | 41.11036984 | -0.762772585 | 0.452689191 | -2.007408599 | 0.04470617 | 0.127414 |
| ENSG00000223960 | CHROMR | 194.45 | -0.7628856 | 0.2701591 | -3.01833 | 0.00254172 | 0.01268 |
| ENSG00000179409 | GEMIN4 | 2738.495 | -0.7629879 | 0.2186673 | -3.637926 | 0.00027484 | 0.001871 |
| ENSG00000165416 | SUGT1 | 2029.765 | -0.7631152 | 0.1825382 | -4.303619 | 1.68E-05 | 0.000157 |
| ENSG00000104804 | TULP2 | 1.899957967 | -0.76398504 | 1.037233128 | -1.681351448 | 0.09269467 | 0.220208 |
| ENSG00000111305 | GSG1 | 2.766204704 | -0.764046888 | 0.861545965 | -1.608960637 | 0.10762495 | 0.245407 |
| ENSG00000260231 | KDM7A-DT | 218.5937 | -0.7642953 | 0.2717718 | -3.006585 | 0.002642 | 0.013103 |
| ENSG00000288033 | | 11.40427 | -0.7649845 | 0.5686283 | -1.764872 | 0.07758532 | 0.192748 |
| ENSG00000086200 | IPO11 | 1634.052266 | -0.765072819 | 0.269201337 | -3.0348208 | 0.00240679 | 0.012109 |
| ENSG00000240707 | LINC01168 | 5.411173 | -0.7652517 | 1.181601 | -1.917783 | 0.0551386 | 0.149396 |
| ENSG00000105939 | ZC3HAV1 | 3904.026649 | -0.765268774 | 0.1607175 | -4.873631031 | 1.10E-06 | 1.35E-05 |
| ENSG00000105993 | DNAJB6 | 4422.500267 | -0.765307853 | 0.191498321 | -4.115228491 | 3.87E-05 | 0.00033 |
| ENSG00000214837 | LINC01347 | 35.5185 | -0.7653729 | 0.3837038 | -2.270778 | 0.0231604 | 0.076374 |
| ENSG00000278463 | H2AC4 | 8.194497 | -0.7654109 | 0.7363587 | -1.613104 | 0.1067218 | 0.243976 |
| ENSG00000267315 | UFM1P2 | 12.42301 | -0.7654837 | 0.4878051 | -1.924055 | 0.0543477 | 0.147902 |
| ENSG00000058804 | NDC1 | 3523.167565 | -0.76569696 | 0.307324229 | -2.710671032 | 0.00671472 | 0.028148 |
| ENSG00000260782 | | 4.8568 | -0.7657507 | 1.087612 | -1.717948 | 0.08580607 | 0.208085 |
| ENSG00000236514 | | 25.08224 | -0.7659839 | 0.4333196 | -2.081554 | 0.03738321 | 0.110939 |
| ENSG00000151303 | | 1.877522 | -0.7662198 | 1.104181 | -1.794959 | 0.0726602 | 0.183718 |
| ENSG00000258314 | | 9.101139 | -0.7663423 | 0.7283319 | -1.634599 | 0.1021332 | 0.23623 |
| ENSG00000267649 | | 18.72484 | -0.7664631 | 0.564498 | -1.784514 | 0.07434023 | 0.18697 |
| ENSG00000286599 | | 70.32875 | -0.7668686 | 0.3679778 | -2.345706 | 0.0189911 | 0.065207 |
| ENSG00000269378 | ITGB1P1 | 1971.261 | -0.766969 | 0.1810247 | -4.369008 | 1.25E-05 | 0.00012 |
| ENSG00000018408 | WWTR1 | 1912.610638 | -0.767316387 | 0.176801358 | -4.467520549 | 7.91E-06 | 8.02E-05 |
| ENSG00000226723 | | 392.4634 | -0.7677083 | 0.2139414 | -3.740595 | 0.00018359 | 0.001314 |
| ENSG00000274987 | | 6.012587 | -0.7686008 | 0.7524087 | -1.615342 | 0.1062366 | 0.243223 |
| ENSG00000286995 | | 3.874742 | -0.768651 | 0.9050748 | -1.583031 | 0.1134144 | 0.254653 |

| | | | | | | | |
|---|---|---|---|---|---|---|---|
| ENSG00000171867 | PRNP | 2793.771 | -0.7691368 | 0.3124484 | -2.716025 | 0.0066071 | 0.027794 |
| ENSG00000224629 | | 3.959316 | -0.7691561 | 0.9812516 | -1.635239 | 0.101999 | 0.236023 |
| ENSG00000230044 | CDK8P1 | 3.194316 | -0.7693101 | 1.107094 | -1.812042 | 0.06997971 | 0.178503 |
| ENSG00000269947 | | 11.27468 | -0.7695643 | 0.6324241 | -1.686173 | 0.09176245 | 0.218565 |
| ENSG00000224812 | TMEM72-AS1 | 3.283651 | -0.7696099 | 0.9322214 | -1.634367 | 0.1021819 | 0.236302 |
| ENSG00000175946 | KLHL38 | 1.437574 | -0.7702923 | 1.163482 | -1.86267 | 0.06250867 | NA |
| ENSG00000183055 | FAM133CP | 2.133704 | -0.7704727 | 1.028617 | -1.633039 | 0.1024609 | 0.236761 |
| ENSG00000279863 | | 45.42334 | -0.7706299 | 0.3805662 | -2.299564 | 0.02147291 | 0.072053 |
| ENSG00000033050 | ABCF2 | 1440.568382 | -0.770749061 | 0.209937215 | -3.819479445 | 0.00013373 | 0.000991 |
| ENSG00000233246 | PHC2-AS1 | 8.065419 | -0.7711542 | 0.6318483 | -1.688627 | 0.09129093 | 0.217867 |
| ENSG00000246375 | PPM1K-DT | 10.25437 | -0.7712325 | 0.6582278 | -1.681036 | 0.09275596 | 0.220255 |
| ENSG00000270587 | | 35.3728 | -0.7712615 | 0.3766292 | -2.315368 | 0.02059278 | 0.069556 |
| ENSG00000166619 | BLCAP | 3291.61 | -0.7713348 | 0.1662811 | -4.781095 | 1.74E-06 | 2.07E-05 |
| ENSG00000237886 | NALT1 | 193.148 | -0.7714602 | 0.3099169 | -2.711404 | 0.00669989 | 0.028099 |
| ENSG00000267076 | CCDC58P3 | 74.13505 | -0.7717761 | 0.3548649 | -2.436241 | 0.01484079 | 0.053474 |
| ENSG00000249661 | TNRC18P1 | 50.93294 | -0.7717922 | 0.4925604 | -1.936264 | 0.05283543 | 0.144735 |
| ENSG00000130813 | SHFL | 1577.765142 | -0.771857279 | 0.336346641 | -2.529116629 | 0.011435 | 0.043316 |
| ENSG00000226421 | SLC25A5P5 | 156.608 | -0.7722513 | 0.2818438 | -2.943576 | 0.00324444 | 0.015507 |
| ENSG00000138166 | DUSP5 | 401.6221537 | -0.772456404 | 0.525220744 | -1.868484155 | 0.06169462 | 0.162406 |
| ENSG00000173272 | MZT2A | 2812.064 | -0.772641 | 0.2171681 | -3.709825 | 0.0002074 | 0.001465 |
| ENSG00000272443 | OR13Z2P | 2.262985 | -0.7726826 | 1.117801 | -1.806237 | 0.07088138 | 0.180301 |
| ENSG00000263916 | | 5.096512 | -0.7727056 | 0.7093612 | -1.63672 | 0.101689 | 0.235614 |
| ENSG00000075336 | TIMM21 | 1314.601084 | -0.77293178 | 0.225333057 | -3.616005677 | 0.00029918 | 0.002017 |
| ENSG00000279124 | | 5.890941 | -0.7730257 | 0.8787523 | -1.593481 | 0.1110523 | 0.251025 |
| ENSG00000101463 | SYNDIG1 | 10.48842179 | -0.773363509 | 0.780456024 | -1.629811327 | 0.10314138 | 0.238003 |
| ENSG00000243107 | | 9.708786 | -0.773499 | 0.6013495 | -1.738921 | 0.0820486 | 0.201251 |
| ENSG00000176472 | ZNF575 | 34.44184 | -0.7738983 | 0.3945401 | -2.246409 | 0.02467783 | 0.080276 |
| ENSG00000096060 | FKBP5 | 1974.210887 | -0.774434834 | 0.316511655 | -2.676472289 | 0.00744017 | 0.030667 |
| ENSG00000255723 | | 1.454931 | -0.7745663 | 1.05836 | -1.668988 | 0.09511984 | NA |
| ENSG00000197208 | SLC22A4 | 275.7752 | -0.7746652 | 0.305637 | -2.757407 | 0.00582618 | 0.02515 |
| ENSG00000149150 | SLC43A1 | 663.3861 | -0.7746819 | 0.7510445 | -1.652905 | 0.0983503 | 0.229831 |
| ENSG00000053702 | NRIP2 | 115.626851 | -0.774941632 | 0.289439736 | -2.884336932 | 0.00392239 | 0.018153 |
| ENSG00000234545 | FAM133B | 546.7904 | -0.7749578 | 0.1817879 | -4.394106 | 1.11E-05 | 0.000109 |

| | | | | | | | |
|---|---|---|---|---|---|---|---|
| ENSG00000182240 | BACE2 | 4511.151 | -0.7752929 | 0.2773205 | -2.989295 | 0.00279622 | 0.013776 |
| ENSG00000286258 | | 18.70535 | -0.7753097 | 0.6585471 | -1.670657 | 0.09478947 | 0.223903 |
| ENSG00000257316 | ASS1P14 | 55.00431 | -0.7762186 | 0.3431682 | -2.509523 | 0.01208942 | 0.045272 |
| ENSG00000274554 | | 6.098656 | -0.7762745 | 0.7908889 | -1.613403 | 0.1066571 | 0.24387 |
| ENSG00000285793 | | 18.2107 | -0.7773773 | 0.6497711 | -1.699473 | 0.08923007 | 0.214335 |
| ENSG00000247934 | | 37.31903 | -0.7776642 | 0.4987236 | -1.925174 | 0.05420755 | 0.147626 |
| ENSG00000248663 | LINC00992 | 439.254 | -0.7778999 | 0.2258858 | -3.604779 | 0.00031242 | 0.002097 |
| ENSG00000280103 | | 1.390885 | -0.7779322 | 1.131764 | -1.747553 | 0.08054153 | NA |
| ENSG00000099957 | P2RX6 | 13.30057615 | -0.778452214 | 0.71047252 | -1.65751116 | 0.09741618 | 0.228287 |
| ENSG00000150556 | LYPD6B | 1360.273 | -0.7785109 | 0.2700467 | -3.077889 | 0.00208473 | 0.01073 |
| ENSG00000229132 | EIF4A1P10 | 3255.997 | -0.7789963 | 0.1984993 | -4.066265 | 4.78E-05 | 0.000398 |
| ENSG00000149328 | GLB1L2 | 971.5523 | -0.7790166 | 0.212792 | -3.832529 | 0.00012683 | 0.000946 |
| ENSG00000273143 | DUSP5-DT | 2.289177 | -0.7791797 | 1.122893 | -1.799459 | 0.07194604 | 0.182311 |
| ENSG00000277224 | H2BC7 | 163.4644 | -0.7792812 | 0.3273498 | -2.625724 | 0.0086465 | 0.034629 |
| ENSG00000176014 | TUBB6 | 2471.899 | -0.7793698 | 0.3901419 | -2.276664 | 0.02280629 | 0.075525 |
| ENSG00000072134 | EPN2 | 1748.041758 | -0.779413115 | 0.203149076 | -3.991323984 | 6.57E-05 | 0.000527 |
| ENSG00000283128 | | 7.666878 | -0.7800335 | 0.5826041 | -1.768548 | 0.07696936 | 0.191659 |
| ENSG00000182648 | LINC01006 | 1182.144 | -0.7805182 | 0.2409758 | -3.420935 | 0.00062406 | 0.003804 |
| ENSG00000234281 | LANCL1-AS1 | 9.647602 | -0.781354 | 0.5648912 | -1.798614 | 0.07207982 | 0.182591 |
| ENSG00000258366 | RTEL1 | 160.9559 | -0.7813986 | 0.2835592 | -2.963058 | 0.00304599 | 0.014725 |
| ENSG00000145916 | RMND5B | 9567.693 | -0.7814709 | 0.1170595 | -6.757565 | 1.40E-11 | 3.95E-10 |
| ENSG00000142002 | DPP9 | 3264.129 | -0.7816824 | 0.1369807 | -5.804059 | 6.47E-09 | 1.22E-07 |
| ENSG00000132109 | TRIM21 | 793.0449368 | -0.782058562 | 0.272444965 | -3.05849691 | 0.0022245 | 0.011325 |
| ENSG00000147117 | ZNF157 | 6.951133 | -0.7820815 | 0.5964737 | -1.759618 | 0.07847266 | 0.19441 |
| ENSG00000060069 | CTDP1 | 1474.482592 | -0.782090713 | 0.162077191 | -4.941842935 | 7.74E-07 | 9.86E-06 |
| ENSG00000205060 | SLC35B4 | 1191.24 | -0.7821098 | 0.1988536 | -4.078118 | 4.54E-05 | 0.000381 |
| ENSG00000118620 | ZNF430 | 242.4900399 | -0.782289172 | 0.28303166 | -2.975124722 | 0.00292869 | 0.014304 |
| ENSG00000262728 | | 10.11303 | -0.7825302 | 0.6204445 | -1.735278 | 0.08269154 | 0.202473 |
| ENSG00000174547 | MRPL11 | 4936.495 | -0.7831227 | 0.316495 | -2.701236 | 0.00690823 | 0.02882 |
| ENSG00000227417 | | 2.198774 | -0.7833191 | 1.158253 | -1.804669 | 0.07112656 | 0.180735 |
| ENSG00000285763 | | 43.37377 | -0.7833647 | 0.4770039 | -1.981099 | 0.04758017 | 0.133434 |
| ENSG00000166250 | CLMP | 13.3447 | -0.7837201 | 0.6002708 | -1.771088 | 0.07654611 | 0.190891 |
| ENSG00000164758 | MED30 | 562.8615 | -0.7839695 | 0.1870537 | -4.324846 | 1.53E-05 | 0.000144 |

| | | | | | | | |
|---|---|---|---|---|---|---|---|
| ENSG00000225026 | | 10.76461 | -0.7839854 | 0.5757108 | -1.791728 | 0.07317652 | 0.184762 |
| ENSG00000252965 | | 3.615441 | -0.7840892 | 0.8268274 | -1.609761 | 0.1074501 | 0.245093 |
| ENSG00000283196 | | 401.6792 | -0.7845663 | 0.2322708 | -3.543724 | 0.00039452 | 0.002562 |
| ENSG00000234764 | E2F6P1 | 159.1008 | -0.7849834 | 0.2178477 | -3.765795 | 0.00016602 | 0.001203 |
| ENSG00000148690 | FRA10AC1 | 1580.895 | -0.7850187 | 0.2030952 | -4.008493 | 6.11E-05 | 0.000495 |
| ENSG00000273759 | | 70.95073 | -0.7853966 | 0.3999592 | -2.262939 | 0.02363944 | 0.077598 |
| ENSG00000230716 | KRT8P7 | 38.29562 | -0.7856059 | 0.3707885 | -2.388943 | 0.01689691 | 0.059336 |
| ENSG00000188089 | PLA2G4E | 2.245226 | -0.7858242 | 1.219039 | -1.953859 | 0.05071786 | 0.140017 |
| ENSG00000275074 | NUDT18 | 549.2742 | -0.7859101 | 0.2069522 | -3.94512 | 7.98E-05 | 0.000627 |
| ENSG00000188342 | GTF2F2 | 2247.709 | -0.7863628 | 0.1489744 | -5.385897 | 7.21E-08 | 1.13E-06 |
| ENSG00000129667 | RHBDF2 | 2149.051914 | -0.786822074 | 0.144652944 | -5.542411013 | 2.98E-08 | 4.96E-07 |
| ENSG00000214077 | GNAQP1 | 81.79456 | -0.786833 | 0.3367734 | -2.584201 | 0.00976049 | 0.038114 |
| ENSG00000141759 | TXNL4A | 3264.39 | -0.7868352 | 0.1949494 | -4.207911 | 2.58E-05 | 0.000229 |
| ENSG00000188078 | | 1.942205 | -0.7871076 | 1.126956 | -1.74986 | 0.08014248 | 0.197524 |
| ENSG00000109674 | NEIL3 | 394.0191816 | -0.78711488 | 0.534091431 | -1.882870891 | 0.05971787 | 0.158458 |
| ENSG00000177791 | MYOZ1 | 244.5294 | -0.7872544 | 0.4896166 | -1.94729 | 0.05149999 | 0.141705 |
| ENSG00000266907 | | 5.166044 | -0.7874345 | 0.8707333 | -1.638001 | 0.1014214 | 0.235198 |
| ENSG00000164221 | CCDC112 | 736.1437 | -0.787536 | 0.1873013 | -4.33737 | 1.44E-05 | 0.000137 |
| ENSG00000196341 | OR8D1 | 9.033085 | -0.7880427 | 0.7070659 | -1.658387 | 0.09723939 | 0.227947 |
| ENSG00000270175 | | 50.0777 | -0.7882984 | 0.3237719 | -2.667276 | 0.00764689 | 0.031368 |
| ENSG00000201944 | | 4.097783 | -0.7883071 | 0.8588032 | -1.634278 | 0.1022005 | 0.236324 |
| ENSG00000048405 | ZNF800 | 1326.601594 | -0.789059978 | 0.191194093 | -4.264982235 | 2.00E-05 | 0.000183 |
| ENSG00000009335 | UBE3C | 7980.547211 | -0.789291703 | 0.16155115 | -4.996736226 | 5.83E-07 | 7.64E-06 |
| ENSG00000230773 | | 14.18586 | -0.7894182 | 0.5366534 | -1.865822 | 0.06206627 | 0.163029 |
| ENSG00000003400 | CASP10 | 37.67902256 | -0.789572228 | 0.611623042 | -1.737401248 | 0.08231637 | 0.201744 |
| ENSG00000067533 | RRP15 | 3753.564157 | -0.789708046 | 0.186188437 | -4.388818894 | 1.14E-05 | 0.000111 |
| ENSG00000164885 | CDK5 | 1755.362 | -0.790371 | 0.2209998 | -3.744811 | 0.00018053 | 0.001296 |
| ENSG00000241685 | ARPC1A | 979.3624 | -0.7903746 | 0.2519818 | -3.327433 | 0.0008765 | 0.005102 |
| ENSG00000271780 | | 173.2563 | -0.7907814 | 0.2567011 | -3.266288 | 0.00108967 | 0.006165 |
| ENSG00000165672 | PRDX3 | 7562.016 | -0.790798 | 0.2252462 | -3.668421 | 0.00024405 | 0.00169 |
| ENSG00000188996 | HUS1B | 2.896653 | -0.7908301 | 1.070757 | -1.687374 | 0.09153142 | 0.218253 |
| ENSG00000174950 | CD164L2 | 38.06062 | -0.7910188 | 0.4051359 | -2.249798 | 0.0244618 | 0.079729 |
| ENSG00000139437 | TCHP | 1727.53 | -0.7910336 | 0.1383067 | -5.820027 | 5.88E-09 | 1.12E-07 |

| | | | | | | | |
|---|---|---|---|---|---|---|---|
| ENSG00000129128 | SPCS3 | 5752.155991 | -0.791075175 | 0.113266684 | -7.081984237 | 1.42E-12 | 4.54E-11 |
| ENSG00000205861 | PCOTH | 1.788279 | -0.7912817 | 1.09845 | -1.717652 | 0.08586017 | 0.208178 |
| ENSG00000188825 | LINC00910 | 347.1794 | -0.7914572 | 0.2112556 | -3.897114 | 9.73E-05 | 0.000748 |
| ENSG00000230534 | | 5.53445 | -0.7917134 | 0.7865323 | -1.628834 | 0.1033482 | 0.238335 |
| ENSG00000164597 | COG5 | 2010.004 | -0.7919546 | 0.2059045 | -3.996409 | 6.43E-05 | 0.000518 |
| ENSG00000251066 | | 27.04831 | -0.7921015 | 0.4191683 | -2.190585 | 0.0284818 | 0.089969 |
| ENSG00000136518 | ACTL6A | 3985.442454 | -0.792103983 | 0.193360733 | -4.258902087 | 2.05E-05 | 0.000187 |
| ENSG00000196954 | CASP4 | 938.2936 | -0.7926913 | 0.38386 | -2.343491 | 0.01910422 | 0.065519 |
| ENSG00000198055 | GRK6 | 5247.133 | -0.7927409 | 0.2192654 | -3.778134 | 0.00015801 | 0.001151 |
| ENSG00000107554 | DNMBP | 927.4490635 | -0.792853223 | 0.281512655 | -3.009877818 | 0.00261353 | 0.012987 |
| ENSG00000252552 | RNU6-307P | 1.534827 | -0.7928545 | 1.084074 | -1.691363 | 0.09076755 | NA |
| ENSG00000186638 | KIF24 | 310.4464 | -0.7928711 | 0.4882647 | -1.99432 | 0.04611711 | 0.130363 |
| ENSG00000221540 | MIR1180 | 2.911632 | -0.7928969 | 0.9515336 | -1.663947 | 0.09612304 | 0.225961 |
| ENSG00000101224 | CDC25B | 7704.81438 | -0.792904532 | 0.232553044 | -3.572081956 | 0.00035416 | 0.002337 |
| ENSG00000151176 | PLBD2 | 2890.096 | -0.7931164 | 0.1815031 | -4.51852 | 6.23E-06 | 6.49E-05 |
| ENSG00000243679 | | 566.9118 | -0.7934025 | 0.2651797 | -3.184675 | 0.00144917 | 0.007879 |
| ENSG00000126001 | CEP250 | 2877.194671 | -0.793458733 | 0.171868707 | -4.744008867 | 2.10E-06 | 2.45E-05 |
| ENSG00000181260 | MTHFD2P7 | 59.08809 | -0.7935002 | 0.45454 | -2.088551 | 0.03674817 | 0.109489 |
| ENSG00000274238 | | 1.45407 | -0.7936098 | 1.299889 | -2.068946 | 0.03855119 | NA |
| ENSG00000254612 | DNAJB6P1 | 1860.1 | -0.7937397 | 0.1421412 | -5.690233 | 1.27E-08 | 2.28E-07 |
| ENSG00000142494 | SLC47A1 | 50.48339 | -0.7938766 | 0.4657474 | -2.055312 | 0.03984885 | 0.116556 |
| ENSG00000119537 | KDSR | 1876.732829 | -0.794057386 | 0.190280035 | -4.319861273 | 1.56E-05 | 0.000147 |
| ENSG00000256995 | | 1.522232 | -0.7941836 | 1.192931 | -1.873946 | 0.06093789 | NA |
| ENSG00000084463 | WBP11 | 3752.229485 | -0.794351642 | 0.221021995 | -3.756590081 | 0.00017224 | 0.001243 |
| ENSG00000267322 | SNHG22 | 12.95574 | -0.7946601 | 0.7421578 | -1.670567 | 0.09480732 | 0.22392 |
| ENSG00000119397 | CNTRL | 551.1378379 | -0.79506837 | 0.288401009 | -2.970138208 | 0.00297666 | 0.014479 |
| ENSG00000265478 | | 4.761286 | -0.7959184 | 0.7434894 | -1.659978 | 0.09691893 | 0.227397 |
| ENSG00000010292 | NCAPD2 | 6604.730774 | -0.796284512 | 0.384894766 | -2.376335944 | 0.01748554 | 0.061015 |
| ENSG00000250950 | | 5.526986 | -0.7964289 | 0.8710299 | -1.632243 | 0.1026284 | 0.237107 |
| ENSG00000163659 | TIPARP | 1880.017 | -0.7966729 | 0.2641726 | -3.202993 | 0.00136007 | 0.007465 |
| ENSG00000265205 | | 63.71182 | -0.7966864 | 0.3841058 | -2.359125 | 0.01831807 | 0.063322 |
| ENSG00000254978 | ALG1L9P | 33.511 | -0.7967675 | 0.3629005 | -2.457911 | 0.0139748 | 0.05089 |
| ENSG00000286833 | | 14.56428 | -0.7967823 | 0.5323725 | -1.90258 | 0.05709539 | 0.153165 |

| | | | | | | | |
|---|---|---|---|---|---|---|---|
| ENSG00000210140 | MT-TC | 14.43839 | -0.7967872 | 0.4937636 | -1.987722 | 0.04684243 | 0.131895 |
| ENSG00000133119 | RFC3 | 3065.373952 | -0.797382926 | 0.418664891 | -2.219598773 | 0.02644602 | 0.084758 |
| ENSG00000207973 | MIR589 | 2.684902 | -0.7974591 | 1.127093 | -1.761695 | 0.07812092 | 0.193727 |
| ENSG00000106617 | PRKAG2 | 878.0985247 | -0.797499454 | 0.170357931 | -4.817722664 | 1.45E-06 | 1.76E-05 |
| ENSG00000278765 | | 22.73171 | -0.7979398 | 0.4891098 | -1.992686 | 0.04629585 | 0.130757 |
| ENSG00000227973 | PIN4P1 | 57.16598 | -0.7980339 | 0.2739103 | -3.109915 | 0.00187141 | 0.00979 |
| ENSG00000226022 | | 3.250558 | -0.7980497 | 1.034641 | -1.72784 | 0.08401695 | 0.204774 |
| ENSG00000062282 | DGAT2 | 842.9378857 | -0.798195781 | 0.275697641 | -3.099785257 | 0.00193661 | 0.010079 |
| ENSG00000232479 | KRT8P52 | 6.520146 | -0.7982872 | 0.8375876 | -1.642525 | 0.1004813 | 0.233609 |
| ENSG00000137547 | MRPL15 | 3743.050445 | -0.798449689 | 0.243878961 | -3.452990829 | 0.00055441 | 0.003434 |
| ENSG00000279573 | | 2.47164 | -0.7985372 | 0.9519797 | -1.627261 | 0.1036817 | 0.238821 |
| ENSG00000171861 | MRM3 | 905.2025 | -0.7986503 | 0.278864 | -3.073224 | 0.00211759 | 0.01088 |
| ENSG00000089177 | KIF16B | 1213.540658 | -0.798901749 | 0.179170439 | -4.585955587 | 4.52E-06 | 4.87E-05 |
| ENSG00000231530 | | 28.6627 | -0.7991774 | 0.3880963 | -2.349943 | 0.01877627 | 0.064587 |
| ENSG00000111641 | NOP2 | 118.2031042 | -0.799332994 | 0.275269263 | -3.10642367 | 0.00189365 | 0.009885 |
| ENSG00000205667 | ARSH | 5.135009 | -0.7993571 | 0.7993106 | -1.650821 | 0.09877521 | 0.230612 |
| ENSG00000181885 | CLDN7 | 6749.789 | -0.7996031 | 0.1985097 | -4.180972 | 2.90E-05 | 0.000255 |
| ENSG00000115919 | KYNU | 3977.752714 | -0.799648365 | 0.352674318 | -2.54189318 | 0.01102539 | 0.042106 |
| ENSG00000276115 | | 4.07852 | -0.7999224 | 0.8888946 | -1.645056 | 0.09995818 | 0.23276 |
| ENSG00000285970 | | 1.623883 | -0.8000127 | 1.297326 | -2.014551 | 0.04395176 | 0.125809 |
| ENSG00000251739 | RNU6-1053P | 7.962269 | -0.8002208 | 0.6879436 | -1.701932 | 0.08876804 | 0.213534 |
| ENSG00000275759 | | 4.958938 | -0.8003392 | 0.7488374 | -1.665611 | 0.09579091 | 0.225447 |
| ENSG00000234997 | | 1.560819 | -0.8006907 | 1.207619 | -1.853063 | 0.06387337 | NA |
| ENSG00000144619 | CNTN4 | 320.1803 | -0.800905 | 0.2864925 | -3.006102 | 0.0026462 | 0.013121 |
| ENSG00000228700 | | 2.436303 | -0.8010886 | 0.9098836 | -1.637801 | 0.1014632 | 0.235275 |
| ENSG00000274523 | RCC1L | 3386.561 | -0.8011334 | 0.2705698 | -3.16943 | 0.00152738 | 0.008236 |
| ENSG00000262171 | | 12.29829 | -0.8014277 | 0.6303038 | -1.765325 | 0.07750924 | 0.192624 |
| ENSG00000203867 | RBM20 | 42.67636 | -0.8018279 | 0.4656348 | -2.080665 | 0.03746461 | 0.111112 |
| ENSG00000065328 | MCM10 | 1674.159189 | -0.801954258 | 0.717899109 | -1.716344595 | 0.08609899 | 0.208491 |
| ENSG00000157045 | NTAN1 | 1216.435 | -0.8020078 | 0.2626399 | -3.244963 | 0.00117466 | 0.006564 |
| ENSG00000155363 | MOV10 | 4738.263 | -0.8023412 | 0.1702294 | -4.84225 | 1.28E-06 | 1.56E-05 |
| ENSG00000125827 | TMX4 | 1819.158652 | -0.802413995 | 0.216027841 | -3.876334195 | 0.00010604 | 0.000806 |
| ENSG00000102024 | PLS3 | 4533.151543 | -0.802920134 | 0.183880441 | -4.501968864 | 6.73E-06 | 6.94E-05 |

| | | | | | | | |
|---|---|---|---|---|---|---|---|
| ENSG00000171428 | NAT1 | 533.3062 | -0.8030667 | 0.2598543 | -3.28202 | 0.00103066 | 0.005878 |
| ENSG00000187720 | THSD4 | 2049.533 | -0.8030814 | 0.5299434 | -1.927237 | 0.05395012 | 0.147106 |
| ENSG00000203814 | H2BC18 | 1314.652 | -0.8033552 | 0.3718434 | -2.436231 | 0.01484119 | 0.053474 |
| ENSG00000181778 | TMEM252 | 2.04137 | -0.8033799 | 1.109786 | -1.731325 | 0.08339389 | 0.203611 |
| ENSG00000185201 | IFITM2 | 3165.918 | -0.8033974 | 0.2446725 | -3.432144 | 0.00059883 | 0.003669 |
| ENSG00000170092 | SPDYE5 | 41.0039 | -0.8039202 | 0.397242 | -2.317135 | 0.02049636 | 0.069301 |
| ENSG00000215032 | GNL3LP1 | 6.912384 | -0.8040456 | 0.6146629 | -1.774395 | 0.0759978 | 0.189898 |
| ENSG00000205336 | ADGRG1 | 8229.258 | -0.8043356 | 0.3600751 | -2.505353 | 0.01223292 | 0.045721 |
| ENSG00000280046 | | 89.44968 | -0.8046344 | 0.3449013 | -2.584114 | 0.00976295 | 0.038118 |
| ENSG00000225766 | DHRS4L1 | 13.19096 | -0.8048532 | 0.4864965 | -2.013911 | 0.0440189 | 0.125946 |
| ENSG00000074266 | EED | 1815.969255 | -0.804858978 | 0.266663694 | -3.191757129 | 0.0014141 | 0.007713 |
| ENSG00000145908 | ZNF300 | 1.336296 | -0.805049 | 1.238531 | -1.978155 | 0.04791121 | NA |
| ENSG00000242294 | STAG3L5P | 663.7115 | -0.8051217 | 0.3120813 | -2.816235 | 0.00485902 | 0.021675 |
| ENSG00000104976 | SNAPC2 | 1093.666747 | -0.805532903 | 0.187005588 | -4.445242934 | 8.78E-06 | 8.78E-05 |
| ENSG00000259123 | | 1.996431 | -0.8056879 | 1.281405 | -1.962055 | 0.04975603 | 0.138021 |
| ENSG00000176971 | FIBIN | 3.684318 | -0.8058603 | 0.8171734 | -1.661176 | 0.09667803 | 0.226932 |
| ENSG00000288086 | | 14.25473 | -0.8063308 | 0.6711471 | -1.731592 | 0.08334619 | 0.203513 |
| ENSG00000244242 | IFITM10 | 138.7759 | -0.8065746 | 0.4531067 | -2.124451 | 0.03363241 | 0.102463 |
| ENSG00000026297 | RNASET2 | 1270.420536 | -0.806644126 | 0.18238076 | -4.564805913 | 5.00E-06 | 5.33E-05 |
| ENSG00000101695 | RNF125 | 267.235492 | -0.806771894 | 0.356683411 | -2.51839976 | 0.01178894 | 0.044397 |
| ENSG00000136943 | CTSV | 174.7585941 | -0.806957843 | 0.573390059 | -1.85989364 | 0.06290058 | 0.164683 |
| ENSG00000039123 | MTREX | 4316.530776 | -0.807636099 | 0.125838535 | -6.49784276 | 8.15E-11 | 2.04E-09 |
| ENSG00000133315 | MACROD1 | 2598.400688 | -0.807883383 | 0.204605569 | -4.108630559 | 3.98E-05 | 0.000338 |
| ENSG00000238078 | LINC01352 | 2.076607 | -0.8079415 | 1.350911 | -2.118045 | 0.03417124 | 0.103736 |
| ENSG00000223956 | LINC01767 | 11.25617 | -0.8080916 | 0.5736655 | -1.839879 | 0.06578601 | 0.170324 |
| ENSG00000087494 | PTHLH | 20.87363464 | -0.808342563 | 0.545314448 | -1.907652184 | 0.05643619 | 0.152013 |
| ENSG00000225920 | RIMKLBP2 | 55.19269 | -0.8084384 | 0.4641958 | -2.092223 | 0.03641857 | 0.108769 |
| ENSG00000248388 | | 3.89824 | -0.8084423 | 0.7910321 | -1.667152 | 0.09548428 | 0.224964 |
| ENSG00000281490 | CICP14 | 1151.838 | -0.8085218 | 0.4065589 | -2.30538 | 0.02114532 | 0.071097 |
| ENSG00000228973 | | 1.144002 | -0.8087975 | 1.371639 | -2.161695 | 0.03064169 | NA |
| ENSG00000204209 | DAXX | 3548.535 | -0.8090916 | 0.2136547 | -3.952717 | 7.73E-05 | 0.00061 |
| ENSG00000012660 | ELOVL5 | 5707.91415 | -0.809171703 | 0.20064603 | -4.183097149 | 2.88E-05 | 0.000253 |
| ENSG00000120675 | DNAJC15 | 2734.189731 | -0.809307833 | 0.204062672 | -4.114824962 | 3.87E-05 | 0.000331 |

| | | | | | | | |
|---|---|---|---|---|---|---|---|
| ENSG00000163040 | CCDC74A | 275.8781 | -0.8093875 | 0.4153663 | -2.267251 | 0.02337491 | 0.076929 |
| ENSG00000226781 | TBCAP1 | 136.2658 | -0.8094078 | 0.2899172 | -3.005557 | 0.00265095 | 0.013138 |
| ENSG00000145730 | PAM | 2687.934 | -0.8096476 | 0.2593673 | -3.310921 | 0.00092989 | 0.005364 |
| ENSG00000137648 | TMPRSS4 | 257.8177979 | -0.809924088 | 0.511404258 | -1.972781329 | 0.04852048 | 0.1354 |
| ENSG00000253463 | HMGB1P19 | 1.940316 | -0.8100141 | 1.241902 | -1.975739 | 0.04818431 | 0.134668 |
| ENSG00000250999 | | 53.34368 | -0.8100721 | 0.3224365 | -2.737968 | 0.006182 | 0.026364 |
| ENSG00000167740 | CYB5D2 | 1193.476 | -0.8101436 | 0.1851682 | -4.501497 | 6.75E-06 | 6.95E-05 |
| ENSG00000279199 | | 14.62388 | -0.8104863 | 0.6420508 | -1.767795 | 0.07709525 | 0.1919 |
| ENSG00000119203 | CPSF3 | 3050.919087 | -0.810695334 | 0.237788931 | -3.585613083 | 0.00033629 | 0.002235 |
| ENSG00000157111 | TMEM171 | 9.416111 | -0.8111732 | 0.6868202 | -1.714765 | 0.08638829 | 0.209039 |
| ENSG00000007202 | KIAA0100 | 18410.97245 | -0.811216101 | 0.158231094 | -5.254247266 | 1.49E-07 | 2.18E-06 |
| ENSG00000152234 | ATP5F1A | 21598.13 | -0.811322 | 0.2197547 | -3.85488 | 0.00011579 | 0.000873 |
| ENSG00000129933 | MAU2 | 2737.447707 | -0.811407263 | 0.165172586 | -5.041739885 | 4.61E-07 | 6.16E-06 |
| ENSG00000255314 | | 2.034115 | -0.8116882 | 1.105065 | -1.718235 | 0.08575373 | 0.207977 |
| ENSG00000276603 | | 16.71728 | -0.8122539 | 0.6098588 | -1.803737 | 0.07127247 | 0.181002 |
| ENSG00000183527 | PSMG1 | 3319.183 | -0.8125506 | 0.2464252 | -3.480495 | 0.00050049 | 0.00314 |
| ENSG00000279602 | | 51.14479 | -0.8131508 | 0.3331118 | -2.689731 | 0.00715096 | 0.029677 |
| ENSG00000272752 | STAG3L5P-PVRIG | 20.24031 | -0.8132941 | 0.4847271 | -2.039894 | 0.04136085 | 0.120054 |
| ENSG00000164818 | DNAAF5 | 4451.361 | -0.8136275 | 0.216126 | -3.927271 | 8.59E-05 | 0.00067 |
| ENSG00000234699 | | 4.734759 | -0.8138376 | 0.8376782 | -1.686234 | 0.09175077 | 0.218565 |
| ENSG00000239306 | RBM14 | 3687.434 | -0.8141755 | 0.1971473 | -4.27539 | 1.91E-05 | 0.000175 |
| ENSG00000003056 | M6PR | 4030.907824 | -0.814913667 | 0.144076207 | -5.735617631 | 9.72E-09 | 1.79E-07 |
| ENSG00000164237 | CMBL | 1549.696 | -0.8151187 | 0.2098697 | -4.042683 | 5.28E-05 | 0.000435 |
| ENSG00000166596 | CFAP52 | 53.40637 | -0.8151792 | 0.3118775 | -2.84789 | 0.00440102 | 0.019944 |
| ENSG00000132386 | SERPINF1 | 103.6422967 | -0.815271272 | 0.317442888 | -2.806230422 | 0.00501248 | 0.022192 |
| ENSG00000134222 | PSRC1 | 1993.630864 | -0.816222504 | 0.388623549 | -2.404653912 | 0.01618779 | 0.057252 |
| ENSG00000169919 | GUSB | 3542.003 | -0.8163876 | 0.212369 | -3.993839 | 6.50E-05 | 0.000522 |
| ENSG00000102362 | SYTL4 | 427.235965 | -0.816924474 | 0.283567124 | -3.093365113 | 0.00197901 | 0.010262 |
| ENSG00000129226 | CD68 | 2.773590557 | -0.817674617 | 1.108139792 | -1.779886487 | 0.07509454 | 0.188225 |
| ENSG00000163710 | PCOLCE2 | 108.3936 | -0.8177118 | 0.4690576 | -2.100093 | 0.03572063 | 0.107287 |
| ENSG00000243406 | MRPS31P5 | 45.22249 | -0.8181324 | 0.3465283 | -2.621613 | 0.00875146 | 0.034981 |
| ENSG00000280347 | | 186.7627 | -0.8184004 | 0.4115852 | -2.316357 | 0.02053878 | 0.0694 |
| ENSG00000115317 | HTRA2 | 1835.380495 | -0.818767418 | 0.157201978 | -5.32182916 | 1.03E-07 | 1.55E-06 |

| | | | | | | | |
|---|---|---|---|---|---|---|---|
| ENSG00000101958 | GLRA2 | 2.190107401 | -0.819186128 | 0.986166484 | -1.705861889 | 0.0880338 | 0.212136 |
| ENSG00000133027 | PEMT | 1065.892425 | -0.819533602 | 0.150837479 | -5.570737117 | 2.54E-08 | 4.28E-07 |
| ENSG00000160993 | ALKBH4 | 812.8907 | -0.8199783 | 0.2146066 | -3.966111 | 7.31E-05 | 0.000579 |
| ENSG00000214106 | PAXIP1-AS2 | 481.4938 | -0.8202434 | 0.2705184 | -3.257658 | 0.00112336 | 0.006327 |
| ENSG00000270557 | | 14.17331 | -0.820498 | 0.5731085 | -1.861579 | 0.06266252 | 0.164254 |
| ENSG00000249628 | LINC00942 | 5.215532 | -0.8214568 | 0.9356235 | -1.637096 | 0.1016105 | 0.235513 |
| ENSG00000197587 | DMBX1 | 86.02372 | -0.8216631 | 0.6404848 | -1.79959 | 0.07192534 | 0.182294 |
| ENSG00000278338 | VWA8-AS1 | 6.521007 | -0.8219477 | 0.7223914 | -1.712876 | 0.08673531 | 0.209726 |
| ENSG00000128581 | IFT22 | 1355.455749 | -0.823102007 | 0.179817765 | -4.697277818 | 2.64E-06 | 3.01E-05 |
| ENSG00000275764 | | 194.4282 | -0.8234629 | 0.2856924 | -3.101776 | 0.00192364 | 0.010019 |
| ENSG00000275485 | | 9.863153 | -0.8237634 | 0.7041671 | -1.730351 | 0.08356763 | 0.203941 |
| ENSG00000196584 | XRCC2 | 1134.14 | -0.8247131 | 0.4348648 | -2.235809 | 0.02536428 | 0.081983 |
| ENSG00000164898 | FMC1 | 58.99068 | -0.8249816 | 0.3074783 | -2.907821 | 0.00363956 | 0.017063 |
| ENSG00000122574 | WIPF3 | 5.65393339 | -0.825433451 | 0.89966445 | -1.678060671 | 0.09333526 | 0.221334 |
| ENSG00000264311 | CCDC58P1 | 18.51207 | -0.8254604 | 0.5157661 | -2.004438 | 0.0450232 | 0.128048 |
| ENSG00000157445 | CACNA2D3 | 14.69983 | -0.8257716 | 0.579898 | -1.871699 | 0.06124822 | 0.161567 |
| ENSG00000223842 | | 9.048048 | -0.8260323 | 0.7652756 | -1.699933 | 0.08914359 | 0.214185 |
| ENSG00000146250 | PRSS35 | 6.217881 | -0.826073 | 0.9825987 | -1.608368 | 0.1077546 | 0.245598 |
| ENSG00000225783 | MIAT | 18.28582 | -0.8264422 | 0.5915322 | -1.848919 | 0.06446944 | 0.167717 |
| ENSG00000229154 | KCNQ5-AS1 | 2.030179 | -0.8265959 | 1.391207 | -2.215622 | 0.02671739 | 0.085458 |
| ENSG00000123360 | PDE1B | 3.396339916 | -0.826627254 | 1.069152336 | -1.699280641 | 0.08926632 | 0.214363 |
| ENSG00000228549 | | 17.23663 | -0.8267669 | 0.4022124 | -2.363715 | 0.01809273 | 0.062723 |
| ENSG00000237594 | | 60.17245 | -0.8268441 | 0.3774608 | -2.480256 | 0.0131288 | 0.048427 |
| ENSG00000229391 | HLA-DRB6 | 11.50034 | -0.8268798 | 0.9398487 | -1.663533 | 0.09620581 | 0.226023 |
| ENSG00000106268 | NUDT1 | 984.4276654 | -0.826980134 | 0.503645277 | -2.035354146 | 0.04181525 | 0.121062 |
| ENSG00000159063 | ALG8 | 2960.291 | -0.8271289 | 0.1178787 | -7.104958 | 1.20E-12 | 3.90E-11 |
| ENSG00000138029 | HADHB | 5683.747839 | -0.827245661 | 0.142527316 | -5.9024859 | 3.58E-09 | 7.07E-08 |
| ENSG00000128203 | ASPHD2 | 77.77782105 | -0.827263524 | 0.292291515 | -3.040783545 | 0.00235963 | 0.011905 |
| ENSG00000133606 | MKRN1 | 8421.116099 | -0.827343038 | 0.115264394 | -7.264017011 | 3.76E-13 | 1.30E-11 |
| ENSG00000268746 | | 1.315079 | -0.8281358 | 1.358889 | -2.095765 | 0.03610303 | NA |
| ENSG00000004478 | FKBP4 | 13212.31399 | -0.828150767 | 0.170588587 | -4.984110926 | 6.22E-07 | 8.10E-06 |
| ENSG00000198443 | KRTAP4-1 | 49.87783 | -0.8283063 | 0.3664886 | -2.540735 | 0.01106196 | 0.042216 |
| ENSG00000271913 | | 38.87146 | -0.8283939 | 0.4661953 | -2.131535 | 0.03304509 | 0.101126 |

| | | | | | | | |
|---|---|---|---|---|---|---|---|
| ENSG00000240790 | | 1.263291 | -0.8284759 | 1.405316 | -2.182851 | 0.02904678 | NA |
| ENSG00000163618 | CADPS | 75.14887 | -0.8293412 | 0.4119445 | -2.331101 | 0.01974802 | 0.067302 |
| ENSG00000141456 | PELP1 | 4343.451 | -0.8294185 | 0.2187829 | -3.946492 | 7.93E-05 | 0.000623 |
| ENSG00000054392 | HHAT | 354.0425059 | -0.829470358 | 0.261493238 | -3.360225539 | 0.00077879 | 0.00461 |
| ENSG00000273374 | | 18.63936 | -0.829471 | 0.710525 | -1.748385 | 0.08039734 | 0.198005 |
| ENSG00000165283 | STOML2 | 6087.709 | -0.8296119 | 0.2277946 | -3.817979 | 0.00013455 | 0.000996 |
| ENSG00000213057 | C1orf220 | 76.22045 | -0.8298078 | 0.4061824 | -2.35248 | 0.01864871 | 0.064214 |
| ENSG00000162851 | TFB2M | 3033.936 | -0.8300686 | 0.2182512 | -3.96545 | 7.33E-05 | 0.000581 |
| ENSG00000145781 | COMMD10 | 1980.734 | -0.8302899 | 0.1949353 | -4.398769 | 1.09E-05 | 0.000106 |
| ENSG00000267698 | | 22.58613 | -0.8306859 | 0.5271732 | -1.987652 | 0.0468502 | 0.131902 |
| ENSG00000090447 | TFAP4 | 1120.376536 | -0.83136869 | 0.172450283 | -4.953318521 | 7.30E-07 | 9.36E-06 |
| ENSG00000230487 | PSMG3-AS1 | 2279.548 | -0.8315545 | 0.2812231 | -3.169115 | 0.00152904 | 0.008244 |
| ENSG00000284713 | SMIM38 | 23.13973 | -0.8316472 | 0.662031 | -1.784965 | 0.07426704 | 0.186838 |
| ENSG00000287299 | | 32.94719 | -0.8317461 | 0.5813476 | -1.891064 | 0.05861579 | 0.156377 |
| ENSG00000066735 | KIF26A | 1166.372651 | -0.831871576 | 0.437222438 | -2.236469861 | 0.02532101 | 0.081894 |
| ENSG00000132518 | GUCY2D | 1.457117619 | -0.831989156 | 1.289011525 | -2.025254583 | 0.04284123 | NA |
| ENSG00000066336 | SPI1 | 2.22827497 | -0.832018396 | 0.961211247 | -1.697052136 | 0.0896868 | 0.215154 |
| ENSG00000151929 | BAG3 | 2977.57 | -0.832274 | 0.2104217 | -4.110812 | 3.94E-05 | 0.000335 |
| ENSG00000113231 | PDE8B | 43.52310412 | -0.832310007 | 0.477120828 | -2.114573975 | 0.03446627 | 0.104476 |
| ENSG00000253731 | PCDHGA6 | 83.90246 | -0.8323881 | 0.3373698 | -2.720983 | 0.00650882 | 0.027458 |
| ENSG00000229207 | SERPINH1P1 | 3.001233 | -0.8327265 | 0.9220389 | -1.685866 | 0.09182166 | 0.218642 |
| ENSG00000115459 | ELMOD3 | 1555.424404 | -0.833490283 | 0.253339936 | -3.481147235 | 0.00049927 | 0.003134 |
| ENSG00000285853 | | 2.987201 | -0.8346327 | 1.210499 | -1.885399 | 0.05937595 | 0.157882 |
| ENSG00000168411 | RFWD3 | 5365.168 | -0.8351447 | 0.2535971 | -3.526497 | 0.0004211 | 0.002704 |
| ENSG00000267390 | | 33.61247 | -0.8351717 | 0.3557173 | -2.618588 | 0.00882945 | 0.035229 |
| ENSG00000172331 | BPGM | 1682.509 | -0.8357023 | 0.3185479 | -2.863921 | 0.00418433 | 0.019168 |
| ENSG00000213397 | HAUS7 | 83.01022 | -0.8358331 | 0.2951116 | -3.055047 | 0.00225026 | 0.011429 |
| ENSG00000231291 | | 1.539324 | -0.8361544 | 1.345329 | -2.039569 | 0.04139324 | NA |
| ENSG00000138316 | ADAMTS14 | 3.917243033 | -0.836406305 | 0.92303083 | -1.712080364 | 0.08688186 | 0.210004 |
| ENSG00000251620 | STPG2-AS1 | 2.001178 | -0.8364682 | 1.312632 | -2.016413 | 0.04375679 | 0.125373 |
| ENSG00000153086 | ACMSD | 6.419275 | -0.8365221 | 0.7345047 | -1.736762 | 0.08242919 | 0.201923 |
| ENSG00000254876 | STRA6LP | 43.24945 | -0.8369942 | 0.5469807 | -1.96088 | 0.04989305 | 0.138272 |
| ENSG00000256862 | PARP11-AS1 | 5.500161 | -0.8375502 | 0.7864887 | -1.701632 | 0.08882443 | 0.213631 |

| | | | | | | | |
|---|---|---|---|---|---|---|---|
| ENSG00000198860 | TSEN15 | 3116.52 | -0.8376477 | 0.161984 | -5.295502 | 1.19E-07 | 1.77E-06 |
| ENSG00000146707 | POMZP3 | 79.49077 | -0.8377264 | 0.3120753 | -2.918075 | 0.003522 | 0.016603 |
| ENSG00000250068 | | 5.855524 | -0.838176 | 0.8129159 | -1.704744 | 0.08824216 | 0.212519 |
| ENSG00000285886 | | 15.39229 | -0.8382471 | 0.6038654 | -1.862594 | 0.06251942 | 0.163992 |
| ENSG00000156017 | CARNMT1 | 778.2428 | -0.838705 | 0.1607056 | -5.343959 | 9.09E-08 | 1.39E-06 |
| ENSG00000283000 | LINC02635 | 12.13445 | -0.8387171 | 0.8194248 | -1.688104 | 0.09139122 | 0.218008 |
| ENSG00000159714 | ZDHHC1 | 13.48502 | -0.8390596 | 0.7456033 | -1.740294 | 0.08180749 | 0.200772 |
| ENSG00000280286 | FRG1KP | 10.76884 | -0.8392473 | 0.5950865 | -1.8758 | 0.06068273 | 0.160349 |
| ENSG00000245534 | RORA-AS1 | 19.5602 | -0.8395025 | 0.4670195 | -2.155563 | 0.03111784 | 0.096515 |
| ENSG00000213211 | KRT18P64 | 12.41471 | -0.839615 | 0.5344526 | -1.981827 | 0.04749859 | 0.133276 |
| ENSG00000279667 | | 11.3535 | -0.839661 | 0.6142502 | -1.856209 | 0.06342379 | 0.165678 |
| ENSG00000272156 | | 8.215073 | -0.8401304 | 0.6863237 | -1.765616 | 0.07746036 | 0.192557 |
| ENSG00000141012 | GALNS | 3816.222 | -0.8403051 | 0.1358171 | -6.289279 | 3.19E-10 | 7.43E-09 |
| ENSG00000100003 | SEC14L2 | 202.0327237 | -0.840399657 | 0.236735288 | -3.728819233 | 0.00019238 | 0.001371 |
| ENSG00000117308 | GALE | 1157.627061 | -0.841022556 | 0.361363005 | -2.600622561 | 0.00930548 | 0.036704 |
| ENSG00000259172 | | 95.75832 | -0.8411 | 0.346536 | -2.688135 | 0.00718524 | 0.029792 |
| ENSG00000229161 | TCP1P1 | 19.4801 | -0.8420535 | 0.530977 | -2.007914 | 0.04465239 | 0.127307 |
| ENSG00000223461 | | 8.330515 | -0.8420625 | 0.7882799 | -1.725446 | 0.0844471 | 0.20554 |
| ENSG00000171681 | ATF7IP | 3498.915 | -0.8421873 | 0.2233283 | -3.942205 | 8.07E-05 | 0.000633 |
| ENSG00000205238 | SPDYE2 | 50.28347 | -0.8424449 | 0.3913789 | -2.452077 | 0.01420342 | 0.051617 |
| ENSG00000111669 | TPI1 | 24791.19877 | -0.84245149 | 0.269254484 | -3.334982035 | 0.00085305 | 0.004981 |
| ENSG00000156587 | UBE2L6 | 3708.255 | -0.842481 | 0.3883774 | -2.469747 | 0.01352086 | 0.049536 |
| ENSG00000285278 | | 79.97814 | -0.8427068 | 0.3150253 | -2.915978 | 0.00354575 | 0.016697 |
| ENSG00000101546 | RBFA | 1170.385588 | -0.842869885 | 0.158628834 | -5.429229644 | 5.66E-08 | 9.04E-07 |
| ENSG00000108064 | TFAM | 4213.854422 | -0.842918075 | 0.206883585 | -4.231304485 | 2.32E-05 | 0.00021 |
| ENSG00000242193 | CRYZL2P | 89.98913 | -0.8430111 | 0.3839884 | -2.491924 | 0.01270533 | 0.047193 |
| ENSG00000233025 | CRYZP1 | 24.00525 | -0.8437832 | 0.5387886 | -1.990254 | 0.04656291 | 0.131316 |
| ENSG00000149656 | LINC00266-1 | 3.441683 | -0.8438441 | 1.122009 | -1.784136 | 0.07440157 | 0.187096 |
| ENSG00000134440 | NARS1 | 11027.179 | -0.844378096 | 0.198829315 | -4.405123226 | 1.06E-05 | 0.000104 |
| ENSG00000117155 | SSX2IP | 1733.066043 | -0.844402724 | 0.189498586 | -4.602344485 | 4.18E-06 | 4.54E-05 |
| ENSG00000163131 | CTSS | 55.0217 | -0.8445649 | 0.7514847 | -1.727014 | 0.08416507 | 0.205023 |
| ENSG00000273723 | SUGT1-DT | 35.99339 | -0.8446404 | 0.4739787 | -2.14931 | 0.03160985 | 0.097724 |
| ENSG00000170439 | METTL7B | 259.8562 | -0.8447278 | 0.3660073 | -2.592624 | 0.00952468 | 0.037347 |

| | | | | | | | |
|---|---|---|---|---|---|---|---|
| ENSG00000167191 | GPRC5B | 195.0713 | -0.8451547 | 0.7559421 | -1.746791 | 0.08067367 | 0.198557 |
| ENSG00000287794 | | 3.453101 | -0.8452491 | 0.9191327 | -1.694725 | 0.09012768 | 0.215827 |
| ENSG00000184898 | RBM43 | 465.1665 | -0.8454307 | 0.1852791 | -4.703201 | 2.56E-06 | 2.94E-05 |
| ENSG00000261170 | | 12.2739 | -0.8455849 | 0.5303323 | -2.005255 | 0.04493574 | 0.127881 |
| ENSG00000115946 | PNO1 | 1836.712571 | -0.8455953 | 0.182449331 | -4.782728532 | 1.73E-06 | 2.06E-05 |
| ENSG00000132792 | CTNNBL1 | 2528.522167 | -0.846458705 | 0.201621568 | -4.354244188 | 1.34E-05 | 0.000127 |
| ENSG00000171530 | TBCA | 4837.209 | -0.8466267 | 0.1688856 | -5.138943 | 2.76E-07 | 3.83E-06 |
| ENSG00000104835 | SARS2 | 135.344904 | -0.846864218 | 0.323934931 | -2.86818518 | 0.00412834 | 0.018951 |
| ENSG00000285844 | | 3.288218 | -0.8472114 | 1.25454 | -1.95962 | 0.05004023 | 0.138578 |
| ENSG00000107371 | EXOSC3 | 1016.92787 | -0.847234679 | 0.18424931 | -4.744086104 | 2.09E-06 | 2.45E-05 |
| ENSG00000196418 | ZNF124 | 754.3635 | -0.847479 | 0.1459962 | -5.911706 | 3.39E-09 | 6.71E-08 |
| ENSG00000004838 | ZMYND10 | 46.97943949 | -0.847914592 | 0.362560972 | -2.624143111 | 0.00868673 | 0.034744 |
| ENSG00000051128 | HOMER3 | 852.4631175 | -0.848268384 | 0.249985615 | -3.584927153 | 0.00033717 | 0.00224 |
| ENSG00000185689 | C6orf201 | 4.132292 | -0.8489953 | 0.7157766 | -1.767241 | 0.07718782 | 0.192059 |
| ENSG00000148680 | HTR7 | 28.12293 | -0.8495693 | 0.6878437 | -1.805719 | 0.07096225 | 0.180421 |
| ENSG00000146963 | LUC7L2 | 1847.796 | -0.8501172 | 0.1794051 | -4.881139 | 1.05E-06 | 1.31E-05 |
| ENSG00000258711 | | 14.98685 | -0.8502033 | 0.8833019 | -1.723144 | 0.0848625 | 0.20631 |
| ENSG00000233223 | | 300.6274 | -0.8503364 | 0.2248636 | -3.948891 | 7.85E-05 | 0.000618 |
| ENSG00000264808 | | 6.959737 | -0.8503793 | 0.7559858 | -1.735321 | 0.08268395 | 0.202473 |
| ENSG00000101812 | H2BW2 | 2.558104673 | -0.850447414 | 1.133525071 | -1.751437711 | 0.07987054 | 0.196966 |
| ENSG00000276988 | | 15.43816 | -0.8504941 | 0.6415876 | -1.847309 | 0.0647024 | 0.168096 |
| ENSG00000168765 | GSTM4 | 2630.534 | -0.8507691 | 0.3466205 | -2.707184 | 0.00678566 | 0.028405 |
| ENSG00000133706 | LARS1 | 10082.22919 | -0.850778388 | 0.187299116 | -4.69304721 | 2.69E-06 | 3.07E-05 |
| ENSG00000105948 | TTC26 | 490.0614768 | -0.850951221 | 0.214662792 | -4.127783161 | 3.66E-05 | 0.000314 |
| ENSG00000197746 | PSAP | 38431.88 | -0.8519878 | 0.2069711 | -4.275126 | 1.91E-05 | 0.000176 |
| ENSG00000134030 | CTIF | 2089.880035 | -0.852294708 | 0.25394196 | -3.538265831 | 0.00040276 | 0.002606 |
| ENSG00000261253 | | 4.608355 | -0.8523827 | 0.8821127 | -1.700773 | 0.08898559 | 0.213887 |
| ENSG00000157388 | CACNA1D | 1342.997 | -0.8523947 | 0.3208932 | -2.906285 | 0.00365748 | 0.017129 |
| ENSG00000286912 | | 75.85089 | -0.8524253 | 0.4135098 | -2.381222 | 0.01725529 | 0.060331 |
| ENSG00000272507 | RNU6-88P | 2.142224 | -0.8525172 | 1.26216 | -1.908707 | 0.05629985 | 0.151707 |
| ENSG00000127129 | EDN2 | 264.480347 | -0.852791223 | 0.452455048 | -2.236420368 | 0.02532425 | 0.081894 |
| ENSG00000196604 | POTEF | 145.0901 | -0.8529624 | 0.5917973 | -1.925307 | 0.05419098 | 0.147611 |
| ENSG00000262089 | | 58.70601 | -0.853012 | 0.3984263 | -2.445765 | 0.01445452 | 0.05235 |

| | | | | | | | |
|---|---|---|---|---|---|---|---|
| ENSG00000145217 | SLC26A1 | 505.2135 | -0.8532077 | 0.2737411 | -3.335673 | 0.00085093 | 0.004972 |
| ENSG00000228952 | LINC02041 | 49.18578 | -0.8533061 | 0.3149258 | -2.946838 | 0.00321042 | 0.015366 |
| ENSG00000180178 | FAR2P1 | 184.6818 | -0.8533655 | 0.3210955 | -2.904695 | 0.00367611 | 0.017183 |
| ENSG00000270983 | | 27.38333 | -0.8534799 | 0.4821999 | -2.14418 | 0.03201849 | 0.098655 |
| ENSG00000124097 | HMGB1P1 | 105.6023201 | -0.853527087 | 0.364824141 | -2.627577153 | 0.00859953 | 0.034472 |
| ENSG00000129116 | PALLD | 3010.880294 | -0.853721038 | 0.300527172 | -3.083265447 | 0.00204742 | 0.010571 |
| ENSG00000105879 | CBLL1 | 1711.892094 | -0.854363573 | 0.17877583 | -4.908465204 | 9.18E-07 | 1.15E-05 |
| ENSG00000101825 | MXRA5 | 31.98850697 | -0.854431208 | 0.697486459 | -1.784116961 | 0.07440466 | 0.187096 |
| ENSG00000284874 | | 22.33939 | -0.8544709 | 0.4502518 | -2.248404 | 0.02455046 | 0.079927 |
| ENSG00000225805 | DEFB131B | 8.553796 | -0.8544861 | 0.6573992 | -1.823114 | 0.06828615 | 0.175243 |
| ENSG00000114054 | PCCB | 5091.501409 | -0.854914371 | 0.247038144 | -3.65842399 | 0.00025377 | 0.001746 |
| ENSG00000180891 | CUEDC1 | 5734.444 | -0.8553584 | 0.2994822 | -3.09602 | 0.00196137 | 0.010184 |
| ENSG00000170296 | GABARAP | 83.78195 | -0.8553795 | 0.3448304 | -2.746569 | 0.00602223 | 0.02582 |
| ENSG00000249242 | TMEM150C | 3782.312 | -0.8554388 | 0.254298 | -3.561626 | 0.00036857 | 0.002416 |
| ENSG00000005075 | POLR2J | 2940.259741 | -0.855450239 | 0.251373905 | -3.563475753 | 0.00036598 | 0.002402 |
| ENSG00000138074 | SLC5A6 | 9558.237738 | -0.855492251 | 0.169139289 | -5.197040415 | 2.02E-07 | 2.90E-06 |
| ENSG00000286619 | | 2102.944 | -0.8556033 | 0.3155846 | -2.959244 | 0.00308395 | 0.014873 |
| ENSG00000204060 | FOXO6 | 369.5896 | -0.8556551 | 0.2163236 | -4.137941 | 3.50E-05 | 0.000302 |
| ENSG00000204899 | MZT1 | 1356.493 | -0.8560737 | 0.2751315 | -3.326333 | 0.00087997 | 0.005119 |
| ENSG00000230562 | FAM133DP | 268.8894 | -0.8561849 | 0.2142908 | -4.162095 | 3.15E-05 | 0.000275 |
| ENSG00000265511 | | 69.23661 | -0.8570397 | 0.3152222 | -2.968239 | 0.00299511 | 0.014543 |
| ENSG00000172428 | COPS9 | 2628.434 | -0.857164 | 0.2621786 | -3.473314 | 0.00051407 | 0.003212 |
| ENSG00000107758 | PPP3CB | 3737.421919 | -0.857285905 | 0.093994348 | -9.192443723 | 3.84E-20 | 2.92E-18 |
| ENSG00000117151 | CTBS | 212.2328012 | -0.85731087 | 0.25442622 | -3.564709349 | 0.00036426 | 0.002393 |
| ENSG00000108590 | MED31 | 277.4896577 | -0.857439174 | 0.179815035 | -4.90286219 | 9.45E-07 | 1.18E-05 |
| ENSG00000241720 | | 268.2954 | -0.8575381 | 0.417369 | -2.376957 | 0.01745614 | 0.060929 |
| ENSG00000003989 | SLC7A2 | 6417.506238 | -0.85778318 | 0.340384908 | -2.791506254 | 0.00524633 | 0.02304 |
| ENSG00000188282 | RUFY4 | 2.576858 | -0.8578765 | 1.328781 | -1.976536 | 0.04809413 | 0.134507 |
| ENSG00000164967 | RPP25L | 1343.629 | -0.8579621 | 0.1944618 | -4.565325 | 4.99E-06 | 5.32E-05 |
| ENSG00000164037 | SLC9B1 | 27.9119 | -0.8582034 | 0.5130901 | -2.07671 | 0.03782833 | 0.11196 |
| ENSG00000237317 | | 2.446104 | -0.8588081 | 1.461155 | -2.202724 | 0.02761418 | 0.087759 |
| ENSG00000122711 | SPINK4 | 19.42017087 | -0.858996107 | 0.742119524 | -1.786322521 | 0.07404705 | 0.186479 |
| ENSG00000236449 | | 277.9404 | -0.8590468 | 0.2542851 | -3.572234 | 0.00035395 | 0.002336 |

| | | | | | | | |
|---|---|---|---|---|---|---|---|
| ENSG00000234828 | IQCM | 3.273508 | -0.8595949 | 1.198935 | -1.90061 | 0.05735317 | 0.153704 |
| ENSG00000218358 | RAET1K | 56.00158 | -0.8598503 | 0.3516268 | -2.70587 | 0.00681256 | 0.028482 |
| ENSG00000270945 | HSPE1P7 | 15.61055 | -0.8599896 | 0.5431582 | -2.007374 | 0.04470986 | 0.127414 |
| ENSG00000111666 | CHPT1 | 1993.874191 | -0.860128936 | 0.231740394 | -3.885912041 | 0.00010195 | 0.000777 |
| ENSG00000104870 | FCGRT | 91.04027305 | -0.861071215 | 0.611434358 | -1.901633705 | 0.05721906 | 0.153421 |
| ENSG00000237217 | | 1.274472 | -0.8612593 | 1.347927 | -2.066156 | 0.03881375 | NA |
| ENSG00000151208 | DLG5 | 6568.921 | -0.8617053 | 0.1978775 | -4.511385 | 6.44E-06 | 6.69E-05 |
| ENSG00000287460 | | 82.27832 | -0.8618005 | 0.3574005 | -2.687694 | 0.00719474 | 0.029826 |
| ENSG00000252494 | RNU6-126P | 3.168027 | -0.8618166 | 1.006576 | -1.731097 | 0.0834344 | 0.203654 |
| ENSG00000143493 | INTS7 | 1439.067 | -0.8622558 | 0.2338964 | -3.862711 | 0.00011214 | 0.000848 |
| ENSG00000166682 | TMPRSS5 | 17.09197 | -0.862264 | 0.5016736 | -2.110871 | 0.03478343 | 0.105138 |
| ENSG00000187260 | WDR86 | 13.56679 | -0.862863 | 0.5963515 | -1.924542 | 0.05428672 | 0.147766 |
| ENSG00000174529 | TMEM81 | 210.0393 | -0.8632455 | 0.2588827 | -3.538526 | 0.00040237 | 0.002604 |
| ENSG00000189089 | RIMKLBP1 | 10.05011 | -0.8634856 | 0.6826664 | -1.816712 | 0.06926125 | 0.177079 |
| ENSG00000282787 | | 15.60037 | -0.8636942 | 0.4952542 | -2.128421 | 0.03330223 | 0.10176 |
| ENSG00000171714 | ANO5 | 82.65791 | -0.8637301 | 0.3799533 | -2.569016 | 0.01019876 | 0.039512 |
| ENSG00000111247 | RAD51AP1 | 679.7372352 | -0.863851363 | 0.55170393 | -2.01758688 | 0.0436343 | 0.125088 |
| ENSG00000119138 | KLF9 | 660.4910526 | -0.863882303 | 0.428290313 | -2.350316297 | 0.01875746 | 0.064547 |
| ENSG00000127152 | BCL11B | 533.1480135 | -0.864549966 | 0.366079289 | -2.649210474 | 0.00806801 | 0.032756 |
| ENSG00000006634 | DBF4 | 1254.769951 | -0.864686951 | 0.375427707 | -2.583618145 | 0.009777 | 0.038161 |
| ENSG00000286753 | | 29.57005 | -0.8653866 | 0.5148048 | -2.08137 | 0.03740008 | 0.110969 |
| ENSG00000239480 | | 3.569072 | -0.8654603 | 0.9396707 | -1.687385 | 0.09152931 | 0.218253 |
| ENSG00000157240 | FZD1 | 702.1755 | -0.8656038 | 0.1968524 | -4.54991 | 5.37E-06 | 5.67E-05 |
| ENSG00000272622 | | 350.1765 | -0.8662297 | 0.4016879 | -2.470031 | 0.01351014 | 0.04951 |
| ENSG00000138185 | ENTPD1 | 762.2344105 | -0.86633339 | 0.189021677 | -4.728372249 | 2.26E-06 | 2.63E-05 |
| ENSG00000225486 | | 57.39376 | -0.8671519 | 0.2995847 | -3.129326 | 0.00175208 | 0.009262 |
| ENSG00000251596 | HADHAP1 | 22.5684 | -0.8679747 | 0.416731 | -2.406885 | 0.01608926 | 0.056982 |
| ENSG00000258759 | | 37.77276 | -0.8686223 | 0.5405335 | -2.048166 | 0.04054372 | 0.118141 |
| ENSG00000133657 | ATP13A3 | 6130.923908 | -0.868629028 | 0.257870564 | -3.581789647 | 0.00034125 | 0.002264 |
| ENSG00000183762 | KREMEN1 | 834.8953 | -0.8689134 | 0.2240737 | -4.048182 | 5.16E-05 | 0.000426 |
| ENSG00000287725 | | 64.71998 | -0.8689661 | 0.6456473 | -1.876831 | 0.06054127 | 0.16012 |
| ENSG00000177602 | HASPIN | 295.1415 | -0.8689753 | 0.4812111 | -2.20191 | 0.02767164 | 0.08787 |
| ENSG00000109771 | LRP2BP | 74.44179676 | -0.869313966 | 0.373733701 | -2.613411323 | 0.00896433 | 0.035639 |

| | | | | | | | |
|---|---|---|---|---|---|---|---|
| ENSG00000287250 | | 5.797734 | -0.8694096 | 0.7587728 | -1.758443 | 0.07867209 | 0.194731 |
| ENSG00000244932 | | 159.5497 | -0.8697522 | 0.3943211 | -2.516903 | 0.01183915 | 0.044545 |
| ENSG00000245614 | DDX11-AS1 | 75.63935 | -0.8702534 | 0.3004149 | -3.133207 | 0.00172907 | 0.009153 |
| ENSG00000230937 | MIR205HG | 80.25316 | -0.8705492 | 0.4240656 | -2.38718 | 0.01697818 | 0.059558 |
| ENSG00000234715 | | 1.394327 | -0.8705567 | 1.442516 | -2.200833 | 0.02774783 | NA |
| ENSG00000261292 | | 6.710918 | -0.8707743 | 0.7121143 | -1.807875 | 0.07062603 | 0.179839 |
| ENSG00000169884 | WNT10B | 145.7624 | -0.871544 | 0.3702614 | -2.650656 | 0.00803357 | 0.032642 |
| ENSG00000246174 | KCTD21-AS1 | 98.30453 | -0.8715711 | 0.3483175 | -2.775649 | 0.00550916 | 0.02398 |
| ENSG00000273148 | LINC00653 | 144.0017 | -0.871754 | 0.2337805 | -3.914972 | 9.04E-05 | 0.000699 |
| ENSG00000272599 | | 58.91924 | -0.8720278 | 0.3750322 | -2.615443 | 0.00891118 | 0.035472 |
| ENSG00000251794 | RNU6-237P | 1.1773 | -0.8727965 | 1.257047 | -1.942326 | 0.05209762 | NA |
| ENSG00000138080 | EMILIN1 | 116.8855988 | -0.873725991 | 0.203993662 | -4.444989673 | 8.79E-06 | 8.78E-05 |
| ENSG00000105887 | MTPN | 9620.972043 | -0.873752743 | 0.141240111 | -6.307158737 | 2.84E-10 | 6.65E-09 |
| ENSG00000223722 | IFITM3P2 | 90.73245 | -0.8738349 | 0.3200892 | -2.982234 | 0.00286153 | 0.014032 |
| ENSG00000186298 | PPP1CC | 8396.586 | -0.8740716 | 0.2079751 | -4.366198 | 1.26E-05 | 0.000121 |
| ENSG00000272791 | | 61.80998 | -0.8740855 | 0.3580326 | -2.7189 | 0.00654994 | 0.027597 |
| ENSG00000156313 | RPGR | 435.8477 | -0.8741716 | 0.1828996 | -4.930787 | 8.19E-07 | 1.04E-05 |
| ENSG00000222041 | CYTOR | 866.7098 | -0.8743957 | 0.2579951 | -3.59433 | 0.00032523 | 0.002172 |
| ENSG00000272906 | | 24.39927 | -0.8743961 | 0.4327481 | -2.353728 | 0.0185862 | 0.064032 |
| ENSG00000127957 | PMS2P3 | 290.3156171 | -0.87446953 | 0.212055001 | -4.290851502 | 1.78E-05 | 0.000165 |
| ENSG00000065427 | KARS1 | 13636.90232 | -0.874516381 | 0.160960304 | -5.581892891 | 2.38E-08 | 4.04E-07 |
| ENSG00000185722 | ANKFY1 | 5001.22 | -0.8745485 | 0.1828998 | -4.928444 | 8.29E-07 | 1.05E-05 |
| ENSG00000286416 | | 33.71561 | -0.874946 | 0.5736766 | -1.991373 | 0.04643989 | 0.131011 |
| ENSG00000107566 | ERLIN1 | 2292.089931 | -0.875473076 | 0.12924237 | -6.874834701 | 6.21E-12 | 1.83E-10 |
| ENSG00000260589 | STAM-AS1 | 48.4709 | -0.8761308 | 0.6117084 | -1.939432 | 0.05244877 | 0.143866 |
| ENSG00000164603 | BMT2 | 579.399 | -0.8761428 | 0.220276 | -4.15533 | 3.25E-05 | 0.000282 |
| ENSG00000106144 | CASP2 | 2610.712224 | -0.87624997 | 0.224342566 | -4.077343048 | 4.56E-05 | 0.000382 |
| ENSG00000188512 | | 9.729333 | -0.8769896 | 0.6960588 | -1.826881 | 0.06771766 | 0.174171 |
| ENSG00000124635 | H2BC11 | 274.203588 | -0.877249724 | 0.573358295 | -2.005932172 | 0.04486349 | 0.127744 |
| ENSG00000103375 | AQP8 | 1.543171292 | -0.87789136 | 1.139107273 | -1.802596517 | 0.0714516 | NA |
| ENSG00000230299 | SNRPEP11 | 9.77016 | -0.8783796 | 0.5408475 | -2.05055 | 0.04031082 | 0.117591 |
| ENSG00000103126 | AXIN1 | 3930.675545 | -0.878940852 | 0.117650982 | -7.563477773 | 3.92E-14 | 1.52E-12 |
| ENSG00000255836 | | 191.6969 | -0.8789523 | 0.281455 | -3.339954 | 0.00083792 | 0.00491 |

| | | | | | | | |
|---|---|---|---|---|---|---|---|
| ENSG00000215126 | CBWD6 | 396.8998 | -0.8789923 | 0.2115183 | -4.323755 | 1.53E-05 | 0.000144 |
| ENSG00000091136 | LAMB1 | 5914.024155 | -0.880003683 | 0.384483927 | -2.59778258 | 0.00938279 | 0.036927 |
| ENSG00000229728 | | 28.98078 | -0.8801253 | 0.488014 | -2.190286 | 0.02850351 | 0.090017 |
| ENSG00000285771 | | 2.359761 | -0.8804693 | 1.189907 | -1.822131 | 0.06843517 | 0.175507 |
| ENSG00000137274 | BPHL | 795.5928582 | -0.881307831 | 0.1656399 | -5.452620488 | 4.96E-08 | 8.00E-07 |
| ENSG00000236703 | MYB-AS1 | 4.488874 | -0.8815104 | 0.9452709 | -1.791737 | 0.07317513 | 0.184762 |
| ENSG00000145757 | SPATA9 | 31.80607 | -0.8820869 | 0.3351766 | -2.892336 | 0.00382389 | 0.017765 |
| ENSG00000205213 | LGR4 | 2079.73 | -0.8823887 | 0.228553 | -4.036547 | 5.42E-05 | 0.000445 |
| ENSG00000204677 | FAM153CP | 335.9105 | -0.8825656 | 0.4150715 | -2.463435 | 0.0137613 | 0.050251 |
| ENSG00000261488 | TBILA | 121.0086 | -0.882636 | 0.6046809 | -1.959051 | 0.05010682 | 0.138705 |
| ENSG00000285812 | | 18.9464 | -0.8827062 | 0.5619216 | -2.021958 | 0.04318066 | 0.124146 |
| ENSG00000139146 | SINHCAF | 1993.683455 | -0.883537364 | 0.205404582 | -4.464543055 | 8.02E-06 | 8.12E-05 |
| ENSG00000276627 | | 2.08195 | -0.8837005 | 1.359695 | -1.998553 | 0.04565676 | 0.129454 |
| ENSG00000073711 | PPP2R3A | 1053.088256 | -0.883722911 | 0.155407412 | -5.787741082 | 7.13E-09 | 1.34E-07 |
| ENSG00000284292 | | 1.724397 | -0.8839403 | 1.376678 | -2.024232 | 0.04294628 | 0.123637 |
| ENSG00000238151 | MLLT10P1 | 7.848135 | -0.8841691 | 0.6540577 | -1.881305 | 0.05993039 | 0.158911 |
| ENSG00000226026 | | 8.295466 | -0.8845016 | 0.6957474 | -1.820419 | 0.06869523 | 0.17597 |
| ENSG00000267249 | | 16.73894 | -0.8845272 | 0.4866307 | -2.196492 | 0.02805672 | 0.088838 |
| ENSG00000144199 | FAHD2B | 342.21 | -0.884696 | 0.1935718 | -4.720221 | 2.36E-06 | 2.73E-05 |
| ENSG00000198839 | ZNF277 | 1015.813 | -0.8849172 | 0.2282704 | -4.060657 | 4.89E-05 | 0.000407 |
| ENSG00000214022 | REPIN1 | 11490.49 | -0.8849234 | 0.1552244 | -5.829611 | 5.56E-09 | 1.06E-07 |
| ENSG00000185875 | THNSL1 | 1224.388 | -0.8853402 | 0.2278702 | -4.07169 | 4.67E-05 | 0.00039 |
| ENSG00000137809 | ITGA11 | 3.81711324 | -0.885348643 | 0.761754123 | -1.787447727 | 0.07386515 | 0.186145 |
| ENSG00000107242 | PIP5K1B | 154.4425461 | -0.885593702 | 0.411940366 | -2.468260833 | 0.01357714 | 0.049701 |
| ENSG00000268362 | | 541.8243 | -0.8855968 | 0.227396 | -4.066556 | 4.77E-05 | 0.000398 |
| ENSG00000184809 | B3GALT5-AS1 | 311.8529 | -0.8857269 | 0.5039827 | -2.180332 | 0.02923285 | 0.091937 |
| ENSG00000212533 | | 4.82369 | -0.8857793 | 0.7410318 | -1.809335 | 0.07039892 | 0.179418 |
| ENSG00000271833 | | 20.65714 | -0.8860389 | 0.5008025 | -2.168447 | 0.03012472 | 0.093975 |
| ENSG00000242159 | ABCF2P1 | 13.52654 | -0.8866295 | 0.6677632 | -1.881687 | 0.05987854 | 0.158821 |
| ENSG00000165376 | CLDN2 | 2.422189 | -0.8868227 | 1.030444 | -1.743124 | 0.08131199 | 0.199838 |
| ENSG00000141664 | ZCCHC2 | 1626.061 | -0.887067 | 0.1971633 | -4.65818 | 3.19E-06 | 3.57E-05 |
| ENSG00000011677 | GABRA3 | 318.9438276 | -0.887316461 | 0.295023386 | -3.239268589 | 0.00119837 | 0.006683 |
| ENSG00000169016 | E2F6 | 1556.478 | -0.8880446 | 0.1278895 | -7.056828 | 1.70E-12 | 5.39E-11 |

| | | | | | | | |
|---|---|---|---|---|---|---|---|
| ENSG00000152778 | IFIT5 | 2038.367 | -0.8888706 | 0.2388827 | -3.909061 | 9.27E-05 | 0.000715 |
| ENSG00000152409 | JMY | 3034.021 | -0.8890478 | 0.2875977 | -3.323366 | 0.00088938 | 0.005166 |
| ENSG00000273297 | | 5.708817 | -0.8895735 | 0.9141819 | -1.767936 | 0.07707164 | 0.19186 |
| ENSG00000166508 | MCM7 | 11800.25 | -0.890008 | 0.4189441 | -2.458447 | 0.01395395 | 0.050842 |
| ENSG00000167522 | ANKRD11 | 10939.97 | -0.8900422 | 0.2605333 | -3.623727 | 0.00029039 | 0.001966 |
| ENSG00000162129 | CLPB | 1565.388 | -0.8903058 | 0.321554 | -3.031726 | 0.0024316 | 0.012206 |
| ENSG00000134330 | IAH1 | 2520.09646 | -0.890359156 | 0.175335117 | -5.223349804 | 1.76E-07 | 2.54E-06 |
| ENSG00000160917 | CPSF4 | 2480.915 | -0.890692 | 0.1834168 | -5.010495 | 5.43E-07 | 7.17E-06 |
| ENSG00000165905 | LARGE2 | 1267.468 | -0.8909951 | 0.2970839 | -3.233525 | 0.00122273 | 0.006806 |
| ENSG00000146776 | ATXN7L1 | 535.7953 | -0.8910541 | 0.21754 | -4.263028 | 2.02E-05 | 0.000184 |
| ENSG00000163029 | SMC6 | 2949.622 | -0.8916655 | 0.171879 | -5.333004 | 9.66E-08 | 1.47E-06 |
| ENSG00000099204 | ABLIM1 | 1232.197542 | -0.892522984 | 0.363032219 | -2.75542481 | 0.0058616 | 0.025257 |
| ENSG00000141665 | FBXO15 | 139.5358 | -0.8925756 | 0.3344456 | -2.933926 | 0.00334704 | 0.015919 |
| ENSG00000143318 | CASQ1 | 2.10421 | -0.8929233 | 1.459084 | -2.119481 | 0.0340498 | 0.103474 |
| ENSG00000261842 | | 1.821612 | -0.8932924 | 1.145326 | -1.782893 | 0.07460367 | 0.187348 |
| ENSG00000287759 | | 6.60863 | -0.8933339 | 0.9194756 | -1.759038 | 0.07857109 | 0.194555 |
| ENSG00000138346 | DNA2 | 1115.366952 | -0.893341347 | 0.391102935 | -2.611463987 | 0.00901555 | 0.035816 |
| ENSG00000272056 | | 3.387896 | -0.8942974 | 0.9839072 | -1.75429 | 0.07938087 | 0.196138 |
| ENSG00000120594 | PLXDC2 | 1951.944104 | -0.894703887 | 0.368919752 | -2.72414372 | 0.00644685 | 0.027236 |
| ENSG00000145246 | ATP10D | 370.2392 | -0.8949172 | 0.2886339 | -3.342243 | 0.00083104 | 0.004874 |
| ENSG00000154839 | SKA1 | 409.6963 | -0.8951862 | 0.5219836 | -2.147356 | 0.03176494 | 0.098078 |
| ENSG00000135632 | SMYD5 | 2930.013177 | -0.895340968 | 0.175838377 | -5.227744135 | 1.72E-07 | 2.49E-06 |
| ENSG00000250802 | ZBED3-AS1 | 219.3986 | -0.8954648 | 0.387306 | -2.622582 | 0.00872663 | 0.034892 |
| ENSG00000286652 | | 8.499177 | -0.8956585 | 0.7999317 | -1.79773 | 0.07221977 | 0.182813 |
| ENSG00000060138 | YBX3 | 6589.976326 | -0.896017668 | 0.149409774 | -6.118781396 | 9.43E-10 | 2.03E-08 |
| ENSG00000134202 | GSTM3 | 14465.50783 | -0.896032359 | 0.248367352 | -3.806426918 | 0.00014099 | 0.001038 |
| ENSG00000232000 | CLCN3P1 | 2.476551 | -0.8960659 | 1.191493 | -1.8004 | 0.07179745 | 0.182126 |
| ENSG00000265018 | AGAP12P | 10.08872 | -0.8961259 | 0.6045473 | -1.97852 | 0.0478701 | 0.134007 |
| ENSG00000277157 | H4C4 | 26.96179 | -0.8967904 | 0.4139371 | -2.491187 | 0.01273171 | 0.047231 |
| ENSG00000150867 | PIP4K2A | 2191.147 | -0.8968974 | 0.2030469 | -4.581387 | 4.62E-06 | 4.97E-05 |
| ENSG00000146701 | MDH2 | 11406.56 | -0.8976308 | 0.207407 | -4.514869 | 6.34E-06 | 6.59E-05 |
| ENSG00000065600 | PACC1 | 634.5700247 | -0.897642 | 0.169357285 | -5.428035906 | 5.70E-08 | 9.09E-07 |
| ENSG00000151353 | TMEM18 | 3182.276 | -0.897703 | 0.1272861 | -7.176901 | 7.13E-13 | 2.37E-11 |

| | | | | | | | |
|---|---|---|---|---|---|---|---|
| ENSG00000164104 | HMGB2 | 10364.14 | -0.8978215 | 0.3936393 | -2.600739 | 0.00930233 | 0.036697 |
| ENSG00000224117 | PTPN2P2 | 5.936581 | -0.8981221 | 0.7484049 | -1.820473 | 0.06868704 | 0.175966 |
| ENSG00000141013 | GAS8 | 2394.126 | -0.8987393 | 0.1231156 | -7.394664 | 1.42E-13 | 5.15E-12 |
| ENSG00000229591 | | 8.909731 | -0.8989174 | 0.6504667 | -1.908773 | 0.05629134 | 0.151699 |
| ENSG00000258857 | | 2.332532 | -0.8991306 | 1.086936 | -1.778572 | 0.07530991 | 0.18868 |
| ENSG00000260097 | SPDYE6 | 60.88177 | -0.8991823 | 0.298903 | -3.241128 | 0.00119058 | 0.006643 |
| ENSG00000080823 | MOK | 695.4231291 | -0.899897989 | 0.188732635 | -4.922142842 | 8.56E-07 | 1.08E-05 |
| ENSG00000197956 | S100A6 | 7598.01 | -0.9001182 | 0.3299674 | -2.994538 | 0.00274861 | 0.013554 |
| ENSG00000050438 | SLC4A8 | 877.4719621 | -0.900139684 | 0.354921668 | -2.82729266 | 0.00469434 | 0.021068 |
| ENSG00000236307 | EEF1E1P1 | 17.28863 | -0.9003759 | 0.5437907 | -2.092789 | 0.03636797 | 0.108713 |
| ENSG00000217275 | | 121.3973 | -0.9006637 | 0.3889717 | -2.626485 | 0.00862718 | 0.034562 |
| ENSG00000259033 | | 28.66595 | -0.9008275 | 0.4040393 | -2.555618 | 0.01059994 | 0.040816 |
| ENSG00000167656 | LY6D | 7.352788 | -0.9016998 | 0.8066891 | -1.813393 | 0.06977129 | 0.178126 |
| ENSG00000267432 | DNAH17-AS1 | 67.14623 | -0.9017108 | 0.3694962 | -2.735353 | 0.00623134 | 0.026524 |
| ENSG00000272950 | | 2.056321 | -0.9018064 | 1.326683 | -1.97585 | 0.0481718 | 0.134668 |
| ENSG00000236567 | TCF3P1 | 1.646454 | -0.902239 | 1.243122 | -1.860783 | 0.06277476 | 0.164467 |
| ENSG00000223459 | TCAF1P1 | 162.7232 | -0.9023621 | 0.293868 | -3.314459 | 0.00091821 | 0.005302 |
| ENSG00000151025 | GPR158 | 319.1189 | -0.9023896 | 0.3102704 | -3.152637 | 0.00161803 | 0.008639 |
| ENSG00000168350 | DEGS2 | 1229.188 | -0.9024582 | 0.3663385 | -2.771059 | 0.00558743 | 0.024269 |
| ENSG00000263648 | | 1.547495 | -0.9024623 | 1.263451 | -1.948207 | 0.05139024 | NA |
| ENSG00000235036 | | 212.0996 | -0.9028057 | 0.4358983 | -2.430405 | 0.01508194 | 0.05415 |
| ENSG00000216368 | | 2.291212 | -0.9028578 | 1.137049 | -1.758997 | 0.07857796 | 0.194555 |
| ENSG00000228444 | | 11.22645 | -0.9028935 | 0.7323308 | -1.834392 | 0.06659578 | 0.171985 |
| ENSG00000272370 | | 10.58848 | -0.9032166 | 0.6976282 | -1.874913 | 0.06080474 | 0.160573 |
| ENSG00000107614 | TRDMT1 | 220.631998 | -0.903443514 | 0.194474356 | -4.790348699 | 1.66E-06 | 1.99E-05 |
| ENSG00000256806 | C17orf100 | 281.6294 | -0.9035526 | 0.2340803 | -4.037562 | 5.40E-05 | 0.000444 |
| ENSG00000230002 | ALMS1-IT1 | 40.5398 | -0.9035854 | 0.5531325 | -2.092902 | 0.03635787 | 0.108713 |
| ENSG00000231686 | ANKRD11P2 | 289.7476 | -0.9037527 | 0.260168 | -3.679686 | 0.00023352 | 0.001626 |
| ENSG00000127418 | FGFRL1 | 2418.528766 | -0.904061167 | 0.189259506 | -4.930561165 | 8.20E-07 | 1.04E-05 |
| ENSG00000234111 | LINC02433 | 2.068847 | -0.905425 | 1.566407 | -2.268015 | 0.02332827 | 0.076814 |
| ENSG00000120694 | HSPH1 | 15053.2329 | -0.90547093 | 0.253568101 | -3.660664531 | 0.00025156 | 0.001732 |
| ENSG00000128524 | ATP6V1F | 3356.804913 | -0.906053179 | 0.21240695 | -4.434754319 | 9.22E-06 | 9.16E-05 |
| ENSG00000136159 | NUDT15 | 1039.46516 | -0.90605673 | 0.4198932 | -2.502431209 | 0.01233436 | 0.04602 |

| | | | | | | | |
|---|---|---|---|---|---|---|---|
| ENSG00000179546 | HTR1D | 73.88172 | -0.9066323 | 0.7462517 | -1.786371 | 0.07403925 | 0.186477 |
| ENSG00000226415 | TPI1P1 | 4667.126 | -0.9066668 | 0.2728203 | -3.543072 | 0.00039549 | 0.002568 |
| ENSG00000287236 | | 20.24771 | -0.9067036 | 0.5684453 | -2.069827 | 0.03846856 | 0.113431 |
| ENSG00000225338 | RPL23AP18 | 11.29985 | -0.907382 | 0.6996487 | -1.887482 | 0.05909553 | 0.157357 |
| ENSG00000196405 | EVL | 6476.888 | -0.9078657 | 0.3402804 | -2.946253 | 0.00321649 | 0.015392 |
| ENSG00000265750 | | 5.955438 | -0.907906 | 0.7920506 | -1.816898 | 0.0692327 | 0.177023 |
| ENSG00000174353 | STAG3L3 | 82.03881 | -0.9081989 | 0.3026738 | -3.241854 | 0.00118755 | 0.006627 |
| ENSG00000102531 | FNDC3A | 2367.412923 | -0.908200562 | 0.245670225 | -3.893078982 | 9.90E-05 | 0.000759 |
| ENSG00000074695 | LMAN1 | 6022.044978 | -0.908558686 | 0.177011591 | -5.273139536 | 1.34E-07 | 1.99E-06 |
| ENSG00000213864 | EEF1B2P2 | 12.81149 | -0.9086226 | 0.5241214 | -2.150265 | 0.03153427 | 0.097525 |
| ENSG00000130600 | H19 | 4602.841825 | -0.908846623 | 1.443818532 | -2.172654925 | NA | NA |
| ENSG00000187372 | PCDHB13 | 128.8149 | -0.9091119 | 0.4578699 | -2.356629 | 0.01844165 | 0.063642 |
| ENSG00000229808 | | 7.986052 | -0.9096611 | 0.780663 | -1.832282 | 0.06690944 | 0.172544 |
| ENSG00000111319 | SCNN1A | 2346.020316 | -0.909856721 | 0.418773522 | -2.517107646 | 0.01183227 | 0.044529 |
| ENSG00000111291 | GPRC5D | 30.94229452 | -0.909976596 | 0.450207262 | -2.385832125 | 0.01704053 | 0.059714 |
| ENSG00000137135 | ARHGEF39 | 378.9372266 | -0.910208543 | 0.474444254 | -2.310295973 | 0.02087177 | 0.070347 |
| ENSG00000270799 | | 1.69363 | -0.9103399 | 1.284554 | -1.908991 | 0.05626329 | 0.151639 |
| ENSG00000179344 | HLA-DQB1 | 320.5359 | -0.9107019 | 0.3970698 | -2.610643 | 0.00903723 | 0.035869 |
| ENSG00000130159 | ECSIT | 1352.209293 | -0.910775247 | 0.200485007 | -4.669433044 | 3.02E-06 | 3.40E-05 |
| ENSG00000049449 | RCN1 | 3255.663747 | -0.910817045 | 0.204828505 | -4.615161639 | 3.93E-06 | 4.29E-05 |
| ENSG00000129673 | AANAT | 15.37068225 | -0.910923156 | 0.644441901 | -1.960456353 | 0.04994248 | 0.138351 |
| ENSG00000182704 | TSKU | 4870.465 | -0.910925 | 0.3698412 | -2.767544 | 0.00564804 | 0.024494 |
| ENSG00000181104 | F2R | 90.35368 | -0.9111009 | 0.476272 | -2.304855 | 0.02117469 | 0.071169 |
| ENSG00000082515 | MRPL22 | 1488.998964 | -0.91122795 | 0.293581146 | -3.343747647 | 0.00082655 | 0.004854 |
| ENSG00000157800 | SLC37A3 | 2184.455 | -0.9119747 | 0.1621823 | -5.756733 | 8.58E-09 | 1.60E-07 |
| ENSG00000222448 | RN7SKP150 | 15.19871 | -0.9120749 | 0.520378 | -2.173842 | 0.02971699 | 0.093064 |
| ENSG00000287676 | | 9.09568 | -0.9128033 | 0.8166842 | -1.81733 | 0.06916655 | 0.176973 |
| ENSG00000247199 | | 9.602354 | -0.913029 | 0.6896222 | -1.892062 | 0.05848272 | 0.156164 |
| ENSG00000243789 | JMJD7 | 22.5499 | -0.9130295 | 0.5260754 | -2.167774 | 0.03017589 | 0.09409 |
| ENSG00000152422 | XRCC4 | 462.0294 | -0.9134496 | 0.2545597 | -3.805231 | 0.00014167 | 0.001042 |
| ENSG00000269765 | | 7.362131 | -0.914122 | 0.7737698 | -1.824924 | 0.06801245 | 0.174727 |
| ENSG00000225891 | DHDDS-AS1 | 10.78756 | -0.9142144 | 0.6367424 | -1.963381 | 0.04960198 | 0.13768 |
| ENSG00000198879 | SFMBT2 | 1046.666 | -0.9142583 | 0.2979077 | -3.322536 | 0.00089203 | 0.005178 |

| | | | | | | | |
|---|---|---|---|---|---|---|---|
| ENSG00000182742 | HOXB4 | 1.840127 | -0.9143268 | 1.371227 | -2.023943 | 0.042976 | 0.123679 |
| ENSG00000132846 | ZBED3 | 927.4846022 | -0.914378846 | 0.315683053 | -3.153870808 | 0.0016112 | 0.008607 |
| ENSG00000275601 | | 25.9161 | -0.9150483 | 0.42482 | -2.494991 | 0.01259604 | 0.046872 |
| ENSG00000228503 | | 79.35894 | -0.9152554 | 0.3246485 | -3.080122 | 0.00206916 | 0.010669 |
| ENSG00000107165 | TYRP1 | 575.2470064 | -0.915781743 | 0.195217237 | -4.846918492 | 1.25E-06 | 1.53E-05 |
| ENSG00000164253 | WDR41 | 2432.418 | -0.9158054 | 0.1581149 | -5.937609 | 2.89E-09 | 5.79E-08 |
| ENSG00000160193 | WDR4 | 1402.548 | -0.9162782 | 0.1556145 | -6.036844 | 1.57E-09 | 3.27E-08 |
| ENSG00000213614 | HEXA | 2379.279 | -0.9163904 | 0.2099535 | -4.53552 | 5.75E-06 | 6.03E-05 |
| ENSG00000285437 | POLR2J3 | 68.35487 | -0.9165501 | 0.3949326 | -2.637573 | 0.00835017 | 0.033666 |
| ENSG00000078043 | PIAS2 | 1189.84837 | -0.916877346 | 0.133938087 | -6.94010788 | 3.92E-12 | 1.20E-10 |
| ENSG00000172164 | SNTB1 | 492.8228 | -0.9172309 | 0.3608754 | -2.831518 | 0.00463277 | 0.020835 |
| ENSG00000228360 | | 2.862428 | -0.9173456 | 1.094025 | -1.77643 | 0.07566203 | 0.189326 |
| ENSG00000152430 | BOLL | 10.59589 | -0.917475 | 0.6782788 | -1.916568 | 0.0552928 | 0.149661 |
| ENSG00000258702 | | 3.333669 | -0.917496 | 0.9669012 | -1.77624 | 0.07569331 | 0.189387 |
| ENSG00000243697 | | 5.89528 | -0.9175989 | 0.9688002 | -1.807783 | 0.07064033 | 0.179843 |
| ENSG00000286389 | | 90.35995 | -0.9184751 | 0.3624345 | -2.827118 | 0.00469691 | 0.021076 |
| ENSG00000132972 | RNF17 | 2.892447592 | -0.919017021 | 1.002976639 | -1.774873841 | 0.07591872 | 0.189721 |
| ENSG00000179715 | PCED1B | 181.853 | -0.9193235 | 0.4056679 | -2.602808 | 0.00924637 | 0.036514 |
| ENSG00000111203 | ITFG2 | 1057.14943 | -0.919399786 | 0.164813661 | -5.71026816 | 1.13E-08 | 2.05E-07 |
| ENSG00000105929 | ATP6V0A4 | 215.0426174 | -0.919416894 | 0.781767579 | -1.852488837 | 0.06395566 | 0.166641 |
| ENSG00000135697 | BCO1 | 13.64513484 | -0.919559613 | 0.697231534 | -1.896409565 | 0.05790591 | 0.15495 |
| ENSG00000226329 | | 1.976051 | -0.9195975 | 1.133064 | -1.757052 | 0.07890892 | 0.19519 |
| ENSG00000275709 | | 2.79451 | -0.9197976 | 1.081252 | -1.766321 | 0.07734204 | 0.192316 |
| ENSG00000136636 | KCTD3 | 11409.68641 | -0.919960477 | 0.204905911 | -4.649286039 | 3.33E-06 | 3.71E-05 |
| ENSG00000224066 | | 10.02377 | -0.9199775 | 0.5837207 | -2.055963 | 0.03978612 | 0.116404 |
| ENSG00000272905 | | 14.28686 | -0.9201892 | 0.6194831 | -1.996706 | 0.04585709 | 0.129822 |
| ENSG00000156876 | SASS6 | 514.0311 | -0.9206849 | 0.3472937 | -2.938063 | 0.0033027 | 0.01574 |
| ENSG00000132000 | PODNL1 | 49.23856788 | -0.920717736 | 0.386472831 | -2.692011724 | 0.00710225 | 0.029507 |
| ENSG00000174871 | CNIH2 | 263.9197 | -0.9216281 | 0.7247532 | -1.923543 | 0.05441188 | 0.148039 |
| ENSG00000255160 | | 7.361962 | -0.9218392 | 0.6538762 | -1.948078 | 0.05140565 | 0.141517 |
| ENSG00000261135 | | 7.472217 | -0.9223086 | 0.6815855 | -1.9126 | 0.05579928 | 0.150702 |
| ENSG00000145911 | N4BP3 | 7461.764 | -0.9228357 | 0.2313731 | -4.177226 | 2.95E-05 | 0.000259 |
| ENSG00000144677 | CTDSPL | 816.6293 | -0.9228784 | 0.1737356 | -5.459702 | 4.77E-08 | 7.71E-07 |

| | | | | | | | |
|---|---|---|---|---|---|---|---|
| ENSG00000164105 | SAP30 | 726.3788 | -0.9229859 | 0.2279493 | -4.236815 | 2.27E-05 | 0.000205 |
| ENSG00000197753 | LHFPL5 | 244.9014 | -0.923751 | 0.3185447 | -3.163102 | 0.00156097 | 0.008389 |
| ENSG00000234645 | YWHAEP5 | 24.04935 | -0.9240407 | 0.4221008 | -2.539135 | 0.01111268 | 0.042378 |
| ENSG00000026559 | KCNG1 | 627.5187546 | -0.925237381 | 0.35494436 | -2.900127949 | 0.0037301 | 0.017405 |
| ENSG00000281026 | N4BP2L2-IT2 | 180.3747 | -0.9255231 | 0.373433 | -2.787363 | 0.00531389 | 0.023271 |
| ENSG00000185040 | SPDYE16 | 4.504544 | -0.925745 | 1.090021 | -1.728361 | 0.0839236 | 0.204603 |
| ENSG00000157350 | ST3GAL2 | 406.391 | -0.9257947 | 0.2504173 | -3.903358 | 9.49E-05 | 0.000731 |
| ENSG00000082212 | ME2 | 1527.789834 | -0.925852177 | 0.184814476 | -5.166458387 | 2.39E-07 | 3.37E-06 |
| ENSG00000249084 | | 3.806186 | -0.9260503 | 1.273 | -1.948821 | 0.05131684 | 0.141333 |
| ENSG00000244676 | | 6.302916 | -0.9262655 | 1.008184 | -1.793259 | 0.07293155 | 0.184264 |
| ENSG00000285918 | | 3.853078 | -0.9266508 | 0.9115904 | -1.782757 | 0.0746258 | 0.187386 |
| ENSG00000188153 | COL4A5 | 1986.766 | -0.9267569 | 0.5121512 | -2.257417 | 0.02398202 | 0.078509 |
| ENSG00000143870 | PDIA6 | 10828.31 | -0.9267868 | 0.1406395 | -6.727227 | 1.73E-11 | 4.79E-10 |
| ENSG00000207088 | SNORA7B | 12.35062 | -0.9270875 | 0.6511101 | -1.966319 | 0.04926178 | 0.137008 |
| ENSG00000229536 | LINC02572 | 10.13845 | -0.9271007 | 0.6592481 | -1.957531 | 0.05028508 | 0.139053 |
| ENSG00000286169 | NA | 426.912 | -0.9278319 | 0.2276866 | -4.275063 | 1.91E-05 | 0.000176 |
| ENSG00000271335 | | 124.8594 | -0.9279505 | 0.3985596 | -2.647543 | 0.00810789 | 0.032883 |
| ENSG00000267270 | PARD6G-AS1 | 40.46627 | -0.9284442 | 0.4295501 | -2.514515 | 0.01191963 | 0.044794 |
| ENSG00000064490 | RFXANK | 1358.160244 | -0.92868734 | 0.196175196 | -4.896893619 | 9.74E-07 | 1.21E-05 |
| ENSG00000072571 | HMMR | 1521.098061 | -0.928835679 | 0.469376987 | -2.371888327 | 0.01769744 | 0.061553 |
| ENSG00000158246 | TENT5B | 251.2676 | -0.9289932 | 0.3449143 | -2.978445 | 0.00289716 | 0.014181 |
| ENSG00000174606 | ANGEL2 | 1437.098 | -0.9293515 | 0.1744891 | -5.472029 | 4.45E-08 | 7.23E-07 |
| ENSG00000197872 | CYRIA | 186.6339 | -0.9294036 | 0.4305974 | -2.510614 | 0.01205214 | 0.045177 |
| ENSG00000162599 | NFIA | 583.9823 | -0.9294245 | 0.3162248 | -3.20082 | 0.00137037 | 0.007513 |
| ENSG00000180616 | SSTR2 | 22.31895 | -0.9296751 | 0.7670127 | -1.88061 | 0.06002503 | 0.15913 |
| ENSG00000075142 | SRI | 2054.329994 | -0.929893832 | 0.193835998 | -4.95433388 | 7.26E-07 | 9.32E-06 |
| ENSG00000106397 | PLOD3 | 5069.672788 | -0.929983775 | 0.123489473 | -7.611986185 | 2.70E-14 | 1.07E-12 |
| ENSG00000198440 | ZNF583 | 54.82319 | -0.9300402 | 0.2848844 | -3.491563 | 0.0004802 | 0.003031 |
| ENSG00000070985 | TRPM5 | 1.562144351 | -0.930172758 | 1.31340507 | -1.920588723 | 0.05478358 | NA |
| ENSG00000279742 | | 58.66822 | -0.9303962 | 0.4649801 | -2.380144 | 0.01730589 | 0.060476 |
| ENSG00000110880 | CORO1C | 4818.160454 | -0.931244888 | 0.203512651 | -4.746155827 | 2.07E-06 | 2.43E-05 |
| ENSG00000172264 | MACROD2 | 765.341 | -0.931899 | 0.2405674 | -4.068114 | 4.74E-05 | 0.000395 |
| ENSG00000162924 | REL | 669.0503 | -0.9320319 | 0.5166595 | -2.244368 | 0.02480877 | 0.0806 |

| | | | | | | | |
|---|---|---|---|---|---|---|---|
| ENSG00000140479 | PCSK6 | 2212.766 | -0.9325131 | 0.2390292 | -4.096791 | 4.19E-05 | 0.000354 |
| ENSG00000226742 | HSBP1L1 | 1601.228 | -0.9330736 | 0.2270627 | -4.29752 | 1.73E-05 | 0.000161 |
| ENSG00000065911 | MTHFD2 | 12099.74934 | -0.933318909 | 0.306845652 | -3.295579136 | 0.00098219 | 0.005637 |
| ENSG00000167515 | TRAPPC2L | 5147.087 | -0.9334391 | 0.1683712 | -5.685948 | 1.30E-08 | 2.33E-07 |
| ENSG00000251661 | | 50.25801 | -0.9336553 | 0.3407231 | -3.005382 | 0.00265248 | 0.013143 |
| ENSG00000182993 | C12orf60 | 24.19666 | -0.93408 | 0.4665854 | -2.38751 | 0.01696295 | 0.05952 |
| ENSG00000121440 | PDZRN3 | 603.9765898 | -0.934194345 | 0.255186438 | -3.879799638 | 0.00010454 | 0.000796 |
| ENSG00000106785 | TRIM14 | 2582.057075 | -0.934500085 | 0.241039603 | -4.09070449 | 4.30E-05 | 0.000363 |
| ENSG00000130675 | MNX1 | 599.3741043 | -0.934714751 | 0.160398094 | -5.957987205 | 2.55E-09 | 5.17E-08 |
| ENSG00000136100 | VPS36 | 2376.220877 | -0.934724173 | 0.165605637 | -5.781609378 | 7.40E-09 | 1.39E-07 |
| ENSG00000102970 | CCL17 | 2.410627182 | -0.934814754 | 1.302855659 | -1.950353819 | 0.05113396 | 0.140944 |
| ENSG00000165282 | PIGO | 2605.822 | -0.9348698 | 0.1464325 | -6.503873 | 7.83E-11 | 1.97E-09 |
| ENSG00000275328 | SPDYE19P | 7.956955 | -0.9359688 | 0.7484271 | -1.879793 | 0.06013633 | 0.159306 |
| ENSG00000233461 | | 96.10106 | -0.9361709 | 0.2396327 | -4.117338 | 3.83E-05 | 0.000328 |
| ENSG00000230359 | TPI1P2 | 58.76652 | -0.9364921 | 0.3305991 | -3.107211 | 0.00188862 | 0.009862 |
| ENSG00000184220 | CMSS1 | 2036.711 | -0.9368391 | 0.2334328 | -4.199401 | 2.68E-05 | 0.000237 |
| ENSG00000086570 | FAT2 | 14.47368041 | -0.937041018 | 0.764233344 | -1.890753056 | 0.05865732 | 0.156457 |
| ENSG00000214248 | | 16.31148 | -0.9373234 | 0.6399577 | -1.998404 | 0.04567288 | 0.129455 |
| ENSG00000106077 | ABHD11 | 4915.328761 | -0.937516817 | 0.235088287 | -4.179065606 | 2.93E-05 | 0.000257 |
| ENSG00000223886 | | 8.374544 | -0.9375943 | 0.6682651 | -1.953738 | 0.05073216 | 0.140042 |
| ENSG00000198576 | ARC | 178.9258 | -0.9376133 | 0.4637868 | -2.417543 | 0.01562568 | 0.055696 |
| ENSG00000231704 | | 8.599237 | -0.9383293 | 0.8286743 | -1.836195 | 0.06632889 | 0.171428 |
| ENSG00000222011 | FAM185A | 565.384 | -0.938832 | 0.1451879 | -6.586152 | 4.51E-11 | 1.17E-09 |
| ENSG00000279641 | | 19.67821 | -0.9394244 | 0.5843138 | -2.08135 | 0.03740185 | 0.110969 |
| ENSG00000155792 | DEPTOR | 762.222 | -0.9395354 | 0.318625 | -3.210629 | 0.00132445 | 0.007314 |
| ENSG00000235859 | | 233.3038 | -0.9395861 | 0.2009465 | -4.841724 | 1.29E-06 | 1.57E-05 |
| ENSG00000188677 | PARVB | 413.8764 | -0.9396519 | 0.2916102 | -3.464244 | 0.00053172 | 0.003312 |
| ENSG00000139629 | GALNT6 | 5384.623 | -0.9397375 | 0.1648029 | -5.858082 | 4.68E-09 | 9.07E-08 |
| ENSG00000285851 | | 5.889432 | -0.9400831 | 0.9958403 | -1.828033 | 0.06754467 | 0.173811 |
| ENSG00000214018 | RRM2P3 | 16.75073 | -0.9401734 | 0.9394481 | -1.849457 | 0.06439185 | 0.167548 |
| ENSG00000259918 | NDUFA5P11 | 98.78025 | -0.9408853 | 0.292504 | -3.457733 | 0.00054474 | 0.003385 |
| ENSG00000181355 | OFCC1 | 24.23631 | -0.9412493 | 0.4627447 | -2.414758 | 0.01574568 | 0.056011 |
| ENSG00000176273 | SLC35G1 | 445.3385 | -0.9421253 | 0.2597296 | -3.844497 | 0.0001208 | 0.000905 |

| | | | | | | | |
|---|---|---|---|---|---|---|---|
| ENSG00000133627 | ACTR3B | 673.6513238 | -0.94245901 | 0.223363021 | -4.397706198 | 1.09E-05 | 0.000107 |
| ENSG00000043039 | BARX2 | 509.7867986 | -0.942621135 | 0.247326264 | -4.011056864 | 6.04E-05 | 0.00049 |
| ENSG00000255843 | | 6.870764 | -0.9430253 | 0.7586324 | -1.883597 | 0.05961954 | 0.158292 |
| ENSG00000153902 | LGI4 | 3.278508 | -0.9430827 | 0.9217828 | -1.793918 | 0.07282632 | 0.184086 |
| ENSG00000226025 | | 3.456677 | -0.9430912 | 1.131026 | -1.831567 | 0.06701589 | 0.172768 |
| ENSG00000226746 | SMCR5 | 2.690647 | -0.9431478 | 1.151884 | -1.87728 | 0.06047977 | 0.160017 |
| ENSG00000273270 | | 835.1737 | -0.9432046 | 0.2313185 | -4.254792 | 2.09E-05 | 0.00019 |
| ENSG00000278058 | | 3.022929 | -0.9432746 | 0.9847104 | -1.829903 | 0.06726441 | 0.173274 |
| ENSG00000143494 | VASH2 | 200.3455 | -0.9436482 | 0.4028481 | -2.694206 | 0.00705566 | 0.029328 |
| ENSG00000117394 | SLC2A1 | 6823.513556 | -0.943729735 | 0.238714774 | -4.15004742 | 3.32E-05 | 0.000288 |
| ENSG00000272338 | | 16.50402 | -0.9438515 | 0.5275691 | -2.219277 | 0.02646786 | 0.084818 |
| ENSG00000230869 | AGAP10P | 14.28722 | -0.9439696 | 0.6482412 | -1.997111 | 0.04581317 | 0.129714 |
| ENSG00000125740 | FOSB | 3148.542024 | -0.943974259 | 1.207147275 | -1.956171777 | 0.05044492 | 0.139437 |
| ENSG00000236809 | SNX25P1 | 70.81866 | -0.9439767 | 0.3327461 | -3.112507 | 0.00185506 | 0.009721 |
| ENSG00000253190 | | 2.039471 | -0.9444785 | 1.196794 | -1.820973 | 0.06861098 | 0.175873 |
| ENSG00000268234 | FKBP4P6 | 24.6435 | -0.9446821 | 0.4058999 | -2.668239 | 0.007625 | 0.031298 |
| ENSG00000102854 | MSLN | 23.91895467 | -0.944695315 | 0.473897926 | -2.391574377 | 0.01677628 | 0.05899 |
| ENSG00000263934 | SNORD3A | 73.82691 | -0.9449474 | 0.4497353 | -2.473586 | 0.01337646 | 0.049109 |
| ENSG00000240898 | | 27.68779 | -0.9450695 | 0.500021 | -2.294158 | 0.02178141 | 0.072858 |
| ENSG00000170464 | DNAJC18 | 502.4955 | -0.9452744 | 0.2285619 | -4.3195 | 1.56E-05 | 0.000147 |
| ENSG00000270332 | SMC2-AS1 | 7.287974 | -0.9455493 | 0.8183584 | -1.851622 | 0.06408012 | 0.166867 |
| ENSG00000170310 | STX8 | 583.2636 | -0.9455606 | 0.21987 | -4.471134 | 7.78E-06 | 7.91E-05 |
| ENSG00000278156 | TSC22D1-AS1 | 58.99397 | -0.9463115 | 0.4837417 | -2.356073 | 0.01846931 | 0.063704 |
| ENSG00000089225 | TBX5 | 3.247158679 | -0.946933017 | 1.624153875 | -2.243835823 | 0.02484297 | 0.080671 |
| ENSG00000207547 | MIR25 | 60.62637 | -0.9470537 | 0.4046599 | -2.686086 | 0.00722945 | 0.029928 |
| ENSG00000235217 | TSPY26P | 164.9913 | -0.9472031 | 0.2736751 | -3.687577 | 0.0002264 | 0.001582 |
| ENSG00000272941 | | 17.66258 | -0.947487 | 0.5742924 | -2.1303 | 0.03314684 | 0.101367 |
| ENSG00000185985 | SLITRK2 | 2.362266 | -0.9476724 | 1.421579 | -2.080388 | 0.03748992 | 0.111144 |
| ENSG00000100092 | SH3BP1 | 212.7814015 | -0.94797907 | 0.465870307 | -2.428053754 | 0.01518009 | 0.054436 |
| ENSG00000238279 | | 20.66596 | -0.9484422 | 0.7649771 | -1.895005 | 0.05809176 | 0.155338 |
| ENSG00000166881 | NEMP1 | 2676.391 | -0.9484681 | 0.2298026 | -4.299066 | 1.72E-05 | 0.00016 |
| ENSG00000196844 | PATE2 | 4.698689 | -0.9490734 | 0.8899113 | -1.826628 | 0.06775575 | 0.174236 |
| ENSG00000204745 | | 88.60926 | -0.949335 | 0.2748276 | -3.685079 | 0.00022863 | 0.001595 |

| | | | | | | | |
|---|---|---|---|---|---|---|---|
| ENSG00000056050 | HPF1 | 826.1180103 | -0.949371249 | 0.222859104 | -4.45400282 | 8.43E-06 | 8.48E-05 |
| ENSG00000078403 | MLLT10 | 3775.13831 | -0.949494367 | 0.216479311 | -4.565247815 | 4.99E-06 | 5.32E-05 |
| ENSG00000215788 | TNFRSF25 | 871.82 | -0.9495226 | 0.2784102 | -3.648444 | 0.00026383 | 0.001809 |
| ENSG00000285999 | | 58.13298 | -0.9496985 | 0.3835717 | -2.791825 | 0.00524117 | 0.023028 |
| ENSG00000225742 | LINC02036 | 2.850639 | -0.9497941 | 1.20014 | -1.821534 | 0.06852576 | 0.175689 |
| ENSG00000279390 | | 10.26062 | -0.9498805 | 0.7742246 | -1.904837 | 0.05680125 | 0.15264 |
| ENSG00000006704 | GTF2IRD1 | 3802.438613 | -0.950126235 | 0.182276404 | -5.36199144 | 8.23E-08 | 1.26E-06 |
| ENSG00000173261 | PLAC8L1 | 15.93767 | -0.9501752 | 0.4985365 | -2.316172 | 0.02054885 | 0.069417 |
| ENSG00000189077 | TMEM120A | 1672.429 | -0.9511863 | 0.1897779 | -5.166782 | 2.38E-07 | 3.37E-06 |
| ENSG00000274943 | | 36.20752 | -0.9512932 | 0.4541076 | -2.469397 | 0.0135341 | 0.049564 |
| ENSG00000147955 | SIGMAR1 | 2160.32 | -0.9519766 | 0.1847891 | -5.304555 | 1.13E-07 | 1.70E-06 |
| ENSG00000108559 | NUP88 | 2781.316315 | -0.952018815 | 0.211184198 | -4.708439499 | 2.50E-06 | 2.87E-05 |
| ENSG00000091527 | CDV3 | 20325.22359 | -0.952226972 | 0.200676448 | -4.948355735 | 7.48E-07 | 9.56E-06 |
| ENSG00000174705 | SH3PXD2B | 712.871 | -0.9523319 | 0.2190682 | -4.528944 | 5.93E-06 | 6.22E-05 |
| ENSG00000056291 | NPFFR2 | 3.945045501 | -0.953347586 | 1.072637032 | -1.800616306 | 0.07176338 | 0.182057 |
| ENSG00000272508 | | 2.735356 | -0.9534455 | 1.245428 | -1.903401 | 0.05698823 | 0.152994 |
| ENSG00000251254 | GTF2F2P1 | 52.38055 | -0.9543639 | 0.2990926 | -3.444431 | 0.00057226 | 0.003528 |
| ENSG00000223646 | | 2.570902 | -0.9545617 | 1.068283 | -1.82671 | 0.06774334 | 0.174221 |
| ENSG00000107807 | TLX1 | 2.431818499 | -0.95490367 | 1.047592562 | -1.848412035 | 0.06454276 | 0.167874 |
| ENSG00000228834 | ATP5MFP2 | 44.8712 | -0.9554455 | 0.4227604 | -2.617036 | 0.0088697 | 0.035347 |
| ENSG00000146001 | PCDHB18P | 22.64679 | -0.9557136 | 0.4107864 | -2.657713 | 0.00786729 | 0.032099 |
| ENSG00000280426 | | 4.934223 | -0.9558812 | 0.9499958 | -1.8481 | 0.0645879 | 0.167901 |
| ENSG00000225264 | ZNRF2P2 | 14.07435 | -0.9560842 | 0.6050924 | -2.093733 | 0.03628376 | 0.108671 |
| ENSG00000185736 | ADARB2 | 15.34261 | -0.9568279 | 0.6525102 | -2.013747 | 0.04403612 | 0.125982 |
| ENSG00000213062 | | 360.4596 | -0.9583244 | 0.3295157 | -3.187399 | 0.00143559 | 0.007819 |
| ENSG00000143476 | DTL | 1085.959 | -0.9590316 | 0.6603794 | -2.031122 | 0.04224261 | 0.122034 |
| ENSG00000107745 | MICU1 | 2950.542734 | -0.960308024 | 0.151841966 | -6.454713227 | 1.08E-10 | 2.69E-09 |
| ENSG00000117528 | ABCD3 | 1720.223251 | -0.960785573 | 0.180273547 | -5.470382089 | 4.49E-08 | 7.30E-07 |
| ENSG00000169682 | SPNS1 | 70.94449 | -0.960953 | 0.3457976 | -3.066749 | 0.00216401 | 0.011082 |
| ENSG00000242670 | RPL22P13 | 11.57794 | -0.9611204 | 0.6604985 | -2.021758 | 0.0432014 | 0.124182 |
| ENSG00000279307 | NDUFAF2P1 | 379.2514 | -0.9617185 | 0.2417812 | -4.170286 | 3.04E-05 | 0.000266 |
| ENSG00000172922 | RNASEH2C | 2882.183 | -0.9618227 | 0.2144331 | -4.666992 | 3.06E-06 | 3.44E-05 |
| ENSG00000164713 | BRI3 | 1745.882 | -0.9618374 | 0.3475935 | -3.059425 | 0.00221763 | 0.011296 |

| | | | | | | | |
|---|---|---|---|---|---|---|---|
| ENSG00000196411 | EPHB4 | 5901.788 | -0.9619047 | 0.1610412 | -6.125467 | 9.04E-10 | 1.96E-08 |
| ENSG00000271978 | | 11.91428 | -0.9634235 | 0.5237731 | -2.271231 | 0.02313302 | 0.076322 |
| ENSG00000084774 | CAD | 7701.597431 | -0.963685593 | 0.158343517 | -6.225273851 | 4.81E-10 | 1.09E-08 |
| ENSG00000227039 | ITGB2-AS1 | 25.3687 | -0.9640052 | 0.502761 | -2.342523 | 0.01915387 | 0.065622 |
| ENSG00000148459 | PDSS1 | 770.5985 | -0.9643558 | 0.3710106 | -2.913336 | 0.0035759 | 0.016813 |
| ENSG00000231434 | HSP90AA3P | 39.19208 | -0.9644113 | 0.3660584 | -2.940275 | 0.00327921 | 0.015642 |
| ENSG00000105793 | GTPBP10 | 946.1467103 | -0.964441788 | 0.152158658 | -6.467289252 | 9.98E-11 | 2.48E-09 |
| ENSG00000087111 | PIGS | 6445.569708 | -0.964657382 | 0.158478328 | -6.226424068 | 4.77E-10 | 1.08E-08 |
| ENSG00000071575 | TRIB2 | 616.802461 | -0.964726897 | 0.259487519 | -3.934976407 | 8.32E-05 | 0.000651 |
| ENSG00000155660 | PDIA4 | 11394.51 | -0.9650106 | 0.2214776 | -4.507068 | 6.57E-06 | 6.80E-05 |
| ENSG00000257752 | | 8.263218 | -0.9652614 | 0.6943398 | -1.987801 | 0.04683372 | 0.131895 |
| ENSG00000142507 | PSMB6 | 4983.173 | -0.9657009 | 0.2704007 | -3.805875 | 0.0001413 | 0.00104 |
| ENSG00000264577 | | 409.9874 | -0.9658049 | 0.1770557 | -5.604493 | 2.09E-08 | 3.58E-07 |
| ENSG00000286975 | | 5.299489 | -0.9661206 | 0.815495 | -1.8801 | 0.0600945 | 0.159261 |
| ENSG00000233947 | ASS1P3 | 156.6868 | -0.9666804 | 0.6077759 | -2.121664 | 0.03386599 | 0.103068 |
| ENSG00000145912 | NHP2 | 5616.243 | -0.9673513 | 0.1898205 | -5.265491 | 1.40E-07 | 2.06E-06 |
| ENSG00000250565 | ATP6V1E2 | 490.6892 | -0.9673786 | 0.1944338 | -5.134021 | 2.84E-07 | 3.92E-06 |
| ENSG00000108179 | PPIF | 6062.625052 | -0.969000319 | 0.340936811 | -3.119175134 | 0.00181358 | 0.009543 |
| ENSG00000257681 | | 7.599207 | -0.9693746 | 0.6939187 | -1.994901 | 0.04605364 | 0.130226 |
| ENSG00000267643 | | 1.511292 | -0.9695999 | 1.460082 | -2.093781 | 0.03627948 | NA |
| ENSG00000072778 | ACADVL | 16923.47324 | -0.96963811 | 0.20094208 | -5.009123984 | 5.47E-07 | 7.21E-06 |
| ENSG00000231793 | DOC2GP | 73.93918 | -0.9703943 | 0.539356 | -2.249024 | 0.02451096 | 0.079854 |
| ENSG00000105988 | NHP2P1 | 1113.637321 | -0.970545728 | 0.219820729 | -4.597336522 | 4.28E-06 | 4.64E-05 |
| ENSG00000205208 | C4orf46 | 1028.556 | -0.9713028 | 0.3042288 | -3.452814 | 0.00055477 | 0.003435 |
| ENSG00000176236 | RPP38-DT | 7.798665 | -0.9713276 | 0.7773707 | -1.922471 | 0.05454655 | 0.148306 |
| ENSG00000162390 | ACOT11 | 71.70428 | -0.9713769 | 0.5493783 | -2.239339 | 0.02513388 | 0.081427 |
| ENSG00000280424 | | 20.30302 | -0.9714813 | 0.669117 | -2.013923 | 0.04401762 | 0.125946 |
| ENSG00000162104 | ADCY9 | 3946.978 | -0.9715748 | 0.2407338 | -4.254956 | 2.09E-05 | 0.00019 |
| ENSG00000235489 | DBF4P1 | 235.9623 | -0.9716662 | 0.414579 | -2.69896 | 0.00695566 | 0.028975 |
| ENSG00000254593 | OR7E126P | 8.730238 | -0.9723751 | 0.8338328 | -1.879213 | 0.06021548 | 0.15946 |
| ENSG00000006047 | YBX2 | 1173.3285 | -0.973630299 | 0.310790417 | -3.396295088 | 0.00068305 | 0.004115 |
| ENSG00000261136 | | 55.0044 | -0.9737481 | 0.4652661 | -2.487362 | 0.01286946 | 0.047689 |
| ENSG00000215067 | ALOX12-AS1 | 236.171 | -0.9738265 | 0.3843338 | -2.860791 | 0.00422586 | 0.019308 |

| | | | | | | | |
|---|---|---|---|---|---|---|---|
| ENSG00000116852 | KIF21B | 624.9747476 | -0.974084535 | 0.580179118 | -2.175764852 | 0.02957285 | 0.092774 |
| ENSG00000164687 | FABP5 | 78.83458 | -0.974719 | 0.4561541 | -2.529495 | 0.01142267 | 0.043288 |
| ENSG00000267710 | EDDM13 | 15.19479 | -0.9753051 | 0.5779734 | -2.181008 | 0.02918279 | 0.091856 |
| ENSG00000245556 | SCAMP1-AS1 | 748.2329 | -0.9754542 | 0.2385941 | -4.285994 | 1.82E-05 | 0.000168 |
| ENSG00000107521 | HPS1 | 4509.257828 | -0.975729507 | 0.242904113 | -4.22942547 | 2.34E-05 | 0.000211 |
| ENSG00000198898 | CAPZA2 | 4781.415 | -0.9760402 | 0.1359647 | -7.312505 | 2.62E-13 | 9.25E-12 |
| ENSG00000236423 | LINC01134 | 23.15775 | -0.9762469 | 0.771927 | -1.940327 | 0.05233998 | 0.1436 |
| ENSG00000184357 | H1-5 | 10.74477 | -0.9764279 | 0.7307799 | -1.964338 | 0.04949092 | 0.137501 |
| ENSG00000105705 | SUGP1 | 1367.024709 | -0.97653213 | 0.15203393 | -6.547276957 | 5.86E-11 | 1.50E-09 |
| ENSG00000248905 | FMN1 | 549.9214 | -0.9765857 | 0.4590721 | -2.524608 | 0.01158275 | 0.043782 |
| ENSG00000158715 | SLC45A3 | 603.4789 | -0.9768818 | 0.2284294 | -4.461148 | 8.15E-06 | 8.23E-05 |
| ENSG00000082516 | GEMIN5 | 4131.306497 | -0.977039409 | 0.259724191 | -3.995482503 | 6.46E-05 | 0.000519 |
| ENSG00000244327 | | 7.299191 | -0.9778352 | 0.8414951 | -1.900004 | 0.05743263 | 0.153885 |
| ENSG00000102302 | FGD1 | 174.0367249 | -0.977934027 | 0.456703721 | -2.534157518 | 0.01127181 | 0.042875 |
| ENSG00000280077 | | 183.2373 | -0.9782455 | 0.3486027 | -3.09912 | 0.00194096 | 0.010096 |
| ENSG00000025156 | HSF2 | 1029.112825 | -0.978347947 | 0.172076933 | -5.827243162 | 5.64E-09 | 1.07E-07 |
| ENSG00000002726 | AOC1 | 1.537549887 | -0.978486065 | 1.449344204 | -2.063752721 | 0.03904117 | NA |
| ENSG00000275084 | SNORD91B | 2.111614 | -0.9806151 | 1.098874 | -1.866818 | 0.06192695 | 0.162813 |
| ENSG00000244161 | FLNB-AS1 | 104.5207 | -0.9808042 | 0.3487742 | -3.107271 | 0.00188823 | 0.009862 |
| ENSG00000086061 | DNAJA1 | 13093.12246 | -0.981006997 | 0.208023929 | -4.901383537 | 9.52E-07 | 1.19E-05 |
| ENSG00000279243 | LINC02736 | 1.357153 | -0.9817236 | 1.465192 | -2.090111 | 0.03660787 | NA |
| ENSG00000103742 | IGDCC4 | 84.05936809 | -0.981901607 | 0.396038577 | -2.814303647 | 0.0048883 | 0.021773 |
| ENSG00000162997 | PRORSD1P | 154.4001 | -0.9833815 | 0.2458334 | -4.208553 | 2.57E-05 | 0.000229 |
| ENSG00000171497 | PPID | 2357.221 | -0.9835038 | 0.156971 | -6.397493 | 1.58E-10 | 3.84E-09 |
| ENSG00000222047 | C10orf55 | 7.771576 | -0.9838443 | 0.8016048 | -1.884275 | 0.05952777 | 0.158148 |
| ENSG00000164323 | CFAP97 | 3167.703 | -0.9840361 | 0.2773159 | -3.78386 | 0.00015442 | 0.001127 |
| ENSG00000122591 | FAM126A | 613.0915911 | -0.984038219 | 0.312575034 | -3.414282065 | 0.0006395 | 0.003885 |
| ENSG00000163517 | HDAC11 | 2170.08 | -0.9843645 | 0.2019746 | -5.044293 | 4.55E-07 | 6.09E-06 |
| ENSG00000239569 | KMT2E-AS1 | 156.1521 | -0.9845347 | 0.2605786 | -3.979202 | 6.91E-05 | 0.000551 |
| ENSG00000273492 | APP-DT | 13.14424 | -0.9845743 | 0.7264461 | -1.986634 | 0.04696292 | 0.132136 |
| ENSG00000230146 | SEPHS1P4 | 52.2678 | -0.9851465 | 0.3736037 | -2.955884 | 0.00311774 | 0.015001 |
| ENSG00000126749 | EMG1 | 2357.121423 | -0.985679608 | 0.246367857 | -4.233075008 | 2.31E-05 | 0.000208 |
| ENSG00000274751 | | 15.47261 | -0.9861466 | 0.6958357 | -2.019623 | 0.0434225 | 0.124636 |

| | | | | | | | |
|---|---|---|---|---|---|---|---|
| ENSG00000120318 | ARAP3 | 55.82513417 | -0.986147844 | 0.514028586 | -2.3779417 | 0.01740958 | 0.060775 |
| ENSG00000262358 | | 2.099703 | -0.9865048 | 1.643528 | -2.206266 | 0.02736539 | 0.087158 |
| ENSG00000254866 | DEFB109D | 9.575813 | -0.9869569 | 0.6836344 | -2.034278 | 0.04192355 | 0.121283 |
| ENSG00000138650 | PCDH10 | 714.2304688 | -0.987266566 | 0.492267082 | -2.432041417 | 0.01501399 | 0.053957 |
| ENSG00000104205 | SGK3 | 172.1350361 | -0.987314403 | 0.302037223 | -3.522517538 | 0.00042747 | 0.002742 |
| ENSG00000180739 | S1PR5 | 100.6676 | -0.9904811 | 0.55393 | -2.272779 | 0.02303952 | 0.076118 |
| ENSG00000227382 | EIF4A2P2 | 4.168577 | -0.9912085 | 0.9874015 | -1.870147 | 0.0614634 | 0.161953 |
| ENSG00000229337 | | 16.92747 | -0.9923765 | 0.527789 | -2.321485 | 0.0202607 | 0.068702 |
| ENSG00000229876 | CASC20 | 1.964633 | -0.9926767 | 1.865654 | -2.411268 | 0.01589714 | 0.056437 |
| ENSG00000196510 | ANAPC7 | 2922.581 | -0.9927563 | 0.2172494 | -4.754046 | 1.99E-06 | 2.34E-05 |
| ENSG00000156869 | FRRS1 | 92.32126 | -0.9930279 | 0.3080101 | -3.4803 | 0.00050085 | 0.003141 |
| ENSG00000159461 | AMFR | 3992.684 | -0.9935767 | 0.2072973 | -4.975248 | 6.52E-07 | 8.45E-06 |
| ENSG00000164627 | KIF6 | 29.75693 | -0.9936945 | 0.3940385 | -2.860486 | 0.00422993 | 0.01932 |
| ENSG00000234604 | | 17.22355 | -0.9937837 | 0.6421811 | -2.112623 | 0.03463302 | 0.104855 |
| ENSG00000115159 | GPD2 | 3584.375946 | -0.99380129 | 0.237342926 | -4.390255553 | 1.13E-05 | 0.00011 |
| ENSG00000261614 | YBX3P1 | 242.3475 | -0.9938183 | 0.2018068 | -5.094938 | 3.49E-07 | 4.74E-06 |
| ENSG00000156502 | SUPV3L1 | 2421.508 | -0.9939545 | 0.1726871 | -5.915955 | 3.30E-09 | 6.56E-08 |
| ENSG00000137691 | CFAP300 | 17.91066989 | -0.994016102 | 0.586678147 | -2.195746185 | 0.02811012 | 0.088965 |
| ENSG00000228121 | PHBP3 | 16.92487 | -0.9941116 | 0.5690852 | -2.237525 | 0.02525207 | 0.081735 |
| ENSG00000272444 | | 39.44683 | -0.9942618 | 0.4431909 | -2.615194 | 0.00891767 | 0.035485 |
| ENSG00000137825 | ITPKA | 175.6938101 | -0.995022671 | 0.624787352 | -2.151955439 | 0.03140086 | 0.097203 |
| ENSG00000254489 | MPPED2-AS1 | 3.846359 | -0.9954518 | 0.9641459 | -1.899149 | 0.05754486 | 0.154124 |
| ENSG00000110013 | SIAE | 1667.895446 | -0.995666946 | 0.205718009 | -5.016491829 | 5.26E-07 | 6.96E-06 |
| ENSG00000140563 | MCTP2 | 344.2167 | -0.9959773 | 0.2569014 | -4.097342 | 4.18E-05 | 0.000354 |
| ENSG00000272109 | | 7.706178 | -0.9960811 | 0.6649278 | -2.052905 | 0.04008183 | 0.117064 |
| ENSG00000215548 | FRG1JP | 71.2175 | -0.9964178 | 0.4196245 | -2.723069 | 0.00646786 | 0.027316 |
| ENSG00000147180 | ZNF711 | 1021.645 | -0.9966634 | 0.2387388 | -4.383832 | 1.17E-05 | 0.000113 |
| ENSG00000265102 | MIR3942 | 2.798971 | -0.9966931 | 1.209163 | -1.843797 | 0.06521277 | 0.169153 |
| ENSG00000129235 | TXNDC17 | 2025.840178 | -0.997304533 | 0.262864907 | -4.038535904 | 5.38E-05 | 0.000442 |
| ENSG00000268379 | | 66.04492 | -0.998559 | 0.3488181 | -3.16069 | 0.00157396 | 0.008446 |
| ENSG00000132300 | PTCD3 | 5765.940877 | -0.998979877 | 0.146727831 | -6.964046114 | 3.31E-12 | 1.02E-10 |
| ENSG00000213694 | S1PR3 | 1171.106 | -0.9992469 | 0.3886942 | -2.912511 | 0.00358536 | 0.016848 |
| ENSG00000250635 | CXXC5-AS1 | 5.428639 | -0.9997955 | 1.026178 | -1.919181 | 0.05496138 | 0.149015 |

| | | | | | | | |
|---|---|---|---|---|---|---|---|
| ENSG00000183691 | NOG | 44.53291 | -1.000054 | 0.5094877 | -2.406735 | 0.01609585 | 0.056997 |
| ENSG00000207146 | | 4.322056 | -1.000128 | 0.7976956 | -1.951544 | 0.05099242 | 0.140585 |
| ENSG00000249650 | | 7.99522 | -1.000275 | 0.7726653 | -1.968768 | 0.04897971 | 0.13638 |
| ENSG00000213977 | TAX1BP3 | 35.26105 | -1.000754 | 0.4043915 | -2.819132 | 0.00481536 | 0.021521 |
| ENSG00000245105 | A2M-AS1 | 52.1836 | -1.001272 | 0.3878058 | -2.910736 | 0.00360579 | 0.016929 |
| ENSG00000186187 | ZNRF1 | 4095.358 | -1.001309 | 0.172893 | -5.945685 | 2.75E-09 | 5.54E-08 |
| ENSG00000227855 | DPY19L2P3 | 63.75834 | -1.001356 | 0.3739184 | -2.994062 | 0.0027529 | 0.013572 |
| ENSG00000118707 | TGIF2 | 2345.051782 | -1.001373575 | 0.143663128 | -7.09653484 | 1.28E-12 | 4.11E-11 |
| ENSG00000105967 | TFEC | 5.810999892 | -1.001445325 | 0.82598132 | -1.929430228 | 0.05367747 | 0.146605 |
| ENSG00000064419 | TNPO3 | 4565.493921 | -1.002072889 | 0.10485971 | -9.637645737 | 5.54E-22 | 5.19E-20 |
| ENSG00000056487 | PHF21B | 275.2823195 | -1.002256843 | 0.376386584 | -2.988217278 | 0.0028061 | 0.01381 |
| ENSG00000275491 | LINC01730 | 33.52338 | -1.002324 | 0.7190101 | -2.037341 | 0.04161592 | 0.120617 |
| ENSG00000154065 | ANKRD29 | 52.16618 | -1.003336 | 0.5069981 | -2.407855 | 0.01604655 | 0.056868 |
| ENSG00000162729 | IGSF8 | 1999.7 | -1.003385 | 0.165881 | -6.192586 | 5.92E-10 | 1.32E-08 |
| ENSG00000178921 | PFAS | 2503.252 | -1.003792 | 0.2100552 | -4.951964 | 7.35E-07 | 9.42E-06 |
| ENSG00000158406 | H4C8 | 1990.037 | -1.003875 | 0.3343342 | -3.310173 | 0.00093239 | 0.005376 |
| ENSG00000170085 | SIMC1 | 948.0913 | -1.003966 | 0.1463422 | -6.983209 | 2.89E-12 | 8.94E-11 |
| ENSG00000276368 | H2AC14 | 7.473147 | -1.004157 | 1.039115 | -1.90167 | 0.05721431 | 0.153422 |
| ENSG00000263883 | EEF1DP7 | 3.439288 | -1.004269 | 1.358593 | -1.931381 | 0.05343595 | 0.146049 |
| ENSG00000287445 | | 2.025547 | -1.00446 | 1.52307 | -2.128254 | 0.03331606 | 0.101791 |
| ENSG00000111196 | MAGOHB | 753.3552635 | -1.004928966 | 0.289452973 | -3.721328803 | 0.00019818 | 0.001406 |
| ENSG00000286532 | PARTICL | 323.1051 | -1.005477 | 0.2156483 | -4.843658 | 1.27E-06 | 1.56E-05 |
| ENSG00000176463 | SLCO3A1 | 477.0436 | -1.005643 | 0.2757627 | -3.894037 | 9.86E-05 | 0.000756 |
| ENSG00000118004 | COLEC11 | 3.137987213 | -1.006545661 | 1.155127494 | -1.86676688 | 0.06193415 | 0.162813 |
| ENSG00000244184 | | 23.9573 | -1.006582 | 0.5360043 | -2.334128 | 0.01958905 | 0.066872 |
| ENSG00000103089 | FA2H | 452.1810845 | -1.006732664 | 0.34796014 | -3.184318568 | 0.00145095 | 0.007885 |
| ENSG00000113356 | POLR3G | 370.058264 | -1.006955858 | 0.374880629 | -3.014449034 | 0.00257447 | 0.012819 |
| ENSG00000224699 | LAMTOR5-AS1 | 178.007 | -1.007064 | 0.2653088 | -4.023488 | 5.73E-05 | 0.000468 |
| ENSG00000169918 | OTUD7A | 195.2812 | -1.007139 | 0.2344128 | -4.497236 | 6.88E-06 | 7.08E-05 |
| ENSG00000120549 | KIAA1217 | 2348.769272 | -1.007169177 | 0.228246713 | -4.618636906 | 3.86E-06 | 4.23E-05 |
| ENSG00000148204 | CRB2 | 8.642526 | -1.007227 | 0.6944725 | -2.054869 | 0.03989164 | 0.116624 |
| ENSG00000272170 | | 2.889734 | -1.00879 | 1.233149 | -1.855967 | 0.06345833 | 0.165703 |
| ENSG00000150627 | WDR17 | 412.7597 | -1.008826 | 0.4080156 | -2.826537 | 0.00470542 | 0.021107 |

| | | | | | | | |
|---|---|---|---|---|---|---|---|
| ENSG00000008083 | JARID2 | 2164.6947 | -1.009164321 | 0.202509906 | -5.148753777 | 2.62E-07 | 3.66E-06 |
| ENSG00000070814 | TCOF1 | 5853.944003 | -1.009378682 | 0.301082387 | -3.607042392 | 0.00030971 | 0.00208 |
| ENSG00000161513 | FDXR | 1846.058 | -1.009937 | 0.1827428 | -5.683451 | 1.32E-08 | 2.35E-07 |
| ENSG00000198795 | ZNF521 | 23.68086 | -1.010195 | 0.4501154 | -2.631869 | 0.00849166 | 0.034112 |
| ENSG00000276564 | | 41.96779 | -1.010388 | 0.547382 | -2.319163 | 0.02038623 | 0.06899 |
| ENSG00000238113 | LINC01410 | 179.8389 | -1.010629 | 0.364039 | -3.095283 | 0.00196625 | 0.010203 |
| ENSG00000234160 | | 75.9392 | -1.010985 | 0.2493297 | -4.26876 | 1.97E-05 | 0.00018 |
| ENSG00000090266 | NDUFB2 | 2684.070526 | -1.011239381 | 0.216950083 | -4.847905656 | 1.25E-06 | 1.53E-05 |
| ENSG00000162643 | DNAI3 | 32.17822 | -1.011264 | 0.4815131 | -2.510528 | 0.01205509 | 0.045182 |
| ENSG00000164136 | IL15 | 55.65434 | -1.011665 | 0.5658732 | -2.266819 | 0.02340131 | 0.076957 |
| ENSG00000151131 | C12orf45 | 1066.338 | -1.011696 | 0.1223581 | -8.374568 | 5.54E-17 | 3.01E-15 |
| ENSG00000124787 | RPP40 | 724.2869936 | -1.011822696 | 0.298090732 | -3.64867286 | 0.0002636 | 0.001808 |
| ENSG00000261430 | | 1.68741 | -1.012633 | 1.489877 | -2.105421 | 0.03525465 | 0.106176 |
| ENSG00000214826 | DDX12P | 546.803 | -1.013961 | 0.4367199 | -2.716659 | 0.00659446 | 0.027745 |
| ENSG00000242612 | DECR2 | 1564.46 | -1.014701 | 0.1908089 | -5.483948 | 4.16E-08 | 6.79E-07 |
| ENSG00000272568 | | 2.777993 | -1.015212 | 1.151615 | -1.910891 | 0.05601853 | 0.151179 |
| ENSG00000181908 | LINC02724 | 6.196359 | -1.015213 | 0.9450296 | -1.888126 | 0.05900909 | 0.157158 |
| ENSG00000271795 | | 8.754351 | -1.015488 | 0.7494694 | -1.99953 | 0.04555109 | 0.129203 |
| ENSG00000166710 | B2M | 16760 | -1.015614 | 0.2490689 | -4.293037 | 1.76E-05 | 0.000163 |
| ENSG00000170423 | KRT78 | 1.680072 | -1.015753 | 1.814123 | -2.356413 | 0.01845238 | 0.06367 |
| ENSG00000204086 | RPA4 | 10.19578 | -1.017714 | 0.6018059 | -2.204039 | 0.02752159 | 0.087571 |
| ENSG00000160182 | TFF1 | 6911.772 | -1.018011 | 0.7192263 | -2.077074 | 0.03779474 | 0.111873 |
| ENSG00000253696 | KBTBD11-OT1 | 5.914344 | -1.019155 | 0.9093105 | -1.909891 | 0.05614723 | 0.151419 |
| ENSG00000270133 | | 10.76884 | -1.019233 | 0.5734264 | -2.281318 | 0.02252966 | 0.074889 |
| ENSG00000260668 | | 70.76769 | -1.019906 | 0.273676 | -3.959334 | 7.52E-05 | 0.000595 |
| ENSG00000075426 | FOSL2 | 4955.122039 | -1.019914964 | 0.326323986 | -3.409849087 | 0.00064999 | 0.003942 |
| ENSG00000165868 | HSPA12A | 614.6384 | -1.020559 | 0.2143434 | -4.948406 | 7.48E-07 | 9.56E-06 |
| ENSG00000136098 | NEK3 | 436.0476698 | -1.020592026 | 0.16876805 | -6.193018439 | 5.90E-10 | 1.32E-08 |
| ENSG00000283297 | TEX52 | 10.009 | -1.020924 | 0.6839344 | -2.090331 | 0.03658806 | 0.109103 |
| ENSG00000277342 | | 6.768117 | -1.021172 | 0.8884307 | -1.952796 | 0.05084376 | 0.140248 |
| ENSG00000138119 | MYOF | 17788.07014 | -1.021937099 | 0.345870822 | -3.25890861 | 0.00111842 | 0.006304 |
| ENSG00000262185 | LINC02861 | 16.34576 | -1.021941 | 0.4639994 | -2.590492 | 0.00958388 | 0.037543 |
| ENSG00000099139 | PCSK5 | 43.14822531 | -1.022529873 | 0.625854286 | -2.173170113 | 0.02976752 | 0.093149 |

| | | | | | | | |
|---|---|---|---|---|---|---|---|
| ENSG00000100167 | SEPTIN3 | 709.7925658 | -1.022666853 | 0.280286628 | -3.88974107 | 0.00010035 | 0.000767 |
| ENSG00000227253 | | 2.820532 | -1.022832 | 1.272243 | -1.885422 | 0.05937289 | 0.157882 |
| ENSG00000204666 | | 15.95619 | -1.022939 | 0.5240844 | -2.388584 | 0.01691345 | 0.059386 |
| ENSG00000129990 | SYT5 | 10.12771649 | -1.023222809 | 0.589519632 | -2.248540979 | 0.02454172 | 0.079917 |
| ENSG00000105865 | DUS4L | 386.9485566 | -1.023489532 | 0.235261632 | -4.554945136 | 5.24E-06 | 5.57E-05 |
| ENSG00000260257 | | 50.83016 | -1.023737 | 0.6015969 | -2.23203 | 0.02561295 | 0.082655 |
| ENSG00000159110 | IFNAR2 | 23.19026 | -1.023762 | 0.4412071 | -2.702676 | 0.00687838 | 0.028726 |
| ENSG00000124256 | ZBP1 | 8.190170102 | -1.023825408 | 1.328635887 | -1.953488614 | 0.05076173 | 0.140109 |
| ENSG00000229781 | RRN3P4 | 1.404357 | -1.023878 | 1.474969 | -2.063903 | 0.0390269 | NA |
| ENSG00000231952 | DPY19L1P2 | 35.59281 | -1.024147 | 0.409215 | -2.854092 | 0.004316 | 0.019639 |
| ENSG00000145700 | ANKRD31 | 58.08322 | -1.024419 | 0.3438818 | -3.265287 | 0.00109353 | 0.006183 |
| ENSG00000242715 | CCDC169 | 44.89464 | -1.024505 | 0.4226744 | -2.790083 | 0.00526945 | 0.02313 |
| ENSG00000232815 | DUX4L50 | 330.8111 | -1.025657 | 0.2722789 | -4.002598 | 6.27E-05 | 0.000506 |
| ENSG00000143502 | SUSD4 | 1147.969 | -1.025679 | 0.3562713 | -3.192994 | 0.00140806 | 0.007685 |
| ENSG00000124784 | RIOK1 | 1336.170939 | -1.026217399 | 0.180961883 | -5.830432872 | 5.53E-09 | 1.06E-07 |
| ENSG00000275763 | ZNF516-DT | 13.19817 | -1.026376 | 0.6950678 | -2.102306 | 0.03552645 | 0.106813 |
| ENSG00000133121 | STARD13 | 3060.011692 | -1.0264374 | 0.273449764 | -3.992450356 | 6.54E-05 | 0.000524 |
| ENSG00000254910 | | 55.08597 | -1.026813 | 0.3742885 | -3.070627 | 0.0021361 | 0.010965 |
| ENSG00000181085 | MAPK15 | 628.5148 | -1.026941 | 0.1805714 | -5.85536 | 4.76E-09 | 9.20E-08 |
| ENSG00000223510 | CDRT15 | 11.23705 | -1.02703 | 0.6977904 | -2.075127 | 0.03797481 | 0.112306 |
| ENSG00000126249 | PDCD2L | 406.6070849 | -1.027165992 | 0.215129898 | -4.962864627 | 6.95E-07 | 8.97E-06 |
| ENSG00000278389 | | 9.63175 | -1.027547 | 0.9662953 | -1.866038 | 0.06203608 | 0.162966 |
| ENSG00000286428 | | 2.642128 | -1.027769 | 1.069877 | -1.925651 | 0.05414797 | 0.147524 |
| ENSG00000205220 | PSMB10 | 668.5469 | -1.028602 | 0.2769366 | -3.952008 | 7.75E-05 | 0.000611 |
| ENSG00000166479 | TMX3 | 2861.058 | -1.028744 | 0.2092494 | -5.100258 | 3.39E-07 | 4.63E-06 |
| ENSG00000075790 | BCAP29 | 34.14174172 | -1.028951538 | 0.412044817 | -2.849826717 | 0.00437431 | 0.019853 |
| ENSG00000235453 | SMIM27 | 141.1781 | -1.029258 | 0.2269621 | -4.730273 | 2.24E-06 | 2.61E-05 |
| ENSG00000245248 | USP2-AS1 | 77.79325 | -1.029347 | 0.3377177 | -3.345478 | 0.00082141 | 0.004833 |
| ENSG00000049541 | RFC2 | 2120.522941 | -1.029567633 | 0.455636461 | -2.662481107 | 0.00775669 | 0.031717 |
| ENSG00000251666 | ZNF346-IT1 | 11.17855 | -1.030134 | 0.6172015 | -2.204903 | 0.02746089 | 0.08741 |
| ENSG00000269971 | | 3.748756 | -1.031209 | 1.138661 | -1.902631 | 0.05708867 | 0.153165 |
| ENSG00000165661 | QSOX2 | 3142.762 | -1.031862 | 0.1340914 | -7.810279 | 5.71E-15 | 2.42E-13 |
| ENSG00000267549 | | 5.042463 | -1.032172 | 0.9253005 | -1.904842 | 0.05680056 | 0.15264 |

| | | | | | | | |
|---|---|---|---|---|---|---|---|
| ENSG00000286058 | | 573.2042 | -1.032232 | 0.3647469 | -3.151055 | 0.00162682 | 0.008669 |
| ENSG00000243024 | | 10.76674 | -1.032459 | 0.6604952 | -2.130921 | 0.03309562 | 0.101234 |
| ENSG00000111231 | GPN3 | 1003.001314 | -1.032462228 | 0.312808956 | -3.580964507 | 0.00034233 | 0.00227 |
| ENSG00000164815 | ORC5 | 1396.334 | -1.032558 | 0.1707591 | -6.197828 | 5.72E-10 | 1.28E-08 |
| ENSG00000240591 | | 17.31481 | -1.032676 | 0.6098441 | -2.231562 | 0.02564391 | 0.082731 |
| ENSG00000212175 | | 8.632019 | -1.032935 | 0.6950909 | -2.084293 | 0.0371335 | 0.11037 |
| ENSG00000214456 | PLIN5 | 390.8991 | -1.033198 | 0.4285699 | -2.788927 | 0.0052883 | 0.023193 |
| ENSG00000258947 | TUBB3 | 241.9629 | -1.034781 | 0.2978621 | -3.74917 | 0.00017742 | 0.001275 |
| ENSG00000169926 | KLF13 | 1974.48 | -1.035001 | 0.2791895 | -3.954168 | 7.68E-05 | 0.000607 |
| ENSG00000231233 | CFAP58-DT | 8.495259 | -1.035206 | 0.9984072 | -1.903803 | 0.05693579 | 0.152878 |
| ENSG00000181031 | RPH3AL | 210.8536 | -1.035268 | 0.2301806 | -4.701192 | 2.59E-06 | 2.96E-05 |
| ENSG00000256571 | | 2.078216 | -1.035941 | 1.315907 | -1.946253 | 0.05162436 | 0.141974 |
| ENSG00000161057 | PSMC2 | 6049.737 | -1.036573 | 0.1717835 | -6.183175 | 6.28E-10 | 1.40E-08 |
| ENSG00000043591 | ADRB1 | 84.33846951 | -1.037103416 | 0.425209809 | -2.812332004 | 0.00491837 | 0.021861 |
| ENSG00000124275 | MTRR | 3267.098691 | -1.03734715 | 0.174189595 | -6.106909905 | 1.02E-09 | 2.18E-08 |
| ENSG00000286482 | | 64.4254 | -1.038614 | 0.4376739 | -2.753348 | 0.00589892 | 0.025387 |
| ENSG00000258743 | LINC02301 | 7.199467 | -1.038906 | 0.7621451 | -2.037534 | 0.04159658 | 0.120574 |
| ENSG00000119457 | SLC46A2 | 7.721360497 | -1.039032767 | 0.965890083 | -1.953222278 | 0.05079326 | 0.140167 |
| ENSG00000150275 | PCDH15 | 5.055794 | -1.039211 | 0.7953201 | -1.989592 | 0.04663591 | 0.13148 |
| ENSG00000267801 | | 9.928992 | -1.039414 | 1.010896 | -1.834801 | 0.06653523 | 0.171862 |
| ENSG00000279212 | | 7.391847 | -1.039784 | 1.035627 | -1.94616 | 0.0516355 | 0.14199 |
| ENSG00000145623 | OSMR | 2530.432 | -1.040453 | 0.3405789 | -3.353569 | 0.00079776 | 0.004708 |
| ENSG00000128609 | NDUFA5 | 2571.082185 | -1.041770798 | 0.153293466 | -6.941799746 | 3.87E-12 | 1.19E-10 |
| ENSG00000265912 | | 9.155187 | -1.041944 | 0.6368787 | -2.18052 | 0.0292189 | 0.091915 |
| ENSG00000129993 | CBFA2T3 | 2885.922376 | -1.042044036 | 0.144634786 | -7.335956222 | 2.20E-13 | 7.84E-12 |
| ENSG00000237651 | C2orf74 | 47.04206 | -1.042147 | 0.3630379 | -3.185907 | 0.00144301 | 0.007852 |
| ENSG00000261251 | | 9.865569 | -1.042714 | 0.7713126 | -2.001252 | 0.04536528 | 0.1288 |
| ENSG00000068001 | HYAL2 | 1068.0577 | -1.042828213 | 0.227262629 | -4.785848873 | 1.70E-06 | 2.03E-05 |
| ENSG00000115271 | GCA | 1984.092478 | -1.042977239 | 0.189910258 | -5.652412473 | 1.58E-08 | 2.78E-07 |
| ENSG00000132259 | CNGA4 | 8.78497648 | -1.043749711 | 0.651842483 | -2.147700931 | 0.03173752 | 0.098015 |
| ENSG00000237529 | | 8.281862 | -1.043827 | 0.7488644 | -2.059661 | 0.03943096 | 0.115629 |
| ENSG00000230450 | NEK2P4 | 16.33441 | -1.043933 | 0.7342801 | -2.075284 | 0.0379602 | 0.112275 |
| ENSG00000162976 | SLC66A3 | 2915.548 | -1.044231 | 0.179596 | -5.971908 | 2.34E-09 | 4.76E-08 |

| | | | | | | | |
|---|---|---|---|---|---|---|---|
| ENSG00000279613 | | 9.177926 | -1.044935 | 0.8421401 | -1.988907 | 0.04671152 | 0.131638 |
| ENSG00000261671 | | 4.064026 | -1.045023 | 1.223562 | -1.961674 | 0.0498004 | 0.138116 |
| ENSG00000124191 | TOX2 | 110.1032926 | -1.045704077 | 0.576993718 | -2.334014635 | 0.01959496 | 0.066883 |
| ENSG00000141682 | PMAIP1 | 704.4515 | -1.04571 | 0.3607803 | -3.206185 | 0.00134508 | 0.007401 |
| ENSG00000171680 | PLEKHG5 | 1480.393 | -1.045729 | 0.1669759 | -6.41064 | 1.45E-10 | 3.54E-09 |
| ENSG00000132170 | PPARG | 1716.661034 | -1.04610924 | 0.659658812 | -2.086604967 | 0.03692386 | 0.109919 |
| ENSG00000285407 | | 61.13183 | -1.047378 | 0.4106574 | -2.912689 | 0.00358332 | 0.016842 |
| ENSG00000117593 | DARS2 | 4195.826607 | -1.047427602 | 0.358225721 | -3.240349122 | 0.00119383 | 0.006659 |
| ENSG00000116667 | C1orf21 | 1089.301459 | -1.048549892 | 0.321428713 | -3.546203771 | 0.00039082 | 0.002544 |
| ENSG00000140950 | MEAK7 | 1999.307 | -1.048616 | 0.1278811 | -8.309235 | 9.63E-17 | 5.11E-15 |
| ENSG00000267419 | ZNF56 | 75.12232 | -1.048667 | 0.3681029 | -3.16655 | 0.00154259 | 0.008303 |
| ENSG00000114757 | PEX5L | 39.42800266 | -1.049149034 | 0.55633167 | -2.384423379 | 0.01710591 | 0.059892 |
| ENSG00000101311 | FERMT1 | 650.2447345 | -1.049206959 | 0.33824207 | -3.403507345 | 0.00066527 | 0.004022 |
| ENSG00000233020 | CHORDC1P5 | 5.972472 | -1.049643 | 1.252432 | -1.943923 | 0.05190472 | 0.142582 |
| ENSG00000253146 | CIR1P1 | 60.79031 | -1.049681 | 0.3522363 | -3.290735 | 0.00099926 | 0.005718 |
| ENSG00000087510 | TFAP2C | 6690.015589 | -1.049806311 | 0.233879782 | -4.716690393 | 2.40E-06 | 2.77E-05 |
| ENSG00000248278 | SUMO2P17 | 73.25877 | -1.049883 | 0.3106359 | -3.644132 | 0.0002683 | 0.001833 |
| ENSG00000003249 | DBNDD1 | 6919.352801 | -1.050970102 | 0.188979879 | -5.725712081 | 1.03E-08 | 1.89E-07 |
| ENSG00000197943 | PLCG2 | 889.1243 | -1.051681 | 0.3371407 | -3.419451 | 0.00062748 | 0.003822 |
| ENSG00000147378 | FATE1 | 16.15824 | -1.051831 | 0.6942343 | -2.150832 | 0.03148942 | 0.097409 |
| ENSG00000118965 | WDR35 | 1015.931242 | -1.052536797 | 0.246896492 | -4.482323317 | 7.38E-06 | 7.55E-05 |
| ENSG00000237004 | ZNRF2P1 | 141.1681 | -1.052578 | 0.3792622 | -3.113589 | 0.00184827 | 0.009695 |
| ENSG00000181192 | DHTKD1 | 4935.403 | -1.053515 | 0.4200188 | -2.882685 | 0.00394301 | 0.018229 |
| ENSG00000156042 | CFAP70 | 78.14281 | -1.053702 | 0.5779811 | -2.340633 | 0.01925108 | 0.065904 |
| ENSG00000143434 | SEMA6C | 395.0596 | -1.054058 | 0.2863991 | -3.933126 | 8.38E-05 | 0.000655 |
| ENSG00000106070 | GRB10 | 1459.374795 | -1.05421475 | 0.463689511 | -2.689157063 | 0.00716327 | 0.029714 |
| ENSG00000232907 | DLGAP4-AS1 | 6.500604 | -1.054296 | 0.8445439 | -1.973435 | 0.04844601 | 0.135235 |
| ENSG00000100399 | CHADL | 23.61368968 | -1.054606701 | 0.388427939 | -3.050761961 | 0.00228262 | 0.011565 |
| ENSG00000010626 | LRRC23 | 266.2605199 | -1.054657134 | 0.212331294 | -5.155527051 | 2.53E-07 | 3.55E-06 |
| ENSG00000250071 | | 8.2992 | -1.054839 | 0.7088503 | -2.113855 | 0.03452769 | 0.104627 |
| ENSG00000157212 | PAXIP1 | 1814.619 | -1.055174 | 0.1820663 | -5.958624 | 2.54E-09 | 5.15E-08 |
| ENSG00000106392 | C1GALT1 | 1527.734593 | -1.055930489 | 0.209676011 | -5.220465485 | 1.78E-07 | 2.57E-06 |
| ENSG00000007264 | MATK | 1170.737227 | -1.056553616 | 0.392541872 | -3.043337779 | 0.0023397 | 0.011817 |

| | | | | | | | |
|---|---|---|---|---|---|---|---|
| ENSG00000143110 | C1orf162 | 40.34935 | -1.057529 | 0.5712102 | -2.35968 | 0.01829072 | 0.063244 |
| ENSG00000091732 | ZC3HC1 | 562.1134412 | -1.057720086 | 0.301209257 | -3.775451572 | 0.00015972 | 0.001162 |
| ENSG00000144843 | ADPRH | 107.8836 | -1.05794 | 0.3124841 | -3.663808 | 0.00024849 | 0.001716 |
| ENSG00000167207 | NOD2 | 82.84109 | -1.058956 | 0.6031506 | -2.30461 | 0.02118842 | 0.071206 |
| ENSG00000276832 | CDRT15P6 | 6.838587 | -1.059063 | 0.6364747 | -2.224004 | 0.02614814 | 0.084004 |
| ENSG00000151743 | AMN1 | 192.8105 | -1.059145 | 0.3195601 | -3.596383 | 0.00032267 | 0.002157 |
| ENSG00000199023 | MIR339 | 7.469 | -1.059358 | 0.7234079 | -2.099415 | 0.03578033 | 0.107418 |
| ENSG00000166321 | NUDT13 | 263.098 | -1.059521 | 0.2319766 | -4.763844 | 1.90E-06 | 2.24E-05 |
| ENSG00000198931 | APRT | 12036.59 | -1.05953 | 0.193285 | -5.669453 | 1.43E-08 | 2.54E-07 |
| ENSG00000272425 | | 18.04432 | -1.059582 | 0.5016362 | -2.548609 | 0.01081534 | 0.041437 |
| ENSG00000251467 | | 2.169789 | -1.059844 | 1.393708 | -2.019018 | 0.04348532 | 0.124769 |
| ENSG00000036672 | USP2 | 444.0171189 | -1.061307391 | 0.287912983 | -3.943109199 | 8.04E-05 | 0.000631 |
| ENSG00000273091 | | 28.87022 | -1.061883 | 0.5995552 | -2.307039 | 0.02105266 | 0.070857 |
| ENSG00000136944 | LMX1B | 2069.892576 | -1.061918122 | 0.204240999 | -5.382906889 | 7.33E-08 | 1.14E-06 |
| ENSG00000272631 | | 307.9888 | -1.061959 | 0.238868 | -4.646067 | 3.38E-06 | 3.76E-05 |
| ENSG00000252202 | | 6.214176 | -1.062683 | 0.7348528 | -2.092732 | 0.03637311 | 0.108713 |
| ENSG00000174028 | FAM3C2P | 729.2471 | -1.062724 | 0.1607102 | -6.759806 | 1.38E-11 | 3.89E-10 |
| ENSG00000112812 | PRSS16 | 360.4660625 | -1.063440871 | 0.292680608 | -3.891613939 | 9.96E-05 | 0.000762 |
| ENSG00000261693 | | 50.18608 | -1.063976 | 0.4150503 | -2.920969 | 0.00348945 | 0.016473 |
| ENSG00000117650 | NEK2 | 1026.476771 | -1.064413696 | 0.54114551 | -2.466359595 | 0.01364942 | 0.049931 |
| ENSG00000105409 | ATP1A3 | 18.23493038 | -1.06466726 | 0.648017152 | -2.208396133 | 0.02721667 | 0.086757 |
| ENSG00000228069 | MTCO3P29 | 4.793066 | -1.064955 | 0.9441895 | -1.946762 | 0.0515633 | 0.141835 |
| ENSG00000152669 | CCNO | 513.3862 | -1.065551 | 0.2201211 | -5.032888 | 4.83E-07 | 6.42E-06 |
| ENSG00000106246 | PTCD1 | 196.6010346 | -1.065859223 | 0.225078172 | -4.929193668 | 8.26E-07 | 1.04E-05 |
| ENSG00000259775 | | 29.27146 | -1.06718 | 0.4269081 | -2.872954 | 0.00406654 | 0.018712 |
| ENSG00000224261 | RPSAP18 | 6.013434 | -1.067236 | 0.8470538 | -2.002966 | 0.04518097 | 0.128417 |
| ENSG00000170827 | CELP | 1.935723 | -1.068052 | 1.479544 | -1.999764 | 0.04552572 | 0.129145 |
| ENSG00000285847 | | 5.486865 | -1.068572 | 0.8112672 | -2.037556 | 0.04159435 | 0.120574 |
| ENSG00000166821 | PEX11A | 1157.641 | -1.069211 | 0.3417997 | -3.427278 | 0.00060967 | 0.003728 |
| ENSG00000268403 | | 65.82255 | -1.069881 | 0.3348468 | -3.488418 | 0.00048589 | 0.003062 |
| ENSG00000178752 | ERFE | 60.40994 | -1.069996 | 0.8361184 | -2.039402 | 0.04140989 | 0.120138 |
| ENSG00000164251 | F2RL1 | 1118.046 | -1.070067 | 0.2833427 | -4.029019 | 5.60E-05 | 0.000458 |
| ENSG00000279069 | | 179.8506 | -1.070114 | 0.1758253 | -6.2454 | 4.23E-10 | 9.67E-09 |

| | | | | | | | |
|---|---|---|---|---|---|---|---|
| ENSG00000285725 | | 80.55085 | -1.070743 | 0.3628789 | -3.270335 | 0.0010742 | 0.006084 |
| ENSG00000132382 | MYBBP1A | 6378.903844 | -1.071121842 | 0.154101919 | -7.086454704 | 1.38E-12 | 4.40E-11 |
| ENSG00000147912 | FBXO10 | 286.6946 | -1.071244 | 0.1965384 | -5.622916 | 1.88E-08 | 3.24E-07 |
| ENSG00000181585 | TMIE | 7.309416 | -1.071613 | 0.7266309 | -2.127556 | 0.03337392 | 0.101933 |
| ENSG00000230306 | BANF1P2 | 2.853566 | -1.071747 | 1.103836 | -1.971005 | 0.04872325 | 0.135809 |
| ENSG00000250444 | CCT5P1 | 54.83385 | -1.071758 | 0.3886809 | -3.109218 | 0.00187584 | 0.009807 |
| ENSG00000237766 | GGTA2P | 10.96539 | -1.071769 | 0.8849988 | -2.012613 | 0.04415534 | 0.126183 |
| ENSG00000168556 | ING2 | 627.3888 | -1.072248 | 0.3307067 | -3.541719 | 0.00039753 | 0.002579 |
| ENSG00000188313 | PLSCR1 | 2958.999 | -1.07236 | 0.362577 | -3.276595 | 0.00105067 | 0.005975 |
| ENSG00000144810 | COL8A1 | 1.738119 | -1.072737 | 1.870072 | -2.268266 | 0.02331301 | 0.076773 |
| ENSG00000120669 | SOHLH2 | 13.08211461 | -1.073657165 | 0.688355592 | -2.176434853 | 0.02952276 | 0.092639 |
| ENSG00000237424 | FOXD2-AS1 | 187.3078 | -1.073983 | 0.4246917 | -2.901083 | 0.00371875 | 0.017355 |
| ENSG00000234230 | ZFX-AS1 | 1.98519 | -1.074514 | 1.359394 | -1.984906 | 0.04715491 | 0.132494 |
| ENSG00000143473 | KCNH1 | 6.356531 | -1.074577 | 0.7966311 | -2.042878 | 0.04106455 | 0.119454 |
| ENSG00000160255 | ITGB2 | 50.34776 | -1.074633 | 0.3930632 | -3.083083 | 0.00204868 | 0.010573 |
| ENSG00000240449 | REPIN1-AS1 | 153.6786 | -1.074755 | 0.1717433 | -6.417061 | 1.39E-10 | 3.41E-09 |
| ENSG00000151725 | CENPU | 2841.976 | -1.075016 | 0.7050652 | -2.192362 | 0.02835337 | 0.089617 |
| ENSG00000161048 | NAPEPLD | 598.2443 | -1.075276 | 0.2541445 | -4.467147 | 7.93E-06 | 8.03E-05 |
| ENSG00000259683 | | 14.62219 | -1.075773 | 0.6903613 | -2.178305 | 0.02938334 | 0.092323 |
| ENSG00000173338 | KCNK7 | 13.39092 | -1.076349 | 0.6975604 | -2.174206 | 0.02968967 | 0.093 |
| ENSG00000282851 | BISPR | 119.3392 | -1.076454 | 0.6074844 | -2.317049 | 0.02050104 | 0.069308 |
| ENSG00000286957 | | 2.595666 | -1.076462 | 1.33921 | -2.00271 | 0.04520844 | 0.128451 |
| ENSG00000179833 | SERTAD2 | 1247.828 | -1.076878 | 0.2347689 | -4.795996 | 1.62E-06 | 1.94E-05 |
| ENSG00000226263 | ISM1-AS1 | 6.152795 | -1.077088 | 0.8204576 | -2.036095 | 0.04174084 | 0.120913 |
| ENSG00000086062 | B4GALT1 | 7589.333903 | -1.077460585 | 0.229016494 | -4.90574864 | 9.31E-07 | 1.17E-05 |
| ENSG00000254691 | | 2.451032 | -1.077508 | 1.107765 | -1.953006 | 0.05081887 | 0.140207 |
| ENSG00000139131 | YARS2 | 774.8800778 | -1.078715764 | 0.190435217 | -5.827389426 | 5.63E-09 | 1.07E-07 |
| ENSG00000263361 | MIR378H | 1.531484 | -1.078861 | 1.990316 | -2.351588 | 0.01869348 | NA |
| ENSG00000232320 | MXRA7P1 | 46.22787 | -1.07928 | 0.3800027 | -3.181498 | 0.00146516 | 0.007942 |
| ENSG00000221838 | AP4M1 | 815.33 | -1.07931 | 0.148112 | -7.419688 | 1.17E-13 | 4.30E-12 |
| ENSG00000277778 | PGM5P2 | 17.30587 | -1.079472 | 0.5620751 | -2.409751 | 0.01596343 | 0.056622 |
| ENSG00000260988 | | 8.443181 | -1.079775 | 0.8229238 | -2.04839 | 0.04052182 | 0.118103 |
| ENSG00000089639 | GMIP | 1145.685652 | -1.080741198 | 0.139353874 | -7.877772279 | 3.33E-15 | 1.46E-13 |

| | | | | | | | |
|---|---|---|---|---|---|---|---|
| ENSG00000260541 | | 8.64147 | -1.082068 | 0.6730091 | -2.17968 | 0.02928122 | 0.092057 |
| ENSG00000264278 | ZNF236-DT | 71.06243 | -1.082299 | 0.3031952 | -3.832701 | 0.00012674 | 0.000946 |
| ENSG00000235748 | SEPTIN14P12 | 8.610441 | -1.082956 | 0.9190932 | -2.011867 | 0.04423399 | 0.126385 |
| ENSG00000120647 | CCDC77 | 327.091827 | -1.084882942 | 0.336606349 | -3.518902285 | 0.00043334 | 0.002773 |
| ENSG00000196776 | CD47 | 14136.34 | -1.085965 | 0.1371397 | -8.050644 | 8.24E-16 | 3.93E-14 |
| ENSG00000287697 | | 12.12164 | -1.086806 | 0.5436156 | -2.471459 | 0.0134563 | 0.049368 |
| ENSG00000278954 | | 1.555335 | -1.087235 | 1.878644 | -2.308664 | 0.02096224 | NA |
| ENSG00000261611 | | 3.296723 | -1.087858 | 0.9657103 | -1.981009 | 0.04759023 | 0.133449 |
| ENSG00000123213 | NLN | 3137.033002 | -1.088027085 | 0.266493496 | -4.333146085 | 1.47E-05 | 0.000139 |
| ENSG00000214900 | LINC01588 | 115.5578 | -1.088812 | 0.3421211 | -3.488415 | 0.00048589 | 0.003062 |
| ENSG00000196503 | ARL9 | 7.714685 | -1.089833 | 0.5646928 | -2.435659 | 0.01486468 | 0.05355 |
| ENSG00000137563 | GGH | 1433.174617 | -1.090084766 | 0.503108108 | -2.625583342 | 0.00865006 | 0.034638 |
| ENSG00000234967 | | 416.1558 | -1.090154 | 0.3039104 | -3.857633 | 0.00011449 | 0.000864 |
| ENSG00000183386 | FHL3 | 348.2565 | -1.090275 | 0.3049533 | -3.849705 | 0.00011826 | 0.00089 |
| ENSG00000120327 | PCDHB14 | 127.8519631 | -1.090508916 | 0.33099995 | -3.588446155 | 0.00033266 | 0.002215 |
| ENSG00000254842 | LINC02551 | 2.502385 | -1.090636 | 1.893694 | -2.292225 | 0.02189264 | 0.073202 |
| ENSG00000254388 | DUTP2 | 12.30245 | -1.090722 | 1.188185 | -2.0309 | 0.04226509 | 0.122072 |
| ENSG00000197150 | ABCB8 | 2099.129 | -1.091458 | 0.1611176 | -6.922368 | 4.44E-12 | 1.33E-10 |
| ENSG00000135747 | ZNF670-ZNF695 | 39.78120785 | -1.091518291 | 0.656809318 | -2.265996717 | 0.02345158 | 0.077067 |
| ENSG00000213420 | GPC2 | 130.3781 | -1.091572 | 0.3492784 | -3.426015 | 0.00061251 | 0.003744 |
| ENSG00000146242 | TPBG | 6365.703 | -1.092326 | 0.3023461 | -3.887061 | 0.00010147 | 0.000775 |
| ENSG00000106028 | SSBP1 | 4316.347832 | -1.092724101 | 0.221268229 | -5.125312406 | 2.97E-07 | 4.10E-06 |
| ENSG00000223959 | AFG3L1P | 2334.593 | -1.093243 | 0.2239412 | -5.086011 | 3.66E-07 | 4.95E-06 |
| ENSG00000095209 | TMEM38B | 1304.100152 | -1.093567219 | 0.200479224 | -5.636396672 | 1.74E-08 | 3.03E-07 |
| ENSG00000171227 | TMEM37 | 163.0459 | -1.093973 | 0.4580244 | -2.795666 | 0.00517929 | 0.022802 |
| ENSG00000134013 | LOXL2 | 258.0626741 | -1.094038142 | 0.451466145 | -2.824996711 | 0.00472811 | 0.021184 |
| ENSG00000249947 | XBP1P1 | 15.35168 | -1.094139 | 0.6405005 | -2.279272 | 0.02265088 | 0.075179 |
| ENSG00000259341 | | 5.738879 | -1.096377 | 0.8229011 | -2.06026 | 0.03937372 | 0.11553 |
| ENSG00000138669 | PRKG2 | 73.54096687 | -1.09651471 | 0.4329442 | -2.92378885 | 0.00345799 | 0.016348 |
| ENSG00000199047 | MIR378A | 3.007371 | -1.097386 | 1.388263 | -1.969262 | 0.04892301 | 0.136251 |
| ENSG00000278356 | | 16.13406 | -1.097543 | 0.5790557 | -2.420344 | 0.01550584 | 0.055351 |
| ENSG00000196743 | GM2A | 1556.314 | -1.098159 | 0.2036929 | -5.576456 | 2.45E-08 | 4.16E-07 |
| ENSG00000111667 | USP5 | 7620.452275 | -1.098553035 | 0.214242382 | -5.341992563 | 9.19E-08 | 1.40E-06 |

| | | | | | | | |
|---|---|---|---|---|---|---|---|
| ENSG00000183011 | NAA38 | 1919.24 | -1.098843 | 0.286316 | -4.108719 | 3.98E-05 | 0.000338 |
| ENSG00000027001 | MIPEP | 973.6081127 | -1.099257135 | 0.188078289 | -6.023048855 | 1.71E-09 | 3.54E-08 |
| ENSG00000237840 | FAM21FP | 62.79823 | -1.099742 | 0.3789108 | -3.23225 | 0.0012282 | 0.006831 |
| ENSG00000128534 | LSM8 | 2134.013531 | -1.099757622 | 0.126673315 | -8.784923637 | 1.56E-18 | 9.95E-17 |
| ENSG00000220517 | ASS1P1 | 473.3007 | -1.099781 | 0.5313741 | -2.540891 | 0.01105705 | 0.042203 |
| ENSG00000013573 | DDX11 | 2244.787544 | -1.099857359 | 0.287594643 | -4.086506329 | 4.38E-05 | 0.000369 |
| ENSG00000124215 | CDH26 | 19.76759011 | -1.099982171 | 0.709422093 | -2.175055653 | 0.02962595 | 0.092877 |
| ENSG00000270953 | | 17.83705 | -1.100139 | 0.5468352 | -2.508282 | 0.012132 | 0.045399 |
| ENSG00000065268 | WDR18 | 4276.740665 | -1.100225106 | 0.232475186 | -4.949738257 | 7.43E-07 | 9.52E-06 |
| ENSG00000254756 | | 8.109325 | -1.100373 | 0.7216273 | -2.173418 | 0.0297489 | 0.09312 |
| ENSG00000286216 | | 27.21021 | -1.100374 | 0.4407471 | -2.886185 | 0.00389943 | 0.018071 |
| ENSG00000131558 | EXOC4 | 3391.784875 | -1.100613598 | 0.136921748 | -8.164590686 | 3.23E-16 | 1.61E-14 |
| ENSG00000287851 | | 19.89116 | -1.101425 | 0.6449635 | -2.282079 | 0.02248469 | 0.074796 |
| ENSG00000204869 | IGFL4 | 5.787893 | -1.102987 | 1.03294 | -1.944734 | 0.051807 | 0.142358 |
| ENSG00000282836 | | 2.347011 | -1.103113 | 1.30899 | -1.988754 | 0.04672834 | 0.131671 |
| ENSG00000134285 | FKBP11 | 916.3917418 | -1.104605468 | 0.181924328 | -6.23285167 | 4.58E-10 | 1.04E-08 |
| ENSG00000288056 | | 12.22683 | -1.10508 | 0.6693086 | -2.256222 | 0.02405674 | 0.0787 |
| ENSG00000049860 | HEXB | 6536.375376 | -1.105168785 | 0.166280549 | -6.79816734 | 1.06E-11 | 3.03E-10 |
| ENSG00000106258 | CYP3A5 | 175.8620045 | -1.105386694 | 0.270716821 | -4.325952356 | 1.52E-05 | 0.000143 |
| ENSG00000244625 | MIATNB | 162.3287 | -1.10729 | 0.2900567 | -4.078905 | 4.52E-05 | 0.00038 |
| ENSG00000183605 | SFXN4 | 2187.808 | -1.107307 | 0.2129543 | -5.393654 | 6.90E-08 | 1.08E-06 |
| ENSG00000215325 | ASS1P10 | 135.6513 | -1.107362 | 0.5220696 | -2.599172 | 0.0093449 | 0.03681 |
| ENSG00000212124 | TAS2R19 | 6.015517 | -1.10977 | 0.8651485 | -2.056905 | 0.03969536 | 0.11623 |
| ENSG00000164440 | TXLNB | 16.38348 | -1.110573 | 0.6699716 | -2.274901 | 0.02291186 | 0.075771 |
| ENSG00000105976 | MET | 2849.184396 | -1.110703746 | 0.250036583 | -4.682573896 | 2.83E-06 | 3.21E-05 |
| ENSG00000177694 | NAALADL2 | 704.4008 | -1.111257 | 0.4683152 | -2.807522 | 0.00499243 | 0.022126 |
| ENSG00000178184 | PARD6G | 410.9021 | -1.111785 | 0.1667948 | -6.825381 | 8.77E-12 | 2.53E-10 |
| ENSG00000198416 | ZNF658B | 52.86123 | -1.112206 | 0.2796351 | -4.21899 | 2.45E-05 | 0.00022 |
| ENSG00000197557 | TTC30A | 337.2411 | -1.112375 | 0.188807 | -6.06636 | 1.31E-09 | 2.76E-08 |
| ENSG00000163638 | ADAMTS9 | 1466.304 | -1.112543 | 0.4346222 | -2.961819 | 0.00305828 | 0.014771 |
| ENSG00000267056 | | 28.79451 | -1.112749 | 0.5357697 | -2.562013 | 0.01040674 | 0.040183 |
| ENSG00000107833 | NPM3 | 2094.267434 | -1.112772031 | 0.276273336 | -4.275940051 | 1.90E-05 | 0.000175 |
| ENSG00000138336 | TET1 | 208.9658128 | -1.113350539 | 0.454932127 | -2.863561387 | 0.00418908 | 0.01918 |

| | | | | | | | |
|---|---|---|---|---|---|---|---|
| ENSG00000279488 | | 7.765367 | -1.113544 | 0.6494999 | -2.263571 | 0.02360053 | 0.077499 |
| ENSG00000111087 | GLI1 | 427.3311534 | -1.113675782 | 0.842503493 | -2.099875567 | 0.03573979 | 0.107332 |
| ENSG00000198420 | TCAF1 | 1845.003 | -1.114052 | 0.1761348 | -6.487121 | 8.75E-11 | 2.19E-09 |
| ENSG00000117479 | SLC19A2 | 3527.704826 | -1.114367274 | 0.346671421 | -3.532258934 | 0.00041203 | 0.002658 |
| ENSG00000128311 | TST | 3486.18547 | -1.114497059 | 0.258085014 | -4.590748196 | 4.42E-06 | 4.78E-05 |
| ENSG00000111788 | | 415.5310489 | -1.114884003 | 0.466791629 | -2.821862807 | 0.00477456 | 0.02136 |
| ENSG00000166886 | NAB2 | 4219.549 | -1.114929 | 0.1815053 | -6.307834 | 2.83E-10 | 6.63E-09 |
| ENSG00000108561 | C1QBP | 7163.140262 | -1.115498354 | 0.223350271 | -5.220674208 | 1.78E-07 | 2.57E-06 |
| ENSG00000175449 | RFESD | 50.58777 | -1.116776 | 0.3492603 | -3.51674 | 0.00043688 | 0.002793 |
| ENSG00000185761 | ADAMTSL5 | 370.8645 | -1.117481 | 0.2756647 | -4.304531 | 1.67E-05 | 0.000156 |
| ENSG00000213066 | CEP43 | 571.0188 | -1.117645 | 0.2282926 | -5.104256 | 3.32E-07 | 4.54E-06 |
| ENSG00000287916 | | 8.325734 | -1.117683 | 0.7412585 | -2.172527 | 0.02981595 | 0.093257 |
| ENSG00000188452 | CERKL | 21.20209 | -1.117689 | 0.5573059 | -2.505339 | 0.01223342 | 0.045721 |
| ENSG00000260389 | WBP11P1 | 4.806657 | -1.118904 | 0.9575216 | -2.003315 | 0.04514352 | 0.128335 |
| ENSG00000185989 | RASA3 | 1133.677 | -1.118989 | 0.2829746 | -4.217968 | 2.47E-05 | 0.000221 |
| ENSG00000197448 | GSTK1 | 3610.764 | -1.119159 | 0.199468 | -5.785119 | 7.25E-09 | 1.36E-07 |
| ENSG00000283511 | | 9.799543 | -1.119199 | 0.7288978 | -2.203708 | 0.02754486 | 0.087624 |
| ENSG00000271551 | | 6.756106 | -1.119513 | 1.250215 | -2.019598 | 0.0434251 | 0.124636 |
| ENSG00000155980 | KIF5A | 140.8 | -1.119693 | 0.2203026 | -5.253974 | 1.49E-07 | 2.18E-06 |
| ENSG00000185499 | MUC1 | 5135.717 | -1.119865 | 0.3604267 | -3.431092 | 0.00060116 | 0.00368 |
| ENSG00000272744 | | 15.10082 | -1.120314 | 0.6713687 | -2.273164 | 0.02301631 | 0.07606 |
| ENSG00000197140 | ADAM32 | 156.4245 | -1.120705 | 0.3163282 | -3.828276 | 0.00012904 | 0.00096 |
| ENSG00000153044 | CENPH | 962.5776 | -1.12113 | 0.4933558 | -2.736619 | 0.00620741 | 0.026447 |
| ENSG00000149218 | ENDOD1 | 1317.253 | -1.12161 | 0.1657184 | -6.918095 | 4.58E-12 | 1.37E-10 |
| ENSG00000214851 | LINC00612 | 13.9731 | -1.121746 | 0.6956143 | -2.221683 | 0.02630475 | 0.084377 |
| ENSG00000287901 | | 2.535446 | -1.123166 | 0.9870132 | -2.019189 | 0.04346756 | 0.124732 |
| ENSG00000168538 | TRAPPC11 | 2201.421 | -1.123227 | 0.1570434 | -7.297777 | 2.93E-13 | 1.03E-11 |
| ENSG00000180881 | CAPS2 | 66.83952 | -1.123625 | 0.4948178 | -2.724867 | 0.00643273 | 0.027181 |
| ENSG00000165181 | SHOC1 | 8.846693 | -1.124022 | 0.773396 | -2.13043 | 0.03313611 | 0.101346 |
| ENSG00000141965 | FEM1A | 112.823 | -1.124295 | 0.3889356 | -3.246941 | 0.00116653 | 0.006539 |
| ENSG00000214253 | FIS1 | 4670.608 | -1.124348 | 0.1895908 | -6.101826 | 1.05E-09 | 2.24E-08 |
| ENSG00000267426 | | 5.572825 | -1.124404 | 0.7110798 | -2.200393 | 0.027779 | 0.088137 |
| ENSG00000115935 | WIPF1 | 173.5264719 | -1.124571963 | 0.456969285 | -2.878892725 | 0.00399074 | 0.018418 |

| | | | | | | | |
|---|---|---|---|---|---|---|---|
| ENSG00000135976 | ANKRD36 | 712.0860062 | -1.124883678 | 0.348038577 | -3.551439562 | 0.00038313 | 0.002502 |
| ENSG00000104833 | TUBB4A | 46.39950707 | -1.125296003 | 0.615551012 | -2.392496918 | 0.01673417 | 0.058865 |
| ENSG00000224138 | | 39.13369 | -1.125802 | 0.3565496 | -3.478337 | 0.00050454 | 0.003161 |
| ENSG00000260763 | | 5.179513 | -1.126647 | 1.092607 | -1.958298 | 0.05019504 | 0.138864 |
| ENSG00000260456 | C16orf95 | 79.37239 | -1.126777 | 0.365872 | -3.409945 | 0.00064976 | 0.003941 |
| ENSG00000223749 | MIR503HG | 1577.001 | -1.128034 | 0.4111519 | -3.129341 | 0.00175199 | 0.009262 |
| ENSG00000145414 | NAF1 | 1002.006 | -1.128575 | 0.2058583 | -5.669645 | 1.43E-08 | 2.54E-07 |
| ENSG00000118246 | FASTKD2 | 2360.509814 | -1.128576409 | 0.210682459 | -5.55200491 | 2.82E-08 | 4.73E-07 |
| ENSG00000130589 | HELZ2 | 7202.026571 | -1.128736986 | 0.285785721 | -4.227776181 | 2.36E-05 | 0.000213 |
| ENSG00000275881 | | 3.057467 | -1.128756 | 1.259501 | -1.976781 | 0.04806634 | 0.134443 |
| ENSG00000135899 | SP110 | 912.4555932 | -1.128812395 | 0.312009488 | -3.904744295 | 9.43E-05 | 0.000727 |
| ENSG00000185614 | INKA1 | 6.658694 | -1.129523 | 0.9850481 | -1.971652 | 0.04864933 | 0.135668 |
| ENSG00000170425 | ADORA2B | 359.9885 | -1.129535 | 0.3575703 | -3.49403 | 0.00047579 | 0.003006 |
| ENSG00000228915 | OR7E128P | 15.1508 | -1.129697 | 0.8869996 | -2.088526 | 0.0367504 | 0.109489 |
| ENSG00000228302 | | 40.22327 | -1.129799 | 0.406286 | -3.150744 | 0.00162855 | 0.008676 |
| ENSG00000100365 | NCF4 | 2.888452563 | -1.12981851 | 1.124043003 | -1.986784622 | 0.04694627 | 0.132126 |
| ENSG00000273069 | | 1.325711 | -1.129917 | 1.996395 | -2.287806 | 0.02214883 | NA |
| ENSG00000279232 | | 2.067804 | -1.12994 | 1.980281 | -2.284513 | 0.02234139 | 0.07445 |
| ENSG00000204790 | | 3.697416 | -1.13039 | 1.287417 | -1.984839 | 0.04716242 | 0.132501 |
| ENSG00000228393 | LINC01004 | 110.7048 | -1.13106 | 0.3689257 | -3.390333 | 0.00069808 | 0.004195 |
| ENSG00000169994 | MYO7B | 4.322643 | -1.131293 | 1.517964 | -2.058739 | 0.03951922 | 0.115816 |
| ENSG00000122859 | NEUROG3 | 2.488368086 | -1.13172925 | 1.706843389 | -2.141943341 | 0.03219804 | 0.099047 |
| ENSG00000106400 | ZNHIT1 | 3929.023289 | -1.131893405 | 0.174838713 | -6.635188889 | 3.24E-11 | 8.61E-10 |
| ENSG00000138035 | PNPT1 | 3726.177044 | -1.132087843 | 0.218033179 | -5.392386952 | 6.95E-08 | 1.09E-06 |
| ENSG00000173320 | STOX2 | 474.5226 | -1.132088 | 0.3299026 | -3.739322 | 0.00018452 | 0.001319 |
| ENSG00000149488 | TMC2 | 8.274456 | -1.132649 | 0.8657133 | -2.065917 | 0.03883635 | 0.114279 |
| ENSG00000278390 | | 25.17827 | -1.132747 | 0.4542279 | -2.903404 | 0.0036913 | 0.017242 |
| ENSG00000101868 | POLA1 | 2273.594299 | -1.133688979 | 0.319257518 | -3.850033991 | 0.0001181 | 0.000889 |
| ENSG00000273391 | | 47.35288 | -1.134266 | 0.3180387 | -3.848769 | 0.00011871 | 0.000892 |
| ENSG00000167491 | GATAD2A | 3274.347 | -1.134823 | 0.1790713 | -6.502315 | 7.91E-11 | 1.99E-09 |
| ENSG00000079112 | CDH17 | 2.83918352 | -1.135063933 | 1.245904717 | -1.974947567 | 0.0482741 | 0.134854 |
| ENSG00000076382 | SPAG5 | 4238.51746 | -1.135275429 | 0.524096397 | -2.66299761 | 0.0077448 | 0.031688 |
| ENSG00000006459 | KDM7A | 1288.428126 | -1.136334126 | 0.385155356 | -3.307298216 | 0.00094201 | 0.005427 |

| | | | | | | | |
|---|---|---|---|---|---|---|---|
| ENSG00000188015 | S100A3 | 7.404784 | -1.136447 | 1.158928 | -1.978402 | 0.04788341 | 0.13403 |
| ENSG00000176170 | SPHK1 | 2043.456 | -1.137201 | 0.3036676 | -4.025128 | 5.69E-05 | 0.000465 |
| ENSG00000184194 | GPR173 | 387.4626 | -1.137424 | 0.3438702 | -3.630925 | 0.00028241 | 0.001918 |
| ENSG00000239486 | | 8.998952 | -1.137689 | 0.8712601 | -2.091202 | 0.03650992 | 0.108971 |
| ENSG00000168763 | CNNM3 | 3502.769 | -1.137733 | 0.1490461 | -7.786779 | 6.87E-15 | 2.90E-13 |
| ENSG00000213252 | | 89.87708 | -1.137735 | 0.3267777 | -3.772273 | 0.00016177 | 0.001176 |
| ENSG00000179082 | C9orf106 | 137.0098 | -1.138988 | 0.360558 | -3.484534 | 0.000493 | 0.003102 |
| ENSG00000229388 | LINC01715 | 41.02333 | -1.139236 | 0.4442203 | -2.963479 | 0.00304183 | 0.01471 |
| ENSG00000141668 | CBLN2 | 1063.036 | -1.139599 | 0.7309941 | -2.242706 | 0.02491581 | 0.080878 |
| ENSG00000233006 | MIR3936HG | 211.1973 | -1.140243 | 0.5373844 | -2.618911 | 0.00882111 | 0.035201 |
| ENSG00000163155 | LYSMD1 | 806.7992 | -1.141339 | 0.2248811 | -5.286814 | 1.24E-07 | 1.85E-06 |
| ENSG00000231927 | | 3.362682 | -1.141654 | 0.9153259 | -2.059966 | 0.03940176 | 0.115586 |
| ENSG00000152076 | CCDC74B | 112.4215 | -1.143745 | 0.4343598 | -3.038588 | 0.0023769 | 0.011981 |
| ENSG00000281398 | SNHG4 | 1664.788 | -1.144974 | 0.235378 | -5.093442 | 3.52E-07 | 4.77E-06 |
| ENSG00000105983 | LMBR1 | 3983.568736 | -1.145115421 | 0.136131854 | -8.537646423 | 1.37E-17 | 8.02E-16 |
| ENSG00000064199 | SPA17 | 115.1126156 | -1.145604694 | 0.222341974 | -5.355330114 | 8.54E-08 | 1.31E-06 |
| ENSG00000163701 | IL17RE | 437.4405 | -1.146381 | 0.3160404 | -3.917027 | 8.96E-05 | 0.000694 |
| ENSG00000206262 | FOXL2NB | 6.897033 | -1.146531 | 0.7654214 | -2.193214 | 0.02829193 | 0.089433 |
| ENSG00000256802 | | 12.85251 | -1.147038 | 0.6877051 | -2.290104 | 0.02201526 | 0.073529 |
| ENSG00000225096 | | 32.99596 | -1.149247 | 0.4964154 | -2.76437 | 0.00570329 | 0.024699 |
| ENSG00000077152 | UBE2T | 3968.734064 | -1.149289989 | 0.366129063 | -3.475495975 | 0.00050991 | 0.00319 |
| ENSG00000213782 | DDX47 | 84.78044 | -1.150518 | 0.3224079 | -3.867827 | 0.00010981 | 0.000831 |
| ENSG00000117597 | UTP25 | 2216.162924 | -1.151229753 | 0.179097303 | -6.59969497 | 4.12E-11 | 1.07E-09 |
| ENSG00000180530 | NRIP1 | 3240.327 | -1.151432 | 0.2929802 | -4.200634 | 2.66E-05 | 0.000236 |
| ENSG00000133597 | ADCK2 | 1771.593458 | -1.151932528 | 0.202814721 | -5.866663371 | 4.45E-09 | 8.65E-08 |
| ENSG00000270923 | TAS2R6P | 5.020052 | -1.152567 | 0.8921694 | -2.082901 | 0.03726026 | 0.110673 |
| ENSG00000128595 | CALU | 7440.768011 | -1.152955095 | 0.148607862 | -7.958731131 | 1.74E-15 | 7.89E-14 |
| ENSG00000238109 | RNF14P3 | 3.073832 | -1.153023 | 1.153908 | -1.980833 | 0.04761006 | 0.133479 |
| ENSG00000135249 | RINT1 | 1135.059405 | -1.153719892 | 0.212620923 | -5.62050618 | 1.90E-08 | 3.29E-07 |
| ENSG00000273062 | RALGPS2-AS1 | 12.53899 | -1.15385 | 0.5555842 | -2.573773 | 0.01005962 | 0.039058 |
| ENSG00000121350 | PYROXD1 | 398.7754912 | -1.154465025 | 0.280596519 | -4.37257298 | 1.23E-05 | 0.000118 |
| ENSG00000255561 | FDXACB1 | 41.68164 | -1.154469 | 0.3660151 | -3.492258 | 0.00047896 | 0.003023 |
| ENSG00000218631 | | 18.2735 | -1.156201 | 0.5353932 | -2.646275 | 0.00813836 | 0.032977 |

| | | | | | | | |
|---|---|---|---|---|---|---|---|
| ENSG00000084444 | FAM234B | 2778.662077 | -1.156692042 | 0.252268305 | -4.819558771 | 1.44E-06 | 1.74E-05 |
| ENSG00000162849 | KIF26B | 19.47872 | -1.157796 | 0.5806861 | -2.519245 | 0.01176068 | 0.044297 |
| ENSG00000236893 | ASS1P7 | 186.3407 | -1.157896 | 0.4815704 | -2.849049 | 0.00438501 | 0.019898 |
| ENSG00000251307 | | 6.386121 | -1.158332 | 0.8500261 | -2.106024 | 0.03520228 | 0.106052 |
| ENSG00000224592 | MIR3659HG | 1.371197 | -1.15945 | 2.160164 | -2.363273 | 0.0181143 | NA |
| ENSG00000263884 | | 19.21366 | -1.159606 | 0.5193939 | -2.694093 | 0.00705805 | 0.029333 |
| ENSG00000163864 | NMNAT3 | 339.8161 | -1.15996 | 0.2687698 | -4.552669 | 5.30E-06 | 5.61E-05 |
| ENSG00000136213 | CHST12 | 1786.070463 | -1.160172634 | 0.197729177 | -6.053668863 | 1.42E-09 | 2.97E-08 |
| ENSG00000229358 | DPY19L1P1 | 260.7781 | -1.160993 | 0.3420088 | -3.706446 | 0.00021019 | 0.001482 |
| ENSG00000267327 | | 11.5896 | -1.16132 | 1.314243 | -2.015289 | 0.04387432 | 0.125655 |
| ENSG00000108641 | B9D1 | 1078.955699 | -1.161971401 | 0.204396975 | -5.874920742 | 4.23E-09 | 8.27E-08 |
| ENSG00000157992 | KRTCAP3 | 390.4811 | -1.162061 | 0.2337665 | -5.187443 | 2.13E-07 | 3.03E-06 |
| ENSG00000197110 | IFNL3 | 6.52033 | -1.16312 | 1.305989 | -2.021924 | 0.04318419 | 0.124146 |
| ENSG00000276900 | | 107.1755 | -1.163208 | 0.3279364 | -3.845871 | 0.00012013 | 0.000901 |
| ENSG00000164989 | CCDC171 | 152.244 | -1.163297 | 0.3649964 | -3.523355 | 0.00042612 | 0.002734 |
| ENSG00000117983 | MUC5B | 75.12204812 | -1.163342821 | 0.638031422 | -2.429762191 | 0.01510873 | 0.054232 |
| ENSG00000161036 | LRWD1 | 1648.389 | -1.164285 | 0.17185 | -6.93399 | 4.09E-12 | 1.24E-10 |
| ENSG00000183098 | GPC6 | 1160.37 | -1.165626 | 0.2166406 | -5.582546 | 2.37E-08 | 4.03E-07 |
| ENSG00000286032 | | 20.68845 | -1.166222 | 0.5288255 | -2.684949 | 0.00725409 | 0.030021 |
| ENSG00000139178 | C1RL | 21.77835896 | -1.166647233 | 0.701346441 | -2.290467952 | 0.0219942 | 0.073468 |
| ENSG00000089327 | FXYD5 | 573.4481429 | -1.168471773 | 0.504008095 | -2.773448418 | 0.00554656 | 0.024131 |
| ENSG00000259405 | ISCA1P4 | 11.71213 | -1.168504 | 0.67993 | -2.341597 | 0.01920142 | 0.06576 |
| ENSG00000234327 | ZNF232-AS1 | 82.68255 | -1.168603 | 0.3661982 | -3.527501 | 0.0004195 | 0.002695 |
| ENSG00000075275 | CELSR1 | 4811.166756 | -1.168760376 | 0.242854069 | -5.098707966 | 3.42E-07 | 4.66E-06 |
| ENSG00000288096 | | 12.58234 | -1.169366 | 0.6207398 | -2.447138 | 0.01439955 | 0.052222 |
| ENSG00000199436 | SNORD9 | 4.723579 | -1.169526 | 1.029215 | -2.01311 | 0.04410302 | 0.126092 |
| ENSG00000230795 | HLA-K | 22.41365 | -1.169569 | 0.5001915 | -2.773184 | 0.00555107 | 0.024142 |
| ENSG00000130707 | ASS1 | 20007.15212 | -1.169790713 | 0.514340661 | -2.758959722 | 0.00579857 | 0.025052 |
| ENSG00000286447 | | 33.04029 | -1.170085 | 0.5146948 | -2.748401 | 0.00598867 | 0.025688 |
| ENSG00000139187 | KLRG1 | 9.039006715 | -1.170154682 | 0.630258133 | -2.417379866 | 0.01563269 | 0.055699 |
| ENSG00000164292 | RHOBTB3 | 9210.416 | -1.170693 | 0.2700463 | -4.586086 | 4.52E-06 | 4.87E-05 |
| ENSG00000183621 | ZNF438 | 265.2159 | -1.171232 | 0.2262607 | -5.38651 | 7.18E-08 | 1.12E-06 |
| ENSG00000224090 | | 3.331942 | -1.17163 | 1.186782 | -2.056772 | 0.03970812 | 0.116241 |

| | | | | | | | |
|---|---|---|---|---|---|---|---|
| ENSG00000225953 | SATB2-AS1 | 372.4144 | -1.172331 | 0.2876501 | -4.334859 | 1.46E-05 | 0.000138 |
| ENSG00000185272 | RBM11 | 97.80063 | -1.172985 | 0.4272388 | -3.129674 | 0.00175001 | 0.009255 |
| ENSG00000111199 | TRPV4 | 70.12890281 | -1.173313242 | 0.519775541 | -2.737419988 | 0.00619232 | 0.026396 |
| ENSG00000178093 | TSSK6 | 60.49677 | -1.173694 | 0.3999388 | -3.30639 | 0.00094506 | 0.00544 |
| ENSG00000151655 | ITIH2 | 9.311491 | -1.174013 | 0.7522736 | -2.240137 | 0.025082 | 0.081328 |
| ENSG00000258128 | MKRN9P | 17.52874 | -1.174251 | 0.5584619 | -2.606108 | 0.00915776 | 0.036256 |
| ENSG00000278192 | | 9.076396 | -1.174457 | 0.8607473 | -2.096592 | 0.03602966 | 0.108044 |
| ENSG00000153094 | BCL2L11 | 2631.873 | -1.174725 | 0.1751063 | -6.873962 | 6.24E-12 | 1.84E-10 |
| ENSG00000261210 | CLEC19A | 2.672051 | -1.175447 | 1.609865 | -2.138295 | 0.03249284 | 0.099723 |
| ENSG00000177359 | | 6.082857 | -1.175581 | 1.024863 | -2.069769 | 0.03847397 | 0.113431 |
| ENSG00000155090 | KLF10 | 6028.686 | -1.175762 | 0.2643866 | -4.693267 | 2.69E-06 | 3.07E-05 |
| ENSG00000275832 | ARHGAP23 | 12.69705 | -1.180549 | 0.6112624 | -2.492584 | 0.01268173 | 0.047151 |
| ENSG00000278107 | | 26.12785 | -1.18068 | 0.4233186 | -3.173285 | 0.00150725 | 0.008141 |
| ENSG00000286240 | | 2.672409 | -1.181417 | 1.488141 | -2.100594 | 0.03567663 | 0.107179 |
| ENSG00000182552 | RWDD4 | 690.4981 | -1.181666 | 0.1571449 | -7.672572 | 1.69E-14 | 6.80E-13 |
| ENSG00000254726 | MEX3A | 3388.852 | -1.182202 | 0.2266548 | -5.432601 | 5.55E-08 | 8.88E-07 |
| ENSG00000225308 | ASS1P11 | 33.63956 | -1.182361 | 0.5745949 | -2.59496 | 0.00946018 | 0.03716 |
| ENSG00000057294 | PKP2 | 1484.299103 | -1.182550406 | 0.254436704 | -4.896415675 | 9.76E-07 | 1.22E-05 |
| ENSG00000102760 | RGCC | 40.79198822 | -1.182844 | 0.570969709 | -2.605609845 | 0.00917109 | 0.036292 |
| ENSG00000004059 | ARF5 | 5981.604057 | -1.183384725 | 0.216301972 | -5.692695199 | 1.25E-08 | 2.25E-07 |
| ENSG00000285755 | | 4.956479 | -1.18365 | 1.034042 | -2.079098 | 0.03760838 | 0.111431 |
| ENSG00000255026 | | 16.81234 | -1.184207 | 0.7519214 | -2.270177 | 0.02319687 | 0.076464 |
| ENSG00000133424 | LARGE1 | 2492.135012 | -1.184531273 | 0.152399568 | -7.91993696 | 2.38E-15 | 1.07E-13 |
| ENSG00000271503 | CCL5 | 109.8345 | -1.185545 | 0.6731878 | -2.374275 | 0.01758344 | 0.061268 |
| ENSG00000228049 | POLR2J2 | 27.76945 | -1.187101 | 0.4107191 | -3.265066 | 0.00109438 | 0.006187 |
| ENSG00000163293 | NIPAL1 | 917.1199 | -1.187306 | 0.3520262 | -3.700691 | 0.00021501 | 0.001512 |
| ENSG00000130045 | NXNL2 | 182.3919428 | -1.187754829 | 0.333559261 | -3.875616562 | 0.00010636 | 0.000808 |
| ENSG00000117868 | ESYT2 | 10248.38688 | -1.188538939 | 0.166479156 | -7.276323497 | 3.43E-13 | 1.20E-11 |
| ENSG00000178660 | ARMC10P1 | 135.3383 | -1.189205 | 0.2585026 | -4.842246 | 1.28E-06 | 1.56E-05 |
| ENSG00000152193 | OBI1 | 385.6019 | -1.191817 | 0.185967 | -6.578422 | 4.75E-11 | 1.23E-09 |
| ENSG00000248698 | LINC01085 | 10.80608 | -1.19275 | 0.8246089 | -2.209177 | 0.02716236 | 0.086609 |
| ENSG00000254343 | | 40.13603 | -1.193038 | 0.5331731 | -2.734797 | 0.00624188 | 0.02656 |
| ENSG00000270055 | | 104.2226 | -1.193169 | 0.2478104 | -5.052528 | 4.36E-07 | 5.84E-06 |

| | | | | | | | |
|---|---|---|---|---|---|---|---|
| ENSG00000249992 | TMEM158 | 24.43321 | -1.193599 | 0.5957682 | -2.538828 | 0.01112244 | 0.042398 |
| ENSG00000230303 | | 10.21149 | -1.195373 | 0.8193595 | -2.209447 | 0.0271436 | 0.086566 |
| ENSG00000279717 | | 2.773893 | -1.198342 | 1.115203 | -2.070802 | 0.03837726 | 0.113208 |
| ENSG00000103544 | VPS35L | 634.0397534 | -1.19989913 | 0.263236367 | -4.803313533 | 1.56E-06 | 1.87E-05 |
| ENSG00000100605 | ITPK1 | 8980.889825 | -1.199906145 | 0.182656172 | -6.661671109 | 2.71E-11 | 7.28E-10 |
| ENSG00000132437 | DDC | 2.891803787 | -1.20029509 | 1.22519587 | -2.066142192 | 0.03881506 | 0.114248 |
| ENSG00000286583 | | 2.089018 | -1.201723 | 1.526179 | -2.120379 | 0.03397408 | 0.103291 |
| ENSG00000217791 | ASS1P9 | 73.65226 | -1.201727 | 0.4889713 | -2.917462 | 0.00352893 | 0.016633 |
| ENSG00000146904 | EPHA1 | 1610.758 | -1.201859 | 0.1431056 | -8.530876 | 1.45E-17 | 8.46E-16 |
| ENSG00000284985 | | 25.71201 | -1.202038 | 0.36822 | -3.593358 | 0.00032644 | 0.002179 |
| ENSG00000174740 | PABPC5 | 24.14474 | -1.202753 | 0.5570502 | -2.675608 | 0.00745939 | 0.030732 |
| ENSG00000272562 | H3-3A-DT | 12.68565 | -1.202904 | 0.5906684 | -2.56071 | 0.01044585 | 0.040299 |
| ENSG00000110328 | GALNT18 | 625.3347834 | -1.203690147 | 0.194304986 | -6.350246733 | 2.15E-10 | 5.15E-09 |
| ENSG00000145945 | FAM50B | 639.8705 | -1.203846 | 0.2725475 | -4.673252 | 2.96E-06 | 3.35E-05 |
| ENSG00000188580 | NKAIN2 | 8.604863 | -1.204325 | 0.7138243 | -2.329901 | 0.01981141 | 0.067466 |
| ENSG00000113248 | PCDHB15 | 23.19946739 | -1.204578401 | 0.61816542 | -2.524559153 | 0.01158435 | 0.043782 |
| ENSG00000156097 | GPR61 | 26.79889 | -1.204619 | 0.4862147 | -2.914812 | 0.00355903 | 0.016746 |
| ENSG00000146192 | FGD2 | 21.51368 | -1.207355 | 1.137273 | -2.005507 | 0.04490891 | 0.127841 |
| ENSG00000215553 | KRT18P3 | 7.844767 | -1.207567 | 0.8606294 | -2.15121 | 0.03145964 | 0.097339 |
| ENSG00000244265 | SIAH2-AS1 | 64.87534 | -1.207765 | 0.418525 | -3.280751 | 0.00103531 | 0.005901 |
| ENSG00000278175 | GLIDR | 188.5904 | -1.208528 | 0.2385528 | -5.286623 | 1.25E-07 | 1.85E-06 |
| ENSG00000185361 | TNFAIP8L1 | 218.7881 | -1.208757 | 0.4278655 | -3.233201 | 0.00122412 | 0.006813 |
| ENSG00000105671 | DDX49 | 2769.801769 | -1.20935867 | 0.202316984 | -6.155479791 | 7.49E-10 | 1.64E-08 |
| ENSG00000111907 | TPD52L1 | 5534.256722 | -1.211582276 | 0.2277028 | -5.565661759 | 2.61E-08 | 4.40E-07 |
| ENSG00000196335 | STK31 | 229.1467 | -1.211754 | 0.2651887 | -4.81766 | 1.45E-06 | 1.76E-05 |
| ENSG00000204442 | FAM155A | 22.19079 | -1.211808 | 0.9387112 | -2.10866 | 0.03497395 | 0.10557 |
| ENSG00000229368 | | 8.643537 | -1.211835 | 0.6783858 | -2.387561 | 0.01696058 | 0.05952 |
| ENSG00000236519 | LINC01424 | 32.82983 | -1.212632 | 0.4920919 | -2.922176 | 0.00347595 | 0.016415 |
| ENSG00000227456 | LINC00310 | 8.668532 | -1.213507 | 0.7958171 | -2.228947 | 0.0258174 | 0.083165 |
| ENSG00000188859 | FAM78B | 102.4285 | -1.213624 | 0.3305839 | -3.97293 | 7.10E-05 | 0.000564 |
| ENSG00000042781 | USH2A | 14.7716234 | -1.213643273 | 0.532052748 | -2.772528827 | 0.00556226 | 0.024179 |
| ENSG00000169550 | MUC15 | 24.61866 | -1.215039 | 1.097388 | -1.917386 | 0.05518887 | 0.149502 |
| ENSG00000267130 | | 7.171883 | -1.215436 | 0.8075712 | -2.194377 | 0.02820829 | 0.089212 |

| | | | | | | | |
|---|---|---|---|---|---|---|---|
| ENSG00000105819 | PMPCB | 4238.323105 | -1.216440118 | 0.122313184 | -10.05805473 | 8.47E-24 | 9.22E-22 |
| ENSG00000255306 | | 33.08679 | -1.216503 | 0.6009607 | -2.56955 | 0.01018306 | 0.039473 |
| ENSG00000250241 | | 32.93748 | -1.216617 | 0.6094593 | -2.569511 | 0.01018423 | 0.039473 |
| ENSG00000274677 | | 20.77619 | -1.216617 | 0.5947343 | -2.597263 | 0.00939698 | 0.036977 |
| ENSG00000169871 | TRIM56 | 4897.031 | -1.218372 | 0.2910121 | -4.468995 | 7.86E-06 | 7.97E-05 |
| ENSG00000168255 | POLR2J3 | 35.14592 | -1.218632 | 0.3312268 | -3.980217 | 6.89E-05 | 0.000549 |
| ENSG00000185615 | PDIA2 | 325.6307 | -1.219346 | 0.2875001 | -4.512074 | 6.42E-06 | 6.67E-05 |
| ENSG00000186889 | TMEM17 | 95.69983 | -1.220856 | 0.26738 | -4.811348 | 1.50E-06 | 1.81E-05 |
| ENSG00000232342 | | 1.843401 | -1.221062 | 1.452312 | -2.030748 | 0.04228054 | 0.122103 |
| ENSG00000155561 | NUP205 | 6570.367 | -1.22191 | 0.2943069 | -4.430628 | 9.40E-06 | 9.32E-05 |
| ENSG00000277386 | | 8.717083 | -1.222506 | 0.6758419 | -2.435618 | 0.01486639 | 0.05355 |
| ENSG00000176490 | DIRAS1 | 197.3235 | -1.223641 | 0.3922564 | -3.488088 | 0.00048649 | 0.003065 |
| ENSG00000273038 | NA | 115.6944 | -1.225124 | 0.3688397 | -3.666067 | 0.00024631 | 0.001703 |
| ENSG00000181350 | LRRC75A | 364.0696 | -1.225227 | 0.178232 | -7.040304 | 1.92E-12 | 6.03E-11 |
| ENSG00000165630 | PRPF18 | 577.548 | -1.225762 | 0.1550859 | -8.054549 | 7.98E-16 | 3.82E-14 |
| ENSG00000278071 | | 10.91302 | -1.227179 | 0.8741155 | -2.177868 | 0.02941584 | 0.09236 |
| ENSG00000249916 | | 56.99039 | -1.229404 | 0.3928603 | -3.50394 | 0.00045843 | 0.00291 |
| ENSG00000091127 | PUS7 | 2161.296176 | -1.229411291 | 0.206999861 | -6.138473895 | 8.33E-10 | 1.81E-08 |
| ENSG00000284052 | | 9.512257 | -1.22951 | 0.7921769 | -2.24192 | 0.02496651 | 0.081023 |
| ENSG00000196092 | PAX5 | 63.2344 | -1.230063 | 0.3818957 | -3.575427 | 0.00034966 | 0.002311 |
| ENSG00000262503 | | 52.22688 | -1.230302 | 0.3908746 | -3.511837 | 0.00044502 | 0.002836 |
| ENSG00000238000 | PSME2P1 | 995.5283 | -1.230402 | 0.2512786 | -5.130301 | 2.89E-07 | 4.00E-06 |
| ENSG00000138622 | HCN4 | 38.33283495 | -1.23088865 | 0.609592091 | -2.595128932 | 0.00945554 | 0.037152 |
| ENSG00000265972 | TXNIP | 30140.56 | -1.231194 | 0.386689 | -3.552284 | 0.0003819 | 0.002496 |
| ENSG00000182575 | NXPH3 | 86.09684 | -1.231223 | 0.6656373 | -2.484093 | 0.01298818 | 0.048028 |
| ENSG00000105669 | COPE | 3462.582854 | -1.231799755 | 0.206971025 | -6.142105399 | 8.14E-10 | 1.77E-08 |
| ENSG00000239389 | PCDHA13 | 49.81706 | -1.232348 | 0.5041883 | -2.916586 | 0.00353886 | 0.016671 |
| ENSG00000265342 | | 5.844449 | -1.232683 | 0.8345928 | -2.244969 | 0.02477011 | 0.080494 |
| ENSG00000189014 | SHLD2P3 | 5.896528 | -1.232839 | 0.7882352 | -2.280566 | 0.02257416 | 0.075 |
| ENSG00000011523 | CEP68 | 1418.523309 | -1.232901245 | 0.142485467 | -8.792184536 | 1.47E-18 | 9.39E-17 |
| ENSG00000220884 | MESTP1 | 158.7529 | -1.232954 | 0.3330522 | -4.005718 | 6.18E-05 | 0.0005 |
| ENSG00000176125 | UFSP1 | 204.4125 | -1.23403 | 0.2250261 | -5.692775 | 1.25E-08 | 2.25E-07 |
| ENSG00000201207 | | 2.904491 | -1.235231 | 1.345963 | -2.11684 | 0.03427341 | 0.104022 |

| | | | | | | | |
|---|---|---|---|---|---|---|---|
| ENSG00000163431 | LMOD1 | 21.10346 | -1.239427 | 0.5938949 | -2.646867 | 0.00812413 | 0.03293 |
| ENSG00000130208 | APOC1 | 29.94959049 | -1.239995332 | 0.612325348 | -2.60736721 | 0.00912415 | 0.036144 |
| ENSG00000004866 | ST7 | 869.6360251 | -1.241200224 | 0.189792839 | -6.707491361 | 1.98E-11 | 5.43E-10 |
| ENSG00000157833 | GAREM2 | 81.30737 | -1.241598 | 0.5285662 | -2.845017 | 0.00444091 | 0.020085 |
| ENSG00000260808 | | 9.786494 | -1.241793 | 0.6925855 | -2.408077 | 0.01603678 | 0.056849 |
| ENSG00000116785 | CFHR3 | 55.64262592 | -1.242537785 | 0.436822064 | -3.260379743 | 0.00111263 | 0.006275 |
| ENSG00000275714 | H3C1 | 19.70112 | -1.243836 | 0.5529598 | -2.746905 | 0.00601605 | 0.025797 |
| ENSG00000261485 | PAN3-AS1 | 82.60522 | -1.244371 | 0.4830227 | -3.027822 | 0.00246323 | 0.012346 |
| ENSG00000273350 | | 4.038264 | -1.24449 | 1.306629 | -2.077453 | 0.03775974 | 0.111794 |
| ENSG00000042493 | CAPG | 1064.871093 | -1.244652202 | 0.357357543 | -3.822289149 | 0.00013222 | 0.00098 |
| ENSG00000132554 | RGS22 | 12.15014541 | -1.244942648 | 1.045742167 | -2.201027094 | 0.02773411 | 0.088016 |
| ENSG00000260852 | FBXL19-AS1 | 703.6199 | -1.244944 | 0.356559 | -3.831745 | 0.00012724 | 0.000948 |
| ENSG00000164307 | ERAP1 | 2663.762 | -1.2456 | 0.2528127 | -5.165494 | 2.40E-07 | 3.38E-06 |
| ENSG00000243083 | LINC00870 | 1.866387 | -1.245833 | 2.18488 | -2.330296 | 0.01979053 | 0.067412 |
| ENSG00000113763 | UNC5A | 69.40278267 | -1.246223071 | 0.532069379 | -2.848995609 | 0.00438575 | 0.019898 |
| ENSG00000107736 | CDH23 | 7.426462817 | -1.24684708 | 0.740126872 | -2.327348435 | 0.01994673 | 0.067753 |
| ENSG00000237330 | RNF223 | 1068.597 | -1.246989 | 0.253873 | -5.151085 | 2.59E-07 | 3.62E-06 |
| ENSG00000225489 | | 99.2289 | -1.247026 | 0.2838318 | -4.66081 | 3.15E-06 | 3.53E-05 |
| ENSG00000272374 | | 34.05205 | -1.247756 | 0.606743 | -2.6181 | 0.0088421 | 0.035258 |
| ENSG00000109063 | MYH3 | 123.8129483 | -1.248420206 | 0.393634485 | -3.540490974 | 0.00039938 | 0.002588 |
| ENSG00000121297 | TSHZ3 | 134.5438201 | -1.248857123 | 0.53051457 | -2.820815568 | 0.00479017 | 0.021419 |
| ENSG00000124249 | KCNK15 | 1431.625013 | -1.250593277 | 0.47967247 | -3.065606522 | 0.00217229 | 0.011108 |
| ENSG00000273816 | | 5.524265 | -1.251017 | 1.171805 | -2.090925 | 0.03653475 | 0.109007 |
| ENSG00000019144 | PHLDB1 | 1000.515507 | -1.251124875 | 0.186401909 | -6.894351181 | 5.41E-12 | 1.61E-10 |
| ENSG00000278022 | | 9.300016 | -1.251437 | 0.7569661 | -2.375921 | 0.01750523 | 0.061054 |
| ENSG00000280159 | | 4.227457 | -1.251905 | 1.134165 | -2.054182 | 0.03995809 | 0.116754 |
| ENSG00000169047 | IRS1 | 5550.251 | -1.252581 | 0.2464177 | -5.318974 | 1.04E-07 | 1.57E-06 |
| ENSG00000227620 | ALG1L8P | 13.43441 | -1.253555 | 0.5043793 | -2.948893 | 0.00318914 | 0.015286 |
| ENSG00000163171 | CDC42EP3 | 2555.786 | -1.254324 | 0.5439445 | -2.817093 | 0.00484604 | 0.021629 |
| ENSG00000196872 | CRACDL | 514.8563 | -1.255164 | 0.3034963 | -4.424299 | 9.68E-06 | 9.57E-05 |
| ENSG00000224511 | LINC00365 | 108.5739 | -1.256292 | 0.5665299 | -2.770623 | 0.00559492 | 0.024289 |
| ENSG00000135097 | MSI1 | 529.2691438 | -1.256731117 | 0.206712173 | -6.297873034 | 3.02E-10 | 7.05E-09 |
| ENSG00000004660 | CAMKK1 | 705.8344758 | -1.257618195 | 0.199465792 | -6.499611532 | 8.05E-11 | 2.02E-09 |

| | | | | | | | |
|---|---|---|---|---|---|---|---|
| ENSG00000186862 | PDZD7 | 181.5035 | -1.257768 | 0.3999366 | -3.528799 | 0.00041745 | 0.002686 |
| ENSG00000279265 | | 20.16676 | -1.258863 | 0.5965992 | -2.663288 | 0.00773811 | 0.031669 |
| ENSG00000254290 | | 43.85421 | -1.259527 | 0.8539394 | -2.298095 | 0.02155638 | 0.072269 |
| ENSG00000168447 | SCNN1B | 79.94239 | -1.259956 | 0.4866305 | -3.02514 | 0.00248518 | 0.012439 |
| ENSG00000178947 | SMIM10L2A | 86.40514 | -1.260296 | 0.4492884 | -3.237753 | 0.00120475 | 0.006716 |
| ENSG00000286305 | | 26.28895 | -1.260587 | 0.4275068 | -3.354367 | 0.00079547 | 0.004697 |
| ENSG00000267385 | | 12.99055 | -1.261454 | 0.6534308 | -2.548141 | 0.01082987 | 0.041485 |
| ENSG00000180694 | TMEM64 | 4974.383 | -1.262106 | 0.2481595 | -5.322504 | 1.02E-07 | 1.55E-06 |
| ENSG00000162840 | MT2P1 | 19.68307 | -1.262144 | 0.938198 | -2.239276 | 0.02513798 | 0.08143 |
| ENSG00000213398 | LCAT | 88.61187 | -1.262356 | 0.3470762 | -3.9486 | 7.86E-05 | 0.000619 |
| ENSG00000011422 | PLAUR | 256.0311119 | -1.262400253 | 0.31517415 | -4.306510431 | 1.66E-05 | 0.000155 |
| ENSG00000105854 | PON2 | 1269.761326 | -1.262462437 | 0.221549889 | -5.922891641 | 3.16E-09 | 6.30E-08 |
| ENSG00000167705 | RILP | 363.2612 | -1.262691 | 0.2915985 | -4.611283 | 4.00E-06 | 4.36E-05 |
| ENSG00000226609 | | 16.87514 | -1.262757 | 0.7244389 | -2.417976 | 0.01560711 | 0.055652 |
| ENSG00000138162 | TACC2 | 2860.14553 | -1.262958653 | 0.298135944 | -4.515870993 | 6.31E-06 | 6.56E-05 |
| ENSG00000170161 | | 174.6881 | -1.264739 | 0.2470817 | -5.340301 | 9.28E-08 | 1.41E-06 |
| ENSG00000255100 | | 28.1322 | -1.265071 | 0.550024 | -2.822256 | 0.0047687 | 0.021337 |
| ENSG00000260167 | | 27.68202 | -1.265493 | 0.3887215 | -3.621236 | 0.0002932 | 0.001982 |
| ENSG00000157600 | TMEM164 | 4225.793 | -1.26564 | 0.2405171 | -5.492607 | 3.96E-08 | 6.48E-07 |
| ENSG00000183833 | CFAP91 | 4.602036 | -1.266186 | 1.057483 | -2.087594 | 0.03683443 | 0.109702 |
| ENSG00000141068 | KSR1 | 590.6689 | -1.266989 | 0.1794369 | -7.232909 | 4.73E-13 | 1.61E-11 |
| ENSG00000279691 | | 9.404759 | -1.268713 | 0.7485215 | -2.365964 | 0.01798321 | 0.062408 |
| ENSG00000047597 | XK | 153.6564484 | -1.268871799 | 0.385606962 | -3.650905428 | 0.00026132 | 0.001795 |
| ENSG00000279479 | | 5.704369 | -1.269219 | 1.209124 | -1.968294 | 0.04903423 | 0.136504 |
| ENSG00000215004 | MESTP4 | 6.099588 | -1.269321 | 0.806752 | -2.28496 | 0.02231517 | 0.074381 |
| ENSG00000272914 | | 5.687226 | -1.27031 | 0.9708706 | -2.211168 | 0.0270242 | 0.086247 |
| ENSG00000285933 | | 13.11995 | -1.271133 | 0.6135398 | -2.628255 | 0.00858241 | 0.034409 |
| ENSG00000118513 | MYB | 2184.589778 | -1.271899966 | 0.439183913 | -3.322487519 | 0.00089219 | 0.005178 |
| ENSG00000153904 | DDAH1 | 1948.366 | -1.272487 | 0.1464907 | -8.81818 | 1.16E-18 | 7.52E-17 |
| ENSG00000262075 | | 3.085693 | -1.272931 | 0.9752781 | -2.179774 | 0.02927425 | 0.092046 |
| ENSG00000163485 | ADORA1 | 208.6966 | -1.273172 | 1.22808 | -2.193906 | 0.0282422 | 0.089298 |
| ENSG00000110400 | NECTIN1 | 2750.196975 | -1.273264013 | 0.235546042 | -5.616837635 | 1.94E-08 | 3.36E-07 |
| ENSG00000286292 | | 6.281923 | -1.273359 | 0.7671236 | -2.345452 | 0.01900401 | 0.065226 |

| | | | | | | | |
|---|---|---|---|---|---|---|---|
| ENSG00000257596 | SCAT2 | 33.73205 | -1.27341 | 0.5486799 | -2.835267 | 0.00457874 | 0.020644 |
| ENSG00000250644 | | 15.96576 | -1.275 | 0.6091712 | -2.660999 | 0.00779091 | 0.031826 |
| ENSG00000213928 | IRF9 | 540.8404 | -1.275162 | 0.41917 | -3.441768 | 0.00057793 | 0.00356 |
| ENSG00000223751 | | 26.40178 | -1.275443 | 0.652603 | -2.58311 | 0.00979142 | 0.038203 |
| ENSG00000232973 | CYP1B1-AS1 | 90.14785 | -1.275686 | 0.4699663 | -3.156042 | 0.00159926 | 0.008556 |
| ENSG00000171811 | CFAP46 | 102.0455 | -1.276248 | 0.3489375 | -3.990232 | 6.60E-05 | 0.000529 |
| ENSG00000224940 | PRRT4 | 113.3988 | -1.276901 | 0.4137859 | -3.483527 | 0.00049485 | 0.003111 |
| ENSG00000258794 | DUX4L27 | 11.54219 | -1.278447 | 0.7389299 | -2.433641 | 0.01494783 | 0.053785 |
| ENSG00000229241 | PNPT1P1 | 19.08317 | -1.278609 | 0.61574 | -2.654485 | 0.00794297 | 0.032353 |
| ENSG00000241913 | RPS29P21 | 8.825516 | -1.280607 | 0.6346097 | -2.593001 | 0.00951426 | 0.037317 |
| ENSG00000146955 | RAB19 | 59.54202 | -1.280657 | 0.4963751 | -3.044396 | 0.00233148 | 0.011786 |
| ENSG00000120915 | EPHX2 | 271.2791586 | -1.280859617 | 0.271331658 | -4.977604552 | 6.44E-07 | 8.36E-06 |
| ENSG00000213513 | IMPDH1P5 | 46.17831 | -1.280951 | 0.3012016 | -4.549285 | 5.38E-06 | 5.69E-05 |
| ENSG00000278416 | PMS2P2 | 9.439034 | -1.281268 | 0.6954307 | -2.475929 | 0.01328898 | 0.048857 |
| ENSG00000261879 | | 80.42935 | -1.281672 | 0.5048594 | -3.016834 | 0.0025543 | 0.012728 |
| ENSG00000239665 | | 549.1003 | -1.282667 | 0.3538945 | -3.958392 | 7.55E-05 | 0.000597 |
| ENSG00000187730 | GABRD | 66.92841 | -1.282722 | 0.4625709 | -3.214605 | 0.00130624 | 0.007224 |
| ENSG00000171791 | BCL2 | 3706.373 | -1.283105 | 0.4661468 | -3.210545 | 0.00132483 | 0.007315 |
| ENSG00000267575 | | 389.9637 | -1.283162 | 0.3247291 | -4.264346 | 2.00E-05 | 0.000183 |
| ENSG00000240207 | | 32.10578 | -1.283651 | 0.6102025 | -2.672148 | 0.00753675 | 0.030993 |
| ENSG00000080224 | EPHA6 | 200.6067688 | -1.284602704 | 0.433446387 | -3.37946939 | 0.00072626 | 0.004336 |
| ENSG00000091140 | DLD | 7023.408227 | -1.286747965 | 0.217402592 | -6.127274051 | 8.94E-10 | 1.94E-08 |
| ENSG00000135925 | WNT10A | 17.53964832 | -1.288311485 | 1.147354107 | -2.124360526 | 0.03364001 | 0.102463 |
| ENSG00000251957 | RNU6-1095P | 3.943783 | -1.288601 | 1.119617 | -2.134465 | 0.03280477 | 0.100529 |
| ENSG00000159217 | IGF2BP1 | 41.02033 | -1.288878 | 0.5250608 | -2.949189 | 0.00318609 | 0.015275 |
| ENSG00000286146 | | 2.902426 | -1.289024 | 1.288877 | -2.112511 | 0.0346426 | 0.104855 |
| ENSG00000202538 | RNU4-2 | 6.780637 | -1.29014 | 1.339524 | -2.133707 | 0.03286679 | 0.100657 |
| ENSG00000123609 | NMI | 1317.441169 | -1.293352126 | 0.249965662 | -5.421766139 | 5.90E-08 | 9.38E-07 |
| ENSG00000123453 | SARDH | 10.90678039 | -1.293835255 | 0.960172068 | -2.268371555 | 0.02330657 | 0.076761 |
| ENSG00000239093 | | 8.537571 | -1.294281 | 0.780201 | -2.357506 | 0.01839815 | 0.063533 |
| ENSG00000156650 | KAT6B | 2327.612 | -1.295807 | 0.2301145 | -5.852406 | 4.85E-09 | 9.36E-08 |
| ENSG00000175741 | RWDD4P2 | 422.3688 | -1.296315 | 0.1773052 | -7.480652 | 7.40E-14 | 2.78E-12 |
| ENSG00000197837 | H4-16 | 337.1887 | -1.296627 | 0.5248841 | -2.9803 | 0.00287966 | 0.014106 |

| | | | | | | | |
|---|---|---|---|---|---|---|---|
| ENSG00000164638 | SLC29A4 | 836.6368 | -1.29748 | 0.3032983 | -4.574106 | 4.78E-06 | 5.12E-05 |
| ENSG00000266904 | LINC00663 | 58.51518 | -1.297787 | 0.4623061 | -3.243176 | 0.00118205 | 0.006599 |
| ENSG00000160867 | FGFR4 | 1937.939 | -1.298363 | 0.4809526 | -3.163316 | 0.00155983 | 0.008386 |
| ENSG00000261664 | TTC39A-AS1 | 69.01636 | -1.298712 | 0.3948939 | -3.672503 | 0.00024019 | 0.001668 |
| ENSG00000231721 | LINC-PINT | 151.5159 | -1.298931 | 0.4810588 | -3.161117 | 0.00157166 | 0.008438 |
| ENSG00000178445 | GLDC | 165.1618 | -1.299222 | 0.5505 | -2.879717 | 0.00398033 | 0.018379 |
| ENSG00000168918 | INPP5D | 18.74339 | -1.29946 | 0.5655688 | -2.832316 | 0.00462122 | 0.020796 |
| ENSG00000064607 | SUGP2 | 5889.692667 | -1.300136183 | 0.208542399 | -6.435945687 | 1.23E-10 | 3.04E-09 |
| ENSG00000197157 | SND1 | 11472.37 | -1.300387 | 0.1349986 | -9.773526 | 1.46E-22 | 1.47E-20 |
| ENSG00000129910 | CDH15 | 14.11388073 | -1.300542829 | 0.67362991 | -2.550730142 | 0.01074975 | 0.041249 |
| ENSG00000109255 | NMU | 85.13909117 | -1.301361424 | 0.628922717 | -2.67461734 | 0.00748146 | 0.030803 |
| ENSG00000106484 | MEST | 3764.80081 | -1.302639662 | 0.236775514 | -5.73544061 | 9.73E-09 | 1.79E-07 |
| ENSG00000158292 | GPR153 | 856.3105 | -1.302882 | 0.2120129 | -6.358253 | 2.04E-10 | 4.90E-09 |
| ENSG00000249740 | OSMR-AS1 | 36.43583 | -1.302891 | 0.5236381 | -2.97646 | 0.00291597 | 0.014251 |
| ENSG00000120324 | PCDHB10 | 59.89614367 | -1.303799581 | 0.515442145 | -3.020040288 | 0.00252741 | 0.01262 |
| ENSG00000183092 | BEGAIN | 179.9481 | -1.304531 | 0.3101497 | -4.508164 | 6.54E-06 | 6.77E-05 |
| ENSG00000117148 | ACTL8 | 23.26656015 | -1.304671464 | 1.326626779 | -2.204557495 | 0.02748516 | 0.087476 |
| ENSG00000248359 | | 2.856652 | -1.307561 | 1.504203 | -2.163032 | 0.03053873 | 0.095054 |
| ENSG00000117114 | ADGRL2 | 1330.052001 | -1.30808121 | 0.277898646 | -4.973418164 | 6.58E-07 | 8.53E-06 |
| ENSG00000279656 | | 18.88071 | -1.309004 | 0.6892785 | -2.547376 | 0.01085365 | 0.041564 |
| ENSG00000272777 | | 11.24761 | -1.309207 | 0.9573624 | -2.251503 | 0.02435373 | 0.079461 |
| ENSG00000153721 | CNKSR3 | 284.0665 | -1.311829 | 0.182967 | -7.344788 | 2.06E-13 | 7.35E-12 |
| ENSG00000284906 | | 4.346626 | -1.312367 | 0.9711953 | -2.219753 | 0.02643553 | 0.084735 |
| ENSG00000250519 | | 42.11506 | -1.315421 | 0.3412994 | -4.172754 | 3.01E-05 | 0.000264 |
| ENSG00000110492 | MDK | 7181.68194 | -1.315922325 | 0.265989285 | -5.203850522 | 1.95E-07 | 2.80E-06 |
| ENSG00000244490 | RWDD4P1 | 41.89241 | -1.317263 | 0.3492793 | -4.094332 | 4.23E-05 | 0.000358 |
| ENSG00000230521 | | 19.97075 | -1.317595 | 0.5299409 | -2.971829 | 0.00296032 | 0.014426 |
| ENSG00000119471 | HSDL2 | 2527.086835 | -1.319672416 | 0.153958227 | -8.722805094 | 2.71E-18 | 1.70E-16 |
| ENSG00000214113 | LYRM4 | 695.2342 | -1.320585 | 0.1946961 | -6.972591 | 3.11E-12 | 9.62E-11 |
| ENSG00000215007 | DNAJA1P3 | 21.29578 | -1.321383 | 0.4797787 | -3.210191 | 0.00132647 | 0.007321 |
| ENSG00000117691 | NENF | 1908.966244 | -1.322007743 | 0.214726216 | -6.368566342 | 1.91E-10 | 4.61E-09 |
| ENSG00000250794 | ALG1L12P | 10.30118 | -1.322706 | 0.6138904 | -2.719962 | 0.00652894 | 0.027517 |
| ENSG00000214063 | TSPAN4 | 2679.084 | -1.322955 | 0.2092779 | -6.52661 | 6.73E-11 | 1.71E-09 |

| | | | | | | | |
|---|---|---|---|---|---|---|---|
| ENSG00000184368 | MAP7D2 | 11.32722 | -1.323097 | 0.5874911 | -2.799094 | 0.00512463 | 0.022602 |
| ENSG00000287630 | | 53.32807 | -1.323891 | 0.4375712 | -3.448908 | 0.00056286 | 0.003476 |
| ENSG00000148832 | PAOX | 5.663618 | -1.324582 | 1.012339 | -2.236372 | 0.02532739 | 0.081894 |
| ENSG00000263712 | MIR4639 | 4.137893 | -1.325744 | 1.155124 | -2.098886 | 0.03582692 | 0.107521 |
| ENSG00000261379 | | 64.29831 | -1.328139 | 0.5784102 | -2.857149 | 0.00427466 | 0.019473 |
| ENSG00000227702 | LINC00111 | 4.391241 | -1.328238 | 1.509909 | -2.105987 | 0.03520546 | 0.106052 |
| ENSG00000175183 | CSRP2 | 104.8668 | -1.328761 | 0.5073894 | -3.120244 | 0.00180702 | 0.009513 |
| ENSG00000140807 | NKD1 | 47.29578 | -1.329099 | 0.4118249 | -3.617513 | 0.00029745 | 0.002007 |
| ENSG00000168961 | LGALS9 | 309.7829 | -1.329665 | 0.6886294 | -2.537352 | 0.01116945 | 0.042534 |
| ENSG00000268401 | | 4.912318 | -1.330317 | 1.019008 | -2.203631 | 0.02755029 | 0.087627 |
| ENSG00000106327 | TFR2 | 3.543611044 | -1.331937126 | 1.164176535 | -2.174399486 | 0.02967515 | 0.092974 |
| ENSG00000259560 | | 2.997785 | -1.332415 | 2.201364 | -2.334517 | 0.01956867 | 0.066811 |
| ENSG00000151729 | SLC25A4 | 3095.924 | -1.333065 | 0.1392432 | -9.708785 | 2.77E-22 | 2.71E-20 |
| ENSG00000242551 | POU5F1P6 | 3.107698 | -1.33311 | 1.070472 | -2.185379 | 0.02886111 | 0.091008 |
| ENSG00000188095 | MESP2 | 7.726522 | -1.333363 | 0.8984471 | -2.314357 | 0.02064814 | 0.069708 |
| ENSG00000171604 | CXXC5 | 6177.987 | -1.334475 | 0.2603422 | -5.482237 | 4.20E-08 | 6.85E-07 |
| ENSG00000183914 | DNAH2 | 18.40917 | -1.33686 | 0.5423323 | -2.971922 | 0.00295942 | 0.014425 |
| ENSG00000259994 | | 88.81773 | -1.337559 | 0.2949748 | -4.814241 | 1.48E-06 | 1.79E-05 |
| ENSG00000284237 | LINC02767 | 6.302093 | -1.340047 | 0.8112842 | -2.38447 | 0.01710373 | 0.059892 |
| ENSG00000136040 | PLXNC1 | 94.10056567 | -1.340069519 | 0.388327776 | -3.828179061 | 0.0001291 | 0.00096 |
| ENSG00000182912 | TSPEAR-AS2 | 24.09588 | -1.340244 | 1.08864 | -2.2549 | 0.02413959 | 0.078918 |
| ENSG00000204852 | TCTN1 | 1502.952 | -1.340465 | 0.1478788 | -9.209871 | 3.27E-20 | 2.50E-18 |
| ENSG00000101180 | HRH3 | 14.73289323 | -1.341238506 | 0.855796829 | -2.356098946 | 0.018468 | 0.063704 |
| ENSG00000124194 | GDAP1L1 | 3.490760836 | -1.341868225 | 1.238910615 | -2.092447608 | 0.03639849 | 0.108745 |
| ENSG00000102879 | CORO1A | 919.1834345 | -1.342251216 | 0.903582671 | -2.342643799 | 0.01914765 | 0.065618 |
| ENSG00000225218 | | 20.79019 | -1.342599 | 0.5757258 | -2.859462 | 0.0042436 | 0.019369 |
| ENSG00000260284 | TPSP2 | 52.01572 | -1.342913 | 0.7809234 | -2.477466 | 0.01323189 | 0.048726 |
| ENSG00000213385 | | 39.13143 | -1.34468 | 0.459687 | -3.356959 | 0.00078805 | 0.004658 |
| ENSG00000090376 | IRAK3 | 1.823785817 | -1.345429573 | 3.676393355 | -2.546140846 | 0.01089212 | 0.041675 |
| ENSG00000153574 | RPIA | 2191.474 | -1.345834 | 0.1777378 | -7.749819 | 9.20E-15 | 3.83E-13 |
| ENSG00000267736 | HMGB2P1 | 4.871491 | -1.345992 | 0.9935987 | -2.25821 | 0.0239326 | 0.078376 |
| ENSG00000241535 | CBX5P1 | 1.754068 | -1.346248 | 2.248159 | -2.293097 | 0.02184241 | 0.073044 |
| ENSG00000117226 | GBP3 | 528.206769 | -1.346592232 | 0.554169279 | -2.98161299 | 0.00286734 | 0.014058 |

| | | | | | | | |
|---|---|---|---|---|---|---|---|
| ENSG00000285905 | BMP8B-AS1 | 6.961658 | -1.348388 | 0.9219776 | -2.283533 | 0.02239899 | 0.074595 |
| ENSG00000287787 | | 8.282781 | -1.349053 | 1.100945 | -2.245723 | 0.02472173 | 0.080396 |
| ENSG00000176387 | HSD11B2 | 226.6637 | -1.349602 | 0.2631541 | -5.380511 | 7.43E-08 | 1.15E-06 |
| ENSG00000271931 | | 1.574376 | -1.350605 | 2.494708 | -2.343543 | 0.01910156 | 0.065519 |
| ENSG00000163126 | ANKRD23 | 100.1074 | -1.350682 | 0.3475467 | -4.215689 | 2.49E-05 | 0.000223 |
| ENSG00000151322 | NPAS3 | 672.1651 | -1.352499 | 0.2729216 | -5.206157 | 1.93E-07 | 2.77E-06 |
| ENSG00000100079 | LGALS2 | 19.46347365 | -1.352782619 | 0.671573155 | -2.635250273 | 0.00840753 | 0.033845 |
| ENSG00000023697 | DERA | 518.0179236 | -1.352786965 | 0.266844546 | -5.332151283 | 9.71E-08 | 1.47E-06 |
| ENSG00000181409 | AATK | 360.6916 | -1.353328 | 0.2066089 | -6.751316 | 1.47E-11 | 4.11E-10 |
| ENSG00000266017 | MIR4477B | 28.7898 | -1.353497 | 0.380697 | -3.910117 | 9.23E-05 | 0.000712 |
| ENSG00000103811 | CTSH | 6930.893375 | -1.353631221 | 0.301901595 | -4.783873479 | 1.72E-06 | 2.05E-05 |
| ENSG00000129255 | MPDU1 | 2852.515706 | -1.353652397 | 0.294635386 | -4.890351916 | 1.01E-06 | 1.25E-05 |
| ENSG00000235381 | | 45.32753 | -1.353725 | 0.3253852 | -4.456687 | 8.32E-06 | 8.38E-05 |
| ENSG00000215156 | | 7.46115 | -1.355112 | 0.703046 | -2.554235 | 0.01064216 | 0.040937 |
| ENSG00000162631 | NTNG1 | 35.60815 | -1.356627 | 0.7463201 | -2.514208 | 0.01193002 | 0.04482 |
| ENSG00000170632 | ARMC10 | 1655.624 | -1.356995 | 0.130489 | -10.526 | 6.56E-26 | 8.98E-24 |
| ENSG00000260236 | | 63.96508 | -1.35726 | 0.351744 | -4.189852 | 2.79E-05 | 0.000247 |
| ENSG00000105708 | ZNF14 | 351.1893021 | -1.357615992 | 0.199394776 | -6.995760224 | 2.64E-12 | 8.21E-11 |
| ENSG00000110719 | TCIRG1 | 1831.762394 | -1.357910236 | 0.251770981 | -5.627290474 | 1.83E-08 | 3.17E-07 |
| ENSG00000166159 | LRTM2 | 7.580267 | -1.357999 | 1.150166 | -2.163952 | 0.03046801 | 0.094868 |
| ENSG00000118402 | ELOVL4 | 468.0335437 | -1.3582504 | 0.247415946 | -5.720721372 | 1.06E-08 | 1.94E-07 |
| ENSG00000173947 | PIFO | 114.2468 | -1.358689 | 0.4967378 | -3.21847 | 0.00128877 | 0.007137 |
| ENSG00000274315 | | 8.594236 | -1.359749 | 0.7609388 | -2.492473 | 0.01268569 | 0.047159 |
| ENSG00000275106 | | 7.575552 | -1.361026 | 1.01911 | -2.249404 | 0.02448679 | 0.079788 |
| ENSG00000275413 | | 19.57258 | -1.361293 | 0.496766 | -3.213946 | 0.00130924 | 0.007236 |
| ENSG00000096968 | JAK2 | 713.0237648 | -1.361886203 | 0.286275228 | -5.041282521 | 4.62E-07 | 6.17E-06 |
| ENSG00000165168 | CYBB | 18.74746 | -1.361955 | 0.5845376 | -2.879512 | 0.00398292 | 0.018388 |
| ENSG00000274422 | | 82.12667 | -1.361994 | 0.5823733 | -2.899005 | 0.00374349 | 0.017452 |
| ENSG00000279926 | | 38.51244 | -1.362237 | 0.4729756 | -3.326446 | 0.00087961 | 0.005118 |
| ENSG00000171903 | CYP4F11 | 280.6084 | -1.363276 | 0.3600509 | -4.143545 | 3.42E-05 | 0.000296 |
| ENSG00000279561 | | 212.9751 | -1.363512 | 0.215281 | -6.538258 | 6.22E-11 | 1.59E-09 |
| ENSG00000179774 | ATOH7 | 4.543568 | -1.364123 | 1.063169 | -2.202904 | 0.0276015 | 0.087737 |
| ENSG00000198088 | NUP62CL | 149.8919 | -1.364581 | 0.4948191 | -3.245116 | 0.00117403 | 0.006563 |

| | | | | | | | |
|---|---|---|---|---|---|---|---|
| ENSG00000225535 | LINC01393 | 12.22819 | -1.365345 | 0.6740203 | -2.609887 | 0.0090572 | 0.035927 |
| ENSG00000204540 | PSORS1C1 | 16.30549 | -1.365586 | 0.6362094 | -2.749577 | 0.00596722 | 0.025609 |
| ENSG00000130856 | ZNF236 | 1413.704752 | -1.365589643 | 0.22726895 | -6.236341585 | 4.48E-10 | 1.02E-08 |
| ENSG00000197417 | SHPK | 243.3057 | -1.366478 | 0.3731337 | -4.029534 | 5.59E-05 | 0.000457 |
| ENSG00000263781 | | 15.42579 | -1.366824 | 0.6133092 | -2.812524 | 0.00491544 | 0.021854 |
| ENSG00000075651 | PLD1 | 553.5149991 | -1.367186001 | 0.514129774 | -3.160421655 | 0.00157541 | 0.008449 |
| ENSG00000229109 | | 11.52743 | -1.368107 | 0.8295846 | -2.432844 | 0.01498076 | 0.053874 |
| ENSG00000272281 | TPTE2P2 | 23.64097 | -1.36811 | 0.538416 | -3.070317 | 0.00213832 | 0.010971 |
| ENSG00000266821 | | 2.285422 | -1.370242 | 1.092839 | -2.169548 | 0.03004108 | 0.093802 |
| ENSG00000106477 | CEP41 | 662.8954154 | -1.370537027 | 0.210496817 | -6.722008046 | 1.79E-11 | 4.95E-10 |
| ENSG00000069482 | GAL | 316.1773778 | -1.371101063 | 1.023441394 | -2.332703952 | 0.01966369 | 0.067086 |
| ENSG00000150873 | C2orf50 | 47.25314 | -1.372143 | 0.4101892 | -3.743678 | 0.00018135 | 0.0013 |
| ENSG00000139211 | AMIGO2 | 1762.505456 | -1.372167915 | 0.49504045 | -3.258737239 | 0.00111909 | 0.006306 |
| ENSG00000171954 | CYP4F22 | 68.31144 | -1.372538 | 0.5965273 | -2.885193 | 0.00391173 | 0.018119 |
| ENSG00000285106 | | 34.32403 | -1.372779 | 0.8558114 | -2.41216 | 0.01585833 | 0.056337 |
| ENSG00000005187 | ACSM3 | 128.9740944 | -1.373403401 | 0.311313231 | -4.706429519 | 2.52E-06 | 2.89E-05 |
| ENSG00000251129 | LINC02506 | 65.18408 | -1.374538 | 0.6028575 | -2.866511 | 0.00415024 | 0.019041 |
| ENSG00000109881 | CCDC34 | 867.7112155 | -1.374870989 | 0.366403715 | -4.103669384 | 4.07E-05 | 0.000345 |
| ENSG00000271151 | | 27.58138 | -1.375116 | 0.6758132 | -2.66804 | 0.00762952 | 0.031311 |
| ENSG00000109536 | FRG1 | 1584.73794 | -1.375594832 | 0.187389719 | -7.526082248 | 5.23E-14 | 2.00E-12 |
| ENSG00000107819 | SFXN3 | 2252.355637 | -1.376526225 | 0.290929838 | -5.00557002 | 5.57E-07 | 7.33E-06 |
| ENSG00000178573 | MAF | 74.27587 | -1.377465 | 0.5770643 | -2.958098 | 0.00309543 | 0.014915 |
| ENSG00000226891 | LINC01359 | 13.17824 | -1.378168 | 0.6021227 | -2.852531 | 0.00433726 | 0.019715 |
| ENSG00000259394 | | 6.10507 | -1.378665 | 0.9635778 | -2.302846 | 0.02128754 | 0.071485 |
| ENSG00000287337 | | 10.76897 | -1.379695 | 0.6131368 | -2.824699 | 0.0047325 | 0.0212 |
| ENSG00000262050 | | 14.02165 | -1.380106 | 0.4897077 | -3.261376 | 0.00110873 | 0.006254 |
| ENSG00000185055 | EFCAB10 | 21.31828 | -1.380664 | 0.508103 | -3.187145 | 0.00143685 | 0.007824 |
| ENSG00000048028 | USP28 | 1274.331298 | -1.381400002 | 0.196162121 | -7.230137267 | 4.83E-13 | 1.64E-11 |
| ENSG00000271889 | | 11.3404 | -1.382736 | 0.7209989 | -2.587095 | 0.00967889 | 0.037845 |
| ENSG00000232445 | EMSLR | 269.6893 | -1.382743 | 0.3954033 | -3.877837 | 0.00010539 | 0.000801 |
| ENSG00000274011 | | 8.306209 | -1.383327 | 1.104843 | -2.196019 | 0.02809059 | 0.088931 |
| ENSG00000234964 | FABP5P7 | 648.1443 | -1.384701 | 0.4125599 | -3.76191 | 0.00016862 | 0.001221 |
| ENSG00000198963 | RORB | 13.18817 | -1.385385 | 1.215597 | -2.040858 | 0.04126495 | 0.119861 |

| | | | | | | | |
|---|---|---|---|---|---|---|---|
| ENSG00000135299 | ANKRD6 | 389.1807824 | -1.385507838 | 0.310241325 | -4.76503404 | 1.89E-06 | 2.23E-05 |
| ENSG00000253276 | CCDC71L | 884.6319 | -1.385592 | 0.2363192 | -6.093573 | 1.10E-09 | 2.36E-08 |
| ENSG00000170608 | FOXA3 | 23.89461 | -1.38564 | 0.6551601 | -2.713984 | 0.00664794 | 0.02793 |
| ENSG00000201684 | RN7SKP239 | 7.543516 | -1.387736 | 0.966178 | -2.321644 | 0.0202521 | 0.068694 |
| ENSG00000206341 | HLA-H | 147.2473 | -1.388204 | 0.4503725 | -3.518811 | 0.00043349 | 0.002773 |
| ENSG00000152229 | PSTPIP2 | 622.8893 | -1.389846 | 0.2192391 | -6.560041 | 5.38E-11 | 1.39E-09 |
| ENSG00000216895 | | 124.3375 | -1.391084 | 0.3519174 | -4.290045 | 1.79E-05 | 0.000165 |
| ENSG00000230148 | HOXB-AS1 | 2.54199 | -1.39245 | 2.716838 | -2.358584 | 0.01834482 | 0.063398 |
| ENSG00000251095 | | 289.4731 | -1.395792 | 0.8997375 | -2.429067 | 0.01513772 | 0.054317 |
| ENSG00000197261 | C6orf141 | 1633.343 | -1.396327 | 0.562438 | -3.05407 | 0.00225759 | 0.01146 |
| ENSG00000272604 | | 56.39828 | -1.396599 | 0.3534347 | -4.287794 | 1.80E-05 | 0.000167 |
| ENSG00000125845 | BMP2 | 28.54349743 | -1.397044005 | 0.706091464 | -2.649913029 | 0.00805125 | 0.032695 |
| ENSG00000162896 | PIGR | 1.888458 | -1.399149 | 2.5259 | -2.338687 | 0.01935163 | 0.06618 |
| ENSG00000267519 | MIR23AHG | 371.134 | -1.400578 | 0.3477457 | -4.373576 | 1.22E-05 | 0.000118 |
| ENSG00000074964 | ARHGEF10L | 1558.197408 | -1.401207206 | 0.208514573 | -6.936237785 | 4.03E-12 | 1.22E-10 |
| ENSG00000169955 | ZNF747 | 163.6377 | -1.401484 | 0.3957947 | -3.921932 | 8.78E-05 | 0.000683 |
| ENSG00000214357 | NEURL1B | 943.5525 | -1.402213 | 0.2705178 | -5.449812 | 5.04E-08 | 8.11E-07 |
| ENSG00000183971 | NPW | 73.4734 | -1.402539 | 0.5118977 | -3.234079 | 0.00122036 | 0.006796 |
| ENSG00000196937 | FAM3C | 972.2074 | -1.404023 | 0.1743976 | -8.222868 | 1.99E-16 | 1.02E-14 |
| ENSG00000267530 | LINC01836 | 304.3828 | -1.404098 | 0.3224138 | -4.669754 | 3.02E-06 | 3.40E-05 |
| ENSG00000270426 | | 14.17927 | -1.405455 | 0.7982479 | -2.484426 | 0.01297604 | 0.04801 |
| ENSG00000162882 | HAAO | 11.43944 | -1.405566 | 1.18885 | -2.294167 | 0.02178089 | 0.072858 |
| ENSG00000226904 | | 8.141628 | -1.406784 | 0.7944172 | -2.511903 | 0.01200822 | 0.045069 |
| ENSG00000270362 | HMGN3-AS1 | 99.13256 | -1.4079 | 0.260696 | -5.64245 | 1.68E-08 | 2.93E-07 |
| ENSG00000163840 | DTX3L | 4578.014 | -1.40844 | 0.3477924 | -4.375957 | 1.21E-05 | 0.000117 |
| ENSG00000163520 | FBLN2 | 4.012619 | -1.408504 | 1.268138 | -2.06247 | 0.03916297 | 0.114975 |
| ENSG00000266852 | MIR4482 | 7.980013 | -1.408828 | 0.8218158 | -2.474379 | 0.01334679 | 0.049034 |
| ENSG00000198853 | RUSC2 | 640.3168 | -1.409347 | 0.233002 | -6.277258 | 3.45E-10 | 7.98E-09 |
| ENSG00000265349 | | 7.650754 | -1.409927 | 0.8093568 | -2.486639 | 0.01289563 | 0.047759 |
| ENSG00000233860 | SHROOM3-AS1 | 7.193701 | -1.412389 | 0.7825622 | -2.507145 | 0.01217107 | 0.04552 |
| ENSG00000160999 | SH2B2 | 311.4347 | -1.412584 | 0.2696924 | -5.505327 | 3.68E-08 | 6.06E-07 |
| ENSG00000274290 | H2BC6 | 122.9305 | -1.413129 | 0.3504782 | -4.371203 | 1.24E-05 | 0.000119 |
| ENSG00000106665 | CLIP2 | 300.6574217 | -1.413888812 | 0.258861169 | -5.710066733 | 1.13E-08 | 2.05E-07 |

| | | | | | | | |
|---|---|---|---|---|---|---|---|
| ENSG00000261324 | | 836.8271 | -1.415681 | 0.4146466 | -3.832173 | 0.00012702 | 0.000947 |
| ENSG00000286116 | | 5.594084 | -1.416145 | 1.128864 | -2.195085 | 0.02815754 | 0.089093 |
| ENSG00000228340 | MIR646HG | 1.62803 | -1.416426 | 4.775824 | -2.592009 | 0.00954173 | 0.037391 |
| ENSG00000281903 | LINC02246 | 7.9507 | -1.41858 | 0.8591941 | -2.430914 | 0.01506078 | 0.054089 |
| ENSG00000128283 | CDC42EP1 | 2679.636546 | -1.418672906 | 0.222136409 | -6.601918702 | 4.06E-11 | 1.06E-09 |
| ENSG00000145439 | CBR4 | 1375.838 | -1.418867 | 0.1438082 | -10.01894 | 1.26E-23 | 1.35E-21 |
| ENSG00000183307 | TMEM121B | 17.1682 | -1.419259 | 0.5624891 | -3.027007 | 0.00246988 | 0.012371 |
| ENSG00000120800 | UTP20 | 3004.430177 | -1.420795174 | 0.239205066 | -6.174308229 | 6.65E-10 | 1.47E-08 |
| ENSG00000260876 | LINC01229 | 2.280913 | -1.422229 | 3.080342 | -2.424009 | 0.01535024 | 0.054913 |
| ENSG00000134533 | RERG | 740.6600772 | -1.423350994 | 0.343814111 | -4.473348101 | 7.70E-06 | 7.84E-05 |
| ENSG00000280924 | LINC00628 | 34.53478 | -1.423383 | 0.5664053 | -3.065955 | 0.00216976 | 0.011105 |
| ENSG00000167646 | DNAAF3 | 143.6209 | -1.423462 | 0.3318207 | -4.61797 | 3.88E-06 | 4.24E-05 |
| ENSG00000255837 | TAS2R20 | 12.83861 | -1.425788 | 0.6002473 | -2.929119 | 0.00339925 | 0.016122 |
| ENSG00000130203 | APOE | 3653.917697 | -1.426598289 | 0.687091819 | -2.808212045 | 0.00498174 | 0.022086 |
| ENSG00000174500 | GCSAM | 133.011 | -1.42765 | 0.5124857 | -3.286512 | 0.00101437 | 0.005793 |
| ENSG00000064652 | SNX24 | 1326.138834 | -1.427681217 | 0.212721926 | -6.919745024 | 4.52E-12 | 1.36E-10 |
| ENSG00000112782 | CLIC5 | 211.7231452 | -1.431380931 | 0.612574914 | -2.908846871 | 0.00362765 | 0.01702 |
| ENSG00000186399 | GOLGA8R | 12.93517 | -1.431517 | 0.7736121 | -2.581689 | 0.00983182 | 0.03833 |
| ENSG00000259687 | LINC01220 | 15.35607 | -1.431795 | 0.5400756 | -3.16114 | 0.00157153 | 0.008438 |
| ENSG00000266783 | | 12.10788 | -1.432003 | 0.6858643 | -2.717106 | 0.00658555 | 0.027716 |
| ENSG00000271387 | | 12.29092 | -1.432084 | 0.6820553 | -2.745894 | 0.00603462 | 0.025848 |
| ENSG00000187608 | ISG15 | 6551.954 | -1.432737 | 0.4304205 | -3.752566 | 0.00017503 | 0.001259 |
| ENSG00000286540 | | 24.48398 | -1.433155 | 0.6080838 | -2.927119 | 0.00342118 | 0.016211 |
| ENSG00000284523 | | 29.96179 | -1.434095 | 0.4807902 | -3.43843 | 0.0005851 | 0.003598 |
| ENSG00000170345 | FOS | 12981.89 | -1.434427 | 0.8691231 | -2.496987 | 0.01252535 | 0.046641 |
| ENSG00000213542 | | 19.95683 | -1.434908 | 0.6089507 | -2.942411 | 0.00325668 | 0.015557 |
| ENSG00000259182 | | 4.160016 | -1.435556 | 1.059327 | -2.290638 | 0.02198437 | 0.073444 |
| ENSG00000250927 | MESTP3 | 11.15585 | -1.437596 | 0.6375286 | -2.858959 | 0.00425034 | 0.019393 |
| ENSG00000105792 | CFAP69 | 119.9743269 | -1.439018796 | 0.4445585 | -3.670252915 | 0.00024231 | 0.00168 |
| ENSG00000227071 | FOCAD-AS1 | 2.901216 | -1.440101 | 1.23569 | -2.170331 | 0.02998182 | 0.093639 |
| ENSG00000169668 | BCRP2 | 2.16697 | -1.440256 | 1.484708 | -2.180542 | 0.02921733 | 0.091915 |
| ENSG00000109861 | CTSC | 930.9298749 | -1.440751625 | 0.253133171 | -5.946761403 | 2.73E-09 | 5.51E-08 |
| ENSG00000251468 | | 4.407906 | -1.441277 | 1.129834 | -2.294654 | 0.02175296 | 0.072799 |

| | | | | | | | |
|---|---|---|---|---|---|---|---|
| ENSG00000287391 | | 12.3151 | -1.441808 | 1.129399 | -2.360462 | 0.0182522 | 0.063152 |
| ENSG00000276384 | | 9.382401 | -1.442115 | 0.7773359 | -2.567076 | 0.010256 | 0.039687 |
| ENSG00000010319 | SEMA3G | 34.55975692 | -1.442398447 | 0.554309541 | -3.140604023 | 0.001686 | 0.008948 |
| ENSG00000038002 | AGA | 1931.828024 | -1.442587038 | 0.32836539 | -4.718974959 | 2.37E-06 | 2.74E-05 |
| ENSG00000125430 | HS3ST3B1 | 105.9660192 | -1.443038654 | 0.462148972 | -3.575711081 | 0.00034928 | 0.00231 |
| ENSG00000263938 | | 5.092809 | -1.443673 | 0.8486749 | -2.481127 | 0.01309677 | 0.048363 |
| ENSG00000174899 | SLC66A1L | 46.40057 | -1.446063 | 0.5043552 | -3.350511 | 0.00080663 | 0.004753 |
| ENSG00000244056 | RN7SL417P | 1.914342 | -1.44724 | 1.599761 | -2.148568 | 0.03166863 | 0.097853 |
| ENSG00000008056 | SYN1 | 7.174105577 | -1.447738227 | 1.314299866 | -2.30318028 | 0.0212687 | 0.071431 |
| ENSG00000106344 | RBM28 | 2773.74402 | -1.448757479 | 0.127277794 | -11.50940908 | 1.18E-30 | 2.42E-28 |
| ENSG00000169599 | NFU1 | 1456.685 | -1.448782 | 0.1700754 | -8.684961 | 3.79E-18 | 2.33E-16 |
| ENSG00000243554 | | 12.47648 | -1.44912 | 0.608678 | -2.928239 | 0.00340888 | 0.016162 |
| ENSG00000226007 | | 31.44691 | -1.453039 | 0.3531198 | -4.445479 | 8.77E-06 | 8.77E-05 |
| ENSG00000212127 | TAS2R14 | 40.43798 | -1.453617 | 0.3365154 | -4.641808 | 3.45E-06 | 3.82E-05 |
| ENSG00000205022 | PABPN1L | 37.25002 | -1.453744 | 0.5639948 | -3.116839 | 0.00182802 | 0.009604 |
| ENSG00000105676 | ARMC6 | 2013.021202 | -1.454134491 | 0.215140326 | -6.979773127 | 2.96E-12 | 9.16E-11 |
| ENSG00000279539 | | 49.37381 | -1.454703 | 0.3778019 | -4.218744 | 2.46E-05 | 0.00022 |
| ENSG00000272719 | | 9.799208 | -1.454965 | 0.8976038 | -2.398191 | 0.01647627 | 0.058123 |
| ENSG00000009950 | MLXIPL | 809.9369174 | -1.455537069 | 0.244534081 | -6.197207536 | 5.75E-10 | 1.29E-08 |
| ENSG00000288029 | | 10.33142 | -1.456472 | 0.7832679 | -2.589752 | 0.00960451 | 0.037599 |
| ENSG00000143942 | CHAC2 | 230.9533 | -1.456734 | 0.4441605 | -3.73368 | 0.0001887 | 0.001346 |
| ENSG00000101331 | CCM2L | 30.00394687 | -1.457247366 | 0.623235139 | -2.954433909 | 0.00313243 | 0.015054 |
| ENSG00000189149 | CRYM-AS1 | 25.66512 | -1.459646 | 0.4805991 | -3.496033 | 0.00047223 | 0.002989 |
| ENSG00000260388 | LINC00562 | 3.55819 | -1.460452 | 1.446925 | -2.235983 | 0.0253529 | 0.081957 |
| ENSG00000139800 | ZIC5 | 119.7243 | -1.46266 | 0.4573771 | -3.644343 | 0.00026808 | 0.001833 |
| ENSG00000189051 | RNF222 | 28.19898 | -1.463193 | 0.6208856 | -2.893358 | 0.00381147 | 0.017716 |
| ENSG00000259755 | | 5.183203 | -1.465324 | 0.8354254 | -2.510932 | 0.01204129 | 0.045149 |
| ENSG00000125148 | MT2A | 1456.350999 | -1.465786595 | 0.773713739 | -2.660929583 | 0.00779253 | 0.031828 |
| ENSG00000273329 | | 405.2905 | -1.466052 | 0.3502279 | -4.522908 | 6.10E-06 | 6.37E-05 |
| ENSG00000216490 | IFI30 | 12.41213 | -1.466569 | 0.7301307 | -2.699312 | 0.0069483 | 0.028954 |
| ENSG00000105717 | PBX4 | 61.34726922 | -1.466575311 | 0.488717488 | -3.478810495 | 0.00050365 | 0.003156 |
| ENSG00000227210 | | 34.9754 | -1.467056 | 0.5031453 | -3.400059 | 0.00067371 | 0.004064 |
| ENSG00000168830 | HTR1E | 45.93544 | -1.467142 | 0.6300178 | -2.945264 | 0.00322679 | 0.015436 |

| | | | | | | | |
|---|---|---|---|---|---|---|---|
| ENSG00000133466 | C1QTNF6 | 2210.416744 | -1.467423395 | 0.306388705 | -5.117248097 | 3.10E-07 | 4.26E-06 |
| ENSG00000150625 | GPM6A | 11.98874 | -1.46757 | 1.052916 | -2.329803 | 0.01981656 | 0.067475 |
| ENSG00000233203 | DHCR24-DT | 11.50871 | -1.470317 | 0.6754182 | -2.81602 | 0.00486226 | 0.021686 |
| ENSG00000105821 | DNAJC2 | 3429.245057 | -1.471825289 | 0.219430694 | -6.911853855 | 4.78E-12 | 1.43E-10 |
| ENSG00000204624 | DISP3 | 88.82896 | -1.4736 | 0.3339351 | -4.743327 | 2.10E-06 | 2.46E-05 |
| ENSG00000196739 | COL27A1 | 2526.283 | -1.474097 | 0.4033412 | -4.055653 | 5.00E-05 | 0.000414 |
| ENSG00000273055 | | 17.69826 | -1.474315 | 0.8017333 | -2.584962 | 0.00973897 | 0.038047 |
| ENSG00000154856 | APCDD1 | 630.3276 | -1.47445 | 0.5233711 | -3.33459 | 0.00085425 | 0.004986 |
| ENSG00000270640 | | 4.178832 | -1.474983 | 1.280797 | -2.276013 | 0.02284521 | 0.075624 |
| ENSG00000227355 | | 8.35676 | -1.475262 | 0.7432835 | -2.66641 | 0.00766662 | 0.031429 |
| ENSG00000196843 | ARID5A | 493.7402 | -1.477878 | 0.2295339 | -6.665404 | 2.64E-11 | 7.12E-10 |
| ENSG00000182534 | MXRA7 | 384.7732 | -1.478734 | 0.3404125 | -4.680852 | 2.86E-06 | 3.24E-05 |
| ENSG00000167601 | AXL | 7.510054 | -1.478957 | 0.7259367 | -2.723436 | 0.00646067 | 0.02729 |
| ENSG00000146909 | NOM1 | 2210.119 | -1.48 | 0.1835476 | -8.243341 | 1.67E-16 | 8.69E-15 |
| ENSG00000214659 | KRT8P26 | 11.86371 | -1.481343 | 0.8544067 | -2.549685 | 0.01078202 | 0.041337 |
| ENSG00000215417 | MIR17HG | 118.1315 | -1.48384 | 0.367012 | -4.403629 | 1.06E-05 | 0.000104 |
| ENSG00000279970 | | 116.9604 | -1.485727 | 0.3081563 | -5.124485 | 2.98E-07 | 4.12E-06 |
| ENSG00000146918 | NCAPG2 | 4592.188 | -1.486197 | 0.7213426 | -2.766659 | 0.00566339 | 0.024555 |
| ENSG00000135083 | CCNJL | 754.913631 | -1.486842855 | 0.273022569 | -5.71989503 | 1.07E-08 | 1.95E-07 |
| ENSG00000072682 | P4HA2 | 1785.837997 | -1.488396102 | 0.170051365 | -8.930246809 | 4.25E-19 | 2.90E-17 |
| ENSG00000145284 | SCD5 | 2017.837 | -1.488427 | 0.169755 | -8.932122 | 4.18E-19 | 2.85E-17 |
| ENSG00000271646 | IRF2-DT | 210.0513 | -1.488494 | 0.2273792 | -6.776739 | 1.23E-11 | 3.49E-10 |
| ENSG00000066382 | MPPED2 | 304.8050891 | -1.489659378 | 0.332405965 | -4.808044793 | 1.52E-06 | 1.83E-05 |
| ENSG00000071243 | ING3 | 691.432735 | -1.489709946 | 0.195594031 | -7.813537426 | 5.56E-15 | 2.36E-13 |
| ENSG00000277072 | STAG3L2 | 459.3867 | -1.490275 | 0.2344877 | -6.587371 | 4.48E-11 | 1.16E-09 |
| ENSG00000119547 | ONECUT2 | 874.3607674 | -1.491079131 | 0.475849518 | -3.603076367 | 0.00031447 | 0.00211 |
| ENSG00000233825 | | 8.823122 | -1.49242 | 0.7483595 | -2.678888 | 0.0073867 | 0.030475 |
| ENSG00000199053 | MIR324 | 32.07717 | -1.493156 | 0.4106663 | -4.019977 | 5.82E-05 | 0.000473 |
| ENSG00000254463 | PPIAP41 | 3.072003 | -1.493993 | 2.130023 | -2.266683 | 0.02340962 | 0.076957 |
| ENSG00000090975 | PITPNM2 | 626.810748 | -1.494251834 | 0.225778063 | -6.849448845 | 7.41E-12 | 2.16E-10 |
| ENSG00000276223 | | 103.2201 | -1.494311 | 0.409365 | -4.047776 | 5.17E-05 | 0.000427 |
| ENSG00000138028 | CGREF1 | 2458.722433 | -1.494744087 | 0.224706647 | -6.887507266 | 5.68E-12 | 1.68E-10 |
| ENSG00000280360 | | 3.586053 | -1.495559 | 1.07515 | -2.363602 | 0.01809825 | 0.062734 |

| | | | | | | | |
|---|---|---|---|---|---|---|---|
| ENSG00000227252 | | 50.51587 | -1.495783 | 0.3221913 | -4.949438 | 7.44E-07 | 9.53E-06 |
| ENSG00000154040 | CABYR | 243.458 | -1.496189 | 0.2551923 | -6.108664 | 1.00E-09 | 2.16E-08 |
| ENSG00000275896 | PRSS2 | 6.120057 | -1.498546 | 2.947635 | -2.384686 | 0.01709368 | 0.059884 |
| ENSG00000253348 | | 21.83095 | -1.498793 | 1.032068 | -2.252905 | 0.02426514 | 0.079235 |
| ENSG00000255353 | ASS1P13 | 44.80103 | -1.499665 | 0.4558943 | -3.73298 | 0.00018923 | 0.00135 |
| ENSG00000273373 | | 298.2084 | -1.499984 | 0.563313 | -3.217938 | 0.00129116 | 0.007148 |
| ENSG00000174799 | CEP135 | 985.6843 | -1.500971 | 0.254691 | -6.145921 | 7.95E-10 | 1.73E-08 |
| ENSG00000229766 | | 4.847679 | -1.502613 | 0.8920318 | -2.481719 | 0.01307503 | 0.048309 |
| ENSG00000168952 | STXBP6 | 167.3828 | -1.503947 | 0.4374054 | -3.860826 | 0.00011301 | 0.000854 |
| ENSG00000161040 | FBXL13 | 18.39456 | -1.505218 | 0.5363419 | -3.320766 | 0.00089771 | 0.005204 |
| ENSG00000019186 | CYP24A1 | 224.5950569 | -1.50667982 | 1.04094193 | -2.410182676 | 0.01594454 | 0.056567 |
| ENSG00000117266 | CDK18 | 5424.222077 | -1.50798505 | 0.281099897 | -5.642128967 | 1.68E-08 | 2.93E-07 |
| ENSG00000273680 | | 24.84856 | -1.509258 | 0.4406402 | -3.844127 | 0.00012098 | 0.000906 |
| ENSG00000117600 | PLPPR4 | 8.417500778 | -1.509624974 | 0.852711531 | -2.557926409 | 0.01052984 | 0.040581 |
| ENSG00000278716 | | 6.700854 | -1.509802 | 1.556008 | -2.348136 | 0.01886765 | 0.064851 |
| ENSG00000287146 | | 7.862983 | -1.512092 | 1.031771 | -2.40252 | 0.01628255 | 0.05752 |
| ENSG00000229116 | | 2.568755 | -1.512189 | 1.4747 | -2.291672 | 0.02192459 | 0.073275 |
| ENSG00000151023 | ENKUR | 15.98743 | -1.512331 | 0.6927854 | -2.825688 | 0.00471793 | 0.021149 |
| ENSG00000281468 | | 84.94713 | -1.513616 | 0.5367126 | -3.343247 | 0.00082804 | 0.004861 |
| ENSG00000255310 | | 94.20496 | -1.513943 | 0.4716579 | -3.648382 | 0.0002639 | 0.001809 |
| ENSG00000133104 | SPART | 1738.033984 | -1.51584339 | 0.2418298 | -6.52883243 | 6.63E-11 | 1.69E-09 |
| ENSG00000282164 | PEG13 | 50.49172 | -1.516072 | 0.4964235 | -3.537342 | 0.00040418 | 0.002613 |
| ENSG00000248734 | | 20.31179 | -1.516176 | 0.7818718 | -2.646082 | 0.008143 | 0.03299 |
| ENSG00000172671 | ZFAND4 | 220.4726 | -1.516771 | 0.2833031 | -5.631195 | 1.79E-08 | 3.11E-07 |
| ENSG00000205583 | STAG3L1 | 12.39866 | -1.517769 | 0.6264674 | -3.00694 | 0.00263892 | 0.013093 |
| ENSG00000234869 | | 76.2531 | -1.519353 | 0.2777282 | -5.740809 | 9.42E-09 | 1.74E-07 |
| ENSG00000286147 | | 2.894065 | -1.519721 | 1.297282 | -2.222894 | 0.02622298 | 0.084165 |
| ENSG00000184497 | TMEM255B | 15.61257 | -1.519739 | 0.6165435 | -3.030717 | 0.00243974 | 0.012242 |
| ENSG00000232124 | | 54.98468 | -1.521011 | 0.5877597 | -3.176626 | 0.00148999 | 0.008063 |
| ENSG00000252412 | | 11.09987 | -1.522081 | 0.7604722 | -2.725024 | 0.00642969 | 0.027172 |
| ENSG00000233874 | MFN1P1 | 5.598399 | -1.522798 | 1.104283 | -2.339941 | 0.01928678 | 0.066009 |
| ENSG00000204959 | ARHGEF34P | 162.1257 | -1.522833 | 0.3108838 | -5.202498 | 1.97E-07 | 2.82E-06 |
| ENSG00000243738 | RN7SL181P | 4.561694 | -1.523413 | 1.075169 | -2.304906 | 0.02117183 | 0.071168 |

| | | | | | | | |
|---|---|---|---|---|---|---|---|
| ENSG00000271631 | | 9.250141 | -1.525941 | 0.8109664 | -2.608775 | 0.00908669 | 0.036023 |
| ENSG00000184523 | PTGER4P2 | 12.87154 | -1.526951 | 0.6791733 | -2.909434 | 0.00362083 | 0.016991 |
| ENSG00000229422 | | 176.9233 | -1.527041 | 0.3524545 | -4.677162 | 2.91E-06 | 3.29E-05 |
| ENSG00000251359 | WWC2-AS2 | 28.95661 | -1.529078 | 0.515741 | -3.436668 | 0.00058892 | 0.003616 |
| ENSG00000266916 | ZNF793-AS1 | 95.32915 | -1.529209 | 0.3426305 | -4.799402 | 1.59E-06 | 1.91E-05 |
| ENSG00000214376 | VSTM5 | 9.969681 | -1.529824 | 0.945534 | -2.500856 | 0.01238936 | 0.046189 |
| ENSG00000243433 | | 13.92471 | -1.531019 | 0.6147487 | -3.07048 | 0.00213715 | 0.010968 |
| ENSG00000286799 | | 6.069751 | -1.53124 | 0.8246318 | -2.606194 | 0.00915545 | 0.036252 |
| ENSG00000227060 | LINC00629 | 25.87773 | -1.531761 | 0.707012 | -2.832705 | 0.0046156 | 0.020787 |
| ENSG00000138030 | KHK | 534.6187422 | -1.536299569 | 0.26595687 | -6.053884726 | 1.41E-09 | 2.97E-08 |
| ENSG00000249210 | GAPDHP38 | 21.80991 | -1.540838 | 0.4531626 | -3.809286 | 0.00013937 | 0.001027 |
| ENSG00000047662 | FAM184B | 43.25573289 | -1.541554192 | 0.531832688 | -3.419892053 | 0.00062646 | 0.003818 |
| ENSG00000184162 | NR2C2AP | 1041.797 | -1.54253 | 0.2610874 | -6.164245 | 7.08E-10 | 1.56E-08 |
| ENSG00000246528 | | 3.817983 | -1.543081 | 0.9894769 | -2.434108 | 0.01492853 | 0.053737 |
| ENSG00000235029 | MNX1-AS2 | 31.26128 | -1.543392 | 0.4492769 | -3.862627 | 0.00011217 | 0.000848 |
| ENSG00000146842 | TMEM209 | 1016.041 | -1.543741 | 0.2443237 | -6.564654 | 5.22E-11 | 1.35E-09 |
| ENSG00000121236 | TRIM6 | 18.02631049 | -1.544144528 | 0.726280512 | -2.780951055 | 0.00541999 | 0.023654 |
| ENSG00000214973 | CHCHD3P3 | 174.822 | -1.544382 | 0.2285097 | -6.989561 | 2.76E-12 | 8.56E-11 |
| ENSG00000142089 | IFITM3 | 11354.9 | -1.545978 | 0.3265859 | -5.057228 | 4.25E-07 | 5.72E-06 |
| ENSG00000168398 | BDKRB2 | 315.3937 | -1.548757 | 0.4206496 | -4.100719 | 4.12E-05 | 0.000349 |
| ENSG00000231711 | LINC00899 | 7.743205 | -1.550509 | 0.7491135 | -2.759415 | 0.0057905 | 0.025032 |
| ENSG00000260710 | | 28.33046 | -1.551819 | 0.8917372 | -2.603065 | 0.00923943 | 0.036501 |
| ENSG00000145779 | TNFAIP8 | 906.8453 | -1.553287 | 0.2920366 | -5.608669 | 2.04E-08 | 3.50E-07 |
| ENSG00000254901 | BORCS8 | 513.6542 | -1.5544 | 0.2119913 | -7.547515 | 4.44E-14 | 1.71E-12 |
| ENSG00000125788 | DEFB126 | 18.3226032 | -1.554408481 | 0.712120849 | -2.86940102 | 0.0041125 | 0.018892 |
| ENSG00000196104 | SPOCK3 | 12.47173 | -1.55912 | 0.6371501 | -3.035059 | 0.00240489 | 0.012106 |
| ENSG00000171798 | KNDC1 | 870.8666 | -1.560584 | 0.4315035 | -4.047392 | 5.18E-05 | 0.000427 |
| ENSG00000154099 | DNAAF1 | 152.496 | -1.562452 | 0.3921064 | -4.371558 | 1.23E-05 | 0.000119 |
| ENSG00000106554 | CHCHD3 | 4356.607059 | -1.563315425 | 0.185896096 | -8.596144966 | 8.24E-18 | 4.91E-16 |
| ENSG00000182578 | CSF1R | 16.26989 | -1.564403 | 0.8856087 | -2.61112 | 0.00902463 | 0.035837 |
| ENSG00000249364 | | 103.0116 | -1.565589 | 0.7295606 | -2.865059 | 0.00416932 | 0.019106 |
| ENSG00000168404 | MLKL | 834.2271 | -1.566708 | 0.2545895 | -6.411718 | 1.44E-10 | 3.52E-09 |
| ENSG00000128573 | FOXP2 | 311.9954908 | -1.567085262 | 0.535539299 | -3.455142851 | 0.00055 | 0.003412 |

| | | | | | | | |
|---|---|---|---|---|---|---|---|
| ENSG00000224043 | CCNT2-AS1 | 155.8432 | -1.567277 | 0.2790219 | -5.8934 | 3.78E-09 | 7.45E-08 |
| ENSG00000272701 | MESTIT1 | 7.527856 | -1.571995 | 0.8540485 | -2.570674 | 0.01015009 | 0.039371 |
| ENSG00000181143 | MUC16 | 34.77424 | -1.572494 | 1.045542 | -2.50964 | 0.01208544 | 0.04527 |
| ENSG00000148154 | UGCG | 2913.472 | -1.574844 | 0.2415554 | -6.76054 | 1.37E-11 | 3.88E-10 |
| ENSG00000123096 | SSPN | 104.6650474 | -1.575329363 | 0.445552891 | -3.976959956 | 6.98E-05 | 0.000556 |
| ENSG00000225439 | BOLA3-AS1 | 45.0924 | -1.576072 | 0.3643382 | -4.679857 | 2.87E-06 | 3.25E-05 |
| ENSG00000261243 | | 8.257839 | -1.57617 | 0.7223682 | -2.847381 | 0.00440806 | 0.019966 |
| ENSG00000277007 | | 18.78315 | -1.576505 | 0.6948835 | -2.929947 | 0.0033902 | 0.016093 |
| ENSG00000259620 | | 3.717821 | -1.57796 | 1.593496 | -2.372567 | 0.01766498 | 0.061472 |
| ENSG00000247853 | | 11.61283 | -1.578037 | 0.7932658 | -2.7089 | 0.00675068 | 0.028281 |
| ENSG00000204022 | LIPJ | 5.462223 | -1.579447 | 0.8540294 | -2.612215 | 0.00899576 | 0.035748 |
| ENSG00000105088 | OLFM2 | 1132.127627 | -1.580230175 | 0.312878896 | -5.373426762 | 7.73E-08 | 1.19E-06 |
| ENSG00000235530 | | 69.40882 | -1.583118 | 0.3750237 | -4.575569 | 4.75E-06 | 5.09E-05 |
| ENSG00000266709 | | 26.75681 | -1.583886 | 0.9927803 | -2.522151 | 0.01166397 | 0.044001 |
| ENSG00000140600 | SH3GL3 | 125.403 | -1.586762 | 0.3114228 | -5.400724 | 6.64E-08 | 1.04E-06 |
| ENSG00000168003 | SLC3A2 | 30182.84 | -1.586888 | 0.3339662 | -5.085082 | 3.67E-07 | 4.97E-06 |
| ENSG00000075886 | TUBA3D | 78.34194263 | -1.58737499 | 0.399065328 | -4.369019959 | 1.25E-05 | 0.00012 |
| ENSG00000064545 | TMEM161A | 1422.744712 | -1.589633472 | 0.176147206 | -9.189977061 | 3.93E-20 | 2.97E-18 |
| ENSG00000143429 | LSP1P4 | 30.58558 | -1.594603 | 0.4633278 | -3.874485 | 0.00010685 | 0.000811 |
| ENSG00000257923 | CUX1 | 9661.675 | -1.596276 | 0.2470384 | -6.735249 | 1.64E-11 | 4.54E-10 |
| ENSG00000236383 | CCDC200 | 4.382558 | -1.596326 | 0.870869 | -2.611333 | 0.00901899 | 0.035824 |
| ENSG00000234290 | NA | 52.04737 | -1.599568 | 0.5175759 | -3.562245 | 0.0003677 | 0.002412 |
| ENSG00000171587 | DSCAM | 1128.663 | -1.599939 | 0.4877701 | -3.768922 | 0.00016395 | 0.001191 |
| ENSG00000276517 | | 37.77324 | -1.600909 | 0.4873731 | -3.767432 | 0.00016494 | 0.001197 |
| ENSG00000214194 | SMIM30 | 1100.588 | -1.602148 | 0.2289637 | -7.225463 | 4.99E-13 | 1.69E-11 |
| ENSG00000286063 | | 5.522834 | -1.602155 | 1.450378 | -2.386981 | 0.01698737 | 0.059582 |
| ENSG00000264365 | | 4.33161 | -1.602587 | 1.072515 | -2.44189 | 0.01461062 | 0.052822 |
| ENSG00000258210 | | 5.596235 | -1.603053 | 0.8831144 | -2.59868 | 0.00935831 | 0.036847 |
| ENSG00000135114 | OASL | 679.1447036 | -1.603573981 | 0.497976423 | -3.712230164 | 0.00020544 | 0.001452 |
| ENSG00000056736 | IL17RB | 932.8690845 | -1.603742246 | 0.295367424 | -5.723653298 | 1.04E-08 | 1.91E-07 |
| ENSG00000157542 | KCNJ6 | 1.471153 | -1.605658 | 2.908824 | -2.409507 | 0.01597411 | NA |
| ENSG00000205221 | VIT | 2.445325 | -1.605792 | 2.949751 | -2.379542 | 0.01733419 | 0.060559 |
| ENSG00000142798 | HSPG2 | 3198.937 | -1.606078 | 0.3853832 | -4.520751 | 6.16E-06 | 6.43E-05 |

| | | | | | | | |
|---|---|---|---|---|---|---|---|
| ENSG00000260976 | LINC01633 | 7.910214 | -1.606365 | 0.7132744 | -2.929358 | 0.00339663 | 0.016115 |
| ENSG00000281357 | ARRDC3-AS1 | 58.03985 | -1.606525 | 0.46248 | -3.921301 | 8.81E-05 | 0.000684 |
| ENSG00000154127 | UBASH3B | 90.74747 | -1.608294 | 0.5206838 | -3.540963 | 0.00039867 | 0.002584 |
| ENSG00000260498 | | 20.07043 | -1.61087 | 1.009109 | -2.560951 | 0.0104386 | 0.040282 |
| ENSG00000063438 | AHRR | 26.66737397 | -1.612812929 | 0.458382925 | -3.943080627 | 8.04E-05 | 0.000631 |
| ENSG00000197536 | IRF1-AS1 | 167.4204 | -1.613563 | 0.4999449 | -3.714277 | 0.00020379 | 0.001442 |
| ENSG00000224977 | LINC02776 | 6.798721 | -1.613755 | 1.137243 | -2.431088 | 0.01505354 | 0.054078 |
| ENSG00000228824 | MIR4500HG | 23.61874 | -1.613999 | 0.419024 | -4.256372 | 2.08E-05 | 0.000189 |
| ENSG00000151388 | ADAMTS12 | 3.351488 | -1.614985 | 1.071283 | -2.475393 | 0.01330898 | 0.048915 |
| ENSG00000151640 | DPYSL4 | 51.7846 | -1.617995 | 0.461178 | -3.950175 | 7.81E-05 | 0.000616 |
| ENSG00000144366 | GULP1 | 1023.477 | -1.618554 | 0.3840643 | -4.600166 | 4.22E-06 | 4.59E-05 |
| ENSG00000082929 | LINC01587 | 18.43995096 | -1.619520028 | 0.711049758 | -2.964541261 | 0.00303135 | 0.014678 |
| ENSG00000178342 | KCNG2 | 9.213434 | -1.624783 | 0.7804656 | -2.777227 | 0.00548249 | 0.023875 |
| ENSG00000246792 | | 100.4565 | -1.625412 | 0.3418708 | -5.094408 | 3.50E-07 | 4.75E-06 |
| ENSG00000115155 | OTOF | 2.451064077 | -1.62609043 | 1.319575982 | -2.340698548 | 0.0192477 | 0.065901 |
| ENSG00000270147 | | 9.214654 | -1.62702 | 0.7472012 | -2.841234 | 0.00449393 | 0.020296 |
| ENSG00000186603 | HPDL | 1525.949 | -1.627467 | 0.454016 | -4.036507 | 5.43E-05 | 0.000445 |
| ENSG00000197043 | ANXA6 | 5980.767 | -1.629061 | 0.3148617 | -5.489218 | 4.04E-08 | 6.60E-07 |
| ENSG00000213014 | VN2R17P | 2.135247 | -1.630858 | 1.739535 | -2.244119 | 0.02482477 | 0.080622 |
| ENSG00000139180 | NDUFA9 | 427.1109424 | -1.631817685 | 0.215783552 | -7.781691905 | 7.16E-15 | 3.02E-13 |
| ENSG00000251583 | MFFP2 | 11.24708 | -1.632126 | 0.8798869 | -2.687248 | 0.00720434 | 0.029848 |
| ENSG00000225213 | | 4.970904 | -1.633845 | 1.036349 | -2.474109 | 0.01335691 | 0.04905 |
| ENSG00000183346 | CABCOCO1 | 13.95824 | -1.634011 | 0.7703622 | -2.848028 | 0.0043991 | 0.019942 |
| ENSG00000189050 | RNFT1 | 1972.388 | -1.636486 | 0.2581568 | -6.605971 | 3.95E-11 | 1.03E-09 |
| ENSG00000126010 | GRPR | 13.00684039 | -1.636809874 | 0.944125821 | -2.644038201 | 0.00819234 | 0.033119 |
| ENSG00000164707 | SLC13A4 | 14.72736 | -1.638111 | 0.8829144 | -2.683136 | 0.00729354 | 0.030156 |
| ENSG00000148541 | FAM13C | 12.73657 | -1.638516 | 1.089569 | -2.544368 | 0.01094757 | 0.041857 |
| ENSG00000277130 | | 6.257404 | -1.640552 | 1.069365 | -2.483662 | 0.01300392 | 0.048073 |
| ENSG00000241362 | RPL36AP43 | 28.7994 | -1.640839 | 0.4692706 | -3.937406 | 8.24E-05 | 0.000645 |
| ENSG00000133131 | MORC4 | 2332.023526 | -1.642022973 | 0.218938336 | -7.725166361 | 1.12E-14 | 4.59E-13 |
| ENSG00000157150 | TIMP4 | 13.52533 | -1.642715 | 1.476578 | -2.248497 | 0.02454449 | 0.079917 |
| ENSG00000138061 | CYP1B1 | 6701.108897 | -1.64393873 | 0.467423718 | -3.981108201 | 6.86E-05 | 0.000547 |
| ENSG00000248988 | PSME2P3 | 23.94903 | -1.644354 | 0.4978525 | -3.758657 | 0.00017083 | 0.001234 |

| | | | | | | | |
|---|---|---|---|---|---|---|---|
| ENSG00000236761 | CTAGE9 | 3.317018 | -1.645558 | 2.131637 | -2.309715 | 0.02090392 | 0.070428 |
| ENSG00000119943 | PYROXD2 | 1974.544772 | -1.645666272 | 0.257092547 | -6.674589349 | 2.48E-11 | 6.71E-10 |
| ENSG00000161133 | USP41 | 8.532261 | -1.646022 | 1.098181 | -2.495742 | 0.0125694 | 0.046786 |
| ENSG00000171695 | LKAAEAR1 | 14.24063 | -1.646908 | 0.7954872 | -2.825302 | 0.00472361 | 0.021171 |
| ENSG00000286181 | | 23.24145 | -1.647328 | 0.5387369 | -3.581869 | 0.00034115 | 0.002264 |
| ENSG00000275191 | | 6.15215 | -1.647603 | 1.274854 | -2.373969 | 0.01759805 | 0.061304 |
| ENSG00000145416 | MARCHF1 | 583.1089 | -1.648855 | 0.1912071 | -8.813714 | 1.21E-18 | 7.79E-17 |
| ENSG00000183508 | TENT5C | 207.7184 | -1.649157 | 0.4288103 | -4.268796 | 1.97E-05 | 0.00018 |
| ENSG00000279277 | | 184.3897 | -1.652287 | 0.3476893 | -5.110691 | 3.21E-07 | 4.40E-06 |
| ENSG00000256913 | | 15.83511 | -1.652319 | 0.6609783 | -3.096935 | 0.00195533 | 0.010161 |
| ENSG00000168785 | TSPAN5 | 1027.534 | -1.652999 | 0.4098983 | -4.452288 | 8.50E-06 | 8.54E-05 |
| ENSG00000234602 | MCIDAS | 223.3465 | -1.653022 | 0.2274435 | -7.503389 | 6.22E-14 | 2.37E-12 |
| ENSG00000151365 | THRSP | 8.627172 | -1.653491 | 1.211051 | -2.49217 | 0.01269651 | 0.047179 |
| ENSG00000271860 | | 7.821727 | -1.654238 | 0.6802365 | -3.078788 | 0.00207845 | 0.010706 |
| ENSG00000184903 | IMMP2L | 415.9464 | -1.655712 | 0.2485664 | -6.919569 | 4.53E-12 | 1.36E-10 |
| ENSG00000232467 | TMA16P2 | 87.30427 | -1.655842 | 0.2275979 | -7.505696 | 6.11E-14 | 2.33E-12 |
| ENSG00000108244 | KRT23 | 87.52858214 | -1.6559632 | 0.758841227 | -2.922980249 | 0.00346699 | 0.016385 |
| ENSG00000106034 | CPED1 | 39.93661923 | -1.656012061 | 0.847270568 | -2.657897665 | 0.00786298 | 0.032086 |
| ENSG00000151474 | FRMD4A | 433.6173 | -1.656544 | 0.2119171 | -8.038743 | 9.08E-16 | 4.30E-14 |
| ENSG00000283199 | C13orf46 | 12.0134 | -1.657117 | 0.6567941 | -3.122023 | 0.00179613 | 0.009465 |
| ENSG00000175773 | ZBTB44-DT | 5.870482 | -1.659633 | 0.9949365 | -2.531906 | 0.01134445 | 0.043071 |
| ENSG00000115604 | IL18R1 | 19.54063331 | -1.660139734 | 0.928107231 | -2.515123053 | 0.01189908 | 0.044729 |
| ENSG00000260115 | | 3.349871 | -1.660245 | 3.892913 | -2.470648 | 0.01348686 | 0.049446 |
| ENSG00000230707 | | 3.081852 | -1.662419 | 1.307929 | -2.389247 | 0.01688294 | 0.059294 |
| ENSG00000215808 | LINC01139 | 186.7822 | -1.66451 | 0.2875211 | -6.066507 | 1.31E-09 | 2.76E-08 |
| ENSG00000231327 | LINC01816 | 40.21799 | -1.664847 | 0.4941497 | -3.845359 | 0.00012038 | 0.000902 |
| ENSG00000267387 | | 7.965915 | -1.66743 | 1.081444 | -2.455954 | 0.01405112 | 0.051147 |
| ENSG00000163545 | NUAK2 | 725.5654 | -1.668159 | 0.2515627 | -6.880214 | 5.98E-12 | 1.76E-10 |
| ENSG00000254039 | | 14.33909 | -1.668372 | 0.5999411 | -3.327504 | 0.00087628 | 0.005102 |
| ENSG00000116014 | KISS1R | 213.7818493 | -1.668695707 | 0.407973928 | -4.492749537 | 7.03E-06 | 7.21E-05 |
| ENSG00000188783 | PRELP | 3.013937 | -1.669002 | 2.910282 | -2.398016 | 0.01648413 | 0.058124 |
| ENSG00000115841 | RMDN2 | 148.0679059 | -1.669623834 | 0.256131498 | -6.770090294 | 1.29E-11 | 3.64E-10 |
| ENSG00000231346 | LINC01160 | 18.73968 | -1.670637 | 0.6925364 | -3.087122 | 0.00202104 | 0.010453 |

| | | | | | | | |
|---|---|---|---|---|---|---|---|
| ENSG00000068079 | IFI35 | 1388.484557 | -1.670639342 | 0.403090642 | -4.55314647 | 5.28E-06 | 5.61E-05 |
| ENSG00000286565 | | 3.692319 | -1.670699 | 1.033886 | -2.51101 | 0.01203863 | 0.045145 |
| ENSG00000258294 | | 4.060092 | -1.67123 | 1.121928 | -2.47085 | 0.01347923 | 0.049438 |
| ENSG00000273265 | CNNM3-DT | 33.26753 | -1.671755 | 0.5190164 | -3.729615 | 0.00019177 | 0.001367 |
| ENSG00000174945 | AMZ1 | 436.8186 | -1.672686 | 0.3902816 | -4.679729 | 2.87E-06 | 3.25E-05 |
| ENSG00000148344 | PTGES | 209.5422 | -1.676258 | 0.5056566 | -3.805741 | 0.00014138 | 0.00104 |
| ENSG00000137975 | CLCA2 | 66.56229154 | -1.676955575 | 0.692448775 | -3.09167118 | 0.00199033 | 0.010308 |
| ENSG00000143851 | PTPN7 | 20.69552 | -1.677758 | 0.8139347 | -2.755172 | 0.00586613 | 0.025273 |
| ENSG00000251453 | HAUS1P1 | 7.210127 | -1.677759 | 0.7709814 | -2.887429 | 0.00388404 | 0.01801 |
| ENSG00000100346 | CACNA1I | 27.21596054 | -1.677852076 | 0.490263764 | -3.904233682 | 9.45E-05 | 0.000728 |
| ENSG00000198756 | COLGALT2 | 66.01111 | -1.680945 | 0.4677512 | -4.050205 | 5.12E-05 | 0.000423 |
| ENSG00000144354 | CDCA7 | 855.5079 | -1.681864 | 0.5567445 | -3.577247 | 0.00034723 | 0.002299 |
| ENSG00000138271 | GPR87 | 16.42522901 | -1.682656755 | 0.597457188 | -3.378312969 | 0.00072932 | 0.004351 |
| ENSG00000270885 | RASL10B | 249.8764 | -1.687622 | 0.4361883 | -4.300584 | 1.70E-05 | 0.000159 |
| ENSG00000104998 | IL27RA | 862.1814845 | -1.688087338 | 0.258770653 | -6.762214567 | 1.36E-11 | 3.84E-10 |
| ENSG00000186891 | TNFRSF18 | 25.76119 | -1.688742 | 0.8358598 | -2.819629 | 0.00480792 | 0.021491 |
| ENSG00000187870 | RNFT1P3 | 14.05934 | -1.691267 | 0.5804845 | -3.469215 | 0.00052198 | 0.003257 |
| ENSG00000231431 | FAR2P4 | 22.56559 | -1.692765 | 0.7736997 | -2.935518 | 0.00332991 | 0.015852 |
| ENSG00000072422 | RHOBTB1 | 563.0456373 | -1.693408601 | 0.29284657 | -6.073125982 | 1.25E-09 | 2.66E-08 |
| ENSG00000179388 | EGR3 | 1458.642 | -1.694029 | 1.828749 | -2.491488 | 0.01272094 | 0.047223 |
| ENSG00000170629 | DPY19L2P2 | 52.26212 | -1.697706 | 0.524719 | -3.75466 | 0.00017358 | 0.001251 |
| ENSG00000115415 | STAT1 | 16537.22509 | -1.699722929 | 0.318311233 | -5.655822522 | 1.55E-08 | 2.73E-07 |
| ENSG00000175175 | PPM1E | 1365.956 | -1.702643 | 0.2292782 | -7.652805 | 1.97E-14 | 7.87E-13 |
| ENSG00000234026 | | 9.335169 | -1.703083 | 0.9298102 | -2.551728 | 0.01071902 | 0.041167 |
| ENSG00000103460 | TOX3 | 21.88287108 | -1.70322494 | 1.094402798 | -2.563531606 | 0.01036133 | 0.040042 |
| ENSG00000280255 | | 6.4976 | -1.703908 | 1.099652 | -2.568389 | 0.01021723 | 0.039572 |
| ENSG00000196569 | LAMA2 | 21.69418 | -1.704295 | 0.5640334 | -3.56282 | 0.00036689 | 0.002407 |
| ENSG00000223935 | LGALSL-DT | 8.160354 | -1.705413 | 0.9531594 | -2.678264 | 0.00740048 | 0.030527 |
| ENSG00000173809 | TDRD12 | 50.73522 | -1.705696 | 0.4753808 | -4.050785 | 5.10E-05 | 0.000422 |
| ENSG00000243056 | EIF4EBP3 | 13.03862 | -1.707414 | 0.6633048 | -3.157887 | 0.00158917 | 0.008511 |
| ENSG00000197122 | SRC | 3083.287 | -1.708801 | 0.181107 | -9.61551 | 6.88E-22 | 6.41E-20 |
| ENSG00000236308 | | 19.30027 | -1.709629 | 0.4521747 | -4.207989 | 2.58E-05 | 0.000229 |
| ENSG00000273014 | | 53.28411 | -1.709695 | 0.3547096 | -5.166743 | 2.38E-07 | 3.37E-06 |

| | | | | | | | |
|---|---|---|---|---|---|---|---|
| ENSG00000218561 | | 2.192418 | -1.712146 | 1.438478 | -2.385881 | 0.01703828 | 0.059714 |
| ENSG00000101098 | RIMS4 | 1361.300967 | -1.713526931 | 0.224247039 | -7.885298816 | 3.14E-15 | 1.39E-13 |
| ENSG00000184271 | POU6F1 | 53.34012 | -1.715157 | 0.4236398 | -4.452492 | 8.49E-06 | 8.54E-05 |
| ENSG00000185668 | POU3F1 | 2.57545 | -1.715676 | 1.771299 | -2.272458 | 0.02305886 | 0.076153 |
| ENSG00000288548 | | 17.54469 | -1.715984 | 0.6309793 | -3.309343 | 0.00093515 | 0.00539 |
| ENSG00000187554 | TLR5 | 226.0728 | -1.718194 | 0.4019251 | -4.684312 | 2.81E-06 | 3.19E-05 |
| ENSG00000188883 | KLRG2 | 166.5687 | -1.721259 | 0.3472164 | -5.303109 | 1.14E-07 | 1.71E-06 |
| ENSG00000112183 | RBM24 | 229.6817357 | -1.724390973 | 0.80618355 | -2.933825344 | 0.00334813 | 0.015919 |
| ENSG00000109794 | FAM149A | 28.34426453 | -1.724705977 | 0.543792095 | -3.685448779 | 0.0002283 | 0.001593 |
| ENSG00000170835 | CEL | 83.32698 | -1.72513 | 0.3444763 | -5.34537 | 9.02E-08 | 1.38E-06 |
| ENSG00000251463 | FKBP4P1 | 1.811993 | -1.72724 | 3.01889 | -2.415802 | 0.01570058 | 0.055888 |
| ENSG00000246596 | | 197.3702 | -1.728198 | 0.4829815 | -4.061641 | 4.87E-05 | 0.000405 |
| ENSG00000175344 | CHRNA7 | 19.4398 | -1.732314 | 0.6033004 | -3.452338 | 0.00055575 | 0.003438 |
| ENSG00000196754 | S100A2 | 40.34081 | -1.734241 | 0.6761353 | -3.222702 | 0.00126988 | 0.007039 |
| ENSG00000286186 | | 3.048246 | -1.734379 | 1.24102 | -2.421274 | 0.01546623 | 0.055239 |
| ENSG00000064692 | SNCAIP | 54.65743827 | -1.735252915 | 0.39901719 | -4.729793849 | 2.25E-06 | 2.61E-05 |
| ENSG00000205847 | OR7E91P | 16.16885 | -1.735321 | 1.066983 | -2.409728 | 0.01596443 | 0.056622 |
| ENSG00000170561 | IRX2 | 3017.256 | -1.736199 | 0.1580551 | -11.13769 | 8.22E-29 | 1.47E-26 |
| ENSG00000272097 | | 17.69004 | -1.737305 | 0.5108625 | -3.877894 | 0.00010537 | 0.000801 |
| ENSG00000254854 | | 24.10748 | -1.738844 | 0.7871934 | -2.970551 | 0.00297266 | 0.014471 |
| ENSG00000055332 | EIF2AK2 | 5956.503602 | -1.739075843 | 0.235783812 | -7.609381054 | 2.75E-14 | 1.09E-12 |
| ENSG00000233847 | ERCC8-AS1 | 3.060183 | -1.739671 | 1.351282 | -2.404617 | 0.01618943 | 0.057252 |
| ENSG00000134470 | IL15RA | 481.0461661 | -1.740241052 | 0.249274446 | -7.218230378 | 5.27E-13 | 1.78E-11 |
| ENSG00000168394 | TAP1 | 2092.197 | -1.741036 | 0.3906181 | -4.841978 | 1.29E-06 | 1.57E-05 |
| ENSG00000261366 | MANEA-DT | 8.952379 | -1.743458 | 0.8960062 | -2.780133 | 0.00543366 | 0.023702 |
| ENSG00000074657 | ZNF532 | 751.3121264 | -1.746849378 | 0.29705067 | -6.180659 | 6.38E-10 | 1.42E-08 |
| ENSG00000213337 | ANKRD39 | 615.5455 | -1.748829 | 0.1900309 | -9.391722 | 5.90E-21 | 4.87E-19 |
| ENSG00000189058 | APOD | 1043.126 | -1.750115 | 0.5572285 | -3.704155 | 0.0002121 | 0.001494 |
| ENSG00000267787 | | 16.36228 | -1.756194 | 0.4765322 | -4.138329 | 3.50E-05 | 0.000302 |
| ENSG00000224184 | MIR3681HG | 3.782323 | -1.756827 | 1.198666 | -2.487239 | 0.01287388 | 0.047692 |
| ENSG00000177189 | RPS6KA3 | 3185.898 | -1.757897 | 0.1868662 | -9.603777 | 7.71E-22 | 7.16E-20 |
| ENSG00000168453 | HR | 1684.19 | -1.761119 | 0.4032533 | -4.76622 | 1.88E-06 | 2.22E-05 |
| ENSG00000180155 | LYNX1 | 46.71256 | -1.764775 | 0.5144647 | -3.923931 | 8.71E-05 | 0.000678 |

| | | | | | | | |
|---|---|---|---|---|---|---|---|
| ENSG00000152240 | HAUS1 | 1248.552 | -1.764891 | 0.3573957 | -5.291572 | 1.21E-07 | 1.81E-06 |
| ENSG00000212464 | SNORA12 | 17.92674 | -1.76537 | 0.7926519 | -2.956003 | 0.00311654 | 0.014998 |
| ENSG00000142623 | PADI1 | 10.79233 | -1.769035 | 1.207194 | -2.517528 | 0.01181816 | 0.044492 |
| ENSG00000174718 | RESF1 | 1526.839 | -1.771483 | 0.2727265 | -6.775089 | 1.24E-11 | 3.53E-10 |
| ENSG00000111801 | BTN3A3 | 50.30026 | -1.772981859 | 0.661978743 | -3.31205941 | 0.00092612 | 0.005344 |
| ENSG00000139190 | VAMP1 | 969.1995999 | -1.773373788 | 0.268838252 | -6.862931923 | 6.75E-12 | 1.98E-10 |
| ENSG00000276903 | H2AC16 | 13.1628 | -1.773921 | 0.7717649 | -3.017832 | 0.0025459 | 0.012696 |
| ENSG00000144837 | PLA1A | 3.392771 | -1.776125 | 1.743401 | -2.378175 | 0.01739858 | 0.060744 |
| ENSG00000172403 | SYNPO2 | 266.4121 | -1.777084 | 0.3900735 | -4.957067 | 7.16E-07 | 9.21E-06 |
| ENSG00000244479 | OR2A1-AS1 | 7.651649 | -1.782415 | 0.8376255 | -2.862357 | 0.00420503 | 0.019229 |
| ENSG00000271857 | | 58.0695 | -1.784111 | 0.4911513 | -4.094365 | 4.23E-05 | 0.000358 |
| ENSG00000168679 | SLC16A4 | 153.1756 | -1.784461 | 0.4876075 | -4.138157 | 3.50E-05 | 0.000302 |
| ENSG00000233871 | DLG5-AS1 | 71.57636 | -1.786427 | 0.3194058 | -5.911882 | 3.38E-09 | 6.71E-08 |
| ENSG00000276337 | | 9.669708 | -1.789036 | 0.9739116 | -2.736094 | 0.00621732 | 0.026477 |
| ENSG00000165379 | LRFN5 | 25.9027 | -1.789277 | 0.8614048 | -2.905827 | 0.00366284 | 0.01715 |
| ENSG00000287875 | | 6.738251 | -1.791261 | 0.7719932 | -3.025082 | 0.00248566 | 0.01244 |
| ENSG00000237527 | | 5.304768 | -1.791326 | 1.112121 | -2.639595 | 0.00830051 | 0.033496 |
| ENSG00000188596 | CFAP54 | 6.712099 | -1.794851 | 1.364196 | -2.44544 | 0.01446755 | 0.052369 |
| ENSG00000225255 | PSLNR | 56.90117 | -1.799269 | 0.5674492 | -3.726577 | 0.0001941 | 0.00138 |
| ENSG00000196154 | S100A4 | 506.21 | -1.800736 | 0.9569998 | -2.800106 | 0.00510858 | 0.022543 |
| ENSG00000138386 | NAB1 | 579.3620353 | -1.801003643 | 0.316527488 | -6.012859099 | 1.82E-09 | 3.75E-08 |
| ENSG00000136840 | ST6GALNAC4 | 600.0788517 | -1.801060737 | 0.261251229 | -7.151678812 | 8.57E-13 | 2.81E-11 |
| ENSG00000203706 | SERTAD4-AS1 | 66.08605 | -1.804437 | 0.3774041 | -5.153542 | 2.56E-07 | 3.58E-06 |
| ENSG00000243415 | | 4.989683 | -1.80754 | 1.222754 | -2.526537 | 0.01151931 | 0.043585 |
| ENSG00000078295 | ADCY2 | 12.55431282 | -1.808392341 | 0.933577183 | -2.679148077 | 0.00738098 | 0.030461 |
| ENSG00000132744 | ACY3 | 50.64487694 | -1.810005371 | 0.550358924 | -3.810401288 | 0.00013874 | 0.001024 |
| ENSG00000249346 | LINC01016 | 16.97671 | -1.813102 | 0.7407155 | -3.1604 | 0.00157553 | 0.008449 |
| ENSG00000206172 | HBA1 | 3.157579 | -1.814275 | 2.064598 | -2.423755 | 0.01536096 | 0.054937 |
| ENSG00000135540 | NHSL1 | 639.5065568 | -1.818219388 | 0.232504848 | -8.045266026 | 8.61E-16 | 4.08E-14 |
| ENSG00000181378 | CFAP65 | 15.11474 | -1.818481 | 0.6911611 | -3.292901 | 0.00099159 | 0.005681 |
| ENSG00000281831 | | 4.360886 | -1.820111 | 1.02824 | -2.652824 | 0.00798215 | 0.032468 |
| ENSG00000111696 | NT5DC3 | 490.7783967 | -1.821006302 | 0.283170273 | -6.715699542 | 1.87E-11 | 5.15E-10 |
| ENSG00000111846 | GCNT2 | 45.30265169 | -1.821806431 | 0.907887827 | -2.813796804 | 0.00489602 | 0.0218 |

| | | | | | | | |
|---|---|---|---|---|---|---|---|
| ENSG00000078237 | TIGAR | 301.7221438 | -1.822183088 | 0.385325103 | -5.106758387 | 3.28E-07 | 4.48E-06 |
| ENSG00000136717 | BIN1 | 102.9744567 | -1.823426318 | 0.816477994 | -2.986239692 | 0.00282431 | 0.013883 |
| ENSG00000134463 | ECHDC3 | 345.5048303 | -1.823773992 | 0.350502614 | -5.546233639 | 2.92E-08 | 4.86E-07 |
| ENSG00000187715 | KBTBD12 | 3.669281 | -1.824355 | 1.21939 | -2.555566 | 0.01060152 | 0.040816 |
| ENSG00000173535 | TNFRSF10C | 108.3874 | -1.824731 | 0.5988285 | -3.623991 | 0.00029009 | 0.001965 |
| ENSG00000277715 | | 27.72436 | -1.826425 | 0.6834764 | -3.31554 | 0.00091466 | 0.005286 |
| ENSG00000205704 | LINC00634 | 22.36419 | -1.826881 | 0.5417672 | -3.882171 | 0.00010353 | 0.000788 |
| ENSG00000157601 | MX1 | 670.5316 | -1.82737 | 0.3358267 | -5.75692 | 8.57E-09 | 1.60E-07 |
| ENSG00000147996 | CBWD5 | 1152.122 | -1.827443 | 0.2215565 | -8.479837 | 2.26E-17 | 1.28E-15 |
| ENSG00000272734 | ADIRF-AS1 | 1097.633 | -1.827633 | 0.2475314 | -7.62681 | 2.41E-14 | 9.53E-13 |
| ENSG00000164741 | DLC1 | 2183.166 | -1.831575 | 0.3805114 | -5.188274 | 2.12E-07 | 3.02E-06 |
| ENSG00000179869 | ABCA13 | 12.54341 | -1.832198 | 1.169859 | -2.635991 | 0.00838919 | 0.033792 |
| ENSG00000286810 | | 20.53747 | -1.83317 | 0.7006796 | -3.278032 | 0.00104534 | 0.005951 |
| ENSG00000150687 | PRSS23 | 3384.286 | -1.833225 | 0.4387109 | -4.6265 | 3.72E-06 | 4.09E-05 |
| ENSG00000137285 | TUBB2B | 27.83271931 | -1.834820466 | 0.625680972 | -3.5485716 | 0.00038733 | 0.002525 |
| ENSG00000196659 | TTC30B | 384.8925 | -1.835889 | 0.1850257 | -10.10222 | 5.40E-24 | 6.05E-22 |
| ENSG00000105726 | ATP13A1 | 3777.970835 | -1.835961803 | 0.144047359 | -12.90970694 | 3.97E-38 | 1.21E-35 |
| ENSG00000069020 | MAST4 | 1459.886442 | -1.836768034 | 0.220026452 | -8.565331424 | 1.08E-17 | 6.38E-16 |
| ENSG00000157657 | ZNF618 | 1590.052 | -1.838304 | 0.239939 | -7.895066 | 2.90E-15 | 1.29E-13 |
| ENSG00000212125 | TAS2R15P | 16.27572 | -1.840701 | 0.6131241 | -3.567042 | 0.00036103 | 0.002377 |
| ENSG00000164308 | ERAP2 | 118.1968 | -1.84075 | 0.5360675 | -3.948521 | 7.86E-05 | 0.000619 |
| ENSG00000135917 | SLC19A3 | 54.83501381 | -1.841256593 | 0.708207936 | -3.155809101 | 0.00160054 | 0.00856 |
| ENSG00000228748 | | 42.48556 | -1.844141 | 0.4588479 | -4.470953 | 7.79E-06 | 7.91E-05 |
| ENSG00000146802 | TMEM168 | 1271.556 | -1.845105 | 0.2285327 | -8.301277 | 1.03E-16 | 5.44E-15 |
| ENSG00000164733 | CTSB | 6965.794 | -1.845597 | 0.2221212 | -8.541497 | 1.32E-17 | 7.77E-16 |
| ENSG00000069702 | TGFBR3 | 93.76517922 | -1.845973395 | 0.352638719 | -5.574716079 | 2.48E-08 | 4.20E-07 |
| ENSG00000248050 | | 8.478458 | -1.846319 | 0.8223394 | -2.97911 | 0.00289087 | 0.014154 |
| ENSG00000171044 | XKR6 | 68.57589 | -1.852157 | 0.503933 | -4.114585 | 3.88E-05 | 0.000331 |
| ENSG00000012171 | SEMA3B | 2331.401638 | -1.853977364 | 0.499270778 | -4.203463189 | 2.63E-05 | 0.000234 |
| ENSG00000232648 | | 29.58669 | -1.855397 | 0.437716 | -4.639177 | 3.50E-06 | 3.87E-05 |
| ENSG00000287120 | | 6.702123 | -1.856815 | 0.8572295 | -2.938316 | 0.0033 | 0.01573 |
| ENSG00000165621 | OXGR1 | 18.02025 | -1.860281 | 0.9233667 | -2.81725 | 0.00484367 | 0.021625 |
| ENSG00000146005 | PSD2 | 30.44995 | -1.860431 | 0.5374351 | -3.969468 | 7.20E-05 | 0.000572 |

| | | | | | | | |
|---|---|---|---|---|---|---|---|
| ENSG00000198498 | TMA16 | 1830.953 | -1.861852 | 0.1380767 | -13.62047 | 3.03E-42 | 1.20E-39 |
| ENSG00000234772 | LINC00412 | 4.294157 | -1.86372 | 1.326127 | -2.503087 | 0.01231152 | 0.045948 |
| ENSG00000231868 | | 31.41744 | -1.865064 | 0.3900569 | -5.150724 | 2.59E-07 | 3.62E-06 |
| ENSG00000229474 | PATL2 | 52.69361 | -1.868925 | 0.6221902 | -3.601618 | 0.00031624 | 0.002121 |
| ENSG00000184788 | SATL1 | 6.377013 | -1.869366 | 0.8285523 | -3.018061 | 0.00254398 | 0.012688 |
| ENSG00000257335 | MGAM | 14.87751 | -1.869824 | 0.864355 | -2.963691 | 0.00303973 | 0.014706 |
| ENSG00000274949 | | 7.943291 | -1.870597 | 1.081166 | -2.706311 | 0.00680352 | 0.028464 |
| ENSG00000185507 | IRF7 | 1897.548 | -1.870823 | 0.4024554 | -5.05249 | 4.36E-07 | 5.84E-06 |
| ENSG00000255176 | | 4.171049 | -1.876031 | 1.196674 | -2.580297 | 0.00987155 | 0.038463 |
| ENSG00000138411 | HECW2 | 476.6203002 | -1.876804337 | 0.351489545 | -5.681825309 | 1.33E-08 | 2.37E-07 |
| ENSG00000183785 | TUBA8 | 11.50195 | -1.877902 | 0.8502568 | -2.96793 | 0.00299813 | 0.014555 |
| ENSG00000140450 | ARRDC4 | 3629.737 | -1.878954 | 0.3321887 | -5.983254 | 2.19E-09 | 4.46E-08 |
| ENSG00000208005 | MIR503 | 12.10033 | -1.883243 | 0.8863145 | -2.879979 | 0.00397702 | 0.018367 |
| ENSG00000237513 | | 5.044704 | -1.883911 | 1.560405 | -2.419064 | 0.01556051 | 0.055516 |
| ENSG00000272256 | | 3.159622 | -1.884886 | 1.436609 | -2.35471 | 0.01853716 | 0.063913 |
| ENSG00000278642 | | 8.883155 | -1.886282 | 1.166358 | -2.599708 | 0.00933031 | 0.036769 |
| ENSG00000169006 | NTSR2 | 5.658264 | -1.888162 | 1.259503 | -2.634944 | 0.00841513 | 0.033866 |
| ENSG00000198721 | ECI2 | 820.6684 | -1.888287 | 0.3104802 | -6.390359 | 1.65E-10 | 4.01E-09 |
| ENSG00000197444 | OGDHL | 491.3483 | -1.894958 | 0.3744856 | -5.431346 | 5.59E-08 | 8.94E-07 |
| ENSG00000273449 | | 68.39374 | -1.895124 | 0.3237879 | -6.168296 | 6.90E-10 | 1.53E-08 |
| ENSG00000152952 | PLOD2 | 4442.058 | -1.897258 | 0.1872124 | -10.31326 | 6.14E-25 | 7.59E-23 |
| ENSG00000167771 | RCOR2 | 249.8442 | -1.900525 | 0.3805545 | -5.364287 | 8.13E-08 | 1.25E-06 |
| ENSG00000106003 | LFNG | 2379.565989 | -1.902052576 | 0.414552605 | -4.994404474 | 5.90E-07 | 7.72E-06 |
| ENSG00000184979 | USP18 | 665.7218 | -1.902358 | 0.4198241 | -4.941947 | 7.73E-07 | 9.86E-06 |
| ENSG00000147642 | SYBU | 860.3792 | -1.903547 | 0.3632798 | -5.583878 | 2.35E-08 | 4.01E-07 |
| ENSG00000262445 | | 3.613648 | -1.903708 | 1.563237 | -2.41693 | 0.01565204 | 0.055753 |
| ENSG00000232303 | DFFBP1 | 9.62872 | -1.906941 | 0.6882374 | -3.373754 | 0.00074151 | 0.004415 |
| ENSG00000281181 | | 1.966417 | -1.909436 | 1.707809 | -2.356725 | 0.01843687 | 0.063633 |
| ENSG00000256650 | RERG-IT1 | 6.136861 | -1.911266 | 0.8462097 | -3.055107 | 0.0022498 | 0.011429 |
| ENSG00000120251 | GRIA2 | 6.091500514 | -1.914401412 | 0.769705414 | -3.146130971 | 0.00165446 | 0.008795 |
| ENSG00000178031 | ADAMTSL1 | 18.85943 | -1.917235 | 0.6587129 | -3.5178 | 0.00043514 | 0.002782 |
| ENSG00000200152 | RNU6-216P | 5.692876 | -1.920825 | 1.077116 | -2.657656 | 0.00786862 | 0.032099 |
| ENSG00000161544 | CYGB | 10.93636 | -1.920927 | 0.7744622 | -3.18422 | 0.00145145 | 0.007886 |

| | | | | | | | |
|---|---|---|---|---|---|---|---|
| ENSG00000166183 | ASPG | 72.60569 | -1.921633 | 0.6267018 | -3.666487 | 0.00024591 | 0.001701 |
| ENSG00000148400 | NOTCH1 | 4071.548 | -1.922253 | 0.3046589 | -6.626338 | 3.44E-11 | 9.10E-10 |
| ENSG00000182179 | UBA7 | 394.8946 | -1.922862 | 0.5275577 | -4.144918 | 3.40E-05 | 0.000294 |
| ENSG00000259803 | SLC22A31 | 43.6588 | -1.924278 | 0.5062681 | -4.295107 | 1.75E-05 | 0.000162 |
| ENSG00000158623 | COPG2 | 1599.665 | -1.924287 | 0.1966657 | -9.977714 | 1.91E-23 | 2.00E-21 |
| ENSG00000124302 | CHST8 | 56.25835402 | -1.924723743 | 0.533098076 | -4.114687153 | 3.88E-05 | 0.000331 |
| ENSG00000115355 | CCDC88A | 23.80629581 | -1.925026536 | 0.493116651 | -4.378698237 | 1.19E-05 | 0.000115 |
| ENSG00000171433 | GLOD5 | 7.28686 | -1.925642 | 0.7826844 | -3.099059 | 0.00194136 | 0.010096 |
| ENSG00000253154 | | 9.225154 | -1.928279 | 0.9944803 | -2.875348 | 0.00403583 | 0.01859 |
| ENSG00000153898 | MCOLN2 | 239.3852 | -1.932012 | 0.2180709 | -9.068652 | 1.20E-19 | 8.61E-18 |
| ENSG00000163082 | SGPP2 | 297.3107 | -1.936612 | 0.365753 | -5.643455 | 1.67E-08 | 2.92E-07 |
| ENSG00000273998 | | 2.252033 | -1.937395 | 1.551825 | -2.368608 | 0.01785515 | 0.06202 |
| ENSG00000141504 | SAT2 | 884.395 | -1.937416 | 0.3096388 | -6.557363 | 5.48E-11 | 1.41E-09 |
| ENSG00000141579 | ZNF750 | 369.373 | -1.937875 | 0.5065042 | -4.317006 | 1.58E-05 | 0.000148 |
| ENSG00000181896 | ZNF101 | 376.1884 | -1.939377 | 0.2139731 | -9.272599 | 1.82E-20 | 1.41E-18 |
| ENSG00000279035 | | 7.594237 | -1.942542 | 0.9776087 | -2.826117 | 0.00471161 | 0.021127 |
| ENSG00000215458 | AATBC | 31.84191 | -1.943224 | 0.4469278 | -4.764602 | 1.89E-06 | 2.23E-05 |
| ENSG00000124429 | POF1B | 836.0552581 | -1.943674967 | 0.326671457 | -6.262046008 | 3.80E-10 | 8.76E-09 |
| ENSG00000286538 | | 8.298071 | -1.944083 | 0.8393104 | -3.063496 | 0.00218767 | 0.011167 |
| ENSG00000173157 | ADAMTS20 | 43.4044 | -1.947462 | 0.599138 | -3.818765 | 0.00013412 | 0.000993 |
| ENSG00000148175 | STOM | 5936.818 | -1.951244 | 0.3745784 | -5.586623 | 2.32E-08 | 3.95E-07 |
| ENSG00000284882 | | 6.830983 | -1.952401 | 1.103378 | -2.722713 | 0.00647483 | 0.027337 |
| ENSG00000077942 | FBLN1 | 3077.031809 | -1.953099702 | 0.563917732 | -4.003931649 | 6.23E-05 | 0.000503 |
| ENSG00000233577 | | 3.407469 | -1.954967 | 1.146359 | -2.663832 | 0.00772561 | 0.031637 |
| ENSG00000174791 | RIN1 | 313.0536 | -1.957635 | 0.3326858 | -6.209711 | 5.31E-10 | 1.19E-08 |
| ENSG00000158321 | AUTS2 | 499.93 | -1.957778 | 0.3213858 | -6.410458 | 1.45E-10 | 3.54E-09 |
| ENSG00000114113 | RBP2 | 6.135453137 | -1.958483061 | 1.077537174 | -2.726141101 | 0.00640796 | 0.027089 |
| ENSG00000231187 | | 43.93966 | -1.960424 | 0.6970295 | -3.38169 | 0.00072042 | 0.004309 |
| ENSG00000243479 | MNX1-AS1 | 439.9785 | -1.963202 | 0.3084155 | -6.660335 | 2.73E-11 | 7.34E-10 |
| ENSG00000081665 | ZNF506 | 49.18832613 | -1.964456866 | 0.413438217 | -5.134559165 | 2.83E-07 | 3.92E-06 |
| ENSG00000185697 | MYBL1 | 1190.266 | -1.964587 | 0.3581196 | -5.831052 | 5.51E-09 | 1.05E-07 |
| ENSG00000147255 | IGSF1 | 142.7484 | -1.966505 | 0.7046139 | -3.474076 | 0.00051262 | 0.003204 |
| ENSG00000119922 | IFIT2 | 727.9652419 | -1.972968318 | 0.5713132 | -4.006415752 | 6.16E-05 | 0.000499 |

| | | | | | | | |
|---|---|---|---|---|---|---|---|
| ENSG00000113273 | ARSB | 441.601589 | -1.974779332 | 0.27366098 | -7.486779244 | 7.06E-14 | 2.66E-12 |
| ENSG00000213144 | | 619.8607 | -1.975541 | 0.2084852 | -9.672472 | 3.95E-22 | 3.79E-20 |
| ENSG00000226419 | SLC16A1-AS1 | 70.74618 | -1.975583 | 0.468872 | -4.66369 | 3.11E-06 | 3.49E-05 |
| ENSG00000154319 | FAM167A | 67.25625 | -1.978491 | 0.4304186 | -5.009555 | 5.46E-07 | 7.20E-06 |
| ENSG00000157303 | SUSD3 | 754.0098 | -1.97885 | 0.5434873 | -4.173042 | 3.01E-05 | 0.000263 |
| ENSG00000256417 | | 58.70769 | -1.979949 | 0.4125308 | -5.191196 | 2.09E-07 | 2.98E-06 |
| ENSG00000064205 | CCN5 | 228.3074661 | -1.982250624 | 0.58195377 | -3.969260643 | 7.21E-05 | 0.000572 |
| ENSG00000274349 | ZNF658 | 147.9203 | -1.982817 | 0.1981041 | -10.19383 | 2.11E-24 | 2.51E-22 |
| ENSG00000171621 | SPSB1 | 752.5825 | -1.983568 | 0.3559744 | -5.908226 | 3.46E-09 | 6.85E-08 |
| ENSG00000122735 | DNAI1 | 16.06352072 | -1.983871391 | 0.686969925 | -3.535129716 | 0.00040758 | 0.002633 |
| ENSG00000235237 | | 1.753285 | -1.988861 | 1.993892 | -2.401675 | 0.01632018 | 0.057623 |
| ENSG00000088881 | EBF4 | 494.5380971 | -1.992495414 | 0.410358388 | -5.257309954 | 1.46E-07 | 2.15E-06 |
| ENSG00000049768 | FOXP3 | 58.50593964 | -1.994544401 | 0.496330372 | -4.50034708 | 6.78E-06 | 6.99E-05 |
| ENSG00000134595 | SOX3 | 400.1792437 | -1.996127914 | 0.316667505 | -6.613656539 | 3.75E-11 | 9.87E-10 |
| ENSG00000265185 | SNORD3B-1 | 25.56995 | -1.998156 | 0.6691494 | -3.597506 | 0.00032128 | 0.00215 |
| ENSG00000115267 | IFIH1 | 1900.355972 | -1.998807405 | 0.359275426 | -5.926887399 | 3.09E-09 | 6.15E-08 |
| ENSG00000124067 | SLC12A4 | 266.6497726 | -2.004307255 | 0.261939577 | -7.893961412 | 2.93E-15 | 1.30E-13 |
| ENSG00000187017 | ESPN | 2770.21 | -2.004585 | 0.2007823 | -10.17743 | 2.50E-24 | 2.93E-22 |
| ENSG00000221261 | MIR1208 | 7.606426 | -2.004802 | 0.9559715 | -2.958745 | 0.00308895 | 0.014887 |
| ENSG00000224842 | | 52.38108 | -2.00851 | 0.4667532 | -4.746608 | 2.07E-06 | 2.43E-05 |
| ENSG00000197479 | PCDHB11 | 29.68194 | -2.009998 | 0.4767562 | -4.659198 | 3.17E-06 | 3.56E-05 |
| ENSG00000169548 | ZNF280A | 7.930284 | -2.010516 | 1.116903 | -2.715643 | 0.00661472 | 0.027817 |
| ENSG00000145934 | TENM2 | 2.473536 | -2.011208 | 1.52646 | -2.41023 | 0.01594245 | 0.056567 |
| ENSG00000150394 | CDH8 | 10.15717 | -2.01222 | 0.7116316 | -3.430606 | 0.00060224 | 0.003685 |
| ENSG00000279884 | | 9.092242 | -2.017436 | 0.8780058 | -3.009061 | 0.00262056 | 0.013019 |
| ENSG00000196132 | MYT1 | 13.38236 | -2.017962 | 0.9472891 | -3.003743 | 0.0026668 | 0.013204 |
| ENSG00000262202 | | 11.09346 | -2.020359 | 1.102108 | -2.806547 | 0.00500756 | 0.022178 |
| ENSG00000247324 | | 2.514199 | -2.0204 | 2.022923 | -2.41782 | 0.0156138 | 0.055669 |
| ENSG00000186352 | ANKRD37 | 98.46129 | -2.022482 | 0.4041128 | -5.394129 | 6.89E-08 | 1.08E-06 |
| ENSG00000084453 | SLCO1A2 | 26.78083547 | -2.022670065 | 0.526429699 | -4.334179585 | 1.46E-05 | 0.000139 |
| ENSG00000196083 | IL1RAP | 328.4855 | -2.023736 | 0.2989018 | -7.055517 | 1.72E-12 | 5.43E-11 |
| ENSG00000166407 | LMO1 | 19.50214 | -2.026854 | 0.9212003 | -3.053439 | 0.00226235 | 0.011482 |
| ENSG00000279798 | | 61.35775 | -2.028299 | 0.4443209 | -4.997046 | 5.82E-07 | 7.63E-06 |

| | | | | | | | |
|---|---|---|---|---|---|---|---|
| ENSG00000130768 | SMPDL3B | 497.0023876 | -2.032889777 | 0.203748787 | -10.1658223 | 2.82E-24 | 3.27E-22 |
| ENSG00000134343 | ANO3 | 18.43968468 | -2.035026259 | 0.579735938 | -4.009878851 | 6.07E-05 | 0.000493 |
| ENSG00000064042 | LIMCH1 | 1905.91127 | -2.035152024 | 0.533163328 | -4.321459273 | 1.55E-05 | 0.000146 |
| ENSG00000198133 | TMEM229B | 1700.238 | -2.035408 | 0.2647652 | -7.952091 | 1.83E-15 | 8.29E-14 |
| ENSG00000179241 | LDLRAD3 | 795.9372 | -2.036031 | 0.2820977 | -7.48669 | 7.06E-14 | 2.66E-12 |
| ENSG00000162496 | DHRS3 | 745.2291 | -2.040155 | 0.4111049 | -5.352189 | 8.69E-08 | 1.33E-06 |
| ENSG00000108961 | RANGRF | 46.56267733 | -2.041523139 | 0.376672504 | -5.77405313 | 7.74E-09 | 1.45E-07 |
| ENSG00000250027 | | 8.113903 | -2.041891 | 0.928292 | -3.013245 | 0.0025847 | 0.012865 |
| ENSG00000265939 | UBE2CP2 | 8.702267 | -2.043909 | 0.7909511 | -3.294941 | 0.00098442 | 0.005645 |
| ENSG00000285730 | | 6.122293 | -2.045053 | 1.046302 | -2.801559 | 0.00508564 | 0.022468 |
| ENSG00000162409 | PRKAA2 | 307.7314 | -2.045259 | 0.4573426 | -4.91368 | 8.94E-07 | 1.12E-05 |
| ENSG00000197977 | ELOVL2 | 932.5243 | -2.047915 | 0.6135836 | -3.940621 | 8.13E-05 | 0.000637 |
| ENSG00000188910 | GJB3 | 294.4799 | -2.049374 | 0.4544066 | -4.968192 | 6.76E-07 | 8.75E-06 |
| ENSG00000160460 | SPTBN4 | 610.3262 | -2.05171 | 0.316121 | -6.795608 | 1.08E-11 | 3.08E-10 |
| ENSG00000142303 | ADAMTS10 | 5.51915 | -2.052995 | 1.467881 | -2.533336 | 0.01129825 | 0.042945 |
| ENSG00000128342 | LIF | 474.9779722 | -2.05579287 | 0.520963503 | -4.449593039 | 8.60E-06 | 8.63E-05 |
| ENSG00000120328 | PCDHB12 | 44.64569193 | -2.05591384 | 0.440796173 | -5.073761237 | 3.90E-07 | 5.26E-06 |
| ENSG00000082175 | PGR | 3388.334107 | -2.05661236 | 1.248569789 | -2.810986048 | 0.00493899 | 0.021929 |
| ENSG00000233593 | LINC02609 | 65.86031 | -2.060066 | 0.5111812 | -4.509463 | 6.50E-06 | 6.74E-05 |
| ENSG00000233191 | | 7.57616 | -2.062267 | 1.271848 | -2.574797 | 0.01002988 | 0.038954 |
| ENSG00000205420 | KRT6A | 4.973118 | -2.065192 | 2.358874 | -2.446228 | 0.01443596 | 0.052319 |
| ENSG00000162174 | ASRGL1 | 194.4632 | -2.065328 | 0.5210319 | -4.462419 | 8.10E-06 | 8.19E-05 |
| ENSG00000109775 | UFSP2 | 1005.264618 | -2.065776973 | 0.181431907 | -11.55851006 | 6.69E-31 | 1.41E-28 |
| ENSG00000272735 | | 11.20528 | -2.076498 | 1.245259 | -2.718389 | 0.00656006 | 0.027631 |
| ENSG00000237732 | CT75 | 44.51281 | -2.079666 | 0.6439388 | -3.825958 | 0.00013027 | 0.000968 |
| ENSG00000111186 | WNT5B | 22.88509914 | -2.079752435 | 0.632273317 | -3.861495532 | 0.0001127 | 0.000852 |
| ENSG00000100739 | BDKRB1 | 2.666852555 | -2.08284345 | 1.663683932 | -2.525579118 | 0.01155078 | 0.04368 |
| ENSG00000109103 | UNC119 | 3767.387447 | -2.08313006 | 0.232366357 | -9.231686772 | 2.66E-20 | 2.05E-18 |
| ENSG00000280916 | FOXCUT | 8.092503 | -2.08676 | 1.172605 | -2.509156 | 0.01210199 | 0.045306 |
| ENSG00000281332 | LINC00997 | 178.4631 | -2.088038 | 0.3614201 | -6.122583 | 9.21E-10 | 1.99E-08 |
| ENSG00000101347 | SAMHD1 | 2145.317835 | -2.088139536 | 0.283708675 | -7.634855298 | 2.26E-14 | 8.99E-13 |
| ENSG00000079393 | DUSP13 | 31.29177928 | -2.088967722 | 0.818710511 | -3.32172448 | 0.00089463 | 0.00519 |
| ENSG00000235044 | PPIAP3 | 4.986636 | -2.09091 | 1.115182 | -2.743582 | 0.00607728 | 0.025989 |

| | | | | | | | |
|---|---|---|---|---|---|---|---|
| ENSG00000156284 | CLDN8 | 17.70191 | -2.091302 | 1.084773 | -2.837688 | 0.00454416 | 0.020498 |
| ENSG00000250305 | TRMT9B | 83.91491 | -2.091536 | 0.5972816 | -4.062493 | 4.86E-05 | 0.000404 |
| ENSG00000229832 | | 2.023527 | -2.091762 | 2.074622 | -2.453588 | 0.01414387 | 0.051443 |
| ENSG00000204176 | SYT15 | 44.51157 | -2.092985 | 0.6680001 | -3.69624 | 0.00021882 | 0.001536 |
| ENSG00000197380 | DACT3 | 60.21793 | -2.099066 | 0.4627139 | -4.957938 | 7.12E-07 | 9.18E-06 |
| ENSG00000240032 | LNCSRLR | 6.084212 | -2.102835 | 0.7841233 | -3.37742 | 0.00073169 | 0.004364 |
| ENSG00000145526 | CDH18 | 93.82166 | -2.104324 | 0.5618992 | -4.283102 | 1.84E-05 | 0.00017 |
| ENSG00000204025 | TRPC5OS | 3.875389 | -2.10493 | 1.329098 | -2.713478 | 0.0066581 | 0.027968 |
| ENSG00000203364 | | 6.406133 | -2.106752 | 0.9785744 | -2.944378 | 0.00323604 | 0.015472 |
| ENSG00000264269 | | 15.47516 | -2.111774 | 0.5375335 | -4.398849 | 1.09E-05 | 0.000106 |
| ENSG00000280208 | GGT4P | 6.576293 | -2.115724 | 1.101227 | -2.712884 | 0.00667005 | 0.027992 |
| ENSG00000280294 | | 7.248032 | -2.116897 | 1.494134 | -2.495255 | 0.01258666 | 0.046843 |
| ENSG00000204632 | HLA-G | 48.70753 | -2.119956 | 0.5347168 | -4.418889 | 9.92E-06 | 9.78E-05 |
| ENSG00000233719 | GOT2P3 | 3.394528 | -2.129417 | 1.208422 | -2.644984 | 0.00816948 | 0.033067 |
| ENSG00000019582 | CD74 | 509.9170701 | -2.133061885 | 0.929773947 | -3.15193712 | 0.00162191 | 0.00865 |
| ENSG00000265943 | | 25.20694 | -2.135079 | 0.4971832 | -4.766664 | 1.87E-06 | 2.22E-05 |
| ENSG00000235121 | | 23.05727 | -2.137313 | 0.6763565 | -3.740569 | 0.0001836 | 0.001314 |
| ENSG00000138496 | PARP9 | 3733.463457 | -2.138874444 | 0.385632664 | -5.904490974 | 3.54E-09 | 6.99E-08 |
| ENSG00000082497 | SERTAD4 | 1227.213568 | -2.141044357 | 0.305056196 | -7.305405695 | 2.76E-13 | 9.71E-12 |
| ENSG00000229321 | | 10.89097 | -2.144007 | 0.6513489 | -3.837529 | 0.00012428 | 0.00093 |
| ENSG00000261770 | | 39.58822 | -2.144324 | 0.6801775 | -3.785723 | 0.00015326 | 0.001119 |
| ENSG00000228784 | LINC00954 | 15.0439 | -2.147018 | 0.6076929 | -4.076272 | 4.58E-05 | 0.000383 |
| ENSG00000265479 | DTX2P1-UPK3BP | 3.452001 | -2.150597 | 1.843332 | -2.466226 | 0.01365451 | 0.049936 |
| ENSG00000251348 | HSPD1P11 | 42.60176 | -2.154442 | 0.5671597 | -4.323141 | 1.54E-05 | 0.000145 |
| ENSG00000159433 | STARD9 | 96.35633 | -2.155394 | 0.3138571 | -7.166399 | 7.70E-13 | 2.54E-11 |
| ENSG00000253978 | | 7.720189 | -2.159254 | 0.8797156 | -3.192983 | 0.00140811 | 0.007685 |
| ENSG00000254911 | SCARNA9 | 10.42614 | -2.159927 | 0.7202487 | -3.618776 | 0.000296 | 0.001999 |
| ENSG00000085831 | TTC39A | 8580.345757 | -2.161594279 | 0.201779861 | -10.91494946 | 9.78E-28 | 1.57E-25 |
| ENSG00000198734 | F5 | 9.90413 | -2.162384 | 0.7203764 | -3.644527 | 0.00026788 | 0.001832 |
| ENSG00000268536 | | 4.575022 | -2.162568 | 1.096732 | -2.84748 | 0.00440669 | 0.019963 |
| ENSG00000274139 | | 3.160796 | -2.166157 | 1.564006 | -2.564736 | 0.01032542 | 0.039932 |
| ENSG00000198223 | CSF2RA | 5.485469 | -2.169216 | 1.110314 | -2.797761 | 0.00514582 | 0.022673 |
| ENSG00000144847 | IGSF11 | 27.80602 | -2.169636 | 0.438698 | -5.352002 | 8.70E-08 | 1.33E-06 |

| | | | | | | | |
|---|---|---|---|---|---|---|---|
| ENSG00000268460 | | 14.1688 | -2.175058 | 0.8480862 | -3.256305 | 0.00112873 | 0.006354 |
| ENSG00000150594 | ADRA2A | 27.92209 | -2.176916 | 0.6365691 | -4.018233 | 5.86E-05 | 0.000477 |
| ENSG00000156140 | ADAMTS3 | 18.99876 | -2.178017 | 0.4809684 | -4.950599 | 7.40E-07 | 9.48E-06 |
| ENSG00000163235 | TGFA | 970.6781 | -2.181432 | 0.3587528 | -6.416577 | 1.39E-10 | 3.42E-09 |
| ENSG00000165406 | MARCHF8 | 1536.307 | -2.18171 | 0.2448539 | -9.138941 | 6.31E-20 | 4.62E-18 |
| ENSG00000288549 | | 3.563391 | -2.183483 | 1.094075 | -2.894122 | 0.00380221 | 0.01768 |
| ENSG00000241106 | HLA-DOB | 8.393976 | -2.184534 | 1.331412 | -2.625033 | 0.00866406 | 0.034684 |
| ENSG00000168758 | SEMA4C | 8166.356 | -2.186636 | 0.1713421 | -12.92188 | 3.39E-38 | 1.05E-35 |
| ENSG00000188886 | ASTL | 18.91459 | -2.190222 | 0.70284 | -3.752197 | 0.00017529 | 0.001261 |
| ENSG00000272462 | | 28.72849 | -2.194182 | 0.4075355 | -5.744879 | 9.20E-09 | 1.71E-07 |
| ENSG00000226942 | IL9RP3 | 21.01769 | -2.194628 | 0.6434201 | -3.981067 | 6.86E-05 | 0.000547 |
| ENSG00000172794 | RAB37 | 109.2758 | -2.195077 | 0.486411 | -4.966108 | 6.83E-07 | 8.84E-06 |
| ENSG00000142619 | PADI3 | 47.20014 | -2.19873 | 0.6947862 | -3.811772 | 0.00013797 | 0.001019 |
| ENSG00000196358 | NTNG2 | 11.18787 | -2.202112 | 1.030527 | -2.960957 | 0.00306685 | 0.0148 |
| ENSG00000224958 | PGM5-AS1 | 13.35896 | -2.203911 | 1.03307 | -2.985613 | 0.00283011 | 0.013907 |
| ENSG00000099998 | GGT5 | 19.35722696 | -2.205340443 | 0.934779155 | -3.203955535 | 0.00135553 | 0.007444 |
| ENSG00000285094 | LINC01488 | 20.97797 | -2.205891 | 0.8023255 | -3.498657 | 0.00046761 | 0.002964 |
| ENSG00000087258 | GNAO1 | 76.36901191 | -2.206301976 | 0.433306325 | -5.497316094 | 3.86E-08 | 6.33E-07 |
| ENSG00000026950 | BTN3A1 | 94.61315665 | -2.207557354 | 0.465083551 | -5.174757416 | 2.28E-07 | 3.23E-06 |
| ENSG00000231566 | LINC02595 | 12.27094 | -2.207942 | 0.9438016 | -3.008676 | 0.00262389 | 0.013033 |
| ENSG00000258499 | LINC02287 | 2.311986 | -2.208425 | 2.679803 | -2.534896 | 0.01124807 | 0.042799 |
| ENSG00000182985 | CADM1 | 1979.529 | -2.208864 | 0.269107 | -8.480629 | 2.24E-17 | 1.27E-15 |
| ENSG00000254226 | LINC01933 | 20.85884 | -2.209198 | 0.7320001 | -3.683685 | 0.00022989 | 0.001603 |
| ENSG00000173156 | RHOD | 4596.085 | -2.210956 | 0.2540871 | -8.941666 | 3.83E-19 | 2.63E-17 |
| ENSG00000188906 | LRRK2 | 54.28871 | -2.21447 | 0.4155689 | -5.71526 | 1.10E-08 | 2.00E-07 |
| ENSG00000250420 | AACSP1 | 35.16718 | -2.216253 | 1.025844 | -3.063463 | NA | NA |
| ENSG00000261051 | | 19.34958 | -2.21852 | 0.546076 | -4.556896 | 5.19E-06 | 5.52E-05 |
| ENSG00000233296 | TMEM18-DT | 15.62686 | -2.219739 | 0.5645512 | -4.440771 | 8.96E-06 | 8.95E-05 |
| ENSG00000143847 | PPFIA4 | 149.6616 | -2.221557 | 0.2982427 | -7.727934 | 1.09E-14 | 4.50E-13 |
| ENSG00000130558 | OLFM1 | 1661.260006 | -2.223952148 | 0.24277779 | -9.393144936 | 5.82E-21 | 4.84E-19 |
| ENSG00000249085 | | 2.313201 | -2.228067 | 1.832186 | -2.523002 | 0.01163577 | 0.043926 |
| ENSG00000285848 | | 2.979096 | -2.230328 | 1.731491 | -2.525019 | 0.01156921 | 0.043743 |
| ENSG00000236545 | | 3.806268 | -2.23152 | 1.029858 | -2.972703 | 0.0029519 | 0.014399 |

| | | | | | | | |
|---|---|---|---|---|---|---|---|
| ENSG00000281383 | | 11.22937 | -2.232775 | 0.8213338 | -3.436651 | 0.00058895 | 0.003616 |
| ENSG00000164647 | STEAP1 | 59.82259 | -2.237364 | 0.7114098 | -3.613835 | 0.0003017 | 0.002032 |
| ENSG00000286427 | | 8.850508 | -2.242577 | 0.8416946 | -3.373564 | 0.00074202 | 0.004417 |
| ENSG00000256115 | LINC02443 | 16.33849 | -2.242718 | 0.8726023 | -3.356718 | 0.00078874 | 0.004661 |
| ENSG00000226380 | | 45.32969 | -2.246023 | 0.6097474 | -4.229087 | 2.35E-05 | 0.000212 |
| ENSG00000243902 | | 3.17371 | -2.249986 | 1.562152 | -2.52232 | 0.01165837 | 0.043986 |
| ENSG00000174885 | NLRP6 | 13.67527 | -2.253751 | 0.7466617 | -3.662624 | 0.00024965 | 0.001721 |
| ENSG00000181458 | TMEM45A | 54.80253 | -2.253844 | 0.7309199 | -3.761152 | 0.00016913 | 0.001223 |
| ENSG00000255794 | RMST | 469.6934 | -2.255158 | 0.3522021 | -6.733845 | 1.65E-11 | 4.58E-10 |
| ENSG00000256546 | | 5.716896 | -2.256438 | 1.007417 | -3.057063 | 0.00223518 | 0.01137 |
| ENSG00000132563 | REEP2 | 207.0231502 | -2.25699201 | 0.524491362 | -4.79291258 | 1.64E-06 | 1.96E-05 |
| ENSG00000196873 | CBWD3 | 236.2606 | -2.263088 | 0.2281848 | -10.12662 | 4.21E-24 | 4.78E-22 |
| ENSG00000268812 | | 26.65847 | -2.264482 | 0.7630244 | -3.631886 | 0.00028136 | 0.001912 |
| ENSG00000111331 | OAS3 | 8264.934946 | -2.265220003 | 0.426463325 | -5.708955255 | 1.14E-08 | 2.06E-07 |
| ENSG00000226965 | | 15.46922 | -2.268007 | 0.6087384 | -4.259247 | 2.05E-05 | 0.000187 |
| ENSG00000224596 | ZMIZ1-AS1 | 13.58286 | -2.269155 | 0.6448424 | -4.080256 | 4.50E-05 | 0.000378 |
| ENSG00000135218 | CD36 | 134.841766 | -2.272164229 | 0.774217748 | -3.555452463 | 0.00037733 | 0.002469 |
| ENSG00000165309 | ARMC3 | 8.023858 | -2.274999 | 0.7602699 | -3.602539 | 0.00031513 | 0.002114 |
| ENSG00000267264 | | 8.248319 | -2.276691 | 0.7351533 | -3.695969 | 0.00021905 | 0.001537 |
| ENSG00000115825 | PRKD3 | 1913.810936 | -2.282175465 | 0.200576457 | -11.5626727 | 6.37E-31 | 1.35E-28 |
| ENSG00000203930 | LINC00632 | 284.6422 | -2.284747 | 0.4617233 | -5.38188 | 7.37E-08 | 1.15E-06 |
| ENSG00000164309 | CMYA5 | 2109.558 | -2.285435 | 0.3986793 | -6.111128 | 9.89E-10 | 2.13E-08 |
| ENSG00000167723 | TRPV3 | 53.90383 | -2.286134 | 0.5059044 | -4.990148 | 6.03E-07 | 7.87E-06 |
| ENSG00000184486 | POU3F2 | 129.4918 | -2.287564 | 0.3518505 | -6.830242 | 8.48E-12 | 2.45E-10 |
| ENSG00000183709 | IFNL2 | 6.574264 | -2.292385 | 1.297311 | -2.601285 | 0.00928753 | 0.036649 |
| ENSG00000128594 | LRRC4 | 29.47895468 | -2.292859354 | 0.672062895 | -4.02135331 | 5.79E-05 | 0.000471 |
| ENSG00000174837 | ADGRE1 | 150.0671 | -2.294401 | 0.5913484 | -4.417524 | 9.98E-06 | 9.84E-05 |
| ENSG00000186115 | CYP4F2 | 3.02795 | -2.298949 | 1.724587 | -2.538724 | 0.01112576 | 0.042404 |
| ENSG00000287777 | | 2.737702 | -2.298987 | 1.719741 | -2.624315 | 0.00868234 | 0.034741 |
| ENSG00000108691 | CCL2 | 466.2966989 | -2.304221843 | 1.322890658 | -2.645976534 | 0.00814555 | 0.032996 |
| ENSG00000106927 | AMBP | 17.55828027 | -2.310299612 | 1.148431334 | -3.034837495 | 0.00240665 | 0.012109 |
| ENSG00000287712 | | 6.111957 | -2.31631 | 1.241401 | -2.624258 | 0.00868381 | 0.034742 |
| ENSG00000214336 | FOXI3 | 29.17327 | -2.316321 | 0.5491881 | -4.697771 | 2.63E-06 | 3.01E-05 |

| | | | | | | | |
|---|---|---|---|---|---|---|---|
| ENSG00000232613 | LINC02576 | 2.535555 | -2.316425 | 2.24601 | -2.54956 | 0.0107859 | 0.04134 |
| ENSG00000162989 | KCNJ3 | 261.0888 | -2.31765 | 0.3929512 | -6.256127 | 3.95E-10 | 9.07E-09 |
| ENSG00000251026 | LINC02163 | 67.7773 | -2.321233 | 0.4378552 | -5.705659 | 1.16E-08 | 2.10E-07 |
| ENSG00000086712 | TXLNG | 4908.699744 | -2.322197987 | 0.177910061 | -13.21763434 | 6.94E-40 | 2.25E-37 |
| ENSG00000132970 | WASF3 | 338.5085994 | -2.322280527 | 0.296482886 | -8.107296183 | 5.18E-16 | 2.53E-14 |
| ENSG00000104267 | CA2 | 326.5204029 | -2.323700605 | 0.485984986 | -5.231616083 | 1.68E-07 | 2.44E-06 |
| ENSG00000264714 | KIAA0895LP1 | 2.854464 | -2.324351 | 1.440052 | -2.668793 | 0.00761244 | 0.031265 |
| ENSG00000120664 | SPART-AS1 | 15.68275888 | -2.330063847 | 0.531787653 | -4.847957833 | 1.25E-06 | 1.53E-05 |
| ENSG00000101955 | SRPX | 23.29994443 | -2.33006405 | 0.513988368 | -4.98296778 | 6.26E-07 | 8.15E-06 |
| ENSG00000102290 | PCDH11X | 157.938037 | -2.332304215 | 0.556865704 | -4.697147698 | 2.64E-06 | 3.01E-05 |
| ENSG00000287665 | | 7.927476 | -2.332407 | 1.355915 | -2.806369 | 0.00501033 | 0.022187 |
| ENSG00000101096 | NFATC2 | 905.9888227 | -2.332422576 | 0.285055267 | -8.44743586 | 2.98E-17 | 1.68E-15 |
| ENSG00000102575 | ACP5 | 19.81633485 | -2.335932252 | 0.574629971 | -4.570415565 | 4.87E-06 | 5.20E-05 |
| ENSG00000133816 | MICAL2 | 1567.921044 | -2.337518937 | 0.277662532 | -8.661482299 | 4.66E-18 | 2.84E-16 |
| ENSG00000206503 | HLA-A | 3328.629 | -2.337651 | 0.4661631 | -5.461142 | 4.73E-08 | 7.66E-07 |
| ENSG00000173193 | PARP14 | 6228.643 | -2.33977 | 0.3666431 | -6.722359 | 1.79E-11 | 4.94E-10 |
| ENSG00000175899 | A2M | 212.6946 | -2.342157 | 1.515566 | -2.791534 | 0.00524589 | 0.02304 |
| ENSG00000170965 | PLAC1 | 153.4123 | -2.34584 | 0.6874904 | -4.044745 | 5.24E-05 | 0.000432 |
| ENSG00000267644 | | 5.068337 | -2.353781 | 1.03339 | -3.017042 | 0.00255254 | 0.012722 |
| ENSG00000134326 | CMPK2 | 472.6638107 | -2.355595247 | 0.592994237 | -4.509876582 | 6.49E-06 | 6.73E-05 |
| ENSG00000276718 | | 26.769 | -2.357413 | 0.5897338 | -4.485885 | 7.26E-06 | 7.43E-05 |
| ENSG00000242173 | ARHGDIG | 317.7616 | -2.357908 | 0.3481776 | -7.094253 | 1.30E-12 | 4.18E-11 |
| ENSG00000286179 | | 20.26156 | -2.364246 | 0.6937827 | -3.976026 | 7.01E-05 | 0.000558 |
| ENSG00000149403 | GRIK4 | 7.219597 | -2.365019 | 0.8913078 | -3.337995 | 0.00084385 | 0.004938 |
| ENSG00000106565 | TMEM176B | 10.23794776 | -2.367662109 | 0.774291914 | -3.723084262 | 0.0001968 | 0.001397 |
| ENSG00000167968 | DNASE1L2 | 352.5261 | -2.372308 | 0.2462094 | -9.855611 | 6.48E-23 | 6.68E-21 |
| ENSG00000161929 | SCIMP | 8.888451 | -2.373898 | 0.8177748 | -3.566361 | 0.00036197 | 0.002383 |
| ENSG00000228649 | SNHG26 | 188.3685 | -2.374069 | 0.3951093 | -6.364862 | 1.95E-10 | 4.71E-09 |
| ENSG00000100298 | APOBEC3H | 31.51267737 | -2.374978318 | 0.688143774 | -3.98682321 | 6.70E-05 | 0.000536 |
| ENSG00000254632 | | 37.42659 | -2.377935 | 0.7142743 | -3.978145 | 6.95E-05 | 0.000554 |
| ENSG00000196839 | ADA | 167.7408 | -2.381223 | 0.5088873 | -5.141781 | 2.72E-07 | 3.78E-06 |
| ENSG00000102313 | ITIH6 | 44.25209996 | -2.390564176 | 0.458673932 | -5.59666785 | 2.19E-08 | 3.73E-07 |
| ENSG00000154330 | PGM5 | 656.0596 | -2.393709 | 0.4007057 | -6.348562 | 2.17E-10 | 5.20E-09 |

| | | | | | | | |
|---|---|---|---|---|---|---|---|
| ENSG00000268879 | IGFL1P1 | 43.80582 | -2.393825 | 1.249885 | -2.66078 | 0.007796 | 0.031837 |
| ENSG00000260910 | LINC00565 | 3.823718 | -2.394937 | 1.306739 | -2.645747 | 0.00815108 | 0.033008 |
| ENSG00000286299 | | 18.50637 | -2.403009 | 0.6356579 | -4.336997 | 1.44E-05 | 0.000137 |
| ENSG00000286707 | | 20.80534 | -2.403835 | 0.6704305 | -4.13339 | 3.57E-05 | 0.000308 |
| ENSG00000260641 | | 43.53598 | -2.403952 | 0.761264 | -3.84495 | 0.00012058 | 0.000903 |
| ENSG00000167780 | SOAT2 | 12.39614 | -2.405025 | 1.480195 | -2.881114 | 0.00396273 | 0.018311 |
| ENSG00000160963 | COL26A1 | 8.643186 | -2.412206 | 0.9090674 | -3.33005 | 0.0008683 | 0.005063 |
| ENSG00000135519 | KCNH3 | 130.6484187 | -2.424794531 | 0.551336878 | -4.905733995 | 9.31E-07 | 1.17E-05 |
| ENSG00000128805 | ARHGAP22 | 16.42463608 | -2.424817475 | 0.727375141 | -3.92616535 | 8.63E-05 | 0.000672 |
| ENSG00000237451 | CDK2AP2P2 | 12.17224 | -2.4288 | 0.8710093 | -3.540278 | 0.00039971 | 0.002589 |
| ENSG00000164171 | ITGA2 | 3566.839 | -2.431151 | 0.4336114 | -6.001252 | 1.96E-09 | 4.02E-08 |
| ENSG00000114737 | CISH | 1073.215454 | -2.437582336 | 0.265961826 | -9.416025835 | 4.68E-21 | 3.95E-19 |
| ENSG00000185760 | KCNQ5 | 98.92768 | -2.44197 | 0.623317 | -4.485735 | 7.27E-06 | 7.43E-05 |
| ENSG00000105695 | MAG | 13.41727541 | -2.447847931 | 1.079347107 | -3.192033225 | 0.00141275 | 0.007707 |
| ENSG00000100867 | DHRS2 | 7142.658497 | -2.448750199 | 0.549979551 | -5.009028146 | 5.47E-07 | 7.21E-06 |
| ENSG00000274825 | | 4.758374 | -2.451661 | 1.337204 | -2.676171 | 0.00744688 | 0.03069 |
| ENSG00000285621 | | 10.45367 | -2.453381 | 0.6963643 | -4.107276 | 4.00E-05 | 0.00034 |
| ENSG00000278598 | MIR6775 | 5.913582 | -2.4539 | 1.163044 | -3.0721 | 0.00212559 | 0.010915 |
| ENSG00000005961 | ITGA2B | 164.1883362 | -2.454486947 | 0.534090321 | -5.082659469 | 3.72E-07 | 5.03E-06 |
| ENSG00000169129 | AFAP1L2 | 568.18 | -2.459635 | 0.4530952 | -5.844228 | 5.09E-09 | 9.81E-08 |
| ENSG00000167617 | CDC42EP5 | 60.13681 | -2.459943 | 0.5902598 | -4.677568 | 2.90E-06 | 3.29E-05 |
| ENSG00000152086 | TUBA3E | 12.79977 | -2.460795 | 0.7765937 | -3.82954 | 0.00012838 | 0.000955 |
| ENSG00000250328 | | 19.39118 | -2.463609 | 1.116279 | -3.152349 | 0.00161963 | 0.008643 |
| ENSG00000106688 | SLC1A1 | 223.7887162 | -2.46423841 | 0.40264061 | -6.487389785 | 8.73E-11 | 2.19E-09 |
| ENSG00000139200 | PIANP | 25.34282287 | -2.467179631 | 0.484024492 | -5.489538149 | 4.03E-08 | 6.59E-07 |
| ENSG00000175920 | DOK7 | 956.158 | -2.470588 | 0.5033492 | -5.362532 | 8.21E-08 | 1.26E-06 |
| ENSG00000240571 | | 7.577372 | -2.470672 | 1.067026 | -3.08127 | 0.0020612 | 0.010634 |
| ENSG00000135272 | MDFIC | 5.759332699 | -2.471154783 | 1.756437998 | -2.632876801 | 0.00846651 | 0.034031 |
| ENSG00000169403 | PTAFR | 47.81649 | -2.472151 | 0.4373806 | -6.047923 | 1.47E-09 | 3.07E-08 |
| ENSG00000135318 | NT5E | 710.6877237 | -2.472665654 | 0.681478238 | -4.213731903 | 2.51E-05 | 0.000224 |
| ENSG00000076706 | MCAM | 74.01735526 | -2.474134091 | 0.634919041 | -4.392078586 | 1.12E-05 | 0.000109 |
| ENSG00000221740 | SNORD93 | 3.701557 | -2.48315 | 1.608833 | -2.661091 | 0.0077888 | 0.031822 |
| ENSG00000287261 | | 21.21606 | -2.489137 | 0.5172818 | -5.248587 | 1.53E-07 | 2.24E-06 |

| | | | | | | | |
|---|---|---|---|---|---|---|---|
| ENSG00000237638 | LINC02245 | 145.2352 | -2.48981 | 0.3147845 | -8.185946 | 2.70E-16 | 1.37E-14 |
| ENSG00000262228 | | 12.72489 | -2.49219 | 0.7394117 | -3.960208 | 7.49E-05 | 0.000593 |
| ENSG00000101773 | RBBP8 | 3212.607925 | -2.4941703 | 0.205444525 | -12.32618439 | 6.55E-35 | 1.63E-32 |
| ENSG00000140538 | NTRK3 | 4.513695 | -2.497971 | 1.176708 | -3.077931 | 0.00208443 | 0.01073 |
| ENSG00000152377 | SPOCK1 | 21.16474 | -2.498265 | 0.7557123 | -3.941497 | 8.10E-05 | 0.000635 |
| ENSG00000169474 | SPRR1A | 4.073968 | -2.502417 | 3.242081 | -2.556308 | 0.01057894 | 0.040753 |
| ENSG00000204525 | HLA-C | 7086.969 | -2.502965 | 0.3499828 | -7.462636 | 8.48E-14 | 3.18E-12 |
| ENSG00000167103 | PIP5KL1 | 106.4651 | -2.509334 | 0.276867 | -9.301371 | 1.39E-20 | 1.09E-18 |
| ENSG00000226953 | NCKAP5-AS2 | 34.69915 | -2.517788 | 0.7660342 | -3.961376 | 7.45E-05 | 0.00059 |
| ENSG00000224294 | PINCR | 2.89086 | -2.520882 | 1.755979 | -2.640141 | 0.00828715 | 0.033452 |
| ENSG00000136883 | KIF12 | 2108.297977 | -2.522724118 | 0.524918279 | -5.286019129 | 1.25E-07 | 1.86E-06 |
| ENSG00000205325 | | 2.795099 | -2.524378 | 3.264756 | -2.555191 | 0.01061295 | 0.040849 |
| ENSG00000102265 | TIMP1 | 750.365526 | -2.525108637 | 0.48716983 | -5.626073367 | 1.84E-08 | 3.19E-07 |
| ENSG00000235890 | TSPEAR-AS1 | 16.89285 | -2.528907 | 0.859705 | -3.660928 | 0.0002513 | 0.001731 |
| ENSG00000003096 | KLHL13 | 155.8602768 | -2.529436938 | 0.412051329 | -6.506748956 | 7.68E-11 | 1.94E-09 |
| ENSG00000121316 | PLBD1 | 427.9989937 | -2.531502816 | 0.427745089 | -6.301025038 | 2.96E-10 | 6.92E-09 |
| ENSG00000103257 | SLC7A5 | 48915.37588 | -2.5364433 | 0.462314496 | -5.901266076 | 3.61E-09 | 7.12E-08 |
| ENSG00000273313 | RBAKDN | 2.289271 | -2.53933 | 1.465813 | -2.683364 | 0.00728856 | 0.03014 |
| ENSG00000255164 | | 5.801365 | -2.545024 | 1.123209 | -3.109796 | 0.00187217 | 0.009792 |
| ENSG00000232044 | SILC1 | 36.61856 | -2.546255 | 0.8995695 | -3.621732 | 0.00029264 | 0.001979 |
| ENSG00000121769 | FABP3 | 78.90721304 | -2.547858993 | 0.447940696 | -6.090066536 | 1.13E-09 | 2.40E-08 |
| ENSG00000227203 | SUB1P1 | 109.1948 | -2.54843 | 0.2486981 | -10.46843 | 1.21E-25 | 1.60E-23 |
| ENSG00000169962 | TAS1R3 | 78.08749 | -2.558572 | 0.2693397 | -9.731396 | 2.22E-22 | 2.19E-20 |
| ENSG00000225087 | | 25.38108 | -2.565329 | 0.4942489 | -5.568896 | 2.56E-08 | 4.33E-07 |
| ENSG00000273108 | | 3.559974 | -2.565634 | 1.77275 | -2.763475 | 0.00571895 | 0.024759 |
| ENSG00000270020 | | 6.439055 | -2.572849 | 1.157259 | -3.072488 | 0.00212282 | 0.010903 |
| ENSG00000286691 | | 15.11054 | -2.57286 | 0.943179 | -3.49615 | 0.00047202 | 0.002988 |
| ENSG00000154556 | SORBS2 | 3.033263 | -2.573143 | 1.598971 | -2.75906 | 0.00579678 | 0.025051 |
| ENSG00000231393 | | 2.539619 | -2.573563 | 2.065927 | -2.615742 | 0.00890338 | 0.035449 |
| ENSG00000119638 | NEK9 | 421.1609024 | -2.58373408 | 0.346103299 | -7.768057031 | 7.97E-15 | 3.34E-13 |
| ENSG00000115596 | WNT6 | 335.702708 | -2.584694528 | 0.610701413 | -4.7556702 | 1.98E-06 | 2.33E-05 |
| ENSG00000149131 | SERPING1 | 60.18158 | -2.588742 | 0.7250847 | -4.171294 | 3.03E-05 | 0.000265 |
| ENSG00000085117 | CD82 | 518.173947 | -2.589218308 | 0.412947392 | -6.639100958 | 3.16E-11 | 8.40E-10 |

| | | | | | | | |
|---|---|---|---|---|---|---|---|
| ENSG00000204682 | MIR1915HG | 431.0411 | -2.591034 | 0.2215431 | -11.87796 | 1.54E-32 | 3.44E-30 |
| ENSG00000223808 | | 857.7648 | -2.593799 | 0.44617 | -6.201648 | 5.59E-10 | 1.25E-08 |
| ENSG00000262966 | | 1.673782 | -2.595508 | 2.335759 | -2.634223 | 0.00843301 | 0.033933 |
| ENSG00000233694 | LINC02579 | 5.560245 | -2.59625 | 0.9744862 | -3.365894 | 0.00076296 | 0.004532 |
| ENSG00000154328 | NEIL2 | 860.3003 | -2.599053 | 0.1713945 | -15.32622 | 5.11E-53 | 3.57E-50 |
| ENSG00000275630 | | 10.67324 | -2.602591 | 0.7459225 | -4.105344 | 4.04E-05 | 0.000343 |
| ENSG00000204778 | CBWD4P | 77.65394 | -2.602871 | 0.338448 | -7.980333 | 1.46E-15 | 6.69E-14 |
| ENSG00000233058 | LINC00884 | 3.720927 | -2.604092 | 1.308654 | -2.899096 | 0.0037424 | 0.01745 |
| ENSG00000113805 | CNTN3 | 159.8303305 | -2.614429435 | 0.636466466 | -4.634241437 | 3.58E-06 | 3.96E-05 |
| ENSG00000059378 | PARP12 | 2120.601517 | -2.617468579 | 0.296591665 | -9.089121178 | 9.98E-20 | 7.17E-18 |
| ENSG00000286570 | | 3.351436 | -2.617569 | 1.140193 | -3.140414 | 0.00168709 | 0.008952 |
| ENSG00000227533 | SLC2A1-AS1 | 39.20345 | -2.619595 | 0.3911166 | -7.019527 | 2.23E-12 | 6.97E-11 |
| ENSG00000188176 | SMTNL2 | 19.94319 | -2.62158 | 1.107854 | -3.305237 | 0.00094896 | 0.005459 |
| ENSG00000100604 | CHGA | 9.386145683 | -2.622106702 | 0.956065966 | -3.547085901 | 0.00038952 | 0.002537 |
| ENSG00000223922 | ASS1P2 | 3.646009 | -2.623514 | 1.599779 | -2.665841 | 0.00767961 | 0.031463 |
| ENSG00000242540 | | 2.672752 | -2.627459 | 2.99853 | -2.603291 | 0.00923334 | 0.036495 |
| ENSG00000005513 | SOX8 | 36.33388508 | -2.630405557 | 1.061316508 | -3.364790172 | 0.00076602 | 0.004546 |
| ENSG00000197635 | DPP4 | 75.65859 | -2.632686 | 0.5141639 | -5.513116 | 3.53E-08 | 5.82E-07 |
| ENSG00000115353 | TACR1 | 5.851994421 | -2.635541014 | 0.967212673 | -3.450934453 | 0.00055865 | 0.003453 |
| ENSG00000287258 | | 4.47923 | -2.636481 | 1.507661 | -2.927322 | 0.00341894 | 0.016204 |
| ENSG00000165188 | RNF183 | 7.206274 | -2.639559 | 0.7504281 | -4.111715 | 3.93E-05 | 0.000334 |
| ENSG00000145063 | | 47.47082 | -2.641991 | 0.3329001 | -8.201408 | 2.38E-16 | 1.21E-14 |
| ENSG00000134917 | ADAMTS8 | 2.645628365 | -2.645410754 | 1.334120266 | -2.794334913 | 0.00520066 | 0.022877 |
| ENSG00000118515 | SGK1 | 397.8351497 | -2.647072878 | 0.450986535 | -6.263708393 | 3.76E-10 | 8.68E-09 |
| ENSG00000240423 | LINC00636 | 17.94084 | -2.654613 | 0.7075055 | -4.345516 | 1.39E-05 | 0.000132 |
| ENSG00000286381 | | 220.7678 | -2.66157 | 0.3113499 | -8.827668 | 1.07E-18 | 6.93E-17 |
| ENSG00000115232 | ITGA4 | 1.845160109 | -2.66275863 | 2.287958261 | -2.66368219 | 0.00772906 | 0.031641 |
| ENSG00000143845 | ETNK2 | 2331.514 | -2.665884 | 0.323616 | -8.518648 | 1.61E-17 | 9.28E-16 |
| ENSG00000111254 | AKAP3 | 16.90204319 | -2.669981305 | 0.690709294 | -4.391984572 | 1.12E-05 | 0.000109 |
| ENSG00000128274 | A4GALT | 1119.363275 | -2.670000184 | 0.381481148 | -7.326669266 | 2.36E-13 | 8.38E-12 |
| ENSG00000187123 | LYPD6 | 1685.181 | -2.670215 | 0.2260977 | -12.06359 | 1.64E-33 | 3.83E-31 |
| ENSG00000104059 | FAM189A1 | 95.82670845 | -2.671185312 | 0.516347157 | -5.513010463 | 3.53E-08 | 5.82E-07 |
| ENSG00000168062 | BATF2 | 204.6901 | -2.672752 | 0.5158211 | -5.607898 | 2.05E-08 | 3.52E-07 |

| | | | | | | | |
|---|---|---|---|---|---|---|---|
| ENSG00000173917 | HOXB2 | 14.95572 | -2.674477 | 1.509485 | -2.755474 | 0.00586072 | 0.025257 |
| ENSG00000162040 | HS3ST6 | 14.79617 | -2.674669 | 1.177516 | -3.111454 | 0.00186168 | 0.009746 |
| ENSG00000256884 | | 3.558395 | -2.680366 | 1.384511 | -2.90508 | 0.00367159 | 0.017171 |
| ENSG00000213949 | ITGA1 | 45.63877 | -2.681391 | 0.5028617 | -5.763696 | 8.23E-09 | 1.54E-07 |
| ENSG00000182393 | IFNL1 | 10.39368 | -2.682513 | 1.235187 | -3.06137 | 0.00220327 | 0.011234 |
| ENSG00000136514 | RTP4 | 109.3291801 | -2.687115942 | 1.112503337 | -3.102602616 | 0.00191827 | 0.009996 |
| ENSG00000275833 | | 5.683101 | -2.688887 | 1.342632 | -2.925344 | 0.00344076 | 0.016287 |
| ENSG00000173581 | CCDC106 | 389.9837 | -2.697428 | 0.2537192 | -10.84355 | 2.14E-27 | 3.38E-25 |
| ENSG00000164294 | GPX8 | 501.3117 | -2.698627 | 0.4084457 | -6.95906 | 3.43E-12 | 1.06E-10 |
| ENSG00000276850 | | 468.7873 | -2.705644 | 1.060265 | -3.341684 | 0.00083272 | 0.004883 |
| ENSG00000134363 | FST | 235.6740038 | -2.705942084 | 0.810744398 | -4.036656971 | 5.42E-05 | 0.000445 |
| ENSG00000108352 | RAPGEFL1 | 3454.605602 | -2.706283701 | 0.29766839 | -9.357463091 | 8.17E-21 | 6.60E-19 |
| ENSG00000230550 | ERLNC1 | 14.42759 | -2.711344 | 0.7929939 | -4.067444 | 4.75E-05 | 0.000396 |
| ENSG00000227394 | RPL11P1 | 4.74416 | -2.711865 | 1.469909 | -2.857753 | 0.00426653 | 0.01945 |
| ENSG00000269918 | | 47.71638 | -2.715225 | 0.5355272 | -5.421776 | 5.90E-08 | 9.38E-07 |
| ENSG00000180592 | SKIDA1 | 129.7377 | -2.720594 | 0.3166193 | -8.865598 | 7.61E-19 | 5.03E-17 |
| ENSG00000236581 | STARD13-AS | 22.9227 | -2.721241 | 0.6544747 | -4.689811 | 2.73E-06 | 3.12E-05 |
| ENSG00000147576 | ADHFE1 | 28.89278 | -2.72762 | 0.7402836 | -4.285499 | 1.82E-05 | 0.000168 |
| ENSG00000243650 | RN7SL834P | 6.458525 | -2.728771 | 1.503665 | -2.751455 | 0.00593312 | 0.025511 |
| ENSG00000267048 | | 2.843239 | -2.730971 | 1.774073 | -2.722706 | 0.00647496 | 0.027337 |
| ENSG00000224177 | LINC00570 | 3.717181 | -2.743346 | 1.417692 | -2.921146 | 0.00348746 | 0.016467 |
| ENSG00000138642 | HERC6 | 1754.527479 | -2.750484111 | 0.449166286 | -6.507724465 | 7.63E-11 | 1.93E-09 |
| ENSG00000258476 | LINC02207 | 8.071505 | -2.752982 | 1.726105 | -2.750134 | 0.0059571 | 0.025586 |
| ENSG00000108830 | RND2 | 189.8407465 | -2.769786577 | 0.476076006 | -6.22071859 | 4.95E-10 | 1.12E-08 |
| ENSG00000113389 | NPR3 | 1149.483943 | -2.774665638 | 0.711095354 | -4.508839732 | 6.52E-06 | 6.76E-05 |
| ENSG00000164756 | SLC30A8 | 4.534431 | -2.776631 | 1.56995 | -2.856718 | 0.00428046 | 0.019491 |
| ENSG00000144868 | TMEM108 | 36.85124 | -2.785156 | 0.611439 | -5.07348 | 3.91E-07 | 5.27E-06 |
| ENSG00000155380 | SLC16A1 | 2091.594 | -2.787102 | 0.2604869 | -10.92251 | 9.00E-28 | 1.47E-25 |
| ENSG00000142959 | BEST4 | 6.589019 | -2.789055 | 0.9757825 | -3.55277 | 0.0003812 | 0.002492 |
| ENSG00000188508 | KRTDAP | 30.38451 | -2.789799 | 0.7427638 | -4.374601 | 1.22E-05 | 0.000117 |
| ENSG00000132357 | CARD6 | 43.24621138 | -2.791652855 | 0.910017075 | -3.719204455 | 0.00019985 | 0.001417 |
| ENSG00000283138 | | 2.382524 | -2.797644 | 2.835639 | -2.635752 | 0.00839511 | 0.033808 |
| ENSG00000101342 | TLDC2 | 2.81873561 | -2.803757356 | 2.349541162 | -2.700752134 | 0.00691829 | 0.028847 |

| | | | | | | | |
|---|---|---|---|---|---|---|---|
| ENSG00000204642 | HLA-F | 433.4935 | -2.809161 | 0.3947508 | -7.449207 | 9.39E-14 | 3.50E-12 |
| ENSG00000150782 | IL18 | 28.34525 | -2.809608 | 0.8162738 | -4.064025 | 4.82E-05 | 0.000401 |
| ENSG00000204779 | FOXD4L5 | 9.563668 | -2.813892 | 0.9043748 | -3.840143 | 0.00012296 | 0.00092 |
| ENSG00000259757 | | 2.021733 | -2.816928 | 4.396449 | -2.605857 | 0.00916448 | 0.036277 |
| ENSG00000272482 | | 3.071488 | -2.817433 | 1.842362 | -2.750483 | 0.00595075 | 0.025563 |
| ENSG00000287346 | | 16.29298 | -2.817447 | 0.779418 | -4.12205 | 3.76E-05 | 0.000321 |
| ENSG00000182175 | RGMA | 154.8884 | -2.817931 | 0.5730743 | -5.370211 | 7.86E-08 | 1.21E-06 |
| ENSG00000100385 | IL2RB | 13.65673471 | -2.818939235 | 0.947698242 | -3.761308234 | 0.00016903 | 0.001223 |
| ENSG00000116711 | PLA2G4A | 7.175211904 | -2.819161611 | 1.177237185 | -3.271093335 | 0.00107133 | 0.006069 |
| ENSG00000111816 | FRK | 306.0943859 | -2.821918869 | 0.464560519 | -6.467483512 | 9.96E-11 | 2.48E-09 |
| ENSG00000230098 | TCERG1L-AS1 | 2.423355 | -2.826695 | 1.920958 | -2.742578 | 0.0060959 | 0.026047 |
| ENSG00000230601 | TEX48 | 3.223362 | -2.831866 | 1.238706 | -3.079483 | 0.0020736 | 0.010687 |
| ENSG00000285697 | | 2.032596 | -2.840633 | 2.161123 | -2.699071 | 0.00695334 | 0.02897 |
| ENSG00000114771 | AADAC | 10.53998239 | -2.841060108 | 1.170835593 | -3.147450792 | 0.00164701 | 0.008763 |
| ENSG00000184611 | KCNH7 | 16.75155 | -2.843366 | 0.7091079 | -4.562261 | 5.06E-06 | 5.39E-05 |
| ENSG00000274114 | ALOX15P1 | 23.13496 | -2.845169 | 0.7998208 | -4.220902 | 2.43E-05 | 0.000218 |
| ENSG00000271553 | | 22.96301 | -2.85045 | 0.7161229 | -4.544732 | 5.50E-06 | 5.80E-05 |
| ENSG00000250510 | GPR162 | 26.015 | -2.854163 | 0.7963439 | -4.224767 | 2.39E-05 | 0.000215 |
| ENSG00000163328 | GPR155 | 379.0821 | -2.85835 | 0.2523793 | -11.53317 | 8.98E-31 | 1.88E-28 |
| ENSG00000170289 | CNGB3 | 37.38085 | -2.860274 | 0.5246568 | -5.886879 | 3.94E-09 | 7.73E-08 |
| ENSG00000203797 | DDO | 5.056908 | -2.865203 | 2.777374 | -2.686339 | 0.00722398 | 0.02992 |
| ENSG00000142156 | COL6A1 | 230.5006 | -2.865439 | 0.4618527 | -6.593186 | 4.30E-11 | 1.12E-09 |
| ENSG00000266256 | LINC00683 | 21.10457 | -2.870892 | 0.6442646 | -4.981772 | 6.30E-07 | 8.19E-06 |
| ENSG00000235811 | | 2.897058 | -2.871187 | 2.530305 | -2.720097 | 0.00652627 | 0.027511 |
| ENSG00000188404 | SELL | 92.85464 | -2.890518 | 1.454218 | -2.838668 | 0.00453023 | 0.020442 |
| ENSG00000134321 | RSAD2 | 288.8287715 | -2.894409651 | 0.655067889 | -4.948915076 | 7.46E-07 | 9.55E-06 |
| ENSG00000163565 | IFI16 | 31.6822 | -2.895615 | 1.172527 | -3.252609 | 0.00114351 | 0.00642 |
| ENSG00000267481 | | 8.212096 | -2.898806 | 0.7990213 | -4.127936 | 3.66E-05 | 0.000314 |
| ENSG00000155850 | SLC26A2 | 3115.639 | -2.902852 | 0.3092784 | -9.641079 | 5.36E-22 | 5.03E-20 |
| ENSG00000287858 | | 13.79446 | -2.903105 | 0.8069354 | -4.133319 | 3.58E-05 | 0.000308 |
| ENSG00000149972 | CNTN5 | 46.90139 | -2.905547 | 0.5374731 | -5.839652 | 5.23E-09 | 1.01E-07 |
| ENSG00000185933 | CALHM1 | 9.049923 | -2.911061 | 0.7541614 | -4.458949 | 8.24E-06 | 8.30E-05 |
| ENSG00000233785 | | 15.3661 | -2.914677 | 0.8291507 | -4.143066 | 3.43E-05 | 0.000296 |

| | | | | | | | |
|---|---|---|---|---|---|---|---|
| ENSG00000166664 | CHRFAM7A | 3.230755 | -2.917711 | 1.913479 | -2.78095 | 0.00542001 | 0.023654 |
| ENSG00000255774 | LINC02747 | 138.1777 | -2.919498 | 0.6372243 | -5.115621 | 3.13E-07 | 4.30E-06 |
| ENSG00000228019 | | 4.481056 | -2.921468 | 1.27199 | -2.980834 | 0.00287465 | 0.014089 |
| ENSG00000270069 | MIR222HG | 68.29918 | -2.923985 | 0.6127784 | -5.191549 | 2.09E-07 | 2.98E-06 |
| ENSG00000197565 | COL4A6 | 57.08459 | -2.926929 | 0.8288854 | -4.21549 | 2.49E-05 | 0.000223 |
| ENSG00000142621 | FHAD1 | 6.111594 | -2.930865 | 1.490508 | -2.924382 | 0.00345141 | 0.016323 |
| ENSG00000286133 | | 20.34577 | -2.934456 | 0.5156435 | -6.083036 | 1.18E-09 | 2.51E-08 |
| ENSG00000256609 | | 10.95451 | -2.939159 | 1.150573 | -3.452176 | 0.00055608 | 0.003439 |
| ENSG00000223865 | HLA-DPB1 | 6.410663 | -2.946376 | 1.759958 | -2.789991 | 0.00527094 | 0.023132 |
| ENSG00000237512 | UNC5B-AS1 | 3.061815 | -2.946967 | 1.376067 | -3.056454 | 0.00223972 | 0.011388 |
| ENSG00000110777 | POU2AF1 | 1.998603956 | -2.949884016 | 1.944793237 | -2.787861105 | 0.00530573 | 0.023247 |
| ENSG00000002822 | MAD1L1 | 23.68158831 | -2.957245752 | 0.522163814 | -6.077589156 | 1.22E-09 | 2.59E-08 |
| ENSG00000116396 | KCNC4 | 190.0082069 | -2.958098364 | 0.428010864 | -7.268208062 | 3.64E-13 | 1.27E-11 |
| ENSG00000139547 | RDH16 | 42.95874 | -2.958243 | 0.5308905 | -5.99618 | 2.02E-09 | 4.14E-08 |
| ENSG00000180638 | SLC47A2 | 7.221042 | -2.962571 | 0.9147416 | -3.812954 | 0.00013732 | 0.001014 |
| ENSG00000108679 | LGALS3BP | 18156.32102 | -2.96666864 | 0.325675958 | -9.376041203 | 6.85E-21 | 5.57E-19 |
| ENSG00000181634 | TNFSF15 | 321.6699 | -2.967848 | 0.8787939 | -4.079368 | 4.52E-05 | 0.000379 |
| ENSG00000196878 | LAMB3 | 610.7322 | -2.972768 | 0.4716175 | -6.679633 | 2.40E-11 | 6.50E-10 |
| ENSG00000171992 | SYNPO | 803.4729 | -2.978191 | 0.509958 | -6.260083 | 3.85E-10 | 8.86E-09 |
| ENSG00000175874 | CREG2 | 24.12934 | -2.979985 | 0.6328918 | -5.173034 | 2.30E-07 | 3.26E-06 |
| ENSG00000111341 | MGP | 528.7022439 | -2.98736726 | 0.78223602 | -4.481250268 | 7.42E-06 | 7.58E-05 |
| ENSG00000144824 | PHLDB2 | 924.6343 | -2.988723 | 0.4727866 | -6.711271 | 1.93E-11 | 5.30E-10 |
| ENSG00000167741 | GGT6 | 467.5665 | -2.989677 | 0.4134309 | -7.579901 | 3.46E-14 | 1.35E-12 |
| ENSG00000225362 | CT62 | 59.23124 | -2.990187 | 0.6294332 | -5.266434 | 1.39E-07 | 2.05E-06 |
| ENSG00000143126 | CELSR2 | 20522.72 | -2.990971 | 0.2749365 | -11.10415 | 1.20E-28 | 2.09E-26 |
| ENSG00000164379 | FOXQ1 | 10.43941 | -2.995753 | 1.437792 | -2.981348 | 0.00286983 | 0.014068 |
| ENSG00000115468 | EFHD1 | 4571.301222 | -3.00172839 | 0.32168769 | -9.597955375 | 8.15E-22 | 7.55E-20 |
| ENSG00000175318 | GRAMD2A | 808.4097 | -3.003805 | 0.8431202 | -4.228657 | 2.35E-05 | 0.000212 |
| ENSG00000272915 | | 2.947955 | -3.006167 | 1.620224 | -2.848553 | 0.00439186 | 0.019923 |
| ENSG00000287900 | | 83.61798 | -3.007935 | 0.4982295 | -6.431308 | 1.27E-10 | 3.12E-09 |
| ENSG00000236751 | LINC01186 | 33.30985 | -3.00961 | 0.5772449 | -5.613815 | 1.98E-08 | 3.41E-07 |
| ENSG00000228261 | ITPRIP-AS1 | 6.401901 | -3.013453 | 1.205215 | -3.30146 | 0.00096183 | 0.005528 |
| ENSG00000139193 | CD27 | 2.745234044 | -3.017722427 | 1.452315006 | -2.93630342 | 0.00332149 | 0.015821 |

| | | | | | | | |
|---|---|---|---|---|---|---|---|
| ENSG00000198670 | LPA | 7.572799 | -3.02804 | 0.9926998 | -3.706961 | 0.00020976 | 0.00148 |
| ENSG00000144857 | BOC | 189.6884 | -3.030082 | 0.2868133 | -10.79852 | 3.50E-27 | 5.40E-25 |
| ENSG00000197888 | UGT2B17 | 6.948655 | -3.030368 | 2.756325 | -2.800167 | 0.00510763 | 0.022542 |
| ENSG00000250320 | EDIL3-DT | 20.43424 | -3.032248 | 0.7429394 | -4.606877 | 4.09E-06 | 4.45E-05 |
| ENSG00000107201 | DDX58 | 1527.824075 | -3.033032793 | 0.408303038 | -7.758039776 | 8.63E-15 | 3.60E-13 |
| ENSG00000165887 | ANKRD2 | 25.95928 | -3.033139 | 0.522619 | -6.165089 | 7.04E-10 | 1.55E-08 |
| ENSG00000130598 | TNNI2 | 6.680285714 | -3.041875502 | 1.325116612 | -3.227816776 | 0.00124739 | 0.006929 |
| ENSG00000123119 | NECAB1 | 160.2766597 | -3.044487214 | 0.59119722 | -5.625757532 | 1.85E-08 | 3.20E-07 |
| ENSG00000175894 | TSPEAR | 47.16517 | -3.045077 | 1.300201 | -3.372943 | 0.00074369 | 0.004426 |
| ENSG00000132329 | RAMP1 | 160.6136656 | -3.050620828 | 0.6854159 | -5.004300201 | 5.61E-07 | 7.37E-06 |
| ENSG00000255710 | | 7.907496 | -3.055432 | 0.8574875 | -4.114054 | 3.89E-05 | 0.000331 |
| ENSG00000178826 | TMEM139 | 334.1035 | -3.055529 | 0.3235831 | -9.683705 | 3.54E-22 | 3.40E-20 |
| ENSG00000161714 | PLCD3 | 4840.326 | -3.057227 | 0.258479 | -12.03506 | 2.32E-33 | 5.37E-31 |
| ENSG00000234745 | HLA-B | 2962.118 | -3.063439 | 0.5174851 | -6.33374 | 2.39E-10 | 5.67E-09 |
| ENSG00000258957 | | 17.22413 | -3.07381 | 0.6398677 | -5.285961 | 1.25E-07 | 1.86E-06 |
| ENSG00000249993 | BFSP2-AS1 | 39.92589 | -3.075099 | 0.5922764 | -5.663626 | 1.48E-08 | 2.62E-07 |
| ENSG00000146151 | HMGCLL1 | 13.40491 | -3.077338 | 0.5800487 | -5.719342 | 1.07E-08 | 1.96E-07 |
| ENSG00000168916 | ZNF608 | 925.5345 | -3.079736 | 0.4500044 | -7.207076 | 5.72E-13 | 1.92E-11 |
| ENSG00000260807 | CEROX1 | 138.5788 | -3.084265 | 0.6253197 | -5.435367 | 5.47E-08 | 8.75E-07 |
| ENSG00000261226 | | 56.33661 | -3.085657 | 0.6064089 | -5.579423 | 2.41E-08 | 4.09E-07 |
| ENSG00000217646 | H2BC16P | 3.861207 | -3.089356 | 2.604314 | -2.770336 | 0.00559986 | 0.024307 |
| ENSG00000171855 | IFNB1 | 7.248939 | -3.091418 | 1.378334 | -3.021657 | 0.00251395 | 0.012567 |
| ENSG00000165949 | IFI27 | 3013.542 | -3.102951 | 0.732716 | -4.81061 | 1.50E-06 | 1.81E-05 |
| ENSG00000264357 | MIR4648 | 1.839634 | -3.108133 | 3.01876 | -2.701213 | 0.00690871 | 0.02882 |
| ENSG00000170231 | FABP6 | 29.42121 | -3.109049 | 0.6546751 | -5.227942 | 1.71E-07 | 2.48E-06 |
| ENSG00000206337 | HCP5 | 74.67321 | -3.110271 | 1.095181 | -3.694709 | 0.00022014 | 0.001543 |
| ENSG00000083454 | P2RX5 | 92.01449295 | -3.117821409 | 0.403166545 | -8.051555867 | 8.17E-16 | 3.91E-14 |
| ENSG00000287670 | | 42.23392 | -3.12275 | 0.6142684 | -5.559224 | 2.71E-08 | 4.55E-07 |
| ENSG00000181418 | DDN | 56.64815 | -3.127954 | 0.7746732 | -4.654301 | 3.25E-06 | 3.63E-05 |
| ENSG00000166402 | TUB | 50.24085 | -3.12998 | 0.6753361 | -5.143163 | 2.70E-07 | 3.76E-06 |
| ENSG00000237356 | | 165.6691 | -3.133425 | 0.2879438 | -11.1063 | 1.17E-28 | 2.06E-26 |
| ENSG00000250337 | PURPL | 29.49784 | -3.134341 | 0.7751321 | -4.613418 | 3.96E-06 | 4.32E-05 |
| ENSG00000183486 | MX2 | 88.55145 | -3.13551 | 0.7694891 | -4.635887 | 3.55E-06 | 3.93E-05 |

| | | | | | | | |
|---|---|---|---|---|---|---|---|
| ENSG00000204267 | TAP2 | 1283.255 | -3.135672 | 0.3299107 | -9.761098 | 1.65E-22 | 1.65E-20 |
| ENSG00000112562 | SMOC2 | 106.0993806 | -3.141247917 | 0.460027004 | -7.177211548 | 7.11E-13 | 2.37E-11 |
| ENSG00000136235 | GPNMB | 1908.364713 | -3.141621488 | 1.30812105 | -3.439026666 | 0.00058381 | 0.003592 |
| ENSG00000287134 | | 26.95391 | -3.141998 | 0.7466522 | -4.786296 | 1.70E-06 | 2.03E-05 |
| ENSG00000204740 | MALRD1 | 1032.84 | -3.155357 | 0.4554093 | -7.289927 | 3.10E-13 | 1.08E-11 |
| ENSG00000132207 | SLX1A | 5.611161964 | -3.159862006 | 1.296671164 | -3.106238933 | 0.00189484 | 0.009888 |
| ENSG00000269899 | NA | 12.87229 | -3.161451 | 0.5878212 | -5.723889 | 1.04E-08 | 1.91E-07 |
| ENSG00000162006 | MSLNL | 17.25256 | -3.163476 | 0.6117979 | -5.629881 | 1.80E-08 | 3.14E-07 |
| ENSG00000152137 | HSPB8 | 8078.554 | -3.163556 | 0.3890851 | -8.439086 | 3.20E-17 | 1.79E-15 |
| ENSG00000095370 | SH2D3C | 11.71180787 | -3.163964378 | 0.69126248 | -5.068179544 | 4.02E-07 | 5.41E-06 |
| ENSG00000184156 | KCNQ3 | 11.05 | -3.166204 | 0.8819663 | -4.120033 | 3.79E-05 | 0.000324 |
| ENSG00000229642 | | 11.24358 | -3.17467 | 0.8118029 | -4.396679 | 1.10E-05 | 0.000107 |
| ENSG00000272046 | | 17.97186 | -3.178474 | 0.7004239 | -5.078784 | 3.80E-07 | 5.13E-06 |
| ENSG00000130303 | BST2 | 2252.231111 | -3.179719368 | 0.819869295 | -4.52534741 | 6.03E-06 | 6.31E-05 |
| ENSG00000255983 | | 23.87878 | -3.183556 | 0.869543 | -4.320264 | 1.56E-05 | 0.000147 |
| ENSG00000254560 | BBOX1-AS1 | 4.383113 | -3.186273 | 1.469977 | -3.019527 | 0.00253169 | 0.012634 |
| ENSG00000238015 | | 1.500786 | -3.192311 | 3.403662 | -2.684658 | 0.00726041 | NA |
| ENSG00000059804 | SLC2A3 | 6.005318823 | -3.195231604 | 1.174830495 | -3.488854972 | 0.00048509 | 0.003059 |
| ENSG00000247796 | | 386.8244 | -3.202542 | 0.2175665 | -14.88322 | 4.24E-50 | 2.56E-47 |
| ENSG00000267034 | NA | 16.64803 | -3.203264 | 0.8266884 | -4.453214 | 8.46E-06 | 8.51E-05 |
| ENSG00000162654 | GBP4 | 21.54251 | -3.204415 | 1.423052 | -3.091947 | 0.00198849 | 0.010301 |
| ENSG00000133321 | PLAAT4 | 133.6191017 | -3.207445626 | 0.56549149 | -6.054133395 | 1.41E-09 | 2.96E-08 |
| ENSG00000272966 | | 4.121466 | -3.208993 | 1.703073 | -2.96165 | 0.00305996 | 0.014774 |
| ENSG00000229124 | VIM-AS1 | 123.2086 | -3.217664 | 0.7775371 | -4.420595 | 9.84E-06 | 9.72E-05 |
| ENSG00000060709 | RIMBP2 | 6.434021505 | -3.227781282 | 2.490857501 | -2.865144254 | 0.00416819 | 0.019104 |
| ENSG00000179133 | C10orf67 | 6.735149 | -3.234736 | 0.9544638 | -4.057662 | 4.96E-05 | 0.000411 |
| ENSG00000260466 | | 15.57619 | -3.240334 | 0.5724366 | -6.059714 | 1.36E-09 | 2.87E-08 |
| ENSG00000183862 | CNGA2 | 10.63426 | -3.241891 | 0.8453754 | -4.440217 | 8.99E-06 | 8.97E-05 |
| ENSG00000163499 | CRYBA2 | 3.134547 | -3.251017 | 1.501161 | -3.038173 | 0.00238017 | 0.011991 |
| ENSG00000123405 | NFE2 | 13.53060833 | -3.256003084 | 0.718539521 | -5.053530106 | 4.34E-07 | 5.82E-06 |
| ENSG00000236347 | | 8.04135 | -3.262544 | 1.388351 | -3.096448 | 0.00195854 | 0.010174 |
| ENSG00000225886 | | 74.98914 | -3.263147 | 0.6101483 | -5.817054 | 5.99E-09 | 1.14E-07 |
| ENSG00000160183 | TMPRSS3 | 294.5929 | -3.263671 | 0.4520558 | -7.578361 | 3.50E-14 | 1.37E-12 |

| | | | | | | | |
|---|---|---|---|---|---|---|---|
| ENSG00000111348 | ARHGDIB | 268.5633808 | -3.268317132 | 0.579859766 | -6.048488365 | 1.46E-09 | 3.06E-08 |
| ENSG00000231389 | HLA-DPA1 | 11.77167 | -3.269112 | 1.436789 | -3.051503 | 0.00227699 | 0.011545 |
| ENSG00000186470 | BTN3A2 | 45.99396 | -3.269955 | 0.5564224 | -6.280619 | 3.37E-10 | 7.84E-09 |
| ENSG00000162595 | DIRAS3 | 12.18898 | -3.273694 | 0.7062184 | -5.155416 | 2.53E-07 | 3.55E-06 |
| ENSG00000167755 | KLK6 | 29.03844 | -3.278903 | 0.9452208 | -3.983607 | 6.79E-05 | 0.000542 |
| ENSG00000242154 | | 14.01376 | -3.283938 | 0.8286469 | -4.43503 | 9.21E-06 | 9.15E-05 |
| ENSG00000100342 | APOL1 | 390.4164747 | -3.288880908 | 0.620426666 | -5.768930336 | 7.98E-09 | 1.49E-07 |
| ENSG00000178882 | RFLNA | 22.09094 | -3.290142 | 0.9497343 | -4.053305 | 5.05E-05 | 0.000418 |
| ENSG00000249942 | | 5.485363 | -3.291598 | 1.027873 | -3.914152 | 9.07E-05 | 0.000701 |
| ENSG00000234511 | C5orf58 | 2.488986 | -3.294118 | 1.546598 | -3.036392 | 0.00239428 | 0.012059 |
| ENSG00000273301 | | 4.205092 | -3.294803 | 1.087767 | -3.694262 | 0.00022053 | 0.001546 |
| ENSG00000126709 | IFI6 | 10065.01238 | -3.304870275 | 0.676680198 | -5.406799516 | 6.42E-08 | 1.01E-06 |
| ENSG00000130287 | NCAN | 1.846018243 | -3.305218201 | 2.949300826 | -2.756425066 | 0.0058437 | 0.025205 |
| ENSG00000161031 | PGLYRP2 | 39.57042 | -3.309579 | 0.9184515 | -4.305433 | 1.67E-05 | 0.000156 |
| ENSG00000258592 | | 13.00898 | -3.317231 | 0.7889242 | -4.751001 | 2.02E-06 | 2.38E-05 |
| ENSG00000166796 | LDHC | 1.612953 | -3.337055 | 3.250799 | -2.729485 | 0.00634333 | 0.026888 |
| ENSG00000267295 | | 3.197427 | -3.340712 | 2.586849 | -2.858864 | 0.00425161 | 0.019396 |
| ENSG00000223573 | TINCR | 3572.04 | -3.343312 | 0.2697072 | -12.61991 | 1.64E-36 | 4.74E-34 |
| ENSG00000204219 | TCEA3 | 399.5726 | -3.345292 | 0.2522873 | -13.47281 | 2.26E-41 | 8.34E-39 |
| ENSG00000287818 | | 3.739079 | -3.345747 | 1.61586 | -3.100784 | 0.00193009 | 0.010049 |
| ENSG00000162894 | FCMR | 276.6695 | -3.347235 | 0.4367824 | -7.996997 | 1.27E-15 | 5.87E-14 |
| ENSG00000248693 | LINC02100 | 9.149213 | -3.350537 | 1.088278 | -3.643624 | 0.00026883 | 0.001836 |
| ENSG00000288557 | | 4.956722 | -3.353286 | 1.760006 | -2.94843 | 0.00319392 | 0.015301 |
| ENSG00000076356 | PLXNA2 | 1521.255911 | -3.354789441 | 0.451410375 | -7.775153654 | 7.54E-15 | 3.17E-13 |
| ENSG00000082482 | KCNK2 | 1762.840437 | -3.359911779 | 0.391476707 | -8.882064758 | 6.56E-19 | 4.38E-17 |
| ENSG00000275004 | ZNF280B | 2.177674 | -3.365318 | 3.650018 | -2.706201 | 0.00680578 | 0.028464 |
| ENSG00000237423 | LINC01522 | 90.33632 | -3.365877 | 0.5171757 | -6.894572 | 5.40E-12 | 1.61E-10 |
| ENSG00000204264 | PSMB8 | 811.213 | -3.365914 | 0.4270295 | -8.195807 | 2.49E-16 | 1.27E-14 |
| ENSG00000111664 | GNB3 | 45.13496062 | -3.369891605 | 0.548955833 | -6.536527394 | 6.30E-11 | 1.61E-09 |
| ENSG00000016402 | IL20RA | 140.7860257 | -3.372006614 | 0.274803738 | -12.47266457 | 1.05E-35 | 2.80E-33 |
| ENSG00000250303 | LINC02762 | 145.5135 | -3.386786 | 0.4399875 | -8.025545 | 1.01E-15 | 4.74E-14 |
| ENSG00000138347 | MYPN | 111.142448 | -3.396989043 | 0.739778026 | -5.135289923 | 2.82E-07 | 3.90E-06 |
| ENSG00000127507 | ADGRE2 | 47.51623996 | -3.39797837 | 0.662138469 | -5.51683713 | 3.45E-08 | 5.71E-07 |

| | | | | | | | |
|---|---|---|---|---|---|---|---|
| ENSG00000242082 | SLC5A4-AS1 | 4.650143 | -3.398888 | 1.323202 | -3.222801 | 0.00126944 | 0.007038 |
| ENSG00000254251 | | 203.8168 | -3.400353 | 0.8879304 | -4.504673 | 6.65E-06 | 6.86E-05 |
| ENSG00000167754 | KLK5 | 3.681283 | -3.402256 | 2.680416 | -2.854655 | 0.00430836 | 0.019611 |
| ENSG00000164172 | MOCS2 | 5640.676 | -3.402569 | 0.1736532 | -19.72406 | 1.34E-86 | 7.12E-83 |
| ENSG00000170214 | ADRA1B | 8.834619 | -3.402697 | 1.144438 | -3.590861 | 0.00032959 | 0.002199 |
| ENSG00000127955 | GNAI1 | 647.3015466 | -3.402750375 | 0.252215368 | -13.6863872 | 1.22E-42 | 5.08E-40 |
| ENSG00000225932 | CTAGE4 | 19.80083 | -3.405179 | 0.7301833 | -5.097849 | 3.44E-07 | 4.68E-06 |
| ENSG00000162944 | RFTN2 | 81.01572 | -3.420717 | 0.4519979 | -7.87892 | 3.30E-15 | 1.46E-13 |
| ENSG00000099960 | SLC7A4 | 6.009533054 | -3.426108377 | 2.202023179 | -2.984043865 | 0.00284466 | 0.01396 |
| ENSG00000133101 | CCNA1 | 122.9046036 | -3.426676665 | 0.513484653 | -7.060131805 | 1.66E-12 | 5.27E-11 |
| ENSG00000266850 | | 14.12257 | -3.426971 | 0.6343472 | -5.856221 | 4.74E-09 | 9.16E-08 |
| ENSG00000272636 | DOC2B | 15.40189 | -3.427008 | 0.6391106 | -5.800184 | 6.62E-09 | 1.25E-07 |
| ENSG00000107954 | NEURL1 | 96.31567248 | -3.427395127 | 0.346761522 | -10.1500152 | 3.31E-24 | 3.83E-22 |
| ENSG00000177614 | PGBD5 | 279.0001 | -3.431114 | 0.3689937 | -9.57894 | 9.80E-22 | 8.86E-20 |
| ENSG00000164120 | HPGD | 833.7167 | -3.434542 | 0.4450589 | -8.047747 | 8.43E-16 | 4.01E-14 |
| ENSG00000238117 | | 123.818 | -3.448325 | 0.8964166 | -4.528713 | 5.93E-06 | 6.22E-05 |
| ENSG00000132429 | POPDC3 | 13.32826151 | -3.453909278 | 0.790504237 | -4.923200912 | 8.51E-07 | 1.07E-05 |
| ENSG00000166206 | GABRB3 | 334.9095 | -3.460434 | 0.3993691 | -8.948121 | 3.62E-19 | 2.48E-17 |
| ENSG00000130775 | THEMIS2 | 50.33204433 | -3.464231597 | 0.460877443 | -7.840255922 | 4.50E-15 | 1.93E-13 |
| ENSG00000169594 | BNC1 | 163.7173 | -3.470453 | 0.4350547 | -8.302784 | 1.02E-16 | 5.38E-15 |
| ENSG00000131773 | KHDRBS3 | 259.4106239 | -3.473663822 | 0.291504248 | -12.14299365 | 6.25E-34 | 1.51E-31 |
| ENSG00000254298 | | 3.222057 | -3.478202 | 1.954787 | -2.956129 | 0.00311527 | 0.014995 |
| ENSG00000142552 | RCN3 | 35.89011 | -3.486707 | 0.8046268 | -4.898243 | 9.67E-07 | 1.21E-05 |
| ENSG00000178597 | PSAPL1 | 2.819603 | -3.487165 | 3.046811 | -2.845299 | 0.00443697 | 0.020076 |
| ENSG00000166220 | TBATA | 5.688563 | -3.498637 | 1.475062 | -3.348208 | 0.00081336 | 0.004788 |
| ENSG00000187094 | CCK | 3.147394 | -3.499098 | 2.818495 | -2.839189 | 0.00452283 | 0.020416 |
| ENSG00000012779 | ALOX5 | 528.7642782 | -3.501324127 | 1.027656239 | -3.667056417 | 0.00024536 | 0.001699 |
| ENSG00000197457 | STMN3 | 2135.626 | -3.503909 | 0.2957502 | -12.06989 | 1.52E-33 | 3.58E-31 |
| ENSG00000279631 | | 50.56167 | -3.50589 | 0.5771405 | -6.482193 | 9.04E-11 | 2.26E-09 |
| ENSG00000157873 | TNFRSF14 | 18.25794 | -3.512554 | 0.8695449 | -4.623664 | 3.77E-06 | 4.14E-05 |
| ENSG00000122786 | CALD1 | 1629.259032 | -3.529673701 | 0.482848818 | -7.670931661 | 1.71E-14 | 6.87E-13 |
| ENSG00000159403 | C1R | 86.56444 | -3.536067 | 0.5555592 | -6.713073 | 1.91E-11 | 5.24E-10 |
| ENSG00000175785 | PRIMA1 | 92.21471 | -3.53739 | 0.7695957 | -5.163361 | 2.43E-07 | 3.41E-06 |

| | | | | | | | |
|---|---|---|---|---|---|---|---|
| ENSG00000051523 | CYBA | 5528.242409 | -3.556138601 | 0.284316082 | -12.72798631 | 4.13E-37 | 1.22E-34 |
| ENSG00000187546 | AGMO | 11.61335 | -3.568318 | 0.8680525 | -4.478876 | 7.50E-06 | 7.65E-05 |
| ENSG00000152213 | ARL11 | 23.47395 | -3.569199 | 0.6632845 | -5.773194 | 7.78E-09 | 1.45E-07 |
| ENSG00000181577 | C6orf223 | 24.2314 | -3.577321 | 2.373617 | -3.064032 | 0.00218376 | 0.011153 |
| ENSG00000273232 | | 9.70146 | -3.585782 | 0.9033186 | -4.554256 | 5.26E-06 | 5.58E-05 |
| ENSG00000101265 | RASSF2 | 193.6971745 | -3.590995852 | 0.548178856 | -6.953431951 | 3.57E-12 | 1.10E-10 |
| ENSG00000169085 | VXN | 31.56556 | -3.593419 | 0.6259829 | -6.182902 | 6.29E-10 | 1.40E-08 |
| ENSG00000071677 | PRLH | 3.380928297 | -3.59424283 | 1.524558986 | -3.199465043 | 0.00137683 | 0.007542 |
| ENSG00000286180 | | 8.403642 | -3.599861 | 1.121091 | -3.778448 | 0.00015781 | 0.00115 |
| ENSG00000265190 | ANXA8 | 59.27903 | -3.601375 | 1.515914 | -3.166647 | 0.00154208 | 0.008303 |
| ENSG00000163220 | S100A9 | 1577.564 | -3.611669 | 1.109385 | -4.059382 | 4.92E-05 | 0.000409 |
| ENSG00000129151 | BBOX1 | 3.962105029 | -3.620613006 | 1.515598833 | -3.284678577 | 0.00102099 | 0.005829 |
| ENSG00000147041 | SYTL5 | 1414.144 | -3.632421 | 0.5387306 | -7.135275 | 9.66E-13 | 3.15E-11 |
| ENSG00000105141 | CASP14 | 4.307461313 | -3.633993898 | 1.305195384 | -3.543867559 | 0.0003943 | 0.002562 |
| ENSG00000186529 | CYP4F3 | 29.29182 | -3.636371 | 0.9161763 | -4.264489 | 2.00E-05 | 0.000183 |
| ENSG00000022267 | FHL1 | 397.8425769 | -3.640462972 | 0.364309052 | -10.2575719 | 1.09E-24 | 1.31E-22 |
| ENSG00000162975 | KCNF1 | 184.3595 | -3.641402 | 0.5156889 | -7.437125 | 1.03E-13 | 3.80E-12 |
| ENSG00000285722 | | 14.84495 | -3.645194 | 2.343323 | -3.120075 | 0.00180805 | 0.009516 |
| ENSG00000089127 | OAS1 | 1647.016867 | -3.648475781 | 0.714298596 | -5.661334358 | 1.50E-08 | 2.65E-07 |
| ENSG00000185686 | PRAME | 395.7588 | -3.65347 | 0.3350824 | -11.132 | 8.76E-29 | 1.55E-26 |
| ENSG00000196364 | PRSS29P | 4.495561 | -3.654218 | 1.479524 | -3.264917 | 0.00109496 | 0.006188 |
| ENSG00000233421 | LINC01783 | 3.611217 | -3.66485 | 1.363316 | -3.342278 | 0.00083094 | 0.004874 |
| ENSG00000138646 | HERC5 | 313.6173931 | -3.677835057 | 0.422033975 | -8.996360162 | 2.33E-19 | 1.63E-17 |
| ENSG00000137672 | TRPC6 | 39.30156339 | -3.684982754 | 0.431245411 | -8.833336099 | 1.02E-18 | 6.60E-17 |
| ENSG00000124143 | ARHGAP40 | 82.41355055 | -3.68942033 | 0.786205777 | -5.238827539 | 1.62E-07 | 2.36E-06 |
| ENSG00000112175 | BMP5 | 3.219634504 | -3.689871531 | 1.59031034 | -3.199432269 | 0.00137699 | 0.007542 |
| ENSG00000162738 | VANGL2 | 11.0819 | -3.691009 | 0.8418697 | -4.839874 | 1.30E-06 | 1.58E-05 |
| ENSG00000079308 | TNS1 | 862.8801494 | -3.69974488 | 0.375409054 | -10.11330274 | 4.82E-24 | 5.45E-22 |
| ENSG00000231652 | | 4.461021 | -3.703461 | 1.09966 | -3.902389 | 9.52E-05 | 0.000733 |
| ENSG00000134538 | SLCO1B1 | 4.61831458 | -3.708766019 | 2.582511216 | -3.005573048 | 0.00265081 | 0.013138 |
| ENSG00000184659 | FOXD4L4 | 6.893022 | -3.710193 | 1.364968 | -3.575923 | 0.000349 | 0.002309 |
| ENSG00000268621 | IGFL2-AS1 | 272.0888 | -3.714632 | 0.8392606 | -4.737042 | 2.17E-06 | 2.53E-05 |
| ENSG00000124491 | F13A1 | 2.773655915 | -3.723177599 | 1.749315655 | -3.172944665 | 0.00150901 | 0.008149 |

| | | | | | | | |
|---|---|---|---|---|---|---|---|
| ENSG00000149043 | SYT8 | 58.45052 | -3.724611 | 0.4856431 | -7.997656 | 1.27E-15 | 5.85E-14 |
| ENSG00000277249 | MIR6784 | 6.115359 | -3.728218 | 0.911161 | -4.507981 | 6.54E-06 | 6.78E-05 |
| ENSG00000065320 | NTN1 | 551.6275028 | -3.736821864 | 0.243369969 | -15.53112805 | 2.14E-54 | 1.72E-51 |
| ENSG00000136960 | ENPP2 | 10.95452939 | -3.760322533 | 1.112162645 | -4.109053248 | 3.97E-05 | 0.000338 |
| ENSG00000064763 | FAR2 | 729.3511081 | -3.763333324 | 0.198883721 | -19.0555628 | 5.91E-81 | 1.74E-77 |
| ENSG00000231367 | LINC02613 | 3.161675 | -3.770346 | 1.622657 | -3.202369 | 0.00136302 | 0.007476 |
| ENSG00000181381 | DDX60L | 1261.693 | -3.800179 | 0.3907162 | -9.995288 | 1.60E-23 | 1.71E-21 |
| ENSG00000286342 | | 63.01429 | -3.81127 | 0.4418665 | -8.919848 | 4.67E-19 | 3.16E-17 |
| ENSG00000168658 | VWA3B | 5.55271 | -3.824422 | 1.290851 | -3.534271 | 0.0004089 | 0.002641 |
| ENSG00000280362 | | 5.697814 | -3.829189 | 0.9925454 | -4.404535 | 1.06E-05 | 0.000104 |
| ENSG00000159164 | SV2A | 104.0946 | -3.838012 | 0.4854893 | -8.233189 | 1.82E-16 | 9.40E-15 |
| ENSG00000257193 | LINC02385 | 23.44507 | -3.840749 | 0.8650773 | -4.769955 | 1.84E-06 | 2.18E-05 |
| ENSG00000173762 | CD7 | 3.770838 | -3.86998 | 1.885698 | -3.152125 | 0.00162087 | 0.008648 |
| ENSG00000231173 | | 4.17812 | -3.873948 | 1.364049 | -3.617505 | 0.00029746 | 0.002007 |
| ENSG00000180176 | TH | 32.96813 | -3.880776 | 0.745959 | -5.712946 | 1.11E-08 | 2.02E-07 |
| ENSG00000215023 | | 5.763499 | -3.883477 | 1.118755 | -4.009824 | 6.08E-05 | 0.000493 |
| ENSG00000137628 | DDX60 | 2070.653438 | -3.88555975 | 0.704203974 | -6.003136159 | 1.94E-09 | 3.97E-08 |
| ENSG00000136689 | IL1RN | 23.48441051 | -3.88586125 | 0.702548602 | -5.990511924 | 2.09E-09 | 4.27E-08 |
| ENSG00000223923 | TNS1-AS1 | 19.13432 | -3.901017 | 0.7918761 | -5.227578 | 1.72E-07 | 2.49E-06 |
| ENSG00000010610 | CD4 | 7.57495072 | -3.905963228 | 1.049028408 | -4.329000515 | 1.50E-05 | 0.000141 |
| ENSG00000106560 | GIMAP2 | 70.74640536 | -3.909027225 | 0.62543306 | -6.639117179 | 3.16E-11 | 8.40E-10 |
| ENSG00000173077 | DELEC1 | 3.910221 | -3.910254 | 1.371857 | -3.482121 | 0.00049746 | 0.003125 |
| ENSG00000254506 | | 10.70736 | -3.917131 | 1.265621 | -3.926843 | 8.61E-05 | 0.00067 |
| ENSG00000178685 | PARP10 | 3305.163 | -3.918289 | 0.4259975 | -9.47957 | 2.55E-21 | 2.23E-19 |
| ENSG00000135549 | PKIB | 2021.30547 | -3.936986927 | 0.364464246 | -11.05024118 | 2.19E-28 | 3.72E-26 |
| ENSG00000112137 | PHACTR1 | 30.59382901 | -3.951583957 | 0.63617609 | -6.602344357 | 4.05E-11 | 1.06E-09 |
| ENSG00000234215 | CT66 | 15.45602 | -3.951675 | 0.8110774 | -5.270427 | 1.36E-07 | 2.01E-06 |
| ENSG00000117322 | CR2 | 9.092434191 | -3.956422806 | 1.032159381 | -4.269966597 | 1.96E-05 | 0.000179 |
| ENSG00000271584 | LINC02550 | 3.084768 | -3.962504 | 1.740626 | -3.275931 | 0.00105314 | 0.005987 |
| ENSG00000143452 | HORMAD1 | 4.46299 | -3.973903 | 1.332084 | -3.729918 | 0.00019154 | 0.001366 |
| ENSG00000205129 | C4orf47 | 5.706489 | -3.983018 | 1.017096 | -4.313614 | 1.61E-05 | 0.000151 |
| ENSG00000233968 | | 17.84345 | -3.988656 | 0.7611774 | -5.72735 | 1.02E-08 | 1.88E-07 |
| ENSG00000162873 | KLHDC8A | 8.822193 | -3.991541 | 0.8723003 | -5.042089 | 4.60E-07 | 6.15E-06 |

| | | | | | | | |
|---|---|---|---|---|---|---|---|
| ENSG00000234210 | | 2.320641 | -3.992021 | 2.635857 | -3.077969 | 0.00208417 | 0.01073 |
| ENSG00000137573 | SULF1 | 605.6036121 | -3.995939844 | 0.421440486 | -9.764413629 | 1.60E-22 | 1.60E-20 |
| ENSG00000273812 | | 7.406532 | -4.010193 | 1.053664 | -4.378858 | 1.19E-05 | 0.000115 |
| ENSG00000106772 | PRUNE2 | 188.5444788 | -4.018845303 | 0.809420118 | -5.51373261 | 3.51E-08 | 5.80E-07 |
| ENSG00000206075 | SERPINB5 | 31.86788 | -4.018953 | 0.7902608 | -5.553307 | 2.80E-08 | 4.70E-07 |
| ENSG00000178662 | CSRNP3 | 8.809916 | -4.03343 | 0.8862238 | -5.008852 | 5.48E-07 | 7.22E-06 |
| ENSG00000153930 | ANKFN1 | 39.82534 | -4.037658 | 0.9387515 | -4.71122 | 2.46E-06 | 2.83E-05 |
| ENSG00000163218 | PGLYRP4 | 7.153241 | -4.038301 | 1.310209 | -3.6396 | 0.00027306 | 0.001862 |
| ENSG00000226674 | TEX41 | 14.57898 | -4.044804 | 1.151079 | -4.142706 | 3.43E-05 | 0.000297 |
| ENSG00000156535 | CD109 | 550.4321 | -4.054155 | 0.4040047 | -10.29822 | 7.18E-25 | 8.71E-23 |
| ENSG00000178586 | OR6B3 | 4.043132 | -4.059059 | 3.080893 | -2.970754 | 0.00297069 | 0.014464 |
| ENSG00000185885 | IFITM1 | 2249.021 | -4.085476 | 0.5166132 | -8.261686 | 1.44E-16 | 7.51E-15 |
| ENSG00000204287 | HLA-DRA | 31.16032 | -4.086753 | 1.153838 | -4.073333 | 4.63E-05 | 0.000387 |
| ENSG00000265763 | ZNF488 | 36.61111 | -4.091258 | 0.6878654 | -6.322217 | 2.58E-10 | 6.08E-09 |
| ENSG00000119917 | IFIT3 | 1455.867579 | -4.097169522 | 0.582242989 | -7.418820315 | 1.18E-13 | 4.32E-12 |
| ENSG00000185745 | IFIT1 | 3477.249 | -4.098059 | 0.6916793 | -6.347098 | 2.19E-10 | 5.24E-09 |
| ENSG00000047617 | ANO2 | 6.719259899 | -4.0988318 | 0.916766228 | -4.861213608 | 1.17E-06 | 1.43E-05 |
| ENSG00000137766 | UNC13C | 4.052949978 | -4.108557608 | 1.440311637 | -3.534933863 | 0.00040788 | 0.002635 |
| ENSG00000215039 | CD27-AS1 | 232.9585 | -4.117572 | 0.2872045 | -14.51217 | 1.01E-47 | 5.50E-45 |
| ENSG00000236182 | | 8.82542 | -4.126532 | 0.8768089 | -5.19074 | 2.09E-07 | 2.99E-06 |
| ENSG00000118971 | CCND2 | 5.232640837 | -4.128879456 | 2.567049487 | -3.180282133 | 0.00147132 | 0.007971 |
| ENSG00000261996 | | 26.17657 | -4.136305 | 0.5345387 | -8.001553 | 1.23E-15 | 5.70E-14 |
| ENSG00000243491 | | 5.669032 | -4.143412 | 1.565395 | -3.545634 | 0.00039167 | 0.002548 |
| ENSG00000164176 | EDIL3 | 471.3861 | -4.144558 | 0.4656416 | -9.205647 | 3.40E-20 | 2.59E-18 |
| ENSG00000108771 | DHX58 | 294.9300506 | -4.145202169 | 0.397576751 | -10.66704914 | 1.45E-26 | 2.13E-24 |
| ENSG00000255790 | | 2.812802 | -4.150907 | 2.014124 | -3.183158 | 0.00145678 | 0.007907 |
| ENSG00000226945 | | 154.7287 | -4.175591 | 0.5875456 | -7.488045 | 6.99E-14 | 2.64E-12 |
| ENSG00000099994 | SUSD2 | 452.2449865 | -4.178745663 | 0.595102689 | -7.407950948 | 1.28E-13 | 4.67E-12 |
| ENSG00000253125 | | 440.2653 | -4.180345 | 0.4643513 | -9.30422 | 1.35E-20 | 1.07E-18 |
| ENSG00000166897 | ELFN2 | 117.0086 | -4.192836 | 0.5728979 | -7.6874 | 1.50E-14 | 6.09E-13 |
| ENSG00000074410 | CA12 | 5392.777333 | -4.192879274 | 0.356604826 | -11.98832172 | 4.09E-33 | 9.29E-31 |
| ENSG00000204983 | PRSS1 | 3.030363 | -4.198201 | 2.931713 | -3.001504 | 0.00268649 | 0.013277 |
| ENSG00000256683 | ZNF350 | 90.71992 | -4.205694 | 0.6232076 | -6.927694 | 4.28E-12 | 1.29E-10 |

| | | | | | | | |
|---|---|---|---|---|---|---|---|
| ENSG00000249715 | FER1L5 | 836.4749 | -4.226001 | 0.3347109 | -12.83151 | 1.09E-37 | 3.30E-35 |
| ENSG00000128284 | APOL3 | 243.4934279 | -4.231002019 | 0.807247562 | -5.753152217 | 8.76E-09 | 1.63E-07 |
| ENSG00000105559 | PLEKHA4 | 693.8455602 | -4.231145038 | 0.737303157 | -6.21396495 | 5.17E-10 | 1.16E-08 |
| ENSG00000233455 | | 3.152801 | -4.235484 | 1.801761 | -3.334814 | 0.00085357 | 0.004983 |
| ENSG00000170819 | BFSP2 | 231.1125 | -4.25208 | 0.8947129 | -5.340707 | 9.26E-08 | 1.41E-06 |
| ENSG00000128285 | MCHR1 | 6.989315191 | -4.269036129 | 1.032837523 | -4.618322277 | 3.87E-06 | 4.23E-05 |
| ENSG00000165816 | VWA2 | 38.39182 | -4.294618 | 0.8760694 | -5.444693 | 5.19E-08 | 8.33E-07 |
| ENSG00000169035 | KLK7 | 1.984221 | -4.297645 | 2.951817 | -3.027083 | 0.00246926 | 0.012371 |
| ENSG00000232487 | RASA3-IT1 | 3.302983 | -4.326568 | 2.21247 | -3.201412 | 0.00136756 | 0.007499 |
| ENSG00000272566 | | 4.122724 | -4.326912 | 2.550457 | -3.293904 | 0.00098806 | 0.005664 |
| ENSG00000142224 | IL19 | 16.66591 | -4.327904 | 0.775473 | -6.031245 | 1.63E-09 | 3.37E-08 |
| ENSG00000186007 | LEMD1 | 4.058191 | -4.328732 | 1.944471 | -3.318887 | 0.00090377 | 0.005233 |
| ENSG00000101336 | HCK | 46.61652508 | -4.358533126 | 0.80650095 | -5.891931167 | 3.82E-09 | 7.51E-08 |
| ENSG00000173227 | SYT12 | 5870.672 | -4.36036 | 0.2720725 | -16.19577 | 5.40E-59 | 6.83E-56 |
| ENSG00000206052 | DOK6 | 56.70155 | -4.362693 | 0.5819123 | -7.829076 | 4.91E-15 | 2.10E-13 |
| ENSG00000248334 | WHAMMP2 | 11.68096 | -4.369891 | 0.8454506 | -5.408101 | 6.37E-08 | 1.01E-06 |
| ENSG00000211448 | DIO2 | 503.4276 | -4.396632 | 0.6096247 | -7.571941 | 3.68E-14 | 1.43E-12 |
| ENSG00000260469 | INSYN1-AS1 | 21.39834 | -4.402129 | 1.05091 | -4.82702 | 1.39E-06 | 1.68E-05 |
| ENSG00000250546 | | 34.72138 | -4.416105 | 0.6285007 | -7.307724 | 2.72E-13 | 9.56E-12 |
| ENSG00000117115 | PADI2 | 752.9029271 | -4.420700054 | 0.795660958 | -6.056367861 | 1.39E-09 | 2.92E-08 |
| ENSG00000205363 | INSYN1 | 278.2683 | -4.42168 | 0.4797632 | -9.495363 | 2.19E-21 | 1.93E-19 |
| ENSG00000235357 | LINC01621 | 19.25823 | -4.430478 | 1.118914 | -4.23428 | 2.29E-05 | 0.000207 |
| ENSG00000259977 | | 33.82245 | -4.432436 | 0.8004608 | -6.013696 | 1.81E-09 | 3.73E-08 |
| ENSG00000250596 | | 2.421992 | -4.447833 | 3.536467 | -2.914524 | 0.00356231 | 0.016758 |
| ENSG00000172987 | HPSE2 | 32.75277 | -4.45404 | 0.7849243 | -6.158349 | 7.35E-10 | 1.62E-08 |
| ENSG00000133863 | TEX15 | 28.65760461 | -4.45719048 | 0.753970202 | -6.32630425 | 2.51E-10 | 5.93E-09 |
| ENSG00000154252 | GAL3ST2 | 52.26599 | -4.468663 | 0.4726857 | -9.702292 | 2.95E-22 | 2.88E-20 |
| ENSG00000221963 | APOL6 | 1770.752 | -4.485895 | 0.7252609 | -6.628214 | 3.40E-11 | 9.00E-10 |
| ENSG00000188001 | TPRG1 | 165.9032 | -4.487491 | 0.4473231 | -10.29435 | 7.47E-25 | 9.02E-23 |
| ENSG00000287048 | | 7.10908 | -4.487504 | 1.043892 | -4.776633 | 1.78E-06 | 2.12E-05 |
| ENSG00000203999 | LINC01270 | 43.36081 | -4.491016 | 0.500932 | -9.161865 | 5.10E-20 | 3.81E-18 |
| ENSG00000225676 | | 7.115514 | -4.493405 | 1.107753 | -4.608741 | 4.05E-06 | 4.41E-05 |
| ENSG00000106538 | RARRES2 | 57.02183055 | -4.494772333 | 0.46417169 | -9.941676284 | 2.74E-23 | 2.86E-21 |

| | | | | | | | |
|---|---|---|---|---|---|---|---|
| ENSG00000143341 | HMCN1 | 356.1597 | -4.508413 | 0.9179675 | -5.483467 | 4.17E-08 | 6.80E-07 |
| ENSG00000124749 | COL21A1 | 53.59949825 | -4.51459508 | 0.795495623 | -6.151547787 | 7.67E-10 | 1.68E-08 |
| ENSG00000134830 | C5AR2 | 1062.671615 | -4.519623909 | 0.321052826 | -14.27097001 | 3.32E-46 | 1.63E-43 |
| ENSG00000131018 | SYNE1 | 1341.608458 | -4.521189837 | 0.312939121 | -14.63768513 | 1.61E-48 | 9.13E-46 |
| ENSG00000123569 | H2BW1 | 12.1954857 | -4.528450005 | 1.049916298 | -4.464699435 | 8.02E-06 | 8.12E-05 |
| ENSG00000189184 | PCDH18 | 66.06687 | -4.530208 | 0.5792416 | -8.168367 | 3.13E-16 | 1.56E-14 |
| ENSG00000150540 | HNMT | 717.5576 | -4.532139 | 0.4189036 | -11.05552 | 2.06E-28 | 3.53E-26 |
| ENSG00000184988 | TMEM106A | 10.38757 | -4.542097 | 0.8769252 | -5.464393 | 4.64E-08 | 7.54E-07 |
| ENSG00000259527 | LINC00052 | 2432.875 | -4.542335 | 0.3318457 | -13.87649 | 8.80E-44 | 3.83E-41 |
| ENSG00000148143 | ZNF462 | 633.5717 | -4.545362 | 0.2571581 | -17.82182 | 4.79E-71 | 8.48E-68 |
| ENSG00000170426 | SDR9C7 | 1.903055 | -4.550124 | 2.970137 | -3.096964 | 0.00195514 | 0.010161 |
| ENSG00000274621 | MIR6867 | 2.100757 | -4.551062 | 2.916753 | -3.128347 | 0.00175793 | 0.009284 |
| ENSG00000196748 | CLPSL2 | 72.7033 | -4.5519 | 1.272118 | -4.372204 | 1.23E-05 | 0.000118 |
| ENSG00000241821 | RN7SL170P | 3.346442 | -4.557395 | 3.617549 | -2.935827 | 0.0033266 | 0.01584 |
| ENSG00000258732 | | 5.373822 | -4.561302 | 2.567498 | -3.390276 | 0.00069822 | 0.004195 |
| ENSG00000204021 | LIPK | 2.291395 | -4.571869 | 3.183551 | -3.025725 | 0.00248038 | 0.012418 |
| ENSG00000205436 | EXOC3L4 | 29.41057 | -4.588527 | 1.230192 | -4.281577 | 1.86E-05 | 0.000171 |
| ENSG00000236313 | VN1R53P | 2.401888 | -4.607887 | 3.662725 | -2.938606 | 0.00329692 | 0.015718 |
| ENSG00000109101 | FOXN1 | 62.84621506 | -4.611752772 | 0.605207466 | -7.965268596 | 1.65E-15 | 7.53E-14 |
| ENSG00000226281 | | 3.96087 | -4.617009 | 1.650286 | -3.674335 | 0.00023847 | 0.001657 |
| ENSG00000258053 | | 11.38303 | -4.618046 | 1.041626 | -4.561739 | 5.07E-06 | 5.40E-05 |
| ENSG00000169071 | ROR2 | 150.509 | -4.677795 | 0.5145796 | -9.381825 | 6.48E-21 | 5.30E-19 |
| ENSG00000167768 | KRT1 | 6.068073 | -4.690747 | 2.502835 | -3.182702 | 0.00145908 | 0.007918 |
| ENSG00000287550 | | 19.88968 | -4.697416 | 0.8186008 | -6.14869 | 7.81E-10 | 1.71E-08 |
| ENSG00000263551 | | 6.534037 | -4.698781 | 1.434452 | -4.01352 | 5.98E-05 | 0.000486 |
| ENSG00000162551 | ALPL | 3.238887 | -4.702255 | 1.830255 | -3.52957 | 0.00041624 | 0.002681 |
| ENSG00000163879 | DNALI1 | 508.864 | -4.704883 | 0.3566056 | -13.40928 | 5.34E-41 | 1.87E-38 |
| ENSG00000272255 | | 3.312802 | -4.717065 | 1.564 | -3.83594 | 0.00012509 | 0.000935 |
| ENSG00000279516 | FAM230C | 41.36792 | -4.718904 | 0.5245993 | -9.168403 | 4.80E-20 | 3.59E-18 |
| ENSG00000276136 | | 3.810546 | -4.731717 | 1.74101 | -3.6445 | 0.00026791 | 0.001832 |
| ENSG00000122707 | RECK | 108.5461541 | -4.741778054 | 0.440475822 | -10.9507421 | 6.59E-28 | 1.09E-25 |
| ENSG00000182492 | BGN | 242.3985 | -4.747317 | 0.8624283 | -6.018292 | 1.76E-09 | 3.63E-08 |
| ENSG00000212312 | RNA5SP109 | 5.548147 | -4.760977 | 1.610336 | -3.71574 | 0.00020261 | 0.001435 |

| | | | | | | | |
|---|---|---|---|---|---|---|---|
| ENSG00000149294 | NCAM1 | 82.41499 | -4.761224 | 0.5036022 | -9.718231 | 2.52E-22 | 2.48E-20 |
| ENSG00000134198 | TSPAN2 | 160.5677293 | -4.778053057 | 0.75120035 | -6.721215532 | 1.80E-11 | 4.97E-10 |
| ENSG00000234261 | | 25.562 | -4.780731 | 0.5168124 | -9.458215 | 3.13E-21 | 2.72E-19 |
| ENSG00000119121 | TRPM6 | 35.35299761 | -4.793963244 | 0.637401751 | -7.675999164 | 1.64E-14 | 6.63E-13 |
| ENSG00000124107 | SLPI | 171.8153336 | -4.823597158 | 0.818769544 | -6.133635934 | 8.59E-10 | 1.86E-08 |
| ENSG00000165863 | C10orf82 | 38.50528 | -4.838104 | 0.7759623 | -6.348827 | 2.17E-10 | 5.19E-09 |
| ENSG00000114450 | GNB4 | 373.1623155 | -4.865859063 | 0.315381152 | -15.61606403 | 5.66E-55 | 5.18E-52 |
| ENSG00000124882 | EREG | 8.546081736 | -4.876155148 | 1.016785327 | -5.220026244 | 1.79E-07 | 2.58E-06 |
| ENSG00000197415 | VEPH1 | 366.8829 | -4.876997 | 0.3982341 | -12.47446 | 1.03E-35 | 2.76E-33 |
| ENSG00000172460 | PRSS30P | 32.58184 | -4.882147 | 0.5330725 | -9.378176 | 6.71E-21 | 5.47E-19 |
| ENSG00000205002 | AARD | 244.6297 | -4.911486 | 0.4667489 | -10.77323 | 4.61E-27 | 7.03E-25 |
| ENSG00000155052 | CNTNAP5 | 37.34433 | -4.91294 | 0.6547046 | -7.682518 | 1.56E-14 | 6.31E-13 |
| ENSG00000069188 | SDK2 | 103.9276858 | -4.942174312 | 0.957952047 | -5.691895314 | 1.26E-08 | 2.26E-07 |
| ENSG00000109321 | AREG | 2857.57042 | -4.967876988 | 0.700565166 | -7.49263121 | 6.75E-14 | 2.56E-12 |
| ENSG00000226080 | RPL6P4 | 9.26045 | -4.984272 | 1.490537 | -4.081112 | 4.48E-05 | 0.000376 |
| ENSG00000137965 | IFI44 | 1244.439251 | -5.011726662 | 0.899939766 | -6.080911946 | 1.20E-09 | 2.54E-08 |
| ENSG00000139192 | TAPBPL | 771.8159514 | -5.020299227 | 0.318015171 | -15.95892415 | 2.47E-57 | 2.62E-54 |
| ENSG00000164128 | NPY1R | 5319.69 | -5.04031 | 0.5732401 | -9.113506 | 7.98E-20 | 5.77E-18 |
| ENSG00000240065 | PSMB9 | 354.1864 | -5.040865 | 0.6174058 | -8.472378 | 2.40E-17 | 1.36E-15 |
| ENSG00000233435 | AGGF1P2 | 13.73176 | -5.050083 | 1.266605 | -4.544019 | 5.52E-06 | 5.81E-05 |
| ENSG00000204140 | CLPSL1 | 29.40454 | -5.053184 | 1.391494 | -4.409116 | 1.04E-05 | 0.000102 |
| ENSG00000264575 | LINC00526 | 16.90079 | -5.055624 | 2.409729 | -3.933812 | 8.36E-05 | 0.000653 |
| ENSG00000287188 | | 8.366605 | -5.065619 | 1.584568 | -3.97463 | 7.05E-05 | 0.000561 |
| ENSG00000255760 | LINC02422 | 2.265978 | -5.070957 | 2.704644 | -3.498562 | 0.00046777 | 0.002964 |
| ENSG00000128422 | KRT17 | 228.90517 | -5.074802732 | 0.558993671 | -9.396561706 | 5.64E-21 | 4.70E-19 |
| ENSG00000164342 | TLR3 | 47.12199 | -5.077336 | 0.9334117 | -5.776736 | 7.62E-09 | 1.43E-07 |
| ENSG00000257258 | | 29.09556 | -5.112375 | 0.7068787 | -7.458363 | 8.76E-14 | 3.28E-12 |
| ENSG00000170190 | SLC16A5 | 543.895 | -5.120701 | 0.4148193 | -12.57712 | 2.82E-36 | 7.81E-34 |
| ENSG00000198947 | DMD | 453.2086 | -5.126376 | 0.2632591 | -19.63163 | 8.30E-86 | 3.68E-82 |
| ENSG00000163359 | COL6A3 | 51.94181 | -5.132529 | 0.6299518 | -8.477456 | 2.30E-17 | 1.30E-15 |
| ENSG00000224272 | | 47.22711 | -5.14951 | 0.5916761 | -8.957554 | 3.32E-19 | 2.29E-17 |
| ENSG00000260177 | | 5.884093 | -5.154109 | 1.342244 | -4.294068 | 1.75E-05 | 0.000163 |
| ENSG00000184828 | ZBTB7C | 33.26678 | -5.168885 | 0.6541959 | -8.150853 | 3.61E-16 | 1.79E-14 |

| | | | | | | | |
|---|---|---|---|---|---|---|---|
| ENSG00000152661 | GJA1 | 140.9156 | -5.178632 | 1.079336 | -5.401589 | 6.61E-08 | 1.04E-06 |
| ENSG00000139988 | RDH12 | 50.79334 | -5.182931 | 0.5054379 | -10.50103 | 8.54E-26 | 1.15E-23 |
| ENSG00000162891 | IL20 | 2047.457 | -5.211214 | 0.3947041 | -13.41319 | 5.06E-41 | 1.79E-38 |
| ENSG00000153404 | PLEKHG4B | 488.9164 | -5.233046 | 0.5919454 | -9.137009 | 6.42E-20 | 4.69E-18 |
| ENSG00000164619 | BMPER | 158.5661 | -5.247236 | 0.5262493 | -10.23951 | 1.32E-24 | 1.57E-22 |
| ENSG00000124935 | SCGB1D2 | 9.696384178 | -5.250245044 | 1.415614668 | -4.339726155 | 1.43E-05 | 0.000135 |
| ENSG00000179314 | WSCD1 | 19.86434 | -5.264347 | 1.341866 | -4.582634 | 4.59E-06 | 4.94E-05 |
| ENSG00000112343 | TRIM38 | 183.9170321 | -5.283953022 | 0.625026119 | -8.52944834 | 1.47E-17 | 8.55E-16 |
| ENSG00000249412 | | 46.1876 | -5.321726 | 0.8495924 | -6.634348 | 3.26E-11 | 8.65E-10 |
| ENSG00000133488 | SEC14L4 | 46.4753984 | -5.327590333 | 0.526245688 | -10.32966474 | 5.17E-25 | 6.48E-23 |
| ENSG00000287692 | | 7.17318 | -5.346908 | 1.502485 | -4.147496 | 3.36E-05 | 0.000291 |
| ENSG00000196208 | GREB1 | 18307.81 | -5.366846 | 0.505928 | -10.87484 | 1.52E-27 | 2.43E-25 |
| ENSG00000264230 | ANXA8L1 | 45.69036 | -5.382565 | 2.470002 | -4.057144 | 4.97E-05 | 0.000412 |
| ENSG00000154274 | C4orf19 | 1135.954 | -5.428886 | 0.3517451 | -15.62348 | 5.04E-55 | 4.78E-52 |
| ENSG00000288046 | | 26.9771 | -5.446562 | 0.7463142 | -7.609448 | 2.75E-14 | 1.09E-12 |
| ENSG00000241469 | LINC00635 | 5.191039 | -5.45562 | 1.847093 | -3.874611 | 0.0001068 | 0.00081 |
| ENSG00000138131 | LOXL4 | 32.23853221 | -5.464344788 | 0.662577943 | -8.488846665 | 2.09E-17 | 1.19E-15 |
| ENSG00000245598 | DACT3-AS1 | 97.53436 | -5.470661 | 0.5130738 | -10.75289 | 5.74E-27 | 8.72E-25 |
| ENSG00000223872 | | 6.913067 | -5.481427 | 2.500745 | -4.063188 | 4.84E-05 | 0.000403 |
| ENSG00000120094 | HOXB1 | 2.241778465 | -5.487035617 | 3.028125485 | -3.386092575 | 0.00070896 | 0.00425 |
| ENSG00000233760 | | 5.885279 | -5.494659 | 1.80979 | -3.930495 | 8.48E-05 | 0.000662 |
| ENSG00000147256 | ARHGAP36 | 114.0563 | -5.500243 | 0.506241 | -11.09876 | 1.27E-28 | 2.21E-26 |
| ENSG00000067066 | SP100 | 1136.720675 | -5.515789797 | 0.53952061 | -10.47988429 | 1.07E-25 | 1.43E-23 |
| ENSG00000204099 | NEU4 | 54.51874 | -5.537752 | 0.622312 | -9.140719 | 6.20E-20 | 4.57E-18 |
| ENSG00000104368 | PLAT | 18.2029395 | -5.541120133 | 1.049353788 | -5.719417613 | 1.07E-08 | 1.96E-07 |
| ENSG00000174827 | PDZK1 | 4614.37 | -5.580062 | 0.5401814 | -10.60807 | 2.73E-26 | 3.86E-24 |
| ENSG00000173114 | LRRN3 | 2.795125 | -5.583792 | 2.682071 | -3.804761 | 0.00014194 | 0.001044 |
| ENSG00000106123 | EPHB6 | 343.09031 | -5.584567172 | 0.6396671 | -9.049826417 | 1.43E-19 | 1.01E-17 |
| ENSG00000241684 | ADAMTS9-AS2 | 4.55614 | -5.628837 | 2.811543 | -3.646871 | 0.00026545 | 0.001819 |
| ENSG00000139044 | B4GALNT3 | 338.065738 | -5.645123371 | 0.502849862 | -11.48444158 | 1.58E-30 | 3.18E-28 |
| ENSG00000179046 | TRIML2 | 9.217378 | -5.672966 | 2.901138 | -3.556344 | 0.00037605 | 0.002462 |
| ENSG00000258754 | LINC01579 | 27.32838 | -5.683483 | 1.319525 | -4.955228 | 7.22E-07 | 9.29E-06 |
| ENSG00000125089 | SH3TC1 | 19.4790702 | -5.69589369 | 1.096342838 | -5.616642601 | 1.95E-08 | 3.36E-07 |

| | | | | | | | |
|---|---|---|---|---|---|---|---|
| ENSG00000106366 | SERPINE1 | 102.3049532 | -5.696502152 | 0.590060485 | -9.929549677 | 3.10E-23 | 3.21E-21 |
| ENSG00000087077 | TRIP6 | 4867.062984 | -5.698176015 | 0.193451632 | -29.54400975 | 7.84E-192 | 2.08E-187 |
| ENSG00000010030 | ETV7 | 56.37880271 | -5.706300677 | 0.693306002 | -8.414232685 | 3.95E-17 | 2.18E-15 |
| ENSG00000135824 | RGS8 | 5.385303204 | -5.707539318 | 1.620784941 | -4.293814477 | 1.76E-05 | 0.000163 |
| ENSG00000263753 | LINC00667 | 130.1032 | -5.717851 | 0.795838 | -7.255014 | 4.02E-13 | 1.38E-11 |
| ENSG00000260289 | | 5.021512 | -5.722416 | 2.598945 | -4.042236 | 5.29E-05 | 0.000436 |
| ENSG00000168306 | ACOX2 | 23.50298 | -5.74288 | 2.470043 | -3.575384 | 0.00034971 | 0.002311 |
| ENSG00000221844 | DPP3P2 | 44.91117 | -5.762479 | 0.5153615 | -11.33707 | 8.60E-30 | 1.67E-27 |
| ENSG00000261390 | MAFTRR | 8.776515 | -5.769591 | 1.466379 | -4.644037 | 3.42E-06 | 3.79E-05 |
| ENSG00000169218 | RSPO1 | 14.62852 | -5.769702 | 1.187314 | -5.27087 | 1.36E-07 | 2.01E-06 |
| ENSG00000132274 | TRIM22 | 201.9432664 | -5.799359673 | 1.371113927 | -4.923312505 | 8.51E-07 | 1.07E-05 |
| ENSG00000099256 | PRTFDC1 | 245.6910894 | -5.80092858 | 0.466822203 | -12.60417826 | 2.00E-36 | 5.72E-34 |
| ENSG00000141293 | SKAP1 | 11.29759 | -5.805298 | 1.290658 | -4.855155 | 1.20E-06 | 1.47E-05 |
| ENSG00000187889 | FYB2 | 161.241 | -5.806587 | 0.6353856 | -9.455985 | 3.20E-21 | 2.77E-19 |
| ENSG00000132915 | PDE6A | 82.58125885 | -5.87733957 | 0.667813233 | -9.090938183 | 9.82E-20 | 7.07E-18 |
| ENSG00000149596 | JPH2 | 224.4206 | -5.888774 | 0.9858248 | -6.165234 | 7.04E-10 | 1.55E-08 |
| ENSG00000233077 | LINC01271 | 2.432738 | -5.913692 | 2.7294 | -3.921577 | 8.80E-05 | 0.000684 |
| ENSG00000110811 | P3H3 | 313.8750371 | -5.92759386 | 0.442279367 | -13.58504835 | 4.91E-42 | 1.89E-39 |
| ENSG00000252845 | RNA5SP101 | 5.823295 | -5.933005 | 1.624767 | -4.435674 | 9.18E-06 | 9.14E-05 |
| ENSG00000068831 | RASGRP2 | 20.45499478 | -5.940273106 | 1.036215745 | -5.938742507 | 2.87E-09 | 5.75E-08 |
| ENSG00000249679 | | 9.554769 | -5.947689 | 1.173334 | -5.255151 | 1.48E-07 | 2.17E-06 |
| ENSG00000259459 | LINC02568 | 217.1739 | -5.956045 | 0.5886172 | -10.38713 | 2.84E-25 | 3.62E-23 |
| ENSG00000204261 | PSMB8-AS1 | 28.78327 | -5.981481 | 1.512611 | -4.658096 | 3.19E-06 | 3.57E-05 |
| ENSG00000286190 | | 9.813895 | -5.991869 | 1.511353 | -4.688984 | 2.75E-06 | 3.13E-05 |
| ENSG00000119660 | DPPA5P4 | 3.060685237 | -5.994388999 | 3.202743681 | -3.450779701 | 0.00055897 | 0.003454 |
| ENSG00000158457 | TSPAN33 | 117.442 | -5.994437 | 0.4330886 | -13.93602 | 3.83E-44 | 1.72E-41 |
| ENSG00000186832 | KRT16 | 108.1166 | -5.998323 | 0.6469397 | -9.503852 | 2.02E-21 | 1.79E-19 |
| ENSG00000136630 | HLX | 53.87067024 | -6.023253584 | 0.533330676 | -11.46848904 | 1.90E-30 | 3.79E-28 |
| ENSG00000141574 | SECTM1 | 977.4418 | -6.047424 | 0.4970015 | -12.40056 | 2.59E-35 | 6.69E-33 |
| ENSG00000101680 | LAMA1 | 4.433313331 | -6.059819378 | 2.687358428 | -4.088362329 | 4.34E-05 | 0.000366 |
| ENSG00000278990 | | 2.631439 | -6.115863 | 2.891742 | -3.790284 | 0.00015048 | 0.0011 |
| ENSG00000111335 | OAS2 | 2204.590715 | -6.124253142 | 0.813629977 | -7.913258002 | 2.51E-15 | 1.13E-13 |
| ENSG00000235688 | SNTG2-AS1 | 9.975958 | -6.148203 | 1.363371 | -5.180877 | 2.21E-07 | 3.13E-06 |

| | | | | | | | |
|---|---|---|---|---|---|---|---|
| ENSG00000251363 | LINC02315 | 14.57831 | -6.157699 | 1.413617 | -5.042099 | 4.60E-07 | 6.15E-06 |
| ENSG00000168754 | FAM178B | 141.079 | -6.164664 | 0.5359182 | -11.708 | 1.16E-31 | 2.55E-29 |
| ENSG00000100842 | EFS | 27.79918314 | -6.175713336 | 0.733117331 | -8.492517386 | 2.02E-17 | 1.16E-15 |
| ENSG00000121690 | DEPDC7 | 26.86178808 | -6.187820992 | 2.543001913 | -4.576177841 | 4.74E-06 | 5.08E-05 |
| ENSG00000147573 | TRIM55 | 71.28823 | -6.195494 | 1.084594 | -6.232074 | 4.60E-10 | 1.05E-08 |
| ENSG00000185640 | KRT79 | 7.450387 | -6.255682 | 1.640099 | -4.59595 | 4.31E-06 | 4.67E-05 |
| ENSG00000285569 | | 7.209666 | -6.256306 | 1.526849 | -4.822438 | 1.42E-06 | 1.72E-05 |
| ENSG00000183715 | OPCML | 7.194904 | -6.259834 | 1.677726 | -4.505001 | 6.64E-06 | 6.86E-05 |
| ENSG00000239899 | RN7SL674P | 39.86076 | -6.287245 | 0.6538861 | -9.784158 | 1.32E-22 | 1.33E-20 |
| ENSG00000157087 | ATP2B2 | 12.57204 | -6.297454 | 1.395307 | -4.92976 | 8.23E-07 | 1.04E-05 |
| ENSG00000284659 | | 3.334552 | -6.297832 | 3.350473 | -3.456748 | 0.00054674 | 0.003395 |
| ENSG00000267107 | PCAT19 | 10.91951 | -6.319875 | 2.65296 | -4.376211 | 1.21E-05 | 0.000117 |
| ENSG00000128815 | WDFY4 | 13.92633543 | -6.340380304 | 1.287016132 | -5.234564336 | 1.65E-07 | 2.40E-06 |
| ENSG00000196167 | COLCA1 | 190.3789 | -6.346494 | 0.7375759 | -8.846836 | 9.00E-19 | 5.91E-17 |
| ENSG00000151224 | MAT1A | 220.3856 | -6.369904 | 1.631498 | -4.717987 | 2.38E-06 | 2.75E-05 |
| ENSG00000106536 | POU6F2 | 69.61965855 | -6.382613756 | 1.316072192 | -5.50952177 | 3.60E-08 | 5.93E-07 |
| ENSG00000052850 | ALX4 | 8.38208999 | -6.39058934 | 1.457018167 | -5.08593251 | 3.66E-07 | 4.95E-06 |
| ENSG00000165626 | BEND7 | 1351.378 | -6.406796 | 0.3523571 | -18.3431 | 3.75E-75 | 8.30E-72 |
| ENSG00000278973 | | 7.568004 | -6.43205 | 1.585092 | -4.796753 | 1.61E-06 | 1.93E-05 |
| ENSG00000203685 | STUM | 48.92052 | -6.436543 | 0.8768581 | -7.473216 | 7.83E-14 | 2.94E-12 |
| ENSG00000177294 | FBXO39 | 5.528144 | -6.437874 | 3.084655 | -3.711571 | 0.00020598 | 0.001455 |
| ENSG00000234464 | | 5.660012 | -6.473777 | 2.625538 | -4.584554 | 4.55E-06 | 4.90E-05 |
| ENSG00000159713 | TPPP3 | 11.89619 | -6.478545 | 1.586325 | -4.834943 | 1.33E-06 | 1.62E-05 |
| ENSG00000205413 | SAMD9 | 427.2583 | -6.570357 | 0.6024012 | -11.14164 | 7.87E-29 | 1.41E-26 |
| ENSG00000287357 | | 4.002747 | -6.572428 | 2.69044 | -4.478373 | 7.52E-06 | 7.67E-05 |
| ENSG00000166106 | ADAMTS15 | 411.3706 | -6.622696 | 0.3481435 | -19.1823 | 5.20E-82 | 1.73E-78 |
| ENSG00000272123 | | 18.8621 | -6.623181 | 0.9776486 | -6.884841 | 5.79E-12 | 1.71E-10 |
| ENSG00000234352 | | 5.877574 | -6.623713 | 2.778426 | -4.302257 | 1.69E-05 | 0.000158 |
| ENSG00000137959 | IFI44L | 1703.729552 | -6.635303498 | 0.954752052 | -7.370178755 | 1.70E-13 | 6.13E-12 |
| ENSG00000177409 | SAMD9L | 77.82612 | -6.636639 | 1.609865 | -4.860131 | 1.17E-06 | 1.44E-05 |
| ENSG00000236499 | LINC00896 | 40.88794 | -6.653614 | 0.6986476 | -9.671504 | 3.98E-22 | 3.81E-20 |
| ENSG00000156689 | GLYATL2 | 113.6256 | -6.657615 | 1.030963 | -6.861193 | 6.83E-12 | 2.00E-10 |
| ENSG00000007216 | SLC13A2 | 301.8876359 | -6.665020551 | 0.506275995 | -13.31293955 | 1.95E-40 | 6.47E-38 |

| | | | | | | | |
|---|---|---|---|---|---|---|---|
| ENSG00000002745 | WNT16 | 51.294075 | -6.679634686 | 0.571442308 | -11.75767517 | 6.45E-32 | 1.43E-29 |
| ENSG00000169181 | GSG1L | 149.9975 | -6.689845 | 0.4667034 | -14.46474 | 2.02E-47 | 1.08E-44 |
| ENSG00000106714 | CNTNAP3 | 15.94548096 | -6.737946337 | 1.079042153 | -6.236192033 | 4.48E-10 | 1.02E-08 |
| ENSG00000287027 | | 4.232835 | -6.746377 | 2.979872 | -4.004059 | 6.23E-05 | 0.000503 |
| ENSG00000120093 | HOXB3 | 17.59843678 | -6.763675844 | 2.682635 | -4.67028979 | 3.01E-06 | 3.39E-05 |
| ENSG00000172927 | MYEOV | 314.2619 | -6.801643 | 0.8706376 | -8.184369 | 2.74E-16 | 1.38E-14 |
| ENSG00000205334 | LINC01460 | 9.597048 | -6.863009 | 1.448869 | -5.442887 | 5.24E-08 | 8.41E-07 |
| ENSG00000101670 | LIPG | 10.16877321 | -6.882439717 | 2.780172405 | -4.494287767 | 6.98E-06 | 7.17E-05 |
| ENSG00000261026 | | 157.4549 | -6.924811 | 1.621944 | -5.013353 | NA | NA |
| ENSG00000231877 | | 6.810504 | -6.929314 | 2.693119 | -4.787368 | 1.69E-06 | 2.02E-05 |
| ENSG00000251165 | F11-AS1 | 9.755691 | -6.945506 | 2.61651 | -5.061473 | 4.16E-07 | 5.60E-06 |
| ENSG00000198074 | AKR1B10 | 284.8494 | -6.956018 | 1.29851 | -5.571078 | 2.53E-08 | 4.28E-07 |
| ENSG00000163221 | S100A12 | 4.076878 | -7.02898 | 2.835483 | -4.463297 | 8.07E-06 | 8.16E-05 |
| ENSG00000218336 | TENM3 | 1908.884 | -7.061518 | 0.3289548 | -21.59667 | 1.93E-103 | 1.71E-99 |
| ENSG00000233633 | | 33.30096 | -7.147702 | 1.363893 | -5.929971 | 3.03E-09 | 6.04E-08 |
| ENSG00000227728 | | 5.679783 | -7.186031 | 2.894138 | -4.43107 | 9.38E-06 | 9.30E-05 |
| ENSG00000224020 | MIR181A2HG | 6.851466 | -7.219302 | 2.739264 | -4.887701 | 1.02E-06 | 1.27E-05 |
| ENSG00000272461 | | 8.171748 | -7.224144 | 2.691718 | -5.091475 | 3.55E-07 | 4.82E-06 |
| ENSG00000132517 | SLC52A1 | 105.3216116 | -7.301509964 | 0.783712756 | -9.261716071 | 2.01E-20 | 1.56E-18 |
| ENSG00000077522 | ACTN2 | 33.10649979 | -7.305730446 | 1.107621524 | -6.572937879 | 4.93E-11 | 1.28E-09 |
| ENSG00000240922 | LSAMP-AS1 | 7.102951 | -7.317025 | 2.861792 | -4.608072 | 4.06E-06 | 4.43E-05 |
| ENSG00000265933 | LINC00668 | 47.78517 | -7.341057 | 1.044102 | -6.90973 | 4.86E-12 | 1.45E-10 |
| ENSG00000132530 | XAF1 | 579.2677542 | -7.429550064 | 1.142703535 | -6.932118215 | 4.15E-12 | 1.25E-10 |
| ENSG00000173262 | SLC2A14 | 23.34592 | -7.460721 | 1.097779 | -6.770935 | 1.28E-11 | 3.62E-10 |
| ENSG00000148513 | ANKRD30A | 90.51831 | -7.512265 | 0.8566211 | -8.9503 | 3.55E-19 | 2.44E-17 |
| ENSG00000284615 | | 137.0426 | -7.518913 | 0.6750691 | -11.17074 | 5.67E-29 | 1.04E-26 |
| ENSG00000069535 | MAOB | 262.6814432 | -7.596733423 | 0.981453616 | -8.108973501 | 5.10E-16 | 2.50E-14 |
| ENSG00000146856 | AGBL3 | 27.75081 | -7.612669 | 1.319146 | -6.491543 | 8.50E-11 | 2.13E-09 |
| ENSG00000168671 | UGT3A2 | 29.96512 | -7.668045 | 2.735457 | -5.402669 | 6.57E-08 | 1.04E-06 |
| ENSG00000273445 | | 129.4237 | -7.687316 | 0.9550804 | -8.370606 | 5.73E-17 | 3.11E-15 |
| ENSG00000237620 | GCNT1P5 | 56.06208 | -7.698548 | 1.324206 | -6.542391 | 6.05E-11 | 1.55E-09 |
| ENSG00000177599 | ZNF491 | 6.865931 | -7.705036 | 2.784151 | -5.226656 | 1.73E-07 | 2.50E-06 |
| ENSG00000280335 | | 6.410263 | -7.732998 | 2.865783 | -4.939486 | 7.83E-07 | 9.95E-06 |

| | | | | | | | |
|---|---|---|---|---|---|---|---|
| ENSG00000144331 | ZNF385B | 744.4702 | -7.841734 | 0.7364335 | -10.91969 | 9.28E-28 | 1.50E-25 |
| ENSG00000204049 | | 175.4813 | -7.901958 | 0.4743051 | -16.66834 | 2.23E-62 | 3.11E-59 |
| ENSG00000174145 | NWD2 | 5.581762 | -8.024917 | 2.8351 | -5.334022 | 9.61E-08 | 1.46E-06 |
| ENSG00000242042 | LINC01743 | 31.0858 | -8.027027 | 1.385659 | -6.527162 | 6.70E-11 | 1.70E-09 |
| ENSG00000169469 | SPRR1B | 7.389309 | -8.065414 | 3.069032 | -4.622071 | 3.80E-06 | 4.17E-05 |
| ENSG00000101194 | SLC17A9 | 216.4512263 | -8.078666684 | 0.898724709 | -9.295606906 | 1.46E-20 | 1.15E-18 |
| ENSG00000225889 | | 6.716589 | -8.106953 | 2.908212 | -5.142853 | 2.71E-07 | 3.77E-06 |
| ENSG00000185156 | MFSD6L | 12.53316 | -8.163944 | 2.756616 | -5.904653 | 3.53E-09 | 6.99E-08 |
| ENSG00000214290 | COLCA2 | 177.8272 | -8.179597 | 1.002291 | -7.933643 | 2.13E-15 | 9.58E-14 |
| ENSG00000214273 | AGGF1P1 | 8.560805 | -8.196096 | 2.900233 | -5.244848 | 1.56E-07 | 2.28E-06 |
| ENSG00000143546 | S100A8 | 149.5605 | -8.285771 | 1.099021 | -7.844134 | 4.36E-15 | 1.87E-13 |
| ENSG00000233854 | POU6F2-AS2 | 36.51179 | -8.313476 | 1.313288 | -7.102934 | 1.22E-12 | 3.95E-11 |
| ENSG00000260877 | | 8.782844 | -8.324662 | 2.85802 | -5.551507 | 2.83E-08 | 4.74E-07 |
| ENSG00000280334 | | 12.59755 | -8.366305 | 2.95003 | -5.21918 | 1.80E-07 | 2.59E-06 |
| ENSG00000172554 | SNTG2 | 8.413854 | -8.369374 | 2.854227 | -5.648156 | 1.62E-08 | 2.84E-07 |
| ENSG00000182667 | NTM | 43.95345 | -8.395347 | 1.174247 | -7.208112 | 5.67E-13 | 1.91E-11 |
| ENSG00000233571 | | 9.420872 | -8.454163 | 2.921738 | -5.418785 | 6.00E-08 | 9.53E-07 |
| ENSG00000164129 | NPY5R | 155.5558 | -8.455494 | 0.6258119 | -13.52916 | 1.05E-41 | 3.94E-39 |
| ENSG00000267313 | | 48.1798 | -8.542361 | 1.364001 | -7.04291 | 1.88E-12 | 5.93E-11 |
| ENSG00000273540 | AGBL1 | 8.332095 | -8.585185 | 2.979361 | -5.322785 | 1.02E-07 | 1.55E-06 |
| ENSG00000018510 | AGPS | 8.432884787 | -8.594943785 | 2.962281645 | -5.38174026 | 7.38E-08 | 1.15E-06 |
| ENSG00000243064 | ABCC13 | 8.13357 | -8.598164 | 2.949965 | -5.459628 | 4.77E-08 | 7.71E-07 |
| ENSG00000115828 | QPCT | 306.699634 | -8.613818771 | 1.322240419 | -7.314719856 | 2.58E-13 | 9.12E-12 |
| ENSG00000256870 | SLC5A8 | 153.4503 | -8.680208 | 0.5897148 | -14.64205 | 1.51E-48 | 8.75E-46 |
| ENSG00000205517 | RGL3 | 288.7608 | -8.680714 | 1.020599 | -8.295065 | 1.09E-16 | 5.71E-15 |
| ENSG00000122691 | TWIST1 | 10.51650589 | -8.891902099 | 2.930259285 | -5.861358239 | 4.59E-09 | 8.92E-08 |
| ENSG00000251350 | LINC02475 | 17.03962 | -8.897808 | 2.932503 | -5.846312 | 5.03E-09 | 9.69E-08 |
| ENSG00000259761 | NA | 209.2061 | -8.941889 | 0.6197841 | -14.40627 | 4.73E-47 | 2.41E-44 |
| ENSG00000162892 | IL24 | 787.7372 | -9.036308 | 0.8195184 | -11.27138 | 1.82E-29 | 3.42E-27 |
| ENSG00000225046 | | 9.430331 | -9.058351 | 2.953944 | -5.931824 | 3.00E-09 | 5.98E-08 |
| ENSG00000235687 | LINC00993 | 64.14978 | -9.182635 | 1.439835 | -7.166365 | 7.70E-13 | 2.54E-11 |
| ENSG00000260220 | CCDC187 | 70.74255 | -9.216238 | 1.011071 | -8.854498 | 8.41E-19 | 5.54E-17 |
| ENSG00000148803 | FUOM | 72.37291 | -9.361983 | 1.33843 | -7.85877 | 3.88E-15 | 1.68E-13 |

| | | | | | | | |
|---|---|---|---|---|---|---|---|
| ENSG00000234918 | | 17.43163 | -9.409704 | 2.978193 | -6.224067 | 4.84E-10 | 1.10E-08 |
| ENSG00000145703 | IQGAP2 | 1119.305 | -9.444624 | 0.4615774 | -20.51705 | 1.52E-93 | 1.01E-89 |
| ENSG00000172137 | CALB2 | 290.7594 | -9.471456 | 1.309041 | -7.370386 | 1.70E-13 | 6.13E-12 |
| ENSG00000246223 | LINC01550 | 20.09543 | -9.547698 | 2.914655 | -6.867451 | 6.54E-12 | 1.92E-10 |
| ENSG00000281406 | BLACAT1 | 63.81521 | -9.556099 | 1.674662 | -6.440451 | 1.19E-10 | 2.95E-09 |
| ENSG00000164778 | EN2 | 65.16964 | -9.667881 | 2.907707 | -7.122582 | 1.06E-12 | 3.44E-11 |
| ENSG00000070748 | CHAT | 14.33958665 | -9.743235404 | 3.043449481 | -6.248624677 | 4.14E-10 | 9.48E-09 |
| ENSG00000172824 | CES4A | 94.83229 | -9.772167 | 1.395048 | -7.867093 | 3.63E-15 | 1.58E-13 |
| ENSG00000088882 | CPXM1 | 155.1743857 | -9.82058076 | 0.905902009 | -10.74222922 | 6.45E-27 | 9.73E-25 |
| ENSG00000160801 | PTH1R | 63.00471 | -9.830244 | 1.425046 | -7.747461 | 9.37E-15 | 3.89E-13 |
| ENSG00000259129 | LINC00648 | 121.7877 | -9.868128 | 1.356445 | -8.171537 | 3.04E-16 | 1.53E-14 |
| ENSG00000006210 | CX3CL1 | 195.0373635 | -9.896848382 | 1.332764429 | -7.71988327 | 1.16E-14 | 4.76E-13 |
| ENSG00000184330 | S100A7A | 71.0086 | -10.01911 | 1.40963 | -7.98365 | 1.42E-15 | 6.53E-14 |
| ENSG00000288073 | | 18.5255 | -10.1226 | 3.030691 | -6.826232 | 8.72E-12 | 2.52E-10 |
| ENSG00000154553 | PDLIM3 | 138.2724 | -10.17159 | 1.05793 | -9.422459 | 4.41E-21 | 3.75E-19 |
| ENSG00000176887 | SOX11 | 322.4224 | -10.17329 | 1.323327 | -8.647711 | 5.25E-18 | 3.17E-16 |
| ENSG00000163873 | GRIK3 | 685.6421 | -10.27396 | 0.6630238 | -15.57138 | 1.14E-54 | 1.01E-51 |
| ENSG00000136574 | GATA4 | 150.6928563 | -10.31168978 | 1.055756576 | -9.544563261 | 1.37E-21 | 1.22E-19 |
| ENSG00000060982 | BCAT1 | 108.3202228 | -10.48777302 | 1.597102516 | -7.367199544 | 1.74E-13 | 6.26E-12 |
| ENSG00000280380 | | 45.18338 | -10.67381 | 3.029692 | -7.77414 | 7.60E-15 | 3.19E-13 |
| ENSG00000221866 | PLXNA4 | 603.3305 | -10.83694 | 0.7067872 | -15.33015 | 4.81E-53 | 3.45E-50 |
| ENSG00000269954 | | 266.0265 | -10.93677 | 1.489962 | -7.498049 | 6.48E-14 | 2.46E-12 |
| ENSG00000168269 | FOXI1 | 53.60166 | -11.3965 | 3.41018 | -6.105829 | 1.02E-09 | 2.19E-08 |
| ENSG00000168491 | CCDC110 | 62.21461 | -12.19435 | 3.199872 | -8.838711 | 9.68E-19 | 6.31E-17 |
| ENSG00000136573 | BLK | 134.2829729 | -13.58840765 | 3.350633459 | -9.820897416 | 9.15E-23 | 9.28E-21 |
| ENSG00000041982 | TNC | 201.9795476 | -13.83745496 | 3.449540452 | -8.843880086 | 9.25E-19 | 6.05E-17 |
| ENSG00000143556 | S100A7 | 202.3806 | -14.0236 | 3.574981 | -8.006589 | 1.18E-15 | 5.49E-14 |
| ENSG00000095627 | TDRD1 | 485.0641476 | -14.60274342 | 3.440785786 | -10.83890494 | 2.25E-27 | 3.52E-25 |
| ENSG00000177822 | TENM3-AS1 | 347.4568 | -14.71457 | 3.461807 | -10.7346 | 7.00E-27 | 1.05E-24 |

al
