## Supplementary Table 10 for "GLYATL1 is associated with metabolic and epigenetic changes and with endocrine resistance in luminal breast cancer"

| | H2Aub | H3 | pH3 | H3.3 | H4 | H3K27ac | H3K27me2 | H3K27me3 | H3K36me2 | H3K36me3 | H3K4me1 |
|---|---|---|---|---|---|---|---|---|---|---|---|
| MCF7_parent | -0.0072737 | 0.00806917 | -0.0642016 | -0.0080692 | 0.19943351 | 0.28713806 | 0.03890029 | -0.289742 | 0.2970825 | 0.04302428 | 0.08191179 |
| MCF7_parent | 0.19936408 | 0.03733567 | 0.15178365 | -0.0373357 | 0.35681028 | 0.41399128 | 0.22916051 | 0.04127017 | 0.40059758 | 0.21439561 | 0.19856399 |
| MCF7_LTED_ | 0.25775964 | 0.03226785 | 0.15390793 | -0.0322679 | 0.00647306 | 0.34130389 | 0.32016184 | -0.0638859 | -0.1340562 | 0.07954582 | -0.0668902 |
| MCF7_LTED_ | 0.63489633 | -0.2225701 | 0.15407279 | 0.22257012 | -0.0499118 | 0.66886712 | 0.65881606 | 0.0056939 | 0.24838448 | -0.0296786 | -0.1242666 |
| MCF7_KO1_1 | 0.46194838 | 0.12135338 | 0.06410288 | -0.1213534 | 0.03142699 | 0.27760052 | 0.20562148 | 0.11314317 | 0.25317208 | 0.12899096 | -0.1153649 |
| MCF7_KO1_2 | 0.22857396 | 0.46982808 | 0.24621765 | -0.4698281 | 0.12149057 | 0.32659066 | 0.13621752 | 0.11165405 | -0.0808177 | 0.10464008 | -0.2253967 |
| MCF7_KO2_1 | 0.21909456 | -0.1325603 | 0.25961478 | 0.13256031 | 0.25388242 | 0.08049215 | -0.1546465 | -0.0035262 | 0.18721852 | 0.37679557 | 0.2728953 |
| MCF7_KO2_2 | -0.5099788 | -0.0763666 | 0.1247409 | 0.07636664 | 0.25954296 | -0.6450174 | -0.7733779 | -0.2323422 | -0.0961698 | 0.37001311 | 0.03402444 |

| H3K4me3 | H3K64ac | H3K9me2 | H3K9me3 | H4K16ac | H4K20me3 | ki-67 | ER | test_0 | test_1 |
|---|---|---|---|---|---|---|---|---|---|
| 0.15950058 | 0.10638891 | 0.06215971 | 0.26045844 | -0.1262686 | 0.05850459 | 0.71316897 | -0.0545142 | NA | NA |
| 0.45206109 | 0.12838221 | 0.25957366 | 0.64897628 | 0.25254382 | 0.38485963 | 0.36210208 | -0.0030363 | NA | NA |
| 0.24903055 | 0.41409068 | -0.0537051 | 0.05403238 | 0.06262933 | 0.11169913 | 0.03796174 | 0.04348294 | NA | NA |
| 0.53030793 | 0.76833169 | 0.00179237 | 0.220543 | 0.26551526 | 0.13383026 | 0.22319143 | -0.0065484 | NA | NA |
| -0.1058841 | 0.18188181 | -0.1700999 | 0.07561778 | 0.03176395 | -0.0177899 | -0.6380714 | 0.4909505 | NA | NA |
| -0.269852 | 0.08475039 | -0.2559106 | -0.0498863 | -0.0702961 | -0.0654075 | -0.5886734 | 0.46952059 | NA | NA |
| -0.055397 | 0.03910219 | 0.16846824 | 0.18756408 | 0.28033532 | 0.10338979 | -0.4986177 | 0.04180373 | NA | NA |
| -0.6264375 | -0.4809823 | -0.0269939 | -0.1021589 | -0.4034157 | -0.206444 | -0.6813679 | 0.00792182 | NA | NA |
