## Supplementary Table 11 for "GLYATL1 is associated with metabolic and epigenetic changes and with endocrine resistance in luminal breast cancer"

| **Antibody_Name** | **target** | **element** | **isotope** | **vendor** | **cat#** |
|---|---|---|---|---|---|
| Ubiquityl-Histone H2A (Lys119) (D27C4) XP® Rabbit mAb (BSA and Azide Free) #45772 | H2Aub | Eu | 153 | Cell Signaling | 45772 |
| Histone H3 Antibody (D1H2) - 115In | H3 | In | 115 | IonPath | 711501 |
| Anti-pHistone H3 [S28] (HTA28)-175Lu | pH3 | Lu | 175 | FLUIDIGM | 3175012A |
| Anti-Histone H3.3 antibody [EPR17899] - ChIP Grade | H3.3 | Gd | 155 | Abcam | ab208690 |
| Histone H4 (D2X4V) Rabbit mAb #13919 | H4 | Tb | 159 | Cell Signaling | 13919 |
| Acetyl-Histone H3 (Lys27) (D5E4) XP Rabbit mAb | H3K27ac | Gd | 160 | Cell Signaling | 8173 |
| Anti-Histone H3 (di methyl K27) antibody - ChIP Grade | H3K27me2 | Nd | 142 | Abcam | ab24684 |
| Histone H3K27me3 antibody (mAb) - 168Er | H3K27me3 | Er | 168 | Active Motif | 61017 |
| Di-Methyl-Histone H3 (Lys36) (C75H12) Rabbit mAb | H3K36me2 | Sm | 149 | Cell Signaling | 2901 |
| Tri-Methyl-Histone H3 (Lys36) (D5A7) XP® Rabbit mAb #4909 | H3K36me3 | Ho | 165 | Cell Signaling | 4909 |
| Mono-Methyl-Histone H3 (Lys4) (D1A9) XP® Rabbit mAb (BSA and Azide Free) | H3K4me1 | Sm | 154 | Cell Signaling | 53138 |
| Tri-Methyl-Histone H3 (Lys4) (C42D8) Rabbit mAb | H3K4me3 | Nd | 145 | Cell Signaling | 9751 |
| Anti-Histone H3 (acetyl K64) antibody [EPR20713] - BSA and Azide free | H3K64ac | Gd | 156 | Abcam | ab251549 |
| Anti-Histone H3 (di methyl K9) antibody [Y49] - BSA and Azide free (ab173325) | H3K9me2 | Eu | 151 | Abcam | ab173325 |
| Tri-Methyl-Histone H3 (Lys9) (D4W1U) Rabbit mAb #13969 | H3K9me3 | Er | 170 | Cell Signaling | 13969 |
| Acetyl-Histone H4 (Lys16) (E2B8W) Rabbit mAb #13534 | H4K16AC | Sm | 152 | Cell Signaling | 13534 |
| Anti-Histone H4 (tri methyl K20) antibody [EPR17001(2)] - BSA and Azide free | H4K20me3 | Dy | 161 | Abcam | ab239410 |
| (3162012B) Anti-Ki-67 (B56)-162Dy | Ki-67 | Yb | 172 | FLUIDIGM | 3162012B |
| Anti-Estrogen Receptor alpha antibody [E115] - Low endotoxin, Azide free | ERa | Dy | 163 | Abcam | ab167611 |
