## Supplementary Table 2 for "GLYATL1 is associated with metabolic and epigenetic changes and with endocrine resistance in luminal breast cancer"

| ChrPos (hg38) | log2FoldChange_LTED_vs_parental | Pvalue_LTED_vs_parental | FDR_LTED_vs_parental |
|---|---|---|---|
| chr15:49838669-49839601 | 6.156277993 | 3.64E-38 | 2.79E-33 |
| chr4:182143759-182144986 | -7.199213268 | 2.91E-33 | 1.11E-28 |
| chr15:49539436-49540460 | 4.316097778 | 3.21E-26 | 8.19E-22 |
| chr15:51366931-51367926 | 4.052288115 | 8.67E-26 | 1.57E-21 |
| chr9:22008695-22009678 | 6.524512945 | 1.03E-25 | 1.57E-21 |
| chr4:182448307-182449165 | -6.317172548 | 4.24E-24 | 5.41E-20 |
| chr15:50115664-50116105 | 4.878544049 | 5.48E-24 | 6.00E-20 |
| chr15:51845174-51845682 | 4.963095667 | 7.84E-24 | 7.51E-20 |
| chr15:49605290-49606088 | 3.905279881 | 8.24E-23 | 7.02E-19 |
| chr15:52902055-52902619 | 4.59481267 | 9.59E-23 | 7.35E-19 |
| chr15:49853029-49853791 | 3.880060346 | 2.88E-22 | 2.00E-18 |
| chr15:49712326-49712731 | 4.476260611 | 3.13E-22 | 2.00E-18 |
| chr15:50239762-50240070 | 4.798315428 | 4.07E-22 | 2.40E-18 |
| chr16:77954477-77954928 | 4.881900607 | 6.81E-22 | 3.69E-18 |
| chr15:49741588-49742358 | 3.887699726 | 7.23E-22 | 3.69E-18 |
| chr15:50287860-50288582 | 3.839566871 | 1.65E-21 | 7.92E-18 |
| chr16:77937349-77938207 | 3.736420873 | 2.40E-21 | 1.08E-17 |
| chr15:50026421-50026809 | 4.400229113 | 7.99E-21 | 3.30E-17 |
| chr15:49730901-49731706 | 3.585458051 | 8.18E-21 | 3.30E-17 |
| chr19:52962415-52963175 | 3.892521324 | 2.25E-20 | 8.11E-17 |
| chr15:51413871-51415182 | 3.379813657 | 2.32E-20 | 8.11E-17 |
| chr15:53455171-53455760 | 3.865900739 | 2.33E-20 | 8.11E-17 |
| chr15:50220110-50220714 | 3.667807719 | 3.14E-20 | 1.05E-16 |
| chr15:49558419-49558913 | 4.022332091 | 4.08E-20 | 1.30E-16 |
| chr15:53454734-53455054 | 4.682828563 | 6.29E-20 | 1.93E-16 |
| chr15:51029798-51031479 | 3.288243147 | 7.11E-20 | 2.10E-16 |
| chr15:52514457-52514739 | 4.94188489 | 1.11E-19 | 3.14E-16 |
| chr15:51180265-51180877 | 3.758153758 | 1.41E-19 | 3.86E-16 |
| chr15:51044836-51045276 | 3.829952261 | 1.53E-19 | 4.05E-16 |
| chr15:53566468-53567136 | 3.911954063 | 1.94E-19 | 4.95E-16 |
| chr15:51442368-51444184 | 3.173461054 | 2.36E-19 | 5.84E-16 |
| chr15:49449120-49450047 | 3.633142173 | 2.82E-19 | 6.75E-16 |
| chr7:64306955-64307774 | 3.599998675 | 4.00E-19 | 9.29E-16 |
| chr15:53038338-53038977 | 3.731893409 | 4.51E-19 | 1.02E-15 |
| chr15:53851563-53852238 | 3.943108387 | 4.99E-19 | 1.09E-15 |
| chr15:52112162-52112684 | 3.657943718 | 5.72E-19 | 1.22E-15 |
| chr15:50786938-50787181 | 4.866085734 | 6.94E-19 | 1.44E-15 |
| chr16:3364709-3365466 | 3.81907448 | 7.34E-19 | 1.48E-15 |
| chr15:49724845-49725417 | 3.60235189 | 8.13E-19 | 1.60E-15 |
| chr15:52875090-52875467 | 4.008156167 | 9.96E-19 | 1.91E-15 |
| chr15:50084146-50085533 | 3.19744341 | 1.06E-18 | 1.98E-15 |
| chr15:51170950-51172027 | 3.255767761 | 1.18E-18 | 2.12E-15 |
| chr13:89163075-89163572 | 4.170405133 | 1.20E-18 | 2.12E-15 |
| chr15:51136895-51137486 | 3.468675244 | 1.21E-18 | 2.12E-15 |
| chr15:51302892-51303154 | 4.32718051 | 1.26E-18 | 2.15E-15 |
| chr15:53540567-53541286 | 3.286160204 | 1.34E-18 | 2.23E-15 |
| chr15:52088049-52088490 | 3.940183945 | 1.49E-18 | 2.42E-15 |
| chr15:50894571-50895048 | 3.724837994 | 1.75E-18 | 2.79E-15 |

| | | | |
|---|---|---|---|
| chr15:51651440-51651680 | 5.033119519 | 1.89E-18 | 2.96E-15 |
| chr9:23826210-23826645 | 4.629479324 | 2.26E-18 | 3.47E-15 |
| chr15:52048832-52049247 | 4.389647646 | 2.46E-18 | 3.69E-15 |
| chr15:50944109-50944443 | 4.624275931 | 2.51E-18 | 3.69E-15 |
| chr15:52041759-52042199 | 3.96386983 | 2.61E-18 | 3.77E-15 |
| chr15:49553424-49554054 | 3.397905495 | 2.78E-18 | 3.94E-15 |
| chr3:75671984-75672783 | 3.693063455 | 3.08E-18 | 4.29E-15 |
| chr15:51707851-51708381 | 3.567585687 | 3.99E-18 | 5.47E-15 |
| chr16:35207401-35207965 | 3.679795198 | 4.19E-18 | 5.63E-15 |
| chr15:52974541-52975003 | 3.487351371 | 5.07E-18 | 6.69E-15 |
| chr15:51482910-51484119 | 3.178067323 | 5.23E-18 | 6.80E-15 |
| chr15:52334477-52335008 | 3.595425351 | 5.46E-18 | 6.95E-15 |
| chr15:49865991-49866736 | 3.171429379 | 5.53E-18 | 6.95E-15 |
| chr7:23844824-23845315 | 4.028711441 | 6.28E-18 | 7.76E-15 |
| chr15:49973078-49973457 | 4.088784214 | 6.62E-18 | 8.05E-15 |
| chr15:49701595-49702052 | 3.714173064 | 6.94E-18 | 8.31E-15 |
| chr15:52077992-52078277 | 4.15624485 | 7.73E-18 | 9.11E-15 |
| chr19:57320238-57320624 | 5.209221989 | 8.68E-18 | 1.01E-14 |
| chr15:53598131-53598491 | 4.558475837 | 9.32E-18 | 1.07E-14 |
| chr15:53134353-53134848 | 4.005370638 | 9.86E-18 | 1.11E-14 |
| chr15:50590329-50590655 | 4.061341446 | 1.15E-17 | 1.28E-14 |
| chr15:52094028-52095031 | 3.177723997 | 1.18E-17 | 1.29E-14 |
| chr15:50194194-50194687 | 3.457784083 | 1.34E-17 | 1.45E-14 |
| chr6:122622455-122622915 | -3.871337324 | 1.68E-17 | 1.78E-14 |
| chr15:49831718-49832053 | 3.863869862 | 1.83E-17 | 1.92E-14 |
| chr11:107293759-107294374 | -3.924639851 | 1.88E-17 | 1.94E-14 |
| chr19:57389612-57390120 | 3.865042164 | 2.16E-17 | 2.21E-14 |
| chr1:208476256-208477240 | -3.189505512 | 2.50E-17 | 2.52E-14 |
| chr15:50417720-50418287 | 3.344373469 | 3.25E-17 | 3.24E-14 |
| chr15:50493511-50493878 | 3.94646223 | 3.46E-17 | 3.40E-14 |
| chr15:51006500-51006846 | 4.009866101 | 3.66E-17 | 3.55E-14 |
| chr15:52392099-52393072 | 3.043655406 | 3.84E-17 | 3.68E-14 |
| chr15:52989568-52989869 | 4.497916403 | 4.32E-17 | 4.08E-14 |
| chr15:49832471-49833059 | 3.165614604 | 4.44E-17 | 4.15E-14 |
| chr15:52464296-52464715 | 3.878791019 | 4.74E-17 | 4.35E-14 |
| chr15:50164496-50164993 | 3.3794712 | 4.77E-17 | 4.35E-14 |
| chr15:50108280-50108746 | 3.431927464 | 4.89E-17 | 4.41E-14 |
| chr15:51335838-51336306 | 3.306437184 | 5.58E-17 | 4.97E-14 |
| chr13:106783760-106784488 | -3.710920349 | 5.76E-17 | 5.07E-14 |
| chr15:50280240-50280498 | 4.055004372 | 6.13E-17 | 5.34E-14 |
| chr15:52148425-52149135 | 3.227136997 | 6.35E-17 | 5.47E-14 |
| chr15:51038040-51039523 | 2.919679287 | 6.42E-17 | 5.47E-14 |
| chr15:52163314-52164012 | 3.190061755 | 6.76E-17 | 5.70E-14 |
| chr15:50163520-50163865 | 3.714279008 | 6.99E-17 | 5.82E-14 |
| chr15:51127600-51127993 | 3.518150303 | 7.75E-17 | 6.39E-14 |
| chr15:50051717-50052551 | 2.996994677 | 8.28E-17 | 6.71E-14 |
| chr15:49984456-49985085 | 3.058796795 | 8.32E-17 | 6.71E-14 |
| chr15:49685026-49685300 | 4.626110266 | 8.53E-17 | 6.77E-14 |
| chr13:71712925-71713519 | -3.567334572 | 8.57E-17 | 6.77E-14 |
| chr15:49557972-49558243 | 4.106729588 | 1.05E-16 | 8.22E-14 |

| | | | |
|---|---|---|---|
| chr15:49793849-49794070 | 4.823163298 | 1.13E-16 | 8.71E-14 |
| chr15:50157061-50157477 | 3.684514186 | 1.31E-16 | 1.00E-13 |
| chr15:49776762-49777036 | 4.303749983 | 1.52E-16 | 1.16E-13 |
| chr15:49734128-49734435 | 3.634846545 | 1.75E-16 | 1.32E-13 |
| chr15:52745405-52745633 | 4.590538339 | 1.77E-16 | 1.32E-13 |
| chr13:59261997-59262673 | -5.443854558 | 2.03E-16 | 1.50E-13 |
| chr15:49795091-49795450 | 3.704709258 | 2.10E-16 | 1.53E-13 |
| chr15:50397550-50398058 | 3.278976744 | 2.43E-16 | 1.76E-13 |
| chrX:31984213-31985005 | -3.898190808 | 2.83E-16 | 2.03E-13 |
| chr15:50980873-50981229 | 3.727202899 | 3.00E-16 | 2.13E-13 |
| chr15:51727214-51727934 | 3.099493409 | 3.34E-16 | 2.34E-13 |
| chr15:49606501-49606816 | 3.773940084 | 3.55E-16 | 2.47E-13 |
| chr15:52545954-52546348 | 3.39817805 | 3.94E-16 | 2.72E-13 |
| chr15:49946712-49947123 | 3.667779467 | 3.97E-16 | 2.72E-13 |
| chr15:50210416-50211398 | 2.854328328 | 5.93E-16 | 4.02E-13 |
| chr15:51121189-51121499 | 3.648445457 | 6.16E-16 | 4.14E-13 |
| chr2:137983481-137984219 | -3.110602574 | 8.89E-16 | 5.88E-13 |
| chr15:49908032-49908241 | 4.342597788 | 8.91E-16 | 5.88E-13 |
| chr15:52710337-52710649 | 3.55135201 | 9.67E-16 | 6.33E-13 |
| chr9:21994282-21994898 | 5.367039006 | 1.03E-15 | 6.70E-13 |
| chr15:50127696-50128653 | 2.852870867 | 1.08E-15 | 6.93E-13 |
| chr15:50410974-50411254 | 3.986932868 | 1.16E-15 | 7.39E-13 |
| chr15:51415277-51415513 | 4.193263205 | 1.18E-15 | 7.47E-13 |
| chr15:52113804-52114030 | 3.97203356 | 1.37E-15 | 8.61E-13 |
| chr15:49965617-49966096 | 3.221697077 | 1.48E-15 | 9.20E-13 |
| chr15:50920797-50921051 | 4.3108905 | 1.59E-15 | 9.82E-13 |
| chr15:50281884-50282247 | 3.453763175 | 1.66E-15 | 1.02E-12 |
| chr1:208309680-208310199 | -3.984044522 | 1.69E-15 | 1.03E-12 |
| chr15:54227460-54227852 | 3.486879335 | 1.83E-15 | 1.11E-12 |
| chr15:50145410-50145719 | 3.195862773 | 1.87E-15 | 1.12E-12 |
| chr15:52104439-52104708 | 3.683735389 | 1.97E-15 | 1.17E-12 |
| chr4:74579665-74580334 | -3.232164637 | 2.22E-15 | 1.31E-12 |
| chr15:49953872-49954197 | 3.683058676 | 2.24E-15 | 1.31E-12 |
| chr15:50956774-50957125 | 3.501484046 | 2.31E-15 | 1.34E-12 |
| chr15:51295249-51295465 | 3.947449873 | 2.38E-15 | 1.37E-12 |
| chr4:178014982-178015700 | -3.581670519 | 2.43E-15 | 1.39E-12 |
| chr11:12802596-12803129 | -3.245627453 | 2.50E-15 | 1.42E-12 |
| chr15:51750868-51752561 | 2.664847908 | 2.86E-15 | 1.60E-12 |
| chr15:51095135-51095376 | 3.667671917 | 2.87E-15 | 1.60E-12 |
| chr15:50256268-50256732 | 2.931454731 | 3.35E-15 | 1.86E-12 |
| chr15:49965023-49965233 | 4.432631047 | 3.57E-15 | 1.97E-12 |
| chr15:49953049-49953429 | 3.478082584 | 3.62E-15 | 1.98E-12 |
| chr15:50074825-50075141 | 3.479806264 | 4.12E-15 | 2.24E-12 |
| chr15:51424660-51425219 | 3.035547521 | 4.52E-15 | 2.44E-12 |
| chrX:36470605-36471395 | -3.993879374 | 4.88E-15 | 2.62E-12 |
| chr15:52492262-52492640 | 3.469867598 | 5.05E-15 | 2.69E-12 |
| chr15:51799132-51799416 | 3.590975786 | 5.11E-15 | 2.70E-12 |
| chr15:50168583-50169154 | 2.965851263 | 6.53E-15 | 3.43E-12 |
| chr15:52054814-52055077 | 3.81380663 | 7.05E-15 | 3.67E-12 |
| chr15:50618720-50618988 | 4.091478695 | 7.43E-15 | 3.85E-12 |

| | | | |
|---|---|---|---|
| chr15:53791543-53791802 | 4.387591353 | 7.55E-15 | 3.88E-12 |
| chr15:51192531-51193050 | 2.801687105 | 7.98E-15 | 4.07E-12 |
| chr15:50232876-50233289 | 3.103215774 | 8.02E-15 | 4.07E-12 |
| chr13:55377192-55377788 | -3.958358974 | 8.13E-15 | 4.10E-12 |
| chr3:17860771-17861344 | 3.794066759 | 8.39E-15 | 4.20E-12 |
| chr15:51280841-51281143 | 3.365704528 | 9.36E-15 | 4.66E-12 |
| chr14:85621963-85622572 | 3.071489516 | 9.99E-15 | 4.94E-12 |
| chr2:102848455-102848815 | -3.724262556 | 1.05E-14 | 5.14E-12 |
| chr7:113944649-113945939 | -2.841755096 | 1.14E-14 | 5.56E-12 |
| chr15:51246180-51246475 | 3.456297299 | 1.20E-14 | 5.83E-12 |
| chr5:23007097-23007659 | -3.087055816 | 1.24E-14 | 5.97E-12 |
| chr15:49710671-49710985 | 3.451206975 | 1.25E-14 | 5.97E-12 |
| chr15:51013991-51014267 | 3.71171191 | 1.26E-14 | 6.01E-12 |
| chr15:49592504-49593259 | 2.87102293 | 1.27E-14 | 6.01E-12 |
| chr15:49742436-49742715 | 3.647403015 | 1.36E-14 | 6.37E-12 |
| chr15:53358681-53359013 | 3.586863477 | 1.46E-14 | 6.83E-12 |
| chr15:50545931-50546728 | 2.831120587 | 1.54E-14 | 7.16E-12 |
| chr7:102079485-102080293 | -3.576425476 | 1.57E-14 | 7.24E-12 |
| chr15:50938074-50938276 | 4.325859903 | 1.70E-14 | 7.78E-12 |
| chr15:53928710-53929028 | 4.338629201 | 1.71E-14 | 7.78E-12 |
| chr15:50193043-50193254 | 3.769563207 | 1.72E-14 | 7.78E-12 |
| chr4:182238234-182238710 | -5.176942902 | 2.01E-14 | 9.04E-12 |
| chr15:50577481-50577804 | 3.360090877 | 2.16E-14 | 9.69E-12 |
| chr15:50116526-50116797 | 3.247815001 | 2.18E-14 | 9.70E-12 |
| chr15:52846760-52847011 | 3.574262261 | 2.29E-14 | 1.01E-11 |
| chr10:59322907-59323829 | -2.857499446 | 2.30E-14 | 1.01E-11 |
| chr5:167952569-167953770 | -2.888112818 | 2.67E-14 | 1.17E-11 |
| chr15:50262040-50262516 | 2.815183194 | 2.72E-14 | 1.18E-11 |
| chr2:197640025-197641025 | -3.120873395 | 2.78E-14 | 1.20E-11 |
| chr13:77760688-77761612 | -3.33894593 | 2.87E-14 | 1.24E-11 |
| chr15:50653281-50653481 | 3.908843502 | 3.12E-14 | 1.34E-11 |
| chr15:52896372-52896705 | 3.812999855 | 3.27E-14 | 1.39E-11 |
| chr15:50279889-50280089 | 3.731650501 | 3.34E-14 | 1.41E-11 |
| chr15:51295498-51295811 | 3.271403359 | 3.34E-14 | 1.41E-11 |
| chr15:49908380-49908603 | 3.8057498 | 3.41E-14 | 1.43E-11 |
| chr15:50938374-50938818 | 3.06701441 | 4.53E-14 | 1.89E-11 |
| chr15:51381108-51381606 | 2.746885119 | 4.56E-14 | 1.89E-11 |
| chr1:210294924-210295339 | -3.349071634 | 4.59E-14 | 1.89E-11 |
| chr15:53478618-53478856 | 4.107806276 | 4.99E-14 | 2.04E-11 |
| chr15:50036348-50036564 | 3.706750757 | 5.19E-14 | 2.11E-11 |
| chr1:177448487-177449399 | -2.717573005 | 5.21E-14 | 2.11E-11 |
| chr15:52951482-52951739 | 3.467493739 | 5.58E-14 | 2.25E-11 |
| chr15:51239982-51240369 | 2.992350131 | 6.34E-14 | 2.55E-11 |
| chr7:114048321-114049145 | -3.265801482 | 6.50E-14 | 2.59E-11 |
| chr7:134986270-134987119 | -2.971038461 | 6.60E-14 | 2.62E-11 |
| chr15:52208324-52208674 | 2.813882489 | 6.63E-14 | 2.62E-11 |
| chr15:53437924-53438341 | 3.156542229 | 6.70E-14 | 2.63E-11 |
| chr15:51037330-51037852 | 2.821973551 | 6.94E-14 | 2.71E-11 |
| chr15:51094653-51094955 | 3.118246362 | 7.01E-14 | 2.73E-11 |
| chr3:13966273-13966656 | 3.501566604 | 7.22E-14 | 2.79E-11 |

| | | | |
|---|---|---|---|
| chr15:51169912-51170112 | 4.241915641 | 7.29E-14 | 2.80E-11 |
| chr15:53519633-53519867 | 4.469437363 | 7.30E-14 | 2.80E-11 |
| chr15:52254204-52254538 | 3.200441951 | 7.60E-14 | 2.90E-11 |
| chr15:52996645-52997098 | 3.19845211 | 9.06E-14 | 3.44E-11 |
| chr7:33867855-33868590 | -2.834284483 | 9.12E-14 | 3.44E-11 |
| chr15:49477445-49477930 | 3.344596477 | 9.69E-14 | 3.64E-11 |
| chr15:49750591-49750811 | 3.546640981 | 1.00E-13 | 3.75E-11 |
| chr10:8702582-8703314 | 2.992588131 | 1.08E-13 | 4.02E-11 |
| chr5:7878302-7878914 | -3.126130676 | 1.15E-13 | 4.24E-11 |
| chr15:50989623-50989936 | 3.525983095 | 1.23E-13 | 4.52E-11 |
| chr15:51977527-51977738 | 3.664177976 | 1.26E-13 | 4.62E-11 |
| chr3:14223700-14224042 | 3.819943474 | 1.27E-13 | 4.62E-11 |
| chr15:53862759-53862962 | 4.684219185 | 1.30E-13 | 4.74E-11 |
| chr13:59750609-59751343 | -2.887250276 | 1.31E-13 | 4.74E-11 |
| chr15:50737629-50737892 | 3.650666753 | 1.33E-13 | 4.76E-11 |
| chr15:51525564-51526122 | 2.982276622 | 1.33E-13 | 4.76E-11 |
| chr15:51162872-51163729 | 2.604958058 | 1.34E-13 | 4.76E-11 |
| chr15:52381161-52381492 | 3.328069004 | 1.36E-13 | 4.82E-11 |
| chr15:50778160-50778387 | 3.379122631 | 1.41E-13 | 4.98E-11 |
| chr15:50315906-50316374 | 2.814800001 | 1.52E-13 | 5.36E-11 |
| chr15:53483671-53483923 | 3.725176479 | 1.57E-13 | 5.48E-11 |
| chr7:140533353-140533903 | -3.717543558 | 1.57E-13 | 5.48E-11 |
| chr2:155256742-155257260 | -3.012049755 | 1.68E-13 | 5.80E-11 |
| chr15:53044979-53045222 | 3.908642465 | 1.68E-13 | 5.80E-11 |
| chr15:52980938-52981216 | 3.403978946 | 1.73E-13 | 5.95E-11 |
| chr15:49870993-49871218 | 3.635591317 | 1.78E-13 | 6.08E-11 |
| chr2:188605716-188606497 | 2.881355835 | 2.10E-13 | 7.14E-11 |
| chr13:41107124-41108267 | -2.564854162 | 2.26E-13 | 7.65E-11 |
| chr15:50476277-50476531 | 3.621464322 | 2.27E-13 | 7.65E-11 |
| chr15:50803327-50803587 | 3.172064939 | 2.38E-13 | 8.01E-11 |
| chr15:49844590-49844790 | 3.485896468 | 2.64E-13 | 8.83E-11 |
| chr15:51758759-51758959 | 3.795142034 | 2.69E-13 | 8.95E-11 |
| chr15:52563652-52563857 | 3.482236294 | 2.85E-13 | 9.46E-11 |
| chr15:51110084-51110384 | 3.080610889 | 3.05E-13 | 1.01E-10 |
| chr13:85738796-85739405 | -3.258717961 | 3.11E-13 | 1.02E-10 |
| chr15:51246609-51246820 | 3.600883957 | 3.26E-13 | 1.07E-10 |
| chr15:53832079-53832373 | 3.257615363 | 3.27E-13 | 1.07E-10 |
| chr15:56886858-56887887 | 2.543024598 | 3.28E-13 | 1.07E-10 |
| chr15:52747819-52748069 | 3.880312555 | 3.37E-13 | 1.09E-10 |
| chr15:52114329-52114538 | 3.531242349 | 3.37E-13 | 1.09E-10 |
| chr15:52466110-52466474 | 3.077835932 | 3.42E-13 | 1.10E-10 |
| chr15:49774949-49775174 | 3.41365478 | 3.48E-13 | 1.11E-10 |
| chr15:52368220-52368743 | 2.901536365 | 3.49E-13 | 1.11E-10 |
| chr10:55467658-55468040 | -3.618267591 | 3.49E-13 | 1.11E-10 |
| chr15:53258356-53258818 | 2.960767403 | 3.51E-13 | 1.11E-10 |
| chr21:9129322-9130001 | 2.836016859 | 3.54E-13 | 1.11E-10 |
| chr15:53409529-53409857 | 3.317463649 | 3.66E-13 | 1.14E-10 |
| chr19:52570076-52570482 | 3.672278286 | 3.67E-13 | 1.14E-10 |
| chr15:50562749-50563012 | 3.671332208 | 3.75E-13 | 1.16E-10 |
| chr15:49620255-49620513 | 2.910656606 | 4.10E-13 | 1.27E-10 |

| | | | |
|---|---|---|---|
| chr15:49908947-49909402 | 2.817606122 | 4.16E-13 | 1.28E-10 |
| chr15:53082764-53082998 | 3.668481693 | 4.43E-13 | 1.36E-10 |
| chrX:78838831-78840072 | 2.902400862 | 4.44E-13 | 1.36E-10 |
| chr4:186500785-186501233 | -3.760713035 | 4.50E-13 | 1.37E-10 |
| chr15:52008476-52008809 | 3.164506377 | 4.73E-13 | 1.43E-10 |
| chr15:50028433-50028633 | 3.581366942 | 4.80E-13 | 1.45E-10 |
| chr15:51146951-51147157 | 3.584731181 | 4.93E-13 | 1.48E-10 |
| chr15:51649919-51650278 | 2.862614178 | 5.03E-13 | 1.51E-10 |
| chr15:51642639-51643055 | 3.053110528 | 5.24E-13 | 1.56E-10 |
| chr15:51580312-51580611 | 3.744252172 | 5.27E-13 | 1.56E-10 |
| chr15:50562224-50562481 | 3.392810965 | 5.30E-13 | 1.57E-10 |
| chr15:54156898-54157323 | 3.221090442 | 5.59E-13 | 1.64E-10 |
| chr6:90739175-90740207 | -2.61331435 | 5.60E-13 | 1.64E-10 |
| chr4:171629435-171630328 | -2.781453129 | 5.67E-13 | 1.66E-10 |
| chr15:50163994-50164330 | 2.930755672 | 5.75E-13 | 1.67E-10 |
| chr5:96718478-96719151 | -2.770835919 | 5.75E-13 | 1.67E-10 |
| chr15:51782441-51782686 | 3.500017664 | 5.91E-13 | 1.71E-10 |
| chr15:56918258-56919984 | 2.467769174 | 6.25E-13 | 1.80E-10 |
| chr15:53020052-53020286 | 3.96936257 | 6.81E-13 | 1.95E-10 |
| chr15:50831627-50831830 | 3.484424115 | 7.42E-13 | 2.12E-10 |
| chr15:51736720-51738288 | 2.39156041 | 7.45E-13 | 2.12E-10 |
| chr18:63259166-63259889 | -2.727073898 | 7.72E-13 | 2.19E-10 |
| chr15:51341193-51342261 | 2.447627465 | 8.39E-13 | 2.37E-10 |
| chr15:54114047-54114273 | 4.095496544 | 8.59E-13 | 2.42E-10 |
| chr15:50228958-50229284 | 2.842675439 | 9.15E-13 | 2.57E-10 |
| chr15:53737021-53737286 | 4.283876613 | 9.47E-13 | 2.65E-10 |
| chr15:49953538-49953781 | 3.411483803 | 9.70E-13 | 2.70E-10 |
| chr8:13156469-13157347 | -2.770362951 | 1.01E-12 | 2.81E-10 |
| chr15:53286219-53286598 | 3.032375377 | 1.13E-12 | 3.13E-10 |
| chr15:49650717-49650962 | 3.250666093 | 1.14E-12 | 3.13E-10 |
| chr12:74871696-74872379 | -2.74569515 | 1.15E-12 | 3.16E-10 |
| chr4:185573776-185574215 | -3.400173475 | 1.31E-12 | 3.59E-10 |
| chr7:64313734-64314267 | 3.22500333 | 1.61E-12 | 4.39E-10 |
| chr4:163304131-163304833 | -2.84654136 | 1.65E-12 | 4.49E-10 |
| chr15:51675436-51675715 | 3.275802964 | 1.76E-12 | 4.76E-10 |
| chr7:146905711-146906458 | -3.307750435 | 1.79E-12 | 4.82E-10 |
| chr15:52848414-52848677 | 3.320816588 | 1.83E-12 | 4.92E-10 |
| chr15:49750143-49750343 | 3.50263811 | 1.86E-12 | 4.98E-10 |
| chr7:11707267-11707995 | 3.054042824 | 1.98E-12 | 5.28E-10 |
| chr15:53024781-53025149 | 3.217775017 | 2.03E-12 | 5.40E-10 |
| chr11:133215415-133216183 | -2.831958276 | 2.05E-12 | 5.43E-10 |
| chr15:51132992-51133212 | 3.081925362 | 2.11E-12 | 5.57E-10 |
| chr15:49547984-49548233 | 3.764787615 | 2.45E-12 | 6.44E-10 |
| chr15:50242526-50242750 | 2.970973651 | 2.48E-12 | 6.50E-10 |
| chr15:51198942-51199144 | 3.21751173 | 2.51E-12 | 6.57E-10 |
| chr15:54481387-54481898 | 3.652537659 | 2.58E-12 | 6.71E-10 |
| chr5:118247437-118248205 | -2.593090779 | 2.72E-12 | 7.06E-10 |
| chr7:124919784-124920261 | -2.961644505 | 2.74E-12 | 7.10E-10 |
| chr15:52100876-52101170 | 2.934152393 | 2.98E-12 | 7.69E-10 |
| chr15:53869115-53869382 | 3.058896748 | 3.03E-12 | 7.80E-10 |

| | | | |
|---|---|---|---|
| chr9:73445352-73445815 | -3.632738175 | 3.06E-12 | 7.84E-10 |
| chr15:51401879-51402270 | 2.568357663 | 3.10E-12 | 7.93E-10 |
| chr15:52594642-52594915 | 3.545232212 | 3.17E-12 | 8.06E-10 |
| chr15:50391591-50391904 | 2.915837504 | 3.23E-12 | 8.19E-10 |
| chr15:52256317-52256667 | 2.609217845 | 3.28E-12 | 8.30E-10 |
| chr15:52861867-52862088 | 3.400209304 | 3.42E-12 | 8.63E-10 |
| chr15:51173180-51173416 | 3.640175632 | 3.54E-12 | 8.87E-10 |
| chr4:74455983-74457097 | -2.631039423 | 3.54E-12 | 8.87E-10 |
| chr12:12776281-12776684 | -3.636721004 | 3.75E-12 | 9.37E-10 |
| chr15:51970287-51970517 | 2.867274516 | 4.02E-12 | 1.00E-09 |
| chr15:52228994-52229262 | 2.881726309 | 4.17E-12 | 1.03E-09 |
| chr15:50777168-50777511 | 2.761427743 | 4.25E-12 | 1.05E-09 |
| chr15:54264852-54265180 | 3.84028676 | 4.28E-12 | 1.06E-09 |
| chr10:52440543-52441239 | -2.743138994 | 4.49E-12 | 1.10E-09 |
| chr15:53684141-53684445 | 3.226212445 | 4.57E-12 | 1.12E-09 |
| chr21:24348664-24349201 | -3.224679698 | 4.71E-12 | 1.15E-09 |
| chr15:51499406-51499761 | 2.664106605 | 4.79E-12 | 1.16E-09 |
| chr15:52860934-52861277 | 2.950614298 | 4.91E-12 | 1.19E-09 |
| chr15:52546446-52546692 | 3.101769369 | 4.96E-12 | 1.20E-09 |
| chr15:54249815-54250041 | 4.001660189 | 5.02E-12 | 1.21E-09 |
| chr15:52093725-52094024 | 3.028092707 | 5.11E-12 | 1.23E-09 |
| chr5:115037557-115038063 | -2.993473126 | 5.31E-12 | 1.27E-09 |
| chr4:37480211-37481108 | -2.58655808 | 5.58E-12 | 1.33E-09 |
| chr4:176691725-176692829 | -2.778526883 | 5.63E-12 | 1.34E-09 |
| chr18:57737670-57738426 | -2.75388863 | 5.89E-12 | 1.40E-09 |
| chr5:39606007-39606449 | 3.969791165 | 5.90E-12 | 1.40E-09 |
| chr15:51403240-51403603 | 2.645133499 | 5.94E-12 | 1.40E-09 |
| chr15:53389210-53389419 | 3.506938211 | 6.31E-12 | 1.48E-09 |
| chr15:51286745-51287198 | 2.487612174 | 6.46E-12 | 1.51E-09 |
| chr15:50242116-50242326 | 3.149459334 | 6.68E-12 | 1.56E-09 |
| chr15:52559178-52559393 | 3.430013941 | 7.25E-12 | 1.69E-09 |
| chr15:51907430-51907953 | 2.45000955 | 7.42E-12 | 1.72E-09 |
| chr15:50298111-50298311 | 3.49448994 | 7.51E-12 | 1.74E-09 |
| chr15:52621955-52622169 | 4.153660387 | 7.88E-12 | 1.82E-09 |
| chr15:51137750-51138009 | 2.972905947 | 8.25E-12 | 1.90E-09 |
| chr15:50094982-50095231 | 3.17780389 | 8.43E-12 | 1.93E-09 |
| chr15:54082893-54083981 | 2.414977288 | 9.43E-12 | 2.16E-09 |
| chr15:51494486-51494713 | 3.486041663 | 1.05E-11 | 2.39E-09 |
| chr15:54197930-54198489 | 2.701820031 | 1.16E-11 | 2.64E-09 |
| chr15:53591975-53592196 | 3.399064618 | 1.19E-11 | 2.70E-09 |
| chr15:50676807-50677309 | 2.58841917 | 1.27E-11 | 2.86E-09 |
| chr15:53472245-53472445 | 4.110074524 | 1.38E-11 | 3.12E-09 |
| chr13:81987892-81988190 | -3.993924866 | 1.45E-11 | 3.25E-09 |
| chr1:207866603-207867200 | -2.527418938 | 1.57E-11 | 3.52E-09 |
| chr6:19538074-19538634 | -2.710375117 | 1.60E-11 | 3.58E-09 |
| chr15:53863079-53863284 | 3.76991337 | 1.63E-11 | 3.63E-09 |
| chr15:51557919-51558248 | 3.301389302 | 1.65E-11 | 3.66E-09 |
| chr15:52765879-52766111 | 2.965033738 | 1.68E-11 | 3.72E-09 |
| chr15:50262714-50262936 | 2.830652311 | 1.79E-11 | 3.95E-09 |
| chr15:50717318-50717761 | 2.679325405 | 1.82E-11 | 4.01E-09 |

| | | | |
|---|---|---|---|
| chr15:50563181-50563412 | 3.137931198 | 2.14E-11 | 4.69E-09 |
| chr4:166197979-166198827 | -2.623687984 | 2.20E-11 | 4.82E-09 |
| chr15:51423474-51423706 | 3.28964804 | 2.22E-11 | 4.84E-09 |
| chr11:34513637-34514046 | 2.972458124 | 2.31E-11 | 5.02E-09 |
| chr15:53175956-53176233 | 3.046153709 | 2.37E-11 | 5.14E-09 |
| chr15:51126474-51126678 | 3.351331583 | 2.39E-11 | 5.17E-09 |
| chr15:54250180-54250385 | 4.072252507 | 2.44E-11 | 5.27E-09 |
| chr15:50876942-50877322 | 2.426018343 | 2.55E-11 | 5.49E-09 |
| chr15:51128071-51128303 | 2.964214301 | 2.67E-11 | 5.72E-09 |
| chr15:50205720-50206113 | 2.492935205 | 2.71E-11 | 5.79E-09 |
| chr15:51786980-51787339 | 2.788384407 | 2.81E-11 | 5.99E-09 |
| chr15:51243814-51244053 | 2.901927925 | 2.84E-11 | 6.05E-09 |
| chr15:52319026-52319226 | 3.501533892 | 2.85E-11 | 6.05E-09 |
| chr7:139588620-139588968 | -3.326060271 | 3.12E-11 | 6.61E-09 |
| chr1:208510207-208511049 | -2.507150809 | 3.14E-11 | 6.63E-09 |
| chr15:51568731-51569648 | 2.327534358 | 3.15E-11 | 6.64E-09 |
| chr4:182306811-182307238 | -4.69668125 | 3.20E-11 | 6.72E-09 |
| chr15:53157663-53157866 | 3.602013843 | 3.26E-11 | 6.83E-09 |
| chr15:53189860-53190199 | 3.098591375 | 3.45E-11 | 7.20E-09 |
| chr7:110522700-110523254 | -2.715538946 | 3.74E-11 | 7.80E-09 |
| chr15:53397994-53398562 | 2.329655975 | 3.82E-11 | 7.93E-09 |
| chr15:51776066-51776512 | 2.456286371 | 3.83E-11 | 7.94E-09 |
| chr7:27346850-27347371 | 2.575510981 | 3.94E-11 | 8.13E-09 |
| chr15:51425702-51426111 | 2.584412731 | 4.00E-11 | 8.23E-09 |
| chr4:84298626-84299401 | -2.854878863 | 4.06E-11 | 8.34E-09 |
| chr15:51759893-51760093 | 3.248410818 | 4.10E-11 | 8.40E-09 |
| chr15:50096197-50096471 | 3.008561832 | 4.23E-11 | 8.65E-09 |
| chr4:305425-305721 | 4.035220865 | 4.29E-11 | 8.73E-09 |
| chr15:51950980-51951228 | 3.088968394 | 4.38E-11 | 8.90E-09 |
| chr15:52798933-52799153 | 3.077055553 | 4.74E-11 | 9.61E-09 |
| chr13:90748680-90749072 | -3.584371547 | 4.76E-11 | 9.61E-09 |
| chr5:64682664-64683060 | -3.683095054 | 4.77E-11 | 9.62E-09 |
| chr15:51600240-51600459 | 3.24217767 | 4.85E-11 | 9.75E-09 |
| chr15:52252876-52253142 | 2.795012797 | 4.97E-11 | 9.97E-09 |
| chr15:51413614-51413814 | 3.033844009 | 5.00E-11 | 1.00E-08 |
| chr15:51982147-51982603 | 2.373973822 | 5.05E-11 | 1.01E-08 |
| chr15:52709549-52710154 | 2.344530058 | 5.20E-11 | 1.03E-08 |
| chr15:54027828-54028028 | 3.572201622 | 5.44E-11 | 1.08E-08 |
| chr15:50641265-50641550 | 2.989108093 | 5.49E-11 | 1.09E-08 |
| chr15:51329727-51329927 | 3.067002002 | 5.66E-11 | 1.12E-08 |
| chr7:23760058-23760443 | 2.993519477 | 5.71E-11 | 1.12E-08 |
| chr15:50944867-50945071 | 3.117902448 | 5.82E-11 | 1.14E-08 |
| chr8:37598434-37599532 | -2.33883111 | 5.93E-11 | 1.16E-08 |
| chr15:49875473-49875718 | 2.733054486 | 6.01E-11 | 1.17E-08 |
| chr15:52139846-52140220 | 2.386395705 | 6.19E-11 | 1.21E-08 |
| chr4:186157771-186158530 | -2.656949805 | 6.26E-11 | 1.22E-08 |
| chr1:246166159-246166808 | 2.665275013 | 6.42E-11 | 1.25E-08 |
| chr15:53976811-53977330 | 2.430707002 | 6.63E-11 | 1.28E-08 |
| chr10:104480840-104481672 | -2.431864004 | 6.84E-11 | 1.32E-08 |
| chr7:16905244-16905685 | 2.631627461 | 6.87E-11 | 1.32E-08 |

| | | | |
|---|---|---|---|
| chr15:52921050-52921323 | 2.763516491 | 7.00E-11 | 1.34E-08 |
| chr15:50149209-50149469 | 2.703448904 | 7.03E-11 | 1.35E-08 |
| chr2:226151164-226151679 | -2.694072925 | 7.46E-11 | 1.42E-08 |
| chr4:101865609-101866088 | 2.980957957 | 7.63E-11 | 1.46E-08 |
| chr2:11530745-11531378 | -2.355449312 | 7.74E-11 | 1.47E-08 |
| chr7:102004112-102004449 | -4.611634895 | 8.55E-11 | 1.62E-08 |
| chr2:1051103-1051899 | -2.587921464 | 8.61E-11 | 1.63E-08 |
| chr15:53282866-53283159 | 2.861677236 | 8.94E-11 | 1.69E-08 |
| chr1:218642124-218643242 | -2.190794655 | 9.28E-11 | 1.75E-08 |
| chr2:177970982-177971754 | -2.633755848 | 9.37E-11 | 1.76E-08 |
| chr15:51800979-51801726 | 2.324630377 | 9.41E-11 | 1.76E-08 |
| chr12:50960306-50960774 | 3.807821589 | 9.64E-11 | 1.80E-08 |
| chr4:156289967-156290697 | -2.747938889 | 1.01E-10 | 1.89E-08 |
| chr4:185534948-185535750 | -2.534763617 | 1.04E-10 | 1.93E-08 |
| chrX:6690163-6690751 | -2.979315473 | 1.05E-10 | 1.94E-08 |
| chr9:1198188-1198645 | 3.774341542 | 1.06E-10 | 1.95E-08 |
| chr2:237558790-237559612 | -2.349478314 | 1.15E-10 | 2.13E-08 |
| chr2:35297354-35297765 | -2.842246396 | 1.16E-10 | 2.14E-08 |
| chr8:102128113-102128627 | 2.614827295 | 1.16E-10 | 2.14E-08 |
| chr2:64273481-64274447 | -2.381937486 | 1.17E-10 | 2.14E-08 |
| chr15:50577878-50578115 | 3.121565246 | 1.17E-10 | 2.14E-08 |
| chr15:50675639-50676020 | 2.568987873 | 1.19E-10 | 2.18E-08 |
| chr15:51720127-51720487 | 2.569206154 | 1.31E-10 | 2.38E-08 |
| chr13:67048138-67048817 | -2.564088789 | 1.34E-10 | 2.43E-08 |
| chr15:53792354-53792554 | 3.955037877 | 1.35E-10 | 2.45E-08 |
| chr15:52018641-52020264 | 2.103415789 | 1.37E-10 | 2.48E-08 |
| chr15:54707702-54708394 | 2.662765661 | 1.41E-10 | 2.55E-08 |
| chr4:89275604-89276048 | 3.065873634 | 1.43E-10 | 2.58E-08 |
| chr8:60583606-60584223 | -2.607588308 | 1.44E-10 | 2.59E-08 |
| chr12:87098339-87099096 | -2.434860512 | 1.45E-10 | 2.60E-08 |
| chr15:52518598-52518807 | 3.113855321 | 1.50E-10 | 2.68E-08 |
| chr15:53549750-53550324 | 2.261076363 | 1.53E-10 | 2.72E-08 |
| chr5:1327043-1327611 | 2.474365067 | 1.73E-10 | 3.08E-08 |
| chr4:37250440-37250970 | 3.095484764 | 1.83E-10 | 3.24E-08 |
| chr15:52336618-52336844 | 2.769690103 | 1.86E-10 | 3.30E-08 |
| chr11:131896998-131897552 | -3.041946761 | 2.03E-10 | 3.58E-08 |
| chr14:42945908-42946378 | 2.854587866 | 2.09E-10 | 3.67E-08 |
| chr12:28191713-28192019 | -3.298742425 | 2.13E-10 | 3.75E-08 |
| chr11:21896812-21897613 | -2.534343112 | 2.20E-10 | 3.85E-08 |
| chr6:24394863-24395326 | -3.462459021 | 2.24E-10 | 3.91E-08 |
| chr7:13641770-13642160 | 3.272146298 | 2.26E-10 | 3.94E-08 |
| chr10:8950816-8952011 | 2.350488258 | 2.35E-10 | 4.10E-08 |
| chr11:38019002-38019889 | -2.329126031 | 2.40E-10 | 4.17E-08 |
| chr4:186472891-186473331 | -3.456761729 | 2.41E-10 | 4.17E-08 |
| chr8:95746164-95746612 | 2.75050402 | 2.44E-10 | 4.22E-08 |
| chr21:36613112-36613479 | 3.272358002 | 2.45E-10 | 4.23E-08 |
| chr7:20441739-20442406 | 2.506668427 | 2.54E-10 | 4.38E-08 |
| chr8:49401237-49401920 | -2.765219625 | 2.56E-10 | 4.39E-08 |
| chr2:5691709-5692438 | -2.488486223 | 2.63E-10 | 4.51E-08 |
| chr18:46113702-46114510 | -2.452220283 | 2.65E-10 | 4.53E-08 |

| | | | |
|---|---|---|---|
| chr1:248067451-248068353 | -2.381849654 | 2.68E-10 | 4.57E-08 |
| chr21:18987419-18987802 | -3.353188471 | 2.76E-10 | 4.70E-08 |
| chr3:41131220-41131676 | 3.071913822 | 2.79E-10 | 4.74E-08 |
| chr13:48258475-48258862 | -3.073966529 | 2.87E-10 | 4.86E-08 |
| chr2:997494-998031 | -3.022698937 | 2.87E-10 | 4.86E-08 |
| chr15:52408924-52409129 | 3.263718121 | 2.97E-10 | 5.01E-08 |
| chr15:53586777-53587169 | 2.477482413 | 3.02E-10 | 5.08E-08 |
| chr18:45016451-45017014 | -2.883229716 | 3.03E-10 | 5.09E-08 |
| chr1:212916535-212917301 | -2.398893564 | 3.06E-10 | 5.14E-08 |
| chr4:182246411-182246835 | -4.526215524 | 3.16E-10 | 5.29E-08 |
| chr14:34700168-34700693 | 2.718665871 | 3.26E-10 | 5.43E-08 |
| chr2:1366482-1366976 | -3.004393854 | 3.45E-10 | 5.74E-08 |
| chr18:72582781-72583530 | -2.741959522 | 3.51E-10 | 5.83E-08 |
| chr8:133620851-133621317 | 2.475529653 | 3.60E-10 | 5.97E-08 |
| chr8:113046116-113047001 | -2.439791383 | 3.88E-10 | 6.43E-08 |
| chr15:49966732-49966949 | 2.864999082 | 3.98E-10 | 6.58E-08 |
| chr4:49161140-49161479 | 3.425204011 | 4.10E-10 | 6.75E-08 |
| chr15:50368842-50369065 | 2.7706798 | 4.12E-10 | 6.78E-08 |
| chr15:51349534-51349833 | 2.689863922 | 4.18E-10 | 6.86E-08 |
| chr6:56728531-56729040 | -2.786228489 | 4.28E-10 | 6.99E-08 |
| chr15:51807098-51807317 | 2.99215336 | 4.28E-10 | 6.99E-08 |
| chr21:31338009-31338322 | 2.862145293 | 4.39E-10 | 7.15E-08 |
| chr7:152737659-152738323 | 2.425092168 | 4.75E-10 | 7.73E-08 |
| chr15:50089527-50089727 | 3.232589032 | 5.01E-10 | 8.13E-08 |
| chr8:66525319-66525692 | -2.519507274 | 5.05E-10 | 8.18E-08 |
| chr7:143882552-143883065 | -2.593976643 | 5.34E-10 | 8.64E-08 |
| chr12:90953642-90954284 | -2.448744353 | 5.38E-10 | 8.67E-08 |
| chr12:52280420-52280762 | 2.78900147 | 5.57E-10 | 8.95E-08 |
| chr15:49675104-49675423 | 2.413949366 | 5.58E-10 | 8.95E-08 |
| chr7:13278009-13278374 | 2.885103981 | 5.59E-10 | 8.95E-08 |
| chr8:8296011-8296776 | -2.742845832 | 5.60E-10 | 8.95E-08 |
| chr7:111499038-111499583 | -2.513208408 | 5.61E-10 | 8.95E-08 |
| chr15:51971059-51972397 | 2.030040773 | 5.67E-10 | 9.03E-08 |
| chr15:51297879-51298079 | 2.860987182 | 5.74E-10 | 9.12E-08 |
| chr13:89162665-89162944 | 3.679291229 | 5.96E-10 | 9.45E-08 |
| chr15:53271753-53272029 | 2.699302192 | 6.01E-10 | 9.50E-08 |
| chr14:84403513-84403899 | 2.898279833 | 6.01E-10 | 9.50E-08 |
| chr15:51688929-51689149 | 2.805778642 | 6.14E-10 | 9.68E-08 |
| chr3:80384671-80385398 | 2.466497391 | 6.16E-10 | 9.69E-08 |
| chr20:19757827-19758533 | 2.193174611 | 6.17E-10 | 9.69E-08 |
| chr15:50148369-50148698 | 2.273016681 | 6.20E-10 | 9.72E-08 |
| chr13:81957075-81957713 | -2.739761713 | 6.21E-10 | 9.72E-08 |
| chr6:28633464-28634241 | -2.686705213 | 6.27E-10 | 9.79E-08 |
| chr15:51777002-51777344 | 2.275621598 | 6.47E-10 | 1.01E-07 |
| chr18:34595714-34596409 | -2.303318586 | 6.49E-10 | 1.01E-07 |
| chr15:52735336-52735826 | 2.240045931 | 6.57E-10 | 1.02E-07 |
| chr15:51985091-51985396 | 2.695113257 | 6.57E-10 | 1.02E-07 |
| chr15:50622387-50622716 | 2.442933874 | 6.61E-10 | 1.02E-07 |
| chr15:52568171-52569540 | 2.019448565 | 6.64E-10 | 1.02E-07 |
| chr7:77007999-77008371 | -3.506370102 | 6.70E-10 | 1.03E-07 |

| | | | |
|---|---|---|---|
| chr15:54228408-54228608 | 3.206546581 | 7.08E-10 | 1.09E-07 |
| chr3:86420996-86421945 | -2.22838846 | 7.10E-10 | 1.09E-07 |
| chr1:59507067-59507596 | -2.873322153 | 7.14E-10 | 1.09E-07 |
| chr4:184310064-184310684 | -2.656631339 | 7.14E-10 | 1.09E-07 |
| chr16:77905346-77905966 | 2.307035365 | 7.16E-10 | 1.09E-07 |
| chr15:54217569-54217780 | 3.300275743 | 7.28E-10 | 1.11E-07 |
| chr15:50390678-50391482 | 2.12222174 | 7.48E-10 | 1.13E-07 |
| chr8:11571774-11572178 | -2.750129844 | 8.08E-10 | 1.22E-07 |
| chr15:50885642-50886238 | 2.090227753 | 8.41E-10 | 1.27E-07 |
| chr4:16034050-16034553 | 2.722723273 | 8.50E-10 | 1.28E-07 |
| chr15:51023481-51023681 | 2.945339251 | 8.75E-10 | 1.32E-07 |
| chr15:51888682-51888894 | 2.854265221 | 8.84E-10 | 1.33E-07 |
| chr2:179646245-179646815 | -2.710010615 | 8.88E-10 | 1.33E-07 |
| chr15:53837251-53837715 | 2.357373815 | 8.89E-10 | 1.33E-07 |
| chr15:53257427-53257642 | 3.04389666 | 8.98E-10 | 1.34E-07 |
| chr6:53464519-53466199 | -2.065956461 | 9.65E-10 | 1.44E-07 |
| chr5:78204092-78204756 | -2.629946972 | 9.68E-10 | 1.44E-07 |
| chr15:49521823-49522192 | 2.276901112 | 9.88E-10 | 1.47E-07 |
| chr15:50003551-50004254 | 2.066752663 | 9.98E-10 | 1.48E-07 |
| chr1:220967043-220967804 | -2.253534658 | 9.99E-10 | 1.48E-07 |
| chr8:136530273-136530911 | 2.433012819 | 1.01E-09 | 1.50E-07 |
| chr14:48285123-48285661 | 2.881601564 | 1.02E-09 | 1.50E-07 |
| chr7:125339912-125340401 | -2.654036648 | 1.03E-09 | 1.52E-07 |
| chr2:38462019-38462798 | -2.288719323 | 1.09E-09 | 1.61E-07 |
| chr15:53012964-53013199 | 3.170585375 | 1.11E-09 | 1.63E-07 |
| chr3:189098855-189099840 | -2.111467991 | 1.14E-09 | 1.67E-07 |
| chr17:15920758-15921157 | 2.978088417 | 1.16E-09 | 1.70E-07 |
| chr7:101126303-101127198 | -2.295037444 | 1.17E-09 | 1.70E-07 |
| chr4:163448561-163449238 | -2.383229523 | 1.19E-09 | 1.73E-07 |
| chr1:206903542-206904201 | -2.130566012 | 1.21E-09 | 1.75E-07 |
| chr15:52789710-52790794 | 2.006829602 | 1.24E-09 | 1.79E-07 |
| chr1:184156326-184157298 | -2.068336384 | 1.24E-09 | 1.79E-07 |
| chr2:175062725-175063385 | -2.439458926 | 1.32E-09 | 1.90E-07 |
| chr15:53384384-53384823 | 2.205853412 | 1.33E-09 | 1.92E-07 |
| chr12:28998456-28999367 | -2.220919667 | 1.33E-09 | 1.92E-07 |
| chr11:131511849-131512227 | -3.359222845 | 1.34E-09 | 1.92E-07 |
| chr3:40396419-40396910 | 2.676257268 | 1.37E-09 | 1.96E-07 |
| chr15:51764603-51764803 | 2.872291159 | 1.39E-09 | 1.99E-07 |
| chr5:107104486-107104992 | -3.015273512 | 1.39E-09 | 1.99E-07 |
| chr15:50907808-50909401 | 1.968586498 | 1.43E-09 | 2.04E-07 |
| chr15:51004403-51004648 | 2.66808764 | 1.44E-09 | 2.04E-07 |
| chr3:95721761-95722329 | -2.266530534 | 1.47E-09 | 2.08E-07 |
| chr13:73368632-73369007 | -2.819892596 | 1.48E-09 | 2.09E-07 |
| chr15:53328393-53328710 | 2.650895536 | 1.52E-09 | 2.14E-07 |
| chr7:142375697-142376321 | -2.24527543 | 1.57E-09 | 2.22E-07 |
| chr15:53992144-53992388 | 3.016416429 | 1.67E-09 | 2.35E-07 |
| chr12:29220685-29221301 | -2.706162348 | 1.69E-09 | 2.37E-07 |
| chr15:52331476-52331974 | 2.111317802 | 1.70E-09 | 2.38E-07 |
| chr2:114797462-114797859 | 2.941809156 | 1.89E-09 | 2.64E-07 |
| chr4:6556243-6556680 | 2.581723775 | 1.94E-09 | 2.71E-07 |

| | | | |
|---|---|---|---|
| chr15:52528004-52529882 | 1.952750692 | 1.95E-09 | 2.72E-07 |
| chr7:11606132-11606473 | 3.749663747 | 1.99E-09 | 2.78E-07 |
| chr11:19978809-19979366 | 2.803596074 | 2.10E-09 | 2.92E-07 |
| chr4:159804244-159805015 | -2.463583167 | 2.11E-09 | 2.92E-07 |
| chr15:52543683-52543897 | 2.884365557 | 2.14E-09 | 2.96E-07 |
| chr15:53218070-53218442 | 2.477034526 | 2.15E-09 | 2.96E-07 |
| chrX:127878030-127878920 | -2.599543358 | 2.15E-09 | 2.96E-07 |
| chr11:9103518-9103929 | 2.851378254 | 2.15E-09 | 2.96E-07 |
| chr15:51681379-51681581 | 2.84117974 | 2.17E-09 | 2.99E-07 |
| chr9:96686694-96687068 | 3.049182791 | 2.24E-09 | 3.08E-07 |
| chr15:52294806-52296099 | 1.961593686 | 2.24E-09 | 3.08E-07 |
| chr7:142323553-142324058 | -2.598293233 | 2.30E-09 | 3.15E-07 |
| chr15:53466551-53466855 | 2.593811631 | 2.35E-09 | 3.20E-07 |
| chr15:50314139-50314745 | 2.068695456 | 2.36E-09 | 3.22E-07 |
| chr2:197198657-197199241 | -2.331048824 | 2.40E-09 | 3.26E-07 |
| chr7:19742295-19742772 | 2.825036257 | 2.40E-09 | 3.26E-07 |
| chr7:114663510-114663824 | -3.194714804 | 2.52E-09 | 3.41E-07 |
| chr6:26020273-26020563 | -2.73238528 | 2.60E-09 | 3.52E-07 |
| chr1:6955571-6955801 | 3.578268867 | 2.67E-09 | 3.61E-07 |
| chr7:135385221-135385566 | -3.029357037 | 2.67E-09 | 3.61E-07 |
| chr12:570464-571691 | -2.12488067 | 2.69E-09 | 3.62E-07 |
| chr1:198657634-198658141 | 2.398680101 | 2.72E-09 | 3.66E-07 |
| chr15:52179271-52180478 | 1.940093746 | 2.79E-09 | 3.74E-07 |
| chr5:53090223-53090564 | -2.474498148 | 2.81E-09 | 3.77E-07 |
| chr12:101219900-101220264 | -2.820979751 | 2.87E-09 | 3.84E-07 |
| chr13:62021163-62021721 | -2.771527052 | 2.88E-09 | 3.85E-07 |
| chr6:143397393-143397852 | -2.434067001 | 3.02E-09 | 4.02E-07 |
| chr7:86282030-86282712 | -2.465191689 | 3.09E-09 | 4.11E-07 |
| chr15:51231830-51232030 | 2.775878938 | 3.13E-09 | 4.16E-07 |
| chr2:189318822-189319837 | -2.153269141 | 3.28E-09 | 4.35E-07 |
| chrX:23803979-23804365 | -2.88319511 | 3.33E-09 | 4.40E-07 |
| chr4:142313856-142314499 | -2.624722012 | 3.46E-09 | 4.57E-07 |
| chr2:81398580-81399213 | -2.412591286 | 3.76E-09 | 4.96E-07 |
| chr2:200300346-200301230 | -2.272156355 | 3.79E-09 | 4.99E-07 |
| chr4:185636887-185637868 | -2.15042389 | 3.80E-09 | 5.00E-07 |
| chr2:70616178-70616695 | -2.319258347 | 3.83E-09 | 5.02E-07 |
| chr9:92097979-92098410 | -2.83032012 | 3.84E-09 | 5.03E-07 |
| chr15:52349169-52349530 | 2.338781219 | 4.15E-09 | 5.42E-07 |
| chr3:79651502-79651814 | 3.015731093 | 4.18E-09 | 5.45E-07 |
| chr15:51281243-51281706 | 2.130416233 | 4.20E-09 | 5.46E-07 |
| chr15:52350446-52350810 | 2.27913483 | 4.20E-09 | 5.46E-07 |
| chr15:50181948-50183041 | 1.934148016 | 4.29E-09 | 5.57E-07 |
| chr4:181957111-181957399 | -4.317322665 | 4.32E-09 | 5.61E-07 |
| chr15:50247856-50248170 | 2.17845746 | 4.54E-09 | 5.87E-07 |
| chr2:205763616-205764209 | -2.237074308 | 4.68E-09 | 6.05E-07 |
| chr7:128122099-128122970 | -2.28673932 | 4.73E-09 | 6.10E-07 |
| chr20:10250767-10251437 | 2.095214196 | 4.74E-09 | 6.11E-07 |
| chr4:64271418-64272191 | -2.062367949 | 4.81E-09 | 6.19E-07 |
| chr10:88474304-88474980 | -2.219872142 | 4.86E-09 | 6.24E-07 |
| chr16:57992874-57993260 | 2.638234907 | 4.94E-09 | 6.33E-07 |

| | | | |
|---|---|---|---|
| chr3:16482703-16483466 | 2.092230285 | 5.05E-09 | 6.46E-07 |
| chr16:77896526-77897396 | 2.106373083 | 5.08E-09 | 6.49E-07 |
| chr12:10100924-10101793 | -2.28920628 | 5.36E-09 | 6.84E-07 |
| chr2:19915953-19917136 | -2.065824469 | 5.46E-09 | 6.94E-07 |
| chr15:52812822-52813038 | 2.819813924 | 5.46E-09 | 6.94E-07 |
| chr14:61153462-61154008 | -2.208997013 | 5.47E-09 | 6.94E-07 |
| chr15:52078876-52079196 | 2.18078228 | 5.50E-09 | 6.96E-07 |
| chr15:51829450-51830469 | 1.914158279 | 5.56E-09 | 7.03E-07 |
| chr7:21076635-21076897 | 2.869136404 | 5.59E-09 | 7.06E-07 |
| chr15:53385081-53385299 | 2.659360195 | 5.66E-09 | 7.13E-07 |
| chr8:53564391-53564756 | 2.601579388 | 5.68E-09 | 7.15E-07 |
| chr7:15625468-15625796 | 3.337264417 | 5.94E-09 | 7.46E-07 |
| chr7:105439039-105439910 | -2.101337709 | 6.00E-09 | 7.52E-07 |
| chr4:26452633-26452991 | 3.049493216 | 6.04E-09 | 7.56E-07 |
| chr11:27423132-27423538 | -3.060654697 | 6.16E-09 | 7.70E-07 |
| chr4:163316814-163317144 | -3.957871333 | 6.31E-09 | 7.87E-07 |
| chr15:49725517-49725717 | 2.61873782 | 6.53E-09 | 8.13E-07 |
| chr1:208184361-208184803 | -2.361978559 | 6.63E-09 | 8.25E-07 |
| chr19:9324046-9324296 | 3.667289391 | 6.70E-09 | 8.32E-07 |
| chr14:71500658-71501251 | -2.565608012 | 6.75E-09 | 8.37E-07 |
| chr10:113958534-113959094 | -2.324750679 | 6.93E-09 | 8.58E-07 |
| chr19:44401417-44401717 | 3.050256782 | 6.97E-09 | 8.61E-07 |
| chr7:95376846-95377486 | -2.326962512 | 7.23E-09 | 8.92E-07 |
| chr15:52824098-52824916 | 1.928333785 | 7.32E-09 | 9.02E-07 |
| chr19:33310471-33311178 | 2.202711899 | 7.40E-09 | 9.10E-07 |
| chr1:209220002-209220539 | -2.756382039 | 7.62E-09 | 9.36E-07 |
| chr5:39459086-39459628 | 2.534477025 | 7.65E-09 | 9.37E-07 |
| chr2:11532200-11532969 | -2.086583979 | 7.73E-09 | 9.46E-07 |
| chr5:119780187-119780925 | -2.458103136 | 7.86E-09 | 9.59E-07 |
| chr11:85844387-85845412 | -1.989015027 | 7.86E-09 | 9.59E-07 |
| chr16:77212013-77213239 | 1.919101085 | 8.03E-09 | 9.78E-07 |
| chr15:53494033-53494233 | 3.039173992 | 8.27E-09 | 1.01E-06 |
| chr19:52963284-52963494 | 4.251483976 | 8.65E-09 | 1.05E-06 |
| chr4:157823729-157824681 | -1.971371866 | 8.80E-09 | 1.07E-06 |
| chr4:76892661-76893017 | 2.727435614 | 8.93E-09 | 1.08E-06 |
| chr3:61064542-61065259 | -2.17906062 | 9.43E-09 | 1.14E-06 |
| chr15:51936216-51936457 | 2.538614106 | 9.56E-09 | 1.15E-06 |
| chr15:52496241-52496472 | 2.666938469 | 9.56E-09 | 1.15E-06 |
| chr2:178144674-178144946 | 4.239295963 | 9.70E-09 | 1.17E-06 |
| chr7:15130736-15131163 | 2.562447143 | 1.00E-08 | 1.20E-06 |
| chr15:53197375-53197896 | 2.050482317 | 1.02E-08 | 1.22E-06 |
| chr7:24733701-24734054 | 2.532018673 | 1.07E-08 | 1.28E-06 |
| chr11:126968965-126969641 | -2.419808771 | 1.10E-08 | 1.31E-06 |
| chr1:91830206-91830587 | -2.88493676 | 1.11E-08 | 1.33E-06 |
| chr15:51991613-51991844 | 2.38017702 | 1.12E-08 | 1.34E-06 |
| chr14:43640040-43640558 | 2.392652493 | 1.13E-08 | 1.34E-06 |
| chr19:57477543-57477828 | 3.006296899 | 1.13E-08 | 1.34E-06 |
| chr1:173122856-173123741 | -2.021403695 | 1.20E-08 | 1.42E-06 |
| chr5:124963307-124964252 | -2.082832205 | 1.20E-08 | 1.42E-06 |
| chr16:80739546-80739918 | -2.431235247 | 1.22E-08 | 1.45E-06 |

| | | | |
|---|---|---|---|
| chr4:137414306-137415105 | -2.116636148 | 1.30E-08 | 1.53E-06 |
| chr14:28801391-28801760 | 2.648495491 | 1.30E-08 | 1.53E-06 |
| chr15:53655538-53655826 | 2.520980467 | 1.30E-08 | 1.54E-06 |
| chr4:171203426-171203961 | -2.361287268 | 1.31E-08 | 1.54E-06 |
| chr15:51621848-51623330 | 1.842426059 | 1.33E-08 | 1.56E-06 |
| chr6:6825118-6825784 | -2.176238402 | 1.34E-08 | 1.57E-06 |
| chr7:22572305-22573496 | 1.903346215 | 1.34E-08 | 1.57E-06 |
| chr7:136419853-136420275 | -3.286427435 | 1.35E-08 | 1.57E-06 |
| chr13:82305620-82306081 | -2.535628087 | 1.35E-08 | 1.58E-06 |
| chr15:51181517-51181809 | 2.287585561 | 1.39E-08 | 1.61E-06 |
| chr14:34929232-34929474 | 3.302476409 | 1.39E-08 | 1.61E-06 |
| chr15:53962857-53963314 | 2.175823686 | 1.39E-08 | 1.61E-06 |
| chr12:50532625-50533162 | 2.345597007 | 1.41E-08 | 1.64E-06 |
| chr14:37616180-37616608 | 2.416132321 | 1.43E-08 | 1.66E-06 |
| chr7:47103819-47104464 | -2.064204327 | 1.46E-08 | 1.69E-06 |
| chr5:1867531-1867817 | 2.985364137 | 1.47E-08 | 1.70E-06 |
| chr3:152021811-152022520 | -2.005680193 | 1.50E-08 | 1.73E-06 |
| chr7:10924450-10924834 | 2.400658718 | 1.51E-08 | 1.73E-06 |
| chr18:65931084-65931524 | -2.395199065 | 1.52E-08 | 1.74E-06 |
| chrX:126459950-126460523 | 2.742090863 | 1.53E-08 | 1.75E-06 |
| chr14:20394659-20394876 | 3.860125049 | 1.54E-08 | 1.76E-06 |
| chr7:80019826-80020836 | -2.20577024 | 1.55E-08 | 1.77E-06 |
| chr13:103732041-103732378 | -2.923584459 | 1.63E-08 | 1.86E-06 |
| chr13:58239011-58239457 | -2.860101006 | 1.65E-08 | 1.89E-06 |
| chr13:87810357-87811424 | -1.949073654 | 1.67E-08 | 1.90E-06 |
| chrX:140491961-140492762 | -2.056456862 | 1.69E-08 | 1.92E-06 |
| chr2:164522486-164523179 | -2.199035682 | 1.70E-08 | 1.93E-06 |
| chr8:101368846-101369165 | 2.727719766 | 1.73E-08 | 1.96E-06 |
| chr16:87925243-87925858 | 2.136857931 | 1.73E-08 | 1.96E-06 |
| chr2:148154335-148155400 | -2.135330617 | 1.74E-08 | 1.97E-06 |
| chr7:24192188-24192568 | 2.90986828 | 1.76E-08 | 1.98E-06 |
| chr15:50828799-50829147 | 2.172191874 | 1.76E-08 | 1.98E-06 |
| chr1:237309362-237310222 | -2.354754909 | 1.78E-08 | 2.00E-06 |
| chr18:34557566-34558457 | -2.231600252 | 1.79E-08 | 2.01E-06 |
| chr3:174147106-174147459 | -2.785633754 | 1.85E-08 | 2.08E-06 |
| chr3:5060041-5060386 | 2.593659333 | 1.86E-08 | 2.08E-06 |
| chr15:49591945-49592145 | 2.611246259 | 1.87E-08 | 2.09E-06 |
| chr7:14654606-14654933 | 3.581242757 | 1.88E-08 | 2.09E-06 |
| chr7:142181270-142181757 | -2.595412732 | 1.92E-08 | 2.14E-06 |
| chrX:82163777-82164235 | -2.506259338 | 1.93E-08 | 2.15E-06 |
| chr7:135653314-135653621 | -3.391301426 | 1.94E-08 | 2.15E-06 |
| chr10:9144865-9145137 | 4.193100017 | 1.95E-08 | 2.17E-06 |
| chr2:12507324-12507653 | -2.967233937 | 1.98E-08 | 2.19E-06 |
| chr2:3348969-3349779 | -2.012150022 | 1.98E-08 | 2.19E-06 |
| chr7:8196337-8196544 | 4.175563253 | 1.99E-08 | 2.20E-06 |
| chr12:31773103-31774253 | -1.996479224 | 1.99E-08 | 2.20E-06 |
| chr11:38994652-38994977 | -3.415736537 | 2.00E-08 | 2.20E-06 |
| chr15:52780558-52780883 | 2.10479558 | 2.02E-08 | 2.22E-06 |
| chr18:9428274-9428701 | 2.603359216 | 2.06E-08 | 2.27E-06 |
| chr15:50797951-50798336 | 1.981177019 | 2.07E-08 | 2.27E-06 |

| | | | |
|---|---|---|---|
| chr16:75044787-75045533 | -1.98109829 | 2.09E-08 | 2.29E-06 |
| chr19:52535625-52536720 | 2.020833196 | 2.12E-08 | 2.32E-06 |
| chr5:5721878-5722267 | -2.673053322 | 2.16E-08 | 2.36E-06 |
| chr13:77478153-77478738 | -2.327476329 | 2.16E-08 | 2.36E-06 |
| chr13:33345676-33346198 | -2.065435916 | 2.17E-08 | 2.37E-06 |
| chr12:25135387-25136228 | -2.019013873 | 2.20E-08 | 2.39E-06 |
| chr5:32785585-32786261 | -2.292626469 | 2.21E-08 | 2.40E-06 |
| chr17:17445762-17447057 | 1.900724678 | 2.22E-08 | 2.40E-06 |
| chr3:108401199-108401584 | 3.224803137 | 2.30E-08 | 2.49E-06 |
| chr2:196607930-196608393 | -2.635483136 | 2.32E-08 | 2.51E-06 |
| chr15:56968797-56969143 | 2.820178285 | 2.34E-08 | 2.53E-06 |
| chr8:86940787-86941089 | 2.339123473 | 2.36E-08 | 2.54E-06 |
| chr10:126940865-126941258 | 2.889286562 | 2.37E-08 | 2.55E-06 |
| chr16:78184533-78185290 | 2.055088373 | 2.42E-08 | 2.61E-06 |
| chr15:51382909-51383109 | 2.298708932 | 2.43E-08 | 2.61E-06 |
| chrX:109270877-109271453 | -2.199493109 | 2.47E-08 | 2.65E-06 |
| chr2:122980683-122981072 | -2.766624622 | 2.47E-08 | 2.65E-06 |
| chr1:33008442-33009055 | 2.12439137 | 2.48E-08 | 2.66E-06 |
| chr13:36509371-36509823 | -2.407236313 | 2.49E-08 | 2.66E-06 |
| chr10:60848678-60849103 | -2.289540507 | 2.50E-08 | 2.67E-06 |
| chr12:29148698-29150328 | -1.880765329 | 2.51E-08 | 2.67E-06 |
| chr3:146052095-146052633 | -2.468775939 | 2.51E-08 | 2.67E-06 |
| chrX:139158330-139158722 | 2.479681758 | 2.51E-08 | 2.67E-06 |
| chr7:118363723-118364380 | -2.274705001 | 2.53E-08 | 2.69E-06 |
| chr4:103443351-103443722 | -2.802637239 | 2.53E-08 | 2.69E-06 |
| chr5:53089303-53089935 | -2.110773874 | 2.57E-08 | 2.72E-06 |
| chr12:124786605-124786834 | 3.035746651 | 2.60E-08 | 2.75E-06 |
| chr6:99802386-99802838 | -2.462925505 | 2.61E-08 | 2.76E-06 |
| chr10:984402-984757 | 2.351160766 | 2.63E-08 | 2.77E-06 |
| chrX:34415111-34415766 | -2.434530964 | 2.66E-08 | 2.80E-06 |
| chr2:226118372-226119437 | -1.954402334 | 2.70E-08 | 2.84E-06 |
| chr11:88522886-88523151 | -4.160573954 | 2.71E-08 | 2.84E-06 |
| chr7:123594587-123595243 | -2.075454185 | 2.72E-08 | 2.85E-06 |
| chr5:170245127-170245847 | -2.233090106 | 2.74E-08 | 2.86E-06 |
| chr7:70788274-70788776 | -2.461718501 | 2.77E-08 | 2.89E-06 |
| chr9:37528446-37529523 | -2.046650281 | 2.80E-08 | 2.92E-06 |
| chr19:35478833-35479324 | 2.304098802 | 2.80E-08 | 2.92E-06 |
| chr10:132575283-132575795 | 2.180031533 | 2.89E-08 | 3.01E-06 |
| chr7:33377487-33377891 | 3.093840744 | 2.90E-08 | 3.01E-06 |
| chr7:13414231-13415154 | 2.001351783 | 2.90E-08 | 3.01E-06 |
| chr14:64093004-64093287 | 2.946896389 | 2.93E-08 | 3.04E-06 |
| chr6:84052142-84052992 | -2.034173109 | 3.02E-08 | 3.12E-06 |
| chr11:14029500-14029813 | -2.459511493 | 3.07E-08 | 3.18E-06 |
| chr7:127833411-127833952 | -2.388768341 | 3.09E-08 | 3.19E-06 |
| chr12:25518507-25518995 | -2.510548926 | 3.10E-08 | 3.19E-06 |
| chr7:127598319-127598736 | -2.328704215 | 3.14E-08 | 3.23E-06 |
| chr12:131315932-131316326 | 2.868321976 | 3.17E-08 | 3.26E-06 |
| chr7:77720770-77721098 | -2.847631278 | 3.19E-08 | 3.28E-06 |
| chr15:93829135-93830146 | -1.92565528 | 3.33E-08 | 3.41E-06 |
| chr15:51561300-51561616 | 2.217243901 | 3.36E-08 | 3.45E-06 |

| | | | |
|---|---|---|---|
| chr15:52241756-52241972 | 2.384874888 | 3.38E-08 | 3.45E-06 |
| chr3:189324548-189325159 | 2.117624082 | 3.45E-08 | 3.52E-06 |
| chr3:189267883-189268700 | -1.990870725 | 3.46E-08 | 3.53E-06 |
| chr15:50854442-50855359 | 1.835524867 | 3.47E-08 | 3.54E-06 |
| chr9:21995258-21995504 | 4.126092149 | 3.49E-08 | 3.55E-06 |
| chr13:104641284-104641899 | -2.167044812 | 3.54E-08 | 3.60E-06 |
| chr2:38515334-38516031 | -2.039897106 | 3.60E-08 | 3.66E-06 |
| chr8:125066620-125067203 | 1.982158625 | 3.67E-08 | 3.72E-06 |
| chr12:97656439-97657223 | -2.478295768 | 3.69E-08 | 3.73E-06 |
| chr7:102064369-102065051 | -2.289820388 | 3.69E-08 | 3.73E-06 |
| chr7:118870281-118870565 | -3.511649643 | 3.75E-08 | 3.78E-06 |
| chr7:22583242-22583624 | 2.15262881 | 3.79E-08 | 3.82E-06 |
| chr10:67230665-67231472 | -2.334291097 | 3.87E-08 | 3.90E-06 |
| chr10:122599297-122599959 | 2.522983615 | 3.89E-08 | 3.91E-06 |
| chr6:163324642-163325082 | 2.847157716 | 3.90E-08 | 3.92E-06 |
| chr13:61507893-61508531 | -2.172834634 | 3.93E-08 | 3.94E-06 |
| chr19:12365500-12365949 | 2.422165983 | 3.94E-08 | 3.95E-06 |
| chr1:96059693-96060181 | -2.18893698 | 4.00E-08 | 4.00E-06 |
| chr11:111735212-111736141 | -2.009507167 | 4.02E-08 | 4.02E-06 |
| chr7:19369625-19370292 | 2.04897564 | 4.04E-08 | 4.03E-06 |
| chr2:35291909-35293026 | -1.937470347 | 4.05E-08 | 4.03E-06 |
| chr2:179574047-179575002 | -1.988249927 | 4.10E-08 | 4.08E-06 |
| chr2:180359551-180360618 | -1.885273547 | 4.15E-08 | 4.12E-06 |
| chr12:39990638-39991053 | -3.114107268 | 4.24E-08 | 4.21E-06 |
| chr11:38971962-38972523 | -2.309089314 | 4.34E-08 | 4.30E-06 |
| chr3:177451829-177452299 | -2.457879916 | 4.48E-08 | 4.43E-06 |
| chr1:210270249-210270815 | -2.09944826 | 4.52E-08 | 4.47E-06 |
| chr7:10780332-10780719 | 3.311454462 | 4.60E-08 | 4.54E-06 |
| chr7:106859026-106859403 | -2.760141239 | 4.67E-08 | 4.60E-06 |
| chr10:19833073-19833524 | -2.384085275 | 4.67E-08 | 4.60E-06 |
| chrX:16832369-16832625 | -2.976824546 | 4.74E-08 | 4.66E-06 |
| chr13:74751699-74752146 | -2.39773496 | 4.78E-08 | 4.70E-06 |
| chr17:17445343-17445609 | 2.546400654 | 4.79E-08 | 4.70E-06 |
| chr19:52637824-52638615 | 1.92802493 | 4.85E-08 | 4.75E-06 |
| chr15:27410826-27411330 | 2.205523252 | 4.90E-08 | 4.79E-06 |
| chr11:3813448-3814000 | 2.216094011 | 4.93E-08 | 4.82E-06 |
| chr14:85587087-85587777 | 1.951420873 | 4.93E-08 | 4.82E-06 |
| chrX:154369322-154369760 | 2.03335543 | 4.96E-08 | 4.84E-06 |
| chr11:124536506-124536962 | -2.22530051 | 5.02E-08 | 4.89E-06 |
| chr7:113170381-113171345 | -2.14881782 | 5.07E-08 | 4.93E-06 |
| chr15:50980555-50980860 | 2.234416265 | 5.15E-08 | 5.00E-06 |
| chr10:60848146-60848569 | -2.393760506 | 5.18E-08 | 5.02E-06 |
| chr10:114819380-114819781 | 2.260908828 | 5.19E-08 | 5.03E-06 |
| chr2:36731974-36732855 | -2.02322932 | 5.23E-08 | 5.06E-06 |
| chr12:100678923-100679355 | -2.238568427 | 5.46E-08 | 5.28E-06 |
| chr17:28536915-28537431 | -2.098386781 | 5.55E-08 | 5.36E-06 |
| chr4:175297930-175298305 | -2.761987239 | 5.56E-08 | 5.36E-06 |
| chr15:52760027-52760310 | 2.191018892 | 5.79E-08 | 5.57E-06 |
| chr15:53684583-53684928 | 2.334722281 | 5.93E-08 | 5.70E-06 |
| chr18:60372422-60373227 | -2.10116418 | 5.94E-08 | 5.70E-06 |

| | | | |
|---|---|---|---|
| chr13:101416587-101417398 | -2.045399588 | 5.95E-08 | 5.71E-06 |
| chr4:183353664-183354563 | -1.985099644 | 5.96E-08 | 5.71E-06 |
| chr19:44051524-44052322 | 1.983107422 | 6.12E-08 | 5.85E-06 |
| chr15:53875194-53875432 | 2.440896772 | 6.18E-08 | 5.91E-06 |
| chr7:78058270-78058773 | -2.742520971 | 6.20E-08 | 5.91E-06 |
| chr16:89430913-89431484 | 2.089448028 | 6.27E-08 | 5.98E-06 |
| chr11:99069804-99070238 | -2.744624595 | 6.37E-08 | 6.06E-06 |
| chr10:62640271-62640770 | -2.460429504 | 6.38E-08 | 6.06E-06 |
| chr1:225677429-225677682 | 3.33066136 | 6.48E-08 | 6.15E-06 |
| chr3:178736315-178736685 | -2.558858085 | 6.51E-08 | 6.18E-06 |
| chr4:94428894-94429424 | 2.176486261 | 6.55E-08 | 6.20E-06 |
| chr4:132540310-132541190 | -1.928751132 | 6.63E-08 | 6.27E-06 |
| chr8:122489665-122490080 | -2.092575175 | 6.68E-08 | 6.31E-06 |
| chr14:77699431-77699876 | 2.081258769 | 6.73E-08 | 6.35E-06 |
| chr4:74620256-74620754 | -2.247680973 | 6.80E-08 | 6.41E-06 |
| chrX:96433726-96434127 | -2.473116325 | 6.83E-08 | 6.43E-06 |
| chr5:87231012-87231407 | -2.58821779 | 6.86E-08 | 6.45E-06 |
| chr4:186067966-186068414 | -2.418895797 | 6.91E-08 | 6.48E-06 |
| chr8:91289139-91289577 | 2.341603776 | 7.04E-08 | 6.61E-06 |
| chrX:78571872-78572206 | 2.56964258 | 7.08E-08 | 6.63E-06 |
| chr4:163478511-163478908 | -2.594804979 | 7.10E-08 | 6.64E-06 |
| chr11:123197804-123198041 | -3.372340588 | 7.17E-08 | 6.70E-06 |
| chr6:156827139-156827754 | -2.078713505 | 7.22E-08 | 6.74E-06 |
| chr4:74545494-74545851 | -2.544304655 | 7.24E-08 | 6.74E-06 |
| chr18:7038104-7038950 | -2.037216725 | 7.31E-08 | 6.80E-06 |
| chr8:66126765-66127234 | -2.155724527 | 7.31E-08 | 6.80E-06 |
| chr19:53622025-53622615 | 2.386258494 | 7.33E-08 | 6.80E-06 |
| chr4:74583124-74583363 | -2.738161231 | 7.33E-08 | 6.80E-06 |
| chr1:225720473-225720887 | 2.396590822 | 7.36E-08 | 6.82E-06 |
| chr3:149388069-149388465 | 2.059968147 | 7.44E-08 | 6.88E-06 |
| chr2:144902916-144903577 | -2.468183138 | 7.62E-08 | 7.04E-06 |
| chr21:40340890-40341342 | -2.539011607 | 7.71E-08 | 7.11E-06 |
| chr2:183348475-183348757 | -3.170890755 | 7.75E-08 | 7.14E-06 |
| chr2:192500713-192501257 | -2.013214708 | 7.76E-08 | 7.15E-06 |
| chr18:43279398-43279751 | -2.43344839 | 8.07E-08 | 7.43E-06 |
| chr2:35298255-35299161 | -1.920304498 | 8.24E-08 | 7.57E-06 |
| chr11:58926244-58926663 | 2.405852126 | 8.27E-08 | 7.59E-06 |
| chr7:75842754-75843326 | -2.297905116 | 8.31E-08 | 7.62E-06 |
| chr8:66534514-66534990 | -2.218271974 | 8.38E-08 | 7.67E-06 |
| chr10:30151714-30152423 | -2.079344923 | 8.43E-08 | 7.71E-06 |
| chr13:75489507-75489801 | -2.864549405 | 8.45E-08 | 7.72E-06 |
| chr19:1114665-1114878 | 3.707685881 | 8.49E-08 | 7.74E-06 |
| chr4:188195799-188196588 | -2.36254786 | 8.51E-08 | 7.75E-06 |
| chr7:21860869-21861165 | 2.927065693 | 8.60E-08 | 7.83E-06 |
| chr1:228059626-228060266 | 1.962505422 | 8.62E-08 | 7.83E-06 |
| chr7:110394572-110394905 | -3.257517441 | 8.62E-08 | 7.83E-06 |
| chr15:51924207-51924410 | 2.367248392 | 8.69E-08 | 7.88E-06 |
| chr22:12097707-12098083 | 2.651717878 | 8.89E-08 | 8.05E-06 |
| chr15:47824176-47824578 | -2.463129834 | 8.91E-08 | 8.06E-06 |
| chr8:109501309-109501570 | 2.85633145 | 8.92E-08 | 8.06E-06 |

| | | | |
|---|---|---|---|
| chr8:127782226-127783014 | 1.84751605 | 8.95E-08 | 8.08E-06 |
| chr8:100345144-100345635 | 2.105961195 | 9.01E-08 | 8.12E-06 |
| chr15:69107145-69107474 | 2.723170576 | 9.05E-08 | 8.14E-06 |
| chr3:180762888-180763375 | -2.347522711 | 9.06E-08 | 8.14E-06 |
| chr7:7529421-7529889 | 2.254523188 | 9.11E-08 | 8.19E-06 |
| chr4:163385499-163385903 | -2.223226983 | 9.18E-08 | 8.24E-06 |
| chrX:23816819-23817522 | -2.185624431 | 9.28E-08 | 8.32E-06 |
| chr5:101091260-101091810 | -2.257567923 | 9.36E-08 | 8.38E-06 |
| chr15:51917526-51917992 | 1.922015207 | 9.41E-08 | 8.41E-06 |
| chr4:49513346-49513957 | 2.250808296 | 9.45E-08 | 8.44E-06 |
| chrX:153059720-153060066 | 2.846915127 | 9.53E-08 | 8.50E-06 |
| chr1:102833195-102833563 | -2.37945595 | 9.54E-08 | 8.50E-06 |
| chr2:35495252-35495685 | -2.526587475 | 9.55E-08 | 8.50E-06 |
| chr5:119355168-119355479 | -2.345856587 | 9.83E-08 | 8.74E-06 |
| chr15:50416052-50416277 | 2.233195625 | 9.91E-08 | 8.80E-06 |
| chr4:101921409-101921726 | 2.664503277 | 9.98E-08 | 8.85E-06 |
| chr7:28409103-28409477 | 2.709415268 | 1.00E-07 | 8.87E-06 |
| chr7:132028533-132028883 | -2.593943694 | 1.00E-07 | 8.88E-06 |
| chr14:28217427-28218188 | 2.146822759 | 1.02E-07 | 9.00E-06 |
| chr4:186021483-186021940 | -2.739944003 | 1.03E-07 | 9.05E-06 |
| chr3:159179123-159179479 | 3.243739079 | 1.03E-07 | 9.11E-06 |
| chr5:100517204-100517738 | -1.91503057 | 1.06E-07 | 9.32E-06 |
| chr11:105289944-105290421 | -2.927461423 | 1.06E-07 | 9.35E-06 |
| chr15:51726334-51726797 | 1.847183507 | 1.07E-07 | 9.44E-06 |
| chr2:21251655-21251880 | -3.304524144 | 1.09E-07 | 9.53E-06 |
| chr15:53130877-53131138 | 2.257301353 | 1.09E-07 | 9.60E-06 |
| chr7:138951947-138952257 | -2.471800478 | 1.11E-07 | 9.70E-06 |
| chr6:158839850-158840133 | 4.013738118 | 1.11E-07 | 9.72E-06 |
| chr15:50840593-50840819 | 2.252035277 | 1.11E-07 | 9.73E-06 |
| chr2:51581694-51582278 | -2.110761246 | 1.12E-07 | 9.74E-06 |
| chrX:116384457-116384893 | 2.542689962 | 1.14E-07 | 9.90E-06 |
| chr21:32495133-32495692 | 2.596051387 | 1.14E-07 | 9.92E-06 |
| chr14:87312118-87312459 | 2.708983893 | 1.16E-07 | 1.01E-05 |
| chr4:174369821-174370488 | -2.332944195 | 1.19E-07 | 1.04E-05 |
| chr15:50353896-50355829 | 1.696699866 | 1.19E-07 | 1.04E-05 |
| chr5:19293201-19293566 | 2.904826541 | 1.20E-07 | 1.04E-05 |
| chr10:84717892-84718139 | 4.038959288 | 1.23E-07 | 1.07E-05 |
| chr15:53808499-53808854 | 2.120568947 | 1.24E-07 | 1.07E-05 |
| chr7:130994571-130995000 | -2.538570203 | 1.24E-07 | 1.07E-05 |
| chr5:100647312-100648167 | -1.865309481 | 1.26E-07 | 1.09E-05 |
| chr7:7586029-7586336 | 2.688732421 | 1.27E-07 | 1.09E-05 |
| chr14:52502295-52502727 | 2.752142567 | 1.27E-07 | 1.10E-05 |
| chr13:75685171-75685757 | -1.95308912 | 1.28E-07 | 1.10E-05 |
| chr15:50764984-50766171 | 1.711652963 | 1.28E-07 | 1.10E-05 |
| chr7:9404112-9404936 | -2.019013436 | 1.28E-07 | 1.10E-05 |
| chr7:119472424-119472970 | -2.460748096 | 1.29E-07 | 1.11E-05 |
| chr14:35747424-35747663 | 3.264480214 | 1.30E-07 | 1.12E-05 |
| chr19:21141850-21142109 | 3.42628547 | 1.31E-07 | 1.12E-05 |
| chr5:103006855-103007310 | -2.171640668 | 1.31E-07 | 1.12E-05 |
| chr4:185304912-185305350 | -2.348090532 | 1.31E-07 | 1.12E-05 |

| | | | |
|---|---|---|---|
| chr1:75861595-75862327 | -2.008732427 | 1.33E-07 | 1.13E-05 |
| chr10:113454561-113455074 | -2.302330905 | 1.33E-07 | 1.13E-05 |
| chr7:101564834-101565107 | -3.092090734 | 1.34E-07 | 1.14E-05 |
| chr15:80854157-80854494 | -2.400339461 | 1.35E-07 | 1.15E-05 |
| chr10:132362268-132362496 | 2.751613698 | 1.37E-07 | 1.16E-05 |
| chr7:138343004-138343316 | -2.868434531 | 1.38E-07 | 1.17E-05 |
| chr3:188964771-188965496 | -2.156917998 | 1.39E-07 | 1.17E-05 |
| chr7:142587667-142587936 | -3.98543456 | 1.39E-07 | 1.17E-05 |
| chr5:45263170-45263700 | 2.740372988 | 1.39E-07 | 1.18E-05 |
| chr7:47409195-47409792 | -1.967638406 | 1.40E-07 | 1.18E-05 |
| chr7:153523781-153524637 | -2.040974259 | 1.41E-07 | 1.19E-05 |
| chr15:57198699-57198943 | 3.438175484 | 1.41E-07 | 1.19E-05 |
| chr7:142438294-142438651 | -2.322099302 | 1.42E-07 | 1.19E-05 |
| chr12:15216011-15216779 | -1.927167279 | 1.42E-07 | 1.19E-05 |
| chr7:30654735-30655056 | 2.884246311 | 1.42E-07 | 1.20E-05 |
| chr7:83629125-83630131 | -2.025179904 | 1.43E-07 | 1.20E-05 |
| chr7:123450482-123451347 | -2.061465807 | 1.44E-07 | 1.21E-05 |
| chr1:210251060-210252045 | -1.814768834 | 1.45E-07 | 1.21E-05 |
| chr11:130533113-130533750 | -2.166708938 | 1.45E-07 | 1.21E-05 |
| chr7:19954116-19954514 | 2.498158043 | 1.47E-07 | 1.23E-05 |
| chr16:80837728-80838226 | 2.15260402 | 1.48E-07 | 1.23E-05 |
| chr10:92384812-92385137 | 2.894115361 | 1.49E-07 | 1.24E-05 |
| chr1:51629058-51629630 | 2.266516193 | 1.50E-07 | 1.25E-05 |
| chr7:11751764-11752044 | 2.689006623 | 1.50E-07 | 1.25E-05 |
| chr3:74985713-74986145 | 2.537049138 | 1.53E-07 | 1.27E-05 |
| chr18:8779511-8779825 | 3.226236891 | 1.55E-07 | 1.28E-05 |
| chr5:17326542-17327090 | 2.132948587 | 1.56E-07 | 1.30E-05 |
| chr11:132858242-132858493 | -3.126462294 | 1.58E-07 | 1.30E-05 |
| chr7:127905595-127905956 | -2.982319144 | 1.60E-07 | 1.32E-05 |
| chr15:53506483-53506683 | 2.678510819 | 1.62E-07 | 1.34E-05 |
| chr8:129883792-129884161 | -2.144341741 | 1.63E-07 | 1.34E-05 |
| chr16:49010413-49010885 | -2.579208465 | 1.63E-07 | 1.34E-05 |
| chr7:142496536-142497224 | -2.001011185 | 1.65E-07 | 1.36E-05 |
| chr16:35811254-35811635 | 2.809679245 | 1.67E-07 | 1.37E-05 |
| chr2:11498502-11499002 | -1.957495797 | 1.68E-07 | 1.38E-05 |
| chr2:11539347-11539877 | -1.92069792 | 1.68E-07 | 1.38E-05 |
| chr10:77872120-77873380 | -1.786580027 | 1.69E-07 | 1.38E-05 |
| chr7:31549640-31549908 | 3.378218132 | 1.75E-07 | 1.43E-05 |
| chr18:69879003-69879396 | -2.816900268 | 1.76E-07 | 1.44E-05 |
| chr8:37243111-37243444 | 2.783943667 | 1.77E-07 | 1.44E-05 |
| chr5:10301378-10302291 | -1.844553416 | 1.78E-07 | 1.45E-05 |
| chr10:58229653-58230477 | -1.956535788 | 1.80E-07 | 1.46E-05 |
| chr7:24492834-24493183 | 2.253414501 | 1.80E-07 | 1.46E-05 |
| chr3:194962241-194962485 | 3.388723758 | 1.81E-07 | 1.48E-05 |
| chr16:78051180-78051488 | 2.444496356 | 1.82E-07 | 1.48E-05 |
| chr7:107880332-107880721 | -2.188061775 | 1.84E-07 | 1.49E-05 |
| chr7:23853781-23854104 | 2.872509765 | 1.90E-07 | 1.54E-05 |
| chr4:128841467-128841843 | 2.545433835 | 1.91E-07 | 1.55E-05 |
| chr11:101441603-101442525 | -1.933761596 | 1.92E-07 | 1.55E-05 |
| chr9:73581161-73581923 | -2.196397139 | 1.93E-07 | 1.56E-05 |

| | | | |
|---|---|---|---|
| chr15:52071611-52071916 | 1.931366959 | 1.94E-07 | 1.57E-05 |
| chr6:122623355-122623671 | -2.463884704 | 1.94E-07 | 1.57E-05 |
| chrX:20361924-20363296 | -1.812642593 | 1.95E-07 | 1.57E-05 |
| chr7:134891166-134891655 | -2.129841773 | 1.95E-07 | 1.57E-05 |
| chr4:185513584-185514323 | -2.042608366 | 1.97E-07 | 1.58E-05 |
| chr10:4280588-4281414 | 1.886533243 | 1.97E-07 | 1.58E-05 |
| chr8:67737589-67737857 | 3.610399838 | 1.97E-07 | 1.58E-05 |
| chr3:116971524-116971869 | -2.559018968 | 1.98E-07 | 1.59E-05 |
| chr7:102063758-102064079 | -2.761867221 | 2.05E-07 | 1.64E-05 |
| chr15:53142102-53142353 | 2.509710225 | 2.07E-07 | 1.65E-05 |
| chr4:81021451-81022199 | -1.967971836 | 2.08E-07 | 1.66E-05 |
| chr7:121313533-121313892 | -3.185849978 | 2.09E-07 | 1.67E-05 |
| chr14:42149591-42149857 | 3.410436159 | 2.12E-07 | 1.69E-05 |
| chr6:164307422-164307812 | -2.925089317 | 2.14E-07 | 1.71E-05 |
| chr20:19775613-19776292 | 1.907179217 | 2.15E-07 | 1.71E-05 |
| chrX:88612917-88613627 | -2.307530006 | 2.15E-07 | 1.71E-05 |
| chr6:135062669-135063177 | -2.003561951 | 2.16E-07 | 1.71E-05 |
| chr9:76379505-76379980 | -1.94935372 | 2.21E-07 | 1.76E-05 |
| chr1:152743196-152743805 | -1.903760066 | 2.26E-07 | 1.79E-05 |
| chr4:86359967-86360253 | -2.601027345 | 2.27E-07 | 1.80E-05 |
| chr3:133206636-133207379 | -1.874258716 | 2.28E-07 | 1.80E-05 |
| chr2:137873878-137874691 | -2.040891573 | 2.29E-07 | 1.81E-05 |
| chr16:27829627-27829895 | 3.224413867 | 2.31E-07 | 1.82E-05 |
| chr5:52943332-52943764 | -2.164427251 | 2.31E-07 | 1.82E-05 |
| chr3:109648066-109648460 | -2.125240978 | 2.32E-07 | 1.83E-05 |
| chr12:26523781-26524088 | -2.612873578 | 2.33E-07 | 1.83E-05 |
| chr19:19105583-19105891 | -2.7106655 | 2.33E-07 | 1.83E-05 |
| chr1:216596309-216596627 | -2.117132208 | 2.39E-07 | 1.87E-05 |
| chr16:76268374-76268602 | 2.858192403 | 2.40E-07 | 1.89E-05 |
| chr12:97500220-97501218 | -1.910082064 | 2.41E-07 | 1.89E-05 |
| chr4:138910572-138911460 | -1.717871289 | 2.43E-07 | 1.91E-05 |
| chr8:11572480-11573200 | -1.975269918 | 2.50E-07 | 1.95E-05 |
| chrX:131813193-131813483 | 2.566099884 | 2.50E-07 | 1.96E-05 |
| chr8:78350400-78351347 | -1.966987964 | 2.51E-07 | 1.96E-05 |
| chr2:227267782-227268129 | -2.697868736 | 2.51E-07 | 1.96E-05 |
| chr8:133899307-133899725 | 2.418033417 | 2.52E-07 | 1.96E-05 |
| chr5:35295013-35295682 | -2.080078081 | 2.60E-07 | 2.02E-05 |
| chr11:65403472-65403682 | 3.392646979 | 2.67E-07 | 2.07E-05 |
| chr3:42179402-42179773 | -2.374744178 | 2.73E-07 | 2.12E-05 |
| chr7:7504898-7505258 | 2.684869802 | 2.77E-07 | 2.15E-05 |
| chr7:18048459-18048833 | 2.634657903 | 2.79E-07 | 2.16E-05 |
| chr2:239181742-239182388 | -2.093344563 | 2.79E-07 | 2.16E-05 |
| chr5:141867666-141868024 | -2.313781328 | 2.82E-07 | 2.18E-05 |
| chr19:57351091-57351423 | 2.525277744 | 2.84E-07 | 2.20E-05 |
| chr4:34922225-34922676 | 2.199526478 | 2.85E-07 | 2.20E-05 |
| chr7:23030612-23030988 | 2.745261009 | 2.86E-07 | 2.20E-05 |
| chr20:52008776-52008995 | 2.749774838 | 2.91E-07 | 2.24E-05 |
| chr11:1100688-1100911 | 3.62677983 | 2.93E-07 | 2.25E-05 |
| chrX:150688865-150689512 | 1.92381165 | 2.93E-07 | 2.25E-05 |
| chr10:60707912-60708188 | -2.778249225 | 2.94E-07 | 2.26E-05 |

| | | | |
|---|---|---|---|
| chr7:134710394-134711233 | -1.959174441 | 2.98E-07 | 2.28E-05 |
| chr9:114156321-114156836 | -2.36462258 | 3.03E-07 | 2.32E-05 |
| chr3:103813971-103814377 | 2.20955688 | 3.17E-07 | 2.42E-05 |
| chr7:24059617-24060209 | 1.922890522 | 3.17E-07 | 2.43E-05 |
| chr1:99648147-99648457 | -2.588060686 | 3.18E-07 | 2.43E-05 |
| chr15:49620613-49621364 | 1.662999637 | 3.19E-07 | 2.43E-05 |
| chr21:31986551-31986834 | 2.743715692 | 3.23E-07 | 2.46E-05 |
| chr14:35384622-35384860 | 2.674377221 | 3.23E-07 | 2.46E-05 |
| chr10:7942898-7943240 | 2.374563977 | 3.24E-07 | 2.46E-05 |
| chr10:120455620-120455963 | -2.437171088 | 3.26E-07 | 2.48E-05 |
| chr12:58302457-58302987 | 2.061340932 | 3.33E-07 | 2.53E-05 |
| chr2:165413087-165413543 | -2.242749295 | 3.34E-07 | 2.53E-05 |
| chr21:28961896-28962283 | 2.568713469 | 3.38E-07 | 2.56E-05 |
| chr12:132055717-132056022 | 2.920537974 | 3.42E-07 | 2.59E-05 |
| chr16:68332319-68332648 | 2.373506736 | 3.43E-07 | 2.59E-05 |
| chr2:36097092-36097383 | -2.74384958 | 3.45E-07 | 2.61E-05 |
| chr3:3708325-3708758 | 2.141271776 | 3.47E-07 | 2.62E-05 |
| chr13:46611125-46611339 | 2.56628045 | 3.47E-07 | 2.62E-05 |
| chr11:103007478-103007912 | 2.206194871 | 3.55E-07 | 2.67E-05 |
| chr11:7538669-7539039 | 2.187549902 | 3.55E-07 | 2.67E-05 |
| chr3:8500627-8501116 | 1.803570967 | 3.63E-07 | 2.73E-05 |
| chr11:129815748-129816322 | 2.18622528 | 3.64E-07 | 2.74E-05 |
| chr7:102317303-102317674 | -2.144182264 | 3.65E-07 | 2.74E-05 |
| chr4:23374019-23374712 | -1.982758608 | 3.70E-07 | 2.77E-05 |
| chr2:104378790-104379207 | 2.239315369 | 3.70E-07 | 2.77E-05 |
| chr7:20607114-20607472 | 2.716939354 | 3.73E-07 | 2.79E-05 |
| chr14:35730187-35730504 | 2.560200316 | 3.76E-07 | 2.81E-05 |
| chr18:48330652-48330930 | -2.679064515 | 3.78E-07 | 2.82E-05 |
| chr15:53978242-53978517 | 2.083088882 | 3.79E-07 | 2.83E-05 |
| chr12:83518461-83518768 | 3.185256119 | 3.81E-07 | 2.84E-05 |
| chr9:107249074-107249627 | -1.952565586 | 3.83E-07 | 2.85E-05 |
| chr6:73028110-73028714 | -1.932948839 | 3.98E-07 | 2.96E-05 |
| chr3:81531001-81532151 | -1.795958911 | 4.05E-07 | 3.01E-05 |
| chr13:39510296-39510546 | -2.906190256 | 4.06E-07 | 3.01E-05 |
| chr19:37210470-37210855 | 2.114735804 | 4.08E-07 | 3.03E-05 |
| chr1:206524862-206525597 | -1.751716679 | 4.10E-07 | 3.04E-05 |
| chr15:49876758-49877078 | 1.871318664 | 4.11E-07 | 3.04E-05 |
| chr5:1326393-1326827 | 2.116616607 | 4.12E-07 | 3.04E-05 |
| chr17:21345316-21345635 | -2.562240117 | 4.12E-07 | 3.04E-05 |
| chr13:81290973-81291272 | -2.607358322 | 4.13E-07 | 3.05E-05 |
| chr7:83677154-83677932 | -1.791406591 | 4.15E-07 | 3.06E-05 |
| chr7:113085697-113086309 | -2.218968259 | 4.19E-07 | 3.09E-05 |
| chr15:53808124-53808365 | 2.143217216 | 4.19E-07 | 3.09E-05 |
| chr11:97573617-97574170 | -2.068733644 | 4.21E-07 | 3.09E-05 |
| chr7:21262218-21262509 | 2.716068849 | 4.21E-07 | 3.09E-05 |
| chrX:32185162-32185666 | -2.337723665 | 4.24E-07 | 3.11E-05 |
| chr4:189090281-189090530 | -3.016628037 | 4.25E-07 | 3.12E-05 |
| chr19:29450373-29450695 | 3.015827553 | 4.26E-07 | 3.12E-05 |
| chr12:29185605-29185987 | -2.203060928 | 4.30E-07 | 3.15E-05 |
| chr7:95491787-95492363 | -2.013167806 | 4.32E-07 | 3.16E-05 |

| | | | |
|---|---|---|---|
| chr12:119171069-119171776 | -1.875201438 | 4.33E-07 | 3.16E-05 |
| chr15:51252826-51253026 | 2.324174098 | 4.35E-07 | 3.17E-05 |
| chr15:51991079-51991545 | 1.771994672 | 4.37E-07 | 3.19E-05 |
| chr14:43604716-43605225 | 2.314062738 | 4.37E-07 | 3.19E-05 |
| chr2:11542397-11542699 | -2.074572663 | 4.39E-07 | 3.19E-05 |
| chr14:43135687-43136225 | 2.253636625 | 4.40E-07 | 3.20E-05 |
| chr4:168587851-168588146 | -2.455034994 | 4.43E-07 | 3.22E-05 |
| chr4:166198914-166199185 | -2.790285955 | 4.47E-07 | 3.24E-05 |
| chr10:9066647-9066929 | 2.597990597 | 4.48E-07 | 3.25E-05 |
| chr10:89922230-89922518 | 2.603009338 | 4.49E-07 | 3.25E-05 |
| chr14:55085097-55085571 | 2.084235034 | 4.50E-07 | 3.25E-05 |
| chr15:56969323-56969596 | 2.658286187 | 4.51E-07 | 3.26E-05 |
| chr8:31497174-31497626 | -2.642633921 | 4.54E-07 | 3.28E-05 |
| chr13:80654596-80655242 | -2.068744215 | 4.55E-07 | 3.28E-05 |
| chrX:138105590-138105990 | 2.450491631 | 4.55E-07 | 3.28E-05 |
| chr6:13455407-13455994 | -2.095742683 | 4.60E-07 | 3.31E-05 |
| chr5:169945642-169946269 | -1.865713072 | 4.60E-07 | 3.31E-05 |
| chr4:148060414-148060998 | -1.935239167 | 4.62E-07 | 3.32E-05 |
| chr7:126987182-126987689 | -2.397089256 | 4.63E-07 | 3.32E-05 |
| chr2:42054659-42055131 | -2.139152871 | 4.71E-07 | 3.38E-05 |
| chr5:1514758-1515099 | 2.066497463 | 4.75E-07 | 3.40E-05 |
| chr12:84069160-84069558 | -2.539004146 | 4.76E-07 | 3.41E-05 |
| chr9:73465807-73466331 | -2.275007069 | 4.77E-07 | 3.41E-05 |
| chr10:34791925-34793196 | -1.749181493 | 4.77E-07 | 3.41E-05 |
| chr4:26110506-26110853 | 2.708938172 | 4.79E-07 | 3.42E-05 |
| chr9:119412355-119412935 | -1.870100746 | 4.86E-07 | 3.46E-05 |
| chr13:49097440-49097879 | -2.304346145 | 4.86E-07 | 3.46E-05 |
| chr8:115930142-115930491 | -2.467336553 | 4.92E-07 | 3.51E-05 |
| chr7:143892475-143892982 | -1.992790849 | 4.99E-07 | 3.55E-05 |
| chr9:107389638-107390215 | -1.902497613 | 4.99E-07 | 3.55E-05 |
| chr4:174474488-174475002 | -1.975742731 | 5.01E-07 | 3.56E-05 |
| chr10:31935715-31935971 | -3.111998034 | 5.02E-07 | 3.56E-05 |
| chr8:54059705-54060211 | -2.063018205 | 5.18E-07 | 3.67E-05 |
| chr12:118107041-118107361 | 2.034984359 | 5.20E-07 | 3.68E-05 |
| chr1:183374811-183375254 | 2.056496395 | 5.21E-07 | 3.69E-05 |
| chr8:10850946-10851241 | -2.983579707 | 5.22E-07 | 3.69E-05 |
| chr13:93233957-93234473 | -1.925482313 | 5.23E-07 | 3.69E-05 |
| chr3:31370782-31371145 | 2.483932173 | 5.28E-07 | 3.72E-05 |
| chr3:21272051-21272349 | 3.270879001 | 5.29E-07 | 3.73E-05 |
| chr12:14416573-14416983 | -2.395919173 | 5.32E-07 | 3.75E-05 |
| chr18:73520255-73520570 | -2.712484454 | 5.37E-07 | 3.78E-05 |
| chr7:153876565-153876966 | -2.420593946 | 5.40E-07 | 3.80E-05 |
| chr12:83232248-83232645 | 2.703322359 | 5.41E-07 | 3.80E-05 |
| chr2:202344190-202344853 | -2.047096377 | 5.47E-07 | 3.83E-05 |
| chr10:65751213-65751687 | -1.97059025 | 5.49E-07 | 3.85E-05 |
| chrX:31470814-31471611 | -2.051979824 | 5.53E-07 | 3.87E-05 |
| chr8:130366265-130366832 | 1.933628887 | 5.54E-07 | 3.88E-05 |
| chr3:929888-930212 | 2.527589017 | 5.63E-07 | 3.93E-05 |
| chr2:64096626-64096907 | -3.335120851 | 5.64E-07 | 3.94E-05 |
| chr16:74487638-74488020 | 2.209840949 | 5.65E-07 | 3.94E-05 |

| | | | |
|---|---|---|---|
| chr12:53685532-53685732 | 3.568986905 | 5.70E-07 | 3.97E-05 |
| chr12:210643-211514 | -1.8632153 | 5.71E-07 | 3.98E-05 |
| chr2:20080440-20081215 | -1.889392102 | 5.73E-07 | 3.98E-05 |
| chr3:116758364-116758630 | -3.865554546 | 5.73E-07 | 3.98E-05 |
| chr7:133617414-133617658 | -3.281749242 | 5.75E-07 | 3.99E-05 |
| chr3:106842527-106843128 | -2.170292136 | 5.78E-07 | 4.01E-05 |
| chr7:159021015-159021362 | -2.34018303 | 5.80E-07 | 4.02E-05 |
| chr2:48016694-48016962 | -2.630074744 | 5.82E-07 | 4.03E-05 |
| chr12:121754984-121755184 | 2.989500466 | 5.85E-07 | 4.05E-05 |
| chrX:35919420-35920286 | -1.705891187 | 5.85E-07 | 4.05E-05 |
| chr4:172411021-172411483 | -2.153310474 | 5.87E-07 | 4.05E-05 |
| chr14:84520744-84521072 | 2.296637922 | 5.88E-07 | 4.06E-05 |
| chr15:52217172-52217560 | 1.774082176 | 5.89E-07 | 4.06E-05 |
| chr7:34814207-34814553 | 2.100244386 | 5.90E-07 | 4.07E-05 |
| chr15:51078342-51078591 | 2.055235739 | 5.91E-07 | 4.07E-05 |
| chr6:102204274-102204793 | -2.092522452 | 5.97E-07 | 4.10E-05 |
| chr14:27805320-27805593 | 3.271255076 | 5.99E-07 | 4.12E-05 |
| chr18:41080681-41081152 | -2.640555951 | 6.04E-07 | 4.15E-05 |
| chr4:172892366-172892683 | -2.57192487 | 6.08E-07 | 4.17E-05 |
| chr1:232529810-232530215 | 2.114187378 | 6.09E-07 | 4.17E-05 |
| chr2:20381299-20381870 | -1.906217383 | 6.10E-07 | 4.17E-05 |
| chr2:102126235-102126597 | 2.35396184 | 6.18E-07 | 4.23E-05 |
| chr2:171763215-171763549 | 2.628912586 | 6.22E-07 | 4.25E-05 |
| chr2:692590-693328 | -1.842239761 | 6.27E-07 | 4.28E-05 |
| chr4:78569266-78569502 | 3.490502967 | 6.32E-07 | 4.31E-05 |
| chr2:11543807-11544095 | -2.258443119 | 6.38E-07 | 4.35E-05 |
| chr8:31833396-31833918 | -2.202509645 | 6.47E-07 | 4.41E-05 |
| chr7:54838220-54838956 | -1.737883437 | 6.52E-07 | 4.44E-05 |
| chr5:88014906-88015677 | -2.109911831 | 6.56E-07 | 4.46E-05 |
| chr2:726197-726885 | -1.811387574 | 6.58E-07 | 4.47E-05 |
| chr2:4672881-4673421 | -2.180709176 | 6.66E-07 | 4.52E-05 |
| chr13:90416478-90417014 | -2.112352991 | 6.66E-07 | 4.52E-05 |
| chr9:82031405-82031947 | -2.127493013 | 6.67E-07 | 4.52E-05 |
| chr12:112149653-112150006 | 2.097334007 | 6.69E-07 | 4.53E-05 |
| chr14:73675178-73675520 | 2.212495232 | 6.79E-07 | 4.59E-05 |
| chr18:69250460-69250903 | -2.421874013 | 6.90E-07 | 4.66E-05 |
| chr5:132378423-132379049 | -1.776700144 | 6.98E-07 | 4.71E-05 |
| chr20:48812526-48813132 | 1.672631953 | 7.01E-07 | 4.72E-05 |
| chr7:8442951-8443260 | 2.499750938 | 7.10E-07 | 4.78E-05 |
| chr2:11484803-11485068 | -2.406749707 | 7.26E-07 | 4.89E-05 |
| chr8:93160207-93160597 | 1.967095724 | 7.29E-07 | 4.90E-05 |
| chrX:147796502-147797105 | 2.000137709 | 7.29E-07 | 4.90E-05 |
| chr10:110802624-110802996 | -2.006122837 | 7.30E-07 | 4.90E-05 |
| chr1:214439467-214440022 | -2.002818397 | 7.37E-07 | 4.95E-05 |
| chr4:174984952-174985263 | -2.677177913 | 7.38E-07 | 4.95E-05 |
| chr7:117791988-117792244 | -2.755588801 | 7.42E-07 | 4.97E-05 |
| chrX:152038773-152039161 | 2.30675318 | 7.43E-07 | 4.97E-05 |
| chr2:60597328-60597736 | -2.178781555 | 7.43E-07 | 4.97E-05 |
| chr7:142898428-142898718 | -2.600484565 | 7.46E-07 | 4.98E-05 |
| chr6:139323658-139324312 | -1.875694801 | 7.49E-07 | 5.00E-05 |

| | | | |
|---|---|---|---|
| chr1:100668254-100669097 | -1.934866678 | 7.53E-07 | 5.02E-05 |
| chr4:5922160-5922410 | 3.489385511 | 7.53E-07 | 5.02E-05 |
| chr13:23212635-23213143 | 1.878642747 | 7.59E-07 | 5.05E-05 |
| chr1:230783489-230783689 | 3.291296424 | 7.59E-07 | 5.05E-05 |
| chr5:107556985-107557725 | -1.741386727 | 7.60E-07 | 5.05E-05 |
| chr2:239061218-239061583 | 2.821607017 | 7.62E-07 | 5.06E-05 |
| chr13:65133165-65133428 | -3.247394454 | 7.63E-07 | 5.06E-05 |
| chr4:168677031-168677422 | -1.961443886 | 7.64E-07 | 5.07E-05 |
| chr3:86922980-86923272 | -2.456934454 | 7.67E-07 | 5.08E-05 |
| chr4:185508190-185508647 | -2.017738065 | 7.73E-07 | 5.12E-05 |
| chr5:97239558-97240436 | -1.753248572 | 7.76E-07 | 5.13E-05 |
| chr2:14191825-14192230 | -2.043104787 | 7.77E-07 | 5.13E-05 |
| chr10:93896956-93897204 | 2.977563459 | 7.84E-07 | 5.17E-05 |
| chr4:38591512-38591876 | -2.19749243 | 7.85E-07 | 5.18E-05 |
| chr7:103660454-103661132 | -1.909752976 | 7.87E-07 | 5.18E-05 |
| chr1:51325905-51326136 | -2.611401686 | 7.88E-07 | 5.19E-05 |
| chr22:46649496-46649816 | 2.675218141 | 7.96E-07 | 5.23E-05 |
| chr6:71489819-71490201 | 2.16082772 | 8.00E-07 | 5.26E-05 |
| chr4:62485385-62485804 | -2.496179745 | 8.05E-07 | 5.28E-05 |
| chr11:101566623-101567402 | -2.018561007 | 8.11E-07 | 5.32E-05 |
| chr12:21214763-21215159 | -2.86158277 | 8.15E-07 | 5.34E-05 |
| chr12:89756621-89757299 | -1.868402555 | 8.16E-07 | 5.34E-05 |
| chr4:172188259-172188990 | -1.760696142 | 8.19E-07 | 5.36E-05 |
| chr2:150376644-150377026 | -2.174486543 | 8.19E-07 | 5.36E-05 |
| chr10:89921424-89921870 | 1.95803501 | 8.21E-07 | 5.36E-05 |
| chr15:55943970-55944368 | 2.370984813 | 8.32E-07 | 5.43E-05 |
| chr6:6758822-6759843 | -1.660880439 | 8.34E-07 | 5.44E-05 |
| chr14:23611411-23611757 | 2.255784382 | 8.39E-07 | 5.47E-05 |
| chr11:38971096-38971519 | -2.065415887 | 8.41E-07 | 5.47E-05 |
| chr7:23287572-23287885 | 2.200332 | 8.43E-07 | 5.48E-05 |
| chr14:29961030-29961391 | 2.40435516 | 8.48E-07 | 5.51E-05 |
| chr5:101658573-101659055 | -2.163582247 | 8.52E-07 | 5.53E-05 |
| chr8:56636978-56637460 | -2.232307217 | 8.56E-07 | 5.55E-05 |
| chrX:22097179-22097677 | 2.310300743 | 8.59E-07 | 5.57E-05 |
| chr16:33798599-33798930 | 2.392571255 | 8.61E-07 | 5.58E-05 |
| chr7:84980977-84981553 | -2.003107679 | 8.70E-07 | 5.63E-05 |
| chr2:99404354-99404743 | 2.131148772 | 8.77E-07 | 5.67E-05 |
| chr5:1864764-1865473 | 1.762762321 | 8.93E-07 | 5.77E-05 |
| chr4:6641065-6641619 | 2.02118441 | 8.95E-07 | 5.78E-05 |
| chr7:20486664-20487103 | 2.211964715 | 9.07E-07 | 5.85E-05 |
| chr7:112459670-112459955 | -2.650189151 | 9.12E-07 | 5.88E-05 |
| chrX:31070024-31070343 | -2.807270167 | 9.23E-07 | 5.95E-05 |
| chr7:19595304-19595621 | 2.581941926 | 9.31E-07 | 5.99E-05 |
| chr1:234702197-234702453 | 2.843436447 | 9.47E-07 | 6.09E-05 |
| chr14:51117442-51117816 | 2.648764717 | 9.53E-07 | 6.12E-05 |
| chr7:142516002-142516321 | -2.153334061 | 9.57E-07 | 6.14E-05 |
| chr14:31667866-31668399 | 2.19388204 | 9.61E-07 | 6.16E-05 |
| chr10:48814916-48815243 | -2.225621919 | 9.63E-07 | 6.17E-05 |
| chr7:47485340-47486152 | -1.768196415 | 9.66E-07 | 6.19E-05 |
| chr18:43876424-43877263 | -1.929314607 | 9.74E-07 | 6.23E-05 |

| | | | |
|---|---|---|---|
| chrX:134670189-134670763 | 1.990245889 | 9.74E-07 | 6.23E-05 |
| chr2:193347307-193348080 | -1.878596926 | 9.77E-07 | 6.24E-05 |
| chr3:23580315-23580731 | 2.258774042 | 9.81E-07 | 6.26E-05 |
| chr5:81906947-81907322 | 2.528789276 | 9.88E-07 | 6.30E-05 |
| chr4:80307784-80308470 | -1.906665841 | 9.89E-07 | 6.30E-05 |
| chr13:33218014-33218480 | -1.858173896 | 9.94E-07 | 6.33E-05 |
| chr13:75968119-75968450 | -2.481336236 | 9.97E-07 | 6.34E-05 |
| chr10:129963555-129964543 | 1.813284769 | 9.99E-07 | 6.35E-05 |
| chr9:75101007-75101545 | -1.970794462 | 1.00E-06 | 6.36E-05 |
| chr7:8597275-8597910 | 1.759173218 | 1.00E-06 | 6.37E-05 |
| chr7:146043159-146043835 | -2.100681696 | 1.01E-06 | 6.40E-05 |
| chr1:69972358-69972580 | -3.315318574 | 1.01E-06 | 6.42E-05 |
| chrX:104386614-104387013 | 2.25785556 | 1.02E-06 | 6.44E-05 |
| chr4:66782770-66783080 | -2.809302241 | 1.02E-06 | 6.45E-05 |
| chr4:165476619-165476973 | 2.03165952 | 1.04E-06 | 6.57E-05 |
| chr10:128451707-128452081 | 2.644408793 | 1.04E-06 | 6.59E-05 |
| chr2:49770906-49771442 | -1.957093937 | 1.05E-06 | 6.61E-05 |
| chr7:19788067-19788448 | 2.318985495 | 1.06E-06 | 6.67E-05 |
| chr14:34361356-34361862 | 1.895145671 | 1.06E-06 | 6.67E-05 |
| chr7:142478470-142479081 | -2.08592911 | 1.06E-06 | 6.67E-05 |
| chr2:77753712-77754565 | -1.853010744 | 1.06E-06 | 6.67E-05 |
| chr5:6858646-6858922 | 2.64602793 | 1.07E-06 | 6.72E-05 |
| chr4:153556357-153556752 | 2.283930098 | 1.07E-06 | 6.73E-05 |
| chr7:128021242-128021821 | -1.771126628 | 1.08E-06 | 6.80E-05 |
| chr2:226215940-226216291 | -2.271200675 | 1.09E-06 | 6.82E-05 |
| chr12:26246515-26246881 | -2.345203562 | 1.09E-06 | 6.82E-05 |
| chr4:185591888-185592110 | -3.4879524 | 1.10E-06 | 6.88E-05 |
| chr1:248810636-248811281 | -1.706131843 | 1.12E-06 | 7.02E-05 |
| chr5:28397555-28397983 | -2.234734851 | 1.13E-06 | 7.06E-05 |
| chr10:60527696-60528017 | -2.576212215 | 1.13E-06 | 7.06E-05 |
| chr3:144008445-144008832 | -1.9797865 | 1.13E-06 | 7.06E-05 |
| chr20:48884983-48885337 | 1.775263318 | 1.14E-06 | 7.12E-05 |
| chr15:78291715-78292093 | 1.993372152 | 1.19E-06 | 7.42E-05 |
| chr2:30604674-30605076 | -2.46899273 | 1.19E-06 | 7.42E-05 |
| chr15:89498223-89498677 | 1.885681951 | 1.20E-06 | 7.45E-05 |
| chr12:7036678-7037270 | -2.098308084 | 1.21E-06 | 7.49E-05 |
| chr7:79386999-79387402 | -2.35288682 | 1.22E-06 | 7.54E-05 |
| chr1:55262822-55263302 | -1.922928555 | 1.22E-06 | 7.54E-05 |
| chr2:33395725-33396039 | -2.407197981 | 1.22E-06 | 7.54E-05 |
| chr12:75982254-75982480 | -2.527035524 | 1.22E-06 | 7.56E-05 |
| chr2:70488965-70489662 | -1.83573212 | 1.23E-06 | 7.59E-05 |
| chr4:159455483-159455893 | -2.512191953 | 1.23E-06 | 7.59E-05 |
| chr3:146049854-146050221 | -2.412318319 | 1.23E-06 | 7.59E-05 |
| chr13:75258230-75258519 | -2.652681858 | 1.23E-06 | 7.60E-05 |
| chr2:163123450-163123949 | -1.835386753 | 1.24E-06 | 7.62E-05 |
| chr1:117226085-117226290 | 2.572029629 | 1.25E-06 | 7.70E-05 |
| chr7:149318056-149318596 | -2.003133522 | 1.25E-06 | 7.70E-05 |
| chr7:129595074-129595370 | -2.440776154 | 1.25E-06 | 7.71E-05 |
| chr3:32890529-32890866 | 2.413442258 | 1.27E-06 | 7.80E-05 |
| chr5:6193289-6193675 | -2.089886207 | 1.27E-06 | 7.80E-05 |

| | | | |
|---|---|---|---|
| chr1:8913653-8913880 | 3.059020944 | 1.27E-06 | 7.82E-05 |
| chr5:126748055-126748410 | -2.171742014 | 1.29E-06 | 7.91E-05 |
| chr14:26427698-26428097 | 2.445872785 | 1.29E-06 | 7.91E-05 |
| chr3:9970039-9970746 | 1.799198663 | 1.30E-06 | 7.94E-05 |
| chr8:96419409-96420127 | 1.696420834 | 1.30E-06 | 7.96E-05 |
| chr3:166788958-166789402 | 2.049138397 | 1.34E-06 | 8.20E-05 |
| chr12:79452744-79453419 | -1.781958707 | 1.35E-06 | 8.21E-05 |
| chr3:40417891-40418459 | 1.906956496 | 1.35E-06 | 8.21E-05 |
| chr10:112992243-112992621 | -2.516114095 | 1.35E-06 | 8.25E-05 |
| chr13:75226058-75226475 | -2.024135786 | 1.36E-06 | 8.31E-05 |
| chr14:84643476-84643857 | 2.35744308 | 1.37E-06 | 8.32E-05 |
| chr1:27968741-27968977 | 2.708924387 | 1.37E-06 | 8.36E-05 |
| chr4:160318943-160319791 | -1.681228078 | 1.38E-06 | 8.41E-05 |
| chr2:167715949-167716313 | -2.362288832 | 1.39E-06 | 8.41E-05 |
| chr6:89298058-89298314 | -2.88557721 | 1.39E-06 | 8.42E-05 |
| chr13:22945885-22946253 | 2.239497881 | 1.40E-06 | 8.48E-05 |
| chr3:81237267-81237524 | -2.89818551 | 1.40E-06 | 8.48E-05 |
| chr10:58515278-58515598 | -2.455876188 | 1.40E-06 | 8.48E-05 |
| chr1:82386541-82387331 | -1.821604425 | 1.40E-06 | 8.48E-05 |
| chr4:149997772-149998488 | -1.822134044 | 1.40E-06 | 8.48E-05 |
| chr14:75222348-75222732 | 2.317552821 | 1.41E-06 | 8.52E-05 |
| chr21:31425227-31425653 | 2.009243153 | 1.42E-06 | 8.54E-05 |
| chr8:89100870-89101266 | 2.323061256 | 1.42E-06 | 8.58E-05 |
| chr2:74897870-74898216 | 2.325289775 | 1.44E-06 | 8.66E-05 |
| chr4:44016565-44016930 | -2.563318097 | 1.44E-06 | 8.66E-05 |
| chr4:133565384-133565890 | -1.870698931 | 1.44E-06 | 8.66E-05 |
| chr7:15494952-15495375 | 2.176184643 | 1.44E-06 | 8.68E-05 |
| chr14:105162437-105162721 | 1.920016225 | 1.45E-06 | 8.69E-05 |
| chr8:93193513-93194055 | 1.992523907 | 1.46E-06 | 8.75E-05 |
| chr6:48994775-48995240 | -1.915264726 | 1.46E-06 | 8.76E-05 |
| chr12:52857382-52857664 | 2.55123283 | 1.47E-06 | 8.83E-05 |
| chr7:158495681-158496089 | -2.122469902 | 1.48E-06 | 8.84E-05 |
| chr5:143659538-143660195 | 2.059201661 | 1.48E-06 | 8.87E-05 |
| chr1:74485183-74485717 | 1.942521945 | 1.49E-06 | 8.90E-05 |
| chr2:18435470-18435910 | -2.359678735 | 1.50E-06 | 8.96E-05 |
| chr21:29785664-29786056 | 2.8777154 | 1.50E-06 | 8.97E-05 |
| chr2:139487878-139488622 | -1.920558437 | 1.51E-06 | 8.99E-05 |
| chr17:60879002-60879282 | 2.316007224 | 1.51E-06 | 8.99E-05 |
| chr2:232563619-232563957 | 2.539774096 | 1.52E-06 | 9.05E-05 |
| chr1:184393108-184393533 | -1.862085737 | 1.52E-06 | 9.06E-05 |
| chr12:6451402-6452156 | -1.874592714 | 1.52E-06 | 9.06E-05 |
| chrX:40721521-40721836 | -3.392195744 | 1.53E-06 | 9.07E-05 |
| chr14:106517630-106517994 | 2.621467088 | 1.53E-06 | 9.07E-05 |
| chr1:177571564-177572086 | -1.931735028 | 1.55E-06 | 9.17E-05 |
| chr13:35714126-35715060 | -1.662133045 | 1.58E-06 | 9.38E-05 |
| chr7:128783155-128783860 | -1.717468226 | 1.58E-06 | 9.38E-05 |
| chr8:81233454-81233662 | 3.413599944 | 1.59E-06 | 9.41E-05 |
| chr12:90090966-90091608 | -2.003850316 | 1.61E-06 | 9.54E-05 |
| chr9:89881115-89881407 | 2.588555163 | 1.62E-06 | 9.55E-05 |
| chr6:142822800-142823534 | -1.741851394 | 1.62E-06 | 9.55E-05 |

| | | | |
|---|---|---|---|
| chr10:16799320-16799674 | -2.501713103 | 1.62E-06 | 9.58E-05 |
| chr4:176785767-176786042 | -2.602551178 | 1.63E-06 | 9.60E-05 |
| chr1:117190562-117190813 | 2.682799504 | 1.64E-06 | 9.68E-05 |
| chr9:17134834-17135197 | 2.658080879 | 1.66E-06 | 9.76E-05 |
| chr15:53252072-53252379 | 1.826045815 | 1.66E-06 | 9.77E-05 |
| chr1:210250686-210250965 | -2.089825841 | 1.66E-06 | 9.77E-05 |
| chr17:39723492-39724005 | -1.938387653 | 1.66E-06 | 9.77E-05 |
| chr13:61508862-61509201 | -2.3472292 | 1.67E-06 | 9.80E-05 |
| chr7:11352786-11353026 | 3.006351594 | 1.69E-06 | 9.89E-05 |
| chr7:16495381-16495659 | 2.219735809 | 1.70E-06 | 9.97E-05 |
| chr16:31404488-31404856 | 2.053694593 | 1.70E-06 | 9.97E-05 |
| chr16:68069923-68070182 | 2.671798139 | 1.72E-06 | 0.000100525 |
| chr7:143372091-143372400 | -2.463038634 | 1.74E-06 | 0.000101936 |
| chr13:65214479-65214892 | -2.335020725 | 1.75E-06 | 0.000102002 |
| chr19:36915916-36916395 | 2.205865048 | 1.75E-06 | 0.000102007 |
| chr13:41610319-41610794 | -2.035358764 | 1.75E-06 | 0.000102256 |
| chr5:56358580-56359465 | -1.836312969 | 1.76E-06 | 0.000102414 |
| chr8:18490849-18491171 | 2.597446623 | 1.77E-06 | 0.000103139 |
| chr7:7737722-7738003 | 2.079674393 | 1.78E-06 | 0.000103667 |
| chr5:72379744-72380363 | -1.88161543 | 1.80E-06 | 0.00010454 |
| chr12:53432648-53433040 | 2.485025628 | 1.80E-06 | 0.000104569 |
| chr3:34174710-34175130 | 1.997969506 | 1.81E-06 | 0.000104786 |
| chr6:38133958-38134305 | 2.082475525 | 1.81E-06 | 0.000105045 |
| chr17:13420925-13421385 | -2.204880027 | 1.82E-06 | 0.000105527 |
| chr11:79059861-79060913 | -1.598497844 | 1.82E-06 | 0.000105626 |
| chr5:2659177-2659739 | -2.046172324 | 1.83E-06 | 0.000106084 |
| chr5:124735233-124735784 | -2.121821944 | 1.84E-06 | 0.000106523 |
| chrX:150739581-150739846 | 2.768734659 | 1.84E-06 | 0.000106523 |
| chr1:235295749-235296403 | 1.678303418 | 1.86E-06 | 0.000107271 |
| chr11:117314315-117314562 | 2.539705589 | 1.87E-06 | 0.000107946 |
| chr1:191513167-191513544 | -2.199787338 | 1.88E-06 | 0.000108183 |
| chr1:119786303-119786856 | 1.830776175 | 1.88E-06 | 0.000108364 |
| chr4:74944198-74944599 | 2.138516515 | 1.89E-06 | 0.000108625 |
| chr10:20330336-20331009 | -1.795809137 | 1.89E-06 | 0.000108795 |
| chr13:95226106-95226444 | 2.296990835 | 1.89E-06 | 0.000108903 |
| chr8:93935990-93936317 | 2.042630205 | 1.91E-06 | 0.000109773 |
| chr2:61970882-61971587 | -1.870722712 | 1.92E-06 | 0.000110126 |
| chr11:85723131-85723558 | -2.12293184 | 1.92E-06 | 0.000110244 |
| chr9:88270893-88271236 | 2.232544663 | 1.93E-06 | 0.00011052 |
| chr12:72520026-72520730 | -1.680753224 | 1.94E-06 | 0.000111219 |
| chr7:18805244-18805549 | 2.582623272 | 1.95E-06 | 0.000111701 |
| chr5:174482600-174483152 | -1.873315129 | 1.96E-06 | 0.000111853 |
| chr11:15618453-15618935 | 1.841123289 | 1.96E-06 | 0.000111889 |
| chr3:66729611-66730271 | -1.802637094 | 1.98E-06 | 0.000112984 |
| chr4:28709711-28710332 | -2.094770791 | 1.98E-06 | 0.000113045 |
| chr11:126610413-126610741 | -2.167034196 | 1.99E-06 | 0.000113255 |
| chr3:117762560-117763050 | 2.051877482 | 1.99E-06 | 0.000113255 |
| chr1:197391866-197392194 | -2.143957937 | 1.99E-06 | 0.000113444 |
| chr7:22667837-22668332 | 1.782385707 | 2.00E-06 | 0.000113741 |
| chr3:60775250-60775574 | -2.360858577 | 2.01E-06 | 0.000113989 |

| | | | |
|---|---|---|---|
| chr4:185654357-185654680 | -2.285054537 | 2.01E-06 | 0.000113989 |
| chrX:35622055-35622316 | -2.996450038 | 2.04E-06 | 0.000115608 |
| chr6:164826931-164827922 | -1.80082967 | 2.04E-06 | 0.000115834 |
| chr14:105138675-105139051 | 2.09023829 | 2.04E-06 | 0.000115834 |
| chr7:116393557-116393962 | -2.838772872 | 2.05E-06 | 0.000115834 |
| chr20:44954246-44954731 | 1.947013596 | 2.05E-06 | 0.000115834 |
| chr15:89150990-89151327 | 2.195275783 | 2.05E-06 | 0.000115834 |
| chr10:3881765-3882175 | 2.106538451 | 2.05E-06 | 0.000115834 |
| chr8:124181392-124181714 | 2.141982124 | 2.05E-06 | 0.000115887 |
| chr1:196622968-196623431 | -2.189399449 | 2.06E-06 | 0.00011632 |
| chr8:116926442-116927498 | -1.662615064 | 2.07E-06 | 0.000116534 |
| chr8:94984630-94984981 | 2.048342327 | 2.07E-06 | 0.000116608 |
| chr8:18491176-18491384 | 3.148544404 | 2.07E-06 | 0.000116639 |
| chr7:19560599-19560803 | 3.720969876 | 2.08E-06 | 0.000116934 |
| chr14:55316609-55316863 | 2.674523439 | 2.10E-06 | 0.000118142 |
| chr15:52227398-52227789 | 1.639566619 | 2.11E-06 | 0.000118735 |
| chr10:1257834-1258126 | 2.748560523 | 2.14E-06 | 0.000120374 |
| chr15:20340945-20341221 | 2.294574279 | 2.16E-06 | 0.000120902 |
| chr7:12221614-12221888 | 2.729960526 | 2.16E-06 | 0.000120902 |
| chr3:28198143-28198404 | 2.74117176 | 2.17E-06 | 0.000121357 |
| chr9:115262412-115262877 | -1.90845836 | 2.17E-06 | 0.000121357 |
| chr15:50685981-50687334 | 1.516593155 | 2.18E-06 | 0.000122083 |
| chr5:68434065-68434584 | -1.859277685 | 2.19E-06 | 0.000122226 |
| chr7:133846434-133846851 | -2.446717691 | 2.19E-06 | 0.000122226 |
| chr4:86977499-86978104 | -1.703861571 | 2.19E-06 | 0.000122437 |
| chr6:3380705-3381023 | 2.164259101 | 2.21E-06 | 0.000123023 |
| chr8:100238118-100238365 | 2.842586082 | 2.23E-06 | 0.00012427 |
| chr5:58085860-58086178 | -2.184438106 | 2.24E-06 | 0.000124455 |
| chr16:64478175-64478755 | 1.996054257 | 2.25E-06 | 0.000124948 |
| chr5:124748705-124749108 | -1.764276039 | 2.25E-06 | 0.000124948 |
| chr8:116195754-116196330 | -1.947828623 | 2.25E-06 | 0.000125229 |
| chr16:87602018-87602343 | 2.319729282 | 2.27E-06 | 0.000125839 |
| chr1:179756695-179756957 | 2.997451523 | 2.27E-06 | 0.000125969 |
| chrX:119552548-119552883 | 2.43635547 | 2.28E-06 | 0.000126625 |
| chr3:169534493-169534930 | 1.905411236 | 2.29E-06 | 0.000126881 |
| chr10:43316859-43317231 | 2.122850266 | 2.29E-06 | 0.000126904 |
| chr4:74374539-74375493 | -1.687152413 | 2.30E-06 | 0.000127311 |
| chr9:98218822-98219185 | 2.169439778 | 2.31E-06 | 0.000127899 |
| chr3:168424294-168424781 | 2.057819136 | 2.33E-06 | 0.000128443 |
| chr16:79677954-79678458 | -1.790927934 | 2.33E-06 | 0.000128571 |
| chr18:55988584-55988981 | -2.584882309 | 2.33E-06 | 0.000128571 |
| chr3:52999246-52999567 | -2.513166497 | 2.33E-06 | 0.000128668 |
| chr13:83780163-83780585 | -2.151938186 | 2.36E-06 | 0.000129891 |
| chr1:15795170-15795695 | 2.040548165 | 2.36E-06 | 0.000129952 |
| chr12:6295058-6295551 | -2.002873069 | 2.36E-06 | 0.000129988 |
| chr19:33447715-33447980 | 2.968716926 | 2.36E-06 | 0.000129988 |
| chr7:139095284-139095649 | -2.378321282 | 2.38E-06 | 0.000130511 |
| chr7:23129179-23129599 | 2.109676493 | 2.38E-06 | 0.000130864 |
| chr10:103490580-103491055 | -1.832888793 | 2.39E-06 | 0.000131165 |
| chr2:17214506-17214997 | -2.061778784 | 2.39E-06 | 0.00013119 |

| | | | |
|---|---|---|---|
| chr2:215749213-215750124 | -1.736796531 | 2.40E-06 | 0.000131558 |
| chr12:7959746-7959956 | -3.009990666 | 2.41E-06 | 0.000131774 |
| chr2:40725020-40725528 | -1.853150187 | 2.41E-06 | 0.000131896 |
| chr7:8305967-8306245 | 2.21376217 | 2.42E-06 | 0.000132209 |
| chr7:19650973-19651284 | 2.34029014 | 2.43E-06 | 0.000132869 |
| chr1:188170016-188170340 | 2.44748511 | 2.45E-06 | 0.000133655 |
| chr4:76536080-76536543 | 2.248243331 | 2.45E-06 | 0.000133737 |
| chr21:36596231-36596525 | 2.832999334 | 2.46E-06 | 0.000134165 |
| chr8:144620501-144620781 | 2.560904048 | 2.47E-06 | 0.000134737 |
| chr2:12456264-12456818 | -1.850377078 | 2.48E-06 | 0.000135104 |
| chr9:73978932-73979608 | -1.881862906 | 2.50E-06 | 0.000135999 |
| chr18:62942548-62942933 | -2.172056045 | 2.50E-06 | 0.000135999 |
| chr2:152466068-152466347 | -2.512041185 | 2.51E-06 | 0.000136279 |
| chr14:105624700-105625622 | 1.655237361 | 2.53E-06 | 0.000137025 |
| chr12:49331633-49332132 | 1.837441405 | 2.53E-06 | 0.00013732 |
| chr6:10201920-10202802 | -1.74538847 | 2.54E-06 | 0.000137831 |
| chr5:61245575-61246438 | -1.650568448 | 2.56E-06 | 0.000138383 |
| chr6:116557846-116558466 | 1.722092028 | 2.57E-06 | 0.000138818 |
| chr8:62690047-62690502 | -2.033394415 | 2.57E-06 | 0.000138947 |
| chr5:11547782-11548144 | -1.988519489 | 2.59E-06 | 0.000139861 |
| chr3:89628913-89629745 | -1.684759442 | 2.59E-06 | 0.000139861 |
| chr5:77620438-77621067 | -1.736602983 | 2.59E-06 | 0.000139861 |
| chr19:15250126-15250517 | -2.400560107 | 2.61E-06 | 0.000140546 |
| chr6:139635570-139636672 | -1.679736194 | 2.61E-06 | 0.000140546 |
| chr7:85577236-85577612 | -2.951149646 | 2.62E-06 | 0.00014102 |
| chr10:93915599-93915885 | -2.521693803 | 2.64E-06 | 0.00014205 |
| chr5:139392652-139392909 | 2.129130893 | 2.65E-06 | 0.000142224 |
| chr19:35569303-35569589 | 2.644549024 | 2.66E-06 | 0.0001429 |
| chr5:166145407-166145809 | -2.328585851 | 2.66E-06 | 0.000142946 |
| chr1:230802770-230803036 | 2.745334035 | 2.67E-06 | 0.000143152 |
| chr8:124166116-124166432 | 2.084597205 | 2.68E-06 | 0.000143717 |
| chr7:113421709-113422175 | -1.769737384 | 2.69E-06 | 0.00014401 |
| chr18:78192042-78192365 | -2.5088089 | 2.69E-06 | 0.00014401 |
| chr8:86714803-86715205 | 2.043532137 | 2.70E-06 | 0.000144351 |
| chr14:87430785-87431075 | 2.559694516 | 2.70E-06 | 0.00014447 |
| chr16:2698297-2698562 | 2.94785918 | 2.71E-06 | 0.000144601 |
| chr1:70587987-70588464 | 1.997171362 | 2.71E-06 | 0.000144601 |
| chr12:107765430-107765630 | 2.395286695 | 2.72E-06 | 0.000144973 |
| chr8:140149643-140150083 | 1.93367347 | 2.73E-06 | 0.00014538 |
| chr13:54614813-54615179 | -2.228963051 | 2.74E-06 | 0.000146164 |
| chr7:18208901-18209425 | 1.864234142 | 2.76E-06 | 0.000146807 |
| chr18:48996947-48997362 | -1.844879663 | 2.79E-06 | 0.000148204 |
| chr15:50742688-50742888 | 2.153763477 | 2.80E-06 | 0.000148737 |
| chr15:54420354-54420672 | 2.63040684 | 2.82E-06 | 0.000149854 |
| chr15:52678131-52679966 | 1.489203417 | 2.83E-06 | 0.000150115 |
| chrX:14409404-14409640 | -3.67313004 | 2.85E-06 | 0.000151169 |
| chr3:116939613-116939900 | -2.563311902 | 2.86E-06 | 0.000151624 |
| chr7:10727153-10727401 | 2.804958204 | 2.86E-06 | 0.000151739 |
| chr7:19279964-19280654 | 2.052099675 | 2.87E-06 | 0.000151822 |
| chr7:122825886-122826120 | -3.326054367 | 2.88E-06 | 0.000152485 |

| | | | |
|---|---|---|---|
| chr7:153835221-153835475 | -3.102324793 | 2.88E-06 | 0.000152485 |
| chr1:7885600-7885882 | 2.490804902 | 2.89E-06 | 0.000152552 |
| chr10:22825839-22826521 | -1.860914052 | 2.89E-06 | 0.000152752 |
| chr3:6769671-6770238 | -1.86372199 | 2.91E-06 | 0.000153392 |
| chr1:223030402-223030758 | 2.080198703 | 2.91E-06 | 0.000153392 |
| chr3:44870471-44870925 | 2.150092437 | 2.92E-06 | 0.000153941 |
| chr7:93117898-93118212 | -2.153247301 | 2.93E-06 | 0.000154498 |
| chr22:42734335-42734815 | -1.823060295 | 2.94E-06 | 0.00015495 |
| chr4:6201120-6201407 | 2.230999101 | 2.95E-06 | 0.000154985 |
| chr3:107992869-107993752 | -1.565846426 | 2.98E-06 | 0.000156401 |
| chrX:145111710-145112094 | 2.330154201 | 2.99E-06 | 0.000156734 |
| chr18:69815019-69815393 | -2.387778638 | 2.99E-06 | 0.000156734 |
| chr13:65065628-65066191 | -1.947330567 | 2.99E-06 | 0.000156734 |
| chr4:41157118-41157795 | -1.82195665 | 2.99E-06 | 0.000156734 |
| chr7:8239629-8240239 | 1.744147364 | 2.99E-06 | 0.000156734 |
| chr12:132204375-132204580 | 2.644172589 | 3.00E-06 | 0.000156891 |
| chr4:132737368-132737673 | -2.327590066 | 3.00E-06 | 0.000157029 |
| chr7:27921211-27921481 | 2.484644147 | 3.02E-06 | 0.000157979 |
| chr18:51614986-51615743 | -1.702217194 | 3.03E-06 | 0.000158064 |
| chr7:131071785-131072418 | -1.706084433 | 3.04E-06 | 0.000158508 |
| chr18:76686683-76688035 | -1.574373871 | 3.04E-06 | 0.000158633 |
| chr5:30099615-30100119 | 1.932845691 | 3.05E-06 | 0.000159074 |
| chrX:36234828-36235826 | -1.769501856 | 3.06E-06 | 0.000159423 |
| chr18:41631712-41632307 | -2.191428651 | 3.06E-06 | 0.000159503 |
| chr7:12328078-12328369 | 2.326770285 | 3.10E-06 | 0.000161162 |
| chr7:115965045-115965499 | -2.313742419 | 3.10E-06 | 0.000161237 |
| chr10:123750952-123751276 | 2.114038414 | 3.11E-06 | 0.000161552 |
| chr6:96104955-96105267 | 2.542378177 | 3.11E-06 | 0.000161552 |
| chr1:246281752-246281956 | 2.443243351 | 3.14E-06 | 0.000162935 |
| chr11:27367830-27368303 | -1.926197563 | 3.15E-06 | 0.000163125 |
| chr4:163747836-163748285 | -2.018259293 | 3.17E-06 | 0.000164262 |
| chr1:211606892-211607182 | -2.833611187 | 3.18E-06 | 0.000164682 |
| chr18:69484833-69485077 | -2.833611187 | 3.18E-06 | 0.000164682 |
| chr15:55882558-55882823 | 2.389426664 | 3.19E-06 | 0.000164682 |
| chr13:28469054-28469604 | 2.109630495 | 3.20E-06 | 0.000165198 |
| chr4:40989632-40990027 | 2.055438702 | 3.20E-06 | 0.000165198 |
| chr6:77368669-77369312 | -1.896831159 | 3.21E-06 | 0.000165881 |
| chr1:151603070-151603484 | 2.036102088 | 3.22E-06 | 0.000166042 |
| chr15:53893387-53893913 | 1.676565575 | 3.22E-06 | 0.000166042 |
| chr8:89558925-89559162 | 2.334587963 | 3.24E-06 | 0.000166996 |
| chr13:104648286-104648571 | -2.571326494 | 3.25E-06 | 0.000167103 |
| chr6:45239594-45240153 | -2.353829074 | 3.26E-06 | 0.0001674 |
| chr6:57792712-57793719 | -1.800379474 | 3.26E-06 | 0.000167554 |
| chr2:190718617-190719464 | -1.607095486 | 3.28E-06 | 0.000168402 |
| chr1:70874095-70874529 | -1.912075659 | 3.28E-06 | 0.000168402 |
| chr3:72255904-72256480 | -1.757463976 | 3.29E-06 | 0.000168987 |
| chr9:78201213-78201764 | -1.891158098 | 3.30E-06 | 0.000169092 |
| chr15:58521608-58522008 | 1.896797181 | 3.31E-06 | 0.000169313 |
| chr8:61767084-61767571 | 1.955943591 | 3.31E-06 | 0.000169335 |
| chr15:52727114-52727476 | 1.76397054 | 3.32E-06 | 0.000169698 |

| | | | |
|---|---|---|---|
| chr2:11389610-11390068 | 2.055346701 | 3.33E-06 | 0.000170249 |
| chr5:115770487-115770831 | -2.120884132 | 3.33E-06 | 0.000170352 |
| chr15:55279884-55280098 | 2.62615516 | 3.36E-06 | 0.000171376 |
| chr3:11653632-11654011 | 2.046184772 | 3.37E-06 | 0.000172032 |
| chr11:30521685-30521989 | -2.242664832 | 3.38E-06 | 0.000172437 |
| chrX:67700357-67700963 | 1.782099199 | 3.40E-06 | 0.000173113 |
| chr6:71121890-71122708 | -1.865478823 | 3.41E-06 | 0.000173362 |
| chr2:22317126-22317429 | -2.486453638 | 3.45E-06 | 0.000175471 |
| chr15:74245362-74245863 | 1.785633641 | 3.46E-06 | 0.000176067 |
| chr19:37334206-37334893 | 1.683577346 | 3.47E-06 | 0.00017611 |
| chr5:129399061-129399373 | -2.324274851 | 3.51E-06 | 0.00017847 |
| chr10:132503983-132504183 | 2.81263931 | 3.52E-06 | 0.000178714 |
| chr6:109679137-109679689 | -1.854840705 | 3.53E-06 | 0.000178714 |
| chr4:166850375-166851146 | -1.679540132 | 3.53E-06 | 0.000178714 |
| chr3:160869789-160870701 | -1.576856753 | 3.53E-06 | 0.000178714 |
| chr6:129801909-129802304 | 1.941320058 | 3.54E-06 | 0.00017916 |
| chr2:199435942-199436157 | -3.33189463 | 3.56E-06 | 0.000179895 |
| chr14:43042425-43042733 | 2.424836559 | 3.56E-06 | 0.000180114 |
| chr3:106685556-106686271 | -1.621669331 | 3.57E-06 | 0.00018022 |
| chr15:53071755-53072566 | 1.517293933 | 3.57E-06 | 0.000180275 |
| chr18:44537595-44537966 | -2.174320867 | 3.59E-06 | 0.00018089 |
| chr15:53634756-53635016 | 1.968248414 | 3.65E-06 | 0.000184184 |
| chr7:54725148-54725593 | -2.224733223 | 3.66E-06 | 0.000184207 |
| chr5:42950344-42950981 | 1.693288308 | 3.66E-06 | 0.000184299 |
| chr6:116229195-116229655 | -1.731130938 | 3.66E-06 | 0.000184299 |
| chr7:80563591-80564569 | -1.616597347 | 3.67E-06 | 0.000184299 |
| chr18:48991304-48992251 | -1.64647161 | 3.69E-06 | 0.000185542 |
| chr13:33208207-33209090 | -1.619335145 | 3.70E-06 | 0.000185542 |
| chr20:2805921-2806567 | -1.781175891 | 3.70E-06 | 0.000185759 |
| chr20:48463800-48464289 | 1.592697489 | 3.71E-06 | 0.000186106 |
| chr22:45048935-45049225 | 3.314153095 | 3.74E-06 | 0.000187325 |
| chr14:35179957-35180289 | 2.461564596 | 3.80E-06 | 0.000190245 |
| chrX:27987702-27988343 | -1.85708942 | 3.83E-06 | 0.000191608 |
| chr8:90485911-90486126 | 2.68820808 | 3.84E-06 | 0.000191854 |
| chr14:44166651-44166886 | 2.932909597 | 3.85E-06 | 0.000192188 |
| chr6:134707346-134707735 | -2.060633709 | 3.86E-06 | 0.000192578 |
| chr13:61685572-61685992 | -1.931732324 | 3.89E-06 | 0.000194306 |
| chr13:38674289-38674664 | -2.021456944 | 3.90E-06 | 0.000194705 |
| chr17:75301996-75302198 | 2.470757318 | 3.93E-06 | 0.00019592 |
| chr8:100692648-100692932 | 2.338134536 | 3.94E-06 | 0.000196114 |
| chr7:139894030-139894334 | -2.450388366 | 3.94E-06 | 0.000196114 |
| chr10:54801087-54801591 | -1.798675076 | 3.94E-06 | 0.000196227 |
| chr4:159677560-159677921 | -2.393030499 | 3.96E-06 | 0.000196835 |
| chr14:31652570-31652842 | 3.081120099 | 3.96E-06 | 0.000196844 |
| chr4:186038013-186038470 | -2.191783321 | 3.97E-06 | 0.000197087 |
| chr10:13567070-13567382 | -2.614788691 | 3.99E-06 | 0.000198082 |
| chr4:167055251-167056214 | -1.631182326 | 4.00E-06 | 0.000198285 |
| chr2:63186361-63186703 | -2.233594769 | 4.00E-06 | 0.000198306 |
| chr15:55882891-55883366 | 1.98863739 | 4.02E-06 | 0.000199065 |
| chr2:142693517-142694106 | -1.793649566 | 4.06E-06 | 0.000200753 |

| | | | |
|---|---|---|---|
| chr1:114033948-114034279 | 1.648029038 | 4.07E-06 | 0.000201162 |
| chr5:155402991-155403905 | -1.665740008 | 4.07E-06 | 0.000201162 |
| chr18:48997844-48998650 | -1.646851642 | 4.08E-06 | 0.000201442 |
| chr16:3983074-3983401 | 1.981768262 | 4.15E-06 | 0.000204682 |
| chr7:16195385-16195644 | 2.663872774 | 4.16E-06 | 0.000205124 |
| chr14:95547299-95547758 | -1.860429755 | 4.20E-06 | 0.000207317 |
| chr15:52287821-52289026 | 1.479507949 | 4.22E-06 | 0.000207969 |
| chr6:137824096-137824706 | -1.888842224 | 4.22E-06 | 0.000207969 |
| chr9:129245465-129245770 | 2.66141675 | 4.24E-06 | 0.000208786 |
| chr4:74583684-74584205 | -1.851399604 | 4.25E-06 | 0.000208862 |
| chr12:17773788-17774449 | -1.970173572 | 4.26E-06 | 0.000209375 |
| chr12:101209801-101210514 | -1.749590825 | 4.26E-06 | 0.000209434 |
| chr3:133551279-133552025 | -1.73830241 | 4.28E-06 | 0.000210167 |
| chr11:65035295-65035539 | 2.689527003 | 4.32E-06 | 0.000211815 |
| chr1:210151173-210151440 | -2.457664072 | 4.34E-06 | 0.000212759 |
| chr7:21024001-21024286 | 2.573071611 | 4.35E-06 | 0.00021293 |
| chr3:169658348-169658809 | -1.703008265 | 4.36E-06 | 0.0002136 |
| chr5:128037004-128037509 | -1.784507175 | 4.39E-06 | 0.000214544 |
| chr3:430071-430519 | 2.096802557 | 4.39E-06 | 0.000214544 |
| chr15:94791774-94792835 | -1.554050034 | 4.41E-06 | 0.000215333 |
| chr7:7795704-7796099 | 1.945745391 | 4.41E-06 | 0.000215366 |
| chr7:98287650-98287995 | -1.894809565 | 4.41E-06 | 0.000215366 |
| chr5:77984583-77985037 | -2.097788406 | 4.42E-06 | 0.000215618 |
| chr5:42931998-42932471 | 1.891967819 | 4.45E-06 | 0.000217056 |
| chr8:66524704-66524980 | -1.905104917 | 4.48E-06 | 0.000218115 |
| chr1:228280353-228280553 | 2.685921823 | 4.48E-06 | 0.000218115 |
| chr11:71555576-71555962 | 2.230434056 | 4.49E-06 | 0.000218227 |
| chr22:44318655-44318961 | 2.584307681 | 4.49E-06 | 0.000218227 |
| chr7:87732383-87733352 | -1.632793746 | 4.49E-06 | 0.000218283 |
| chr13:60144341-60144541 | -3.164635425 | 4.50E-06 | 0.000218515 |
| chr3:52359611-52359862 | 2.664141914 | 4.52E-06 | 0.000219423 |
| chr4:177367318-177367700 | -1.90868502 | 4.54E-06 | 0.000219941 |
| chr1:23699827-23700102 | -2.534773998 | 4.55E-06 | 0.000220383 |
| chr6:11805270-11805593 | -2.025038641 | 4.59E-06 | 0.000222318 |
| chr7:13929827-13930087 | 2.887445345 | 4.60E-06 | 0.000222497 |
| chr2:163755026-163755340 | 2.178316638 | 4.60E-06 | 0.000222497 |
| chr3:8035527-8035831 | 2.578942894 | 4.61E-06 | 0.000223005 |
| chr1:93779568-93780238 | -1.737142633 | 4.63E-06 | 0.000223498 |
| chr14:105766321-105767097 | 1.670280137 | 4.64E-06 | 0.000224151 |
| chr1:191363267-191363968 | -1.584467265 | 4.65E-06 | 0.000224151 |
| chr4:40238886-40239221 | 1.930483344 | 4.67E-06 | 0.000225149 |
| chr7:15117573-15117917 | 2.3697041 | 4.71E-06 | 0.000227193 |
| chr5:156835963-156836285 | -2.204667595 | 4.73E-06 | 0.000227578 |
| chr7:51324505-51324823 | -2.042602703 | 4.73E-06 | 0.000227637 |
| chr12:25377517-25377890 | -2.257306369 | 4.74E-06 | 0.000227933 |
| chr11:66686886-66687151 | 2.185718162 | 4.75E-06 | 0.000228244 |
| chr7:24510946-24511246 | 1.994092594 | 4.78E-06 | 0.000229709 |
| chr1:213464899-213465484 | -2.024444746 | 4.79E-06 | 0.000229964 |
| chr19:35327769-35328027 | 2.269864503 | 4.81E-06 | 0.000230753 |
| chr7:127383094-127383653 | -1.91670984 | 4.82E-06 | 0.000231346 |

| | | | |
|---|---|---|---|
| chr5:56319249-56319702 | -1.942161871 | 4.83E-06 | 0.00023136 |
| chr5:173497702-173497949 | 2.304483999 | 4.83E-06 | 0.00023136 |
| chr1:246797614-246798000 | -1.779406948 | 4.85E-06 | 0.000232089 |
| chr1:210272740-210273116 | -1.925137632 | 4.85E-06 | 0.000232119 |
| chr7:21914581-21914822 | 2.161726373 | 4.87E-06 | 0.000232588 |
| chr5:76773-77013 | 2.446859019 | 4.89E-06 | 0.000233522 |
| chr16:21539329-21539611 | 2.765123594 | 4.92E-06 | 0.000234943 |
| chr13:73194139-73194681 | -1.954143681 | 4.92E-06 | 0.00023495 |
| chr9:130549177-130549442 | 2.03754146 | 4.93E-06 | 0.000235151 |
| chr7:111445548-111445830 | -2.231281507 | 4.95E-06 | 0.000235682 |
| chr10:6423255-6423664 | 1.981601122 | 4.99E-06 | 0.000237753 |
| chr3:175787439-175788041 | 2.091370366 | 5.01E-06 | 0.000238258 |
| chr12:62022595-62022916 | 2.52623691 | 5.01E-06 | 0.000238352 |
| chr7:131546030-131546424 | -1.946767249 | 5.03E-06 | 0.000238918 |
| chr7:138776183-138776831 | -1.727056515 | 5.04E-06 | 0.000239336 |
| chr8:96702084-96702339 | 2.385565947 | 5.07E-06 | 0.000240515 |
| chr2:80149943-80150231 | 2.568284251 | 5.07E-06 | 0.0002406 |
| chr2:187986176-187986825 | -1.776087069 | 5.08E-06 | 0.0002406 |
| chr3:175136025-175137043 | -1.668939243 | 5.08E-06 | 0.0002406 |
| chr16:12024499-12024808 | 2.24275962 | 5.09E-06 | 0.000240932 |
| chr15:51576856-51577101 | 1.828164274 | 5.09E-06 | 0.000240932 |
| chr5:96648673-96649069 | -1.811846349 | 5.13E-06 | 0.000242579 |
| chr7:121328677-121330082 | -1.534186506 | 5.16E-06 | 0.00024383 |
| chr9:1101053-1101382 | 2.062260028 | 5.16E-06 | 0.000243966 |
| chr7:125833896-125834195 | -2.469445478 | 5.18E-06 | 0.000244727 |
| chr5:17312222-17312697 | 1.769763212 | 5.19E-06 | 0.000244978 |
| chr4:94430458-94430812 | 2.175777643 | 5.20E-06 | 0.000244978 |
| chr8:106176157-106176383 | 3.619102959 | 5.20E-06 | 0.000244978 |
| chr14:37903561-37903771 | 2.510499763 | 5.24E-06 | 0.000246812 |
| chr17:56173992-56174643 | -1.652567292 | 5.25E-06 | 0.000247014 |
| chr11:37650888-37651188 | -2.788458268 | 5.26E-06 | 0.000247565 |
| chr3:183966285-183966871 | 1.750692985 | 5.30E-06 | 0.000249115 |
| chr18:76553977-76554272 | -2.51156554 | 5.31E-06 | 0.000249397 |
| chr15:54486135-54486380 | 2.56694605 | 5.31E-06 | 0.000249397 |
| chr3:131601445-131601811 | 2.139715136 | 5.32E-06 | 0.000249397 |
| chr13:59247270-59247575 | -2.386806436 | 5.35E-06 | 0.000250669 |
| chr1:220404657-220405017 | 2.448930755 | 5.38E-06 | 0.000252317 |
| chr12:106793109-106793951 | -1.832615438 | 5.39E-06 | 0.000252625 |
| chr18:76619621-76620592 | -1.56964453 | 5.41E-06 | 0.000253218 |
| chr6:82569802-82570486 | -1.612249778 | 5.43E-06 | 0.000253914 |
| chr7:7791219-7791438 | 2.409922386 | 5.44E-06 | 0.000254288 |
| chr8:140588589-140589557 | -1.582793749 | 5.44E-06 | 0.000254359 |
| chr16:76871403-76871832 | 1.869134027 | 5.48E-06 | 0.000255872 |
| chr7:21330043-21330272 | 2.564377639 | 5.49E-06 | 0.000256306 |
| chr19:54179862-54180510 | -1.610925259 | 5.50E-06 | 0.000256609 |
| chr13:53556482-53556887 | -2.107238945 | 5.51E-06 | 0.000256609 |
| chr7:51123698-51124077 | -1.902334477 | 5.52E-06 | 0.000257017 |
| chr14:77699027-77699312 | 2.293525035 | 5.52E-06 | 0.000257017 |
| chr2:186561283-186561826 | -1.864484727 | 5.53E-06 | 0.000257038 |
| chr15:90095527-90096159 | 1.647055345 | 5.55E-06 | 0.000258115 |

| | | | |
|---|---|---|---|
| chr15:54605601-54605976 | 1.858430456 | 5.60E-06 | 0.00026011 |
| chr3:84906030-84906430 | 2.11695414 | 5.62E-06 | 0.000260929 |
| chr8:85418009-85418253 | 2.868918324 | 5.63E-06 | 0.000261106 |
| chr21:21048068-21048590 | -2.06359477 | 5.63E-06 | 0.000261106 |
| chr7:33532116-33532414 | 2.15281162 | 5.64E-06 | 0.00026125 |
| chr1:113993626-113994080 | 1.620094864 | 5.67E-06 | 0.000262494 |
| chr7:24079807-24080075 | 2.560721727 | 5.67E-06 | 0.000262494 |
| chr7:17692364-17692609 | 2.56020633 | 5.72E-06 | 0.000264571 |
| chr18:48152538-48153055 | -2.078894594 | 5.77E-06 | 0.000266718 |
| chr2:27752726-27753358 | -1.740460594 | 5.78E-06 | 0.000266944 |
| chr5:122382912-122383666 | -1.832621347 | 5.81E-06 | 0.000268465 |
| chr1:76177614-76178870 | -1.69879098 | 5.82E-06 | 0.000268465 |
| chr5:119360938-119361447 | -1.785694828 | 5.82E-06 | 0.000268542 |
| chr11:88095994-88096341 | -2.445324706 | 5.82E-06 | 0.000268544 |
| chr1:199127067-199127267 | 3.077857936 | 5.86E-06 | 0.00026989 |
| chr7:24492570-24492825 | 2.140635483 | 5.90E-06 | 0.000271699 |
| chr3:79291030-79291396 | 2.654158499 | 5.94E-06 | 0.00027334 |
| chr6:20366656-20366939 | -2.631985658 | 5.94E-06 | 0.00027334 |
| chr16:77677171-77677490 | 1.90926872 | 5.95E-06 | 0.00027334 |
| chr6:12485189-12485636 | 2.017843543 | 5.98E-06 | 0.000274714 |
| chr2:201357234-201357562 | -1.908992977 | 5.98E-06 | 0.000274714 |
| chr18:66155720-66155991 | -2.90578644 | 5.99E-06 | 0.000274714 |
| chr4:166169175-166169485 | -2.094118822 | 6.02E-06 | 0.000276096 |
| chr13:61665146-61665900 | -1.604824799 | 6.03E-06 | 0.00027625 |
| chr8:86165711-86165972 | 2.202826886 | 6.03E-06 | 0.00027635 |
| chr3:8483260-8483704 | 1.916188656 | 6.06E-06 | 0.000277381 |
| chr15:54460210-54460561 | 2.377388156 | 6.07E-06 | 0.000277728 |
| chr10:61592433-61592918 | -1.897400402 | 6.08E-06 | 0.000278058 |
| chrX:23807160-23808042 | -1.544153947 | 6.09E-06 | 0.000278058 |
| chr2:148594190-148594477 | 2.483849477 | 6.10E-06 | 0.000278325 |
| chr3:27559851-27560368 | -1.707481981 | 6.13E-06 | 0.000279528 |
| chr1:208090349-208090953 | -1.668754497 | 6.24E-06 | 0.000284692 |
| chr12:53796585-53796861 | 2.897710725 | 6.26E-06 | 0.000285565 |
| chr4:108264692-108264964 | 2.110927761 | 6.27E-06 | 0.000285625 |
| chr14:97016366-97016695 | 2.198650254 | 6.27E-06 | 0.000285629 |
| chr7:17474220-17474499 | 2.626100742 | 6.29E-06 | 0.000286388 |
| chr7:14761664-14762050 | 2.195762314 | 6.33E-06 | 0.000287621 |
| chr1:151885529-151885791 | 2.114609728 | 6.34E-06 | 0.0002881 |
| chr19:52601515-52602205 | 1.819723479 | 6.34E-06 | 0.0002881 |
| chr7:47786478-47787094 | -1.748840629 | 6.35E-06 | 0.000288197 |
| chr10:68288940-68289256 | -2.428822587 | 6.36E-06 | 0.000288667 |
| chr7:113118060-113119202 | -1.557374134 | 6.40E-06 | 0.000290345 |
| chr7:153518179-153518932 | -1.722668905 | 6.41E-06 | 0.000290452 |
| chr17:62807882-62808920 | -1.509922399 | 6.42E-06 | 0.000290738 |
| chr2:168056242-168056765 | -1.744992663 | 6.46E-06 | 0.000292571 |
| chr3:194370438-194370731 | 2.312311595 | 6.50E-06 | 0.000293814 |
| chr1:188432454-188433020 | -1.768562598 | 6.54E-06 | 0.000295556 |
| chr11:70664228-70664428 | 2.570606761 | 6.56E-06 | 0.000296365 |
| chr10:19232451-19232783 | -2.365290234 | 6.60E-06 | 0.00029806 |
| chr7:139603245-139603742 | -1.886622102 | 6.62E-06 | 0.000298801 |

| | | | |
|---|---|---|---|
| chr10:96432508-96432713 | 2.901567136 | 6.64E-06 | 0.000299664 |
| chr5:124888162-124888431 | -2.615057467 | 6.67E-06 | 0.000300608 |
| chr18:63890247-63890689 | -2.0222189 | 6.68E-06 | 0.000300698 |
| chr7:8304210-8304467 | 2.093945212 | 6.69E-06 | 0.000301052 |
| chr2:79105323-79105611 | -2.424702181 | 6.70E-06 | 0.000301284 |
| chr3:106699109-106699605 | -1.821377236 | 6.70E-06 | 0.000301284 |
| chr13:78605258-78605677 | -1.825505952 | 6.75E-06 | 0.000303422 |
| chr15:88439401-88439655 | 2.563925261 | 6.76E-06 | 0.000303743 |
| chr4:76724017-76724246 | -2.881331698 | 6.77E-06 | 0.000303998 |
| chr4:39568898-39569210 | 2.405903873 | 6.80E-06 | 0.000305196 |
| chr5:13945750-13946889 | -1.560392786 | 6.82E-06 | 0.000305824 |
| chr5:39712943-39713166 | 2.146537149 | 6.88E-06 | 0.000308264 |
| chr7:121995003-121995309 | -2.204312544 | 6.91E-06 | 0.00030964 |
| chr15:50614310-50614510 | 2.051102654 | 6.92E-06 | 0.00030964 |
| chr7:116511479-116512461 | -1.673952239 | 6.92E-06 | 0.00030964 |
| chr10:119309184-119309425 | 2.544671947 | 6.95E-06 | 0.000310235 |
| chr15:72170170-72170826 | -1.641671441 | 6.95E-06 | 0.000310235 |
| chr6:9810672-9811394 | -1.908620472 | 6.95E-06 | 0.000310235 |
| chr8:89208464-89208699 | 2.524870822 | 6.95E-06 | 0.000310235 |
| chr6:113763298-113763572 | 2.735714741 | 6.96E-06 | 0.000310235 |
| chr1:245372706-245373004 | 2.483645466 | 7.01E-06 | 0.000312314 |
| chr1:196674799-196675551 | -1.61950236 | 7.04E-06 | 0.000313589 |
| chr17:57016686-57017174 | 1.64863084 | 7.05E-06 | 0.00031397 |
| chr16:89772158-89772358 | 2.757581234 | 7.10E-06 | 0.000315799 |
| chr1:100006365-100006765 | -2.032025411 | 7.18E-06 | 0.00031909 |
| chr7:7903637-7903837 | 2.842480921 | 7.18E-06 | 0.000319174 |
| chr10:56622390-56622833 | -2.223946525 | 7.22E-06 | 0.000320849 |
| chr17:82972925-82973324 | 1.891779325 | 7.25E-06 | 0.000321856 |
| chr12:125161064-125161302 | 2.739731421 | 7.28E-06 | 0.000322894 |
| chr5:105616537-105617074 | -1.788158367 | 7.30E-06 | 0.000323698 |
| chr7:34004695-34005116 | 2.021835265 | 7.31E-06 | 0.000323787 |
| chr10:98314438-98314913 | -1.787267155 | 7.31E-06 | 0.000323787 |
| chr15:82597863-82598217 | 1.943921179 | 7.33E-06 | 0.000324277 |
| chr2:240946390-240947215 | -1.576605507 | 7.33E-06 | 0.000324277 |
| chrX:25404478-25404724 | -3.233313279 | 7.34E-06 | 0.000324277 |
| chr2:159362662-159362927 | 2.861747441 | 7.34E-06 | 0.000324277 |
| chr19:38768815-38769015 | 3.241705467 | 7.35E-06 | 0.000324329 |
| chr13:40102772-40103304 | -1.682996052 | 7.35E-06 | 0.000324329 |
| chr7:16509724-16510389 | 1.696527437 | 7.37E-06 | 0.00032505 |
| chr11:106552845-106553084 | -3.01509907 | 7.37E-06 | 0.00032505 |
| chr1:87423906-87424204 | -2.898636757 | 7.39E-06 | 0.000325537 |
| chr7:126803110-126803395 | -2.237239571 | 7.41E-06 | 0.000325944 |
| chrX:25288737-25289112 | -2.237239571 | 7.41E-06 | 0.000325944 |
| chr12:9454044-9454363 | -2.255375614 | 7.41E-06 | 0.00032614 |
| chr5:149887747-149888194 | -1.924341821 | 7.42E-06 | 0.00032626 |
| chr18:3230354-3230870 | -1.608317677 | 7.44E-06 | 0.000326749 |
| chr1:248171958-248172495 | -1.988861787 | 7.44E-06 | 0.000326749 |
| chr7:158647177-158647571 | -1.799936509 | 7.46E-06 | 0.00032743 |
| chr13:67382745-67383007 | -2.897364219 | 7.47E-06 | 0.00032743 |
| chr12:124594954-124595154 | 2.636297635 | 7.53E-06 | 0.000329884 |

| | | | |
|---|---|---|---|
| chr6:48658909-48659451 | -1.878498686 | 7.54E-06 | 0.000329884 |
| chr8:88678824-88679072 | 2.258746416 | 7.54E-06 | 0.000329884 |
| chr11:25625158-25625519 | 2.186326346 | 7.54E-06 | 0.000329884 |
| chr7:109905308-109905587 | -2.836828923 | 7.56E-06 | 0.000330616 |
| chr14:56755026-56755347 | 2.035272551 | 7.59E-06 | 0.000331722 |
| chr7:115231196-115231518 | -2.604459314 | 7.61E-06 | 0.000332515 |
| chr15:29064392-29064731 | 2.341867668 | 7.62E-06 | 0.000332515 |
| chr3:115658757-115659218 | -1.863801692 | 7.64E-06 | 0.000333305 |
| chr14:96065564-96066203 | 1.551276 | 7.67E-06 | 0.000334357 |
| chr4:52840979-52841273 | 1.995483151 | 7.76E-06 | 0.000338272 |
| chr8:11631140-11631587 | -2.157563346 | 7.77E-06 | 0.000338272 |
| chr3:20597583-20598053 | 1.874958224 | 7.77E-06 | 0.000338307 |
| chr11:25971199-25971399 | -3.061714242 | 7.83E-06 | 0.000340637 |
| chr4:185235742-185236247 | -1.827562743 | 7.84E-06 | 0.000340863 |
| chr1:8763876-8764385 | -1.889089352 | 7.85E-06 | 0.000341384 |
| chr9:119855377-119855662 | 2.524927383 | 7.88E-06 | 0.000342239 |
| chr13:92573474-92573801 | -2.100932302 | 7.89E-06 | 0.000342286 |
| chr3:172292917-172293217 | 2.300258423 | 7.89E-06 | 0.000342286 |
| chr3:20400272-20400626 | -1.849230109 | 7.91E-06 | 0.00034301 |
| chr22:32765745-32766001 | 3.009503856 | 7.93E-06 | 0.000343687 |
| chr6:57800216-57800480 | -3.242803843 | 7.99E-06 | 0.000345955 |
| chr13:90145291-90145876 | -1.933509309 | 8.02E-06 | 0.000347032 |
| chr12:64816439-64816724 | 1.945699697 | 8.02E-06 | 0.000347143 |
| chr10:62048901-62049484 | -1.624396066 | 8.03E-06 | 0.000347178 |
| chr3:13233479-13233757 | 1.992232547 | 8.05E-06 | 0.000347771 |
| chr1:150139253-150139596 | 2.034097272 | 8.08E-06 | 0.000348869 |
| chr10:43574255-43574778 | 1.688679621 | 8.08E-06 | 0.000348869 |
| chr1:53982507-53982812 | -2.029262464 | 8.10E-06 | 0.000349396 |
| chr6:148885677-148885976 | 2.714061702 | 8.12E-06 | 0.000350189 |
| chr9:75890425-75890761 | -2.119507185 | 8.21E-06 | 0.000353811 |
| chr1:81097888-81098367 | -2.020671429 | 8.22E-06 | 0.000354216 |
| chr7:71251705-71252104 | -1.976819107 | 8.24E-06 | 0.000354651 |
| chr12:114813147-114813715 | 1.646810351 | 8.27E-06 | 0.000355784 |
| chr22:43343266-43343530 | -2.711536566 | 8.28E-06 | 0.000356145 |
| chr3:50605154-50605545 | -1.993865133 | 8.32E-06 | 0.000357531 |
| chr4:8794110-8794458 | 2.282481382 | 8.36E-06 | 0.000358906 |
| chr7:10649237-10649712 | 2.023497201 | 8.39E-06 | 0.000360214 |
| chr8:116937911-116938944 | -1.501426499 | 8.40E-06 | 0.000360214 |
| chr8:13365084-13365575 | -1.933831427 | 8.41E-06 | 0.000360477 |
| chr12:15243058-15243927 | -1.695852131 | 8.41E-06 | 0.000360477 |
| chr7:50069616-50070233 | -1.682728015 | 8.42E-06 | 0.000360477 |
| chr7:101805053-101805518 | -1.883177661 | 8.42E-06 | 0.000360477 |
| chr5:123438888-123439418 | -1.831732728 | 8.43E-06 | 0.000360775 |
| chr3:19826996-19827315 | 2.719376158 | 8.50E-06 | 0.000363596 |
| chr5:79193204-79194386 | -1.556715293 | 8.54E-06 | 0.000364937 |
| chr4:83172794-83173047 | 2.823014529 | 8.54E-06 | 0.000364937 |
| chr13:108728369-108728722 | -2.094723953 | 8.57E-06 | 0.000365934 |
| chr3:150149722-150150138 | 1.802845883 | 8.58E-06 | 0.000365934 |
| chr6:104602681-104602913 | 2.513755502 | 8.59E-06 | 0.000366026 |
| chr8:103723516-103723942 | 1.845745316 | 8.59E-06 | 0.000366026 |

| | | | |
|---|---|---|---|
| chr6:148307376-148307786 | 1.811730048 | 8.65E-06 | 0.000368431 |
| chr7:35345916-35346179 | 2.60470951 | 8.67E-06 | 0.000369218 |
| chr14:29513004-29513237 | 2.511111047 | 8.78E-06 | 0.00037366 |
| chr3:189293138-189293527 | -1.850776722 | 8.83E-06 | 0.000375412 |
| chr19:37990420-37990648 | 2.142172349 | 8.85E-06 | 0.00037594 |
| chr7:113628997-113629761 | -1.656763144 | 8.86E-06 | 0.000376227 |
| chr18:31684766-31686069 | 1.516204666 | 8.86E-06 | 0.000376227 |
| chr2:227781087-227781555 | -2.043867931 | 8.87E-06 | 0.000376227 |
| chr7:21542755-21543873 | 1.524677303 | 8.88E-06 | 0.000376514 |
| chr1:81254523-81254850 | -1.89252531 | 8.89E-06 | 0.000376834 |
| chr7:142391274-142391508 | -2.417433179 | 8.91E-06 | 0.000377191 |
| chr13:95226604-95226842 | 2.45596939 | 8.91E-06 | 0.000377191 |
| chr10:132458566-132458810 | 1.829994246 | 8.93E-06 | 0.000377494 |
| chr2:175233475-175233769 | 2.290176242 | 8.93E-06 | 0.000377494 |
| chr1:25210351-25210634 | 2.728391283 | 8.94E-06 | 0.000377978 |
| chr2:231421387-231421723 | 2.327781196 | 8.99E-06 | 0.00037992 |
| chr15:52171493-52171714 | 1.852464225 | 9.06E-06 | 0.000382362 |
| chr3:101801313-101801596 | 2.383135133 | 9.07E-06 | 0.000382774 |
| chr14:62501188-62501433 | 2.6074107 | 9.08E-06 | 0.000382919 |
| chr15:52921403-52921756 | 1.812620539 | 9.10E-06 | 0.000383345 |
| chr14:105084430-105084664 | 2.197810266 | 9.17E-06 | 0.00038617 |
| chr14:97163652-97164296 | 1.677159663 | 9.20E-06 | 0.000387203 |
| chr12:58034388-58035315 | 1.630031274 | 9.34E-06 | 0.000393153 |
| chr10:132359393-132359593 | 2.319742826 | 9.40E-06 | 0.000395389 |
| chr14:59886381-59886651 | 2.522333007 | 9.42E-06 | 0.000395745 |
| chr8:118849958-118850562 | -1.734863569 | 9.42E-06 | 0.000395764 |
| chr2:240095621-240096670 | -1.637540703 | 9.45E-06 | 0.000396627 |
| chr14:34112352-34112622 | 2.159238676 | 9.52E-06 | 0.000399589 |
| chr4:7805599-7805866 | 2.121285985 | 9.53E-06 | 0.000399589 |
| chr1:117043726-117043926 | 2.732581517 | 9.54E-06 | 0.000399701 |
| chr5:161399866-161400415 | -2.054808197 | 9.54E-06 | 0.000399701 |
| chr15:55062438-55062834 | 2.152708534 | 9.60E-06 | 0.00040214 |
| chr2:179665508-179666029 | -1.999300091 | 9.63E-06 | 0.000403191 |
| chr5:126846783-126847097 | -1.976683921 | 9.65E-06 | 0.000403812 |
| chr11:42979299-42979922 | -1.672014499 | 9.66E-06 | 0.000403995 |
| chr7:141231514-141232007 | -1.830650613 | 9.75E-06 | 0.000407519 |
| chr2:89029180-89029509 | -2.19185897 | 9.79E-06 | 0.000408632 |
| chr7:101124810-101125135 | -2.001391906 | 9.81E-06 | 0.000409362 |
| chr6:105608860-105609741 | 1.674777988 | 9.81E-06 | 0.000409362 |
| chr19:35666163-35666534 | 1.926922644 | 9.84E-06 | 0.000410065 |
| chr7:19171292-19171530 | 2.982678984 | 9.84E-06 | 0.000410081 |
| chr14:27069784-27069984 | 2.987431579 | 9.87E-06 | 0.000410877 |
| chrX:98414931-98415220 | 2.368595716 | 9.95E-06 | 0.000414017 |
| chr13:29693619-29693908 | -2.029547709 | 9.97E-06 | 0.000414899 |
| chr2:9708830-9709547 | -1.703121221 | 1.00E-05 | 0.000415849 |
| chr9:106862887-106863248 | -2.131347154 | 1.00E-05 | 0.000417272 |
| chr10:25682200-25682472 | -2.389953161 | 1.01E-05 | 0.000418694 |
| chr2:213279104-213279426 | 2.501079181 | 1.01E-05 | 0.00041961 |
| chr15:52008077-52008307 | 1.928894713 | 1.01E-05 | 0.000420918 |
| chr6:95758281-95759039 | -1.590403194 | 1.02E-05 | 0.000421877 |

| | | | |
|---|---|---|---|
| chr3:8307577-8308014 | -2.110820343 | 1.02E-05 | 0.000422821 |
| chr10:80471650-80472110 | -1.727846783 | 1.02E-05 | 0.000422821 |
| chr16:76716188-76716778 | 1.743982502 | 1.02E-05 | 0.000422891 |
| chr6:134123908-134124287 | -1.73901336 | 1.02E-05 | 0.000423514 |
| chr4:188109556-188109952 | -2.177942512 | 1.03E-05 | 0.000424989 |
| chr14:31263669-31263994 | 1.852789916 | 1.03E-05 | 0.000427271 |
| chr18:71398432-71398919 | -1.964634878 | 1.04E-05 | 0.000427625 |
| chr15:53717015-53717245 | 1.905613003 | 1.04E-05 | 0.000428397 |
| chr15:96344333-96344581 | 2.38834071 | 1.04E-05 | 0.000429859 |
| chr10:11446672-11446882 | 2.822946628 | 1.05E-05 | 0.000431133 |
| chr17:14266341-14266635 | -2.445515381 | 1.05E-05 | 0.000431538 |
| chr5:175355588-175355909 | 2.328545231 | 1.05E-05 | 0.000432088 |
| chr8:96796758-96797052 | 1.931777033 | 1.05E-05 | 0.000432121 |
| chr3:25854196-25854964 | 1.601975832 | 1.06E-05 | 0.000436678 |
| chr7:12943388-12943729 | 2.007848459 | 1.06E-05 | 0.000437959 |
| chr8:141154065-141154490 | 1.640059416 | 1.07E-05 | 0.000438345 |
| chr6:129440753-129441013 | -2.257358707 | 1.07E-05 | 0.000438508 |
| chr4:171308742-171309035 | -2.436042529 | 1.07E-05 | 0.00043901 |
| chr3:10551802-10552403 | -1.69006792 | 1.07E-05 | 0.000440596 |
| chr10:29277313-29277670 | 1.991791559 | 1.08E-05 | 0.000441569 |
| chrX:8342185-8342565 | -2.128249483 | 1.08E-05 | 0.000442889 |
| chr6:125867636-125868051 | -1.771234819 | 1.08E-05 | 0.000443126 |
| chr10:100561963-100562355 | 1.898709823 | 1.08E-05 | 0.000443798 |
| chr3:17136002-17136229 | 2.50204265 | 1.09E-05 | 0.000445905 |
| chr9:106879124-106879332 | -3.187279249 | 1.09E-05 | 0.000447696 |
| chrX:30580661-30580998 | -2.415897048 | 1.09E-05 | 0.000447696 |
| chr7:21944671-21944926 | 1.691591066 | 1.10E-05 | 0.000448519 |
| chr2:200266057-200266461 | -2.443385013 | 1.10E-05 | 0.000448771 |
| chr6:53382654-53383327 | -1.758061404 | 1.10E-05 | 0.000450064 |
| chr8:81439586-81439803 | 3.520935915 | 1.10E-05 | 0.000450296 |
| chr1:107512618-107512869 | 2.828118021 | 1.11E-05 | 0.000451158 |
| chr13:106376735-106377195 | -1.856173303 | 1.11E-05 | 0.000453372 |
| chr8:90751011-90751340 | 1.885458811 | 1.11E-05 | 0.000453544 |
| chr5:102628658-102629150 | -2.062401999 | 1.12E-05 | 0.000454353 |
| chr10:44628471-44628837 | 2.352347609 | 1.12E-05 | 0.000457271 |
| chrX:126190803-126191342 | -1.962583555 | 1.13E-05 | 0.000460271 |
| chr16:35429140-35429603 | 2.255551042 | 1.14E-05 | 0.000461957 |
| chr15:50466605-50466805 | 2.022041256 | 1.14E-05 | 0.000462232 |
| chr6:68575985-68576556 | -1.967899446 | 1.14E-05 | 0.000462777 |
| chr4:127579451-127579918 | 2.024794783 | 1.14E-05 | 0.000462904 |
| chr14:37801502-37801721 | 2.429282983 | 1.14E-05 | 0.000463052 |
| chr10:24466072-24466674 | -1.661993184 | 1.14E-05 | 0.000463933 |
| chr5:18356422-18356832 | 2.099943716 | 1.14E-05 | 0.000463933 |
| chr8:122269666-122270280 | -1.635771297 | 1.15E-05 | 0.000464763 |
| chr6:72522663-72523225 | -1.631247653 | 1.15E-05 | 0.000465122 |
| chr10:121324678-121325202 | 1.695536593 | 1.15E-05 | 0.000465352 |
| chr11:46797633-46797833 | 2.700076614 | 1.15E-05 | 0.000465419 |
| chr13:75617147-75617619 | -1.899143834 | 1.15E-05 | 0.000466198 |
| chr10:72784631-72784908 | -2.324805837 | 1.16E-05 | 0.000468927 |
| chrX:17470179-17470588 | -2.324512199 | 1.16E-05 | 0.000469312 |

| | | | |
|---|---|---|---|
| chrX:95464877-95465274 | 2.047744096 | 1.16E-05 | 0.000469312 |
| chr7:11752080-11752281 | 2.563849857 | 1.17E-05 | 0.000470268 |
| chr7:19320742-19320943 | 3.173239748 | 1.17E-05 | 0.000472904 |
| chr8:91958172-91958416 | 2.573176295 | 1.17E-05 | 0.000472904 |
| chr2:133581368-133581855 | -2.097224567 | 1.18E-05 | 0.00047425 |
| chr5:156543320-156543682 | -2.030123213 | 1.18E-05 | 0.000476114 |
| chr12:91381360-91381774 | -1.743144282 | 1.19E-05 | 0.000477235 |
| chr8:91205204-91205645 | 1.684916646 | 1.19E-05 | 0.000477594 |
| chr12:21419143-21419762 | -1.948623974 | 1.19E-05 | 0.000479494 |
| chr3:8001828-8002028 | 2.676804263 | 1.20E-05 | 0.000480388 |
| chr4:168304874-168305405 | -1.678057877 | 1.20E-05 | 0.000480514 |
| chr2:128519646-128519959 | 2.367085206 | 1.20E-05 | 0.000481731 |
| chr20:48599193-48599558 | 1.531768437 | 1.20E-05 | 0.000482546 |
| chr5:73083073-73083403 | -2.202697072 | 1.20E-05 | 0.000482834 |
| chr14:97281738-97282069 | 2.212418236 | 1.21E-05 | 0.000484194 |
| chr14:71696069-71696300 | 2.83451719 | 1.21E-05 | 0.000484399 |
| chr19:47336734-47337252 | -1.638438143 | 1.23E-05 | 0.000490346 |
| chr2:186748091-186748825 | -1.575102177 | 1.23E-05 | 0.000493458 |
| chr1:96060419-96061110 | -1.666918569 | 1.24E-05 | 0.000495873 |
| chr2:212640475-212640803 | -1.980426186 | 1.25E-05 | 0.000497382 |
| chr2:65123935-65124300 | -2.364829596 | 1.25E-05 | 0.000497565 |
| chr3:9149577-9149901 | 1.974266244 | 1.25E-05 | 0.000499676 |
| chr7:139041366-139041791 | -2.110324573 | 1.26E-05 | 0.000502808 |
| chr6:166330641-166331033 | -2.055402688 | 1.26E-05 | 0.000503392 |
| chr7:112572031-112572501 | -1.737316783 | 1.26E-05 | 0.000503392 |
| chr15:86863955-86864369 | -1.910430761 | 1.27E-05 | 0.000504023 |
| chr12:77027119-77027400 | 2.287091317 | 1.27E-05 | 0.000506627 |
| chr2:118142872-118143144 | -2.421604719 | 1.27E-05 | 0.000506627 |
| chr9:12740703-12741190 | -1.993882134 | 1.27E-05 | 0.000506648 |
| chr21:45513291-45513735 | 1.699264353 | 1.27E-05 | 0.000506648 |
| chr9:131744644-131745265 | 1.57176895 | 1.28E-05 | 0.000506726 |
| chr2:36433294-36433734 | -1.618544509 | 1.28E-05 | 0.000506726 |
| chr16:76277145-76277688 | 1.700395978 | 1.28E-05 | 0.000506726 |
| chr1:229445189-229445535 | 2.360080604 | 1.28E-05 | 0.000507029 |
| chr10:13679597-13680149 | -1.878131483 | 1.28E-05 | 0.000508462 |
| chr1:10890783-10891117 | 1.835007493 | 1.29E-05 | 0.000510987 |
| chr12:53336043-53336247 | 2.487514626 | 1.30E-05 | 0.00051303 |
| chr19:752463-752952 | 1.717972136 | 1.30E-05 | 0.000514928 |
| chr16:46380722-46381159 | 1.752220928 | 1.30E-05 | 0.000514928 |
| chr12:42266890-42267090 | 2.68169958 | 1.30E-05 | 0.000514928 |
| chr9:125567975-125568262 | 2.549507406 | 1.30E-05 | 0.000514928 |
| chr3:102032422-102032697 | -2.344434232 | 1.30E-05 | 0.00051501 |
| chr1:231726350-231726581 | 2.802282866 | 1.31E-05 | 0.000516762 |
| chr7:113763318-113763567 | -2.783235231 | 1.31E-05 | 0.000516869 |
| chr11:131696701-13169711C | -1.923062458 | 1.32E-05 | 0.000520573 |
| chr2:130720129-130720374 | -2.506605206 | 1.33E-05 | 0.000524077 |
| chr12:53804352-53804880 | 1.531073639 | 1.33E-05 | 0.00052501 |
| chr15:69561309-69561639 | 2.002858027 | 1.34E-05 | 0.000526554 |
| chr6:39270042-39270247 | 2.781669144 | 1.34E-05 | 0.000526631 |
| chr3:88747248-88747534 | 2.35883858 | 1.35E-05 | 0.000529358 |

| | | | |
|---|---|---|---|
| chr15:97977088-97977657 | -1.657602009 | 1.35E-05 | 0.000529926 |
| chr7:83475453-83475727 | -2.188123369 | 1.35E-05 | 0.000529926 |
| chr4:76570191-76570734 | 1.64074364 | 1.35E-05 | 0.000530143 |
| chr5:115857899-115858328 | -1.837446142 | 1.36E-05 | 0.000533594 |
| chr13:73369103-73369458 | -2.027974263 | 1.36E-05 | 0.000533594 |
| chr21:43729905-43730115 | 2.479949466 | 1.36E-05 | 0.000533639 |
| chr1:92589061-92589326 | -2.817882396 | 1.37E-05 | 0.000536023 |
| chr19:11988482-11988701 | 2.569759567 | 1.37E-05 | 0.000537741 |
| chr3:184008081-184008408 | 1.965082839 | 1.37E-05 | 0.000538111 |
| chr2:186575349-186575617 | -1.853075864 | 1.38E-05 | 0.000538857 |
| chr7:12942642-12943182 | 1.658718594 | 1.38E-05 | 0.000539997 |
| chr8:20878904-20879479 | -2.133023282 | 1.38E-05 | 0.000541243 |
| chr10:8273085-8273285 | 2.816391565 | 1.39E-05 | 0.000542855 |
| chr11:75326089-75326739 | -1.556781876 | 1.39E-05 | 0.000544599 |
| chr1:81513901-81514408 | -1.962390402 | 1.40E-05 | 0.000544599 |
| chr7:21335371-21335571 | 2.546363039 | 1.40E-05 | 0.000547612 |
| chr4:172666794-172667344 | -1.696524697 | 1.41E-05 | 0.000548006 |
| chr2:39466309-39466660 | -1.893601965 | 1.41E-05 | 0.000549869 |
| chr1:31390009-31390422 | 1.716652333 | 1.41E-05 | 0.000550155 |
| chr3:190265797-190266054 | -2.284423716 | 1.41E-05 | 0.000550155 |
| chr12:19572933-19573415 | -1.995491065 | 1.42E-05 | 0.000550663 |
| chr3:180306616-180307119 | -1.681286953 | 1.42E-05 | 0.000551876 |
| chr16:73065090-73065336 | 2.28296139 | 1.42E-05 | 0.000551876 |
| chr14:37575388-37575747 | 1.613732117 | 1.42E-05 | 0.000551876 |
| chr10:65754158-65754478 | -2.16217784 | 1.42E-05 | 0.000553105 |
| chr12:60215148-60215646 | -1.838046439 | 1.43E-05 | 0.000553197 |
| chr13:91208319-91208602 | -2.673384953 | 1.43E-05 | 0.000553209 |
| chr7:134612151-134612473 | -2.27677201 | 1.43E-05 | 0.000553436 |
| chr14:34930174-34930395 | 2.417377436 | 1.44E-05 | 0.000556048 |
| chr2:155190511-155190763 | -2.49782837 | 1.44E-05 | 0.000556048 |
| chr10:33264220-33264657 | -1.764237792 | 1.44E-05 | 0.000556321 |
| chrX:7086408-7086962 | -1.877669435 | 1.44E-05 | 0.000556452 |
| chr16:76796516-76796800 | 1.929387964 | 1.44E-05 | 0.000556559 |
| chr16:77269488-77269826 | 1.986215079 | 1.44E-05 | 0.000556559 |
| chr3:80016311-80016589 | 2.538172893 | 1.44E-05 | 0.000557418 |
| chr3:64965960-64966167 | 1.79090746 | 1.45E-05 | 0.000558782 |
| chr7:10769763-10770018 | 2.405051787 | 1.45E-05 | 0.000559441 |
| chr15:51213661-51213861 | 1.936271438 | 1.45E-05 | 0.000559441 |
| chr1:195345230-195345551 | -1.825483663 | 1.45E-05 | 0.000559441 |
| chr5:123819403-123820373 | 1.515863254 | 1.46E-05 | 0.000563051 |
| chr4:152278318-152279236 | -1.560932944 | 1.46E-05 | 0.000563051 |
| chr14:105738176-105738691 | 1.678389386 | 1.46E-05 | 0.000563887 |
| chr18:66724699-66725018 | -2.056754087 | 1.47E-05 | 0.000563967 |
| chr8:19421575-19421949 | 1.84697552 | 1.47E-05 | 0.000565266 |
| chrX:118652783-118653162 | 1.893841993 | 1.47E-05 | 0.000565266 |
| chr3:138889694-138889999 | 2.655153517 | 1.47E-05 | 0.000565266 |
| chr5:178585597-178585797 | 2.7732339 | 1.47E-05 | 0.000565632 |
| chr5:150790346-150790820 | -2.074597209 | 1.48E-05 | 0.000566454 |
| chr3:156637553-156638075 | -1.689409855 | 1.48E-05 | 0.000567709 |
| chr4:73080625-73081174 | -1.761112895 | 1.48E-05 | 0.000567709 |

| | | | |
|---|---|---|---|
| chr1:177571193-177571491 | -1.933741511 | 1.48E-05 | 0.000567709 |
| chr7:24166329-24166570 | 2.234870645 | 1.48E-05 | 0.000568299 |
| chr5:154395638-154396106 | -1.66784857 | 1.49E-05 | 0.00056915 |
| chr10:96596603-96597023 | 1.839300267 | 1.49E-05 | 0.00057101 |
| chr5:96651966-96652277 | -1.840691307 | 1.49E-05 | 0.000571796 |
| chr18:34709026-34709543 | -1.821913687 | 1.50E-05 | 0.000574435 |
| chr10:60663879-60664234 | -1.803324759 | 1.50E-05 | 0.000574435 |
| chr11:80281169-80281511 | 2.10954012 | 1.50E-05 | 0.00057453 |
| chr12:50202001-50202570 | 1.653271526 | 1.51E-05 | 0.00057453 |
| chr6:142361489-142362219 | -1.563402925 | 1.51E-05 | 0.00057453 |
| chr15:56261575-56261790 | 2.556188073 | 1.51E-05 | 0.00057568 |
| chr1:203643057-203643686 | -1.516056301 | 1.51E-05 | 0.00057733 |
| chr20:58741363-58741587 | 1.957461438 | 1.52E-05 | 0.000577846 |
| chr10:128626621-128627060 | -1.903576061 | 1.52E-05 | 0.000578404 |
| chrX:147148015-147148231 | 3.144338626 | 1.52E-05 | 0.000579278 |
| chrX:118207822-118208117 | 2.540211008 | 1.53E-05 | 0.00058311 |
| chr10:53517874-53518177 | -2.191529082 | 1.54E-05 | 0.000583867 |
| chr10:53284737-53285154 | -2.013858781 | 1.54E-05 | 0.000584029 |
| chr15:87471051-87471425 | -2.186529724 | 1.55E-05 | 0.00058941 |
| chr3:14243764-14244112 | 1.668258413 | 1.55E-05 | 0.000590035 |
| chr7:114533058-114533578 | -1.615936286 | 1.56E-05 | 0.000593434 |
| chr2:206783812-206784050 | -2.338892065 | 1.56E-05 | 0.000593434 |
| chr16:60553530-60554228 | -1.77092574 | 1.57E-05 | 0.000593676 |
| chr13:79869298-79869730 | -1.734383351 | 1.57E-05 | 0.000596261 |
| chr7:132580344-132580594 | -2.095112356 | 1.57E-05 | 0.000596261 |
| chr13:25148620-25149000 | 2.179989735 | 1.58E-05 | 0.000597543 |
| chr10:114526538-114527281 | -1.551292276 | 1.59E-05 | 0.000599809 |
| chr7:24506193-24506393 | 2.531971496 | 1.59E-05 | 0.000599809 |
| chr11:106152004-106152448 | -1.925533506 | 1.59E-05 | 0.000600374 |
| chr6:157469330-157469866 | -1.758859586 | 1.59E-05 | 0.000602359 |
| chr1:185001290-185002136 | -1.619082256 | 1.60E-05 | 0.000602394 |
| chr14:48937645-48938168 | -1.719143093 | 1.60E-05 | 0.000602394 |
| chr8:128141727-128142553 | -1.403353512 | 1.60E-05 | 0.000602683 |
| chr12:121878299-121878501 | 2.931048247 | 1.61E-05 | 0.000605663 |
| chrX:153722888-153723134 | 1.839943909 | 1.61E-05 | 0.000605918 |
| chr20:41603744-41604292 | 1.651557599 | 1.61E-05 | 0.000606492 |
| chr16:76619425-76619641 | 2.217838259 | 1.63E-05 | 0.000612513 |
| chr13:80399643-80400168 | -1.938273223 | 1.63E-05 | 0.000614717 |
| chr8:100471784-100472675 | 1.484708329 | 1.64E-05 | 0.000617216 |
| chr12:82634755-82635629 | -1.486315046 | 1.65E-05 | 0.000618643 |
| chr8:123661893-123662566 | -1.520641087 | 1.65E-05 | 0.000619222 |
| chr12:49674349-49674622 | 2.10254593 | 1.65E-05 | 0.000621251 |
| chr15:55552880-55553123 | 2.45869156 | 1.65E-05 | 0.000621251 |
| chr5:102154904-102155571 | 1.684573019 | 1.66E-05 | 0.000621887 |
| chr9:112841196-112841645 | 1.950315734 | 1.66E-05 | 0.000622858 |
| chr5:93622276-93622635 | -1.718561684 | 1.68E-05 | 0.00062995 |
| chr9:84620703-84620993 | 2.748810228 | 1.69E-05 | 0.000632059 |
| chr5:32330577-32330891 | 1.845315399 | 1.69E-05 | 0.000632827 |
| chr10:32360354-32360691 | 2.039132223 | 1.69E-05 | 0.00063391 |
| chr4:185470809-185472051 | -1.438014449 | 1.70E-05 | 0.000634522 |

| | | | |
|---|---|---|---|
| chr16:316750-317331 | -1.662476356 | 1.70E-05 | 0.000634522 |
| chr13:101875914-101876463 | -1.849869489 | 1.70E-05 | 0.000636031 |
| chr1:197783278-197783932 | 1.627852301 | 1.70E-05 | 0.000636842 |
| chr9:79664371-79664844 | -2.196231409 | 1.71E-05 | 0.000636842 |
| chr4:162727192-162727506 | -2.066705309 | 1.71E-05 | 0.000638652 |
| chr14:27348714-27349005 | 2.548702872 | 1.71E-05 | 0.000638652 |
| chr8:124872252-124872546 | 2.101928367 | 1.71E-05 | 0.000638766 |
| chr2:191415763-191416224 | 1.748621865 | 1.72E-05 | 0.000640113 |
| chr10:306719-307150 | 1.631917997 | 1.72E-05 | 0.000640113 |
| chr1:107337009-107337354 | 1.623540395 | 1.72E-05 | 0.000640742 |
| chr6:125759047-125759499 | -1.70648327 | 1.72E-05 | 0.000640973 |
| chr3:164194829-164195169 | -2.024024899 | 1.73E-05 | 0.000644172 |
| chr6:11805700-11805949 | -2.37699962 | 1.74E-05 | 0.000646404 |
| chr5:26942547-26942847 | 2.027716003 | 1.74E-05 | 0.000646404 |
| chr5:5891479-5891707 | -2.269568048 | 1.74E-05 | 0.000646404 |
| chr13:71290900-71291647 | -1.512828916 | 1.74E-05 | 0.000646404 |
| chr8:86444552-86445380 | 1.603111039 | 1.74E-05 | 0.000646404 |
| chr14:65027533-65027841 | 1.986098327 | 1.74E-05 | 0.000646713 |
| chr11:124538086-124538477 | -1.956043096 | 1.75E-05 | 0.000648854 |
| chr7:19795887-19796201 | 2.216508105 | 1.76E-05 | 0.000650262 |
| chr2:45752448-45753310 | -1.551554699 | 1.76E-05 | 0.000650555 |
| chr10:42665062-42665857 | -1.545853704 | 1.76E-05 | 0.000650555 |
| chr5:129177455-129178069 | -1.461774098 | 1.76E-05 | 0.00065111 |
| chr17:11520511-11520755 | 2.783256814 | 1.76E-05 | 0.00065111 |
| chr19:4732604-4732965 | 1.852599459 | 1.77E-05 | 0.000653109 |
| chr1:207867286-207867542 | -1.864475702 | 1.77E-05 | 0.000653806 |
| chr1:116853699-116854157 | 1.529138527 | 1.77E-05 | 0.000654159 |
| chr2:186572760-186573065 | -1.954090764 | 1.77E-05 | 0.000654908 |
| chr16:87806602-87807538 | -1.433879298 | 1.78E-05 | 0.000655962 |
| chr8:96409974-96410306 | 2.127630549 | 1.78E-05 | 0.000656789 |
| chr17:72412694-72412898 | 2.454988076 | 1.79E-05 | 0.000658451 |
| chr9:120043477-120043924 | -1.826766184 | 1.79E-05 | 0.000661013 |
| chr14:94450031-94450444 | 1.793958339 | 1.80E-05 | 0.000661013 |
| chr1:244353755-244354211 | -1.754596804 | 1.80E-05 | 0.000661338 |
| chr2:156175830-156176435 | 1.617369079 | 1.80E-05 | 0.000661338 |
| chr4:108367961-108368257 | 2.26348982 | 1.80E-05 | 0.000661871 |
| chr10:90407143-90407594 | 1.699810412 | 1.80E-05 | 0.000661997 |
| chr3:189842680-189843537 | -1.534959773 | 1.81E-05 | 0.000664741 |
| chr7:140443986-140444374 | -1.797141881 | 1.82E-05 | 0.000667137 |
| chr4:186587481-186587739 | -2.632534321 | 1.82E-05 | 0.000667295 |
| chr13:62209658-62209870 | -2.8004893 | 1.82E-05 | 0.000668061 |
| chr15:58323380-58323645 | 2.541726356 | 1.83E-05 | 0.00066909 |
| chr14:26234245-26234500 | 2.554955321 | 1.83E-05 | 0.00066909 |
| chr8:102535724-102536067 | -1.853815417 | 1.83E-05 | 0.000669148 |
| chr9:104928099-104928418 | 2.133313979 | 1.84E-05 | 0.000671907 |
| chr1:39829961-39830510 | 1.660631561 | 1.84E-05 | 0.000672002 |
| chr22:31179278-31179800 | 1.658388237 | 1.85E-05 | 0.000676513 |
| chr1:74614447-74614707 | -2.455252785 | 1.85E-05 | 0.000676935 |
| chr1:106480741-106480991 | 1.715480565 | 1.86E-05 | 0.000678889 |
| chr5:1180924-1181275 | 1.923702002 | 1.86E-05 | 0.000680812 |

| | | | |
|---|---|---|---|
| chr6:142313321-142313620 | -2.525183004 | 1.87E-05 | 0.000681861 |
| chr5:129584354-129585282 | -1.621796298 | 1.88E-05 | 0.000686207 |
| chr19:35216930-35217203 | 2.310148702 | 1.89E-05 | 0.000687964 |
| chr2:16828899-16829320 | -2.12540832 | 1.90E-05 | 0.000692068 |
| chr18:61639961-61640248 | -2.160658805 | 1.90E-05 | 0.000693584 |
| chr2:85713217-85713699 | -1.626671368 | 1.91E-05 | 0.00069533 |
| chr1:177487504-177487855 | -1.873921331 | 1.91E-05 | 0.000695807 |
| chr4:15149890-15150466 | 1.635597839 | 1.91E-05 | 0.000695807 |
| chr3:123360196-123360554 | 2.187069792 | 1.91E-05 | 0.000695826 |
| chr15:97680550-97681173 | -1.674547707 | 1.92E-05 | 0.000698667 |
| chr8:93478282-93478482 | 2.737519606 | 1.93E-05 | 0.000700421 |
| chr17:44987382-44987600 | 2.379652776 | 1.94E-05 | 0.000704394 |
| chr8:22986915-22987541 | 1.596184282 | 1.94E-05 | 0.000705248 |
| chr19:2922882-2923251 | 2.359587458 | 1.95E-05 | 0.000706603 |
| chr16:82045897-82046210 | 2.15740626 | 1.95E-05 | 0.000707219 |
| chr5:124747341-124747607 | -1.94171225 | 1.95E-05 | 0.000707329 |
| chr3:124774310-124774564 | 1.983901788 | 1.96E-05 | 0.000709581 |
| chr7:146018970-146019218 | -2.432835271 | 1.97E-05 | 0.000712076 |
| chr8:105319042-105319520 | 1.784284806 | 1.97E-05 | 0.000712076 |
| chr5:14077626-14077982 | 2.126708262 | 1.97E-05 | 0.000712586 |
| chrX:16832039-16832239 | -2.524000327 | 1.97E-05 | 0.000712833 |
| chrX:84267505-84267929 | -2.296046436 | 1.97E-05 | 0.000713312 |
| chr9:83611556-83611922 | -1.911215343 | 1.98E-05 | 0.000714658 |
| chr12:131404138-131404341 | 2.520861523 | 1.98E-05 | 0.000714835 |
| chr8:93663536-93663740 | 2.158961925 | 1.98E-05 | 0.000715815 |
| chr1:109239469-109241148 | -1.369720983 | 1.99E-05 | 0.000716123 |
| chr2:77086468-77086895 | -2.075909062 | 1.99E-05 | 0.000716123 |
| chr7:127708572-127709222 | -1.67186661 | 1.99E-05 | 0.000717512 |
| chr20:63361711-63361936 | 2.158486379 | 1.99E-05 | 0.000717512 |
| chr13:23790864-23791110 | 2.543685706 | 2.00E-05 | 0.000718576 |
| chr8:6258519-6258948 | -2.034783584 | 2.01E-05 | 0.000721798 |
| chr18:44198214-44198470 | -2.545594615 | 2.01E-05 | 0.000722856 |
| chr10:132434730-132434964 | 1.884226505 | 2.01E-05 | 0.000723356 |
| chr1:240769030-240769260 | -2.456788684 | 2.01E-05 | 0.000723356 |
| chr2:20203594-20203940 | -1.763196874 | 2.02E-05 | 0.000723951 |
| chr2:71379223-71379570 | -2.25434042 | 2.02E-05 | 0.000724803 |
| chr1:177256225-177256769 | -1.559645242 | 2.03E-05 | 0.000727103 |
| chr13:81291406-81291643 | -2.90428362 | 2.03E-05 | 0.000727227 |
| chr15:89067183-89067665 | 1.621944728 | 2.03E-05 | 0.000729014 |
| chr15:39622970-39623245 | 2.439958467 | 2.04E-05 | 0.000729014 |
| chr2:226124542-226124953 | -1.857220934 | 2.04E-05 | 0.000729014 |
| chr7:8474779-8475030 | 2.133080008 | 2.04E-05 | 0.000729014 |
| chr15:56979346-56979610 | 2.253280088 | 2.04E-05 | 0.000729744 |
| chr15:50423798-50424806 | 1.366882093 | 2.04E-05 | 0.000729744 |
| chr19:57947555-57948004 | 1.681407756 | 2.04E-05 | 0.000729744 |
| chr8:43099610-43100073 | 1.918846745 | 2.04E-05 | 0.000730522 |
| chr14:78523103-78523390 | 2.440999208 | 2.05E-05 | 0.000733291 |
| chr17:57286495-57287045 | -1.542017659 | 2.06E-05 | 0.000733776 |
| chr5:136788187-136788807 | -1.898580893 | 2.06E-05 | 0.00073423 |
| chr14:34599208-34599457 | 2.122742367 | 2.06E-05 | 0.00073423 |

| | | | |
|---|---|---|---|
| chr12:25500765-25501315 | -1.663610087 | 2.07E-05 | 0.000737398 |
| chr19:35903659-35904052 | 1.679616668 | 2.07E-05 | 0.000737657 |
| chr3:34367603-34367935 | 2.060442414 | 2.08E-05 | 0.000742407 |
| chr7:80940622-80941025 | -2.067569535 | 2.10E-05 | 0.000746554 |
| chr14:75150409-75150762 | 2.002735052 | 2.10E-05 | 0.000748719 |
| chr16:4538726-4538969 | 2.149503293 | 2.11E-05 | 0.000748719 |
| chr2:81408350-81408939 | -1.839500236 | 2.11E-05 | 0.000748719 |
| chr9:113884906-113885321 | -1.700370567 | 2.11E-05 | 0.000750194 |
| chr4:164731915-164732187 | -2.236507194 | 2.11E-05 | 0.000750194 |
| chr3:12627092-12627318 | 2.354628539 | 2.12E-05 | 0.000752081 |
| chr8:69161505-69162235 | -1.476774151 | 2.12E-05 | 0.000753441 |
| chr7:112782533-112783101 | -1.911606485 | 2.13E-05 | 0.000754868 |
| chr7:14562214-14562460 | 2.353859531 | 2.13E-05 | 0.000755287 |
| chr4:76200979-76201233 | -2.296622996 | 2.14E-05 | 0.000756672 |
| chr18:13542251-13542665 | 1.926215656 | 2.14E-05 | 0.000759782 |
| chr11:134385671-134386064 | 1.795487696 | 2.15E-05 | 0.000761316 |
| chr14:29586871-29587095 | 2.523392432 | 2.15E-05 | 0.000761546 |
| chr12:67519828-67520135 | 1.903706926 | 2.17E-05 | 0.000767022 |
| chr12:49734413-49734641 | 2.623776691 | 2.17E-05 | 0.000768435 |
| chr10:62844246-62844638 | -1.969589182 | 2.18E-05 | 0.000769712 |
| chr7:254247-254633 | 1.617847523 | 2.18E-05 | 0.000770137 |
| chr4:2743293-2743604 | 1.876043202 | 2.19E-05 | 0.000772002 |
| chr8:93829987-93830382 | 1.600629226 | 2.19E-05 | 0.000772373 |
| chrX:138886810-138887173 | 1.920908194 | 2.19E-05 | 0.000772373 |
| chr7:14162210-14162410 | 3.103911 | 2.20E-05 | 0.000774221 |
| chr8:111738927-111739217 | 2.187103545 | 2.20E-05 | 0.000775517 |
| chr5:17418066-17418395 | 2.168336553 | 2.20E-05 | 0.000775517 |
| chr8:37142765-37143166 | -1.664396978 | 2.20E-05 | 0.000776006 |
| chr5:122390253-122390540 | -2.283326413 | 2.20E-05 | 0.000776006 |
| chr10:132306928-132307280 | 1.559548429 | 2.22E-05 | 0.000779267 |
| chr21:40074775-40075123 | -1.706902674 | 2.22E-05 | 0.000780997 |
| chr10:8829653-8830068 | 2.018906288 | 2.23E-05 | 0.000782718 |
| chr2:146734089-146734570 | -1.916838302 | 2.23E-05 | 0.000783601 |
| chr8:137070116-137070407 | 2.594088065 | 2.24E-05 | 0.000785993 |
| chr10:27319693-27320317 | -1.839732604 | 2.24E-05 | 0.000787904 |
| chr7:19508318-19508580 | 2.506335344 | 2.25E-05 | 0.000788604 |
| chr16:81302713-81302937 | 2.26635572 | 2.25E-05 | 0.000788604 |
| chr5:115389859-115390145 | -1.894275376 | 2.25E-05 | 0.000788771 |
| chr2:64606339-64607369 | -1.40949779 | 2.25E-05 | 0.000789085 |
| chr7:127486714-127487080 | -2.137951562 | 2.25E-05 | 0.000789085 |
| chr8:65283220-65283610 | 2.362866066 | 2.26E-05 | 0.000790059 |
| chr20:38346405-38346848 | 2.04580605 | 2.26E-05 | 0.000790799 |
| chr20:56132823-56133127 | 2.04580605 | 2.26E-05 | 0.000790799 |
| chr17:56394040-56394299 | 2.147558494 | 2.27E-05 | 0.000792916 |
| chr11:90240023-90240917 | -1.487319122 | 2.28E-05 | 0.000794647 |
| chr10:5547926-5548492 | 1.595878285 | 2.28E-05 | 0.000794647 |
| chr7:19129529-19129829 | 2.507455436 | 2.28E-05 | 0.000795574 |
| chr7:12220868-12221116 | 2.113281552 | 2.28E-05 | 0.000795574 |
| chr6:165691517-165691996 | -1.925192091 | 2.29E-05 | 0.000798528 |
| chr12:43471189-43471473 | -2.538135029 | 2.29E-05 | 0.000799904 |

| | | | |
|---|---|---|---|
| chr7:103065674-103066108 | -1.713634701 | 2.31E-05 | 0.00080404 |
| chr8:22738793-22739619 | -1.421247873 | 2.31E-05 | 0.00080404 |
| chr8:102078706-102078995 | 2.237492694 | 2.31E-05 | 0.00080404 |
| chr7:142159035-142159539 | -1.865211137 | 2.31E-05 | 0.00080404 |
| chr6:133745649-133746500 | -1.497707271 | 2.31E-05 | 0.00080404 |
| chr2:125224791-125225452 | -1.659045339 | 2.31E-05 | 0.00080404 |
| chr1:152959128-152959353 | 2.627211312 | 2.32E-05 | 0.000806244 |
| chr2:200184987-200185332 | -1.95640167 | 2.32E-05 | 0.000806821 |
| chr15:95319218-95319597 | 2.180892246 | 2.34E-05 | 0.000811064 |
| chr3:107289392-107289662 | 2.044377918 | 2.35E-05 | 0.000813918 |
| chr7:19988468-19988767 | 2.593112747 | 2.37E-05 | 0.000821038 |
| chr16:34594971-34595343 | 1.559000907 | 2.38E-05 | 0.000823453 |
| chr17:82127691-82127935 | 2.425610824 | 2.39E-05 | 0.000826996 |
| chr1:207668768-207669054 | 2.078322788 | 2.39E-05 | 0.00082953 |
| chr7:158818259-158818998 | -1.513721202 | 2.41E-05 | 0.000835814 |
| chr9:137407492-137407819 | 2.278860084 | 2.42E-05 | 0.000837252 |
| chr7:118637405-118637874 | -1.987792286 | 2.42E-05 | 0.000837252 |
| chr7:48291495-48292098 | -1.648126197 | 2.43E-05 | 0.00084077 |
| chr12:50273973-50274173 | 2.915788582 | 2.43E-05 | 0.000841071 |
| chr12:52280917-52281142 | 2.094901578 | 2.44E-05 | 0.00084165 |
| chr10:52701672-52702040 | -1.820595433 | 2.44E-05 | 0.000841812 |
| chr2:96350684-96350924 | 2.72874502 | 2.44E-05 | 0.000842835 |
| chr4:19455833-19456160 | 1.95243916 | 2.44E-05 | 0.000843492 |
| chr8:92974272-92974660 | 2.06587203 | 2.45E-05 | 0.000845081 |
| chr3:1583070-1583513 | 1.934861599 | 2.45E-05 | 0.000845425 |
| chr7:110442298-110442679 | -1.731953156 | 2.46E-05 | 0.00084826 |
| chr10:64379819-64380234 | -1.725537995 | 2.47E-05 | 0.000850577 |
| chr8:133216379-133216779 | 1.753849909 | 2.47E-05 | 0.000851839 |
| chr7:18788226-18788581 | 2.190765929 | 2.48E-05 | 0.00085367 |
| chr14:46691750-46692187 | 1.662676894 | 2.49E-05 | 0.000855653 |
| chr2:186570508-186571096 | -1.545757221 | 2.50E-05 | 0.000859815 |
| chr10:79779351-79779647 | -2.349038532 | 2.50E-05 | 0.000860128 |
| chr6:152115762-152115981 | -2.214255975 | 2.51E-05 | 0.000860347 |
| chr7:114530031-114530454 | -1.873349145 | 2.51E-05 | 0.000861474 |
| chr5:119355678-119356171 | -1.647466373 | 2.51E-05 | 0.000862211 |
| chr7:50353396-50354120 | 1.750137802 | 2.51E-05 | 0.000862211 |
| chr1:80441710-80442023 | -2.179689937 | 2.53E-05 | 0.000865965 |
| chr7:124785239-124785515 | -2.047827649 | 2.53E-05 | 0.000865965 |
| chr7:12943863-12944090 | 2.06910445 | 2.54E-05 | 0.000869306 |
| chr3:23626171-23626486 | 2.579623081 | 2.54E-05 | 0.00086994 |
| chr3:14551392-14551674 | 1.802075527 | 2.55E-05 | 0.000871379 |
| chr4:173209349-173210363 | -1.417179629 | 2.55E-05 | 0.00087247 |
| chr17:15334486-15334742 | 2.51988625 | 2.55E-05 | 0.000873424 |
| chr7:23285596-23285944 | 1.800992789 | 2.56E-05 | 0.000876439 |
| chr4:1222461-1222714 | 2.335448937 | 2.57E-05 | 0.000876568 |
| chr7:25073028-25073248 | 2.486366628 | 2.57E-05 | 0.000876755 |
| chr13:76242911-76243670 | -1.570393913 | 2.58E-05 | 0.00088029 |
| chrX:31352940-31353392 | -1.897833472 | 2.58E-05 | 0.00088029 |
| chr7:123148857-123149311 | -1.869356271 | 2.58E-05 | 0.000881364 |
| chr7:116801098-116801649 | -1.650955508 | 2.59E-05 | 0.00088242 |

| | | | |
|---|---|---|---|
| chr4:123547029-123547799 | 1.456425348 | 2.60E-05 | 0.000886252 |
| chr5:102450303-102450653 | -1.860934032 | 2.60E-05 | 0.000886847 |
| chr15:80953523-80954755 | -1.43476422 | 2.61E-05 | 0.000887434 |
| chr1:194214193-194215093 | 1.528714009 | 2.61E-05 | 0.000887434 |
| chrX:105328781-105329256 | -1.797281522 | 2.62E-05 | 0.000890934 |
| chr7:122217089-122217423 | -2.264877613 | 2.62E-05 | 0.000891973 |
| chr1:113837667-113837966 | 1.64136537 | 2.63E-05 | 0.000892537 |
| chr3:97493742-97494128 | 2.266361557 | 2.63E-05 | 0.000892804 |
| chr12:29428038-29428310 | -2.190443951 | 2.63E-05 | 0.000892953 |
| chr4:2930680-2930880 | 2.062020963 | 2.64E-05 | 0.000894704 |
| chr16:327072-327744 | -1.559747953 | 2.64E-05 | 0.000896226 |
| chr16:79776306-79776984 | -1.621092612 | 2.64E-05 | 0.000896296 |
| chr10:121597543-121598801 | 1.445712869 | 2.65E-05 | 0.000896563 |
| chr5:17365673-17366252 | 1.460570063 | 2.65E-05 | 0.000897667 |
| chr7:129958653-129958931 | -2.081807588 | 2.65E-05 | 0.000897667 |
| chr4:183388792-183389026 | -2.503168584 | 2.65E-05 | 0.000898492 |
| chr7:4172211-4172464 | 1.981042777 | 2.66E-05 | 0.000899364 |
| chrX:40578133-40578371 | 2.283018981 | 2.67E-05 | 0.000903338 |
| chr14:89519751-89519951 | 2.735482658 | 2.67E-05 | 0.000903338 |
| chr9:114558140-114558686 | 1.54757641 | 2.69E-05 | 0.000907897 |
| chr7:108033195-108033703 | -1.756456972 | 2.69E-05 | 0.000907897 |
| chr16:77588879-77589085 | 1.939160625 | 2.69E-05 | 0.000907897 |
| chr15:56582501-56582744 | 2.487285339 | 2.69E-05 | 0.000907897 |
| chr13:112131897-112132504 | -1.761141932 | 2.69E-05 | 0.000908435 |
| chr8:93461668-93461968 | 2.313783956 | 2.70E-05 | 0.00090981 |
| chr7:21345675-21345980 | 1.967018109 | 2.71E-05 | 0.000911478 |
| chr9:114539097-114539423 | 2.213069449 | 2.71E-05 | 0.000913429 |
| chr10:2717117-2717341 | 2.594760924 | 2.72E-05 | 0.00091455 |
| chr8:86695804-86696013 | 2.32999495 | 2.72E-05 | 0.000916128 |
| chr15:52727806-52728046 | 1.700449697 | 2.73E-05 | 0.000916492 |
| chr2:145845511-145845878 | -2.220034427 | 2.73E-05 | 0.000916492 |
| chr9:113137372-113137857 | 1.777991939 | 2.73E-05 | 0.000916492 |
| chr16:77519846-77520065 | 2.097977955 | 2.73E-05 | 0.000918721 |
| chr1:148424220-148424607 | 1.833463168 | 2.74E-05 | 0.000920836 |
| chr10:77865574-77866323 | -1.461400106 | 2.74E-05 | 0.000920836 |
| chr22:43144898-43145370 | 1.838368641 | 2.75E-05 | 0.000922126 |
| chr14:91290727-91291419 | 1.497894598 | 2.75E-05 | 0.000922126 |
| chr12:86725318-86725618 | -2.226977226 | 2.76E-05 | 0.000923452 |
| chr4:14225736-14226035 | 2.095736251 | 2.77E-05 | 0.000926418 |
| chr4:87553259-87553586 | 1.961425232 | 2.77E-05 | 0.000927069 |
| chr3:120719140-120719947 | -1.467332797 | 2.77E-05 | 0.000927557 |
| chr7:24435171-24435374 | 2.326868487 | 2.78E-05 | 0.000930143 |
| chr8:103908319-103908519 | 2.699584265 | 2.78E-05 | 0.000930143 |
| chr8:130559274-130559611 | 2.388949341 | 2.78E-05 | 0.000930473 |
| chr3:156812445-156813001 | -1.398156175 | 2.79E-05 | 0.00093071 |
| chrX:153886955-153887360 | 1.723709513 | 2.79E-05 | 0.000931251 |
| chr14:105629526-105629771 | 1.881951065 | 2.79E-05 | 0.00093189 |
| chr2:41755176-41755396 | -2.511967977 | 2.79E-05 | 0.00093189 |
| chr2:124362031-124362309 | -2.415611271 | 2.80E-05 | 0.000933205 |
| chr14:76690764-76691121 | 1.75528436 | 2.81E-05 | 0.000937714 |

| | | | |
|---|---|---|---|
| chr16:54088681-54089067 | -2.050290339 | 2.81E-05 | 0.000938019 |
| chr5:71748393-71748813 | -1.907983784 | 2.82E-05 | 0.000939248 |
| chr10:108453247-108453701 | 1.906335341 | 2.83E-05 | 0.000943924 |
| chr2:73020944-73021216 | 2.062376602 | 2.86E-05 | 0.000950318 |
| chr14:34113162-34113476 | 1.804724364 | 2.86E-05 | 0.000951516 |
| chr8:133725153-133725463 | 1.858996951 | 2.86E-05 | 0.000951739 |
| chr4:142777352-142777933 | -1.827197438 | 2.86E-05 | 0.000951739 |
| chr7:127629279-127629749 | -1.588469033 | 2.87E-05 | 0.000954924 |
| chr20:48421197-48421438 | 1.578488213 | 2.89E-05 | 0.000958054 |
| chr7:25259646-25259938 | 2.08710345 | 2.89E-05 | 0.000958054 |
| chr15:53666016-53666216 | 1.886111125 | 2.89E-05 | 0.000959252 |
| chr18:48336422-48336700 | -2.341838001 | 2.90E-05 | 0.00096271 |
| chr2:200638082-200638403 | 2.387735261 | 2.90E-05 | 0.00096271 |
| chr13:92175396-92175713 | -1.96056013 | 2.90E-05 | 0.00096271 |
| chr5:156561874-156562398 | -1.864482873 | 2.92E-05 | 0.000968405 |
| chr6:52856067-52856532 | -1.666465605 | 2.93E-05 | 0.000968741 |
| chr17:28533142-28533513 | -1.728094798 | 2.93E-05 | 0.000969622 |
| chr1:153415157-153415701 | -1.579903026 | 2.93E-05 | 0.000969763 |
| chr5:62699357-62699726 | 1.859826017 | 2.94E-05 | 0.000970922 |
| chr3:78823792-78824281 | -1.608036374 | 2.94E-05 | 0.000973104 |
| chr17:74960702-74961078 | -1.6678492 | 2.95E-05 | 0.00097428 |
| chr7:23742223-23742484 | 1.920253186 | 2.96E-05 | 0.000977849 |
| chr7:115868473-115868707 | -2.847247857 | 2.97E-05 | 0.000979018 |
| chr13:110625521-110625930 | 1.962873022 | 2.97E-05 | 0.000980211 |
| chr13:71379545-71379909 | -2.123587977 | 2.98E-05 | 0.000982019 |
| chr15:87576564-87577503 | -1.44175057 | 2.98E-05 | 0.000982347 |
| chr7:136169760-136169980 | -2.846485548 | 2.98E-05 | 0.000982347 |
| chr6:140463380-140463671 | -1.842616505 | 2.98E-05 | 0.000982347 |
| chr14:29621657-29622212 | 1.766884552 | 2.98E-05 | 0.000982347 |
| chr8:127035437-127036168 | -1.461401557 | 2.99E-05 | 0.000982347 |
| chr10:6059802-6060059 | 2.084586113 | 2.99E-05 | 0.000982347 |
| chr7:121297953-121298253 | -2.397251005 | 2.99E-05 | 0.000982347 |
| chr16:5665347-5665752 | 2.132671563 | 2.99E-05 | 0.000982347 |
| chr9:91856784-91856984 | 2.842003657 | 2.99E-05 | 0.000982347 |
| chr15:73365890-73366229 | 1.814110085 | 2.99E-05 | 0.000982347 |
| chr4:174538080-174538595 | -1.677886612 | 2.99E-05 | 0.000982347 |
| chr14:50965456-50965837 | 1.992061076 | 3.01E-05 | 0.000986235 |
| chr13:28545572-28546070 | 1.683466779 | 3.01E-05 | 0.000988124 |
| chrX:111823119-111823406 | 1.939660848 | 3.02E-05 | 0.000989268 |
| chr6:83090078-83090528 | -1.861033022 | 3.03E-05 | 0.000992876 |
| chr7:13115192-13115619 | 1.737693856 | 3.04E-05 | 0.000996661 |
| chr1:153461472-153461851 | -1.954240596 | 3.05E-05 | 0.000999904 |
| chr8:37566751-37567834 | -1.515781593 | 3.05E-05 | 0.000999904 |
| chr2:171553855-171554275 | -1.561400683 | 3.07E-05 | 0.001004865 |
| chr1:114540252-114541197 | 1.368859287 | 3.07E-05 | 0.001004865 |
| chr15:56766881-56767131 | 2.38540681 | 3.08E-05 | 0.001006642 |
| chr4:181786357-181787252 | -1.792585085 | 3.09E-05 | 0.001009925 |
| chr16:34585782-34586044 | 1.564692635 | 3.10E-05 | 0.001010903 |
| chr11:84947991-84948504 | -1.781962141 | 3.10E-05 | 0.001011055 |
| chr14:37113095-37113780 | 1.550770025 | 3.11E-05 | 0.001015499 |

| | | | |
|---|---|---|---|
| chr8:93177067-93177357 | 1.75778479 | 3.12E-05 | 0.001016392 |
| chr16:74910297-74910585 | 2.261957537 | 3.12E-05 | 0.001016955 |
| chr1:218336369-218336804 | -1.517486001 | 3.13E-05 | 0.001020499 |
| chr20:45448403-45448772 | 1.718849812 | 3.14E-05 | 0.0010221 |
| chr11:85757920-85758373 | 1.601590644 | 3.15E-05 | 0.001027184 |
| chr20:48514626-48514921 | 1.655227968 | 3.16E-05 | 0.00102916 |
| chr3:13124884-13125116 | 1.95097747 | 3.17E-05 | 0.001029916 |
| chr7:32668557-32668856 | 2.110031108 | 3.17E-05 | 0.001029916 |
| chr7:21794513-21794728 | 2.456500586 | 3.17E-05 | 0.001029916 |
| chr7:241673-242184 | 1.711192782 | 3.19E-05 | 0.001036993 |
| chr11:70915571-70915892 | 2.166156621 | 3.19E-05 | 0.001036993 |
| chr2:226020270-226020526 | -2.591838075 | 3.20E-05 | 0.001036993 |
| chr2:38926007-38926306 | -1.840270427 | 3.20E-05 | 0.001036993 |
| chr3:164683599-164683950 | -1.914593996 | 3.20E-05 | 0.001037594 |
| chr3:72821114-72821396 | 2.676577424 | 3.21E-05 | 0.001041655 |
| chr1:11531125-11531999 | -1.474036808 | 3.22E-05 | 0.001042607 |
| chr10:65780147-65780466 | -2.276416915 | 3.22E-05 | 0.001042661 |
| chr17:82781762-82782256 | 1.58295852 | 3.22E-05 | 0.001043941 |
| chr16:78075308-78075508 | 2.145258727 | 3.23E-05 | 0.001045387 |
| chr3:178705916-178706500 | -1.75340672 | 3.24E-05 | 0.001048743 |
| chr2:6225977-6226340 | -1.836100554 | 3.25E-05 | 0.001050773 |
| chr13:80086160-80086777 | -1.597647217 | 3.25E-05 | 0.001051517 |
| chr1:148409235-148409445 | 2.334796422 | 3.26E-05 | 0.001053974 |
| chr10:124065306-124065678 | 1.722281917 | 3.26E-05 | 0.001054546 |
| chr9:98455797-98456297 | -1.75877542 | 3.28E-05 | 0.001059014 |
| chr13:33613270-33613571 | -2.000051483 | 3.29E-05 | 0.001059898 |
| chr3:30081885-30082297 | -1.737943119 | 3.29E-05 | 0.001059898 |
| chr7:93862793-93863063 | -2.237665152 | 3.29E-05 | 0.001059898 |
| chr21:38913093-38913804 | -1.422071576 | 3.29E-05 | 0.001060949 |
| chr14:73997453-73997653 | 2.385819219 | 3.29E-05 | 0.001061504 |
| chr17:7477376-7477821 | 1.732387163 | 3.30E-05 | 0.001062312 |
| chr2:15912175-15912684 | -1.517976157 | 3.30E-05 | 0.001062769 |
| chr15:71747515-71747971 | 1.674240079 | 3.30E-05 | 0.001063409 |
| chr8:76843517-76843734 | 2.681330697 | 3.32E-05 | 0.001067596 |
| chr7:42336677-42337095 | -1.79873727 | 3.32E-05 | 0.001067596 |
| chr4:33456325-33456656 | -2.001335806 | 3.32E-05 | 0.001067596 |
| chr3:53518734-53519280 | 1.749882188 | 3.34E-05 | 0.001071617 |
| chr11:14029926-14030609 | -1.447392334 | 3.34E-05 | 0.001072655 |
| chr3:143357368-143357642 | 2.140841619 | 3.34E-05 | 0.001072655 |
| chr7:38273379-38273918 | 1.764836213 | 3.35E-05 | 0.001075601 |
| chr5:125461484-125462161 | -1.61948422 | 3.36E-05 | 0.001077115 |
| chr1:211592186-211592581 | -1.941764839 | 3.36E-05 | 0.001078471 |
| chr4:8352994-8353219 | 2.374607136 | 3.37E-05 | 0.001078514 |
| chr2:236178129-236178397 | 2.014229185 | 3.37E-05 | 0.001079549 |
| chr12:125471898-125472185 | 2.157383463 | 3.38E-05 | 0.001081563 |
| chr1:116879882-116880093 | 2.043879216 | 3.38E-05 | 0.001081809 |
| chr10:24542444-24543250 | -1.49906418 | 3.38E-05 | 0.001081809 |
| chr4:188417304-188417504 | -3.40030853 | 3.38E-05 | 0.001081809 |
| chr7:7556760-7557091 | 1.772962068 | 3.39E-05 | 0.00108466 |
| chr8:133620226-133620560 | 1.708383143 | 3.40E-05 | 0.001086095 |

| | | | |
|---|---|---|---|
| chr15:88727588-88727835 | 2.316424318 | 3.41E-05 | 0.00109006 |
| chr5:14316680-14316881 | 2.468865814 | 3.44E-05 | 0.001096993 |
| chr8:99696902-99697245 | 2.071857822 | 3.44E-05 | 0.001098812 |
| chr2:5474734-5474995 | 2.14770585 | 3.45E-05 | 0.001099023 |
| chr1:222893241-222893819 | -1.62920567 | 3.46E-05 | 0.001101942 |
| chr3:179037282-179037754 | -1.52020834 | 3.46E-05 | 0.001104061 |
| chr14:43778628-43779211 | 1.55556224 | 3.47E-05 | 0.001106437 |
| chr12:69468236-69468695 | -1.684525065 | 3.48E-05 | 0.001107958 |
| chr8:56890022-56890297 | -2.237997679 | 3.49E-05 | 0.001109822 |
| chr5:104062544-104063197 | -1.495723549 | 3.49E-05 | 0.001111414 |
| chr2:43130771-43131787 | -1.418750777 | 3.51E-05 | 0.001115044 |
| chr20:58294568-58294936 | 1.586886873 | 3.51E-05 | 0.001115255 |
| chr3:14232636-14232911 | 1.894395801 | 3.51E-05 | 0.001115255 |
| chr6:140062203-140063145 | -1.632604398 | 3.51E-05 | 0.001115784 |
| chr1:172359713-172360006 | 1.880125151 | 3.52E-05 | 0.001118654 |
| chr9:79103261-79104199 | -1.518317542 | 3.54E-05 | 0.00112219 |
| chr13:42985918-42986278 | -1.65578562 | 3.54E-05 | 0.001122685 |
| chr18:43661602-43661857 | -2.150226713 | 3.54E-05 | 0.001123839 |
| chr1:222252780-222253026 | 2.393011476 | 3.55E-05 | 0.001123869 |
| chr8:122683389-122683806 | 1.578506088 | 3.55E-05 | 0.001123869 |
| chr13:81131955-81132155 | -2.932040833 | 3.56E-05 | 0.001126221 |
| chr13:61286189-61286608 | -1.78762279 | 3.56E-05 | 0.001128301 |
| chr3:194512758-194513107 | -1.864574675 | 3.57E-05 | 0.001128301 |
| chr13:107069287-107069685 | -1.886784712 | 3.57E-05 | 0.001128301 |
| chr9:21194944-21195414 | -2.012259103 | 3.57E-05 | 0.001128301 |
| chr14:45679974-45680226 | 2.299904956 | 3.57E-05 | 0.001128301 |
| chr16:51994670-51994945 | -2.25733156 | 3.57E-05 | 0.001128301 |
| chr18:3623596-3624442 | -1.54830449 | 3.59E-05 | 0.001133224 |
| chr10:99767952-99768424 | 1.937172644 | 3.59E-05 | 0.001133746 |
| chr5:64939411-64940550 | -1.365764826 | 3.59E-05 | 0.001133746 |
| chr2:15471681-15471947 | -2.186308796 | 3.60E-05 | 0.001133803 |
| chr20:60162105-60162517 | 1.889155615 | 3.60E-05 | 0.001133803 |
| chr18:69685968-69686243 | -2.727551223 | 3.60E-05 | 0.001133803 |
| chr3:107987038-107987587 | -1.514971357 | 3.61E-05 | 0.001136516 |
| chr5:81592180-81592520 | -1.956658597 | 3.61E-05 | 0.001136751 |
| chr13:40341555-40341947 | -1.76703242 | 3.61E-05 | 0.001137137 |
| chr3:8619598-8619976 | 1.563892722 | 3.61E-05 | 0.001137137 |
| chr20:15490665-15490941 | 2.237400027 | 3.62E-05 | 0.001137718 |
| chr14:30914098-30914307 | 2.834153156 | 3.62E-05 | 0.001138694 |
| chr10:49701053-49701316 | -2.148760133 | 3.63E-05 | 0.001141257 |
| chr7:40717282-40717802 | -1.577991317 | 3.64E-05 | 0.00114363 |
| chr5:100506393-100506632 | -2.248607865 | 3.66E-05 | 0.00114818 |
| chr7:156929895-156930431 | -1.738335234 | 3.67E-05 | 0.001151933 |
| chr18:77590973-77591331 | -1.931385665 | 3.67E-05 | 0.001152907 |
| chr19:10712242-10712466 | 2.457637882 | 3.68E-05 | 0.001153155 |
| chr12:29242234-29242612 | -1.825784589 | 3.69E-05 | 0.001157563 |
| chr5:169445443-169445650 | 2.822363386 | 3.70E-05 | 0.001158336 |
| chr14:73017753-73017985 | 2.295510076 | 3.70E-05 | 0.001159344 |
| chr6:145755911-145756646 | -1.549024862 | 3.70E-05 | 0.001159344 |
| chr20:6976422-6976944 | -1.745831686 | 3.70E-05 | 0.001159344 |

| | | | |
|---|---|---|---|
| chr1:107996549-107996797 | 2.250642123 | 3.71E-05 | 0.001159344 |
| chr11:75024471-75024671 | 2.318583521 | 3.71E-05 | 0.001159886 |
| chr2:179646885-179647245 | -1.714605279 | 3.71E-05 | 0.001159886 |
| chr5:50303163-50303373 | 2.445665662 | 3.71E-05 | 0.001159886 |
| chr2:37344446-37345178 | -1.532098464 | 3.71E-05 | 0.001159886 |
| chr8:133723056-133723321 | 1.950477232 | 3.72E-05 | 0.001162583 |
| chr5:17216097-17218371 | 1.324917979 | 3.73E-05 | 0.001162696 |
| chr13:56712455-56712816 | -2.196218315 | 3.73E-05 | 0.001162835 |
| chr13:40564501-40565007 | -1.60808949 | 3.73E-05 | 0.001163662 |
| chr14:64720318-64720608 | 1.590753152 | 3.73E-05 | 0.001164222 |
| chr20:54579394-54579683 | 1.508027262 | 3.75E-05 | 0.001167539 |
| chr11:65471903-65472501 | 1.486230333 | 3.75E-05 | 0.001167539 |
| chr6:34544921-34545491 | -1.459965592 | 3.76E-05 | 0.001171051 |
| chr10:59362171-59363412 | -1.375533577 | 3.77E-05 | 0.001171907 |
| chr11:111097178-111097586 | -1.843911411 | 3.77E-05 | 0.001173627 |
| chr10:53886489-53887290 | -1.575646632 | 3.78E-05 | 0.001174222 |
| chr2:233486882-233487388 | 1.800460116 | 3.79E-05 | 0.001177025 |
| chrX:9269191-9269704 | -1.72532264 | 3.80E-05 | 0.001180069 |
| chr7:16803912-16804357 | 1.550675885 | 3.82E-05 | 0.001185503 |
| chr13:93244130-93244805 | -1.511685402 | 3.82E-05 | 0.001185595 |
| chr14:90264376-90264597 | 2.195136855 | 3.82E-05 | 0.001185595 |
| chr21:41112978-41113847 | -1.4217929 | 3.82E-05 | 0.001185979 |
| chr7:17863025-17863417 | 1.773539797 | 3.83E-05 | 0.001187866 |
| chr7:148270673-148270919 | 2.059532844 | 3.85E-05 | 0.0011928 |
| chr2:20071389-20072372 | -1.352800589 | 3.86E-05 | 0.001196161 |
| chr1:238809804-238810052 | 2.293585985 | 3.87E-05 | 0.001199488 |
| chr3:19772546-19772834 | 2.370807845 | 3.88E-05 | 0.001201157 |
| chr14:54596536-54596851 | 2.309856852 | 3.88E-05 | 0.001201157 |
| chr5:56763993-56764523 | -1.603163315 | 3.88E-05 | 0.001201498 |
| chr7:157851810-157852041 | -2.287593095 | 3.89E-05 | 0.001201742 |
| chr11:104385487-104385946 | -2.248303986 | 3.89E-05 | 0.001201742 |
| chr3:12753696-12753983 | 1.805479595 | 3.89E-05 | 0.001201742 |
| chr14:52027134-52027404 | 2.15989606 | 3.90E-05 | 0.001204843 |
| chr18:57581314-57581982 | -1.540062377 | 3.93E-05 | 0.001212647 |
| chr4:827694-828199 | 1.68647302 | 3.94E-05 | 0.001214874 |
| chr10:132065698-132066093 | 1.62875567 | 3.94E-05 | 0.001215692 |
| chr7:130084136-130084445 | -2.043768324 | 3.94E-05 | 0.001215692 |
| chr7:84303035-84303269 | -2.532841186 | 3.95E-05 | 0.001216984 |
| chrX:35694825-35695334 | -1.995655686 | 3.97E-05 | 0.001223028 |
| chr10:64441828-64442567 | -1.49888012 | 3.98E-05 | 0.0012253 |
| chr20:60397413-60397667 | 2.299104982 | 3.99E-05 | 0.001228881 |
| chr1:246647106-246647473 | -1.682740494 | 3.99E-05 | 0.001228998 |
| chr1:248252702-248253071 | -2.021498704 | 4.02E-05 | 0.001236056 |
| chr9:38195595-38196172 | -1.556815521 | 4.03E-05 | 0.001237827 |
| chr14:63663416-63664436 | -1.355433407 | 4.03E-05 | 0.001237827 |
| chr8:90253329-90253625 | 1.95939276 | 4.03E-05 | 0.001237827 |
| chr17:44628077-44628277 | 2.687898141 | 4.03E-05 | 0.001237827 |
| chr5:28374392-28375061 | -1.457081505 | 4.03E-05 | 0.001238652 |
| chr7:142353021-142353221 | -2.657641411 | 4.04E-05 | 0.001240049 |
| chr10:122135251-122135580 | 2.072371448 | 4.05E-05 | 0.001242207 |

| | | | |
|---|---|---|---|
| chr10:1703215-1703503 | 2.248549216 | 4.06E-05 | 0.001244805 |
| chr6:124348661-124348906 | -2.547778325 | 4.07E-05 | 0.001245975 |
| chr2:155966180-155966577 | -2.083995534 | 4.07E-05 | 0.001245975 |
| chr12:32934741-32935046 | 2.455809515 | 4.07E-05 | 0.001245975 |
| chr19:47805384-47805605 | 2.6666965 | 4.07E-05 | 0.001246999 |
| chr6:111564663-111565079 | -1.646321763 | 4.08E-05 | 0.001247308 |
| chr2:126991006-126991591 | -1.753587142 | 4.08E-05 | 0.001247897 |
| chr4:183798002-183798853 | -1.557867793 | 4.08E-05 | 0.001248765 |
| chrX:153610474-153611083 | 1.588298862 | 4.09E-05 | 0.001249972 |
| chr4:164215210-164215703 | -1.799750769 | 4.09E-05 | 0.001250018 |
| chr3:107812684-107813323 | -1.583050403 | 4.09E-05 | 0.001250492 |
| chr15:54364552-54364766 | 2.538295319 | 4.10E-05 | 0.001250632 |
| chr13:37640576-37640789 | -2.381232753 | 4.10E-05 | 0.001251756 |
| chr14:32413637-32413896 | 2.371312328 | 4.10E-05 | 0.001251756 |
| chr4:139469877-139470174 | 1.968163436 | 4.11E-05 | 0.001252296 |
| chr2:64906909-64907617 | -1.518570748 | 4.11E-05 | 0.001253589 |
| chr20:22824130-22824449 | -1.833610355 | 4.13E-05 | 0.001258604 |
| chr4:165493295-165493727 | -1.634592492 | 4.14E-05 | 0.001259588 |
| chr14:36751190-36751436 | 1.933907016 | 4.14E-05 | 0.001260653 |
| chr5:32230786-32231078 | 1.908964531 | 4.14E-05 | 0.001261273 |
| chr15:68748818-68749105 | 2.457369111 | 4.15E-05 | 0.001263804 |
| chrX:28205994-28206274 | -2.03098411 | 4.16E-05 | 0.001265808 |
| chr5:129135511-129136013 | -1.521725669 | 4.16E-05 | 0.001265808 |
| chr1:197620649-197621272 | -1.429183366 | 4.17E-05 | 0.001266779 |
| chr13:70715015-70715554 | -1.876991248 | 4.18E-05 | 0.001269663 |
| chr7:142783302-142783695 | -1.499544398 | 4.19E-05 | 0.001270657 |
| chr13:20171378-20171812 | 1.62580675 | 4.20E-05 | 0.001274227 |
| chr5:157828898-157829345 | -1.72934728 | 4.20E-05 | 0.001274227 |
| chr20:21725928-21726244 | 2.645884311 | 4.20E-05 | 0.001274227 |
| chr7:110396879-110397140 | -2.419901541 | 4.20E-05 | 0.001274307 |
| chr19:35484945-35485178 | 2.654961054 | 4.22E-05 | 0.001276601 |
| chr19:56165102-56165311 | 2.654961054 | 4.22E-05 | 0.001276601 |
| chr13:79560939-79561319 | -2.243528663 | 4.22E-05 | 0.001278076 |
| chr3:118531859-118532286 | 1.711733758 | 4.25E-05 | 0.001285804 |
| chr7:139844779-139845144 | -1.879847597 | 4.25E-05 | 0.001285804 |
| chr8:109186653-109187003 | 1.82777647 | 4.26E-05 | 0.001286849 |
| chr19:32844152-32844501 | 2.08278736 | 4.26E-05 | 0.001287968 |
| chr5:134439295-134439726 | -1.541977373 | 4.27E-05 | 0.001291419 |
| chr13:36091813-36092180 | -1.978953403 | 4.28E-05 | 0.001292595 |
| chr1:215425189-215425608 | -1.707503255 | 4.30E-05 | 0.001297213 |
| chr12:125890984-125891251 | 2.232152213 | 4.30E-05 | 0.001298424 |
| chr20:48447400-48447799 | 1.438052907 | 4.31E-05 | 0.001301086 |
| chr2:204452433-204452977 | -1.680779351 | 4.32E-05 | 0.00130366 |
| chr5:28077411-28077750 | 1.804965117 | 4.33E-05 | 0.001304568 |
| chr16:22844788-22844988 | 2.679458372 | 4.34E-05 | 0.001306957 |
| chr7:141975949-141976767 | -1.427111325 | 4.34E-05 | 0.001306957 |
| chr5:159331394-159331883 | -1.7891931 | 4.35E-05 | 0.001308271 |
| chr20:48492115-48492468 | 1.469826608 | 4.35E-05 | 0.001308662 |
| chr12:72932010-72932247 | 2.182975281 | 4.35E-05 | 0.001308662 |
| chr1:216822472-216822888 | -1.540772706 | 4.36E-05 | 0.001311209 |

| | | | |
|---|---|---|---|
| chr1:172728012-172728398 | -1.530017335 | 4.39E-05 | 0.00131967 |
| chr1:219204673-219205185 | -1.558183528 | 4.39E-05 | 0.001320425 |
| chr1:208231856-208232244 | -1.893149611 | 4.40E-05 | 0.00132239 |
| chr8:88759503-88759895 | 1.999773178 | 4.41E-05 | 0.001323425 |
| chr14:38030668-38030947 | 2.051788622 | 4.42E-05 | 0.001324864 |
| chr12:82650609-82650957 | -1.606488288 | 4.42E-05 | 0.001324864 |
| chr10:118112721-118113060 | -2.185073796 | 4.42E-05 | 0.001324864 |
| chr7:145582865-145583846 | -1.585462126 | 4.44E-05 | 0.001330036 |
| chr7:104944816-104945495 | -1.390836539 | 4.44E-05 | 0.001330036 |
| chr21:45381124-45381383 | 2.046552109 | 4.44E-05 | 0.001330036 |
| chr12:94275019-94275363 | 1.906494301 | 4.44E-05 | 0.001330036 |
| chr10:115997751-115998031 | 2.222344312 | 4.44E-05 | 0.001330036 |
| chr16:87487399-87487926 | -1.554347105 | 4.45E-05 | 0.001331138 |
| chr9:133690169-133690605 | 1.691604586 | 4.46E-05 | 0.001334474 |
| chr19:33032431-33032780 | 1.786910808 | 4.47E-05 | 0.001336329 |
| chr13:93299766-93300213 | -1.90135692 | 4.48E-05 | 0.001339102 |
| chr6:41577905-41578332 | 1.683815395 | 4.50E-05 | 0.001343024 |
| chr5:98180008-98181206 | -1.450872141 | 4.50E-05 | 0.001343899 |
| chr8:80873187-80873553 | 1.83875546 | 4.50E-05 | 0.001343899 |
| chr7:120856715-120857430 | -1.451915433 | 4.51E-05 | 0.001345245 |
| chr18:63260175-63260559 | -1.661862271 | 4.51E-05 | 0.001345558 |
| chr8:134794101-134794403 | 1.947173561 | 4.52E-05 | 0.001346197 |
| chr20:19767111-19767441 | 1.724289498 | 4.52E-05 | 0.001348337 |
| chr3:15520759-15520975 | 2.105736707 | 4.53E-05 | 0.001348337 |
| chr20:41203106-41203319 | 2.166228102 | 4.53E-05 | 0.001350166 |
| chr2:150154036-150155006 | -1.439661246 | 4.54E-05 | 0.001350541 |
| chr12:132210275-132210698 | 1.775807134 | 4.54E-05 | 0.00135124 |
| chr2:205716153-205716487 | -1.667414054 | 4.55E-05 | 0.001352304 |
| chr18:76856943-76857799 | -1.481183717 | 4.55E-05 | 0.001353771 |
| chr20:63509924-63510162 | 2.283238521 | 4.56E-05 | 0.001354359 |
| chr8:8600942-8601329 | -2.214262309 | 4.56E-05 | 0.001354359 |
| chr1:68881165-68882117 | -1.430098379 | 4.56E-05 | 0.001355594 |
| chr14:65851031-65851367 | 1.969301553 | 4.58E-05 | 0.001360439 |
| chr2:196601375-196602169 | -1.443443269 | 4.59E-05 | 0.001361566 |
| chr4:163343588-163345004 | -1.396511935 | 4.60E-05 | 0.001364099 |
| chr3:171173659-171174198 | -1.468355274 | 4.60E-05 | 0.001364508 |
| chr11:763087-763454 | 1.782771165 | 4.60E-05 | 0.00136476 |
| chr4:610793-611209 | 1.832627291 | 4.62E-05 | 0.001368386 |
| chr7:69780346-69780736 | -1.799331318 | 4.62E-05 | 0.001368513 |
| chr20:32606963-32607205 | 2.218082835 | 4.63E-05 | 0.001369076 |
| chr13:104024409-104024794 | -1.945364527 | 4.63E-05 | 0.001369076 |
| chr7:127408007-127408276 | -2.201820826 | 4.64E-05 | 0.00137207 |
| chr19:38282058-38282322 | -2.131542578 | 4.66E-05 | 0.001376793 |
| chr18:22871700-22872113 | -1.840056786 | 4.67E-05 | 0.001380063 |
| chr7:138958603-138958894 | -2.10582265 | 4.67E-05 | 0.001380063 |
| chr3:76572420-76572627 | 2.51200337 | 4.67E-05 | 0.001380565 |
| chr4:74457166-74457809 | -1.568366484 | 4.68E-05 | 0.001380918 |
| chr6:144726152-144726488 | -1.754281405 | 4.68E-05 | 0.001380918 |
| chr1:215425841-215426041 | -2.643640236 | 4.69E-05 | 0.00138381 |
| chr18:51152065-51152480 | -1.758633366 | 4.70E-05 | 0.001385875 |

| | | | |
|---|---|---|---|
| chr6:138577478-138577970 | -1.576502019 | 4.70E-05 | 0.001385875 |
| chr1:190020222-190020777 | -1.463053117 | 4.70E-05 | 0.001385875 |
| chr16:77971558-77972183 | 1.414765338 | 4.71E-05 | 0.001387505 |
| chr10:44752940-44753281 | 1.850484772 | 4.71E-05 | 0.001388319 |
| chr3:133461318-133461646 | -1.510510997 | 4.73E-05 | 0.00139178 |
| chr19:42655811-42656055 | 2.119773221 | 4.73E-05 | 0.00139178 |
| chr4:86300579-86300911 | 1.943833787 | 4.74E-05 | 0.001392788 |
| chr3:172541289-172541785 | -1.714689917 | 4.74E-05 | 0.001393598 |
| chr9:69207540-69207840 | -1.819637355 | 4.76E-05 | 0.001397292 |
| chr7:98321327-98321809 | -1.796888135 | 4.76E-05 | 0.001397292 |
| chr3:15986518-15986780 | 2.351336847 | 4.76E-05 | 0.001398474 |
| chr1:19920898-19921133 | 2.542507639 | 4.77E-05 | 0.001399384 |
| chr8:103898019-103898335 | 1.906510974 | 4.77E-05 | 0.001399384 |
| chr13:51683619-51684103 | -1.857048964 | 4.78E-05 | 0.001402452 |
| chr3:128020727-128020979 | 2.349608138 | 4.79E-05 | 0.001404894 |
| chr14:28952755-28953076 | 1.84633498 | 4.81E-05 | 0.001408471 |
| chrX:26852036-26852300 | -2.147341054 | 4.81E-05 | 0.001408765 |
| chr13:39833837-39834108 | 2.279444032 | 4.81E-05 | 0.001408765 |
| chr12:112199066-112199309 | 2.077753994 | 4.81E-05 | 0.001409681 |
| chr16:11497415-11497685 | 1.939880653 | 4.82E-05 | 0.001410641 |
| chr8:80066104-80066310 | 2.413177111 | 4.83E-05 | 0.001414205 |
| chr6:83803558-83803882 | -2.054156222 | 4.86E-05 | 0.001420698 |
| chr3:161890110-161890879 | -1.364839117 | 4.86E-05 | 0.001420702 |
| chr19:38290679-38291166 | 1.658338293 | 4.87E-05 | 0.001421701 |
| chr17:38761732-38761992 | 2.266530616 | 4.87E-05 | 0.001421847 |
| chr4:19615061-19615625 | -1.781312718 | 4.87E-05 | 0.001423411 |
| chr2:149080943-149081379 | -1.557910536 | 4.89E-05 | 0.001426798 |
| chr7:13255526-13255779 | 2.192011514 | 4.90E-05 | 0.001429073 |
| chr16:28603543-28603948 | 1.756346271 | 4.90E-05 | 0.001429073 |
| chr2:165902937-165903336 | -1.786168687 | 4.90E-05 | 0.001429073 |
| chr7:130941493-130941799 | -1.808891191 | 4.90E-05 | 0.001429073 |
| chr5:130897424-130898275 | -1.360031735 | 4.94E-05 | 0.001440454 |
| chr9:83596946-83597415 | -1.687199081 | 4.95E-05 | 0.001440501 |
| chr1:202084739-202085696 | -1.331608374 | 4.97E-05 | 0.001445436 |
| chr3:97232452-97233103 | -1.412164727 | 4.98E-05 | 0.001448695 |
| chr6:113672776-113673190 | 1.793861754 | 4.99E-05 | 0.001450522 |
| chr1:164581844-164582254 | 1.805273569 | 4.99E-05 | 0.001450522 |
| chr10:107340276-107340750 | -1.859204388 | 4.99E-05 | 0.001451146 |
| chr3:75329838-75330038 | 2.15810586 | 5.00E-05 | 0.001453432 |
| chr7:13489521-13489736 | 2.350097112 | 5.00E-05 | 0.001453432 |
| chr10:52475641-52475864 | 2.204701335 | 5.01E-05 | 0.001453648 |
| chr14:36807000-36807267 | 2.283544645 | 5.02E-05 | 0.001456844 |
| chr14:40008355-40008634 | 2.15308645 | 5.03E-05 | 0.001458621 |
| chr19:41776814-41777092 | 2.038760627 | 5.03E-05 | 0.001459163 |
| chr1:114182002-114182338 | 1.536755113 | 5.04E-05 | 0.001460415 |
| chr20:49592244-49592470 | 2.108161273 | 5.04E-05 | 0.001460415 |
| chr11:37370829-37371138 | 1.923881815 | 5.04E-05 | 0.001460415 |
| chr16:57797616-57798298 | 1.52677092 | 5.04E-05 | 0.0014608 |
| chr15:101503009-101503516 | -1.56095454 | 5.05E-05 | 0.001462277 |
| chr20:23235440-23235685 | -2.07238438 | 5.06E-05 | 0.001462888 |

| | | | |
|---|---|---|---|
| chr8:96945715-96946189 | -1.555469588 | 5.06E-05 | 0.001464802 |
| chr20:39849377-39849960 | -1.835377559 | 5.09E-05 | 0.001470684 |
| chr15:40342647-40342902 | 1.714156475 | 5.09E-05 | 0.001470818 |
| chr13:56046307-56046749 | -1.986905795 | 5.09E-05 | 0.001470818 |
| chr12:41966010-41966384 | -1.77822479 | 5.09E-05 | 0.001471131 |
| chr11:106990300-106990635 | -1.998041175 | 5.10E-05 | 0.001471131 |
| chr18:834148-834349 | 2.422783873 | 5.10E-05 | 0.00147216 |
| chr7:81956601-81956827 | -2.415305548 | 5.11E-05 | 0.001473385 |
| chr5:99037717-99038473 | -1.62326429 | 5.14E-05 | 0.001481443 |
| chr18:76941350-76941639 | -2.023516584 | 5.14E-05 | 0.001481443 |
| chr14:72905038-72905309 | 1.786190844 | 5.15E-05 | 0.001483896 |
| chr19:45938194-45938394 | 2.532815727 | 5.16E-05 | 0.001485616 |
| chr14:67479925-67480880 | -1.354907281 | 5.17E-05 | 0.001489721 |
| chr1:235246399-235246852 | -2.111808497 | 5.18E-05 | 0.001489795 |
| chr13:38711151-38711381 | -2.223867437 | 5.18E-05 | 0.001489795 |
| chr10:102583849-102584225 | 1.799228404 | 5.18E-05 | 0.001489989 |
| chr2:190497411-190497881 | -1.697807416 | 5.19E-05 | 0.001490864 |
| chr8:104273276-104273720 | 1.616751135 | 5.19E-05 | 0.001492375 |
| chr15:85852948-85853168 | 2.163257449 | 5.19E-05 | 0.001492375 |
| chr17:75993520-75993774 | 1.900083416 | 5.20E-05 | 0.001493191 |
| chr14:20848640-20849280 | 1.525909949 | 5.21E-05 | 0.001494555 |
| chr14:105241797-105242055 | 1.911790472 | 5.21E-05 | 0.001495121 |
| chr3:192825684-192826303 | -1.531079501 | 5.23E-05 | 0.001499461 |
| chr7:192102-192650 | 1.460882796 | 5.23E-05 | 0.001499809 |
| chr9:1337076-1337598 | 1.520466356 | 5.24E-05 | 0.001501869 |
| chr5:79011560-79011826 | -2.197214679 | 5.25E-05 | 0.001503302 |
| chr8:136793669-136793894 | 2.207641052 | 5.25E-05 | 0.001503302 |
| chr4:70019440-70019859 | -1.81360501 | 5.25E-05 | 0.001503886 |
| chr4:189722435-189722719 | -2.779921925 | 5.26E-05 | 0.001506954 |
| chr11:121281135-121281432 | 1.937480451 | 5.28E-05 | 0.00151003 |
| chr7:7514852-7515052 | 2.196748328 | 5.29E-05 | 0.00151411 |
| chr14:20924610-20924994 | -2.214775664 | 5.34E-05 | 0.001527061 |
| chr3:50314865-50315172 | 2.113838903 | 5.37E-05 | 0.001533217 |
| chr6:80537300-80537793 | -1.718692971 | 5.37E-05 | 0.001533217 |
| chr14:106061988-106062586 | 1.617880858 | 5.37E-05 | 0.001534308 |
| chr16:46400529-46400797 | 1.54233091 | 5.38E-05 | 0.00153446 |
| chr11:78708242-78708442 | 2.356739768 | 5.38E-05 | 0.00153446 |
| chr10:132397651-132398126 | 1.466311988 | 5.40E-05 | 0.001540133 |
| chr9:81320908-81321527 | -1.411524225 | 5.40E-05 | 0.001540133 |
| chr2:240893583-240893820 | -1.953779851 | 5.40E-05 | 0.001540232 |
| chr3:144843156-144843605 | -1.875524179 | 5.41E-05 | 0.001540232 |
| chr20:641567-641862 | -2.33061245 | 5.41E-05 | 0.001541403 |
| chr5:116032111-116032449 | -1.793811896 | 5.42E-05 | 0.00154212 |
| chr21:8455775-8456015 | 1.765757113 | 5.42E-05 | 0.001543003 |
| chr19:38799554-38799770 | 2.261882738 | 5.43E-05 | 0.00154396 |
| chr20:56804272-56805142 | 1.31581144 | 5.43E-05 | 0.001545675 |
| chr3:10285918-10286247 | 1.686906236 | 5.44E-05 | 0.001546177 |
| chr9:69505893-69506261 | -1.885959685 | 5.44E-05 | 0.001546177 |
| chr9:21836373-21836907 | -1.591827885 | 5.44E-05 | 0.001546177 |
| chr8:22685468-22685931 | -1.749236699 | 5.45E-05 | 0.001546825 |

| | | | |
|---|---|---|---|
| chr1:223513284-223513715 | -1.618842094 | 5.45E-05 | 0.001547954 |
| chr6:46748726-46748942 | 2.778193836 | 5.46E-05 | 0.001549096 |
| chr3:71384389-71384812 | -1.868556413 | 5.46E-05 | 0.001549096 |
| chr13:84865981-84866355 | -2.017462956 | 5.46E-05 | 0.001549096 |
| chr14:83974281-83974492 | 2.340166721 | 5.48E-05 | 0.001554752 |
| chr14:105748282-105748563 | 1.880721164 | 5.50E-05 | 0.001558313 |
| chr3:16141748-16141992 | 2.404521446 | 5.50E-05 | 0.001558407 |
| chr14:104932389-104932589 | 2.095741786 | 5.53E-05 | 0.001564997 |
| chr12:124130718-124131105 | 1.696128053 | 5.55E-05 | 0.001570764 |
| chr1:50976620-50976820 | 2.418333962 | 5.56E-05 | 0.001572282 |
| chr5:52922100-52922848 | -1.401828332 | 5.56E-05 | 0.001572282 |
| chr3:33030171-33030415 | 2.330777248 | 5.56E-05 | 0.001572319 |
| chr4:24349260-24349485 | 2.504040034 | 5.56E-05 | 0.001572549 |
| chr12:42756923-42757332 | 1.599886886 | 5.57E-05 | 0.001573414 |
| chr13:71751179-71751436 | -2.273048415 | 5.58E-05 | 0.001575618 |
| chr7:116930478-116930961 | -1.546162155 | 5.58E-05 | 0.001575794 |
| chr3:28788590-28789067 | 1.685005965 | 5.59E-05 | 0.001575794 |
| chr17:82436995-82437197 | 2.066755109 | 5.59E-05 | 0.001575794 |
| chr2:7802626-7802932 | 2.204238449 | 5.60E-05 | 0.001577898 |
| chr15:52653281-52653988 | 1.372631584 | 5.62E-05 | 0.001583386 |
| chr1:181446107-181446487 | 1.675994847 | 5.63E-05 | 0.001585648 |
| chr4:7156846-7157150 | 1.839950329 | 5.64E-05 | 0.001587848 |
| chr5:117418110-117418670 | -1.996713455 | 5.64E-05 | 0.001587848 |
| chrX:135665165-135665599 | 1.859653653 | 5.64E-05 | 0.001587848 |
| chr2:164069502-164069880 | -1.912464141 | 5.65E-05 | 0.001589575 |
| chr4:96701919-96702399 | -1.840881268 | 5.66E-05 | 0.001591898 |
| chr1:218377270-218377682 | 1.491113065 | 5.66E-05 | 0.001592568 |
| chr9:76403341-76403697 | -1.817354596 | 5.70E-05 | 0.001600543 |
| chr8:121610799-121611248 | -1.507837182 | 5.70E-05 | 0.001600543 |
| chr1:206895108-206896233 | -1.296505264 | 5.72E-05 | 0.001605893 |
| chr7:16804631-16805424 | 1.399841504 | 5.72E-05 | 0.001605893 |
| chr3:131778027-131778470 | 1.808930875 | 5.74E-05 | 0.001610845 |
| chr18:60892891-60893235 | -1.616190746 | 5.76E-05 | 0.001614532 |
| chr2:189541349-189542049 | -1.476457578 | 5.76E-05 | 0.001614532 |
| chr1:190084545-190085070 | -1.567787254 | 5.76E-05 | 0.001614652 |
| chr16:30911086-30911309 | 2.153675056 | 5.77E-05 | 0.001615901 |
| chr20:49270263-49270540 | 1.993858057 | 5.80E-05 | 0.001623622 |
| chr14:66843206-66843452 | 2.397908726 | 5.80E-05 | 0.001624559 |
| chr3:186255747-186256398 | -1.473885532 | 5.81E-05 | 0.001625516 |
| chr18:37166984-37167304 | -1.891125366 | 5.82E-05 | 0.001627399 |
| chr12:109119107-109119312 | 2.133174194 | 5.82E-05 | 0.001627399 |
| chr4:104490823-104491701 | -1.408853778 | 5.82E-05 | 0.001627696 |
| chr4:132530876-132531174 | -2.090868868 | 5.83E-05 | 0.001631021 |
| chr4:163007644-163007984 | -1.859637334 | 5.85E-05 | 0.001635637 |
| chr2:101566715-101567205 | 1.628788753 | 5.86E-05 | 0.001637447 |
| chr5:17114224-17114471 | 1.906732217 | 5.88E-05 | 0.001640947 |
| chr14:36582506-36582762 | 1.785530224 | 5.89E-05 | 0.001642963 |
| chr4:1329125-1329450 | 1.974898986 | 5.89E-05 | 0.001642963 |
| chr21:15189905-15190229 | -2.33424202 | 5.89E-05 | 0.001642963 |
| chr2:32976840-32977719 | -1.419990691 | 5.89E-05 | 0.001642963 |

| | | | |
|---|---|---|---|
| chr2:28377770-28378333 | -1.455932339 | 5.90E-05 | 0.001642963 |
| chr10:22651334-22651850 | -1.595580785 | 5.90E-05 | 0.001642963 |
| chr2:201984048-201984464 | 1.787439403 | 5.90E-05 | 0.001642963 |
| chr16:64715567-64715858 | 2.134059142 | 5.90E-05 | 0.001643977 |
| chr7:16098068-16098489 | 1.535530195 | 5.94E-05 | 0.001652036 |
| chr16:10932914-10933166 | 2.184717689 | 5.94E-05 | 0.001653889 |
| chr5:115366134-115366734 | -1.407133431 | 5.95E-05 | 0.00165414 |
| chr14:25945586-25946021 | 1.751582163 | 5.96E-05 | 0.001656498 |
| chr8:51794042-51794283 | 2.608214106 | 5.97E-05 | 0.001657339 |
| chr11:67648248-67648494 | 1.822028787 | 5.97E-05 | 0.001657339 |
| chr7:128469580-128469828 | -2.394670837 | 5.97E-05 | 0.00165882 |
| chr5:105868135-105868750 | -1.634593479 | 5.98E-05 | 0.001659294 |
| chr2:185628954-185629461 | -1.728390972 | 5.98E-05 | 0.001659294 |
| chr3:133447595-133448353 | -1.424298645 | 5.98E-05 | 0.001659419 |
| chr2:7791965-7792733 | -1.358014827 | 5.99E-05 | 0.001660364 |
| chr9:78312138-78312443 | 2.080886762 | 6.00E-05 | 0.001662799 |
| chr19:2430953-2431153 | 2.255588241 | 6.01E-05 | 0.001665704 |
| chr5:81171744-81172126 | 1.770590468 | 6.03E-05 | 0.001669486 |
| chr4:176789966-176790283 | -1.671142404 | 6.04E-05 | 0.001672822 |
| chr3:11482446-11482646 | 2.401225904 | 6.07E-05 | 0.001679399 |
| chr1:209207673-209207993 | -1.801879438 | 6.07E-05 | 0.001679399 |
| chr15:89108110-89108364 | 2.004523251 | 6.08E-05 | 0.001682846 |
| chr4:8330708-8330908 | 2.494218681 | 6.09E-05 | 0.00168365 |
| chrX:123814928-123815135 | 2.185069356 | 6.10E-05 | 0.001684487 |
| chr4:184825575-184826841 | -1.421521421 | 6.10E-05 | 0.001684487 |
| chr10:22573740-22574297 | -1.8584563 | 6.10E-05 | 0.001684487 |
| chr9:94987699-94987938 | 2.482364328 | 6.11E-05 | 0.0016875 |
| chr16:46387816-46388058 | 1.460948573 | 6.11E-05 | 0.0016875 |
| chr14:65907351-65907551 | 2.507903531 | 6.12E-05 | 0.001688067 |
| chr14:83954865-83955080 | 2.507903531 | 6.12E-05 | 0.001688067 |
| chr14:84316040-84316550 | 1.657141956 | 6.12E-05 | 0.001688067 |
| chr3:124282887-124283322 | 1.795913 | 6.12E-05 | 0.001688067 |
| chr3:190585805-190587023 | -1.335638588 | 6.15E-05 | 0.001693504 |
| chr7:138773606-138774205 | -1.4494908 | 6.16E-05 | 0.001695775 |
| chr1:189336475-189336764 | 2.093858503 | 6.16E-05 | 0.001695775 |
| chr4:4968398-4968678 | 2.076367887 | 6.17E-05 | 0.001698574 |
| chr15:91942716-91943462 | -1.391442588 | 6.18E-05 | 0.00170031 |
| chr4:125638696-125639710 | -1.411807174 | 6.20E-05 | 0.001705163 |
| chr2:40844260-40844892 | -1.474858817 | 6.20E-05 | 0.001705163 |
| chr8:134495406-134495725 | 1.952557957 | 6.21E-05 | 0.001705594 |
| chr20:48680251-48680575 | 1.388745604 | 6.21E-05 | 0.001706643 |
| chr5:93807897-93808250 | 1.638654325 | 6.23E-05 | 0.00170925 |
| chr13:74330045-74330350 | -2.259771083 | 6.23E-05 | 0.00170925 |
| chr19:58209767-58210032 | 2.088062733 | 6.23E-05 | 0.00170925 |
| chrX:150486056-150486299 | 2.187469755 | 6.23E-05 | 0.001709877 |
| chr7:48311437-48311868 | -1.759101163 | 6.24E-05 | 0.001712605 |
| chr16:4565122-4565354 | 2.086187203 | 6.26E-05 | 0.001715087 |
| chr8:134467690-134468127 | 1.599293794 | 6.26E-05 | 0.001716794 |
| chr20:723797-723997 | -2.646843172 | 6.27E-05 | 0.001718558 |
| chr2:162490015-162490667 | -1.456924475 | 6.28E-05 | 0.001720234 |

| | | | |
|---|---|---|---|
| chr8:107399889-107400197 | 2.294792862 | 6.29E-05 | 0.001722383 |
| chr8:99696216-99696741 | 1.632526863 | 6.31E-05 | 0.00172581 |
| chr1:218025362-218025845 | 1.468404845 | 6.31E-05 | 0.001726314 |
| chr14:29370212-29370492 | 2.089245645 | 6.33E-05 | 0.001731939 |
| chr6:143896093-143897042 | -1.416632916 | 6.34E-05 | 0.00173239 |
| chr12:54053434-54053779 | 1.946758019 | 6.34E-05 | 0.001733131 |
| chr10:67531032-67531354 | -1.740392496 | 6.38E-05 | 0.001742167 |
| chr8:123851991-123852240 | 2.115203266 | 6.38E-05 | 0.001742167 |
| chr5:111984069-111984465 | -1.851700671 | 6.40E-05 | 0.00174718 |
| chr19:29691057-29691260 | 1.982185894 | 6.41E-05 | 0.001749859 |
| chr4:3437069-3437419 | 1.688629527 | 6.42E-05 | 0.001749883 |
| chr11:131033140-131033668 | -1.74902224 | 6.42E-05 | 0.001749945 |
| chr12:14961046-14961602 | -1.768170475 | 6.43E-05 | 0.001751496 |
| chr5:168282184-168282589 | -1.495900107 | 6.43E-05 | 0.001752315 |
| chr2:156675965-156676364 | -1.730863312 | 6.43E-05 | 0.001752315 |
| chrX:149351283-149351601 | 2.255158207 | 6.44E-05 | 0.001753957 |
| chr8:85835939-85836514 | -1.481770709 | 6.44E-05 | 0.001753957 |
| chr2:27171451-27171700 | -2.092146676 | 6.46E-05 | 0.001757884 |
| chr6:55758547-55759086 | -1.591151956 | 6.47E-05 | 0.00176 |
| chr14:71583875-71584137 | 2.502204684 | 6.47E-05 | 0.00176 |
| chr1:214660583-214660787 | -1.973249941 | 6.49E-05 | 0.001764448 |
| chr4:187606550-187606842 | -1.96726518 | 6.52E-05 | 0.001771216 |
| chr4:165398479-165398964 | -1.529079476 | 6.56E-05 | 0.001783019 |
| chr18:48028610-48029064 | -1.633220764 | 6.57E-05 | 0.001783506 |
| chr5:15271094-15272220 | -1.407369426 | 6.59E-05 | 0.00178951 |
| chr8:104101934-104102235 | 1.80488302 | 6.60E-05 | 0.001791142 |
| chr7:26365016-26365373 | 1.727804663 | 6.61E-05 | 0.001792993 |
| chr7:94289070-94289358 | -1.928106182 | 6.62E-05 | 0.001793849 |
| chr5:100789852-100790582 | -1.551125203 | 6.62E-05 | 0.001794663 |
| chr7:130901333-130901748 | -1.489447142 | 6.62E-05 | 0.001794663 |
| chr4:1810015-1810509 | 1.571992235 | 6.64E-05 | 0.001799301 |
| chr4:170853502-170853769 | -1.968194619 | 6.65E-05 | 0.001800757 |
| chr5:119501611-119502159 | 1.645231225 | 6.65E-05 | 0.001800757 |
| chr5:38830519-38831690 | -1.376106581 | 6.66E-05 | 0.001802438 |
| chr2:180337140-180337559 | -1.733672418 | 6.68E-05 | 0.001807129 |
| chr12:113725874-113726177 | 2.074566871 | 6.69E-05 | 0.001809855 |
| chr10:32379404-32379616 | -2.186062853 | 6.70E-05 | 0.001810544 |
| chr9:107260552-107261146 | -1.702484633 | 6.71E-05 | 0.001812843 |
| chr16:77739826-77740068 | 1.825632537 | 6.71E-05 | 0.001813317 |
| chr2:133435343-133435650 | -1.810440006 | 6.73E-05 | 0.001816998 |
| chr19:45540737-45540942 | 2.603105318 | 6.74E-05 | 0.00181975 |
| chr6:26172585-26172864 | 2.076956547 | 6.74E-05 | 0.00181975 |
| chr6:30799181-30799608 | 1.615028553 | 6.77E-05 | 0.001824553 |
| chr16:80324970-80325370 | 1.864983282 | 6.81E-05 | 0.001835504 |
| chr21:39211229-39211594 | 1.864983282 | 6.81E-05 | 0.001835504 |
| chr12:90978059-90978500 | -1.690512066 | 6.81E-05 | 0.001835504 |
| chr1:160159947-160160364 | 1.840370947 | 6.82E-05 | 0.001837654 |
| chr8:101012589-101012914 | 1.57231702 | 6.83E-05 | 0.001837917 |
| chr7:117012740-117012968 | -2.060978126 | 6.83E-05 | 0.001839219 |
| chr15:89756924-89757239 | 1.63881406 | 6.85E-05 | 0.001843923 |

| | | | |
|---|---|---|---|
| chr3:23315769-23316045 | 2.166943416 | 6.86E-05 | 0.001844437 |
| chr21:32403232-32403468 | 1.923926587 | 6.87E-05 | 0.001845484 |
| chrX:144006859-144007093 | 2.235754944 | 6.87E-05 | 0.001845484 |
| chr14:96453635-96453903 | 2.084175995 | 6.88E-05 | 0.001848559 |
| chr20:48790870-48791425 | 1.332151555 | 6.89E-05 | 0.00185126 |
| chr9:109506159-109506424 | 2.224369346 | 6.91E-05 | 0.001855275 |
| chr1:227695863-227696130 | 2.391931315 | 6.94E-05 | 0.00186203 |
| chr7:85628008-85628527 | -1.794613704 | 6.94E-05 | 0.00186203 |
| chr3:130094306-130094919 | -1.583273676 | 6.95E-05 | 0.001864608 |
| chr4:186622239-186622619 | -1.982976553 | 6.96E-05 | 0.001864608 |
| chr20:31068071-31068935 | 1.360242857 | 6.96E-05 | 0.001864608 |
| chr7:119901030-119901405 | -2.065559229 | 6.98E-05 | 0.001869079 |
| chr8:52969616-52969957 | 2.365874761 | 6.98E-05 | 0.00187013 |
| chr2:54755679-54756243 | -1.67956274 | 6.99E-05 | 0.001871814 |
| chr1:59057171-59057821 | -1.40787181 | 7.02E-05 | 0.00187878 |
| chr7:157018952-157019620 | -1.556254275 | 7.05E-05 | 0.001886438 |
| chr7:141754966-141755260 | -1.701014132 | 7.06E-05 | 0.001886758 |
| chr3:25690852-25691560 | 1.446925717 | 7.06E-05 | 0.001886758 |
| chr1:208445724-208446092 | -1.910901225 | 7.06E-05 | 0.00188815 |
| chr7:142856802-142857060 | -1.805283501 | 7.07E-05 | 0.001889046 |
| chr15:94029325-94030034 | -1.455785596 | 7.07E-05 | 0.001889499 |
| chr10:60942774-60944879 | -1.282402659 | 7.08E-05 | 0.001889924 |
| chr14:31401428-31401682 | 2.163091807 | 7.08E-05 | 0.001890656 |
| chr1:153407238-153407533 | 1.935030211 | 7.09E-05 | 0.001890688 |
| chr8:91627001-91627249 | 2.075567189 | 7.09E-05 | 0.001890742 |
| chr4:48728859-48729362 | -1.561493799 | 7.10E-05 | 0.001891577 |
| chr12:113457221-113457439 | 2.236591776 | 7.11E-05 | 0.001894955 |
| chr7:7852371-7853290 | 1.359395717 | 7.11E-05 | 0.001894955 |
| chr15:33866002-33866681 | 1.434059026 | 7.12E-05 | 0.001894955 |
| chr19:52857331-52857880 | 1.490961517 | 7.12E-05 | 0.001894955 |
| chr3:169083957-169084208 | -2.068850296 | 7.14E-05 | 0.001899366 |
| chr4:74661598-74661948 | -1.91955559 | 7.16E-05 | 0.001905471 |
| chr1:117226427-117226637 | 2.115597094 | 7.16E-05 | 0.001905471 |
| chr4:64875-65201 | 1.660589936 | 7.19E-05 | 0.001911598 |
| chr11:129014091-129014429 | 1.662747497 | 7.19E-05 | 0.001911638 |
| chr7:104548930-104549239 | -1.913028718 | 7.21E-05 | 0.001914803 |
| chrX:105635756-105636228 | -1.492990457 | 7.22E-05 | 0.001917493 |
| chr20:8703441-8703921 | 1.846216033 | 7.26E-05 | 0.001927908 |
| chr10:107545071-107546683 | -1.326540724 | 7.26E-05 | 0.001927908 |
| chr21:8401578-8401827 | 1.92551495 | 7.33E-05 | 0.001944932 |
| chr22:20887355-20887811 | -1.625921471 | 7.34E-05 | 0.001946365 |
| chr3:72255300-72255595 | -1.889130465 | 7.34E-05 | 0.001946365 |
| chr1:170614553-170614815 | 2.18108804 | 7.34E-05 | 0.001946365 |
| chr8:69488695-69489483 | -1.418308075 | 7.35E-05 | 0.001946964 |
| chr5:96714976-96715185 | -1.951827199 | 7.37E-05 | 0.001950861 |
| chr7:154183474-154183887 | -1.941059789 | 7.40E-05 | 0.001959066 |
| chr12:113828622-113828829 | 2.289633747 | 7.42E-05 | 0.00196293 |
| chr14:83671385-83671616 | 2.481261821 | 7.43E-05 | 0.001964545 |
| chr8:37144687-37145214 | -1.419183092 | 7.43E-05 | 0.0019649 |
| chr4:93980013-93980244 | 2.473582906 | 7.45E-05 | 0.001969689 |

| | | | |
|---|---|---|---|
| chr4:5038265-5038508 | 2.123597374 | 7.46E-05 | 0.001972008 |
| chr14:54565118-54566228 | 1.298772733 | 7.47E-05 | 0.001973897 |
| chr5:107619652-107620093 | -2.008237669 | 7.49E-05 | 0.001978586 |
| chr8:129932214-129932422 | 2.070731891 | 7.49E-05 | 0.001978586 |
| chr12:22275655-22276265 | -1.432435864 | 7.51E-05 | 0.001982777 |
| chr1:84847182-84848062 | -1.358654181 | 7.53E-05 | 0.001986706 |
| chr21:35256460-35256939 | 1.590152699 | 7.54E-05 | 0.001988765 |
| chr7:15088580-15089284 | -1.595690771 | 7.54E-05 | 0.001989254 |
| chr7:114021128-114021617 | -1.643417437 | 7.55E-05 | 0.001990192 |
| chr4:6453413-6453683 | 2.212303572 | 7.55E-05 | 0.001990192 |
| chr17:81794250-81794877 | 1.619857464 | 7.56E-05 | 0.001990192 |
| chr1:163953212-163953442 | 2.391131902 | 7.56E-05 | 0.001990312 |
| chr13:66037855-66038600 | -1.358312152 | 7.56E-05 | 0.001990312 |
| chr13:82757992-82758459 | -1.633027404 | 7.57E-05 | 0.001991022 |
| chr2:2083257-2083783 | -1.455172627 | 7.57E-05 | 0.001991022 |
| chr13:28623950-28624724 | -1.465079856 | 7.60E-05 | 0.001997502 |
| chr4:172809126-172809728 | -1.474760446 | 7.62E-05 | 0.002002667 |
| chr6:147382990-147383268 | -2.454048099 | 7.63E-05 | 0.002003721 |
| chr3:133448559-133449028 | -1.508518301 | 7.64E-05 | 0.002006572 |
| chr15:56049699-56049997 | 1.793409428 | 7.65E-05 | 0.002008733 |
| chr21:39625019-39625227 | 2.942886036 | 7.65E-05 | 0.002009062 |
| chr5:22991854-22992230 | 1.904788482 | 7.68E-05 | 0.002014908 |
| chr18:35583549-35583852 | 2.297767173 | 7.68E-05 | 0.002015255 |
| chr7:117212908-117213299 | -1.632501276 | 7.69E-05 | 0.002016182 |
| chr4:58817834-58818397 | -1.554099566 | 7.70E-05 | 0.002017525 |
| chr2:154901038-154901532 | -1.681729166 | 7.71E-05 | 0.002019814 |
| chr10:117609069-117609476 | 1.566346704 | 7.71E-05 | 0.002020689 |
| chr2:156803194-156803746 | -1.574329957 | 7.72E-05 | 0.002021839 |
| chr3:148990204-148990410 | -2.057563769 | 7.73E-05 | 0.002022629 |
| chrX:83634885-83635581 | -1.503773662 | 7.74E-05 | 0.002024285 |
| chr7:32259669-32259910 | 1.871140207 | 7.75E-05 | 0.002026271 |
| chr7:83648420-83649316 | -1.440162545 | 7.75E-05 | 0.002027988 |
| chr1:119907000-119907200 | 2.227435239 | 7.79E-05 | 0.00203542 |
| chr19:55210140-55210432 | 1.875285176 | 7.81E-05 | 0.0020403 |
| chr1:162263352-162263552 | 2.487316021 | 7.81E-05 | 0.0020403 |
| chr12:51074162-51074409 | 2.297127333 | 7.81E-05 | 0.002040743 |
| chr17:20821295-20821530 | 2.39172445 | 7.82E-05 | 0.002042467 |
| chr4:61437750-61438120 | -2.059694126 | 7.85E-05 | 0.002048446 |
| chr1:171714727-171714945 | 2.374786871 | 7.86E-05 | 0.002051711 |
| chr12:32060638-32060904 | -1.988576973 | 7.87E-05 | 0.002051823 |
| chr7:116723827-116724573 | -1.374164539 | 7.89E-05 | 0.002056371 |
| chr18:69845609-69846247 | -1.588744314 | 7.89E-05 | 0.002056371 |
| chr8:21611114-21611483 | -1.849952039 | 7.89E-05 | 0.002056371 |
| chr7:79361277-79361509 | -2.363506606 | 7.90E-05 | 0.002056822 |
| chr14:105646570-105647140 | 1.489273957 | 7.90E-05 | 0.002056822 |
| chr1:6058703-6058937 | 2.038857973 | 7.90E-05 | 0.002056822 |
| chr8:98590491-98590722 | 2.357811333 | 7.92E-05 | 0.002059104 |
| chr11:18669286-18670195 | 1.344749919 | 7.92E-05 | 0.002059104 |
| chr10:21525460-21526810 | -1.308278989 | 7.93E-05 | 0.002060675 |
| chr16:76648676-76648876 | 2.059179429 | 7.93E-05 | 0.00206175 |

| | | | |
|---|---|---|---|
| chr8:107697062-107697262 | 2.296158295 | 7.95E-05 | 0.00206642 |
| chr1:195103555-195103841 | 2.105341685 | 7.96E-05 | 0.00206642 |
| chr3:133986302-133986603 | 1.735646347 | 7.96E-05 | 0.002066954 |
| chr1:215113738-215114635 | -1.348988396 | 7.96E-05 | 0.002066954 |
| chr21:39908265-39908888 | -1.393193896 | 7.97E-05 | 0.002067002 |
| chr7:17139282-17139482 | 2.730626414 | 7.97E-05 | 0.002068408 |
| chr4:177546814-177547067 | -2.116179292 | 7.98E-05 | 0.002069423 |
| chr20:15496547-15496776 | 2.223562002 | 7.98E-05 | 0.002069423 |
| chr14:42735521-42735820 | 2.289387202 | 7.99E-05 | 0.002071464 |
| chr16:85465945-85466375 | -1.468049664 | 8.03E-05 | 0.002078567 |
| chr12:123095602-123095828 | -2.076539113 | 8.03E-05 | 0.002078567 |
| chr22:30313051-30313282 | 2.281540068 | 8.04E-05 | 0.002082067 |
| chr7:81073583-81073954 | 1.586281509 | 8.07E-05 | 0.002088426 |
| chr13:45168240-45168660 | -1.656871069 | 8.10E-05 | 0.0020958 |
| chr5:155291078-155291518 | -1.587927247 | 8.11E-05 | 0.00209604 |
| chr1:208496513-208496920 | -1.865021585 | 8.11E-05 | 0.00209604 |
| chr21:41933561-41934105 | -1.642842735 | 8.11E-05 | 0.00209604 |
| chr11:61268750-61269022 | 2.157484379 | 8.12E-05 | 0.002098572 |
| chr3:156879699-156880107 | 1.840858301 | 8.13E-05 | 0.002100417 |
| chr14:86536571-86536771 | 2.476499396 | 8.14E-05 | 0.002100417 |
| chr8:88049662-88049890 | 2.015677385 | 8.14E-05 | 0.002101619 |
| chr14:39853447-39853647 | 2.166875549 | 8.17E-05 | 0.00210724 |
| chr2:165469459-165469848 | -1.532076865 | 8.17E-05 | 0.00210777 |
| chr22:40410276-40410522 | -2.229955283 | 8.19E-05 | 0.002110258 |
| chr14:92780713-92780954 | 2.16804291 | 8.21E-05 | 0.00211507 |
| chr1:117577961-117578259 | 1.783511245 | 8.21E-05 | 0.00211507 |
| chr7:119197616-119198264 | -1.442626119 | 8.21E-05 | 0.00211507 |
| chrX:129139906-129140106 | 2.728360012 | 8.22E-05 | 0.00211507 |
| chr1:110771318-110771556 | 2.162080548 | 8.22E-05 | 0.00211507 |
| chrX:106124236-106124557 | -2.089015559 | 8.22E-05 | 0.00211507 |
| chr10:130036910-130037167 | 2.097088535 | 8.23E-05 | 0.002117445 |
| chr10:129689654-129690164 | 1.495282073 | 8.25E-05 | 0.002120926 |
| chr2:173044837-173045091 | 1.978289717 | 8.27E-05 | 0.002124486 |
| chr1:231606789-231607048 | 1.934830252 | 8.27E-05 | 0.002125311 |
| chr13:41178265-41178532 | -2.498436714 | 8.30E-05 | 0.002132086 |
| chr7:142632566-142632864 | -1.84253279 | 8.30E-05 | 0.002132086 |
| chr18:37299426-37300122 | -1.361323016 | 8.31E-05 | 0.002132498 |
| chr1:220881352-220881560 | -2.16418354 | 8.31E-05 | 0.00213266 |
| chr1:16061368-16061573 | 2.355583802 | 8.32E-05 | 0.002133528 |
| chr8:69780977-69781341 | -2.084397456 | 8.32E-05 | 0.002133528 |
| chr3:158869353-158869612 | 2.025200457 | 8.35E-05 | 0.002140245 |
| chr1:199931730-199932419 | -1.481134976 | 8.35E-05 | 0.002140245 |
| chr3:106284729-106285067 | -1.755540898 | 8.37E-05 | 0.002144746 |
| chr10:784495-785303 | 1.342819939 | 8.38E-05 | 0.00214487 |
| chr7:140517761-140518381 | -1.393827246 | 8.38E-05 | 0.002145239 |
| chr13:47091740-47092033 | 1.779754449 | 8.39E-05 | 0.00214658 |
| chr19:40920868-40921695 | 1.349683269 | 8.39E-05 | 0.002147446 |
| chr16:23257706-23258009 | 1.895214091 | 8.41E-05 | 0.002150469 |
| chr7:117157663-117157871 | -2.205024934 | 8.41E-05 | 0.002150469 |
| chr7:24361972-24362378 | 1.710244113 | 8.41E-05 | 0.002150469 |

| | | | |
|---|---|---|---|
| chr2:8493430-8493667 | 1.953999254 | 8.44E-05 | 0.002155442 |
| chr3:33286542-33287069 | -1.714880634 | 8.45E-05 | 0.002157744 |
| chr17:19364637-19364964 | -1.622765618 | 8.45E-05 | 0.002157871 |
| chr20:47918926-47919195 | 1.42680191 | 8.46E-05 | 0.00215861 |
| chr7:148747322-148747695 | -1.657127908 | 8.47E-05 | 0.002161866 |
| chr9:92729376-92729627 | 2.014083909 | 8.49E-05 | 0.002165165 |
| chr6:36302369-36302855 | 1.55939381 | 8.49E-05 | 0.002165165 |
| chr9:135816576-135816923 | 1.924042332 | 8.49E-05 | 0.002165165 |
| chr7:83876658-83876889 | -2.366091827 | 8.50E-05 | 0.002166303 |
| chr14:99004185-99004483 | 1.668023477 | 8.51E-05 | 0.002166925 |
| chr17:32512442-32512728 | 2.463123644 | 8.53E-05 | 0.002171123 |
| chr14:70130363-70130585 | 2.101275958 | 8.53E-05 | 0.002171123 |
| chr11:111357655-111357969 | -2.064899137 | 8.53E-05 | 0.002171123 |
| chr1:238974650-238975232 | -1.702058439 | 8.53E-05 | 0.002171123 |
| chr14:80215681-80216133 | -1.553827061 | 8.56E-05 | 0.002175326 |
| chr1:196608297-196609644 | -1.281759354 | 8.56E-05 | 0.002175326 |
| chr14:76667458-76667658 | 2.369146137 | 8.56E-05 | 0.002176263 |
| chr18:23257816-23258057 | -1.745044364 | 8.57E-05 | 0.002177758 |
| chr7:36303122-36303813 | -1.352617587 | 8.58E-05 | 0.002179186 |
| chr7:253737-254141 | 1.522510539 | 8.59E-05 | 0.002181645 |
| chr5:64074590-64074896 | -1.806895291 | 8.60E-05 | 0.002183842 |
| chr3:27935580-27935820 | 2.452199177 | 8.63E-05 | 0.00218868 |
| chr2:216620116-216620506 | 1.702134512 | 8.63E-05 | 0.00218868 |
| chr14:22822912-22823298 | 1.637356887 | 8.63E-05 | 0.002189164 |
| chr7:100866434-100866744 | -1.657429905 | 8.69E-05 | 0.002202236 |
| chr20:56530097-56530297 | 1.924789847 | 8.71E-05 | 0.002206858 |
| chr2:165068134-165068391 | -1.937174671 | 8.71E-05 | 0.002206858 |
| chr20:50423220-50423447 | 1.791021317 | 8.72E-05 | 0.002206858 |
| chr4:175253169-175253782 | -1.523278607 | 8.72E-05 | 0.002206858 |
| chr2:202288749-202288957 | -2.621514194 | 8.73E-05 | 0.002209721 |
| chr14:39289505-39289806 | 2.165902577 | 8.74E-05 | 0.002209721 |
| chr11:4531520-4532025 | -1.69802966 | 8.74E-05 | 0.002211317 |
| chr3:172496971-172497305 | 2.147018655 | 8.76E-05 | 0.002215667 |
| chr20:48875748-48876102 | 1.365249375 | 8.78E-05 | 0.00221825 |
| chr14:42421396-42421683 | 2.019291237 | 8.79E-05 | 0.002221055 |
| chr13:89695449-89696057 | -1.528195903 | 8.79E-05 | 0.002221055 |
| chr20:57770369-57770642 | 1.645028333 | 8.83E-05 | 0.002230112 |
| chr18:63063161-63063627 | -1.619098785 | 8.84E-05 | 0.002230268 |
| chr13:22983858-22984397 | 1.390505908 | 8.84E-05 | 0.002230302 |
| chr6:113980934-113981286 | -1.760783077 | 8.86E-05 | 0.002234633 |
| chr1:83818837-83819505 | -1.531191617 | 8.86E-05 | 0.002234633 |
| chr7:12124895-12125283 | 1.709056017 | 8.88E-05 | 0.002238988 |
| chr7:125833543-125833783 | -2.711774907 | 8.90E-05 | 0.002242842 |
| chr14:95297748-95297993 | 1.866592639 | 8.91E-05 | 0.00224355 |
| chr1:44425390-44425598 | 2.562121713 | 8.92E-05 | 0.002244943 |
| chr15:48890081-48890281 | 2.463038574 | 8.93E-05 | 0.002248326 |
| chr2:21403981-21404682 | -1.464760034 | 8.93E-05 | 0.002248326 |
| chr4:99188841-99189238 | -2.197106264 | 8.94E-05 | 0.002249919 |
| chr10:22997156-22997547 | -1.602574758 | 8.99E-05 | 0.002259786 |
| chr16:15857895-15858279 | 1.829593292 | 9.00E-05 | 0.002262646 |

| | | | |
|---|---|---|---|
| chr14:39939645-39939865 | 2.359675722 | 9.02E-05 | 0.002266092 |
| chr10:3831007-3831275 | 1.861830701 | 9.05E-05 | 0.002274229 |
| chr1:244223095-244223379 | 2.08525183 | 9.06E-05 | 0.002274349 |
| chr10:102769424-102769683 | 1.967345189 | 9.06E-05 | 0.002274349 |
| chr16:4774922-4775404 | -1.655030994 | 9.07E-05 | 0.002275759 |
| chr18:57519701-57520034 | -2.430016512 | 9.07E-05 | 0.002275759 |
| chr16:78211648-78212295 | 1.346255453 | 9.09E-05 | 0.002280358 |
| chr12:16076570-16076865 | -1.883686385 | 9.10E-05 | 0.002281146 |
| chr1:32981749-32982027 | 1.99007548 | 9.11E-05 | 0.00228423 |
| chr8:43100409-43100691 | 2.018657218 | 9.12E-05 | 0.002286285 |
| chr8:98809035-98809366 | 1.8406926 | 9.14E-05 | 0.002290217 |
| chr10:53256546-53257088 | -1.582254266 | 9.16E-05 | 0.002292904 |
| chr2:128638965-128639178 | 2.061654343 | 9.17E-05 | 0.00229631 |
| chr14:105569168-105569454 | 1.741162359 | 9.19E-05 | 0.002298516 |
| chr5:88889356-88889848 | -1.778666961 | 9.19E-05 | 0.002298912 |
| chr21:31630197-31630452 | 1.911994048 | 9.20E-05 | 0.002298912 |
| chr7:70716582-70716886 | -1.994331545 | 9.20E-05 | 0.002298912 |
| chr20:48889853-48890257 | 1.352448462 | 9.20E-05 | 0.002298912 |
| chr2:51031811-51032691 | -1.386398294 | 9.20E-05 | 0.002298912 |
| chr7:101590335-101590536 | -2.264680608 | 9.22E-05 | 0.002302785 |
| chr12:48999329-48999625 | -1.824383464 | 9.24E-05 | 0.002307979 |
| chr9:96777633-96778336 | 1.434458064 | 9.25E-05 | 0.002309027 |
| chr8:100667622-100667848 | 1.784660409 | 9.27E-05 | 0.002313006 |
| chr4:10174326-10174747 | 1.47020309 | 9.31E-05 | 0.002321289 |
| chr18:55405759-55406149 | -2.27737018 | 9.32E-05 | 0.002323575 |
| chr11:67062042-67062987 | -1.331455071 | 9.33E-05 | 0.002324973 |
| chr9:118057934-118058617 | -1.65892156 | 9.37E-05 | 0.002336 |
| chr4:7391437-7391663 | 2.550576596 | 9.38E-05 | 0.002336755 |
| chr4:119080706-119081514 | 1.450180162 | 9.39E-05 | 0.002336755 |
| chr8:107764518-107764802 | 2.345187387 | 9.39E-05 | 0.002336755 |
| chr11:101033970-101034439 | -1.712702574 | 9.44E-05 | 0.002348585 |
| chr2:168120902-168121106 | -2.2974985 | 9.45E-05 | 0.002348585 |
| chr3:7264150-7264414 | 2.088726493 | 9.45E-05 | 0.002348585 |
| chr1:64072389-64072632 | 2.143890325 | 9.45E-05 | 0.002348585 |
| chr12:101594750-101595141 | -1.758410455 | 9.45E-05 | 0.002348585 |
| chr13:41098376-41098829 | 1.802958467 | 9.45E-05 | 0.002348585 |
| chr7:34190382-34190618 | 1.937379263 | 9.46E-05 | 0.002348585 |
| chr1:179787476-179787676 | 2.20513116 | 9.49E-05 | 0.002355496 |
| chr3:42079711-42080099 | -1.815963223 | 9.50E-05 | 0.00235844 |
| chr2:188319897-188320186 | -2.045441014 | 9.50E-05 | 0.00235844 |
| chr7:116952891-116954598 | -1.268579618 | 9.55E-05 | 0.00236889 |
| chr14:35385594-35385835 | 1.753924885 | 9.59E-05 | 0.00237813 |
| chr1:187038778-187039207 | -1.652795971 | 9.59E-05 | 0.00237813 |
| chr14:20400968-20401339 | 1.764619245 | 9.64E-05 | 0.002389097 |
| chr14:61104558-61104800 | 2.132032244 | 9.64E-05 | 0.002389097 |
| chr19:19643423-19644173 | -1.422574254 | 9.67E-05 | 0.002394966 |
| chr7:1892280-1892574 | 1.677672934 | 9.70E-05 | 0.00240019 |
| chr19:35362351-35362628 | 2.340567328 | 9.71E-05 | 0.002401959 |
| chr5:18506881-18507460 | 1.394182268 | 9.73E-05 | 0.002406313 |
| chr3:107986462-107986917 | -1.44339035 | 9.73E-05 | 0.002406313 |

| | | | |
|---|---|---|---|
| chr14:77956441-77956829 | 1.642616299 | 9.73E-05 | 0.002406313 |
| chr20:56859702-56860398 | -1.322168801 | 9.75E-05 | 0.002410808 |
| chr7:30988532-30989093 | -1.407065388 | 9.77E-05 | 0.002414689 |
| chr12:130992956-130993183 | 2.352663178 | 9.78E-05 | 0.002416141 |
| chr6:29604401-29604689 | 2.010107701 | 9.80E-05 | 0.002418775 |
| chr9:101790301-101790784 | 1.639135852 | 9.83E-05 | 0.002427617 |
| chr6:43798795-43799379 | 1.425423213 | 9.84E-05 | 0.002427869 |
| chr7:140460867-140461067 | -2.566421304 | 9.85E-05 | 0.002429088 |
| chrX:120824076-120824507 | 1.686889448 | 9.86E-05 | 0.002431147 |
| chr21:41715519-41716239 | -1.402140536 | 9.87E-05 | 0.002432072 |
| chr10:1233570-1233834 | 2.12942447 | 9.87E-05 | 0.002432072 |
| chr1:116901996-116902748 | 1.382081616 | 9.88E-05 | 0.002433679 |
| chr1:147072762-147073065 | 1.839601789 | 9.88E-05 | 0.002434578 |
| chr6:18405664-18405948 | 1.988309138 | 9.90E-05 | 0.002437406 |
| chrX:37912176-37912718 | -1.723889924 | 9.93E-05 | 0.002444746 |
| chr6:87706772-87707128 | 1.758775931 | 9.96E-05 | 0.002450747 |
| chr12:114635949-114636378 | 1.649314687 | 9.96E-05 | 0.002450747 |
| chr1:229845757-229845957 | 2.154571241 | 9.97E-05 | 0.00245115 |
| chr7:74581841-74582434 | -1.393825369 | 0.000100204 | 0.002463218 |
| chr7:102008964-102009285 | -1.829734802 | 0.000100418 | 0.002467093 |
| chr5:102545239-102545575 | -1.573824355 | 0.000100426 | 0.002467093 |
| chr18:38427261-38427584 | -1.893516641 | 0.000100843 | 0.002476546 |
| chr4:10327454-10327838 | 2.063069283 | 0.000101008 | 0.002479803 |
| chr8:32143231-32144145 | -1.569117793 | 0.000101172 | 0.002483023 |
| chr10:24465550-24465992 | -1.597552663 | 0.000101274 | 0.002484731 |
| chr1:247390086-247390550 | -1.317378036 | 0.000101375 | 0.002486408 |
| chr6:82567735-82568044 | -1.761020552 | 0.000101726 | 0.002493524 |
| chr15:91074915-91075214 | 1.996425374 | 0.00010173 | 0.002493524 |
| chr4:168589240-168589560 | -1.814485026 | 0.000101954 | 0.002498207 |
| chr4:74400433-74400753 | -1.884029683 | 0.000102352 | 0.002506632 |
| chr7:23246590-23246925 | 1.501524112 | 0.000102392 | 0.002506632 |
| chr5:170173252-170173609 | -1.722612445 | 0.000102396 | 0.002506632 |
| chr3:43879097-43879760 | -1.541739526 | 0.000102439 | 0.002506897 |
| chr1:198656605-198656953 | 1.599002814 | 0.000102582 | 0.002509595 |
| chr4:153761889-153762170 | 2.198694149 | 0.000102649 | 0.002510437 |
| chr12:107764723-107765010 | 1.564515555 | 0.000102861 | 0.002514343 |
| chr1:247936589-247937709 | -1.276982139 | 0.000102875 | 0.002514343 |
| chr7:142338610-142339169 | -1.549210626 | 0.000102908 | 0.002514352 |
| chr10:56012025-56012545 | -1.468437453 | 0.00010337 | 0.002524323 |
| chr4:182865567-182865998 | -1.634358764 | 0.000103382 | 0.002524323 |
| chr14:31753407-31753619 | 2.137103934 | 0.000103596 | 0.002528743 |
| chr18:78713824-78714151 | -1.99523075 | 0.000103738 | 0.002531399 |
| chr5:111694032-111694502 | -1.810065832 | 0.000103899 | 0.00253453 |
| chr2:147989854-147990117 | -2.03859507 | 0.000104001 | 0.002536205 |
| chr7:109551833-109552258 | -2.066448813 | 0.000104434 | 0.00254589 |
| chr9:93886922-93887192 | -2.180693545 | 0.000104464 | 0.00254589 |
| chr4:184968780-184969140 | -1.764604112 | 0.000104658 | 0.00254979 |
| chr1:190019752-190020139 | -1.418396401 | 0.000104743 | 0.002551063 |
| chr16:14308026-14308375 | 1.601225886 | 0.000104787 | 0.002551307 |
| chr14:72893104-72893774 | 1.439640681 | 0.000104991 | 0.00255547 |

| | | | |
|---|---|---|---|
| chr4:3047433-3048085 | 1.412143343 | 0.000105156 | 0.002558679 |
| chr5:125447031-125447575 | 1.608447246 | 0.000105323 | 0.002561747 |
| chr3:189043677-189043989 | -1.805137488 | 0.000105349 | 0.002561747 |
| chr12:79415778-79416109 | 1.60374204 | 0.000105437 | 0.002562451 |
| chr2:221124896-221125334 | -1.890209817 | 0.000105445 | 0.002562451 |
| chr8:111526514-111526784 | 2.247260693 | 0.000105578 | 0.002564859 |
| chr16:28605399-28605744 | 1.895357459 | 0.000106048 | 0.002575465 |
| chr1:58183361-58183655 | 1.988252928 | 0.000106157 | 0.002577309 |
| chr20:48766575-48766797 | 1.439339956 | 0.000106593 | 0.002587066 |
| chr11:130315745-13031600 | -1.856786331 | 0.000106647 | 0.002587557 |
| chr7:19249007-19249226 | 2.318576233 | 0.000106823 | 0.002591003 |
| chr13:66543826-66544097 | -1.813876842 | 0.000106928 | 0.002592723 |
| chr3:51933106-51933356 | 2.243771374 | 0.00010734 | 0.002601908 |
| chr1:240876183-240876383 | 2.44132051 | 0.000107471 | 0.002604249 |
| chr4:140659873-140660763 | -1.339043687 | 0.000107703 | 0.002609038 |
| chr14:25901241-25901442 | 2.435808597 | 0.000108242 | 0.002621287 |
| chr12:75850797-75851538 | -1.461528649 | 0.000108354 | 0.002623166 |
| chr2:241777234-241777576 | -1.639689726 | 0.000108573 | 0.002627417 |
| chr10:53140495-53140927 | -1.615329477 | 0.000108598 | 0.002627417 |
| chr15:88648673-88648994 | 1.67783089 | 0.000108785 | 0.002630604 |
| chr7:1939569-1940746 | -1.296709577 | 0.000108799 | 0.002630604 |
| chr15:49783752-49784073 | 1.554427628 | 0.000108874 | 0.002631587 |
| chr10:21527069-21527637 | -1.490610466 | 0.000109152 | 0.002637468 |
| chr1:188713231-188713679 | 1.579660937 | 0.000109294 | 0.002640078 |
| chr10:13706828-13707271 | -1.675069796 | 0.000109373 | 0.002641145 |
| chr7:102917653-102917917 | -2.302537522 | 0.000109526 | 0.002644013 |
| chr5:53959741-53960139 | -1.870943327 | 0.000109571 | 0.002644271 |
| chr20:50014300-50014582 | 1.625669796 | 0.000109633 | 0.002644928 |
| chr14:34078297-34078508 | 1.873944108 | 0.000109714 | 0.002646045 |
| chr8:98110546-98110746 | 2.178565791 | 0.00010992 | 0.002650197 |
| chr16:21554669-21555141 | 1.448139398 | 0.000110013 | 0.002651446 |
| chr2:110732453-110732935 | 1.619974541 | 0.000110077 | 0.002651446 |
| chr21:36193242-36193641 | 1.486199902 | 0.00011012 | 0.002651446 |
| chr9:120386509-120387222 | -1.379356184 | 0.000110123 | 0.002651446 |
| chr7:53031784-53032491 | -1.420585385 | 0.000110145 | 0.002651446 |
| chrX:137929926-137930360 | 1.583716953 | 0.000110346 | 0.002655445 |
| chr7:18824339-18824610 | 2.059551719 | 0.000110489 | 0.002658041 |
| chr3:157494261-157494960 | -1.359246357 | 0.000110525 | 0.002658084 |
| chr20:33138852-33139085 | 2.536293641 | 0.00011082 | 0.002664345 |
| chr15:56549181-56549422 | 2.325266168 | 0.000111228 | 0.002673314 |
| chr18:72653061-72653283 | -2.200899424 | 0.000111277 | 0.002673647 |
| chr15:57496922-57497373 | 1.596389303 | 0.000111353 | 0.002674632 |
| chr18:78431580-78431856 | -2.196326993 | 0.000111472 | 0.002676667 |
| chr7:156545014-156545616 | -1.542366803 | 0.00011152 | 0.00267698 |
| chr1:247272859-247273535 | -1.453034095 | 0.000111585 | 0.0026777 |
| chr3:172156306-172156688 | -1.547327131 | 0.000111695 | 0.002679488 |
| chr3:76949505-76950011 | 1.61853354 | 0.000111734 | 0.002679559 |
| chr12:131863222-131863592 | -1.587858355 | 0.000111768 | 0.002679559 |
| chr10:84279820-84280609 | -1.435231556 | 0.000111895 | 0.002681759 |
| chr14:35347088-35347509 | 1.558734919 | 0.000112086 | 0.002685508 |

| | | | |
|---|---|---|---|
| chr10:26869284-26869893 | -1.520203834 | 0.000112504 | 0.002694677 |
| chr2:45664019-45664320 | -2.134129027 | 0.000112663 | 0.002697652 |
| chr6:87452249-87452548 | 1.816651556 | 0.000112813 | 0.0027004 |
| chrX:79325165-79325700 | -1.578104554 | 0.000112872 | 0.002700964 |
| chr14:61152819-61153264 | -1.560127516 | 0.000112921 | 0.002701297 |
| chr20:34345421-34345762 | 2.050853644 | 0.000113599 | 0.002716662 |
| chr14:105723623-105724207 | 1.580746144 | 0.000113684 | 0.002717848 |
| chr13:41587534-41587928 | -1.604527862 | 0.000113772 | 0.002719096 |
| chr1:173682226-173682456 | 1.979426766 | 0.000114019 | 0.002724166 |
| chr6:82347952-82348507 | -1.491676334 | 0.000114112 | 0.002725535 |
| chr16:89395153-89395666 | -1.439074366 | 0.00011423 | 0.002727502 |
| chr11:101079584-101079948 | -1.917366393 | 0.000114436 | 0.002730978 |
| chr7:103504490-103505402 | -1.443713588 | 0.000114447 | 0.002730978 |
| chr12:131702406-131702639 | 2.030265907 | 0.0001146 | 0.002733781 |
| chr8:117671950-117672350 | -1.742360827 | 0.000114721 | 0.002734714 |
| chr18:62452197-62452597 | -1.747736859 | 0.000114733 | 0.002734714 |
| chr3:10553575-10553963 | -1.553671275 | 0.000114746 | 0.002734714 |
| chr4:70019028-70019344 | -1.869198286 | 0.000114797 | 0.002735028 |
| chrX:145600232-145600507 | 2.309489096 | 0.000114831 | 0.002735028 |
| chr12:70796985-70797314 | 1.75808496 | 0.000115116 | 0.002740431 |
| chr17:18235655-18236336 | -1.37639844 | 0.000115129 | 0.002740431 |
| chr3:7106760-7107025 | 1.889158857 | 0.000115174 | 0.002740647 |
| chr8:11730665-11731049 | -1.727221301 | 0.000115509 | 0.002747769 |
| chr7:17752921-17753283 | 1.728225011 | 0.000115598 | 0.002749018 |
| chr1:31955133-31955349 | 2.247156902 | 0.00011582 | 0.002752679 |
| chr3:135926941-135927565 | -1.527911036 | 0.000115851 | 0.002752679 |
| chr11:77141012-77141335 | 1.606668242 | 0.00011586 | 0.002752679 |
| chrX:28586325-28586590 | -2.052596812 | 0.000116103 | 0.002756344 |
| chr1:220878898-220879805 | -1.392931267 | 0.000116138 | 0.002756344 |
| chr5:80247082-80247921 | -1.310938341 | 0.000116151 | 0.002756344 |
| chr14:71471591-71472232 | 1.391871519 | 0.000116158 | 0.002756344 |
| chr10:9038154-9038852 | -1.350432942 | 0.000116757 | 0.0027697 |
| chr14:32820658-32820939 | 1.883832819 | 0.000116909 | 0.002772466 |
| chr2:156986312-156986666 | -1.77520019 | 0.000117129 | 0.002776811 |
| chr2:11121462-11122371 | -1.363556728 | 0.00011718 | 0.002777161 |
| chr1:187655553-187655826 | 2.252913202 | 0.000117295 | 0.002779026 |
| chr7:131859175-131859530 | -1.578812471 | 0.000117396 | 0.00278056 |
| chr5:112182993-112183687 | -1.467140467 | 0.000117887 | 0.002790753 |
| chr4:186201583-186202236 | -1.407134813 | 0.000117955 | 0.002790753 |
| chr9:113096698-113096928 | 2.314584575 | 0.000117975 | 0.002790753 |
| chr22:33400902-33401174 | 2.41144174 | 0.000118 | 0.002790753 |
| chr7:119320336-119320578 | -2.422310391 | 0.000118038 | 0.002790753 |
| chr22:20720992-20721227 | 2.323401098 | 0.000118045 | 0.002790753 |
| chr4:182925285-182925750 | -1.51153659 | 0.000118421 | 0.002798492 |
| chr13:37918763-37919335 | -1.593609696 | 0.000118445 | 0.002798492 |
| chr11:62500256-62500545 | 1.700548686 | 0.000118627 | 0.00280182 |
| chr15:100889403-100889746 | 1.684909173 | 0.000118659 | 0.00280182 |
| chr1:145232819-145233206 | 1.980462297 | 0.000118707 | 0.002802093 |
| chr10:69890582-69890802 | 1.910480156 | 0.000119001 | 0.002808169 |
| chr15:93117204-93117418 | 2.187334938 | 0.000119138 | 0.002810533 |

| | | | |
|---|---|---|---|
| chr4:169033604-169034055 | -1.412788174 | 0.000119189 | 0.002810864 |
| chr4:555743-555947 | 2.522572533 | 0.000119424 | 0.00281554 |
| chr1:91745005-91745205 | -2.618017974 | 0.000120089 | 0.002829684 |
| chr7:112789534-112790846 | -1.258045452 | 0.00012011 | 0.002829684 |
| chr19:54432085-54432438 | 1.526774484 | 0.000120135 | 0.002829684 |
| chr2:219968606-219969153 | -1.420658176 | 0.000120302 | 0.002832543 |
| chr8:125278214-125278562 | -1.48771371 | 0.00012033 | 0.002832543 |
| chr15:38034924-38035166 | 2.127612042 | 0.000120987 | 0.002847133 |
| chr14:35669799-35670247 | 1.408663575 | 0.000121419 | 0.002856419 |
| chr10:110845514-110846458 | -1.337826416 | 0.000121557 | 0.002858801 |
| chr5:20622042-20622257 | 2.316425994 | 0.000121756 | 0.00286118 |
| chr15:62549511-62549761 | 2.115836631 | 0.000121758 | 0.00286118 |
| chr2:10407896-10408263 | -1.50935493 | 0.000121771 | 0.00286118 |
| chr1:14940160-14940401 | 2.329408745 | 0.000122051 | 0.0028669 |
| chr4:183960070-183960379 | -1.894548397 | 0.000122152 | 0.002868392 |
| chr2:212956355-212957227 | -1.361033689 | 0.000122267 | 0.002870205 |
| chr10:3869169-3869415 | 2.114328278 | 0.000122659 | 0.002878528 |
| chr5:74017130-74017405 | -1.897051549 | 0.000122791 | 0.002880732 |
| chr7:121994606-121994874 | -1.854678489 | 0.00012338 | 0.00289366 |
| chr4:186554418-186555389 | -1.382212779 | 0.00012383 | 0.002903327 |
| chr8:116802832-116803293 | 1.402570569 | 0.000124054 | 0.002907696 |
| chr19:44625998-44626281 | 1.930196575 | 0.00012425 | 0.002911401 |
| chr7:87316847-87317532 | -1.447115582 | 0.000124336 | 0.002912519 |
| chr8:128133938-128134368 | 1.389760836 | 0.000124851 | 0.002922696 |
| chr20:19784583-19785219 | 1.347770097 | 0.000124874 | 0.002922696 |
| chr13:48078149-48078548 | -1.682302702 | 0.000124885 | 0.002922696 |
| chr7:10727921-10728168 | 2.163437045 | 0.000124966 | 0.002923281 |
| chr9:1173081-1173564 | 1.482640088 | 0.000124986 | 0.002923281 |
| chr12:100952352-100952902 | -1.452930939 | 0.000125061 | 0.002923837 |
| chr17:82522090-82522503 | -1.553303201 | 0.000125086 | 0.002923837 |
| chr7:102318506-102319212 | -1.301816108 | 0.000125452 | 0.002931501 |
| chr11:68260700-68260962 | 1.846260595 | 0.000125904 | 0.002941164 |
| chr5:116546117-116546378 | -1.93612439 | 0.000126174 | 0.002946566 |
| chr19:41948804-41949004 | 2.240297401 | 0.000126236 | 0.002947133 |
| chr10:52472266-52472872 | -1.388017183 | 0.000126303 | 0.002947294 |
| chr16:81866819-81867077 | 1.979064072 | 0.00012632 | 0.002947294 |
| chr7:106365303-106365740 | -1.520783684 | 0.000126442 | 0.002949234 |
| chr2:58428810-58429332 | -1.608121047 | 0.000126838 | 0.002957583 |
| chr12:125890538-125890824 | 1.91254764 | 0.000126963 | 0.002959578 |
| chr15:83278184-83278423 | -2.257240883 | 0.000127041 | 0.002960497 |
| chr5:104079713-104080447 | -1.382108189 | 0.000127145 | 0.002962027 |
| chrX:119401490-119401964 | 1.432712249 | 0.000127265 | 0.002963925 |
| chr20:48710999-48711203 | 1.415529299 | 0.000127656 | 0.002972119 |
| chr3:195769580-195769780 | 2.167058417 | 0.000127782 | 0.002974154 |
| chr3:108114339-108115032 | -1.426435906 | 0.000128058 | 0.002979393 |
| chr7:21938582-21938832 | 1.721044016 | 0.000128085 | 0.002979393 |
| chr7:130577921-130578137 | -2.680305777 | 0.000128183 | 0.002980257 |
| chr13:70611682-70611934 | -2.727776936 | 0.0001282 | 0.002980257 |
| chr4:183566340-183566713 | -1.860545324 | 0.000128315 | 0.002982035 |
| chr12:25457975-25458671 | -1.363291448 | 0.00012847 | 0.002984738 |

| | | | |
|---|---|---|---|
| chr7:140624175-140624767 | -1.386796293 | 0.000128567 | 0.002986069 |
| chr13:79307768-79308296 | -1.633945256 | 0.000129138 | 0.002998435 |
| chr12:68236586-68237096 | -1.612369791 | 0.000129219 | 0.002999391 |
| chr9:80481800-80482695 | -1.363255981 | 0.00012926 | 0.002999441 |
| chr7:77537990-77538533 | -1.380837998 | 0.000129542 | 0.003004436 |
| chr8:107606636-107607047 | 1.470328612 | 0.000129554 | 0.003004436 |
| chr6:119289773-119289974 | 2.102156961 | 0.000130071 | 0.003015529 |
| chr7:12898275-12898497 | 2.150065305 | 0.000130205 | 0.003017719 |
| chr1:148086270-148086517 | -2.14262084 | 0.000130505 | 0.003023747 |
| chr7:8284071-8284343 | 1.70145413 | 0.00013066 | 0.00302608 |
| chr13:50209004-50209204 | -2.442492117 | 0.000130684 | 0.00302608 |
| chr7:21681992-21682239 | 2.210873459 | 0.00013074 | 0.003026448 |
| chr16:78240490-78240923 | 1.388840513 | 0.000130794 | 0.003026784 |
| chr7:141649871-141650528 | -1.356601548 | 0.000131201 | 0.003035281 |
| chrX:29478115-29478997 | -1.363614864 | 0.000131285 | 0.003036311 |
| chr18:41391605-41391914 | -1.890799686 | 0.000131427 | 0.003038676 |
| chr13:114077990-114078260 | -1.895185285 | 0.000131631 | 0.003042479 |
| chr1:186523880-186524144 | 2.223319489 | 0.000131722 | 0.003043208 |
| chr7:12647471-12648069 | 1.371251982 | 0.000131742 | 0.003043208 |
| chr8:89253989-89254395 | 1.55256287 | 0.000131805 | 0.003043754 |
| chr7:23759759-23760012 | 1.868830515 | 0.00013191 | 0.00304526 |
| chr6:137044060-137045670 | -1.271491077 | 0.000132071 | 0.003048058 |
| chr7:127477654-127478266 | -1.540095263 | 0.000132187 | 0.003049812 |
| chr8:103624418-103624720 | 1.93201989 | 0.000132914 | 0.003065673 |
| chr5:160685352-160685985 | -1.552002661 | 0.000133085 | 0.003068693 |
| chr7:107768334-107768664 | -1.830541906 | 0.000133186 | 0.003069585 |
| chr14:46551320-46551545 | -2.479707573 | 0.000133231 | 0.003069585 |
| chr14:32134582-32135068 | 1.499591733 | 0.000133244 | 0.003069585 |
| chr21:41520618-41520818 | 2.30347422 | 0.000133501 | 0.003074582 |
| chr18:32308825-32309025 | -2.47204747 | 0.000133848 | 0.003081352 |
| chr1:223148837-223149544 | -1.462813427 | 0.000133876 | 0.003081352 |
| chr2:200741482-200742217 | 1.32524715 | 0.000133958 | 0.003082315 |
| chr3:8556434-8556776 | -1.756111889 | 0.000134085 | 0.003084306 |
| chr13:76813931-76814269 | -1.862976315 | 0.000134151 | 0.003084903 |
| chr1:238581354-238581767 | -1.903350498 | 0.000134225 | 0.003085696 |
| chr19:46491789-46492020 | 1.846812026 | 0.000134301 | 0.003086499 |
| chr5:766112-766421 | 1.750301216 | 0.000134401 | 0.003087626 |
| chr11:130450050-130450408 | -1.64363956 | 0.00013443 | 0.003087626 |
| chr7:18085908-18087051 | 1.296509038 | 0.000134875 | 0.003096901 |
| chr8:106206164-106206534 | 1.520624023 | 0.000134975 | 0.003098281 |
| chr4:107690148-107690587 | 1.591011874 | 0.000135046 | 0.003098928 |
| chr7:29456304-29456601 | 2.097065411 | 0.000135086 | 0.003098928 |
| chr1:35298611-35298899 | 1.775591237 | 0.000135125 | 0.003098928 |
| chr2:9754155-9754584 | -1.575641123 | 0.000135245 | 0.003100754 |
| chr3:157070600-157071046 | -1.475744124 | 0.000135885 | 0.003114088 |
| chr15:90344998-90345260 | 1.898358276 | 0.000135908 | 0.003114088 |
| chr11:67372942-67373382 | 1.40143265 | 0.000136092 | 0.003117182 |
| chr4:185115925-185116242 | -1.800514877 | 0.000136144 | 0.003117182 |
| chr4:86970547-86971002 | -1.498328641 | 0.000136165 | 0.003117182 |
| chr18:74234346-74234569 | -2.56542221 | 0.000136517 | 0.003124318 |

| | | | |
|---|---|---|---|
| chr8:115866809-115868110 | -1.228401929 | 0.000136644 | 0.003126284 |
| chrX:85289348-85289809 | -1.871565561 | 0.000136725 | 0.003126788 |
| chr7:105673335-105673990 | -1.543908333 | 0.000136748 | 0.003126788 |
| chr14:60233249-60233553 | -1.806167904 | 0.000137029 | 0.003132285 |
| chr6:37307828-37308054 | 1.964636282 | 0.000137459 | 0.003141171 |
| chr15:56711683-56712222 | 1.616196444 | 0.0001376 | 0.003142955 |
| chr10:490973-491260 | 1.583702 | 0.000137619 | 0.003142955 |
| chr4:14879558-14880115 | 1.414741055 | 0.000137867 | 0.003147679 |
| chr13:23187042-23187319 | 1.90073444 | 0.000137999 | 0.003149755 |
| chr7:82559343-82559752 | -1.797183215 | 0.000138227 | 0.003154029 |
| chr2:21818558-21818917 | -2.204197416 | 0.000138387 | 0.003156751 |
| chr7:143884902-143885566 | -1.35731119 | 0.000138516 | 0.003158751 |
| chr4:183722219-183722570 | -1.714394501 | 0.000138577 | 0.003159199 |
| chr13:80882140-80882497 | -1.722834832 | 0.000138697 | 0.003160989 |
| chr2:219785255-219785523 | 2.06070499 | 0.000138786 | 0.003162085 |
| chr2:12050407-12050785 | -1.672152613 | 0.000138897 | 0.003163656 |
| chr3:161569371-161569795 | 1.562563172 | 0.000139014 | 0.00316539 |
| chr17:78374253-78374540 | 1.581471802 | 0.000139088 | 0.003166133 |
| chr14:85596189-85596428 | 2.232382663 | 0.000139263 | 0.003168933 |
| chr7:24970789-24971035 | 1.969975602 | 0.000139326 | 0.003168933 |
| chr7:158817026-158817283 | -1.822712108 | 0.000139335 | 0.003168933 |
| chr5:19734751-19735451 | 1.395120156 | 0.000139704 | 0.003176377 |
| chr5:149877868-149878717 | -1.350057791 | 0.000140088 | 0.003184172 |
| chr5:9299566-9299816 | 2.286636251 | 0.000140427 | 0.003190935 |
| chr3:176633563-176633863 | -1.78307549 | 0.00014049 | 0.003191407 |
| chr5:17316755-17317391 | 1.373727884 | 0.000140617 | 0.003193351 |
| chr3:98919469-98919718 | -2.052455219 | 0.00014087 | 0.003198145 |
| chr10:22428155-22428462 | 1.840368458 | 0.0001411 | 0.003201467 |
| chr7:21641777-21641977 | 2.500883822 | 0.0001411 | 0.003201467 |
| chr10:22655146-22655388 | -2.11288505 | 0.000141478 | 0.003209085 |
| chr3:40588368-40588731 | 1.659896288 | 0.000141796 | 0.003215361 |
| chr16:383707-383925 | 1.932515598 | 0.000141874 | 0.003216163 |
| chr3:101036257-101036645 | 1.639214702 | 0.000142161 | 0.003221721 |
| chr7:24217612-24217812 | 2.203868123 | 0.000142209 | 0.003221857 |
| chr18:74082425-74082905 | -1.431998726 | 0.000142357 | 0.003224269 |
| chr6:9463125-9463648 | -1.516032021 | 0.000142455 | 0.003225526 |
| chr18:39647880-39648484 | -1.591492383 | 0.00014272 | 0.003230574 |
| chr10:124665692-124665991 | 1.680582124 | 0.000142894 | 0.003233564 |
| chrX:26783500-26783877 | -1.794280345 | 0.000142992 | 0.003234808 |
| chr1:247431393-247432118 | -1.415832025 | 0.0001431 | 0.003236304 |
| chrX:154428380-154428785 | 1.395929807 | 0.000143152 | 0.003236517 |
| chr10:6126577-6126780 | 2.218336309 | 0.000143241 | 0.003237592 |
| chr1:104648237-104648660 | -1.718780604 | 0.000143719 | 0.003247442 |
| chr5:96108498-96108883 | -1.793812928 | 0.000143774 | 0.003247719 |
| chr14:69954736-69954936 | 2.155596559 | 0.000143968 | 0.003251148 |
| chr3:120438999-120439395 | 1.571139178 | 0.000144219 | 0.003255848 |
| chr5:119255692-119256437 | -1.363115975 | 0.000144418 | 0.003259382 |
| chr7:4724662-4725952 | -1.268640484 | 0.000144615 | 0.003262872 |
| chr8:14305704-14306057 | -1.836569727 | 0.000145066 | 0.003272081 |
| chr2:28325924-28326978 | -1.326791577 | 0.000145119 | 0.003272309 |

| | | | |
|---|---|---|---|
| chr17:14737863-14738607 | -1.316726412 | 0.000145176 | 0.003272641 |
| chr3:168480268-168480571 | 2.212884796 | 0.000145397 | 0.003276651 |
| chrX:16187097-16187630 | 1.81317506 | 0.000145534 | 0.00327877 |
| chr1:114657663-114658014 | 1.388688738 | 0.000145603 | 0.003279354 |
| chr7:128596399-128597121 | -1.365574281 | 0.000145832 | 0.003283549 |
| chr9:30264348-30264574 | 2.495018138 | 0.000146249 | 0.003290931 |
| chr7:17186445-17186661 | 2.037624474 | 0.000146294 | 0.003290931 |
| chr3:1250394-1250672 | 1.994521387 | 0.00014633 | 0.003290931 |
| chr6:165708433-165708780 | -1.85255719 | 0.00014636 | 0.003290931 |
| chr3:160653376-160653608 | 2.096637588 | 0.000146374 | 0.003290931 |
| chr2:206921220-206921659 | 1.954570894 | 0.000146613 | 0.003295319 |
| chr14:47919202-47919402 | 2.40838703 | 0.000146966 | 0.003302293 |
| chr16:29710271-29710661 | 1.6602786 | 0.000147092 | 0.003304167 |
| chr1:244273919-244274277 | 1.733300116 | 0.0001474 | 0.003310111 |
| chr5:84287997-84288308 | -1.843892689 | 0.000147781 | 0.003317683 |
| chr1:237288224-237288487 | 2.047333639 | 0.000147867 | 0.003318657 |
| chr2:216884174-216884419 | 2.041067149 | 0.000147964 | 0.003319849 |
| chr8:97476061-97476438 | 1.798032876 | 0.000148086 | 0.003321628 |
| chr5:101390211-101390621 | -1.608060475 | 0.000148231 | 0.00332389 |
| chr19:19146031-19146270 | -2.04977648 | 0.000148286 | 0.003324168 |
| chr20:48386138-48386411 | 1.365423355 | 0.000148516 | 0.00332835 |
| chr17:47824338-47824618 | 1.772561908 | 0.000148823 | 0.003334254 |
| chr10:132447832-132448090 | 1.490118582 | 0.000149184 | 0.00334077 |
| chr14:39423983-39424232 | 1.761845314 | 0.000149201 | 0.00334077 |
| chr3:127052518-127052760 | 1.886853962 | 0.000149278 | 0.003341508 |
| chr15:50815948-50816583 | 1.257472909 | 0.000149686 | 0.003349374 |
| chr3:27360339-27360962 | -1.366297413 | 0.000149745 | 0.003349374 |
| chr20:48355519-48355854 | 1.367603533 | 0.000149803 | 0.003349374 |
| chr15:28222664-28222959 | 1.988240433 | 0.000149804 | 0.003349374 |
| chr5:160287036-160288110 | -1.321221014 | 0.00015058 | 0.003365727 |
| chr12:19138541-19138869 | -2.092428984 | 0.000150693 | 0.003367271 |
| chr4:160307533-160307780 | -2.306027036 | 0.000150803 | 0.003367829 |
| chr11:8597274-8597692 | -1.823333022 | 0.000150806 | 0.003367829 |
| chr3:154852402-154852698 | 1.830001947 | 0.000151023 | 0.003371692 |
| chr14:39393362-39393621 | 1.805431441 | 0.000151276 | 0.003376377 |
| chr17:74176713-74177040 | 1.510850231 | 0.000151361 | 0.003377279 |
| chr8:107265484-107265805 | 1.619466407 | 0.000151708 | 0.003383535 |
| chr8:125025030-125025409 | 1.574899645 | 0.00015173 | 0.003383535 |
| chr2:43158295-43159086 | -1.322358215 | 0.000152373 | 0.003395912 |
| chr2:112692440-112692935 | 1.420570358 | 0.000152373 | 0.003395912 |
| chr2:160932914-160933392 | -1.431466335 | 0.000152657 | 0.003400157 |
| chr6:111711454-111711756 | -1.810440191 | 0.000152661 | 0.003400157 |
| chr5:20555091-20555338 | -2.087198981 | 0.000152697 | 0.003400157 |
| chr14:105225157-105225398 | 1.691937384 | 0.000152807 | 0.003401624 |
| chr7:142181896-142182302 | -1.651808441 | 0.000152923 | 0.003403207 |
| chr8:86428821-86429056 | 1.804195702 | 0.00015325 | 0.003409039 |
| chr7:19612389-19612642 | 2.379037977 | 0.000153274 | 0.003409039 |
| chr5:68178600-68178924 | -1.749139405 | 0.000153342 | 0.003409569 |
| chr14:95685030-95685282 | 1.912219719 | 0.000153495 | 0.003411515 |
| chr3:75906380-75907591 | -1.381299459 | 0.000153545 | 0.003411515 |

| | | | |
|---|---|---|---|
| chr13:28548902-28549158 | 1.787063915 | 0.000153563 | 0.003411515 |
| chr2:132068730-132068986 | 2.04132835 | 0.000153668 | 0.003412858 |
| chr13:78324691-78324936 | -2.042083075 | 0.000154299 | 0.003425879 |
| chr12:87795739-87795969 | 1.985049756 | 0.000154371 | 0.003425899 |
| chr14:105169081-105169407 | 1.474211146 | 0.000154389 | 0.003425899 |
| chr4:115340053-115340426 | -1.696544728 | 0.000154484 | 0.003426454 |
| chr10:11859981-11860312 | 1.886809778 | 0.000154504 | 0.003426454 |
| chr3:102205941-102206199 | 1.952831985 | 0.000154554 | 0.003426582 |
| chr3:33218538-33218781 | 2.381812776 | 0.000154885 | 0.003431558 |
| chr5:108437305-108437910 | -1.491114955 | 0.00015491 | 0.003431558 |
| chr20:48774170-48774406 | 1.400878442 | 0.000154913 | 0.003431558 |
| chr1:79361138-79361532 | -1.674695927 | 0.000155047 | 0.003432624 |
| chr1:184425463-184426328 | -1.323824592 | 0.000155051 | 0.003432624 |
| chr13:66657147-66657866 | -1.400725106 | 0.0001557 | 0.003445998 |
| chr6:27495484-27496366 | -1.374819546 | 0.000155931 | 0.003450119 |
| chr7:99210286-99210560 | -1.97451817 | 0.000156325 | 0.003457171 |
| chr5:149937487-149938013 | -1.479044236 | 0.00015634 | 0.003457171 |
| chr14:90757005-90757230 | -1.845095159 | 0.000156404 | 0.003457581 |
| chr9:107341114-107341571 | -1.442808919 | 0.000156964 | 0.003468482 |
| chr3:186772857-186773133 | -1.990038345 | 0.000156987 | 0.003468482 |
| chr3:100426144-100426448 | -1.867642064 | 0.000157048 | 0.003468808 |
| chr21:31780921-31781248 | 1.527638799 | 0.000157197 | 0.003470352 |
| chr16:77875933-77876208 | 1.721689616 | 0.000157208 | 0.003470352 |
| chr13:35855533-35856033 | -1.519548326 | 0.000157495 | 0.003475644 |
| chr14:105682936-105683437 | 1.472220475 | 0.000157538 | 0.003475644 |
| chr6:109527983-109528270 | -1.900500196 | 0.00015867 | 0.003499604 |
| chr9:74770499-74771261 | -1.382208274 | 0.000158756 | 0.003499721 |
| chrX:100745770-100746027 | -1.927520883 | 0.000158767 | 0.003499721 |
| chr2:210197094-210197405 | 2.201996749 | 0.0001589 | 0.003501641 |
| chrX:123678697-123679017 | 2.182323499 | 0.000159021 | 0.003502672 |
| chr1:209427795-209428142 | -1.605207092 | 0.000159038 | 0.003502672 |
| chr5:4028279-4029304 | -1.360229652 | 0.000159116 | 0.003503393 |
| chr10:21732187-21732512 | -1.60708951 | 0.000159193 | 0.00350408 |
| chr17:57535295-57535617 | 1.467038786 | 0.000159318 | 0.00350582 |
| chr7:131654731-131655217 | -1.557904428 | 0.000159612 | 0.003511283 |
| chr6:37759224-37759700 | 1.408474818 | 0.000159819 | 0.003513924 |
| chr1:245345108-245345863 | -1.404026483 | 0.000159825 | 0.003513924 |
| chr7:138449633-138450253 | -1.736917133 | 0.00015987 | 0.003513924 |
| chr7:107147519-107147827 | -1.638049686 | 0.000160146 | 0.003518983 |
| chr17:80900747-80901292 | -1.443432929 | 0.000160196 | 0.003519073 |
| chr8:68853237-68854824 | -1.222681232 | 0.000160647 | 0.003527966 |
| chr6:38253402-38253617 | 2.19662265 | 0.000160939 | 0.003533363 |
| chr13:79361298-79361670 | -1.709052597 | 0.000161075 | 0.003535347 |
| chr14:42241665-42241905 | 2.285457055 | 0.000161149 | 0.003535959 |
| chr1:107008514-107009007 | 1.345933514 | 0.000161497 | 0.003542585 |
| chr5:113090630-113091470 | -1.343126811 | 0.000161796 | 0.003548118 |
| chr3:172129886-172130701 | -1.263014517 | 0.00016192 | 0.003549818 |
| chr7:19272888-19273149 | 2.279579956 | 0.000162169 | 0.003554274 |
| chr2:237731738-237732002 | -2.027993096 | 0.000162262 | 0.003555284 |
| chr7:96135181-96135683 | -1.681259903 | 0.000162377 | 0.003556788 |

| | | | |
|---|---|---|---|
| chr8:54405450-54405746 | -1.976448409 | 0.000162641 | 0.003561558 |
| chr7:31671656-31671861 | 1.995279342 | 0.000163033 | 0.003569127 |
| chr14:82740733-82741079 | 1.666236991 | 0.000163254 | 0.003572931 |
| chr14:97977970-97978196 | -2.036976336 | 0.000163601 | 0.003579517 |
| chr1:114088846-114089065 | 1.451937678 | 0.000163663 | 0.003579845 |
| chr7:25031436-25031638 | 1.945247331 | 0.000164736 | 0.003602284 |
| chr8:122803001-122804002 | -1.268009267 | 0.000164934 | 0.003604717 |
| chr1:175655879-175656109 | 2.198659002 | 0.000164941 | 0.003604717 |
| chr1:108721347-108721642 | -1.844290799 | 0.000165508 | 0.003616069 |
| chr14:47103803-47104121 | 2.067813913 | 0.000165627 | 0.003617629 |
| chr22:11618106-11618381 | 1.83377601 | 0.000165801 | 0.003620232 |
| chr7:51376418-51376799 | 1.575756282 | 0.00016584 | 0.003620232 |
| chr6:78646680-78647469 | -1.392295547 | 0.000166168 | 0.003626352 |
| chr7:127823061-127823438 | -1.568526065 | 0.000166511 | 0.003632805 |
| chr7:133182840-133183708 | -1.382795076 | 0.000167013 | 0.003642721 |
| chr6:13454690-13455299 | -1.427762595 | 0.000167083 | 0.003643198 |
| chr1:214213210-214213570 | 1.790738551 | 0.000167158 | 0.003643806 |
| chr10:19038427-19038847 | -1.607540753 | 0.000167223 | 0.003644185 |
| chr4:150591358-150591870 | -1.442342451 | 0.000167423 | 0.003647506 |
| chr4:129544974-129545174 | 2.487331646 | 0.000167747 | 0.003653528 |
| chr4:188427262-188427646 | -1.869492628 | 0.000168121 | 0.003660641 |
| chr7:22726335-22726727 | 1.361302916 | 0.000168421 | 0.003666129 |
| chr9:115385743-115386044 | 1.796252509 | 0.000168484 | 0.003666454 |
| chr14:83625089-83625290 | 2.201209378 | 0.000168798 | 0.003672233 |
| chr12:81475660-81476294 | -1.515740496 | 0.000168987 | 0.003675321 |
| chr13:114151675-114151963 | -1.896790886 | 0.000169432 | 0.003683941 |
| chr3:57544260-57544553 | -2.19954872 | 0.00016961 | 0.003686765 |
| chr3:111210384-111210751 | -1.777249962 | 0.00016995 | 0.00369207 |
| chr3:195030783-195031146 | 1.58901923 | 0.00016995 | 0.00369207 |
| chr17:75834512-75834745 | 2.019281106 | 0.000170162 | 0.003695614 |
| chr9:747246-747845 | 1.452120594 | 0.000170239 | 0.003696254 |
| chr1:208988725-208989214 | -1.65272933 | 0.000170629 | 0.003703654 |
| chr5:25889130-25889330 | 2.476650831 | 0.000170786 | 0.003706028 |
| chr5:13060608-13060851 | 2.205148449 | 0.000170908 | 0.003707628 |
| chr7:106065006-106065672 | -1.426880449 | 0.000171088 | 0.003710482 |
| chr11:34631695-34632418 | 1.37076278 | 0.000171207 | 0.003712011 |
| chr4:159047970-159048587 | -1.539950833 | 0.000172058 | 0.003727776 |
| chr4:8192760-8192985 | 1.81202135 | 0.000172062 | 0.003727776 |
| chr12:130098020-130098493 | -1.775360792 | 0.00017208 | 0.003727776 |
| chr13:94659284-94659493 | -2.516746485 | 0.00017216 | 0.00372814 |
| chr7:22623447-22623647 | 2.27085911 | 0.000172195 | 0.00372814 |
| chrX:132498951-132499172 | 2.473300609 | 0.000172488 | 0.003733443 |
| chr7:123622676-123622933 | -1.930197033 | 0.000172999 | 0.003742101 |
| chr3:150515639-150515860 | 2.026304823 | 0.000173018 | 0.003742101 |
| chr18:35874282-35874640 | 1.828613793 | 0.000173035 | 0.003742101 |
| chr19:29225725-29226039 | 1.846110244 | 0.000173598 | 0.003752569 |
| chr2:23381200-23381424 | -1.867293293 | 0.000173617 | 0.003752569 |
| chr20:55856406-55856762 | 1.788979728 | 0.000173732 | 0.003754012 |
| chr6:108684769-108685369 | -1.415095329 | 0.000173819 | 0.003754834 |
| chr8:57210722-57210974 | 1.819400405 | 0.000174008 | 0.003757843 |

| | | | |
|---|---|---|---|
| chr4:118194699-118194923 | 2.194149604 | 0.000174065 | 0.00375795 |
| chr12:14260093-14260293 | -2.033227333 | 0.000174111 | 0.00375795 |
| chr10:99320650-99320850 | 2.13790415 | 0.000174319 | 0.003759509 |
| chr10:132566951-132567153 | 1.910774437 | 0.000174358 | 0.003759509 |
| chr14:20224222-20224422 | 2.135592204 | 0.00017438 | 0.003759509 |
| chr17:74137506-74137827 | 1.701477594 | 0.000174404 | 0.003759509 |
| chr4:89509785-89510086 | 2.029970908 | 0.000174432 | 0.003759509 |
| chr8:11862509-11862913 | -1.421479731 | 0.000174478 | 0.003759509 |
| chr7:100866942-100867421 | -1.432197514 | 0.000174728 | 0.003763839 |
| chr4:177047124-177047694 | -1.400904558 | 0.000174935 | 0.003767246 |
| chr7:110186321-110186721 | -1.627841337 | 0.000175354 | 0.003775202 |
| chr1:58288394-58288615 | 2.02858186 | 0.00017561 | 0.003779111 |
| chr1:91806865-91807275 | -1.591944786 | 0.000175634 | 0.003779111 |
| chr4:106525061-106525883 | -1.586067315 | 0.000175929 | 0.003784405 |
| chr4:90946706-90947213 | 1.633166321 | 0.000176132 | 0.003786937 |
| chr13:61011368-61011646 | -1.898773671 | 0.000176146 | 0.003786937 |
| chr8:124180208-124180700 | 1.395310312 | 0.00017652 | 0.003793925 |
| chr18:57698438-57699222 | -1.4375063 | 0.000176843 | 0.003799791 |
| chr10:50511547-50512497 | -1.263394185 | 0.000177103 | 0.003804303 |
| chr4:189980347-189980629 | -1.928137064 | 0.0001774 | 0.003809624 |
| chr12:71469267-71469974 | -1.38135302 | 0.000177593 | 0.003812708 |
| chr12:15208160-15208448 | -2.026458974 | 0.000177911 | 0.003817753 |
| chr1:150732428-150732812 | 1.760612886 | 0.000177928 | 0.003817753 |
| chr10:96487832-96488032 | 2.270688699 | 0.000178341 | 0.003825544 |
| chr14:53700047-53700337 | 1.867273039 | 0.000179251 | 0.003843791 |
| chr4:75546633-75546883 | -2.133732056 | 0.000179292 | 0.003843791 |
| chr22:29795713-29796117 | 1.496323875 | 0.000179575 | 0.003848715 |
| chr12:3097501-3097813 | -1.802812602 | 0.00017966 | 0.003848715 |
| chr16:5059797-5060141 | 1.908125278 | 0.000179672 | 0.003848715 |
| chr6:87574160-87574568 | -1.596890828 | 0.000180411 | 0.003863463 |
| chr15:60387041-60387257 | 2.075955026 | 0.000180481 | 0.003863869 |
| chr2:45909636-45910058 | -1.510218768 | 0.000180789 | 0.003869397 |
| chr7:56105501-56105797 | 1.684844388 | 0.000181087 | 0.003874689 |
| chr5:147391870-147392175 | -1.651675413 | 0.000181139 | 0.003874716 |
| chr2:142868212-142868555 | -1.560054676 | 0.000181307 | 0.003876767 |
| chr1:143246914-143247199 | 1.586023742 | 0.000181336 | 0.003876767 |
| chr14:74945718-74946064 | 1.432222075 | 0.000181905 | 0.003887842 |
| chr20:50779175-50779436 | 1.319651788 | 0.000181972 | 0.003888199 |
| chr1:168836569-168836864 | 2.366390494 | 0.000182532 | 0.003899064 |
| chr14:28141696-28141896 | 2.185457539 | 0.000183156 | 0.003911314 |
| chr20:37657700-37657962 | 1.896875882 | 0.000183256 | 0.003911727 |
| chr9:83093910-83094212 | 2.172180413 | 0.000183278 | 0.003911727 |
| chr7:39016482-39017118 | -1.295723585 | 0.000183473 | 0.003914802 |
| chr3:5563197-5563440 | 1.893288513 | 0.000183555 | 0.003915467 |
| chr22:17528955-17529622 | -1.463733435 | 0.000183802 | 0.003919647 |
| chr1:201582714-201583260 | 1.380310521 | 0.000184338 | 0.003929975 |
| chr7:112689438-112689893 | -1.739391146 | 0.000184646 | 0.003935459 |
| chr10:87568109-87568672 | 1.656350941 | 0.000184736 | 0.003935488 |
| chr5:143928910-143929422 | -1.556373882 | 0.00018475 | 0.003935488 |
| chr7:125563657-125564342 | -1.3804644 | 0.000184989 | 0.003939483 |

| | | | |
|---|---|---|---|
| chr3:139522555-139522793 | -1.973139366 | 0.000185332 | 0.003945674 |
| chr14:62650085-62650405 | 1.791146133 | 0.000185547 | 0.003949155 |
| chr2:45889124-45889497 | -1.54857428 | 0.000185654 | 0.003950338 |
| chr8:23010741-23010945 | 2.827517735 | 0.000185752 | 0.003951328 |
| chr5:68396709-68397148 | -1.510792439 | 0.000185809 | 0.00395144 |
| chr6:37515862-37517130 | 1.265830404 | 0.000185882 | 0.003951904 |
| chr18:21242171-21242712 | 1.508418958 | 0.000186094 | 0.003955317 |
| chr20:49413326-49413548 | 1.819731322 | 0.000186204 | 0.003956558 |
| chr16:76854291-76854658 | 1.491626437 | 0.000186419 | 0.003960019 |
| chr16:81289724-81290019 | 1.639670795 | 0.000186938 | 0.003969943 |
| chr20:55806234-55806434 | 2.257295488 | 0.000187181 | 0.00397401 |
| chr8:86238949-86239167 | 2.111951602 | 0.000187261 | 0.003974608 |
| chr14:77303484-77304140 | 1.292951142 | 0.000187384 | 0.003976115 |
| chr5:116947780-116948350 | -1.331235027 | 0.000187524 | 0.003977988 |
| chr5:137142350-137142990 | -1.400137824 | 0.000187653 | 0.00397962 |
| chr14:85817704-85817918 | 2.278441699 | 0.000188658 | 0.003999817 |
| chr1:170758450-170758661 | 2.199440099 | 0.000188767 | 0.004001023 |
| chr14:75851754-75851966 | 2.269333319 | 0.000188969 | 0.00400419 |
| chr15:26629013-26629470 | -1.529230721 | 0.000189033 | 0.004004448 |
| chr4:121226376-121226693 | 1.718401922 | 0.000189154 | 0.004004454 |
| chr9:124779304-124779647 | 1.546060868 | 0.000189206 | 0.004004454 |
| chr16:31393399-31393625 | 2.069936464 | 0.000189231 | 0.004004454 |
| chr4:63262070-63262701 | -1.60876991 | 0.000189242 | 0.004004454 |
| chr2:48545268-48545632 | -1.608308543 | 0.000189372 | 0.004006096 |
| chr5:34191232-34191865 | 1.32132308 | 0.000189775 | 0.00401351 |
| chr5:99368747-99369076 | -1.926761 | 0.000190011 | 0.004017397 |
| chr5:57094869-57095774 | -1.253805201 | 0.000190163 | 0.0040195 |
| chr13:73060964-73061193 | -1.717764901 | 0.000190266 | 0.00402057 |
| chr16:68973714-68974085 | 1.656808446 | 0.000190572 | 0.004025926 |
| chr18:48782483-48782958 | -1.607773382 | 0.000191227 | 0.004038645 |
| chr7:40232543-40232807 | -1.922142621 | 0.000191635 | 0.00404615 |
| chrX:119663222-119663422 | 2.614074034 | 0.000191712 | 0.004046659 |
| chr9:98142493-98142798 | -1.630734201 | 0.000192014 | 0.004051917 |
| chr15:62447994-62448194 | 2.196880903 | 0.000192095 | 0.004052513 |
| chr8:126349109-126349341 | 1.929485812 | 0.000192551 | 0.004061016 |
| chr13:30002528-30002825 | -1.888815785 | 0.000193033 | 0.004070052 |
| chr3:127006155-127006687 | -1.378175612 | 0.000193822 | 0.004085161 |
| chr11:22128477-22128779 | -2.127286783 | 0.000193856 | 0.004085161 |
| chr16:72841643-72841983 | 1.556113918 | 0.000193962 | 0.004086265 |
| chr7:129796325-129796823 | -1.470693335 | 0.000194143 | 0.004088948 |
| chr7:104231324-104231861 | -1.46633707 | 0.00019449 | 0.00409513 |
| chr6:27405855-27406316 | -1.841695294 | 0.000194633 | 0.004097016 |
| chr1:19681884-19682112 | 2.200574519 | 0.000194687 | 0.004097038 |
| chr8:48571419-48571667 | 2.058965499 | 0.000194833 | 0.004097993 |
| chr7:120110181-120110557 | -1.628141397 | 0.000194839 | 0.004097993 |
| chr6:67066523-67066844 | -1.608616505 | 0.000194924 | 0.004098656 |
| chr1:53264462-53264713 | 1.851916793 | 0.000195389 | 0.004107302 |
| chr5:7603143-7603437 | 1.927542624 | 0.000195876 | 0.004116401 |
| chr15:55318105-55318621 | 1.420801216 | 0.000196458 | 0.004127345 |
| chr16:80027025-80027510 | -1.333405606 | 0.000196504 | 0.004127345 |

| | | | |
|---|---|---|---|
| chr7:55429908-55430186 | -1.735571971 | 0.000196971 | 0.004136007 |
| chr1:93673186-93673555 | 1.691049232 | 0.000197142 | 0.004136148 |
| chr17:57293104-57293463 | -1.414311778 | 0.000197188 | 0.004136148 |
| chr6:3664676-3665121 | 1.421136533 | 0.000197191 | 0.004136148 |
| chr4:48058603-48059011 | -1.821799975 | 0.000197219 | 0.004136148 |
| chr1:150906426-150906999 | 1.331529692 | 0.000197247 | 0.004136148 |
| chr11:129284763-129285027 | -2.198208409 | 0.000197578 | 0.00414195 |
| chr2:172241276-172241476 | 2.35483798 | 0.000197769 | 0.004144817 |
| chr12:26017502-26017883 | -1.977141868 | 0.00019788 | 0.004145333 |
| chr4:3302823-3303043 | 2.262570371 | 0.000197902 | 0.004145333 |
| chr8:62035336-62035828 | 1.414083835 | 0.00019796 | 0.004145423 |
| chr3:108664367-108664779 | 1.722681846 | 0.000198342 | 0.004152287 |
| chr13:56666207-56666537 | -2.196990916 | 0.000199043 | 0.00416583 |
| chr3:34284998-34285272 | 1.719766355 | 0.000199107 | 0.004166018 |
| chr1:238238704-238239706 | -1.351233339 | 0.000199254 | 0.004167959 |
| chr7:7745057-7745348 | 1.768022375 | 0.000199434 | 0.004170591 |
| chr10:51581243-51581539 | 1.751175361 | 0.000199621 | 0.004173365 |
| chr2:35035918-35036189 | -1.700690763 | 0.000200035 | 0.004180875 |
| chr1:200483487-200483838 | 1.533044225 | 0.000200342 | 0.004185498 |
| chr12:117738275-117738857 | 1.431212069 | 0.000200409 | 0.004185498 |
| chr20:58285458-58285697 | 1.778914565 | 0.000200441 | 0.004185498 |
| chr3:175258698-175259281 | -1.434837732 | 0.000200475 | 0.004185498 |
| chrX:103460059-103460292 | 2.001592789 | 0.000200615 | 0.004187276 |
| chr12:6080195-6080725 | -1.461765833 | 0.000200808 | 0.00419018 |
| chr5:90073059-90073936 | -1.359871782 | 0.000201165 | 0.00419648 |
| chr3:49222137-49222412 | 1.962377867 | 0.00020144 | 0.004200141 |
| chr7:57450007-57450313 | 1.920540694 | 0.000201494 | 0.004200141 |
| chr12:110948022-110948419 | 1.631312307 | 0.000201505 | 0.004200141 |
| chr5:157024243-157024584 | 1.742981809 | 0.000201625 | 0.004201493 |
| chr8:100401694-100401976 | 2.048883524 | 0.000202571 | 0.004220066 |
| chr17:59454521-59454881 | 1.426319011 | 0.000202794 | 0.004223565 |
| chr10:114179200-114179405 | -1.886659155 | 0.000203923 | 0.004245928 |
| chr3:42900777-42901329 | 1.540749758 | 0.000203987 | 0.004246109 |
| chr1:248054272-248054622 | -1.998973046 | 0.000204171 | 0.004248768 |
| chr14:64148692-64148892 | 2.361056696 | 0.000204322 | 0.004250766 |
| chr7:37346736-37347041 | 1.561044516 | 0.000204696 | 0.004257391 |
| chr12:79835035-79835300 | 2.109639608 | 0.000204841 | 0.004259239 |
| chr8:143095083-143095469 | 1.45800802 | 0.000204992 | 0.004261231 |
| chr22:39349683-39349994 | 1.823604574 | 0.000205256 | 0.004265553 |
| chr4:3292156-3292406 | 1.650454058 | 0.000205418 | 0.004267765 |
| chr4:163672847-163673118 | -1.961015466 | 0.000205896 | 0.004276228 |
| chr2:222499116-222499643 | -1.7849094 | 0.000205946 | 0.004276228 |
| chr9:69105102-69105619 | -1.337388584 | 0.000205992 | 0.004276228 |
| chr8:110790944-110791279 | 1.686644136 | 0.000206102 | 0.004277342 |
| chr7:23211547-23212165 | 1.296385587 | 0.000206226 | 0.00427876 |
| chr20:56682912-56683939 | 1.218017031 | 0.000206334 | 0.004279399 |
| chr8:95700687-95700995 | 1.769861796 | 0.000206389 | 0.004279399 |
| chr4:169185688-169185989 | -1.817507018 | 0.000206425 | 0.004279399 |
| chr3:23222782-23222982 | 2.248230048 | 0.000206526 | 0.004280336 |
| chr19:44435466-44435778 | 1.850428827 | 0.000206916 | 0.004287267 |

| | | | |
|---|---|---|---|
| chr14:23973017-23973312 | 2.063819778 | 0.000207254 | 0.004293101 |
| chr9:122070247-122070572 | 2.163585095 | 0.000207573 | 0.004298553 |
| chr15:88162970-88163271 | 1.809038086 | 0.000208124 | 0.004308803 |
| chr9:11980418-11980663 | -2.103692644 | 0.000208352 | 0.004312362 |
| chr2:48767366-48768014 | -1.441576441 | 0.000208783 | 0.004320119 |
| chr5:113990648-113990859 | -2.100082951 | 0.000208889 | 0.004321138 |
| chr13:66900172-66900482 | -2.004645185 | 0.000209209 | 0.004326589 |
| chr20:54402659-54402930 | 1.378623337 | 0.000209353 | 0.00432839 |
| chr19:45551754-45552004 | 2.254000573 | 0.000209672 | 0.00433383 |
| chr7:14125626-14125880 | 2.104270983 | 0.000210745 | 0.004354818 |
| chr13:34729411-34729705 | -2.055012939 | 0.000211002 | 0.004358954 |
| chr19:34341817-34342042 | 2.100975085 | 0.000211201 | 0.004361717 |
| chr2:190551906-190552222 | -1.848521626 | 0.000211249 | 0.004361717 |
| chr10:9111938-9112754 | 1.307121003 | 0.000211691 | 0.004369171 |
| chr16:31487709-31488262 | 1.367541342 | 0.000211724 | 0.004369171 |
| chrY:56833269-56833493 | 1.874603902 | 0.000212277 | 0.004379402 |
| chr1:150227321-150227521 | 1.952477245 | 0.000212374 | 0.004380215 |
| chr5:500402-500840 | 1.597561656 | 0.000212533 | 0.004382318 |
| chr14:43651054-43652142 | 1.244474139 | 0.000213072 | 0.00439225 |
| chr3:64393167-64393367 | 1.402123419 | 0.000213375 | 0.004397313 |
| chr7:23299015-23299950 | 1.216163347 | 0.000213584 | 0.004400445 |
| chr16:69181106-69181337 | 2.121607669 | 0.00021368 | 0.004401233 |
| chr5:16673372-16673616 | 1.830045012 | 0.000213856 | 0.004403684 |
| chr7:116972019-116972241 | -1.988235056 | 0.000214552 | 0.004416825 |
| chr15:46634886-46635183 | -2.058694973 | 0.00021485 | 0.004421768 |
| chr14:38345881-38346483 | 1.327031955 | 0.000215099 | 0.004425713 |
| chr14:44014003-44014281 | 1.827688811 | 0.000215351 | 0.004429703 |
| chr2:51424493-51424741 | -2.185143154 | 0.000215682 | 0.004435321 |
| chr14:104976232-104976490 | 1.467580694 | 0.000215797 | 0.004436484 |
| chr14:30746867-30747123 | 1.8434151 | 0.00021591 | 0.00443763 |
| chr1:206898636-206898976 | -1.351175775 | 0.00021656 | 0.004448619 |
| chr15:82889613-82889886 | 2.037559464 | 0.000216561 | 0.004448619 |
| chr4:16044216-16044565 | 1.612025464 | 0.000216923 | 0.004454024 |
| chr19:53365501-53365818 | 1.574676254 | 0.000216982 | 0.004454024 |
| chr10:19220892-19221109 | -2.229981781 | 0.000217053 | 0.004454024 |
| chr2:11391501-11392031 | -1.348800164 | 0.000217057 | 0.004454024 |
| chr14:28659447-28659694 | 2.057136184 | 0.000217281 | 0.004457427 |
| chr8:101158979-101159186 | 1.815415074 | 0.000217347 | 0.004457473 |
| chr4:189940173-189941295 | -1.225690569 | 0.000217399 | 0.004457473 |
| chr6:106119546-106119881 | 1.684573801 | 0.000217773 | 0.004463946 |
| chr12:101924644-101924950 | -1.832890645 | 0.000218009 | 0.004466723 |
| chr7:15948122-15948367 | 2.228075825 | 0.000218025 | 0.004466723 |
| chr7:33640951-33641217 | 1.771478019 | 0.000218362 | 0.004472432 |
| chr1:246998711-246999010 | 1.771992988 | 0.000218484 | 0.004473717 |
| chrX:31985483-31985776 | -1.820805301 | 0.000218806 | 0.004479129 |
| chr8:130483038-130483786 | 1.263792233 | 0.000219127 | 0.00448449 |
| chr15:66842204-66842689 | 1.440543075 | 0.000219241 | 0.004485628 |
| chr2:106723592-106723837 | 2.049722847 | 0.000219359 | 0.004486836 |
| chrX:129327224-129327797 | 1.377057691 | 0.000219814 | 0.004494951 |
| chr11:30407336-30407818 | -1.539560836 | 0.000220362 | 0.004504965 |

| | | | |
|---|---|---|---|
| chr10:121988731-121989348 | -1.42781166 | 0.000220677 | 0.004509118 |
| chr20:29883856-29884101 | 2.092176219 | 0.000220683 | 0.004509118 |
| chr7:105580035-105580684 | -1.397030716 | 0.000220875 | 0.004511661 |
| chr11:131025671-131025946 | -2.005826278 | 0.000220925 | 0.004511661 |
| chr7:105680529-105681013 | -1.513786164 | 0.000221205 | 0.004516169 |
| chr10:42367923-42368207 | 1.707158003 | 0.000221368 | 0.00451757 |
| chr13:83914583-83914792 | -2.602825851 | 0.000221419 | 0.00451757 |
| chr18:56563664-56563868 | -2.12678614 | 0.000221451 | 0.00451757 |
| chrX:88786964-88787202 | 1.9926147 | 0.00022152 | 0.004517784 |
| chr2:36403573-36404240 | -1.298925422 | 0.000222605 | 0.004538694 |
| chr2:70617578-70618104 | -1.382672585 | 0.000222713 | 0.004539695 |
| chr1:246219671-246220046 | -1.609132092 | 0.00022295 | 0.004543317 |
| chr4:118997366-118997622 | 2.04694131 | 0.000223063 | 0.004544419 |
| chr1:84724388-84724735 | -1.542180432 | 0.000223493 | 0.004551953 |
| chr10:88037001-88037201 | 2.350509734 | 0.000223611 | 0.004552847 |
| chr8:126715305-126715630 | 1.623542162 | 0.000223655 | 0.004552847 |
| chr7:68813664-68814140 | -1.604376016 | 0.000223835 | 0.004554584 |
| chr16:80299487-80299689 | -2.257138286 | 0.00022386 | 0.004554584 |
| chr21:40445162-40445687 | -1.549815189 | 0.000223968 | 0.004555588 |
| chr7:142326580-142327072 | -1.66074118 | 0.000224223 | 0.004559567 |
| chr4:75708388-75708710 | -1.872124205 | 0.000224573 | 0.004565454 |
| chr13:33349306-33349764 | -1.332806092 | 0.000224777 | 0.004566375 |
| chr3:57596530-57596832 | 1.450362868 | 0.000224781 | 0.004566375 |
| chr14:42456985-42457187 | 1.958269775 | 0.000224797 | 0.004566375 |
| chr7:106568567-106569368 | -1.299497981 | 0.000224899 | 0.004567247 |
| chr5:167030532-167030887 | -1.653067031 | 0.000224987 | 0.004567824 |
| chr1:230824325-230824525 | 2.035928339 | 0.000225373 | 0.004574454 |
| chr7:36404959-36405373 | 1.515810321 | 0.000225777 | 0.004581423 |
| chr17:6831632-6832146 | -1.546259185 | 0.000226014 | 0.004585033 |
| chr13:92074647-92074991 | -1.712826152 | 0.000226115 | 0.004585853 |
| chr4:168877829-168878041 | 2.157687107 | 0.00022632 | 0.004588813 |
| chr4:13640217-13640618 | 1.694907009 | 0.000227399 | 0.004609462 |
| chr1:5876216-5876434 | 2.324630564 | 0.000227763 | 0.004615617 |
| chr11:41784913-41785264 | 1.66257722 | 0.000228036 | 0.004619934 |
| chr8:96736590-96736890 | 1.881722576 | 0.000228423 | 0.004626546 |
| chr2:198973197-198973528 | -1.869969793 | 0.000228509 | 0.004627066 |
| chr16:88832695-88832895 | 1.90802369 | 0.000228778 | 0.004631298 |
| chr18:49021192-49022176 | -1.212911203 | 0.00022892 | 0.004632942 |
| chr1:84759855-84760085 | -2.000122532 | 0.000229301 | 0.004639422 |
| chr7:23357472-23357953 | 1.464113108 | 0.000229419 | 0.004640589 |
| chr18:40529563-40529793 | -2.184140076 | 0.000229885 | 0.004648339 |
| chr14:34797899-34798122 | 2.042432008 | 0.000229924 | 0.004648339 |
| chr7:157411418-157411928 | -1.457014145 | 0.000230161 | 0.004651919 |
| chr2:240629842-240630267 | 1.596889153 | 0.000230387 | 0.004655252 |
| chr3:28459033-28459461 | -1.538685206 | 0.000230522 | 0.004656758 |
| chr14:96741409-96741743 | 1.485682231 | 0.00023082 | 0.004661549 |
| chr7:138797836-138798234 | -1.624380124 | 0.000230972 | 0.004663389 |
| chr19:7294276-7294627 | 1.462230583 | 0.000231643 | 0.004675695 |
| chr6:14316644-14317022 | 1.647587766 | 0.000232085 | 0.004683381 |
| chr8:26437696-26438528 | -1.389106721 | 0.000232684 | 0.004694245 |

| | | | |
|---|---|---|---|
| chr9:18711172-18711510 | -1.882126765 | 0.00023286 | 0.00469655 |
| chr8:101975406-101975613 | 1.87048331 | 0.000233319 | 0.004704353 |
| chr1:40636412-40636612 | 2.039816534 | 0.00023337 | 0.004704353 |
| chrX:105683933-105684672 | -1.457208115 | 0.000233653 | 0.004708836 |
| chr2:34031149-34031485 | -1.827398351 | 0.000234375 | 0.004722127 |
| chr10:110680326-110680811 | -1.453346934 | 0.000234615 | 0.004725733 |
| chr9:121599229-121599792 | 1.313690184 | 0.000235208 | 0.004736427 |
| chr4:185005041-185005547 | -1.417337012 | 0.000235384 | 0.004738729 |
| chr7:13981118-13981553 | 1.410717058 | 0.000235593 | 0.004740756 |
| chr16:46398792-46399042 | 1.353496192 | 0.000235609 | 0.004740756 |
| chr21:46002886-46003306 | 1.631183366 | 0.000235746 | 0.004741074 |
| chrX:141128415-141128749 | 1.548544135 | 0.000235769 | 0.004741074 |
| chr15:71266198-71266417 | 2.160006963 | 0.00023581 | 0.004741074 |
| chr20:46199243-46199492 | 1.690475773 | 0.000236191 | 0.004747491 |
| chr20:48886327-48886572 | 1.35408962 | 0.000236412 | 0.004750681 |
| chr5:113447677-113447895 | -2.152862304 | 0.000236585 | 0.004752923 |
| chr5:157385064-157385370 | 1.736323272 | 0.000236706 | 0.004754096 |
| chr12:115872139-115872473 | 1.476319082 | 0.000237028 | 0.004758218 |
| chr2:125710595-125711130 | 1.395050661 | 0.000237035 | 0.004758218 |
| chr3:13791810-13792093 | 1.588544383 | 0.000237338 | 0.00476305 |
| chr14:88488184-88488522 | -1.506908608 | 0.0002376 | 0.004765222 |
| chr11:94832719-94833250 | 1.475166741 | 0.000237608 | 0.004765222 |
| chr11:127074141-127074457 | -1.812915146 | 0.000237633 | 0.004765222 |
| chr2:218315972-218316287 | 1.636067551 | 0.000238224 | 0.004774914 |
| chr7:150016549-150017158 | -1.492963198 | 0.000238241 | 0.004774914 |
| chr11:129861717-129862457 | 1.289624051 | 0.000238353 | 0.004775918 |
| chr20:46808070-46808277 | 1.872571018 | 0.000238673 | 0.004781069 |
| chr15:64053031-64053288 | 1.901733036 | 0.000238935 | 0.004785071 |
| chr10:895342-895701 | 1.500051122 | 0.000239235 | 0.004789825 |
| chr8:99685939-99686387 | 1.608991806 | 0.000239554 | 0.004794954 |
| chr14:105617084-105617823 | 1.248385344 | 0.000239894 | 0.004800509 |
| chr16:57265903-57266291 | 1.740297666 | 0.000240069 | 0.004802749 |
| chr9:24579853-24580636 | -1.476605072 | 0.000240559 | 0.004811311 |
| chr4:176123585-176124050 | -1.490081536 | 0.000240668 | 0.004812239 |
| chr7:158205644-158206081 | -1.850487193 | 0.000240774 | 0.00481309 |
| chr17:4560551-4560784 | -1.893896333 | 0.000240872 | 0.004813787 |
| chr14:94398176-94398400 | 2.031423125 | 0.000241157 | 0.004818231 |
| chr13:82647409-82647813 | -1.826392601 | 0.000241431 | 0.004822453 |
| chr18:45829380-45829849 | -1.371231123 | 0.000241554 | 0.004823649 |
| chr2:138122324-138123217 | -1.386631356 | 0.000241943 | 0.004829401 |
| chr10:19847464-19847984 | -1.538399839 | 0.000241968 | 0.004829401 |
| chr12:42865378-42865644 | 1.752421324 | 0.000242057 | 0.004829925 |
| chr16:88650482-88651288 | -1.296276108 | 0.000242698 | 0.00484145 |
| chr7:108146074-108146736 | -1.458057203 | 0.00024296 | 0.004845409 |
| chr6:15021754-15022190 | -1.510410107 | 0.00024315 | 0.004847948 |
| chr11:34483423-34483686 | 1.98080454 | 0.000243608 | 0.004855809 |
| chr6:125918792-125919499 | -1.370038739 | 0.000243751 | 0.004856462 |
| chr15:39455593-39455934 | 1.546216805 | 0.000243798 | 0.004856462 |
| chr10:58305624-58305941 | -1.835700978 | 0.000243831 | 0.004856462 |
| chr6:103122853-103123455 | -1.379237864 | 0.000244115 | 0.004860851 |

| | | | |
|---|---|---|---|
| chr8:56270650-56271007 | -1.849632767 | 0.000244196 | 0.004860936 |
| chr4:26003079-26003305 | 2.312727517 | 0.000244246 | 0.004860936 |
| chr7:23618031-23618282 | 1.954625536 | 0.000244506 | 0.004864849 |
| chr3:149236323-149236528 | 1.937234026 | 0.000244972 | 0.004872857 |
| chr3:18492879-18493162 | 1.781109987 | 0.000245436 | 0.004880829 |
| chr5:90161503-90161882 | -1.671869323 | 0.000245579 | 0.004882376 |
| chr12:94822802-94823440 | -1.46493453 | 0.000245642 | 0.004882376 |
| chr7:142153450-142153715 | -1.757210008 | 0.000246559 | 0.004899331 |
| chrX:30212209-30212470 | -2.082435237 | 0.00024666 | 0.004900064 |
| chr1:68050176-68050649 | -1.596576199 | 0.000246878 | 0.004902536 |
| chr13:25733998-25734198 | 2.09584298 | 0.000246912 | 0.004902536 |
| chr1:198655631-198655853 | 1.743525874 | 0.000247133 | 0.004905648 |
| chr15:31398561-31399123 | -1.301538465 | 0.000247879 | 0.004919187 |
| chr8:23984422-23984897 | -1.571413692 | 0.000247989 | 0.004920091 |
| chr21:30076042-30076277 | 2.211249689 | 0.000248506 | 0.004929084 |
| chr7:136803013-136803385 | -1.669172925 | 0.000248787 | 0.004933371 |
| chr11:30518098-30518399 | -1.600232616 | 0.000248882 | 0.004933472 |
| chr2:8573433-8574117 | -1.237069135 | 0.000248921 | 0.004933472 |
| chr10:132448416-132448801 | 1.287611844 | 0.000249039 | 0.004934544 |
| chr8:134753880-134754540 | 1.394334802 | 0.000249221 | 0.004936863 |
| chr6:37381494-37381905 | 1.400841974 | 0.000249332 | 0.004937788 |
| chr5:115634707-115635029 | -1.488736034 | 0.000249992 | 0.004949579 |
| chr8:31828411-31828723 | -2.014493165 | 0.000251702 | 0.004982155 |
| chr7:142459509-142460045 | -1.378040826 | 0.000251987 | 0.004986514 |
| chr15:31351366-31351806 | -1.400277049 | 0.000252319 | 0.004991787 |
| chr22:31055526-31055939 | 1.394045911 | 0.000252645 | 0.004996949 |
| chr19:39457218-39457427 | 1.90229304 | 0.000253144 | 0.005005531 |
| chr4:104896650-104897044 | -1.570547372 | 0.000253334 | 0.005007993 |
| chr3:835632-835837 | 2.232507824 | 0.000253722 | 0.005014363 |
| chr7:10501496-10501696 | 2.028881453 | 0.000254199 | 0.005022494 |
| chr3:64186891-64187528 | 1.190643867 | 0.000254727 | 0.00503163 |
| chr12:51980395-51980598 | 1.932146958 | 0.000254948 | 0.005034232 |
| chr7:120926899-120927469 | -1.545604223 | 0.00025499 | 0.005034232 |
| chr16:68756213-68756564 | 1.570517755 | 0.000255608 | 0.005045132 |
| chr1:236399490-236400106 | 1.311002811 | 0.000255757 | 0.005046066 |
| chr7:134696282-134696541 | -1.840859505 | 0.000255787 | 0.005046066 |
| chr20:40687699-40688104 | 1.419078995 | 0.00025595 | 0.005047976 |
| chr7:30844459-30844702 | 1.931233328 | 0.000256192 | 0.005051448 |
| chr14:105585339-105585539 | 2.032325704 | 0.000256621 | 0.005057401 |
| chr7:27739920-27741061 | 1.202154669 | 0.000256631 | 0.005057401 |
| chr2:14524182-14524382 | 2.314578374 | 0.000256692 | 0.005057401 |
| chr1:151135402-151135860 | 1.319781886 | 0.000256794 | 0.005058112 |
| chr14:29906176-29906570 | 1.686824118 | 0.00025711 | 0.005062676 |
| chr18:54931169-54931444 | -2.031601143 | 0.000257158 | 0.005062676 |
| chr19:3384902-3385275 | -1.611580123 | 0.000257997 | 0.005076441 |
| chr22:23908247-23908482 | 1.826038593 | 0.000258021 | 0.005076441 |
| chr11:76604266-76604579 | 1.669178064 | 0.000258166 | 0.005076441 |
| chr8:100258993-100259277 | 1.569620872 | 0.000258169 | 0.005076441 |
| chr2:65388076-65388684 | -1.314876042 | 0.000258204 | 0.005076441 |
| chr12:78878989-78879279 | 2.077064853 | 0.000258254 | 0.005076441 |

| | | | |
|---|---|---|---|
| chr4:124869612-124869982 | 1.970927722 | 0.000259188 | 0.005093482 |
| chr7:112743831-112744204 | -1.422480523 | 0.000259614 | 0.005100556 |
| chr7:138343791-138344165 | -1.666667193 | 0.000260248 | 0.005111689 |
| chr2:27082616-27082826 | -1.762360151 | 0.000260413 | 0.00511362 |
| chr19:50844326-50844564 | 1.890676754 | 0.000260651 | 0.005116992 |
| chr2:121699431-121699724 | 1.683036385 | 0.000261435 | 0.005131071 |
| chr9:73132305-73132736 | -1.66087896 | 0.000261535 | 0.00513171 |
| chrX:144845613-144846280 | 1.350279061 | 0.000261861 | 0.005135714 |
| chr13:50975750-50976218 | -1.852495688 | 0.000261873 | 0.005135714 |
| chr5:166598554-166598816 | -1.732645113 | 0.000262031 | 0.005137506 |
| chr1:112280492-112280701 | 2.031038538 | 0.000262267 | 0.005140809 |
| chr8:116103013-116103344 | -1.531566287 | 0.000262892 | 0.00515175 |
| chr1:17229071-17229351 | -1.984781849 | 0.000263483 | 0.005162012 |
| chr2:149422352-149423315 | -1.247824525 | 0.000264307 | 0.005176832 |
| chr2:54698616-54698963 | -1.970328996 | 0.000264523 | 0.005178854 |
| chr22:29716853-29717143 | 1.597333203 | 0.000264547 | 0.005178854 |
| chr15:89268004-89268264 | 1.728678584 | 0.000264613 | 0.005178854 |
| chr1:227751141-227751341 | 2.074399935 | 0.000265114 | 0.005187328 |
| chr13:90404884-90405174 | -1.892946233 | 0.000266422 | 0.005211596 |
| chr1:110390884-110391279 | -1.505925097 | 0.000267275 | 0.005225833 |
| chr3:193836974-193837669 | -1.356299606 | 0.00026734 | 0.005225833 |
| chrX:118773884-118774114 | 2.203352823 | 0.000267355 | 0.005225833 |
| chr1:109246175-109246576 | -1.394735463 | 0.000267685 | 0.005230957 |
| chr8:107151404-107151679 | 1.719480225 | 0.000267872 | 0.005233277 |
| chr9:77449245-77450014 | 1.270971231 | 0.00026824 | 0.005237955 |
| chr7:3108824-3109270 | -1.46938505 | 0.000268248 | 0.005237955 |
| chr13:23859172-23859415 | 2.022432895 | 0.00026843 | 0.005239685 |
| chr5:143594916-143596165 | 1.200529621 | 0.000268474 | 0.005239685 |
| chr1:179330191-179330391 | 2.148782034 | 0.000268575 | 0.005240325 |
| chr2:65377566-65378156 | -1.268803309 | 0.000269129 | 0.005249791 |
| chr7:113159820-113160100 | -1.617574453 | 0.000269206 | 0.005249961 |
| chr19:18725635-18725891 | 2.224682885 | 0.000269572 | 0.005255759 |
| chr3:73542163-73542537 | -1.503450963 | 0.000269805 | 0.005258968 |
| chr10:110456621-110457333 | -1.305916773 | 0.000270109 | 0.005262298 |
| chr1:80918456-80918677 | -2.169495944 | 0.000270113 | 0.005262298 |
| chr7:141014521-141015489 | -1.189691753 | 0.000271125 | 0.005280669 |
| chr2:84300023-84300744 | -1.435690208 | 0.000271535 | 0.005287298 |
| chr10:67817980-67818870 | -1.251636915 | 0.000271741 | 0.00528922 |
| chr14:42928674-42928890 | 1.820835665 | 0.000271771 | 0.00528922 |
| chr14:84144800-84145038 | 2.217281179 | 0.000271906 | 0.005289493 |
| chr14:28157682-28158018 | 1.68866486 | 0.000271975 | 0.005289493 |
| chrX:30618190-30618834 | -1.337936816 | 0.000271992 | 0.005289493 |
| chr2:195559404-195559608 | -2.154322626 | 0.000272727 | 0.005302424 |
| chr1:202411512-202411996 | -1.475275667 | 0.000272796 | 0.005302424 |
| chr6:107620124-107620448 | 1.84968408 | 0.000273004 | 0.00530401 |
| chr1:44850064-44850264 | 2.222717604 | 0.000273016 | 0.00530401 |
| chr19:42377077-42377284 | 2.304499535 | 0.000273121 | 0.005304709 |
| chr6:130282248-130282610 | -1.639123098 | 0.000273308 | 0.005306819 |
| chr8:80543056-80543399 | -1.57956906 | 0.000273368 | 0.005306819 |
| chr1:61642367-61643242 | -1.225031444 | 0.000273518 | 0.005308373 |

| | | | |
|---|---|---|---|
| chr11:47924260-47924618 | 1.558526398 | 0.000273718 | 0.005310918 |
| chr11:3021264-3021505 | 1.846272774 | 0.000273913 | 0.005313364 |
| chr6:92023619-92023894 | -2.073292769 | 0.000274027 | 0.005314216 |
| chr4:173416031-173416608 | -1.358340187 | 0.000274218 | 0.00531659 |
| chr2:46127936-46128136 | 2.312570955 | 0.000274321 | 0.005317234 |
| chr4:114615955-114616841 | -1.241467242 | 0.000274593 | 0.005319918 |
| chr1:116276953-116277186 | 1.753700725 | 0.000274598 | 0.005319918 |
| chr6:135162731-135163020 | -1.777649788 | 0.000275091 | 0.005328115 |
| chr7:92584844-92585263 | -1.435803612 | 0.000275776 | 0.00533882 |
| chr2:3692377-3692754 | -1.536511154 | 0.000275783 | 0.00533882 |
| chr3:154454238-154454459 | 2.067051203 | 0.000275939 | 0.005340494 |
| chr11:1224457-1224753 | -2.028425271 | 0.000276287 | 0.005345159 |
| chr3:189071052-189071806 | -1.431672581 | 0.00027634 | 0.005345159 |
| chr1:162002170-162002370 | 2.225931744 | 0.00027639 | 0.005345159 |
| chr14:72189384-72189584 | 2.228029825 | 0.000277171 | 0.005358918 |
| chr4:89482297-89482572 | -1.915986304 | 0.0002774 | 0.005361986 |
| chr3:36908390-36908692 | -1.633568186 | 0.000278527 | 0.005382418 |
| chr13:37827687-37828328 | -1.453629875 | 0.00027864 | 0.005382677 |
| chr17:47140162-47140399 | 2.078217868 | 0.000278681 | 0.005382677 |
| chr8:102600399-102601736 | -1.165066901 | 0.000278896 | 0.005385477 |
| chr3:49158220-49158432 | 1.919899642 | 0.000279104 | 0.005388136 |
| chr5:67779948-67780205 | 1.924465274 | 0.000279252 | 0.00538964 |
| chr1:114300552-114300770 | 1.421517672 | 0.000279786 | 0.005398589 |
| chr18:72235388-72235853 | -1.514451598 | 0.000279874 | 0.005398642 |
| chr17:81566601-81567136 | 1.40683616 | 0.000279932 | 0.005398642 |
| chr3:63303298-63303583 | 1.48580589 | 0.000280069 | 0.005398642 |
| chr5:174483374-174483792 | -1.474201987 | 0.000280071 | 0.005398642 |
| chr14:75794964-75795164 | 2.034352721 | 0.000280448 | 0.005403566 |
| chr12:31116987-31117431 | -1.526511047 | 0.000280527 | 0.005403566 |
| chr8:101051336-101052384 | 1.19511921 | 0.000280538 | 0.005403566 |
| chr7:22488892-22489267 | 1.379642324 | 0.000280715 | 0.005404677 |
| chr11:133821564-133822015 | -1.665495961 | 0.000280737 | 0.005404677 |
| chr7:158001461-158002064 | -1.364989374 | 0.000283197 | 0.00545067 |
| chr13:84951226-84951478 | -2.079882282 | 0.000284252 | 0.005469081 |
| chr17:59367146-59367417 | 1.46427886 | 0.000284296 | 0.005469081 |
| chr6:86146717-86147256 | 1.447128558 | 0.000284397 | 0.005469531 |
| chr3:111027270-111027620 | 1.976190522 | 0.000284462 | 0.005469531 |
| chrX:109713041-109713332 | -1.721105008 | 0.000285437 | 0.005486142 |
| chr7:101203921-101204285 | -1.624747968 | 0.00028547 | 0.005486142 |
| chr8:86965265-86965465 | 1.969326565 | 0.0002856 | 0.00548728 |
| chr2:45851678-45851929 | -1.922186698 | 0.000286423 | 0.005501705 |
| chr16:46920559-46920822 | 1.757023123 | 0.000286817 | 0.005507895 |
| chr10:59809382-59809730 | -1.439792768 | 0.00028699 | 0.00550983 |
| chr19:9914677-9915022 | 1.629472559 | 0.000287242 | 0.00551329 |
| chr14:48970910-48971224 | 1.768172537 | 0.000287409 | 0.005515123 |
| chr4:37528077-37528792 | -1.30685721 | 0.000288084 | 0.00552668 |
| chr15:99199476-99199679 | 2.205983815 | 0.000288303 | 0.005529495 |
| chr12:132189514-132189954 | 1.355235377 | 0.000288458 | 0.005531098 |
| chr8:100416823-100417722 | 1.228162003 | 0.000289034 | 0.005540751 |
| chr12:90945103-90945407 | -1.6379679 | 0.00028946 | 0.005547521 |

| | | | |
|---|---|---|---|
| chr7:31849535-31849778 | 2.005352525 | 0.000290043 | 0.005554585 |
| chrX:154502854-154503136 | 2.005352525 | 0.000290043 | 0.005554585 |
| chr7:24227774-24228039 | 1.656057116 | 0.000290046 | 0.005554585 |
| chr16:76551980-76552294 | 1.618363834 | 0.00029033 | 0.005558642 |
| chr13:75506847-75507843 | -1.324036964 | 0.000290523 | 0.00556095 |
| chr9:83060616-83060901 | -1.442451628 | 0.000290648 | 0.005561941 |
| chr9:135384417-135384617 | 2.010094391 | 0.000290724 | 0.005562004 |
| chrX:80799760-80800129 | -1.791381129 | 0.000291075 | 0.005567337 |
| chr1:116632350-116633042 | 1.318980811 | 0.000291184 | 0.005568023 |
| chr7:20304271-20304526 | 1.71771643 | 0.000292198 | 0.005586024 |
| chr4:47692784-47693004 | 1.959610973 | 0.0002923 | 0.005586579 |
| chr7:4734404-4734619 | 1.845723527 | 0.000292426 | 0.00558759 |
| chr2:240840478-240840821 | 1.51593404 | 0.000292846 | 0.005594223 |
| chr5:150615132-150615584 | -1.356208252 | 0.000293768 | 0.005610449 |
| chr7:83468136-83468790 | -1.359295207 | 0.000294338 | 0.005619919 |
| chr7:74059932-74060547 | -1.492763758 | 0.000294585 | 0.005621754 |
| chr16:32168642-32168889 | 1.918028469 | 0.000294594 | 0.005621754 |
| chr14:89185997-89186598 | 1.374251988 | 0.000294654 | 0.005621754 |
| chr7:108347725-108348061 | -1.530739405 | 0.000295467 | 0.005635861 |
| chrX:107155384-107156425 | -1.324911055 | 0.00029568 | 0.005638518 |
| chrX:34426352-34426705 | -1.714952446 | 0.000295931 | 0.005641666 |
| chr14:84904480-84904716 | 2.083546166 | 0.000295992 | 0.005641666 |
| chr13:95535190-95535770 | -1.371223901 | 0.00029654 | 0.005649363 |
| chr13:22451282-22451591 | -1.840519675 | 0.000296543 | 0.005649363 |
| chr6:155185616-155186020 | -1.529670641 | 0.000297149 | 0.005657986 |
| chr3:148450764-148451227 | -1.65686343 | 0.000297288 | 0.005657986 |
| chr2:179428806-179429802 | -1.318925947 | 0.000297293 | 0.005657986 |
| chr7:32415880-32416208 | 1.6462905 | 0.000297302 | 0.005657986 |
| chr1:55191300-55191533 | 1.919779286 | 0.000297365 | 0.005657986 |
| chr4:94076208-94076418 | -2.208646935 | 0.000297911 | 0.005666967 |
| chr15:48855496-48855767 | -1.924020419 | 0.000298522 | 0.005676671 |
| chr2:63614534-63614953 | 1.408871005 | 0.000298569 | 0.005676671 |
| chr10:133181972-133182333 | 1.668731511 | 0.000299487 | 0.005692708 |
| chr3:22328631-22329103 | -1.51929724 | 0.000300042 | 0.005701848 |
| chr4:180550179-180550482 | -1.632973561 | 0.000300305 | 0.005705384 |
| chr7:17424513-17424713 | 2.281378939 | 0.000300385 | 0.005705384 |
| chr7:156942971-156944079 | -1.244973787 | 0.000300452 | 0.005705384 |
| chr7:23190940-23191326 | 1.399822827 | 0.000300539 | 0.005705617 |
| chr2:106172611-106172852 | 2.005864421 | 0.000300964 | 0.005712283 |
| chr5:67165932-67166523 | -1.367903754 | 0.00030161 | 0.005723112 |
| chr1:113855212-113855419 | 1.461561938 | 0.000301787 | 0.005723909 |
| chr20:33308992-33309263 | 1.967968701 | 0.00030184 | 0.005723909 |
| chr7:14788709-14789023 | 1.841556792 | 0.000301876 | 0.005723909 |
| chr3:152639203-152639579 | -1.799869453 | 0.000303094 | 0.005744879 |
| chr3:150190793-150191062 | 2.050101334 | 0.000303148 | 0.005744879 |
| chr4:7034674-7035078 | 1.419454504 | 0.000303207 | 0.005744879 |
| chr8:69893553-69894027 | 1.671256895 | 0.000303335 | 0.005745893 |
| chr4:1566420-1566644 | 1.66803093 | 0.000303538 | 0.005748319 |
| chr2:190277321-190277949 | -1.328690712 | 0.000303856 | 0.005751595 |
| chr6:27236268-27236574 | 1.784220671 | 0.000303861 | 0.005751595 |

| | | | |
|---|---|---|---|
| chr2:188050741-188051097 | -1.682716734 | 0.00030462 | 0.005764496 |
| chr5:58541485-58542011 | -1.488024354 | 0.000304693 | 0.005764496 |
| chr4:119730212-119730577 | 1.773867803 | 0.000305447 | 0.005777333 |
| chr7:108088850-108089055 | -2.052895042 | 0.000305667 | 0.005779911 |
| chr10:90836144-90836431 | 1.825860881 | 0.000305734 | 0.005779911 |
| chr3:179464172-179465255 | -1.229683323 | 0.00030603 | 0.005784071 |
| chr7:120733205-120733468 | -1.858545693 | 0.000306279 | 0.005787349 |
| chr14:34527263-34527463 | 2.058578376 | 0.000306869 | 0.005797075 |
| chr9:136599135-136599382 | 1.705787782 | 0.000306999 | 0.005798089 |
| chr8:22694847-22695937 | -1.187867819 | 0.000307225 | 0.005800935 |
| chr8:101388196-101388532 | 1.373832827 | 0.000308812 | 0.005829461 |
| chr1:245045485-245045862 | -1.45141678 | 0.000309086 | 0.005833204 |
| chr17:62780338-62780663 | -1.384131093 | 0.000309251 | 0.005834879 |
| chr6:49542081-49542294 | -2.110271277 | 0.000309372 | 0.005835724 |
| chr4:59994161-59994361 | -2.426162628 | 0.000310644 | 0.005858278 |
| chr7:125181361-125182266 | -1.218055637 | 0.000311186 | 0.005867048 |
| chr2:24362290-24362516 | 1.728382762 | 0.000311526 | 0.005871778 |
| chr20:48898967-48899384 | 1.225469348 | 0.00031159 | 0.005871778 |
| chr13:77137014-77137436 | -1.716677391 | 0.000312314 | 0.005883972 |
| chr10:3126670-3127405 | -1.253020793 | 0.000312957 | 0.005894639 |
| chr20:48598061-48598275 | 1.316486871 | 0.000314011 | 0.005911959 |
| chr17:57582985-57583260 | 1.475817508 | 0.000314031 | 0.005911959 |
| chr12:52061141-52061348 | 1.705196275 | 0.000314281 | 0.005915221 |
| chr17:48241079-48241381 | -1.808457631 | 0.000314497 | 0.005917825 |
| chr14:77910690-77910890 | 1.799153535 | 0.000314852 | 0.005923042 |
| chr17:59462017-59462217 | 1.665768642 | 0.000315669 | 0.005936965 |
| chr7:140403824-140404729 | -1.266074332 | 0.000316083 | 0.005943299 |
| chr11:76770996-76771329 | -1.344011148 | 0.000316217 | 0.00594358 |
| chr3:120586476-120586685 | 2.060670346 | 0.000316254 | 0.00594358 |
| chr17:53824540-53824776 | 1.994514714 | 0.000316766 | 0.005951743 |
| chr9:98146053-98146779 | -1.254308359 | 0.00031693 | 0.005953369 |
| chr1:31185545-31185949 | 1.488092152 | 0.000317127 | 0.005954883 |
| chrX:154489944-154490393 | 1.354854999 | 0.000317166 | 0.005954883 |
| chr16:82281232-82281468 | 2.059753422 | 0.000317382 | 0.005956566 |
| chr7:121099502-121099925 | -1.598622583 | 0.000317456 | 0.005956566 |
| chr6:55522385-55522731 | -1.987640041 | 0.000317489 | 0.005956566 |
| chrX:154369029-154369229 | 1.575098173 | 0.000317696 | 0.00595784 |
| chr2:233331363-233331563 | 2.396818165 | 0.000317787 | 0.00595784 |
| chr8:96663340-96663559 | 2.181739156 | 0.00031779 | 0.00595784 |
| chr21:36499982-36500417 | 1.482951624 | 0.000318574 | 0.005969672 |
| chr19:35352888-35353232 | 1.703473441 | 0.000318577 | 0.005969672 |
| chr2:65614159-65614359 | 2.409386111 | 0.000318711 | 0.005970731 |
| chr3:73638060-73638422 | -1.786898489 | 0.000319202 | 0.005977736 |
| chr11:88523482-88523730 | -2.314639843 | 0.000319241 | 0.005977736 |
| chr6:282909-283109 | 2.01529888 | 0.000319428 | 0.005979764 |
| chr6:4491556-4491918 | 1.524859367 | 0.000320266 | 0.005993997 |
| chr7:117480952-117481309 | -1.713358599 | 0.000320482 | 0.005996326 |
| chr12:24679014-24679487 | -1.769687455 | 0.000320547 | 0.005996326 |
| chr1:155695199-155695405 | 1.871444278 | 0.000320988 | 0.006001922 |
| chr12:128190256-128190949 | -1.441850868 | 0.000321003 | 0.006001922 |

| | | | |
|---|---|---|---|
| chr12:70418074-70418537 | -1.434478978 | 0.000321315 | 0.006006292 |
| chr7:15742414-15742623 | 1.932536757 | 0.000322412 | 0.006025333 |
| chr6:149080270-149080493 | 1.944764006 | 0.000323636 | 0.006045322 |
| chr7:14002719-14003416 | -1.433659667 | 0.00032364 | 0.006045322 |
| chr2:9226100-9226458 | 1.72719204 | 0.000323828 | 0.006047363 |
| chr3:78857529-78857936 | -1.479349906 | 0.000324114 | 0.006051238 |
| chr1:113826059-113826378 | 1.334987305 | 0.000324195 | 0.006051278 |
| chr1:230218674-230218911 | 2.068782818 | 0.000324622 | 0.006057773 |
| chr12:94744407-94744626 | -2.218935129 | 0.000324945 | 0.006061259 |
| chr6:157756732-157757016 | 1.657920991 | 0.000324994 | 0.006061259 |
| chr5:140828412-140828698 | 1.686649276 | 0.000325078 | 0.006061259 |
| chr8:130498069-130498302 | 1.812367513 | 0.000325126 | 0.006061259 |
| chr5:152230092-152230727 | -1.381401041 | 0.000325235 | 0.006061822 |
| chr7:127322662-127323210 | -1.510057277 | 0.000325577 | 0.006066726 |
| chr14:28318744-28318944 | 2.127463038 | 0.000325738 | 0.006068238 |
| chr7:12524871-12525149 | 1.598001188 | 0.000326836 | 0.006085922 |
| chr4:8941267-8941596 | 1.596037586 | 0.000326846 | 0.006085922 |
| chr6:170340330-170340640 | 1.733246289 | 0.000327211 | 0.006090721 |
| chr12:111733588-111733788 | 2.06891985 | 0.000327262 | 0.006090721 |
| chr2:142181030-142181525 | -1.432568494 | 0.000327415 | 0.006092088 |
| chr4:167857703-167858583 | -1.219690235 | 0.00032833 | 0.006105205 |
| chr2:191703351-191703898 | -1.487666756 | 0.000328345 | 0.006105205 |
| chr4:163048545-163048776 | -1.792911855 | 0.000328369 | 0.006105205 |
| chr3:175126928-175127738 | -1.211168054 | 0.000328439 | 0.006105205 |
| chr16:88094139-88094371 | 2.005477355 | 0.000328767 | 0.006109817 |
| chr5:4555907-4556107 | 2.274212982 | 0.000329912 | 0.00612962 |
| chr5:93410603-93410823 | 2.047027524 | 0.000330035 | 0.00613028 |
| chr3:16513522-16513971 | 1.405638193 | 0.000330173 | 0.00613028 |
| chr3:3245598-3245838 | 1.986281794 | 0.000330188 | 0.00613028 |
| chrX:9530887-9531195 | -1.719692016 | 0.000330537 | 0.006134442 |
| chr6:100454076-100454282 | 1.907653693 | 0.000330572 | 0.006134442 |
| chr13:67821744-67821949 | -2.226456503 | 0.000331602 | 0.006150573 |
| chr13:79807860-79808060 | -2.226456503 | 0.000331602 | 0.006150573 |
| chr7:128322048-128322390 | -1.527457837 | 0.000332081 | 0.006157964 |
| chr5:57364510-57364774 | -1.931797507 | 0.000332388 | 0.00616216 |
| chr16:71578032-71578285 | 2.059993534 | 0.000332662 | 0.006165757 |
| chr2:69756075-69756457 | -1.606218426 | 0.000334269 | 0.006194039 |
| chr7:116422328-116422547 | -2.115415983 | 0.000334908 | 0.006204376 |
| chr20:52437432-52437634 | 2.115666858 | 0.000335113 | 0.006206672 |
| chr7:52541311-52541691 | -1.674938667 | 0.000335236 | 0.006207456 |
| chr1:177585153-177586194 | -1.177179788 | 0.00033532 | 0.006207507 |
| chr2:76197430-76198323 | -1.302400507 | 0.000335699 | 0.006213033 |
| chr1:106920443-106920750 | 1.406479685 | 0.000336485 | 0.006226068 |
| chr2:164568458-164568827 | -1.910505048 | 0.000336574 | 0.006226219 |
| chr4:61541018-61541245 | -2.203853588 | 0.000336746 | 0.006227623 |
| chr2:41848071-41848394 | -1.725123474 | 0.000336813 | 0.006227623 |
| chr1:1272409-1272874 | 1.370515614 | 0.000338291 | 0.006253281 |
| chr12:40484183-40484625 | 1.742981623 | 0.000338364 | 0.006253281 |
| chr2:1251506-1251958 | 1.612692629 | 0.000338472 | 0.00625378 |
| chr8:87374464-87374681 | 1.952116193 | 0.000338732 | 0.006257066 |

| | | | |
|---|---|---|---|
| chr3:172546152-172546366 | 2.11127069 | 0.000338871 | 0.006258131 |
| chr11:133383252-133384063 | -1.409426865 | 0.000338972 | 0.006258483 |
| chr22:30706001-30706243 | 2.006427081 | 0.000339816 | 0.006271352 |
| chr7:23277462-23277689 | 1.484207718 | 0.000339833 | 0.006271352 |
| chr1:31862167-31862367 | 2.30515851 | 0.000340239 | 0.006276496 |
| chr9:70601566-70601887 | -1.581004225 | 0.000340322 | 0.006276496 |
| chr12:27303374-27303703 | -1.767138659 | 0.000340438 | 0.006276496 |
| chr5:31593484-31594119 | -1.35237172 | 0.000340484 | 0.006276496 |
| chr3:184081289-184081524 | 1.946102711 | 0.000340568 | 0.006276496 |
| chr18:79752062-79752355 | -1.947349775 | 0.000340603 | 0.006276496 |
| chr8:41306375-41306666 | -1.979223985 | 0.000340897 | 0.006280403 |
| chr14:95384598-95384836 | 1.83044732 | 0.000341602 | 0.006291884 |
| chr8:24955066-24955517 | -1.67558661 | 0.000341758 | 0.006293247 |
| chr14:31678169-31678386 | 1.90308642 | 0.000342008 | 0.006296335 |
| chr10:97197058-97197450 | 1.484351352 | 0.000342406 | 0.00630215 |
| chr17:68618952-68619152 | 1.829453138 | 0.000343173 | 0.00631368 |
| chr1:196786075-196787102 | -1.181971377 | 0.000343197 | 0.00631368 |
| chr7:48140036-48140540 | -1.38568957 | 0.000343954 | 0.006324216 |
| chr3:71395453-71395705 | -2.0595337 | 0.000343959 | 0.006324216 |
| chr10:7282170-7282410 | 2.060105046 | 0.000344018 | 0.006324216 |
| chr21:22885209-22885565 | 1.802712815 | 0.000344671 | 0.006334701 |
| chr8:62735951-62736778 | -1.284120262 | 0.000345166 | 0.006342282 |
| chr11:70259171-70259410 | -1.941822082 | 0.00034604 | 0.006355685 |
| chr3:170233476-170233779 | -1.659425771 | 0.000346061 | 0.006355685 |
| chr11:65020022-65020402 | 1.630761838 | 0.000346155 | 0.006355889 |
| chr4:11614407-11614783 | 1.413841264 | 0.000346434 | 0.006359486 |
| chr3:172465424-172465666 | 1.941692834 | 0.000346983 | 0.006368038 |
| chr4:189202304-189202566 | -1.75146489 | 0.000347698 | 0.006379622 |
| chr1:96164191-96164966 | 1.31890091 | 0.000348707 | 0.006396612 |
| chr21:41147696-41148002 | -1.535381022 | 0.000348876 | 0.006398182 |
| chr4:186079089-186079626 | -1.544697115 | 0.000349118 | 0.006401077 |
| chr7:11810327-11810586 | 1.966572183 | 0.000349317 | 0.006402868 |
| chr5:73522947-73523276 | -1.663995913 | 0.000349399 | 0.006402868 |
| chr10:42756871-42757151 | 1.860933336 | 0.000349466 | 0.006402868 |
| chr12:43277541-43277882 | 1.770890978 | 0.000349686 | 0.006403839 |
| chr19:30510651-30510895 | 1.770890978 | 0.000349686 | 0.006403839 |
| chr9:105088488-105088844 | -1.67198528 | 0.00035062 | 0.006419408 |
| chr12:53167121-53167375 | 1.821781377 | 0.000351058 | 0.006425882 |
| chr7:142384295-142384906 | -1.372328301 | 0.000351227 | 0.006427448 |
| chrX:31589447-31589918 | -1.766365006 | 0.000351497 | 0.006430852 |
| chr1:11061920-11062243 | 1.753622136 | 0.000351673 | 0.006430864 |
| chr13:114019466-114019732 | 1.565140167 | 0.000351696 | 0.006430864 |
| chr4:1130683-1130895 | 1.935987292 | 0.00035175 | 0.006430864 |
| chr4:173013314-173013788 | -1.460522563 | 0.00035217 | 0.006437014 |
| chr12:111763793-111764058 | 1.952285424 | 0.000353312 | 0.006456353 |
| chr14:67068865-67069065 | 2.284629959 | 0.000353921 | 0.006465933 |
| chr16:75820681-75821550 | -1.304684178 | 0.000354166 | 0.006467643 |
| chr10:103458849-103459231 | -1.407456222 | 0.000354183 | 0.006467643 |
| chr20:50656709-50657144 | 1.244219181 | 0.000354414 | 0.006470318 |
| chr2:136082622-136082926 | 1.892264319 | 0.000354978 | 0.006479069 |

| | | | |
|---|---|---|---|
| chr3:14962313-14962604 | 1.707476794 | 0.000355081 | 0.006479413 |
| chr12:32674866-32675135 | 1.723847625 | 0.000355318 | 0.006482192 |
| chr4:83093135-83093526 | -1.684861948 | 0.000357037 | 0.006511514 |
| chr12:57280199-57280615 | -1.385524076 | 0.000357096 | 0.006511514 |
| chr5:92432141-92432468 | -1.645688887 | 0.000357592 | 0.00651902 |
| chr14:104805267-104805645 | 1.317534551 | 0.000358028 | 0.006525417 |
| chr5:8430480-8430710 | 2.106499115 | 0.000358482 | 0.006532134 |
| chr6:63671135-63671430 | 1.69193558 | 0.000358817 | 0.006536682 |
| chr4:160746333-160746604 | -2.19119766 | 0.000358929 | 0.006537171 |
| chr3:129106409-129106682 | 1.56266558 | 0.000359368 | 0.006543615 |
| chr13:22665581-22665796 | -2.060435061 | 0.000360101 | 0.006555399 |
| chr6:120005391-120005767 | -1.749131891 | 0.000360359 | 0.006558537 |
| chr16:14529106-14529407 | 1.61856589 | 0.000360519 | 0.006559893 |
| chr5:40895309-40896012 | 1.33626705 | 0.000360647 | 0.006560667 |
| chr7:143284668-143285048 | -1.288104987 | 0.000360777 | 0.006561473 |
| chr3:65331830-65332095 | 1.683395524 | 0.000361139 | 0.006566485 |
| chr18:73891526-73891838 | -1.599819697 | 0.000361344 | 0.006568653 |
| chr9:127545341-127545884 | 1.256951488 | 0.000361738 | 0.006573517 |
| chr7:82534654-82535108 | -1.651277172 | 0.000361828 | 0.006573517 |
| chr9:106794250-106794683 | -1.495422642 | 0.000361873 | 0.006573517 |
| chr20:11799106-11799800 | -1.345393088 | 0.000361954 | 0.006573517 |
| chr7:152574924-152575685 | -1.332817331 | 0.000362507 | 0.006581996 |
| chr13:95740152-95740897 | -1.459490087 | 0.000362684 | 0.006583652 |
| chr5:11359427-11359758 | 1.61782763 | 0.000362776 | 0.006583755 |
| chr13:79441663-79442054 | -1.653337637 | 0.000362988 | 0.006586049 |
| chr14:81965351-81965551 | 2.111530417 | 0.000363376 | 0.00659153 |
| chr14:20373490-20373690 | 1.859490879 | 0.000364301 | 0.006606732 |
| chr22:36465624-36466014 | 1.547299622 | 0.000364595 | 0.006610511 |
| chr13:95420631-95420961 | -1.720472191 | 0.000365161 | 0.006619208 |
| chr8:22678402-22678611 | -1.833638854 | 0.00036533 | 0.006620694 |
| chr8:128201726-128202196 | -1.297890548 | 0.000365748 | 0.006626709 |
| chr7:129948853-129949455 | -1.204020205 | 0.000366049 | 0.006630596 |
| chr3:157581969-157582279 | -1.617267859 | 0.000367541 | 0.00665605 |
| chrX:7068838-7069536 | -1.337519945 | 0.000367728 | 0.006657263 |
| chr15:29812225-29812635 | 1.525669461 | 0.000367782 | 0.006657263 |
| chr7:45485050-45485343 | 1.882352459 | 0.00036836 | 0.006666149 |
| chr2:163407452-163407697 | 1.815264322 | 0.000368744 | 0.006671528 |
| chr16:56951322-56951604 | 1.603039603 | 0.000368832 | 0.006671535 |
| chr18:65938617-65938974 | -1.500652313 | 0.000369083 | 0.006674502 |
| chr14:37778109-37778383 | 1.65844074 | 0.000369463 | 0.006679807 |
| chr7:92924869-92925197 | -1.773688769 | 0.000369645 | 0.006681508 |
| chr12:27549773-27550078 | -1.651254038 | 0.000369915 | 0.006684821 |
| chr3:114412353-114412644 | -1.770652445 | 0.000370588 | 0.0066954 |
| chr4:147274462-147274740 | 2.097800959 | 0.000370887 | 0.006699224 |
| chr3:154987664-154988031 | 1.636644616 | 0.000371179 | 0.006702913 |
| chr8:98305957-98306810 | 1.246589206 | 0.000371734 | 0.006711365 |
| chr3:87145574-87145941 | -1.434434439 | 0.000371882 | 0.006712452 |
| chr16:77782802-77783053 | 1.438732712 | 0.000372059 | 0.006714058 |
| chr13:114078459-114079116 | -1.269329464 | 0.000372479 | 0.006720058 |
| chr4:106763048-106763465 | -1.649443917 | 0.000372657 | 0.006721682 |

| | | | |
|---|---|---|---|
| chr4:118757679-118758249 | -1.289475854 | 0.000372803 | 0.006721958 |
| chr11:126333679-126333900 | 1.987575079 | 0.000372848 | 0.006721958 |
| chr6:7672671-7673551 | -1.247687916 | 0.000372971 | 0.006722607 |
| chr7:157490540-157490838 | -1.913360593 | 0.000373176 | 0.006724723 |
| chr16:49948359-49948657 | 1.684537557 | 0.000373417 | 0.006727468 |
| chr14:43822620-43822822 | 1.889814378 | 0.000373622 | 0.006728674 |
| chr14:93545008-93545263 | 2.10042998 | 0.000373664 | 0.006728674 |
| chr11:64335107-64335499 | -1.371685616 | 0.000373747 | 0.006728674 |
| chr16:46401294-46401710 | 1.240205326 | 0.000374874 | 0.006747376 |
| chr5:88279539-88279877 | -1.874710551 | 0.00037521 | 0.006751844 |
| chr6:57288131-57288563 | 1.40032437 | 0.000375519 | 0.006755809 |
| chr9:109422687-109422968 | 1.80856042 | 0.000377406 | 0.006786885 |
| chr4:168674900-168675204 | -1.514968073 | 0.000377423 | 0.006786885 |
| chr1:174093812-174094118 | -1.651291824 | 0.000377701 | 0.006790295 |
| chr7:104883512-104884223 | -1.247864538 | 0.000378189 | 0.006797461 |
| chrX:129042852-129043058 | 2.363351783 | 0.000379056 | 0.006811446 |
| chr7:51295479-51295775 | -1.580087675 | 0.000379808 | 0.006823367 |
| chr14:42149068-42149268 | 2.28759116 | 0.000379994 | 0.006823811 |
| chr4:94488682-94489051 | -1.712615411 | 0.000380011 | 0.006823811 |
| chr1:174848693-174849498 | -1.366848349 | 0.000380437 | 0.006829863 |
| chr8:106045882-106046335 | 1.419530575 | 0.000380917 | 0.006836369 |
| chr13:48783407-48783663 | -1.720699111 | 0.000380978 | 0.006836369 |
| chr7:34341360-34341630 | -1.797068711 | 0.000381248 | 0.00683922 |
| chrX:32233089-32233437 | -1.915231273 | 0.000381315 | 0.00683922 |
| chr1:242057487-242057735 | 1.788280779 | 0.000381436 | 0.006839782 |
| chr8:86993995-86994411 | 1.410166687 | 0.000381736 | 0.006842745 |
| chr5:148006166-148006405 | -1.971803753 | 0.00038178 | 0.006842745 |
| chr4:70120435-70120659 | -2.039703232 | 0.000381952 | 0.006844223 |
| chr15:93196022-93196418 | -1.637241416 | 0.000382042 | 0.006844238 |
| chr2:96809162-96809581 | 1.622885453 | 0.000382166 | 0.006844855 |
| chr19:1565975-1566265 | 1.844716785 | 0.00038226 | 0.006844944 |
| chr20:48808032-48808922 | 1.141943636 | 0.000382699 | 0.006849853 |
| chr1:200684090-200684314 | 1.755245256 | 0.000382713 | 0.006849853 |
| chr1:24294832-24295140 | -1.813068816 | 0.000383848 | 0.006868563 |
| chr4:125432143-125432533 | 1.538214533 | 0.000384003 | 0.006869543 |
| chr21:36510338-36510538 | 1.971344665 | 0.000384082 | 0.006869543 |
| chr15:77135032-77135377 | 1.749798126 | 0.000384731 | 0.00687954 |
| chr10:56175262-56175580 | -1.789538112 | 0.000385044 | 0.006883536 |
| chr12:94389498-94390422 | -1.168313864 | 0.00038565 | 0.00689276 |
| chr2:102227685-102228111 | 1.361826461 | 0.000386463 | 0.006905683 |
| chr12:52050647-52051613 | 1.157207199 | 0.000386837 | 0.006910763 |
| chr3:173089440-173089646 | 2.358761099 | 0.00038734 | 0.006918134 |
| chr5:75696096-75696359 | -1.86990038 | 0.000387457 | 0.00691861 |
| chr7:103406785-103407271 | -1.290271102 | 0.000388163 | 0.006929601 |
| chr13:90866709-90867672 | -1.25440522 | 0.000388424 | 0.006932636 |
| chr14:104093818-104094127 | 1.452675289 | 0.000389749 | 0.006954676 |
| chr5:147675249-147675506 | -1.739145672 | 0.000389948 | 0.00695476 |
| chr3:43185618-43186291 | 1.507340817 | 0.000389996 | 0.00695476 |
| chr13:69326599-69326925 | -2.037384064 | 0.000390026 | 0.00695476 |
| chr14:89359550-89359806 | 1.697730209 | 0.000390685 | 0.006964885 |

| | | | |
|---|---|---|---|
| chr7:33745118-33745346 | 1.6818559 | 0.000390897 | 0.006967044 |
| chr4:20138048-20138278 | 2.03828149 | 0.000391246 | 0.006970143 |
| chr14:39212013-39212577 | 1.431472699 | 0.000391302 | 0.006970143 |
| chr14:45543408-45543619 | 2.042766315 | 0.000391358 | 0.006970143 |
| chr1:197788880-197789175 | 1.497440076 | 0.000391435 | 0.006970143 |
| chr22:28772492-28773653 | 1.187886077 | 0.000391546 | 0.006970508 |
| chr6:33813492-33813692 | 2.092304317 | 0.000391885 | 0.006974922 |
| chr18:24472503-24472882 | 1.696548352 | 0.000392521 | 0.006984477 |
| chr14:105562599-105562845 | 1.687057646 | 0.000392605 | 0.006984477 |
| chr5:126845796-126846078 | -1.584859159 | 0.000392752 | 0.006985484 |
| chr2:37585290-37586057 | -1.306547639 | 0.000393163 | 0.006991041 |
| chr1:248262650-248262950 | -2.051312309 | 0.000393272 | 0.006991041 |
| chr19:4267080-4267349 | 1.690567502 | 0.000393338 | 0.006991041 |
| chr4:107862756-107863082 | -1.664821991 | 0.000393672 | 0.006995353 |
| chr14:61595465-61595740 | 1.678646933 | 0.000393988 | 0.00699934 |
| chr1:237056169-237056388 | 1.973676808 | 0.000394256 | 0.007002485 |
| chr13:92806262-92806514 | -2.047045842 | 0.000394447 | 0.007003389 |
| chr7:42222179-42222385 | 1.971933842 | 0.000394502 | 0.007003389 |
| chrX:133414839-133416088 | 1.20280033 | 0.000394582 | 0.007003389 |
| chr10:24464113-24464321 | -1.852238672 | 0.000394788 | 0.007005435 |
| chr1:210252606-210252969 | -1.365455613 | 0.000395078 | 0.007008958 |
| chr4:2949075-2949371 | 1.699478374 | 0.00039523 | 0.007010034 |
| chr2:39665004-39665467 | -1.691070632 | 0.000395501 | 0.007013205 |
| chr17:83160871-83161227 | 1.451015441 | 0.000395616 | 0.007013621 |
| chr9:106863360-106863922 | -1.393526517 | 0.000395837 | 0.007015922 |
| chr3:11221959-11222307 | 1.60319485 | 0.000396089 | 0.007018268 |
| chr16:70714909-70715602 | 1.208863669 | 0.00039618 | 0.007018268 |
| chr7:135704178-135704417 | -1.918718053 | 0.000396244 | 0.007018268 |
| chr8:95294635-95294835 | 1.877065576 | 0.000396513 | 0.007020781 |
| chr7:11892373-11892635 | 1.949712866 | 0.000396569 | 0.007020781 |
| chr8:108067842-108068323 | 1.478884255 | 0.000397254 | 0.007029587 |
| chr3:24591702-24592103 | -1.510407817 | 0.000397361 | 0.007029587 |
| chr5:153476836-153477057 | -1.967662445 | 0.000397418 | 0.007029587 |
| chr10:53894224-53894440 | -2.120306084 | 0.000397434 | 0.007029587 |
| chr1:22131914-22132161 | 1.765766411 | 0.00039779 | 0.007034263 |
| chr10:55263348-55263702 | -1.482551601 | 0.00039831 | 0.007040943 |
| chr11:97683681-97684135 | -1.488848301 | 0.000398351 | 0.007040943 |
| chr13:59196345-59196587 | -2.037498928 | 0.000399641 | 0.007062104 |
| chr6:113490391-113490683 | -1.603395339 | 0.000399994 | 0.007066716 |
| chr4:4248050-4248849 | 1.227918203 | 0.00040078 | 0.007078976 |
| chr5:128271959-128272410 | -1.472334425 | 0.000400894 | 0.007079355 |
| chr7:17742531-17742838 | 1.59760741 | 0.000401906 | 0.007095584 |
| chr6:35726863-35727201 | 1.482561941 | 0.00040284 | 0.007109013 |
| chr13:74290409-74290788 | -1.31706035 | 0.000402852 | 0.007109013 |
| chr15:56052830-56053059 | 1.609962155 | 0.000403067 | 0.007111163 |
| chr20:57893187-57893387 | 1.601053908 | 0.000403309 | 0.007113803 |
| chr20:48500451-48500726 | 1.335113598 | 0.000404231 | 0.007125305 |
| chr21:14327350-14327722 | -1.576470933 | 0.000404236 | 0.007125305 |
| chr20:48491129-48491546 | 1.213106446 | 0.00040424 | 0.007125305 |
| chr13:67416155-67416757 | -1.349989838 | 0.000405291 | 0.00714219 |

| | | | |
|---|---|---|---|
| chr12:66823888-66824171 | 1.654350349 | 0.000405419 | 0.007142789 |
| chr18:75802764-75803046 | 1.640879199 | 0.000405573 | 0.007143868 |
| chr7:134562219-134562474 | -1.865715479 | 0.000406025 | 0.007150188 |
| chr14:51898908-51899118 | 1.921347869 | 0.000406334 | 0.007153988 |
| chr8:17504819-17505242 | -1.790402616 | 0.000406889 | 0.007162102 |
| chr16:34586732-34586998 | 1.254293861 | 0.000408169 | 0.007182991 |
| chr12:128163018-128163320 | -1.740898463 | 0.000408424 | 0.007185821 |
| chr20:48642151-48642382 | 1.276562649 | 0.00040893 | 0.007193082 |
| chr2:118296418-118296687 | 1.93154149 | 0.000409136 | 0.007195042 |
| chr8:124026419-124026729 | 1.82505094 | 0.000409891 | 0.0072053 |
| chr4:180028584-180029034 | -1.469927627 | 0.000409907 | 0.0072053 |
| chr14:74206580-74206825 | 1.773703961 | 0.000410899 | 0.007221087 |
| chr5:176834930-176835219 | 1.614810558 | 0.000411128 | 0.007223447 |
| chr5:102554790-102555077 | -1.76925665 | 0.000411504 | 0.007226847 |
| chr12:117507561-117507864 | 1.545569005 | 0.00041151 | 0.007226847 |
| chr10:110822175-110823336 | -1.171334488 | 0.000412107 | 0.007235665 |
| chr12:124231853-124232377 | 1.407051654 | 0.000412333 | 0.007237973 |
| chr5:102780493-102780780 | -1.617759658 | 0.000413473 | 0.007254906 |
| chr7:127252804-127254241 | -1.169897783 | 0.000413877 | 0.007254906 |
| chr7:112391857-112392208 | -1.616079316 | 0.000413974 | 0.007254906 |
| chr8:108459170-108459493 | 1.813595399 | 0.000414048 | 0.007254906 |
| chr7:144726928-144727134 | -2.090650814 | 0.000414067 | 0.007254906 |
| chr1:247357754-247358057 | -1.506640407 | 0.000414092 | 0.007254906 |
| chr16:16098806-16099064 | 1.871491936 | 0.000414189 | 0.007254906 |
| chr8:107886122-107886415 | 1.693081779 | 0.00041423 | 0.007254906 |
| chr4:8586648-8586927 | 1.503702911 | 0.000414277 | 0.007254906 |
| chr14:64786031-64786231 | 1.807428749 | 0.000414291 | 0.007254906 |
| chr13:72311916-72312445 | -1.639614419 | 0.000414339 | 0.007254906 |
| chr13:99552994-99553321 | -1.876175645 | 0.000414957 | 0.00726407 |
| chr11:17026802-17027049 | 1.978332289 | 0.000415093 | 0.007264797 |
| chr2:189226041-189226293 | 1.735393997 | 0.000416362 | 0.007285338 |
| chr14:52596022-52596294 | 2.091440681 | 0.000416487 | 0.007285854 |
| chr7:118352940-118353195 | -1.959191945 | 0.000417495 | 0.007301825 |
| chr13:33017491-33017961 | -1.356999899 | 0.000418195 | 0.007311475 |
| chr14:46490247-46490919 | -1.314901002 | 0.000418237 | 0.007311475 |
| chr8:134288163-134288679 | 1.434529107 | 0.000418686 | 0.007317639 |
| chr1:68342495-68343149 | -1.264315178 | 0.000419202 | 0.00732499 |
| chr7:133955397-133955597 | -2.695420672 | 0.000419656 | 0.007331249 |
| chr4:173231964-173232277 | -1.548387143 | 0.000419786 | 0.007331855 |
| chr5:895590-896144 | 1.313075273 | 0.000419886 | 0.007331926 |
| chr12:102730983-102731351 | -1.539550647 | 0.000420148 | 0.007334829 |
| chr14:26391353-26391553 | 1.927816177 | 0.000420251 | 0.007334953 |
| chr12:97279828-97280273 | -1.406382107 | 0.000420595 | 0.00733906 |
| chr18:68172899-68173580 | -1.330839567 | 0.000420678 | 0.00733906 |
| chr3:108383367-108383627 | 1.950247966 | 0.000421361 | 0.007349306 |
| chr7:155333343-155333584 | -2.01200547 | 0.000422165 | 0.007361659 |
| chr5:148482464-148483373 | -1.249563902 | 0.000422436 | 0.007364708 |
| chr11:15967157-15967440 | -1.637398558 | 0.000422847 | 0.007370196 |
| chr13:101918205-101918692 | -1.562053687 | 0.000423397 | 0.007378101 |
| chr13:92968015-92968691 | -1.40871993 | 0.000424106 | 0.007388782 |

| | | | |
|---|---|---|---|
| chr13:75173639-75174416 | -1.221404908 | 0.000425504 | 0.007411458 |
| chr10:61164436-61164799 | -1.502511098 | 0.000425764 | 0.007414294 |
| chr14:52457959-52458173 | 2.080979679 | 0.000427037 | 0.007434764 |
| chr14:33950251-33951314 | 1.165978222 | 0.00042737 | 0.007438885 |
| chrX:149571909-149572221 | 1.670435331 | 0.000427722 | 0.007443022 |
| chr7:130904087-130904691 | -1.269882902 | 0.000427802 | 0.007443022 |
| chr8:11717696-11717896 | -2.148292086 | 0.000428215 | 0.007448507 |
| chr6:452787-453120 | 1.731225352 | 0.000428544 | 0.007452535 |
| chr3:127196927-127197154 | 1.90602685 | 0.000428785 | 0.007455038 |
| chr2:219480649-219480849 | 1.745132643 | 0.00042931 | 0.007458465 |
| chr7:131007263-131007807 | -1.299732245 | 0.000429315 | 0.007458465 |
| chr6:50381087-50381378 | -1.820761555 | 0.000429338 | 0.007458465 |
| chr1:163290062-163290597 | -1.360753158 | 0.000429371 | 0.007458465 |
| chr14:93037107-93037808 | -1.273978902 | 0.000429966 | 0.007467106 |
| chr20:34143210-34143463 | 2.152629565 | 0.000430409 | 0.007471721 |
| chr4:188110067-188110328 | -1.825405218 | 0.000430427 | 0.007471721 |
| chr1:90762073-90762408 | -1.733978109 | 0.000430912 | 0.00747844 |
| chr5:5369586-5369854 | 1.941446002 | 0.000431249 | 0.007481422 |
| chr7:140992411-140992842 | -1.37055549 | 0.000431279 | 0.007481422 |
| chr3:171537462-171537729 | 1.957255102 | 0.000431414 | 0.007482068 |
| chr20:57426186-57426639 | 1.278268596 | 0.000431659 | 0.007484634 |
| chr8:123000768-123001143 | 1.476089729 | 0.00043194 | 0.007487801 |
| chr3:4504751-4504983 | 1.647152536 | 0.000432689 | 0.007499097 |
| chr14:92540925-92541158 | 1.964554963 | 0.000433248 | 0.00750708 |
| chr8:103871924-103872124 | 1.699945704 | 0.000434104 | 0.007520208 |
| chr17:77389372-77389946 | -1.355580774 | 0.000434226 | 0.007520632 |
| chr5:57157076-57157593 | -1.591311202 | 0.00043489 | 0.007530428 |
| chr13:32468574-32469121 | -1.357538653 | 0.000435218 | 0.007534399 |
| chr4:38724437-38724641 | 2.072176679 | 0.000435748 | 0.007541868 |
| chr10:31582604-31583120 | -1.297473801 | 0.00043598 | 0.00754418 |
| chr4:25299625-25300381 | 1.266653086 | 0.000436563 | 0.007552566 |
| chr21:9734055-9734340 | 1.841028949 | 0.000436728 | 0.007553468 |
| chr10:28884118-28884783 | 1.354744591 | 0.000436881 | 0.007553468 |
| chr2:123045792-123046049 | -2.186717964 | 0.000436911 | 0.007553468 |
| chr13:23178134-23178334 | 2.088577149 | 0.000437928 | 0.007567825 |
| chr12:52771868-52772068 | 1.963220871 | 0.000437939 | 0.007567825 |
| chr3:194730416-194730744 | -1.755949692 | 0.000438081 | 0.007568585 |
| chr2:216873697-216873947 | 1.817713392 | 0.000438903 | 0.007579963 |
| chr7:83547455-83548146 | -1.257595828 | 0.000438938 | 0.007579963 |
| chr11:7671202-7671659 | 1.570392121 | 0.000439045 | 0.007580106 |
| chr20:48891757-48892096 | 1.228886691 | 0.000439371 | 0.00758274 |
| chr11:132623533-132623781 | -2.095618346 | 0.000439396 | 0.00758274 |
| chr1:7407371-7407632 | 1.839345845 | 0.000439742 | 0.007587014 |
| chr6:122028888-122029439 | -1.364098187 | 0.000439844 | 0.007587055 |
| chr18:34951800-34952057 | -2.030435975 | 0.000440475 | 0.007595449 |
| chr5:1017816-1018087 | 1.565517195 | 0.000440661 | 0.007595449 |
| chr5:31226943-31227326 | 1.66142455 | 0.000440679 | 0.007595449 |
| chr8:95376069-95376286 | 1.825918673 | 0.000440727 | 0.007595449 |
| chr21:38474360-38474632 | 1.685601164 | 0.000441055 | 0.007599401 |
| chr4:108940708-108941019 | 1.639340818 | 0.000441293 | 0.007601784 |

| | | | |
|---|---|---|---|
| chr3:133494530-133494756 | -1.821951237 | 0.000441451 | 0.007602796 |
| chr6:134529858-134530265 | -1.49857773 | 0.000441569 | 0.007603119 |
| chr20:38117618-38117818 | 2.148795641 | 0.000442061 | 0.007608366 |
| chr7:83936152-83936429 | -1.696633027 | 0.000442072 | 0.007608366 |
| chr4:6674737-6675005 | 1.435949646 | 0.00044258 | 0.007615396 |
| chr1:221040908-221041270 | -1.731851343 | 0.000442751 | 0.007616621 |
| chr9:304200-305098 | -1.368132956 | 0.000443327 | 0.007623162 |
| chr4:64864530-64865465 | -1.256984178 | 0.00044333 | 0.007623162 |
| chr7:128266787-128267816 | -1.189484639 | 0.000444134 | 0.00763527 |
| chr7:86911778-86911996 | -1.950196544 | 0.000444599 | 0.00764156 |
| chr7:123517012-123517309 | -2.073486288 | 0.000445256 | 0.00765113 |
| chr4:176964113-176964664 | -1.520627386 | 0.00044558 | 0.00765499 |
| chr21:45330135-45330571 | 1.300873031 | 0.000446563 | 0.00766978 |
| chr9:124267365-124267565 | 1.739749799 | 0.000446641 | 0.00766978 |
| chr2:19003603-19003899 | 1.725909558 | 0.000446888 | 0.007672303 |
| chr5:111063682-111064050 | -1.489714338 | 0.000447653 | 0.007683714 |
| chr1:41862800-41863097 | 1.551712833 | 0.000448726 | 0.007700408 |
| chr3:1404272-1404646 | 1.415833375 | 0.000448994 | 0.00770327 |
| chr3:39982998-39983303 | 1.709157116 | 0.000449295 | 0.007706719 |
| chr7:17094874-17095477 | 1.232937406 | 0.000449547 | 0.007709286 |
| chr4:170497108-170497662 | -1.433896136 | 0.000449653 | 0.007709286 |
| chr11:13332682-13333022 | -1.496018182 | 0.000449834 | 0.007709286 |
| chr3:5081374-5081588 | 1.86481972 | 0.000449848 | 0.007709286 |
| chr7:135562760-135563089 | -1.368021864 | 0.000450142 | 0.007712607 |
| chr20:33908198-33908845 | 1.433704337 | 0.000451689 | 0.00773587 |
| chr1:6034007-6034357 | -1.49369341 | 0.000451702 | 0.00773587 |
| chr1:209076776-209076993 | -2.02641338 | 0.000453695 | 0.007768266 |
| chr7:142556778-142557130 | -1.499398305 | 0.000454177 | 0.007773183 |
| chr10:47165664-47166060 | -1.524144335 | 0.000454198 | 0.007773183 |
| chr5:64900642-64900988 | -1.434636329 | 0.000454343 | 0.007773183 |
| chr5:128538272-128538555 | 1.825670345 | 0.000454408 | 0.007773183 |
| chr7:137181676-137181886 | -2.232275832 | 0.000454565 | 0.007773183 |
| chr16:5391339-5391621 | 1.902746589 | 0.00045459 | 0.007773183 |
| chr10:132048299-132048504 | 1.826274707 | 0.000454898 | 0.007776699 |
| chr7:24786926-24787181 | 1.821522637 | 0.000455057 | 0.007777691 |
| chr1:204199273-204199515 | 1.545692002 | 0.000456238 | 0.007796052 |
| chr7:152571283-152571499 | 2.083806203 | 0.000456335 | 0.007796052 |
| chr1:235843439-235843679 | -1.873768957 | 0.000456873 | 0.007801834 |
| chr14:99296543-99296833 | -1.786030052 | 0.000456877 | 0.007801834 |
| chr2:47034231-47034822 | -1.267027988 | 0.000457684 | 0.007812884 |
| chr10:103238918-103239263 | 1.538889827 | 0.000457728 | 0.007812884 |
| chr3:22491141-22491349 | 1.996369002 | 0.000458059 | 0.00781679 |
| chrX:154540992-154541581 | 1.216004259 | 0.000458379 | 0.00782051 |
| chr5:149951285-149951637 | -1.693093897 | 0.000458592 | 0.007821052 |
| chr4:165464856-165465296 | -1.577410989 | 0.000458615 | 0.007821052 |
| chrX:153886138-153886338 | 1.793825815 | 0.000459825 | 0.007839939 |
| chr2:122748398-122749295 | -1.248186511 | 0.000460021 | 0.007841547 |
| chr6:77707618-77707924 | -1.711062473 | 0.000460667 | 0.007850815 |
| chr11:76772562-76773053 | -1.299626552 | 0.000460823 | 0.007851716 |
| chr13:72255958-72256238 | -1.611115935 | 0.00046151 | 0.007861681 |

| | | | |
|---|---|---|---|
| chr17:61213177-61213468 | 1.246675157 | 0.0004625 | 0.007876797 |
| chr1:107283241-107283450 | 1.368344292 | 0.00046294 | 0.007882529 |
| chr7:157000430-157000923 | -1.471857959 | 0.000463627 | 0.007892475 |
| chr7:82605296-82605617 | -1.66452487 | 0.000464645 | 0.007908044 |
| chr7:123580814-123581211 | -1.432198561 | 0.000465023 | 0.007911468 |
| chr12:19223099-19223365 | -1.917560362 | 0.000465115 | 0.007911468 |
| chr7:12738704-12739025 | 1.654871422 | 0.00046521 | 0.007911468 |
| chr2:137951676-137951933 | -1.789373974 | 0.00046533 | 0.007911468 |
| chr8:102897982-102898527 | -1.240520503 | 0.000465362 | 0.007911468 |
| chr9:112764489-112765235 | -1.244797692 | 0.000465786 | 0.007916307 |
| chr14:36544491-36544691 | 2.016781427 | 0.000465853 | 0.007916307 |
| chr7:139133269-139134141 | -1.203342397 | 0.000466063 | 0.007918104 |
| chr1:108846022-108846412 | 1.496013824 | 0.000466466 | 0.007923198 |
| chr4:144338933-144339187 | 1.701671449 | 0.000466633 | 0.007924284 |
| chr4:129822716-129823004 | 1.754142025 | 0.000466963 | 0.007927845 |
| chr1:198234502-198234979 | -1.432048585 | 0.00046705 | 0.007927845 |
| chr3:112097408-112097808 | 1.370461037 | 0.000467596 | 0.007934659 |
| chr14:56022162-56022453 | 1.817813381 | 0.000467658 | 0.007934659 |
| chr20:48339904-48340365 | 1.188524012 | 0.000468234 | 0.007942662 |
| chr7:126608166-126608923 | -1.247761066 | 0.000468458 | 0.007944713 |
| chr1:116919919-116920174 | 1.610927693 | 0.000468752 | 0.007947927 |
| chr13:42155734-42155934 | 1.912073767 | 0.000469809 | 0.007964093 |
| chr4:183422338-183422677 | -1.692697892 | 0.000471121 | 0.00798456 |
| chr18:57029836-57030130 | -1.753358668 | 0.000471456 | 0.007986806 |
| chr16:19095768-19095974 | 1.867541765 | 0.000471462 | 0.007986806 |
| chr15:55742403-55743681 | 1.160783313 | 0.00047253 | 0.008001726 |
| chr7:76054060-76054540 | -1.384154239 | 0.000472649 | 0.008001726 |
| chr3:150905890-150906247 | 1.506570233 | 0.000472783 | 0.008001726 |
| chr7:27235319-27235649 | 1.555118388 | 0.000472858 | 0.008001726 |
| chr20:22828034-22828234 | 2.008455727 | 0.000472864 | 0.008001726 |
| chr9:30539523-30540081 | -1.53941308 | 0.0004736 | 0.008012398 |
| chr7:148501286-148502073 | -1.227160431 | 0.000474461 | 0.008023452 |
| chr7:13107951-13108192 | 1.790378438 | 0.000474462 | 0.008023452 |
| chr12:15152549-15152756 | -2.169120712 | 0.000474826 | 0.00802782 |
| chr4:82744718-82744954 | -1.690195489 | 0.000475215 | 0.008032639 |
| chr6:90982157-90982363 | -2.150319332 | 0.000475464 | 0.008035073 |
| chr3:5596801-5597001 | 2.007048582 | 0.000475588 | 0.008035399 |
| chr4:74545081-74545338 | -1.664627132 | 0.000476393 | 0.008047215 |
| chr16:11506247-11506754 | 1.333729925 | 0.00047672 | 0.008050964 |
| chr6:5036287-5036577 | -1.711880636 | 0.000477217 | 0.00805624 |
| chr6:150535410-150535614 | 2.070257464 | 0.000477242 | 0.00805624 |
| chr5:147098228-147098813 | -1.360858573 | 0.000477613 | 0.008060724 |
| chr7:138161550-138162301 | -1.259676392 | 0.000478227 | 0.008069299 |
| chr20:55811790-55812057 | 1.823471689 | 0.000479227 | 0.008084398 |
| chr4:8697858-8698075 | 1.784618679 | 0.000479335 | 0.008084438 |
| chr3:64242412-64242839 | 1.174068112 | 0.00047969 | 0.008088648 |
| chrX:7217006-7217230 | 2.22021765 | 0.000480027 | 0.008092546 |
| chr12:111034561-111034842 | 1.517323943 | 0.00048022 | 0.008094013 |
| chr20:18077203-18077403 | 1.997822699 | 0.000480703 | 0.008100381 |
| chr19:34994476-34994733 | 1.818203174 | 0.000480904 | 0.008101983 |

| | | | |
|---|---|---|---|
| chr3:19913998-19914373 | 1.787059931 | 0.000481623 | 0.008112307 |
| chr2:183105564-183106003 | 1.656019282 | 0.000482681 | 0.008128344 |
| chr15:78194376-78194857 | -1.601984934 | 0.000482833 | 0.008129124 |
| chr4:73599032-73599566 | -1.501010639 | 0.000482957 | 0.008129426 |
| chr2:137963997-137964833 | -1.283266986 | 0.000483145 | 0.00813067 |
| chr7:12544607-12545157 | 1.353797369 | 0.000483243 | 0.00813067 |
| chr6:83905651-83906068 | 1.638120152 | 0.000484047 | 0.008141529 |
| chr1:20166181-20166381 | -2.257978648 | 0.00048424 | 0.008141529 |
| chr1:227379637-227379955 | 1.865633714 | 0.000484295 | 0.008141529 |
| chr12:3863867-3864124 | 1.816902937 | 0.000484314 | 0.008141529 |
| chr8:60499416-60499843 | 1.471230722 | 0.000485936 | 0.008167003 |
| chr5:66223669-66223963 | -1.720709121 | 0.000487361 | 0.008189157 |
| chr8:87459467-87459765 | 1.616166553 | 0.000487603 | 0.00819143 |
| chr12:76137844-76138136 | 1.817757185 | 0.000487882 | 0.008194315 |
| chr5:124749307-124749507 | -1.936884954 | 0.000488409 | 0.008200094 |
| chr12:120136248-120136770 | 1.314219488 | 0.00048844 | 0.008200094 |
| chr1:116714275-116714475 | 1.905338193 | 0.000488858 | 0.008205321 |
| chr16:87582655-87582899 | 1.704361956 | 0.000489438 | 0.008213261 |
| chr1:114301711-114302060 | 1.265467641 | 0.000489672 | 0.008215387 |
| chr13:90732084-90732382 | -1.566432585 | 0.000490072 | 0.008220297 |
| chr19:12840591-12840908 | 1.444051331 | 0.000490226 | 0.008221073 |
| chr7:20797306-20798571 | 1.1362792 | 0.000490872 | 0.008230116 |
| chr14:54634409-54634627 | 1.774034591 | 0.00049133 | 0.008235991 |
| chr1:51990619-51990861 | 1.653263351 | 0.000491644 | 0.008236509 |
| chr1:164719741-164720033 | 1.81156502 | 0.000491665 | 0.008236509 |
| chr5:139643056-139643847 | -1.15947232 | 0.000491684 | 0.008236509 |
| chr6:91708549-91708906 | -1.539948222 | 0.000492192 | 0.008243215 |
| chr8:128824502-128824702 | 1.946291156 | 0.000492651 | 0.008249106 |
| chr11:114055079-114055300 | -2.17406325 | 0.000492966 | 0.008252571 |
| chr2:84247505-84248004 | -1.358056048 | 0.000493422 | 0.00825841 |
| chr11:131246720-131246933 | -2.124532715 | 0.000493975 | 0.008265851 |
| chr5:175027186-175027855 | -1.339984155 | 0.000494114 | 0.008266378 |
| chr18:66770963-66771238 | -2.002150318 | 0.000494368 | 0.008268658 |
| chr1:166479071-166479360 | -1.752320616 | 0.000494466 | 0.008268658 |
| chr14:41785679-41786030 | 1.813888872 | 0.000495144 | 0.008277809 |
| chr1:181230222-181230551 | 1.549553859 | 0.000495293 | 0.008277809 |
| chr7:133853820-133854190 | -1.336409221 | 0.000495338 | 0.008277809 |
| chr8:77037537-77037807 | -1.880487301 | 0.000495798 | 0.0082837 |
| chr11:80436611-80436854 | -2.1729951 | 0.000496351 | 0.008291137 |
| chr13:83709801-83710098 | -1.818618637 | 0.000497233 | 0.008304048 |
| chr14:37641174-37641530 | 1.284886639 | 0.000498293 | 0.00831994 |
| chr3:152258439-152258777 | -1.567175122 | 0.000498586 | 0.008323026 |
| chr7:2522886-2523869 | -1.145478145 | 0.000498981 | 0.008327806 |
| chr8:68620152-68620372 | 2.318496601 | 0.000499523 | 0.008335035 |
| chr5:72611177-72611487 | -1.6369904 | 0.000500668 | 0.008352313 |
| chr7:32471042-32471272 | 1.717672531 | 0.000500952 | 0.008352687 |
| chr11:110065256-110065524 | -1.855786732 | 0.000501114 | 0.008352687 |
| chr6:154554384-154554584 | 1.896863182 | 0.000501116 | 0.008352687 |
| chr11:27040278-27041086 | -1.182399511 | 0.000501126 | 0.008352687 |
| chr3:196792860-196793150 | 1.482193015 | 0.000501462 | 0.008356463 |

| | | | |
|---|---|---|---|
| chr17:11608364-11608765 | -1.620736197 | 0.000501869 | 0.008361427 |
| chr3:12271783-12272033 | 1.879332405 | 0.000501978 | 0.008361428 |
| chr21:10384473-10384737 | 1.774879468 | 0.000502343 | 0.008363627 |
| chr4:37068674-37068935 | -1.855302055 | 0.000502366 | 0.008363627 |
| chr10:80491314-80492122 | -1.209964094 | 0.000502438 | 0.008363627 |
| chr15:90067795-90068020 | 1.718630682 | 0.000502828 | 0.008368299 |
| chr8:19326961-19327299 | 1.649889614 | 0.00050331 | 0.008374512 |
| chr8:66455585-66455812 | 2.324066928 | 0.000506243 | 0.008419674 |
| chr21:22363427-22363855 | -1.528047541 | 0.000506244 | 0.008419674 |
| chr2:96951924-96952242 | -1.442886061 | 0.000506668 | 0.008424887 |
| chr14:37793419-37793640 | 1.785449625 | 0.000507439 | 0.008435882 |
| chr7:118689443-118689679 | -1.996334972 | 0.000508189 | 0.008446518 |
| chr6:109678701-109679077 | -1.550704038 | 0.000509028 | 0.008458632 |
| chrX:24995479-24995781 | -1.879599775 | 0.000509388 | 0.008462784 |
| chr4:16140715-16141267 | 1.322612772 | 0.000509811 | 0.008467968 |
| chr4:5888909-5889163 | 1.814044386 | 0.000509983 | 0.008468988 |
| chr8:94092522-94093030 | -1.351154443 | 0.000512362 | 0.008506647 |
| chr6:166006077-166006420 | 1.420994953 | 0.000513186 | 0.008518487 |
| chr2:44068341-44068695 | 1.590578681 | 0.000513396 | 0.008520125 |
| chr13:62629193-62629620 | -1.836518111 | 0.000513631 | 0.008522177 |
| chr16:23540408-23540608 | 1.894483084 | 0.000513857 | 0.00852409 |
| chr14:34059802-34060815 | 1.169497482 | 0.000514531 | 0.008533423 |
| chr4:185142651-185143943 | -1.155560557 | 0.000515912 | 0.008554473 |
| chr5:71865893-71866321 | -1.46235775 | 0.000517248 | 0.008574772 |
| chr20:43384620-43384934 | -1.709801631 | 0.000517612 | 0.00857768 |
| chr6:32046493-32046693 | 1.952111708 | 0.000517648 | 0.00857768 |
| chr20:50703356-50703694 | 1.16492428 | 0.00051793 | 0.008580503 |
| chr11:1573564-1573890 | 1.447636362 | 0.000518579 | 0.008589399 |
| chrX:22196346-22196801 | -1.672017187 | 0.000519589 | 0.008604269 |
| chr11:110479208-110479749 | -1.438309631 | 0.000519943 | 0.008608274 |
| chr13:85313685-85314096 | -1.469901939 | 0.000520219 | 0.008609407 |
| chr22:41863729-41864012 | 2.124014936 | 0.000520236 | 0.008609407 |
| chr4:80566573-80566901 | -1.739504613 | 0.000521167 | 0.008622944 |
| chr17:41812757-41812958 | 1.64856681 | 0.000521715 | 0.008630155 |
| chr3:62574331-62574549 | 1.386991169 | 0.000522103 | 0.008633816 |
| chr10:80399327-80399619 | -1.713832502 | 0.000522162 | 0.008633816 |
| chr21:42376261-42377064 | -1.17514739 | 0.000522777 | 0.008642125 |
| chr11:117095910-117096116 | -1.895007029 | 0.000522995 | 0.008643865 |
| chr8:102179390-102179693 | 1.506105445 | 0.000523216 | 0.008644132 |
| chr3:132297540-132297773 | 2.00203821 | 0.000523237 | 0.008644132 |
| chr4:186473661-186473861 | -1.999575885 | 0.000524345 | 0.008660573 |
| chr5:172637020-172637592 | -1.230896844 | 0.000524989 | 0.008669335 |
| chr15:62341891-62342124 | 1.949186272 | 0.0005254 | 0.008674027 |
| chr5:1707457-1707802 | 1.51031794 | 0.000525512 | 0.008674027 |
| chr6:39173067-39173321 | 1.99612748 | 0.000525613 | 0.008674027 |
| chr8:88418428-88418672 | 1.556754347 | 0.000525751 | 0.00867444 |
| chr19:1566472-1566804 | 1.510441104 | 0.0005262 | 0.008679979 |
| chr4:184814941-184815416 | -1.291659442 | 0.000526471 | 0.008682578 |
| chr17:65064807-65065030 | 1.531194655 | 0.000526792 | 0.008686011 |
| chr10:121511982-121512372 | 1.384831241 | 0.00052694 | 0.008686573 |

| | | | |
|---|---|---|---|
| chr12:47979284-47979484 | 1.772102021 | 0.000527794 | 0.008698785 |
| chr1:96596250-96596598 | -1.506684483 | 0.000528239 | 0.008704238 |
| chr22:36465177-36465508 | 1.440169695 | 0.000529979 | 0.008731038 |
| chr3:19395026-19395321 | -1.76377985 | 0.0005304 | 0.0087345 |
| chr3:147712361-147713375 | -1.272189205 | 0.000530417 | 0.0087345 |
| chr1:100928209-100928440 | -2.151342816 | 0.000530765 | 0.008737795 |
| chr2:63043962-63044202 | -1.844911277 | 0.000530845 | 0.008737795 |
| chr14:35390878-35391410 | 1.261935861 | 0.000531195 | 0.008741671 |
| chr10:23499276-23499658 | -1.708304167 | 0.000532006 | 0.008753144 |
| chr11:13781107-13781526 | -1.403833695 | 0.000532209 | 0.008754602 |
| chr8:100244354-100244554 | 1.929637872 | 0.000532746 | 0.008760992 |
| chr17:53791993-53792256 | 1.845951186 | 0.000532826 | 0.008760992 |
| chr11:12746551-12747158 | -1.266094684 | 0.000533128 | 0.008763754 |
| chr16:15996533-15996771 | 1.822486603 | 0.000533226 | 0.008763754 |
| chr7:157318396-157318632 | -1.787565588 | 0.000533369 | 0.008763754 |
| chr7:154071930-154072220 | -1.666516592 | 0.000533452 | 0.008763754 |
| chr4:734873-735073 | 1.93406544 | 0.000534452 | 0.0087783 |
| chr9:14261925-14262414 | -1.503919169 | 0.000534696 | 0.008780432 |
| chr1:220906497-220907286 | -1.239999241 | 0.000535049 | 0.008784338 |
| chr16:29302556-29302809 | 1.723056226 | 0.000535568 | 0.008790976 |
| chr3:158744010-158744350 | 1.493748797 | 0.000535789 | 0.008792438 |
| chr13:62512246-62512926 | -1.328884239 | 0.000535949 | 0.008792438 |
| chr11:21753417-21753672 | -2.149813817 | 0.000536001 | 0.008792438 |
| chr4:174632270-174632625 | -1.709749002 | 0.000536253 | 0.008794686 |
| chr10:79685155-79685749 | -1.294032736 | 0.000537689 | 0.008816343 |
| chr1:243580203-243580403 | 2.064947102 | 0.000538176 | 0.00882245 |
| chr7:21923223-21923423 | 1.849517093 | 0.000538409 | 0.008824372 |
| chr11:2892875-2893141 | 1.559476176 | 0.000538651 | 0.008826453 |
| chr8:111858209-111858543 | 1.454082787 | 0.000539166 | 0.008832289 |
| chr11:64787712-64787922 | 1.725799376 | 0.000539237 | 0.008832289 |
| chr6:3193076-3193276 | 2.119915426 | 0.000541369 | 0.008865154 |
| chr10:34427210-34427747 | -1.245305917 | 0.000541475 | 0.008865154 |
| chr21:42396010-42396385 | -1.562620027 | 0.000541713 | 0.008867157 |
| chr10:4849558-4850033 | 1.217153621 | 0.000541913 | 0.008868524 |
| chr12:32956346-32956881 | 1.363189939 | 0.000542156 | 0.008870615 |
| chr20:48875189-48875654 | 1.17116555 | 0.000542405 | 0.008871476 |
| chr4:57233476-57233853 | 1.323838354 | 0.00054244 | 0.008871476 |
| chr5:95780951-95781615 | -1.23621047 | 0.000542836 | 0.008876044 |
| chr8:76152180-76152436 | 1.984651293 | 0.000543617 | 0.008883758 |
| chr4:74451148-74451439 | -1.59006967 | 0.000543659 | 0.008883758 |
| chr5:57004810-57005158 | -1.735686543 | 0.000543731 | 0.008883758 |
| chr16:82152595-82153258 | 1.216891988 | 0.000543771 | 0.008883758 |
| chr9:74281705-74281998 | -1.726430472 | 0.000544358 | 0.008891447 |
| chr3:106345386-106345945 | -1.556534512 | 0.00054486 | 0.008897755 |
| chr12:109937636-109937836 | -2.014830741 | 0.000545088 | 0.008897876 |
| chr21:9715680-9715880 | 2.212197367 | 0.000545155 | 0.008897876 |
| chr13:61125557-61125908 | -1.417898587 | 0.000545216 | 0.008897876 |
| chr1:169439052-169439388 | 1.528872412 | 0.000545553 | 0.00890148 |
| chrX:153970917-153971464 | 1.203640486 | 0.000546603 | 0.008914507 |
| chr3:18425323-18425532 | 1.661472542 | 0.000546673 | 0.008914507 |

| | | | |
|---|---|---|---|
| chr8:100026264-100026493 | 1.809382252 | 0.000546817 | 0.008914507 |
| chr8:103438458-103438720 | 1.809382252 | 0.000546817 | 0.008914507 |
| chr9:104791864-104792203 | -1.7815324 | 0.000547029 | 0.008916069 |
| chr19:58098242-58098462 | 1.93158524 | 0.000547319 | 0.008918898 |
| chr12:79452037-79452397 | -1.387879315 | 0.000548198 | 0.008931323 |
| chr1:40078637-40078882 | 1.651646161 | 0.000549395 | 0.008948922 |
| chr8:66453477-66453842 | -1.531645756 | 0.000549706 | 0.008952085 |
| chr6:116261404-116261604 | 2.054758436 | 0.000550222 | 0.008958592 |
| chrY:11323847-11324069 | 1.930525166 | 0.000551169 | 0.008972092 |
| chr2:46967163-46967719 | -1.205515705 | 0.000551399 | 0.008973929 |
| chr2:46967853-46968499 | -1.183538121 | 0.000551709 | 0.008977068 |
| chr7:108592782-108593046 | -1.534086961 | 0.000552199 | 0.008983138 |
| chr17:49278107-49278322 | 1.878547874 | 0.000552808 | 0.008991131 |
| chr2:35783720-35784112 | -1.381013178 | 0.000552937 | 0.008991323 |
| chr1:19338191-19338687 | -1.446034095 | 0.000553072 | 0.008991609 |
| chr7:23811657-23811903 | 1.769389775 | 0.000553425 | 0.008995453 |
| chr14:27311315-27311607 | 1.70731882 | 0.000555132 | 0.009019863 |
| chr5:82365697-82366145 | -1.436519189 | 0.000555163 | 0.009019863 |
| chr4:185383493-185384171 | -1.24109452 | 0.000555579 | 0.009024613 |
| chr18:23143274-23143795 | -1.36980816 | 0.000555691 | 0.009024613 |
| chr1:146020045-146020305 | 1.445489849 | 0.000556614 | 0.00903769 |
| chr3:138129079-138129450 | 1.707759123 | 0.000556767 | 0.009038267 |
| chr7:139017054-139017455 | -1.373371015 | 0.000558206 | 0.009056825 |
| chr5:68691670-68692332 | -1.353816156 | 0.000558332 | 0.009056825 |
| chr14:44745327-44745698 | 1.529941514 | 0.000558372 | 0.009056825 |
| chr11:35380599-35380949 | -1.476575952 | 0.000558383 | 0.009056825 |
| chr3:189043217-189043603 | -1.534745242 | 0.000558821 | 0.009062001 |
| chr7:142320053-142320320 | -1.567304655 | 0.00055987 | 0.009074744 |
| chr3:23475937-23476188 | 1.922663825 | 0.000560027 | 0.009074744 |
| chr14:59324544-59324758 | 2.061300929 | 0.000560071 | 0.009074744 |
| chr6:127781752-127782334 | -1.295882129 | 0.00056008 | 0.009074744 |
| chr5:87048113-87048525 | -1.447095187 | 0.000560217 | 0.009074767 |
| chr9:20885053-20885327 | -1.976190644 | 0.000560319 | 0.009074767 |
| chr8:100982444-100982738 | 1.601718439 | 0.000560575 | 0.009077004 |
| chr13:46080597-46080869 | 1.734941068 | 0.000560731 | 0.009077232 |
| chr16:1486402-1486763 | 1.604005593 | 0.000560861 | 0.009077232 |
| chr1:223720655-223721469 | -1.214954198 | 0.000560955 | 0.009077232 |
| chr8:89282389-89282613 | 1.726635122 | 0.000561063 | 0.009077232 |
| chr7:119482066-119482604 | -1.351375919 | 0.000561455 | 0.009081654 |
| chr21:43929707-43930025 | 1.446742959 | 0.000561733 | 0.009083306 |
| chr14:31350988-31351235 | 1.988925456 | 0.000561794 | 0.009083306 |
| chr15:98749210-98749463 | -1.622751546 | 0.000561935 | 0.009083663 |
| chr2:112661885-112662143 | 1.844576743 | 0.00056211 | 0.009084587 |
| chr1:168939534-168939857 | -1.465781909 | 0.000562588 | 0.009088424 |
| chr5:1175201-1175439 | 1.740002387 | 0.000562607 | 0.009088424 |
| chr3:155124788-155125073 | -1.498587761 | 0.000562704 | 0.009088424 |
| chr1:31471555-31471793 | 1.804929444 | 0.000563666 | 0.009102044 |
| chr9:130446142-130446723 | -1.216961056 | 0.000564417 | 0.00911225 |
| chr4:12115997-12116463 | 1.358958763 | 0.000564994 | 0.009119642 |
| chr7:41136436-41136688 | 1.980815974 | 0.000565565 | 0.009126939 |

| | | | |
|---|---|---|---|
| chr10:1629511-1630152 | -1.230843807 | 0.000565797 | 0.009128768 |
| chr8:127409006-127409479 | 1.353733319 | 0.000566292 | 0.00913483 |
| chr7:54687556-54688026 | -1.39520202 | 0.000566477 | 0.009135886 |
| chr14:67892435-67892918 | 1.412114622 | 0.000566637 | 0.009136543 |
| chr20:56525582-56525880 | 1.524314127 | 0.000566954 | 0.009139729 |
| chr2:217951464-217951910 | -1.352716443 | 0.000567131 | 0.009140666 |
| chr12:113371517-113371738 | 1.918311787 | 0.00056728 | 0.009141146 |
| chr15:79104991-79105198 | 1.921494474 | 0.000567504 | 0.009142827 |
| chr14:37845618-37845818 | 1.803606077 | 0.000568282 | 0.009153438 |
| chr13:47409106-47409378 | 1.476236157 | 0.000568464 | 0.009154443 |
| chr12:33795497-33795768 | 1.804593501 | 0.000569439 | 0.009168217 |
| chr1:117912861-117913221 | 1.719239784 | 0.00057011 | 0.009177101 |
| chr4:168421678-168422069 | -1.555130077 | 0.000570609 | 0.009181933 |
| chr7:156160842-156161056 | -1.97352189 | 0.00057065 | 0.009181933 |
| chr2:58325381-58325878 | -1.453876843 | 0.000571735 | 0.009195539 |
| chr7:131627623-131628164 | -1.340293231 | 0.000571736 | 0.009195539 |
| chr22:20056041-20056425 | 1.611840114 | 0.000572459 | 0.009205238 |
| chr8:46013669-46013964 | 1.830578863 | 0.000574017 | 0.009228361 |
| chr4:187172045-187172473 | -1.728873874 | 0.000574488 | 0.009233989 |
| chr13:51990891-51991142 | -1.92523046 | 0.000575418 | 0.009245113 |
| chr8:86258513-86258908 | 1.265912935 | 0.000575421 | 0.009245113 |
| chr7:22452245-22452809 | 1.224892613 | 0.00057586 | 0.009248275 |
| chr8:22732980-22733431 | -1.396283492 | 0.00057586 | 0.009248275 |
| chr2:117412006-117412313 | -1.621176864 | 0.000576171 | 0.009249516 |
| chr2:131831903-131832180 | -1.545103913 | 0.000576178 | 0.009249516 |
| chr7:32088545-32088790 | 1.657205293 | 0.000576531 | 0.009253241 |
| chr12:104790410-104790867 | -1.462936136 | 0.000576831 | 0.00925611 |
| chr3:153779506-153779780 | 1.876654872 | 0.000577057 | 0.009257797 |
| chr11:84687334-84687850 | -1.267014496 | 0.000577541 | 0.009263626 |
| chr16:2692498-2692918 | 1.537255292 | 0.00057784 | 0.009266478 |
| chr7:31892119-31892383 | 1.90032777 | 0.000578107 | 0.009268831 |
| chr13:103471717-103472285 | -1.338333148 | 0.000578681 | 0.009276093 |
| chr12:64147347-64147725 | 1.517540249 | 0.00057905 | 0.009280061 |
| chr13:59357922-59358335 | -1.669781333 | 0.000579233 | 0.009281057 |
| chr5:53477039-53477239 | 1.929819213 | 0.000579719 | 0.009286894 |
| chr4:8534017-8534791 | 1.191812953 | 0.000580267 | 0.009293725 |
| chr7:136050036-136050326 | 1.819388702 | 0.000580562 | 0.009295563 |
| chr7:116730584-116730970 | -1.57617664 | 0.000580624 | 0.009295563 |
| chr5:180616634-180616862 | 1.775460592 | 0.000580818 | 0.009296736 |
| chr6:118940852-118941076 | 1.73266447 | 0.000581184 | 0.009300152 |
| chr1:110098257-110098655 | -1.359193027 | 0.000581275 | 0.009300152 |
| chr4:170102463-170102834 | -1.44821536 | 0.000581646 | 0.009304157 |
| chr11:61601627-61602046 | 1.370084687 | 0.000582143 | 0.00931015 |
| chr6:20550217-20550771 | -1.345099568 | 0.000583146 | 0.009324256 |
| chr7:81684946-81685369 | -1.342410751 | 0.000584672 | 0.009346701 |
| chr8:93104599-93105137 | 1.29284276 | 0.00058498 | 0.009349663 |
| chr18:54246791-54247449 | -1.291153288 | 0.000585101 | 0.009349663 |
| chr16:2799687-2800286 | 1.328656828 | 0.000585988 | 0.009361875 |
| chr5:176822233-176822735 | -1.267783679 | 0.000586272 | 0.009364466 |
| chr2:33411532-33412225 | 1.192920691 | 0.000586679 | 0.009369018 |

| | | | |
|---|---|---|---|
| chr8:80407257-80407707 | 1.499234946 | 0.000587789 | 0.009383399 |
| chr1:31932564-31932768 | 1.734810002 | 0.000587825 | 0.009383399 |
| chr6:57029119-57029647 | -1.261933826 | 0.000588032 | 0.009384745 |
| chr10:123864095-123864414 | 1.675371757 | 0.000589117 | 0.009400116 |
| chr7:106779998-106780289 | -1.719702359 | 0.000589751 | 0.009408265 |
| chr4:39548754-39549049 | 1.849141028 | 0.000589989 | 0.009408777 |
| chr5:124755700-124755953 | -1.698658787 | 0.000590029 | 0.009408777 |
| chr7:16978395-16978667 | 1.532936373 | 0.000590201 | 0.009409561 |
| chr4:183157338-183157538 | -2.34006913 | 0.000590369 | 0.009410282 |
| chr13:72166139-72166765 | -1.257724376 | 0.000590511 | 0.009410598 |
| chr9:110241829-110242456 | -1.295539459 | 0.000591487 | 0.009424183 |
| chr3:21839534-21839997 | -1.300413642 | 0.000592036 | 0.009430966 |
| chr7:35906460-35906858 | 1.381087391 | 0.000592196 | 0.009431563 |
| chr4:17707685-17708133 | 1.453522354 | 0.000592901 | 0.009438907 |
| chr10:80429655-80430036 | -1.363058572 | 0.000592904 | 0.009438907 |
| chr7:23596993-23598169 | 1.112854374 | 0.000593372 | 0.009444407 |
| chr7:112970844-112971110 | -1.613463029 | 0.00059358 | 0.009445755 |
| chr10:129703825-129704125 | 1.52031486 | 0.000595115 | 0.009468209 |
| chr7:128168658-128169035 | -1.333391813 | 0.000595425 | 0.00947117 |
| chr2:127217983-127218241 | -1.691517639 | 0.000596413 | 0.009484337 |
| chr4:123478733-123479399 | 1.197017599 | 0.0005965 | 0.009484337 |
| chr12:63928921-63929152 | 1.772612841 | 0.000597561 | 0.0094974 |
| chr10:50653080-50653515 | -1.554184371 | 0.000597569 | 0.0094974 |
| chr8:39032184-39032871 | -1.329008591 | 0.000598104 | 0.009503932 |
| chr8:126041966-126042179 | 1.826515388 | 0.000599414 | 0.009522763 |
| chr12:132638799-132639023 | 1.595517323 | 0.00059975 | 0.009525724 |
| chr8:87799089-87799294 | 2.027831815 | 0.000599849 | 0.009525724 |
| chr2:85589960-85590336 | 1.359169399 | 0.000600624 | 0.009536062 |
| chr3:161358580-161358780 | 1.915187616 | 0.00060152 | 0.0095483 |
| chr10:2393063-2393492 | 1.496429924 | 0.00060232 | 0.009559015 |
| chr8:135324482-135324756 | 1.978449695 | 0.000602738 | 0.009563683 |
| chr7:2825239-2825439 | 1.833548342 | 0.000603912 | 0.009580317 |
| chr7:130440670-130441472 | -1.135672514 | 0.000604062 | 0.009580722 |
| chr1:173506495-173506777 | -1.500955829 | 0.000604468 | 0.009585168 |
| chr4:168318160-168318966 | -1.164286738 | 0.000604809 | 0.009588595 |
| chr14:74040840-74041040 | 2.028987847 | 0.000605763 | 0.009601725 |
| chr2:162091406-162092097 | -1.295816939 | 0.000606 | 0.009603496 |
| chr8:58721659-58721970 | -1.733715499 | 0.000606128 | 0.009603541 |
| chr12:82634406-82634698 | -1.515207659 | 0.00060676 | 0.009611569 |
| chr16:34581196-34581758 | 1.148921514 | 0.000606986 | 0.009613154 |
| chr6:41734721-41735096 | 1.356041735 | 0.000607111 | 0.009613154 |
| chr15:91279434-91279703 | 1.786096014 | 0.000607986 | 0.009625022 |
| chr20:4171983-4172770 | 1.26699781 | 0.000608322 | 0.00962748 |
| chr4:3488224-3488424 | 1.727342478 | 0.000608394 | 0.00962748 |
| chr12:119695212-119695435 | 1.917256604 | 0.000608644 | 0.00962748 |
| chr14:20271200-20271406 | 1.917256604 | 0.000608644 | 0.00962748 |
| chr4:26133536-26133751 | 1.880768246 | 0.000609387 | 0.009637245 |
| chr18:43932190-43932490 | -1.790900918 | 0.000609684 | 0.009639946 |
| chr18:51614058-51614330 | -1.579563619 | 0.000609831 | 0.00964028 |
| chr5:146327214-146328301 | -1.191003028 | 0.000610323 | 0.009646076 |

| | | | |
|---|---|---|---|
| chr11:117915396-117915596 | 1.730478222 | 0.000610937 | 0.009653788 |
| chr2:14255002-14255694 | -1.169258981 | 0.000611223 | 0.009656319 |
| chr18:76525253-76525713 | -1.313505299 | 0.000612468 | 0.00967308 |
| chr7:78903166-78903605 | -1.654282485 | 0.000612537 | 0.00967308 |
| chr15:54282424-54282624 | 1.890699184 | 0.000613391 | 0.009683975 |
| chr12:88781896-88782282 | -1.365758779 | 0.000613479 | 0.009683975 |
| chr12:94649772-94650077 | -1.639774629 | 0.00061371 | 0.009685619 |
| chr2:84996894-84997233 | 1.507992532 | 0.000614348 | 0.00969369 |
| chr4:106987289-106987672 | -1.563702545 | 0.00061486 | 0.009699782 |
| chr20:48296809-48297036 | 1.271450661 | 0.000617542 | 0.009740084 |
| chr4:74370890-74371158 | -1.453700811 | 0.000618267 | 0.009749504 |
| chr8:80964975-80965241 | 1.64807041 | 0.000618414 | 0.009749814 |
| chr2:196051489-196052195 | -1.227983642 | 0.000619463 | 0.009764344 |
| chr8:85959097-85959297 | 2.02701381 | 0.000620687 | 0.009781433 |
| chr9:22342608-22343247 | 1.333739063 | 0.000620941 | 0.009781433 |
| chr12:67377561-67377936 | -1.661442168 | 0.000620996 | 0.009781433 |
| chr1:228782877-228783077 | 2.036792545 | 0.000621145 | 0.009781433 |
| chr3:87206049-87206562 | -1.388776887 | 0.000621185 | 0.009781433 |
| chr13:40116044-40116422 | -1.513834996 | 0.000621629 | 0.009785928 |
| chr1:113206414-113206799 | -1.326514827 | 0.0006218 | 0.009785928 |
| chr20:50298695-50299271 | -1.226785332 | 0.000621854 | 0.009785928 |
| chr15:69468324-69469136 | 1.177005898 | 0.000622619 | 0.009795964 |
| chr10:119254040-119254367 | 1.753260398 | 0.000623353 | 0.009802199 |
| chr6:135270767-135271705 | -1.187954914 | 0.000623392 | 0.009802199 |
| chr16:54540867-54541360 | 1.389913391 | 0.000623566 | 0.009802199 |
| chr3:63960223-63960728 | -1.273697828 | 0.000623772 | 0.009802199 |
| chr5:175015050-175015552 | -1.306806564 | 0.000623777 | 0.009802199 |
| chr10:5532105-5532357 | 1.71530171 | 0.000623885 | 0.009802199 |
| chr1:28865567-28865767 | 1.97831212 | 0.000623911 | 0.009802199 |
| chr20:55581203-55581643 | 1.149768279 | 0.000624048 | 0.009802335 |
| chr4:79652235-79652676 | 1.446314637 | 0.000624316 | 0.009804536 |
| chr15:89399877-89400148 | 1.367526884 | 0.000624465 | 0.009804871 |
| chrX:154097406-154098151 | 1.171817725 | 0.000625445 | 0.009818241 |
| chr14:43686426-43686663 | 1.823543472 | 0.000626828 | 0.009837945 |
| chr2:52109923-52110268 | -1.754513351 | 0.000627 | 0.00983863 |
| chr2:119715696-119715929 | 1.904654665 | 0.000627232 | 0.009840245 |
| chr15:88081323-88081527 | 1.786741236 | 0.000628002 | 0.009850317 |
| chr7:77481598-77482336 | -1.209779028 | 0.000628769 | 0.009858452 |
| chr13:48542813-48543247 | -1.512948053 | 0.000628778 | 0.009858452 |
| chr11:75748844-75749236 | -1.752960843 | 0.000629595 | 0.009869243 |
| chr4:69170591-69170861 | -1.687308696 | 0.000629821 | 0.009870767 |
| chr1:214050462-214050919 | -1.417677209 | 0.000630131 | 0.0098736 |
| chr14:28375296-28375612 | 1.467363683 | 0.000630833 | 0.009882579 |
| chr1:56032259-56032540 | -2.047837517 | 0.000631163 | 0.009885733 |
| chr2:10762693-10763249 | -1.244634674 | 0.000632067 | 0.009897869 |
| chr15:44347953-44348153 | 2.206719129 | 0.000633073 | 0.009911582 |
| chr5:29023435-29023907 | 1.364092982 | 0.000634745 | 0.009934303 |
| chr12:52655783-52656096 | 1.475577322 | 0.000634783 | 0.009934303 |
| chr1:74352321-74352773 | -1.504401972 | 0.000635472 | 0.009943046 |
| chr22:35452608-35452855 | 1.661639928 | 0.00063787 | 0.009978542 |

| | | | |
|---|---|---|---|
| chr2:36279665-36279871 | -2.054780842 | 0.000638105 | 0.009979239 |
| chr22:39617198-39617588 | -1.662652649 | 0.000638175 | 0.009979239 |
| chr3:184124410-184124631 | -1.926016705 | 0.000638376 | 0.009980334 |
| chr9:135512894-135513347 | 1.547734762 | 0.000639205 | 0.009991265 |
| chr18:28421932-28422258 | -1.721402713 | 0.000639866 | 0.00999822 |
| chr4:95084654-95085011 | 1.485873615 | 0.000640028 | 0.00999822 |
| chrX:39857778-39858221 | -1.472481242 | 0.000640042 | 0.00999822 |
| chr5:61525632-61526150 | -1.331479606 | 0.00064068 | 0.01000616 |
| chr5:14588842-14589537 | 1.194124873 | 0.00064108 | 0.0100097 |
| chr10:66627172-66627498 | -1.451820414 | 0.000641168 | 0.0100097 |
| chr5:148397968-148398219 | -1.626072115 | 0.00064143 | 0.010011741 |
| chr7:87747395-87747624 | -1.907685993 | 0.000641916 | 0.010017288 |
| chr6:20279610-20279924 | -1.861718471 | 0.00064219 | 0.010017339 |
| chr7:126790769-126791138 | -1.549582884 | 0.0006422 | 0.010017339 |
| chr5:2757733-2758410 | -1.163849653 | 0.000642311 | 0.010017339 |
| chr3:144252652-144253064 | -1.577327705 | 0.000642881 | 0.010024179 |
| chr2:151777555-151777817 | -1.606909235 | 0.0006433 | 0.010028156 |
| chr19:41848829-41849365 | 1.391094032 | 0.0006435 | 0.010028156 |
| chr18:49510720-49510998 | -1.716552219 | 0.000643529 | 0.010028156 |
| chr7:87767661-87768322 | -1.271574701 | 0.00064388 | 0.010031591 |
| chr7:10778891-10779091 | 2.100019582 | 0.000644505 | 0.010039291 |
| chr7:26491328-26491603 | 1.672500633 | 0.000644713 | 0.010040489 |
| chr15:83286220-83286544 | -1.520893987 | 0.000644888 | 0.010041166 |
| chr14:72516895-72517143 | 1.530397688 | 0.000645026 | 0.010041274 |
| chr7:16075188-16075396 | 1.909502032 | 0.000647413 | 0.010076391 |
| chr5:60260620-60260989 | -1.856805127 | 0.000648357 | 0.010089028 |
| chr8:61273382-61273604 | -1.850210481 | 0.000649127 | 0.010098966 |
| chr1:29783142-29783386 | 1.924221575 | 0.000649619 | 0.010102728 |
| chr7:12189893-12190214 | 1.75250724 | 0.000649632 | 0.010102728 |
| chr14:20959319-20959586 | 1.789284783 | 0.00065017 | 0.010109034 |
| chr11:65308800-65309008 | 1.737992665 | 0.000650817 | 0.010117046 |
| chr4:171186787-171187055 | -1.638732152 | 0.000651032 | 0.0101175 |
| chr14:72311190-72311409 | 2.035040846 | 0.000651111 | 0.0101175 |
| chr7:103279935-103281015 | -1.197699515 | 0.000652185 | 0.010132138 |
| chr4:74659816-74660117 | -1.681931395 | 0.000654197 | 0.010160141 |
| chr18:13800875-13801451 | 1.268709651 | 0.000654253 | 0.010160141 |
| chr16:1356985-1357215 | 1.647689737 | 0.000654416 | 0.010160622 |
| chr17:81065580-81065901 | 1.575176101 | 0.000654709 | 0.010163112 |
| chr2:196206791-196207217 | -1.404708881 | 0.000655424 | 0.010171642 |
| chr12:51229035-51229236 | 1.960437453 | 0.000655524 | 0.010171642 |
| chr20:48819145-48819398 | 1.237044755 | 0.000655723 | 0.010172676 |
| chr4:183289006-183289370 | -1.540622564 | 0.000656135 | 0.010176994 |
| chr1:156823699-156823905 | 2.021992308 | 0.000656613 | 0.010182352 |
| chr15:91974439-91974882 | -1.463382539 | 0.000657031 | 0.01018677 |
| chr14:65849507-65849707 | 1.827112595 | 0.000657314 | 0.010187081 |
| chr14:35535275-35535496 | 1.596590946 | 0.000657317 | 0.010187081 |
| chr6:142002841-142003322 | -1.766800926 | 0.000657664 | 0.010188483 |
| chr4:3830282-3830482 | 1.809795497 | 0.000657824 | 0.010188483 |
| chr1:67298804-67299070 | -1.755447487 | 0.000657916 | 0.010188483 |
| chr6:161287061-161287506 | -1.60388229 | 0.000657939 | 0.010188483 |

| | | | |
|---|---|---|---|
| chr5:139696481-139697728 | -1.11260061 | 0.000658119 | 0.010189204 |
| chr22:33342327-33342651 | 1.889353538 | 0.000658837 | 0.010197933 |
| chr5:111991491-111992103 | 1.334176225 | 0.000658949 | 0.010197933 |
| chr20:45218621-45219003 | 1.704412297 | 0.0006594 | 0.010202859 |
| chr5:134279594-134280058 | -1.28312932 | 0.000659568 | 0.010203403 |
| chr6:100633147-100633474 | -1.488402794 | 0.00065979 | 0.010204769 |
| chrX:28587326-28587547 | -1.844086405 | 0.00066025 | 0.010209828 |
| chr6:125821901-125822200 | -1.514837001 | 0.000661742 | 0.010230833 |
| chr10:88428485-88429053 | -1.50619309 | 0.000662809 | 0.010245262 |
| chr17:3976057-3976472 | -1.563751856 | 0.000663079 | 0.010247368 |
| chr6:98827678-98828403 | -1.29514382 | 0.000663405 | 0.010250334 |
| chr3:20250388-20250588 | 1.825832095 | 0.000663794 | 0.010254285 |
| chr10:11827085-11827285 | 1.978290661 | 0.000664178 | 0.010258141 |
| chr9:72478004-72478759 | -1.274762594 | 0.000664774 | 0.010265285 |
| chr3:48268338-48269544 | -1.212330077 | 0.000666106 | 0.010283782 |
| chr15:56821829-56822206 | 1.605810463 | 0.000666542 | 0.010288429 |
| chr7:70725531-70725906 | -1.597130376 | 0.000667019 | 0.010293727 |
| chr8:39852694-39853096 | -1.436810708 | 0.000667219 | 0.010294742 |
| chr13:30907355-30907600 | -1.912916455 | 0.000668125 | 0.010306631 |
| chr7:54491640-54491968 | -1.709797327 | 0.000669114 | 0.010317928 |
| chr3:171404654-171404987 | 1.447587198 | 0.000669131 | 0.010317928 |
| chr9:69089346-69089813 | -1.579136766 | 0.000669261 | 0.010317928 |
| chr8:92264413-92264637 | 2.00836014 | 0.000669793 | 0.010324061 |
| chr1:145579676-145579983 | 1.634445629 | 0.000670563 | 0.010333845 |
| chr4:8817534-8817959 | 1.524791632 | 0.000670936 | 0.010337516 |
| chr7:2720812-2721344 | -1.197861824 | 0.000671323 | 0.010341403 |
| chr6:170438383-170438851 | -1.365044144 | 0.000671904 | 0.010348269 |
| chr2:77772722-77772958 | -2.050441512 | 0.000672169 | 0.010350264 |
| chr5:42931224-42931894 | 1.235404842 | 0.000672749 | 0.010357118 |
| chr8:105740908-105741136 | 1.681882661 | 0.000673615 | 0.010368367 |
| chr3:30188755-30189082 | -1.454920161 | 0.000673961 | 0.010371602 |
| chr5:1111769-1112160 | 1.35830985 | 0.000674101 | 0.010371682 |
| chr20:44663015-44663221 | 1.74252242 | 0.000674781 | 0.010378276 |
| chr20:54084639-54085485 | 1.086677449 | 0.000674801 | 0.010378276 |
| chr4:96228457-96228938 | -1.284654962 | 0.000676093 | 0.010396062 |
| chr8:144691347-144691627 | 1.675381396 | 0.00067666 | 0.010401685 |
| chr7:22356016-22357415 | 1.110203126 | 0.000676958 | 0.010401685 |
| chr6:157688978-157689535 | -1.463868354 | 0.000677072 | 0.010401685 |
| chr2:220198904-220199464 | -1.35401712 | 0.000677103 | 0.010401685 |
| chr13:29908888-29909659 | -1.226875143 | 0.000677137 | 0.010401685 |
| chr18:29712040-29712241 | 1.859050853 | 0.000677447 | 0.010404002 |
| chr20:58315446-58315714 | 1.407264069 | 0.000677631 | 0.010404002 |
| chr17:55635271-55635725 | -1.464298876 | 0.000677789 | 0.010404002 |
| chr2:133431239-133431620 | -1.56421438 | 0.000677831 | 0.010404002 |
| chrX:25524220-25524512 | -1.699120095 | 0.000678846 | 0.01041749 |
| chr19:38417541-38417875 | 1.506532977 | 0.000679461 | 0.010424166 |
| chr5:61304949-61305357 | -1.268597181 | 0.000679553 | 0.010424166 |
| chr8:80965373-80965582 | 1.774820369 | 0.00067969 | 0.010424175 |
| chr7:110283017-110283472 | -1.414942245 | 0.000680162 | 0.010425615 |
| chr10:77650117-77650317 | 1.960279982 | 0.000680191 | 0.010425615 |

| | | | |
|---|---|---|---|
| chr9:98453800-98454247 | -1.450606853 | 0.000680192 | 0.010425615 |
| chr16:89530908-89531401 | 1.468993549 | 0.000680335 | 0.010425733 |
| chr12:107912249-107912981 | -1.247324141 | 0.000680566 | 0.010426498 |
| chr1:222472702-222473088 | 1.72789714 | 0.000680658 | 0.010426498 |
| chr8:100617003-100617217 | 1.6301465 | 0.000681277 | 0.010432389 |
| chr7:85750041-85750730 | -1.274236546 | 0.000681314 | 0.010432389 |
| chr4:184961692-184961995 | -1.53128474 | 0.000681786 | 0.010437517 |
| chr13:32080311-32080723 | -1.712696151 | 0.000681939 | 0.010437739 |
| chr1:114081617-114081848 | 1.274254003 | 0.000682072 | 0.010437739 |
| chr19:35692016-35692371 | 1.528875023 | 0.000682897 | 0.010448276 |
| chr18:74657600-74657915 | -1.580536113 | 0.000683149 | 0.0104488 |
| chr5:84059642-84059929 | -1.807117217 | 0.000683204 | 0.0104488 |
| chr18:51537483-51537788 | -1.584571178 | 0.000685057 | 0.010473896 |
| chr6:73697361-73697638 | -1.550282507 | 0.000685171 | 0.010473896 |
| chr9:65675472-65676033 | -1.272107869 | 0.000685255 | 0.010473896 |
| chr11:58845231-58845586 | -1.814101679 | 0.000686531 | 0.010491306 |
| chr6:27814487-27815303 | -1.160235205 | 0.000686687 | 0.010491587 |
| chr8:87568868-87569097 | 1.713779469 | 0.000687973 | 0.010507184 |
| chr10:30626899-30627099 | 2.028136832 | 0.000687982 | 0.010507184 |
| chr3:133466781-133467035 | -1.464425732 | 0.000688465 | 0.010512474 |
| chr12:94141137-94141441 | 1.682349205 | 0.000688678 | 0.010513622 |
| chr1:172895746-172896116 | -1.588084066 | 0.000689247 | 0.010520211 |
| chr4:116829223-116829670 | -1.568676007 | 0.000689521 | 0.010522307 |
| chr10:23702915-23703449 | -1.230725916 | 0.000690531 | 0.01053562 |
| chr1:192998196-192998434 | 1.654754152 | 0.000690705 | 0.010536175 |
| chr10:77582748-77583096 | -1.398538983 | 0.000691234 | 0.010542149 |
| chr18:7839110-7839310 | 1.964533246 | 0.000691409 | 0.010542715 |
| chr12:39849417-39849647 | 1.91076214 | 0.000691797 | 0.010544438 |
| chr17:78288617-78288817 | 1.91076214 | 0.000691797 | 0.010544438 |
| chr11:2885886-2886125 | 1.640355086 | 0.000692161 | 0.010546985 |
| chr8:91134176-91134379 | 2.00336333 | 0.00069224 | 0.010546985 |
| chr2:44715519-44715995 | -1.314621596 | 0.000693161 | 0.010558926 |
| chr6:151563335-151563751 | 1.386534525 | 0.000693517 | 0.010561192 |
| chr20:48567967-48568291 | 1.189941685 | 0.000693586 | 0.010561192 |
| chr11:80718663-80719109 | 1.296557309 | 0.00069386 | 0.010563268 |
| chr5:67215476-67216326 | -1.238506184 | 0.000694696 | 0.010573899 |
| chr17:78204412-78204617 | 1.898142152 | 0.000695331 | 0.010581467 |
| chr14:45212141-45212347 | 1.61478232 | 0.000695827 | 0.010586902 |
| chr10:98598246-98598568 | 1.66660076 | 0.000696784 | 0.010599366 |
| chr8:109378844-109379099 | 1.600805142 | 0.00069707 | 0.010600199 |
| chr14:32252093-32252380 | 1.382249517 | 0.000697116 | 0.010600199 |
| chr7:19747808-19748041 | 1.592305426 | 0.000697394 | 0.01060233 |
| chr17:20291968-20292233 | 1.760371673 | 0.0006977 | 0.01060389 |
| chr1:165383107-165383369 | 1.720253115 | 0.000697774 | 0.01060389 |
| chr15:97448424-97449433 | -1.211278549 | 0.000698001 | 0.010605246 |
| chr5:84018085-84019058 | -1.188732853 | 0.00069833 | 0.010608137 |
| chr8:128451689-128451889 | 1.90739769 | 0.000699792 | 0.010628238 |
| chr21:42571036-42571466 | -1.386581678 | 0.000700495 | 0.010636811 |
| chr13:49018964-49019756 | -1.172071825 | 0.000700737 | 0.01063837 |
| chr17:2111611-2111889 | 1.741846011 | 0.000701613 | 0.010649569 |

| | | | |
|---|---|---|---|
| chr3:106140446-106140798 | 1.502617348 | 0.000701924 | 0.010650663 |
| chr13:61954170-61954475 | -1.67310636 | 0.000701964 | 0.010650663 |
| chr1:151492186-151492512 | 1.33181162 | 0.000702416 | 0.010655415 |
| chr7:143285332-143285553 | -1.402977253 | 0.000703777 | 0.010673064 |
| chr3:189563911-189564457 | -1.467206467 | 0.000703858 | 0.010673064 |
| chr12:29155094-29155475 | -1.449741534 | 0.00070575 | 0.010699634 |
| chr3:134471390-134471683 | 1.651741428 | 0.000705958 | 0.01070063 |
| chr7:135557549-135558798 | -1.108973194 | 0.000706095 | 0.01070063 |
| chr5:149623998-149624258 | 1.741121015 | 0.000706318 | 0.010701893 |
| chr16:77010413-77010613 | 1.761499322 | 0.000707056 | 0.010707582 |
| chr6:28826826-28827058 | 1.798645871 | 0.000707298 | 0.010707582 |
| chr14:55876249-55876487 | 1.718189186 | 0.000707448 | 0.010707582 |
| chr7:116710737-116710991 | -1.837492578 | 0.000707455 | 0.010707582 |
| chr11:21948867-21949274 | -1.276130233 | 0.000707463 | 0.010707582 |
| chr13:29572664-29572897 | 1.778796703 | 0.000707532 | 0.010707582 |
| chr8:108608362-108608804 | 1.298382488 | 0.000707841 | 0.010708824 |
| chr14:93683339-93683576 | 1.681705498 | 0.000707893 | 0.010708824 |
| chr11:64128900-64129107 | 1.737137386 | 0.000708421 | 0.010714689 |
| chr1:217493379-217493594 | -1.772221148 | 0.000709555 | 0.010729722 |
| chr10:13769808-13770131 | -1.767396427 | 0.000709978 | 0.010734004 |
| chr22:24775802-24776085 | 1.579468535 | 0.000710228 | 0.01073566 |
| chr8:104943748-104943997 | 1.851476833 | 0.000710418 | 0.010736425 |
| chr11:112780794-112781059 | -1.856352047 | 0.000710677 | 0.010738223 |
| chr16:76492109-76492317 | 1.847386209 | 0.000711677 | 0.010751202 |
| chr4:183438770-183439116 | -1.593149547 | 0.000712012 | 0.010754141 |
| chr3:140725216-140725460 | 1.829497793 | 0.000712822 | 0.010763042 |
| chr2:79670194-79670575 | -1.534159757 | 0.000712964 | 0.010763042 |
| chr8:37302098-37303039 | 1.156305648 | 0.000713022 | 0.010763042 |
| chr19:46115475-46115743 | -1.737493317 | 0.000714079 | 0.010774265 |
| chr19:45750535-45750801 | 1.74242035 | 0.000714184 | 0.010774265 |
| chr15:36904308-36904508 | -2.103263001 | 0.000714238 | 0.010774265 |
| chr7:40726434-40727041 | -1.312769517 | 0.000714328 | 0.010774265 |
| chr3:175150713-175151427 | -1.157072122 | 0.000714692 | 0.010777636 |
| chr2:2868735-2868953 | 1.846991875 | 0.000714935 | 0.01077917 |
| chr1:114197076-114197517 | 1.188002406 | 0.000715966 | 0.010792594 |
| chr12:108495049-108495355 | -1.538651157 | 0.00071651 | 0.010798662 |
| chr20:19212842-19213529 | 1.214245736 | 0.000716743 | 0.010800051 |
| chr8:52765960-52766328 | 1.666272748 | 0.000717003 | 0.010800484 |
| chr3:146750709-146750960 | 2.271747165 | 0.000717053 | 0.010800484 |
| chr5:17375486-17375795 | 1.374991802 | 0.000717927 | 0.010811514 |
| chr14:102187575-102187984 | 1.39401075 | 0.00071909 | 0.010826905 |
| chr10:88703334-88703542 | -2.007061419 | 0.000719267 | 0.010827442 |
| chr2:56557125-56557702 | 1.355444247 | 0.000720618 | 0.010845641 |
| chr2:160492634-160492916 | 1.541813115 | 0.000720772 | 0.010845831 |
| chr7:95084819-95085080 | -2.165036754 | 0.000721728 | 0.010858081 |
| chr12:56689223-56689425 | -1.646921002 | 0.000722258 | 0.01086393 |
| chr3:164192670-164192972 | 1.644961511 | 0.000722614 | 0.010867152 |
| chr20:3046772-3047155 | 1.293327254 | 0.000723678 | 0.010881014 |
| chr13:29369178-29369422 | -1.603088341 | 0.000724263 | 0.010885699 |
| chr13:41442238-41442478 | 1.679638092 | 0.000724274 | 0.010885699 |

| | | | |
|---|---|---|---|
| chr8:108139667-108140232 | -1.206485585 | 0.000724494 | 0.010886872 |
| chr13:51695489-51696056 | -1.397964705 | 0.000725665 | 0.010902328 |
| chr7:16386068-16386389 | 1.506095996 | 0.00072619 | 0.010906001 |
| chr15:78263292-78263545 | 1.586765216 | 0.000726329 | 0.010906001 |
| chr4:185597649-185597914 | -1.801029001 | 0.000726336 | 0.010906001 |
| chr13:71835091-71835384 | -1.706728769 | 0.00072681 | 0.0109107 |
| chr15:78764028-78764285 | 1.652986976 | 0.000726934 | 0.0109107 |
| chr16:83803914-83804140 | 1.422309014 | 0.000727465 | 0.010916531 |
| chr12:52036778-52037160 | 1.254781038 | 0.000727653 | 0.010916983 |
| chr5:153445634-153446569 | -1.124150687 | 0.00072778 | 0.010916983 |
| chr9:79776795-79777013 | 1.773795435 | 0.000727997 | 0.01091811 |
| chr12:116280533-116280801 | 1.49803948 | 0.000728197 | 0.010918958 |
| chr8:144331893-144332204 | 1.301625337 | 0.000728648 | 0.010923586 |
| chr3:175368518-175368737 | 2.008720181 | 0.000729213 | 0.010929189 |
| chr13:78660117-78660679 | -1.27243945 | 0.000729307 | 0.010929189 |
| chr8:137335401-137335765 | -1.608696212 | 0.00072994 | 0.010936539 |
| chr11:84688496-84688949 | -1.312717415 | 0.000730293 | 0.010939687 |
| chr7:110407322-110407583 | -1.791924882 | 0.000731729 | 0.010959065 |
| chr16:23021504-23021704 | 1.949787331 | 0.000732132 | 0.010962946 |
| chr3:111033650-111033979 | 1.602931972 | 0.000732296 | 0.010963267 |
| chr7:141674770-141675090 | -1.463728781 | 0.000732675 | 0.010966312 |
| chr14:41520897-41521142 | 1.80605663 | 0.000732786 | 0.010966312 |
| chr7:121438255-121438810 | -1.325323675 | 0.000733502 | 0.010974881 |
| chr17:61213822-61214179 | 1.145887004 | 0.000734006 | 0.010980278 |
| chr13:90336403-90336991 | -1.272678801 | 0.000734532 | 0.010984946 |
| chr12:62620262-62620703 | 1.459897417 | 0.000734604 | 0.010984946 |
| chr2:62362556-62362944 | -1.415491486 | 0.000735086 | 0.010989998 |
| chr11:129853902-129854393 | 1.285702592 | 0.00073584 | 0.010999136 |
| chr5:58094022-58094569 | -1.222228504 | 0.000736907 | 0.011011099 |
| chr15:39630429-39630632 | 1.900760142 | 0.000736928 | 0.011011099 |
| chr18:73296714-73297077 | -1.454783017 | 0.000737233 | 0.01101351 |
| chr1:225657535-225657853 | 1.496258471 | 0.000738977 | 0.011037405 |
| chr17:81161946-81162185 | 1.550535878 | 0.000739568 | 0.011044079 |
| chr4:6783213-6783884 | 1.164643301 | 0.000740756 | 0.011057706 |
| chr13:40146515-40146820 | -1.575009145 | 0.000740769 | 0.011057706 |
| chr1:236420966-236421372 | 1.238139814 | 0.000742217 | 0.011077155 |
| chr7:65235664-65235955 | -1.721909057 | 0.000743073 | 0.011087778 |
| chr3:164244018-164244481 | -1.424132164 | 0.000743935 | 0.011098474 |
| chr1:168752190-168752450 | 1.583567229 | 0.000744578 | 0.011105906 |
| chr2:179745705-179746072 | -1.580158768 | 0.000744988 | 0.011109856 |
| chr1:45348830-45349063 | 1.958585008 | 0.000745347 | 0.011112396 |
| chr12:109130031-109130283 | 1.592175045 | 0.000745448 | 0.011112396 |
| chr8:107647403-107647703 | 1.546626132 | 0.000745848 | 0.01111619 |
| chr6:27548465-27548799 | -1.31187027 | 0.000746577 | 0.011124896 |
| chr8:105873376-105873600 | 1.945019367 | 0.000747316 | 0.011129788 |
| chr21:31872848-31873330 | 1.317186605 | 0.000747333 | 0.011129788 |
| chr12:57795444-57795766 | -1.697436623 | 0.000747341 | 0.011129788 |
| chr19:10292282-10292548 | 1.461327931 | 0.000748462 | 0.011144323 |
| chr9:38115133-38115557 | -1.408759051 | 0.000748964 | 0.011146578 |
| chrX:102958874-102959282 | -1.459458322 | 0.000748998 | 0.011146578 |

| | | | |
|---|---|---|---|
| chr5:62790402-62790965 | -1.251079087 | 0.000749179 | 0.011146578 |
| chr7:80247727-80248247 | -1.617123287 | 0.000749196 | 0.011146578 |
| chr21:9330192-9330455 | 1.597645685 | 0.000749833 | 0.011153895 |
| chr18:31157290-31157738 | 1.355934084 | 0.000751351 | 0.011174307 |
| chr15:89139813-89140043 | 1.557412418 | 0.000751884 | 0.011180055 |
| chr1:150206173-150206529 | 1.381494006 | 0.000752822 | 0.011191837 |
| chr13:112214044-112214528 | 1.338542484 | 0.000753251 | 0.011196037 |
| chr18:56709968-56710348 | -1.844763076 | 0.000754641 | 0.011214524 |
| chr19:49011814-49012123 | 1.610585127 | 0.000754847 | 0.011215408 |
| chr3:42734665-42734865 | 1.896820758 | 0.000755397 | 0.011221405 |
| chr13:92035048-92035279 | -2.019190998 | 0.000756163 | 0.011230614 |
| chr15:88804301-88804501 | 1.684048189 | 0.000756476 | 0.011233078 |
| chr7:41274416-41274616 | 2.073401796 | 0.000756758 | 0.011235086 |
| chr18:46264282-46264604 | -1.662986683 | 0.00075705 | 0.011237256 |
| chr12:103852700-103852961 | -1.737562801 | 0.000757384 | 0.011240031 |
| chr1:114863165-114863367 | 1.833174319 | 0.000757641 | 0.011241666 |
| chr7:134527476-134527676 | -2.086234503 | 0.000758336 | 0.011249805 |
| chr17:51487957-51488315 | -1.30570846 | 0.000760984 | 0.011286904 |
| chr16:65813422-65813908 | 1.286737114 | 0.000762051 | 0.011298434 |
| chr15:88157512-88157834 | 1.552648757 | 0.000762057 | 0.011298434 |
| chr10:132407267-132408299 | 1.092690347 | 0.000762514 | 0.011302751 |
| chr7:115002797-115002997 | -2.085298787 | 0.000762787 | 0.011302751 |
| chr6:143034906-143035274 | -1.529830128 | 0.00076279 | 0.011302751 |
| chr4:185244818-185245165 | -1.496857009 | 0.000764339 | 0.011323505 |
| chr6:37172249-37172556 | 1.24899178 | 0.000765261 | 0.011334969 |
| chr12:25135094-25135294 | -1.906458892 | 0.000765585 | 0.011336778 |
| chr4:151602250-151602472 | 1.787738307 | 0.000765679 | 0.011336778 |
| chr1:157162371-157162598 | 1.628887031 | 0.000768249 | 0.011372641 |
| chr6:75524303-75524983 | 1.199887504 | 0.000768449 | 0.011373407 |
| chr1:60156799-60157265 | -1.274045894 | 0.000768781 | 0.011376115 |
| chr12:72583930-72584175 | -1.953428819 | 0.00076906 | 0.011378043 |
| chr13:71683221-71683736 | -1.499837399 | 0.000770495 | 0.011397076 |
| chr1:108334384-108334586 | 1.94086305 | 0.000771105 | 0.011401562 |
| chr6:48746881-48747842 | -1.174834555 | 0.000771286 | 0.011401562 |
| chr1:223331280-223331629 | -1.693139962 | 0.00077132 | 0.011401562 |
| chr2:145853572-145853919 | -1.595776566 | 0.000771533 | 0.011401562 |
| chr6:158475417-158475719 | 1.697616707 | 0.000771593 | 0.011401562 |
| chr19:19668423-19669048 | -1.218383925 | 0.000771691 | 0.011401562 |
| chr5:25713118-25713786 | -1.280477197 | 0.000772968 | 0.011415467 |
| chr15:71790015-71790716 | 1.339771592 | 0.000773194 | 0.011415467 |
| chr7:143172420-143172708 | -1.644507661 | 0.00077322 | 0.011415467 |
| chr14:72725446-72725712 | 1.600983287 | 0.000773228 | 0.011415467 |
| chr3:4752777-4753072 | 1.520554282 | 0.000773425 | 0.01141617 |
| chr15:99450524-99450794 | -1.633641533 | 0.000774116 | 0.011424179 |
| chr2:214310937-214311432 | -1.249961443 | 0.000774517 | 0.011427897 |
| chr8:121833654-121833891 | 1.563464084 | 0.00077476 | 0.011429279 |
| chr15:71096596-71097028 | -1.409225577 | 0.000775172 | 0.011433158 |
| chr1:183366975-183367196 | 1.899201667 | 0.000775952 | 0.011442461 |
| chr17:59768666-59768904 | 1.519945332 | 0.000777603 | 0.011464591 |
| chr20:25227139-25227586 | 1.388674257 | 0.000778633 | 0.01147732 |

| | | | |
|---|---|---|---|
| chr6:22071738-22071996 | -2.268891413 | 0.000778766 | 0.01147732 |
| chr12:25347588-25348239 | -1.292032307 | 0.000781533 | 0.011515887 |
| chr4:163372113-163372503 | -1.680387584 | 0.000781707 | 0.011516242 |
| chr6:98308230-98308984 | -1.215960278 | 0.000781876 | 0.011516507 |
| chr20:40925855-40926090 | 1.726119485 | 0.00078269 | 0.011526288 |
| chr19:34319255-34319522 | 1.559602696 | 0.000784619 | 0.011551213 |
| chr20:47719709-47719916 | 1.176832893 | 0.000784757 | 0.011551213 |
| chr11:115869854-11587015 | -1.663710492 | 0.000784841 | 0.011551213 |
| chr4:185425109-185426522 | -1.093085749 | 0.000784986 | 0.011551213 |
| chr5:157865890-157866140 | 1.806468057 | 0.000785173 | 0.01155175 |
| chr17:39077943-39078190 | 1.679807818 | 0.000785538 | 0.011554898 |
| chrX:134717897-134718761 | 1.231570425 | 0.000787214 | 0.011575154 |
| chr14:105660989-105661267 | 1.408695745 | 0.000787294 | 0.011575154 |
| chr4:7539696-7540075 | 1.335492578 | 0.000787378 | 0.011575154 |
| chr14:105690651-10569130 | 1.212343088 | 0.000787652 | 0.011575154 |
| chr7:135672829-135673165 | -1.359744685 | 0.000787709 | 0.011575154 |
| chr2:168856685-168856951 | 1.699667614 | 0.000787821 | 0.011575154 |
| chr9:21316404-21316714 | -1.877426517 | 0.000788114 | 0.01157724 |
| chr2:139773393-139773987 | -1.301254241 | 0.000788284 | 0.01157751 |
| chr7:154151492-154151893 | -1.627443735 | 0.000788757 | 0.011582243 |
| chr5:103224431-103225270 | -1.11716677 | 0.000789132 | 0.011585535 |
| chr11:30960449-30960720 | -1.588170887 | 0.000789622 | 0.011590494 |
| chrY:56873684-56873904 | 1.778173816 | 0.000790218 | 0.011597027 |
| chr1:85884749-85884986 | -1.904794569 | 0.000790407 | 0.01159758 |
| chr14:44820746-44820956 | 1.888360261 | 0.000792048 | 0.011619437 |
| chr2:224679123-224679683 | -1.369244697 | 0.000792856 | 0.01162906 |
| chr1:172449125-172449390 | 1.38374633 | 0.000794497 | 0.011650896 |
| chr11:48097115-48097592 | 1.323355795 | 0.000794671 | 0.011651218 |
| chr16:68406947-68407199 | 1.698690682 | 0.000795396 | 0.011659614 |
| chr14:76227610-76227852 | 1.799995304 | 0.000796052 | 0.011664358 |
| chr8:126024454-126024769 | 1.65436633 | 0.000796162 | 0.011664358 |
| chr5:138724441-138725747 | -1.07697612 | 0.000796176 | 0.011664358 |
| chr5:140232100-140232334 | 2.059409653 | 0.000796737 | 0.011670342 |
| chr11:111303403-11130388 | -1.317431001 | 0.000799345 | 0.011706313 |
| chr4:567117-567317 | 2.05845206 | 0.00080003 | 0.011714106 |
| chr9:98779078-98779448 | 1.374791308 | 0.000801048 | 0.011726769 |
| chr8:97147893-97148093 | 1.920306082 | 0.000801544 | 0.011731782 |
| chr13:33642561-33643261 | -1.137749211 | 0.000803851 | 0.011763312 |
| chr6:64999354-65000243 | -1.18390721 | 0.000804102 | 0.011764734 |
| chr7:138097365-138097743 | -1.340363832 | 0.000804757 | 0.011772062 |
| chr3:98291612-98291812 | 2.000261042 | 0.00080586 | 0.011785955 |
| chr7:11345583-11345805 | 1.646310796 | 0.000807101 | 0.01179742 |
| chr12:53129034-53129360 | 1.590383223 | 0.000807133 | 0.01179742 |
| chr1:233543620-233544064 | -1.327610916 | 0.000807137 | 0.01179742 |
| chr12:49869074-49869638 | 1.20458936 | 0.00080726 | 0.01179742 |
| chr21:7917958-7918189 | 1.839508578 | 0.000807623 | 0.01180048 |
| chr1:234939843-234940081 | 1.892919626 | 0.000809337 | 0.011821027 |
| chr14:32365542-32365842 | 1.443104116 | 0.000809338 | 0.011821027 |
| chr2:191404492-191404697 | 1.992096326 | 0.000810578 | 0.011835471 |
| chr1:59856907-59857140 | 1.714342728 | 0.000810636 | 0.011835471 |

| | | | |
|---|---|---|---|
| chr17:50324202-50324422 | 1.786535451 | 0.000810995 | 0.011838454 |
| chr20:47248661-47249034 | 1.113845878 | 0.000811612 | 0.011845212 |
| chr16:18983720-18984351 | 1.259544349 | 0.000812796 | 0.011860223 |
| chr1:217712867-217713561 | -1.156881949 | 0.000812952 | 0.011860247 |
| chr10:12214874-12215151 | -1.799357739 | 0.000816144 | 0.011903279 |
| chr7:11950393-11950659 | 1.58613295 | 0.000816212 | 0.011903279 |
| chr20:48092463-48092695 | 1.218031159 | 0.000816615 | 0.01190688 |
| chr13:54536254-54536486 | -1.754110027 | 0.000817548 | 0.011918225 |
| chr17:36183429-36183779 | 1.54160761 | 0.000817997 | 0.011922498 |
| chr8:83447112-83447426 | -1.872401194 | 0.000818304 | 0.011924703 |
| chr9:15837083-15837374 | -1.534463455 | 0.000819445 | 0.011939054 |
| chr19:5553794-5554156 | -1.552464016 | 0.000820195 | 0.011947721 |
| chr7:101399887-101400168 | -1.777663038 | 0.0008205 | 0.011949501 |
| chr1:106364337-106364556 | 1.363905437 | 0.000820629 | 0.011949501 |
| chr5:39131724-39132009 | 1.930119016 | 0.000821191 | 0.011955404 |
| chr3:126023633-126023967 | 1.549076851 | 0.000821471 | 0.011957204 |
| chr3:61065506-61065787 | -1.622014363 | 0.000821746 | 0.011958934 |
| chr1:54755407-54755611 | 1.930761606 | 0.000822106 | 0.011961914 |
| chr3:32346421-32346720 | 1.805194087 | 0.000822588 | 0.011964163 |
| chr4:175380886-175381401 | -1.276138357 | 0.000822689 | 0.011964163 |
| chr13:86922861-86923273 | -1.478997802 | 0.000822729 | 0.011964163 |
| chr7:98086171-98086694 | 1.209863831 | 0.000824114 | 0.011981546 |
| chrX:154371180-154371561 | 1.194181216 | 0.000824238 | 0.011981546 |
| chr3:176508118-176508318 | 2.066349048 | 0.000824483 | 0.011982846 |
| chr19:27718049-27718256 | -2.01326374 | 0.000824867 | 0.011986144 |
| chr5:148320403-148320838 | -1.525035512 | 0.000825418 | 0.01199172 |
| chr11:108200019-10820048€ | -1.447368932 | 0.000825647 | 0.01199172 |
| chr11:69179845-69180065 | 1.662584536 | 0.00082581 | 0.01199172 |
| chr7:116426376-116426722 | -1.551687527 | 0.000825876 | 0.01199172 |
| chr3:14566183-14566383 | 1.751984343 | 0.000826254 | 0.011994299 |
| chr1:218020649-218021006 | -1.62999564 | 0.000826367 | 0.011994299 |
| chr8:123233181-123233388 | 1.73650094 | 0.000827027 | 0.012000244 |
| chr10:86752365-86752732 | -1.229351766 | 0.00082709 | 0.012000244 |
| chr14:36613562-36613762 | 1.798260486 | 0.000827748 | 0.012007525 |
| chr7:3937048-3937563 | 1.404902989 | 0.000828272 | 0.012010928 |
| chr7:41955835-41956265 | -1.260679132 | 0.000828297 | 0.012010928 |
| chr7:111019034-111019574 | -1.565319749 | 0.000829158 | 0.012021144 |
| chr12:117433013-117433255 | 1.931956662 | 0.000829672 | 0.01202632 |
| chr4:74098932-74099512 | -1.378006562 | 0.000830149 | 0.012028898 |
| chr20:46782486-46782710 | 1.497572455 | 0.000830164 | 0.012028898 |
| chr5:99057460-99057810 | -1.493655866 | 0.000830654 | 0.012033725 |
| chr12:20220370-20220630 | -1.712503736 | 0.000831672 | 0.012046191 |
| chr7:125316386-125317048 | -1.262386365 | 0.00083286 | 0.01206112 |
| chr2:19009344-19009992 | 1.226159027 | 0.000833289 | 0.012063298 |
| chr5:79255739-79256941 | -1.113564427 | 0.000833334 | 0.012063298 |
| chr1:209849594-209849859 | -2.021898873 | 0.000833483 | 0.012063298 |
| chr3:104279260-104279760 | -1.370528284 | 0.000833896 | 0.012065509 |
| chr13:72251771-72252176 | -1.603669903 | 0.000834289 | 0.012065509 |
| chr12:12916251-12916551 | -1.381267737 | 0.000834363 | 0.012065509 |
| chr10:62492118-62492404 | -1.606842059 | 0.000834405 | 0.012065509 |

| | | | |
|---|---|---|---|
| chr3:159486639-159486868 | -1.947530229 | 0.000834423 | 0.012065509 |
| chr22:27718142-27718640 | 1.375915547 | 0.000835722 | 0.012082022 |
| chr5:17634739-17634992 | 1.760130221 | 0.000836461 | 0.012090417 |
| chr12:6069490-6069690 | -1.895810995 | 0.000838244 | 0.012113913 |
| chr7:24812384-24812681 | 1.447815817 | 0.000838994 | 0.012120994 |
| chr14:51792949-51793149 | 1.829185801 | 0.000839051 | 0.012120994 |
| chr9:26833906-26834191 | -1.921467175 | 0.000839278 | 0.012121989 |
| chr14:38600683-38600904 | 1.686275781 | 0.000840503 | 0.012137398 |
| chr6:126262257-126262565 | 1.706850471 | 0.000840877 | 0.012140509 |
| chr19:28074699-28075057 | -1.603076347 | 0.000842064 | 0.012155355 |
| chr2:30938104-30938304 | -2.09503915 | 0.000842473 | 0.012158974 |
| chr4:186474050-186474446 | -1.618549398 | 0.000842861 | 0.012162283 |
| chr7:107103458-107103755 | -1.817895251 | 0.000843622 | 0.012170001 |
| chr7:140973035-140973536 | -1.300770087 | 0.000843804 | 0.012170001 |
| chr7:84013690-84013986 | -1.858949321 | 0.000843873 | 0.012170001 |
| chr10:80047928-80048243 | 1.609252685 | 0.000844629 | 0.012178619 |
| chr1:216531337-216531608 | -1.516439412 | 0.000845073 | 0.0121823 |
| chr7:112481794-112482215 | -1.261100583 | 0.000845355 | 0.0121823 |
| chr1:244887341-244887641 | 1.455881534 | 0.000845362 | 0.0121823 |
| chr7:15230801-15231218 | 1.486723366 | 0.000847443 | 0.01221 |
| chr7:15198767-15199221 | 1.389157513 | 0.000849023 | 0.012228254 |
| chr20:25354198-25354403 | 1.719558263 | 0.000849029 | 0.012228254 |
| chr18:77055021-77055430 | -1.451588799 | 0.000849191 | 0.012228282 |
| chr11:66012901-66013191 | 1.470963482 | 0.000850943 | 0.012251206 |
| chr2:138814541-138814820 | -1.576060446 | 0.000851296 | 0.012253989 |
| chr15:90087256-90088074 | -1.128346434 | 0.000852166 | 0.012264219 |
| chr3:183942999-183943207 | 1.922133282 | 0.000853249 | 0.012277495 |
| chrX:39953998-39954239 | 1.68851418 | 0.000853947 | 0.012285234 |
| chr5:91864878-91865354 | -1.520821114 | 0.000854714 | 0.012292157 |
| chr6:119237308-119237544 | 1.756231077 | 0.000854749 | 0.012292157 |
| chr7:46058946-46059320 | 1.729174961 | 0.000855368 | 0.012297181 |
| chr20:31710643-31710943 | 1.572364855 | 0.00085542 | 0.012297181 |
| chr6:31403336-31403856 | 1.224125913 | 0.000855698 | 0.012298879 |
| chr7:113281165-113281777 | -1.318797366 | 0.00085658 | 0.012309246 |
| chr5:15880186-15880458 | 1.867703873 | 0.000856754 | 0.012309426 |
| chr8:128564098-128564308 | 1.825798199 | 0.000857791 | 0.012322022 |
| chr21:7956436-7956689 | 1.568973974 | 0.000858512 | 0.012326785 |
| chr1:60924910-60925321 | -1.453901538 | 0.000858648 | 0.012326785 |
| chr13:77317199-77317478 | -1.713907941 | 0.000858698 | 0.012326785 |
| chr18:36411817-36412083 | -1.781561916 | 0.000858766 | 0.012326785 |
| chr5:139711195-139711684 | -1.255854931 | 0.000859736 | 0.012338393 |
| chr1:234746807-234747046 | 1.730870655 | 0.000862634 | 0.012377673 |
| chr18:48050219-48050720 | -1.334796067 | 0.000863043 | 0.012381221 |
| chr6:25720888-25721116 | -1.817690188 | 0.00086352 | 0.012385735 |
| chr18:55096303-55096678 | -1.474179882 | 0.000864869 | 0.012402007 |
| chr13:91782083-91782439 | -1.5592839 | 0.000864978 | 0.012402007 |
| chr8:93734304-93734720 | 1.363732418 | 0.000865255 | 0.012403665 |
| chr3:175270137-175270846 | -1.205185918 | 0.00086542 | 0.012403708 |
| chr14:105700496-105700738 | 1.566986647 | 0.000865927 | 0.012408656 |
| chr1:151491739-151492023 | 1.375258996 | 0.000866471 | 0.012414126 |

| | | | |
|---|---|---|---|
| chr9:90878571-90878897 | 1.586259628 | 0.000866937 | 0.012417667 |
| chr3:127849223-127849430 | 1.921888499 | 0.000867042 | 0.012417667 |
| chr2:240688400-240688855 | 1.400485791 | 0.000868549 | 0.012434677 |
| chr20:19026177-19026484 | 1.630533047 | 0.000868555 | 0.012434677 |
| chr5:119243952-119244285 | -1.48569476 | 0.000869984 | 0.01245214 |
| chr6:135173731-135174376 | -1.228227609 | 0.000870099 | 0.01245214 |
| chr17:55026457-55026742 | -1.6506066 | 0.000870727 | 0.012458791 |
| chr1:232642109-232642330 | 1.829529468 | 0.000870904 | 0.012459 |
| chr2:125313688-125314144 | -1.38872962 | 0.000871977 | 0.012472022 |
| chr3:165225641-165225957 | -1.408498419 | 0.00087215 | 0.012472162 |
| chr10:118835651-118836296 | -1.191473814 | 0.000874092 | 0.012497603 |
| chrX:149630658-149631239 | 1.203285927 | 0.000874606 | 0.012500757 |
| chr11:26316568-26316804 | 1.817680844 | 0.000874639 | 0.012500757 |
| chr17:74147546-74147746 | 1.711167805 | 0.000875242 | 0.012507047 |
| chr12:23950624-23951036 | -1.497480475 | 0.000875568 | 0.012509079 |
| chr3:19682154-19682385 | 1.920932522 | 0.000875711 | 0.012509079 |
| chr14:51175960-51176185 | 1.88063274 | 0.000876974 | 0.012524789 |
| chr11:47234629-47234893 | 1.444028634 | 0.000877486 | 0.012529761 |
| chr14:64872199-64872449 | 1.827973718 | 0.000878272 | 0.01253865 |
| chr1:117108443-117108815 | 1.438404864 | 0.000879096 | 0.012548077 |
| chr2:206764717-206766162 | -1.078493573 | 0.000881248 | 0.012576453 |
| chr10:26081052-26081385 | -1.496525939 | 0.000882041 | 0.012585431 |
| chr10:13700486-13701016 | -1.240157512 | 0.000882534 | 0.012590121 |
| chr2:103361769-103362148 | 1.637543528 | 0.000883105 | 0.012595925 |
| chr5:31626684-31626994 | 1.680270023 | 0.000883892 | 0.01260451 |
| chr10:132452994-132453528 | 1.146877198 | 0.000884173 | 0.01260451 |
| chr7:90928476-90929427 | -1.127457275 | 0.000884201 | 0.01260451 |
| chr16:29613328-29613966 | 1.180747315 | 0.000885529 | 0.012618933 |
| chr4:8260630-8261161 | 1.227275207 | 0.000885542 | 0.012618933 |
| chrX:119421442-119421655 | 2.047540397 | 0.000886653 | 0.012629793 |
| chr4:144170263-144170508 | 1.863698895 | 0.000886977 | 0.012629793 |
| chr7:29978306-29978609 | -1.72498345 | 0.000887024 | 0.012629793 |
| chr3:154995884-154996258 | 1.692673695 | 0.000887027 | 0.012629793 |
| chr3:190534970-190535371 | -1.402812056 | 0.000887128 | 0.012629793 |
| chr13:73210691-73211059 | -1.680198696 | 0.000887639 | 0.012634715 |
| chr1:154133176-154133376 | 1.993877643 | 0.000888759 | 0.012648315 |
| chr3:75642885-75643273 | -1.497441055 | 0.000888941 | 0.012648549 |
| chr7:12863916-12864175 | 1.588543361 | 0.000889752 | 0.012657257 |
| chr13:73290569-73291116 | -1.425826729 | 0.000889883 | 0.012657257 |
| chr5:169139795-169140069 | 1.669851606 | 0.000890188 | 0.012659237 |
| chr2:25310970-25311955 | -1.118805311 | 0.00089086 | 0.012664063 |
| chr9:38046832-38047047 | 1.662068961 | 0.000890876 | 0.012664063 |
| chr6:15026763-15027251 | -1.228425389 | 0.000891093 | 0.012664063 |
| chr16:17316079-17316310 | 1.636106996 | 0.000891188 | 0.012664063 |
| chr7:118183620-118184487 | -1.092235939 | 0.000891656 | 0.012666726 |
| chr4:184733222-184733606 | -1.239942945 | 0.000891706 | 0.012666726 |
| chr16:52565283-52565530 | 1.550575622 | 0.00089222 | 0.012671676 |
| chr6:49787216-49787560 | -1.44512541 | 0.000892736 | 0.012676648 |
| chr7:28062126-28062595 | 1.286516905 | 0.000893451 | 0.012684451 |
| chr1:113877716-113877949 | 1.322171044 | 0.000893718 | 0.012685172 |

| | | | |
|---|---|---|---|
| chr17:48752288-48752488 | 1.913973217 | 0.000893833 | 0.012685172 |
| chr14:101318248-101318563 | 1.406850261 | 0.00089424 | 0.0126886 |
| chr1:151945652-151946198 | -1.121674775 | 0.00089587 | 0.012709371 |
| chr14:91701625-91701825 | 1.781563069 | 0.000896822 | 0.012720523 |
| chr9:133513018-133513301 | 1.659815528 | 0.000899036 | 0.012749574 |
| chr7:111098614-111098970 | -1.569637377 | 0.000899364 | 0.012751866 |
| chr14:30733603-30733921 | 1.775687424 | 0.00090258 | 0.012795101 |
| chr19:13884735-13885543 | -1.217856508 | 0.000903191 | 0.012799131 |
| chr11:118956043-118956900 | 1.113651719 | 0.000903265 | 0.012799131 |
| chr18:73891927-73892224 | -1.495127022 | 0.000903366 | 0.012799131 |
| chr7:140062180-140063341 | -1.14370935 | 0.000903561 | 0.012799524 |
| chr6:143337623-143337950 | -1.375911467 | 0.000903961 | 0.012801199 |
| chr16:29721686-29721886 | 1.692431968 | 0.000904013 | 0.012801199 |
| chr1:114256648-114257077 | 1.141752244 | 0.00090496 | 0.012812234 |
| chr3:22730048-22730311 | 1.515656999 | 0.000905609 | 0.012816881 |
| chr4:139892781-139893290 | -1.431636683 | 0.000905622 | 0.012816881 |
| chr2:207810923-207811299 | -1.490055452 | 0.000906698 | 0.012829728 |
| chr11:126082906-126083375 | -1.351406038 | 0.000908615 | 0.01285449 |
| chr5:39034536-39035135 | -1.29336053 | 0.000908818 | 0.012854983 |
| chr1:55428988-55429629 | 1.146578359 | 0.000909566 | 0.012863184 |
| chr2:153213032-153213688 | 1.146993981 | 0.000911075 | 0.012882148 |
| chr8:74197613-74197889 | 1.713513214 | 0.000911362 | 0.012883829 |
| chr1:218451354-218451554 | 1.738019301 | 0.000911865 | 0.012888571 |
| chr19:41792563-41792770 | 1.770683018 | 0.000912644 | 0.012897201 |
| chr4:185007568-185007789 | -1.686731646 | 0.000913494 | 0.012906836 |
| chr8:62523231-62523513 | 1.555050063 | 0.000914638 | 0.012918542 |
| chr3:168649404-168649664 | 1.814489828 | 0.00091466 | 0.012918542 |
| chr20:41764533-41764733 | 2.043420921 | 0.000916252 | 0.01293215 |
| chr21:37965141-37965341 | 2.043420921 | 0.000916252 | 0.01293215 |
| chr12:107579-108375 | -1.214528515 | 0.000916278 | 0.01293215 |
| chr15:45574032-45574411 | -1.36873151 | 0.000916361 | 0.01293215 |
| chr13:29567470-29567745 | -1.519410536 | 0.000916468 | 0.01293215 |
| chr7:111191848-111192179 | -1.524300156 | 0.000917851 | 0.012947813 |
| chr20:59357504-59357761 | 1.436844139 | 0.000917916 | 0.012947813 |
| chr4:159966475-159966718 | -1.874543001 | 0.00091831 | 0.012950989 |
| chr10:121510136-121510705 | 1.263321438 | 0.000918688 | 0.012953946 |
| chr1:75707195-75707637 | -1.34948977 | 0.000921006 | 0.012984242 |
| chr3:65332315-65332566 | 1.807153188 | 0.000921715 | 0.01299131 |
| chr1:106662426-106662932 | 1.120924401 | 0.000921847 | 0.01299131 |
| chr9:31909101-31909377 | -1.801020487 | 0.000922327 | 0.012995683 |
| chr2:160469243-160469609 | -1.535089937 | 0.000923039 | 0.013003323 |
| chr14:36217047-36217303 | 1.695888956 | 0.000923336 | 0.013005115 |
| chr3:129751418-129751618 | 1.91270197 | 0.000924344 | 0.01301693 |
| chr3:14170965-14171243 | 1.467051208 | 0.000924992 | 0.013023655 |
| chr4:105158874-105159405 | -1.217224635 | 0.000926099 | 0.013035366 |
| chr2:217610432-217611073 | -1.30037882 | 0.000926164 | 0.013035366 |
| chr2:60677143-60677955 | -1.180642738 | 0.000926406 | 0.013036385 |
| chr7:22559892-22560166 | 1.283181653 | 0.000926793 | 0.013039434 |
| chr17:41547426-41548271 | -1.169567655 | 0.000927525 | 0.013046432 |
| chr2:191607133-191607422 | -1.703007507 | 0.000927631 | 0.013046432 |

| | | | |
|---|---|---|---|
| chr7:17559506-17559730 | 1.537658793 | 0.000927988 | 0.013049052 |
| chr8:127994747-127995164 | 1.2333264 | 0.000929067 | 0.013059995 |
| chr7:23895851-23896051 | 1.661790983 | 0.000929107 | 0.013059995 |
| chr4:147608670-147608980 | 1.630937009 | 0.000930386 | 0.013073663 |
| chr14:66189571-66189782 | 1.974991351 | 0.000930421 | 0.013073663 |
| chr8:61308347-61308672 | -1.543317318 | 0.000931492 | 0.01308632 |
| chr2:80054364-80054708 | -1.510537461 | 0.00093168 | 0.013086562 |
| chr3:66473638-66473979 | 1.429732838 | 0.000932667 | 0.013097132 |
| chr19:5563957-5564354 | -1.399897235 | 0.000932787 | 0.013097132 |
| chr20:29878160-29878726 | 1.223938056 | 0.00093307 | 0.013097132 |
| chr5:19044597-19045794 | -1.111173623 | 0.000933306 | 0.013097132 |
| chr12:100163611-100164264 | -1.146774965 | 0.000933313 | 0.013097132 |
| chr7:138089575-138089834 | -1.686238955 | 0.000933458 | 0.013097132 |
| chr12:102939851-102940116 | -1.675956553 | 0.000934453 | 0.013108689 |
| chr6:120671539-120671931 | -1.422008436 | 0.000935219 | 0.013117031 |
| chr20:31060254-31060790 | 1.261354002 | 0.000935658 | 0.013120792 |
| chr3:160935378-160936008 | -1.230402728 | 0.000936495 | 0.013130117 |
| chr13:25971429-25971700 | 1.598007357 | 0.00093698 | 0.013133715 |
| chr3:11514454-11514681 | 1.603101438 | 0.000937094 | 0.013133715 |
| chr18:58223703-58224365 | -1.353206918 | 0.000937721 | 0.013140098 |
| chr8:48581972-48582172 | 2.12520148 | 0.000939131 | 0.013157456 |
| chr10:71713209-71713939 | -1.203785757 | 0.000939768 | 0.013163967 |
| chr1:50189820-50190096 | 1.654326075 | 0.000941027 | 0.013179192 |
| chr17:71402554-71403395 | -1.155169227 | 0.000941447 | 0.013182455 |
| chr6:121838831-121839167 | -1.659288569 | 0.000941604 | 0.013182455 |
| chr8:53523720-53524237 | 1.235090327 | 0.000941884 | 0.013183974 |
| chr10:44172475-44172832 | -1.57431223 | 0.00094213 | 0.013184996 |
| chr10:52805473-52805952 | -1.459487699 | 0.000942498 | 0.013187744 |
| chr3:12780097-12780477 | 1.606173902 | 0.000943691 | 0.013202032 |
| chr16:11584092-11584365 | 1.466201879 | 0.000944244 | 0.013207355 |
| chr9:5879962-5880173 | -1.796703877 | 0.000945611 | 0.013221533 |
| chrX:154347462-154347662 | 1.855237683 | 0.000945648 | 0.013221533 |
| chr4:41475405-41475822 | -1.318559049 | 0.000945776 | 0.013221533 |
| chr5:142011725-142012000 | -1.606764883 | 0.000947416 | 0.013242045 |
| chrX:107716440-107716677 | 1.475217273 | 0.000947722 | 0.013243905 |
| chr7:2883620-2883867 | 1.523240438 | 0.000947948 | 0.013244654 |
| chr14:90257015-90257351 | 1.341111924 | 0.000948205 | 0.013245823 |
| chr7:144391818-144392303 | -1.328481221 | 0.000948941 | 0.013253697 |
| chr3:63240614-63241178 | 1.155375949 | 0.000949253 | 0.013255629 |
| chr4:48109536-48109948 | 1.586517566 | 0.000950069 | 0.013264613 |
| chr16:71339543-71340205 | -1.250681585 | 0.0009516 | 0.01328115 |
| chr4:130421168-130421533 | -1.585921884 | 0.0009516 | 0.01328115 |
| chr2:84919851-84920528 | -1.194959485 | 0.000951872 | 0.013282075 |
| chr19:44680660-44680930 | 1.767779433 | 0.000952013 | 0.013282075 |
| chr7:102306879-102307307 | -1.36282839 | 0.00095243 | 0.013285467 |
| chr11:112881385-112881820 | -1.3114848 | 0.000953157 | 0.013293198 |
| chr6:79453568-79453869 | -1.543055656 | 0.000954425 | 0.013308458 |
| chr1:115194342-115195019 | 1.163672757 | 0.000954727 | 0.013310245 |
| chr2:217147893-217148294 | 1.585664867 | 0.000955553 | 0.013319328 |
| chr13:42415950-42416169 | 1.749208507 | 0.000956057 | 0.013323932 |

| | | | |
|---|---|---|---|
| chr2:189698362-189698872 | -1.310177071 | 0.000956638 | 0.013327758 |
| chr9:108552792-108553032 | -1.758293547 | 0.000956983 | 0.013327758 |
| chr2:48778996-48779415 | -1.419835725 | 0.000957175 | 0.013327758 |
| chr13:85653793-85654066 | -1.92642438 | 0.000957178 | 0.013327758 |
| chr15:55479691-55479906 | 1.596805384 | 0.000957201 | 0.013327758 |
| chr17:66052411-66052661 | 1.610420912 | 0.000957655 | 0.013330121 |
| chr8:132495860-132496178 | 1.627400462 | 0.000957734 | 0.013330121 |
| chr7:110677411-110677693 | 1.634594389 | 0.000957893 | 0.013330121 |
| chr21:6358938-6359257 | 1.564906928 | 0.0009584 | 0.013334754 |
| chr12:75010976-75011498 | -1.275837436 | 0.00095868 | 0.013335145 |
| chr12:46143346-46143609 | 1.824546691 | 0.000958776 | 0.013335145 |
| chr8:99061749-99062037 | 1.684920723 | 0.000959962 | 0.013349216 |
| chr14:100012278-100012803 | -1.425643626 | 0.000960363 | 0.013350979 |
| chr3:149245659-149246321 | 1.234286437 | 0.000960437 | 0.013350979 |
| chr10:62107151-62107468 | -1.496946519 | 0.000961671 | 0.013365715 |
| chr2:148936953-148937231 | -1.609653338 | 0.000962488 | 0.013373218 |
| chr1:8026715-8027110 | 1.476801396 | 0.00096256 | 0.013373218 |
| chr2:55021787-55022290 | -1.481690915 | 0.000963622 | 0.013385535 |
| chr4:74671139-74671465 | -1.359258777 | 0.000964892 | 0.013400752 |
| chr22:19605814-19606014 | 2.038069198 | 0.000965685 | 0.013409331 |
| chr3:67518312-67518584 | -1.720459572 | 0.000966039 | 0.013411825 |
| chr19:46648096-46648296 | 1.919587203 | 0.000966514 | 0.013415991 |
| chr20:41364349-41364660 | 1.321858708 | 0.000968066 | 0.013435089 |
| chr15:85047261-85047549 | 1.539120196 | 0.000968517 | 0.013438919 |
| chr1:113940343-113940614 | 1.236890562 | 0.000969388 | 0.013448574 |
| chr7:16616571-16616838 | 1.699831395 | 0.000970102 | 0.013455014 |
| chr10:115664727-115664971 | -1.924329558 | 0.000970204 | 0.013455014 |
| chr14:36341972-36342172 | 1.974063514 | 0.000971401 | 0.01346918 |
| chr14:80877603-80877921 | 1.537562731 | 0.000971672 | 0.013469231 |
| chr18:3319978-3320189 | 1.864532836 | 0.000971756 | 0.013469231 |
| chr13:74036369-74036811 | -1.169747924 | 0.000972598 | 0.013478465 |
| chr19:49201962-49202293 | 1.80555486 | 0.000973757 | 0.013490434 |
| chr21:40054330-40054880 | -1.151534797 | 0.000973814 | 0.013490434 |
| chr15:84390975-84391260 | 1.742403838 | 0.000974094 | 0.013491872 |
| chr7:118251002-118251207 | -2.236402687 | 0.000975087 | 0.013500738 |
| chr7:136911381-136911628 | -2.236402687 | 0.000975087 | 0.013500738 |
| chr3:75887274-75887476 | 2.118299027 | 0.000976091 | 0.013512209 |
| chr17:72974708-72975018 | 1.631977131 | 0.000977271 | 0.013526093 |
| chr12:17704946-17705242 | -1.636011534 | 0.000977882 | 0.013532105 |
| chr11:65932095-65932344 | 1.604320619 | 0.000978945 | 0.013542995 |
| chr12:40727837-40728185 | -1.428775276 | 0.000979022 | 0.013542995 |
| chr14:29446464-29446664 | 1.910255594 | 0.000979999 | 0.013554055 |
| chr10:59043206-59043911 | -1.18296861 | 0.000980421 | 0.013557454 |
| chr7:145398790-145399226 | -1.414962739 | 0.000981935 | 0.013575931 |
| chr14:22614214-22614559 | 1.374717168 | 0.000982295 | 0.013578463 |
| chr5:58317024-58317270 | 1.867988908 | 0.000984384 | 0.013604878 |
| chr12:65666324-65666572 | 1.774036466 | 0.000984635 | 0.013605895 |
| chr7:117171131-117171431 | -1.432068827 | 0.000984834 | 0.013606193 |
| chr14:91638994-91639250 | 1.922944909 | 0.000985302 | 0.013609079 |
| chr10:77431199-77431643 | 1.258690481 | 0.000985398 | 0.013609079 |

| | | | |
|---|---|---|---|
| chr14:105930275-105930523 | 1.683728273 | 0.000986325 | 0.013619422 |
| chr7:33869925-33870159 | 1.702263627 | 0.000986932 | 0.013625354 |
| chr5:40165507-40165741 | 1.613680675 | 0.000987738 | 0.013634023 |
| chr7:137899132-137899764 | -1.220536995 | 0.000988657 | 0.013644248 |
| chr2:209368438-209368747 | 1.347080054 | 0.000989284 | 0.013650449 |
| chr1:114290467-114290675 | 1.247226306 | 0.000989526 | 0.013651322 |
| chr4:142936143-142936351 | 1.967258982 | 0.000989945 | 0.013654645 |
| chr3:8196263-8196534 | 1.549921582 | 0.000990659 | 0.013662033 |
| chr20:62836236-62836649 | 1.220065051 | 0.000992066 | 0.013678978 |
| chr8:61273705-61274144 | -1.455994314 | 0.000992431 | 0.013681545 |
| chr8:11867556-11867808 | 1.571155738 | 0.000992806 | 0.013684261 |
| chr14:75794644-75794844 | 1.772376668 | 0.000993932 | 0.013695059 |
| chr8:52168987-52169390 | -1.515680619 | 0.000994126 | 0.013695059 |
| chr8:63265364-63265665 | -1.515680619 | 0.000994126 | 0.013695059 |
| chr20:57047905-57048632 | -1.082659143 | 0.000994871 | 0.013702616 |
| chr12:70304473-70305517 | -1.128085563 | 0.000995032 | 0.013702616 |
| chr14:44700860-44701114 | 1.76637609 | 0.000997973 | 0.013739301 |
| chr10:47687406-47687693 | -1.727847093 | 0.000998055 | 0.013739301 |
| chr13:47226539-47226739 | 1.808156223 | 0.000999493 | 0.013756631 |
| chr6:152943451-152943823 | -1.483993064 | 0.001000413 | 0.013766813 |
| chr7:104934328-104934599 | -1.788493909 | 0.00100163 | 0.013779833 |
| chr12:52243385-52243740 | 1.27661924 | 0.001001849 | 0.013779833 |
| chr5:168226216-168226600 | 1.562138525 | 0.001001898 | 0.013779833 |
| chr14:51627361-51627659 | 1.694452919 | 0.001002327 | 0.01378325 |
| chr13:27960263-27961238 | -1.122814807 | 0.001002668 | 0.013785466 |
| chr13:109088058-109088395 | -1.603191852 | 0.001003858 | 0.013798565 |
| chrX:16662077-16663389 | -1.095143238 | 0.001003981 | 0.013798565 |
| chr6:16711625-16711851 | -1.806316481 | 0.001004389 | 0.013801701 |
| chr3:4580495-4581052 | -1.168643387 | 0.001005244 | 0.013810972 |
| chr1:163766837-163767248 | 1.475703607 | 0.001006789 | 0.013824581 |
| chr3:162370036-162370275 | 1.811246061 | 0.001006841 | 0.013824581 |
| chr2:223563517-223563746 | -1.865775682 | 0.001006913 | 0.013824581 |
| chr8:54191644-54192165 | -1.28562086 | 0.001006956 | 0.013824581 |
| chr1:114216528-114216950 | 1.125418567 | 0.00100809 | 0.013837665 |
| chr10:21637024-21637524 | 1.466697075 | 0.00100892 | 0.013845339 |
| chr3:12853246-12853625 | 1.329890105 | 0.00100901 | 0.013845339 |
| chr13:73741477-73741750 | -1.863625422 | 0.001009506 | 0.013849664 |
| chr4:153492532-153493134 | 1.263286331 | 0.001010368 | 0.013859012 |
| chr7:116591475-116591675 | -2.116523601 | 0.001011919 | 0.01387558 |
| chr7:39810172-39811726 | -1.072673911 | 0.001011938 | 0.01387558 |
| chr5:124906432-124907210 | -1.263426653 | 0.001012585 | 0.013881963 |
| chr20:59491906-59492152 | 1.526526778 | 0.001013912 | 0.013897672 |
| chr3:19375968-19376756 | -1.25609362 | 0.001015052 | 0.013910799 |
| chr13:21202723-21202997 | 1.534226162 | 0.001015458 | 0.013913878 |
| chr3:142975394-142975690 | -1.390451891 | 0.001015931 | 0.013915593 |
| chr4:111336690-111337078 | -1.685593833 | 0.001015946 | 0.013915593 |
| chr3:184775684-184775948 | 1.741771687 | 0.001016321 | 0.01391824 |
| chr8:136009791-136010026 | 1.88924557 | 0.001016579 | 0.01391928 |
| chr14:40025461-40025822 | 1.562874106 | 0.001016818 | 0.013920071 |
| chr2:137888943-137889250 | -1.430870753 | 0.001017715 | 0.013929859 |

| | | | |
|---|---|---|---|
| chr3:169321367-169321787 | 1.265626767 | 0.00101885 | 0.013942526 |
| chr14:42564668-42564968 | 1.563296349 | 0.001019004 | 0.013942526 |
| chr10:79956105-79956846 | -1.169589543 | 0.001019632 | 0.013948624 |
| chr1:213061339-213061573 | -1.861998406 | 0.001020379 | 0.013956349 |
| chr2:19176866-19177321 | 1.385576757 | 0.001020602 | 0.013956913 |
| chr19:50136048-50136478 | 1.402200035 | 0.001022106 | 0.013974979 |
| chr8:100748033-100748409 | 1.205850875 | 0.001022746 | 0.013981237 |
| chr9:76586539-76586772 | 1.730370246 | 0.001023464 | 0.013988553 |
| chr4:175770584-175771549 | -1.19048248 | 0.001024852 | 0.014005026 |
| chr19:52601056-52601413 | 1.321778495 | 0.001027104 | 0.01403218 |
| chr1:204130151-204130581 | -1.151101185 | 0.001027205 | 0.01403218 |
| chr18:10413136-10413336 | -1.760025458 | 0.001028482 | 0.014047125 |
| chr2:190467210-190467430 | -1.692479544 | 0.001028936 | 0.014050812 |
| chr15:48324939-48325327 | 1.293994431 | 0.001029342 | 0.0140521 |
| chr19:47608009-47608398 | 1.260672443 | 0.001029397 | 0.0140521 |
| chr7:20432262-20432484 | 1.848195307 | 0.001030336 | 0.014062423 |
| chr12:79546503-79546703 | 1.42263808 | 0.00103098 | 0.014068697 |
| chr2:45821441-45822589 | -1.108748446 | 0.001031517 | 0.014073528 |
| chr6:134690087-134690389 | -1.551631638 | 0.001031762 | 0.014074355 |
| chr8:56093605-56094133 | -1.288484531 | 0.001032054 | 0.01407583 |
| chr17:62259212-62259567 | 1.481692788 | 0.001032509 | 0.014079537 |
| chr9:137351486-137351686 | 1.795157118 | 0.001033455 | 0.014089926 |
| chr9:124453351-124453843 | 1.217578907 | 0.001034535 | 0.014102148 |
| chr2:7053770-7054047 | -1.517221829 | 0.001034963 | 0.014105463 |
| chr11:12798747-12799004 | -1.463686709 | 0.001035404 | 0.014108969 |
| chr19:29405090-29405296 | 1.593084196 | 0.001035896 | 0.014111728 |
| chr12:131264687-131265142 | 1.324848692 | 0.001035975 | 0.014111728 |
| chr2:120480981-120481336 | 1.523226083 | 0.001036475 | 0.014116036 |
| chr20:47603706-47603957 | 1.156605047 | 0.00103715 | 0.014122715 |
| chr16:12901298-12901659 | 1.379139418 | 0.001038363 | 0.014136719 |
| chr12:100279170-100279441 | -1.695347308 | 0.001039277 | 0.014146644 |
| chr1:102550695-102551133 | -1.212631187 | 0.001039816 | 0.014151477 |
| chr5:20043403-20043603 | 1.959272458 | 0.001040297 | 0.014155501 |
| chr8:109940619-109940987 | 1.473824648 | 0.001041714 | 0.014172269 |
| chr4:159764381-159764706 | -1.44854906 | 0.001042208 | 0.014176466 |
| chr8:109229330-109229771 | 1.381794363 | 0.001043541 | 0.014192087 |
| chr10:6477269-6477469 | 1.986851896 | 0.001045434 | 0.01421531 |
| chr5:5650232-5650476 | -1.659398418 | 0.001045962 | 0.014219957 |
| chr13:51246880-51247107 | -1.765053731 | 0.001046529 | 0.014225143 |
| chr3:108818436-108818764 | -1.355451491 | 0.001046746 | 0.014225575 |
| chr16:86232676-86232928 | 1.812549707 | 0.001046964 | 0.014226016 |
| chr14:31775643-31775959 | 1.653302174 | 0.001047501 | 0.014229225 |
| chr3:102162890-102164083 | -1.06100954 | 0.001047572 | 0.014229225 |
| chr1:207797381-207797821 | -1.243496276 | 0.001048622 | 0.014240967 |
| chr13:91438539-91439349 | -1.114529958 | 0.001050156 | 0.014259268 |
| chr20:48317551-48317862 | 1.161647954 | 0.001050937 | 0.01426735 |
| chr19:50460343-50460543 | 1.758564271 | 0.001051442 | 0.01426994 |
| chr18:647164-647435 | 2.050276809 | 0.001051545 | 0.01426994 |
| chrX:105063756-105064081 | -1.449218904 | 0.001051687 | 0.01426994 |
| chr3:15364119-15364439 | 1.342244203 | 0.001051995 | 0.014271593 |

| | | | |
|---|---|---|---|
| chr4:186132942-186133219 | -1.4564538 | 0.001053804 | 0.014293603 |
| chr20:46118589-46118789 | 1.688259043 | 0.001055275 | 0.014308912 |
| chr2:2230047-2230722 | -1.324322943 | 0.001055306 | 0.014308912 |
| chr18:63254799-63255075 | -1.463551172 | 0.001056441 | 0.014321763 |
| chr8:91359627-91359994 | 1.546587784 | 0.00105708 | 0.014325347 |
| chr1:6959606-6959806 | 1.443976143 | 0.00105724 | 0.014325347 |
| chr22:30771353-30771596 | 1.692336675 | 0.001057266 | 0.014325347 |
| chr6:138493496-138494233 | -1.290294477 | 0.001057646 | 0.014327965 |
| chr1:106542601-106543119 | 1.111848423 | 0.001057938 | 0.014329384 |
| chr7:50984999-50985199 | 1.966265349 | 0.001058906 | 0.014337562 |
| chr6:90938134-90938472 | -1.507764443 | 0.001058916 | 0.014337562 |
| chr17:39642834-39643132 | 1.641611941 | 0.001060682 | 0.014358939 |
| chr11:85620124-85620360 | 1.904051031 | 0.001064247 | 0.014404648 |
| chr3:119768635-119768839 | 1.850512952 | 0.00106532 | 0.014416626 |
| chr14:102055346-102055806 | -1.338033323 | 0.001066208 | 0.014426098 |
| chr19:43486321-43486743 | 1.534574859 | 0.0010686 | 0.014455914 |
| chr5:57989491-57989730 | -1.803630934 | 0.001069477 | 0.014465214 |
| chr1:33280569-33280769 | 1.896414231 | 0.00106983 | 0.014467437 |
| chr12:112926519-112926899 | 1.323029496 | 0.001070319 | 0.014471498 |
| chr8:108278207-108278474 | 1.675297815 | 0.001071457 | 0.014480897 |
| chr5:166382013-166382361 | -1.618944627 | 0.001071579 | 0.014480897 |
| chr13:33992967-33993207 | -1.952317608 | 0.001071581 | 0.014480897 |
| chr17:59360833-59361075 | 1.476955032 | 0.001072548 | 0.014491402 |
| chr7:105580928-105581819 | -1.090724207 | 0.001074964 | 0.01452149 |
| chr2:241093917-241094198 | 1.576976881 | 0.001075468 | 0.014525311 |
| chr7:26083330-26083530 | 1.718144633 | 0.001075751 | 0.014525311 |
| chr19:45517967-45518167 | 1.902151717 | 0.001075816 | 0.014525311 |
| chr7:24701737-24701937 | 1.597820725 | 0.001076801 | 0.014536058 |
| chr13:79446320-79446557 | -1.75498063 | 0.001077671 | 0.014545237 |
| chr15:86863319-86863545 | -1.756657175 | 0.001080047 | 0.014574735 |
| chr14:80564752-80565580 | -1.191224804 | 0.001080385 | 0.01457673 |
| chr4:185183736-185183944 | -1.895511525 | 0.001080997 | 0.014582427 |
| chr14:50879041-50879253 | 1.791060601 | 0.00108228 | 0.014597161 |
| chr10:51893071-51893645 | -1.263468523 | 0.001084496 | 0.014622665 |
| chr12:123765668-123765876 | 1.794795256 | 0.001084553 | 0.014622665 |
| chr1:116714607-116714933 | 1.516875798 | 0.001085374 | 0.014631168 |
| chr5:136066828-136067244 | -1.347852808 | 0.001086223 | 0.014640036 |
| chr2:117525821-117526101 | -1.801254763 | 0.001086631 | 0.01464152 |
| chr3:143707879-143708395 | -1.275979344 | 0.001086716 | 0.01464152 |
| chr2:121689324-121689968 | -1.203783858 | 0.00108727 | 0.014646413 |
| chr20:55218220-55218437 | 1.764046205 | 0.001088405 | 0.014657432 |
| chr5:41889743-41889943 | 2.216213564 | 0.001088471 | 0.014657432 |
| chr8:96746443-96746643 | 2.01050581 | 0.00109027 | 0.014677198 |
| chr8:29773447-29774440 | -1.132227744 | 0.00109046 | 0.014677198 |
| chr4:35520053-35520363 | -1.481339198 | 0.001090513 | 0.014677198 |
| chr13:85261637-85262044 | -1.721847671 | 0.001090777 | 0.014678175 |
| chrX:55268204-55268407 | 1.899302249 | 0.001091002 | 0.014678619 |
| chr14:105702355-105702645 | 1.372137349 | 0.001091621 | 0.014682198 |
| chr16:46924540-46924740 | 1.962191922 | 0.001091824 | 0.014682198 |
| chr12:116703254-116703550 | 1.422697254 | 0.001091843 | 0.014682198 |

| | | | |
|---|---|---|---|
| chr4:156933964-156934340 | -1.428988373 | 0.001092249 | 0.014685089 |
| chr6:64504512-64504989 | -1.204659291 | 0.001092606 | 0.014687315 |
| chr18:55234507-55235055 | -1.240784395 | 0.001093087 | 0.014688467 |
| chr4:40475685-40475986 | -1.508080984 | 0.001093159 | 0.014688467 |
| chr5:1077157-1077357 | 1.650940415 | 0.001093267 | 0.014688467 |
| chr1:165456176-165456573 | 1.387800979 | 0.001093961 | 0.014695209 |
| chr4:164924898-164925208 | -1.385216119 | 0.0010943 | 0.014697181 |
| chr6:122568745-122569530 | -1.158152241 | 0.001095493 | 0.014710634 |
| chr7:108340659-108341442 | -1.204889554 | 0.001095797 | 0.014712136 |
| chrX:154051310-154051529 | 1.741995204 | 0.001096017 | 0.014712506 |
| chr21:31351463-31351795 | 1.300377443 | 0.001098419 | 0.014742178 |
| chr5:75536656-75536930 | -1.715133733 | 0.001099924 | 0.014758394 |
| chr3:23821105-23821305 | 1.798323373 | 0.001100206 | 0.014758394 |
| chr4:74191117-74191321 | 1.798323373 | 0.001100206 | 0.014758394 |
| chr7:142412577-142413108 | -1.250022678 | 0.001100663 | 0.014759822 |
| chr16:34573953-34574415 | 1.147583611 | 0.00110076 | 0.014759822 |
| chr3:10191942-10192336 | 1.278758257 | 0.00110089 | 0.014759822 |
| chr7:37375563-37375780 | 1.836387275 | 0.001101208 | 0.0147615 |
| chr11:119859748-11985995 | 1.752430891 | 0.001101796 | 0.014766798 |
| chr7:23181584-23182743 | 1.057258646 | 0.001102296 | 0.014770919 |
| chrX:16771047-16772218 | -1.071509123 | 0.001103363 | 0.014782631 |
| chr4:70531982-70532299 | -1.407498578 | 0.001103859 | 0.014786687 |
| chr1:106726496-106726856 | 1.170693086 | 0.001104791 | 0.014795266 |
| chr15:94110730-94110991 | 1.588421245 | 0.001104885 | 0.014795266 |
| chr20:54886759-54887133 | 1.143299642 | 0.001106269 | 0.014811213 |
| chr7:81478381-81479156 | -1.089646 | 0.001107029 | 0.014816156 |
| chr5:127752385-127752847 | -1.331827976 | 0.001107095 | 0.014816156 |
| chr14:53901048-53901282 | 1.750928405 | 0.001107219 | 0.014816156 |
| chr7:135281501-135281821 | -1.296327084 | 0.001107546 | 0.014817943 |
| chr12:53251171-53252809 | 1.051568111 | 0.001108759 | 0.014831586 |
| chr7:122166728-122166928 | 2.104039096 | 0.001110165 | 0.0148478 |
| chr20:48154393-48154918 | 1.09140822 | 0.001110636 | 0.014851511 |
| chr3:193870550-193871394 | -1.173191365 | 0.001112445 | 0.014873104 |
| chr7:124702133-124702352 | -1.878785581 | 0.001112833 | 0.014875694 |
| chr1:242226239-242226444 | 1.750310809 | 0.001113147 | 0.014877292 |
| chr5:121052962-121053730 | -1.200799695 | 0.001114254 | 0.014889427 |
| chr10:34157063-34157279 | -1.983855337 | 0.001114651 | 0.014889427 |
| chr4:106287375-106287738 | -1.457374943 | 0.001114801 | 0.014889427 |
| chr6:140388201-140388773 | -1.328212974 | 0.001114832 | 0.014889427 |
| chr10:77512362-77512621 | 1.837099167 | 0.001115478 | 0.014894856 |
| chr4:82692590-82692790 | 1.957905312 | 0.001115627 | 0.014894856 |
| chr7:40986329-40986840 | 1.28820748 | 0.001116816 | 0.014908132 |
| chr20:56936499-56937127 | -1.127105433 | 0.001117994 | 0.01492083 |
| chr14:55794525-55794958 | 1.336446448 | 0.001118159 | 0.01492083 |
| chr14:101917143-10191750 | 1.427415458 | 0.001118351 | 0.01492083 |
| chr2:162545323-162545616 | -1.562015601 | 0.001118839 | 0.01492473 |
| chr20:47269780-47270066 | 1.108050952 | 0.001119118 | 0.014925855 |
| chr7:33589057-33589294 | 1.753871739 | 0.001119942 | 0.014934252 |
| chr7:155239510-155240319 | -1.179209667 | 0.001120158 | 0.014934524 |
| chr5:42834510-42834935 | 1.245980538 | 0.001120572 | 0.014937451 |

| | | | |
|---|---|---|---|
| chr20:49899839-49900128 | 1.334918178 | 0.001121163 | 0.014942729 |
| chr21:42375494-42375709 | -1.504358376 | 0.001122283 | 0.014955057 |
| chr1:119906631-119906843 | 1.71303135 | 0.001124666 | 0.0149842 |
| chr8:70615031-70615454 | -1.299803627 | 0.001125781 | 0.014993848 |
| chr14:45391606-45391980 | 1.414042464 | 0.001125781 | 0.014993848 |
| chr7:49001533-49002077 | -1.218640492 | 0.001127132 | 0.015008382 |
| chr21:40318362-40319030 | -1.136088479 | 0.001127702 | 0.015008382 |
| chr14:95461386-95461636 | 1.674569984 | 0.001127797 | 0.015008382 |
| chr11:111069191-111069430 | -1.550680305 | 0.001127798 | 0.015008382 |
| chr14:21287794-21287994 | 1.896012324 | 0.001127852 | 0.015008382 |
| chrX:112306456-112306711 | 1.882042648 | 0.001128269 | 0.015009618 |
| chr1:234535884-234536139 | 1.612226727 | 0.001128337 | 0.015009618 |
| chr2:192878890-192879117 | -1.965338024 | 0.001129177 | 0.015016279 |
| chr6:10133947-10134691 | -1.262060475 | 0.001129229 | 0.015016279 |
| chr19:1183678-1183878 | 1.683577232 | 0.00113052 | 0.015030831 |
| chr20:59596825-59597056 | 1.476986499 | 0.001131666 | 0.01504303 |
| chr3:61847264-61847599 | 1.209210193 | 0.001131923 | 0.01504303 |
| chr14:75240515-75240759 | 1.630005944 | 0.001132026 | 0.01504303 |
| chr14:90778091-90778307 | 1.608517644 | 0.001132288 | 0.015043072 |
| chr12:111597841-111598062 | 1.456179769 | 0.001132473 | 0.015043072 |
| chr1:80634051-80634732 | -1.230859474 | 0.001132618 | 0.015043072 |
| chr7:134254380-134254643 | -1.741671164 | 0.001133295 | 0.015049448 |
| chr2:88934015-88934383 | -1.282985917 | 0.001134371 | 0.015061124 |
| chr13:36965937-36966146 | 1.788335143 | 0.00113506 | 0.015067663 |
| chr14:104537247-104537700 | 1.347560455 | 0.001135415 | 0.015069771 |
| chr16:1992230-1992482 | 1.375637552 | 0.001136535 | 0.015082018 |
| chr7:133117868-133118135 | -1.701711556 | 0.001136988 | 0.015085418 |
| chr3:2134574-2134827 | 1.827312741 | 0.00113771 | 0.015092387 |
| chr9:124569012-124569247 | 1.77580062 | 0.001138004 | 0.015093673 |
| chr7:108611253-108611652 | -1.207670427 | 0.001138751 | 0.015100968 |
| chr2:16620620-16620989 | -1.255709936 | 0.001139154 | 0.015103693 |
| chr7:54839271-54839890 | -1.165848192 | 0.001140255 | 0.015115674 |
| chr2:204051525-204051902 | -1.351913919 | 0.001141358 | 0.015126771 |
| chr7:17238087-17238292 | 1.942391477 | 0.001141487 | 0.015126771 |
| chr10:15585270-15585580 | -1.50146803 | 0.001141886 | 0.015129422 |
| chr1:159141977-159142311 | -1.479060658 | 0.001142082 | 0.015129422 |
| chr2:94870859-94871429 | -1.192241031 | 0.001142529 | 0.015132731 |
| chr1:222452017-222452232 | 1.832585385 | 0.001143743 | 0.015146193 |
| chr5:153352641-153353158 | -1.489742367 | 0.001144532 | 0.015152016 |
| chr20:47516006-47516295 | 1.132438264 | 0.001144578 | 0.015152016 |
| chr3:141122470-141122888 | -1.360297208 | 0.001144883 | 0.015153437 |
| chr21:9341274-9341538 | 1.570562756 | 0.001145267 | 0.015155226 |
| chr5:76787281-76787630 | -1.350438617 | 0.001145414 | 0.015155226 |
| chr20:47559707-47559983 | 1.130195655 | 0.001146455 | 0.015165897 |
| chr7:12449699-12449906 | 1.676442989 | 0.001146689 | 0.015165897 |
| chr7:147101618-147101854 | 1.888549173 | 0.001146856 | 0.015165897 |
| chr6:98416255-98416710 | 1.367712549 | 0.001147012 | 0.015165897 |
| chr1:107107862-107108070 | 1.242069611 | 0.001147545 | 0.015170324 |
| chr7:151918058-151918310 | 1.942704758 | 0.001148138 | 0.015173292 |
| chr14:27082344-27082556 | 1.791812826 | 0.001148166 | 0.015173292 |

| | | | |
|---|---|---|---|
| chr18:60053834-60054191 | -1.623546296 | 0.001149079 | 0.015179746 |
| chr2:46898971-46899236 | -1.904788322 | 0.001149126 | 0.015179746 |
| chr8:130497782-130497989 | 1.579067657 | 0.001149248 | 0.015179746 |
| chr2:210072953-210073189 | -1.677330935 | 0.00114966 | 0.015182564 |
| chr12:112759847-112760047 | 1.953553647 | 0.001151107 | 0.015199054 |
| chr10:77732156-77732455 | -1.712368868 | 0.001151562 | 0.01519981 |
| chr4:147937620-147937827 | 1.938922237 | 0.001151588 | 0.01519981 |
| chr5:126032092-126032405 | -1.731620159 | 0.001151759 | 0.01519981 |
| chr1:210257879-210258299 | -1.378593533 | 0.001152295 | 0.015204258 |
| chr17:61253426-61253693 | 1.117095951 | 0.001153093 | 0.015210165 |
| chr5:83366655-83366964 | 1.88107226 | 0.001153338 | 0.015210165 |
| chr4:1290712-1290944 | 1.505513918 | 0.001153338 | 0.015210165 |
| chr13:49098244-49098461 | -1.842605196 | 0.001153932 | 0.015215375 |
| chr10:24469984-24470494 | 1.194316507 | 0.001154255 | 0.01521702 |
| chr6:42444463-42444884 | 1.363865743 | 0.001154804 | 0.015220568 |
| chr18:43596187-43596406 | -1.79093059 | 0.001155102 | 0.015220568 |
| chr12:14554224-14554770 | -1.316244623 | 0.001155254 | 0.015220568 |
| chr16:53061710-53062170 | -1.461735421 | 0.001155319 | 0.015220568 |
| chr10:58435812-58436052 | -1.708232685 | 0.001155555 | 0.015221067 |
| chr2:38685074-38685938 | -1.080850592 | 0.001156992 | 0.015234934 |
| chr18:78306320-78306546 | -1.961300476 | 0.001157006 | 0.015234934 |
| chr7:133252771-133253804 | -1.08495 | 0.001157313 | 0.015236369 |
| chr16:72179965-72180324 | 1.575362518 | 0.00115783 | 0.015240546 |
| chrX:133297323-133297824 | 1.357232404 | 0.001158252 | 0.015242719 |
| chr6:114543758-114544169 | -1.426029974 | 0.001158392 | 0.015242719 |
| chr1:154681217-154681514 | 1.739048023 | 0.001158911 | 0.015244653 |
| chr4:138308418-138308685 | -1.791481507 | 0.00115899 | 0.015244653 |
| chr2:133942285-133942746 | 1.440891238 | 0.001159196 | 0.015244653 |
| chr18:69469398-69469632 | -1.97583829 | 0.001159335 | 0.015244653 |
| chr14:44104238-44104440 | 1.939569282 | 0.001160039 | 0.015251285 |
| chr16:57818662-57818883 | 1.618895348 | 0.001160321 | 0.01525238 |
| chr8:122017081-122017855 | -1.149874089 | 0.001163124 | 0.0152866 |
| chr12:82666457-82666803 | -1.492745251 | 0.001164587 | 0.015300702 |
| chr20:47833131-47834260 | 1.076327786 | 0.001164596 | 0.015300702 |
| chr4:125522790-125523361 | -1.167505136 | 0.001165669 | 0.015312164 |
| chr14:105229309-105229547 | 1.580944334 | 0.001166042 | 0.01531445 |
| chr14:54935793-54936141 | 1.527721354 | 0.001167005 | 0.015324465 |
| chr13:31183742-31183964 | 1.77934086 | 0.00116801 | 0.01533475 |
| chr20:48555553-48555784 | 1.186379659 | 0.001168251 | 0.01533475 |
| chr16:21631077-21631279 | 1.800545012 | 0.001168584 | 0.01533475 |
| chr16:14013779-14014337 | -1.308092418 | 0.001168589 | 0.01533475 |
| chr13:96993757-96994421 | -1.348567625 | 0.001169242 | 0.015336854 |
| chr2:37324396-37325524 | -1.100599976 | 0.001169257 | 0.015336854 |
| chr18:48838893-48839114 | -1.612749821 | 0.00116935 | 0.015336854 |
| chr4:167265220-167265684 | -1.278900924 | 0.001169826 | 0.015340474 |
| chr14:29078901-29079117 | 1.648857662 | 0.001170062 | 0.015340951 |
| chr20:48827117-48828725 | 1.024422835 | 0.001170861 | 0.015348801 |
| chr16:72961906-72962143 | -1.497465575 | 0.001171646 | 0.015356464 |
| chr2:35338784-35339155 | -1.558244343 | 0.001173051 | 0.01537224 |
| chr1:244280896-244281275 | -1.558107475 | 0.001174308 | 0.015386087 |

| | | | |
|---|---|---|---|
| chr20:29445217-29445915 | 1.130155398 | 0.001177606 | 0.015426659 |
| chr2:18232790-18233429 | -1.297332585 | 0.001178213 | 0.015431976 |
| chr16:77788188-77788784 | 1.13859811 | 0.001178798 | 0.015436999 |
| chr16:35084020-35084291 | 1.639721352 | 0.001179095 | 0.01543824 |
| chr1:158272733-158273151 | 1.492259587 | 0.001179568 | 0.0154418 |
| chr7:121611372-121611645 | -1.816964856 | 0.001180368 | 0.015449635 |
| chr7:21713046-21713290 | 1.481774967 | 0.001181831 | 0.015466145 |
| chr1:183349193-183349498 | 1.405902443 | 0.001183279 | 0.01548117 |
| chrX:154478408-154479454 | 1.070748618 | 0.001183383 | 0.01548117 |
| chr14:50347568-50347830 | 1.526417001 | 0.001183869 | 0.015484298 |
| chr3:14839201-14839665 | 1.231300919 | 0.001184027 | 0.015484298 |
| chr13:78767741-78767952 | -1.840573273 | 0.00118552 | 0.01550118 |
| chr9:114804584-114804838 | 1.578206588 | 0.001186626 | 0.015512999 |
| chr4:111960163-111960516 | -1.349345559 | 0.001187247 | 0.015515827 |
| chr1:208219619-208220145 | -1.278552108 | 0.001187248 | 0.015515827 |
| chr7:140656490-140657305 | -1.123599305 | 0.001188269 | 0.015526528 |
| chr1:162037044-162037316 | 1.459104864 | 0.001188531 | 0.0155273 |
| chr2:76599680-76600285 | -1.182761748 | 0.001190962 | 0.015552446 |
| chr11:133934754-133935030 | -1.5045699 | 0.001191044 | 0.015552446 |
| chr9:131773934-131774171 | 1.645262221 | 0.001191065 | 0.015552446 |
| chr17:55705801-55706161 | -1.355376992 | 0.001191544 | 0.015555217 |
| chr6:26020672-26020944 | -1.336572521 | 0.001191683 | 0.015555217 |
| chr14:54578068-54578275 | 1.671017739 | 0.001192537 | 0.015563712 |
| chr22:41971141-41971360 | 1.744635746 | 0.001192877 | 0.015565497 |
| chr12:17589079-17589279 | -2.0366422 | 0.001193142 | 0.015566309 |
| chr6:122696801-122697368 | -1.107302959 | 0.001196423 | 0.015605664 |
| chr1:3762696-3762919 | 1.739534337 | 0.001196995 | 0.015605664 |
| chr17:52597524-52598109 | -1.261277872 | 0.001197112 | 0.015605664 |
| chr4:176195229-176196019 | -1.095652082 | 0.001197149 | 0.015605664 |
| chr10:32501691-32502484 | -1.247840611 | 0.001197255 | 0.015605664 |
| chr8:101670793-101671101 | 1.501037343 | 0.001197381 | 0.015605664 |
| chr4:188738300-188738668 | -1.636570547 | 0.001197785 | 0.015607035 |
| chr1:240359829-240360037 | 1.872592459 | 0.001197893 | 0.015607035 |
| chr2:200288679-200289417 | -1.181131015 | 0.001198309 | 0.015609795 |
| chr8:99492882-99493115 | 1.652469077 | 0.001198867 | 0.015611362 |
| chr18:43285885-43286636 | -1.298999489 | 0.001198931 | 0.015611362 |
| chr7:135232751-135233047 | -1.487950108 | 0.00119904 | 0.015611362 |
| chr10:61628730-61629175 | -1.372802381 | 0.001199674 | 0.015611398 |
| chr4:141549219-141549588 | -1.449012495 | 0.001199834 | 0.015611398 |
| chr2:84515949-84516881 | -1.068116053 | 0.001199841 | 0.015611398 |
| chr7:45978045-45978628 | -1.113124121 | 0.001199858 | 0.015611398 |
| chr11:9075664-9076477 | -1.213696607 | 0.001200617 | 0.015617245 |
| chr8:123745382-123745886 | 1.262798645 | 0.001200715 | 0.015617245 |
| chr5:40853492-40853803 | 1.566970199 | 0.001202173 | 0.015630862 |
| chr3:186977306-186977718 | -1.372606079 | 0.001202231 | 0.015630862 |
| chr5:113686592-113686918 | -1.53258071 | 0.001202374 | 0.015630862 |
| chr17:58560791-58561001 | 1.272481067 | 0.001204148 | 0.015648955 |
| chr14:100274013-100274218 | -1.892202672 | 0.001204174 | 0.015648955 |
| chr5:95392690-95393071 | -1.477547693 | 0.001204412 | 0.015649385 |
| chr16:82667351-82667591 | 1.739437798 | 0.001204848 | 0.015652407 |

| | | | |
|---|---|---|---|
| chr2:202546917-202547186 | -1.784386261 | 0.001205597 | 0.015659472 |
| chr2:174547961-174548312 | -1.495095627 | 0.00120608 | 0.015663093 |
| chr22:15761244-15761477 | 1.678717973 | 0.001207107 | 0.015672126 |
| chr6:161618829-161619098 | -1.657269503 | 0.00120731 | 0.015672126 |
| chr14:94323663-94323872 | 1.711064203 | 0.001207389 | 0.015672126 |
| chr17:62778852-62779375 | -1.118853596 | 0.001208168 | 0.015679279 |
| chr11:27205201-27205532 | 1.413764985 | 0.001208349 | 0.015679279 |
| chr3:42572352-42572704 | -1.409880676 | 0.001210081 | 0.015697147 |
| chr4:168077081-168077587 | -1.292254258 | 0.001210154 | 0.015697147 |
| chr17:74152716-74153100 | 1.280734001 | 0.001210537 | 0.015697147 |
| chr22:17396668-17397036 | 1.401506344 | 0.001210546 | 0.015697147 |
| chr1:225865599-225865799 | 1.733738346 | 0.001211355 | 0.015704978 |
| chr18:69959749-69960016 | -1.790430061 | 0.001213364 | 0.015728372 |
| chr8:88713015-88713336 | 1.363624532 | 0.001214097 | 0.015733277 |
| chr16:4152120-4152805 | -1.128152682 | 0.001214153 | 0.015733277 |
| chr20:46894261-46894897 | 1.165107344 | 0.001214866 | 0.015739847 |
| chr8:135792512-135792841 | 1.518193298 | 0.001215833 | 0.015749718 |
| chr19:37979934-37980135 | 1.778013354 | 0.001216479 | 0.01575542 |
| chr6:102907332-102907638 | -1.696586791 | 0.001217425 | 0.015765002 |
| chrX:9206762-9207091 | -1.682676436 | 0.001218292 | 0.015773569 |
| chr11:62505231-62505455 | 1.781749963 | 0.001218667 | 0.01577575 |
| chrX:109732955-109733894 | 1.144591665 | 0.001219555 | 0.015784589 |
| chr7:43808199-43808897 | -1.153637575 | 0.001220562 | 0.015794953 |
| chr11:124509869-124510732 | -1.285115277 | 0.001221473 | 0.015804066 |
| chr11:76302681-76302881 | 1.742823037 | 0.001222051 | 0.015808879 |
| chr1:153647554-153647792 | 1.580210936 | 0.001222924 | 0.015817498 |
| chr6:55528687-55528949 | -1.59470913 | 0.001225267 | 0.015845138 |
| chr4:40975449-40975803 | 1.279292648 | 0.001225975 | 0.015849994 |
| chr8:91392698-91393084 | 1.497969114 | 0.001226057 | 0.015849994 |
| chr3:102098742-102098987 | 1.670948295 | 0.001226694 | 0.015855563 |
| chr11:75231760-75232041 | 1.469292499 | 0.001227363 | 0.01586139 |
| chr16:2163278-2163498 | 1.658635313 | 0.001227559 | 0.01586139 |
| chr15:39630004-39630246 | 1.607708679 | 0.001228093 | 0.015865606 |
| chr12:6647330-6647541 | -1.903193353 | 0.00122897 | 0.015874268 |
| chr14:58122706-58123033 | 1.515831836 | 0.001229258 | 0.015875305 |
| chr20:58665190-58665500 | 1.44144404 | 0.001230415 | 0.015887571 |
| chrX:6690996-6691250 | -1.813743168 | 0.001230909 | 0.01589127 |
| chr13:79117970-79118213 | -1.673030034 | 0.001231175 | 0.015892033 |
| chr16:29783563-29783810 | 1.385827848 | 0.001231728 | 0.015892219 |
| chr3:177574243-177574641 | -1.278114251 | 0.001231794 | 0.015892219 |
| chr4:42713418-42713751 | -1.386977407 | 0.001231944 | 0.015892219 |
| chr16:67000499-67000808 | -1.67146302 | 0.001232019 | 0.015892219 |
| chr17:51667215-51667424 | 1.735246644 | 0.001232818 | 0.01589984 |
| chr17:77720448-77720648 | 1.840262336 | 0.001233408 | 0.015904775 |
| chr8:100445174-100445374 | 1.739650633 | 0.001236257 | 0.015938826 |
| chr4:184894290-184894511 | -1.780447407 | 0.001237462 | 0.015951691 |
| chr1:241219463-241219804 | -1.421637704 | 0.001237796 | 0.015953304 |
| chr7:47771275-47771576 | -1.508002239 | 0.001238371 | 0.015958026 |
| chr7:22821864-22822328 | 1.2046068 | 0.001239229 | 0.015961425 |
| chr5:178078177-178078952 | -1.073968271 | 0.001239247 | 0.015961425 |

| | | | |
|---|---|---|---|
| chr12:113135256-113135486 | 1.451631208 | 0.001239779 | 0.015961425 |
| chr1:117224878-117225196 | 1.490444641 | 0.001239806 | 0.015961425 |
| chr21:45447790-45448219 | -1.283521216 | 0.001239854 | 0.015961425 |
| chr3:62860802-62861002 | 1.354114513 | 0.001240079 | 0.015961425 |
| chr6:127366285-127367005 | -1.24608236 | 0.001240092 | 0.015961425 |
| chr6:122438936-122439176 | -1.861786007 | 0.001240466 | 0.015963549 |
| chr2:65300153-65300481 | -1.388213099 | 0.001241125 | 0.015969356 |
| chr7:88111343-88111636 | -1.682725605 | 0.00124227 | 0.015981395 |
| chr16:27364530-27364978 | 1.247624698 | 0.001244471 | 0.016007021 |
| chr9:101465179-101465710 | -1.183566979 | 0.001245771 | 0.016021062 |
| chr2:226048672-226049006 | -1.724602435 | 0.001246022 | 0.016021591 |
| chr7:111828022-111828328 | -1.816954058 | 0.001246874 | 0.016029867 |
| chrX:92260079-92260538 | -1.325099438 | 0.001248225 | 0.016044535 |
| chr17:35795641-35795874 | -1.744301017 | 0.001248832 | 0.016045147 |
| chr7:112516059-112516275 | -1.723575388 | 0.001248895 | 0.016045147 |
| chr9:22103989-22104264 | 1.448647882 | 0.001248901 | 0.016045147 |
| chr18:41955719-41955957 | -1.509527534 | 0.001249562 | 0.016047075 |
| chr21:25626793-25627219 | 1.312513555 | 0.001249712 | 0.016047075 |
| chr1:106793789-106794009 | 1.256244331 | 0.001249805 | 0.016047075 |
| chr2:160125921-160126223 | -1.633729788 | 0.001249888 | 0.016047075 |
| chr16:75208388-75208588 | 1.634724747 | 0.001250331 | 0.016047487 |
| chr20:47295001-47295281 | 1.101451304 | 0.001250339 | 0.016047487 |
| chr1:240435986-240436682 | -1.206608525 | 0.001250735 | 0.016048872 |
| chr13:101101645-101101848 | -1.836911096 | 0.001250866 | 0.016048872 |
| chr7:134127191-134127741 | -1.156023384 | 0.00125129 | 0.01604927 |
| chrX:129877103-129877318 | 1.715063098 | 0.001251316 | 0.01604927 |
| chr15:46280819-46281082 | -1.638649513 | 0.001251812 | 0.016052949 |
| chr13:21367185-21367385 | 1.933894363 | 0.001253991 | 0.016078201 |
| chr13:102402395-102402595 | -2.079652736 | 0.001254495 | 0.016081585 |
| chr10:11612603-11613156 | -1.142911424 | 0.001254675 | 0.016081585 |
| chr7:156554534-156554948 | -1.554309812 | 0.001255538 | 0.016081999 |
| chr6:41162232-41162432 | 2.087498372 | 0.001255706 | 0.016081999 |
| chr7:128218364-128218632 | -1.545641924 | 0.001255818 | 0.016081999 |
| chr15:71801195-71801480 | 1.37415082 | 0.001255996 | 0.016081999 |
| chr16:30775902-30776196 | 1.26253078 | 0.001256173 | 0.016081999 |
| chr9:96279946-96280303 | 1.335905688 | 0.001256323 | 0.016081999 |
| chr7:55572375-55572742 | 1.308764545 | 0.001256371 | 0.016081999 |
| chr7:121395482-121396772 | -1.040311241 | 0.001256432 | 0.016081999 |
| chr1:150952276-150952485 | 1.626981182 | 0.001256596 | 0.016081999 |
| chr20:48195349-48195641 | 1.11898765 | 0.001256927 | 0.016083543 |
| chr6:47180694-47181013 | -1.430669898 | 0.001257368 | 0.016086499 |
| chr3:142372315-142372515 | 1.870366623 | 0.001258031 | 0.016092292 |
| chr2:9769298-9770727 | -1.049220482 | 0.001259982 | 0.016114566 |
| chr3:165883920-165884643 | -1.225424698 | 0.001261276 | 0.016128423 |
| chr1:27416454-27416732 | 1.617911257 | 0.00126239 | 0.01613997 |
| chr3:186843007-186843343 | 1.434024585 | 0.001264099 | 0.016158828 |
| chr14:75364677-75364956 | 1.562333896 | 0.001264287 | 0.016158828 |
| chr11:84558484-84558854 | -1.479997217 | 0.001266429 | 0.016183505 |
| chr14:105632005-105632613 | 1.167150384 | 0.001269143 | 0.016215487 |
| chr1:106571462-106571665 | 1.224887611 | 0.001269914 | 0.016222627 |

| | | | |
|---|---|---|---|
| chr3:52688275-52688680 | 1.360776936 | 0.00127067 | 0.016227595 |
| chr1:27864600-27864800 | 1.661932294 | 0.001270863 | 0.016227595 |
| chr3:150180474-150180674 | 1.665783644 | 0.001270938 | 0.016227595 |
| chr7:132650609-132650818 | -1.862577106 | 0.001271518 | 0.016232303 |
| chr14:71956780-71957312 | 1.31583549 | 0.001272069 | 0.016236628 |
| chr14:30790949-30791262 | 1.599211078 | 0.001273409 | 0.01624771 |
| chr8:101410762-101410980 | 1.480567496 | 0.001273552 | 0.01624771 |
| chr5:143246024-143246427 | -1.228593945 | 0.001273573 | 0.01624771 |
| chr7:135169133-135169389 | -1.347177411 | 0.001274999 | 0.01626319 |
| chr13:52004533-52004900 | -1.37281205 | 0.001275638 | 0.01626863 |
| chr5:49601471-49601741 | 1.439426528 | 0.001276057 | 0.016271268 |
| chr1:191126798-191127192 | -1.441972581 | 0.001276659 | 0.016276236 |
| chr5:47397-47606 | 1.870613646 | 0.001277549 | 0.016284867 |
| chr14:49641960-49642193 | 1.581337871 | 0.001278077 | 0.016288898 |
| chr8:16214644-16215362 | -1.351657365 | 0.001279378 | 0.016302767 |
| chr20:47378097-47378371 | 1.08811118 | 0.001279623 | 0.016303173 |
| chr12:20681044-20681415 | -1.549808331 | 0.00128022 | 0.016303862 |
| chr2:202107979-202108191 | 1.921210477 | 0.001280243 | 0.016303862 |
| chr2:144389539-144389964 | -1.440176969 | 0.001280316 | 0.016303862 |
| chr19:51799264-51799723 | 1.310849667 | 0.001280799 | 0.016304499 |
| chr19:58009524-58010042 | 1.181366977 | 0.001280866 | 0.016304499 |
| chr22:24945282-24945702 | 1.310631953 | 0.001281004 | 0.016304499 |
| chr4:37463541-37463784 | -1.734739918 | 0.001282121 | 0.016315773 |
| chr4:118968046-118968420 | -1.33066088 | 0.001282316 | 0.016315773 |
| chr3:39716895-39717194 | -1.693481072 | 0.001282767 | 0.016318809 |
| chr6:18406020-18406220 | 1.922322974 | 0.001283015 | 0.016319251 |
| chr20:37279244-37279607 | -1.423667629 | 0.001284956 | 0.016341222 |
| chrX:137654576-137654787 | 1.826789787 | 0.001285319 | 0.016341814 |
| chr4:61070346-61070843 | -1.394098829 | 0.001285429 | 0.016341814 |
| chr7:67943644-67943935 | -1.805601978 | 0.001285842 | 0.016343741 |
| chr5:64347432-64347645 | -1.781081145 | 0.001286174 | 0.016343741 |
| chr10:125836617-125837259 | 1.115217784 | 0.00128622 | 0.016343741 |
| chr15:79102133-79102592 | -1.299186773 | 0.001287976 | 0.016363345 |
| chr3:24679840-24680040 | 1.87351435 | 0.001288376 | 0.016365704 |
| chr9:73079324-73079628 | -1.524706947 | 0.001288806 | 0.016368458 |
| chr9:33211998-33212209 | -1.865953237 | 0.001289181 | 0.016370502 |
| chr21:17612470-17613154 | 1.149713202 | 0.001289801 | 0.016375669 |
| chr8:128554929-128555256 | 1.400377511 | 0.001290069 | 0.016376358 |
| chr7:105876160-105877596 | -1.050113494 | 0.001290777 | 0.016382623 |
| chr7:112769957-112770456 | -1.203242661 | 0.001291002 | 0.016382762 |
| chr18:58091174-58091638 | -1.381582524 | 0.001291529 | 0.016386514 |
| chr8:65450393-65450723 | -1.41276632 | 0.001291725 | 0.016386514 |
| chr12:62983541-62983899 | 1.370467582 | 0.001291959 | 0.016386776 |
| chr20:47749024-47749254 | 1.125824926 | 0.001292466 | 0.016390492 |
| chr11:83681128-83681616 | -1.404244318 | 0.001293042 | 0.016395083 |
| chr9:6628769-6629063 | -1.745780969 | 0.001293347 | 0.016396228 |
| chr9:114305654-114306165 | 1.142475463 | 0.001294112 | 0.01640322 |
| chr19:33428383-33428781 | 1.424803635 | 0.00129456 | 0.016406178 |
| chr18:39800160-39800476 | -1.296330728 | 0.0012949 | 0.016407781 |
| chr7:92432317-92432901 | -1.281036733 | 0.001295234 | 0.01640929 |

| | | | |
|---|---|---|---|
| chr19:57681584-57682304 | 1.128805434 | 0.001296075 | 0.016415691 |
| chr14:35229496-35229705 | 1.700258801 | 0.001296167 | 0.016415691 |
| chr8:22743719-22744040 | -1.223246255 | 0.00129673 | 0.016418843 |
| chr20:42572515-42572962 | -1.151477224 | 0.001296845 | 0.016418843 |
| chr5:91390327-91390649 | -1.623867046 | 0.001298188 | 0.016433136 |
| chr4:187150687-187151007 | -1.690346858 | 0.001298529 | 0.016434735 |
| chr14:104774464-104774871 | 1.136419472 | 0.001298924 | 0.016436888 |
| chr15:55588010-55588542 | 1.200852756 | 0.001299128 | 0.016436888 |
| chr4:168587326-168587657 | -1.251355939 | 0.001300001 | 0.016445223 |
| chr10:64648979-64649246 | -1.62941281 | 0.001301389 | 0.01645783 |
| chr1:217889451-217889723 | -1.319790075 | 0.001301428 | 0.01645783 |
| chr3:53292413-53292649 | 1.994829953 | 0.001303955 | 0.016487074 |
| chr10:78756652-78757071 | -1.31098044 | 0.001304296 | 0.016488657 |
| chr16:85015805-85016138 | 1.273632689 | 0.001307078 | 0.016521109 |
| chr20:48287879-48288083 | 1.219999866 | 0.001307491 | 0.016523603 |
| chr4:123315592-123316027 | 1.260179656 | 0.001308256 | 0.016530536 |
| chr5:126539097-126539523 | -1.253549415 | 0.001308763 | 0.016533433 |
| chr6:28710423-28710765 | 1.321480766 | 0.001308917 | 0.016533433 |
| chr11:126115770-126116408 | -1.119775701 | 0.001310403 | 0.016549482 |
| chr10:81041680-81041926 | 1.689921065 | 0.001311356 | 0.016558787 |
| chr13:78806532-78806784 | -1.775829787 | 0.001311873 | 0.016562586 |
| chr16:77918753-77919096 | 1.28713034 | 0.001312975 | 0.016573661 |
| chr3:20787092-20787354 | 1.604718466 | 0.001313183 | 0.016573661 |
| chr2:68110182-68110527 | -1.355102414 | 0.001313777 | 0.01657746 |
| chr1:175534163-175534488 | 1.620515529 | 0.00131416 | 0.01657746 |
| chr17:67018582-67018840 | 1.349899979 | 0.00131431 | 0.01657746 |
| chr20:56922513-56922852 | 1.253085962 | 0.001314554 | 0.01657746 |
| chr14:72115884-72116140 | 1.657021959 | 0.001314566 | 0.01657746 |
| chr8:60448281-60448840 | -1.178589811 | 0.001315509 | 0.016586624 |
| chr17:7560447-7560664 | -1.484233922 | 0.001315925 | 0.016589142 |
| chr20:14142190-14142517 | -1.469077443 | 0.001316387 | 0.016592234 |
| chr10:85435864-85436064 | 1.817026565 | 0.001317043 | 0.016597777 |
| chrX:39855739-39856006 | -1.482815427 | 0.001320322 | 0.016635117 |
| chr11:128506980-128507377 | -1.489720152 | 0.00132044 | 0.016635117 |
| chr16:53732562-53732767 | 1.728868793 | 0.001321471 | 0.016645361 |
| chr7:26662699-26662934 | 1.709745924 | 0.001322219 | 0.016651152 |
| chr14:29737628-29737828 | 1.870073303 | 0.001322365 | 0.016651152 |
| chr14:31090823-31091048 | 1.469033863 | 0.001323518 | 0.016662935 |
| chr10:29099013-29099408 | 1.272138375 | 0.001324024 | 0.016663999 |
| chr9:66900479-66901040 | -1.244203098 | 0.001324038 | 0.016663999 |
| chr13:79377909-79378295 | -1.368072799 | 0.001324602 | 0.016668364 |
| chr1:227729051-227729284 | 1.365778403 | 0.001325933 | 0.016682373 |
| chr3:167020577-167020827 | 1.923700482 | 0.001326556 | 0.01668473 |
| chr7:15961310-15961549 | 1.923700482 | 0.001326556 | 0.01668473 |
| chr9:129268759-129268959 | 1.716676843 | 0.001327049 | 0.016688189 |
| chr7:34874887-34875101 | 1.696301771 | 0.001332276 | 0.016748425 |
| chr8:96569304-96569504 | 1.696301771 | 0.001332276 | 0.016748425 |
| chr15:74316459-74316659 | 1.722904159 | 0.00133518 | 0.016782185 |
| chr14:33938393-33938593 | 1.552934995 | 0.001335918 | 0.016788701 |
| chr17:80280797-80281049 | 1.480462588 | 0.001337164 | 0.016801608 |

| | | | |
|---|---|---|---|
| chr3:175089131-175089355 | 1.732364337 | 0.00133776 | 0.016806338 |
| chr5:15284736-15285035 | 1.584405231 | 0.001339486 | 0.016825268 |
| chr16:78117049-78117295 | 1.613846168 | 0.001341096 | 0.016842721 |
| chr5:119578937-119579333 | -1.186721645 | 0.001341915 | 0.016850244 |
| chr14:79388667-79388867 | 1.773874489 | 0.001342649 | 0.0168567 |
| chr16:77603746-77604345 | 1.124361364 | 0.001343209 | 0.016860965 |
| chr5:120817896-120818289 | -1.62775021 | 0.001344452 | 0.016871451 |
| chr7:98802039-98802468 | -1.292704876 | 0.001344484 | 0.016871451 |
| chr3:21117376-21117871 | -1.357695644 | 0.001344954 | 0.016874581 |
| chr16:15154922-15155179 | -1.65792014 | 0.001347284 | 0.016901047 |
| chr18:8389656-8389917 | -1.597830233 | 0.001347529 | 0.016901355 |
| chr12:96254209-96255208 | -1.134742023 | 0.001348782 | 0.016914306 |
| chr9:88544177-88544748 | -1.339324564 | 0.001349257 | 0.016917486 |
| chr3:14573297-14573516 | 1.531026069 | 0.001349772 | 0.016921173 |
| chr5:143124573-143125018 | -1.259547186 | 0.001351009 | 0.016932093 |
| chr13:51844969-51845503 | -1.213829208 | 0.001351085 | 0.016932093 |
| chr20:48816443-48816757 | 1.119156856 | 0.001352506 | 0.016947138 |
| chrX:20514533-20514808 | -1.674555722 | 0.001352903 | 0.016949339 |
| chr10:77773767-77773972 | -1.729825191 | 0.001354218 | 0.016963039 |
| chr4:101954027-101954305 | 1.535750566 | 0.001354682 | 0.016966074 |
| chr8:134665669-134666163 | 1.326543889 | 0.00135709 | 0.016993188 |
| chr3:187955100-187955300 | 1.92225649 | 0.001357296 | 0.016993188 |
| chr1:106936281-106936713 | 1.172160201 | 0.001357591 | 0.016993188 |
| chr11:126992170-126992382 | -1.877867439 | 0.001357734 | 0.016993188 |
| chr5:181099301-181100240 | 1.153407167 | 0.001358599 | 0.017001239 |
| chr2:449536-449792 | -1.694213146 | 0.001358991 | 0.017001847 |
| chr5:72378662-72379270 | -1.262145053 | 0.001359092 | 0.017001847 |
| chr5:90128589-90128909 | -1.618577319 | 0.001360055 | 0.01701112 |
| chr1:107298946-107299221 | 1.205735953 | 0.001361982 | 0.01703244 |
| chr4:163748456-163748809 | -1.396619183 | 0.001362246 | 0.01703297 |
| chr10:121824019-121824302 | 1.462899543 | 0.001362797 | 0.01703707 |
| chr8:38714383-38714665 | 1.426674565 | 0.001363818 | 0.017047055 |
| chr18:50051416-50051632 | -1.877075192 | 0.001364466 | 0.017052375 |
| chr5:123700597-123700953 | -1.621821485 | 0.001365037 | 0.017056734 |
| chr2:179440202-179440438 | -1.766390934 | 0.001365758 | 0.017060812 |
| chr1:170113058-170113844 | -1.115351944 | 0.001365809 | 0.017060812 |
| chr17:78386411-78386611 | 1.724753262 | 0.001366124 | 0.017061967 |
| chr14:31620519-31620767 | 1.634142421 | 0.001366868 | 0.017066324 |
| chr7:86598366-86598613 | -1.613579718 | 0.001366919 | 0.017066324 |
| chr20:47567888-47568291 | 1.058975155 | 0.001367225 | 0.017067235 |
| chr7:150631082-150631297 | 1.688396868 | 0.001367607 | 0.017067235 |
| chr3:83934227-83934896 | -1.12792047 | 0.00136766 | 0.017067235 |
| chr1:226652840-226653040 | 1.63844147 | 0.001369146 | 0.017082995 |
| chr21:35217458-35217904 | 1.335461205 | 0.001369956 | 0.017090319 |
| chrX:149909511-149909711 | 1.764292693 | 0.001370236 | 0.01709103 |
| chr11:84637037-84637451 | -1.398701502 | 0.001370552 | 0.017092191 |
| chr15:90149220-90149582 | 1.340678994 | 0.001371165 | 0.017095221 |
| chr10:106577405-106577634 | -1.624121601 | 0.001371241 | 0.017095221 |
| chr5:88145592-88145903 | 1.275958656 | 0.001371853 | 0.017099064 |
| chr7:139758450-139758822 | -1.406572021 | 0.001372128 | 0.017099064 |

| | | | |
|---|---|---|---|
| chr7:116236070-116236323 | -1.668220623 | 0.001372219 | 0.017099064 |
| chr13:33892533-33892923 | -1.396566383 | 0.001372664 | 0.017101836 |
| chr19:36170800-36171043 | 1.654534633 | 0.001373559 | 0.017110199 |
| chr22:31436243-31436542 | 1.654669707 | 0.001373862 | 0.017111189 |
| chr22:31530541-31530746 | 1.803827245 | 0.00137449 | 0.017115209 |
| chr3:9507673-9507873 | 1.530295335 | 0.001374631 | 0.017115209 |
| chr2:28074828-28075028 | -2.1000153 | 0.001375059 | 0.017115748 |
| chr16:23486510-23486892 | 1.252964614 | 0.001375227 | 0.017115748 |
| chr8:100238485-100238692 | 1.606225075 | 0.001375345 | 0.017115748 |
| chr1:46549653-46549884 | -1.546109479 | 0.001377064 | 0.017130462 |
| chr1:217315583-217315792 | -1.546109479 | 0.001377064 | 0.017130462 |
| chr8:136082679-136083375 | -1.229081363 | 0.001377198 | 0.017130462 |
| chr1:228185479-228185812 | 1.476693266 | 0.001377852 | 0.017135816 |
| chr4:828962-829248 | 1.367320459 | 0.001378785 | 0.017144634 |
| chr7:23055693-23055946 | 1.371936502 | 0.001380459 | 0.017162675 |
| chr7:47002275-47002475 | 2.061765983 | 0.0013814 | 0.017169785 |
| chr12:31820882-31821220 | -1.542859025 | 0.00138148 | 0.017169785 |
| chr20:50283860-50284626 | -1.076389552 | 0.001382011 | 0.017173608 |
| chr14:95432251-95432491 | 1.424466885 | 0.001382325 | 0.017174727 |
| chr20:47610915-47611215 | 1.089993953 | 0.001382912 | 0.017179231 |
| chr14:32549689-32549894 | 1.722512403 | 0.001383719 | 0.017186465 |
| chr20:41670845-41671045 | 1.685051856 | 0.001386263 | 0.017215275 |
| chr3:27507940-27508214 | 1.465821832 | 0.001387477 | 0.017227555 |
| chr1:116888137-116888373 | 1.498467668 | 0.001388186 | 0.017233566 |
| chr13:46866111-46866600 | -1.095080962 | 0.00138869 | 0.01723703 |
| chr15:65466562-65466872 | 1.607612934 | 0.001389696 | 0.01724672 |
| chr10:5090326-5090596 | 1.597584278 | 0.001391451 | 0.017265712 |
| chr3:129936894-129937149 | -1.502435081 | 0.001393137 | 0.01728383 |
| chr19:2014391-2014665 | 1.685289838 | 0.001393905 | 0.017289628 |
| chr7:7667039-7667244 | 1.656849529 | 0.001394077 | 0.017289628 |
| chr16:70288123-70288461 | 1.265505061 | 0.001394281 | 0.017289628 |
| chr6:96450140-96450838 | -1.143495069 | 0.001395166 | 0.017297807 |
| chr7:11786041-11786589 | 1.122370267 | 0.00139689 | 0.017316372 |
| chr2:163878804-163879059 | -1.496899599 | 0.001397232 | 0.017317816 |
| chr1:155044918-155045421 | 1.172044333 | 0.001398995 | 0.017336854 |
| chr6:10451871-10452232 | -1.500895242 | 0.001399656 | 0.017342251 |
| chr10:43293212-43293486 | -1.483756093 | 0.001400702 | 0.017352393 |
| chr6:166949642-166949903 | -1.704293183 | 0.001401855 | 0.017363878 |
| chr5:171451176-171451376 | 1.656114909 | 0.001402392 | 0.017367717 |
| chr20:47542750-47543410 | 1.036320667 | 0.001403095 | 0.017373612 |
| chr3:19688480-19688835 | 1.397356987 | 0.001403563 | 0.017376603 |
| chr18:76380829-76381201 | -1.2247274 | 0.001404065 | 0.017380008 |
| chr12:9372340-9372563 | -1.726586165 | 0.001404667 | 0.017384659 |
| chr17:62853767-62853996 | 1.476647804 | 0.001405279 | 0.017389423 |
| chr3:140910895-140911248 | -1.455114517 | 0.001405748 | 0.0173911 |
| chr6:154284625-154284907 | -1.852097813 | 0.001405869 | 0.0173911 |
| chr7:22594042-22594437 | 1.255696981 | 0.00140842 | 0.017419845 |
| chr14:24433080-24433345 | 1.224362813 | 0.001408675 | 0.017420185 |
| chr15:78209379-78209685 | 1.518707296 | 0.001410068 | 0.017434602 |
| chr8:110651216-110651430 | 1.561920997 | 0.00141047 | 0.017436762 |

| | | | |
|---|---|---|---|
| chr1:65550684-65550948 | -1.770759657 | 0.001410705 | 0.017436852 |
| chr2:25135182-25135433 | 1.622003323 | 0.001411861 | 0.017448326 |
| chr12:109951711-109952153 | -1.261223169 | 0.001412132 | 0.017448864 |
| chr21:31236062-31236479 | 1.297041853 | 0.001412374 | 0.017449038 |
| chr15:54699050-54699421 | 1.421679584 | 0.001412836 | 0.017451933 |
| chr15:91328687-91328887 | 1.858089844 | 0.001415284 | 0.017477062 |
| chr12:72699384-72699755 | -1.392896011 | 0.001415512 | 0.017477062 |
| chr2:55824598-55824992 | -1.517494895 | 0.001415555 | 0.017477062 |
| chr12:102051425-102051846 | 1.199663566 | 0.001416461 | 0.017484571 |
| chr10:59633144-59633374 | -1.737370364 | 0.001416619 | 0.017484571 |
| chr4:124884438-124884833 | -1.427033911 | 0.001417283 | 0.017489942 |
| chr10:59830669-59831036 | -1.289621888 | 0.001418054 | 0.017496646 |
| chr12:52926091-52926579 | 1.15649483 | 0.00142187 | 0.0175409 |
| chr1:6625825-6626045 | 1.605480983 | 0.001422456 | 0.017545303 |
| chr7:20365549-20365784 | 1.74560709 | 0.001423273 | 0.017552557 |
| chr7:132830759-132831186 | -1.524632778 | 0.001423555 | 0.01755265 |
| chr15:75709345-75709588 | 1.410683045 | 0.001423739 | 0.01755265 |
| chr1:154908839-154909223 | 1.431503861 | 0.00142441 | 0.017558097 |
| chr7:19464331-19464577 | 1.634970757 | 0.001425826 | 0.01757273 |
| chr4:163702021-163702294 | -1.744532832 | 0.001426287 | 0.017575585 |
| chr12:118102083-118102305 | 1.476915979 | 0.001426617 | 0.017575896 |
| chr17:61723074-61723619 | 1.037248495 | 0.001426771 | 0.017575896 |
| chr2:189412431-189412715 | -1.637237948 | 0.001427739 | 0.017584077 |
| chr1:212027815-212028015 | -2.179443733 | 0.001428109 | 0.017584077 |
| chr1:64996901-64997217 | -1.565316022 | 0.001428304 | 0.017584077 |
| chr21:8987983-8988357 | -1.165750177 | 0.001428488 | 0.017584077 |
| chr4:53702353-53702863 | -1.29917527 | 0.001428775 | 0.017584077 |
| chr1:151865093-151865299 | 1.446962739 | 0.001428812 | 0.017584077 |
| chr5:16700951-16701327 | 1.426988915 | 0.001429685 | 0.017591988 |
| chr8:98799505-98799831 | 1.442642883 | 0.001430297 | 0.017593996 |
| chr4:117311240-117311440 | 1.755287941 | 0.001430307 | 0.017593996 |
| chr7:125057969-125058245 | -1.707291513 | 0.001430818 | 0.017597458 |
| chr9:12666924-12667127 | -1.916329032 | 0.001432268 | 0.017612465 |
| chr14:74663350-74663633 | 1.583128023 | 0.001435009 | 0.01764333 |
| chr15:71279150-71279563 | 1.326882426 | 0.001436793 | 0.01766118 |
| chr17:7324207-7324438 | -1.565725294 | 0.001436921 | 0.01766118 |
| chr14:105281977-105282242 | 1.488185443 | 0.001437597 | 0.01766665 |
| chr3:149222217-149222450 | 1.617988618 | 0.001438061 | 0.017669523 |
| chr10:44339226-44339656 | -1.29918814 | 0.001438406 | 0.017670921 |
| chr6:10314513-10314879 | 1.299068482 | 0.001442782 | 0.017720868 |
| chr11:35927756-35928125 | -1.341111194 | 0.001442964 | 0.017720868 |
| chr5:122129408-122129661 | -1.641473476 | 0.001443394 | 0.017720868 |
| chr2:241522416-241522658 | 1.750517245 | 0.001443397 | 0.017720868 |
| chrX:153353247-153353578 | 1.598435739 | 0.001445591 | 0.017744962 |
| chr20:42982211-42982761 | -1.297889599 | 0.001445994 | 0.017747067 |
| chr16:46389614-46390095 | 1.053756734 | 0.001447345 | 0.017760801 |
| chr4:41358678-41358918 | 1.665205884 | 0.001448335 | 0.017762446 |
| chr14:71727287-71727492 | 1.621167874 | 0.001448491 | 0.017762446 |
| chr14:34180470-34180714 | 1.533344068 | 0.00144854 | 0.017762446 |
| chr8:22748172-22748730 | -1.156814079 | 0.001448751 | 0.017762446 |

| | | | |
|---|---|---|---|
| chr1:107091609-107091809 | 1.265150228 | 0.00144882 | 0.017762446 |
| chr7:134632766-134633301 | -1.479217111 | 0.00144887 | 0.017762446 |
| chr3:16148976-16149176 | 1.715280129 | 0.001450479 | 0.017778315 |
| chr14:52373489-52373848 | -1.369510561 | 0.001450628 | 0.017778315 |
| chr22:30802612-30803250 | 1.145169849 | 0.001451334 | 0.017784118 |
| chr7:101085039-101086046 | -1.063868344 | 0.001452177 | 0.017788487 |
| chr3:182226702-182227326 | -1.489525198 | 0.001452359 | 0.017788487 |
| chr19:57544266-57544466 | 1.84687464 | 0.001452572 | 0.017788487 |
| chr1:108520123-108521045 | -1.090469356 | 0.00145296 | 0.017788487 |
| chr14:105589385-105589660 | 1.601311994 | 0.001452978 | 0.017788487 |
| chr1:168631531-168632071 | 1.204814385 | 0.001453086 | 0.017788487 |
| chr1:106453798-106454131 | 1.191802682 | 0.001453315 | 0.017788487 |
| chr8:106966219-106966419 | 1.973383361 | 0.001453614 | 0.017789295 |
| chr1:212760180-212760516 | -1.406483959 | 0.001454748 | 0.017800332 |
| chr16:56610111-56610702 | -1.197249314 | 0.001455995 | 0.017812697 |
| chr5:66765012-66765252 | -1.650826191 | 0.001456223 | 0.017812697 |
| chr21:44836521-44836941 | 1.224875809 | 0.001457384 | 0.017824056 |
| chr7:135976694-135977860 | -1.035000602 | 0.001458714 | 0.017837465 |
| chr14:34228052-34228344 | 1.227191059 | 0.001459389 | 0.01784288 |
| chr20:45771239-45771500 | 1.339991104 | 0.001460633 | 0.017855233 |
| chr7:3482313-3482822 | -1.288152337 | 0.001461188 | 0.017858761 |
| chr20:45910632-45911321 | 1.072576454 | 0.001461485 | 0.017858761 |
| chr1:236336175-236336502 | 1.316494563 | 0.001461621 | 0.017858761 |
| chr20:19783896-19784450 | 1.090690155 | 0.001462877 | 0.017871259 |
| chr20:57024585-57024868 | 1.301267726 | 0.001465782 | 0.017902527 |
| chr20:48794223-48794465 | 1.136378076 | 0.001465903 | 0.017902527 |
| chr2:188396336-188396687 | -1.350917314 | 0.001466314 | 0.017904682 |
| chrX:33725596-33726518 | -1.145420299 | 0.001468228 | 0.017925198 |
| chrX:80184874-80185156 | -1.628769425 | 0.001469468 | 0.017937478 |
| chr21:34959842-34960042 | 1.839797817 | 0.001469931 | 0.017938484 |
| chr3:120573275-120573664 | -1.468443956 | 0.001470018 | 0.017938484 |
| chr15:79983125-79983438 | 1.506075932 | 0.001470518 | 0.017941721 |
| chr5:57037374-57037730 | -1.324464181 | 0.001473016 | 0.01796934 |
| chr12:117142919-117143539 | 1.102137203 | 0.001473473 | 0.017970422 |
| chr2:158184727-158185346 | 1.123936672 | 0.001473574 | 0.017970422 |
| chr7:122902711-122903083 | -1.245650421 | 0.001474983 | 0.017984743 |
| chr7:48459069-48459589 | -1.16209176 | 0.001477132 | 0.018008084 |
| chr1:3127301-3127501 | 1.677199752 | 0.00147775 | 0.018012746 |
| chr3:10758589-10759095 | 1.101727382 | 0.001478175 | 0.018015069 |
| chrX:140933122-140933436 | 1.438292603 | 0.001478548 | 0.01801675 |
| chr8:62164635-62165615 | -1.093140251 | 0.00147937 | 0.018023755 |
| chr3:62574629-62574832 | 1.271474028 | 0.001479594 | 0.018023755 |
| chr16:1689269-1689609 | 1.47890288 | 0.001481369 | 0.018042514 |
| chr12:13255738-13256080 | -1.247295295 | 0.00148205 | 0.018045124 |
| chr17:38492116-38492340 | 1.652203156 | 0.001482167 | 0.018045124 |
| chr4:2914716-2914967 | 1.969894213 | 0.001482324 | 0.018045124 |
| chr3:18767856-18768247 | 1.388349497 | 0.001482525 | 0.018045124 |
| chr17:68627851-68628436 | -1.123727597 | 0.001484025 | 0.018056512 |
| chr7:18778641-18778846 | 1.643239374 | 0.001484133 | 0.018056512 |
| chr15:56505832-56506034 | 1.669147051 | 0.001484168 | 0.018056512 |

| | | | |
|---|---|---|---|
| chr8:129973100-129973481 | 1.337628201 | 0.001486817 | 0.018085865 |
| chr6:34202376-34203174 | -1.183938956 | 0.001487512 | 0.018091449 |
| chr14:75410446-75410678 | 1.613591108 | 0.001488798 | 0.01810422 |
| chr5:122330909-122331205 | -1.642503367 | 0.001489087 | 0.018104856 |
| chr5:53874114-53874844 | -1.243928224 | 0.001489422 | 0.018106055 |
| chr6:168667068-168667577 | -1.447930464 | 0.00149009 | 0.018111312 |
| chr2:127128456-127129015 | -1.240513041 | 0.001490397 | 0.018112162 |
| chr3:14405953-14406191 | 1.207819685 | 0.001490676 | 0.018112677 |
| chr3:181868585-181868894 | 1.410882808 | 0.001492279 | 0.018129285 |
| chr18:63306381-63306654 | -1.638842751 | 0.001493006 | 0.018135236 |
| chr5:4700272-4700490 | -1.796031403 | 0.001493503 | 0.018138405 |
| chr10:37007912-37008800 | -1.142565162 | 0.001494196 | 0.018143947 |
| chr12:112630666-112630960 | 1.525799909 | 0.001494561 | 0.018145504 |
| chr2:160583146-160583447 | 1.482569995 | 0.001495171 | 0.018150035 |
| chr22:45757018-45757442 | -1.579942895 | 0.001495704 | 0.018153048 |
| chr11:47916543-47916867 | 1.343751236 | 0.001495893 | 0.018153048 |
| chr12:124381668-124382115 | -1.259498848 | 0.001496744 | 0.018159848 |
| chr5:179888475-179888695 | 1.500140759 | 0.001496928 | 0.018159848 |
| chr8:128721704-128722202 | 1.371635555 | 0.001497573 | 0.018162326 |
| chrX:150694300-150694614 | 1.208554798 | 0.001497606 | 0.018162326 |
| chr8:79351014-79351308 | 1.542406408 | 0.001499876 | 0.018186978 |
| chr3:115512465-115513412 | -1.144933233 | 0.001502307 | 0.018213566 |
| chr16:27324772-27324981 | 1.721617146 | 0.001502573 | 0.018213918 |
| chr5:181233836-181234097 | 1.325661916 | 0.001502994 | 0.018215803 |
| chr7:101321257-101322360 | -1.109374956 | 0.001503204 | 0.018215803 |
| chr18:66523293-66524077 | -1.112574927 | 0.00150419 | 0.018224862 |
| chr14:72702155-72702355 | 1.673576005 | 0.001505799 | 0.018241481 |
| chr2:241101477-241101706 | 1.510952993 | 0.00150637 | 0.018245513 |
| chr1:219517152-219517418 | -1.383973824 | 0.00150817 | 0.018264427 |
| chr10:55410611-55410943 | -1.535239344 | 0.001509622 | 0.018279113 |
| chr8:115495239-115495439 | -2.066178457 | 0.001510327 | 0.018284764 |
| chr10:12690668-12691174 | -1.403296952 | 0.001510907 | 0.018288901 |
| chr8:85836641-85837219 | -1.150197371 | 0.001511888 | 0.018297877 |
| chr4:113328624-113328832 | -1.987102794 | 0.001513 | 0.018308443 |
| chr10:34739805-34740234 | -1.261458381 | 0.001513631 | 0.018313195 |
| chr8:108740111-108740339 | 1.742352637 | 0.001516085 | 0.018339982 |
| chr4:36824772-36825021 | -1.854159145 | 0.001516661 | 0.018344061 |
| chr12:87086265-87086475 | 1.724403108 | 0.001517199 | 0.018347663 |
| chr7:46695126-46695446 | 1.310735311 | 0.001519694 | 0.01837494 |
| chr13:24651809-24652154 | 1.310482675 | 0.001520207 | 0.018378238 |
| chr20:18943381-18943672 | 1.615158554 | 0.001520963 | 0.018384483 |
| chr17:50290125-50290345 | 1.540348519 | 0.00152232 | 0.018397984 |
| chr2:35724785-35725123 | -1.576576653 | 0.001522756 | 0.018400351 |
| chr14:19918042-19918470 | -1.364341588 | 0.001523552 | 0.018407067 |
| chr16:25119593-25120008 | 1.210800962 | 0.001525273 | 0.018421335 |
| chr12:89237069-89237455 | 1.376561996 | 0.001525324 | 0.018421335 |
| chr1:187862528-187862946 | -1.28713286 | 0.001525454 | 0.018421335 |
| chrX:123779994-123780305 | 1.267179704 | 0.001526026 | 0.018425336 |
| chr18:34809765-34810150 | -1.320906078 | 0.001527199 | 0.018436593 |
| chr7:105531452-105531821 | -1.165931784 | 0.001527778 | 0.018440674 |

| | | | |
|---|---|---|---|
| chr13:57716339-57716625 | -1.537322663 | 0.001528585 | 0.018447508 |
| chr6:111788761-111788961 | -2.072495849 | 0.001529053 | 0.018450246 |
| chr7:47991673-47992428 | -1.098249286 | 0.001530677 | 0.01846694 |
| chr1:199814435-199814692 | 1.701966009 | 0.001531352 | 0.01847218 |
| chr14:42994150-42994509 | 1.358947146 | 0.00153217 | 0.01847913 |
| chr2:9594743-9595024 | -1.683270428 | 0.001533131 | 0.018484803 |
| chr19:34677202-34677931 | 1.08980538 | 0.001533291 | 0.018484803 |
| chr3:64817110-64817343 | 1.154364747 | 0.001533587 | 0.018484803 |
| chr7:101017021-101017321 | -1.542004329 | 0.001533605 | 0.018484803 |
| chr3:84586416-84586844 | -1.503167001 | 0.001535879 | 0.018509296 |
| chr10:28180707-28181037 | -1.440746412 | 0.00153625 | 0.01851086 |
| chr17:62002550-62003533 | 1.022250944 | 0.001536997 | 0.018510931 |
| chr14:76640971-76641282 | 1.413084395 | 0.001537469 | 0.018510931 |
| chr5:111830465-111830764 | -1.440492458 | 0.001537621 | 0.018510931 |
| chr14:36136256-36136613 | 1.518167792 | 0.001537775 | 0.018510931 |
| chr4:82525360-82526096 | -1.116144075 | 0.00153778 | 0.018510931 |
| chr11:94301147-94301359 | -1.857143801 | 0.001538535 | 0.018510931 |
| chr14:80585221-80585730 | -1.284798191 | 0.001538867 | 0.018510931 |
| chr17:59756386-59756793 | 1.164440076 | 0.001539003 | 0.018510931 |
| chr5:104652344-104652832 | -1.333213821 | 0.001539022 | 0.018510931 |
| chr13:93325522-93326810 | -1.110065283 | 0.001539034 | 0.018510931 |
| chr11:30366040-30366367 | -1.670217058 | 0.001539127 | 0.018510931 |
| chr7:123720630-123720830 | -1.966828728 | 0.001539155 | 0.018510931 |
| chr7:131069272-131069526 | -1.402616147 | 0.001540181 | 0.01851759 |
| chr19:52296825-52297299 | 1.262272935 | 0.001540192 | 0.01851759 |
| chr2:101341490-101342113 | -1.204519869 | 0.001541293 | 0.018527915 |
| chr7:140316082-140316298 | -1.884881167 | 0.001541619 | 0.018528927 |
| chr2:78115804-78116032 | -1.762619425 | 0.001542874 | 0.018541112 |
| chr5:93899079-93899351 | -1.573258494 | 0.001544608 | 0.018559034 |
| chr14:67547252-67547688 | 1.16575736 | 0.001545597 | 0.018566451 |
| chr9:137899643-137899923 | 1.664075923 | 0.00154571 | 0.018566451 |
| chr5:101101765-101101965 | -1.911011582 | 0.001546213 | 0.018569288 |
| chr2:86556502-86556728 | 1.900745233 | 0.001546431 | 0.018569288 |
| chr4:187944166-187944398 | -1.985774849 | 0.001549245 | 0.01860016 |
| chr7:127550362-127550562 | -1.897480022 | 0.001550503 | 0.018612356 |
| chr14:39627172-39627470 | 1.466828577 | 0.00155106 | 0.01861612 |
| chr12:122910079-122910441 | -1.669611323 | 0.001552458 | 0.018628137 |
| chr8:104575319-104575580 | 1.53753322 | 0.001552547 | 0.018628137 |
| chr1:99659723-99660228 | -1.295629005 | 0.001553041 | 0.018631142 |
| chr2:41442003-41442286 | -1.462652451 | 0.001554359 | 0.018644029 |
| chr5:43455197-43455397 | 1.849165533 | 0.001554655 | 0.018644672 |
| chr7:29609821-29610070 | -1.909932131 | 0.001556273 | 0.018661152 |
| chr10:50310770-50311159 | -1.231711279 | 0.001556664 | 0.018662913 |
| chr7:16098683-16098896 | 1.480087602 | 0.001556916 | 0.018663015 |
| chr7:140924077-140925357 | -1.031084595 | 0.001559022 | 0.018685272 |
| chr9:129873791-129873991 | 1.755417872 | 0.00155926 | 0.018685272 |
| chr3:167927973-167928348 | -1.21397015 | 0.001561076 | 0.018704103 |
| chr1:174897478-174897678 | 1.855466815 | 0.001562534 | 0.018718653 |
| chr21:9246280-9248665 | 1.01329408 | 0.001565679 | 0.018753393 |
| chr4:1400934-1401182 | 1.729510838 | 0.001566131 | 0.018753604 |

| | | | |
|---|---|---|---|
| chr16:29779471-29779776 | 1.255510178 | 0.001566186 | 0.018753604 |
| chr4:173536924-173537207 | -1.419177784 | 0.001569481 | 0.018790123 |
| chr2:105888628-105889319 | -1.163150886 | 0.001570146 | 0.018795146 |
| chr9:107527179-107527531 | 1.564790086 | 0.001571033 | 0.018802831 |
| chr7:24201316-24201516 | 1.620226601 | 0.001572066 | 0.018812254 |
| chr5:66666914-66667137 | -1.676750224 | 0.001575335 | 0.018845984 |
| chr13:54573223-54573715 | -1.21319133 | 0.001575377 | 0.018845984 |
| chr9:114619408-114619655 | 1.451468578 | 0.001575959 | 0.018848474 |
| chr15:92771965-92772166 | 1.703112687 | 0.001576077 | 0.018848474 |
| chr13:50912066-50912367 | -1.350356195 | 0.001578657 | 0.018876387 |
| chr7:96303024-96304048 | -1.116652103 | 0.001579409 | 0.018882423 |
| chr7:140645208-140645426 | -1.685725663 | 0.001580355 | 0.018888976 |
| chr5:135139921-135140309 | -1.395927136 | 0.00158045 | 0.018888976 |
| chr1:210265037-210265312 | -1.358013912 | 0.001586188 | 0.018952799 |
| chr15:53910820-53911174 | 1.174406417 | 0.001586285 | 0.018952799 |
| chrX:155612375-155613118 | 1.102705481 | 0.001586773 | 0.018955683 |
| chr1:19485431-19486132 | 1.104408437 | 0.001587335 | 0.018959433 |
| chr20:48485203-48485516 | 1.123936106 | 0.001588403 | 0.018969237 |
| chr4:86973103-86973399 | -1.249886268 | 0.001589203 | 0.018975826 |
| chrX:150544289-150544489 | 1.608295229 | 0.001589927 | 0.018981514 |
| chr4:56681152-56681636 | 1.151618753 | 0.001590603 | 0.018986627 |
| chr7:91394808-91395543 | -1.210059133 | 0.00159125 | 0.018991394 |
| chr7:13047121-13047321 | 1.581179574 | 0.001593691 | 0.019017564 |
| chr3:59654182-59654397 | 1.733824793 | 0.001594076 | 0.019019199 |
| chr3:66442301-66442650 | -1.485952533 | 0.00159533 | 0.019031195 |
| chr4:114012589-114013527 | -1.098698083 | 0.001596406 | 0.019041072 |
| chrX:153598532-153598852 | 1.271942591 | 0.0015968 | 0.019042803 |
| chr16:78060355-78060560 | 1.56270318 | 0.001597832 | 0.019052144 |
| chr2:44115222-44115422 | 1.898354029 | 0.001598111 | 0.019052505 |
| chr22:18087577-18087819 | 1.83764448 | 0.00159935 | 0.019064308 |
| chr2:36373249-36373465 | -1.527187112 | 0.001600567 | 0.019075857 |
| chr14:105564159-105564471 | 1.510555811 | 0.00160094 | 0.01907733 |
| chr16:48489381-48489581 | 1.907123011 | 0.001601748 | 0.019083103 |
| chr2:39920142-39920727 | -1.231201156 | 0.001601923 | 0.019083103 |
| chr5:174831982-174832317 | 1.636777813 | 0.001602727 | 0.019089715 |
| chr13:85799100-85799730 | -1.091299379 | 0.001603688 | 0.019098194 |
| chr5:155085658-155086057 | 1.199795811 | 0.001604144 | 0.019100658 |
| chr12:29073716-29073993 | -1.524826733 | 0.001607796 | 0.019141172 |
| chr7:158802925-158803547 | -1.201316184 | 0.001609591 | 0.019159553 |
| chr4:125096598-125097010 | -1.496266572 | 0.001611932 | 0.019184448 |
| chr2:35428587-35428886 | -1.403896953 | 0.001613441 | 0.019199422 |
| chr10:94123351-94123662 | 1.370707929 | 0.001614188 | 0.019204185 |
| chr5:103041108-103041355 | -1.514735776 | 0.001614343 | 0.019204185 |
| chrX:153667854-153668254 | 1.183849241 | 0.001615044 | 0.019209547 |
| chr11:101436584-101436802 | -1.758827231 | 0.001616669 | 0.019225893 |
| chr9:121147508-121147899 | 1.514743843 | 0.001617322 | 0.019230669 |
| chr5:181150209-181150409 | 1.736092552 | 0.001617655 | 0.019231651 |
| chr1:35589283-35589483 | 1.671850571 | 0.001618901 | 0.019240954 |
| chr16:68417621-68418305 | 1.149728382 | 0.00161894 | 0.019240954 |
| chr2:205629478-205629791 | -1.520370664 | 0.001619982 | 0.019250355 |

| | | | |
|---|---|---|---|
| chr21:45329615-45330025 | 1.156415379 | 0.001620622 | 0.019250577 |
| chr20:48576983-48577236 | 1.116960041 | 0.001620652 | 0.019250577 |
| chr7:108109342-108109891 | -1.209749561 | 0.001620755 | 0.019250577 |
| chr1:175621082-175621301 | 1.733948534 | 0.001623367 | 0.019278611 |
| chr20:46013236-46013472 | 1.438116127 | 0.001624152 | 0.019284951 |
| chr13:48873697-48873906 | -1.796990596 | 0.001624979 | 0.019291783 |
| chr3:64317120-64317493 | 1.089178363 | 0.001626517 | 0.019307043 |
| chr7:51225130-51225609 | -1.189446092 | 0.00162726 | 0.019312873 |
| chr3:75717982-75718182 | 1.643202798 | 0.00162828 | 0.019321983 |
| chr12:122087914-122088114 | 1.835858352 | 0.00162867 | 0.019323618 |
| chr10:52637689-52638042 | -1.640778452 | 0.001629115 | 0.019325911 |
| chr20:48898248-48898448 | 1.161018789 | 0.001630444 | 0.019338685 |
| chr7:101699008-101699218 | -1.736586513 | 0.001631157 | 0.019344141 |
| chr13:68304400-68304819 | -1.498801611 | 0.001631564 | 0.019345807 |
| chr3:72591592-72592118 | -1.24071297 | 0.001631802 | 0.019345807 |
| chr7:111192554-111192833 | -1.620707874 | 0.001632884 | 0.019355634 |
| chr7:70727016-70727314 | -1.496985167 | 0.00163452 | 0.019372027 |
| chr5:86567823-86568175 | -1.450272459 | 0.001635391 | 0.019379362 |
| chr18:77634607-77634807 | -2.079986623 | 0.001635956 | 0.019383058 |
| chr2:99636649-99637021 | -1.424787311 | 0.001637726 | 0.019401024 |
| chr2:27973704-27973904 | -1.841645691 | 0.001638504 | 0.019407243 |
| chr7:15785432-15785641 | 1.707860435 | 0.001638891 | 0.019408825 |
| chr3:16068140-16068515 | 1.260654708 | 0.001639237 | 0.019409921 |
| chr9:135480471-135480860 | 1.249358571 | 0.001640312 | 0.019419655 |
| chr4:133584712-133585101 | -1.413469157 | 0.001640623 | 0.019420329 |
| chr5:32942536-32942820 | -1.595333701 | 0.001641301 | 0.019425356 |
| chr13:74575017-74575276 | -1.598117064 | 0.00164243 | 0.019435716 |
| chr7:103074143-103075926 | -1.001777599 | 0.001643091 | 0.019440543 |
| chr7:139732153-139732570 | -1.191466196 | 0.001646793 | 0.019481326 |
| chr1:113255386-113256018 | 1.166134224 | 0.001648932 | 0.019503619 |
| chr7:100082587-100082854 | -1.470268594 | 0.001650536 | 0.019519587 |
| chr1:54854632-54854870 | 1.471801157 | 0.001651896 | 0.01953011 |
| chr7:32676248-32676450 | 1.60205935 | 0.001651936 | 0.01953011 |
| chr10:15725520-15726031 | -1.194524372 | 0.001654433 | 0.01955661 |
| chr8:80366763-80367714 | -1.044286574 | 0.001655231 | 0.019563024 |
| chr7:135335650-135336061 | -1.138480583 | 0.001655848 | 0.019567297 |
| chr9:120393708-120394053 | -1.323609599 | 0.001656284 | 0.019569439 |
| chr10:30645323-30645771 | -1.363653716 | 0.001658848 | 0.019595457 |
| chr2:26777879-26778129 | 1.553768198 | 0.001659254 | 0.019595457 |
| chr3:129015213-129015413 | 1.553768198 | 0.001659254 | 0.019595457 |
| chr15:41589659-41589862 | 1.800689759 | 0.001659775 | 0.019597694 |
| chr12:103575270-103576069 | -1.065712206 | 0.001659954 | 0.019597694 |
| chr3:189944252-189945022 | -1.068671684 | 0.001660441 | 0.019600423 |
| chr14:35877378-35877606 | 1.567681204 | 0.001662917 | 0.019626623 |
| chr20:51833288-51833761 | 1.070130604 | 0.001663735 | 0.019633257 |
| chr8:132893388-132894340 | -1.098111714 | 0.001666226 | 0.019658589 |
| chr22:46704968-46705198 | 1.699127785 | 0.001666395 | 0.019658589 |
| chr7:134845543-134845865 | -1.408043384 | 0.00166774 | 0.019671428 |
| chr12:109116252-109116522 | 1.371737667 | 0.001668029 | 0.019671809 |
| chr3:141287361-141287578 | 1.828474256 | 0.001668359 | 0.019672665 |

| | | | |
|---|---|---|---|
| chr18:27860123-27860418 | 1.540448503 | 0.001668976 | 0.019676763 |
| chr3:159045000-159045916 | -1.056897016 | 0.00166922 | 0.019676763 |
| chr14:44795797-44796016 | 1.577158645 | 0.001672232 | 0.019708636 |
| chr7:16615266-16615503 | 1.59675036 | 0.001672756 | 0.019708636 |
| chr7:116354697-116355389 | -1.159874699 | 0.001672908 | 0.019708636 |
| chr3:189937280-189938131 | -1.1347681 | 0.001672953 | 0.019708636 |
| chr3:15117647-15117847 | 1.563621116 | 0.001673399 | 0.019710866 |
| chr7:139340943-139341919 | -1.013037783 | 0.001673757 | 0.019712051 |
| chr17:77056278-77056559 | 1.3575149 | 0.001674404 | 0.01971664 |
| chr14:43679985-43680226 | 1.715574813 | 0.001677951 | 0.019755371 |
| chr9:136305927-136306351 | 1.28319314 | 0.00167856 | 0.019759502 |
| chr10:87589561-87589854 | -1.568601221 | 0.001679474 | 0.019765603 |
| chr8:121724190-121724894 | -1.062149295 | 0.001679594 | 0.019765603 |
| chrX:47781793-47782078 | -1.883929334 | 0.001680714 | 0.019775742 |
| chr3:121099236-121099483 | 1.778250741 | 0.001681011 | 0.019776209 |
| chr11:76903380-76903683 | -1.463122526 | 0.001682712 | 0.019790164 |
| chr5:5271125-5271557 | -1.249246003 | 0.001682714 | 0.019790164 |
| chr18:74541942-74542760 | -1.144606942 | 0.001683994 | 0.019801406 |
| chr8:8289212-8289748 | -1.367318723 | 0.001684187 | 0.019801406 |
| chr8:101409817-101410573 | 1.084836329 | 0.001684568 | 0.019802845 |
| chr9:893742-894319 | 1.142253885 | 0.001686147 | 0.019816368 |
| chr1:198672194-198672498 | 1.396636082 | 0.001686235 | 0.019816368 |
| chr20:48043204-48043427 | 1.145749275 | 0.00168777 | 0.019831174 |
| chr14:79317490-79317690 | 1.897079717 | 0.001688013 | 0.019831174 |
| chr7:134141250-134141593 | -1.435720045 | 0.001688562 | 0.019834583 |
| chr18:9694860-9695286 | 1.36453294 | 0.001689625 | 0.019844035 |
| chr17:57603976-57604514 | 1.106740422 | 0.001690521 | 0.019851514 |
| chr15:91954092-91954864 | -1.084280647 | 0.001691617 | 0.019861336 |
| chr14:88483518-88483908 | 1.374967972 | 0.001694034 | 0.019886671 |
| chr9:32743530-32744239 | -1.21011491 | 0.001694407 | 0.019887997 |
| chr21:15706712-15706987 | 1.434292681 | 0.00169587 | 0.019902119 |
| chr8:141698491-141698730 | 1.486023069 | 0.001696865 | 0.019910749 |
| chr3:61619860-61620350 | 1.133246984 | 0.00169815 | 0.019922777 |
| chr15:89866380-89866650 | 1.82857176 | 0.001700054 | 0.019942055 |
| chr5:42078619-42078985 | -1.430888782 | 0.001702067 | 0.019962616 |
| chrX:20056708-20057123 | -1.237790513 | 0.001704023 | 0.019982504 |
| chr11:70805876-70806331 | -1.224029106 | 0.001704923 | 0.019989991 |
| chr7:140696433-140697622 | -1.015963325 | 0.001705797 | 0.019997182 |
| chr4:92051452-92051896 | 1.225012664 | 0.001706445 | 0.020001723 |
| chr3:160685157-160685434 | -1.479294338 | 0.001706853 | 0.020003441 |
| chr16:27597507-27597783 | 1.646722968 | 0.001707123 | 0.020003545 |
| chr5:122295179-122295496 | -1.431805176 | 0.001707755 | 0.020007885 |
| chr1:219317796-219318118 | -1.191240618 | 0.00170906 | 0.020020123 |
| chr21:31052896-31053108 | 1.824213347 | 0.00170991 | 0.020026693 |
| chr15:84665987-84666393 | 1.392918162 | 0.001710144 | 0.020026693 |
| chr4:40576803-40577519 | -1.108142558 | 0.001710887 | 0.020029394 |
| chr7:117505324-117505572 | -1.61357428 | 0.001710897 | 0.020029394 |
| chr15:98727503-98727934 | 1.169759852 | 0.001712417 | 0.020044122 |
| chr6:27553171-27553803 | -1.170996606 | 0.001713352 | 0.020051831 |
| chr5:42733926-42734258 | 1.478773974 | 0.001713599 | 0.020051831 |

| | | | |
|---|---|---|---|
| chr4:75930835-75931073 | 1.694412473 | 0.001715766 | 0.020073311 |
| chr7:54817150-54817584 | -1.434776311 | 0.001715959 | 0.020073311 |
| chr10:1359043-1359728 | 1.106359709 | 0.001716531 | 0.020076944 |
| chr6:161273709-161274332 | -1.145856535 | 0.001717332 | 0.020083249 |
| chr21:40173817-40174342 | -1.261954934 | 0.001719781 | 0.020107724 |
| chr6:26240057-26240477 | -1.29647922 | 0.00171995 | 0.020107724 |
| chr11:13741069-13741308 | 1.653820094 | 0.001721691 | 0.020125006 |
| chr6:21987463-21987757 | -1.533643782 | 0.00172216 | 0.020125087 |
| chr14:105853613-105853848 | 1.411519793 | 0.001722223 | 0.020125087 |
| chr3:116939131-116939570 | -1.304227616 | 0.001722597 | 0.020126386 |
| chr3:57170080-57170515 | -1.266367878 | 0.001723225 | 0.020130651 |
| chr1:170518019-170518223 | -1.731433557 | 0.001724729 | 0.020141396 |
| chr13:75372466-75372738 | -1.51645252 | 0.001724906 | 0.020141396 |
| chr7:32936723-32936979 | 1.327894477 | 0.001724933 | 0.020141396 |
| chr17:65531560-65531778 | 1.349217439 | 0.00172541 | 0.020143896 |
| chr1:115019418-115020128 | 1.150672798 | 0.001726087 | 0.020146232 |
| chr17:26619877-26620276 | 1.164643687 | 0.001726286 | 0.020146232 |
| chr17:55146734-55146995 | 1.589218078 | 0.001726533 | 0.020146232 |
| chr1:186428168-186428371 | 1.832163122 | 0.001726854 | 0.020146232 |
| chr18:976261-976461 | -1.981878513 | 0.001727182 | 0.020146232 |
| chr13:50969739-50970066 | -1.561102917 | 0.001727188 | 0.020146232 |
| chr1:217687440-217687851 | -1.196768988 | 0.001727716 | 0.020149323 |
| chr3:61684395-61684615 | 1.237480751 | 0.001728589 | 0.020156434 |
| chr3:123409603-123410425 | -1.061141516 | 0.001729683 | 0.020166127 |
| chr8:66522016-66523168 | -1.015921694 | 0.001731202 | 0.020178759 |
| chr1:63847329-63847661 | 1.395448874 | 0.001731439 | 0.020178759 |
| chr15:80294196-80294396 | 1.832393362 | 0.001731557 | 0.020178759 |
| chr3:62476521-62476825 | 1.163883512 | 0.001732964 | 0.020192093 |
| chr1:84619378-84619622 | -1.56635893 | 0.001735379 | 0.02021407 |
| chr20:45927363-45927650 | 1.395599418 | 0.001735528 | 0.02021407 |
| chr16:23679381-23679782 | 1.208901064 | 0.001735642 | 0.02021407 |
| chr9:126532175-126532567 | 1.35479095 | 0.001736027 | 0.020215481 |
| chr14:69619868-69620458 | 1.115342889 | 0.001736329 | 0.02021593 |
| chr2:48586373-48586869 | -1.247031858 | 0.001739333 | 0.020247818 |
| chr3:19933928-19934540 | -1.097112474 | 0.001739743 | 0.020249522 |
| chr1:155952323-155952523 | 1.739110417 | 0.001741406 | 0.020265797 |
| chr7:23272897-23273586 | 1.069707287 | 0.001741717 | 0.020266342 |
| chr8:57379097-57379638 | 1.306504526 | 0.001742266 | 0.020269651 |
| chr1:96972113-96972433 | 1.517595445 | 0.00174267 | 0.020270815 |
| chr6:4856851-4857051 | -2.278639684 | 0.001742895 | 0.020270815 |
| chr7:144127740-144128107 | -1.142096431 | 0.001743964 | 0.020280162 |
| chr14:45453315-45453625 | 1.571014279 | 0.00174519 | 0.020291338 |
| chr7:104979474-104979785 | -1.289861153 | 0.001747543 | 0.020315616 |
| chr11:29269426-29269646 | -1.790201892 | 0.001748003 | 0.020317885 |
| chr10:23375305-23375783 | -1.244703004 | 0.001749133 | 0.020327931 |
| chr5:62291828-62292068 | -1.617520771 | 0.001750157 | 0.020335628 |
| chr1:145707463-145708608 | -1.016287955 | 0.001750659 | 0.020335628 |
| chr1:97044033-97044426 | 1.473792127 | 0.001750812 | 0.020335628 |
| chr7:115173191-115173852 | -1.225564255 | 0.001750857 | 0.020335628 |
| chr14:58661812-58662068 | 1.570413109 | 0.001751876 | 0.020342103 |

| | | | |
|---|---|---|---|
| chr16:11296962-11297438 | 1.193750532 | 0.001751945 | 0.020342103 |
| chr10:97270642-97270864 | -1.852640094 | 0.001753426 | 0.020351643 |
| chrX:112574946-112575243 | 1.536384529 | 0.00175344 | 0.020351643 |
| chr5:119418211-119418411 | -1.589387958 | 0.001753564 | 0.020351643 |
| chr2:200296825-200297075 | -1.550392033 | 0.001757017 | 0.020388632 |
| chr5:174642879-174643322 | 1.248824297 | 0.00175747 | 0.0203908 |
| chr20:41810236-41810490 | 1.393826724 | 0.001758714 | 0.020402142 |
| chrX:115696916-115697116 | 1.822099285 | 0.001759158 | 0.020404208 |
| chr8:108352055-108352291 | 1.408638469 | 0.001760294 | 0.020412322 |
| chr17:54639237-54639842 | -1.186625775 | 0.00176039 | 0.020412322 |
| chr7:33846281-33846582 | -1.352673825 | 0.001760857 | 0.020414645 |
| chr8:124215414-124215663 | 1.364968045 | 0.001761291 | 0.020416591 |
| chr1:114207582-114207901 | 1.127059301 | 0.001762292 | 0.020425101 |
| chr16:3166649-3166862 | 1.537806385 | 0.00176373 | 0.020438674 |
| chr16:85555320-85555757 | -1.160615445 | 0.001766252 | 0.02046481 |
| chr6:108117578-108117778 | -1.881390621 | 0.00176708 | 0.020471302 |
| chr6:149079921-149080121 | 1.882672125 | 0.001768896 | 0.02048924 |
| chr15:55367982-55368221 | 1.725074175 | 0.001769914 | 0.020497937 |
| chr9:22100749-22100968 | -1.719130933 | 0.001770991 | 0.020507311 |
| chr3:24143680-24143880 | 2.024088196 | 0.001771563 | 0.020508977 |
| chr4:168692048-168692488 | -1.312082187 | 0.00177167 | 0.020508977 |
| chr7:21058987-21059359 | 1.376981653 | 0.001772278 | 0.020512909 |
| chr1:53102638-53103016 | 1.435005423 | 0.001772956 | 0.020517313 |
| chr20:47887275-47887553 | 1.095535769 | 0.001773194 | 0.020517313 |
| chr14:22391917-22392236 | 1.28414209 | 0.001774091 | 0.020524599 |
| chr2:135181786-135181986 | 1.767355233 | 0.001775803 | 0.020540049 |
| chr1:11241832-11242041 | 1.693968519 | 0.001775963 | 0.020540049 |
| chr1:186348133-186348732 | -1.137654449 | 0.001776264 | 0.020540432 |
| chr14:56013096-56013333 | 1.625943886 | 0.001776678 | 0.020542118 |
| chrX:74179523-74179723 | 1.693775944 | 0.001779021 | 0.020563898 |
| chr1:182334322-182334522 | -1.795806841 | 0.001779098 | 0.020563898 |
| chr4:119026850-119027050 | 1.820380484 | 0.001779934 | 0.020570457 |
| chr7:29921484-29921760 | 1.394167913 | 0.001781857 | 0.020589568 |
| chr14:49429648-49429991 | 1.534046613 | 0.00178224 | 0.020590894 |
| chr3:122504357-122504557 | 1.827302295 | 0.001782747 | 0.020593647 |
| chr14:80434464-80435287 | -1.119603352 | 0.001783317 | 0.020597118 |
| chr12:89464896-89465431 | -1.223888347 | 0.001784374 | 0.020603131 |
| chr8:101903887-101904087 | 1.677655757 | 0.001784375 | 0.020603131 |
| chr3:64178502-64178840 | 1.088999363 | 0.001786408 | 0.020623499 |
| chr5:100900500-100901241 | -1.062523042 | 0.001788352 | 0.020642831 |
| chr2:23524364-23524791 | -1.201084252 | 0.001788673 | 0.020643424 |
| chr13:37215605-37216055 | -1.344005287 | 0.001790418 | 0.020660454 |
| chr3:42683902-42684430 | -1.211571219 | 0.001791012 | 0.0206642 |
| chr7:112132864-112133225 | -1.76166079 | 0.001791476 | 0.020666433 |
| chr16:85720166-85720508 | 1.159877959 | 0.001793302 | 0.020684393 |
| chr7:77982718-77983168 | 1.283935335 | 0.00179442 | 0.020694166 |
| chr18:67200869-67201241 | -1.642170231 | 0.001795517 | 0.020703702 |
| chr19:2579243-2579623 | 1.331402944 | 0.001796732 | 0.020713211 |
| chr4:153294851-153295731 | -1.069873508 | 0.001796882 | 0.020713211 |
| chr13:31784875-31785075 | 1.773363409 | 0.00179756 | 0.020716321 |

| | | | |
|---|---|---|---|
| chr7:108054615-108054879 | -1.58217278 | 0.001797693 | 0.020716321 |
| chr18:65415248-65415723 | -1.400924416 | 0.001798819 | 0.02072618 |
| chr5:147397572-147397928 | -1.296395263 | 0.00179969 | 0.020733102 |
| chr20:48794484-48794797 | 1.082164452 | 0.001800986 | 0.020744104 |
| chr7:121081335-121081675 | -1.325892151 | 0.001801348 | 0.020744104 |
| chr12:78641908-78642353 | -1.381056539 | 0.001801725 | 0.020744104 |
| chr8:96653355-96653578 | 1.623266955 | 0.001801728 | 0.020744104 |
| chr19:29415929-29416158 | 1.610565944 | 0.001802877 | 0.020754214 |
| chr17:40322048-40322698 | -1.089904471 | 0.001803154 | 0.020754285 |
| chr14:79337974-79338211 | -1.586574195 | 0.001803647 | 0.020754826 |
| chr7:16667821-16668685 | 1.021456063 | 0.001803743 | 0.020754826 |
| chr20:42191306-42191857 | -1.029367725 | 0.001804903 | 0.020765052 |
| chr5:139746209-139746799 | -1.101107449 | 0.001806035 | 0.020774966 |
| chr10:80495141-80495515 | -1.255255601 | 0.001806382 | 0.020775829 |
| chr2:144326738-144327331 | -1.266831736 | 0.00180883 | 0.02080087 |
| chr20:51230079-51230348 | 1.353051701 | 0.00180945 | 0.020804877 |
| chr2:97067316-97067797 | 1.238520893 | 0.001810173 | 0.020810062 |
| chr13:59727079-59727387 | -1.555011541 | 0.001814153 | 0.020852692 |
| chr12:69868743-69868979 | 1.621690466 | 0.001815994 | 0.020870717 |
| chr5:59799104-59799779 | 1.159187475 | 0.001816788 | 0.020872447 |
| chr19:52942472-52942818 | 1.247622859 | 0.001816993 | 0.020872447 |
| chr10:30553910-30554236 | -1.39822968 | 0.001817345 | 0.020872447 |
| chrX:128914726-128914949 | 1.87796988 | 0.001817467 | 0.020872447 |
| chr2:41774052-41774828 | -1.097622494 | 0.001817506 | 0.020872447 |
| chr2:15920004-15920353 | -1.307714546 | 0.001820308 | 0.020901488 |
| chr10:95266701-95267054 | -1.408529264 | 0.001820645 | 0.020902226 |
| chr10:26335076-26335750 | -1.150252099 | 0.001820951 | 0.020902604 |
| chr8:121037196-121037396 | 1.759825648 | 0.001822123 | 0.020912931 |
| chr7:157905711-157906106 | -1.440875233 | 0.001822845 | 0.02091808 |
| chr4:24598009-24598519 | 1.122644809 | 0.001823472 | 0.020922143 |
| chr1:234997236-234997843 | 1.177049411 | 0.001824759 | 0.020933777 |
| chr9:86741293-86742013 | -1.152733824 | 0.001825565 | 0.020939882 |
| chr9:127721354-127722411 | -1.046473577 | 0.001826293 | 0.020945102 |
| chr2:180240807-180241007 | -1.778707679 | 0.001828819 | 0.020970933 |
| chr7:142483410-142483636 | -1.569939401 | 0.001829529 | 0.020973136 |
| chr12:52907785-52908370 | 1.062652937 | 0.001829559 | 0.020973136 |
| chr3:81490456-81490849 | -1.201689419 | 0.001830777 | 0.020983958 |
| chr15:74828906-74829216 | 1.269208215 | 0.001833961 | 0.021017308 |
| chr6:141077546-141078069 | -1.1683254 | 0.001835689 | 0.021033974 |
| chr2:27421882-27422328 | 1.260069956 | 0.001837269 | 0.021048922 |
| chr14:33964504-33964727 | 1.648326701 | 0.001838716 | 0.021062361 |
| chr15:87015752-87016401 | -1.134902975 | 0.001839647 | 0.021069874 |
| chr7:8483403-8483603 | 1.825273864 | 0.001840885 | 0.021080902 |
| chr8:121260872-121261445 | -1.070226493 | 0.00184178 | 0.021085894 |
| chr13:67343569-67344309 | -1.106785752 | 0.001842363 | 0.021085894 |
| chr17:4513353-4514533 | -1.045647851 | 0.001842419 | 0.021085894 |
| chr4:44147756-44148021 | 1.573935176 | 0.001842422 | 0.021085894 |
| chr4:102380045-102380984 | -1.120877141 | 0.001844831 | 0.021110308 |
| chr1:109892449-109892832 | -1.264401253 | 0.001845237 | 0.021111802 |
| chr3:9130434-9130634 | 1.623552804 | 0.001850145 | 0.021164794 |

| | | | |
|---|---|---|---|
| chr7:35955673-35955951 | 1.728839197 | 0.001852014 | 0.021183012 |
| chr4:115655655-115655951 | 1.512441159 | 0.001852347 | 0.021183656 |
| chr1:54314555-54314857 | 1.558339072 | 0.001853069 | 0.021188759 |
| chr7:156907268-156907540 | -1.492396908 | 0.001856481 | 0.021224602 |
| chr22:48370095-48370567 | -1.383175561 | 0.001859211 | 0.021252639 |
| chr17:7703628-7703981 | -1.367013096 | 0.001860956 | 0.021269411 |
| chr7:17120151-17120426 | 1.545455282 | 0.001864388 | 0.021303051 |
| chr14:50944322-50945310 | 1.040927603 | 0.0018645 | 0.021303051 |
| chr8:124351761-124352241 | -1.15676004 | 0.001864949 | 0.021303051 |
| chr6:27209714-27210170 | -1.184020916 | 0.001865011 | 0.021303051 |
| chr10:1232022-1232222 | 1.581109482 | 0.001865826 | 0.021309181 |
| chr16:75501823-75502165 | -1.319555486 | 0.001873031 | 0.021388285 |
| chr8:69771688-69771898 | 1.818126536 | 0.001874849 | 0.021404516 |
| chr4:25400501-25400933 | 1.326181192 | 0.001875285 | 0.021404516 |
| chr16:30913398-30913600 | 1.676588064 | 0.001875291 | 0.021404516 |
| chr12:62116241-62116524 | 1.596493392 | 0.001877733 | 0.021429202 |
| chr7:19982893-19983682 | 1.104837412 | 0.00187883 | 0.021438528 |
| chr16:87020551-87020831 | 1.64556473 | 0.001879579 | 0.021443886 |
| chr18:46433471-46433821 | -1.389445981 | 0.00188058 | 0.021452101 |
| chr1:160726855-160727110 | 1.415720689 | 0.001880983 | 0.021453511 |
| chr4:115090496-115090978 | -1.429359401 | 0.001881525 | 0.021456501 |
| chr19:39307312-39307676 | 1.212564262 | 0.001882427 | 0.021462392 |
| chr5:26482437-26483127 | -1.102591391 | 0.001882863 | 0.021462392 |
| chr5:53993605-53993847 | -1.547533816 | 0.001882882 | 0.021462392 |
| chr2:133331621-133331833 | -1.763807256 | 0.001884246 | 0.021474743 |
| chr2:19021008-19021352 | -1.329447261 | 0.001886876 | 0.021500239 |
| chr2:147414683-147414967 | 1.417274596 | 0.001887044 | 0.021500239 |
| chr10:12621077-12621277 | 1.818222906 | 0.001888398 | 0.021512465 |
| chrX:67792650-67792953 | 1.378390615 | 0.00188888 | 0.02151476 |
| chr7:16298865-16299207 | 1.369062696 | 0.001889417 | 0.021517669 |
| chr3:3082540-3082740 | 1.765314344 | 0.001890734 | 0.02152947 |
| chr21:36188054-36188609 | 1.094929131 | 0.001894827 | 0.021572877 |
| chr1:239386256-239386991 | -1.112241019 | 0.001896295 | 0.02158638 |
| chr5:42168230-42168430 | 1.803779767 | 0.001897817 | 0.021600492 |
| chr1:12049674-12049947 | 1.502276185 | 0.00190012 | 0.021623494 |
| chr1:113973167-113973417 | 1.150624903 | 0.001901813 | 0.021639553 |
| chr12:43795907-43796363 | -1.225032962 | 0.001904231 | 0.021663849 |
| chr3:11252566-11252766 | 1.647819016 | 0.0019064 | 0.021685297 |
| chr10:125905404-125905716 | 1.558455092 | 0.001908037 | 0.021700704 |
| chr4:28278100-28278358 | 1.441022933 | 0.001909427 | 0.02171328 |
| chr1:21634134-21634334 | 1.566435226 | 0.001912107 | 0.021740537 |
| chr7:91852233-91852738 | -1.29389642 | 0.001913727 | 0.021755725 |
| chr11:48055657-48056074 | 1.167560839 | 0.001914502 | 0.021761305 |
| chr9:1009116-1009606 | 1.16603421 | 0.001916365 | 0.021779254 |
| chr3:137946453-137946707 | -1.481891979 | 0.001917315 | 0.021785628 |
| chr2:64876179-64876635 | -1.328797681 | 0.001917495 | 0.021785628 |
| chr15:87520650-87521065 | -1.269936371 | 0.001917859 | 0.021786534 |
| chr7:12508709-12509030 | 1.433155451 | 0.001919185 | 0.021798366 |
| chr1:191427669-191428014 | -1.447437661 | 0.001920397 | 0.021808902 |
| chr3:175338652-175339563 | -1.044077749 | 0.001921281 | 0.021815702 |

| | | | |
|---|---|---|---|
| chr5:178553843-178554153 | 1.546939573 | 0.001922032 | 0.021817814 |
| chr5:42483783-42484076 | 1.386161308 | 0.001922036 | 0.021817814 |
| chr4:151737958-151738211 | 1.613597203 | 0.001922992 | 0.021825201 |
| chr14:33709764-33710141 | -1.213580196 | 0.001923256 | 0.021825201 |
| chr7:157418496-157419143 | -1.183664712 | 0.001923596 | 0.021825816 |
| chr7:16377723-16377963 | 1.565131246 | 0.001924527 | 0.021831173 |
| chr10:132223848-132224048 | 1.623034123 | 0.001924915 | 0.021831173 |
| chr5:168974151-168974351 | 1.870727806 | 0.001924922 | 0.021831173 |
| chr11:102391164-102391395 | -1.682758085 | 0.001926389 | 0.021844578 |
| chr20:47603332-47603610 | 1.074315382 | 0.001932442 | 0.021909969 |
| chr16:84396530-84397034 | 1.171007069 | 0.001933495 | 0.021916123 |
| chr15:77569133-77569413 | 1.423705699 | 0.001933557 | 0.021916123 |
| chr17:79289962-79290252 | 1.452582042 | 0.001933915 | 0.021916941 |
| chr10:99573868-99574068 | 1.734135415 | 0.001934202 | 0.02191695 |
| chr10:45362041-45362291 | -1.773509592 | 0.001934931 | 0.021921974 |
| chr2:20208076-20208519 | -1.135241321 | 0.001936695 | 0.02193667 |
| chr4:8267250-8267479 | 1.418953301 | 0.001936801 | 0.02193667 |
| chr12:23949709-23950007 | -1.478013081 | 0.001940285 | 0.021972886 |
| chr3:23475497-23475697 | 1.752362231 | 0.001943205 | 0.02199928 |
| chr4:170750556-170750879 | -1.444308149 | 0.001943466 | 0.02199928 |
| chr11:102306069-102306515 | -1.422042959 | 0.001944315 | 0.02199928 |
| chr2:65486351-65486985 | -1.135497626 | 0.001944336 | 0.02199928 |
| chr2:179898908-179899426 | -1.207187976 | 0.001944342 | 0.02199928 |
| chr4:126214023-126214332 | -1.477282926 | 0.001944412 | 0.02199928 |
| chr12:54387473-54388071 | 1.06775476 | 0.001944626 | 0.02199928 |
| chr1:158999638-159000253 | -1.210573696 | 0.001945744 | 0.022006048 |
| chr20:10469588-10470097 | -1.230752328 | 0.001945798 | 0.022006048 |
| chr3:63499604-63499888 | 1.320006028 | 0.001946421 | 0.022007278 |
| chr20:18856011-18856211 | 1.875840036 | 0.001946482 | 0.022007278 |
| chr3:153140474-153140708 | 1.444496954 | 0.0019475 | 0.022015548 |
| chr7:135335337-135335596 | -1.493051575 | 0.001948428 | 0.022019872 |
| chr2:174990080-174990652 | -1.170810535 | 0.001948458 | 0.022019872 |
| chr20:41997970-41998288 | 1.425632918 | 0.001950829 | 0.022043418 |
| chr6:163170257-163170653 | -1.284917838 | 0.001951432 | 0.022046983 |
| chr2:226932502-226932756 | -1.760267109 | 0.001953057 | 0.022059639 |
| chrX:150363114-150363370 | 1.306904468 | 0.001953128 | 0.022059639 |
| chr5:148280985-148281678 | -1.243571483 | 0.001954147 | 0.022067896 |
| chr20:45770962-45771162 | 1.616122541 | 0.001956522 | 0.022091459 |
| chr6:104907920-104909056 | -1.054519626 | 0.001957105 | 0.022094782 |
| chr3:20917542-20917787 | 1.642812648 | 0.001963479 | 0.022163476 |
| chr4:1554850-1555283 | 1.197452306 | 0.001965182 | 0.022177082 |
| chr2:127906118-127906510 | -1.262946237 | 0.001965263 | 0.022177082 |
| chr13:75876356-75876990 | -1.178504697 | 0.001966689 | 0.022187555 |
| chr5:127867325-127867689 | -1.29577995 | 0.00196677 | 0.022187555 |
| chr10:57202452-57202710 | -1.646596514 | 0.001967082 | 0.022187804 |
| chr21:36909935-36910511 | 1.250032139 | 0.001967714 | 0.022191667 |
| chr10:61896937-61897929 | -1.043077254 | 0.001970381 | 0.022218471 |
| chr15:89601698-89601945 | 1.713602479 | 0.001971163 | 0.022224023 |
| chr16:67535902-67536151 | 1.498385673 | 0.001971795 | 0.022227875 |
| chr5:16889690-16889941 | 1.241503225 | 0.001973648 | 0.022245488 |

| | | | |
|---|---|---|---|
| chr14:63659869-63660069 | -1.476196684 | 0.001973993 | 0.022246111 |
| chr7:153103183-153103551 | -1.335216648 | 0.001974966 | 0.022253803 |
| chr13:75182083-75182500 | -1.437116739 | 0.001975583 | 0.022257485 |
| chr19:38004315-38004895 | -1.167942563 | 0.001977273 | 0.022273247 |
| chr14:34664770-34664986 | 1.614645795 | 0.00197816 | 0.022279963 |
| chr8:125743038-125743242 | 1.486940578 | 0.001978522 | 0.022280762 |
| chr4:188177929-188178265 | -1.8086891 | 0.00197892 | 0.022281974 |
| chr3:72625644-72626079 | -1.250731439 | 0.001981543 | 0.022308226 |
| chr1:200161868-200162335 | 1.168200868 | 0.001982352 | 0.022314059 |
| chr1:217590577-217591064 | -1.254757901 | 0.001983095 | 0.022319139 |
| chr20:32731439-32731820 | 1.298360475 | 0.001983634 | 0.022321927 |
| chr16:46394439-46395046 | 1.003175435 | 0.001987211 | 0.022358898 |
| chr11:66749082-66749410 | 1.199864709 | 0.001987956 | 0.022363996 |
| chr13:33692030-33692325 | -1.571037918 | 0.001988646 | 0.022367831 |
| chrX:120497835-120498344 | 1.243181023 | 0.001988881 | 0.022367831 |
| chr3:171904961-171905310 | 1.593158868 | 0.001991259 | 0.022391293 |
| chr5:173781507-173781758 | 1.359973161 | 0.001992109 | 0.022394348 |
| chr6:134117318-134117814 | -1.182622505 | 0.001992188 | 0.022394348 |
| chr12:12017087-12017540 | -1.313354336 | 0.001992408 | 0.022394348 |
| chr14:99654280-99655141 | 1.064239079 | 0.001996082 | 0.022432355 |
| chr11:130535485-130535839 | -1.286082846 | 0.001996797 | 0.022437104 |
| chr15:88982816-88983089 | 1.490629223 | 0.001999491 | 0.022464082 |
| chr18:26225329-26225633 | -1.305150365 | 0.002000984 | 0.022477554 |
| chr1:147357706-147357912 | 1.603959669 | 0.002005922 | 0.02252972 |
| chr22:42532925-42533172 | -1.544211303 | 0.00200625 | 0.022530099 |
| chr7:4515157-4515394 | 1.696714099 | 0.002007497 | 0.022540806 |
| chr1:212782447-212782685 | -1.767240158 | 0.002008159 | 0.022541935 |
| chr15:87745614-87745881 | 1.511818648 | 0.002008186 | 0.022541935 |
| chr7:130485720-130487098 | -1.00723016 | 0.002011532 | 0.022576179 |
| chr3:118575015-118575261 | 1.543303469 | 0.002012502 | 0.022583764 |
| chr14:33285507-33285707 | 1.761187484 | 0.002014279 | 0.022600395 |
| chr7:17871542-17871742 | 1.674873775 | 0.002015342 | 0.022609014 |
| chr5:155529276-155529796 | -1.228807231 | 0.002016579 | 0.022618753 |
| chr10:22727710-22727982 | -1.573358683 | 0.002016801 | 0.022618753 |
| chr14:21070139-21070490 | 1.192395546 | 0.00201904 | 0.022640549 |
| chr4:149835243-149835769 | -1.403218054 | 0.002019713 | 0.022644785 |
| chr1:77417122-77417398 | -1.42449784 | 0.00202107 | 0.022656684 |
| chr3:63058-63403 | 1.482645419 | 0.002021837 | 0.022661968 |
| chr17:27552459-27552721 | -1.453124814 | 0.002022144 | 0.022662094 |
| chr10:88616773-88617035 | 1.749055263 | 0.002023122 | 0.022669731 |
| chr15:53910193-53910670 | 1.056649739 | 0.002024465 | 0.022680979 |
| chr4:48360264-48360892 | -1.34292451 | 0.002024717 | 0.022680979 |
| chr22:28818216-28818907 | 1.035151024 | 0.002025179 | 0.022682837 |
| chr19:43519542-43519761 | 1.528880607 | 0.002025764 | 0.022686073 |
| chr14:24483609-24483853 | 1.573114764 | 0.002027085 | 0.022697541 |
| chr16:77993662-77993881 | 1.34687684 | 0.002027618 | 0.022699133 |
| chr20:48405892-48406211 | 1.069021236 | 0.002027889 | 0.022699133 |
| chr4:175110011-175110254 | -1.714286922 | 0.00202859 | 0.022699133 |
| chr14:36784559-36784806 | 1.500901179 | 0.002028694 | 0.022699133 |
| chr1:1758686-1759112 | 1.179275331 | 0.002028708 | 0.022699133 |

| | | | |
|---|---|---|---|
| chr12:124686053-124686350 | 1.342877373 | 0.002029195 | 0.022700249 |
| chr9:87119317-87119714 | 1.20493674 | 0.0020294 | 0.022700249 |
| chr10:63637845-63638185 | -1.302078449 | 0.002030166 | 0.0227055 |
| chr10:73906665-73907224 | -1.134217167 | 0.002032036 | 0.022721624 |
| chr12:62519763-62520017 | -1.477268113 | 0.002032201 | 0.022721624 |
| chr16:15907894-15908583 | -1.066079279 | 0.002032568 | 0.02272241 |
| chr12:69476225-69476517 | -1.549457075 | 0.002032975 | 0.022723647 |
| chr5:9996982-9997182 | 1.861171286 | 0.002034855 | 0.02274135 |
| chr4:2817854-2818315 | 1.208015208 | 0.00203724 | 0.022764674 |
| chr1:159076547-159076912 | -1.411330365 | 0.002038301 | 0.022769095 |
| chr10:60955898-60956470 | -1.089433335 | 0.002038533 | 0.022769095 |
| chr7:24040060-24040463 | 1.22788705 | 0.002038793 | 0.022769095 |
| chr17:36920017-36920333 | 1.277941103 | 0.002038824 | 0.022769095 |
| chr7:113831551-113831829 | -1.74277445 | 0.002039654 | 0.022775044 |
| chr3:173932739-173932954 | -1.713144096 | 0.002040152 | 0.02277726 |
| chr1:11080271-11080849 | 1.210874566 | 0.002040447 | 0.02277726 |
| chr13:65638436-65638974 | -1.219008304 | 0.002041359 | 0.022784126 |
| chr14:69457466-69457869 | 1.219871107 | 0.002043473 | 0.022802349 |
| chr17:14792818-14793054 | -1.600353395 | 0.002043587 | 0.022802349 |
| chr9:35045378-35045578 | 1.758297066 | 0.002044195 | 0.022805814 |
| chr4:186097064-186097295 | -1.595626065 | 0.002044757 | 0.022808759 |
| chr18:67217583-67217838 | -1.717435585 | 0.00204693 | 0.022829672 |
| chr11:111334553-111335276 | -1.082998932 | 0.002047273 | 0.022830173 |
| chr11:17413570-17414013 | 1.150193979 | 0.002049338 | 0.022848753 |
| chr14:97432719-97432919 | 1.718032316 | 0.002049535 | 0.022848753 |
| chr7:22725718-22725952 | 1.322271971 | 0.002055072 | 0.022907143 |
| chr11:69217945-69218181 | 1.556148965 | 0.002055958 | 0.022909989 |
| chr20:47583694-47583983 | 1.055388663 | 0.002056098 | 0.022909989 |
| chr10:131943453-131943769 | 1.334705108 | 0.002056224 | 0.022909989 |
| chr3:109018901-109019191 | -1.372525777 | 0.002060737 | 0.022956932 |
| chr7:11486350-11486550 | 1.569505372 | 0.002063726 | 0.022986893 |
| chr4:48491357-48491557 | 1.8055273 | 0.002066502 | 0.023014459 |
| chr20:59445720-59445971 | 1.297031 | 0.002067024 | 0.023016936 |
| chr1:145545883-145546112 | 1.547514674 | 0.002069507 | 0.023041228 |
| chr10:63491641-63492170 | -1.303760503 | 0.002070492 | 0.02304775 |
| chr2:177812786-177813141 | -1.337031286 | 0.002070694 | 0.02304775 |
| chr16:25049718-25049973 | 1.598172843 | 0.002071683 | 0.023052263 |
| chr10:74573309-74574224 | -1.108020207 | 0.002071701 | 0.023052263 |
| chr8:108628311-108628511 | 1.704576246 | 0.002072062 | 0.023052934 |
| chr11:15988667-15989317 | -1.118764198 | 0.002074184 | 0.023072438 |
| chr20:44885957-44886239 | 1.19307373 | 0.002074418 | 0.023072438 |
| chr13:33787268-33787641 | -1.349751092 | 0.002075598 | 0.023082221 |
| chr14:39766334-39766534 | 1.529724369 | 0.002077169 | 0.02309131 |
| chr17:78717463-78717867 | -1.365729299 | 0.002077247 | 0.02309131 |
| chr17:78934130-78934461 | 1.277575514 | 0.002077408 | 0.02309131 |
| chr12:52236008-52236469 | 1.105116159 | 0.002077858 | 0.02309131 |
| chr14:102731777-102731987 | 1.644057692 | 0.002077923 | 0.02309131 |
| chr7:5071737-5072688 | -1.049854763 | 0.002079072 | 0.023100327 |
| chr8:125679510-125679784 | 1.424430599 | 0.002079337 | 0.023100327 |
| chr11:103325943-103326418 | -1.222439719 | 0.002081998 | 0.023126535 |

| | | | |
|---|---|---|---|
| chr1:31838206-31838408 | 1.708900738 | 0.002082393 | 0.023127574 |
| chr2:122352032-122352491 | -1.275277307 | 0.002083448 | 0.02313594 |
| chr4:185395827-185397407 | -0.984633873 | 0.002085442 | 0.023154723 |
| chr1:161179789-161180004 | 1.399312477 | 0.002086685 | 0.023163683 |
| chr15:57138329-57138529 | 1.861081868 | 0.002086854 | 0.023163683 |
| chr3:104242101-104242301 | 1.924934989 | 0.002087726 | 0.023170005 |
| chr12:79614658-79615001 | 1.194256708 | 0.002088704 | 0.023177505 |
| chr10:29590163-29590494 | 1.396551511 | 0.002089161 | 0.023179217 |
| chr1:151611283-151611490 | 1.282714038 | 0.002090682 | 0.023192739 |
| chr7:20576001-20576201 | 1.663184616 | 0.002091237 | 0.023193234 |
| chr22:21926710-21926946 | 1.571735104 | 0.002091332 | 0.023193234 |
| chr14:105623220-105623450 | 1.425768683 | 0.002092648 | 0.023204469 |
| chr16:67900699-67900987 | 1.518374939 | 0.002094187 | 0.023218168 |
| chrX:8136623-8136854 | -1.795257213 | 0.002097476 | 0.023251279 |
| chr15:96449257-96449623 | 1.533868244 | 0.002098994 | 0.023264733 |
| chr14:46063893-46064257 | -1.293279855 | 0.002099467 | 0.023266617 |
| chr3:189773030-189773230 | -1.949399192 | 0.002101363 | 0.023284258 |
| chr7:103857415-103857722 | -1.56284117 | 0.002102265 | 0.023290879 |
| chr9:124658136-124658713 | -1.146922563 | 0.002102784 | 0.023293268 |
| chr7:117938037-117938473 | -1.272867124 | 0.002103738 | 0.023300466 |
| chr2:15142144-15142870 | -1.105276918 | 0.00210409 | 0.023301 |
| chr4:12903833-12904076 | -1.637453484 | 0.002104571 | 0.023302958 |
| chr3:179476268-179476749 | -1.235995874 | 0.002108155 | 0.02333927 |
| chr8:103788652-103789441 | 1.088885176 | 0.002108986 | 0.023345091 |
| chr20:45104236-45104451 | 1.662555254 | 0.002110325 | 0.023356539 |
| chr2:40244664-40244961 | -1.2917805 | 0.002112945 | 0.023379979 |
| chr3:64789317-64789601 | 1.105756292 | 0.002113306 | 0.023379979 |
| chr18:57557839-57558231 | -1.192799488 | 0.002113358 | 0.023379979 |
| chr19:15187368-15187828 | 1.331057152 | 0.002116062 | 0.023406512 |
| chr8:92894463-92895000 | 1.191818307 | 0.002118003 | 0.023424603 |
| chr17:59367739-59367954 | 1.28476515 | 0.002118931 | 0.023430323 |
| chr18:71784691-71785147 | -1.25413862 | 0.002119435 | 0.023430323 |
| chr8:33066079-33066368 | -1.745589638 | 0.002119438 | 0.023430323 |
| chr10:5283506-5283816 | 1.216789976 | 0.002119759 | 0.023430494 |
| chr13:75733119-75733857 | -1.060836829 | 0.002121547 | 0.023446872 |
| chr3:168782307-168782586 | 1.608298784 | 0.002124293 | 0.023470914 |
| chr1:167109136-167109429 | 1.283146794 | 0.002124335 | 0.023470914 |
| chr1:180732599-180732862 | 1.695729084 | 0.002125644 | 0.023481997 |
| chr11:33331201-33331457 | 1.577175139 | 0.002126502 | 0.023488082 |
| chr18:24502697-24503001 | 1.39183167 | 0.002127612 | 0.023496952 |
| chr7:79308256-79308506 | -1.692503052 | 0.002128729 | 0.023505902 |
| chr1:195745684-195745972 | -1.250319658 | 0.002129452 | 0.0235105 |
| chr21:43804074-43804274 | 1.672501713 | 0.002132267 | 0.023537426 |
| chr6:130018466-130018669 | -1.785112658 | 0.002132675 | 0.023537426 |
| chr3:192273398-192273951 | 1.143682422 | 0.002132812 | 0.023537426 |
| chr19:45130188-45130391 | 1.514853242 | 0.00213373 | 0.023544165 |
| chr1:6519968-6520281 | 1.307888394 | 0.002134962 | 0.023549199 |
| chr13:49215032-49215380 | -1.326051533 | 0.002135318 | 0.023549199 |
| chr1:223479946-223480211 | 1.541639803 | 0.002135396 | 0.023549199 |
| chr19:29671158-29672019 | 1.049487066 | 0.002135416 | 0.023549199 |

| | | | |
|---|---|---|---|
| chr11:12178800-12179139 | 1.463214066 | 0.002137461 | 0.023568357 |
| chr2:156803827-156804027 | -1.768944165 | 0.002138071 | 0.023571699 |
| chr5:88172644-88173456 | -1.110444681 | 0.002138598 | 0.023574115 |
| chr12:123436214-123437052 | 1.090961018 | 0.002139409 | 0.023579663 |
| chr13:112711499-112712152 | -1.125541158 | 0.00214095 | 0.023593251 |
| chr20:59857606-59858022 | 1.217218438 | 0.002141931 | 0.023600664 |
| chr5:157290219-157290457 | 1.537835924 | 0.002145722 | 0.023639034 |
| chr3:136676999-136677293 | 1.702701716 | 0.002146186 | 0.023640748 |
| chr14:33986907-33987123 | 1.571083691 | 0.002148499 | 0.023660737 |
| chr16:74279888-74280245 | 1.349850999 | 0.002148618 | 0.023660737 |
| chr13:68816774-68817090 | -1.490839122 | 0.002148955 | 0.023661046 |
| chr4:48329327-48329588 | 1.513521554 | 0.002151321 | 0.023683693 |
| chr3:180457067-180457675 | -1.233247773 | 0.002152604 | 0.02369442 |
| chr3:127412870-127413100 | 1.540900447 | 0.002153183 | 0.023697384 |
| chr4:110287011-110287474 | -1.366733421 | 0.002154041 | 0.023703425 |
| chr10:3852285-3852717 | 1.138135484 | 0.002154936 | 0.023709869 |
| chr14:90443461-90444637 | -0.987597505 | 0.00215529 | 0.023710355 |
| chr2:186895458-186895904 | 1.29523022 | 0.002159088 | 0.023748725 |
| chr6:123171417-123171918 | -1.420692821 | 0.002159492 | 0.023749767 |
| chr20:24691969-24692293 | 1.468188347 | 0.002160773 | 0.023760439 |
| chr3:14268018-14268302 | 1.234064621 | 0.002162276 | 0.023773556 |
| chr13:55086554-55086955 | -1.367514039 | 0.002162647 | 0.023774223 |
| chr16:79293893-79294107 | -1.80761367 | 0.002163512 | 0.023780318 |
| chr7:21617033-21618369 | 1.003823743 | 0.002164847 | 0.023791578 |
| chr4:45621832-45622306 | -1.183810605 | 0.002166355 | 0.023804739 |
| chr7:70446790-70447236 | -1.279678841 | 0.002168458 | 0.023819992 |
| chrX:118603917-118604237 | 1.342057173 | 0.002168632 | 0.023819992 |
| chr6:53679477-53679783 | -1.366558426 | 0.002168676 | 0.023819992 |
| chr10:132036552-132037060 | 1.237269153 | 0.002169181 | 0.023822129 |
| chr5:62931644-62931882 | 1.509063166 | 0.002171732 | 0.023846724 |
| chr10:103605774-103606183 | -1.281923004 | 0.002172367 | 0.02384953 |
| chr7:17557859-17558078 | 1.442857907 | 0.00217261 | 0.02384953 |
| chr7:140646873-140647380 | -1.125669038 | 0.002173276 | 0.023853425 |
| chr12:63276831-63277258 | 1.250748645 | 0.002174565 | 0.023864148 |
| chr14:63174733-63175104 | -1.330715381 | 0.002175526 | 0.023871242 |
| chrX:74015446-74015697 | 1.489839974 | 0.002175834 | 0.023871242 |
| chr12:128111966-128112371 | -1.320653841 | 0.002176809 | 0.023878519 |
| chr3:64272880-64273136 | 1.080623827 | 0.002177897 | 0.023887031 |
| chr4:8719753-8719953 | 1.709116341 | 0.002178382 | 0.02388893 |
| chr3:159417897-159418164 | -1.557803402 | 0.002179149 | 0.023893615 |
| chr16:28539000-28539200 | 1.595056364 | 0.002179432 | 0.023893615 |
| chr8:93102462-93102947 | 1.084126989 | 0.002180761 | 0.023904758 |
| chr5:90803423-90803706 | -1.529566181 | 0.002185294 | 0.023951018 |
| chr16:84423677-84424029 | 1.388089466 | 0.002187118 | 0.02396758 |
| chr1:211352946-211353229 | -1.394820026 | 0.002187972 | 0.023972837 |
| chr10:4736253-4736473 | 1.56126648 | 0.002188283 | 0.023972837 |
| chr15:55439713-55440005 | 1.261848767 | 0.002188536 | 0.023972837 |
| chr20:56574799-56574999 | 1.463276316 | 0.002191276 | 0.023998826 |
| chr2:94897914-94898216 | 1.527819298 | 0.002191535 | 0.023998826 |
| chr1:31422713-31422958 | 1.424807689 | 0.00219236 | 0.024004434 |

| | | | |
|---|---|---|---|
| chr8:124856626-124857288 | 1.021972283 | 0.002192963 | 0.0240076 |
| chr8:124128930-124129166 | 1.550518811 | 0.002193759 | 0.024012882 |
| chr6:165641642-165642025 | -1.462749413 | 0.002195249 | 0.024025231 |
| chr4:86730319-86730632 | -1.482281762 | 0.002195514 | 0.024025231 |
| chr4:182916753-182918231 | -1.015874668 | 0.002197689 | 0.024045596 |
| chr5:2415739-2416000 | -1.523495642 | 0.002199624 | 0.024063337 |
| chr2:120423228-120423434 | 1.498405701 | 0.002200101 | 0.024065114 |
| chr4:189124259-189124472 | -1.858282577 | 0.002201361 | 0.024075466 |
| chr7:124482312-124482556 | -1.83513584 | 0.002202071 | 0.024079788 |
| chr2:85037055-85037509 | -1.479164551 | 0.002202548 | 0.024081573 |
| chr4:102512190-102512417 | -1.620040458 | 0.002203686 | 0.024087445 |
| chr1:171541352-171541564 | 1.652047867 | 0.002203714 | 0.024087445 |
| chr1:194909248-194909569 | -1.5305686 | 0.002204372 | 0.024090254 |
| chr9:15403185-15403440 | -1.519354734 | 0.0022046 | 0.024090254 |
| chr20:48319905-48320268 | 1.085738979 | 0.002206863 | 0.024111539 |
| chr1:227961565-227961773 | 1.700758604 | 0.002207834 | 0.024118712 |
| chr20:41588197-41588599 | 1.235974769 | 0.002209983 | 0.024133253 |
| chr8:33245003-33245559 | -1.185358664 | 0.002210048 | 0.024133253 |
| chr18:62485678-62486141 | -1.139497192 | 0.00221011 | 0.024133253 |
| chr13:41040544-41041023 | 1.259394535 | 0.002210628 | 0.024135468 |
| chr1:215539745-215540693 | -1.016010337 | 0.002210945 | 0.024135489 |
| chr12:112808997-112809302 | 1.694703542 | 0.002213036 | 0.024153159 |
| chr3:136643452-136643653 | 1.516926587 | 0.002213194 | 0.024153159 |
| chr7:222970-223180 | 1.484636467 | 0.002213954 | 0.024158015 |
| chr5:26329675-26329877 | 1.619273201 | 0.00221482 | 0.024161052 |
| chr14:81684766-81684999 | 1.431945317 | 0.002215112 | 0.024161052 |
| chrX:133620886-133621143 | 1.44008026 | 0.002215476 | 0.024161052 |
| chr10:58412749-58413056 | -1.261275204 | 0.002215494 | 0.024161052 |
| chr2:150614259-150614498 | 1.659768123 | 0.002217393 | 0.024178321 |
| chr1:16630076-16630567 | 1.153200962 | 0.002219597 | 0.024198918 |
| chr13:73542259-73542797 | -1.268775644 | 0.002220353 | 0.02420371 |
| chr1:166490083-166490549 | -1.229856926 | 0.002223473 | 0.024234277 |
| chr17:40958124-40958425 | 1.330347592 | 0.002224506 | 0.024242084 |
| chr7:105664907-105665386 | -1.206439868 | 0.002224837 | 0.024242248 |
| chr8:114921694-114922520 | -1.046976753 | 0.002225989 | 0.024251353 |
| chr13:45669497-45670283 | -1.127961586 | 0.002230587 | 0.024297992 |
| chr5:155068647-155069284 | -1.15627108 | 0.002231176 | 0.024300945 |
| chr15:56338312-56338599 | 1.523185309 | 0.002232002 | 0.024306493 |
| chr21:30857563-30857851 | 1.582436077 | 0.002232745 | 0.024310681 |
| chr14:42759526-42759738 | 1.676403833 | 0.002233021 | 0.024310681 |
| chrX:129433529-129433783 | 1.622746766 | 0.002233727 | 0.024314906 |
| chr6:8481773-8482041 | -1.693020771 | 0.002234587 | 0.024320813 |
| chr3:155019653-155020275 | -1.062732939 | 0.00223686 | 0.024342098 |
| chr2:192749669-192750051 | -1.478325603 | 0.00223853 | 0.024356531 |
| chr5:53482381-53483330 | -1.049022184 | 0.002238822 | 0.024356531 |
| chr20:18984273-18984588 | -1.241310676 | 0.002241375 | 0.024380844 |
| chr3:157626686-157627211 | -1.096817734 | 0.002250365 | 0.024475041 |
| chr15:101625665-101625992 | 1.394031952 | 0.002250674 | 0.024475041 |
| chr8:142710311-142710565 | 1.46818727 | 0.00225233 | 0.024489573 |
| chr15:76418751-76419202 | 1.252916722 | 0.00225309 | 0.024494357 |

| | | | |
|---|---|---|---|
| chr16:72635180-72635583 | -1.391954418 | 0.002254649 | 0.024502068 |
| chr12:34065405-34065632 | 1.68670972 | 0.002254679 | 0.024502068 |
| chr16:48524532-48524791 | 1.580577187 | 0.002254858 | 0.024502068 |
| chr16:24045590-24045827 | 1.510752516 | 0.002255078 | 0.024502068 |
| chr11:17921737-17921937 | 1.80114403 | 0.002255642 | 0.024504717 |
| chr17:62963833-62964035 | 1.110398793 | 0.002257087 | 0.02451694 |
| chr5:76601528-76601775 | -1.49846633 | 0.002258338 | 0.024527054 |
| chr1:208166857-208167109 | -1.502839566 | 0.002259906 | 0.024540605 |
| chr19:39052555-39052758 | 1.546092718 | 0.002261931 | 0.024557942 |
| chr14:90819952-90820263 | 1.189618342 | 0.002262143 | 0.024557942 |
| chr13:72194616-72194936 | -1.37966236 | 0.002262478 | 0.024558093 |
| chr16:87867497-87867774 | 1.240923949 | 0.002263408 | 0.024561197 |
| chr2:224971190-224971643 | -1.265774098 | 0.002263471 | 0.024561197 |
| chr10:92216641-92216994 | -1.198695202 | 0.002263726 | 0.024561197 |
| chr13:53782047-53782249 | -1.754862172 | 0.00226538 | 0.024575673 |
| chr14:63468376-63468576 | 1.742003548 | 0.002266064 | 0.024579611 |
| chr13:44172229-44172736 | -1.273713628 | 0.002267038 | 0.024585685 |
| chr14:71204806-71205006 | 1.593826794 | 0.002267266 | 0.024585685 |
| chr4:6955651-6955866 | 1.516395991 | 0.002268247 | 0.024592838 |
| chr12:87274651-87274916 | -1.533350297 | 0.002269499 | 0.024602938 |
| chr13:36431668-36432619 | -1.010716024 | 0.002269971 | 0.024604569 |
| chr12:122841586-122841814 | 1.407000903 | 0.00227036 | 0.024605304 |
| chr2:20796482-20796787 | -1.407452224 | 0.002271196 | 0.024610882 |
| chr7:11831486-11832301 | 1.035776955 | 0.002276542 | 0.02466533 |
| chr16:69853842-69854103 | 1.592061964 | 0.002276873 | 0.024665426 |
| chr5:66494510-66494770 | -1.39437861 | 0.002277546 | 0.024669226 |
| chr21:37484660-37484930 | 1.621191994 | 0.002278116 | 0.024671914 |
| chr12:93991021-93991395 | -1.336323193 | 0.002279965 | 0.024686112 |
| chr11:68643671-68643937 | 1.492609334 | 0.002280071 | 0.024686112 |
| chr9:72038972-72039303 | -1.244372666 | 0.002282994 | 0.024714269 |
| chr14:70972361-70972875 | 1.183029911 | 0.002285725 | 0.02474033 |
| chr10:44864097-44864374 | -1.240087631 | 0.002286832 | 0.024748823 |
| chr19:38428883-38429221 | 1.515513522 | 0.002287204 | 0.024749346 |
| chr15:39095639-39095851 | 1.532548524 | 0.002288475 | 0.024754058 |
| chr2:169393135-169393586 | -1.305250191 | 0.002288594 | 0.024754058 |
| chr4:170776860-170777334 | -1.553126037 | 0.002288608 | 0.024754058 |
| chr11:13354342-13354743 | -1.289060811 | 0.002290769 | 0.024772073 |
| chr9:77687722-77688105 | -1.171393789 | 0.00229092 | 0.024772073 |
| chr6:82875727-82876061 | -1.723853305 | 0.002291885 | 0.024779002 |
| chr3:162697091-162697293 | 1.690941872 | 0.00229418 | 0.02480032 |
| chr3:71022603-71022919 | -1.43280356 | 0.002295157 | 0.024807381 |
| chr19:2952990-2953308 | 1.252666072 | 0.002296775 | 0.024821366 |
| chr8:124961381-124961864 | 1.05950078 | 0.002297267 | 0.024823187 |
| chr20:17297307-17297517 | 1.532065025 | 0.00229777 | 0.024825119 |
| chr3:150323286-150323544 | 1.527687299 | 0.002298918 | 0.024834019 |
| chr4:92599023-92599406 | -1.273460089 | 0.002299622 | 0.024835762 |
| chr5:60463172-60463852 | -1.222710936 | 0.002299728 | 0.024835762 |
| chr8:124907033-124907304 | 1.30647628 | 0.002301306 | 0.024849305 |
| chr7:137169204-137169583 | -1.263687348 | 0.00230207 | 0.024854055 |
| chr12:112799568-112799768 | 1.74044632 | 0.002302684 | 0.02485718 |

| | | | |
|---|---|---|---|
| chr2:109690672-109691165 | 1.145222146 | 0.002303467 | 0.024858925 |
| chr1:214660951-214661232 | -1.217300833 | 0.002303495 | 0.024858925 |
| chr17:57730180-57730452 | 1.286166239 | 0.002306371 | 0.024886465 |
| chr7:112939329-112940876 | -0.968640307 | 0.002309322 | 0.024914794 |
| chr1:153478264-153479021 | -1.157821706 | 0.00231074 | 0.024926586 |
| chr1:223639285-223639673 | 1.2920594 | 0.002318873 | 0.025007904 |
| chr14:24566597-24566852 | 1.542062728 | 0.00231911 | 0.025007904 |
| chr2:20677657-20677874 | -1.45184411 | 0.002319258 | 0.025007904 |
| chr13:96983961-96984572 | -1.111612373 | 0.00232171 | 0.025027456 |
| chr5:176823997-176824197 | 1.556675665 | 0.002322192 | 0.025027456 |
| chr20:41477038-41477261 | 1.444782267 | 0.002322228 | 0.025027456 |
| chr7:143841713-143841975 | -1.566987426 | 0.002322378 | 0.025027456 |
| chr2:101751649-101752042 | 1.546043691 | 0.002323089 | 0.025029747 |
| chr4:175671529-175671959 | -1.271599529 | 0.002323243 | 0.025029747 |
| chr18:48356950-48357498 | -1.27889964 | 0.002324245 | 0.025037021 |
| chr2:159083712-159083998 | 1.616263593 | 0.002326083 | 0.025051565 |
| chr1:107074891-107075122 | 1.167555528 | 0.002326249 | 0.025051565 |
| chr10:5938383-5938683 | 1.401575589 | 0.002327598 | 0.025062563 |
| chr1:213983986-213984217 | -1.529441945 | 0.002332828 | 0.025115349 |
| chr1:151548733-151549262 | 1.035338037 | 0.002333881 | 0.025120162 |
| chr9:34560248-34560556 | 1.396030786 | 0.002333931 | 0.025120162 |
| chr6:139133173-139133643 | -1.112281313 | 0.002334669 | 0.025124579 |
| chr5:71778779-71779047 | -1.503783166 | 0.002336079 | 0.025136224 |
| chr12:53190158-53190509 | 1.127354258 | 0.002338569 | 0.025158568 |
| chr2:182068798-182069336 | -1.308873447 | 0.002338812 | 0.025158568 |
| chr12:108768640-108769101 | -1.274248413 | 0.002341067 | 0.025176494 |
| chr18:58785377-58785843 | 1.236777407 | 0.002341136 | 0.025176494 |
| chr7:13920047-13920722 | 1.012575098 | 0.002342355 | 0.025179679 |
| chr7:29449845-29450225 | 1.293896756 | 0.002342686 | 0.025179679 |
| chr8:10756697-10757199 | -1.174826241 | 0.002342954 | 0.025179679 |
| chr22:28971504-28971865 | 1.42594738 | 0.002342988 | 0.025179679 |
| chr5:164517796-164518851 | -1.06936807 | 0.002343185 | 0.025179679 |
| chr20:47851593-47852210 | 1.000784086 | 0.002343404 | 0.025179679 |
| chr17:66394604-66394804 | 1.545108542 | 0.002344838 | 0.025191556 |
| chr15:40253387-40253601 | 1.539100434 | 0.002346432 | 0.025205148 |
| chr2:189172298-189172498 | -2.014411023 | 0.002347873 | 0.025217093 |
| chr7:42832181-42833023 | -1.09858878 | 0.002348876 | 0.02522433 |
| chr9:105912674-105913473 | -1.101813407 | 0.002350593 | 0.025237742 |
| chr17:56110752-56111055 | -1.29996727 | 0.002351098 | 0.025237742 |
| chr7:27662262-27663272 | 1.0005312 | 0.002351332 | 0.025237742 |
| chr4:27094716-27095150 | 1.195743405 | 0.002351443 | 0.025237742 |
| chr5:101306972-101307196 | -1.833322759 | 0.002351832 | 0.02523839 |
| chr2:215930896-215931209 | 1.203655618 | 0.002353452 | 0.025252233 |
| chr1:182340169-182341073 | -1.036225384 | 0.002355005 | 0.025265363 |
| chr3:17026542-17026795 | 1.678711641 | 0.002355733 | 0.025269632 |
| chr7:108114135-108114472 | -1.233728798 | 0.002357081 | 0.025280552 |
| chr7:151795355-151795730 | -1.30392973 | 0.00235954 | 0.025303382 |
| chr5:1831717-1832004 | 1.252630265 | 0.002360197 | 0.025305472 |
| chr10:125344561-125345228 | -1.235509414 | 0.002360395 | 0.025305472 |
| chr7:142811203-142811481 | -1.426556566 | 0.002361786 | 0.025316837 |

| | | | |
|---|---|---|---|
| chr15:100721143-100721435 | 1.336114416 | 0.002362899 | 0.025324209 |
| chr1:106581452-106581734 | 1.114186934 | 0.002363135 | 0.025324209 |
| chr19:6161233-6161768 | 1.12348442 | 0.002363526 | 0.025324863 |
| chr2:96964697-96965064 | -1.144456372 | 0.002364482 | 0.025330222 |
| chr8:128176558-128177853 | -0.964446431 | 0.002365044 | 0.025330222 |
| chr20:47645979-47646439 | 1.005432016 | 0.002365079 | 0.025330222 |
| chr12:80079948-80080925 | -1.084661319 | 0.002365349 | 0.025330222 |
| chr2:142857440-142857655 | -1.61234457 | 0.002367484 | 0.025349551 |
| chr22:45290284-45290922 | 1.174181948 | 0.002369499 | 0.025367576 |
| chr5:20328417-20328698 | 1.618919807 | 0.00237001 | 0.025369503 |
| chr6:1418598-1418810 | 1.778463915 | 0.002372727 | 0.025393677 |
| chr6:94673256-94673676 | 1.17201392 | 0.002372931 | 0.025393677 |
| chr11:37135672-37136295 | 1.143364353 | 0.00237521 | 0.025414508 |
| chr8:102032818-102033018 | 1.553112246 | 0.002375541 | 0.025414508 |
| chr1:48600177-48600670 | -1.253390722 | 0.002377017 | 0.025425291 |
| chr14:33466492-33466818 | 1.383664361 | 0.002377213 | 0.025425291 |
| chr14:44463515-44463740 | 1.699918673 | 0.002381841 | 0.025471239 |
| chr1:152982814-152983171 | 1.560954426 | 0.002383926 | 0.025489981 |
| chr20:51952087-51952312 | 1.408083831 | 0.002384678 | 0.025494462 |
| chr10:21336546-21337135 | -1.168381703 | 0.00238579 | 0.025497982 |
| chr13:68552031-68552344 | -1.543954479 | 0.002386018 | 0.025497982 |
| chr2:216754476-216755089 | -1.187295359 | 0.002386037 | 0.025497982 |
| chr9:94300577-94301087 | -1.158036116 | 0.002386609 | 0.025497982 |
| chr22:32411443-32411701 | 1.46973981 | 0.002386671 | 0.025497982 |
| chr20:11627956-11628202 | -1.660072361 | 0.002387213 | 0.02550021 |
| chr10:8102916-8103439 | -1.115820436 | 0.002387926 | 0.025504273 |
| chr13:62014333-62014593 | -1.738037012 | 0.002390364 | 0.025526754 |
| chr11:61513841-61514159 | -1.262743058 | 0.002391561 | 0.025535981 |
| chr1:16718712-16718913 | 1.688551695 | 0.002393526 | 0.02554874 |
| chr2:200184735-200184979 | -1.54369178 | 0.002393648 | 0.02554874 |
| chr1:52867518-52867760 | 1.538101247 | 0.002393756 | 0.02554874 |
| chr16:4392527-4393522 | -1.029092445 | 0.002395096 | 0.025559478 |
| chr18:42030681-42030888 | -1.732667551 | 0.002396809 | 0.025574196 |
| chr1:186730509-186730773 | 1.584217655 | 0.002397319 | 0.025576071 |
| chr20:2851979-2852256 | -1.550566224 | 0.002398545 | 0.025585394 |
| chr1:59831498-59832013 | -1.248260194 | 0.002399077 | 0.025585394 |
| chr2:47193044-47193480 | -1.179135373 | 0.002399194 | 0.025585394 |
| chr1:67459215-67459415 | 1.732730575 | 0.002400339 | 0.02559404 |
| chr7:107890590-107891708 | -0.987647997 | 0.002401444 | 0.025602256 |
| chr2:237421461-237421993 | -1.283905985 | 0.002403298 | 0.025618456 |
| chr2:124804056-124804302 | 1.486260351 | 0.002403871 | 0.025621004 |
| chr1:209823159-209823558 | -1.209353699 | 0.002404989 | 0.025629356 |
| chr5:55547060-55547312 | -1.373123438 | 0.002406977 | 0.025646972 |
| chr1:169308384-169308638 | 1.412332105 | 0.002407714 | 0.025650799 |
| chr13:41352893-41353284 | -1.309037003 | 0.002408259 | 0.025650799 |
| chr1:227953959-227954198 | 1.424903375 | 0.00240834 | 0.025650799 |
| chr1:54137379-54137726 | 1.291735704 | 0.002409425 | 0.025658318 |
| chr11:19370304-19370509 | 1.610650712 | 0.002409716 | 0.025658318 |
| chr1:210292274-210292855 | -1.034571059 | 0.002410085 | 0.025658685 |
| chr10:119377632-119378225 | -1.217975331 | 0.00241142 | 0.025666557 |

| | | | |
|---|---|---|---|
| chr13:83730590-83731044 | -1.253624187 | 0.002411666 | 0.025666557 |
| chr1:27299370-27300003 | 1.118023059 | 0.00241183 | 0.025666557 |
| chr2:156537056-156537473 | -1.329111038 | 0.002413038 | 0.025675855 |
| chr8:136296354-136296747 | -1.442203428 | 0.002414164 | 0.02568427 |
| chr1:6199430-6199882 | 1.116672616 | 0.002415068 | 0.025690313 |
| chr5:157312472-157312737 | -1.378029124 | 0.002415801 | 0.025691408 |
| chr18:55781729-55781940 | -1.731368844 | 0.002415841 | 0.025691408 |
| chr17:182141-182394 | -1.496390551 | 0.002416748 | 0.02569748 |
| chr5:141889177-141889499 | -1.549458208 | 0.002417973 | 0.025706945 |
| chr1:161725869-161727303 | 0.987832684 | 0.002420582 | 0.025728595 |
| chr4:127975211-127975546 | 1.460617054 | 0.002420774 | 0.025728595 |
| chr4:95178245-95178575 | 1.503196428 | 0.002421024 | 0.025728595 |
| chr4:100444305-100444586 | 1.520683222 | 0.002421353 | 0.025728595 |
| chr5:10323808-10324166 | 1.230747516 | 0.002422226 | 0.025734304 |
| chr16:77387850-77388381 | 1.068283084 | 0.002423567 | 0.025743312 |
| chr7:122834035-122835179 | -1.013136288 | 0.002423746 | 0.025743312 |
| chr5:78065709-78066112 | -1.203899516 | 0.002426827 | 0.025772464 |
| chr4:59553372-59553673 | -1.418509499 | 0.002427337 | 0.025774313 |
| chrX:47297880-47298145 | 1.545133076 | 0.002428834 | 0.025786627 |
| chr20:48325330-48325620 | 1.070347383 | 0.002429261 | 0.025787595 |
| chrX:64176934-64177174 | -1.563548024 | 0.00243007 | 0.025792605 |
| chr6:41767170-41767632 | 1.110632913 | 0.002430465 | 0.025793227 |
| chr1:236459178-236459432 | 1.639667418 | 0.002432322 | 0.025809363 |
| chr9:935324-935711 | 1.200518516 | 0.002433615 | 0.025819502 |
| chr4:76526621-76526894 | 1.297556017 | 0.002434581 | 0.025826178 |
| chr7:28960665-28960887 | 1.581740229 | 0.002435596 | 0.025833371 |
| chr7:118041463-118041987 | -1.1854304 | 0.002440125 | 0.025877822 |
| chr11:111565198-111565410 | -1.690849367 | 0.002441114 | 0.025883363 |
| chr3:61754143-61754460 | 1.11374104 | 0.002441491 | 0.025883363 |
| chr5:23832145-23832460 | 1.429598071 | 0.002441661 | 0.025883363 |
| chr14:90337072-90337272 | 1.462744213 | 0.002444558 | 0.025910485 |
| chr1:150163156-150163524 | 1.082125539 | 0.002446099 | 0.025923235 |
| chr14:36855958-36856786 | 1.023309742 | 0.002446768 | 0.02592674 |
| chr16:88325388-88325588 | 1.551803784 | 0.002448816 | 0.025944363 |
| chr3:156045288-156045520 | 1.526159053 | 0.002449108 | 0.025944363 |
| chr4:186723680-186724063 | -1.123617924 | 0.002450051 | 0.025950763 |
| chr7:19442241-19442742 | 1.14984211 | 0.002452009 | 0.025967907 |
| chr7:114422565-114422810 | -1.559876275 | 0.002452599 | 0.025968193 |
| chr2:45796875-45797250 | 1.459443119 | 0.002452826 | 0.025968193 |
| chr12:23472885-23473313 | -1.51685355 | 0.002453052 | 0.025968193 |
| chr5:167587853-167588094 | -1.466538585 | 0.002454996 | 0.025985174 |
| chr18:38545162-38545408 | -1.49406981 | 0.002455695 | 0.025988983 |
| chr11:71566039-71566570 | -1.149824585 | 0.002456757 | 0.025996635 |
| chr22:36454947-36455434 | 1.096871658 | 0.002458629 | 0.026012846 |
| chr1:109168286-109168703 | -1.215585357 | 0.002458977 | 0.026012942 |
| chr12:41932763-41933056 | -1.519117065 | 0.002459779 | 0.026014496 |
| chr1:154758329-154758670 | 1.378711803 | 0.002459803 | 0.026014496 |
| chr12:113512498-113512710 | 1.6839241 | 0.002463687 | 0.026051979 |
| chr3:133700103-133700467 | 1.365697086 | 0.002464516 | 0.026056753 |
| chr12:128449205-128450100 | -1.077490395 | 0.002464958 | 0.026056753 |

| | | | |
|---|---|---|---|
| chr3:50184325-50184530 | 1.464840714 | 0.00246527 | 0.026056753 |
| chr9:79165918-79166380 | -1.218527594 | 0.002465499 | 0.026056753 |
| chr11:12517897-12518337 | 1.166811733 | 0.002465941 | 0.026057833 |
| chr4:89915985-89916354 | 1.209509233 | 0.002467423 | 0.026069894 |
| chr12:67017763-67017963 | 1.779418012 | 0.002468166 | 0.026074151 |
| chr16:370460-370690 | 1.322770266 | 0.002473281 | 0.026124582 |
| chr18:10221458-10221856 | -1.288651587 | 0.002474866 | 0.026137718 |
| chr7:153323018-153323623 | -1.311646356 | 0.002475674 | 0.026142651 |
| chr16:77037697-77038210 | 1.08552325 | 0.002479112 | 0.026175354 |
| chr1:207848218-207848508 | -1.318857736 | 0.00248031 | 0.026184392 |
| chr4:74544720-74544944 | -1.512485787 | 0.00248135 | 0.026186612 |
| chr16:46399170-46399904 | 0.991540062 | 0.002481521 | 0.026186612 |
| chr5:56350349-56351146 | -1.063787968 | 0.002481546 | 0.026186612 |
| chr8:29529495-29530147 | -1.232280251 | 0.002481922 | 0.026186977 |
| chr8:11713280-11713550 | -1.626888483 | 0.002483444 | 0.026199432 |
| chr21:36698890-36699433 | 1.140421874 | 0.002485467 | 0.026217161 |
| chr10:112798053-11279833 | 1.578402758 | 0.002486231 | 0.026221606 |
| chr17:72463333-72463651 | -1.1779077 | 0.00248715 | 0.026227688 |
| chr2:165018323-165018601 | -1.497788256 | 0.002490693 | 0.026258652 |
| chr13:40504043-40504399 | -1.284548499 | 0.002490771 | 0.026258652 |
| chr1:165783923-165784364 | 1.361493375 | 0.002492772 | 0.026276126 |
| chr10:35858626-35859321 | -1.161339355 | 0.002493365 | 0.026278765 |
| chr7:42373379-42373714 | -1.24353023 | 0.002497171 | 0.026315258 |
| chr7:107951933-107952293 | -1.278724752 | 0.002497694 | 0.026317147 |
| chr4:157405193-157405534 | -1.387132254 | 0.002499602 | 0.02633363 |
| chr7:154892535-154892840 | -1.444041796 | 0.002505354 | 0.026390604 |
| chr2:1010563-1010932 | -1.466066431 | 0.002506544 | 0.026392226 |
| chr8:60444435-60445005 | -1.171555706 | 0.002506558 | 0.026392226 |
| chr7:129501886-129503216 | -1.022642978 | 0.002506708 | 0.026392226 |
| chr13:23739728-23739928 | 1.730777189 | 0.002507506 | 0.026392226 |
| chr4:40395836-40396163 | -1.348431323 | 0.002507759 | 0.026392226 |
| chr2:25966511-25966897 | 1.216461295 | 0.00250781 | 0.026392226 |
| chr5:9775326-9775619 | 1.36337825 | 0.002507991 | 0.026392226 |
| chr2:133266108-133266812 | -1.119558182 | 0.002508264 | 0.026392226 |
| chr20:59960660-59960860 | 1.561858129 | 0.002508877 | 0.026393731 |
| chr1:236947216-236947452 | 1.644625416 | 0.002509096 | 0.026393731 |
| chr1:226737889-226738820 | 1.068333898 | 0.002510888 | 0.026406945 |
| chr2:7899547-7900116 | -1.320527148 | 0.002511252 | 0.026406945 |
| chr1:28532409-28532740 | -1.544448797 | 0.002511386 | 0.026406945 |
| chr7:110590505-110590761 | -1.651310158 | 0.002513931 | 0.026430081 |
| chr1:113630083-113630323 | 1.372049619 | 0.002514864 | 0.026436264 |
| chr9:36037372-36037635 | -1.560934848 | 0.002517254 | 0.02645738 |
| chr8:61850402-61851204 | -1.154637205 | 0.002517564 | 0.02645738 |
| chr13:37495103-37495541 | -1.208047719 | 0.002519799 | 0.026476353 |
| chr9:98686904-98687604 | 1.080002163 | 0.002520197 | 0.026476353 |
| chr1:53992147-53992405 | -1.468533801 | 0.002520406 | 0.026476353 |
| chr13:77306526-77306835 | -1.645395701 | 0.002521366 | 0.026482811 |
| chr5:102090732-102091109 | -1.18996303 | 0.002522056 | 0.026486432 |
| chr8:88356302-88356548 | 1.506481206 | 0.002522968 | 0.026492375 |
| chr6:27211466-27211703 | -1.373280174 | 0.002523893 | 0.02649846 |

| | | | |
|---|---|---|---|
| chr9:37668358-37668635 | -1.591497953 | 0.002524586 | 0.026501576 |
| chr1:223394139-223394539 | -1.465377714 | 0.002524882 | 0.026501576 |
| chr8:56892897-56893199 | -1.346890452 | 0.002525939 | 0.026509043 |
| chr5:567472-567752 | 1.581374031 | 0.002527247 | 0.026519136 |
| chr5:92688048-92688346 | -1.626178426 | 0.002527843 | 0.026520513 |
| chr16:2919767-2920057 | 1.30362789 | 0.002528071 | 0.026520513 |
| chr5:159951432-159951714 | -1.36863646 | 0.002529439 | 0.026531233 |
| chr7:128259240-128260489 | -0.991794318 | 0.00253069 | 0.026540721 |
| chr4:183658547-183659641 | -1.008187654 | 0.002533616 | 0.026567774 |
| chr18:42114392-42114648 | -1.567132562 | 0.002534957 | 0.026569965 |
| chr10:110375108-110375328 | 1.643767338 | 0.002535039 | 0.026569965 |
| chr1:108208908-108209971 | -1.012091529 | 0.002535225 | 0.026569965 |
| chr4:66306414-66306639 | 1.580079513 | 0.002535436 | 0.026569965 |
| chr20:48758206-48758461 | 1.07377412 | 0.002535559 | 0.026569965 |
| chr19:1914124-1914580 | -1.247375282 | 0.002536236 | 0.026573426 |
| chr1:209267095-209267295 | -1.889622148 | 0.002537584 | 0.026583879 |
| chr5:61010850-61011144 | -1.457401487 | 0.002538261 | 0.026583879 |
| chr11:117183961-117184303 | 1.164646607 | 0.002538555 | 0.026583879 |
| chr18:65788428-65788763 | -1.271484686 | 0.002538621 | 0.026583879 |
| chr11:101129424-101129951 | -1.08856144 | 0.002540319 | 0.026598018 |
| chr11:116787860-116788587 | 1.058621443 | 0.002541901 | 0.026610952 |
| chr15:89575737-89576234 | 1.07487895 | 0.002542913 | 0.026617908 |
| chr16:10257197-10257398 | 1.602789583 | 0.002545173 | 0.026637923 |
| chr4:189793033-189793363 | -1.30120015 | 0.002546485 | 0.026648018 |
| chr7:81447421-81447693 | -1.414770136 | 0.002547096 | 0.026650768 |
| chr10:48814004-48814366 | -1.311323954 | 0.00254748 | 0.02665115 |
| chr5:32331239-32331527 | 1.211780534 | 0.002547968 | 0.026652614 |
| chr14:95946433-95946633 | 1.904306089 | 0.002548771 | 0.026657374 |
| chr10:25875638-25875909 | -1.463856556 | 0.002549239 | 0.026658632 |
| chr13:22744714-22745501 | -1.091990002 | 0.0025521 | 0.02667868 |
| chr9:121222553-121223237 | -1.127908181 | 0.002552172 | 0.02667868 |
| chr1:6601138-6601396 | 1.334535135 | 0.002552201 | 0.02667868 |
| chrX:131677561-131678000 | 1.182999085 | 0.002553597 | 0.026689631 |
| chr2:226031027-226031570 | -1.133582593 | 0.002555152 | 0.026702247 |
| chr16:30798024-30798268 | 1.601246393 | 0.002556674 | 0.026714504 |
| chr4:7110835-7111035 | 1.66876548 | 0.002557763 | 0.026722237 |
| chr22:38921214-38921414 | 1.636990339 | 0.002561324 | 0.026755801 |
| chr17:49290402-49290648 | 1.602376794 | 0.002562417 | 0.026763566 |
| chr9:76906045-76906942 | -1.015907856 | 0.002563205 | 0.026768142 |
| chr18:46677661-46677960 | -1.460529786 | 0.002563882 | 0.026771564 |
| chr4:168153657-168154232 | -1.13185363 | 0.002564522 | 0.026774604 |
| chr7:20277055-20277255 | 1.722817578 | 0.002565072 | 0.0267767 |
| chr7:13742130-13742475 | 1.231588484 | 0.0025656 | 0.02677811 |
| chr20:48866514-48866825 | 1.065057794 | 0.002566238 | 0.02677811 |
| chr18:51278078-51278397 | -1.400662022 | 0.002566256 | 0.02677811 |
| chr1:46206706-46206906 | 1.641489136 | 0.00256705 | 0.02678275 |
| chr1:217695860-217696624 | -0.993151095 | 0.0025684 | 0.026793182 |
| chr8:105398506-105398706 | 1.681895963 | 0.002569269 | 0.026798603 |
| chr12:99959600-99960279 | -1.167242706 | 0.002569918 | 0.026801721 |
| chr15:88657427-88657733 | 1.190013339 | 0.002572828 | 0.026828423 |

| | | | |
|---|---|---|---|
| chr7:121351755-121352069 | -1.708396863 | 0.002573203 | 0.026828676 |
| chr6:142633475-142633700 | -1.713424493 | 0.002575057 | 0.026844352 |
| chr1:168621395-168621661 | 1.49371723 | 0.002575921 | 0.026846103 |
| chr4:176563656-176563885 | -1.720502724 | 0.002575989 | 0.026846103 |
| chr8:103693813-103694066 | 1.317606544 | 0.002576604 | 0.026846103 |
| chr21:39756740-39757032 | 1.507417009 | 0.002576626 | 0.026846103 |
| chr4:25865735-25865943 | 1.521497246 | 0.002578522 | 0.026862204 |
| chr1:100502189-100502722 | -1.124153918 | 0.00257936 | 0.026867277 |
| chr1:160743348-160743792 | 1.377846807 | 0.002579889 | 0.026869135 |
| chr8:122489198-122489544 | -1.218526146 | 0.002580345 | 0.026870239 |
| chr8:90726268-90726495 | 1.465504517 | 0.002581671 | 0.026880386 |
| chr14:37611349-37611581 | 1.305269502 | 0.002585018 | 0.026911582 |
| chr1:145891184-145891512 | 1.08015243 | 0.002586143 | 0.026919636 |
| chr14:65095821-65096128 | 1.244547848 | 0.002587536 | 0.026930482 |
| chr4:120000987-120001355 | -1.437095548 | 0.002588534 | 0.026937209 |
| chr7:20652080-20652289 | 1.489968271 | 0.002590948 | 0.026958667 |
| chr12:88834397-88834597 | -1.856332359 | 0.002591885 | 0.02696475 |
| chrX:81238158-81238830 | -1.216227815 | 0.002593986 | 0.026982943 |
| chr3:76824728-76825099 | 1.486302744 | 0.002598534 | 0.027023488 |
| chr3:14410709-14411022 | 1.197310583 | 0.0025987 | 0.027023488 |
| chrX:18099671-18100294 | 1.053707432 | 0.002598941 | 0.027023488 |
| chr12:13362231-13362635 | -1.272358888 | 0.002601233 | 0.027043649 |
| chr1:219518650-219519028 | -1.153381111 | 0.002601776 | 0.027045625 |
| chr7:20496231-20496525 | 1.395716759 | 0.00260319 | 0.02705665 |
| chr6:92918164-92918460 | 1.444405232 | 0.002603733 | 0.027058619 |
| chr1:216175744-216176020 | -1.363032586 | 0.002605906 | 0.027077532 |
| chr17:83024627-83024922 | 1.255206483 | 0.002608524 | 0.027100876 |
| chr20:48528225-48528430 | 1.130144452 | 0.00260886 | 0.027100876 |
| chrX:117407807-117408099 | 1.638208391 | 0.002611147 | 0.027120958 |
| chr7:120857560-120858552 | -1.003285412 | 0.002612034 | 0.027126021 |
| chr5:31692690-31692898 | 1.626550594 | 0.002612343 | 0.027126021 |
| chr6:98315530-98315823 | -1.761679478 | 0.002615605 | 0.027156214 |
| chr11:34685360-34685704 | -1.385487898 | 0.002617118 | 0.027168247 |
| chr6:145042687-145042946 | -1.663392328 | 0.002618546 | 0.027179388 |
| chrX:150363561-150364258 | 1.043193891 | 0.002619033 | 0.027180761 |
| chr17:69246820-69247119 | -1.339939944 | 0.002623118 | 0.027219472 |
| chr1:31476832-31477899 | 1.028164109 | 0.002628256 | 0.027267097 |
| chr11:34223199-34223399 | 1.597717546 | 0.00262842 | 0.027267097 |
| chr12:103742398-103742881 | -1.216822969 | 0.002629409 | 0.027273662 |
| chr8:22694422-22694789 | -1.159913075 | 0.002630265 | 0.027278851 |
| chr7:148051770-148052194 | -1.489955208 | 0.002632366 | 0.027296947 |
| chr10:132516731-132517591 | 1.003663539 | 0.002634058 | 0.027307206 |
| chr9:116661152-116661423 | 1.474465085 | 0.002634068 | 0.027307206 |
| chr10:44460292-44460678 | 1.471252849 | 0.002635507 | 0.027318427 |
| chr11:83021485-83021765 | -1.329349756 | 0.002636506 | 0.027325091 |
| chr2:28436437-28437834 | -0.988633713 | 0.002637869 | 0.027335516 |
| chr14:105676130-105676372 | 1.671448806 | 0.002638262 | 0.02733589 |
| chr15:90316211-90316502 | 1.234285639 | 0.002639614 | 0.027346206 |
| chr2:27008667-27009106 | -1.441664705 | 0.002640054 | 0.027347061 |
| chr7:116249985-116250484 | -1.14300355 | 0.00264234 | 0.027367042 |

| | | | |
|---|---|---|---|
| chr20:47651957-47652222 | 1.029633281 | 0.002643852 | 0.027379003 |
| chr4:132809586-132809992 | -1.327323553 | 0.002644253 | 0.027379458 |
| chr8:40143895-40144257 | -1.524253617 | 0.002644832 | 0.027381745 |
| chr3:111616559-111616907 | 1.171231451 | 0.002646149 | 0.027391678 |
| chr5:152038943-152039247 | -1.45530732 | 0.002647072 | 0.027397536 |
| chr17:61293559-61293907 | 1.013799259 | 0.00264923 | 0.02741617 |
| chr12:109291423-109291825 | -1.214140182 | 0.002649702 | 0.027416211 |
| chr17:45309309-45309509 | 1.671362166 | 0.00264995 | 0.027416211 |
| chrX:9419569-9419827 | -1.754288665 | 0.002651333 | 0.027426818 |
| chr2:212567621-212568336 | -1.184820123 | 0.002652228 | 0.027432376 |
| chr7:7438855-7439274 | 1.178401479 | 0.002652959 | 0.027436234 |
| chr16:75156914-75157197 | -1.540094701 | 0.002654527 | 0.027448744 |
| chr2:72115237-72116476 | -0.982625734 | 0.002655257 | 0.027452581 |
| chr13:82285365-82285600 | -1.72760104 | 0.002655622 | 0.027452652 |
| chr4:156367203-156368065 | -1.102302539 | 0.00265791 | 0.0274726 |
| chr11:103738838-103739610 | -1.073531573 | 0.002658289 | 0.027472816 |
| chr11:105016232-105016915 | -1.238396403 | 0.002665252 | 0.02754106 |
| chr16:50698229-50698457 | -1.64022783 | 0.002666317 | 0.027548345 |
| chr2:100887739-100888032 | 1.377404829 | 0.002667766 | 0.027559605 |
| chr7:112868669-112869520 | -1.135621662 | 0.002668534 | 0.027563817 |
| chr12:94684787-94685028 | 1.600868863 | 0.00267093 | 0.027581356 |
| chr13:27536441-27536650 | -1.470887849 | 0.002670952 | 0.027581356 |
| chr3:183951684-183952212 | 1.226637127 | 0.002672006 | 0.027588527 |
| chr2:177430008-177430234 | 1.635270155 | 0.002672392 | 0.027588791 |
| chr3:123877857-123878137 | 1.440550817 | 0.002674346 | 0.027605244 |
| chr6:139485855-139486084 | -1.752588368 | 0.002678143 | 0.027640719 |
| chr8:1735500-1735761 | -1.473368712 | 0.002678663 | 0.027642127 |
| chr12:130850045-130850273 | 1.508660009 | 0.002679001 | 0.027642127 |
| chr17:56926692-56927597 | 1.015734776 | 0.002679617 | 0.02764476 |
| chr10:58148513-58149228 | -1.103129937 | 0.002680009 | 0.027645074 |
| chr3:152211564-152212077 | 1.141319477 | 0.002681139 | 0.027653017 |
| chr14:94465819-94466542 | 1.067055834 | 0.002682478 | 0.027663104 |
| chr21:43287777-43288358 | 1.110131472 | 0.002685903 | 0.027694688 |
| chr13:79384225-79384470 | -1.633268664 | 0.002686528 | 0.027697412 |
| chr20:24629733-24630061 | 1.468223879 | 0.002687725 | 0.027706019 |
| chr10:6053898-6054381 | -1.178316974 | 0.002688148 | 0.02770666 |
| chr7:7339238-7339510 | 1.445253348 | 0.002689219 | 0.027713967 |
| chr3:14118553-14118916 | 1.184675515 | 0.002689622 | 0.027714395 |
| chr13:94712848-94713116 | -1.599146823 | 0.002690128 | 0.027715883 |
| chrX:150716899-150717144 | 1.415153661 | 0.002691216 | 0.027723365 |
| chr5:141668474-141668847 | 1.329634618 | 0.002692203 | 0.027729798 |
| chr4:6773205-6773492 | 1.269097684 | 0.002693006 | 0.027734339 |
| chr16:89637995-89638388 | 1.203966357 | 0.00269427 | 0.027738752 |
| chr7:156840281-156840517 | -1.885591008 | 0.002694332 | 0.027738752 |
| chr3:11759947-11760349 | 1.168403843 | 0.00269452 | 0.027738752 |
| chr2:163191431-163191684 | -1.681050397 | 0.002698122 | 0.0277721 |
| chr4:54356594-54356794 | 1.82128814 | 0.002698509 | 0.027772347 |
| chr4:24137617-24137817 | 1.759971182 | 0.002698957 | 0.027773229 |
| chrX:123720332-123720911 | 1.068857546 | 0.002699466 | 0.027774738 |
| chr8:128889394-128889657 | 1.44849879 | 0.00270068 | 0.0277835 |

| | | | |
|---|---|---|---|
| chr20:51446786-51447745 | -1.005413067 | 0.002702524 | 0.027798733 |
| chr8:129551210-129551716 | -1.094580931 | 0.002704419 | 0.027814494 |
| chr2:9897410-9897725 | -1.364758505 | 0.002707552 | 0.027842403 |
| chr6:88520323-88520716 | -1.363736752 | 0.002707859 | 0.027842403 |
| chr18:46152506-46153485 | -1.044424026 | 0.002708934 | 0.02784972 |
| chr3:14822869-14823424 | 1.081011625 | 0.002710129 | 0.02785826 |
| chr8:111169156-111169356 | 1.672747079 | 0.002710718 | 0.027860577 |
| chr3:177284344-177284861 | -1.239305512 | 0.002715738 | 0.027908429 |
| chr8:116173422-116174072 | -1.108451264 | 0.002716541 | 0.02791294 |
| chr14:97365242-97365504 | 1.552567684 | 0.002717901 | 0.027923175 |
| chr7:111357664-111358123 | -1.231424558 | 0.002721461 | 0.027955997 |
| chr14:43342327-43342699 | 1.3675415 | 0.002722775 | 0.027965744 |
| chr12:49967083-49968127 | 0.988802887 | 0.002724206 | 0.027974699 |
| chr6:4937862-4938062 | -1.761151309 | 0.002724377 | 0.027974699 |
| chr16:25014540-25015064 | 1.133613488 | 0.002724743 | 0.027974706 |
| chr5:654241-654560 | 1.539954525 | 0.002729678 | 0.028021622 |
| chrX:30712968-30713381 | -1.219648946 | 0.002730646 | 0.028026698 |
| chr7:134358288-134358527 | -1.515291865 | 0.00273143 | 0.028026698 |
| chr10:63872312-63872619 | -1.337907226 | 0.002731692 | 0.028026698 |
| chr7:141785978-141786313 | -1.278502314 | 0.002731877 | 0.028026698 |
| chr2:68448562-68449150 | -1.308114381 | 0.002732001 | 0.028026698 |
| chr2:7431350-7431740 | -1.200615387 | 0.002735095 | 0.028052081 |
| chr5:776993-777280 | 1.269616882 | 0.002735216 | 0.028052081 |
| chr1:183060374-183061057 | -1.042259192 | 0.002735574 | 0.028052081 |
| chr20:54505762-54506104 | 1.026007371 | 0.002736662 | 0.028056453 |
| chr9:6715702-6716723 | -0.992009128 | 0.002736733 | 0.028056453 |
| chr8:117405081-117405324 | 1.568583935 | 0.002737546 | 0.028061034 |
| chr4:92482913-92483787 | -1.051187441 | 0.002741128 | 0.028093992 |
| chr5:163104573-163104847 | -1.431203377 | 0.002744366 | 0.02812342 |
| chr5:96868691-96869431 | -1.050893187 | 0.002746293 | 0.028139399 |
| chr3:105154747-105154955 | 1.600790839 | 0.002746869 | 0.028140853 |
| chr11:69768178-69768596 | 1.29088009 | 0.002747169 | 0.028140853 |
| chr3:160460370-160460629 | 1.463664042 | 0.002747861 | 0.028142656 |
| chr14:82855893-82856131 | -1.418087792 | 0.00274808 | 0.028142656 |
| chr15:49421962-49422246 | -1.314548608 | 0.002748571 | 0.028143922 |
| chr13:72221138-72221705 | -1.091079761 | 0.002751694 | 0.028172136 |
| chr2:48749726-48750464 | -1.088614177 | 0.002752181 | 0.028173362 |
| chr2:21281724-21282483 | -1.038272121 | 0.002752613 | 0.028174015 |
| chr21:14273725-14274053 | -1.797514189 | 0.002753363 | 0.028177932 |
| chr8:101004215-101004440 | 1.404475677 | 0.002754318 | 0.028183941 |
| chr1:204455573-204455822 | 1.303125043 | 0.002755089 | 0.028186306 |
| chr2:132383791-132384178 | 1.284745695 | 0.002755285 | 0.028186306 |
| chr20:48528746-48528952 | 1.117738409 | 0.002755916 | 0.028188999 |
| chr5:78278694-78278971 | -1.758970203 | 0.002759335 | 0.028207991 |
| chr9:93466852-93467052 | 1.675536604 | 0.002759382 | 0.028207991 |
| chr16:78227820-78228217 | 1.091699465 | 0.002759473 | 0.028207991 |
| chr2:124869308-124869541 | 1.656258915 | 0.002759716 | 0.028207991 |
| chr16:87608467-87608667 | 1.414953277 | 0.002759776 | 0.028207991 |
| chr2:151289309-151289867 | -1.154103328 | 0.002759982 | 0.028207991 |
| chr12:40238495-40238758 | -1.675003918 | 0.002761471 | 0.028212887 |

| | | | |
|---|---|---|---|
| chr3:166725759-166725986 | 1.805434203 | 0.002761529 | 0.028212887 |
| chr2:54965915-54966699 | -1.061447594 | 0.002761565 | 0.028212887 |
| chr2:195520604-195521168 | 1.093589353 | 0.002762214 | 0.028214041 |
| chr8:119655131-119655470 | -1.296896661 | 0.002762415 | 0.028214041 |
| chr12:118071644-118071917 | 1.620376774 | 0.002764281 | 0.02822934 |
| chr2:151928377-151928786 | -1.170638847 | 0.002765265 | 0.028230763 |
| chr15:69813917-69814193 | 1.392134949 | 0.002765442 | 0.028230763 |
| chr11:67022218-67023315 | -0.982716131 | 0.002765722 | 0.028230763 |
| chrX:154457666-154458104 | 1.122346669 | 0.002765894 | 0.028230763 |
| chr3:155125304-155125513 | -1.703442348 | 0.002769374 | 0.028262513 |
| chr11:20707832-20708552 | -1.130511526 | 0.002774806 | 0.02831418 |
| chr7:139644220-139644506 | -1.464135845 | 0.002775943 | 0.028321987 |
| chr5:176438890-176439242 | 1.241512509 | 0.00277631 | 0.028321987 |
| chr13:38656748-38657042 | -1.285150758 | 0.002776716 | 0.02832235 |
| chr1:232322258-232322586 | 1.249604609 | 0.00278042 | 0.028356365 |
| chr14:50943843-50944251 | 1.170702603 | 0.00278132 | 0.028361763 |
| chr7:134646472-134647378 | -0.970659879 | 0.002782301 | 0.028362064 |
| chr1:234544195-234544883 | 1.064351242 | 0.002782355 | 0.028362064 |
| chr15:98763143-98763716 | -1.093873594 | 0.00278246 | 0.028362064 |
| chr20:48159508-48159808 | 1.053258465 | 0.002782845 | 0.028362219 |
| chrX:100746224-100746522 | -1.457077727 | 0.002784832 | 0.028378696 |
| chr7:123527633-123528405 | -1.108131096 | 0.002786125 | 0.028388097 |
| chr10:25682601-25682880 | -1.478678853 | 0.002787546 | 0.028398796 |
| chr8:79310230-79310646 | 1.544917046 | 0.002792044 | 0.028439735 |
| chr20:50700482-50700765 | 1.008873392 | 0.002792307 | 0.028439735 |
| chr15:54923738-54923938 | 1.773539437 | 0.002793314 | 0.028446208 |
| chr9:87570553-87570886 | 1.619739236 | 0.002795216 | 0.028459747 |
| chr17:61796827-61797066 | 1.050921024 | 0.002795386 | 0.028459747 |
| chr10:60414736-60415238 | -1.107331054 | 0.00279933 | 0.028496113 |
| chr2:451630-451830 | -1.779282964 | 0.002804427 | 0.028544209 |
| chr9:911026-911377 | 1.203459895 | 0.002804958 | 0.028545824 |
| chr2:219167979-219168234 | 1.465880105 | 0.00280541 | 0.028546325 |
| chrX:107729564-107730007 | -1.106885947 | 0.002805753 | 0.028546325 |
| chr7:34198830-34199097 | 1.420893774 | 0.002807765 | 0.028563003 |
| chr3:123621760-123622205 | 1.057648727 | 0.002808441 | 0.028565494 |
| chr14:28676430-28676760 | 1.374692136 | 0.002808755 | 0.028565494 |
| chr20:37279703-37280346 | -1.16576566 | 0.002813145 | 0.02860634 |
| chr4:37765841-37766077 | 1.484730616 | 0.002815509 | 0.028626585 |
| chr12:77179201-77179497 | -1.5020101 | 0.002817529 | 0.028643321 |
| chr1:196567923-196568813 | -1.040513767 | 0.002819218 | 0.028656686 |
| chr6:24788320-24788570 | 1.568069268 | 0.002821471 | 0.028675787 |
| chr1:113954111-113954311 | 1.193028353 | 0.002823341 | 0.028690747 |
| chr8:81569199-81569841 | 1.127022048 | 0.002823692 | 0.028690747 |
| chr11:21674316-21675385 | -1.034620962 | 0.002825011 | 0.02870034 |
| chr8:90544362-90544599 | 1.47373707 | 0.002827008 | 0.02871682 |
| chr7:68321053-68321506 | -1.295221491 | 0.002827829 | 0.028718404 |
| chr2:190679928-190680684 | -1.010784299 | 0.002827913 | 0.028718404 |
| chr7:118081044-118081353 | -1.331934824 | 0.002829013 | 0.028725763 |
| chrX:42888417-42888669 | -1.576249443 | 0.002829603 | 0.028727955 |
| chr1:103106401-103107309 | -1.112668546 | 0.002830461 | 0.028732856 |

| | | | |
|---|---|---|---|
| chr1:114892176-114892543 | 1.140891742 | 0.002832778 | 0.028752563 |
| chr16:11320120-11320325 | 1.51125763 | 0.00283419 | 0.028763086 |
| chr7:103929841-103930074 | -1.748108749 | 0.002835477 | 0.02877115 |
| chr10:58436164-58436486 | -1.270224987 | 0.002836229 | 0.02877115 |
| chr8:108184324-108184630 | 1.379209123 | 0.002836384 | 0.02877115 |
| chr7:11918199-11918462 | 1.504252005 | 0.002836486 | 0.02877115 |
| chr2:8939227-8939427 | 1.59562478 | 0.002837014 | 0.028772696 |
| chr1:71461506-71461748 | -1.498564249 | 0.002838569 | 0.028784657 |
| chr10:18104325-18104634 | -1.378561193 | 0.002840666 | 0.028802109 |
| chr20:47309977-47310322 | 0.995521152 | 0.002843087 | 0.028822843 |
| chr2:179622720-179623199 | -1.33136627 | 0.002843486 | 0.028823072 |
| chr7:14473194-14473452 | 1.416527742 | 0.002844536 | 0.028826965 |
| chr8:134129042-134129287 | 1.698033943 | 0.002844623 | 0.028826965 |
| chr18:59996429-59996653 | -1.838442664 | 0.00284725 | 0.028849774 |
| chr11:80155460-80155749 | -1.358059393 | 0.002848229 | 0.028855876 |
| chr13:32267812-32268400 | -1.048278915 | 0.002848642 | 0.028856249 |
| chr12:68738841-68739117 | 1.620161694 | 0.002850587 | 0.028871595 |
| chrX:31266273-31267249 | -1.03308191 | 0.002850911 | 0.028871595 |
| chr7:4234460-4234735 | 1.519686092 | 0.002851558 | 0.028874332 |
| chr13:79393466-79394340 | -1.011374209 | 0.002853693 | 0.028892134 |
| chr7:34348785-34349008 | 1.49017751 | 0.002856071 | 0.028912393 |
| chr7:117835025-117835307 | -1.368181548 | 0.00285796 | 0.028927685 |
| chr16:76770442-76770642 | 1.457318713 | 0.002860126 | 0.028945789 |
| chr1:192739313-192740102 | -1.122616816 | 0.002861084 | 0.028951659 |
| chr19:44689235-44689435 | 1.548497644 | 0.00286273 | 0.028964493 |
| chr5:17375152-17375352 | 1.570912442 | 0.002864271 | 0.028976259 |
| chr20:59925798-59926464 | 1.011934846 | 0.00286715 | 0.029001556 |
| chrX:40518138-40518847 | 1.052902758 | 0.002869285 | 0.029014032 |
| chr5:93062617-93062878 | -1.473186813 | 0.002869574 | 0.029014032 |
| chr1:175573556-175573915 | -1.307305263 | 0.002869673 | 0.029014032 |
| chr16:81852632-81852832 | 1.49849485 | 0.002869898 | 0.029014032 |
| chr16:48384590-48384867 | 1.455722634 | 0.002871769 | 0.02902911 |
| chr1:43421408-43421662 | 1.551416039 | 0.002877182 | 0.029079994 |
| chr19:10333867-10334124 | 1.219506528 | 0.002881165 | 0.029114728 |
| chr15:91456171-91456566 | 1.201022882 | 0.002881378 | 0.029114728 |
| chr9:15697086-15697286 | -1.880642042 | 0.002882244 | 0.029119635 |
| chr13:110017517-110017777 | -1.45182147 | 0.002884391 | 0.029137482 |
| chr5:64680389-64680679 | -1.469985348 | 0.002885337 | 0.029143192 |
| chr2:2662598-2663311 | -1.067861518 | 0.002886396 | 0.029150052 |
| chr10:95807428-95807883 | -1.413511493 | 0.002888885 | 0.029171343 |
| chr7:14433027-14433227 | 1.697605726 | 0.00289046 | 0.029181036 |
| chr14:68748151-68748530 | 1.287881058 | 0.002890607 | 0.029181036 |
| chr5:107472554-107473147 | -1.045259482 | 0.002892279 | 0.02919067 |
| chr17:27551661-27552213 | -1.131539134 | 0.002892326 | 0.02919067 |
| chr8:100663575-100663894 | 1.228330324 | 0.002892704 | 0.02919067 |
| chr4:104571375-104571886 | -1.200133606 | 0.002896741 | 0.029222385 |
| chr12:102896108-102896441 | -1.335356724 | 0.002897081 | 0.029222385 |
| chr10:53598935-53599353 | -1.41099658 | 0.002897133 | 0.029222385 |
| chr1:248686877-248687416 | -1.09801826 | 0.002897372 | 0.029222385 |
| chr14:43665710-43665971 | 1.483771345 | 0.002902686 | 0.029272126 |

| | | | |
|---|---|---|---|
| chr7:115989277-115989975 | -1.023444083 | 0.002905046 | 0.029292074 |
| chr7:37320172-37320769 | -1.248627097 | 0.002906052 | 0.029298356 |
| chr7:131625778-131626152 | -1.188387341 | 0.002906721 | 0.029301242 |
| chr1:230267425-230267625 | 1.522162484 | 0.002907282 | 0.029303047 |
| chr7:152564834-152565034 | 1.647213092 | 0.002907782 | 0.029304227 |
| chr10:75994325-75994708 | -1.474961516 | 0.002909308 | 0.029313461 |
| chr2:38153015-38153230 | 1.559532492 | 0.002909463 | 0.029313461 |
| chr4:106329076-106329354 | -1.407042042 | 0.002913351 | 0.029348775 |
| chrX:23796431-23797966 | -0.978870357 | 0.002913813 | 0.029349569 |
| chr7:116671686-116672653 | -1.002931861 | 0.00291548 | 0.029361964 |
| chr11:84558162-84558415 | -1.39666271 | 0.00291581 | 0.029361964 |
| chr1:146400497-146400825 | 1.357594648 | 0.002917934 | 0.029379492 |
| chr2:110796807-110797060 | 1.37471058 | 0.002918528 | 0.029381611 |
| chr7:112428103-112428463 | -1.516946998 | 0.002920056 | 0.029393133 |
| chr2:22994335-22994669 | 1.453153344 | 0.002920864 | 0.029395649 |
| chr5:163978970-163979277 | -1.386592018 | 0.002921073 | 0.029395649 |
| chr8:10850437-10850822 | -1.311700599 | 0.002922502 | 0.029406168 |
| chr17:38511563-38511776 | 1.694219487 | 0.002927172 | 0.029449289 |
| chr16:84994797-84995211 | 1.138738172 | 0.002928085 | 0.029454607 |
| chr16:87789676-87790968 | -0.964187318 | 0.002929255 | 0.029460238 |
| chr3:108764601-108765303 | -1.033989956 | 0.002929414 | 0.029460238 |
| chr1:209398590-209398918 | -1.345440602 | 0.002930526 | 0.029467555 |
| chr19:13051378-13051722 | 1.366797612 | 0.002931704 | 0.029475533 |
| chr13:34011756-34012065 | -1.413333507 | 0.002936508 | 0.029516778 |
| chr12:51817210-51817459 | 1.371306 | 0.002936577 | 0.029516778 |
| chr10:96124331-96125022 | -1.122285472 | 0.002937499 | 0.029522174 |
| chr10:59350273-59350915 | -1.111808149 | 0.00294125 | 0.029555997 |
| chr14:104794061-104794457 | 1.057751778 | 0.00294319 | 0.029569286 |
| chr8:2733721-2733947 | -1.790703586 | 0.002943345 | 0.029569286 |
| chr7:21767272-21767687 | 1.048281296 | 0.002944253 | 0.029574539 |
| chr16:83113390-83113854 | -1.257708344 | 0.002946693 | 0.02959457 |
| chr1:245527970-245528362 | -1.333990304 | 0.002947033 | 0.02959457 |
| chr8:120784999-120785307 | 1.262883803 | 0.002947406 | 0.02959457 |
| chr13:65011476-65011738 | -1.623826725 | 0.002947919 | 0.029595836 |
| chr10:80413593-80413904 | -1.295392309 | 0.002948479 | 0.029596366 |
| chr14:71004531-71004797 | 1.579444122 | 0.002948744 | 0.029596366 |
| chr3:118843175-118843399 | 1.619212119 | 0.002950698 | 0.029612102 |
| chr2:191494534-191494879 | 1.356143171 | 0.002951228 | 0.029613544 |
| chr7:120740430-120740665 | -1.814125082 | 0.002953091 | 0.029625096 |
| chr3:63239546-63239806 | 1.133737804 | 0.002953153 | 0.029625096 |
| chrX:150693130-150693477 | 1.228089011 | 0.002954413 | 0.029633856 |
| chr3:190238455-190239590 | -1.044270154 | 0.002955147 | 0.029637343 |
| chr1:197841204-197841635 | -1.243299794 | 0.002957082 | 0.029652863 |
| chr2:215218671-215218923 | -1.422397636 | 0.002962802 | 0.029703283 |
| chr11:85182773-85183106 | -1.208351852 | 0.002962885 | 0.029703283 |
| chr7:135710901-135711139 | -1.430580248 | 0.00296403 | 0.029710741 |
| chr13:25406460-25406674 | 1.34492316 | 0.002964653 | 0.029710741 |
| chr7:82760346-82760653 | -1.723433727 | 0.002964792 | 0.029710741 |
| chr6:32189406-32189606 | 1.504029652 | 0.00296763 | 0.029735287 |
| chr14:23830150-23830358 | 1.514998764 | 0.002968574 | 0.029740855 |

| | | | |
|---|---|---|---|
| chr6:7431713-7432510 | -1.145800335 | 0.002972135 | 0.029772638 |
| chr22:46996645-46996902 | -1.488992978 | 0.002976044 | 0.029806002 |
| chr12:104255419-104255653 | -1.588137469 | 0.002976518 | 0.029806002 |
| chr2:194319241-194319585 | -1.274776202 | 0.002976632 | 0.029806002 |
| chr13:60347746-60348035 | -1.492455302 | 0.002980465 | 0.029840477 |
| chr4:6927086-6927352 | 1.421976261 | 0.002982671 | 0.029858667 |
| chr10:25638462-25638816 | -1.307934578 | 0.002983662 | 0.029864687 |
| chr9:135689485-135689710 | 1.533897525 | 0.002985149 | 0.029875664 |
| chr1:101769591-101769830 | -1.717993109 | 0.002985677 | 0.02987705 |
| chr19:47096202-47096725 | 1.074648362 | 0.002988072 | 0.029897104 |
| chr1:9842863-9843068 | 1.586245762 | 0.002990655 | 0.029919048 |
| chr4:186190732-186192471 | -0.966416037 | 0.002993547 | 0.029941657 |
| chr7:95246897-95247189 | -1.474757124 | 0.002993697 | 0.029941657 |
| chr2:206274668-206274882 | 1.616350489 | 0.002994292 | 0.029942209 |
| chr14:91829509-91829849 | 1.281513595 | 0.002994534 | 0.029942209 |
| chr4:182745146-182745728 | -1.039502035 | 0.002996143 | 0.029951852 |
| chr5:141664990-141665314 | -1.207378897 | 0.00299628 | 0.029951852 |
| chr7:23334527-23335012 | 1.037987906 | 0.002996973 | 0.029954869 |
| chr2:83694787-83695619 | -1.179138919 | 0.002997541 | 0.029956648 |
| chr9:111663307-111663601 | -1.299280867 | 0.002998028 | 0.02995719 |
| chr7:116167207-116167486 | -1.436197195 | 0.00299851 | 0.02995719 |
| chr7:56078327-56078527 | 1.558789038 | 0.002998798 | 0.02995719 |
| chr18:57794425-57794665 | -1.706590507 | 0.00299916 | 0.02995719 |
| chr7:7363927-7364171 | 1.423613582 | 0.00300014 | 0.029961599 |
| chr20:25360698-25360965 | 1.29732793 | 0.003000743 | 0.029961599 |
| chr19:1914902-1915156 | -1.415326108 | 0.003000774 | 0.029961599 |
| chr14:37936968-37937351 | 1.217688037 | 0.003001858 | 0.02996466 |
| chrX:129039477-129039677 | 1.549445291 | 0.003002002 | 0.02996466 |
| chr15:62269215-62269953 | 1.145881111 | 0.003002254 | 0.02996466 |
| chr16:21541076-21541404 | 1.300611627 | 0.00300356 | 0.029973788 |
| chr2:26018085-26018437 | 1.172115987 | 0.003004315 | 0.029977421 |
| chr7:23105170-23106699 | 0.943367488 | 0.003006483 | 0.029991763 |
| chr7:8418723-8418961 | 1.271906546 | 0.003006684 | 0.029991763 |
| chrX:151905246-151905446 | 1.80127099 | 0.003007039 | 0.029991763 |
| chrX:116520055-116520683 | -1.205105474 | 0.003007318 | 0.029991763 |
| chr11:74344244-74344460 | 1.752829677 | 0.003008458 | 0.029995681 |
| chr16:33816448-33817008 | 1.089326137 | 0.003008494 | 0.029995681 |
| chr3:9715844-9716127 | 1.193094729 | 0.003009817 | 0.030003953 |
| chr4:3462619-3462924 | -1.333701369 | 0.003010336 | 0.030003953 |
| chr13:29732503-29732722 | 1.575523443 | 0.003010635 | 0.030003953 |
| chr13:102401250-102401552 | -1.622413706 | 0.003011093 | 0.030003953 |
| chr7:101856452-101857366 | -0.977497234 | 0.003011281 | 0.030003953 |
| chr1:107195894-107196174 | 1.094950279 | 0.003014257 | 0.030029693 |
| chr20:50812874-50813690 | 0.954794874 | 0.003015716 | 0.030040325 |
| chr7:131513490-131513854 | -1.145991326 | 0.003016116 | 0.030040406 |
| chr5:21145113-21145369 | 1.744051607 | 0.003018725 | 0.03006248 |
| chrX:136346540-136346773 | 1.590145411 | 0.003022296 | 0.030094137 |
| chr22:43993010-43993245 | -1.705255654 | 0.003022915 | 0.03009639 |
| chr18:58135835-58136038 | -1.751433576 | 0.003023876 | 0.030101407 |
| chr6:13725599-13725878 | 1.339978989 | 0.003024443 | 0.030101407 |

| | | | |
|---|---|---|---|
| chr2:64807319-64807761 | -1.122599843 | 0.003024899 | 0.030101407 |
| chr2:18685698-18685986 | 1.656445699 | 0.003024991 | 0.030101407 |
| chr7:4057122-4057372 | 1.607064606 | 0.00302721 | 0.03011958 |
| chr1:193507841-193508101 | 1.418960787 | 0.003027901 | 0.03012254 |
| chr6:158374067-158374334 | -1.476422866 | 0.003028785 | 0.030127423 |
| chr3:104103420-104103675 | 1.590174241 | 0.003032593 | 0.030160038 |
| chr9:76818265-76818629 | -1.282128891 | 0.003032851 | 0.030160038 |
| chr2:32617626-32618104 | -1.237553106 | 0.003033507 | 0.030162649 |
| chr1:116706186-116706590 | 1.161418579 | 0.003034583 | 0.03016943 |
| chr5:107248137-107248402 | -1.416946893 | 0.003036806 | 0.03018762 |
| chr4:11428484-11429389 | 1.023758485 | 0.003037417 | 0.030189774 |
| chr1:144077274-144077643 | 1.447018197 | 0.003038497 | 0.030194538 |
| chr4:165351470-165351722 | -1.412924038 | 0.003038685 | 0.030194538 |
| chr7:19659350-19659550 | 1.719319222 | 0.003039344 | 0.030197173 |
| chr9:70420816-70421073 | -1.125725255 | 0.003040322 | 0.030199379 |
| chr15:45445255-45446042 | -1.137996414 | 0.003040629 | 0.030199379 |
| chr1:216031742-216031942 | -1.665925481 | 0.003040748 | 0.030199379 |
| chr4:13701824-13702293 | -1.164758998 | 0.003043534 | 0.030223125 |
| chr14:53946744-53947202 | 1.177624772 | 0.003044129 | 0.030225118 |
| chr3:170062762-170063018 | -1.299904795 | 0.003047519 | 0.030254856 |
| chr3:74290194-74290565 | 1.469531167 | 0.003048399 | 0.030259677 |
| chr14:40180634-40180888 | 1.605091957 | 0.003050078 | 0.030272422 |
| chr1:106850806-106851060 | 1.095225893 | 0.003053025 | 0.030297749 |
| chr11:31787148-31787664 | -1.181889718 | 0.003058937 | 0.030348924 |
| chr8:85659819-85660389 | 1.100714805 | 0.003058974 | 0.030348924 |
| chr21:15199431-15200477 | -1.033009502 | 0.003059391 | 0.030349125 |
| chr13:48400973-48401220 | -1.571897029 | 0.00306 | 0.030350742 |
| chrX:67698153-67698691 | 1.054104267 | 0.003060346 | 0.030350742 |
| chr14:65216731-65216939 | 1.383754015 | 0.003065224 | 0.030395187 |
| chr1:197404207-197404430 | -1.464443058 | 0.003068206 | 0.030418845 |
| chr19:46304044-46304401 | 1.173682261 | 0.003068404 | 0.030418845 |
| chr2:39913634-39914057 | -1.284264568 | 0.003069832 | 0.030429068 |
| chr17:78854102-78854302 | 1.495157572 | 0.003072762 | 0.030447324 |
| chr20:48194395-48194601 | 1.087261708 | 0.003072765 | 0.030447324 |
| chr7:16652846-16653244 | 1.023114381 | 0.003072866 | 0.030447324 |
| chr8:144661244-144661504 | 1.367713537 | 0.003073439 | 0.030449065 |
| chr5:125696837-125697148 | -1.335726678 | 0.003076722 | 0.030474785 |
| chr18:63265023-63265223 | -1.575680342 | 0.003076831 | 0.030474785 |
| chr5:61679023-61679308 | -1.299685156 | 0.003077949 | 0.030481924 |
| chr1:34825102-34825310 | -1.663821764 | 0.003079425 | 0.030492594 |
| chr10:132375682-132375882 | 1.432702134 | 0.003081803 | 0.030508564 |
| chr14:37660462-37660662 | 1.362505967 | 0.003081834 | 0.030508564 |
| chr10:6202303-6203764 | 0.950081912 | 0.003086651 | 0.030552304 |
| chr8:76431281-76431567 | 1.339604541 | 0.003087826 | 0.030554536 |
| chr3:58657863-58658063 | 1.611719847 | 0.003088073 | 0.030554536 |
| chr3:115254832-115255073 | 1.611719847 | 0.003088073 | 0.030554536 |
| chr5:100927773-100928017 | -1.684998951 | 0.003090138 | 0.030571016 |
| chr7:143120819-143121442 | -1.36759411 | 0.003091504 | 0.030580585 |
| chr9:102514575-102515001 | -1.283288124 | 0.003093042 | 0.030591845 |
| chr11:106706302-106706877 | -1.154873868 | 0.003095085 | 0.030606411 |

| | | | |
|---|---|---|---|
| chr1:210477353-210477819 | -1.194799456 | 0.003095313 | 0.030606411 |
| chr11:63567151-63567509 | -1.442249998 | 0.003096195 | 0.030611179 |
| chrX:79809086-79809295 | 1.544457358 | 0.003097655 | 0.030621669 |
| chr15:89058334-89058585 | 1.40749233 | 0.003101578 | 0.030656488 |
| chr10:94371466-94371705 | 1.605784936 | 0.003103799 | 0.030674483 |
| chr4:176217665-176217865 | -1.960186294 | 0.003104343 | 0.030675909 |
| chr7:42195017-42195217 | -1.875436386 | 0.003105271 | 0.030679852 |
| chr4:170805300-170805756 | -1.141957603 | 0.003105543 | 0.030679852 |
| chr20:47894704-47896196 | 0.936495555 | 0.003107687 | 0.030697076 |
| chr7:24874317-24874529 | 1.473320909 | 0.00310936 | 0.030709641 |
| chr1:48619433-48619916 | -1.204536384 | 0.003110535 | 0.030717055 |
| chr8:99138554-99138846 | 1.539316673 | 0.003110913 | 0.030717055 |
| chr18:56650938-56651847 | -0.980866175 | 0.003112185 | 0.030725495 |
| chr3:119588216-119588528 | -1.320714452 | 0.003112577 | 0.030725495 |
| chr5:147721188-147722031 | -1.03723253 | 0.00311297 | 0.030725495 |
| chr18:62436651-62436964 | -1.616317221 | 0.003117365 | 0.030764912 |
| chr20:63846091-63846886 | -0.991496623 | 0.003117887 | 0.030766094 |
| chr11:63999177-63999779 | -1.062748816 | 0.003119062 | 0.03077373 |
| chr9:35102967-35103755 | -1.005855405 | 0.003120128 | 0.030778138 |
| chr3:37595115-37595758 | 1.05446206 | 0.003120673 | 0.030778138 |
| chr11:94838000-94838597 | 1.101016135 | 0.003120714 | 0.030778138 |
| chr2:154914676-154914940 | -1.516069119 | 0.003122614 | 0.030792916 |
| chr5:1882432-1883112 | 1.005712684 | 0.003126705 | 0.030826083 |
| chr14:104800847-104801262 | 1.040175371 | 0.003127063 | 0.030826083 |
| chr4:185742893-185743098 | -1.876665616 | 0.003127185 | 0.030826083 |
| chr7:101733959-101734169 | -1.686517055 | 0.003128006 | 0.030830209 |
| chr8:98541167-98541377 | 1.602375214 | 0.003131306 | 0.030858766 |
| chr12:92973602-92974017 | -1.289289037 | 0.003132819 | 0.030868209 |
| chrX:87728065-87728911 | -1.025717263 | 0.00313307 | 0.030868209 |
| chr16:86844714-86845095 | -1.166090407 | 0.00313437 | 0.030872944 |
| chr11:38824670-38824874 | -1.706820562 | 0.003134818 | 0.030872944 |
| chr5:100479015-100479486 | -1.257123896 | 0.003134939 | 0.030872944 |
| chr4:35079743-35079943 | -1.958291114 | 0.003135162 | 0.030872944 |
| chr5:163473692-163473959 | -1.474457951 | 0.003135925 | 0.030874939 |
| chr2:222673585-222674127 | -1.317842769 | 0.00313617 | 0.030874939 |
| chr6:135182654-135183432 | -1.015507477 | 0.003137662 | 0.030884384 |
| chr5:180792804-180793018 | 1.528697475 | 0.003137936 | 0.030884384 |
| chr15:57600195-57600548 | -1.346363273 | 0.003140607 | 0.03090479 |
| chr10:13219967-13220234 | -1.419066173 | 0.003140816 | 0.03090479 |
| chr17:6421883-6422356 | -1.228339289 | 0.003142381 | 0.030916219 |
| chr12:22934931-22935342 | -1.568260652 | 0.003145468 | 0.030942622 |
| chrX:23318119-23318516 | -1.161268178 | 0.003145961 | 0.030943494 |
| chr14:34794543-34794752 | 1.569372974 | 0.003148182 | 0.030961363 |
| chr3:194091164-194091590 | -1.164370694 | 0.003151802 | 0.030992992 |
| chr10:66180738-66181084 | -1.234777273 | 0.003154662 | 0.031017132 |
| chr7:107874660-107874993 | -1.21177141 | 0.003155237 | 0.031018808 |
| chr1:193519857-193520274 | -1.167920513 | 0.00315611 | 0.031023403 |
| chr19:4706901-4707271 | -1.416429738 | 0.003156995 | 0.031028125 |
| chr3:69403452-69404032 | 1.106343383 | 0.003159819 | 0.031051896 |
| chr10:76128212-76128872 | -1.148981581 | 0.00316089 | 0.03105844 |

| | | | |
|---|---|---|---|
| chr5:68534336-68534818 | -1.182329142 | 0.003161536 | 0.031060801 |
| chr11:2094183-2094401 | -1.5862586 | 0.003161949 | 0.031060878 |
| chr10:3853756-3853976 | 1.32939616 | 0.003164582 | 0.031082667 |
| chr12:13101001-13101913 | -0.999433188 | 0.003165098 | 0.031082667 |
| chr13:40159760-40160190 | 1.189302329 | 0.003165384 | 0.031082667 |
| chr2:172998493-172999071 | 1.054640122 | 0.003168282 | 0.031104829 |
| chr4:170589657-170590012 | -1.126085194 | 0.003168622 | 0.031104829 |
| chr13:77779542-77779841 | -1.434951322 | 0.003168859 | 0.031104829 |
| chr3:179450936-179452033 | -1.017864709 | 0.003169348 | 0.031105648 |
| chr7:73612788-73613174 | -1.313262848 | 0.003170522 | 0.031113182 |
| chr6:163634456-163635130 | -1.065775415 | 0.003172698 | 0.031130553 |
| chr1:195068705-195068999 | 1.441124174 | 0.00317369 | 0.031135166 |
| chr3:67647471-67647903 | 1.287819738 | 0.003173981 | 0.031135166 |
| chr2:231716225-231716469 | -1.33922688 | 0.003174826 | 0.031139466 |
| chr7:132387644-132388091 | -1.369696201 | 0.003176916 | 0.031155973 |
| chr5:56763754-56763961 | -1.593624202 | 0.003177366 | 0.031156406 |
| chr10:71138526-71138955 | 1.311382342 | 0.003178479 | 0.031163333 |
| chr3:112751760-112752009 | -1.691070359 | 0.003179534 | 0.031169688 |
| chr4:185559696-185560121 | -1.307109109 | 0.003182036 | 0.031189464 |
| chr20:54358354-54358609 | 1.045547175 | 0.003182366 | 0.031189464 |
| chr4:170400422-170400690 | -1.439144811 | 0.003185157 | 0.031212827 |
| chr2:142874129-142874865 | -0.967584 | 0.003186351 | 0.031220531 |
| chr7:151776274-151776521 | 1.535726915 | 0.003186819 | 0.031221128 |
| chr8:130323633-130323908 | 1.394352026 | 0.003187844 | 0.031227178 |
| chr1:205449885-205450256 | -1.052836621 | 0.003190857 | 0.031251218 |
| chr17:69277746-69278090 | -1.399161599 | 0.003191114 | 0.031251218 |
| chr1:209590511-209590802 | -1.204305243 | 0.003192874 | 0.031264462 |
| chr16:73673671-73674038 | 1.487950867 | 0.003196289 | 0.031293904 |
| chr3:180063821-180064101 | -1.449527317 | 0.003197199 | 0.031296077 |
| chr1:78544027-78544365 | -1.301356605 | 0.003197328 | 0.031296077 |
| chr1:240958756-240958956 | 1.683604694 | 0.003198514 | 0.03130297 |
| chr2:128269432-128269655 | 1.454564352 | 0.00319885 | 0.03130297 |
| chr18:67418084-67418843 | -1.103647628 | 0.003199734 | 0.031307622 |
| chr1:217421719-217421940 | 1.444151793 | 0.003200635 | 0.031312438 |
| chrX:111402274-111402653 | -1.374644086 | 0.00320155 | 0.031315055 |
| chr14:42586665-42586873 | 1.571090772 | 0.003201719 | 0.031315055 |
| chr6:49622712-49623807 | -1.03159876 | 0.003202278 | 0.031316526 |
| chr7:154789482-154789747 | -1.454401956 | 0.003205625 | 0.031345255 |
| chr20:33009832-33010051 | 1.725577701 | 0.003206189 | 0.031346763 |
| chrX:31137556-31137784 | -1.595031042 | 0.003208016 | 0.031360626 |
| chr3:158766432-158766656 | 1.54793916 | 0.003209434 | 0.031370489 |
| chr3:153958042-153958303 | -1.638077965 | 0.003210522 | 0.031377119 |
| chr17:58909321-58909547 | 1.135154201 | 0.003212207 | 0.031389579 |
| chr10:90712434-90713346 | -1.05942075 | 0.003213913 | 0.031402251 |
| chr3:161558446-161558656 | 1.464878471 | 0.003215888 | 0.031417538 |
| chr1:8314373-8314611 | 1.407049749 | 0.003216322 | 0.031417777 |
| chr17:55324568-55324768 | -1.651652533 | 0.003217197 | 0.031422313 |
| chr1:200090391-200090967 | -1.034115078 | 0.003217859 | 0.03142478 |
| chr9:107438206-107439051 | -1.05196366 | 0.003218986 | 0.031428873 |
| chr5:137622809-137623144 | -1.264783063 | 0.003219125 | 0.031428873 |

| | | | |
|---|---|---|---|
| chr3:129095350-129095654 | 1.454053818 | 0.003219509 | 0.031428873 |
| chr7:92048465-92048733 | -1.493856679 | 0.003220854 | 0.031437995 |
| chr1:72693349-72693815 | 1.133828265 | 0.003222487 | 0.03144618 |
| chr1:65264477-65264809 | -1.383710909 | 0.003222513 | 0.03144618 |
| chr2:189614715-189614989 | -1.544173808 | 0.003226139 | 0.031477548 |
| chr20:48259474-48259676 | 1.106982414 | 0.003226823 | 0.031480216 |
| chr1:168939989-168940340 | -1.456972016 | 0.00322837 | 0.031491298 |
| chr1:114181308-114181738 | 1.035304972 | 0.003229413 | 0.031493618 |
| chr16:8776141-8776383 | 1.477603708 | 0.00322943 | 0.031493618 |
| chr3:135788379-135788711 | 1.424651674 | 0.003230517 | 0.031495516 |
| chr10:31940825-31941193 | -1.175980853 | 0.003230853 | 0.031495516 |
| chr19:663117-663734 | 1.05165003 | 0.003230857 | 0.031495516 |
| chr1:114687821-114688086 | 1.070409452 | 0.003233972 | 0.031521865 |
| chr5:78678379-78679305 | -1.045411178 | 0.00323599 | 0.031534823 |
| chr5:150915192-150915458 | -1.732501454 | 0.003236124 | 0.031534823 |
| chr4:76208353-76208714 | -1.16135564 | 0.003237392 | 0.031543165 |
| chr1:58942932-58943312 | 1.184857889 | 0.003240185 | 0.031566366 |
| chr12:24563245-24563479 | -1.434746042 | 0.003240773 | 0.031568078 |
| chr1:108845262-108845462 | 1.522379612 | 0.003241512 | 0.031571266 |
| chr7:24572508-24574411 | 0.939333352 | 0.003242686 | 0.031578389 |
| chr4:109332995-109333195 | -1.798899573 | 0.003243068 | 0.031578389 |
| chr12:3999346-3999880 | -1.133669258 | 0.003244462 | 0.031587952 |
| chr5:633907-634157 | 1.428060127 | 0.003245029 | 0.031589457 |
| chr16:14372352-14372552 | 1.603445259 | 0.003248677 | 0.031620954 |
| chrX:115892971-115893171 | 1.521609585 | 0.003249746 | 0.031627334 |
| chr7:12490163-12490477 | 1.140093475 | 0.003250844 | 0.031634007 |
| chr20:48585129-48585365 | 1.082815031 | 0.003253636 | 0.031657156 |
| chr7:140238682-140239120 | -1.216104245 | 0.003255717 | 0.03167236 |
| chr12:122355454-122355654 | 1.733016536 | 0.003256025 | 0.03167236 |
| chr6:118776940-118777490 | -1.203483909 | 0.003258437 | 0.031691797 |
| chr2:65867897-65868477 | -1.331038319 | 0.003260677 | 0.031704097 |
| chr2:101410556-101411141 | -1.126194827 | 0.003260817 | 0.031704097 |
| chr15:31338963-31339217 | -1.379638103 | 0.003260943 | 0.031704097 |
| chr11:92211752-92211984 | -1.648555441 | 0.003264168 | 0.031729024 |
| chr6:108966879-108967548 | -1.05911533 | 0.00326438 | 0.031729024 |
| chr2:40732816-40733180 | -1.43594369 | 0.003264956 | 0.031729024 |
| chr8:124638388-124638618 | 1.425053817 | 0.003265242 | 0.031729024 |
| chr10:11805805-11806005 | 1.68707891 | 0.003265578 | 0.031729024 |
| chr7:50921119-50921481 | -1.164311216 | 0.003266454 | 0.031732458 |
| chr10:109820357-109820566 | 1.641696724 | 0.003267173 | 0.031732458 |
| chr17:38743923-38744123 | 1.641696724 | 0.003267173 | 0.031732458 |
| chr8:96972257-96972457 | 1.722955093 | 0.003274731 | 0.031801828 |
| chr12:14936192-14936444 | -1.655592308 | 0.0032788 | 0.031837311 |
| chr2:104947352-104947804 | 1.15018335 | 0.003280106 | 0.031845957 |
| chr4:161252492-161252826 | -1.231505907 | 0.003281855 | 0.031856955 |
| chr1:201511948-201512530 | 0.998959302 | 0.00328207 | 0.031856955 |
| chr8:108225276-108225617 | 1.184725653 | 0.00328434 | 0.031871289 |
| chrX:140712538-140713329 | -1.100631528 | 0.003284526 | 0.031871289 |
| chr10:70920391-70920895 | -1.09863504 | 0.003284795 | 0.031871289 |
| chr17:56652725-56652929 | 1.343754365 | 0.003288695 | 0.031902713 |

| | | | |
|---|---|---|---|
| chr11:93065371-93065665 | -1.363419622 | 0.003288866 | 0.031902713 |
| chr6:165725296-165725545 | -1.674228907 | 0.003289703 | 0.031906785 |
| chr12:47773051-47773904 | 1.029694744 | 0.003290721 | 0.03191078 |
| chr14:40601023-40601235 | 1.744036786 | 0.003290948 | 0.03191078 |
| chr11:101760059-101760429 | -1.292424225 | 0.003291676 | 0.031913809 |
| chr3:10708625-10708970 | 1.301820989 | 0.003295788 | 0.031947945 |
| chr21:38755635-38756241 | -1.075876032 | 0.003296031 | 0.031947945 |
| chr1:67468272-67468679 | -1.180091883 | 0.003297637 | 0.031953728 |
| chr16:66081838-66082340 | -1.17833656 | 0.003297751 | 0.031953728 |
| chr2:232925265-232925486 | 1.565166857 | 0.003297968 | 0.031953728 |
| chr20:44058093-44058443 | 1.315850206 | 0.003298562 | 0.031953728 |
| chr3:174639929-174640736 | -1.025679605 | 0.003298713 | 0.031953728 |
| chr7:7541446-7541730 | 1.190744852 | 0.003299574 | 0.031956136 |
| chr16:445754-446061 | 1.450538015 | 0.003300183 | 0.031956136 |
| chr13:45490238-45490488 | -1.684838693 | 0.003300213 | 0.031956136 |
| chr4:75180720-75181516 | -1.087923209 | 0.003300978 | 0.031959506 |
| chr7:12494661-12495007 | 1.136911758 | 0.003306226 | 0.032006272 |
| chr17:44095694-44095974 | 1.252163639 | 0.003308604 | 0.032023961 |
| chr17:35587116-35587813 | 1.068726939 | 0.003308889 | 0.032023961 |
| chr1:107208642-107208847 | 1.116691366 | 0.003310947 | 0.032038229 |
| chr7:108805865-108806167 | -1.36381003 | 0.0033112 | 0.032038229 |
| chr3:43638642-43639088 | -1.386454281 | 0.003314379 | 0.032064946 |
| chr1:119997076-119997280 | 1.638465222 | 0.00331512 | 0.032065651 |
| chr1:204867453-204867668 | 1.307234997 | 0.003315289 | 0.032065651 |
| chr2:628641-629247 | -1.169155569 | 0.003317535 | 0.032083318 |
| chr1:214719300-214719517 | -1.474829457 | 0.003319093 | 0.032094343 |
| chr4:186946722-186947021 | -1.214460418 | 0.003320263 | 0.032097177 |
| chr10:81570912-81571193 | 1.412278891 | 0.003320523 | 0.032097177 |
| chr2:228681618-228681819 | -1.644980739 | 0.003320967 | 0.032097177 |
| chr7:134432299-134433115 | -0.979382256 | 0.003321463 | 0.032097177 |
| chr2:164536891-164537157 | -1.535164968 | 0.003321481 | 0.032097177 |
| chr19:58228276-58229106 | 1.036773947 | 0.003322763 | 0.032105513 |
| chr8:61656181-61656615 | -1.345184984 | 0.003324767 | 0.032119554 |
| chr1:152215740-152216003 | 1.316280531 | 0.003325054 | 0.032119554 |
| chr2:120319117-120319405 | 1.331531168 | 0.003327028 | 0.032134571 |
| chr1:85502627-85503120 | -1.086849011 | 0.003329443 | 0.032153841 |
| chr5:85638475-85639086 | -1.160438425 | 0.003330772 | 0.032162619 |
| chr10:64120615-64121066 | -1.212964537 | 0.003332229 | 0.032172639 |
| chr7:120337357-120337586 | -1.784672586 | 0.003334543 | 0.032190925 |
| chr3:70982830-70983185 | 1.17353757 | 0.003335864 | 0.032196509 |
| chr5:135843736-135844101 | 1.390465202 | 0.003335962 | 0.032196509 |
| chr1:107237874-107238188 | 1.043172344 | 0.003337224 | 0.032204634 |
| chr9:106892700-106893243 | -1.128630242 | 0.003338233 | 0.032210308 |
| chr6:156925838-156926377 | -1.218485752 | 0.00334085 | 0.032231507 |
| chr9:38187054-38187258 | -1.711695386 | 0.003341469 | 0.032233418 |
| chr6:20200488-20201135 | -1.159947951 | 0.003342298 | 0.032234151 |
| chr1:197883938-197884790 | -0.992797704 | 0.003342386 | 0.032234151 |
| chr17:74758008-74758579 | -1.035927418 | 0.003343702 | 0.032242784 |
| chr8:80232054-80232336 | 1.18544445 | 0.003346829 | 0.032265091 |
| chr18:61581798-61582252 | -1.122430227 | 0.003347345 | 0.032265091 |

| | | | |
|---|---|---|---|
| chr7:14049483-14050148 | 1.045126613 | 0.003347761 | 0.032265091 |
| chr11:36334037-36334251 | 1.682471083 | 0.003348174 | 0.032265091 |
| chr11:39267852-39268207 | -1.429761793 | 0.003348406 | 0.032265091 |
| chr4:157204595-157205055 | -1.136491307 | 0.003348542 | 0.032265091 |
| chr20:48663728-48664119 | 0.982540926 | 0.003349322 | 0.032268542 |
| chr7:21086180-21086381 | 1.395591558 | 0.003349798 | 0.032269075 |
| chr10:131453525-131453753 | 1.682015577 | 0.003350908 | 0.032275706 |
| chr3:11021980-11022180 | 1.411099265 | 0.003352749 | 0.032289388 |
| chr4:52848370-52848615 | 1.352325263 | 0.003354364 | 0.032300876 |
| chr17:66717714-66718163 | -1.093353879 | 0.003354878 | 0.032301771 |
| chr17:61629990-61630290 | 1.002608532 | 0.003362422 | 0.032368185 |
| chr7:142960974-142961295 | -1.300063088 | 0.003362621 | 0.032368185 |
| chr6:102795101-102795468 | -1.292048057 | 0.003366309 | 0.032399611 |
| chr14:90791156-90791375 | 1.478120204 | 0.003368078 | 0.032412567 |
| chr2:85106670-85107195 | 1.160574262 | 0.003368647 | 0.032413976 |
| chr7:80078098-80078425 | -1.526230227 | 0.003380263 | 0.032521662 |
| chr17:41872007-41872249 | 1.635983127 | 0.003380882 | 0.032523534 |
| chr3:73637150-73637372 | -1.572493318 | 0.00338213 | 0.03253145 |
| chr5:139709061-139709536 | -1.105219515 | 0.003382981 | 0.032533547 |
| chr10:113552089-113552407 | 1.3014637 | 0.003383197 | 0.032533547 |
| chr7:130996825-130997025 | -1.545957328 | 0.003385369 | 0.032550354 |
| chr3:114144990-114145723 | -1.014945906 | 0.003392345 | 0.032609588 |
| chr7:18573529-18573729 | 1.601899836 | 0.003392472 | 0.032609588 |
| chr1:33218800-33219181 | 1.231297734 | 0.003392807 | 0.032609588 |
| chr12:82931231-82932011 | 1.021075236 | 0.003393676 | 0.032611902 |
| chr10:5873505-5873805 | 1.157645166 | 0.003393899 | 0.032611902 |
| chr5:76180962-76181481 | -1.168520888 | 0.003394495 | 0.032611921 |
| chr1:219416429-219417011 | -1.031244919 | 0.003394916 | 0.032611921 |
| chr10:84613258-84614051 | -0.964717086 | 0.003395194 | 0.032611921 |
| chr6:99547638-99548045 | -1.276951541 | 0.003395603 | 0.032611921 |
| chr8:88474304-88474997 | 1.091109469 | 0.003398374 | 0.032634439 |
| chr6:30198351-30198624 | 1.543176014 | 0.003401315 | 0.032658587 |
| chr2:218310926-218311138 | 1.420781108 | 0.00340189 | 0.032660017 |
| chr7:14787985-14788261 | 1.343511751 | 0.003403807 | 0.032672421 |
| chr5:124806430-124806953 | -1.265559762 | 0.003404035 | 0.032672421 |
| chr3:149924841-149925153 | 1.393186475 | 0.003404775 | 0.032675434 |
| chr17:59390794-59391179 | 1.114664341 | 0.003406028 | 0.032683367 |
| chr13:47182208-47182427 | 1.373903206 | 0.003407925 | 0.032697473 |
| chr8:84237026-84237252 | 1.779009243 | 0.00340935 | 0.032707048 |
| chr12:49707384-49707853 | 1.04573894 | 0.003411685 | 0.032725357 |
| chr10:43076937-43077138 | 1.392638891 | 0.003413337 | 0.032733988 |
| chr1:11052177-11052421 | 1.504632461 | 0.003413483 | 0.032733988 |
| chr19:53333133-53333390 | 1.448241849 | 0.003414778 | 0.032733988 |
| chr10:1241189-1241457 | 1.43394714 | 0.003414893 | 0.032733988 |
| chr2:15269061-15269371 | -1.432184869 | 0.003415081 | 0.032733988 |
| chr3:62587363-62587579 | 1.163865705 | 0.003415148 | 0.032733988 |
| chr4:2402030-2402379 | 1.178517703 | 0.003415624 | 0.032734449 |
| chr11:104969477-104969763 | -1.470360832 | 0.00341696 | 0.032743161 |
| chr5:38782990-38783333 | 1.360006153 | 0.003418176 | 0.032745933 |
| chr17:61518047-61518489 | 0.966720575 | 0.003418505 | 0.032745933 |

| | | | |
|---|---|---|---|
| chr14:101191134-101191445 | 1.346267552 | 0.003418532 | 0.032745933 |
| chr20:48088934-48089160 | 1.064117538 | 0.003420909 | 0.032764615 |
| chr17:20997254-20997533 | 1.263753983 | 0.003422411 | 0.032774902 |
| chr7:117435469-117435809 | -1.346553321 | 0.003424181 | 0.032787756 |
| chr1:247961727-247962048 | 1.520440241 | 0.00342553 | 0.032793331 |
| chr3:69086132-69086737 | -1.021808599 | 0.00342562 | 0.032793331 |
| chr10:80320185-80320851 | -1.019326191 | 0.003426308 | 0.032795821 |
| chr4:103567262-103567466 | -1.729425612 | 0.003427645 | 0.032804525 |
| chr9:32525180-32525479 | -1.264604601 | 0.003428196 | 0.032805699 |
| chr5:102562012-102562495 | -1.146877548 | 0.003428901 | 0.032808351 |
| chr21:37071858-37072336 | 1.049659764 | 0.003430115 | 0.032815863 |
| chr3:115967902-115968106 | 1.714212588 | 0.003430772 | 0.032818059 |
| chr6:6320312-6320761 | -1.284577934 | 0.003432637 | 0.032831791 |
| chr5:170358142-170358827 | -1.109312759 | 0.003433088 | 0.03283201 |
| chr7:34638505-34638751 | 1.543023493 | 0.003433876 | 0.032835019 |
| chr9:83451319-83451682 | -1.380446739 | 0.00343426 | 0.032835019 |
| chr10:956191-956409 | 1.44742203 | 0.003438473 | 0.032871199 |
| chr1:246597382-246597732 | -1.211496936 | 0.003440011 | 0.032881805 |
| chr1:184838647-184839043 | -1.134172958 | 0.003440655 | 0.032882942 |
| chr1:205805108-205805675 | -1.028234998 | 0.003440989 | 0.032882942 |
| chrX:104408745-104409082 | -1.401675679 | 0.003444149 | 0.032902665 |
| chr7:21128559-21128781 | 1.364015783 | 0.003444357 | 0.032902665 |
| chr11:46237776-46238023 | 1.304428452 | 0.003444737 | 0.032902665 |
| chr20:11716190-11716545 | -1.204872049 | 0.003445512 | 0.032902665 |
| chr3:189159997-189160197 | 1.780097626 | 0.003445629 | 0.032902665 |
| chr6:53264230-53264430 | 1.780097626 | 0.003445629 | 0.032902665 |
| chr1:154182175-154182484 | 1.153111322 | 0.003446255 | 0.03290454 |
| chr4:3531926-3532155 | 1.379534131 | 0.003446783 | 0.03290548 |
| chr9:38025884-38026157 | 1.393875044 | 0.003451143 | 0.032941613 |
| chr17:25919975-25920432 | 0.991079147 | 0.003451485 | 0.032941613 |
| chr9:30732110-30732586 | -1.222505251 | 0.003451857 | 0.032941613 |
| chr2:48707478-48707840 | -1.212271524 | 0.003459464 | 0.033010095 |
| chr3:4034692-4034914 | 1.558289829 | 0.003461939 | 0.033029595 |
| chrX:134542090-134542319 | 1.675844452 | 0.003463699 | 0.033041466 |
| chr9:98306251-98306452 | 1.541326147 | 0.003464127 | 0.033041466 |
| chr12:119902100-119902364 | 1.285235024 | 0.003464728 | 0.033041466 |
| chr11:70747089-70747340 | 1.456303991 | 0.003464908 | 0.033041466 |
| chr8:84621310-84621683 | 1.279682886 | 0.003465771 | 0.033042262 |
| chr7:29988921-29989962 | 1.001010489 | 0.003466215 | 0.033042262 |
| chr9:110842212-110842456 | -1.365884305 | 0.003466285 | 0.033042262 |
| chr7:99314393-99314767 | -1.362988948 | 0.003467298 | 0.033047801 |
| chr2:118242272-118242512 | 1.424945083 | 0.00346841 | 0.033054289 |
| chrX:123884909-123885112 | 1.59722119 | 0.003469822 | 0.033063638 |
| chr7:22372853-22373206 | 1.20206941 | 0.003476295 | 0.033121194 |
| chr19:58002507-58003010 | 1.152213929 | 0.003478091 | 0.03313265 |
| chr6:24389133-24389778 | -1.114283314 | 0.003478362 | 0.03313265 |
| chr12:75921941-75922188 | 1.324975475 | 0.003479279 | 0.033137267 |
| chr1:161676962-161677186 | 1.591285088 | 0.003481741 | 0.033154364 |
| chrX:136768510-136768785 | 1.325034745 | 0.003481939 | 0.033154364 |
| chr5:34706703-34707228 | 1.138248504 | 0.00348242 | 0.033154821 |

| | | | |
|---|---|---|---|
| chr21:42125712-42126342 | -1.046490847 | 0.00348374 | 0.033163271 |
| chr13:81061046-81061953 | -1.059070662 | 0.003485346 | 0.03317443 |
| chr6:108712926-108713387 | -1.086108168 | 0.00348608 | 0.033177299 |
| chr14:66717743-66718037 | 1.464163228 | 0.003486892 | 0.033180903 |
| chr15:22795330-22795869 | -1.245022923 | 0.003490708 | 0.033213089 |
| chr3:109900686-109901126 | -1.16719701 | 0.003492956 | 0.033230355 |
| chr15:39580720-39581418 | 0.985543404 | 0.00349516 | 0.033247197 |
| chr7:40117282-40117519 | -1.55164264 | 0.00349664 | 0.033257146 |
| chr18:50611468-50612126 | 1.055346703 | 0.003497319 | 0.03325859 |
| chr4:97589567-97590659 | -0.977968895 | 0.00349766 | 0.03325859 |
| chr22:41543804-41544232 | 1.088729056 | 0.003498441 | 0.033261887 |
| chr1:151547141-151548205 | 0.968115855 | 0.003499763 | 0.033270327 |
| chr5:171629141-171629367 | 1.391796048 | 0.003500401 | 0.033272078 |
| chr7:123701566-123701852 | -1.265490705 | 0.003500816 | 0.033272078 |
| chr8:86836719-86837032 | -1.235929666 | 0.003502035 | 0.033279544 |
| chrX:119870654-119872285 | 0.953047775 | 0.003502825 | 0.033282921 |
| chr2:23213012-23213536 | 1.180675813 | 0.003503569 | 0.033285495 |
| chr17:54603659-54603944 | 1.458152574 | 0.003503965 | 0.033285495 |
| chr8:61655193-61655639 | -1.251108935 | 0.003506578 | 0.033306184 |
| chr11:90134423-90134767 | -1.496846529 | 0.003508974 | 0.033324816 |
| chr16:4315618-4316129 | 1.055931804 | 0.003511427 | 0.033343974 |
| chr7:143494722-143494922 | -1.755465727 | 0.003512199 | 0.03334718 |
| chr4:168479979-168480855 | -0.975841052 | 0.003512845 | 0.033347721 |
| chr1:205449351-205449685 | -1.097830684 | 0.003513127 | 0.033347721 |
| chr20:48376320-48376635 | 1.028865884 | 0.003513606 | 0.033348142 |
| chr14:92603553-92603817 | 1.279252469 | 0.003521763 | 0.033421051 |
| chr20:58556613-58556905 | 1.135420411 | 0.003522161 | 0.033421051 |
| chr2:68403198-68403538 | -1.447584501 | 0.003523386 | 0.033428544 |
| chr15:69467652-69468130 | 1.036065934 | 0.003523858 | 0.033428878 |
| chr17:79349853-79350123 | 1.360074977 | 0.003527011 | 0.033454651 |
| chr8:123930809-123931020 | 1.454381296 | 0.003528615 | 0.033465717 |
| chr9:117957279-117957537 | -1.618223565 | 0.003531695 | 0.033489861 |
| chr19:52690215-52690803 | 1.06125074 | 0.003532138 | 0.033489861 |
| chr12:16189925-16190187 | -1.505426583 | 0.003532472 | 0.033489861 |
| chr7:16794107-16794453 | 1.245798112 | 0.003533054 | 0.033491234 |
| chr16:17064460-17064937 | 1.113753505 | 0.003536463 | 0.033519406 |
| chr7:157875506-157875774 | -1.351470168 | 0.003536921 | 0.033519603 |
| chr2:73256666-73256868 | -1.419235131 | 0.003538885 | 0.033534068 |
| chr14:30683595-30683882 | 1.465855275 | 0.003540306 | 0.033543379 |
| chrX:151713918-151714166 | 1.262448046 | 0.003546498 | 0.033597897 |
| chr14:53226574-53227643 | -1.011886514 | 0.003547535 | 0.033603565 |
| chr14:38809160-38809414 | 1.45664994 | 0.003548227 | 0.033605965 |
| chr14:50924339-50924580 | 1.58857383 | 0.003548921 | 0.033608385 |
| chr7:17038871-17039103 | 1.427318772 | 0.003552374 | 0.033636922 |
| chr20:50806332-50806583 | 1.00966919 | 0.003555399 | 0.033661407 |
| chr6:13383499-13383893 | -1.281192221 | 0.003561206 | 0.033708505 |
| chr16:53511137-53511732 | 1.054737356 | 0.003561253 | 0.033708505 |
| chr16:24989049-24989332 | 1.213263512 | 0.003562194 | 0.033713242 |
| chr9:11254458-11254740 | 1.380618252 | 0.003563164 | 0.033718258 |
| chrX:84724527-84724824 | -1.411521373 | 0.003563712 | 0.033719284 |

| | | | |
|---|---|---|---|
| chr9:81371883-81372164 | -1.371349633 | 0.003565574 | 0.033732729 |
| chr5:361455-361660 | 1.593435749 | 0.00356659 | 0.033738183 |
| chr7:117993358-117993771 | -1.095314472 | 0.003567376 | 0.033741451 |
| chr20:33789586-33789834 | 1.30671964 | 0.003568177 | 0.033744861 |
| chr11:79566129-79566546 | 1.269272556 | 0.003572798 | 0.033782745 |
| chr2:36889749-36890108 | -1.384008489 | 0.00357321 | 0.033782745 |
| chr4:170991023-170991511 | -1.184228754 | 0.003573506 | 0.033782745 |
| chr6:97288290-97288560 | -1.397809503 | 0.00357405 | 0.033783728 |
| chrX:119692443-119692851 | 1.144699795 | 0.003574856 | 0.033787178 |
| chrX:154490514-154491069 | 0.997866401 | 0.003578042 | 0.033813115 |
| chr8:134457208-134457535 | -1.226350162 | 0.003578781 | 0.033815927 |
| chr2:48622813-48623020 | -1.683028322 | 0.003579249 | 0.033816177 |
| chr7:122679191-122679707 | -1.012004726 | 0.003582308 | 0.033840642 |
| chr12:14418578-14418881 | -1.464281833 | 0.003582722 | 0.033840642 |
| chr9:120738193-120738393 | 1.768340402 | 0.003583456 | 0.033843404 |
| chr10:6401527-6402117 | 1.017439607 | 0.003585384 | 0.033857439 |
| chr5:148687392-148687829 | -1.093957737 | 0.003587046 | 0.033868966 |
| chr4:137732819-137733681 | -0.99850034 | 0.003588276 | 0.033876406 |
| chr3:101725472-101725833 | -1.155327486 | 0.003590847 | 0.033893471 |
| chr9:23829651-23830120 | -1.277467 | 0.003591177 | 0.033893471 |
| chr1:204868304-204868565 | -1.338126489 | 0.003591411 | 0.033893471 |
| chr7:108497178-108497432 | -1.394485552 | 0.003594148 | 0.033915124 |
| chr8:127035019-127035349 | -1.192722556 | 0.003595416 | 0.033922911 |
| chr4:124908954-124909235 | -1.259501351 | 0.003599857 | 0.033960631 |
| chr7:20599140-20599375 | 1.340097534 | 0.003602024 | 0.033971227 |
| chrX:130216448-130216740 | -1.429113801 | 0.003602188 | 0.033971227 |
| chr19:4008295-4008593 | 1.152397971 | 0.00360231 | 0.033971227 |
| chr20:48457607-48457941 | 0.993739238 | 0.003603104 | 0.03397195 |
| chr7:105652393-105652699 | -1.159790131 | 0.003603278 | 0.03397195 |
| chr17:75078447-75079280 | -1.009176037 | 0.003603717 | 0.03397195 |
| chr22:31290578-31290965 | 1.076165123 | 0.003604797 | 0.033977951 |
| chr13:81875417-81875632 | -1.636490295 | 0.003606832 | 0.03399295 |
| chr3:136095414-136095629 | 1.591143214 | 0.003607708 | 0.033997026 |
| chr17:48169308-48169587 | 1.402470335 | 0.003610541 | 0.03401954 |
| chrX:17690206-17690406 | 1.707900686 | 0.003611333 | 0.034022817 |
| chr10:58435436-58435636 | -1.529929124 | 0.003612101 | 0.034025871 |
| chr4:167688365-167689122 | -1.033377697 | 0.003615678 | 0.034055372 |
| chr6:35776418-35777078 | -1.102462162 | 0.003620457 | 0.034096195 |
| chr17:74430563-74431278 | 0.98757724 | 0.003621213 | 0.034098416 |
| chr11:47139122-47139521 | -1.144150086 | 0.003621995 | 0.034098416 |
| chr16:88662431-88662725 | 1.190148147 | 0.003622028 | 0.034098416 |
| chr12:100567238-100567503 | -1.640067996 | 0.00362292 | 0.034102003 |
| chr2:16218699-16219047 | -1.183056481 | 0.003623299 | 0.034102003 |
| chr16:65113442-65113642 | 1.590625667 | 0.003627097 | 0.034133553 |
| chr2:187541772-187542023 | 1.374585769 | 0.003629849 | 0.034154207 |
| chr10:1570655-1570992 | 1.246780844 | 0.003630284 | 0.034154207 |
| chr3:25582241-25582654 | 1.045154164 | 0.00363079 | 0.034154207 |
| chr1:147223876-147224139 | 1.347051983 | 0.003631074 | 0.034154207 |
| chr4:49512652-49512879 | 1.391818512 | 0.003632272 | 0.034160104 |
| chr10:51947470-51947798 | -1.4207443 | 0.003632741 | 0.034160104 |

| | | | |
|---|---|---|---|
| chr13:65217474-65217952 | -1.11945926 | 0.003633039 | 0.034160104 |
| chr7:104922876-104923236 | -1.122384093 | 0.003633516 | 0.034160397 |
| chr14:103146757-103146968 | 1.519341206 | 0.003636164 | 0.03417769 |
| chr14:67313207-67313407 | 1.453729433 | 0.003636247 | 0.03417769 |
| chr4:107380886-107381362 | -1.123880729 | 0.003638832 | 0.034197786 |
| chr5:98905916-98906172 | -1.412116267 | 0.00363969 | 0.034200216 |
| chr8:90666166-90666370 | 1.372114819 | 0.003639983 | 0.034200216 |
| chr1:116909734-116910710 | 0.961018835 | 0.003644044 | 0.034234172 |
| chr3:129385403-129385943 | -1.123720809 | 0.003645054 | 0.034239335 |
| chr1:51324246-51324633 | -1.161723056 | 0.003645487 | 0.034239335 |
| chr11:117717639-117718330 | 1.017149023 | 0.003646267 | 0.034242467 |
| chr3:186974786-186975301 | -1.072340678 | 0.003647639 | 0.034251149 |
| chr2:75311949-75312344 | -1.121555287 | 0.003649571 | 0.034265097 |
| chr6:152110974-152111535 | -1.080373231 | 0.003651115 | 0.034275394 |
| chr14:87653961-87654254 | 1.424627859 | 0.003652369 | 0.034282959 |
| chr7:73587635-73588221 | -1.118659067 | 0.003652993 | 0.034284616 |
| chr22:11931245-11932569 | 0.980488821 | 0.003656287 | 0.034309094 |
| chr7:1511277-1511477 | 1.305793416 | 0.003656496 | 0.034309094 |
| chr7:24966169-24966640 | 1.113681647 | 0.003657382 | 0.034313205 |
| chr1:155977691-155977952 | 1.232117002 | 0.003659317 | 0.034325438 |
| chr10:56116751-56116981 | -1.468303361 | 0.003659582 | 0.034325438 |
| chr6:5034808-5035112 | 1.282721122 | 0.003660999 | 0.034334525 |
| chr2:40168014-40168505 | -1.143464177 | 0.003661531 | 0.034335277 |
| chr2:222583024-222583242 | -1.67897067 | 0.003662099 | 0.034335277 |
| chr10:110840477-110840897 | -1.205732072 | 0.003662424 | 0.034335277 |
| chr8:125947149-125947459 | 1.335079667 | 0.003670364 | 0.03440551 |
| chr7:129054439-129055558 | -0.948785765 | 0.003671352 | 0.034408098 |
| chr8:97528804-97529004 | 1.51798892 | 0.003671577 | 0.034408098 |
| chr16:28609343-28609591 | 1.367395398 | 0.0036724 | 0.034408098 |
| chr3:111582032-111582413 | 1.114607762 | 0.003672436 | 0.034408098 |
| chr14:75427452-75428116 | 1.009594729 | 0.003676198 | 0.034439127 |
| chr18:63155243-63156082 | -0.983547357 | 0.003678516 | 0.034456633 |
| chr9:5440495-5440738 | 1.314890327 | 0.003681622 | 0.034481507 |
| chr8:95100410-95100999 | 0.991476692 | 0.003682459 | 0.034485134 |
| chr5:137833454-137833799 | -1.221190214 | 0.003683016 | 0.03448614 |
| chr10:125406897-125407853 | -0.979723917 | 0.00368569 | 0.034503831 |
| chr4:175509346-175509709 | -1.182714296 | 0.00368674 | 0.034503831 |
| chr1:84606774-84607239 | -1.188951473 | 0.003687284 | 0.034503831 |
| chr1:209692773-209693194 | -1.254025036 | 0.003687304 | 0.034503831 |
| chr20:47839167-47840200 | 0.937988059 | 0.003687726 | 0.034503831 |
| chr1:227491443-227491816 | 1.281575837 | 0.003687944 | 0.034503831 |
| chr7:82044573-82044821 | -1.57843522 | 0.003688058 | 0.034503831 |
| chr6:122405483-122405756 | -1.455813371 | 0.003689163 | 0.034508053 |
| chr18:48142350-48142915 | -1.11852341 | 0.00368941 | 0.034508053 |
| chr5:65481838-65482131 | -1.182022402 | 0.003690465 | 0.034513706 |
| chrX:122363062-122364428 | -0.949083586 | 0.003693099 | 0.034534124 |
| chr2:205047508-205047793 | -1.439420578 | 0.003694443 | 0.034542477 |
| chr12:82857315-82857558 | 1.547798472 | 0.003697012 | 0.034562217 |
| chr8:93856417-93856763 | 1.290565342 | 0.003697456 | 0.034562217 |
| chr8:25660076-25660847 | -1.092693595 | 0.003698361 | 0.034566457 |

| | | | |
|---|---|---|---|
| chr7:107506065-107507163 | -0.937945278 | 0.003700715 | 0.034582012 |
| chr14:51035635-51036267 | 1.097665548 | 0.003700928 | 0.034582012 |
| chr22:20313678-20313946 | 1.492693123 | 0.003707549 | 0.034639656 |
| chr10:32378623-32378989 | -1.136765466 | 0.00370839 | 0.034643289 |
| chr16:88922409-88922783 | -1.074336821 | 0.003715647 | 0.034705393 |
| chr17:49995025-49995445 | 1.040396615 | 0.003715944 | 0.034705393 |
| chr12:132445971-132446464 | 1.040836764 | 0.00371656 | 0.034706921 |
| chr1:152697901-152698285 | 1.358672864 | 0.003720159 | 0.034732196 |
| chr14:89988110-89988327 | 1.44934338 | 0.003720173 | 0.034732196 |
| chr8:86910628-86911229 | 1.024203013 | 0.003721659 | 0.034741833 |
| chr2:137173359-137173819 | -1.339900842 | 0.003723878 | 0.034758312 |
| chr11:105315371-105315618 | -1.590950137 | 0.00372446 | 0.034759508 |
| chr6:50192886-50193101 | -1.712961129 | 0.003728172 | 0.034789912 |
| chr14:69475209-69475582 | 1.133412765 | 0.00373089 | 0.034811042 |
| chr11:9530948-9531163 | 1.585753002 | 0.003733282 | 0.034829118 |
| chr2:20378700-20379506 | -0.971705868 | 0.003735181 | 0.034842587 |
| chr4:82388483-82388718 | 1.57938157 | 0.003738245 | 0.034866928 |
| chr8:7026763-7026988 | -1.611089369 | 0.003739243 | 0.034871994 |
| chr10:2928288-2928689 | 1.152193402 | 0.003740728 | 0.034878122 |
| chr17:61218460-61218682 | 1.008498062 | 0.003740811 | 0.034878122 |
| chr15:85831564-85831881 | 1.153888899 | 0.00374237 | 0.034886013 |
| chr5:34190061-34190278 | 1.241364094 | 0.003742591 | 0.034886013 |
| chrX:85243756-85245215 | -0.957289519 | 0.003743023 | 0.034886013 |
| chr3:111151534-111151932 | 1.264041991 | 0.003744129 | 0.034892073 |
| chr9:69023677-69024124 | -1.131787139 | 0.003744845 | 0.034894508 |
| chr1:228830676-228830932 | 1.486508157 | 0.00375071 | 0.034944904 |
| chr10:58291375-58291922 | -1.113311457 | 0.003752099 | 0.034953595 |
| chr4:165124388-165124877 | -1.120048236 | 0.00375325 | 0.034960067 |
| chr1:159959385-159959953 | 1.090317113 | 0.003755209 | 0.034973267 |
| chr1:108816141-108816866 | -1.018058805 | 0.00375558 | 0.034973267 |
| chr14:43275419-43275722 | 1.34953813 | 0.00375636 | 0.034976282 |
| chr10:116044729-116045211 | -1.255609966 | 0.003757609 | 0.034983663 |
| chr7:41107437-41108399 | -0.988405832 | 0.003760672 | 0.035007924 |
| chr14:101796522-101796722 | 1.623231328 | 0.003764026 | 0.035034891 |
| chr8:100457636-100457867 | 1.372650941 | 0.003765568 | 0.03504324 |
| chr5:103014068-103014834 | -1.049898428 | 0.003765838 | 0.03504324 |
| chr7:129830071-129830281 | -1.38380742 | 0.003766646 | 0.035046503 |
| chr3:49903857-49904188 | 1.197621724 | 0.003768664 | 0.035061019 |
| chr1:104706762-104707143 | -1.18067666 | 0.003770278 | 0.035071775 |
| chr9:124110353-124110668 | 1.252515915 | 0.003771417 | 0.03507701 |
| chr1:79198901-79199548 | -1.262944825 | 0.003771756 | 0.03507701 |
| chr14:55122586-55122967 | 1.052614583 | 0.003772728 | 0.035081796 |
| chr19:42438498-42438698 | 1.502900032 | 0.003773832 | 0.035087803 |
| chr13:70339794-70340136 | -1.215825149 | 0.003775248 | 0.035096707 |
| chr20:48326865-48327192 | 1.017430569 | 0.003776317 | 0.03509967 |
| chr13:38265143-38265376 | -1.549226277 | 0.003776483 | 0.03509967 |
| chr1:192736957-192737315 | -1.357109451 | 0.003777804 | 0.035107692 |
| chr21:10147312-10147515 | 1.623248226 | 0.003778874 | 0.035113372 |
| chr17:49566060-49566408 | -1.139788228 | 0.003779336 | 0.035113407 |
| chr16:19116800-19117000 | 1.412231185 | 0.003781934 | 0.035133282 |

| | | | |
|---|---|---|---|
| chr11:128453601-128453801 | -1.708585175 | 0.003783533 | 0.035141424 |
| chr16:69713065-69713564 | -1.126816526 | 0.003783727 | 0.035141424 |
| chr4:183262469-183262967 | -1.137631173 | 0.003784485 | 0.035144205 |
| chr4:157220188-157220912 | -1.018825015 | 0.003785129 | 0.03514592 |
| chr2:177375725-177376014 | -1.456615745 | 0.003786152 | 0.035147262 |
| chr1:198686916-198687140 | 1.484675759 | 0.003786495 | 0.035147262 |
| chr1:106531932-106532191 | 1.081511182 | 0.003786649 | 0.035147262 |
| chr14:66771121-66771497 | 1.389612615 | 0.003787223 | 0.035148333 |
| chrX:66437731-66437958 | 1.536234329 | 0.003788472 | 0.03515566 |
| chr2:58634065-58634447 | -1.451803981 | 0.003789587 | 0.035161753 |
| chr14:74763607-74764192 | 1.05140806 | 0.003792665 | 0.03518605 |
| chr1:23792538-23792749 | 1.53139567 | 0.00379374 | 0.035191761 |
| chr2:221448076-221448520 | -1.274279231 | 0.003795032 | 0.035199487 |
| chr3:130828169-130828369 | 1.759703238 | 0.003795532 | 0.03519986 |
| chr13:78137509-78138200 | -1.056445404 | 0.003797634 | 0.035215097 |
| chr13:51080281-51080967 | -1.071804891 | 0.003798991 | 0.035223417 |
| chr1:115364773-115365112 | 1.277074205 | 0.00380033 | 0.035225565 |
| chrX:129768262-129768462 | 1.823384584 | 0.003800409 | 0.035225565 |
| chr7:151405465-151405665 | -1.609696015 | 0.003800602 | 0.035225565 |
| chr20:47257252-47257742 | 0.939104708 | 0.003801426 | 0.035228939 |
| chr14:26762243-26762443 | 1.666984187 | 0.003805415 | 0.035258565 |
| chr5:112738320-112738604 | -1.196142616 | 0.003805543 | 0.035258565 |
| chr2:222436011-222436431 | -1.247205207 | 0.003807699 | 0.035271447 |
| chr12:124465008-124465347 | 1.270718538 | 0.003807854 | 0.035271447 |
| chr16:27258489-27258689 | 1.516180615 | 0.003809853 | 0.035283091 |
| chr2:106560970-106561183 | 1.516028548 | 0.003810173 | 0.035283091 |
| chr18:76376644-76377205 | -1.120710806 | 0.003810493 | 0.035283091 |
| chr13:94640669-94640978 | -1.458246626 | 0.003812932 | 0.035298747 |
| chr19:50649484-50649819 | 1.266648314 | 0.00381332 | 0.035298747 |
| chr15:80710109-80710376 | 1.454504915 | 0.003813566 | 0.035298747 |
| chr1:170094664-170095255 | -1.095717559 | 0.003814923 | 0.035302749 |
| chr4:7067932-7068711 | 0.949545297 | 0.003815071 | 0.035302749 |
| chr17:61600537-61600889 | 0.977444658 | 0.00381538 | 0.035302749 |
| chr1:120133501-120133911 | 1.174103945 | 0.003818797 | 0.0353292 |
| chr8:70456374-70456749 | 1.173224265 | 0.003819161 | 0.0353292 |
| chr6:80527457-80528493 | -0.953474224 | 0.00382093 | 0.035337087 |
| chr6:47701309-47701697 | -1.172734014 | 0.003820936 | 0.035337087 |
| chr8:70770668-70771214 | -1.190543065 | 0.003821508 | 0.035337646 |
| chr9:124045331-124045760 | 1.153507989 | 0.003821919 | 0.035337646 |
| chr20:54226428-54226705 | 1.024307285 | 0.00382305 | 0.035343838 |
| chr6:120077432-120077878 | -1.105250019 | 0.003824383 | 0.035348802 |
| chr5:159913148-159913373 | -1.566415582 | 0.003824509 | 0.035348802 |
| chr10:45250434-45250647 | -1.587498199 | 0.003827708 | 0.035374101 |
| chr5:136249767-136250073 | -1.405993493 | 0.003830917 | 0.035398732 |
| chr7:135089755-135089980 | -1.766644364 | 0.003831298 | 0.035398732 |
| chr16:50368628-50369172 | 1.111619911 | 0.003833218 | 0.035410213 |
| chr7:126148929-126149451 | -1.080613787 | 0.003833465 | 0.035410213 |
| chr11:90248108-90248381 | -1.456056511 | 0.003835829 | 0.035427787 |
| chr4:186169963-186170445 | -1.212104358 | 0.003836687 | 0.035431436 |
| chr18:10417088-10417628 | -1.049601022 | 0.00383773 | 0.035435252 |

| | | | |
|---|---|---|---|
| chr1:106602705-106602931 | 1.075152607 | 0.003838025 | 0.035435252 |
| chr9:124399521-124399721 | 1.822842669 | 0.003839355 | 0.035443259 |
| chr20:41717363-41717828 | 1.075403244 | 0.003840765 | 0.035447938 |
| chr6:138572249-138572799 | -0.999655472 | 0.003840787 | 0.035447938 |
| chr9:120161773-120162282 | -1.160597041 | 0.003843472 | 0.03546441 |
| chr5:178927845-178928065 | 1.613470933 | 0.003843497 | 0.03546441 |
| chr6:56024984-56025314 | -1.488225418 | 0.003844592 | 0.035469516 |
| chr6:131792326-131792556 | -1.607841613 | 0.003844977 | 0.035469516 |
| chr4:16057162-16057616 | 1.321036917 | 0.003845655 | 0.035471502 |
| chr20:44089990-44090338 | 1.154695421 | 0.003849102 | 0.035499026 |
| chr5:92042288-92042647 | -1.14937908 | 0.003851409 | 0.035516027 |
| chr7:32688714-32688936 | 1.456132967 | 0.003856372 | 0.035554482 |
| chr11:60285671-60286052 | -1.351551254 | 0.003856507 | 0.035554482 |
| chr3:140585412-140585750 | 1.412771374 | 0.003856972 | 0.03555449 |
| chr3:65810819-65811019 | 1.710860827 | 0.003857514 | 0.035555207 |
| chr15:64700054-64700257 | 1.266954995 | 0.00385799 | 0.035555321 |
| chr11:47981183-47981527 | 1.120890549 | 0.003859204 | 0.035562229 |
| chr9:91423741-91424794 | 0.986094476 | 0.003863624 | 0.035596013 |
| chr5:137660287-137660678 | -1.180865373 | 0.003863929 | 0.035596013 |
| chr20:47598757-47599262 | 0.95200362 | 0.003864264 | 0.035596013 |
| chr14:43914177-43914377 | 1.540453808 | 0.003864897 | 0.035597562 |
| chr14:91872600-91872946 | 1.089485195 | 0.00386637 | 0.035606853 |
| chr1:81414415-81414750 | -1.166783988 | 0.003867257 | 0.035610742 |
| chr4:90039335-90039731 | 1.357373074 | 0.003867781 | 0.035611284 |
| chr15:73127366-73127617 | 1.517885361 | 0.003870412 | 0.035625334 |
| chr15:80895057-80895281 | 1.517885361 | 0.003870412 | 0.035625334 |
| chr4:173125137-173125667 | -1.118255072 | 0.003870702 | 0.035625334 |
| chr16:24844562-24844762 | 1.408538699 | 0.003871889 | 0.035631983 |
| chr7:136086799-136087124 | -1.222483842 | 0.00387284 | 0.03563645 |
| chr1:206843642-206844221 | -1.00087733 | 0.003876311 | 0.03566411 |
| chr19:57916403-57916888 | 1.065541877 | 0.003877006 | 0.035666224 |
| chr3:151631082-151631420 | -1.223032097 | 0.003878006 | 0.035669212 |
| chr7:107579623-107580774 | -0.952545733 | 0.003878262 | 0.035669212 |
| chr2:71064080-71064368 | -1.146360791 | 0.003880785 | 0.03568813 |
| chr4:6225485-6225717 | 1.543933261 | 0.003881348 | 0.035689027 |
| chr6:2908435-2908676 | 1.450000151 | 0.003882991 | 0.035697404 |
| chr7:37547484-37547684 | 1.656435074 | 0.003883456 | 0.035697404 |
| chr8:1755476-1755978 | 1.03211563 | 0.003883866 | 0.035697404 |
| chr11:14453240-14453542 | 1.463884481 | 0.003884123 | 0.035697404 |
| chr20:47296681-47296916 | 0.991181492 | 0.003888664 | 0.035734856 |
| chr15:91136309-91136575 | 1.410585114 | 0.003890844 | 0.035748717 |
| chr19:40744429-40745057 | 1.037532563 | 0.003891398 | 0.035748717 |
| chr7:158863205-158864217 | -0.989087687 | 0.003891911 | 0.035748717 |
| chr6:132937424-132937897 | -1.117863803 | 0.003892039 | 0.035748717 |
| chr11:69274573-69274925 | -1.103256709 | 0.003900072 | 0.035818212 |
| chr1:189330758-189331490 | -1.014459257 | 0.003904159 | 0.035851444 |
| chr1:208107769-208108468 | -1.112873605 | 0.003904764 | 0.035852708 |
| chr1:39270087-39270339 | 1.611324298 | 0.003906695 | 0.035862885 |
| chr21:33930996-33931196 | -1.850255008 | 0.003907066 | 0.035862885 |
| chr7:106959871-106960098 | -1.440010096 | 0.003907277 | 0.035862885 |

| | | | |
|---|---|---|---|
| chr20:48740274-48740506 | 1.044919099 | 0.003908036 | 0.035864271 |
| chr2:48759776-48760092 | -1.543544997 | 0.003908364 | 0.035864271 |
| chr9:97737588-97737828 | 1.396412789 | 0.003911127 | 0.035885331 |
| chr4:1581181-1581394 | 1.255226839 | 0.003913543 | 0.035902988 |
| chr7:40604685-40605314 | -1.097087723 | 0.003914308 | 0.035902988 |
| chr6:1838608-1839016 | -1.303892381 | 0.003914546 | 0.035902988 |
| chr1:106555202-106556229 | 0.916514217 | 0.003914926 | 0.035902988 |
| chr14:95575637-95575978 | 1.387383112 | 0.003915425 | 0.03590327 |
| chr2:164600016-164600723 | -1.024588535 | 0.003916452 | 0.03590839 |
| chr2:45218986-45219454 | 1.079475436 | 0.003917431 | 0.035913068 |
| chr3:135373460-135373691 | 1.548212809 | 0.003918898 | 0.035922218 |
| chr2:191219107-191219932 | 0.97281823 | 0.003920292 | 0.035930693 |
| chr13:38824493-38824741 | -1.51011719 | 0.003920826 | 0.035931291 |
| chr1:190863749-190864067 | -1.282796957 | 0.003922361 | 0.035937357 |
| chr1:210222258-210222604 | -1.357153284 | 0.003922426 | 0.035937357 |
| chr4:2907961-2908476 | 1.084359488 | 0.003923524 | 0.035938892 |
| chr5:175778805-175779394 | -1.070507451 | 0.003923532 | 0.035938892 |
| chr1:50181072-50181680 | -1.066359184 | 0.003926183 | 0.035956723 |
| chr5:43338428-43338777 | 1.398841807 | 0.003926417 | 0.035956723 |
| chr13:47897358-47898074 | -1.047558715 | 0.003927607 | 0.035962703 |
| chr1:44813525-44813966 | 1.266988415 | 0.003928008 | 0.035962703 |
| chr21:44299611-44300149 | 1.005128792 | 0.003933082 | 0.036004849 |
| chr18:65567448-65567660 | -1.583119907 | 0.003935143 | 0.036016529 |
| chr2:153451661-153451886 | -1.660623679 | 0.003935768 | 0.036016529 |
| chr17:47310460-47310660 | -1.660623679 | 0.003935768 | 0.036016529 |
| chr16:70162153-70162392 | 1.373992307 | 0.003936614 | 0.036019968 |
| chr7:138835177-138835453 | -1.558090738 | 0.003938816 | 0.036035812 |
| chr4:82491370-82491570 | 1.812628191 | 0.003939935 | 0.036041749 |
| chr6:114975761-114976061 | -1.346241664 | 0.003941811 | 0.036050622 |
| chr3:179750784-179751053 | -1.383488438 | 0.003941846 | 0.036050622 |
| chr12:47828893-47829556 | -1.006173185 | 0.003943739 | 0.036063636 |
| chr2:233711454-233711662 | 1.440665537 | 0.003944341 | 0.036064834 |
| chr17:74368778-74368988 | 1.285160413 | 0.003947579 | 0.036085528 |
| chr22:29377994-29378404 | 1.059416555 | 0.003947757 | 0.036085528 |
| chr6:168377478-168378051 | -1.29801608 | 0.003948443 | 0.036085528 |
| chr4:174521761-174522920 | -0.947560066 | 0.003948723 | 0.036085528 |
| chr1:212011967-212012245 | -1.352650061 | 0.003949415 | 0.036085528 |
| chr7:114405487-114405765 | -1.362500266 | 0.00394943 | 0.036085528 |
| chr8:120672936-120673136 | 1.561247981 | 0.003952341 | 0.036107818 |
| chr14:59778821-59779167 | 1.409100182 | 0.003954552 | 0.036120413 |
| chr10:100743654-100743942 | -1.287348875 | 0.003954662 | 0.036120413 |
| chr15:77935121-77935377 | 1.611638336 | 0.003955226 | 0.036121252 |
| chr7:84100899-84101157 | -1.397694827 | 0.00395592 | 0.036123287 |
| chr14:65359682-65359926 | 1.349407128 | 0.003957393 | 0.036132434 |
| chrX:39726203-39726553 | -1.323276781 | 0.003958252 | 0.036135969 |
| chr1:35524337-35524606 | 1.414447244 | 0.003960045 | 0.036148028 |
| chr10:95368750-95368972 | 1.615126787 | 0.003960933 | 0.03615183 |
| chr12:93244181-93244518 | -1.208587943 | 0.003965493 | 0.036189141 |
| chr3:135757469-135757669 | -1.660607795 | 0.003966955 | 0.036198167 |
| chr5:97400682-97400968 | -1.444552786 | 0.003968396 | 0.036205253 |

| | | | |
|---|---|---|---|
| chr18:61322200-61322623 | -1.135499685 | 0.003968676 | 0.036205253 |
| chr7:22255889-22256148 | 1.422590004 | 0.003971797 | 0.036229096 |
| chr20:58006188-58006404 | 1.21494986 | 0.003972702 | 0.036229096 |
| chr18:54922237-54922513 | -1.382424147 | 0.003972708 | 0.036229096 |
| chr11:121258014-121258370 | 1.177070995 | 0.003975482 | 0.036250073 |
| chr5:170151360-170151836 | -1.155574212 | 0.00398269 | 0.036311475 |
| chr14:73752345-73752703 | 1.110347717 | 0.003984691 | 0.036321699 |
| chr14:76117161-76117366 | 1.434605475 | 0.003985013 | 0.036321699 |
| chr9:26250083-26250372 | -1.559412305 | 0.003985233 | 0.036321699 |
| chr2:158973722-158973969 | -1.363888793 | 0.003989442 | 0.036355737 |
| chr5:150211333-150211628 | -1.243339932 | 0.003991811 | 0.036372994 |
| chr20:45833537-45834639 | 0.947361539 | 0.003993407 | 0.036383214 |
| chr9:127910581-127910994 | 1.240505598 | 0.003994243 | 0.036386503 |
| chr8:130349480-130349747 | 1.480622735 | 0.003996383 | 0.036401669 |
| chr10:82150151-82150447 | -1.403090041 | 0.003998004 | 0.036412108 |
| chr8:60458679-60458923 | -1.441760983 | 0.004000723 | 0.036432538 |
| chr1:182003893-182004499 | -1.096143443 | 0.004002915 | 0.03644817 |
| chr7:14834018-14834253 | 1.574360389 | 0.004005178 | 0.036464444 |
| chr5:54415787-54416757 | -0.965586053 | 0.004005703 | 0.036464894 |
| chr10:35563275-35563638 | -1.254896129 | 0.004007705 | 0.036478777 |
| chr12:64924174-64924452 | 1.259807484 | 0.004008501 | 0.036479879 |
| chr4:175280866-175281124 | -1.453484835 | 0.004009143 | 0.036479879 |
| chr2:177234137-177234337 | 1.70024217 | 0.004009254 | 0.036479879 |
| chr3:9955082-9955537 | 1.025113673 | 0.004009908 | 0.036481496 |
| chr6:157428755-157429090 | 1.208258514 | 0.004011057 | 0.036487619 |
| chr3:126067986-126068186 | 1.700203379 | 0.004013188 | 0.036502667 |
| chr13:82275585-82275854 | -1.618086545 | 0.004014581 | 0.036511009 |
| chr8:131884200-131884400 | 1.647923459 | 0.004015752 | 0.036517322 |
| chr7:114085630-114086771 | -0.931241793 | 0.004016846 | 0.036522931 |
| chr2:161576518-161576793 | -1.617572021 | 0.004019588 | 0.036543527 |
| chr4:108060224-108060461 | 1.294427953 | 0.004021458 | 0.036554059 |
| chr5:14646225-14646437 | 1.549392635 | 0.00402251 | 0.036554059 |
| chr7:22727239-22727441 | 1.341097267 | 0.004022574 | 0.036554059 |
| chr1:16566495-16566837 | 1.243685865 | 0.004022654 | 0.036554059 |
| chr1:247401319-247401581 | -1.402139471 | 0.004025274 | 0.036573527 |
| chr20:38471374-38471574 | 1.555202713 | 0.004026173 | 0.036577361 |
| chrX:37348731-37350213 | -0.93470439 | 0.004027379 | 0.036583941 |
| chr14:66003762-66004044 | 1.412733371 | 0.004027853 | 0.036583941 |
| chr5:115574616-115574907 | -1.375004435 | 0.004030821 | 0.036603081 |
| chr2:190418746-190419171 | -1.068513262 | 0.004030915 | 0.036603081 |
| chr11:70233338-70234015 | 0.999100626 | 0.004033387 | 0.036621187 |
| chr2:188395908-188396188 | -1.336886973 | 0.004034868 | 0.036630293 |
| chr2:180520272-180520542 | 1.345938153 | 0.004037574 | 0.036650512 |
| chr22:46803471-46803708 | -1.611372351 | 0.004039476 | 0.036659896 |
| chr5:66443165-66444070 | -1.067765822 | 0.00403995 | 0.036659896 |
| chr1:180144618-180145000 | 1.176284868 | 0.004040395 | 0.036659896 |
| chr8:88556224-88556450 | 1.328979971 | 0.004040521 | 0.036659896 |
| chr7:88226866-88227644 | -0.959282256 | 0.00404215 | 0.036670333 |
| chr4:5365791-5366108 | 1.220923426 | 0.004046822 | 0.036708368 |
| chr5:65382701-65382901 | 1.655943274 | 0.004048438 | 0.036718684 |

| | | | |
|---|---|---|---|
| chr8:100409936-100410194 | 1.373984674 | 0.004049195 | 0.0367212 |
| chr20:63817811-63818076 | 1.301170631 | 0.004051463 | 0.036737416 |
| chr8:119437417-119437713 | -1.173945477 | 0.004052355 | 0.036741159 |
| chr3:127746930-127747172 | 1.394000492 | 0.00405341 | 0.036746377 |
| chr18:58161737-58162021 | -1.247880405 | 0.004054584 | 0.03675103 |
| chr3:189584624-189585422 | -1.065819159 | 0.004054883 | 0.03675103 |
| chr2:67733459-67733964 | -1.112703472 | 0.004055691 | 0.036754011 |
| chr7:120674163-120674477 | -1.375271763 | 0.004056444 | 0.036756483 |
| chr2:73851332-73851567 | -1.506045213 | 0.004057208 | 0.036759065 |
| chr7:98013390-98013809 | -1.137439775 | 0.004058291 | 0.03676361 |
| chr16:77851082-77851382 | 1.19490989 | 0.00405867 | 0.03676361 |
| chr4:135670611-135670897 | -1.395569814 | 0.004059473 | 0.036766545 |
| chr10:97418815-97419487 | -1.076346639 | 0.004062618 | 0.036789523 |
| chr18:75752769-75753281 | -1.167921391 | 0.004062971 | 0.036789523 |
| chr5:41951523-41951859 | 1.272978783 | 0.004063512 | 0.036790074 |
| chr4:166257727-166258100 | -1.312032881 | 0.004064881 | 0.036798051 |
| chr1:150895908-150896108 | 1.43192886 | 0.004065468 | 0.036798051 |
| chr7:128031065-128031809 | -1.048454109 | 0.004065834 | 0.036798051 |
| chr7:82050135-82050576 | -1.189663553 | 0.004066669 | 0.036801269 |
| chr17:17781016-17781750 | -1.000218642 | 0.004068194 | 0.036810718 |
| chr1:212485846-212486495 | -1.051885686 | 0.004070896 | 0.036830819 |
| chr10:124308176-124308434 | -1.575765184 | 0.004071569 | 0.036832558 |
| chr7:95004593-95004932 | -1.366396126 | 0.004072069 | 0.036832735 |
| chr16:50548126-50548587 | 1.16188317 | 0.004074556 | 0.036850873 |
| chr12:110993064-110993416 | 1.196817104 | 0.00407563 | 0.036856237 |
| chr19:17747467-17747778 | 1.225876297 | 0.004077094 | 0.03686513 |
| chr1:200477809-200478200 | 1.08629833 | 0.004080699 | 0.036893372 |
| chr2:42558783-42559234 | -1.000296252 | 0.004081601 | 0.036897172 |
| chr19:46796148-46796706 | -1.056828922 | 0.004082469 | 0.036900662 |
| chr7:107563228-107564630 | -0.941710858 | 0.004083792 | 0.036908268 |
| chr2:85394789-85395348 | -1.027422917 | 0.004088665 | 0.036947952 |
| chr2:219672985-219673248 | 1.446939169 | 0.004091962 | 0.036973385 |
| chrX:151396653-151397880 | 0.941561832 | 0.00409519 | 0.036996787 |
| chr1:67736013-67736567 | -1.196681453 | 0.004095518 | 0.036996787 |
| chr1:41574495-41574743 | 1.430574207 | 0.004098067 | 0.037010477 |
| chr1:16565793-16566322 | 1.096923848 | 0.004098255 | 0.037010477 |
| chr1:169115979-169116477 | -1.104010219 | 0.004098482 | 0.037010477 |
| chr3:172307191-172307594 | -1.262304675 | 0.004098972 | 0.037010535 |
| chr9:84349669-84350119 | -1.136200391 | 0.004101582 | 0.037029738 |
| chr7:15525960-15526188 | 1.447366844 | 0.004103929 | 0.037046565 |
| chr16:77599843-77600125 | 1.11636853 | 0.004105355 | 0.037055076 |
| chr4:7917678-7917976 | 1.289184345 | 0.004108648 | 0.037080431 |
| chr14:72924224-72924811 | 0.955082375 | 0.004112792 | 0.03711346 |
| chr9:131678620-131678820 | -1.805306426 | 0.004114399 | 0.037123587 |
| chr14:62732639-62732873 | 1.64141102 | 0.004115802 | 0.037131872 |
| chr11:68411488-68411836 | 1.263874152 | 0.004118219 | 0.037149299 |
| chr9:82410353-82410748 | -1.122319009 | 0.004119514 | 0.037156611 |
| chr6:130021061-130021412 | -1.17451846 | 0.004121002 | 0.037165655 |
| chr7:19416785-19417019 | 1.529064525 | 0.004121633 | 0.037166976 |
| chr5:9775643-9775867 | 1.457948647 | 0.004122233 | 0.037168008 |

| | | | |
|---|---|---|---|
| chr16:34572531-34573293 | 0.976398063 | 0.004123145 | 0.037171441 |
| chr7:144835148-144836434 | -0.93483703 | 0.004123584 | 0.037171441 |
| chr9:135393664-135394249 | -1.040033806 | 0.004126458 | 0.037192974 |
| chr1:113519598-113519929 | 1.263471219 | 0.00413232 | 0.037241428 |
| chr1:83692648-83693357 | -1.121585055 | 0.004133213 | 0.03724384 |
| chr3:170049548-170050131 | 1.02152039 | 0.00413356 | 0.03724384 |
| chr22:32649476-32649823 | 1.419450909 | 0.004134946 | 0.037251952 |
| chr7:8261287-8262766 | 0.904843575 | 0.004137908 | 0.037273028 |
| chr1:210232406-210233947 | -0.924296377 | 0.004138487 | 0.037273028 |
| chr20:61980988-61981327 | -1.143841626 | 0.004138745 | 0.037273028 |
| chr1:148276709-148276909 | 1.457193883 | 0.00414099 | 0.037288862 |
| chr5:80679688-80679937 | -1.521689125 | 0.004144161 | 0.037308715 |
| chr19:797640-797985 | 1.085707118 | 0.004144654 | 0.037308715 |
| chr13:62358185-62358456 | -1.253596567 | 0.004144655 | 0.037308715 |
| chr19:41994127-41994402 | 1.230564854 | 0.004146021 | 0.037316622 |
| chr2:241093292-241093528 | 1.379220657 | 0.004146718 | 0.037318518 |
| chr8:109579011-109579211 | 1.401215406 | 0.004147398 | 0.037320252 |
| chr8:121951323-121951546 | 1.511460431 | 0.004150353 | 0.037342456 |
| chr12:96061867-96062187 | -1.233152039 | 0.004152552 | 0.037357855 |
| chr16:46400013-46400271 | 1.039133531 | 0.004153618 | 0.037363053 |
| chr5:141483329-141483587 | 1.53789333 | 0.004157461 | 0.037393239 |
| chr13:39016763-39017165 | -1.124226299 | 0.004159428 | 0.037406535 |
| chr2:154426060-154426528 | -1.287048986 | 0.004160082 | 0.03740803 |
| chr6:37484192-37484497 | 1.129984993 | 0.004161999 | 0.037420875 |
| chr1:114684498-114684698 | 1.120959707 | 0.004162633 | 0.037421774 |
| chr10:4982807-4983024 | 1.482663534 | 0.004163076 | 0.037421774 |
| chr2:179647338-179647538 | -1.568166732 | 0.004167366 | 0.037455939 |
| chr1:93579569-93580168 | 1.088937004 | 0.004168429 | 0.037461106 |
| chr8:90888679-90888968 | -1.248468994 | 0.004172571 | 0.037493681 |
| chr9:71397861-71398269 | -1.251034065 | 0.004173033 | 0.037493681 |
| chr4:742658-742983 | 1.19818894 | 0.004178022 | 0.037534105 |
| chr1:184502285-184502547 | -1.576841967 | 0.004179636 | 0.037540601 |
| chr2:207146209-207146409 | -1.779765218 | 0.004179851 | 0.037540601 |
| chr12:46546900-46547215 | -1.272942405 | 0.004180215 | 0.037540601 |
| chr13:85592354-85592783 | -1.239685659 | 0.004181356 | 0.037546446 |
| chr1:202106299-202108106 | -0.909148234 | 0.004182472 | 0.037552071 |
| chr4:38267541-38267786 | 1.531811129 | 0.004185706 | 0.037576707 |
| chr3:63891573-63892069 | 0.979356793 | 0.004188691 | 0.037595005 |
| chr10:50891027-50891497 | -1.234313878 | 0.004188726 | 0.037595005 |
| chr12:15931584-15931809 | -1.499261302 | 0.004190122 | 0.037603128 |
| chr4:183398364-183399006 | -1.093621947 | 0.004194188 | 0.037635211 |
| chr7:18059706-18059906 | 1.545696346 | 0.004195401 | 0.037641686 |
| chr6:34908078-34908473 | -1.352095029 | 0.004199721 | 0.037676036 |
| chr2:210068712-210068912 | 1.645334093 | 0.004200899 | 0.037681238 |
| chr3:24963851-24964058 | 1.641934849 | 0.004201284 | 0.037681238 |
| chr7:140315169-140315414 | -1.687244661 | 0.004202737 | 0.037686175 |
| chr10:1598650-1598896 | 1.392775956 | 0.004202819 | 0.037686175 |
| chr6:6906969-6907254 | -1.641359255 | 0.004205234 | 0.037702601 |
| chr3:150268785-150269336 | 1.050843417 | 0.00420575 | 0.037702601 |
| chr3:7637716-7637980 | 1.450663607 | 0.004206127 | 0.037702601 |

| | | | |
|---|---|---|---|
| chr10:10834170-10834400 | 1.655458362 | 0.004207188 | 0.037707701 |
| chr20:46055407-46055670 | 1.356459288 | 0.00421165 | 0.037743282 |
| chr6:166512574-166512885 | -1.305119584 | 0.004212281 | 0.037744525 |
| chr5:61790671-61791095 | -1.072802134 | 0.004213426 | 0.037750364 |
| chr5:149005080-149005432 | -1.32538712 | 0.004214929 | 0.037759417 |
| chr21:38414356-38415235 | -1.059203555 | 0.004220487 | 0.037804785 |
| chr10:52735829-52736598 | -1.047531939 | 0.004221012 | 0.03780507 |
| chr5:62233168-62233662 | 1.181979983 | 0.004222333 | 0.037812485 |
| chr12:92827909-92828264 | -1.280920411 | 0.004224061 | 0.03782354 |
| chr15:90035288-90035818 | 1.032195404 | 0.004227353 | 0.037848587 |
| chr6:14711145-14711368 | 1.503106501 | 0.004231047 | 0.037875389 |
| chr5:171911309-171911525 | 1.619922048 | 0.004231335 | 0.037875389 |
| chr1:232629298-232629615 | 1.142299538 | 0.004233029 | 0.037882728 |
| chr8:22565561-22566064 | 1.10616324 | 0.004233167 | 0.037882728 |
| chr16:29807690-29808092 | 0.996298709 | 0.004233638 | 0.037882728 |
| chr7:149840770-149841059 | -1.165582198 | 0.00423793 | 0.037916703 |
| chr5:42108480-42108910 | 1.038740445 | 0.00424095 | 0.0379393 |
| chr3:94961237-94962076 | -0.953148804 | 0.00424268 | 0.03794687 |
| chr8:124661555-124661792 | 1.246096826 | 0.004242787 | 0.03794687 |
| chrX:30864982-30865401 | -1.079482686 | 0.004244414 | 0.037954204 |
| chr4:72900935-72901307 | -1.328149811 | 0.004244598 | 0.037954204 |
| chr8:53486631-53487073 | 1.117146633 | 0.004246503 | 0.037966811 |
| chr5:748543-748812 | 1.371783378 | 0.004248597 | 0.037976249 |
| chr19:18785180-18785520 | 1.110775045 | 0.004249023 | 0.037976249 |
| chr11:68380423-68380795 | 1.084311182 | 0.004249046 | 0.037976249 |
| chr8:90726572-90726906 | 1.220409855 | 0.004249965 | 0.037980036 |
| chr11:113885883-11388633s | -1.311589879 | 0.004251985 | 0.037993656 |
| chr5:125647044-125647514 | -1.2740596 | 0.004254737 | 0.038013439 |
| chr9:135392533-135393055 | -1.074618821 | 0.004255191 | 0.038013439 |
| chr5:5129437-5129871 | 1.259717765 | 0.004256056 | 0.03801673 |
| chr19:3968575-3969087 | 0.987307346 | 0.004257994 | 0.03802961 |
| chr4:158578573-158578820 | -1.436889442 | 0.004258986 | 0.038034032 |
| chr8:107254652-107255031 | 1.138865477 | 0.004260323 | 0.038041544 |
| chr1:106226420-106226794 | 1.100421259 | 0.004261565 | 0.038048201 |
| chr1:97174103-97174626 | -1.106706874 | 0.004265077 | 0.038075114 |
| chr2:19389382-19390089 | -1.070352564 | 0.004267608 | 0.038093274 |
| chr12:50075180-50075572 | 1.025220996 | 0.004269673 | 0.038107266 |
| chr15:62896421-62897584 | 0.922921074 | 0.004272062 | 0.038124149 |
| chr14:105928181-105928412 | 1.473083018 | 0.004274228 | 0.038139038 |
| chr5:178086239-178086706 | -1.011025841 | 0.004275931 | 0.038149784 |
| chr16:28210080-28210280 | 1.382741786 | 0.004276833 | 0.038153382 |
| chr7:107952522-107953118 | -1.038518966 | 0.00427733 | 0.038153382 |
| chr20:48341341-48341649 | 0.990259453 | 0.004278234 | 0.038153464 |
| chr2:74560461-74561188 | -1.028439041 | 0.004278611 | 0.038153464 |
| chr17:21117970-21118203 | 1.543798129 | 0.004278833 | 0.038153464 |
| chr13:49421307-49421868 | -1.092823222 | 0.004279392 | 0.03815401 |
| chr1:150813184-150813396 | 1.285454321 | 0.004280489 | 0.03815662 |
| chr1:63655675-63655937 | 1.581011892 | 0.004280681 | 0.03815662 |
| chr2:135245074-135245407 | -1.466128626 | 0.004281757 | 0.038161778 |
| chr6:80509264-80509981 | -1.063572377 | 0.00428241 | 0.038163155 |

| | | | |
|---|---|---|---|
| chr4:139600036-139600355 | 1.180182073 | 0.00428479 | 0.038178101 |
| chr2:129558568-129558852 | 1.347509637 | 0.004285083 | 0.038178101 |
| chr5:137141882-137142263 | -1.20950079 | 0.004287211 | 0.038192611 |
| chr11:69131191-69131410 | 1.42353018 | 0.004290475 | 0.038217245 |
| chr6:27236820-27237031 | 1.388552999 | 0.004305456 | 0.03834343 |
| chr10:50659736-50660684 | -0.990140605 | 0.004305828 | 0.03834343 |
| chr3:66010647-66010990 | 1.262158847 | 0.004306167 | 0.03834343 |
| chr18:50203777-50204348 | -1.142477374 | 0.004306733 | 0.03834343 |
| chr4:98573790-98574535 | 0.993864623 | 0.004307357 | 0.03834343 |
| chr8:51121703-51122241 | -1.110187899 | 0.004307643 | 0.03834343 |
| chrX:28177232-28177595 | -1.221573447 | 0.004308874 | 0.038349931 |
| chr12:69553650-69554277 | -1.051102035 | 0.004313453 | 0.038382783 |
| chr10:60684911-60685189 | -1.271694871 | 0.004313567 | 0.038382783 |
| chr7:136857669-136857983 | -1.391512863 | 0.004314189 | 0.038383861 |
| chr3:63945636-63946002 | 1.001655351 | 0.004315344 | 0.038389673 |
| chr11:2269010-2269432 | 1.122604668 | 0.004321707 | 0.038440477 |
| chr12:67388845-67389232 | -1.247612523 | 0.004322058 | 0.038440477 |
| chr11:2168826-2169733 | -1.025522323 | 0.00432257 | 0.038440574 |
| chr8:81098177-81098439 | 1.566645235 | 0.004323179 | 0.038441527 |
| chr7:111364415-111364648 | -1.636593569 | 0.004324705 | 0.038450631 |
| chr9:39791395-39791652 | -1.606607201 | 0.004330359 | 0.038496431 |
| chr5:23007872-23008074 | -1.599293904 | 0.004332079 | 0.03850726 |
| chr3:24237942-24238363 | 1.092192553 | 0.004333508 | 0.038515487 |
| chr3:177358270-177358924 | -0.965077771 | 0.004334251 | 0.038517631 |
| chr5:35644620-35644958 | -1.387388282 | 0.004335938 | 0.038528152 |
| chr2:72945364-72945609 | 1.40411107 | 0.004339883 | 0.038558732 |
| chr18:63257230-63257781 | -1.005760242 | 0.004340963 | 0.038559441 |
| chrX:153032656-153032999 | 1.195922996 | 0.004340969 | 0.038559441 |
| chr16:34593005-34593542 | 0.96270846 | 0.004342414 | 0.038567801 |
| chr5:149485613-149486613 | -0.942727114 | 0.004344225 | 0.038579419 |
| chr8:123911898-123912127 | 1.293614423 | 0.004344758 | 0.038579681 |
| chr8:144619837-144620236 | 1.129738487 | 0.004349772 | 0.038619721 |
| chr14:45112807-45113832 | 0.960110158 | 0.004350993 | 0.038626095 |
| chr7:112572756-112573403 | -1.0843926 | 0.004353393 | 0.03864292 |
| chr7:102450326-102450657 | -1.145055737 | 0.004354279 | 0.038646308 |
| chr4:168679371-168680249 | -0.967875577 | 0.004358796 | 0.03868192 |
| chr3:96106773-96106973 | 1.559861715 | 0.004361284 | 0.038691804 |
| chr11:62556201-62556997 | -0.970285771 | 0.00436141 | 0.038691804 |
| chr14:68864246-68864595 | 1.169976966 | 0.004361425 | 0.038691804 |
| chr1:192747989-192748289 | -1.438108904 | 0.004364498 | 0.038714584 |
| chr13:39469399-39469643 | 1.529317873 | 0.004368099 | 0.038737558 |
| chr14:83794246-83794461 | 1.529317873 | 0.004368099 | 0.038737558 |
| chr11:19912763-19913043 | 1.403389767 | 0.004368766 | 0.038738988 |
| chr2:201232215-201232595 | -1.061300364 | 0.004375297 | 0.038792407 |
| chr4:185811701-185811947 | -1.516902317 | 0.004376794 | 0.038800453 |
| chr7:139230616-139231730 | -0.919911467 | 0.004377217 | 0.038800453 |
| chr3:53391371-53391729 | -1.410765039 | 0.00438639 | 0.038877266 |
| chr8:124498166-124498400 | 1.366854848 | 0.004387413 | 0.038881836 |
| chr9:38708001-38708296 | -1.346382172 | 0.004390616 | 0.038905719 |
| chr8:128139685-128140313 | -0.935127886 | 0.004391881 | 0.038912429 |

| | | | |
|---|---|---|---|
| chr13:54572073-54572392 | -1.118618506 | 0.004394335 | 0.038927871 |
| chr16:77998455-77999085 | 0.97764435 | 0.00439464 | 0.038927871 |
| chr10:122101923-122102123 | 1.492835891 | 0.00439743 | 0.038939811 |
| chr9:5590104-5590304 | 1.638000696 | 0.004398026 | 0.038939811 |
| chr2:120291412-120291943 | 1.082560056 | 0.004398369 | 0.038939811 |
| chr3:52470986-52471255 | 1.343755492 | 0.004399419 | 0.038939811 |
| chr1:101025247-101026445 | -0.942713062 | 0.004399528 | 0.038939811 |
| chr14:95442577-95442841 | 1.124660862 | 0.004399579 | 0.038939811 |
| chr7:111730579-111730890 | -1.179376601 | 0.004399797 | 0.038939811 |
| chr14:32667775-32668202 | 1.126264199 | 0.004400053 | 0.038939811 |
| chr8:79900850-79901314 | -1.054423159 | 0.00440211 | 0.038953193 |
| chr12:93467121-93468269 | -0.931227549 | 0.004402582 | 0.038953193 |
| chr2:96897478-96897783 | -1.162112087 | 0.004403573 | 0.038957464 |
| chr8:108120222-108120759 | 1.121184984 | 0.004405097 | 0.038966442 |
| chr12:69232094-69232335 | 1.484177303 | 0.004405671 | 0.038967025 |
| chr2:213806184-213806453 | -1.483544963 | 0.004407042 | 0.038974654 |
| chrX:104909672-104910531 | -0.988650048 | 0.004407561 | 0.03897474 |
| chr11:29962350-29962621 | -1.396507963 | 0.004408309 | 0.038976855 |
| chr2:175525655-175526116 | -1.373584017 | 0.004409256 | 0.03898073 |
| chr20:50810863-50812203 | 0.900510298 | 0.004413486 | 0.039013631 |
| chr3:95521503-95521768 | -1.602246262 | 0.004414971 | 0.039019765 |
| chr15:57376290-57376609 | 1.205169591 | 0.004415199 | 0.039019765 |
| chr17:54435873-54436604 | -1.069259219 | 0.004415916 | 0.039021598 |
| chr8:94504609-94504869 | 1.45506938 | 0.004416928 | 0.03902604 |
| chr1:209627687-209628022 | -1.134867027 | 0.004420781 | 0.039050483 |
| chr15:70834008-70834208 | 1.497066256 | 0.004421033 | 0.039050483 |
| chr7:127338097-127338346 | -1.719492061 | 0.004421223 | 0.039050483 |
| chr6:53302085-53302334 | -1.430531326 | 0.004425956 | 0.03908247 |
| chr3:170883971-170884212 | -1.43402398 | 0.004426748 | 0.03908247 |
| chr13:75357854-75358307 | -1.084718075 | 0.004426856 | 0.03908247 |
| chr8:130409402-130409602 | 1.562473641 | 0.004426885 | 0.03908247 |
| chr4:10157800-10158058 | 1.317583735 | 0.004428315 | 0.03909059 |
| chr3:146115345-146115967 | -1.043978431 | 0.004429646 | 0.039097835 |
| chr6:6856613-6857573 | -1.038020217 | 0.004435465 | 0.039144688 |
| chr18:28365476-28365721 | 1.500609157 | 0.004437262 | 0.03915604 |
| chr2:64360103-64360595 | -1.150126774 | 0.004442044 | 0.039193724 |
| chr6:133240614-133241091 | -1.17347007 | 0.004447527 | 0.039237587 |
| chr19:53102712-53103059 | 1.183896931 | 0.00444805 | 0.039237678 |
| chr2:214587545-214587783 | -1.640274294 | 0.004449625 | 0.039247056 |
| chr5:38433114-38433371 | 1.414408499 | 0.004452297 | 0.039266104 |
| chr9:1018191-1018453 | 1.421475779 | 0.004454485 | 0.039280882 |
| chr22:37186033-37186599 | -1.077661634 | 0.004457145 | 0.039299739 |
| chr11:47234096-47234487 | 1.128276199 | 0.00445765 | 0.039299739 |
| chr2:9168254-9168509 | 1.331123134 | 0.00445958 | 0.039312238 |
| chr8:74073157-74073559 | 1.110586484 | 0.004463121 | 0.039338921 |
| chr5:3139532-3139888 | -1.287766691 | 0.004466006 | 0.039359826 |
| chr12:82517803-82518192 | -1.032770051 | 0.004470483 | 0.039394752 |
| chr1:114374236-114374679 | 0.9606738 | 0.00447244 | 0.039407465 |
| chr1:6431029-6431402 | 1.130450119 | 0.004473898 | 0.039410456 |
| chr4:6750942-6751286 | 1.089321836 | 0.004474148 | 0.039410456 |

| | | | |
|---|---|---|---|
| chr12:79424081-79424633 | 0.998827437 | 0.004474359 | 0.039410456 |
| chr4:143095390-143095651 | 1.525106967 | 0.004474837 | 0.039410456 |
| chr8:97451326-97451553 | 1.341185211 | 0.004476949 | 0.039421148 |
| chr7:114087202-114087409 | -1.315303923 | 0.00447708 | 0.039421148 |
| chr10:35042806-35043086 | -1.412361995 | 0.004481713 | 0.039455504 |
| chrX:28206736-28207050 | -1.544033342 | 0.004482012 | 0.039455504 |
| chr1:114606868-114607115 | 1.022064587 | 0.004483837 | 0.039462662 |
| chr15:71103154-71103611 | -1.142211076 | 0.004484162 | 0.039462662 |
| chr1:203821140-203821377 | 1.3469363 | 0.00448437 | 0.039462662 |
| chr6:142355760-142356144 | -1.363905854 | 0.00448554 | 0.039468421 |
| chr16:5065022-5065320 | 1.25186845 | 0.004486424 | 0.039471669 |
| chr1:245203511-245203799 | 1.488576937 | 0.0044896 | 0.039490254 |
| chr7:127504794-127505068 | -1.506400676 | 0.004490057 | 0.039490254 |
| chr2:113587281-113587556 | -1.348562822 | 0.004490082 | 0.039490254 |
| chr2:21267672-21267975 | -1.330557526 | 0.004490841 | 0.039492389 |
| chr12:130948634-130949036 | -1.369402914 | 0.004492174 | 0.039499582 |
| chr1:219907653-219907853 | 1.637582673 | 0.004498698 | 0.039552411 |
| chr5:112649845-112650312 | -1.026693912 | 0.004501362 | 0.039571289 |
| chr17:62298677-62298905 | 1.469036447 | 0.004502207 | 0.039574182 |
| chr8:60659028-60659315 | 1.474451762 | 0.004504642 | 0.039591037 |
| chr9:112065960-112066184 | 1.327585874 | 0.004505604 | 0.039594955 |
| chrX:117743720-117744170 | 1.157494496 | 0.004513085 | 0.039654669 |
| chr1:200432514-200432874 | 1.123501243 | 0.004513434 | 0.039654669 |
| chr16:84841869-84842167 | -1.242337235 | 0.004515891 | 0.039671703 |
| chr1:234875720-234875994 | 1.24670458 | 0.004516923 | 0.039676224 |
| chr16:87968634-87968834 | -1.604069847 | 0.004519203 | 0.039691694 |
| chr8:90098091-90098446 | 1.123987698 | 0.004522022 | 0.039711903 |
| chr5:56454138-56454791 | -1.117316883 | 0.004523492 | 0.039716979 |
| chr18:37778327-37778847 | -1.14396177 | 0.004523637 | 0.039716979 |
| chr6:156980419-156980902 | -1.143700488 | 0.004525132 | 0.039725557 |
| chr13:26742675-26742921 | 1.518986673 | 0.004526046 | 0.039729028 |
| chrX:19904781-19905165 | -1.13874096 | 0.004527377 | 0.03973616 |
| chr3:196022112-196022400 | -1.347961575 | 0.004529648 | 0.039751544 |
| chr4:186699339-186699587 | -1.684844387 | 0.00453523 | 0.039795969 |
| chr20:36632647-36633104 | 1.177771744 | 0.004539335 | 0.039827428 |
| chr7:80594282-80594771 | -1.250196602 | 0.004540838 | 0.039836054 |
| chr9:68568060-68568466 | -1.149792316 | 0.004542236 | 0.039843757 |
| chr1:164623069-164623291 | 1.423007604 | 0.004544077 | 0.039855342 |
| chr14:99460706-99461253 | -1.077350069 | 0.004545145 | 0.039860151 |
| chr9:120532801-120533032 | 1.551915378 | 0.004546491 | 0.03986739 |
| chr22:24303069-24303831 | 1.027717985 | 0.004547947 | 0.039875595 |
| chr1:221709027-221709388 | -1.189208933 | 0.004550752 | 0.039895616 |
| chr1:200130625-200131135 | -1.021022009 | 0.004551747 | 0.039899779 |
| chr11:35381100-35381383 | -1.460584843 | 0.004553547 | 0.039906927 |
| chr14:26191218-26191664 | 1.170894395 | 0.004553604 | 0.039906927 |
| chr6:41222932-41223185 | 1.486893025 | 0.004554964 | 0.039914278 |
| chr8:112500743-112501138 | 1.11429952 | 0.004557782 | 0.039931943 |
| chr14:105640745-105641810 | 0.975609639 | 0.004558022 | 0.039931943 |
| chr2:66080142-66080429 | -1.491563522 | 0.004559594 | 0.039941151 |
| chr3:62568151-62568445 | 1.041465525 | 0.004564119 | 0.039976214 |

| | | | |
|---|---|---|---|
| chr15:95652648-95652882 | 1.523711774 | 0.004566457 | 0.039992118 |
| chr6:14464072-14464465 | -1.346743813 | 0.004566986 | 0.039992186 |
| chr1:81541312-81541649 | -1.27898815 | 0.004568725 | 0.040002839 |
| chr14:59880138-59880338 | 1.596268223 | 0.004574155 | 0.040045807 |
| chr8:133303409-133303698 | 1.315723172 | 0.004576166 | 0.040058834 |
| chrX:24093619-24094362 | -1.013765074 | 0.004578472 | 0.040070578 |
| chr9:93053135-93053630 | -1.103974631 | 0.004578553 | 0.040070578 |
| chr20:41377464-41377693 | 1.224185293 | 0.004580079 | 0.040076633 |
| chr10:74207725-74208152 | -1.302037245 | 0.004580291 | 0.040076633 |
| chr10:21494757-21494957 | -1.402993432 | 0.004581807 | 0.040085315 |
| chr4:168923744-168924111 | -1.130260973 | 0.004585447 | 0.04011258 |
| chr6:142827490-142827732 | -1.39363853 | 0.004588214 | 0.040132205 |
| chr14:105761374-105762099 | 1.046824102 | 0.004589075 | 0.040135156 |
| chr7:116862214-116863348 | -0.908589242 | 0.004592421 | 0.040159839 |
| chr5:115856360-115856659 | -1.428457022 | 0.004594289 | 0.040171583 |
| chr11:119134501-119134701 | 1.448743132 | 0.004595407 | 0.040176775 |
| chr4:158722265-158722732 | -1.07597169 | 0.004598017 | 0.040193171 |
| chr9:2027671-2028254 | -1.12301939 | 0.00459879 | 0.040193171 |
| chr1:247487168-247487480 | -1.090012651 | 0.004598856 | 0.040193171 |
| chr20:43388510-43388710 | -1.626545418 | 0.0046053 | 0.040244902 |
| chr11:85737718-85738149 | -1.158048867 | 0.004608857 | 0.040271396 |
| chr3:127802447-127802668 | 1.614666664 | 0.004610781 | 0.040279014 |
| chr8:138907694-138907909 | 1.614666664 | 0.004610781 | 0.040279014 |
| chr7:109456156-109456454 | -1.401328035 | 0.004611477 | 0.040280502 |
| chr17:59280368-59280603 | 1.27949188 | 0.004612791 | 0.040287388 |
| chr5:88729176-88729920 | -1.041271807 | 0.004614503 | 0.040296033 |
| chr20:33796463-33797214 | 1.01741332 | 0.004614832 | 0.040296033 |
| chr11:71055965-71056346 | 1.294875447 | 0.004616878 | 0.040309304 |
| chr4:82542904-82543579 | 1.016890309 | 0.004618434 | 0.040316314 |
| chr2:19819415-19819931 | -1.102378626 | 0.004618734 | 0.040316314 |
| chr14:44505542-44505837 | 1.398049077 | 0.00462047 | 0.040326873 |
| chr14:105666368-105666768 | 1.111759491 | 0.004625916 | 0.040366865 |
| chr4:163331724-163333056 | -0.915252796 | 0.004626105 | 0.040366865 |
| chr7:105981882-105982171 | -1.337274054 | 0.004631361 | 0.040408127 |
| chr19:34999737-35000017 | 1.146076293 | 0.004635796 | 0.040442218 |
| chr2:216192912-216193147 | -1.547827574 | 0.004637205 | 0.040449896 |
| chr14:72962857-72963347 | -1.016000454 | 0.004638221 | 0.040454158 |
| chr20:55631055-55631323 | 1.325201807 | 0.004639183 | 0.040456323 |
| chr16:57816264-57816752 | 1.042260559 | 0.004639655 | 0.040456323 |
| chr1:86992915-86993130 | 1.459774002 | 0.004640053 | 0.040456323 |
| chr20:24208701-24208972 | 1.27366166 | 0.004646371 | 0.040506795 |
| chr1:179164546-179164903 | 1.340513282 | 0.004651163 | 0.040543957 |
| chr1:108106657-108106876 | 1.383579722 | 0.004653463 | 0.040559392 |
| chr3:153506298-153506530 | -1.403358521 | 0.004657344 | 0.040588606 |
| chr4:113116371-113117257 | -0.984688143 | 0.0046612 | 0.040617592 |
| chr13:63636160-63636518 | -1.227355736 | 0.004668531 | 0.040676849 |
| chr3:106963425-106963707 | -1.382474975 | 0.004671736 | 0.04070014 |
| chr22:44085730-44086047 | -1.446883904 | 0.004674025 | 0.040715452 |
| chr21:39765550-39765754 | 1.414819992 | 0.004674832 | 0.040717857 |
| chr3:171783259-171783459 | -1.590631649 | 0.00467662 | 0.040728802 |

| | | | |
|---|---|---|---|
| chr6:132734717-132735265 | -1.133725483 | 0.004678185 | 0.040737795 |
| chr20:47897363-47898142 | 0.927575494 | 0.004683239 | 0.040774369 |
| chr8:141147185-141147385 | 1.447193631 | 0.004683449 | 0.040774369 |
| chr9:124320320-124320592 | 1.166728682 | 0.004684337 | 0.040777468 |
| chr17:7797466-7797720 | 1.513232401 | 0.004686628 | 0.040788319 |
| chr11:78030320-78030954 | -1.039854042 | 0.004686648 | 0.040788319 |
| chr10:35228212-35228427 | -1.487050445 | 0.004687452 | 0.04079068 |
| chrX:25404133-25404348 | -1.708871933 | 0.004688466 | 0.040790827 |
| chr5:136424847-136425223 | -1.258709907 | 0.004688534 | 0.040790827 |
| chr14:24630585-24631490 | 0.988642797 | 0.004689474 | 0.040794375 |
| chr3:89955236-89955523 | 1.350785924 | 0.00469351 | 0.040824854 |
| chr20:59208176-59208397 | 1.247928902 | 0.004695451 | 0.0408371 |
| chr2:216294362-216294694 | -1.356414151 | 0.004697842 | 0.040850174 |
| chr17:62671469-62672647 | -0.908948249 | 0.004698021 | 0.040850174 |
| chr2:150166059-150166728 | -1.017868085 | 0.004699732 | 0.040860417 |
| chr6:95923593-95923882 | -1.270787329 | 0.004702926 | 0.040879868 |
| chr11:6357329-6357602 | -1.517906795 | 0.004703036 | 0.040879868 |
| chr14:37642160-37642394 | 1.163891079 | 0.004704764 | 0.040890247 |
| chr2:84103800-84104596 | -0.942631673 | 0.004707025 | 0.040905263 |
| chr4:170465206-170465646 | -1.223123722 | 0.004709856 | 0.040925217 |
| chr13:72648772-72649227 | -1.213888905 | 0.004711133 | 0.040931676 |
| chr11:6812432-6812759 | 1.189261362 | 0.004715158 | 0.040961999 |
| chr2:34183862-34184134 | -1.427738835 | 0.004717737 | 0.04097976 |
| chr1:28196981-28197367 | -1.180836532 | 0.004719829 | 0.04098912 |
| chr14:94890512-94890765 | 1.283672546 | 0.004719885 | 0.04098912 |
| chr4:158624970-158625241 | -1.341522152 | 0.004720429 | 0.040989197 |
| chr5:163649610-163650070 | -1.178352661 | 0.004721137 | 0.040990703 |
| chr7:111312991-111313339 | -1.24102663 | 0.004722877 | 0.041001161 |
| chr17:61245899-61246219 | 0.956329496 | 0.004725175 | 0.04101647 |
| chr3:61642501-61642737 | 1.093213105 | 0.004727045 | 0.041026072 |
| chr14:71931984-71932840 | 0.967389021 | 0.004727352 | 0.041026072 |
| chrX:16605019-16605277 | -1.534941635 | 0.004729428 | 0.041035948 |
| chr3:149993445-149993737 | 1.305609236 | 0.004729561 | 0.041035948 |
| chr20:47872883-47873342 | 0.948622541 | 0.004730758 | 0.041041684 |
| chr2:4258469-4258797 | -1.474840498 | 0.004732735 | 0.041054186 |
| chr10:19209276-19209693 | -1.097955724 | 0.004737988 | 0.041095104 |
| chr20:56946294-56946494 | 1.270738081 | 0.004740801 | 0.041100517 |
| chr7:105817079-105817319 | -1.669611037 | 0.004740856 | 0.041100517 |
| chr1:244322649-244323265 | 1.060746322 | 0.004741008 | 0.041100517 |
| chr16:46757827-46758036 | 1.556062128 | 0.004741032 | 0.041100517 |
| chr5:11596719-11596919 | 1.431342708 | 0.004741295 | 0.041100517 |
| chr11:19344772-19345014 | 1.472356475 | 0.004742155 | 0.041103323 |
| chr14:59367532-59367732 | 1.735821682 | 0.004742834 | 0.041104558 |
| chr7:43439352-43440202 | -0.983662285 | 0.004743937 | 0.041109473 |
| chr11:22152374-22153108 | -1.029570067 | 0.00474634 | 0.041122394 |
| chr1:162753496-162753732 | 1.393817375 | 0.004746895 | 0.041122394 |
| chr14:59275678-59276071 | -1.146472531 | 0.004747541 | 0.041122394 |
| chr3:146148403-146148668 | -1.317053623 | 0.004747575 | 0.041122394 |
| chr2:168480811-168481473 | -1.099089494 | 0.004748255 | 0.041123638 |
| chr17:60185541-60185793 | 1.016280615 | 0.004748806 | 0.041123755 |

| | | | |
|---|---|---|---|
| chr8:124913493-124913738 | 1.189692734 | 0.004750974 | 0.04113788 |
| chr2:231192478-231192708 | 1.456361844 | 0.00475749 | 0.04118862 |
| chr1:37434428-37434656 | 1.252217482 | 0.004757909 | 0.04118862 |
| chr10:1232374-1232898 | 1.093481058 | 0.004759369 | 0.041196607 |
| chr9:93071606-93072793 | -0.95175347 | 0.004761052 | 0.041206523 |
| chr8:135852824-135853525 | 1.017578267 | 0.004762476 | 0.041211885 |
| chr2:74206127-74206396 | -1.292384534 | 0.004762748 | 0.041211885 |
| chr20:19081917-19082230 | 1.302060445 | 0.00476472 | 0.041224301 |
| chr11:105368888-105369088 | -1.751699032 | 0.004767711 | 0.041241001 |
| chr8:79932659-79932859 | 1.58894518 | 0.004767727 | 0.041241001 |
| chr7:123624633-123625098 | -1.203445104 | 0.004768953 | 0.041244076 |
| chr7:143299868-143300220 | -1.120176288 | 0.004769159 | 0.041244076 |
| chr14:65413334-65413540 | 1.133873206 | 0.004772619 | 0.04126776 |
| chr1:68010214-68010480 | 1.306887731 | 0.004772975 | 0.04126776 |
| chr14:81289441-81289643 | -1.489915902 | 0.004773827 | 0.041270466 |
| chr10:56782577-56782828 | -1.647979629 | 0.004779488 | 0.041312383 |
| chr10:63866458-63866756 | -1.096150652 | 0.004779816 | 0.041312383 |
| chr20:12065135-12065790 | -1.073656826 | 0.004780293 | 0.041312383 |
| chr8:98978112-98978433 | 1.230982452 | 0.00478353 | 0.041335058 |
| chr5:2530475-2530675 | 1.555167133 | 0.004783995 | 0.041335058 |
| chr12:110245433-110245779 | -1.31914289 | 0.004786429 | 0.041351424 |
| chr19:39845721-39845921 | 1.455599569 | 0.004791562 | 0.0413911 |
| chr7:145372627-145373138 | -1.137923069 | 0.004792228 | 0.041392184 |
| chr3:64701111-64701330 | 1.02243468 | 0.004793991 | 0.041402747 |
| chr17:75356663-75357063 | 1.19132532 | 0.004795728 | 0.041413084 |
| chr4:115113589-115114378 | -1.010623809 | 0.004796428 | 0.041414455 |
| chr5:149666049-149666440 | 1.255738603 | 0.004799478 | 0.041436123 |
| chr14:21319834-21320501 | 1.014800006 | 0.004800611 | 0.04144124 |
| chr2:62802190-62802444 | -1.344161966 | 0.004801234 | 0.041441947 |
| chr8:124961137-124961337 | 1.212174303 | 0.004804455 | 0.04146508 |
| chr11:116836033-116836593 | -1.033416555 | 0.00480752 | 0.041486856 |
| chr6:3324233-3324471 | 1.459092546 | 0.004812739 | 0.041523587 |
| chr2:58594253-58594659 | -1.264402074 | 0.004813124 | 0.041523587 |
| chr14:97802180-97802380 | 1.543206321 | 0.004813402 | 0.041523587 |
| chr10:84540393-84541030 | -0.98992447 | 0.004816245 | 0.041543436 |
| chr9:106642944-106643622 | -1.08717918 | 0.004821632 | 0.041582258 |
| chr2:148209667-148210016 | -1.144981733 | 0.004822016 | 0.041582258 |
| chr1:215888222-215888532 | -1.065308127 | 0.004822374 | 0.041582258 |
| chr5:139660128-139661554 | -0.923414547 | 0.004822961 | 0.041582643 |
| chr18:64924042-64924245 | -1.587661806 | 0.004824634 | 0.041592385 |
| chr21:36112820-36113630 | 0.987856714 | 0.004828759 | 0.041623257 |
| chr3:16621275-16621481 | 1.610860968 | 0.00483519 | 0.041674005 |
| chr14:101978756-101979034 | 1.40817506 | 0.004836332 | 0.041675673 |
| chr6:137237488-137237743 | -1.568814677 | 0.004836954 | 0.041675673 |
| chr7:106050100-106050524 | -1.133450462 | 0.004837015 | 0.041675673 |
| chr3:8629438-8629858 | -1.013400383 | 0.004845277 | 0.041742168 |
| chr17:43941272-43941472 | 1.408290894 | 0.004848027 | 0.041756665 |
| chr19:42177144-42177438 | 1.244791196 | 0.00484805 | 0.041756665 |
| chr14:41070574-41070806 | 1.445372064 | 0.004849543 | 0.041761235 |
| chr5:18744978-18745301 | -1.142001364 | 0.004849671 | 0.041761235 |

| | | | |
|---|---|---|---|
| chr7:80499315-80499684 | -1.249255757 | 0.004851146 | 0.041769245 |
| chr2:10340389-10340711 | -1.260962575 | 0.004853537 | 0.041785132 |
| chr8:13159236-13159613 | -1.126914404 | 0.004855094 | 0.041793841 |
| chr15:88614485-88614867 | 1.049566462 | 0.00485737 | 0.041808741 |
| chr2:48584563-48585077 | -1.046399061 | 0.004858412 | 0.041813009 |
| chr7:29254721-29255024 | 1.31101319 | 0.004862959 | 0.041847443 |
| chr13:70842317-70842610 | -1.23196615 | 0.004863885 | 0.041850711 |
| chr2:151043501-151043895 | -1.094611808 | 0.004864705 | 0.041853069 |
| chr7:17158349-17158573 | 1.400374232 | 0.00486897 | 0.041885055 |
| chr12:101607907-101608190 | -1.280654095 | 0.004872704 | 0.041905139 |
| chr1:39625693-39626040 | 1.266677102 | 0.004872793 | 0.041905139 |
| chr6:104252131-104252635 | -1.105291381 | 0.004872945 | 0.041905139 |
| chr2:5673527-5673727 | -1.636904133 | 0.004873952 | 0.041909092 |
| chr7:112422871-112423269 | -1.149884003 | 0.004875947 | 0.04191405 |
| chr19:49916934-49917263 | 1.144945739 | 0.004876006 | 0.04191405 |
| chr7:143408163-143409484 | -0.921356036 | 0.004876169 | 0.04191405 |
| chr1:215165790-215166637 | -0.989686855 | 0.004876789 | 0.041914672 |
| chr7:122919630-122920202 | -1.139220112 | 0.004879334 | 0.041927743 |
| chr4:136220077-136220277 | 1.546212049 | 0.004879404 | 0.041927743 |
| chr3:40467943-40468466 | 1.023960604 | 0.004880967 | 0.041934403 |
| chr2:164542547-164543091 | -1.058966535 | 0.004881274 | 0.041934403 |
| chr7:127107547-127108094 | -1.029926204 | 0.004885953 | 0.041969897 |
| chrX:56026271-56026509 | -1.496712146 | 0.004892251 | 0.042019143 |
| chr12:97608771-97609231 | -1.269773351 | 0.004892783 | 0.042019143 |
| chr13:100836796-100837046 | -1.589887705 | 0.004893845 | 0.04202356 |
| chr4:8440555-8441303 | 0.964504985 | 0.004896151 | 0.042038643 |
| chr14:71036440-71036715 | 1.281028014 | 0.004900225 | 0.042067551 |
| chr5:113900925-113901144 | -1.616234086 | 0.004900615 | 0.042067551 |
| chr4:37471969-37472300 | -1.22905386 | 0.004903954 | 0.042091492 |
| chr20:48124050-48124271 | 1.023774196 | 0.004906365 | 0.04210747 |
| chr7:29562923-29563357 | 1.076522412 | 0.004908215 | 0.042117709 |
| chr2:23421013-23421438 | 1.191269844 | 0.004908657 | 0.042117709 |
| chr3:153553004-153553921 | -0.947169059 | 0.004909867 | 0.042123372 |
| chr2:9339720-9340111 | 1.093235308 | 0.0049114 | 0.04213181 |
| chr5:10555405-10555679 | 1.168007814 | 0.004912989 | 0.042140717 |
| chr2:186975234-186975444 | 1.470229792 | 0.004913992 | 0.042144601 |
| chr9:1049983-1050690 | 0.922114055 | 0.004914851 | 0.042147253 |
| chr18:24108672-24109254 | -1.175520819 | 0.004916024 | 0.042152598 |
| chr6:44068689-44068919 | 1.302444487 | 0.004917135 | 0.042154871 |
| chr8:47554158-47554438 | 1.459838025 | 0.00491739 | 0.042154871 |
| chr3:72722913-72723282 | 1.085325918 | 0.00491971 | 0.042165721 |
| chr9:88256227-88256522 | 1.199052704 | 0.004919756 | 0.042165721 |
| chr12:62021840-62022159 | 1.277591051 | 0.004922683 | 0.04218498 |
| chr5:91027470-91027757 | 1.358645554 | 0.004923104 | 0.04218498 |
| chr20:48705164-48705580 | 0.938789989 | 0.004923775 | 0.042185614 |
| chr9:116968323-116968654 | 1.361494173 | 0.004924448 | 0.042185614 |
| chr12:14417772-14418019 | -1.421473301 | 0.004925373 | 0.042185614 |
| chr6:113530790-113531504 | -1.082435176 | 0.004925419 | 0.042185614 |
| chr14:95516043-95516387 | 1.090611057 | 0.004925931 | 0.042185614 |
| chr8:89824368-89824748 | 1.153713048 | 0.004928512 | 0.04219604 |

| | | | |
|---|---|---|---|
| chr19:50461257-50461618 | 1.102025131 | 0.004928542 | 0.04219604 |
| chr3:171615773-171615975 | -1.473083147 | 0.004928801 | 0.04219604 |
| chr7:11331196-11331456 | 1.169287185 | 0.00493252 | 0.042223163 |
| chr2:18244870-18245127 | -1.520661834 | 0.004942541 | 0.042304223 |
| chr13:30514617-30514817 | 1.620180937 | 0.004943328 | 0.042306232 |
| chr2:60435663-60435952 | 1.491312838 | 0.004945476 | 0.042319641 |
| chrX:154411387-154412282 | 0.966775017 | 0.004946 | 0.042319641 |
| chr7:139775803-139776127 | -1.119039365 | 0.004946592 | 0.042319984 |
| chr3:88509390-88509887 | 1.133239921 | 0.00494766 | 0.042324398 |
| chr13:23969684-23969929 | 1.357521288 | 0.004948688 | 0.042328462 |
| chr11:10899742-10900236 | 1.074521499 | 0.00495222 | 0.042353948 |
| chr20:45428254-45428454 | 1.66035563 | 0.004956289 | 0.042375302 |
| chr2:193583987-193584391 | -1.102814019 | 0.004956351 | 0.042375302 |
| chr12:64010103-64010527 | 1.112229792 | 0.004957238 | 0.042375302 |
| chr13:29841457-29842339 | -0.991110872 | 0.004957348 | 0.042375302 |
| chr3:84032013-84032309 | 1.332424041 | 0.004957482 | 0.042375302 |
| chr12:62807960-62808311 | 1.28419251 | 0.004961626 | 0.042405444 |
| chr1:221435469-221435724 | -1.452761853 | 0.004962251 | 0.042405444 |
| chr12:31238947-31239147 | -1.449454971 | 0.004963196 | 0.042405444 |
| chr21:38408900-38409266 | 1.252800983 | 0.004963222 | 0.042405444 |
| chr1:32742416-32742789 | 1.19246659 | 0.004966197 | 0.042426127 |
| chr12:12009039-12009729 | -1.084561097 | 0.004967133 | 0.042429398 |
| chr13:73287958-73288249 | -1.367565431 | 0.004971095 | 0.042458502 |
| chr2:241952265-241952495 | 1.386033971 | 0.004976048 | 0.042496067 |
| chr5:26569133-26570133 | -1.001686814 | 0.00498009 | 0.042523335 |
| chr19:9620926-9621664 | 1.026415635 | 0.00498035 | 0.042523335 |
| chr1:18464755-18465064 | 1.28656259 | 0.004981452 | 0.042528002 |
| chr6:37807791-37808590 | 0.959319581 | 0.004983451 | 0.042534095 |
| chr2:40725931-40726416 | -1.120148401 | 0.004983668 | 0.042534095 |
| chr18:79290144-79290344 | -1.813833699 | 0.004983831 | 0.042534095 |
| chr14:92652887-92653108 | 1.300850267 | 0.004986211 | 0.042543396 |
| chr1:57072859-57073071 | -1.475239521 | 0.004986494 | 0.042543396 |
| chr14:75457928-75458135 | 1.323841317 | 0.004986587 | 0.042543396 |
| chrX:103082799-103083044 | 1.448360956 | 0.004998382 | 0.042639286 |
| chr7:117337226-117337647 | -1.411298586 | 0.005001788 | 0.042663586 |
| chr8:27603130-27603542 | 1.168618332 | 0.005003949 | 0.042677266 |
| chr1:215606694-215606901 | -1.427043152 | 0.005008021 | 0.042707243 |
| chr4:48967741-48967972 | 1.612892449 | 0.005008903 | 0.04271001 |
| chr4:163484853-163485260 | -1.153734551 | 0.005009798 | 0.042712892 |
| chr11:24637531-24638055 | -1.073858892 | 0.00501456 | 0.042748043 |
| chr17:60223871-60224437 | 0.940976646 | 0.005015037 | 0.042748043 |
| chr19:47581606-47581938 | 1.168181615 | 0.005016152 | 0.042752794 |
| chr14:34629373-34630568 | 0.906942191 | 0.005018565 | 0.042767108 |
| chr4:75058289-75058575 | 1.403362053 | 0.005018948 | 0.042767108 |
| chr7:82594653-82595070 | -1.329008268 | 0.005020604 | 0.042775133 |
| chr13:114047416-114047861 | -1.157795981 | 0.005021006 | 0.042775133 |
| chr2:83347870-83348166 | -1.313424218 | 0.005024742 | 0.042799848 |
| chr18:60676778-60677145 | -1.260271771 | 0.005025025 | 0.042799848 |
| chr14:22100296-22100605 | 1.147570882 | 0.005027299 | 0.042814459 |
| chr11:131911140-131911743 | -1.213628313 | 0.005029161 | 0.04282556 |

| | | | |
|---|---|---|---|
| chr12:124636250-124636511 | 1.197195883 | 0.005033336 | 0.042856346 |
| chr20:57871252-57871470 | 1.234622372 | 0.005034134 | 0.042858383 |
| chr5:135559426-135559654 | -1.529998943 | 0.005035769 | 0.042867534 |
| chr1:214261535-214262146 | -1.027219533 | 0.00503655 | 0.042869425 |
| chr16:30409442-30409784 | 1.086318339 | 0.005037913 | 0.042876257 |
| chr6:120141561-120142456 | -1.018316757 | 0.005042448 | 0.042910086 |
| chr14:29822143-29822569 | 1.184275315 | 0.005044715 | 0.042924614 |
| chr4:157180087-157181059 | -0.972280802 | 0.005045986 | 0.042930665 |
| chr16:25413994-25414315 | 1.320502076 | 0.005046652 | 0.042930734 |
| chr13:33418180-33418673 | -1.090187161 | 0.005047115 | 0.042930734 |
| chr4:68420273-68421015 | -1.05873682 | 0.005051659 | 0.042964613 |
| chr14:54652323-54652553 | 1.339707032 | 0.005057253 | 0.04300742 |
| chr2:31197218-31197539 | 1.261163722 | 0.005060732 | 0.043032229 |
| chr1:106595219-106595539 | 0.995693747 | 0.005063087 | 0.043039493 |
| chr5:158659704-158660242 | -1.230969517 | 0.005063376 | 0.043039493 |
| chr12:124422148-124423353 | -0.930356299 | 0.005063723 | 0.043039493 |
| chr1:48630465-48630769 | 1.387494934 | 0.005064354 | 0.043039493 |
| chr19:17282673-17282996 | 1.24097118 | 0.005064445 | 0.043039493 |
| chr4:76105801-76106145 | -1.281650259 | 0.005064957 | 0.043039493 |
| chr6:145491183-145491673 | -1.120522492 | 0.005067371 | 0.043055238 |
| chr14:60998798-60998998 | 1.494482441 | 0.005071463 | 0.043082081 |
| chr2:196208777-196209151 | 1.176202943 | 0.005071655 | 0.043082081 |
| chr11:17263778-17264106 | -1.441828755 | 0.005072371 | 0.043083381 |
| chr15:77976797-77977054 | 1.469093352 | 0.005074443 | 0.043096209 |
| chr17:62017272-62018518 | 0.897817628 | 0.005075761 | 0.043102622 |
| chr11:70815245-70815638 | 1.114243196 | 0.005081335 | 0.043141514 |
| chr3:162945621-162945859 | 1.584386443 | 0.005081467 | 0.043141514 |
| chr17:61259101-61259346 | 0.97954806 | 0.005083883 | 0.043155353 |
| chr22:12175808-12176605 | 0.951511342 | 0.005084224 | 0.043155353 |
| chr15:97977732-97978185 | -1.022689453 | 0.005090668 | 0.043205267 |
| chr10:79538326-79538593 | -1.688528689 | 0.005096951 | 0.043250207 |
| chr9:12817706-12818118 | -1.108832113 | 0.00509753 | 0.043250207 |
| chr3:30112869-30113171 | 1.326957292 | 0.005097656 | 0.043250207 |
| chr13:36910648-36910887 | -1.369370809 | 0.00509993 | 0.043264711 |
| chr19:39746938-39747496 | -1.112529119 | 0.00510227 | 0.043279763 |
| chr20:47366832-47367039 | 0.973485018 | 0.005102981 | 0.043281004 |
| chr21:44370300-44370718 | 1.047856378 | 0.005104136 | 0.043286011 |
| chr1:68454596-68454863 | -1.39293129 | 0.005105718 | 0.043294636 |
| chr12:27732369-27732595 | -1.626907755 | 0.005110181 | 0.043327688 |
| chr8:81288062-81288357 | 1.346911779 | 0.005112672 | 0.043344011 |
| chrX:106397804-106398051 | -1.380644593 | 0.005115122 | 0.043359983 |
| chr8:120193639-120193839 | 1.65501572 | 0.005116374 | 0.043365794 |
| chr2:30696670-30696886 | 1.532213534 | 0.005119481 | 0.043387329 |
| chr19:34172754-34172955 | 1.284794485 | 0.005123496 | 0.043416561 |
| chr2:84888558-84888931 | -1.328674196 | 0.005126083 | 0.043433568 |
| chr11:33829515-33830065 | 1.161347897 | 0.005126637 | 0.043433568 |
| chr2:212124538-212124983 | -1.211731073 | 0.005128296 | 0.043442821 |
| chr18:62510779-62511149 | -1.369580565 | 0.005129844 | 0.043451131 |
| chr11:120137815-120138358 | 1.101641963 | 0.005131232 | 0.043454513 |
| chr14:91115702-91116007 | 1.21158905 | 0.005131378 | 0.043454513 |

| | | | |
|---|---|---|---|
| chr20:47640054-47640260 | 0.983627792 | 0.005133519 | 0.043463081 |
| chr9:38227714-38227981 | -1.21076247 | 0.005133524 | 0.043463081 |
| chr20:48598742-48598955 | 1.018131985 | 0.005134598 | 0.043467369 |
| chr3:39177759-39178227 | 1.008397802 | 0.005136604 | 0.043476199 |
| chr11:12556136-12556529 | -1.221909274 | 0.005136994 | 0.043476199 |
| chr6:55882769-55882976 | -1.779826282 | 0.005137343 | 0.043476199 |
| chr1:218259803-218260275 | -1.040832153 | 0.005142434 | 0.043513761 |
| chr7:134532305-134532595 | -1.325641762 | 0.005142917 | 0.043513761 |
| chr14:43871031-43871231 | 1.310453372 | 0.005144899 | 0.043521106 |
| chr6:154172292-154172906 | -1.037362234 | 0.005144922 | 0.043521106 |
| chr20:47340055-47340400 | 0.930289546 | 0.00514734 | 0.043534538 |
| chr8:103957422-103957672 | 1.351819741 | 0.005147646 | 0.043534538 |
| chr7:115030490-115030747 | -1.520837248 | 0.005148504 | 0.043536992 |
| chr14:90797464-90798076 | 0.994654881 | 0.005151432 | 0.04355694 |
| chr18:109264-109611 | 1.066188325 | 0.005153196 | 0.043563385 |
| chr4:88369918-88370136 | 1.574850118 | 0.005153331 | 0.043563385 |
| chr1:36754692-36754917 | 1.475792324 | 0.005154029 | 0.043564482 |
| chr20:54577614-54577814 | 1.058013003 | 0.005155822 | 0.043570587 |
| chr16:30183387-30183662 | 1.103680231 | 0.005155889 | 0.043570587 |
| chr16:72013368-72013679 | -1.107802623 | 0.005159873 | 0.043599444 |
| chr10:116183800-116184172 | 1.210727664 | 0.005161802 | 0.043610258 |
| chr12:96070986-96071353 | -1.136824726 | 0.005162623 | 0.043610258 |
| chr12:101108197-101108469 | -1.343165019 | 0.00516286 | 0.043610258 |
| chr6:36120987-36121200 | 1.451214732 | 0.005164684 | 0.043620853 |
| chr5:141840666-141840993 | -1.274494632 | 0.005167043 | 0.043635971 |
| chr16:73126502-73126804 | -1.303137063 | 0.005168214 | 0.043641049 |
| chr5:65037853-65038162 | -1.07282346 | 0.005169527 | 0.043646784 |
| chr12:21501188-21502391 | -0.90655322 | 0.005170032 | 0.043646784 |
| chr5:103808525-103808881 | -1.180857351 | 0.005170632 | 0.043647039 |
| chr19:7643715-7644392 | 1.006034571 | 0.005175681 | 0.043673011 |
| chr4:186144613-186145356 | -1.043454953 | 0.005175749 | 0.043673011 |
| chr13:46252874-46253074 | 1.525724757 | 0.005175765 | 0.043673011 |
| chrX:23211088-23211476 | -1.325199859 | 0.005175989 | 0.043673011 |
| chr8:88866177-88866412 | 1.425113603 | 0.005177115 | 0.043677703 |
| chr12:74376365-74376571 | -1.735516647 | 0.00517852 | 0.043684748 |
| chr2:177521043-177521413 | -1.337250035 | 0.00518196 | 0.043702839 |
| chr2:18404809-18405142 | -1.291632728 | 0.005182225 | 0.043702839 |
| chr20:55865771-55866173 | 1.178676875 | 0.005182376 | 0.043702839 |
| chr3:134419679-134419922 | 1.442451376 | 0.005183816 | 0.043710173 |
| chr5:57494649-57495010 | -1.252213222 | 0.005186831 | 0.043730787 |
| chr1:247131781-247132154 | -1.243227195 | 0.005188848 | 0.04374298 |
| chr3:128945882-128946082 | 1.479912203 | 0.005194746 | 0.043782233 |
| chr5:129126418-129127168 | -0.92715798 | 0.005194777 | 0.043782233 |
| chr3:139088946-139089155 | 1.47023186 | 0.005195417 | 0.043782233 |
| chr16:1379005-1379314 | 1.122922551 | 0.00519579 | 0.043782233 |
| chr8:64376638-64377299 | -1.087792605 | 0.005197007 | 0.04378767 |
| chr14:75884020-75884346 | 1.389435383 | 0.005198333 | 0.043794026 |
| chr8:123589154-123589355 | 1.406411157 | 0.005199991 | 0.043803175 |
| chr4:158722942-158724016 | -0.918355247 | 0.005201201 | 0.043805945 |
| chr4:138682804-138683229 | -1.198434164 | 0.005201463 | 0.043805945 |

| | | | |
|---|---|---|---|
| chr6:96542806-96543025 | -1.717020814 | 0.005203495 | 0.04381824 |
| chr9:128218693-128219189 | 0.973571191 | 0.005204134 | 0.043818808 |
| chr20:47265979-47266181 | 0.97226735 | 0.005209551 | 0.043859604 |
| chr17:37488052-37488510 | 1.096537938 | 0.005210774 | 0.043863262 |
| chr5:149151788-149152087 | -1.240123421 | 0.005211131 | 0.043863262 |
| chr11:102047190-102047787 | -1.06092291 | 0.00521234 | 0.043868616 |
| chr9:37798598-37799059 | -1.11798013 | 0.005214042 | 0.043878123 |
| chr13:60990127-60990387 | -1.50042214 | 0.005216148 | 0.04389103 |
| chr13:108734709-108735039 | -1.312833265 | 0.005219294 | 0.043908029 |
| chr8:23068474-23069413 | 0.944556941 | 0.005219315 | 0.043908029 |
| chr5:111680672-111681567 | -0.936151207 | 0.005221339 | 0.043920236 |
| chr1:113918297-113918497 | 1.042025922 | 0.005223125 | 0.043930439 |
| chr9:35646269-35646586 | -1.155376743 | 0.005231195 | 0.043992897 |
| chr3:174414998-174415244 | -1.492651514 | 0.005231758 | 0.043992897 |
| chrX:141278297-141278604 | 1.236737289 | 0.005232274 | 0.043992897 |
| chr12:124188523-124188997 | 1.057115446 | 0.005236075 | 0.044020031 |
| chr6:121885373-121886041 | -1.033782127 | 0.005237415 | 0.044026466 |
| chr7:18321008-18321363 | 1.086649991 | 0.005239165 | 0.044036346 |
| chr10:59181730-59182003 | -1.282026316 | 0.005240258 | 0.044040702 |
| chr2:189485957-189486225 | -1.234273941 | 0.00524231 | 0.044053112 |
| chr8:47669069-47669558 | 1.069421144 | 0.005244983 | 0.044067722 |
| chr5:119299521-119300321 | -0.959878223 | 0.005245581 | 0.044067722 |
| chr4:188085629-188085854 | -1.715485513 | 0.005245774 | 0.044067722 |
| chr4:53908-54321 | 0.969863773 | 0.00524654 | 0.044069328 |
| chrX:153554795-153555113 | 1.137380473 | 0.005251508 | 0.04410435 |
| chr15:39185093-39185374 | -1.296137208 | 0.005251861 | 0.04410435 |
| chr1:113885275-113885508 | 1.069976535 | 0.005252592 | 0.044105655 |
| chr9:97501421-97501739 | 1.202873548 | 0.00526346 | 0.044192069 |
| chr7:130095872-130096120 | -1.485086129 | 0.005264696 | 0.044193166 |
| chr13:88005205-88005629 | -1.254975407 | 0.005265298 | 0.044193166 |
| chr7:150839352-150839593 | -1.458453066 | 0.005265321 | 0.044193166 |
| chr1:216721879-216722252 | -1.082787842 | 0.005266457 | 0.04419786 |
| chr13:65425572-65425772 | -1.621440313 | 0.00526749 | 0.044201687 |
| chr10:80177461-80177765 | 1.306279895 | 0.005268895 | 0.04420572 |
| chr20:56566325-56567043 | 0.940387238 | 0.005269124 | 0.04420572 |
| chr6:65671895-65672389 | -1.052402338 | 0.005269718 | 0.044205857 |
| chr1:106548463-106548903 | 0.955742602 | 0.005272477 | 0.044221782 |
| chr14:96936038-96936238 | 1.361162304 | 0.00527277 | 0.044221782 |
| chr1:226512362-226512573 | 1.493269684 | 0.005277945 | 0.044260341 |
| chr2:238098533-238098804 | 1.248027332 | 0.005280271 | 0.044274999 |
| chr16:1979400-1979787 | 1.062922378 | 0.005281368 | 0.04427808 |
| chr7:121366919-121367119 | -1.714179564 | 0.005281989 | 0.04427808 |
| chr14:105194363-105194608 | 1.126743345 | 0.005282372 | 0.04427808 |
| chr20:63573377-63573801 | 1.04914119 | 0.005284824 | 0.044293781 |
| chr13:52599536-52600561 | -0.958735686 | 0.005286005 | 0.044296844 |
| chr5:58127898-58128137 | 1.547729827 | 0.005287429 | 0.044296844 |
| chr3:63332287-63332620 | 1.092347534 | 0.005287461 | 0.044296844 |
| chr1:106407460-106407747 | 1.098564235 | 0.005287501 | 0.044296844 |
| chr11:7541004-7541447 | 1.134179887 | 0.005289129 | 0.044305634 |
| chr16:63387802-63388733 | -1.007669167 | 0.005291451 | 0.044320241 |

| | | | |
|---|---|---|---|
| chr3:27768098-27768729 | 0.999439102 | 0.005292875 | 0.044325321 |
| chr7:102412857-102413125 | -1.324451738 | 0.005293285 | 0.044325321 |
| chrX:105260704-105261017 | -1.32106821 | 0.005293793 | 0.044325321 |
| chr20:59988163-59988442 | 1.109915883 | 0.00529548 | 0.044330692 |
| chr7:42407825-42408176 | -1.176727664 | 0.005295592 | 0.044330692 |
| chr7:135509594-135510715 | -0.889330071 | 0.005296788 | 0.044335859 |
| chr16:15765921-15766203 | 1.186721172 | 0.005298367 | 0.044344239 |
| chr14:35591231-35591431 | 1.564306962 | 0.005299625 | 0.044345075 |
| chr16:28255322-28255522 | 1.564306962 | 0.005299625 | 0.044345075 |
| chr16:86806809-86807166 | -1.210914044 | 0.005300717 | 0.044349372 |
| chr5:38196139-38196407 | 1.188200837 | 0.005308076 | 0.044406093 |
| chr10:133123938-133124172 | 1.327681305 | 0.00530871 | 0.044406549 |
| chr16:54175259-54175672 | -1.115523682 | 0.005309869 | 0.044411398 |
| chr7:21147125-21147401 | 1.171882212 | 0.005312219 | 0.044426199 |
| chr11:4406262-4406611 | -1.13915161 | 0.005314204 | 0.04443795 |
| chr18:50308200-50308518 | -1.148547415 | 0.005314791 | 0.044438011 |
| chr10:43350017-43350853 | 0.9728404 | 0.005315822 | 0.044441781 |
| chr2:241848989-241849338 | -1.198417355 | 0.005318036 | 0.044452099 |
| chr14:96219637-96219845 | 1.519117329 | 0.005318217 | 0.044452099 |
| chr4:28562977-28563263 | -1.405149414 | 0.005319666 | 0.044455122 |
| chr5:96624744-96625212 | -1.238810864 | 0.005319739 | 0.044455122 |
| chr2:5111233-5111995 | -0.989300715 | 0.005320998 | 0.044458905 |
| chr3:31288744-31289055 | 1.261744864 | 0.005321352 | 0.044458905 |
| chr2:162490767-162491236 | -1.016509665 | 0.005322317 | 0.04446212 |
| chr8:141206294-141206841 | 1.005404553 | 0.005322932 | 0.044462408 |
| chr16:72836189-72836502 | 1.143538173 | 0.005325771 | 0.044481274 |
| chr7:33122666-33122965 | 1.1927403 | 0.005326697 | 0.044484163 |
| chr3:171943649-171943981 | 1.371610332 | 0.005332789 | 0.044530184 |
| chr1:207838584-207838809 | -1.535158392 | 0.005334356 | 0.044538417 |
| chr3:129967715-129967951 | 1.349127639 | 0.005335526 | 0.044538625 |
| chr14:99003769-99004058 | 1.304390722 | 0.005335544 | 0.044538625 |
| chrX:104950238-104950942 | -0.963759018 | 0.005337827 | 0.044552828 |
| chr15:91141974-91142392 | 1.181190908 | 0.005340677 | 0.044568317 |
| chr7:20598508-20598855 | 1.052569409 | 0.005340846 | 0.044568317 |
| chr16:89383027-89384030 | -0.943645947 | 0.00534227 | 0.04457535 |
| chr4:174283003-174284526 | -0.89032496 | 0.005345985 | 0.04460149 |
| chr10:113665932-113666439 | 0.976793682 | 0.005352745 | 0.044653028 |
| chr20:30281677-30281969 | 1.445716524 | 0.005355253 | 0.044669083 |
| chr7:94492547-94492920 | 1.231329173 | 0.005356836 | 0.044677427 |
| chr1:109728913-109729309 | -1.019363752 | 0.005358795 | 0.044688899 |
| chr9:74184331-74184591 | -1.552068877 | 0.005360818 | 0.044700907 |
| chr6:138582527-138583091 | -0.989499211 | 0.005361921 | 0.044705234 |
| chr14:29805757-29806025 | 1.416117496 | 0.005363741 | 0.044715546 |
| chr18:72114835-72115319 | -1.187930026 | 0.005369475 | 0.044758473 |
| chr1:159141504-159141875 | -1.090730568 | 0.005370749 | 0.044764222 |
| chr15:71114824-71115575 | -0.956744079 | 0.0053722 | 0.044771447 |
| chr4:6603352-6603652 | 1.375004576 | 0.005373052 | 0.044773678 |
| chr7:157832036-157832286 | -1.396374716 | 0.00537634 | 0.044792299 |
| chr10:91936601-91936809 | 1.604705486 | 0.005376472 | 0.044792299 |
| chr7:30892850-30893060 | 1.554811064 | 0.00537704 | 0.044792299 |

| | | | |
|---|---|---|---|
| chr3:89474857-89475075 | 1.374458134 | 0.005380742 | 0.044818264 |
| chr3:163929325-163929577 | -1.286178608 | 0.00538501 | 0.044848941 |
| chr13:91017398-91018059 | -1.016963043 | 0.005387069 | 0.044856307 |
| chr14:74793866-74794082 | 1.328153484 | 0.005388008 | 0.044856307 |
| chr6:11241741-11242023 | -1.423766178 | 0.005388241 | 0.044856307 |
| chr3:154743050-154743252 | 1.634673074 | 0.00538829 | 0.044856307 |
| chr3:177635706-177635934 | -1.568086348 | 0.005388822 | 0.044856307 |
| chr17:56195567-56195918 | 1.285185953 | 0.00539162 | 0.044874727 |
| chr7:33519102-33519303 | 1.43983832 | 0.005393885 | 0.044888419 |
| chr6:93165945-93166145 | 1.660771582 | 0.005394437 | 0.044888419 |
| chr5:133065573-133065867 | 1.16515417 | 0.005398489 | 0.044914227 |
| chrX:114975029-114975254 | 1.410808716 | 0.005398711 | 0.044914227 |
| chr16:34587146-34587450 | 0.98450288 | 0.005399851 | 0.044918836 |
| chr10:102226060-102226410 | 1.062648563 | 0.005401735 | 0.044929625 |
| chr14:94466656-94467016 | 1.145065334 | 0.005402526 | 0.044931332 |
| chr2:120820954-120821254 | -1.273791688 | 0.005403457 | 0.044934195 |
| chr16:31457990-31458371 | 1.041633116 | 0.005404917 | 0.044941458 |
| chr20:55888715-55888955 | 1.274345935 | 0.005406429 | 0.044949157 |
| chr7:106294424-106294709 | -1.33062157 | 0.005408221 | 0.044959177 |
| chr4:25922420-25922758 | 1.331951674 | 0.005410304 | 0.044971611 |
| chr7:158482918-158483137 | -1.514075684 | 0.005411275 | 0.044974806 |
| chr4:170025638-170026020 | -1.087497673 | 0.005413055 | 0.044984718 |
| chr3:63748258-63748458 | 1.030939967 | 0.005415348 | 0.044994827 |
| chr2:72013053-72013272 | 1.425506592 | 0.005415446 | 0.044994827 |
| chr6:122580800-122581111 | -1.224937114 | 0.005418195 | 0.045012789 |
| chr7:105387853-105389558 | -0.901145556 | 0.005419832 | 0.045017327 |
| chr3:128245611-128245831 | 1.485254632 | 0.005419917 | 0.045017327 |
| chr8:94001977-94002215 | 1.418110286 | 0.005421815 | 0.045028217 |
| chr18:65762618-65762882 | -1.426537008 | 0.005423654 | 0.04503861 |
| chr12:112387751-112388101 | 1.212920508 | 0.005429421 | 0.04508161 |
| chr2:138421869-138422336 | -1.044232114 | 0.005433733 | 0.045111786 |
| chr1:35574947-35575429 | 1.05999639 | 0.005434233 | 0.045111786 |
| chr20:42206575-42206937 | -0.965466358 | 0.005437527 | 0.045134239 |
| chr10:5518135-5518444 | 1.141744877 | 0.005438548 | 0.045134841 |
| chr5:65163706-65164240 | -1.171262742 | 0.005438777 | 0.045134841 |
| chr2:12514727-12515305 | -1.096964607 | 0.005441979 | 0.045156518 |
| chr3:64521752-64522248 | 0.954616475 | 0.005442871 | 0.04515903 |
| chr20:8903795-8904024 | 1.50860482 | 0.005443838 | 0.045162169 |
| chr20:47520577-47521186 | 0.899445729 | 0.00544756 | 0.045188153 |
| chr14:65818335-65818574 | 1.204938332 | 0.005453459 | 0.04523219 |
| chr19:40323762-40324003 | 1.336709706 | 0.005454732 | 0.045237846 |
| chr3:50264591-50265261 | -0.979450664 | 0.005456366 | 0.045242369 |
| chr17:57940325-57941130 | -0.934219037 | 0.005456458 | 0.045242369 |
| chr8:92119727-92119928 | 1.434151612 | 0.005459242 | 0.045260558 |
| chr16:48384005-48384205 | 1.470724709 | 0.005460607 | 0.045266973 |
| chr4:84159857-84160057 | -1.765536714 | 0.005461927 | 0.045269802 |
| chr16:22615231-22615464 | 1.487228204 | 0.00546213 | 0.045269802 |
| chr13:47211308-47211560 | 1.259812336 | 0.005463042 | 0.045272466 |
| chr7:8371796-8372026 | 1.252511671 | 0.005464311 | 0.045278082 |
| chr11:82682236-82682896 | 0.974053102 | 0.005465459 | 0.045282699 |

| | | | |
|---|---|---|---|
| chr16:76729564-76729872 | 1.17171863 | 0.005469393 | 0.04531039 |
| chr18:62399759-62400351 | -1.056649715 | 0.005476605 | 0.04536506 |
| chr14:64184167-64184367 | 1.521249474 | 0.005477176 | 0.04536506 |
| chr10:97612530-97612774 | 1.188726297 | 0.005478183 | 0.045368497 |
| chr10:124892949-124893351 | 1.108284409 | 0.005479266 | 0.045371406 |
| chr4:57132084-57132585 | 1.093620569 | 0.005479718 | 0.045371406 |
| chr7:123557315-123558375 | -0.898951439 | 0.005481141 | 0.045378281 |
| chr7:83769442-83769982 | -1.225759159 | 0.005482153 | 0.045381756 |
| chr6:151702940-151703553 | 0.953404907 | 0.005483213 | 0.045385625 |
| chr10:3056513-3056770 | 1.34616196 | 0.005484139 | 0.045386953 |
| chr8:91076623-91076936 | 1.133969237 | 0.005484558 | 0.045386953 |
| chr3:97720267-97720868 | -1.00898858 | 0.005485207 | 0.045387423 |
| chr11:84233346-84233626 | -1.491555633 | 0.005488325 | 0.045408321 |
| chr1:164644747-164645306 | -1.094601583 | 0.005489422 | 0.045412489 |
| chr3:120003758-120004663 | -0.99223728 | 0.0054946 | 0.045450422 |
| chr9:109205658-109205863 | -1.593970604 | 0.005498281 | 0.045475956 |
| chr1:26404737-26405013 | 1.36189599 | 0.005498971 | 0.045476068 |
| chr19:29777472-29777849 | 1.252955726 | 0.005499481 | 0.045476068 |
| chr20:47236157-47236933 | 0.886723251 | 0.005512703 | 0.045567914 |
| chr8:81050626-81050946 | 1.19010199 | 0.005512834 | 0.045567914 |
| chr22:46525680-46526218 | -1.223480298 | 0.005512954 | 0.045567914 |
| chr3:190424440-190425114 | -0.935731889 | 0.005513003 | 0.045567914 |
| chr9:89767111-89767311 | 1.524647328 | 0.005513562 | 0.045567914 |
| chr8:613909-614318 | 1.17214694 | 0.00552112 | 0.045625461 |
| chr20:16589677-16589913 | -1.614819969 | 0.005526355 | 0.045663792 |
| chr10:130127775-130128121 | 1.054429782 | 0.005528318 | 0.045675092 |
| chr21:33848497-33848865 | -1.217779782 | 0.00553101 | 0.0456924 |
| chr6:144322001-144323079 | -0.967665586 | 0.005532206 | 0.045693587 |
| chr6:98117977-98118177 | -1.843277424 | 0.005532346 | 0.045693587 |
| chr20:41399525-41399945 | 0.990726149 | 0.005533375 | 0.045697158 |
| chrX:26653615-26654187 | -1.018615284 | 0.005534142 | 0.045698 |
| chr2:137866017-137866263 | -1.405985867 | 0.005534669 | 0.045698 |
| chr2:39593583-39594122 | -0.993965286 | 0.005535619 | 0.045700913 |
| chrX:14508163-14508530 | -1.304409912 | 0.0055367 | 0.045704918 |
| chr5:17401703-17402108 | 1.047491383 | 0.005538544 | 0.045715216 |
| chrX:126723646-126724495 | -1.048785336 | 0.005542553 | 0.045743372 |
| chr1:84632896-84633486 | -0.976090908 | 0.005544869 | 0.045757562 |
| chr16:4415560-4416045 | 1.0510908 | 0.005553257 | 0.045821843 |
| chr17:76586707-76586997 | 1.280455458 | 0.005556209 | 0.045841268 |
| chr1:211447669-211447916 | -1.536605877 | 0.005559348 | 0.045862226 |
| chr13:64723650-64724159 | -1.005984853 | 0.005560525 | 0.045863271 |
| chr20:51871819-51872237 | 1.045022903 | 0.005560672 | 0.045863271 |
| chr8:67733786-67734018 | -1.597550621 | 0.005561576 | 0.045865796 |
| chr8:134002128-134002384 | 1.382612954 | 0.005567078 | 0.045906229 |
| chr17:21764718-21764918 | 1.599065924 | 0.005572665 | 0.045947353 |
| chr20:38180719-38181160 | -1.174562604 | 0.005576036 | 0.045968068 |
| chr1:116724780-116725036 | 1.148500988 | 0.005576377 | 0.045968068 |
| chr3:150094391-150094995 | -1.019834219 | 0.005578529 | 0.045980853 |
| chr11:118530897-118531283 | 1.096957026 | 0.005579969 | 0.045987776 |
| chr7:17489554-17489754 | 1.434385272 | 0.005583002 | 0.046007826 |

| | | | |
|---|---|---|---|
| chrX:155264063-155264645 | 1.037861503 | 0.005585176 | 0.046017814 |
| chr7:143240346-143240912 | -1.053160232 | 0.005585415 | 0.046017814 |
| chr5:151232679-151232940 | 1.348580661 | 0.005586233 | 0.046019602 |
| chr22:46375250-46375450 | 1.417297177 | 0.005587217 | 0.046022764 |
| chrX:151083022-151083222 | 1.476341961 | 0.005589365 | 0.046033691 |
| chr11:119067017-119067285 | 1.17464137 | 0.005589948 | 0.046033691 |
| chr5:95664493-95664828 | -1.2390985 | 0.005590346 | 0.046033691 |
| chr2:8301980-8302342 | -1.215987759 | 0.00559328 | 0.046052904 |
| chr2:211423153-211423400 | -1.482473167 | 0.005597065 | 0.046079112 |
| chr5:68884972-68885197 | -1.521590694 | 0.005600546 | 0.046097627 |
| chr5:66746656-66747138 | -1.179272953 | 0.005601084 | 0.046097627 |
| chr1:245108920-245109199 | 1.153098773 | 0.005601119 | 0.046097627 |
| chr14:66758072-66758317 | 1.466837862 | 0.005602496 | 0.046101708 |
| chr12:5022489-5022783 | -1.33548333 | 0.005602818 | 0.046101708 |
| chr19:14505859-14506213 | 1.164466018 | 0.005605343 | 0.046117533 |
| chr1:9128487-9128941 | 1.053455377 | 0.005606432 | 0.046121541 |
| chr2:159516521-159516749 | -1.520294122 | 0.005607605 | 0.046122675 |
| chr12:52163843-52164336 | 0.992272154 | 0.005607774 | 0.046122675 |
| chr14:39317248-39317448 | 1.493360255 | 0.005608597 | 0.046124496 |
| chr10:59432622-59432991 | 1.216092927 | 0.005610007 | 0.04612662 |
| chr17:42763350-42763686 | 1.174147109 | 0.005610059 | 0.04612662 |
| chr7:124029501-124029817 | 1.271587691 | 0.005612843 | 0.046144559 |
| chr5:78584553-78585061 | -1.018069406 | 0.00561533 | 0.046160052 |
| chr20:47867284-47867921 | 0.905149502 | 0.005617028 | 0.04616905 |
| chr16:83813910-83814574 | -1.032681916 | 0.005618023 | 0.046172277 |
| chr2:151261193-151262069 | -0.922982124 | 0.005619675 | 0.046180905 |
| chrX:25020545-25021460 | -0.98037549 | 0.005622169 | 0.046196439 |
| chr20:48005466-48006065 | 0.907161163 | 0.005627814 | 0.046237869 |
| chr12:5503842-5504138 | -1.303828013 | 0.005632336 | 0.046270057 |
| chr1:43254182-43254387 | 1.596116976 | 0.005635736 | 0.046283204 |
| chr20:47524202-47524431 | 0.959989817 | 0.005635743 | 0.046283204 |
| chr14:24491473-24492349 | -1.023285723 | 0.005635872 | 0.046283204 |
| chrX:129747602-129747802 | 1.757254205 | 0.005636352 | 0.046283204 |
| chr7:137964186-137964674 | -1.086108769 | 0.005641155 | 0.046312837 |
| chr13:42595054-42595309 | -1.334437721 | 0.00564117 | 0.046312837 |
| chr12:724320-724695 | -1.319996254 | 0.005644072 | 0.046331699 |
| chr7:9217255-9217758 | -1.176482991 | 0.005645056 | 0.046333737 |
| chr14:61154177-61154448 | -1.231870313 | 0.00564553 | 0.046333737 |
| chr7:129110550-129111059 | -1.094524477 | 0.005647633 | 0.046346037 |
| chr21:41121777-41122299 | -1.014351891 | 0.005649705 | 0.04635807 |
| chr2:192660573-192660923 | -1.187387929 | 0.005651552 | 0.046364375 |
| chr7:50970180-50970437 | -1.36902557 | 0.005652134 | 0.046364375 |
| chr1:247576540-247576787 | -1.338504941 | 0.005652288 | 0.046364375 |
| chr17:61708262-61708489 | 0.972665594 | 0.005654036 | 0.046373748 |
| chr16:4354145-4354360 | 1.247369185 | 0.00565494 | 0.046376199 |
| chr3:184011115-184011586 | 1.012928778 | 0.00565619 | 0.046381485 |
| chr4:27100732-27101355 | 1.033988552 | 0.005658417 | 0.046392044 |
| chr17:76940767-76941113 | 1.15998892 | 0.005658689 | 0.046392044 |
| chr3:20124524-20124724 | 1.691638983 | 0.005659899 | 0.046395427 |
| chr10:73110896-73111420 | -0.996807654 | 0.005660312 | 0.046395427 |

| | | | |
|---|---|---|---|
| chr2:160142505-160142714 | 1.485134244 | 0.005663324 | 0.046413038 |
| chr4:88240138-88240560 | -1.091550992 | 0.005663907 | 0.046413038 |
| chr1:206864943-206865879 | -0.898446345 | 0.005664278 | 0.046413038 |
| chr17:58561666-58562013 | 0.984198553 | 0.00566885 | 0.04644553 |
| chr9:12794130-12794825 | -1.101647309 | 0.005672884 | 0.046473614 |
| chr3:52606639-52606967 | 1.256983199 | 0.005675462 | 0.04648473 |
| chr15:86877078-86877717 | -1.001690525 | 0.005675626 | 0.04648473 |
| chr4:122703098-122703455 | -1.181891741 | 0.005676575 | 0.04648473 |
| chr7:93890041-93891127 | -0.943357155 | 0.005676668 | 0.04648473 |
| chrX:153609923-153610281 | 1.105667581 | 0.005678144 | 0.04649114 |
| chrX:138945363-138945563 | 1.689658076 | 0.005678664 | 0.04649114 |
| chr5:40477821-40478260 | -1.132910333 | 0.005685091 | 0.046538783 |
| chr12:52191803-52192511 | 0.912050313 | 0.005686253 | 0.046543328 |
| chrX:153636336-153636656 | 1.116097603 | 0.005687507 | 0.046548616 |
| chr19:36055138-36055379 | 1.090113222 | 0.005690293 | 0.046563217 |
| chr14:97236373-97236573 | 1.557988112 | 0.005690506 | 0.046563217 |
| chr5:108481437-108481718 | -1.344158987 | 0.005692961 | 0.046578327 |
| chr22:37335348-37335649 | 1.246388429 | 0.005695016 | 0.046590166 |
| chr13:55999772-56000010 | -1.557258782 | 0.005696632 | 0.046598415 |
| chr3:16503417-16503747 | 1.18671895 | 0.005698472 | 0.046608487 |
| chr12:87086487-87086766 | 1.263754581 | 0.005699191 | 0.046609393 |
| chr6:28810083-28810489 | 1.134451868 | 0.005703601 | 0.046640483 |
| chr3:113685565-113685868 | -1.19077454 | 0.005706781 | 0.04666151 |
| chr4:39648767-39649228 | -1.187947762 | 0.005708475 | 0.046670378 |
| chr15:100211941-100212172 | 1.46942708 | 0.005709212 | 0.046671422 |
| chr5:7843123-7843411 | 1.467595573 | 0.005713115 | 0.046698345 |
| chr14:20855647-20856182 | 0.990716439 | 0.005714824 | 0.046707335 |
| chr10:88401788-88402276 | -1.285818467 | 0.005716465 | 0.04671576 |
| chr10:121553834-121554223 | 1.117362494 | 0.005717863 | 0.046722206 |
| chr13:62243085-62243366 | -1.556562225 | 0.005719117 | 0.046727467 |
| chr10:67920177-67920444 | -1.605904879 | 0.005722423 | 0.046749494 |
| chr5:112146765-112147273 | -1.123550382 | 0.005724309 | 0.046759915 |
| chr20:17883089-17883289 | 1.434303473 | 0.005724973 | 0.04676035 |
| chr3:160077951-160078186 | 1.439905367 | 0.00572857 | 0.04678475 |
| chr5:136724256-136724535 | 1.366929119 | 0.005729249 | 0.046784815 |
| chr7:101538837-101539118 | -1.237840352 | 0.0057298 | 0.046784815 |
| chr3:127439520-127439929 | 1.263260057 | 0.005735725 | 0.046825087 |
| chr11:2377169-2377453 | 1.290761945 | 0.005735954 | 0.046825087 |
| chr20:40648754-40649008 | 1.348640938 | 0.005737225 | 0.046830476 |
| chr5:68395924-68396430 | -1.123362347 | 0.005742935 | 0.046872085 |
| chr20:32527484-32527873 | 1.24054398 | 0.005744273 | 0.046878018 |
| chr1:32872626-32872943 | 1.115591951 | 0.005745751 | 0.046885083 |
| chr3:89474601-89474805 | 1.483564826 | 0.005748388 | 0.046901607 |
| chr19:8050631-8050848 | 1.325567756 | 0.005751331 | 0.046920626 |
| chr6:153104986-153105315 | -1.392312278 | 0.005752787 | 0.046927504 |
| chr8:57579173-57579375 | 1.596045755 | 0.005753553 | 0.046928758 |
| chr4:6909220-6910265 | 0.922373362 | 0.005757414 | 0.046955253 |
| chr1:182969375-182969725 | 1.166190757 | 0.005759021 | 0.046963359 |
| chr17:19581663-19581890 | 1.412290875 | 0.00575974 | 0.046964223 |
| chrX:16808046-16808338 | -1.2386753 | 0.005761924 | 0.04697703 |

| | | | |
|---|---|---|---|
| chr10:28939249-28940190 | -0.935251493 | 0.005764748 | 0.046991111 |
| chr6:167116138-167116564 | -1.286726406 | 0.005764878 | 0.046991111 |
| chr14:33908663-33908863 | 1.516900037 | 0.005766688 | 0.047000872 |
| chr1:168744433-168744665 | 1.521638939 | 0.005770589 | 0.047027661 |
| chr1:932987-933304 | 1.244360536 | 0.005771309 | 0.047028526 |
| chr5:87099524-87099883 | -1.159281055 | 0.005775138 | 0.047051312 |
| chr1:223319471-223319747 | 1.232763592 | 0.005775698 | 0.047051312 |
| chr2:61144757-61145248 | -0.96494689 | 0.005776139 | 0.047051312 |
| chr4:168547254-168547486 | -1.503706682 | 0.005776628 | 0.047051312 |
| chr15:76848967-76849188 | 1.432874819 | 0.005777176 | 0.047051312 |
| chr1:220810198-220810920 | -0.973556424 | 0.00577779 | 0.047051312 |
| chr10:53844116-53844373 | -1.400419768 | 0.005779096 | 0.04705247 |
| chr21:38622454-38622762 | 1.265329098 | 0.005779455 | 0.04705247 |
| chr8:57225361-57225691 | 1.171894566 | 0.005780325 | 0.04705247 |
| chr11:45823500-45823709 | 1.342434958 | 0.005780388 | 0.04705247 |
| chr5:42099739-42100279 | 0.984765314 | 0.005781125 | 0.047053468 |
| chr16:57576483-57576714 | 1.399586255 | 0.005784834 | 0.04707366 |
| chr3:165751723-165751923 | 1.399586255 | 0.005784834 | 0.04707366 |
| chr12:32605430-32605631 | 1.44790983 | 0.005785495 | 0.047074038 |
| chr21:43376254-43376454 | 1.460985642 | 0.005787174 | 0.047081431 |
| chr8:95996684-95996887 | 1.629130182 | 0.005787633 | 0.047081431 |
| chr5:180534025-180534225 | 1.587847776 | 0.005789776 | 0.047093864 |
| chr19:1470567-1470891 | 1.135618464 | 0.005796592 | 0.047144299 |
| chr9:131545255-131546983 | 0.899811345 | 0.005799349 | 0.047161721 |
| chr3:174529326-174529801 | -1.078151142 | 0.005801578 | 0.047174836 |
| chr11:65553358-65553630 | -1.182048394 | 0.005809134 | 0.047231267 |
| chr2:232885397-232885597 | 1.455645961 | 0.005813752 | 0.0472638 |
| chr4:147741784-147741984 | 1.361819893 | 0.005816555 | 0.04728157 |
| chr2:88191461-88191791 | -1.105977235 | 0.005818258 | 0.047289484 |
| chr15:90342287-90342553 | -1.435011079 | 0.005819338 | 0.047289484 |
| chr7:143914990-143915304 | -1.258171818 | 0.005819516 | 0.047289484 |
| chr4:107863854-107864152 | -1.181393545 | 0.005820186 | 0.047289484 |
| chr10:142517-142880 | 1.334960208 | 0.005820614 | 0.047289484 |
| chr20:47221165-47221561 | 0.9083442 | 0.005823788 | 0.047309505 |
| chr16:56608222-56608739 | 0.977029204 | 0.005824314 | 0.047309505 |
| chr3:107697404-107698283 | -0.959169795 | 0.005825905 | 0.047312834 |
| chr7:49579261-49580319 | -0.96712952 | 0.005825959 | 0.047312834 |
| chr18:75249335-75249759 | 1.208803763 | 0.005827456 | 0.047313059 |
| chr13:92355310-92355572 | -1.393901762 | 0.005827839 | 0.047313059 |
| chr2:2134084-2134361 | -1.393901762 | 0.005827839 | 0.047313059 |
| chr3:124747443-124747719 | 1.257294332 | 0.00582955 | 0.047318362 |
| chr7:143416621-143416850 | -1.405891493 | 0.005829727 | 0.047318362 |
| chr5:115599576-115599794 | -1.367929213 | 0.005834167 | 0.047347686 |
| chr19:1064884-1065248 | 1.150133165 | 0.00583482 | 0.047347686 |
| chr3:107401491-107401753 | 1.381927566 | 0.005835194 | 0.047347686 |
| chr5:103888835-103889763 | -0.946746674 | 0.005837642 | 0.04736254 |
| chrX:153900919-153901119 | 1.409895175 | 0.005841132 | 0.047385832 |
| chr22:37887630-37887916 | 1.214318757 | 0.005843771 | 0.047396481 |
| chr7:110758480-110758708 | 1.409622059 | 0.00584407 | 0.047396481 |
| chr3:196135650-196136237 | 1.043587229 | 0.0058443 | 0.047396481 |

| | | | |
|---|---|---|---|
| chr2:53688098-53688419 | 1.19175188 | 0.005846462 | 0.047408996 |
| chr3:154904353-154904859 | -1.045430886 | 0.005847834 | 0.047415107 |
| chr14:51638127-51638576 | -1.101766822 | 0.005848757 | 0.047417401 |
| chr4:137468882-137469438 | -1.130620296 | 0.005849355 | 0.047417401 |
| chr10:61923187-61923941 | -1.001023511 | 0.005851288 | 0.047428056 |
| chr7:12297705-12298096 | 0.99204209 | 0.005856493 | 0.047465222 |
| chr4:169133801-169134483 | -1.003075219 | 0.005858834 | 0.047479175 |
| chr12:116434112-116435046 | -0.951941412 | 0.005859951 | 0.047483201 |
| chr7:44224561-44225246 | -0.956393091 | 0.005861589 | 0.047491454 |
| chr20:52053974-52054198 | 1.340647982 | 0.00586662 | 0.047527189 |
| chr12:132479192-132479524 | 1.138993523 | 0.005868214 | 0.047535078 |
| chr14:59303152-59303585 | -1.097861938 | 0.005869846 | 0.04754327 |
| chr13:60226586-60226881 | -1.347218867 | 0.005873814 | 0.04756805 |
| chr12:111034207-111034464 | 1.397801721 | 0.005874757 | 0.04756805 |
| chr15:20340387-20340789 | 1.106855468 | 0.005874768 | 0.04756805 |
| chr5:108196332-108196600 | -1.474385173 | 0.00587801 | 0.047585282 |
| chr4:114827543-114827934 | -1.551420989 | 0.005878595 | 0.047585282 |
| chr8:93105345-93105826 | 0.956544516 | 0.005878759 | 0.047585282 |
| chr6:170398822-170399237 | 1.115873179 | 0.005880631 | 0.047595404 |
| chr5:10090139-10090383 | -1.424311888 | 0.005882429 | 0.04760057 |
| chr7:8380406-8380725 | 1.259445538 | 0.005882511 | 0.04760057 |
| chr21:18865229-18865429 | -1.701589578 | 0.005886154 | 0.047619984 |
| chr3:166034561-166034761 | -1.701589578 | 0.005886154 | 0.047619984 |
| chr5:12272411-12272784 | -1.153065512 | 0.005889525 | 0.047639036 |
| chrX:131501645-131501845 | 1.48923382 | 0.005889752 | 0.047639036 |
| chr3:164673722-164674050 | -1.274821034 | 0.005890741 | 0.047640265 |
| chr1:115992849-115993068 | -1.42250341 | 0.005891148 | 0.047640265 |
| chr1:153677715-153677962 | 1.194174905 | 0.0058921 | 0.047642936 |
| chr2:62306580-62307463 | -0.908120325 | 0.005896665 | 0.047669912 |
| chr2:216108419-216108719 | -1.140705685 | 0.00589668 | 0.047669912 |
| chr1:225768712-225768950 | 1.229656112 | 0.005899178 | 0.047685078 |
| chr18:73942154-73942647 | -1.021556513 | 0.005900646 | 0.047691906 |
| chr4:83381043-83381466 | 1.183526514 | 0.005902461 | 0.047697501 |
| chr7:131544854-131545321 | -1.124086729 | 0.005902717 | 0.047697501 |
| chr7:51315467-51317256 | -0.88524719 | 0.005903205 | 0.047697501 |
| chr17:8056747-8057144 | 1.258505561 | 0.005904071 | 0.047699469 |
| chr3:53219932-53220270 | 1.075895398 | 0.005906595 | 0.047714828 |
| chr17:61118209-61118640 | 1.075801989 | 0.005910797 | 0.047743738 |
| chr17:80979256-80979560 | 1.3331316 | 0.005911961 | 0.047748103 |
| chr1:245258390-245258779 | 1.25294615 | 0.005915131 | 0.047768671 |
| chr3:177420432-177421033 | 1.01485232 | 0.00592022 | 0.047801698 |
| chr4:168569131-168569692 | -1.012925764 | 0.005920468 | 0.047801698 |
| chr14:94474457-94474739 | 1.277119189 | 0.005922163 | 0.047810347 |
| chr8:26151347-26151636 | 1.332963218 | 0.005925534 | 0.047832515 |
| chr2:215389755-215390067 | -1.330713701 | 0.005928618 | 0.047844974 |
| chr11:33266509-33266790 | -1.291103547 | 0.00592866 | 0.047844974 |
| chr19:55027418-55027680 | 1.31919334 | 0.00592895 | 0.047844974 |
| chr1:192983412-192983649 | -1.372476751 | 0.005929663 | 0.047845682 |
| chr7:23463054-23463259 | 1.266184289 | 0.005931665 | 0.047856803 |
| chr16:11698256-11698456 | 1.550173494 | 0.005935233 | 0.047880547 |

| | | | |
|---|---|---|---|
| chr13:85716400-85716822 | -1.411754933 | 0.005935931 | 0.047881135 |
| chr16:74485965-74486206 | 1.357957504 | 0.005939483 | 0.047904743 |
| chr5:18969135-18969350 | 1.517571995 | 0.005942007 | 0.04792006 |
| chr7:141663043-141663791 | -0.918944512 | 0.005943853 | 0.047927276 |
| chr9:136282254-136282836 | -1.025786027 | 0.005944445 | 0.047927276 |
| chr10:132493452-132493716 | 1.194711322 | 0.005944779 | 0.047927276 |
| chr1:107387566-107388063 | 0.943953055 | 0.005950496 | 0.047968322 |
| chr12:90103123-90103377 | -1.35179495 | 0.005954371 | 0.047994512 |
| chr18:46518488-46518718 | -1.398849173 | 0.005955448 | 0.047998146 |
| chr1:156126251-156126562 | -1.164297117 | 0.005956643 | 0.047998315 |
| chr2:216596370-216596570 | 1.511234373 | 0.005957265 | 0.047998315 |
| chr14:94080512-94080757 | 1.157051871 | 0.005957349 | 0.047998315 |
| chr9:83219132-83220039 | -0.913677095 | 0.00596042 | 0.048016783 |
| chr9:111851563-111851929 | -1.079769956 | 0.005961222 | 0.048016783 |
| chr21:34163060-34163330 | 1.281411695 | 0.005961521 | 0.048016783 |
| chr2:129827571-129827771 | 1.591775951 | 0.005962844 | 0.048022395 |
| chr14:49659865-49660411 | -1.007084576 | 0.005964755 | 0.048027759 |
| chr13:47500536-47500759 | 1.228685557 | 0.005964764 | 0.048027759 |
| chr9:2292061-2292676 | 1.073924359 | 0.005968349 | 0.048051573 |
| chr2:190046064-190046403 | 1.335586404 | 0.005969217 | 0.04805351 |
| chr7:128123315-128123751 | -1.088953716 | 0.005973502 | 0.048082959 |
| chr7:16524422-16524675 | 1.337660687 | 0.005975376 | 0.048092989 |
| chr17:76924337-76924586 | 1.331357597 | 0.005977342 | 0.048103764 |
| chr2:113039080-113039280 | 1.582128081 | 0.005980057 | 0.048120553 |
| chr4:143828997-143829461 | -1.26250913 | 0.005984638 | 0.048152362 |
| chr2:101900402-101900602 | 1.554711122 | 0.005994136 | 0.04822372 |
| chr5:113064824-113065024 | 1.431908299 | 0.005996882 | 0.048238283 |
| chr8:114585582-114587039 | -0.89467049 | 0.005997206 | 0.048238283 |
| chr1:180315863-180316099 | 1.590040436 | 0.00599917 | 0.048249022 |
| chr18:66714773-66715817 | -0.977817452 | 0.006006675 | 0.048301898 |
| chr6:107880279-107880668 | 1.172829881 | 0.006007509 | 0.048301898 |
| chr21:34361909-34362109 | 1.381689272 | 0.006008773 | 0.048301898 |
| chr7:10950468-10950785 | 1.272031296 | 0.006008964 | 0.048301898 |
| chr1:208652343-208652644 | -1.212535212 | 0.00600909 | 0.048301898 |
| chr6:43771331-43771673 | 1.035036876 | 0.006010033 | 0.048301898 |
| chr2:62157347-62157718 | -1.429901593 | 0.006010158 | 0.048301898 |
| chr16:4115309-4115566 | 1.264048076 | 0.006012446 | 0.048315221 |
| chr1:207868211-207868600 | -1.001240813 | 0.006013312 | 0.048317116 |
| chr16:74068105-74068337 | 1.479836473 | 0.006018282 | 0.048351977 |
| chr2:216750737-216751131 | 1.17713511 | 0.006022912 | 0.048384101 |
| chr8:143187065-143187265 | 1.200253701 | 0.006023633 | 0.048384817 |
| chr7:21568853-21569126 | 1.255867078 | 0.006026991 | 0.048406717 |
| chr15:78666701-78666938 | 1.46734438 | 0.006028432 | 0.048411515 |
| chr1:175481261-175481526 | 1.199961783 | 0.006028852 | 0.048411515 |
| chr20:47263817-47264093 | 0.927587402 | 0.006032082 | 0.048429847 |
| chr1:206449614-206449979 | -1.087099654 | 0.006032399 | 0.048429847 |
| chr20:48911178-48911433 | 0.961637481 | 0.006033112 | 0.048430498 |
| chr3:161454654-161454898 | 1.245586958 | 0.006036409 | 0.048451889 |
| chr7:21409939-21410156 | 1.312255814 | 0.006038976 | 0.048467414 |
| chr10:66931505-66931708 | -1.603933819 | 0.00604215 | 0.048487808 |

| | | | |
|---|---|---|---|
| chr20:47816969-47817291 | 0.945950015 | 0.006046467 | 0.048517221 |
| chr12:96720396-96720714 | -1.278294619 | 0.006047081 | 0.048517221 |
| chr4:149796367-149796684 | 1.230587199 | 0.006047833 | 0.048518175 |
| chr7:157461906-157462169 | -1.301083051 | 0.006049444 | 0.048526019 |
| chr14:49241232-49241432 | 1.546671028 | 0.006052628 | 0.048546475 |
| chr6:165692108-165692478 | -1.33484765 | 0.006055247 | 0.048562399 |
| chr15:87212028-87212467 | -1.153005069 | 0.006057726 | 0.048577196 |
| chr18:57446406-57446649 | -1.323466876 | 0.006058524 | 0.048578507 |
| chr10:60434511-60434773 | -1.453392507 | 0.006059496 | 0.048581221 |
| chr20:38113506-38113870 | 1.22989798 | 0.006060773 | 0.048586372 |
| chr4:36235247-36235658 | -1.079321974 | 0.006065162 | 0.048616471 |
| chr10:21084007-21084235 | 1.387975313 | 0.006067139 | 0.048618009 |
| chr2:127085155-127085666 | -0.983472682 | 0.006067178 | 0.048618009 |
| chr4:164022950-164023483 | -1.216024868 | 0.006067257 | 0.048618009 |
| chr21:39373669-39373917 | 1.127270684 | 0.00606795 | 0.048618471 |
| chr17:40909357-40909597 | 1.323932532 | 0.006070263 | 0.048631919 |
| chr19:53333487-53334117 | 0.983414354 | 0.006075946 | 0.048669138 |
| chr11:131668884-131669099 | -1.550777644 | 0.006076179 | 0.048669138 |
| chr7:139479644-139479844 | -1.400859243 | 0.006081651 | 0.048707874 |
| chr1:207864991-207865323 | -1.02508446 | 0.006087682 | 0.048751087 |
| chr1:15229762-15230122 | -1.329797434 | 0.006088443 | 0.048752081 |
| chr6:42512752-42512987 | 1.383478898 | 0.006092397 | 0.048778642 |
| chr5:17304994-17305407 | 1.010573627 | 0.006095506 | 0.048798441 |
| chr15:89057521-89057818 | 1.118448719 | 0.006097649 | 0.048810492 |
| chr16:56382873-56383472 | -0.977515448 | 0.006103513 | 0.04885233 |
| chr17:61437935-61438280 | 0.922159645 | 0.006111332 | 0.048907454 |
| chr6:101417906-101418235 | -1.176158185 | 0.006111676 | 0.048907454 |
| chr3:64813797-64813997 | 0.996922906 | 0.006113154 | 0.048914171 |
| chr3:120604835-120605219 | 1.167243097 | 0.006117033 | 0.048940097 |
| chr10:7288086-7288408 | 1.098399731 | 0.006122948 | 0.048982302 |
| chr9:763982-764454 | 0.969239392 | 0.006127592 | 0.049010747 |
| chr3:86050106-86050381 | 1.389743375 | 0.006127783 | 0.049010747 |
| chr3:78988376-78988634 | -1.477739511 | 0.006129461 | 0.049015417 |
| chr10:114087790-114088037 | -1.34872722 | 0.006129646 | 0.049015417 |
| chr18:12407636-12408247 | 0.951455072 | 0.0061312 | 0.04902273 |
| chr1:211675133-211676205 | -0.885292844 | 0.006134057 | 0.049040454 |
| chr4:133150834-133152606 | -0.889775497 | 0.00613881 | 0.049073334 |
| chr20:36422017-36422240 | 1.620660161 | 0.006140739 | 0.049083632 |
| chr12:96617221-96617424 | -1.558410104 | 0.006142242 | 0.049090525 |
| chr11:29343729-29344080 | -1.117178682 | 0.006144889 | 0.049106558 |
| chr21:33066767-33067024 | 1.534966918 | 0.006147372 | 0.049121276 |
| chr5:108212943-108213522 | -0.982130865 | 0.006148275 | 0.049123075 |
| chr8:109865307-109865594 | 1.283188068 | 0.006148879 | 0.049123075 |
| chr2:176796716-176797163 | -1.089655387 | 0.006150739 | 0.04913281 |
| chr19:18919291-18919874 | -0.973510634 | 0.006153825 | 0.049151054 |
| chr8:101233942-101234147 | 1.288865952 | 0.006154893 | 0.049151054 |
| chr11:77216710-77216910 | 1.471779325 | 0.006154947 | 0.049151054 |
| chr3:31695493-31696062 | 1.09504594 | 0.006156074 | 0.04915493 |
| chr4:15457681-15458083 | 1.187845267 | 0.006159515 | 0.049177276 |
| chr20:19751740-19751995 | 1.342172567 | 0.006164834 | 0.049214617 |

| | | | |
|---|---|---|---|
| chr6:6960567-6960813 | 1.422178739 | 0.006170245 | 0.049251159 |
| chr6:53011879-53012352 | 1.078219961 | 0.006170697 | 0.049251159 |
| chr12:109069720-109069937 | 1.462159408 | 0.006171539 | 0.049252754 |
| chr6:83008676-83009030 | -1.228651401 | 0.006175048 | 0.049275623 |
| chr17:38869250-38869539 | 1.030892892 | 0.006176394 | 0.049279296 |
| chr22:28923733-28924285 | 0.998172094 | 0.006176795 | 0.049279296 |
| chr1:180778829-180779286 | -1.062000742 | 0.006178462 | 0.049287464 |
| chr7:118153621-118153881 | -1.396276868 | 0.006182568 | 0.04931509 |
| chr7:142357909-142358272 | -1.060439586 | 0.00618436 | 0.049324246 |
| chr13:71252222-71252480 | -1.387305359 | 0.006185259 | 0.04932628 |
| chr3:73718426-73718687 | 1.313990273 | 0.006188372 | 0.049345973 |
| chr12:46455679-46456312 | 0.950675371 | 0.006192341 | 0.049372482 |
| chr2:45881714-45882053 | 1.355709226 | 0.006197722 | 0.049410244 |
| chr6:156688074-156688380 | -1.09239333 | 0.006199903 | 0.049422491 |
| chr7:143007949-143008522 | -1.04123821 | 0.006200823 | 0.049424678 |
| chr22:28829435-28829695 | 1.104675534 | 0.006201967 | 0.049428654 |
| chr7:22586829-22587109 | 1.051506191 | 0.006205429 | 0.049451104 |
| chr5:128350521-128351111 | -1.017479428 | 0.006206731 | 0.049456334 |
| chr21:43815395-43815660 | 1.211727223 | 0.006207857 | 0.049460167 |
| chr11:64153086-64153404 | 1.080420164 | 0.006215813 | 0.049518146 |
| chr12:40225354-40225554 | -1.553754999 | 0.006216427 | 0.049518146 |
| chr7:116809738-116810733 | -0.953819528 | 0.006217581 | 0.049522191 |
| chr11:75125681-75125965 | 1.498214246 | 0.006221835 | 0.049550926 |
| chr13:22945310-22945620 | 1.278919849 | 0.006224074 | 0.049563602 |
| chr6:144634103-144634303 | 1.675843748 | 0.006229899 | 0.049604831 |
| chr12:48350243-48350620 | 1.054127131 | 0.006230835 | 0.049607132 |
| chr10:7121307-7121596 | 1.110117159 | 0.006232593 | 0.049615338 |
| chr15:97847580-97848050 | 1.31724622 | 0.006233161 | 0.049615338 |
| chr15:84962948-84963406 | 0.975860295 | 0.006235004 | 0.049622843 |
| chr7:45347757-45348407 | -0.989341691 | 0.006235399 | 0.049622843 |
| chr1:152467881-152468221 | 1.18968084 | 0.006237792 | 0.04963673 |
| chr8:37179877-37180130 | 1.350308553 | 0.006239715 | 0.049639397 |
| chr20:10186051-10186488 | 1.165738851 | 0.006240068 | 0.049639397 |
| chr16:49019427-49019627 | -1.584178005 | 0.00624007 | 0.049639397 |
| chr12:131796170-131796472 | 1.13187159 | 0.006241343 | 0.049644363 |
| chr2:24021362-24021737 | -1.045286133 | 0.006246651 | 0.049680836 |
| chr1:190019324-190019590 | -1.226779824 | 0.006247225 | 0.049680836 |
| chr5:21483660-21484292 | 0.925676323 | 0.006249263 | 0.049691885 |
| chrX:153539487-153539768 | 1.136950937 | 0.006253927 | 0.04972381 |
| chr5:79495311-79495726 | -1.203947469 | 0.00626042 | 0.049770271 |
| chr9:4334439-4334739 | -1.427603473 | 0.006261269 | 0.049771859 |
| chr10:64041163-64041821 | -0.943783659 | 0.006265405 | 0.049797426 |
| chr5:105644615-105644815 | -1.829449285 | 0.006265785 | 0.049797426 |
| chr8:124749589-124749879 | 1.154703444 | 0.006267049 | 0.049798802 |
| chrX:115966918-115967299 | -1.186126558 | 0.006267258 | 0.049798802 |
| chr1:106607943-106608304 | 0.956251424 | 0.006268213 | 0.049801225 |
| chr7:97065023-97065307 | -1.326608976 | 0.006273177 | 0.049835494 |
| chr20:55578577-55579789 | 0.872523825 | 0.006276762 | 0.04985881 |
| chr20:59134173-59134472 | 1.148918208 | 0.006279612 | 0.049876279 |
| chr14:21557410-21557936 | -1.042978684 | 0.006283428 | 0.049901416 |

| | | | |
|---|---|---|---|
| chr5:167049933-167050808 | -0.944947881 | 0.006288195 | 0.049934096 |
| chr4:108360964-108361594 | 1.007895803 | 0.006289018 | 0.049935451 |
| chr1:114299718-114299980 | 1.014492913 | 0.006292824 | 0.049960495 |
| chr3:170492733-170493548 | -0.956400497 | 0.006296191 | 0.04998205 |
| chr1:34984621-34984903 | 1.233927223 | 0.006298658 | 0.049996451 |
| chr2:187530064-187530370 | -1.139330447 | 0.00630017 | 0.050000429 |
| chr7:33204666-33204866 | 1.436860591 | 0.00630181 | 0.050000429 |
| chr2:185006541-185006845 | -1.291172811 | 0.006301996 | 0.050000429 |
| chr19:54979644-54979853 | 1.533490938 | 0.006302068 | 0.050000429 |
| chr3:20032125-20032357 | 1.410559896 | 0.006302422 | 0.050000429 |
| chr17:25389752-25390079 | 1.10658374 | 0.006303493 | 0.050003715 |
| chr20:58611637-58611837 | 1.095507647 | 0.006304141 | 0.050003715 |
| chr7:88243328-88243856 | -1.077779105 | 0.006306002 | 0.0500133 |
| chr9:137711658-137711878 | 1.434665116 | 0.006307 | 0.050014641 |
| chr6:10520940-10521545 | -0.935262665 | 0.006307567 | 0.050014641 |
| chr6:11492556-11492809 | -1.332890034 | 0.006308129 | 0.050014641 |
| chr13:76016842-76017168 | -1.146744313 | 0.006310142 | 0.050025426 |
| chr5:181072897-181073097 | 1.451831209 | 0.006310897 | 0.050025777 |
| chr4:99825172-99825594 | -1.251097243 | 0.006311493 | 0.050025777 |
| chr15:73379906-73380106 | 1.341559573 | 0.006314399 | 0.050043633 |
| chr10:67855186-67855935 | -0.944145522 | 0.006317237 | 0.050060948 |
| chr11:17513496-17513696 | 1.624008289 | 0.006324748 | 0.050115288 |
| chr1:38951158-38951477 | 1.191190578 | 0.006325569 | 0.050115662 |
| chr1:119954924-119955153 | 1.281855557 | 0.006326103 | 0.050115662 |
| chr2:194845519-194846127 | -1.056022272 | 0.006328164 | 0.050121836 |
| chr7:49522011-49522521 | -1.032908038 | 0.006328191 | 0.050121836 |
| chr8:108948843-108949143 | 1.238936618 | 0.006331135 | 0.050139972 |
| chr17:61617412-61617730 | 0.919167249 | 0.006333647 | 0.050154683 |
| chr4:159315842-159316263 | -1.091809769 | 0.006339085 | 0.050192554 |
| chr8:106725797-106726281 | -0.921398427 | 0.00634154 | 0.050206807 |
| chr3:196110479-196110744 | 1.20062626 | 0.00634278 | 0.050208899 |
| chr3:181702190-181702390 | -1.503584939 | 0.006343115 | 0.050208899 |
| chr10:3468927-3469454 | 1.001762255 | 0.006344498 | 0.050214662 |
| chr15:20341326-20341639 | 1.194759894 | 0.006347377 | 0.050232257 |
| chr3:137862149-137862349 | 1.505571878 | 0.006348782 | 0.050238185 |
| chr1:185800061-185800270 | 1.38059208 | 0.006350686 | 0.050248068 |
| chr9:98773039-98773285 | 1.221494221 | 0.006352017 | 0.050253405 |
| chr1:247372668-247373251 | -0.923327478 | 0.006356912 | 0.050286943 |
| chr20:47957305-47957505 | 0.979099209 | 0.006359119 | 0.050299208 |
| chr5:92463996-92464430 | -1.227698309 | 0.006366432 | 0.050351855 |
| chr17:78617676-78618023 | -1.271250601 | 0.006369624 | 0.050369373 |
| chr14:105665620-105666109 | 1.03883792 | 0.006369962 | 0.050369373 |
| chr1:43836451-43836814 | -1.145604122 | 0.00637419 | 0.050397605 |
| chr12:4286341-4286655 | -1.340762547 | 0.006378254 | 0.050424531 |
| chr7:57404436-57405123 | 0.933088955 | 0.006381653 | 0.050446197 |
| chr13:32391020-32391623 | -1.010708196 | 0.006387048 | 0.050478831 |
| chr4:158824462-158824795 | -1.411345639 | 0.006387099 | 0.050478831 |
| chr4:46124558-46125306 | -1.011799469 | 0.006387764 | 0.050478884 |
| chr10:62924252-62924854 | -0.971260981 | 0.006389631 | 0.050488429 |
| chr7:81470000-81470416 | -1.1011034 | 0.006390821 | 0.050489612 |

| | | | |
|---|---|---|---|
| chr12:116495188-116495709 | -1.021814977 | 0.006391098 | 0.050489612 |
| chr3:183845162-183845986 | 0.942235401 | 0.006391803 | 0.050489973 |
| chr4:39540126-39540702 | -0.98920997 | 0.006393941 | 0.050496628 |
| chr11:72454481-72454808 | 1.346842809 | 0.006393964 | 0.050496628 |
| chr4:84259136-84259637 | -1.146128741 | 0.006397748 | 0.050518117 |
| chr2:219127047-219127247 | 1.510145778 | 0.006398345 | 0.050518117 |
| chr13:40233384-40233626 | -1.521251443 | 0.006398957 | 0.050518117 |
| chr7:155356577-155357085 | -1.086366059 | 0.006399322 | 0.050518117 |
| chr6:149854928-149855288 | 1.080025762 | 0.006400441 | 0.050521743 |
| chr5:7686901-7687101 | -1.64357064 | 0.006401605 | 0.050525728 |
| chr15:56149326-56149563 | 1.403146004 | 0.006403987 | 0.050535117 |
| chr19:16570020-16570233 | 1.367202128 | 0.006404113 | 0.050535117 |
| chr20:56263497-56263708 | 1.270976826 | 0.006405856 | 0.050543659 |
| chr20:47242206-47242684 | 0.888847716 | 0.006409569 | 0.050567754 |
| chr11:64226098-64226640 | 0.96981085 | 0.006410684 | 0.050571342 |
| chr5:92749969-92750206 | -1.436843443 | 0.006412176 | 0.0505779 |
| chr8:127952098-127952513 | 0.943620701 | 0.006415348 | 0.050596188 |
| chr2:19989583-19990531 | -0.897026239 | 0.006415815 | 0.050596188 |
| chr7:52857339-52857722 | -1.234817636 | 0.006416753 | 0.050598382 |
| chr15:63561265-63561548 | 1.386818752 | 0.006422488 | 0.050638387 |
| chr14:24254719-24254926 | 1.330087881 | 0.006426767 | 0.050666889 |
| chr10:74743339-74743944 | -0.994875518 | 0.006427583 | 0.050666889 |
| chr7:78587534-78587769 | -1.525004839 | 0.006428192 | 0.050666889 |
| chr7:28041505-28041772 | 1.30794025 | 0.006428748 | 0.050666889 |
| chr8:93780059-93780303 | 1.308889175 | 0.006429728 | 0.050669407 |
| chr1:115829312-115829588 | -1.382844364 | 0.006432212 | 0.050683767 |
| chr4:13654110-13654458 | 1.040730703 | 0.00643565 | 0.050705645 |
| chr10:86793837-86794061 | 1.356146833 | 0.006438197 | 0.050720497 |
| chr21:9256969-9257221 | 1.230977373 | 0.006440268 | 0.050731591 |
| chr12:7832161-7832483 | -1.231503368 | 0.006441574 | 0.050736668 |
| chr7:131692038-131692443 | -1.054158801 | 0.006442584 | 0.050739402 |
| chr5:88189360-88190733 | -0.897339498 | 0.006446652 | 0.050766226 |
| chr5:110620570-110620860 | -1.174719928 | 0.006448923 | 0.050778888 |
| chr7:128862526-128863171 | -0.899215305 | 0.006453883 | 0.050812726 |
| chr20:41233971-41234272 | 1.177767326 | 0.006459078 | 0.050847514 |
| chr7:158589966-158590318 | -1.151374566 | 0.006459629 | 0.050847514 |
| chrX:104972599-104973127 | -1.071709927 | 0.006463635 | 0.050873821 |
| chr20:54499133-54499462 | 0.944681967 | 0.006466301 | 0.050889578 |
| chr1:33007235-33007563 | 1.036640354 | 0.006467802 | 0.050895255 |
| chr3:150631475-150631753 | 1.332359326 | 0.006468351 | 0.050895255 |
| chr4:120007553-120007852 | -1.193689393 | 0.00647362 | 0.050931479 |
| chrX:107445568-107445973 | -1.146267497 | 0.006474454 | 0.050932811 |
| chr10:61190404-61190887 | -1.063390269 | 0.006477316 | 0.0509501 |
| chr20:47572999-47573199 | 0.962265989 | 0.006480701 | 0.050971491 |
| chr1:236503026-236503622 | -0.946700729 | 0.006482558 | 0.050980867 |
| chr1:246805046-246805585 | -0.947914848 | 0.006483994 | 0.050986921 |
| chr1:182812451-182812732 | 1.280620265 | 0.006492189 | 0.051046129 |
| chrX:33434496-33434849 | -1.128284175 | 0.006494239 | 0.051057003 |
| chr14:105538397-105538649 | 1.217750549 | 0.006495673 | 0.051063039 |
| chr20:54557424-54557920 | 0.902382683 | 0.006503267 | 0.051117492 |

| | | | |
|---|---|---|---|
| chr1:202085853-202086342 | -0.91422592 | 0.006506811 | 0.051125357 |
| chr12:16939837-16940498 | -0.966327834 | 0.006506917 | 0.051125357 |
| chr12:93551096-93551458 | -1.27881217 | 0.006506929 | 0.051125357 |
| chr12:107622081-107622284 | -1.47497494 | 0.006506936 | 0.051125357 |
| chr8:128204321-128204676 | 1.049038176 | 0.006513549 | 0.051163861 |
| chr10:132518616-132518889 | 0.991919348 | 0.006513684 | 0.051163861 |
| chr2:91571137-91571389 | -1.46869114 | 0.00651384 | 0.051163861 |
| chr3:16016962-16017213 | 1.334002996 | 0.006516597 | 0.051180267 |
| chr11:86470008-86470208 | 1.477635252 | 0.006517679 | 0.051183519 |
| chr7:13032638-13032851 | 1.343148482 | 0.00651968 | 0.05118923 |
| chr2:2333153-2333518 | -1.288400353 | 0.006519742 | 0.05118923 |
| chr5:149264543-149265233 | -0.947127225 | 0.006528454 | 0.051252375 |
| chr18:69326116-69326434 | -1.234956788 | 0.006536266 | 0.051308452 |
| chr15:68335808-68336017 | 1.386763797 | 0.006540185 | 0.051333957 |
| chr3:64198971-64199235 | 0.984585789 | 0.006541803 | 0.051341392 |
| chr14:105687981-105688242 | 1.193828776 | 0.006542638 | 0.051342691 |
| chr15:80447423-80447726 | 1.225516227 | 0.006549509 | 0.051391342 |
| chr17:51941170-51941370 | 1.494581506 | 0.006551965 | 0.051405352 |
| chr2:6675725-6675925 | -1.639702813 | 0.006554524 | 0.051420166 |
| chr1:54935001-54935201 | 1.445303223 | 0.006556213 | 0.051428149 |
| chr5:121108482-121109200 | -0.97749344 | 0.006557244 | 0.051430971 |
| chr20:54571655-54571873 | 0.99468152 | 0.00655836 | 0.051434115 |
| chr3:149523795-149524045 | 1.434833671 | 0.006558987 | 0.051434115 |
| chr6:51679145-51679604 | -1.208404924 | 0.006562469 | 0.051456154 |
| chr4:26787985-26788373 | 1.105420015 | 0.006563556 | 0.051459407 |
| chr17:58772844-58773094 | 1.0783041 | 0.006568282 | 0.051491195 |
| chr12:16385973-16386625 | -1.021664893 | 0.006570545 | 0.051503666 |
| chr12:93091053-93091253 | 1.537637978 | 0.006574271 | 0.051522324 |
| chr19:32701409-32701618 | 1.537637978 | 0.006574271 | 0.051522324 |
| chr16:29611547-29612043 | 1.02720335 | 0.006576869 | 0.051537418 |
| chr10:90658870-90659100 | -1.38185021 | 0.006581962 | 0.051572054 |
| chr13:84455174-84455457 | -1.434638147 | 0.006583602 | 0.051577431 |
| chr5:119744055-119744349 | -1.155935176 | 0.006583995 | 0.051577431 |
| chr3:164550902-164551181 | -1.260387219 | 0.006589693 | 0.051616795 |
| chr17:22977540-22977823 | 1.455678295 | 0.006590954 | 0.051621389 |
| chr2:155157797-155158229 | -1.08458634 | 0.006593877 | 0.05163901 |
| chr3:31445817-31446070 | 1.269112015 | 0.00659487 | 0.051641503 |
| chr4:89681210-89681597 | -1.210975451 | 0.006595947 | 0.051644665 |
| chr1:79353219-79353638 | -1.102764479 | 0.006602696 | 0.05168344 |
| chr13:81927357-81927639 | -1.207313304 | 0.006603842 | 0.05168344 |
| chr1:30034915-30035133 | 1.428660384 | 0.006604661 | 0.05168344 |
| chr16:34592713-34592913 | 1.121878288 | 0.006604747 | 0.05168344 |
| chrX:119779769-119780008 | 1.350148218 | 0.00660481 | 0.05168344 |
| chr7:45686074-45686274 | 1.49861823 | 0.006604947 | 0.05168344 |
| chr18:48990741-48991244 | -1.01954704 | 0.006606875 | 0.051688632 |
| chr21:36966045-36967136 | 0.910161209 | 0.00660696 | 0.051688632 |
| chr9:70418836-70420648 | -0.858299164 | 0.006608803 | 0.051697774 |
| chr4:161304902-161305160 | -1.536319273 | 0.006617772 | 0.051755568 |
| chr15:55648447-55648688 | 1.215004114 | 0.006617993 | 0.051755568 |
| chr12:53370841-53371641 | 0.892194267 | 0.006619395 | 0.051755568 |

| | | | |
|---|---|---|---|
| chr12:81143043-81143510 | -1.003189927 | 0.006620195 | 0.051755568 |
| chr1:117284587-117284938 | 1.049767087 | 0.006621285 | 0.051755568 |
| chr12:45290238-45290586 | -1.170813785 | 0.006621656 | 0.051755568 |
| chr1:248905928-248907037 | 0.874580461 | 0.006621857 | 0.051755568 |
| chr7:115593813-115594149 | -1.392146509 | 0.006621922 | 0.051755568 |
| chr14:55427429-55427766 | 1.136852815 | 0.00662227 | 0.051755568 |
| chr20:54824072-54824419 | 0.935999258 | 0.006625104 | 0.051772432 |
| chr7:95866052-95866315 | -1.37121599 | 0.006626418 | 0.051777421 |
| chr6:56394055-56394349 | -1.266580566 | 0.006629215 | 0.051793179 |
| chr16:16974345-16974554 | 1.529616695 | 0.006631022 | 0.051793179 |
| chr10:123743887-123744419 | 0.972771704 | 0.00663109 | 0.051793179 |
| chr4:167234354-167234822 | -1.094124443 | 0.006631138 | 0.051793179 |
| chr1:168130160-168130374 | 1.409603161 | 0.006632588 | 0.051799218 |
| chr5:58080419-58080948 | -1.051337501 | 0.006643022 | 0.051874893 |
| chr7:138460725-138461301 | -0.953421476 | 0.006644024 | 0.051874893 |
| chr3:64966200-64966414 | 1.071793245 | 0.006644308 | 0.051874893 |
| chr14:94001767-94002174 | 0.972456392 | 0.006646021 | 0.051882974 |
| chr2:31274091-31274357 | 1.373017391 | 0.006648476 | 0.051896853 |
| chr7:93231727-93232926 | -0.868932884 | 0.006656152 | 0.05194818 |
| chr11:27396022-27396222 | -1.541076011 | 0.006656407 | 0.05194818 |
| chr5:123455794-123456039 | -1.348288681 | 0.006657691 | 0.051951803 |
| chr1:167473889-167474359 | -1.161762854 | 0.006658228 | 0.051951803 |
| chr5:118752624-118752884 | -1.430339278 | 0.006660067 | 0.051960861 |
| chr3:188052363-188052716 | -1.107940562 | 0.00666094 | 0.051962385 |
| chr2:65901506-65901789 | 1.497989918 | 0.006663102 | 0.051973962 |
| chr7:41189940-41190141 | -1.522282949 | 0.006666171 | 0.051992603 |
| chr9:95726821-95727070 | 1.323369749 | 0.006670887 | 0.052024092 |
| chr5:14929624-14929850 | 1.477800266 | 0.006671811 | 0.052026004 |
| chr2:2766241-2766465 | 1.456625244 | 0.006674629 | 0.052042683 |
| chr4:172344403-172344966 | -1.146273491 | 0.006681174 | 0.052085506 |
| chr10:86642648-86642857 | 1.194792253 | 0.006681481 | 0.052085506 |
| chr5:59724550-59724829 | -1.192205493 | 0.006682905 | 0.052087951 |
| chr19:8209647-8209847 | 1.431827625 | 0.006683154 | 0.052087951 |
| chr3:171277741-171277995 | 1.301908864 | 0.00668991 | 0.052134182 |
| chr17:61877270-61877599 | 0.917692853 | 0.006690447 | 0.052134182 |
| chr6:35305170-35305440 | 1.255137102 | 0.00669514 | 0.052165449 |
| chr5:143589171-143589605 | 1.179576768 | 0.006698718 | 0.052188017 |
| chr2:139377647-139377939 | -1.586154512 | 0.006703237 | 0.052217916 |
| chr8:2511251-2511565 | -1.383678525 | 0.006706635 | 0.052239079 |
| chr3:159067393-159067685 | -1.403871879 | 0.006707408 | 0.052239785 |
| chr6:42555928-42556239 | -1.365297648 | 0.006715287 | 0.052295839 |
| chr12:63869735-63870014 | -1.265737046 | 0.006717457 | 0.052303289 |
| chr8:43050837-43051042 | 1.412162829 | 0.006717643 | 0.052303289 |
| chr4:117203268-117203595 | -1.222426229 | 0.00672019 | 0.052303289 |
| chr6:130136200-130136484 | 1.345491608 | 0.006720243 | 0.052303289 |
| chr15:22979969-22981252 | 0.915830027 | 0.006720307 | 0.052303289 |
| chr20:1427812-1428012 | 1.57048557 | 0.00672034 | 0.052303289 |
| chr2:204145233-204146013 | -0.945819407 | 0.006722179 | 0.052312292 |
| chr11:22331903-22332124 | -1.465469721 | 0.006724832 | 0.052327619 |
| chr17:64815285-64816180 | -0.901332967 | 0.006733033 | 0.052386112 |

| | | | |
|---|---|---|---|
| chr12:94756857-94757115 | 1.381757373 | 0.006739657 | 0.052422111 |
| chr1:8903672-8903877 | 1.461077001 | 0.006740395 | 0.052422111 |
| chr14:41600631-41600845 | 1.341877506 | 0.006740464 | 0.052422111 |
| chr8:116351613-116352494 | -0.974482012 | 0.006740629 | 0.052422111 |
| chr8:95033734-95034019 | 1.215118191 | 0.00674108 | 0.052422111 |
| chr1:173914307-173914887 | -0.964865263 | 0.0067438 | 0.052437937 |
| chr17:61259420-61259629 | 0.952317383 | 0.006745081 | 0.052442574 |
| chr3:169173219-169174014 | 0.934948197 | 0.006746207 | 0.052446006 |
| chr6:73134576-73134844 | -1.319918142 | 0.006752156 | 0.052486932 |
| chr8:140881483-140881683 | 1.554858093 | 0.006755637 | 0.052508665 |
| chr14:24378433-24378720 | 1.088206211 | 0.006761239 | 0.052546877 |
| chr7:87627624-87629282 | -0.875106479 | 0.006762193 | 0.052548963 |
| chr20:47926802-47927002 | 1.009836564 | 0.006767865 | 0.052587112 |
| chr7:16915498-16915743 | 1.213904563 | 0.006768475 | 0.052587112 |
| chr13:78658327-78659322 | -0.927774195 | 0.006770823 | 0.05260002 |
| chr10:121272389-121272589 | 1.568500159 | 0.006775739 | 0.052632044 |
| chr16:1192970-1193599 | -0.983150844 | 0.006776597 | 0.052632044 |
| chr14:90418533-90418785 | 1.341056526 | 0.006777332 | 0.052632044 |
| chr14:58387531-58387801 | 1.343599636 | 0.006777693 | 0.052632044 |
| chrX:134951319-134951549 | 1.340356321 | 0.006783282 | 0.052670112 |
| chr8:104737323-104737523 | 1.340233173 | 0.006788009 | 0.052700988 |
| chr7:135621253-135621656 | -1.166484878 | 0.006788634 | 0.052700988 |
| chr14:52835508-52835896 | -1.132773064 | 0.006790801 | 0.052712463 |
| chr1:212147367-212148045 | -0.950328677 | 0.006792991 | 0.052724123 |
| chr5:140827821-140828335 | 1.094679583 | 0.006794123 | 0.052727345 |
| chr7:108222665-108222974 | -1.445805422 | 0.006794782 | 0.052727345 |
| chr19:35731366-35731666 | 1.152055897 | 0.006796352 | 0.052734188 |
| chr2:121334964-121335274 | 1.282568963 | 0.006798198 | 0.052740808 |
| chr7:149635570-149636038 | -1.064914573 | 0.006798582 | 0.052740808 |
| chr8:11589376-11589769 | -1.08487824 | 0.006807112 | 0.052801636 |
| chr5:115054286-115054889 | -1.052657797 | 0.006811276 | 0.052828582 |
| chr11:20337655-20337885 | -1.428478962 | 0.006817606 | 0.052872329 |
| chr13:45527879-45528289 | -1.144227329 | 0.006830323 | 0.052965591 |
| chr22:35474079-35474420 | 1.330667323 | 0.006831776 | 0.052971497 |
| chrX:11787622-11787909 | -1.440618286 | 0.006833091 | 0.052976336 |
| chr14:31805473-31805719 | 1.492080896 | 0.006837065 | 0.053001781 |
| chr15:22786002-22787673 | 0.90040388 | 0.006841946 | 0.053034254 |
| chr13:59176793-59177029 | -1.399002286 | 0.006846215 | 0.05305527 |
| chr4:73853146-73853817 | -1.05292457 | 0.006846299 | 0.05305527 |
| chr6:15089603-15090584 | -0.89978948 | 0.006846735 | 0.05305527 |
| chr20:47974485-47974741 | 0.944656046 | 0.006847591 | 0.053056543 |
| chr11:27397658-27397962 | -1.153255215 | 0.006852243 | 0.053087218 |
| chr1:77447228-77448319 | -0.911980487 | 0.006853059 | 0.053088168 |
| chr7:142626538-142626865 | -1.359126751 | 0.006854496 | 0.053089395 |
| chr7:105401589-105402125 | -1.025767352 | 0.006855071 | 0.053089395 |
| chr7:127447417-127447726 | -1.185667228 | 0.006855296 | 0.053089395 |
| chr11:68763045-68763674 | 0.898668229 | 0.006858451 | 0.053108459 |
| chr2:37546394-37546650 | -1.391348161 | 0.006862142 | 0.05313167 |
| chr20:48326488-48326769 | 0.952882655 | 0.006864372 | 0.053141837 |
| chrX:154486525-154487231 | 0.921052748 | 0.006864842 | 0.053141837 |

| | | | |
|---|---|---|---|
| chr1:231049408-231049777 | 1.220893894 | 0.006866686 | 0.053150747 |
| chr13:32125412-32125630 | -1.623395265 | 0.006870963 | 0.05317311 |
| chr17:27472335-27472535 | -1.623395265 | 0.006870963 | 0.05317311 |
| chr21:36587497-36587756 | 1.367409858 | 0.006873039 | 0.053173562 |
| chr6:161353173-161353677 | -1.082181458 | 0.006873095 | 0.053173562 |
| chr1:10899950-10900970 | -0.922050402 | 0.006873104 | 0.053173562 |
| chr20:25328819-25329054 | 1.355683322 | 0.006874592 | 0.053179706 |
| chr19:5298433-5298876 | 1.067042466 | 0.006876635 | 0.053190143 |
| chr6:119284968-119285197 | -1.48399689 | 0.006878506 | 0.053194113 |
| chr4:168585564-168585844 | -1.111441586 | 0.00687915 | 0.053194113 |
| chr4:97656112-97656336 | -1.487452122 | 0.006879539 | 0.053194113 |
| chrX:87485697-87486028 | 1.183178691 | 0.006879926 | 0.053194113 |
| chr19:49097554-49098278 | 0.953072004 | 0.006883687 | 0.053217822 |
| chr6:6699732-6700358 | -0.970215817 | 0.006885245 | 0.0532245 |
| chr6:125245316-125246251 | -0.892423749 | 0.006889963 | 0.053255599 |
| chr14:77208779-77208979 | 1.248221272 | 0.006892902 | 0.053269299 |
| chr11:103055211-103055462 | -1.483770872 | 0.006893126 | 0.053269299 |
| chr5:141910224-141910684 | -1.098954639 | 0.00689753 | 0.053296451 |
| chr20:44714762-44714976 | 1.157940757 | 0.006898031 | 0.053296451 |
| chr1:201034732-201034976 | 1.206573308 | 0.006901119 | 0.053310387 |
| chr22:28784386-28784595 | 1.431596575 | 0.006901838 | 0.053310387 |
| chr22:23849342-23850017 | 0.962781559 | 0.006902324 | 0.053310387 |
| chr14:78848302-78848575 | 1.148319374 | 0.006902617 | 0.053310387 |
| chr5:173642327-173642760 | -0.999657365 | 0.006903819 | 0.05331429 |
| chrX:139204917-139205520 | 0.951972643 | 0.006911605 | 0.053369039 |
| chr18:64244600-64244952 | -1.2286332 | 0.00691367 | 0.053379611 |
| chr5:126228154-126228357 | 1.608926964 | 0.006922126 | 0.053435694 |
| chr17:74431694-74432446 | 0.905546673 | 0.006922388 | 0.053435694 |
| chr2:144449028-144449337 | -1.213189803 | 0.006923084 | 0.053435694 |
| chr7:131040501-131040927 | -1.115123873 | 0.006923724 | 0.053435694 |
| chr7:18215418-18215660 | 1.268248967 | 0.006930603 | 0.053483401 |
| chr12:3122951-3123271 | -1.288128403 | 0.006933587 | 0.053501036 |
| chr16:85458364-85458583 | 1.176339841 | 0.006934333 | 0.053501403 |
| chr4:109441589-109442176 | -1.091873177 | 0.006937953 | 0.053523947 |
| chr5:83738165-83738570 | 1.316784056 | 0.006941705 | 0.053547503 |
| chr15:89655340-89655540 | 1.241622138 | 0.006942845 | 0.053550904 |
| chr15:85044353-85044614 | 1.179396468 | 0.006945109 | 0.053558705 |
| chr14:38466925-38467174 | 1.446185169 | 0.006945255 | 0.053558705 |
| chr1:145761610-145761857 | 1.179883658 | 0.006948044 | 0.053574824 |
| chr10:132418605-132419239 | 0.905367498 | 0.006951915 | 0.053599276 |
| chr20:47592908-47593190 | 0.9332129 | 0.006952828 | 0.053600927 |
| chr14:99596130-99596407 | 1.165191259 | 0.006953773 | 0.053602818 |
| chr15:72167229-72167429 | 1.461286262 | 0.006955415 | 0.053607334 |
| chr14:68473076-68473336 | -1.276922034 | 0.006955758 | 0.053607334 |
| chr11:118171738-118171938 | 1.291603966 | 0.006959349 | 0.053629614 |
| chr7:135170149-135171088 | -0.882751944 | 0.006964097 | 0.053660804 |
| chrX:116285419-116285985 | -1.05102209 | 0.006968996 | 0.053693151 |
| chr14:23063744-23063944 | 1.218710815 | 0.006969782 | 0.053693813 |
| chr2:85954716-85955622 | 0.919098371 | 0.006971207 | 0.053699389 |
| chr2:69263878-69264356 | -1.257739451 | 0.006972434 | 0.053700203 |

| | | | |
|---|---|---|---|
| chr1:154501226-154501627 | 1.081090921 | 0.006972714 | 0.053700203 |
| chr14:89560905-89561143 | 1.371825456 | 0.006982267 | 0.053768371 |
| chrX:153334381-153334627 | 1.295420429 | 0.006984281 | 0.053778475 |
| chr22:32080948-32081210 | 1.173368256 | 0.0069864 | 0.053789381 |
| chr16:70778065-70778393 | -1.170437882 | 0.006989128 | 0.053804983 |
| chr5:118684444-118684682 | 1.479052162 | 0.006990808 | 0.053812506 |
| chr13:41973292-41973714 | -1.091423457 | 0.006995049 | 0.053839742 |
| chr7:130668052-130669284 | -0.889039955 | 0.007004692 | 0.053908552 |
| chrX:50629917-50630141 | 1.335029984 | 0.007005803 | 0.053911378 |
| chr9:37342437-37342733 | -1.382514035 | 0.007006467 | 0.053911378 |
| chr1:102790162-102790423 | -1.62639975 | 0.007009997 | 0.053933128 |
| chr1:209566118-209566415 | -1.076768709 | 0.007012019 | 0.053942393 |
| chr2:86599028-86599427 | -1.418569002 | 0.00701261 | 0.053942393 |
| chr5:148851395-148851600 | 1.388590857 | 0.007014327 | 0.05395019 |
| chr8:22918470-22918750 | 1.204364164 | 0.007015876 | 0.053952922 |
| chr13:84645763-84646068 | -1.308108528 | 0.007016337 | 0.053952922 |
| chr11:2892142-2892736 | 0.987212099 | 0.007016795 | 0.053952922 |
| chr15:74397044-74397244 | 1.324625747 | 0.007017553 | 0.05395334 |
| chr3:52210795-52211002 | 1.383878989 | 0.007020405 | 0.05396985 |
| chr5:78066277-78066532 | -1.288881352 | 0.007022677 | 0.053981897 |
| chr9:122383438-122383667 | -1.508820589 | 0.007029746 | 0.054030817 |
| chr15:87045712-87046262 | -1.044415763 | 0.007030769 | 0.05403326 |
| chr1:114651610-114652401 | 0.8837789 | 0.007036731 | 0.054073655 |
| chr9:98135545-98135935 | 1.199663016 | 0.007043201 | 0.054117942 |
| chr2:75627957-75628503 | -1.101176707 | 0.007045562 | 0.054129806 |
| chr8:17836779-17837005 | 1.472902174 | 0.007046158 | 0.054129806 |
| chr14:106638933-106639139 | -1.574560359 | 0.007047699 | 0.054136223 |
| chr8:103641389-103641620 | 1.39509068 | 0.007052648 | 0.054163748 |
| chr10:3777098-3777325 | 1.122771647 | 0.007052697 | 0.054163748 |
| chr6:72182865-72183288 | 1.04788437 | 0.007053673 | 0.05416582 |
| chr11:85481446-85481843 | 1.215008744 | 0.007055095 | 0.054171309 |
| chr17:57609363-57609636 | 1.135115391 | 0.007057246 | 0.054182396 |
| chr1:225898536-225898792 | 1.247207819 | 0.007061385 | 0.054204722 |
| chr10:106711425-106711654 | -1.487753789 | 0.007062952 | 0.054204722 |
| chr1:208238942-208239363 | -1.180743738 | 0.007063129 | 0.054204722 |
| chr16:25138261-25138495 | 1.413213624 | 0.007063653 | 0.054204722 |
| chr10:109650160-109650720 | 0.953720209 | 0.007063691 | 0.054204722 |
| chr20:47211903-47212103 | 0.934463187 | 0.007064429 | 0.054204953 |
| chr7:105648835-105649379 | -0.99489792 | 0.0070666 | 0.054216184 |
| chr2:163173652-163173914 | -1.192380253 | 0.007067813 | 0.054220061 |
| chr1:106278461-106278661 | 1.1389972 | 0.007071849 | 0.054245593 |
| chr3:62859830-62860154 | 0.991700434 | 0.007074979 | 0.054264166 |
| chr8:122689661-122689918 | 1.126424877 | 0.007076314 | 0.054268972 |
| chr7:130025994-130026493 | -1.015403844 | 0.007079647 | 0.054289101 |
| chr5:74315995-74316780 | -0.921763842 | 0.007082282 | 0.054295818 |
| chr1:154104566-154105058 | 1.08374815 | 0.00708241 | 0.054295818 |
| chr1:72899131-72899389 | -1.490726541 | 0.007082649 | 0.054295818 |
| chr7:103447543-103448008 | -0.986466527 | 0.007090864 | 0.054353356 |
| chr14:95273302-95273603 | 1.164279938 | 0.007099064 | 0.054406168 |
| chr4:8108916-8109165 | 1.350195019 | 0.007099173 | 0.054406168 |

| | | | |
|---|---|---|---|
| chr11:79393670-79393942 | -1.162261908 | 0.007100211 | 0.054408677 |
| chr7:124489818-124490266 | -1.161356894 | 0.007101315 | 0.054411695 |
| chr14:24396252-24396490 | 1.173040141 | 0.007107024 | 0.054449998 |
| chr3:158722935-158723789 | 0.877645275 | 0.007108441 | 0.054455406 |
| chr7:37693987-37695207 | -0.879858823 | 0.007110913 | 0.054468897 |
| chr4:173317074-173317361 | -1.125780781 | 0.007113739 | 0.054485095 |
| chr9:79759479-79759746 | 1.24889757 | 0.007121011 | 0.054535344 |
| chr6:135770849-135771254 | -1.277706675 | 0.007122369 | 0.054540294 |
| chr7:96414980-96415219 | -1.388900121 | 0.007124261 | 0.054549326 |
| chr1:107728025-107728254 | 1.187527881 | 0.007126675 | 0.054561909 |
| chr13:70248269-70248615 | -1.338924233 | 0.007127328 | 0.054561909 |
| chr6:40648908-40649308 | 1.077361047 | 0.007132397 | 0.054589031 |
| chr15:66670757-66670983 | 1.312805037 | 0.007132432 | 0.054589031 |
| chr20:46614514-46614738 | 1.314143631 | 0.007133009 | 0.054589031 |
| chr4:173752624-173752842 | -1.522289946 | 0.007134858 | 0.054596272 |
| chr1:100249179-100250277 | -0.8778285 | 0.00713538 | 0.054596272 |
| chr16:76345238-76345438 | 1.411262051 | 0.007136539 | 0.054599687 |
| chr1:228861313-228861587 | 1.21748435 | 0.007146685 | 0.054671855 |
| chr22:29344143-29344492 | 1.03796241 | 0.007149284 | 0.054686278 |
| chr17:80024117-80024317 | 1.33950965 | 0.007150291 | 0.054688523 |
| chr4:79492134-79492374 | 1.291609387 | 0.007152103 | 0.054696923 |
| chr15:55217647-55217920 | 1.090590841 | 0.007153141 | 0.054699396 |
| chr5:103434144-103434623 | -0.9210287 | 0.007155169 | 0.054704005 |
| chr5:80181305-80181541 | -1.236509928 | 0.007155171 | 0.054704005 |
| chr7:88306757-88307154 | -1.053495237 | 0.007158132 | 0.054717913 |
| chr21:41360239-41360595 | -1.117475318 | 0.007158419 | 0.054717913 |
| chr11:96070540-96070950 | -1.264919222 | 0.007159724 | 0.054722429 |
| chr5:178467901-178468170 | -1.248994412 | 0.00716119 | 0.054728177 |
| chr3:62765723-62765972 | 1.161099426 | 0.007162284 | 0.054731081 |
| chr4:7554903-7555294 | 1.104755315 | 0.007163085 | 0.054731744 |
| chr11:60943254-60943556 | 1.07472903 | 0.007167 | 0.054756193 |
| chr21:8465245-8465548 | 0.947598445 | 0.007168346 | 0.054761016 |
| chr2:194326106-194326560 | -1.078186733 | 0.007169583 | 0.054765011 |
| chr3:103654042-103654306 | 1.315068959 | 0.007177782 | 0.054821484 |
| chr16:75550136-75550612 | -0.982287577 | 0.007178408 | 0.054821484 |
| chr11:12342264-12342588 | 1.376017922 | 0.007183864 | 0.054857688 |
| chr2:20649591-20649857 | -1.07523209 | 0.007185132 | 0.054860803 |
| chr7:142385506-142385925 | -1.063019141 | 0.007185704 | 0.054860803 |
| chr7:128208509-128208770 | -1.088777792 | 0.007187978 | 0.054871474 |
| chr7:12770233-12770467 | 1.209811916 | 0.007188534 | 0.054871474 |
| chr7:106071377-106071841 | -0.999637681 | 0.00719101 | 0.054884906 |
| chr8:92896340-92896587 | 1.271647422 | 0.007192385 | 0.054889935 |
| chr12:100363542-100363807 | -1.391888604 | 0.007193976 | 0.054896608 |
| chr2:139488910-139489284 | -1.229295541 | 0.007194843 | 0.054897753 |
| chr8:71052450-71052912 | -1.195461606 | 0.007196532 | 0.054905175 |
| chr2:44974088-44974525 | -1.065702284 | 0.007202179 | 0.054937325 |
| chr9:31908311-31908673 | -1.203252769 | 0.00720218 | 0.054937325 |
| chr2:31980738-31981311 | -1.009321562 | 0.007204828 | 0.054952055 |
| chr7:10638262-10638462 | 1.418462175 | 0.007213781 | 0.055006868 |
| chr6:139587568-139587865 | 1.223376664 | 0.007214105 | 0.055006868 |

| | | | |
|---|---|---|---|
| chr15:85047667-85047913 | 1.254581329 | 0.007214168 | 0.055006868 |
| chr6:148191642-148192159 | 0.983860227 | 0.007217544 | 0.055027128 |
| chr3:66316015-66316318 | -1.483199457 | 0.007218745 | 0.055030812 |
| chr5:149011016-149011317 | -1.143830429 | 0.007220627 | 0.055037002 |
| chr9:504058-505251 | 0.890255497 | 0.007220994 | 0.055037002 |
| chr5:1522924-1523384 | 0.96874198 | 0.007223373 | 0.055049657 |
| chr7:19846368-19846598 | 1.439800353 | 0.007227979 | 0.055079285 |
| chr7:132580727-132581087 | -1.05138807 | 0.007232203 | 0.055103965 |
| chr12:10213140-10213607 | -1.041919146 | 0.007232656 | 0.055103965 |
| chr2:119518341-119518562 | 1.326138869 | 0.007233418 | 0.055104291 |
| chr16:3582281-3582607 | 1.260245134 | 0.007235618 | 0.055115569 |
| chr8:22999829-23000183 | 1.158029106 | 0.007236674 | 0.055118132 |
| chr7:74475610-74475962 | 1.091010522 | 0.007240393 | 0.055140982 |
| chr18:62486287-62486766 | -0.990402812 | 0.007243202 | 0.055156893 |
| chr11:70577872-70578169 | 1.25745364 | 0.007244865 | 0.055164071 |
| chr20:34986687-34986907 | 1.319612258 | 0.007245626 | 0.055164382 |
| chr12:7439622-7440447 | -0.968111026 | 0.00725111 | 0.055200649 |
| chr5:88683533-88684088 | -0.932806533 | 0.007254435 | 0.055216186 |
| chr2:214379286-214379784 | -1.011381995 | 0.007254592 | 0.055216186 |
| chr3:168864963-168865215 | -1.319209597 | 0.007256776 | 0.055227325 |
| chr6:56246581-56246824 | -1.357477834 | 0.007262307 | 0.05526393 |
| chr10:73944-74248 | 1.216634114 | 0.007264324 | 0.055273788 |
| chr8:100900462-100901011 | 0.948609268 | 0.007266202 | 0.055282583 |
| chr19:46890797-46891088 | 1.251243555 | 0.007269629 | 0.055303165 |
| chr14:32254905-32255374 | 1.02130214 | 0.007271066 | 0.055308611 |
| chr10:54063452-54063654 | -1.487659631 | 0.007276699 | 0.05534596 |
| chr1:238485059-238486181 | -0.893403088 | 0.007292453 | 0.055456569 |
| chr19:4707367-4707666 | -1.481219286 | 0.007292689 | 0.055456569 |
| chr2:143334903-143335103 | 1.398291984 | 0.007294217 | 0.055461444 |
| chr12:66403817-66404123 | 1.17648602 | 0.007294777 | 0.055461444 |
| chr3:102053397-102053597 | -1.619829478 | 0.007297247 | 0.055474717 |
| chr16:31135601-31135955 | 0.991476681 | 0.007298711 | 0.05548034 |
| chr7:349153-349476 | 1.170418204 | 0.00730563 | 0.055527428 |
| chr15:93899935-93900185 | -1.321313633 | 0.007306458 | 0.055528207 |
| chr7:98733096-98733379 | -1.229181245 | 0.007310074 | 0.055545661 |
| chr12:15706115-15706360 | -1.393502199 | 0.007310933 | 0.055545661 |
| chr20:48846471-48846840 | 0.931074321 | 0.007312207 | 0.055545661 |
| chr2:120564335-120564683 | 1.130061675 | 0.007312299 | 0.055545661 |
| chr14:82642425-82642947 | -0.964999969 | 0.007312794 | 0.055545661 |
| chr1:88877972-88878253 | -1.416399684 | 0.007313104 | 0.055545661 |
| chr3:156801767-156802201 | -0.978141003 | 0.007317134 | 0.055567032 |
| chrX:39612311-39612592 | -1.261723485 | 0.007318356 | 0.055567032 |
| chr5:991631-991831 | 1.37828976 | 0.007319291 | 0.055567032 |
| chr9:72527312-72527611 | 1.395269378 | 0.007319375 | 0.055567032 |
| chr3:63208381-63208669 | 1.034757985 | 0.00731969 | 0.055567032 |
| chr14:25821496-25821864 | 1.083316989 | 0.00732224 | 0.055567032 |
| chr16:11916566-11916920 | 0.968261007 | 0.007322392 | 0.055567032 |
| chr20:38604845-38605275 | 1.023619674 | 0.007322706 | 0.055567032 |
| chr12:70138636-70138891 | 1.41135375 | 0.007323086 | 0.055567032 |
| chr16:79632393-79632617 | -1.32002722 | 0.007323779 | 0.055567032 |

| | | | |
|---|---|---|---|
| chr16:450639-450988 | -1.18704504 | 0.007324898 | 0.055567032 |
| chr1:106972910-106973134 | 1.064690169 | 0.007325621 | 0.055567032 |
| chr3:64239022-64239222 | 0.973773856 | 0.007326009 | 0.055567032 |
| chr17:9672697-9673150 | 1.079426197 | 0.00732607 | 0.055567032 |
| chr11:47206232-47206492 | 1.36515441 | 0.007327857 | 0.055575081 |
| chr1:242239443-242239831 | 0.998246953 | 0.007328762 | 0.055576447 |
| chr20:49385020-49385305 | 1.128236946 | 0.007338031 | 0.055641228 |
| chr1:51314326-51314745 | 1.065294931 | 0.007341197 | 0.055659729 |
| chr1:146894443-146894741 | 1.376621085 | 0.007343447 | 0.055671278 |
| chr2:201612458-201613208 | -0.982264953 | 0.007345437 | 0.055680854 |
| chr12:112136931-112137471 | 0.967138308 | 0.007348083 | 0.055688935 |
| chr1:15617893-15618249 | 1.077893204 | 0.007348577 | 0.055688935 |
| chr7:8423173-8423425 | 1.244009628 | 0.007348683 | 0.055688935 |
| chr9:83642744-83643162 | -1.061067412 | 0.007352065 | 0.055709054 |
| chr5:149904306-149904763 | -0.991671538 | 0.007354402 | 0.05572125 |
| chr7:1980211-1980411 | 1.352024921 | 0.007355276 | 0.055722363 |
| chr13:65966957-65967489 | -1.03986582 | 0.007361207 | 0.055761778 |
| chr2:102111590-102111950 | 1.288040592 | 0.007362799 | 0.055768322 |
| chr4:184331224-184331739 | -1.046652254 | 0.007366513 | 0.055790943 |
| chr4:137474950-137475196 | -1.353303594 | 0.007372792 | 0.055832976 |
| chr5:107869679-107869922 | 1.245928411 | 0.007373793 | 0.055835043 |
| chr4:6470499-6470700 | 1.292632532 | 0.007375773 | 0.055844515 |
| chr20:8305465-8305958 | -1.075765384 | 0.007377101 | 0.055849052 |
| chr21:9810266-9810511 | 1.329912587 | 0.007385498 | 0.055900586 |
| chr22:20494934-20495306 | 1.123210124 | 0.007385586 | 0.055900586 |
| chr14:90036494-90036726 | 1.402895214 | 0.007386569 | 0.055900586 |
| chr1:1763967-1764614 | -1.003727447 | 0.007386827 | 0.055900586 |
| chr18:56869194-56869474 | -1.358460725 | 0.007387645 | 0.055901261 |
| chr10:21335096-21335343 | -1.252135223 | 0.007392108 | 0.055924999 |
| chr2:163610997-163611565 | -0.983444246 | 0.007392425 | 0.055924999 |
| chr15:62248608-62248987 | -1.124990563 | 0.007392972 | 0.055924999 |
| chr3:150667422-150667778 | 1.168794209 | 0.007393808 | 0.055925801 |
| chr14:93359880-93360184 | 1.36249534 | 0.007397597 | 0.055948938 |
| chr17:62444558-62444894 | 1.114902844 | 0.007402647 | 0.055981608 |
| chr13:20089892-20090092 | 1.515647964 | 0.007408112 | 0.056017405 |
| chr12:56206594-56206878 | 1.103502864 | 0.007409765 | 0.056024378 |
| chr2:121238995-121239241 | -1.417163622 | 0.00741569 | 0.056063644 |
| chr8:74337415-74337863 | -1.156036089 | 0.007418528 | 0.056078755 |
| chr4:119788671-119788871 | 1.55806964 | 0.007419785 | 0.056078755 |
| chr3:88031730-88032167 | -1.142157574 | 0.007419942 | 0.056078755 |
| chr10:67114726-67115391 | -0.996191073 | 0.007420617 | 0.056078755 |
| chr6:158183429-158183682 | -1.426300973 | 0.007422315 | 0.056083517 |
| chr12:88052692-88053060 | -1.058348949 | 0.007422711 | 0.056083517 |
| chr3:195582958-195583347 | -1.072713128 | 0.007426487 | 0.056106518 |
| chr2:137517635-137517836 | 1.443796274 | 0.007428507 | 0.056116241 |
| chr6:133837728-133838395 | -0.922959495 | 0.007430196 | 0.056123473 |
| chr19:7197099-7197554 | 0.990792391 | 0.00743357 | 0.056143422 |
| chr12:10747744-10748237 | -1.081787785 | 0.007443497 | 0.056211474 |
| chr3:71532717-71533018 | -1.331523786 | 0.007444048 | 0.056211474 |
| chr5:87210820-87211356 | -1.003290583 | 0.007445129 | 0.0562141 |

| | | | |
|---|---|---|---|
| chr4:186725877-186726153 | -1.027458054 | 0.007446717 | 0.05622055 |
| chr10:117677348-117677602 | -1.360447401 | 0.007448681 | 0.056224466 |
| chr6:129671800-129672153 | -1.283054731 | 0.007448703 | 0.056224466 |
| chr5:113123673-113123899 | -1.318971475 | 0.007450842 | 0.05623507 |
| chr9:127786375-127786815 | 0.990857473 | 0.007455322 | 0.056263343 |
| chr16:89047686-89047965 | -1.220752888 | 0.007463594 | 0.056320219 |
| chr2:201258224-201258621 | -1.106386813 | 0.007467622 | 0.056345063 |
| chr17:74199754-74199954 | 1.135678248 | 0.007470929 | 0.056360324 |
| chr10:101032757-101033330 | -0.918203866 | 0.007472223 | 0.056360324 |
| chrX:119398931-119399754 | 0.917591555 | 0.00747253 | 0.056360324 |
| chr1:10722611-10723226 | -0.983038875 | 0.007473653 | 0.056360324 |
| chr7:106598714-106599276 | -0.96522465 | 0.007474868 | 0.056360324 |
| chr2:197287364-197287653 | 1.197096513 | 0.007474957 | 0.056360324 |
| chr17:61562188-61562388 | 0.950533238 | 0.007475666 | 0.056360324 |
| chr17:80538641-80538845 | 1.448531804 | 0.007475956 | 0.056360324 |
| chr18:31241490-31241732 | 1.293967746 | 0.007476842 | 0.056360324 |
| chr6:122850677-122850998 | 1.213858737 | 0.007477 | 0.056360324 |
| chr13:79391419-79391667 | -1.205314864 | 0.007478132 | 0.056362909 |
| chr5:98913975-98914208 | -1.407423815 | 0.007478814 | 0.056362909 |
| chr2:49026354-49026645 | 1.311763983 | 0.007479642 | 0.056363603 |
| chr22:19629835-19630168 | 1.190554058 | 0.007483753 | 0.056389038 |
| chr5:154850256-154850847 | -0.955618914 | 0.007486556 | 0.056402983 |
| chr15:73265519-73265949 | 1.050270087 | 0.007487076 | 0.056402983 |
| chr1:223212567-223212916 | -1.157869832 | 0.007496381 | 0.056467525 |
| chr4:83074037-83074342 | -1.214672064 | 0.007498266 | 0.05646939 |
| chr10:11658800-11659238 | 1.063236535 | 0.007498653 | 0.05646939 |
| chr1:2311567-2311833 | 1.147786471 | 0.00749905 | 0.05646939 |
| chr4:182809766-182810045 | -1.147856585 | 0.007499909 | 0.05646939 |
| chr1:193186460-193186740 | -1.191892561 | 0.007500313 | 0.05646939 |
| chr9:110213447-110213975 | -1.065777786 | 0.007503022 | 0.056479745 |
| chr4:3827932-3828266 | 1.163000068 | 0.007503163 | 0.056479745 |
| chr13:114036742-114037232 | -1.121851921 | 0.007504934 | 0.05648533 |
| chr11:1573894-1574204 | 1.070405783 | 0.007505379 | 0.05648533 |
| chr19:50922286-50922587 | 1.17568684 | 0.007508698 | 0.056504757 |
| chr16:57772546-57772746 | 1.591835312 | 0.007512152 | 0.056525197 |
| chr21:29433691-29434002 | 1.139449416 | 0.00751412 | 0.056534458 |
| chr17:59984731-59985015 | 1.103919633 | 0.007519902 | 0.05656485 |
| chr6:135743492-135744015 | -1.157691679 | 0.007520219 | 0.05656485 |
| chr1:35476067-35476567 | -1.138858785 | 0.007520374 | 0.05656485 |
| chr2:164103296-164103685 | -1.269757457 | 0.007521951 | 0.056571158 |
| chr8:90436803-90437081 | 1.32381304 | 0.007524885 | 0.056587664 |
| chr1:37395306-37395567 | -1.319049227 | 0.007527737 | 0.056603563 |
| chr4:8503792-8503992 | 1.323574055 | 0.007529465 | 0.056609208 |
| chr1:218139004-218139204 | -1.416195651 | 0.007529966 | 0.056609208 |
| chrX:114611078-114611400 | -1.180054733 | 0.007532283 | 0.056621069 |
| chr10:50554915-50555196 | -1.318811407 | 0.007534855 | 0.056634852 |
| chr17:42929206-42929432 | 1.367217001 | 0.0075382 | 0.056654437 |
| chr12:114355868-114356093 | -1.488034347 | 0.007541732 | 0.056671735 |
| chr2:20008718-20009006 | 1.151735014 | 0.007541981 | 0.056671735 |
| chr20:50284750-50285111 | -1.020637465 | 0.007547134 | 0.056704892 |

| | | | |
|---|---|---|---|
| chr4:10694191-10694826 | -1.033155172 | 0.007549679 | 0.056718454 |
| chr10:688728-690253 | 0.875879107 | 0.007552958 | 0.056737523 |
| chr20:56905182-56905427 | 1.237357813 | 0.007553878 | 0.056738875 |
| chr3:112731350-112731632 | 1.17690332 | 0.007555488 | 0.056745404 |
| chr5:42381930-42382427 | 1.063286611 | 0.007557397 | 0.056750837 |
| chr1:198745916-198746208 | 1.294192091 | 0.007557693 | 0.056750837 |
| chr7:34036956-34037529 | -0.920682315 | 0.007559358 | 0.056757779 |
| chr7:38146996-38147316 | -1.288372368 | 0.007561503 | 0.05676832 |
| chr9:27487953-27488385 | 1.098489712 | 0.007566254 | 0.056798426 |
| chr12:111024200-111024538 | 1.067390259 | 0.007567429 | 0.056801676 |
| chr3:39151036-39151906 | 0.885434989 | 0.007568452 | 0.05680379 |
| chr10:60692974-60693483 | -1.213476093 | 0.007570053 | 0.056810243 |
| chr3:62593169-62593388 | 1.072147061 | 0.007571978 | 0.05681913 |
| chr16:74362584-74362796 | 1.330227006 | 0.007574408 | 0.056830833 |
| chr12:88181875-88182495 | -0.949413952 | 0.007575022 | 0.056830833 |
| chr16:21156794-21156994 | 1.298960925 | 0.007578323 | 0.056850035 |
| chr7:2643924-2644213 | 1.109116195 | 0.007580286 | 0.056859195 |
| chr18:59249223-59249564 | -1.181984456 | 0.007584912 | 0.056878686 |
| chr12:103632543-103632954 | -0.979817946 | 0.007584922 | 0.056878686 |
| chr5:135553931-135554132 | 1.396794152 | 0.007585112 | 0.056878686 |
| chr13:110809934-110810371 | 1.07658781 | 0.007587304 | 0.056889559 |
| chr18:976546-977011 | -1.153833755 | 0.007589291 | 0.05689889 |
| chr1:207649645-207649893 | 1.256485085 | 0.00759197 | 0.056913405 |
| chr4:158836024-158836285 | -1.500857095 | 0.007595203 | 0.05693207 |
| chr5:122759770-122760819 | -0.932021633 | 0.007601004 | 0.056969979 |
| chr1:238951854-238952180 | -1.175908114 | 0.007602755 | 0.056977535 |
| chr12:106644855-106645154 | 1.243052011 | 0.007624183 | 0.05713253 |
| chr17:2709362-2709786 | 0.997326869 | 0.007627647 | 0.057152703 |
| chr14:99508701-99508949 | 1.267589597 | 0.007628367 | 0.057152703 |
| chr5:112927711-112928257 | -0.959748097 | 0.007629914 | 0.057158707 |
| chr15:70437866-70438175 | -1.164638863 | 0.007632263 | 0.057170716 |
| chr5:127752872-127753301 | -1.096229402 | 0.00763305 | 0.05717102 |
| chr5:91787722-91788104 | 1.16390331 | 0.007633956 | 0.057172218 |
| chr9:114154632-114154862 | -1.342868518 | 0.007635676 | 0.057179513 |
| chr7:25116816-25117139 | -1.215535829 | 0.007638378 | 0.057189933 |
| chr6:89657490-89658015 | 1.019381638 | 0.00763856 | 0.057189933 |
| chr5:40775404-40775679 | 1.258131978 | 0.007643365 | 0.057220313 |
| chr13:52616788-52617853 | -0.872275607 | 0.007644114 | 0.057220332 |
| chr8:100889178-100889723 | 0.978520349 | 0.007646268 | 0.057230864 |
| chr11:106601530-106602179 | -1.021710549 | 0.007650033 | 0.057249413 |
| chr16:56372409-56372668 | -1.346281389 | 0.007650241 | 0.057249413 |
| chr3:169879911-169880115 | 1.308173352 | 0.0076523 | 0.057252315 |
| chr15:46502685-46502982 | -1.327349104 | 0.007652833 | 0.057252315 |
| chr4:117798992-117799229 | -1.478159021 | 0.00765287 | 0.057252315 |
| chr13:36299641-36299943 | 1.179182857 | 0.007654469 | 0.057258686 |
| chr1:51690720-51691216 | 1.05638739 | 0.007656565 | 0.057268775 |
| chr12:24069909-24070238 | -1.313572416 | 0.007658149 | 0.057275028 |
| chr9:12908966-12909556 | 1.010177073 | 0.007666634 | 0.057332895 |
| chr2:210337229-210337815 | -1.106519121 | 0.007675081 | 0.057386877 |
| chr10:46551146-46551354 | 1.207114156 | 0.007675693 | 0.057386877 |

| | | | |
|---|---|---|---|
| chr13:93473429-93473638 | -1.50764894 | 0.0076761 | 0.057386877 |
| chr5:142362804-142363048 | 1.313244792 | 0.007682408 | 0.057428439 |
| chr5:56388591-56389026 | -1.176533991 | 0.007684757 | 0.057440389 |
| chr4:152709779-152710031 | 1.226344085 | 0.007687639 | 0.057448505 |
| chr14:100334966-100335225 | -1.225498715 | 0.007687883 | 0.057448505 |
| chr11:46361326-46361551 | -1.169925937 | 0.007688092 | 0.057448505 |
| chr7:123650588-123650907 | -1.297640954 | 0.007689049 | 0.057450056 |
| chr22:31857430-31857652 | -1.432234461 | 0.007694137 | 0.057482468 |
| chr2:215136766-215137510 | -0.95722099 | 0.007699572 | 0.05751746 |
| chr20:6746796-6747624 | -0.898289072 | 0.007705114 | 0.057552238 |
| chr4:145703902-145704138 | 1.26913816 | 0.00770573 | 0.057552238 |
| chr15:40060777-40060977 | 1.502729882 | 0.007709508 | 0.057574845 |
| chr18:78985058-78985321 | -1.254748831 | 0.007711228 | 0.057582078 |
| chr16:4614787-4615162 | 0.93993937 | 0.007712281 | 0.057584329 |
| chrX:40074381-40074581 | 1.64176444 | 0.007715633 | 0.057603747 |
| chr2:212711980-212712180 | -1.609901762 | 0.007721732 | 0.057643661 |
| chr19:45528216-45529288 | 0.889878691 | 0.007726076 | 0.057668032 |
| chr3:175787124-175787372 | 1.384009702 | 0.007726502 | 0.057668032 |
| chr13:23488431-23488845 | -1.175995636 | 0.007729167 | 0.057682304 |
| chr17:61672679-61672889 | 0.942078012 | 0.007740573 | 0.057761804 |
| chr1:226548437-226548637 | 1.392865398 | 0.007744039 | 0.057782037 |
| chr12:92340615-92340817 | -1.474225414 | 0.007751056 | 0.057828766 |
| chr6:140953043-140953342 | -1.441506328 | 0.007753349 | 0.057834895 |
| chr17:61543648-61544485 | 0.854516105 | 0.007753387 | 0.057834895 |
| chr17:80490881-80491334 | 1.060650913 | 0.007755936 | 0.057844525 |
| chr1:223886675-223886875 | 1.41533159 | 0.007756188 | 0.057844525 |
| chr2:64978933-64979143 | -1.410812785 | 0.007757509 | 0.057848744 |
| chr17:68924803-68925042 | -1.304029915 | 0.00776256 | 0.057880779 |
| chr2:218169877-218170082 | 1.507030325 | 0.007765522 | 0.057892516 |
| chr19:47447456-47447890 | 1.013253223 | 0.007766624 | 0.057892516 |
| chr20:33477270-33477806 | -1.044901515 | 0.007767374 | 0.057892516 |
| chr8:91419736-91420022 | 1.295771848 | 0.007767658 | 0.057892516 |
| chr2:214504826-214505195 | -1.357930354 | 0.007767912 | 0.057892516 |
| chr9:792776-792980 | 1.436587962 | 0.007771318 | 0.057909134 |
| chrX:31993475-31994007 | -1.174278174 | 0.007771653 | 0.057909134 |
| chr10:12909199-12909675 | 1.093434959 | 0.007775177 | 0.057929757 |
| chr2:120203963-120204335 | 1.085748073 | 0.007776354 | 0.057932895 |
| chr12:79112112-79112384 | 1.252575801 | 0.007777249 | 0.057933923 |
| chr15:55288973-55289252 | 1.064599633 | 0.007779132 | 0.057942324 |
| chr3:107334119-107334607 | -1.111346332 | 0.007782628 | 0.057962725 |
| chr1:152467308-152467558 | 1.462582299 | 0.007785395 | 0.057974534 |
| chr2:168964208-168964987 | -0.90591127 | 0.00778643 | 0.057974534 |
| chr4:8603636-8603836 | 1.392226701 | 0.007786483 | 0.057974534 |
| chr3:79417190-79417417 | 1.506474635 | 0.007789029 | 0.057981576 |
| chr1:165162411-165163291 | -0.91280425 | 0.00779017 | 0.057981576 |
| chr7:11212863-11213297 | 0.935520068 | 0.007790335 | 0.057981576 |
| chr1:114190965-114191384 | 0.915149669 | 0.007790456 | 0.057981576 |
| chr14:28982266-28982628 | 1.199473705 | 0.007793374 | 0.057997663 |
| chr21:40841230-40841430 | -1.330777653 | 0.007799096 | 0.058034602 |
| chr3:128698666-128699302 | 0.991262333 | 0.007801969 | 0.058050349 |

| | | | |
|---|---|---|---|
| chr17:79728971-79729250 | 1.236927326 | 0.007803534 | 0.058051286 |
| chr5:100774773-100775457 | -0.954675055 | 0.007803611 | 0.058051286 |
| chr19:4053725-4054012 | -1.518604122 | 0.007805195 | 0.058057435 |
| chr1:171213976-171214372 | -1.102777354 | 0.007812843 | 0.058108683 |
| chr12:26465449-26465731 | -1.409541162 | 0.007814545 | 0.058115701 |
| chr17:7558082-7558417 | -1.120527144 | 0.007815663 | 0.058118378 |
| chr3:64331609-64331872 | 0.946318051 | 0.007823238 | 0.058169061 |
| chr2:73255019-73255219 | -1.323331797 | 0.007828196 | 0.058185192 |
| chr10:97533384-97533601 | 1.316897965 | 0.007828576 | 0.058185192 |
| chr10:106506549-106506847 | -1.309773688 | 0.007828585 | 0.058185192 |
| chr20:44433569-44433840 | 1.356715607 | 0.007828869 | 0.058185192 |
| chr3:19972255-19972590 | 1.196359322 | 0.007829205 | 0.058185192 |
| chr8:123653678-123654036 | -1.029340976 | 0.007830369 | 0.058188203 |
| chrX:120418327-120418528 | 1.507132936 | 0.007835575 | 0.058221242 |
| chrX:105636587-105637136 | -1.016491705 | 0.007836391 | 0.058221661 |
| chr1:31172127-31172652 | 0.939650735 | 0.007841209 | 0.058251808 |
| chr15:95875357-95875787 | -1.019927856 | 0.007844353 | 0.05826951 |
| chr2:39229888-39230387 | -0.94387207 | 0.007845174 | 0.058269965 |
| chrX:30343286-30343528 | -1.340345418 | 0.007854499 | 0.058333571 |
| chr13:53197336-53197746 | -1.126650845 | 0.007859456 | 0.058364727 |
| chr12:49358715-49358915 | 1.320097337 | 0.007861253 | 0.058372413 |
| chr1:107210800-107211140 | 0.945072308 | 0.007869945 | 0.058431292 |
| chr8:138500447-138500787 | -1.124508379 | 0.007874866 | 0.058462164 |
| chr14:76678753-76678958 | 1.467846061 | 0.007877252 | 0.058467888 |
| chr2:65256105-65256321 | -1.472843863 | 0.007877926 | 0.058467888 |
| chr2:227601499-227601755 | -1.472843863 | 0.007877926 | 0.058467888 |
| chr12:69407905-69408221 | -1.358515289 | 0.00787944 | 0.058473458 |
| chr20:48516151-48516535 | 0.952514866 | 0.007881172 | 0.058480649 |
| chr5:154461542-154461895 | -1.117667662 | 0.007883779 | 0.058494327 |
| chr11:61891139-61891339 | 1.293093116 | 0.0078853 | 0.058499948 |
| chr1:220984306-220984516 | -1.552733084 | 0.007892628 | 0.05854864 |
| chr19:37172615-37173115 | 0.984190288 | 0.007896758 | 0.058573612 |
| chr17:62644550-62644870 | -1.000904578 | 0.007897526 | 0.058573633 |
| chr2:177029476-177029855 | -1.058764483 | 0.007898343 | 0.058574028 |
| chr8:143982470-143982674 | 1.20741839 | 0.007900992 | 0.058587998 |
| chr17:35764094-35764920 | 0.927907791 | 0.007905674 | 0.058617045 |
| chr12:25385436-25385640 | -1.180732361 | 0.007910895 | 0.058650078 |
| chr1:33298712-33299171 | 1.021468697 | 0.007915104 | 0.05867561 |
| chr4:6679417-6679617 | 1.33300666 | 0.007917589 | 0.058688352 |
| chr17:61302931-61304039 | 0.841133164 | 0.007919608 | 0.058697641 |
| chr20:47326819-47327077 | 0.895781983 | 0.007922902 | 0.058716373 |
| chr3:109664803-109665368 | -1.041920641 | 0.007930998 | 0.058767373 |
| chr5:91397997-91398203 | -1.495965218 | 0.007931317 | 0.058767373 |
| chr16:76157673-76158099 | -1.156780813 | 0.007934295 | 0.058783754 |
| chr3:100595422-100595757 | -1.327287594 | 0.007937066 | 0.058798595 |
| chr5:112737474-112738242 | -0.884199208 | 0.007944768 | 0.058849966 |
| chr20:49594586-49594786 | 1.327843117 | 0.007953019 | 0.058905388 |
| chr20:48545020-48545415 | 0.892048916 | 0.007953925 | 0.0589064 |
| chrX:142873287-142873554 | 1.412477478 | 0.007956564 | 0.058920251 |
| chr15:70427681-70428125 | -1.088505864 | 0.007957652 | 0.058922615 |

| | | | |
|---|---|---|---|
| chrX:106119633-106119910 | 1.213209802 | 0.007959187 | 0.058925416 |
| chr1:149845413-149845687 | 1.083327313 | 0.007959569 | 0.058925416 |
| chr8:89151269-89151751 | 1.036452886 | 0.007961041 | 0.058930625 |
| chrX:86171417-86171765 | -1.087157785 | 0.007969707 | 0.058989076 |
| chr19:18763928-18764499 | -1.086409301 | 0.007972298 | 0.05900255 |
| chr1:150650127-150650730 | 0.88277848 | 0.007976248 | 0.059026084 |
| chr20:54280794-54281001 | 0.954401846 | 0.007983432 | 0.059073542 |
| chr3:184636723-184637095 | -1.114253914 | 0.007988672 | 0.059106606 |
| chr7:106542502-106542798 | -1.223829918 | 0.008000643 | 0.059189463 |
| chr10:24457557-24458450 | -0.909345063 | 0.008004386 | 0.059211436 |
| chr7:80134302-80135536 | -0.89253895 | 0.008010113 | 0.059248082 |
| chr5:95664215-95664421 | -1.495225003 | 0.008011874 | 0.059255388 |
| chr1:214808894-214809201 | -1.02848533 | 0.008016004 | 0.059276455 |
| chr8:134510799-134511233 | 1.14815893 | 0.00801627 | 0.059276455 |
| chr5:27140548-27140809 | 1.373639223 | 0.008019161 | 0.059292111 |
| chr7:102478881-102479135 | -1.38534221 | 0.008023338 | 0.059317269 |
| chr6:122692013-122692514 | -0.924857151 | 0.008031253 | 0.059370061 |
| chr13:106530333-106530767 | 0.99886519 | 0.008034558 | 0.059384894 |
| chr7:149664661-149665030 | -0.992004078 | 0.00803481 | 0.059384894 |
| chr7:115707906-115708118 | -1.493738393 | 0.0080356 | 0.059385005 |
| chr14:65334896-65335121 | 1.2220216 | 0.008044114 | 0.059442191 |
| chr13:39012760-39013166 | -1.085506536 | 0.00805109 | 0.059488005 |
| chr15:40070044-40070459 | 1.129509699 | 0.008052575 | 0.059493238 |
| chr5:78367422-78367801 | -1.215974604 | 0.008057153 | 0.059521322 |
| chr8:142257970-142258176 | 1.296263212 | 0.008060167 | 0.05953019 |
| chr7:138981216-138982205 | -0.920091342 | 0.00806066 | 0.05953019 |
| chr5:171437129-171437329 | 1.262497359 | 0.008060684 | 0.05953019 |
| chr1:208310420-208310657 | -1.492401163 | 0.008066341 | 0.059566231 |
| chr8:88935779-88936041 | 1.286755611 | 0.008072398 | 0.059600693 |
| chr8:81286709-81287019 | -1.192142689 | 0.008072564 | 0.059600693 |
| chr20:47476728-47477065 | 0.885663115 | 0.008076648 | 0.059625103 |
| chr11:14022728-14023159 | -1.067890319 | 0.008080232 | 0.059645812 |
| chr6:27545534-27546077 | -0.945230772 | 0.008084489 | 0.059671485 |
| chr15:56496606-56496873 | 1.098601896 | 0.008087281 | 0.059680607 |
| chr20:63498049-63498377 | 0.993641042 | 0.008087282 | 0.059680607 |
| chr3:133463304-133464112 | -0.881659972 | 0.008088253 | 0.059682025 |
| chr5:148825925-148826689 | -0.887866446 | 0.008090243 | 0.059686994 |
| chr4:69485159-69485478 | -1.449994791 | 0.008090485 | 0.059686994 |
| chr2:177519985-177520345 | -1.236904905 | 0.008092036 | 0.059692689 |
| chr2:190469857-190470128 | -1.340386938 | 0.008096901 | 0.059722825 |
| chr16:73000612-73001057 | -0.941537018 | 0.008099002 | 0.059732575 |
| chr12:74039084-74039540 | -0.937275316 | 0.008103878 | 0.059762789 |
| chr10:34905368-34905568 | 1.398481135 | 0.008106017 | 0.059770124 |
| chr9:21828970-21829578 | -0.992033477 | 0.008107882 | 0.059770124 |
| chr1:216464442-216464866 | 1.07783132 | 0.008108228 | 0.059770124 |
| chr3:115986575-115986926 | -1.170655295 | 0.008108425 | 0.059770124 |
| chr10:8390094-8390296 | 1.487415446 | 0.008108773 | 0.059770124 |
| chr1:120475350-120475603 | 1.344401603 | 0.008117059 | 0.059819892 |
| chr7:113099181-113099521 | -1.279401311 | 0.008117087 | 0.059819892 |
| chr8:143491062-143491409 | 1.104708887 | 0.008118866 | 0.059827251 |

| | | | |
|---|---|---|---|
| chr5:6875351-6875750 | -1.141152031 | 0.008123633 | 0.059851186 |
| chr10:55899604-55900297 | -0.946035967 | 0.008123676 | 0.059851186 |
| chr7:37064305-37064577 | 1.360004307 | 0.008125247 | 0.059857001 |
| chr12:101861745-101862032 | -1.129439969 | 0.008128389 | 0.059874392 |
| chr16:88316233-88316454 | 1.32793359 | 0.008130724 | 0.059885833 |
| chr11:94746790-94746991 | -1.433081852 | 0.008131546 | 0.059886131 |
| chr1:113790703-113791044 | 1.123871185 | 0.008134899 | 0.05990181 |
| chr12:48882186-48882807 | 0.987233664 | 0.008135238 | 0.05990181 |
| chr22:24772457-24772985 | 1.003546265 | 0.008137391 | 0.059911903 |
| chr7:141158375-141158884 | 1.144297122 | 0.008139649 | 0.059922771 |
| chr2:74560043-74560302 | -1.195653744 | 0.008142537 | 0.059938271 |
| chr6:135183860-135184154 | -1.014503526 | 0.008143438 | 0.059939143 |
| chr3:180036635-180037246 | -1.068427265 | 0.008149045 | 0.059974654 |
| chr12:50057239-50057812 | 0.935013683 | 0.008153316 | 0.059995069 |
| chr7:17848421-17848621 | 1.386054534 | 0.008153385 | 0.059995069 |
| chr8:17538785-17539052 | 1.259987276 | 0.008155404 | 0.060004163 |
| chr8:100487820-100488254 | 0.921131987 | 0.00815692 | 0.060004201 |
| chr7:130149595-130149847 | -1.356056469 | 0.008156975 | 0.060004201 |
| chr11:35389712-35390287 | -1.000902733 | 0.008157916 | 0.060005361 |
| chr2:45257425-45257692 | 1.331811807 | 0.008161954 | 0.060029296 |
| chr3:158570812-158571865 | -0.855862738 | 0.008166506 | 0.060057017 |
| chr7:51350112-51350788 | -0.904136382 | 0.008168622 | 0.060066806 |
| chr4:35407720-35408009 | 1.130614759 | 0.008170815 | 0.060077171 |
| chr1:153749467-153749716 | 1.074437702 | 0.008171957 | 0.060079798 |
| chr19:48608869-48609158 | 1.078834324 | 0.008174517 | 0.060092856 |
| chr3:185120196-185120439 | 1.313280848 | 0.008175651 | 0.060095427 |
| chr3:114691875-114692696 | -0.93989088 | 0.008176483 | 0.06009578 |
| chr20:54587545-54587854 | 0.923523719 | 0.008178553 | 0.060105229 |
| chr14:24081365-24081940 | 0.931153592 | 0.008187861 | 0.060167857 |
| chr18:71638531-71638825 | -1.525438421 | 0.008189428 | 0.060171793 |
| chr15:43285611-43285916 | 1.136507868 | 0.008189967 | 0.060171793 |
| chr4:163679208-163679560 | -1.157812568 | 0.008198584 | 0.060229331 |
| chr9:128098765-128098984 | 1.141414436 | 0.008202347 | 0.060244263 |
| chr12:3141931-3142131 | -1.501929577 | 0.008202569 | 0.060244263 |
| chr1:99773648-99774085 | -1.131281449 | 0.00820333 | 0.060244263 |
| chr14:52934827-52935097 | 1.416742518 | 0.008204863 | 0.060244263 |
| chr1:64982987-64983661 | -0.899151685 | 0.008205131 | 0.060244263 |
| chr8:105568823-105569033 | 1.316544542 | 0.008205334 | 0.060244263 |
| chr7:96595896-96596413 | -1.048985241 | 0.008206539 | 0.060247332 |
| chr16:30437433-30437694 | 1.196293971 | 0.008208621 | 0.060256844 |
| chr13:61414937-61415722 | -0.886377482 | 0.008210949 | 0.060267078 |
| chr4:80322751-80323153 | -1.095003118 | 0.008212375 | 0.060267078 |
| chr9:126851732-126852012 | 1.338243763 | 0.008212833 | 0.060267078 |
| chr11:19419924-19420259 | -1.154242453 | 0.008213161 | 0.060267078 |
| chr7:18831277-18831514 | 1.455753683 | 0.008216892 | 0.06028868 |
| chr1:195105163-195105552 | 1.16928318 | 0.008219226 | 0.060295939 |
| chr5:82375822-82376086 | -1.398258544 | 0.008220484 | 0.060295939 |
| chr10:99132954-99133304 | -1.18871034 | 0.008221077 | 0.060295939 |
| chr7:125383305-125383630 | -1.386549737 | 0.008221726 | 0.060295939 |
| chr3:64341675-64341934 | 0.945034902 | 0.008221816 | 0.060295939 |

| | | | |
|---|---|---|---|
| chr16:2010249-2010537 | 1.140996622 | 0.00822275 | 0.06029702 |
| chr7:158704422-158705476 | -0.844249432 | 0.008225061 | 0.060308197 |
| chr6:95680067-95680388 | -1.27179693 | 0.008231929 | 0.060349975 |
| chr18:50115153-50115481 | -1.32225408 | 0.008232335 | 0.060349975 |
| chr6:109486152-109486362 | 1.286915832 | 0.008234283 | 0.060355517 |
| chr14:41640153-41640497 | 1.098588472 | 0.008234666 | 0.060355517 |
| chr20:32484361-32484592 | 1.220199943 | 0.008240088 | 0.060389481 |
| chr1:110230497-110230877 | -1.052439092 | 0.008242505 | 0.06040142 |
| chrX:104343525-104344101 | -0.937578864 | 0.008247865 | 0.060434606 |
| chr9:104093588-104095008 | -0.846787593 | 0.008248612 | 0.060434606 |
| chr3:3109900-3110600 | -0.956046099 | 0.008251909 | 0.06045211 |
| chr9:131716837-131717037 | 1.522941789 | 0.008252579 | 0.06045211 |
| chr2:48760717-48761102 | -1.297435978 | 0.008257917 | 0.060485433 |
| chr7:128360502-128360756 | -1.285047774 | 0.008260834 | 0.060501015 |
| chr7:128361129-128362371 | -0.885360111 | 0.008265757 | 0.060526356 |
| chr14:84848718-84848918 | 1.579506808 | 0.008265874 | 0.060526356 |
| chr20:38165262-38165653 | 1.027647313 | 0.00827477 | 0.060585709 |
| chr4:61594573-61594773 | 1.527005376 | 0.008277495 | 0.060599871 |
| chr16:89476328-89476982 | -0.929802191 | 0.008280373 | 0.060615144 |
| chr1:216064664-216064886 | 1.359510771 | 0.008284712 | 0.060641113 |
| chr3:173556630-173556856 | -1.463524653 | 0.008285987 | 0.060644658 |
| chr6:44058568-44058841 | 1.260157855 | 0.008288397 | 0.060656501 |
| chr4:166091952-166092185 | -1.338590641 | 0.008302953 | 0.060753205 |
| chr7:142885507-142886186 | -0.916730104 | 0.008303197 | 0.060753205 |
| chr5:130871510-130871782 | -1.218808936 | 0.008309193 | 0.060791271 |
| chr18:72013381-72013799 | -1.215168028 | 0.008310248 | 0.060793182 |
| chr7:138785799-138786169 | -1.150184458 | 0.008313062 | 0.060805131 |
| chr1:101004148-101004484 | -1.121247928 | 0.008313468 | 0.060805131 |
| chr5:35331162-35331368 | 1.495764965 | 0.008318099 | 0.060833197 |
| chr12:74641755-74642010 | -1.342788318 | 0.008326307 | 0.060884718 |
| chr8:141023489-141023692 | 1.328944851 | 0.008326733 | 0.060884718 |
| chr10:43904189-43904460 | -1.345130859 | 0.008330636 | 0.060902614 |
| chr5:106241104-106242000 | -0.898332542 | 0.00833077 | 0.060902614 |
| chr21:36980545-36980927 | 1.115699557 | 0.008334216 | 0.06092199 |
| chr13:90055159-90055532 | -1.012701071 | 0.00833663 | 0.060932501 |
| chr10:55601739-55602519 | -0.989963902 | 0.008338445 | 0.060932501 |
| chr7:110534197-110535101 | -0.926037859 | 0.008338791 | 0.060932501 |
| chr7:15721002-15721204 | 1.451775391 | 0.008339286 | 0.060932501 |
| chr17:57766591-57766826 | 1.111493453 | 0.00833963 | 0.060932501 |
| chr4:2008549-2009680 | 0.883959401 | 0.00834372 | 0.060956573 |
| chr13:43058221-43058710 | -0.929454978 | 0.008348905 | 0.060988638 |
| chr3:160941783-160942251 | -0.994506093 | 0.00835206 | 0.061005865 |
| chr7:112424970-112425177 | -1.570230546 | 0.00835487 | 0.061020575 |
| chr21:20739541-20739788 | 1.338353467 | 0.008360594 | 0.061056563 |
| chr5:124979216-124979475 | -1.365607117 | 0.00836314 | 0.061069333 |
| chr19:16615006-16615206 | 1.417611616 | 0.008363955 | 0.061069462 |
| chr17:15248005-15248422 | -1.092273892 | 0.008365494 | 0.061072744 |
| chr7:127975951-127976151 | -1.523465807 | 0.008365998 | 0.061072744 |
| chr17:74368291-74368633 | 0.993148043 | 0.00836876 | 0.061087083 |
| chr7:101217908-101218436 | -0.910344285 | 0.008371014 | 0.061097719 |

| | | | |
|---|---|---|---|
| chr4:94076560-94076761 | -1.461596723 | 0.008373922 | 0.061108469 |
| chr17:65633899-65634865 | -0.878298274 | 0.008374658 | 0.061108469 |
| chr10:104888-105585 | -0.998532539 | 0.00837488 | 0.061108469 |
| chr20:17611753-17611999 | 1.246007126 | 0.008379726 | 0.061135655 |
| chr8:104067030-104067242 | 1.442119384 | 0.008380201 | 0.061135655 |
| chr1:12186166-12186507 | 1.334159569 | 0.008381553 | 0.061139696 |
| chr5:95975025-95975413 | -1.149519052 | 0.008384126 | 0.061152641 |
| chr16:81715732-81715992 | 1.231019579 | 0.008386446 | 0.061163743 |
| chr18:50218822-50219088 | -1.397145554 | 0.008388904 | 0.061175846 |
| chr17:28655331-28655560 | 1.396953725 | 0.008392805 | 0.061198466 |
| chr5:84431363-84431633 | -1.18635073 | 0.008396505 | 0.061216662 |
| chr14:50735397-50735603 | 1.348981174 | 0.008396898 | 0.061216662 |
| chr3:157205187-157205401 | 1.237623282 | 0.008403866 | 0.061261635 |
| chr4:108192035-108192413 | -1.133613492 | 0.00840594 | 0.061268251 |
| chr14:29926973-29928536 | 0.855966194 | 0.008406373 | 0.061268251 |
| chr10:56370765-56371028 | -1.548783796 | 0.008407218 | 0.061268581 |
| chr3:160983972-160984291 | -1.241884878 | 0.008412748 | 0.061303053 |
| chr12:97404555-97404877 | -1.071791693 | 0.008415122 | 0.061314517 |
| chr6:141421369-141421682 | -1.332908612 | 0.008419233 | 0.061338641 |
| chr9:76215116-76215486 | -1.088861362 | 0.008421805 | 0.06134929 |
| chr14:58638640-58638840 | 1.265769825 | 0.008422372 | 0.06134929 |
| chr7:705353-705773 | 1.006190113 | 0.008423643 | 0.06134929 |
| chr2:169141818-169142146 | -1.096541203 | 0.008423898 | 0.06134929 |
| chr1:200755627-200755886 | 1.157293942 | 0.008427481 | 0.061369556 |
| chr4:170421151-170421444 | -1.297889818 | 0.008431635 | 0.061393972 |
| chr13:43799180-43799557 | 1.086826458 | 0.008433783 | 0.061403773 |
| chr19:53196846-53197381 | 0.945312617 | 0.008436179 | 0.061413279 |
| chr17:40694664-40694930 | 1.219062245 | 0.008436691 | 0.061413279 |
| chr2:161273213-161273413 | -1.458235617 | 0.008437994 | 0.061416925 |
| chr7:134514608-134514921 | -1.147549964 | 0.0084422 | 0.0614417 |
| chr16:2868060-2868752 | 0.926622936 | 0.008443004 | 0.06144172 |
| chr1:148439070-148439270 | 1.292431712 | 0.008445683 | 0.061455379 |
| chr4:141853133-141853860 | -0.898916197 | 0.008447732 | 0.061464448 |
| chr11:131025100-131025326 | -1.547829338 | 0.008449034 | 0.061468086 |
| chr2:173448933-173449543 | -0.968752404 | 0.008450408 | 0.061472249 |
| chr18:76565227-76565698 | -0.96501916 | 0.008453191 | 0.061486655 |
| chr7:92425417-92425926 | -0.961173644 | 0.008458781 | 0.061517745 |
| chr2:61707965-61708340 | 1.056109846 | 0.008459071 | 0.061517745 |
| chr4:185203411-185204347 | -0.885896783 | 0.008461036 | 0.061526193 |
| chr9:120384564-120384810 | -1.108356226 | 0.008462261 | 0.061526646 |
| chr3:150938852-150939145 | 1.320997636 | 0.008462704 | 0.061526646 |
| chr3:20057302-20057502 | 1.307444334 | 0.008464625 | 0.061534769 |
| chr9:111839407-111839810 | 1.134978404 | 0.008474547 | 0.061601056 |
| chr7:123054587-123055123 | -1.016605516 | 0.008477639 | 0.061617686 |
| chr13:79631264-79631644 | -1.010763288 | 0.008481227 | 0.061637922 |
| chrX:129842826-129843076 | 1.163486131 | 0.008482551 | 0.061639984 |
| chr15:89751808-89752204 | 0.976034794 | 0.00848312 | 0.061639984 |
| chr5:36139066-36139676 | -1.078121492 | 0.008487631 | 0.061661614 |
| chr3:138609457-138609660 | 1.180224735 | 0.008488246 | 0.061661614 |
| chr18:58195414-58195792 | -1.061775285 | 0.008488511 | 0.061661614 |

| | | | |
|---|---|---|---|
| chr7:68474095-68474295 | 1.569093073 | 0.008492789 | 0.061686842 |
| chr17:79008708-79008908 | 1.27069827 | 0.008493645 | 0.061687208 |
| chr1:150400986-150401219 | 1.313214427 | 0.008494974 | 0.061691014 |
| chr3:62399358-62399621 | 1.044104001 | 0.008498638 | 0.061706681 |
| chr1:45575835-45576106 | 1.319655164 | 0.008498742 | 0.061706681 |
| chr4:42767811-42768030 | -1.540650084 | 0.008501234 | 0.061711115 |
| chr4:92735109-92735325 | -1.540650084 | 0.008501234 | 0.061711115 |
| chr5:93850363-93850572 | -1.583007439 | 0.008501769 | 0.061711115 |
| chr9:110575980-110576352 | -1.280429894 | 0.008505356 | 0.061731303 |
| chr7:128209790-128210148 | -0.974980603 | 0.008509962 | 0.061754409 |
| chr19:41774541-41774823 | 1.144579061 | 0.008510151 | 0.061754409 |
| chr5:108607378-108607722 | -1.294708216 | 0.008511718 | 0.061759927 |
| chr12:92340869-92341299 | -1.088854929 | 0.008513223 | 0.061765 |
| chr1:163922898-163923163 | -1.463077722 | 0.008528736 | 0.061871692 |
| chr20:56954694-56954908 | 1.100048947 | 0.008533791 | 0.061896338 |
| chr3:63777173-63777536 | 0.945962274 | 0.008534478 | 0.061896338 |
| chr2:25340092-25340847 | -0.924952877 | 0.008534557 | 0.061896338 |
| chr2:191758833-191759263 | -0.947968837 | 0.008536652 | 0.061905677 |
| chr3:164737320-164737520 | 1.490664399 | 0.008539148 | 0.061914474 |
| chr14:102421458-102421708 | -1.240835151 | 0.008539482 | 0.061914474 |
| chr5:130817596-130818547 | -0.876476723 | 0.008540633 | 0.061916154 |
| chr13:44191443-44192006 | -0.956890341 | 0.008541329 | 0.061916154 |
| chr13:102556061-102556314 | -1.417258037 | 0.008547663 | 0.061956207 |
| chr1:201467887-201469693 | 0.837723335 | 0.008549989 | 0.061963246 |
| chr2:163174296-163174498 | -1.366406186 | 0.008550252 | 0.061963246 |
| chr2:126506911-126507361 | -1.246809532 | 0.008553846 | 0.061983427 |
| chr15:90098008-90098451 | 0.932406327 | 0.008557821 | 0.06200637 |
| chr5:16588572-16588870 | 1.092179763 | 0.008559601 | 0.062013399 |
| chr2:158381227-158381480 | -1.319021101 | 0.008560871 | 0.062016737 |
| chr3:141928307-141928517 | 1.392902013 | 0.008563981 | 0.062033403 |
| chr9:70484576-70484853 | -1.152820516 | 0.008571363 | 0.062081007 |
| chr1:117013218-117013774 | 0.932033709 | 0.008573136 | 0.062087979 |
| chr11:12066854-12067062 | 1.445845945 | 0.00857467 | 0.062091648 |
| chr4:171238166-171238534 | -1.274365234 | 0.008575616 | 0.062091648 |
| chr20:54546804-54547089 | 0.929726501 | 0.008576074 | 0.062091648 |
| chr5:123814153-123814553 | -1.048597962 | 0.008579317 | 0.062106432 |
| chr8:100974932-100975399 | 1.028632752 | 0.008579737 | 0.062106432 |
| chr3:25603781-25604087 | -1.089689943 | 0.008581862 | 0.062115952 |
| chr3:111561675-111562372 | -0.996361146 | 0.008587996 | 0.062154478 |
| chr7:33214002-33214260 | 1.268886563 | 0.008592721 | 0.062182799 |
| chr4:175536920-175537193 | -1.492593619 | 0.008597486 | 0.062211406 |
| chr10:24360609-24360855 | -1.421474602 | 0.008600998 | 0.062230944 |
| chr1:231040159-231040631 | 0.969125031 | 0.008603957 | 0.062246471 |
| chr14:84578757-84579061 | 1.366016805 | 0.00860605 | 0.062255737 |
| chr11:44862354-44862600 | 1.276331175 | 0.008607147 | 0.062257794 |
| chr5:158725251-158725522 | -1.221630066 | 0.008614237 | 0.062298913 |
| chr19:43316734-43316968 | 1.292595693 | 0.008614458 | 0.062298913 |
| chr16:87363911-87364166 | 1.223919961 | 0.00861948 | 0.062329351 |
| chr5:80475004-80475294 | -1.239816303 | 0.008626786 | 0.062376293 |
| chr20:11363344-11363810 | -1.156452862 | 0.008630142 | 0.062394674 |

| | | | |
|---|---|---|---|
| chr5:129767895-129768305 | -0.971494697 | 0.008634384 | 0.06241945 |
| chr11:133809551-133810135 | -1.065158761 | 0.008638542 | 0.062439394 |
| chr2:16608621-16609383 | -0.890637691 | 0.008638772 | 0.062439394 |
| chr12:96489311-96490120 | -0.888006929 | 0.008640983 | 0.062440866 |
| chr18:79062743-79063114 | 1.085492416 | 0.008641105 | 0.062440866 |
| chr7:108211581-108212270 | -1.013139528 | 0.008641421 | 0.062440866 |
| chr3:111870141-111870426 | 1.264322233 | 0.008647899 | 0.062476031 |
| chr6:44156532-44156846 | 1.08924962 | 0.008647918 | 0.062476031 |
| chr1:237055814-237056014 | 1.396158078 | 0.008649729 | 0.062480901 |
| chr3:24373155-24373643 | 1.040014429 | 0.008650223 | 0.062480901 |
| chr9:105294890-105295090 | -1.700188608 | 0.008651044 | 0.062480942 |
| chr10:1289696-1290399 | -0.925935861 | 0.008658621 | 0.062529769 |
| chr18:63161799-63162065 | -1.196025839 | 0.008663341 | 0.062557962 |
| chr21:45070056-45070292 | 1.176500961 | 0.008670312 | 0.062598698 |
| chr13:71477652-71478072 | -1.292387584 | 0.008670617 | 0.062598698 |
| chr5:17655479-17655745 | 1.288961117 | 0.008671797 | 0.062599703 |
| chr7:123492510-123492836 | -1.192991621 | 0.00867239 | 0.062599703 |
| chr4:7102900-7103660 | 0.899351604 | 0.008687853 | 0.062705415 |
| chr7:6564739-6564958 | 1.26041807 | 0.008691828 | 0.062722284 |
| chr8:110040505-110040832 | 1.26041807 | 0.008691828 | 0.062722284 |
| chr10:29064278-29064669 | -1.17266212 | 0.008694787 | 0.062737729 |
| chr6:143194785-143194985 | 1.485788254 | 0.008698323 | 0.062757333 |
| chr4:27075605-27075907 | -1.354686897 | 0.008700747 | 0.062764882 |
| chr15:78457203-78457437 | 1.259197364 | 0.008701007 | 0.062764882 |
| chr7:79741799-79742016 | -1.443911321 | 0.008702908 | 0.062767687 |
| chr10:6201587-6201853 | 1.016209952 | 0.008703034 | 0.062767687 |
| chr11:65932496-65932806 | 1.154327886 | 0.008703873 | 0.06276783 |
| chr20:45995326-45995867 | 0.912859858 | 0.008705701 | 0.0627751 |
| chrX:132940720-132941191 | 1.029574694 | 0.008707035 | 0.062778811 |
| chr7:128109072-128109272 | -1.303842234 | 0.008709953 | 0.06279394 |
| chr14:37681401-37681601 | 1.276215171 | 0.0087175 | 0.062842438 |
| chr1:193338130-193338493 | -1.170000262 | 0.008719453 | 0.062850606 |
| chr12:58247747-58248099 | 1.200868892 | 0.008726508 | 0.062886571 |
| chr1:82795817-82796103 | -1.276027365 | 0.008726559 | 0.062886571 |
| chr3:63787615-63787849 | 1.045441232 | 0.008727574 | 0.062886571 |
| chr6:119141188-119141504 | -1.151970449 | 0.008727726 | 0.062886571 |
| chr5:65640186-65640540 | -1.263402785 | 0.008730676 | 0.062901914 |
| chr12:82089767-82090051 | 1.286075798 | 0.008732425 | 0.0629086 |
| chr4:98284559-98285082 | -1.003327758 | 0.008735218 | 0.062922801 |
| chr1:33342056-33342635 | 0.978685937 | 0.008737284 | 0.062931768 |
| chr4:64144999-64145286 | 1.155410798 | 0.00874018 | 0.062943473 |
| chr8:118439727-118440123 | 1.124060021 | 0.008740552 | 0.062943473 |
| chr3:14672303-14672503 | 1.38901457 | 0.008744463 | 0.06296572 |
| chr10:53808576-53809207 | -1.026274194 | 0.008746993 | 0.062978022 |
| chr15:71071087-71071376 | -1.22367462 | 0.008750952 | 0.063000602 |
| chr15:63093493-63093892 | 1.007190765 | 0.008752541 | 0.063006128 |
| chr1:225839722-225839922 | 1.252029319 | 0.008754734 | 0.063006507 |
| chrX:154373372-154373831 | 0.929077708 | 0.008755562 | 0.063006507 |
| chr3:145735585-145736095 | -0.975918602 | 0.008755777 | 0.063006507 |
| chr6:24574852-24575330 | -1.284286849 | 0.008755883 | 0.063006507 |

| | | | |
|---|---|---|---|
| chr17:1736436-1737072 | -1.013711516 | 0.008760237 | 0.063028902 |
| chr12:100987888-100988138 | -1.272739381 | 0.008760641 | 0.063028902 |
| chr16:86658747-86658987 | 1.310367211 | 0.008769433 | 0.063086232 |
| chr7:134016643-134017030 | -1.206704253 | 0.008777588 | 0.063138972 |
| chr1:215241532-215241769 | -1.278098139 | 0.008782073 | 0.063165305 |
| chr4:57430768-57431078 | 1.129649398 | 0.008783495 | 0.063169606 |
| chr8:73185019-73185219 | 1.474072483 | 0.008785818 | 0.063180382 |
| chr1:242243193-242243395 | 1.408767991 | 0.008789156 | 0.06319845 |
| chrX:152895653-152896244 | 0.903869189 | 0.008793494 | 0.063223709 |
| chr7:116984864-116985151 | -1.065553403 | 0.008794947 | 0.063226255 |
| chr13:81142303-81143057 | -0.915656551 | 0.00879559 | 0.063226255 |
| chr5:122076679-122078429 | -0.846221233 | 0.008796323 | 0.063226255 |
| chr7:137592264-137592640 | -1.102937382 | 0.008800075 | 0.063247292 |
| chr17:61779208-61779412 | 0.913542239 | 0.00880157 | 0.063250871 |
| chr15:25237111-25237356 | 1.418401318 | 0.00880301 | 0.063250871 |
| chr7:133139322-133139725 | -1.040847284 | 0.00880305 | 0.063250871 |
| chr4:145333825-145334533 | 1.000444109 | 0.008804135 | 0.063252735 |
| chr1:104878569-104878784 | -1.453307431 | 0.008805622 | 0.063257489 |
| chr19:50099756-50100005 | 1.388301926 | 0.008809905 | 0.063274138 |
| chr2:55699353-55699664 | -1.358030215 | 0.008810504 | 0.063274138 |
| chr17:56946854-56947269 | -1.0424374 | 0.008810624 | 0.063274138 |
| chr5:22289799-22290009 | 1.456650065 | 0.008811243 | 0.063274138 |
| chr4:157818960-157820089 | -0.846492335 | 0.008816297 | 0.063301103 |
| chr21:40852596-40852852 | -1.20466788 | 0.00881665 | 0.063301103 |
| chr10:64014016-64014334 | -1.262940962 | 0.008823035 | 0.063335 |
| chr13:27965704-27965921 | 1.242242118 | 0.008823794 | 0.063335 |
| chr12:52800959-52801159 | 1.223663662 | 0.008824464 | 0.063335 |
| chr3:182597148-182598087 | 0.909640315 | 0.008824678 | 0.063335 |
| chr14:52863040-52863274 | -1.423839534 | 0.008833827 | 0.063394726 |
| chr15:101458071-101458292 | 1.382844233 | 0.008837831 | 0.063417519 |
| chr17:61643604-61644021 | 0.869124337 | 0.008842039 | 0.063441777 |
| chr4:68463480-68464290 | -0.950784645 | 0.008845864 | 0.06346292 |
| chr3:146476655-146476858 | 1.41778471 | 0.008846643 | 0.06346292 |
| chr5:109618895-109619192 | -1.375126377 | 0.008848455 | 0.063469979 |
| chr1:209747576-209748389 | -0.854931543 | 0.008850761 | 0.063480578 |
| chr4:175891854-175892208 | -1.162348866 | 0.008854106 | 0.063498623 |
| chr1:184379377-184379825 | -1.016257417 | 0.008857312 | 0.063510304 |
| chr2:223619536-223619756 | 1.45092558 | 0.008857392 | 0.063510304 |
| chr20:47559212-47559519 | 0.902134854 | 0.008858234 | 0.063510395 |
| chr2:231412352-231412697 | 0.987865966 | 0.008859679 | 0.063514812 |
| chr4:33089857-33090151 | -1.175338827 | 0.008863115 | 0.063533504 |
| chr6:161922652-161923242 | -1.049446216 | 0.008864161 | 0.063535058 |
| chr2:9283940-9284152 | -1.541911054 | 0.008868881 | 0.063562939 |
| chr22:23670709-23671017 | 1.274283214 | 0.00887747 | 0.063618549 |
| chr2:203543992-203544367 | 1.105086384 | 0.0088829 | 0.063651508 |
| chr14:34480989-34481377 | 1.010744506 | 0.008887753 | 0.063680329 |
| chr14:87133575-87133869 | 1.221978235 | 0.008890114 | 0.06368343 |
| chr2:25982417-25982908 | 0.987871476 | 0.008890226 | 0.06368343 |
| chr2:30960989-30961189 | 1.487929495 | 0.00889068 | 0.06368343 |
| chr6:135258960-135259502 | -0.973088698 | 0.008893351 | 0.063691158 |

| | | | |
|---|---|---|---|
| chr1:87598500-87598752 | -1.416919461 | 0.008893421 | 0.063691158 |
| chr7:115129836-115130399 | -1.103708094 | 0.008897212 | 0.063709451 |
| chr6:158233449-158233718 | -1.40565287 | 0.008897638 | 0.063709451 |
| chr17:75249668-75250380 | -0.965847455 | 0.008899907 | 0.063719743 |
| chr14:105228571-105228785 | 1.241098211 | 0.008901505 | 0.063720059 |
| chr4:2377824-2378065 | 1.43501508 | 0.008901615 | 0.063720059 |
| chr11:15769125-15769556 | 0.974989591 | 0.008903509 | 0.063723775 |
| chr2:101106652-101106867 | 1.44432914 | 0.008903797 | 0.063723775 |
| chrX:23964247-23964593 | -1.256920696 | 0.008910011 | 0.06376024 |
| chr9:94914258-94914594 | 1.034458381 | 0.008910556 | 0.06376024 |
| chr10:113927189-113927760 | 0.992104192 | 0.008912346 | 0.063767091 |
| chrX:108945955-108946186 | -1.437841126 | 0.008915226 | 0.063781743 |
| chr10:63166867-63167148 | -1.107180612 | 0.00891616 | 0.06378247 |
| chr5:62621304-62621677 | -1.022112006 | 0.008917603 | 0.063786834 |
| chr11:5380785-5381043 | -1.35774081 | 0.008923707 | 0.063824537 |
| chr17:82675432-82675632 | 1.406252783 | 0.008926239 | 0.063836692 |
| chr5:1140253-1140641 | 0.958736907 | 0.008927497 | 0.06383973 |
| chr4:119676088-119676347 | 1.302733008 | 0.00893 | 0.063851664 |
| chr22:38202437-38202983 | 0.958431788 | 0.008932105 | 0.063860758 |
| chr13:91739723-91740267 | -1.051853682 | 0.008934651 | 0.063873002 |
| chr4:158367855-158368810 | -0.87269919 | 0.008936524 | 0.063880433 |
| chr12:6044238-6044613 | -1.174788359 | 0.008937837 | 0.063883856 |
| chr13:48126893-48127226 | -1.147651763 | 0.008949411 | 0.06396062 |
| chr9:83620758-83621119 | -0.962102167 | 0.00896117 | 0.064031859 |
| chr20:48272134-48272480 | 0.908438774 | 0.008962025 | 0.064031859 |
| chr1:175864285-175864509 | 1.278171549 | 0.008962692 | 0.064031859 |
| chr5:146291496-146291725 | -1.263866041 | 0.008962722 | 0.064031859 |
| chrX:96424092-96424340 | -1.404445371 | 0.008964721 | 0.064040171 |
| chr12:50353679-50353940 | 1.409237402 | 0.008968436 | 0.064060734 |
| chr3:135895193-135895478 | 1.136666312 | 0.008974266 | 0.0640964 |
| chr11:30914211-30914478 | -1.150213603 | 0.008976362 | 0.064099483 |
| chr21:39137139-39137503 | -1.006649775 | 0.00897637 | 0.064099483 |
| chr4:105088166-105088505 | -1.169942692 | 0.008981105 | 0.064127319 |
| chr20:48716695-48716909 | 0.943859485 | 0.008988222 | 0.064167921 |
| chr16:33767980-33768180 | 1.385019664 | 0.008989031 | 0.064167921 |
| chr9:113111510-113111960 | 1.136480569 | 0.008989304 | 0.064167921 |
| chr15:36591783-36591983 | -1.529101606 | 0.008991838 | 0.064177673 |
| chr19:55376400-55376628 | -1.377810583 | 0.008992345 | 0.064177673 |
| chr16:87986317-87986574 | 1.346099566 | 0.008993792 | 0.064182019 |
| chr12:125314110-125314375 | -1.268581151 | 0.008996191 | 0.064188945 |
| chr18:49447868-49448085 | 1.388877677 | 0.008996534 | 0.064188945 |
| chr14:82622137-82622398 | -1.268997924 | 0.008997654 | 0.064188945 |
| chr5:180248777-180249113 | -1.305975296 | 0.008998114 | 0.064188945 |
| chr1:202845212-202845542 | 1.007129127 | 0.009000861 | 0.064202567 |
| chr2:69816998-69817260 | -1.180310746 | 0.009002078 | 0.064205272 |
| chr16:1719887-1720171 | 1.217932564 | 0.009003553 | 0.064209817 |
| chr16:87093261-87093684 | 1.146904086 | 0.009006198 | 0.064222701 |
| chr5:166488167-166488972 | -0.940853541 | 0.009007829 | 0.06422835 |
| chr1:185383920-185384817 | -0.945439667 | 0.009010848 | 0.0642439 |
| chr5:176006062-176006373 | -1.217145567 | 0.009014971 | 0.064263505 |

| | | | |
|---|---|---|---|
| chr5:173380429-173380720 | 1.217807031 | 0.009015275 | 0.064263505 |
| chr20:48705828-48706042 | 0.949724162 | 0.00901668 | 0.064266599 |
| chr11:129315589-129316424 | 0.891596975 | 0.009017387 | 0.064266599 |
| chr14:73208365-73208613 | 1.260827586 | 0.009023154 | 0.064301722 |
| chr20:47156915-47159668 | 0.816442758 | 0.009026148 | 0.064317076 |
| chr1:16173448-16173875 | 1.072441053 | 0.009031777 | 0.064351199 |
| chr4:101885597-101885952 | 1.047999218 | 0.009035359 | 0.064367805 |
| chr21:40299634-40299871 | 1.383657706 | 0.009035787 | 0.064367805 |
| chr16:54094618-54095078 | -1.036255431 | 0.009039825 | 0.06439058 |
| chr17:29588594-29588928 | 1.013092348 | 0.009041708 | 0.064398008 |
| chr18:41532441-41532780 | -1.174680562 | 0.009045657 | 0.064420142 |
| chr2:224484910-224485894 | -0.895761436 | 0.009048874 | 0.064437065 |
| chr2:11483054-11483826 | -0.871577449 | 0.009051313 | 0.064448446 |
| chr1:106363361-106363629 | 0.986944083 | 0.009056172 | 0.064475896 |
| chr2:79512363-79513116 | -0.890169533 | 0.009056933 | 0.064475896 |
| chr16:61395486-61395978 | -1.118163052 | 0.009057693 | 0.064475896 |
| chr1:100982711-100983121 | -1.115068759 | 0.009058825 | 0.064477962 |
| chr14:40213496-40214431 | 0.898400651 | 0.009061498 | 0.064490997 |
| chr9:135525109-135525341 | 1.368062148 | 0.009063511 | 0.06449859 |
| chr4:185208993-185210726 | -0.842950259 | 0.009064248 | 0.06449859 |
| chr3:189144746-189144950 | -1.354042109 | 0.00906573 | 0.064503145 |
| chr4:94147835-94148647 | -0.878216365 | 0.009067601 | 0.064510466 |
| chr1:154813222-154813662 | -1.000491222 | 0.009069279 | 0.064514187 |
| chr9:10612888-10613414 | 0.958731678 | 0.009070118 | 0.064514187 |
| chr10:33290929-33291654 | -0.961585634 | 0.00907065 | 0.064514187 |
| chr14:74457273-74457489 | 1.232728352 | 0.009076865 | 0.064552397 |
| chr5:172340144-172340382 | 1.314174482 | 0.009086342 | 0.064613802 |
| chr7:48035163-48036203 | -0.852411221 | 0.009088569 | 0.064623638 |
| chr3:86733050-86733544 | -1.067914283 | 0.009090174 | 0.064629057 |
| chr20:11585777-11586103 | -1.170159021 | 0.009091885 | 0.064629834 |
| chr15:78368923-78369123 | 1.44585561 | 0.009091971 | 0.064629834 |
| chr19:488814-489334 | 0.97220118 | 0.009095575 | 0.064645926 |
| chr6:110688634-110689623 | -0.884018167 | 0.009096712 | 0.064645926 |
| chr5:81842842-81843221 | 1.095026843 | 0.009096766 | 0.064645926 |
| chr3:183912024-183912476 | 0.987504019 | 0.00910045 | 0.064663796 |
| chr3:185645481-185646312 | -0.94402365 | 0.00910145 | 0.064663796 |
| chr16:35188661-35188876 | 1.218209412 | 0.009101812 | 0.064663796 |
| chrX:16882673-16883191 | -0.94621311 | 0.009106393 | 0.064690344 |
| chr4:56505283-56505492 | -1.483071436 | 0.009110411 | 0.064712889 |
| chr16:3305185-3305864 | 0.926177839 | 0.00911547 | 0.064742819 |
| chr10:57898326-57898688 | -1.121681289 | 0.009118472 | 0.064755175 |
| chr1:242240100-242240323 | 1.406644375 | 0.0091189 | 0.064755175 |
| chr13:60009516-60009805 | -1.265991078 | 0.009131167 | 0.064836278 |
| chr3:61210406-61210640 | -1.533219888 | 0.009143234 | 0.064913528 |
| chr6:24504400-24505069 | -0.915734089 | 0.009143741 | 0.064913528 |
| chr4:2041723-2042120 | 1.009793998 | 0.009144603 | 0.064913635 |
| chr5:168365039-168366080 | -0.855130618 | 0.009150348 | 0.064948396 |
| chr15:40582720-40583232 | -1.014409663 | 0.009151513 | 0.064950648 |
| chr16:89211034-89211238 | -1.439783511 | 0.009158627 | 0.064990176 |
| chr11:13415191-13415578 | -1.097677987 | 0.009159343 | 0.064990176 |

| | | | |
|---|---|---|---|
| chr6:135195758-135196009 | -1.558208339 | 0.009161088 | 0.064990176 |
| chr7:109844474-109844705 | -1.601967032 | 0.009161286 | 0.064990176 |
| chr10:98664538-98664804 | -1.255338226 | 0.009161851 | 0.064990176 |
| chr1:208198599-208198830 | -1.394613332 | 0.009162171 | 0.064990176 |
| chr1:147114216-147114617 | 0.999912115 | 0.009166314 | 0.065004106 |
| chr16:1706520-1706756 | 1.137319132 | 0.009166461 | 0.065004106 |
| chr6:6881778-6882163 | -1.253498114 | 0.00916668 | 0.065004106 |
| chr4:109989067-109989420 | -1.232602109 | 0.009167797 | 0.065006007 |
| chr13:73248712-73249014 | -1.100051258 | 0.0091712 | 0.065020278 |
| chr6:54137605-54137805 | 1.422162838 | 0.009171506 | 0.065020278 |
| chr8:75305341-75305597 | -1.256617015 | 0.009174369 | 0.065034552 |
| chr21:35273277-35273665 | 1.106294547 | 0.009178001 | 0.065049631 |
| chrX:106182188-106182388 | 1.462701234 | 0.009178194 | 0.065049631 |
| chr12:53452443-53452869 | 0.942797858 | 0.009180046 | 0.065056738 |
| chr1:62241637-62241941 | 1.196064948 | 0.009181956 | 0.065058371 |
| chr2:152755020-152755468 | -1.025765643 | 0.009182708 | 0.065058371 |
| chr20:54521170-54521379 | 0.945468496 | 0.009182823 | 0.065058371 |
| chr20:46485492-46485692 | 1.380790723 | 0.009186608 | 0.065079169 |
| chr4:2468684-2469230 | 0.913578614 | 0.009189821 | 0.065095913 |
| chr4:68271867-68272379 | -1.013022319 | 0.009196988 | 0.065137278 |
| chr1:214910576-214911024 | -1.091719942 | 0.009197361 | 0.065137278 |
| chr3:102238301-102238540 | 1.254503022 | 0.009200574 | 0.065154007 |
| chr2:98869241-98870245 | -0.877247335 | 0.009206606 | 0.065189779 |
| chr1:125173925-125174205 | 1.014131578 | 0.009207327 | 0.065189779 |
| chr4:6687956-6688357 | 0.908461758 | 0.00921107 | 0.065210254 |
| chr6:129432774-129433103 | -1.129843185 | 0.009212093 | 0.065211476 |
| chr5:122704279-122704521 | -1.28005342 | 0.009216371 | 0.065235729 |
| chr10:116511489-116511689 | -1.580921872 | 0.009218082 | 0.065241813 |
| chr5:62692462-62692679 | 1.22853483 | 0.009222383 | 0.065266227 |
| chr16:57720255-57720479 | 1.380154311 | 0.00923237 | 0.065327672 |
| chr3:19075173-19075373 | 1.443282729 | 0.009232771 | 0.065327672 |
| chr1:24594204-24594572 | 1.022386784 | 0.009233938 | 0.065329158 |
| chr11:69115160-69115406 | 1.1714089 | 0.009234686 | 0.065329158 |
| chr1:65267767-65267977 | -1.383686807 | 0.00923885 | 0.065349154 |
| chr13:39676366-39676644 | -1.252704922 | 0.009239317 | 0.065349154 |
| chr5:139896724-139897177 | -1.104384919 | 0.009240071 | 0.065349154 |
| chr5:132370468-132370676 | -1.16133947 | 0.009243329 | 0.065366164 |
| chr15:71136462-71136717 | -1.324150616 | 0.00924774 | 0.065391322 |
| chr3:53927155-53927865 | -0.966041614 | 0.009251489 | 0.06541179 |
| chr14:46405185-46405589 | -1.105775246 | 0.009253403 | 0.065414755 |
| chr1:205477378-205478429 | -0.845703844 | 0.009253616 | 0.065414755 |
| chr8:125017401-125017626 | 1.200131885 | 0.009255068 | 0.065418987 |
| chr12:80868357-80869232 | -0.883113515 | 0.009267331 | 0.065499627 |
| chr12:49568895-49569126 | 1.144732434 | 0.009269118 | 0.065506212 |
| chr16:33491368-33491886 | 1.000815804 | 0.00927652 | 0.06555248 |
| chr13:76039463-76039808 | -1.284860544 | 0.009279614 | 0.065561958 |
| chr2:1719510-1719814 | -1.249273433 | 0.009279776 | 0.065561958 |
| chr8:11466422-11466813 | 1.038990264 | 0.009280967 | 0.065561958 |
| chr14:94637093-94637379 | 1.101816627 | 0.009281284 | 0.065561958 |
| chr7:77903701-77903946 | -1.263411145 | 0.009282539 | 0.065564778 |

| | | | |
|---|---|---|---|
| chr3:87026349-87026573 | 1.332102414 | 0.00929461 | 0.065638677 |
| chr22:42553685-42553948 | 1.175079315 | 0.009294715 | 0.065638677 |
| chr20:47167976-47169357 | 0.819934059 | 0.00929882 | 0.065661093 |
| chr13:94362390-94362716 | -1.212613005 | 0.009300116 | 0.065661093 |
| chr19:5046389-5046621 | 1.214779648 | 0.00930046 | 0.065661093 |
| chr13:77053267-77053519 | 1.330851506 | 0.009308534 | 0.065712038 |
| chr1:187185458-187185783 | -1.176737785 | 0.009317455 | 0.065760322 |
| chr10:93363854-93364178 | -1.168783099 | 0.009317864 | 0.065760322 |
| chr11:8290163-8290558 | -1.224298376 | 0.009318284 | 0.065760322 |
| chr5:59496479-59497117 | -1.031584533 | 0.009318806 | 0.065760322 |
| chr5:41139867-41140189 | 1.30130388 | 0.009320063 | 0.065763134 |
| chr1:221853630-221853839 | 1.442124884 | 0.009331096 | 0.06583327 |
| chr17:57068220-57068620 | -0.953253254 | 0.009331721 | 0.06583327 |
| chr7:156999740-156999940 | -1.311331201 | 0.00933338 | 0.065838907 |
| chr3:106566348-106566625 | -1.25961176 | 0.009335188 | 0.065845605 |
| chr10:104277886-104278090 | 1.355109746 | 0.009343436 | 0.065897716 |
| chr4:136591131-136591384 | 1.394217221 | 0.009344566 | 0.065899618 |
| chr17:61431728-61431928 | 0.914867132 | 0.009351524 | 0.065942618 |
| chr10:91643170-91643428 | 1.383058903 | 0.009354834 | 0.065959356 |
| chr7:17939685-17940908 | 0.82873328 | 0.00935562 | 0.065959356 |
| chr15:78230089-78230396 | 1.195305227 | 0.009356856 | 0.065962003 |
| chr9:124390865-124391123 | 1.200508069 | 0.009369365 | 0.066044109 |
| chr17:61907119-61907859 | 0.840262694 | 0.009371246 | 0.066051296 |
| chr17:61790645-61790949 | 0.883462153 | 0.009373969 | 0.06606441 |
| chr13:79237974-79238333 | -1.186431048 | 0.009375302 | 0.066067723 |
| chr14:105659928-105660188 | 1.120073674 | 0.009378871 | 0.066086796 |
| chr5:164341175-164341545 | -1.01211517 | 0.00938367 | 0.066110237 |
| chr19:1193048-1193291 | 1.386637844 | 0.009383923 | 0.066110237 |
| chr3:31300980-31301214 | 1.359025211 | 0.009385059 | 0.066112163 |
| chr12:30851128-30851433 | -1.054162908 | 0.009391273 | 0.066149858 |
| chr15:59247216-59247677 | 1.090154465 | 0.009397882 | 0.066190324 |
| chr6:70280381-70280882 | -1.073429316 | 0.009399851 | 0.066194243 |
| chr3:180306246-180306446 | -1.412139838 | 0.009400567 | 0.066194243 |
| chr14:60664856-60665460 | 0.940422423 | 0.009402427 | 0.066194243 |
| chr1:200150582-200151346 | -0.860709806 | 0.009402484 | 0.066194243 |
| chr8:104801349-104801577 | 1.32990957 | 0.009402758 | 0.066194243 |
| chr8:72881737-72881937 | -1.559925547 | 0.009404751 | 0.066202186 |
| chr14:65528450-65528666 | 1.402129193 | 0.009411163 | 0.06624124 |
| chr17:13024486-13024787 | -1.017188458 | 0.009415142 | 0.066261412 |
| chr8:115950116-115950406 | -1.181828585 | 0.009415759 | 0.066261412 |
| chr5:10635815-10636052 | 1.197728593 | 0.00941836 | 0.066273634 |
| chr6:110693150-110693770 | -0.94905651 | 0.009428362 | 0.066337919 |
| chr20:47141475-47143243 | 0.817224527 | 0.00943148 | 0.066353767 |
| chr1:23574379-23574611 | 1.337271663 | 0.00943384 | 0.066364278 |
| chr5:2779477-2779785 | -1.068535864 | 0.009437886 | 0.066386643 |
| chr14:92720028-92720253 | 1.320845605 | 0.009448827 | 0.066456711 |
| chr18:2385174-2385977 | -0.974172636 | 0.009449582 | 0.066456711 |
| chr10:63935738-63936208 | -1.082388506 | 0.009461604 | 0.066535149 |
| chr3:63218799-63219322 | 0.907410321 | 0.009465432 | 0.066550091 |
| chr7:17600351-17600608 | 1.117224194 | 0.009465826 | 0.066550091 |

| | | | |
|---|---|---|---|
| chr11:114160099-114160331 | -1.220103453 | 0.009466334 | 0.066550091 |
| chr1:219702614-219702876 | 1.174816315 | 0.009467996 | 0.066555665 |
| chr12:52301348-52301581 | 1.320495012 | 0.009474706 | 0.066596729 |
| chr14:37874486-37875036 | 0.900293249 | 0.009479324 | 0.066623074 |
| chr14:36960638-36960838 | 1.415552844 | 0.009481409 | 0.066631619 |
| chr16:47143835-47144137 | 1.075992635 | 0.009491562 | 0.066696375 |
| chr5:137205950-137206473 | -1.035219784 | 0.009492365 | 0.066696375 |
| chr21:39523492-39524259 | -0.882975967 | 0.00949475 | 0.066703853 |
| chr20:24134813-24135188 | 1.126485228 | 0.00949517 | 0.066703853 |
| chr1:32292552-32292930 | 0.993897216 | 0.009496852 | 0.066709556 |
| chr12:15942840-15943049 | -1.439020721 | 0.009502914 | 0.066746014 |
| chr8:120868674-120869193 | -0.901898529 | 0.009506113 | 0.066762361 |
| chr4:75889312-75889931 | -1.075548071 | 0.009509623 | 0.066768106 |
| chr10:59580242-59580555 | -1.24285815 | 0.009509767 | 0.066768106 |
| chr8:88591992-88592207 | 1.238295892 | 0.009510008 | 0.066768106 |
| chr1:82778677-82778894 | -1.245139473 | 0.009511044 | 0.066768106 |
| chr5:100401121-100401835 | -0.958916313 | 0.009512098 | 0.066768106 |
| chr15:46636259-46636648 | -1.107660329 | 0.009512159 | 0.066768106 |
| chr9:20611758-20612390 | -1.069400251 | 0.009514623 | 0.0667788 |
| chr11:116742841-116743149 | 1.210869268 | 0.009515426 | 0.0667788 |
| chr2:101350172-101350530 | 1.229448884 | 0.009519902 | 0.066804097 |
| chr11:122981764-122982038 | -1.226568893 | 0.009524573 | 0.066829655 |
| chr5:113249331-113249817 | -1.185346782 | 0.009526121 | 0.066829655 |
| chr3:183428041-183429063 | 0.878220881 | 0.009526161 | 0.066829655 |
| chr1:34781686-34782389 | 0.902402397 | 0.009529001 | 0.066843457 |
| chr3:180631339-180631618 | -1.190350203 | 0.009531059 | 0.066851134 |
| chr16:84556035-84556235 | 1.316275293 | 0.00953184 | 0.066851134 |
| chr20:31566260-31566481 | -1.230019133 | 0.009532832 | 0.066851976 |
| chr5:16913471-16913851 | -1.032455995 | 0.009535297 | 0.066863138 |
| chr2:155223851-155224349 | -1.068058317 | 0.009548035 | 0.066942905 |
| chr3:153685971-153686175 | -1.436424332 | 0.00954842 | 0.066942905 |
| chr5:22853233-22853564 | -1.127328043 | 0.009550032 | 0.066948082 |
| chr10:88253769-88254073 | -1.280287635 | 0.009552371 | 0.066958356 |
| chr14:76975996-76976574 | -0.914544793 | 0.00955712 | 0.066985517 |
| chr1:191851759-191852037 | -1.285456561 | 0.009561269 | 0.067001309 |
| chr17:27475172-27475637 | -1.03713901 | 0.009562917 | 0.067001309 |
| chr7:22846148-22846477 | 1.142081055 | 0.009563437 | 0.067001309 |
| chr7:74708569-74708841 | -1.502470968 | 0.009563602 | 0.067001309 |
| chr17:60773625-60773970 | 0.876500493 | 0.009563746 | 0.067001309 |
| chr1:11654636-11655023 | 1.033806778 | 0.009570042 | 0.067039289 |
| chr15:95872543-95872760 | 1.265022139 | 0.009575482 | 0.067071268 |
| chr5:1853976-1854221 | 1.175341463 | 0.009578359 | 0.067077004 |
| chr2:65367483-65367927 | -0.964295699 | 0.009578466 | 0.067077004 |
| chr7:123355644-123356187 | -0.920287877 | 0.009578927 | 0.067077004 |
| chr14:96083561-96084305 | 0.865767278 | 0.009580195 | 0.067079749 |
| chr14:26905326-26905604 | 1.230385038 | 0.0095811 | 0.067079958 |
| chr9:108855935-108856180 | 1.261558756 | 0.009586347 | 0.067110561 |
| chr1:240459252-240459603 | -1.213687177 | 0.009590468 | 0.067133279 |
| chr1:42943171-42943627 | 0.93251288 | 0.009592762 | 0.067143202 |
| chr16:79639546-79640414 | -0.871445895 | 0.009595634 | 0.067150534 |

| | | | |
|---|---|---|---|
| chr16:85436911-85437888 | -0.86120566 | 0.009596556 | 0.067150534 |
| chr5:64417464-64417889 | -1.07819964 | 0.009597271 | 0.067150534 |
| chr21:39579324-39580104 | -0.893409894 | 0.009597938 | 0.067150534 |
| chr3:83567450-83568310 | -0.990131778 | 0.00959887 | 0.067150534 |
| chr13:79541487-79541766 | -1.373093452 | 0.009599127 | 0.067150534 |
| chr14:86217486-86217805 | -1.183454524 | 0.009599945 | 0.067150534 |
| chr16:74743545-74744296 | -0.868044787 | 0.009602282 | 0.067160756 |
| chr19:44512366-44512599 | 1.306949309 | 0.009605954 | 0.067180305 |
| chr19:17466283-17466483 | 1.302917537 | 0.00961154 | 0.067213234 |
| chr10:3295001-3295435 | 0.981893436 | 0.00961474 | 0.067224236 |
| chr5:83473042-83474776 | -0.830219321 | 0.009615039 | 0.067224236 |
| chr19:10119397-10120253 | 0.886980951 | 0.009615745 | 0.067224236 |
| chr1:248243215-248243623 | -1.050911709 | 0.009622709 | 0.067266785 |
| chrY:56837040-56837342 | 1.11285222 | 0.009632928 | 0.067332072 |
| chr1:195385178-195385378 | 1.286007333 | 0.009635437 | 0.067343467 |
| chr2:190354369-190354733 | -1.15761802 | 0.009640908 | 0.067375558 |
| chr1:176216661-176216962 | -1.277931074 | 0.009642344 | 0.067379448 |
| chr5:151134275-151134679 | 1.006526769 | 0.009650087 | 0.067427412 |
| chr4:175266627-175266858 | -1.471874832 | 0.00965291 | 0.067440983 |
| chr20:59482441-59482683 | 1.212318502 | 0.009663553 | 0.067509185 |
| chr19:2624596-2624995 | -1.14867922 | 0.009664711 | 0.067511118 |
| chr5:173917825-173918500 | -0.927383548 | 0.00966635 | 0.067516414 |
| chr3:100682970-100683289 | 1.193609736 | 0.00967023 | 0.067537362 |
| chr12:52992962-52993235 | 1.20323339 | 0.009671115 | 0.067537387 |
| chr5:140633071-140633606 | -0.933434341 | 0.009673199 | 0.067545783 |
| chr3:155512378-155512656 | 1.412375474 | 0.009679717 | 0.067585135 |
| chr2:130836679-130836953 | 1.129019018 | 0.009685148 | 0.067616894 |
| chr4:175954007-175954491 | -1.086465518 | 0.009686162 | 0.067617813 |
| chr2:109691238-109691507 | 1.179975057 | 0.009695256 | 0.067675127 |
| chr3:22032474-22032685 | 1.366759643 | 0.009696571 | 0.067678142 |
| chr20:47345864-47346608 | 0.826571742 | 0.009702495 | 0.06771332 |
| chr2:29583100-29583313 | -1.40755082 | 0.009712188 | 0.0677748 |
| chr3:8607561-8607877 | 0.988777297 | 0.00971341 | 0.067777154 |
| chr19:29771791-29772225 | 1.036149279 | 0.009719175 | 0.067807016 |
| chr3:151770083-151770401 | -1.429684656 | 0.009720262 | 0.067807016 |
| chr7:131327228-131328538 | -0.860646258 | 0.009720345 | 0.067807016 |
| chrX:145858372-145858923 | -1.094184415 | 0.009727033 | 0.067847494 |
| chr5:68661317-68661571 | -1.363660975 | 0.009728693 | 0.067852895 |
| chr8:38786990-38787263 | 1.126468597 | 0.009734509 | 0.067885953 |
| chr5:141843245-141843572 | 1.048358135 | 0.009735962 | 0.067885953 |
| chr20:48686099-48686860 | 0.832575032 | 0.009737486 | 0.067885953 |
| chr10:34305158-34305795 | -0.921488986 | 0.009737493 | 0.067885953 |
| chr10:87701523-87701879 | -1.238586758 | 0.009738429 | 0.067885953 |
| chr9:70477995-70478251 | -1.307130942 | 0.009738749 | 0.067885953 |
| chr12:31952927-31953254 | -1.166627395 | 0.009739777 | 0.067886947 |
| chr7:53793514-53793870 | -1.154458272 | 0.009742191 | 0.067897592 |
| chr16:57372235-57372435 | -1.439720022 | 0.009744338 | 0.067900921 |
| chr1:113506754-113507079 | 1.172095774 | 0.009745626 | 0.067900921 |
| chr4:148562494-148562698 | 1.470795044 | 0.009745756 | 0.067900921 |
| chr15:58332172-58332575 | 0.937435835 | 0.009746213 | 0.067900921 |

| | | | |
|---|---|---|---|
| chr13:61880868-61881224 | -1.093668377 | 0.009750249 | 0.067922866 |
| chr9:127556032-127556371 | -1.125881754 | 0.009751436 | 0.067924958 |
| chr3:148246217-148247110 | -0.972618181 | 0.00975651 | 0.067954124 |
| chr8:127668477-127669487 | -0.826414362 | 0.009758572 | 0.067958269 |
| chr7:159015383-159016085 | -0.894244631 | 0.009758879 | 0.067958269 |
| chr11:93476832-93477155 | -1.152696829 | 0.009759916 | 0.067959315 |
| chr12:27081620-27081890 | -1.276577085 | 0.009761174 | 0.067961898 |
| chr9:71678264-71678495 | -1.453229208 | 0.009762473 | 0.067964769 |
| chr7:121446528-121446861 | -1.033784537 | 0.009765188 | 0.067977489 |
| chr4:102246493-102246738 | 1.20979668 | 0.009768055 | 0.067991272 |
| chr13:92103719-92104205 | 1.074984249 | 0.009769977 | 0.067996264 |
| chr1:216566879-216567135 | -1.085802393 | 0.009770547 | 0.067996264 |
| chr17:55838321-55838531 | 1.315126434 | 0.00977318 | 0.068008407 |
| chr3:152323588-152323833 | 1.431858332 | 0.009776596 | 0.068026005 |
| chr4:149056667-149056942 | 1.178913548 | 0.009785854 | 0.068084236 |
| chr7:131303203-131303461 | -1.231999584 | 0.009788152 | 0.068094039 |
| chr1:200433207-200433517 | 1.130695 | 0.009794867 | 0.068134423 |
| chr1:208669054-208669442 | -1.155350467 | 0.009795735 | 0.068134423 |
| chr20:47328861-47329163 | 0.865540866 | 0.009798518 | 0.0681413 |
| chr17:721320-721617 | -1.295886062 | 0.009798693 | 0.0681413 |
| chr4:6695671-6696578 | 0.862684403 | 0.009799392 | 0.0681413 |
| chr16:87622274-87622516 | 1.139390537 | 0.009803475 | 0.068157865 |
| chr6:53573331-53573648 | -1.150717581 | 0.009803553 | 0.068157865 |
| chr1:102497883-102498129 | -1.432955321 | 0.009806319 | 0.068167654 |
| chr7:150869530-150869870 | -1.075618477 | 0.00980674 | 0.068167654 |
| chr8:38553680-38554651 | -0.879207477 | 0.009807888 | 0.06816945 |
| chr9:135204989-135205383 | 1.16674739 | 0.009810175 | 0.06817159 |
| chr6:157592462-157592822 | -1.186222278 | 0.00981066 | 0.06817159 |
| chr10:133689320-133689564 | 1.123779566 | 0.009811095 | 0.06817159 |
| chr17:80047182-80047609 | -1.020345541 | 0.009811755 | 0.06817159 |
| chr4:168487472-168487972 | -1.071045106 | 0.009816197 | 0.068196271 |
| chr1:186806533-186806859 | -1.046447658 | 0.009820809 | 0.068222125 |
| chr11:130653775-130654220 | -1.099393243 | 0.009826302 | 0.068249826 |
| chr18:74819745-74820115 | -1.000557997 | 0.009828467 | 0.068249826 |
| chr2:36333688-36333942 | -1.165688737 | 0.00982902 | 0.068249826 |
| chr10:84541272-84541496 | -1.103006225 | 0.009829283 | 0.068249826 |
| chr17:51336924-51337165 | 1.227037948 | 0.009829654 | 0.068249826 |
| chr8:144060638-144060838 | 1.159998106 | 0.009830243 | 0.068249826 |
| chr20:25308824-25309058 | 1.256969358 | 0.009831032 | 0.068249826 |
| chr5:33550018-33550374 | 1.076068047 | 0.009832629 | 0.068254726 |
| chr4:174461494-174462053 | -1.081071137 | 0.009833839 | 0.068256944 |
| chr1:229094781-229095039 | -1.205503056 | 0.00983474 | 0.068257017 |
| chr10:24552324-24552742 | -0.90996004 | 0.00983703 | 0.068266727 |
| chr4:143166383-143167327 | -0.850517372 | 0.009840738 | 0.068286276 |
| chr3:175383586-175383797 | 1.356007913 | 0.00984248 | 0.068292177 |
| chr12:12736751-12737009 | -1.403985031 | 0.009846926 | 0.068316838 |
| chr7:130888127-130888332 | -1.391990878 | 0.009849921 | 0.068329738 |
| chr11:68750503-68750774 | -1.088268981 | 0.009850569 | 0.068329738 |
| chr7:111873157-111873553 | -1.200060678 | 0.009853201 | 0.068333553 |
| chr4:162839862-162840142 | -1.236641051 | 0.009853566 | 0.068333553 |

| | | | |
|---|---|---|---|
| chr15:43649562-43649810 | 1.329956454 | 0.009853989 | 0.068333553 |
| chr1:174892062-174892318 | 1.36350545 | 0.009855425 | 0.068333553 |
| chr17:61304388-61304659 | 0.886530751 | 0.009855578 | 0.068333553 |
| chr10:112811495-112812500 | 0.867279788 | 0.009867447 | 0.068400297 |
| chr20:19552174-19552499 | -1.1537258 | 0.009867486 | 0.068400297 |
| chr14:84780026-84780226 | 1.503434415 | 0.009867882 | 0.068400297 |
| chr20:47296024-47296488 | 0.847446248 | 0.009874992 | 0.068443384 |
| chr2:187002349-187002661 | 1.219508702 | 0.009877167 | 0.06845227 |
| chr16:8911902-8912115 | 1.294904914 | 0.009880748 | 0.068470896 |
| chr17:17063584-17064168 | -0.978290481 | 0.009885826 | 0.068499887 |
| chr10:26891723-26891923 | -1.401543204 | 0.009886726 | 0.068499927 |
| chr19:57559517-57560033 | 1.015456389 | 0.009889463 | 0.0685127 |
| chr14:28583676-28583942 | 1.313113352 | 0.009893453 | 0.068534141 |
| chr3:64655261-64655664 | 0.887165964 | 0.009897965 | 0.068556478 |
| chr7:14170880-14171175 | 1.144369376 | 0.009898467 | 0.068556478 |
| chrX:24395830-24396119 | -1.148552345 | 0.009904329 | 0.068585676 |
| chr3:129197485-129197717 | 1.189111903 | 0.009904473 | 0.068585676 |
| chr14:72404085-72404659 | 0.924898915 | 0.009906914 | 0.068596381 |
| chr9:33221327-33221692 | -1.074654631 | 0.009909482 | 0.068606702 |
| chrX:154918008-154918232 | 1.371795173 | 0.009910195 | 0.068606702 |
| chr4:184983936-184984255 | -1.023953666 | 0.009915637 | 0.068634286 |
| chr6:18493470-18493982 | 0.957045247 | 0.009915971 | 0.068634286 |
| chr5:1883631-1883960 | 0.951295663 | 0.009929753 | 0.06872347 |
| chr5:180021790-180022210 | -1.140341698 | 0.009937848 | 0.068767772 |
| chr20:50671896-50673435 | 0.808623376 | 0.009938604 | 0.068767772 |
| chrX:151753400-151753756 | 1.003197112 | 0.009938924 | 0.068767772 |
| chr3:192896839-192897398 | 0.936983663 | 0.009939744 | 0.068767772 |
| chr3:9314942-9315173 | 1.137485341 | 0.009951741 | 0.068836261 |
| chr7:234681-235528 | 0.860711087 | 0.009952012 | 0.068836261 |
| chr1:222009203-222009410 | -1.435786595 | 0.009952339 | 0.068836261 |
| chr2:199692470-199693018 | -0.999162675 | 0.009953779 | 0.068840009 |
| chr6:75563021-75563429 | -1.011560494 | 0.009956581 | 0.068853172 |
| chr22:32204982-32205190 | 1.190594662 | 0.009959552 | 0.068867505 |
| chr9:134125778-134125978 | 1.26135193 | 0.009960926 | 0.068870791 |
| chr15:75145050-75145924 | -0.879167589 | 0.009963138 | 0.068879864 |
| chr10:64091849-64092219 | -1.157457405 | 0.009965228 | 0.068888097 |
| chr7:143263111-143264103 | -0.867933534 | 0.009966807 | 0.068892803 |
| chr5:92185555-92186086 | -0.975817441 | 0.009967735 | 0.068892996 |
| chr7:93601409-93601894 | -1.102739548 | 0.009972941 | 0.068922766 |
| chr7:19587401-19587601 | 1.353551318 | 0.009976524 | 0.068941235 |
| chr1:210361271-210361554 | -1.31521026 | 0.009977413 | 0.068941235 |
| chr4:175457989-175458322 | 1.038174592 | 0.009978661 | 0.068943642 |
| chr6:112145303-112145563 | -1.422933827 | 0.009983809 | 0.068969197 |
| chr7:115840590-115841290 | -0.907326595 | 0.00998416 | 0.068969197 |
| chr5:100940304-100940504 | -1.603529572 | 0.009988683 | 0.068994221 |
| chr1:186722795-186723028 | -1.339153986 | 0.009990711 | 0.069002007 |
| chr20:58192551-58192758 | 1.051886646 | 0.009992698 | 0.069009509 |
| chr6:25175245-25175646 | -1.225570917 | 0.009999905 | 0.069053055 |
| chr14:104882308-104882709 | 0.905752939 | 0.010002573 | 0.069065254 |
| chr16:87834518-87834941 | -0.935110379 | 0.010009057 | 0.069095007 |

| | | | |
|---|---|---|---|
| chr2:215974081-215974371 | -1.075021068 | 0.010009232 | 0.069095007 |
| chr9:38641881-38642854 | -0.915552845 | 0.010009587 | 0.069095007 |
| chr22:47102733-47103039 | 1.175327272 | 0.010011886 | 0.069099637 |
| chr6:27157171-27157384 | 1.322916095 | 0.010012061 | 0.069099637 |
| chr7:101671380-101671591 | -1.360511782 | 0.010014055 | 0.069107174 |
| chr7:119175943-119176292 | -1.197139504 | 0.010019852 | 0.069140951 |
| chr17:62800407-62801071 | -0.85029797 | 0.010024165 | 0.069164482 |
| chr18:63046256-63046835 | -1.000115842 | 0.010025354 | 0.06916646 |
| chr16:21518151-21518746 | 0.901198324 | 0.010027485 | 0.069174934 |
| chr10:95147667-95148082 | -1.121679837 | 0.010044538 | 0.06928633 |
| chr18:59697269-59697833 | -0.98747374 | 0.010048104 | 0.069304689 |
| chr14:57057889-57058724 | -0.888251672 | 0.010049244 | 0.069306316 |
| chr7:146019343-146019543 | -1.601950686 | 0.010065372 | 0.069407947 |
| chr17:61718483-61718683 | 0.909560625 | 0.010066028 | 0.069407947 |
| chr18:76528795-76529044 | -1.227059117 | 0.010066698 | 0.069407947 |
| chr16:69567117-69567439 | 1.091210727 | 0.010068089 | 0.069411293 |
| chr7:85829883-85830148 | -1.304390187 | 0.010069143 | 0.069412313 |
| chr6:73803841-73804148 | -1.086112554 | 0.010074809 | 0.069445128 |
| chr13:77002655-77002892 | -1.309383514 | 0.010077097 | 0.069454651 |
| chr4:142033878-142034381 | 1.040371133 | 0.010081089 | 0.069469826 |
| chr2:150133688-150133951 | -1.359325663 | 0.010081112 | 0.069469826 |
| chr11:24755326-24755739 | 1.110900267 | 0.010083939 | 0.06948246 |
| chr14:47040530-47040886 | -1.051112729 | 0.010084759 | 0.06948246 |
| chr2:77104578-77104852 | -1.256259713 | 0.010089093 | 0.06950607 |
| chr14:37436557-37436831 | 1.114091909 | 0.010091571 | 0.06951689 |
| chr8:79684223-79684490 | 1.291303204 | 0.010095544 | 0.06953766 |
| chr5:127515853-127516287 | -0.999854484 | 0.01009699 | 0.06953766 |
| chr5:10832953-10833224 | 1.186372908 | 0.010097475 | 0.06953766 |
| chr13:75704883-75705634 | -0.893236738 | 0.010098216 | 0.06953766 |
| chr1:114275485-114275702 | 0.957008112 | 0.010099885 | 0.0695429 |
| chr13:47287458-47287658 | 1.241188241 | 0.01010433 | 0.069567255 |
| chr6:108995361-108995687 | 1.043216882 | 0.010106451 | 0.069575609 |
| chr20:59310526-59310775 | 1.18575102 | 0.010112423 | 0.069608971 |
| chr11:30333454-30333737 | -1.1524453 | 0.010113521 | 0.069608971 |
| chrX:150536764-150536965 | 1.312801587 | 0.01011413 | 0.069608971 |
| chr3:49461728-49461972 | 1.21606781 | 0.010114932 | 0.069608971 |
| chr7:105325698-105325933 | -1.379929209 | 0.010116954 | 0.069616637 |
| chr15:74462009-74462218 | 1.281538608 | 0.010121955 | 0.069644795 |
| chr1:33137974-33138297 | 1.120238407 | 0.010124824 | 0.06965828 |
| chr3:194494394-194495359 | -0.921814693 | 0.010128159 | 0.069674969 |
| chr16:73477545-73478591 | -0.864546183 | 0.010131082 | 0.069688822 |
| chr17:44981659-44981961 | 1.183109979 | 0.010132565 | 0.069692764 |
| chr12:51875309-51875528 | 1.230567278 | 0.010138249 | 0.06972132 |
| chr12:55742925-55743379 | 0.967497021 | 0.010138537 | 0.06972132 |
| chr6:109455667-109456213 | 0.925610151 | 0.010142687 | 0.069743606 |
| chr7:36195848-36196264 | 1.031398998 | 0.010147667 | 0.069771582 |
| chr13:100646288-100646619 | -1.105699828 | 0.010150974 | 0.069784658 |
| chr17:58742359-58742587 | 0.964858571 | 0.0101532 | 0.069784658 |
| chr15:90706417-90706714 | -1.180300205 | 0.010153251 | 0.069784658 |
| chr7:124929436-124930287 | -0.823975884 | 0.010154398 | 0.069784658 |

| | | | |
|---|---|---|---|
| chr14:76985303-76985683 | 1.061651418 | 0.010154667 | 0.069784658 |
| chr18:41323852-41324052 | -1.46800599 | 0.010155109 | 0.069784658 |
| chr12:82694369-82694643 | -1.234000489 | 0.010156167 | 0.069784658 |
| chr10:225722-225922 | 1.455053159 | 0.010158205 | 0.069784658 |
| chr16:50879331-50879629 | 1.306511223 | 0.010158588 | 0.069784658 |
| chr10:56296729-56296957 | -1.276031102 | 0.010158676 | 0.069784658 |
| chr8:144351544-144351757 | 1.129101295 | 0.010159605 | 0.069784781 |
| chr4:166501885-166502393 | -0.953881887 | 0.010163542 | 0.069805565 |
| chr12:64569175-64569607 | 0.996726352 | 0.010173384 | 0.069866905 |
| chr5:170611846-170612153 | 1.124922163 | 0.010176539 | 0.069882308 |
| chr1:183899708-183899931 | 1.255386488 | 0.010178118 | 0.069886885 |
| chr14:73787032-73787439 | 0.949171865 | 0.010183524 | 0.06991774 |
| chr20:47840306-47840843 | 0.855622858 | 0.010187088 | 0.069932374 |
| chr8:130421408-130421720 | 1.103114668 | 0.010187481 | 0.069932374 |
| chr7:104252458-104252776 | -1.059953381 | 0.01018883 | 0.069935366 |
| chr6:4253786-4254164 | 1.112764869 | 0.010191094 | 0.069944642 |
| chr13:34627946-34628674 | 0.925741286 | 0.010192786 | 0.069949992 |
| chr3:133547439-133547879 | -0.982758599 | 0.010194717 | 0.069956975 |
| chr2:207289262-207289659 | 1.022494948 | 0.010196999 | 0.069964531 |
| chr17:61908289-61908699 | 0.855909696 | 0.010197644 | 0.069964531 |
| chrX:26286618-26287469 | -0.893723593 | 0.010199108 | 0.06996831 |
| chr3:79467497-79468097 | -0.945948212 | 0.010204159 | 0.069996693 |
| chr1:198881065-198881534 | -0.981459202 | 0.010207202 | 0.070011302 |
| chr7:96548749-96549065 | -1.173289789 | 0.010210175 | 0.070025426 |
| chr1:113798982-113799273 | 1.043568845 | 0.010211431 | 0.070027772 |
| chr2:42141333-42142016 | -0.988489507 | 0.010218532 | 0.070070197 |
| chr4:705441-706169 | 0.845448958 | 0.010230263 | 0.070144362 |
| chr6:90459363-90459768 | 0.992151057 | 0.010232347 | 0.070150849 |
| chr1:81007643-81008106 | -1.020783914 | 0.01023355 | 0.070150849 |
| chr8:85774078-85774286 | 1.521130231 | 0.010233956 | 0.070150849 |
| chr5:178247063-178247268 | 1.271550691 | 0.01023739 | 0.070168113 |
| chr20:36153946-36154205 | 1.097683967 | 0.010239422 | 0.070175757 |
| chr6:129659886-129660457 | -0.959524992 | 0.010243483 | 0.070197313 |
| chr17:50858968-50859216 | -1.165383444 | 0.010245752 | 0.070206586 |
| chr22:30279397-30279842 | -1.097365531 | 0.010247768 | 0.07021214 |
| chr13:91313157-91313381 | -1.428327439 | 0.010248515 | 0.07021214 |
| chr5:119439106-119439866 | -0.853241827 | 0.010249312 | 0.07021214 |
| chr3:62614871-62615071 | 1.024244051 | 0.010254783 | 0.070243339 |
| chr4:79937429-79937904 | -1.097438655 | 0.010256277 | 0.070247293 |
| chr2:68732076-68732344 | -1.434107275 | 0.010258999 | 0.070259652 |
| chr7:121530473-121530722 | -1.377372803 | 0.010260987 | 0.070262726 |
| chr6:6741385-6741596 | -1.448279427 | 0.010261518 | 0.070262726 |
| chr1:209428209-209428922 | -0.876033746 | 0.010262198 | 0.070262726 |
| chr7:91880339-91881472 | -0.835961485 | 0.010264582 | 0.070272769 |
| chr3:141360941-141361619 | 0.933643316 | 0.010267369 | 0.070275985 |
| chr8:141507814-141508160 | 1.081475584 | 0.010267559 | 0.070275985 |
| chr10:92387257-92387581 | -1.264164755 | 0.010267804 | 0.070275985 |
| chr17:73570544-73570833 | -1.246426526 | 0.010269591 | 0.070281941 |
| chr5:14929285-14929552 | 1.247890095 | 0.010274442 | 0.070308861 |
| chr4:49286953-49287721 | 0.943224435 | 0.010277127 | 0.070320951 |

| | | | |
|---|---|---|---|
| chr4:166622921-166623122 | -1.448636302 | 0.010280432 | 0.070337286 |
| chr7:84025459-84025884 | -1.098010241 | 0.010290999 | 0.070403296 |
| chr4:114780277-114780732 | -1.075363452 | 0.010292053 | 0.070404222 |
| chr6:75905178-75905447 | -1.327131143 | 0.010294926 | 0.070417591 |
| chr5:39005633-39005888 | -1.235148179 | 0.01029756 | 0.070429318 |
| chr1:34884836-34885220 | 1.049246899 | 0.010304772 | 0.070467241 |
| chr7:113015064-113015290 | -1.445966163 | 0.010304944 | 0.070467241 |
| chr7:128431235-128431559 | -0.983790027 | 0.010308072 | 0.07048234 |
| chr5:132506208-132506424 | 1.198443284 | 0.010311264 | 0.070493841 |
| chr9:137900338-137900589 | 1.398038417 | 0.010311594 | 0.070493841 |
| chr9:70449578-70449834 | -1.375327647 | 0.010314197 | 0.070499621 |
| chr18:63783275-63783720 | -1.156569365 | 0.01031428 | 0.070499621 |
| chr5:162068485-162068875 | -0.939071655 | 0.010317345 | 0.070505378 |
| chr5:29860004-29860253 | 1.321479122 | 0.010318774 | 0.070505378 |
| chr8:61495969-61496443 | 0.974139157 | 0.010319276 | 0.070505378 |
| chr11:21026900-21027145 | 1.423362281 | 0.010319579 | 0.070505378 |
| chr6:2902656-2903075 | 1.01042222 | 0.010319723 | 0.070505378 |
| chr2:207632721-207633541 | -0.875646553 | 0.010323421 | 0.070518482 |
| chr10:8026863-8027134 | 1.10894688 | 0.010323854 | 0.070518482 |
| chr9:108473288-108473689 | -1.149216902 | 0.010324402 | 0.070518482 |
| chr15:86104124-86104722 | 1.042973761 | 0.010326325 | 0.070520884 |
| chr6:154457811-154458313 | -0.958265201 | 0.010326594 | 0.070520884 |
| chr1:36436956-36437265 | 1.059247886 | 0.010331674 | 0.070549283 |
| chr17:78986678-78987047 | -1.186463534 | 0.010336754 | 0.070577134 |
| chr5:115674453-115674736 | -1.245590214 | 0.010337595 | 0.070577134 |
| chr6:105121983-105122278 | -1.218566407 | 0.010339225 | 0.070581976 |
| chr3:134597718-134597967 | 1.070004221 | 0.010341461 | 0.070590948 |
| chr19:53808761-53808970 | 1.30242262 | 0.010343955 | 0.070601685 |
| chr3:76045101-76045373 | -1.356829315 | 0.010346387 | 0.070605117 |
| chr3:169998679-169998906 | 1.248609991 | 0.010346446 | 0.070605117 |
| chr10:50992371-50992571 | -1.36433487 | 0.010347222 | 0.070605117 |
| chr13:27131255-27131455 | -1.435207271 | 0.010355009 | 0.070651962 |
| chr12:105199483-105199811 | -1.138109035 | 0.010356856 | 0.070658268 |
| chr5:21807830-21808372 | -0.920501151 | 0.010360893 | 0.070679515 |
| chr7:22351131-22351333 | 1.240495382 | 0.010364558 | 0.0706871 |
| chr7:35714836-35715294 | 0.900023756 | 0.010364745 | 0.0706871 |
| chr12:64608337-64608562 | 1.05428265 | 0.010364772 | 0.0706871 |
| chr14:35133841-35134099 | 1.211910137 | 0.010368394 | 0.07070551 |
| chr15:79204761-79205032 | 1.120433491 | 0.010372651 | 0.070728245 |
| chr2:235107165-235107467 | 1.239984624 | 0.010374171 | 0.07073231 |
| chr14:26917102-26917456 | 1.162185262 | 0.010379731 | 0.070761271 |
| chr1:46756749-46758239 | -0.836027649 | 0.010380265 | 0.070761271 |
| chr11:101299376-101299662 | -1.28630304 | 0.01038136 | 0.07076244 |
| chr5:175378494-175378871 | 1.019120222 | 0.010387219 | 0.07078735 |
| chrX:153971560-153972973 | 0.82030918 | 0.010387232 | 0.07078735 |
| chr11:126573288-126573494 | -1.507239903 | 0.010387786 | 0.07078735 |
| chr15:91941172-91941432 | -1.129612624 | 0.010395542 | 0.070830142 |
| chr10:96789725-96789925 | 1.354182843 | 0.010396766 | 0.070830142 |
| chr7:139620829-139621138 | -1.140160136 | 0.010396839 | 0.070830142 |
| chr15:90221288-90221628 | 0.957185634 | 0.010398448 | 0.070831442 |

| | | | |
|---|---|---|---|
| chr8:119439116-119439454 | -1.055222168 | 0.010398879 | 0.070831442 |
| chr4:180131019-180131496 | -1.14812513 | 0.010400465 | 0.070835946 |
| chr1:235906532-235906867 | 1.170888646 | 0.010411924 | 0.070899156 |
| chr8:42692534-42692821 | -1.153185469 | 0.010412958 | 0.070899156 |
| chr2:33809988-33810514 | -1.114255776 | 0.010413191 | 0.070899156 |
| chr7:86340140-86340893 | -0.947001144 | 0.010413447 | 0.070899156 |
| chr8:100746200-100746787 | 0.893964425 | 0.010422195 | 0.070951361 |
| chr13:80527529-80527893 | -1.292026558 | 0.010424078 | 0.070951361 |
| chr14:71754496-71754948 | -0.961348522 | 0.010424607 | 0.070951361 |
| chr16:53041608-53041845 | 1.398873109 | 0.010425475 | 0.070951361 |
| chr11:12223992-12224267 | 1.205074977 | 0.010427066 | 0.070951361 |
| chr5:42500203-42500469 | 1.171204908 | 0.01042756 | 0.070951361 |
| chr7:101017521-101017755 | -1.25297444 | 0.010427968 | 0.070951361 |
| chr1:224482614-224483069 | -1.012768247 | 0.010428522 | 0.070951361 |
| chr8:72773129-72773523 | 1.031882777 | 0.01043449 | 0.070981872 |
| chr2:110811791-110812052 | 1.127008157 | 0.010435726 | 0.070981872 |
| chr8:2086656-2086963 | -1.167541776 | 0.010435786 | 0.070981872 |
| chr14:96051796-96052071 | 1.079185951 | 0.010437217 | 0.070985305 |
| chr4:23373294-23373668 | -1.127556722 | 0.010438737 | 0.070986883 |
| chr16:66651001-66651282 | -1.158032717 | 0.010439302 | 0.070986883 |
| chrX:154419741-154420353 | 0.972075768 | 0.010444025 | 0.071009848 |
| chr1:165539341-165539803 | -1.09113525 | 0.010445297 | 0.071009848 |
| chr3:194002957-194003999 | -0.835781938 | 0.010445459 | 0.071009848 |
| chr16:19899206-19899620 | 1.039422016 | 0.010450029 | 0.07103352 |
| chr4:147596988-147597245 | 1.199401567 | 0.010450827 | 0.07103352 |
| chr16:75336098-75336355 | -1.26735158 | 0.010451723 | 0.07103352 |
| chr8:21409473-21409865 | -1.154361887 | 0.010458191 | 0.071071178 |
| chr3:171261300-171261896 | -0.933491146 | 0.010461748 | 0.071084947 |
| chr7:122238390-122239178 | -0.909758275 | 0.010462407 | 0.071084947 |
| chr4:100880065-100880346 | -1.189979857 | 0.010463001 | 0.071084947 |
| chr11:108417628-108417874 | -1.336363981 | 0.010464104 | 0.071086141 |
| chr2:95315107-95315344 | 1.013252557 | 0.010468042 | 0.071106589 |
| chr10:110407873-110408774 | -0.833437605 | 0.010471486 | 0.07112016 |
| chr16:2223889-2224255 | 0.977043474 | 0.010471896 | 0.07112016 |
| chr1:114310711-114311070 | 0.893112687 | 0.010486564 | 0.071213462 |
| chr7:106041112-106041581 | -0.963754064 | 0.010492805 | 0.071249528 |
| chr20:47346783-47347065 | 0.866577028 | 0.010496663 | 0.07126941 |
| chr1:246545304-246545513 | 1.380934911 | 0.010506724 | 0.071331403 |
| chr20:37758357-37758646 | 1.207230863 | 0.010511532 | 0.071357721 |
| chr8:28351410-28352279 | -0.925920748 | 0.01051296 | 0.071361091 |
| chr12:86660828-86661106 | -1.164876111 | 0.010516557 | 0.071379186 |
| chr17:63460717-63460964 | 1.046768586 | 0.010521024 | 0.071403178 |
| chr9:124659066-124659616 | -1.001875868 | 0.010524985 | 0.071414353 |
| chr15:66774302-66774704 | 0.961568839 | 0.010525263 | 0.071414353 |
| chr4:170122270-170122658 | -0.956953592 | 0.010525466 | 0.071414353 |
| chr20:55577010-55578510 | 0.808647232 | 0.010527668 | 0.071418271 |
| chr1:162877873-162878307 | 1.022337075 | 0.010530104 | 0.071418271 |
| chr7:25602966-25603740 | -1.015056887 | 0.010531067 | 0.071418271 |
| chr20:50699368-50699992 | 0.823109344 | 0.010532549 | 0.071418271 |
| chr11:1772574-1772781 | -1.398943526 | 0.010532978 | 0.071418271 |

| | | | |
|---|---|---|---|
| chr13:29454868-29455479 | -0.914502562 | 0.010533144 | 0.071418271 |
| chr8:104063731-104063967 | 1.321330962 | 0.01053346 | 0.071418271 |
| chr2:153345292-153345580 | -1.394379127 | 0.010533501 | 0.071418271 |
| chr17:80030776-80031158 | 0.960851626 | 0.010537846 | 0.071438218 |
| chr12:22046069-22047057 | -0.855801537 | 0.010538307 | 0.071438218 |
| chr6:96196562-96197076 | 0.96087532 | 0.010539865 | 0.071442459 |
| chr14:53325030-53325230 | 1.385518036 | 0.010542398 | 0.071453306 |
| chrX:74250973-74251364 | -0.977886337 | 0.010543793 | 0.071454613 |
| chr4:174778409-174779080 | -1.008805568 | 0.010544456 | 0.071454613 |
| chr14:29257197-29257436 | 1.403497326 | 0.010547448 | 0.071468565 |
| chr9:120383603-120383876 | -1.075060567 | 0.01055146 | 0.07148943 |
| chr1:32900641-32900924 | 1.075570021 | 0.010554785 | 0.071505508 |
| chrX:129939939-129940336 | 1.230981213 | 0.010555699 | 0.071505508 |
| chr1:29124605-29124970 | 1.035165348 | 0.010556865 | 0.071507081 |
| chr1:247172273-247172477 | -1.309170148 | 0.010559432 | 0.071518149 |
| chr2:54103720-54104174 | -0.948017722 | 0.010561384 | 0.071525043 |
| chr3:73093974-73094619 | -0.899524302 | 0.01056802 | 0.071560489 |
| chr20:47224100-47224816 | 0.816292395 | 0.010569171 | 0.071560489 |
| chr3:165404323-165404936 | -0.907696526 | 0.010569419 | 0.071560489 |
| chr2:177164519-177164890 | -1.047974938 | 0.010573771 | 0.071583629 |
| chr18:568729-569286 | 0.965046253 | 0.010576297 | 0.071594399 |
| chr7:122885651-122887112 | -0.817208654 | 0.010578626 | 0.071603843 |
| chr4:172707618-172708239 | -1.025066029 | 0.010579922 | 0.071606288 |
| chr22:21952089-21952382 | 1.079743676 | 0.010585501 | 0.071636918 |
| chr20:45293741-45294078 | 0.988186732 | 0.010586318 | 0.071636918 |
| chr5:122849766-122850099 | -0.952423845 | 0.010587978 | 0.071641825 |
| chr10:3584901-3585746 | 0.895052225 | 0.010595081 | 0.07168356 |
| chr6:158661017-158661467 | 1.047997045 | 0.010596384 | 0.071686044 |
| chr5:172236112-172236312 | -1.354669532 | 0.010601393 | 0.071707137 |
| chr4:25021203-25021614 | 1.083743815 | 0.01060177 | 0.071707137 |
| chr2:184527166-184527413 | -1.282329756 | 0.010602309 | 0.071707137 |
| chr7:112916300-112917157 | -0.878954932 | 0.010604147 | 0.071709244 |
| chr10:22513593-22514373 | -0.921217516 | 0.010604493 | 0.071709244 |
| chr19:19377763-19378128 | -1.075504823 | 0.01060607 | 0.071713582 |
| chr5:162323425-162323900 | -0.879791605 | 0.010610055 | 0.071734193 |
| chr3:18843103-18843817 | 0.860363113 | 0.010615683 | 0.071764858 |
| chr5:119244413-119244673 | -1.188108814 | 0.010616463 | 0.071764858 |
| chr3:154071433-154071633 | 1.422786176 | 0.010620082 | 0.071773101 |
| chr15:87804946-87805441 | 1.012747762 | 0.010620108 | 0.071773101 |
| chr12:127773608-127773916 | -1.117111548 | 0.010621259 | 0.071773101 |
| chr13:73065902-73066367 | -0.961604782 | 0.010622193 | 0.071773101 |
| chr9:38186486-38187029 | -1.0361254 | 0.010623072 | 0.071773101 |
| chr15:98873418-98874277 | -0.84379879 | 0.010624173 | 0.071773101 |
| chr17:36602222-36602468 | 1.139515629 | 0.01062424 | 0.071773101 |
| chr10:125029809-125030028 | 1.328236 | 0.010631836 | 0.071818084 |
| chr7:136687740-136688009 | -1.270065442 | 0.010633692 | 0.071824288 |
| chr12:123946479-123946719 | 1.323854618 | 0.010635051 | 0.071827138 |
| chr2:200259227-200260159 | -0.853571813 | 0.010642262 | 0.071869506 |
| chr10:104120920-104121351 | -0.912558994 | 0.01064656 | 0.071884081 |
| chr14:105685037-105685297 | 1.101292225 | 0.01064714 | 0.071884081 |

| | | | |
|---|---|---|---|
| chr15:41894774-41895172 | -1.049633728 | 0.010647235 | 0.071884081 |
| chr14:82313886-82314161 | 1.280341798 | 0.010655701 | 0.071934904 |
| chr20:19757432-19757756 | 1.02791201 | 0.010663208 | 0.071979237 |
| chr10:97070043-97070391 | -1.042301281 | 0.010664535 | 0.071981853 |
| chr6:13710499-13710707 | 1.075974335 | 0.010667536 | 0.07199577 |
| chr2:205972145-205972398 | 1.272033381 | 0.010669763 | 0.072004269 |
| chr10:125370626-125371329 | -0.977732347 | 0.010671253 | 0.072004269 |
| chr12:48925641-48925912 | -1.228804108 | 0.010671615 | 0.072004269 |
| chr8:99477434-99477788 | 1.251606126 | 0.010679173 | 0.072048427 |
| chr7:39400963-39401226 | -1.410306646 | 0.01068004 | 0.072048427 |
| chr3:9951959-9952568 | 0.868974346 | 0.010684529 | 0.072072362 |
| chr13:80572222-80572628 | -1.149530577 | 0.0106867 | 0.072080663 |
| chr3:195970213-195970480 | 1.26908903 | 0.010688278 | 0.072084959 |
| chr2:48047207-48047491 | -1.289301485 | 0.010690472 | 0.072093411 |
| chr4:177632515-177632715 | -1.545002807 | 0.010691485 | 0.072093899 |
| chr2:16553320-16553533 | -1.416019477 | 0.010705361 | 0.072175318 |
| chr13:49280942-49281750 | -0.918136078 | 0.010705443 | 0.072175318 |
| chr19:12333501-12333971 | 0.956307823 | 0.010707455 | 0.072182526 |
| chr6:34077341-34077541 | 1.33996462 | 0.010713702 | 0.072213088 |
| chr8:27490921-27491757 | -0.882885307 | 0.010714796 | 0.072213088 |
| chr7:141551177-141551789 | -0.919426441 | 0.010714816 | 0.072213088 |
| chr10:132331511-132332482 | 0.819400316 | 0.010719803 | 0.07224035 |
| chr16:89502199-89502501 | 1.111714878 | 0.010724638 | 0.072266571 |
| chr14:105773119-105773346 | 1.348518731 | 0.010727623 | 0.072280333 |
| chr5:56534042-56534307 | -1.448958371 | 0.010731495 | 0.072288073 |
| chr13:34156759-34157410 | -0.859024881 | 0.010731559 | 0.072288073 |
| chr14:105669447-105669730 | 1.163771546 | 0.010731602 | 0.072288073 |
| chr5:59665927-59666127 | -1.59877003 | 0.010732754 | 0.072289478 |
| chr7:70694944-70695310 | -1.293594831 | 0.010737418 | 0.072314536 |
| chr14:95340512-95340712 | 1.306322917 | 0.010740271 | 0.07232513 |
| chr3:163011274-163012026 | -0.905345868 | 0.010740879 | 0.07232513 |
| chr2:142857867-142858104 | -1.193243627 | 0.010746213 | 0.072354685 |
| chr7:16042098-16042382 | 1.22836917 | 0.010751598 | 0.072384582 |
| chr5:170632355-170632646 | -1.062054726 | 0.010758553 | 0.072425044 |
| chr3:32794565-32795055 | 1.00700913 | 0.010762163 | 0.072442983 |
| chr5:103120792-103121060 | -1.029977847 | 0.010769237 | 0.07248423 |
| chr17:43983316-43983662 | 1.212129073 | 0.010774205 | 0.072511301 |
| chr14:45305190-45305390 | 1.309779258 | 0.010781453 | 0.072553703 |
| chr14:84958734-84959042 | 1.088310978 | 0.010786214 | 0.072579373 |
| chrX:120316639-120316990 | 1.054267559 | 0.0107874 | 0.072580976 |
| chr13:69226184-69226384 | -1.605659561 | 0.01080071 | 0.072664147 |
| chr4:88334997-88335654 | -0.873022984 | 0.010803347 | 0.072675511 |
| chr4:183722685-183723022 | -1.039072193 | 0.010804309 | 0.072675603 |
| chr6:118769018-118769275 | 1.309489825 | 0.010812685 | 0.072725557 |
| chr1:214466785-214467477 | -0.916557149 | 0.010816307 | 0.072743536 |
| chr17:58578518-58578940 | 0.878700448 | 0.010821628 | 0.072772934 |
| chr3:10378553-10378753 | 1.220335798 | 0.010824737 | 0.07278745 |
| chr7:34622121-34622417 | 1.147322869 | 0.0108295 | 0.072813087 |
| chr6:62802967-62803578 | 0.919366601 | 0.01083291 | 0.072829625 |
| chr9:113987095-113987345 | 1.304464924 | 0.010834702 | 0.07283528 |

| | | | |
|---|---|---|---|
| chr2:15964950-15965290 | 1.035083362 | 0.010837865 | 0.072850153 |
| chr7:42574778-42574978 | -1.531320388 | 0.010841041 | 0.072865107 |
| chr1:204562743-204562981 | 1.053519829 | 0.010845272 | 0.072887152 |
| chr15:89040833-89041215 | 1.015659006 | 0.010846427 | 0.07288852 |
| chr12:94262209-94262821 | -0.939052998 | 0.010848181 | 0.072893918 |
| chr3:104198231-104198431 | 1.385580183 | 0.01085125 | 0.072906061 |
| chr7:24820220-24820589 | 0.928359517 | 0.010851892 | 0.072906061 |
| chr3:172550507-172550898 | -0.983389588 | 0.010856598 | 0.072931286 |
| chr16:55623263-55623512 | -1.32287016 | 0.010864539 | 0.072975706 |
| chr19:19618639-19619004 | -1.070643058 | 0.010865873 | 0.072975706 |
| chr7:31773191-31773417 | 1.30072456 | 0.010866689 | 0.072975706 |
| chr6:85660106-85660657 | -0.907829563 | 0.010867978 | 0.072975706 |
| chr13:33279825-33280310 | -0.995325827 | 0.01086813 | 0.072975706 |
| chr10:73920075-73920448 | -1.047414652 | 0.010868925 | 0.072975706 |
| chr12:106365887-106366087 | -1.497468289 | 0.010871223 | 0.072984739 |
| chr12:79512105-79512589 | -1.016098347 | 0.010878364 | 0.073014175 |
| chr1:116674724-116675147 | 0.915686461 | 0.01087849 | 0.073014175 |
| chr14:105664718-105664972 | 1.111524223 | 0.010879043 | 0.073014175 |
| chr13:66765631-66765924 | -1.121447926 | 0.010879419 | 0.073014175 |
| chr7:128738782-128740000 | -0.823402896 | 0.010881284 | 0.073020299 |
| chr2:6930973-6931177 | -1.539216451 | 0.010888427 | 0.073059545 |
| chr6:41846724-41847076 | 1.023159086 | 0.01088904 | 0.073059545 |
| chr8:137040479-137041186 | -0.927209279 | 0.010899839 | 0.07312148 |
| chr13:30769889-30770145 | 1.189366728 | 0.010900179 | 0.07312148 |
| chr7:8014267-8014712 | 1.002125214 | 0.010902724 | 0.07312799 |
| chr13:106619944-106620281 | -1.055615032 | 0.010903059 | 0.07312799 |
| chr21:29499685-29500155 | -0.92448824 | 0.010904464 | 0.073131014 |
| chr7:33722334-33723046 | -0.905540789 | 0.01090571 | 0.073132971 |
| chr10:45672170-45673171 | -0.837070542 | 0.010908605 | 0.073145984 |
| chr12:126094010-126094210 | 1.354702833 | 0.010909943 | 0.073148553 |
| chr1:174497369-174497900 | -1.048266797 | 0.010913512 | 0.073165609 |
| chr4:132642219-132642497 | -1.151100675 | 0.010914397 | 0.073165609 |
| chr20:4761037-4761517 | 1.166844307 | 0.010915639 | 0.073167534 |
| chr8:98745713-98745913 | 1.445395179 | 0.010918175 | 0.073173752 |
| chr10:97861971-97862238 | 1.077698005 | 0.010918476 | 0.073173752 |
| chr3:147147100-147147334 | 1.285396242 | 0.010925159 | 0.073212131 |
| chr9:70418165-70418680 | -0.850122149 | 0.010929549 | 0.073233485 |
| chr10:94133382-94133721 | 1.232303387 | 0.010930257 | 0.073233485 |
| chr1:26538075-26538323 | 1.249567415 | 0.010934055 | 0.07325253 |
| chr11:68405987-68406282 | 1.199983862 | 0.010935021 | 0.073252592 |
| chr3:127396366-127396721 | -1.047054868 | 0.010938881 | 0.073270042 |
| chr1:63557445-63557699 | -1.208349307 | 0.010939538 | 0.073270042 |
| chr15:89750739-89751275 | 0.889093765 | 0.010942801 | 0.073285491 |
| chr3:64776146-64776462 | 0.891813478 | 0.010949558 | 0.073324337 |
| chr8:108527242-108527638 | 1.093532139 | 0.0109593 | 0.07338316 |
| chr5:83509425-83509903 | -0.978021658 | 0.010961195 | 0.073389437 |
| chr5:108681363-108681643 | -1.090003962 | 0.010965719 | 0.073409469 |
| chr15:56732930-56733841 | 0.832977249 | 0.010966104 | 0.073409469 |
| chr1:188715631-188715916 | -1.14980166 | 0.010967864 | 0.073414841 |
| chr4:692891-693198 | 1.149370364 | 0.010970606 | 0.073426781 |

| | | | |
|---|---|---|---|
| chr7:129144059-129145032 | -0.857770384 | 0.010971634 | 0.073427249 |
| chr14:70908232-70908606 | 1.018762359 | 0.010973966 | 0.073436436 |
| chr7:123748267-123749803 | -0.811083036 | 0.010975018 | 0.073437067 |
| chr8:128220802-128221149 | 1.010658316 | 0.010978158 | 0.073445509 |
| chr17:81372557-81372870 | 1.117140673 | 0.010978197 | 0.073445509 |
| chr11:69106465-69106926 | 0.929795232 | 0.010983413 | 0.073473988 |
| chr8:140116749-140116970 | 1.202383874 | 0.01098522 | 0.073479665 |
| chr21:42269444-42269833 | -1.016970695 | 0.010994602 | 0.073535996 |
| chr4:8472146-8472426 | 1.208963528 | 0.010999199 | 0.073560323 |
| chr1:114702043-114702243 | 1.009557875 | 0.011003211 | 0.073580733 |
| chr8:140950342-140950656 | -1.063540403 | 0.01101235 | 0.073635419 |
| chr7:102363456-102364425 | -0.82869988 | 0.011016178 | 0.073654589 |
| chr20:55160029-55160229 | 1.321601594 | 0.011021928 | 0.073677191 |
| chr13:80030538-80030862 | -1.077909359 | 0.011023037 | 0.073677191 |
| chr10:77305697-77305903 | 1.283394042 | 0.011023167 | 0.073677191 |
| chrX:149881061-149881529 | 1.053486756 | 0.011023405 | 0.073677191 |
| chr14:105738851-105739807 | 0.896089415 | 0.011032227 | 0.073729722 |
| chr3:188349598-188349877 | -1.23387162 | 0.01103997 | 0.073775039 |
| chr13:24133587-24133856 | 1.378324032 | 0.011041144 | 0.073776445 |
| chr5:1394451-1394651 | 1.181937652 | 0.011043285 | 0.073784322 |
| chr4:148652042-148652496 | -1.138784838 | 0.011046176 | 0.073797202 |
| chr5:110785000-110785721 | -0.989836339 | 0.011050298 | 0.073818301 |
| chr3:11029236-11029436 | 1.330177337 | 0.01105339 | 0.073831489 |
| chr3:102180226-102181106 | -0.869013796 | 0.011054199 | 0.073831489 |
| chr3:107868791-107869209 | -1.117151825 | 0.011055553 | 0.073834099 |
| chr7:90298212-90298525 | -1.135208104 | 0.011060853 | 0.07386065 |
| chr4:182938185-182938447 | -1.2485314 | 0.011061457 | 0.07386065 |
| chr16:14116711-14117060 | 1.223162008 | 0.011063322 | 0.073866666 |
| chr17:61653698-61653936 | 0.886847646 | 0.011066487 | 0.073881361 |
| chr6:158213996-158214540 | -0.971080766 | 0.011070293 | 0.073900333 |
| chr2:85132814-85133619 | 0.84915203 | 0.01107322 | 0.073907754 |
| chr7:47591660-47592034 | 1.100046393 | 0.011073334 | 0.073907754 |
| chr11:86741940-86742164 | 1.187826289 | 0.011074398 | 0.073908416 |
| chr1:198890656-198891010 | -0.960211907 | 0.011079398 | 0.073935345 |
| chr3:15831956-15832234 | 1.236971518 | 0.011080382 | 0.073935472 |
| chr16:9160212-9160412 | 1.279091995 | 0.011082608 | 0.073943886 |
| chr7:138563768-138564740 | -0.851349551 | 0.011085598 | 0.073957397 |
| chr7:120089991-120090285 | -1.098239569 | 0.011098716 | 0.074038466 |
| chrX:25034922-25035230 | -1.100923185 | 0.01110123 | 0.074048789 |
| chr7:120800332-120800541 | -1.47461256 | 0.011102271 | 0.074049287 |
| chr6:138188383-138188949 | 0.921491175 | 0.0111072 | 0.07406448 |
| chr14:62769102-62769382 | 1.246586422 | 0.01110741 | 0.07406448 |
| chr3:62845664-62845917 | 1.138623918 | 0.011107449 | 0.07406448 |
| chrX:35243881-35244264 | -1.295192886 | 0.011112025 | 0.074088545 |
| chr17:61440108-61440584 | 0.828112231 | 0.011115865 | 0.074107702 |
| chr8:62860874-62861128 | 1.25801066 | 0.011122219 | 0.074138243 |
| chr11:73359399-73359818 | -0.987637423 | 0.011122381 | 0.074138243 |
| chr6:89951545-89951812 | -1.283193251 | 0.011125862 | 0.074147421 |
| chr1:107148703-107149114 | 0.88784844 | 0.011126078 | 0.074147421 |
| chr4:71184292-71184509 | 1.310168302 | 0.011126661 | 0.074147421 |

| | | | |
|---|---|---|---|
| chr12:93127742-93128639 | -0.89429224 | 0.011132304 | 0.074167189 |
| chr8:104678695-104679137 | 0.964673498 | 0.011132401 | 0.074167189 |
| chr16:82435261-82435542 | -1.452124238 | 0.011132532 | 0.074167189 |
| chr7:32512183-32512484 | 1.18064479 | 0.011139971 | 0.074208521 |
| chr1:157585889-157586089 | 1.381097698 | 0.01114222 | 0.074208521 |
| chr11:21552716-21553222 | 0.980114309 | 0.011142448 | 0.074208521 |
| chr3:2523495-2523700 | 1.350933019 | 0.01114261 | 0.074208521 |
| chr3:40327668-40327918 | 1.183180771 | 0.011146696 | 0.074229283 |
| chr10:24383703-24384023 | -1.075158586 | 0.011150684 | 0.074246655 |
| chr16:30417756-30418449 | 0.863861345 | 0.011151243 | 0.074246655 |
| chr6:36269741-36270473 | -0.943825566 | 0.011152469 | 0.074248367 |
| chr12:14260581-14260998 | -0.957537209 | 0.011160134 | 0.074292945 |
| chr2:112666273-112666483 | 1.478647069 | 0.011164376 | 0.074314729 |
| chr1:234487498-234487760 | 1.152880821 | 0.011166142 | 0.074318342 |
| chr12:110043180-110043413 | -1.203176857 | 0.01116708 | 0.074318342 |
| chr11:68890052-68891184 | -0.855436607 | 0.011167829 | 0.074318342 |
| chr10:61000947-61001865 | -0.874245968 | 0.011171762 | 0.074338061 |
| chr13:79683790-79684038 | -1.344191049 | 0.01117311 | 0.074340569 |
| chr15:41791546-41791758 | 1.380916228 | 0.011174556 | 0.074343738 |
| chr7:953525-953840 | 0.958866016 | 0.011180426 | 0.074376331 |
| chr11:98361972-98362172 | -1.491883723 | 0.01118209 | 0.074376744 |
| chr17:61210403-61211804 | 0.80042914 | 0.011182429 | 0.074376744 |
| chr5:113989657-113990024 | -1.123430517 | 0.011185118 | 0.07438817 |
| chr1:215566962-215568198 | -0.804290381 | 0.011187932 | 0.074400426 |
| chr17:55327715-55328099 | -1.040630469 | 0.011189165 | 0.07440217 |
| chr4:170708775-170709079 | -1.269673108 | 0.011190447 | 0.07440424 |
| chr2:70984483-70984683 | -1.321980323 | 0.011192182 | 0.074409317 |
| chrX:41984606-41985091 | -0.980767514 | 0.011196235 | 0.074429802 |
| chr4:177511359-177512011 | -0.859661627 | 0.011199172 | 0.074436581 |
| chr12:87266347-87266610 | -1.295846744 | 0.011200073 | 0.074436581 |
| chrX:37036638-37037176 | -0.883010984 | 0.011200858 | 0.074436581 |
| chr5:132389643-132390019 | -0.934975541 | 0.01120114 | 0.074436581 |
| chr1:56570875-56571415 | 0.905101734 | 0.011202455 | 0.074437153 |
| chr4:2962772-2963987 | 0.835786026 | 0.011203169 | 0.074437153 |
| chr2:207594387-207594613 | -1.457623851 | 0.011204281 | 0.074438083 |
| chr4:110516352-110516763 | -0.941937417 | 0.011205897 | 0.074441217 |
| chr19:40464406-40464606 | 1.082770208 | 0.01120701 | 0.074441217 |
| chr7:145083005-145083247 | -1.334165692 | 0.011207667 | 0.074441217 |
| chr3:71142514-71142718 | -1.384967735 | 0.011211264 | 0.074458649 |
| chr7:69067416-69067668 | -1.436190723 | 0.011213901 | 0.074469713 |
| chr2:238105780-238106050 | -1.2010684 | 0.011223318 | 0.074523243 |
| chr15:59154186-59154386 | 1.406328547 | 0.011223907 | 0.074523243 |
| chr2:162075400-162075600 | 1.392528675 | 0.01122815 | 0.074544955 |
| chr15:70494891-70495154 | 1.097800806 | 0.011233435 | 0.07457358 |
| chr1:176316111-176316440 | -1.276908094 | 0.011240077 | 0.074611205 |
| chr10:110181984-110182378 | 1.08721865 | 0.011241442 | 0.074613805 |
| chr1:147819079-147819279 | 1.252990399 | 0.011249742 | 0.074662427 |
| chr2:15189360-15189582 | -1.380024587 | 0.011265278 | 0.074759065 |
| chr17:74750389-74750630 | 1.05134959 | 0.011267349 | 0.07476633 |
| chr2:26299719-26299996 | -1.316219618 | 0.011268714 | 0.074768911 |

| | | | |
|---|---|---|---|
| chr3:52471591-52471791 | -1.383877047 | 0.011276952 | 0.074817095 |
| chr1:242225593-242226114 | 0.897884229 | 0.011279613 | 0.074828266 |
| chr9:76810345-76810693 | -1.114933305 | 0.011282329 | 0.07483219 |
| chr20:54462998-54463270 | 0.894858593 | 0.011282677 | 0.07483219 |
| chr7:80883499-80883905 | -0.995823816 | 0.011283134 | 0.07483219 |
| chrX:23931215-23931645 | -1.163573155 | 0.011287725 | 0.07485616 |
| chr9:1005113-1005351 | 1.112804874 | 0.011291598 | 0.074875364 |
| chr8:131028773-131029079 | -1.07462846 | 0.011295373 | 0.074891593 |
| chr15:43903791-43904081 | -1.158672796 | 0.011296 | 0.074891593 |
| chr2:183137911-183138112 | -1.45335678 | 0.011301713 | 0.074922982 |
| chr16:85497946-85498146 | 1.209535812 | 0.011305134 | 0.074939178 |
| chr13:75219584-75220074 | -0.976633164 | 0.01131009 | 0.074965547 |
| chr1:114391777-114391984 | 1.018689024 | 0.011314182 | 0.074986183 |
| chr12:3803850-3804130 | -1.352188425 | 0.011315824 | 0.074990581 |
| chr14:31865211-31865417 | 1.275570389 | 0.01132895 | 0.075071069 |
| chr4:6134676-6135051 | 1.06160691 | 0.011343992 | 0.075161837 |
| chr14:86892406-86892683 | 1.251395163 | 0.011344764 | 0.075161837 |
| chr5:75668899-75669424 | -1.056248745 | 0.01134559 | 0.075161837 |
| chr1:109057150-109057367 | 1.212373865 | 0.011353864 | 0.075210147 |
| chr18:13216147-13216355 | 1.193979538 | 0.01135676 | 0.07522283 |
| chr5:106779873-106781101 | -0.834198705 | 0.011358038 | 0.075224787 |
| chr13:69556609-69557102 | -1.065585698 | 0.01136046 | 0.075234328 |
| chr4:6673744-6674194 | 0.88346606 | 0.011363781 | 0.075249818 |
| chr5:17517610-17517855 | 1.128945613 | 0.011365394 | 0.075253995 |
| chr7:155298841-155299450 | -0.859746956 | 0.011372191 | 0.075292494 |
| chrX:132489261-132489585 | 1.033581327 | 0.011381978 | 0.075344358 |
| chr7:48144383-48144740 | -1.090464503 | 0.011381991 | 0.075344358 |
| chr7:134768020-134768305 | -1.250091666 | 0.011390954 | 0.075397172 |
| chr20:48839574-48839841 | 0.906043278 | 0.011392612 | 0.075401633 |
| chr2:174578018-174578245 | -1.346497831 | 0.011404374 | 0.075472961 |
| chr20:47338526-47338726 | 0.878709558 | 0.011410353 | 0.075506011 |
| chr5:81856926-81857923 | -0.859187185 | 0.011418243 | 0.075551696 |
| chr7:85464376-85464621 | -1.393070907 | 0.011421202 | 0.075561098 |
| chr7:108002828-108003518 | -0.926643629 | 0.01142219 | 0.075561098 |
| chr5:66807801-66808094 | -1.181980784 | 0.011422623 | 0.075561098 |
| chr13:79300006-79300729 | -0.905869398 | 0.011424068 | 0.075564137 |
| chr19:45873001-45873261 | 1.224991179 | 0.011426537 | 0.075573939 |
| chr10:48805553-48805912 | -1.114631833 | 0.011428797 | 0.075582368 |
| chr4:163251175-163251464 | -1.202807034 | 0.01143431 | 0.075612301 |
| chrX:54640757-54641320 | -0.885757564 | 0.011442473 | 0.075659748 |
| chr14:39627504-39627707 | 1.324250663 | 0.011444675 | 0.075667781 |
| chr14:35331651-35331921 | 1.007264804 | 0.011447955 | 0.075682931 |
| chr13:35164325-35164688 | -1.073296663 | 0.011453476 | 0.075712901 |
| chr12:57139063-57139282 | 1.244360901 | 0.01145506 | 0.075716842 |
| chr11:39306701-39307130 | -1.049369562 | 0.011456856 | 0.075722179 |
| chr17:8176242-8176614 | -1.026266386 | 0.011463921 | 0.075762341 |
| chr21:38343694-38343988 | 1.123831739 | 0.011465517 | 0.075766352 |
| chr9:115118496-115119205 | -0.948198047 | 0.011472833 | 0.075808157 |
| chr3:141345041-141345384 | 1.296026765 | 0.011475614 | 0.075819993 |
| chr9:125511804-125512015 | 1.226985509 | 0.011481675 | 0.075853498 |

| | | | |
|---|---|---|---|
| chr8:69073441-69073648 | -1.39577942 | 0.011482784 | 0.075854286 |
| chr1:113279694-113280484 | -0.944994135 | 0.011485933 | 0.075868549 |
| chr6:52962400-52962675 | 1.322465515 | 0.011491515 | 0.075894521 |
| chr12:106136250-106136455 | -1.298422042 | 0.011491846 | 0.075894521 |
| chr11:44900022-44900293 | 1.086353082 | 0.011502097 | 0.075955674 |
| chr1:228021456-228021679 | 1.139803844 | 0.011503548 | 0.075958708 |
| chr5:56480720-56480920 | -1.368729213 | 0.011509515 | 0.075991563 |
| chr5:122653493-122653914 | 0.977538982 | 0.011511743 | 0.075999721 |
| chr14:105760655-105760912 | 1.183452794 | 0.011517431 | 0.076030726 |
| chr16:85347258-85347517 | -1.060845679 | 0.011528462 | 0.076086088 |
| chr5:180753522-180753931 | 1.051873763 | 0.011528492 | 0.076086088 |
| chr9:38172574-38172973 | -1.254727976 | 0.011529524 | 0.076086088 |
| chr19:1169921-1170372 | -0.99996842 | 0.01152979 | 0.076086088 |
| chr10:132307530-132308677 | 0.815209897 | 0.011536304 | 0.07612252 |
| chr5:132253037-132253424 | -1.113172382 | 0.011538502 | 0.076129866 |
| chr5:98839694-98840349 | -0.917862569 | 0.011539405 | 0.076129866 |
| chr10:97196251-97196878 | 0.894575586 | 0.01154042 | 0.076130013 |
| chr5:17833111-17833355 | 1.354813242 | 0.011542221 | 0.076135338 |
| chr14:75554522-75554887 | 0.939356652 | 0.011544642 | 0.076144751 |
| chr4:83509102-83509548 | -1.029749692 | 0.011548074 | 0.076160827 |
| chr3:130649443-130649643 | 1.561515968 | 0.011553487 | 0.076189974 |
| chr4:143441783-143441984 | -1.527033021 | 0.011555982 | 0.076199869 |
| chr4:100935116-100935362 | -1.243892167 | 0.011557729 | 0.076202276 |
| chr7:7817245-7817485 | 1.16447998 | 0.011558336 | 0.076202276 |
| chr2:144004733-144005148 | -1.106208819 | 0.011562816 | 0.076222564 |
| chr17:60016540-60017019 | 0.860862364 | 0.011563779 | 0.076222564 |
| chr18:48983060-48983317 | -1.312583645 | 0.011564398 | 0.076222564 |
| chr6:111645110-111645328 | 1.209105543 | 0.011571778 | 0.076264647 |
| chr9:99905893-99906233 | 1.005842661 | 0.011572955 | 0.076265845 |
| chr3:26865480-26865714 | 1.3786899 | 0.011575321 | 0.076274878 |
| chr8:19013620-19014029 | 0.984483908 | 0.011577836 | 0.076284888 |
| chr6:53293769-53293969 | 1.443587336 | 0.011578902 | 0.076285349 |
| chr7:22822463-22823101 | 0.829239648 | 0.011583065 | 0.076306216 |
| chr13:110524652-110524929 | -1.200884511 | 0.011586473 | 0.076322105 |
| chr15:101167742-101168566 | -0.863286097 | 0.011588974 | 0.076332016 |
| chr1:181361-181603 | -1.292716634 | 0.011593358 | 0.076354331 |
| chr3:84836772-84837470 | -0.87099273 | 0.011597941 | 0.076372099 |
| chr2:66575320-66575629 | -1.042795215 | 0.01159805 | 0.076372099 |
| chr4:91734745-91734954 | -1.485294374 | 0.011603439 | 0.076401025 |
| chr7:16926289-16926567 | 1.052055328 | 0.011607659 | 0.076422239 |
| chr10:102422689-102422930 | 0.983483845 | 0.01161449 | 0.076460642 |
| chr8:115938982-115939421 | -0.959539915 | 0.01161704 | 0.076470863 |
| chr3:64133163-64133554 | 0.859122241 | 0.011620966 | 0.076490131 |
| chr12:11580417-11580845 | -1.010399585 | 0.011626125 | 0.076517519 |
| chr4:9703521-9703805 | 1.192899727 | 0.011628742 | 0.076522047 |
| chr8:29738439-29738737 | -1.236892229 | 0.011629708 | 0.076522047 |
| chr1:164591905-164592111 | 1.121122724 | 0.011629809 | 0.076522047 |
| chr13:66618778-66619021 | -1.177060784 | 0.011631778 | 0.076524385 |
| chr4:141549662-141549944 | -1.117321631 | 0.011632162 | 0.076524385 |
| chr17:21030442-21030725 | 1.399267896 | 0.011633998 | 0.076529895 |

| | | | |
|---|---|---|---|
| chr5:87541140-87541381 | -1.311469274 | 0.011636946 | 0.076542716 |
| chr3:86089133-86089355 | 1.4263974 | 0.011642349 | 0.076559255 |
| chr3:167978015-167978457 | 0.977719829 | 0.011642399 | 0.076559255 |
| chr1:54519759-54519982 | -1.316048513 | 0.011642458 | 0.076559255 |
| chr19:48216949-48217199 | -1.405536047 | 0.011644184 | 0.076564033 |
| chr20:63651854-63652353 | -0.904294364 | 0.011651047 | 0.076601094 |
| chr1:198451136-198451437 | -1.27366099 | 0.011652548 | 0.076601094 |
| chr1:26853200-26853409 | 1.135194925 | 0.01165282 | 0.076601094 |
| chr19:33416299-33416618 | 1.022869083 | 0.011655189 | 0.076610096 |
| chr10:67849288-67849914 | -0.885010541 | 0.011657707 | 0.076620075 |
| chr20:48501999-48503331 | 0.803698148 | 0.011665436 | 0.076662279 |
| chr12:55656879-55657081 | 1.346625882 | 0.011666204 | 0.076662279 |
| chr12:57935830-57936393 | -0.893107421 | 0.01166713 | 0.076662279 |
| chr4:57308314-57308550 | 1.223479942 | 0.011681567 | 0.076745514 |
| chr4:82421163-82421465 | -1.230971245 | 0.011681801 | 0.076745514 |
| chr8:66902837-66903200 | -1.200986229 | 0.011685949 | 0.076766182 |
| chr5:50651942-50652352 | 0.993224073 | 0.011690781 | 0.076786773 |
| chrX:107331033-107331283 | 1.299116427 | 0.01169167 | 0.076786773 |
| chr3:40782134-40782397 | 1.220058145 | 0.01169209 | 0.076786773 |
| chr4:128601465-128601687 | 1.287089605 | 0.011700262 | 0.076833859 |
| chr6:42916080-42916304 | 1.29557352 | 0.011706839 | 0.076867776 |
| chr12:74871218-74871520 | -1.12265916 | 0.011707433 | 0.076867776 |
| chr2:102661754-102662184 | -1.077073097 | 0.011714804 | 0.076909582 |
| chr11:59866433-59866658 | -1.377047482 | 0.01171587 | 0.076909991 |
| chr12:46342625-46343700 | 0.872502643 | 0.011719975 | 0.0769252 |
| chrX:148500839-148501204 | 1.157050272 | 0.011720195 | 0.0769252 |
| chr17:61834224-61834535 | 0.853166464 | 0.011728116 | 0.076970595 |
| chrX:119501720-119502077 | 1.101691578 | 0.011735105 | 0.077004186 |
| chr15:60392630-60392847 | -1.181029931 | 0.011735244 | 0.077004186 |
| chr5:168371275-168371549 | -1.242370152 | 0.011739283 | 0.077024089 |
| chrX:105701188-105701982 | -0.931939236 | 0.011742443 | 0.077032048 |
| chr1:153649507-153649986 | 0.950272141 | 0.011742506 | 0.077032048 |
| chr4:111943757-111944109 | -1.004968318 | 0.011748752 | 0.077066424 |
| chr17:61518807-61519187 | 0.837292543 | 0.011754107 | 0.077085258 |
| chr3:85253995-85254248 | 1.290586968 | 0.011754271 | 0.077085258 |
| chr10:35328115-35328334 | -1.376423137 | 0.011755069 | 0.077085258 |
| chr2:234422284-234422617 | 1.188391106 | 0.011755648 | 0.077085258 |
| chr20:47278091-47278352 | 0.858073915 | 0.011761778 | 0.077118858 |
| chr21:43262519-43262830 | 1.064212506 | 0.011778224 | 0.07722008 |
| chr3:18987923-18988164 | 1.267946375 | 0.011780616 | 0.077229157 |
| chr20:62088412-62088851 | 0.919112697 | 0.011782562 | 0.077235305 |
| chr4:7323851-7324280 | 0.980665131 | 0.011786466 | 0.077248111 |
| chr3:38154870-38155185 | -1.060213206 | 0.011786532 | 0.077248111 |
| chr3:42013414-42013685 | 1.085863749 | 0.011787724 | 0.077249316 |
| chr20:48468357-48468783 | 0.85374999 | 0.011789543 | 0.077254628 |
| chr20:54521460-54521711 | 0.895256357 | 0.011794787 | 0.077282379 |
| chr1:174325751-174325951 | 1.391628116 | 0.011797324 | 0.077288756 |
| chr2:117958266-117958681 | 1.033498394 | 0.011797777 | 0.077288756 |
| chr13:43553527-43553752 | -1.375724539 | 0.01179907 | 0.07729062 |
| chr1:114093569-114093849 | 0.877009107 | 0.011800881 | 0.077295872 |

| | | | |
|---|---|---|---|
| chr20:51715110-51715459 | 0.962865446 | 0.011804145 | 0.07731064 |
| chr7:134316321-134317514 | -0.817730721 | 0.01180617 | 0.077317297 |
| chr20:41233101-41233337 | 1.151099685 | 0.01181175 | 0.077347227 |
| chr18:33201460-33202124 | -0.959825869 | 0.011812838 | 0.077347741 |
| chr12:33588118-33588597 | -1.035498595 | 0.01182001 | 0.077388088 |
| chr18:45793045-45793912 | -0.878337104 | 0.011822006 | 0.077394542 |
| chr1:33140817-33141090 | 1.21492046 | 0.011828071 | 0.077427634 |
| chr12:89579603-89579858 | -1.209542273 | 0.011829252 | 0.077428752 |
| chr17:7627496-7628091 | -0.938547452 | 0.011831307 | 0.077435588 |
| chr14:44578538-44578795 | 1.161345472 | 0.011834114 | 0.077447345 |
| chr7:1071684-1072118 | 0.966881513 | 0.011837935 | 0.077465731 |
| chr5:64959514-64959714 | -1.482433356 | 0.011840211 | 0.077474009 |
| chr11:47153072-47153450 | 0.931558902 | 0.011842255 | 0.077480766 |
| chr9:109213153-109213386 | 1.344566629 | 0.01184525 | 0.077490292 |
| chr3:112699762-112700076 | -1.277465797 | 0.011845734 | 0.077490292 |
| chr12:91382052-91382545 | -0.965924641 | 0.011847576 | 0.077495729 |
| chr1:179660400-179660644 | -1.374937604 | 0.011848817 | 0.077497227 |
| chr5:11182699-11183049 | -1.021402874 | 0.011851809 | 0.07749913 |
| chr20:45138805-45139047 | 1.274350245 | 0.011852036 | 0.07749913 |
| chr19:12789827-12790383 | 0.884264878 | 0.011852142 | 0.07749913 |
| chr7:13042750-13043114 | 1.134299849 | 0.011853528 | 0.07749974 |
| chr6:149024380-149024620 | -1.357863911 | 0.011854941 | 0.07749974 |
| chr10:67492428-67492691 | -1.26688373 | 0.011855269 | 0.07749974 |
| chr12:94764824-94765154 | -1.14924286 | 0.011858073 | 0.077511456 |
| chr6:167767341-167767646 | -1.138751568 | 0.011859837 | 0.077516374 |
| chr4:65176548-65176804 | -1.260866726 | 0.011862406 | 0.077524272 |
| chr4:38160925-38161251 | -1.065202444 | 0.011863069 | 0.077524272 |
| chr12:113796761-113796961 | 1.308569877 | 0.011866286 | 0.077537943 |
| chr5:25331554-25331837 | 1.125729401 | 0.011868071 | 0.077537943 |
| chr12:9750817-9751081 | -1.224509479 | 0.011868197 | 0.077537943 |
| chr17:63001503-63002550 | 0.803786597 | 0.011873701 | 0.077567285 |
| chr8:127851322-127851714 | -0.891764393 | 0.011877745 | 0.077575882 |
| chr16:90077230-90078108 | -0.823986395 | 0.011878772 | 0.077575882 |
| chr17:65941970-65942253 | -1.132261899 | 0.011879299 | 0.077575882 |
| chr5:85663508-85663940 | -1.075981053 | 0.011880496 | 0.077575882 |
| chr10:21724720-21725101 | -1.039567966 | 0.011881076 | 0.077575882 |
| chr19:29442828-29443076 | 1.368144994 | 0.011882104 | 0.077575882 |
| chr20:20816137-20816354 | 1.368144994 | 0.011882104 | 0.077575882 |
| chr7:101814951-101816121 | -0.843657703 | 0.01188512 | 0.077588964 |
| chr5:62108381-62108661 | -1.373861315 | 0.011891541 | 0.077624265 |
| chr14:34139206-34139408 | 1.115750535 | 0.011895163 | 0.077641297 |
| chr8:23487082-23487411 | -1.249235759 | 0.011899115 | 0.077660475 |
| chr3:118176797-118177066 | -1.283468441 | 0.011905174 | 0.077691768 |
| chr4:70522530-70522757 | 1.253553162 | 0.011905938 | 0.077691768 |
| chr3:78480666-78481497 | -0.906520961 | 0.011917371 | 0.077759752 |
| chr1:169868992-169869368 | -0.960553549 | 0.01192428 | 0.077791864 |
| chr3:141782426-141782783 | -1.033226416 | 0.011924824 | 0.077791864 |
| chr8:93588186-93588442 | 1.169854125 | 0.011925338 | 0.077791864 |
| chr6:53744205-53744908 | 0.867619817 | 0.011935324 | 0.077850379 |
| chr2:193102246-193102623 | -1.153392969 | 0.011948251 | 0.077928063 |

| | | | |
|---|---|---|---|
| chr7:152588283-152588863 | -0.97324513 | 0.011962354 | 0.078013401 |
| chr10:30423409-30423872 | -0.982285542 | 0.011965877 | 0.078029737 |
| chr9:72229297-72229596 | -1.301830186 | 0.011970502 | 0.078052597 |
| chr3:153506764-153507133 | -0.940932467 | 0.01197142 | 0.078052597 |
| chr14:104639938-104640528 | 0.898855986 | 0.01197365 | 0.078060495 |
| chr11:77654741-77655084 | -1.106277478 | 0.011984513 | 0.078124672 |
| chr3:81295711-81296753 | -0.842856367 | 0.01198975 | 0.078152159 |
| chr10:78927219-78927668 | -0.910497838 | 0.011990883 | 0.078152894 |
| chr10:21021802-21022390 | -0.947036284 | 0.012002542 | 0.078222232 |
| chr1:195030247-195030610 | 1.055945751 | 0.012006973 | 0.078240769 |
| chr11:103433520-103433729 | -1.399694925 | 0.012007429 | 0.078240769 |
| chr13:78767362-78767580 | -1.485717268 | 0.012013558 | 0.078269394 |
| chr11:17760396-17760698 | 0.976297903 | 0.012014836 | 0.078269394 |
| chr20:48657669-48658647 | 0.797863726 | 0.012015313 | 0.078269394 |
| chr13:49976681-49976905 | 1.314472231 | 0.012015908 | 0.078269394 |
| chr2:189486291-189486806 | -0.879272193 | 0.012019955 | 0.0782891 |
| chr2:143558517-143558888 | -1.096866919 | 0.012025232 | 0.078316817 |
| chr3:134645025-134645390 | -0.985552792 | 0.012027137 | 0.078322563 |
| chr15:63770456-63770658 | 1.341573737 | 0.012030254 | 0.078332608 |
| chr5:125357657-125358173 | -1.003135909 | 0.012030724 | 0.078332608 |
| chr5:128274613-128274878 | -1.306804942 | 0.0120326 | 0.078338167 |
| chr21:8228602-8228893 | 0.934527517 | 0.012037038 | 0.078360403 |
| chr4:169069760-169070701 | -0.809528993 | 0.012042889 | 0.078391828 |
| chr17:13415851-13416409 | -1.046279024 | 0.012049412 | 0.078421543 |
| chr21:41704260-41704587 | -1.030786852 | 0.012049501 | 0.078421543 |
| chr10:26734171-26734536 | 1.053869796 | 0.012052525 | 0.078429831 |
| chr19:54426188-54426640 | 0.972402176 | 0.012052821 | 0.078429831 |
| chr10:21144097-21144341 | -1.243171189 | 0.012057471 | 0.078453425 |
| chr15:89139157-89139357 | 1.198659698 | 0.012059234 | 0.078458235 |
| chr6:79610580-79610911 | 1.039480068 | 0.012061472 | 0.078466129 |
| chr11:131674007-131674302 | -1.159142824 | 0.012063876 | 0.078469137 |
| chr6:137881249-137881506 | 1.346244427 | 0.012063982 | 0.078469137 |
| chr19:44784228-44784814 | 0.977575265 | 0.012066917 | 0.078481565 |
| chr2:47215894-47216321 | -0.99359813 | 0.012071182 | 0.078501431 |
| chr1:55041022-55041418 | -1.063032972 | 0.012072021 | 0.078501431 |
| chr6:35249677-35249877 | 1.424154632 | 0.01207631 | 0.07852266 |
| chrX:16836464-16836760 | -1.014309476 | 0.012077683 | 0.078524921 |
| chr7:55407060-55407295 | 1.275890444 | 0.012083524 | 0.078556234 |
| chr10:45470417-45470861 | -1.046533953 | 0.012085187 | 0.07856038 |
| chr5:12146901-12147292 | -1.310402446 | 0.012087843 | 0.078567134 |
| chr12:14514000-14514578 | -0.906125359 | 0.012088306 | 0.078567134 |
| chr10:92343116-92343329 | 1.365920654 | 0.012090435 | 0.078567134 |
| chr3:190038098-190038437 | -1.21594765 | 0.012090685 | 0.078567134 |
| chr10:9766785-9767035 | 1.288786496 | 0.012091822 | 0.078567134 |
| chr3:105368618-105369304 | -0.823038809 | 0.012092622 | 0.078567134 |
| chr1:223258635-223259243 | -0.932182518 | 0.012094884 | 0.078567134 |
| chr16:2585367-2585567 | 1.428675694 | 0.012095455 | 0.078567134 |
| chr3:104909270-104909470 | 1.428675694 | 0.012095455 | 0.078567134 |
| chr8:8461263-8462011 | 0.873457147 | 0.012097778 | 0.078575568 |
| chr2:160875966-160876356 | -1.004489143 | 0.012100639 | 0.078587487 |

| | | | |
|---|---|---|---|
| chr1:28153248-28153448 | 1.335902855 | 0.012102921 | 0.078595642 |
| chr9:347767-347992 | 1.335264636 | 0.012105977 | 0.078604148 |
| chr2:42752165-42752574 | -0.997708866 | 0.012106282 | 0.078604148 |
| chr20:2768300-2768685 | 1.014653538 | 0.012108837 | 0.078611482 |
| chr2:20411356-20411820 | -0.879974219 | 0.012109464 | 0.078611482 |
| chr20:54881153-54881353 | 0.977725104 | 0.01211282 | 0.07862661 |
| chr6:28648124-28648838 | -0.909691187 | 0.012114303 | 0.078627865 |
| chr3:190384289-190384641 | -1.014470711 | 0.012116103 | 0.078627865 |
| chr1:92485840-92486178 | -1.021001194 | 0.012116259 | 0.078627865 |
| chr15:70008940-70009186 | 1.100376797 | 0.012117118 | 0.078627865 |
| chr5:60688055-60688629 | -0.866306049 | 0.012123568 | 0.078663057 |
| chr21:43712209-43712535 | 0.945791935 | 0.012127253 | 0.078680302 |
| chr9:90517089-90517791 | 0.9113327 | 0.012133668 | 0.078715257 |
| chr17:38799363-38800249 | 0.841320265 | 0.012136106 | 0.078724407 |
| chr4:174222309-174222547 | -1.25865396 | 0.012137484 | 0.078726681 |
| chr16:2474236-2474548 | 1.001665957 | 0.012144818 | 0.078767584 |
| chr7:27200211-27200541 | 1.05950366 | 0.012148702 | 0.078786108 |
| chr11:37003549-37004030 | -0.952499415 | 0.012151096 | 0.078794963 |
| chr1:106543950-106544160 | 0.96873365 | 0.012152226 | 0.078795622 |
| chr13:49313759-49314485 | -0.892886418 | 0.012154781 | 0.078805522 |
| chr4:997720-998586 | -0.883796982 | 0.012158623 | 0.078823758 |
| chr17:63428298-63428596 | 0.910550031 | 0.01216194 | 0.078838595 |
| chr5:146515871-146516235 | -1.124698889 | 0.012163403 | 0.078841404 |
| chr1:43235682-43236193 | -0.926056444 | 0.012165922 | 0.078848639 |
| chr2:211378567-211378905 | -1.105505951 | 0.012167763 | 0.078848639 |
| chr1:70257743-70258018 | -1.476196957 | 0.012168635 | 0.078848639 |
| chr17:5593699-5593899 | -1.476196957 | 0.012168635 | 0.078848639 |
| chr1:41667441-41667641 | 1.287924237 | 0.012174559 | 0.078880353 |
| chr16:1678688-1678986 | 0.982570527 | 0.012175939 | 0.078882623 |
| chr2:15748770-15749057 | -1.067687938 | 0.012180826 | 0.078907612 |
| chr12:22138-22413 | 1.122525443 | 0.012183509 | 0.078912746 |
| chr8:79914332-79914549 | 1.227355267 | 0.012183678 | 0.078912746 |
| chr2:21333690-21334028 | -1.135431971 | 0.012188733 | 0.078938813 |
| chr16:66478759-66479065 | 0.995192919 | 0.012190455 | 0.078943294 |
| chr12:113245653-113246881 | 0.811685427 | 0.0121946 | 0.078963458 |
| chr7:156819619-156820000 | -1.176649759 | 0.012196595 | 0.078964157 |
| chr2:149146952-149147182 | -1.394748511 | 0.012196768 | 0.078964157 |
| chr2:40820429-40820632 | -1.432868065 | 0.012204549 | 0.079000569 |
| chr7:157302451-157302804 | -1.096988507 | 0.012206459 | 0.079000569 |
| chr7:142813171-142813819 | -0.93018165 | 0.012206478 | 0.079000569 |
| chr2:75379961-75380220 | -1.361947576 | 0.012206517 | 0.079000569 |
| chr6:6629492-6629790 | 1.273206902 | 0.012208796 | 0.079008645 |
| chr9:72843942-72844615 | -0.908179231 | 0.012215967 | 0.079048378 |
| chr10:43255571-43255775 | -1.183881932 | 0.012222396 | 0.079083297 |
| chr1:99766334-99766928 | -0.929068045 | 0.012228723 | 0.079111655 |
| chr8:144241242-144241518 | 1.093642227 | 0.012228843 | 0.079111655 |
| chr12:25194779-25195659 | -0.832233369 | 0.012233741 | 0.079136656 |
| chr6:36914251-36914729 | -1.014176513 | 0.012236048 | 0.079144901 |
| chr6:116317116-116317428 | -1.208742713 | 0.012250307 | 0.079230441 |
| chr4:164847719-164848022 | -1.305504193 | 0.012253511 | 0.079241504 |

| | | | |
|---|---|---|---|
| chr15:90201014-90201880 | 0.837831672 | 0.012254086 | 0.079241504 |
| chr16:84964622-84964835 | 1.252522877 | 0.012256032 | 0.079243186 |
| chr10:5325991-5326213 | 1.275221371 | 0.012256415 | 0.079243186 |
| chr3:117628749-117629470 | -0.86074844 | 0.012261059 | 0.079266525 |
| chr7:18120099-18120323 | 1.175718281 | 0.012264489 | 0.079282014 |
| chr1:83931736-83931997 | -1.173862399 | 0.012270048 | 0.079311258 |
| chr8:99078731-99079160 | 1.026786309 | 0.012272407 | 0.07931981 |
| chr12:27779423-27780854 | -0.825119461 | 0.012276636 | 0.079340454 |
| chr1:114180661-114181084 | 0.87905852 | 0.012279692 | 0.079348347 |
| chr1:170067227-170067512 | -1.262237137 | 0.012279929 | 0.079348347 |
| chr4:31800705-31801029 | 1.028857503 | 0.012282074 | 0.079352566 |
| chr6:104201643-104201936 | -1.132304929 | 0.012282653 | 0.079352566 |
| chr4:139530859-139531119 | -1.155686554 | 0.012285854 | 0.079366553 |
| chr17:17496212-17496676 | 0.877107737 | 0.012288491 | 0.079376897 |
| chr18:34773895-34774343 | -1.087652077 | 0.012289565 | 0.079377141 |
| chr8:132097527-132097874 | -1.123673099 | 0.012292183 | 0.079387361 |
| chr13:35789385-35789655 | -1.15748882 | 0.012296563 | 0.079408953 |
| chr1:212432689-212433705 | -0.840876453 | 0.01230197 | 0.079437175 |
| chr4:180456921-180457419 | -0.976282668 | 0.012311755 | 0.079493666 |
| chr17:61842688-61843103 | 0.828881786 | 0.012313367 | 0.079497375 |
| chr12:60684768-60685286 | 0.956857599 | 0.012314858 | 0.079500299 |
| chr3:100776722-100777266 | 1.00433087 | 0.01231894 | 0.079519954 |
| chr8:92801447-92801680 | 1.2601173 | 0.012323225 | 0.079524522 |
| chr14:76034589-76034789 | 1.357311733 | 0.012323585 | 0.079524522 |
| chr21:40887066-40887326 | -1.235409713 | 0.012323723 | 0.079524522 |
| chr3:4720228-4720556 | -0.953096085 | 0.0123238 | 0.079524522 |
| chr5:114177247-114177531 | -1.37996096 | 0.012327394 | 0.079541019 |
| chr3:98343802-98344276 | -0.979307505 | 0.012339726 | 0.079596617 |
| chr1:195738946-195739617 | -0.885848786 | 0.012340626 | 0.079596617 |
| chr11:30046636-30046931 | -1.26347591 | 0.012341405 | 0.079596617 |
| chr5:96434177-96434676 | -0.936285262 | 0.012341678 | 0.079596617 |
| chr20:14036064-14036267 | -1.47409934 | 0.012342309 | 0.079596617 |
| chr5:63227894-63228393 | -1.225768926 | 0.012343227 | 0.079596617 |
| chr17:45415651-45415851 | 1.294467303 | 0.012344104 | 0.079596617 |
| chr7:130875376-130875605 | -1.2879818 | 0.012344321 | 0.079596617 |
| chr2:22626734-22626967 | -1.361273858 | 0.012345649 | 0.079597192 |
| chr3:156681042-156681399 | -0.983094085 | 0.012346488 | 0.079597192 |
| chrX:77390425-77390710 | -1.141174095 | 0.012350466 | 0.07961614 |
| chr6:120406023-120406265 | -1.251082349 | 0.012358681 | 0.07966239 |
| chr6:125192067-125192727 | -0.854890951 | 0.012360173 | 0.079664353 |
| chr3:146870990-146871190 | 1.464717872 | 0.012361065 | 0.079664353 |
| chr1:145877972-145878298 | 0.933571777 | 0.012364916 | 0.079682471 |
| chr4:49323020-49323268 | 1.355541641 | 0.012380262 | 0.079774654 |
| chr5:91219094-91219720 | -0.918224837 | 0.012383344 | 0.079787803 |
| chr1:85613118-85613388 | -1.176958304 | 0.01238715 | 0.07980456 |
| chr17:57995276-57995953 | -0.836440194 | 0.012388378 | 0.07980456 |
| chr10:112949884-112950426 | -0.877427219 | 0.012389069 | 0.07980456 |
| chr3:43155570-43156042 | -1.055839475 | 0.012390208 | 0.079805188 |
| chr15:80679514-80679796 | 1.100163157 | 0.012391902 | 0.079809393 |
| chr16:31333706-31334049 | 1.009454874 | 0.012393417 | 0.079812442 |

| | | | |
|---|---|---|---|
| chr14:67440532-67440882 | 0.980086884 | 0.012395951 | 0.07981831 |
| chr14:98903745-98904253 | -0.949578607 | 0.012396412 | 0.07981831 |
| chr7:157397776-157398298 | -0.886762726 | 0.012398022 | 0.079821966 |
| chr4:76897080-76898497 | 0.817356726 | 0.012405839 | 0.079865586 |
| chr18:44098498-44099299 | -0.899388781 | 0.012407706 | 0.079870897 |
| chr2:170714468-170714720 | 1.167871315 | 0.012416 | 0.07991757 |
| chr11:102794369-102794630 | -1.284371681 | 0.012427599 | 0.079980451 |
| chr6:36848695-36848962 | -1.086268202 | 0.012427857 | 0.079980451 |
| chr20:15236863-15237102 | -1.381531666 | 0.012430269 | 0.079989254 |
| chr1:114589179-114589409 | 0.985380414 | 0.012431396 | 0.079989791 |
| chr1:75053279-75053599 | -1.109023639 | 0.012435962 | 0.080012449 |
| chr6:106808024-106808259 | -1.354216419 | 0.012437664 | 0.080016683 |
| chr20:54904103-54904326 | 0.93683132 | 0.012440517 | 0.08002832 |
| chr7:16604098-16604452 | 0.970799479 | 0.012443316 | 0.080039607 |
| chr10:93435051-93435481 | -0.947541395 | 0.012451371 | 0.08008469 |
| chr2:14781230-14781538 | 1.0514726 | 0.012457045 | 0.080114463 |
| chr1:184155911-184156249 | -0.959385088 | 0.012459227 | 0.080121768 |
| chr10:26269832-26270081 | -1.267056048 | 0.012463421 | 0.080142012 |
| chr20:50776713-50776915 | 0.877522849 | 0.012469022 | 0.0801713 |
| chrX:141814999-141815221 | 1.387302423 | 0.012473395 | 0.080192691 |
| chr7:102404588-102404902 | -0.971118736 | 0.012476023 | 0.080202853 |
| chr4:49577868-49578068 | 1.391750428 | 0.012480755 | 0.080218836 |
| chr7:24820947-24821160 | 1.108492536 | 0.012480977 | 0.080218836 |
| chr8:74574225-74574517 | 1.051673307 | 0.01248165 | 0.080218836 |
| chr20:47906787-47907628 | 0.808871389 | 0.012488237 | 0.080254441 |
| chr17:13024077-13024354 | -1.044802013 | 0.012492387 | 0.080274377 |
| chr6:57162133-57162356 | 1.229547867 | 0.012496088 | 0.080291428 |
| chr20:48019976-48020431 | 0.835204579 | 0.012498403 | 0.080294816 |
| chr7:102134398-102134822 | -0.980839374 | 0.012498711 | 0.080294816 |
| chr12:99852651-99853179 | -0.934011731 | 0.012502402 | 0.080311788 |
| chr4:59892377-59892618 | 1.089980309 | 0.012510621 | 0.080357854 |
| chr11:77515517-77515884 | 1.056592658 | 0.012514368 | 0.080370788 |
| chr13:69565464-69565859 | -0.975379773 | 0.012514733 | 0.080370788 |
| chr4:16043853-16044053 | 1.269054189 | 0.012516151 | 0.080373159 |
| chr2:50344527-50344875 | -1.12887011 | 0.012519592 | 0.080382899 |
| chr19:4556077-4557008 | 0.853277414 | 0.012520719 | 0.080382899 |
| chr11:31986457-31986768 | -1.201441012 | 0.012520815 | 0.080382899 |
| chr15:74763516-74763913 | 1.023844376 | 0.012523088 | 0.080384818 |
| chr22:17299098-17299377 | 1.226535905 | 0.012523212 | 0.080384818 |
| chr2:121811328-121811623 | 1.019886763 | 0.012524542 | 0.080386617 |
| chr6:34145872-34146091 | 1.215233417 | 0.012525684 | 0.080387217 |
| chr8:120153752-120154154 | -1.079782023 | 0.012540197 | 0.080473615 |
| chr10:71658553-71658911 | 1.113020922 | 0.012543478 | 0.080487932 |
| chr9:124078779-124078979 | 1.372588836 | 0.012545214 | 0.080492327 |
| chr3:134755191-134755453 | 1.035707862 | 0.012548965 | 0.080506668 |
| chr17:80244052-80244282 | 1.20788731 | 0.01254955 | 0.080506668 |
| chr3:179441096-179441475 | -1.053084874 | 0.012551338 | 0.080509579 |
| chr5:125719721-125720313 | -0.909561633 | 0.012552105 | 0.080509579 |
| chr9:76375159-76375487 | -0.927234306 | 0.012569481 | 0.080614276 |
| chr3:11246069-11246288 | 1.211652773 | 0.012572912 | 0.080629535 |

| | | | |
|---|---|---|---|
| chr11:62908924-62909802 | 0.838563877 | 0.012575019 | 0.080636293 |
| chr8:123228982-123229373 | 1.004426923 | 0.01258275 | 0.08067912 |
| chr3:148207785-148208056 | -1.192631276 | 0.012586962 | 0.080697045 |
| chr5:76483560-76483779 | -1.253064716 | 0.012587652 | 0.080697045 |
| chr7:37584345-37584922 | 0.921504794 | 0.012593816 | 0.080727992 |
| chr4:97496230-97497005 | -0.882187563 | 0.012594587 | 0.080727992 |
| chr4:55948076-55948280 | -1.177747082 | 0.012595673 | 0.080728201 |
| chr18:497045-497245 | -1.432746078 | 0.012602128 | 0.080762821 |
| chrX:26446592-26446880 | -1.267966635 | 0.012603837 | 0.080767013 |
| chr13:23753500-23753805 | -1.106649153 | 0.012607211 | 0.08078188 |
| chr19:17419651-17420136 | 0.902693451 | 0.012613133 | 0.080813066 |
| chr4:112086192-112087109 | -0.967471036 | 0.012616015 | 0.080824106 |
| chr4:37452953-37454048 | -0.813384852 | 0.012617102 | 0.080824106 |
| chr11:69984811-69985080 | 1.138735005 | 0.01261802 | 0.080824106 |
| chr8:108673063-108673610 | 0.90075038 | 0.012619566 | 0.080827248 |
| chr17:77037143-77037357 | -1.254855188 | 0.012622413 | 0.08083557 |
| chr7:128343535-128344335 | -0.835354737 | 0.012622975 | 0.08083557 |
| chr1:165129573-165130096 | -0.952908587 | 0.012626584 | 0.080851925 |
| chr9:137611737-137612588 | 0.828449452 | 0.012629617 | 0.080861154 |
| chr1:245825210-245825425 | -1.308100227 | 0.012630136 | 0.080861154 |
| chr9:82524533-82524959 | -1.207561305 | 0.012641161 | 0.080914043 |
| chr1:27166389-27166682 | 1.281380423 | 0.012641538 | 0.080914043 |
| chrX:123906564-123906782 | 1.29509665 | 0.012642463 | 0.080914043 |
| chr3:189842362-189842570 | -1.232171929 | 0.012642621 | 0.080914043 |
| chr7:846716-846916 | 1.271905059 | 0.012651381 | 0.080963344 |
| chr12:131830152-131830779 | 0.858466644 | 0.012653702 | 0.080971432 |
| chr1:202105841-202106089 | -0.934172795 | 0.01265813 | 0.080993008 |
| chr10:38578915-38579146 | 1.378164116 | 0.012673058 | 0.081077842 |
| chr2:64362621-64362893 | -1.130479556 | 0.012673505 | 0.081077842 |
| chr19:11479557-11479820 | -1.319929863 | 0.012680666 | 0.081116883 |
| chr5:30428671-30429555 | -0.84901075 | 0.012683724 | 0.081129668 |
| chr5:120073793-120074028 | -1.315671712 | 0.012687924 | 0.081148104 |
| chr20:50772825-50773025 | 0.874237577 | 0.012688724 | 0.081148104 |
| chr8:125273812-125274223 | -0.871968807 | 0.012690289 | 0.081150816 |
| chr12:49297402-49297846 | 0.87099008 | 0.012691267 | 0.081150816 |
| chr11:67382470-67382865 | 0.948688862 | 0.01269312 | 0.081155896 |
| chr7:151062301-151063305 | -0.806507715 | 0.012698545 | 0.081183805 |
| chr4:100081590-100081902 | -1.18291299 | 0.012702404 | 0.081201704 |
| chr9:32136745-32137383 | -0.907307816 | 0.012703805 | 0.081203885 |
| chr16:70471244-70471453 | 1.234761871 | 0.012711847 | 0.081243935 |
| chr15:89171572-89171800 | 1.169178539 | 0.012712939 | 0.081243935 |
| chr10:100728973-100729376 | 0.94686067 | 0.012713252 | 0.081243935 |
| chr22:33513962-33514340 | -1.089937884 | 0.012719277 | 0.081275662 |
| chr12:76172445-76172993 | -0.977838573 | 0.012723977 | 0.081298915 |
| chr14:32939697-32940026 | 0.947138751 | 0.012726537 | 0.081304943 |
| chr20:48725516-48725870 | 0.841694089 | 0.012727481 | 0.081304943 |
| chr7:3592402-3593057 | -0.944693256 | 0.012728104 | 0.081304943 |
| chr11:96343051-96343644 | -0.883677399 | 0.012733379 | 0.081331855 |
| chr1:228735905-228736140 | 1.123069681 | 0.012740041 | 0.081367627 |
| chr5:160165123-160165518 | 1.008069519 | 0.012750903 | 0.08143021 |

| | | | |
|---|---|---|---|
| chr21:40166403-40166887 | -0.962107803 | 0.012752973 | 0.081436647 |
| chr11:71864038-71864568 | -0.990640778 | 0.012759994 | 0.081474687 |
| chr7:94003778-94004983 | -0.831961858 | 0.012768751 | 0.081523811 |
| chr2:131157716-131157991 | -1.178637209 | 0.012773801 | 0.081549257 |
| chr10:79264034-79264307 | -1.017411158 | 0.012776373 | 0.081558879 |
| chr13:73169120-73169443 | -1.111171629 | 0.01277832 | 0.081564516 |
| chr14:93160306-93160609 | 1.04290778 | 0.012781133 | 0.08156844 |
| chr5:62206227-62206633 | -1.053200223 | 0.012782414 | 0.08156844 |
| chrX:143634830-143635194 | 1.040319837 | 0.012782862 | 0.08156844 |
| chr13:23770379-23770617 | -1.252497283 | 0.012783193 | 0.08156844 |
| chr4:170464650-170464857 | -1.423377633 | 0.012792881 | 0.081623459 |
| chr15:90664959-90665188 | 1.026124248 | 0.012793954 | 0.081623512 |
| chr14:70394698-70394974 | 1.106926488 | 0.012798318 | 0.081644554 |
| chr7:13117185-13117416 | 1.142950664 | 0.01280162 | 0.081657305 |
| chr7:22082640-22083753 | 0.795341502 | 0.012802448 | 0.081657305 |
| chr13:91347040-91349445 | -0.785304783 | 0.012804609 | 0.081664285 |
| chr17:62004167-62004896 | 0.807631008 | 0.012809783 | 0.081690489 |
| chr10:8978184-8978398 | -1.249343627 | 0.012816571 | 0.081721874 |
| chr1:107688383-107688761 | -0.895063939 | 0.012817567 | 0.081721874 |
| chr11:24773320-24773794 | -0.972636991 | 0.012817905 | 0.081721874 |
| chr20:54846136-54846426 | 0.872473611 | 0.012834956 | 0.081823779 |
| chr10:119371077-119371404 | -1.105789988 | 0.012841174 | 0.081856607 |
| chr15:78951752-78951956 | 1.188704456 | 0.012843078 | 0.081861932 |
| chr8:96467152-96467427 | 1.031102514 | 0.012846221 | 0.081869346 |
| chr1:211258352-211258591 | -0.998840876 | 0.012847162 | 0.081869346 |
| chr2:53433489-53433792 | 1.221129412 | 0.012847446 | 0.081869346 |
| chr12:53046904-53048014 | 0.807293224 | 0.012851566 | 0.081886384 |
| chr20:62958856-62959094 | 1.117800019 | 0.012852512 | 0.081886384 |
| chr2:27481124-27481418 | -1.200906653 | 0.012853326 | 0.081886384 |
| chr2:209788978-209789200 | -1.421287309 | 0.012854406 | 0.081886458 |
| chr5:16631836-16632094 | 1.27211141 | 0.012857207 | 0.081897489 |
| chr7:111752208-111752521 | -1.00158345 | 0.012864604 | 0.081937797 |
| chr6:18997393-18997728 | -1.09864867 | 0.012866157 | 0.081940873 |
| chr1:40879891-40880211 | 1.210968572 | 0.012873023 | 0.081977789 |
| chr7:15546388-15546640 | 1.224248905 | 0.012879852 | 0.082014464 |
| chr14:68838397-68838681 | -1.254663147 | 0.012883533 | 0.082024591 |
| chr1:36408475-36408864 | 1.003133391 | 0.012884259 | 0.082024591 |
| chr11:79918099-79918299 | 1.422592802 | 0.012884794 | 0.082024591 |
| chr16:78235228-78235737 | 0.884802525 | 0.012885882 | 0.082024591 |
| chr2:43273550-43274225 | -0.940310892 | 0.012886795 | 0.082024591 |
| chr9:133410144-133410389 | 1.173047379 | 0.01288894 | 0.082031428 |
| chr2:31180272-31180546 | -1.329688272 | 0.012893608 | 0.08204873 |
| chr13:23451297-23451519 | 1.319317528 | 0.0128938 | 0.08204873 |
| chr2:200748026-200748602 | 0.898518246 | 0.012905943 | 0.082119179 |
| chr19:38683514-38684314 | 0.854847164 | 0.012911802 | 0.08214964 |
| chr6:40320670-40321197 | 0.996383429 | 0.012914279 | 0.082158578 |
| chr13:42364342-42364648 | -1.152891629 | 0.012932923 | 0.082267541 |
| chr1:116777868-116778101 | 1.175524174 | 0.012935407 | 0.082267541 |
| chr12:54080711-54081073 | -1.220638668 | 0.012935505 | 0.082267541 |
| chr19:2524936-2525358 | -0.984240389 | 0.012935702 | 0.082267541 |

| | | | |
|---|---|---|---|
| chr5:44028036-44028236 | 1.497906938 | 0.012939679 | 0.082286005 |
| chr2:18210248-18210656 | -0.980419705 | 0.012941038 | 0.082287816 |
| chr4:101864721-101864921 | 1.357817989 | 0.012951538 | 0.08234529 |
| chr7:24419156-24419413 | 1.110468529 | 0.012952226 | 0.08234529 |
| chr11:79063007-79063536 | -0.887403072 | 0.012966257 | 0.082427659 |
| chr3:31444559-31444772 | 1.190144774 | 0.012971567 | 0.082448325 |
| chr1:168387173-168387754 | -0.867081169 | 0.01297166 | 0.082448325 |
| chr17:55735876-55736076 | -1.390891077 | 0.012975935 | 0.082463611 |
| chr4:22029813-22030238 | -1.097487653 | 0.012976218 | 0.082463611 |
| chr16:28879599-28880170 | 0.888119925 | 0.012980679 | 0.082485119 |
| chr4:2255535-2256149 | -0.95345521 | 0.012985379 | 0.082507421 |
| chr20:23412161-23412443 | -1.159955872 | 0.012988737 | 0.082507421 |
| chr3:12987003-12987345 | 0.910486212 | 0.012989113 | 0.082507421 |
| chr15:44719747-44720024 | -1.277320004 | 0.012989526 | 0.082507421 |
| chr1:216528814-216529267 | -0.899208839 | 0.012990155 | 0.082507421 |
| chr15:90106621-90107112 | 0.861762217 | 0.012990649 | 0.082507421 |
| chr12:90828726-90828931 | -1.357066953 | 0.012998982 | 0.082553503 |
| chr14:76876049-76876363 | 0.945844142 | 0.013013555 | 0.082639203 |
| chr7:138890702-138891428 | -0.871677303 | 0.013017444 | 0.082656336 |
| chr17:45740566-45740825 | -1.180812441 | 0.013018411 | 0.082656336 |
| chr8:98036930-98037408 | 0.88929006 | 0.013019524 | 0.082656554 |
| chr8:89198326-89198531 | 1.23080068 | 0.013022383 | 0.082667857 |
| chr5:78510301-78510685 | -1.05422763 | 0.013023798 | 0.082669988 |
| chr1:168766592-168767168 | 0.937579172 | 0.013026499 | 0.082674823 |
| chr7:139108593-139110166 | -0.802347655 | 0.013026717 | 0.082674823 |
| chr7:8254634-8254952 | 0.960612037 | 0.013030032 | 0.082689012 |
| chr2:38465790-38466080 | -1.019581273 | 0.013032005 | 0.08269014 |
| chr4:163550292-163550644 | -0.947790824 | 0.013032368 | 0.08269014 |
| chr2:10659956-10660405 | 0.991432213 | 0.013042577 | 0.082748061 |
| chr7:21914999-21915217 | 1.191482294 | 0.013045937 | 0.082762525 |
| chr4:176660618-176660975 | -1.121549725 | 0.013047919 | 0.082763021 |
| chr7:124276030-124276379 | -1.012687234 | 0.013048175 | 0.082763021 |
| chr14:105649221-105649679 | 1.013409036 | 0.013052611 | 0.082784304 |
| chr14:69999804-70000131 | 1.21135358 | 0.013056433 | 0.082801691 |
| chr10:11472194-11472527 | -1.024605307 | 0.013060241 | 0.082818984 |
| chr20:47941543-47941818 | 0.858557901 | 0.013061377 | 0.082819334 |
| chr3:2452369-2452935 | -1.014942058 | 0.013066846 | 0.082838778 |
| chr14:30874111-30875033 | 0.837903157 | 0.013067378 | 0.082838778 |
| chr7:15143909-15144265 | 0.988022329 | 0.013067687 | 0.082838778 |
| chr1:114473920-114474136 | 0.959588928 | 0.013069118 | 0.082840996 |
| chr1:202988819-202989091 | 1.052989202 | 0.013077498 | 0.082887256 |
| chr5:20032910-20033110 | 1.304139519 | 0.013083269 | 0.08290614 |
| chr20:54823089-54823383 | 0.880527646 | 0.013083568 | 0.08290614 |
| chr12:53061093-53061316 | 1.013455707 | 0.013083723 | 0.08290614 |
| chr7:54483584-54484065 | -0.973682966 | 0.013089166 | 0.082933769 |
| chr8:108930299-108931094 | 0.894051813 | 0.013091949 | 0.082944547 |
| chrX:154096990-154097295 | 0.96777856 | 0.013094151 | 0.082951524 |
| chr9:34126245-34127110 | -0.851039149 | 0.013095216 | 0.082951524 |
| chr5:92222480-92222835 | -1.002290321 | 0.013097944 | 0.082961948 |
| chr15:35018663-35019368 | -0.977846354 | 0.013111776 | 0.083042694 |

| | | | |
|---|---|---|---|
| chr16:48116728-48117001 | -1.282376817 | 0.013114104 | 0.08305057 |
| chr11:101336289-101336793 | -1.13735168 | 0.013117522 | 0.083064221 |
| chr8:126736173-126736618 | 0.968064514 | 0.013118427 | 0.083064221 |
| chr13:67229504-67229843 | -0.964186159 | 0.01312069 | 0.083071681 |
| chr12:131863853-131864297 | -0.993420535 | 0.01312251 | 0.083076342 |
| chr14:40214579-40214779 | 1.381911326 | 0.013125361 | 0.083084674 |
| chr12:69254034-69254257 | -1.190369541 | 0.013126162 | 0.083084674 |
| chr5:148844689-148844967 | -1.344310231 | 0.013127079 | 0.083084674 |
| chr21:14700112-14700357 | 1.293986181 | 0.013128754 | 0.083088412 |
| chr1:3619067-3619280 | 1.218156281 | 0.013133236 | 0.08310814 |
| chr20:63610662-63610922 | 1.184768638 | 0.013134041 | 0.08310814 |
| chr8:131916199-131917011 | -0.88213194 | 0.013138893 | 0.083131976 |
| chr17:77728315-77728517 | -1.24851011 | 0.01314044 | 0.083134901 |
| chr7:150398642-150398893 | -1.33273655 | 0.013145499 | 0.083156594 |
| chr16:17037505-17037874 | 1.003247837 | 0.013146039 | 0.083156594 |
| chr1:208284885-208285124 | -1.354910257 | 0.013148036 | 0.083162359 |
| chr10:61140786-61141105 | -1.050048031 | 0.013150269 | 0.083169615 |
| chr18:63132901-63133300 | -1.013183102 | 0.013151916 | 0.083170202 |
| chr8:46029473-46029706 | 1.122892619 | 0.013152532 | 0.083170202 |
| chr5:165395938-165396264 | -1.189026423 | 0.013154081 | 0.083173133 |
| chr5:76428052-76428252 | 1.329640323 | 0.013155986 | 0.08317831 |
| chr13:92187416-92187698 | -1.142037028 | 0.01316711 | 0.083241777 |
| chr1:106713167-106713395 | 0.935884589 | 0.013168712 | 0.083245036 |
| chr13:54572595-54572927 | -1.045632092 | 0.013179127 | 0.083298701 |
| chr17:73507671-73508052 | 1.031830955 | 0.013179376 | 0.083298701 |
| chr14:77303021-77303262 | 1.050790892 | 0.01319536 | 0.083392845 |
| chr7:114088279-114088541 | -1.06813946 | 0.013201046 | 0.083421902 |
| chr14:70671403-70671637 | 1.265915348 | 0.01320815 | 0.083459912 |
| chr6:160810310-160810536 | 1.281692941 | 0.013213717 | 0.083488203 |
| chr20:54329557-54330101 | 0.81884279 | 0.013216951 | 0.08349403 |
| chr4:183443766-183444928 | -0.805065713 | 0.013217086 | 0.08349403 |
| chr17:48576793-48577077 | -1.189200572 | 0.013217908 | 0.08349403 |
| chr7:155472818-155473125 | -1.128366318 | 0.01322294 | 0.083518928 |
| chr6:146028067-146028586 | -0.928373471 | 0.013232499 | 0.083572413 |
| chr20:47233531-47233937 | 0.811305546 | 0.013235569 | 0.083584917 |
| chr8:93672295-93673276 | 0.876525315 | 0.01323887 | 0.083598868 |
| chr4:152265107-152265562 | 1.011779098 | 0.013241944 | 0.083611392 |
| chr1:221612349-221612851 | -0.960763908 | 0.01325082 | 0.08365851 |
| chr1:223628277-223628511 | 1.177264577 | 0.01325159 | 0.08365851 |
| chr4:128565621-128565856 | 1.157437374 | 0.013254025 | 0.083666988 |
| chr1:34859250-34860025 | 0.838598219 | 0.01325967 | 0.083695727 |
| chr15:49146130-49146417 | -1.22150987 | 0.013260782 | 0.083695854 |
| chr5:122746535-122746773 | -1.248583681 | 0.013264899 | 0.083714938 |
| chr2:25078637-25079062 | 1.120966269 | 0.013266643 | 0.083719052 |
| chr3:826546-826746 | 1.458057917 | 0.013268766 | 0.08372555 |
| chr2:102620390-102620724 | 0.972856836 | 0.013271848 | 0.083738102 |
| chr8:12951290-12952471 | -0.845456966 | 0.013273797 | 0.083743503 |
| chr5:43414117-43414355 | 1.28159113 | 0.013278084 | 0.083763656 |
| chr10:13299885-13300421 | 0.86732113 | 0.013285493 | 0.083803493 |
| chr1:218671546-218671983 | -0.973502521 | 0.013289725 | 0.083823291 |

| | | | |
|---|---|---|---|
| chr9:113150742-113151179 | 0.88071131 | 0.013292546 | 0.083832866 |
| chr12:101608438-101609113 | -0.859192339 | 0.013293631 | 0.083832866 |
| chr7:235640-235956 | 0.954032733 | 0.013294526 | 0.083832866 |
| chr1:196071463-196071750 | -1.067942182 | 0.013306542 | 0.083901735 |
| chr1:156678059-156678381 | 0.956427163 | 0.013307773 | 0.083902592 |
| chr8:75421720-75421920 | 1.357846814 | 0.013311273 | 0.083917755 |
| chr6:147806768-147807428 | -0.888901253 | 0.013313496 | 0.08392486 |
| chr21:44778403-44778609 | 1.295895534 | 0.013326852 | 0.083998494 |
| chr5:25105963-25106243 | -1.282831295 | 0.013327369 | 0.083998494 |
| chr18:33097016-33097411 | -1.189476086 | 0.013329156 | 0.084002842 |
| chr1:201461029-201462226 | 0.794129073 | 0.013330615 | 0.084005128 |
| chr12:50258231-50258500 | 1.185805804 | 0.013335401 | 0.084028379 |
| chr12:46392641-46393005 | -0.958821917 | 0.0133386 | 0.084041623 |
| chr4:6709422-6710105 | 0.860395457 | 0.013343172 | 0.084063518 |
| chr11:38115873-38116098 | 1.229466794 | 0.013355666 | 0.084134797 |
| chr15:96209156-96209822 | -0.874792395 | 0.013356682 | 0.084134797 |
| chr18:45359292-45359784 | -0.985391791 | 0.013362572 | 0.08416498 |
| chr20:58227715-58228048 | 0.958866268 | 0.013372282 | 0.084219217 |
| chr2:223944371-223944876 | 0.953852013 | 0.01337397 | 0.084222926 |
| chr9:78004522-78004879 | -0.99957617 | 0.013375427 | 0.084225178 |
| chr13:72180620-72180820 | -1.50487788 | 0.01337661 | 0.084225708 |
| chrX:99757750-99758229 | -0.968900514 | 0.013378005 | 0.084227567 |
| chr9:80391993-80392246 | 1.200380764 | 0.013381998 | 0.084245783 |
| chr18:45212761-45213110 | -1.045355439 | 0.013383577 | 0.084248802 |
| chr15:63473889-63474488 | -0.860420736 | 0.013392874 | 0.084297969 |
| chr8:100672784-100673031 | 1.083027212 | 0.013393588 | 0.084297969 |
| chr12:121110417-121110689 | 1.142498823 | 0.013398967 | 0.084324898 |
| chr3:133012801-133013050 | -1.417184678 | 0.013405214 | 0.084357284 |
| chr8:98705573-98705812 | 1.023904287 | 0.013407762 | 0.084366388 |
| chr8:14051510-14051781 | -1.303792783 | 0.013410445 | 0.084376342 |
| chr1:6343571-6343899 | 1.024838606 | 0.013414269 | 0.084393471 |
| chr17:12522780-12523243 | 1.033751443 | 0.013423615 | 0.08444534 |
| chr2:215274695-215275150 | -1.045171626 | 0.013424817 | 0.084445969 |
| chr7:158638117-158638421 | -1.300352219 | 0.013428411 | 0.084461641 |
| chr3:86578488-86578720 | 1.34009981 | 0.013430354 | 0.084466929 |
| chr4:177505113-177505567 | -0.980857481 | 0.01343152 | 0.084467326 |
| chr9:137881939-137882139 | 1.235842886 | 0.013436713 | 0.084492628 |
| chr1:193395491-193396578 | -0.823067407 | 0.013437749 | 0.084492628 |
| chr4:105950942-105951252 | 1.252689005 | 0.013443059 | 0.084519086 |
| chr6:13711463-13712409 | 0.80464942 | 0.013445484 | 0.084527393 |
| chr16:78133953-78134224 | 0.96877259 | 0.013447791 | 0.084534963 |
| chr6:28684208-28684510 | 1.05113483 | 0.013449604 | 0.084537375 |
| chr17:62955604-62956459 | -0.84913578 | 0.013450382 | 0.084537375 |
| chr19:49390407-49390621 | 1.303242477 | 0.013453831 | 0.084552117 |
| chr20:19935034-19935563 | -0.978358282 | 0.013455129 | 0.084553341 |
| chr20:47546649-47547008 | 0.82346758 | 0.013460339 | 0.084579147 |
| chr3:18294918-18295809 | -0.828759802 | 0.013464914 | 0.084600952 |
| chr18:2302524-2302883 | -1.319102211 | 0.013466544 | 0.084604259 |
| chr10:11916046-11916468 | -0.959342699 | 0.013472273 | 0.084633311 |
| chr8:17832070-17832347 | -1.233251374 | 0.013474874 | 0.08463522 |

| | | | |
|---|---|---|---|
| chr20:47304183-47304957 | 0.782221284 | 0.013476596 | 0.08463522 |
| chr8:124262743-124263150 | -0.907124125 | 0.013477948 | 0.08463522 |
| chr8:12436404-12436624 | -1.44679878 | 0.013477989 | 0.08463522 |
| chr2:48601359-48601906 | -0.899044042 | 0.0134781 | 0.08463522 |
| chr10:32939392-32939917 | -0.976471586 | 0.013481396 | 0.084638949 |
| chr4:13528047-13528247 | -1.455933311 | 0.013482668 | 0.084638949 |
| chr4:184537201-184538361 | -0.814687128 | 0.013482689 | 0.084638949 |
| chr3:172860738-172861076 | 1.217027086 | 0.01348361 | 0.084638949 |
| chr4:105088645-105088888 | -1.265218887 | 0.013484217 | 0.084638949 |
| chr11:64395778-64396047 | 1.209645898 | 0.013485665 | 0.084641108 |
| chr17:77508208-77508455 | 1.189561921 | 0.013499215 | 0.084708654 |
| chr15:37926202-37926483 | 1.201549076 | 0.013499566 | 0.084708654 |
| chr4:100558945-100559150 | 1.449927019 | 0.013499744 | 0.084708654 |
| chr5:87013099-87013376 | -1.16072558 | 0.013504636 | 0.084725306 |
| chr13:49495458-49496708 | -0.826995588 | 0.013505541 | 0.084725306 |
| chr13:26837910-26838363 | -0.908240919 | 0.013505715 | 0.084725306 |
| chr19:940454-941357 | -0.873290953 | 0.013514949 | 0.084776296 |
| chr13:88979903-88980134 | -1.269132554 | 0.01351992 | 0.084800534 |
| chr2:132651157-132651511 | -1.170769651 | 0.013521431 | 0.08480307 |
| chr2:67608987-67609349 | -1.321253755 | 0.013526119 | 0.084825527 |
| chr11:103036037-103036292 | -1.247226935 | 0.013529634 | 0.084831805 |
| chr7:75527418-75527751 | -1.056677926 | 0.013530031 | 0.084831805 |
| chr20:47643234-47643522 | 0.836129861 | 0.013530442 | 0.084831805 |
| chr13:84870051-84870580 | -0.919297904 | 0.013531924 | 0.084831875 |
| chr10:90921032-90921634 | 0.962854039 | 0.013533039 | 0.084831875 |
| chr18:1905722-1906097 | -1.264355931 | 0.013533774 | 0.084831875 |
| chr16:20149717-20150257 | 0.925986154 | 0.013536491 | 0.084841966 |
| chr4:98828815-98829151 | -1.106154972 | 0.013538702 | 0.084845664 |
| chr5:59548696-59549335 | -0.865980782 | 0.013539296 | 0.084845664 |
| chr10:80952785-80953160 | -1.283366409 | 0.013541401 | 0.084851915 |
| chr22:31291126-31291326 | 1.288486254 | 0.013547224 | 0.084881462 |
| chr8:1745144-1745892 | 0.905469501 | 0.013556826 | 0.084934681 |
| chr2:108823332-108823619 | 1.179575355 | 0.013558584 | 0.084938747 |
| chr17:82097264-82097622 | 0.904396447 | 0.013566632 | 0.084982214 |
| chr8:129727638-129727871 | 1.17959631 | 0.013573155 | 0.085015571 |
| chr8:99313407-99313684 | 1.113540984 | 0.013574176 | 0.085015571 |
| chr17:3959342-3959792 | 0.914013524 | 0.013578051 | 0.085032893 |
| chr3:37860385-37860728 | 0.985870768 | 0.01357943 | 0.085034578 |
| chr15:88467037-88467940 | 0.820687241 | 0.013580994 | 0.085036585 |
| chr11:76783797-76784063 | -1.009898825 | 0.01358197 | 0.085036585 |
| chrX:110238272-110238602 | -1.176993318 | 0.013589845 | 0.085072249 |
| chr13:76384160-76384431 | -1.286865638 | 0.013589887 | 0.085072249 |
| chr1:206935276-206936012 | -0.828775503 | 0.013591184 | 0.085073419 |
| chr10:28668651-28669040 | 0.925453787 | 0.013593397 | 0.085080321 |
| chr12:32036591-32036798 | -1.388660776 | 0.013602479 | 0.085130209 |
| chr16:85464936-85465252 | -0.9673478 | 0.013604449 | 0.085135582 |
| chr1:8061306-8061557 | 1.254612924 | 0.013614485 | 0.085191428 |
| chr3:182412286-182412529 | 1.40156733 | 0.01361608 | 0.085191833 |
| chrX:20266847-20267598 | -0.824086098 | 0.013616773 | 0.085191833 |
| chr1:196360820-196361725 | -0.804778047 | 0.013627319 | 0.085250853 |

| | | | |
|---|---|---|---|
| chr1:201877910-201878216 | 1.022466263 | 0.013630362 | 0.085262927 |
| chr2:144015111-144015473 | -1.120753378 | 0.013641397 | 0.08532499 |
| chr1:35545060-35545398 | 1.016774678 | 0.013644266 | 0.085335969 |
| chr7:42876518-42876770 | -1.129026618 | 0.013648079 | 0.085352851 |
| chr20:47479110-47479364 | 0.837005714 | 0.01364999 | 0.085357836 |
| chr7:725633-726040 | -0.898262253 | 0.013658858 | 0.08540632 |
| chr10:103168524-103169025 | -1.131101473 | 0.013665294 | 0.085439589 |
| chr1:195779540-195779905 | -1.235363218 | 0.013666993 | 0.085443239 |
| chr17:66818888-66819097 | 1.271746802 | 0.013670161 | 0.085454896 |
| chr14:71314568-71314768 | 1.246015844 | 0.013672405 | 0.085454896 |
| chr1:105419186-105419809 | -0.946080098 | 0.013673513 | 0.085454896 |
| chr1:45802788-45803225 | 0.942840933 | 0.013673712 | 0.085454896 |
| chr3:62935180-62935403 | 0.939900591 | 0.013674434 | 0.085454896 |
| chr17:61606979-61607214 | 0.853628741 | 0.013681812 | 0.085491518 |
| chr1:118117215-118117415 | 1.295705935 | 0.013682526 | 0.085491518 |
| chr1:109371315-109371709 | 1.051051519 | 0.013684868 | 0.085492912 |
| chr3:64965341-64965925 | 0.828483112 | 0.01368498 | 0.085492912 |
| chr2:190216864-190217338 | -0.994991882 | 0.013687092 | 0.085498038 |
| chr6:152910054-152910254 | -1.545123802 | 0.013688033 | 0.085498038 |
| chr4:186735911-186736198 | -1.085998995 | 0.013690994 | 0.085503516 |
| chrX:3550171-3550625 | -0.965860388 | 0.013691141 | 0.085503516 |
| chr12:97462657-97463207 | -0.875917887 | 0.013694173 | 0.085510905 |
| chr1:106318581-106318833 | 0.961907692 | 0.013694557 | 0.085510905 |
| chr10:22690756-22691383 | -1.020694586 | 0.0136975 | 0.085515783 |
| chr1:50333679-50333938 | 1.101252409 | 0.013697819 | 0.085515783 |
| chr4:152328640-152328923 | -1.311609089 | 0.013698686 | 0.085515783 |
| chr13:71849870-71850122 | 1.275062549 | 0.013700782 | 0.085521901 |
| chr17:10800470-10800797 | 0.983595791 | 0.013705533 | 0.085544586 |
| chr11:59261242-59261527 | -1.129615202 | 0.01370762 | 0.085549866 |
| chr10:67837957-67838354 | -0.925776741 | 0.013708612 | 0.085549866 |
| chr13:45327666-45328135 | -0.915668501 | 0.013710864 | 0.085556953 |
| chr5:168851983-168852255 | -1.269711488 | 0.0137144 | 0.085572046 |
| chr1:96473871-96474337 | -0.999336748 | 0.013715739 | 0.085573433 |
| chr10:34217520-34217904 | 1.03818053 | 0.01372414 | 0.08561888 |
| chrX:132181180-132181812 | -0.863083327 | 0.013726115 | 0.085624228 |
| chr13:28442956-28443268 | -1.163008154 | 0.013727675 | 0.08562699 |
| chr3:346176-346487 | 1.025804791 | 0.013733722 | 0.085657735 |
| chr2:155815838-155816285 | -0.92183019 | 0.013739307 | 0.085685594 |
| chr1:39701883-39702147 | 1.122227249 | 0.013745877 | 0.085719586 |
| chr20:10034691-10035495 | 0.814039961 | 0.013747734 | 0.085724195 |
| chr20:61113611-61113852 | 1.142984076 | 0.01375648 | 0.085771746 |
| chr1:197131417-197131757 | 1.030424076 | 0.013762092 | 0.085799756 |
| chr12:95649263-95649611 | -1.056057399 | 0.013767099 | 0.08582399 |
| chr11:116217039-116217369 | -1.311501323 | 0.013769227 | 0.085830273 |
| chr8:9949978-9950179 | 1.378578149 | 0.013770402 | 0.085830616 |
| chr1:154669652-154669989 | 1.029271932 | 0.013784415 | 0.085904373 |
| chr12:9814142-9814738 | -0.943034697 | 0.013784478 | 0.085904373 |
| chr5:132979562-132979814 | 1.167387704 | 0.013789005 | 0.085925597 |
| chrX:47185982-47186272 | 1.269359465 | 0.013791227 | 0.085928608 |
| chr8:101805403-101805920 | 1.01063634 | 0.013792802 | 0.085928608 |

| | | | |
|---|---|---|---|
| chr14:67868048-67868411 | 1.099109906 | 0.013792853 | 0.085928608 |
| chr1:119809026-119809289 | 1.046861979 | 0.0137972 | 0.085948706 |
| chr1:246875034-246875258 | -1.2632921 | 0.013798979 | 0.0859528 |
| chr16:11357607-11357977 | 0.907856212 | 0.013804804 | 0.085982095 |
| chr5:100904204-100904537 | -0.885207378 | 0.013806924 | 0.085985094 |
| chr10:89695290-89695559 | 1.197929891 | 0.01380753 | 0.085985094 |
| chr13:86277025-86277389 | -1.117129919 | 0.013814544 | 0.086021777 |
| chr6:110542498-110542846 | -1.025563101 | 0.013818432 | 0.086030768 |
| chr1:1059030-1059757 | 0.872939723 | 0.013819442 | 0.086030768 |
| chr11:104968463-104968762 | -1.15502903 | 0.013820012 | 0.086030768 |
| chr15:88636129-88636390 | 0.969274204 | 0.013820479 | 0.086030768 |
| chr3:66197660-66197896 | 1.293965436 | 0.013827062 | 0.086057765 |
| chr5:14307607-14307807 | 1.293965436 | 0.013827062 | 0.086057765 |
| chr10:63268844-63269504 | -0.833929088 | 0.013833766 | 0.086088587 |
| chr6:26402087-26402625 | -0.924652763 | 0.013834261 | 0.086088587 |
| chr10:104563486-104563811 | -1.241886242 | 0.013838584 | 0.086108493 |
| chr10:45308861-45309079 | -1.376438852 | 0.013844266 | 0.086135415 |
| chr10:4722091-4722321 | 1.30134043 | 0.013845366 | 0.086135415 |
| chr7:39353606-39353814 | -1.437349512 | 0.013846283 | 0.086135415 |
| chr10:76036420-76036743 | -1.070909692 | 0.013849032 | 0.086145521 |
| chr12:121812432-121812724 | 0.91597882 | 0.013854591 | 0.086160525 |
| chr2:234981364-234981674 | 1.155068653 | 0.013854596 | 0.086160525 |
| chr8:3283991-3284345 | -1.046513004 | 0.013854817 | 0.086160525 |
| chr1:244583407-244583607 | 1.371578755 | 0.013860242 | 0.08618027 |
| chr3:18782606-18782806 | 1.371578755 | 0.013860242 | 0.08618027 |
| chr21:38459572-38459772 | 1.230097466 | 0.013862787 | 0.086189099 |
| chr5:149606009-149606397 | -1.041114688 | 0.013869429 | 0.086221673 |
| chr3:107285260-107285529 | 1.184665132 | 0.013870277 | 0.086221673 |
| chr7:158828615-158830336 | -0.778585444 | 0.013871633 | 0.086223111 |
| chr3:163785723-163786249 | -0.899933997 | 0.013875055 | 0.086233091 |
| chr18:63319642-63321177 | -0.795755238 | 0.013875707 | 0.086233091 |
| chr2:156191532-156191830 | -1.260222952 | 0.013877056 | 0.086233091 |
| chr2:29748269-29748583 | -1.216607514 | 0.013877741 | 0.086233091 |
| chr16:1363512-1363798 | 1.058248309 | 0.013879478 | 0.086236895 |
| chr20:54620735-54620935 | 0.896455474 | 0.013886347 | 0.086265968 |
| chr1:234498899-234499445 | 0.892831396 | 0.013886409 | 0.086265968 |
| chrX:107655033-107655691 | -0.853437505 | 0.013888492 | 0.086271915 |
| chr7:131027260-131027497 | -1.31623663 | 0.013891576 | 0.086284071 |
| chr18:61551573-61551821 | -1.137689784 | 0.013904033 | 0.086354448 |
| chr1:192539761-192540079 | -1.04284252 | 0.013908002 | 0.086372094 |
| chr2:162333963-162334305 | -1.086386383 | 0.013918567 | 0.086427309 |
| chr12:87568994-87569232 | -1.375388957 | 0.013919148 | 0.086427309 |
| chr10:101779996-101780228 | -1.258719274 | 0.013924262 | 0.086449171 |
| chr7:49483233-49483499 | -1.212070392 | 0.013924926 | 0.086449171 |
| chr8:32038657-32038924 | -1.324414795 | 0.013928608 | 0.086457913 |
| chr2:205720882-205721603 | -0.876895317 | 0.013929115 | 0.086457913 |
| chr11:108616661-108616956 | -1.255687068 | 0.013931992 | 0.086457913 |
| chr7:55588561-55588926 | 1.227123433 | 0.013932425 | 0.086457913 |
| chr8:80874339-80875021 | 0.8841252 | 0.01393262 | 0.086457913 |
| chr3:161875533-161875782 | 1.187522265 | 0.013934733 | 0.086457913 |

| | | | |
|---|---|---|---|
| chr10:49550424-49551043 | 0.924237893 | 0.013934811 | 0.086457913 |
| chr16:88083402-88083602 | 1.309135884 | 0.013935361 | 0.086457913 |
| chr2:231222962-231223264 | 1.083669756 | 0.013940356 | 0.086481902 |
| chr16:34582109-34582515 | 0.863648904 | 0.013942823 | 0.086490201 |
| chr9:83299650-83299889 | -1.249077993 | 0.013944031 | 0.086490692 |
| chr10:114724218-114724864 | -0.92145223 | 0.013945759 | 0.086494409 |
| chr20:46699750-46700099 | 0.9942891 | 0.013950017 | 0.086509592 |
| chr20:36847091-36847354 | 1.074662006 | 0.013950465 | 0.086509592 |
| chr11:74138542-74138882 | 0.988713418 | 0.013960905 | 0.086567325 |
| chr6:4280727-4280932 | 1.425585858 | 0.013963138 | 0.086574167 |
| chr16:9053080-9053462 | 0.901200827 | 0.013969085 | 0.08660403 |
| chr1:2890396-2890678 | 1.241952912 | 0.013972686 | 0.086619345 |
| chr5:101085779-101086307 | -1.054219919 | 0.01397734 | 0.086641191 |
| chr3:8873387-8873645 | 1.190428516 | 0.013982361 | 0.086665301 |
| chr11:37811450-37811732 | -1.282598494 | 0.013985458 | 0.086677485 |
| chr8:136058686-136058898 | 1.291118272 | 0.013994089 | 0.086723961 |
| chr8:126275331-126275586 | 1.023286589 | 0.013998961 | 0.086747138 |
| chr1:181228150-181228396 | 1.058273133 | 0.014008523 | 0.086788591 |
| chr11:65429896-65430173 | 1.031423842 | 0.014008976 | 0.086788591 |
| chr8:94545661-94545954 | 1.052159795 | 0.014009048 | 0.086788591 |
| chr5:42215658-42215872 | 1.256314002 | 0.014012936 | 0.086805655 |
| chr8:134639940-134640140 | 1.255524941 | 0.014016288 | 0.086819403 |
| chr7:108525382-108526796 | -0.77444318 | 0.014017616 | 0.08682061 |
| chr11:63622375-63622751 | 1.046256271 | 0.014019752 | 0.086822771 |
| chr15:72210870-72211099 | 1.16244317 | 0.014020231 | 0.086822771 |
| chrX:115690603-115690901 | 1.064046587 | 0.014025511 | 0.086848447 |
| chr4:98839397-98839643 | -1.411655412 | 0.01403052 | 0.08686794 |
| chr12:22443914-22444128 | -1.342297894 | 0.014030926 | 0.08686794 |
| chr12:20368066-20368640 | -0.925755113 | 0.014035152 | 0.086887083 |
| chr7:23679885-23681314 | 0.783859484 | 0.014038258 | 0.086899288 |
| chr4:182784410-182785375 | -0.874712709 | 0.014040498 | 0.086906133 |
| chr18:37383112-37383365 | -1.183107747 | 0.014048905 | 0.086951147 |
| chr7:157851358-157851739 | -1.087966127 | 0.014052292 | 0.086961368 |
| chr19:17655012-17655348 | -1.095523311 | 0.014052826 | 0.086961368 |
| chr3:5787873-5788449 | -0.920555801 | 0.014055731 | 0.086972319 |
| chr13:79405508-79406882 | -0.796925237 | 0.01405754 | 0.08697649 |
| chr14:64410596-64410956 | 1.054964716 | 0.01406094 | 0.086990502 |
| chr9:112971425-112971710 | 1.084246632 | 0.01406653 | 0.08701806 |
| chr9:76916109-76916581 | -0.984823317 | 0.01407276 | 0.087049569 |
| chr1:211236687-211237158 | -1.009232263 | 0.014077558 | 0.087071363 |
| chr7:40052210-40052839 | -0.986873401 | 0.014078556 | 0.087071363 |
| chr2:138072269-138072643 | -1.082070907 | 0.01409213 | 0.087148282 |
| chr5:162324046-162324838 | -0.839832772 | 0.014095846 | 0.087164226 |
| chr2:74236824-74237047 | -1.412129716 | 0.014097523 | 0.08716756 |
| chr6:76779101-76779352 | -1.357371879 | 0.014099839 | 0.087174843 |
| chr11:42951298-42951568 | 1.087236391 | 0.014111634 | 0.08724073 |
| chr2:208489671-208490116 | -0.935901612 | 0.01411295 | 0.087241825 |
| chr2:208552884-208553160 | -1.276364399 | 0.014115979 | 0.087253516 |
| chr7:141737829-141738750 | -0.791710376 | 0.014119354 | 0.087267336 |
| chrX:40097811-40098144 | 0.940150257 | 0.014120643 | 0.087268259 |

| | | | |
|---|---|---|---|
| chr4:38060886-38061269 | -1.126824812 | 0.014125192 | 0.08728771 |
| chr5:149932737-149933059 | -1.069343194 | 0.014126068 | 0.08728771 |
| chr11:536755-537640 | -0.838421632 | 0.014128076 | 0.087293074 |
| chr18:23872712-23873177 | -0.988389828 | 0.014130628 | 0.087301804 |
| chr17:51947304-51947632 | 1.154454356 | 0.014132076 | 0.087303709 |
| chr13:55410483-55410766 | -1.333933306 | 0.014136882 | 0.08732636 |
| chr2:238883891-238884193 | 1.301647584 | 0.014139198 | 0.087333628 |
| chr18:60414232-60414447 | -1.335453921 | 0.01414055 | 0.087334936 |
| chr3:176926089-176926289 | -1.444806936 | 0.014143788 | 0.087342383 |
| chr11:32987820-32988161 | 1.092981113 | 0.01414461 | 0.087342383 |
| chr1:168084798-168085081 | -1.031002943 | 0.014145175 | 0.087342383 |
| chr20:47419828-47420466 | 0.784935047 | 0.014149046 | 0.087359245 |
| chr3:101827347-101828135 | -0.852497744 | 0.014150889 | 0.087363489 |
| chr7:34628250-34628450 | 1.289520513 | 0.014153139 | 0.087363489 |
| chr10:11875837-11876128 | -1.043486757 | 0.014153154 | 0.087363489 |
| chr2:219398838-219399247 | 0.914461359 | 0.014159762 | 0.087397234 |
| chr2:25992052-25992324 | -1.159851803 | 0.014163724 | 0.087414647 |
| chr3:4533647-4534160 | 0.921239999 | 0.014169132 | 0.087440979 |
| chr20:54777756-54778479 | 0.798316438 | 0.014173051 | 0.087456478 |
| chr4:31846718-31847161 | -1.0688017 | 0.014174773 | 0.087456478 |
| chr17:61279249-61280024 | 0.786745395 | 0.014175067 | 0.087456478 |
| chr2:167953805-167954006 | -1.338576586 | 0.014178872 | 0.087472907 |
| chr3:134473871-134474259 | 0.930102369 | 0.014180088 | 0.087473369 |
| chr3:39153514-39153867 | 0.865040473 | 0.01419578 | 0.087563121 |
| chr1:107087117-107087341 | 0.939625237 | 0.014198922 | 0.08757545 |
| chr12:51210787-51211044 | 1.184241867 | 0.014201631 | 0.087582287 |
| chr1:203847593-203847807 | 1.235182333 | 0.014203837 | 0.087582287 |
| chr2:168160756-168161314 | -0.912961061 | 0.014204062 | 0.087582287 |
| chr22:25134099-25134325 | -1.410605068 | 0.014204603 | 0.087582287 |
| chr5:119453172-119453405 | -1.057957162 | 0.014212927 | 0.087626559 |
| chr1:108211509-108211924 | 1.073088391 | 0.014215917 | 0.087637946 |
| chr1:39954961-39955231 | 1.031910155 | 0.014218936 | 0.087649504 |
| chr19:51777900-51778106 | 1.340222137 | 0.014220801 | 0.087653945 |
| chr3:127443710-127443933 | 1.222771026 | 0.01422696 | 0.087684855 |
| chr20:46814593-46815241 | 0.853718779 | 0.014228958 | 0.087690117 |
| chr5:96687909-96688183 | -1.039359282 | 0.014243884 | 0.087775041 |
| chr21:32599056-32599277 | 1.310841262 | 0.014246727 | 0.087785504 |
| chr13:60755405-60755624 | -1.274670274 | 0.014248154 | 0.087787238 |
| chr21:40923437-40923740 | -1.131693241 | 0.014252368 | 0.087801197 |
| chr7:1425693-1426049 | 0.93071177 | 0.014253282 | 0.087801197 |
| chr20:32695227-32695428 | 1.264450024 | 0.01425394 | 0.087801197 |
| chr11:67027569-67028391 | -0.812294798 | 0.014255964 | 0.087801197 |
| chr21:44498861-44499106 | 1.186562593 | 0.014256149 | 0.087801197 |
| chr5:171123207-171123407 | 1.322364214 | 0.014257748 | 0.087803986 |
| chr17:79965954-79966221 | 1.176242358 | 0.014264867 | 0.087840762 |
| chr2:150526825-150527217 | -0.997900747 | 0.014267415 | 0.087846899 |
| chr14:65164212-65164458 | 1.045026072 | 0.014269098 | 0.087846899 |
| chr13:45540246-45540503 | 1.130420209 | 0.014269303 | 0.087846899 |
| chr4:151540742-151541121 | 0.940964527 | 0.014270726 | 0.087848603 |
| chr2:191713850-191714197 | -1.082133864 | 0.014273914 | 0.087860668 |

| | | | |
|---|---|---|---|
| chr2:190417134-190417486 | -1.007285336 | 0.014274979 | 0.087860668 |
| chr7:5396546-5397806 | -0.790699421 | 0.014279158 | 0.087879328 |
| chr10:42070434-42070700 | 1.092961794 | 0.014280915 | 0.087880383 |
| chr16:2692928-2693273 | 1.082040612 | 0.014281623 | 0.087880383 |
| chr3:180295853-180296347 | -0.988646417 | 0.014287477 | 0.087909347 |
| chr12:131912626-131912885 | -1.119957401 | 0.014289155 | 0.087912611 |
| chr3:50195435-50195696 | -1.276734705 | 0.014291388 | 0.087919287 |
| chr8:107255122-107255322 | 1.263744688 | 0.014292985 | 0.087922051 |
| chr7:122971346-122971672 | -1.08664968 | 0.014296939 | 0.087930266 |
| chr20:47413008-47413210 | 0.851652976 | 0.01429699 | 0.087930266 |
| chr17:61284834-61285037 | 0.846721021 | 0.014297997 | 0.087930266 |
| chr5:111686024-111686313 | -1.063226344 | 0.01429891 | 0.087930266 |
| chr20:31914687-31914942 | 1.12226863 | 0.01430764 | 0.087970717 |
| chr14:52775348-52775667 | 1.282161369 | 0.014308036 | 0.087970717 |
| chr1:177373452-177373661 | 1.274653726 | 0.014308933 | 0.087970717 |
| chr2:71049738-71050270 | -0.926606569 | 0.014314413 | 0.087993694 |
| chr7:123534082-123535356 | -0.772318759 | 0.014314967 | 0.087993694 |
| chr11:126386660-126387090 | -0.9624014 | 0.014317792 | 0.087998544 |
| chr14:73786709-73786909 | 1.06212516 | 0.014318053 | 0.087998544 |
| chr17:66897859-66898137 | 1.019798885 | 0.014322355 | 0.088017925 |
| chr16:27867512-27867989 | 0.853769884 | 0.014326284 | 0.088035012 |
| chr2:156794182-156794463 | -1.276425175 | 0.014331956 | 0.0880628 |
| chr17:79074561-79075134 | 0.844437389 | 0.014333706 | 0.088065502 |
| chr11:17239393-17239662 | 1.082558355 | 0.014334694 | 0.088065502 |
| chr3:129673813-129674431 | 0.845752703 | 0.01433789 | 0.088068187 |
| chr7:123070758-123071058 | -1.154156627 | 0.0143392 | 0.088068187 |
| chr20:44355214-44355414 | 1.236889 | 0.014339659 | 0.088068187 |
| chr14:106565289-106565537 | 1.304683394 | 0.014340878 | 0.088068187 |
| chr15:96165749-96165986 | 1.304683394 | 0.014340878 | 0.088068187 |
| chr7:12623925-12624228 | 0.934664608 | 0.014345267 | 0.088088081 |
| chr5:96595531-96595764 | -1.17353921 | 0.014347755 | 0.088092585 |
| chr19:49403838-49404078 | 1.155465052 | 0.0143483 | 0.088092585 |
| chr13:53217258-53217530 | -1.339361759 | 0.014352937 | 0.088108766 |
| chr20:50710705-50710926 | 0.83872993 | 0.014353236 | 0.088108766 |
| chr4:87189015-87189341 | -1.076953895 | 0.014360118 | 0.088143954 |
| chr17:8345776-8346197 | 1.06012731 | 0.014364883 | 0.088166138 |
| chr11:48080567-48080894 | 1.201470156 | 0.014366686 | 0.088170139 |
| chr12:124964784-124965014 | 1.21370734 | 0.014369398 | 0.088179722 |
| chr7:100889359-100890591 | -0.789571506 | 0.014381466 | 0.088246713 |
| chr6:73180751-73181365 | -0.901742598 | 0.014390749 | 0.088292653 |
| chr1:217917505-217917826 | -0.934469332 | 0.014391258 | 0.088292653 |
| chr4:10093409-10093624 | 1.171676734 | 0.014393993 | 0.088302366 |
| chr4:164768554-164769008 | -1.072214344 | 0.014398251 | 0.088321416 |
| chr15:65201863-65202067 | 1.175493622 | 0.014400186 | 0.088326215 |
| chr20:62704467-62704861 | 0.880792264 | 0.014401973 | 0.088330106 |
| chr18:61651126-61651522 | -0.891253336 | 0.014406849 | 0.088352939 |
| chr16:8565529-8565765 | 1.135341356 | 0.014412172 | 0.088378507 |
| chr10:35614404-35614627 | -1.149898533 | 0.014425929 | 0.088452258 |
| chr3:63779222-63779514 | 0.899053148 | 0.014426508 | 0.088452258 |
| chr20:57717828-57718136 | 0.885113068 | 0.014436746 | 0.088507949 |

| | | | |
|---|---|---|---|
| chr17:41738084-41738576 | 0.927719546 | 0.014441053 | 0.088523887 |
| chr18:54357385-54358628 | -0.802681178 | 0.014442736 | 0.088523887 |
| chr2:209970294-209970501 | 1.349140275 | 0.0144434 | 0.088523887 |
| chr3:174859167-174859395 | -1.004343775 | 0.014444202 | 0.088523887 |
| chr8:47059918-47060179 | 1.231964394 | 0.014445122 | 0.088523887 |
| chr4:116180106-116180327 | -1.276047228 | 0.014448785 | 0.088539249 |
| chr2:41698835-41699286 | -1.033970428 | 0.014450246 | 0.088541125 |
| chr22:39150795-39151063 | 1.18689897 | 0.014455245 | 0.088564671 |
| chr3:167379914-167380597 | -0.801351581 | 0.014469846 | 0.088647041 |
| chr2:208040275-208040521 | -1.120965773 | 0.01447196 | 0.088652903 |
| chr14:53871122-53871322 | 1.400430997 | 0.014484508 | 0.08872034 |
| chr20:47392895-47393185 | 0.813539232 | 0.014485284 | 0.08872034 |
| chr6:164005014-164005232 | -1.478554209 | 0.014487635 | 0.088720569 |
| chr22:46638968-46639254 | 1.231185548 | 0.014488392 | 0.088720569 |
| chr11:76973415-76973638 | 1.239534228 | 0.014488795 | 0.088720569 |
| chr14:29895555-29896077 | 0.919439913 | 0.014490852 | 0.088723118 |
| chr5:162209022-162209538 | -0.908611127 | 0.014491527 | 0.088723118 |
| chr13:44243128-44243631 | -0.897406264 | 0.014495891 | 0.088742742 |
| chr8:3646683-3646883 | -1.426970542 | 0.01450181 | 0.088768233 |
| chr11:44936494-44936893 | 1.043969847 | 0.014503163 | 0.088768233 |
| chr14:104553061-104553291 | 1.109429006 | 0.014504418 | 0.088768233 |
| chr9:63823469-63824067 | 0.888197982 | 0.014504689 | 0.088768233 |
| chr14:43596680-43596986 | 1.271249338 | 0.014506197 | 0.088770369 |
| chr12:101522874-101523103 | -1.306377003 | 0.014511177 | 0.088786665 |
| chr16:62213852-62214101 | -1.306377003 | 0.014511177 | 0.088786665 |
| chr1:27659849-27660358 | 0.889424766 | 0.01451471 | 0.088792609 |
| chr6:138382521-138382737 | -1.275243146 | 0.014515124 | 0.088792609 |
| chr1:183623010-183623588 | -0.88962287 | 0.014515625 | 0.088792609 |
| chr6:26224944-26225400 | -1.036393323 | 0.0145179 | 0.088799434 |
| chrX:80405281-80405899 | -0.975348211 | 0.014531788 | 0.088877285 |
| chr13:23915333-23915681 | 1.007804989 | 0.014534898 | 0.088882433 |
| chr6:156762504-156762862 | -1.148374054 | 0.014534949 | 0.088882433 |
| chr5:2695418-2695851 | -0.912670919 | 0.014548828 | 0.088954487 |
| chr5:67214969-67215378 | -0.939189768 | 0.014549054 | 0.088954487 |
| chr15:72628350-72628550 | -1.372294103 | 0.014551296 | 0.088961094 |
| chr17:3976646-3976891 | -1.305768601 | 0.014558286 | 0.088996726 |
| chrX:29617604-29618127 | -1.119437616 | 0.014560544 | 0.089003428 |
| chr8:140149091-140149313 | 1.122052458 | 0.014562887 | 0.089010654 |
| chr7:55964228-55965214 | 0.845441431 | 0.014566193 | 0.089023756 |
| chr2:5932528-5933053 | -0.932972149 | 0.014567848 | 0.089026771 |
| chr14:73740503-73741153 | 0.835349653 | 0.014569326 | 0.089028704 |
| chr12:93020341-93020628 | 1.053458553 | 0.014576587 | 0.089064322 |
| chr3:82568942-82569142 | 1.26881986 | 0.01457748 | 0.089064322 |
| chr6:125974281-125974570 | -1.161476943 | 0.014582291 | 0.089086613 |
| chrY:11305265-11305672 | 0.996284949 | 0.01459092 | 0.089132224 |
| chr6:163423123-163423504 | -1.185418467 | 0.014594902 | 0.089145612 |
| chrX:29044898-29045098 | -1.476993297 | 0.014595439 | 0.089145612 |
| chr8:47542292-47542556 | 1.091295568 | 0.014601029 | 0.089161715 |
| chr12:69992271-69992534 | 1.283953791 | 0.01460123 | 0.089161715 |
| chr1:6245126-6245328 | 1.135802065 | 0.014601566 | 0.089161715 |

| | | | |
|---|---|---|---|
| chr5:124700975-124701457 | -1.013913201 | 0.014604072 | 0.089169909 |
| chr10:59290593-59290793 | -1.305087864 | 0.01461116 | 0.089206078 |
| chr18:336376-336688 | 1.260109326 | 0.014613333 | 0.089212236 |
| chr14:68512542-68512781 | 1.157224142 | 0.014628846 | 0.089299829 |
| chr3:51967431-51967631 | 1.304281753 | 0.014632234 | 0.089313394 |
| chr1:171696104-171696481 | -1.079400568 | 0.014635912 | 0.089328729 |
| chr10:32210336-32210620 | -1.034927063 | 0.014645586 | 0.089380651 |
| chr19:7853254-7853520 | 1.154814412 | 0.014647731 | 0.089384067 |
| chr7:33824422-33824695 | 1.032695357 | 0.014648479 | 0.089384067 |
| chr5:163261613-163261995 | -0.953245156 | 0.014652168 | 0.089399461 |
| chr7:100053258-100053562 | -1.008436323 | 0.01466764 | 0.089483744 |
| chr1:216221036-216221771 | -0.827882215 | 0.014668317 | 0.089483744 |
| chr4:68498435-68498733 | -1.066039117 | 0.01466949 | 0.089483775 |
| chr7:82699676-82700331 | -0.830858609 | 0.014675163 | 0.089507566 |
| chr20:48063466-48063739 | 0.850836177 | 0.014675727 | 0.089507566 |
| chr1:172662807-172663059 | -1.195979742 | 0.014678923 | 0.089519933 |
| chr19:50252423-50252649 | 1.171248316 | 0.014682245 | 0.089533069 |
| chr3:63179798-63180471 | 0.830266093 | 0.014686474 | 0.08955173 |
| chr3:158570571-158570771 | -1.101624547 | 0.014689899 | 0.089565482 |
| chr12:47944994-47945375 | -0.964525701 | 0.014692909 | 0.089573276 |
| chr3:174477575-174477910 | -1.154169558 | 0.014693515 | 0.089573276 |
| chr15:56619788-56619988 | 1.29895768 | 0.014699439 | 0.08960226 |
| chr19:8364562-8364762 | 1.358890106 | 0.014702594 | 0.089614364 |
| chr17:59975011-59975399 | 0.86300366 | 0.014707906 | 0.089639609 |
| chrX:119512504-119512768 | 1.07528897 | 0.01471069 | 0.089649447 |
| chr5:170473828-170474095 | 1.261866586 | 0.014714101 | 0.089656666 |
| chrX:24090797-24091054 | 1.287906201 | 0.014714215 | 0.089656666 |
| chr7:140653442-140653733 | -0.96348991 | 0.014720243 | 0.089686264 |
| chr1:39788459-39788659 | 1.227497501 | 0.014723571 | 0.089699408 |
| chr9:83544250-83544793 | -0.872825795 | 0.014728244 | 0.089720741 |
| chr9:82466985-82467216 | -1.321513085 | 0.014736222 | 0.08976221 |
| chr1:204479623-204480210 | -0.799419808 | 0.014739602 | 0.089771429 |
| chr3:20254322-20255128 | -0.841725454 | 0.014740989 | 0.089771429 |
| chr11:119156216-119156445 | 1.057520308 | 0.014742058 | 0.089771429 |
| chr2:179842279-179842817 | -1.043358964 | 0.014742422 | 0.089771429 |
| chr7:139845155-139845450 | -1.186962339 | 0.01475331 | 0.089822088 |
| chr16:84369999-84370202 | 1.02638207 | 0.014754092 | 0.089822088 |
| chr9:134299878-134300078 | 1.170940953 | 0.014754259 | 0.089822088 |
| chr11:15569717-15569966 | -1.30803942 | 0.01475666 | 0.089824142 |
| chr7:130204770-130205893 | -0.791437931 | 0.01475694 | 0.089824142 |
| chr1:114215290-114215500 | 0.910374771 | 0.01476046 | 0.089838426 |
| chr1:107378452-107379063 | 0.802328615 | 0.014764883 | 0.08985821 |
| chr2:159755816-159756137 | -0.991545457 | 0.014769127 | 0.089872805 |
| chr1:177476762-177477231 | -1.008328083 | 0.014769627 | 0.089872805 |
| chr18:22170115-22170548 | 0.926341636 | 0.014774968 | 0.089898164 |
| chr13:96104339-96105014 | 0.887630654 | 0.014779601 | 0.089915414 |
| chr12:69602984-69603255 | -1.156791913 | 0.01478015 | 0.089915414 |
| chr3:61621206-61621407 | 0.989594976 | 0.014783019 | 0.089925732 |
| chr1:114053897-114054492 | 0.80969704 | 0.014789015 | 0.089955058 |
| chr11:120815728-120816012 | 1.138706458 | 0.014797477 | 0.089993023 |

| | | | |
|---|---|---|---|
| chr15:89522233-89522454 | 1.227646487 | 0.014797605 | 0.089993023 |
| chr1:114269932-114270178 | 0.880411889 | 0.01480235 | 0.090004736 |
| chr16:3696922-3697128 | 1.243160769 | 0.014802921 | 0.090004736 |
| chr3:147930518-147930763 | -1.271615025 | 0.014804195 | 0.090004736 |
| chr7:92133628-92135388 | -0.770411106 | 0.01480423 | 0.090004736 |
| chr15:91315793-91316129 | -1.194675787 | 0.014806591 | 0.090011952 |
| chr18:54606955-54607919 | -0.862465036 | 0.014809772 | 0.090024143 |
| chr2:217396211-217396504 | 1.116471888 | 0.014815532 | 0.090052014 |
| chr1:112464456-112464951 | 0.867431521 | 0.014817697 | 0.090058027 |
| chr10:89027972-89028172 | 1.357796972 | 0.014820706 | 0.090069171 |
| chr12:14885755-14886195 | -1.024394169 | 0.014826379 | 0.090082303 |
| chr3:72636727-72636951 | 1.285646494 | 0.014826393 | 0.090082303 |
| chr3:84031629-84031829 | 1.285646494 | 0.014826393 | 0.090082303 |
| chr14:85934260-85935083 | -0.849109517 | 0.014829598 | 0.090094627 |
| chr14:104933303-104933666 | 0.892984532 | 0.014831601 | 0.090099652 |
| chr6:98633826-98634097 | -1.347656351 | 0.014840772 | 0.090148219 |
| chr20:19254995-19255337 | -0.997035199 | 0.014843689 | 0.090158791 |
| chr16:87305672-87306367 | -0.873165703 | 0.014848682 | 0.090161664 |
| chr7:2333446-2333694 | 1.143927026 | 0.01484939 | 0.090161664 |
| chr1:209937649-209938099 | -1.011242985 | 0.014849731 | 0.090161664 |
| chr1:192363243-192363516 | -1.175986694 | 0.014850142 | 0.090161664 |
| chr3:164124238-164124525 | 1.206627319 | 0.014850561 | 0.090161664 |
| chr4:85335356-85335697 | 1.008629527 | 0.014851222 | 0.090161664 |
| chr7:139604021-139604247 | -1.131036588 | 0.01485765 | 0.090186436 |
| chr2:15799422-15799681 | -1.004899585 | 0.014857657 | 0.090186436 |
| chr4:40191474-40191837 | 0.950705543 | 0.014862949 | 0.090211412 |
| chr20:41344589-41344814 | 0.899086727 | 0.014881556 | 0.090317192 |
| chr16:53433525-53433968 | 0.873947742 | 0.014882781 | 0.090317478 |
| chr7:128135634-128136380 | -0.84506611 | 0.014894846 | 0.090383534 |
| chr10:81547590-81547876 | -1.338267891 | 0.014896401 | 0.090385811 |
| chr18:56565775-56566032 | 1.137937308 | 0.014899022 | 0.090388809 |
| chr8:52483960-52484246 | -1.1859654 | 0.014900527 | 0.090388809 |
| chr7:22673044-22673779 | 0.819110366 | 0.014901543 | 0.090388809 |
| chr4:5916239-5916493 | 1.226428354 | 0.014901994 | 0.090388809 |
| chr11:95093016-95093265 | -1.175015471 | 0.014903845 | 0.090388809 |
| chr17:29369850-29370112 | 1.075195216 | 0.014903973 | 0.090388809 |
| chr9:69223570-69223857 | -1.108445246 | 0.014916285 | 0.090456317 |
| chr13:76231747-76232107 | -1.105994932 | 0.014918815 | 0.090464502 |
| chr4:91791397-91792039 | -0.898906261 | 0.014920814 | 0.090467516 |
| chr1:34360878-34361116 | 1.302197063 | 0.014921674 | 0.090467516 |
| chr6:33633709-33633925 | 1.181266845 | 0.014926957 | 0.090489276 |
| chr20:56407637-56407837 | 1.237539061 | 0.014927624 | 0.090489276 |
| chr13:46937084-46937285 | 1.099697875 | 0.014934363 | 0.090522961 |
| chr1:115665233-115666560 | -0.789858556 | 0.014938367 | 0.090540071 |
| chr5:12557077-12557336 | 1.117813831 | 0.014940368 | 0.09054352 |
| chr12:81758386-81759590 | -0.842609607 | 0.0149413 | 0.09054352 |
| chr1:232378167-232379022 | 0.850975448 | 0.014944176 | 0.090553787 |
| chr2:96950943-96951533 | -0.85681868 | 0.014946662 | 0.090561693 |
| chr21:36747202-36748098 | 0.88771104 | 0.014949534 | 0.090571931 |
| chr15:55289355-55290816 | 0.781194585 | 0.014955551 | 0.090601222 |

| | | | |
|---|---|---|---|
| chr1:213984459-213985312 | -0.869865211 | 0.014960277 | 0.090621063 |
| chr18:14178628-14179562 | 0.831170405 | 0.014961192 | 0.090621063 |
| chr1:114686330-114686579 | 0.888436462 | 0.014965363 | 0.090639161 |
| chr1:209601899-209602399 | -0.929507958 | 0.014966649 | 0.090639788 |
| chr17:77450207-77450628 | -0.992479691 | 0.014969145 | 0.090645648 |
| chr10:34025175-34025436 | -1.36093694 | 0.014969983 | 0.090645648 |
| chr2:212919295-212919604 | -1.07537676 | 0.014972294 | 0.090651608 |
| chr5:10057469-10057761 | 1.171898634 | 0.014973333 | 0.090651608 |
| chr5:133552917-133553261 | -1.103514097 | 0.01498918 | 0.090740376 |
| chr6:133952251-133952843 | -0.832180056 | 0.014998843 | 0.090791699 |
| chr2:36952798-36953029 | 1.304353248 | 0.01500113 | 0.090798374 |
| chr4:142874830-142875462 | -0.952730508 | 0.015003156 | 0.090798451 |
| chr10:29104986-29105420 | 0.901203859 | 0.015003513 | 0.090798451 |
| chr7:130989321-130989807 | -0.93093814 | 0.015009063 | 0.090819307 |
| chr12:102702646-102703035 | -1.138990304 | 0.01500933 | 0.090819307 |
| chr13:90968586-90968847 | -1.208492641 | 0.015022224 | 0.090890148 |
| chr4:173301532-173301875 | -1.158526933 | 0.015023948 | 0.0908934 |
| chr7:51162689-51163309 | -0.842759014 | 0.015034232 | 0.09094844 |
| chr15:67446700-67446974 | 1.249698953 | 0.015049382 | 0.091032897 |
| chr5:101346815-101347254 | -0.974798866 | 0.015053204 | 0.091048833 |
| chr4:177033254-177033458 | -1.360400747 | 0.015054525 | 0.091049634 |
| chr15:87500235-87500588 | -0.987495134 | 0.015073421 | 0.091156722 |
| chrX:153764011-153764707 | 0.84791497 | 0.015081125 | 0.091196115 |
| chr3:78154819-78155019 | 1.268806542 | 0.015086426 | 0.091220967 |
| chr5:68736695-68736946 | 1.322209365 | 0.015092648 | 0.091251393 |
| chr14:94458199-94458402 | 1.268405742 | 0.015101339 | 0.091296735 |
| chr3:172194740-172195038 | -1.089932399 | 0.0151095 | 0.091336541 |
| chr2:83823185-83823453 | -1.248492165 | 0.01511142 | 0.091336541 |
| chr7:22350865-22351088 | 1.127663165 | 0.0151115 | 0.091336541 |
| chr2:62724893-62725837 | -0.822642052 | 0.015113055 | 0.091338738 |
| chr6:15517805-15518005 | 1.366517728 | 0.015117693 | 0.091359561 |
| chrX:135433324-135433524 | 1.257292545 | 0.015124221 | 0.091391801 |
| chr1:173504806-173505006 | 1.268204183 | 0.015132234 | 0.091426217 |
| chr11:86967662-86967914 | -1.335222973 | 0.015132303 | 0.091426217 |
| chr1:210311092-210311625 | -0.843447299 | 0.015139093 | 0.091460034 |
| chr11:38989755-38990506 | -0.882654004 | 0.015141324 | 0.091465393 |
| chr17:15892705-15893313 | -0.925015798 | 0.015142923 | 0.091465393 |
| chr7:81244253-81244549 | -1.135966076 | 0.015143562 | 0.091465393 |
| chr7:105531922-105532752 | -0.783740714 | 0.01514909 | 0.091491573 |
| chr4:156800551-156800778 | -1.35913635 | 0.01515107 | 0.091496318 |
| chr10:50501938-50502485 | -0.863966614 | 0.015156677 | 0.091517725 |
| chr4:38179675-38180124 | -1.020603841 | 0.015157927 | 0.091517725 |
| chr20:47372599-47373417 | 0.770059257 | 0.015158784 | 0.091517725 |
| chr8:126281093-126281361 | 1.148682061 | 0.015159392 | 0.091517725 |
| chr8:47967302-47967600 | 1.085089524 | 0.015165173 | 0.091545409 |
| chr11:102869457-102869657 | 1.32698412 | 0.01516955 | 0.091564621 |
| chr6:71831672-71832610 | -0.886584088 | 0.015170942 | 0.091565804 |
| chr2:88500486-88500954 | -1.036855656 | 0.015177308 | 0.091597016 |
| chr5:157131559-157132332 | -0.845618348 | 0.015181124 | 0.091612827 |
| chr10:30186742-30186989 | -1.211513365 | 0.015183771 | 0.091621584 |

| | | | |
|---|---|---|---|
| chr2:152782132-152782442 | -1.189110848 | 0.015186653 | 0.091631763 |
| chr14:104794699-104794907 | 1.008945863 | 0.015192111 | 0.091657475 |
| chr3:64392523-64392739 | 0.8768001 | 0.015194275 | 0.091663311 |
| chr12:70719718-70720149 | -1.031810532 | 0.015207177 | 0.091733927 |
| chr8:126449774-126450002 | 1.162396915 | 0.015220533 | 0.091807265 |
| chr1:114239934-114240352 | 0.83179748 | 0.015224264 | 0.091822539 |
| chr1:155117719-155118076 | 0.977334019 | 0.015229704 | 0.091848123 |
| chr1:177612740-177612970 | 1.229254047 | 0.015231506 | 0.091851761 |
| chr14:87522498-87522902 | -0.931040338 | 0.015237176 | 0.09187872 |
| chr5:66436821-66437063 | -1.221623554 | 0.015246493 | 0.091927669 |
| chr8:140480679-140480895 | 1.251402473 | 0.015253807 | 0.091964531 |
| chr5:74624528-74624943 | -0.954052259 | 0.015257729 | 0.09198094 |
| chr12:118376171-118376575 | 0.862961261 | 0.015259413 | 0.091983854 |
| chr1:150604553-150605131 | 0.845599386 | 0.015268297 | 0.092030164 |
| chr13:20479893-20480132 | 1.316688906 | 0.015277987 | 0.092081331 |
| chr3:86388886-86389169 | -1.095205638 | 0.015279213 | 0.092081471 |
| chr19:38774145-38774365 | 1.255085111 | 0.015281325 | 0.09208696 |
| chr11:61899144-61899479 | 0.911284496 | 0.015286621 | 0.092111631 |
| chr3:57756742-57757099 | 0.96382229 | 0.015292154 | 0.092137723 |
| chr15:55993027-55994432 | 0.774512563 | 0.015293856 | 0.092140734 |
| chr1:68443100-68443471 | -0.982808467 | 0.015296973 | 0.092150768 |
| chr1:168907048-168907769 | 0.831792851 | 0.015297927 | 0.092150768 |
| chr3:156587612-156587967 | 1.091728432 | 0.015300517 | 0.09215831 |
| chr21:36385005-36385925 | 0.821194031 | 0.015301585 | 0.09215831 |
| chr7:31122705-31122905 | 1.306655025 | 0.015303722 | 0.092163937 |
| chr5:145636247-145636762 | -0.990165478 | 0.015305638 | 0.092168232 |
| chr2:5433827-5434177 | 1.100321891 | 0.015307974 | 0.092175055 |
| chr6:104825312-104825512 | -1.460494574 | 0.015315851 | 0.092210539 |
| chr3:173794001-173794362 | -0.972293551 | 0.015316274 | 0.092210539 |
| chr13:98106001-98106201 | 1.351368712 | 0.015317741 | 0.092212124 |
| chr2:199330460-199330775 | -1.238261649 | 0.015322995 | 0.092236507 |
| chr13:47146105-47146584 | 0.920226124 | 0.015324977 | 0.092241195 |
| chr20:21239166-21239480 | 1.163705453 | 0.015326494 | 0.09224308 |
| chr7:84536219-84536509 | -1.21678089 | 0.015330004 | 0.092252924 |
| chr15:40507094-40507422 | 1.000721196 | 0.015330538 | 0.092252924 |
| chr3:164649707-164649964 | -1.248552758 | 0.015334998 | 0.092268813 |
| chr4:148203600-148203821 | 1.158454119 | 0.015335587 | 0.092268813 |
| chr3:162964491-162964696 | 1.450873037 | 0.015337696 | 0.092272465 |
| chr11:82094129-82094363 | -1.273002745 | 0.015339507 | 0.092272465 |
| chr1:61076807-61077486 | -0.802875503 | 0.015339807 | 0.092272465 |
| chr1:236748533-236748801 | 1.052942377 | 0.015346826 | 0.092307441 |
| chr3:14261243-14261447 | 1.099932489 | 0.015352306 | 0.092333157 |
| chr10:3067342-3067740 | 0.885420789 | 0.015355283 | 0.092343811 |
| chr8:14404725-14405004 | -1.285424079 | 0.015362264 | 0.092378545 |
| chr17:42067321-42067741 | -1.011993338 | 0.015364995 | 0.092387714 |
| chr18:26919918-26920156 | 1.290609857 | 0.015370999 | 0.092416563 |
| chr10:60916056-60916285 | -1.324567853 | 0.01537409 | 0.092427893 |
| chr20:56277285-56277540 | 1.120062293 | 0.015379209 | 0.092451417 |
| chr2:61910627-61910885 | -1.167247659 | 0.015381266 | 0.09245653 |
| chr5:132465046-132465317 | -1.087447434 | 0.015387003 | 0.092474833 |

| | | | |
|---|---|---|---|
| chr2:20196544-20196970 | 0.948861895 | 0.015388547 | 0.092474833 |
| chr9:124011368-124011957 | 0.82680373 | 0.015388766 | 0.092474833 |
| chr3:158313421-158313743 | -1.048053461 | 0.015389139 | 0.092474833 |
| chr14:41825693-41826091 | -1.060672108 | 0.015391173 | 0.092479804 |
| chr21:33122050-33122388 | -1.148769275 | 0.015399507 | 0.092520394 |
| chr2:70108616-70109222 | -0.840749115 | 0.015400343 | 0.092520394 |
| chr17:794899-795531 | -0.974929358 | 0.015402904 | 0.092523696 |
| chr4:79141557-79141889 | 0.986334543 | 0.015403376 | 0.092523696 |
| chr17:47673437-47673690 | 1.071904231 | 0.015404516 | 0.092523696 |
| chr1:117216087-117216347 | 1.095502376 | 0.015406391 | 0.092527705 |
| chr4:16001009-16001232 | 1.208040152 | 0.015413085 | 0.092560657 |
| chr13:58869717-58869930 | -1.329823028 | 0.015419608 | 0.09259257 |
| chr1:156294799-156295009 | 1.103363506 | 0.015432091 | 0.092659063 |
| chr2:17171474-17171813 | -0.914360719 | 0.0154331 | 0.092659063 |
| chr8:13276016-13276907 | -0.838639083 | 0.015438662 | 0.092685194 |
| chr1:82290808-82291060 | -1.22243083 | 0.015450427 | 0.09274856 |
| chr1:162587286-162587723 | 0.910268208 | 0.015452968 | 0.092756546 |
| chr4:2761572-2761785 | 1.175104519 | 0.015455304 | 0.092763303 |
| chr15:98850930-98851267 | 0.853343973 | 0.015457559 | 0.092769571 |
| chr8:143156642-143156900 | 0.999237333 | 0.015461434 | 0.092785559 |
| chr21:43884515-43884833 | 1.047997553 | 0.015463538 | 0.092790918 |
| chr15:48962865-48963161 | 1.045779347 | 0.015467111 | 0.092801258 |
| chr14:54493420-54493653 | -1.147828725 | 0.015467684 | 0.092801258 |
| chr1:195710860-195711224 | -1.123117435 | 0.015470176 | 0.092804095 |
| chr19:45079010-45079383 | 0.93994162 | 0.015470579 | 0.092804095 |
| chr9:114538271-114538771 | 0.93779938 | 0.0154782 | 0.092842542 |
| chr3:156230284-156230773 | -0.960857557 | 0.015485106 | 0.092876697 |
| chr10:37276409-37276864 | -1.061994138 | 0.015491274 | 0.092906417 |
| chr7:17099565-17099830 | 1.079965566 | 0.015497725 | 0.092937832 |
| chr13:65985580-65986057 | -0.901410764 | 0.015508151 | 0.092993079 |
| chr7:135210959-135212110 | -0.777178992 | 0.015514563 | 0.093023352 |
| chr12:31659028-31659892 | -0.809989395 | 0.015515628 | 0.093023352 |
| chr1:107384985-107385637 | 0.802077506 | 0.015518646 | 0.093029579 |
| chr8:97918451-97918901 | 0.934959052 | 0.015519095 | 0.093029579 |
| chr5:82669778-82670073 | -1.258595996 | 0.015522177 | 0.093034518 |
| chr14:26553424-26553624 | 1.355232249 | 0.015522347 | 0.093034518 |
| chr17:77524512-77524712 | -1.433263772 | 0.015528743 | 0.093065572 |
| chr7:142483972-142484214 | -1.228118454 | 0.015533738 | 0.093088226 |
| chr1:204623378-204623806 | -0.850075502 | 0.015537103 | 0.093101113 |
| chr11:12087482-12087741 | -1.161560992 | 0.015539855 | 0.093110322 |
| chr2:176768182-176768619 | -0.950732172 | 0.01555324 | 0.093183234 |
| chr5:138955989-138956315 | 1.168890747 | 0.015557671 | 0.093198582 |
| chr10:14572117-14572418 | 1.08282255 | 0.015558235 | 0.093198582 |
| chr6:32609844-32610110 | -1.148395452 | 0.015560213 | 0.093203144 |
| chr3:61783950-61784208 | 0.928126374 | 0.015572505 | 0.093269481 |
| chr1:188463938-188464194 | -1.203581807 | 0.015574634 | 0.093274943 |
| chr2:162301216-162301487 | -1.108225098 | 0.015576729 | 0.093280197 |
| chr10:78691290-78691672 | -1.021542912 | 0.015578988 | 0.093283329 |
| chr4:169659921-169660717 | -0.800846513 | 0.015579687 | 0.093283329 |
| chr16:10579623-10579897 | 1.015620526 | 0.015586644 | 0.093317474 |

| | | | |
|---|---|---|---|
| chr5:176237954-176238490 | -0.991814619 | 0.015588512 | 0.093317474 |
| chr2:166424889-166425173 | -1.348716689 | 0.015589203 | 0.093317474 |
| chr10:119025334-119025632 | 0.974527407 | 0.015590261 | 0.093317474 |
| chr4:108635083-108635528 | -0.995168624 | 0.015594174 | 0.093333605 |
| chr12:29380440-29381507 | -0.782592074 | 0.015597248 | 0.093344713 |
| chr17:59482299-59482623 | 0.977996974 | 0.015598616 | 0.093345605 |
| chr10:113753148-113753348 | -1.35825155 | 0.015600538 | 0.093349821 |
| chr15:55846208-55846434 | 1.097741942 | 0.015602335 | 0.093353282 |
| chr8:103498540-103499875 | 0.769629664 | 0.015605249 | 0.093363425 |
| chr13:109275902-109276136 | -1.35399791 | 0.01561575 | 0.093418955 |
| chr2:41989836-41990151 | -1.029760453 | 0.015632057 | 0.09350921 |
| chr7:153989869-153990071 | -1.371661677 | 0.015635581 | 0.093520492 |
| chr13:45851452-45851652 | 1.233113699 | 0.015636384 | 0.093520492 |
| chr14:58465474-58465757 | 1.066066374 | 0.015638878 | 0.093528107 |
| chr10:133378789-133379111 | 1.030719168 | 0.015652432 | 0.093601859 |
| chr5:173045488-173045794 | -1.013981786 | 0.015659156 | 0.093634761 |
| chr12:45388659-45388871 | -1.326778289 | 0.015663112 | 0.093651106 |
| chr7:86939219-86939549 | -0.989883607 | 0.015665306 | 0.09365692 |
| chr21:42850822-42851127 | -0.947961922 | 0.015670436 | 0.09367169 |
| chr18:69664540-69664747 | -1.389996992 | 0.015671444 | 0.09367169 |
| chr18:67200352-67200566 | -1.389996992 | 0.015671444 | 0.09367169 |
| chr12:116910772-116911102 | 0.954512595 | 0.015676321 | 0.09369255 |
| chr6:38797527-38797872 | 1.094160039 | 0.01567738 | 0.09369255 |
| chr2:142124-142697 | -0.901296488 | 0.015678975 | 0.093693978 |
| chr20:51023062-51023370 | 0.898693468 | 0.015680069 | 0.093693978 |
| chr20:54312526-54312791 | 0.844544999 | 0.015681639 | 0.093693978 |
| chr2:179939301-179939552 | -1.24147031 | 0.015682991 | 0.093693978 |
| chr2:189521594-189522344 | -0.887106824 | 0.015683733 | 0.093693978 |
| chr4:18367054-18367333 | 1.000358639 | 0.01568961 | 0.09372178 |
| chrX:84046418-84046791 | -0.972600784 | 0.015694708 | 0.093744929 |
| chr10:78459036-78459322 | 1.156937131 | 0.015696499 | 0.093748315 |
| chr12:19862012-19862595 | -0.985509642 | 0.015706719 | 0.093802042 |
| chr10:31468231-31468588 | -1.014654156 | 0.015711325 | 0.093819081 |
| chr20:54254325-54254560 | 0.860724649 | 0.015712021 | 0.093819081 |
| chr13:54210476-54211119 | -0.806664489 | 0.015718977 | 0.093853303 |
| chr4:41909137-41909445 | -1.120419051 | 0.01572188 | 0.093863322 |
| chr3:185558445-185558731 | 1.111884053 | 0.015724751 | 0.093873151 |
| chr1:147875674-147875944 | 1.118283737 | 0.015735015 | 0.093927107 |
| chrX:36568989-36569215 | -1.37131248 | 0.015737246 | 0.093928888 |
| chr7:143901387-143902427 | -0.774155455 | 0.015737765 | 0.093928888 |
| chr11:6468990-6469474 | -0.871805206 | 0.01574323 | 0.093954187 |
| chr12:101909310-101909570 | 1.156282145 | 0.015750233 | 0.093988656 |
| chr7:14686903-14687103 | 1.223519995 | 0.015753178 | 0.093998909 |
| chr18:34659664-34659993 | -1.104656094 | 0.015757189 | 0.094015521 |
| chr22:11814668-11814953 | 1.026747714 | 0.015761108 | 0.094027465 |
| chr20:54536736-54537080 | 0.824512122 | 0.015761645 | 0.094027465 |
| chrX:119482456-119483183 | 0.841187501 | 0.015766906 | 0.094051526 |
| chr5:122493932-122494515 | -0.948919502 | 0.015771364 | 0.094061363 |
| chr8:89561332-89561550 | 1.200405448 | 0.015771458 | 0.094061363 |
| chr17:62950019-62950485 | -0.862920217 | 0.015772238 | 0.094061363 |

| | | | |
|---|---|---|---|
| chr2:204453895-204454214 | -1.083171379 | 0.015777874 | 0.094087654 |
| chr3:67384785-67385716 | -0.867222315 | 0.015780164 | 0.094093989 |
| chr7:116006261-116006583 | -1.257286648 | 0.015785942 | 0.094121112 |
| chr3:24365248-24365573 | 1.03035411 | 0.015790471 | 0.094126933 |
| chr2:47088458-47088943 | 1.036172149 | 0.015790908 | 0.094126933 |
| chr8:143571465-143572176 | -0.839015696 | 0.015791428 | 0.094126933 |
| chr7:108016806-108017119 | -1.128479793 | 0.015791832 | 0.094126933 |
| chr4:183505989-183506511 | -0.823646928 | 0.015796483 | 0.094140253 |
| chr7:38722082-38722313 | 1.249409823 | 0.015796524 | 0.094140253 |
| chr9:15824351-15824553 | -1.397114941 | 0.015799171 | 0.094148709 |
| chr13:25198518-25198836 | 1.147747073 | 0.015805834 | 0.09418109 |
| chr1:19762304-19762553 | -1.272377657 | 0.015807683 | 0.094184779 |
| chr4:159947070-159947290 | -1.291973722 | 0.015813405 | 0.094211545 |
| chr20:49831147-49831362 | 1.178131247 | 0.015816058 | 0.09421658 |
| chr5:90084205-90084673 | -1.066036496 | 0.015816709 | 0.09421658 |
| chr14:34203475-34203675 | 1.203389297 | 0.015828681 | 0.094272099 |
| chr4:86168429-86168731 | -1.125428249 | 0.015828897 | 0.094272099 |
| chr6:32439557-32439964 | -1.015001081 | 0.015830095 | 0.094272099 |
| chr17:76880605-76880840 | 1.152844348 | 0.015830951 | 0.094272099 |
| chrX:110401968-110402994 | -0.830551649 | 0.015834377 | 0.094285175 |
| chr8:118565297-118565582 | -1.104317206 | 0.015839318 | 0.094307268 |
| chr5:73596760-73596996 | -1.324202546 | 0.015840616 | 0.094307665 |
| chr7:55144363-55144740 | -0.959759987 | 0.015843069 | 0.094314946 |
| chr7:22924538-22924738 | 1.252919967 | 0.015851339 | 0.094356845 |
| chr2:129056302-129056789 | -0.912921327 | 0.015856894 | 0.094382579 |
| chr4:68504859-68505250 | -0.921250727 | 0.015860066 | 0.094394127 |
| chr2:58447992-58448438 | -0.86293835 | 0.01586408 | 0.094410681 |
| chr1:106628021-106628269 | 0.86204993 | 0.01587781 | 0.094485057 |
| chr2:177552415-177553635 | -0.783661792 | 0.015879234 | 0.09448619 |
| chr2:158010360-158010605 | -1.214986729 | 0.015881576 | 0.094487218 |
| chr4:6664639-6664896 | 1.037408236 | 0.015882309 | 0.094487218 |
| chr5:775659-776140 | 0.883419096 | 0.015883106 | 0.094487218 |
| chr22:37560110-37560607 | -0.964499861 | 0.01588831 | 0.0945041 |
| chr20:47441161-47441620 | 0.784499791 | 0.015888411 | 0.0945041 |
| chr5:135566229-135566472 | -1.103267194 | 0.015889659 | 0.094504186 |
| chr12:130701788-130702009 | 1.321551676 | 0.015899352 | 0.09454888 |
| chr21:37220670-37221545 | 0.808262422 | 0.015899641 | 0.09454888 |
| chr7:154720143-154720548 | -1.126483221 | 0.015903307 | 0.094563338 |
| chr20:46752689-46752997 | 0.935757395 | 0.015913282 | 0.094615309 |
| chr5:43997919-43998189 | 1.064504484 | 0.015916228 | 0.094624801 |
| chr4:173025119-173025597 | -0.855385921 | 0.015917348 | 0.094624801 |
| chr12:120640450-120640687 | 1.134142006 | 0.015920298 | 0.094634995 |
| chr9:77589358-77589877 | -0.830708239 | 0.015926746 | 0.094665979 |
| chr4:173371852-173372112 | -0.910451034 | 0.015928268 | 0.094667682 |
| chr1:83882853-83883512 | -0.852233657 | 0.015932313 | 0.094684379 |
| chr1:246413423-246413684 | 1.285802187 | 0.015946353 | 0.094753694 |
| chr7:128869117-128869845 | -0.831654248 | 0.01594645 | 0.094753694 |
| chr5:4644158-4644512 | 1.154290864 | 0.015948855 | 0.094760639 |
| chr22:23415534-23415734 | 1.247669864 | 0.015951033 | 0.094766229 |
| chr3:912813-913120 | 1.058398759 | 0.015952939 | 0.094770204 |

| | | | |
|---|---|---|---|
| chr11:27166232-27166495 | -1.197788925 | 0.015958532 | 0.094796082 |
| chr2:62605100-62605883 | -0.911995671 | 0.015962672 | 0.094813324 |
| chr12:14886375-14886681 | -1.076182031 | 0.015976554 | 0.094888423 |
| chr12:15801814-15802064 | -1.316723314 | 0.015991723 | 0.09497115 |
| chr10:117239650-117239883 | 1.287145383 | 0.015996847 | 0.094994217 |
| chr7:130278070-130278306 | -1.205030409 | 0.015999748 | 0.095004086 |
| chr7:149209617-149209863 | -1.06556643 | 0.016003062 | 0.095011666 |
| chr5:137822161-137822501 | -1.029499299 | 0.016003505 | 0.095011666 |
| chr5:93579968-93580393 | -0.93135884 | 0.016005869 | 0.095018339 |
| chr14:34564011-34564594 | 0.927041856 | 0.016016528 | 0.095072549 |
| chr5:128436771-128437126 | -1.115591522 | 0.016017482 | 0.095072549 |
| chr6:123380311-123380538 | -1.231662036 | 0.016027216 | 0.095122955 |
| chr2:32937951-32938399 | -0.855362944 | 0.016033466 | 0.095152678 |
| chr7:94395458-94395736 | -1.047179253 | 0.016034995 | 0.095154385 |
| chr1:33220922-33221328 | -0.977552395 | 0.016040163 | 0.095172394 |
| chr19:17260908-17261190 | 0.993836225 | 0.016040514 | 0.095172394 |
| chr4:179867296-179867754 | -0.880247122 | 0.016045984 | 0.095197476 |
| chr8:95192967-95193406 | 0.829514602 | 0.016053098 | 0.095232307 |
| chr15:61750278-61750993 | 0.811261407 | 0.016056295 | 0.095243898 |
| chr13:108902680-108903075 | -1.158306696 | 0.016065409 | 0.095284968 |
| chr20:21518461-21518693 | 1.075758716 | 0.016066971 | 0.095284968 |
| chr17:17783297-17783638 | -0.944284611 | 0.016067068 | 0.095284968 |
| chr19:44304934-44305201 | 1.079651168 | 0.016068193 | 0.095284968 |
| chr3:53708368-53708597 | -1.354293868 | 0.016074172 | 0.095310211 |
| chr2:34353810-34354086 | -1.178322633 | 0.016074938 | 0.095310211 |
| chr6:166747742-166748185 | -1.140533485 | 0.016076558 | 0.095312443 |
| chr1:180726013-180726436 | -1.051350543 | 0.016080048 | 0.09532576 |
| chr2:66391017-66391449 | -1.019775522 | 0.016082407 | 0.095329253 |
| chr20:48089355-48089678 | 0.826180204 | 0.016083126 | 0.095329253 |
| chr13:82323177-82323435 | -1.076053512 | 0.016085629 | 0.095336714 |
| chr6:44312845-44313051 | 1.192108712 | 0.016089544 | 0.095352544 |
| chr6:152618448-152618798 | -1.281534038 | 0.016094958 | 0.095377251 |
| chr10:125035129-125035340 | 1.26481863 | 0.016098537 | 0.095391085 |
| chr8:142911992-142912242 | 1.060822386 | 0.016109494 | 0.095448627 |
| chr1:114531432-114531673 | 0.868285605 | 0.016113676 | 0.095466026 |
| chr2:8934467-8934771 | -1.223056383 | 0.016115258 | 0.095468014 |
| chr1:114735605-114736012 | 0.857717883 | 0.016116678 | 0.095469046 |
| chr6:135227508-135227755 | -1.300533797 | 0.016121766 | 0.095491802 |
| chr2:193103069-193103417 | -1.102857288 | 0.016138965 | 0.095586286 |
| chr16:90047429-90048088 | -0.8629172 | 0.016147573 | 0.095625977 |
| chr1:106563995-106564195 | 0.912068857 | 0.016148163 | 0.095625977 |
| chr2:35410292-35410727 | -1.049289841 | 0.016156647 | 0.095668827 |
| chrX:153793876-153794770 | 0.793130992 | 0.016166799 | 0.095721542 |
| chr7:134420409-134420885 | -0.912628559 | 0.016172759 | 0.095749434 |
| chr11:606159-606636 | -0.944986813 | 0.016176103 | 0.095761832 |
| chr13:48532330-48533228 | -0.784369412 | 0.016185354 | 0.095797647 |
| chr8:98218681-98218902 | 1.184120836 | 0.01618606 | 0.095797647 |
| chr8:95366382-95366733 | 1.003531192 | 0.016186864 | 0.095797647 |
| chr2:31489225-31489748 | -0.870530362 | 0.016187154 | 0.095797647 |
| chr14:30610582-30611302 | 0.836665472 | 0.016191021 | 0.09581313 |

| | | | |
|---|---|---|---|
| chr5:178081608-178082087 | -0.821292657 | 0.016192656 | 0.095815406 |
| chr3:156475143-156475369 | -1.17343095 | 0.016200772 | 0.095856025 |
| chr4:128517728-128518047 | -1.085542939 | 0.01621207 | 0.095914161 |
| chr1:53919076-53919382 | -1.103849552 | 0.016213101 | 0.095914161 |
| chr7:87218962-87219214 | -1.012504572 | 0.016214986 | 0.095917908 |
| chr14:64704677-64705508 | 0.80997781 | 0.016216574 | 0.095919896 |
| chr2:202339560-202339945 | -0.946395218 | 0.016223611 | 0.095954111 |
| chr10:46092446-46092667 | 1.227485258 | 0.016230613 | 0.095988118 |
| chr1:100894653-100895189 | -0.813578089 | 0.016240888 | 0.096041472 |
| chr2:18753060-18753382 | -1.078860258 | 0.016242595 | 0.096044149 |
| chr1:200714689-200714950 | -1.056880377 | 0.016250555 | 0.096083807 |
| chr1:246963992-246964216 | 1.156754965 | 0.016252769 | 0.096089478 |
| chr3:151107938-151108258 | 1.091982554 | 0.016261804 | 0.096135477 |
| chr3:107999176-107999975 | -0.804094984 | 0.016273403 | 0.096196626 |
| chr5:95365730-95366057 | 1.019113601 | 0.016299782 | 0.096337841 |
| chr19:9768503-9769081 | 0.874730535 | 0.016299806 | 0.096337841 |
| chr3:129514727-129514983 | 1.043615572 | 0.016313689 | 0.09641128 |
| chr6:127879658-127880056 | -0.990898288 | 0.016314748 | 0.09641128 |
| chr4:44157581-44158319 | 0.845993257 | 0.016316854 | 0.096416284 |
| chr10:6144524-6145462 | 0.785822063 | 0.016318985 | 0.096421445 |
| chrX:115030550-115031188 | 0.882338203 | 0.016327272 | 0.096462967 |
| chr2:241762928-241763386 | -0.96734177 | 0.016330714 | 0.096475865 |
| chr9:40991654-40992780 | -0.784906959 | 0.016332708 | 0.096480208 |
| chr6:42429313-42430191 | -0.881305462 | 0.016335337 | 0.096488297 |
| chr11:12376529-12376729 | 1.283801136 | 0.01634007 | 0.096508053 |
| chr1:65170499-65170854 | 0.962263521 | 0.016342266 | 0.096508053 |
| chr2:25967069-25967287 | 1.204469375 | 0.01634246 | 0.096508053 |
| chr4:176001540-176002614 | -0.824282894 | 0.016346717 | 0.096525751 |
| chr4:156558187-156559036 | -0.84607762 | 0.016350791 | 0.096536181 |
| chr17:76789024-76789400 | 1.056202364 | 0.016351996 | 0.096536181 |
| chr1:200229258-200229458 | 1.200530196 | 0.016352263 | 0.096536181 |
| chr1:222622815-222623065 | 1.147692202 | 0.016357356 | 0.096558809 |
| chr2:227415597-227415847 | -1.190275389 | 0.016360991 | 0.096572826 |
| chrX:129655612-129655843 | 1.038546424 | 0.01637218 | 0.096631423 |
| chr8:100239052-100239281 | 1.243517699 | 0.016374052 | 0.096635033 |
| chr9:94094496-94094914 | -1.152051174 | 0.016385133 | 0.096688893 |
| chr3:193834595-193834879 | -1.029904533 | 0.016385702 | 0.096688893 |
| chr19:11060787-11061048 | 1.208114318 | 0.016391806 | 0.096717463 |
| chr19:57799178-57799385 | -1.317355993 | 0.016398789 | 0.096751216 |
| chr10:5148285-5148498 | 1.265658299 | 0.016403342 | 0.096769334 |
| chr13:29902360-29902623 | -1.17762442 | 0.016404386 | 0.096769334 |
| chr5:25119657-25120023 | -0.99066469 | 0.016405988 | 0.096769484 |
| chr14:90103945-90104192 | 1.102492925 | 0.016407025 | 0.096769484 |
| chr5:143186029-143186250 | -1.208803748 | 0.016409305 | 0.096769484 |
| chr3:102673546-102674043 | -0.911820863 | 0.016409463 | 0.096769484 |
| chrX:155026518-155027537 | 0.78014531 | 0.016411902 | 0.096776415 |
| chr3:136804038-136804692 | 0.825264456 | 0.016415722 | 0.096791492 |
| chr6:78460800-78461098 | 1.072305927 | 0.016418039 | 0.096797704 |
| chr20:9912948-9913224 | -1.070291373 | 0.016423726 | 0.096823785 |
| chr14:55705624-55706578 | 0.806877341 | 0.016425341 | 0.096825853 |

| | | | |
|---|---|---|---|
| chr15:71252392-71252733 | 1.011108067 | 0.016435398 | 0.096877691 |
| chr7:134676773-134677091 | -1.293231492 | 0.016437108 | 0.096880313 |
| chr1:173489191-173489501 | 0.948054643 | 0.016441913 | 0.096901183 |
| chr2:196526887-196527647 | -0.829211534 | 0.016447889 | 0.09692339 |
| chr17:42554500-42555248 | 0.845827316 | 0.016448211 | 0.09692339 |
| chr8:123922025-123922798 | -0.815967842 | 0.016461226 | 0.096992624 |
| chr8:89572957-89573260 | 1.080036615 | 0.016467826 | 0.097024052 |
| chr15:91430735-91431889 | 0.786745159 | 0.016470865 | 0.09703449 |
| chr2:557863-558325 | -0.998275985 | 0.016484731 | 0.097108713 |
| chr2:60656051-60656251 | -1.287784975 | 0.016494033 | 0.097152505 |
| chr13:71390058-71390533 | -0.993741577 | 0.0164947 | 0.097152505 |
| chr2:13223997-13224259 | -1.250535468 | 0.016499364 | 0.097172503 |
| chr7:79317690-79317994 | -1.053449366 | 0.016503821 | 0.097191284 |
| chr7:17405860-17406194 | 1.010647322 | 0.016508246 | 0.097209868 |
| chr17:78359879-78361091 | 0.7751401 | 0.01651487 | 0.097241403 |
| chr19:8362023-8362423 | 0.943070451 | 0.016519346 | 0.097260285 |
| chr12:54399352-54399610 | 1.010479559 | 0.016521843 | 0.09726751 |
| chr7:98315656-98316047 | -0.981134518 | 0.016535566 | 0.097340824 |
| chr9:95810469-95810700 | 1.232192968 | 0.016537876 | 0.097346941 |
| chr8:128108189-128108427 | -0.951184367 | 0.016539669 | 0.097350015 |
| chr5:114803403-114803680 | -1.132027215 | 0.016543607 | 0.097361774 |
| chr11:78961212-78961532 | -0.956549279 | 0.016544208 | 0.097361774 |
| chr18:76874882-76875660 | -0.914084636 | 0.016551038 | 0.097394486 |
| chr2:24421804-24422009 | -1.287075721 | 0.016554879 | 0.09740961 |
| chr13:33497444-33497644 | -1.419727696 | 0.016558351 | 0.097422559 |
| chr10:48555533-48555813 | -1.24344547 | 0.016562114 | 0.097437217 |
| chr11:3747684-3747985 | -1.114895091 | 0.016563711 | 0.097439128 |
| chr7:99033965-99034292 | 1.13616787 | 0.016566668 | 0.097441864 |
| chr8:128766236-128766526 | 1.168354622 | 0.016566719 | 0.097441864 |
| chr4:186712765-186712978 | -1.305464131 | 0.016576229 | 0.097490317 |
| chr7:73979204-73979444 | 1.095998278 | 0.016578482 | 0.097496081 |
| chr10:11716062-11716422 | -1.004116866 | 0.016584776 | 0.097519218 |
| chr1:78544570-78544872 | -1.140744818 | 0.016584962 | 0.097519218 |
| chrX:92690362-92690565 | -1.451391548 | 0.016588046 | 0.097524323 |
| chr10:54369058-54369320 | -1.124023217 | 0.016588375 | 0.097524323 |
| chr17:21705179-21705419 | 1.244804757 | 0.016592818 | 0.097542955 |
| chr21:36297991-36298553 | 0.905250787 | 0.016598418 | 0.097562002 |
| chr17:15910913-15911155 | 1.150100526 | 0.016598604 | 0.097562002 |
| chr14:56470229-56470429 | 1.30916988 | 0.016600166 | 0.097563699 |
| chr20:48945701-48946418 | 0.810544792 | 0.016604312 | 0.097580579 |
| chr17:41655339-41656156 | -0.834707651 | 0.016608772 | 0.097590598 |
| chr7:132004453-132004925 | -0.996819432 | 0.016609333 | 0.097590598 |
| chr3:187319723-187320026 | -0.969971716 | 0.01660993 | 0.097590598 |
| chr5:145719069-145719293 | -1.370892533 | 0.016611111 | 0.097590598 |
| chr3:194089703-194090011 | -1.109431108 | 0.016616927 | 0.097617281 |
| chr7:22442183-22442749 | 0.852711518 | 0.016627195 | 0.097670114 |
| chrX:36961670-36961925 | -1.121843676 | 0.016629033 | 0.097673422 |
| chr6:41528816-41529016 | 1.284898045 | 0.016631291 | 0.097679195 |
| chr8:127933836-127934036 | 1.093053508 | 0.016637848 | 0.097704619 |
| chrX:31279301-31279626 | -1.012371618 | 0.016638355 | 0.097704619 |

| | | | |
|---|---|---|---|
| chr6:6724374-6725507 | -0.803501316 | 0.016639445 | 0.097704619 |
| chr14:105537690-105538375 | 0.832155918 | 0.016646195 | 0.097736764 |
| chr8:92335440-92335921 | 0.934652342 | 0.016649476 | 0.097748538 |
| chr6:9481862-9482230 | -1.303752932 | 0.016653657 | 0.097765593 |
| chr14:46761046-46761446 | 1.067659516 | 0.016658887 | 0.097788804 |
| chr13:97256463-97256871 | -0.995550847 | 0.01667247 | 0.097861041 |
| chr10:7828200-7828440 | 1.20972529 | 0.016676721 | 0.097878491 |
| chr14:87049914-87050761 | -0.81088894 | 0.016678311 | 0.097880324 |
| chr12:53295343-53295736 | 0.859172803 | 0.016685586 | 0.097915523 |
| chr5:160649383-160649741 | -1.082178884 | 0.016692078 | 0.097946116 |
| chr10:118332406-118332629 | -1.312562222 | 0.016698176 | 0.097974399 |
| chr13:20165892-20166092 | 1.209293697 | 0.016714873 | 0.098064855 |
| chr5:102295737-102297152 | -0.762849391 | 0.016716173 | 0.098064975 |
| chr12:72639912-72640426 | -0.999152359 | 0.016717516 | 0.098065346 |
| chr17:64471397-64471722 | 0.996421518 | 0.016723043 | 0.098086126 |
| chr9:128250150-128250741 | 0.822561074 | 0.016724505 | 0.098086126 |
| chr2:47128315-47128707 | -1.069137441 | 0.016725707 | 0.098086126 |
| chr16:22436598-22437730 | 0.813271872 | 0.016726179 | 0.098086126 |
| chr8:81111368-81112545 | 0.784308317 | 0.016729058 | 0.0980955 |
| chr18:23282797-23283090 | 1.155642794 | 0.016732064 | 0.098105617 |
| chr21:39468267-39468476 | 1.121531729 | 0.016736201 | 0.098122367 |
| chr11:129694515-129694930 | -0.953556152 | 0.016741706 | 0.098146767 |
| chr4:169009435-169010976 | -0.760179917 | 0.016742925 | 0.098146767 |
| chr20:41441274-41441644 | 0.986905331 | 0.016747378 | 0.098161791 |
| chr11:132312688-132312972 | -1.173960103 | 0.016748897 | 0.098161791 |
| chr11:75422035-75422652 | 0.816834338 | 0.016749331 | 0.098161791 |
| chr9:119020440-119020694 | -1.262993519 | 0.016759118 | 0.098211635 |
| chr1:247653052-247653289 | -1.381699442 | 0.01676215 | 0.098219076 |
| chr14:52700729-52700929 | 1.158752531 | 0.016762951 | 0.098219076 |
| chr20:3716242-3716671 | 0.994067237 | 0.016766467 | 0.098226025 |
| chr2:61917656-61918139 | -0.983436269 | 0.016766701 | 0.098226025 |
| chr17:49267549-49268078 | 0.894864923 | 0.016770116 | 0.098237646 |
| chr13:31524380-31524622 | 1.225743354 | 0.016771249 | 0.098237646 |
| chr5:26415803-26416075 | 1.196405886 | 0.016775259 | 0.098246069 |
| chr2:218699005-218699310 | 1.142591819 | 0.016775926 | 0.098246069 |
| chr2:173943706-173944166 | 0.83831281 | 0.016777357 | 0.098246069 |
| chr8:97172280-97172499 | 1.23351719 | 0.016778137 | 0.098246069 |
| chr3:9503287-9503548 | 0.937598863 | 0.016779098 | 0.098246069 |
| chr10:27483075-27483328 | -1.156462845 | 0.016781226 | 0.098251018 |
| chr2:81283027-81283260 | -1.303987535 | 0.016784856 | 0.098260847 |
| chr8:104316321-104316558 | 1.094235417 | 0.016785469 | 0.098260847 |
| chr16:4142267-4142776 | -0.924909454 | 0.016789789 | 0.098278623 |
| chr7:8607811-8608090 | 1.018344567 | 0.01679651 | 0.098307871 |
| chr4:177548106-177548306 | -1.375853105 | 0.016797351 | 0.098307871 |
| chr6:164013800-164014209 | -0.999456848 | 0.01680711 | 0.098357471 |
| chr11:112355753-112355986 | -1.311221016 | 0.016812897 | 0.098369465 |
| chr19:19532131-19532572 | -1.083368869 | 0.016813488 | 0.098369465 |
| chr3:142866818-142867360 | -0.797252239 | 0.016813707 | 0.098369465 |
| chrX:154805386-154805654 | 1.031123167 | 0.016814863 | 0.098369465 |
| chr8:101424231-101424452 | 1.125589492 | 0.016815579 | 0.098369465 |

| | | | |
|---|---|---|---|
| chr5:175809709-175810065 | 0.888087197 | 0.016827268 | 0.098430332 |
| chr17:82592128-82592641 | 0.884899686 | 0.016830024 | 0.098438937 |
| chr11:47185248-47186006 | 0.810781333 | 0.016833438 | 0.098451392 |
| chr7:157289297-157290097 | -0.7842024 | 0.016838291 | 0.098469591 |
| chr17:14785063-14785313 | -1.132397222 | 0.01683912 | 0.098469591 |
| chr1:151858262-151858483 | 0.984037398 | 0.016860653 | 0.098577509 |
| chr2:102142512-102143100 | 0.898153364 | 0.016861748 | 0.098577509 |
| chr1:114218642-114219128 | 0.796338538 | 0.016862523 | 0.098577509 |
| chr9:10268850-10269111 | -1.119482206 | 0.016862764 | 0.098577509 |
| chr9:336063-336385 | 0.998582829 | 0.016864008 | 0.098577509 |
| chr14:54566596-54567127 | 0.832451525 | 0.016866541 | 0.098584797 |
| chr7:12941550-12942041 | 0.86165869 | 0.016874157 | 0.098615465 |
| chr12:53451701-53452348 | 0.792051393 | 0.016874362 | 0.098615465 |
| chr9:127904982-127905308 | 1.077662369 | 0.016879857 | 0.098640055 |
| chr17:44376151-44376412 | -1.108440619 | 0.016888159 | 0.098671365 |
| chr7:106042847-106043133 | -1.057164438 | 0.016888979 | 0.098671365 |
| chr15:57505885-57506644 | 0.79985031 | 0.016889078 | 0.098671365 |
| chr9:81587569-81587824 | 1.225111066 | 0.01689163 | 0.098674386 |
| chr2:179020199-179020528 | -0.993914134 | 0.016892248 | 0.098674386 |
| chr20:58162075-58162378 | 0.962566149 | 0.016893513 | 0.098674386 |
| chr8:96160845-96161045 | 1.132708609 | 0.016894746 | 0.098674386 |
| chr5:115875432-115875740 | -1.222147217 | 0.016897199 | 0.098675739 |
| chr7:148501021-148501232 | -1.215409399 | 0.016897554 | 0.098675739 |
| chr1:15886285-15886720 | -1.012645567 | 0.016915999 | 0.098775922 |
| chr7:89718962-89719436 | -0.886247603 | 0.016918652 | 0.098783889 |
| chr13:35988023-35988964 | -0.795088112 | 0.016924031 | 0.098807767 |
| chr20:47620601-47620823 | 0.850343779 | 0.016929315 | 0.098813601 |
| chr13:91417595-91417803 | -1.379648684 | 0.016930035 | 0.098813601 |
| chr19:47790495-47790748 | 1.098946378 | 0.016930575 | 0.098813601 |
| chr4:3532317-3532907 | 0.837829871 | 0.016931168 | 0.098813601 |
| chr14:44803325-44803594 | 0.983290075 | 0.016931758 | 0.098813601 |
| chr1:110215059-110215306 | -1.090037648 | 0.016933116 | 0.098813601 |
| chr5:150616038-150616336 | -1.004474445 | 0.016934058 | 0.098813601 |
| chr1:108709030-108709458 | -0.896232797 | 0.016939557 | 0.098831586 |
| chr2:86337897-86338235 | 0.966736019 | 0.01693972 | 0.098831586 |
| chrX:78024345-78025079 | -0.836991371 | 0.016944802 | 0.098848635 |
| chrX:102934645-102934951 | 1.022480211 | 0.016945222 | 0.098848635 |
| chr20:56233892-56234256 | 1.032245062 | 0.01696133 | 0.098935069 |
| chr4:133081255-133081512 | -1.220908573 | 0.01696456 | 0.098946376 |
| chr12:64931509-64931709 | 1.232011941 | 0.016975735 | 0.099004016 |
| chrX:135421497-135422169 | 0.832729108 | 0.016987483 | 0.099064992 |
| chr18:61635870-61636579 | -0.848653442 | 0.016989694 | 0.099070346 |
| chr9:130908505-130908726 | 1.179254995 | 0.016995045 | 0.09909199 |
| chr12:33108212-33108412 | 1.281329634 | 0.016997024 | 0.09909199 |
| chr7:30134561-30135374 | 0.803945202 | 0.016997286 | 0.09909199 |
| chr2:85471051-85471358 | -1.189317632 | 0.017002283 | 0.099113583 |
| chr15:66017076-66017276 | 1.201385226 | 0.017007465 | 0.09913625 |
| chr6:88730624-88731069 | -0.92820506 | 0.017013775 | 0.099159574 |
| chr10:73358206-73358997 | -0.806922584 | 0.017014817 | 0.099159574 |
| chr7:155412520-155413046 | -0.878582816 | 0.017015349 | 0.099159574 |

| | | | |
|---|---|---|---|
| chr7:127620890-127621090 | -1.270451727 | 0.017019787 | 0.099177892 |
| chr18:35507764-35508284 | 0.908005483 | 0.017021721 | 0.099181619 |
| chr2:30915579-30915838 | -1.155123832 | 0.017023994 | 0.099187324 |
| chr4:2469410-2469652 | 1.055656753 | 0.017025502 | 0.099188565 |
| chr12:104358414-104358685 | -1.023041894 | 0.017026919 | 0.099189281 |
| chr20:47689301-47689510 | 0.82118043 | 0.017028594 | 0.099191493 |
| chr4:134698839-134699045 | -1.360911896 | 0.017034598 | 0.09921641 |
| chr14:52400249-52400531 | 1.058290161 | 0.017035461 | 0.09921641 |
| chr3:163622418-163622793 | -0.891627025 | 0.01703764 | 0.099221561 |
| chr9:126611607-126611916 | 0.941385058 | 0.017043626 | 0.099248877 |
| chr3:11498737-11498969 | 1.110125452 | 0.017048808 | 0.099271509 |
| chr22:29470158-29470780 | 0.832916014 | 0.017052622 | 0.099286173 |
| chr11:71252272-71252952 | 0.827815169 | 0.017057917 | 0.099308925 |
| chr13:57624612-57625355 | -0.858622014 | 0.017059553 | 0.099308925 |
| chr1:64184996-64185241 | -1.254074149 | 0.01706119 | 0.099308925 |
| chr5:88116438-88116804 | -1.002974677 | 0.017061714 | 0.099308925 |
| chr12:100063295-100063591 | -1.167986036 | 0.017065893 | 0.099325701 |
| chr13:51906408-51906927 | -0.892711962 | 0.017069429 | 0.099338732 |
| chr3:194604964-194605165 | -1.274553732 | 0.017078852 | 0.099382521 |
| chr1:114955626-114955826 | 1.069925912 | 0.017079547 | 0.099382521 |
| chr12:79560266-79560861 | 0.889943213 | 0.017082472 | 0.099391549 |
| chr7:26423585-26423786 | 1.258553894 | 0.017083693 | 0.099391549 |
| chr20:48912075-48912684 | 0.777548722 | 0.017088191 | 0.09941017 |
| chr2:160166028-160166411 | 0.908696464 | 0.017095652 | 0.099446026 |
| chr12:15181725-15182069 | -1.039241084 | 0.017101232 | 0.099470931 |
| chr19:8396720-8397743 | -0.85311314 | 0.017107164 | 0.099497886 |
| chr5:87424879-87425379 | -0.92224006 | 0.017116562 | 0.099540732 |
| chr11:65780077-65780355 | 0.917940722 | 0.017117129 | 0.099540732 |
| chr3:154334968-154335454 | -0.971071676 | 0.017124489 | 0.099575974 |
| chr7:114141477-114141677 | -1.358055233 | 0.017131699 | 0.099610338 |
| chr1:174413615-174414093 | -0.95407951 | 0.017141084 | 0.099657344 |
| chrX:104270922-104271368 | -0.887718579 | 0.01714518 | 0.099673598 |
| chr12:8544225-8544523 | -1.202954137 | 0.017151236 | 0.099701239 |
| chr8:117329046-117329304 | -1.074769467 | 0.01716056 | 0.099747869 |
| chr11:78104928-78105183 | -1.272932082 | 0.017161878 | 0.099747962 |
| chr1:41546541-41546755 | 1.2384687 | 0.017163481 | 0.099749714 |
| chr4:62709730-62709980 | -1.180370841 | 0.017169043 | 0.099769686 |
| chr22:21673739-21674053 | 1.055408048 | 0.017169522 | 0.099769686 |
| chr1:191993652-191993946 | -1.147178454 | 0.017172966 | 0.099782134 |
| chr18:36572121-36572458 | -1.20611945 | 0.017177908 | 0.09980328 |
| chr12:132100004-132100240 | 1.221406494 | 0.017194769 | 0.099893666 |
| chr9:15431006-15431655 | -0.916736894 | 0.017196521 | 0.099896271 |
| chr7:152759360-152760522 | -0.784717177 | 0.017199014 | 0.099903181 |
| chr14:39884273-39884508 | 1.098703593 | 0.017200579 | 0.099904696 |
| chr20:54236209-54236498 | 0.834671236 | 0.017203598 | 0.099908461 |
| chr1:107266949-107267296 | 0.852625446 | 0.017203835 | 0.099908461 |
| chr8:103597376-103597642 | 1.05745571 | 0.017213657 | 0.099952752 |
| chr8:29350333-29351116 | 0.81513266 | 0.017214071 | 0.099952752 |
| chr3:53145270-53145716 | 0.868930147 | 0.017218538 | 0.099971116 |
| chr7:115910568-115910819 | -1.150396788 | 0.017223138 | 0.099983326 |

| | | | |
|---|---|---|---|
| chr8:93960972-93961350 | 0.883838194 | 0.017223814 | 0.099983326 |
| chr3:101609805-101610041 | 1.224361021 | 0.017224867 | 0.099983326 |
| chr11:29448623-29448835 | -1.278431027 | 0.017225861 | 0.099983326 |
| chr2:37323929-37324219 | -1.051451628 | 0.017228119 | 0.099988858 |
| chr11:134639930-134640320 | -0.962219425 | 0.017229714 | 0.099990546 |
| chr3:151156658-151156953 | -0.980839339 | 0.017232097 | 0.099996798 |
| chr18:76559045-76559748 | -0.862324691 | 0.017234245 | 0.100001525 |
| chr8:109928056-109928409 | 1.049753079 | 0.017235522 | 0.100001525 |
| chr1:125176873-125177181 | 0.850871942 | 0.017239294 | 0.100015841 |
| chr2:157497649-157497953 | -1.036657635 | 0.017249205 | 0.10006576 |
| chr2:157125293-157125574 | -1.206412092 | 0.017253522 | 0.100076727 |
| chr14:39316310-39316510 | 1.328278142 | 0.017253707 | 0.100076727 |
| chr7:155738728-155738971 | -1.285346878 | 0.017256249 | 0.100083893 |
| chr2:48106141-48106905 | -0.817319435 | 0.017265275 | 0.100128666 |
| chr7:105012520-105014345 | -0.747947496 | 0.017271205 | 0.100155471 |
| chr2:9608609-9608827 | 1.197988496 | 0.017275278 | 0.100171513 |
| chr16:88925300-88925562 | -0.905293462 | 0.017281321 | 0.100198969 |
| chr11:48184520-48184781 | 1.15561355 | 0.017285472 | 0.100209862 |
| chr19:42253478-42253769 | 0.932978191 | 0.017285815 | 0.100209862 |
| chr4:186227029-186227255 | -1.274798081 | 0.017288002 | 0.100214958 |
| chr1:6489902-6490740 | 0.828126869 | 0.017291923 | 0.100230101 |
| chr22:23956611-23956846 | 1.161801489 | 0.017296406 | 0.100243132 |
| chr15:31598583-31599103 | -0.889781325 | 0.017296788 | 0.100243132 |
| chr6:153412415-153412673 | 1.131296248 | 0.017303692 | 0.100275561 |
| chr1:199029679-199029879 | 1.210590192 | 0.017306732 | 0.100280866 |
| chr5:100311914-100312121 | -1.300468397 | 0.017307225 | 0.100280866 |
| chr3:64344857-64345060 | 0.867933361 | 0.017316787 | 0.10032732 |
| chr16:31472548-31473191 | 0.810180542 | 0.017317861 | 0.10032732 |
| chr11:76648304-76648531 | 1.300728356 | 0.017322043 | 0.100339655 |
| chr7:106947289-106947489 | -1.287703754 | 0.017323972 | 0.100339655 |
| chr7:142853908-142854399 | -0.853647609 | 0.017324138 | 0.100339655 |
| chr2:2755803-2756016 | 1.175785195 | 0.017325228 | 0.100339655 |
| chr16:28974054-28974358 | 0.971903877 | 0.017328388 | 0.100350368 |
| chr4:180057443-180057707 | -0.969902155 | 0.017333066 | 0.100369877 |
| chr17:47127682-47127946 | -1.376198128 | 0.017336307 | 0.100381058 |
| chr6:21935646-21935890 | -1.145878466 | 0.017340214 | 0.100396093 |
| chr19:38736912-38737145 | 1.055766168 | 0.017346899 | 0.100424291 |
| chr15:90233230-90234491 | 0.760962025 | 0.017347706 | 0.100424291 |
| chr2:109480563-109480763 | 1.295446233 | 0.01735492 | 0.100454018 |
| chr6:36396064-36396282 | 1.291037878 | 0.017355463 | 0.100454018 |
| chr3:185152810-185153280 | 0.960643822 | 0.017358148 | 0.100460706 |
| chr7:11341341-11341617 | 0.97895204 | 0.017359241 | 0.100460706 |
| chr1:240122810-240123401 | -0.817638952 | 0.017363165 | 0.10047583 |
| chrX:154459140-154459550 | 0.932251569 | 0.01736559 | 0.100482273 |
| chr12:125683746-125684085 | -1.043364504 | 0.017376119 | 0.100535603 |
| chr14:93345021-93345221 | 1.361611142 | 0.017379849 | 0.100549589 |
| chr13:47082048-47082506 | 0.914168249 | 0.01738717 | 0.100584352 |
| chr6:50025164-50025698 | -0.881042741 | 0.017392389 | 0.100600301 |
| chr20:50684297-50684773 | 0.772931232 | 0.017393587 | 0.100600301 |
| chr10:48444396-48444603 | -1.138430894 | 0.017394195 | 0.100600301 |

| | | | |
|---|---|---|---|
| chr11:12497436-12497651 | -1.338615089 | 0.017395179 | 0.100600301 |
| chr6:38212494-38212834 | 0.977440497 | 0.017402248 | 0.10063359 |
| chr6:106477806-106478072 | -1.209673513 | 0.017404306 | 0.100637892 |
| chr7:2498430-2498644 | 1.048036092 | 0.017413741 | 0.10068485 |
| chrX:37060909-37061380 | -0.936324223 | 0.017420646 | 0.100717176 |
| chr4:162163779-162164646 | -0.784341083 | 0.017436124 | 0.100799054 |
| chr4:148117012-148117212 | -1.446348164 | 0.017438227 | 0.100803609 |
| chr7:50449604-50450791 | -0.778117104 | 0.017441822 | 0.100808425 |
| chr1:174194045-174194267 | -1.249838972 | 0.017442508 | 0.100808425 |
| chr3:145017983-145018203 | -1.394930254 | 0.017443007 | 0.100808425 |
| chr17:21253390-21253590 | 1.177543882 | 0.01745934 | 0.100886858 |
| chr11:128848907-128849128 | -1.417447733 | 0.017460478 | 0.100886858 |
| chr12:109927659-109927861 | -1.205272104 | 0.017460529 | 0.100886858 |
| chr5:155700336-155700630 | -1.050751305 | 0.017470489 | 0.100936795 |
| chr18:3602827-3603151 | -1.067691165 | 0.017473417 | 0.100946104 |
| chr20:47825047-47826354 | 0.755170318 | 0.017475341 | 0.100949604 |
| chr7:16695661-16695976 | 0.887837521 | 0.017477729 | 0.100955791 |
| chr7:20084178-20084422 | 1.242654588 | 0.017481772 | 0.100971534 |
| chr12:10673807-10674561 | -0.819807972 | 0.017484772 | 0.100981249 |
| chr11:12269103-12269415 | -1.013172098 | 0.017486826 | 0.100985497 |
| chr7:143305392-143305803 | -0.894448115 | 0.017495089 | 0.101025603 |
| chr12:123075707-123076130 | 0.920343543 | 0.017496863 | 0.101028234 |
| chr3:123884007-123884702 | 0.82752969 | 0.017501089 | 0.101045019 |
| chr12:20212747-20213177 | -0.883283387 | 0.017509568 | 0.101086355 |
| chr14:29222733-29222933 | 1.181169919 | 0.017512033 | 0.101092969 |
| chr9:124220917-124221257 | 0.977268451 | 0.017514962 | 0.101102262 |
| chr3:97662609-97663019 | -0.948054165 | 0.017522253 | 0.101136729 |
| chr6:69974263-69974526 | -1.177659198 | 0.017526267 | 0.101152278 |
| chr7:121581916-121582209 | -1.254799584 | 0.017534121 | 0.101189984 |
| chr7:128672996-128673311 | -0.994030108 | 0.017539418 | 0.101199147 |
| chr6:79544895-79545471 | -0.803433066 | 0.017539458 | 0.101199147 |
| chr9:73916031-73916558 | -0.939210604 | 0.017540594 | 0.101199147 |
| chr2:96872841-96873090 | -0.958231599 | 0.017540992 | 0.101199147 |
| chr22:36065778-36066200 | 0.952194151 | 0.017546687 | 0.101224385 |
| chr13:110444367-110444650 | 1.078058342 | 0.017550344 | 0.101230442 |
| chr13:33350004-33350837 | -0.793493156 | 0.01755038 | 0.101230442 |
| chr5:6408146-6408413 | 1.187328398 | 0.017569095 | 0.101330763 |
| chr4:102263082-102263361 | 1.102291197 | 0.017571838 | 0.101330864 |
| chr16:81889971-81890333 | -0.995765975 | 0.017572146 | 0.101330864 |
| chr5:91463954-91464446 | -0.929257513 | 0.01757308 | 0.101330864 |
| chr6:80510549-80510800 | 1.10772689 | 0.017587007 | 0.101392191 |
| chr2:216034041-216034297 | -1.07526919 | 0.017587468 | 0.101392191 |
| chr11:128905075-128905302 | 1.15884951 | 0.017587685 | 0.101392191 |
| chr12:53538239-53538692 | 0.936833113 | 0.017603909 | 0.101478088 |
| chr17:51217894-51218094 | -1.304622435 | 0.017607406 | 0.10149061 |
| chr4:140602831-140603084 | 1.159382819 | 0.017612023 | 0.101509587 |
| chr10:116143182-116143674 | -1.004070367 | 0.017625168 | 0.10157771 |
| chr7:37230330-37230591 | -1.075551821 | 0.017632996 | 0.101615178 |
| chr9:126642893-126643313 | -0.906066102 | 0.017634358 | 0.101615387 |
| chr8:60736089-60736473 | -1.137156715 | 0.017637426 | 0.101625422 |

| | | | |
|---|---|---|---|
| chr13:48094572-48095516 | -0.800236384 | 0.01764066 | 0.101636414 |
| chr2:2120817-2121345 | -0.901955275 | 0.017654079 | 0.101706075 |
| chr11:68804715-68805076 | 1.019192285 | 0.017662453 | 0.101746669 |
| chr7:57404024-57404363 | 0.838230142 | 0.017663961 | 0.101747707 |
| chr8:81803311-81803596 | -1.215423078 | 0.017669315 | 0.10176702 |
| chr10:85125680-85126397 | -0.84214873 | 0.017671265 | 0.10176702 |
| chr11:118573151-118573392 | 1.066355043 | 0.017671298 | 0.10176702 |
| chr7:16678792-16679473 | 0.801252524 | 0.017692918 | 0.101876941 |
| chr17:82848062-82848536 | 0.891128469 | 0.017693044 | 0.101876941 |
| chr15:98761086-98761325 | 1.103280305 | 0.017694842 | 0.101879636 |
| chr12:48001409-48001923 | 0.862120241 | 0.017701294 | 0.101903709 |
| chr4:3467002-3467268 | -1.06826244 | 0.017701683 | 0.101903709 |
| chr13:27361953-27363014 | -0.791881264 | 0.01770723 | 0.101927983 |
| chr14:101749791-101750131 | 1.020348765 | 0.01771036 | 0.101934711 |
| chr4:163548515-163548799 | -0.98298775 | 0.017711339 | 0.101934711 |
| chr7:117127620-117127914 | -1.017335156 | 0.01771239 | 0.101934711 |
| chr3:150183381-150183643 | 1.045470476 | 0.017715026 | 0.101942226 |
| chr3:184345365-184345565 | 1.139893508 | 0.017721419 | 0.10196387 |
| chr7:135661885-135663139 | -0.76305675 | 0.017721449 | 0.10196387 |
| chr12:117100360-117100579 | 1.085169692 | 0.017728083 | 0.10199438 |
| chr14:102481844-102482212 | 0.90641148 | 0.017730968 | 0.101994607 |
| chr11:110208757-110209706 | -0.830275973 | 0.017731008 | 0.101994607 |
| chr7:106153755-106154183 | -0.946920712 | 0.01773288 | 0.101994607 |
| chr5:61275557-61275871 | -1.231815044 | 0.017734691 | 0.101994607 |
| chr7:138738667-138739073 | -0.941708627 | 0.017735818 | 0.101994607 |
| chr6:164130001-164130448 | -0.988172139 | 0.017736109 | 0.101994607 |
| chr6:26743297-26743508 | 1.151784598 | 0.017742965 | 0.102021165 |
| chr8:104849995-104850329 | 1.007037652 | 0.01774339 | 0.102021165 |
| chr7:132249746-132250008 | -1.07435765 | 0.017744787 | 0.102021541 |
| chr12:51590360-51591013 | 0.900409937 | 0.017750982 | 0.102047994 |
| chr19:57362911-57363845 | 0.793749848 | 0.017753557 | 0.102047994 |
| chr14:100666449-100666649 | 1.147981874 | 0.017754212 | 0.102047994 |
| chr10:106508957-106509571 | -0.897077365 | 0.017754715 | 0.102047994 |
| chr11:78209119-78209326 | 1.088414513 | 0.017757694 | 0.102051517 |
| chr14:34331684-34332489 | 0.781311014 | 0.017758691 | 0.102051517 |
| chr5:11279199-11279412 | 1.251438446 | 0.017759324 | 0.102051517 |
| chr1:34825572-34826074 | -1.014119285 | 0.017760943 | 0.102053168 |
| chr1:209993627-209993884 | -1.2043445 | 0.017762783 | 0.102056084 |
| chr1:197245801-197246246 | 0.947087629 | 0.01776921 | 0.102077907 |
| chr1:32335179-32335435 | 0.934329878 | 0.017769245 | 0.102077907 |
| chr12:132560861-132561460 | 0.822399057 | 0.01777631 | 0.102110835 |
| chr17:41773100-41773421 | 0.933873222 | 0.017777921 | 0.102112432 |
| chr6:125769888-125770797 | -0.811068741 | 0.01778721 | 0.102153817 |
| chr2:241507361-241507773 | 0.901224924 | 0.017787792 | 0.102153817 |
| chr5:132460204-132460519 | -0.943498314 | 0.017793684 | 0.102160013 |
| chrX:132875520-132875935 | 1.164143124 | 0.017793837 | 0.102160013 |
| chr17:71889389-71889589 | -1.461729739 | 0.017794205 | 0.102160013 |
| chr18:45282494-45282694 | -1.461729739 | 0.017794205 | 0.102160013 |
| chr8:107609047-107609259 | 1.01623281 | 0.017798744 | 0.102178422 |
| chr5:74541342-74541803 | -0.997868714 | 0.017800536 | 0.102181049 |

| | | | |
|---|---|---|---|
| chr5:162096559-162096763 | -1.395653066 | 0.017803432 | 0.102190022 |
| chr4:177441801-177442840 | -0.763103105 | 0.017805462 | 0.102190142 |
| chr11:820013-820597 | 0.855855006 | 0.017806121 | 0.102190142 |
| chr14:75522357-75522763 | 0.89818378 | 0.017808147 | 0.102191835 |
| chr4:1841576-1841776 | 1.348084645 | 0.017809083 | 0.102191835 |
| chr1:86792744-86793319 | -0.877756168 | 0.017817291 | 0.102231275 |
| chr7:140326150-140326861 | -0.851027961 | 0.017824423 | 0.102264541 |
| chr13:81480718-81480986 | -1.196211113 | 0.01783332 | 0.102304339 |
| chr11:22634927-22635127 | -1.293764765 | 0.01783403 | 0.102304339 |
| chr1:87318271-87318611 | -1.100796542 | 0.017841721 | 0.102329961 |
| chr7:20776956-20777180 | 1.150173728 | 0.017843209 | 0.102329961 |
| chr11:60370505-60371318 | 0.839697412 | 0.017843656 | 0.102329961 |
| chr7:133198392-133198646 | -1.281012024 | 0.017843839 | 0.102329961 |
| chr1:172497181-172497659 | -0.83691681 | 0.017849006 | 0.102351934 |
| chr12:124774722-124775056 | -1.085224235 | 0.017854694 | 0.10237689 |
| chr8:82440942-82441302 | 1.029235275 | 0.017859383 | 0.102396111 |
| chr3:112437627-112437988 | -0.965390338 | 0.017864327 | 0.102415959 |
| chr2:83196149-83196616 | -0.999914667 | 0.017865518 | 0.102415959 |
| chr6:56387477-56388021 | -1.000970151 | 0.017871741 | 0.10244397 |
| chr8:94798947-94799335 | 1.027853443 | 0.017879465 | 0.102480576 |
| chr17:59190636-59190976 | 0.940409803 | 0.017883459 | 0.102494322 |
| chr8:101438315-101438840 | 0.80131762 | 0.017884538 | 0.102494322 |
| chr10:103232349-103232570 | 1.155823507 | 0.017887561 | 0.102503979 |
| chr11:40514201-40514417 | -1.196219742 | 0.017892734 | 0.102525955 |
| chr21:8466651-8467004 | 0.860354952 | 0.017899305 | 0.102555939 |
| chr6:108200611-108201132 | -0.900035632 | 0.017901177 | 0.102558993 |
| chr10:3482156-3482760 | 0.805675871 | 0.01790314 | 0.10256257 |
| chr7:77247300-77247530 | -1.280747055 | 0.017906146 | 0.102572125 |
| chr4:48954380-48954635 | 1.285302107 | 0.017907718 | 0.102573461 |
| chr14:53452389-53452619 | 1.26569131 | 0.01790982 | 0.102577829 |
| chr14:80567970-80568511 | -0.910471046 | 0.017920638 | 0.102632117 |
| chr12:113152805-113153636 | 0.80203372 | 0.017930829 | 0.102682807 |
| chr5:129129118-129129632 | -0.808203137 | 0.017933001 | 0.10268757 |
| chr7:129091507-129091845 | -0.975510316 | 0.017944829 | 0.102747622 |
| chr16:84407775-84407975 | 1.134236231 | 0.017949973 | 0.102761892 |
| chr8:68621927-68622844 | -0.827454162 | 0.017950003 | 0.102761892 |
| chr4:74670669-74670980 | -1.029952203 | 0.01795215 | 0.102766502 |
| chr5:172144414-172144747 | -0.980603747 | 0.017962822 | 0.102819912 |
| chr4:165016910-165017299 | -0.903250028 | 0.0179658 | 0.102829275 |
| chr16:27052933-27053133 | 1.28379453 | 0.017969905 | 0.102845092 |
| chr15:98904616-98904819 | 1.082719994 | 0.017977227 | 0.10287284 |
| chr20:38685976-38686176 | 1.324682834 | 0.017978659 | 0.10287284 |
| chr21:38831534-38832078 | -0.855304521 | 0.017978923 | 0.10287284 |
| chr3:169545673-169545947 | -1.088739998 | 0.017980124 | 0.10287284 |
| chr12:99842365-99842664 | -1.150673771 | 0.017990097 | 0.102918253 |
| chr7:50821800-50822051 | 1.111064668 | 0.017990748 | 0.102918253 |
| chr3:25724995-25725661 | 0.807351913 | 0.018004764 | 0.102983865 |
| chr6:154590697-154591255 | -0.877967588 | 0.018004913 | 0.102983865 |
| chr14:38329369-38329721 | 0.961265693 | 0.018006249 | 0.102983865 |
| chr16:68085678-68085878 | 1.11354826 | 0.018015317 | 0.103028035 |

| | | | |
|---|---|---|---|
| chr5:181182016-181182236 | 0.996967785 | 0.018024838 | 0.103064831 |
| chr2:142951176-142951460 | -1.175488939 | 0.018025904 | 0.103064831 |
| chr9:96166561-96166769 | 1.319840263 | 0.018026041 | 0.103064831 |
| chr3:185544968-185545183 | 1.298315882 | 0.018027131 | 0.103064831 |
| chr2:83115600-83116056 | -0.891021247 | 0.018029052 | 0.103065173 |
| chr15:22262553-22263087 | -0.928993819 | 0.018030541 | 0.103065173 |
| chr1:61322073-61323207 | -0.801755176 | 0.018031227 | 0.103065173 |
| chr8:85655741-85655975 | 1.187155462 | 0.018036615 | 0.103086321 |
| chr14:40890959-40891212 | 1.171598999 | 0.018037617 | 0.103086321 |
| chr4:26061293-26061541 | 1.107669497 | 0.01804105 | 0.103097524 |
| chr7:101743388-101743599 | -1.251928008 | 0.018043171 | 0.103097524 |
| chr3:51539424-51539819 | -1.009725905 | 0.018043614 | 0.103097524 |
| chr1:189719671-189720039 | -1.095003568 | 0.018051343 | 0.103133997 |
| chr16:85750196-85750609 | 0.884276805 | 0.018053986 | 0.103141404 |
| chr3:134300600-134300838 | 1.105748511 | 0.018055459 | 0.103142127 |
| chr14:101857307-101857554 | -1.291171902 | 0.018065312 | 0.103190723 |
| chr7:101000072-101000364 | 1.073268604 | 0.01807468 | 0.103236534 |
| chr4:148587551-148587846 | -1.215101979 | 0.018079335 | 0.103255427 |
| chr15:82921213-82921549 | 0.889780007 | 0.018082699 | 0.103263731 |
| chr17:57823797-57824083 | 1.021183195 | 0.018083484 | 0.103263731 |
| chrX:149504029-149504591 | -0.802839058 | 0.0180873 | 0.103277825 |
| chrX:130098512-130098745 | 1.126869195 | 0.018095026 | 0.103314237 |
| chr16:80948888-80949324 | -0.920851326 | 0.01809669 | 0.103316043 |
| chr10:47384068-47384902 | -0.78661928 | 0.018098832 | 0.103320571 |
| chr6:134657336-134657632 | 1.186322434 | 0.018105727 | 0.10335223 |
| chr4:17586138-17586409 | 1.144510732 | 0.018107811 | 0.103356426 |
| chr5:168748293-168748657 | -0.906652407 | 0.018109839 | 0.103360303 |
| chr1:115975931-115977098 | 0.787441969 | 0.0181116 | 0.103362655 |
| chr12:97198553-97198843 | -1.241065352 | 0.018118093 | 0.10339201 |
| chr17:54553961-54554224 | 1.234116557 | 0.018120381 | 0.103397366 |
| chr1:202583591-202583801 | 1.078802483 | 0.018138651 | 0.10349391 |
| chr21:15290398-15290870 | -1.021757275 | 0.018150893 | 0.10355605 |
| chr3:108001498-108001798 | 0.935024789 | 0.018157561 | 0.10357964 |
| chr1:217048119-217048653 | -0.86077594 | 0.018157732 | 0.10357964 |
| chr1:177451255-177451543 | -0.942086233 | 0.018167524 | 0.103620604 |
| chr14:55739771-55739971 | 1.160403238 | 0.018168055 | 0.103620604 |
| chr5:129322832-129323185 | -0.99135791 | 0.01816897 | 0.103620604 |
| chr1:201741484-201742016 | -0.809637089 | 0.018172976 | 0.103635736 |
| chr6:12331900-12332458 | 0.814574222 | 0.018192205 | 0.103733484 |
| chr13:77326402-77327494 | -0.75992098 | 0.018192824 | 0.103733484 |
| chr19:38095730-38096011 | 1.013411245 | 0.018197489 | 0.103752366 |
| chr13:107819040-107819278 | 1.28199984 | 0.018201238 | 0.103766016 |
| chr1:15247009-15247686 | -0.877570517 | 0.018205175 | 0.103780739 |
| chr19:35000072-35000585 | 0.838110306 | 0.018209544 | 0.103797924 |
| chr12:44961009-44961243 | -1.213780865 | 0.018213554 | 0.10381306 |
| chr14:67311893-67312250 | 0.935515481 | 0.018217146 | 0.103825807 |
| chr17:62135705-62136211 | 0.78088673 | 0.018234756 | 0.103900973 |
| chr1:155176677-155176950 | 0.851776286 | 0.01823543 | 0.103900973 |
| chr12:131047418-131047654 | 1.13970161 | 0.018235484 | 0.103900973 |
| chr1:111030504-111030819 | -1.049288967 | 0.018235758 | 0.103900973 |

| | | | |
|---|---|---|---|
| chr8:108864602-108865059 | -0.90403945 | 0.018238443 | 0.10390854 |
| chr7:7996476-7997107 | 0.800795669 | 0.018245141 | 0.103938971 |
| chr17:59988348-59988900 | 0.847569526 | 0.018251464 | 0.103967265 |
| chr4:77044555-77044868 | 1.04943203 | 0.018260476 | 0.104010871 |
| chr5:15077652-15077852 | 1.218041097 | 0.018263206 | 0.104018682 |
| chr9:131348659-131348891 | 1.139653439 | 0.018277442 | 0.104092031 |
| chr10:45362517-45362815 | -1.124885983 | 0.01828113 | 0.104104563 |
| chr2:6139118-6139346 | -1.292635729 | 0.01828236 | 0.104104563 |
| chr3:127822741-127822970 | 1.050827476 | 0.018291556 | 0.104149189 |
| chr2:23902510-23902769 | -1.132399731 | 0.018293566 | 0.104152894 |
| chr5:159441462-159441703 | -1.109170242 | 0.018300266 | 0.104182242 |
| chr1:214871295-214871544 | 1.132749119 | 0.018302291 | 0.104182242 |
| chr2:182240862-182241333 | -1.013381346 | 0.0183028 | 0.104182242 |
| chr2:65452229-65452943 | -0.819173454 | 0.018306718 | 0.104195391 |
| chr7:98348774-98349155 | -1.027319447 | 0.018308142 | 0.104195391 |
| chr11:65988941-65989544 | 0.82152958 | 0.01830919 | 0.104195391 |
| chr14:73759346-73759557 | 0.926773571 | 0.018312746 | 0.104205682 |
| chr13:51520448-51520932 | -0.882720505 | 0.018313718 | 0.104205682 |
| chr7:101144006-101144431 | -0.949480127 | 0.018321844 | 0.104244178 |
| chr10:124035715-124036247 | -0.930123763 | 0.018332813 | 0.104298842 |
| chr5:181173362-181174250 | -0.774259777 | 0.0183352 | 0.104304676 |
| chr17:1087219-1088241 | -0.78879835 | 0.018344343 | 0.104348944 |
| chr2:7248550-7248815 | 1.235600617 | 0.018352477 | 0.104381508 |
| chr3:128280468-128280767 | 1.044505073 | 0.018352793 | 0.104381508 |
| chr20:62257115-62257315 | -1.22023605 | 0.018354562 | 0.104383367 |
| chrX:8724384-8724592 | -1.309834386 | 0.018356348 | 0.104383367 |
| chr8:63241486-63241863 | -0.960647977 | 0.018357206 | 0.104383367 |
| chr17:63891242-63891493 | 1.179101036 | 0.018368443 | 0.10443951 |
| chr12:92602635-92603602 | 0.799251548 | 0.018372315 | 0.104453776 |
| chr14:102508441-102508815 | 0.850448989 | 0.018381267 | 0.104496915 |
| chr3:154552211-154552700 | -0.937435982 | 0.018393583 | 0.104559174 |
| chr10:1268925-1269207 | -0.988049454 | 0.018403079 | 0.104605396 |
| chr14:75303550-75303826 | 0.952739359 | 0.018408738 | 0.104629797 |
| chr1:178379927-178380135 | -1.413009061 | 0.01841291 | 0.104645751 |
| chr10:2448289-2448538 | -1.294263104 | 0.018419313 | 0.104674377 |
| chr13:113538696-113538930 | 1.106527292 | 0.018424521 | 0.104685714 |
| chr1:209651188-209651572 | -0.921788574 | 0.018424721 | 0.104685714 |
| chr10:82683868-82684301 | -1.033608215 | 0.018425407 | 0.104685714 |
| chr3:146189353-146189930 | -0.821527087 | 0.018432271 | 0.104702632 |
| chr9:73667746-73668101 | -0.94675524 | 0.018433126 | 0.104702632 |
| chr10:89392235-89392703 | -0.865524562 | 0.01843371 | 0.104702632 |
| chr2:70887693-70888158 | 0.918204043 | 0.018433851 | 0.104702632 |
| chr20:48638082-48638420 | 0.796876398 | 0.018437052 | 0.10471305 |
| chr2:41414465-41415250 | -0.853193267 | 0.018441265 | 0.104729216 |
| chr1:114237752-114238061 | 0.842148225 | 0.018448233 | 0.104761027 |
| chr8:69942205-69942860 | -0.856518344 | 0.018455525 | 0.104794664 |
| chr8:47826537-47826917 | -0.938975159 | 0.018460158 | 0.104813204 |
| chr6:87293528-87293832 | -1.091380934 | 0.018465714 | 0.104836986 |
| chr5:372828-374058 | -0.771939 | 0.018467835 | 0.104841259 |
| chr3:185825670-185825870 | 1.270612461 | 0.018470816 | 0.104849725 |

| | | | |
|---|---|---|---|
| chr3:59446698-59446957 | -1.259334229 | 0.018472063 | 0.104849725 |
| chr2:187500703-187501413 | 0.782749936 | 0.018473637 | 0.104850891 |
| chr11:111327877-111328131 | -1.361113014 | 0.01847816 | 0.104868791 |
| chr1:229221104-229222005 | 0.793089473 | 0.018482543 | 0.104873153 |
| chr5:166102829-166103460 | -0.851844185 | 0.018482783 | 0.104873153 |
| chr2:34025192-34025561 | -1.047873528 | 0.018483034 | 0.104873153 |
| chr17:38825008-38825729 | 0.821427499 | 0.018486978 | 0.104887764 |
| chr1:209900047-209900306 | -1.193912375 | 0.018496041 | 0.104931411 |
| chr9:33106850-33107188 | -1.033251633 | 0.018503415 | 0.104965474 |
| chrX:20516190-20516585 | -1.106084212 | 0.018507761 | 0.104982356 |
| chr18:63936154-63936701 | -0.866155814 | 0.018512214 | 0.104999844 |
| chr3:128867189-128867640 | -0.92241979 | 0.018517461 | 0.10502183 |
| chr5:179194663-179194863 | 1.182134793 | 0.018520044 | 0.105028702 |
| chr2:162317912-162319235 | -0.755452195 | 0.018539263 | 0.105129917 |
| chr6:75368086-75368681 | 0.833540604 | 0.018549085 | 0.105177828 |
| chr2:222655430-222656746 | -0.756797903 | 0.018552459 | 0.105189173 |
| chr11:67859213-67859456 | 1.274658373 | 0.018558983 | 0.105211656 |
| chr2:231569179-231569418 | 1.193404975 | 0.01855917 | 0.105211656 |
| chr10:57795569-57796139 | -0.913041942 | 0.018564758 | 0.105235546 |
| chrX:100928548-100929702 | -0.779601269 | 0.018570499 | 0.105260306 |
| chr1:27851737-27851980 | 1.249405365 | 0.018582738 | 0.105321375 |
| chr2:217644448-217644780 | 1.116687835 | 0.018584023 | 0.105321375 |
| chr7:159197056-159197501 | -0.967725491 | 0.018587673 | 0.105334269 |
| chr4:165346080-165346290 | -1.341827555 | 0.018609567 | 0.105438477 |
| chr4:166900063-166900269 | -1.341827555 | 0.018609567 | 0.105438477 |
| chr15:80660132-80660467 | 0.92412168 | 0.018611346 | 0.105438477 |
| chr18:45920561-45920798 | 1.246038026 | 0.018611566 | 0.105438477 |
| chr13:51733820-51734020 | -1.353552558 | 0.018616606 | 0.105459231 |
| chr3:6936098-6936325 | 1.133137198 | 0.018622156 | 0.105472778 |
| chr4:124213-125018 | 0.799308204 | 0.018623171 | 0.105472778 |
| chr5:90709396-90709664 | -1.022707787 | 0.018623284 | 0.105472778 |
| chr5:10375749-10376116 | -1.036592843 | 0.018624503 | 0.105472778 |
| chr7:20959264-20959642 | 0.873558412 | 0.018630798 | 0.105500631 |
| chr1:37346599-37346803 | 1.280814786 | 0.018645603 | 0.105576661 |
| chr16:46388579-46388832 | 0.810445689 | 0.018649912 | 0.105593255 |
| chr6:109229556-109229959 | -0.934332112 | 0.018651778 | 0.105596022 |
| chr1:230744973-230745173 | 1.139391943 | 0.018658985 | 0.10562902 |
| chr7:141032710-141033243 | -0.983492705 | 0.018674265 | 0.105707707 |
| chr14:51751976-51752287 | 0.989908092 | 0.01868179 | 0.105725027 |
| chr8:82562009-82562369 | 0.985292585 | 0.018681912 | 0.105725027 |
| chr3:133595261-133595663 | -0.923176997 | 0.018681927 | 0.105725027 |
| chr21:38458594-38458794 | 1.123973018 | 0.018682844 | 0.105725027 |
| chr3:124063498-124063698 | 1.333082791 | 0.018688589 | 0.105743131 |
| chr10:132142476-132142746 | 0.964847963 | 0.018689666 | 0.105743131 |
| chr9:72822474-72822898 | -1.04701607 | 0.018690183 | 0.105743131 |
| chr6:32048108-32048480 | 0.906321584 | 0.018692889 | 0.105750631 |
| chr13:109257061-109257311 | -1.235354893 | 0.01870108 | 0.105789159 |
| chr7:21254117-21254374 | 1.036453562 | 0.018713995 | 0.105854404 |
| chr7:22109669-22110046 | 0.890253869 | 0.018718589 | 0.105872573 |
| chr3:11867602-11867906 | 1.151742199 | 0.018724594 | 0.105898716 |

| | | | |
|---|---|---|---|
| chr6:11043488-11043688 | -1.256770989 | 0.018726846 | 0.105903636 |
| chr5:6772133-6772378 | 1.187984441 | 0.018734588 | 0.105939603 |
| chr4:77994142-77994791 | -0.883376139 | 0.018738683 | 0.105953932 |
| chr20:56451296-56451716 | 0.88485891 | 0.018740481 | 0.105953932 |
| chr1:204813316-204813735 | 0.879602259 | 0.018741271 | 0.105953932 |
| chr15:41660200-41660977 | 0.787110319 | 0.018742896 | 0.105955304 |
| chr1:216243155-216243426 | 1.175163045 | 0.018754057 | 0.106010577 |
| chr7:23261870-23262099 | 1.027469696 | 0.018759496 | 0.106026402 |
| chr5:157918784-157919398 | 0.902080122 | 0.018759624 | 0.106026402 |
| chr13:48037243-48038327 | -0.755790137 | 0.018762227 | 0.106033292 |
| chr6:79617798-79618003 | 1.240797062 | 0.018766533 | 0.106049803 |
| chr10:106401258-106401799 | -1.040016748 | 0.018773383 | 0.106069927 |
| chr1:114191525-114191888 | 0.836361148 | 0.018774026 | 0.106069927 |
| chr10:110789680-110790324 | -0.896657231 | 0.018774571 | 0.106069927 |
| chr4:171846294-171846560 | -0.99782097 | 0.018775631 | 0.106069927 |
| chr15:92797129-92797625 | 0.842611947 | 0.018782502 | 0.106098653 |
| chr20:23200937-23201312 | -0.977828796 | 0.018783485 | 0.106098653 |
| chr2:190554420-190554714 | -1.070052079 | 0.018792251 | 0.10614034 |
| chr4:42282022-42282555 | -0.915009208 | 0.018798444 | 0.106167491 |
| chr1:151836478-151837312 | 0.751835061 | 0.018807341 | 0.106209912 |
| chr19:5557869-5558069 | -1.289901389 | 0.018809529 | 0.10621444 |
| chr12:126493292-126493937 | -0.936951091 | 0.018816227 | 0.106244433 |
| chr4:141860852-141861315 | -0.987858217 | 0.018827331 | 0.106299296 |
| chr9:36419597-36420323 | -0.864748011 | 0.018832947 | 0.106323171 |
| chr22:29394977-29395294 | 0.960336573 | 0.018836236 | 0.106333902 |
| chr11:78606723-78606995 | 1.193999154 | 0.018839548 | 0.106344764 |
| chr15:23878048-23878282 | 1.074638248 | 0.018842316 | 0.106352554 |
| chr7:51838748-51839596 | -0.789950493 | 0.018847356 | 0.106373165 |
| chr10:95807117-95807344 | -1.26073709 | 0.018850017 | 0.106380348 |
| chr2:235766239-235766571 | -0.959706138 | 0.018858102 | 0.10640726 |
| chr10:908201-908401 | 1.287445684 | 0.018858328 | 0.10640726 |
| chr7:150110494-150110960 | -0.899987464 | 0.018860618 | 0.10640726 |
| chr17:61267858-61268068 | 0.828567255 | 0.018861282 | 0.10640726 |
| chr2:174585114-174585821 | 0.810653371 | 0.018861874 | 0.10640726 |
| chr19:5048057-5048654 | -0.855853021 | 0.018863118 | 0.10640726 |
| chr3:141255945-141256343 | -0.894436691 | 0.018866254 | 0.106417119 |
| chr1:222543881-222544208 | -0.993663409 | 0.018869075 | 0.106425193 |
| chr3:145939561-145939831 | -1.164868025 | 0.018872741 | 0.106438034 |
| chr6:151475515-151476187 | -0.814238057 | 0.018879166 | 0.106459836 |
| chr7:46590972-46591809 | -0.825584377 | 0.018879385 | 0.106459836 |
| chr16:72961014-72961793 | -0.807210428 | 0.01888341 | 0.106474695 |
| chr1:106268972-106269195 | 0.941743477 | 0.018884907 | 0.106475304 |
| chr6:112337228-112337500 | -1.0520986 | 0.018893702 | 0.106517053 |
| chr8:77957096-77957647 | -0.925064569 | 0.018895122 | 0.10651722 |
| chr17:27488995-27489344 | -1.060019279 | 0.018898786 | 0.106524991 |
| chr6:140366888-140367356 | -0.846022352 | 0.018899281 | 0.106524991 |
| chr13:77810511-77810788 | -1.009324731 | 0.018900934 | 0.106526421 |
| chr17:61323847-61324607 | 0.749275677 | 0.018902316 | 0.106526421 |
| chr7:149007612-149007817 | -1.145550076 | 0.018906202 | 0.106540488 |
| chr7:141654533-141654832 | -0.926187152 | 0.01891015 | 0.106548516 |

| | | | |
|---|---|---|---|
| chr10:70566099-70566309 | 1.214166046 | 0.018912542 | 0.106548516 |
| chr13:39765922-39766122 | -1.283811888 | 0.018913725 | 0.106548516 |
| chr1:116329619-116329900 | 0.976679634 | 0.018915122 | 0.106548516 |
| chr18:22466528-22467035 | 0.841670943 | 0.018916135 | 0.106548516 |
| chr12:124389294-124389511 | 1.017423961 | 0.018917209 | 0.106548516 |
| chr2:105650524-105650889 | -0.973876667 | 0.018917361 | 0.106548516 |
| chr8:48270987-48271294 | 1.01380514 | 0.018920418 | 0.106557903 |
| chr12:122421864-122422621 | 0.819585623 | 0.018924618 | 0.106573722 |
| chr1:83053907-83054174 | -1.074947779 | 0.018930634 | 0.106599767 |
| chr22:23402308-23403219 | 0.800715886 | 0.01893321 | 0.106606436 |
| chr10:80171185-80171681 | 0.827243081 | 0.018939174 | 0.106632183 |
| chrX:24959330-24959728 | -1.092084959 | 0.018949452 | 0.106682214 |
| chrX:121197590-121197875 | -1.257105123 | 0.018952649 | 0.106692371 |
| chr21:40487646-40487860 | -1.232177735 | 0.018955701 | 0.106701715 |
| chr1:226988230-226988468 | 1.029297686 | 0.01895855 | 0.10670991 |
| chr7:108358544-108359049 | -0.875889982 | 0.018966294 | 0.10673933 |
| chr7:1689996-1690208 | 1.266695235 | 0.018968069 | 0.10673933 |
| chr17:56305148-56305383 | -1.210005948 | 0.018968453 | 0.10673933 |
| chr2:189982834-189983117 | -1.088677699 | 0.018969349 | 0.10673933 |
| chr1:119388483-119388767 | -1.23362235 | 0.018973053 | 0.106752334 |
| chr20:54345525-54345777 | 0.829678389 | 0.018979293 | 0.106779602 |
| chr20:33400841-33401703 | 0.835625194 | 0.018984334 | 0.106800121 |
| chr10:124565770-124566297 | -0.921510918 | 0.018985932 | 0.106801269 |
| chr21:41778054-41778601 | -0.838955225 | 0.018993327 | 0.106832664 |
| chr1:107071565-107072004 | 0.823325384 | 0.018994302 | 0.106832664 |
| chr20:61423100-61423406 | -0.915359565 | 0.018999269 | 0.106852757 |
| chr7:5291277-5291489 | 1.068678718 | 0.019005937 | 0.106882415 |
| chr7:117385897-117386184 | -1.041953224 | 0.019011226 | 0.106904312 |
| chr12:69985598-69986018 | -0.901997177 | 0.019017159 | 0.106929826 |
| chr10:132396325-132396525 | 1.112517021 | 0.019042046 | 0.107060247 |
| chr5:118792065-118792396 | -0.895627884 | 0.019043148 | 0.107060247 |
| chr3:102374208-102374459 | 1.184873234 | 0.01904765 | 0.107077697 |
| chr20:31566594-31567489 | -0.784864111 | 0.019056859 | 0.107121607 |
| chr5:165029489-165029866 | -0.853386281 | 0.01905909 | 0.107126289 |
| chr14:97088674-97088956 | 1.17638849 | 0.019063408 | 0.107142702 |
| chr17:16294827-16295056 | 1.233072124 | 0.019069654 | 0.107169943 |
| chr12:9331201-9331483 | -1.047356778 | 0.019081637 | 0.107229421 |
| chr15:69848635-69848862 | -1.288377069 | 0.019083393 | 0.107231425 |
| chr1:61083436-61083786 | -1.022540878 | 0.019086525 | 0.107241164 |
| chr17:46958175-46958377 | 1.228111128 | 0.019090017 | 0.107252788 |
| chr11:9983235-9983517 | 1.055824399 | 0.019094008 | 0.107252788 |
| chr20:63485492-63485692 | 1.081495586 | 0.01909691 | 0.107252788 |
| chr1:43689242-43689552 | 1.148375166 | 0.019096948 | 0.107252788 |
| chr7:11193153-11193866 | 0.801083766 | 0.019099199 | 0.107252788 |
| chr2:118680397-118680725 | 1.143924857 | 0.019099225 | 0.107252788 |
| chr12:97471068-97471779 | -0.882430587 | 0.019099704 | 0.107252788 |
| chr3:169604597-169604919 | -0.926991061 | 0.019099792 | 0.107252788 |
| chr15:55407631-55408964 | 0.748309428 | 0.019102046 | 0.10725591 |
| chr1:64378848-64379106 | -1.085243015 | 0.019105636 | 0.10725591 |
| chr17:68034154-68034475 | -0.837879766 | 0.019106519 | 0.10725591 |

| | | | |
|---|---|---|---|
| chr8:99959968-99960254 | 1.203998176 | 0.0191071 | 0.10725591 |
| chr4:82695016-82695250 | -1.354740185 | 0.019107592 | 0.10725591 |
| chr3:25397163-25397381 | 1.228273451 | 0.01910977 | 0.10725591 |
| chr1:228130923-228131489 | 0.818180752 | 0.01911072 | 0.10725591 |
| chr16:25106859-25107379 | 0.816412124 | 0.019111547 | 0.10725591 |
| chr19:51712704-51712972 | 1.032677724 | 0.019115541 | 0.107266721 |
| chr1:36751530-36751812 | 1.065727925 | 0.019116273 | 0.107266721 |
| chr7:17181558-17181812 | 1.131255436 | 0.019118544 | 0.107271611 |
| chr2:212987607-212987837 | -1.203930075 | 0.019127798 | 0.107315674 |
| chr17:61538669-61539342 | 0.75571647 | 0.019131909 | 0.10733088 |
| chr21:39048090-39048422 | -0.970138463 | 0.019138709 | 0.107361168 |
| chr9:78213793-78214017 | -1.271574363 | 0.0191427 | 0.10737569 |
| chr10:59536689-59537134 | -1.003905869 | 0.019146953 | 0.107391686 |
| chr20:36100937-36101207 | 1.239233596 | 0.019153023 | 0.107417871 |
| chr17:50062369-50062827 | -0.861309693 | 0.019154656 | 0.107419167 |
| chr11:81822316-81822535 | 1.09966681 | 0.019157682 | 0.107421236 |
| chr4:154507021-154507489 | 0.848000135 | 0.019157829 | 0.107421236 |
| chr12:93081901-93082228 | 1.009348525 | 0.019168029 | 0.107463914 |
| chr10:101839804-101840011 | 1.007441898 | 0.019168246 | 0.107463914 |
| chr13:109704842-109705525 | -0.875215883 | 0.019177844 | 0.107509858 |
| chr18:37309053-37309323 | -1.150136454 | 0.019179973 | 0.107513927 |
| chr8:118548382-118549281 | -0.779167833 | 0.019185155 | 0.107531167 |
| chr1:192546157-192546570 | -1.017739738 | 0.019185855 | 0.107531167 |
| chr13:41019193-41019491 | -0.975067364 | 0.019187793 | 0.107534161 |
| chr5:135462267-135462688 | -0.861938402 | 0.019189493 | 0.107535131 |
| chrY:56855238-56855519 | 1.058742221 | 0.019190773 | 0.107535131 |
| chr22:37805071-37805271 | 1.156731216 | 0.019192582 | 0.107537407 |
| chr13:37869875-37870744 | -0.815979258 | 0.019197371 | 0.107555155 |
| chr12:53867513-53867966 | 0.914634989 | 0.019198585 | 0.107555155 |
| chr4:183098832-183099600 | -0.809892106 | 0.019199961 | 0.107555155 |
| chr8:90865258-90865458 | 1.280440322 | 0.01920591 | 0.107580617 |
| chr8:56441672-56441937 | -1.297401774 | 0.019209719 | 0.107594085 |
| chr4:164935451-164935793 | -1.012826201 | 0.019218609 | 0.107636009 |
| chr13:30176121-30176383 | -1.152137124 | 0.019221839 | 0.107646229 |
| chr14:67392055-67392407 | 0.889318426 | 0.019223577 | 0.107648097 |
| chr5:170721296-170721507 | 1.252037212 | 0.019228628 | 0.107668515 |
| chr2:131120683-131121263 | 0.854175764 | 0.019230146 | 0.107669144 |
| chr3:134126260-134126460 | 1.38134397 | 0.019232663 | 0.107675368 |
| chr5:17654684-17654918 | 1.099328411 | 0.019242647 | 0.107723397 |
| chrX:151802068-151802390 | 1.055867525 | 0.019250474 | 0.10775934 |
| chr1:217029138-217029662 | -0.902704193 | 0.019253234 | 0.107761485 |
| chr8:121426680-121427136 | 0.852628219 | 0.01925367 | 0.107761485 |
| chr1:29202646-29203188 | 0.861187618 | 0.019255445 | 0.107763545 |
| chr2:65326328-65326649 | -0.994916385 | 0.019267489 | 0.107823077 |
| chr13:57581958-57582335 | -0.928966346 | 0.019272624 | 0.107841424 |
| chr20:54161547-54162042 | 0.76919232 | 0.019273583 | 0.107841424 |
| chr4:3443314-3443714 | 0.929800289 | 0.019277515 | 0.107855551 |
| chr19:48788232-48788477 | 1.07075305 | 0.019281091 | 0.107856003 |
| chr2:166493991-166494572 | -0.855900852 | 0.019281759 | 0.107856003 |
| chr8:144233595-144233803 | 1.13047611 | 0.019281819 | 0.107856003 |

| | | | |
|---|---|---|---|
| chr14:105526440-105526700 | 0.966767897 | 0.019289561 | 0.107884986 |
| chr6:64469397-64469763 | -0.950642826 | 0.019289816 | 0.107884986 |
| chr15:96264246-96264842 | -0.861829228 | 0.019306604 | 0.107970994 |
| chr4:174538973-174539238 | -1.115363808 | 0.019309374 | 0.107974052 |
| chr3:73438782-73438990 | 1.255824438 | 0.019309969 | 0.107974052 |
| chr2:189581603-189582125 | -0.927406174 | 0.019311606 | 0.107975325 |
| chr7:95899968-95900237 | -1.304049647 | 0.019319103 | 0.108009363 |
| chr14:40724801-40725182 | 0.999321579 | 0.019323941 | 0.108028527 |
| chr17:72488503-72489311 | -0.845366912 | 0.019325401 | 0.108028807 |
| chr19:53467487-53467874 | 0.873132132 | 0.019328459 | 0.108038023 |
| chr5:61319739-61320015 | -1.086048051 | 0.019330927 | 0.108043934 |
| chr12:120534734-120535013 | 1.093245706 | 0.019335061 | 0.108059156 |
| chr14:79342112-79342469 | 0.974989536 | 0.019338808 | 0.108072217 |
| chr9:83962196-83962917 | -0.825005874 | 0.019344617 | 0.108090148 |
| chr19:44847615-44848008 | 0.863559537 | 0.019344838 | 0.108090148 |
| chr15:97013054-97013624 | 0.907992777 | 0.019347474 | 0.108096991 |
| chr12:53938305-53939636 | 0.744060361 | 0.019355128 | 0.10812703 |
| chr19:12676407-12676635 | 1.032739616 | 0.019355673 | 0.10812703 |
| chrX:102042726-102043139 | -0.925886849 | 0.019360221 | 0.10813847 |
| chr17:75087574-75087937 | -1.002771753 | 0.019360543 | 0.10813847 |
| chr7:34130503-34130888 | -0.912614562 | 0.019364789 | 0.108147909 |
| chr17:40072028-40072902 | -0.810945584 | 0.019365056 | 0.108147909 |
| chr7:3119650-3119850 | 1.289536126 | 0.019385726 | 0.108252497 |
| chr7:42292848-42293622 | -0.796388585 | 0.019386609 | 0.108252497 |
| chr2:5735076-5735320 | -1.380611469 | 0.019393501 | 0.10828309 |
| chr4:149645734-149646162 | -1.01230116 | 0.019397752 | 0.108298933 |
| chr2:86831936-86832167 | 1.286949011 | 0.019405936 | 0.108335112 |
| chr7:129611325-129612364 | -0.78080735 | 0.01940706 | 0.108335112 |
| chr19:42051900-42052100 | 1.122904566 | 0.019415518 | 0.108358037 |
| chr10:109938318-109938554 | 1.207900414 | 0.01941664 | 0.108358037 |
| chr17:5033179-5033517 | 1.084920814 | 0.019417501 | 0.108358037 |
| chr5:154405189-154405575 | -0.860076583 | 0.019417843 | 0.108358037 |
| chrX:30036509-30036796 | -1.120166882 | 0.019418238 | 0.108358037 |
| chr2:55855750-55855986 | 1.158025265 | 0.019421862 | 0.108370371 |
| chr8:130422081-130422643 | -0.907239682 | 0.019426576 | 0.108388782 |
| chr7:4768660-4768866 | 1.180983283 | 0.019429658 | 0.108398082 |
| chr19:14062009-14062255 | 1.05841992 | 0.019434625 | 0.108417896 |
| chr12:53701683-53702267 | 0.843210028 | 0.019436154 | 0.108418535 |
| chr5:125198609-125198940 | -1.102251333 | 0.019437752 | 0.108419556 |
| chr10:21372902-21373204 | -1.020632424 | 0.019445434 | 0.108454509 |
| chr1:96376685-96377205 | 0.876178496 | 0.019458002 | 0.108510077 |
| chr11:35943692-35944627 | -0.784383586 | 0.019458229 | 0.108510077 |
| chr21:42621225-42621642 | -0.879806598 | 0.019469354 | 0.108564215 |
| chr7:143003492-143003708 | -1.217205081 | 0.019476022 | 0.108593493 |
| chr17:60028199-60028404 | 1.338091824 | 0.019486402 | 0.10863228 |
| chr6:53869757-53870405 | 0.811783596 | 0.019486594 | 0.10863228 |
| chr8:133242886-133243367 | 0.849033492 | 0.019487232 | 0.10863228 |
| chr18:76573237-76573637 | -1.002836273 | 0.019494314 | 0.108663853 |
| chr7:131753680-131754049 | -1.175244908 | 0.01949798 | 0.108676383 |
| chr3:48300886-48301779 | 0.821280114 | 0.01950744 | 0.1087212 |

| | | | |
|---|---|---|---|
| chr2:199455639-199456286 | -0.796945454 | 0.019513199 | 0.108738937 |
| chr12:100128545-100128842 | -0.981665447 | 0.019513461 | 0.108738937 |
| chr10:59595645-59595894 | -1.186891736 | 0.019519951 | 0.108767194 |
| chr6:148790368-148790737 | 1.125237983 | 0.019522198 | 0.108771807 |
| chr4:72959216-72959960 | -0.858437225 | 0.019525755 | 0.108780489 |
| chr2:136179181-136179543 | 0.993903872 | 0.019526596 | 0.108780489 |
| chr15:85854462-85854722 | 0.978199162 | 0.019530146 | 0.108792356 |
| chr3:61980935-61981297 | 0.872192484 | 0.019532072 | 0.108795176 |
| chr6:30818569-30819027 | 0.918166188 | 0.01953617 | 0.108803146 |
| chr18:48937499-48937908 | -0.901381102 | 0.019536343 | 0.108803146 |
| chr13:32650828-32651071 | -1.085684914 | 0.019544095 | 0.108838407 |
| chr3:9800587-9800860 | 1.014511935 | 0.019550779 | 0.108864843 |
| chr10:35195408-35195618 | -1.231508556 | 0.019552901 | 0.108864843 |
| chr3:170160147-170160359 | 1.121521426 | 0.019553104 | 0.108864843 |
| chr7:158957882-158958500 | -0.805663946 | 0.019558557 | 0.108882314 |
| chr5:88502224-88502566 | -1.159235492 | 0.019560015 | 0.108882314 |
| chr9:33785967-33786203 | -1.29198178 | 0.019560506 | 0.108882314 |
| chr1:53407156-53407411 | -1.157565647 | 0.019566957 | 0.108910313 |
| chr7:157432550-157433261 | -0.805535902 | 0.019568941 | 0.108913444 |
| chr11:121452142-121452647 | 0.870776293 | 0.01957438 | 0.108935801 |
| chr5:61599794-61600205 | -0.856383704 | 0.01957708 | 0.108942917 |
| chr16:87835064-87835460 | -0.882832248 | 0.01957907 | 0.108946079 |
| chr1:246552160-246552501 | 0.975656181 | 0.019591935 | 0.109009746 |
| chr18:12278294-12278545 | 1.164145968 | 0.019593537 | 0.109010744 |
| chr9:80605209-80605409 | -1.420686488 | 0.019598914 | 0.109032744 |
| chr7:47337687-47337952 | -1.105032819 | 0.019600586 | 0.109034129 |
| chr1:85224884-85225843 | -0.797280361 | 0.019602652 | 0.109037704 |
| chr1:185627219-185627544 | -1.130191005 | 0.019611184 | 0.109076759 |
| chr1:216723195-216723746 | -0.855592678 | 0.01961252 | 0.109076759 |
| chr17:54186903-54187103 | -1.231640014 | 0.019618447 | 0.109101802 |
| chr1:214602798-214603632 | -0.759042812 | 0.019621656 | 0.109111732 |
| chr5:126875139-126875883 | -0.830569855 | 0.019625115 | 0.109122519 |
| chr16:2849918-2850150 | 1.240046401 | 0.019626445 | 0.109122519 |
| chr9:32525963-32526789 | -0.777167068 | 0.01964011 | 0.109183395 |
| chr14:92986286-92986657 | 0.877429294 | 0.019640244 | 0.109183395 |
| chr2:73284189-73285214 | -0.77708039 | 0.01964691 | 0.10921253 |
| chr4:184474426-184475386 | -0.776453651 | 0.019651449 | 0.109229839 |
| chr17:61689783-61690202 | 0.772363751 | 0.019655926 | 0.109246799 |
| chr8:128001172-128001418 | 1.005816118 | 0.019680603 | 0.109361812 |
| chr20:56752051-56752549 | 0.832784107 | 0.019680685 | 0.109361812 |
| chr7:128531093-128531868 | -0.79853257 | 0.019680901 | 0.109361812 |
| chr7:44512705-44512946 | 1.163279248 | 0.019690604 | 0.109407791 |
| chr20:62087333-62087535 | 1.075701161 | 0.019697153 | 0.109436247 |
| chr4:107877336-107877714 | 0.902743276 | 0.019699211 | 0.109439742 |
| chr13:38470129-38470567 | -1.053343526 | 0.019706259 | 0.109470964 |
| chr17:78592238-78592876 | -0.866418421 | 0.019712774 | 0.109499214 |
| chr20:47280257-47280946 | 0.745328484 | 0.019718268 | 0.10951471 |
| chr1:194160002-194160377 | -0.95333882 | 0.019719624 | 0.10951471 |
| chr17:74431377-74431577 | 1.006386585 | 0.019719851 | 0.10951471 |
| chr20:57071226-57071884 | 0.768067382 | 0.01972258 | 0.109521923 |

| | | | |
|---|---|---|---|
| chr12:89461815-89462109 | -0.934776295 | 0.0197309 | 0.109560186 |
| chr16:66924002-66924291 | 1.151548164 | 0.019733508 | 0.109566726 |
| chr3:87771293-87771612 | -0.930389132 | 0.019735602 | 0.109570418 |
| chr16:74145934-74146134 | 1.186073975 | 0.019737085 | 0.109570709 |
| chr14:97177048-97177309 | 1.018810027 | 0.019744969 | 0.109606538 |
| chr1:198109295-198109690 | 0.959739752 | 0.019761467 | 0.109688671 |
| chr7:157336482-157337574 | -0.760391618 | 0.019762628 | 0.109688671 |
| chr2:22904242-22904475 | -1.178171895 | 0.01976412 | 0.109689005 |
| chr5:11311969-11312213 | 1.048082278 | 0.019766942 | 0.109692173 |
| chr7:102287377-102288048 | -0.84153626 | 0.019767554 | 0.109692173 |
| chr10:132850372-132850608 | 1.198284335 | 0.019769752 | 0.109696426 |
| chr9:84266114-84266315 | -1.373324662 | 0.019771949 | 0.109700671 |
| chr2:91571723-91571923 | -1.348503175 | 0.019775232 | 0.10971094 |
| chr7:152411479-152411741 | -1.16696648 | 0.019778028 | 0.109718508 |
| chr4:121050314-121050612 | -1.134243842 | 0.019783785 | 0.109742499 |
| chr10:10911378-10911802 | -1.017933078 | 0.019804105 | 0.109843737 |
| chr17:61740164-61740375 | 0.819333642 | 0.019804902 | 0.109843737 |
| chr20:52738020-52738294 | -1.219199653 | 0.019809037 | 0.109844734 |
| chr6:43770646-43771149 | 0.823898594 | 0.019809978 | 0.109844734 |
| chr19:49391705-49392203 | 0.862478388 | 0.019810639 | 0.109844734 |
| chr21:37072434-37073554 | 0.752498085 | 0.019810816 | 0.109844734 |
| chr16:22007483-22008115 | 0.839570937 | 0.019815281 | 0.109857825 |
| chr19:19628704-19628909 | -1.200510503 | 0.019816045 | 0.109857825 |
| chr4:169922679-169922879 | -1.426264128 | 0.01982262 | 0.109886325 |
| chr1:153572543-153572867 | 0.881735959 | 0.019826164 | 0.109898018 |
| chr4:30720193-30720449 | 0.884390305 | 0.019828943 | 0.109905471 |
| chr5:174295288-174295551 | -1.245511385 | 0.019831599 | 0.109907945 |
| chr9:83322477-83322677 | -1.333283528 | 0.019832258 | 0.109907945 |
| chr12:55728348-55728608 | 0.90128126 | 0.019840796 | 0.10994731 |
| chr3:171752985-171753452 | 0.901166579 | 0.019849971 | 0.109984662 |
| chr3:23308607-23308893 | 1.209846059 | 0.019850407 | 0.109984662 |
| chr5:105840713-105841099 | -1.113730156 | 0.019854222 | 0.109997844 |
| chr17:35578292-35578738 | 0.939927884 | 0.019858041 | 0.110011051 |
| chr9:84868732-84868932 | -1.417756851 | 0.01986112 | 0.110020152 |
| chr7:93870132-93870460 | 1.00184133 | 0.019865797 | 0.110038105 |
| chr1:46557518-46557769 | -1.043959454 | 0.019869403 | 0.11005012 |
| chr4:148557905-148558252 | -1.118151484 | 0.019875143 | 0.110073959 |
| chr10:44173506-44173877 | -0.906847088 | 0.019879548 | 0.110090397 |
| chr7:87039563-87040447 | -0.780196345 | 0.01988295 | 0.110101281 |
| chr20:19290100-19290313 | -1.200592118 | 0.019897865 | 0.110175906 |
| chr12:110431369-110432006 | -0.880556536 | 0.019908064 | 0.110217079 |
| chr1:118184746-118185774 | -0.776806704 | 0.019908177 | 0.110217079 |
| chr12:69101540-69101869 | -0.931099425 | 0.019913786 | 0.110238254 |
| chr12:79071605-79071805 | -1.347023022 | 0.01991488 | 0.110238254 |
| chrX:153927841-153928770 | 0.756976144 | 0.019917536 | 0.110244992 |
| chr5:66814242-66814689 | -0.995296098 | 0.019919287 | 0.110246719 |
| chr1:198970849-198971289 | -0.831168837 | 0.01993805 | 0.110342595 |
| chr6:100228862-100229306 | 0.82394005 | 0.019939783 | 0.110344216 |
| chr5:131270981-131271187 | -1.37197946 | 0.019946216 | 0.110371848 |
| chr2:155250831-155251134 | -1.03556491 | 0.019948374 | 0.110375818 |

| | | | |
|---|---|---|---|
| chr1:109997676-109997957 | -1.202406941 | 0.01994982 | 0.110375846 |
| chr1:145394146-145394356 | 1.143538892 | 0.019962776 | 0.110438019 |
| chr14:61940661-61941341 | 0.846539964 | 0.01996394 | 0.110438019 |
| chr20:50856160-50856454 | 0.888611845 | 0.019966175 | 0.110442412 |
| chr3:9820985-9821247 | 0.932070911 | 0.01996985 | 0.110454762 |
| chr5:102295049-102295268 | -1.041479585 | 0.019974159 | 0.110470625 |
| chr11:3581145-3581516 | 0.933751339 | 0.019976382 | 0.110473425 |
| chr1:187383447-187383911 | -1.045907929 | 0.019977549 | 0.110473425 |
| chr5:137325754-137326004 | -1.294317979 | 0.019985096 | 0.110507121 |
| chr20:41110883-41111212 | 0.86566997 | 0.019986527 | 0.110507121 |
| chr3:64761933-64762204 | 0.815857895 | 0.019990717 | 0.110516052 |
| chrX:35786242-35786612 | -1.164908287 | 0.019991027 | 0.110516052 |
| chr7:741261-741461 | 1.240356713 | 0.019993987 | 0.110524441 |
| chr3:38479807-38480125 | 0.968102981 | 0.019996646 | 0.110531168 |
| chr14:34873251-34875575 | 0.736063858 | 0.020000597 | 0.110545031 |
| chr5:61817800-61818052 | -1.120492616 | 0.020013962 | 0.110610921 |
| chr12:10153199-10153811 | -0.87662897 | 0.020017954 | 0.110625005 |
| chr20:22807683-22807995 | -1.082485087 | 0.020020726 | 0.110632344 |
| chr7:10680659-10680947 | 1.097643392 | 0.020022967 | 0.110636748 |
| chr1:51252667-51252906 | 1.228233856 | 0.020036267 | 0.110694272 |
| chr1:189132343-189132554 | 1.228233856 | 0.020036267 | 0.110694272 |
| chr5:112479530-112479818 | 1.036080899 | 0.020038136 | 0.110696616 |
| chr1:248218458-248218787 | -1.171755733 | 0.020042053 | 0.110710273 |
| chr16:19337883-19338083 | 1.282045837 | 0.020045438 | 0.11072099 |
| chr3:10595022-10595257 | 0.970896314 | 0.020050706 | 0.1107421 |
| chr20:56316591-56316841 | 1.060321963 | 0.020056833 | 0.110767957 |
| chr6:55175278-55175560 | -1.261983613 | 0.02005925 | 0.110773324 |
| chr12:82752208-82752670 | -0.902274297 | 0.02006468 | 0.110795325 |
| chr14:69611319-69611959 | 0.778717358 | 0.020068758 | 0.110809855 |
| chr6:36782892-36783157 | -1.08925575 | 0.020075719 | 0.110836373 |
| chr13:40635655-40635855 | -1.247046387 | 0.020077763 | 0.110836373 |
| chr11:36227486-36227691 | 1.243415607 | 0.0200779 | 0.110836373 |
| chr1:25820502-25821013 | 0.847537332 | 0.020086488 | 0.110865526 |
| chr10:32207545-32207797 | -1.049643915 | 0.020087767 | 0.110865526 |
| chr5:132225520-132225769 | -1.077321713 | 0.020087824 | 0.110865526 |
| chr5:42571235-42571559 | 1.113681807 | 0.020089681 | 0.110865526 |
| chr1:162168198-162168589 | 0.863168746 | 0.020090416 | 0.110865526 |
| chr10:13586673-13587686 | -0.767998149 | 0.02009889 | 0.1109043 |
| chr1:36323570-36323816 | 0.965844823 | 0.020105663 | 0.110931618 |
| chr5:142922166-142922451 | -1.118763622 | 0.02010921 | 0.110931618 |
| chr6:10759316-10759721 | -0.900001733 | 0.020111102 | 0.110931618 |
| chr19:29515486-29515686 | 1.231886619 | 0.020111147 | 0.110931618 |
| chr4:162497630-162497917 | -1.05510179 | 0.020112874 | 0.110931618 |
| chr12:124829199-124829616 | 0.911484034 | 0.020113157 | 0.110931618 |
| chr6:127118792-127119136 | 1.056961319 | 0.02011492 | 0.110931618 |
| chr1:164579416-164579660 | 1.110478798 | 0.020115423 | 0.110931618 |
| chr3:64302663-64302871 | 0.875721755 | 0.020119319 | 0.110940131 |
| chr17:48275419-48275656 | -1.278643449 | 0.020119862 | 0.110940131 |
| chr1:191142481-191142681 | -1.350688965 | 0.020134759 | 0.111002451 |
| chr8:22768276-22768885 | -0.865194609 | 0.020136309 | 0.111002451 |

| | | | |
|---|---|---|---|
| chr20:31610612-31611075 | -0.816861486 | 0.020136966 | 0.111002451 |
| chr16:31094558-31095171 | 0.782308952 | 0.020138002 | 0.111002451 |
| chr11:34359005-34359258 | 0.970061152 | 0.020138408 | 0.111002451 |
| chr2:192029139-192029439 | -1.069248062 | 0.020150482 | 0.111061018 |
| chr1:106731588-106731829 | 0.894759866 | 0.020159658 | 0.111103597 |
| chr17:54373835-54374247 | -0.939639208 | 0.020164601 | 0.111122676 |
| chr20:48329756-48330167 | 0.787609809 | 0.02016602 | 0.111122676 |
| chr7:112160572-112160886 | -1.058645418 | 0.020170609 | 0.111139969 |
| chr20:47540375-47540735 | 0.773792037 | 0.020172275 | 0.111141157 |
| chr21:38965897-38966497 | -0.792820048 | 0.020177607 | 0.111162541 |
| chr3:98612557-98613199 | -0.830348449 | 0.02017938 | 0.111164313 |
| chr13:24220122-24220455 | 0.880780347 | 0.020183739 | 0.111174651 |
| chr12:105444054-105444328 | 0.992214978 | 0.020185417 | 0.111174651 |
| chr6:21102595-21102895 | -1.191783982 | 0.020185609 | 0.111174651 |
| chr4:183407105-183407490 | -0.922920348 | 0.020189825 | 0.111189876 |
| chr5:84383697-84385285 | -0.761984646 | 0.020191922 | 0.111193432 |
| chr19:55212478-55212804 | -0.985233425 | 0.020210245 | 0.111286337 |
| chr3:144304473-144304829 | 1.003203172 | 0.020214307 | 0.111300704 |
| chr5:93078442-93078956 | -0.89144205 | 0.020218402 | 0.111313429 |
| chr2:102235499-102235799 | -0.972852461 | 0.020219523 | 0.111313429 |
| chr13:51781106-51781789 | -0.926461223 | 0.020223131 | 0.111318829 |
| chr6:165835348-165835631 | -1.101821122 | 0.02022341 | 0.111318829 |
| chr18:72632624-72632883 | -1.169960583 | 0.020232554 | 0.11135609 |
| chr9:70049712-70049912 | -1.291095431 | 0.020234393 | 0.11135609 |
| chr4:186669552-186669761 | -1.306381571 | 0.020235951 | 0.11135609 |
| chr8:129552113-129552362 | -1.076068834 | 0.020236802 | 0.11135609 |
| chr10:13548387-13548624 | -1.207905926 | 0.020237499 | 0.11135609 |
| chr1:153494933-153495221 | 0.950251038 | 0.020238899 | 0.11135609 |
| chr1:114488815-114489186 | 0.810899784 | 0.02024048 | 0.111356794 |
| chrX:68454548-68454784 | -1.324968625 | 0.020245798 | 0.111378054 |
| chr14:102235392-102235613 | 1.043227048 | 0.020250044 | 0.111393057 |
| chr19:35449181-35449981 | 0.8023968 | 0.020251433 | 0.111393057 |
| chr13:87999874-88000161 | -1.127004929 | 0.020253357 | 0.111395645 |
| chr16:78263283-78263595 | 0.990204939 | 0.020258436 | 0.11141558 |
| chr10:67763527-67763750 | -0.940220379 | 0.020262956 | 0.111424443 |
| chr11:78822991-78823484 | -0.940220379 | 0.020262956 | 0.111424443 |
| chr1:186828603-186829166 | -0.891093863 | 0.020270731 | 0.111459201 |
| chr10:73602011-73602211 | 1.335566438 | 0.020272618 | 0.111461579 |
| chr4:114386742-114386953 | -1.248173519 | 0.020277963 | 0.111482964 |
| chr1:113642202-113642410 | -1.087273004 | 0.020279432 | 0.111483042 |
| chr20:47271206-47271752 | 0.747179317 | 0.02029159 | 0.111541876 |
| chr13:84605846-84606209 | -1.196025856 | 0.020294436 | 0.111549049 |
| chr14:37760308-37760508 | 1.100943125 | 0.020295836 | 0.111549049 |
| chr8:100881940-100882220 | 0.998447137 | 0.020297262 | 0.111549049 |
| chr6:55617338-55617600 | -1.115860677 | 0.020304073 | 0.111578474 |
| chr7:19051837-19052106 | 1.172512365 | 0.020307953 | 0.111591794 |
| chr1:247613848-247614048 | -1.280370137 | 0.020316394 | 0.11163017 |
| chr12:8038522-8039343 | -0.794267176 | 0.020321846 | 0.11165212 |
| chr14:45392384-45392584 | 1.205728213 | 0.020324252 | 0.111657335 |
| chr8:88249096-88249296 | 1.197343888 | 0.020327211 | 0.111664552 |

| | | | |
|---|---|---|---|
| chr14:36450013-36450213 | 1.205525944 | 0.02032848 | 0.111664552 |
| chr1:119779807-119780319 | 0.793425387 | 0.020330939 | 0.111670052 |
| chr9:34430846-34431423 | 0.92934814 | 0.02033569 | 0.111688142 |
| chr3:23479501-23479886 | -1.136178839 | 0.020341288 | 0.111710876 |
| chr20:61942435-61942662 | 1.164358103 | 0.020343358 | 0.111714241 |
| chr7:36037051-36037741 | -0.761064752 | 0.020355568 | 0.111773279 |
| chr13:51066743-51066943 | -1.302441562 | 0.020361716 | 0.111799028 |
| chrX:94862311-94863082 | -0.785810309 | 0.020370983 | 0.111841891 |
| chr10:99140891-99141453 | -0.872796452 | 0.020384274 | 0.111906843 |
| chr7:123570378-123570858 | -0.98421723 | 0.02039117 | 0.111936686 |
| chr11:22192256-22193365 | -0.771808958 | 0.020397334 | 0.111962499 |
| chr5:32627931-32628225 | 1.033063068 | 0.020399479 | 0.111966251 |
| chr8:94567477-94567919 | 0.84586157 | 0.020400982 | 0.111966479 |
| chr5:175079966-175080205 | 1.237293858 | 0.020410534 | 0.112009538 |
| chr2:235027696-235027918 | 1.145348633 | 0.020412285 | 0.112009538 |
| chr1:180830614-180830912 | 1.171993592 | 0.020413213 | 0.112009538 |
| chr9:119429230-119429509 | -1.288727278 | 0.020415561 | 0.112014403 |
| chr6:156933176-156933447 | -1.086893952 | 0.020422849 | 0.112046363 |
| chr8:11807953-11808643 | 0.831447897 | 0.020442224 | 0.112138609 |
| chr18:45691317-45691566 | -1.080026104 | 0.020442589 | 0.112138609 |
| chr1:211735682-211736108 | -0.847895507 | 0.020452076 | 0.112176237 |
| chr10:789453-789766 | 0.834004391 | 0.020452377 | 0.112176237 |
| chr2:169258126-169258326 | -1.348376649 | 0.020457109 | 0.112194159 |
| chr1:58767235-58767463 | -1.278842804 | 0.020470363 | 0.112242746 |
| chr1:162699448-162699648 | -1.278842804 | 0.020470363 | 0.112242746 |
| chr10:25280165-25280399 | -1.278842804 | 0.020470363 | 0.112242746 |
| chr1:41901415-41901639 | 1.167306952 | 0.020475275 | 0.112261647 |
| chr17:59815200-59815498 | 0.986891612 | 0.020479856 | 0.112273038 |
| chr18:824281-824482 | -1.311242927 | 0.020480283 | 0.112273038 |
| chr11:27464219-27464753 | -0.850701492 | 0.020484104 | 0.112285952 |
| chr10:65941292-65941594 | -1.125120932 | 0.020490199 | 0.112311326 |
| chr12:130958997-130959197 | 1.272724968 | 0.020508471 | 0.11240344 |
| chr15:84579443-84579753 | 1.051415725 | 0.020510138 | 0.112404533 |
| chr6:113375941-113376180 | -1.185366006 | 0.020513337 | 0.112414024 |
| chr4:162350371-162350663 | 0.996680375 | 0.020519776 | 0.112439764 |
| chr11:88292561-88292829 | -1.30428682 | 0.020520969 | 0.112439764 |
| chr8:81104365-81104671 | 0.941500491 | 0.020530073 | 0.112481607 |
| chr2:155559383-155559665 | 1.203539387 | 0.020533256 | 0.112491 |
| chr3:87832610-87833048 | 0.926941971 | 0.020536482 | 0.112496397 |
| chr2:15993360-15993632 | -1.15458538 | 0.020538506 | 0.112496397 |
| chr12:89247569-89247853 | -1.137627452 | 0.020538645 | 0.112496397 |
| chr15:90154921-90155394 | 0.854102908 | 0.02054385 | 0.112516859 |
| chr4:56836694-56836959 | 1.031893213 | 0.020550796 | 0.112546857 |
| chr3:107065053-107065596 | -0.87860479 | 0.020552781 | 0.112549687 |
| chrX:123859383-123860707 | 0.737185591 | 0.020557875 | 0.112569536 |
| chr14:66213141-66213423 | -1.307720981 | 0.020559412 | 0.112569759 |
| chr8:55934755-55935010 | 1.174204828 | 0.020561542 | 0.112569759 |
| chr3:7285539-7285739 | 1.190392135 | 0.020562324 | 0.112569759 |
| chr12:88554416-88554628 | -1.194917433 | 0.020571255 | 0.112601941 |
| chr4:162874181-162874529 | -1.132443222 | 0.020572256 | 0.112601941 |

| | | | |
|---|---|---|---|
| chr12:53221428-53221882 | 0.833015997 | 0.020572611 | 0.112601941 |
| chr14:28633865-28634088 | 1.125108862 | 0.020578663 | 0.112627025 |
| chr1:79497244-79497546 | -1.050527019 | 0.020585795 | 0.11265801 |
| chr14:51533287-51533506 | 1.133229392 | 0.020605293 | 0.112756662 |
| chr15:58951656-58951970 | 0.955562726 | 0.020613182 | 0.112791775 |
| chr17:27494263-27494588 | -1.0310734 | 0.020619 | 0.112815553 |
| chr3:133676688-133676890 | 1.271124367 | 0.02063231 | 0.112876062 |
| chr13:97133991-97134738 | -0.833869727 | 0.020633005 | 0.112876062 |
| chr5:179135865-179136194 | 0.985541394 | 0.020641383 | 0.112913832 |
| chr5:181216084-181216529 | -0.831844816 | 0.020650192 | 0.112953955 |
| chr2:152861072-152861727 | 0.8383324 | 0.020664911 | 0.113026401 |
| chr1:85830688-85831211 | -0.958392422 | 0.020669489 | 0.113043366 |
| chr7:50372676-50372946 | 1.14700629 | 0.020681267 | 0.113099709 |
| chr17:67490536-67491610 | -0.775918538 | 0.020684101 | 0.113107136 |
| chr15:76235220-76235581 | 0.979508528 | 0.020691412 | 0.113139038 |
| chr6:49870413-49870696 | -1.204880044 | 0.020694827 | 0.11314443 |
| chr10:121753842-121754043 | -1.307809128 | 0.020695351 | 0.11314443 |
| chr14:20530973-20531253 | 1.00859043 | 0.020700526 | 0.113164649 |
| chr3:128965794-128966088 | 1.134535901 | 0.020706356 | 0.113188442 |
| chr14:91149390-91149683 | 0.967098301 | 0.020712828 | 0.113215746 |
| chr14:99579160-99579524 | 0.833289468 | 0.020726339 | 0.113275044 |
| chr20:775802-776033 | 1.129531562 | 0.020726634 | 0.113275044 |
| chr6:30100806-30101149 | 0.928184182 | 0.020730882 | 0.113290179 |
| chr5:17897938-17898176 | -1.236762595 | 0.020734725 | 0.113303101 |
| chr1:161738750-161739000 | 1.179387628 | 0.0207502 | 0.113379579 |
| chr3:127124208-127124457 | 1.204737167 | 0.020759399 | 0.113421753 |
| chr16:23434434-23434636 | 1.206626914 | 0.020767532 | 0.113458099 |
| chr12:66115669-66115891 | 1.232058217 | 0.020778786 | 0.11351149 |
| chr5:151223189-151223418 | 1.044859743 | 0.020782725 | 0.113524915 |
| chr9:137436092-137436356 | 0.996999669 | 0.020786543 | 0.113537676 |
| chr7:105939993-105940786 | -0.826776594 | 0.020788381 | 0.113539621 |
| chr8:133488423-133488875 | 0.832811112 | 0.020792933 | 0.113550667 |
| chr4:59904465-59904736 | -1.171496373 | 0.020793368 | 0.113550667 |
| chr13:34421408-34421608 | -1.301472106 | 0.020803086 | 0.113592551 |
| chr2:84022848-84023158 | -1.039923169 | 0.020804002 | 0.113592551 |
| chr18:63676663-63676881 | -1.261101377 | 0.02081989 | 0.113665735 |
| chr5:96733834-96734493 | -0.774687496 | 0.020821688 | 0.113665735 |
| chr20:44521540-44522506 | 0.757790275 | 0.020821856 | 0.113665735 |
| chr5:158770439-158770844 | -0.945929427 | 0.020829031 | 0.113696803 |
| chr16:75733743-75734221 | -0.947840402 | 0.020832863 | 0.113704604 |
| chr5:22068149-22068349 | 1.196974766 | 0.020833428 | 0.113704604 |
| chr11:12813668-12813964 | -1.054080041 | 0.020849966 | 0.11376673 |
| chr17:77214751-77215049 | 0.930640134 | 0.020850428 | 0.11376673 |
| chr1:226671411-226671777 | 1.025781256 | 0.020850733 | 0.11376673 |
| chr14:53946443-53946739 | 1.045734716 | 0.020850991 | 0.11376673 |
| chr16:3009631-3010417 | 0.774552151 | 0.020854241 | 0.11376673 |
| chr14:64823276-64823613 | 0.878971312 | 0.02085429 | 0.11376673 |
| chr1:199545797-199546122 | -0.896068972 | 0.02085661 | 0.11376673 |
| chrX:101595748-101595948 | 1.245512454 | 0.020860007 | 0.11376673 |
| chr13:49560418-49561005 | -0.953421949 | 0.020860416 | 0.11376673 |

| | | | |
|---|---|---|---|
| chr11:65416363-65416595 | 0.928582166 | 0.020860441 | 0.11376673 |
| chr22:37020979-37021263 | -1.192321553 | 0.020861144 | 0.11376673 |
| chr1:50620596-50621355 | -0.823048307 | 0.020862805 | 0.113767693 |
| chr10:74948193-74948400 | -1.361120288 | 0.020869157 | 0.113794234 |
| chr3:60775983-60776299 | -0.993199728 | 0.020873717 | 0.113810995 |
| chr8:115127832-115128263 | -0.889371527 | 0.020883003 | 0.11385117 |
| chr2:143575360-143575605 | -1.072225017 | 0.020884057 | 0.11385117 |
| chr7:23247342-23247542 | 1.134723521 | 0.020887091 | 0.113859611 |
| chr8:13275629-13275960 | -0.9857602 | 0.020912464 | 0.113987447 |
| chr21:8452002-8452568 | 0.768149049 | 0.020913517 | 0.113987447 |
| chr3:32781238-32781819 | 0.797914696 | 0.020916347 | 0.113991614 |
| chr5:51841378-51841685 | -1.060644757 | 0.020917257 | 0.113991614 |
| chr5:175921350-175921618 | 1.074830867 | 0.020920135 | 0.113999188 |
| chr1:203893203-203893498 | 0.989775641 | 0.020923085 | 0.114007155 |
| chr2:227717717-227718152 | -1.10973641 | 0.020925794 | 0.114010817 |
| chr12:43654172-43654374 | -1.300240856 | 0.020927582 | 0.114010817 |
| chr17:14048215-14048438 | -1.268315209 | 0.020928221 | 0.114010817 |
| chr17:67834030-67834230 | -1.239947186 | 0.020930075 | 0.114012812 |
| chr8:64262665-64262974 | -0.951943563 | 0.020933935 | 0.11401934 |
| chr11:88338421-88338630 | -1.190621929 | 0.020934704 | 0.11401934 |
| chr11:33508020-33508271 | -1.113988926 | 0.020935738 | 0.11401934 |
| chr4:123997104-123997422 | -1.112165172 | 0.020942062 | 0.114041212 |
| chr4:153576111-153576395 | -1.208242169 | 0.02094273 | 0.114041212 |
| chr3:95751604-95752582 | -0.817776289 | 0.020951025 | 0.114078273 |
| chr21:39951237-39951470 | 1.112171701 | 0.02095959 | 0.114116801 |
| chr2:104733596-104733984 | -0.914444856 | 0.020975834 | 0.114197128 |
| chr20:41738672-41738904 | 0.92546268 | 0.020980517 | 0.114211827 |
| chr10:87903258-87903742 | -0.918594849 | 0.020981515 | 0.114211827 |
| chr15:64653870-64654070 | 1.14364787 | 0.020989157 | 0.114221335 |
| chr1:78673157-78673375 | -1.169884455 | 0.020990442 | 0.114221335 |
| chr10:105383237-105383446 | -1.169884455 | 0.020990442 | 0.114221335 |
| chr3:168002729-168003423 | -0.823244681 | 0.020990522 | 0.114221335 |
| chr19:33498629-33498831 | 1.191193419 | 0.020990716 | 0.114221335 |
| chr20:50959297-50959542 | 0.946980652 | 0.020992389 | 0.11422233 |
| chr20:54574146-54574591 | 0.778335056 | 0.020995867 | 0.114233141 |
| chr10:106189260-106189561 | -1.088203745 | 0.020997771 | 0.114235389 |
| chr18:74196089-74196661 | -0.889845725 | 0.02100006 | 0.114237056 |
| chr7:16709051-16709436 | 0.963598609 | 0.021003414 | 0.114237056 |
| chr5:131417138-131417605 | -0.999906324 | 0.021003479 | 0.114237056 |
| chr15:87672463-87672683 | 1.089561629 | 0.021004041 | 0.114237056 |
| chr8:29635993-29636254 | -1.183758933 | 0.021008811 | 0.114247473 |
| chr4:23369467-23369779 | -1.094015273 | 0.021008939 | 0.114247473 |
| chr1:106704029-106704229 | 0.878948321 | 0.021011376 | 0.114252619 |
| chr10:114876998-114877372 | -0.901676301 | 0.021027099 | 0.114330003 |
| chr1:161020083-161020347 | 0.928270687 | 0.021028669 | 0.114330423 |
| chr21:41024054-41025285 | -0.771267775 | 0.021032138 | 0.114341169 |
| chr20:54759279-54759513 | 0.842804995 | 0.021039425 | 0.114369984 |
| chr2:142112978-142113212 | -1.272915604 | 0.021040423 | 0.114369984 |
| chrX:15619104-15619327 | -1.316853484 | 0.021048264 | 0.11439715 |
| chr15:79464123-79464463 | 1.116593064 | 0.021048407 | 0.11439715 |

| | | | |
|---|---|---|---|
| chr14:37571278-37571478 | 0.993927429 | 0.021050728 | 0.114401649 |
| chr2:10773734-10774721 | -0.7785151 | 0.021066746 | 0.114480583 |
| chr20:48460702-48461010 | 0.794918401 | 0.021071085 | 0.114496042 |
| chrX:25968842-25969276 | -1.010200144 | 0.021086635 | 0.114572411 |
| chr1:234975253-234975453 | 1.029443285 | 0.021094969 | 0.114609566 |
| chr17:44587321-44587695 | 0.912182705 | 0.021100176 | 0.114628144 |
| chr1:23559422-23559711 | 0.906332358 | 0.02110138 | 0.114628144 |
| chr6:43396350-43396713 | 0.938140773 | 0.02110786 | 0.114655214 |
| chr7:121046935-121047202 | -1.157092212 | 0.021112162 | 0.11467045 |
| chr14:77314018-77314370 | 0.887094648 | 0.021117907 | 0.114688399 |
| chr7:130373725-130374169 | 0.914511783 | 0.02111846 | 0.114688399 |
| chr1:195517354-195517641 | 1.113035592 | 0.021124353 | 0.114705589 |
| chr11:66070059-66070480 | 0.877936708 | 0.02112604 | 0.114705589 |
| chr14:24297669-24297882 | 0.915662213 | 0.021126116 | 0.114705589 |
| chr5:155049669-155050191 | -0.878271919 | 0.021134242 | 0.114741577 |
| chr11:16893491-16894246 | 0.801592105 | 0.021138374 | 0.114755125 |
| chr7:42869098-42870006 | -0.775664944 | 0.021139733 | 0.114755125 |
| chr11:78870172-78870372 | 1.25599807 | 0.021144947 | 0.114767676 |
| chr20:47560027-47560323 | 0.78165547 | 0.02114504 | 0.114767676 |
| chr7:28854016-28854531 | 0.873311909 | 0.021147462 | 0.114770672 |
| chr13:29894083-29894324 | -1.134773989 | 0.021148588 | 0.114770672 |
| chr16:2842502-2842921 | -0.991084729 | 0.021150795 | 0.114774521 |
| chr7:140176878-140177780 | -0.768324803 | 0.021154045 | 0.11478216 |
| chr8:106950498-106950832 | 0.97048178 | 0.021155199 | 0.11478216 |
| chr20:54499634-54499850 | 0.831173836 | 0.021159724 | 0.114791871 |
| chr7:149238007-149238284 | 1.029209803 | 0.021160557 | 0.114791871 |
| chr3:197599895-197600331 | 0.9017755 | 0.02116376 | 0.114791871 |
| chr4:163166114-163167349 | -0.741507628 | 0.0211649 | 0.114791871 |
| chr18:74353246-74353505 | -1.171460945 | 0.021165348 | 0.114791871 |
| chr17:39705738-39706293 | 0.850078346 | 0.021165978 | 0.114791871 |
| chr12:26845181-26845573 | -1.020913642 | 0.021193057 | 0.114930599 |
| chr15:55064594-55064794 | 1.291490058 | 0.021198068 | 0.114941549 |
| chr2:134491604-134492425 | -0.784328026 | 0.021198077 | 0.114941549 |
| chr1:160205636-160206309 | 0.804765225 | 0.021204178 | 0.114966499 |
| chr16:56452127-56452407 | 0.991165562 | 0.021206189 | 0.114969265 |
| chr1:206713748-206714509 | 0.773827423 | 0.021211278 | 0.114982484 |
| chr2:100777457-100777686 | -1.297486378 | 0.021211629 | 0.114982484 |
| chr16:88525111-88525331 | 1.032640201 | 0.021214271 | 0.114988673 |
| chr1:115811124-115811388 | 0.959145971 | 0.021223524 | 0.115030691 |
| chr19:12836269-12836492 | 1.037799208 | 0.021229296 | 0.115051111 |
| chr11:65900359-65900813 | 0.866635802 | 0.021230295 | 0.115051111 |
| chr11:65588789-65589031 | 0.955031567 | 0.021238872 | 0.115089453 |
| chr2:45874426-45874711 | -1.079297235 | 0.021243396 | 0.115105824 |
| chr20:51131130-51131334 | 1.056531763 | 0.021256983 | 0.115168309 |
| chr5:132674687-132674975 | -1.162265427 | 0.021259047 | 0.115168309 |
| chr4:90174471-90175275 | 0.760374679 | 0.021260563 | 0.115168309 |
| chr17:61563386-61563591 | 0.817464121 | 0.02126094 | 0.115168309 |
| chrX:153662253-153662537 | 1.002562299 | 0.021265567 | 0.115182598 |
| chr5:131304023-131304349 | -1.201464635 | 0.021267591 | 0.115182598 |
| chr11:12270151-12270388 | -1.111336647 | 0.021271178 | 0.115182598 |

| | | | |
|---|---|---|---|
| chr11:25982085-25982320 | -1.168370049 | 0.021271422 | 0.115182598 |
| chr13:43116304-43116527 | -1.168370049 | 0.021271422 | 0.115182598 |
| chr7:92527646-92529126 | -0.731725527 | 0.021272597 | 0.115182598 |
| chr11:100919080-100919391 | -1.163906102 | 0.021276757 | 0.115193742 |
| chr8:131903844-131905027 | -0.748528257 | 0.021277662 | 0.115193742 |
| chr21:42044917-42045221 | 0.987480421 | 0.021280273 | 0.115193987 |
| chr4:170259489-170260359 | -0.785895869 | 0.021280714 | 0.115193987 |
| chr3:192907071-192907517 | -0.854078029 | 0.021283713 | 0.115202082 |
| chr13:91406716-91406999 | -1.073307954 | 0.021286995 | 0.115208288 |
| chr8:120654481-120654707 | 1.145189649 | 0.021287867 | 0.115208288 |
| chr11:79054754-79055072 | -0.945660253 | 0.021294659 | 0.115236905 |
| chr13:80342635-80343160 | -0.831456454 | 0.021296517 | 0.11523882 |
| chr7:112449898-112451359 | -0.724849428 | 0.021300888 | 0.115254337 |
| chr2:212912465-212913095 | -0.809282201 | 0.021304013 | 0.115263106 |
| chr2:217092887-217093099 | -1.302655401 | 0.021307155 | 0.115271964 |
| chr5:168359838-168360265 | 0.827003144 | 0.021310552 | 0.115282201 |
| chr14:95197546-95197940 | 0.859667248 | 0.021316373 | 0.115305549 |
| chr14:90508817-90509281 | 0.782234253 | 0.021322875 | 0.115332578 |
| chr8:126609391-126609921 | 0.82474161 | 0.021328288 | 0.115353714 |
| chr7:111561869-111562929 | -0.73980113 | 0.021336572 | 0.115390373 |
| chr17:81019614-81019874 | 1.044543659 | 0.02134603 | 0.115431744 |
| chr9:38672351-38672740 | -0.944220517 | 0.021347235 | 0.115431744 |
| chr11:64379581-64379781 | 1.184417964 | 0.021351294 | 0.115445546 |
| chr3:175312637-175313261 | -0.851407637 | 0.021353904 | 0.115451514 |
| chr11:8192597-8192846 | 1.015958161 | 0.021360027 | 0.115469626 |
| chr2:52369235-52369460 | -1.29587146 | 0.021360268 | 0.115469626 |
| chr4:112296915-112297806 | -0.791604009 | 0.021364369 | 0.1154785 |
| chr10:77837633-77837917 | -0.93518618 | 0.021364924 | 0.1154785 |
| chr16:66277972-66278970 | -0.754989736 | 0.021370091 | 0.115490267 |
| chr3:87069809-87070418 | 0.81624723 | 0.021370116 | 0.115490267 |
| chr3:118949471-118949892 | -1.043049551 | 0.021374886 | 0.115507899 |
| chr7:2348060-2348310 | 0.97794058 | 0.021377477 | 0.115509046 |
| chr8:48503963-48504508 | 0.845980095 | 0.021378113 | 0.115509046 |
| chrX:57821770-57822271 | -0.913749391 | 0.021382126 | 0.115522582 |
| chr14:103099122-103099590 | -0.842688524 | 0.021384459 | 0.115527042 |
| chr1:151992425-151993032 | 0.752000941 | 0.021391417 | 0.115556484 |
| chr15:66842749-66843119 | 0.9345532 | 0.021395513 | 0.115565846 |
| chr12:80763928-80764144 | -1.240689755 | 0.021396167 | 0.115565846 |
| chr16:3939363-3939563 | 1.191756988 | 0.021401943 | 0.11558645 |
| chr17:6676188-6676473 | -1.129576913 | 0.021403421 | 0.11558645 |
| chr3:116118535-116118775 | -1.209685744 | 0.021404507 | 0.11558645 |
| chr14:47917361-47917607 | -1.169231311 | 0.021417181 | 0.11564674 |
| chr12:39587287-39587629 | -1.113492093 | 0.021419355 | 0.115650328 |
| chr17:65093993-65094407 | 0.900406381 | 0.021424484 | 0.115669872 |
| chr7:120950390-120951478 | -0.730885214 | 0.021428036 | 0.115680898 |
| chr1:183276370-183276719 | -0.971435166 | 0.021446569 | 0.115772793 |
| chr18:63890914-63891247 | -0.965691674 | 0.021449742 | 0.115781766 |
| chr2:131833007-131833207 | -1.268838789 | 0.021458176 | 0.115812377 |
| chr2:96869793-96870395 | -0.793363024 | 0.021458436 | 0.115812377 |
| chr4:90328595-90328834 | 1.142904989 | 0.021464567 | 0.115837303 |

| | | | |
|---|---|---|---|
| chr12:118103275-118104543 | 0.748624726 | 0.021466312 | 0.115838562 |
| chr20:59194600-59194800 | 1.01211427 | 0.021478563 | 0.115896512 |
| chr20:47149716-47150520 | 0.739560579 | 0.021494248 | 0.115972978 |
| chr4:165355820-165356313 | -0.915557666 | 0.021497117 | 0.115980289 |
| chr6:84227466-84228292 | -0.763891624 | 0.021504304 | 0.116010896 |
| chr14:59044732-59045047 | -1.098557665 | 0.021508447 | 0.116016054 |
| chr2:58973763-58974330 | -0.855962258 | 0.02150917 | 0.116016054 |
| chr1:109191360-109191638 | -0.931217302 | 0.021511576 | 0.116016054 |
| chr7:159030468-159031110 | -0.818973567 | 0.02151225 | 0.116016054 |
| chrX:9207223-9207490 | -1.174237195 | 0.021513839 | 0.116016054 |
| chr7:19984112-19984380 | 0.977544843 | 0.021515684 | 0.116016054 |
| chr4:160525007-160525233 | -1.268152557 | 0.021515859 | 0.116016054 |
| chr1:8211815-8212302 | -0.993097441 | 0.02152023 | 0.11603146 |
| chr19:1566926-1567126 | 1.194636286 | 0.021547268 | 0.116164328 |
| chr6:27290562-27290836 | -1.077261549 | 0.021547905 | 0.116164328 |
| chr7:148339243-148340007 | -0.783121101 | 0.021550239 | 0.116168734 |
| chr4:173177259-173177470 | -1.258810036 | 0.021561746 | 0.116220741 |
| chr1:96865755-96866303 | -0.935413122 | 0.02156292 | 0.116220741 |
| chr20:47429443-47429643 | 0.791451057 | 0.021567682 | 0.116238233 |
| chr2:230067838-230068612 | 0.828012011 | 0.021572637 | 0.116251127 |
| chr20:48578391-48578687 | 0.787190792 | 0.021573109 | 0.116251127 |
| chr14:45706104-45706550 | 0.895392162 | 0.021577785 | 0.116268148 |
| chr12:91355430-91355756 | -0.926932541 | 0.021581643 | 0.116280758 |
| chr8:135422431-135422656 | 1.198411295 | 0.021584338 | 0.1162871 |
| chr19:55606282-55606559 | 0.885471382 | 0.021587533 | 0.116296137 |
| chrX:155070687-155072108 | 0.748230487 | 0.021594072 | 0.116321127 |
| chr6:118796543-118796942 | -0.897508495 | 0.021595208 | 0.116321127 |
| chr12:52583028-52583274 | 0.974005059 | 0.021597603 | 0.116325848 |
| chr8:120772546-120772786 | 1.101270061 | 0.021609768 | 0.116383189 |
| chr20:55106186-55106386 | 1.138763631 | 0.021620369 | 0.11642605 |
| chr2:200199942-200200201 | -1.192759411 | 0.021620765 | 0.11642605 |
| chr4:49120949-49121154 | 1.273099378 | 0.021634806 | 0.116493473 |
| chr7:135774325-135774608 | -1.172652661 | 0.021639479 | 0.116505639 |
| chr4:176065376-176066320 | -0.740039443 | 0.021640174 | 0.116505639 |
| chr3:188296761-188297347 | -0.947536966 | 0.021642604 | 0.116505639 |
| chr8:687066-687733 | 0.865409798 | 0.021643148 | 0.116505639 |
| chr18:68660359-68660650 | -1.163708572 | 0.02165489 | 0.116560657 |
| chr2:121004984-121005184 | 1.185142973 | 0.021657457 | 0.116566285 |
| chr17:64582987-64583268 | 1.105595775 | 0.021662428 | 0.116584851 |
| chr10:88387153-88387459 | -1.013694351 | 0.021672925 | 0.116621413 |
| chr17:74369335-74369554 | 1.007337812 | 0.02167373 | 0.116621413 |
| chr3:127959974-127960250 | 0.991759534 | 0.021673787 | 0.116621413 |
| chr1:164321349-164321568 | 1.086803141 | 0.021677724 | 0.116634404 |
| chr17:60997022-60997275 | 0.994978615 | 0.021682418 | 0.116651471 |
| chr10:9699425-9699625 | -1.359532953 | 0.021693288 | 0.116701754 |
| chr2:240069787-240070252 | -0.891965322 | 0.021695948 | 0.11670787 |
| chr10:132421512-132421805 | 0.875768535 | 0.021700148 | 0.116722272 |
| chr2:155952981-155953245 | -1.251392221 | 0.021701955 | 0.1167236 |
| chr12:52156843-52157126 | 0.940864002 | 0.021703442 | 0.1167236 |
| chr4:95362897-95363176 | -1.093778532 | 0.021705807 | 0.116728125 |

| | | | |
|---|---|---|---|
| chr7:38306302-38306554 | 1.18848257 | 0.021711011 | 0.116742726 |
| chr5:24644826-24645276 | -0.907543783 | 0.021711569 | 0.116742726 |
| chr14:53953128-53954226 | 0.751024357 | 0.021717472 | 0.116766272 |
| chr8:51851019-51851570 | -0.858989865 | 0.021719759 | 0.116770375 |
| chr8:111114203-111114431 | 1.048036327 | 0.021728551 | 0.116809446 |
| chrX:152897838-152898074 | 0.975250304 | 0.021730223 | 0.116810238 |
| chr15:89016737-89017094 | -0.941831077 | 0.02173689 | 0.116835686 |
| chr5:587096-587341 | 1.052593456 | 0.021738007 | 0.116835686 |
| chr19:12035395-12035985 | 0.833650098 | 0.021741837 | 0.116848075 |
| chr11:5808451-5808765 | -0.955280011 | 0.021747389 | 0.116869718 |
| chr6:158085848-158086130 | 1.17151119 | 0.021753174 | 0.116892609 |
| chr1:218619092-218619376 | -1.037859303 | 0.021757943 | 0.116909181 |
| chr1:119516743-119517015 | -1.185722209 | 0.02175931 | 0.116909181 |
| chr10:95290588-95291145 | 0.838490584 | 0.021761443 | 0.116912443 |
| chr4:171903090-171903327 | -1.23157679 | 0.021764105 | 0.116913833 |
| chr10:15245684-15245884 | -1.237284758 | 0.021764806 | 0.116913833 |
| chr5:147628051-147628434 | -0.866316521 | 0.021766279 | 0.116913833 |
| chr2:127023490-127023706 | -1.191380363 | 0.021775304 | 0.116954112 |
| chr14:86791082-86791439 | 1.031522606 | 0.021779112 | 0.116965916 |
| chrX:14848312-14849076 | -0.841837322 | 0.02178092 | 0.116965916 |
| chrX:32561218-32561432 | 1.167220173 | 0.021782082 | 0.116965916 |
| chr8:95519228-95519601 | -0.948994657 | 0.021796682 | 0.117031756 |
| chr16:35525853-35526559 | 0.816185264 | 0.021797398 | 0.117031756 |
| chr8:95853901-95854101 | 1.211466396 | 0.021801728 | 0.117046807 |
| chr14:93736313-93736659 | 0.916545214 | 0.021812986 | 0.117099042 |
| chr18:2766249-2766549 | -1.179399703 | 0.021822724 | 0.117134342 |
| chr20:54503927-54504319 | 0.769677177 | 0.021823081 | 0.117134342 |
| chr10:93386702-93386902 | 1.248562261 | 0.021824148 | 0.117134342 |
| chr18:61799094-61799379 | -1.08513138 | 0.021826948 | 0.117141166 |
| chr3:64139544-64139772 | 0.838451848 | 0.021837489 | 0.117189526 |
| chr3:165056532-165056847 | -1.010346463 | 0.021843726 | 0.11721479 |
| chr10:51882737-51883079 | -1.168283205 | 0.021851381 | 0.117240054 |
| chr1:173454561-173454785 | 1.229806142 | 0.021851495 | 0.117240054 |
| chr2:68063719-68064382 | -0.805057806 | 0.02187923 | 0.117377718 |
| chr5:138065106-138065481 | -1.024330203 | 0.021880216 | 0.117377718 |
| chr4:39384139-39384371 | 1.183377305 | 0.021883353 | 0.117386324 |
| chr20:50824213-50824618 | -0.931556323 | 0.021888095 | 0.117403544 |
| chr2:119492082-119492442 | 0.902448785 | 0.021893915 | 0.117421304 |
| chr3:154105289-154105498 | -1.205215982 | 0.02189494 | 0.117421304 |
| chr15:41001560-41001760 | 1.154161969 | 0.021896004 | 0.117421304 |
| chr5:171670597-171670798 | 1.275533693 | 0.021898816 | 0.117428168 |
| chr3:168671120-168671465 | 0.958339204 | 0.02190158 | 0.117431625 |
| chr6:71365677-71366183 | 0.886913302 | 0.021902526 | 0.117431625 |
| chr8:127776680-127776880 | 1.089732219 | 0.021907074 | 0.117447792 |
| chr17:61335861-61336388 | 0.73584255 | 0.02191684 | 0.117491928 |
| chr2:153871617-153871902 | 1.118970899 | 0.021928887 | 0.117548283 |
| chr7:34900500-34900737 | 1.060226149 | 0.021931635 | 0.117554789 |
| chr7:22433193-22434033 | 0.783304902 | 0.02193415 | 0.117560045 |
| chr13:67366634-67366895 | -1.26311513 | 0.021940559 | 0.117586175 |
| chr10:52644737-52645067 | -0.930280578 | 0.021943428 | 0.117593325 |

| | | | |
|---|---|---|---|
| chr2:30072275-30072589 | -0.98884003 | 0.021947356 | 0.117601243 |
| chr16:57619631-57620823 | 0.76231382 | 0.021947976 | 0.117601243 |
| chr8:69329671-69329955 | -1.041831271 | 0.021950312 | 0.117602053 |
| chr7:116293265-116293505 | -1.17798656 | 0.021951196 | 0.117602053 |
| chr2:173049265-173049630 | 0.935075936 | 0.021957213 | 0.117626063 |
| chr12:12560867-12561331 | 0.857647576 | 0.021965434 | 0.117661876 |
| chr10:67558887-67559229 | -1.086278029 | 0.021967921 | 0.117666974 |
| chr20:53344966-53345202 | 1.074262621 | 0.021978373 | 0.11771473 |
| chr7:39742092-39742624 | -0.798789862 | 0.021988192 | 0.117759088 |
| chr4:126583334-126583962 | -0.834150272 | 0.021998587 | 0.117806524 |
| chr3:167885333-167885643 | -0.961351145 | 0.0220059 | 0.117837448 |
| chr17:29567785-29568150 | 0.832303445 | 0.022027929 | 0.117947168 |
| chr12:52674830-52675030 | 1.282284071 | 0.022032161 | 0.117961586 |
| chr16:28936522-28936854 | 0.868954106 | 0.022036024 | 0.117974026 |
| chr7:17741643-17741843 | 1.100041249 | 0.022049393 | 0.118034073 |
| chr5:153786551-153786761 | -1.317769683 | 0.022050321 | 0.118034073 |
| chr3:32144155-32144444 | 0.974269661 | 0.022056416 | 0.118058448 |
| chr3:192907788-192908551 | -0.772703092 | 0.022063814 | 0.118089799 |
| chr14:99551985-99552194 | 1.103125209 | 0.022073287 | 0.11813225 |
| chr13:20391550-20391800 | 0.96146904 | 0.022081285 | 0.11816105 |
| chr3:15805736-15805970 | -1.163414602 | 0.022082221 | 0.11816105 |
| chr14:32711238-32711502 | 1.026763952 | 0.022083908 | 0.11816105 |
| chr11:116772669-116773532 | 0.787024192 | 0.022085585 | 0.11816105 |
| chr13:71303176-71303444 | -1.114151124 | 0.022089173 | 0.11816105 |
| chr11:117648417-117648617 | 1.157634968 | 0.022089324 | 0.11816105 |
| chr4:175560640-175560956 | -1.02004812 | 0.022090098 | 0.11816105 |
| chr7:128537542-128537742 | -1.433249507 | 0.022091547 | 0.11816105 |
| chrX:24294632-24294950 | -1.13012626 | 0.022092548 | 0.11816105 |
| chr15:90510320-90510544 | 1.156603147 | 0.02209512 | 0.11816656 |
| chr7:130063039-130063243 | -1.272765741 | 0.022115383 | 0.11826667 |
| chr3:23959111-23959699 | -0.840678653 | 0.022119093 | 0.118275578 |
| chr4:24488513-24488943 | 0.974254014 | 0.022120355 | 0.118275578 |
| chr10:35613904-35614111 | -1.184971058 | 0.022122347 | 0.118275578 |
| chr14:52131702-52132040 | 0.913028742 | 0.022123223 | 0.118275578 |
| chr6:30415698-30416087 | 1.04335526 | 0.022126908 | 0.118282126 |
| chr10:59287772-59287975 | -1.235027785 | 0.022127535 | 0.118282126 |
| chr1:109498134-109498695 | -0.81384439 | 0.022130611 | 0.118288629 |
| chr8:74173926-74174252 | 1.015897904 | 0.022131839 | 0.118288629 |
| chr7:129635353-129635863 | -0.969425148 | 0.022133447 | 0.11828897 |
| chr11:130434423-130434782 | 0.887924236 | 0.022135028 | 0.11828917 |
| chr10:129522439-129522642 | 1.141569597 | 0.022136581 | 0.11828922 |
| chr1:94884097-94884435 | -0.987542691 | 0.022139902 | 0.118298717 |
| chr2:180715024-180715465 | -0.910586017 | 0.022142195 | 0.118302718 |
| chr3:20511822-20512317 | 0.825998696 | 0.022147675 | 0.118323747 |
| chr8:126131429-126131662 | 1.026247388 | 0.022155809 | 0.118351812 |
| chr6:53410708-53410992 | -1.06219306 | 0.022156018 | 0.118351812 |
| chr1:95043344-95043722 | -0.960295051 | 0.022162957 | 0.118375904 |
| chr5:158075288-158075643 | -0.928463383 | 0.022163618 | 0.118375904 |
| chr2:9336816-9337361 | -0.848820397 | 0.022165393 | 0.118377135 |
| chr12:53983666-53983937 | 0.989390155 | 0.022169879 | 0.118388166 |

| | | | |
|---|---|---|---|
| chr10:67862226-67862552 | -0.983673639 | 0.022171011 | 0.118388166 |
| chr8:142389651-142390042 | 0.962980697 | 0.022172094 | 0.118388166 |
| chr13:38905781-38906207 | -0.894469769 | 0.022174624 | 0.11838835 |
| chr12:30695310-30696330 | -0.766944127 | 0.022175219 | 0.11838835 |
| chr3:21171010-21171335 | -0.970037723 | 0.022179481 | 0.118402858 |
| chr5:77619769-77619971 | -1.197865388 | 0.022186609 | 0.118432655 |
| chr7:127287078-127287278 | -1.393102387 | 0.022195127 | 0.118469873 |
| chr14:104077972-104078346 | 0.834228297 | 0.022200249 | 0.118488955 |
| chr6:29887271-29887754 | -0.998694717 | 0.022203765 | 0.11849947 |
| chr6:49667910-49668825 | 0.764461945 | 0.022206073 | 0.118503535 |
| chr1:33181589-33182290 | 0.76664651 | 0.022209477 | 0.118513443 |
| chr14:31449945-31450364 | 0.839986391 | 0.022212984 | 0.118518222 |
| chr1:172727395-172727648 | -1.144556306 | 0.022213466 | 0.118518222 |
| chr1:89808406-89808934 | -0.824548297 | 0.022215392 | 0.118520247 |
| chr11:97269552-97269894 | -1.003825528 | 0.022218877 | 0.118530582 |
| chr5:102782194-102782429 | -1.311048596 | 0.022226213 | 0.118561468 |
| chr15:87154860-87155153 | -1.063235893 | 0.0222343 | 0.118596347 |
| chr12:29783393-29784617 | -0.792812847 | 0.022247648 | 0.118659282 |
| chr7:38107510-38107813 | -0.928173653 | 0.02225183 | 0.118673329 |
| chr1:17837947-17838205 | 1.255057138 | 0.022254874 | 0.118678328 |
| chr20:52504102-52504317 | 1.222881663 | 0.022255865 | 0.118678328 |
| chr7:108435939-108436188 | -1.190927678 | 0.022265709 | 0.118722559 |
| chr17:64311395-64311801 | 0.772592165 | 0.02226841 | 0.118728698 |
| chr3:110752414-110752614 | 1.170858465 | 0.022271956 | 0.118739341 |
| chr12:93314489-93315517 | -0.73167476 | 0.022276923 | 0.118753638 |
| chr5:130545326-130545895 | -0.819567841 | 0.022277737 | 0.118753638 |
| chrX:34365561-34365782 | -1.249423424 | 0.022282182 | 0.118760529 |
| chr19:50399068-50399299 | 1.104041349 | 0.022282495 | 0.118760529 |
| chr5:141094439-141095194 | -0.845847014 | 0.022285384 | 0.118760529 |
| chrX:50997029-50997233 | -1.240530886 | 0.022285838 | 0.118760529 |
| chr17:43251574-43251849 | 0.96748525 | 0.02228678 | 0.118760529 |
| chr3:138686777-138687048 | 1.054022882 | 0.022290667 | 0.118772983 |
| chr4:163544407-163544682 | -0.953910576 | 0.022308558 | 0.118860048 |
| chr8:69072805-69073197 | -0.982238119 | 0.02231321 | 0.118876565 |
| chr20:43583282-43583522 | 1.139857734 | 0.022318409 | 0.118889352 |
| chr16:642276-642476 | 0.992167024 | 0.022318713 | 0.118889352 |
| chr7:128381246-128381455 | -1.15799873 | 0.022323915 | 0.118908792 |
| chr18:79528843-79529165 | -0.973723964 | 0.022328275 | 0.11892375 |
| chr9:78167376-78167749 | -0.868986611 | 0.022332395 | 0.118928358 |
| chr20:45319587-45319834 | 0.955215595 | 0.022333097 | 0.118928358 |
| chr2:146412646-146412988 | -0.995533081 | 0.022333796 | 0.118928358 |
| chr6:10355236-10355692 | -0.948623994 | 0.02234031 | 0.118954774 |
| chrX:107205774-107206959 | -0.738159579 | 0.022352003 | 0.11900632 |
| chr7:48311025-48311375 | -1.112645251 | 0.022353097 | 0.11900632 |
| chr20:60270270-60270832 | 0.759875179 | 0.022358197 | 0.119025207 |
| chr11:31641583-31641812 | -1.231724082 | 0.022360751 | 0.119030531 |
| chr4:35525185-35526048 | -0.856058873 | 0.022368952 | 0.119065913 |
| chr17:50164002-50164424 | -0.855575699 | 0.022377108 | 0.119101054 |
| chr5:13668848-13669078 | 1.131494397 | 0.022390459 | 0.119163836 |
| chr4:8352106-8352599 | -0.972271834 | 0.022392949 | 0.119168809 |

| | | | |
|---|---|---|---|
| chr10:52740854-52741247 | -0.874928828 | 0.022399892 | 0.119197482 |
| chr2:199485845-199486267 | -0.910775609 | 0.022406348 | 0.119220038 |
| chr2:200635392-200635682 | -1.181997972 | 0.022407243 | 0.119220038 |
| chr16:15509037-15509494 | -0.927346224 | 0.022414828 | 0.119252115 |
| chr12:104137295-104138590 | -0.735391192 | 0.022420802 | 0.119271646 |
| chr7:39453802-39454114 | -1.061972474 | 0.022421612 | 0.119271646 |
| chr2:22984219-22984517 | -1.260813181 | 0.022440648 | 0.11936462 |
| chr2:109613762-109614676 | 0.743268544 | 0.022449974 | 0.119405936 |
| chr16:4918710-4919385 | 0.807679463 | 0.022453017 | 0.119413829 |
| chr5:2592064-2592497 | -0.940355774 | 0.022457593 | 0.119429879 |
| chr4:164997393-164997855 | -0.880939033 | 0.022460564 | 0.119435569 |
| chr1:221886931-221887580 | -0.787147371 | 0.022462759 | 0.119435569 |
| chr15:74482751-74482970 | 1.096920815 | 0.022465039 | 0.119435569 |
| chr10:106808625-106809034 | -1.000280455 | 0.022466278 | 0.119435569 |
| chr16:31476943-31477185 | 0.898867104 | 0.022466457 | 0.119435569 |
| chr4:176348561-176349102 | -0.893517864 | 0.022471644 | 0.11945085 |
| chr3:64327431-64327647 | 0.84344815 | 0.022472449 | 0.11945085 |
| chr10:75512485-75512751 | 1.052510786 | 0.02247891 | 0.119476904 |
| chr5:110590289-110590543 | -1.037652063 | 0.022486906 | 0.119511112 |
| chr2:17830553-17831214 | -0.759383605 | 0.022493793 | 0.119532891 |
| chr7:13742757-13743032 | 0.923852794 | 0.022494124 | 0.119532891 |
| chr10:3538373-3539128 | 0.788420251 | 0.022503605 | 0.119574983 |
| chr12:130197169-130197396 | 1.134846168 | 0.022505896 | 0.119578861 |
| chr8:99340183-99340429 | 1.138376574 | 0.022510174 | 0.119593298 |
| chr15:91185514-91185832 | 1.093020228 | 0.022511843 | 0.119593872 |
| chr7:122887260-122887600 | -0.852770575 | 0.022519009 | 0.119621555 |
| chr12:51521662-51521887 | 1.066498588 | 0.022520176 | 0.119621555 |
| chr10:12549409-12549748 | -1.022737582 | 0.022523214 | 0.119625557 |
| chr17:17072087-17072359 | -1.060910745 | 0.022524052 | 0.119625557 |
| chr2:98829000-98829200 | 1.160870443 | 0.022532716 | 0.119663275 |
| chr20:54429113-54429425 | 0.781326192 | 0.022537155 | 0.119674195 |
| chr4:136831594-136831930 | -0.917560254 | 0.022538654 | 0.119674195 |
| chr6:12008460-12009194 | -0.774283187 | 0.022540743 | 0.119674195 |
| chr8:123096339-123096813 | 0.855722098 | 0.022541382 | 0.119674195 |
| chr2:131335371-131335649 | -1.176843081 | 0.022544037 | 0.119674195 |
| chr4:88514926-88515405 | 0.798377974 | 0.022544143 | 0.119674195 |
| chr1:12229755-12230315 | 0.869584828 | 0.022547258 | 0.119680123 |
| chr21:41902975-41903183 | 1.167841837 | 0.022548384 | 0.119680123 |
| chr12:53973458-53974029 | 0.818212043 | 0.02255215 | 0.119691821 |
| chr7:42198356-42198684 | -0.954736639 | 0.022555391 | 0.119700735 |
| chr5:74016435-74016736 | -0.986730643 | 0.022566562 | 0.119748165 |
| chr2:104046527-104046837 | 1.002600727 | 0.022567455 | 0.119748165 |
| chr7:143345361-143346182 | -0.806367285 | 0.022575411 | 0.11978209 |
| chr14:69168633-69168897 | -1.18325798 | 0.022577652 | 0.119785686 |
| chr21:39666830-39667212 | -0.91046735 | 0.02257965 | 0.11978799 |
| chr6:143941875-143942622 | -0.780846598 | 0.022584152 | 0.119800673 |
| chr7:86926644-86927011 | -0.980767712 | 0.022585196 | 0.119800673 |
| chr9:68560084-68560440 | -1.033694881 | 0.022586731 | 0.119800673 |
| chr2:73254574-73254779 | 1.095252385 | 0.022593135 | 0.119823308 |
| chr1:195685894-195686210 | -0.979396893 | 0.022594126 | 0.119823308 |

| | | | |
|---|---|---|---|
| chr1:43206701-43207137 | 0.862452852 | 0.022596936 | 0.119829915 |
| chr11:76836341-76837038 | -0.836135068 | 0.022599247 | 0.119833876 |
| chr15:48855873-48856199 | -0.972314384 | 0.022610483 | 0.119878995 |
| chr7:131347045-131347504 | -0.969622935 | 0.022611609 | 0.119878995 |
| chr5:152116889-152117282 | -1.008954805 | 0.02261245 | 0.119878995 |
| chr11:15006203-15006406 | -1.288696482 | 0.022638401 | 0.120008272 |
| chr21:45140312-45140701 | 0.827505058 | 0.022651152 | 0.120067559 |
| chr5:148781298-148781761 | -1.01762485 | 0.022657859 | 0.120083455 |
| chr4:165623495-165623994 | -0.867467974 | 0.022659544 | 0.120083455 |
| chr3:17238170-17238416 | -1.055950919 | 0.022661871 | 0.120083455 |
| chr3:127074988-127075352 | 0.841293287 | 0.022664388 | 0.120083455 |
| chr2:229408292-229408812 | -0.8174531 | 0.022664443 | 0.120083455 |
| chr10:123451689-123451939 | -1.158836677 | 0.022665121 | 0.120083455 |
| chr12:93047203-93047473 | -1.158836677 | 0.022665121 | 0.120083455 |
| chr1:48498546-48498861 | -0.898911811 | 0.022670401 | 0.120096579 |
| chr3:173140994-173141219 | -1.317034291 | 0.022670733 | 0.120096579 |
| chr10:126387148-126387348 | 1.263066372 | 0.022676006 | 0.120109437 |
| chr20:54314433-54314750 | 0.78388548 | 0.022676295 | 0.120109437 |
| chr4:15001438-15003254 | 0.721994635 | 0.022681281 | 0.120127537 |
| chr4:158805630-158806005 | -0.93158606 | 0.022686847 | 0.120148715 |
| chr3:153140885-153141110 | 1.058074812 | 0.022689447 | 0.120154177 |
| chr22:19171486-19172012 | 0.799573788 | 0.022698071 | 0.120191543 |
| chr5:44913347-44913553 | 1.256945163 | 0.022704581 | 0.120217704 |
| chr14:28294955-28295156 | 1.050774571 | 0.022706178 | 0.120217854 |
| chr2:129532289-129532505 | -1.201179173 | 0.022710257 | 0.12023114 |
| chr18:74331409-74331742 | -0.956347621 | 0.022716996 | 0.120231307 |
| chr2:132346939-132347792 | -0.762889396 | 0.022717228 | 0.120231307 |
| chr6:38012644-38012902 | 1.055737682 | 0.022718395 | 0.120231307 |
| chr7:102715762-102716111 | -1.132844561 | 0.022718565 | 0.120231307 |
| chr1:158087331-158087891 | -0.830365482 | 0.022719329 | 0.120231307 |
| chr16:85751054-85751285 | 0.921793519 | 0.022719703 | 0.120231307 |
| chr1:107066602-107066971 | 0.809602894 | 0.022729406 | 0.120254625 |
| chr12:2433344-2433544 | 1.303781978 | 0.022729547 | 0.120254625 |
| chr4:6715832-6716854 | 0.762313678 | 0.022729596 | 0.120254625 |
| chr7:123797911-123798562 | -0.806283256 | 0.022730677 | 0.120254625 |
| chr1:202199957-202200655 | -0.751650646 | 0.022731957 | 0.120254625 |
| chr5:176996559-176996776 | 1.223634243 | 0.02273942 | 0.120285804 |
| chr17:58462908-58463197 | 1.075542196 | 0.022746614 | 0.120308579 |
| chr2:75425019-75425351 | -1.173515699 | 0.022746866 | 0.120308579 |
| chr8:119144058-119144347 | 1.041880633 | 0.022750621 | 0.120320131 |
| chr17:74749897-74750115 | 0.952178285 | 0.022755851 | 0.120339488 |
| chr18:24215592-24215954 | 0.929708198 | 0.022760466 | 0.120355586 |
| chr2:68185599-68185893 | -0.99257578 | 0.022767244 | 0.120375432 |
| chr15:77704115-77704315 | 1.074563606 | 0.022767361 | 0.120375432 |
| chr1:217477008-217477333 | 0.971106503 | 0.022770306 | 0.120382694 |
| chr7:17558982-17559400 | 0.868565919 | 0.022780862 | 0.12043019 |
| chr6:139713583-139713897 | -1.150659365 | 0.022788976 | 0.120456558 |
| chr7:47571268-47571614 | -0.851559925 | 0.022788994 | 0.120456558 |
| chr14:74805332-74805602 | 1.099253328 | 0.022793478 | 0.120464577 |
| chr8:106958804-106959080 | 1.082577406 | 0.022793655 | 0.120464577 |

| | | | |
|---|---|---|---|
| chr2:120630211-120630713 | 0.847124979 | 0.022803783 | 0.12050979 |
| chr1:178310669-178311060 | -0.911551975 | 0.022820089 | 0.120587647 |
| chr14:103317235-103317435 | 1.09962449 | 0.022825967 | 0.120603106 |
| chr15:95988329-95988814 | -0.924311779 | 0.022827459 | 0.120603106 |
| chr2:219495391-219495688 | 0.856796791 | 0.022827737 | 0.120603106 |
| chr7:29988003-29988314 | 0.930650524 | 0.022829894 | 0.120606186 |
| chr7:18345747-18346001 | 0.977663099 | 0.022832498 | 0.12061163 |
| chr10:62264687-62265050 | -0.881125119 | 0.022838455 | 0.120634777 |
| chr3:133272169-133272369 | 1.349155092 | 0.022841659 | 0.120637163 |
| chr3:64546929-64547201 | 0.812345633 | 0.022842055 | 0.120637163 |
| chr2:183356885-183357244 | -1.034453143 | 0.022851694 | 0.120679749 |
| chr10:121077385-121077607 | 1.197147927 | 0.022856209 | 0.120695276 |
| chr1:182953471-182953693 | -1.122783807 | 0.022859698 | 0.12070538 |
| chr1:165816232-165816469 | -1.288019319 | 0.022865274 | 0.120726504 |
| chr8:3783092-3783584 | -0.833688124 | 0.022873795 | 0.120753937 |
| chr17:7444946-7445156 | -1.248664243 | 0.022875212 | 0.120753937 |
| chr15:87015368-87015668 | -0.972947235 | 0.022876091 | 0.120753937 |
| chr3:113685940-113686698 | -0.79804439 | 0.022876774 | 0.120753937 |
| chr5:850523-850892 | 0.839953076 | 0.022882947 | 0.120778201 |
| chr4:186726329-186727062 | -0.780569752 | 0.022894657 | 0.120831685 |
| chr8:22864848-22865139 | -1.009566334 | 0.022898024 | 0.120841133 |
| chr7:17706783-17707079 | 0.900647595 | 0.022903829 | 0.120863441 |
| chr11:19072600-19072865 | 0.991923058 | 0.022905678 | 0.120864877 |
| chr10:87578941-87579422 | -0.910514942 | 0.022916163 | 0.120910759 |
| chr4:102737133-102737527 | 0.826409707 | 0.02291753 | 0.120910759 |
| chr14:22200978-22201292 | 0.944095989 | 0.022919441 | 0.120912519 |
| chr15:74642214-74642696 | 0.851738354 | 0.02292334 | 0.120924761 |
| chr10:86970940-86971632 | -0.763844459 | 0.022926637 | 0.120933829 |
| chr20:55580047-55580253 | 0.827704864 | 0.022929099 | 0.120938487 |
| chrX:138745347-138745547 | 1.205513614 | 0.022934171 | 0.120956913 |
| chr11:86781577-86781858 | -1.014685548 | 0.022952405 | 0.121044749 |
| chr1:160219627-160219902 | 0.997920682 | 0.022954374 | 0.121046802 |
| chr6:30686690-30687579 | 0.756992315 | 0.022957941 | 0.121057283 |
| chr7:131222302-131222513 | -1.273611183 | 0.022961534 | 0.121067895 |
| chr7:95840943-95841187 | -1.189196835 | 0.022966933 | 0.12108803 |
| chr6:30728442-30728664 | 1.090206916 | 0.022974898 | 0.121121691 |
| chrX:145817409-145817725 | -1.318441388 | 0.022982743 | 0.121154713 |
| chr4:96786459-96786857 | -0.987465411 | 0.022985692 | 0.121161921 |
| chr1:3587219-3587802 | 0.888844827 | 0.022988287 | 0.121165643 |
| chr5:142816548-142816996 | -0.90311429 | 0.0229897 | 0.121165643 |
| chr5:118611190-118611713 | -0.882769954 | 0.022991142 | 0.121165643 |
| chr4:16226402-16226873 | 0.855617096 | 0.022997792 | 0.121178322 |
| chr2:16425530-16426138 | -0.854664484 | 0.022997862 | 0.121178322 |
| chr3:192965341-192965692 | -0.923036957 | 0.022999315 | 0.121178322 |
| chr2:185197017-185197289 | -1.199649848 | 0.022999874 | 0.121178322 |
| chr4:56821382-56821993 | 0.80403927 | 0.023010286 | 0.121224843 |
| chr2:241733200-241733711 | -0.845763672 | 0.023023393 | 0.121283678 |
| chr1:2085152-2085493 | -1.025221754 | 0.023025007 | 0.121283678 |
| chr7:148940754-148941703 | -0.751418445 | 0.023026202 | 0.121283678 |
| chr2:157327492-157327892 | -0.838378775 | 0.023045572 | 0.12137736 |

| | | | |
|---|---|---|---|
| chr13:82867370-82867570 | -1.284906249 | 0.023050823 | 0.121396669 |
| chr15:69010459-69010770 | 1.060860573 | 0.02305285 | 0.121396754 |
| chr17:41811640-41812570 | 0.761829181 | 0.023054008 | 0.121396754 |
| chr5:95846997-95847333 | -0.843724026 | 0.023065473 | 0.12144878 |
| chr5:59543821-59544337 | -0.888686945 | 0.023073438 | 0.121482375 |
| chr17:41421226-41421426 | 1.230391821 | 0.023075094 | 0.121482743 |
| chr9:105662348-105662669 | -1.065432799 | 0.023077893 | 0.12148436 |
| chr2:20145865-20146065 | -1.171656165 | 0.023078572 | 0.12148436 |
| chr3:121072300-121072500 | -1.340989201 | 0.023084655 | 0.121506129 |
| chr20:3470770-3471184 | 0.867211146 | 0.023085879 | 0.121506129 |
| chr2:233448314-233448559 | -1.215147605 | 0.023102673 | 0.121586165 |
| chr10:124093204-124093857 | 0.768709741 | 0.023105256 | 0.121591409 |
| chr14:99361078-99361278 | 1.167646132 | 0.02310764 | 0.121595605 |
| chr6:33490377-33490577 | 1.247829291 | 0.023110706 | 0.12159602 |
| chr16:22206974-22207218 | 0.968669233 | 0.023110893 | 0.12159602 |
| chr2:196516625-196517058 | -0.977035168 | 0.023120775 | 0.121639659 |
| chr12:21437482-21438484 | -0.755491003 | 0.023127313 | 0.121649507 |
| chr15:79304428-79304683 | 1.1563323 | 0.023127889 | 0.121649507 |
| chr5:138789724-138790205 | -0.922875363 | 0.023128988 | 0.121649507 |
| chr1:219089042-219089281 | -1.108367213 | 0.023128997 | 0.121649507 |
| chr7:90346122-90347115 | -0.740755124 | 0.023138818 | 0.121692809 |
| chr7:124764798-124766332 | -0.725303516 | 0.02314067 | 0.121694196 |
| chr12:62505363-62505563 | 1.214897493 | 0.023153992 | 0.121755898 |
| chr10:19832687-19832906 | -1.092955541 | 0.023158069 | 0.121766953 |
| chr1:201177051-201177345 | 0.946778463 | 0.023159273 | 0.121766953 |
| chr5:67621628-67621932 | -1.039137899 | 0.023165573 | 0.121791723 |
| chr17:4533512-4533730 | -1.097155822 | 0.023169548 | 0.121799646 |
| chr12:110355117-110355395 | -1.06380934 | 0.023170383 | 0.121799646 |
| chr4:4165733-4165983 | 1.134965482 | 0.023171849 | 0.121799646 |
| chr21:9031830-9032297 | 0.97149383 | 0.023185128 | 0.121853299 |
| chr12:79532820-79533217 | 0.867917216 | 0.023185237 | 0.121853299 |
| chr13:87670915-87672301 | -0.731284888 | 0.023196247 | 0.121899552 |
| chr12:40032459-40032786 | 1.049200414 | 0.02319722 | 0.121899552 |
| chrX:129726341-129726734 | 0.917744125 | 0.023200526 | 0.121904357 |
| chr17:16881019-16881621 | 0.90656977 | 0.02320245 | 0.121904357 |
| chr9:69168908-69169224 | -1.055605664 | 0.023202907 | 0.121904357 |
| chr2:29165663-29165863 | 1.229231363 | 0.023208471 | 0.121925228 |
| chr2:70212515-70212858 | -1.169981985 | 0.023224849 | 0.122002905 |
| chr2:231000403-231000737 | 1.001491539 | 0.023236544 | 0.122055972 |
| chr1:99965624-99966318 | -0.77303594 | 0.02324578 | 0.122091496 |
| chr20:52426848-52427156 | -0.99627966 | 0.023246494 | 0.122091496 |
| chr11:67055015-67055685 | -0.73885071 | 0.023256096 | 0.122130241 |
| chrX:100642338-100642594 | -1.11516946 | 0.023257059 | 0.122130241 |
| chr19:51627084-51627284 | 1.278310417 | 0.023265146 | 0.122164336 |
| chr2:207734300-207734571 | 0.997162741 | 0.02326957 | 0.122179193 |
| chr3:112293361-112294346 | -0.766414821 | 0.023271832 | 0.122182701 |
| chr2:234366963-234367209 | 1.166337794 | 0.023279989 | 0.122217152 |
| chr8:85474716-85475085 | 0.883121877 | 0.023282728 | 0.122223156 |
| chr1:152291576-152291776 | 1.157772803 | 0.023289713 | 0.122246963 |
| chr3:67154393-67154737 | -1.018941015 | 0.023290454 | 0.122246963 |

| | | | |
|---|---|---|---|
| chr4:103425409-103425692 | -0.996393804 | 0.023295033 | 0.122258719 |
| chr5:62752273-62752592 | -0.975018107 | 0.023295885 | 0.122258719 |
| chr12:6272581-6273229 | -0.858487731 | 0.023299846 | 0.122271134 |
| chr9:69346666-69346939 | 1.068113239 | 0.023302457 | 0.12227646 |
| chr15:75862992-75863217 | 1.245767409 | 0.02331125 | 0.122296716 |
| chr2:62293359-62294145 | -0.765833182 | 0.02331204 | 0.122296716 |
| chr5:179803089-179803865 | -0.760713674 | 0.023313713 | 0.122296716 |
| chr8:132193864-132194125 | 1.021897931 | 0.023315946 | 0.122296716 |
| chr1:106986571-106987185 | 0.775104297 | 0.023317702 | 0.122296716 |
| chr4:116664163-116664517 | -0.923396163 | 0.023317794 | 0.122296716 |
| chr2:208657547-208657811 | -1.030981428 | 0.023318866 | 0.122296716 |
| chr10:53298539-53298739 | -1.31862846 | 0.023319086 | 0.122296716 |
| chr2:190023753-190024153 | -1.146863036 | 0.023323822 | 0.122313179 |
| chr9:136398989-136399311 | 1.054143189 | 0.023326165 | 0.122317094 |
| chr3:147008478-147008805 | 1.042290799 | 0.023348367 | 0.122425139 |
| chr5:168401867-168402736 | -0.757504517 | 0.023354463 | 0.122448723 |
| chr8:98345433-98345702 | 1.001211359 | 0.023358774 | 0.122456513 |
| chr10:125294220-125294516 | 0.938167082 | 0.023359145 | 0.122456513 |
| chr20:41192084-41192545 | 0.897231735 | 0.023362009 | 0.122458314 |
| chr8:144413439-144413780 | 0.807041879 | 0.023363911 | 0.122458314 |
| chr14:76593321-76593555 | 1.107058988 | 0.023364283 | 0.122458314 |
| chr7:51733031-51733250 | -1.192393784 | 0.023369494 | 0.122477249 |
| chr12:42442296-42442640 | -1.071100177 | 0.023382505 | 0.122537057 |
| chr11:15629748-15629977 | -1.102314829 | 0.02339042 | 0.12257015 |
| chr1:215082565-215084009 | -0.73499997 | 0.023394203 | 0.122573499 |
| chr21:41973915-41974375 | -0.843158616 | 0.023394258 | 0.122573499 |
| chr20:47536089-47536415 | 0.757976605 | 0.023397547 | 0.122582346 |
| chr9:134644752-134645002 | 1.04579451 | 0.023403952 | 0.122607521 |
| chr3:21988558-21989273 | -0.837910327 | 0.023419037 | 0.122678159 |
| chr14:73569109-73569643 | 0.863794098 | 0.023421867 | 0.122684599 |
| chr17:46182958-46183158 | 1.288294783 | 0.023433644 | 0.122737895 |
| chr7:30889362-30889571 | 1.201725817 | 0.023436616 | 0.12274507 |
| chr7:100559823-100560105 | -0.959514787 | 0.023450755 | 0.122810729 |
| chr20:53582431-53582726 | 0.765095572 | 0.023454384 | 0.122821335 |
| chr21:10413154-10413914 | 0.762606592 | 0.023461935 | 0.122852483 |
| chr4:127963980-127964321 | 0.805243657 | 0.023466987 | 0.122870541 |
| chr20:58150047-58150283 | 0.968834828 | 0.023471863 | 0.122887673 |
| chr5:79878735-79879064 | -0.956101919 | 0.023477765 | 0.122910175 |
| chr5:175967951-175968434 | 0.894991382 | 0.023483446 | 0.122931515 |
| chr12:76092381-76092768 | -0.9259073 | 0.023491466 | 0.122965099 |
| chr5:133287650-133287875 | 1.086090973 | 0.023497107 | 0.122986223 |
| chr7:127347529-127348341 | -0.775624132 | 0.02349929 | 0.122989249 |
| chr8:124464450-124464803 | -0.892232053 | 0.023501473 | 0.122992271 |
| chr11:47259222-47259425 | 1.024549148 | 0.023504311 | 0.122998725 |
| chr12:25959743-25959943 | -1.152155252 | 0.023519941 | 0.123072109 |
| chr9:72203254-72203546 | -1.036747096 | 0.023523851 | 0.123079932 |
| chr7:138262654-138263075 | -0.961044755 | 0.023524648 | 0.123079932 |
| chr2:184753919-184754298 | -0.925747254 | 0.023528004 | 0.123083538 |
| chr8:91846075-91846315 | 1.266870385 | 0.02352855 | 0.123083538 |
| chr5:52778839-52779090 | -1.191010742 | 0.023534063 | 0.123096124 |

| | | | |
|---|---|---|---|
| chr14:34029961-34030307 | -0.889113081 | 0.023535265 | 0.123096124 |
| chr14:53458352-53458622 | 1.106744933 | 0.023535776 | 0.123096124 |
| chr9:94301152-94301387 | -1.076866339 | 0.023538839 | 0.123099534 |
| chr5:176135376-176135732 | 1.013097497 | 0.023541235 | 0.123099534 |
| chr7:130051248-130051632 | -0.882320319 | 0.023541851 | 0.123099534 |
| chr5:23984646-23984968 | -1.147172106 | 0.023543117 | 0.123099534 |
| chr15:62773128-62773374 | -1.313222525 | 0.023544461 | 0.123099534 |
| chr4:136810526-136810921 | -1.153862153 | 0.023554213 | 0.123142121 |
| chr2:54838580-54838909 | -0.939548661 | 0.023559339 | 0.12315728 |
| chr1:218521427-218521924 | -0.830196647 | 0.023560328 | 0.12315728 |
| chr10:123614966-123615202 | 1.120217332 | 0.023562087 | 0.123158077 |
| chr13:28693550-28694411 | -0.782849052 | 0.023568171 | 0.123174506 |
| chr8:22881059-22881259 | 1.260822163 | 0.023568445 | 0.123174506 |
| chr19:52095653-52096112 | 0.809048427 | 0.02357174 | 0.123183324 |
| chr8:3062249-3062464 | -1.341578133 | 0.023580828 | 0.123215127 |
| chr17:61900168-61900604 | 0.742720715 | 0.023581042 | 0.123215127 |
| chr20:48421534-48421845 | 0.78394905 | 0.023585372 | 0.123227683 |
| chr17:58893702-58893911 | 0.90846053 | 0.023586662 | 0.123227683 |
| chr14:23057457-23058387 | 0.741902496 | 0.0235903 | 0.123233228 |
| chr7:20539261-20539504 | 1.042143748 | 0.02359094 | 0.123233228 |
| chr3:64964746-64965274 | 0.759469211 | 0.023596374 | 0.123253212 |
| chr2:169131935-169132177 | 1.068398198 | 0.023608315 | 0.123304821 |
| chr3:153073733-153073987 | 1.010423839 | 0.023610251 | 0.123304821 |
| chr1:228378169-228378921 | 0.77860033 | 0.023611082 | 0.123304821 |
| chr12:94101478-94101712 | 0.969665604 | 0.023624412 | 0.123366027 |
| chr2:189762289-189762990 | -0.746168451 | 0.023633008 | 0.123402505 |
| chr1:113912784-113913261 | 0.75781459 | 0.023636315 | 0.123411359 |
| chr3:81242935-81243190 | -1.052603539 | 0.023644665 | 0.123446548 |
| chrX:46792054-46792309 | 1.177579377 | 0.023647017 | 0.123450413 |
| chrX:87583709-87583973 | -1.179130093 | 0.023649619 | 0.123455588 |
| chr16:4176886-4177302 | -0.927183281 | 0.023651422 | 0.12345659 |
| chr10:112721123-112721396 | 0.996516311 | 0.023672189 | 0.123556572 |
| chr7:157404968-157405570 | -0.768596591 | 0.023679756 | 0.123587646 |
| chr8:23008698-23008898 | -1.30141618 | 0.023687252 | 0.12361835 |
| chr12:96215229-96215607 | -0.818166419 | 0.023699576 | 0.123672181 |
| chr15:31397081-31397652 | -0.782984605 | 0.023702169 | 0.123672181 |
| chr17:62000981-62002499 | 0.714149469 | 0.023702601 | 0.123672181 |
| chr1:156907305-156907505 | 0.967685665 | 0.023704023 | 0.123672181 |
| chr12:52139582-52139783 | 0.9691493 | 0.023716896 | 0.123730917 |
| chr13:106423531-106424000 | -0.979532459 | 0.023720317 | 0.123740341 |
| chr7:106659897-106661450 | -0.730389599 | 0.02372758 | 0.123766847 |
| chr2:17851780-17852246 | -0.886383945 | 0.023728629 | 0.123766847 |
| chr15:25438789-25439352 | 0.804536017 | 0.023736625 | 0.123800127 |
| chr7:116341624-116341824 | -1.340013898 | 0.023748496 | 0.123850538 |
| chr4:3461262-3462130 | -0.760416992 | 0.023749523 | 0.123850538 |
| chr1:168709964-168710390 | 0.831798321 | 0.023757858 | 0.123879781 |
| chr3:22978292-22978494 | -1.247670601 | 0.023758364 | 0.123879781 |
| chr9:13063989-13064539 | -0.932143133 | 0.023761613 | 0.123888291 |
| chr6:28657754-28658404 | -0.794051723 | 0.023766799 | 0.123903963 |
| chr12:75891206-75891450 | 1.000041595 | 0.023767853 | 0.123903963 |

| | | | |
|---|---|---|---|
| chr3:8579413-8579613 | 1.219197456 | 0.023771295 | 0.123913475 |
| chr3:128559600-128559800 | 1.107684916 | 0.023777704 | 0.123938449 |
| chr7:54831712-54831941 | -1.213796788 | 0.023785762 | 0.123967628 |
| chr6:321187-321432 | 0.985392194 | 0.023786537 | 0.123967628 |
| chr10:79054168-79054431 | 1.056767049 | 0.023789771 | 0.123976049 |
| chr20:17557643-17558183 | 0.831660862 | 0.023803817 | 0.124040807 |
| chr1:114363920-114364130 | 0.896322171 | 0.023808749 | 0.124058075 |
| chr11:95512633-95512920 | 1.096999778 | 0.023811809 | 0.124059872 |
| chr11:70145073-70145273 | 1.113700227 | 0.023812332 | 0.124059872 |
| chr10:88951589-88951825 | -1.099012005 | 0.023815634 | 0.124068639 |
| chr6:15510611-15510811 | 1.208261066 | 0.023830763 | 0.124124288 |
| chr14:43724769-43725005 | 1.06260058 | 0.023830934 | 0.124124288 |
| chr5:142965271-142965594 | -1.004351939 | 0.023831176 | 0.124124288 |
| chr1:209827519-209828364 | -0.751613713 | 0.023833766 | 0.124129337 |
| chr10:42769628-42770048 | 0.882612491 | 0.023842231 | 0.124164983 |
| chr17:50153961-50154385 | -0.832996949 | 0.023852801 | 0.124211588 |
| chr3:121938149-121938895 | -0.785150297 | 0.023874174 | 0.124299107 |
| chr12:106112168-106112636 | -0.966088123 | 0.023874333 | 0.124299107 |
| chr1:149959399-149959667 | 1.031146937 | 0.023874474 | 0.124299107 |
| chr6:136526556-136526891 | -1.105412463 | 0.023886487 | 0.124353201 |
| chr5:82597533-82597733 | -1.299592013 | 0.023889253 | 0.124358367 |
| chr3:119171192-119171392 | 1.235736265 | 0.023891679 | 0.124358367 |
| chr5:13695099-13695480 | -0.86784458 | 0.023892348 | 0.124358367 |
| chr2:57418648-57419349 | -0.913402738 | 0.023894957 | 0.124363498 |
| chr7:22103696-22104008 | 0.858350105 | 0.023910103 | 0.124433874 |
| chr4:6427810-6428048 | 1.126411518 | 0.023930468 | 0.124531398 |
| chr1:71916955-71917181 | -1.246237356 | 0.023934414 | 0.124535346 |
| chr3:32340169-32340423 | -1.09002577 | 0.023934477 | 0.124535346 |
| chr2:105187098-105187427 | 0.97519894 | 0.023947078 | 0.12459245 |
| chr19:35545378-35546059 | 0.793856263 | 0.02395338 | 0.124616776 |
| chr5:132239009-132239209 | 1.115780296 | 0.023959726 | 0.124641329 |
| chr5:178818208-178818425 | -1.212286045 | 0.023964519 | 0.1246578 |
| chr15:66946698-66946908 | 1.093611005 | 0.023977085 | 0.124706398 |
| chr11:61638334-61638719 | 0.880772546 | 0.023977117 | 0.124706398 |
| chr8:101360792-101361297 | 0.773711939 | 0.023984829 | 0.124738039 |
| chr6:76702185-76702599 | -1.106008689 | 0.0239936 | 0.124775188 |
| chr7:42888087-42888459 | 1.070408153 | 0.023995348 | 0.124775807 |
| chr11:95158833-95159042 | -1.191877637 | 0.023997518 | 0.124778628 |
| chr8:140176262-140176518 | -1.099298167 | 0.023999548 | 0.124780712 |
| chr16:642558-642806 | 0.981826169 | 0.024009878 | 0.124809721 |
| chr14:23129676-23130315 | 0.786280139 | 0.024009964 | 0.124809721 |
| chr11:26964651-26965373 | -0.835429272 | 0.024011019 | 0.124809721 |
| chr5:115033854-115034245 | -0.923544044 | 0.024011643 | 0.124809721 |
| chr17:61573790-61574030 | 0.772748317 | 0.024015113 | 0.12481929 |
| chr16:5616179-5616379 | 1.152307931 | 0.024018522 | 0.124828542 |
| chr16:89990635-89990835 | -1.167886225 | 0.02402187 | 0.124837472 |
| chr20:50801618-50802586 | 0.712959264 | 0.024032134 | 0.124873816 |
| chr13:76019394-76019872 | -0.865335076 | 0.024032961 | 0.124873816 |
| chr7:22001415-22001710 | 0.881492844 | 0.024034538 | 0.124873816 |
| chr2:210430507-210430907 | 0.939053218 | 0.024035382 | 0.124873816 |

| | | | |
|---|---|---|---|
| chr8:95304338-95304538 | 1.081737752 | 0.024042286 | 0.124899079 |
| chr3:123799724-123799967 | 1.136940743 | 0.024043505 | 0.124899079 |
| chr2:11370577-11371135 | -0.814048783 | 0.024056531 | 0.124958273 |
| chr19:37294336-37294584 | 1.18738337 | 0.024058579 | 0.124960441 |
| chr9:135972015-135972236 | 0.977836563 | 0.024065403 | 0.124987413 |
| chr10:48626911-48627259 | -1.274539993 | 0.024067093 | 0.124987721 |
| chr4:180146135-180146750 | -0.873463043 | 0.024072268 | 0.125002735 |
| chr6:85537129-85537410 | -1.132657321 | 0.024073247 | 0.125002735 |
| chr4:155418650-155419498 | -0.845804375 | 0.024085491 | 0.125057838 |
| chr4:183504158-183505731 | -0.718956881 | 0.024090054 | 0.125073053 |
| chr10:96864148-96864575 | 0.858878411 | 0.024092815 | 0.125075883 |
| chr1:32361756-32362075 | 0.915915148 | 0.024093864 | 0.125075883 |
| chr3:152828600-152828843 | 1.14747413 | 0.024095943 | 0.1250782 |
| chr5:75203063-75203316 | -1.073981218 | 0.024116829 | 0.125169658 |
| chr5:83418426-83418809 | -1.073981218 | 0.024116829 | 0.125169658 |
| chr1:56811453-56811663 | -1.147871722 | 0.024123322 | 0.125194877 |
| chr1:144523619-144524449 | 0.764111729 | 0.024130569 | 0.125224006 |
| chr9:89395047-89395247 | 1.259775017 | 0.024141368 | 0.125270945 |
| chr11:67055829-67056183 | -0.8089689 | 0.024142884 | 0.125270945 |
| chr21:33931400-33931835 | -0.972397737 | 0.024145589 | 0.125276496 |
| chr13:109971666-109972064 | -0.996474654 | 0.024162029 | 0.125352435 |
| chr18:22170688-22170888 | 1.154858327 | 0.024163497 | 0.125352435 |
| chr12:12398475-12398708 | -1.190534861 | 0.024167369 | 0.125357646 |
| chr2:30446977-30447978 | -0.760327806 | 0.024167774 | 0.125357646 |
| chr3:13475832-13476476 | -0.74388007 | 0.024170235 | 0.125361929 |
| chr2:121036230-121036517 | 0.990943324 | 0.024173792 | 0.12537189 |
| chr15:88913029-88913578 | 0.790907234 | 0.024177191 | 0.125381031 |
| chr20:57498847-57499235 | 0.79833921 | 0.024181302 | 0.125393862 |
| chr20:51311980-51312193 | 0.930357783 | 0.024197257 | 0.125468109 |
| chr8:122324797-122325081 | 1.031266277 | 0.024206764 | 0.12550891 |
| chr4:65670234-65670972 | -0.786725876 | 0.024209474 | 0.125514468 |
| chr17:61701809-61702019 | 0.798903584 | 0.024212022 | 0.125519187 |
| chr10:72980968-72981200 | -1.219086152 | 0.024214228 | 0.125522127 |
| chr1:31214642-31215023 | 0.986468283 | 0.024222722 | 0.125557665 |
| chr14:77123384-77124066 | -0.802558521 | 0.024230285 | 0.12558187 |
| chr19:10074168-10074519 | 0.898328528 | 0.02423067 | 0.12558187 |
| chr1:215992683-215992977 | -0.969938737 | 0.024233221 | 0.125585603 |
| chr11:69915506-69915746 | 0.95418675 | 0.024234668 | 0.125585603 |
| chr6:83431783-83432098 | 0.87386763 | 0.024244762 | 0.125626321 |
| chr3:106360745-106361328 | 0.819941836 | 0.024246413 | 0.125626321 |
| chr7:113281954-113282180 | -1.142224089 | 0.024248025 | 0.125626321 |
| chr20:58944803-58945003 | 1.110246377 | 0.024249084 | 0.125626321 |
| chrX:115345418-115345653 | 1.053734011 | 0.024260609 | 0.125669923 |
| chr17:64481041-64481249 | 1.142999191 | 0.02426078 | 0.125669923 |
| chr1:219385885-219386206 | -0.837272845 | 0.024265351 | 0.125685102 |
| chr8:13122724-13123371 | -0.888849999 | 0.024270098 | 0.125698612 |
| chr18:48777153-48777577 | -0.91844935 | 0.024273424 | 0.125698612 |
| chr14:50525088-50525316 | 1.065871117 | 0.024273783 | 0.125698612 |
| chr2:237483055-237483700 | -0.799516238 | 0.024275412 | 0.125698612 |
| chr5:116572377-116573001 | -0.766889688 | 0.024276162 | 0.125698612 |

| | | | |
|---|---|---|---|
| chr15:96072875-96073129 | -1.027836487 | 0.024279043 | 0.125700563 |
| chrX:40581208-40581550 | 0.917097383 | 0.02427982 | 0.125700563 |
| chrX:112003950-112004470 | -0.893048667 | 0.024281629 | 0.125701435 |
| chr13:43054834-43055034 | -1.101987697 | 0.02429393 | 0.125752063 |
| chr7:90153846-90154805 | -0.763672391 | 0.024294691 | 0.125752063 |
| chr14:71647092-71647428 | 0.878415361 | 0.024302535 | 0.125784169 |
| chr1:209850448-209850776 | -0.874883905 | 0.024311511 | 0.125822128 |
| chr13:50377112-50377418 | -1.123614493 | 0.02432965 | 0.125907499 |
| chr14:105733472-105733884 | 0.872433476 | 0.024335923 | 0.125918014 |
| chr6:43454356-43454793 | 0.866977729 | 0.024336201 | 0.125918014 |
| chr3:64776814-64777164 | 0.777403033 | 0.024336612 | 0.125918014 |
| chr4:37091678-37092125 | -1.118512024 | 0.024346752 | 0.125960424 |
| chr2:135819302-135819711 | -0.859659266 | 0.02434836 | 0.125960424 |
| chr3:135816910-135817110 | 1.184797556 | 0.024351336 | 0.125960424 |
| chr8:133496454-133496790 | -0.899469796 | 0.024351384 | 0.125960424 |
| chr2:5204469-5205026 | -0.848121461 | 0.024355565 | 0.12596768 |
| chr11:132402136-132402345 | -1.188976968 | 0.024357646 | 0.12596768 |
| chr4:170420299-170420727 | -0.999418284 | 0.024357719 | 0.12596768 |
| chr8:126570365-126571055 | 0.753611347 | 0.024365791 | 0.126000919 |
| chr19:47663514-47663796 | 0.983366848 | 0.024369559 | 0.126011902 |
| chr20:50292572-50293026 | -0.810812212 | 0.024387116 | 0.126094179 |
| chr7:106176659-106176989 | -0.925893489 | 0.024391371 | 0.126107668 |
| chr5:166399904-166400131 | 1.084402388 | 0.024394102 | 0.126113275 |
| chr12:11650040-11650323 | -1.051061777 | 0.024396445 | 0.126116881 |
| chr12:77685207-77685475 | -1.015252509 | 0.024398726 | 0.126117044 |
| chr2:200844502-200844846 | 0.985268521 | 0.024399769 | 0.126117044 |
| chr19:6476133-6476468 | 0.876444572 | 0.024405712 | 0.126139258 |
| chr13:45380870-45381182 | -1.025122767 | 0.024408488 | 0.126145096 |
| chr9:25517614-25517860 | -1.328530277 | 0.024412406 | 0.126154891 |
| chr12:52884879-52885318 | 0.813519846 | 0.024413676 | 0.126154891 |
| chr21:36989701-36990564 | 0.769407512 | 0.024423384 | 0.12619654 |
| chr18:54775786-54775986 | -1.26748712 | 0.024442754 | 0.126288112 |
| chr16:8678691-8678904 | 0.995408418 | 0.024446038 | 0.12628935 |
| chr3:57078828-57079089 | 1.023799661 | 0.02444629 | 0.12628935 |
| chr2:65049241-65049718 | -0.858384681 | 0.024448537 | 0.126292444 |
| chr17:61621373-61622176 | 0.719399725 | 0.024462432 | 0.126355702 |
| chr10:91251535-91251748 | -1.213308366 | 0.024471514 | 0.126387115 |
| chr13:31424170-31424528 | -0.858603012 | 0.024472115 | 0.126387115 |
| chr13:50244940-50245440 | -0.890027377 | 0.024473462 | 0.126387115 |
| chr3:155449822-155450022 | -1.247271602 | 0.024477702 | 0.126400492 |
| chr3:129512589-129512886 | 0.999902527 | 0.024493163 | 0.126471407 |
| chr17:61343885-61344158 | 0.758684242 | 0.024495921 | 0.126471407 |
| chr10:65358590-65358863 | -1.034997422 | 0.024496387 | 0.126471407 |
| chr3:177345640-177346034 | -0.939702796 | 0.024503196 | 0.126498041 |
| chr2:11132428-11133307 | -0.742003084 | 0.02451051 | 0.126527273 |
| chr13:33090716-33091610 | 0.767188662 | 0.024513069 | 0.1265293 |
| chr13:69857342-69857557 | -1.143768124 | 0.024515448 | 0.1265293 |
| chr18:35231021-35231314 | -0.997776677 | 0.024515857 | 0.1265293 |
| chrX:107496802-107497146 | -0.912620589 | 0.024520859 | 0.126544112 |
| chr4:153271861-153272140 | -1.163046612 | 0.02452203 | 0.126544112 |

| | | | |
|---|---|---|---|
| chr6:131828225-131828506 | 1.026472218 | 0.024525552 | 0.126546361 |
| chr1:43858269-43858824 | 0.804385511 | 0.024525769 | 0.126546361 |
| chr5:94181992-94182268 | -1.006847993 | 0.024528157 | 0.12655016 |
| chr18:71114948-71115583 | -0.887799297 | 0.024531272 | 0.12655771 |
| chr6:109165550-109165829 | 1.144169199 | 0.024538439 | 0.126586163 |
| chr6:170461973-170462366 | 0.951991099 | 0.024541335 | 0.126592577 |
| chr13:77329668-77329986 | -0.8651544 | 0.024549118 | 0.126616224 |
| chr1:147327123-147327598 | 0.892344406 | 0.024549224 | 0.126616224 |
| chr7:123191058-123191893 | -0.752958026 | 0.024554574 | 0.126635292 |
| chr10:43854150-43854462 | 0.959893858 | 0.024568226 | 0.126697172 |
| chr5:2642102-2642302 | 1.160538312 | 0.024574763 | 0.12671442 |
| chr10:69425648-69426118 | -0.915423787 | 0.024574878 | 0.12671442 |
| chr5:72835795-72835995 | -1.308225627 | 0.024582626 | 0.126745843 |
| chr7:137893813-137894198 | -1.19528268 | 0.024589422 | 0.126763542 |
| chr10:15878100-15878439 | -1.034001794 | 0.024589731 | 0.126763542 |
| chr11:30128392-30128622 | -1.186063781 | 0.024591022 | 0.126763542 |
| chr11:1051116-1051490 | 0.880705407 | 0.024602122 | 0.126812231 |
| chr12:120178818-120179139 | 1.043614066 | 0.024608643 | 0.126833616 |
| chr1:195152348-195152651 | 1.063495167 | 0.024609582 | 0.126833616 |
| chr1:143214576-143214943 | 0.901376512 | 0.024611853 | 0.126836793 |
| chr8:94905341-94905699 | 0.852379508 | 0.024621375 | 0.126877329 |
| chr5:63962846-63963422 | -0.839398874 | 0.024623065 | 0.126877503 |
| chr14:59507768-59507986 | 1.085328263 | 0.024627134 | 0.126883068 |
| chr10:96118601-96119347 | -0.76959389 | 0.024627457 | 0.126883068 |
| chr9:32217084-32217564 | -0.976044741 | 0.024631147 | 0.126893549 |
| chr3:101926750-101927114 | 1.03520767 | 0.0246358 | 0.126908988 |
| chr3:126322416-126322867 | -0.968644071 | 0.024648343 | 0.126957339 |
| chr16:75070543-75070917 | 0.818682412 | 0.0246485 | 0.126957339 |
| chr5:148374688-148375324 | -0.803612549 | 0.024653065 | 0.126972318 |
| chr5:135466201-135466454 | -0.965996327 | 0.024655288 | 0.12697523 |
| chr10:86968302-86968699 | -0.81743684 | 0.024664386 | 0.127013548 |
| chr5:178090801-178091254 | -0.813683718 | 0.024667064 | 0.127018802 |
| chr10:131885267-131885712 | 0.83668225 | 0.024669577 | 0.127023207 |
| chr6:40999568-40999824 | -1.060949405 | 0.024685794 | 0.127090273 |
| chr21:17327413-17327618 | -1.271346388 | 0.02468592 | 0.127090273 |
| chr18:54223630-54224120 | -0.844980179 | 0.024693494 | 0.127120725 |
| chr22:40396956-40397316 | 0.917795324 | 0.024697917 | 0.127129893 |
| chr8:125434642-125434975 | -0.753696939 | 0.024698651 | 0.127129893 |
| chr1:51622535-51622801 | 0.962422511 | 0.024700252 | 0.127129893 |
| chr13:97221991-97222666 | -0.784167427 | 0.024702789 | 0.127134412 |
| chr12:113135555-113136068 | 0.77350082 | 0.024709044 | 0.127158063 |
| chr12:27940806-27941074 | -1.033559548 | 0.024710748 | 0.127158291 |
| chr20:50697931-50698327 | 0.734471585 | 0.024720304 | 0.127197314 |
| chr4:71419272-71419545 | -1.180451943 | 0.024721652 | 0.127197314 |
| chr11:66624660-66624887 | 1.084394059 | 0.024729465 | 0.127228973 |
| chr5:148850911-148851308 | 0.882621646 | 0.024736027 | 0.127254187 |
| chr15:34062015-34062244 | 1.035761861 | 0.024740804 | 0.127270215 |
| chr8:64931133-64931333 | 1.244386155 | 0.024754644 | 0.127332866 |
| chr2:206004111-206004638 | -0.839450866 | 0.024761967 | 0.127359757 |
| chr10:18436999-18437212 | 1.084208892 | 0.024763197 | 0.127359757 |

| | | | |
|---|---|---|---|
| chr5:46434666-46434936 | 1.04740271 | 0.0247658 | 0.127364597 |
| chr6:108707630-108707846 | -1.149903482 | 0.024767505 | 0.127364815 |
| chr1:144103446-144104727 | 0.724377931 | 0.024769651 | 0.127367306 |
| chr12:22616-23037 | 0.90731009 | 0.024780265 | 0.127410771 |
| chr10:12631853-12632088 | 1.05951459 | 0.024785308 | 0.127410771 |
| chr14:79045290-79045580 | 1.05951459 | 0.024785308 | 0.127410771 |
| chr1:207672493-207672858 | -0.847025288 | 0.024785495 | 0.127410771 |
| chr18:80230368-80230600 | -1.10007884 | 0.024786418 | 0.127410771 |
| chr7:51867232-51867503 | 1.087713004 | 0.024788208 | 0.127411423 |
| chr20:51494977-51495223 | 0.959716759 | 0.024794542 | 0.127435429 |
| chr1:37038168-37038967 | -0.785759201 | 0.024805559 | 0.127483503 |
| chr12:103840764-103842289 | -0.713407139 | 0.024807504 | 0.127484044 |
| chr3:184443466-184444169 | 0.819624146 | 0.024810467 | 0.127484044 |
| chr10:132172649-132172856 | 1.075322964 | 0.02481108 | 0.127484044 |
| chr6:52087803-52088031 | -1.1347328 | 0.02481232 | 0.127484044 |
| chr13:77329122-77329587 | -0.783816337 | 0.024817161 | 0.127500367 |
| chr11:117105931-117106355 | 0.941997112 | 0.024820659 | 0.127509794 |
| chr16:85741443-85741693 | 1.087369593 | 0.024829867 | 0.127548542 |
| chr17:67884101-67884501 | -0.998808868 | 0.024836487 | 0.127573998 |
| chr18:61838589-61838834 | -1.239571329 | 0.024847926 | 0.127624196 |
| chr7:128398601-128398930 | -0.86046148 | 0.024851375 | 0.127633359 |
| chr5:23909756-23910003 | 1.191225374 | 0.024853672 | 0.127636601 |
| chr21:19544522-19544755 | 1.166452037 | 0.024860165 | 0.127661386 |
| chr14:41420761-41421131 | -1.07109564 | 0.024868324 | 0.127694728 |
| chr14:81302568-81303151 | -0.80939865 | 0.024874255 | 0.127702813 |
| chr10:24483687-24484074 | -0.875761274 | 0.024874261 | 0.127702813 |
| chr7:120987635-120987954 | -0.859949309 | 0.024874899 | 0.127702813 |
| chr17:61888160-61888679 | 0.728350955 | 0.02487838 | 0.127712128 |
| chr11:34002467-34002753 | 0.916044848 | 0.024888507 | 0.127755556 |
| chr14:82691151-82691363 | 1.165689314 | 0.024893842 | 0.12777438 |
| chr13:72199804-72200015 | -1.208606276 | 0.024903223 | 0.127813969 |
| chr2:190328859-190329059 | -1.297720701 | 0.024912007 | 0.127850487 |
| chr11:78062003-78062251 | -1.03403797 | 0.024913863 | 0.127851452 |
| chr22:19881213-19881446 | 0.949245491 | 0.02491944 | 0.127866992 |
| chr6:168102652-168103135 | -0.925583931 | 0.024921416 | 0.127866992 |
| chr7:136406031-136406698 | -0.820729907 | 0.024921898 | 0.127866992 |
| chr18:26487547-26487788 | -1.094382896 | 0.024929529 | 0.127897582 |
| chr5:81736179-81736408 | -1.243375676 | 0.024936755 | 0.127923114 |
| chr2:64000665-64000964 | -0.904985743 | 0.024937845 | 0.127923114 |
| chr2:85421393-85421712 | 0.847452672 | 0.02494583 | 0.127955508 |
| chr8:89401992-89402233 | 1.034415827 | 0.024948354 | 0.12795816 |
| chr4:40702528-40703186 | 0.854949612 | 0.024949687 | 0.12795816 |
| chr12:104603870-104604765 | -0.789861002 | 0.024962533 | 0.128006626 |
| chr1:248611972-248612331 | 0.959117781 | 0.02496272 | 0.128006626 |
| chr12:89566449-89567234 | -0.731269471 | 0.024964149 | 0.128006626 |
| chr9:92657869-92658069 | 1.129277044 | 0.024971024 | 0.128031108 |
| chr2:61426286-61426640 | -0.824070402 | 0.024972265 | 0.128031108 |
| chr20:36587343-36587568 | -1.0816218 | 0.024976665 | 0.128036886 |
| chr6:26364967-26365610 | -0.795396598 | 0.024976734 | 0.128036886 |
| chr6:143299676-143300039 | -1.068840728 | 0.02498833 | 0.128087757 |

| | | | |
|---|---|---|---|
| chr3:102471067-102471320 | -1.104627412 | 0.024991343 | 0.128094636 |
| chr1:204307227-204307934 | -0.744857708 | 0.024998992 | 0.128125266 |
| chr10:53842080-53842435 | -1.022762538 | 0.025009089 | 0.128168443 |
| chr10:47406720-47407417 | -0.756603288 | 0.025022824 | 0.12823026 |
| chr6:35782458-35782836 | -0.89042697 | 0.025026611 | 0.128241088 |
| chr3:81910321-81910592 | -0.986010968 | 0.025033272 | 0.128259407 |
| chr10:63866866-63867239 | -0.850034322 | 0.025034418 | 0.128259407 |
| chr12:79547073-79548207 | 0.715716758 | 0.025035208 | 0.128259407 |
| chr13:97127783-97128131 | 0.955417351 | 0.025038042 | 0.128265351 |
| chr2:27250010-27250536 | -0.871543382 | 0.025046243 | 0.128293524 |
| chr4:47972793-47973059 | 1.110410024 | 0.025046987 | 0.128293524 |
| chr19:6530825-6531142 | 0.96536387 | 0.025050569 | 0.128293524 |
| chr1:203956756-203956963 | 1.097895552 | 0.025051716 | 0.128293524 |
| chrX:81201669-81202278 | -0.803435926 | 0.025051913 | 0.128293524 |
| chr17:77914734-77915094 | 0.926853952 | 0.025059354 | 0.128319541 |
| chr1:174023556-174023776 | -0.934896607 | 0.025060343 | 0.128319541 |
| chr4:68369275-68369803 | 0.860372824 | 0.025062151 | 0.128320221 |
| chr9:93967200-93967400 | 1.173817171 | 0.025064418 | 0.128323253 |
| chr13:108897031-108897305 | -1.023978045 | 0.02508387 | 0.128391977 |
| chr6:16277757-16278089 | 1.035741328 | 0.025083881 | 0.128391977 |
| chr2:66021330-66021636 | -1.001389772 | 0.025084149 | 0.128391977 |
| chr6:138087562-138087900 | -0.992169786 | 0.025084544 | 0.128391977 |
| chr3:4071174-4071819 | -0.863318266 | 0.025091696 | 0.128420009 |
| chr12:122421400-122421766 | 0.903875167 | 0.025110157 | 0.12849309 |
| chr14:39367350-39367579 | 1.077252029 | 0.025110702 | 0.12849309 |
| chr3:14268464-14268767 | 0.938181016 | 0.025114102 | 0.12849309 |
| chr3:126171230-126171473 | 1.213759672 | 0.025115794 | 0.12849309 |
| chr2:17646136-17646416 | -0.920944426 | 0.025118899 | 0.12849309 |
| chr20:47747831-47748418 | 0.720593131 | 0.025119731 | 0.12849309 |
| chr6:2875848-2876335 | -0.895130494 | 0.025119802 | 0.12849309 |
| chr5:52685480-52685938 | -0.91520803 | 0.02512091 | 0.12849309 |
| chr1:28737612-28738036 | 0.89308886 | 0.025121068 | 0.12849309 |
| chr6:67086133-67086806 | -0.855676625 | 0.025132487 | 0.128542918 |
| chr16:3020766-3021620 | 0.734402027 | 0.025135972 | 0.128552158 |
| chr2:157291035-157291625 | 0.823058635 | 0.02514157 | 0.128572205 |
| chr4:28225045-28225324 | -0.998333262 | 0.025155393 | 0.128634312 |
| chr3:175685535-175686056 | -0.967031699 | 0.025161057 | 0.128648577 |
| chr3:150751737-150752277 | 0.770887999 | 0.025161541 | 0.128648577 |
| chr10:36899654-36899959 | -0.927457365 | 0.025177188 | 0.128719989 |
| chr2:173568151-173568362 | -1.206087434 | 0.025184649 | 0.128749544 |
| chr16:71600515-71600878 | 1.043435924 | 0.025188484 | 0.128760559 |
| chr22:39152579-39153048 | 0.884751075 | 0.02519138 | 0.128766772 |
| chr1:42682777-42683283 | 0.790768692 | 0.025196788 | 0.128785824 |
| chr3:155934250-155934551 | -0.957056454 | 0.025208216 | 0.128825064 |
| chr6:100220526-100221028 | 0.89792668 | 0.025209264 | 0.128825064 |
| chr10:117185866-117186198 | -0.931545774 | 0.025210612 | 0.128825064 |
| chr16:35297882-35298147 | 1.192888593 | 0.025212316 | 0.128825064 |
| chr1:6220256-6220609 | 0.876265117 | 0.025212872 | 0.128825064 |
| chr4:83582269-83582794 | -0.827725498 | 0.025220183 | 0.128848641 |
| chr4:3346020-3346370 | 1.023948016 | 0.025220849 | 0.128848641 |

| | | | |
|---|---|---|---|
| chr5:168848185-168848461 | 1.121627826 | 0.025223756 | 0.128854899 |
| chr10:15069290-15069595 | -1.04127775 | 0.025226311 | 0.128857071 |
| chr12:77675635-77675902 | 1.206880349 | 0.025227545 | 0.128857071 |
| chr2:55419537-55420427 | -0.742748981 | 0.02523131 | 0.128867714 |
| chr10:22616215-22616436 | -1.16067314 | 0.025233211 | 0.128868011 |
| chr14:29846535-29846802 | 1.127722413 | 0.025234732 | 0.128868011 |
| chr4:156198469-156199226 | -0.862460645 | 0.025240931 | 0.128891077 |
| chr1:220511849-220512150 | -1.040768218 | 0.025251005 | 0.128933929 |
| chr16:11359102-11359837 | 0.742493066 | 0.025263038 | 0.128986773 |
| chr12:96249472-96249860 | 0.973416451 | 0.025265418 | 0.128990329 |
| chr13:74287827-74288045 | -1.102293238 | 0.025268313 | 0.128996514 |
| chr2:67215259-67215572 | -0.914530447 | 0.025275623 | 0.129019307 |
| chr13:81442974-81443357 | -0.982875936 | 0.025276145 | 0.129019307 |
| chrX:131746282-131746961 | 0.806415683 | 0.025284456 | 0.129053132 |
| chr3:156075364-156075582 | 1.157799884 | 0.025293976 | 0.129086013 |
| chr1:216587768-216588250 | -0.905024686 | 0.025294595 | 0.129086013 |
| chr12:120231384-120231783 | 0.788440514 | 0.025297298 | 0.129086013 |
| chr7:4384498-4384744 | 1.13211516 | 0.025297637 | 0.129086013 |
| chr2:195498222-195498562 | -0.994364972 | 0.025307915 | 0.129129855 |
| chr14:90419179-90419666 | -0.825402888 | 0.025310266 | 0.129133255 |
| chr1:207751720-207753040 | -0.713303828 | 0.02532363 | 0.129190131 |
| chr8:108458429-108458629 | 1.069662574 | 0.025324786 | 0.129190131 |
| chr5:169099795-169100047 | -1.228791235 | 0.025349844 | 0.129309351 |
| chr4:85418143-85418484 | -0.928641094 | 0.025355746 | 0.129325684 |
| chr8:74948100-74948727 | -0.872815738 | 0.025356422 | 0.129325684 |
| chr7:128409656-128410704 | -0.722868054 | 0.025359129 | 0.129330488 |
| chr13:46951632-46951886 | 0.961509757 | 0.02536074 | 0.129330488 |
| chr4:60938758-60938971 | -1.211028256 | 0.02536274 | 0.129332082 |
| chr13:49225758-49226013 | -1.121359409 | 0.02537964 | 0.129407793 |
| chr3:15462456-15462697 | 1.086270181 | 0.025380965 | 0.129407793 |
| chr20:47422437-47422940 | 0.721219861 | 0.025395346 | 0.1294725 |
| chr4:145891714-145892512 | -0.846265532 | 0.025398306 | 0.129478975 |
| chr6:43096553-43097164 | 0.824627452 | 0.025401505 | 0.129486668 |
| chr3:64209881-64210554 | 0.723933738 | 0.025410226 | 0.129522507 |
| chr7:149780983-149781244 | -1.004362304 | 0.025412493 | 0.129525448 |
| chr2:53989360-53989913 | -0.82616692 | 0.025419725 | 0.129553688 |
| chr12:49827512-49828028 | 0.797809454 | 0.025424368 | 0.12956873 |
| chr9:14238508-14238876 | -0.911971828 | 0.025426058 | 0.12956873 |
| chr7:54758684-54758954 | -0.88609432 | 0.025430916 | 0.129584867 |
| chr13:49775313-49775606 | -1.153089372 | 0.025452727 | 0.129687379 |
| chr5:168038599-168038894 | 1.022295083 | 0.025460021 | 0.12971592 |
| chr20:12156359-12156598 | -1.184102362 | 0.025462196 | 0.129718378 |
| chr7:42696705-42696951 | 1.166372558 | 0.025466615 | 0.129732262 |
| chr19:39557818-39558018 | 1.125416242 | 0.025471529 | 0.12974867 |
| chr8:100858452-100858722 | 0.930941601 | 0.025479324 | 0.129779749 |
| chr3:101865988-101866476 | -0.82024962 | 0.025482366 | 0.129786614 |
| chr3:21017036-21017839 | -0.754267054 | 0.025486328 | 0.129798166 |
| chr5:112492379-112492704 | -0.900705464 | 0.025508494 | 0.129889529 |
| chr18:37335343-37335726 | -0.883938641 | 0.025508632 | 0.129889529 |
| chr3:74848284-74848515 | 1.121922344 | 0.025509353 | 0.129889529 |

| | | | |
|---|---|---|---|
| chr3:14096797-14097081 | 0.96949421 | 0.025516602 | 0.129917807 |
| chr7:6283066-6283339 | 1.08547044 | 0.025518573 | 0.129919209 |
| chr8:94574161-94574456 | 0.900475051 | 0.025525479 | 0.129945734 |
| chr8:18957535-18957846 | 0.94757675 | 0.025530824 | 0.12996431 |
| chr13:104015676-104015876 | -1.378191523 | 0.025536364 | 0.129975239 |
| chr2:80122658-80122858 | -1.378191523 | 0.025536364 | 0.129975239 |
| chr1:113638724-113639591 | -0.762659422 | 0.025550332 | 0.130037696 |
| chr8:129518423-129518749 | 1.132575201 | 0.025557677 | 0.130066442 |
| chr1:246705313-246705781 | 0.830289737 | 0.02556681 | 0.130104278 |
| chr14:95056543-95056844 | 0.863480429 | 0.025568935 | 0.130106451 |
| chr4:70133593-70133878 | -1.06870814 | 0.025581141 | 0.130152762 |
| chr2:185208319-185208674 | -0.877125399 | 0.025584417 | 0.130152762 |
| chr1:205556238-205556438 | 1.097450285 | 0.025585169 | 0.130152762 |
| chr20:48648481-48648953 | 0.730694523 | 0.025585763 | 0.130152762 |
| chr2:1743049-1743334 | 0.9087138 | 0.025587552 | 0.130152762 |
| chr17:67901882-67902189 | 0.942393846 | 0.025589072 | 0.130152762 |
| chr5:80672208-80672478 | -1.090410989 | 0.025589927 | 0.130152762 |
| chr5:92641105-92641633 | -0.841897911 | 0.025593112 | 0.130160323 |
| chr6:45500993-45501433 | -0.861550807 | 0.025596804 | 0.130170459 |
| chr14:69299370-69299570 | 1.093685595 | 0.02559924 | 0.130174206 |
| chr6:33732045-33732344 | 0.956479659 | 0.025602593 | 0.13018262 |
| chr3:158963810-158964483 | -0.762991777 | 0.025611454 | 0.130215308 |
| chr14:28774395-28774778 | 0.8316542 | 0.025612421 | 0.130215308 |
| chr1:96962987-96963221 | 1.043913994 | 0.025617874 | 0.130234393 |
| chr7:23696973-23697328 | 0.839830597 | 0.02562369 | 0.130253384 |
| chr1:189678860-189679083 | -1.086196751 | 0.02562501 | 0.130253384 |
| chr1:189369003-189369557 | -0.845529159 | 0.025628215 | 0.130261034 |
| chr1:112955342-112956965 | -0.718011944 | 0.025631553 | 0.130269359 |
| chr17:61783868-61784139 | 0.775921667 | 0.025633536 | 0.130270795 |
| chr19:17916016-17916243 | 1.151637977 | 0.02564201 | 0.130305221 |
| chr3:12994625-12995331 | 0.731147546 | 0.025644296 | 0.130308191 |
| chr2:28377245-28377491 | -1.040366523 | 0.025655053 | 0.130354209 |
| chr5:68549192-68549694 | -0.823756303 | 0.025658759 | 0.130364393 |
| chr15:42490652-42491373 | 0.79506487 | 0.025666681 | 0.130395997 |
| chr4:51885450-51885753 | -1.000447494 | 0.025669092 | 0.130399602 |
| chr5:150656263-150657064 | 0.771383359 | 0.025677688 | 0.13042743 |
| chr6:10838008-10839002 | -0.741735984 | 0.025677975 | 0.13042743 |
| chr5:21865436-21865636 | 1.187775686 | 0.025686341 | 0.130450952 |
| chr8:62963278-62963490 | 1.12757495 | 0.025687421 | 0.130450952 |
| chr1:91736898-91737329 | -0.976329414 | 0.025687713 | 0.130450952 |
| chr2:189624633-189625258 | -0.847385275 | 0.025693273 | 0.130466283 |
| chr11:72785898-72786161 | -0.891981736 | 0.025694373 | 0.130466283 |
| chr2:41456973-41457428 | -0.857450753 | 0.025696789 | 0.130466283 |
| chr1:218284585-218285649 | -0.709350227 | 0.025697543 | 0.130466283 |
| chr3:8632649-8632906 | -0.961568055 | 0.025702057 | 0.130469361 |
| chr20:47416726-47417010 | 0.745854136 | 0.025703051 | 0.130469361 |
| chr3:63733550-63733766 | 0.829629691 | 0.025704882 | 0.130469361 |
| chr22:22207862-22208167 | 1.023372858 | 0.025705912 | 0.130469361 |
| chr7:30148973-30149607 | -0.865402376 | 0.025706663 | 0.130469361 |
| chr15:31366984-31367252 | -1.024722496 | 0.025711037 | 0.130482917 |

| | | | |
|---|---|---|---|
| chr7:108569225-108570680 | -0.704782512 | 0.025731682 | 0.130571776 |
| chr20:48686902-48687102 | 0.798561036 | 0.025732133 | 0.130571776 |
| chr12:52949610-52950090 | 0.756828782 | 0.025733659 | 0.130571776 |
| chr7:128300836-128301134 | -1.076527739 | 0.025740262 | 0.130596631 |
| chr7:95357515-95357836 | -1.145220633 | 0.025744066 | 0.130607287 |
| chr7:142854884-142855145 | -0.940232127 | 0.025752721 | 0.130642543 |
| chr7:20960556-20960758 | 1.187002641 | 0.025755786 | 0.130649442 |
| chr11:66134387-66134862 | 0.876107858 | 0.025766546 | 0.130677069 |
| chr20:47329818-47330234 | 0.729423111 | 0.025767382 | 0.130677069 |
| chr19:1415470-1415710 | 1.052327256 | 0.025768667 | 0.130677069 |
| chr10:541687-542042 | 0.964252421 | 0.025769412 | 0.130677069 |
| chr16:11537484-11537728 | 1.222242394 | 0.02576976 | 0.130677069 |
| chr1:114640334-114640534 | 0.868270988 | 0.025772923 | 0.130684462 |
| chr16:391789-392067 | 0.98407588 | 0.025780929 | 0.130712285 |
| chr1:245344499-245344766 | 1.137617425 | 0.025781822 | 0.130712285 |
| chr4:148061845-148062314 | -0.890228619 | 0.025784105 | 0.130715212 |
| chr11:75241652-75242023 | 0.808548551 | 0.025791012 | 0.130741576 |
| chr5:37661302-37661873 | 0.824160816 | 0.025794568 | 0.13075095 |
| chr17:57375507-57376329 | -0.75771619 | 0.025799596 | 0.130767787 |
| chr8:94059378-94059789 | 0.860336305 | 0.025802166 | 0.130772163 |
| chr2:215314947-215315238 | 0.924676377 | 0.025804527 | 0.130775477 |
| chr2:11416-11989 | 0.798998634 | 0.02580726 | 0.130780679 |
| chr5:95456250-95456560 | -1.076592107 | 0.025811586 | 0.130784387 |
| chr9:76064813-76065260 | -1.049265086 | 0.025811849 | 0.130784387 |
| chr22:29799252-29799669 | 0.809666509 | 0.025813112 | 0.130784387 |
| chr12:69425440-69425943 | -0.89002998 | 0.025817666 | 0.130793865 |
| chr13:51280695-51280897 | 1.256374562 | 0.025818397 | 0.130793865 |
| chr15:100559247-100559593 | 0.996553495 | 0.025838376 | 0.13088 |
| chrX:16619149-16619426 | -1.212100902 | 0.025838816 | 0.13088 |
| chr4:158193229-158193487 | -1.038445331 | 0.025857736 | 0.130967175 |
| chr10:50739582-50740352 | -0.768043534 | 0.025867419 | 0.130999208 |
| chr13:73708411-73708678 | -1.149933428 | 0.025867479 | 0.130999208 |
| chr6:58784767-58785072 | 0.812580142 | 0.025874508 | 0.13102614 |
| chr20:58038242-58038631 | 0.818482229 | 0.025884915 | 0.131070182 |
| chr3:150738380-150738671 | -0.88271059 | 0.025886962 | 0.131071882 |
| chr11:111977173-111977610 | 0.873324584 | 0.025899433 | 0.131126362 |
| chr4:9102494-9103001 | 0.809549203 | 0.02590514 | 0.131146592 |
| chr3:178823589-178823888 | -0.980167597 | 0.025911185 | 0.131163522 |
| chr17:61563120-61563320 | 0.790022497 | 0.025912005 | 0.131163522 |
| chr3:98645032-98645415 | -0.931962085 | 0.02591362 | 0.131163522 |
| chr6:26033143-26033859 | -0.765905035 | 0.025919499 | 0.131180663 |
| chr8:69994621-69994949 | 0.938366346 | 0.025920431 | 0.131180663 |
| chr3:119892037-119892499 | -0.876803164 | 0.025922915 | 0.131184573 |
| chr8:139989245-139989511 | 1.11471792 | 0.025925145 | 0.131187192 |
| chr1:39785528-39785761 | -1.028216226 | 0.025948533 | 0.131292651 |
| chr15:87490055-87490286 | 1.133560727 | 0.025950166 | 0.131292651 |
| chr7:123769451-123769651 | -1.171276826 | 0.025951126 | 0.131292651 |
| chr7:150956907-150957145 | 1.032811093 | 0.025956151 | 0.131309402 |
| chr7:3272739-3272948 | 1.082149384 | 0.025958643 | 0.131313341 |
| chr11:117458008-117458248 | 1.117665768 | 0.025961589 | 0.131319571 |

| | | | |
|---|---|---|---|
| chr7:42372624-42373135 | -0.920479463 | 0.025970053 | 0.131353713 |
| chr3:40631383-40632009 | 0.81475777 | 0.025974659 | 0.13136834 |
| chr3:79996164-79996478 | 0.993935804 | 0.025976437 | 0.13136866 |
| chr15:84658136-84658763 | 0.744430687 | 0.025984875 | 0.131401544 |
| chr4:139676973-139677173 | 1.144985451 | 0.025986369 | 0.131401544 |
| chr17:59505328-59505734 | 0.823233545 | 0.025994335 | 0.131432276 |
| chr5:10453566-10453937 | 0.852387621 | 0.025995878 | 0.131432276 |
| chr1:53176578-53177038 | 0.897896503 | 0.026000779 | 0.131448384 |
| chr5:127313920-127314279 | -0.909389678 | 0.0260059 | 0.131465596 |
| chr20:47481769-47482151 | 0.729351012 | 0.026017388 | 0.131514993 |
| chr9:94819323-94819791 | 0.795641156 | 0.02602448 | 0.131542165 |
| chrX:147174659-147174946 | 0.890234214 | 0.026026317 | 0.131542773 |
| chr11:83241360-83241733 | -0.830238789 | 0.026029893 | 0.131547943 |
| chr7:155204628-155204833 | 1.128757786 | 0.026030773 | 0.131547943 |
| chr10:86556554-86557229 | 0.740285406 | 0.026035285 | 0.131562064 |
| chr22:36978082-36978349 | 0.955140314 | 0.026042144 | 0.131585463 |
| chr5:7170621-7170920 | -1.090063215 | 0.026045341 | 0.131585463 |
| chr6:21462241-21462832 | -0.873042602 | 0.026045365 | 0.131585463 |
| chr6:10133496-10133728 | -1.195292877 | 0.026047358 | 0.131585463 |
| chr10:132799684-132799950 | 1.093367164 | 0.026048502 | 0.131585463 |
| chr15:93386874-93387074 | 1.217981982 | 0.026052797 | 0.131598484 |
| chr6:80922124-80922324 | -1.280770661 | 0.026062285 | 0.131637731 |
| chr1:112881452-112882048 | -0.884576357 | 0.026067148 | 0.131651859 |
| chr13:30108365-30109317 | -0.734249246 | 0.026068518 | 0.131651859 |
| chr2:42931700-42931900 | -1.039322371 | 0.026078013 | 0.131682801 |
| chr14:32201055-32201473 | 0.746407454 | 0.026078082 | 0.131682801 |
| chr6:106746133-106747015 | 0.764745747 | 0.026100191 | 0.131785753 |
| chr2:158723580-158724867 | -0.711887778 | 0.026104785 | 0.131800265 |
| chr18:57141639-57142044 | -0.883800056 | 0.026111182 | 0.13181829 |
| chr8:17535196-17535472 | -1.13944421 | 0.026111843 | 0.13181829 |
| chr1:107031002-107031266 | 0.837340253 | 0.026114772 | 0.13181829 |
| chr3:141728626-141728853 | -1.173058769 | 0.026115236 | 0.13181829 |
| chr17:9162535-9162885 | -1.030790801 | 0.026127401 | 0.131871003 |
| chr20:48905090-48905328 | 0.776544625 | 0.026130525 | 0.131878083 |
| chr12:25501485-25502020 | -0.792651623 | 0.026139598 | 0.131908235 |
| chr9:120935446-120935748 | 0.946605289 | 0.026139942 | 0.131908235 |
| chr3:149115225-149115427 | 1.102177393 | 0.02614373 | 0.131918661 |
| chr15:60592474-60592706 | 1.042278665 | 0.026149259 | 0.131937871 |
| chr17:7021929-7022194 | -1.016635408 | 0.026151865 | 0.131942333 |
| chr20:57235068-57235420 | 0.729482591 | 0.026156435 | 0.131948622 |
| chr14:69739546-69739805 | 0.949320166 | 0.026156556 | 0.131948622 |
| chr3:190261802-190262067 | -1.127463022 | 0.026161961 | 0.131967199 |
| chr13:75549193-75550534 | -0.711299239 | 0.026164699 | 0.131970297 |
| chr20:54399867-54400091 | 0.797871007 | 0.02616602 | 0.131970297 |
| chr3:31448370-31448974 | 0.841702121 | 0.026168981 | 0.131976548 |
| chr10:69800214-69800469 | -0.941574221 | 0.026174232 | 0.131994342 |
| chr4:617655-618215 | 0.889672046 | 0.02617926 | 0.131996251 |
| chr16:2027533-2027733 | 0.996675115 | 0.026180216 | 0.131996251 |
| chr7:107879458-107879706 | -1.003090917 | 0.026182459 | 0.131996251 |
| chr6:22906285-22906494 | -1.127685976 | 0.026182567 | 0.131996251 |

| | | | |
|---|---|---|---|
| chr20:51803224-51803429 | 0.93400719 | 0.026183224 | 0.131996251 |
| chr5:14597671-14597931 | 1.052765562 | 0.026191194 | 0.132023121 |
| chr8:102927706-102927957 | 0.971413509 | 0.026192001 | 0.132023121 |
| chr8:129948414-129948656 | -1.080627622 | 0.026200156 | 0.132049447 |
| chr11:87005216-87005711 | 0.822731523 | 0.02620067 | 0.132049447 |
| chr2:44150468-44150939 | -1.00297361 | 0.026202622 | 0.132050599 |
| chr17:61903341-61903561 | 0.770727941 | 0.026209514 | 0.132076642 |
| chr3:81173338-81173880 | -0.890035465 | 0.026224267 | 0.132142037 |
| chr2:55793460-55794336 | -0.736294248 | 0.026227294 | 0.132142037 |
| chr4:6876134-6876398 | 1.124781117 | 0.026234441 | 0.132142037 |
| chr20:54717411-54718069 | 0.724863978 | 0.02623782 | 0.132142037 |
| chr1:180994387-180994724 | 0.973032521 | 0.02623834 | 0.132142037 |
| chr9:73271126-73271398 | -1.053901427 | 0.026238864 | 0.132142037 |
| chr20:47512976-47513237 | 0.765136631 | 0.026239821 | 0.132142037 |
| chr20:47259588-47259867 | 0.743291629 | 0.026240948 | 0.132142037 |
| chr13:28528874-28529074 | 1.198877923 | 0.026241393 | 0.132142037 |
| chr16:72498408-72498608 | 1.198877923 | 0.026241393 | 0.132142037 |
| chr10:110425962-110426519 | 0.827697546 | 0.026241461 | 0.132142037 |
| chr17:50021614-50021865 | 1.170138577 | 0.026247647 | 0.132164502 |
| chr5:74766380-74766662 | -0.92211109 | 0.026256105 | 0.132195451 |
| chr13:43023213-43024146 | -0.733480307 | 0.026257244 | 0.132195451 |
| chr20:47187734-47188229 | 0.720312693 | 0.0262605 | 0.132203154 |
| chr2:85490980-85491317 | 1.139319042 | 0.026270004 | 0.132238107 |
| chr4:76215194-76215421 | 1.006303265 | 0.026272481 | 0.132238107 |
| chr2:212863201-212863556 | -1.02759428 | 0.026275158 | 0.132238107 |
| chr3:48436080-48436359 | 0.97806699 | 0.026275559 | 0.132238107 |
| chr9:129494482-129494737 | -0.903130075 | 0.026276072 | 0.132238107 |
| chr8:124395599-124395968 | 0.87785135 | 0.026283959 | 0.132261407 |
| chr2:175002595-175002837 | 0.96617701 | 0.026284154 | 0.132261407 |
| chr7:135238075-135238515 | -0.859144643 | 0.026298908 | 0.132318858 |
| chr6:75795053-75795335 | -1.216978675 | 0.026299025 | 0.132318858 |
| chr5:65480394-65480811 | -0.906963323 | 0.026308703 | 0.132358859 |
| chr22:39145575-39145829 | 0.966112743 | 0.026315427 | 0.13238091 |
| chr19:11798407-11798607 | -1.255976948 | 0.026316541 | 0.13238091 |
| chr17:61993501-61993701 | 1.010315188 | 0.026321543 | 0.132397379 |
| chr12:88466528-88467282 | -0.772829501 | 0.026328992 | 0.132426156 |
| chr8:144950597-144951078 | 0.851756592 | 0.026333311 | 0.132429341 |
| chr3:69179417-69179694 | -1.094357155 | 0.026334811 | 0.132429341 |
| chr3:110917181-110917523 | -1.094357155 | 0.026334811 | 0.132429341 |
| chr21:36059858-36061026 | 0.72919136 | 0.026337736 | 0.132435361 |
| chr7:124101230-124101491 | -1.069765535 | 0.026358075 | 0.132528932 |
| chr10:86614866-86615193 | 0.941225253 | 0.026367359 | 0.13256515 |
| chr14:92995639-92996132 | -0.842091554 | 0.026368738 | 0.13256515 |
| chr11:60925070-60925300 | 0.982122153 | 0.02637417 | 0.132583759 |
| chr11:61160909-61162044 | 0.717153952 | 0.026376398 | 0.132586259 |
| chr3:153174141-153174439 | -0.867356055 | 0.026380651 | 0.13259894 |
| chr3:33203258-33203487 | 1.150242005 | 0.02638897 | 0.132632056 |
| chr13:86542117-86542523 | -0.935270125 | 0.02639191 | 0.132636621 |
| chr1:202964696-202965015 | -0.811278769 | 0.026393396 | 0.132636621 |
| chr17:28326068-28326495 | 0.839810618 | 0.026395072 | 0.132636621 |

| | | | |
|---|---|---|---|
| chr10:97138251-97138479 | 1.0505584 | 0.026400369 | 0.132654542 |
| chr5:26148508-26148716 | 1.189904444 | 0.02641138 | 0.132701166 |
| chr7:76063163-76063807 | -0.862946541 | 0.026415426 | 0.132712792 |
| chr8:102423919-102424169 | 1.06419729 | 0.026417163 | 0.132712815 |
| chr3:71423202-71423808 | -0.797786476 | 0.026426846 | 0.132752755 |
| chr3:18444966-18445926 | 0.753029892 | 0.026432035 | 0.132770121 |
| chr4:184648441-184650167 | -0.717530895 | 0.026437611 | 0.132789421 |
| chr13:28035260-28035498 | -1.096025023 | 0.026443466 | 0.132810124 |
| chr9:89605403-89605646 | 0.925486927 | 0.026447776 | 0.132821575 |
| chr17:49932918-49933208 | 1.021837723 | 0.026449213 | 0.132821575 |
| chr3:64881149-64881427 | 0.786335132 | 0.026459582 | 0.132864936 |
| chr8:105026063-105026552 | 0.856914771 | 0.026462715 | 0.132871964 |
| chr6:29628600-29629137 | -0.78495257 | 0.026468763 | 0.132885401 |
| chr9:34603717-34603981 | -1.00313297 | 0.02646886 | 0.132885401 |
| chr7:44522377-44522747 | 0.993745146 | 0.026478585 | 0.132915069 |
| chr1:109570563-109570885 | 0.900102337 | 0.026479379 | 0.132915069 |
| chr4:77471876-77472122 | 1.035278173 | 0.026479974 | 0.132915069 |
| chr1:116842818-116843018 | 1.018040073 | 0.026483114 | 0.132921865 |
| chr2:28956352-28956672 | -0.856890659 | 0.026484797 | 0.132921865 |
| chr7:128904514-128904966 | -0.828597117 | 0.026498874 | 0.132969816 |
| chr19:4830843-4831089 | -1.176609472 | 0.026498976 | 0.132969816 |
| chr10:533290-533490 | 1.028687009 | 0.026499557 | 0.132969816 |
| chr7:116591808-116592051 | -1.040513712 | 0.026501704 | 0.132971877 |
| chr16:78079210-78079474 | 0.964320212 | 0.026512829 | 0.133018988 |
| chr4:82201473-82201688 | 1.142968113 | 0.026519938 | 0.133045944 |
| chr3:1387962-1388205 | 1.068633509 | 0.026532206 | 0.133091026 |
| chr8:100416272-100416472 | 0.975035932 | 0.026532398 | 0.133091026 |
| chr7:116499191-116500097 | -0.755092178 | 0.026548575 | 0.133158021 |
| chr5:42078146-42078603 | -0.897840804 | 0.02654923 | 0.133158021 |
| chrX:153598966-153599603 | 0.756638234 | 0.026551464 | 0.133160509 |
| chr21:39071772-39072245 | -0.848794474 | 0.026565034 | 0.133219849 |
| chr12:10364541-10365481 | -0.803709035 | 0.02656935 | 0.133232769 |
| chr3:129971196-129971790 | -0.841923172 | 0.026572837 | 0.133238134 |
| chr8:120737618-120737818 | 1.139406576 | 0.026573897 | 0.133238134 |
| chr2:151437545-151437856 | -0.996224547 | 0.026582943 | 0.133274767 |
| chr4:1870137-1871271 | 0.728574182 | 0.026593122 | 0.133309103 |
| chr1:160607160-160607394 | 1.152494244 | 0.026593271 | 0.133309103 |
| chr17:215345-215588 | -1.178412495 | 0.02659736 | 0.133312308 |
| chr20:47487597-47487946 | 0.729565904 | 0.02659739 | 0.133312308 |
| chr6:24451634-24452038 | -0.920007377 | 0.026601194 | 0.133322649 |
| chr15:80970095-80970560 | 0.890471067 | 0.026619943 | 0.133407892 |
| chr10:59719379-59719679 | -1.001645773 | 0.026625805 | 0.133424597 |
| chr10:28334329-28334735 | -0.882523142 | 0.026626759 | 0.133424597 |
| chr10:127196352-12719721ς | 0.772019732 | 0.026638387 | 0.133474137 |
| chr9:128026726-128027205 | -0.853495934 | 0.026650666 | 0.13351182 |
| chr12:45655229-45655589 | -0.876325299 | 0.026650934 | 0.13351182 |
| chr6:113763899-113764100 | 1.175299091 | 0.026651135 | 0.13351182 |
| chr18:13165714-13166045 | 1.010020909 | 0.026656929 | 0.133532113 |
| chr17:64331524-64331965 | 0.730452017 | 0.026667012 | 0.133573889 |
| chr3:107394498-107394763 | 0.96055286 | 0.026684012 | 0.133650154 |

| | | | |
|---|---|---|---|
| chr4:169942471-169942777 | -0.897973653 | 0.026685726 | 0.133650154 |
| chr5:16813039-16813317 | 0.966221149 | 0.026693297 | 0.133679332 |
| chr22:12692332-12692574 | 1.013751872 | 0.026703574 | 0.133722059 |
| chr7:4003425-4003742 | 0.946573308 | 0.026705333 | 0.133722129 |
| chr5:106269207-106269602 | -0.942161151 | 0.026711425 | 0.133743893 |
| chr12:46259097-46259459 | 0.807084289 | 0.026714213 | 0.133749115 |
| chr1:211323528-211323777 | -0.959923544 | 0.026716466 | 0.133751654 |
| chr5:58446114-58446410 | -1.030832227 | 0.026720314 | 0.133756411 |
| chr2:207230266-207230527 | -1.084507254 | 0.026720908 | 0.133756411 |
| chr14:75488828-75489076 | 0.944519398 | 0.026722752 | 0.133756905 |
| chr17:4743390-4743647 | 1.20092019 | 0.026734764 | 0.133808288 |
| chr3:18666711-18666996 | 0.999355955 | 0.026739581 | 0.133823656 |
| chr18:23667148-23667402 | 1.099481193 | 0.026742719 | 0.133830619 |
| chr3:28342647-28342921 | 1.050956498 | 0.026756489 | 0.133890784 |
| chr18:54444373-54444591 | -1.250266359 | 0.026760495 | 0.133902088 |
| chr14:21783278-21783551 | 1.105317673 | 0.026765084 | 0.133912517 |
| chr10:99297652-99297884 | 1.192582676 | 0.026766075 | 0.133912517 |
| chr11:117143872-117144811 | 0.740034537 | 0.026771914 | 0.133932983 |
| chr13:29443535-29443795 | 1.127152286 | 0.026779936 | 0.133964368 |
| chr2:72367752-72368180 | -0.89721936 | 0.02678947 | 0.133992807 |
| chr1:115147785-115148453 | -0.847891271 | 0.026790945 | 0.133992807 |
| chr9:2424685-2424885 | -1.307369036 | 0.026791934 | 0.133992807 |
| chr6:367404-367647 | 1.0387046 | 0.026792616 | 0.133992807 |
| chr16:53434742-53435014 | 0.928220601 | 0.026802778 | 0.134034883 |
| chr3:83946359-83947084 | -0.841022203 | 0.026804562 | 0.134035053 |
| chr11:4016531-4016966 | -0.907115426 | 0.026820046 | 0.134095856 |
| chr7:141220412-141220618 | -1.103415318 | 0.026820221 | 0.134095856 |
| chr8:6852450-6852814 | -0.960812743 | 0.026823904 | 0.134099146 |
| chr13:114109453-114110082 | -0.888281998 | 0.02682438 | 0.134099146 |
| chr5:121630401-121631232 | -0.82898127 | 0.026831196 | 0.13412447 |
| chr20:54157914-54158158 | 0.775505341 | 0.02683565 | 0.134137985 |
| chr20:41894117-41894441 | 0.87865266 | 0.026850614 | 0.134204028 |
| chr4:18593626-18593826 | 1.130591464 | 0.026857073 | 0.134227557 |
| chr13:23159249-23160950 | 0.703379914 | 0.026863105 | 0.134245634 |
| chr1:118122335-118122631 | -0.946180084 | 0.026864194 | 0.134245634 |
| chr2:164093150-164093373 | -1.221863535 | 0.026867833 | 0.134255059 |
| chr7:127477374-127477574 | -1.349894659 | 0.026886869 | 0.134341422 |
| chr5:165497520-165498063 | -0.885523981 | 0.026895858 | 0.134377571 |
| chr13:44853493-44854099 | 0.83231499 | 0.026914321 | 0.134453658 |
| chr14:104977741-104977959 | 0.898668392 | 0.026914596 | 0.134453658 |
| chr22:45317647-45318039 | -0.933853681 | 0.026920756 | 0.134475662 |
| chr1:170115493-170115707 | 1.203910414 | 0.026936649 | 0.134546279 |
| chr16:86368970-86369361 | 0.956766646 | 0.026943255 | 0.134570503 |
| chr3:21405609-21406133 | -0.933459852 | 0.026950146 | 0.134588907 |
| chr10:95079009-95079256 | -1.225940719 | 0.026950453 | 0.134588907 |
| chr17:38869616-38870237 | 0.762068798 | 0.026957365 | 0.134612038 |
| chr5:77076875-77078078 | -0.735736683 | 0.026958598 | 0.134612038 |
| chr5:112733878-112734158 | 1.133966353 | 0.026963798 | 0.134623929 |
| chr17:44103034-44103536 | 0.803815184 | 0.02696485 | 0.134623929 |
| chr12:52217155-52217615 | 0.772499939 | 0.026966251 | 0.134623929 |

| | | | |
|---|---|---|---|
| chr16:31073161-31073576 | 0.780331898 | 0.026969933 | 0.134633542 |
| chr5:80346094-80346441 | -0.923522358 | 0.026972423 | 0.134637197 |
| chrX:42715257-42715687 | -0.968124665 | 0.026975287 | 0.134642722 |
| chr12:127293600-127293864 | -1.079907023 | 0.02697964 | 0.134655595 |
| chr5:83799006-83799297 | -1.141469251 | 0.026981381 | 0.134655595 |
| chr3:170118251-170118465 | 1.083471196 | 0.026985754 | 0.134665553 |
| chr12:107992745-107993207 | -0.85523705 | 0.026987425 | 0.134665553 |
| chr13:112906218-112906529 | 1.003663366 | 0.026988649 | 0.134665553 |
| chr8:94914342-94914583 | 1.006237373 | 0.026998114 | 0.134704011 |
| chr5:2252135-2252454 | 0.994093201 | 0.027006762 | 0.134738385 |
| chr10:133378141-133378684 | 0.799530518 | 0.027016853 | 0.134779956 |
| chr5:168114420-168114779 | -0.822576876 | 0.027029183 | 0.134832685 |
| chr2:195322421-195322706 | -1.048465743 | 0.027032775 | 0.134841537 |
| chr5:145962065-145962286 | -1.205588651 | 0.027034477 | 0.134841537 |
| chrX:110229465-110230220 | -0.783158044 | 0.027036353 | 0.134842118 |
| chr15:62325680-62326857 | -0.748571571 | 0.027043989 | 0.134862473 |
| chrX:153910378-153911301 | 0.741774557 | 0.027046054 | 0.134862473 |
| chr2:35101386-35101663 | -1.123816452 | 0.027047934 | 0.134862473 |
| chr2:237685471-237685715 | 1.106445027 | 0.027048774 | 0.134862473 |
| chr4:189143210-189143697 | -0.794259626 | 0.027049235 | 0.134862473 |
| chr2:198968215-198968505 | -1.098436223 | 0.027052586 | 0.134870403 |
| chr7:98294769-98295387 | -0.781366909 | 0.027061213 | 0.13489181 |
| chr8:143597108-143597549 | 0.766852226 | 0.027061848 | 0.13489181 |
| chr12:104986082-104986958 | -0.727183595 | 0.027062161 | 0.13489181 |
| chr15:90648455-90648851 | 0.852251602 | 0.027069809 | 0.134913932 |
| chr2:94874233-94874493 | 0.911833474 | 0.027070121 | 0.134913932 |
| chr10:48614826-48615288 | -0.95397339 | 0.027079923 | 0.134944021 |
| chr1:6205753-6206438 | 0.77856983 | 0.02708037 | 0.134944021 |
| chr1:193183047-193183574 | -0.833945774 | 0.027081824 | 0.134944021 |
| chr6:42978552-42978774 | 1.033643792 | 0.027083203 | 0.134944021 |
| chr2:100291018-100292465 | -0.71063888 | 0.027087366 | 0.13495599 |
| chr4:131209187-131209932 | -0.826017098 | 0.027099816 | 0.135009238 |
| chr7:22480619-22480904 | 0.922251593 | 0.027103623 | 0.135012796 |
| chr2:58480225-58480607 | -0.957346729 | 0.027104431 | 0.135012796 |
| chrX:38175172-38175650 | -0.927794113 | 0.027106885 | 0.135012796 |
| chr10:114087362-114087664 | -1.069154138 | 0.027107578 | 0.135012796 |
| chr3:63980786-63981079 | 0.829929287 | 0.027110897 | 0.135020549 |
| chr16:631255-631459 | 0.93698306 | 0.027116375 | 0.135031145 |
| chr4:168028573-168029518 | -0.760156715 | 0.027116549 | 0.135031145 |
| chr3:64728537-64728816 | 0.765227806 | 0.027136166 | 0.13512005 |
| chr16:14003171-14003619 | -0.826970254 | 0.027140638 | 0.135133536 |
| chr3:197676599-197676861 | 1.024775598 | 0.027143903 | 0.135141008 |
| chr16:70114020-70114479 | 0.796769549 | 0.027146575 | 0.135145533 |
| chr21:44199974-44200230 | 1.109082942 | 0.027152155 | 0.13516453 |
| chr15:34159276-34159528 | -1.219506671 | 0.027160858 | 0.135178164 |
| chr14:50763403-50763603 | 1.163175802 | 0.02716248 | 0.135178164 |
| chr12:15011267-15011488 | -1.216378849 | 0.027164384 | 0.135178164 |
| chr10:125145098-125145507 | -0.8649325 | 0.027164617 | 0.135178164 |
| chr2:392287-392604 | 1.077368923 | 0.027165193 | 0.135178164 |
| chr4:24988767-24988973 | 1.062441875 | 0.027165479 | 0.135178164 |

| | | | |
|---|---|---|---|
| chr4:164956665-164957564 | -0.727022835 | 0.027171128 | 0.135193413 |
| chr7:76513013-76513413 | -0.895279761 | 0.027172073 | 0.135193413 |
| chr12:97465089-97465301 | -1.047507 | 0.027175366 | 0.135201019 |
| chr15:95267503-95267816 | 1.01481611 | 0.027181004 | 0.135220287 |
| chr5:77643082-77643320 | -1.169289859 | 0.027186355 | 0.135227487 |
| chr10:60883127-60883563 | -0.849133291 | 0.027187942 | 0.135227487 |
| chr7:39085831-39086087 | -1.098133706 | 0.027188765 | 0.135227487 |
| chr6:20644869-20645309 | -0.99799494 | 0.02718951 | 0.135227487 |
| chr9:90517799-90518073 | 1.059277312 | 0.027193325 | 0.135237679 |
| chrX:153723815-153725156 | 0.707146688 | 0.027203127 | 0.135271429 |
| chr17:68888691-68888919 | -1.164531421 | 0.027205953 | 0.135271429 |
| chr17:65100030-65101088 | -0.716866031 | 0.027206561 | 0.135271429 |
| chr5:64407173-64407760 | -0.811095757 | 0.027208715 | 0.135271429 |
| chr21:38397930-38398130 | 1.176013283 | 0.027208938 | 0.135271429 |
| chr8:123374061-123374971 | -0.741759125 | 0.027210831 | 0.135272064 |
| chr14:94549369-94549699 | 0.927764742 | 0.027219448 | 0.135306121 |
| chr16:86364750-86365126 | 1.002008449 | 0.027226349 | 0.135331647 |
| chr5:113846944-113847398 | -0.79606997 | 0.027233255 | 0.135353688 |
| chr4:124997179-124997443 | -1.05896204 | 0.027234946 | 0.135353688 |
| chrX:28586770-28587200 | -0.889026787 | 0.027238616 | 0.135353688 |
| chr6:35777376-35777684 | -0.999675047 | 0.027239766 | 0.135353688 |
| chr12:11431992-11432270 | -1.09025265 | 0.027240693 | 0.135353688 |
| chr7:41794475-41794721 | 1.098664822 | 0.027241383 | 0.135353688 |
| chr7:2733211-2733526 | 0.893440337 | 0.027243913 | 0.135357483 |
| chrX:153333855-153334315 | 0.798624923 | 0.027250326 | 0.135377318 |
| chr3:66428776-66429289 | -0.81476791 | 0.027253393 | 0.135377318 |
| chr3:32276031-32276956 | -0.771515176 | 0.0272541 | 0.135377318 |
| chr19:42313308-42313559 | 1.029959693 | 0.027254973 | 0.135377318 |
| chr4:24583544-24584988 | 0.706366938 | 0.027269392 | 0.13544016 |
| chr11:30984303-30984859 | -0.937542016 | 0.027281093 | 0.135489495 |
| chr4:122602397-122603553 | -0.751827605 | 0.027283539 | 0.135492858 |
| chr10:3313668-3314003 | 1.026919099 | 0.027287863 | 0.135505548 |
| chr5:7627378-7627635 | 1.147496142 | 0.027299068 | 0.135544849 |
| chr8:122678857-122679337 | -0.769879513 | 0.027299315 | 0.135544849 |
| chr18:37270118-37270354 | -0.993142267 | 0.027313932 | 0.135608639 |
| chrX:153649668-153649981 | 0.872613595 | 0.027323757 | 0.135640993 |
| chr4:1855911-1856907 | 0.735050903 | 0.02732486 | 0.135640993 |
| chr3:8156909-8157207 | 1.138439676 | 0.02732576 | 0.135640993 |
| chr6:151689877-151690112 | 0.915614342 | 0.027330988 | 0.135649973 |
| chr4:94475822-94476294 | -0.824681022 | 0.02733111 | 0.135649973 |
| chr3:64642815-64643143 | 0.771605889 | 0.027334838 | 0.135659692 |
| chr1:85605871-85606425 | -0.843364198 | 0.027344972 | 0.135701195 |
| chr10:64430798-64431087 | -0.982199668 | 0.02735673 | 0.135742028 |
| chr10:1036287-1036545 | 1.001212701 | 0.027356744 | 0.135742028 |
| chrX:106894377-106895409 | -0.740418162 | 0.027360403 | 0.135751396 |
| chr21:15722613-15723003 | 0.845039247 | 0.027366563 | 0.135773167 |
| chr7:115707185-115707606 | -0.868836062 | 0.027371129 | 0.135787029 |
| chr5:16314831-16315031 | 1.212490935 | 0.027376238 | 0.135803579 |
| chr4:74649235-74649481 | -1.191884237 | 0.027379729 | 0.135812106 |
| chr2:26145054-26145371 | 0.913709248 | 0.027386825 | 0.135838509 |

| | | | |
|---|---|---|---|
| chr5:102583803-102584046 | -1.119716335 | 0.027402201 | 0.135905977 |
| chr5:34842563-34842955 | 0.820809486 | 0.027404167 | 0.13590693 |
| chr5:127097275-127097622 | -1.098814997 | 0.027407391 | 0.135914121 |
| chr6:26295896-26296576 | -0.777665747 | 0.027413889 | 0.135924573 |
| chr7:108385147-108385428 | -0.978735038 | 0.027414725 | 0.135924573 |
| chr14:68397732-68398115 | -0.843980499 | 0.02741482 | 0.135924573 |
| chr21:42474051-42474695 | -0.778566832 | 0.027418418 | 0.135933617 |
| chr10:11930638-11930838 | -1.290405695 | 0.027420951 | 0.135937379 |
| chr10:83373729-83374039 | -0.95080189 | 0.027440507 | 0.136025524 |
| chr12:76455728-76456079 | 0.852813669 | 0.027446701 | 0.136047427 |
| chr10:99560554-99560872 | 0.893888307 | 0.027454627 | 0.136070007 |
| chr7:138501983-138502336 | -1.084355215 | 0.027454808 | 0.136070007 |
| chr6:133198800-133199045 | -1.180320664 | 0.02746418 | 0.136107652 |
| chr3:146682260-146682512 | -1.008069757 | 0.02746888 | 0.136122138 |
| chr2:100786360-100786596 | 1.188724332 | 0.027475471 | 0.136145996 |
| chr19:12722916-12723139 | 1.080616423 | 0.027481689 | 0.136167999 |
| chr8:37691132-37691801 | -0.757587773 | 0.027497699 | 0.13623852 |
| chr19:6236862-6237157 | 0.988393155 | 0.027501724 | 0.136240788 |
| chr1:26396350-26396568 | -1.253464471 | 0.0275034 | 0.136240788 |
| chr9:134778953-134779451 | 0.840789629 | 0.027504296 | 0.136240788 |
| chr14:34147472-34147704 | 1.121747812 | 0.027505269 | 0.136240788 |
| chr8:23853976-23854500 | -0.840432303 | 0.027509141 | 0.136251156 |
| chr17:37373542-37373784 | 1.11771357 | 0.027513769 | 0.136265269 |
| chr14:34202214-34202525 | 0.944299416 | 0.027519214 | 0.13628343 |
| chr13:51858932-51859216 | 1.049295442 | 0.027522575 | 0.136291263 |
| chr2:218898264-218898464 | -1.182008554 | 0.027539497 | 0.136366249 |
| chr3:141051022-141051620 | 0.789947369 | 0.027551485 | 0.136416795 |
| chr2:108271980-108272210 | -1.047498657 | 0.027560509 | 0.136445469 |
| chr5:124842181-124842574 | -0.993963733 | 0.027560838 | 0.136445469 |
| chr11:74312403-74312603 | -1.166568488 | 0.027567852 | 0.136461584 |
| chr10:37489027-37489684 | 0.76661184 | 0.02756873 | 0.136461584 |
| chr14:67837075-67837278 | 1.122023334 | 0.027569436 | 0.136461584 |
| chr1:215334016-215334216 | -1.187443557 | 0.027577539 | 0.136492875 |
| chr2:55518940-55520580 | -0.696708653 | 0.0275896 | 0.13654375 |
| chr17:4413231-4413436 | 1.109538647 | 0.027595957 | 0.136566392 |
| chr5:61145876-61146244 | -1.120111481 | 0.027605257 | 0.13660359 |
| chr7:130357140-130357352 | -1.100092684 | 0.027608883 | 0.13661271 |
| chr11:120801863-120802092 | -1.143236449 | 0.02764545 | 0.136780124 |
| chr7:4020430-4020668 | 1.095541177 | 0.027646287 | 0.136780124 |
| chr4:40255662-40256344 | 0.802898009 | 0.027649429 | 0.136786838 |
| chr13:52405859-52406150 | -1.042794247 | 0.027681349 | 0.136934188 |
| chr16:89732540-89732797 | 1.017363057 | 0.027682788 | 0.136934188 |
| chr7:44973806-44974105 | -0.883748135 | 0.027688953 | 0.136944704 |
| chr1:155255060-155255816 | 0.752633017 | 0.027689232 | 0.136944704 |
| chr5:160526683-160527238 | -0.872167062 | 0.027690919 | 0.136944704 |
| chr3:10164703-10165017 | 0.873175824 | 0.027692063 | 0.136944704 |
| chr4:77881040-77881262 | -1.210920859 | 0.027695042 | 0.136950599 |
| chr5:58235329-58235670 | -0.964733697 | 0.027697355 | 0.136953195 |
| chr19:11324402-11324701 | 0.950131533 | 0.027712695 | 0.137020207 |
| chr7:139483451-139484127 | -0.748842969 | 0.027744747 | 0.137169829 |

| | | | |
|---|---|---|---|
| chr7:2437231-2437462 | -0.953332911 | 0.027758822 | 0.13723056 |
| chr14:99029643-99029853 | 1.034223368 | 0.027763983 | 0.137247218 |
| chr20:32521810-32522212 | 0.931549358 | 0.027781406 | 0.13732449 |
| chr14:76387124-76387525 | 0.804376454 | 0.027785177 | 0.137334271 |
| chr21:41134004-41134228 | -1.096123439 | 0.027795537 | 0.137376613 |
| chr8:118712465-118712755 | -1.00423879 | 0.027806021 | 0.137412356 |
| chr6:7112841-7113158 | -0.984819282 | 0.027806356 | 0.137412356 |
| chr8:91566647-91566896 | 1.12753304 | 0.027813217 | 0.137437397 |
| chr20:47678557-47678860 | 0.728341619 | 0.027817912 | 0.137447022 |
| chr5:178644630-178644897 | 1.208906983 | 0.027818752 | 0.137447022 |
| chr12:76158778-76159209 | -0.825953239 | 0.027830158 | 0.13749451 |
| chr17:61231301-61231802 | 0.709614397 | 0.027839576 | 0.137532174 |
| chr5:58542146-58542450 | -1.061690961 | 0.027846542 | 0.137557718 |
| chr6:10248680-10248883 | -1.182533072 | 0.027855617 | 0.137585665 |
| chr13:99191708-99192400 | 0.778815734 | 0.027856651 | 0.137585665 |
| chr3:133499226-133499573 | -0.924607468 | 0.027857587 | 0.137585665 |
| chr10:132468001-132468210 | 0.997979372 | 0.027861445 | 0.137595851 |
| chr1:200327401-200327601 | 1.000463525 | 0.027863281 | 0.137596051 |
| chr2:158497474-158497870 | -0.888263828 | 0.027872454 | 0.137632477 |
| chr12:108618105-108618426 | -0.879700322 | 0.027888777 | 0.137704207 |
| chr2:86633112-86633365 | -1.120855587 | 0.027893255 | 0.137717441 |
| chr14:99793134-99793529 | 0.830836355 | 0.027903698 | 0.137753791 |
| chr7:12475423-12475623 | 1.130760684 | 0.027904952 | 0.137753791 |
| chr11:96207408-96207690 | -1.034620994 | 0.027906037 | 0.137753791 |
| chr7:123154260-123154673 | -0.817631125 | 0.027910324 | 0.137753791 |
| chr5:55897829-55898242 | -0.976163585 | 0.027911053 | 0.137753791 |
| chr9:89484246-89484580 | 0.992850702 | 0.027911404 | 0.137753791 |
| chr13:21060889-21061910 | 0.718421163 | 0.027915236 | 0.13776383 |
| chrX:147255106-147255343 | 1.080768136 | 0.027918552 | 0.137771321 |
| chr12:52051692-52051958 | 0.845455394 | 0.027928437 | 0.137811224 |
| chr1:8816833-8817418 | 0.835821106 | 0.02793207 | 0.137820276 |
| chr20:14845498-14845846 | -0.928713778 | 0.027937055 | 0.137835994 |
| chr6:12324088-12324633 | 0.761953604 | 0.027940136 | 0.137842324 |
| chr3:128590965-128591270 | -0.937008659 | 0.027943352 | 0.137849313 |
| chr7:1290339-1290685 | 0.843834416 | 0.027952391 | 0.137885026 |
| chr20:57119867-57120190 | 0.72270362 | 0.027965084 | 0.137938493 |
| chr3:108153260-108153829 | -0.789166255 | 0.027966831 | 0.137938493 |
| chr6:55711392-55711615 | -1.237979743 | 0.027970114 | 0.137945809 |
| chr5:87400380-87400698 | -0.850669337 | 0.027975425 | 0.137958691 |
| chr4:163385004-163385271 | -0.96691178 | 0.027976327 | 0.137958691 |
| chr8:92759258-92759458 | 1.196132524 | 0.027979509 | 0.1379655 |
| chr16:78169913-78170238 | 0.916881158 | 0.027985533 | 0.137986327 |
| chr17:72151576-72152043 | 0.805090235 | 0.027990035 | 0.137999645 |
| chr5:132830515-132830752 | 0.954010949 | 0.027998626 | 0.138033118 |
| chr7:25069872-25070342 | 0.845503458 | 0.028014359 | 0.138101797 |
| chr4:173237166-173237514 | -0.976622189 | 0.028020795 | 0.138124635 |
| chr5:2074710-2075017 | 1.163782389 | 0.028032245 | 0.138172187 |
| chr12:132219013-132219222 | 1.064531998 | 0.028037516 | 0.138185269 |
| chr20:47748466-47748948 | 0.715224805 | 0.028038506 | 0.138185269 |
| chr20:35781770-35781981 | -1.19006586 | 0.028047866 | 0.138222511 |

| | | | |
|---|---|---|---|
| chr3:72289258-72289769 | -0.90029636 | 0.028052968 | 0.13823876 |
| chr1:201650388-201650681 | 0.826360518 | 0.028058733 | 0.138258279 |
| chr2:177724806-177725038 | -1.065515556 | 0.028062816 | 0.138269507 |
| chr17:62447787-62448248 | 0.820748748 | 0.028066778 | 0.138280136 |
| chr2:231662036-231662735 | 0.728187594 | 0.028082201 | 0.138339335 |
| chr10:115799480-115799698 | 1.124389456 | 0.028083335 | 0.138339335 |
| chr1:966105-966617 | 0.848378957 | 0.028085159 | 0.138339335 |
| chr19:46542650-46543034 | 0.866435457 | 0.028086016 | 0.138339335 |
| chr17:61317477-61318138 | 0.702228584 | 0.028097048 | 0.138384777 |
| chr22:47452250-47452535 | -1.043826284 | 0.028102586 | 0.138403161 |
| chr4:183987212-183988440 | -0.742593608 | 0.028116479 | 0.138460959 |
| chr13:28751124-28751359 | -1.03882329 | 0.028117936 | 0.138460959 |
| chr1:32972830-32973291 | 0.858023053 | 0.028124112 | 0.138482469 |
| chr20:58907760-58908147 | -0.789819973 | 0.028129947 | 0.138502301 |
| chr2:120879268-120880220 | -0.787365793 | 0.028138677 | 0.138536381 |
| chr7:104096741-104096983 | -1.13092559 | 0.028140917 | 0.138538509 |
| chr17:55466623-55466836 | 1.137476861 | 0.02814538 | 0.138551578 |
| chr9:105881591-105882038 | -0.910029083 | 0.028158649 | 0.138607992 |
| chr8:90412322-90412636 | 1.004248687 | 0.028163312 | 0.138622041 |
| chr10:32893602-32893819 | -1.285885309 | 0.028172294 | 0.138648435 |
| chr12:100868609-100868836 | -1.285885309 | 0.028172294 | 0.138648435 |
| chr7:87464849-87465691 | -0.753949477 | 0.028183102 | 0.138692719 |
| chr1:149944127-149944468 | 0.878954265 | 0.02818572 | 0.138696695 |
| chr3:107897535-107898014 | -0.923392607 | 0.028196735 | 0.138741988 |
| chr10:86647790-86648015 | 0.911993227 | 0.028204453 | 0.138771052 |
| chrX:154019875-154020088 | 0.988726114 | 0.028206721 | 0.138773299 |
| chrX:47482419-47483356 | 0.728738257 | 0.028224866 | 0.138853655 |
| chr13:24307415-24307626 | 1.066121056 | 0.028236543 | 0.138901555 |
| chr14:75711742-75712229 | 0.835057273 | 0.028238228 | 0.138901555 |
| chr7:30166159-30166587 | 0.838690443 | 0.028244407 | 0.138916093 |
| chr7:157676162-157676371 | -1.149843517 | 0.02824481 | 0.138916093 |
| chr11:122580146-122580413 | -1.002949043 | 0.028250511 | 0.138935215 |
| chr2:73647642-73648004 | -0.928105345 | 0.028256656 | 0.13895391 |
| chr8:90392449-90392649 | 1.249491869 | 0.028257939 | 0.13895391 |
| chr2:26449800-26450000 | 1.128262094 | 0.028275461 | 0.139031146 |
| chr8:58411007-58411934 | -0.745590784 | 0.02828115 | 0.139045768 |
| chr1:231550366-231550692 | 0.978643695 | 0.028282064 | 0.139045768 |
| chr5:20601672-20602324 | -0.803128406 | 0.028285886 | 0.139054154 |
| chr4:145234154-145234457 | -0.90866568 | 0.028287399 | 0.139054154 |
| chr1:218459893-218460788 | -0.718883912 | 0.028303455 | 0.139124156 |
| chr17:61485649-61485862 | 0.745638934 | 0.028310492 | 0.139149819 |
| chr20:37907487-37907764 | 0.963915148 | 0.028320183 | 0.13918852 |
| chr2:100991547-100991755 | 1.09787246 | 0.028325854 | 0.139207462 |
| chr12:12600019-12600230 | 1.178218614 | 0.028328753 | 0.139211447 |
| chr7:96949785-96950042 | -1.170350089 | 0.028330298 | 0.139211447 |
| chr7:152711332-152711592 | -0.974826051 | 0.028337702 | 0.139238902 |
| chr17:64954059-64954463 | 0.966446679 | 0.028344008 | 0.139260954 |
| chr1:154926327-154926528 | 1.0169004 | 0.028347685 | 0.13927009 |
| chr8:64177863-64178140 | -1.092690075 | 0.028353325 | 0.139275982 |
| chr19:52779697-52779897 | 1.290734053 | 0.028354754 | 0.139275982 |

| | | | |
|---|---|---|---|
| chr20:37406325-37406538 | 1.10190219 | 0.028355897 | 0.139275982 |
| chr5:50885609-50885809 | 1.216796563 | 0.028356155 | 0.139275982 |
| chr22:35631687-35631888 | 1.133990944 | 0.02836535 | 0.139306974 |
| chr10:89898579-89898848 | 1.127175621 | 0.028366101 | 0.139306974 |
| chr14:56676072-56676407 | 0.916371859 | 0.028370606 | 0.13932017 |
| chrX:130192289-130192805 | 0.831788001 | 0.028377063 | 0.139342944 |
| chr15:89785103-89785420 | -0.972839009 | 0.028386312 | 0.13937943 |
| chr1:113597555-113597774 | 1.065363508 | 0.028404675 | 0.139460658 |
| chr7:157387019-157387513 | -0.871534338 | 0.028411296 | 0.139484227 |
| chr9:135255315-135255525 | 1.055972539 | 0.028420092 | 0.13951847 |
| chr22:24270454-24271475 | 0.714880562 | 0.028424358 | 0.139530475 |
| chr1:48216893-48217213 | 0.958098979 | 0.028432335 | 0.13956069 |
| chr8:98983228-98983428 | 1.135857133 | 0.028439603 | 0.139586598 |
| chr3:8456244-8457096 | 0.716077218 | 0.028441257 | 0.139586598 |
| chr7:98280764-98281919 | -0.713021611 | 0.028450163 | 0.139621367 |
| chr7:30420666-30420959 | 0.998864343 | 0.028454154 | 0.139624798 |
| chr12:107057085-107057379 | 0.958531165 | 0.028454507 | 0.139624798 |
| chr12:71455156-71455702 | -0.841211204 | 0.0284566 | 0.139626128 |
| chr3:149463530-149463753 | 1.199179905 | 0.028460138 | 0.139634546 |
| chr12:124512665-124512974 | -0.897056525 | 0.028464625 | 0.13964762 |
| chr1:7704483-7704914 | 0.845870553 | 0.028470031 | 0.139665197 |
| chr20:47258721-47259077 | 0.722866062 | 0.028475543 | 0.139683292 |
| chr22:46538497-46538825 | -0.91730253 | 0.028481503 | 0.139701788 |
| chr1:168637403-168637707 | -0.854571317 | 0.028484451 | 0.139701788 |
| chr4:55841449-55842567 | -0.756729915 | 0.028484783 | 0.139701788 |
| chr5:150228371-150228571 | 1.122945912 | 0.028487488 | 0.139706113 |
| chr4:152273398-152273658 | -1.028888928 | 0.028499665 | 0.139756885 |
| chr8:139340852-139341248 | -0.881812604 | 0.028513166 | 0.139814145 |
| chr11:83207104-83207485 | -0.955880918 | 0.028524545 | 0.139853135 |
| chr6:7050971-7051438 | -0.968658286 | 0.028525852 | 0.139853135 |
| chr4:3292467-3293060 | 0.734941941 | 0.028526594 | 0.139853135 |
| chr14:35932921-35933175 | 0.964222761 | 0.028531865 | 0.139870029 |
| chr2:28269219-28269612 | -0.915950036 | 0.028534731 | 0.139875131 |
| chr20:43689776-43689976 | 1.109123098 | 0.028545905 | 0.13991958 |
| chr1:216915752-216915952 | -1.137719557 | 0.028547451 | 0.13991958 |
| chr7:88219659-88220741 | -0.713826395 | 0.028555957 | 0.139952319 |
| chr17:61755262-61755632 | 0.725640254 | 0.028559644 | 0.139958232 |
| chr1:32637023-32637454 | 0.895949914 | 0.028560817 | 0.139958232 |
| chr2:46876791-46877419 | -0.771930523 | 0.028566689 | 0.139978055 |
| chr6:125858680-125859130 | -0.798721086 | 0.028582907 | 0.140048568 |
| chr20:47171522-47171722 | 0.756774165 | 0.028586997 | 0.140054673 |
| chr17:83089295-83089925 | 0.737539527 | 0.028587809 | 0.140054673 |
| chr20:42662281-42662502 | 1.07231491 | 0.028592165 | 0.140067057 |
| chr19:8586093-8586319 | 1.12584105 | 0.028599063 | 0.140091892 |
| chr5:9515054-9515279 | 1.162030226 | 0.028607819 | 0.140125828 |
| chr7:130560990-130561329 | -1.069250364 | 0.028612503 | 0.14013981 |
| chr3:137248273-137248567 | 1.023456582 | 0.028621558 | 0.140175201 |
| chr6:53326171-53327301 | -0.727483794 | 0.028625548 | 0.14018578 |
| chr2:174395235-174396607 | -0.712777919 | 0.028628814 | 0.140192815 |
| chr16:73015552-73016123 | -0.737853612 | 0.028634968 | 0.140213991 |

| | | | |
|---|---|---|---|
| chr2:48531038-48531581 | 0.761378641 | 0.028638175 | 0.140220544 |
| chr7:94907220-94908957 | -0.715850878 | 0.028642804 | 0.140220544 |
| chr17:45791953-45792290 | 0.948488203 | 0.02864356 | 0.140220544 |
| chr10:83274048-83274348 | -1.10391253 | 0.028643627 | 0.140220544 |
| chr17:83202114-83202468 | 0.831940903 | 0.028657262 | 0.140278331 |
| chr1:195856192-195856411 | 1.092000004 | 0.028665066 | 0.140307571 |
| chr16:70789349-70789727 | -0.885960953 | 0.028676755 | 0.140348484 |
| chrX:17291505-17291944 | 0.805484312 | 0.028677088 | 0.140348484 |
| chr8:91854869-91855111 | 1.011785507 | 0.02868577 | 0.140382008 |
| chr7:801912-802445 | -0.822361087 | 0.028689855 | 0.140393031 |
| chr8:132383473-132383696 | 1.142200854 | 0.028694463 | 0.140397272 |
| chr1:114248711-114249193 | 0.740887376 | 0.028695109 | 0.140397272 |
| chr7:148084097-148084429 | -0.936771509 | 0.028696219 | 0.140397272 |
| chr10:18140331-18140962 | 0.829056746 | 0.028702447 | 0.140417624 |
| chr18:78305731-78306070 | -1.091257033 | 0.028704044 | 0.140417624 |
| chr3:9422662-9423453 | 0.738052584 | 0.02870757 | 0.140425909 |
| chr2:236745880-236746152 | 0.987350169 | 0.028718109 | 0.1404656 |
| chr10:120979452-120979678 | 1.051492007 | 0.028719351 | 0.1404656 |
| chr6:170400074-170400274 | 1.07966277 | 0.028723593 | 0.140468202 |
| chr8:54134813-54135444 | -0.743050709 | 0.028724242 | 0.140468202 |
| chr11:48052217-48052618 | 0.776198603 | 0.028725383 | 0.140468202 |
| chr4:5998371-5998843 | 0.837253114 | 0.028734881 | 0.140505685 |
| chr7:110204524-110205098 | -0.855821237 | 0.0287388 | 0.140515878 |
| chr12:113466551-113466752 | 1.069943615 | 0.028742099 | 0.140523041 |
| chr16:23182359-23182854 | 0.754813623 | 0.028745378 | 0.140530106 |
| chr11:82676550-82676783 | -1.027305574 | 0.028751877 | 0.140552911 |
| chr19:18940742-18941433 | -0.9635602 | 0.028755299 | 0.1405604 |
| chr3:167983113-167983551 | 0.90364496 | 0.028757078 | 0.1405604 |
| chr11:45983014-45983664 | -0.841682453 | 0.028759519 | 0.140563365 |
| chr19:45577605-45577858 | 1.005002044 | 0.028765965 | 0.140585902 |
| chr17:9968971-9969203 | 1.053691944 | 0.028774138 | 0.14060826 |
| chr17:61734324-61734596 | 0.751118989 | 0.028774252 | 0.14060826 |
| chr14:88535167-88535531 | -0.905258415 | 0.028776045 | 0.14060826 |
| chr12:45268932-45269577 | -0.842809226 | 0.028779436 | 0.140615862 |
| chr5:100903110-100903851 | -0.717946038 | 0.028787247 | 0.140645056 |
| chr8:140347120-140347356 | 1.040898549 | 0.028792097 | 0.140651571 |
| chr16:37389144-37389369 | 1.170579799 | 0.028792251 | 0.140651571 |
| chrX:116667326-116667691 | -0.913483384 | 0.02879978 | 0.14067386 |
| chr3:177664382-177665364 | -0.75062672 | 0.028800486 | 0.14067386 |
| chr21:36134858-36136292 | 0.717006034 | 0.028804181 | 0.140680377 |
| chr11:75151281-75151481 | 1.101453716 | 0.028805492 | 0.140680377 |
| chr17:17682677-17682952 | 0.874245324 | 0.028817495 | 0.140730024 |
| chr12:79414224-79414640 | 0.836369178 | 0.028826945 | 0.140767202 |
| chr5:171850592-171850912 | 0.850413225 | 0.028843092 | 0.140837076 |
| chr1:155324939-155325401 | 0.750522467 | 0.028850394 | 0.140856803 |
| chr2:181239482-181239767 | -1.039294715 | 0.028851947 | 0.140856803 |
| chr7:128459389-128459599 | -0.961889171 | 0.028852647 | 0.140856803 |
| chr8:11016557-11016757 | -1.200348673 | 0.028856676 | 0.140867499 |
| chr12:118061258-118061727 | 0.824774334 | 0.028871502 | 0.140927134 |
| chr17:58953012-58953253 | 0.809200143 | 0.028872571 | 0.140927134 |

| | | | |
|---|---|---|---|
| chr7:130069954-130070757 | -0.719624883 | 0.028879029 | 0.140949675 |
| chr4:127417790-127418100 | 1.116985242 | 0.028883224 | 0.140961174 |
| chr14:36374061-36374417 | 0.789580491 | 0.028886073 | 0.140966095 |
| chr16:30615425-30615715 | -0.968523064 | 0.028893096 | 0.140991388 |
| chr9:113944872-113945150 | 1.104927897 | 0.028903729 | 0.141034297 |
| chr9:134189962-134190187 | 1.05561654 | 0.028909641 | 0.141054159 |
| chr1:106571042-106571391 | 0.760437564 | 0.028920997 | 0.141100582 |
| chr6:45608382-45608869 | -0.854690537 | 0.028923849 | 0.14110551 |
| chr15:66673463-66673902 | -0.857805643 | 0.028936017 | 0.141152559 |
| chr8:128460134-128460693 | 0.812949022 | 0.028937177 | 0.141152559 |
| chr3:169880276-169880485 | 1.09137723 | 0.028941754 | 0.141165899 |
| chr2:120669512-120669801 | -0.983000445 | 0.028945131 | 0.141173382 |
| chr15:58361137-58361531 | 0.921363826 | 0.028951622 | 0.1411934 |
| chr12:122726149-122726377 | 1.090375512 | 0.028954877 | 0.1411934 |
| chr11:26276842-26277128 | -0.961202697 | 0.028956654 | 0.1411934 |
| chr3:129016398-129016753 | 0.810993809 | 0.028958328 | 0.1411934 |
| chr2:160062162-160063024 | -0.768561152 | 0.028958449 | 0.1411934 |
| chr10:132446760-132447135 | 0.756531268 | 0.02896767 | 0.141229372 |
| chr4:70601420-70601791 | -0.848800531 | 0.028969945 | 0.141231475 |
| chr3:23862671-23862969 | 1.156107704 | 0.028975759 | 0.141241133 |
| chr2:122883281-122883532 | -1.270839369 | 0.028975901 | 0.141241133 |
| chr4:3880643-3880868 | 1.158008448 | 0.028977456 | 0.141241133 |
| chr13:41455418-41456158 | -0.737314638 | 0.028990088 | 0.14129366 |
| chr3:43565748-43565956 | -1.271593058 | 0.02899192 | 0.14129366 |
| chr17:77131999-77132255 | -1.023509927 | 0.029002534 | 0.141336398 |
| chr12:55707833-55708140 | 0.904501473 | 0.029012515 | 0.141360496 |
| chr14:43449235-43449644 | 0.899167236 | 0.029012887 | 0.141360496 |
| chr1:211630192-211630964 | -0.797894375 | 0.029013014 | 0.141360496 |
| chr6:64330076-64330354 | -1.25725609 | 0.029026754 | 0.141408794 |
| chr7:139753067-139753443 | -0.94306305 | 0.029028217 | 0.141408794 |
| chr4:2413348-2413613 | 0.885377299 | 0.029028463 | 0.141408794 |
| chr3:72446576-72446804 | 1.018955592 | 0.029037272 | 0.14143456 |
| chr5:137091561-137091775 | -1.165878159 | 0.029037444 | 0.14143456 |
| chr1:226646004-226646228 | 0.949789964 | 0.02904095 | 0.141435223 |
| chr3:171115826-171116026 | 1.182791465 | 0.029041272 | 0.141435223 |
| chr11:107849809-107850140 | -0.938424253 | 0.029046844 | 0.14145337 |
| chr17:38351461-38352118 | 0.774212731 | 0.029061557 | 0.141516023 |
| chr8:95056957-95057170 | 0.96970215 | 0.029063901 | 0.141518444 |
| chr5:125409948-125410426 | -0.83806829 | 0.029078537 | 0.141567537 |
| chr10:13383506-13383732 | -1.183720939 | 0.029078957 | 0.141567537 |
| chr5:62589584-62589904 | -1.056801716 | 0.029079526 | 0.141567537 |
| chr20:47226395-47226595 | 0.75644791 | 0.029084628 | 0.141583379 |
| chr10:112810885-112811219 | 0.869044932 | 0.029087917 | 0.141585006 |
| chr15:38991247-38991504 | 1.092589049 | 0.029088658 | 0.141585006 |
| chr14:36604915-36605308 | 0.810257519 | 0.029093466 | 0.141591938 |
| chr1:35718012-35718439 | 0.849946 | 0.029093778 | 0.141591938 |
| chr12:130014036-130014327 | -0.873637927 | 0.029101903 | 0.141602205 |
| chr10:70549903-70550314 | 0.865130531 | 0.029102015 | 0.141602205 |
| chrX:39734614-39734822 | 1.149883247 | 0.029104871 | 0.141602205 |
| chr7:18178372-18178572 | 1.030232608 | 0.029105022 | 0.141602205 |

| | | | |
|---|---|---|---|
| chr10:9068887-9069484 | -0.817664198 | 0.029105128 | 0.141602205 |
| chrX:45515586-45515930 | -0.961950474 | 0.029108678 | 0.141610482 |
| chr5:113294476-113295142 | -0.754542895 | 0.02911974 | 0.14165269 |
| chr7:126501562-126501782 | 1.072574641 | 0.029121051 | 0.14165269 |
| chr3:61627816-61628223 | 0.749486363 | 0.029125907 | 0.141667312 |
| chr8:89801208-89802073 | 0.734579857 | 0.029127755 | 0.141667312 |
| chr16:1331687-1332120 | 0.845653395 | 0.029135447 | 0.141695731 |
| chr7:107420489-107420883 | -1.081629245 | 0.029140623 | 0.141711908 |
| chr3:159451087-159451386 | -0.949192904 | 0.029152172 | 0.141759076 |
| chr4:185202390-185202697 | -0.977055727 | 0.029155661 | 0.141767041 |
| chr2:21396223-21396647 | -0.87671713 | 0.029163061 | 0.141792438 |
| chr2:31783528-31784007 | -0.873138087 | 0.029164585 | 0.141792438 |
| chr7:106957799-106958098 | -0.955159364 | 0.029170494 | 0.141812169 |
| chr5:39052318-39052518 | 1.194913844 | 0.029175901 | 0.141825056 |
| chr19:55069047-55069919 | 0.736850694 | 0.029176847 | 0.141825056 |
| chr6:138291094-138291296 | 1.133866254 | 0.029180086 | 0.141831803 |
| chr6:116500896-116501493 | -0.766630912 | 0.029187823 | 0.141860409 |
| chrX:72126618-72126884 | 1.002465957 | 0.029206291 | 0.141934427 |
| chr15:65333321-65333542 | 1.013954383 | 0.029206757 | 0.141934427 |
| chr7:143002172-143002372 | -1.293439528 | 0.029241839 | 0.142072829 |
| chr18:50268172-50268761 | -0.786345492 | 0.029242501 | 0.142072829 |
| chr18:59899400-59901077 | -0.698120606 | 0.02924324 | 0.142072829 |
| chr11:62880698-62881921 | 0.699135457 | 0.02924503 | 0.142072829 |
| chr17:19003987-19004360 | 0.833810629 | 0.029245762 | 0.142072829 |
| chr1:74352907-74353107 | -1.182521035 | 0.029246362 | 0.142072829 |
| chr8:126796579-126796844 | 0.99980388 | 0.029257527 | 0.142118056 |
| chr7:144019490-144019987 | -0.836155682 | 0.02927141 | 0.142167868 |
| chr11:859739-859951 | 1.012902544 | 0.029271492 | 0.142167868 |
| chr3:50365179-50365601 | 0.835155567 | 0.029276116 | 0.142181314 |
| chr17:61330543-61331040 | 0.704404787 | 0.02928097 | 0.142195872 |
| chr4:172392793-172393558 | -0.75486776 | 0.029307662 | 0.142316474 |
| chr8:40848990-40849428 | -0.885694648 | 0.02931356 | 0.142336098 |
| chr21:46519067-46519274 | 1.147072456 | 0.029327686 | 0.142395663 |
| chr12:124637564-124637928 | 0.869356748 | 0.029340523 | 0.142443404 |
| chr5:111143010-111143260 | -1.090115115 | 0.029341237 | 0.142443404 |
| chr5:120447918-120448185 | -1.085980804 | 0.029345863 | 0.142456839 |
| chr16:23500568-23500979 | -0.960646373 | 0.029349873 | 0.142467278 |
| chr10:35581554-35581948 | -0.964451529 | 0.029356534 | 0.142490583 |
| chrX:24081591-24082122 | -0.850474429 | 0.029365178 | 0.142523511 |
| chr2:111887798-111888151 | 0.809060053 | 0.02937049 | 0.142540266 |
| chr6:18324038-18324472 | 0.967764107 | 0.029374562 | 0.142550997 |
| chr6:104333492-104333714 | 1.08009089 | 0.029386432 | 0.14259518 |
| chr5:170896771-170897131 | -0.915893322 | 0.029387389 | 0.14259518 |
| chr19:19192481-19192948 | -0.779802768 | 0.029392087 | 0.142608948 |
| chr14:91398708-91399046 | 0.885525608 | 0.029400217 | 0.142639362 |
| chr19:55141103-55141411 | 0.95300709 | 0.029405338 | 0.142655174 |
| chr17:76506445-76506898 | 0.804637647 | 0.029411477 | 0.142675921 |
| chr1:209652832-209653435 | -0.742290864 | 0.029414853 | 0.142683267 |
| chr1:196372974-196373307 | -0.867084549 | 0.029431529 | 0.142750775 |
| chr6:100176573-100176964 | 0.843189499 | 0.029432496 | 0.142750775 |

| | | | |
|---|---|---|---|
| chrX:19585746-19585982 | 1.047087398 | 0.029443227 | 0.142793779 |
| chr11:64871639-64871846 | 1.170014562 | 0.029460223 | 0.142857971 |
| chr8:103635335-103635852 | 0.840159906 | 0.029460393 | 0.142857971 |
| chr1:44729293-44729493 | 0.980632886 | 0.029462056 | 0.142857971 |
| chr6:24064146-24064418 | -1.004108134 | 0.029467365 | 0.142874672 |
| chr9:37430874-37431220 | -0.96268135 | 0.029490487 | 0.142973704 |
| chrX:150693672-150694112 | 0.779575799 | 0.029491522 | 0.142973704 |
| chr4:184805661-184805861 | -1.266711449 | 0.029494121 | 0.142977259 |
| chr17:35119320-35120123 | 0.748829368 | 0.02951039 | 0.143041769 |
| chr5:68972418-68972775 | -1.197399594 | 0.029511162 | 0.143041769 |
| chr3:23685730-23686149 | 0.787358703 | 0.029514313 | 0.143047993 |
| chr8:134700446-134700646 | 1.123774518 | 0.029516306 | 0.1430486 |
| chr5:145531426-145531793 | 0.838458537 | 0.029519093 | 0.143053063 |
| chr20:53929396-53930022 | 0.69726188 | 0.029534178 | 0.143117113 |
| chr4:142443573-142443801 | -1.148177475 | 0.029538876 | 0.143130828 |
| chr6:156822718-156823088 | -0.922340501 | 0.029544421 | 0.143148642 |
| chr2:231557839-231558153 | 0.898859693 | 0.02954847 | 0.143159206 |
| chr1:118132054-118132360 | -1.048209028 | 0.029551808 | 0.143165205 |
| chr14:34081884-34082088 | 1.020623766 | 0.029553445 | 0.143165205 |
| chr8:66870449-66870960 | 0.820511913 | 0.029556677 | 0.143171812 |
| chr7:101077616-101077953 | -0.836317573 | 0.02955856 | 0.143171881 |
| chr4:69689591-69689793 | -1.231452564 | 0.029562871 | 0.14318371 |
| chr22:23971100-23971303 | 1.111197232 | 0.029572028 | 0.143219007 |
| chr12:3214981-3215397 | 0.799162976 | 0.029579886 | 0.143248009 |
| chr5:64946858-64947512 | -0.76161925 | 0.029593927 | 0.143306951 |
| chr20:54600909-54601184 | 0.755482441 | 0.029603888 | 0.143346127 |
| chr2:53455030-53455416 | -0.930115987 | 0.02961024 | 0.143367823 |
| chr5:131312217-131313351 | -0.71939455 | 0.029619232 | 0.143402301 |
| chr12:46585940-46586190 | 1.050613881 | 0.029631291 | 0.143451621 |
| chr6:49540616-49540980 | -0.863506497 | 0.029640849 | 0.143488827 |
| chr5:23711675-23711875 | 1.161552067 | 0.029645524 | 0.143502391 |
| chr5:119411710-119412417 | -0.751611952 | 0.029649997 | 0.143514977 |
| chr4:6688844-6690255 | 0.700154696 | 0.029660983 | 0.143559083 |
| chr2:38527986-38528386 | -0.909156901 | 0.029668486 | 0.143584166 |
| chr3:53551767-53552011 | -1.151953305 | 0.029671696 | 0.143584166 |
| chr16:54929582-54929999 | 0.752949664 | 0.029671787 | 0.143584166 |
| chr8:23345350-23345671 | -1.038500909 | 0.029682824 | 0.143628504 |
| chr19:42742029-42742242 | 1.11049998 | 0.029695756 | 0.143682004 |
| chr11:33015588-33016402 | -0.821150056 | 0.029704918 | 0.143717257 |
| chr5:112109339-112109689 | -0.996424348 | 0.029718407 | 0.143773442 |
| chr6:29531545-29531893 | 0.988041508 | 0.029727174 | 0.143806775 |
| chr15:77527175-77527565 | 0.858490773 | 0.029736856 | 0.14384453 |
| chr14:58151783-58152378 | 0.752847353 | 0.029741013 | 0.143855557 |
| chr2:172545759-172546075 | -1.0115555 | 0.029745177 | 0.143866619 |
| chr14:41925491-41925761 | 0.967976687 | 0.029759005 | 0.143924198 |
| chrX:123588830-123589105 | 1.016674509 | 0.029760839 | 0.143924198 |
| chr4:2220005-2220300 | 0.988778281 | 0.029779476 | 0.143992556 |
| chr6:81806828-81807386 | 0.817974198 | 0.029780443 | 0.143992556 |
| chr2:27717100-27717435 | -0.82544093 | 0.029780612 | 0.143992556 |
| chr9:109441360-109441640 | -0.929057659 | 0.029790878 | 0.144033108 |

| | | | |
|---|---|---|---|
| chr1:106659637-106660215 | 0.715480045 | 0.02979522 | 0.144042363 |
| chr12:123384611-123384811 | 0.926100645 | 0.029796553 | 0.144042363 |
| chr1:112727642-112728073 | 0.867846963 | 0.029801711 | 0.14405821 |
| chr4:112938523-112938787 | -1.066733849 | 0.029805245 | 0.14406147 |
| chr2:66495791-66496424 | -0.786926401 | 0.029806146 | 0.14406147 |
| chr11:75351130-75351520 | 0.791911391 | 0.02981016 | 0.144065931 |
| chr1:217630670-217631671 | -0.690645532 | 0.029810829 | 0.144065931 |
| chr16:77675552-77676057 | 0.755185731 | 0.029822266 | 0.144112111 |
| chr10:5888857-5889416 | 0.733523133 | 0.029824778 | 0.144115162 |
| chr4:781523-782787 | 0.691366644 | 0.02983311 | 0.144139917 |
| chr19:10260122-10260572 | -0.847026232 | 0.029833663 | 0.144139917 |
| chr2:10432296-10433272 | -0.719436359 | 0.029835607 | 0.144140219 |
| chr20:54892658-54893014 | 0.737538171 | 0.029841324 | 0.144158752 |
| chr16:11333675-11334158 | 0.774347819 | 0.029843876 | 0.144161989 |
| chr3:50445949-50446195 | 1.087890345 | 0.029851671 | 0.144188561 |
| chr11:131748189-131748389 | -1.302936385 | 0.029853446 | 0.144188561 |
| chr6:27866544-27866756 | -0.981757768 | 0.029855022 | 0.144188561 |
| chr19:40862431-40862717 | 1.090249832 | 0.029875594 | 0.144278583 |
| chr7:5606958-5607199 | 0.998434987 | 0.029879154 | 0.144278583 |
| chr9:69122710-69123115 | -0.77542728 | 0.029879311 | 0.144278583 |
| chr10:112804114-112804324 | 1.101586214 | 0.02988744 | 0.144308744 |
| chr14:59788240-59788440 | 1.229885878 | 0.02989188 | 0.144321084 |
| chr5:133154745-133155001 | 0.998190709 | 0.029897619 | 0.1443397 |
| chr9:76692060-76692488 | -0.930956342 | 0.029913577 | 0.144399288 |
| chr3:188362506-188362794 | 0.95007725 | 0.029914736 | 0.144399288 |
| chr16:87909412-87909684 | -0.981395344 | 0.029915616 | 0.144399288 |
| chr6:142071955-142072434 | -0.752460306 | 0.02992405 | 0.144430902 |
| chr1:210234086-210234534 | -0.799761216 | 0.029931136 | 0.144456005 |
| chr20:10316696-10316917 | -1.170913101 | 0.029934831 | 0.144464735 |
| chr1:198582649-198583017 | 0.888102616 | 0.029941855 | 0.14448336 |
| chr15:83753768-83754478 | 0.74806038 | 0.029943819 | 0.14448336 |
| chr1:174954110-174954345 | -1.071460166 | 0.029944347 | 0.14448336 |
| chr10:49615711-49616180 | -0.816671539 | 0.029957587 | 0.144531022 |
| chr7:111865090-111865367 | -1.114822303 | 0.029960166 | 0.144531022 |
| chr7:131843931-131844241 | 0.975997833 | 0.029960316 | 0.144531022 |
| chr4:40455545-40455843 | -0.935799306 | 0.02996177 | 0.144531022 |
| chr1:196523105-196523332 | 1.162173196 | 0.029975385 | 0.144580251 |
| chr19:36797250-36797651 | 0.806566774 | 0.029975749 | 0.144580251 |
| chr8:85379236-85379574 | 0.879136969 | 0.029982222 | 0.144602368 |
| chr5:178097135-178097362 | 1.043591651 | 0.029987486 | 0.144618651 |
| chr3:188164104-188164770 | -0.779418058 | 0.029989833 | 0.144620868 |
| chr3:118472727-118472970 | 1.027218675 | 0.029994849 | 0.144635953 |
| chr12:31653281-31653608 | -0.952764552 | 0.030016498 | 0.144727001 |
| chr21:25970631-25970904 | 0.970571929 | 0.030017508 | 0.144727001 |
| chr14:105737767-105738076 | 0.926502555 | 0.030027069 | 0.144762531 |
| chr1:171251370-171252025 | -0.767239368 | 0.030028656 | 0.144762531 |
| chr4:86831225-86832090 | -0.720598804 | 0.030035796 | 0.144787841 |
| chr11:33329759-33329972 | 1.032611759 | 0.030039759 | 0.144797164 |
| chr11:60376625-60376970 | 0.949279752 | 0.030041509 | 0.144797164 |
| chr7:77696034-77697161 | -0.699322316 | 0.030047004 | 0.144806136 |

| | | | |
|---|---|---|---|
| chr15:70462584-70463589 | -0.709527843 | 0.03004715 | 0.144806136 |
| chr1:155127658-155127978 | 0.778620517 | 0.030050802 | 0.144814627 |
| chr17:69058315-69058587 | -0.896917238 | 0.030060476 | 0.144843234 |
| chr15:35000500-35000783 | -1.056987689 | 0.030060519 | 0.144843234 |
| chr13:77697096-77698052 | -0.718821221 | 0.030064493 | 0.144853272 |
| chr17:59989530-59989859 | 0.878724246 | 0.030071801 | 0.144879374 |
| chr1:242459879-242460221 | 0.941451196 | 0.030078873 | 0.144904333 |
| chr13:41980330-41980534 | 1.166071247 | 0.030082944 | 0.144912959 |
| chr11:120802204-120802605 | -0.868564789 | 0.030084446 | 0.144912959 |
| chr13:33375612-33375889 | -0.986423438 | 0.030089202 | 0.144924908 |
| chr9:126513404-126513842 | 0.848981514 | 0.03009071 | 0.144924908 |
| chr13:98537372-98537707 | 1.0746932 | 0.030103644 | 0.144978088 |
| chr10:62454765-62454965 | -1.233276005 | 0.030115977 | 0.145028371 |
| chr12:46209975-46210480 | -0.752810584 | 0.030118005 | 0.14502902 |
| chrX:85003044-85004240 | -0.713299177 | 0.030127381 | 0.145060069 |
| chr6:133614885-133615085 | -1.331353231 | 0.030128239 | 0.145060069 |
| chr3:100893670-100893976 | -1.026863982 | 0.030146474 | 0.145138747 |
| chr16:78659298-78659712 | -0.941683109 | 0.030148727 | 0.145140475 |
| chr13:67224257-67224549 | -0.849766992 | 0.030156013 | 0.145166426 |
| chr12:109613128-109613332 | 0.991246074 | 0.030158521 | 0.145169379 |
| chr2:172427965-172428165 | 1.065727892 | 0.0301857 | 0.145291078 |
| chrX:150201090-150201329 | 0.9862769 | 0.03019686 | 0.145331349 |
| chr4:64174386-64174685 | 1.095324075 | 0.03019786 | 0.145331349 |
| chr9:95004483-95004952 | 0.847987949 | 0.030206458 | 0.1453636 |
| chr5:6526027-6526270 | 0.948392389 | 0.030208897 | 0.145366207 |
| chr20:48864874-48865449 | 0.719514498 | 0.030212491 | 0.145374372 |
| chr13:23979466-23980048 | 0.785193629 | 0.030215385 | 0.145379167 |
| chr2:160492991-160493733 | 0.762369083 | 0.030220006 | 0.145392159 |
| chr20:653141-653770 | 0.782754554 | 0.03022188 | 0.145392159 |
| chr13:113291935-113292270 | -1.008330186 | 0.03022783 | 0.145404131 |
| chr16:31873608-31874447 | 0.747447267 | 0.030228164 | 0.145404131 |
| chr12:103751139-103751507 | -0.909709219 | 0.030234245 | 0.145424252 |
| chr3:56557592-56557857 | -0.91633561 | 0.030247809 | 0.145480359 |
| chr7:142939467-142939943 | -0.827341519 | 0.030251774 | 0.145490298 |
| chr8:141267299-141267499 | 1.119199819 | 0.03025978 | 0.145512037 |
| chr3:5389028-5389463 | 0.8047733 | 0.030261084 | 0.145512037 |
| chrX:53482093-53482378 | 0.962602222 | 0.030262699 | 0.145512037 |
| chr20:44204150-44204640 | 0.77868104 | 0.030264164 | 0.145512037 |
| chr17:45124445-45124923 | -0.821340876 | 0.03026579 | 0.145512037 |
| chr5:129241565-129242377 | -0.737992448 | 0.030270297 | 0.145518914 |
| chr2:49210137-49210666 | -0.789215671 | 0.030272436 | 0.145518914 |
| chr20:54761047-54761516 | 0.731799429 | 0.030274208 | 0.145518914 |
| chr4:128574711-128574974 | 0.935324482 | 0.030274817 | 0.145518914 |
| chr1:175518221-175518455 | 1.084227439 | 0.03029456 | 0.145604676 |
| chr2:18387411-18388115 | -0.76781928 | 0.030297887 | 0.145611533 |
| chr1:17475806-17476190 | 0.910443235 | 0.030302243 | 0.145623335 |
| chr8:107020327-107020660 | 0.891471453 | 0.030306967 | 0.145636904 |
| chr2:158848189-158848755 | -0.846890435 | 0.030313115 | 0.14565719 |
| chr1:87025268-87025486 | -1.106602184 | 0.030314991 | 0.14565719 |
| chr17:65700002-65700264 | -1.036415472 | 0.030322466 | 0.145683972 |

| | | | |
|---|---|---|---|
| chr16:83643968-83644174 | 1.254179082 | 0.030335626 | 0.145738059 |
| chr1:44660214-44660460 | -1.232874923 | 0.030353213 | 0.145806703 |
| chr4:28686711-28686955 | 1.076985278 | 0.03035372 | 0.145806703 |
| chr1:193019525-193019830 | -0.956960557 | 0.030361216 | 0.145830413 |
| chr5:10517231-10517629 | 0.975772193 | 0.030362462 | 0.145830413 |
| chr7:76373203-76373518 | 0.903585399 | 0.030371569 | 0.145860975 |
| chr18:72543499-72543760 | -1.039939623 | 0.030372633 | 0.145860975 |
| chr8:100835435-100835792 | 0.824024098 | 0.030386247 | 0.145916332 |
| chr13:102457166-102457732 | -0.817395753 | 0.030387968 | 0.145916332 |
| chr1:153348460-153349052 | -0.816025098 | 0.030396635 | 0.145943104 |
| chr10:7354569-7354890 | 1.131244075 | 0.030397353 | 0.145943104 |
| chr5:129461427-129461972 | -0.72603034 | 0.030408077 | 0.145985443 |
| chr15:74787815-74788168 | -0.798912851 | 0.030412019 | 0.145995219 |
| chr6:116442605-116442866 | 1.145082858 | 0.03041902 | 0.146019682 |
| chr5:135191171-135191903 | -0.760398089 | 0.03042318 | 0.146030501 |
| chr12:111994591-111995044 | 0.820913527 | 0.030426662 | 0.146038066 |
| chr13:59766299-59766630 | -1.013354511 | 0.030450756 | 0.146144557 |
| chr2:130288476-130289006 | 0.806180638 | 0.030457668 | 0.146168571 |
| chrX:78357675-78358259 | -0.766744226 | 0.030463463 | 0.146184114 |
| chr20:54586551-54586968 | 0.725457939 | 0.030466566 | 0.146184114 |
| chr4:499009-499511 | 0.785578207 | 0.03046663 | 0.146184114 |
| chr7:18693488-18693788 | 0.913521684 | 0.030469593 | 0.146189178 |
| chr12:124865823-124866023 | 0.963215764 | 0.030496168 | 0.14630096 |
| chr6:108074107-108075221 | 0.717575698 | 0.03049671 | 0.14630096 |
| chr12:74816578-74817135 | -0.825236306 | 0.030504163 | 0.14632755 |
| chr1:86434455-86434745 | -1.024620444 | 0.030519641 | 0.146392633 |
| chr15:80804905-80805521 | 0.763624028 | 0.030539611 | 0.146479255 |
| chr17:77734448-77736086 | -0.710608705 | 0.030548258 | 0.146502867 |
| chr6:33987556-33988677 | 0.705400762 | 0.030548358 | 0.146502867 |
| chr8:119867132-119867432 | -0.893479887 | 0.030558495 | 0.146542308 |
| chr9:129096540-129096740 | 1.021121293 | 0.030566041 | 0.146569322 |
| chr21:46244618-46244921 | 0.890664941 | 0.030568508 | 0.146571978 |
| chr1:48719143-48719389 | -1.115767117 | 0.030578801 | 0.146612158 |
| chr2:119310858-119311211 | -0.954867396 | 0.030581938 | 0.146618023 |
| chr1:116290351-116290578 | 0.934884089 | 0.030585235 | 0.146624658 |
| chr1:230917303-230917504 | -1.198527511 | 0.030589008 | 0.1466265 |
| chr11:32173453-32173849 | 0.855728271 | 0.030589447 | 0.1466265 |
| chr17:16171597-16172039 | 0.851373509 | 0.030592485 | 0.146631889 |
| chr11:46826689-46827083 | 0.815507147 | 0.030598136 | 0.146642229 |
| chr8:130727003-130727265 | -1.032626619 | 0.03059847 | 0.146642229 |
| chr1:216399288-216399518 | -1.130065395 | 0.030607156 | 0.146662342 |
| chrX:84118826-84119200 | -0.897348409 | 0.030607792 | 0.146662342 |
| chr11:78907588-78907856 | 0.912073111 | 0.030610231 | 0.146662342 |
| chr5:12100602-12100861 | -1.064708884 | 0.030611702 | 0.146662342 |
| chr2:21422117-21422476 | -0.859471023 | 0.030613884 | 0.146662342 |
| chr4:1684255-1684741 | 0.797024694 | 0.030614151 | 0.146662342 |
| chr3:122021374-122021574 | 1.069709317 | 0.030624548 | 0.146702976 |
| chr21:31908369-31908701 | 0.863540297 | 0.030633947 | 0.14672457 |
| chr17:60684279-60684510 | 0.779719233 | 0.030634242 | 0.14672457 |
| chr6:10954562-10954834 | -1.076405238 | 0.030635417 | 0.14672457 |

| | | | |
|---|---|---|---|
| chr14:67820531-67820807 | 0.943863498 | 0.030636998 | 0.14672457 |
| chr16:58027043-58027243 | 1.013353929 | 0.03063863 | 0.14672457 |
| chr15:93419957-93420473 | -0.790764159 | 0.030649859 | 0.146769171 |
| chr18:32470533-32471638 | 0.695816948 | 0.030667481 | 0.146842116 |
| chr20:54292298-54292513 | 0.771060849 | 0.030668925 | 0.146842116 |
| chr13:22965109-22965658 | -0.840169153 | 0.030678587 | 0.1468792 |
| chr14:68519212-68519426 | -1.084780824 | 0.030694334 | 0.14694541 |
| chr6:83815660-83815972 | -1.055249256 | 0.030697767 | 0.146952661 |
| chr3:99925590-99925897 | 0.938269366 | 0.03070039 | 0.146956037 |
| chr4:48954777-48955113 | 0.854083273 | 0.030711492 | 0.146989245 |
| chr1:231584171-231584490 | 0.947490581 | 0.030711833 | 0.146989245 |
| chr19:42108262-42108462 | 1.187849699 | 0.030713083 | 0.146989245 |
| chr7:99498562-99498808 | -0.981747578 | 0.030719061 | 0.147008675 |
| chr6:123529881-123530159 | -1.217502315 | 0.030724042 | 0.147023331 |
| chr1:199272704-199272988 | -0.973612615 | 0.030731667 | 0.147049775 |
| chr1:220369999-220370889 | -0.75684655 | 0.030733407 | 0.147049775 |
| chr7:6928155-6928573 | -0.939259177 | 0.030738479 | 0.147064861 |
| chrX:124911453-124911861 | 0.876084573 | 0.030757285 | 0.14713916 |
| chr7:5088474-5089323 | -0.728755625 | 0.030757849 | 0.14713916 |
| chr1:202268021-202268429 | -0.795065729 | 0.030761013 | 0.147145109 |
| chr5:99192130-99192367 | -1.084108461 | 0.030763012 | 0.147145483 |
| chr5:65483797-65483997 | -1.060930842 | 0.030766562 | 0.147153276 |
| chr17:58988350-58988665 | 0.749242004 | 0.030772147 | 0.147170805 |
| chr20:47878387-47878590 | 0.78005693 | 0.030781584 | 0.147206748 |
| chr2:46164941-46165223 | 0.907086031 | 0.030796372 | 0.147264361 |
| chr8:128658165-128658380 | 0.990422453 | 0.030797524 | 0.147264361 |
| chr15:90219680-90219978 | 0.897567685 | 0.030799397 | 0.147264361 |
| chr17:56880060-56880333 | 0.930408597 | 0.030805311 | 0.147283449 |
| chr16:84186375-84186587 | 0.923381003 | 0.030808815 | 0.147287092 |
| chr1:51297560-51298070 | -0.802755602 | 0.030809918 | 0.147287092 |
| chr1:173691021-173691459 | -0.904362618 | 0.030823465 | 0.147330722 |
| chr19:42602023-42602381 | -1.020516082 | 0.030823647 | 0.147330722 |
| chr5:56269607-56269866 | 1.016679128 | 0.030824813 | 0.147330722 |
| chr14:50677220-50677432 | 1.146071905 | 0.030834793 | 0.147369231 |
| chr20:47621294-47621494 | 0.775753468 | 0.030850871 | 0.147436876 |
| chr1:102826062-102826645 | -0.885384983 | 0.030856033 | 0.147452348 |
| chr4:142901111-142901362 | -1.146581837 | 0.030863363 | 0.147478181 |
| chr15:68228727-68228977 | 0.891704456 | 0.030869682 | 0.147499177 |
| chr3:128435081-128435569 | 0.776509493 | 0.030873464 | 0.147508047 |
| chr9:21253060-21253551 | -0.957605797 | 0.030876604 | 0.147508691 |
| chr4:1899269-1899480 | 0.987888273 | 0.030877449 | 0.147508691 |
| chr1:35748269-35748758 | 0.84257028 | 0.030880918 | 0.147516067 |
| chr1:180213457-180213757 | -0.89826099 | 0.030891833 | 0.147559006 |
| chrX:100447573-100448118 | -0.788873094 | 0.030920452 | 0.1476865 |
| chr5:79513417-79515017 | -0.684622735 | 0.030922439 | 0.147686786 |
| chr1:237901349-237901683 | -0.893847603 | 0.030939383 | 0.147737546 |
| chr20:64037711-64038821 | 0.705899279 | 0.030939861 | 0.147737546 |
| chr5:132808969-132809198 | 1.091069646 | 0.03094033 | 0.147737546 |
| chr10:104473371-104473705 | -0.993595031 | 0.030942511 | 0.147737546 |
| chr12:112163343-112163560 | 1.098604183 | 0.030942707 | 0.147737546 |

| | | | |
|---|---|---|---|
| chr11:3110099-3110466 | 0.831755627 | 0.030947163 | 0.147746821 |
| chr2:98731152-98731640 | 0.823936266 | 0.030948507 | 0.147746821 |
| chr21:32096508-32096747 | -1.082452973 | 0.030950998 | 0.147749509 |
| chr19:6279989-6280319 | 0.901586299 | 0.030959365 | 0.147772276 |
| chr3:193910894-193911560 | -0.766777462 | 0.030959624 | 0.147772276 |
| chr14:28751457-28751817 | 0.807898857 | 0.030962578 | 0.147777167 |
| chr9:26955249-26955595 | -1.062990079 | 0.030984312 | 0.147871687 |
| chr1:212455911-212456533 | -0.78421219 | 0.030989056 | 0.147885117 |
| chr8:108811638-108811859 | 1.1138033 | 0.03099628 | 0.14791038 |
| chrX:119753546-119754007 | 0.847633353 | 0.031001469 | 0.147925267 |
| chr14:104825878-104826903 | 0.702441089 | 0.031003261 | 0.147925267 |
| chr16:24409101-24409341 | 1.054093135 | 0.03100741 | 0.147935849 |
| chr9:78115196-78115428 | -1.152254697 | 0.031012756 | 0.147952146 |
| chr17:14805437-14805637 | -1.228071165 | 0.031021612 | 0.147983628 |
| chr20:56003066-56003956 | -0.737156744 | 0.031023218 | 0.147983628 |
| chr3:8681227-8681535 | 1.087930581 | 0.031036545 | 0.14803798 |
| chr12:107990321-107990640 | -1.005296409 | 0.031047824 | 0.148082561 |
| chr3:171113700-171113970 | -1.05769318 | 0.031056468 | 0.14810143 |
| chr12:53232163-53233152 | 0.698095009 | 0.031057865 | 0.14810143 |
| chr16:30055035-30055256 | 0.976045313 | 0.031058917 | 0.14810143 |
| chr1:115993443-115993855 | -0.938550888 | 0.031059511 | 0.14810143 |
| chr15:70588107-70588390 | 1.00820613 | 0.031069588 | 0.148140259 |
| chr1:215095565-215095940 | -0.854897774 | 0.03107285 | 0.148146593 |
| chr6:155268133-155268418 | -1.068237892 | 0.031077555 | 0.148159807 |
| chr4:473631-474484 | 0.710376453 | 0.03107976 | 0.1481611 |
| chr5:71887794-71888029 | -1.108086201 | 0.031086212 | 0.148182639 |
| chr1:182806260-182806648 | 0.786397716 | 0.031105222 | 0.148264034 |
| chr6:161370876-161371207 | -0.943208584 | 0.031111416 | 0.148280323 |
| chr2:1680328-1680575 | 1.020737577 | 0.03111251 | 0.148280323 |
| chr1:114324024-114324488 | 0.743124967 | 0.031118394 | 0.14829914 |
| chr22:45919010-45919848 | 0.748841803 | 0.031120346 | 0.148299219 |
| chrX:150982668-150983447 | 0.722296707 | 0.03114887 | 0.148425917 |
| chr10:546905-547591 | 0.727479743 | 0.031156857 | 0.148454743 |
| chr11:73178089-73178435 | 0.802757382 | 0.031159231 | 0.148456821 |
| chr11:67031718-67032007 | -0.813005401 | 0.031161831 | 0.148456926 |
| chr9:120344244-120344559 | -0.904758064 | 0.031164724 | 0.148456926 |
| chr20:6691951-6692265 | -1.103940408 | 0.031165237 | 0.148456926 |
| chr11:72025149-72025591 | 0.777291337 | 0.031167003 | 0.148456926 |
| chr2:199485039-199485239 | -1.167756971 | 0.031174648 | 0.148484109 |
| chr1:168111168-168111515 | -0.978140841 | 0.031185275 | 0.148525494 |
| chr17:60186881-60187277 | 0.744474242 | 0.031203431 | 0.148602008 |
| chr1:36204201-36204401 | -1.104020488 | 0.031205219 | 0.148602008 |
| chr17:65733246-65733446 | 1.138898244 | 0.031217682 | 0.148652119 |
| chr10:31774369-31774625 | 1.055444968 | 0.03122651 | 0.148684913 |
| chr17:43833116-43833316 | 1.019945211 | 0.031232348 | 0.148703473 |
| chr16:30628798-30628998 | 1.103905131 | 0.031234496 | 0.148704458 |
| chr15:94168312-94168535 | -1.055565991 | 0.031249433 | 0.148764975 |
| chr19:30425469-30425740 | 1.179800883 | 0.031251219 | 0.148764975 |
| chr17:26594751-26595037 | 0.845713809 | 0.031253032 | 0.148764975 |
| chrX:135031814-135032694 | 0.710358881 | 0.031260847 | 0.148792931 |

| | | | |
|---|---|---|---|
| chr2:135997358-135997702 | -0.842547639 | 0.031266581 | 0.148810982 |
| chr7:45182459-45182659 | -1.106500869 | 0.031274696 | 0.148840357 |
| chr18:41954942-41955579 | -0.725895654 | 0.031288151 | 0.148890134 |
| chr5:119452212-119453075 | -0.694308377 | 0.031290116 | 0.148890134 |
| chr15:97881777-97882094 | 1.064583407 | 0.03129251 | 0.148890134 |
| chr18:62239590-62239793 | -1.166958223 | 0.031292928 | 0.148890134 |
| chr22:45934560-45935270 | -0.767811372 | 0.031307651 | 0.148942129 |
| chr1:85641697-85642161 | -0.893225324 | 0.031307744 | 0.148942129 |
| chr19:54114518-54114722 | 0.9796649 | 0.031311356 | 0.148948942 |
| chr6:117467404-117467604 | 1.171879726 | 0.031313064 | 0.148948942 |
| chr19:45092656-45092971 | 0.811297216 | 0.031316813 | 0.148957529 |
| chr15:74438737-74438999 | 0.861659765 | 0.031324315 | 0.148983962 |
| chr17:61745451-61745709 | 0.745923079 | 0.031327687 | 0.148990752 |
| chr17:60017901-60018159 | 0.819687249 | 0.031337392 | 0.14902352 |
| chr7:123824684-123825050 | -0.804490529 | 0.031338466 | 0.14902352 |
| chr18:11733009-11733325 | 0.902814058 | 0.03135275 | 0.149082189 |
| chr17:50112930-50113172 | -0.902769183 | 0.031359536 | 0.149101168 |
| chr18:80229748-80230102 | -0.922495678 | 0.031362472 | 0.149101168 |
| chr17:61763169-61763661 | 0.704032571 | 0.031362579 | 0.149101168 |
| chr2:216006241-216006722 | -0.777144774 | 0.031366263 | 0.149109432 |
| chr17:62520974-62521205 | 0.993206154 | 0.031376261 | 0.149147705 |
| chr6:73382493-73382722 | -1.113241224 | 0.031382207 | 0.149166716 |
| chr12:63901320-63901770 | 0.762697518 | 0.031388207 | 0.149175066 |
| chr4:150952137-150952355 | -1.327924101 | 0.031389119 | 0.149175066 |
| chr4:68368846-68369113 | 0.989257221 | 0.031389804 | 0.149175066 |
| chr12:73935831-73936167 | -0.890767462 | 0.031392632 | 0.149175803 |
| chr7:103811983-103812219 | -1.114610298 | 0.031397655 | 0.149175803 |
| chr15:20293772-20293975 | 1.166400745 | 0.03140043 | 0.149175803 |
| chr18:74282345-74282545 | 1.166400745 | 0.03140043 | 0.149175803 |
| chr5:56029451-56030161 | -0.8398628 | 0.031401534 | 0.149175803 |
| chr10:11204403-11204603 | -1.193964878 | 0.031402963 | 0.149175803 |
| chr1:23019941-23020187 | 1.035059008 | 0.031403588 | 0.149175803 |
| chr7:139524737-139524957 | -1.083525683 | 0.031412614 | 0.149200431 |
| chr6:132675052-132675358 | -0.940166438 | 0.031412667 | 0.149200431 |
| chr2:104254110-104254341 | -1.192437342 | 0.031417792 | 0.149215526 |
| chr14:58864914-58865124 | -1.166051637 | 0.031427657 | 0.14924914 |
| chr1:219391230-219391430 | 1.106948534 | 0.031428765 | 0.14924914 |
| chr3:113626794-113627197 | -0.804247126 | 0.03143525 | 0.149264211 |
| chr8:131993254-131993560 | -1.088078833 | 0.031435835 | 0.149264211 |
| chr7:129145167-129145367 | -1.174509147 | 0.031451028 | 0.149327096 |
| chr5:151984495-151984753 | 1.006121096 | 0.031475065 | 0.14943196 |
| chr16:72891649-72891910 | 1.016391535 | 0.031483047 | 0.149458382 |
| chr13:27383142-27383624 | -0.768861765 | 0.031486058 | 0.149458382 |
| chr3:13420160-13420812 | 0.732715549 | 0.031486482 | 0.149458382 |
| chr9:134992918-134993119 | -1.128739651 | 0.031499235 | 0.149509658 |
| chr8:38429543-38429883 | -0.978448203 | 0.031511213 | 0.149557247 |
| chr2:3533867-3534175 | -0.899658728 | 0.031520381 | 0.149591493 |
| chr19:57614046-57614368 | 0.849152759 | 0.031523093 | 0.149595098 |
| chr18:39799085-39799299 | -1.101190489 | 0.031533251 | 0.149634036 |
| chr10:17512466-17512784 | -1.020600706 | 0.031548618 | 0.149690722 |

| | | | |
|---|---|---|---|
| chr7:129952774-129953178 | -0.737011705 | 0.031549308 | 0.149690722 |
| chr6:79758244-79758542 | -1.07165434 | 0.031551058 | 0.149690722 |
| chr12:120725627-120726176 | 0.744491053 | 0.031555338 | 0.149701757 |
| chr1:209576552-209577074 | -0.766169686 | 0.031571889 | 0.149771007 |
| chr13:35321255-35321993 | -0.826473481 | 0.031575076 | 0.14977685 |
| chr12:78584730-78585072 | -1.010033301 | 0.031579801 | 0.14978999 |
| chr11:111983492-111983791 | 0.977611217 | 0.031605593 | 0.149903051 |
| chr5:146025056-146025334 | -1.092242019 | 0.031614397 | 0.149930127 |
| chr1:222618872-222619159 | -0.918393204 | 0.031615216 | 0.149930127 |
| chr1:227234988-227235229 | 0.97018273 | 0.03163512 | 0.150015236 |
| chr14:50067803-50068003 | 0.996397343 | 0.031638108 | 0.15002012 |
| chr2:189366417-189366705 | -1.073812832 | 0.031641295 | 0.150025094 |
| chr2:156804035-156804293 | -1.096347101 | 0.031643073 | 0.150025094 |
| chr20:54449971-54450221 | 0.753104684 | 0.031647234 | 0.150035536 |
| chr13:43963770-43964121 | -0.930365525 | 0.031658924 | 0.150081672 |
| chr2:167331422-167331893 | -0.842353075 | 0.031686029 | 0.150194109 |
| chr17:61377072-61377392 | 0.712031988 | 0.031686562 | 0.150194109 |
| chr19:17539691-17540105 | 0.793108901 | 0.031693779 | 0.150219022 |
| chr8:124924679-124925017 | 0.806423517 | 0.031696877 | 0.150224412 |
| chr1:167854202-167854467 | 0.919841629 | 0.031703142 | 0.150244813 |
| chr7:131105888-131106671 | -0.72741321 | 0.031706788 | 0.150252797 |
| chr5:150272582-150272791 | 1.00475117 | 0.031709812 | 0.150257837 |
| chr14:104865700-104866203 | 0.748099102 | 0.031718065 | 0.150287649 |
| chr4:111337416-111337657 | -1.185668895 | 0.031724572 | 0.150309186 |
| chr16:29700491-29700835 | 0.82521244 | 0.031727099 | 0.150311862 |
| chr7:149883405-149883906 | -0.798251988 | 0.031729759 | 0.150315173 |
| chr15:99704437-99704695 | 0.913560143 | 0.03173279 | 0.150320238 |
| chr2:121888277-121888604 | 0.870957936 | 0.03173662 | 0.150329086 |
| chr4:115664156-115664757 | 0.798376701 | 0.0317568 | 0.150415375 |
| chr4:12845090-12845540 | 0.883309211 | 0.031761112 | 0.150426502 |
| chr12:110613673-110614490 | -0.743073553 | 0.031765583 | 0.150438378 |
| chr3:175940719-175941734 | -0.729791069 | 0.031768012 | 0.150440104 |
| chr1:32611991-32612979 | 0.727535872 | 0.031769875 | 0.150440104 |
| chr8:102231424-102231624 | 1.018921495 | 0.031795819 | 0.150553656 |
| chr14:68137706-68138150 | -0.794437404 | 0.031798245 | 0.150555836 |
| chr8:107482446-107482708 | 1.067758558 | 0.031804847 | 0.15057779 |
| chr4:173196648-173197103 | -0.82266857 | 0.031820311 | 0.150634718 |
| chr10:89185688-89185991 | -0.94884898 | 0.031820803 | 0.150634718 |
| chr3:97139248-97139453 | -1.138150523 | 0.031824415 | 0.150642512 |
| chr3:168165684-168165976 | -0.879854362 | 0.031827618 | 0.150648367 |
| chrX:143115264-143116038 | 0.732746976 | 0.031837126 | 0.150684063 |
| chr7:143287964-143288817 | -0.690766476 | 0.031849136 | 0.150731595 |
| chr3:175188539-175188901 | 0.991822576 | 0.031854031 | 0.150745448 |
| chr6:52904567-52904896 | -0.919137454 | 0.031857904 | 0.150754468 |
| chr1:219543940-219544481 | -0.844841864 | 0.031864624 | 0.150776953 |
| chr10:71964096-71964536 | 0.848157116 | 0.031869457 | 0.15079051 |
| chr9:106864123-106864365 | -1.014169128 | 0.031877754 | 0.150820456 |
| chrX:30593885-30594133 | -0.987913135 | 0.031883208 | 0.150825608 |
| chr14:63388852-63389127 | -0.982884508 | 0.031883539 | 0.150825608 |
| chr16:32865432-32865832 | 0.85996426 | 0.031884748 | 0.150825608 |

| | | | |
|---|---|---|---|
| chr13:73133468-73133985 | -0.824979539 | 0.031889287 | 0.150837766 |
| chr12:93154794-93155101 | -1.044443492 | 0.031895881 | 0.150859642 |
| chr10:29791775-29792092 | 0.950778191 | 0.031900623 | 0.150872757 |
| chr17:82578010-82578252 | 1.209867606 | 0.031910897 | 0.150912035 |
| chrX:36241560-36241760 | -1.275051127 | 0.03191292 | 0.150912285 |
| chr15:28123424-28123715 | 0.970348888 | 0.031916563 | 0.150918215 |
| chr5:17218539-17218739 | 0.887462412 | 0.031918113 | 0.150918215 |
| chr6:95767888-95768353 | -0.915523251 | 0.031923732 | 0.150935469 |
| chr2:202149404-202149787 | -1.024600182 | 0.031928834 | 0.150950275 |
| chr2:235793370-235793570 | 1.120244865 | 0.03193701 | 0.150979615 |
| chr14:105701944-105702260 | 0.891138384 | 0.031940517 | 0.15098688 |
| chr19:35673203-35673705 | 0.797623258 | 0.031943003 | 0.150989315 |
| chr22:20320601-20320831 | 0.963998004 | 0.031949505 | 0.15101073 |
| chr11:35337426-35337728 | 1.050559701 | 0.031953738 | 0.151017493 |
| chr6:20839639-20839911 | -1.174981504 | 0.031954877 | 0.151017493 |
| chr2:85872794-85873126 | -0.886206838 | 0.031957001 | 0.151018214 |
| chr7:152076294-152076826 | -0.79517961 | 0.031960152 | 0.15102379 |
| chr8:100395761-100396523 | 0.729489063 | 0.031962167 | 0.151023998 |
| chr20:17605256-17605633 | 0.824868073 | 0.031964663 | 0.151026481 |
| chr11:72039794-72040051 | 0.889693998 | 0.031968704 | 0.151036258 |
| chrX:88425981-88426750 | -0.795090468 | 0.0319726 | 0.151038373 |
| chr3:71684795-71685413 | -0.858816831 | 0.031973094 | 0.151038373 |
| chr2:64138050-64138250 | 1.124282071 | 0.031977791 | 0.151044286 |
| chr17:48036962-48037767 | 0.720018449 | 0.031978288 | 0.151044286 |
| chr12:109853162-109853383 | -1.05926364 | 0.03198252 | 0.151054962 |
| chr5:113032692-113032939 | -1.017739513 | 0.032000359 | 0.151118915 |
| chr1:146970692-146970984 | 1.047129471 | 0.032001347 | 0.151118915 |
| chr4:168501215-168501797 | -0.783442529 | 0.032002965 | 0.151118915 |
| chr11:34825654-34825875 | 1.102890238 | 0.03200395 | 0.151118915 |
| chr14:52835098-52835370 | -1.062707267 | 0.032013852 | 0.151152732 |
| chr18:63074734-63075450 | -0.760930172 | 0.032015239 | 0.151152732 |
| chr1:8409182-8409496 | -1.132098458 | 0.03201703 | 0.151152732 |
| chr5:77751348-77752035 | -0.801685728 | 0.032041389 | 0.151258413 |
| chr10:99412082-99412287 | 1.107303877 | 0.032050431 | 0.151291777 |
| chr12:47882766-47883169 | -0.856378333 | 0.032060367 | 0.151329358 |
| chr1:207876219-207876674 | -0.861835645 | 0.032074987 | 0.151383911 |
| chr17:40513705-40513955 | 1.047417895 | 0.032077191 | 0.151383911 |
| chr20:19137203-19137529 | 1.035736958 | 0.032077852 | 0.151383911 |
| chr2:154697404-154698679 | -0.714810063 | 0.032080935 | 0.151389134 |
| chr8:96365442-96365678 | 0.915078624 | 0.03208882 | 0.151417021 |
| chr4:183721772-183722035 | -0.91064707 | 0.032091172 | 0.151418792 |
| chr5:733159-733515 | 0.890813276 | 0.032095219 | 0.15142053 |
| chr19:10617573-10617797 | 1.065975092 | 0.032095493 | 0.15142053 |
| chrX:107429163-107429462 | 0.899751564 | 0.032112536 | 0.151491611 |
| chr2:71277536-71277949 | -0.795847615 | 0.032116039 | 0.151498806 |
| chr7:94970348-94970761 | -1.135128269 | 0.032122119 | 0.15151816 |
| chr1:6235869-6236312 | 0.761390137 | 0.032124896 | 0.151521932 |
| chr8:131903527-131903739 | -0.988484885 | 0.03212944 | 0.151531364 |
| chr3:126321798-126322173 | 0.949170717 | 0.03213328 | 0.151531364 |
| chr1:154567834-154568068 | 0.93946519 | 0.032134242 | 0.151531364 |

| | | | |
|---|---|---|---|
| chr19:44807988-44808495 | 0.772382654 | 0.032134806 | 0.151531364 |
| chr3:63619189-63619471 | 0.816461487 | 0.032140193 | 0.151547437 |
| chr4:108502455-108502791 | -1.000709003 | 0.032155717 | 0.151611305 |
| chr14:33710294-33710703 | -0.894996065 | 0.032160913 | 0.151626476 |
| chr11:43162071-43162280 | 1.049687307 | 0.032180228 | 0.151690318 |
| chr1:108034213-108034413 | 1.049687307 | 0.032180228 | 0.151690318 |
| chr3:146035410-146035650 | -1.092978021 | 0.032180394 | 0.151690318 |
| chr3:135705458-135705724 | 1.005739051 | 0.03218492 | 0.151702322 |
| chr5:174699033-174699269 | 0.996133236 | 0.032189809 | 0.151716033 |
| chr8:101437011-101437844 | 0.693175887 | 0.032193283 | 0.151723072 |
| chr13:108269926-108270249 | -0.956976155 | 0.032196892 | 0.151730752 |
| chr6:73129751-73130479 | -0.735458458 | 0.032198918 | 0.151730965 |
| chr22:28669378-28669755 | 0.895431635 | 0.032217259 | 0.151797018 |
| chr12:79300000-79300663 | 0.746920859 | 0.032218002 | 0.151797018 |
| chr3:32003075-32003288 | 1.194395736 | 0.032219407 | 0.151797018 |
| chr5:165805675-165806132 | -0.852157632 | 0.03222086 | 0.151797018 |
| chr4:112664739-112664961 | 1.059385187 | 0.032225469 | 0.1518094 |
| chr14:94981324-94981534 | 1.131807711 | 0.032234758 | 0.151836145 |
| chr6:55875268-55875655 | -1.081770213 | 0.03223511 | 0.151836145 |
| chr16:17476734-17477121 | 0.859193104 | 0.032239714 | 0.151848497 |
| chr1:216193859-216194419 | -0.735900149 | 0.032244081 | 0.151857552 |
| chr20:10671522-10671824 | -0.813502273 | 0.032245687 | 0.151857552 |
| chr13:35630512-35631056 | -0.774089991 | 0.032247582 | 0.151857552 |
| chr5:108084156-108084445 | -0.988033173 | 0.032251975 | 0.151867201 |
| chr4:188160239-188160477 | -1.120507473 | 0.032253595 | 0.151867201 |
| chr3:23162935-23163271 | 0.882281831 | 0.032256402 | 0.151869188 |
| chr3:158771603-158772261 | -0.773592855 | 0.032257981 | 0.151869188 |
| chr3:32110350-32110669 | 0.796596393 | 0.032266987 | 0.151902253 |
| chr3:169505837-169506050 | -1.091860538 | 0.032272039 | 0.151914378 |
| chr8:85842517-85842925 | 0.802209569 | 0.032273528 | 0.151914378 |
| chr9:32218093-32218394 | -1.100408925 | 0.032283248 | 0.151950799 |
| chr18:69838671-69839544 | -0.681774164 | 0.032286369 | 0.151956152 |
| chr10:14759323-14759542 | -1.096321177 | 0.032292475 | 0.151972572 |
| chr11:17353376-17353860 | 0.776650795 | 0.032293825 | 0.151972572 |
| chr5:154868442-154869028 | -0.787349225 | 0.032297741 | 0.15198167 |
| chr10:102818937-102819489 | 0.864852657 | 0.032299979 | 0.151982866 |
| chr2:149313492-149313976 | -0.776854954 | 0.032308182 | 0.152012131 |
| chr4:176883254-176883454 | -1.253797123 | 0.032318514 | 0.152045769 |
| chr3:64678600-64678921 | 0.739831674 | 0.0323193 | 0.152045769 |
| chr1:218480168-218481589 | -0.685305283 | 0.032323576 | 0.152056549 |
| chr10:33376136-33376402 | -1.13896772 | 0.032332619 | 0.152080228 |
| chr7:119269940-119270520 | -0.765971964 | 0.03233344 | 0.152080228 |
| chr5:24547160-24547903 | -0.803902419 | 0.032334564 | 0.152080228 |
| chr20:48400759-48400959 | 0.776472869 | 0.032337433 | 0.152084384 |
| chr13:30892825-30893341 | -0.804058024 | 0.032341365 | 0.152093543 |
| chr14:92641610-92641846 | 0.97333993 | 0.032354547 | 0.152146194 |
| chr2:47724356-47724691 | -1.048137265 | 0.032356888 | 0.152147867 |
| chr3:24829054-24829948 | -0.766645483 | 0.03236856 | 0.152185134 |
| chr3:107824871-107825149 | 0.920064361 | 0.032368786 | 0.152185134 |
| chrX:119235941-119236916 | 0.715237298 | 0.032370782 | 0.152185179 |

| | | | |
|---|---|---|---|
| chr1:1658928-1659353 | 0.80906355 | 0.032373317 | 0.152187761 |
| chr1:192732879-192733088 | -1.134038821 | 0.032380726 | 0.152213252 |
| chr9:89437228-89437580 | -0.833259755 | 0.032388104 | 0.152238594 |
| chr16:53429178-53429607 | 0.855395121 | 0.03240239 | 0.152296402 |
| chr1:234573653-234573853 | -1.214224964 | 0.032409058 | 0.152318397 |
| chr9:121598049-121598799 | 0.726283467 | 0.032434539 | 0.152428804 |
| chr7:16432162-16432462 | 0.835147359 | 0.03244041 | 0.152447049 |
| chr16:3443296-3444398 | 0.713149196 | 0.03244332 | 0.152451372 |
| chr4:175469442-175469694 | -1.138208337 | 0.032447126 | 0.152459907 |
| chr15:90529180-90531039 | 0.690113907 | 0.032451897 | 0.152472974 |
| chr3:176734120-176734355 | -1.133398685 | 0.032458692 | 0.152486735 |
| chr15:100693488-100693888 | 0.789744901 | 0.032459602 | 0.152486735 |
| chr4:95191067-95191809 | -0.790351256 | 0.032460796 | 0.152486735 |
| chr1:106853066-106854118 | 0.688748186 | 0.032465393 | 0.152498977 |
| chr19:7871387-7871833 | 0.773533938 | 0.032470041 | 0.152511462 |
| chr20:59933485-59934474 | 0.685333291 | 0.032472955 | 0.1525158 |
| chrX:119410281-119410863 | 0.80165724 | 0.032484232 | 0.152559411 |
| chr10:5654550-5654780 | 1.133077713 | 0.032487186 | 0.152563935 |
| chr8:110329874-110330329 | 0.794270022 | 0.032490332 | 0.152569358 |
| chr7:139340052-139340866 | -0.68404389 | 0.032493318 | 0.152574032 |
| chr11:78474496-78474806 | 0.919256653 | 0.032496041 | 0.152577465 |
| chr17:80219811-80220527 | 0.735819996 | 0.032504425 | 0.152607479 |
| chr15:64461723-64461972 | 0.945151285 | 0.032515946 | 0.152652218 |
| chr2:156649507-156649838 | -1.026480165 | 0.032521223 | 0.152667638 |
| chr14:85876680-85876936 | -1.209440438 | 0.032531848 | 0.152701784 |
| chr20:10611380-10611690 | -0.895342634 | 0.032532483 | 0.152701784 |
| chr17:61714159-61714583 | 0.703955751 | 0.032539922 | 0.152721122 |
| chr5:69330244-69330444 | 1.002395246 | 0.032540589 | 0.152721122 |
| chr2:88987024-88987293 | -1.09469738 | 0.032543555 | 0.152725686 |
| chr7:111785013-111785222 | 1.06362313 | 0.032553158 | 0.152761395 |
| chr9:36913980-36914180 | -1.269519069 | 0.032572262 | 0.152841685 |
| chr2:240703199-240703483 | 1.005321637 | 0.032576886 | 0.15285402 |
| chr4:9691359-9692019 | 0.727616801 | 0.032581105 | 0.152864456 |
| chr19:773137-773398 | 1.116653251 | 0.032586489 | 0.152880354 |
| chr3:62461164-62461469 | 0.786643847 | 0.032589254 | 0.152883964 |
| chr10:123583742-123584378 | 0.786854664 | 0.0325917 | 0.152886079 |
| chrX:36908854-36909085 | -1.165525481 | 0.032608567 | 0.15295584 |
| chr4:166840163-166840376 | -1.087909499 | 0.032612381 | 0.152964365 |
| chr14:48805139-48805712 | 0.800849068 | 0.032618831 | 0.152985252 |
| chr9:127162070-127162316 | 0.951028849 | 0.032621683 | 0.152989263 |
| chr13:69935094-69935328 | -1.034475906 | 0.032633061 | 0.153027401 |
| chr5:57879085-57879330 | -1.071163211 | 0.032633809 | 0.153027401 |
| chr11:32090685-32091859 | -0.70919363 | 0.03263757 | 0.153035671 |
| chr17:10053626-10053954 | -1.110302188 | 0.032640538 | 0.153040221 |
| chr19:17511308-17511763 | 0.755083362 | 0.032649824 | 0.153074395 |
| chr1:38065187-38065524 | 0.963991526 | 0.032653239 | 0.153081039 |
| chr12:20646350-20646597 | -1.058878317 | 0.032672579 | 0.153162332 |
| chr10:9209085-9209519 | 0.77381965 | 0.032683696 | 0.153205073 |
| chr17:61602691-61603421 | 0.683198033 | 0.032688726 | 0.15321928 |
| chr9:124010986-124011238 | 0.904231747 | 0.032695217 | 0.153240332 |

| | | | |
|---|---|---|---|
| chr6:63307860-63308120 | -0.957619909 | 0.032731773 | 0.153402281 |
| chr7:28020383-28020606 | 1.049557426 | 0.032737765 | 0.153417017 |
| chr17:61763906-61764540 | 0.686444791 | 0.032738922 | 0.153417017 |
| chr17:61593437-61593813 | 0.703807505 | 0.032746848 | 0.153439209 |
| chr2:199979911-199980325 | -0.931758099 | 0.032747662 | 0.153439209 |
| chr1:196910237-196910553 | -0.912590352 | 0.032749851 | 0.15344008 |
| chr1:168386572-168386772 | 1.052773989 | 0.032752633 | 0.153443734 |
| chr12:52965110-52965590 | -0.815833756 | 0.032761919 | 0.153477855 |
| chr2:97748502-97748824 | -0.908919108 | 0.032767347 | 0.153490577 |
| chrX:112642410-112642624 | 1.110454257 | 0.032768642 | 0.153490577 |
| chr17:78497215-78497415 | 0.952303107 | 0.032776352 | 0.153517306 |
| chr1:215517421-215517940 | -0.739692139 | 0.03278044 | 0.15352707 |
| chr17:32486285-32486657 | 0.846402231 | 0.032783459 | 0.153531826 |
| chr1:112880952-112881439 | -0.835412495 | 0.032786132 | 0.153534961 |
| chr8:93075425-93075732 | -1.032526703 | 0.032836589 | 0.153752612 |
| chr17:72634612-72635196 | -0.773897992 | 0.032836623 | 0.153752612 |
| chr11:3641966-3642392 | 0.801398543 | 0.032858179 | 0.153844142 |
| chr7:146772663-146772904 | -0.984435158 | 0.032860603 | 0.15384609 |
| chr18:58097326-58097716 | -0.939133451 | 0.032868878 | 0.153875428 |
| chr3:149487640-149487937 | -0.883066603 | 0.032873151 | 0.153885984 |
| chr1:43494060-43494325 | 1.057627255 | 0.032875149 | 0.153885984 |
| chr1:152952381-152952769 | 0.977559996 | 0.032878362 | 0.153891623 |
| chr10:79293049-79293281 | -0.977383255 | 0.032884382 | 0.153910398 |
| chr7:106626850-106627337 | -0.956915753 | 0.032893991 | 0.15394414 |
| chr21:41859394-41859611 | 1.019992607 | 0.03289561 | 0.15394414 |
| chr20:48082322-48082644 | 0.744808374 | 0.032900158 | 0.153956022 |
| chr3:8710191-8710588 | 0.829316255 | 0.032909027 | 0.153984662 |
| chrY:56846022-56846412 | 0.888821149 | 0.032913042 | 0.153984662 |
| chr15:83955878-83956227 | 1.101270066 | 0.03291346 | 0.153984662 |
| chr20:59561123-59561397 | -0.90944796 | 0.032915448 | 0.153984662 |
| chr3:75314925-75315213 | 1.024250706 | 0.032916327 | 0.153984662 |
| chr2:227774352-227774610 | -1.129563151 | 0.032918849 | 0.153987062 |
| chr12:113297270-113297704 | 0.886453194 | 0.032924474 | 0.153998749 |
| chr11:71112627-71112832 | -1.089962914 | 0.032925368 | 0.153998749 |
| chr10:37789649-37789886 | 1.113268161 | 0.032931552 | 0.154018273 |
| chr1:53101633-53101955 | 0.937183647 | 0.032939411 | 0.154045627 |
| chr15:34582754-34583331 | 0.775413996 | 0.032943312 | 0.154054467 |
| chrX:80875128-80875413 | -0.992522078 | 0.032947499 | 0.154057983 |
| chr1:11484046-11484251 | -1.109933567 | 0.032949476 | 0.154057983 |
| chr13:109644011-109644454 | 0.828256367 | 0.03295196 | 0.154057983 |
| chr4:76580205-76580413 | -1.108832295 | 0.032952106 | 0.154057983 |
| chrX:11532333-11532591 | -0.903707781 | 0.032955002 | 0.154062119 |
| chr10:53998027-53998239 | -1.130276443 | 0.03296393 | 0.154085212 |
| chr21:36509979-36510256 | 0.939208527 | 0.032963963 | 0.154085212 |
| chr5:77491796-77492908 | -0.697781768 | 0.032967483 | 0.154092265 |
| chr11:83361923-83362287 | -0.944446253 | 0.032969575 | 0.154092641 |
| chr10:99484623-99485003 | 0.831032202 | 0.032977396 | 0.154119794 |
| chr6:55046700-55047147 | -0.921879151 | 0.032985952 | 0.154150377 |
| chr3:172142396-172142784 | -0.828608644 | 0.033006288 | 0.154236006 |
| chr7:6026269-6026775 | -0.772733668 | 0.033011127 | 0.15424921 |

| | | | |
|---|---|---|---|
| chr3:194158845-194159676 | -0.71488081 | 0.03302712 | 0.154309724 |
| chr8:101994239-101994726 | 0.867593454 | 0.033028105 | 0.154309724 |
| chr16:31451712-31452108 | 0.784766834 | 0.033030932 | 0.154313521 |
| chr1:177200390-177200613 | -1.093113158 | 0.033041514 | 0.154345074 |
| chr9:98502200-98502400 | 1.197488584 | 0.033041714 | 0.154345074 |
| chr21:36141440-36141882 | 0.773566536 | 0.033068873 | 0.154455931 |
| chr4:3304824-3305024 | -1.28079121 | 0.033069478 | 0.154455931 |
| chr20:54930999-54931727 | 0.695269639 | 0.033085984 | 0.154502713 |
| chr14:77817365-77817763 | 0.838230344 | 0.033087707 | 0.154502713 |
| chr7:51161108-51161329 | -0.991764816 | 0.033088174 | 0.154502713 |
| chr13:29909734-29910647 | -0.716910522 | 0.033090171 | 0.154502713 |
| chr20:53851907-53852410 | 0.685281159 | 0.033091559 | 0.154502713 |
| chr12:14629953-14630471 | -0.82611335 | 0.033091593 | 0.154502713 |
| chr19:52335856-52336358 | 0.838194595 | 0.033096573 | 0.154508194 |
| chr10:48506680-48506928 | -1.029231163 | 0.0330968 | 0.154508194 |
| chr6:152232857-152233373 | -0.890364039 | 0.033101922 | 0.154522691 |
| chr13:31043900-31044140 | -0.943292402 | 0.03312292 | 0.154611292 |
| chr13:45524126-45524518 | 0.904609977 | 0.033129853 | 0.154634235 |
| chr20:48282833-48283123 | 0.74825981 | 0.033134441 | 0.154646228 |
| chr6:52352462-52352792 | 0.855841469 | 0.03315699 | 0.154742047 |
| chr2:172248964-172249258 | -0.997137549 | 0.033159811 | 0.154745782 |
| chr20:38434981-38435622 | 0.748343939 | 0.033167838 | 0.154769331 |
| chr7:21945398-21946540 | 0.676352486 | 0.033168896 | 0.154769331 |
| chr3:75580130-75580568 | 0.77851936 | 0.033187463 | 0.154846537 |
| chr7:133049838-133050169 | -1.112997576 | 0.033189519 | 0.154846699 |
| chr17:2084557-2084757 | -1.155350227 | 0.033207823 | 0.154918275 |
| chr7:50197625-50197825 | -1.227807443 | 0.033208904 | 0.154918275 |
| chr6:88629846-88630138 | -1.036375111 | 0.033219765 | 0.154959507 |
| chr3:5004787-5004987 | 1.024442853 | 0.033224051 | 0.154970061 |
| chr1:229254913-229255192 | 1.035741747 | 0.033226606 | 0.154972547 |
| chr3:24149133-24149405 | -1.066245661 | 0.033236849 | 0.155010887 |
| chr1:57835729-57836038 | 0.978548832 | 0.033251986 | 0.155072043 |
| chr5:11334479-11334704 | -1.02205267 | 0.033266404 | 0.15512984 |
| chr12:42652127-42652452 | -0.955883696 | 0.033277708 | 0.155173112 |
| chr11:61580985-61581742 | 0.719681643 | 0.033291609 | 0.155226576 |
| chr7:17408669-17408869 | 1.083956397 | 0.033293226 | 0.155226576 |
| chr16:29790143-29791060 | 0.688542606 | 0.033297149 | 0.155229286 |
| chr8:138385149-138385520 | -0.971813693 | 0.033297859 | 0.155229286 |
| chrX:139126561-139126860 | 0.887030847 | 0.033304702 | 0.155245455 |
| chr20:33685705-33686015 | 0.821003058 | 0.03330538 | 0.155245455 |
| chr17:36608580-36609487 | -0.763790464 | 0.033311169 | 0.15526148 |
| chr2:229930787-229931030 | 0.997968849 | 0.03331287 | 0.15526148 |
| chr13:33643413-33643919 | -0.7757009 | 0.033329148 | 0.155327899 |
| chr19:39032581-39032855 | 0.86500617 | 0.033336046 | 0.155350598 |
| chr6:94785076-94785338 | -1.106461884 | 0.033344461 | 0.155380363 |
| chr1:154221344-154221601 | 0.817368288 | 0.033348971 | 0.155386169 |
| chr6:148856185-148856663 | 0.860099283 | 0.033349763 | 0.155386169 |
| chr17:61250660-61251161 | 0.693664631 | 0.033354526 | 0.155394795 |
| chr2:46326323-46326608 | -1.003574498 | 0.033355671 | 0.155394795 |
| chr1:210214938-210215244 | -1.105786948 | 0.033363395 | 0.155413514 |

| | | | |
|---|---|---|---|
| chr10:103542867-103543355 | -0.828230818 | 0.033363745 | 0.155413514 |
| chr3:112932365-112932747 | 0.905415233 | 0.033373078 | 0.155447536 |
| chr14:70995686-70995974 | -0.957346178 | 0.033380938 | 0.155463575 |
| chr4:174714454-174714732 | -0.907121399 | 0.033381544 | 0.155463575 |
| chr17:72764137-72764337 | 1.199600529 | 0.033382608 | 0.155463575 |
| chr20:35632715-35633023 | 1.094220232 | 0.033386865 | 0.155473951 |
| chr4:119683661-119683997 | 0.871619561 | 0.033395653 | 0.155505424 |
| chr7:80771676-80771911 | -1.193972504 | 0.033403872 | 0.155534241 |
| chr21:31457798-31458085 | 0.942728518 | 0.033407864 | 0.155543378 |
| chr6:37742042-37742286 | 0.889305873 | 0.033414177 | 0.155554093 |
| chr3:77296633-77296881 | -1.088845626 | 0.033415996 | 0.155554093 |
| chr1:154519695-154520006 | -0.948529169 | 0.033416256 | 0.155554093 |
| chr17:61655284-61655567 | 0.725278891 | 0.033423655 | 0.155579083 |
| chr17:50918087-50918337 | 0.849493007 | 0.033430001 | 0.155598013 |
| chr6:139881723-139882120 | -0.894013468 | 0.033431783 | 0.155598013 |
| chr3:62134064-62134572 | -0.728999638 | 0.03343575 | 0.155607023 |
| chr12:117062677-117062964 | 0.887331225 | 0.033447111 | 0.155650444 |
| chr3:45183222-45183485 | -0.991823965 | 0.033470609 | 0.155742217 |
| chr3:64817447-64818188 | 0.689909872 | 0.033472626 | 0.155742217 |
| chr3:162696682-162697014 | 0.856097715 | 0.03347293 | 0.155742217 |
| chr1:191163594-191164057 | -0.836569454 | 0.033475287 | 0.155743727 |
| chr7:18649420-18649812 | 0.876172291 | 0.033477467 | 0.155744413 |
| chr14:32938435-32938635 | 0.909221335 | 0.033481371 | 0.15575312 |
| chr6:83708669-83709189 | 0.857149299 | 0.033501731 | 0.155838369 |
| chr10:88810284-88810822 | -0.82117365 | 0.033505949 | 0.155848529 |
| chr10:4270084-4270403 | 0.902612586 | 0.033514419 | 0.155857039 |
| chr13:28000581-28001109 | -0.814678274 | 0.033515577 | 0.155857039 |
| chr2:238107309-238107833 | -0.832723767 | 0.03351647 | 0.155857039 |
| chr1:191615503-191615769 | 0.952816722 | 0.033516795 | 0.155857039 |
| chr17:74125158-74125481 | 0.815985589 | 0.033518267 | 0.155857039 |
| chr11:116808907-116809189 | -1.035773798 | 0.033519983 | 0.155857039 |
| chr11:61588664-61588968 | -0.86154729 | 0.033525122 | 0.155871475 |
| chr20:48128728-48128928 | 0.788356218 | 0.033552374 | 0.155988712 |
| chrX:53832001-53832212 | 1.016218685 | 0.03355757 | 0.156003403 |
| chr15:99910017-99910217 | 1.033236038 | 0.033563792 | 0.156011083 |
| chr4:38591959-38592236 | -1.030796057 | 0.03356518 | 0.156011083 |
| chr1:161487402-161487770 | -0.907560728 | 0.033566697 | 0.156011083 |
| chr8:124989695-124989900 | 0.985399507 | 0.033567366 | 0.156011083 |
| chr6:88465866-88466102 | -1.230145439 | 0.033587595 | 0.156095635 |
| chr12:13223148-13223649 | -0.784382773 | 0.033591833 | 0.156105862 |
| chr2:130853554-130853770 | 1.041067283 | 0.033603706 | 0.156151566 |
| chr17:61376273-61376632 | 0.697754995 | 0.033608805 | 0.156165791 |
| chr8:80091610-80092045 | 0.777065453 | 0.033619659 | 0.156206748 |
| chr7:99208064-99208484 | -0.916178286 | 0.03362881 | 0.15623536 |
| chr2:67954120-67954362 | -1.08846051 | 0.033629895 | 0.15623536 |
| chr11:71438385-71438633 | -0.982272062 | 0.033635265 | 0.156250835 |
| chr14:26262443-26262920 | 0.870047368 | 0.033650534 | 0.156312289 |
| chr4:161147916-161148641 | -0.773505514 | 0.033658657 | 0.156340546 |
| chr15:54880836-54881052 | 1.131797037 | 0.033661118 | 0.156342496 |
| chr2:138827447-138827770 | -0.904518427 | 0.033665571 | 0.156345642 |

| | | | |
|---|---|---|---|
| chr4:90127074-90128829 | 0.669720301 | 0.033665876 | 0.156345642 |
| chr4:177250085-177250554 | -0.756381723 | 0.033678712 | 0.156394321 |
| chr11:60334001-60334341 | -0.901813494 | 0.03368044 | 0.156394321 |
| chr2:211766570-211767194 | -0.722154214 | 0.03370109 | 0.156480726 |
| chr7:134605943-134606266 | -0.985646098 | 0.03371007 | 0.156512937 |
| chr12:99983921-99985338 | -0.695132303 | 0.03371747 | 0.156532611 |
| chr6:92126079-92126491 | -0.86608749 | 0.033718393 | 0.156532611 |
| chr3:50503595-50503805 | 1.046901602 | 0.033730218 | 0.156578016 |
| chr4:161356525-161356728 | -1.274960508 | 0.03373805 | 0.156604888 |
| chr15:59161186-59161571 | -0.799907173 | 0.033742758 | 0.156617255 |
| chr9:4866240-4866643 | -0.840568488 | 0.033751319 | 0.1566475 |
| chr22:48586184-48586422 | 1.017708299 | 0.033771935 | 0.15673369 |
| chr7:20835163-20835369 | 1.020691603 | 0.033774897 | 0.156737943 |
| chr1:34788130-34788425 | 0.858469292 | 0.033778128 | 0.156743444 |
| chr17:82397851-82398398 | -0.803434051 | 0.033784664 | 0.156750015 |
| chr12:11648682-11649990 | -0.713139829 | 0.033786071 | 0.156750015 |
| chr8:71811263-71811463 | -1.191458147 | 0.033787208 | 0.156750015 |
| chr1:156025946-156026274 | 0.801395455 | 0.033787727 | 0.156750015 |
| chr4:184733764-184734576 | -0.693648738 | 0.033794994 | 0.156756368 |
| chr17:41864713-41865682 | 0.735675141 | 0.033795491 | 0.156756368 |
| chr8:6331352-6331587 | -1.042026084 | 0.033796735 | 0.156756368 |
| chr3:13346447-13346968 | -0.738996636 | 0.03379728 | 0.156756368 |
| chr6:25720479-25720804 | -0.952013416 | 0.033800696 | 0.156762722 |
| chr11:1072126-1072584 | -0.856897784 | 0.033805423 | 0.156775156 |
| chr4:136634133-136634440 | -0.991384683 | 0.033813334 | 0.156800136 |
| chr21:40049254-40049454 | 1.092681853 | 0.033814902 | 0.156800136 |
| chr21:23016371-23016922 | -0.838142839 | 0.033821729 | 0.156815524 |
| chr11:73838436-73838691 | -1.111832677 | 0.033822904 | 0.156815524 |
| chr1:114751702-114751933 | 0.766973072 | 0.033824361 | 0.156815524 |
| chr10:80966245-80966944 | -0.70885802 | 0.033830738 | 0.1568356 |
| chr9:128035202-128035641 | 0.785594954 | 0.033853065 | 0.156929612 |
| chrX:72045904-72046305 | 0.851652836 | 0.033855543 | 0.156931603 |
| chr2:20675776-20676124 | -0.767708433 | 0.033863072 | 0.156948684 |
| chr18:55737737-55738124 | -0.872743897 | 0.033863812 | 0.156948684 |
| chr12:55955862-55956472 | 0.737914055 | 0.033866964 | 0.156948684 |
| chr3:146160238-146161694 | -0.670146648 | 0.033868884 | 0.156948684 |
| chr20:49104205-49104558 | 0.940269955 | 0.033869469 | 0.156948684 |
| chrX:103489789-103490075 | -0.953304171 | 0.033878324 | 0.156980219 |
| chr12:25276269-25276518 | 1.062275251 | 0.033884369 | 0.156998739 |
| chr5:112491332-112491535 | -1.127686015 | 0.033893605 | 0.157032034 |
| chr7:117375513-117375810 | -1.074723832 | 0.033903002 | 0.15705989 |
| chr1:214329814-214330101 | 1.030412528 | 0.033903717 | 0.15705989 |
| chr13:60459476-60459905 | -0.937762977 | 0.033923137 | 0.157140354 |
| chr4:102715114-102715315 | -1.206917072 | 0.033927945 | 0.157153125 |
| chr1:209784219-209785132 | -0.688127425 | 0.033942849 | 0.157212655 |
| chr8:116730166-116730382 | 1.002829626 | 0.033960207 | 0.157271528 |
| chr16:9151119-9151426 | -0.938940452 | 0.033960766 | 0.157271528 |
| chr5:96876918-96877172 | -0.918468 | 0.033961717 | 0.157271528 |
| chr11:61815153-61815800 | 0.73799451 | 0.033987653 | 0.157382122 |
| chr8:120633773-120633991 | 1.042602377 | 0.034004029 | 0.157440762 |

| | | | |
|---|---|---|---|
| chr12:79505195-79505423 | 1.102831649 | 0.034004427 | 0.157440762 |
| chr2:191377280-191377590 | -0.943514463 | 0.03401054 | 0.157459553 |
| chr18:46067488-46067774 | -0.930918567 | 0.034016179 | 0.157476145 |
| chr5:1174341-1174782 | 0.773602874 | 0.034021633 | 0.15747931 |
| chr7:127792646-127793081 | -0.823958476 | 0.034022522 | 0.15747931 |
| chr3:8683013-8683240 | -1.126883645 | 0.034023029 | 0.15747931 |
| chr3:182426463-182426857 | 0.86292345 | 0.034025176 | 0.157479739 |
| chr19:40592259-40592532 | -0.950724851 | 0.034029224 | 0.157488958 |
| chr1:197944221-197944421 | -1.153215876 | 0.034043608 | 0.157546013 |
| chr17:63753239-63753463 | 1.01898169 | 0.034047727 | 0.15755556 |
| chr16:65889460-65889686 | 1.027167037 | 0.034050802 | 0.157560271 |
| chr8:142677329-142677545 | 0.909652295 | 0.034059695 | 0.157591907 |
| chr3:107825354-107826331 | -0.711995709 | 0.034070073 | 0.157630404 |
| chr8:140486805-140487108 | 0.993584454 | 0.034081539 | 0.157665541 |
| chr6:39923453-39923818 | 0.836125653 | 0.034081783 | 0.157665541 |
| chr14:88078824-88079124 | 1.008578871 | 0.034083863 | 0.157665647 |
| chr20:24992115-24992384 | 0.830378834 | 0.034095973 | 0.157712142 |
| chr1:159197972-159198282 | 0.854848423 | 0.034106772 | 0.15775257 |
| chr5:83085203-83085455 | 1.057176757 | 0.034115865 | 0.1577795 |
| chr8:121803791-121804023 | -0.997936809 | 0.034117714 | 0.1577795 |
| chr5:65482771-65483012 | -0.948763145 | 0.034118772 | 0.1577795 |
| chrX:126832885-126833120 | 1.051810554 | 0.0341232 | 0.157790456 |
| chr7:74365324-74365897 | -0.763812477 | 0.034129145 | 0.157801109 |
| chr18:58080726-58081025 | -1.063872173 | 0.034129623 | 0.157801109 |
| chr22:37525423-37525678 | -0.972390129 | 0.034139251 | 0.157836101 |
| chr16:79770109-79770908 | -0.719579173 | 0.034150407 | 0.15787815 |
| chr15:39188990-39189305 | -0.943414176 | 0.034154258 | 0.157881446 |
| chr20:54220710-54221071 | 0.717551923 | 0.034155747 | 0.157881446 |
| chr8:115099366-115099720 | -0.889152994 | 0.034159223 | 0.157881446 |
| chr10:77606813-77607163 | -0.806556277 | 0.034160752 | 0.157881446 |
| chr4:167190150-167190359 | -1.20536933 | 0.034162332 | 0.157881446 |
| chr5:35172365-35172574 | 1.050767596 | 0.034163483 | 0.157881446 |
| chr6:132893868-132894385 | 0.85878392 | 0.034167955 | 0.157892589 |
| chr7:111763543-111763823 | -0.946478741 | 0.034178344 | 0.157931075 |
| chr3:154746029-154746239 | -1.098399151 | 0.034188873 | 0.1579702 |
| chr4:177309440-177310464 | -0.679802648 | 0.034196647 | 0.157996592 |
| chr8:94522554-94522877 | 0.908379579 | 0.034215686 | 0.158075025 |
| chr9:135393190-135393627 | -0.826245372 | 0.034220901 | 0.158089585 |
| chr7:94764454-94764872 | -0.891509306 | 0.034224783 | 0.158097989 |
| chr22:44710398-44711165 | -0.793366655 | 0.034229357 | 0.158109586 |
| chr21:36155557-36157156 | 0.667987619 | 0.034233339 | 0.158118445 |
| chr15:89633388-89633604 | 1.059997737 | 0.034241208 | 0.158145258 |
| chr8:73651511-73651979 | 0.816859544 | 0.034245008 | 0.158153278 |
| chr8:20435336-20435853 | -0.826204937 | 0.034252893 | 0.158180158 |
| chr17:53516693-53516943 | -1.083864161 | 0.034257907 | 0.158193776 |
| chr12:69290783-69291320 | -0.767976301 | 0.034263755 | 0.158211247 |
| chr20:33002468-33002668 | 1.060167443 | 0.034280647 | 0.158279707 |
| chr14:91383420-91384011 | 0.728112641 | 0.03428505 | 0.158284109 |
| chr14:40628632-40629588 | -0.725265949 | 0.034285732 | 0.158284109 |
| chr12:64625308-64626006 | 0.735785923 | 0.034292618 | 0.15830636 |

| | | | |
|---|---|---|---|
| chr11:1762465-1762874 | 0.840161977 | 0.03429988 | 0.158330344 |
| chr12:73383662-73384351 | -0.701207189 | 0.034320355 | 0.158415315 |
| chr20:10783833-10784306 | -0.937279579 | 0.034323976 | 0.158422487 |
| chr2:14743522-14744173 | 0.776063976 | 0.034336297 | 0.158469806 |
| chr5:123090541-123091001 | 0.798038131 | 0.034340204 | 0.158478294 |
| chr2:241891340-241891540 | 0.992156623 | 0.034344347 | 0.158487869 |
| chr3:154301567-154301999 | -0.743648603 | 0.034350652 | 0.158498417 |
| chr4:128725542-128725792 | 0.950809884 | 0.034352331 | 0.158498417 |
| chr8:123783963-123784275 | 0.95150743 | 0.034352839 | 0.158498417 |
| chr5:36666506-36666745 | 1.180187519 | 0.034358434 | 0.158514687 |
| chr15:95895502-95896084 | -0.736728394 | 0.034378942 | 0.158590257 |
| chr2:65320385-65320599 | -1.045516764 | 0.03437986 | 0.158590257 |
| chr21:21458104-21458426 | -1.121410197 | 0.034382837 | 0.158590257 |
| chr5:27365214-27365607 | 1.005623406 | 0.034383093 | 0.158590257 |
| chrX:51148931-51149197 | -1.030942594 | 0.034388567 | 0.15860356 |
| chr8:42492375-42492846 | 0.8329349 | 0.034390719 | 0.15860356 |
| chr11:17351542-17352081 | 0.776345192 | 0.034392187 | 0.15860356 |
| chr4:45453291-45453566 | -0.9083309 | 0.034396608 | 0.158614403 |
| chrX:35142260-35142480 | -1.220294098 | 0.034402962 | 0.15862525 |
| chr20:55859114-55859330 | 1.080935841 | 0.034405334 | 0.15862525 |
| chr11:66070741-66071381 | 0.748682343 | 0.034407053 | 0.15862525 |
| chr7:128320954-128321410 | -0.86227666 | 0.034407241 | 0.15862525 |
| chr19:58182759-58183988 | 0.690979532 | 0.034421802 | 0.15868283 |
| chr4:4289574-4290572 | 0.714032372 | 0.034432441 | 0.158701466 |
| chr7:22110254-22110617 | 0.749588409 | 0.034432992 | 0.158701466 |
| chr2:21351575-21351977 | -0.901692884 | 0.034433245 | 0.158701466 |
| chr11:14844285-14844983 | -0.785166948 | 0.03443413 | 0.158701466 |
| chr7:129607889-129608089 | -1.014553722 | 0.034447861 | 0.158755201 |
| chr5:102865702-102866529 | -0.777173631 | 0.03445469 | 0.158771351 |
| chr10:68093995-68094254 | -1.004351945 | 0.034459322 | 0.158771351 |
| chr11:33817344-33817590 | 1.167972459 | 0.034460895 | 0.158771351 |
| chr14:74549501-74549807 | 0.843589887 | 0.034461029 | 0.158771351 |
| chr4:116676976-116677190 | -1.168815549 | 0.034461726 | 0.158771351 |
| chr8:74547913-74548160 | -1.035429959 | 0.034469246 | 0.158796449 |
| chr2:193738729-193739038 | -0.984408147 | 0.034478699 | 0.158830449 |
| chr17:61776711-61777088 | 0.706720341 | 0.034492287 | 0.158882687 |
| chr10:24488753-24489114 | -0.991796566 | 0.034494186 | 0.158882687 |
| chr5:142655756-142655992 | 0.945696356 | 0.034525779 | 0.15901865 |
| chr5:166145105-166145305 | -1.261390137 | 0.034534865 | 0.159050935 |
| chr12:74259994-74260902 | -0.719305745 | 0.034543609 | 0.159081643 |
| chr11:11842497-11842697 | 1.07618667 | 0.03454872 | 0.159095622 |
| chr18:63969948-63971221 | -0.679862539 | 0.034561025 | 0.159142721 |
| chr17:61325132-61325692 | 0.681177866 | 0.034564407 | 0.159148727 |
| chr14:105532865-105533065 | 0.912071135 | 0.034567422 | 0.159153047 |
| chr3:53163030-53163293 | 0.87677168 | 0.034571652 | 0.1591537 |
| chr13:41402702-41403187 | -0.828526562 | 0.034571718 | 0.1591537 |
| chr12:132236847-132237204 | 0.896083191 | 0.034581373 | 0.159188585 |
| chr5:77030138-77031129 | -0.699041646 | 0.034592482 | 0.159221887 |
| chr3:63048575-63049139 | 0.71696532 | 0.034594354 | 0.159221887 |
| chr14:54773150-54773761 | 0.694119223 | 0.034595943 | 0.159221887 |

| | | | |
|---|---|---|---|
| chr16:58025377-58025831 | 0.771350523 | 0.034598264 | 0.159221887 |
| chr5:137535497-137535846 | -0.912683016 | 0.034598998 | 0.159221887 |
| chr20:46021811-46022413 | 0.695374961 | 0.034602611 | 0.159228953 |
| chr17:38572937-38573204 | 1.011985203 | 0.034606875 | 0.159239008 |
| chr17:61735081-61735813 | 0.67623991 | 0.034616541 | 0.159273922 |
| chr19:19034362-19034626 | -0.926757553 | 0.034626608 | 0.159310672 |
| chr13:88069044-88069321 | -0.936934612 | 0.034641065 | 0.159361946 |
| chr14:54446504-54446733 | -0.96795522 | 0.034641912 | 0.159361946 |
| chr16:76000309-76000641 | -1.066539323 | 0.034644525 | 0.159364399 |
| chr21:41612667-41613262 | -0.751534476 | 0.034656097 | 0.159408059 |
| chr13:37155716-37156088 | 0.983374461 | 0.034676757 | 0.15948818 |
| chrX:123770480-123770773 | 0.886888276 | 0.034677678 | 0.15948818 |
| chr3:138013392-138013630 | -1.135172619 | 0.034680671 | 0.15949237 |
| chr14:105724931-105725155 | 1.054329956 | 0.034693062 | 0.159530411 |
| chr7:26466517-26466930 | 1.025642032 | 0.034693107 | 0.159530411 |
| chr17:60043750-60044217 | 0.698575365 | 0.034717945 | 0.159635044 |
| chr17:61828035-61828294 | 0.741670189 | 0.034720905 | 0.159639076 |
| chr20:48604082-48604346 | 0.739412796 | 0.034741112 | 0.159701842 |
| chr1:27496785-27497123 | 0.943979143 | 0.034742039 | 0.159701842 |
| chr14:82366851-82367220 | -0.798201963 | 0.034743729 | 0.159701842 |
| chr1:114215527-114215937 | 0.716044283 | 0.03474492 | 0.159701842 |
| chr5:35516809-35517095 | 1.03158241 | 0.034744978 | 0.159701842 |
| chr8:70471657-70472265 | -0.760131128 | 0.034755047 | 0.159738539 |
| chr1:114488543-114488752 | 0.813518749 | 0.034758008 | 0.159742566 |
| chr5:110738536-110739661 | -0.677246794 | 0.034781165 | 0.159839407 |
| chr7:149854597-149855101 | -0.795635112 | 0.034789838 | 0.159869674 |
| chr7:118104905-118105129 | -1.114854553 | 0.034796319 | 0.159889867 |
| chr20:370307-370627 | 0.854795816 | 0.034800365 | 0.15989887 |
| chr11:26297218-26297523 | -0.884955393 | 0.034802637 | 0.159899721 |
| chr4:161419730-161420201 | -0.835794207 | 0.034808312 | 0.159916204 |
| chr19:37293688-37293907 | 1.09835631 | 0.0348183 | 0.159952503 |
| chrX:71179008-71179289 | 0.902374389 | 0.034825552 | 0.159976227 |
| chr10:110487319-110487973 | -0.781498972 | 0.034827932 | 0.159977569 |
| chr3:64528349-64528600 | 0.754609428 | 0.034847558 | 0.160058122 |
| chrX:112941488-112941756 | -1.035548208 | 0.03486049 | 0.160107923 |
| chr22:25362249-25362504 | 0.903614303 | 0.03486878 | 0.160136398 |
| chr1:48093706-48094463 | -0.73123764 | 0.034877306 | 0.160165955 |
| chr20:55533884-55534203 | 0.924209966 | 0.03489093 | 0.160218915 |
| chr3:93865150-93865394 | 1.081048352 | 0.034893132 | 0.160219424 |
| chr9:38443255-38443475 | 1.085463749 | 0.034912501 | 0.160298756 |
| chr13:24683881-24684165 | 0.828935222 | 0.034916947 | 0.160309566 |
| chr1:198788499-198789064 | -0.770552073 | 0.034921343 | 0.160320141 |
| chrX:37302654-37302935 | -0.976760679 | 0.034934669 | 0.160359295 |
| chr5:69094356-69094786 | -0.774500605 | 0.034934697 | 0.160359295 |
| chrX:102715003-102715268 | -0.979636926 | 0.03493615 | 0.160359295 |
| chr2:22795344-22795927 | -0.79029557 | 0.034938504 | 0.16036049 |
| chr2:112726662-112727089 | 0.765181284 | 0.03494528 | 0.160381987 |
| chr11:118641822-118642022 | 0.936695447 | 0.034949543 | 0.160384103 |
| chr7:10995254-10995454 | 1.011145909 | 0.034950022 | 0.160384103 |
| chr2:178397042-178397345 | -0.954882395 | 0.034952021 | 0.160384103 |

| | | | |
|---|---|---|---|
| chr12:49945604-49945804 | 0.998011064 | 0.034955963 | 0.160386263 |
| chr1:113630610-113631011 | 0.820407636 | 0.034956678 | 0.160386263 |
| chr16:74298113-74298395 | 0.809509584 | 0.034972187 | 0.160447812 |
| chr12:53324766-53325179 | 0.757348865 | 0.034976675 | 0.160458795 |
| chr16:10729681-10730020 | -0.899076407 | 0.034979987 | 0.160464383 |
| chr17:26885638-26885957 | 0.763654079 | 0.034994644 | 0.160522009 |
| chr17:26619478-26619762 | 0.977786007 | 0.035001416 | 0.160538452 |
| chr11:47768293-47768513 | -0.857230205 | 0.035002419 | 0.160538452 |
| chr3:14805186-14805406 | 1.007886495 | 0.03501047 | 0.16055804 |
| chr2:111659861-111660069 | 1.073817816 | 0.035010881 | 0.16055804 |
| chr20:54809024-54809229 | 0.786536773 | 0.035018675 | 0.16058417 |
| chr7:7201375-7202068 | -0.76689267 | 0.035030378 | 0.160628222 |
| chr12:49336850-49337376 | 0.745546131 | 0.03503446 | 0.16063733 |
| chr8:22164511-22164764 | 0.926010203 | 0.035037881 | 0.160643402 |
| chr7:22732829-22733079 | 0.901423174 | 0.035042864 | 0.160656634 |
| chr4:175309184-175309384 | -1.164136811 | 0.03504753 | 0.160662076 |
| chr8:82092185-82092474 | 0.89245021 | 0.035048245 | 0.160662076 |
| chrX:40411420-40411620 | 1.153044833 | 0.035059043 | 0.160697379 |
| chr8:81035356-81035556 | 1.082004096 | 0.03506014 | 0.160697379 |
| chr14:50542910-50543200 | 0.84078498 | 0.035071571 | 0.160730871 |
| chr8:72882032-72882456 | -0.945437986 | 0.035071643 | 0.160730871 |
| chr1:216034573-216034868 | -0.816361243 | 0.035075135 | 0.160737259 |
| chr1:36373846-36374064 | 0.965214858 | 0.035082075 | 0.160751326 |
| chr7:117837913-117838152 | -1.070989918 | 0.035083127 | 0.160751326 |
| chr5:96807291-96808645 | -0.670203194 | 0.035084498 | 0.160751326 |
| chr2:39431504-39432042 | -0.839629639 | 0.035093999 | 0.160785244 |
| chr6:137172263-137172696 | -0.945175843 | 0.035102599 | 0.160815029 |
| chr8:11361775-11362054 | 0.958712272 | 0.035105712 | 0.160819673 |
| chr4:107354411-107354617 | -1.2213071 | 0.035118759 | 0.160864405 |
| chr9:11064922-11065361 | 0.747230899 | 0.035119675 | 0.160864405 |
| chr10:67976952-67977221 | -1.009754387 | 0.035130258 | 0.160899145 |
| chr1:211928978-211929313 | -0.874657962 | 0.035132269 | 0.160899145 |
| chr13:41310937-41312039 | -0.677525862 | 0.03513356 | 0.160899145 |
| chr13:33628537-33628737 | -1.234592524 | 0.035135683 | 0.160899253 |
| chr9:72557545-72557922 | -0.838590211 | 0.035142863 | 0.160922515 |
| chr2:73893383-73893656 | -1.136130288 | 0.035157529 | 0.16098005 |
| chr13:34172234-34172494 | 1.05950516 | 0.035162822 | 0.160994668 |
| chr8:100888336-100888795 | 0.803122138 | 0.035171488 | 0.161024723 |
| chrX:78019626-78020183 | -0.803192864 | 0.035176535 | 0.161034646 |
| chr17:28292984-28293277 | -1.146487158 | 0.035177859 | 0.161034646 |
| chr2:191690390-191690616 | -1.095287048 | 0.035181263 | 0.161040609 |
| chr9:131555820-131556208 | 0.889249184 | 0.035200057 | 0.161117011 |
| chr11:102374581-102374781 | -1.272323007 | 0.035207047 | 0.161139381 |
| chr11:75803245-75803551 | 0.792405612 | 0.035212917 | 0.161156621 |
| chr4:150256898-150257262 | -0.905533688 | 0.035218533 | 0.161172696 |
| chr5:110588569-110588825 | 0.930966689 | 0.035226375 | 0.161182896 |
| chr1:60343685-60344006 | -0.978712254 | 0.035227189 | 0.161182896 |
| chr4:158856911-158857207 | -0.91359421 | 0.035229004 | 0.161182896 |
| chr19:55291941-55292230 | -0.944262365 | 0.035229931 | 0.161182896 |
| chr14:61713689-61713889 | 1.10362896 | 0.03523128 | 0.161182896 |

| | | | |
|---|---|---|---|
| chr1:37752536-37752858 | 0.927043846 | 0.035235117 | 0.161190825 |
| chr21:40642007-40642412 | -0.88187221 | 0.035237266 | 0.161191031 |
| chr13:33817892-33818761 | -0.688665458 | 0.035243377 | 0.161209364 |
| chr12:103768788-103769548 | -0.767748314 | 0.035249502 | 0.161227753 |
| chr4:122945092-122945969 | -0.691861784 | 0.035257416 | 0.161244171 |
| chr5:111835251-111835474 | -1.106018514 | 0.035259811 | 0.161244171 |
| chr6:3748334-3748629 | 0.812071589 | 0.035260933 | 0.161244171 |
| chr1:113993186-113993390 | 0.798065004 | 0.035261509 | 0.161244171 |
| chr1:8412281-8412835 | 0.793134042 | 0.035265623 | 0.161253361 |
| chr14:36332334-36332668 | 0.975942527 | 0.035268949 | 0.161258946 |
| chr12:51341514-51341735 | 0.972823786 | 0.035274717 | 0.161273075 |
| chr14:90345779-90346312 | 0.812044017 | 0.035276249 | 0.161273075 |
| chr20:45963053-45963441 | 0.839787628 | 0.035283885 | 0.161298364 |
| chr14:22002587-22002993 | -0.993534228 | 0.035295027 | 0.161339673 |
| chr19:54430913-54431208 | 0.84193795 | 0.035305404 | 0.161368281 |
| chr14:82872560-82872763 | 1.047113547 | 0.035305498 | 0.161368281 |
| chr17:63065990-63066394 | 0.827929161 | 0.035317835 | 0.161415041 |
| chr20:20360329-20360714 | 0.811714976 | 0.035327611 | 0.161441351 |
| chr2:235567912-235568198 | -0.97480294 | 0.035327806 | 0.161441351 |
| chr9:77759467-77759676 | -1.136121642 | 0.035330022 | 0.161441852 |
| chr1:114250243-114250622 | 0.726355769 | 0.035335225 | 0.161455995 |
| chr13:58239593-58239793 | -1.173918949 | 0.035349803 | 0.161512978 |
| chr8:80682915-80683254 | 0.832550156 | 0.035354483 | 0.161520293 |
| chr19:44807128-44807471 | 0.790990661 | 0.035355621 | 0.161520293 |
| chr1:37603183-37603383 | 1.063990841 | 0.035365439 | 0.161543577 |
| chr10:9931297-9931602 | -1.063495767 | 0.035366593 | 0.161543577 |
| chr8:144189496-144189900 | -0.8478645 | 0.035367042 | 0.161543577 |
| chr3:94525065-94525731 | -0.757372469 | 0.03537078 | 0.161551021 |
| chr10:91381030-91381537 | -0.840620444 | 0.035376741 | 0.161568614 |
| chr15:60393058-60393453 | -0.752451254 | 0.035380098 | 0.161574318 |
| chr10:67763822-67765382 | -0.668789906 | 0.03538726 | 0.161597392 |
| chr11:120236511-120237162 | -0.722991859 | 0.035397669 | 0.161632313 |
| chr5:88622726-88623486 | -0.719304409 | 0.035399126 | 0.161632313 |
| chr10:90319582-90319816 | -1.045783119 | 0.035401395 | 0.161633038 |
| chr10:12150232-12150485 | -1.041606114 | 0.035407248 | 0.161650131 |
| chr4:6746777-6747437 | 0.722115745 | 0.035416293 | 0.161670896 |
| chr4:126018601-126018831 | -1.139808908 | 0.03541727 | 0.161670896 |
| chr1:70132765-70133510 | -0.745609244 | 0.035418126 | 0.161670896 |
| chr10:1242562-1242803 | 1.090229115 | 0.03543257 | 0.161727192 |
| chr2:98800698-98800937 | 1.019276885 | 0.035439806 | 0.1617419 |
| chr22:25462570-25462851 | 0.874910585 | 0.035440014 | 0.1617419 |
| chr5:172731427-172731690 | 0.918583454 | 0.035443331 | 0.161746644 |
| chr6:26171859-26172231 | 0.84100364 | 0.035445275 | 0.161746644 |
| chr10:92800004-92800338 | -1.030331108 | 0.035456611 | 0.161776573 |
| chr5:141040030-141040504 | -0.770052969 | 0.035456784 | 0.161776573 |
| chr20:47746123-47746552 | 0.686551606 | 0.035458168 | 0.161776573 |
| chr3:141804980-141805180 | 1.172843644 | 0.035460724 | 0.161778602 |
| chr10:88728460-88728864 | -0.854497979 | 0.035475801 | 0.161837748 |
| chr11:46127121-46127620 | -0.746549802 | 0.035485012 | 0.161870133 |
| chr5:150500634-150501181 | -0.770935098 | 0.035498246 | 0.161920862 |

| | | | |
|---|---|---|---|
| chr14:96056654-96056993 | 0.799786676 | 0.035509313 | 0.161961701 |
| chr3:62058398-62058694 | 0.798393569 | 0.035515229 | 0.161978356 |
| chr16:1078644-1078885 | 0.909102951 | 0.035517193 | 0.161978356 |
| chr16:80105034-80105518 | -0.863518255 | 0.035520362 | 0.161983169 |
| chr4:102217159-102217746 | 0.799682064 | 0.035538703 | 0.162057161 |
| chr20:47908984-47909568 | 0.693121006 | 0.035547934 | 0.162089611 |
| chr10:72340466-72340737 | -0.843109936 | 0.035553285 | 0.162104363 |
| chr11:34600438-34600669 | -1.061642747 | 0.035573788 | 0.162188195 |
| chr18:55699333-55699533 | -1.191042348 | 0.035576625 | 0.162190706 |
| chr12:101908394-101908817 | 0.727401441 | 0.035580683 | 0.162190706 |
| chr2:174039746-174040232 | -0.820494445 | 0.035580689 | 0.162190706 |
| chr4:57405374-57405727 | 0.838897966 | 0.035592239 | 0.162233704 |
| chrX:93420137-93420974 | -0.770266385 | 0.035595835 | 0.162240443 |
| chr2:15923647-15923896 | -0.956743384 | 0.035600904 | 0.162253894 |
| chr3:125062924-125063572 | 0.7293333 | 0.035621385 | 0.162337582 |
| chr11:9363379-9363720 | 0.851089831 | 0.035632047 | 0.162376205 |
| chr20:62717372-62717612 | -0.936128674 | 0.035634098 | 0.162376205 |
| chr17:61731312-61731720 | 0.690747548 | 0.03564201 | 0.1624026 |
| chr2:84829459-84829688 | 1.073221495 | 0.03565778 | 0.162445357 |
| chr1:58290751-58290951 | -1.278725769 | 0.035658237 | 0.162445357 |
| chr3:10224938-10225211 | 0.847812675 | 0.035659584 | 0.162445357 |
| chr1:247184168-247184572 | -0.905624163 | 0.035661803 | 0.162445357 |
| chr8:52965493-52965719 | -1.092437762 | 0.035661995 | 0.162445357 |
| chr2:191382691-191383240 | 0.73177029 | 0.035666687 | 0.162457075 |
| chr8:144354408-144354747 | 0.764238839 | 0.035676253 | 0.162490985 |
| chr4:20760093-20760324 | 1.151747996 | 0.035705189 | 0.16261311 |
| chr14:35687388-35687622 | 0.922481411 | 0.03570753 | 0.162614106 |
| chr7:116802930-116803166 | -0.933787721 | 0.035711248 | 0.162621373 |
| chr19:46193212-46193425 | 1.017006259 | 0.035720162 | 0.162644115 |
| chr3:45617482-45617839 | -0.93818927 | 0.035720844 | 0.162644115 |
| chr16:73055596-73055935 | 0.797800536 | 0.035723581 | 0.162644115 |
| chr2:121219067-121219290 | 1.036091895 | 0.035724732 | 0.162644115 |
| chr21:42164420-42164868 | 0.828928622 | 0.035744715 | 0.162711311 |
| chr2:10775338-10775545 | 1.006621711 | 0.035747774 | 0.162711311 |
| chr3:63583042-63583299 | 0.848228327 | 0.035747952 | 0.162711311 |
| chr3:190201684-190202125 | -0.925362908 | 0.035749984 | 0.162711311 |
| chr7:92608592-92609228 | -0.730048888 | 0.03575011 | 0.162711311 |
| chr15:87287189-87287519 | -0.850457637 | 0.035758284 | 0.162738847 |
| chr10:5951122-5951410 | -1.024351614 | 0.035760859 | 0.162740902 |
| chr3:71453467-71454265 | 0.752300821 | 0.035765961 | 0.162749482 |
| chr1:102434548-102434753 | -1.118580748 | 0.035766993 | 0.162749482 |
| chr1:216459728-216459993 | -0.939529256 | 0.0357725 | 0.162764876 |
| chr7:731954-732520 | -0.764193479 | 0.035794057 | 0.162837006 |
| chr7:134589370-134590026 | -0.73347741 | 0.035795006 | 0.162837006 |
| chr10:112116429-112116705 | -1.102315118 | 0.035796366 | 0.162837006 |
| chr5:157342750-157343141 | -0.851310319 | 0.035796854 | 0.162837006 |
| chr7:24587812-24588184 | 0.793574044 | 0.035810511 | 0.162889459 |
| chr7:35241959-35242283 | 0.862049867 | 0.035824324 | 0.162942618 |
| chr4:74608638-74608962 | -0.936456115 | 0.035829397 | 0.162956021 |
| chr3:5250988-5251233 | -0.941900956 | 0.035841104 | 0.16299959 |

| | | | |
|---|---|---|---|
| chr1:23651137-23651505 | 0.852017697 | 0.035843774 | 0.163002055 |
| chr2:62215255-62215837 | -0.74062465 | 0.035854529 | 0.16303942 |
| chr7:56106208-56106834 | 0.704908131 | 0.035856246 | 0.16303942 |
| chr5:76774595-76774936 | -0.846919612 | 0.035869945 | 0.163092031 |
| chr11:12800124-12800488 | -0.779303689 | 0.035873879 | 0.163100239 |
| chr1:152189894-152190094 | 0.892515247 | 0.03589919 | 0.163205632 |
| chr1:217953940-217954299 | -0.730179089 | 0.035909518 | 0.163238222 |
| chr16:16858068-16858307 | 1.005686761 | 0.035910619 | 0.163238222 |
| chr9:86228345-86228598 | 1.122040302 | 0.035933934 | 0.163334515 |
| chr15:67731136-67731346 | 1.133740705 | 0.03594413 | 0.163370182 |
| chr9:115015085-115015302 | 0.974816149 | 0.035946046 | 0.163370182 |
| chr14:83981795-83982149 | 1.002948492 | 0.035951401 | 0.163384831 |
| chr6:128911057-128911765 | -0.776157088 | 0.035956215 | 0.163397017 |
| chr19:32718292-32718641 | 0.79965901 | 0.035970297 | 0.163436036 |
| chr4:71713677-71713893 | -1.115046188 | 0.035970767 | 0.163436036 |
| chr5:172932425-172932700 | 0.829829912 | 0.035972157 | 0.163436036 |
| chr20:55584262-55586236 | 0.658063309 | 0.035973946 | 0.163436036 |
| chr1:202091130-202091438 | 0.792314678 | 0.035975466 | 0.163436036 |
| chr15:94230937-94232060 | -0.681037791 | 0.035986955 | 0.163478537 |
| chr8:49795374-49795835 | -0.810719977 | 0.036008254 | 0.163565595 |
| chr7:133081514-133082449 | -0.680998768 | 0.036020757 | 0.163612687 |
| chr15:63496982-63497337 | 0.806645141 | 0.036038887 | 0.163677554 |
| chr14:24351410-24351832 | -0.914942803 | 0.03603931 | 0.163677554 |
| chr21:38669677-38669877 | -1.14155246 | 0.036043397 | 0.163686413 |
| chr7:73827610-73827929 | -0.793939225 | 0.036047135 | 0.163693685 |
| chr2:210222527-210222727 | 1.12169214 | 0.036055658 | 0.163722686 |
| chr9:121499600-121499876 | 0.862260035 | 0.036066109 | 0.163760441 |
| chr4:65416890-65417356 | -0.841919352 | 0.036072609 | 0.163780249 |
| chr22:21029041-21029313 | 0.982783962 | 0.036080123 | 0.163804656 |
| chr3:120907642-120908788 | -0.669490886 | 0.03608318 | 0.163808828 |
| chrX:56053072-56053344 | -0.946558063 | 0.036086507 | 0.163814225 |
| chr20:47679082-47679430 | 0.684951642 | 0.036094854 | 0.163842411 |
| chr8:40148601-40148852 | -1.232568528 | 0.036125816 | 0.163970722 |
| chr7:14381024-14381224 | 1.074331853 | 0.036127401 | 0.163970722 |
| chr18:58067758-58067967 | -1.132734628 | 0.036143841 | 0.164027152 |
| chr20:42205558-42205805 | 0.801188906 | 0.036144116 | 0.164027152 |
| chr21:31531000-31531479 | -0.791328856 | 0.036149229 | 0.16403682 |
| chr14:48262967-48263530 | 0.844278172 | 0.036150528 | 0.16403682 |
| chr3:167493230-167493642 | -0.78247699 | 0.036161703 | 0.16407781 |
| chr2:189763054-189763361 | -0.781261226 | 0.036174029 | 0.164124021 |
| chr10:35336986-35337600 | -0.757664871 | 0.036179236 | 0.164133888 |
| chr6:109397129-109397630 | -0.801329516 | 0.036180488 | 0.164133888 |
| chr14:49479115-49479877 | 0.716832036 | 0.036185252 | 0.164145783 |
| chr8:93955665-93955933 | 0.880825475 | 0.036189858 | 0.164156958 |
| chr22:39662449-39662649 | 1.049463412 | 0.036195008 | 0.164170599 |
| chr3:109731749-109732185 | 0.819404906 | 0.036205837 | 0.164209994 |
| chr19:41449481-41449726 | 0.904097652 | 0.036210294 | 0.164220488 |
| chr21:37257738-37258721 | 0.710716974 | 0.036219301 | 0.164251616 |
| chr6:81785458-81785762 | 0.947732767 | 0.036228278 | 0.164272721 |
| chr4:13891841-13892441 | 0.785360144 | 0.03622915 | 0.164272721 |

| | | | |
|---|---|---|---|
| chr9:18393336-18394072 | -0.761412863 | 0.036230387 | 0.164272721 |
| chrX:85499437-85499702 | -0.956338505 | 0.036234614 | 0.164282166 |
| chr10:81116210-81116712 | 0.768843841 | 0.036238131 | 0.16428839 |
| chr1:188873049-188873330 | 1.090703393 | 0.03624148 | 0.164293852 |
| chr5:74182033-74182247 | -1.131812295 | 0.036252671 | 0.164326082 |
| chr19:13747576-13748091 | 0.750656556 | 0.036252879 | 0.164326082 |
| chr4:22129499-22129837 | 0.917348471 | 0.036259029 | 0.164344238 |
| chr5:94347154-94347387 | -1.161455559 | 0.0362747 | 0.164405539 |
| chr12:112916468-112916725 | -1.076080042 | 0.036277746 | 0.164409623 |
| chr3:146222573-146223086 | -0.789913332 | 0.036285269 | 0.164433988 |
| chr14:58939808-58940057 | 1.053149225 | 0.036287448 | 0.164434137 |
| chr5:138214010-138214210 | -0.94244127 | 0.036298166 | 0.16447298 |
| chrX:73862351-73862551 | 1.097571683 | 0.0363012 | 0.164476999 |
| chr7:116780827-116781411 | -0.708455491 | 0.036306843 | 0.164492842 |
| chr4:158210074-158210777 | -0.676078565 | 0.036317424 | 0.164531051 |
| chr10:5629885-5630363 | 0.766542121 | 0.03632169 | 0.164540647 |
| chr11:46718280-46718537 | 0.898354014 | 0.036330654 | 0.164571527 |
| chr4:83552129-83552516 | -0.890690154 | 0.036333864 | 0.164576338 |
| chr5:119435504-119435887 | -0.753748061 | 0.036338848 | 0.164589183 |
| chr20:51743075-51743311 | -0.813507368 | 0.036353579 | 0.16464617 |
| chr13:48731969-48732361 | -0.869858506 | 0.036360476 | 0.164667673 |
| chr3:160784795-160785136 | 0.854266709 | 0.036368182 | 0.164692837 |
| chr2:190769688-190770240 | -0.876384602 | 0.036380605 | 0.164733057 |
| chr13:20706674-20706912 | -0.873150689 | 0.036381363 | 0.164733057 |
| chr2:55015037-55015290 | -0.901878016 | 0.036385704 | 0.164734991 |
| chr3:8185396-8185799 | 0.992558075 | 0.036387259 | 0.164734991 |
| chr3:166304595-166304928 | -0.971429089 | 0.036388383 | 0.164734991 |
| chr4:168273914-168274224 | -0.912239166 | 0.03639039 | 0.164734991 |
| chr1:77858709-77859013 | 1.020533697 | 0.036398662 | 0.16474808 |
| chr5:179023579-179024183 | -0.725087164 | 0.036401239 | 0.16474808 |
| chr1:27556805-27557735 | -0.71179289 | 0.036401483 | 0.16474808 |
| chr12:94833001-94833213 | -1.056704065 | 0.036401882 | 0.16474808 |
| chr8:125768838-125769038 | 1.081010245 | 0.036405981 | 0.164754596 |
| chr9:74379801-74380096 | -0.916281101 | 0.036408356 | 0.164754596 |
| chr3:11169821-11170021 | 1.032186437 | 0.036409772 | 0.164754596 |
| chr10:12597290-12597553 | -1.196439745 | 0.036412068 | 0.164755253 |
| chr1:91166162-91166406 | -1.01951393 | 0.036421917 | 0.164790086 |
| chr4:87220991-87221356 | -0.782029925 | 0.036426815 | 0.164802517 |
| chr7:128444028-128444706 | -0.713612615 | 0.036430348 | 0.164808769 |
| chr5:35171170-35171375 | 1.166940605 | 0.036436579 | 0.164827228 |
| chr15:84804273-84804694 | 0.789848327 | 0.036447502 | 0.164866906 |
| chr20:44911492-44911867 | 0.741042556 | 0.036452858 | 0.164877594 |
| chr12:113130295-113130852 | -0.822305707 | 0.036455144 | 0.164877594 |
| chr3:156886656-156887030 | 0.949527306 | 0.036457616 | 0.164877594 |
| chr20:50283042-50283242 | 0.891487085 | 0.036458472 | 0.164877594 |
| chr8:144358233-144359060 | 0.700187712 | 0.03646543 | 0.164899325 |
| chr17:61768927-61769517 | 0.676407138 | 0.036469087 | 0.164906128 |
| chr10:98414620-98415741 | -0.678260402 | 0.036473424 | 0.164916008 |
| chr1:10519209-10519466 | -1.11620282 | 0.036483507 | 0.164948318 |
| chrX:10062192-10062786 | -0.765727047 | 0.036486604 | 0.164948318 |

| | | | |
|---|---|---|---|
| chr2:55843175-55843446 | 0.930812322 | 0.036487028 | 0.164948318 |
| chr3:143700256-143700542 | 1.013457149 | 0.036491335 | 0.164958058 |
| chr20:48137112-48137686 | 0.689774399 | 0.036497771 | 0.164969855 |
| chr20:48066930-48067149 | 0.74777632 | 0.036498251 | 0.164969855 |
| chr14:94937862-94938192 | 0.901970608 | 0.03650543 | 0.164992416 |
| chr9:36980129-36980504 | -0.871892243 | 0.036507549 | 0.164992416 |
| chr3:143851672-143851904 | -1.13537636 | 0.036511296 | 0.164999618 |
| chr12:81627444-81627898 | -0.858210339 | 0.036518025 | 0.165020295 |
| chr5:10457511-10458484 | -0.68139958 | 0.036531112 | 0.165060455 |
| chr1:89022360-89022940 | -0.898899389 | 0.036531221 | 0.165060455 |
| chr19:14574246-14574533 | 0.876696364 | 0.036534592 | 0.165065199 |
| chrX:47582142-47582674 | -0.844500414 | 0.036536579 | 0.165065199 |
| chr7:103148926-103149837 | -0.682662775 | 0.036548624 | 0.165103592 |
| chr11:1628838-1629058 | 1.053725659 | 0.036550305 | 0.165103592 |
| chr7:54018455-54018778 | -1.130415356 | 0.036551542 | 0.165103592 |
| chr1:218214390-218214920 | -0.813150442 | 0.036564553 | 0.165152626 |
| chr2:179422088-179422384 | -0.936468313 | 0.03658174 | 0.165220517 |
| chr4:183905670-183905988 | -0.965972705 | 0.036586885 | 0.165234014 |
| chr7:52547579-52547911 | -0.918260633 | 0.036589863 | 0.165237725 |
| chr7:132225714-132225918 | -1.045197482 | 0.036593578 | 0.165244763 |
| chr17:63412232-63412491 | 0.858621583 | 0.03659644 | 0.165247947 |
| chr15:60134469-60134856 | -0.946727686 | 0.036604321 | 0.165273796 |
| chr1:45011977-45012445 | 0.722090038 | 0.036611691 | 0.16529733 |
| chr5:168353190-168353516 | 0.851895072 | 0.036625021 | 0.16534456 |
| chr20:53979048-53979268 | 0.722670039 | 0.036627594 | 0.16534456 |
| chr9:136695437-136695743 | 0.808080837 | 0.036629748 | 0.16534456 |
| chr20:46979531-46979755 | 1.063359081 | 0.036632195 | 0.16534456 |
| chr2:235026987-235027317 | 0.758025606 | 0.036632942 | 0.16534456 |
| chr10:61271479-61271679 | -1.195725147 | 0.03664723 | 0.165393284 |
| chr5:125190786-125191122 | -0.924222699 | 0.036648054 | 0.165393284 |
| chr7:55153486-55153769 | 1.044430145 | 0.036653763 | 0.165407244 |
| chr17:10749247-10749496 | 1.00825899 | 0.036655465 | 0.165407244 |
| chr6:43038898-43039170 | 0.905497298 | 0.036669232 | 0.165449414 |
| chr2:109208133-109208418 | -1.023181782 | 0.03667192 | 0.165449414 |
| chr1:146966458-146966730 | -1.018176448 | 0.036672841 | 0.165449414 |
| chr17:61789998-61790226 | 0.728027054 | 0.036673447 | 0.165449414 |
| chr1:28526899-28527353 | 0.775342342 | 0.036680814 | 0.165472907 |
| chr6:138662435-138662812 | 0.835318241 | 0.036700432 | 0.165551662 |
| chr20:54897863-54899152 | 0.661352755 | 0.036704852 | 0.165561851 |
| chr10:92530059-92530259 | -1.230701513 | 0.036713926 | 0.165593034 |
| chr7:35731492-35731999 | 0.830813191 | 0.036719464 | 0.165608261 |
| chr7:134943881-134944151 | -0.976279939 | 0.036723051 | 0.165614693 |
| chr12:59843369-59843656 | 0.923076273 | 0.036763267 | 0.165776543 |
| chr14:34322995-34323211 | 0.923076273 | 0.036763267 | 0.165776543 |
| chrX:153947333-153947681 | 0.799161154 | 0.036767298 | 0.165784967 |
| chr1:177567494-177567737 | -0.880808126 | 0.036781648 | 0.165839911 |
| chr2:95313433-95313702 | -0.904359669 | 0.036787177 | 0.165848528 |
| chr7:103244965-103245211 | -1.076580788 | 0.036787888 | 0.165848528 |
| chr8:80767830-80768406 | -0.768363235 | 0.036790538 | 0.165850715 |
| chr10:88791938-88792312 | 0.858760153 | 0.036796328 | 0.16585999 |

| | | | |
|---|---|---|---|
| chr3:149339947-149340210 | 0.880823634 | 0.036796967 | 0.16585999 |
| chr5:11390731-11391070 | -0.918519618 | 0.036799089 | 0.16585999 |
| chr18:13451748-13451997 | 1.004643843 | 0.036810736 | 0.165902723 |
| chr4:36386802-36387084 | 1.064544063 | 0.036826266 | 0.165962955 |
| chr4:97910078-97910338 | -1.005681222 | 0.036832155 | 0.165979731 |
| chr1:47173716-47174029 | 0.93141556 | 0.036835438 | 0.165984765 |
| chr9:33102436-33102709 | 1.038670187 | 0.036842033 | 0.166004722 |
| chr16:32939918-32940129 | 1.070285882 | 0.036847263 | 0.166014779 |
| chr12:46461142-46461356 | 1.107671944 | 0.036848599 | 0.166014779 |
| chr6:119148565-119148880 | 0.814114554 | 0.036850894 | 0.166015357 |
| chr9:89425698-89425915 | 1.059020964 | 0.036855695 | 0.166018111 |
| chr17:48659996-48660250 | 1.152163891 | 0.036855839 | 0.166018111 |
| chr4:177596075-177596340 | -0.987743806 | 0.036858981 | 0.166022506 |
| chr19:55643404-55643725 | 0.778672783 | 0.036866664 | 0.166039032 |
| chr1:33470539-33470817 | 0.808962775 | 0.036866984 | 0.166039032 |
| chr3:46878746-46879234 | -0.840837651 | 0.036873183 | 0.166057189 |
| chr1:245488386-245488715 | -0.985181521 | 0.036879379 | 0.166075332 |
| chr7:116210251-116211655 | -0.669781271 | 0.036886086 | 0.166095775 |
| chr2:9190929-9191129 | 1.073833264 | 0.036890125 | 0.1661042 |
| chr1:82759106-82759394 | -0.922832346 | 0.036899074 | 0.16613062 |
| chr2:55692801-55694100 | -0.672691052 | 0.036900329 | 0.16613062 |
| chr20:53585885-53586127 | 0.716844225 | 0.03691115 | 0.166169574 |
| chr12:57452984-57453209 | -0.932163446 | 0.03691642 | 0.166183533 |
| chr3:62379396-62379661 | 0.817647221 | 0.036919134 | 0.166185989 |
| chr7:149377395-149377773 | -0.821557383 | 0.036926962 | 0.166211462 |
| chr14:76956466-76957173 | 0.69541346 | 0.036929848 | 0.166214688 |
| chr8:116473225-116473829 | 0.726580709 | 0.036953408 | 0.166310955 |
| chr2:219399916-219400725 | 0.715758792 | 0.036955673 | 0.166311381 |
| chr1:56024202-56024596 | -0.965763285 | 0.036960197 | 0.166321971 |
| chr6:121593048-121593517 | -0.858971804 | 0.036971213 | 0.166358794 |
| chr15:80202567-80202773 | 1.022636749 | 0.036972722 | 0.166358794 |
| chr12:124992863-124993175 | 0.786519323 | 0.036981325 | 0.1663791 |
| chr2:185289748-185290527 | -0.698873445 | 0.036986659 | 0.1663791 |
| chr20:63969236-63969477 | 0.887567044 | 0.036989502 | 0.1663791 |
| chr2:41869777-41870036 | -0.930498244 | 0.03699065 | 0.1663791 |
| chrX:41010029-41010812 | -0.770444299 | 0.036990783 | 0.1663791 |
| chr10:55989085-55989468 | -0.999841977 | 0.036992278 | 0.1663791 |
| chrX:150568193-150569330 | 0.668089416 | 0.036992435 | 0.1663791 |
| chr2:38230652-38230897 | 1.081164512 | 0.036997468 | 0.166391972 |
| chr8:16204200-16204751 | -0.901014532 | 0.037002502 | 0.166404845 |
| chr9:136631416-136631702 | 0.821183644 | 0.037007804 | 0.166418919 |
| chr15:40282132-40282776 | 0.737551214 | 0.03701355 | 0.16643499 |
| chr5:125696462-125696730 | -1.009510495 | 0.037024969 | 0.166476568 |
| chr6:45504876-45505326 | -0.802674916 | 0.037030156 | 0.16649012 |
| chr14:74612479-74612863 | 0.773962492 | 0.037036851 | 0.166510451 |
| chr12:116503046-116503703 | -0.782135532 | 0.037043569 | 0.166530883 |
| chr12:88579203-88579436 | -0.850851146 | 0.037047553 | 0.16653902 |
| chr3:155336093-155336421 | 0.90859968 | 0.037051672 | 0.166547766 |
| chr4:118772162-118772428 | 0.952984155 | 0.03705845 | 0.166568461 |
| chr20:23586066-23586356 | 0.981091245 | 0.037073355 | 0.166617498 |

| | | | |
|---|---|---|---|
| chr5:32713785-32714396 | -0.68591048 | 0.037073709 | 0.166617498 |
| chr5:84411893-84412093 | -1.156598727 | 0.037078001 | 0.166625997 |
| chr14:79299212-79299508 | 0.902578453 | 0.037080049 | 0.166625997 |
| chr6:34553709-34554131 | -0.742955935 | 0.037082124 | 0.166625997 |
| chr11:121460732-121461202 | 0.841343989 | 0.037094343 | 0.166667796 |
| chr5:155499285-155499702 | -0.889964603 | 0.037095777 | 0.166667796 |
| chr5:140926477-140926677 | 1.088888786 | 0.037103497 | 0.166692708 |
| chr13:73722180-73722492 | -0.998225379 | 0.037124285 | 0.166776321 |
| chr16:85147351-85147756 | 0.817616706 | 0.037132975 | 0.166805582 |
| chr9:122356724-122356948 | -1.028270537 | 0.03713948 | 0.166825023 |
| chr4:48121115-48121497 | 0.778039801 | 0.037147303 | 0.166850382 |
| chr6:76633346-76633657 | -0.909926567 | 0.037149544 | 0.166850668 |
| chr7:122143796-122144607 | -0.691646428 | 0.037167027 | 0.166919403 |
| chr5:33641862-33642577 | -0.745725454 | 0.037170917 | 0.166922172 |
| chr18:64721119-64721332 | -1.25187656 | 0.037172 | 0.166922172 |
| chr3:167901217-167901417 | -1.185342188 | 0.03717859 | 0.166941977 |
| chr7:158350525-158350841 | -1.045031349 | 0.037181378 | 0.166944713 |
| chr5:13478908-13479216 | 0.823531831 | 0.037191771 | 0.166981595 |
| chr5:66856146-66856370 | 1.024096518 | 0.037199772 | 0.166994382 |
| chr5:175746963-175747255 | 0.881820014 | 0.037200919 | 0.166994382 |
| chr7:144854917-144855198 | -0.934776898 | 0.037201158 | 0.166994382 |
| chr5:125412188-125412389 | -1.028735646 | 0.037211831 | 0.167026611 |
| chr2:42906941-42907198 | -0.865301772 | 0.037214434 | 0.167026611 |
| chr10:95675787-95675987 | 1.134980157 | 0.037217544 | 0.167026611 |
| chr1:166354496-166354875 | -0.946688109 | 0.037218789 | 0.167026611 |
| chr10:114635987-114636204 | 1.019263059 | 0.037220189 | 0.167026611 |
| chr19:35509909-35510351 | 0.774011861 | 0.037221417 | 0.167026611 |
| chr9:108743840-108744419 | -0.768698212 | 0.037227448 | 0.167043893 |
| chr1:146013093-146013336 | 0.827556679 | 0.037233305 | 0.16705278 |
| chr1:84603967-84604283 | -0.909088846 | 0.037233789 | 0.16705278 |
| chr1:232542643-232542907 | 0.954950212 | 0.037237264 | 0.16705859 |
| chr3:64930955-64931442 | 0.690479325 | 0.037246253 | 0.16707442 |
| chr20:47906263-47906488 | 0.775844374 | 0.037246459 | 0.16707442 |
| chr7:140397838-140398720 | -0.718961028 | 0.037247334 | 0.16707442 |
| chr1:111513462-111513889 | -0.828794068 | 0.037252707 | 0.167088738 |
| chr8:85351495-85351955 | 0.78125973 | 0.037264685 | 0.167132681 |
| chr1:56645144-56646161 | -0.736442569 | 0.037268807 | 0.167141383 |
| chr16:56789430-56789703 | 0.959885878 | 0.037276543 | 0.167166294 |
| chr14:36581262-36581865 | 0.752357457 | 0.037285317 | 0.167195855 |
| chr20:14919835-14920076 | -1.07094106 | 0.037288016 | 0.167198173 |
| chr6:118903250-118903538 | -0.989748711 | 0.03729076 | 0.167200692 |
| chr3:166028944-166029152 | 1.032913139 | 0.037299461 | 0.167221852 |
| chr6:119147177-119147662 | 0.776960086 | 0.037300043 | 0.167221852 |
| chr9:85511157-85511413 | 1.094252619 | 0.037302134 | 0.167221852 |
| chr14:45201795-45202252 | 0.776507184 | 0.037304209 | 0.167221852 |
| chr20:32608013-32608226 | -1.18079097 | 0.037308306 | 0.167230433 |
| chr4:105990842-105991374 | 0.759147689 | 0.03732009 | 0.167260317 |
| chr6:14066883-14067186 | -1.042753044 | 0.037320784 | 0.167260317 |
| chr5:52948524-52948861 | -0.825052503 | 0.037321522 | 0.167260317 |
| chr6:161498365-161498784 | -0.873005795 | 0.037337223 | 0.167320896 |

| | | | |
|---|---|---|---|
| chr21:43210517-43210717 | 1.088300878 | 0.03734634 | 0.16734899 |
| chr2:37343516-37343767 | -0.948064959 | 0.03734786 | 0.16734899 |
| chr5:90182732-90183076 | -0.841019915 | 0.037354002 | 0.167366724 |
| chr2:69893784-69894299 | -0.765889375 | 0.037363706 | 0.167399133 |
| chr7:144777138-144777840 | -0.748616724 | 0.037365605 | 0.167399133 |
| chr6:358337-358591 | 0.90622919 | 0.037380316 | 0.16745525 |
| chr7:14902585-14903182 | -0.824701716 | 0.037385461 | 0.167461757 |
| chr10:1347779-1347979 | 0.985363918 | 0.037386597 | 0.167461757 |
| chr2:68251108-68251382 | -0.782916551 | 0.037389429 | 0.167461757 |
| chr10:9853475-9853768 | 1.026036001 | 0.037390511 | 0.167461757 |
| chr2:21095142-21095342 | -1.250330645 | 0.037413898 | 0.167556705 |
| chr5:58445600-58445800 | 1.133375972 | 0.037429714 | 0.167617742 |
| chr4:163269949-163270149 | -1.206772045 | 0.037433706 | 0.1676206 |
| chr19:50710353-50710553 | 1.118720839 | 0.037434728 | 0.1676206 |
| chr7:95434617-95435277 | -0.718327911 | 0.037441038 | 0.167639058 |
| chr13:97257560-97258425 | -0.701716325 | 0.037451595 | 0.167676528 |
| chr5:132281946-132282146 | 1.09544237 | 0.037455087 | 0.167682366 |
| chr14:29548925-29549190 | 0.859335923 | 0.037459849 | 0.167685618 |
| chr19:39131129-39131435 | 0.855788948 | 0.03746019 | 0.167685618 |
| chr10:131981378-131982294 | 0.683137769 | 0.037467507 | 0.167694884 |
| chr5:121988254-121988553 | -1.144494296 | 0.037469412 | 0.167694884 |
| chr4:87866063-87866331 | 1.062944364 | 0.03747033 | 0.167694884 |
| chr6:16770968-16771253 | 0.888133489 | 0.037473012 | 0.167694884 |
| chr20:48035296-48035686 | 0.695347688 | 0.037474931 | 0.167694884 |
| chr8:82578639-82578920 | 0.954365076 | 0.037476056 | 0.167694884 |
| chr12:52147435-52147637 | 0.881929084 | 0.037479323 | 0.167694884 |
| chr6:158040390-158040718 | 0.916196179 | 0.037481553 | 0.167694884 |
| chr2:150485143-150485902 | -0.694934083 | 0.037481958 | 0.167694884 |
| chr9:137534848-137535170 | 0.807176687 | 0.037486838 | 0.167706924 |
| chr11:67119615-67119871 | 0.874506543 | 0.037503305 | 0.167763827 |
| chr4:164350617-164350980 | -0.892031716 | 0.037503936 | 0.167763827 |
| chr7:105589459-105589855 | -0.77038217 | 0.037523447 | 0.167841307 |
| chr9:76704566-76704925 | -0.88210899 | 0.037545399 | 0.167929693 |
| chr14:27203417-27203762 | 1.08605952 | 0.037551138 | 0.167945559 |
| chrX:7159196-7159593 | -0.970428764 | 0.037574035 | 0.168034457 |
| chr5:5025044-5025378 | -1.011361359 | 0.03757612 | 0.168034457 |
| chr7:69551078-69551278 | -1.204519386 | 0.03757966 | 0.168034457 |
| chr16:11133280-11133539 | 1.051506667 | 0.037581216 | 0.168034457 |
| chr5:133944383-133944723 | -0.897808317 | 0.037582262 | 0.168034457 |
| chr2:237512771-237513182 | -0.823348568 | 0.03758534 | 0.168034457 |
| chr7:129824812-129826163 | -0.662950892 | 0.037586366 | 0.168034457 |
| chr14:61482624-61482982 | 0.804047002 | 0.037593807 | 0.168057917 |
| chr1:31394642-31395197 | 0.732439505 | 0.037597717 | 0.168065591 |
| chr6:73714248-73714486 | -1.172438543 | 0.037600042 | 0.168066179 |
| chr10:43105479-43105842 | -0.823720759 | 0.037608792 | 0.168095485 |
| chr6:169655242-169655442 | 1.02549313 | 0.037623567 | 0.168145899 |
| chr1:162846724-162847173 | 0.791396927 | 0.037624461 | 0.168145899 |
| chr8:125362638-125362838 | 0.978117947 | 0.037634839 | 0.168176399 |
| chr7:155805024-155805863 | -0.719909564 | 0.037635675 | 0.168176399 |
| chr20:51221702-51221988 | 0.890568472 | 0.037649306 | 0.168227499 |

| | | | |
|---|---|---|---|
| chr3:169634685-169635218 | -0.757117179 | 0.037660203 | 0.168256006 |
| chr1:170417888-170418088 | 1.147264206 | 0.037660316 | 0.168256006 |
| chr8:60447869-60448149 | -0.952470394 | 0.037662274 | 0.168256006 |
| chr12:53492771-53493526 | 0.680949471 | 0.037674389 | 0.168300319 |
| chr1:188371487-188371757 | -1.067529317 | 0.037690854 | 0.168364052 |
| chr4:53522389-53522631 | 1.018337091 | 0.037697294 | 0.168383006 |
| chr16:25015141-25015623 | 0.747167513 | 0.037726093 | 0.168498837 |
| chr9:137550334-137550581 | 0.813318488 | 0.037727624 | 0.168498837 |
| chr11:986694-986972 | 0.910726558 | 0.037732246 | 0.16850226 |
| chr8:26142984-26143662 | -0.746834219 | 0.037734368 | 0.16850226 |
| chr14:106694719-106695036 | -1.084135287 | 0.037734988 | 0.16850226 |
| chr12:112286198-112286439 | 1.012937248 | 0.037743806 | 0.168531814 |
| chr3:193816677-193816983 | -0.935330719 | 0.037751616 | 0.168556863 |
| chr7:107743175-107744692 | -0.660346873 | 0.037759486 | 0.168582179 |
| chr11:76799078-76799638 | -0.751619093 | 0.037766945 | 0.168598811 |
| chr3:16399452-16399821 | -0.871134 | 0.037767612 | 0.168598811 |
| chr6:84626637-84627169 | -0.805615009 | 0.037786883 | 0.168673687 |
| chr2:219552336-219552869 | 0.723027789 | 0.037790978 | 0.168673687 |
| chr12:132622872-132623845 | 0.693517227 | 0.037790989 | 0.168673687 |
| chr5:129878821-129879042 | -1.14248253 | 0.037813226 | 0.168761302 |
| chr6:140427911-140428251 | -0.92218834 | 0.037815024 | 0.168761302 |
| chr7:91565842-91566076 | -1.078072587 | 0.037819528 | 0.168765036 |
| chr1:185505684-185505982 | -1.005767206 | 0.037820266 | 0.168765036 |
| chr2:172985684-172985907 | 0.920245712 | 0.037822942 | 0.168767149 |
| chr8:108703890-108704100 | 0.924479504 | 0.037835151 | 0.168808911 |
| chr1:107276294-107276515 | 0.772158779 | 0.037836708 | 0.168808911 |
| chr7:20280360-20280808 | 0.82011142 | 0.03784121 | 0.168819168 |
| chr1:219581506-219581715 | -1.012327392 | 0.037845582 | 0.168828841 |
| chr2:182642318-182642750 | -0.72800328 | 0.037851943 | 0.168847387 |
| chr1:114660301-114660604 | 0.74945872 | 0.037857278 | 0.168861354 |
| chr3:151387738-151387938 | 1.101814651 | 0.037870352 | 0.168909839 |
| chr7:117119559-117119778 | -1.125299737 | 0.037874104 | 0.168916634 |
| chr13:25988681-25989065 | 0.832017547 | 0.037876285 | 0.168916634 |
| chr8:63007541-63008285 | -0.686779772 | 0.037879882 | 0.168922843 |
| chr21:41955160-41955367 | -0.956797943 | 0.037887348 | 0.168946308 |
| chr11:35082338-35082992 | -0.721833185 | 0.037891012 | 0.168952814 |
| chr20:45456693-45456932 | 0.930443913 | 0.037894021 | 0.168956397 |
| chr20:53665896-53666575 | 0.659898454 | 0.037898074 | 0.168964638 |
| chr19:52734683-52735362 | 0.734102769 | 0.037900979 | 0.168967756 |
| chr10:28410923-28411321 | 0.859474745 | 0.037910225 | 0.168999141 |
| chr9:101705074-101705634 | -0.756975165 | 0.037913694 | 0.169004773 |
| chr8:95890603-95890853 | 0.882942568 | 0.037922761 | 0.16902856 |
| chr17:8832665-8832887 | -1.083091794 | 0.037923442 | 0.16902856 |
| chr3:64931451-64931825 | 0.699483702 | 0.037927407 | 0.169036399 |
| chr17:17056283-17056588 | -0.935832936 | 0.037930864 | 0.169038365 |
| chr8:102227764-102227969 | 0.973459036 | 0.03793226 | 0.169038365 |
| chr15:78315759-78315991 | 0.905978228 | 0.037937461 | 0.16904774 |
| chr14:60514410-60514656 | -1.079045394 | 0.037942098 | 0.16904774 |
| chr17:40314291-40314527 | -0.941398229 | 0.037943068 | 0.16904774 |
| chr1:145545146-145545398 | 0.985712504 | 0.037943189 | 0.16904774 |

| | | | |
|---|---|---|---|
| chr10:2083623-2083840 | -1.156676944 | 0.037955159 | 0.169091238 |
| chr19:1672060-1672304 | 0.969904407 | 0.037967976 | 0.169138504 |
| chr5:173724453-173724727 | 0.999173691 | 0.037972408 | 0.169148414 |
| chr1:154546519-154546744 | 0.929837929 | 0.037980671 | 0.169175386 |
| chr12:15910888-15911719 | -0.695972793 | 0.037990875 | 0.169208896 |
| chr6:75201869-75202556 | 0.750220576 | 0.037992611 | 0.169208896 |
| chr11:62424734-62425025 | 0.913085066 | 0.038010756 | 0.169279867 |
| chr5:142280198-142280453 | 0.946519831 | 0.03802187 | 0.169319525 |
| chr1:29271749-29272094 | 0.822222401 | 0.038038301 | 0.169375062 |
| chr21:43611898-43612098 | 0.958787038 | 0.038041228 | 0.169375062 |
| chr4:74444675-74445870 | -0.673409965 | 0.038041856 | 0.169375062 |
| chr2:190126906-190127106 | -1.29030796 | 0.038043184 | 0.169375062 |
| chr8:126817082-126817399 | -0.807749652 | 0.038053684 | 0.169411968 |
| chr14:59213488-59213688 | 1.067846332 | 0.038066358 | 0.169454335 |
| chr3:93949671-93950575 | -0.675863652 | 0.038067624 | 0.169454335 |
| chr5:159917054-159917335 | 0.98764951 | 0.038070581 | 0.169457653 |
| chr2:36777668-36777941 | -1.124749741 | 0.0380791 | 0.169485726 |
| chr5:112419375-112420671 | 0.682089897 | 0.038082466 | 0.169490865 |
| chr10:23338183-23338539 | -0.843625729 | 0.038089028 | 0.169510222 |
| chr14:25132676-25133169 | -0.730663132 | 0.038091815 | 0.16951278 |
| chr6:108914196-108914571 | -0.905574395 | 0.038105774 | 0.169565049 |
| chr2:86764719-86765015 | -0.924840987 | 0.038117027 | 0.169605275 |
| chr10:14185154-14185998 | -0.725444265 | 0.038130105 | 0.169653614 |
| chr8:123693808-123694118 | 0.786576708 | 0.038147024 | 0.169708535 |
| chrX:119516276-119516592 | 0.827278893 | 0.038149412 | 0.169708535 |
| chr4:134762114-134762353 | -1.089469933 | 0.038149836 | 0.169708535 |
| chr7:38932543-38932844 | -0.909461436 | 0.038151308 | 0.169708535 |
| chr13:59846902-59847133 | -1.081800171 | 0.038157849 | 0.169727778 |
| chr5:94613636-94613883 | -0.853789091 | 0.038201428 | 0.169893258 |
| chr7:106178801-106179001 | -1.077968882 | 0.038201482 | 0.169893258 |
| chr20:675559-675904 | 0.944986609 | 0.038201704 | 0.169893258 |
| chr15:96325920-96326222 | 0.766634001 | 0.038204531 | 0.169895972 |
| chr18:74148174-74148844 | -0.687206816 | 0.038210311 | 0.169911813 |
| chr19:16172952-16173302 | 0.912127664 | 0.038213221 | 0.169914894 |
| chr1:32695367-32695578 | 1.055398565 | 0.038216588 | 0.169920001 |
| chr14:97776930-97777382 | 0.742906376 | 0.038224952 | 0.16994733 |
| chr7:101232176-101232960 | -0.696304887 | 0.038232704 | 0.169966634 |
| chr10:61188723-61188967 | -1.024841368 | 0.038233731 | 0.169966634 |
| chr3:61669700-61669963 | 0.773579012 | 0.038236109 | 0.169967345 |
| chr1:182687209-182687705 | -0.785867371 | 0.038245019 | 0.169997091 |
| chr10:86668224-86668547 | 0.745087118 | 0.038254023 | 0.170015835 |
| chr6:27547289-27547928 | -0.72355655 | 0.038255862 | 0.170015835 |
| chr6:13796847-13797149 | 0.793397171 | 0.038255893 | 0.170015835 |
| chr16:1257624-1257885 | 1.018611377 | 0.038261742 | 0.170031969 |
| chr14:33727834-33728352 | -0.752226654 | 0.038268303 | 0.17005126 |
| chr15:23073853-23074114 | -1.061283019 | 0.03828872 | 0.170122395 |
| chr9:3740886-3741153 | -0.94706665 | 0.038288751 | 0.170122395 |
| chr13:98761854-98762054 | -1.179506022 | 0.038298784 | 0.17014867 |
| chr3:151918924-151919169 | 1.055755753 | 0.038299941 | 0.17014867 |
| chr10:23795803-23796081 | -0.981212059 | 0.038301327 | 0.17014867 |

| | | | |
|---|---|---|---|
| chr17:41521815-41522633 | -0.697220303 | 0.03830648 | 0.170158509 |
| chr3:156024975-156025256 | -0.937651056 | 0.03831001 | 0.170158509 |
| chr15:74239961-74240487 | 0.71919118 | 0.038311805 | 0.170158509 |
| chr3:121883530-121883832 | -0.957440796 | 0.038312425 | 0.170158509 |
| chr14:77498430-77499491 | 0.707740432 | 0.038317608 | 0.170171668 |
| chr13:82490269-82490617 | -1.054015599 | 0.038320272 | 0.170173635 |
| chr4:62644449-62644714 | -0.992002606 | 0.038324868 | 0.170184179 |
| chr4:1192102-1192302 | 1.155473708 | 0.038329164 | 0.170186451 |
| chr3:102124270-102124610 | -1.004873931 | 0.038329822 | 0.170186451 |
| chr6:138934116-138934360 | -1.169231009 | 0.038336321 | 0.170200415 |
| chr4:24472704-24473311 | 0.716049117 | 0.038339521 | 0.170200415 |
| chr20:54817432-54818464 | 0.660094746 | 0.03833963 | 0.170200415 |
| chr3:12025692-12025892 | 1.179833039 | 0.038343811 | 0.170209114 |
| chr3:160112547-160112751 | -1.094279376 | 0.038366662 | 0.170300682 |
| chr17:19937618-19937861 | 0.917403507 | 0.038370727 | 0.170308862 |
| chr19:2602393-2602666 | 0.876625488 | 0.038380667 | 0.170335326 |
| chr8:125696691-125697123 | -0.72390091 | 0.038381136 | 0.170335326 |
| chr7:129664476-129664837 | -0.90095845 | 0.03838408 | 0.170338524 |
| chr7:7181733-7183424 | -0.651973703 | 0.038392534 | 0.170366177 |
| chr6:35241132-35241440 | 0.815286251 | 0.038406755 | 0.170419413 |
| chr7:38846215-38846866 | -0.795162159 | 0.038413635 | 0.170440068 |
| chr7:23013728-23014497 | 0.662173621 | 0.038416447 | 0.170442676 |
| chr4:163535740-163535961 | -1.026202769 | 0.03843631 | 0.17052093 |
| chr16:85514669-85515800 | -0.667439855 | 0.038440831 | 0.170523588 |
| chr16:63289892-63290170 | -0.950327098 | 0.038442003 | 0.170523588 |
| chr1:975832-976731 | 0.703776519 | 0.038443586 | 0.170523588 |
| chr13:39684604-39684876 | -0.989211801 | 0.03844638 | 0.170526108 |
| chr4:98584298-98584657 | 0.859928211 | 0.03845016 | 0.170533004 |
| chrX:154019066-154019500 | 0.783686955 | 0.038467087 | 0.170598204 |
| chr7:65874012-65874212 | 1.113091057 | 0.038474867 | 0.17061314 |
| chr5:176132726-176133011 | 0.913189521 | 0.038474909 | 0.17061314 |
| chr3:64810093-64810353 | 0.73420447 | 0.038477214 | 0.170613487 |
| chr13:48595719-48595968 | -0.978725026 | 0.038482331 | 0.170626208 |
| chr2:241102166-241102672 | 0.743880314 | 0.038484536 | 0.170626208 |
| chr20:2651814-2652771 | 0.704118867 | 0.038495619 | 0.170665469 |
| chr3:9098636-9099028 | 0.74041639 | 0.038509041 | 0.170705317 |
| chr20:36154282-36154650 | 0.807019081 | 0.038509063 | 0.170705317 |
| chr3:127667258-127668044 | -0.745843321 | 0.038519355 | 0.170740241 |
| chr22:20131068-20131324 | 0.860849352 | 0.038521398 | 0.170740241 |
| chr2:162994840-162995040 | -1.213234747 | 0.038531676 | 0.170775918 |
| chr16:23596109-23596585 | 0.73550306 | 0.038546177 | 0.170822618 |
| chr13:59391689-59391943 | -0.924396138 | 0.038546672 | 0.170822618 |
| chr20:47889399-47889617 | 0.743555785 | 0.038552482 | 0.170838488 |
| chr1:6960505-6960708 | 0.862053971 | 0.038557435 | 0.170846504 |
| chr10:14186233-14186658 | -0.855277589 | 0.038558751 | 0.170846504 |
| chr2:22156382-22157076 | -0.777294895 | 0.038574491 | 0.170906363 |
| chr11:65498470-65498670 | -0.946182264 | 0.038584544 | 0.170935244 |
| chr4:142615126-142615811 | -0.710976994 | 0.038590222 | 0.170935244 |
| chr7:18352402-18352627 | 0.886171568 | 0.038590273 | 0.170935244 |
| chr4:6200495-6200764 | 0.860486279 | 0.038593951 | 0.170935244 |

| | | | |
|---|---|---|---|
| chr18:79380558-79381221 | -0.761309508 | 0.038595297 | 0.170935244 |
| chr14:81459799-81459999 | 1.065498908 | 0.038598536 | 0.170935244 |
| chr17:54603308-54603588 | 1.031542062 | 0.038598779 | 0.170935244 |
| chr9:113688037-113688489 | -0.799636088 | 0.038598857 | 0.170935244 |
| chr10:69516308-69516583 | -0.874878942 | 0.038603249 | 0.170944818 |
| chr21:45088736-45088936 | 1.097785382 | 0.03860646 | 0.170949157 |
| chrX:149491810-149492277 | -0.838102678 | 0.038609057 | 0.170950773 |
| chr5:113336944-113337267 | -0.868253128 | 0.038616784 | 0.170975108 |
| chr6:102162169-102162369 | -1.284987025 | 0.038621518 | 0.170986189 |
| chr3:9600450-9600876 | 0.895332382 | 0.038634687 | 0.171032781 |
| chr9:79154291-79154596 | -1.020680258 | 0.038636506 | 0.171032781 |
| chr17:15262515-15262750 | 0.935682216 | 0.038641249 | 0.17104081 |
| chr19:54162246-54162731 | 0.733248396 | 0.038643472 | 0.17104081 |
| chr16:2167944-2168204 | 0.947637708 | 0.038645017 | 0.17104081 |
| chr13:81171082-81171320 | -1.101820269 | 0.038649363 | 0.171050163 |
| chr12:51028131-51028426 | 0.800248467 | 0.038656103 | 0.171070114 |
| chr14:35332084-35332565 | 0.68846442 | 0.038662752 | 0.171089657 |
| chr7:8638115-8638382 | 0.920387386 | 0.038674074 | 0.171129875 |
| chr15:93043577-93043833 | -0.927902615 | 0.038678676 | 0.171140355 |
| chr21:43659255-43659589 | 0.844552697 | 0.038714029 | 0.17128689 |
| chr1:108953437-108953795 | -0.85291069 | 0.03872338 | 0.171318367 |
| chr21:31343585-31344560 | 0.69714922 | 0.038727392 | 0.17131851 |
| chr12:78447479-78447688 | -1.193766147 | 0.038727884 | 0.17131851 |
| chr2:220478539-220479286 | 0.750378694 | 0.038739909 | 0.171361811 |
| chr6:28837392-28837670 | 0.78035773 | 0.038751685 | 0.171394605 |
| chr6:58840323-58840586 | 0.879375426 | 0.038755593 | 0.171394605 |
| chr17:47987326-47987587 | 0.876118765 | 0.038757932 | 0.171394605 |
| chr2:162261473-162261915 | -0.883684864 | 0.038759203 | 0.171394605 |
| chr17:75720691-75721586 | 0.678747904 | 0.038760256 | 0.171394605 |
| chr5:72280795-72281246 | -0.87765236 | 0.038761512 | 0.171394605 |
| chr3:177377133-177377572 | -0.772849477 | 0.038762981 | 0.171394605 |
| chr17:58093854-58094054 | 0.978506556 | 0.038767274 | 0.171403697 |
| chr2:222959256-222959692 | -0.906172317 | 0.038777119 | 0.171437335 |
| chr16:57808112-57808462 | 0.83879946 | 0.038784617 | 0.171460591 |
| chr10:43110796-43111140 | -0.813881683 | 0.038787864 | 0.171465051 |
| chr2:154482976-154483273 | -0.973067197 | 0.038794897 | 0.171486248 |
| chr3:66938358-66938683 | -0.916663162 | 0.038803828 | 0.171515828 |
| chrX:33808514-33808772 | 1.059289887 | 0.038808968 | 0.171522586 |
| chr16:22232406-22233251 | 0.709636114 | 0.038809834 | 0.171522586 |
| chr11:84993452-84993743 | -0.912613896 | 0.038813296 | 0.171527994 |
| chr17:40123303-40123538 | 0.898219444 | 0.038815846 | 0.171529369 |
| chr15:80513075-80513275 | 1.146951094 | 0.03882313 | 0.171551666 |
| chr2:9323933-9324157 | -1.019692767 | 0.038830595 | 0.171574754 |
| chr9:122390024-122390590 | -0.813953332 | 0.038834497 | 0.171582103 |
| chr7:134547403-134547865 | -0.758759775 | 0.038837032 | 0.171583408 |
| chr7:128236970-128237274 | -0.954959285 | 0.038839909 | 0.171585925 |
| chr12:39806992-39807416 | -0.796322954 | 0.03884208 | 0.171585925 |
| chr18:52670997-52671362 | -0.969566556 | 0.038846788 | 0.171596827 |
| chr3:33882197-33882456 | 0.983807265 | 0.038849361 | 0.1715983 |
| chr3:50519525-50519768 | 0.993524656 | 0.038851652 | 0.171598529 |

| | | | |
|---|---|---|---|
| chr15:60006293-60006759 | -0.876039719 | 0.03885843 | 0.171608818 |
| chr20:20549692-20550227 | 0.737564542 | 0.038858461 | 0.171608818 |
| chr6:103273078-103273330 | -1.07542787 | 0.038863968 | 0.171623245 |
| chr2:160994487-160994715 | 0.971339853 | 0.038868097 | 0.171626295 |
| chr21:30408621-30409299 | 0.697362573 | 0.038870208 | 0.171626295 |
| chr14:61461807-61462027 | 0.89990159 | 0.038871378 | 0.171626295 |
| chr2:16635891-16636361 | -0.826523659 | 0.038880802 | 0.17165486 |
| chr10:35388000-35388210 | 1.044469858 | 0.038884121 | 0.17165486 |
| chr6:147204456-147204713 | -0.889421501 | 0.038884568 | 0.17165486 |
| chr8:6840547-6840978 | -0.84163345 | 0.038900929 | 0.171717191 |
| chr12:118165080-118165407 | 0.943582223 | 0.038916425 | 0.171775696 |
| chr12:96416166-96416526 | -1.006393495 | 0.038921893 | 0.171783717 |
| chrX:80403573-80403789 | -1.08809773 | 0.038923017 | 0.171783717 |
| chr19:55526290-55526638 | 0.860351269 | 0.038927286 | 0.171783717 |
| chr10:116826893-116827392 | -0.824795642 | 0.038929526 | 0.171783717 |
| chr4:163494091-163495298 | -0.653564957 | 0.038929649 | 0.171783717 |
| chr7:104643645-104643845 | -1.263701278 | 0.038931694 | 0.171783717 |
| chr3:163327206-163327504 | 0.901871001 | 0.038939715 | 0.171809216 |
| chr5:111092076-111092898 | -0.678918538 | 0.038955689 | 0.1718698 |
| chr2:78546791-78547095 | -1.102539522 | 0.038966486 | 0.171905635 |
| chr3:75502740-75503295 | -0.809059141 | 0.038968299 | 0.171905635 |
| chr2:13006884-13007257 | -0.838889757 | 0.03897071 | 0.171906377 |
| chr12:79044977-79045942 | -0.66662974 | 0.038973449 | 0.171908559 |
| chr13:51542597-51542929 | -0.972389461 | 0.038992851 | 0.171984242 |
| chr14:72547795-72548095 | 0.998951615 | 0.039000092 | 0.172006278 |
| chr8:102511715-102511915 | 0.946137794 | 0.039003286 | 0.172010464 |
| chr8:129053041-129053283 | -1.073840067 | 0.039007191 | 0.172011519 |
| chr5:170497663-170498097 | -0.804306413 | 0.039008564 | 0.172011519 |
| chr17:61382976-61383515 | 0.65967853 | 0.03901026 | 0.172011519 |
| chr2:173667537-173667766 | -1.095094741 | 0.039016472 | 0.17202901 |
| chr7:140977810-140978067 | -1.064234513 | 0.039019556 | 0.172032709 |
| chr2:75153310-75153613 | 0.808752892 | 0.039029355 | 0.172051113 |
| chr11:30241112-30241443 | -1.099705921 | 0.039032122 | 0.172051113 |
| chr12:89180041-89180250 | -1.099705921 | 0.039032122 | 0.172051113 |
| chr17:61377558-61377784 | 0.703849762 | 0.039032712 | 0.172051113 |
| chr3:64292710-64293259 | 0.682785379 | 0.039038509 | 0.172062246 |
| chr7:7761229-7761616 | 0.798019751 | 0.039039729 | 0.172062246 |
| chr8:92102745-92103903 | 0.673858717 | 0.039050237 | 0.172098659 |
| chr3:98699272-98699528 | 1.054300191 | 0.03905296 | 0.172100759 |
| chr7:42097751-42098154 | -0.792290538 | 0.039058708 | 0.172107932 |
| chr22:33038144-33038604 | -0.846170332 | 0.03905908 | 0.172107932 |
| chr2:206120831-206121200 | 1.027150769 | 0.039074013 | 0.172163832 |
| chr8:132712210-132712750 | 0.786161238 | 0.039078958 | 0.172175718 |
| chr1:246709899-246710336 | 0.775025606 | 0.039082265 | 0.172180387 |
| chr13:73684311-73684652 | -0.900615669 | 0.039086893 | 0.172190878 |
| chr1:206550515-206550870 | -0.798682092 | 0.039097477 | 0.1722276 |
| chr10:19246267-19246654 | -0.888095723 | 0.039109107 | 0.172268927 |
| chr8:82129980-82130222 | 1.039721316 | 0.039123436 | 0.172322139 |
| chrX:30735667-30736133 | -0.862012685 | 0.039138545 | 0.17237878 |
| chr18:2541858-2542235 | -1.050158208 | 0.039153853 | 0.172436286 |

| | | | |
|---|---|---|---|
| chr1:200643151-200643514 | 0.793508258 | 0.039159738 | 0.172449378 |
| chr5:148491367-148491807 | -0.880551873 | 0.039161327 | 0.172449378 |
| chr10:68980916-68981313 | -1.021334109 | 0.039165332 | 0.172457102 |
| chr17:1332653-1332909 | -1.056557461 | 0.039169176 | 0.172464118 |
| chr1:32464946-32465197 | 0.940246706 | 0.039172404 | 0.172468421 |
| chr20:561777-562004 | 0.984938151 | 0.039174958 | 0.172469754 |
| chr2:215828501-215828827 | -0.817252503 | 0.039185387 | 0.172505757 |
| chr3:10136294-10136500 | 1.00219615 | 0.03918965 | 0.172510752 |
| chr19:54471036-54471474 | 0.865563811 | 0.039191024 | 0.172510752 |
| chr1:246809575-246809820 | -0.875478502 | 0.039200215 | 0.172541294 |
| chr4:37383647-37383847 | 1.118249869 | 0.039214809 | 0.172595617 |
| chr9:78130073-78130416 | -0.831481813 | 0.039220906 | 0.172612539 |
| chr21:41411230-41411629 | -0.872087778 | 0.039229383 | 0.172625572 |
| chr5:66717753-66718087 | -0.901453965 | 0.039230652 | 0.172625572 |
| chr11:79315371-79315636 | -0.895459512 | 0.039232204 | 0.172625572 |
| chr9:129166822-129167306 | 0.76058006 | 0.03923288 | 0.172625572 |
| chr2:50402381-50402685 | -0.948302655 | 0.039246596 | 0.172676009 |
| chr3:40192805-40193005 | 1.076955504 | 0.039250314 | 0.172682452 |
| chr13:54733624-54733954 | -0.91684585 | 0.039259119 | 0.172711273 |
| chr1:29136594-29136932 | -0.868433 | 0.039289395 | 0.17283016 |
| chr17:58706797-58706997 | 0.782943542 | 0.039290655 | 0.17283016 |
| chr3:15003707-15003907 | 1.22798573 | 0.039295023 | 0.172839451 |
| chr16:10616511-10617700 | -0.664488614 | 0.039305276 | 0.172871999 |
| chr7:155316442-155317430 | -0.687644189 | 0.039306935 | 0.172871999 |
| chr20:32035114-32035362 | 0.876522343 | 0.039320397 | 0.172920462 |
| chr5:103066404-103066711 | -1.01417067 | 0.039322468 | 0.172920462 |
| chr7:31866935-31867135 | 1.050780234 | 0.039346253 | 0.173015127 |
| chr17:18156450-18156668 | -0.886773822 | 0.039350905 | 0.17301594 |
| chr7:90403216-90404175 | -0.664418713 | 0.039350954 | 0.17301594 |
| chr1:222470327-222470689 | -0.854105951 | 0.039369732 | 0.173088569 |
| chr1:186079169-186079498 | -0.993655391 | 0.039374976 | 0.173101695 |
| chr21:32359793-32360015 | 0.998251032 | 0.039410908 | 0.17324972 |
| chr7:50492620-50493405 | -0.706159778 | 0.039425992 | 0.173296836 |
| chr3:62249797-62249999 | 0.87928637 | 0.03942615 | 0.173296836 |
| chr10:110431030-110431324 | 0.835230841 | 0.039434805 | 0.173324939 |
| chr17:61424700-61425118 | 0.670766307 | 0.0394454 | 0.173361559 |
| chr4:133146381-133147146 | -0.716758715 | 0.039451492 | 0.173378387 |
| chr4:175252702-175252930 | -0.988473093 | 0.039455579 | 0.173386407 |
| chr3:160547727-160547949 | 1.024647149 | 0.039459167 | 0.173392227 |
| chr9:96868838-96869038 | 1.012617852 | 0.039467147 | 0.173417348 |
| chr6:119727055-119728654 | -0.663295613 | 0.039473522 | 0.173435416 |
| chr5:33997387-33997714 | -0.850208873 | 0.039481881 | 0.173462197 |
| chr10:8013934-8014149 | 1.169786244 | 0.03948572 | 0.173469117 |
| chr7:139505648-139505888 | -0.973834428 | 0.039497212 | 0.173509652 |
| chr3:52880083-52880403 | -0.935351997 | 0.039502988 | 0.173525078 |
| chr10:13074808-13075154 | 0.862777472 | 0.03951534 | 0.173559545 |
| chr21:45736886-45737239 | 0.872739362 | 0.039515487 | 0.173559545 |
| chr2:642183-642383 | 1.09768522 | 0.039518296 | 0.173559545 |
| chr5:66498777-66499152 | -0.841432445 | 0.039519895 | 0.173559545 |
| chr12:47361338-47361708 | -0.822505074 | 0.039524471 | 0.173569696 |

| | | | |
|---|---|---|---|
| chr14:69612072-69612308 | 0.826813642 | 0.039527142 | 0.173571475 |
| chr21:37009873-37010204 | 0.847937378 | 0.039531346 | 0.173575801 |
| chr5:139373810-139374025 | -0.99739552 | 0.039532658 | 0.173575801 |
| chr19:46925264-46925628 | 0.852589727 | 0.039539702 | 0.173596784 |
| chr17:1053807-1054036 | -1.177940588 | 0.039543834 | 0.173604974 |
| chr3:127961902-127962129 | 0.976949347 | 0.03956138 | 0.173659174 |
| chr20:15491651-15491858 | 1.030798815 | 0.039561477 | 0.173659174 |
| chr8:80565752-80566201 | -0.816183741 | 0.039565555 | 0.173659174 |
| chr10:68097241-68097503 | -0.9567892 | 0.039566954 | 0.173659174 |
| chr13:75480687-75482782 | -0.65096246 | 0.039567512 | 0.173659174 |
| chr5:32793871-32794460 | 0.773984701 | 0.039573824 | 0.173676931 |
| chr15:66940876-66941091 | 1.066584206 | 0.039581289 | 0.173694752 |
| chr15:92636720-92637506 | -0.672682616 | 0.039584321 | 0.173694752 |
| chr3:52409733-52410893 | 0.661804295 | 0.039584685 | 0.173694752 |
| chr11:57277873-57278487 | -0.751350955 | 0.039596231 | 0.173735463 |
| chr7:49424147-49424441 | -0.952918868 | 0.039603874 | 0.173752245 |
| chr4:168816633-168817109 | -0.745503079 | 0.039604591 | 0.173752245 |
| chr16:88540178-88540380 | 0.911896089 | 0.039612626 | 0.173777545 |
| chr9:77291728-77292115 | -0.806431032 | 0.039623252 | 0.173814212 |
| chr20:54008273-54008601 | 0.685858972 | 0.039642838 | 0.173888233 |
| chr5:91438927-91439236 | -0.891270705 | 0.039646357 | 0.173888233 |
| chr17:21505385-21505953 | -0.88329231 | 0.039646935 | 0.173888233 |
| chr13:24605830-24606325 | -0.824494262 | 0.039660969 | 0.173939112 |
| chr2:193800909-193801180 | -0.965739121 | 0.039665042 | 0.173939112 |
| chr17:62130148-62131610 | 0.648214795 | 0.039667283 | 0.173939112 |
| chr12:53224536-53224818 | 0.780899174 | 0.039669501 | 0.173939112 |
| chr13:32164671-32165013 | -0.806222528 | 0.039670627 | 0.173939112 |
| chr17:61505945-61506182 | 0.712063921 | 0.039672607 | 0.173939112 |
| chr6:106771957-106772194 | -1.141492647 | 0.039674426 | 0.173939112 |
| chr5:149228978-149229333 | -0.792700736 | 0.039682611 | 0.173965044 |
| chr11:57903479-57904076 | -0.737453089 | 0.039692709 | 0.173989059 |
| chrX:63560605-63561273 | 0.729355641 | 0.03969773 | 0.173989059 |
| chr22:37828342-37828581 | 1.027437745 | 0.039699237 | 0.173989059 |
| chr19:27793065-27794533 | -0.658256336 | 0.039700704 | 0.173989059 |
| chr6:150741829-150742086 | -0.972858497 | 0.039703339 | 0.173989059 |
| chr6:146905330-146905893 | -0.723781976 | 0.039707915 | 0.173989059 |
| chr17:4389128-4389381 | -1.036317372 | 0.039710018 | 0.173989059 |
| chr6:41536917-41537218 | 0.805079585 | 0.0397108 | 0.173989059 |
| chr1:2082820-2083072 | 1.09096194 | 0.039715021 | 0.173989059 |
| chr3:196867217-196868681 | -0.674651209 | 0.039715663 | 0.173989059 |
| chr15:71296319-71296568 | 0.969589987 | 0.039716387 | 0.173989059 |
| chr2:159787728-159788150 | -0.897705462 | 0.039717676 | 0.173989059 |
| chr14:96424258-96424696 | 0.76470703 | 0.039718095 | 0.173989059 |
| chrX:152958746-152959368 | 0.732826455 | 0.039720148 | 0.173989059 |
| chr7:83119004-83119412 | -0.846240015 | 0.03972215 | 0.173989059 |
| chr7:22550241-22550546 | 0.80158975 | 0.039729887 | 0.174012998 |
| chr11:47523253-47523656 | 0.945308439 | 0.039734216 | 0.174022011 |
| chr17:61659031-61659263 | 0.712956552 | 0.039743739 | 0.174053772 |
| chr10:63720029-63720375 | -0.815679268 | 0.039756032 | 0.174091009 |
| chr16:75555164-75556680 | -0.652602203 | 0.039757295 | 0.174091009 |

| | | | |
|---|---|---|---|
| chr2:215418615-215418910 | -0.917289145 | 0.039759079 | 0.174091009 |
| chr15:76312823-76313169 | 0.781543761 | 0.03976133 | 0.174091009 |
| chr7:148698265-148700236 | -0.656305619 | 0.039772262 | 0.174118434 |
| chr14:100010345-100010623 | -0.953906466 | 0.039772479 | 0.174118434 |
| chr5:121628915-121629174 | -1.095145562 | 0.039774411 | 0.174118434 |
| chr11:72189666-72189868 | 1.050764345 | 0.039780548 | 0.174135351 |
| chr9:128295332-128295626 | 0.804608651 | 0.039790845 | 0.174170472 |
| chr1:13749075-13750000 | 0.701723716 | 0.039812374 | 0.174254754 |
| chr14:97771533-97772103 | 0.722300584 | 0.039814967 | 0.174256148 |
| chr11:82546617-82547099 | -0.839317584 | 0.039820209 | 0.174269137 |
| chr15:91522696-91522896 | 1.06158276 | 0.039823002 | 0.174271405 |
| chr5:169350786-169351150 | -0.889183815 | 0.039827299 | 0.174278948 |
| chr9:19128016-19128497 | -0.82769167 | 0.039829274 | 0.174278948 |
| chr17:66275762-66275994 | 0.893974925 | 0.039837693 | 0.174305832 |
| chr6:106888258-106888769 | 0.765644421 | 0.039847313 | 0.174337966 |
| chr19:34556726-34556941 | -1.078252745 | 0.039850543 | 0.174342143 |
| chr2:96824599-96824830 | -0.928430379 | 0.039855938 | 0.174355793 |
| chr7:80248319-80248661 | -0.950143693 | 0.039859104 | 0.174359686 |
| chr1:61865826-61866026 | 1.087124344 | 0.039863914 | 0.17436101 |
| chr1:202365704-202365909 | 0.963859461 | 0.039863958 | 0.17436101 |
| chr12:103558977-103559280 | 0.985409059 | 0.039881261 | 0.174426735 |
| chr7:7321716-7322113 | 0.731632031 | 0.039902432 | 0.174496342 |
| chr8:76736857-76737200 | 0.906549986 | 0.039903531 | 0.174496342 |
| chr20:8935172-8935420 | -1.009758219 | 0.039904008 | 0.174496342 |
| chr9:20658122-20658882 | -0.711724973 | 0.03991683 | 0.174533836 |
| chr5:93478000-93478279 | -1.196207635 | 0.039917138 | 0.174533836 |
| chr14:103679392-103679715 | 0.896428213 | 0.039920915 | 0.174537209 |
| chr12:57632460-57633442 | 0.686485452 | 0.039925946 | 0.174537209 |
| chr8:143142739-143142964 | 0.912415851 | 0.039927837 | 0.174537209 |
| chr4:159102579-159104159 | -0.655163268 | 0.039928349 | 0.174537209 |
| chr6:95596102-95596376 | 0.950618716 | 0.039929299 | 0.174537209 |
| chr20:45844083-45844418 | 0.919998503 | 0.039933621 | 0.174546143 |
| chr7:74462652-74462911 | -1.01375102 | 0.039936319 | 0.17454798 |
| chr5:144196711-144197004 | -0.973136721 | 0.039940367 | 0.174552256 |
| chr11:127106211-127106436 | -1.118937194 | 0.039941853 | 0.174552256 |
| chr9:122347398-122347639 | 0.936823756 | 0.039945182 | 0.174553778 |
| chr3:64304331-64305240 | 0.663178916 | 0.039946758 | 0.174553778 |
| chr4:40825518-40825803 | 1.157174037 | 0.039967545 | 0.174627364 |
| chr6:56388157-56388403 | -1.052874345 | 0.039968156 | 0.174627364 |
| chr6:107185721-107185953 | 0.968907381 | 0.03998096 | 0.174666476 |
| chr12:109462388-109462798 | -0.768353401 | 0.039981667 | 0.174666476 |
| chr2:135792432-135792637 | -0.932238701 | 0.039985825 | 0.17467468 |
| chr12:12737192-12737433 | -1.01835934 | 0.039995975 | 0.1746991 |
| chr13:101514779-101515060 | -1.01835934 | 0.039995975 | 0.1746991 |
| chr14:105108332-105108678 | 0.982216638 | 0.040014891 | 0.174764958 |
| chr2:131486525-131487119 | -0.737960016 | 0.040018088 | 0.174764958 |
| chr6:139858468-139858756 | -0.9337036 | 0.040019703 | 0.174764958 |
| chr12:96578569-96578826 | -0.992079482 | 0.040020176 | 0.174764958 |
| chr20:20860319-20860519 | 1.032219191 | 0.040037997 | 0.174824936 |
| chr2:20143356-20143681 | 0.787665099 | 0.040038474 | 0.174824936 |

| | | | |
|---|---|---|---|
| chr17:68351432-68351738 | -0.775427792 | 0.040042386 | 0.174827134 |
| chr12:65602534-65602883 | -0.840111655 | 0.040043541 | 0.174827134 |
| chr5:166787659-166787880 | -1.013941497 | 0.040049344 | 0.174835031 |
| chr3:53136339-53136645 | 0.932358794 | 0.040049913 | 0.174835031 |
| chr13:21290903-21291108 | 1.052777193 | 0.040052447 | 0.174835273 |
| chr11:115582107-115582387 | -0.919779891 | 0.040054532 | 0.174835273 |
| chr10:100433650-100434161 | 0.754408965 | 0.040060229 | 0.174843485 |
| chr4:52816339-52816605 | -1.049173163 | 0.040060978 | 0.174843485 |
| chr3:154140076-154140503 | 0.895041163 | 0.040077383 | 0.174905122 |
| chr5:174875503-174875840 | 0.951987283 | 0.040101159 | 0.17499892 |
| chr9:72524955-72525588 | -0.751145975 | 0.04011642 | 0.175055544 |
| chr14:39243856-39244079 | 1.006227864 | 0.04013719 | 0.175130385 |
| chr22:18157150-18157484 | 0.843995429 | 0.040138142 | 0.175130385 |
| chr12:53213329-53213566 | 0.851003863 | 0.040143857 | 0.175142234 |
| chr1:177334375-177334636 | -0.939401778 | 0.04014631 | 0.175142234 |
| chr19:10450892-10451175 | 0.933286213 | 0.040147715 | 0.175142234 |
| chr20:54682630-54682871 | 0.712172243 | 0.040158445 | 0.175179069 |
| chr10:132403847-132404067 | 0.822328336 | 0.040168633 | 0.175210622 |
| chr3:154262616-154262842 | 0.967392044 | 0.040170251 | 0.175210622 |
| chr6:134052097-134052878 | -0.709743631 | 0.04018174 | 0.175238528 |
| chr14:69717177-69717377 | 1.046028676 | 0.040182409 | 0.175238528 |
| chr5:143361455-143361771 | -1.04490471 | 0.040184587 | 0.175238528 |
| chr11:79493559-79494143 | -0.74660857 | 0.040185798 | 0.175238528 |
| chr7:34088091-34088805 | -0.711336371 | 0.040190651 | 0.175242091 |
| chr3:158010521-158010757 | 1.041640882 | 0.040191189 | 0.175242091 |
| chr1:154418292-154418628 | 0.817665065 | 0.04019758 | 0.175259983 |
| chr5:774984-775184 | 0.954240775 | 0.040204269 | 0.175271681 |
| chr7:157815089-157815322 | -1.012943926 | 0.040204838 | 0.175271681 |
| chr7:50531186-50531957 | -0.701400646 | 0.040210801 | 0.17528767 |
| chr17:61668386-61669223 | 0.651175986 | 0.040213081 | 0.17528767 |
| chr1:247400802-247401053 | -1.059585615 | 0.040224209 | 0.175326203 |
| chr13:95246399-95246635 | 1.034198909 | 0.040236103 | 0.175368069 |
| chr2:48915284-48915596 | -0.882881856 | 0.04025105 | 0.175415402 |
| chr17:10357823-10358094 | -0.922076503 | 0.040251541 | 0.175415402 |
| chr2:205885503-205886165 | -0.716139704 | 0.040264524 | 0.175454369 |
| chr2:152200584-152200991 | -0.738102608 | 0.040265062 | 0.175454369 |
| chr17:69092503-69092703 | 1.133758677 | 0.040276061 | 0.175492314 |
| chr9:73667189-73667551 | -0.958898528 | 0.040283462 | 0.175510813 |
| chr15:80749475-80750165 | 0.680978737 | 0.040284888 | 0.175510813 |
| chr5:177301525-177302359 | 0.671376388 | 0.040295179 | 0.175545666 |
| chr6:135523262-135523545 | -1.011637606 | 0.040302521 | 0.175567672 |
| chr14:56135128-56135347 | 1.033538111 | 0.040306528 | 0.175575145 |
| chr18:69400226-69400898 | -0.757493684 | 0.040315455 | 0.175604048 |
| chr22:29787325-29787697 | 0.811179446 | 0.040322455 | 0.175621575 |
| chr2:186460481-186461001 | -0.833006329 | 0.040327547 | 0.175621575 |
| chr5:112333352-112333745 | 0.836665535 | 0.040330624 | 0.175621575 |
| chr8:40373396-40373716 | 0.854231251 | 0.04033091 | 0.175621575 |
| chr1:177112819-177113042 | 1.055149152 | 0.040330939 | 0.175621575 |
| chr1:154405884-154406501 | 0.725770232 | 0.040350074 | 0.175694911 |
| chr6:31712306-31712579 | 0.891499473 | 0.040366884 | 0.175752646 |

| | | | |
|---|---|---|---|
| chr5:157447227-157447690 | -0.827007465 | 0.040367921 | 0.175752646 |
| chr17:38584394-38584932 | 0.727172626 | 0.04037204 | 0.17575284 |
| chr2:55742333-55742533 | -1.298657737 | 0.040372553 | 0.17575284 |
| chr5:141478380-141478659 | 0.803069649 | 0.040385454 | 0.175785609 |
| chr10:101141201-101141665 | -0.804913728 | 0.040385852 | 0.175785609 |
| chr4:3765363-3765830 | 0.772432133 | 0.040386963 | 0.175785609 |
| chr16:87420646-87420902 | 0.894812562 | 0.040392324 | 0.175798959 |
| chr21:39082605-39082939 | -0.774968505 | 0.040399222 | 0.175818991 |
| chr17:62855934-62856259 | -0.767137845 | 0.040409837 | 0.175855203 |
| chr9:126398560-126398760 | 0.986165661 | 0.04041493 | 0.175867375 |
| chrX:87409465-87409846 | -0.801379822 | 0.040420596 | 0.175882046 |
| chr1:221263807-221264084 | -0.972976555 | 0.04042647 | 0.175897614 |
| chr18:22466090-22466371 | 0.837177879 | 0.040434715 | 0.175919352 |
| chr7:42407262-42407462 | -1.102544319 | 0.040437054 | 0.175919352 |
| chr14:21024768-21025235 | 0.7716262 | 0.04043905 | 0.175919352 |
| chr1:219564151-219564491 | -0.8295933 | 0.04044065 | 0.175919352 |
| chr20:47962805-47963364 | 0.671136836 | 0.040448269 | 0.175942506 |
| chr13:39982661-39982941 | 0.966242129 | 0.040450621 | 0.175942749 |
| chr1:227999226-227999669 | 0.762600605 | 0.04046504 | 0.175995476 |
| chr8:87565707-87566041 | -0.950678964 | 0.040476505 | 0.17603535 |
| chr6:35786636-35787380 | -0.74348262 | 0.04048877 | 0.176078694 |
| chr1:115568015-115568248 | 1.047905576 | 0.040497409 | 0.176106268 |
| chr8:103500281-103500650 | 0.725281687 | 0.040506512 | 0.176132553 |
| chr5:59562874-59563414 | -0.758014237 | 0.040509081 | 0.176132553 |
| chr3:64150568-64151413 | 0.649416307 | 0.040510349 | 0.176132553 |
| chr12:98998758-98999233 | -0.84235057 | 0.040518452 | 0.176150031 |
| chr2:158753729-158754322 | -0.772346129 | 0.040518993 | 0.176150031 |
| chr16:72856868-72857130 | 0.87262776 | 0.040521266 | 0.176150031 |
| chr1:106419096-106419647 | 0.696943426 | 0.040532567 | 0.176177028 |
| chr18:58764222-58764589 | -0.822710156 | 0.040535796 | 0.176177028 |
| chr17:8894974-8895286 | 0.882206033 | 0.040536865 | 0.176177028 |
| chr2:46158195-46158432 | 1.01018952 | 0.040538855 | 0.176177028 |
| chr5:104179100-104179392 | -0.915188257 | 0.040540352 | 0.176177028 |
| chr10:110151435-110151783 | -0.889698077 | 0.040541272 | 0.176177028 |
| chr12:53220643-53221378 | 0.681997493 | 0.040548038 | 0.17618563 |
| chr5:672752-673116 | -0.775851233 | 0.040548976 | 0.17618563 |
| chr8:139758317-139758615 | 0.858470378 | 0.040552656 | 0.17618563 |
| chr3:62433234-62433525 | 0.814809984 | 0.040554171 | 0.17618563 |
| chr5:87019625-87020470 | -0.710627083 | 0.040555303 | 0.17618563 |
| chr19:10864516-10864740 | 0.98494893 | 0.040557048 | 0.17618563 |
| chr15:84735090-84735804 | -0.718649953 | 0.040563405 | 0.176199618 |
| chr2:137886476-137886804 | -0.836259171 | 0.040564867 | 0.176199618 |
| chr12:25839540-25839758 | -1.142545036 | 0.040574271 | 0.176230473 |
| chr6:10314991-10315292 | 0.82056661 | 0.040579394 | 0.176242736 |
| chr12:23472279-23472730 | -0.89353835 | 0.040615159 | 0.176388068 |
| chr9:19925957-19926506 | -0.77132507 | 0.040626054 | 0.176425385 |
| chr21:45513864-45514129 | 0.844406674 | 0.040635827 | 0.176457825 |
| chr5:108260588-108260910 | -0.818243785 | 0.040641744 | 0.176471311 |
| chr8:99904727-99905339 | -0.7013078 | 0.040646208 | 0.176471311 |
| chr6:81767341-81767902 | 0.750851299 | 0.040646833 | 0.176471311 |

| | | | |
|---|---|---|---|
| chr7:142957042-142957328 | -0.98049605 | 0.040648145 | 0.176471311 |
| chr7:83819964-83820248 | -1.107068009 | 0.040666985 | 0.176543101 |
| chr11:65468903-65469180 | 0.827967889 | 0.040669538 | 0.176544183 |
| chr1:200654264-200654554 | -0.864571776 | 0.040676764 | 0.176565544 |
| chr10:133420798-133421034 | 1.006216784 | 0.040681694 | 0.176576942 |
| chr5:161071086-161072160 | -0.718670261 | 0.040684058 | 0.176577199 |
| chr6:37325776-37326051 | 0.831897356 | 0.040699612 | 0.1766347 |
| chr3:53161029-53161238 | 0.856539226 | 0.040703731 | 0.176638691 |
| chrX:104113124-104113583 | -0.778489582 | 0.040705247 | 0.176638691 |
| chr1:6245924-6246217 | 0.838061587 | 0.040707447 | 0.176638691 |
| chr6:63882308-63882590 | -0.984153598 | 0.040716275 | 0.176666993 |
| chr9:82734404-82734710 | 1.064279363 | 0.040734714 | 0.176736991 |
| chr10:19251349-19251587 | -1.094248294 | 0.040760007 | 0.176826876 |
| chr3:61696668-61696868 | 0.832939408 | 0.040760047 | 0.176826876 |
| chr21:14063805-14064144 | 0.838055576 | 0.04076397 | 0.176833884 |
| chr17:38559387-38559934 | 0.759980127 | 0.040789455 | 0.17693442 |
| chr15:91412664-91412892 | 1.032535704 | 0.040821739 | 0.177063759 |
| chr6:13407796-13408015 | -1.158332692 | 0.040825595 | 0.177063759 |
| chr14:94342379-94342800 | 0.847273854 | 0.040827466 | 0.177063759 |
| chr9:107755358-107755579 | 1.068342765 | 0.040828585 | 0.177063759 |
| chr2:17309698-17310077 | -0.805974758 | 0.040830826 | 0.177063759 |
| chr7:24290262-24290612 | 0.765462768 | 0.040835688 | 0.177074819 |
| chr3:96192955-96193560 | -0.788304694 | 0.040838232 | 0.177075833 |
| chr11:76919888-76920480 | -0.743531658 | 0.040843103 | 0.177086931 |
| chr3:167740969-167741324 | -0.753540926 | 0.040853694 | 0.177122829 |
| chr1:99685542-99686117 | -0.789143819 | 0.0408656 | 0.177164424 |
| chr12:72279001-72279240 | -1.046537941 | 0.040883894 | 0.177233706 |
| chr6:73339337-73339769 | -0.819922368 | 0.040914767 | 0.177357506 |
| chr8:133033146-133033779 | -0.737870704 | 0.040921204 | 0.177375372 |
| chr17:68953301-68953526 | -0.971505722 | 0.040929976 | 0.177400864 |
| chr1:234433450-234433867 | -0.93136537 | 0.040933699 | 0.177400864 |
| chr10:23981950-23982224 | -1.047813982 | 0.040935963 | 0.177400864 |
| chr15:40450023-40450418 | -0.804195618 | 0.040936346 | 0.177400864 |
| chr18:63848146-63848476 | -0.871703234 | 0.040939944 | 0.177405489 |
| chr6:166032701-166033204 | -0.778405861 | 0.040943783 | 0.177405489 |
| chr10:37137347-37137856 | -0.866776332 | 0.040944932 | 0.177405489 |
| chr7:106239351-106239733 | -0.819827163 | 0.040946674 | 0.177405489 |
| chr4:38017648-38017911 | -1.025209691 | 0.040958652 | 0.177447349 |
| chr6:118705893-118706110 | -1.108050132 | 0.040962015 | 0.177451885 |
| chr1:827342-827770 | 0.800619192 | 0.040971487 | 0.177482884 |
| chr5:41869603-41870935 | 0.653574345 | 0.040974846 | 0.1774874 |
| chr8:106447447-106448141 | -0.713370508 | 0.040982758 | 0.177511637 |
| chr14:24123331-24123547 | 0.976082133 | 0.0409947 | 0.17755264 |
| chr3:137970973-137971608 | -0.716307805 | 0.040998316 | 0.17755264 |
| chr9:21316904-21317194 | -1.003679052 | 0.040999176 | 0.17755264 |
| chr3:122415338-122415673 | -0.728498024 | 0.041005107 | 0.177565228 |
| chr13:75537357-75538436 | -0.661896151 | 0.041008154 | 0.177565228 |
| chr12:99847921-99848476 | -0.71602736 | 0.041009035 | 0.177565228 |
| chr1:69305716-69305916 | 1.108496969 | 0.041013269 | 0.177567351 |
| chr15:37764631-37764922 | -1.084155907 | 0.041015769 | 0.177567351 |

| | | | |
|---|---|---|---|
| chr17:38320490-38320895 | 0.806847813 | 0.041016478 | 0.177567351 |
| chr3:149180067-149180393 | -0.840299598 | 0.041023509 | 0.177587759 |
| chr12:122937715-122937915 | 1.088623222 | 0.041089365 | 0.177838255 |
| chr11:22976035-22976235 | -1.20612854 | 0.04109061 | 0.177838255 |
| chr12:106301710-106303101 | -0.664093638 | 0.041091746 | 0.177838255 |
| chr12:121232962-121233538 | 0.730982856 | 0.041093716 | 0.177838255 |
| chr4:24471065-24471525 | 0.769701844 | 0.041093782 | 0.177838255 |
| chr3:63015133-63015413 | 0.804877422 | 0.041095301 | 0.177838255 |
| chr10:46033778-46033978 | -0.98924985 | 0.041107055 | 0.177875771 |
| chr20:44728713-44729408 | 0.678880098 | 0.041108613 | 0.177875771 |
| chr8:90031728-90031986 | 0.943855178 | 0.041113635 | 0.177887457 |
| chr1:207997854-207998191 | -0.838197886 | 0.041127137 | 0.177935826 |
| chr7:7271117-7271434 | -1.026302419 | 0.041134615 | 0.177958132 |
| chr9:757631-757831 | 0.98558603 | 0.041141444 | 0.177968006 |
| chr1:207818856-207819205 | -0.786981951 | 0.041141543 | 0.177968006 |
| chr8:64669949-64670292 | -0.837107647 | 0.041174328 | 0.178099774 |
| chr7:142420601-142420896 | -0.909513383 | 0.041182011 | 0.178117149 |
| chr9:112174673-112175136 | 0.796356627 | 0.041182995 | 0.178117149 |
| chr16:85878896-85879109 | 1.058459114 | 0.041191134 | 0.178129338 |
| chr18:54119538-54120072 | -0.755076581 | 0.041191477 | 0.178129338 |
| chr2:186595268-186595474 | -0.99494171 | 0.041194875 | 0.178129338 |
| chr7:10889349-10889639 | 0.788895072 | 0.041195112 | 0.178129338 |
| chrX:111164816-111165058 | 0.959927746 | 0.041211645 | 0.178182718 |
| chr5:86749908-86750218 | -0.8621999 | 0.041213393 | 0.178182718 |
| chr3:195454409-195454663 | -1.082996179 | 0.041214433 | 0.178182718 |
| chr16:68265259-68265661 | 0.755199615 | 0.041225411 | 0.178220124 |
| chr7:47950867-47951096 | -0.878787348 | 0.041230526 | 0.178232179 |
| chr6:17580106-17581489 | -0.672065914 | 0.04124465 | 0.178283175 |
| chr6:135069730-135070049 | -1.055329761 | 0.041254852 | 0.178317216 |
| chr4:138560463-138560750 | -0.925778815 | 0.041260112 | 0.178329892 |
| chr5:493960-494160 | 0.948732524 | 0.041271095 | 0.178367298 |
| chr2:142286480-142286992 | -0.77560767 | 0.041276399 | 0.178375067 |
| chr14:63504623-63504990 | 0.771087806 | 0.041277548 | 0.178375067 |
| chr10:30414136-30414398 | -0.984239957 | 0.041280688 | 0.178378572 |
| chr10:89398068-89398298 | -1.070053368 | 0.041299869 | 0.178451396 |
| chr1:218489131-218489331 | 1.009630722 | 0.041309044 | 0.178480971 |
| chr7:140638506-140638761 | -0.929521618 | 0.041314135 | 0.178492905 |
| chr7:80098944-80099144 | -1.225312363 | 0.04132527 | 0.178530943 |
| chr2:85446712-85447036 | 0.965199987 | 0.04133503 | 0.178551241 |
| chr4:1712132-1713427 | 0.657484402 | 0.041337451 | 0.178551241 |
| chr12:41983916-41984178 | -1.032099101 | 0.041338714 | 0.178551241 |
| chr7:13525122-13525417 | 0.834624315 | 0.041339289 | 0.178551241 |
| chr19:33195238-33195717 | 0.742565397 | 0.041344751 | 0.178564766 |
| chr20:57171239-57171524 | 0.673388486 | 0.041348932 | 0.17856795 |
| chr18:45523196-45523503 | -1.027712501 | 0.041350149 | 0.17856795 |
| chr17:36534093-36535806 | 0.660722524 | 0.041376349 | 0.178660806 |
| chr1:26357319-26357546 | -1.02955478 | 0.041380557 | 0.178660806 |
| chr10:112191190-112191474 | 1.038556431 | 0.04138069 | 0.178660806 |
| chr17:42779992-42780262 | 0.997982012 | 0.041380978 | 0.178660806 |
| chr14:23551071-23551379 | 0.756586966 | 0.041384866 | 0.178667525 |

| | | | |
|---|---|---|---|
| chr8:136280232-136280522 | 0.952477802 | 0.0413967 | 0.178708543 |
| chr6:138726483-138726755 | -0.992252548 | 0.041405934 | 0.178738337 |
| chr2:68909875-68910120 | -0.957008616 | 0.04142199 | 0.17879324 |
| chr1:218124102-218124302 | -1.070654414 | 0.04142332 | 0.17879324 |
| chrX:23901115-23901395 | -0.881122604 | 0.041428937 | 0.178798301 |
| chr11:21752885-21753160 | -0.98807448 | 0.041429159 | 0.178798301 |
| chr20:47652409-47653220 | 0.650879979 | 0.041461924 | 0.178929627 |
| chr13:28324195-28324532 | -0.794517394 | 0.041471 | 0.178958716 |
| chr18:45665172-45665385 | -0.998554415 | 0.041480631 | 0.178990196 |
| chr3:149307641-149307862 | 1.11803471 | 0.041489972 | 0.179018911 |
| chr7:43764858-43765416 | -0.704693086 | 0.041493704 | 0.179018911 |
| chr14:83329376-83329612 | 1.03510983 | 0.041495561 | 0.179018911 |
| chr6:105374014-105374393 | 0.793109898 | 0.041496631 | 0.179018911 |
| chr1:158527240-158527525 | -1.000253058 | 0.041509691 | 0.179056503 |
| chr14:23149785-23150039 | 0.895340164 | 0.041510019 | 0.179056503 |
| chr21:40467027-40467227 | -1.233682452 | 0.041512695 | 0.179057968 |
| chr4:26263178-26263463 | 0.827558059 | 0.041523895 | 0.179066358 |
| chr3:136994927-136995246 | -0.866678022 | 0.041529681 | 0.179066358 |
| chr2:3059334-3059534 | 1.069852163 | 0.041529955 | 0.179066358 |
| chr9:70204711-70205171 | -0.793584457 | 0.0415317 | 0.179066358 |
| chr1:235630209-235630550 | 0.867217104 | 0.041533485 | 0.179066358 |
| chr10:63865255-63865544 | -0.82831778 | 0.041533622 | 0.179066358 |
| chrX:119757821-119758784 | 0.683091691 | 0.04153404 | 0.179066358 |
| chr3:121811877-121812113 | 1.005156436 | 0.041535497 | 0.179066358 |
| chr4:11228662-11228927 | 0.889882002 | 0.041535674 | 0.179066358 |
| chr3:120868746-120868976 | 1.107916665 | 0.041548924 | 0.179109301 |
| chr10:110897011-110897411 | -0.99905131 | 0.04155031 | 0.179109301 |
| chr2:174670404-174670719 | -1.040240022 | 0.041562622 | 0.179148791 |
| chr17:31805045-31805418 | 0.818528187 | 0.041567376 | 0.179148791 |
| chrX:115960751-115960997 | 1.057917756 | 0.041567414 | 0.179148791 |
| chr2:14178829-14179111 | -0.990102369 | 0.041569612 | 0.179148791 |
| chr2:212254384-212254992 | -0.744592193 | 0.041572083 | 0.179148791 |
| chr2:152473698-152474099 | 0.83568859 | 0.041574274 | 0.179148791 |
| chr1:195608899-195609231 | -0.828659337 | 0.041575837 | 0.179148791 |
| chr10:17217275-17217578 | -1.019972961 | 0.041578373 | 0.179149642 |
| chr13:48303176-48304251 | -0.671454789 | 0.041584846 | 0.179165886 |
| chr1:40107374-40108052 | 0.735000118 | 0.041587686 | 0.179165886 |
| chr10:75108377-75108948 | -0.729211286 | 0.041589158 | 0.179165886 |
| chr3:8971074-8971274 | 0.952133588 | 0.041598489 | 0.17919601 |
| chr3:23039352-23039615 | -1.02657783 | 0.041620635 | 0.179274561 |
| chr7:123285098-123285383 | -0.902293476 | 0.041621403 | 0.179274561 |
| chr11:65133360-65133588 | 0.848546578 | 0.041630366 | 0.179303085 |
| chr6:66258775-66259632 | -0.692054704 | 0.04164743 | 0.179366497 |
| chr1:66436317-66436876 | -0.768436614 | 0.041673524 | 0.179468792 |
| chr6:169789112-169789356 | -0.956006626 | 0.041687281 | 0.179509706 |
| chr10:96640867-96641198 | 0.821735781 | 0.04168771 | 0.179509706 |
| chr1:21176570-21177372 | -0.721255577 | 0.041693997 | 0.179519437 |
| chr1:186754599-186755111 | -0.81121403 | 0.041695403 | 0.179519437 |
| chr9:127579580-127579780 | 1.007844344 | 0.041696999 | 0.179519437 |
| chr4:164100589-164101391 | -0.732758096 | 0.041710081 | 0.17956567 |

| | | | |
|---|---|---|---|
| chr5:151881842-151882069 | -1.159549178 | 0.041720834 | 0.179601275 |
| chr20:47757076-47757276 | 0.745572206 | 0.041723039 | 0.179601275 |
| chr4:122700118-122700718 | -0.746163521 | 0.041739209 | 0.179660788 |
| chr20:47278658-47279143 | 0.656971985 | 0.041754393 | 0.179714981 |
| chrX:108704995-108705195 | 1.093628206 | 0.04175649 | 0.179714981 |
| chr3:157700473-157700774 | -0.851376603 | 0.041766461 | 0.17973821 |
| chr20:47816560-47816774 | 0.73104703 | 0.041766606 | 0.17973821 |
| chr16:27972513-27972881 | 0.803149476 | 0.041768925 | 0.17973821 |
| chr9:76444375-76444575 | 1.16568332 | 0.041777201 | 0.179763729 |
| chr4:103009148-103009415 | -0.942762051 | 0.041780379 | 0.179767308 |
| chr3:172623660-172624615 | -0.676972394 | 0.041783679 | 0.179771409 |
| chr4:179945212-179945583 | -1.02896153 | 0.041802094 | 0.179840541 |
| chr2:74553295-74553518 | -0.926976201 | 0.041808564 | 0.179858278 |
| chr18:14431004-14431618 | 0.75020507 | 0.041830385 | 0.179942048 |
| chr7:100929577-100929777 | 1.113640694 | 0.041859309 | 0.18005636 |
| chr2:30208456-30208746 | 0.876474294 | 0.041864703 | 0.180069455 |
| chr13:23466329-23466996 | 0.72357125 | 0.041885903 | 0.180150526 |
| chr12:66103364-66103789 | -0.934143895 | 0.041890548 | 0.180160394 |
| chr5:3236974-3237174 | 1.084616315 | 0.041898547 | 0.180184683 |
| chr5:141604307-141604617 | -0.977334508 | 0.041907517 | 0.180204901 |
| chr4:175622960-175623280 | -0.972832763 | 0.041907952 | 0.180204901 |
| chr10:22087165-22087480 | -0.940783084 | 0.041921173 | 0.180251635 |
| chr17:83144926-83145246 | 0.820086321 | 0.041927191 | 0.180267393 |
| chr2:196502618-196502960 | -0.861722014 | 0.041933407 | 0.180284005 |
| chr7:141701282-141701889 | -0.731221292 | 0.041937177 | 0.180290098 |
| chr12:70553631-70553941 | 1.040036637 | 0.041944971 | 0.180313485 |
| chr1:168082238-168082625 | -0.765875366 | 0.041953557 | 0.180340278 |
| chr18:63342297-63342506 | -1.165147272 | 0.04196695 | 0.180381966 |
| chr4:1579241-1579632 | 0.809593968 | 0.041967964 | 0.180381966 |
| chr6:41941110-41941323 | 0.899198862 | 0.041970798 | 0.18038403 |
| chr3:184379018-184379221 | 0.975227406 | 0.041977557 | 0.180402959 |
| chr2:43389794-43390181 | 0.882870315 | 0.04198278 | 0.180415286 |
| chr7:100827203-100828042 | -0.689370866 | 0.041990221 | 0.180437146 |
| chr10:17229856-17230963 | -0.665506857 | 0.041993468 | 0.180440977 |
| chr1:65215651-65215979 | -0.8361101 | 0.042001175 | 0.180461976 |
| chr18:73908347-73908604 | -1.042722583 | 0.042003066 | 0.180461976 |
| chr17:45371179-45371458 | -0.943623512 | 0.042012281 | 0.18049145 |
| chr8:66713827-66714420 | -0.741133904 | 0.042020089 | 0.180513735 |
| chr7:127391675-127393436 | -0.646560817 | 0.04202347 | 0.180513735 |
| chr12:65163386-65163632 | -1.01140176 | 0.042024536 | 0.180513735 |
| chr20:61194865-61195094 | 0.929716208 | 0.042032654 | 0.180538483 |
| chr20:58888775-58889202 | 0.730848469 | 0.042038662 | 0.180549744 |
| chr2:100268854-100269112 | -0.905182277 | 0.042039988 | 0.180549744 |
| chr14:50667693-50668821 | 0.665247941 | 0.042044633 | 0.180559571 |
| chr4:13907277-13907608 | 0.919960608 | 0.042061957 | 0.180623843 |
| chr5:154937566-154938625 | -0.655166707 | 0.042064434 | 0.18062436 |
| chr9:132406755-132406996 | 0.900473808 | 0.042068422 | 0.18062582 |
| chr12:21527619-21528169 | -0.84325081 | 0.042069489 | 0.18062582 |
| chr5:143825180-143825453 | 0.883903571 | 0.042074388 | 0.180636732 |
| chr1:218498186-218498481 | 0.800191694 | 0.042090201 | 0.180692303 |

| | | | |
|---|---|---|---|
| chr16:30896029-30897332 | 0.658310915 | 0.042092048 | 0.180692303 |
| chr20:44765589-44766000 | -0.755274445 | 0.042102779 | 0.180728241 |
| chr13:22107531-22107809 | 0.946813158 | 0.04211599 | 0.180770301 |
| chr4:133909887-133910773 | 0.693962326 | 0.042118531 | 0.180770301 |
| chr13:40664829-40665197 | -0.767006838 | 0.042119655 | 0.180770301 |
| chr2:237590307-237591868 | -0.64984412 | 0.042123329 | 0.180775944 |
| chr7:102395701-102396980 | -0.660248734 | 0.042132819 | 0.180806545 |
| chr6:96435999-96436206 | 1.054186064 | 0.042138525 | 0.18081603 |
| chr11:96411041-96411714 | -0.763529705 | 0.042142482 | 0.18081603 |
| chr5:3344575-3344830 | 1.018620379 | 0.042143751 | 0.18081603 |
| chr6:7418251-7418918 | -0.734525259 | 0.042146362 | 0.18081603 |
| chr1:226550269-226550652 | 0.799955475 | 0.042146829 | 0.18081603 |
| chr1:226912667-226913006 | 0.827831587 | 0.0421503 | 0.180820799 |
| chr14:45610314-45610644 | 0.858477096 | 0.042155682 | 0.180833762 |
| chr12:100671808-100672378 | -0.720718764 | 0.042164058 | 0.180859566 |
| chr16:31386697-31386962 | -0.967650776 | 0.042170046 | 0.180867044 |
| chr5:122415547-122415925 | -0.77768281 | 0.042170522 | 0.180867044 |
| chr17:65082599-65082836 | 0.888935137 | 0.042191171 | 0.180940025 |
| chr2:237692754-237693065 | 0.831062336 | 0.04219442 | 0.180940025 |
| chr13:66617544-66618155 | -0.746727472 | 0.042194623 | 0.180940025 |
| chr3:140597752-140598142 | -0.97274024 | 0.042200766 | 0.180956241 |
| chr2:201345010-201345291 | -0.887978282 | 0.04221333 | 0.180999989 |
| chr6:43758586-43758803 | 0.939017017 | 0.042220179 | 0.181019224 |
| chr22:20021866-20022144 | 0.851530162 | 0.042226522 | 0.18103629 |
| chr18:31684214-31684744 | 0.767314062 | 0.042243187 | 0.181097605 |
| chr17:55207194-55207399 | 1.041720121 | 0.042253965 | 0.181119224 |
| chr10:94183612-94183864 | 1.074167787 | 0.042255322 | 0.181119224 |
| chr16:27355787-27356018 | 1.074167787 | 0.042255322 | 0.181119224 |
| chr10:62437480-62437832 | -0.935919858 | 0.042262281 | 0.181138921 |
| chr8:108505600-108506534 | -0.67124525 | 0.042284033 | 0.181222015 |
| chr7:127458246-127458648 | -0.991213258 | 0.042291912 | 0.181245645 |
| chr17:48566169-48566369 | 0.975770424 | 0.042315349 | 0.181335945 |
| chr10:46048213-46048622 | -0.805636228 | 0.04232563 | 0.181369856 |
| chr14:94722846-94723119 | 1.023785378 | 0.042331651 | 0.181381821 |
| chr8:29555358-29556058 | 0.728475948 | 0.042335517 | 0.181381821 |
| chr17:81733751-81734006 | 0.784927898 | 0.042335524 | 0.181381821 |
| chr5:69042835-69043414 | -0.743698518 | 0.042342243 | 0.181400465 |
| chr10:24415782-24416181 | -0.829711729 | 0.042352556 | 0.181434503 |
| chr9:126545981-126546189 | 1.097529349 | 0.042356921 | 0.181443059 |
| chr13:77026227-77027649 | -0.648897795 | 0.042360734 | 0.18144925 |
| chr9:134681403-134681611 | 1.044910541 | 0.042373704 | 0.181494657 |
| chr20:54134584-54134935 | 0.676937304 | 0.042377843 | 0.181502241 |
| chr17:57899383-57900184 | -0.673193298 | 0.042383818 | 0.181517684 |
| chr22:26645034-26645682 | 0.697878325 | 0.042404481 | 0.181591025 |
| chr8:87536742-87537023 | 1.026371752 | 0.042405683 | 0.181591025 |
| chr6:141561179-141561440 | 0.905351094 | 0.042410021 | 0.181599454 |
| chr14:96167406-96167817 | 0.770877513 | 0.042417444 | 0.181621089 |
| chr5:131382344-131382791 | -0.74482509 | 0.042422671 | 0.181633321 |
| chr10:23035550-23035884 | -0.863862629 | 0.042426932 | 0.181641416 |
| chr2:181715055-181715356 | 0.865976064 | 0.042434689 | 0.181657319 |

| | | | |
|---|---|---|---|
| chr1:116453498-116454305 | 0.674016837 | 0.042435389 | 0.181657319 |
| chr10:55629776-55630106 | -0.745589432 | 0.042448416 | 0.181702933 |
| chr5:473701-474167 | 0.717201674 | 0.042453553 | 0.181714774 |
| chr4:56181304-56181546 | -1.08345997 | 0.042468278 | 0.181767647 |
| chr12:64836849-64837178 | 0.811169699 | 0.042478847 | 0.181802728 |
| chrX:11759035-11759237 | -0.948164424 | 0.042489943 | 0.181840059 |
| chr1:17441633-17441833 | 0.978373093 | 0.042525325 | 0.181981315 |
| chr3:146640868-146641218 | -0.74615458 | 0.042528633 | 0.181984329 |
| chr2:1047584-1047785 | -1.025062736 | 0.042530779 | 0.181984329 |
| chr3:175026810-175027236 | -0.828460523 | 0.042534903 | 0.181991809 |
| chr4:140195917-140196208 | -1.017056495 | 0.042539251 | 0.182000251 |
| chr7:18698319-18698697 | 0.804147434 | 0.042543499 | 0.182008263 |
| chr9:122546032-122546568 | -0.793579815 | 0.042550606 | 0.182028504 |
| chr1:108526540-108526856 | -0.910338309 | 0.042552995 | 0.182028559 |
| chr7:153192851-153193576 | -0.736231715 | 0.042557936 | 0.182039533 |
| chr15:63041869-63043125 | 0.659787251 | 0.042561902 | 0.182046333 |
| chr3:87309765-87310064 | 1.047322745 | 0.042575037 | 0.182088257 |
| chr2:172735830-172736090 | -1.081424273 | 0.042576456 | 0.182088257 |
| chr6:168530677-168530934 | -0.97360067 | 0.042587124 | 0.182123714 |
| chr4:183130660-183131020 | -1.016030011 | 0.042595359 | 0.182143983 |
| chrX:126190515-126190720 | -1.175812213 | 0.042596618 | 0.182143983 |
| chr3:12158846-12159133 | 0.863882546 | 0.042612363 | 0.182201141 |
| chr7:148297271-148297616 | -0.957200634 | 0.042619501 | 0.182221496 |
| chr5:94707517-94707832 | -0.991223617 | 0.042626197 | 0.182239954 |
| chr6:23659232-23659479 | -0.988172588 | 0.042629432 | 0.182243615 |
| chr4:169024822-169025137 | -0.841331871 | 0.042633114 | 0.182249188 |
| chr4:9898522-9898897 | 0.915241533 | 0.04265403 | 0.18232843 |
| chr4:113901958-113902391 | -0.742663331 | 0.042670021 | 0.182386611 |
| chr14:24035863-24036096 | 0.905541879 | 0.042673228 | 0.18239014 |
| chr3:155247118-155247332 | 1.086694523 | 0.042688242 | 0.182432044 |
| chr8:110530629-110530829 | 1.117030091 | 0.042688704 | 0.182432044 |
| chr16:77422291-77422491 | 1.011458143 | 0.042690174 | 0.182432044 |
| chr2:1854809-1855071 | -1.02985039 | 0.042695077 | 0.18244282 |
| chr10:22927616-22928362 | -0.696537514 | 0.042714429 | 0.182505694 |
| chr1:33149748-33150515 | 0.706162906 | 0.042714555 | 0.182505694 |
| chr3:11799284-11799783 | 0.781656199 | 0.042717648 | 0.182508734 |
| chr20:47239294-47239729 | 0.662325998 | 0.042723618 | 0.182524062 |
| chr8:79873373-79873882 | -0.792312198 | 0.042730775 | 0.18254446 |
| chr1:114448802-114449031 | 0.797493049 | 0.042745688 | 0.182597989 |
| chr5:77168249-77169222 | -0.688023027 | 0.042749026 | 0.182602066 |
| chr10:10719197-10719465 | 0.955791799 | 0.042757038 | 0.182626108 |
| chr15:98853214-98853520 | 0.716931882 | 0.04276165 | 0.182635627 |
| chr3:123725750-123725950 | 1.002515252 | 0.04277758 | 0.182693479 |
| chr5:154849819-154850019 | -0.963012799 | 0.042794372 | 0.182755011 |
| chr3:31934859-31935098 | 0.950655981 | 0.042806158 | 0.182795152 |
| chr19:50050965-50051297 | 0.760218991 | 0.042810522 | 0.182803601 |
| chr3:192964983-192965295 | -0.828444667 | 0.042819802 | 0.182833038 |
| chr11:112321894-112322229 | -0.861407735 | 0.042835864 | 0.182891428 |
| chr11:113289550-113289897 | -0.891622429 | 0.042847659 | 0.182930104 |
| chr2:23384608-23385223 | -0.729963646 | 0.042849697 | 0.182930104 |

| | | | |
|---|---|---|---|
| chr10:121245771-121246269 | 0.766061881 | 0.0428533 | 0.182934004 |
| chr14:77622532-77622990 | -0.747072966 | 0.042859514 | 0.182934004 |
| chr14:30085975-30086185 | 1.062803847 | 0.042860522 | 0.182934004 |
| chr20:29044167-29044536 | 0.895095205 | 0.042861425 | 0.182934004 |
| chr3:103406145-103406606 | -0.88687205 | 0.042862548 | 0.182934004 |
| chr2:97355854-97356174 | 0.905437314 | 0.042866598 | 0.182941096 |
| chr19:39160320-39160806 | 0.714932629 | 0.042880864 | 0.182989859 |
| chr10:3970439-3970689 | 0.961305993 | 0.0428828 | 0.182989859 |
| chr20:53840521-53840730 | 0.69648604 | 0.042886489 | 0.182995411 |
| chr2:36322187-36322467 | -0.963666269 | 0.042908621 | 0.183079649 |
| chr8:98913423-98913658 | 0.903228945 | 0.042921117 | 0.183122769 |
| chr4:139294851-139295183 | -0.763767511 | 0.042925489 | 0.183131227 |
| chr15:40108125-40108344 | 0.844573732 | 0.042936265 | 0.183167001 |
| chr6:75855151-75855382 | -1.089257011 | 0.042950776 | 0.183215129 |
| chr15:66931632-66932352 | 0.692498181 | 0.042952329 | 0.183215129 |
| chr12:50283053-50283903 | 0.678041823 | 0.042959052 | 0.183233606 |
| chr5:175820649-175820868 | 0.958407491 | 0.042964505 | 0.183246663 |
| chr15:68372557-68372789 | 0.902702626 | 0.042971404 | 0.183253418 |
| chr10:23158436-23158737 | -0.825836945 | 0.04297218 | 0.183253418 |
| chr13:98562979-98563356 | -0.901843951 | 0.042973264 | 0.183253418 |
| chr2:35220491-35220888 | -0.897831363 | 0.042977024 | 0.183259256 |
| chr15:99708629-99708947 | -0.816743859 | 0.042983577 | 0.183270045 |
| chr12:82517082-82517374 | -0.829002539 | 0.042985368 | 0.183270045 |
| chr1:223097387-223097587 | 1.097808693 | 0.042989223 | 0.183270045 |
| chr10:78745786-78745986 | 1.097808693 | 0.042989223 | 0.183270045 |
| chr6:79946764-79948058 | -0.650595052 | 0.042991514 | 0.183270045 |
| chr10:122703905-122704197 | 0.896663423 | 0.042999351 | 0.183293257 |
| chr9:82774584-82775438 | -0.722014115 | 0.043004755 | 0.183306092 |
| chr1:195733919-195734382 | -0.842605858 | 0.043027316 | 0.183392059 |
| chr6:122869484-122869765 | -0.844593152 | 0.043036989 | 0.183423083 |
| chr9:104386466-104386696 | -1.141585198 | 0.043061713 | 0.183518246 |
| chr16:87571125-87571798 | 0.704261233 | 0.043076399 | 0.183570626 |
| chr4:8176833-8177111 | 0.833178023 | 0.0430836 | 0.183591104 |
| chr8:134843688-134844006 | -0.806454032 | 0.043098247 | 0.183643302 |
| chr9:121286938-121287640 | -0.710674769 | 0.043109664 | 0.183681736 |
| chr20:45987337-45987620 | 0.815187132 | 0.043115471 | 0.183688276 |
| chr1:20212957-20213244 | -1.057886575 | 0.043115993 | 0.183688276 |
| chr12:131895794-131896859 | 0.657316154 | 0.04312184 | 0.183698789 |
| chr14:90716049-90716491 | 0.730590833 | 0.043123256 | 0.183698789 |
| chr4:8183172-8183817 | 0.703739924 | 0.043129325 | 0.183703785 |
| chr20:47428769-47429288 | 0.650116399 | 0.043130215 | 0.183703785 |
| chr3:161069787-161070240 | -0.80177479 | 0.043131621 | 0.183703785 |
| chr18:63241382-63241658 | -0.830792149 | 0.043137766 | 0.183719744 |
| chr4:24594852-24595287 | 0.720179407 | 0.043156085 | 0.183787545 |
| chr11:86151114-86151368 | 0.99828555 | 0.043186094 | 0.183905123 |
| chr4:8580243-8580821 | 0.705435447 | 0.043193924 | 0.183928245 |
| chrX:116284364-116284660 | -0.918860915 | 0.043196798 | 0.183930262 |
| chr17:69084919-69085273 | -0.760437696 | 0.043223645 | 0.184033032 |
| chr6:157126402-157126607 | -1.177394971 | 0.043227534 | 0.184033032 |
| chr16:31221670-31221877 | 1.041018706 | 0.04322814 | 0.184033032 |

| | | | |
|---|---|---|---|
| chr3:72339136-72339340 | 0.972015899 | 0.04324757 | 0.184105522 |
| chr16:76769791-76769991 | 1.03714286 | 0.04325398 | 0.18412258 |
| chr7:30413162-30413369 | 0.931526991 | 0.043265824 | 0.184162767 |
| chr2:11582783-11583027 | -0.79744011 | 0.043281363 | 0.184218676 |
| chr3:174409457-174409770 | -0.931537007 | 0.04329704 | 0.184275164 |
| chr1:80779339-80779584 | -1.00817405 | 0.043299934 | 0.184277245 |
| chr16:25040465-25040753 | 0.964900866 | 0.043309942 | 0.1843096 |
| chr3:181363265-181363482 | -1.057418286 | 0.043315339 | 0.18431387 |
| chr5:175533996-175534464 | 0.747392463 | 0.043315983 | 0.18431387 |
| chr3:136703356-136703603 | 0.98233722 | 0.043318778 | 0.18431387 |
| chr14:77152181-77152527 | 0.767539092 | 0.043320567 | 0.18431387 |
| chr3:12676326-12676636 | -0.863727709 | 0.043336362 | 0.184366673 |
| chr13:71218999-71219250 | -1.005258246 | 0.04333779 | 0.184366673 |
| chr10:49612913-49613150 | -0.904029683 | 0.043342742 | 0.184377501 |
| chr1:173628618-173628897 | -1.024990382 | 0.043346495 | 0.18438323 |
| chr5:73615198-73615549 | -0.997879772 | 0.043351804 | 0.184395578 |
| chr11:47373941-47374211 | 0.795242902 | 0.043354836 | 0.184398239 |
| chr14:90513362-90513998 | -0.682913985 | 0.043358421 | 0.184398612 |
| chr1:158329906-158330146 | 0.958723693 | 0.043359737 | 0.184398612 |
| chr1:78145072-78145321 | -1.035555895 | 0.043366077 | 0.184415337 |
| chr7:142782971-142783181 | -0.957870159 | 0.043375125 | 0.184442479 |
| chr6:116680290-116680623 | -0.767193739 | 0.043377274 | 0.184442479 |
| chr3:119490168-119490459 | -0.866125335 | 0.043384461 | 0.184458326 |
| chr14:103127036-103127296 | 0.826144145 | 0.043386745 | 0.184458326 |
| chr20:50711015-50711536 | 0.648768342 | 0.04339023 | 0.184458326 |
| chr1:170457281-170458450 | -0.665151098 | 0.04339063 | 0.184458326 |
| chr8:89233439-89233679 | 0.949950962 | 0.043402521 | 0.184491437 |
| chr17:61510257-61510594 | 0.670846912 | 0.043403235 | 0.184491437 |
| chr1:234564611-234565047 | 0.784847708 | 0.043418047 | 0.184544163 |
| chr3:15272979-15273203 | 0.878105608 | 0.043424351 | 0.184560718 |
| chr1:70510104-70510562 | -0.819631345 | 0.043431434 | 0.184568055 |
| chr18:73980373-73980723 | -0.86111365 | 0.043434598 | 0.184568055 |
| chr17:60465894-60466097 | 0.737010751 | 0.043435669 | 0.184568055 |
| chr16:72987529-72987976 | -0.751460226 | 0.043435712 | 0.184568055 |
| chr4:154314341-154314564 | -1.056792993 | 0.043441309 | 0.184571364 |
| chr4:165343816-165344016 | -1.056792993 | 0.043441309 | 0.184571364 |
| chr11:96389242-96390493 | -0.659503305 | 0.043453422 | 0.184612591 |
| chr2:29114091-29114338 | -1.002517587 | 0.043459229 | 0.184627026 |
| chr19:55640746-55641290 | 0.707758681 | 0.043470408 | 0.184664278 |
| chr1:205476801-205477028 | 0.818517399 | 0.043481215 | 0.184699949 |
| chr14:55541024-55541627 | 0.721317096 | 0.04349092 | 0.184730931 |
| chr7:43674559-43674831 | -1.048328112 | 0.043504283 | 0.184777448 |
| chr7:156560164-156560467 | -1.015042415 | 0.043515123 | 0.184808085 |
| chr9:114611074-114612027 | 0.668685264 | 0.04351632 | 0.184808085 |
| chr14:71403707-71404009 | 0.847393954 | 0.043531589 | 0.184862685 |
| chrX:85309704-85310152 | -0.840398695 | 0.04354552 | 0.184903195 |
| chr17:66841388-66841700 | -0.759918929 | 0.043545955 | 0.184903195 |
| chr22:24996663-24996868 | 0.951915975 | 0.043566877 | 0.18497261 |
| chr22:16774733-16774940 | -1.05686852 | 0.043569545 | 0.18497261 |
| chr3:168137077-168137320 | -1.05686852 | 0.043569545 | 0.18497261 |

| | | | |
|---|---|---|---|
| chr2:197850019-197850219 | -1.183474202 | 0.043572007 | 0.184972815 |
| chr7:156640073-156641253 | -0.646789082 | 0.043582567 | 0.185007393 |
| chr5:111127529-111127826 | -0.848089443 | 0.043587957 | 0.185012245 |
| chr1:215076180-215076768 | -0.7030945 | 0.043590404 | 0.185012245 |
| chr17:35141778-35142507 | 0.734244441 | 0.043590954 | 0.185012245 |
| chr10:119198610-119198834 | 0.990454043 | 0.043594735 | 0.185015623 |
| chr8:102123590-102124334 | 0.706475912 | 0.043596579 | 0.185015623 |
| chr11:63961638-63962067 | -0.736718503 | 0.043608536 | 0.185056117 |
| chr17:59945593-59945843 | 0.741296638 | 0.043616779 | 0.185080848 |
| chr6:16776745-16776945 | -1.182290629 | 0.043621426 | 0.185083114 |
| chr17:75896387-75896788 | 0.757231114 | 0.043625942 | 0.185083114 |
| chr13:33800078-33800505 | 0.814977 | 0.043625988 | 0.185083114 |
| chr2:74205393-74205985 | -0.735111897 | 0.043628821 | 0.185083114 |
| chr18:54157652-54157984 | -1.009413328 | 0.043630854 | 0.185083114 |
| chr8:117083010-117083522 | -0.767792882 | 0.043632444 | 0.185083114 |
| chr3:9552341-9552587 | 0.90415355 | 0.043634222 | 0.185083114 |
| chr1:170597797-170598018 | -1.038816093 | 0.04364731 | 0.185125088 |
| chr19:37991976-37992176 | 0.992042973 | 0.04364895 | 0.185125088 |
| chr11:18853126-18853687 | -0.755064557 | 0.04365804 | 0.185153392 |
| chr12:64889713-64890399 | 0.659543469 | 0.043668623 | 0.18517051 |
| chr18:58310498-58310915 | -0.873407092 | 0.04366885 | 0.18517051 |
| chr6:51453841-51454196 | -0.873545636 | 0.043669326 | 0.18517051 |
| chr14:101817556-101817782 | 1.067639733 | 0.043672353 | 0.185173098 |
| chr6:149544882-149545160 | -0.81577707 | 0.043677114 | 0.185176941 |
| chrX:47193549-47193896 | 0.7418865 | 0.043678093 | 0.185176941 |
| chr5:76818600-76819823 | -0.653665925 | 0.043685822 | 0.185199459 |
| chr16:29290910-29291248 | 0.781832316 | 0.043691798 | 0.185208661 |
| chrX:41734638-41734838 | -1.169596179 | 0.043693014 | 0.185208661 |
| chr2:13581545-13581803 | -1.002952843 | 0.043700218 | 0.185208661 |
| chr1:201302605-201302895 | -0.78552764 | 0.043701411 | 0.185208661 |
| chr12:96985454-96985950 | -0.777981564 | 0.043703808 | 0.185208661 |
| chr19:17021656-17022132 | -0.846636756 | 0.043703923 | 0.185208661 |
| chr4:88515657-88515981 | 0.758029922 | 0.043704912 | 0.185208661 |
| chr8:76344566-76344885 | 0.883979207 | 0.043719872 | 0.18526181 |
| chr16:73055079-73055396 | 0.746661674 | 0.043761513 | 0.185428008 |
| chr19:10942087-10942387 | 0.821670989 | 0.043768392 | 0.185446901 |
| chr17:78689143-78689354 | 0.965881734 | 0.043787358 | 0.185513863 |
| chr3:71652832-71653152 | -0.873416655 | 0.043789039 | 0.185513863 |
| chr11:267908-268227 | 0.990937779 | 0.043797219 | 0.185524428 |
| chr20:20832706-20832994 | 0.941967757 | 0.043797534 | 0.185524428 |
| chr1:32817002-32818422 | 0.64373372 | 0.043798796 | 0.185524428 |
| chr10:59865030-59865503 | -0.762897832 | 0.043802133 | 0.185528305 |
| chr3:75658253-75659133 | 0.709079507 | 0.043810627 | 0.185554023 |
| chr17:74008226-74008568 | 0.839728486 | 0.043815786 | 0.185556352 |
| chr14:41606879-41607133 | 0.835940407 | 0.04381602 | 0.185556352 |
| chr1:32712229-32712630 | 0.735392931 | 0.043818837 | 0.185558027 |
| chr1:180157414-180158106 | 0.674879071 | 0.043832278 | 0.185604688 |
| chr16:81765041-81765615 | 0.742430444 | 0.043852921 | 0.185681838 |
| chr1:145711117-145711691 | 0.690154848 | 0.043875844 | 0.185768632 |
| chr7:24381955-24382318 | 0.760420807 | 0.043882141 | 0.185785026 |

| | | | |
|---|---|---|---|
| chr3:134029442-134029758 | 0.809403252 | 0.043886175 | 0.185791841 |
| chr14:23109322-23109522 | 0.916449539 | 0.043907221 | 0.185870667 |
| chr19:52527960-52528265 | 0.811037647 | 0.043912808 | 0.185884047 |
| chr1:177256874-177257095 | -0.965978969 | 0.043923989 | 0.185921108 |
| chr2:187843339-187843788 | -0.805268073 | 0.043966718 | 0.186090858 |
| chr2:119837790-119838053 | -0.998551612 | 0.04396895 | 0.186090858 |
| chr10:33508958-33509903 | -0.688306747 | 0.043978535 | 0.186121142 |
| chr5:143222709-143223211 | -0.710950388 | 0.043982498 | 0.186127636 |
| chr19:34173223-34173597 | 0.809849286 | 0.043985943 | 0.186131932 |
| chr9:74887444-74887955 | -0.822545857 | 0.043989574 | 0.186133188 |
| chr3:31892102-31892356 | 1.025920913 | 0.043991098 | 0.186133188 |
| chr2:12931052-12931624 | -0.784602458 | 0.043995676 | 0.186134773 |
| chr20:54222375-54223568 | 0.639186792 | 0.043996331 | 0.186134773 |
| chr9:100601212-100601508 | -0.814507 | 0.044002755 | 0.1861461 |
| chr4:169088101-169088301 | -1.153157433 | 0.044006294 | 0.1861461 |
| chr2:17300196-17300454 | -1.055849054 | 0.044006297 | 0.1861461 |
| chr10:2657230-2657450 | -1.101602777 | 0.044019161 | 0.186190238 |
| chr3:151976302-151976578 | 1.002268742 | 0.044022328 | 0.186193353 |
| chr12:82896992-82897504 | 0.776101463 | 0.044030856 | 0.186219143 |
| chr8:48271517-48271824 | -1.072895051 | 0.044042868 | 0.186259665 |
| chr21:42449895-42450305 | -0.721527347 | 0.044052837 | 0.186281313 |
| chr1:209587627-209588169 | -0.708937891 | 0.044052849 | 0.186281313 |
| chr8:99492362-99492582 | 1.081300633 | 0.044069449 | 0.186341223 |
| chr19:47707183-47707648 | 0.751902146 | 0.044079901 | 0.186375135 |
| chr20:2840550-2840897 | 0.782297561 | 0.044088599 | 0.186401195 |
| chr12:125006563-125006787 | 1.032455672 | 0.044092179 | 0.186401195 |
| chr7:4286328-4286694 | -0.87055012 | 0.044093363 | 0.186401195 |
| chr20:56818981-56819573 | -0.710958565 | 0.044097079 | 0.186406619 |
| chr1:204898517-204898776 | 0.935262993 | 0.044112282 | 0.186460597 |
| chr10:80740385-80740629 | -1.035114071 | 0.044133232 | 0.186538861 |
| chr11:57434616-57434898 | -1.057524227 | 0.044139349 | 0.186554426 |
| chr4:8242305-8242526 | 0.988497292 | 0.044144869 | 0.186567464 |
| chrX:151871400-151871600 | 0.984749182 | 0.044158599 | 0.186615201 |
| chr12:51391027-51391993 | 0.663589091 | 0.044169273 | 0.186650015 |
| chr7:115679935-115680231 | -0.84511955 | 0.044173559 | 0.18665783 |
| chr11:17735016-17735216 | 0.897553339 | 0.044178941 | 0.186670277 |
| chr13:73417044-73417583 | -0.722339101 | 0.04419264 | 0.186717865 |
| chr7:76392911-76393995 | -0.648447072 | 0.044195367 | 0.186719089 |
| chr2:168211162-168211603 | -0.741588116 | 0.044243169 | 0.186910741 |
| chrX:119880826-119881179 | 0.747483012 | 0.044246169 | 0.186913108 |
| chr1:34696264-34696551 | -1.013722933 | 0.044256578 | 0.186946773 |
| chr6:74277760-74278191 | -0.806434292 | 0.044262619 | 0.186955465 |
| chr1:246723884-246725166 | -0.643722149 | 0.044263516 | 0.186955465 |
| chr5:19278283-19278502 | 0.983856008 | 0.044277996 | 0.186996049 |
| chr13:102091238-102091645 | -0.792679219 | 0.044278005 | 0.186996049 |
| chr5:132280315-132280557 | -1.023365161 | 0.044284159 | 0.187011729 |
| chr7:101817537-101817815 | -0.893146356 | 0.044288126 | 0.187018173 |
| chr11:10394693-10395193 | 0.744315951 | 0.044296229 | 0.187042086 |
| chr7:93622047-93622308 | -0.803965895 | 0.044300585 | 0.187050168 |
| chr3:14265538-14265960 | 0.733865852 | 0.044305872 | 0.187062185 |

| | | | |
|---|---|---|---|
| chr13:23147842-23148231 | -0.88611228 | 0.044311184 | 0.187074306 |
| chr5:20572627-20572956 | -0.912216135 | 0.044317031 | 0.187076866 |
| chr1:39491237-39491712 | 0.750168667 | 0.044320178 | 0.187076866 |
| chr1:147216059-147216361 | 0.893152624 | 0.044321504 | 0.187076866 |
| chr12:91423400-91423628 | -0.997499444 | 0.044321557 | 0.187076866 |
| chr14:104131055-104131330 | 0.794817307 | 0.044330514 | 0.187104365 |
| chr6:7129379-7129678 | -0.921882975 | 0.044337962 | 0.187109754 |
| chr13:40273262-40273519 | -1.003691752 | 0.044339064 | 0.187109754 |
| chr19:37293359-37293623 | 0.83060888 | 0.044339117 | 0.187109754 |
| chr17:61368266-61368592 | 0.664407345 | 0.044345153 | 0.187124921 |
| chr14:48568751-48569155 | 0.875855346 | 0.044351023 | 0.187139386 |
| chr1:24150286-24150758 | 0.809253372 | 0.044367455 | 0.187198411 |
| chr19:42313571-42313876 | 0.83609262 | 0.04437667 | 0.187226979 |
| chr3:108552640-108552996 | 0.838745657 | 0.044384806 | 0.187250999 |
| chr10:68348529-68348782 | -0.940354343 | 0.044389821 | 0.187261842 |
| chr1:242425331-242425576 | -1.033838392 | 0.044399898 | 0.187287716 |
| chrX:150364365-150364895 | 0.744909077 | 0.044400843 | 0.187287716 |
| chr14:21768196-21768617 | 0.811013418 | 0.044421729 | 0.187360558 |
| chr1:191320213-191320439 | -1.015892558 | 0.044423002 | 0.187360558 |
| chr6:32861160-32861543 | -0.817727949 | 0.044425516 | 0.18736085 |
| chr17:45680435-45680635 | 1.020643297 | 0.044428943 | 0.187364988 |
| chr4:185039852-185040062 | -1.01268566 | 0.044439587 | 0.187399562 |
| chr7:105521388-105522761 | -0.641772961 | 0.044454676 | 0.187452876 |
| chr3:122572858-122573456 | -0.7704342 | 0.044460195 | 0.187465831 |
| chr11:2193315-2193597 | -0.861315894 | 0.044472051 | 0.187505503 |
| chr17:68577088-68577511 | -0.798961526 | 0.044477528 | 0.187518277 |
| chr15:85330944-85331604 | 0.709573303 | 0.044488753 | 0.187551364 |
| chr2:138877546-138877797 | -1.075584848 | 0.044491728 | 0.187551364 |
| chr14:32211156-32211358 | 0.959399684 | 0.044494974 | 0.187551364 |
| chr6:143878342-143878710 | -0.824336949 | 0.044498431 | 0.187551364 |
| chr1:236099617-236099992 | -0.768822963 | 0.044499108 | 0.187551364 |
| chr9:134356607-134357423 | -0.664132195 | 0.044500765 | 0.187551364 |
| chr20:47195550-47195799 | 0.678995433 | 0.04450251 | 0.187551364 |
| chr2:95292783-95293274 | -0.747165815 | 0.04451147 | 0.187578807 |
| chr11:21018191-21018435 | -1.093549577 | 0.044520894 | 0.187608201 |
| chr14:33792765-33792965 | 0.947522158 | 0.044544989 | 0.187699416 |
| chr22:36462175-36462430 | 0.889711737 | 0.044557849 | 0.18773561 |
| chr7:106768362-106768611 | -1.138223769 | 0.044559605 | 0.18773561 |
| chr6:104710562-104710876 | -0.812531455 | 0.04456093 | 0.18773561 |
| chr16:30856518-30857243 | 0.706514426 | 0.044565699 | 0.18774538 |
| chr8:130032781-130033750 | 0.667711413 | 0.044579261 | 0.187776248 |
| chr13:47171566-47171829 | 0.935887917 | 0.044580205 | 0.187776248 |
| chr12:51372387-51372709 | 0.973361094 | 0.044580378 | 0.187776248 |
| chrX:9267426-9267843 | -0.784799903 | 0.044589895 | 0.187797724 |
| chr2:203394621-203394902 | -0.905952107 | 0.044592167 | 0.187797724 |
| chr5:142888590-142888816 | -1.021177479 | 0.044592829 | 0.187797724 |
| chr9:119555894-119556833 | -0.698877192 | 0.044598034 | 0.187809318 |
| chr2:191571056-191571313 | -1.00937473 | 0.044601649 | 0.18781422 |
| chr10:31759162-31760568 | -0.63918329 | 0.044612479 | 0.187849502 |
| chr2:217672448-217672900 | 0.740891976 | 0.044621821 | 0.18787001 |

| | | | |
|---|---|---|---|
| chr16:87471568-87471780 | 0.986461924 | 0.044622254 | 0.18787001 |
| chr1:246748915-246749234 | -0.832241454 | 0.04462473 | 0.187870114 |
| chr21:37563723-37565087 | 0.6453575 | 0.044631279 | 0.187878421 |
| chr5:67244255-67244567 | -0.956907561 | 0.044631607 | 0.187878421 |
| chr1:150548702-150549510 | 0.668573887 | 0.044643839 | 0.187919586 |
| chr1:174144092-174144644 | -0.800698512 | 0.044655979 | 0.187960361 |
| chr8:23854761-23854990 | -0.984128722 | 0.044659846 | 0.187966314 |
| chrX:27683830-27684085 | -0.99161396 | 0.044665104 | 0.187978115 |
| chr1:215599463-215599715 | -0.901488818 | 0.04467098 | 0.187992521 |
| chr1:36516409-36516613 | 0.917637299 | 0.044677842 | 0.188008561 |
| chr13:51330868-51331129 | -1.03558942 | 0.044682153 | 0.188008561 |
| chr21:38296902-38297168 | -1.03558942 | 0.044682153 | 0.188008561 |
| chr5:22242635-22242841 | -1.116299351 | 0.044691349 | 0.188036425 |
| chr14:39338961-39339365 | -0.83097792 | 0.044693683 | 0.188036425 |
| chr5:1799342-1799584 | 0.790988846 | 0.044703993 | 0.188063928 |
| chr3:170832441-170832655 | 1.076006543 | 0.044705129 | 0.188063928 |
| chr17:62110908-62113308 | 0.62588677 | 0.044711256 | 0.188079376 |
| chr9:111524953-111526165 | -0.658478887 | 0.044733678 | 0.188163367 |
| chr2:765772-766051 | -0.983735601 | 0.044738265 | 0.188172332 |
| chr15:74307882-74308337 | 0.763429164 | 0.044750358 | 0.188212864 |
| chr15:33756549-33756749 | 1.088816276 | 0.044761874 | 0.188250962 |
| chr7:128060105-128060353 | -1.003532209 | 0.044772251 | 0.188284273 |
| chr13:34560630-34560901 | 1.017464387 | 0.044776624 | 0.188292327 |
| chr13:29354460-29354742 | 0.903261486 | 0.044779978 | 0.188296097 |
| chr6:112116272-112116611 | -0.831795256 | 0.044796903 | 0.188356927 |
| chr12:115440446-11544077 | 1.001704919 | 0.044805386 | 0.188382259 |
| chr4:86223476-86223866 | -0.766312612 | 0.044821634 | 0.188423557 |
| chr5:83799469-83799742 | -0.991316599 | 0.044823929 | 0.188423557 |
| chr7:39732913-39733764 | -0.641673865 | 0.04482578 | 0.188423557 |
| chrX:53048333-53048787 | -0.816100343 | 0.044826217 | 0.188423557 |
| chr10:60901886-60902086 | -1.051418724 | 0.044827504 | 0.188423557 |
| chr2:71416797-71417094 | -0.94173288 | 0.04483197 | 0.188431993 |
| chr7:8823540-8823748 | -1.047525457 | 0.044835589 | 0.188436864 |
| chr9:105296927-105297212 | -0.997915118 | 0.044846216 | 0.188471193 |
| chr1:11262087-11262298 | -1.073739935 | 0.044876328 | 0.188577761 |
| chr8:65041649-65042093 | -0.714310939 | 0.044876496 | 0.188577761 |
| chr20:47434110-47434582 | 0.644332442 | 0.044887491 | 0.188613618 |
| chr2:102020661-102021094 | 0.757568547 | 0.04489962 | 0.188654236 |
| chr15:66876119-66876366 | 1.033260846 | 0.044903103 | 0.188658528 |
| chr9:130432457-130432742 | -0.855350097 | 0.044909021 | 0.188673048 |
| chr2:43840904-43841182 | 0.93192061 | 0.044925796 | 0.188709076 |
| chr17:76102960-76103166 | 0.911695447 | 0.044926809 | 0.188709076 |
| chr3:27570765-27571000 | -0.977412932 | 0.044929381 | 0.188709076 |
| chr6:37625009-37625243 | 0.888635576 | 0.044929494 | 0.188709076 |
| chr20:52937795-52938172 | -0.79252522 | 0.044929911 | 0.188709076 |
| chr20:645532-645874 | 0.855673372 | 0.044932398 | 0.188709176 |
| chr4:28614-29478 | 0.691723549 | 0.044939071 | 0.188726858 |
| chr1:206001476-206001747 | -0.788916323 | 0.044955236 | 0.188784398 |
| chr3:114928997-114929205 | 1.059594032 | 0.044961199 | 0.188799089 |
| chr13:29918213-29919098 | -0.669517494 | 0.044966765 | 0.188812116 |

| | | | |
|---|---|---|---|
| chr1:15424256-15424597 | -0.977391181 | 0.044975323 | 0.188837702 |
| chr10:9331593-9331848 | 0.869956172 | 0.044993029 | 0.188901691 |
| chr7:116446847-116447048 | -1.046720928 | 0.045002727 | 0.188930294 |
| chr6:6820916-6821247 | -0.884040479 | 0.045004773 | 0.188930294 |
| chr15:85413493-85413715 | 1.09221885 | 0.045008478 | 0.188935494 |
| chr6:145267100-145267374 | -1.042782398 | 0.04504702 | 0.189086929 |
| chr1:20732077-20732334 | 1.024786718 | 0.045050696 | 0.189091999 |
| chr1:7042850-7043121 | 0.997538714 | 0.04507334 | 0.189176679 |
| chr19:50266683-50267360 | 0.69163157 | 0.045076691 | 0.189180382 |
| chr2:188513928-188514268 | -0.814839978 | 0.045084336 | 0.189202103 |
| chr1:27542579-27542824 | 0.915763312 | 0.045090281 | 0.189216686 |
| chr22:43232734-43233035 | 0.959977307 | 0.045099843 | 0.189234937 |
| chr2:166181355-166181879 | -0.722658885 | 0.04510742 | 0.189234937 |
| chr7:6489649-6489887 | -0.900591043 | 0.045108938 | 0.189234937 |
| chr7:10637127-10637383 | 0.906680582 | 0.045110753 | 0.189234937 |
| chr9:105159670-105159988 | -0.850706079 | 0.045111754 | 0.189234937 |
| chr9:33044092-33045019 | -0.67685891 | 0.045113565 | 0.189234937 |
| chr7:111486641-111486856 | -1.042655445 | 0.045113755 | 0.189234937 |
| chr4:6110346-6110600 | 0.852215452 | 0.045114387 | 0.189234937 |
| chr14:90397409-90398137 | 0.672268151 | 0.045122103 | 0.18925694 |
| chr4:139214077-139214436 | -0.846368707 | 0.045128287 | 0.189272518 |
| chr1:218306252-218306524 | 0.880000548 | 0.045140445 | 0.189311239 |
| chr19:37469143-37469671 | -0.801098393 | 0.045142461 | 0.189311239 |
| chr7:96314366-96314774 | -0.804935813 | 0.04514853 | 0.189326327 |
| chr12:131711764-131712101 | 0.834420687 | 0.045160426 | 0.189365851 |
| chr1:201888575-201888775 | 0.881796961 | 0.045167084 | 0.189383403 |
| chr11:61513225-61513765 | -0.703872836 | 0.045176633 | 0.189409886 |
| chr1:211698493-211698908 | -0.767000023 | 0.045178887 | 0.189409886 |
| chr20:53988059-53988299 | 0.690739073 | 0.045181948 | 0.189409886 |
| chr13:36858213-36858707 | -0.733639037 | 0.045185169 | 0.189409886 |
| chr17:45430830-45432200 | 0.64620855 | 0.04518576 | 0.189409886 |
| chr2:27753914-27754387 | -0.788788939 | 0.045194079 | 0.18942433 |
| chr10:20531173-20531923 | -0.701105095 | 0.04519765 | 0.18942433 |
| chr21:45111794-45111994 | -1.180809095 | 0.045197761 | 0.18942433 |
| chr3:185677165-185677541 | 0.793158808 | 0.045199095 | 0.18942433 |
| chr14:42756130-42756438 | -0.932334142 | 0.045203551 | 0.189432643 |
| chr10:87038431-87038671 | 0.922790275 | 0.045208921 | 0.189444787 |
| chr19:2491380-2491597 | 0.933551732 | 0.04521228 | 0.189448504 |
| chr6:82688551-82688823 | 0.973671681 | 0.045218384 | 0.18946372 |
| chr7:127588016-127589137 | -0.641880855 | 0.045226103 | 0.189485619 |
| chr6:108589327-108589786 | -0.802498334 | 0.045228557 | 0.189485619 |
| chr14:30375311-30375683 | 0.748308681 | 0.045231353 | 0.189486974 |
| chr22:33861370-33861691 | -0.945810797 | 0.04523742 | 0.189502028 |
| chr7:88835195-88835395 | -1.203001113 | 0.045262576 | 0.189597042 |
| chr14:74302106-74302458 | 0.743454325 | 0.045266606 | 0.189603558 |
| chr17:61589263-61590336 | 0.632012943 | 0.045269111 | 0.189603684 |
| chrX:119488849-119489123 | 0.910568086 | 0.04527236 | 0.189606929 |
| chr1:236141996-236142871 | -0.680750162 | 0.045284313 | 0.189646622 |
| chr10:97718597-97719141 | -0.685145262 | 0.045287445 | 0.189649375 |
| chr7:112257164-112257364 | -1.196778277 | 0.0452919 | 0.189657665 |

| | | | |
|---|---|---|---|
| chr6:29014726-29015191 | 0.771534853 | 0.045302352 | 0.189691066 |
| chr10:58189267-58189725 | -0.820044408 | 0.045316134 | 0.189738406 |
| chr16:29594681-29594881 | 1.038420586 | 0.045321616 | 0.18975099 |
| chr3:185519176-185519384 | -1.068267112 | 0.045339468 | 0.189806791 |
| chr7:73762276-73762646 | -0.961792865 | 0.045339898 | 0.189806791 |
| chr4:56762573-56762773 | 1.006401364 | 0.045343897 | 0.18981316 |
| chr1:101002664-101002872 | -1.089406214 | 0.045352268 | 0.189837831 |
| chr12:51401186-51401550 | -0.907578358 | 0.045369995 | 0.18990166 |
| chr7:150738856-150739273 | -1.031013027 | 0.045382981 | 0.189945636 |
| chr16:30786371-30786728 | 0.769794534 | 0.045391695 | 0.189971732 |
| chr11:16520504-16521369 | -0.685800336 | 0.045410712 | 0.190040939 |
| chr11:10405986-10406188 | 1.133035125 | 0.045413904 | 0.190043919 |
| chr12:123270594-123270967 | -0.794183967 | 0.045428096 | 0.190092926 |
| chr6:21738261-21739160 | -0.653943156 | 0.045435513 | 0.19011358 |
| chr16:29995989-29996592 | 0.695024967 | 0.045453977 | 0.190180454 |
| chr8:57209606-57210511 | 0.673668784 | 0.045463462 | 0.19020975 |
| chr3:150808349-150808616 | 0.990322031 | 0.045471246 | 0.190230047 |
| chr6:142551383-142551633 | -0.989822698 | 0.045473383 | 0.190230047 |
| chr12:32678947-32679932 | -0.650330408 | 0.045475761 | 0.190230047 |
| chr9:68746888-68747270 | -0.873340558 | 0.045480583 | 0.190239831 |
| chr6:69179654-69180152 | -0.713153934 | 0.045488796 | 0.190253432 |
| chr4:3106625-3106888 | 0.886297329 | 0.045488801 | 0.190253432 |
| chr21:8216799-8217159 | 0.669630826 | 0.045494329 | 0.190266167 |
| chr12:96315767-96316498 | -0.706243765 | 0.045500729 | 0.190282551 |
| chr8:140821548-140821748 | 1.075354157 | 0.045516407 | 0.190337724 |
| chr1:16351931-16353077 | 0.67334237 | 0.045521829 | 0.190350008 |
| chr11:68732412-68732612 | 1.018890741 | 0.045537183 | 0.190395587 |
| chr7:105006448-105007086 | -0.70796933 | 0.04553799 | 0.190395587 |
| chr4:158768518-158769136 | -0.751060276 | 0.045540183 | 0.190395587 |
| chr7:98520557-98520940 | -0.929988699 | 0.045542982 | 0.190396901 |
| chr6:57216173-57216394 | -1.065425913 | 0.045556139 | 0.190441511 |
| chr4:8315485-8316238 | 0.695382037 | 0.045567061 | 0.190470962 |
| chr9:32551685-32551939 | -0.822581088 | 0.045568155 | 0.190470962 |
| chr8:118344007-118344369 | -0.785187086 | 0.04558122 | 0.190515178 |
| chr17:54937470-54937868 | -0.931969905 | 0.045592026 | 0.19054995 |
| chr17:2346441-2346906 | 0.736090522 | 0.045599096 | 0.190569102 |
| chr4:124974044-124974625 | -0.768291023 | 0.04561273 | 0.190615687 |
| chr12:122023350-122023654 | 0.806485669 | 0.045615817 | 0.190618188 |
| chr3:14791368-14791657 | 0.753034523 | 0.045636655 | 0.190694869 |
| chr3:123515422-123515622 | 1.036867268 | 0.045639691 | 0.190697153 |
| chr7:142445087-142445339 | -1.054195145 | 0.045644756 | 0.19069752 |
| chr7:143256437-143256676 | -1.054195145 | 0.045644756 | 0.19069752 |
| chr4:2009886-2010106 | 0.939482332 | 0.045656391 | 0.190735729 |
| chr3:141698945-141699188 | 0.971553859 | 0.045661636 | 0.190744485 |
| chr5:88602599-88602997 | -0.839295607 | 0.045663466 | 0.190744485 |
| chr9:111760169-111760369 | -1.13245917 | 0.045674613 | 0.190780645 |
| chr18:61901373-61901914 | -0.746486989 | 0.045684282 | 0.190802512 |
| chr2:567479-568162 | -0.719925979 | 0.045684828 | 0.190802512 |
| chr8:100564045-100564302 | 0.765150029 | 0.045694171 | 0.190831131 |
| chr7:16644948-16646750 | 0.625814172 | 0.045699592 | 0.190843368 |

| | | | |
|---|---|---|---|
| chr4:33316629-33316923 | -0.949348011 | 0.045703242 | 0.190848209 |
| chr1:201463058-201463347 | 0.748419778 | 0.045710792 | 0.190869334 |
| chr5:153147990-153148729 | -0.704168829 | 0.045725041 | 0.190918431 |
| chr5:15912731-15912931 | 1.149271186 | 0.045729295 | 0.190918641 |
| chr20:29079974-29080424 | 0.757349925 | 0.045730829 | 0.190918641 |
| chr4:120648713-120649098 | 0.798822237 | 0.045732567 | 0.190918641 |
| chr7:123916474-123916732 | -0.919316522 | 0.045743062 | 0.190952051 |
| chr8:122995300-122995566 | 0.854202197 | 0.045763655 | 0.19102761 |
| chr10:97185678-97186133 | 0.755600699 | 0.04576953 | 0.19102961 |
| chr1:157234998-157235259 | 0.897090621 | 0.045771931 | 0.19102961 |
| chr3:161415584-161416222 | 0.66290428 | 0.045773979 | 0.19102961 |
| chr1:192429637-192429856 | 1.028032984 | 0.04577542 | 0.19102961 |
| chr14:23649800-23650171 | -0.841625504 | 0.0457766 | 0.19102961 |
| chr2:32940975-32941372 | -0.760818233 | 0.045783497 | 0.191041682 |
| chr20:14337507-14337860 | 0.788602457 | 0.04578448 | 0.191041682 |
| chr13:93375520-93375724 | -1.118423976 | 0.04579175 | 0.191061615 |
| chr2:10386902-10387337 | 0.768299059 | 0.045799716 | 0.191084445 |
| chr1:110291439-110291673 | -1.093406591 | 0.04580395 | 0.191091704 |
| chr16:24523364-24523686 | 0.901439479 | 0.045813308 | 0.191120342 |
| chr1:29324727-29325172 | -0.773199886 | 0.045827195 | 0.191167863 |
| chr3:170488848-170489169 | -0.788516056 | 0.045833725 | 0.191184696 |
| chr18:76349140-76349352 | -0.87318816 | 0.04584021 | 0.191201337 |
| chr5:83714737-83715152 | -0.72000503 | 0.045857502 | 0.19126305 |
| chr2:26724156-26724381 | -1.006127507 | 0.045862693 | 0.191272762 |
| chr3:71258101-71258585 | -0.824936753 | 0.045864823 | 0.191272762 |
| chr5:18745410-18745994 | -0.692483367 | 0.045867465 | 0.19127337 |
| chr16:75323401-75323601 | -1.114162607 | 0.045885221 | 0.191337 |
| chr1:215432820-215433480 | -0.725820215 | 0.0458927 | 0.191357776 |
| chr15:68554068-68554353 | 0.800882456 | 0.04589638 | 0.191362705 |
| chr17:10618166-10618521 | -0.76002011 | 0.045903189 | 0.191372611 |
| chr5:18984496-18985415 | -0.69342253 | 0.045904226 | 0.191372611 |
| chr7:55255512-55256083 | -0.688194447 | 0.045906249 | 0.191372611 |
| chr2:6772459-6772836 | -0.8087467 | 0.045916488 | 0.191404884 |
| chr8:67902093-67902527 | -0.859732977 | 0.045922547 | 0.191413692 |
| chr9:28127049-28127327 | -1.074957771 | 0.045923868 | 0.191413692 |
| chr4:67711777-67712591 | -0.680437289 | 0.045926096 | 0.191413692 |
| chr13:99617446-99617660 | -1.042415426 | 0.04593362 | 0.191423996 |
| chr3:186644755-186645042 | -0.936443298 | 0.045936181 | 0.191423996 |
| chr12:110894433-110894740 | 0.787253152 | 0.045937924 | 0.191423996 |
| chrX:9709058-9709339 | 0.90980065 | 0.045940965 | 0.191423996 |
| chr7:13990424-13990870 | 0.68741122 | 0.04594106 | 0.191423996 |
| chr10:1368867-1369179 | 0.790339268 | 0.045947771 | 0.1914381 |
| chr9:5879493-5879831 | -0.858650876 | 0.045949441 | 0.1914381 |
| chr11:5624646-5625262 | -0.749062522 | 0.04595331 | 0.191440999 |
| chr4:133483646-133483961 | -0.812550186 | 0.045955227 | 0.191440999 |
| chr5:158695558-158695821 | -0.884552308 | 0.045959623 | 0.191440999 |
| chr5:181086017-181086252 | 0.988241709 | 0.045960131 | 0.191440999 |
| chr4:63717512-63717712 | 1.108181936 | 0.045981674 | 0.191519602 |
| chr13:44815732-44816064 | -0.757997646 | 0.045984001 | 0.191519602 |
| chrX:12814537-12814739 | -1.127071787 | 0.045992395 | 0.191544151 |

| | | | |
|---|---|---|---|
| chr6:26064623-26064823 | -1.039280159 | 0.0460034 | 0.191577863 |
| chr11:61891494-61892008 | 0.705587287 | 0.04600549 | 0.191577863 |
| chr15:98338608-98338811 | 1.050913494 | 0.046017349 | 0.191616832 |
| chr5:132227302-132228423 | -0.6616584 | 0.046023888 | 0.191633646 |
| chr1:191528282-191528680 | -0.756796006 | 0.046027585 | 0.191638625 |
| chr5:38533714-38534035 | -1.021914128 | 0.046047937 | 0.191712944 |
| chr7:37549821-37550112 | -0.922369081 | 0.046051689 | 0.191718147 |
| chr20:38599431-38599969 | 0.723137853 | 0.046069636 | 0.191772089 |
| chr8:15238286-15238600 | -0.861084353 | 0.04606996 | 0.191772089 |
| chr8:140048938-140049138 | 0.941151229 | 0.046075136 | 0.191772089 |
| chr12:101414630-101414874 | -0.943918811 | 0.046075735 | 0.191772089 |
| chr15:40928877-40929505 | 0.681419735 | 0.04607716 | 0.191772089 |
| chr10:112769594-112769918 | -0.88327227 | 0.046082006 | 0.191781841 |
| chr5:95440882-95441095 | -1.061321419 | 0.046087414 | 0.19179393 |
| chr1:146771415-146771780 | -0.856135735 | 0.046094298 | 0.191794621 |
| chr4:152995742-152996507 | -0.677204646 | 0.04609638 | 0.191794621 |
| chr3:95348077-95348277 | 1.067816697 | 0.046098044 | 0.191794621 |
| chr7:153190572-153190811 | -0.972878907 | 0.046098289 | 0.191794621 |
| chr7:101944997-101945328 | -0.815670233 | 0.046100096 | 0.191794621 |
| chr1:207001943-207002157 | -0.937503513 | 0.046108576 | 0.19181597 |
| chr18:864847-865088 | -0.939758236 | 0.046110234 | 0.19181597 |
| chr11:113906209-113906498 | 0.885687232 | 0.046117031 | 0.19183383 |
| chr8:108582536-108583049 | -0.706917437 | 0.046123863 | 0.191851833 |
| chr20:32079736-32080070 | -0.886579435 | 0.046143091 | 0.191921393 |
| chr5:165757765-165758203 | -0.876823515 | 0.046159825 | 0.191980573 |
| chr9:88984622-88984879 | 0.962152323 | 0.046165618 | 0.191994247 |
| chr11:34242334-34242700 | 0.916562669 | 0.046172092 | 0.192010748 |
| chr11:73342599-73342799 | 0.921169037 | 0.046189346 | 0.19205992 |
| chr6:7347480-7348123 | -0.768359292 | 0.046194128 | 0.19205992 |
| chr2:37671551-37672932 | -0.660369049 | 0.046194344 | 0.19205992 |
| chr13:77698200-77698627 | -0.756513957 | 0.046196183 | 0.19205992 |
| chr6:128303033-128303503 | -0.849164422 | 0.046196449 | 0.19205992 |
| chr13:28329152-28329387 | 1.029816484 | 0.046203719 | 0.192078382 |
| chr20:4085252-4085853 | 0.721093675 | 0.046205904 | 0.192078382 |
| chr20:16222866-16223104 | 1.026058185 | 0.046214689 | 0.192099315 |
| chr2:173532671-173533412 | -0.753291085 | 0.046215953 | 0.192099315 |
| chr1:11736391-11736759 | 0.798698425 | 0.046219707 | 0.192104498 |
| chr8:106769374-106769586 | 0.879346432 | 0.04622333 | 0.192109134 |
| chr12:61090305-61090544 | 1.065006509 | 0.046238589 | 0.192155196 |
| chr2:37890366-37890756 | -0.942480925 | 0.046239429 | 0.192155196 |
| chr1:113750012-113750232 | 0.87018309 | 0.046243992 | 0.192163737 |
| chr5:72920509-72921016 | -0.830694033 | 0.046258779 | 0.192214759 |
| chr19:46704331-46704923 | -0.755956635 | 0.046262248 | 0.192218752 |
| chr14:66316292-66316512 | 0.983750644 | 0.046271431 | 0.192246478 |
| chr9:9914238-9914491 | -1.013163491 | 0.046279997 | 0.192270116 |
| chr2:21413198-21413436 | 1.012305953 | 0.046282139 | 0.192270116 |
| chr15:70132183-70132402 | 0.975057405 | 0.046303612 | 0.192348896 |
| chr2:14945140-14945400 | -1.042888548 | 0.046308962 | 0.19236069 |
| chr9:136439515-136440070 | 0.714431994 | 0.046313554 | 0.192369335 |
| chr14:40005117-40005496 | -0.784104325 | 0.04631915 | 0.192372888 |

| | | | |
|---|---|---|---|
| chr6:130320898-130321190 | 0.895255965 | 0.046319431 | 0.192372888 |
| chr12:25405018-25405302 | -0.877786926 | 0.046327873 | 0.19239752 |
| chr6:163352342-163352807 | 0.759548541 | 0.046346668 | 0.19245589 |
| chr4:6625205-6625547 | 0.948418069 | 0.046348815 | 0.19245589 |
| chr7:139118461-139118661 | -1.125269197 | 0.046349463 | 0.19245589 |
| chr17:16887438-16887671 | -1.111830122 | 0.046352258 | 0.192457067 |
| chr13:45728762-45729367 | -0.69034627 | 0.046364592 | 0.192497847 |
| chr18:63949112-63949734 | -0.733390536 | 0.046369301 | 0.192504943 |
| chr2:66024549-66024749 | -1.088005613 | 0.046371683 | 0.192504943 |
| chr20:54316198-54316429 | 0.712658075 | 0.046373838 | 0.192504943 |
| chr10:72109832-72110032 | -1.090585093 | 0.046378263 | 0.192512882 |
| chr11:24310497-24310841 | -1.010618817 | 0.04638809 | 0.192543241 |
| chr8:57113776-57114033 | -0.854987983 | 0.046396565 | 0.192567985 |
| chr6:26556458-26556919 | -0.680842432 | 0.046405659 | 0.192589412 |
| chr16:85637835-85638119 | -0.791407909 | 0.046406754 | 0.192589412 |
| chr20:46021512-46021729 | 0.781716242 | 0.046414367 | 0.192610572 |
| chr8:28799862-28800222 | -0.86022969 | 0.046424392 | 0.192635958 |
| chr14:57182914-57183114 | 1.04224674 | 0.046425512 | 0.192635958 |
| chr3:108305570-108305834 | 0.942962408 | 0.046439779 | 0.192684721 |
| chr15:58346247-58346659 | 0.789450741 | 0.046449187 | 0.192713322 |
| chr2:63698874-63699373 | -0.743490051 | 0.046467218 | 0.19277769 |
| chr5:123937256-123937527 | -0.924040327 | 0.046482482 | 0.192830578 |
| chr16:3023363-3024461 | 0.640589713 | 0.046487242 | 0.192833791 |
| chr3:197541190-197541406 | 0.871002441 | 0.04648829 | 0.192833791 |
| chr19:2051129-2051526 | 0.731861778 | 0.046494945 | 0.192843385 |
| chrX:44675390-44675703 | -0.953558749 | 0.046495637 | 0.192843385 |
| chr2:165698457-165698931 | -0.80727917 | 0.046499554 | 0.192845818 |
| chr15:56491162-56491432 | 0.800254964 | 0.046507368 | 0.192845818 |
| chr1:206872143-206872459 | -0.776969113 | 0.046508241 | 0.192845818 |
| chr1:246699072-246699533 | -0.657545987 | 0.046508796 | 0.192845818 |
| chr10:63629431-63631021 | -0.626417605 | 0.046509989 | 0.192845818 |
| chr18:61882618-61882897 | -1.044681468 | 0.046513798 | 0.192845818 |
| chrX:106118780-106119342 | 0.718625976 | 0.04651401 | 0.192845818 |
| chr1:113917430-113918193 | 0.649894219 | 0.046516358 | 0.192845818 |
| chr7:101163531-101163842 | 0.74616864 | 0.046521483 | 0.19285663 |
| chr14:20841123-20841335 | 1.056805706 | 0.046525558 | 0.192858177 |
| chr4:158170520-158171468 | -0.658746869 | 0.04653095 | 0.192858177 |
| chr14:91555704-91555918 | 0.963654777 | 0.046533054 | 0.192858177 |
| chr13:38647264-38647743 | -0.737723066 | 0.046533787 | 0.192858177 |
| chr7:142421024-142421320 | -0.873173848 | 0.046536956 | 0.192858177 |
| chr3:23501445-23501727 | -0.971449726 | 0.046538365 | 0.192858177 |
| chr12:119716483-119716933 | 0.765204109 | 0.046539476 | 0.192858177 |
| chr20:56628431-56628830 | 0.675153916 | 0.046544709 | 0.192867332 |
| chr17:73290387-73290840 | -0.747228658 | 0.046546719 | 0.192867332 |
| chr9:70258450-70259648 | -0.636713584 | 0.046559233 | 0.192903216 |
| chr6:43076086-43076379 | 0.852414467 | 0.046560415 | 0.192903216 |
| chr17:16486474-16487045 | -0.74846086 | 0.046567534 | 0.192922281 |
| chr17:78791900-78792172 | 0.874080535 | 0.046596414 | 0.193031491 |
| chr4:101410966-101411832 | -0.699447857 | 0.046601513 | 0.193042177 |
| chr1:216310798-216311255 | -0.728344292 | 0.046608591 | 0.19306106 |

| | | | |
|---|---|---|---|
| chr3:176868059-176868686 | -0.707747865 | 0.046614281 | 0.193074191 |
| chr12:26043196-26043546 | -0.939700126 | 0.046619575 | 0.193084515 |
| chr2:25590230-25590572 | -0.942159137 | 0.046621814 | 0.193084515 |
| chr1:153229317-153229595 | 0.856677797 | 0.04663574 | 0.193127082 |
| chrX:30060830-30061096 | -0.902567684 | 0.04663759 | 0.193127082 |
| chr17:74296622-74296822 | 0.938314638 | 0.046639654 | 0.193127082 |
| chr11:10304662-10305172 | 0.721273271 | 0.046644304 | 0.193135902 |
| chr14:47899183-47899458 | -0.972595384 | 0.046647767 | 0.193139802 |
| chr7:34158817-34159103 | -0.859492815 | 0.046652834 | 0.193150345 |
| chr6:141677693-141678091 | -0.943269215 | 0.046662434 | 0.193158677 |
| chr17:52828630-52829000 | -0.719709572 | 0.046662689 | 0.193158677 |
| chr7:145653148-145653708 | -0.785905235 | 0.046664722 | 0.193158677 |
| chr10:60807394-60807859 | -0.82372958 | 0.04666493 | 0.193158677 |
| chr9:104968133-104969193 | -0.663568652 | 0.046678462 | 0.193204254 |
| chr3:169769269-169769729 | -0.668261463 | 0.046682685 | 0.193211294 |
| chr9:37582631-37583016 | -0.967705327 | 0.046707502 | 0.193303566 |
| chr4:82360567-82361025 | -0.812700507 | 0.046716459 | 0.193326373 |
| chr15:39668475-39668805 | -0.862811351 | 0.046718113 | 0.193326373 |
| chr14:101743752-101743977 | 1.031491719 | 0.046720582 | 0.193326373 |
| chr10:7475629-7475829 | 1.039646197 | 0.046725302 | 0.193333291 |
| chr3:48232715-48232988 | 0.877982039 | 0.046727833 | 0.193333291 |
| chr2:9473506-9475131 | -0.638274004 | 0.046729824 | 0.193333291 |
| chr5:175460148-175460444 | 0.822608952 | 0.046735438 | 0.19334608 |
| chr12:10901526-10901789 | -0.950334874 | 0.046739057 | 0.193350611 |
| chr9:33925248-33925516 | -0.938013759 | 0.04674535 | 0.193350811 |
| chr8:13535943-13536234 | -1.03223565 | 0.046746041 | 0.193350811 |
| chr11:86592939-86593167 | 0.932171606 | 0.046746675 | 0.193350811 |
| chr8:88249459-88249833 | 0.78995608 | 0.046759002 | 0.193391357 |
| chr1:39901814-39902577 | 0.663549382 | 0.046769175 | 0.193422988 |
| chr9:130243287-130243487 | 0.959374232 | 0.04678317 | 0.193459997 |
| chrX:33434160-33434383 | -1.095403163 | 0.046783173 | 0.193459997 |
| chr17:49508126-49508735 | -0.696691653 | 0.046788599 | 0.193471995 |
| chr10:118459661-118459976 | -0.925144463 | 0.046815298 | 0.193571949 |
| chr8:138260153-138260605 | -0.842017284 | 0.046822509 | 0.193591319 |
| chr10:52949952-52950185 | -0.907087703 | 0.046829406 | 0.193609387 |
| chr12:87509494-87509710 | -1.041866849 | 0.046833452 | 0.193615668 |
| chr10:52817964-52818389 | -0.774907988 | 0.046837674 | 0.193622674 |
| chr20:47204285-47204767 | 0.635729736 | 0.046843453 | 0.193636117 |
| chr20:25206988-25207577 | -0.747481796 | 0.046850045 | 0.193652919 |
| chr12:71041614-71041853 | 1.038473996 | 0.04685436 | 0.193656677 |
| chr7:1570897-1571170 | -0.789049813 | 0.046856009 | 0.193656677 |
| chr16:47460791-47461690 | 0.676429663 | 0.046863894 | 0.193678817 |
| chr18:46249613-46249929 | -1.004614249 | 0.046871781 | 0.193700965 |
| chr8:143953724-143953933 | 0.78307312 | 0.046882326 | 0.193728392 |
| chr12:11730237-11730659 | -0.788407721 | 0.046883474 | 0.193728392 |
| chr16:77951810-77952133 | 0.756497666 | 0.046886026 | 0.193728489 |
| chrX:9351743-9352226 | -0.761073581 | 0.046888901 | 0.193729922 |
| chr4:7042412-7043383 | 0.652178173 | 0.046898474 | 0.193748935 |
| chr19:19203302-19203604 | -0.846175036 | 0.046898561 | 0.193748935 |
| chr6:26440256-26440861 | -0.787290699 | 0.046904141 | 0.193761542 |

| | | | |
|---|---|---|---|
| chr17:59016752-59016994 | 0.714644561 | 0.046907449 | 0.193764761 |
| chr14:75634746-75635255 | 0.704021281 | 0.04691257 | 0.19377547 |
| chr21:39385754-39386074 | -0.755228836 | 0.046924472 | 0.19381418 |
| chr7:138663678-138664715 | -0.675485203 | 0.046928554 | 0.193820596 |
| chr8:87127940-87128168 | 0.972074061 | 0.046931714 | 0.193823198 |
| chr16:34587915-34588454 | 0.643521627 | 0.046938464 | 0.193839951 |
| chr11:31487929-31488177 | 0.887952389 | 0.046941018 | 0.193839951 |
| chr4:128379909-128380190 | 0.796108793 | 0.04694336 | 0.193839951 |
| chr7:104862712-104863082 | -0.774827626 | 0.046946046 | 0.193840598 |
| chr1:239976474-239976877 | -0.870849416 | 0.046972421 | 0.193939046 |
| chr5:133848177-133848584 | 0.799732237 | 0.04698262 | 0.193970704 |
| chr5:160311996-160312954 | -0.667165659 | 0.047003429 | 0.194037461 |
| chr10:107230964-107231357 | -0.87421251 | 0.047004997 | 0.194037461 |
| chr16:89921813-89922030 | -0.788517129 | 0.047006387 | 0.194037461 |
| chr8:142728332-142728656 | 0.742256148 | 0.047009484 | 0.194039793 |
| chr6:76276278-76276533 | -1.019678023 | 0.047017318 | 0.194061677 |
| chr10:102451283-102451621 | 0.776543381 | 0.047023446 | 0.194076514 |
| chr5:95834622-95835276 | -0.669765865 | 0.047045581 | 0.194157411 |
| chr10:66272369-66272666 | -1.007329114 | 0.047048404 | 0.194158604 |
| chr20:41003071-41003914 | 0.648196632 | 0.047057499 | 0.194185678 |
| chr15:71295076-71295933 | 0.689183041 | 0.047065728 | 0.194197178 |
| chr7:140343190-140343471 | -0.93059362 | 0.047066954 | 0.194197178 |
| chr12:95942467-95943715 | -0.64583618 | 0.047067889 | 0.194197178 |
| chr7:152030180-152030400 | -0.878135622 | 0.047082892 | 0.194248621 |
| chr4:2583575-2583977 | 0.778589061 | 0.047100731 | 0.194311755 |
| chr2:168336555-168337217 | -0.660330829 | 0.047116483 | 0.194366274 |
| chr12:84856705-84856936 | 1.008173489 | 0.047132072 | 0.194420115 |
| chr11:71343571-71343892 | 0.782920099 | 0.047140929 | 0.194446179 |
| chr3:142270461-142270834 | 0.88899177 | 0.047145546 | 0.194447884 |
| chr19:32580566-32581685 | 0.644530167 | 0.047146417 | 0.194447884 |
| chr22:30290906-30291259 | 0.758838846 | 0.047157972 | 0.194478439 |
| chr5:62791128-62791364 | -0.888640588 | 0.047158902 | 0.194478439 |
| chr3:8741557-8741943 | -0.746501153 | 0.047169391 | 0.194506226 |
| chr11:47143324-47143794 | 0.725864665 | 0.047170717 | 0.194506226 |
| chr16:4307410-4308183 | 0.649980943 | 0.047186947 | 0.194562678 |
| chr14:104815462-104816424 | 0.646294972 | 0.047223211 | 0.194701532 |
| chr16:80792550-80792956 | 0.869914744 | 0.047225705 | 0.194701532 |
| chr20:48027248-48027460 | 0.719267677 | 0.047236362 | 0.194734989 |
| chr9:31800026-31800266 | -1.072013818 | 0.047255495 | 0.194803385 |
| chr12:6038164-6038431 | -0.89135398 | 0.047262347 | 0.19482115 |
| chr3:62325880-62326280 | 0.71924145 | 0.047266942 | 0.194829612 |
| chr10:95756349-95756560 | -0.989073709 | 0.047288615 | 0.194904448 |
| chr1:39816848-39817078 | 0.871302232 | 0.047290185 | 0.194904448 |
| chr8:15656615-15657220 | -0.872936827 | 0.047305827 | 0.194958427 |
| chr7:23570602-23571168 | 0.684850221 | 0.04731497 | 0.194985622 |
| chr19:35958195-35958463 | 0.909325656 | 0.047320357 | 0.194997334 |
| chr2:20509947-20510536 | -0.686533091 | 0.047327817 | 0.195017585 |
| chr8:141041528-141041783 | -0.912645138 | 0.047335 | 0.195036693 |
| chr5:103562160-103563214 | -0.641594057 | 0.047344217 | 0.195064183 |
| chr11:34632495-34632722 | 0.911090593 | 0.047349193 | 0.195074192 |

| | | | |
|---|---|---|---|
| chr2:95158850-95159575 | 0.696937898 | 0.047355192 | 0.195088421 |
| chr17:60006058-60006645 | 0.64916806 | 0.047358286 | 0.195090675 |
| chr12:99284314-99284514 | 1.000937978 | 0.047362979 | 0.195099518 |
| chr14:105273007-105273439 | 0.744814802 | 0.047366282 | 0.195102635 |
| chr11:99693918-99694146 | -1.040084932 | 0.047371101 | 0.195112 |
| chr1:218958172-218958488 | -0.841851143 | 0.047389606 | 0.195177724 |
| chr9:131096258-131096575 | 0.780503246 | 0.047394498 | 0.19518738 |
| chr10:116670197-116670607 | -0.826341977 | 0.047405585 | 0.195222547 |
| chr14:104865124-104865454 | 0.715999489 | 0.047424418 | 0.195289611 |
| chr9:123929376-123930319 | 0.663384754 | 0.047442866 | 0.195355078 |
| chr8:124965783-124966120 | 0.701179239 | 0.047456021 | 0.195398744 |
| chr12:124388753-124389121 | 0.709558936 | 0.047469413 | 0.195443384 |
| chr14:72925946-72927176 | 0.62969855 | 0.047474625 | 0.195454338 |
| chr8:89877297-89877575 | 0.757753171 | 0.047483655 | 0.195481011 |
| chr1:84936062-84936580 | -0.784502001 | 0.047491894 | 0.195495867 |
| chr6:75687625-75687881 | 0.882136552 | 0.047492366 | 0.195495867 |
| chr3:97257888-97258470 | -0.716196299 | 0.047514782 | 0.195570719 |
| chr3:10141553-10142320 | 0.647876558 | 0.047515655 | 0.195570719 |
| chr12:116916133-116916396 | 0.824083494 | 0.047523642 | 0.195580892 |
| chr3:98770382-98770844 | -0.841581407 | 0.047526651 | 0.195580892 |
| chr3:61783483-61783706 | 0.749893815 | 0.047531218 | 0.195580892 |
| chr2:166631619-166631861 | -0.973497424 | 0.04753123 | 0.195580892 |
| chr13:37074745-37074994 | -0.879154869 | 0.047532059 | 0.195580892 |
| chr18:63963668-63964003 | -0.823130756 | 0.047533741 | 0.195580892 |
| chr12:84910930-84912156 | -0.651665989 | 0.047535994 | 0.195580892 |
| chr10:59981514-59981944 | 0.875668762 | 0.047541064 | 0.195591249 |
| chr11:68298921-68299121 | 0.992749232 | 0.047555008 | 0.195638111 |
| chr17:59919704-59919904 | 0.729743313 | 0.047563706 | 0.195663388 |
| chr13:41905633-41905935 | -0.823721545 | 0.047576771 | 0.195706626 |
| chr6:95133612-95134136 | -0.811316147 | 0.047581994 | 0.195717603 |
| chr19:7872000-7872405 | 0.696663576 | 0.047593235 | 0.195753332 |
| chr7:140478725-140478998 | -0.795235118 | 0.047598745 | 0.195760034 |
| chr5:93953526-93953881 | -0.873177393 | 0.047599974 | 0.195760034 |
| chr13:45841587-45841792 | 0.958996957 | 0.047607739 | 0.195781462 |
| chr21:41634275-41634612 | -0.811196375 | 0.047612804 | 0.195784814 |
| chr6:154824227-154824477 | 1.008939385 | 0.047616499 | 0.195784814 |
| chr12:105245827-105246504 | -0.772080528 | 0.04761942 | 0.195784814 |
| chr12:101172283-101172710 | -0.763542799 | 0.047620833 | 0.195784814 |
| chr3:62686584-62686802 | -0.835873954 | 0.04762133 | 0.195784814 |
| chr10:58952508-58952976 | -0.808817653 | 0.047625148 | 0.195790002 |
| chr8:143684304-143684636 | 0.765088555 | 0.047651294 | 0.195886982 |
| chr3:71533195-71533415 | -0.979626054 | 0.04765873 | 0.195897243 |
| chr2:168534901-168535120 | 1.055830792 | 0.047658994 | 0.195897243 |
| chr12:70534533-70534831 | 0.992344669 | 0.04766146 | 0.195897243 |
| chr3:9353178-9353505 | 0.734867137 | 0.047669005 | 0.195917743 |
| chr5:124043456-124043831 | -0.746609834 | 0.047683236 | 0.195965721 |
| chrX:96684232-96685961 | -0.630494884 | 0.047685867 | 0.195966023 |
| chr8:116324732-116325171 | -0.735832563 | 0.047689085 | 0.195968735 |
| chr12:76738703-76738956 | 0.939339637 | 0.047699501 | 0.196001029 |
| chr3:130972021-130972288 | 0.975762717 | 0.047703513 | 0.196007004 |

| | | | |
|---|---|---|---|
| chr1:61082322-61082621 | -0.787573125 | 0.047728397 | 0.196098732 |
| chr9:97470183-97470383 | 1.047562908 | 0.04773422 | 0.196112141 |
| chr1:166700725-166700925 | -1.167653916 | 0.047754452 | 0.196184743 |
| chr13:59364821-59365139 | -0.997392691 | 0.047762928 | 0.196201311 |
| chr6:129709857-129710566 | -0.65434061 | 0.047763606 | 0.196201311 |
| chr9:20621747-20622787 | -0.642881826 | 0.047778241 | 0.196250906 |
| chr8:108734045-108734507 | 0.726101201 | 0.04778935 | 0.196286014 |
| chr7:99274237-99274457 | -1.00888983 | 0.047792933 | 0.196290209 |
| chr6:134400938-134401288 | 0.788763099 | 0.04780903 | 0.196345797 |
| chr2:152025628-152026264 | -0.706728683 | 0.047812618 | 0.196350008 |
| chr14:59679902-59680215 | -0.939516351 | 0.047824783 | 0.19638944 |
| chr13:45301228-45301670 | -0.715319225 | 0.047838179 | 0.196433921 |
| chr11:35526326-35526823 | -0.705209986 | 0.047851954 | 0.196479952 |
| chr1:44762018-44762218 | 1.132964355 | 0.047862647 | 0.196506147 |
| chr1:216600714-216601691 | -0.633509964 | 0.047864766 | 0.196506147 |
| chr17:60735852-60736294 | 0.66252207 | 0.047866027 | 0.196506147 |
| chr1:109893569-109893769 | 0.965544506 | 0.047877576 | 0.196535049 |
| chr11:79973810-79974072 | -0.984688564 | 0.047878197 | 0.196535049 |
| chr1:203649813-203650050 | 0.856912294 | 0.047890815 | 0.196576314 |
| chr13:93226672-93228727 | -0.627994886 | 0.047903232 | 0.19660928 |
| chr18:48848585-48849132 | -0.75059512 | 0.047906394 | 0.19660928 |
| chr1:201353241-201353763 | -0.701125161 | 0.047906544 | 0.19660928 |
| chr9:130701170-130701370 | 0.968216852 | 0.04792065 | 0.196646659 |
| chr17:48374661-48374962 | 0.842530138 | 0.047920785 | 0.196646659 |
| chr2:156366800-156367126 | -0.905588168 | 0.047926315 | 0.196658816 |
| chr8:48254023-48254387 | 0.809137991 | 0.047929533 | 0.19666149 |
| chr7:90244786-90245845 | -0.659718078 | 0.047944694 | 0.196713166 |
| chr2:35461840-35462234 | -0.812557943 | 0.047960426 | 0.196767174 |
| chr17:10327629-10327921 | 0.832784349 | 0.047979662 | 0.196824174 |
| chr20:54301545-54302357 | 0.632200821 | 0.047981384 | 0.196824174 |
| chr8:103120201-103120507 | -0.857816705 | 0.047982025 | 0.196824174 |
| chr8:91343643-91343876 | 0.935176555 | 0.047994129 | 0.196863286 |
| chr2:164620827-164622082 | -0.634408048 | 0.048002797 | 0.196888301 |
| chr20:44116731-44116961 | 0.835406857 | 0.048006562 | 0.196893201 |
| chr8:94417772-94418064 | 0.852920827 | 0.048026515 | 0.196961372 |
| chr12:110039028-110039427 | 0.741491919 | 0.048028324 | 0.196961372 |
| chr13:39942437-39943263 | -0.664530334 | 0.048032876 | 0.196969496 |
| chr1:113520577-113520851 | -0.892588636 | 0.048036757 | 0.19697487 |
| chr13:39496693-39496893 | 0.975186931 | 0.048041999 | 0.19698582 |
| chr20:51160494-51160811 | 0.775949544 | 0.048044797 | 0.196986755 |
| chr3:153421919-153422217 | -0.943593795 | 0.048071013 | 0.197083695 |
| chr18:54958656-54958994 | -0.901926766 | 0.048075406 | 0.197091158 |
| chr8:133453354-133453604 | 0.765520697 | 0.048082368 | 0.197106271 |
| chr7:112206109-112207322 | -0.629682576 | 0.048084793 | 0.197106271 |
| chr3:16069244-16069452 | 0.929604842 | 0.04808681 | 0.197106271 |
| chr17:64311961-64312258 | 0.682421746 | 0.048099978 | 0.197142634 |
| chr6:117207001-117207201 | -1.092736873 | 0.04810129 | 0.197142634 |
| chr1:109792214-109792508 | -0.824971833 | 0.0481034 | 0.197142634 |
| chr1:6553706-6554346 | 0.769999044 | 0.048115953 | 0.197171256 |
| chr12:79059598-79059909 | -0.814230726 | 0.048117924 | 0.197171256 |

| | | | |
|---|---|---|---|
| chrX:40014714-40015009 | 0.770083915 | 0.048118103 | 0.197171256 |
| chr8:129819916-129820416 | -0.75596034 | 0.048121505 | 0.197174649 |
| chr11:267572-267894 | -0.890358665 | 0.048124098 | 0.19717473 |
| chr12:106551407-106551923 | -0.76602946 | 0.048156172 | 0.197295596 |
| chr20:57736427-57736627 | 0.808303915 | 0.048176813 | 0.197369607 |
| chr3:39447902-39448385 | 0.769768081 | 0.048183955 | 0.197388315 |
| chr15:40323189-40323715 | 0.703754706 | 0.048196806 | 0.197424969 |
| chr7:157619852-157620088 | -1.024974508 | 0.048198056 | 0.197424969 |
| chr11:68726128-68726357 | 1.127941608 | 0.048204831 | 0.197426893 |
| chr8:137725767-137726285 | -0.810205452 | 0.048205839 | 0.197426893 |
| chr2:219369382-219369887 | -0.734413693 | 0.048206256 | 0.197426893 |
| chr4:49150670-49150908 | 1.103278298 | 0.048209002 | 0.197427587 |
| chr17:62835867-62836107 | 0.860893262 | 0.048231157 | 0.197507339 |
| chr7:16846819-16847956 | 0.639632064 | 0.048233632 | 0.197507339 |
| chr10:84845574-84845972 | -0.753749615 | 0.048258945 | 0.197600432 |
| chr18:74269074-74269588 | -0.752357392 | 0.048269633 | 0.197630945 |
| chr5:90585534-90585734 | -1.152249934 | 0.048272061 | 0.197630945 |
| chr17:61219180-61219894 | 0.627463114 | 0.048278313 | 0.197630945 |
| chr19:15452106-15452644 | 0.689820506 | 0.048279045 | 0.197630945 |
| chr3:17993213-17993413 | 1.096980674 | 0.048279294 | 0.197630945 |
| chr1:168145863-168146213 | -0.783702602 | 0.048292026 | 0.197669993 |
| chr13:101751535-101751824 | -1.03501876 | 0.048293992 | 0.197669993 |
| chr4:80989973-80990437 | -0.820429053 | 0.048300192 | 0.197675432 |
| chr17:38444133-38444333 | 0.966219367 | 0.048300481 | 0.197675432 |
| chr8:19003654-19003939 | 0.889254062 | 0.048304916 | 0.197683026 |
| chr6:125953528-125953891 | -0.713271019 | 0.048312977 | 0.197705455 |
| chr1:53172898-53173115 | 0.974073742 | 0.048321585 | 0.197730118 |
| chr21:8453783-8454207 | 0.65912469 | 0.048335204 | 0.197775287 |
| chr16:85461937-85462294 | -0.712337853 | 0.04837128 | 0.19791233 |
| chr5:104709778-104710003 | -1.119321293 | 0.048379299 | 0.197934571 |
| chr20:53905269-53905555 | 0.663980719 | 0.048394914 | 0.197987886 |
| chr10:81126580-81126805 | 1.012285687 | 0.048405886 | 0.19800693 |
| chr1:226132935-226133546 | -0.661789314 | 0.048407258 | 0.19800693 |
| chr15:73934736-73934996 | 0.864380585 | 0.048407421 | 0.19800693 |
| chr1:197137492-197137812 | -0.868268077 | 0.048409906 | 0.19800693 |
| chr17:57445020-57445422 | 0.718046725 | 0.048421219 | 0.198033242 |
| chr1:228635472-228635680 | 1.016217203 | 0.048424909 | 0.198033242 |
| chr1:190634169-190634524 | 0.766243502 | 0.048426018 | 0.198033242 |
| chr15:99961102-99961356 | -0.903592011 | 0.048426677 | 0.198033242 |
| chr18:37343385-37343585 | -0.982093965 | 0.048453548 | 0.198132552 |
| chr9:72353083-72353353 | -1.048123394 | 0.048459585 | 0.198146662 |
| chr12:42446686-42447162 | -0.769620381 | 0.048463679 | 0.19815283 |
| chr2:492213-492692 | -0.781292221 | 0.048474887 | 0.198188078 |
| chr19:38319602-38320035 | -0.737770975 | 0.048480776 | 0.198198444 |
| chrX:69158430-69158910 | 0.783783692 | 0.048482596 | 0.198198444 |
| chr15:98881616-98881869 | 0.809197066 | 0.048487588 | 0.198208277 |
| chr1:112466047-112466506 | 0.70553932 | 0.048507394 | 0.198278664 |
| chr1:196891035-196891283 | -0.921421638 | 0.04851391 | 0.198294719 |
| chr6:149846147-149846678 | -0.750776211 | 0.048520348 | 0.198310454 |
| chr16:82917252-82917498 | -0.982048057 | 0.048525011 | 0.198318935 |

| | | | |
|---|---|---|---|
| chr16:1225024-1225443 | 0.785219624 | 0.048530889 | 0.198332378 |
| chr14:104863992-104864202 | 0.811241601 | 0.04853445 | 0.198332554 |
| chr17:79393300-79393641 | 0.788705898 | 0.048536109 | 0.198332554 |
| chr10:34974449-34974674 | -0.912727693 | 0.048543559 | 0.198352422 |
| chr5:103232449-103232749 | -0.813807163 | 0.048548706 | 0.198362187 |
| chr9:107282745-107284212 | -0.624415445 | 0.048551127 | 0.198362187 |
| chr20:57755644-57755914 | 0.770609341 | 0.048574366 | 0.198446553 |
| chr6:51118811-51119458 | 0.698549639 | 0.048584571 | 0.19847766 |
| chr20:7692470-7692834 | -0.920559722 | 0.048607151 | 0.198559316 |
| chr3:13431532-13431796 | -0.955602439 | 0.0486177 | 0.198591822 |
| chr10:95456524-95456858 | -1.000603196 | 0.048627834 | 0.198622629 |
| chr14:34375183-34375383 | 0.991557883 | 0.048638619 | 0.198656021 |
| chr19:50537484-50538212 | 0.715047112 | 0.048641195 | 0.198656021 |
| chr1:99645691-99646360 | -0.730389823 | 0.048650356 | 0.198674893 |
| chr16:19886244-19886502 | 0.857259745 | 0.048651001 | 0.198674893 |
| chr5:124663107-124663307 | 1.135840909 | 0.04867771 | 0.198773368 |
| chr20:21916189-21916507 | -0.877536787 | 0.048694792 | 0.198832526 |
| chr10:31947650-31947924 | -0.858564416 | 0.048712391 | 0.198893787 |
| chr7:138459620-138460585 | -0.62896298 | 0.048720384 | 0.198906402 |
| chr19:34171866-34172684 | 0.67319978 | 0.048721869 | 0.198906402 |
| chr5:170505327-170505527 | -1.030511307 | 0.048723268 | 0.198906402 |
| chr5:150295222-150295571 | -0.778999371 | 0.048727002 | 0.198911045 |
| chr1:20554422-20554759 | -0.886390941 | 0.048734078 | 0.198929333 |
| chr3:152444119-152444348 | -0.996425764 | 0.04874492 | 0.198962992 |
| chr7:76074259-76074489 | 1.00450597 | 0.048749167 | 0.198969726 |
| chr2:188367174-188367575 | -0.794114255 | 0.048756061 | 0.19898263 |
| chr11:40274118-40274382 | -0.968902553 | 0.048757522 | 0.19898263 |
| chr8:125589526-125589745 | 0.822187364 | 0.048767246 | 0.199011714 |
| chr3:181947456-181947716 | -1.007905014 | 0.048779327 | 0.199050411 |
| chr9:35518974-35519394 | 0.795064607 | 0.048798215 | 0.199116882 |
| chr4:140667240-140667664 | -0.868419455 | 0.048803805 | 0.199129089 |
| chr3:72152653-72153147 | 0.796782997 | 0.048821541 | 0.199190849 |
| chr19:18594108-18594592 | 0.793775245 | 0.048836652 | 0.19924014 |
| chr10:24432831-24433090 | -1.10439657 | 0.048838823 | 0.19924014 |
| chr12:79121537-79121789 | -0.891611484 | 0.048847675 | 0.199265643 |
| chr7:129224421-129225833 | -0.626076329 | 0.048854347 | 0.199275214 |
| chr7:8201053-8201261 | 1.037736319 | 0.048856745 | 0.199275214 |
| chr7:42736944-42737328 | -0.738005706 | 0.048857824 | 0.199275214 |
| chr2:134207635-134207922 | -0.939738389 | 0.048862337 | 0.199283012 |
| chr14:64703962-64704579 | 0.662403948 | 0.048868714 | 0.199298414 |
| chr20:48295557-48296000 | 0.654884419 | 0.048872653 | 0.199303871 |
| chr10:84214619-84214926 | -0.784561753 | 0.048885162 | 0.199339849 |
| chr8:9045163-9045494 | 0.834854834 | 0.048886679 | 0.199339849 |
| chr2:30115611-30116025 | -0.769160685 | 0.04889318 | 0.199347409 |
| chr20:54373892-54374296 | 0.663197995 | 0.048893736 | 0.199347409 |
| chr1:33455838-33456145 | 0.813232468 | 0.048899941 | 0.19935896 |
| chr3:130480763-130481096 | -0.821074577 | 0.048901773 | 0.19935896 |
| chr13:45073881-45074147 | -0.916003857 | 0.048904895 | 0.199361082 |
| chr3:163916777-163917461 | -0.665608953 | 0.048910642 | 0.199373902 |
| chr15:98915393-98915606 | 0.868486616 | 0.048929998 | 0.199442193 |

| | | | |
|---|---|---|---|
| chr7:134562965-134563314 | -0.933961981 | 0.048945641 | 0.199495339 |
| chr16:18998645-18999650 | 0.677539775 | 0.048954454 | 0.199520647 |
| chr15:63396664-63396923 | -0.978743347 | 0.048973458 | 0.199582205 |
| chr10:68231125-68231929 | -0.675477767 | 0.048974767 | 0.199582205 |
| chr17:62725640-62726319 | 0.664225729 | 0.048982786 | 0.199601469 |
| chr2:207847704-207847993 | 0.98331014 | 0.048986364 | 0.199601469 |
| chr1:176375039-176375283 | 0.851035515 | 0.048987309 | 0.199601469 |
| chr17:42061709-42062225 | -0.74967093 | 0.048996896 | 0.199621952 |
| chr21:38527012-38527368 | -0.900155712 | 0.048997547 | 0.199621952 |
| chr2:227546018-227546235 | -1.072227741 | 0.049026361 | 0.199728722 |
| chr20:54611128-54611465 | 0.669467285 | 0.049033476 | 0.199734714 |
| chr2:53767372-53768394 | -0.634819893 | 0.049036341 | 0.199734714 |
| chr7:143387792-143388097 | -0.829476787 | 0.049038307 | 0.199734714 |
| chr5:181159287-181159784 | 0.676297948 | 0.049040113 | 0.199734714 |
| chr8:102444177-102444377 | 0.981288578 | 0.049043025 | 0.199734714 |
| chrX:30302107-30302517 | -0.840862274 | 0.049043472 | 0.199734714 |
| chr1:22414255-22414572 | 0.883936314 | 0.049052836 | 0.199762234 |
| chr8:19171850-19172050 | 1.077487484 | 0.049057593 | 0.19977071 |
| chr17:8152373-8152633 | -0.774715424 | 0.049060132 | 0.19977071 |
| chr14:76566411-76566702 | 0.931348474 | 0.049071891 | 0.199807972 |
| chr8:60707233-60707576 | 0.844471965 | 0.049081811 | 0.199827688 |
| chr20:40688973-40690345 | 0.632054077 | 0.049081949 | 0.199827688 |
| chr1:228213000-228213250 | 0.787493862 | 0.049095265 | 0.199870254 |
| chr10:53836316-53836516 | -1.209010133 | 0.049097621 | 0.199870254 |
| chr13:32829791-32829991 | 0.982708277 | 0.049108039 | 0.199902044 |
| chr5:61363045-61363245 | -1.134153319 | 0.049111033 | 0.199902674 |
| chr2:95550845-95551707 | 0.668358548 | 0.049113412 | 0.199902674 |
| chr3:18005187-18005868 | -0.79473103 | 0.049141016 | 0.200004405 |
| chr8:87955834-87956038 | 1.019840069 | 0.04915026 | 0.200031401 |
| chr4:160835763-160836189 | -0.714483216 | 0.049154804 | 0.200039273 |
| chr8:52100851-52101427 | 0.749481557 | 0.049161991 | 0.200057894 |
| chr4:96013053-96013450 | 0.862898064 | 0.049172198 | 0.200073805 |
| chr4:27168655-27168887 | -1.026208574 | 0.049172475 | 0.200073805 |
| chr20:48097520-48097946 | 0.652531693 | 0.049173735 | 0.200073805 |
| chr11:72136605-72136815 | 1.043562199 | 0.049183687 | 0.200103673 |
| chr6:39189984-39190283 | 0.807868026 | 0.049190698 | 0.20012157 |
| chr1:210584427-210584852 | -0.757097569 | 0.04919995 | 0.200146672 |
| chr14:100645735-100646043 | 0.756005029 | 0.049202092 | 0.200146672 |
| chr6:119314433-119314652 | -1.169226466 | 0.049206069 | 0.200152222 |
| chr6:152356732-152357050 | -0.866983012 | 0.049211379 | 0.200163195 |
| chr3:150221337-150221589 | 0.778868888 | 0.049225382 | 0.200209524 |
| chr5:104646269-104646949 | -0.721801142 | 0.049232448 | 0.200219209 |
| chr3:170244684-170245138 | 0.791713881 | 0.049233702 | 0.200219209 |
| chr22:22687663-22687868 | 0.945932548 | 0.049235602 | 0.200219209 |
| chr13:114030542-114030802 | 0.930490419 | 0.049240555 | 0.200221296 |
| chrX:106302048-106302757 | 0.661263591 | 0.049241342 | 0.200221296 |
| chr3:125116536-125116811 | 0.927258403 | 0.049254286 | 0.200263303 |
| chr2:210548923-210549209 | -0.925648651 | 0.049264132 | 0.200292705 |
| chr8:61054077-61054978 | -0.649458519 | 0.049269528 | 0.200304016 |
| chr11:19903647-19904129 | -0.80177218 | 0.049281921 | 0.200343768 |

| | | | |
|---|---|---|---|
| chr5:123209642-123209909 | -1.024634714 | 0.049286739 | 0.200352726 |
| chr14:21558033-21558301 | -0.926127907 | 0.049290929 | 0.200359128 |
| chr1:234657317-234657550 | 0.924359594 | 0.049299873 | 0.200378115 |
| chr14:34498792-34499135 | 0.717851868 | 0.04930083 | 0.200378115 |
| chr20:62782781-62782981 | 0.926957209 | 0.049319043 | 0.200439657 |
| chr3:152298551-152299562 | -0.63517576 | 0.049321204 | 0.200439657 |
| chr2:27378189-27378389 | 0.901095844 | 0.049327894 | 0.200451023 |
| chr8:34289901-34290403 | -0.871196361 | 0.049329233 | 0.200451023 |
| chr6:15816986-15817562 | -0.702299278 | 0.049339178 | 0.200480803 |
| chr1:156571911-156572111 | 0.929453873 | 0.049347997 | 0.200498254 |
| chr2:54455896-54456728 | 0.667868472 | 0.049348706 | 0.200498254 |
| chr18:3238915-3239115 | 0.886870053 | 0.049355 | 0.200507754 |
| chr16:28610376-28610597 | 0.942917476 | 0.049356278 | 0.200507754 |
| chr16:3234522-3235595 | 0.654982372 | 0.04936237 | 0.20052187 |
| chr12:81329146-81329502 | -0.764971629 | 0.049372025 | 0.200550459 |
| chr13:34151953-34152294 | 0.818822176 | 0.049383786 | 0.200587597 |
| chr12:93198262-93198474 | -1.000539233 | 0.049387093 | 0.200590396 |
| chr5:37668452-37668792 | 0.800838448 | 0.049390132 | 0.200592106 |
| chr11:33997850-33998050 | -0.993869589 | 0.049403973 | 0.200637686 |
| chr12:104201193-104201559 | -0.771053354 | 0.04940968 | 0.200644811 |
| chr4:158553514-158553929 | -0.898087924 | 0.049410965 | 0.200644811 |
| chr4:94207356-94208320 | -0.641277183 | 0.049416427 | 0.200656358 |
| chr5:165045918-165046470 | -0.72371569 | 0.049423111 | 0.200663341 |
| chr12:53344731-53345521 | 0.678368385 | 0.049423891 | 0.200663341 |
| chr14:31251345-31251545 | 0.981590523 | 0.049429008 | 0.200663341 |
| chr5:28014923-28015123 | 1.005246482 | 0.049431171 | 0.200663341 |
| chr5:95620408-95620719 | -0.931518257 | 0.049431241 | 0.200663341 |
| chr20:48698910-48699199 | 0.662941523 | 0.049440777 | 0.200677103 |
| chr3:64121812-64122139 | 0.676521827 | 0.049445621 | 0.200677103 |
| chr20:24034785-24035227 | 0.779587742 | 0.049447186 | 0.200677103 |
| chr17:81727263-81727579 | 0.752352412 | 0.049447796 | 0.200677103 |
| chr20:53836998-53837589 | 0.63026004 | 0.049449537 | 0.200677103 |
| chr15:33007371-33007581 | -1.04507525 | 0.049450346 | 0.200677103 |
| chr9:137295379-137295837 | 0.771897544 | 0.049463583 | 0.20071677 |
| chr17:58771985-58772185 | 0.775620252 | 0.049467855 | 0.20071677 |
| chr2:3222574-3223485 | -0.648559228 | 0.049467979 | 0.20071677 |
| chr7:73916507-73916820 | 0.830011888 | 0.049487484 | 0.200785278 |
| chr5:90221912-90222271 | 0.850792612 | 0.049496874 | 0.200806148 |
| chr3:63371642-63371842 | 0.83138905 | 0.049497869 | 0.200806148 |
| chr17:2401488-2402219 | -0.650663576 | 0.049505205 | 0.20081958 |
| chr1:201285845-201286225 | 0.690666788 | 0.0495076 | 0.20081958 |
| chr2:237848955-237849486 | -0.858863155 | 0.049511727 | 0.20081958 |
| chr3:72061307-72061575 | 0.882810455 | 0.049514961 | 0.20081958 |
| chr20:53813286-53813530 | 0.66560189 | 0.04951646 | 0.20081958 |
| chr4:170027198-170027748 | -0.664769962 | 0.049518193 | 0.20081958 |
| chr3:23976296-23976509 | 1.039345853 | 0.049519526 | 0.20081958 |
| chr22:38924425-38924837 | 0.780949548 | 0.049524154 | 0.200827716 |
| chr1:99707623-99707874 | -1.031518002 | 0.049534161 | 0.200857665 |
| chr9:100714707-100715088 | 0.815722965 | 0.049551231 | 0.200916252 |
| chr7:38359043-38359377 | -0.805226013 | 0.049570623 | 0.200984245 |

| | | | |
|---|---|---|---|
| chr6:33995869-33996077 | 1.045484761 | 0.049579147 | 0.201008169 |
| chr1:167319108-167319716 | -0.79492246 | 0.049590692 | 0.20104434 |
| chr8:93404277-93405050 | 0.656247456 | 0.049600058 | 0.201071672 |
| chr2:117349562-117349959 | -0.874306857 | 0.049605615 | 0.20108356 |
| chr20:5528924-5529214 | -0.854342794 | 0.049608506 | 0.201084642 |
| chr19:40605242-40605911 | 0.658690336 | 0.049621207 | 0.201125481 |
| chr6:10140969-10141169 | 1.079964732 | 0.049637444 | 0.201152107 |
| chr3:64352239-64352670 | 0.660143668 | 0.049639594 | 0.201152107 |
| chr11:133064129-133064836 | -0.766282551 | 0.049641497 | 0.201152107 |
| chr9:70318092-70318370 | -1.060909797 | 0.049641745 | 0.201152107 |
| chr12:52069392-52070310 | 0.638923302 | 0.049643118 | 0.201152107 |
| chr17:10757104-10757403 | 0.796897881 | 0.049643527 | 0.201152107 |
| chr1:165738061-165738378 | -0.936804561 | 0.049649512 | 0.201155083 |
| chr1:172969629-172969883 | -0.936804561 | 0.049649512 | 0.201155083 |
| chr6:69164925-69165227 | -0.825018869 | 0.049655297 | 0.201167883 |
| chr14:74217742-74217942 | 0.871628561 | 0.049658621 | 0.201170713 |
| chr16:31117340-31118182 | 0.637669489 | 0.049665145 | 0.201186505 |
| chr3:174302798-174303142 | -0.872306113 | 0.049671816 | 0.201202892 |
| chr19:34964005-34964453 | 0.790821199 | 0.049689375 | 0.201263375 |
| chr1:247330934-247332594 | -0.617478252 | 0.04969256 | 0.201265636 |
| chr18:55477831-55478061 | -1.130176873 | 0.049697422 | 0.201271621 |
| chr17:59017215-59017453 | 0.711742925 | 0.049699935 | 0.201271621 |
| chr11:64996903-64997185 | 0.843422638 | 0.049701918 | 0.201271621 |
| chr8:66123857-66124618 | -0.668266269 | 0.049725407 | 0.201348998 |
| chr1:36802107-36802364 | 0.989207392 | 0.049726281 | 0.201348998 |
| chr14:24080910-24081337 | 0.708991074 | 0.049738667 | 0.201388508 |
| chr10:91627835-91628042 | 0.967988788 | 0.049745228 | 0.201401447 |
| chr5:122458268-122458598 | -0.822937362 | 0.049747119 | 0.201401447 |
| chr8:53879522-53880408 | 0.671952289 | 0.049751126 | 0.201407026 |
| chr10:60335799-60336034 | -0.970470053 | 0.049767298 | 0.201461851 |
| chr10:7818051-7819433 | 0.628788347 | 0.049774082 | 0.201478669 |
| chr5:132526544-132527058 | -0.730526815 | 0.04980971 | 0.201612233 |
| chr11:75350714-75350927 | 0.797095704 | 0.049815168 | 0.201623674 |
| chr3:111781773-111782154 | -0.837741144 | 0.049822623 | 0.201643197 |
| chr2:114610473-114611083 | -0.774260146 | 0.049826978 | 0.201650173 |
| chr3:184491165-184491365 | 0.999547339 | 0.049834127 | 0.201668454 |
| chr3:83501394-83501698 | 0.91188668 | 0.04985089 | 0.201720615 |
| chr10:11210597-11210797 | -1.099587033 | 0.049852282 | 0.201720615 |
| chr12:43848096-43848437 | -0.823621827 | 0.049863362 | 0.201754794 |
| chr12:45745970-45746262 | 0.858918104 | 0.049869463 | 0.201768824 |
| chr3:33915775-33916115 | -0.774159605 | 0.049884882 | 0.201810716 |
| chr1:182244485-182244833 | -0.790918873 | 0.049885085 | 0.201810716 |
| chr1:215957913-215958277 | 0.939765622 | 0.049903256 | 0.201873568 |
| chr5:133981683-133981971 | -0.9971793 | 0.049908953 | 0.201881051 |
| chr16:75287738-75288449 | -0.692115708 | 0.049910375 | 0.201881051 |
| chrX:140918340-140918952 | -0.667662858 | 0.049920505 | 0.201901671 |
| chrX:132488952-132489152 | 0.993572536 | 0.049920743 | 0.201901671 |
| chr10:113425668-113426054 | -0.885914471 | 0.049925062 | 0.201903647 |
| chr7:69597468-69598483 | -0.681079572 | 0.04992831 | 0.201903647 |
| chr14:104768512-104769130 | 0.668077024 | 0.049930111 | 0.201903647 |

| | | | |
|---|---|---|---|
| chr4:16148268-16148496 | 1.019959977 | 0.049931772 | 0.201903647 |
| chr6:123415917-123416124 | -1.116453626 | 0.049952077 | 0.20197027 |
| chr1:41919935-41920307 | 0.799884078 | 0.04995352 | 0.20197027 |
| chr7:106056949-106057261 | -0.733005743 | 0.049972516 | 0.20203434 |
| chr14:36070474-36070759 | 0.771637911 | 0.04997464 | 0.20203434 |
| chr3:193922401-193922766 | -0.753864185 | 0.049984076 | 0.202061828 |
| chr6:122381559-122381852 | -0.791905154 | 0.049994822 | 0.202082182 |
| chr1:108058274-108058474 | 1.00526494 | 0.049997301 | 0.202082182 |
| chr3:108914912-108915444 | -0.702830142 | 0.050000133 | 0.202082182 |
| chr1:218619523-218619776 | -0.896034065 | 0.050001068 | 0.202082182 |
| chr7:116138239-116138531 | -0.838277268 | 0.050002298 | 0.202082182 |
| chr1:33050691-33050964 | 0.812768969 | 0.050005813 | 0.202085729 |
| chrX:3345301-3345571 | 0.84096549 | 0.050009209 | 0.202088794 |
| chr6:20933168-20933708 | -0.872819504 | 0.050019932 | 0.202121467 |
| chr5:180135112-180135542 | -0.720728599 | 0.050033643 | 0.202166207 |
| chr4:74475105-74475305 | -1.065182719 | 0.050039072 | 0.202177481 |
| chr12:85951890-85952176 | -0.962932713 | 0.050055515 | 0.202228476 |
| chr1:10803988-10804764 | -0.697576532 | 0.050056972 | 0.202228476 |
| chr11:80699703-80700006 | 0.891751917 | 0.050061192 | 0.202234865 |
| chr3:195771891-195772120 | 0.99124457 | 0.050074184 | 0.202276685 |
| chr5:111511944-111513358 | -0.619517421 | 0.050081586 | 0.20229493 |
| chr7:38304961-38305185 | -1.087491644 | 0.050085256 | 0.20229493 |
| chr13:39274034-39274332 | -0.840903302 | 0.050086622 | 0.20229493 |
| chr11:67253757-67254175 | -0.805618168 | 0.050107231 | 0.202367502 |
| chr19:48752434-48753315 | 0.640453074 | 0.050113536 | 0.202382298 |
| chr1:106995209-106995409 | 0.816943028 | 0.050138604 | 0.202468499 |
| chr3:54322158-54322696 | -0.777594529 | 0.050140166 | 0.202468499 |
| chr10:3468191-3468482 | 0.835829342 | 0.050155439 | 0.202519501 |
| chr2:36418005-36418205 | -1.049400776 | 0.050168322 | 0.202560847 |
| chr6:139655593-139655815 | -0.928896977 | 0.050179133 | 0.202593822 |
| chr5:132661252-132661539 | -0.893367169 | 0.050203972 | 0.202683426 |
| chr4:139243517-139243869 | -0.989862848 | 0.050208001 | 0.202689013 |
| chr1:216687116-216687489 | -0.835907858 | 0.050231774 | 0.202771537 |
| chr17:64312966-64313255 | 0.67446188 | 0.050233736 | 0.202771537 |
| chr21:46378122-46378322 | 1.012038098 | 0.050245953 | 0.202799964 |
| chr15:97652890-97653172 | -0.893684732 | 0.050246072 | 0.202799964 |
| chr18:51491682-51491927 | -0.974028389 | 0.050256492 | 0.202831339 |
| chr17:50918946-50920348 | -0.623544814 | 0.050274719 | 0.202894211 |
| chr11:13772306-13772720 | -0.837933262 | 0.050291393 | 0.202942517 |
| chr9:707001-707501 | 0.740019945 | 0.050291986 | 0.202942517 |
| chr14:94355575-94355776 | 0.934977809 | 0.050295759 | 0.202947056 |
| chr7:91249610-91250116 | -0.917321798 | 0.050314613 | 0.203005334 |
| chr2:9703758-9704026 | 0.857828802 | 0.050315501 | 0.203005334 |
| chr1:42462813-42463780 | -0.64533922 | 0.050324471 | 0.203030835 |
| chr14:69151400-69152162 | 0.655355778 | 0.050330081 | 0.203042778 |
| chr10:18627410-18628116 | -0.673488803 | 0.050337316 | 0.203061273 |
| chr5:20679168-20679414 | 0.977387927 | 0.050343907 | 0.20307717 |
| chr6:86192083-86192375 | -0.84647185 | 0.050353612 | 0.203105625 |
| chr7:148941826-148942200 | -0.685641717 | 0.050362336 | 0.203128194 |
| chr14:30980113-30980423 | 0.76308916 | 0.050364509 | 0.203128194 |

| | | | |
|---|---|---|---|
| chr2:63424500-63425174 | -0.7192949 | 0.050382207 | 0.203188875 |
| chr11:29260984-29261410 | -0.795590904 | 0.050388189 | 0.203202307 |
| chr1:16727183-16727848 | 0.687295285 | 0.050398052 | 0.203231384 |
| chr20:57100534-57101183 | 0.625324978 | 0.050403409 | 0.203242292 |
| chr16:81913190-81913409 | 0.954360129 | 0.050406449 | 0.203243853 |
| chr1:218299413-218299695 | -0.77239091 | 0.050422909 | 0.203299521 |
| chr1:169741933-169742380 | -0.866582915 | 0.050428717 | 0.203306223 |
| chr3:170117215-170117481 | -0.841375154 | 0.050429924 | 0.203306223 |
| chr8:108468931-108469131 | 0.908838242 | 0.050432531 | 0.203306223 |
| chr5:107258008-107258332 | -0.871525976 | 0.050444038 | 0.20332508 |
| chr17:39730165-39730739 | 0.687835981 | 0.050447826 | 0.20332508 |
| chr18:46345910-46346181 | 0.956334019 | 0.050448084 | 0.20332508 |
| chr2:18144652-18144869 | -1.067356399 | 0.050449782 | 0.20332508 |
| chr1:230773166-230773382 | 0.889535517 | 0.050450477 | 0.20332508 |
| chr1:248622526-248622984 | -0.737499633 | 0.050455369 | 0.203334101 |
| chr1:215146169-215146826 | -0.688397699 | 0.050489473 | 0.203460839 |
| chr5:142680952-142681202 | -0.857988423 | 0.050515755 | 0.203556042 |
| chr14:23358184-23358505 | 0.814493654 | 0.05051953 | 0.203560551 |
| chr2:186980070-186980280 | 0.908603707 | 0.050524726 | 0.203562749 |
| chr16:88003081-88003292 | 1.061110346 | 0.050527615 | 0.203562749 |
| chr2:231482733-231483382 | 0.731087844 | 0.050529757 | 0.203562749 |
| chr13:100695024-100695357 | -0.827128989 | 0.050530703 | 0.203562749 |
| chr3:32037101-32037395 | 0.855394965 | 0.050535553 | 0.203571585 |
| chr6:103675838-103676166 | -0.840884473 | 0.050542042 | 0.203587022 |
| chr1:97063456-97063904 | -0.787083292 | 0.050551866 | 0.203615888 |
| chr1:169308798-169308998 | 1.070409641 | 0.050558246 | 0.203617446 |
| chr1:28260533-28260739 | 0.88663543 | 0.050559412 | 0.203617446 |
| chr1:180939077-180939277 | 0.880092534 | 0.050560225 | 0.203617446 |
| chr18:47175911-47176554 | -0.699105772 | 0.050567175 | 0.203634733 |
| chr11:118869843-118870981 | 0.646719716 | 0.050580668 | 0.203678366 |
| chr3:15552517-15552717 | 0.936714817 | 0.050589551 | 0.203703431 |
| chr2:15920687-15921092 | -0.767123215 | 0.050597682 | 0.203724201 |
| chr8:125546960-125547480 | -0.68961752 | 0.050600027 | 0.203724201 |
| chr7:16355750-16355950 | 1.075279342 | 0.050621556 | 0.203797414 |
| chr7:42932838-42933119 | -0.764917757 | 0.050623531 | 0.203797414 |
| chr20:54389520-54389721 | 0.718718971 | 0.050628297 | 0.203805893 |
| chr10:77744334-77744576 | -0.95301201 | 0.050648133 | 0.203868247 |
| chr17:67663551-67663751 | 0.881856824 | 0.050649108 | 0.203868247 |
| chr7:1873296-1874187 | 0.68138883 | 0.050660886 | 0.203904943 |
| chr6:120108140-120108722 | -0.797148825 | 0.050667123 | 0.203918027 |
| chr1:6058968-6059411 | 0.80139117 | 0.050669459 | 0.203918027 |
| chr17:61341331-61341585 | 0.660970436 | 0.050710083 | 0.204070798 |
| chr12:101279374-101280483 | -0.631122027 | 0.05072194 | 0.204107791 |
| chr13:29937926-29938195 | 0.918123588 | 0.050735316 | 0.20413272 |
| chr2:43478420-43478714 | 0.94261059 | 0.050736127 | 0.20413272 |
| chr19:41004601-41004914 | 0.94261059 | 0.050736127 | 0.20413272 |
| chr16:75491795-75492416 | -0.665533307 | 0.050747245 | 0.204148697 |
| chr3:130854462-130854805 | 0.882070739 | 0.050747417 | 0.204148697 |
| chr7:11427791-11428196 | 0.825370601 | 0.050748091 | 0.204148697 |
| chr20:40860710-40860948 | 0.966914309 | 0.050755591 | 0.204159174 |

| | | | |
|---|---|---|---|
| chr22:31122756-31122979 | 0.886124962 | 0.050756025 | 0.204159174 |
| chr20:48384174-48384535 | 0.650498914 | 0.050761503 | 0.204170493 |
| chr3:137054873-137055177 | -1.071417034 | 0.050769857 | 0.204193374 |
| chr6:73942608-73942974 | -0.771410881 | 0.050780012 | 0.204223499 |
| chr19:12294665-12295138 | 0.714456704 | 0.050784889 | 0.204232393 |
| chr3:174125325-174125639 | -0.807501639 | 0.050788227 | 0.204235096 |
| chr12:131860520-131861294 | -0.657741141 | 0.050800916 | 0.204266949 |
| chr5:152592504-152592741 | -0.999516716 | 0.05080148 | 0.204266949 |
| chr3:64156909-64158193 | 0.612812395 | 0.050806696 | 0.204277205 |
| chrX:135098605-135099221 | 0.638967308 | 0.05081808 | 0.204307519 |
| chr1:162023585-162024607 | 0.654949241 | 0.050823202 | 0.204307519 |
| chr3:111591755-111591955 | 1.050498471 | 0.050824262 | 0.204307519 |
| chr5:66197677-66198765 | -0.666821708 | 0.050825648 | 0.204307519 |
| chr8:124798365-124798565 | 0.885832157 | 0.050827568 | 0.204307519 |
| chr18:13531028-13531248 | 0.995095679 | 0.050838443 | 0.204329792 |
| chr19:10085875-10086079 | 0.995095679 | 0.050838443 | 0.204329792 |
| chr2:94587806-94588042 | 0.977934319 | 0.050844099 | 0.204340648 |
| chr2:120067891-120068652 | -0.743852393 | 0.050853349 | 0.204340648 |
| chr3:124359787-124360040 | 1.013453714 | 0.050855414 | 0.204340648 |
| chr14:105414933-105415380 | 0.703217715 | 0.050855898 | 0.204340648 |
| chr6:98054572-98054833 | -1.059163373 | 0.050856387 | 0.204340648 |
| chr4:140238432-140238893 | -0.794434045 | 0.050857145 | 0.204340648 |
| chrY:56846975-56847241 | 0.890253054 | 0.050860627 | 0.204341319 |
| chrX:110450129-110450568 | 0.758792523 | 0.050862646 | 0.204341319 |
| chr4:17614155-17615279 | 0.637493394 | 0.050872434 | 0.20436993 |
| chr16:16143154-16143424 | 0.901835674 | 0.050883728 | 0.204404581 |
| chr3:154314238-154314495 | -1.065614743 | 0.050889861 | 0.204418502 |
| chr1:223345828-223346121 | -0.922654363 | 0.050892573 | 0.204418679 |
| chr6:148345300-148345559 | 0.827480491 | 0.050901584 | 0.204437512 |
| chr10:131875004-131875254 | 0.837147465 | 0.050902598 | 0.204437512 |
| chr10:5551734-5552365 | -0.684655253 | 0.050905871 | 0.20443994 |
| chr9:108270132-108270376 | 0.902096271 | 0.05090939 | 0.204443357 |
| chr15:40436317-40436737 | -0.758986867 | 0.050913821 | 0.204447722 |
| chr6:165987661-165988471 | 0.684987886 | 0.050915813 | 0.204447722 |
| chr6:111565876-111566221 | -0.734909635 | 0.050928312 | 0.204487196 |
| chr7:78019748-78020846 | -0.649758473 | 0.050951442 | 0.204569344 |
| chr1:111848200-111848553 | 0.684038528 | 0.050958919 | 0.204588645 |
| chr6:33702434-33702690 | 0.852032388 | 0.050976313 | 0.204647754 |
| chr12:100279927-100280154 | -0.970243597 | 0.050987787 | 0.204683092 |
| chr2:71177065-71177265 | -1.072743072 | 0.051004111 | 0.204737899 |
| chr12:67424896-67425110 | -0.974180352 | 0.051031463 | 0.204836961 |
| chr3:71658171-71658581 | -0.744415092 | 0.05104461 | 0.204879001 |
| chr5:134245371-134245583 | -1.03583141 | 0.051056878 | 0.204905485 |
| chr1:197063312-197063545 | -0.904878817 | 0.051062792 | 0.204905485 |
| chr8:17200721-17201042 | -0.795164363 | 0.051064853 | 0.204905485 |
| chr16:30429842-30430664 | 0.640446376 | 0.051066291 | 0.204905485 |
| chr19:18097585-18097904 | 0.779995221 | 0.051066665 | 0.204905485 |
| chr1:47449751-47450260 | -0.763017999 | 0.051068108 | 0.204905485 |
| chr6:126312115-126312570 | -0.789047172 | 0.05107028 | 0.204905485 |
| chrX:123740123-123740373 | 0.858943145 | 0.051073225 | 0.204905485 |

| | | | |
|---|---|---|---|
| chr13:64764005-64764444 | -0.730885099 | 0.051075966 | 0.204905485 |
| chr12:94562184-94562420 | -0.931566098 | 0.051078059 | 0.204905485 |
| chr9:129052548-129052784 | 0.938762252 | 0.051080625 | 0.204905485 |
| chr5:101590482-101590711 | -1.003039644 | 0.051085559 | 0.204914547 |
| chr7:128235366-128235604 | -1.039608272 | 0.051097628 | 0.204937389 |
| chr10:51010964-51011247 | -0.865304837 | 0.051097801 | 0.204937389 |
| chr13:57631208-57632562 | -0.659286878 | 0.051100013 | 0.204937389 |
| chr7:120555017-120555286 | -0.890096331 | 0.051101952 | 0.204937389 |
| chr12:56468265-56468673 | 0.720600587 | 0.051116516 | 0.204985066 |
| chr7:119572070-119572404 | -0.966306184 | 0.051128052 | 0.205020598 |
| chr1:2049483-2049788 | 0.73823208 | 0.051131968 | 0.205025572 |
| chr7:132654279-132654479 | -1.018772163 | 0.051135328 | 0.205028315 |
| chr15:87752678-87752878 | 1.003960904 | 0.051152058 | 0.205084663 |
| chr20:50783348-50783585 | 0.667209164 | 0.051168905 | 0.205134721 |
| chr3:118863803-118864372 | -0.756620231 | 0.051169898 | 0.205134721 |
| chr10:123559401-123559648 | 0.843025116 | 0.051182808 | 0.205175742 |
| chr9:125296886-125297263 | 0.804213868 | 0.051189063 | 0.205190081 |
| chr5:103444236-103444975 | -0.651252697 | 0.051196974 | 0.205201703 |
| chr11:97610934-97611208 | -0.944451656 | 0.051197319 | 0.205201703 |
| chr5:1641608-1641861 | 0.83800235 | 0.051203153 | 0.205214352 |
| chr10:18233903-18234165 | -0.893820825 | 0.051215054 | 0.205251312 |
| chr11:10751654-10751967 | 0.919223658 | 0.051233336 | 0.205313844 |
| chr7:20777673-20778287 | 0.721202093 | 0.051242955 | 0.205339552 |
| chr5:102670279-102670525 | -0.951079054 | 0.051245111 | 0.205339552 |
| chr2:182784756-182785054 | -0.872766674 | 0.051265803 | 0.205411722 |
| chr1:234326595-234326991 | -0.767387981 | 0.051274782 | 0.205436957 |
| chr2:6654330-6654674 | -0.874153404 | 0.051284115 | 0.205463606 |
| chr5:95673104-95673482 | -0.824787495 | 0.051289754 | 0.205475453 |
| chr12:26195387-26195749 | -0.867698602 | 0.05129585 | 0.205489131 |
| chr12:120641285-120641650 | 0.781752568 | 0.051310198 | 0.205535865 |
| chr13:41696596-41697011 | -0.867675307 | 0.051323815 | 0.205579663 |
| chr5:174777154-174777515 | 0.736692514 | 0.051332227 | 0.20560261 |
| chr6:84549998-84550402 | -0.950911583 | 0.051343698 | 0.205637803 |
| chr11:15903326-15903535 | -0.989611125 | 0.051352108 | 0.205655051 |
| chr5:68950079-68950456 | -0.839074958 | 0.051353372 | 0.205655051 |
| chr1:197042942-197043326 | -0.813043607 | 0.051362879 | 0.205679813 |
| chr10:104340018-104340221 | 0.92789922 | 0.051365138 | 0.205679813 |
| chr12:52213501-52213780 | 0.748969866 | 0.051367609 | 0.205679813 |
| chr2:27118301-27118789 | -0.735476082 | 0.051377003 | 0.205702879 |
| chr10:17200888-17201877 | -0.645616762 | 0.051378738 | 0.205702879 |
| chr16:47143451-47143651 | 1.043037624 | 0.051404881 | 0.205796791 |
| chr20:48522867-48523509 | 0.635615931 | 0.051411361 | 0.205811978 |
| chr3:192812703-192813011 | 0.879523381 | 0.051424583 | 0.205854154 |
| chr21:40324378-40325039 | -0.677796161 | 0.051430949 | 0.205868884 |
| chr5:12078305-12078639 | -0.817293164 | 0.051448433 | 0.205928109 |
| chr10:62781363-62781701 | -1.070022 | 0.051453731 | 0.205938561 |
| chr5:44935668-44935900 | 0.992568038 | 0.051457643 | 0.205943459 |
| chr1:6157802-6158217 | 0.730904623 | 0.051463309 | 0.205955379 |
| chr16:80415760-80416009 | 1.038193734 | 0.05146675 | 0.205958392 |
| chr3:151314683-151314912 | -0.939807493 | 0.051470444 | 0.205962416 |

| | | | |
|---|---|---|---|
| chr9:121498167-121498486 | 0.744546218 | 0.051480863 | 0.205993352 |
| chr10:1953075-1953342 | 0.96935594 | 0.051489162 | 0.206015801 |
| chr20:47545751-47546551 | 0.621414403 | 0.051505614 | 0.20606336 |
| chr6:40695142-40695462 | -0.817631468 | 0.051506427 | 0.20606336 |
| chr1:214331920-214332163 | -0.867786622 | 0.05151371 | 0.206081736 |
| chr10:133357940-133358165 | -0.96897141 | 0.051517684 | 0.206086764 |
| chr5:9060317-9060532 | 1.006076949 | 0.051520346 | 0.206086764 |
| chr20:54412242-54412552 | 0.670395175 | 0.051534843 | 0.206127556 |
| chr3:151789913-151790674 | -0.691063659 | 0.051535924 | 0.206127556 |
| chr9:138053486-138053697 | 0.942245802 | 0.051543201 | 0.206135557 |
| chr7:22234268-22234622 | 0.827634529 | 0.051545415 | 0.206135557 |
| chr13:99774747-99774988 | 1.021454957 | 0.051546419 | 0.206135557 |
| chr14:104972228-104972515 | 0.712842277 | 0.051548686 | 0.206135557 |
| chr1:21324609-21324889 | -0.935858782 | 0.051553416 | 0.206143713 |
| chr10:12534879-12535125 | 0.926963906 | 0.051556835 | 0.206146629 |
| chr2:25228761-25229285 | -0.693002558 | 0.051562126 | 0.206157024 |
| chr9:99184649-99184906 | 0.843252569 | 0.051567705 | 0.206168573 |
| chr15:55196394-55197391 | 0.620551582 | 0.051587288 | 0.206236105 |
| chr20:32007884-32008084 | 0.988290226 | 0.051593093 | 0.206236895 |
| chr5:139482240-139482503 | 0.899441809 | 0.051599037 | 0.206236895 |
| chr11:47979408-47979648 | 0.83094949 | 0.051599052 | 0.206236895 |
| chr9:74886768-74887310 | -0.75853579 | 0.051601344 | 0.206236895 |
| chr2:19746690-19747029 | -0.799996786 | 0.051603032 | 0.206236895 |
| chr1:161192536-161192842 | 0.774694585 | 0.051603635 | 0.206236895 |
| chr3:78675103-78675973 | -0.631520653 | 0.051608811 | 0.206246821 |
| chr3:30135279-30135491 | -1.035232177 | 0.051622486 | 0.206290204 |
| chrX:129678146-129678569 | 0.711255529 | 0.051625051 | 0.206290204 |
| chr4:165827134-165827415 | -0.992920354 | 0.051629123 | 0.206295717 |
| chr2:84257942-84258185 | -1.001940729 | 0.051638802 | 0.206323631 |
| chr2:8189419-8189638 | -1.064606418 | 0.051641494 | 0.206323631 |
| chr17:52410469-52410761 | 0.889323323 | 0.051654012 | 0.206362881 |
| chr7:76177939-76178573 | -0.687879147 | 0.051687475 | 0.206463145 |
| chr14:54472418-54472722 | 0.842808618 | 0.051687556 | 0.206463145 |
| chr8:89611943-89612143 | 0.862183794 | 0.051688859 | 0.206463145 |
| chr20:54369234-54369473 | 0.692902123 | 0.051689887 | 0.206463145 |
| chr19:27851455-27851872 | -0.911400263 | 0.051692727 | 0.206463726 |
| chr17:68389107-68389403 | 0.806967819 | 0.051700562 | 0.20648354 |
| chr17:59974335-59974549 | 0.72660559 | 0.051703077 | 0.20648354 |
| chr16:78241186-78241612 | 0.748530549 | 0.051715516 | 0.20652245 |
| chr4:61372679-61373193 | -0.769816778 | 0.05174964 | 0.206636939 |
| chr11:47186321-47186692 | 0.709212544 | 0.051749888 | 0.206636939 |
| chr7:50795171-50796065 | -0.662540732 | 0.051754224 | 0.206636939 |
| chr7:138008735-138009161 | -0.734911144 | 0.051754972 | 0.206636939 |
| chr12:100148979-100149595 | -0.7106738 | 0.051761027 | 0.206650345 |
| chr2:127014284-127014580 | 0.852455267 | 0.051764511 | 0.206653488 |
| chr12:45173193-45173604 | -0.828238158 | 0.051770356 | 0.206666053 |
| chr10:129703412-129703612 | 0.981897236 | 0.051790274 | 0.206734796 |
| chr6:162727164-162728271 | -0.622030863 | 0.051794319 | 0.206740171 |
| chr16:21278078-21278547 | 0.73216366 | 0.051797782 | 0.206743222 |
| chr3:9976996-9977333 | 0.759019372 | 0.051807071 | 0.206769527 |

| | | | |
|---|---|---|---|
| chr1:106345567-106345777 | 0.756282126 | 0.051812609 | 0.20678086 |
| chr5:153467165-153467433 | -1.029531752 | 0.051816172 | 0.20678431 |
| chr19:33505713-33505945 | 0.935456962 | 0.051829011 | 0.206814743 |
| chr11:15027084-15027362 | 0.935113576 | 0.05183129 | 0.206814743 |
| chr17:65931463-65931690 | 1.005806597 | 0.051831896 | 0.206814743 |
| chr3:187913553-187914001 | -0.747709453 | 0.05183598 | 0.206816277 |
| chr1:155651252-155651771 | 0.708566683 | 0.051837679 | 0.206816277 |
| chr22:35257358-35257873 | 0.754265426 | 0.051848283 | 0.206840005 |
| chr2:138846280-138846773 | -0.772164463 | 0.051849025 | 0.206840005 |
| chr4:182903908-182904108 | -1.18451402 | 0.051882971 | 0.206964649 |
| chr2:45675167-45675885 | -0.695036814 | 0.05189234 | 0.206991246 |
| chr14:35949959-35950264 | -0.909653335 | 0.051896406 | 0.206996691 |
| chr3:158532167-158532416 | -0.955764472 | 0.051910306 | 0.207032675 |
| chr4:120648237-120648492 | -0.966469549 | 0.051913741 | 0.207032675 |
| chr14:74502061-74502478 | 0.71308228 | 0.051914293 | 0.207032675 |
| chr6:114051116-114051532 | 0.752178001 | 0.051916236 | 0.207032675 |
| chr21:37060394-37060594 | 1.01990615 | 0.051920584 | 0.207038218 |
| chr5:24804839-24805098 | -1.059081273 | 0.051925176 | 0.207038218 |
| chr15:58604180-58604959 | 0.674267541 | 0.051925732 | 0.207038218 |
| chr17:61699685-61700129 | 0.640087544 | 0.051931849 | 0.207051835 |
| chrX:3348255-3348507 | 0.82955283 | 0.051948079 | 0.207102416 |
| chr3:132231700-132231960 | -0.833298364 | 0.051951028 | 0.207102416 |
| chr4:67871934-67872190 | -0.948466105 | 0.051954117 | 0.207102416 |
| chr2:102100368-102100653 | 1.01589902 | 0.051955348 | 0.207102416 |
| chr3:73469318-73469651 | -0.914477103 | 0.051965703 | 0.207130133 |
| chr20:47969521-47969755 | 0.680085811 | 0.051967707 | 0.207130133 |
| chrX:94361007-94361548 | 0.75520986 | 0.051971524 | 0.20713457 |
| chr12:10721749-10722809 | -0.633559357 | 0.051974849 | 0.207137047 |
| chr6:39432696-39432930 | 0.98659718 | 0.051979967 | 0.20714667 |
| chr3:99499299-99499540 | -0.931767456 | 0.051986868 | 0.207163397 |
| chr7:23531205-23532528 | 0.615237889 | 0.052003952 | 0.207220698 |
| chr10:88851525-88852054 | -0.696207033 | 0.052008876 | 0.207223352 |
| chr1:3651978-3653242 | 0.665006623 | 0.052010664 | 0.207223352 |
| chr16:66695937-66696511 | 0.727728491 | 0.052017535 | 0.207223352 |
| chr20:47735483-47735863 | 0.639311037 | 0.052017705 | 0.207223352 |
| chr1:190373664-190374133 | -0.71926903 | 0.05201814 | 0.207223352 |
| chr7:74269834-74270253 | -0.925514278 | 0.052038991 | 0.207295635 |
| chr5:52845977-52846183 | -1.084115023 | 0.052052522 | 0.207337767 |
| chr18:923183-923444 | 0.927742923 | 0.052054979 | 0.207337767 |
| chr2:209771425-209772133 | -0.685919743 | 0.052061338 | 0.207343863 |
| chr3:123328881-123329142 | 0.775051277 | 0.052064618 | 0.207343863 |
| chr6:10505491-10505819 | -0.787824006 | 0.052065988 | 0.207343863 |
| chr6:12815886-12816602 | -0.685534594 | 0.052068857 | 0.207343863 |
| chr8:59643895-59644095 | -1.138339913 | 0.05207004 | 0.207343863 |
| chr2:72638244-72638502 | -1.063693894 | 0.052091991 | 0.207413863 |
| chr12:53984862-53985159 | 0.750828957 | 0.052093033 | 0.207413863 |
| chr8:120431443-120431815 | -0.833103922 | 0.052102507 | 0.207440804 |
| chr13:60369199-60369421 | -1.035267255 | 0.052115874 | 0.207483242 |
| chr12:89648607-89648910 | -0.943944589 | 0.052121075 | 0.207493169 |
| chr1:208132045-208132535 | -0.713613767 | 0.052138182 | 0.207550489 |

| | | | |
|---|---|---|---|
| chr10:97117499-97117718 | 0.953296633 | 0.052158178 | 0.207612332 |
| chr5:60210864-60211221 | -0.886529198 | 0.052159137 | 0.207612332 |
| chr10:13562528-13562746 | -1.006949015 | 0.052164679 | 0.207623607 |
| chrX:40174139-40174420 | 0.790348945 | 0.052169 | 0.20762759 |
| chr12:53352640-53352929 | 0.769615312 | 0.052171099 | 0.20762759 |
| chr5:103414631-103414933 | -0.901298242 | 0.052174726 | 0.207631239 |
| chr15:68285361-68285657 | -1.057382889 | 0.052177467 | 0.20763136 |
| chr19:45326165-45326584 | 0.740590661 | 0.052190041 | 0.207670615 |
| chr20:47845677-47846022 | 0.661121832 | 0.05219436 | 0.207677015 |
| chr9:133658561-133658842 | 0.870326559 | 0.052200652 | 0.207691266 |
| chr16:84688827-84689027 | 1.035349588 | 0.052205328 | 0.207699083 |
| chr5:109395467-109395734 | 0.876118436 | 0.052215557 | 0.207728994 |
| chr7:15643042-15643242 | 0.976749111 | 0.052221978 | 0.207740211 |
| chr1:20613601-20614189 | 0.734272886 | 0.052223799 | 0.207740211 |
| chr5:115169342-115170792 | -0.612210611 | 0.052227472 | 0.207744034 |
| chr9:132575174-132575541 | 0.844055869 | 0.0522389 | 0.207778708 |
| chr6:138816099-138816304 | -0.947296212 | 0.052248437 | 0.207805853 |
| chr5:158855513-158855730 | 0.98551901 | 0.052268438 | 0.207871611 |
| chr6:63817158-63817616 | -0.765992313 | 0.052270397 | 0.207871611 |
| chr10:12785479-12785902 | 0.735627909 | 0.052275332 | 0.20788045 |
| chr5:66645047-66645357 | -0.868912783 | 0.052303493 | 0.207981642 |
| chr2:43169129-43169643 | -0.742523843 | 0.05232287 | 0.208037361 |
| chr15:70488067-70488369 | 0.774917833 | 0.052322936 | 0.208037361 |
| chr13:94641674-94641970 | -0.911250126 | 0.052336923 | 0.208071615 |
| chr5:129771150-129771691 | -0.695519344 | 0.052336982 | 0.208071615 |
| chr4:46147077-46147472 | -0.876932384 | 0.052343276 | 0.208085839 |
| chr19:43640338-43640545 | 0.971939714 | 0.052346404 | 0.20808748 |
| chr1:80656095-80656438 | -0.794591661 | 0.052366398 | 0.208142617 |
| chr2:216683892-216684207 | -0.760626303 | 0.052366632 | 0.208142617 |
| chr12:102879179-102879518 | -0.781765345 | 0.052368424 | 0.208142617 |
| chr10:16827574-16828603 | -0.656123826 | 0.05237623 | 0.208162844 |
| chrX:142076535-142076993 | 0.755891056 | 0.052382102 | 0.208172271 |
| chr7:66537524-66538156 | -0.734609463 | 0.052384035 | 0.208172271 |
| chr4:162461390-162461699 | -0.919181534 | 0.052395843 | 0.208208396 |
| chr4:94758041-94758646 | 0.724934008 | 0.052404259 | 0.208215694 |
| chr15:67945911-67946646 | -0.685447986 | 0.052405831 | 0.208215694 |
| chr7:16989775-16990348 | 0.648407956 | 0.052407757 | 0.208215694 |
| chr1:35726857-35727178 | 0.857217345 | 0.052408549 | 0.208215694 |
| chr10:76103285-76103599 | -0.874866142 | 0.052414503 | 0.208220541 |
| chr3:195895805-195896204 | 0.729380456 | 0.052416597 | 0.208220541 |
| chr7:11261402-11262193 | 0.627744404 | 0.052417922 | 0.208220541 |
| chr11:118790441-118791418 | 0.640807888 | 0.05243103 | 0.208261812 |
| chr7:128888788-128888999 | -0.883432248 | 0.052435456 | 0.208268599 |
| chr5:105882914-105883437 | -0.701385945 | 0.052463947 | 0.208370959 |
| chr3:156217761-156218714 | 0.665575952 | 0.0524706 | 0.208385677 |
| chr3:150791560-150791765 | 0.939055888 | 0.052473092 | 0.208385677 |
| chr8:97540940-97541168 | 0.877535506 | 0.052488872 | 0.208435198 |
| chr5:105593070-105593510 | -0.912743436 | 0.052491002 | 0.208435198 |
| chr1:107375022-107375258 | 0.742969731 | 0.052497982 | 0.208452112 |
| chr10:52409186-52409822 | -0.740907152 | 0.052508551 | 0.208464606 |

| | | | |
|---|---|---|---|
| chr20:8114414-8115027 | -0.717095835 | 0.052508677 | 0.208464606 |
| chr3:24264778-24265164 | 0.774338781 | 0.052509291 | 0.208464606 |
| chr1:64273388-64274275 | -0.675105774 | 0.052520517 | 0.208498369 |
| chr2:64622697-64622924 | -0.965676609 | 0.052568998 | 0.208680021 |
| chr1:53014732-53015103 | 0.781952857 | 0.052579895 | 0.208709344 |
| chr1:164787360-164787663 | -0.802473622 | 0.052582918 | 0.208709344 |
| chr15:79744492-79744844 | -0.804885554 | 0.052584557 | 0.208709344 |
| chr8:123310667-123311147 | -0.685442154 | 0.052587648 | 0.208710804 |
| chr2:191758471-191758737 | -0.920369828 | 0.052593884 | 0.208724739 |
| chr12:25385755-25386729 | -0.633478148 | 0.052599226 | 0.208735129 |
| chr5:142880556-142880820 | -0.880436289 | 0.052610501 | 0.20876906 |
| chr14:80213570-80213887 | -0.738280402 | 0.052614769 | 0.208775186 |
| chr6:142537729-142538243 | -0.750997093 | 0.05262083 | 0.208788422 |
| chr20:51533778-51533978 | 0.919603543 | 0.052624016 | 0.208790252 |
| chr12:26831985-26832336 | -0.77164634 | 0.052632778 | 0.208814203 |
| chr20:48053086-48053286 | 0.69633473 | 0.052637511 | 0.208822169 |
| chr16:88383804-88384055 | 0.819719042 | 0.052650377 | 0.208862396 |
| chr2:185291081-185291375 | -0.752484757 | 0.052654961 | 0.208869767 |
| chr5:3997182-3997598 | -0.897366981 | 0.052674688 | 0.208937201 |
| chr16:1573290-1573749 | -0.727029984 | 0.05267759 | 0.208937898 |
| chr9:135499582-135500094 | 0.677478432 | 0.052682924 | 0.208948239 |
| chr7:74788742-74788942 | -1.135399075 | 0.052693481 | 0.208979289 |
| chr15:32994879-32995184 | -0.94117018 | 0.05270022 | 0.208995202 |
| chr1:189094053-189094493 | -0.842309402 | 0.052718009 | 0.209049033 |
| chr13:71320545-71320824 | -0.912921775 | 0.052719251 | 0.209049033 |
| chr21:46545371-46545819 | -0.791118165 | 0.052726106 | 0.209065393 |
| chrX:40922452-40922673 | -1.025512625 | 0.052738624 | 0.209100684 |
| chr12:6817564-6817778 | 0.999797405 | 0.052740464 | 0.209100684 |
| chr21:40325113-40325365 | 0.822188025 | 0.052750685 | 0.209120764 |
| chr6:154292859-154293450 | 0.695236406 | 0.052750987 | 0.209120764 |
| chr6:26183498-26184063 | -0.686597424 | 0.052758441 | 0.209138354 |
| chr7:139602197-139602443 | -0.906123137 | 0.052760883 | 0.209138354 |
| chr11:128859859-128860099 | -0.947784847 | 0.052765478 | 0.209145748 |
| chr12:33427086-33427580 | -0.74497826 | 0.052789222 | 0.209229037 |
| chr7:32427600-32428026 | -0.688691521 | 0.052806849 | 0.209288076 |
| chr4:1346549-1347341 | 0.624874746 | 0.052822619 | 0.209339748 |
| chr18:6929611-6930353 | -0.696212995 | 0.052825745 | 0.209341312 |
| chr5:132450624-132451034 | 0.72992311 | 0.052830841 | 0.209350678 |
| chr2:208015462-208015753 | 0.861557798 | 0.052837055 | 0.209364473 |
| chr20:40471312-40471512 | 0.944746923 | 0.052840839 | 0.209368641 |
| chr17:73311523-73311737 | 0.74374527 | 0.052844494 | 0.209370283 |
| chr3:102113420-102113723 | 0.852191777 | 0.052846719 | 0.209370283 |
| chr2:155472483-155472792 | -0.89251763 | 0.052854655 | 0.209390897 |
| chr9:80146609-80146875 | -1.025295255 | 0.052865426 | 0.209413263 |
| chr10:60010405-60010605 | 1.032654292 | 0.052865766 | 0.209413263 |
| chr5:65102764-65103812 | -0.657535803 | 0.052883995 | 0.209474643 |
| chr10:23869434-23869972 | 0.721202534 | 0.052887666 | 0.209478353 |
| chr7:143967373-143967876 | -0.665446434 | 0.05290217 | 0.209524971 |
| chr8:142447541-142447941 | 0.719547914 | 0.052908693 | 0.209539975 |
| chr20:44136153-44136395 | 0.955118796 | 0.052930263 | 0.209614566 |

| | | | |
|---|---|---|---|
| chr3:110984920-110985329 | -0.812738275 | 0.052939945 | 0.209642074 |
| chr3:125350741-125351098 | -0.937433219 | 0.05294964 | 0.209669628 |
| chr9:34408792-34409239 | -0.906312275 | 0.05295448 | 0.20967796 |
| chr7:106111714-106112820 | -0.622819574 | 0.052959705 | 0.209686346 |
| chr22:42995101-42995488 | 0.782731902 | 0.052963413 | 0.209686346 |
| chr16:15874742-15875099 | 0.867285307 | 0.052964808 | 0.209686346 |
| chr9:133892204-133892714 | -0.757361367 | 0.05297095 | 0.209696118 |
| chr21:9016139-9016679 | 0.762530464 | 0.05297275 | 0.209696118 |
| chr5:78294222-78295425 | -0.622624703 | 0.052977004 | 0.209702122 |
| chr4:102257593-102258017 | -0.743742821 | 0.0529963 | 0.209767666 |
| chr17:61899539-61899762 | 0.66990199 | 0.053013551 | 0.209822247 |
| chr16:57341888-57342145 | 0.986051601 | 0.053016257 | 0.209822247 |
| chr7:22325192-22325424 | 0.939829866 | 0.05301854 | 0.209822247 |
| chr10:8581772-8582311 | 0.715605828 | 0.05302366 | 0.209822247 |
| chrX:131753068-131753520 | 0.699900603 | 0.053023781 | 0.209822247 |
| chr10:13687633-13687886 | -1.04465932 | 0.053035135 | 0.209856336 |
| chr1:206803529-206803729 | -0.918431688 | 0.053037934 | 0.209856576 |
| chr4:169758462-169758747 | -0.829815599 | 0.053044447 | 0.209871509 |
| chr19:43651994-43652631 | 0.689445176 | 0.053051603 | 0.209881537 |
| chr14:70585325-70585583 | -0.998486101 | 0.05305246 | 0.209881537 |
| chr11:9535226-9535510 | -0.920497517 | 0.053062478 | 0.209910332 |
| chr19:38837580-38838139 | 0.662655564 | 0.0530666 | 0.209915801 |
| chrX:102691240-102692024 | -0.665784922 | 0.053083375 | 0.209971315 |
| chr4:172313895-172314095 | -1.064441457 | 0.05308981 | 0.209982647 |
| chr3:157129022-157129491 | 0.706526574 | 0.053091721 | 0.209982647 |
| chr5:39137868-39138077 | -1.036980487 | 0.053118533 | 0.210063316 |
| chr7:68547896-68548199 | -0.905685959 | 0.053119995 | 0.210063316 |
| chr13:85897037-85897414 | -0.830287411 | 0.053122393 | 0.210063316 |
| chrX:2889792-2889992 | -1.20405954 | 0.053123083 | 0.210063316 |
| chr17:74780936-74781187 | 0.755475999 | 0.053127091 | 0.210068322 |
| chrX:129773930-129774430 | 0.757735363 | 0.053134298 | 0.210068751 |
| chr14:88978425-88979412 | -0.632065048 | 0.053135134 | 0.210068751 |
| chr2:29583664-29583882 | 1.014564655 | 0.053135424 | 0.210068751 |
| chr14:66610131-66610398 | -0.980286236 | 0.05315055 | 0.210116512 |
| chr1:23800553-23801111 | -0.690286342 | 0.053152989 | 0.210116512 |
| chr10:52215823-52216023 | -1.039509623 | 0.053163443 | 0.210146994 |
| chr17:56884471-56884787 | -0.817229251 | 0.053172574 | 0.210172245 |
| chrX:100750980-100751242 | -0.94875748 | 0.053177675 | 0.210181565 |
| chr2:152048099-152048299 | -1.108366486 | 0.053189522 | 0.210217546 |
| chr3:62308885-62309093 | 0.847697995 | 0.053196251 | 0.210233296 |
| chr13:52751247-52751455 | -1.05615503 | 0.053202235 | 0.210246102 |
| chr2:32553921-32554125 | -1.108359945 | 0.053208054 | 0.210258252 |
| chr14:106605844-10660620[illegible] | -0.759060314 | 0.053221289 | 0.210299705 |
| chr16:2235597-2236084 | -0.733436316 | 0.053240447 | 0.21036456 |
| chr1:63981028-63981463 | 0.709370541 | 0.053244104 | 0.210367753 |
| chr12:53097875-53098177 | 0.794771324 | 0.053247528 | 0.210367753 |
| chr20:57991203-57991809 | -0.659888381 | 0.053249492 | 0.210367753 |
| chr14:73836693-73837138 | 0.713702067 | 0.053256776 | 0.210375478 |
| chr13:80151799-80152687 | -0.6707548 | 0.053256939 | 0.210375478 |
| chr7:112495877-112496214 | -0.804711308 | 0.053262889 | 0.210388137 |

| | | | |
|---|---|---|---|
| chr17:61230966-61231232 | 0.650377217 | 0.053267143 | 0.210394092 |
| chr1:179292443-179292710 | 0.738877093 | 0.053278232 | 0.210427047 |
| chr4:169492873-169493301 | -0.740243441 | 0.053287935 | 0.210454521 |
| chr3:46705049-46705319 | 0.801523752 | 0.053290951 | 0.210455583 |
| chr15:81370146-81370355 | 1.091016006 | 0.05330747 | 0.210509969 |
| chrX:94493931-94494131 | 1.097366227 | 0.053313454 | 0.210521409 |
| chrX:154121827-154122309 | 0.708491077 | 0.053315862 | 0.210521409 |
| chr19:53842187-53842422 | 0.894865493 | 0.053332 | 0.21057428 |
| chr5:101897933-101898197 | -0.969401932 | 0.053344928 | 0.210614472 |
| chr6:116533562-116533828 | -0.942626169 | 0.053371899 | 0.210710103 |
| chr6:76708747-76709131 | 0.854316565 | 0.053386513 | 0.210756936 |
| chr15:67148212-67148604 | 0.725427732 | 0.05340198 | 0.210799854 |
| chr1:184370919-184371218 | -0.820094103 | 0.053402886 | 0.210799854 |
| chr4:169015845-169016097 | -0.892571571 | 0.053411657 | 0.210823612 |
| chr11:69730552-69730752 | 0.909196764 | 0.053418923 | 0.210841432 |
| chr15:20342046-20342246 | 0.974692956 | 0.053439002 | 0.21090859 |
| chr7:29563607-29564523 | 0.651248807 | 0.053441444 | 0.21090859 |
| chr20:50770843-50771225 | 0.630808727 | 0.053444791 | 0.210910938 |
| chr4:173834110-173834421 | -0.902393631 | 0.053456648 | 0.210946864 |
| chr11:13009111-13009406 | 0.844559456 | 0.053475027 | 0.211008525 |
| chr20:54515898-54516220 | 0.661575174 | 0.053495812 | 0.211077476 |
| chr1:106820359-106820776 | 0.668578374 | 0.053498011 | 0.211077476 |
| chr2:66576441-66577660 | -0.649634877 | 0.053504771 | 0.211093279 |
| chr15:90202000-90202200 | 0.829820113 | 0.053507963 | 0.211095004 |
| chr16:57759397-57759853 | -0.727999961 | 0.05351197 | 0.211099942 |
| chr15:47621144-47621816 | 0.691444906 | 0.053524354 | 0.211137924 |
| chr5:34803023-34803223 | 1.066447838 | 0.053539629 | 0.211184339 |
| chr1:93666753-93667009 | 0.967531916 | 0.053541641 | 0.211184339 |
| chr8:73419757-73420701 | 0.645767052 | 0.053549292 | 0.211184339 |
| chr7:128842030-128842357 | -0.757671265 | 0.053549643 | 0.211184339 |
| chr17:60199911-60200422 | 0.65441984 | 0.053549901 | 0.211184339 |
| chrX:12988770-12989216 | 0.706705478 | 0.053552976 | 0.211185597 |
| chr7:64998002-64998374 | 0.750135523 | 0.053557435 | 0.211192312 |
| chr9:137054009-137054263 | 0.781577909 | 0.053566437 | 0.211209311 |
| chr12:106282426-106282696 | -0.850896396 | 0.053568691 | 0.211209311 |
| chr19:50658777-50659300 | 0.652088163 | 0.053575986 | 0.211209311 |
| chr7:38311454-38311886 | 0.777930586 | 0.053576153 | 0.211209311 |
| chr17:69414335-69414922 | -0.668872884 | 0.053577564 | 0.211209311 |
| chr2:30481022-30481383 | -0.834995772 | 0.053578285 | 0.211209311 |
| chr9:106333533-106333915 | -0.785868622 | 0.053589886 | 0.211239224 |
| chr7:17206286-17206840 | 0.698216291 | 0.053591387 | 0.211239224 |
| chr3:132445927-132446152 | -1.03268723 | 0.053597414 | 0.21124957 |
| chr11:12245740-12246024 | -0.830969706 | 0.053599526 | 0.21124957 |
| chr20:53692512-53692822 | 0.635607252 | 0.053635188 | 0.211379249 |
| chr7:139035081-139036398 | -0.632543979 | 0.053639634 | 0.211385898 |
| chr6:139124901-139125259 | -0.750325739 | 0.053655691 | 0.211438302 |
| chr1:220086317-220086517 | -0.916070579 | 0.053660234 | 0.211445332 |
| chr4:140220309-140220672 | -1.068554216 | 0.053666121 | 0.211457653 |
| chr1:48673812-48674116 | -0.838960396 | 0.053675047 | 0.21148195 |
| chr15:59179672-59180055 | -0.812193989 | 0.053689788 | 0.211506733 |

| | | | |
|---|---|---|---|
| chr12:119178539-119179096 | -0.663642966 | 0.053689849 | 0.211506733 |
| chr12:57611338-57611859 | 0.705700937 | 0.053691493 | 0.211506733 |
| chr5:25362795-25363228 | 0.742748155 | 0.053692379 | 0.211506733 |
| chr19:35135414-35135700 | 0.768868685 | 0.053697291 | 0.211512481 |
| chr5:159262872-159263795 | -0.637637414 | 0.053700828 | 0.211512481 |
| chr13:48000680-48001583 | -0.642931927 | 0.053704862 | 0.211512481 |
| chr13:41903438-41903781 | 0.853051739 | 0.05370488 | 0.211512481 |
| chr4:111977539-111977752 | -1.035088367 | 0.053723831 | 0.211576241 |
| chr1:191629895-191630296 | -0.750737546 | 0.053738162 | 0.211611952 |
| chr5:126492912-126493112 | -1.029213701 | 0.053738422 | 0.211611952 |
| chr2:58571202-58571593 | -0.706306944 | 0.053747689 | 0.211637569 |
| chr19:34265131-34265478 | -0.813943706 | 0.053768427 | 0.211708347 |
| chr8:93983040-93983522 | 0.697803032 | 0.053782816 | 0.211736362 |
| chr4:71270871-71271097 | -1.088331386 | 0.053783386 | 0.211736362 |
| chr19:9435346-9435783 | 0.716354149 | 0.053783832 | 0.211736362 |
| chr5:164504108-164504494 | -0.797015309 | 0.053790289 | 0.211747369 |
| chr6:146880938-146881344 | -0.912225288 | 0.053792156 | 0.211747369 |
| chr2:70864568-70864924 | -0.799163103 | 0.053795197 | 0.211748462 |
| chr5:147188689-147189139 | -0.825108393 | 0.053830882 | 0.211878042 |
| chr8:86044184-86044442 | 0.780760535 | 0.053846919 | 0.211930276 |
| chr8:63489696-63490566 | -0.719250035 | 0.053853972 | 0.211947148 |
| chr11:417043-417664 | 0.698837416 | 0.053877049 | 0.21202708 |
| chr21:32455693-32455961 | 0.846779103 | 0.053891145 | 0.212071662 |
| chr22:42801603-42801803 | -1.034359761 | 0.05389532 | 0.212077199 |
| chr2:188692963-188693724 | -0.649547383 | 0.053903297 | 0.212097695 |
| chr5:105350337-105350707 | -0.918874385 | 0.053917981 | 0.212140699 |
| chr18:63511340-63511748 | -0.731425591 | 0.053919763 | 0.212140699 |
| chr2:68751700-68752067 | -0.732561815 | 0.053937201 | 0.212198408 |
| chr8:110026541-110026838 | -0.797073258 | 0.05395665 | 0.212260194 |
| chr9:113341712-113342050 | 0.718971035 | 0.053959861 | 0.212260194 |
| chr4:28167368-28167965 | -0.693563783 | 0.053961216 | 0.212260194 |
| chr7:144452121-144452358 | -0.945488127 | 0.053974 | 0.212297103 |
| chr20:53768303-53768703 | 0.6275083 | 0.053976141 | 0.212297103 |
| chr5:93607691-93608193 | -0.703453676 | 0.053993069 | 0.212352783 |
| chr17:67224041-67224249 | -0.961742004 | 0.054000754 | 0.212372106 |
| chr5:147835370-147835767 | 0.783192952 | 0.054015037 | 0.212417378 |
| chr6:30722575-30722870 | 0.736656385 | 0.054017912 | 0.21241778 |
| chr17:31654925-31655418 | 0.723051758 | 0.054028011 | 0.212446589 |
| chr1:27155412-27155612 | 0.920200653 | 0.054037414 | 0.212472662 |
| chr8:96928498-96928712 | 0.876181034 | 0.054041047 | 0.212476044 |
| chr2:148201064-148201856 | -0.703294578 | 0.054051168 | 0.212504931 |
| chr2:217713471-217713785 | 0.790132874 | 0.054056514 | 0.212510372 |
| chr5:96256859-96257101 | -0.917680485 | 0.054059714 | 0.212510372 |
| chr1:207329023-207329223 | 0.900604109 | 0.054060872 | 0.212510372 |
| chr4:169152824-169153229 | -0.803118278 | 0.054065186 | 0.212516427 |
| chr5:168316205-168316752 | -0.67005089 | 0.054082896 | 0.212575132 |
| chr2:136176950-136177167 | 0.923217532 | 0.054099628 | 0.21262858 |
| chr1:107235366-107235778 | 0.654563646 | 0.054102044 | 0.21262858 |
| chr5:12805199-12805775 | -0.742077374 | 0.05410483 | 0.212628623 |
| chr8:61789950-61790303 | -0.809777242 | 0.054125036 | 0.21269273 |

| | | | |
|---|---|---|---|
| chr18:61088742-61089097 | -0.856170845 | 0.054126694 | 0.21269273 |
| chr2:207930769-207931476 | -0.720804438 | 0.054152433 | 0.212782962 |
| chr5:14727389-14727745 | 0.74144116 | 0.054167381 | 0.212830779 |
| chr12:53757251-53757659 | 0.748824382 | 0.054170585 | 0.212832458 |
| chr22:22520034-22520734 | -0.78833424 | 0.054189326 | 0.212892408 |
| chr6:55636387-55636866 | -0.718704942 | 0.054191546 | 0.212892408 |
| chr1:34407138-34407529 | -0.770805265 | 0.054195985 | 0.212892408 |
| chr9:74496904-74497437 | -0.733953108 | 0.054196958 | 0.212892408 |
| chr8:123260387-123260611 | 0.89458808 | 0.054203532 | 0.212907317 |
| chr6:16683753-16684016 | -0.965887222 | 0.054208288 | 0.212915085 |
| chr6:96089277-96089732 | -0.90393568 | 0.054212142 | 0.212916228 |
| chr2:89842356-89842556 | -1.014742364 | 0.054214137 | 0.212916228 |
| chr3:62486620-62486874 | 0.74157951 | 0.054243147 | 0.213014945 |
| chr13:45300335-45300924 | -0.678554899 | 0.054244833 | 0.213014945 |
| chr10:28263215-28263890 | -0.700399537 | 0.05427534 | 0.21312382 |
| chr7:138245792-138246001 | -0.99962727 | 0.054280624 | 0.213128475 |
| chr18:74147568-74148090 | -0.638424565 | 0.054282089 | 0.213128475 |
| chr2:150548594-150548912 | -0.849767638 | 0.054286683 | 0.213135593 |
| chr5:893531-893901 | -0.804242515 | 0.054296388 | 0.213160864 |
| chr4:44583501-44583755 | -0.903457318 | 0.054298684 | 0.213160864 |
| chr6:96436322-96436596 | 0.884561575 | 0.054321969 | 0.213233608 |
| chr19:1445257-1445474 | 0.856382213 | 0.054325603 | 0.213233608 |
| chr1:6358681-6359007 | -0.973279508 | 0.054326141 | 0.213233608 |
| chr10:34732917-34733161 | -0.935305328 | 0.054328346 | 0.213233608 |
| chr7:41188866-41189190 | 0.772801608 | 0.054338881 | 0.213264034 |
| chr2:25141177-25141449 | -0.842416717 | 0.054342268 | 0.213266402 |
| chr10:98831180-98831453 | 1.067701659 | 0.054388073 | 0.213435232 |
| chr2:54763688-54763991 | -0.892973285 | 0.054402379 | 0.213464209 |
| chr3:188525954-188526428 | -0.843474794 | 0.054403305 | 0.213464209 |
| chr6:109925778-109926042 | -1.029574775 | 0.054403815 | 0.213464209 |
| chr13:44758580-44759228 | -0.683791392 | 0.054422812 | 0.213527816 |
| chr11:44301892-44302219 | -0.876190433 | 0.054427479 | 0.21353097 |
| chr6:37059378-37060019 | -0.719555068 | 0.05442919 | 0.21353097 |
| chr10:23095981-23096562 | -0.686325835 | 0.054438348 | 0.213555964 |
| chr8:55545132-55545341 | -1.02212805 | 0.054447005 | 0.213578992 |
| chr4:184625790-184626171 | -0.79972447 | 0.054454672 | 0.213598131 |
| chr4:112765629-112765973 | -0.74680435 | 0.054458269 | 0.213601305 |
| chr7:44560009-44560277 | 0.924514068 | 0.054464667 | 0.213615463 |
| chr3:62255569-62255803 | 0.83452968 | 0.054486325 | 0.213689471 |
| chr8:96520595-96520938 | 0.864763359 | 0.054493504 | 0.213706687 |
| chrX:128781181-128781919 | 0.687501673 | 0.054501183 | 0.213725862 |
| chr1:172055696-172056481 | -0.668470121 | 0.054507705 | 0.2137405 |
| chr19:43701416-43701717 | 0.879005149 | 0.054515587 | 0.213760467 |
| chr10:7049626-7049826 | 1.058410186 | 0.054523813 | 0.213781784 |
| chr1:79363619-79363869 | -0.991578064 | 0.054529914 | 0.213786406 |
| chr15:36803977-36804389 | -0.886691383 | 0.054530572 | 0.213786406 |
| chr2:180006700-180007552 | -0.626407203 | 0.054533954 | 0.213788726 |
| chr1:106845185-106845559 | 0.670084149 | 0.054541123 | 0.21380589 |
| chr6:90369003-90369554 | 0.676019288 | 0.054551456 | 0.213835456 |
| chr5:167964610-167964910 | -0.865759141 | 0.054560701 | 0.213860754 |

| | | | |
|---|---|---|---|
| chr5:164274-164506 | 0.895901299 | 0.054568922 | 0.213882038 |
| chr22:22883998-22884419 | 0.763441955 | 0.054601611 | 0.213999215 |
| chr4:128220171-128220644 | -0.778618276 | 0.054607936 | 0.214004399 |
| chr2:160278458-160278733 | -0.870304073 | 0.05460852 | 0.214004399 |
| chr5:3522340-3522555 | 1.034126511 | 0.054611349 | 0.214004542 |
| chr1:89632692-89634102 | 0.636518541 | 0.054618388 | 0.214021179 |
| chr7:128037543-128037748 | -0.966616712 | 0.054631605 | 0.214062022 |
| chr11:1693985-1694351 | 0.757098106 | 0.054641844 | 0.214091192 |
| chr2:181178869-181179152 | -0.886089886 | 0.0546497 | 0.214111026 |
| chr17:61720528-61720820 | 0.641874119 | 0.054661491 | 0.214113816 |
| chr12:13390445-13390858 | -0.849147293 | 0.054662623 | 0.214113816 |
| chr17:61753595-61754183 | 0.618101818 | 0.054663827 | 0.214113816 |
| chr9:70106591-70106824 | -1.063284146 | 0.054668408 | 0.214113816 |
| chrX:135688203-135688459 | -1.063284146 | 0.054668408 | 0.214113816 |
| chr11:64383426-64383732 | -1.012304003 | 0.054668634 | 0.214113816 |
| chr3:194300371-194300739 | -0.740245666 | 0.054670136 | 0.214113816 |
| chr2:61853682-61854312 | -0.631593704 | 0.054672768 | 0.214113816 |
| chr7:70898991-70899191 | -1.162367685 | 0.054679862 | 0.214130654 |
| chr10:102431816-102432024 | 0.876065154 | 0.054700641 | 0.214198338 |
| chr17:38867546-38867754 | 0.799184148 | 0.054702736 | 0.214198338 |
| chr1:202330033-202330451 | -0.740792091 | 0.054711477 | 0.214221617 |
| chr17:13230537-13230812 | -0.90352408 | 0.054717525 | 0.214234349 |
| chr3:175269313-175269569 | 0.860561988 | 0.054721542 | 0.214239128 |
| chr20:25133280-25133480 | 0.993402588 | 0.054731239 | 0.214266146 |
| chr9:88534003-88535630 | 0.614380178 | 0.054738728 | 0.214284515 |
| chr21:21063206-21063444 | 0.902566326 | 0.054743777 | 0.214293331 |
| chr3:123732516-123732716 | 0.888008628 | 0.054750201 | 0.214295087 |
| chr10:14603951-14604787 | 0.655619132 | 0.054750843 | 0.214295087 |
| chr20:38173868-38174089 | -1.052722772 | 0.054752616 | 0.214295087 |
| chr9:131329118-131329500 | -0.821294403 | 0.054759802 | 0.214312265 |
| chr12:130761726-130762079 | 0.708111976 | 0.054772947 | 0.214331262 |
| chr1:2546810-2547228 | 0.680960238 | 0.054775641 | 0.214331262 |
| chr3:127536457-127536880 | -0.731677863 | 0.054775753 | 0.214331262 |
| chr4:123397213-123397875 | 0.665910851 | 0.054775845 | 0.214331262 |
| chr3:71710999-71711234 | 0.939912685 | 0.054782868 | 0.214347798 |
| chr13:47773949-47774677 | -0.697277255 | 0.054793778 | 0.214379536 |
| chr4:17817028-17817350 | 0.831750421 | 0.054803265 | 0.214400556 |
| chr8:119461635-119461971 | -0.804570579 | 0.054804747 | 0.214400556 |
| chr4:140635810-140636096 | -0.894804295 | 0.05481008 | 0.214410473 |
| chrX:31679879-31680118 | -0.999799978 | 0.054815774 | 0.214419687 |
| chr2:58634598-58635304 | -0.681385179 | 0.054818032 | 0.214419687 |
| chr13:74040240-74040560 | -0.719028627 | 0.054826376 | 0.214441377 |
| chr9:69027742-69028041 | -0.852400992 | 0.054836675 | 0.214470712 |
| chr11:40293725-40294644 | -0.632195796 | 0.054855056 | 0.214525933 |
| chr3:123602365-123602700 | 0.743551116 | 0.054856394 | 0.214525933 |
| chr8:61095906-61096426 | -0.728247345 | 0.054861764 | 0.214535505 |
| chrX:104803232-104803570 | -0.843208865 | 0.054865678 | 0.214535505 |
| chr3:158259997-158260292 | 0.932823173 | 0.054867241 | 0.214535505 |
| chr1:145446147-145446433 | 1.008963097 | 0.054888811 | 0.214600036 |
| chr20:52033349-52033827 | -0.691186545 | 0.054889639 | 0.214600036 |

| | | | |
|---|---|---|---|
| chr3:72164907-72165235 | 0.892394841 | 0.05489607 | 0.214600036 |
| chr22:42800734-42801173 | -0.752971568 | 0.054896532 | 0.214600036 |
| chr16:30775306-30775805 | 0.676739774 | 0.054897749 | 0.214600036 |
| chrX:56233755-56234087 | -0.800193109 | 0.054909245 | 0.214634024 |
| chr5:80181670-80182258 | -0.671333422 | 0.054914279 | 0.214642754 |
| chr1:155636773-155637003 | 1.00311491 | 0.05493208 | 0.214701381 |
| chr2:11692959-11693252 | -0.727128632 | 0.054953938 | 0.214775854 |
| chr10:125514931-125515208 | -0.970420603 | 0.054959326 | 0.214785956 |
| chr22:46596890-46597330 | 0.728495883 | 0.054962477 | 0.214787317 |
| chr5:134626444-134626866 | 0.798356643 | 0.054972729 | 0.214816424 |
| chr9:87812899-87813257 | 0.808079248 | 0.054990959 | 0.214867168 |
| chr17:38853379-38854041 | 0.6778404 | 0.054991323 | 0.214867168 |
| chr20:56106502-56106806 | 0.882340267 | 0.055013975 | 0.214922177 |
| chr20:37519630-37519894 | 0.909794084 | 0.055014178 | 0.214922177 |
| chr1:186679982-186680878 | -0.637100404 | 0.055016452 | 0.214922177 |
| chr2:191318838-191319156 | -0.781946688 | 0.055016622 | 0.214922177 |
| chr16:4840018-4840383 | 0.823554011 | 0.055024706 | 0.214942799 |
| chr18:74040815-74041172 | -0.776028403 | 0.055033834 | 0.214967497 |
| chr17:36486408-36486986 | 0.681169496 | 0.055041357 | 0.214982159 |
| chr11:8722165-8722460 | 0.83099393 | 0.055043199 | 0.214982159 |
| chr4:155953132-155954390 | -0.629131992 | 0.05504641 | 0.214983741 |
| chr4:187994016-187994228 | -1.100584554 | 0.055051816 | 0.214989043 |
| chr10:69230742-69231057 | -0.952269124 | 0.055053379 | 0.214989043 |
| chr17:61784587-61784899 | 0.643831216 | 0.055067307 | 0.215032474 |
| chr1:86502161-86502513 | -0.790622936 | 0.055073186 | 0.215044471 |
| chr11:128956491-128956830 | -0.769379814 | 0.055092513 | 0.215108977 |
| chr1:117537541-117537888 | -0.682390799 | 0.055112952 | 0.215174666 |
| chr7:124497278-124497534 | -1.005829871 | 0.055116825 | 0.215174666 |
| chr15:62831835-62832035 | -1.051171986 | 0.055117762 | 0.215174666 |
| chr2:171069649-171069849 | -1.105812074 | 0.055127449 | 0.215199059 |
| chr15:55457309-55457802 | 0.780485438 | 0.055129735 | 0.215199059 |
| chr3:50245849-50246489 | -0.726617171 | 0.055132501 | 0.215199059 |
| chr20:17533467-17533786 | -0.714778136 | 0.055135245 | 0.215199059 |
| chr13:23375124-23376102 | 0.622697739 | 0.055145659 | 0.215221946 |
| chr9:5799322-5799528 | -1.118728719 | 0.055146726 | 0.215221946 |
| chr9:118829242-118829644 | -0.974750892 | 0.05516285 | 0.215273907 |
| chr2:133844288-133844612 | -0.776535439 | 0.055168435 | 0.215284736 |
| chr10:49379639-49379940 | -0.769080723 | 0.05518581 | 0.215341572 |
| chr14:95319249-95319449 | 0.895841739 | 0.055201088 | 0.21539022 |
| chr1:85282335-85282810 | -0.69329849 | 0.055210816 | 0.215417211 |
| chr7:140810945-140811446 | -0.75599655 | 0.055215794 | 0.215425661 |
| chr20:49508636-49509143 | 0.693170608 | 0.05521913 | 0.215427706 |
| chr7:103344048-103344990 | -0.629523567 | 0.055234699 | 0.215477478 |
| chr5:14569993-14570240 | 0.863047047 | 0.055247311 | 0.215515706 |
| chr2:189094370-189094637 | -0.957145813 | 0.055253179 | 0.215517741 |
| chr7:56074349-56074659 | 0.775266263 | 0.055253459 | 0.215517741 |
| chr9:114914199-114914413 | 0.914971646 | 0.05526396 | 0.215547728 |
| chr3:18818251-18818622 | 0.739771024 | 0.055288538 | 0.215632615 |
| chr7:17674590-17675437 | 0.656081051 | 0.055292101 | 0.215635537 |
| chr13:57794721-57795070 | -0.865318647 | 0.05529596 | 0.215639609 |

| | | | |
|---|---|---|---|
| chr4:70584354-70584968 | -0.686723738 | 0.055300586 | 0.215646675 |
| chr7:48154803-48155391 | -0.683354039 | 0.055304789 | 0.215652089 |
| chr5:173165784-173165998 | 0.924798298 | 0.055311131 | 0.215665841 |
| chr15:91312180-91312423 | 0.944187663 | 0.055317205 | 0.21567855 |
| chr6:75852839-75853096 | 0.880548127 | 0.055322512 | 0.215688269 |
| chr17:66508738-66509450 | 0.663807596 | 0.055335845 | 0.215729273 |
| chr1:12595808-12596194 | 0.768218188 | 0.055381985 | 0.215898168 |
| chr5:175803744-175803954 | -1.000957056 | 0.055390192 | 0.215919174 |
| chr13:40921219-40922227 | -0.614141755 | 0.055395172 | 0.2159276 |
| chrX:153507522-153507943 | 0.696844784 | 0.055401404 | 0.21594091 |
| chr16:2521240-2521519 | 0.782402113 | 0.055409156 | 0.215960139 |
| chr12:27588934-27589192 | 0.930057151 | 0.055413551 | 0.215966284 |
| chr1:107254386-107254616 | 0.708056775 | 0.055417803 | 0.215968972 |
| chr4:20313639-20314007 | -0.772399224 | 0.055421159 | 0.215968972 |
| chr12:47870206-47870470 | -0.909488802 | 0.055422697 | 0.215968972 |
| chr2:140586409-140586766 | -0.733938979 | 0.055427824 | 0.215977966 |
| chr17:61523940-61524810 | 0.608307269 | 0.05543544 | 0.215996658 |
| chr7:112898736-112899025 | -0.796929548 | 0.055447236 | 0.216031632 |
| chr2:4992273-4992524 | -0.875831633 | 0.055463873 | 0.216075494 |
| chr8:104237320-104237520 | 0.959925918 | 0.055464133 | 0.216075494 |
| chrX:108892580-108892905 | -0.811338483 | 0.055475705 | 0.216109586 |
| chr4:177381856-177382059 | -1.039608769 | 0.055479787 | 0.216114501 |
| chr1:45499926-45500691 | 0.638044329 | 0.05548367 | 0.21611801 |
| chr3:112665039-112665562 | -0.770062558 | 0.055487517 | 0.21611801 |
| chr20:31678580-31679141 | 0.68863675 | 0.055489149 | 0.21611801 |
| chr8:141073043-141073320 | 0.770875463 | 0.055505529 | 0.216170815 |
| chr20:54323787-54324327 | 0.635849247 | 0.055512176 | 0.216185715 |
| chr17:80387074-80387313 | 0.969117056 | 0.055517614 | 0.216195906 |
| chr15:47717056-47717559 | 0.748709574 | 0.05553243 | 0.216242609 |
| chr9:109183571-109183882 | -0.79102809 | 0.055539641 | 0.21625154 |
| chr15:80403713-80404157 | 0.707034359 | 0.055540368 | 0.21625154 |
| chr13:95604207-95604413 | -1.136458191 | 0.055549846 | 0.216277453 |
| chr2:235449652-235450200 | -0.823149957 | 0.055587246 | 0.21641207 |
| chr9:82961555-82961840 | -0.797157782 | 0.055606179 | 0.216474782 |
| chr7:77536501-77537755 | -0.610887501 | 0.055613995 | 0.216494209 |
| chr1:35505954-35506250 | 0.811368728 | 0.055618267 | 0.216499839 |
| chr3:46960804-46961011 | 0.966676913 | 0.055630701 | 0.216536968 |
| chr2:49490586-49491165 | -0.656379563 | 0.055633457 | 0.216536968 |
| chr12:85317613-85318531 | -0.647217921 | 0.055638737 | 0.216546519 |
| chr4:173538622-173538840 | 0.972008615 | 0.055646672 | 0.216566402 |
| chr15:30172992-30173232 | 0.978950093 | 0.055654142 | 0.21658447 |
| chr17:32157163-32157638 | 0.718109162 | 0.055659865 | 0.216589486 |
| chr8:24906073-24906273 | -1.042481582 | 0.055661084 | 0.216589486 |
| chr17:78137888-78138230 | 0.700959446 | 0.055677219 | 0.216626992 |
| chr14:39368379-39368617 | 0.902161746 | 0.055679536 | 0.216626992 |
| chr10:13660075-13660314 | -0.926983939 | 0.05567973 | 0.216626992 |
| chr14:48792632-48792939 | 0.821090242 | 0.055682551 | 0.216626992 |
| chr3:144295010-144295319 | -1.000739916 | 0.055684858 | 0.216626992 |
| chr3:189572306-189572759 | -0.797827885 | 0.055695781 | 0.216658481 |
| chr6:36579758-36580133 | -0.861176595 | 0.055701996 | 0.21667166 |

| | | | |
|---|---|---|---|
| chr3:169696556-169696790 | -1.007405914 | 0.055731906 | 0.216774853 |
| chr3:15063279-15063565 | 0.715693022 | 0.055734184 | 0.216774853 |
| chr7:132930950-132931156 | -1.121426933 | 0.055743542 | 0.216800246 |
| chr3:129506330-129506841 | -0.673483179 | 0.055751876 | 0.216808245 |
| chr7:19602948-19603148 | 0.971845054 | 0.055752661 | 0.216808245 |
| chr8:11867215-11867423 | 0.923142921 | 0.055754087 | 0.216808245 |
| chr9:137300929-137301129 | 0.851260332 | 0.0557631 | 0.216827422 |
| chr2:70645815-70646304 | -0.765078572 | 0.055764678 | 0.216827422 |
| chr7:12638944-12639202 | 0.81623701 | 0.055779386 | 0.216867624 |
| chr7:101562720-101562920 | -1.003137225 | 0.055780678 | 0.216867624 |
| chr17:82234843-82235094 | 0.716850279 | 0.055784756 | 0.216872474 |
| chr22:20063774-20064046 | 0.861089968 | 0.055794745 | 0.216900302 |
| chr1:29390391-29390665 | 1.006260655 | 0.05580214 | 0.216918044 |
| chr10:126350531-126350928 | -0.831055975 | 0.055805256 | 0.216919153 |
| chr3:62246150-62246555 | -0.690512171 | 0.055810713 | 0.21692936 |
| chrX:31036820-31037203 | -0.830018362 | 0.055820192 | 0.216955197 |
| chr5:136686336-136686628 | -0.836111925 | 0.055830688 | 0.216984983 |
| chrX:16695189-16695425 | -0.93130926 | 0.055836028 | 0.216986479 |
| chr10:95806796-95806996 | -1.101638836 | 0.055836736 | 0.216986479 |
| chr3:189481694-189481894 | -1.10124645 | 0.055850557 | 0.21702918 |
| chr5:42989229-42989562 | -0.811150024 | 0.055855028 | 0.217035545 |
| chr3:165953995-165954195 | 0.996647386 | 0.055860351 | 0.217045225 |
| chrX:103666560-103666760 | 1.021143355 | 0.055863267 | 0.217045548 |
| chr7:12942150-12942350 | 0.916465533 | 0.055868494 | 0.217054849 |
| chr22:31630741-31631143 | 0.702087956 | 0.055872762 | 0.217060425 |
| chr4:156595164-156595414 | 1.039134358 | 0.055876945 | 0.217065671 |
| chr9:39463809-39464032 | -1.03691567 | 0.055885972 | 0.217089733 |
| chr4:4260245-4260554 | 0.823584429 | 0.055892782 | 0.217095675 |
| chr2:240565831-240566058 | -0.800866507 | 0.055893169 | 0.217095675 |
| chr18:24016719-24016977 | -0.91040054 | 0.055901046 | 0.217115266 |
| chr14:31336730-31337107 | 0.71523717 | 0.055908765 | 0.217119139 |
| chr11:62171210-62171504 | 0.826346587 | 0.055909353 | 0.217119139 |
| chr7:111750182-111750522 | -0.757176183 | 0.05591157 | 0.217119139 |
| chr5:110330047-110330421 | -0.910294956 | 0.055915116 | 0.217119139 |
| chr7:89549629-89550206 | -0.712914057 | 0.055917085 | 0.217119139 |
| chr21:32412866-32413140 | 0.784072835 | 0.055920338 | 0.217119139 |
| chr4:185019895-185020150 | -0.998991615 | 0.055925543 | 0.217119139 |
| chr13:42720437-42720732 | -0.822009764 | 0.055927458 | 0.217119139 |
| chr15:92869135-92870639 | 0.612463608 | 0.055927546 | 0.217119139 |
| chr1:203154176-203154722 | -0.669358072 | 0.055936417 | 0.217132348 |
| chr15:87520341-87520577 | -0.929203645 | 0.055936616 | 0.217132348 |
| chr13:77698790-77699030 | -0.858758523 | 0.055950803 | 0.217176415 |
| chr16:2964538-2964805 | 0.750770182 | 0.055977348 | 0.217267273 |
| chr5:62719949-62720310 | -0.718146711 | 0.055982413 | 0.217267273 |
| chr1:103525367-103526428 | -0.620867916 | 0.055983635 | 0.217267273 |
| chr16:18946250-18946626 | -0.801361244 | 0.055986185 | 0.217267273 |
| chr7:89731636-89731884 | -1.08474981 | 0.055988389 | 0.217267273 |
| chr16:30948151-30949370 | 0.618354034 | 0.056012902 | 0.217351391 |
| chr1:210328500-210329775 | -0.635046966 | 0.056020002 | 0.217364207 |
| chr6:119493966-119494444 | -0.804197724 | 0.056021878 | 0.217364207 |

| | | | |
|---|---|---|---|
| chr5:125277499-125277860 | -0.809405284 | 0.056028195 | 0.217377707 |
| chr14:34139841-34140151 | 0.744367071 | 0.056031889 | 0.217381034 |
| chr18:48475917-48476266 | -0.814571981 | 0.056072645 | 0.217525435 |
| chr3:23651056-23651340 | 0.919550395 | 0.056074788 | 0.217525435 |
| chr2:40827642-40827842 | -1.068024335 | 0.056092404 | 0.217568303 |
| chr15:65824158-65824457 | 0.78043937 | 0.056092811 | 0.217568303 |
| chr3:53664870-53665098 | 0.912712149 | 0.056096103 | 0.217568303 |
| chr4:85474715-85475652 | -0.632207345 | 0.056097497 | 0.217568303 |
| chr4:88195899-88196110 | 0.99171981 | 0.056100036 | 0.217568303 |
| chr1:175963239-175963439 | 1.096835677 | 0.056123209 | 0.217647155 |
| chr17:55315187-55315410 | 0.974154394 | 0.056129992 | 0.217662443 |
| chr20:50770097-50770545 | 0.618841602 | 0.056134408 | 0.217668552 |
| chr18:70598855-70599162 | -0.844910603 | 0.056139788 | 0.217678399 |
| chr5:77775590-77776602 | -0.618187919 | 0.056148803 | 0.217702336 |
| chr11:68764002-68764226 | 0.809472647 | 0.056158837 | 0.217709203 |
| chr4:169791715-169792063 | -0.753398503 | 0.056160882 | 0.217709203 |
| chr5:140302777-140303523 | -0.64113592 | 0.056164432 | 0.217709203 |
| chr12:120079374-120080088 | 0.664834923 | 0.056165281 | 0.217709203 |
| chr15:93287489-93287694 | 1.006927501 | 0.056166586 | 0.217709203 |
| chr1:64686777-64687051 | -0.866306576 | 0.05617029 | 0.217709203 |
| chr2:183543808-183544094 | -0.915835684 | 0.056170463 | 0.217709203 |
| chr9:78439532-78439819 | -0.813266124 | 0.056179522 | 0.2177333 |
| chr6:158910467-158910735 | 0.870270796 | 0.056200158 | 0.217802262 |
| chr20:53769688-53769961 | 0.63357459 | 0.05621197 | 0.21783702 |
| chr20:47377264-47377464 | 0.658900632 | 0.05624083 | 0.217937837 |
| chr1:100132560-100133472 | -0.619443991 | 0.056247889 | 0.217947035 |
| chr1:41789924-41790293 | 0.764789409 | 0.056248892 | 0.217947035 |
| chr16:78081323-78081523 | 0.999274277 | 0.056255496 | 0.217952072 |
| chr12:131829081-131829371 | 0.746466613 | 0.056257905 | 0.217952072 |
| chr21:40521073-40521290 | -1.020805684 | 0.056259695 | 0.217952072 |
| chr9:127887850-127888115 | 0.820701106 | 0.05626157 | 0.217952072 |
| chr11:76824978-76825549 | -0.693083106 | 0.056268653 | 0.217968491 |
| chr19:38831148-38832160 | 0.622607757 | 0.056280021 | 0.218001503 |
| chr9:134413763-134414077 | 0.745001483 | 0.056285993 | 0.218012279 |
| chr11:83922363-83922580 | -1.076018206 | 0.056288493 | 0.218012279 |
| chr1:107315709-107315934 | 0.711451565 | 0.056298389 | 0.218039584 |
| chr4:81405862-81406338 | 0.756679938 | 0.056301457 | 0.218040447 |
| chr2:25568999-25569230 | -1.009414272 | 0.056307043 | 0.218051058 |
| chr7:106588951-106589330 | 0.785096264 | 0.056320874 | 0.218093598 |
| chr6:139656048-139656448 | -0.692116359 | 0.056344697 | 0.21815546 |
| chrX:153469811-153471186 | 0.608490897 | 0.056344784 | 0.21815546 |
| chr2:35114804-35115078 | -0.859253915 | 0.056345391 | 0.21815546 |
| chrX:134459653-134460785 | 0.621794568 | 0.056355032 | 0.21817932 |
| chr19:19385250-19385998 | -0.645134756 | 0.056358632 | 0.21817932 |
| chr1:225611634-225612330 | 0.725221661 | 0.056360096 | 0.21817932 |
| chr1:100463756-100463995 | -0.848174915 | 0.05636451 | 0.218185384 |
| chr6:31558450-31558686 | 1.00109948 | 0.056372575 | 0.218199521 |
| chr12:130764684-130764903 | -0.887769318 | 0.056373858 | 0.218199521 |
| chr5:31820120-31820320 | -1.066072122 | 0.056386646 | 0.218230044 |
| chr9:83937845-83938234 | -0.768046131 | 0.056388981 | 0.218230044 |

| | | | |
|---|---|---|---|
| chr12:52025410-52025657 | 0.748883437 | 0.056390288 | 0.218230044 |
| chr2:131818598-131818902 | -0.862024596 | 0.056406918 | 0.218273021 |
| chr20:53569825-53570183 | 0.628678722 | 0.05640709 | 0.218273021 |
| chr3:107855735-107856146 | -0.678002076 | 0.05641637 | 0.218297905 |
| chr2:33421853-33422153 | 0.846995949 | 0.056458034 | 0.218448088 |
| chr10:77380538-77380782 | -1.017097852 | 0.056467017 | 0.218471811 |
| chr1:106535676-106536206 | 0.64029619 | 0.056478325 | 0.218498021 |
| chr1:207203102-207203625 | 0.721716701 | 0.056479494 | 0.218498021 |
| chr18:5003910-5004858 | -0.672303516 | 0.056484672 | 0.218507019 |
| chr8:128889992-128890192 | 0.963191996 | 0.056488419 | 0.218510484 |
| chr1:184465729-184466080 | -0.876709895 | 0.056504787 | 0.218562763 |
| chr14:73526429-73527079 | 0.677696606 | 0.056509507 | 0.218569989 |
| chr12:27821218-27821514 | -0.862857523 | 0.056515211 | 0.218581016 |
| chr14:54025785-54025995 | 1.008911566 | 0.056519713 | 0.218586311 |
| chr1:156368222-156368579 | 0.704174026 | 0.056522286 | 0.218586311 |
| chr7:104403624-104403824 | -1.117943075 | 0.056525572 | 0.218587989 |
| chr18:59323893-59324180 | -1.011602172 | 0.056531698 | 0.218600643 |
| chrX:30303023-30303293 | -0.811126153 | 0.056561289 | 0.218704031 |
| chr7:122014796-122015179 | -0.908600426 | 0.056572498 | 0.218734505 |
| chr15:79635302-79635531 | 0.950404266 | 0.056574879 | 0.218734505 |
| chr7:91242243-91242443 | -1.055145818 | 0.056586177 | 0.218767144 |
| chr5:10962150-10962608 | -0.727570928 | 0.056594987 | 0.218774389 |
| chr5:108685824-108686029 | -0.940495485 | 0.056596016 | 0.218774389 |
| chr11:106156273-106156608 | 0.833539587 | 0.056596616 | 0.218774389 |
| chr6:132715263-132715605 | -0.752117408 | 0.056610296 | 0.218816228 |
| chr7:42236794-42237792 | -0.630628555 | 0.05661648 | 0.218827601 |
| chr7:25069106-25069306 | 0.939322331 | 0.05661895 | 0.218827601 |
| chr2:62047823-62048060 | 1.03805329 | 0.056630281 | 0.218860355 |
| chr6:127299843-127300144 | 0.796484918 | 0.05664018 | 0.218882876 |
| chr3:154811879-154812087 | 1.017318964 | 0.056641822 | 0.218882876 |
| chr22:45597517-45597871 | 0.8303454 | 0.056647032 | 0.218890672 |
| chr1:21659178-21659577 | 0.759479947 | 0.056649553 | 0.218890672 |
| chr1:200974194-200974404 | 0.771820301 | 0.056657986 | 0.21891222 |
| chr2:195307825-195308271 | 0.743771174 | 0.05668261 | 0.218982468 |
| chr20:23420131-23420338 | 0.892889524 | 0.056684394 | 0.218982468 |
| chr4:87126076-87126315 | -1.063971263 | 0.056684742 | 0.218982468 |
| chr18:35376796-35377514 | 0.676575975 | 0.056691779 | 0.218998615 |
| chrX:105745872-105746288 | -0.751806622 | 0.056704622 | 0.219031776 |
| chr20:53381903-53382309 | -0.825512612 | 0.05670824 | 0.219031776 |
| chr20:58693069-58693739 | 0.63843589 | 0.05670894 | 0.219031776 |
| chr10:79238578-79238892 | -0.708807834 | 0.056723084 | 0.219069832 |
| chr15:22979219-22979513 | 0.799867376 | 0.056724511 | 0.219069832 |
| chr5:38667073-38667300 | 0.840498849 | 0.056741347 | 0.21912381 |
| chr6:116111990-116112277 | -0.824285932 | 0.056747288 | 0.219135706 |
| chr2:215137818-215138387 | -0.721611214 | 0.056763481 | 0.219187193 |
| chr8:58137835-58138094 | 0.842195677 | 0.056770879 | 0.219204299 |
| chr17:78352722-78353579 | 0.637288384 | 0.056773633 | 0.219204299 |
| chr1:23539777-23540172 | 0.71430649 | 0.056791964 | 0.219250074 |
| chr8:101891896-101892099 | 0.922159101 | 0.056792393 | 0.219250074 |
| chr7:148963583-148964016 | -0.690671996 | 0.056794073 | 0.219250074 |

| | | | |
|---|---|---|---|
| chr3:78855101-78855471 | -0.864305295 | 0.05679754 | 0.219252411 |
| chr3:27253777-27254200 | -0.689250718 | 0.056814195 | 0.219305656 |
| chr4:21412788-21413204 | -0.792313282 | 0.056851071 | 0.219436943 |
| chr7:11343900-11344353 | 0.751837226 | 0.056869635 | 0.219494447 |
| chr6:94942005-94942800 | -0.665649981 | 0.056871698 | 0.219494447 |
| chr13:26668443-26668709 | -0.970081272 | 0.056875969 | 0.219499872 |
| chr20:24497965-24498165 | 0.96741758 | 0.056893797 | 0.219537378 |
| chr6:72316834-72317069 | -0.977433785 | 0.056900847 | 0.219537378 |
| chr11:107840622-107841108 | -0.734759763 | 0.056902406 | 0.219537378 |
| chr12:84508335-84508900 | -0.673926921 | 0.056903631 | 0.219537378 |
| chr15:60414170-60414401 | 0.90057601 | 0.056904234 | 0.219537378 |
| chr14:23154277-23154820 | -0.690342375 | 0.05690538 | 0.219537378 |
| chr3:48465548-48465962 | 0.710153093 | 0.056905743 | 0.219537378 |
| chr1:78839737-78839990 | -1.020774125 | 0.056916957 | 0.219569585 |
| chr5:178000301-178000518 | 0.912389312 | 0.056924604 | 0.219588029 |
| chr2:11829793-11830348 | -0.687110012 | 0.056930669 | 0.219591503 |
| chr20:23161674-23161941 | -0.916602019 | 0.056931237 | 0.219591503 |
| chr1:207981974-207982347 | -0.789774587 | 0.056934463 | 0.219592892 |
| chr12:8860536-8860760 | -1.024209441 | 0.056956476 | 0.21966674 |
| chr14:34554260-34555735 | 0.622349226 | 0.056968108 | 0.21969452 |
| chr4:170138636-170138924 | -0.783774234 | 0.056969414 | 0.21969452 |
| chr10:35120324-35120598 | -0.856792132 | 0.056972747 | 0.219696318 |
| chr20:51231228-51231428 | 0.830929374 | 0.056977356 | 0.219703032 |
| chr20:41029387-41029616 | 0.701416671 | 0.056988701 | 0.21973069 |
| chr14:51534507-51535127 | 0.652628645 | 0.056990264 | 0.21973069 |
| chr7:94888412-94888858 | -0.772798093 | 0.057003635 | 0.219771185 |
| chr12:113149520-113150048 | -0.691525178 | 0.057016946 | 0.219811445 |
| chr6:36360491-36360886 | 0.728164329 | 0.05704095 | 0.21989292 |
| chr6:28981589-28981835 | 0.724849632 | 0.057043881 | 0.219893156 |
| chr3:146646272-146646484 | -0.953411742 | 0.057047472 | 0.219895934 |
| chr12:3120180-3120587 | -0.814788029 | 0.057050572 | 0.219896822 |
| chr18:32333815-32334139 | 0.867585874 | 0.057053836 | 0.219898342 |
| chr3:64412810-64413010 | 0.716305305 | 0.057088739 | 0.220000388 |
| chr11:408747-408962 | 0.902813404 | 0.057088915 | 0.220000388 |
| chr10:16545143-16545466 | -0.814618675 | 0.057088926 | 0.220000388 |
| chr5:168987454-168988286 | -0.663081417 | 0.057096239 | 0.220017503 |
| chr4:148215242-148215562 | -0.77700889 | 0.057099993 | 0.220020903 |
| chr7:20152473-20153311 | 0.64496442 | 0.057106087 | 0.22003332 |
| chr19:1978563-1978835 | 0.84850056 | 0.057149342 | 0.220188911 |
| chr8:129825324-129825585 | 0.817697728 | 0.057158441 | 0.220212896 |
| chr2:27414274-27414523 | -0.915356106 | 0.057163874 | 0.22021895 |
| chr1:48039245-48039686 | -0.802889108 | 0.05716613 | 0.22021895 |
| chr2:135785086-135785307 | -0.897237797 | 0.057168635 | 0.22021895 |
| chr7:150405254-150405502 | -0.791129099 | 0.05717634 | 0.22023756 |
| chr7:112480380-112480646 | -0.816805849 | 0.05718458 | 0.220258225 |
| chrX:68209076-68209417 | -0.947399104 | 0.057195196 | 0.22028804 |
| chr20:48006750-48007070 | 0.643916667 | 0.057228486 | 0.220405179 |
| chr15:80911803-80912120 | -0.866900402 | 0.057239098 | 0.220430793 |
| chr2:199606448-199606648 | -1.122706338 | 0.057244774 | 0.220430793 |
| chr7:39703266-39703494 | -0.906152763 | 0.057245666 | 0.220430793 |

| | | | |
|---|---|---|---|
| chr2:36371969-36372404 | -0.731090201 | 0.057246644 | 0.220430793 |
| chr16:3922396-3922610 | 0.913632126 | 0.05725763 | 0.220454874 |
| chr20:41387422-41387882 | -0.683288839 | 0.057258764 | 0.220454874 |
| chr20:49887376-49887827 | -0.676992291 | 0.057261529 | 0.220454874 |
| chr14:68191699-68192144 | -0.798366498 | 0.057271738 | 0.220483097 |
| chr7:100465845-100466124 | -0.877437028 | 0.057276945 | 0.220487124 |
| chr15:78360405-78360627 | 0.90783617 | 0.057278539 | 0.220487124 |
| chr1:244326896-244327112 | 0.917017974 | 0.057283358 | 0.220494599 |
| chr12:130636476-130636676 | 1.018725239 | 0.057288792 | 0.220504435 |
| chr3:73192040-73192548 | -0.732905911 | 0.057303814 | 0.220551179 |
| chr15:64760026-64760247 | 0.898764031 | 0.057307429 | 0.220554014 |
| chr4:3073883-3074818 | 0.614178066 | 0.057317189 | 0.220580494 |
| chr7:17130254-17130778 | 0.660052327 | 0.057329383 | 0.220607973 |
| chr17:68075154-68075609 | -0.865799856 | 0.057331752 | 0.220607973 |
| chr12:33021365-33021602 | 0.91051076 | 0.057332967 | 0.220607973 |
| chr15:92652647-92652913 | -0.814980194 | 0.05734339 | 0.220637001 |
| chrX:74344023-74344354 | 0.814064444 | 0.057365121 | 0.220709529 |
| chr7:134565569-134565826 | -0.941091743 | 0.057374434 | 0.220734279 |
| chr14:94769765-94770013 | 0.88037684 | 0.057381646 | 0.220750939 |
| chr4:84952101-84952543 | -0.753883902 | 0.057389419 | 0.220769759 |
| chr13:22983152-22983508 | 0.704280963 | 0.057406197 | 0.220823214 |
| chr14:54870626-54871146 | 0.711780857 | 0.057411012 | 0.220830651 |
| chrX:118495510-118495914 | 0.824532598 | 0.057420812 | 0.22085726 |
| chr11:33629935-33630767 | -0.683272499 | 0.057438624 | 0.22091468 |
| chr8:46917762-46918030 | 0.809449949 | 0.05745932 | 0.220979063 |
| chr20:32976363-32977126 | 0.657913586 | 0.057461132 | 0.220979063 |
| chr3:140715146-140715426 | 0.829201754 | 0.057473095 | 0.220996766 |
| chr3:53255409-53256273 | 0.64552277 | 0.057474308 | 0.220996766 |
| chr15:79004405-79004777 | -0.725696016 | 0.057474388 | 0.220996766 |
| chr2:191638493-191638855 | 0.817930696 | 0.057518636 | 0.221155808 |
| chrX:130060127-130061134 | 0.621142466 | 0.057530526 | 0.221190424 |
| chr7:107169370-107170009 | -0.617842905 | 0.057534827 | 0.221195861 |
| chr4:112285398-112286556 | -0.616236533 | 0.057543098 | 0.221216558 |
| chr7:70459539-70459868 | -0.812097441 | 0.057549975 | 0.221231896 |
| chr1:224733406-224733649 | 0.895625163 | 0.057565616 | 0.221261922 |
| chr8:42805295-42805771 | 0.727033487 | 0.057565985 | 0.221261922 |
| chr20:25247121-25248815 | 0.608177391 | 0.057566448 | 0.221261922 |
| chr1:17250672-17251069 | -0.776725959 | 0.057575611 | 0.221278883 |
| chr7:51263536-51264167 | -0.632471957 | 0.057576637 | 0.221278883 |
| chr12:93881034-93881298 | -0.977437472 | 0.057587448 | 0.22130933 |
| chr1:171927984-171928246 | 0.922690469 | 0.057593258 | 0.221320557 |
| chr2:238529311-238529658 | 0.807647647 | 0.057601096 | 0.221336794 |
| chr20:63657324-63658424 | 0.604791851 | 0.05760326 | 0.221336794 |
| chr7:102748411-102749419 | -0.609334866 | 0.057613192 | 0.221363856 |
| chr16:80751683-80751913 | -0.996099628 | 0.057618932 | 0.221364889 |
| chr22:21555886-21556427 | 0.724834013 | 0.057619888 | 0.221364889 |
| chr8:99523528-99523728 | 0.971891128 | 0.057622128 | 0.221364889 |
| chr7:149826421-149826825 | -0.722909693 | 0.057629537 | 0.22137861 |
| chr17:78307880-78308223 | 0.764103036 | 0.057632692 | 0.22137861 |
| chr3:25725792-25726126 | 0.73210368 | 0.057634368 | 0.22137861 |

| | | | |
|---|---|---|---|
| chr3:70308768-70308974 | -1.036807238 | 0.057642861 | 0.221388969 |
| chr5:158594338-158594615 | 0.759842017 | 0.057643315 | 0.221388969 |
| chr1:202674465-202674679 | 0.892702694 | 0.057645733 | 0.221388969 |
| chr1:200184175-200184649 | -0.648619946 | 0.057656746 | 0.221410635 |
| chrX:73510973-73511687 | -0.69783748 | 0.057661887 | 0.221410635 |
| chr22:32412045-32412362 | 0.783959425 | 0.057664519 | 0.221410635 |
| chrX:65196874-65197320 | -0.769099034 | 0.057664561 | 0.221410635 |
| chr17:48505836-48506154 | -0.799060707 | 0.057665822 | 0.221410635 |
| chr8:11564565-11564934 | -0.776763231 | 0.057672951 | 0.221423328 |
| chr3:180617052-180617537 | -0.893630195 | 0.057674908 | 0.221423328 |
| chr1:88922235-88922555 | -0.827204911 | 0.057690352 | 0.221471525 |
| chr22:27990815-27991064 | -1.001262719 | 0.057703391 | 0.221510483 |
| chr7:1418503-1418829 | 0.717435767 | 0.057713943 | 0.221539892 |
| chr20:56689075-56689805 | -0.62595967 | 0.057724337 | 0.221568686 |
| chr7:134724644-134724932 | -0.858104001 | 0.057739277 | 0.22161493 |
| chr19:53612744-53612944 | 0.951641975 | 0.057751887 | 0.221643256 |
| chr20:3866051-3866958 | -0.649052065 | 0.057752442 | 0.221643256 |
| chr22:41941332-41941584 | 0.939508317 | 0.057763788 | 0.221675699 |
| chr4:52933-53834 | 0.605130518 | 0.057767832 | 0.221678354 |
| chr7:87091505-87091894 | -0.724912403 | 0.057770267 | 0.221678354 |
| chr6:43990742-43991114 | -0.787253615 | 0.057777982 | 0.221694759 |
| chr1:41418951-41419716 | -0.651053393 | 0.057780329 | 0.221694759 |
| chr5:154117131-154117348 | 0.989143988 | 0.057793595 | 0.221734557 |
| chr7:37234927-37235134 | -1.011398732 | 0.057802075 | 0.221744593 |
| chr8:41421160-41421381 | -1.081334217 | 0.057803699 | 0.221744593 |
| chr9:88173463-88173686 | 0.957896482 | 0.057804893 | 0.221744593 |
| chr18:66616630-66617030 | -0.760699209 | 0.057812813 | 0.221755295 |
| chr11:102746757-102747619 | -0.685982004 | 0.057813471 | 0.221755295 |
| chr7:83994700-83995498 | -0.738712002 | 0.057816435 | 0.221755565 |
| chr12:20624685-20624952 | -1.039131334 | 0.057821452 | 0.221763704 |
| chr14:21614382-21614586 | 0.984321142 | 0.05782915 | 0.221782129 |
| chr9:34633936-34634202 | -0.774911949 | 0.057839314 | 0.221810006 |
| chr4:88908620-88908921 | 0.942757354 | 0.057870547 | 0.221918676 |
| chr4:6415218-6415683 | -0.782808125 | 0.057873783 | 0.221919978 |
| chr1:175041032-175041232 | 1.01144479 | 0.057889339 | 0.221960395 |
| chr8:109954267-109954512 | 0.847807801 | 0.057890117 | 0.221960395 |
| chrX:16711988-16713061 | -0.625255077 | 0.057894385 | 0.221965653 |
| chr2:140891880-140892113 | 0.930960448 | 0.057899956 | 0.221975904 |
| chr7:5418411-5419247 | 0.625003089 | 0.057908686 | 0.221998264 |
| chr2:70984975-70985183 | -0.922058046 | 0.057933657 | 0.222077839 |
| chr5:104910883-104911762 | -0.635614073 | 0.05793524 | 0.222077839 |
| chr13:46614785-46615104 | -0.856272651 | 0.057946928 | 0.2220866 |
| chr16:75764175-75764505 | -0.706427948 | 0.057948621 | 0.2220866 |
| chr1:156608127-156608429 | 0.739980358 | 0.057951641 | 0.2220866 |
| chr1:196794234-196794443 | -1.020206928 | 0.057951652 | 0.2220866 |
| chr4:37408211-37408895 | -0.685830881 | 0.057952018 | 0.2220866 |
| chr22:23856724-23858130 | 0.610256721 | 0.057965941 | 0.222128848 |
| chrX:38008722-38009019 | -0.956361907 | 0.057973446 | 0.2221465 |
| chr2:173402429-173402764 | -0.753778908 | 0.057978821 | 0.222155983 |
| chr7:7424249-7424809 | 0.666436921 | 0.05798491 | 0.222168206 |

| | | | |
|---|---|---|---|
| chr6:92767124-92767497 | -0.829965294 | 0.058018156 | 0.222284471 |
| chr8:145065981-145066264 | -0.92500811 | 0.058023577 | 0.222285995 |
| chrX:25175804-25176150 | -0.848818643 | 0.058024356 | 0.222285995 |
| chr3:100875277-100875477 | -1.034515622 | 0.058038044 | 0.222322107 |
| chr16:1872144-1872541 | 0.789601606 | 0.058041924 | 0.222322107 |
| chr4:3607574-3607793 | 0.96270014 | 0.058042487 | 0.222322107 |
| chr14:57329989-57330216 | 0.953115294 | 0.058048574 | 0.222334307 |
| chr1:32240204-32240823 | 0.695596768 | 0.058052463 | 0.22233809 |
| chr7:26291581-26292406 | 0.656919619 | 0.05806725 | 0.222383605 |
| chr11:43546822-43547385 | 0.740977159 | 0.058093085 | 0.22247143 |
| chr10:52443820-52444217 | 0.785795349 | 0.058114421 | 0.222542015 |
| chr8:124080804-124081221 | 0.726072273 | 0.058130401 | 0.222592083 |
| chr19:1248435-1249143 | 0.640073275 | 0.058148354 | 0.222641123 |
| chr10:135201-136069 | 0.620073582 | 0.058149019 | 0.222641123 |
| chr3:48898016-48899135 | 0.651399592 | 0.058155 | 0.22264243 |
| chr16:31168965-31169185 | 0.960255868 | 0.058155172 | 0.22264243 |
| chr1:115494716-115495120 | -0.789508914 | 0.058182613 | 0.222736355 |
| chr12:576110-576375 | -0.839991102 | 0.058188007 | 0.222745878 |
| chr1:151946314-151946666 | -0.729206771 | 0.058205989 | 0.22280358 |
| chr7:158856052-158857087 | -0.610425936 | 0.058214494 | 0.222813819 |
| chr4:16001370-16001822 | 0.685415937 | 0.05821521 | 0.222813819 |
| chr10:103967560-103967760 | 0.887877267 | 0.058218532 | 0.222813819 |
| chr13:29754478-29754741 | -0.836406321 | 0.058220295 | 0.222813819 |
| chr8:56557922-56558205 | -0.89086887 | 0.058243062 | 0.222881838 |
| chr16:17304187-17304851 | 0.665912841 | 0.058245882 | 0.222881838 |
| chr1:190563267-190563676 | 0.726738009 | 0.058246795 | 0.222881838 |
| chr13:91923276-91923514 | -0.97490188 | 0.058251162 | 0.222887417 |
| chr7:19040401-19040601 | 1.095007343 | 0.058261789 | 0.222916947 |
| chr5:14108540-14108820 | -0.733879586 | 0.05827012 | 0.222937691 |
| chr17:40619935-40620360 | 0.725043618 | 0.058286532 | 0.222989346 |
| chr20:55845893-55846111 | -0.995605482 | 0.058302296 | 0.223038521 |
| chr6:13229178-13229470 | -0.915858691 | 0.058311207 | 0.223061472 |
| chr2:41791606-41791851 | -0.902442463 | 0.058318196 | 0.223067932 |
| chr1:155129721-155130111 | 0.674915278 | 0.058318718 | 0.223067932 |
| chr9:120755888-120756088 | 0.972645462 | 0.058322025 | 0.223069446 |
| chr3:102465927-102466296 | -0.904173339 | 0.058331519 | 0.22309462 |
| chr5:139137701-139137988 | -0.806240568 | 0.058353755 | 0.223164224 |
| chr19:13817905-13818598 | 0.669584432 | 0.058359695 | 0.223164224 |
| chr19:11265750-11266042 | 0.805180479 | 0.0583606 | 0.223164224 |
| chr6:85472914-85473164 | 0.82972439 | 0.058361368 | 0.223164224 |
| chr1:229843871-229844071 | 0.898234661 | 0.058365531 | 0.223169008 |
| chr6:133241193-133241693 | -0.779598515 | 0.058369199 | 0.223171895 |
| chr7:35151762-35152397 | 0.700125167 | 0.058373781 | 0.223178276 |
| chr5:139459343-139459543 | 1.001995762 | 0.058386744 | 0.223216438 |
| chr20:41300971-41301178 | 0.844241518 | 0.058392057 | 0.223216438 |
| chr11:46273989-46274314 | 0.751050017 | 0.058393166 | 0.223216438 |
| chr7:54074782-54074988 | 1.011232862 | 0.058397073 | 0.223216438 |
| chr19:42529246-42529446 | 0.91632395 | 0.058398328 | 0.223216438 |
| chr1:171655335-171655805 | -0.69249967 | 0.058401534 | 0.223217557 |
| chr13:38684692-38685030 | -0.694707885 | 0.058408634 | 0.223226341 |

| | | | |
|---|---|---|---|
| chr12:65602978-65603204 | -0.920086903 | 0.058409747 | 0.223226341 |
| chr11:535566-535891 | 0.719903815 | 0.058412573 | 0.223226341 |
| chr8:142295408-142295708 | 0.752617613 | 0.058425359 | 0.223264071 |
| chr7:17304957-17305699 | 0.668447021 | 0.058433235 | 0.22328303 |
| chr1:183051687-183051953 | 0.884022927 | 0.05847221 | 0.223420818 |
| chr12:83870901-83871826 | -0.630979464 | 0.058485734 | 0.223461351 |
| chr2:16763576-16763832 | -0.882173902 | 0.058498277 | 0.223497536 |
| chr1:148010906-148011132 | 0.879173903 | 0.058502428 | 0.223497536 |
| chr14:93115785-93116124 | 0.691329093 | 0.058503955 | 0.223497536 |
| chr7:107661516-107662207 | -0.661835321 | 0.058510666 | 0.223512029 |
| chr3:51719151-51719406 | -0.973804033 | 0.058525377 | 0.223557078 |
| chr16:2009741-2010038 | 0.739672638 | 0.058545767 | 0.223623818 |
| chr3:196026119-196026377 | -0.951995562 | 0.05855886 | 0.223655696 |
| chr10:74875979-74876319 | -0.811122955 | 0.058559951 | 0.223655696 |
| chr14:60266470-60266729 | 0.876107444 | 0.058569018 | 0.223679174 |
| chr20:54006848-54007281 | 0.619817981 | 0.058585623 | 0.22373144 |
| chr7:149820749-149821336 | -0.68479255 | 0.058596868 | 0.223763229 |
| chr3:50123473-50123742 | -0.837090236 | 0.058618341 | 0.223834072 |
| chr17:42577322-42577980 | 0.66304939 | 0.058632386 | 0.223876547 |
| chr4:6574728-6575451 | 0.646493681 | 0.058642205 | 0.223895924 |
| chr8:104369107-104369307 | 0.874485307 | 0.058643995 | 0.223895924 |
| chr3:65030906-65031488 | -0.753206949 | 0.058646227 | 0.223895924 |
| chr8:144372754-144373231 | 0.670663073 | 0.058651091 | 0.223903338 |
| chr7:14458494-14458954 | 0.812736926 | 0.058665458 | 0.223947027 |
| chr10:97511406-97511808 | 0.808107409 | 0.058671516 | 0.223958994 |
| chr2:197443981-197444260 | -0.983029752 | 0.058681818 | 0.223975859 |
| chr7:14903288-14903695 | -0.84260898 | 0.058683997 | 0.223975859 |
| chr8:129999180-129999394 | 0.80834043 | 0.058684703 | 0.223975859 |
| chr7:144657041-144657241 | -1.054858937 | 0.058696706 | 0.223997774 |
| chr1:188887637-188887859 | 1.071439088 | 0.058696735 | 0.223997774 |
| chr5:113679009-113679252 | -0.898557381 | 0.058699216 | 0.223997774 |
| chr8:143382628-143383026 | 0.669596746 | 0.058716369 | 0.224048429 |
| chr6:70369344-70369669 | -0.763292384 | 0.058718338 | 0.224048429 |
| chr3:9351797-9351997 | 0.840906032 | 0.058727087 | 0.224070653 |
| chr7:101138284-101138803 | -0.704555633 | 0.05874139 | 0.224095914 |
| chr20:47865969-47866217 | 0.683170881 | 0.058746672 | 0.224095914 |
| chr6:135180496-135180731 | -0.748076645 | 0.058747749 | 0.224095914 |
| chr20:16654898-16655231 | 0.805094691 | 0.058748271 | 0.224095914 |
| chr10:31318281-31319312 | -0.633441726 | 0.05874887 | 0.224095914 |
| chr7:42493528-42494527 | -0.637551334 | 0.058752613 | 0.224095914 |
| chr8:121227377-121227602 | -1.001162712 | 0.058754181 | 0.224095914 |
| chr22:30386116-30386316 | 0.959693583 | 0.058765089 | 0.224126366 |
| chr14:75831928-75832342 | 0.702460927 | 0.058769093 | 0.224130479 |
| chr8:3536551-3536809 | -0.898081407 | 0.058775752 | 0.224138171 |
| chr7:111569133-111569363 | -0.967641205 | 0.05877696 | 0.224138171 |
| chr3:85262867-85263569 | -0.753090992 | 0.058782756 | 0.224149117 |
| chr4:166594172-166594668 | -0.732929467 | 0.058786439 | 0.224152003 |
| chr1:53592622-53593044 | 0.703961016 | 0.058799622 | 0.224181136 |
| chr11:69293861-69294500 | -0.640352537 | 0.058799931 | 0.224181136 |
| chr4:118664701-118664942 | -0.991247037 | 0.058809764 | 0.224207472 |

| | | | |
|---|---|---|---|
| chr5:129459539-129460551 | -0.610467958 | 0.058827669 | 0.224260508 |
| chr1:190997668-190998011 | 0.921416459 | 0.05882953 | 0.224260508 |
| chr5:168698943-168699425 | -0.731718914 | 0.058836172 | 0.224274671 |
| chr3:56811085-56811680 | -0.700460182 | 0.058842057 | 0.224285947 |
| chr1:153957907-153958363 | 0.658508028 | 0.058845083 | 0.224286324 |
| chr17:81959674-81959890 | 0.788201035 | 0.058848949 | 0.224289901 |
| chr11:119481305-119481606 | 0.727297764 | 0.058864946 | 0.22433971 |
| chr5:94617863-94619247 | -0.601904645 | 0.058882773 | 0.224396492 |
| chr21:21897510-21897898 | 0.779943707 | 0.058889122 | 0.224405448 |
| chr12:45324414-45324668 | -0.958868772 | 0.058890981 | 0.224405448 |
| chr3:150779632-150779886 | 0.799412184 | 0.058895143 | 0.224410147 |
| chr8:143794628-143794893 | 0.772733937 | 0.058913039 | 0.224467175 |
| chr14:66782907-66783199 | -0.893268622 | 0.05893534 | 0.224534697 |
| chr20:53835379-53835818 | 0.611981371 | 0.058936622 | 0.224534697 |
| chr5:443435-443902 | 0.66007505 | 0.058964109 | 0.224628247 |
| chr14:73560183-73560400 | 0.934887724 | 0.058992889 | 0.224726715 |
| chr5:40841104-40841509 | -0.83342664 | 0.059003996 | 0.22473592 |
| chr18:62186528-62187627 | -0.604515887 | 0.059005191 | 0.22473592 |
| chr7:21551130-21551561 | 0.662761463 | 0.059006799 | 0.22473592 |
| chr8:66531536-66531854 | -0.821883941 | 0.059007038 | 0.22473592 |
| chr4:25862779-25863009 | 0.916733771 | 0.05901495 | 0.224745683 |
| chr10:5659810-5660187 | 0.735522036 | 0.059015467 | 0.224745683 |
| chr7:87220073-87220808 | -0.626699649 | 0.05902104 | 0.224755674 |
| chr1:204573424-204573778 | 0.710594357 | 0.059031233 | 0.224755674 |
| chr2:163169563-163169794 | -0.799869435 | 0.05903392 | 0.224755674 |
| chr3:113222494-113222709 | -0.994426418 | 0.059035068 | 0.224755674 |
| chr9:29212014-29212525 | -0.708828597 | 0.059035552 | 0.224755674 |
| chr8:47945342-47945630 | 0.945120997 | 0.059035691 | 0.224755674 |
| chr4:154399487-154400016 | -0.719056624 | 0.059063998 | 0.22485227 |
| chr1:27300882-27301146 | 0.825973202 | 0.059069248 | 0.224861085 |
| chr18:45738736-45738991 | -0.849743995 | 0.059087063 | 0.224917728 |
| chr14:54001421-54001846 | -0.751573096 | 0.059092346 | 0.224926662 |
| chr1:183022398-183022599 | 0.858036984 | 0.059096244 | 0.224930326 |
| chr10:61902315-61902577 | -0.826258391 | 0.059100715 | 0.224936169 |
| chr10:22633043-22634001 | -0.634136488 | 0.059103741 | 0.224936513 |
| chr6:152897843-152898190 | -0.737581015 | 0.059113492 | 0.224953911 |
| chr2:137314769-137315626 | -0.66442748 | 0.059116752 | 0.224953911 |
| chr8:55520947-55521215 | 0.860410713 | 0.05911712 | 0.224953911 |
| chr2:196131317-196131736 | -0.780595563 | 0.059130482 | 0.224993584 |
| chrX:32246729-32247030 | -0.886401984 | 0.059136445 | 0.2250051 |
| chr6:36132104-36132304 | 0.913577996 | 0.059140421 | 0.225009054 |
| chr2:14927016-14927469 | 0.701095183 | 0.059152121 | 0.225031866 |
| chr4:121711585-121712113 | -0.707351898 | 0.059152291 | 0.225031866 |
| chr4:168541585-168541840 | -1.02448777 | 0.059164692 | 0.225064364 |
| chr1:217281803-217282029 | -0.994838511 | 0.059166708 | 0.225064364 |
| chr5:55451285-55451591 | -0.877078274 | 0.059173962 | 0.225080784 |
| chr5:108060612-108060896 | -1.065933815 | 0.05919317 | 0.22514267 |
| chr6:43724944-43725532 | 0.691146396 | 0.059202014 | 0.225165132 |
| chr5:125408528-125408832 | -0.765391059 | 0.059221092 | 0.225226511 |
| chr17:61502886-61503207 | 0.638317798 | 0.05923624 | 0.225272938 |

| | | | |
|---|---|---|---|
| chr2:119517487-119517898 | 0.748605685 | 0.059243147 | 0.225288024 |
| chr6:11434694-11435238 | 0.6977443 | 0.059248747 | 0.22529814 |
| chr20:57115979-57116249 | 0.6230962 | 0.059270508 | 0.2253697 |
| chr4:69830559-69831339 | -0.624327081 | 0.059277874 | 0.225386526 |
| chr1:220337141-220337916 | -0.634578902 | 0.059283989 | 0.225398592 |
| chr3:169907585-169907822 | -0.887689234 | 0.059288307 | 0.225403822 |
| chr1:169124443-169124767 | -0.741918855 | 0.059302723 | 0.225447444 |
| chr3:87367868-87368294 | -0.816931164 | 0.059312618 | 0.225455591 |
| chr15:71126077-71126388 | -0.754706893 | 0.059313338 | 0.225455591 |
| chr3:165206585-165207332 | -0.630933915 | 0.059314664 | 0.225455591 |
| chr12:106126436-106126850 | 0.746656973 | 0.059316636 | 0.225455591 |
| chr10:30583796-30584155 | -0.76317577 | 0.059323794 | 0.225471614 |
| chr1:230681671-230681871 | 0.930850695 | 0.059342846 | 0.225532837 |
| chr8:129996617-129997100 | 0.676181907 | 0.059355593 | 0.225570095 |
| chr17:1187012-1187510 | -0.716691668 | 0.059361564 | 0.225581598 |
| chr14:74251584-74252042 | 0.678160623 | 0.05936935 | 0.225599996 |
| chr14:41521971-41522275 | 0.756987025 | 0.059375453 | 0.225603778 |
| chr9:135438658-135439099 | -0.775556256 | 0.059376234 | 0.225603778 |
| chr14:105732693-105733161 | 0.726337663 | 0.059384604 | 0.225624391 |
| chr17:73956642-73956975 | 0.718138502 | 0.059392207 | 0.22564209 |
| chr12:122021783-122022059 | 0.759749435 | 0.059396561 | 0.225645912 |
| chr1:38858787-38860458 | 0.604146048 | 0.059399103 | 0.225645912 |
| chr20:6122739-6123612 | -0.660271255 | 0.059410645 | 0.22567857 |
| chr14:64060617-64061057 | -0.679737338 | 0.059417304 | 0.225685931 |
| chrX:42176823-42177030 | -0.985782631 | 0.05941908 | 0.225685931 |
| chr2:12684386-12684809 | -0.778738184 | 0.059421419 | 0.225685931 |
| chr1:31511022-31511518 | 0.674469776 | 0.059431907 | 0.225714575 |
| chr13:79480849-79481980 | -0.605679066 | 0.059458317 | 0.225803687 |
| chr3:184417718-184418186 | 0.721067778 | 0.05949097 | 0.225916495 |
| chr3:189958527-189958906 | -0.780304574 | 0.059509469 | 0.225975545 |
| chr10:28334988-28335402 | -0.795128453 | 0.059522858 | 0.226015185 |
| chr19:7870712-7870974 | 0.732871112 | 0.059529488 | 0.226018866 |
| chr12:107299579-107299856 | -0.9600193 | 0.059529727 | 0.226018866 |
| chr12:52865329-52865611 | 0.731432013 | 0.059551014 | 0.226088483 |
| chr10:1048356-1049719 | 0.59263202 | 0.059557779 | 0.226102964 |
| chr2:18117675-18117936 | -0.897101312 | 0.059562765 | 0.226110691 |
| chr14:75041097-75041312 | 0.97385035 | 0.059571589 | 0.226132984 |
| chr15:91939026-91939226 | 0.960798966 | 0.059595521 | 0.226205581 |
| chr13:73287134-73287527 | -0.788148976 | 0.059599164 | 0.226205581 |
| chr6:167005468-167005668 | -1.140063185 | 0.059602522 | 0.226205581 |
| chr8:65010333-65010533 | -1.140063185 | 0.059602522 | 0.226205581 |
| chr1:116589337-116589817 | 0.685786104 | 0.059608489 | 0.226206828 |
| chr5:5494570-5495034 | 0.654393834 | 0.059608755 | 0.226206828 |
| chr20:53566533-53566908 | 0.627301217 | 0.059619546 | 0.226221319 |
| chr3:149547926-149548400 | -0.725202297 | 0.059619546 | 0.226221319 |
| chrX:10014225-10015349 | 0.611003503 | 0.059621431 | 0.226221319 |
| chr10:115963951-115964151 | -1.036951742 | 0.059641355 | 0.226274471 |
| chr1:172358793-172359353 | 0.669607391 | 0.059643745 | 0.226274471 |
| chr14:105119020-105119303 | 0.722873573 | 0.059647694 | 0.226274471 |
| chr11:23113499-23113934 | -0.769023265 | 0.059648415 | 0.226274471 |

| | | | |
|---|---|---|---|
| chr3:151816015-151816543 | -0.673354993 | 0.059654417 | 0.226274471 |
| chr14:93043442-93043957 | -0.669424447 | 0.059655358 | 0.226274471 |
| chr12:50504492-50505701 | 0.620631356 | 0.059661725 | 0.226274471 |
| chr7:77538710-77538932 | -0.766118619 | 0.059661837 | 0.226274471 |
| chr13:27858042-27858494 | -0.763947192 | 0.059662018 | 0.226274471 |
| chr7:64665451-64665679 | -0.909674663 | 0.059692129 | 0.226377466 |
| chr1:109452893-109453234 | -0.877569652 | 0.059704415 | 0.226412852 |
| chr17:61802148-61802440 | 0.646002796 | 0.05971251 | 0.226425532 |
| chr7:136184067-136184297 | -1.026302753 | 0.059715569 | 0.226425532 |
| chr10:17376030-17376312 | -0.817902458 | 0.059716624 | 0.226425532 |
| chr14:93880162-93880426 | -0.942190594 | 0.059730538 | 0.226467083 |
| chr17:37608833-37609080 | 0.877294862 | 0.059737553 | 0.22646992 |
| chr2:153295863-153296103 | -0.923284514 | 0.059737999 | 0.22646992 |
| chr9:79764217-79765028 | -0.678163453 | 0.059740153 | 0.22646992 |
| chr6:91274731-91275037 | 0.899024214 | 0.05975452 | 0.226501751 |
| chr4:82363973-82364194 | -1.067617448 | 0.05975656 | 0.226501751 |
| chr13:71949374-71949747 | -0.913645068 | 0.059757813 | 0.226501751 |
| chr1:167510176-167510610 | -0.78635297 | 0.059760374 | 0.226501751 |
| chr19:49031720-49031959 | 0.901647238 | 0.059774174 | 0.226542849 |
| chr3:54354050-54354357 | 0.830267067 | 0.0597791 | 0.226542943 |
| chr2:38720690-38721236 | -0.712667043 | 0.059780112 | 0.226542943 |
| chr14:92404446-92404672 | 0.871851732 | 0.059800911 | 0.226591704 |
| chr14:94770136-94770336 | 0.927563682 | 0.059801226 | 0.226591704 |
| chr2:90398370-90398796 | -0.75752334 | 0.059801851 | 0.226591704 |
| chr19:3387869-3388161 | -0.860756744 | 0.059807621 | 0.226602362 |
| chr18:77251221-77251567 | -0.812736656 | 0.059815502 | 0.226616333 |
| chr1:174289873-174290339 | -0.832265197 | 0.059817224 | 0.226616333 |
| chr3:157493724-157493927 | -0.85233353 | 0.059831915 | 0.226660782 |
| chr1:54886374-54886976 | -0.642805552 | 0.059843095 | 0.226691929 |
| chr4:73222889-73223294 | -0.685237646 | 0.059880648 | 0.226822968 |
| chr6:30913917-30914562 | 0.642329947 | 0.059887836 | 0.22683767 |
| chr4:169264019-169264219 | -1.112360537 | 0.05989045 | 0.22683767 |
| chr7:93092839-93093064 | -0.96593221 | 0.059900093 | 0.22686298 |
| chr10:88221468-88221778 | -0.960866355 | 0.059904394 | 0.226868056 |
| chr6:40353557-40354292 | -0.707623639 | 0.059916478 | 0.226900772 |
| chr15:42207865-42208532 | -0.668276174 | 0.059920997 | 0.226900772 |
| chr5:31706250-31706510 | 0.84102772 | 0.059921917 | 0.226900772 |
| chr3:8219087-8219314 | 0.872333567 | 0.059936476 | 0.226944685 |
| chr7:117673959-117674192 | -1.042128696 | 0.059950077 | 0.226984969 |
| chr1:201503499-201503881 | 0.652007809 | 0.059959046 | 0.227007708 |
| chr20:47734022-47734242 | 0.644493069 | 0.059965944 | 0.227022608 |
| chr11:125164499-125165191 | 0.68180351 | 0.059977773 | 0.227031525 |
| chr6:52361931-52362872 | 0.640232368 | 0.059977963 | 0.227031525 |
| chr14:95994438-95994647 | 0.868794125 | 0.059979189 | 0.227031525 |
| chr5:160289742-160289973 | -0.839206633 | 0.059980151 | 0.227031525 |
| chr9:104682641-104683029 | -0.791025542 | 0.059984894 | 0.227038262 |
| chr5:119537790-119537990 | 0.97253593 | 0.059988861 | 0.22704206 |
| chr7:135177562-135177971 | -0.867616881 | 0.060002591 | 0.227082807 |
| chr3:32239579-32239856 | 0.757968632 | 0.060015216 | 0.227119371 |
| chr2:179268364-179268805 | -0.714180115 | 0.060021765 | 0.227132936 |

| | | | |
|---|---|---|---|
| chr11:66336303-66337178 | -0.628196763 | 0.060026791 | 0.227137272 |
| chr2:20291048-20291537 | -0.681723486 | 0.06002884 | 0.227137272 |
| chr1:209248720-209248988 | -0.814499741 | 0.060036514 | 0.22715375 |
| chr6:155154286-155155012 | -0.722247551 | 0.060040899 | 0.22715375 |
| chr6:31042967-31043233 | 0.871349791 | 0.060042088 | 0.22715375 |
| chr2:237482547-237482876 | -0.712331666 | 0.060045781 | 0.227156504 |
| chr15:69490735-69490937 | 1.02536357 | 0.060052879 | 0.227172138 |
| chr1:85146520-85147365 | 0.638472832 | 0.060069106 | 0.22720046 |
| chr4:4541887-4542472 | 0.674756428 | 0.060071358 | 0.22720046 |
| chr11:61293826-61295077 | 0.615803569 | 0.060072874 | 0.22720046 |
| chr12:89529850-89530054 | -0.852987549 | 0.060077132 | 0.22720046 |
| chr12:22262327-22262599 | -0.990027595 | 0.06007794 | 0.22720046 |
| chr3:175821423-175821676 | -0.906473889 | 0.060078157 | 0.22720046 |
| chr8:127038657-127039043 | -0.704377243 | 0.060085 | 0.227207915 |
| chrX:16826349-16826750 | -0.768658756 | 0.060086074 | 0.227207915 |
| chr1:107172454-107172676 | 0.724860836 | 0.060089024 | 0.227207915 |
| chr7:157760508-157760708 | -0.999506826 | 0.060092344 | 0.227209255 |
| chr1:211407545-211408173 | -0.700673264 | 0.060097007 | 0.227215675 |
| chr11:64924008-64924723 | 0.63906876 | 0.060125519 | 0.227312258 |
| chr19:11271088-11271329 | -1.033832749 | 0.060143742 | 0.227369933 |
| chr12:51430245-51430445 | 0.917292911 | 0.060148163 | 0.227375429 |
| chr9:6049897-6050137 | -0.876526555 | 0.060171409 | 0.22745208 |
| chr5:100534908-100536054 | -0.603448263 | 0.060182382 | 0.227482339 |
| chr16:29816840-29817129 | 0.696972307 | 0.060186608 | 0.227487037 |
| chr14:66031910-66032227 | 0.821803598 | 0.060189563 | 0.227487037 |
| chr3:124995516-124995733 | 0.834355412 | 0.060211405 | 0.227558363 |
| chr4:168848149-168848484 | -0.911807543 | 0.060220546 | 0.227579074 |
| chr1:214505057-214505579 | -0.716827172 | 0.060222825 | 0.227579074 |
| chr2:88941210-88941481 | -0.835443888 | 0.060243214 | 0.227644894 |
| chr8:123443156-123443501 | 0.733275863 | 0.060257072 | 0.227686033 |
| chr1:188372794-188373204 | -0.683447288 | 0.060264399 | 0.227702491 |
| chr17:40113148-40113897 | 0.641027689 | 0.060283095 | 0.227761898 |
| chr12:87117552-87117839 | -0.816388187 | 0.06029201 | 0.227784349 |
| chr1:33180669-33181330 | 0.623769329 | 0.060310731 | 0.227843843 |
| chr7:9083651-9084244 | -0.66292488 | 0.060347883 | 0.227972956 |
| chr8:90637895-90638095 | 0.933182636 | 0.060358434 | 0.228001005 |
| chr7:16365045-16365597 | 0.720314507 | 0.060363908 | 0.228001005 |
| chr5:97208273-97208796 | 0.763297363 | 0.060364235 | 0.228001005 |
| chr22:23205809-23206054 | -0.897114742 | 0.060370425 | 0.228013148 |
| chr3:122560731-122560957 | -0.833283327 | 0.060377197 | 0.228027485 |
| chr8:102073960-102074160 | 0.955995419 | 0.060385049 | 0.228035496 |
| chr11:132028555-132028963 | -0.864427795 | 0.060386624 | 0.228035496 |
| chr1:190416281-190416763 | -0.752039068 | 0.060389606 | 0.228035496 |
| chr17:39699874-39700085 | 0.902306368 | 0.060392035 | 0.228035496 |
| chr12:53066027-53066271 | 0.714247365 | 0.060395084 | 0.228035496 |
| chr3:114275027-114275238 | 0.978668718 | 0.060397175 | 0.228035496 |
| chr8:3284618-3284873 | -0.879613191 | 0.060401041 | 0.228038855 |
| chr5:17658245-17658638 | -0.933272688 | 0.060430112 | 0.22813737 |
| chr16:81940827-81941042 | -0.887808131 | 0.060435673 | 0.228147122 |
| chr6:153503777-153504151 | -0.79422931 | 0.060443387 | 0.228165001 |

| | | | |
|---|---|---|---|
| chr5:15034871-15035071 | 1.053201321 | 0.060455086 | 0.228196557 |
| chr3:153087840-153088167 | 0.776609542 | 0.060457703 | 0.228196557 |
| chr11:9545751-9545951 | 1.028629462 | 0.060466437 | 0.228210134 |
| chr12:30989813-30990192 | -0.727745512 | 0.060467257 | 0.228210134 |
| chr4:109589028-109589972 | -0.623625532 | 0.060493833 | 0.228289979 |
| chr8:138616571-138616843 | 0.89945872 | 0.060494372 | 0.228289979 |
| chr1:58652145-58652877 | 0.676170731 | 0.060515775 | 0.228359502 |
| chr18:76491448-76492507 | -0.618494908 | 0.060527018 | 0.228390683 |
| chr4:14667590-14667917 | 0.780638925 | 0.060545987 | 0.228451006 |
| chr4:42655864-42656147 | 0.735880156 | 0.060579338 | 0.22856559 |
| chr12:94459529-94460657 | -0.607527863 | 0.060588718 | 0.228589725 |
| chr5:107472146-107472346 | -0.980850324 | 0.060592646 | 0.228593289 |
| chr3:9500570-9500880 | 0.680792451 | 0.060606779 | 0.228635351 |
| chr3:138947698-138947915 | 0.879314648 | 0.060620604 | 0.228676248 |
| chr20:44884977-44885817 | 0.609261512 | 0.060636257 | 0.228718204 |
| chr10:66455328-66455714 | -0.737135734 | 0.060637697 | 0.228718204 |
| chr5:80083326-80083862 | 0.672946412 | 0.060654302 | 0.228769574 |
| chr7:20172793-20173052 | 0.781465975 | 0.060663167 | 0.228791149 |
| chr1:16440184-16441074 | -0.645152929 | 0.060670743 | 0.228791149 |
| chr9:116860548-116860842 | 0.800686203 | 0.060671286 | 0.228791149 |
| chr1:232880453-232880721 | 0.827557738 | 0.060671966 | 0.228791149 |
| chr12:32051590-32051929 | -0.902125477 | 0.060682705 | 0.228820384 |
| chr3:2699903-2700183 | -1.0517633 | 0.060689098 | 0.228833229 |
| chr11:130603447-130603693 | -0.984096188 | 0.06070151 | 0.228862929 |
| chr5:25255176-25255536 | 0.722747298 | 0.060702948 | 0.228862929 |
| chr2:74740698-74741046 | -0.775285759 | 0.060711084 | 0.228873662 |
| chr1:114428807-114429019 | 0.736877664 | 0.060711769 | 0.228873662 |
| chr20:28308003-28308314 | 0.707849024 | 0.060716467 | 0.228880109 |
| chr2:240898442-240898654 | 0.832572275 | 0.060750301 | 0.228996388 |
| chr5:117454656-117456228 | -0.595737109 | 0.060760911 | 0.229018724 |
| chr4:602656-602953 | 0.756656073 | 0.060762205 | 0.229018724 |
| chr20:23998485-23998839 | 0.856865806 | 0.060768977 | 0.229032985 |
| chr8:120036353-120036615 | -0.915349265 | 0.060784628 | 0.229074642 |
| chr4:163384683-163384883 | -0.997799283 | 0.060786009 | 0.229074642 |
| chr6:170153814-170154029 | 0.85920184 | 0.060806399 | 0.22914021 |
| chr18:31102267-31102566 | 0.780846255 | 0.06081405 | 0.229150174 |
| chr1:1021235-1021531 | 0.756448656 | 0.060815562 | 0.229150174 |
| chr7:25568419-25569175 | 0.654979199 | 0.060818015 | 0.229150174 |
| chr1:10776601-10776832 | 0.9074782 | 0.060825809 | 0.229168273 |
| chr2:120509093-120509293 | 0.977400434 | 0.060839894 | 0.229210067 |
| chr2:8525281-8526284 | -0.630147226 | 0.060847054 | 0.229225773 |
| chr2:41775662-41776049 | -0.708179978 | 0.060876559 | 0.229325651 |
| chr11:130861726-130862352 | -0.694508276 | 0.060886886 | 0.2293449 |
| chr8:68927416-68927816 | -0.825081259 | 0.060890375 | 0.2293449 |
| chr2:15546550-15546938 | -0.787078403 | 0.060890649 | 0.2293449 |
| chr21:40321042-40321242 | -0.833484114 | 0.060917107 | 0.229432302 |
| chr1:206455740-206456325 | -0.65806115 | 0.060919843 | 0.229432302 |
| chr3:14678193-14678393 | 0.880740087 | 0.060932312 | 0.229460296 |
| chr2:14633049-14633249 | 0.787958311 | 0.060933265 | 0.229460296 |
| chr13:79382958-79383375 | -0.723260901 | 0.06093768 | 0.229465643 |

| | | | |
|---|---|---|---|
| chr11:87924235-87924438 | -1.114990793 | 0.060946855 | 0.229488916 |
| chr10:29102536-29102902 | -0.784770434 | 0.060950875 | 0.22949214 |
| chr5:123941793-123942017 | 0.900812539 | 0.060953702 | 0.22949214 |
| chr11:64234065-64234265 | 0.89467161 | 0.060964005 | 0.229519654 |
| chr2:96145294-96145755 | 0.659723116 | 0.060981497 | 0.229574229 |
| chr20:47272451-47272690 | 0.635456761 | 0.061009237 | 0.229667375 |
| chr22:31927061-31927277 | -1.057460667 | 0.061016425 | 0.22968315 |
| chr5:58183511-58183894 | -0.734438979 | 0.06103639 | 0.229738841 |
| chr7:92461620-92461986 | -0.76429247 | 0.061037216 | 0.229738841 |
| chr17:50297420-50297737 | -0.720463324 | 0.061040588 | 0.229740247 |
| chr8:123185078-123185403 | -0.715143343 | 0.061091909 | 0.22992211 |
| chr12:47105758-47106216 | -0.745588875 | 0.061106611 | 0.229966149 |
| chr6:13486028-13486243 | -0.826973811 | 0.061130643 | 0.23004529 |
| chr5:173782159-173782433 | 0.775447286 | 0.061136587 | 0.230056357 |
| chr9:7536406-7536641 | -0.975953544 | 0.061150066 | 0.230088769 |
| chr15:83305339-83306308 | -0.663769733 | 0.061151206 | 0.230088769 |
| chr2:143110825-143111189 | -0.768510147 | 0.061162701 | 0.23012072 |
| chr3:69691135-69691696 | -0.712029998 | 0.061166833 | 0.230124967 |
| chr15:49395940-49396682 | -0.69180635 | 0.061179902 | 0.230162835 |
| chrX:132171525-132171813 | 0.801778641 | 0.061186992 | 0.230178206 |
| chr2:121075795-121076158 | -0.723905052 | 0.061200856 | 0.230219057 |
| chr1:170427007-170427207 | -1.059451458 | 0.061207123 | 0.230231328 |
| chr18:61651589-61652219 | -0.652640237 | 0.061216322 | 0.230245055 |
| chr9:121538299-121538544 | 0.838847688 | 0.061217267 | 0.230245055 |
| chr7:151832359-151832559 | -1.097929253 | 0.061219787 | 0.230245055 |
| chr19:28633776-28634067 | 0.81695429 | 0.06122735 | 0.230250926 |
| chr1:157138046-157138931 | 0.607653955 | 0.061227358 | 0.230250926 |
| chr17:732462-733241 | -0.685108196 | 0.061231077 | 0.230253612 |
| chr16:67642742-67642945 | -1.079597513 | 0.061238199 | 0.230269091 |
| chr5:154713709-154714123 | 0.757886884 | 0.061258058 | 0.230328191 |
| chrX:141176578-141177623 | 0.605131516 | 0.061259928 | 0.230328191 |
| chr3:171447370-171447658 | 0.838390911 | 0.061285482 | 0.230412964 |
| chr1:70685254-70685828 | -0.664179893 | 0.061296211 | 0.230441995 |
| chr5:168219806-168220186 | -0.719652463 | 0.061299392 | 0.230442648 |
| chr1:223696425-223696752 | -0.782954467 | 0.061307991 | 0.230463666 |
| chr17:61383909-61384218 | 0.617656767 | 0.0613141 | 0.230475322 |
| chr9:89604826-89605110 | 0.721065187 | 0.061319598 | 0.230484682 |
| chr2:15891197-15891406 | -0.932642983 | 0.061328136 | 0.230505465 |
| chr1:32901012-32901589 | 0.670466052 | 0.061338446 | 0.230532908 |
| chr8:87055148-87055777 | 0.652646063 | 0.061349034 | 0.230561394 |
| chr9:30943889-30944266 | -0.878683223 | 0.061365894 | 0.230613443 |
| chr6:4334118-4334744 | -0.683577573 | 0.061370525 | 0.230619536 |
| chr7:101192510-101192838 | -0.758708439 | 0.061374671 | 0.230623081 |
| chr6:89509729-89510103 | 0.869917663 | 0.061382467 | 0.230623081 |
| chr11:101775035-101775433 | -0.864398132 | 0.061386005 | 0.230623081 |
| chr5:180869013-180869414 | 0.717435202 | 0.061386905 | 0.230623081 |
| chr7:101363617-101364243 | -0.682679825 | 0.061388434 | 0.230623081 |
| chr1:234125342-234125638 | -0.78828013 | 0.061389527 | 0.230623081 |
| chr2:70686879-70687380 | -0.716178712 | 0.061421106 | 0.23072876 |
| chr9:85408523-85408782 | 0.994578798 | 0.06142368 | 0.23072876 |

| | | | |
|---|---|---|---|
| chr7:19665458-19665703 | 0.844061577 | 0.061439568 | 0.230737847 |
| chr2:178414640-178415116 | -0.635179927 | 0.061440487 | 0.230737847 |
| chr4:7675970-7676183 | 1.015890561 | 0.061442379 | 0.230737847 |
| chrX:24498284-24498484 | -1.015583871 | 0.061443993 | 0.230737847 |
| chr2:186641730-186642088 | -0.736564276 | 0.061444249 | 0.230737847 |
| chr13:41457154-41457917 | -0.625661328 | 0.061446786 | 0.230737847 |
| chr17:37358770-37359570 | 0.644861606 | 0.061447179 | 0.230737847 |
| chr8:102400275-102400737 | 0.700166262 | 0.061452151 | 0.230744208 |
| chr16:46390185-46390918 | 0.5908939 | 0.061458613 | 0.230744208 |
| chr11:19467859-19468101 | 0.929472619 | 0.061459462 | 0.230744208 |
| chr22:19896472-19896845 | -0.820403682 | 0.061460919 | 0.230744208 |
| chr17:17714418-17714647 | 0.828585799 | 0.061467959 | 0.230759331 |
| chr2:119190311-119190669 | -0.81660648 | 0.061472572 | 0.230765344 |
| chr11:63804395-63804595 | 0.918148772 | 0.061504179 | 0.230861371 |
| chr16:3065387-3065600 | 0.918148772 | 0.061504179 | 0.230861371 |
| chr15:84633698-84634107 | 0.677464866 | 0.061507317 | 0.23086184 |
| chr14:47044849-47045176 | 0.750773047 | 0.061534664 | 0.230949444 |
| chr18:43587711-43587922 | -1.00062684 | 0.061536685 | 0.230949444 |
| chr1:169109812-169110042 | -0.753634101 | 0.061556509 | 0.231012532 |
| chr11:66956754-66957262 | -0.66393345 | 0.061566903 | 0.231034426 |
| chr14:105300632-105301733 | 0.593169326 | 0.061568374 | 0.231034426 |
| chr7:97024682-97025006 | 0.751669418 | 0.061585509 | 0.231078337 |
| chr1:21856051-21856414 | -0.798099578 | 0.061587967 | 0.231078337 |
| chr12:48818387-48818735 | 0.717954927 | 0.061589398 | 0.231078337 |
| chr2:39730749-39731041 | -0.818413896 | 0.061592139 | 0.231078337 |
| chr3:186827235-186827920 | -0.64505903 | 0.061597203 | 0.231086021 |
| chr7:35015318-35015796 | -0.745136168 | 0.061604985 | 0.231103899 |
| chr14:70255190-70255735 | -0.689084218 | 0.0616196 | 0.23114741 |
| chr16:28491340-28491540 | 0.839611796 | 0.061624696 | 0.23115521 |
| chr1:151125635-151126322 | -0.668299536 | 0.061634332 | 0.231180038 |
| chr9:78821212-78821463 | -0.86216673 | 0.061641426 | 0.231195327 |
| chr7:127838074-127838274 | -1.043049332 | 0.061647412 | 0.231197701 |
| chr11:71615039-71615239 | 0.883925308 | 0.061648094 | 0.231197701 |
| chr11:121478234-121479301 | 0.637629701 | 0.061671112 | 0.231272707 |
| chr10:113853863-113855646 | 0.605033888 | 0.061678204 | 0.231287984 |
| chr12:101945898-101946150 | -0.910527893 | 0.061687942 | 0.231313177 |
| chr6:82951348-82951868 | -0.702691594 | 0.061697881 | 0.231339128 |
| chr13:30022879-30023256 | 0.735382359 | 0.061713574 | 0.231386644 |
| chrX:67543802-67544229 | 0.675242181 | 0.061723951 | 0.231406165 |
| chr3:194204020-194204647 | 0.705693326 | 0.061724821 | 0.231406165 |
| chr21:40840745-40841063 | -0.81308929 | 0.061732943 | 0.231425293 |
| chr1:155177170-155177548 | 0.64220454 | 0.061737578 | 0.231431344 |
| chr1:27671927-27672686 | -0.639060351 | 0.061743327 | 0.231441575 |
| chr1:173210297-173210536 | 0.892426445 | 0.061750822 | 0.231458345 |
| chr20:25384725-25384925 | 1.028603641 | 0.061762539 | 0.231468913 |
| chr13:69264188-69265053 | -0.643980124 | 0.061762618 | 0.231468913 |
| chr2:38060714-38060927 | -1.005166631 | 0.061762704 | 0.231468913 |
| chr7:50555833-50556123 | -0.863477424 | 0.061784892 | 0.231540743 |
| chr5:160869915-160870129 | -1.048080096 | 0.061790739 | 0.231551328 |
| chr17:59138377-59138760 | -0.711099484 | 0.061795024 | 0.231556062 |

| | | | |
|---|---|---|---|
| chr12:14240141-14240747 | -0.636140436 | 0.061807491 | 0.23159145 |
| chr21:38126632-38127098 | -0.648043728 | 0.061822558 | 0.231623321 |
| chrX:17361728-17361955 | -1.064919041 | 0.061824015 | 0.231623321 |
| chr7:138246572-138246907 | -0.913909388 | 0.061827441 | 0.231623321 |
| chr9:130766378-130766578 | 0.969385063 | 0.061828088 | 0.231623321 |
| chr2:10362453-10362776 | -0.767130993 | 0.061870435 | 0.231761291 |
| chr5:145811879-145812533 | -0.711418417 | 0.061870967 | 0.231761291 |
| chr17:61368634-61369850 | 0.586012799 | 0.061880135 | 0.23177386 |
| chr5:154976267-154976492 | 0.999245298 | 0.061880372 | 0.23177386 |
| chr4:134953243-134953494 | 0.894636714 | 0.061890018 | 0.231782476 |
| chrX:45824486-45824890 | -0.795518546 | 0.061891244 | 0.231782476 |
| chr13:26601808-26602080 | -0.930454566 | 0.061891747 | 0.231782476 |
| chr2:46134932-46135166 | -0.921090782 | 0.061899117 | 0.231788812 |
| chr15:48336986-48337220 | -0.815011519 | 0.061899489 | 0.231788812 |
| chr5:101678431-101678955 | -0.68205072 | 0.061907364 | 0.231800194 |
| chr20:56809764-56810087 | -0.722517995 | 0.061908579 | 0.231800194 |
| chr3:127672466-127673376 | 0.614252729 | 0.061926551 | 0.231854082 |
| chr4:108018560-108018898 | -0.806442818 | 0.061929024 | 0.231854082 |
| chr14:95895775-95895994 | 0.823356948 | 0.061942565 | 0.231875775 |
| chrX:87227305-87227616 | -0.872504373 | 0.061943391 | 0.231875775 |
| chr9:83048085-83048599 | 0.760629261 | 0.061943897 | 0.231875775 |
| chr17:81901667-81901913 | 0.821582253 | 0.061958118 | 0.23191768 |
| chr12:52215624-52216020 | 0.690681866 | 0.061964016 | 0.231928428 |
| chr20:47351973-47352844 | 0.589095663 | 0.061983136 | 0.231988658 |
| chr10:27240263-27241629 | -0.60308038 | 0.061993501 | 0.232016121 |
| chr7:155380164-155380613 | -0.706970219 | 0.061999852 | 0.232028557 |
| chr12:54587119-54587566 | -0.77731021 | 0.062020985 | 0.23209631 |
| chr1:202586117-202586370 | 0.77609207 | 0.062024198 | 0.232096996 |
| chr10:96511281-96511535 | 0.741218613 | 0.06203006 | 0.232106738 |
| chr7:82740330-82740615 | -0.9268746 | 0.06203286 | 0.232106738 |
| chr8:30422916-30423593 | -0.681867193 | 0.062039613 | 0.232120672 |
| chr7:139499951-139500342 | -0.680894458 | 0.062048688 | 0.232137939 |
| chr8:92053761-92053961 | 0.946284677 | 0.062050288 | 0.232137939 |
| chr8:46956799-46957416 | 0.676515906 | 0.062056582 | 0.232143228 |
| chr10:5692478-5692808 | 0.871853844 | 0.062057761 | 0.232143228 |
| chr2:33674026-33674226 | -0.964217911 | 0.062084593 | 0.232230008 |
| chr10:59944295-59944858 | -0.689881374 | 0.062087021 | 0.232230008 |
| chr1:214694404-214695271 | -0.647682302 | 0.0621113 | 0.232277172 |
| chr1:25503895-25504111 | -0.984292725 | 0.06211411 | 0.232277172 |
| chr6:44151854-44152201 | 0.711697761 | 0.062118407 | 0.232277172 |
| chr6:148263003-148263461 | -0.805551433 | 0.06212188 | 0.232277172 |
| chr15:41513947-41514261 | 0.879500083 | 0.062122765 | 0.232277172 |
| chr17:1455970-1456629 | -0.629494788 | 0.062123602 | 0.232277172 |
| chr8:144390840-144391201 | 0.718724835 | 0.062123716 | 0.232277172 |
| chr7:147547310-147547595 | -0.846820151 | 0.062123882 | 0.232277172 |
| chr3:21205493-21205693 | 1.026605759 | 0.062136372 | 0.232312536 |
| chr9:119961813-119962187 | -0.727256093 | 0.062151075 | 0.232356167 |
| chr4:80183706-80184378 | -0.6652455 | 0.062158317 | 0.232361149 |
| chr2:39354384-39354751 | -0.91473687 | 0.062158472 | 0.232361149 |
| chr5:112614328-112614635 | 0.804725409 | 0.062163539 | 0.232361705 |

| | | | |
|---|---|---|---|
| chr4:111954480-111954975 | -0.768050867 | 0.062167127 | 0.232361705 |
| chr12:13196513-13196732 | -1.003636154 | 0.062167719 | 0.232361705 |
| chr19:48741580-48741784 | 0.804683219 | 0.062177881 | 0.232385436 |
| chr1:147716961-147717565 | 0.613411971 | 0.062182406 | 0.232385436 |
| chr19:18588624-18589288 | 0.669776569 | 0.062183263 | 0.232385436 |
| chr13:43045548-43045752 | -1.004329622 | 0.062186199 | 0.232385436 |
| chr4:163038012-163038583 | -0.714158321 | 0.062201572 | 0.232431547 |
| chr17:59642605-59642851 | 0.721496607 | 0.062209161 | 0.232442392 |
| chr7:105304523-105304960 | -0.785348782 | 0.062210542 | 0.232442392 |
| chr21:37700734-37700984 | 0.835618674 | 0.062220402 | 0.232467898 |
| chr14:20998098-20998508 | 0.732575798 | 0.062233387 | 0.232504818 |
| chr12:16347321-16347830 | -0.663533703 | 0.062236353 | 0.232504818 |
| chr22:37018763-37019261 | 0.756088225 | 0.062253809 | 0.232555853 |
| chr7:128476298-128477404 | -0.607413274 | 0.062256084 | 0.232555853 |
| chr10:43384829-43385029 | 0.978333343 | 0.062260359 | 0.232560485 |
| chr3:177816750-177817212 | -0.80665386 | 0.062267083 | 0.232564388 |
| chr2:100606935-100607182 | 0.871324236 | 0.06226985 | 0.232564388 |
| chr11:73269650-73270002 | 0.696172073 | 0.06227051 | 0.232564388 |
| chr2:208399229-208399654 | -0.735378842 | 0.062285045 | 0.232607336 |
| chr11:76257075-76257398 | 0.816943111 | 0.0622941 | 0.232629814 |
| chr10:128095343-128095735 | -0.758413945 | 0.062297242 | 0.232630208 |
| chr10:72337223-72337481 | -0.77181395 | 0.062315 | 0.232685182 |
| chrX:66913874-66914323 | -0.821736733 | 0.062325846 | 0.232704747 |
| chr6:54846151-54847333 | -0.601097678 | 0.062329216 | 0.232704747 |
| chr4:82744290-82744643 | -0.715031186 | 0.062329351 | 0.232704747 |
| chr2:233854288-233854993 | -0.642517809 | 0.062338443 | 0.232727354 |
| chr15:69059132-69059412 | 0.776209647 | 0.062343405 | 0.232734537 |
| chr10:28652785-28652985 | -1.0045458 | 0.062347144 | 0.232737157 |
| chr12:52395780-52396172 | 0.695504004 | 0.06236895 | 0.232799835 |
| chr14:64294746-64295156 | 0.709414035 | 0.062370011 | 0.232799835 |
| chr16:80607153-80607638 | -0.705192947 | 0.062375745 | 0.232809894 |
| chr12:53048409-53048634 | 0.70580524 | 0.062392283 | 0.232860279 |
| chr19:588131-588448 | 0.728617774 | 0.062400567 | 0.232879852 |
| chrX:71095250-71095538 | 0.841027205 | 0.062404924 | 0.23288477 |
| chr18:13140088-13140288 | 0.982198154 | 0.062411769 | 0.23289897 |
| chr3:192851734-192852470 | -0.628309461 | 0.062415436 | 0.23290131 |
| chr3:101695871-101696105 | -0.849733693 | 0.062421896 | 0.232914076 |
| chr15:101501234-101501434 | -0.863151585 | 0.062425731 | 0.23291704 |
| chr3:64925671-64925936 | 0.649248536 | 0.06243343 | 0.232923244 |
| chr14:75069167-75070695 | -0.597643479 | 0.062433473 | 0.232923244 |
| chr8:63145650-63146106 | -0.732966693 | 0.062440003 | 0.232932914 |
| chr22:35231565-35231839 | -0.861019361 | 0.062443101 | 0.232932914 |
| chr2:237567049-237567271 | -0.935940069 | 0.062449655 | 0.232932914 |
| chr7:111089904-111090128 | -0.977421679 | 0.062451627 | 0.232932914 |
| chr16:705758-706166 | -0.737874691 | 0.062452499 | 0.232932914 |
| chr5:74201457-74201716 | -0.899184777 | 0.062457026 | 0.232932914 |
| chr5:163437143-163438255 | -0.592619093 | 0.062457345 | 0.232932914 |
| chr3:39414421-39414756 | -0.799046178 | 0.062465038 | 0.232950265 |
| chr8:78553206-78553518 | 0.889791035 | 0.062469301 | 0.232954825 |
| chr6:44079243-44079482 | 0.756237392 | 0.062479529 | 0.232981628 |

| | | | |
|---|---|---|---|
| chr1:26353965-26354368 | 0.684266858 | 0.062484042 | 0.232987118 |
| chr17:59289771-59290487 | 0.629553041 | 0.062501486 | 0.233035004 |
| chr14:68774696-68775212 | 0.657391873 | 0.062502967 | 0.233035004 |
| chr14:59576232-59577220 | -0.609599707 | 0.062510658 | 0.23305081 |
| chr19:53772949-53773149 | 0.981836582 | 0.06251329 | 0.23305081 |
| chr20:63955843-63956140 | -0.698962461 | 0.062564496 | 0.233216997 |
| chr7:32680235-32681229 | -0.614221316 | 0.062566645 | 0.233216997 |
| chr7:8285053-8285397 | 0.72368329 | 0.062566999 | 0.233216997 |
| chr17:80252772-80253188 | 0.702931094 | 0.062584484 | 0.233270826 |
| chr10:33132738-33133019 | -0.851062806 | 0.062594167 | 0.233295566 |
| chr22:29813647-29813900 | -0.885149703 | 0.062599544 | 0.233304258 |
| chrX:123732325-123733327 | 0.612749593 | 0.062609152 | 0.233319106 |
| chr12:26113887-26114893 | -0.602368351 | 0.062609618 | 0.233319106 |
| chr7:6977402-6977704 | -0.820727706 | 0.062613001 | 0.233320364 |
| chr14:28629337-28629901 | 0.655331717 | 0.062621549 | 0.233340872 |
| chr17:39036006-39036330 | 0.809612726 | 0.062630724 | 0.233363708 |
| chr5:6196590-6196872 | 0.956160804 | 0.062640887 | 0.233386682 |
| chr8:54165908-54166459 | -0.715649902 | 0.062645527 | 0.233386682 |
| chr6:92892679-92892902 | 0.962319674 | 0.062649558 | 0.233386682 |
| chr8:64611704-64611940 | -1.031366094 | 0.062652712 | 0.233386682 |
| chr12:47768391-47768666 | 0.83559114 | 0.062652873 | 0.233386682 |
| chr15:39579816-39580041 | 0.766679113 | 0.062655361 | 0.233386682 |
| chr2:1660153-1660398 | 0.937307695 | 0.062658211 | 0.233386682 |
| chr14:90060231-90060950 | 0.621986686 | 0.062668947 | 0.233415324 |
| chr7:32732912-32733155 | -0.791771589 | 0.062674779 | 0.233425698 |
| chr13:40974656-40974997 | -0.727126605 | 0.062689766 | 0.23347017 |
| chr11:126091826-126092560 | -0.708646004 | 0.062694125 | 0.233475055 |
| chr3:16927525-16927856 | 0.771876389 | 0.062697651 | 0.233476839 |
| chr12:90901227-90901502 | -1.026757591 | 0.062700925 | 0.233477681 |
| chr19:2236213-2237132 | 0.615321726 | 0.06271358 | 0.233507863 |
| chr20:53890931-53891536 | 0.596669709 | 0.062715125 | 0.233507863 |
| chr17:82735468-82735823 | 0.750895732 | 0.062729258 | 0.23354356 |
| chr1:207711304-207711723 | -0.705723618 | 0.062732246 | 0.23354356 |
| chr3:139823322-139823568 | 0.860030678 | 0.062734154 | 0.23354356 |
| chr20:52448520-52448834 | -0.851125867 | 0.062736905 | 0.23354356 |
| chr17:82889462-82889987 | 0.639344165 | 0.062750358 | 0.233582295 |
| chr7:30143903-30144184 | 0.840044307 | 0.062757008 | 0.2335957 |
| chr5:61161952-61162812 | -0.609461279 | 0.062794995 | 0.23372574 |
| chr4:99056086-99056341 | -0.85308478 | 0.062801445 | 0.233738393 |
| chr20:53793893-53794132 | 0.624698913 | 0.062808066 | 0.233751684 |
| chr2:222424936-222425323 | -0.805905424 | 0.062815135 | 0.233763591 |
| chr1:56797411-56797651 | -0.967542872 | 0.062817368 | 0.233763591 |
| chr8:126425628-126426007 | -0.739913321 | 0.062820964 | 0.233765622 |
| chr4:166462865-166463073 | -0.992217432 | 0.062831599 | 0.233793842 |
| chr18:42298826-42299101 | -0.902231239 | 0.062840686 | 0.233816299 |
| chr10:114714437-114714779 | 0.733522851 | 0.062859248 | 0.233874007 |
| chr5:179760183-179760618 | -0.702081347 | 0.062870447 | 0.233904316 |
| chr4:37566787-37567080 | -0.81097457 | 0.062875251 | 0.23391083 |
| chr14:103520715-103521766 | 0.602005414 | 0.062882222 | 0.233917249 |
| chr1:55385028-55385277 | -0.964779505 | 0.062883554 | 0.233917249 |

| | | | |
|---|---|---|---|
| chr2:219457979-219458356 | -0.728932849 | 0.062888159 | 0.233917249 |
| chr3:108203038-108203316 | -0.844044975 | 0.062889188 | 0.233917249 |
| chr3:159765065-159765461 | 0.671951524 | 0.062916165 | 0.233995616 |
| chr6:145813789-145815467 | -0.593532466 | 0.062916365 | 0.233995616 |
| chr7:140479060-140480027 | -0.601641331 | 0.06292179 | 0.234004435 |
| chr6:72952884-72953213 | -0.810598358 | 0.062930446 | 0.234025267 |
| chr2:17518142-17518774 | -0.60777628 | 0.062946636 | 0.234074114 |
| chr7:6172214-6172421 | -0.890290115 | 0.062950009 | 0.234075297 |
| chr9:37840011-37840322 | -0.985443241 | 0.062961074 | 0.234099362 |
| chr13:113647520-113647836 | 0.814863102 | 0.062965203 | 0.234099362 |
| chr13:67736580-67736855 | -0.975326162 | 0.062965646 | 0.234099362 |
| chr8:80240571-80241010 | 0.649033239 | 0.062983034 | 0.234152645 |
| chr3:117275-117769 | 0.723035444 | 0.063007538 | 0.234232381 |
| chr12:34218269-34219105 | -0.623251087 | 0.063027955 | 0.234296914 |
| chr6:108549806-108550010 | -1.013358839 | 0.063031365 | 0.234298224 |
| chr14:53231314-53231654 | -0.798066061 | 0.063036022 | 0.234304164 |
| chr16:23508782-23509025 | 0.797253508 | 0.063045071 | 0.234326432 |
| chr17:28384476-28384723 | -0.816512914 | 0.063062528 | 0.234344739 |
| chr5:169641247-169641682 | -0.681658486 | 0.063063011 | 0.234344739 |
| chr5:140547416-140547873 | -0.653559817 | 0.063063837 | 0.234344739 |
| chr18:35756532-35756732 | 0.980011646 | 0.063064472 | 0.234344739 |
| chr16:19167655-19167969 | 0.733808365 | 0.063065288 | 0.234344739 |
| chr2:42047727-42047994 | 0.838716909 | 0.063069095 | 0.234347521 |
| chr4:183256892-183257092 | -1.078684906 | 0.063075102 | 0.234350265 |
| chr2:37148330-37148727 | 0.681513519 | 0.063075951 | 0.234350265 |
| chr7:113404301-113404540 | -0.928046692 | 0.063103601 | 0.234430262 |
| chr8:130828397-130828640 | -0.928046692 | 0.063103601 | 0.234430262 |
| chr19:54439439-54439869 | 0.712122658 | 0.063119546 | 0.234473802 |
| chr10:109866504-109866768 | 0.814348318 | 0.063121442 | 0.234473802 |
| chr19:52297327-52297784 | 0.734352843 | 0.063129877 | 0.234493768 |
| chr19:49850347-49851239 | 0.615231091 | 0.063147981 | 0.234549644 |
| chr7:72405274-72405692 | -0.792489188 | 0.063157079 | 0.234572066 |
| chr12:119176321-119176677 | -0.684197613 | 0.063175142 | 0.234627779 |
| chr7:97027718-97027995 | -0.839725752 | 0.063193998 | 0.234686435 |
| chr12:62837742-62837951 | 1.00412069 | 0.063201871 | 0.234696718 |
| chr15:64515167-64515568 | 0.749473427 | 0.063203785 | 0.234696718 |
| chr8:101080352-101081339 | 0.606795935 | 0.063210162 | 0.234696718 |
| chr12:7859961-7860214 | -0.906002826 | 0.063211318 | 0.234696718 |
| chr10:90930207-90930785 | 0.685905276 | 0.063212082 | 0.234696718 |
| chr10:1465518-1465911 | -0.713475084 | 0.063223283 | 0.234726932 |
| chr8:144412917-144413351 | 0.62240657 | 0.063249426 | 0.234812611 |
| chr7:126856724-126856956 | -1.005397742 | 0.063266527 | 0.234858794 |
| chr6:116616715-116617135 | -0.686899313 | 0.063268209 | 0.234858794 |
| chr22:50732279-50732479 | 0.926357307 | 0.063272077 | 0.234858794 |
| chr6:4835455-4836309 | -0.643225616 | 0.063274126 | 0.234858794 |
| chr3:12875952-12876427 | 0.685060299 | 0.063279 | 0.234865508 |
| chr12:56932887-56933303 | -0.715857562 | 0.063287602 | 0.234886057 |
| chr12:107079256-107079509 | -0.998798901 | 0.063299042 | 0.234917136 |
| chr12:120925736-120926169 | -0.741633903 | 0.063306173 | 0.234932223 |
| chr12:128154838-128155066 | -0.967371781 | 0.063313856 | 0.234949356 |

| | | | |
|---|---|---|---|
| chr3:64038476-64038906 | 0.619306185 | 0.063339656 | 0.235024677 |
| chr17:77247067-77247657 | 0.668609453 | 0.063340605 | 0.235024677 |
| chr16:31012064-31012292 | 0.867658194 | 0.063345772 | 0.235024677 |
| chr22:11817765-11818026 | 0.750921208 | 0.063352654 | 0.235024677 |
| chr7:39643229-39643496 | -0.937550254 | 0.063354197 | 0.235024677 |
| chr6:2791276-2791533 | 0.83373335 | 0.063356725 | 0.235024677 |
| chr12:114399581-114400140 | -0.735902158 | 0.063358761 | 0.235024677 |
| chr11:68841009-68841384 | 0.707221904 | 0.063360235 | 0.235024677 |
| chr5:3575656-3575930 | 0.951926208 | 0.063364573 | 0.235024677 |
| chr12:68040453-68040927 | -0.724442963 | 0.063364827 | 0.235024677 |
| chr3:64859087-64859423 | 0.631619247 | 0.063376549 | 0.235056777 |
| chr7:51941358-51941940 | -0.671071983 | 0.063397602 | 0.235123479 |
| chr4:104671503-104671765 | -0.912933411 | 0.063405625 | 0.235137587 |
| chr17:2960212-2960438 | 0.886709172 | 0.063407544 | 0.235137587 |
| chr10:7233285-7234199 | 0.598129439 | 0.063410839 | 0.235138426 |
| chr1:125183464-125184284 | 0.58917253 | 0.063421343 | 0.235165997 |
| chr1:207828556-207828895 | -0.672702868 | 0.063433597 | 0.235187203 |
| chr19:40541238-40541721 | 0.732039842 | 0.06343377 | 0.235187203 |
| chr3:196843886-196844182 | 0.860022585 | 0.063436271 | 0.235187203 |
| chr8:89071710-89072175 | 0.688645285 | 0.063453801 | 0.235235805 |
| chr1:215005222-215005422 | -0.903602635 | 0.06345552 | 0.235235805 |
| chr3:22977060-22977478 | 0.777909452 | 0.063468422 | 0.235254453 |
| chr12:95721181-95721818 | -0.693953305 | 0.063468533 | 0.235254453 |
| chrX:24024769-24026136 | -0.590875432 | 0.063469761 | 0.235254453 |
| chr2:12057062-12057294 | -1.04856572 | 0.063499086 | 0.235351762 |
| chr19:38658622-38658946 | 0.74012433 | 0.063515979 | 0.235392102 |
| chr10:121994359-121995598 | 0.610331968 | 0.063516114 | 0.235392102 |
| chr19:55642943-55643362 | -0.658295648 | 0.063523922 | 0.23540965 |
| chr7:76293694-76293992 | 0.707867479 | 0.063530419 | 0.235422341 |
| chr12:56987484-56987924 | 0.726781692 | 0.063533585 | 0.23542269 |
| chr6:72983993-72984781 | -0.657205848 | 0.063549776 | 0.235471296 |
| chr5:168047380-168047885 | -0.681877905 | 0.063561892 | 0.23549819 |
| chr17:81429203-81429575 | 0.717839951 | 0.063563181 | 0.23549819 |
| chr14:105248202-105248657 | 0.652346521 | 0.063567164 | 0.235501559 |
| chr12:90854099-90854338 | -0.867079148 | 0.06357092 | 0.235504086 |
| chr3:17095102-17095302 | 0.949782995 | 0.063574755 | 0.235506906 |
| chr11:17238351-17238817 | 0.669594374 | 0.063591396 | 0.235557164 |
| chr19:34773020-34773423 | 0.682436049 | 0.063612642 | 0.235624163 |
| chr8:121454366-121455024 | -0.646318877 | 0.063615633 | 0.235624163 |
| chr4:171812348-171813491 | -0.611509679 | 0.063621546 | 0.235634672 |
| chr3:112473107-112473469 | 0.77974635 | 0.063640537 | 0.235685746 |
| chr2:136327162-136327871 | 0.61356881 | 0.063641488 | 0.235685746 |
| chr16:30052927-30053244 | 0.719393486 | 0.063655867 | 0.235724818 |
| chr3:166775385-166775624 | -0.895489069 | 0.063660146 | 0.235724818 |
| chr18:36420245-36420585 | -0.803429664 | 0.063663949 | 0.235724818 |
| chr4:170374722-170375180 | -0.699267311 | 0.063664344 | 0.235724818 |
| chr12:47744010-47744365 | 0.666962352 | 0.063678462 | 0.235765698 |
| chr13:52551485-52552140 | -0.6919735 | 0.06369271 | 0.235797214 |
| chrX:23782597-23783592 | -0.600109443 | 0.063694838 | 0.235797214 |
| chr9:34074243-34074556 | -0.780721338 | 0.0636972 | 0.235797214 |

| | | | |
|---|---|---|---|
| chr1:224644627-224644860 | 0.849628765 | 0.063700578 | 0.235797214 |
| chr17:65588840-65589040 | 0.901199325 | 0.063702361 | 0.235797214 |
| chr20:35463759-35463973 | -0.888237141 | 0.063710715 | 0.235816745 |
| chr9:107628610-107628868 | -0.893374161 | 0.063718569 | 0.235819764 |
| chr15:92839820-92840527 | 0.664930657 | 0.063722206 | 0.235819764 |
| chr4:138911989-138912209 | -0.748947388 | 0.063722665 | 0.235819764 |
| chr6:78867154-78868101 | -0.604965438 | 0.063724499 | 0.235819764 |
| chr7:32130940-32131177 | 0.897721565 | 0.06372692 | 0.235819764 |
| chr5:179638445-179638953 | -0.833251866 | 0.063736053 | 0.235842171 |
| chr1:183657934-183658728 | -0.626958513 | 0.063744676 | 0.235862691 |
| chr2:10546820-10547276 | 0.798303919 | 0.063771321 | 0.235949883 |
| chr7:42843239-42843543 | -0.755727959 | 0.06379408 | 0.236022696 |
| chr1:191206878-191207152 | -0.927171183 | 0.063807473 | 0.236049449 |
| chr10:8977792-8978016 | -0.927171183 | 0.063807473 | 0.236049449 |
| chr5:1516078-1516622 | 0.614051459 | 0.063816267 | 0.236060613 |
| chr9:128094324-128094743 | -0.684052588 | 0.063816652 | 0.236060613 |
| chr4:173257798-173258071 | -0.814562851 | 0.063821019 | 0.236065368 |
| chr2:226524357-226524905 | -0.728264056 | 0.063826597 | 0.236074606 |
| chr5:177339365-177339582 | -0.876886814 | 0.06383231 | 0.236084338 |
| chr10:123180539-123181007 | 0.770587161 | 0.063848087 | 0.236131292 |
| chr4:128323128-128323380 | 0.866010864 | 0.063868518 | 0.236192877 |
| chr3:119635866-119636128 | -0.81983571 | 0.063870965 | 0.236192877 |
| chr4:56743437-56744074 | 0.673136167 | 0.06387607 | 0.236192877 |
| chr2:108288652-108288869 | 0.941109388 | 0.063877069 | 0.236192877 |
| chr10:15106450-15106768 | 0.700360005 | 0.063884376 | 0.236208494 |
| chr14:31819309-31819640 | 0.794810849 | 0.063890778 | 0.236220769 |
| chr14:95241223-95241490 | 0.751282058 | 0.063900449 | 0.236245123 |
| chr11:85206890-85207412 | -0.729661999 | 0.063904532 | 0.23624882 |
| chr2:129297726-129297973 | 0.99348073 | 0.063908724 | 0.236252918 |
| chr13:41213826-41214115 | 0.847078143 | 0.063920022 | 0.236283285 |
| chr7:117944198-117944614 | -0.886603532 | 0.063940165 | 0.236346343 |
| chr2:174495405-174495951 | -0.709737837 | 0.063964402 | 0.236424526 |
| chr2:153287142-153287722 | -0.686969949 | 0.063969751 | 0.236428404 |
| chr1:174005182-174005410 | -0.941610803 | 0.063971623 | 0.236428404 |
| chr4:130403249-130403609 | 0.786795863 | 0.063988879 | 0.236480773 |
| chr22:43014562-43014865 | -0.842216015 | 0.063999399 | 0.236508245 |
| chr14:52568535-52568837 | -0.825147346 | 0.064018984 | 0.236569211 |
| chr9:130834658-130835426 | 0.662932784 | 0.064024169 | 0.236576963 |
| chr11:129846273-129846975 | 0.663344196 | 0.064034261 | 0.236602841 |
| chr5:79200648-79200872 | -0.940058098 | 0.064038111 | 0.236605657 |
| chr14:98907880-98908200 | -0.82754524 | 0.064058921 | 0.236671135 |
| chr9:75566911-75567258 | -0.856089397 | 0.064062901 | 0.236674425 |
| chr5:42219009-42219209 | 0.91740956 | 0.064068696 | 0.236684423 |
| chr12:113439576-113439860 | 0.770040735 | 0.064077283 | 0.236701125 |
| chr3:172666079-172667219 | -0.597493694 | 0.064079395 | 0.236701125 |
| chr8:106004315-106004735 | 0.702774422 | 0.064084096 | 0.236707077 |
| chr5:12918864-12919109 | 0.940206619 | 0.064088418 | 0.236711629 |
| chr10:68471449-68472261 | -0.618175867 | 0.064126742 | 0.236841763 |
| chr3:177394079-177394473 | -0.796120561 | 0.064136582 | 0.236866689 |
| chr1:92960407-92960748 | 0.733730572 | 0.064142079 | 0.236875573 |

| | | | |
|---|---|---|---|
| chr12:110124023-110124888 | -0.61811603 | 0.064153606 | 0.236894874 |
| chr14:69483880-69484722 | 0.628043415 | 0.064158421 | 0.236894874 |
| chr3:161701160-161701366 | 0.953047585 | 0.064162152 | 0.236894874 |
| chr17:58851566-58851980 | 0.652228771 | 0.064164918 | 0.236894874 |
| chr16:29815859-29816558 | 0.598523608 | 0.064168153 | 0.236894874 |
| chr2:63188939-63189308 | -0.710482827 | 0.064168767 | 0.236894874 |
| chr2:85827427-85827886 | 0.724452966 | 0.064168948 | 0.236894874 |
| chr13:87274306-87274589 | -0.79685851 | 0.064177639 | 0.236915544 |
| chr11:82858420-82858637 | -0.960567864 | 0.064194611 | 0.236966783 |
| chr12:31073520-31074874 | -0.602069253 | 0.064231313 | 0.237069888 |
| chr7:114426881-114427192 | -0.836952333 | 0.064231771 | 0.237069888 |
| chr6:79455526-79455771 | -0.936856081 | 0.064231825 | 0.237069888 |
| chr7:87344987-87346100 | -0.594203433 | 0.064235053 | 0.237070382 |
| chr1:85134934-85135801 | -0.637476044 | 0.064251449 | 0.237111911 |
| chr2:88659242-88659974 | -0.634922384 | 0.064258452 | 0.237111911 |
| chr1:228406096-228406306 | 0.793587766 | 0.064261166 | 0.237111911 |
| chr11:118929035-118929357 | 0.661806026 | 0.064262365 | 0.237111911 |
| chr1:113714864-113715123 | 0.746512885 | 0.064264157 | 0.237111911 |
| chr20:31608317-31608862 | 0.617611325 | 0.064266506 | 0.237111911 |
| chr5:18048484-18048778 | 0.766257684 | 0.064268418 | 0.237111911 |
| chr3:103446875-103447221 | 0.913988024 | 0.064271062 | 0.237111911 |
| chr7:55549826-55550352 | 0.733422381 | 0.0642855 | 0.237138828 |
| chr13:60163366-60164556 | -0.593267766 | 0.064287403 | 0.237138828 |
| chr2:189142964-189143512 | -0.66300212 | 0.064287642 | 0.237138828 |
| chr3:189316168-189317047 | 0.61035939 | 0.064295112 | 0.237154964 |
| chr14:75566308-75566718 | 0.658335276 | 0.064307886 | 0.237190663 |
| chr5:151684665-151685336 | -0.672971736 | 0.064323426 | 0.237226706 |
| chr9:16131362-16131572 | -0.96482041 | 0.064326526 | 0.237226706 |
| chr17:31558862-31559345 | 0.702084245 | 0.064326946 | 0.237226706 |
| chr5:78359806-78360750 | -0.602194086 | 0.064340142 | 0.237263951 |
| chr8:22367809-22368147 | 0.746266111 | 0.064366428 | 0.237339139 |
| chr1:114010900-114011316 | 0.648112911 | 0.064366726 | 0.237339139 |
| chr17:39152917-39153172 | -0.807481203 | 0.064391036 | 0.237417349 |
| chr6:149696665-149697027 | -0.90232999 | 0.064411629 | 0.237481853 |
| chr15:45167288-45167851 | 0.630686038 | 0.064417245 | 0.237491131 |
| chr10:133143943-133144222 | 0.734553193 | 0.064427616 | 0.237517938 |
| chr10:122353919-122354141 | 0.992807199 | 0.064433045 | 0.237526523 |
| chr10:123009412-123009708 | 0.720367323 | 0.064439349 | 0.237538332 |
| chr3:109528732-109529213 | -0.745458248 | 0.06446525 | 0.237622378 |
| chr10:69650195-69650536 | -0.758173712 | 0.064481068 | 0.237669251 |
| chr20:10511217-10511482 | -0.795083422 | 0.064487323 | 0.237680873 |
| chr12:52327541-52327962 | 0.692044468 | 0.064490473 | 0.237681048 |
| chr10:16809408-16809634 | -0.894302431 | 0.064494728 | 0.237685298 |
| chr2:16775006-16775368 | -0.746695071 | 0.064507547 | 0.237721109 |
| chr2:83347400-83347741 | -0.815861061 | 0.064513139 | 0.237730281 |
| chr9:123401985-123402250 | 0.77973226 | 0.064522398 | 0.237739966 |
| chr12:100898943-100899235 | 0.914465684 | 0.064523233 | 0.237739966 |
| chr18:73842878-73843083 | -0.990246471 | 0.064525075 | 0.237739966 |
| chr1:156627532-156627884 | 0.731934488 | 0.064530018 | 0.237746744 |
| chr17:79814398-79815151 | -0.630511777 | 0.064566761 | 0.237862072 |

| | | | |
|---|---|---|---|
| chr8:90097628-90097956 | 0.779618964 | 0.064567529 | 0.237862072 |
| chr7:4744670-4745553 | -0.60193535 | 0.06458179 | 0.237903168 |
| chr11:17812307-17812997 | 0.648786595 | 0.064588547 | 0.237916621 |
| chr19:57849599-57850184 | 0.664169862 | 0.064591948 | 0.237917712 |
| chr16:85618075-85618312 | 0.732501159 | 0.064596542 | 0.237920531 |
| chr1:100884056-100884412 | -0.86067944 | 0.064603112 | 0.237920531 |
| chr17:1123877-1124077 | -0.976557453 | 0.064613826 | 0.237920531 |
| chr10:125023765-125024310 | 0.685304435 | 0.064616136 | 0.237920531 |
| chr18:48901973-48902278 | -0.889187772 | 0.064616149 | 0.237920531 |
| chr2:44549126-44549406 | -0.844998486 | 0.06461644 | 0.237920531 |
| chr21:8436274-8436474 | 0.698941418 | 0.064618054 | 0.237920531 |
| chr4:17728328-17728974 | -0.692658772 | 0.0646227 | 0.237920531 |
| chr5:54821797-54822244 | -0.810919937 | 0.064625396 | 0.237920531 |
| chr3:127637577-127637923 | -0.719828845 | 0.064626692 | 0.237920531 |
| chr1:228279443-228279645 | 0.88032394 | 0.06462687 | 0.237920531 |
| chr2:64259441-64259643 | -1.017263078 | 0.064661086 | 0.238035061 |
| chr3:166829854-166830468 | -0.674287058 | 0.064674862 | 0.238074333 |
| chr2:155308441-155308767 | 0.782998709 | 0.064680377 | 0.238083199 |
| chr17:82384614-82384905 | 0.713023365 | 0.064693945 | 0.238121702 |
| chr15:91965604-91966194 | -0.68280456 | 0.064699054 | 0.238129067 |
| chr8:58801510-58801928 | -0.917592755 | 0.064716115 | 0.238180421 |
| chr1:3772384-3772884 | 0.663127999 | 0.064721548 | 0.238188974 |
| chr1:47997294-47997911 | 0.685362804 | 0.064730239 | 0.238209518 |
| chr17:60311868-60312075 | 0.662471841 | 0.064736798 | 0.238222214 |
| chr17:14813680-14813982 | -0.838207951 | 0.064754641 | 0.23827643 |
| chr16:1562987-1563224 | 0.855805486 | 0.06476264 | 0.238294421 |
| chr12:47964232-47964508 | 0.791097708 | 0.064778045 | 0.238338203 |
| chr4:94472927-94473286 | -0.800425583 | 0.06478076 | 0.238338203 |
| chr3:107752628-107752988 | -0.720970846 | 0.064784538 | 0.238340656 |
| chr2:161625638-161625862 | -0.853508271 | 0.064800363 | 0.238387433 |
| chr20:54462728-54462976 | 0.652608435 | 0.064806032 | 0.238396842 |
| chr13:76822750-76823080 | -0.823838362 | 0.064811859 | 0.238406828 |
| chr5:64165368-64166541 | -0.610497114 | 0.064850359 | 0.23852261 |
| chr5:172276027-172276282 | 0.959843633 | 0.064852198 | 0.23852261 |
| chr1:10901074-10901774 | -0.655014541 | 0.064854627 | 0.23852261 |
| chr13:77489940-77490172 | -0.954319134 | 0.064855786 | 0.23852261 |
| chr6:122772716-122773348 | -0.66374073 | 0.064860125 | 0.238527118 |
| chr6:116087497-116087750 | -0.918055421 | 0.064872646 | 0.238561716 |
| chr7:47326254-47326585 | -0.740448489 | 0.064878327 | 0.238571157 |
| chr11:59182509-59182740 | -0.897594387 | 0.064888018 | 0.238588998 |
| chr5:146390507-146390812 | -0.774141864 | 0.064889407 | 0.238588998 |
| chr5:6448982-6449262 | 0.810772022 | 0.064909602 | 0.238647557 |
| chr10:32981069-32981376 | -0.729224644 | 0.064911562 | 0.238647557 |
| chr19:46697250-46697764 | -0.672728046 | 0.064915886 | 0.238652001 |
| chr14:59374085-59374342 | -0.988133165 | 0.064933266 | 0.238704442 |
| chr22:35914603-35914808 | 0.884241175 | 0.064957429 | 0.238781815 |
| chr2:12399580-12399808 | -1.016085789 | 0.064979299 | 0.238850749 |
| chr16:25276751-25276951 | 0.940962156 | 0.064983671 | 0.238855059 |
| chr12:14752703-14753110 | -0.770277285 | 0.06499158 | 0.238855059 |
| chr3:134813633-134813860 | -0.953490691 | 0.064992698 | 0.238855059 |

| | | | |
|---|---|---|---|
| chr1:200372126-200372524 | 0.651234941 | 0.064992941 | 0.238855059 |
| chr4:160283018-160283316 | -0.832218804 | 0.065002762 | 0.238879694 |
| chr3:87321528-87321728 | 1.035993091 | 0.065009371 | 0.238890133 |
| chr19:373680-373945 | -0.919753778 | 0.065014844 | 0.238890133 |
| chr3:15064126-15065558 | 0.591234337 | 0.065017684 | 0.238890133 |
| chr6:75747421-75747757 | 0.66471941 | 0.065019943 | 0.238890133 |
| chr16:28975391-28975591 | 0.828471123 | 0.065024721 | 0.238890133 |
| chr5:133316361-133316585 | -0.944829977 | 0.065025436 | 0.238890133 |
| chr5:79689245-79690544 | -0.608030216 | 0.065027427 | 0.238890133 |
| chr6:154166161-154166435 | -0.800189943 | 0.065037682 | 0.238916354 |
| chr18:79962253-79962541 | -0.76751202 | 0.065060845 | 0.238975819 |
| chr4:169311653-169311994 | -0.737964402 | 0.065061471 | 0.238975819 |
| chr4:107747849-107748593 | -0.704817713 | 0.065063227 | 0.238975819 |
| chr2:10086985-10087326 | -0.817533407 | 0.065070404 | 0.238990724 |
| chr5:80991027-80991227 | 0.935382239 | 0.065086161 | 0.23902882 |
| chr7:36272661-36273392 | -0.636104906 | 0.065087016 | 0.23902882 |
| chr19:38229688-38230002 | 0.729054059 | 0.065099944 | 0.239064841 |
| chr8:122845987-122846539 | -0.654765895 | 0.06511838 | 0.239121081 |
| chrX:46566973-46567227 | 0.874993018 | 0.065134588 | 0.239169136 |
| chr7:27235724-27236016 | 0.795671137 | 0.065141839 | 0.239184302 |
| chr4:145732831-145733824 | -0.621225989 | 0.06517871 | 0.239308216 |
| chr11:60757120-60757763 | -0.712892257 | 0.06518452 | 0.239318077 |
| chr3:141123856-141124126 | 0.847360724 | 0.065192524 | 0.239335995 |
| chr10:88280778-88281241 | 0.816464283 | 0.065242302 | 0.239491935 |
| chr6:87848550-87848972 | 0.724969809 | 0.065243942 | 0.239491935 |
| chr2:46408672-46409541 | -0.644501077 | 0.065244377 | 0.239491935 |
| chr5:94026127-94026371 | 0.859218446 | 0.065265787 | 0.239556362 |
| chr10:4650779-4651025 | 0.969980734 | 0.065268906 | 0.239556362 |
| chr5:91357303-91357570 | -0.882414093 | 0.065271308 | 0.239556362 |
| chr2:130372926-130373136 | 0.788260382 | 0.065278583 | 0.239571588 |
| chr10:114002449-114002755 | -0.836250922 | 0.065295642 | 0.239622717 |
| chr11:87832350-87832727 | -0.896222952 | 0.065305263 | 0.239646545 |
| chr4:141509364-141510099 | -0.696693668 | 0.06532161 | 0.239695054 |
| chr19:1652602-1653265 | 0.630685347 | 0.065338755 | 0.239746486 |
| chr17:61562532-61562755 | 0.635546931 | 0.065344518 | 0.23975615 |
| chr6:112879697-112880383 | -0.64104056 | 0.065350805 | 0.239767735 |
| chr6:34154796-34155018 | 0.807569495 | 0.06536134 | 0.239794904 |
| chr20:43657184-43658288 | 0.602107174 | 0.065366071 | 0.239800778 |
| chr10:69385354-69386095 | 0.637632393 | 0.065372288 | 0.239807174 |
| chr20:54217256-54217468 | 0.66122829 | 0.065374073 | 0.239807174 |
| chr4:135418368-135418698 | -0.76358816 | 0.065385158 | 0.239834004 |
| chr2:223560895-223561362 | -0.842923714 | 0.065387648 | 0.239834004 |
| chr1:196779165-196779450 | -0.804096587 | 0.065407146 | 0.239894038 |
| chr6:119312989-119313419 | -0.832477523 | 0.065426219 | 0.239952504 |
| chr2:71548708-71548912 | -0.855153028 | 0.065432088 | 0.239962546 |
| chr12:28202757-28203119 | -0.814662484 | 0.065437424 | 0.239970628 |
| chr11:129308621-129308822 | -1.014306532 | 0.065459579 | 0.240040386 |
| chr11:85464349-85464591 | -0.877163915 | 0.065467212 | 0.240056889 |
| chr3:105827943-105828261 | -0.769751882 | 0.065479837 | 0.240091691 |
| chr2:65539441-65539641 | -1.038768641 | 0.065496917 | 0.240142827 |

| | | | |
|---|---|---|---|
| chr2:47008787-47009320 | -0.700303067 | 0.06550477 | 0.240160128 |
| chr18:10243428-10243628 | 0.901770109 | 0.06552295 | 0.240215284 |
| chr8:88685432-88685663 | 0.904491906 | 0.065541453 | 0.240271624 |
| chrX:134964901-134965722 | 0.661579698 | 0.06556655 | 0.240352128 |
| chr7:106009936-106010184 | -0.891397308 | 0.065581326 | 0.240394794 |
| chr11:36376114-36376629 | 0.709415433 | 0.065599372 | 0.240440978 |
| chr17:60510358-60510635 | 0.645835945 | 0.065600202 | 0.240440978 |
| chr2:10493541-10493864 | 0.74104879 | 0.065605634 | 0.240449389 |
| chr17:58409748-58409977 | 0.93129071 | 0.065612626 | 0.240463511 |
| chr17:28497867-28498485 | -0.636662756 | 0.065627651 | 0.240507071 |
| chr3:63733183-63733389 | 0.701089709 | 0.065640175 | 0.240539455 |
| chr21:34030101-34030456 | 0.704821945 | 0.065642766 | 0.240539455 |
| chr5:55601410-55601610 | -0.991941453 | 0.065648919 | 0.240550498 |
| chr5:85691-85944 | 0.800340533 | 0.065659536 | 0.240577897 |
| chr3:64273546-64273787 | 0.667339403 | 0.065681009 | 0.240644009 |
| chr4:112348435-112349106 | -0.630856497 | 0.065683861 | 0.240644009 |
| chr14:96362000-96362336 | 0.671562814 | 0.065692963 | 0.240665847 |
| chr2:57497565-57497862 | 0.737473776 | 0.065702666 | 0.240689886 |
| chr1:114564477-114564965 | 0.628002643 | 0.065713677 | 0.240718714 |
| chr5:140363125-140363534 | -0.692484736 | 0.065726281 | 0.240753376 |
| chr8:127191221-127191421 | 0.96030471 | 0.065738947 | 0.240788259 |
| chr3:71186897-71187214 | -0.93116553 | 0.065765542 | 0.240874155 |
| chr5:173766371-173766987 | 0.642774921 | 0.065780625 | 0.240917884 |
| chr22:17234155-17234417 | -0.846344736 | 0.065787662 | 0.24093214 |
| chr1:81550787-81551055 | -0.825536594 | 0.065811046 | 0.241006259 |
| chr16:1581148-1581348 | 0.955156097 | 0.065825287 | 0.24104689 |
| chr8:135212939-135213158 | -0.978492124 | 0.06583028 | 0.241047888 |
| chr5:101260719-101261263 | -0.697154073 | 0.065831851 | 0.241047888 |
| chr20:48224651-48225102 | 0.616936236 | 0.065839482 | 0.241052893 |
| chr1:166859700-166859900 | 0.914021276 | 0.06583951 | 0.241052893 |
| chr8:77037949-77038238 | -0.911744882 | 0.065844242 | 0.241058699 |
| chr8:90593238-90593573 | 0.68099243 | 0.065851558 | 0.241062465 |
| chr2:19955868-19956152 | -0.756397612 | 0.065851563 | 0.241062465 |
| chr1:67514407-67514670 | -0.933927066 | 0.06585969 | 0.241080697 |
| chr20:45405947-45406720 | 0.642157169 | 0.065880239 | 0.241140589 |
| chr10:84222452-84222719 | -0.834303674 | 0.065882346 | 0.241140589 |
| chr1:112268436-112268676 | 0.881163719 | 0.065904244 | 0.241209219 |
| chr6:106804450-106804902 | -0.695259477 | 0.065911307 | 0.241223548 |
| chr13:24727999-24728361 | 0.707459545 | 0.065922601 | 0.24125202 |
| chr17:59033469-59033686 | 0.667649569 | 0.065925384 | 0.24125202 |
| chr17:73380848-73381155 | 0.781880174 | 0.065930466 | 0.241259094 |
| chr11:28098321-28098559 | 0.958227155 | 0.065950487 | 0.24132083 |
| chr1:1859252-1859859 | 0.680756441 | 0.065954304 | 0.241323274 |
| chrX:144335488-144335699 | 0.844879726 | 0.065962074 | 0.241331677 |
| chr2:97709911-97710236 | -0.706952034 | 0.065966232 | 0.241331677 |
| chr5:163489008-163489394 | -0.709293445 | 0.065969197 | 0.241331677 |
| chr15:65401375-65401695 | 0.693102874 | 0.065971069 | 0.241331677 |
| chr4:169015404-169015684 | -0.730622694 | 0.065972349 | 0.241331677 |
| chr20:42483698-42484043 | -0.65257446 | 0.065987883 | 0.24137698 |
| chr5:126594474-126595712 | -0.589534831 | 0.065994295 | 0.24138891 |

| | | | |
|---|---|---|---|
| chr7:115004232-115004432 | -0.999269239 | 0.065999592 | 0.241393131 |
| chr4:103424787-103425283 | -0.632540043 | 0.06600175 | 0.241393131 |
| chr11:60915233-60915696 | 0.669516409 | 0.066025679 | 0.241469122 |
| chr7:109699509-109700035 | -0.829431549 | 0.066039625 | 0.241508596 |
| chr17:8057706-8058014 | -0.795110956 | 0.066046373 | 0.241521747 |
| chr1:199774865-199775086 | -0.9570008 | 0.066052028 | 0.241530899 |
| chr4:125719997-125720381 | -0.731593414 | 0.066055562 | 0.241532295 |
| chr19:55418164-55418578 | 0.732450272 | 0.066061339 | 0.241541892 |
| chr12:12449603-12449953 | 0.778875679 | 0.066080493 | 0.241600399 |
| chr7:3627778-3628185 | -0.806430484 | 0.066102083 | 0.241665115 |
| chr2:9753734-9754035 | -0.789215246 | 0.066104502 | 0.241665115 |
| chr5:140402444-140402660 | -0.740887262 | 0.066129071 | 0.241743401 |
| chr19:41883905-41884556 | 0.61360682 | 0.066139275 | 0.241768963 |
| chr10:119199352-119199763 | -0.688649524 | 0.066142374 | 0.241768963 |
| chr11:1203055-1203279 | 0.861053698 | 0.066152247 | 0.241793515 |
| chr8:102814674-102814902 | 0.689993969 | 0.066163949 | 0.241824754 |
| chr4:180557321-180557719 | -0.767038724 | 0.066179388 | 0.241869642 |
| chr12:119066534-119066923 | -0.808596883 | 0.066195696 | 0.241917705 |
| chr1:109649424-109649939 | -0.74856528 | 0.066202201 | 0.241929941 |
| chr10:147838-148288 | 0.688524112 | 0.066208236 | 0.241933562 |
| chr20:38961944-38962947 | 0.602281894 | 0.066209507 | 0.241933562 |
| chr3:48446531-48446918 | 0.711902319 | 0.066228509 | 0.241991456 |
| chr1:11099294-11099601 | 0.740986565 | 0.06623941 | 0.242012483 |
| chr13:46835023-46835308 | 0.737566244 | 0.066240581 | 0.242012483 |
| chr6:53793346-53793886 | -0.629750832 | 0.066245294 | 0.242018163 |
| chr11:35304047-35304709 | -0.706709193 | 0.066250651 | 0.242019438 |
| chrX:154397419-154398937 | 0.586669409 | 0.066251961 | 0.242019438 |
| chr15:79283670-79283904 | 0.848640521 | 0.066262166 | 0.242045179 |
| chr22:24592995-24593422 | 0.686348389 | 0.0662672 | 0.242052029 |
| chr5:129481683-129481917 | -0.945650469 | 0.066288336 | 0.242117689 |
| chr3:15325270-15325867 | 0.651719823 | 0.066299876 | 0.242148297 |
| chr4:1683750-1683950 | 0.891727031 | 0.066312589 | 0.242183184 |
| chrX:103817528-103817784 | -0.896958352 | 0.066339246 | 0.242268991 |
| chr20:48635801-48636334 | 0.593618028 | 0.066348175 | 0.242281823 |
| chr2:7510414-7510704 | 0.891732309 | 0.066349084 | 0.242281823 |
| chr14:67411998-67412495 | 0.661920431 | 0.066367661 | 0.242320262 |
| chr8:140098525-140099716 | -0.587273 | 0.066368174 | 0.242320262 |
| chr12:102950311-102950887 | -0.662215922 | 0.066369098 | 0.242320262 |
| chr3:50601068-50601868 | -0.638137951 | 0.066375643 | 0.242332612 |
| chrX:119468060-119469566 | 0.586589612 | 0.066387707 | 0.24235937 |
| chr3:64122736-64122966 | 0.656102526 | 0.066389298 | 0.24235937 |
| chr1:65687447-65688218 | -0.661284533 | 0.066398562 | 0.242381641 |
| chr3:164558761-164558961 | -1.012192725 | 0.066403113 | 0.242385077 |
| chr6:129667625-129667933 | -0.801290062 | 0.066408991 | 0.242385077 |
| chr5:42036410-42036687 | -0.969183259 | 0.066408994 | 0.242385077 |
| chr1:37513597-37513960 | 0.753658209 | 0.066421746 | 0.242420075 |
| chr17:72465328-72465907 | -0.614835763 | 0.066455103 | 0.24252802 |
| chr18:78953575-78953833 | -0.944946079 | 0.066460155 | 0.24252802 |
| chr14:64487479-64487742 | 0.898173948 | 0.066463095 | 0.24252802 |
| chr1:217911794-217912409 | -0.646872657 | 0.066468911 | 0.24252802 |

| | | | |
|---|---|---|---|
| chr17:58728663-58728863 | 0.695515731 | 0.06646967 | 0.24252802 |
| chr16:84190850-84191050 | -0.87943534 | 0.066470314 | 0.24252802 |
| chr8:28411703-28412180 | 0.720221798 | 0.066477237 | 0.242535709 |
| chr1:99788954-99789233 | -0.84973724 | 0.066478752 | 0.242535709 |
| chr20:32762041-32762712 | 0.637017081 | 0.06649416 | 0.24258037 |
| chr3:64097633-64097958 | 0.619572468 | 0.066503096 | 0.24260142 |
| chr11:70948950-70949806 | -0.629243063 | 0.066518497 | 0.242643207 |
| chr17:83109743-83110199 | 0.633093583 | 0.066522244 | 0.242643207 |
| chr12:97961093-97961390 | -0.825834448 | 0.066524778 | 0.242643207 |
| chr18:21184460-21184717 | 0.827983867 | 0.066527218 | 0.242643207 |
| chr2:231463782-231465353 | 0.582935802 | 0.066537365 | 0.242668666 |
| chr5:78472578-78472781 | -1.009782658 | 0.066545729 | 0.242687619 |
| chr5:55546706-55546911 | -0.987091905 | 0.066550312 | 0.242692779 |
| chr20:54485469-54485788 | 0.631324035 | 0.066557585 | 0.242698938 |
| chr20:57134866-57135086 | 0.625801958 | 0.066558336 | 0.242698938 |
| chr14:67118176-67118804 | 0.651631071 | 0.0665681 | 0.242721773 |
| chr16:11636264-11636710 | 0.644057272 | 0.066573643 | 0.242721773 |
| chr5:96661863-96663548 | -0.578665769 | 0.066574101 | 0.242721773 |
| chr11:17846780-17847114 | 0.732534226 | 0.066600144 | 0.242803072 |
| chr4:159484862-159485197 | -0.752052467 | 0.066602738 | 0.242803072 |
| chr5:31270459-31270678 | -0.95038787 | 0.066613374 | 0.242816698 |
| chr1:203487162-203488307 | -0.584823053 | 0.066614601 | 0.242816698 |
| chr6:47134799-47135009 | 0.870278807 | 0.066615982 | 0.242816698 |
| chr7:35823494-35823802 | 0.764087283 | 0.066628072 | 0.242849211 |
| chr6:50842492-50842746 | -0.786146862 | 0.06663435 | 0.242860542 |
| chr5:170216350-170216618 | -0.805908493 | 0.066639658 | 0.242868335 |
| chr5:147453319-147454403 | -0.632445548 | 0.066646918 | 0.242871451 |
| chrX:85109291-85109531 | -0.925492692 | 0.066649642 | 0.242871451 |
| chr20:47693702-47694223 | 0.590202821 | 0.066650022 | 0.242871451 |
| chr11:80505863-80506309 | -0.767382191 | 0.066656043 | 0.24288184 |
| chr14:105745619-105745931 | 0.784114311 | 0.06666174 | 0.242891049 |
| chr2:199624846-199625350 | 0.665324117 | 0.066668628 | 0.242893791 |
| chr8:132875976-132876176 | -1.051068962 | 0.06667186 | 0.242893791 |
| chr2:26730685-26730889 | 0.938379605 | 0.066672003 | 0.242893791 |
| chr17:62961992-62963625 | 0.574495421 | 0.066681262 | 0.242904929 |
| chr8:61563234-61563779 | -0.682344086 | 0.066681401 | 0.242904929 |
| chr4:115905885-115906328 | -0.72176676 | 0.06668856 | 0.242919461 |
| chr12:46219733-46219999 | 0.825602851 | 0.066708381 | 0.242980109 |
| chr1:148371480-148371840 | 0.665379821 | 0.066723931 | 0.243025194 |
| chr3:153752640-153752840 | 0.96022541 | 0.066730632 | 0.243038048 |
| chr8:98471157-98471382 | 0.794096566 | 0.066761126 | 0.243126582 |
| chrX:154378476-154379505 | 0.582843354 | 0.066761286 | 0.243126582 |
| chr20:43743924-43744168 | 0.840191532 | 0.066767706 | 0.243138406 |
| chr1:26997719-26998118 | 0.75184218 | 0.066780826 | 0.243174626 |
| chr3:142117015-142117454 | -0.769855264 | 0.066799592 | 0.2432314 |
| chr7:124007128-124007371 | -0.947375702 | 0.066812875 | 0.243268206 |
| chr1:61226754-61227232 | -0.676239925 | 0.066850752 | 0.243394549 |
| chr9:22103245-22103599 | 0.724193471 | 0.066863034 | 0.2434277 |
| chr18:54209532-54209802 | -0.903490061 | 0.066868604 | 0.243429305 |
| chr15:43168786-43169078 | -0.843010708 | 0.06687225 | 0.243429305 |

| | | | |
|---|---|---|---|
| chr8:143904659-143905068 | 0.679485893 | 0.066875202 | 0.243429305 |
| chr15:73908320-73908709 | 0.681822698 | 0.066877135 | 0.243429305 |
| chr7:80145820-80146089 | -0.925398504 | 0.06687936 | 0.243429305 |
| chr13:88429719-88430181 | -0.74509596 | 0.066890857 | 0.243459586 |
| chr14:53020751-53020951 | 1.013693425 | 0.066894726 | 0.243462106 |
| chr1:17233803-17234115 | -0.752893553 | 0.066906054 | 0.243491767 |
| chr12:50763760-50764969 | 0.594239566 | 0.066911696 | 0.243500734 |
| chr4:145557639-145558083 | -0.68081917 | 0.066918222 | 0.243512919 |
| chr16:25111365-25112085 | 0.607016926 | 0.066922872 | 0.243518275 |
| chr2:204625196-204625452 | 0.896719275 | 0.066928997 | 0.243528995 |
| chr2:79235316-79235614 | -0.95248281 | 0.066934896 | 0.243538895 |
| chr20:43187891-43188463 | -0.740449324 | 0.066938332 | 0.243539832 |
| chr6:37489637-37490229 | 0.681970942 | 0.066944521 | 0.243550785 |
| chr16:4936243-4936754 | 0.671663059 | 0.066949739 | 0.243558206 |
| chr5:133245584-133245879 | -0.766445951 | 0.066958503 | 0.243578522 |
| chr20:48357181-48357400 | 0.664564858 | 0.066970363 | 0.243610101 |
| chr11:66910080-66910304 | -0.985927143 | 0.066977077 | 0.243617372 |
| chr11:67582610-67582867 | 0.755291555 | 0.066978721 | 0.243617372 |
| chr10:63682334-63682568 | -0.950142132 | 0.066985582 | 0.243625099 |
| chr12:93145178-93145500 | 0.809620339 | 0.066987205 | 0.243625099 |
| chr2:104078867-104079140 | 0.892631381 | 0.066997577 | 0.243651256 |
| chr11:67033253-67033480 | -0.812745255 | 0.067003286 | 0.243660453 |
| chr13:81017407-81017870 | -0.804508937 | 0.067024731 | 0.243726872 |
| chr6:25963525-25963795 | -1.028447646 | 0.067036168 | 0.243754286 |
| chr1:99969659-99970659 | -0.595667326 | 0.067038632 | 0.243754286 |
| chr10:90227249-90227497 | -0.917167843 | 0.067046638 | 0.243771828 |
| chr8:131804289-131804498 | 0.855958704 | 0.067051225 | 0.243776939 |
| chr12:90959638-90959869 | -0.917662118 | 0.06705981 | 0.243796581 |
| chr12:76180464-76180791 | 0.781184706 | 0.067063606 | 0.243798815 |
| chrX:55167193-55167402 | 0.954285226 | 0.067070072 | 0.243800449 |
| chr1:173478121-173478392 | -0.68870085 | 0.067070419 | 0.243800449 |
| chr2:68385627-68385827 | -1.032829976 | 0.06707813 | 0.243816911 |
| chr10:11642212-11642495 | -0.931323889 | 0.067107451 | 0.243901678 |
| chr5:91226097-91226439 | -0.733303542 | 0.067107817 | 0.243901678 |
| chr6:166112500-166112838 | -0.973651647 | 0.067115424 | 0.243917755 |
| chr7:73971828-73972198 | -0.712479067 | 0.067120039 | 0.243922959 |
| chr13:46818265-46818835 | 0.668931762 | 0.067135893 | 0.243958548 |
| chr9:68685831-68686095 | -0.884317485 | 0.0671362 | 0.243958548 |
| chr19:43984482-43984863 | 0.768409983 | 0.067142493 | 0.243967194 |
| chr5:178859683-178860426 | -0.617600295 | 0.067144948 | 0.243967194 |
| chr6:41653177-41653520 | 0.671820458 | 0.067156669 | 0.243977219 |
| chr2:196751048-196751563 | -0.668144848 | 0.067156672 | 0.243977219 |
| chr1:114571341-114571562 | 0.689032106 | 0.067157259 | 0.243977219 |
| chr6:148375727-148375927 | -1.049197691 | 0.067167421 | 0.244002567 |
| chr1:151762955-151763687 | 0.605857824 | 0.067176377 | 0.244023531 |
| chrX:123777571-123778128 | 0.63633254 | 0.067180097 | 0.244025475 |
| chr1:2525571-2525782 | 0.908821219 | 0.067194105 | 0.244064787 |
| chr3:62852671-62853101 | 0.735837707 | 0.067202247 | 0.24408279 |
| chr19:50492313-50492553 | 0.792749375 | 0.067205816 | 0.244084184 |
| chr13:97018032-97018556 | -0.720302371 | 0.067218575 | 0.244118952 |

| | | | |
|---|---|---|---|
| chr2:89145333-89145635 | -0.871584751 | 0.067224267 | 0.244121383 |
| chr6:69232859-69233428 | -0.675983939 | 0.067225617 | 0.244121383 |
| chr7:6938281-6938647 | -0.712075368 | 0.067234974 | 0.244140743 |
| chr2:11482705-11482948 | -0.740709666 | 0.067237915 | 0.244140743 |
| chrX:131830477-131831214 | 0.594436434 | 0.067240507 | 0.244140743 |
| chr5:75020069-75020335 | -0.980085597 | 0.067247443 | 0.244148402 |
| chr14:47626210-47627324 | -0.609419957 | 0.067248989 | 0.244148402 |
| chr19:4352678-4353112 | 0.771873654 | 0.067255696 | 0.244161183 |
| chr15:80778959-80779762 | 0.619904744 | 0.067258935 | 0.244161374 |
| chr7:16564927-16565415 | 0.651516921 | 0.067264171 | 0.244166402 |
| chrX:103404616-103405109 | 0.691690583 | 0.067266693 | 0.244166402 |
| chr1:9824436-9824676 | 0.788192909 | 0.067280729 | 0.244205782 |
| chr11:116219086-116219484 | -0.931295725 | 0.067285699 | 0.24420762 |
| chr17:76789416-76789781 | 0.732328238 | 0.067287719 | 0.24420762 |
| chr18:76565890-76566108 | -0.916357403 | 0.067292519 | 0.24420762 |
| chr1:214328757-214329171 | -0.671124191 | 0.067294928 | 0.24420762 |
| chr1:172620694-172620931 | -0.919891117 | 0.067297172 | 0.24420762 |
| chr7:123865573-123866082 | -0.696106885 | 0.067306089 | 0.244212607 |
| chr15:64248024-64248276 | -0.904953174 | 0.067306282 | 0.244212607 |
| chr12:41190375-41190744 | -0.782004426 | 0.067308108 | 0.244212607 |
| chr13:20525356-20526538 | 0.602374341 | 0.067315079 | 0.244226337 |
| chr1:244273057-244273298 | 0.834002486 | 0.067344589 | 0.244317729 |
| chr6:2158665-2158956 | -0.835156553 | 0.067346647 | 0.244317729 |
| chr4:2843043-2844125 | 0.614680967 | 0.067354738 | 0.244330592 |
| chr5:157290575-157290917 | 0.795284303 | 0.06736018 | 0.244330592 |
| chr11:73295254-73295744 | 0.662101231 | 0.067362887 | 0.244330592 |
| chr13:25468245-25468445 | 0.962401761 | 0.067362947 | 0.244330592 |
| chr8:53751017-53751276 | -0.821679763 | 0.067372791 | 0.244354729 |
| chr14:58072540-58072941 | 0.685129708 | 0.067383785 | 0.244383036 |
| chr20:38601654-38601857 | -0.959761427 | 0.067396437 | 0.244417352 |
| chr11:20068774-20069007 | -0.932625183 | 0.06742679 | 0.244515854 |
| chr12:130214778-130215030 | -0.902248373 | 0.067435648 | 0.244533726 |
| chr15:43266891-43267095 | -0.963665945 | 0.067438101 | 0.244533726 |
| chr17:76626092-76626292 | 0.920249209 | 0.067455078 | 0.244583712 |
| chr12:49264447-49265749 | 0.584886665 | 0.067490635 | 0.244701058 |
| chr3:150773644-150774199 | -0.648296459 | 0.067494756 | 0.244703891 |
| chr7:44264159-44264798 | -0.664542636 | 0.067497804 | 0.244703891 |
| chr4:154416265-154417385 | -0.622777757 | 0.067506275 | 0.244712456 |
| chr10:11424434-11424819 | 0.755069816 | 0.067506554 | 0.244712456 |
| chr12:12975764-12976014 | -0.856345157 | 0.067534995 | 0.244803972 |
| chr3:30721099-30721420 | 0.794506567 | 0.067547703 | 0.244838456 |
| chr6:141090135-141090364 | -1.042922195 | 0.067583799 | 0.244938745 |
| chr7:44415644-44415844 | -1.042922195 | 0.067583799 | 0.244938745 |
| chr2:69958692-69958913 | -0.984346139 | 0.067584962 | 0.244938745 |
| chr14:69878527-69878727 | 0.939819954 | 0.067605799 | 0.245002672 |
| chr22:39240620-39241396 | 0.634176177 | 0.06762755 | 0.245069908 |
| chr17:29567017-29567249 | 0.732176536 | 0.067646409 | 0.245126657 |
| chr12:990406-991732 | -0.601003958 | 0.067652067 | 0.245135566 |
| chr14:34672634-34672834 | 0.930102414 | 0.067655808 | 0.245137529 |
| chr12:123772064-123772301 | -0.914511597 | 0.067682101 | 0.245221199 |

| | | | |
|---|---|---|---|
| chr16:77406714-77407000 | 0.752291655 | 0.067711071 | 0.245312248 |
| chr1:216940200-216940409 | 0.924184257 | 0.067716794 | 0.245312248 |
| chr12:84874601-84874829 | -0.849621704 | 0.067716835 | 0.245312248 |
| chr7:76291313-76292326 | -0.593512132 | 0.067746284 | 0.245393193 |
| chr1:172746317-172746599 | 0.779205967 | 0.067751559 | 0.245393193 |
| chr15:72475343-72475543 | 0.800369383 | 0.067753912 | 0.245393193 |
| chr17:61854049-61854252 | 0.63563969 | 0.067754553 | 0.245393193 |
| chr10:79335832-79336144 | 0.815066589 | 0.067755193 | 0.245393193 |
| chr11:76768491-76768706 | -0.750445163 | 0.067782308 | 0.245479796 |
| chr17:40013755-40014058 | 0.729335979 | 0.067804914 | 0.245545103 |
| chr13:65456264-65456575 | -0.858942809 | 0.067806751 | 0.245545103 |
| chr20:54877839-54878174 | 0.624999383 | 0.067811356 | 0.245550175 |
| chr1:48552475-48552796 | -0.826622506 | 0.067825586 | 0.245590098 |
| chr20:40251343-40251645 | -0.838229277 | 0.067836435 | 0.245617774 |
| chr12:113312554-113312833 | 0.755935275 | 0.067844151 | 0.245634103 |
| chr12:48000999-48001325 | 0.704802477 | 0.067853145 | 0.245655059 |
| chr3:166263525-166263752 | -0.884330904 | 0.067865595 | 0.245657393 |
| chr6:12957508-12958001 | -0.770821402 | 0.067867257 | 0.245657393 |
| chr9:125513628-125513828 | 0.88705645 | 0.067868436 | 0.245657393 |
| chr9:135441140-135441350 | -0.934208974 | 0.067874318 | 0.245657393 |
| chr3:176274246-176274446 | -1.00019007 | 0.067875027 | 0.245657393 |
| chr4:73143057-73143299 | -1.00019007 | 0.067875027 | 0.245657393 |
| chr12:64633992-64634220 | 0.83207699 | 0.067876232 | 0.245657393 |
| chr5:145854749-145854949 | -0.978891626 | 0.067882617 | 0.245668896 |
| chr7:621971-622312 | 0.699140047 | 0.067889414 | 0.24568189 |
| chr2:168518729-168518964 | 0.93800234 | 0.067910759 | 0.245744382 |
| chr5:159095893-159096154 | -0.957985675 | 0.067913097 | 0.245744382 |
| chr15:85825703-85826181 | 0.646561711 | 0.067917009 | 0.245746932 |
| chr3:71930618-71930939 | 0.735691808 | 0.067937343 | 0.245808901 |
| chr13:47511229-47511452 | 0.845550425 | 0.067943693 | 0.245820269 |
| chr7:151727569-151728130 | -0.66226076 | 0.067950766 | 0.245834249 |
| chr4:140037420-140037830 | 0.694227834 | 0.067968087 | 0.245873433 |
| chr2:186693646-186694827 | -0.587612423 | 0.067972406 | 0.245873433 |
| chr20:15296486-15296790 | 0.746573269 | 0.067976691 | 0.245873433 |
| chr5:132668227-132668432 | 0.951630636 | 0.067979254 | 0.245873433 |
| chr19:11380954-11381895 | 0.599387407 | 0.067979631 | 0.245873433 |
| chr5:15915801-15916233 | -0.796957828 | 0.06798085 | 0.245873433 |
| chr2:124283177-124283541 | 0.737960741 | 0.068037054 | 0.246065098 |
| chr6:4357640-4358017 | -0.688343577 | 0.068055138 | 0.246111584 |
| chr1:42380245-42380594 | 0.744415087 | 0.068056332 | 0.246111584 |
| chr19:10158229-10158599 | 0.802985114 | 0.068071141 | 0.24615352 |
| chr13:75760198-75760890 | -0.649184037 | 0.068078291 | 0.246167758 |
| chr22:49971279-49971975 | -0.629836585 | 0.068087116 | 0.246180116 |
| chr7:17531105-17531305 | 0.903937337 | 0.068088134 | 0.246180116 |
| chr15:89358540-89359183 | 0.604344861 | 0.068107126 | 0.246237165 |
| chr14:75711176-75711436 | 0.806513461 | 0.068149276 | 0.246362135 |
| chr16:10883027-10883382 | 0.707425064 | 0.068149908 | 0.246362135 |
| chr2:55442471-55442876 | -0.731093764 | 0.068151338 | 0.246362135 |
| chr2:177572194-177572448 | -0.924167349 | 0.068160305 | 0.246382926 |
| chr10:61942664-61943429 | -0.684360539 | 0.068164374 | 0.24638601 |

| | | | |
|---|---|---|---|
| chr13:45394075-45394295 | -0.862547966 | 0.06816959 | 0.246393243 |
| chr14:23640619-23640915 | 0.720779639 | 0.068181958 | 0.24642632 |
| chr17:81937030-81937710 | 0.611540002 | 0.068199658 | 0.246473173 |
| chr3:57582832-57583168 | 0.722840003 | 0.06820218 | 0.246473173 |
| chr4:105594660-105595043 | 0.735896252 | 0.068204572 | 0.246473173 |
| chr2:205679754-205680416 | -0.626127947 | 0.068210285 | 0.246478535 |
| chr2:238158760-238159016 | 0.851728811 | 0.068215289 | 0.246478535 |
| chr10:62816289-62816933 | -0.594253414 | 0.068220771 | 0.246478535 |
| chr15:88152768-88152975 | -0.873325203 | 0.068220941 | 0.246478535 |
| chr7:17489891-17490362 | 0.688287268 | 0.068222139 | 0.246478535 |
| chr10:83691724-83692648 | -0.636777367 | 0.068227076 | 0.24648135 |
| chr20:37564347-37564949 | 0.69655351 | 0.068229933 | 0.24648135 |
| chr15:33216305-33216621 | -0.860584371 | 0.068232569 | 0.24648135 |
| chr6:151658001-151658515 | -0.632939998 | 0.068240924 | 0.246499908 |
| chr4:84664925-84665178 | -0.898484941 | 0.068246274 | 0.246507615 |
| chr6:150720908-150721311 | 0.765282412 | 0.068264489 | 0.246561125 |
| chr10:85492890-85493165 | -0.887484717 | 0.068270481 | 0.246561125 |
| chr14:41957345-41957757 | 0.744510909 | 0.068270742 | 0.246561125 |
| chr19:44356464-44356941 | 0.684162285 | 0.068281973 | 0.246590063 |
| chr1:77832187-77832803 | -0.670516766 | 0.068297377 | 0.246634068 |
| chr10:12264214-12264775 | -0.649743315 | 0.068301438 | 0.246637108 |
| chr2:44103760-44103998 | -0.925815268 | 0.068322289 | 0.246700774 |
| chr14:88775348-88776181 | 0.627506389 | 0.068326895 | 0.246705779 |
| chr1:190261880-190262318 | 0.786637719 | 0.068342957 | 0.246752146 |
| chr18:54828214-54828902 | -0.610511169 | 0.068354047 | 0.246780559 |
| chr5:106601602-106602048 | -0.666899006 | 0.06836801 | 0.246819339 |
| chr5:95554405-95555543 | -0.577746009 | 0.068382512 | 0.246850883 |
| chr2:191566290-191566542 | 0.772601643 | 0.068383191 | 0.246850883 |
| chr17:66649290-66649490 | 0.907846978 | 0.068391986 | 0.246871002 |
| chr12:55646177-55646649 | -0.657708996 | 0.068432943 | 0.247007207 |
| chr14:42754608-42754864 | -1.114553314 | 0.068466395 | 0.247116309 |
| chr19:44071718-44072463 | 0.63865745 | 0.068492253 | 0.247197995 |
| chr1:179517067-179517342 | 0.891539719 | 0.068499392 | 0.247212115 |
| chr2:196510332-196510552 | -0.943339439 | 0.068524879 | 0.247292448 |
| chr22:18905759-18906640 | 0.615818086 | 0.068534373 | 0.247307436 |
| chr1:158095774-158096033 | -0.871687281 | 0.068535487 | 0.247307436 |
| chr11:68296516-68296719 | 0.869806894 | 0.068562428 | 0.247393001 |
| chr10:99042771-99042971 | -1.028450624 | 0.068571104 | 0.247412656 |
| chr1:29415452-29415736 | 0.75313098 | 0.068580166 | 0.247433699 |
| chr10:10426876-10427187 | -0.896057465 | 0.068585949 | 0.247442912 |
| chr1:227947495-227948083 | 0.65131286 | 0.068597884 | 0.247474318 |
| chr18:46469416-46469874 | -0.682802538 | 0.068613716 | 0.24751978 |
| chr2:97997868-97998172 | 0.729537132 | 0.068626536 | 0.247554372 |
| chr12:67658209-67658597 | -0.873247797 | 0.068639583 | 0.24758978 |
| chr19:35976587-35976787 | 0.861262368 | 0.068658278 | 0.247644299 |
| chr13:40777209-40777856 | -0.607861063 | 0.068661161 | 0.247644299 |
| chr7:127628551-127628806 | -0.867935506 | 0.068681585 | 0.247706305 |
| chr17:74981917-74982940 | 0.598236227 | 0.068694504 | 0.247741237 |
| chr2:65573172-65573628 | -0.666944525 | 0.068707999 | 0.247778241 |
| chr3:16489965-16490544 | 0.626431635 | 0.068737518 | 0.24787303 |

| | | | |
|---|---|---|---|
| chr17:66638977-66639247 | -0.853066381 | 0.068749761 | 0.247905509 |
| chr4:118989263-118989781 | 0.674605256 | 0.068762276 | 0.24792399 |
| chr16:89240452-89241583 | -0.595686302 | 0.068767181 | 0.24792399 |
| chr17:63579604-63579942 | 0.685134175 | 0.068767527 | 0.24792399 |
| chr2:142582369-142582630 | -0.875547655 | 0.068767828 | 0.24792399 |
| chr7:140294505-140294764 | -0.989432231 | 0.068778546 | 0.247950963 |
| chr10:37290179-37290467 | -0.783643967 | 0.068813401 | 0.24806238 |
| chr11:2436977-2437177 | 1.021834207 | 0.068815927 | 0.24806238 |
| chr14:105374485-105374819 | 0.725827977 | 0.068819834 | 0.248064795 |
| chr2:85298770-85299096 | -0.743112361 | 0.068832476 | 0.24809869 |
| chr1:217135365-217135909 | -0.636646803 | 0.068846058 | 0.248135973 |
| chr17:75471874-75472535 | -0.61448945 | 0.068851449 | 0.248140521 |
| chr2:215983245-215983515 | -0.733681272 | 0.068853797 | 0.248140521 |
| chr8:135442815-135443253 | -0.825757665 | 0.068857178 | 0.248141036 |
| chr21:42614755-42615363 | -0.621121831 | 0.068862607 | 0.248141238 |
| chr17:61747926-61748379 | 0.595176016 | 0.068867518 | 0.248141238 |
| chr6:3500714-3500914 | 0.95700264 | 0.068868049 | 0.248141238 |
| chr2:215736745-215736982 | -0.942383345 | 0.068873706 | 0.248141238 |
| chr7:81159406-81159744 | -0.788223453 | 0.06887499 | 0.248141238 |
| chr6:152218883-152219239 | -0.767605468 | 0.068876665 | 0.248141238 |
| chr8:143814897-143815281 | 0.664344894 | 0.068887243 | 0.248167677 |
| chr2:81901083-81901337 | -0.884934422 | 0.068895297 | 0.248185025 |
| chr17:66675904-66676202 | -0.695627524 | 0.068900521 | 0.248192172 |
| chr1:174986998-174987638 | -0.61878526 | 0.068926101 | 0.248272645 |
| chr1:30816595-30816814 | -0.960601907 | 0.068933318 | 0.248281953 |
| chr16:75677377-75677623 | -0.981070535 | 0.068935166 | 0.248281953 |
| chrX:47365972-47366604 | -0.627249265 | 0.068945771 | 0.248305949 |
| chr7:86848432-86848632 | -1.011723454 | 0.06894831 | 0.248305949 |
| chr3:6066898-6067512 | -0.661257944 | 0.068967952 | 0.248365015 |
| chr14:76904716-76905371 | 0.613716875 | 0.068978951 | 0.248375089 |
| chr6:71383496-71383812 | -0.720947683 | 0.068979712 | 0.248375089 |
| chr7:140672611-140674004 | -0.583766145 | 0.068980474 | 0.248375089 |
| chr7:142557999-142558322 | -0.73822857 | 0.06900001 | 0.248433754 |
| chr21:42607186-42607386 | -1.017278218 | 0.069005166 | 0.248434899 |
| chr17:61713000-61713535 | 0.588721066 | 0.069006812 | 0.248434899 |
| chr4:183721186-183721651 | -0.653421798 | 0.069014139 | 0.248449602 |
| chrX:119237188-119237502 | 0.751870505 | 0.069019798 | 0.248458303 |
| chr15:86572137-86572629 | 0.713581468 | 0.069026339 | 0.248470177 |
| chr14:41749501-41749844 | -0.800337851 | 0.069031962 | 0.248478744 |
| chr13:87677501-87678246 | -0.625479088 | 0.069052919 | 0.248542502 |
| chr12:21751473-21751913 | -0.768146263 | 0.069069942 | 0.248592096 |
| chr11:71299269-71299634 | 0.680846047 | 0.069073307 | 0.248592528 |
| chr1:114642591-114642791 | 0.706229419 | 0.069081151 | 0.248609081 |
| chr7:91759266-91759825 | -0.696886272 | 0.06908705 | 0.248618632 |
| chr8:89922732-89922938 | 0.788955289 | 0.069101383 | 0.248625788 |
| chr1:174990369-174990672 | -0.743562831 | 0.069101912 | 0.248625788 |
| chr18:74495675-74496982 | -0.575808511 | 0.069102346 | 0.248625788 |
| chr10:121974325-121975497 | 0.597741719 | 0.069103775 | 0.248625788 |
| chr9:132114222-132114422 | 0.893232707 | 0.069105262 | 0.248625788 |
| chr1:41003663-41004103 | 0.669799127 | 0.069120973 | 0.248670635 |

| | | | |
|---|---|---|---|
| chr12:107885119-107885586 | 0.730662362 | 0.069130487 | 0.248693186 |
| chr4:1289303-1290063 | 0.610242678 | 0.069137465 | 0.248706612 |
| chr20:54572811-54573236 | 0.609824487 | 0.069140821 | 0.248707009 |
| chrX:125378297-125378513 | 0.971151751 | 0.069148793 | 0.24872401 |
| chr3:40580580-40580780 | 0.899123514 | 0.069176341 | 0.248811415 |
| chr17:62136302-62138312 | 0.566964205 | 0.069187354 | 0.248839348 |
| chr17:48671664-48672229 | -0.651853497 | 0.069207264 | 0.248899271 |
| chr8:90333569-90333865 | 0.770271376 | 0.069216309 | 0.248920116 |
| chr10:70405033-70405387 | -0.744284012 | 0.069220107 | 0.248922093 |
| chr3:184688486-184688952 | -0.705092172 | 0.069229998 | 0.248927099 |
| chr4:125070863-125071182 | -0.740518351 | 0.069230738 | 0.248927099 |
| chr13:21060215-21060750 | 0.66418234 | 0.069231245 | 0.248927099 |
| chr1:117590632-117590912 | 0.741197068 | 0.069243481 | 0.248959412 |
| chr15:36887315-36887618 | -0.896738215 | 0.069256734 | 0.248977194 |
| chr7:107462949-107463406 | -0.792068613 | 0.069257038 | 0.248977194 |
| chr5:122480187-122480435 | -0.906114615 | 0.069258175 | 0.248977194 |
| chr11:118607431-118608248 | 0.618585739 | 0.069287848 | 0.249058782 |
| chr7:68731028-68731421 | -0.734080329 | 0.069290211 | 0.249058782 |
| chr3:195720915-195721190 | 0.811982331 | 0.069290913 | 0.249058782 |
| chr5:149586735-149586935 | 0.937812394 | 0.069293873 | 0.249058782 |
| chr1:215644345-215644545 | 0.892845332 | 0.06930838 | 0.249099239 |
| chr1:3695674-3695972 | 0.732677243 | 0.069315202 | 0.249112073 |
| chr13:113002130-113002427 | 0.742899319 | 0.069322832 | 0.24912781 |
| chr14:49302945-49303145 | 0.917183441 | 0.069336764 | 0.24916619 |
| chr3:33884007-33884478 | -0.718057092 | 0.069351688 | 0.249200958 |
| chr16:57764773-57765049 | 0.807932406 | 0.069352943 | 0.249200958 |
| chr16:67170334-67170857 | 0.626576627 | 0.069389724 | 0.249321428 |
| chr13:105979506-105979903 | -0.767489403 | 0.069397997 | 0.249339458 |
| chr10:91749592-91749909 | -0.786702832 | 0.069405748 | 0.249355614 |
| chr2:186602413-186602844 | -0.734391211 | 0.069415374 | 0.249378507 |
| chr4:77473570-77473826 | -0.938255078 | 0.069442567 | 0.249456655 |
| chr2:137508759-137509162 | -0.661018442 | 0.069445106 | 0.249456655 |
| chr7:91830040-91830322 | -0.829260534 | 0.0694481 | 0.249456655 |
| chr11:88103452-88104062 | -0.648368443 | 0.069453027 | 0.249456655 |
| chr5:167535993-167536336 | -1.010136846 | 0.069453405 | 0.249456655 |
| chr12:45543865-45544114 | 0.80116254 | 0.069465481 | 0.249488333 |
| chr13:52850365-52851884 | -0.585830399 | 0.069473816 | 0.24950007 |
| chr10:118592967-118593293 | -0.813026731 | 0.069475262 | 0.24950007 |
| chr3:187914330-187914595 | 0.769548617 | 0.069479082 | 0.249502095 |
| chr1:202130211-202130674 | -0.626692473 | 0.069488082 | 0.249522721 |
| chr7:38908614-38909384 | -0.599485778 | 0.069492515 | 0.249526947 |
| chr1:47031017-47031478 | -0.705672826 | 0.069498129 | 0.249535409 |
| chr10:94546150-94546542 | -0.701174783 | 0.06950149 | 0.249535783 |
| chr5:179452796-179453031 | 0.878890073 | 0.069512428 | 0.249563362 |
| chr7:113289965-113290206 | -0.946596633 | 0.06953333 | 0.2496195 |
| chr5:149550943-149551156 | -0.75581868 | 0.06953458 | 0.2496195 |
| chr19:13833358-13834056 | 0.646997216 | 0.069543571 | 0.24964008 |
| chr12:57213380-57213622 | 0.747360481 | 0.069575438 | 0.249738164 |
| chr3:32570075-32571414 | 0.580886429 | 0.069577413 | 0.249738164 |
| chr17:15659716-15660001 | 0.838693685 | 0.069580954 | 0.249739173 |

| | | | |
|---|---|---|---|
| chr1:1318287-1318548 | -0.907806688 | 0.069585194 | 0.249742692 |
| chr17:81444490-81445289 | -0.626523793 | 0.069600701 | 0.249776901 |
| chr6:140462741-140463301 | -0.631834703 | 0.06960441 | 0.249776901 |
| chr11:126282879-126283851 | 0.603627885 | 0.069604505 | 0.249776901 |
| chr6:32153513-32154419 | 0.606593366 | 0.069615003 | 0.249799564 |
| chr10:92517113-92517551 | -0.738121734 | 0.069618571 | 0.249799564 |
| chr19:45059489-45059804 | 0.736467033 | 0.069621422 | 0.249799564 |
| chr2:231715182-231715544 | -0.69484068 | 0.069623861 | 0.249799564 |
| chr2:7014398-7014640 | -0.939573469 | 0.06963797 | 0.249838485 |
| chr10:85928122-85928451 | -0.847671136 | 0.069660988 | 0.249908013 |
| chr18:6920024-6920433 | -0.706426329 | 0.069663873 | 0.249908013 |
| chr3:138300231-138300431 | -0.93856702 | 0.069670105 | 0.249908847 |
| chr14:51335635-51335835 | 0.942918369 | 0.069670628 | 0.249908847 |
| chr10:61795529-61795754 | -0.85236143 | 0.069690877 | 0.249969777 |
| chr1:158078157-158078407 | 0.908803933 | 0.069703907 | 0.250004809 |
| chr1:213476090-213476313 | -1.053116734 | 0.069716348 | 0.250018164 |
| chr19:33064291-33065127 | 0.605262086 | 0.069716671 | 0.250018164 |
| chr10:30384768-30385273 | -0.676202981 | 0.069717419 | 0.250018164 |
| chr9:109330149-109330385 | 0.840301736 | 0.069748452 | 0.250117747 |
| chr7:11415156-11415449 | 0.743661257 | 0.069758663 | 0.250142654 |
| chr10:48672292-48672811 | -0.784698886 | 0.069775493 | 0.250191296 |
| chr2:121480148-121480478 | -0.734755971 | 0.0697821 | 0.2501997 |
| chr8:100425000-100425200 | 0.865670335 | 0.069784367 | 0.2501997 |
| chr7:44913351-44913615 | 0.798411845 | 0.069795221 | 0.250215694 |
| chr2:127278592-127278968 | -0.784323344 | 0.069795359 | 0.250215694 |
| chr21:10402910-10403324 | 0.729754199 | 0.069804629 | 0.250237218 |
| chr7:93442399-93443106 | -0.696887065 | 0.06982134 | 0.250285415 |
| chr3:106628439-106628681 | -0.948487362 | 0.069825765 | 0.250289568 |
| chr21:14926979-14927228 | 0.846944849 | 0.069835464 | 0.250312009 |
| chr13:38814285-38814678 | -0.897131049 | 0.06983856 | 0.250312009 |
| chr1:173429471-173429805 | -0.668146479 | 0.069851854 | 0.25033615 |
| chr16:12350954-12351201 | 0.91563731 | 0.069853307 | 0.25033615 |
| chr5:14984887-14985091 | 0.865900185 | 0.069855097 | 0.25033615 |
| chr3:102324528-102324832 | -0.879880434 | 0.069884212 | 0.250413312 |
| chr3:71584779-71584980 | -0.82431896 | 0.069884746 | 0.250413312 |
| chr12:71706695-71706907 | 0.936568137 | 0.069892661 | 0.250413312 |
| chr10:59895704-59895909 | -0.884089042 | 0.069895024 | 0.250413312 |
| chr4:57156314-57156573 | -0.816335664 | 0.069897992 | 0.250413312 |
| chr10:21292813-21293204 | -0.819070197 | 0.069898088 | 0.250413312 |
| chr19:35995367-35995995 | 0.642573014 | 0.069899506 | 0.250413312 |
| chr16:19432497-19432777 | -0.901282358 | 0.069907172 | 0.25042907 |
| chr11:71253012-71253324 | 0.704263195 | 0.069920986 | 0.250466844 |
| chr7:148151788-148152147 | -0.794899785 | 0.069932738 | 0.25049723 |
| chr3:64853944-64854286 | 0.625950206 | 0.069953218 | 0.250556555 |
| chr5:69188931-69190232 | -0.588028309 | 0.06995584 | 0.250556555 |
| chr3:100144277-100144531 | -0.8489946 | 0.069985006 | 0.2506493 |
| chr17:56172448-56172978 | -0.627208906 | 0.069989956 | 0.250655312 |
| chr5:72767629-72767983 | -0.77948766 | 0.070000095 | 0.250679905 |
| chr3:14248810-14249473 | 0.6187295 | 0.070005635 | 0.250686208 |
| chr22:23141903-23142207 | -0.734025842 | 0.070010954 | 0.250686208 |

| | | | |
|---|---|---|---|
| chr8:107755487-107755854 | -0.743784186 | 0.07001167 | 0.250686208 |
| chr11:22633596-22633868 | -0.837126786 | 0.070018647 | 0.250693698 |
| chrX:85561203-85561448 | -0.988169848 | 0.070020305 | 0.250693698 |
| chr1:53682123-53682398 | 0.766367638 | 0.070042411 | 0.250761125 |
| chr7:9835254-9835550 | -0.780558821 | 0.070060895 | 0.250815583 |
| chr3:131193673-131194242 | -0.716156759 | 0.070068285 | 0.25083032 |
| chr6:144375512-144375851 | -0.888850392 | 0.070113212 | 0.250965121 |
| chr2:23041502-23041836 | -0.840963283 | 0.07011543 | 0.250965121 |
| chr1:228102355-228102856 | 0.637260107 | 0.070120493 | 0.250965121 |
| chr11:27429426-27430359 | -0.643771817 | 0.070124021 | 0.250965121 |
| chr2:121128869-121129274 | 0.727239675 | 0.070127882 | 0.250965121 |
| chr10:58384828-58386085 | -0.577652353 | 0.07013341 | 0.250965121 |
| chr20:54788471-54788854 | 0.623627305 | 0.070140875 | 0.250965121 |
| chr11:64331813-64332013 | 0.827721152 | 0.070142591 | 0.250965121 |
| chr5:179208082-179208414 | 0.726495276 | 0.070147259 | 0.250965121 |
| chr15:77870487-77870775 | 0.723922752 | 0.07014837 | 0.250965121 |
| chr3:130901486-130901997 | -0.760589787 | 0.070152757 | 0.250965121 |
| chr11:16782169-16782552 | 0.704770101 | 0.070156525 | 0.250965121 |
| chr7:35410378-35410612 | -0.99683787 | 0.070156681 | 0.250965121 |
| chr16:4506419-4506748 | 0.744789925 | 0.07015774 | 0.250965121 |
| chr6:121977693-121977912 | -0.930468443 | 0.070157945 | 0.250965121 |
| chr9:62801329-62801605 | -0.818911206 | 0.070158347 | 0.250965121 |
| chr11:20754918-20755221 | -0.809414838 | 0.070163441 | 0.250971625 |
| chr16:90081993-90082898 | -0.604887445 | 0.070171347 | 0.250988186 |
| chr6:145110171-145110525 | -0.823289273 | 0.070182649 | 0.251016895 |
| chr13:86762092-86762389 | -0.956163573 | 0.070190997 | 0.251029269 |
| chr2:28300808-28301124 | -0.910719085 | 0.070193255 | 0.251029269 |
| chr2:10901234-10901449 | -0.891591493 | 0.070195938 | 0.251029269 |
| chr6:121603538-121604049 | -0.723065844 | 0.070209854 | 0.251041788 |
| chr6:158758781-158758988 | 0.970268291 | 0.070215145 | 0.251041788 |
| chr16:22092335-22092535 | 0.87022774 | 0.070216204 | 0.251041788 |
| chr7:17474935-17475504 | 0.671841135 | 0.070217487 | 0.251041788 |
| chr5:179658669-179658897 | 0.903270182 | 0.070217981 | 0.251041788 |
| chr1:227946605-227946885 | 0.701093222 | 0.070224256 | 0.251041788 |
| chr11:106172333-106172977 | -0.65461169 | 0.070224335 | 0.251041788 |
| chr3:139117060-139117547 | 0.648266388 | 0.070225649 | 0.251041788 |
| chr4:15955731-15955981 | 0.791929987 | 0.07024716 | 0.251106969 |
| chr19:47608507-47609265 | 0.601643444 | 0.070255377 | 0.251124625 |
| chr7:42237930-42238299 | -0.670554874 | 0.070271058 | 0.25116232 |
| chr5:134367022-134367352 | -0.669554581 | 0.070272478 | 0.25116232 |
| chr19:55146744-55147240 | 0.65959343 | 0.070300227 | 0.251245447 |
| chr3:63947676-63947898 | 0.683526596 | 0.070302294 | 0.251245447 |
| chr12:123533762-123534624 | 0.624091224 | 0.070321016 | 0.251300635 |
| chr17:61701445-61701766 | 0.611925996 | 0.070334047 | 0.251333908 |
| chr3:16476633-16476966 | 0.759464064 | 0.070336887 | 0.251333908 |
| chr5:110865738-110866237 | -0.659332345 | 0.070354018 | 0.251370054 |
| chr9:3240733-3241030 | -0.776046359 | 0.070354543 | 0.251370054 |
| chr8:58641420-58641773 | -0.761948491 | 0.070356845 | 0.251370054 |
| chr6:31598535-31598885 | 0.713722249 | 0.070362105 | 0.251377126 |
| chr5:98768174-98768380 | -0.763866427 | 0.070371165 | 0.251397772 |

| | | | |
|---|---|---|---|
| chr17:69607215-69607478 | -0.749317638 | 0.070377546 | 0.251407041 |
| chr12:7265335-7265608 | -0.907137162 | 0.070380539 | 0.251407041 |
| chr15:99435107-99435597 | -0.656089347 | 0.070383603 | 0.251407041 |
| chr14:34179750-34179973 | 0.86096921 | 0.070388258 | 0.251411947 |
| chr10:116878078-116878523 | -0.707422364 | 0.070417129 | 0.251503345 |
| chr20:18496034-18496371 | -0.738724858 | 0.070443663 | 0.251576799 |
| chr19:9827335-9828032 | 0.630447449 | 0.070444262 | 0.251576799 |
| chrX:9862380-9862882 | -0.695358874 | 0.070455547 | 0.251605373 |
| chr13:44818449-44818716 | -0.75153627 | 0.070467409 | 0.251636009 |
| chr3:64962255-64963185 | 0.580044353 | 0.070477505 | 0.251660331 |
| chr4:130563256-130564242 | -0.618333112 | 0.070488317 | 0.25168721 |
| chr7:27179764-27180067 | 0.766724688 | 0.070493928 | 0.251695516 |
| chr2:60296665-60296865 | 1.010555841 | 0.070532739 | 0.251822354 |
| chr2:6772978-6773498 | -0.640965748 | 0.070544481 | 0.251852539 |
| chr12:48817941-48818141 | 0.764234512 | 0.070548481 | 0.251855086 |
| chr12:12586783-12587240 | 0.722063424 | 0.070557643 | 0.251876059 |
| chr7:140049153-140049545 | -0.83776096 | 0.070561795 | 0.251879147 |
| chr1:207499513-207499838 | -0.671461015 | 0.070575917 | 0.251917818 |
| chr19:10780482-10780735 | -0.982526902 | 0.070581425 | 0.251925745 |
| chr7:126937285-126937510 | 0.968492999 | 0.070587876 | 0.251937034 |
| chr12:52948710-52949280 | 0.593247653 | 0.070596067 | 0.251947639 |
| chrX:68827825-68828134 | 0.687711537 | 0.070600711 | 0.251947639 |
| chr10:98361054-98361357 | -0.7171935 | 0.070601514 | 0.251947639 |
| chr10:3239569-3239863 | 0.744599127 | 0.070604519 | 0.251947639 |
| chr6:64774878-64775146 | 0.768982081 | 0.070607289 | 0.251947639 |
| chr1:106772875-106773675 | 0.587638585 | 0.070614132 | 0.251960326 |
| chr18:65725430-65725766 | -0.760117793 | 0.070617628 | 0.251961065 |
| chr1:200802933-200803333 | -0.646112589 | 0.070631004 | 0.251987052 |
| chr7:10811199-10811703 | 0.693821184 | 0.070631489 | 0.251987052 |
| chr11:61149970-61150189 | 0.817480862 | 0.070638881 | 0.252001693 |
| chr5:91032-91321 | 0.758809725 | 0.070657525 | 0.252056469 |
| chr2:150204523-150204822 | -0.863366066 | 0.070664673 | 0.252070231 |
| chr4:162959118-162959526 | -0.72008223 | 0.070674752 | 0.25209445 |
| chr19:45123397-45123928 | 0.645030556 | 0.070693088 | 0.252148114 |
| chr9:126661469-126662236 | -0.596801762 | 0.07070157 | 0.25216382 |
| chr1:114596006-114596212 | 0.701048304 | 0.070704073 | 0.25216382 |
| chr5:150623307-150623507 | -0.889793981 | 0.070713995 | 0.252169589 |
| chr3:112640409-112641774 | -0.585293511 | 0.070714681 | 0.252169589 |
| chr4:140689693-140690102 | -0.753273164 | 0.070715564 | 0.252169589 |
| chr4:179660776-179661203 | -0.777413625 | 0.070722475 | 0.252182498 |
| chr6:74091457-74091711 | -0.928177204 | 0.070732182 | 0.252202082 |
| chr4:56506010-56506374 | -0.872614956 | 0.070737045 | 0.252202082 |
| chr6:166586781-166587137 | -0.776257929 | 0.070737842 | 0.252202082 |
| chr2:48344193-48344402 | -0.954273487 | 0.070757014 | 0.252258697 |
| chr14:30283853-30284217 | 0.722354084 | 0.070761801 | 0.252262605 |
| chr14:42110195-42110519 | 0.701402779 | 0.070767572 | 0.252262605 |
| chr21:44428281-44428955 | 0.595655493 | 0.070768433 | 0.252262605 |
| chr10:7756495-7756695 | 0.880518671 | 0.070773721 | 0.252262605 |
| chr10:53458086-53458429 | -0.708983185 | 0.070774571 | 0.252262605 |
| chr9:105901517-105902135 | -0.627225667 | 0.070812973 | 0.252365474 |

| | | | |
|---|---|---|---|
| chr3:39125299-39126399 | 0.600792109 | 0.070813188 | 0.252365474 |
| chrX:134237066-134237304 | 0.880705607 | 0.070813313 | 0.252365474 |
| chr6:80214470-80214791 | -0.748682481 | 0.070819828 | 0.252376953 |
| chr5:60671924-60672124 | -0.991085576 | 0.070828826 | 0.252397279 |
| chr12:77996773-77997055 | -0.889575658 | 0.070841829 | 0.252431876 |
| chr2:156935685-156935996 | 0.821101345 | 0.070855677 | 0.252463999 |
| chrX:40085219-40085665 | 0.713092447 | 0.070857434 | 0.252463999 |
| chr11:441812-442212 | 0.750193052 | 0.070865433 | 0.252480758 |
| chr13:71374497-71374799 | -0.938112574 | 0.070877454 | 0.252511848 |
| chr3:14857532-14857787 | 0.731440624 | 0.070881564 | 0.252514747 |
| chr7:5412941-5413476 | 0.66618397 | 0.070895858 | 0.252553927 |
| chr5:73152737-73152969 | -0.933452772 | 0.070902769 | 0.252560532 |
| chr11:60486000-60486414 | -0.717402736 | 0.070905644 | 0.252560532 |
| chr7:39565827-39566934 | -0.588586686 | 0.070910391 | 0.252560532 |
| chr15:56364965-56365902 | 0.593932016 | 0.070910897 | 0.252560532 |
| chr3:153796719-153797656 | -0.589369916 | 0.070932543 | 0.252625887 |
| chr12:125186183-125186444 | 0.690363145 | 0.070965122 | 0.252721024 |
| chr9:69324555-69325319 | 0.65530593 | 0.070967031 | 0.252721024 |
| chr10:114166060-114166376 | 0.817079617 | 0.070973928 | 0.252721024 |
| chr1:176809865-176810238 | 0.793604593 | 0.070974711 | 0.252721024 |
| chr6:157227729-157227929 | -0.984947948 | 0.070975747 | 0.252721024 |
| chr6:146041321-146041627 | -0.804456724 | 0.070981984 | 0.252731485 |
| chr7:132251300-132251812 | -0.654738761 | 0.070987111 | 0.252737997 |
| chr19:44577087-44577295 | 0.880315878 | 0.071000403 | 0.252763454 |
| chr6:82365458-82365847 | 0.650887075 | 0.071000859 | 0.252763454 |
| chr13:88744322-88744637 | -0.888186014 | 0.071007277 | 0.252774559 |
| chr13:24085849-24086049 | 0.949846934 | 0.071032793 | 0.252853644 |
| chr20:48550667-48551079 | 0.605628247 | 0.071040531 | 0.25286944 |
| chr20:48153573-48153838 | 0.631934785 | 0.071046152 | 0.2528777 |
| chr18:46555776-46556014 | -0.932110241 | 0.071057382 | 0.252905925 |
| chr16:30756084-30756338 | -0.89550189 | 0.071062681 | 0.252913037 |
| chr6:43020987-43021909 | 0.622281519 | 0.071066729 | 0.252915697 |
| chr1:16216177-16216486 | 0.699487058 | 0.07108358 | 0.252950277 |
| chr14:102539871-102540281 | -0.684131433 | 0.071086487 | 0.252950277 |
| chr11:129695675-129695875 | -0.994010317 | 0.071089651 | 0.252950277 |
| chr2:178282120-178282328 | -0.994010317 | 0.071089651 | 0.252950277 |
| chr8:128767081-128767304 | 0.848429597 | 0.071099343 | 0.252973015 |
| chr22:50293591-50293978 | 0.712453097 | 0.071117876 | 0.253023758 |
| chr5:11841853-11842063 | 1.030835554 | 0.071122929 | 0.253023758 |
| chr12:12696533-12696865 | -0.692493595 | 0.071123512 | 0.253023758 |
| chr19:5156230-5156493 | -0.932871065 | 0.07113762 | 0.2530622 |
| chr7:48299642-48299986 | -0.685948106 | 0.071149351 | 0.253092181 |
| chr16:77298471-77299122 | 0.671481759 | 0.071157241 | 0.253108497 |
| chr19:41958322-41958830 | 0.631607238 | 0.071162496 | 0.253115436 |
| chr22:43185916-43186304 | 0.722935006 | 0.071166418 | 0.253117639 |
| chr11:68127982-68128948 | 0.603147089 | 0.071171145 | 0.2531227 |
| chr12:33012434-33012808 | 0.764753092 | 0.071184793 | 0.25315949 |
| chr4:78125828-78126435 | 0.641700723 | 0.071192055 | 0.253168228 |
| chr1:209879553-209879824 | -0.828768678 | 0.071193858 | 0.253168228 |
| chr3:71208510-71209164 | -0.6215793 | 0.0712116 | 0.253212735 |

| | | | |
|---|---|---|---|
| chr6:34179982-34180309 | 0.746124197 | 0.071212984 | 0.253212735 |
| chr6:137939971-137940253 | -0.917637837 | 0.071221439 | 0.253231048 |
| chrX:153934973-153935702 | 0.617677785 | 0.071231629 | 0.253255526 |
| chr3:164726075-164726466 | 0.7526317 | 0.071269862 | 0.25334474 |
| chr2:5236439-5236812 | -0.887456583 | 0.071271217 | 0.25334474 |
| chr2:1665767-1666180 | -0.734264927 | 0.071275885 | 0.25334474 |
| chr14:103561414-103562002 | 0.644175198 | 0.071277271 | 0.25334474 |
| chr5:156647292-156647525 | -0.950962323 | 0.071278105 | 0.25334474 |
| chr3:104510285-104510863 | -0.66461553 | 0.071281236 | 0.25334474 |
| chr6:125153558-125154064 | -0.613856844 | 0.071283099 | 0.25334474 |
| chr21:40320709-40320933 | -0.756516867 | 0.071283172 | 0.25334474 |
| chr2:69714505-69715012 | -0.647464961 | 0.071288842 | 0.253353138 |
| chr3:23859012-23859360 | -0.750215306 | 0.071304604 | 0.253397403 |
| chr14:105944969-105945272 | 0.707043402 | 0.071316835 | 0.253429113 |
| chr7:152871111-152871369 | -0.913711106 | 0.071320373 | 0.25342993 |
| chr22:41426816-41427051 | 0.892395582 | 0.071336535 | 0.253475607 |
| chr2:21296569-21296890 | -0.881804922 | 0.071355213 | 0.253525487 |
| chr10:8225259-8225559 | -0.807392379 | 0.071357191 | 0.253525487 |
| chr1:40848590-40848891 | 0.760451755 | 0.071370628 | 0.253561471 |
| chr17:58971869-58972256 | 0.62264066 | 0.071374679 | 0.253564108 |
| chr2:126876525-126876974 | -0.833493943 | 0.071388882 | 0.253589064 |
| chr3:87390449-87390775 | -0.783395877 | 0.071390836 | 0.253589064 |
| chr7:108027981-108028434 | -0.738483072 | 0.071391633 | 0.253589064 |
| chr14:65226847-65227676 | 0.580448307 | 0.07142838 | 0.253707831 |
| chr1:236147767-236148025 | 0.757730473 | 0.07145351 | 0.253782701 |
| chr7:75728217-75728683 | -0.714260371 | 0.071456083 | 0.253782701 |
| chr7:106030037-106030268 | -0.820035701 | 0.071468765 | 0.253815979 |
| chr6:100881974-100882204 | -0.764809659 | 0.071486221 | 0.253866206 |
| chr11:34512902-34513355 | 0.678052355 | 0.071495464 | 0.253867538 |
| chr13:25468594-25468993 | 0.729764219 | 0.071499663 | 0.253867538 |
| chr11:72222503-72222965 | -0.664381081 | 0.071502459 | 0.253867538 |
| chr5:115295847-115297021 | -0.580584357 | 0.071502563 | 0.253867538 |
| chr10:67788241-67788835 | -0.645226158 | 0.071504066 | 0.253867538 |
| chr14:51565004-51565496 | 0.639957645 | 0.071506476 | 0.253867538 |
| chr1:17529407-17529795 | -0.751346634 | 0.071516981 | 0.253893069 |
| chr1:904268-904468 | 0.958714287 | 0.071524862 | 0.253909282 |
| chr21:40386338-40386621 | -0.676033259 | 0.071541586 | 0.253956886 |
| chr9:114551780-114552318 | 0.624091595 | 0.071550731 | 0.253966017 |
| chr17:64312268-64312597 | 0.616097056 | 0.071550787 | 0.253966017 |
| chr7:32140646-32140873 | 0.789671901 | 0.071555359 | 0.253970479 |
| chr20:59770222-59770481 | 0.74411252 | 0.071569648 | 0.254000572 |
| chr6:75997564-75997798 | -0.915060791 | 0.071570467 | 0.254000572 |
| chr3:136661318-136661627 | -0.74460351 | 0.071578718 | 0.254014007 |
| chr3:165282767-165282967 | -1.046048608 | 0.071580883 | 0.254014007 |
| chr13:40963750-40964468 | -0.640020685 | 0.071586938 | 0.254023727 |
| chr3:43738224-43738606 | -0.784364405 | 0.071595348 | 0.254038068 |
| chr2:185700957-185701189 | -0.93132373 | 0.071600147 | 0.254038068 |
| chr1:26035284-26035641 | 0.671566836 | 0.071601196 | 0.254038068 |
| chr14:96098693-96099010 | 0.650507924 | 0.071604241 | 0.254038068 |
| chr16:72952764-72953599 | 0.624642858 | 0.071622407 | 0.254080294 |

| | | | |
|---|---|---|---|
| chr10:21184319-21184550 | 0.901731444 | 0.071622775 | 0.254080294 |
| chr2:164504266-164504664 | -0.722504551 | 0.071626439 | 0.254081529 |
| chr3:24048121-24048334 | -0.920135856 | 0.071641593 | 0.254107062 |
| chr2:148006312-148006592 | -0.809805559 | 0.071645384 | 0.254107062 |
| chr9:73269438-73269762 | -0.839645866 | 0.071645415 | 0.254107062 |
| chr10:101012781-101013426 | -0.631298923 | 0.071646903 | 0.254107062 |
| chr6:26568682-26569012 | -0.648140774 | 0.071667356 | 0.25416784 |
| chr11:12248322-12248959 | -0.602433079 | 0.071687267 | 0.254225484 |
| chr14:41419729-41419988 | -0.862334297 | 0.071691862 | 0.254225484 |
| chr7:139777941-139778825 | -0.628633656 | 0.07169604 | 0.254225484 |
| chr17:46977384-46977730 | -0.677306378 | 0.071696882 | 0.254225484 |
| chr8:90472752-90473088 | 0.683124364 | 0.071701266 | 0.254229264 |
| chr16:1705163-1705504 | -0.78044053 | 0.071736126 | 0.254341096 |
| chrX:101658387-101658952 | -0.636305684 | 0.07175601 | 0.254399824 |
| chr17:61340718-61340936 | 0.613842141 | 0.07176925 | 0.254434992 |
| chr1:114323114-114323334 | 0.674165573 | 0.071804213 | 0.254547163 |
| chr15:92306747-92307096 | -0.767786027 | 0.071814379 | 0.254571423 |
| chr10:71975531-71975776 | -0.973692064 | 0.07182566 | 0.254594688 |
| chr13:21557992-21558354 | 0.722206055 | 0.071827587 | 0.254594688 |
| chr12:53061430-53062040 | 0.598161605 | 0.071840099 | 0.254618881 |
| chr11:85685812-85686483 | -0.646167952 | 0.071841059 | 0.254618881 |
| chr12:126856957-126857250 | -0.857060128 | 0.071866887 | 0.254691804 |
| chr3:124463334-124463588 | -0.890077956 | 0.071870983 | 0.254691804 |
| chr7:151833733-151833941 | -0.92203064 | 0.071871606 | 0.254691804 |
| chr6:16332132-16332418 | -0.810739777 | 0.071898602 | 0.254775688 |
| chr11:56509443-56509778 | -0.699754725 | 0.071924068 | 0.254846853 |
| chr3:32106942-32107321 | 0.678839105 | 0.071925337 | 0.254846853 |
| chr1:234182359-234182573 | -0.917101308 | 0.071943033 | 0.254874627 |
| chr4:90137709-90137909 | 0.95271011 | 0.071945019 | 0.254874627 |
| chr5:58452517-58452832 | -0.832008174 | 0.071945563 | 0.254874627 |
| chr14:105370834-105371134 | 0.701634171 | 0.071946482 | 0.254874627 |
| chr18:48453131-48453331 | -0.89075073 | 0.071965282 | 0.254918735 |
| chr6:6781167-6781596 | -0.815665632 | 0.071965586 | 0.254918735 |
| chr7:401679-401998 | 0.739058809 | 0.071972162 | 0.254930241 |
| chr7:16660992-16661447 | 0.673098451 | 0.071988807 | 0.254977414 |
| chr11:129457163-129457719 | 0.640513179 | 0.072016089 | 0.255048375 |
| chr20:53599526-53600019 | 0.578693412 | 0.072018134 | 0.255048375 |
| chr13:29428270-29429077 | -0.613249521 | 0.072018828 | 0.255048375 |
| chr4:87804865-87805206 | -0.861905459 | 0.072035147 | 0.255094379 |
| chr4:183905116-183905508 | -0.704345664 | 0.072065531 | 0.255184083 |
| chr2:40451034-40451432 | -0.739803394 | 0.07206714 | 0.255184083 |
| chr13:110137743-110138006 | -0.751698915 | 0.072081806 | 0.255212551 |
| chrX:12984592-12984873 | 0.858880317 | 0.072081841 | 0.255212551 |
| chr20:12617024-12617232 | 0.94680703 | 0.072087567 | 0.255221033 |
| chr16:28292331-28292617 | 0.709308291 | 0.072097909 | 0.255245854 |
| chr3:195974155-195974414 | 0.853193707 | 0.07210596 | 0.25525534 |
| chr11:108341014-108341386 | -0.729998368 | 0.072109207 | 0.25525534 |
| chr11:2930774-2931080 | 0.650639363 | 0.072115833 | 0.25525534 |
| chr5:119274339-119274813 | -0.628011012 | 0.072115899 | 0.25525534 |
| chr5:180670206-180670474 | 0.856326846 | 0.072117246 | 0.25525534 |

| | | | |
|---|---|---|---|
| chr2:218250793-218250993 | -0.99206179 | 0.072125604 | 0.255273133 |
| chr18:46104144-46104992 | -0.604441403 | 0.07215231 | 0.255355857 |
| chr10:15209891-15210314 | 0.717143902 | 0.072171288 | 0.255404872 |
| chr1:1073302-1073504 | -0.772163638 | 0.072172826 | 0.255404872 |
| chr6:78467216-78467462 | -1.061148542 | 0.072199982 | 0.25548917 |
| chr5:78607809-78608083 | -0.94449841 | 0.072204231 | 0.255492406 |
| chr14:93013395-93013595 | -0.861207737 | 0.0722352 | 0.255590186 |
| chr10:86899768-86899996 | 0.79453383 | 0.07225134 | 0.255625507 |
| chr3:8421343-8421618 | 0.726968226 | 0.072251855 | 0.255625507 |
| chr2:118112772-118113369 | -0.652111667 | 0.072287887 | 0.25574118 |
| chr18:50143489-50143734 | -0.935015252 | 0.072310484 | 0.255809314 |
| chr6:122361535-122361852 | -0.8274531 | 0.072319624 | 0.255824128 |
| chr3:68873139-68873461 | -0.825251697 | 0.072321349 | 0.255824128 |
| chr19:28692299-28692588 | -0.973316838 | 0.072327187 | 0.255832968 |
| chr4:123506003-123506234 | 0.794845657 | 0.072335977 | 0.255838197 |
| chr3:170734457-170734724 | -0.774042773 | 0.072336649 | 0.255838197 |
| chr9:70412769-70413201 | -0.653283528 | 0.072343886 | 0.255838197 |
| chr20:48137980-48138311 | 0.61209389 | 0.072347832 | 0.255838197 |
| chr12:86484448-86484663 | -0.97193642 | 0.072348611 | 0.255838197 |
| chr22:42498568-42498966 | 0.725063962 | 0.072348699 | 0.255838197 |
| chr10:11684024-11684321 | 0.678214519 | 0.072355363 | 0.255849951 |
| chr18:57802328-57802682 | -0.701396269 | 0.072367461 | 0.255880441 |
| chr6:76697636-76697931 | -0.896376192 | 0.072370664 | 0.255880441 |
| chr7:25895411-25895646 | -0.90735903 | 0.07238086 | 0.25590468 |
| chr17:15341473-15342017 | -0.678150201 | 0.072396082 | 0.25594669 |
| chr22:31080756-31081424 | 0.601296316 | 0.072415667 | 0.255980147 |
| chr13:85865106-85865316 | -0.988514459 | 0.072416473 | 0.255980147 |
| chr18:26764702-26764940 | -0.966739836 | 0.072426772 | 0.255980147 |
| chr14:30558723-30559570 | 0.582026833 | 0.072436836 | 0.255980147 |
| chr7:39557175-39557447 | -0.884665984 | 0.072437714 | 0.255980147 |
| chr3:151034366-151034621 | 0.8253497 | 0.072439821 | 0.255980147 |
| chr12:47838643-47838906 | -0.770549482 | 0.072440422 | 0.255980147 |
| chr12:57491430-57491740 | -0.770549482 | 0.072440422 | 0.255980147 |
| chr17:69212976-69213352 | -0.735102234 | 0.07244093 | 0.255980147 |
| chr4:81130545-81131395 | -0.65398128 | 0.072442134 | 0.255980147 |
| chr1:185386220-185386420 | 1.035946394 | 0.072442295 | 0.255980147 |
| chr12:51531935-51532194 | 0.73275154 | 0.072467431 | 0.25605716 |
| chr2:26741970-26742436 | 0.740792511 | 0.072501942 | 0.256167286 |
| chr6:91330212-91330581 | -0.70799875 | 0.072513845 | 0.256192105 |
| chr7:106284472-106285829 | -0.567157866 | 0.072515653 | 0.256192105 |
| chr2:163838966-163839346 | -0.873519263 | 0.072523488 | 0.256199978 |
| chr10:47340478-47340823 | 0.757419802 | 0.072524569 | 0.256199978 |
| chr1:217802363-217802626 | 0.795653572 | 0.072534821 | 0.256215327 |
| chr3:105752797-105753160 | -0.886179684 | 0.072535602 | 0.256215327 |
| chr19:50098889-50099508 | 0.643219495 | 0.072540582 | 0.256221106 |
| chr1:45512688-45513021 | -0.781587879 | 0.072553587 | 0.25625523 |
| chr6:73028841-73029248 | -0.736424949 | 0.072565654 | 0.256286034 |
| chr14:99065389-99065868 | -0.718316957 | 0.072578936 | 0.256321131 |
| chr13:85508786-85509322 | -0.679710351 | 0.072588076 | 0.256341594 |
| chr16:4021089-4021548 | -0.677091923 | 0.072608165 | 0.256400719 |

| | | | |
|---|---|---|---|
| chr20:48123534-48123748 | 0.640385916 | 0.072614159 | 0.256410069 |
| chr20:48060708-48061036 | 0.607032081 | 0.072631702 | 0.256460198 |
| chr20:54742807-54743031 | 0.658094941 | 0.072640466 | 0.256479322 |
| chr7:128863498-128863834 | -0.64187313 | 0.072652884 | 0.256511348 |
| chr1:90850971-90851870 | -0.615207226 | 0.072670371 | 0.256561003 |
| chr14:44657787-44657987 | 0.944201992 | 0.072681417 | 0.256561003 |
| chr16:15899666-15900158 | 0.639662618 | 0.072681587 | 0.256561003 |
| chr1:54681334-54681697 | -0.758312386 | 0.072684306 | 0.256561003 |
| chr16:4402498-4402993 | 0.63900689 | 0.072687857 | 0.256561003 |
| chr3:61814313-61814513 | 0.697977236 | 0.072689895 | 0.256561003 |
| chr18:8623810-8624045 | -0.987630279 | 0.07269455 | 0.256561003 |
| chr8:54135703-54135912 | -0.833572313 | 0.072695753 | 0.256561003 |
| chr15:55168236-55168924 | 0.606641925 | 0.072697084 | 0.256561003 |
| chr4:158671382-158672858 | -0.563783623 | 0.072709668 | 0.256593596 |
| chr10:95496948-95497334 | -0.767537844 | 0.072715878 | 0.256603693 |
| chr1:226107501-226107738 | 0.714270469 | 0.072722572 | 0.256615499 |
| chr2:64784496-64785028 | -0.698666162 | 0.072738073 | 0.25665302 |
| chr12:121811594-121812360 | 0.608451987 | 0.072739905 | 0.25665302 |
| chr20:19212029-19212295 | 0.757901152 | 0.072755761 | 0.256697147 |
| chr1:185646211-185646651 | -0.631140104 | 0.072759799 | 0.256699574 |
| chr7:4931529-4932292 | -0.628081478 | 0.072768786 | 0.256719459 |
| chr20:44553371-44553872 | 0.683626121 | 0.072811817 | 0.256859441 |
| chr9:41273624-41274271 | -0.643311233 | 0.072823155 | 0.256872936 |
| chr16:86865847-86866130 | -0.730632882 | 0.072827193 | 0.256872936 |
| chr1:20137122-20137356 | 0.874695603 | 0.072829025 | 0.256872936 |
| chr5:128084385-128084730 | -0.671658484 | 0.072835411 | 0.256872936 |
| chr7:34134250-34134535 | 0.679844646 | 0.07283575 | 0.256872936 |
| chr19:7394883-7395205 | 0.725406746 | 0.072835757 | 0.256872936 |
| chr17:10843637-10844188 | -0.67482292 | 0.072848459 | 0.256905906 |
| chr3:58433394-58434276 | -0.593409361 | 0.07286536 | 0.256950495 |
| chr4:4995152-4995453 | 0.773991356 | 0.07286781 | 0.256950495 |
| chr15:90886065-90886307 | 0.729174543 | 0.072875588 | 0.256966097 |
| chr10:104580826-104581305 | -0.667321884 | 0.07289444 | 0.257020744 |
| chr4:92600684-92600963 | -0.924533896 | 0.072910663 | 0.257054921 |
| chr12:114668143-114668476 | 0.726698365 | 0.072910843 | 0.257054921 |
| chr2:55827493-55827693 | 0.939294639 | 0.07292807 | 0.257085292 |
| chr2:60807587-60807787 | 0.939294639 | 0.07292807 | 0.257085292 |
| chr12:95487048-95487278 | -0.971970168 | 0.072933567 | 0.257085292 |
| chr4:174523013-174523230 | -0.822319194 | 0.072937968 | 0.257085292 |
| chr6:108739483-108739945 | 0.655245417 | 0.072938883 | 0.257085292 |
| chr12:54033789-54034166 | 0.704419537 | 0.072939589 | 0.257085292 |
| chr1:211625080-211625385 | -0.746279974 | 0.072968574 | 0.257175625 |
| chr4:153649518-153649808 | 0.737181267 | 0.072988624 | 0.257226232 |
| chr6:31680658-31681043 | 0.680077462 | 0.072989647 | 0.257226232 |
| chr3:41277865-41278341 | -0.76771023 | 0.072996874 | 0.25723987 |
| chr2:141467483-141467683 | 0.943435891 | 0.073008412 | 0.257268698 |
| chr20:32483444-32484180 | 0.638906831 | 0.073024023 | 0.257311874 |
| chr13:25620000-25620369 | 0.780645689 | 0.073035728 | 0.257341284 |
| chr19:3104063-3104405 | -0.797046407 | 0.073061579 | 0.257420531 |
| chr14:26597689-26598816 | 0.574667489 | 0.073068232 | 0.257432135 |

| | | | |
|---|---|---|---|
| chr10:1432106-1432465 | 0.742439505 | 0.073082965 | 0.257472202 |
| chr3:10320778-10321523 | 0.588665134 | 0.073087838 | 0.257477531 |
| chr10:103794339-103794539 | 0.902057571 | 0.073093647 | 0.257486158 |
| chr12:49636929-49637252 | 0.70804777 | 0.073109818 | 0.257531285 |
| chr11:61446289-61446515 | 0.799273806 | 0.073118737 | 0.257550861 |
| chr2:220277452-220277980 | 0.668010359 | 0.073133254 | 0.257590154 |
| chr5:151033752-151033974 | 0.916420951 | 0.073137656 | 0.257593817 |
| chr3:87771690-87772066 | -0.6798841 | 0.073149413 | 0.257610438 |
| chr1:50329200-50329434 | -0.973198597 | 0.07315151 | 0.257610438 |
| chr4:115657953-115658361 | 0.792514932 | 0.07315543 | 0.257610438 |
| chr7:115864778-115865105 | -0.858589527 | 0.073155824 | 0.257610438 |
| chr1:31715051-31715945 | 0.590597853 | 0.073167312 | 0.257628693 |
| chr10:6151454-6151737 | 0.675352209 | 0.073172921 | 0.257628693 |
| chr8:30600806-30601006 | -1.057614987 | 0.073173545 | 0.257628693 |
| chr7:152859715-152860025 | -0.748834041 | 0.073177102 | 0.257628693 |
| chr5:95404083-95404288 | -0.944069592 | 0.073177819 | 0.257628693 |
| chr17:78971394-78971640 | 0.728022607 | 0.07320389 | 0.257708636 |
| chr1:6245529-6245799 | 0.767010842 | 0.073241925 | 0.25783069 |
| chr6:163181192-163181502 | -0.837329202 | 0.073251113 | 0.257851186 |
| chr7:34025413-34026295 | -0.640585417 | 0.073254782 | 0.257852258 |
| chr1:108311583-108311796 | 0.972353885 | 0.073269469 | 0.257892104 |
| chr3:127006839-127007052 | -0.808037714 | 0.073297291 | 0.257978184 |
| chr6:122082027-122082379 | -0.794968641 | 0.073302553 | 0.257984852 |
| chr1:28113154-28113388 | -0.914500177 | 0.073308409 | 0.257985922 |
| chr19:52028096-52028760 | 0.660134277 | 0.073315551 | 0.257985922 |
| chr2:96679927-96680389 | -0.763661329 | 0.073315729 | 0.257985922 |
| chr7:29684353-29685398 | 0.615269654 | 0.073316569 | 0.257985922 |
| chr3:160163226-160163487 | -0.814465991 | 0.073319692 | 0.257985922 |
| chr12:12779558-12779866 | -0.794691906 | 0.073327702 | 0.258002259 |
| chr1:215129474-215129674 | -0.993394373 | 0.073363714 | 0.258117114 |
| chr7:30594093-30595449 | 0.584555722 | 0.07337621 | 0.258149226 |
| chr4:88697581-88698092 | 0.704722303 | 0.073380449 | 0.258152285 |
| chr2:197158829-197159043 | 0.88232523 | 0.073400531 | 0.258211077 |
| chr20:47506369-47506771 | 0.574052782 | 0.073410725 | 0.258235084 |
| chr10:30831487-30831832 | -0.740345584 | 0.073436143 | 0.258312636 |
| chr8:85171753-85172331 | -0.628348329 | 0.073471698 | 0.258414383 |
| chr4:121941773-121941973 | 0.997230844 | 0.073472789 | 0.258414383 |
| chr8:76367864-76368096 | 0.907528677 | 0.073480936 | 0.258414383 |
| chr20:53883212-53883800 | 0.574800083 | 0.073481437 | 0.258414383 |
| chr22:20430971-20431212 | 0.774948104 | 0.073481932 | 0.258414383 |
| chr22:24803133-24803338 | 0.912723051 | 0.073493618 | 0.258443618 |
| chr15:42395725-42396108 | -0.817024839 | 0.073499278 | 0.258447898 |
| chr12:107014007-107014431 | -0.667646679 | 0.073501581 | 0.258447898 |
| chr2:15860859-15861102 | -0.830830373 | 0.073516213 | 0.258487485 |
| chr9:99413653-99413970 | -0.810749133 | 0.073532397 | 0.258532525 |
| chr2:46296552-46296845 | -0.787605669 | 0.07355826 | 0.258602224 |
| chr5:11267121-11267585 | -0.703368171 | 0.073559764 | 0.258602224 |
| chr12:17641839-17642297 | -0.815495915 | 0.073562346 | 0.258602224 |
| chr5:56415396-56415713 | -0.818381722 | 0.073583938 | 0.258666262 |
| chr1:193755099-193755373 | -0.832471118 | 0.073587854 | 0.258668161 |

| | | | |
|---|---|---|---|
| chr6:151436271-151436894 | -0.620636201 | 0.073598856 | 0.258677455 |
| chr4:25159580-25161121 | 0.571945212 | 0.073599274 | 0.258677455 |
| chr19:38298776-38299318 | 0.664981798 | 0.07360199 | 0.258677455 |
| chr2:169767710-169767979 | 0.809137286 | 0.073604002 | 0.258677455 |
| chr2:37832632-37833024 | -0.706438283 | 0.073621281 | 0.258725643 |
| chr11:706039-706239 | 0.778016063 | 0.073624467 | 0.258725643 |
| chr1:32142810-32143124 | -0.826603523 | 0.07364034 | 0.258769554 |
| chr8:133570268-133570804 | 0.604873334 | 0.073658927 | 0.258823 |
| chr10:116774764-116775252 | -0.65151265 | 0.073697371 | 0.258937618 |
| chr21:42463903-42464407 | -0.665023432 | 0.073698305 | 0.258937618 |
| chrX:99302028-99302320 | -0.816662134 | 0.073718195 | 0.258995624 |
| chr10:115653523-115653895 | -0.828413503 | 0.073728128 | 0.258998053 |
| chrX:154546559-154547978 | 0.570135927 | 0.073728335 | 0.258998053 |
| chr2:155471917-155472291 | -0.710090259 | 0.073729027 | 0.258998053 |
| chr11:76577228-76577496 | 0.774618626 | 0.073737134 | 0.259014657 |
| chr3:88508498-88509225 | 0.640506178 | 0.073757193 | 0.259073238 |
| chr13:113275392-113275607 | 0.840529322 | 0.073766898 | 0.259095452 |
| chr7:5624533-5624758 | 0.859254074 | 0.073773223 | 0.25910579 |
| chr17:27892970-27893614 | -0.621927926 | 0.073781002 | 0.259113967 |
| chr1:35707463-35707734 | 0.755501457 | 0.073784732 | 0.259113967 |
| chr6:43575291-43576471 | 0.579755826 | 0.073785697 | 0.259113967 |
| chr14:29111520-29111912 | 0.673027533 | 0.073799832 | 0.259144381 |
| chr5:63648426-63648678 | -0.877920237 | 0.073803263 | 0.259144381 |
| chr14:105570290-105570555 | 0.862587376 | 0.073804504 | 0.259144381 |
| chr10:59967029-59967434 | -0.636908215 | 0.073809331 | 0.259149455 |
| chr1:172742635-172742979 | -0.689989761 | 0.07381861 | 0.259167885 |
| chr6:139436511-139437343 | -0.609604768 | 0.073825313 | 0.259167885 |
| chr4:85796667-85797395 | -0.635139259 | 0.07382648 | 0.259167885 |
| chr3:1387397-1387779 | 0.705042644 | 0.07382811 | 0.259167885 |
| chr10:49585027-49585533 | -0.774446428 | 0.073833676 | 0.259175551 |
| chr6:75735610-75736219 | -0.670788988 | 0.073845248 | 0.259204297 |
| chr17:59043971-59044469 | 0.592650154 | 0.073862715 | 0.259253731 |
| chr13:72405469-72405782 | -0.894728996 | 0.073883486 | 0.259314757 |
| chr9:93142027-93142227 | 0.859180737 | 0.073902627 | 0.259359255 |
| chr5:38537192-38537418 | 0.94450323 | 0.073903795 | 0.259359255 |
| chr1:207240755-207241290 | -0.704308698 | 0.073906319 | 0.259359255 |
| chr4:155758703-155760131 | -0.57260865 | 0.073913868 | 0.259367674 |
| chr11:101129022-101129277 | -0.812617757 | 0.073915488 | 0.259367674 |
| chr6:127208214-127208474 | -1.002859391 | 0.073957027 | 0.25950155 |
| chr19:44800402-44800758 | 0.674542916 | 0.073960742 | 0.259502699 |
| chr4:170227991-170228249 | -0.879571392 | 0.073973083 | 0.259534115 |
| chr6:19477611-19477918 | 0.781160775 | 0.073983055 | 0.259557217 |
| chr1:151036436-151036941 | 0.626521565 | 0.073987222 | 0.259559181 |
| chr7:101282962-101283365 | -0.69523052 | 0.073991729 | 0.259559181 |
| chr3:174303588-174303788 | -0.957339603 | 0.073993777 | 0.259559181 |
| chr6:152863897-152864201 | -0.768686271 | 0.07400398 | 0.259583088 |
| chr1:247281280-247281490 | -0.932896763 | 0.074019229 | 0.259624689 |
| chr4:82380281-82381082 | -0.587948846 | 0.074024976 | 0.259632962 |
| chr3:31561887-31562185 | -0.767269399 | 0.07402899 | 0.259635158 |
| chr10:81876267-81876520 | 0.844993757 | 0.074040467 | 0.259658583 |

| | | | |
|---|---|---|---|
| chr1:204746817-204747152 | -0.648481354 | 0.074042447 | 0.259658583 |
| chr12:112112518-112112725 | 0.769235908 | 0.074080853 | 0.259781379 |
| chr11:78149717-78149950 | 0.822633916 | 0.074084485 | 0.259782225 |
| chr3:126715442-126715642 | 0.865181955 | 0.07409312 | 0.259800613 |
| chr16:8661037-8661237 | 0.855822347 | 0.074114975 | 0.259860673 |
| chr5:33329079-33329302 | 0.956646572 | 0.074119498 | 0.259860673 |
| chr6:27115055-27115255 | 1.014195448 | 0.074123685 | 0.259860673 |
| chr4:168481019-168481266 | -0.749159953 | 0.074123814 | 0.259860673 |
| chr1:102837549-102837883 | -0.722896342 | 0.074128047 | 0.259860949 |
| chr19:12550993-12551982 | 0.59377111 | 0.074130676 | 0.259860949 |
| chr4:1580209-1580877 | 0.599938725 | 0.074159512 | 0.25993845 |
| chr5:79133485-79134195 | -0.648746062 | 0.074159569 | 0.25993845 |
| chr7:38349828-38350241 | -0.660317226 | 0.07416842 | 0.259957582 |
| chr7:101882030-101882365 | -0.743542224 | 0.074174472 | 0.259966901 |
| chr6:27505472-27506069 | -0.593564015 | 0.074198308 | 0.260035817 |
| chr4:76724722-76725332 | -0.749871919 | 0.074205743 | 0.260035817 |
| chr3:64949613-64949821 | 0.631668349 | 0.07420885 | 0.260035817 |
| chr8:109690239-109690528 | -0.71679102 | 0.074210352 | 0.260035817 |
| chr3:37213095-37213385 | 0.71296573 | 0.074211999 | 0.260035817 |
| chr20:45971634-45972714 | 0.564902838 | 0.074215151 | 0.260035817 |
| chr21:37176736-37176978 | 0.833838099 | 0.074217891 | 0.260035817 |
| chr17:64801348-64801573 | 0.814044438 | 0.074227454 | 0.260057429 |
| chr19:18173637-18173857 | 0.809765949 | 0.074231328 | 0.260059112 |
| chr1:197363856-197364091 | -0.932202288 | 0.074237106 | 0.260067463 |
| chr13:114226822-114227329 | -0.704209408 | 0.074241557 | 0.260071165 |
| chr19:55186095-55186358 | 0.880311036 | 0.074247008 | 0.260078368 |
| chrX:72686194-72686489 | 0.798095311 | 0.074269305 | 0.260144581 |
| chr7:87318846-87319615 | -0.624404167 | 0.074297392 | 0.2602242 |
| chr1:23772163-23772363 | 0.975323294 | 0.074298828 | 0.2602242 |
| chr7:133235365-133235720 | -0.823361606 | 0.074317643 | 0.2602782 |
| chr9:77688176-77688472 | -0.752247306 | 0.074323648 | 0.260287332 |
| chr20:50678073-50678660 | 0.569667462 | 0.074330133 | 0.260293426 |
| chr7:4766003-4766373 | -0.735763549 | 0.074332182 | 0.260293426 |
| chr11:107598685-107598885 | -0.982034047 | 0.074345386 | 0.260327766 |
| chr3:11185645-11186255 | 0.62346504 | 0.074358536 | 0.260361914 |
| chr20:54642892-54644216 | 0.561194746 | 0.074365648 | 0.260374916 |
| chr8:57597737-57598001 | 0.803341775 | 0.074375993 | 0.26039924 |
| chr12:45628763-45629059 | -0.928557625 | 0.07439896 | 0.260467745 |
| chr11:37088291-37088986 | -0.679190053 | 0.074406834 | 0.260483413 |
| chr14:34723006-34723278 | 0.758900099 | 0.074420763 | 0.260508438 |
| chr10:34474056-34474439 | -0.758896509 | 0.074420783 | 0.260508438 |
| chr12:91178108-91178487 | -0.778996467 | 0.074437469 | 0.26054824 |
| chr11:63024105-63024390 | 0.743955982 | 0.074438954 | 0.26054824 |
| chr6:27837167-27837571 | 0.665711408 | 0.074454126 | 0.260589442 |
| chr6:52697863-52698234 | -0.777836309 | 0.074477687 | 0.260659997 |
| chr20:53839518-53839921 | 0.5792504 | 0.074483582 | 0.260668722 |
| chr5:36553198-36553398 | -1.03499649 | 0.074487675 | 0.260671142 |
| chr12:104957998-104959342 | -0.573070542 | 0.074497862 | 0.260687436 |
| chr17:58121950-58122204 | 0.767066702 | 0.074499136 | 0.260687436 |
| chr20:31719323-31719670 | 0.632837079 | 0.074510129 | 0.260713999 |

| | | | |
|---|---|---|---|
| chr1:33367801-33368383 | 0.6504943 | 0.074514164 | 0.260716209 |
| chr7:104595905-104596157 | -0.9786495 | 0.074550876 | 0.260832751 |
| chr3:164310807-164311247 | -0.827718189 | 0.074586186 | 0.260944374 |
| chr17:49408005-49408499 | -0.638072757 | 0.074595847 | 0.260966257 |
| chr17:60045006-60045407 | 0.599771202 | 0.074641061 | 0.261112514 |
| chr5:77086555-77087696 | -0.579120695 | 0.074656025 | 0.261152938 |
| chr14:77068226-77068544 | 0.671458107 | 0.074669629 | 0.261188601 |
| chr6:37657556-37657789 | 0.800532354 | 0.074680411 | 0.261214393 |
| chr20:11892276-11892531 | -0.732357168 | 0.074688029 | 0.261229111 |
| chr14:79544061-79544573 | -0.66126916 | 0.074695527 | 0.261233984 |
| chr11:67847977-67848879 | -0.647699493 | 0.074696241 | 0.261233984 |
| chr13:31945258-31945958 | -0.654093252 | 0.074703029 | 0.2612458 |
| chr15:87605030-87605230 | 0.907388226 | 0.074711017 | 0.261261813 |
| chr2:125696205-125697102 | 0.582483667 | 0.074721032 | 0.261284907 |
| chr19:43455266-43455996 | 0.629402907 | 0.074739807 | 0.261338635 |
| chr12:6093183-6093428 | -0.944009668 | 0.0747677 | 0.261424235 |
| chr6:154615505-154616042 | -0.619084497 | 0.074775951 | 0.261441154 |
| chr5:52915159-52915804 | -0.611050639 | 0.074787429 | 0.261469356 |
| chr2:37588326-37588526 | -0.939062387 | 0.0748006 | 0.261503471 |
| chr13:109576268-109576862 | 0.615158155 | 0.07485119 | 0.261668394 |
| chr18:52339713-52340392 | -0.658992536 | 0.074867479 | 0.2617134 |
| chr18:67104678-67105024 | -0.70274319 | 0.074883127 | 0.261756155 |
| chr7:158123778-158124122 | -0.813481395 | 0.074905325 | 0.261815255 |
| chr8:93635922-93636167 | 0.746522242 | 0.074909979 | 0.261815255 |
| chr10:129045156-129045499 | 0.913970742 | 0.074910285 | 0.261815255 |
| chr5:154811954-154812212 | -0.734796807 | 0.074916529 | 0.261825135 |
| chr6:138035026-138035346 | -0.840365618 | 0.074925657 | 0.261845096 |
| chr7:100803254-100803830 | -0.662783866 | 0.074933053 | 0.261848058 |
| chr5:150637674-150637882 | 0.811771627 | 0.074934675 | 0.261848058 |
| chr14:105581622-105581859 | 0.877631958 | 0.074940132 | 0.261848058 |
| chr19:54266124-54266393 | 0.818133297 | 0.074940175 | 0.261848058 |
| chr19:17337303-17337663 | 0.647954349 | 0.074948693 | 0.261856977 |
| chr11:129593146-129593519 | -0.788861684 | 0.074949562 | 0.261856977 |
| chr3:67653952-67655105 | -0.581955746 | 0.074953305 | 0.261858114 |
| chr20:53966753-53967025 | 0.605373494 | 0.074960449 | 0.261871132 |
| chr5:150497634-150497926 | -0.723631271 | 0.07497255 | 0.261901465 |
| chr16:1474885-1475431 | 0.632877119 | 0.07497796 | 0.261908421 |
| chr17:60766631-60767128 | 0.5773175 | 0.074985191 | 0.261910894 |
| chr19:1104386-1104782 | 0.677693338 | 0.074985504 | 0.261910894 |
| chr1:82775448-82775784 | -0.684148778 | 0.075002113 | 0.261956964 |
| chr11:126990325-126990680 | -0.739008133 | 0.075007145 | 0.261962598 |
| chr1:209534549-209534808 | -0.883593744 | 0.075017764 | 0.261987743 |
| chr11:34357830-34358308 | 0.60762747 | 0.075028918 | 0.262008692 |
| chr10:91088834-91089119 | -0.819503941 | 0.075030601 | 0.262008692 |
| chr3:37779620-37779820 | -1.033071978 | 0.075037352 | 0.262015874 |
| chr6:150514965-150515496 | -0.656991596 | 0.075039497 | 0.262015874 |
| chr1:210730057-210730341 | -0.892296297 | 0.075049374 | 0.262037095 |
| chr1:182198076-182198366 | -0.811097616 | 0.075052414 | 0.262037095 |
| chr22:23182605-23182862 | 0.761184997 | 0.075070906 | 0.262078676 |
| chr10:59880361-59880640 | -0.84776433 | 0.075071165 | 0.262078676 |

| | | | |
|---|---|---|---|
| chr2:150207958-150208338 | -0.774716386 | 0.075090234 | 0.262133304 |
| chr4:138836224-138836522 | -0.955956036 | 0.075108397 | 0.262175813 |
| chr10:102393922-102394891 | 0.584423083 | 0.075109254 | 0.262175813 |
| chr5:155004336-155004670 | -0.79464845 | 0.075116168 | 0.262188002 |
| chr5:88675717-88676424 | -0.585093687 | 0.075123723 | 0.262190512 |
| chr17:30831812-30832072 | 0.80110003 | 0.075123731 | 0.262190512 |
| chr2:138336098-138336440 | 0.818112201 | 0.075137841 | 0.26222642 |
| chr11:102374995-102375342 | -0.843797661 | 0.075140864 | 0.26222642 |
| chr14:27686067-27686390 | 0.750537831 | 0.075148655 | 0.262241663 |
| chr9:81588193-81588526 | 0.733029928 | 0.075168513 | 0.262299016 |
| chr17:58962799-58963062 | 0.649730124 | 0.075175124 | 0.262310138 |
| chr7:123637254-123637534 | -0.873541768 | 0.07519432 | 0.262365171 |
| chr17:58615440-58615686 | 0.641549033 | 0.075218534 | 0.262437709 |
| chr1:58841529-58841850 | -0.784436103 | 0.075226745 | 0.262454405 |
| chr8:115647988-115648631 | -0.633388358 | 0.075234394 | 0.262469141 |
| chr5:118967146-118967426 | -0.772176179 | 0.075242025 | 0.262483811 |
| chr4:165459891-165460277 | 0.642079103 | 0.075254081 | 0.262513918 |
| chr7:150876741-150877874 | -0.608132856 | 0.075279046 | 0.262581642 |
| chr20:60550525-60550872 | -0.789623363 | 0.075282261 | 0.262581642 |
| chr1:218156467-218157046 | -0.603714616 | 0.075285833 | 0.262581642 |
| chr1:20152335-20152623 | 0.771851539 | 0.075287203 | 0.262581642 |
| chr1:99457634-99458056 | -0.779038342 | 0.07529751 | 0.262605635 |
| chr5:5600723-5600964 | 0.93346333 | 0.07531132 | 0.262634651 |
| chr7:16891393-16891697 | 0.663803686 | 0.075312685 | 0.262634651 |
| chr19:10334831-10335180 | 0.64188046 | 0.075324454 | 0.262663738 |
| chr19:2041822-2042416 | 0.643999399 | 0.075330233 | 0.262671937 |
| chr8:122937046-122937540 | 0.636446119 | 0.075352386 | 0.262737225 |
| chr3:14144348-14145043 | -0.587796649 | 0.075361063 | 0.262755145 |
| chr11:67791605-67792103 | -0.706087717 | 0.075364384 | 0.262755145 |
| chr6:111615209-111615553 | -0.688668344 | 0.075376163 | 0.262780467 |
| chr7:99376018-99376302 | -0.7406352 | 0.075378506 | 0.262780467 |
| chr21:8234108-8234916 | 0.56516515 | 0.075382568 | 0.262782672 |
| chr22:25475531-25475869 | 0.833323158 | 0.075397546 | 0.262813244 |
| chrX:107716807-107717247 | 0.627274075 | 0.075398198 | 0.262813244 |
| chr2:86072712-86073011 | 0.802311746 | 0.075427745 | 0.262904275 |
| chr9:96619479-96619679 | 0.876445525 | 0.075466329 | 0.263026795 |
| chr11:82775729-82776537 | 0.614639302 | 0.075472768 | 0.26303436 |
| chr6:44313216-44313537 | 0.68170795 | 0.075475365 | 0.26303436 |
| chr13:22978926-22979665 | 0.586654708 | 0.075490818 | 0.263076249 |
| chr13:45821250-45821490 | 0.85756009 | 0.075511375 | 0.263135918 |
| chr4:47485047-47485717 | -0.592336722 | 0.075552756 | 0.263268145 |
| chr1:110548370-110548695 | -0.756756485 | 0.075562229 | 0.263289184 |
| chr17:68506292-68506492 | -0.875079954 | 0.075573697 | 0.26331373 |
| chr12:15227101-15227387 | -0.920437411 | 0.075576147 | 0.26331373 |
| chr12:130863258-130863648 | 0.721944838 | 0.075589386 | 0.263347881 |
| chr8:91940356-91940556 | 1.02165599 | 0.075602597 | 0.263362976 |
| chr16:12271111-12271448 | -0.741850209 | 0.075602707 | 0.263362976 |
| chr9:134367885-134368087 | 0.736979058 | 0.075609154 | 0.263362976 |
| chrX:30904301-30904548 | -0.915352242 | 0.075612273 | 0.263362976 |
| chr8:72388593-72389065 | -0.737444235 | 0.075612495 | 0.263362976 |

| | | | |
|---|---|---|---|
| chr1:182909714-182909914 | 0.863120964 | 0.075614342 | 0.263362976 |
| chr13:33119327-33119551 | 0.782319962 | 0.075618313 | 0.263364834 |
| chr1:195981739-195982030 | 0.83508615 | 0.075629032 | 0.263390195 |
| chr21:21063915-21064166 | -0.885992643 | 0.075635789 | 0.263396672 |
| chr3:152471770-152472315 | -0.620816615 | 0.075637767 | 0.263396672 |
| chr8:11846779-11846979 | -0.929319418 | 0.075649676 | 0.263426172 |
| chr5:91380021-91381250 | -0.56009803 | 0.075663319 | 0.263461704 |
| chr2:121489323-121489634 | 0.768240478 | 0.075670748 | 0.263463093 |
| chrX:103157553-103157894 | 0.775500917 | 0.075675297 | 0.263463093 |
| chr7:158572210-158572718 | -0.653551811 | 0.075676956 | 0.263463093 |
| chr3:64831696-64831918 | 0.634589884 | 0.075677471 | 0.263463093 |
| chr5:38650509-38650753 | -0.855591474 | 0.075683104 | 0.263470732 |
| chr6:47135767-47136110 | 0.78001101 | 0.075702393 | 0.263514695 |
| chr11:86422143-86422671 | -0.696233325 | 0.075702611 | 0.263514695 |
| chr8:57205699-57206284 | 0.618802135 | 0.075706246 | 0.263515375 |
| chr18:56571083-56571283 | -1.026645909 | 0.075717637 | 0.263543055 |
| chr4:85210300-85210762 | -0.708926099 | 0.075741354 | 0.263588132 |
| chr5:36180522-36180802 | 0.811513971 | 0.07574382 | 0.263588132 |
| chr5:127197524-127198087 | -0.703922057 | 0.075746107 | 0.263588132 |
| chr1:229844496-229844929 | 0.690710178 | 0.075748321 | 0.263588132 |
| chr6:145886214-145886620 | 0.694504621 | 0.075748481 | 0.263588132 |
| chr20:7740754-7741061 | 0.876062517 | 0.075751229 | 0.263588132 |
| chr1:108533334-108534175 | -0.596144926 | 0.075762034 | 0.263613758 |
| chr11:68853932-68854132 | 0.878352206 | 0.075799044 | 0.263729385 |
| chr1:106671926-106672509 | 0.592723537 | 0.075806797 | 0.263729385 |
| chr4:84531706-84532122 | -0.769408032 | 0.075807179 | 0.263729385 |
| chr3:140855022-140855275 | 0.813846425 | 0.075809033 | 0.263729385 |
| chr9:133992727-133993266 | 0.611584297 | 0.07581997 | 0.263755459 |
| chr2:119744430-119744683 | -0.802897024 | 0.075831798 | 0.26378463 |
| chr1:78333510-78333947 | -0.785824936 | 0.075838325 | 0.263795357 |
| chr1:40592229-40592499 | 0.753884914 | 0.075860422 | 0.263850425 |
| chr8:109378471-109378671 | 0.845064431 | 0.075861851 | 0.263850425 |
| chr10:69449779-69450237 | -0.628341905 | 0.075864487 | 0.263850425 |
| chr11:30539217-30539449 | -0.906388851 | 0.07587628 | 0.263879462 |
| chr10:85317787-85318088 | 0.847921692 | 0.075894865 | 0.263931697 |
| chr11:827753-828004 | 0.809342812 | 0.075898189 | 0.263931697 |
| chr22:43955270-43955753 | 0.695916712 | 0.075919833 | 0.263982874 |
| chr1:2135469-2135688 | -0.87711052 | 0.075922829 | 0.263982874 |
| chrX:84173654-84173924 | -0.911656482 | 0.075923241 | 0.263982874 |
| chr20:51372271-51372678 | 0.614012692 | 0.075941571 | 0.264021246 |
| chr10:123573183-123573559 | 0.647581864 | 0.075942588 | 0.264021246 |
| chr2:96508101-96508351 | 0.727805308 | 0.075944615 | 0.264021246 |
| chr8:35899318-35899651 | 0.748738687 | 0.075951535 | 0.264033324 |
| chr8:124129530-124129730 | 0.791573633 | 0.075957776 | 0.264040537 |
| chr22:30755524-30755742 | 0.837141964 | 0.075960502 | 0.264040537 |
| chr2:183295070-183295540 | -0.665331366 | 0.075981502 | 0.264101554 |
| chr11:9076669-9076992 | -0.739342053 | 0.076000941 | 0.264143442 |
| chrX:40148051-40148390 | 0.789683673 | 0.076001078 | 0.264143442 |
| chr6:132669668-132670009 | -0.702259374 | 0.076004124 | 0.264143442 |
| chr8:20789730-20789978 | -0.911864788 | 0.076010087 | 0.264143442 |

| | | | |
|---|---|---|---|
| chr1:114403158-114403469 | 0.647685612 | 0.07601079 | 0.264143442 |
| chr7:17617917-17618156 | 0.767249705 | 0.076019291 | 0.264144222 |
| chr10:59965118-59965878 | -0.598363707 | 0.076020823 | 0.264144222 |
| chr3:17982531-17982827 | 0.750784504 | 0.076021357 | 0.264144222 |
| chr3:82463093-82463581 | -0.690434819 | 0.076028775 | 0.264158018 |
| chr8:130538178-130538392 | 0.888383008 | 0.076037745 | 0.264177205 |
| chrX:119852574-119853395 | 0.574863509 | 0.076051613 | 0.264213406 |
| chr8:119638790-119639095 | 0.734862371 | 0.076060294 | 0.264231585 |
| chr21:42363367-42363968 | -0.609410137 | 0.076076316 | 0.264273237 |
| chr11:22875954-22876231 | 0.739889467 | 0.076079182 | 0.264273237 |
| chr2:109644299-109645282 | 0.607293289 | 0.07609662 | 0.26431819 |
| chr5:176352248-176352743 | -0.712834239 | 0.076099022 | 0.26431819 |
| chr13:44436686-44437382 | -0.574821187 | 0.076106859 | 0.264333429 |
| chr3:74613510-74613795 | 0.793375959 | 0.076111816 | 0.264334463 |
| chr11:48049952-48050378 | 0.624607518 | 0.076114057 | 0.264334463 |
| chr1:216778405-216778738 | -0.779474911 | 0.076125655 | 0.264359159 |
| chr7:30905975-30906393 | -0.738844034 | 0.076128068 | 0.264359159 |
| chr20:58652003-58652203 | 0.696741663 | 0.076143657 | 0.264401308 |
| chr12:54033189-54033656 | 0.629488654 | 0.076162447 | 0.264454569 |
| chr13:33723824-33724169 | -0.666142088 | 0.076168936 | 0.264465117 |
| chr16:2033501-2033750 | -0.825236166 | 0.076178775 | 0.264487293 |
| chr11:47980210-47980432 | 0.76944103 | 0.076190089 | 0.264514589 |
| chr8:39107309-39107989 | -0.650707683 | 0.076199077 | 0.264533808 |
| chr2:127208379-127208833 | -0.692803966 | 0.076219802 | 0.264593768 |
| chr9:128635190-128636309 | -0.573627932 | 0.076232055 | 0.264624315 |
| chr7:90455278-90455706 | -0.727816354 | 0.076250292 | 0.264675632 |
| chr5:36178023-36178299 | 0.757790726 | 0.076269287 | 0.264729571 |
| chr17:54103060-54103306 | 0.905064802 | 0.076282355 | 0.264752963 |
| chr1:172177671-172178231 | 0.621113684 | 0.076282937 | 0.264752963 |
| chr1:77240277-77240537 | -0.906658465 | 0.07629025 | 0.264766351 |
| chr13:21721183-21721441 | 0.840785507 | 0.076304044 | 0.264802231 |
| chr7:51488217-51488471 | -1.036948333 | 0.076318709 | 0.264841129 |
| chr1:184597803-184598003 | -0.980752003 | 0.076332009 | 0.264875287 |
| chr8:615493-615769 | 0.816488257 | 0.076394681 | 0.265072775 |
| chr1:156609092-156609368 | 0.708157192 | 0.076404138 | 0.265072775 |
| chrX:123960065-123960915 | 0.575685855 | 0.076405397 | 0.265072775 |
| chr6:106885746-106886006 | 0.807837476 | 0.076406457 | 0.265072775 |
| chr4:149914582-149914831 | -0.929971689 | 0.076408137 | 0.265072775 |
| chr5:84435739-84436329 | -0.623728819 | 0.076409678 | 0.265072775 |
| chr7:21261795-21262037 | 0.768772229 | 0.076465962 | 0.265256018 |
| chr12:34022006-34023006 | -0.603142056 | 0.076479534 | 0.265284721 |
| chr6:8707986-8708443 | -0.75263703 | 0.076481161 | 0.265284721 |
| chr9:133238287-133239453 | -0.582626427 | 0.076516374 | 0.26539485 |
| chr1:170765766-170766014 | -0.824275934 | 0.076520926 | 0.265398625 |
| chr1:117570930-117571511 | 0.5976549 | 0.076539755 | 0.265446675 |
| chr9:36205192-36205498 | -0.783417798 | 0.076541743 | 0.265446675 |
| chr14:25842208-25843005 | -0.618282773 | 0.076545173 | 0.265446675 |
| chr18:76983206-76983614 | -0.653317533 | 0.076552616 | 0.265448973 |
| chr5:143569512-143569731 | 0.880171666 | 0.076555501 | 0.265448973 |
| chr5:68456710-68457063 | -0.700228764 | 0.076556229 | 0.265448973 |

| | | | |
|---|---|---|---|
| chr1:201584909-201585125 | -0.824354005 | 0.076565678 | 0.265450969 |
| chr7:19078932-19079303 | 0.716217243 | 0.076566656 | 0.265450969 |
| chr4:88698160-88698360 | 0.937999052 | 0.076567198 | 0.265450969 |
| chr18:62451001-62451386 | -0.740995974 | 0.076578855 | 0.265479369 |
| chr6:19246790-19247587 | -0.660637283 | 0.076584842 | 0.265488112 |
| chr15:38696268-38696561 | -0.856807487 | 0.07659504 | 0.265511452 |
| chr6:142377252-142377533 | -0.755196379 | 0.076606086 | 0.265537731 |
| chr4:184264205-184264471 | -0.838031531 | 0.076611625 | 0.265544917 |
| chr4:45371638-45372040 | -0.869273078 | 0.076638506 | 0.265622812 |
| chr9:26880316-26880563 | -0.897332666 | 0.076641032 | 0.265622812 |
| chr2:47225534-47225927 | -0.791151342 | 0.076660586 | 0.265678565 |
| chr7:98021341-98022241 | -0.651812894 | 0.076665757 | 0.26568447 |
| chr4:143358893-143359282 | 0.651917631 | 0.076686789 | 0.265745336 |
| chr5:22649155-22649523 | -0.718810512 | 0.076697565 | 0.265770659 |
| chr4:30256141-30256516 | -0.767115466 | 0.076706051 | 0.265784062 |
| chr11:16632198-16632704 | -0.735329733 | 0.076710981 | 0.265784062 |
| chr19:36572920-36574152 | 0.594711168 | 0.076711839 | 0.265784062 |
| chr11:65720244-65720944 | 0.61123014 | 0.076716523 | 0.265788271 |
| chr3:25857032-25857247 | 0.825067066 | 0.076738285 | 0.265851648 |
| chr1:84783157-84783387 | 0.837238891 | 0.076742838 | 0.265855401 |
| chr3:134039610-134039831 | 0.829460303 | 0.076761921 | 0.265909487 |
| chr19:2426978-2427217 | 0.856269732 | 0.076794638 | 0.266010639 |
| chr3:78022301-78022501 | 0.869379126 | 0.076798065 | 0.266010639 |
| chr2:38146975-38147351 | 0.769329138 | 0.076811107 | 0.266043787 |
| chr10:60140614-60141216 | -0.609165597 | 0.076834326 | 0.266112177 |
| chr1:45570101-45570379 | 0.71047152 | 0.07686459 | 0.266204965 |
| chr1:101238697-101239116 | -0.671389601 | 0.07687119 | 0.266215788 |
| chr1:149927126-149927364 | 0.697067437 | 0.07688031 | 0.266235338 |
| chr6:30751580-30751909 | -0.719142487 | 0.076893616 | 0.266269382 |
| chr20:57779398-57779812 | 0.604848673 | 0.07691521 | 0.266332124 |
| chr14:76050951-76051151 | 0.889187514 | 0.076931579 | 0.266376765 |
| chr3:42901-43163 | -0.886717922 | 0.076961443 | 0.266468129 |
| chr4:98166313-98166974 | -0.655134047 | 0.076971944 | 0.266483942 |
| chr17:44350824-44351024 | 0.771865522 | 0.076972966 | 0.266483942 |
| chr11:78259764-78260467 | -0.617492958 | 0.076994515 | 0.266546503 |
| chr2:64488025-64488300 | -0.726763382 | 0.077009443 | 0.266586136 |
| chr4:116001319-116001579 | 0.828256046 | 0.077021746 | 0.266604913 |
| chr11:65180959-65181488 | 0.653206041 | 0.077021826 | 0.266604913 |
| chr8:144580496-144580963 | 0.618239628 | 0.077031931 | 0.26662637 |
| chr12:50166646-50167838 | 0.567089986 | 0.077034984 | 0.26662637 |
| chr9:20683894-20684916 | -0.568318326 | 0.07705832 | 0.266695091 |
| chr10:116809459-116809928 | -0.70797961 | 0.07706507 | 0.266706403 |
| chr2:213083175-213083471 | -0.850615421 | 0.07706876 | 0.266707128 |
| chr13:36628959-36629384 | 0.716820367 | 0.077080314 | 0.266735065 |
| chr5:55159748-55160294 | -0.666532921 | 0.077087217 | 0.266746907 |
| chr22:35589553-35590060 | -0.686854886 | 0.077093315 | 0.266755962 |
| chr20:1466548-1467192 | 0.634570996 | 0.077097507 | 0.266758419 |
| chr9:110583159-110583403 | -0.881059575 | 0.077108281 | 0.26678365 |
| chr21:42411418-42411782 | -0.688728335 | 0.07712215 | 0.266819587 |
| chr13:109743787-109744156 | -0.748167412 | 0.077128691 | 0.266830071 |

| | | | |
|---|---|---|---|
| chr8:133234735-133235055 | -0.717832827 | 0.077132145 | 0.266830071 |
| chr1:246200200-246200742 | -0.674446536 | 0.077144868 | 0.266862037 |
| chr7:15291677-15291991 | -0.872717557 | 0.077172722 | 0.266946336 |
| chr22:42962555-42962829 | -0.930354282 | 0.077189567 | 0.26699255 |
| chr16:65480104-65480430 | 0.71223406 | 0.077194131 | 0.266996286 |
| chr3:19348959-19349451 | 0.680302651 | 0.077213237 | 0.267050313 |
| chr3:131306674-131307010 | -0.884034313 | 0.077221391 | 0.267066459 |
| chr1:59290026-59290702 | -0.63791928 | 0.077239665 | 0.267117604 |
| chr7:18590825-18591071 | 0.780903841 | 0.077254622 | 0.267157271 |
| chr4:151714238-151714600 | -0.808564111 | 0.077274202 | 0.26721282 |
| chr19:1028058-1028394 | 0.686388285 | 0.077277848 | 0.26721282 |
| chr20:57175749-57175988 | 0.589541612 | 0.077281148 | 0.26721282 |
| chr17:80278598-80278818 | 0.893202369 | 0.077300656 | 0.267268214 |
| chr5:90983746-90984053 | -0.833771298 | 0.077313924 | 0.267302025 |
| chr11:16613430-16614253 | -0.595784998 | 0.077322532 | 0.267319725 |
| chr1:107830659-107830859 | 0.884425844 | 0.077339555 | 0.26734093 |
| chr10:84179122-84179732 | -0.640291083 | 0.07734673 | 0.26734093 |
| chr12:88179538-88179738 | -0.825394494 | 0.077347494 | 0.26734093 |
| chr5:173329438-173330220 | 0.579370275 | 0.077350447 | 0.26734093 |
| chr8:27837739-27838100 | -0.771603758 | 0.077350745 | 0.26734093 |
| chr16:70771119-70771419 | 0.728297695 | 0.077353737 | 0.26734093 |
| chr14:58677582-58677799 | 0.914058368 | 0.077358188 | 0.26734093 |
| chr7:17009095-17009295 | 0.858285065 | 0.077360068 | 0.26734093 |
| chr8:48755799-48756027 | 0.858285065 | 0.077360068 | 0.26734093 |
| chr18:5155228-5155533 | -0.811379648 | 0.077378567 | 0.267384026 |
| chr15:90479285-90480099 | 0.591730541 | 0.077383955 | 0.267384026 |
| chr5:66646595-66646985 | -0.679177408 | 0.077384163 | 0.267384026 |
| chr4:27075987-27076277 | -0.861517308 | 0.077390521 | 0.267384026 |
| chrX:129961076-129961649 | 0.608093237 | 0.077393677 | 0.267384026 |
| chr20:53671411-53671747 | 0.578147818 | 0.077394892 | 0.267384026 |
| chr4:105140987-105141192 | 0.860698751 | 0.077397082 | 0.267384026 |
| chr10:77125310-77125623 | -0.812963737 | 0.077400455 | 0.267384026 |
| chr2:158357728-158358106 | 0.690530773 | 0.077407934 | 0.267397807 |
| chr8:6841114-6841572 | -0.728862815 | 0.07742116 | 0.267431436 |
| chr12:103833975-103834194 | -0.916484294 | 0.077439776 | 0.26748368 |
| chr16:81737810-81738348 | -0.621232137 | 0.077456915 | 0.267529013 |
| chr22:37634565-37634917 | -0.69254286 | 0.077459883 | 0.267529013 |
| chr20:9567031-9567235 | -0.93055369 | 0.077463391 | 0.267529068 |
| chr2:64859165-64860197 | -0.585714575 | 0.077478463 | 0.267569064 |
| chr5:53803005-53803434 | -0.689851056 | 0.077485094 | 0.267579903 |
| chr5:69083442-69083788 | -0.656879122 | 0.077494881 | 0.26760164 |
| chr2:212945615-212945873 | -0.859950341 | 0.077500065 | 0.26760748 |
| chr12:52004156-52004366 | 0.789356812 | 0.077513322 | 0.267632233 |
| chr15:78811469-78811693 | 0.77510446 | 0.077514219 | 0.267632233 |
| chr9:33617610-33617876 | -0.855086507 | 0.077518795 | 0.267635971 |
| chr10:47552788-47553120 | -0.720935786 | 0.077531217 | 0.267666798 |
| chr6:31736317-31736620 | 0.687740265 | 0.07754548 | 0.267703978 |
| chr6:48928906-48929107 | -0.987607091 | 0.077555715 | 0.267716507 |
| chr5:109468795-109469160 | -0.66001211 | 0.077556098 | 0.267716507 |
| chr1:67524213-67524441 | 0.860228865 | 0.077561689 | 0.267723748 |

| | | | |
|---|---|---|---|
| chr2:68908488-68908688 | 0.907220343 | 0.077571735 | 0.267746362 |
| chr8:69356647-69357497 | -0.62488281 | 0.077577299 | 0.267753504 |
| chr1:110258801-110259021 | -0.889711065 | 0.077596967 | 0.267809324 |
| chr7:44146091-44146443 | 0.678391647 | 0.077603978 | 0.267821456 |
| chr22:36202998-36203217 | 0.873235472 | 0.077608145 | 0.267823774 |
| chr16:48542177-48542391 | 0.88368356 | 0.077612821 | 0.26782785 |
| chr15:85729785-85730169 | 0.731705046 | 0.077621181 | 0.267844634 |
| chr2:70525254-70525474 | 0.814923121 | 0.077636673 | 0.267886027 |
| chr16:19113553-19114141 | 0.613514444 | 0.077676398 | 0.267994794 |
| chr2:96858752-96859043 | -0.663901804 | 0.077679419 | 0.267994794 |
| chr14:35040564-35040870 | 0.739743316 | 0.077679968 | 0.267994794 |
| chr12:62873165-62874051 | 0.606126007 | 0.077682185 | 0.267994794 |
| chr18:74291130-74292365 | -0.5627436 | 0.077692366 | 0.268017851 |
| chr11:117178611-117179004 | 0.64456899 | 0.077697793 | 0.268024504 |
| chr14:32201590-32201943 | 0.616016943 | 0.077708129 | 0.268048093 |
| chr14:37778490-37778697 | 0.876954609 | 0.077722195 | 0.268084542 |
| chr2:23292083-23292364 | -0.887311694 | 0.077729803 | 0.268093186 |
| chr4:3954684-3954943 | 0.691656115 | 0.077731699 | 0.268093186 |
| chr12:21774440-21774717 | -0.899131453 | 0.077746296 | 0.268131462 |
| chr5:60883348-60883548 | -1.017784402 | 0.077758834 | 0.268155682 |
| chr16:58756395-58756672 | 0.761107371 | 0.077760318 | 0.268155682 |
| chr1:223756053-223756302 | 0.769756756 | 0.077775178 | 0.268188436 |
| chr4:53328671-53329352 | -0.666383227 | 0.077777544 | 0.268188436 |
| chr9:4666111-4666597 | -0.681275616 | 0.077780317 | 0.268188436 |
| chr13:78876171-78876519 | -0.730455186 | 0.077785421 | 0.268193967 |
| chr20:44966142-44967054 | 0.573538957 | 0.077798385 | 0.268226595 |
| chr14:22598889-22599185 | 0.657244883 | 0.077829962 | 0.26832339 |
| chr20:44959758-44960803 | 0.578066362 | 0.077837074 | 0.268327331 |
| chr10:21329962-21330391 | -0.673121413 | 0.077838109 | 0.268327331 |
| chr10:113799647-113799924 | 0.819149191 | 0.077842366 | 0.268329935 |
| chr4:174873985-174874258 | -0.801517415 | 0.077851934 | 0.268350841 |
| chr14:86654686-86654908 | 0.879437876 | 0.077858304 | 0.268360729 |
| chr3:38118020-38118422 | 0.640719247 | 0.077871371 | 0.268393692 |
| chr3:140745327-140745527 | 0.933894548 | 0.077885587 | 0.268430617 |
| chrX:55488497-55490060 | 0.584772751 | 0.077897799 | 0.268460629 |
| chr2:182980109-182980321 | -0.92919243 | 0.07790446 | 0.26847151 |
| chr5:122419187-122419585 | 0.70801058 | 0.0779241 | 0.268527113 |
| chr1:11059935-11060454 | 0.634449879 | 0.077932313 | 0.268541514 |
| chr5:172437457-172437726 | 0.728940928 | 0.077935288 | 0.268541514 |
| chr10:86400840-86401507 | 0.600973413 | 0.077947716 | 0.268572261 |
| chr15:34224632-34225400 | 0.581804364 | 0.077973746 | 0.268642319 |
| chr10:70232941-70233976 | -0.570777556 | 0.077975062 | 0.268642319 |
| chr7:15326163-15326364 | 0.893658183 | 0.077988466 | 0.268676419 |
| chr19:11227518-11227818 | -0.81497478 | 0.077997243 | 0.268684757 |
| chr9:78295791-78295991 | -0.905266636 | 0.078000005 | 0.268684757 |
| chr5:163408272-163408843 | -0.644910206 | 0.078004323 | 0.268684757 |
| chr4:184717307-184717588 | -0.898214088 | 0.078004913 | 0.268684757 |
| chr1:184580233-184580442 | -1.001556219 | 0.078014867 | 0.268694887 |
| chr12:89783364-89783564 | -1.001556219 | 0.078014867 | 0.268694887 |
| chr3:85230247-85230447 | -1.001144491 | 0.078047073 | 0.268782882 |

| | | | |
|---|---|---|---|
| chr17:57309584-57309859 | -0.744728367 | 0.078047432 | 0.268782882 |
| chr2:64599460-64599952 | -0.630835976 | 0.078054856 | 0.2687839 |
| chr14:75088647-75088991 | 0.717074072 | 0.078055984 | 0.2687839 |
| chr17:82613933-82614196 | -0.749620266 | 0.078058252 | 0.2687839 |
| chr3:58525994-58526348 | -0.875427146 | 0.078061875 | 0.2687843 |
| chr2:235779527-235779766 | 0.849953538 | 0.078068714 | 0.268795769 |
| chr20:48679663-48679948 | 0.596006876 | 0.078075567 | 0.268807283 |
| chr1:209190682-209191216 | -0.693925056 | 0.078080474 | 0.268812099 |
| chr5:171640914-171641229 | 0.791270315 | 0.078096742 | 0.268853362 |
| chr10:60099166-60099445 | -0.855246601 | 0.078102318 | 0.268853362 |
| chr2:148989954-148990227 | -0.774252225 | 0.078102986 | 0.268853362 |
| chr12:103706107-103706353 | -0.953735121 | 0.078114406 | 0.268880594 |
| chr1:161766113-161766702 | 0.608521589 | 0.078121249 | 0.268892069 |
| chr20:57278993-57279749 | 0.559769309 | 0.078139724 | 0.268943578 |
| chr3:197906555-197906900 | -0.763382246 | 0.078159541 | 0.268999703 |
| chr3:10460122-10460812 | 0.606597004 | 0.078166923 | 0.269002802 |
| chr8:48621796-48622186 | 0.660177357 | 0.078167463 | 0.269002802 |
| chr9:137029397-137029638 | 0.692709591 | 0.078188343 | 0.269058858 |
| chr21:37266858-37268295 | 0.557717789 | 0.078191934 | 0.269058858 |
| chr15:45660228-45660835 | 0.615974461 | 0.078194287 | 0.269058858 |
| chr1:246904290-246904490 | -0.89878073 | 0.078230079 | 0.269138058 |
| chr15:85443761-85444319 | -0.66612778 | 0.078230771 | 0.269138058 |
| chr15:101264747-101264998 | 0.760037714 | 0.078233078 | 0.269138058 |
| chrX:18886418-18887151 | 0.6028434 | 0.078233817 | 0.269138058 |
| chr6:90479249-90479657 | -0.631668071 | 0.07823792 | 0.269138058 |
| chr10:63504115-63504384 | -0.810702948 | 0.078238379 | 0.269138058 |
| chr9:127916933-127917362 | 0.608462049 | 0.078245379 | 0.269150052 |
| chr17:40476196-40476485 | 0.714825601 | 0.078250008 | 0.269150765 |
| chr4:83599221-83599538 | -0.787717889 | 0.078252611 | 0.269150765 |
| chr2:190534191-190535147 | -0.589698221 | 0.078269692 | 0.269197428 |
| chr2:11829275-11829509 | -0.817484863 | 0.078274143 | 0.269200654 |
| chr7:107168356-107169130 | -0.559865816 | 0.07829962 | 0.269266351 |
| chr9:93600132-93600348 | 0.861565932 | 0.078300274 | 0.269266351 |
| chr8:142355684-142355906 | -0.835198981 | 0.078326291 | 0.269343734 |
| chr12:3132751-3132951 | -0.949644031 | 0.078365642 | 0.269466959 |
| chr13:44159570-44159997 | -0.698893402 | 0.078375536 | 0.269488887 |
| chr1:155250508-155251174 | 0.601646431 | 0.078387748 | 0.269518781 |
| chr10:82221825-82222076 | -0.857464257 | 0.078397982 | 0.269534139 |
| chr17:74738337-74738649 | -0.761164109 | 0.07839925 | 0.269534139 |
| chr16:89986742-89986974 | -0.906616656 | 0.078403013 | 0.269534981 |
| chr18:34122786-34123045 | -0.810268723 | 0.078406646 | 0.269535376 |
| chr3:133384442-133384774 | -0.687147894 | 0.078414134 | 0.269549024 |
| chr11:116615215-116615468 | -0.882285725 | 0.078425287 | 0.269575271 |
| chr10:99279537-99280121 | 0.612808432 | 0.078438218 | 0.269607623 |
| chr17:63435019-63436190 | -0.561676998 | 0.07844937 | 0.269633861 |
| chr4:148563264-148563627 | -0.84869877 | 0.078462184 | 0.269665806 |
| chr15:38564033-38565334 | 0.588190431 | 0.078468476 | 0.269670186 |
| chr22:43114930-43115294 | 0.706739415 | 0.078470986 | 0.269670186 |
| chr6:82385789-82386289 | -0.692614742 | 0.078474017 | 0.269670186 |
| chr16:83826125-83826598 | -0.620951998 | 0.078506593 | 0.269763541 |

| | | | |
|---|---|---|---|
| chr15:70196301-70197072 | 0.579271542 | 0.078508225 | 0.269763541 |
| chr21:34889156-34889404 | 0.746906592 | 0.078512672 | 0.269766725 |
| chr11:45881960-45882187 | -0.749605676 | 0.078516238 | 0.269766881 |
| chr4:185269533-185269733 | -0.999551261 | 0.078528894 | 0.269798267 |
| chr1:114298698-114298922 | 0.653001303 | 0.078559675 | 0.269891918 |
| chr2:82950234-82950434 | 0.940495033 | 0.078586029 | 0.269960511 |
| chr6:140436083-140436344 | -0.920756548 | 0.078588412 | 0.269960511 |
| chr10:43569725-43569931 | 0.848600266 | 0.078590211 | 0.269960511 |
| chr6:35327484-35327684 | 0.873739097 | 0.078612956 | 0.270026534 |
| chr3:61680172-61680372 | 0.694374159 | 0.078617508 | 0.270030064 |
| chr16:70746064-70747037 | -0.592479358 | 0.078637971 | 0.270088244 |
| chr5:87411958-87413453 | -0.556998548 | 0.078642542 | 0.270091834 |
| chr3:189812874-189813075 | 0.880335891 | 0.078692062 | 0.270249794 |
| chr2:161662560-161662865 | -0.946467777 | 0.078711054 | 0.270302903 |
| chr10:87622672-87622917 | -0.797904059 | 0.078728095 | 0.270349309 |
| chr1:105137485-105138139 | -0.644393901 | 0.078738458 | 0.270372777 |
| chr13:23960585-23960870 | 0.754339055 | 0.078754998 | 0.270417452 |
| chr5:178279987-178280278 | 0.671404095 | 0.078787333 | 0.270516357 |
| chrX:123764921-123765477 | 0.59626524 | 0.078796337 | 0.27053515 |
| chr1:35682355-35682658 | 0.905211866 | 0.078801324 | 0.270540148 |
| chr6:142885149-142885802 | -0.640617501 | 0.078806966 | 0.270547398 |
| chr17:42312235-42312747 | -0.708184744 | 0.07882033 | 0.270581154 |
| chr7:141472308-141472560 | 0.999998276 | 0.078867882 | 0.270732265 |
| chr10:109922864-109923764 | 0.590372192 | 0.078872268 | 0.270735192 |
| chr5:141397770-141398055 | -0.754872475 | 0.078876704 | 0.270738288 |
| chr7:69412647-69413000 | -0.714656563 | 0.078885833 | 0.270757496 |
| chr5:18043400-18043686 | 0.741015086 | 0.078892541 | 0.270768389 |
| chr3:195785961-195786348 | 0.711501571 | 0.078903152 | 0.270792678 |
| chr1:51977052-51977368 | 0.794672218 | 0.078908595 | 0.27079923 |
| chr4:147292600-147292905 | -0.841582935 | 0.078913066 | 0.270802444 |
| chr2:47965008-47965359 | -0.719741195 | 0.078918932 | 0.270810445 |
| chr11:82927303-82927776 | -0.701215685 | 0.078923442 | 0.270813793 |
| chr5:75510797-75512232 | -0.559676003 | 0.07893307 | 0.270834701 |
| chr17:60480650-60481375 | 0.562549482 | 0.078940693 | 0.270848727 |
| chr1:34985187-34985690 | 0.665337713 | 0.078954702 | 0.270884662 |
| chr1:202410898-202411227 | -0.685365887 | 0.078964563 | 0.270906365 |
| chr9:119963834-119964160 | -0.715534994 | 0.078970826 | 0.270915722 |
| chrX:10579046-10579544 | 0.704792753 | 0.078981347 | 0.270939682 |
| chr3:122005079-122005318 | 0.930325797 | 0.07898718 | 0.270947563 |
| chr9:133403900-133404172 | 0.681946218 | 0.078993105 | 0.270955755 |
| chr8:17766421-17766638 | -0.986151625 | 0.078998351 | 0.270961622 |
| chr20:3045563-3046621 | 0.578421971 | 0.079015929 | 0.27100978 |
| chr7:42072022-42072245 | -0.890231468 | 0.079021582 | 0.271017039 |
| chr12:132275169-132275435 | 0.684849441 | 0.079028887 | 0.271022216 |
| chr21:41678341-41679384 | -0.56196829 | 0.079030166 | 0.271022216 |
| chr18:21704005-21705103 | 0.586099419 | 0.079045082 | 0.271061238 |
| chr17:70808603-70809048 | -0.674692787 | 0.079056885 | 0.271083355 |
| chr20:14879129-14879780 | -0.653733041 | 0.079058608 | 0.271083355 |
| chr16:17277998-17278409 | 0.741920933 | 0.079080264 | 0.271145476 |
| chrX:34332386-34332663 | -0.785407783 | 0.079117105 | 0.271259656 |

| | | | |
|---|---|---|---|
| chr3:179316057-179316346 | -0.776709944 | 0.079140498 | 0.271321672 |
| chrX:136528713-136528991 | -0.79692773 | 0.079142275 | 0.271321672 |
| chr1:48291253-48291752 | -0.758373084 | 0.079160795 | 0.271373023 |
| chr14:80587116-80587358 | -0.763062932 | 0.079165232 | 0.271376091 |
| chr15:74322607-74323104 | 0.648094526 | 0.079181695 | 0.271420384 |
| chr20:44210417-44211437 | 0.557807504 | 0.079192281 | 0.271444528 |
| chr20:47371072-47371372 | 0.5823176 | 0.079200574 | 0.271451431 |
| chr7:44856566-44857322 | -0.589666188 | 0.079202591 | 0.271451431 |
| chr11:65121967-65122448 | 0.601382801 | 0.079204924 | 0.271451431 |
| chr19:55036244-55036582 | 0.859491427 | 0.079210996 | 0.271460102 |
| chr16:4639289-4639607 | -0.847428277 | 0.079218538 | 0.271473806 |
| chr20:45791475-45792630 | 0.574586 | 0.079225463 | 0.271482976 |
| chr1:43935735-43935983 | 0.731432335 | 0.0792283 | 0.271482976 |
| chr16:70251044-70252243 | 0.57393826 | 0.079243124 | 0.271518806 |
| chr2:32627412-32628591 | -0.563499449 | 0.079245844 | 0.271518806 |
| chr4:6640088-6640961 | 0.589478898 | 0.079256497 | 0.271543165 |
| chr2:134466217-134466417 | -0.901212979 | 0.079287133 | 0.271635979 |
| chr7:42941024-42941618 | -0.660230726 | 0.079302686 | 0.271671477 |
| chr19:11924568-11925331 | 0.602862122 | 0.079304585 | 0.271671477 |
| chr12:58765939-58766552 | -0.647070014 | 0.07930921 | 0.271675175 |
| chr10:93334689-93334960 | -0.914842597 | 0.079316853 | 0.271684546 |
| chr14:75943203-75943476 | 0.882921734 | 0.079319037 | 0.271684546 |
| chr4:985151-985430 | 0.700669859 | 0.07932714 | 0.271700153 |
| chr5:149141278-149142232 | -0.591000187 | 0.079333641 | 0.271710272 |
| chr4:173168027-173170245 | -0.557506583 | 0.07937019 | 0.271823301 |
| chr4:155796114-155796443 | -0.711797762 | 0.079381618 | 0.271850287 |
| chr1:160767610-160767848 | -0.936913833 | 0.079387508 | 0.271858308 |
| chr3:100125687-100125887 | -0.965958287 | 0.079401397 | 0.271881566 |
| chr3:119351069-119351309 | -0.965958287 | 0.079401397 | 0.271881566 |
| chr9:101549585-101549785 | -0.984833416 | 0.079406686 | 0.271887528 |
| chr18:37019157-37019901 | -0.616207467 | 0.079411382 | 0.271891458 |
| chr10:80281033-80281280 | 0.736429194 | 0.079418438 | 0.271900835 |
| chr11:29527659-29527925 | -0.812598585 | 0.079422406 | 0.271900835 |
| chr3:86772234-86772581 | 0.764058387 | 0.079424767 | 0.271900835 |
| chr6:37791161-37791800 | 0.589490608 | 0.079429108 | 0.271903548 |
| chr3:176953762-176954153 | -0.705581683 | 0.079452315 | 0.271960406 |
| chr20:9884075-9884404 | -0.729762321 | 0.079452816 | 0.271960406 |
| chr4:65190814-65191168 | -0.674110426 | 0.079459004 | 0.271962735 |
| chr8:100827024-100827343 | 0.635158012 | 0.079460596 | 0.271962735 |
| chr6:35737597-35737845 | 0.76094302 | 0.079464634 | 0.271964408 |
| chr16:67430764-67431311 | 0.634536049 | 0.079472247 | 0.271978317 |
| chr22:37710582-37711079 | -0.767201533 | 0.079495276 | 0.272044977 |
| chr12:102362940-102363229 | -0.917906851 | 0.079512885 | 0.272084403 |
| chr10:79430904-79431114 | 0.858025828 | 0.079513899 | 0.272084403 |
| chr6:64774312-64774572 | 0.791279626 | 0.079528357 | 0.272121726 |
| chr2:66212754-66213383 | -0.615942815 | 0.079537364 | 0.272140391 |
| chr9:76447459-76448209 | -0.621138616 | 0.079554877 | 0.272177103 |
| chr15:94164296-94164533 | 0.75267576 | 0.079555198 | 0.272177103 |
| chr9:79382564-79382838 | -0.918011463 | 0.079562235 | 0.272189025 |
| chr15:67803148-67803352 | 0.822571614 | 0.079583057 | 0.272248101 |

| | | | |
|---|---|---|---|
| chr9:100700653-100700915 | 0.807611538 | 0.079611523 | 0.272333323 |
| chr5:80055779-80056007 | -0.909717501 | 0.079618473 | 0.272344938 |
| chr20:48843235-48843435 | 0.656879808 | 0.079623431 | 0.272349742 |
| chr18:63578261-63578701 | -0.824809651 | 0.079627516 | 0.272351438 |
| chr3:32684766-32685942 | 0.566804053 | 0.079631162 | 0.272351438 |
| chr12:5796155-5796691 | -0.709047338 | 0.079637628 | 0.272351438 |
| chr17:40489975-40490199 | 0.890050848 | 0.079641651 | 0.272351438 |
| chr6:11427308-11427592 | 0.915245595 | 0.079646751 | 0.272351438 |
| chr7:51213233-51213433 | 0.885684245 | 0.079648513 | 0.272351438 |
| chr4:147843156-147843556 | -0.631908527 | 0.079648809 | 0.272351438 |
| chr1:36150848-36151156 | 0.64036319 | 0.079660651 | 0.272373936 |
| chr2:10992851-10993190 | -0.83907059 | 0.079662498 | 0.272373936 |
| chr2:112690935-112691390 | -0.664658607 | 0.079682705 | 0.27242984 |
| chr1:246931031-246932600 | -0.550245312 | 0.079685959 | 0.27242984 |
| chr16:83802042-83802517 | -0.616599591 | 0.079695215 | 0.272449326 |
| chr2:31192318-31192521 | -0.898267405 | 0.079701374 | 0.272458225 |
| chr15:40928559-40928840 | 0.70063496 | 0.079709836 | 0.272474998 |
| chr7:129401453-129401795 | -0.727315311 | 0.079725599 | 0.272516723 |
| chr17:50079676-50080290 | -0.595262785 | 0.079733807 | 0.272532622 |
| chr7:56287438-56288298 | 0.610491589 | 0.079752446 | 0.272581653 |
| chr7:82247915-82248115 | -0.960197073 | 0.079755267 | 0.272581653 |
| chr19:8505771-8506457 | 0.641029465 | 0.079758915 | 0.272581963 |
| chr7:149421802-149422915 | -0.574737338 | 0.079795709 | 0.272695544 |
| chr13:43388358-43388589 | 0.809450674 | 0.079801152 | 0.272701982 |
| chr8:108599493-108599892 | 0.676161034 | 0.079805982 | 0.272706327 |
| chr3:109655432-109655730 | -0.843555225 | 0.07983705 | 0.272787917 |
| chr12:65306566-65306885 | 0.725358948 | 0.07983729 | 0.272787917 |
| chr15:55917879-55918138 | 0.762855766 | 0.079843976 | 0.272787917 |
| chr2:177763323-177764019 | -0.613600037 | 0.0798441 | 0.272787917 |
| chr2:186734267-186734602 | -0.798277278 | 0.07985731 | 0.272820885 |
| chr3:143699576-143699851 | -0.901677749 | 0.079876891 | 0.272875613 |
| chr13:31431719-31431965 | -0.81332473 | 0.079902202 | 0.27294627 |
| chr3:137035678-137036034 | -0.641866168 | 0.079904698 | 0.27294627 |
| chr5:59893806-59894477 | -0.626794144 | 0.079914558 | 0.272961622 |
| chr2:77977530-77977982 | -0.76017269 | 0.079920622 | 0.272961622 |
| chr1:165114305-165114583 | 0.777760775 | 0.079923189 | 0.272961622 |
| chr8:95085393-95086486 | 0.56565753 | 0.079923442 | 0.272961622 |
| chr2:198968522-198968852 | -0.896272091 | 0.079928974 | 0.272968349 |
| chr12:84305054-84305295 | 0.917088072 | 0.079938707 | 0.272989419 |
| chr17:76659870-76660102 | 0.800456102 | 0.079985675 | 0.273126025 |
| chr3:160115196-160115605 | -0.759603223 | 0.079985838 | 0.273126025 |
| chr5:167934781-167935003 | -0.875432354 | 0.08000072 | 0.273153852 |
| chr1:2652913-2653163 | 0.842170227 | 0.080001117 | 0.273153852 |
| chr3:15572157-15572520 | -0.717294723 | 0.08000732 | 0.273162857 |
| chr15:40471821-40472382 | 0.604418845 | 0.080014497 | 0.27317519 |
| chr11:116148147-116148398 | -0.946156142 | 0.080029155 | 0.273202723 |
| chr17:48522060-48522628 | -0.621912565 | 0.080030829 | 0.273202723 |
| chr2:193963605-193963849 | -0.787013102 | 0.080033258 | 0.273202723 |
| chr12:132600464-132600667 | 0.796260262 | 0.080047657 | 0.2732397 |
| chr1:196355195-196355558 | -0.721487155 | 0.080069376 | 0.273301664 |

| | | | |
|---|---|---|---|
| chr16:21188541-21189048 | 0.668583195 | 0.08007368 | 0.273304179 |
| chr10:74213748-74213948 | -1.047140005 | 0.080094276 | 0.2733623 |
| chr2:51012953-51013429 | -0.59961228 | 0.080125248 | 0.273455827 |
| chr13:34156316-34156554 | -0.723952495 | 0.080136028 | 0.273480437 |
| chr2:64410731-64410960 | -0.827850903 | 0.080141075 | 0.273485478 |
| chr13:108003591-108004417 | -0.604273814 | 0.08014482 | 0.273486077 |
| chr3:177443427-177443812 | -0.745537499 | 0.080150956 | 0.273494836 |
| chr16:58027336-58027883 | 0.635799449 | 0.080170925 | 0.273548931 |
| chrX:37905253-37906222 | -0.628583867 | 0.080175809 | 0.273548931 |
| chr9:117073330-117073836 | -0.724702866 | 0.080177519 | 0.273548931 |
| chr20:48172485-48172685 | 0.643224489 | 0.080200364 | 0.273605103 |
| chr11:762554-762836 | 0.768128567 | 0.080201125 | 0.273605103 |
| chr3:16601284-16601514 | 0.80408335 | 0.080218715 | 0.273652926 |
| chr10:96025571-96025917 | -0.726072863 | 0.080240702 | 0.273715746 |
| chr2:23401606-23402028 | -0.739307598 | 0.080247105 | 0.273725401 |
| chr11:120325037-120325859 | 0.616129076 | 0.080260296 | 0.273738329 |
| chr3:143951406-143951692 | -0.747101363 | 0.080263559 | 0.273738329 |
| chr4:160175132-160175607 | -0.67071884 | 0.080264286 | 0.273738329 |
| chr1:40509071-40509688 | 0.681377597 | 0.080266508 | 0.273738329 |
| chr10:96229575-96229846 | 0.763329086 | 0.080268758 | 0.273738329 |
| chr14:46717262-46717467 | 0.908384657 | 0.0802765 | 0.273752549 |
| chr3:105970423-105970623 | -0.982531698 | 0.080287073 | 0.273776418 |
| chr16:81756130-81756387 | 0.838335039 | 0.080295076 | 0.273791523 |
| chr5:142097752-142097957 | -0.977835857 | 0.080304088 | 0.273809855 |
| chr10:77587116-77587395 | -0.730827011 | 0.080307599 | 0.273809855 |
| chr11:27717422-27717622 | -0.866166633 | 0.080322888 | 0.273836847 |
| chr10:109223496-109223913 | -0.68045839 | 0.080325305 | 0.273836847 |
| chr4:132871723-132871984 | 0.749041695 | 0.080326238 | 0.273836847 |
| chr4:31997051-31997581 | -0.686483422 | 0.080332592 | 0.273846327 |
| chr15:45118083-45118435 | 0.617783073 | 0.080349212 | 0.273890797 |
| chr1:1000686-1001008 | 0.655387972 | 0.080358632 | 0.273910721 |
| chr2:114101845-114102309 | -0.743963891 | 0.080376083 | 0.273958017 |
| chr2:177354659-177354859 | -0.968956425 | 0.080384358 | 0.273965029 |
| chr5:78899849-78900209 | -0.767009658 | 0.080385291 | 0.273965029 |
| chr8:64283761-64284036 | -0.891209904 | 0.080393182 | 0.273967187 |
| chr6:116059919-116061242 | -0.574389311 | 0.080395982 | 0.273967187 |
| chr1:116619147-116619368 | 0.821396661 | 0.08039904 | 0.273967187 |
| chr15:77155856-77156596 | -0.625251239 | 0.080400227 | 0.273967187 |
| chr16:69329900-69331025 | 0.572037421 | 0.080404272 | 0.273968789 |
| chr1:198493382-198493582 | 0.856267649 | 0.080412158 | 0.273975013 |
| chr3:97432968-97433298 | 0.673160453 | 0.080415632 | 0.273975013 |
| chr6:159460036-159460830 | -0.684107484 | 0.080416826 | 0.273975013 |
| chr3:164250986-164251291 | -0.770762601 | 0.080434525 | 0.274023127 |
| chr2:149064543-149064818 | 0.829413199 | 0.080438612 | 0.274024867 |
| chr2:109613430-109613646 | 0.80726613 | 0.080447748 | 0.274043806 |
| chr4:156584045-156584305 | -0.90469265 | 0.080462736 | 0.274082677 |
| chr3:55526425-55526635 | 0.890021852 | 0.080472532 | 0.274103861 |
| chr13:71979531-71980002 | -0.703126664 | 0.080481738 | 0.274123032 |
| chr3:165403827-165404259 | -0.723066982 | 0.08049394 | 0.274152406 |
| chr2:156589734-156590058 | -0.843747269 | 0.080498191 | 0.274154696 |

| | | | |
|---|---|---|---|
| chr4:94290400-94290600 | -1.032661307 | 0.080511052 | 0.274186312 |
| chr21:36069686-36070153 | 0.624159206 | 0.080524617 | 0.274220322 |
| chr2:42931483-42931683 | -0.786762074 | 0.080530907 | 0.274229553 |
| chr6:106773507-106773913 | -0.674155203 | 0.08054638 | 0.274269123 |
| chr10:102596541-102596832 | 0.715826622 | 0.080549687 | 0.274269123 |
| chrX:17374689-17375566 | -0.591114043 | 0.080568272 | 0.274319904 |
| chr8:141386775-141387019 | 0.663642342 | 0.080571761 | 0.274319904 |
| chr5:163306521-163306793 | -0.787228038 | 0.080580518 | 0.274337531 |
| chr1:41862439-41862639 | 0.781006863 | 0.080604748 | 0.274407828 |
| chr4:52911784-52912031 | -0.888870666 | 0.080608619 | 0.274408813 |
| chr7:8296025-8296457 | 0.598898623 | 0.080613736 | 0.274414043 |
| chr8:121829869-121830397 | -0.61297956 | 0.080621615 | 0.27442867 |
| chr1:215824276-215824770 | -0.629794159 | 0.080627955 | 0.274438059 |
| chr3:182710612-182710977 | -0.775700443 | 0.080634579 | 0.274448415 |
| chr2:21372703-21372996 | -0.743209156 | 0.080640795 | 0.274457378 |
| chr5:111278848-111279189 | -0.71570951 | 0.080650205 | 0.274477214 |
| chr15:57867971-57868214 | 0.966323746 | 0.080660053 | 0.274492717 |
| chr1:109066089-109066889 | -0.586896877 | 0.080661925 | 0.274492717 |
| chr3:19485376-19485576 | 0.8581218 | 0.080671343 | 0.274512574 |
| chr1:35877105-35877305 | 0.956459086 | 0.080679512 | 0.274528179 |
| chr4:6269492-6270076 | 0.58659003 | 0.080694263 | 0.274566182 |
| chr6:87483817-87484397 | -0.613456598 | 0.080717766 | 0.274625836 |
| chr12:117189949-117190701 | 0.590118726 | 0.080718964 | 0.274625836 |
| chr2:73085681-73085943 | -0.71259831 | 0.080726703 | 0.274639973 |
| chr3:172987306-172987585 | -0.748208045 | 0.080732193 | 0.274646455 |
| chr20:54340289-54341011 | 0.56306287 | 0.080739396 | 0.274658763 |
| chr6:29722968-29723958 | -0.591586451 | 0.080746443 | 0.274670543 |
| chr1:83055383-83055701 | -0.675881264 | 0.080755858 | 0.274690374 |
| chr11:75342655-75343308 | 0.586967585 | 0.080760675 | 0.274694563 |
| chr12:56629544-56629793 | 0.753434082 | 0.080780303 | 0.274749128 |
| chr13:98142753-98143607 | 0.600375736 | 0.080822696 | 0.274881113 |
| chr4:98728398-98728608 | -0.935717928 | 0.080833167 | 0.274904523 |
| chr2:6877579-6878339 | -0.619125878 | 0.080864329 | 0.274998298 |
| chr10:6163342-6164176 | 0.566657701 | 0.080873278 | 0.275016525 |
| chr17:7688040-7688649 | -0.591947048 | 0.08092425 | 0.275177647 |
| chr13:112506138-112506338 | -1.013694611 | 0.080928664 | 0.275178724 |
| chr10:119233663-119233885 | 0.914602572 | 0.08093175 | 0.275178724 |
| chr20:20161552-20161799 | 0.940731116 | 0.080940927 | 0.275197716 |
| chr2:102664440-102664859 | -0.633698528 | 0.080946417 | 0.2752 |
| chr9:20941128-20941416 | -0.759612513 | 0.080948782 | 0.2752 |
| chr9:105064485-105064867 | 0.728098435 | 0.080957776 | 0.275203457 |
| chr12:117671726-117672064 | 0.700304364 | 0.080958464 | 0.275203457 |
| chr13:32607724-32607955 | -0.905914179 | 0.080962519 | 0.275203457 |
| chr4:110622183-110622539 | 0.645362314 | 0.080964978 | 0.275203457 |
| chr18:11947244-11947614 | -0.805528953 | 0.080967758 | 0.275203457 |
| chr3:64873424-64873624 | 0.639703953 | 0.080975717 | 0.275216766 |
| chr6:43817216-43817439 | 0.820401182 | 0.080978857 | 0.275216766 |
| chr9:12819836-12820101 | -0.785612558 | 0.080992647 | 0.275251423 |
| chr21:41142119-41142662 | -0.630434298 | 0.080997861 | 0.275256934 |
| chrX:31293528-31293744 | -0.912912759 | 0.081007565 | 0.275277703 |

| | | | |
|---|---|---|---|
| chr14:96503034-96503234 | 0.72695606 | 0.081015696 | 0.275293125 |
| chrX:30141965-30142315 | -0.72419715 | 0.081022291 | 0.275303326 |
| chr20:19802676-19803204 | 0.614155544 | 0.081044588 | 0.275366876 |
| chr9:107484221-107484457 | -0.805258506 | 0.081053232 | 0.275384034 |
| chr16:67198965-67199346 | 0.635355065 | 0.081056855 | 0.275384133 |
| chr5:73513084-73513322 | 0.9486107 | 0.081071507 | 0.275417949 |
| chr1:207659617-207660384 | -0.578255351 | 0.081073998 | 0.275417949 |
| chr18:63979496-63979745 | -0.812054055 | 0.081078636 | 0.275421495 |
| chr4:82524572-82525202 | -0.611635966 | 0.081092975 | 0.275438182 |
| chr1:209646686-209646901 | -0.80494279 | 0.081093348 | 0.275438182 |
| chr10:59805083-59805299 | -0.844329973 | 0.081094332 | 0.275438182 |
| chr13:24680255-24680611 | 0.744310852 | 0.081104606 | 0.275449722 |
| chr16:30053443-30053899 | 0.673731191 | 0.08110492 | 0.275449722 |
| chr5:132293250-132295163 | -0.549703433 | 0.081150874 | 0.275593575 |
| chr2:64228234-64228591 | -0.701939701 | 0.081157365 | 0.275603404 |
| chr4:75432748-75432995 | -0.891610152 | 0.081162686 | 0.27560926 |
| chr13:35649364-35649815 | -0.806354661 | 0.081181637 | 0.275661396 |
| chr2:183172515-183172867 | -0.729752944 | 0.081189849 | 0.275677062 |
| chr15:70569071-70569466 | 0.694681987 | 0.081217353 | 0.275739625 |
| chr12:48023138-48023507 | 0.685060866 | 0.081219518 | 0.275739625 |
| chr7:64685829-64686029 | 0.838558144 | 0.081221198 | 0.275739625 |
| chr1:232629883-232631025 | 0.561901976 | 0.081222669 | 0.275739625 |
| chr7:23863264-23863523 | 0.721127372 | 0.081230764 | 0.275748059 |
| chr18:46193140-46193340 | -0.962513264 | 0.081232351 | 0.275748059 |
| chr7:86846804-86847185 | -0.653547447 | 0.081248012 | 0.275782396 |
| chr10:79237877-79238272 | -0.615763605 | 0.081249665 | 0.275782396 |
| chr10:3068068-3069305 | 0.56298825 | 0.081258397 | 0.275799819 |
| chr8:72316570-72316876 | -0.774262669 | 0.081290296 | 0.275895864 |
| chr12:5944667-5944992 | -0.74683205 | 0.081303137 | 0.275927224 |
| chr6:158339246-158339540 | -0.846338655 | 0.081333492 | 0.276018019 |
| chrX:24149200-24150603 | -0.562681359 | 0.081339947 | 0.276027699 |
| chr12:52907036-52907482 | 0.594004797 | 0.08139836 | 0.276201332 |
| chr6:30844113-30844328 | 0.892222337 | 0.081400068 | 0.276201332 |
| chr12:88579650-88580917 | -0.558302974 | 0.081402403 | 0.276201332 |
| chr13:81930064-81930273 | -0.904604071 | 0.081410039 | 0.276201332 |
| chr1:207579183-207579965 | -0.590788083 | 0.08141055 | 0.276201332 |
| chr12:77010617-77010887 | -0.862785761 | 0.081412742 | 0.276201332 |
| chr3:165414828-165415221 | -0.804161632 | 0.081416988 | 0.27620351 |
| chr1:107282558-107283061 | 0.591355679 | 0.081450326 | 0.276273252 |
| chr11:22304328-22304700 | -0.728119858 | 0.081451269 | 0.276273252 |
| chr5:168324848-168325568 | -0.60233249 | 0.081451418 | 0.276273252 |
| chr5:83721856-83722863 | -0.569355693 | 0.08145453 | 0.276273252 |
| chr10:55630571-55631804 | -0.564031604 | 0.08145809 | 0.276273252 |
| chrX:110985454-110985793 | 0.701393396 | 0.081462237 | 0.276273252 |
| chr1:111747282-111747573 | -0.698837365 | 0.081462786 | 0.276273252 |
| chr19:12484678-12485006 | 0.700842414 | 0.081466522 | 0.276273695 |
| chr14:100313808-100314285 | -0.675263425 | 0.081476732 | 0.27629609 |
| chr8:127398175-127398581 | 0.62486579 | 0.081488562 | 0.276323979 |
| chr3:148737150-148737635 | -0.684238473 | 0.081493144 | 0.276327286 |
| chr7:107147152-107147362 | -0.872108607 | 0.081536119 | 0.27646077 |

| | | | |
|---|---|---|---|
| chr4:152156504-152156816 | -0.786211439 | 0.081571576 | 0.276568756 |
| chr10:101110959-101111869 | -0.602689326 | 0.081593977 | 0.276632466 |
| chr1:6471473-6471821 | 0.699245519 | 0.081607398 | 0.276651225 |
| chr10:22079982-22080280 | -0.889312392 | 0.081609861 | 0.276651225 |
| chr3:176663918-176664197 | -0.768153511 | 0.081610342 | 0.276651225 |
| chr17:52085146-52085416 | 0.924000099 | 0.081615431 | 0.276656237 |
| chr11:65786744-65787080 | 0.625284967 | 0.08162203 | 0.276666366 |
| chr17:38703728-38704232 | 0.60860768 | 0.081638302 | 0.276694769 |
| chr20:53857693-53858286 | 0.5582151 | 0.081640246 | 0.276694769 |
| chr7:92424393-92424898 | -0.59727583 | 0.081641243 | 0.276694769 |
| chr14:21917925-21918209 | 0.744160756 | 0.08164575 | 0.276697804 |
| chr10:103696459-103696763 | 0.738027052 | 0.081680857 | 0.276801592 |
| chr7:114312492-114312970 | -0.743486151 | 0.0816836 | 0.276801592 |
| chr7:49639210-49639463 | 0.804207693 | 0.081694649 | 0.276826791 |
| chr5:135033118-135033462 | 0.704027454 | 0.081716953 | 0.276870775 |
| chr1:29703637-29703893 | 0.730803179 | 0.081717442 | 0.276870775 |
| chr10:80596452-80596991 | -0.619155085 | 0.08171847 | 0.276870775 |
| chr4:180058184-180059562 | -0.572030266 | 0.081727631 | 0.27688957 |
| chr1:203046330-203046620 | -0.67208814 | 0.081736834 | 0.276903052 |
| chr17:82957462-82957767 | 0.686476784 | 0.081738838 | 0.276903052 |
| chr12:14394220-14394716 | -0.758559042 | 0.081756357 | 0.276950158 |
| chr20:54431009-54431529 | 0.570102659 | 0.081760724 | 0.276952704 |
| chr19:7534680-7534890 | 0.769350464 | 0.081772986 | 0.276981995 |
| chr19:34254026-34254758 | 0.600171193 | 0.081806723 | 0.27708402 |
| chr7:116598356-116598561 | -0.941142403 | 0.081825916 | 0.277136779 |
| chr17:68474381-68474743 | -0.686145852 | 0.081830367 | 0.277139604 |
| chr18:48008230-48008685 | -0.663245283 | 0.081836334 | 0.277147563 |
| chr6:52589071-52589271 | 0.848007871 | 0.081854205 | 0.277195832 |
| chr20:36093575-36093926 | 0.71260967 | 0.081868703 | 0.277232677 |
| chr3:122942983-122943255 | 0.682777111 | 0.081872611 | 0.277233656 |
| chr10:132536850-132537893 | 0.551126365 | 0.081893903 | 0.277293503 |
| chr5:129400348-129400716 | -0.719707169 | 0.081914435 | 0.277327315 |
| chr5:76388293-76388567 | -0.777943971 | 0.081916487 | 0.277327315 |
| chr10:30348797-30349664 | -0.60347734 | 0.081916844 | 0.277327315 |
| chr6:142474427-142475034 | -0.631338178 | 0.081918367 | 0.277327315 |
| chr15:45037131-45037434 | 0.69315476 | 0.081929002 | 0.277351064 |
| chr19:48810932-48811187 | 0.698663187 | 0.081939777 | 0.277362866 |
| chr22:21960413-21960703 | 0.793128987 | 0.081945735 | 0.277362866 |
| chr4:127788591-127788917 | -0.679802273 | 0.081945935 | 0.277362866 |
| chr2:239265259-239265520 | 0.836182466 | 0.081947684 | 0.277362866 |
| chrX:25427883-25428167 | -0.849716833 | 0.081950588 | 0.277362866 |
| chr2:132669418-132669751 | 0.725593358 | 0.081958938 | 0.277375158 |
| chr18:72651171-72651376 | -0.963311002 | 0.08196146 | 0.277375158 |
| chr14:34547424-34548406 | 0.588004981 | 0.081981183 | 0.277429653 |
| chr7:157243431-157243653 | -0.949379519 | 0.081991269 | 0.277440332 |
| chr2:71802079-71802289 | 0.788978639 | 0.081992298 | 0.277440332 |
| chr8:120757098-120757416 | 0.682547958 | 0.081995278 | 0.277440332 |
| chr21:41612091-41612535 | -0.620551205 | 0.081998822 | 0.277440332 |
| chr2:168178699-168178941 | 0.948477767 | 0.08200461 | 0.277447664 |
| chr6:149709018-149709305 | 0.705099614 | 0.082010927 | 0.277448117 |

| | | | |
|---|---|---|---|
| chr12:33499240-33499516 | -0.872130473 | 0.082011986 | 0.277448117 |
| chr3:113179618-113179822 | 0.860733831 | 0.082025544 | 0.277481734 |
| chr1:167650547-167650871 | 0.825113902 | 0.082031529 | 0.277489728 |
| chr12:49342509-49342799 | 0.678001105 | 0.082052795 | 0.27754941 |
| chr19:4182314-4182764 | 0.626072497 | 0.08206116 | 0.277554405 |
| chr1:194889187-194889662 | 0.689915316 | 0.082061516 | 0.277554405 |
| chr7:126460554-126460856 | -0.878691762 | 0.082075756 | 0.277590314 |
| chr12:22101508-22101977 | -0.757061767 | 0.082113598 | 0.277694544 |
| chr1:187513227-187513507 | -0.785948145 | 0.082113822 | 0.277694544 |
| chr16:88437430-88437687 | 0.715312859 | 0.0821189 | 0.277699459 |
| chr11:35326271-35326752 | -0.715266524 | 0.082126451 | 0.27771107 |
| chr8:41542686-41543046 | -0.743701629 | 0.082130946 | 0.27771107 |
| chr14:56817836-56818122 | 0.820423509 | 0.082133207 | 0.27771107 |
| chr3:5023459-5023666 | -0.745893577 | 0.082145392 | 0.277740016 |
| chr7:36161944-36162527 | -0.593053146 | 0.082156158 | 0.27776416 |
| chr9:123434667-123434951 | 0.714037092 | 0.082173501 | 0.277810535 |
| chr7:80150901-80151119 | -0.899463651 | 0.082183914 | 0.277833482 |
| chr8:6560425-6560678 | 0.890228751 | 0.082191322 | 0.277846267 |
| chr3:128426331-128426609 | 0.688841021 | 0.082230805 | 0.277967475 |
| chr15:74250917-74251117 | 0.851344556 | 0.082241287 | 0.277990642 |
| chr9:85698374-85698993 | -0.671599921 | 0.082261977 | 0.278048312 |
| chr4:150254874-150255176 | 0.900927856 | 0.082278157 | 0.278090734 |
| chr16:53349886-53350243 | -0.702672272 | 0.082295098 | 0.278135724 |
| chr5:150474236-150474543 | 0.701079167 | 0.0823028 | 0.278149485 |
| chr19:31572785-31573308 | 0.628264659 | 0.082314662 | 0.278177306 |
| chr7:39580218-39580772 | -0.615085577 | 0.082323447 | 0.278190153 |
| chr5:83430455-83430744 | -0.730739034 | 0.082326981 | 0.278190153 |
| chr6:84095059-84095362 | 0.777152194 | 0.082329356 | 0.278190153 |
| chr1:64989546-64989870 | -0.720277823 | 0.082350116 | 0.2782407 |
| chr1:155323082-155323560 | 0.576850527 | 0.082351578 | 0.2782407 |
| chr3:62131679-62131898 | -0.706852854 | 0.082360773 | 0.278259496 |
| chr1:110746650-110747003 | 0.750820135 | 0.082367049 | 0.27826843 |
| chr10:61662724-61663274 | -0.640827315 | 0.082393116 | 0.278344223 |
| chr2:105454785-105455124 | -0.701969036 | 0.082427498 | 0.278435964 |
| chr5:174636052-174636639 | 0.647708895 | 0.08242754 | 0.278435964 |
| chr5:88720407-88720607 | -0.980742619 | 0.082441052 | 0.27846933 |
| chr5:142915645-142916157 | -0.664503372 | 0.082472338 | 0.278562729 |
| chr7:12951535-12951833 | 0.708239539 | 0.082481009 | 0.278579735 |
| chr5:149638079-149638356 | -0.836279798 | 0.082486229 | 0.278585085 |
| chr3:65907785-65907993 | -0.941203771 | 0.082497237 | 0.278599772 |
| chr21:8461944-8462401 | 0.607594385 | 0.08249785 | 0.278599772 |
| chr20:54276704-54276942 | 0.608766061 | 0.082539803 | 0.278729168 |
| chr7:8351519-8352367 | -0.585643963 | 0.082560265 | 0.278785978 |
| chr20:57142066-57142391 | 0.570115828 | 0.082572742 | 0.278815823 |
| chr10:84334052-84334502 | -0.693121478 | 0.082580521 | 0.278829803 |
| chr16:75139547-75139777 | -0.904064968 | 0.082602731 | 0.278892502 |
| chr1:170636290-170636607 | -0.873587021 | 0.082615922 | 0.278918739 |
| chr2:240686137-240686413 | 0.862295067 | 0.082617782 | 0.278918739 |
| chr3:107235857-107236205 | -0.643265858 | 0.082629682 | 0.278946623 |
| chr1:114369936-114370139 | 0.646902506 | 0.082635189 | 0.278952922 |

| | | | |
|---|---|---|---|
| chr1:100895339-100896954 | -0.550534099 | 0.082643466 | 0.278968573 |
| chr1:202131951-202133286 | -0.552005399 | 0.082648328 | 0.278972697 |
| chr1:83182192-83182465 | -0.891447747 | 0.082652615 | 0.278974878 |
| chr5:129664326-129664687 | -0.740071267 | 0.082708912 | 0.279152596 |
| chr5:171063730-171063935 | 0.92750016 | 0.08272469 | 0.279193551 |
| chr9:22119473-22119934 | -0.73355351 | 0.082743734 | 0.279231481 |
| chr4:61223432-61223817 | -0.75400983 | 0.082744186 | 0.279231481 |
| chr20:53647448-53647708 | 0.586272563 | 0.082748884 | 0.279231481 |
| chr4:164351161-164351371 | -0.93834533 | 0.082750506 | 0.279231481 |
| chr1:210373108-210374685 | -0.551281789 | 0.082765156 | 0.279268619 |
| chr12:131400800-131401097 | -0.826146857 | 0.082792501 | 0.279341628 |
| chr6:27864909-27865728 | -0.572574773 | 0.082794085 | 0.279341628 |
| chr3:149388756-149389062 | 0.639484531 | 0.082800307 | 0.279349625 |
| chr2:232633062-232633672 | -0.642247368 | 0.082806028 | 0.279349625 |
| chr14:58639980-58640215 | 0.736501334 | 0.082807393 | 0.279349625 |
| chr7:37448316-37448552 | -0.851953299 | 0.082811421 | 0.279350917 |
| chr3:133894992-133895237 | 0.765135943 | 0.082820845 | 0.279370405 |
| chr22:37807509-37808076 | 0.618790319 | 0.082828041 | 0.27938238 |
| chr10:103507466-103507672 | 0.807964689 | 0.082857926 | 0.279457057 |
| chr2:74919791-74920305 | 0.632505428 | 0.082858087 | 0.279457057 |
| chr15:91975592-91975938 | -0.684729518 | 0.082861122 | 0.279457057 |
| chr18:36688095-36688341 | -0.824714566 | 0.082875802 | 0.279494263 |
| chr2:218448984-218449263 | -0.720753 | 0.082891783 | 0.279535857 |
| chr1:15834822-15835289 | 0.621319969 | 0.082898926 | 0.279540829 |
| chr12:60494299-60494703 | -0.719076487 | 0.082900554 | 0.279540829 |
| chr4:1384623-1384823 | 0.942957489 | 0.082906297 | 0.279547892 |
| chr22:29080686-29080920 | -1.013761607 | 0.082912943 | 0.279549238 |
| chr2:150588331-150588716 | 0.749501021 | 0.082916583 | 0.279549238 |
| chr22:23281864-23282141 | 0.709310416 | 0.082923001 | 0.279549238 |
| chr5:90802711-90803003 | -0.708870878 | 0.082923074 | 0.279549238 |
| chr15:101174429-101174659 | 0.888222987 | 0.082928192 | 0.279549238 |
| chr17:61906311-61906745 | 0.569035823 | 0.082931986 | 0.279549238 |
| chr1:6385625-6386067 | 0.689202312 | 0.082932235 | 0.279549238 |
| chr2:69026022-69026503 | -0.706263509 | 0.082938598 | 0.279558388 |
| chr8:63063794-63064070 | -0.787024277 | 0.082950147 | 0.279585015 |
| chr22:43171741-43171941 | -0.918201051 | 0.082976294 | 0.279660844 |
| chr8:88982997-88983487 | 0.613242068 | 0.082982836 | 0.279670588 |
| chr17:72430735-72431389 | -0.618433768 | 0.083030127 | 0.279804698 |
| chr15:100803081-100803494 | 0.752474483 | 0.083036605 | 0.279804698 |
| chr2:139790937-139791146 | -0.920135834 | 0.083037161 | 0.279804698 |
| chr22:34105386-34105940 | -0.664155376 | 0.083037235 | 0.279804698 |
| chr13:82681657-82681891 | -0.885159387 | 0.083051778 | 0.279841394 |
| chr14:23120829-23121126 | 0.646003142 | 0.083060353 | 0.279857982 |
| chr20:56577558-56577881 | 0.610628757 | 0.083068624 | 0.279870117 |
| chr12:71511783-71512126 | 0.761557374 | 0.08307126 | 0.279870117 |
| chr3:137121703-137122038 | 0.685445889 | 0.083115121 | 0.280005574 |
| chr16:70707029-70707556 | 0.602526221 | 0.083132404 | 0.2800448 |
| chr17:70684253-70684522 | -0.910061678 | 0.083135938 | 0.2800448 |
| chr3:34277173-34277407 | 0.833513734 | 0.083137729 | 0.2800448 |
| chr2:69559660-69560039 | -0.669937712 | 0.083150843 | 0.28007666 |

| | | | |
|---|---|---|---|
| chr10:91671902-91672644 | -0.609313369 | 0.083155493 | 0.280080012 |
| chr6:154448424-154448746 | -0.697075814 | 0.083160892 | 0.280085885 |
| chr8:107888932-107889171 | 0.800280023 | 0.083178255 | 0.280132048 |
| chr16:75224424-75225066 | 0.610416518 | 0.083182359 | 0.280133316 |
| chr12:132489837-132490224 | 0.602573778 | 0.083189211 | 0.280133316 |
| chr3:156831850-156832186 | -0.712037129 | 0.083189599 | 0.280133316 |
| chr9:37485382-37486853 | -0.557505679 | 0.083210637 | 0.280191845 |
| chrX:69991571-69991979 | -0.815469267 | 0.083222697 | 0.280220141 |
| chr20:49604511-49604786 | 0.655700702 | 0.083252108 | 0.280306407 |
| chr16:29253170-29253464 | -0.86141324 | 0.083258986 | 0.280306407 |
| chr1:2586298-2586993 | 0.575191181 | 0.083259293 | 0.280306407 |
| chr2:47191820-47192681 | -0.597770869 | 0.083289579 | 0.280381345 |
| chr5:180126835-180127412 | -0.638319999 | 0.083293817 | 0.280381345 |
| chr2:151238900-151239158 | -0.822920558 | 0.083294606 | 0.280381345 |
| chr2:203782301-203782665 | -0.682791305 | 0.083299636 | 0.280381345 |
| chr9:110206723-110207717 | -0.590084446 | 0.083299848 | 0.280381345 |
| chr1:237799769-237800200 | -0.588961667 | 0.083318587 | 0.2804321 |
| chr10:117264861-117265129 | 0.827068867 | 0.083330929 | 0.280456969 |
| chr6:125197756-125198302 | -0.61121903 | 0.083333756 | 0.280456969 |
| chr20:35576074-35576554 | 0.68287738 | 0.083336956 | 0.280456969 |
| chr2:220120074-220120436 | -0.77042907 | 0.083355926 | 0.280508488 |
| chr7:102432746-102433690 | -0.585121118 | 0.083374886 | 0.280559971 |
| chr13:44218683-44218961 | 0.777759364 | 0.083380181 | 0.280565466 |
| chr2:151999053-151999324 | -0.762160399 | 0.083389727 | 0.280585264 |
| chr5:10288625-10289341 | -0.560516231 | 0.083394917 | 0.280590407 |
| chr8:142693942-142694533 | 0.697734945 | 0.083400807 | 0.280597903 |
| chr6:29702342-29702809 | -0.718684934 | 0.083410775 | 0.280619118 |
| chr1:150972055-150972461 | 0.602818525 | 0.083415761 | 0.280623571 |
| chr1:197774688-197776387 | -0.544732706 | 0.083422757 | 0.280634785 |
| chr6:104711006-104711786 | -0.593252826 | 0.083440454 | 0.280663678 |
| chr4:121071401-121071665 | 0.791133157 | 0.083441409 | 0.280663678 |
| chr11:117316058-117316755 | 0.609985982 | 0.083442335 | 0.280663678 |
| chr9:133881049-133881527 | 0.632447306 | 0.083466147 | 0.280729757 |
| chr2:4714925-4715610 | -0.651436666 | 0.083469308 | 0.280729757 |
| chr7:148966199-148967338 | -0.560584714 | 0.083499748 | 0.28081981 |
| chr3:106824861-106825080 | -0.877175655 | 0.083538131 | 0.28090763 |
| chr15:29269501-29270006 | 0.661033967 | 0.083539407 | 0.28090763 |
| chr1:36323153-36323353 | 0.770017532 | 0.08354052 | 0.28090763 |
| chr1:211643415-211644170 | -0.607592293 | 0.083540526 | 0.28090763 |
| chr4:185283065-185283332 | -0.854772832 | 0.083574298 | 0.281008859 |
| chr5:167861199-167861432 | -0.865944724 | 0.083578352 | 0.281010156 |
| chr22:23147661-23147919 | 0.742346318 | 0.08360756 | 0.281096028 |
| chr1:118483316-118483593 | 0.803722364 | 0.083697846 | 0.28138723 |
| chr16:24499301-24499507 | 0.852520928 | 0.083704037 | 0.281395697 |
| chr5:73995130-73995330 | -0.931920921 | 0.083709052 | 0.281400211 |
| chr9:88344554-88345036 | 0.664283746 | 0.083726773 | 0.281447432 |
| chr7:156660044-156660285 | -0.891687832 | 0.083739632 | 0.281476858 |
| chr8:22909039-22909316 | 0.74913957 | 0.083742874 | 0.281476858 |
| chr11:77074133-77074467 | 0.708439481 | 0.083752133 | 0.281492984 |
| chr1:36386438-36386977 | 0.580878705 | 0.083755019 | 0.281492984 |

| | | | |
|---|---|---|---|
| chr2:173021968-173022345 | 0.673775514 | 0.083768281 | 0.281525207 |
| chr3:101070297-101070607 | -0.677533349 | 0.083793149 | 0.281589413 |
| chr5:141629899-141630099 | 0.962917947 | 0.083794735 | 0.281589413 |
| chr5:132446126-132446326 | -0.912335398 | 0.083805932 | 0.281610213 |
| chr3:176430340-176430554 | -0.857750859 | 0.083812889 | 0.281610213 |
| chr10:53403687-53403898 | -0.957621743 | 0.083814506 | 0.281610213 |
| chr6:50353637-50353959 | -0.786590697 | 0.083815626 | 0.281610213 |
| chr11:27039946-27040180 | -0.786156541 | 0.083830752 | 0.281648684 |
| chr16:88049598-88049851 | 0.799141529 | 0.083839658 | 0.281666256 |
| chr5:58604396-58604629 | 0.923574913 | 0.08387548 | 0.281774247 |
| chr1:234486940-234487230 | 0.728871534 | 0.083883126 | 0.281787577 |
| chr2:234702865-234703282 | -0.714917216 | 0.083887183 | 0.281788854 |
| chr2:113157093-113157976 | 0.571830239 | 0.083897379 | 0.281810362 |
| chr1:171237839-171238039 | 0.905344758 | 0.083901629 | 0.281810362 |
| chr10:31646128-31646365 | -0.92326427 | 0.083905305 | 0.281810362 |
| chr1:217990662-217990934 | 0.760592721 | 0.083910472 | 0.281810362 |
| chr6:151676645-151676845 | 0.822687258 | 0.083911976 | 0.281810362 |
| chr7:141718314-141718557 | -0.864572796 | 0.083922552 | 0.281832555 |
| chr16:78045823-78046337 | 0.59484026 | 0.083932048 | 0.281832555 |
| chr17:59907588-59907836 | 0.615878582 | 0.083932391 | 0.281832555 |
| chr21:41854874-41855261 | -0.734549744 | 0.083933297 | 0.281832555 |
| chr10:18400868-18401126 | -0.790772904 | 0.083961482 | 0.281914841 |
| chr22:37088170-37088370 | -0.934681973 | 0.083974658 | 0.281946728 |
| chr6:80133006-80133218 | -0.848898036 | 0.083978416 | 0.281946989 |
| chr17:44014841-44015168 | 0.68425241 | 0.083988816 | 0.281969551 |
| chr19:53431363-53432198 | 0.575972192 | 0.084008359 | 0.282022804 |
| chrX:152915673-152916025 | 0.666884327 | 0.084019096 | 0.282035848 |
| chr14:61438602-61439744 | 0.56243479 | 0.084019606 | 0.282035848 |
| chr9:129382997-129383439 | -0.630303905 | 0.084030624 | 0.282060477 |
| chr5:39424647-39425465 | 0.596807802 | 0.084038082 | 0.282069124 |
| chr1:77605077-77605717 | -0.628359178 | 0.084040563 | 0.282069124 |
| chr15:99113626-99113826 | 0.873066521 | 0.084047931 | 0.282072812 |
| chr18:45014216-45014477 | -0.841105823 | 0.084049025 | 0.282072812 |
| chr21:39763374-39763852 | -0.654514676 | 0.084054392 | 0.28207847 |
| chr3:132284795-132284995 | -0.978051466 | 0.084062586 | 0.282093612 |
| chr5:76867759-76868412 | -0.609540458 | 0.084077412 | 0.282131009 |
| chr22:43777270-43777521 | -0.900530894 | 0.084084184 | 0.282141377 |
| chr4:124996671-124997090 | -0.76423773 | 0.084094698 | 0.282159648 |
| chr7:133990952-133991256 | -0.782250888 | 0.084096994 | 0.282159648 |
| chr16:53130082-53131115 | 0.559258031 | 0.084112313 | 0.282198687 |
| chr14:39593644-39593880 | 0.88316737 | 0.084117805 | 0.282204758 |
| chrX:118068718-118069206 | -0.66438987 | 0.084130379 | 0.282228603 |
| chr4:89158937-89159208 | 0.753919762 | 0.08413228 | 0.282228603 |
| chr7:101495916-101496151 | -0.894132487 | 0.084145271 | 0.282259826 |
| chr6:127026254-127026560 | 0.738275941 | 0.084152892 | 0.282273032 |
| chr13:38577083-38577327 | -0.857960318 | 0.084163227 | 0.282295341 |
| chr1:2204726-2205406 | 0.605983085 | 0.084194117 | 0.282376543 |
| chr20:54579830-54580067 | 0.618406854 | 0.084194807 | 0.282376543 |
| chr10:72260712-72261117 | -0.628256661 | 0.084223263 | 0.282448379 |
| chr2:72973630-72973925 | -0.675772959 | 0.084223599 | 0.282448379 |

| | | | |
|---|---|---|---|
| chr1:45854700-45855100 | 0.675076144 | 0.084263017 | 0.282554659 |
| chr6:28910819-28911210 | -0.755716079 | 0.084264031 | 0.282554659 |
| chr12:108457254-108457586 | -0.725013912 | 0.084269608 | 0.282554659 |
| chr12:57765137-57765701 | 0.591252988 | 0.084270041 | 0.282554659 |
| chr16:30935215-30935510 | 0.710171359 | 0.084274204 | 0.282556252 |
| chr16:31382594-31382853 | 0.779431127 | 0.084292533 | 0.282605341 |
| chr3:174316438-174316739 | -0.895435583 | 0.084308279 | 0.282645766 |
| chr7:145848784-145849048 | -0.861028689 | 0.084319214 | 0.282662482 |
| chr7:16071331-16071590 | 0.758322006 | 0.08432788 | 0.282662482 |
| chr1:119613816-119614085 | 0.742096632 | 0.084331051 | 0.282662482 |
| chr20:57189788-57190316 | 0.551550414 | 0.084335452 | 0.282662482 |
| chr7:127651527-127652524 | -0.549035496 | 0.084336747 | 0.282662482 |
| chr12:53543518-53543732 | 0.791051723 | 0.084343842 | 0.282662482 |
| chr1:183209573-183209931 | -0.778686209 | 0.084345134 | 0.282662482 |
| chr3:15121161-15121388 | 0.73935107 | 0.084348644 | 0.282662482 |
| chr16:27235054-27235292 | 0.750776312 | 0.084350046 | 0.282662482 |
| chr3:105891265-105891494 | -0.853757266 | 0.084350156 | 0.282662482 |
| chr2:222708289-222708498 | 0.883054673 | 0.08435743 | 0.282668087 |
| chr12:6914215-6914522 | 0.742024623 | 0.084359207 | 0.282668087 |
| chr10:118778748-118779361 | -0.661650839 | 0.084373467 | 0.282703505 |
| chrX:138034261-138034499 | -0.874958439 | 0.084394566 | 0.282753596 |
| chr2:215361744-215362223 | -0.612173345 | 0.084396413 | 0.282753596 |
| chr6:10528191-10528494 | -0.893857396 | 0.084401951 | 0.282753596 |
| chr3:89098813-89099350 | -0.662387624 | 0.084405835 | 0.282753596 |
| chr22:28792510-28793361 | -0.597581568 | 0.084406868 | 0.282753596 |
| chr5:6159598-6160004 | -0.755155145 | 0.084412944 | 0.282756194 |
| chr3:128488286-128488684 | 0.64718564 | 0.084415024 | 0.282756194 |
| chr5:171770045-171770461 | 0.687033166 | 0.084425895 | 0.28277588 |
| chr4:112726138-112726442 | -0.862764863 | 0.084428282 | 0.28277588 |
| chr14:31456799-31457727 | 0.555981346 | 0.084434766 | 0.282785237 |
| chr10:7787836-7788933 | 0.551457403 | 0.08443958 | 0.282788999 |
| chr20:60272732-60273005 | -0.699395746 | 0.084457303 | 0.282835991 |
| chr14:98808560-98808787 | 0.765882226 | 0.084465745 | 0.28284721 |
| chr5:53318365-53318697 | -0.774857532 | 0.08447386 | 0.28284721 |
| chr17:49371208-49371794 | -0.650428845 | 0.084479825 | 0.28284721 |
| chr12:80622358-80622604 | -0.831797287 | 0.084480029 | 0.28284721 |
| chr2:32165154-32166222 | -0.555858694 | 0.084481365 | 0.28284721 |
| chr12:54432312-54432764 | -0.669509697 | 0.084482802 | 0.28284721 |
| chr1:143264525-143264849 | 0.638627582 | 0.084489121 | 0.282856008 |
| chr10:59966363-59966563 | -0.85545172 | 0.084493514 | 0.282858355 |
| chr10:26157741-26157968 | -0.962718751 | 0.08450301 | 0.282877785 |
| chr4:60944084-60944426 | -0.758076147 | 0.084508483 | 0.282883749 |
| chr5:61768736-61769077 | -0.630856188 | 0.084521242 | 0.282906638 |
| chr1:37314828-37315114 | -0.898914904 | 0.084522706 | 0.282906638 |
| chr13:24511943-24513046 | 0.565297203 | 0.08452973 | 0.282917789 |
| chr8:108666776-108667359 | -0.595710689 | 0.084537302 | 0.282930774 |
| chr12:42229909-42230173 | -0.915886915 | 0.084544579 | 0.282935788 |
| chr13:46184285-46184752 | -0.649460868 | 0.084546185 | 0.282935788 |
| chr12:53053588-53054683 | 0.546031196 | 0.084557802 | 0.282954122 |
| chr1:86104756-86105489 | 0.590054617 | 0.084560195 | 0.282954122 |

| | | | |
|---|---|---|---|
| chr13:60473618-60474321 | -0.601050128 | 0.08456318 | 0.282954122 |
| chr10:61284362-61284574 | -0.917618622 | 0.084566435 | 0.282954122 |
| chr10:47553159-47553765 | -0.611601646 | 0.084583791 | 0.282999836 |
| chr1:248696267-248696824 | -0.620399701 | 0.08458982 | 0.28300765 |
| chr19:13093893-13094143 | 0.801166444 | 0.084597437 | 0.283020774 |
| chr6:107144759-107145167 | -0.8101138 | 0.084612221 | 0.283057875 |
| chr21:30227647-30228051 | 0.699888354 | 0.084638199 | 0.283131264 |
| chr5:88438118-88438488 | -0.808285422 | 0.084641549 | 0.283131264 |
| chr10:59893919-59894496 | -0.610142778 | 0.084674256 | 0.283225332 |
| chr12:109433797-109434006 | -0.787320827 | 0.084679552 | 0.283225332 |
| chr12:107027669-107027878 | 0.871235119 | 0.084680759 | 0.283225332 |
| chr5:162684072-162684481 | 0.684002546 | 0.084705204 | 0.28328889 |
| chr4:154626711-154627344 | -0.611357739 | 0.084707157 | 0.28328889 |
| chr12:49322691-49323542 | 0.560829341 | 0.084727895 | 0.283345879 |
| chr7:23086053-23086353 | 0.669221145 | 0.084742517 | 0.28338241 |
| chr2:158954402-158954693 | -0.791875419 | 0.084756958 | 0.283402529 |
| chr21:36854598-36854813 | -0.932289872 | 0.084761705 | 0.283402529 |
| chr7:455315-455565 | 0.69552237 | 0.084762688 | 0.283402529 |
| chr7:713375-713669 | 0.707032188 | 0.084765745 | 0.283402529 |
| chr15:45253179-45253738 | -0.663714934 | 0.084767027 | 0.283402529 |
| chr2:212005126-212005399 | -0.787135681 | 0.084772857 | 0.283409653 |
| chr19:42396434-42397374 | 0.561364924 | 0.084788034 | 0.28343883 |
| chr2:71452620-71452938 | -0.684556726 | 0.084788983 | 0.28343883 |
| chr14:64716672-64717135 | 0.604021433 | 0.084798408 | 0.28345797 |
| chr5:23419114-23419486 | -0.644487889 | 0.084813193 | 0.28349016 |
| chr6:51613626-51613974 | 0.693715568 | 0.08481605 | 0.28349016 |
| chr10:79242759-79243125 | 0.631447975 | 0.084821222 | 0.28349016 |
| chr15:40069181-40069490 | 0.711620129 | 0.084822837 | 0.28349016 |
| chr7:104207447-104208474 | -0.572725236 | 0.084848212 | 0.283551299 |
| chr1:212566736-212566936 | -0.936537704 | 0.084848532 | 0.283551299 |
| chr8:125273022-125273553 | -0.593112578 | 0.084874926 | 0.283627136 |
| chr14:50215469-50215757 | -0.768953756 | 0.084883867 | 0.283644644 |
| chr14:68951040-68951451 | -0.645286207 | 0.08490169 | 0.283688078 |
| chr8:124953752-124954076 | 0.635078352 | 0.084910501 | 0.283688078 |
| chr2:38302674-38302947 | 0.864408375 | 0.084911262 | 0.283688078 |
| chr20:50981448-50981736 | 0.663780755 | 0.084911675 | 0.283688078 |
| chr12:98754136-98754336 | 0.885471436 | 0.084927777 | 0.283729501 |
| chr16:31134685-31134915 | 0.706560774 | 0.084940088 | 0.28375826 |
| chr2:88454811-88455103 | -0.742239415 | 0.084945183 | 0.283762906 |
| chr5:55864278-55864879 | -0.666024455 | 0.084957212 | 0.283790012 |
| chr8:89759291-89759526 | 0.625975972 | 0.084960704 | 0.283790012 |
| chr20:57939557-57939776 | 0.69026553 | 0.084975766 | 0.283827949 |
| chr11:79116598-79117026 | -0.623795169 | 0.084987209 | 0.283853795 |
| chr1:107081667-107081872 | 0.662439953 | 0.085000505 | 0.283885828 |
| chr7:75905899-75906230 | -0.901987719 | 0.085017973 | 0.283931794 |
| chr10:84004198-84004439 | -0.916254537 | 0.085037165 | 0.283983511 |
| chr21:42580122-42580322 | -0.884994334 | 0.085042116 | 0.283984874 |
| chr3:189383596-189383962 | -0.71635139 | 0.085044986 | 0.283984874 |
| chr2:15430985-15431200 | -0.990039177 | 0.085055878 | 0.283992066 |
| chr15:86906445-86906791 | -0.761631725 | 0.08505872 | 0.283992066 |

| | | | |
|---|---|---|---|
| chr4:103306881-103307646 | 0.62027055 | 0.08505968 | 0.283992066 |
| chrX:115950080-115950358 | 0.750414702 | 0.085064224 | 0.283992066 |
| chr19:6460544-6460744 | 0.777390817 | 0.085065672 | 0.283992066 |
| chr6:89298484-89298719 | -0.905085516 | 0.085069445 | 0.283992288 |
| chr14:26008363-26008597 | 0.833383477 | 0.085078895 | 0.284011462 |
| chr11:34837504-34837754 | -0.881267342 | 0.085093784 | 0.28404594 |
| chr3:141273846-141274144 | -0.728686657 | 0.085096637 | 0.28404594 |
| chr4:2536076-2536776 | 0.571285755 | 0.085121237 | 0.284115674 |
| chr1:48817611-48817968 | -0.71246585 | 0.085127749 | 0.284125034 |
| chr2:179440732-179440933 | -0.890339842 | 0.085137257 | 0.284132113 |
| chr6:125154133-125155394 | -0.550025187 | 0.085137287 | 0.284132113 |
| chr1:203914370-203914668 | 0.719123704 | 0.085145058 | 0.284141636 |
| chr3:167573859-167574124 | -0.83836974 | 0.085147557 | 0.284141636 |
| chr5:73565081-73566111 | -0.557536877 | 0.085159639 | 0.284169579 |
| chr12:80710489-80710790 | -0.754389246 | 0.085163694 | 0.284170734 |
| chr1:32677610-32677810 | 0.953954785 | 0.085194695 | 0.284261797 |
| chr14:91325797-91326257 | 0.621728994 | 0.085203268 | 0.284271899 |
| chr12:6764052-6764662 | -0.574455957 | 0.085205143 | 0.284271899 |
| chr5:180154789-180154989 | 0.881576518 | 0.08521302 | 0.284285802 |
| chr1:64039522-64039956 | 0.644973002 | 0.085218115 | 0.284290421 |
| chrX:48083265-48083520 | 0.809027071 | 0.085246478 | 0.284372661 |
| chr1:161083092-161083895 | 0.562350076 | 0.085272834 | 0.284432245 |
| chr1:171373436-171374193 | -0.586568235 | 0.085275378 | 0.284432245 |
| chr19:44723962-44724165 | 0.754313321 | 0.085276355 | 0.284432245 |
| chr9:92152047-92152395 | 0.68351947 | 0.085279188 | 0.284432245 |
| chr17:19371696-19371928 | 0.88958097 | 0.085286335 | 0.284443699 |
| chrX:4830848-4831343 | 0.63579725 | 0.085290341 | 0.28444468 |
| chr4:178296567-178297003 | -0.734127462 | 0.085297406 | 0.284455861 |
| chr4:185011527-185011765 | -0.814052733 | 0.085323434 | 0.284530276 |
| chr7:17044908-17045147 | 0.801586057 | 0.08537112 | 0.284676907 |
| chr2:231925102-231925479 | 0.732962626 | 0.085378878 | 0.284690387 |
| chr11:62082102-62082429 | 0.706272485 | 0.085383804 | 0.284694421 |
| chr15:94680858-94681087 | -0.934698951 | 0.085393786 | 0.284703088 |
| chr17:40433801-40434026 | 0.824317958 | 0.085393834 | 0.284703088 |
| chr3:113194760-113195357 | -0.678062249 | 0.085415951 | 0.284764434 |
| chr14:74955421-74955758 | 0.661077767 | 0.085435486 | 0.284817167 |
| chr9:120650388-120650606 | -0.961964441 | 0.085443809 | 0.284832522 |
| chr1:218612968-218613483 | -0.642100694 | 0.085466429 | 0.28488187 |
| chr10:19008636-19008936 | -0.896215749 | 0.085467283 | 0.28488187 |
| chr10:3888270-3888545 | 0.864504097 | 0.085469766 | 0.28488187 |
| chr3:152283667-152283872 | 0.854953717 | 0.085484141 | 0.284917389 |
| chr4:175934838-175935461 | -0.597605574 | 0.085500832 | 0.284960625 |
| chr16:88905598-88905865 | -0.749528511 | 0.085509311 | 0.284968354 |
| chr11:123072254-123072797 | 0.619867218 | 0.08551059 | 0.284968354 |
| chr2:88024475-88024699 | -0.925782339 | 0.085540016 | 0.28502317 |
| chr21:29196332-29196597 | 0.800865357 | 0.08554877 | 0.28502317 |
| chr9:121268382-121268791 | 0.656267819 | 0.085548896 | 0.28502317 |
| chr1:207775887-207776115 | -0.902214783 | 0.085551809 | 0.28502317 |
| chr11:3808351-3808865 | 0.623707766 | 0.085552439 | 0.28502317 |
| chr7:132318779-132319128 | -0.665141396 | 0.085552744 | 0.28502317 |

| | | | |
|---|---|---|---|
| chr14:64891859-64892086 | 0.803668588 | 0.085553077 | 0.28502317 |
| chr4:124851814-124852169 | 0.637220139 | 0.085560833 | 0.285033602 |
| chr2:14240591-14240828 | -0.876900949 | 0.085569327 | 0.285033602 |
| chr4:48127789-48128233 | -0.640025577 | 0.085570712 | 0.285033602 |
| chr7:35835438-35835682 | 0.924998909 | 0.085571089 | 0.285033602 |
| chr1:37793922-37794267 | 0.67039245 | 0.085587151 | 0.2850642 |
| chrX:10119306-10119929 | 0.599961473 | 0.085591027 | 0.2850642 |
| chr4:8422333-8423046 | 0.60346075 | 0.085594655 | 0.2850642 |
| chr15:64046418-64046893 | 0.62498772 | 0.085596747 | 0.2850642 |
| chr4:78650026-78650435 | 0.696213947 | 0.085601671 | 0.2850642 |
| chr6:129730561-129730844 | -0.846634273 | 0.085606178 | 0.2850642 |
| chr17:19513920-19514166 | -0.86219251 | 0.085608335 | 0.2850642 |
| chr12:114637234-114637524 | 0.812604965 | 0.085610621 | 0.2850642 |
| chr20:45347984-45348184 | 0.729934216 | 0.085613758 | 0.2850642 |
| chr18:79497218-79497518 | -0.83240035 | 0.085621285 | 0.28507168 |
| chr20:54851794-54852125 | 0.583759195 | 0.085626405 | 0.28507168 |
| chr2:163345984-163346244 | -0.768815206 | 0.085627166 | 0.28507168 |
| chr4:174828639-174829996 | -0.548918667 | 0.085645214 | 0.285098915 |
| chr15:76335876-76336122 | -0.756606592 | 0.085645508 | 0.285098915 |
| chr10:46565648-46565917 | -0.764388947 | 0.085647576 | 0.285098915 |
| chr20:8454171-8454703 | -0.654360624 | 0.08565023 | 0.285098915 |
| chr4:94540417-94541067 | -0.608647412 | 0.08565617 | 0.285106302 |
| chr13:42333509-42334714 | -0.557992205 | 0.085696381 | 0.285227752 |
| chr12:90779295-90779560 | -0.929913443 | 0.085707027 | 0.285235818 |
| chr6:44921834-44922086 | -0.889855073 | 0.085708169 | 0.285235818 |
| chr19:29084365-29084621 | 0.779165411 | 0.085709972 | 0.285235818 |
| chr4:125013900-125014200 | -0.866592377 | 0.085759898 | 0.285389572 |
| chr6:134123610-134123817 | -0.886118391 | 0.085777448 | 0.285435578 |
| chr3:64467698-64468026 | 0.591756061 | 0.085781521 | 0.285436734 |
| chr5:108345606-108345931 | -0.710698931 | 0.085793734 | 0.285464978 |
| chr17:57243165-57243803 | -0.586148626 | 0.085812354 | 0.285514028 |
| chr13:74286990-74287384 | -0.631988199 | 0.085817669 | 0.285514028 |
| chr11:123640965-123641238 | -0.758972117 | 0.085822514 | 0.285514028 |
| chr6:47034934-47035300 | -0.696760223 | 0.085823381 | 0.285514028 |
| chr16:30786906-30787366 | 0.620742728 | 0.085830807 | 0.285526335 |
| chrX:72018559-72019053 | 0.64096581 | 0.085851592 | 0.28556405 |
| chr20:40286893-40287575 | 0.591849069 | 0.085855048 | 0.28556405 |
| chr7:54664199-54665340 | -0.564805735 | 0.085855072 | 0.28556405 |
| chr6:113936140-113936450 | -0.709051616 | 0.085857052 | 0.28556405 |
| chr1:143244921-143245123 | 0.929651245 | 0.085863542 | 0.285573239 |
| chr4:114597827-114598762 | -0.59736203 | 0.085869977 | 0.285582246 |
| chrX:110992714-110992914 | 0.883259683 | 0.085879943 | 0.285602994 |
| chr15:39530945-39531222 | -0.853512369 | 0.085891237 | 0.285628156 |
| chr10:73252298-73253483 | -0.559721557 | 0.085905025 | 0.285652023 |
| chr2:38396109-38396437 | 0.74388999 | 0.085906344 | 0.285652023 |
| chr6:90296034-90296585 | 0.623567994 | 0.085909598 | 0.285652023 |
| chrX:95071619-95072161 | -0.655861157 | 0.085922178 | 0.285656419 |
| chrX:27336537-27336763 | -0.941508189 | 0.085923607 | 0.285656419 |
| chr5:131171489-131171803 | -0.6337435 | 0.085925533 | 0.285656419 |
| chr1:169809528-169809910 | 0.70430486 | 0.085925833 | 0.285656419 |

| | | | |
|---|---|---|---|
| chr12:42569342-42569599 | -0.880918311 | 0.085936267 | 0.285678713 |
| chr7:157010427-157011565 | -0.560645594 | 0.085961514 | 0.285740586 |
| chr9:30967598-30967804 | -0.985752855 | 0.085968299 | 0.285740586 |
| chr14:105411514-105411714 | 0.733191504 | 0.085969521 | 0.285740586 |
| chr17:58839866-58840666 | 0.566621224 | 0.085969796 | 0.285740586 |
| chr17:61777294-61778060 | 0.547817802 | 0.085974514 | 0.285743872 |
| chr5:161544282-161544609 | -0.666772931 | 0.085982249 | 0.285757184 |
| chr22:35376984-35377224 | -0.845997717 | 0.085997319 | 0.285794871 |
| chr17:48593967-48594177 | -0.802779659 | 0.086011674 | 0.285828793 |
| chr17:62206609-62206926 | 0.6826318 | 0.086017811 | 0.285828793 |
| chr1:196225081-196225717 | -0.615803373 | 0.086018717 | 0.285828793 |
| chr8:128167969-128169030 | -0.553574383 | 0.086047141 | 0.28590457 |
| chr6:43229100-43229703 | 0.61461887 | 0.086048985 | 0.28590457 |
| chr6:76684987-76685290 | -0.790295482 | 0.086059051 | 0.285925618 |
| chr12:53508601-53508932 | 0.638696197 | 0.086088799 | 0.286012051 |
| chr12:26402693-26402971 | -0.837647955 | 0.086107599 | 0.286062108 |
| chr19:45338535-45339031 | 0.626758316 | 0.086130353 | 0.286125295 |
| chr7:27372393-27372997 | 0.689932459 | 0.086137285 | 0.286127062 |
| chr16:77584254-77584538 | 0.653572467 | 0.086141298 | 0.286127062 |
| chr13:45362995-45363290 | -0.703789494 | 0.08614212 | 0.286127062 |
| chr5:171838158-171838419 | -0.793567996 | 0.086145822 | 0.286127062 |
| chr20:53707379-53707660 | 0.566214271 | 0.086163998 | 0.286175026 |
| chr10:32881964-32882345 | -0.848993859 | 0.086170866 | 0.286185432 |
| chr9:109451662-109451941 | 0.738989748 | 0.086186608 | 0.286225308 |
| chr1:29245808-29246065 | 0.741716577 | 0.086192209 | 0.2862315 |
| chr6:17505028-17505577 | -0.742588885 | 0.08620012 | 0.286236829 |
| chr10:109451866-109452234 | -0.82406173 | 0.086201285 | 0.286236829 |
| chr6:153728047-153728247 | 0.894266854 | 0.086211233 | 0.286257458 |
| chrX:23907259-23908328 | -0.553266687 | 0.086232754 | 0.28631651 |
| chr9:92669381-92669763 | 0.645944721 | 0.086254309 | 0.286363298 |
| chr3:42773135-42773404 | -0.832900005 | 0.086254321 | 0.286363298 |
| chr13:106859441-106859895 | -0.707437172 | 0.086259098 | 0.28636675 |
| chr3:131026279-131027242 | -0.552178208 | 0.086267958 | 0.286383756 |
| chr12:82314377-82314920 | -0.610781255 | 0.086289481 | 0.286442797 |
| chr10:21675217-21675455 | -0.829482699 | 0.086297579 | 0.286452401 |
| chr12:65607395-65607953 | 0.606339638 | 0.086299852 | 0.286452401 |
| chr5:93597031-93597712 | -0.57522345 | 0.08630438 | 0.286455023 |
| chr16:74243284-74243507 | 0.890956322 | 0.086310196 | 0.286461917 |
| chr14:90256379-90256926 | 0.607409259 | 0.086314417 | 0.286463518 |
| chr2:134228540-134228804 | -0.79980086 | 0.086320217 | 0.286466947 |
| chr5:82124-82449 | 0.688697406 | 0.086322927 | 0.286466947 |
| chr15:62782022-62782851 | 0.580186212 | 0.086331174 | 0.286481905 |
| chr12:7091950-7092301 | -0.79199157 | 0.086340334 | 0.286499895 |
| chr11:47998774-47999214 | 0.65853868 | 0.086350945 | 0.286522695 |
| chr8:119054108-119054342 | 0.848465813 | 0.086360485 | 0.286541943 |
| chr20:21640611-21640881 | 0.756270815 | 0.086396196 | 0.286648017 |
| chr6:130872314-130872609 | -0.789871871 | 0.086406847 | 0.286665809 |
| chr1:212567099-212567339 | -0.836857024 | 0.086409041 | 0.286665809 |
| chr17:59199138-59199417 | 0.709132041 | 0.086423066 | 0.286699924 |
| chr9:35914384-35914717 | -0.781834923 | 0.086437787 | 0.286731158 |

| | | | |
|---|---|---|---|
| chr1:220918593-220918981 | -0.710771301 | 0.086443341 | 0.286731158 |
| chrX:34656593-34657976 | -0.558463761 | 0.086445649 | 0.286731158 |
| chr1:100459599-100459855 | -0.761069481 | 0.08644745 | 0.286731158 |
| chr16:27109816-27110016 | -0.92696009 | 0.086452258 | 0.286734694 |
| chr7:134328025-134328237 | -0.936399986 | 0.086470398 | 0.286756568 |
| chrX:134384399-134384632 | -0.936399986 | 0.086470398 | 0.286756568 |
| chr17:3003962-3004698 | 0.599783616 | 0.086471639 | 0.286756568 |
| chr12:48021747-48022257 | 0.60689068 | 0.086473823 | 0.286756568 |
| chr2:71643803-71644416 | -0.609680565 | 0.086483431 | 0.28677602 |
| chrX:130110072-130110801 | 0.595221501 | 0.086493233 | 0.286787005 |
| chr3:128433940-128434207 | 0.743012429 | 0.08649423 | 0.286787005 |
| chr12:19129558-19130378 | -0.58458914 | 0.086522623 | 0.286861118 |
| chr5:61311907-61312171 | -0.773921789 | 0.08652407 | 0.286861118 |
| chr15:40098784-40099190 | 0.601141331 | 0.086544654 | 0.286881127 |
| chr11:117535227-117535441 | 0.756230573 | 0.08655064 | 0.286881127 |
| chr16:28092729-28093313 | -0.671750479 | 0.086551131 | 0.286881127 |
| chr6:66986767-66987488 | -0.580680861 | 0.086553612 | 0.286881127 |
| chr21:40857507-40857766 | 0.857967304 | 0.086554645 | 0.286881127 |
| chr4:140764970-140765170 | 0.888848798 | 0.08655676 | 0.286881127 |
| chr6:51163914-51164114 | 0.888848798 | 0.08655676 | 0.286881127 |
| chr19:55580549-55581140 | 0.616354612 | 0.086560655 | 0.286881127 |
| chr1:145156028-145156328 | 0.700953712 | 0.086563802 | 0.286881127 |
| chr2:34366073-34366313 | -0.859370524 | 0.086579402 | 0.286920416 |
| chr1:61077837-61078266 | -0.589225651 | 0.086589076 | 0.286940064 |
| chr10:44978538-44979216 | -0.572219503 | 0.086599879 | 0.28695495 |
| chr19:12880630-12881014 | -0.713878109 | 0.086601058 | 0.28695495 |
| chr20:45891144-45891506 | 0.599725508 | 0.086612979 | 0.286982041 |
| chr11:47257240-47258002 | -0.605827102 | 0.086647093 | 0.287065627 |
| chr4:2241474-2242621 | 0.554597091 | 0.086650079 | 0.287065627 |
| chr2:223725630-223725889 | -0.8032983 | 0.086650969 | 0.287065627 |
| chr8:66512210-66513158 | -0.561920793 | 0.086653192 | 0.287065627 |
| chr3:13895836-13896186 | 0.628090078 | 0.086679711 | 0.287127486 |
| chr7:76728579-76729084 | -0.68216945 | 0.086680008 | 0.287127486 |
| chr13:97567964-97568271 | -0.788815347 | 0.086684876 | 0.287127486 |
| chr17:76793868-76794068 | 0.835158692 | 0.086686854 | 0.287127486 |
| chr7:55486379-55486630 | 0.838749422 | 0.086692808 | 0.287134794 |
| chr1:237806806-237807495 | -0.620174019 | 0.086699936 | 0.287145007 |
| chr1:204149157-204150131 | -0.555223568 | 0.086703387 | 0.287145007 |
| chr21:38733961-38734226 | -0.707786341 | 0.086740972 | 0.287251639 |
| chr1:9292743-9293002 | 0.817849189 | 0.086743082 | 0.287251639 |
| chr5:38899795-38900073 | 0.802844621 | 0.086764928 | 0.287296756 |
| chr11:78127483-78127763 | -0.812269515 | 0.086768266 | 0.287296756 |
| chr13:103730445-103730750 | -0.812269515 | 0.086768266 | 0.287296756 |
| chr15:74818687-74818903 | 0.743165996 | 0.086771704 | 0.287296756 |
| chr6:1296146-1296420 | 0.826271117 | 0.086781727 | 0.287317526 |
| chr16:71938755-71939223 | -0.678369466 | 0.086786262 | 0.287320126 |
| chr3:155158370-155158927 | 0.638361414 | 0.086791892 | 0.28732635 |
| chr11:107740996-107741297 | -0.820244073 | 0.086796621 | 0.287329591 |
| chr6:41538596-41538947 | -0.749543013 | 0.08681501 | 0.287373647 |
| chr7:18064294-18064579 | -0.735913521 | 0.086817431 | 0.287373647 |

| | | | |
|---|---|---|---|
| chr8:96282758-96283265 | -0.601189903 | 0.086836237 | 0.287411319 |
| chr10:84534196-84534589 | -0.658039021 | 0.086836314 | 0.287411319 |
| chr3:164658357-164658594 | -0.82009971 | 0.08692105 | 0.287679354 |
| chr1:156115291-156115677 | 0.623464258 | 0.08692678 | 0.287685889 |
| chr20:47747380-47747695 | 0.568660161 | 0.086932593 | 0.287692702 |
| chr8:62901773-62902207 | -0.651377251 | 0.086966669 | 0.287793042 |
| chr4:134785966-134786172 | 0.857204425 | 0.086991926 | 0.287864191 |
| chr11:75376284-75376484 | -0.853097776 | 0.087003688 | 0.287890681 |
| chr6:123715340-123715771 | -0.732010963 | 0.08702367 | 0.287944366 |
| chr19:58380509-58381488 | 0.548088679 | 0.08703294 | 0.287956022 |
| chrX:107626527-107627276 | -0.573313864 | 0.087034818 | 0.287956022 |
| chr9:69180662-69181027 | -0.686804362 | 0.087041948 | 0.287956022 |
| chr8:103152071-103152386 | 0.646308627 | 0.087042226 | 0.287956022 |
| chr17:49861178-49861472 | -0.710359482 | 0.087049824 | 0.287968725 |
| chr17:29243826-29244065 | 0.833780352 | 0.087058692 | 0.287972917 |
| chr20:290068-290626 | 0.618297817 | 0.087061214 | 0.287972917 |
| chr1:206726332-206726649 | -0.611681817 | 0.087062366 | 0.287972917 |
| chr15:91445724-91445957 | -0.927791012 | 0.087072072 | 0.287986708 |
| chr5:164597344-164597637 | -0.74044357 | 0.087077653 | 0.287986708 |
| chr5:43017505-43017705 | 0.70279891 | 0.087077811 | 0.287986708 |
| chr5:151239355-151239582 | -0.766524764 | 0.087097028 | 0.288037829 |
| chr9:113465493-113465886 | 0.629111106 | 0.087112103 | 0.288059246 |
| chr17:17355085-17355307 | 0.764967133 | 0.087116906 | 0.288059246 |
| chr7:134546973-134547198 | -0.761396959 | 0.087118171 | 0.288059246 |
| chr12:87995634-87996010 | -0.655044097 | 0.087118542 | 0.288059246 |
| chr3:58008181-58009273 | 0.577676523 | 0.087135213 | 0.288090844 |
| chr13:83731121-83731441 | -0.704423984 | 0.087144602 | 0.288090844 |
| chrX:99717751-99717951 | 0.939779506 | 0.087147559 | 0.288090844 |
| chr15:78278218-78278786 | 0.594208015 | 0.087149272 | 0.288090844 |
| chr1:10796099-10796300 | -0.828164597 | 0.087151469 | 0.288090844 |
| chr1:32892879-32893637 | 0.589604836 | 0.087154071 | 0.288090844 |
| chr15:84079322-84079704 | -0.798365618 | 0.087154417 | 0.288090844 |
| chr1:3910807-3911155 | 0.680912542 | 0.087170018 | 0.288129983 |
| chr17:8332613-8332876 | 0.78216029 | 0.087195209 | 0.288200815 |
| chr6:127592937-127593903 | -0.581663316 | 0.087207798 | 0.288225417 |
| chr6:158704500-158704817 | 0.759198809 | 0.087210175 | 0.288225417 |
| chr10:86584059-86584390 | 0.638638671 | 0.087219128 | 0.288242573 |
| chr15:84747551-84749036 | 0.548827253 | 0.087224997 | 0.288248257 |
| chr6:137845460-137845731 | 0.73003839 | 0.087228372 | 0.288248257 |
| chr20:24150069-24150309 | 0.739324566 | 0.087243151 | 0.288266768 |
| chr5:162068107-162068311 | -0.882305557 | 0.087246629 | 0.288266768 |
| chr11:57644876-57645148 | -0.734770811 | 0.08724704 | 0.288266768 |
| chr15:84633300-84633666 | 0.658975646 | 0.087249023 | 0.288266768 |
| chr14:24332249-24332473 | 0.681432332 | 0.087259611 | 0.288289319 |
| chr16:68434785-68435171 | -0.716592729 | 0.087267586 | 0.288303238 |
| chrX:95057040-95057280 | -0.934040952 | 0.087279684 | 0.288324278 |
| chr15:94859000-94859200 | -0.906728004 | 0.087281481 | 0.288324278 |
| chr15:61702578-61702826 | 0.759147574 | 0.087289388 | 0.288336565 |
| chr8:122008151-122008412 | 0.762616417 | 0.08729674 | 0.288336565 |
| chr9:137422625-137423700 | 0.564494116 | 0.087312642 | 0.288336565 |

| | | | |
|---|---|---|---|
| chr12:66693965-66694240 | -0.718289185 | 0.087314231 | 0.288336565 |
| chr16:68842951-68843195 | -0.771615587 | 0.08732173 | 0.288336565 |
| chr10:32980341-32981013 | -0.572959046 | 0.087321731 | 0.288336565 |
| chr5:38614098-38614545 | 0.606841849 | 0.08732179 | 0.288336565 |
| chr2:148812573-148812782 | -0.88810677 | 0.087321858 | 0.288336565 |
| chr15:91258371-91258686 | 0.708021519 | 0.087322586 | 0.288336565 |
| chr2:180979999-180981131 | -0.549919906 | 0.087327349 | 0.288336565 |
| chr20:57450703-57450970 | 0.70508434 | 0.08732877 | 0.288336565 |
| chr14:47873011-47873273 | -0.86049664 | 0.087330358 | 0.288336565 |
| chr6:29748950-29749280 | 0.624912021 | 0.08734836 | 0.288374862 |
| chr1:147855077-147855352 | -0.774032486 | 0.087349484 | 0.288374862 |
| chr3:142366365-142366744 | -0.724605629 | 0.08736169 | 0.288402731 |
| chr16:79162562-79162789 | -0.903418543 | 0.087372975 | 0.288427561 |
| chr5:71900893-71901184 | 0.863996005 | 0.087377423 | 0.288429815 |
| chr2:120722350-120722588 | 0.920564277 | 0.087385987 | 0.288445658 |
| chr3:162717173-162717521 | 0.701941787 | 0.087395656 | 0.288465149 |
| chr13:44425633-44425833 | -0.891762397 | 0.087417201 | 0.288523834 |
| chr3:87321232-87321494 | 0.819015602 | 0.087438262 | 0.288580914 |
| chr3:173553653-173553974 | -0.765582471 | 0.087449739 | 0.288606361 |
| chr12:120078886-120079174 | 0.692965195 | 0.087461786 | 0.288625864 |
| chr1:155050459-155051452 | 0.5590932 | 0.087463182 | 0.288625864 |
| chr3:172836507-172836961 | -0.712789049 | 0.087472898 | 0.28864011 |
| chr2:69724642-69725060 | -0.695274906 | 0.087475033 | 0.28864011 |
| chr3:144636899-144637290 | -0.682011429 | 0.087481976 | 0.288650589 |
| chr7:8408992-8409257 | 0.743045602 | 0.087488549 | 0.288659846 |
| chr5:401335-401546 | 0.80108289 | 0.087507116 | 0.288703035 |
| chr6:141980047-141980654 | -0.691137826 | 0.087509175 | 0.288703035 |
| chr6:1604361-1604663 | 0.725468917 | 0.087526807 | 0.288748773 |
| chr20:53797745-53798202 | 0.550019051 | 0.087533469 | 0.288758318 |
| chr15:64854939-64855139 | 0.805574685 | 0.087542326 | 0.288775103 |
| chr2:98592994-98593308 | 0.756763287 | 0.087546134 | 0.288775234 |
| chr14:50578419-50578650 | 0.891530466 | 0.087554406 | 0.288790085 |
| chr9:81604909-81605184 | -0.755848696 | 0.087577064 | 0.288852387 |
| chr14:73728752-73729036 | 0.649743261 | 0.087589748 | 0.288881787 |
| chr17:59366851-59367060 | 0.700705817 | 0.087617112 | 0.288959597 |
| chr3:32947703-32948082 | 0.666075319 | 0.087626833 | 0.288979221 |
| chr19:4471131-4471371 | 0.746085994 | 0.087646336 | 0.289027779 |
| chr17:71587833-71588033 | -0.921368191 | 0.087651523 | 0.289027779 |
| chr3:196968681-196969025 | 0.663691737 | 0.087652874 | 0.289027779 |
| chr13:37341909-37342161 | -0.874211091 | 0.087661053 | 0.289037186 |
| chr10:97336936-97337711 | -0.563462562 | 0.087663375 | 0.289037186 |
| chr2:37475572-37475928 | -0.874028853 | 0.087667043 | 0.289037186 |
| chr5:160438392-160439548 | -0.55746443 | 0.087672731 | 0.2890435 |
| chr17:8226501-8227155 | -0.563689746 | 0.087681986 | 0.289061574 |
| chr14:79466726-79466934 | 0.950221911 | 0.087689112 | 0.289068869 |
| chr4:6782319-6782862 | 0.59028176 | 0.087692232 | 0.289068869 |
| chr1:153797272-153797472 | 0.753422778 | 0.087695516 | 0.289068869 |
| chr13:20043962-20044362 | -0.75296759 | 0.087710066 | 0.289097397 |
| chr6:3991341-3991565 | 0.770701902 | 0.087718233 | 0.289097397 |
| chr6:24282993-24283935 | -0.588397537 | 0.08771922 | 0.289097397 |

| | | | |
|---|---|---|---|
| chr2:168639663-168639905 | -0.886448229 | 0.087719263 | 0.289097397 |
| chr19:38304992-38305338 | 0.632470422 | 0.087734506 | 0.289127575 |
| chr3:149352478-149353030 | -0.57956487 | 0.087735967 | 0.289127575 |
| chr5:103445781-103446011 | -0.808226001 | 0.087747677 | 0.289153729 |
| chr2:71923856-71924306 | -0.630556783 | 0.087803254 | 0.289324427 |
| chr9:91503621-91503944 | -0.932716526 | 0.08780838 | 0.289328876 |
| chr4:98794691-98794929 | -0.921262307 | 0.087821779 | 0.289360583 |
| chr10:89328007-89328437 | -0.773060386 | 0.087857366 | 0.289459643 |
| chrX:132219359-132219648 | 0.649608789 | 0.087862986 | 0.289459643 |
| chr17:36544207-36545814 | 0.550372778 | 0.087863177 | 0.289459643 |
| chr3:42118810-42119182 | 0.676424629 | 0.087914465 | 0.289616155 |
| chr12:109134204-109134435 | 0.804567773 | 0.087928249 | 0.289649111 |
| chr18:20921071-20921655 | 0.644835254 | 0.087932239 | 0.289649799 |
| chr2:112868046-112868666 | -0.639195634 | 0.087938181 | 0.289656921 |
| chr14:38105185-38105504 | 0.736179514 | 0.087943106 | 0.28966069 |
| chr10:76148209-76148428 | -0.89027121 | 0.087956115 | 0.289691086 |
| chr5:9118464-9118844 | -0.723377955 | 0.087967472 | 0.289716038 |
| chr7:95511785-95512288 | -0.678894824 | 0.087978394 | 0.289724956 |
| chr10:64332593-64332793 | -0.969709959 | 0.0879875 | 0.289724956 |
| chr12:61857997-61858197 | -0.969709959 | 0.0879875 | 0.289724956 |
| chr13:97028861-97029061 | -0.969709959 | 0.0879875 | 0.289724956 |
| chr8:51230862-51231062 | -0.954362493 | 0.087989086 | 0.289724956 |
| chr7:103447144-103447495 | -0.627073603 | 0.088006125 | 0.28976861 |
| chr1:154351465-154351881 | 0.65720327 | 0.08801332 | 0.289778755 |
| chr17:14062131-14062665 | 0.627922278 | 0.08801677 | 0.289778755 |
| chr22:37784890-37785090 | 0.873738251 | 0.088032855 | 0.289819257 |
| chr18:76801786-76802539 | -0.589546335 | 0.088052166 | 0.289870378 |
| chr17:61667939-61668304 | 0.563968259 | 0.088061512 | 0.289888691 |
| chr5:85921637-85921900 | -0.831041006 | 0.088075073 | 0.289920875 |
| chr11:27485998-27486314 | -0.73599933 | 0.088082916 | 0.289934239 |
| chr5:149761351-149761824 | -0.661958614 | 0.088098984 | 0.289974671 |
| chr1:215102713-215102913 | -0.876109151 | 0.088122165 | 0.290038509 |
| chr3:11319556-11319849 | -0.854694394 | 0.088139778 | 0.29008402 |
| chr6:50842862-50843279 | -0.685242552 | 0.088164174 | 0.29014011 |
| chr5:103139128-103139612 | -0.657412259 | 0.088167795 | 0.29014011 |
| chr7:3426988-3427308 | 0.881626168 | 0.08816818 | 0.29014011 |
| chr1:147051665-147051868 | 0.876219748 | 0.088177156 | 0.290157185 |
| chr10:19210770-19211203 | -0.666832741 | 0.088193162 | 0.290197393 |
| chr1:63322143-63322826 | 0.589725157 | 0.08819852 | 0.290202561 |
| chr12:87866578-87867028 | -0.674483032 | 0.088202463 | 0.29020307 |
| chr12:98743473-98743673 | 0.846750672 | 0.088237334 | 0.290305336 |
| chr7:104834725-104835015 | -1.013651107 | 0.0882458 | 0.290320725 |
| chr10:132965164-132965600 | 0.65942855 | 0.088262179 | 0.290362142 |
| chr2:186601974-186602174 | -0.905021498 | 0.088267513 | 0.290367222 |
| chr7:50679209-50679702 | -0.627325147 | 0.088311069 | 0.290498034 |
| chr2:238278579-238278849 | -0.854123518 | 0.088318885 | 0.290507551 |
| chr14:72789066-72789299 | 0.819373008 | 0.088321545 | 0.290507551 |
| chr11:100753615-100753854 | -0.872254536 | 0.088329407 | 0.290520939 |
| chr2:153967608-153967808 | 0.911297021 | 0.088338405 | 0.290538063 |
| chr2:129737725-129737925 | 0.901444868 | 0.088349888 | 0.290563359 |

| | | | |
|---|---|---|---|
| chr17:38674145-38675191 | 0.592820321 | 0.088357618 | 0.290576308 |
| chr2:18599-18963 | -0.784715704 | 0.088372397 | 0.290612438 |
| chr3:112582257-112582519 | -0.812351559 | 0.088377758 | 0.290617596 |
| chr18:48913247-48913666 | -0.693148154 | 0.088398877 | 0.290674568 |
| chr3:162411289-162412013 | -0.594452526 | 0.088428968 | 0.290761034 |
| chr1:175742957-175743337 | 0.64803655 | 0.088434626 | 0.290767161 |
| chr14:59361037-59361857 | -0.564973058 | 0.088442635 | 0.290781016 |
| chr15:73710161-73710521 | -0.666621081 | 0.088451786 | 0.290798624 |
| chr5:172350582-172350927 | 0.694265971 | 0.088465777 | 0.290832142 |
| chr1:193069839-193070374 | -0.621609211 | 0.088477963 | 0.290859725 |
| chr6:157187100-157187789 | -0.592871513 | 0.088490689 | 0.290889081 |
| chr8:32223615-32223829 | -0.9136443 | 0.088494873 | 0.290890355 |
| chr4:161586609-161586866 | -0.833818832 | 0.088503845 | 0.290897896 |
| chr11:82733566-82733955 | 0.695569894 | 0.088507437 | 0.290897896 |
| chr5:171912186-171912396 | 0.769168558 | 0.088510884 | 0.290897896 |
| chr1:184982061-184982478 | -0.645221356 | 0.088512354 | 0.290897896 |
| chr5:122946752-122946994 | -0.957587882 | 0.088538818 | 0.290972392 |
| chr16:81939222-81939773 | -0.629873998 | 0.088577138 | 0.291067992 |
| chr8:37916013-37916571 | -0.644842675 | 0.088581117 | 0.291067992 |
| chr8:34267025-34267397 | -0.653865746 | 0.088582912 | 0.291067992 |
| chr6:408622-409166 | -0.646905046 | 0.088583103 | 0.291067992 |
| chr17:61815881-61816430 | 0.550795681 | 0.088615376 | 0.2911543 |
| chr13:42078327-42078645 | -0.829489281 | 0.088616969 | 0.2911543 |
| chr1:196477262-196477607 | -0.730404254 | 0.088644662 | 0.291222245 |
| chr14:68736730-68736930 | 0.802763888 | 0.088645355 | 0.291222245 |
| chr17:63116192-63116501 | -0.674307154 | 0.088649368 | 0.291222245 |
| chr10:73956659-73956859 | -0.888348294 | 0.088655121 | 0.291222245 |
| chr21:38636636-38637069 | -0.701169578 | 0.088658787 | 0.291222245 |
| chr10:91923341-91924459 | -0.551265603 | 0.088660454 | 0.291222245 |
| chr4:171969180-171969487 | -0.796222913 | 0.088694751 | 0.291322412 |
| chr1:231139673-231140570 | 0.586912702 | 0.088716464 | 0.291381239 |
| chr20:53377096-53377296 | 0.888005709 | 0.088723594 | 0.291391673 |
| chr14:89027109-89027599 | 0.612852555 | 0.088727247 | 0.291391673 |
| chr8:60762824-60763183 | -0.677637517 | 0.088772207 | 0.291526832 |
| chr18:35345224-35345689 | 0.601038844 | 0.088779398 | 0.291537953 |
| chr12:132559862-132560062 | 0.733274796 | 0.088789278 | 0.2915579 |
| chrX:30971758-30972073 | -0.664167516 | 0.088827792 | 0.291671595 |
| chr1:204841791-204842166 | 0.623454199 | 0.088834032 | 0.291671595 |
| chr3:72746264-72746814 | 0.640541774 | 0.088841417 | 0.291671595 |
| chr14:43613031-43613326 | 0.765274262 | 0.088842249 | 0.291671595 |
| chr2:100903935-100904227 | 0.699003655 | 0.088845062 | 0.291671595 |
| chr20:47369201-47369411 | 0.585575547 | 0.088846742 | 0.291671595 |
| chr3:171789082-171789390 | -0.694764151 | 0.088857865 | 0.291683567 |
| chr10:6297581-6298186 | 0.622214154 | 0.088859919 | 0.291683567 |
| chr13:97109812-97110389 | 0.627530476 | 0.088864133 | 0.291683567 |
| chrX:106802155-106803393 | -0.550202935 | 0.088865615 | 0.291683567 |
| chr6:27091642-27092359 | -0.578435729 | 0.088877358 | 0.291699805 |
| chr6:117167599-117168321 | -0.630835087 | 0.088883098 | 0.291699805 |
| chr12:71707450-71707650 | 0.855111563 | 0.08888528 | 0.291699805 |
| chr11:75503860-75504100 | 0.818818703 | 0.088889406 | 0.291699805 |

| | | | |
|---|---|---|---|
| chr20:53638071-53638271 | 0.584114237 | 0.088893499 | 0.291699805 |
| chr17:15893576-15893776 | -0.957346581 | 0.088895464 | 0.291699805 |
| chr17:63860076-63860336 | 0.737180861 | 0.088899804 | 0.291699805 |
| chrX:79641170-79641496 | -0.809248135 | 0.088901019 | 0.291699805 |
| chr15:73448266-73448626 | 0.750265871 | 0.088905787 | 0.29170296 |
| chr17:72183131-72183700 | 0.600748956 | 0.088923613 | 0.291748953 |
| chr13:34933555-34933851 | 0.756915419 | 0.088929426 | 0.291755535 |
| chr12:108939724-108939931 | 0.773217173 | 0.088939891 | 0.291776732 |
| chr22:17972326-17972570 | 0.841584998 | 0.088943503 | 0.291776732 |
| chr6:16754922-16755201 | 0.80211813 | 0.088965126 | 0.291829675 |
| chr1:31170670-31171216 | 0.56586634 | 0.08896726 | 0.291829675 |
| chr12:113048746-113048950 | 0.812101124 | 0.088994725 | 0.291907269 |
| chr2:86250042-86250357 | -0.718803541 | 0.089011761 | 0.291944959 |
| chr5:174610920-174611302 | 0.686284456 | 0.089013836 | 0.291944959 |
| chr8:80889997-80890311 | 0.765565132 | 0.089032172 | 0.291992599 |
| chr2:182014323-182014649 | -0.724564407 | 0.089037335 | 0.291997033 |
| chr1:151768413-151768626 | 0.718255434 | 0.089043509 | 0.292004782 |
| chr17:73929920-73930309 | -0.740440713 | 0.089058072 | 0.292027616 |
| chr3:106361400-106361854 | -0.723594765 | 0.089058094 | 0.292027616 |
| chr1:222124683-222124937 | -0.830398162 | 0.089065519 | 0.292029686 |
| chr5:442836-443226 | 0.620938286 | 0.089066348 | 0.292029686 |
| chr12:69920736-69921001 | -0.823125758 | 0.089071323 | 0.2920335 |
| chr11:108647789-108648616 | -0.600240082 | 0.089092131 | 0.292089225 |
| chr11:119692788-119693073 | -0.879873836 | 0.08910594 | 0.292121999 |
| chr11:36149634-36150286 | -0.597873671 | 0.089111429 | 0.292127492 |
| chr21:20443660-20443869 | -0.922421542 | 0.089127471 | 0.292167584 |
| chr11:63508076-63508276 | 0.801574531 | 0.0891427 | 0.292205005 |
| chr2:60555143-60555432 | -0.675997943 | 0.089157978 | 0.292242582 |
| chr2:111984267-111984479 | 0.887212994 | 0.089171661 | 0.292251794 |
| chr17:75436025-75436474 | -0.696607373 | 0.089180359 | 0.292251794 |
| chr17:38870413-38870674 | 0.706168383 | 0.089180707 | 0.292251794 |
| chr7:104967808-104968008 | -0.951373266 | 0.089187461 | 0.292251794 |
| chr3:155323089-155323388 | -0.710558267 | 0.08918843 | 0.292251794 |
| chr10:33150733-33151238 | -0.637650569 | 0.089188916 | 0.292251794 |
| chr8:91028447-91028752 | 0.737600008 | 0.089189967 | 0.292251794 |
| chr8:132899075-132899351 | 0.701096072 | 0.089191302 | 0.292251794 |
| chr7:127585262-127586017 | -0.554165871 | 0.089205319 | 0.292285224 |
| chr1:205563720-205564441 | -0.565363941 | 0.089215255 | 0.292305278 |
| chr11:84236513-84236762 | -0.870476048 | 0.089221111 | 0.292311968 |
| chr18:75161723-75162071 | -0.663704038 | 0.089248756 | 0.292352943 |
| chr13:60627236-60627436 | -0.899872666 | 0.089249577 | 0.292352943 |
| chr5:122500807-122501294 | -0.667914401 | 0.089251027 | 0.292352943 |
| chr13:36088088-36088327 | -0.864833454 | 0.089251903 | 0.292352943 |
| chr1:194813171-194813723 | -0.670547367 | 0.089252696 | 0.292352943 |
| chr1:236753396-236754255 | -0.582595619 | 0.089262994 | 0.29236899 |
| chr2:153497990-153498620 | -0.608800401 | 0.08926862 | 0.29236899 |
| chr20:47205331-47206164 | 0.533358816 | 0.089269042 | 0.29236899 |
| chr6:87213386-87213791 | -0.657760678 | 0.089282155 | 0.292399439 |
| chr10:37538350-37538685 | -0.72600483 | 0.089297622 | 0.292428703 |
| chr9:137482197-137482463 | 0.728038933 | 0.08930088 | 0.292428703 |

| | | | |
|---|---|---|---|
| chr14:72500502-72500830 | 0.680129635 | 0.08930254 | 0.292428703 |
| chr8:70273514-70273881 | -0.720994239 | 0.089335482 | 0.292524059 |
| chr3:40642047-40642459 | 0.792221502 | 0.089339295 | 0.292524059 |
| chr1:214270111-214270403 | -0.806967268 | 0.089357048 | 0.292552671 |
| chr7:102127094-102127374 | -0.97568376 | 0.089359488 | 0.292552671 |
| chr7:117750684-117750889 | -0.97568376 | 0.089359488 | 0.292552671 |
| chr14:49621446-49621836 | 0.615319802 | 0.089399066 | 0.29266974 |
| chr17:78769380-78769688 | 0.800773929 | 0.08941555 | 0.292711196 |
| chr22:42169006-42169284 | 0.742989618 | 0.089444732 | 0.292794216 |
| chr10:77866661-77866901 | -0.694520564 | 0.089451015 | 0.292802274 |
| chr1:195020010-195020272 | -0.872461954 | 0.089455182 | 0.292803406 |
| chr12:120474532-120474937 | 0.598188207 | 0.089460306 | 0.292807606 |
| chr19:4139746-4140060 | 0.688589108 | 0.089464108 | 0.292807606 |
| chr11:62906019-62906284 | 0.704150626 | 0.089476571 | 0.292835888 |
| chr15:35038728-35038975 | -0.854720266 | 0.089503711 | 0.292912201 |
| chr6:82614630-82614935 | -0.910508404 | 0.089542991 | 0.293025756 |
| chr1:208560313-208560565 | -0.892764823 | 0.089546058 | 0.293025756 |
| chr7:98878238-98879282 | -0.563336256 | 0.089574405 | 0.293105998 |
| chr4:167506953-167507153 | -0.960395006 | 0.089578845 | 0.293108008 |
| chr10:124693436-124693662 | -0.75373685 | 0.089584378 | 0.293113595 |
| chr6:131864051-131864260 | -0.895237176 | 0.089593292 | 0.293130247 |
| chr15:75578414-75578749 | 0.72809215 | 0.089598411 | 0.293134478 |
| chr5:33859274-33859743 | -0.745555413 | 0.089623642 | 0.293204504 |
| chr13:33285346-33286143 | -0.55246192 | 0.089631643 | 0.293218162 |
| chr9:89549136-89549371 | 0.803182589 | 0.089645071 | 0.293249567 |
| chr18:65206672-65206876 | -0.868440347 | 0.089689877 | 0.293382468 |
| chr9:107158373-107158703 | -0.729962361 | 0.089693356 | 0.293382468 |
| chr5:126600090-126601490 | -0.545108731 | 0.089743213 | 0.293521839 |
| chr9:76423923-76424148 | -0.817446924 | 0.089743626 | 0.293521839 |
| chr2:149587143-149588398 | -0.544776518 | 0.089749578 | 0.293528777 |
| chr6:37698987-37699226 | 0.755174398 | 0.089756939 | 0.29354032 |
| chr11:65471420-65471678 | 0.685223106 | 0.089763536 | 0.293549365 |
| chr7:30420288-30420498 | 0.860794132 | 0.08978533 | 0.293596846 |
| chr3:170730265-170730467 | 0.845021113 | 0.089785718 | 0.293596846 |
| chr8:117885499-117885740 | -0.875372899 | 0.089831196 | 0.293733021 |
| chr3:135935191-135935783 | -0.624290374 | 0.089849028 | 0.293778793 |
| chr9:105694154-105695470 | -0.545481583 | 0.089876596 | 0.293856392 |
| chr20:49773954-49774154 | 0.852810749 | 0.089893201 | 0.293898143 |
| chr17:61687702-61688542 | 0.539218103 | 0.089930947 | 0.294009004 |
| chr1:31937467-31938079 | 0.58725929 | 0.089950325 | 0.294054077 |
| chr7:76260270-76260668 | -0.649272635 | 0.089952409 | 0.294054077 |
| chr8:139832271-139832712 | 0.667234719 | 0.089964894 | 0.294082343 |
| chr14:87992412-87993696 | 0.551066639 | 0.089994383 | 0.294139559 |
| chr4:156294597-156294949 | -0.70150584 | 0.089994721 | 0.294139559 |
| chr4:132332986-132333243 | -0.813982495 | 0.089997978 | 0.294139559 |
| chr2:54864206-54864448 | -0.86921546 | 0.090000789 | 0.294139559 |
| chr8:82662462-82662905 | 0.678691863 | 0.090001592 | 0.294139559 |
| chr15:68888280-68888706 | 0.699057747 | 0.090011095 | 0.294158071 |
| chr9:14238043-14238383 | -0.726307408 | 0.090024873 | 0.294180174 |
| chr1:169070662-169070862 | 0.859806689 | 0.090027545 | 0.294180174 |

| | | | |
|---|---|---|---|
| chr2:85053878-85054131 | -0.771144862 | 0.090029377 | 0.294180174 |
| chr7:149753664-149753992 | -0.672292986 | 0.090043541 | 0.294213909 |
| chr9:78358349-78358637 | -0.812200714 | 0.090051176 | 0.294214979 |
| chr6:132728684-132728893 | -0.897457723 | 0.090051548 | 0.294214979 |
| chr15:90412668-90413154 | 0.639810848 | 0.090059666 | 0.294225207 |
| chr10:98536625-98536861 | 0.791831544 | 0.090062358 | 0.294225207 |
| chr8:71059050-71059763 | -0.598311993 | 0.090079231 | 0.294267782 |
| chr14:95560639-95561434 | 0.579872807 | 0.09010716 | 0.294308067 |
| chr2:101419116-101419365 | -0.766138607 | 0.090109773 | 0.294308067 |
| chr6:139027566-139028078 | -0.620559981 | 0.090112107 | 0.294308067 |
| chr14:93509711-93509911 | 0.963247346 | 0.090112997 | 0.294308067 |
| chr16:27314028-27314600 | 0.639964568 | 0.090113567 | 0.294308067 |
| chr2:175111479-175111726 | -0.742820813 | 0.090114609 | 0.294308067 |
| chr15:82647460-82647712 | 0.780846574 | 0.090120912 | 0.294316106 |
| chr2:106030605-106030887 | 0.73692243 | 0.090125276 | 0.294317816 |
| chr11:64246844-64247099 | 0.66774008 | 0.090155064 | 0.294402544 |
| chr1:152410203-152410605 | 0.637866706 | 0.090169275 | 0.294436402 |
| chr10:92689733-92690742 | 0.597002283 | 0.090179044 | 0.294452327 |
| chrX:77937403-77937629 | 0.838686562 | 0.090181838 | 0.294452327 |
| chr12:132300799-13230103€ | 0.853165723 | 0.090193169 | 0.294476776 |
| chr13:28684209-28684938 | -0.579949289 | 0.090201066 | 0.294490012 |
| chr2:61016057-61016257 | 0.724710754 | 0.090218158 | 0.294523326 |
| chr17:80160122-80160322 | 0.932116434 | 0.090218958 | 0.294523326 |
| chr13:49628285-49628648 | -0.673725062 | 0.090223848 | 0.294526741 |
| chr14:50550379-50550605 | 0.798815021 | 0.090232116 | 0.294541184 |
| chr6:131652544-131652947 | 0.703865697 | 0.090241639 | 0.294552513 |
| chr4:165112195-165113498 | -0.533974146 | 0.090245409 | 0.294552513 |
| chr18:10726304-10726912 | -0.712361033 | 0.090252886 | 0.294552513 |
| chr12:48924609-48925281 | -0.604341264 | 0.090253987 | 0.294552513 |
| chr11:84217739-84217962 | -0.929097398 | 0.090255911 | 0.294552513 |
| chr5:66742180-66742434 | -0.71596916 | 0.090258652 | 0.294552513 |
| chr5:55173155-55173675 | -0.614064631 | 0.090273975 | 0.294589972 |
| chr17:17810053-17810743 | -0.589554215 | 0.090279206 | 0.294594493 |
| chr2:152925647-152926421 | -0.58780634 | 0.090292353 | 0.294624848 |
| chr20:50998969-50999253 | 0.742789505 | 0.090306542 | 0.294658599 |
| chr1:177460516-177460785 | 0.798483182 | 0.090320623 | 0.294691993 |
| chr10:101307801-10130806€ | -0.779992716 | 0.090325383 | 0.294694975 |
| chr20:47624413-47624671 | 0.57792623 | 0.090342104 | 0.294736979 |
| chr16:4945585-4946178 | 0.567855965 | 0.090353555 | 0.294761786 |
| chr13:39262977-39263294 | -0.643997686 | 0.090366361 | 0.294791013 |
| chr14:38056265-38056543 | 0.798884024 | 0.090384684 | 0.294828016 |
| chr7:34057760-34058262 | -0.61516271 | 0.0903854 | 0.294828016 |
| chr16:4299183-4299675 | 0.616882851 | 0.090406225 | 0.294872824 |
| chr6:35952001-35952226 | 0.891069171 | 0.090410206 | 0.294872824 |
| chr10:44761162-44761388 | 0.879525086 | 0.090410682 | 0.294872824 |
| chrX:74133475-74133693 | 0.847967151 | 0.090422692 | 0.294899442 |
| chr17:1966596-1967671 | -0.570073094 | 0.090433387 | 0.294921771 |
| chr6:72953954-72954433 | -0.665005973 | 0.09044127 | 0.294934923 |
| chr3:193190422-193190813 | -0.636437407 | 0.09046319 | 0.294993852 |
| chr11:7986715-7987549 | -0.556340791 | 0.090472319 | 0.295011067 |

| | | | |
|---|---|---|---|
| chr7:22351339-22351646 | 0.677016865 | 0.090480058 | 0.295023746 |
| chr2:237660916-237661127 | 0.882955605 | 0.090508234 | 0.295054509 |
| chr8:101493714-101494531 | 0.543887282 | 0.090509428 | 0.295054509 |
| chr15:43492801-43493534 | -0.594013361 | 0.090511852 | 0.295054509 |
| chr7:142739992-142740192 | -1.000646103 | 0.090512445 | 0.295054509 |
| chr12:124518177-124519145 | 0.568192157 | 0.090512604 | 0.295054509 |
| chr3:51706821-51707100 | 0.699890888 | 0.090513262 | 0.295054509 |
| chrX:135112211-135112437 | 0.746772288 | 0.090516448 | 0.295054509 |
| chr1:108962667-108962868 | 0.73025585 | 0.090522252 | 0.295060876 |
| chr11:120303058-120303284 | 0.749541282 | 0.090527449 | 0.295065261 |
| chr16:56729847-56730357 | 0.596283908 | 0.090546349 | 0.295114313 |
| chr3:154126706-154127129 | -0.61003914 | 0.090576228 | 0.295191289 |
| chr7:130897002-130897301 | -0.676788861 | 0.090580087 | 0.295191289 |
| chr2:11777766-11778088 | 0.760019623 | 0.090581525 | 0.295191289 |
| chr7:65251883-65252581 | 0.619430272 | 0.09060801 | 0.295262413 |
| chr1:209875579-209875831 | -0.747568073 | 0.090611057 | 0.295262413 |
| chr5:161611744-161612147 | -0.758260359 | 0.090625294 | 0.295296248 |
| chr20:13538650-13538912 | -0.937202998 | 0.090631563 | 0.295304118 |
| chr4:135319768-135319997 | -0.850182672 | 0.090642333 | 0.295326651 |
| chr1:147358148-147358416 | 0.771747236 | 0.090659409 | 0.295358087 |
| chr4:169070816-169071148 | -0.59145492 | 0.09066065 | 0.295358087 |
| chr2:106997369-106997646 | 0.795551707 | 0.090663546 | 0.295358087 |
| chr14:32013140-32013340 | 0.877504104 | 0.090672925 | 0.295376083 |
| chr2:16718890-16719348 | -0.68008532 | 0.09070363 | 0.295463546 |
| chr8:107419252-107419452 | 0.833985643 | 0.090716672 | 0.295488735 |
| chr10:90871310-90872659 | -0.5461708 | 0.090719075 | 0.295488735 |
| chrX:151082124-151082903 | 0.590597613 | 0.090725194 | 0.295496102 |
| chr5:73030504-73030840 | 0.710003627 | 0.090734678 | 0.295514431 |
| chr1:84604403-84604791 | -0.721643851 | 0.090740834 | 0.295521918 |
| chr5:91101774-91102131 | -0.64194278 | 0.090758575 | 0.295567135 |
| chr17:82973535-82973874 | 0.676903997 | 0.090784983 | 0.29564057 |
| chr1:60573117-60573437 | 0.748894122 | 0.090792538 | 0.295652609 |
| chr5:139651503-139651703 | -0.723700998 | 0.090804049 | 0.295660198 |
| chr16:2148079-2148694 | 0.580303009 | 0.090807989 | 0.295660198 |
| chr20:58236668-58237374 | 0.564132692 | 0.090808725 | 0.295660198 |
| chr13:26067554-26067754 | 0.839893017 | 0.090811346 | 0.295660198 |
| chrX:103710792-103711845 | -0.55475625 | 0.090814162 | 0.295660198 |
| chr1:198699259-198699939 | -0.569519024 | 0.090821809 | 0.295672528 |
| chr2:72893626-72894169 | -0.595443313 | 0.090835075 | 0.295703155 |
| chr7:143737059-143737368 | -0.714817273 | 0.090841406 | 0.2957112 |
| chr3:159763527-159764974 | 0.545563488 | 0.090866443 | 0.295780135 |
| chr1:5992065-5993457 | 0.544289507 | 0.090875798 | 0.295798022 |
| chr2:62305033-62305343 | -0.652259757 | 0.09088514 | 0.295815862 |
| chr1:154878526-154878786 | 0.700273676 | 0.090892074 | 0.295825864 |
| chr4:184287638-184288155 | -0.676592725 | 0.090906382 | 0.295859864 |
| chr20:53444397-53444606 | 0.736633275 | 0.090928603 | 0.295919616 |
| chr6:135211129-135211749 | -0.629522628 | 0.090941249 | 0.2959482 |
| chr16:75059493-75059760 | 0.739119176 | 0.090947959 | 0.295957467 |
| chr15:60127667-60127959 | -0.758616167 | 0.090954891 | 0.295960278 |
| chr1:205659403-205659748 | 0.617264933 | 0.09095752 | 0.295960278 |

| | | | |
|---|---|---|---|
| chrX:138026030-138026252 | 0.836685557 | 0.090960411 | 0.295960278 |
| chr16:3690489-3690778 | 0.672232183 | 0.090968557 | 0.295974217 |
| chr12:106178590-106178968 | -0.671926819 | 0.09097554 | 0.295984368 |
| chr6:69812735-69813358 | 0.63003559 | 0.09100635 | 0.296072034 |
| chr7:12403217-12404750 | 0.532621177 | 0.091019779 | 0.296103151 |
| chr9:119289733-119290184 | -0.688211097 | 0.091037784 | 0.296138335 |
| chr11:62786472-62786864 | 0.590488306 | 0.091038324 | 0.296138335 |
| chr18:35122648-35123195 | -0.602994722 | 0.091045657 | 0.296149614 |
| chr14:73633482-73633989 | 0.610013667 | 0.09109842 | 0.29630866 |
| chr14:77603854-77604272 | -0.614175651 | 0.091135269 | 0.296415936 |
| chr11:26649828-26650109 | -0.771430231 | 0.091150715 | 0.296453588 |
| chr6:43667278-43667715 | 0.627918729 | 0.091164625 | 0.296482657 |
| chr8:131936262-131936626 | -0.737752653 | 0.091167392 | 0.296482657 |
| chr10:1800963-1801210 | 0.857271246 | 0.091177499 | 0.296502942 |
| chr15:72237402-72237651 | 0.767786001 | 0.091181762 | 0.296504222 |
| chr8:89321296-89321496 | 0.849474894 | 0.091188213 | 0.296512616 |
| chr20:57143100-57143670 | 0.538797404 | 0.091196691 | 0.2965276 |
| chr13:31234772-31235104 | -0.693061172 | 0.091205879 | 0.296542262 |
| chr19:5132216-5132668 | 0.614239436 | 0.091208941 | 0.296542262 |
| chr2:173027531-173027809 | 0.731861678 | 0.091227332 | 0.296589471 |
| chr10:93751937-93752153 | -0.822513105 | 0.091238824 | 0.296614245 |
| chr17:56392189-56392826 | -0.634549318 | 0.091244337 | 0.296617979 |
| chr5:107542941-107543233 | -0.753716245 | 0.091247715 | 0.296617979 |
| chr2:75321704-75321904 | -0.935551453 | 0.091266936 | 0.296652135 |
| chr14:64511122-64511477 | 0.624558618 | 0.091270188 | 0.296652135 |
| chr5:127572897-127573291 | -0.70339661 | 0.091271435 | 0.296652135 |
| chr2:109449494-109449736 | 0.737111078 | 0.091273708 | 0.296652135 |
| chr11:112163918-112164210 | -0.882829519 | 0.091285257 | 0.296666452 |
| chr17:80706419-80706662 | 0.782116379 | 0.091285857 | 0.296666452 |
| chr3:193554672-193554981 | -0.824697312 | 0.091300593 | 0.296701756 |
| chr19:29276299-29276633 | 0.868059499 | 0.091307999 | 0.296713239 |
| chr8:85583052-85583323 | 0.720538342 | 0.091323459 | 0.296750895 |
| chr14:56339936-56340405 | 0.648478347 | 0.091328954 | 0.296756163 |
| chr20:63069909-63070130 | 0.709842521 | 0.091337257 | 0.296770559 |
| chr7:116625573-116625802 | -0.960634611 | 0.091345471 | 0.296784663 |
| chr1:119968082-119968321 | 0.707769818 | 0.091375762 | 0.296852069 |
| chr19:45862531-45862731 | 0.734093041 | 0.091376136 | 0.296852069 |
| chr1:84122897-84123097 | 0.972778468 | 0.091379309 | 0.296852069 |
| chr10:34836292-34836564 | -0.82813626 | 0.091381715 | 0.296852069 |
| chr17:61640962-61641221 | 0.57622422 | 0.091398561 | 0.296894205 |
| chr17:8965430-8965875 | -0.686684024 | 0.091427268 | 0.296945669 |
| chr14:42911492-42912009 | 0.637175223 | 0.09142808 | 0.296945669 |
| chr12:120265706-120266178 | 0.595327567 | 0.091436377 | 0.296945669 |
| chr1:189662329-189662575 | -0.800693278 | 0.091436618 | 0.296945669 |
| chr10:30611879-30612273 | -0.717545159 | 0.09143675 | 0.296945669 |
| chr17:58096999-58097203 | -0.785849961 | 0.091437657 | 0.296945669 |
| chr1:112242826-112243140 | -0.831995684 | 0.091453228 | 0.296981547 |
| chr14:69107492-69107745 | 0.721997477 | 0.091459918 | 0.296981547 |
| chr8:107302242-107302932 | 0.605654741 | 0.091460332 | 0.296981547 |
| chr8:67777406-67777772 | -0.772453318 | 0.091466481 | 0.296986857 |

| | | | |
|---|---|---|---|
| chr1:204460327-204460583 | 0.696167758 | 0.091469719 | 0.296986857 |
| chr5:166710368-166710638 | -0.822111639 | 0.091479689 | 0.297006643 |
| chr17:55470698-55470938 | -0.826882933 | 0.091492234 | 0.297034785 |
| chr10:18594652-18595163 | -0.652661053 | 0.091499814 | 0.297035822 |
| chr2:14555376-14555638 | 0.754045195 | 0.091500307 | 0.297035822 |
| chr10:79726695-79726975 | -0.836915734 | 0.091505499 | 0.297037065 |
| chr8:143665071-143665315 | 0.832223912 | 0.091508443 | 0.297037065 |
| chr3:124298889-124299272 | 0.668122835 | 0.091517267 | 0.297046076 |
| chr5:145107245-145107548 | -0.889736899 | 0.091518973 | 0.297046076 |
| chr1:236452044-236452360 | -0.680181796 | 0.091526358 | 0.297057465 |
| chr16:78223516-78223716 | 0.84269843 | 0.091533666 | 0.297068599 |
| chr6:151616688-151617248 | -0.636892338 | 0.091548593 | 0.297088783 |
| chr20:47668297-47668816 | 0.53986011 | 0.091551416 | 0.297088783 |
| chr1:30369194-30369534 | 0.794643375 | 0.091553378 | 0.297088783 |
| chr3:17478698-17478959 | 0.768428062 | 0.091555395 | 0.297088783 |
| chr3:64867515-64868094 | 0.557640207 | 0.091569645 | 0.297122443 |
| chr1:15703099-15703380 | -0.814742637 | 0.091583164 | 0.297153725 |
| chr7:139430815-139431412 | -0.596966664 | 0.091616398 | 0.29724135 |
| chr15:90319995-90320425 | 0.588821153 | 0.091617929 | 0.29724135 |
| chr17:61525118-61525474 | 0.561412273 | 0.091621935 | 0.297241761 |
| chr1:234316879-234317184 | 0.786448286 | 0.091635798 | 0.297268358 |
| chr12:125183173-125183663 | 0.602547582 | 0.091637893 | 0.297268358 |
| chr3:63149913-63150480 | 0.586217612 | 0.091652645 | 0.297303626 |
| chr5:154467540-154468199 | -0.609496386 | 0.091675366 | 0.297352467 |
| chr2:130262302-130262737 | -0.67199338 | 0.091675463 | 0.297352467 |
| chr1:235253187-235253523 | 0.656754121 | 0.09170413 | 0.29742054 |
| chr6:119110548-119111079 | -0.642197201 | 0.091704214 | 0.29742054 |
| chr1:227787890-227788648 | 0.545272974 | 0.091713938 | 0.297439489 |
| chr7:36746925-36747268 | -0.786710878 | 0.091770079 | 0.297599054 |
| chr10:80283166-80283393 | -0.82417844 | 0.09177454 | 0.297599054 |
| chr10:42975602-42975962 | -0.654669697 | 0.09177841 | 0.297599054 |
| chr14:47333009-47333254 | -0.881501419 | 0.091780793 | 0.297599054 |
| chr5:1801294-1801553 | 0.656228964 | 0.09178932 | 0.297599054 |
| chr15:29348395-29348595 | -0.934065683 | 0.091791763 | 0.297599054 |
| chr11:21814434-21814643 | -0.921097139 | 0.091792453 | 0.297599054 |
| chr17:60262535-60262854 | 0.592254973 | 0.091794385 | 0.297599054 |
| chr9:92624662-92624999 | -0.801949478 | 0.091798095 | 0.297599054 |
| chr5:143051627-143051923 | -0.809955497 | 0.091844287 | 0.297722682 |
| chr2:232550368-232551212 | 0.565593778 | 0.091846799 | 0.297722682 |
| chr16:56332697-56333022 | 0.680311428 | 0.091847886 | 0.297722682 |
| chr19:48858664-48858910 | 0.753366153 | 0.091868649 | 0.297777385 |
| chr4:174870878-174871152 | -0.825245433 | 0.091885514 | 0.297812838 |
| chr4:74038517-74039333 | 0.593988828 | 0.091887997 | 0.297812838 |
| chr19:46514969-46515174 | -0.863199171 | 0.091891247 | 0.297812838 |
| chr10:98361465-98361679 | -0.826751569 | 0.091918917 | 0.297889915 |
| chr13:54162129-54162400 | -0.801503348 | 0.091936445 | 0.297934119 |
| chr12:91618378-91618679 | -0.745859621 | 0.091943937 | 0.297945795 |
| chr1:114795490-114795766 | 0.641281698 | 0.091972719 | 0.29802646 |
| chr12:104592169-104592574 | -0.823991746 | 0.091992076 | 0.298064783 |
| chr2:23351497-23351896 | -0.621654159 | 0.091992326 | 0.298064783 |

| | | | |
|---|---|---|---|
| chr10:118522648-118523098 | -0.678445686 | 0.092003175 | 0.29808733 |
| chr8:29655338-29655752 | 0.652443911 | 0.092013299 | 0.298101749 |
| chr2:29233467-29233840 | 0.660069682 | 0.092015406 | 0.298101749 |
| chr1:33349697-33350219 | 0.578589228 | 0.092023387 | 0.298114998 |
| chr10:109800919-109801511 | -0.681278074 | 0.092032647 | 0.298132394 |
| chr10:106565342-106565919 | -0.610794198 | 0.092038995 | 0.298134136 |
| chr10:8054251-8054873 | 0.546103559 | 0.092041527 | 0.298134136 |
| chr20:23420395-23420711 | 0.658806335 | 0.092044861 | 0.298134136 |
| chr7:24715817-24716150 | 0.615555858 | 0.092051247 | 0.298134136 |
| chrX:149503536-149503935 | -0.598525881 | 0.09205264 | 0.298134136 |
| chr19:11035957-11036248 | 0.676545819 | 0.092062556 | 0.29813753 |
| chr5:34192243-34192646 | 0.627225052 | 0.092068928 | 0.29813753 |
| chr8:67761317-67761641 | -0.686015207 | 0.092069946 | 0.29813753 |
| chr14:68146854-68147126 | -0.896901708 | 0.09207136 | 0.29813753 |
| chr5:17330490-17330717 | 0.736757193 | 0.092073143 | 0.29813753 |
| chr13:98984174-98984427 | 0.827488023 | 0.092083746 | 0.298159263 |
| chr13:52967981-52968609 | -0.587865329 | 0.09210609 | 0.298219009 |
| chr10:72261238-72261438 | -0.834413116 | 0.092117901 | 0.298222092 |
| chr11:34155671-34155875 | -0.834413116 | 0.092117901 | 0.298222092 |
| chr14:104800287-104800655 | 0.613401495 | 0.092118719 | 0.298222092 |
| chr5:131263199-131264792 | -0.536128034 | 0.0921361 | 0.298254175 |
| chr20:54306908-54307129 | 0.603179841 | 0.092139423 | 0.298254175 |
| chr19:34712746-34712983 | 0.892167415 | 0.092140307 | 0.298254175 |
| chr6:126339843-126340647 | -0.553804983 | 0.092170133 | 0.298338119 |
| chr11:61792042-61793188 | 0.548896502 | 0.092198921 | 0.298418692 |
| chr19:29714745-29715639 | 0.566217545 | 0.092205539 | 0.298423142 |
| chr2:36658547-36658779 | -0.845254894 | 0.092208636 | 0.298423142 |
| chr2:50476266-50476499 | -0.87470541 | 0.09221198 | 0.298423142 |
| chr20:57135920-57136120 | 0.576937639 | 0.092264621 | 0.298574948 |
| chr8:123258106-123258383 | -0.739339089 | 0.092266681 | 0.298574948 |
| chr2:69565369-69565653 | 0.691325673 | 0.092279668 | 0.298599491 |
| chr19:52527092-52527881 | 0.560444398 | 0.092282059 | 0.298599491 |
| chr3:194263918-194264226 | -0.74882271 | 0.092287522 | 0.298604557 |
| chr1:30119031-30119284 | 0.931586152 | 0.092317707 | 0.29868236 |
| chr9:2621805-2622560 | 0.613052605 | 0.092327336 | 0.29868236 |
| chr20:53691220-53692173 | 0.528490547 | 0.092328957 | 0.29868236 |
| chr4:82395037-82395237 | -0.950882622 | 0.092331339 | 0.29868236 |
| chr9:87336928-87337149 | 0.796448837 | 0.092336627 | 0.29868236 |
| chrX:130165318-130166489 | 0.536856618 | 0.092338971 | 0.29868236 |
| chr10:11782877-11783144 | 0.757518237 | 0.092341018 | 0.29868236 |
| chr20:53462860-53463096 | 0.806778546 | 0.092342753 | 0.29868236 |
| chr13:58141907-58142135 | -1.026760436 | 0.09236226 | 0.298732843 |
| chr10:112756158-112756490 | 0.707406479 | 0.092368894 | 0.298741691 |
| chr1:113773571-113773837 | 0.745332655 | 0.092399763 | 0.298828916 |
| chr1:24201965-24202395 | 0.712178728 | 0.092414417 | 0.298855393 |
| chr8:106245683-106246013 | -0.755600424 | 0.092416447 | 0.298855393 |
| chr12:15881830-15883028 | -0.540464776 | 0.092419652 | 0.298855393 |
| chr3:132161687-132161887 | -0.934017055 | 0.092434318 | 0.298882373 |
| chr3:2810866-2811066 | 0.894872545 | 0.092435796 | 0.298882373 |
| chr12:22437080-22437794 | -0.62681877 | 0.09246514 | 0.298964387 |

| | | | |
|---|---|---|---|
| chr19:40614608-40615174 | 0.568671758 | 0.092469769 | 0.298964387 |
| chr1:225917520-225917796 | -0.759213014 | 0.092472867 | 0.298964387 |
| chr14:32307918-32308561 | 0.581353558 | 0.092492778 | 0.299016144 |
| chr12:31728476-31729528 | -0.542262848 | 0.0925146 | 0.299074072 |
| chr3:186540549-186540884 | -0.708436824 | 0.092529418 | 0.299109355 |
| chr5:9971644-9971912 | -0.832460648 | 0.09255069 | 0.299165498 |
| chrX:39900841-39901112 | -0.817146591 | 0.092558348 | 0.299177632 |
| chr16:30984943-30985291 | 0.628528277 | 0.092574768 | 0.299218084 |
| chr1:107187513-107187869 | 0.590024195 | 0.0925938 | 0.29926566 |
| chrX:107493265-107493471 | -0.818968605 | 0.092597299 | 0.29926566 |
| chr11:126925758-126926087 | -0.742894752 | 0.09260678 | 0.299283676 |
| chr7:128106873-128107155 | -0.696564324 | 0.092614571 | 0.299287757 |
| chr7:1851639-1851954 | 0.627656209 | 0.092615855 | 0.299287757 |
| chr17:35567774-35568242 | 0.590146114 | 0.092629207 | 0.299318281 |
| chr7:110774162-110774397 | -0.830921222 | 0.092636299 | 0.299328576 |
| chr12:31326000-31326493 | -0.575676185 | 0.092650536 | 0.299361952 |
| chr12:10212140-10213040 | -0.557455017 | 0.092655662 | 0.299365891 |
| chr19:18417460-18417833 | 0.686000949 | 0.09266306 | 0.299377169 |
| chr9:78704248-78704506 | -0.823414375 | 0.092682114 | 0.299426104 |
| chr15:39273841-39274494 | 0.564467346 | 0.092688415 | 0.299433836 |
| chr1:190860083-190860540 | -0.627304345 | 0.09269303 | 0.299436122 |
| chr1:101258007-101258277 | -0.813109777 | 0.09270594 | 0.299457134 |
| chr16:27742432-27742713 | 0.719831099 | 0.092714413 | 0.299457134 |
| chr1:58809245-58809621 | -0.676765153 | 0.092714561 | 0.299457134 |
| chr15:44591444-44591644 | 0.903918558 | 0.092718333 | 0.299457134 |
| chr6:118861919-118862145 | 0.905484636 | 0.092719076 | 0.299457134 |
| chr20:47860312-47860916 | 0.54659496 | 0.09273396 | 0.299470321 |
| chr19:33373115-33374217 | 0.554725195 | 0.092737627 | 0.299470321 |
| chr4:159192088-159192350 | -0.828279854 | 0.092737659 | 0.299470321 |
| chr9:137263270-137263632 | -0.774274119 | 0.092738926 | 0.299470321 |
| chr7:87875817-87877203 | -0.535326568 | 0.092742701 | 0.299470321 |
| chr10:33010088-33010779 | -0.596091726 | 0.092747597 | 0.299473509 |
| chr6:62150015-62150344 | -0.737708971 | 0.092784765 | 0.299579503 |
| chr3:19919064-19919370 | -0.838571958 | 0.092791254 | 0.299579503 |
| chr10:46786676-46787163 | -0.635390345 | 0.092792153 | 0.299579503 |
| chr3:52005571-52005845 | -0.799425508 | 0.092803182 | 0.299602488 |
| chr17:71764527-71764883 | -0.655906326 | 0.092811468 | 0.299615277 |
| chr8:127775829-127776040 | 0.764324847 | 0.092814965 | 0.299615277 |
| chr13:33761850-33762079 | -0.8484903 | 0.092830748 | 0.299653604 |
| chr14:100271302-10027160C | 0.762622212 | 0.092838978 | 0.29966423 |
| chr13:18874867-18875236 | -0.66063612 | 0.092842139 | 0.29966423 |
| chr8:73330587-73330916 | 0.744398689 | 0.092848407 | 0.29966423 |
| chr19:45799061-45799450 | 0.650219534 | 0.092849684 | 0.29966423 |
| chr19:1812240-1813226 | 0.547377317 | 0.092881258 | 0.299753506 |
| chr12:25690054-25690536 | -0.681769203 | 0.092885808 | 0.299755564 |
| chr12:124906743-124907098 | 0.629006254 | 0.092904151 | 0.299802132 |
| chr5:156584736-156585029 | -0.823846553 | 0.09291833 | 0.299835261 |
| chr7:141073910-141074306 | 0.638253563 | 0.092926235 | 0.299845385 |
| chr3:53846186-53846831 | 0.594958294 | 0.092931305 | 0.299845385 |
| chr16:78002956-78003423 | 0.566752034 | 0.0929358 | 0.299845385 |

| | | | |
|---|---|---|---|
| chr14:76947053-76947835 | 0.560572707 | 0.09293712 | 0.299845385 |
| chr2:112475037-112475278 | 0.79781665 | 0.092952408 | 0.299880778 |
| chr17:65633627-65633827 | -0.767489019 | 0.092955918 | 0.299880778 |
| chr20:54622967-54623643 | 0.543374763 | 0.0929676 | 0.299905836 |
| chr6:125906230-125906766 | -0.622237904 | 0.093003783 | 0.300009928 |
| chr20:50278488-50279287 | -0.56705904 | 0.093010838 | 0.300020058 |
| chr3:115168104-115168416 | 0.737605935 | 0.093029402 | 0.300067305 |
| chr6:35461332-35461566 | 0.749553579 | 0.093044573 | 0.300103607 |
| chr7:76423293-76423919 | -0.59344798 | 0.093050309 | 0.300109474 |
| chr2:216792478-216792695 | 0.810192353 | 0.09305692 | 0.300118164 |
| chr8:103694424-103694688 | 0.728852911 | 0.093072902 | 0.300157074 |
| chr4:159483840-159484040 | -0.856768429 | 0.093115432 | 0.30027651 |
| chr21:42691805-42692005 | 0.724652002 | 0.093117775 | 0.30027651 |
| chr4:35541696-35542394 | -0.643482883 | 0.09312582 | 0.300289815 |
| chr17:62084118-62084380 | 0.702669493 | 0.093132262 | 0.30029795 |
| chr6:75449767-75450224 | 0.690371422 | 0.093141666 | 0.300315633 |
| chr6:142146911-142147647 | -0.545409738 | 0.09319459 | 0.300473631 |
| chr8:69466542-69467127 | -0.634429161 | 0.093218893 | 0.300539342 |
| chr3:156435089-156435297 | -0.859123494 | 0.0932382 | 0.300588942 |
| chr5:179139213-179139415 | -0.920772475 | 0.093256104 | 0.300634011 |
| chr12:120529610-120529897 | 0.716344784 | 0.093285152 | 0.30071046 |
| chr5:42723483-42723710 | 0.910378479 | 0.093287667 | 0.30071046 |
| chr2:187658960-187659450 | -0.632087249 | 0.093303656 | 0.300749348 |
| chr5:43073755-43074226 | -0.616428377 | 0.093310179 | 0.300757721 |
| chr10:80684277-80684605 | -0.622034388 | 0.09331606 | 0.300764025 |
| chr20:33933530-33933730 | 0.854435661 | 0.09332572 | 0.300771696 |
| chr3:100525596-100526303 | -0.58834932 | 0.093326291 | 0.300771696 |
| chr8:144020542-144020886 | 0.661156919 | 0.09334207 | 0.300799063 |
| chr21:36959448-36959787 | -0.733860808 | 0.093342634 | 0.300799063 |
| chr7:151539121-151539334 | 0.848261678 | 0.093353092 | 0.300820113 |
| chr4:153222265-153223375 | 0.564355072 | 0.093362039 | 0.300830262 |
| chr5:175464691-175465040 | -0.657537806 | 0.093364094 | 0.300830262 |
| chr17:62629375-62629617 | 0.629148443 | 0.093372204 | 0.300843743 |
| chr12:83584302-83584571 | -0.92864406 | 0.093394533 | 0.300903032 |
| chr3:169754762-169755086 | -0.718835265 | 0.093399608 | 0.300906729 |
| chr6:35415078-35415627 | 0.593301381 | 0.093447088 | 0.301047038 |
| chr5:154287147-154287481 | 0.660307425 | 0.093463399 | 0.301086926 |
| chr1:213374366-213374807 | -0.699612334 | 0.093474323 | 0.301102743 |
| chr1:188355142-188355388 | -0.82625514 | 0.093476168 | 0.301102743 |
| chr3:144323842-144324418 | -0.582672453 | 0.093492846 | 0.301131324 |
| chr5:115047271-115047471 | -0.942650908 | 0.093492901 | 0.301131324 |
| chr1:42206170-42206526 | 0.673401219 | 0.093515207 | 0.301183478 |
| chr17:9729081-9729318 | -0.786331452 | 0.093516955 | 0.301183478 |
| chr2:161761217-161761519 | 0.779318195 | 0.093521326 | 0.301184895 |
| chr14:40494570-40494790 | 0.826791576 | 0.093529745 | 0.301199348 |
| chr21:44939800-44940247 | 0.606749516 | 0.093543467 | 0.301230878 |
| chr1:107689248-107689567 | -0.623890844 | 0.09355238 | 0.301234117 |
| chr2:42844352-42844619 | 0.75020858 | 0.093553704 | 0.301234117 |
| chr15:61834172-61834395 | 0.756908843 | 0.093556267 | 0.301234117 |
| chr13:25509937-25510137 | 0.87493788 | 0.093565877 | 0.301252401 |

| | | | |
|---|---|---|---|
| chrX:40474086-40474470 | 0.603710636 | 0.093574588 | 0.301267787 |
| chr7:16299940-16300140 | 0.83279664 | 0.093588609 | 0.30130027 |
| chr2:201233871-201234183 | -0.672114953 | 0.093617084 | 0.30137928 |
| chr9:69120480-69122511 | -0.536890224 | 0.093646943 | 0.301461469 |
| chr2:224849372-224849672 | -0.768856792 | 0.093653517 | 0.301461469 |
| chr5:132556607-132557349 | -0.556980549 | 0.093654418 | 0.301461469 |
| chr1:101290866-101291066 | -0.848499354 | 0.093661208 | 0.301470661 |
| chr14:66198504-66198723 | 0.906646982 | 0.093690147 | 0.301533038 |
| chr7:130356335-130356711 | -0.700780584 | 0.093690836 | 0.301533038 |
| chr17:60767790-60768371 | 0.541489368 | 0.093692393 | 0.301533038 |
| chr8:81551669-81551930 | -0.742885978 | 0.093723267 | 0.30161973 |
| chr7:12067213-12067851 | 0.591679851 | 0.093733994 | 0.301635114 |
| chr18:68152791-68153140 | -0.659298034 | 0.09373592 | 0.301635114 |
| chr11:123560601-123560885 | 0.792633017 | 0.093741101 | 0.301639117 |
| chr8:94357657-94357998 | 0.645817884 | 0.093778571 | 0.301743963 |
| chr9:133782576-133782776 | 0.857258742 | 0.093785123 | 0.301743963 |
| chr20:53718178-53718440 | 0.560406874 | 0.093785499 | 0.301743963 |
| chr11:9364213-9364846 | 0.569559358 | 0.093795856 | 0.301757905 |
| chr5:2493786-2494012 | 0.860819122 | 0.093797708 | 0.301757905 |
| chr21:9367209-9367739 | 0.61021058 | 0.093804995 | 0.301768678 |
| chr7:43778565-43779210 | -0.586195843 | 0.093829278 | 0.301834122 |
| chr18:26655925-26656426 | 0.645951708 | 0.093837983 | 0.301849451 |
| chr2:11506134-11506382 | -0.800616409 | 0.093845498 | 0.301860954 |
| chr20:54362252-54362489 | 0.596586671 | 0.093862462 | 0.301902843 |
| chrX:30652535-30653881 | -0.53115607 | 0.093875324 | 0.301917024 |
| chr2:201680755-201681079 | -0.72413269 | 0.093875372 | 0.301917024 |
| chr4:169757146-169758220 | -0.547070323 | 0.093878692 | 0.301917024 |
| chr5:34526785-34527234 | -0.651616052 | 0.093883277 | 0.301919097 |
| chr22:23893873-23895676 | 0.532177461 | 0.093910248 | 0.301971882 |
| chr4:26320470-26321065 | 0.555013472 | 0.093910755 | 0.301971882 |
| chr4:73671390-73671671 | -0.794458793 | 0.093916802 | 0.301971882 |
| chr1:110356453-110356804 | 0.677631069 | 0.093916842 | 0.301971882 |
| chr7:40680651-40680979 | -0.664251684 | 0.093919396 | 0.301971882 |
| chr8:69619996-69620294 | -0.728040444 | 0.093931153 | 0.301997013 |
| chr4:113411491-113411902 | -0.678030894 | 0.093954273 | 0.302058672 |
| chr14:68937806-68938029 | 0.690769099 | 0.093965646 | 0.302078358 |
| chr3:144904825-144905056 | 0.775736004 | 0.093968281 | 0.302078358 |
| chr6:53734971-53735367 | -0.696945043 | 0.093998039 | 0.302161341 |
| chr4:56544593-56545108 | 0.600217172 | 0.094005626 | 0.302172074 |
| chr5:134591377-134591642 | -0.677276518 | 0.094009265 | 0.302172074 |
| chr12:102816881-102817342 | -0.673044002 | 0.094013392 | 0.302172664 |
| chr7:151091463-151091703 | -0.761709597 | 0.094029657 | 0.302212263 |
| chr3:158991203-158991505 | -0.66172287 | 0.094038293 | 0.302219088 |
| chr14:67964429-67964726 | 0.755771332 | 0.094040013 | 0.302219088 |
| chr4:177376138-177376382 | -0.854370722 | 0.094045544 | 0.302219088 |
| chr19:39474211-39474444 | 0.736720174 | 0.094047557 | 0.302219088 |
| chr14:60514788-60515239 | -0.680728615 | 0.094072771 | 0.302287434 |
| chr9:100537530-100537868 | -0.77604059 | 0.094100869 | 0.302353313 |
| chr12:80632833-80633141 | -0.723570793 | 0.094103878 | 0.302353313 |
| chr1:172499549-172500226 | -0.55730847 | 0.094105111 | 0.302353313 |

| | | | |
|---|---|---|---|
| chr1:223683898-223684487 | 0.623476697 | 0.094115283 | 0.302373317 |
| chr14:35404524-35405119 | 0.549371121 | 0.094155602 | 0.302490169 |
| chr10:48295520-48296004 | 0.669940643 | 0.094169811 | 0.302519734 |
| chr20:53853044-53853516 | 0.541292673 | 0.094175241 | 0.302519734 |
| chr15:72319659-72320613 | 0.5639392 | 0.094176649 | 0.302519734 |
| chr3:4429764-4429964 | -0.953255364 | 0.094184861 | 0.302533429 |
| chr5:13935793-13936040 | 0.801994377 | 0.09419785 | 0.302562468 |
| chr6:101114738-101114974 | -0.795979436 | 0.094252968 | 0.302726816 |
| chr1:209805478-209806582 | -0.539494487 | 0.094259573 | 0.30273534 |
| chr3:120558827-120559410 | -0.572502646 | 0.094267248 | 0.3027473 |
| chr16:29805037-29806228 | 0.531644256 | 0.094282149 | 0.302782465 |
| chr3:80696256-80696722 | 0.574239149 | 0.09429458 | 0.302794032 |
| chr8:22048850-22049223 | 0.732490074 | 0.094299907 | 0.302794032 |
| chr1:51622079-51622289 | 0.845002783 | 0.094300025 | 0.302794032 |
| chr5:154045731-154045978 | -0.868861705 | 0.094301558 | 0.302794032 |
| chr8:81928838-81929038 | 0.902442223 | 0.094310086 | 0.302794198 |
| chr7:108455745-108456759 | -0.555971816 | 0.094319402 | 0.302794198 |
| chr12:80329325-80329743 | -0.738992525 | 0.094321274 | 0.302794198 |
| chr12:131097741-131098046 | 0.784334919 | 0.094321914 | 0.302794198 |
| chr10:613013-614275 | 0.548079058 | 0.094323531 | 0.302794198 |
| chr8:136369834-136370276 | -0.65299221 | 0.09432582 | 0.302794198 |
| chr7:99142475-99142740 | -0.638841182 | 0.094330351 | 0.302794198 |
| chr12:105214176-105215871 | 0.534421995 | 0.094333224 | 0.302794198 |
| chr17:52516979-52517504 | 0.661927611 | 0.094339834 | 0.302802729 |
| chr9:69972002-69972271 | -0.917650755 | 0.094369709 | 0.302885932 |
| chr3:173529061-173529900 | -0.568628111 | 0.094373984 | 0.302886965 |
| chr17:75517524-75517730 | 0.632749075 | 0.094395038 | 0.30293917 |
| chr5:178593129-178593380 | 0.810542431 | 0.094400758 | 0.30293917 |
| chr7:96075468-96076262 | -0.598888575 | 0.094405671 | 0.30293917 |
| chr8:97267558-97267787 | 0.710285532 | 0.094406065 | 0.30293917 |
| chr5:155943883-155944142 | -0.727300705 | 0.094411351 | 0.302943445 |
| chr6:149974931-149975185 | -0.871963339 | 0.094444781 | 0.303037963 |
| chr20:57120273-57120538 | 0.555094922 | 0.094448717 | 0.303037963 |
| chr9:70413711-70414518 | -0.548940012 | 0.094469208 | 0.303082696 |
| chr4:74320490-74321151 | -0.577566133 | 0.09447057 | 0.303082696 |
| chr6:28758134-28758627 | -0.59847989 | 0.094488128 | 0.303106149 |
| chr7:44921598-44922437 | -0.561315354 | 0.094489518 | 0.303106149 |
| chr10:132898581-132899105 | 0.65312059 | 0.094489748 | 0.303106149 |
| chr1:182396759-182397015 | 0.718688274 | 0.094500555 | 0.303127293 |
| chr3:908935-909592 | 0.624325548 | 0.094504252 | 0.303127293 |
| chr13:26629275-26629482 | 0.837928203 | 0.094520303 | 0.303166087 |
| chr4:173334099-173335640 | -0.523063 | 0.094547067 | 0.303227691 |
| chr17:61318437-61318666 | 0.570763351 | 0.094552111 | 0.303227691 |
| chr7:35624331-35624930 | -0.566440561 | 0.094553901 | 0.303227691 |
| chr21:30147001-30147355 | 0.702691893 | 0.094556525 | 0.303227691 |
| chr1:30977294-30977511 | -0.844411026 | 0.094560698 | 0.303227691 |
| chr10:132408515-132408942 | 0.598125616 | 0.094563254 | 0.303227691 |
| chr20:45362796-45363813 | 0.534666041 | 0.094587018 | 0.303280342 |
| chr15:69044949-69045233 | -0.805558144 | 0.09458759 | 0.303280342 |
| chr12:34186604-34186854 | 0.737575647 | 0.094609056 | 0.303336476 |

| | | | |
|---|---|---|---|
| chr1:16167508-16168135 | 0.619174224 | 0.094623545 | 0.303370234 |
| chr10:70107337-70107559 | -0.817390519 | 0.094632495 | 0.303386237 |
| chr2:27408985-27410121 | -0.545326913 | 0.094644987 | 0.303398022 |
| chr1:79646910-79647325 | 0.64777386 | 0.094645119 | 0.303398022 |
| chr5:173690253-173690580 | 0.778591855 | 0.094648051 | 0.303398022 |
| chr16:19885339-19885598 | -0.803200636 | 0.094656961 | 0.303413893 |
| chr21:45615348-45615597 | -0.830101469 | 0.094675956 | 0.303436925 |
| chr7:111741773-111742108 | -0.608777681 | 0.094680382 | 0.303436925 |
| chr11:4636941-4637236 | -0.717985672 | 0.09468223 | 0.303436925 |
| chr13:37404104-37404525 | 0.599142812 | 0.09468224 | 0.303436925 |
| chr14:32382753-32383019 | 0.700674328 | 0.094683948 | 0.303436925 |
| chr11:119107117-119107528 | 0.570296087 | 0.094719676 | 0.303538728 |
| chr1:144002502-144002730 | -0.874853496 | 0.094733272 | 0.303569603 |
| chr3:73255548-73255942 | -0.716705686 | 0.094738739 | 0.303574424 |
| chr3:190017801-190018062 | -0.867485547 | 0.094755074 | 0.303614071 |
| chr10:100338498-100338761 | 0.69681669 | 0.094766238 | 0.303637146 |
| chr2:27715094-27715294 | -0.856735946 | 0.094775977 | 0.303645776 |
| chr20:20192857-20193158 | 0.716095842 | 0.094776857 | 0.303645776 |
| chr14:105314302-105314762 | 0.594315587 | 0.094807317 | 0.303730663 |
| chr13:79736346-79736681 | -0.755398876 | 0.094813382 | 0.303737392 |
| chr2:222613801-222614027 | -0.827483537 | 0.094825797 | 0.303764465 |
| chr1:241706914-241707406 | 0.603663937 | 0.094835112 | 0.303772899 |
| chr3:167308692-167308892 | 0.913630916 | 0.094839315 | 0.303772899 |
| chr11:113773108-113774218 | 0.554889352 | 0.094840324 | 0.303772899 |
| chr13:36006016-36007318 | -0.534730291 | 0.094847467 | 0.303783079 |
| chr4:94158883-94159249 | -0.723059317 | 0.094860104 | 0.303803219 |
| chr8:119982508-119983537 | -0.544652173 | 0.094861685 | 0.303803219 |
| chr7:103297207-103297903 | -0.557946265 | 0.094877459 | 0.303837387 |
| chr17:10210103-10210692 | 0.568317508 | 0.094881121 | 0.303837387 |
| chr7:110090183-110090383 | -0.943189009 | 0.09488425 | 0.303837387 |
| chr13:94703650-94703850 | -1.00604357 | 0.094911604 | 0.303912278 |
| chr10:9061761-9062390 | -0.614262843 | 0.094918303 | 0.303921029 |
| chr15:80711242-80711451 | 0.913516111 | 0.094938437 | 0.303972793 |
| chr1:180168737-180169181 | 0.570399173 | 0.094946839 | 0.303986993 |
| chr1:233294962-233295888 | 0.571820565 | 0.094958205 | 0.304010678 |
| chr6:122755986-122756209 | -0.861477616 | 0.095006678 | 0.304153158 |
| chr7:148457593-148457805 | 0.858570486 | 0.095019426 | 0.304181261 |
| chr1:151735053-151735377 | 0.586969961 | 0.095033596 | 0.304213913 |
| chr20:53758620-53759031 | 0.540621488 | 0.095041831 | 0.304227562 |
| chr2:160241213-160242172 | -0.572915659 | 0.095047157 | 0.304231901 |
| chr2:100902989-100903189 | 0.8360777 | 0.095067529 | 0.304284398 |
| chr1:236516949-236517173 | 0.825488639 | 0.095084785 | 0.304320082 |
| chr10:17434703-17435029 | -0.788676654 | 0.095086621 | 0.304320082 |
| chr6:108560610-108561048 | 0.579812075 | 0.095105149 | 0.304358284 |
| chr3:163829360-163829648 | 0.773647304 | 0.095106502 | 0.304358284 |
| chr5:54061645-54061949 | -0.742519082 | 0.095115266 | 0.304367414 |
| chrX:49235253-49235794 | 0.602856215 | 0.0951173 | 0.304367414 |
| chr12:132560356-132560556 | 0.755299728 | 0.095129494 | 0.304393722 |
| chr21:45915376-45915615 | 0.736314439 | 0.095140533 | 0.304416333 |
| chr13:101149976-101150282 | -0.756207571 | 0.095175701 | 0.304516141 |

| | | | |
|---|---|---|---|
| chr13:68893703-68894071 | 0.725230573 | 0.095195183 | 0.304565757 |
| chr10:91344096-91344408 | 0.717123817 | 0.095220846 | 0.304635142 |
| chr7:45086327-45086563 | 0.775445899 | 0.095244122 | 0.304688249 |
| chr22:46150778-46151021 | -0.842674081 | 0.095245399 | 0.304688249 |
| chr8:116887348-116887859 | 0.589919 | 0.095258873 | 0.304718632 |
| chr5:61172304-61172517 | -0.948226535 | 0.095272501 | 0.304749501 |
| chr12:125237108-125237447 | 0.628827264 | 0.095277832 | 0.304753831 |
| chr1:173775675-173775943 | -0.819748435 | 0.095284542 | 0.304762572 |
| chr12:54892010-54892573 | 0.578606459 | 0.095292689 | 0.304775909 |
| chr19:57363848-57364054 | 0.796787201 | 0.095307022 | 0.304809024 |
| chr13:70668322-70668662 | -0.636840617 | 0.095346831 | 0.30491018 |
| chr17:59871918-59872160 | 0.580824244 | 0.095346841 | 0.30491018 |
| chr1:225454184-225454509 | 0.670683901 | 0.095350589 | 0.30491018 |
| chr4:45971162-45971372 | -0.883204271 | 0.095366319 | 0.30493025 |
| chr17:57366330-57367039 | -0.56545242 | 0.095366592 | 0.30493025 |
| chr1:226002740-226003073 | -0.620179381 | 0.095368804 | 0.30493025 |
| chr13:80978655-80978937 | -0.853334068 | 0.095419013 | 0.305078057 |
| chr1:106818184-106818651 | 0.55690208 | 0.095424171 | 0.305081817 |
| chr4:25312282-25313135 | 0.556036943 | 0.095438899 | 0.305116172 |
| chr16:88035436-88035699 | -0.787317634 | 0.095456504 | 0.305159725 |
| chr6:142156496-142156779 | -0.785384124 | 0.0954667 | 0.305179585 |
| chr9:97237988-97238497 | 0.626715559 | 0.095493177 | 0.305251489 |
| chr15:85080415-85080676 | 0.730551625 | 0.095510486 | 0.305280147 |
| chr13:33846758-33847306 | -0.68351326 | 0.095512691 | 0.305280147 |
| chr19:49426163-49426455 | 0.713686666 | 0.095514095 | 0.305280147 |
| chr17:74361440-74361795 | 0.672589776 | 0.095525621 | 0.305304251 |
| chr1:15617073-15617780 | 0.565644854 | 0.095562817 | 0.305410392 |
| chr3:171049836-171050300 | -0.597156692 | 0.095585027 | 0.305464246 |
| chr6:76275765-76276134 | -0.629654948 | 0.095587641 | 0.305464246 |
| chr6:135668210-135668548 | -0.712980483 | 0.095610335 | 0.305524026 |
| chr18:70205522-70206341 | -0.543683619 | 0.095642283 | 0.30561337 |
| chr7:77798018-77799315 | -0.526965768 | 0.095662576 | 0.305665468 |
| chr21:21003785-21004095 | 0.754236447 | 0.095682796 | 0.305717326 |
| chr12:69785335-69785535 | 0.848463376 | 0.09568987 | 0.305727179 |
| chr5:109740273-109740675 | -0.81451132 | 0.095716517 | 0.305799564 |
| chr4:123508508-123509092 | -0.548687523 | 0.095730472 | 0.305818389 |
| chr22:30763824-30764082 | 0.743477079 | 0.095730879 | 0.305818389 |
| chr1:61056883-61057406 | -0.59780845 | 0.095734383 | 0.305818389 |
| chr7:138001496-138002560 | -0.528509004 | 0.09574695 | 0.305844523 |
| chr1:184418670-184418909 | 0.739314716 | 0.095756315 | 0.305844523 |
| chr6:87926972-87927336 | 0.734864078 | 0.095758009 | 0.305844523 |
| chr7:82810694-82810934 | -0.811620377 | 0.09575853 | 0.305844523 |
| chr13:30136007-30136211 | -0.822358229 | 0.095798035 | 0.305932999 |
| chr12:33180984-33181333 | -0.664120078 | 0.095800624 | 0.305932999 |
| chr1:168471164-168471461 | 0.755794573 | 0.095805041 | 0.305932999 |
| chr3:17213150-17213429 | -0.780536423 | 0.095810117 | 0.305932999 |
| chr3:24662695-24662927 | 0.831205816 | 0.095812282 | 0.305932999 |
| chr10:69426824-69427336 | -0.662220445 | 0.095814014 | 0.305932999 |
| chr5:175293505-175293714 | 0.882496228 | 0.095814181 | 0.305932999 |
| chr1:116825813-116826052 | 0.75968966 | 0.09582637 | 0.305959168 |

| | | | |
|---|---|---|---|
| chr11:83132548-83132748 | 0.906678735 | 0.095854725 | 0.306036949 |
| chr20:11544127-11544386 | -0.770618591 | 0.095862283 | 0.306048327 |
| chr1:219173622-219174550 | -0.534977869 | 0.095867374 | 0.3060486 |
| chr5:97192585-97193782 | -0.535104523 | 0.095870357 | 0.3060486 |
| chr3:21406268-21406581 | -0.839818044 | 0.095880764 | 0.306060755 |
| chr7:73742270-73743373 | -0.534390424 | 0.095882153 | 0.306060755 |
| chr2:108189327-108189671 | -0.698832151 | 0.095893256 | 0.306083444 |
| chr10:132886525-132886740 | 0.759086021 | 0.09591091 | 0.30611975 |
| chr11:80180985-80181185 | 0.923259285 | 0.095912621 | 0.30611975 |
| chr4:119786015-119786222 | -0.922166737 | 0.095929289 | 0.30615719 |
| chr12:2794514-2795124 | 0.582187123 | 0.095932343 | 0.30615719 |
| chr10:1242891-1243126 | 0.845061951 | 0.09594812 | 0.306194786 |
| chr3:141913263-141913494 | 0.811310857 | 0.095969662 | 0.306241977 |
| chr1:2189408-2190006 | 0.575905821 | 0.095970901 | 0.306241977 |
| chrX:154762455-154763439 | 0.538125704 | 0.095985835 | 0.306276877 |
| chr10:13440229-13440793 | -0.631133456 | 0.09600076 | 0.306311743 |
| chr10:32210736-32211120 | -0.638528761 | 0.096040613 | 0.306426145 |
| chr8:84020224-84020884 | 0.589417884 | 0.096045861 | 0.306430128 |
| chr12:130014549-130014952 | -0.676982398 | 0.096050496 | 0.306432156 |
| chr12:65556687-65557092 | -0.590348476 | 0.0960625 | 0.306457693 |
| chr11:23113077-23113497 | 0.662194629 | 0.096068574 | 0.306464311 |
| chr18:47754546-47754974 | -0.650428105 | 0.096073585 | 0.306467536 |
| chr3:5236296-5236513 | -0.797305897 | 0.096078657 | 0.306470958 |
| chr18:48541096-48541439 | 0.686620873 | 0.096088669 | 0.306490133 |
| chr13:24221158-24221365 | 0.777573406 | 0.096095006 | 0.306497586 |
| chr11:80727426-80728155 | -0.586322002 | 0.096100752 | 0.306503157 |
| chr16:19127608-19127808 | 0.867083906 | 0.096109354 | 0.306517832 |
| chr22:38398766-38399035 | 0.773310138 | 0.096118525 | 0.306534323 |
| chr14:61491291-61491522 | 0.74614883 | 0.096139361 | 0.306577716 |
| chr1:109112630-109113037 | -0.5714469 | 0.096141405 | 0.306577716 |
| chr6:158008268-158008548 | -0.789306864 | 0.096144135 | 0.306577716 |
| chr14:105511654-105512180 | 0.612665181 | 0.096160712 | 0.306617738 |
| chr20:54292778-54293025 | 0.589524952 | 0.09616469 | 0.306617738 |
| chr13:42115580-42115827 | -0.832930482 | 0.096174245 | 0.306635444 |
| chr5:55226428-55227738 | -0.540782563 | 0.0961831 | 0.30664034 |
| chr2:179580624-179581031 | -0.688555913 | 0.096183784 | 0.30664034 |
| chr6:104414710-104415096 | -0.630157745 | 0.096195183 | 0.306663919 |
| chr15:49215938-49216822 | -0.560866751 | 0.09622148 | 0.306734991 |
| chr2:84511021-84511632 | -0.572474895 | 0.09623783 | 0.306770329 |
| chr14:91253120-91253686 | 0.591507773 | 0.096240573 | 0.306770329 |
| chr2:70249434-70249638 | -0.758202589 | 0.09626134 | 0.306799129 |
| chr9:104961288-104961508 | -0.851434792 | 0.096261624 | 0.306799129 |
| chr5:143860066-143860623 | -0.622920755 | 0.096265443 | 0.306799129 |
| chrX:108312934-108313259 | -0.723336726 | 0.096266131 | 0.306799129 |
| chr17:75755688-75755923 | -0.682238504 | 0.096269628 | 0.306799129 |
| chr14:32934080-32934734 | 0.582987952 | 0.096278282 | 0.306813946 |
| chr1:199522823-199523042 | 0.821642907 | 0.096291398 | 0.306835909 |
| chr1:65527183-65527515 | -0.724877162 | 0.096293183 | 0.306835909 |
| chr20:48098076-48098462 | 0.550780405 | 0.096300939 | 0.306847862 |
| chr7:142866378-142866639 | -0.678636549 | 0.09632927 | 0.30692537 |

| | | | |
|---|---|---|---|
| chr10:87008406-87008679 | -0.770646244 | 0.096360831 | 0.307013165 |
| chr3:187745358-187745853 | -0.573455823 | 0.096388898 | 0.307086052 |
| chr3:39180533-39181249 | 0.554255579 | 0.096391724 | 0.307086052 |
| chr3:122092545-122092865 | -0.663619299 | 0.09641614 | 0.307151069 |
| chr16:69398370-69398627 | -0.838751157 | 0.096422835 | 0.307159624 |
| chr12:101552420-101552664 | -0.783159592 | 0.096427812 | 0.30716271 |
| chr6:401326-401725 | 0.743262386 | 0.096449803 | 0.307197948 |
| chr9:89661947-89662297 | 0.617493507 | 0.096450349 | 0.307197948 |
| chr2:122390407-122390688 | -0.793008166 | 0.096450902 | 0.307197948 |
| chr1:183605930-183606188 | 0.718491394 | 0.096462866 | 0.307206491 |
| chr4:87092741-87093248 | -0.616798011 | 0.096464105 | 0.307206491 |
| chr1:215488979-215489208 | 0.706911224 | 0.096467582 | 0.307206491 |
| chr11:42770504-42770741 | -0.825181157 | 0.096473401 | 0.307206491 |
| chr18:15006410-15006610 | 0.908920563 | 0.096473631 | 0.307206491 |
| chr14:48398720-48399047 | -0.666952735 | 0.096488576 | 0.307236356 |
| chr4:6986480-6987635 | 0.548366886 | 0.09649103 | 0.307236356 |
| chr12:17476467-17476702 | -0.824150073 | 0.096502949 | 0.307255249 |
| chr4:85127638-85128150 | -0.605352704 | 0.096504983 | 0.307255249 |
| chr1:105072539-105072757 | -0.839455921 | 0.096529961 | 0.307318393 |
| chr17:75320536-75320758 | 0.788725216 | 0.096532838 | 0.307318393 |
| chr10:120253155-120253561 | 0.658091822 | 0.09659296 | 0.307497019 |
| chr12:132153827-132154250 | 0.609698413 | 0.096601488 | 0.307511392 |
| chr3:17468056-17468380 | 0.714184411 | 0.096635503 | 0.307606893 |
| chr2:36819370-36819674 | -0.721211447 | 0.096642518 | 0.307616442 |
| chr1:248825532-248826849 | -0.522488076 | 0.096649228 | 0.307621345 |
| chr5:145888024-145888291 | -0.721220977 | 0.09665808 | 0.307621345 |
| chr19:17719982-17720289 | -0.718133533 | 0.096658706 | 0.307621345 |
| chr14:69129315-69129810 | 0.645045526 | 0.096660117 | 0.307621345 |
| chr12:66087427-66087688 | 0.769988169 | 0.096672579 | 0.307648227 |
| chr3:187319343-187319578 | -0.821453032 | 0.09667902 | 0.307652191 |
| chr10:80662704-80663196 | -0.600464751 | 0.096681855 | 0.307652191 |
| chr3:101925396-101925633 | 0.801046201 | 0.096723888 | 0.307771828 |
| chr18:48951993-48952449 | -0.596281457 | 0.096727485 | 0.307771828 |
| chr11:61860349-61860673 | -0.67899602 | 0.09673729 | 0.307790242 |
| chrX:12901376-12901770 | -0.806912694 | 0.096744931 | 0.307801775 |
| chr2:128250338-128250602 | 0.801054146 | 0.096761505 | 0.307841722 |
| chrX:120251479-120251883 | 0.657025366 | 0.096776842 | 0.307877731 |
| chr10:21173419-21173619 | 0.73244982 | 0.09679157 | 0.307911804 |
| chr7:117951788-117952017 | -0.831248022 | 0.096803234 | 0.30792706 |
| chr18:57881866-57882169 | -0.718743211 | 0.096807085 | 0.30792706 |
| chr10:76632713-76633018 | -0.685483272 | 0.096809414 | 0.30792706 |
| chr10:59626060-59626335 | -0.665179475 | 0.096812441 | 0.30792706 |
| chr4:130712021-130712343 | 0.809596773 | 0.096818983 | 0.307935083 |
| chr2:166900683-166900971 | -0.718948512 | 0.096824548 | 0.307940002 |
| chr16:84629704-84629982 | 0.734331201 | 0.09683551 | 0.307962084 |
| chr10:131840114-131840417 | 0.692500169 | 0.096845495 | 0.307973397 |
| chr5:26992121-26992342 | 0.849878758 | 0.096847107 | 0.307973397 |
| chr10:88565176-88565430 | -0.80641003 | 0.09685232 | 0.307977194 |
| chr8:142679777-142680046 | 0.648607844 | 0.096861782 | 0.3079945 |
| chr12:107131815-107132104 | -0.706512816 | 0.096900759 | 0.308105651 |

| | | | |
|---|---|---|---|
| chr13:60396743-60397986 | -0.530717473 | 0.096908916 | 0.308118502 |
| chr2:20437136-20437356 | -0.80874603 | 0.096912844 | 0.308118502 |
| chr16:3611077-3612119 | 0.554126662 | 0.096922783 | 0.308137317 |
| chr19:48812252-48812547 | 0.641270078 | 0.096936579 | 0.30816839 |
| chr10:19168664-19168926 | -0.749054937 | 0.096944906 | 0.308182076 |
| chr12:56291462-56291852 | 0.623914702 | 0.096960785 | 0.308219766 |
| chr7:114402625-114402856 | -0.890914964 | 0.096978019 | 0.308261762 |
| chr1:51569299-51569615 | 0.659709225 | 0.097001363 | 0.308314766 |
| chr14:104650340-104650658 | 0.624007472 | 0.097002741 | 0.308314766 |
| chr3:183479868-183480310 | -0.633612652 | 0.097063086 | 0.30846941 |
| chr4:41374787-41375003 | -0.902197521 | 0.097063471 | 0.30846941 |
| chr16:279047-279247 | 0.787236876 | 0.097063474 | 0.30846941 |
| chr2:185676318-185676728 | -0.640120428 | 0.097068144 | 0.308471457 |
| chr21:41956265-41956492 | 0.671907265 | 0.097090753 | 0.308529937 |
| chr4:186555558-186555764 | -0.783602491 | 0.097094599 | 0.308529937 |
| chr1:29697981-29698342 | 0.641338802 | 0.0971151 | 0.308582283 |
| chr1:1890546-1890866 | -0.737927533 | 0.09712007 | 0.308585268 |
| chr7:142231724-142232052 | -0.657860716 | 0.097124294 | 0.308585268 |
| chr11:82771813-82772013 | 0.841527299 | 0.097133828 | 0.308585268 |
| chr19:10420197-10420402 | -0.784140045 | 0.097134615 | 0.308585268 |
| chr19:57240184-57241019 | 0.562200255 | 0.097136176 | 0.308585268 |
| chr3:36969872-36970201 | 0.742120437 | 0.097164617 | 0.308651629 |
| chr2:132352490-132353827 | -0.540663536 | 0.097165121 | 0.308651629 |
| chr8:81633170-81633613 | -0.616891803 | 0.097170136 | 0.30865286 |
| chr1:68600050-68600432 | -0.750012617 | 0.097173566 | 0.30865286 |
| chr1:78608106-78608349 | -0.817216128 | 0.097178915 | 0.308657057 |
| chr1:108705145-108705442 | 0.754091889 | 0.09718904 | 0.308676419 |
| chr1:152869885-152870133 | 0.733597752 | 0.097212029 | 0.30872675 |
| chr2:127415899-127416424 | 0.593300132 | 0.097212945 | 0.30872675 |
| chr1:66262282-66262556 | -0.737830253 | 0.097217672 | 0.308728966 |
| chr13:46853100-46853319 | 0.68302364 | 0.0972461 | 0.308806444 |
| chrX:142484033-142484472 | 0.619669944 | 0.097251142 | 0.308809658 |
| chr16:89560398-89560978 | -0.559076936 | 0.097286443 | 0.308908948 |
| chr2:98086486-98087521 | -0.547834009 | 0.097311993 | 0.308977271 |
| chr5:95712298-95712787 | -0.647385834 | 0.097322206 | 0.308995295 |
| chr8:70315673-70315992 | -0.752589465 | 0.097334995 | 0.308995295 |
| chr2:197378480-197378934 | -0.642910634 | 0.097337755 | 0.308995295 |
| chr7:85635024-85635275 | -0.763851927 | 0.097338604 | 0.308995295 |
| chr1:55893213-55893524 | -0.776984034 | 0.09734092 | 0.308995295 |
| chr11:73334034-73334310 | 0.744434304 | 0.097341866 | 0.308995295 |
| chr1:180663599-180663838 | 0.789128626 | 0.097349879 | 0.30899941 |
| chr7:139232197-139232493 | -0.657793304 | 0.097351227 | 0.30899941 |
| chr2:96874135-96874379 | -0.660783234 | 0.097369538 | 0.309043378 |
| chr1:31413498-31413923 | 0.620554324 | 0.097373157 | 0.309043378 |
| chr13:49847756-49848312 | -0.617289318 | 0.09737718 | 0.309043378 |
| chr10:119608382-119608661 | 0.682645125 | 0.097391194 | 0.309075053 |
| chr3:150545499-150545720 | 0.63832279 | 0.09739785 | 0.309083375 |
| chrX:18654985-18655276 | 0.759663947 | 0.097424089 | 0.309153837 |
| chr16:48609395-48609595 | 0.864641338 | 0.097429173 | 0.309155251 |
| chr5:1644961-1645333 | 0.648648709 | 0.097432604 | 0.309155251 |

| | | | |
|---|---|---|---|
| chr9:130866476-130866818 | 0.626044127 | 0.097442631 | 0.309174263 |
| chr7:16420314-16421920 | 0.521911568 | 0.097458551 | 0.30921197 |
| chr7:131507962-131508324 | -0.659932445 | 0.097468603 | 0.309231058 |
| chr4:154023908-154024416 | -0.620066328 | 0.09747745 | 0.309242308 |
| chrX:123961267-123962026 | 0.543670424 | 0.097487609 | 0.309242308 |
| chr14:59965589-59965883 | 0.740584408 | 0.097487609 | 0.309242308 |
| chr8:25615789-25616052 | -0.794529299 | 0.097488293 | 0.309242308 |
| chr6:28598067-28598310 | -0.798973286 | 0.097502608 | 0.309274915 |
| chr5:54093283-54093577 | -0.848150374 | 0.097523808 | 0.309319197 |
| chr7:87507507-87507787 | -0.744187594 | 0.097524643 | 0.309319197 |
| chr5:87825705-87825936 | -0.863843641 | 0.097539214 | 0.309352608 |
| chr19:11197412-11198035 | 0.601801755 | 0.097549161 | 0.309371351 |
| chr15:63387742-63388072 | -0.664389776 | 0.097559912 | 0.309392639 |
| chr12:73656486-73656692 | -0.88533224 | 0.097564053 | 0.309392967 |
| chr10:60732725-60733745 | -0.541219567 | 0.097569538 | 0.309397557 |
| chr15:29668082-29668335 | 0.720358906 | 0.097581299 | 0.30941508 |
| chrX:131633588-131633848 | 0.716240019 | 0.097583141 | 0.30941508 |
| chr1:245042732-245043345 | 0.602247042 | 0.09759909 | 0.309452846 |
| chr1:8817488-8818149 | 0.595278627 | 0.097619924 | 0.309506096 |
| chr1:236543132-236543363 | 0.7858693 | 0.097641344 | 0.309561198 |
| chrX:134520832-134521417 | 0.63669814 | 0.097649208 | 0.309573323 |
| chr5:151923959-151924383 | -0.709536688 | 0.097657383 | 0.30958643 |
| chr13:72187020-72187363 | -0.688365324 | 0.097686039 | 0.30966446 |
| chr8:108986401-108986885 | -0.718294959 | 0.097723301 | 0.309769765 |
| chr2:96067893-96068110 | -0.871209683 | 0.097729044 | 0.309775155 |
| chr4:86218899-86219109 | 0.84008493 | 0.097737614 | 0.309789503 |
| chr1:175234849-175235073 | 0.912086357 | 0.097764619 | 0.309862282 |
| chr6:134090110-134090440 | -0.818876649 | 0.097787693 | 0.309922593 |
| chr6:30828312-30828996 | 0.591198215 | 0.097803714 | 0.309960548 |
| chr21:36939487-36939748 | 0.749308669 | 0.097820069 | 0.309999559 |
| chr3:161096802-161097121 | -0.676972747 | 0.097824362 | 0.310000343 |
| chr17:74987897-74988560 | 0.572947905 | 0.097842905 | 0.310036048 |
| chr2:215212263-215212507 | 0.749434213 | 0.097851334 | 0.310036048 |
| chr20:58168288-58168649 | 0.645412226 | 0.0978515 | 0.310036048 |
| chr19:57769424-57770026 | 0.577051985 | 0.097852815 | 0.310036048 |
| chr19:4084132-4084342 | 0.773396818 | 0.097858351 | 0.310036048 |
| chr9:546222-546440 | 0.927888597 | 0.097859907 | 0.310036048 |
| chr5:123155338-123156031 | 0.575524174 | 0.097874122 | 0.310068262 |
| chr1:218490268-218490606 | 0.658724443 | 0.097879941 | 0.310073877 |
| chr2:29117757-29118325 | 0.60744979 | 0.097887927 | 0.310086355 |
| chr13:38223099-38223463 | -0.700679534 | 0.097920838 | 0.31016883 |
| chr16:70696716-70697535 | 0.543278645 | 0.097922059 | 0.31016883 |
| chr14:26281634-26281915 | 0.698027054 | 0.097931656 | 0.31017665 |
| chr5:7762276-7762726 | -0.589453075 | 0.09793501 | 0.31017665 |
| chr10:71570181-71570524 | 0.690363199 | 0.097936672 | 0.31017665 |
| chr2:71282506-71282754 | 0.702750604 | 0.097944462 | 0.310187482 |
| chr5:85459035-85459257 | 0.880916775 | 0.097948189 | 0.310187482 |
| chr6:128731241-128731465 | 0.778123899 | 0.097958292 | 0.310204538 |
| chr1:28518488-28519049 | 0.590328557 | 0.097961672 | 0.310204538 |
| chr2:165306460-165306673 | -0.891314995 | 0.097972949 | 0.31022666 |

| | | | |
|---|---|---|---|
| chr4:170198826-170199026 | -0.899772695 | 0.097976755 | 0.31022666 |
| chr6:26595775-26596131 | -0.637114927 | 0.097984607 | 0.310238702 |
| chr2:131672481-131673799 | -0.536739726 | 0.098001999 | 0.310280411 |
| chr20:19409011-19409296 | 0.712530611 | 0.09800588 | 0.310280411 |
| chr7:47253538-47254735 | -0.545977931 | 0.098035015 | 0.310359829 |
| chr14:76653919-76654416 | 0.603761608 | 0.098091349 | 0.310505854 |
| chr7:7662615-7662829 | 0.73619643 | 0.09809273 | 0.310505854 |
| chr12:110468026-110469469 | -0.532355205 | 0.098093298 | 0.310505854 |
| chr4:173494044-173494717 | -0.564886789 | 0.098110258 | 0.310546709 |
| chr5:72955377-72956829 | -0.533584357 | 0.098138515 | 0.310607167 |
| chr4:157978815-157979110 | -0.669058947 | 0.098139697 | 0.310607167 |
| chr17:80261363-80261764 | 0.614688537 | 0.09814152 | 0.310607167 |
| chr13:21297066-21297328 | 0.685903927 | 0.098148733 | 0.310617166 |
| chr1:212013639-212013952 | -0.673153968 | 0.09816737 | 0.310663318 |
| chr16:87575105-87575336 | -0.748053122 | 0.09817223 | 0.310664645 |
| chr20:1440134-1440422 | -0.794885575 | 0.098175899 | 0.310664645 |
| chr5:123422874-123424075 | -0.52717104 | 0.098192208 | 0.310703421 |
| chr7:6271617-6272092 | -0.59515589 | 0.098202579 | 0.310723406 |
| chr6:32061772-32062027 | 0.900353837 | 0.098247607 | 0.310853044 |
| chr3:17645157-17645490 | 0.649687699 | 0.098266136 | 0.310898829 |
| chr1:66253350-66253623 | 0.789696964 | 0.098273517 | 0.310906525 |
| chr5:21686842-21687155 | 0.713992311 | 0.098278193 | 0.310906525 |
| chr4:147578038-147578270 | 0.77041697 | 0.09828331 | 0.310906525 |
| chr1:171360105-171360476 | -0.654753163 | 0.09828826 | 0.310906525 |
| chr10:31133664-31133967 | -0.724619795 | 0.098291375 | 0.310906525 |
| chr12:53299267-53300115 | 0.5486411 | 0.098293793 | 0.310906525 |
| chr5:174271420-174271620 | -0.929455969 | 0.098296972 | 0.310906525 |
| chr2:196849946-196850325 | -0.630789107 | 0.098342817 | 0.31103869 |
| chr5:166788028-166788649 | -0.617181767 | 0.098356447 | 0.311067207 |
| chr4:153467173-153467912 | 0.547478325 | 0.098361158 | 0.311067207 |
| chr7:79293655-79294081 | -0.663192017 | 0.09836439 | 0.311067207 |
| chr18:34224790-34225225 | -0.636562971 | 0.098368731 | 0.311067207 |
| chr3:62971911-62972145 | 0.656543982 | 0.098372132 | 0.311067207 |
| chr11:113781615-113781849 | 0.871634673 | 0.09837986 | 0.311078807 |
| chr5:112266405-112266769 | -0.708201472 | 0.09840823 | 0.311155672 |
| chr15:90102010-90102951 | 0.539816686 | 0.098425581 | 0.311197692 |
| chr13:39612143-39612391 | -0.747226045 | 0.098433524 | 0.311209965 |
| chr4:108030340-108030691 | -0.667424279 | 0.098441738 | 0.311223091 |
| chr21:43298993-43299216 | 0.802517038 | 0.098497725 | 0.311377701 |
| chr6:41693998-41694327 | -0.814240522 | 0.09849877 | 0.311377701 |
| chr5:106367355-106367686 | -0.744117407 | 0.098511403 | 0.311404789 |
| chr17:77002110-77002310 | 0.853872552 | 0.098540521 | 0.311483985 |
| chr1:30708069-30708410 | 0.648500771 | 0.098560063 | 0.311524811 |
| chr20:54416911-54417111 | 0.600962323 | 0.098561568 | 0.311524811 |
| chr12:89424914-89425114 | -0.921125789 | 0.098579396 | 0.311559635 |
| chr12:53005988-53007296 | 0.523241585 | 0.098580718 | 0.311559635 |
| chr5:96702462-96703511 | -0.528869399 | 0.098616872 | 0.311661042 |
| chr21:41147230-41147430 | -0.806693897 | 0.098621631 | 0.311663229 |
| chr7:128380210-128380466 | 0.830365008 | 0.098628743 | 0.311672847 |
| chr3:170071118-170071396 | -0.807098807 | 0.098644495 | 0.311709769 |

| | | | |
|---|---|---|---|
| chr2:96760459-96761082 | -0.549395363 | 0.098680528 | 0.311810774 |
| chr11:33723006-33723633 | 0.616821748 | 0.098692663 | 0.311836257 |
| chr5:173187353-173187758 | -0.609680521 | 0.098711353 | 0.31188245 |
| chrX:115085870-115086249 | 0.734724977 | 0.098725842 | 0.311915367 |
| chrX:74420291-74420918 | 0.577316606 | 0.098737543 | 0.311939473 |
| chr5:127228834-127229905 | -0.543582702 | 0.098767296 | 0.312020605 |
| chr21:40382526-40383491 | -0.538940655 | 0.09879705 | 0.312090006 |
| chr21:39562533-39563407 | -0.559868866 | 0.09879741 | 0.312090006 |
| chr1:184110617-184110931 | 0.69384957 | 0.098804162 | 0.312098467 |
| chr4:176615038-176615358 | -0.804671684 | 0.098809496 | 0.312102448 |
| chr22:21651738-21652574 | 0.564965803 | 0.09884183 | 0.312174023 |
| chr2:216839785-216840228 | -0.713294375 | 0.098843071 | 0.312174023 |
| chr7:55271114-55271380 | -0.70344172 | 0.098845242 | 0.312174023 |
| chr3:184005967-184006447 | 0.579530724 | 0.098848453 | 0.312174023 |
| chrX:116546490-116546734 | 0.856200587 | 0.098860439 | 0.312199009 |
| chr4:16360574-16361658 | 0.56659485 | 0.098891485 | 0.31226969 |
| chr7:4801050-4801486 | 0.604351734 | 0.098894611 | 0.31226969 |
| chr15:66504499-66505663 | 0.534485039 | 0.098895047 | 0.31226969 |
| chr7:139618976-139619215 | -0.735953669 | 0.098906512 | 0.312293022 |
| chr17:81926465-81927258 | 0.553336199 | 0.098928946 | 0.312350985 |
| chr20:53517157-53517522 | 0.636814984 | 0.098935097 | 0.312355825 |
| chr19:58326714-58327819 | 0.529264489 | 0.098938632 | 0.312355825 |
| chr9:89365372-89365572 | 0.807437015 | 0.098966236 | 0.312430098 |
| chr5:34957977-34958445 | -0.656366673 | 0.098975595 | 0.312443003 |
| chr7:38215770-38216070 | 0.649504913 | 0.098978479 | 0.312443003 |
| chr1:111346089-111346944 | -0.544412649 | 0.098989593 | 0.312464143 |
| chr12:80837402-80837868 | -0.625049006 | 0.09899672 | 0.312464143 |
| chr5:65213929-65214315 | -0.627067233 | 0.09899741 | 0.312464143 |
| chr7:33932837-33933180 | -0.59144938 | 0.099033733 | 0.312559843 |
| chr9:37472394-37472909 | -0.65832328 | 0.099035889 | 0.312559843 |
| chr10:133140403-133140639 | 0.844771936 | 0.09904274 | 0.31256656 |
| chr2:69795984-69796379 | -0.773724394 | 0.099046176 | 0.31256656 |
| chr5:134411439-134412656 | -0.549890789 | 0.099063276 | 0.312601852 |
| chr17:143888-144123 | -0.728620772 | 0.099065519 | 0.312601852 |
| chr9:22508365-22509145 | -0.567969941 | 0.099073054 | 0.312612756 |
| chr1:84573857-84574854 | -0.533842137 | 0.099082619 | 0.3126166 |
| chr13:38763780-38764169 | -0.62931354 | 0.099085181 | 0.3126166 |
| chr12:91422863-91423313 | -0.638432911 | 0.099086512 | 0.3126166 |
| chr6:158405497-158405713 | 0.813187883 | 0.099092484 | 0.312622568 |
| chr17:79897651-79898011 | 0.703136742 | 0.099100602 | 0.312635308 |
| chr19:39167532-39167851 | 0.625943936 | 0.099142487 | 0.31274337 |
| chr5:92653840-92654329 | -0.758093298 | 0.099145937 | 0.31274337 |
| chr7:25894919-25895261 | 0.774212597 | 0.099147894 | 0.31274337 |
| chr19:49355197-49355459 | -0.827206087 | 0.099151183 | 0.31274337 |
| chr19:6744956-6745347 | 0.635437884 | 0.099179042 | 0.312805499 |
| chr6:37538044-37538440 | 0.634737012 | 0.099183865 | 0.312805499 |
| chr12:120534201-120534533 | 0.626697629 | 0.099186159 | 0.312805499 |
| chr3:119258057-119258328 | -0.668360965 | 0.09918721 | 0.312805499 |
| chr6:166956071-166956330 | -0.694418877 | 0.099220404 | 0.312890747 |
| chr8:138505580-138505808 | 0.881933386 | 0.099222408 | 0.312890747 |

| | | | |
|---|---|---|---|
| chr12:104286543-104287501 | -0.534006848 | 0.099233621 | 0.312913228 |
| chr5:4615370-4615666 | -0.83249418 | 0.099256156 | 0.312971408 |
| chr6:122399099-122400335 | -0.522773169 | 0.09926598 | 0.312982374 |
| chrX:111100187-111100526 | 0.628732051 | 0.099271865 | 0.312982374 |
| chr10:794544-794924 | 0.571370431 | 0.099271888 | 0.312982374 |
| chr17:61788688-61789186 | 0.537948846 | 0.099283143 | 0.312993478 |
| chr9:131096909-131097260 | 0.633037919 | 0.09928358 | 0.312993478 |
| chr1:211680757-211681050 | -0.615377936 | 0.099293326 | 0.313011324 |
| chr4:151106879-151107090 | 0.777251249 | 0.0993094 | 0.313049116 |
| chr11:118251954-118252554 | -0.57101332 | 0.099356267 | 0.313168045 |
| chr4:189557560-189557760 | -0.957453487 | 0.099359742 | 0.313168045 |
| chr12:12655928-12656214 | -0.741236647 | 0.099361545 | 0.313168045 |
| chr5:29016183-29016414 | -0.861419845 | 0.099371213 | 0.313168045 |
| chr5:41414720-41414972 | -0.878504028 | 0.099374475 | 0.313168045 |
| chr1:97820517-97820741 | -0.829576045 | 0.099375739 | 0.313168045 |
| chr11:75728941-75729181 | -0.829576045 | 0.099375739 | 0.313168045 |
| chrX:128156701-128156958 | 0.825126559 | 0.099389342 | 0.313187549 |
| chr9:125428092-125428292 | 0.812525764 | 0.099390669 | 0.313187549 |
| chr10:8049567-8049860 | 0.598036026 | 0.09939419 | 0.313187549 |
| chr20:32584147-32584471 | 0.651242471 | 0.099411621 | 0.31322931 |
| chr3:123471-123811 | -0.887046064 | 0.09941562 | 0.31322931 |
| chr7:20217270-20218006 | 0.527915142 | 0.09942641 | 0.313244081 |
| chr2:76681353-76681564 | -0.919676212 | 0.099428484 | 0.313244081 |
| chr19:46571594-46571794 | 0.87180446 | 0.099441531 | 0.313271846 |
| chr17:56520047-56520362 | 0.713533677 | 0.099445474 | 0.313271846 |
| chr3:169122450-169122695 | 0.815345859 | 0.099456906 | 0.313278921 |
| chr14:72154844-72155107 | 0.720357526 | 0.099458169 | 0.313278921 |
| chr12:47819725-47820904 | 0.536977909 | 0.099459986 | 0.313278921 |
| chr7:101182396-101182970 | -0.549239069 | 0.099479236 | 0.31330759 |
| chr2:213929103-213929541 | -0.638899884 | 0.099480837 | 0.31330759 |
| chr14:91331302-91331512 | 0.695071592 | 0.099481355 | 0.31330759 |
| chr11:74417882-74418160 | 0.763110482 | 0.099491178 | 0.313325648 |
| chr5:61934080-61934341 | -0.728067101 | 0.099498825 | 0.313336852 |
| chr1:203932021-203932221 | -0.760137131 | 0.099509251 | 0.313348441 |
| chrX:47463907-47464224 | -0.719352678 | 0.099510684 | 0.313348441 |
| chr1:202943749-202943949 | 0.760882398 | 0.099541461 | 0.313432476 |
| chr6:44655484-44655904 | -0.643729647 | 0.099553466 | 0.313452203 |
| chr1:113119373-113119611 | 0.860675822 | 0.099555908 | 0.313452203 |
| chr3:98188568-98188890 | -0.748327504 | 0.099567998 | 0.313477385 |
| chr19:2456984-2457324 | 0.672760547 | 0.09958857 | 0.313529273 |
| chr19:53879308-53879683 | 0.596580972 | 0.099601921 | 0.313533408 |
| chr20:58881115-58881346 | 0.719262964 | 0.099603228 | 0.313533408 |
| chr11:121319339-121319554 | 0.858317663 | 0.099603983 | 0.313533408 |
| chr17:44385177-44385872 | 0.568801709 | 0.099606252 | 0.313533408 |
| chr8:134521786-134522107 | 0.736083134 | 0.099617845 | 0.313557018 |
| chr8:10587316-10587607 | -0.693518553 | 0.099625332 | 0.313567705 |
| chr5:177600652-177600929 | 0.678129469 | 0.099646241 | 0.313620633 |
| chr12:69768606-69768846 | 0.788936941 | 0.099658779 | 0.313647209 |
| chr1:3624630-3625154 | 0.570995789 | 0.099666774 | 0.313659488 |
| chr2:126372464-126372713 | 0.90358993 | 0.09971618 | 0.313802083 |

| | | | |
|---|---|---|---|
| chr2:210202903-210203433 | -0.693217046 | 0.099726892 | 0.313821264 |
| chr17:51861013-51861275 | -0.728576361 | 0.099733171 | 0.313821264 |
| chr16:67393335-67393705 | -0.65978475 | 0.099734562 | 0.313821264 |
| chr14:71206876-71207088 | -0.80439555 | 0.099740391 | 0.313822333 |
| chr3:121610106-121610351 | 0.85503669 | 0.099746504 | 0.313822333 |
| chr13:57511591-57512150 | -0.636139448 | 0.099747189 | 0.313822333 |
| chrX:13747925-13748125 | 0.846248427 | 0.099752054 | 0.313824754 |
| chr14:68863270-68863480 | 0.788684549 | 0.099757714 | 0.313827877 |
| chr12:78146398-78146598 | -0.941865269 | 0.099761238 | 0.313827877 |
| chr2:205025121-205025755 | -0.647615247 | 0.099790897 | 0.313908289 |
| chr5:138449707-138450443 | -0.564970982 | 0.099799745 | 0.313913115 |
| chr4:6632949-6633890 | 0.549950883 | 0.099800625 | 0.313913115 |
| chr7:101862331-101862557 | -0.883720722 | 0.099813172 | 0.313918408 |
| chr6:26197265-26197479 | 0.678721573 | 0.099816089 | 0.313918408 |
| chr5:50710896-50711607 | -0.600738422 | 0.099820222 | 0.313918408 |
| chr19:37982079-37982442 | 0.653687842 | 0.099822043 | 0.313918408 |
| chr7:105368423-105368755 | -0.829183708 | 0.099825978 | 0.313918408 |
| chr1:88998773-88999022 | 0.834079959 | 0.09982689 | 0.313918408 |
| chr22:20456377-20456622 | -0.738853841 | 0.099833742 | 0.313924407 |
| chr14:35438803-35439096 | 0.718727567 | 0.099840104 | 0.313924407 |
| chr5:118956829-118957029 | -0.858561129 | 0.099841088 | 0.313924407 |
| chr9:98795997-98796294 | 0.670656048 | 0.099860148 | 0.31397145 |
| chr17:56662022-56662259 | 0.685601908 | 0.099876703 | 0.314010617 |
| chr17:9021130-9021742 | -0.634745861 | 0.099892729 | 0.314048116 |
| chr3:134373629-134373938 | 0.585495279 | 0.099910815 | 0.314092087 |
| chr5:131109809-131110319 | -0.624756711 | 0.0999211 | 0.314110431 |
| chr14:86022257-86022530 | 0.823829618 | 0.099924849 | 0.314110431 |
| chr6:36912668-36912887 | 0.791033936 | 0.099943125 | 0.314154993 |
| chr18:57771570-57771947 | 0.643224469 | 0.099952014 | 0.314170048 |
| chr6:108145688-108145888 | -0.894072151 | 0.099965378 | 0.314199163 |
| chr12:109910928-109911189 | -0.712142084 | 0.099980821 | 0.314234811 |
| chr5:180328575-180329023 | 0.599295827 | 0.099986535 | 0.314239882 |
| chr6:127606112-127606570 | -0.670671012 | 0.100021569 | 0.314337092 |
| chr10:52004404-52004631 | -0.819296813 | 0.100028677 | 0.314342002 |
| chr9:113143537-113143737 | 0.804951176 | 0.100031336 | 0.314342002 |
| chr14:49294774-49295061 | 0.693734639 | 0.100038276 | 0.314350919 |
| chr22:37561594-37562074 | -0.621042274 | 0.100043699 | 0.314355068 |
| chr17:14736559-14737123 | -0.632839492 | 0.100067174 | 0.314392439 |
| chr19:9791565-9792962 | 0.536800556 | 0.100070123 | 0.314392439 |
| chr5:30040590-30040850 | 0.720962093 | 0.100074168 | 0.314392439 |
| chr12:7757064-7757286 | -0.843300682 | 0.10007693 | 0.314392439 |
| chr4:40526035-40526257 | -0.891825162 | 0.100078481 | 0.314392439 |
| chr5:134554499-134554797 | 0.698110104 | 0.100080212 | 0.314392439 |
| chr8:95066847-95067972 | 0.524329473 | 0.100092259 | 0.314417394 |
| chr12:40970399-40970647 | -0.89735899 | 0.100100712 | 0.314431057 |
| chr4:126005010-126005251 | -0.861067064 | 0.100107728 | 0.314440204 |
| chr1:54568030-54568285 | -0.912420915 | 0.100112324 | 0.31444175 |
| chr7:129417148-129417413 | -0.798848032 | 0.100116562 | 0.314442172 |
| chr1:235679678-235679978 | -0.751321398 | 0.100138069 | 0.314485737 |
| chr11:44621168-44621406 | 0.725142079 | 0.100142345 | 0.314485737 |

| | | | |
|---|---|---|---|
| chr2:218357234-218357551 | -0.69968337 | 0.100142746 | 0.314485737 |
| chr10:28738885-28739120 | -0.929086007 | 0.100154457 | 0.314497111 |
| chr3:43644502-43644825 | -0.700606398 | 0.100157932 | 0.314497111 |
| chr4:83306951-83307395 | -0.727659953 | 0.10016332 | 0.314497111 |
| chr14:32560296-32560509 | 0.757667101 | 0.100166649 | 0.314497111 |
| chr5:86679445-86679881 | -0.61630053 | 0.100169875 | 0.314497111 |
| chr4:164779824-164780277 | -0.666380474 | 0.100170995 | 0.314497111 |
| chr5:161535770-161536133 | -0.666188187 | 0.100189879 | 0.314536463 |
| chr3:58358358-58358558 | 0.869565083 | 0.100191982 | 0.314536463 |
| chr4:499698-500059 | 0.670660203 | 0.100198314 | 0.314536463 |
| chr20:54385488-54385702 | 0.583816685 | 0.10019995 | 0.314536463 |
| chr2:27955109-27955325 | -0.889524942 | 0.100208952 | 0.314541419 |
| chr8:29784823-29785138 | 0.77540573 | 0.100209738 | 0.314541419 |
| chr13:41848459-41848731 | -0.776193075 | 0.100235598 | 0.314609699 |
| chr20:35306837-35307314 | 0.608469785 | 0.100244364 | 0.314624325 |
| chr10:70888515-70888734 | 0.736555804 | 0.100251675 | 0.314631127 |
| chr1:177490298-177490548 | -0.754491857 | 0.100254744 | 0.314631127 |
| chr1:24112429-24112865 | 0.66105535 | 0.100262805 | 0.314633235 |
| chr17:44066552-44066834 | 0.638237448 | 0.100263628 | 0.314633235 |
| chr5:170644801-170645087 | 0.805850863 | 0.100290181 | 0.314702519 |
| chr16:58814396-58814617 | -0.814881672 | 0.100299422 | 0.314702519 |
| chr1:210400633-210400976 | -0.633738184 | 0.100300474 | 0.314702519 |
| chr16:4768534-4768992 | 0.55945079 | 0.100302135 | 0.314702519 |
| chr4:146241972-146242753 | -0.561422697 | 0.100310738 | 0.314707785 |
| chr20:54398226-54398442 | 0.592534632 | 0.100312028 | 0.314707785 |
| chr19:6022322-6022620 | 0.736121531 | 0.100326666 | 0.314740822 |
| chr16:88926876-88927330 | -0.566662394 | 0.100334321 | 0.314751949 |
| chr19:10861235-10861449 | -0.80714534 | 0.100352543 | 0.314796223 |
| chr3:4426624-4427058 | -0.681247444 | 0.100373393 | 0.314835957 |
| chrX:85004367-85004592 | 0.741593795 | 0.100373427 | 0.314835957 |
| chr8:117615193-117615643 | -0.684547173 | 0.100389846 | 0.314874567 |
| chr15:55498068-55498648 | 0.561311656 | 0.10044345 | 0.315029799 |
| chr8:115115749-115115949 | -0.847250599 | 0.100453689 | 0.315049018 |
| chr8:94968040-94968903 | 0.559124583 | 0.100458909 | 0.315052493 |
| chr11:57402198-57402461 | -0.739202122 | 0.10047094 | 0.315065841 |
| chr15:85176977-85177527 | 0.646565091 | 0.100471389 | 0.315065841 |
| chr3:170627038-170627238 | 0.806905757 | 0.100479268 | 0.315077655 |
| chr14:30906899-30907102 | 0.806409215 | 0.100505175 | 0.315126972 |
| chr12:118061967-118062774 | 0.563654765 | 0.100507023 | 0.315126972 |
| chr1:176609586-176609845 | -0.844743193 | 0.100510898 | 0.315126972 |
| chr17:60274869-60275182 | 0.571936167 | 0.100511447 | 0.315126972 |
| chr7:25093287-25093719 | -0.593998683 | 0.100533213 | 0.315182319 |
| chr6:64724105-64724359 | -0.990287085 | 0.100543722 | 0.315202366 |
| chr21:44897719-44898086 | 0.632471485 | 0.100549507 | 0.315207607 |
| chr22:11829455-11829757 | -0.807106953 | 0.100558617 | 0.315223269 |
| chr2:200335744-200335996 | -0.81510306 | 0.100568212 | 0.315240448 |
| chr9:21255728-21256264 | -0.696482358 | 0.100586253 | 0.315278931 |
| chr14:89231837-89232107 | 0.784608654 | 0.100592051 | 0.315278931 |
| chr7:155984066-155984348 | -0.863589016 | 0.100592833 | 0.315278931 |
| chr16:56344597-56345375 | -0.554378081 | 0.100611335 | 0.315312839 |

| | | | |
|---|---|---|---|
| chr1:211382540-211383346 | -0.541977917 | 0.100611882 | 0.315312839 |
| chr12:122190910-122191205 | 0.71109505 | 0.100627842 | 0.315349961 |
| chr2:55442044-55442350 | -0.68082124 | 0.100639041 | 0.315372158 |
| chr13:46089291-46089496 | -0.791220389 | 0.100651946 | 0.315399698 |
| chr12:71731926-71732646 | -0.561600797 | 0.100677712 | 0.315456986 |
| chr11:64887898-64888195 | 0.658672624 | 0.100678587 | 0.315456986 |
| chr14:89104473-89104757 | -0.773312018 | 0.100682579 | 0.315456986 |
| chr2:23939391-23940026 | -0.586174202 | 0.100706039 | 0.315505742 |
| chr8:74338225-74338505 | 0.690233382 | 0.100706376 | 0.315505742 |
| chr1:215419017-215419217 | -0.807149751 | 0.100738198 | 0.315592536 |
| chr7:132710034-132710301 | -0.900726111 | 0.100761613 | 0.315652984 |
| chr17:36182975-36183195 | 0.803071009 | 0.10077449 | 0.315675229 |
| chr3:127100831-127101041 | -0.857580983 | 0.100776954 | 0.315675229 |
| chr11:13344154-13344505 | -0.618769169 | 0.100785415 | 0.315688827 |
| chr2:231780735-231781797 | 0.539917753 | 0.100791837 | 0.315696037 |
| chr15:63896601-63897131 | -0.629913241 | 0.100799203 | 0.315706203 |
| chr9:92988658-92988858 | 0.828617282 | 0.100808579 | 0.315721139 |
| chr2:34373895-34374191 | -0.793983581 | 0.100812213 | 0.315721139 |
| chr10:112307898-112308152 | -0.751193719 | 0.100829103 | 0.315734758 |
| chr11:75800630-75801018 | 0.640046426 | 0.100829399 | 0.315734758 |
| chr17:21201341-21201547 | -0.841458057 | 0.100831926 | 0.315734758 |
| chr15:75639716-75640014 | 0.637075192 | 0.100835049 | 0.315734758 |
| chr5:8864110-8864422 | 0.681539943 | 0.100837165 | 0.315734758 |
| chr2:124810173-124810552 | -0.676850521 | 0.100848558 | 0.315757528 |
| chr5:386406-387207 | 0.549033631 | 0.100855938 | 0.315767731 |
| chr3:113532329-113532578 | 0.638536598 | 0.100878369 | 0.315825056 |
| chr7:101244220-101245436 | -0.526394257 | 0.100884607 | 0.315830242 |
| chr8:73747931-73748223 | -0.700163463 | 0.10088827 | 0.315830242 |
| chr19:45160923-45161164 | 0.674529512 | 0.100905767 | 0.315852822 |
| chr14:96065106-96065329 | 0.697672871 | 0.100906589 | 0.315852822 |
| chr21:37366182-37367259 | 0.52928589 | 0.100907849 | 0.315852822 |
| chr20:47984031-47984285 | 0.575285995 | 0.100914554 | 0.315860904 |
| chr1:116111338-116111577 | 0.7092442 | 0.100933397 | 0.315894808 |
| chr16:84754390-84754670 | 0.658959641 | 0.100936738 | 0.315894808 |
| chr11:32487367-32487610 | 0.735055349 | 0.100937754 | 0.315894808 |
| chr7:111446045-111446335 | -0.738614111 | 0.100954173 | 0.315923709 |
| chr18:50878412-50879821 | -0.537984985 | 0.100955235 | 0.315923709 |
| chr5:135162981-135163181 | -0.922650461 | 0.100965028 | 0.315941452 |
| chr5:132272647-132272973 | 0.690778107 | 0.100985569 | 0.315992822 |
| chr13:33057513-33057851 | -0.662242508 | 0.10102787 | 0.316089417 |
| chr9:121686893-121687582 | 0.581849803 | 0.101027994 | 0.316089417 |
| chr4:70197929-70198138 | 0.814895204 | 0.101028814 | 0.316089417 |
| chr20:15336398-15336883 | -0.767017498 | 0.101054351 | 0.316156405 |
| chr10:70387269-70387918 | -0.553967219 | 0.101068148 | 0.316186658 |
| chr4:177339246-177339467 | -0.868439349 | 0.101085118 | 0.316226837 |
| chr7:47282171-47282539 | 0.679896251 | 0.101099519 | 0.316258144 |
| chr17:6420126-6420350 | -0.857783339 | 0.101103381 | 0.316258144 |
| chr9:20198376-20198639 | 0.694929783 | 0.101124328 | 0.31630546 |
| chr9:106755683-106755987 | -0.862415857 | 0.101126763 | 0.31630546 |
| chr15:100383811-100384263 | 0.624511889 | 0.101131284 | 0.316306687 |

| | | | |
|---|---|---|---|
| chr1:222096207-222096426 | -0.860464215 | 0.10115721 | 0.316371773 |
| chr22:31630362-31630586 | 0.664714468 | 0.101162483 | 0.316371773 |
| chr5:151825416-151825701 | 0.677024823 | 0.101165792 | 0.316371773 |
| chr3:94002103-94002324 | -0.839410868 | 0.101168609 | 0.316371773 |
| chr14:31206860-31208411 | 0.517956268 | 0.101189887 | 0.316425398 |
| chr9:113593871-113594203 | -0.647810328 | 0.101212568 | 0.31647524 |
| chr9:111621943-111622211 | -0.789582722 | 0.101214087 | 0.31647524 |
| chr1:28877190-28877524 | 0.64897224 | 0.101223219 | 0.316476338 |
| chr11:58286347-58286590 | 0.739376691 | 0.101227763 | 0.316476338 |
| chr3:104612133-104612478 | -0.785281066 | 0.10122803 | 0.316476338 |
| chr21:44083101-44083443 | 0.631071749 | 0.101234506 | 0.316476338 |
| chr2:209795950-209796202 | -0.777602135 | 0.10123509 | 0.316476338 |
| chr1:229557992-229559311 | 0.523815644 | 0.101239611 | 0.31647756 |
| chr3:64955050-64955593 | 0.542884117 | 0.101309311 | 0.316682523 |
| chr17:68377999-68379192 | -0.519569331 | 0.101364859 | 0.316826721 |
| chr18:12023370-12023828 | -0.707225312 | 0.101367168 | 0.316826721 |
| chr13:23140633-23140859 | 0.747016157 | 0.101369952 | 0.316826721 |
| chr19:40055743-40056388 | 0.609935913 | 0.101371981 | 0.316826721 |
| chr7:66051056-66051369 | -0.718151263 | 0.101384261 | 0.316852177 |
| chr6:107560540-107560874 | -0.815000381 | 0.101391721 | 0.316862568 |
| chr3:130328488-130329126 | 0.602074797 | 0.101398465 | 0.316870721 |
| chr17:67559489-67559702 | -0.713395652 | 0.101405713 | 0.316880446 |
| chr3:159039428-159039905 | -0.702766543 | 0.101430717 | 0.316937103 |
| chr17:2828447-2828750 | -0.716314218 | 0.101432117 | 0.316937103 |
| chr10:125912016-125912349 | 0.664995944 | 0.101451709 | 0.316976409 |
| chr9:6054531-6054815 | 0.752553845 | 0.10145297 | 0.316976409 |
| chr6:128202326-128202611 | -0.898657779 | 0.101474418 | 0.317028176 |
| chr7:77990274-77990642 | -0.735135639 | 0.101477814 | 0.317028176 |
| chr17:39174960-39175222 | -0.890190886 | 0.101492752 | 0.317061917 |
| chr6:138411241-138411441 | -0.853943487 | 0.101498453 | 0.317066799 |
| chr20:23441872-23442195 | 0.680177954 | 0.10150374 | 0.317070388 |
| chr11:18802486-18803101 | -0.629249619 | 0.10155261 | 0.317210115 |
| chr1:217089477-217090422 | -0.535938037 | 0.10158299 | 0.317291192 |
| chr17:63679281-63679538 | -0.637137606 | 0.101586849 | 0.317291192 |
| chr11:36594241-36595040 | -0.537038361 | 0.101594308 | 0.317301557 |
| chr19:41836007-41836229 | 0.740917263 | 0.10160239 | 0.317304982 |
| chr3:122660612-122660972 | -0.679014664 | 0.101603688 | 0.317304982 |
| chr12:12986060-12986337 | -0.693757557 | 0.101615287 | 0.317325691 |
| chr4:27684553-27684866 | 0.675227595 | 0.101621706 | 0.317325691 |
| chr18:108320-108572 | 0.642992483 | 0.101622743 | 0.317325691 |
| chr13:29989404-29989605 | -0.867747718 | 0.101627077 | 0.317326291 |
| chr22:23716885-23717593 | -0.585928596 | 0.101654475 | 0.317367377 |
| chr9:20423290-20423490 | -0.920413428 | 0.101655864 | 0.317367377 |
| chr8:98166200-98166429 | -0.702700175 | 0.101659393 | 0.317367377 |
| chr3:128095163-128095585 | 0.58926379 | 0.101659482 | 0.317367377 |
| chr21:42654342-42654605 | -0.684402677 | 0.101660945 | 0.317367377 |
| chr16:85311382-85311626 | -0.700792494 | 0.101666178 | 0.317370783 |
| chr3:2135195-2135395 | 0.937276255 | 0.101679984 | 0.317387691 |
| chr5:168840962-168841335 | -0.760423751 | 0.101682163 | 0.317387691 |
| chr8:114170858-114171097 | -0.867111092 | 0.101684021 | 0.317387691 |

| | | | |
|---|---|---|---|
| chr3:116140131-116140423 | -0.717170997 | 0.101707507 | 0.317448068 |
| chr14:95241718-95241920 | 0.80433344 | 0.10172044 | 0.317475501 |
| chr22:23521415-23522339 | 0.544449876 | 0.101732178 | 0.317499203 |
| chr2:54906422-54906644 | -0.908559272 | 0.101746008 | 0.317529433 |
| chr2:47016510-47016729 | 0.781402012 | 0.10176315 | 0.317569993 |
| chr9:137129618-137130457 | 0.550045995 | 0.101787439 | 0.317632857 |
| chr10:34628444-34628710 | -0.786428194 | 0.101827354 | 0.317744471 |
| chr16:50706337-50706605 | 0.756118101 | 0.101849017 | 0.317790061 |
| chr1:27608694-27608908 | 0.726189408 | 0.101850259 | 0.317790061 |
| chr7:121605293-121605577 | -0.721435743 | 0.101865514 | 0.317824718 |
| chr5:142758070-142758328 | -0.867257603 | 0.10187379 | 0.317837596 |
| chr20:42507465-42507722 | 0.756061797 | 0.101878338 | 0.317838846 |
| chr10:70559119-70559748 | -0.564829876 | 0.101890457 | 0.317863712 |
| chr2:119223404-119224235 | 0.554399557 | 0.101897965 | 0.317874192 |
| chr6:94272343-94272631 | -0.79027379 | 0.101908789 | 0.317885674 |
| chr15:68541451-68541729 | 0.756603644 | 0.101909943 | 0.317885674 |
| chr2:74130032-74130461 | -0.69337231 | 0.10193303 | 0.317936019 |
| chr2:235201862-235202177 | -0.700016704 | 0.101934382 | 0.317936019 |
| chr7:2031574-2031877 | 0.736863083 | 0.10196285 | 0.317986682 |
| chr7:155643909-155644986 | -0.540531996 | 0.101962927 | 0.317986682 |
| chr7:51263060-51263413 | -0.642357953 | 0.101963948 | 0.317986682 |
| chr12:124771855-124772360 | 0.593761783 | 0.101967765 | 0.317986682 |
| chr10:64219802-64220003 | -0.942783828 | 0.101971376 | 0.317986682 |
| chr6:35972564-35972764 | -0.972779516 | 0.101983804 | 0.318012494 |
| chr16:28505064-28505470 | 0.613748236 | 0.101998796 | 0.318046299 |
| chr1:202871369-202871683 | 0.634224464 | 0.102037343 | 0.31815355 |
| chr2:229273693-229274064 | -0.601370097 | 0.102062569 | 0.318181752 |
| chr4:87241368-87241728 | -0.69732853 | 0.102068941 | 0.318181752 |
| chr19:11738786-11739371 | 0.561922874 | 0.102073068 | 0.318181752 |
| chr3:17730394-17730659 | 0.701477103 | 0.102077581 | 0.318181752 |
| chr17:51461529-51462006 | -0.564281284 | 0.102079536 | 0.318181752 |
| chr6:139549496-139549988 | -0.643268588 | 0.102079706 | 0.318181752 |
| chr19:44989261-44989561 | 0.690705018 | 0.102084532 | 0.318181752 |
| chr4:155582959-155583220 | -0.763312003 | 0.102086526 | 0.318181752 |
| chr6:95866846-95867072 | -0.811202757 | 0.102089007 | 0.318181752 |
| chr6:53492732-53493303 | -0.634460795 | 0.1020908 | 0.318181752 |
| chr5:150993906-150994254 | 0.665350297 | 0.102092067 | 0.318181752 |
| chr6:115544902-115545883 | -0.558673073 | 0.102102251 | 0.318200549 |
| chr1:203271074-203271316 | 0.613404803 | 0.102112638 | 0.318211168 |
| chr19:17606212-17606627 | -0.637131923 | 0.102124025 | 0.318211168 |
| chr12:126874491-126874749 | -0.791489926 | 0.102124093 | 0.318211168 |
| chr20:34129714-34129947 | 0.751146045 | 0.102124405 | 0.318211168 |
| chr14:55107900-55108310 | -0.661403424 | 0.102126424 | 0.318211168 |
| chr2:1502183-1502437 | 0.785659832 | 0.102144382 | 0.318254181 |
| chr4:187380101-187380785 | -0.596201701 | 0.102158123 | 0.318284052 |
| chr7:149355023-149356091 | -0.519328894 | 0.102183624 | 0.318350559 |
| chr17:9728614-9728814 | 0.79105651 | 0.102192491 | 0.318365239 |
| chr12:12786775-12787237 | -0.611282543 | 0.102213893 | 0.318418968 |
| chr14:22559561-22559848 | 0.683012381 | 0.102231509 | 0.318460896 |
| chr14:55881940-55882140 | 0.812148394 | 0.102254964 | 0.318508062 |

| | | | |
|---|---|---|---|
| chr17:17891617-17891817 | 0.812148394 | 0.102254964 | 0.318508062 |
| chr9:37944913-37945338 | -0.669661506 | 0.10227309 | 0.318551572 |
| chr2:134692554-134692849 | -0.726977988 | 0.102278192 | 0.318554514 |
| chr4:161254508-161254736 | -0.707115144 | 0.102294632 | 0.318583416 |
| chr15:75201650-75202842 | 0.528196382 | 0.10229694 | 0.318583416 |
| chr10:77890185-77890450 | -0.670878386 | 0.102299945 | 0.318583416 |
| chr19:33509793-33510023 | 0.771219943 | 0.102313181 | 0.318611686 |
| chr3:150280583-150280783 | 0.898022439 | 0.10233266 | 0.318659395 |
| chr8:90549321-90549569 | 0.737423948 | 0.102344886 | 0.318684514 |
| chr2:213119729-213120055 | -0.738850927 | 0.102355762 | 0.31870543 |
| chr19:19663586-19664258 | -0.553187311 | 0.102365931 | 0.318724139 |
| chr1:173062219-173062549 | -0.685405375 | 0.10237466 | 0.318736649 |
| chr14:83714208-83714473 | 0.793585316 | 0.102378268 | 0.318736649 |
| chr3:195556433-195556688 | -0.754002952 | 0.102405311 | 0.318807888 |
| chr14:96592638-96592979 | 0.617290358 | 0.102424292 | 0.318843773 |
| chr2:71100526-71100852 | -0.688871765 | 0.102428686 | 0.318843773 |
| chr6:158759246-158759446 | -0.862031829 | 0.102429322 | 0.318843773 |
| chr14:33475564-33475979 | -0.639071911 | 0.102475109 | 0.318973343 |
| chr18:70454756-70454999 | -0.795323999 | 0.102489804 | 0.318993165 |
| chr4:46932048-46932265 | -0.795323999 | 0.102489804 | 0.318993165 |
| chr12:43776457-43776835 | -0.763850872 | 0.102504086 | 0.319024659 |
| chr12:128969459-128969980 | -0.601331696 | 0.102510293 | 0.319028749 |
| chr10:5949352-5949702 | 0.611831083 | 0.102513728 | 0.319028749 |
| chr1:115458835-115459081 | 0.677279088 | 0.102535434 | 0.31908334 |
| chr14:77007814-77008445 | 0.532547207 | 0.102568948 | 0.319168706 |
| chr4:138871087-138871399 | -0.633580716 | 0.102571196 | 0.319168706 |
| chr1:206030683-206031085 | -0.569188929 | 0.102607185 | 0.319262692 |
| chr20:40268342-40268622 | 0.694432243 | 0.102610384 | 0.319262692 |
| chr3:116920657-116921471 | -0.592092021 | 0.102613901 | 0.319262692 |
| chr17:62819817-62820213 | 0.575528687 | 0.10262172 | 0.319262981 |
| chr10:56327210-56327586 | -0.784909899 | 0.102622327 | 0.319262981 |
| chr4:127379826-127380070 | -0.889577254 | 0.102645286 | 0.319321441 |
| chr6:145734694-145736119 | -0.516060434 | 0.10265123 | 0.31932435 |
| chr10:93746767-93747118 | -0.63089054 | 0.102654556 | 0.31932435 |
| chr1:160941410-160941793 | -0.721117162 | 0.102682374 | 0.319397916 |
| chr11:70784813-70785038 | -0.858008984 | 0.10270416 | 0.319441481 |
| chr7:99751729-99752138 | -0.819565477 | 0.102707482 | 0.319441481 |
| chr3:150620276-150620476 | 0.814515248 | 0.102708887 | 0.319441481 |
| chr11:71466425-71466670 | -0.745250294 | 0.102723571 | 0.319474185 |
| chrX:119887662-119888464 | 0.550123751 | 0.10273104 | 0.31947781 |
| chr12:111100105-111100305 | 0.879959987 | 0.102741979 | 0.31947781 |
| chr1:212286229-212286503 | -0.640759199 | 0.10274443 | 0.31947781 |
| chr15:31034198-31034502 | 0.680854832 | 0.102744556 | 0.31947781 |
| chr3:132230266-132230557 | -0.726218645 | 0.102745585 | 0.31947781 |
| chr16:86517729-86518043 | 0.620036092 | 0.102777169 | 0.319535421 |
| chr5:115802017-115802612 | -0.598215629 | 0.102786373 | 0.319535421 |
| chr4:133921414-133921636 | 0.778993664 | 0.102794946 | 0.319535421 |
| chr5:166562414-166562614 | -0.942607016 | 0.102795785 | 0.319535421 |
| chr12:69108106-69108410 | 0.65535256 | 0.102796159 | 0.319535421 |
| chr17:879673-879919 | -0.821403186 | 0.102796412 | 0.319535421 |

| | | | |
|---|---|---|---|
| chr17:61294775-61295230 | 0.530078148 | 0.102799272 | 0.319535421 |
| chr16:77861770-77861970 | 0.789765276 | 0.102805241 | 0.319535421 |
| chr15:97652044-97652431 | -0.626308867 | 0.102812063 | 0.319535421 |
| chr12:100234205-100234551 | 0.649395912 | 0.102812459 | 0.319535421 |
| chr10:31083723-31084095 | -0.652873277 | 0.102813052 | 0.319535421 |
| chr9:87675495-87675963 | 0.619544406 | 0.102815925 | 0.319535421 |
| chr7:80019222-80019519 | -0.715222365 | 0.102818326 | 0.319535421 |
| chr5:39648807-39649111 | 0.722608573 | 0.102824378 | 0.319541268 |
| chr17:17681200-17681601 | 0.604974951 | 0.102868838 | 0.319666467 |
| chr10:77371843-77372154 | 0.692625883 | 0.102882769 | 0.319696794 |
| chr21:34086726-34087162 | 0.698550179 | 0.102888694 | 0.319702238 |
| chr9:6413985-6414340 | 0.68306756 | 0.102898892 | 0.31972096 |
| chr4:68630408-68630731 | -0.73767738 | 0.102906743 | 0.319732389 |
| chr12:84842443-84842686 | -0.853870342 | 0.102912028 | 0.319735845 |
| chr11:101127312-101128502 | -0.54899076 | 0.102922728 | 0.319756122 |
| chr8:40897674-40898011 | 0.636625617 | 0.102942008 | 0.319793249 |
| chr13:51814991-51815442 | -0.613015405 | 0.102943025 | 0.319793249 |
| chr7:117715873-117716126 | -0.729195866 | 0.102985994 | 0.319886713 |
| chr13:30465641-30466542 | -0.523471749 | 0.102990598 | 0.319886713 |
| chr7:2158758-2159001 | 0.820911323 | 0.102991362 | 0.319886713 |
| chr5:172978794-172979038 | -0.70592711 | 0.102992016 | 0.319886713 |
| chr19:54391846-54392093 | 0.753205911 | 0.102993986 | 0.319886713 |
| chr1:217696781-217697180 | -0.629738633 | 0.103013221 | 0.319933485 |
| chr3:12444709-12445045 | 0.703981629 | 0.103018883 | 0.319938101 |
| chr14:59583707-59584131 | 0.671571186 | 0.103051086 | 0.320025142 |
| chr7:4006752-4007588 | -0.562501325 | 0.103068755 | 0.320064188 |
| chr6:154981562-154981805 | -0.850954353 | 0.103072234 | 0.320064188 |
| chr3:166425994-166426216 | -0.794111336 | 0.103076191 | 0.320064188 |
| chr6:122799942-122800255 | -0.689816826 | 0.103081164 | 0.320066659 |
| chr9:81549508-81550196 | -0.599320924 | 0.10309702 | 0.320102921 |
| chr5:175444034-175444281 | 0.751375793 | 0.103108643 | 0.320118452 |
| chr1:99737022-99737327 | -0.678223742 | 0.103110378 | 0.320118452 |
| chr20:54540018-54540705 | 0.526227329 | 0.103117863 | 0.320121982 |
| chr12:121351666-121352909 | 0.527055867 | 0.103119871 | 0.320121982 |
| chr14:36450590-36450902 | -0.685827951 | 0.103130645 | 0.320125364 |
| chr7:52018157-52018804 | -0.578889458 | 0.103133005 | 0.320125364 |
| chr3:154583920-154584342 | -0.721524411 | 0.103133494 | 0.320125364 |
| chr3:59313456-59313661 | -0.83450398 | 0.103148434 | 0.320158766 |
| chr10:88974659-88974999 | -0.748391334 | 0.103162838 | 0.320190504 |
| chr9:135506908-135507108 | 0.797294736 | 0.103213572 | 0.320334994 |
| chr4:40325478-40325724 | 0.786306572 | 0.103238465 | 0.320399272 |
| chr15:44584238-44584466 | 0.732224474 | 0.103255521 | 0.320431669 |
| chr2:111122866-111123330 | 0.593352008 | 0.103257267 | 0.320431669 |
| chr4:84639220-84639532 | -0.818376767 | 0.103274255 | 0.320471407 |
| chr19:8145924-8146199 | -0.741600343 | 0.103279878 | 0.320475875 |
| chr3:109334223-109334445 | 0.826356681 | 0.103292908 | 0.320503329 |
| chr17:5485858-5487041 | -0.53032086 | 0.103302087 | 0.320514932 |
| chr1:52071702-52071902 | 0.902842869 | 0.103307153 | 0.320514932 |
| chr22:19479094-19480362 | 0.528283205 | 0.103309197 | 0.320514932 |
| chr7:90708853-90709081 | -0.843644825 | 0.103313391 | 0.320514968 |

| | | | |
|---|---|---|---|
| chr3:120137530-120137739 | 0.77824789 | 0.103325071 | 0.320538225 |
| chr17:36592749-36592992 | 0.770576182 | 0.103329312 | 0.320538401 |
| chr6:53112354-53113304 | -0.544343275 | 0.103339865 | 0.320558162 |
| chr5:163677167-163677435 | -0.847075586 | 0.103349475 | 0.320572436 |
| chr2:50536163-50536638 | -0.684128242 | 0.103352834 | 0.320572436 |
| chr2:78934751-78934961 | -0.883037203 | 0.103370082 | 0.320609764 |
| chr9:71604296-71604764 | -0.686980836 | 0.103373238 | 0.320609764 |
| chr5:472985-473399 | 0.570922794 | 0.103379791 | 0.320617111 |
| chr7:56064082-56064641 | 0.565708606 | 0.103385596 | 0.320622136 |
| chr3:27888619-27888883 | 0.740520356 | 0.10339823 | 0.320648339 |
| chr1:206555634-206555895 | -0.646427742 | 0.103414263 | 0.32068508 |
| chr16:1651888-1652088 | 0.881529128 | 0.103425822 | 0.320695191 |
| chr5:134020194-134020829 | -0.56003381 | 0.103425894 | 0.320695191 |
| chr22:29524502-29524735 | 0.794007916 | 0.103439337 | 0.320723894 |
| chr10:99450907-99451250 | -0.688245682 | 0.10344706 | 0.320734861 |
| chr14:89767845-89768045 | 0.844810611 | 0.103452499 | 0.320738745 |
| chr2:1891242-1891709 | -0.637850906 | 0.103470662 | 0.320773482 |
| chrX:85379237-85379910 | -0.567617887 | 0.103472076 | 0.320773482 |
| chr5:110115498-110115805 | -0.736326643 | 0.103477971 | 0.320778778 |
| chr17:64344405-64345041 | 0.524431091 | 0.103497485 | 0.320826292 |
| chr10:30359678-30360261 | -0.550277353 | 0.10350698 | 0.320842745 |
| chr1:64343142-64344181 | -0.525571013 | 0.103515076 | 0.320854861 |
| chr4:170875897-170876265 | -0.678331873 | 0.103549667 | 0.320949094 |
| chr2:48566504-48566896 | -0.650802511 | 0.10356849 | 0.320994453 |
| chr13:75432191-75432410 | -0.88482081 | 0.103590043 | 0.321035402 |
| chr5:32898977-32899247 | -0.837805589 | 0.103590082 | 0.321035402 |
| chr10:15371256-15371699 | -0.672736323 | 0.103602326 | 0.32106036 |
| chr8:74576209-74576913 | -0.582253191 | 0.103614244 | 0.321076227 |
| chr17:81097997-81098274 | -0.674923418 | 0.103615826 | 0.321076227 |
| chr6:33621151-33622214 | 0.546850092 | 0.103623556 | 0.321087192 |
| chr3:79835658-79835912 | 0.724460255 | 0.103649469 | 0.321141967 |
| chr6:25637916-25638264 | 0.70937899 | 0.103649616 | 0.321141967 |
| chr2:75064815-75065275 | -0.623213077 | 0.103678149 | 0.321211418 |
| chr4:109574359-109574874 | -0.616532894 | 0.103680415 | 0.321211418 |
| chr13:102726532-102726748 | -0.823662767 | 0.103689197 | 0.321215593 |
| chr4:158418711-158419048 | -0.811086607 | 0.103693268 | 0.321215593 |
| chr11:81644277-81644699 | -0.616897976 | 0.10369434 | 0.321215593 |
| chr8:92073151-92073567 | 0.652102156 | 0.103714454 | 0.321264914 |
| chr7:77439594-77439968 | -0.62033232 | 0.103722058 | 0.321275478 |
| chr5:61641835-61642575 | -0.587420146 | 0.103729615 | 0.321285899 |
| chr6:18343155-18343641 | 0.603648991 | 0.103772873 | 0.321406891 |
| chr2:25933556-25933757 | -0.951613998 | 0.103811908 | 0.321505042 |
| chr16:16194933-16195190 | 0.712121405 | 0.103812955 | 0.321505042 |
| chr22:24554920-24556081 | 0.518918471 | 0.103859018 | 0.321616612 |
| chr1:85544524-85545230 | -0.609627596 | 0.103859139 | 0.321616612 |
| chr12:87557067-87557498 | -0.715279975 | 0.103861573 | 0.321616612 |
| chr8:122157871-122158177 | 0.700773444 | 0.10386618 | 0.321617881 |
| chr3:4563065-4563266 | 0.802081865 | 0.103877443 | 0.321639757 |
| chr16:55966013-55966404 | 0.709955033 | 0.103888756 | 0.321658525 |
| chr14:105091361-105091999 | 0.542307892 | 0.1038919 | 0.321658525 |

| | | | |
|---|---|---|---|
| chr3:188268475-188268707 | -0.830456883 | 0.103933448 | 0.321749079 |
| chr4:101033998-101034337 | -0.830456883 | 0.103933448 | 0.321749079 |
| chr9:105401734-105402152 | -0.813908723 | 0.103933746 | 0.321749079 |
| chr18:48230581-48230818 | -0.775402377 | 0.103965671 | 0.321834909 |
| chr9:114434850-114435124 | 0.696611069 | 0.103986953 | 0.321870684 |
| chr19:1181694-1182065 | 0.631683775 | 0.103988358 | 0.321870684 |
| chr12:116590595-116590895 | 0.712423565 | 0.10398983 | 0.321870684 |
| chr6:169933401-169933622 | 0.71258137 | 0.103995704 | 0.321875862 |
| chr12:118182320-118182633 | 0.65679294 | 0.104012614 | 0.321915197 |
| chr5:133405851-133406051 | 0.865413705 | 0.104021535 | 0.321929803 |
| chr10:75099427-75099628 | 0.768843193 | 0.104040003 | 0.321969047 |
| chr12:92543380-92543827 | 0.597186841 | 0.10404621 | 0.321969047 |
| chr5:57887981-57888203 | 0.776638718 | 0.104046822 | 0.321969047 |
| chr1:152013678-152014001 | 0.5920031 | 0.104051318 | 0.321969957 |
| chr10:60778004-60779120 | -0.518763674 | 0.104060407 | 0.321985078 |
| chr3:64874038-64874647 | 0.532317104 | 0.104072245 | 0.322006091 |
| chr1:9181311-9181733 | 0.679353653 | 0.104075603 | 0.322006091 |
| chr2:96073173-96073432 | 0.798961639 | 0.104088554 | 0.322033158 |
| chr10:45594211-45595141 | -0.544552873 | 0.104097824 | 0.322043649 |
| chr15:45377938-45378491 | -0.631253435 | 0.104108481 | 0.322043649 |
| chr21:46228274-46229314 | 0.524186491 | 0.104109069 | 0.322043649 |
| chr1:205013822-205014226 | -0.603205503 | 0.104111498 | 0.322043649 |
| chr20:41643674-41643874 | 0.728299122 | 0.10411296 | 0.322043649 |
| chr6:163814488-163814924 | -0.674077736 | 0.104124474 | 0.322062204 |
| chr17:82397190-82397632 | 0.63120468 | 0.104127365 | 0.322062204 |
| chr2:54979686-54979985 | -0.713437007 | 0.104163138 | 0.322152954 |
| chr7:13179154-13179361 | 0.78963954 | 0.104165115 | 0.322152954 |
| chr3:189292659-189293027 | -0.668779986 | 0.104173375 | 0.322165496 |
| chr12:27922548-27922808 | -0.78519039 | 0.104185467 | 0.32217248 |
| chr8:38064100-38064477 | 0.633708479 | 0.10418558 | 0.32217248 |
| chr2:862591-863005 | -0.67096238 | 0.104188247 | 0.32217248 |
| chr11:61610045-61610391 | 0.628064467 | 0.104206279 | 0.322215235 |
| chr7:38398782-38399056 | 0.658163529 | 0.104217947 | 0.322238309 |
| chr14:70475560-70475840 | 0.752571029 | 0.10425983 | 0.322354801 |
| chr21:9839232-9839469 | 0.795020705 | 0.104264581 | 0.322356483 |
| chr13:99956928-99957272 | 0.675404704 | 0.104281397 | 0.322392449 |
| chr3:104635960-104636187 | 0.733184809 | 0.104284629 | 0.322392449 |
| chr1:156123258-156124049 | 0.539654326 | 0.104305191 | 0.322443004 |
| chr6:2841224-2842133 | -0.533269986 | 0.104314473 | 0.322458689 |
| chr16:69112770-69112970 | 0.748456989 | 0.104325808 | 0.32248072 |
| chr7:1130474-1131089 | -0.562603689 | 0.104332919 | 0.322489689 |
| chr3:99367169-99367480 | 0.775443697 | 0.104368398 | 0.322572414 |
| chr14:31356027-31356369 | 0.653012556 | 0.104372217 | 0.322572414 |
| chr5:162436218-162436474 | 0.768212633 | 0.104372312 | 0.322572414 |
| chr11:4607778-4608467 | -0.58689283 | 0.104379618 | 0.322577433 |
| chr1:1746278-1746626 | 0.648028654 | 0.104382356 | 0.322577433 |
| chr1:101797761-101798087 | -0.665323075 | 0.104417306 | 0.32266969 |
| chr2:96899056-96899256 | 0.797634801 | 0.104420631 | 0.32266969 |
| chr11:102390703-102391028 | -0.636908744 | 0.104439565 | 0.32271518 |
| chr1:248873079-248873393 | -0.54709852 | 0.104454263 | 0.322738278 |

| | | | |
|---|---|---|---|
| chr7:100091742-100091942 | -0.826065002 | 0.104461251 | 0.322738278 |
| chr8:72479036-72479701 | -0.607203231 | 0.1044641 | 0.322738278 |
| chr5:139451567-139451788 | 0.800669863 | 0.104465743 | 0.322738278 |
| chr2:9756581-9756808 | 0.756702471 | 0.1044681 | 0.322738278 |
| chr9:8857383-8857734 | 0.742635078 | 0.10447417 | 0.322744018 |
| chr3:183291705-183292036 | 0.63856717 | 0.104480055 | 0.322748291 |
| chrX:24259642-24259923 | -0.749705874 | 0.104486823 | 0.322748291 |
| chr10:6365325-6365888 | 0.601244325 | 0.10448819 | 0.322748291 |
| chr4:103719059-103719843 | -0.601610348 | 0.104506072 | 0.322772788 |
| chr3:182705046-182705246 | -0.878440108 | 0.104510036 | 0.322772788 |
| chr13:39696689-39696953 | -0.81885938 | 0.104512507 | 0.322772788 |
| chr3:116129199-116129532 | -0.727326395 | 0.104512971 | 0.322772788 |
| chr19:1244488-1244688 | 0.711441763 | 0.104533647 | 0.322816883 |
| chr14:73569749-73570102 | 0.641068799 | 0.104537334 | 0.322816883 |
| chr8:121710936-121711213 | 0.66778822 | 0.104539888 | 0.322816883 |
| chr2:195534310-195534685 | -0.702778764 | 0.104549069 | 0.322832223 |
| chr7:150346565-150346886 | -0.777911778 | 0.104559086 | 0.322849877 |
| chr5:58239716-58240154 | 0.625483118 | 0.104563214 | 0.322849877 |
| chr10:84125965-84126455 | -0.638072296 | 0.104579805 | 0.322888095 |
| chr5:71586506-71587700 | -0.516978471 | 0.10461016 | 0.322956924 |
| chr12:109777629-109777899 | -0.654839951 | 0.104613104 | 0.322956924 |
| chr11:69697005-69697238 | 0.751786837 | 0.104614743 | 0.322956924 |
| chr5:160243130-160243522 | -0.631956257 | 0.104620959 | 0.322960667 |
| chr2:153354484-153354734 | -0.900790825 | 0.104624563 | 0.322960667 |
| chr8:109953725-109954239 | -0.618732724 | 0.104628601 | 0.322960667 |
| chr16:70594633-70594958 | 0.669043392 | 0.104640942 | 0.322979092 |
| chr11:1876404-1876604 | 0.803132661 | 0.104643035 | 0.322979092 |
| chr4:98745457-98745695 | 0.80987386 | 0.104647216 | 0.322979092 |
| chr5:109701853-109702071 | 0.854497676 | 0.104678304 | 0.323060672 |
| chr2:70381888-70382219 | -0.716946964 | 0.10468208 | 0.323060672 |
| chr8:9150409-9151417 | 0.545412182 | 0.104687402 | 0.323064083 |
| chr4:179553220-179553459 | -0.845966298 | 0.104737001 | 0.323199935 |
| chr17:71438687-71438966 | 0.785077643 | 0.104741589 | 0.323199935 |
| chr12:108582275-108582568 | 0.739734145 | 0.104744079 | 0.323199935 |
| chr16:80319276-80319549 | -0.713236291 | 0.10475281 | 0.323201017 |
| chr5:180117122-180117448 | 0.612705189 | 0.10475746 | 0.323201017 |
| chr11:120684746-120685150 | 0.677130719 | 0.104758916 | 0.323201017 |
| chr11:43567478-43567708 | 0.777177285 | 0.104761302 | 0.323201017 |
| chr14:95516513-95516749 | 0.715358124 | 0.104773226 | 0.323224791 |
| chr1:247681299-247681644 | -0.602072604 | 0.104778931 | 0.323229375 |
| chr7:2921876-2922102 | -0.720308827 | 0.104812634 | 0.323320327 |
| chr19:47713178-47713403 | 0.734273726 | 0.10481709 | 0.323321058 |
| chr21:17370291-17370749 | 0.634792759 | 0.104860321 | 0.323441388 |
| chr18:57629838-57630865 | -0.544687674 | 0.104880272 | 0.323489903 |
| chr5:127079898-127080571 | -0.563739363 | 0.104885536 | 0.32349312 |
| chr1:187491081-187491304 | 0.873414096 | 0.104897518 | 0.323517052 |
| chr1:182818608-182818808 | 0.849915121 | 0.104922046 | 0.323573621 |
| chr12:131069084-131069352 | -0.739029921 | 0.104924306 | 0.323573621 |
| chr14:58501346-58501569 | 0.721543574 | 0.104931453 | 0.323582637 |
| chr11:93986592-93987074 | -0.618794341 | 0.104945736 | 0.323608575 |

| | | | |
|---|---|---|---|
| chr6:107115785-107116422 | -0.577027973 | 0.104950019 | 0.323608575 |
| chr14:78280714-78280957 | 0.802378699 | 0.104952534 | 0.323608575 |
| chr16:28218159-28218403 | 0.759778994 | 0.104979036 | 0.32367009 |
| chr1:209335174-209335475 | -0.745593949 | 0.104981444 | 0.32367009 |
| chr7:92916502-92917198 | -0.587576913 | 0.104985158 | 0.32367009 |
| chr2:127136641-127136907 | 0.818443314 | 0.104990231 | 0.323672708 |
| chr8:61336046-61336293 | -0.839935522 | 0.105002406 | 0.323697218 |
| chr14:79540442-79540748 | 0.644558609 | 0.105007623 | 0.323700276 |
| chr5:102681442-102681898 | -0.642719659 | 0.105014164 | 0.323707419 |
| chr7:120452956-120453233 | -0.816883649 | 0.105022394 | 0.323719764 |
| chr3:167684410-167684748 | -0.645909166 | 0.105030529 | 0.323731817 |
| chr1:1407067-1407482 | -0.641714171 | 0.105045573 | 0.323762287 |
| chr10:18121729-18122074 | -0.803273317 | 0.105048866 | 0.323762287 |
| chr10:125896232-125896799 | 0.560678704 | 0.105057363 | 0.323775453 |
| chr2:158359519-158360180 | -0.605123535 | 0.105065867 | 0.323788635 |
| chr2:48111016-48112250 | -0.526795988 | 0.1050866 | 0.323839506 |
| chr3:184261352-184262213 | 0.541643754 | 0.10509521 | 0.323853013 |
| chrX:155215754-155217068 | 0.528700296 | 0.105112664 | 0.323893772 |
| chr7:129779447-129780745 | -0.516773942 | 0.105137084 | 0.323955992 |
| chr11:17563454-17563675 | 0.740968561 | 0.105162456 | 0.323997849 |
| chr2:11835517-11835861 | -0.6573549 | 0.105163015 | 0.323997849 |
| chr19:4598048-4598281 | -0.831540227 | 0.105163354 | 0.323997849 |
| chr2:58532666-58533106 | -0.62600286 | 0.105174106 | 0.324017947 |
| chr12:82840918-82841281 | -0.860795128 | 0.105188634 | 0.324049675 |
| chr1:215872703-215873186 | -0.646092891 | 0.10520457 | 0.324085738 |
| chr12:113092802-113093015 | 0.700325085 | 0.105210768 | 0.324091802 |
| chr20:47698282-47698482 | 0.554723918 | 0.10521816 | 0.32410154 |
| chr9:137188446-137189465 | 0.534072124 | 0.105225189 | 0.324110163 |
| chr12:117729204-117729449 | 0.900431052 | 0.105273319 | 0.324233272 |
| chr6:31877630-31877876 | 0.697798104 | 0.10527362 | 0.324233272 |
| chr1:213196499-213196833 | -0.767514514 | 0.105278374 | 0.324234879 |
| chr1:152277620-152277930 | 0.666305878 | 0.105300703 | 0.324290613 |
| chr11:67978838-67979101 | 0.687054584 | 0.105318501 | 0.32433239 |
| chr7:137337912-137338166 | -0.791535797 | 0.105333725 | 0.324366237 |
| chr8:141227939-141228179 | -0.719567189 | 0.105341321 | 0.324376591 |
| chr3:171283691-171283921 | -0.834731848 | 0.105350635 | 0.324392235 |
| chr19:8412942-8413371 | 0.58467894 | 0.105357747 | 0.324401097 |
| chr20:48489144-48489449 | 0.562969918 | 0.105363636 | 0.324406191 |
| chr2:17697888-17698142 | -0.905574105 | 0.105379761 | 0.324442803 |
| chr1:106659235-106659478 | 0.575574353 | 0.105389696 | 0.324460354 |
| chr2:174741934-174742232 | 0.820722267 | 0.105408024 | 0.32450374 |
| chr22:20799313-20799872 | -0.588814106 | 0.105429722 | 0.3245575 |
| chr13:45310958-45311355 | -0.612118204 | 0.105439345 | 0.324574083 |
| chr3:58215043-58215346 | -0.739471247 | 0.105476596 | 0.324666695 |
| chr1:6612795-6613851 | 0.529541997 | 0.105477905 | 0.324666695 |
| chr7:5332316-5332516 | 0.75927575 | 0.105485634 | 0.324677442 |
| chr22:23192238-23192645 | 0.594358533 | 0.105501983 | 0.324690021 |
| chr1:106265034-106265278 | 0.611401052 | 0.105502429 | 0.324690021 |
| chr2:127906593-127906862 | -0.722869111 | 0.105502434 | 0.324690021 |
| chr18:2994605-2994902 | -0.735391421 | 0.105509127 | 0.32469758 |

| | | | |
|---|---|---|---|
| chr14:96039932-96040149 | 0.64238871 | 0.105540353 | 0.32478063 |
| chr5:12016070-12016307 | 0.813989964 | 0.105599851 | 0.324950674 |
| chr11:63593178-63593385 | 0.768221577 | 0.10561523 | 0.324971895 |
| chr14:89900032-89900241 | 0.768221577 | 0.10561523 | 0.324971895 |
| chrX:130393453-130393760 | 0.708273875 | 0.105626932 | 0.324994489 |
| chr3:97230563-97230796 | 0.713770541 | 0.105631056 | 0.324994489 |
| chr4:18850738-18851470 | -0.626331661 | 0.105645393 | 0.325025549 |
| chr10:78533393-78533593 | -0.921326776 | 0.105664168 | 0.325070259 |
| chr8:124728365-124728715 | 0.586390679 | 0.105669808 | 0.325074558 |
| chr20:1225357-1226072 | 0.572398002 | 0.105677699 | 0.325085782 |
| chr19:12872889-12873511 | 0.561467958 | 0.105734062 | 0.32523672 |
| chr5:104800480-104800896 | -0.717272966 | 0.105735255 | 0.32523672 |
| chr17:68049049-68049386 | -0.632095623 | 0.105742987 | 0.325247447 |
| chr12:79555494-79555797 | 0.664315664 | 0.105749999 | 0.325247823 |
| chr6:149560651-149560997 | -0.601817726 | 0.105751599 | 0.325247823 |
| chr3:14224923-14225620 | 0.562244116 | 0.105761689 | 0.325265798 |
| chr5:177474398-177474598 | 0.741048385 | 0.105807865 | 0.325386638 |
| chr5:102536426-102536676 | -0.911156225 | 0.105809474 | 0.325386638 |
| chr5:76715705-76716411 | -0.581412836 | 0.105814805 | 0.325389974 |
| chr22:41998574-41998995 | 0.642334182 | 0.105819356 | 0.325390909 |
| chrX:40472026-40472405 | 0.621194408 | 0.105828916 | 0.325404548 |
| chr2:162840653-162841003 | -0.631534346 | 0.105839948 | 0.325404548 |
| chr11:69670315-69670535 | -0.761948776 | 0.105840779 | 0.325404548 |
| chr13:44591515-44591753 | -0.761948776 | 0.105840779 | 0.325404548 |
| chr1:189720085-189720316 | -0.864917227 | 0.105851282 | 0.325412323 |
| chr3:84547896-84549074 | -0.511839949 | 0.105851802 | 0.325412323 |
| chr10:114786482-114786811 | -0.644893563 | 0.105858853 | 0.32541583 |
| chr5:122092025-122092455 | -0.658034778 | 0.105862644 | 0.32541583 |
| chr6:45310252-45310547 | -0.797703094 | 0.105866686 | 0.32541583 |
| chr19:4375030-4375964 | 0.539408757 | 0.10586993 | 0.32541583 |
| chr22:45009015-45009245 | 0.823010657 | 0.105883454 | 0.325426978 |
| chr20:50141845-50142534 | 0.544314344 | 0.105887936 | 0.325426978 |
| chr18:76709665-76710080 | -0.657858624 | 0.105888754 | 0.325426978 |
| chr20:53813759-53814266 | 0.519570421 | 0.105890546 | 0.325426978 |
| chr1:96220634-96220950 | -0.787735914 | 0.105921743 | 0.325502206 |
| chr2:68151631-68151960 | -0.808295764 | 0.105923521 | 0.325502206 |
| chr12:105331067-105331333 | -0.671360259 | 0.105946633 | 0.325560174 |
| chr8:144148488-144148805 | 0.583702247 | 0.105960407 | 0.325583442 |
| chr8:23922716-23922977 | -0.779390916 | 0.105962704 | 0.325583442 |
| chr13:41668069-41668547 | -0.572532622 | 0.105969997 | 0.325592794 |
| chr11:124762087-124762907 | 0.571946653 | 0.105979318 | 0.325608377 |
| chr2:69012893-69013366 | -0.62300635 | 0.1059858 | 0.325614979 |
| chr14:54420526-54421045 | -0.594536022 | 0.105989966 | 0.325614979 |
| chrX:38561162-38561639 | -0.598314589 | 0.106000002 | 0.325618516 |
| chr20:54563018-54563236 | 0.572509929 | 0.106002801 | 0.325618516 |
| chr7:129210628-129210881 | -0.728379837 | 0.106004469 | 0.325618516 |
| chr5:31895702-31895944 | 0.822257838 | 0.10601365 | 0.325618516 |
| chr10:102119599-102119898 | 0.605177569 | 0.10601434 | 0.325618516 |
| chr5:141692119-141692543 | -0.613165921 | 0.106016616 | 0.325618516 |
| chr7:156951665-156951882 | -0.724522066 | 0.106034841 | 0.325653026 |

| | | | |
|---|---|---|---|
| chr21:44979484-44979717 | 0.744822699 | 0.106036352 | 0.325653026 |
| chr20:45080962-45081201 | 0.726202838 | 0.106046631 | 0.325671542 |
| chr19:57279781-57280107 | 0.591936673 | 0.106061235 | 0.325703335 |
| chr6:52598197-52598594 | 0.584745012 | 0.106078989 | 0.325744801 |
| chr7:75762494-75762800 | -0.689656329 | 0.106087054 | 0.325756512 |
| chr3:49786001-49786201 | 0.757943629 | 0.106103491 | 0.325793927 |
| chr21:36028519-36028821 | 0.69467639 | 0.106114136 | 0.325813559 |
| chr17:27424013-27424213 | 0.86014506 | 0.106119861 | 0.32581808 |
| chr1:231357860-231358109 | 0.727848444 | 0.106143237 | 0.325876793 |
| chr4:149264205-149264510 | -0.690769377 | 0.106152558 | 0.325882425 |
| chr12:14773696-14774675 | -0.521886018 | 0.106153578 | 0.325882425 |
| chr1:229768158-229768585 | 0.633643757 | 0.106202114 | 0.326018364 |
| chr3:33717637-33718609 | 0.533524335 | 0.106213777 | 0.326026204 |
| chr3:63726582-63727531 | 0.514136391 | 0.106213858 | 0.326026204 |
| chr1:187413870-187414257 | -0.736230768 | 0.106217433 | 0.326026204 |
| chr8:140100418-140100999 | -0.549670955 | 0.106230842 | 0.32604429 |
| chr18:56645453-56645763 | -0.7662973 | 0.106231836 | 0.32604429 |
| chr10:112289-112553 | 0.717053305 | 0.106263066 | 0.326127079 |
| chr2:68595271-68595522 | -0.757050987 | 0.10629266 | 0.326204839 |
| chr6:70317526-70318000 | -0.70973267 | 0.106312325 | 0.326240942 |
| chr7:102434112-102434483 | -0.62723928 | 0.106314454 | 0.326240942 |
| chr12:53078742-53079696 | 0.516233684 | 0.106317198 | 0.326240942 |
| chr9:113508130-113508330 | 0.784704356 | 0.106327567 | 0.326259693 |
| chr2:36459866-36460172 | -0.641898826 | 0.106342558 | 0.326269912 |
| chr1:202177578-202177870 | 0.608753983 | 0.10634349 | 0.326269912 |
| chr5:9766575-9766830 | 0.730606791 | 0.106343671 | 0.326269912 |
| chr9:114869890-114870261 | -0.683614345 | 0.106353889 | 0.326275982 |
| chr17:72560978-72561325 | -0.588850122 | 0.106354166 | 0.326275982 |
| chr12:75944263-75944702 | 0.671648498 | 0.106368346 | 0.326306419 |
| chr3:141962514-141962901 | -0.614714494 | 0.106389119 | 0.326333948 |
| chr3:12867983-12868329 | 0.555266678 | 0.106391935 | 0.326333948 |
| chr16:70561590-70561877 | 0.70341186 | 0.106392736 | 0.326333948 |
| chr12:95820192-95820403 | -0.794495472 | 0.106394356 | 0.326333948 |
| chr9:137046422-137046804 | 0.592830515 | 0.106410526 | 0.32637048 |
| chr12:103495442-103496485 | -0.544825792 | 0.106420985 | 0.326382375 |
| chr14:22829667-22830389 | 0.545001981 | 0.106422924 | 0.326382375 |
| chr12:122637786-122638204 | 0.634248886 | 0.106434914 | 0.326406082 |
| chr2:34965667-34965881 | -0.899656366 | 0.106455312 | 0.326455571 |
| chrX:35076156-35076365 | -0.889870018 | 0.106480175 | 0.326518749 |
| chr19:14072573-14072850 | 0.66138847 | 0.106508538 | 0.326592653 |
| chr1:184548087-184548287 | 0.872586573 | 0.106518216 | 0.326609259 |
| chr8:54558961-54559319 | -0.656908536 | 0.106525609 | 0.326617856 |
| chr1:228213561-228213848 | 0.610906728 | 0.106529545 | 0.326617856 |
| chr19:45294293-45294601 | -0.69527236 | 0.106536729 | 0.32662681 |
| chr8:142545455-142545911 | 0.588357833 | 0.106542796 | 0.326632342 |
| chr6:43927123-43927574 | 0.600932074 | 0.106554433 | 0.326654948 |
| chr1:212699056-212699483 | -0.649195418 | 0.106569095 | 0.326686825 |
| chrX:105765798-105766074 | -0.719240628 | 0.106578866 | 0.326703708 |
| chr7:74051781-74052072 | -0.769742493 | 0.1066132 | 0.326787294 |
| chr19:35045791-35046022 | 0.700405398 | 0.106614664 | 0.326787294 |

| | | | |
|---|---|---|---|
| chr6:87731794-87732421 | -0.55743952 | 0.106623036 | 0.326799884 |
| chr10:25015886-25017212 | -0.509613805 | 0.106631562 | 0.326805538 |
| chr5:135566520-135566791 | -0.788539612 | 0.106635751 | 0.326805538 |
| chr9:35664778-35665439 | -0.57497166 | 0.106637677 | 0.326805538 |
| chr20:59455516-59456084 | 0.570311733 | 0.106644893 | 0.326814582 |
| chr20:53565285-53566390 | 0.509984177 | 0.106671765 | 0.3268545 |
| chr1:36397217-36398196 | 0.532861694 | 0.106674408 | 0.3268545 |
| chr1:110524247-110524832 | -0.573842533 | 0.106675375 | 0.3268545 |
| chr1:37497873-37498104 | -0.845751099 | 0.106675958 | 0.3268545 |
| chr18:49813465-49814514 | -0.539457408 | 0.10668337 | 0.3268545 |
| chr19:2034907-2035290 | 0.702046419 | 0.106683514 | 0.3268545 |
| chr3:78311440-78311640 | 0.893918366 | 0.106702313 | 0.326899025 |
| chr9:113222377-113222601 | 0.660922227 | 0.106737908 | 0.326995002 |
| chr8:103207930-103208272 | 0.606444245 | 0.106744837 | 0.327003154 |
| chr10:132964692-132964892 | 0.818138349 | 0.106752738 | 0.327014284 |
| chr2:105005571-105005982 | 0.636703958 | 0.10676987 | 0.32705369 |
| chr19:29834995-29835351 | 0.613999649 | 0.106779048 | 0.327068728 |
| chr14:33780982-33781182 | -0.814806038 | 0.106793919 | 0.3271012 |
| chr3:156561592-156561824 | 0.778009915 | 0.106816313 | 0.327146475 |
| chr4:147782529-147782830 | 0.681251493 | 0.106817239 | 0.327146475 |
| chr13:43805144-43806161 | -0.561356371 | 0.106825337 | 0.327158199 |
| chr15:97126615-97127210 | -0.60840296 | 0.106837789 | 0.327183257 |
| chr15:41971578-41971861 | -0.655544461 | 0.106846414 | 0.327196592 |
| chr1:220527995-220528747 | 0.57702611 | 0.106898368 | 0.32734261 |
| chr1:143264143-143264380 | 0.786013664 | 0.106904888 | 0.327349492 |
| chr12:131851306-131851545 | 0.666959529 | 0.106909357 | 0.327350094 |
| chr3:126355359-126355559 | -0.790030148 | 0.106926268 | 0.327353933 |
| chr5:79611814-79612989 | -0.525459704 | 0.106927525 | 0.327353933 |
| chrX:99885544-99885744 | -0.888214351 | 0.106927842 | 0.327353933 |
| chr15:23040180-23040380 | -0.817773991 | 0.106931405 | 0.327353933 |
| chr6:124759446-124759718 | -0.778807406 | 0.106933248 | 0.327353933 |
| chr14:49692972-49693594 | 0.570499101 | 0.106938762 | 0.327353933 |
| chr14:101965184-101965623 | 0.56593552 | 0.106942211 | 0.327353933 |
| chr21:34526767-34527215 | 0.615238517 | 0.106944789 | 0.327353933 |
| chr12:53099466-53099666 | 0.714467797 | 0.106956931 | 0.32737802 |
| chr13:26942402-26942666 | 0.770320568 | 0.10696343 | 0.327384835 |
| chr8:9831106-9831306 | 0.828619499 | 0.106975506 | 0.327408716 |
| chr7:56064782-56065350 | 0.615346675 | 0.106994478 | 0.327453702 |
| chr14:58238729-58240078 | -0.518872962 | 0.107013703 | 0.327499458 |
| chr1:223500741-223500941 | -0.815413573 | 0.107020199 | 0.327506259 |
| chr7:137326028-137326232 | -0.90662676 | 0.107035272 | 0.327539303 |
| chr1:40477674-40478055 | -0.662605073 | 0.107050453 | 0.327560374 |
| chr2:149087586-149088215 | -0.601645753 | 0.107050708 | 0.327560374 |
| chr4:74480250-74480844 | -0.567535719 | 0.107055902 | 0.327563188 |
| chr12:53727357-53728184 | 0.522932065 | 0.107062132 | 0.327569167 |
| chr10:116398333-116398740 | -0.710123732 | 0.107082505 | 0.327618419 |
| chr20:53713381-53713927 | 0.514369387 | 0.107095644 | 0.327645536 |
| chr12:102567194-102567526 | 0.717432425 | 0.107115872 | 0.327694335 |
| chr22:31107278-31108138 | 0.536880648 | 0.107125194 | 0.327709772 |
| chr12:47949509-47949792 | -0.642093282 | 0.107145234 | 0.327749623 |

| | | | |
|---|---|---|---|
| chr10:91062155-91062439 | -0.843708374 | 0.107146777 | 0.327749623 |
| chr12:101523237-101523622 | -0.69923059 | 0.107156808 | 0.327767224 |
| chr5:159772880-159773091 | -0.811142867 | 0.107166393 | 0.327783458 |
| chr15:38702454-38702970 | -0.624883512 | 0.107186997 | 0.327833389 |
| chr4:105894922-105896674 | 0.515839254 | 0.107196671 | 0.327849892 |
| chr22:35380956-35381192 | 0.755103426 | 0.107209034 | 0.327866299 |
| chr10:68156497-68156703 | -0.858952648 | 0.107210594 | 0.327866299 |
| chr11:124195203-124195409 | 0.813225385 | 0.107219277 | 0.327879767 |
| chr2:192035792-192036357 | -0.619285052 | 0.10722707 | 0.327890513 |
| chr1:154870177-154870618 | 0.55656424 | 0.107240791 | 0.327913384 |
| chr5:76337940-76338326 | -0.609410642 | 0.107243109 | 0.327913384 |
| chr9:78369687-78370028 | -0.646877653 | 0.107257594 | 0.327931797 |
| chr11:77020165-77020583 | -0.619024987 | 0.10725769 | 0.327931797 |
| chr6:163579772-163580196 | -0.728738976 | 0.107270056 | 0.327956517 |
| chr17:35588529-35588745 | 0.903220529 | 0.107309479 | 0.328063955 |
| chr3:166060625-166061548 | -0.574618286 | 0.107323943 | 0.328095083 |
| chr20:54164412-54164644 | 0.558639293 | 0.107332738 | 0.32810888 |
| chr21:41280467-41280777 | -0.764754078 | 0.107339795 | 0.328117362 |
| chr4:160701788-160702081 | 0.796736572 | 0.107352084 | 0.328126114 |
| chr8:87832088-87832428 | 0.581848117 | 0.107352313 | 0.328126114 |
| chr5:121961537-121962450 | -0.517449543 | 0.107355506 | 0.328126114 |
| chr13:26707550-26708160 | -0.586151108 | 0.107366052 | 0.32814526 |
| chr1:207885089-207885382 | -0.637830698 | 0.107372595 | 0.328152166 |
| chr17:28604836-28605124 | 0.827898282 | 0.107387737 | 0.328177801 |
| chr7:84106267-84106578 | -0.782421162 | 0.107389549 | 0.328177801 |
| chrX:94854073-94855028 | -0.533724134 | 0.107402249 | 0.328193673 |
| chr2:125399114-125399578 | -0.602950385 | 0.107403309 | 0.328193673 |
| chr7:130732820-130733131 | -0.751621076 | 0.107410816 | 0.328203524 |
| chrX:118727006-118728035 | 0.527657355 | 0.107435551 | 0.328266013 |
| chr5:111133369-111133662 | -0.749775046 | 0.107462303 | 0.32833466 |
| chr17:56319513-56319872 | -0.616834848 | 0.107475837 | 0.328342585 |
| chr6:140486149-140486395 | -0.821555672 | 0.107477753 | 0.328342585 |
| chr7:4226425-4226674 | -0.821555672 | 0.107477753 | 0.328342585 |
| chr2:230219792-230220297 | -0.576727528 | 0.107492809 | 0.328375488 |
| chr15:93955104-93955313 | -0.858232254 | 0.10751001 | 0.328414419 |
| chr12:98397234-98398109 | -0.532909452 | 0.107514125 | 0.328414419 |
| chr13:42490974-42491339 | -0.731094609 | 0.107524586 | 0.328414438 |
| chr7:11748980-11749214 | 0.70566888 | 0.107526764 | 0.328414438 |
| chr21:41420105-41420944 | -0.540929071 | 0.107527767 | 0.328414438 |
| chr5:170744191-170744816 | -0.569152303 | 0.107531276 | 0.328414438 |
| chr2:172172478-172172822 | 0.645490061 | 0.10754284 | 0.328436663 |
| chr13:35720757-35721307 | -0.556775409 | 0.107548073 | 0.328439557 |
| chr4:2462795-2463116 | 0.717912005 | 0.107553805 | 0.328443969 |
| chr1:169529022-169529851 | -0.53384611 | 0.107596823 | 0.328562242 |
| chr12:22624700-22626153 | -0.520841317 | 0.107612587 | 0.328597284 |
| chr6:151633470-151634119 | 0.549366567 | 0.107619355 | 0.328604855 |
| chr5:1443105-1443576 | 0.604566625 | 0.107629338 | 0.32862224 |
| chr12:48925331-48925625 | -0.697728817 | 0.107637897 | 0.328635279 |
| chr15:90648098-90648383 | 0.633510913 | 0.10764412 | 0.328641184 |
| chr14:65959354-65959625 | -0.665026935 | 0.107661732 | 0.328681856 |

| | | | |
|---|---|---|---|
| chr6:145187381-145187970 | -0.565544692 | 0.107679726 | 0.328714422 |
| chr2:177618255-177619262 | -0.523539867 | 0.107680979 | 0.328714422 |
| chrX:153520993-153521285 | -0.741288849 | 0.107694748 | 0.328727126 |
| chr18:8612262-8612604 | 0.604013258 | 0.107696319 | 0.328727126 |
| chr2:240983191-240983401 | 0.827309411 | 0.107703262 | 0.328727126 |
| chr14:105622479-105622957 | 0.575066345 | 0.107706504 | 0.328727126 |
| chr4:15396233-15396523 | 0.66346456 | 0.107715287 | 0.328727126 |
| chrX:25523703-25523903 | -0.883244125 | 0.107715295 | 0.328727126 |
| chr22:22397118-22397583 | -0.656529238 | 0.107718252 | 0.328727126 |
| chrX:4670718-4670963 | 0.790005096 | 0.107719463 | 0.328727126 |
| chr7:616083-616283 | 0.768038436 | 0.107737339 | 0.328768584 |
| chr15:70825114-70825644 | 0.589543915 | 0.107766802 | 0.328845395 |
| chr11:84899777-84900081 | -0.670743828 | 0.10778416 | 0.328885267 |
| chr20:54894034-54894416 | 0.540319634 | 0.107794333 | 0.328903208 |
| chr5:170380380-170380601 | 0.756380497 | 0.10780364 | 0.32891851 |
| chr2:23872453-23872783 | -0.80952877 | 0.107825199 | 0.328971187 |
| chr15:101017396-101017662 | -0.691132167 | 0.107832428 | 0.328974441 |
| chr9:90420280-90420611 | 0.699583498 | 0.107834852 | 0.328974441 |
| chr8:104746470-104746689 | 0.727553562 | 0.107844428 | 0.328977535 |
| chr9:93313049-93313295 | 0.715633197 | 0.107844453 | 0.328977535 |
| chr2:35112367-35112729 | -0.645716056 | 0.107860657 | 0.329013865 |
| chr5:115839078-115839306 | -0.64505576 | 0.10786832 | 0.329018067 |
| chr20:47180740-47181079 | 0.524102485 | 0.107870623 | 0.329018067 |
| chr2:53442859-53443294 | -0.645833869 | 0.107910846 | 0.329127652 |
| chr2:120245289-120245804 | 0.567161063 | 0.107917389 | 0.32913022 |
| chr2:199973732-199974104 | -0.661778108 | 0.10792269 | 0.32913022 |
| chr5:34181895-34182225 | 0.70362896 | 0.107924575 | 0.32913022 |
| chr1:94579856-94580482 | -0.612802392 | 0.107943505 | 0.329174848 |
| chr8:133375303-133376018 | -0.532695559 | 0.107980355 | 0.329271018 |
| chr13:29771309-29771537 | -0.817439642 | 0.107983636 | 0.329271018 |
| chr10:98748439-98748659 | -0.876278948 | 0.107994076 | 0.32927528 |
| chr13:86567162-86567437 | -0.85330115 | 0.107996412 | 0.32927528 |
| chr8:143793800-143794000 | 0.704671964 | 0.108000232 | 0.32927528 |
| chr1:80218171-80218436 | -0.741862099 | 0.108003324 | 0.32927528 |
| chr3:146745437-146745719 | 0.75106756 | 0.108006521 | 0.32927528 |
| chr20:57184204-57184672 | 0.515001661 | 0.10803624 | 0.329352782 |
| chr7:121658165-121658424 | -0.781576735 | 0.108042312 | 0.329357972 |
| chr21:8420006-8420651 | 0.514088662 | 0.10804654 | 0.329357972 |
| chr12:53845291-53845796 | 0.581371465 | 0.108062603 | 0.329393833 |
| chr1:145599477-145599717 | 0.729306848 | 0.10807033 | 0.329404282 |
| chr11:130827794-130828128 | -0.639369995 | 0.108076292 | 0.329409348 |
| chr9:90801241-90802235 | 0.563244685 | 0.108091491 | 0.32944257 |
| chr4:183098241-183098748 | -0.594716722 | 0.108113487 | 0.329484891 |
| chr2:152123520-152123771 | 0.873132104 | 0.108113977 | 0.329484891 |
| chr10:117663829-117664109 | -0.786866886 | 0.108133567 | 0.329531486 |
| chr20:37342849-37343054 | -0.820580787 | 0.108140416 | 0.329533979 |
| chr1:85707818-85708803 | -0.524842262 | 0.108152577 | 0.329533979 |
| chr6:41073582-41074016 | 0.552522629 | 0.108156571 | 0.329533979 |
| chr11:33311152-33311485 | -0.778901269 | 0.108157496 | 0.329533979 |
| chr13:30976687-30976958 | -0.778901269 | 0.108157496 | 0.329533979 |

| | | | |
|---|---|---|---|
| chr3:65051853-65052386 | -0.670986815 | 0.108160945 | 0.329533979 |
| chr8:17001508-17001762 | -0.903758555 | 0.108168559 | 0.329533979 |
| chr1:218031900-218032159 | -0.718963062 | 0.108168791 | 0.329533979 |
| chr3:97896722-97896948 | -0.910216014 | 0.108196833 | 0.329594489 |
| chr6:14046539-14046771 | -0.855240766 | 0.108200242 | 0.329594489 |
| chr5:95892811-95893114 | -0.659144031 | 0.108201558 | 0.329594489 |
| chr6:16781284-16781703 | 0.646625248 | 0.10823619 | 0.329674902 |
| chr19:54431517-54431908 | 0.61178059 | 0.108236931 | 0.329674902 |
| chr14:90658701-90658943 | 0.75972894 | 0.108251551 | 0.329674902 |
| chr8:119226406-119226621 | -0.772106572 | 0.10825239 | 0.329674902 |
| chr15:66655678-66656310 | -0.573420814 | 0.108257276 | 0.329674902 |
| chr16:87705742-87706039 | 0.632401399 | 0.108261143 | 0.329674902 |
| chr17:19469451-19469651 | 0.762228755 | 0.108261727 | 0.329674902 |
| chr11:86524031-86524573 | 0.602679364 | 0.108262378 | 0.329674902 |
| chr4:26022416-26022616 | 0.83321449 | 0.108269006 | 0.329681984 |
| chr4:76579581-76580100 | -0.622087037 | 0.108277721 | 0.329695421 |
| chr8:86356723-86356967 | 0.672556477 | 0.108283176 | 0.329698927 |
| chr12:6200920-6201492 | 0.571986439 | 0.108306985 | 0.329758317 |
| chr1:11099713-11100091 | 0.58077393 | 0.108318739 | 0.329780999 |
| chr19:50705533-50705793 | 0.693301126 | 0.108327705 | 0.329795192 |
| chr3:107241219-107241621 | -0.590070324 | 0.108351671 | 0.329855049 |
| chr2:2172280-2172480 | -0.88898662 | 0.108367848 | 0.32987816 |
| chr1:32335989-32336435 | 0.561421981 | 0.108367873 | 0.32987816 |
| chr14:57112265-57112797 | -0.677101499 | 0.108373396 | 0.329881866 |
| chr2:211455288-211455525 | -0.788911313 | 0.108397506 | 0.329941471 |
| chr6:31729214-31730385 | 0.523360134 | 0.10840377 | 0.329941471 |
| chr5:23446533-23446848 | -0.73518823 | 0.108405896 | 0.329941471 |
| chr1:184373274-184373477 | -0.82523521 | 0.108427101 | 0.329992905 |
| chr9:133861876-133862076 | -0.940200861 | 0.10843337 | 0.329998875 |
| chr7:155768429-155768633 | -0.803728914 | 0.108441532 | 0.330010607 |
| chr12:110338886-110339105 | -0.854203342 | 0.108457272 | 0.330045398 |
| chr21:43690641-43690898 | -0.779780639 | 0.108468877 | 0.330064056 |
| chr7:55133901-55134140 | -0.8509503 | 0.108475529 | 0.330064056 |
| chr16:11257312-11257572 | 0.592297484 | 0.108476326 | 0.330064056 |
| chr12:128329918-128330304 | -0.73319246 | 0.108499271 | 0.330116343 |
| chr5:72245016-72245224 | -0.813247532 | 0.108502127 | 0.330116343 |
| chr10:44043343-44043696 | 0.727729101 | 0.108526493 | 0.330177364 |
| chr20:53332033-53332320 | 0.764803155 | 0.108533613 | 0.330185918 |
| chr7:140175200-140175808 | -0.540725832 | 0.108555731 | 0.330240093 |
| chr1:16366929-16367989 | 0.533984323 | 0.108581253 | 0.330301609 |
| chr6:68634567-68635655 | -0.525960737 | 0.108585424 | 0.330301609 |
| chr22:49962399-49962661 | 0.662162008 | 0.108592552 | 0.330301609 |
| chr6:126550369-126550820 | -0.595904301 | 0.108593196 | 0.330301609 |
| chr2:189892884-189893194 | -0.756116388 | 0.108609328 | 0.330309496 |
| chr14:38835626-38835912 | 0.764113724 | 0.108609399 | 0.330309496 |
| chr5:59587511-59587846 | -0.617954681 | 0.108611337 | 0.330309496 |
| chr2:50746357-50746952 | -0.613840244 | 0.108613032 | 0.330309496 |
| chr7:5555883-5556263 | 0.554863117 | 0.108621935 | 0.330323461 |
| chr1:51841242-51841453 | 0.824643648 | 0.108631643 | 0.330339871 |
| chr2:21395424-21395691 | -0.667913867 | 0.108637452 | 0.330344425 |

| | | | |
|---|---|---|---|
| chr18:3236475-3236825 | 0.600058035 | 0.108658936 | 0.330368925 |
| chr1:115714528-115715324 | -0.518866642 | 0.108660835 | 0.330368925 |
| chr4:93710176-93710548 | 0.630544338 | 0.108661553 | 0.330368925 |
| chr1:106632887-106633416 | 0.531920797 | 0.108662755 | 0.330368925 |
| chr11:82038295-82038527 | -0.827904516 | 0.108680932 | 0.330411078 |
| chr8:139545538-139545836 | -0.79519331 | 0.108700724 | 0.330438563 |
| chr21:36070222-36070586 | 0.648107362 | 0.108702153 | 0.330438563 |
| chr1:187941767-187942013 | -0.843863979 | 0.108704181 | 0.330438563 |
| chr20:41078429-41078668 | 0.671200573 | 0.10871122 | 0.330438563 |
| chr20:33992727-33992927 | -0.678360277 | 0.108711536 | 0.330438563 |
| chr19:41204287-41204571 | -0.731401541 | 0.108722593 | 0.330456956 |
| chr10:125064651-125065131 | 0.563279734 | 0.108726213 | 0.330456956 |
| chr4:6313239-6313612 | 0.61277301 | 0.108733881 | 0.330467154 |
| chr9:119375951-119376255 | -0.701591831 | 0.10875949 | 0.330531875 |
| chr9:113808270-113808476 | -0.864408023 | 0.108808087 | 0.330666452 |
| chr3:149395798-149396153 | 0.602869673 | 0.108815119 | 0.330674706 |
| chr2:102488149-102488520 | 0.649233596 | 0.108840053 | 0.33073696 |
| chr1:160084028-160084363 | 0.605248227 | 0.108844238 | 0.33073696 |
| chr5:180289537-180289844 | -0.578159138 | 0.108857985 | 0.330758144 |
| chr10:116787850-116788085 | 0.758427719 | 0.108859843 | 0.330758144 |
| chr2:61854381-61855414 | -0.512294063 | 0.108901129 | 0.330839221 |
| chr3:56042397-56042651 | 0.7182007 | 0.108905926 | 0.330839221 |
| chr2:30707058-30707361 | 0.60308831 | 0.108905928 | 0.330839221 |
| chr9:109253969-109254303 | -0.696943712 | 0.108906674 | 0.330839221 |
| chr1:218323511-218323762 | -0.703032808 | 0.108911188 | 0.330839221 |
| chr1:88683649-88685159 | -0.511588949 | 0.108914997 | 0.330839221 |
| chr4:141132511-141132985 | 0.595212603 | 0.108919195 | 0.330839221 |
| chr6:135909788-135910137 | -0.647669064 | 0.108921069 | 0.330839221 |
| chr7:109092214-109092556 | -0.689068609 | 0.108961081 | 0.330947633 |
| chr3:157582427-157582627 | -0.812901134 | 0.108969942 | 0.330952057 |
| chr5:125478584-125479050 | -0.599839208 | 0.108971176 | 0.330952057 |
| chr18:70781800-70782220 | -0.643401811 | 0.108990266 | 0.330996915 |
| chr2:96116867-96117348 | 0.575034049 | 0.108997047 | 0.331002783 |
| chr12:101818032-101818241 | -0.841598705 | 0.109008151 | 0.331002783 |
| chr8:115375481-115375717 | 0.809490523 | 0.109008594 | 0.331002783 |
| chr7:12389090-12389438 | 0.605361695 | 0.109010661 | 0.331002783 |
| chr17:58352895-58353147 | -0.621634373 | 0.109013798 | 0.331002783 |
| chr3:111667333-111667558 | 0.760749902 | 0.109030272 | 0.331039685 |
| chr6:142385499-142385775 | 0.69981385 | 0.109038033 | 0.331050132 |
| chr3:94121126-94121368 | -0.821805406 | 0.109050878 | 0.33105978 |
| chr6:96939377-96939611 | -0.821805406 | 0.109050878 | 0.33105978 |
| chr8:128361901-128362280 | -0.648195183 | 0.109061554 | 0.33105978 |
| chr5:21688681-21688881 | 0.758247789 | 0.109062808 | 0.33105978 |
| chr3:195859346-195859616 | 0.695677171 | 0.109065521 | 0.33105978 |
| chr19:39166560-39167490 | 0.528750433 | 0.109067135 | 0.33105978 |
| chrX:11759243-11759694 | -0.562213336 | 0.109095203 | 0.33113186 |
| chr11:76782961-76783670 | -0.529617153 | 0.109117864 | 0.331187521 |
| chr4:24911966-24913189 | 0.521687781 | 0.109130199 | 0.331211838 |
| chr7:76317706-76319007 | -0.514039626 | 0.109135012 | 0.331213329 |
| chr6:21433704-21433992 | -0.714198269 | 0.10914876 | 0.331241932 |

| | | | |
|---|---|---|---|
| chr1:81802786-81803045 | -0.707944095 | 0.109163997 | 0.33127505 |
| chr20:46791790-46792015 | 0.714769119 | 0.109176792 | 0.33130076 |
| chr3:9916805-9917391 | 0.531778575 | 0.109186318 | 0.331307193 |
| chr12:26032803-26033003 | -0.854226003 | 0.10918756 | 0.331307193 |
| chr7:1059377-1060018 | -0.555643462 | 0.109199839 | 0.331323431 |
| chr1:120850379-120851478 | 0.54165207 | 0.10920156 | 0.331323431 |
| chr12:64883619-64883954 | -0.660601412 | 0.109217906 | 0.331337885 |
| chr2:204108294-204108556 | -0.669963966 | 0.109218999 | 0.331337885 |
| chr3:129205068-129205363 | 0.722869441 | 0.109219297 | 0.331337885 |
| chr4:147160973-147161233 | 0.726800346 | 0.109226776 | 0.331347457 |
| chr12:53767522-53768047 | 0.586317019 | 0.109246274 | 0.331390713 |
| chr4:55125975-55126273 | -0.825550917 | 0.109249713 | 0.331390713 |
| chr1:214516931-214517131 | -0.798869688 | 0.109254011 | 0.331390713 |
| chr22:43537052-43537408 | -0.724114984 | 0.10926154 | 0.331400433 |
| chr20:10659754-10660386 | -0.57850604 | 0.109273648 | 0.331424036 |
| chr17:69394341-69395052 | -0.536129832 | 0.109291775 | 0.331458798 |
| chr14:43795744-43795968 | 0.717402884 | 0.109293761 | 0.331458798 |
| chr5:105695899-105696191 | -0.756358817 | 0.109302256 | 0.331471442 |
| chr12:132768419-132768744 | -0.683302287 | 0.109309922 | 0.33148157 |
| chr17:56840333-56840545 | 0.799275529 | 0.109331104 | 0.331532683 |
| chrX:8731595-8732487 | 0.551448055 | 0.109335697 | 0.331533491 |
| chr9:70392256-70392501 | -0.828895452 | 0.1093444 | 0.331533639 |
| chr9:133866947-133867179 | -0.828895452 | 0.1093444 | 0.331533639 |
| chr4:88211988-88212361 | 0.637470312 | 0.10935282 | 0.331539462 |
| chr3:175186603-175186850 | -0.800433983 | 0.109356758 | 0.331539462 |
| chr16:49857700-49858023 | 0.685829157 | 0.109359301 | 0.331539462 |
| chr1:248752113-248752520 | -0.606239601 | 0.109368391 | 0.331553902 |
| chr1:28314083-28314300 | 0.798893123 | 0.109380398 | 0.331566753 |
| chr14:35356191-35356395 | 0.72959208 | 0.109381285 | 0.331566753 |
| chr5:66717419-66717641 | -0.868001939 | 0.109391973 | 0.331582199 |
| chr11:78179529-78179989 | -0.626778328 | 0.109395035 | 0.331582199 |
| chr7:140336104-140336341 | -0.862986189 | 0.109399973 | 0.331584049 |
| chr3:185253815-185254444 | -0.592643524 | 0.10942584 | 0.331649331 |
| chr6:39092276-39092503 | 0.801836411 | 0.109431678 | 0.331653905 |
| chr12:55019829-55020193 | -0.636256119 | 0.109452075 | 0.331702603 |
| chr1:199243318-199243816 | -0.547933725 | 0.109464318 | 0.331713029 |
| chr5:143355726-143356104 | -0.588747148 | 0.109465331 | 0.331713029 |
| chr5:39692078-39692305 | 0.70326188 | 0.109468503 | 0.331713029 |
| chr1:153986720-153987286 | 0.532882453 | 0.109491151 | 0.331768535 |
| chr1:197081773-197081973 | -0.82370984 | 0.109512072 | 0.3318117 |
| chr20:59302731-59302975 | 0.665843987 | 0.109514057 | 0.3318117 |
| chr5:69407590-69407835 | -0.817624553 | 0.109540592 | 0.331878974 |
| chr9:129174835-129175035 | 0.681238901 | 0.109552055 | 0.331900581 |
| chr2:42493887-42495296 | -0.513900505 | 0.109574093 | 0.33195422 |
| chr5:69364478-69364741 | -0.696931655 | 0.109593859 | 0.332000975 |
| chr7:139321111-139321350 | -0.761240814 | 0.109607824 | 0.332030154 |
| chr11:79186318-79186524 | 0.883352687 | 0.109613166 | 0.332033209 |
| chr1:186315783-186315983 | -0.899046947 | 0.109619777 | 0.332040108 |
| chr16:740788-741585 | 0.539202694 | 0.109710364 | 0.332301359 |
| chr1:204129081-204129975 | -0.510943095 | 0.109723283 | 0.332327353 |

| | | | |
|---|---|---|---|
| chr13:102591977-102592512 | -0.575681671 | 0.109730376 | 0.332335699 |
| chr12:104295360-104295661 | -0.63626424 | 0.109741061 | 0.332354925 |
| chr3:58176870-58177131 | -0.793325671 | 0.109766008 | 0.332417338 |
| chr4:125741830-125742187 | 0.643650204 | 0.109773699 | 0.33242749 |
| chrX:144334677-144334877 | 0.77425349 | 0.109787819 | 0.332457109 |
| chr5:9801796-9802009 | 0.763800397 | 0.109798117 | 0.332475156 |
| chr4:60836741-60836941 | 0.85155045 | 0.109832743 | 0.332566862 |
| chr17:82103999-82104342 | 0.574761624 | 0.10984006 | 0.33257306 |
| chr16:67173794-67174621 | 0.521965484 | 0.109843471 | 0.33257306 |
| chr12:46131822-46132056 | 0.782811831 | 0.109849669 | 0.332578683 |
| chr5:8340758-8341041 | -0.732300485 | 0.109863345 | 0.332606948 |
| chr4:25233558-25235023 | 0.515336764 | 0.10988024 | 0.332634325 |
| chr18:76351934-76352501 | -0.548954371 | 0.109881071 | 0.332634325 |
| chr1:19484874-19485079 | 0.740945663 | 0.109932135 | 0.33276955 |
| chr1:171644869-171645505 | -0.52724672 | 0.10993446 | 0.33276955 |
| chr22:25333938-25334409 | 0.652024706 | 0.109938769 | 0.33276955 |
| chr15:80971294-80971502 | 0.79898022 | 0.109954005 | 0.332802519 |
| chr13:32415101-32415350 | -0.69855074 | 0.109965213 | 0.332823296 |
| chr20:46520996-46521353 | 0.601355672 | 0.11003006 | 0.333006408 |
| chr11:33682396-33682808 | -0.639797283 | 0.110039958 | 0.33302321 |
| chr10:73957247-73957688 | -0.650347004 | 0.110060543 | 0.333072353 |
| chr1:203601227-203601475 | 0.643198846 | 0.110086502 | 0.333137754 |
| chr10:62594037-62594378 | -0.633556353 | 0.110091598 | 0.333140018 |
| chr3:141212174-141212454 | 0.709383964 | 0.110108157 | 0.33317697 |
| chr8:60847527-60847733 | -0.898990038 | 0.110149298 | 0.333288295 |
| chr4:165327193-165328251 | -0.504771673 | 0.110158778 | 0.333297057 |
| chr14:81294264-81294464 | 0.876733846 | 0.110160893 | 0.333297057 |
| chr14:52313892-52314483 | 0.559312333 | 0.110193636 | 0.333382958 |
| chr9:128420140-128420505 | 0.591972341 | 0.110213752 | 0.333429416 |
| chr2:71714810-71715010 | -0.851690034 | 0.11022409 | 0.333429416 |
| chr11:121649817-121650098 | -0.729937564 | 0.110225876 | 0.333429416 |
| chr3:179410429-179410667 | -0.778311124 | 0.110226399 | 0.333429416 |
| chr12:121918303-121919073 | -0.579996034 | 0.110235195 | 0.333437275 |
| chr16:88811313-88812888 | -0.504512046 | 0.110238909 | 0.333437275 |
| chr12:122961470-122961994 | -0.582347048 | 0.110242052 | 0.333437275 |
| chr10:60008934-60009304 | -0.666370742 | 0.11025932 | 0.33347634 |
| chr4:138897749-138898511 | -0.536686464 | 0.110280291 | 0.333526602 |
| chr5:150161189-150161953 | -0.527255085 | 0.110297271 | 0.33356479 |
| chr19:11339570-11339982 | 0.657772387 | 0.110334236 | 0.33366341 |
| chr3:187739990-187740241 | -0.638306588 | 0.110359018 | 0.333725182 |
| chr15:75749854-75750063 | 0.836323843 | 0.110373949 | 0.333757161 |
| chr15:95026206-95026508 | 0.654734548 | 0.110395554 | 0.333809318 |
| chr3:102063254-102063454 | 0.868640481 | 0.110416283 | 0.333858724 |
| chr1:59426934-59427590 | -0.546278512 | 0.110422779 | 0.333858724 |
| chr17:73506437-73506704 | 0.690070196 | 0.110424965 | 0.333858724 |
| chr8:51809170-51809637 | -0.626671302 | 0.110446536 | 0.333910768 |
| chr9:78296168-78296531 | -0.684758999 | 0.110460738 | 0.333927325 |
| chr2:215529334-215529757 | -0.664082053 | 0.110460797 | 0.333927325 |
| chr2:149020243-149020874 | 0.557228497 | 0.110465087 | 0.333927325 |
| chr2:12177014-12177263 | 0.720080916 | 0.110483159 | 0.333967486 |

| | | | |
|---|---|---|---|
| chr19:16363340-16363540 | -0.939556182 | 0.110487695 | 0.333967486 |
| chr1:106879337-106880494 | 0.504035008 | 0.110492574 | 0.333967486 |
| chr12:12926949-12927922 | -0.52754166 | 0.110495807 | 0.333967486 |
| chr14:100239888-100240152 | 0.617590852 | 0.110540169 | 0.334088389 |
| chr14:41609784-41610215 | 0.561455113 | 0.110547261 | 0.334091623 |
| chr5:97142499-97142983 | -0.611594942 | 0.11054996 | 0.334091623 |
| chr3:64507996-64509177 | 0.505103333 | 0.110573425 | 0.334149356 |
| chr18:34647301-34647597 | -0.744842916 | 0.110579565 | 0.334154733 |
| chr15:79314876-79315160 | 0.625761793 | 0.110594277 | 0.334171962 |
| chr5:177939256-177939626 | 0.634507291 | 0.110597079 | 0.334171962 |
| chr2:144890826-144891196 | -0.617224658 | 0.110602387 | 0.334171962 |
| chr12:107405060-107405353 | 0.655615321 | 0.110608131 | 0.334171962 |
| chr13:43132151-43132625 | -0.576658049 | 0.110610503 | 0.334171962 |
| chr6:149840614-149841018 | -0.628230949 | 0.110623103 | 0.334171962 |
| chr6:97947794-97948290 | -0.610494738 | 0.110624034 | 0.334171962 |
| chr2:138250226-138250442 | -0.838441647 | 0.110624422 | 0.334171962 |
| chr19:40523226-40523611 | 0.618686661 | 0.110627332 | 0.334171962 |
| chr12:93085563-93085820 | -0.796178716 | 0.110628879 | 0.334171962 |
| chr15:94236742-94237006 | -0.766147001 | 0.110636637 | 0.334182222 |
| chr17:17849138-17849472 | -0.64311019 | 0.110672757 | 0.334278146 |
| chr2:178280318-178280579 | -0.720189296 | 0.110690332 | 0.334318049 |
| chr20:55798893-55799102 | 0.759648819 | 0.110701133 | 0.334324413 |
| chr5:55172047-55172398 | -0.586326 | 0.110701165 | 0.334324413 |
| chr12:123583938-123584989 | 0.531136958 | 0.110729391 | 0.334396476 |
| chr14:69002111-69002376 | 0.644400978 | 0.110739317 | 0.334408468 |
| chr6:96391806-96392069 | -0.755191922 | 0.110742091 | 0.334408468 |
| chr8:78615357-78615633 | -0.702190554 | 0.110750112 | 0.334419512 |
| chr1:174165602-174165812 | -0.805928155 | 0.11075663 | 0.334426012 |
| chr12:27751668-27751985 | -0.808446759 | 0.11078002 | 0.334483457 |
| chr8:91008787-91009100 | -0.817342442 | 0.110805646 | 0.334525342 |
| chr13:112640651-112641055 | -0.702722752 | 0.110810069 | 0.334525342 |
| chr14:100377807-100378137 | 0.57101403 | 0.110810854 | 0.334525342 |
| chr22:35299590-35300762 | 0.515854023 | 0.110814715 | 0.334525342 |
| chr17:69098911-69099158 | 0.801935933 | 0.110815722 | 0.334525342 |
| chr6:156661974-156662484 | -0.62108345 | 0.11082476 | 0.334528925 |
| chrX:131675678-131675965 | 0.674285139 | 0.11082564 | 0.334528925 |
| chr19:48551941-48552375 | 0.558456044 | 0.110839454 | 0.334551553 |
| chr8:94025095-94025295 | 0.827874302 | 0.11084187 | 0.334551553 |
| chr20:54134116-54134350 | 0.547219686 | 0.110853227 | 0.334572653 |
| chr14:45364481-45364771 | 0.650779373 | 0.110895954 | 0.334688426 |
| chr10:34938109-34938309 | -0.911521012 | 0.110904443 | 0.334689006 |
| chr5:122192929-122193326 | -0.610093156 | 0.110913834 | 0.334689006 |
| chr17:60374210-60374635 | 0.540014982 | 0.110913888 | 0.334689006 |
| chr4:173360726-173360958 | -0.802860522 | 0.110915192 | 0.334689006 |
| chr5:50856516-50856758 | -0.884405187 | 0.11092121 | 0.334689006 |
| chr10:77919634-77920183 | -0.539006194 | 0.110927355 | 0.334689006 |
| chr5:21920290-21920542 | 0.838515066 | 0.110929091 | 0.334689006 |
| chrX:12771317-12771788 | -0.636805821 | 0.11093109 | 0.334689006 |
| chr2:35199319-35199707 | -0.61701199 | 0.110941131 | 0.334692186 |
| chr14:39417093-39417293 | 0.771642179 | 0.110949638 | 0.334692186 |

| | | | |
|---|---|---|---|
| chr2:137928572-137928838 | -0.821177393 | 0.110950401 | 0.334692186 |
| chr5:161546246-161546828 | -0.586233388 | 0.110951209 | 0.334692186 |
| chr18:8704355-8705664 | 0.515829846 | 0.110953985 | 0.334692186 |
| chr2:98608160-98608917 | 0.539624373 | 0.11097276 | 0.334735642 |
| chr17:10114466-10114818 | -0.660974065 | 0.110985579 | 0.334761131 |
| chr18:3447531-3447749 | -0.667193468 | 0.111002805 | 0.33479991 |
| chr15:80072313-80072648 | -0.647265749 | 0.111020105 | 0.334838909 |
| chr17:80270916-80271189 | 0.645201473 | 0.111047488 | 0.334898068 |
| chr19:34404019-34404948 | 0.532892987 | 0.111048461 | 0.334898068 |
| chr10:122279719-122279999 | 0.680102985 | 0.111063637 | 0.334930652 |
| chr15:90665274-90666039 | 0.533558634 | 0.111085558 | 0.334972201 |
| chr5:78504693-78505417 | -0.550218866 | 0.111086158 | 0.334972201 |
| chr17:68646936-68647174 | 0.720746594 | 0.111095335 | 0.334975447 |
| chr7:41045555-41045927 | 0.594010591 | 0.111095978 | 0.334975447 |
| chr20:56933379-56933704 | -0.624981993 | 0.111110093 | 0.335004822 |
| chr2:152730596-152731348 | -0.534986371 | 0.111129044 | 0.335048776 |
| chr3:150447714-150448393 | 0.614067753 | 0.111168606 | 0.335141594 |
| chr20:20825026-20825286 | 0.670176252 | 0.111172421 | 0.335141594 |
| chr1:151300737-151301182 | 0.555138553 | 0.111172952 | 0.335141594 |
| chr7:93575483-93576068 | -0.622296797 | 0.111185389 | 0.3351659 |
| chr7:117781752-117782030 | -0.741974488 | 0.111219406 | 0.335255254 |
| chr11:3605867-3606067 | 0.82633649 | 0.111225629 | 0.335260825 |
| chr11:124115298-124115567 | -0.718935965 | 0.111258293 | 0.335346088 |
| chr10:136093-136869 | 0.525663588 | 0.111278058 | 0.335369592 |
| chr17:50396695-50397132 | 0.571156479 | 0.11127826 | 0.335369592 |
| chr2:18559959-18561024 | -0.51100007 | 0.111282756 | 0.335369592 |
| chr9:126522352-126522596 | 0.866475618 | 0.111290507 | 0.335369592 |
| chr14:77458340-77458658 | 0.575933979 | 0.111292075 | 0.335369592 |
| chr4:109978726-109978967 | -0.830633249 | 0.111292352 | 0.335369592 |
| chr17:7295306-7295506 | -0.750468439 | 0.111304572 | 0.335393225 |
| chr1:45003955-45004155 | 0.794065672 | 0.11134926 | 0.335514688 |
| chr2:189633397-189633871 | -0.595781036 | 0.111370438 | 0.33555906 |
| chr13:41557913-41558114 | -0.884126451 | 0.111372745 | 0.33555906 |
| chr22:18023500-18024129 | 0.56080836 | 0.111399443 | 0.335606334 |
| chr20:47173790-47175213 | 0.497015671 | 0.111400107 | 0.335606334 |
| chr3:146407747-146407997 | 0.765619448 | 0.111401575 | 0.335606334 |
| chr17:72502757-72503216 | 0.619034467 | 0.111423766 | 0.335659989 |
| chr14:64806454-64806703 | 0.650325314 | 0.111429816 | 0.335665016 |
| chr3:61807948-61808148 | 0.616419683 | 0.111435222 | 0.335668106 |
| chr18:66886832-66887052 | -0.784678977 | 0.111467582 | 0.335733713 |
| chr7:148556881-148557538 | -0.54290024 | 0.111467746 | 0.335733713 |
| chr10:114623834-114624231 | 0.598390198 | 0.111470148 | 0.335733713 |
| chr8:91596972-91597226 | 0.737390841 | 0.111486656 | 0.335770236 |
| chr11:313740-313994 | -0.832341078 | 0.111494596 | 0.335780951 |
| chr10:29735150-29737166 | 0.508912649 | 0.111510603 | 0.335815958 |
| chr19:18064757-18064957 | 0.814135101 | 0.111527785 | 0.335854501 |
| chr22:19544008-19544703 | -0.596511655 | 0.111535381 | 0.335864176 |
| chr3:23430151-23430566 | -0.686069619 | 0.111542682 | 0.335872961 |
| chr5:109380783-109380983 | -0.850490898 | 0.11155199 | 0.335877823 |
| chr10:90652555-90652855 | -0.695424029 | 0.111553325 | 0.335877823 |

| | | | |
|---|---|---|---|
| chr3:94928992-94929316 | 0.625013966 | 0.11155987 | 0.335877823 |
| chr5:120457842-120458622 | -0.532165145 | 0.111561831 | 0.335877823 |
| chr7:98354445-98355270 | -0.551073158 | 0.111584921 | 0.335934142 |
| chr5:97182984-97184016 | -0.504631043 | 0.111590905 | 0.335938957 |
| chr4:56799896-56800492 | -0.593257023 | 0.111625909 | 0.3360153 |
| chr10:82349667-82350058 | 0.662072042 | 0.111628235 | 0.3360153 |
| chr16:15482133-15482664 | -0.596652823 | 0.11162942 | 0.3360153 |
| chr1:91226564-91227087 | -0.647922615 | 0.111673367 | 0.336134379 |
| chr9:117466254-117466466 | 0.772934443 | 0.111688529 | 0.33615141 |
| chr13:98217929-98218129 | 0.914068862 | 0.111690663 | 0.33615141 |
| chr21:9373204-9373409 | 0.855737011 | 0.111692187 | 0.33615141 |
| chr4:109422462-109423167 | -0.55794255 | 0.111718923 | 0.336206977 |
| chr1:10555570-10555875 | -0.819014598 | 0.111729152 | 0.336206977 |
| chr8:66142215-66142538 | -0.768752795 | 0.111730293 | 0.336206977 |
| chr1:248873535-248874723 | -0.49723363 | 0.111733988 | 0.336206977 |
| chr6:4210679-4210986 | -0.671884 | 0.111736642 | 0.336206977 |
| chr17:409879-410530 | -0.5169534 | 0.111736977 | 0.336206977 |
| chr20:18491440-18491640 | -0.91007722 | 0.111745028 | 0.336210911 |
| chr17:65941590-65941811 | -0.817645768 | 0.11174706 | 0.336210911 |
| chr11:113690631-113691007 | -0.672705228 | 0.111766953 | 0.336257556 |
| chr16:4426068-4426448 | 0.602898181 | 0.111783859 | 0.336295215 |
| chr5:133256314-133256514 | 0.764759098 | 0.111795679 | 0.336308998 |
| chr11:15852266-15852511 | -0.829552777 | 0.111797219 | 0.336308998 |
| chr3:163556923-163557152 | 0.764867427 | 0.111815985 | 0.336341666 |
| chr9:21802098-21803326 | -0.507076271 | 0.111817139 | 0.336341666 |
| chr10:69789017-69789331 | -0.751744688 | 0.111821247 | 0.336341666 |
| chr10:128268117-128268347 | 0.729044133 | 0.111825862 | 0.336342344 |
| chr7:98872427-98872678 | -0.732389046 | 0.111838173 | 0.336366169 |
| chr13:39655455-39656404 | -0.517765675 | 0.111858328 | 0.336413581 |
| chr17:61791928-61792259 | 0.531691531 | 0.111884704 | 0.3364797 |
| chr5:73201651-73202418 | -0.540529438 | 0.111898042 | 0.336506604 |
| chr4:26975626-26976426 | -0.556961446 | 0.111906784 | 0.336519685 |
| chr11:47248485-47249395 | 0.525786342 | 0.111912157 | 0.336522635 |
| chr17:4575637-4576029 | 0.69654914 | 0.111917814 | 0.33652644 |
| chr14:43471455-43471758 | -0.834979884 | 0.111923422 | 0.336530096 |
| chr4:117134495-117134695 | -0.881949212 | 0.111929989 | 0.336536636 |
| chr6:4256771-4257313 | -0.62156391 | 0.111941736 | 0.33655238 |
| chr12:48890735-48891070 | -0.663819934 | 0.11194401 | 0.33655238 |
| chr6:44247710-44247966 | 0.613525287 | 0.111955041 | 0.336572336 |
| chr8:115934827-115935157 | -0.652844884 | 0.111967439 | 0.33658336 |
| chr1:40612295-40612951 | -0.608469628 | 0.111967493 | 0.33658336 |
| chr15:98823200-98823494 | 0.725281771 | 0.111981669 | 0.3366078 |
| chr3:132262740-132263350 | -0.62699622 | 0.111984409 | 0.3366078 |
| chr7:40186631-40186868 | -0.843005251 | 0.111990264 | 0.336612194 |
| chr5:161966595-161966857 | -0.699346371 | 0.112010181 | 0.336635718 |
| chr1:198567673-198568169 | -0.663093131 | 0.11201057 | 0.336635718 |
| chr11:94732116-94732405 | -0.820434135 | 0.112011271 | 0.336635718 |
| chr8:85515027-85515307 | 0.648685447 | 0.11202043 | 0.33665004 |
| chr1:212466948-212467184 | -0.78193387 | 0.112025776 | 0.336652901 |
| chr11:117861228-117861526 | 0.660875898 | 0.112034674 | 0.336666436 |

| | | | |
|---|---|---|---|
| chr8:17614354-17614616 | 0.72643395 | 0.112041609 | 0.336669285 |
| chr18:77089043-77089486 | -0.583844635 | 0.112045018 | 0.336669285 |
| chr8:24891849-24892122 | -0.811303652 | 0.112048804 | 0.336669285 |
| chr3:184645416-184645742 | -0.735479162 | 0.112075263 | 0.336735326 |
| chr7:30504268-30504592 | 0.633170305 | 0.112079573 | 0.336735326 |
| chr14:55557047-55557636 | 0.555135973 | 0.112090652 | 0.336749711 |
| chr20:23186522-23186759 | -0.800891228 | 0.11209531 | 0.336749711 |
| chr3:185537431-185537873 | 0.659800355 | 0.112105221 | 0.336749711 |
| chr20:54451447-54451880 | 0.524907495 | 0.112108666 | 0.336749711 |
| chr1:113575384-113575738 | 0.69514555 | 0.112109041 | 0.336749711 |
| chr2:241774775-241774976 | -0.736531355 | 0.112110961 | 0.336749711 |
| chr5:10538989-10539319 | 0.592514675 | 0.112115125 | 0.336749711 |
| chr9:130172070-130172457 | 0.626796682 | 0.112121382 | 0.336755304 |
| chr2:130720490-130720795 | -0.841672913 | 0.112135418 | 0.336784258 |
| chr8:42894232-42894710 | 0.559988691 | 0.112145831 | 0.336784461 |
| chr17:8200117-8200868 | -0.543376376 | 0.112149914 | 0.336784461 |
| chr19:46211172-46211568 | 0.600086393 | 0.112156012 | 0.336784461 |
| chr10:93893616-93894348 | -0.57093578 | 0.112156393 | 0.336784461 |
| chr1:113207045-113207354 | -0.634781627 | 0.112157462 | 0.336784461 |
| chr4:159345481-159345820 | -0.818914495 | 0.112162238 | 0.336785604 |
| chr12:54549297-54549650 | 0.65503783 | 0.11218006 | 0.336825915 |
| chr9:135440783-135441011 | -0.855784391 | 0.112189318 | 0.336827319 |
| chr3:8248538-8248773 | -0.865966839 | 0.112195164 | 0.336827319 |
| chr10:77287706-77288175 | -0.610903283 | 0.112196484 | 0.336827319 |
| chr14:24298930-24300186 | 0.51017603 | 0.112199047 | 0.336827319 |
| chr9:131478512-131478801 | 0.636816139 | 0.112203648 | 0.336827319 |
| chr5:32249767-32249979 | 0.774743521 | 0.112206903 | 0.336827319 |
| chr1:43414549-43414971 | 0.652822487 | 0.112290325 | 0.337047023 |
| chr11:2202138-2202776 | 0.570207354 | 0.112292249 | 0.337047023 |
| chr3:57692501-57693368 | 0.525375384 | 0.112293289 | 0.337047023 |
| chr7:73867612-73867917 | -0.767639141 | 0.112313509 | 0.337094507 |
| chr10:7918278-7918562 | 0.7448835 | 0.112338823 | 0.337157278 |
| chr2:210363295-210363577 | -0.656440872 | 0.112345394 | 0.337163791 |
| chr11:118488527-118488785 | 0.737862247 | 0.112364384 | 0.337195424 |
| chr14:94396094-94396341 | 0.694506831 | 0.112365117 | 0.337195424 |
| chr20:56091831-56092077 | 0.690236118 | 0.112369136 | 0.337195424 |
| chr15:68691648-68692223 | 0.551026639 | 0.112384979 | 0.337229758 |
| chr4:139323539-139323818 | 0.792279124 | 0.112399765 | 0.337260917 |
| chr15:91123588-91123817 | 0.782838325 | 0.112410572 | 0.337280136 |
| chrX:97587082-97587553 | 0.58082551 | 0.11242488 | 0.337309859 |
| chr2:70132330-70132683 | 0.689082086 | 0.112438175 | 0.337326277 |
| chr15:40764597-40764822 | 0.694961183 | 0.112439157 | 0.337326277 |
| chr17:47687913-47688113 | 0.682373165 | 0.112447656 | 0.337329026 |
| chr15:58485128-58485409 | 0.763085503 | 0.112448879 | 0.337329026 |
| chr12:42145647-42145868 | -0.692282172 | 0.112464827 | 0.337363661 |
| chr14:23562762-23563014 | 0.638190251 | 0.112485814 | 0.337413406 |
| chr5:28505032-28505664 | -0.557175203 | 0.112503053 | 0.337451905 |
| chr7:11212129-11212659 | 0.548387114 | 0.112507496 | 0.337452021 |
| chr5:36739445-36739971 | -0.575929478 | 0.112525989 | 0.337494278 |
| chr8:40054560-40055332 | -0.571263675 | 0.112536112 | 0.337511428 |

| | | | |
|---|---|---|---|
| chr5:17860734-17861065 | 0.628343004 | 0.112541846 | 0.337515414 |
| chr5:110441012-110441362 | -0.632178231 | 0.112555276 | 0.337542478 |
| chr3:62298620-62298822 | -0.769632469 | 0.112567981 | 0.337558231 |
| chr10:132637894-132638224 | 0.743663661 | 0.112571646 | 0.337558231 |
| chr20:50712872-50713130 | 0.532450194 | 0.112576226 | 0.337558231 |
| chr11:119056442-119057790 | 0.504215305 | 0.11257815 | 0.337558231 |
| chr3:53804804-53805309 | -0.655840435 | 0.112588739 | 0.337576771 |
| chr3:149170709-149170957 | 0.831193529 | 0.11261826 | 0.337632667 |
| chr16:28609792-28610116 | 0.67489497 | 0.112619362 | 0.337632667 |
| chr11:59775934-59776134 | -0.872037455 | 0.112620601 | 0.337632667 |
| chr12:17665786-17666014 | -0.848627817 | 0.11263663 | 0.337667509 |
| chr7:22853769-22855238 | 0.509850681 | 0.112645881 | 0.33768203 |
| chr1:205851987-205852456 | 0.638541764 | 0.112659704 | 0.337710256 |
| chr15:75672903-75673192 | 0.626654424 | 0.112672427 | 0.337735181 |
| chr18:67515950-67517349 | -0.502812877 | 0.112689001 | 0.337767808 |
| chr1:203626456-203627149 | -0.52125368 | 0.112695211 | 0.337767808 |
| chr5:122751372-122751886 | -0.647324408 | 0.112696536 | 0.337767808 |
| chr12:53174858-53175544 | 0.558756251 | 0.112725483 | 0.33784135 |
| chr3:4977614-4977837 | -0.575464216 | 0.112733293 | 0.337851544 |
| chr8:141290335-141290638 | -0.713567179 | 0.112751091 | 0.337869952 |
| chr11:44978627-44979458 | -0.526860349 | 0.11275154 | 0.337869952 |
| chr18:64104102-64104379 | -0.721977458 | 0.112752665 | 0.337869952 |
| chr3:100316594-100317024 | -0.620848711 | 0.112757386 | 0.337870885 |
| chr15:74815217-74815417 | 0.705594893 | 0.11276547 | 0.33787861 |
| chr5:74483534-74483814 | -0.770459993 | 0.112774478 | 0.33787861 |
| chr11:86371274-86371565 | -0.766533872 | 0.11277574 | 0.33787861 |
| chr10:110398653-110398920 | -0.775409914 | 0.112781722 | 0.33787861 |
| chr5:104986353-104986772 | -0.626911722 | 0.112782012 | 0.33787861 |
| chr10:31862365-31862792 | -0.670428994 | 0.11280168 | 0.337920991 |
| chr7:101154105-101155113 | -0.527042059 | 0.112807783 | 0.337920991 |
| chr8:30605019-30605286 | -0.857931359 | 0.112817945 | 0.337920991 |
| chr5:60944904-60946251 | -0.506846868 | 0.112821563 | 0.337920991 |
| chr14:53265004-53265403 | 0.666095763 | 0.112821829 | 0.337920991 |
| chr5:133878782-133879011 | 0.823583331 | 0.11282262 | 0.337920991 |
| chr3:101387088-101387384 | 0.752775928 | 0.112838726 | 0.337956019 |
| chr22:36470311-36470518 | 0.750244038 | 0.112862074 | 0.338009028 |
| chr9:133161012-133161292 | -0.663651616 | 0.112865247 | 0.338009028 |
| chr18:37476264-37476502 | -0.800814552 | 0.112906741 | 0.338120078 |
| chr3:161411743-161412486 | 0.51845001 | 0.11292458 | 0.338160284 |
| chr1:11663965-11664320 | 0.659598544 | 0.112962446 | 0.338253488 |
| chr4:47836588-47837336 | -0.554145423 | 0.112968443 | 0.338253488 |
| chr4:173369480-173370263 | -0.515937341 | 0.112971076 | 0.338253488 |
| chr7:44075342-44075567 | 0.637550865 | 0.112973363 | 0.338253488 |
| chr20:57526592-57526994 | 0.575169861 | 0.112978013 | 0.338254192 |
| chr10:113553573-113553773 | 0.91472374 | 0.112985466 | 0.338263289 |
| chr1:220296222-220296604 | -0.600853731 | 0.113001914 | 0.338299315 |
| chr5:172315206-172315505 | 0.736709378 | 0.113019348 | 0.338326395 |
| chr9:134025737-134025950 | 0.676824447 | 0.113020698 | 0.338326395 |
| chr17:27654992-27655210 | 0.696158853 | 0.113024206 | 0.338326395 |
| chr15:58193446-58193882 | -0.730087682 | 0.113033049 | 0.338339646 |

| | | | |
|---|---|---|---|
| chr6:36679405-36679613 | 0.677763505 | 0.113038656 | 0.338342726 |
| chr1:217945155-217945418 | -0.6635825 | 0.113042909 | 0.338342726 |
| chr3:131123860-131124069 | 0.894490802 | 0.113068741 | 0.338406822 |
| chr7:33586867-33587240 | 0.661505096 | 0.113077283 | 0.338407002 |
| chr11:125874574-125874832 | -0.719621539 | 0.113077634 | 0.338407002 |
| chr8:2030614-2030892 | -0.684937197 | 0.1130929 | 0.33843947 |
| chr1:155903776-155904016 | -0.878978583 | 0.113097823 | 0.338440984 |
| chr19:45444245-45444531 | -0.663338223 | 0.113104975 | 0.338449169 |
| chr2:217837238-217837510 | -0.727066086 | 0.113115691 | 0.338468015 |
| chr4:58247891-58248125 | -0.780935327 | 0.113138033 | 0.338521649 |
| chr15:55526534-55526834 | 0.617083863 | 0.113168438 | 0.338599399 |
| chr7:155285576-155285920 | -0.716161865 | 0.113181962 | 0.338616154 |
| chr16:66297150-66297713 | 0.594084628 | 0.113182876 | 0.338616154 |
| chr15:33068596-33069076 | -0.543326236 | 0.113193731 | 0.33862607 |
| chr11:127345747-127346120 | -0.813035851 | 0.11319846 | 0.33862607 |
| chr1:224204739-224205108 | 0.585511732 | 0.113199449 | 0.33862607 |
| chr14:25516703-25517251 | 0.581524365 | 0.113218892 | 0.338659183 |
| chr5:66988021-66988266 | -0.773854349 | 0.113219358 | 0.338659183 |
| chr1:162854230-162854537 | 0.614538004 | 0.113235851 | 0.338695295 |
| chr2:232981439-232981690 | 0.67724253 | 0.11324387 | 0.338706058 |
| chr3:63349673-63349887 | 0.647782706 | 0.113256496 | 0.338730601 |
| chr2:66030551-66030886 | -0.734636289 | 0.113271174 | 0.338761276 |
| chr7:130598010-130598356 | -0.77414517 | 0.113282465 | 0.338771406 |
| chr17:48878227-48878481 | 0.655826195 | 0.113283404 | 0.338771406 |
| chr3:10010770-10011693 | 0.509860485 | 0.113316172 | 0.338856175 |
| chr14:55354598-55354954 | 0.585272812 | 0.113321367 | 0.338858484 |
| chr3:189233292-189233637 | -0.672082567 | 0.113341745 | 0.338906192 |
| chr16:67676606-67676813 | 0.697334997 | 0.113355415 | 0.338927515 |
| chr18:57437499-57438227 | -0.533474863 | 0.113360708 | 0.338927515 |
| chr11:69524222-69524789 | -0.561084103 | 0.113362146 | 0.338927515 |
| chr19:47743200-47743972 | 0.543395176 | 0.113379437 | 0.338965983 |
| chr1:182588715-182589706 | -0.521153845 | 0.113406375 | 0.339033292 |
| chr5:19891562-19892066 | -0.610882905 | 0.113412823 | 0.339039341 |
| chr10:98363218-98363604 | 0.566715263 | 0.11341927 | 0.339045386 |
| chr7:41942873-41943532 | -0.553011216 | 0.113433393 | 0.339074375 |
| chr17:74446663-74447018 | -0.62948439 | 0.11344679 | 0.339101191 |
| chr16:31130841-31131262 | 0.54885379 | 0.113452344 | 0.339104564 |
| chr22:42990244-42990557 | 0.780776407 | 0.113471474 | 0.339137951 |
| chr1:60765299-60765529 | -0.728068433 | 0.113476236 | 0.339137951 |
| chr7:123747634-123747897 | -0.60859561 | 0.113476793 | 0.339137951 |
| chr9:38436376-38436655 | -0.842506853 | 0.113482606 | 0.339142098 |
| chr7:156764000-156764387 | -0.635752315 | 0.113487413 | 0.339143236 |
| chr5:2780284-2780981 | -0.518355575 | 0.11349546 | 0.339154055 |
| chr12:18141716-18141922 | -0.846624251 | 0.113501186 | 0.339157938 |
| chr5:115069406-115069727 | -0.643964895 | 0.113525008 | 0.339215894 |
| chr10:62268042-62269249 | -0.511583846 | 0.113534178 | 0.339230064 |
| chr1:26169547-26170217 | 0.571854164 | 0.113539202 | 0.339231847 |
| chr7:134532773-134533061 | -0.797403808 | 0.113587779 | 0.339337194 |
| chr6:31497523-31498169 | 0.561042966 | 0.113590311 | 0.339337194 |
| chr7:140239629-140239868 | -0.78498426 | 0.113591449 | 0.339337194 |

| | | | |
|---|---|---|---|
| chr5:98928009-98929893 | -0.495105415 | 0.113592176 | 0.339337194 |
| chr22:39135162-39135409 | 0.743980321 | 0.113602032 | 0.33934382 |
| chr3:18153594-18153908 | 0.796165264 | 0.113603252 | 0.33934382 |
| chr15:62135758-62136027 | -0.716994149 | 0.113629545 | 0.339389789 |
| chr8:123065500-123066045 | -0.538152526 | 0.113634517 | 0.339389789 |
| chr7:45768472-45769484 | -0.51399204 | 0.113636377 | 0.339389789 |
| chr15:99385016-99385336 | 0.664380133 | 0.11363655 | 0.339389789 |
| chr1:27643672-27644139 | -0.651022954 | 0.113640788 | 0.339389789 |
| chr2:152635688-152636041 | 0.632879597 | 0.113662192 | 0.339440482 |
| chr2:46174653-46174853 | -0.799243333 | 0.113703141 | 0.33954954 |
| chr17:27241280-27241971 | 0.536260914 | 0.113731964 | 0.339612767 |
| chr4:151553875-151554109 | 0.777314977 | 0.11373525 | 0.339612767 |
| chr3:29609422-29609801 | -0.659781498 | 0.113741968 | 0.339612767 |
| chr3:161151168-161151412 | 0.753406343 | 0.113742043 | 0.339612767 |
| chr20:53635220-53635592 | 0.514830749 | 0.113750343 | 0.339613694 |
| chr6:57317400-57317995 | -0.530226833 | 0.113754699 | 0.339613694 |
| chr22:31661842-31662706 | 0.518021451 | 0.113755651 | 0.339613694 |
| chr3:168877828-168878374 | -0.534655041 | 0.113769138 | 0.339633162 |
| chr18:68714469-68715564 | -0.508974025 | 0.113773064 | 0.339633162 |
| chr7:155007977-155008260 | -0.681885948 | 0.113775469 | 0.339633162 |
| chr14:25050012-25050454 | 0.61498836 | 0.113785744 | 0.339650602 |
| chr15:79074360-79074560 | 0.760706385 | 0.11379043 | 0.339651356 |
| chr13:87280243-87280636 | -0.688326268 | 0.11380439 | 0.339679794 |
| chr19:35107728-35107928 | 0.851774962 | 0.113809699 | 0.339680299 |
| chr6:13813454-13814845 | 0.506686643 | 0.113818261 | 0.339680299 |
| chr6:133130954-133131296 | -0.671443896 | 0.113821444 | 0.339680299 |
| chr1:189149465-189150344 | -0.506006476 | 0.113822292 | 0.339680299 |
| chr1:3900020-3901094 | 0.505953086 | 0.113844971 | 0.339720609 |
| chr3:131794820-131795347 | -0.613837706 | 0.113846228 | 0.339720609 |
| chr1:150852203-150852436 | 0.68663634 | 0.113850885 | 0.339720609 |
| chr1:229199341-229199636 | 0.620400445 | 0.113853535 | 0.339720609 |
| chr20:53646574-53646880 | 0.52514852 | 0.113879314 | 0.3397843 |
| chr12:27022295-27022954 | -0.551784144 | 0.113894726 | 0.33981705 |
| chr1:117366593-117367639 | 0.507478275 | 0.113900659 | 0.339821521 |
| chr2:128978099-128978500 | 0.639568614 | 0.113909194 | 0.339833753 |
| chr8:85029429-85029735 | -0.759955004 | 0.113920136 | 0.339853164 |
| chr5:77316550-77316853 | -0.812545159 | 0.113937628 | 0.339892114 |
| chr9:79571161-79572052 | -0.50220392 | 0.113947908 | 0.339909546 |
| chr3:123552193-123552407 | -0.814846252 | 0.11396007 | 0.33992274 |
| chr6:85580421-85581249 | -0.537968334 | 0.113963885 | 0.33992274 |
| chr11:67202224-67202450 | 0.81294781 | 0.113970401 | 0.33992274 |
| chr8:122244317-122245068 | -0.528545418 | 0.113971294 | 0.33992274 |
| chr16:58501210-58501573 | 0.62280249 | 0.113979375 | 0.33992274 |
| chr2:95682589-95682850 | 0.700232704 | 0.113986227 | 0.33992274 |
| chr1:114192340-114192694 | 0.542489769 | 0.113988277 | 0.33992274 |
| chr11:78987826-78988031 | -0.78869086 | 0.113994042 | 0.33992274 |
| chr13:98842955-98843202 | -0.73133015 | 0.113995857 | 0.33992274 |
| chr10:14174120-14174320 | -0.885295079 | 0.113996694 | 0.33992274 |
| chr3:13419203-13420063 | 0.523694066 | 0.114001168 | 0.339922851 |
| chr17:60535659-60535895 | 0.559738009 | 0.11401843 | 0.339961091 |

| | | | |
|---|---|---|---|
| chr16:30112918-30113134 | 0.736747548 | 0.114022935 | 0.339961297 |
| chr8:121512889-121513121 | 0.785217719 | 0.11404558 | 0.340015581 |
| chr8:100212734-100213725 | 0.509382473 | 0.114058645 | 0.340041302 |
| chr2:74548444-74549781 | -0.51118403 | 0.114075916 | 0.340079559 |
| chr12:119659326-119659581 | -0.776267968 | 0.114105011 | 0.340132285 |
| chr8:140668325-140668601 | 0.7031957 | 0.114113359 | 0.340132285 |
| chr3:52447039-52447344 | 0.593741359 | 0.114113834 | 0.340132285 |
| chr14:75981836-75982152 | 0.56317614 | 0.114115713 | 0.340132285 |
| chr14:106842668-106843013 | 0.610746282 | 0.114115798 | 0.340132285 |
| chr1:161071898-161072112 | 0.730502305 | 0.114129593 | 0.34015284 |
| chr2:186952069-186952310 | 0.714811466 | 0.114132229 | 0.34015284 |
| chr1:173628050-173628507 | -0.652490527 | 0.114137781 | 0.34015284 |
| chr12:64486600-64486820 | 0.812063212 | 0.11414728 | 0.34015284 |
| chr11:118126384-118126810 | -0.650215527 | 0.114149246 | 0.34015284 |
| chr11:44584941-44585168 | 0.764793462 | 0.114153769 | 0.34015284 |
| chr11:62862716-62862916 | 0.764793462 | 0.114153769 | 0.34015284 |
| chr4:28481646-28481908 | 0.718102304 | 0.114185532 | 0.340234254 |
| chr1:28193811-28194099 | -0.667134957 | 0.114234007 | 0.340353569 |
| chr20:34255639-34255937 | 0.693696601 | 0.114237057 | 0.340353569 |
| chr8:129390041-129390241 | -0.913015996 | 0.114239333 | 0.340353569 |
| chr19:43503682-43504981 | 0.508908644 | 0.114244744 | 0.340353569 |
| chr5:24652279-24653015 | -0.548055478 | 0.114247785 | 0.340353569 |
| chr9:133387234-133388193 | 0.530546543 | 0.114259955 | 0.340376591 |
| chr2:237691325-237692332 | 0.518819716 | 0.114270122 | 0.340393642 |
| chr17:61533213-61533544 | 0.523708323 | 0.114284183 | 0.340422294 |
| chr19:57435041-57435944 | 0.522150241 | 0.114289969 | 0.340426296 |
| chr3:147945946-147946554 | -0.576823648 | 0.114295648 | 0.340429977 |
| chr3:95705928-95706304 | -0.748916599 | 0.114312042 | 0.34046557 |
| chr5:132396639-132396858 | -0.803245166 | 0.114324553 | 0.340489599 |
| chr10:63976555-63976895 | -0.597458693 | 0.114367631 | 0.340604658 |
| chrX:8455563-8455862 | -0.745798951 | 0.114376554 | 0.340612352 |
| chr17:41734457-41734983 | 0.578845092 | 0.114383707 | 0.340612352 |
| chr6:91080955-91081233 | -0.76347111 | 0.114388781 | 0.340612352 |
| chr2:9790844-9791059 | 0.702937052 | 0.114392413 | 0.340612352 |
| chr3:161340215-161340446 | 0.69534272 | 0.114393328 | 0.340612352 |
| chr21:41648270-41648472 | 0.793985646 | 0.114398778 | 0.340612352 |
| chr3:160707513-160707740 | 0.704606077 | 0.114403795 | 0.340612352 |
| chr13:112723007-112723365 | -0.695596576 | 0.114409443 | 0.340612352 |
| chr2:239094163-239094432 | -0.829065007 | 0.114414568 | 0.340612352 |
| chr3:123451259-123451523 | 0.621579097 | 0.114414668 | 0.340612352 |
| chr12:51082878-51084025 | 0.520384466 | 0.114419715 | 0.340614143 |
| chr1:171800448-171800672 | -0.712982701 | 0.114427397 | 0.340621858 |
| chr4:134073123-134073611 | -0.599039371 | 0.114431198 | 0.340621858 |
| chr1:187396316-187396596 | -0.748445173 | 0.114445716 | 0.340645755 |
| chr17:47362284-47362545 | -0.786941698 | 0.114448118 | 0.340645755 |
| chr4:79591826-79592639 | 0.536291136 | 0.114457997 | 0.340661926 |
| chr1:172431628-172432414 | -0.53054335 | 0.114490535 | 0.340745534 |
| chr4:17549648-17549940 | 0.697784357 | 0.114508209 | 0.340773673 |
| chr18:23452801-23453697 | 0.533472379 | 0.11451212 | 0.340773673 |
| chr7:52925424-52925694 | -0.696054647 | 0.114513332 | 0.340773673 |

| | | | |
|---|---|---|---|
| chr20:57101656-57103016 | 0.493714327 | 0.114526298 | 0.340795571 |
| chr13:29702992-29703256 | -0.811704928 | 0.114532456 | 0.340795571 |
| chr10:46325141-46325377 | -0.782701 | 0.114534546 | 0.340795571 |
| chr1:207963260-207964162 | -0.52682591 | 0.114538482 | 0.340795571 |
| chr3:175037774-175037974 | -0.824774608 | 0.114544106 | 0.340799071 |
| chr6:29752985-29753367 | 0.591553902 | 0.114563454 | 0.340825238 |
| chr7:101108887-101109274 | -0.687189318 | 0.11456361 | 0.340825238 |
| chr9:71725988-71726291 | 0.671683323 | 0.114569445 | 0.340825238 |
| chr13:36275195-36275539 | -0.625275561 | 0.114570693 | 0.340825238 |
| chr1:7975298-7975596 | 0.683548006 | 0.114584837 | 0.340845682 |
| chr2:12759198-12759703 | -0.637254929 | 0.114587804 | 0.340845682 |
| chr13:73090456-73090755 | -0.745272375 | 0.114592282 | 0.340845682 |
| chr8:61725128-61725404 | 0.729641699 | 0.114598483 | 0.340845682 |
| chr4:37602566-37602894 | -0.725981395 | 0.114604885 | 0.340845682 |
| chr8:60792963-60793163 | -0.820338291 | 0.114608495 | 0.340845682 |
| chr13:73058563-73059937 | -0.50408051 | 0.114608705 | 0.340845682 |
| chr15:89785507-89785707 | -0.843403905 | 0.11462489 | 0.34087184 |
| chr20:17530772-17531641 | 0.538220444 | 0.114631034 | 0.34087184 |
| chr10:12455292-12455492 | 0.863686951 | 0.114634962 | 0.34087184 |
| chr13:113331292-113331799 | 0.599729328 | 0.114635295 | 0.34087184 |
| chr3:172289643-172289931 | -0.63656943 | 0.114654714 | 0.340916352 |
| chr2:241890731-241891221 | 0.560058248 | 0.114681248 | 0.340982018 |
| chr13:98507326-98507678 | -0.730093931 | 0.114686911 | 0.340985624 |
| chr6:71791004-71791461 | 0.635732133 | 0.114694444 | 0.340994788 |
| chr17:15261274-15261498 | 0.674409351 | 0.114707893 | 0.341021542 |
| chr3:184040036-184040269 | 0.737441576 | 0.114716038 | 0.341024693 |
| chr10:3197299-3197734 | 0.577135275 | 0.114717855 | 0.341024693 |
| chr2:54945671-54946184 | -0.573548566 | 0.114728043 | 0.341041747 |
| chr2:231533810-231534012 | -0.792180039 | 0.114735797 | 0.341051566 |
| chr5:78627382-78627608 | -0.748980409 | 0.114743741 | 0.34106195 |
| chr8:118122627-118122909 | -0.748697233 | 0.114753419 | 0.341077486 |
| chr1:209860364-209860634 | -0.726391226 | 0.114780446 | 0.341144583 |
| chr2:112891370-112891683 | -0.698157961 | 0.114798931 | 0.341180995 |
| chr17:5265064-5265284 | -0.855190304 | 0.114801603 | 0.341180995 |
| chr8:112459861-112460222 | 0.667429425 | 0.11481409 | 0.341204871 |
| chr2:85091885-85092305 | 0.601054017 | 0.114866533 | 0.341347483 |
| chr2:2613224-2613976 | -0.527631578 | 0.114886201 | 0.341392691 |
| chr9:127700588-127700848 | -0.734548303 | 0.114898826 | 0.341416964 |
| chr3:133428216-133428933 | -0.529215939 | 0.114937049 | 0.341491068 |
| chr13:60395986-60396664 | -0.512903029 | 0.114940296 | 0.341491068 |
| chr5:18267547-18267764 | 0.74146682 | 0.114941499 | 0.341491068 |
| chr14:85683111-85683320 | 0.800154771 | 0.114944051 | 0.341491068 |
| chr5:14029978-14030189 | 0.80569246 | 0.114946049 | 0.341491068 |
| chr11:46885998-46886248 | 0.738735201 | 0.114964635 | 0.341523529 |
| chr11:17352947-17353159 | 0.689985781 | 0.114967293 | 0.341523529 |
| chr22:25488311-25488620 | -0.672921749 | 0.114972426 | 0.341523529 |
| chr1:107247802-107248036 | 0.578167131 | 0.114974804 | 0.341523529 |
| chr11:86360830-86361061 | -0.734498816 | 0.114985589 | 0.341542323 |
| chr5:76818167-76818426 | -0.616484132 | 0.115007432 | 0.341593964 |
| chr5:178161744-178162005 | 0.650064759 | 0.115011896 | 0.341593981 |

| | | | |
|---|---|---|---|
| chr6:3209164-3209557 | 0.587839414 | 0.115049175 | 0.341691456 |
| chr12:122054417-122054647 | 0.698510792 | 0.115086565 | 0.341789255 |
| chr7:40536340-40536715 | -0.767931776 | 0.115123837 | 0.341886695 |
| chr10:114879734-114879973 | -0.713985373 | 0.115168876 | 0.342007194 |
| chr7:43764476-43764681 | -0.729362051 | 0.115196817 | 0.34207691 |
| chr12:132614950-132615189 | 0.685350828 | 0.115204899 | 0.34208138 |
| chr2:171181602-171181827 | -0.879849395 | 0.115207251 | 0.34208138 |
| chr17:79699602-79699900 | 0.647107446 | 0.115218987 | 0.34210297 |
| chr5:170030732-170030948 | -0.83116401 | 0.11522541 | 0.342108783 |
| chr20:58692268-58692964 | 0.522733791 | 0.115254834 | 0.342166237 |
| chr19:58409193-58409455 | 0.632719621 | 0.115258183 | 0.342166237 |
| chr11:117554618-117554952 | 0.613136831 | 0.115260217 | 0.342166237 |
| chr3:12993183-12993452 | 0.594922056 | 0.115262624 | 0.342166237 |
| chr1:111945844-111946098 | 0.609008809 | 0.115268668 | 0.342166609 |
| chr15:89379886-89380174 | 0.57679174 | 0.11527168 | 0.342166609 |
| chr6:164665426-164665658 | -0.890471151 | 0.115276315 | 0.342167112 |
| chr14:68819391-68819667 | -0.638967721 | 0.115285987 | 0.342182563 |
| chrX:119565085-119565630 | 0.554499313 | 0.115293672 | 0.342192117 |
| chr12:56966349-56966831 | 0.593364749 | 0.115307735 | 0.342220602 |
| chr12:82326996-82327438 | 0.66256968 | 0.115322937 | 0.342245118 |
| chr7:24665706-24665962 | 0.712583884 | 0.115324929 | 0.342245118 |
| chr19:35869191-35869416 | 0.674929837 | 0.11533303 | 0.342255903 |
| chr2:212539157-212539469 | 0.619345538 | 0.115350026 | 0.342281644 |
| chr1:24319238-24319470 | 0.716476115 | 0.115350638 | 0.342281644 |
| chr3:176841003-176841261 | -0.778177097 | 0.115357342 | 0.342288279 |
| chr20:54973586-54973847 | 0.705658148 | 0.115366993 | 0.342303662 |
| chr19:3665850-3666072 | 0.773028861 | 0.115378149 | 0.342323504 |
| chr1:106929617-106929820 | 0.650405382 | 0.115391939 | 0.342351164 |
| chr18:12060243-12060443 | 0.765143314 | 0.11539738 | 0.342352173 |
| chr10:102645460-102645700 | 0.612817681 | 0.115401215 | 0.342352173 |
| chr3:122564099-122564912 | -0.525502517 | 0.115415904 | 0.342382493 |
| chr17:27247335-27247993 | -0.554461351 | 0.115423537 | 0.342391878 |
| chr10:68955450-68956896 | -0.504465607 | 0.115453668 | 0.342468002 |
| chr21:25881826-25882104 | 0.683142063 | 0.11546377 | 0.342484707 |
| chr14:40677030-40677393 | 0.660475782 | 0.115472079 | 0.342496096 |
| chr19:40378596-40378817 | 0.754419009 | 0.115480339 | 0.342507337 |
| chr10:32367952-32368450 | -0.605931287 | 0.115487731 | 0.342516003 |
| chr3:114152458-114153131 | -0.527137901 | 0.115492861 | 0.34251796 |
| chr1:26531076-26531445 | 0.614947315 | 0.115508684 | 0.342538527 |
| chr4:158255826-158256038 | -0.76083758 | 0.115512161 | 0.342538527 |
| chr6:65259150-65259410 | -0.771322165 | 0.115526335 | 0.342538527 |
| chr13:50387078-50387388 | -0.674872691 | 0.115528581 | 0.342538527 |
| chr7:150878169-150878408 | 0.754156076 | 0.115528931 | 0.342538527 |
| chr3:176939037-176939397 | 0.600452253 | 0.115533233 | 0.342538527 |
| chr10:68796229-68796490 | -0.855817968 | 0.115535272 | 0.342538527 |
| chr7:97024309-97024509 | 0.794929063 | 0.115538906 | 0.342538527 |
| chr2:153819506-153819858 | -0.655745025 | 0.115543421 | 0.342538527 |
| chr19:19776141-19776938 | -0.531976065 | 0.115546321 | 0.342538527 |
| chr1:209513386-209513622 | -0.695293415 | 0.115554551 | 0.342538527 |
| chr5:140169422-140169698 | 0.706014976 | 0.115557589 | 0.342538527 |

| | | | |
|---|---|---|---|
| chr11:14041185-14041389 | -0.842325736 | 0.115561127 | 0.342538527 |
| chrX:120250468-120251213 | 0.519613007 | 0.115562383 | 0.342538527 |
| chr6:28673693-28674131 | -0.569113666 | 0.11557085 | 0.342550372 |
| chr15:99053333-99053943 | -0.558172045 | 0.115589794 | 0.342593268 |
| chr1:154328355-154328803 | 0.555342278 | 0.115604731 | 0.342624287 |
| chr4:103032453-103032892 | -0.60321161 | 0.115614051 | 0.342638656 |
| chrX:118839300-118839542 | 0.731120398 | 0.115639239 | 0.342700048 |
| chr12:76037865-76038181 | -0.671183743 | 0.115697835 | 0.342860438 |
| chrX:25726615-25726878 | -0.815217828 | 0.115726408 | 0.342931848 |
| chr8:110485397-110485745 | 0.667885029 | 0.115737811 | 0.342952377 |
| chr10:93542459-93542673 | 0.737513985 | 0.115754882 | 0.342978315 |
| chr15:34708391-34708879 | -0.619170066 | 0.115755517 | 0.342978315 |
| chr7:77678999-77679361 | -0.713349334 | 0.115784089 | 0.343044947 |
| chr13:89066938-89067214 | -0.784543241 | 0.115788386 | 0.343044947 |
| chr20:59673688-59673914 | 0.6376716 | 0.115791437 | 0.343044947 |
| chr5:154072336-154072635 | -0.630509948 | 0.115818779 | 0.343112684 |
| chr5:28240043-28240413 | 0.68918928 | 0.115833275 | 0.343142363 |
| chr12:89328857-89329084 | 0.711623902 | 0.115875172 | 0.343224643 |
| chr19:40662847-40663763 | -0.529276267 | 0.115878559 | 0.343224643 |
| chr2:9723336-9723536 | 0.784442532 | 0.115880716 | 0.343224643 |
| chr1:177779627-177780454 | -0.530792718 | 0.115883445 | 0.343224643 |
| chr13:98349981-98350256 | -0.681919306 | 0.115887752 | 0.343224643 |
| chr13:76886051-76886998 | -0.517698431 | 0.115892164 | 0.343224643 |
| chr10:47254605-47254877 | -0.735853095 | 0.115892406 | 0.343224643 |
| chr7:27160691-27161315 | 0.54099535 | 0.115916719 | 0.343283379 |
| chr4:7101998-7102334 | 0.592680454 | 0.115927471 | 0.343296873 |
| chr9:21326592-21326898 | -0.711366802 | 0.115930237 | 0.343296873 |
| chr11:117833379-117833928 | 0.549251576 | 0.115940366 | 0.343306744 |
| chr11:61628070-61628502 | 0.583374185 | 0.115944088 | 0.343306744 |
| chr9:88197909-88198178 | 0.784371579 | 0.115950731 | 0.343306744 |
| chr16:970228-971242 | 0.518934104 | 0.115951492 | 0.343306744 |
| chr1:89889947-89890681 | -0.519193504 | 0.115979769 | 0.343368027 |
| chr17:7294533-7295150 | -0.537827966 | 0.115981153 | 0.343368027 |
| chr8:97090009-97090412 | -0.581201303 | 0.115995324 | 0.343396713 |
| chr1:75908649-75909120 | 0.614715004 | 0.116009825 | 0.343426374 |
| chr19:19192105-19192324 | -0.725678574 | 0.116024985 | 0.343449575 |
| chr11:70755506-70755858 | 0.609223537 | 0.116026627 | 0.343449575 |
| chr2:182870264-182870566 | -0.644073705 | 0.116035101 | 0.343456484 |
| chr20:25339502-25339702 | 0.80787361 | 0.116039591 | 0.343456484 |
| chr9:83166266-83166920 | -0.513945653 | 0.116042409 | 0.343456484 |
| chr7:44325479-44325730 | -0.694640536 | 0.116053932 | 0.343463893 |
| chr8:99905472-99905811 | -0.613830952 | 0.116056144 | 0.343463893 |
| chr6:127413264-127413473 | 0.761146152 | 0.11605836 | 0.343463893 |
| chr2:16281892-16282364 | -0.601489596 | 0.116073589 | 0.343480238 |
| chr4:24687427-24687663 | 0.755825854 | 0.116076122 | 0.343480238 |
| chr12:57590803-57591386 | 0.53692973 | 0.116078211 | 0.343480238 |
| chr6:113211086-113211600 | 0.607059403 | 0.116084369 | 0.343480238 |
| chr4:95761560-95761825 | 0.696535886 | 0.116095325 | 0.343480238 |
| chr3:179346763-179348254 | -0.500707683 | 0.116095773 | 0.343480238 |
| chr6:3511463-3511873 | -0.602911301 | 0.116099532 | 0.343480238 |

| | | | |
|---|---|---|---|
| chr6:37169568-37170302 | 0.511009302 | 0.116100584 | 0.343480238 |
| chr16:31507644-31508624 | 0.531743334 | 0.116104228 | 0.343480238 |
| chrX:129925437-129925968 | 0.613983581 | 0.116127282 | 0.34353324 |
| chrX:48511619-48511819 | 0.732542513 | 0.116138764 | 0.34353324 |
| chr2:43178743-43179378 | -0.575259936 | 0.116138821 | 0.34353324 |
| chr1:147721772-147722073 | -0.640772804 | 0.116145823 | 0.34353324 |
| chr15:70324270-70324610 | 0.593113521 | 0.116150926 | 0.34353324 |
| chr10:26438095-26438390 | -0.796115611 | 0.116154514 | 0.34353324 |
| chr20:37710865-37711307 | 0.626678736 | 0.116155816 | 0.34353324 |
| chr10:29142498-29143524 | -0.531083216 | 0.116162509 | 0.34353324 |
| chr8:97304806-97305082 | 0.758252055 | 0.116163608 | 0.34353324 |
| chr3:8239792-8240185 | 0.586712468 | 0.116166979 | 0.34353324 |
| chr1:243263415-243263736 | 0.725974448 | 0.116177061 | 0.343539306 |
| chr7:90412769-90413210 | -0.597084571 | 0.116177997 | 0.343539306 |
| chr3:156136002-156136222 | -0.769865853 | 0.116221704 | 0.343655284 |
| chr2:33064376-33064668 | -0.77360698 | 0.11624552 | 0.343712442 |
| chr3:149393471-149393717 | 0.681833021 | 0.116271095 | 0.343769034 |
| chr1:210407288-210407505 | -0.785397438 | 0.116273633 | 0.343769034 |
| chr14:81655489-81656072 | -0.549062949 | 0.116285379 | 0.343790496 |
| chr10:115441988-115442225 | -0.837842673 | 0.116295223 | 0.343806335 |
| chr8:120073188-120073501 | -0.695598326 | 0.116300883 | 0.343809803 |
| chr20:57115569-57115821 | 0.52057786 | 0.116309662 | 0.343822488 |
| chr13:111429123-111429591 | -0.642296786 | 0.116349117 | 0.343916661 |
| chr2:26330077-26330375 | 0.631604711 | 0.116350496 | 0.343916661 |
| chr12:52945528-52945816 | 0.573938401 | 0.116369928 | 0.34396083 |
| chr5:139543389-139543661 | -0.718726509 | 0.116388938 | 0.343999795 |
| chrX:106317873-106318090 | -0.866048152 | 0.11639209 | 0.343999795 |
| chr5:139391760-139392025 | 0.637457789 | 0.116402539 | 0.344016758 |
| chr8:68872592-68872987 | -0.658955699 | 0.116406809 | 0.344016758 |
| chr8:89678795-89679069 | 0.684113302 | 0.116434231 | 0.344066882 |
| chr6:139349486-139349786 | 0.662591253 | 0.116435347 | 0.344066882 |
| chr15:73487617-73488067 | 0.651997913 | 0.116437241 | 0.344066882 |
| chr16:12899414-12900073 | 0.542678626 | 0.116459429 | 0.344115017 |
| chr2:199405286-199405594 | -0.778950297 | 0.116462512 | 0.344115017 |
| chr21:44510402-44510826 | 0.572819147 | 0.116470417 | 0.344125103 |
| chr3:64049400-64049600 | 0.565862073 | 0.11647951 | 0.344128846 |
| chr12:107092020-107092514 | -0.528411956 | 0.116480666 | 0.344128846 |
| chr17:40582237-40582574 | -0.760465793 | 0.116487449 | 0.344135615 |
| chr19:38515046-38515388 | 0.633032266 | 0.116508405 | 0.344171745 |
| chr12:121795003-121795242 | 0.596591922 | 0.116508662 | 0.344171745 |
| chr2:35455006-35455326 | -0.651730871 | 0.116523955 | 0.344203649 |
| chr2:21351304-21351518 | -0.805623456 | 0.116535738 | 0.344216293 |
| chr4:73127884-73128370 | -0.645289068 | 0.116542541 | 0.344216293 |
| chr12:98475197-98475690 | 0.588957894 | 0.116546999 | 0.344216293 |
| chr17:60886659-60887621 | -0.515687358 | 0.116547245 | 0.344216293 |
| chr11:126405396-126405736 | 0.613768057 | 0.116562028 | 0.344216293 |
| chr5:31937815-31938015 | 0.759682236 | 0.116566257 | 0.344216293 |
| chr4:71067402-71067648 | 0.789738078 | 0.116571349 | 0.344216293 |
| chr7:87236885-87237248 | -0.651862437 | 0.116571492 | 0.344216293 |
| chr3:180859057-180859312 | 0.692883084 | 0.116571564 | 0.344216293 |

| | | | |
|---|---|---|---|
| chr3:4350861-4351141 | -0.659623154 | 0.116573489 | 0.344216293 |
| chr14:75984809-75985062 | 0.612100677 | 0.116577651 | 0.344216293 |
| chr4:119822185-119822432 | 0.786393519 | 0.116582263 | 0.344216646 |
| chr6:79048698-79048960 | -0.754360111 | 0.116605887 | 0.34427313 |
| chrX:14029123-14030258 | -0.509667081 | 0.116634929 | 0.344336734 |
| chr17:14098746-14098963 | 0.742208867 | 0.116638024 | 0.344336734 |
| chr18:10234949-10235157 | -0.799211808 | 0.116640912 | 0.344336734 |
| chr8:65143358-65143681 | -0.666127069 | 0.116649996 | 0.344350286 |
| chr7:15144453-15145057 | 0.57847783 | 0.116681107 | 0.344418974 |
| chr14:31419845-31421055 | 0.502537364 | 0.116682831 | 0.344418974 |
| chr4:163754944-163755366 | -0.671576132 | 0.11668675 | 0.344418974 |
| chr2:175031045-175031321 | -0.643078512 | 0.116705671 | 0.344450156 |
| chr22:24937182-24937705 | 0.574793885 | 0.116710233 | 0.344450156 |
| chr2:56077581-56077804 | -0.81127824 | 0.1167108 | 0.344450156 |
| chr20:56554996-56555219 | 0.623273135 | 0.116740908 | 0.344520579 |
| chr20:48733199-48733603 | 0.516945747 | 0.116746791 | 0.344520579 |
| chr4:190122510-190123073 | -0.614585043 | 0.116748151 | 0.344520579 |
| chr6:21632631-21632925 | -0.831689443 | 0.11676739 | 0.344541704 |
| chr20:54050048-54050592 | 0.509335465 | 0.116768769 | 0.344541704 |
| chr8:128317055-128317418 | -0.592832265 | 0.116776338 | 0.344541704 |
| chr9:114291012-114291394 | 0.585681526 | 0.116782375 | 0.344541704 |
| chr3:147383733-147383951 | -0.667750978 | 0.11678502 | 0.344541704 |
| chr6:79180298-79180498 | 0.799331061 | 0.116787279 | 0.344541704 |
| chr16:68264713-68265075 | 0.59333011 | 0.116790746 | 0.344541704 |
| chr3:39877869-39878560 | -0.597134983 | 0.116791283 | 0.344541704 |
| chr2:179751089-179751451 | -0.699118943 | 0.116805718 | 0.344571023 |
| chr7:148437621-148437822 | 0.829184035 | 0.116816137 | 0.34458849 |
| chr6:34609584-34609943 | -0.628817796 | 0.116861699 | 0.344709621 |
| chr8:120568791-120569024 | -0.77753618 | 0.116896569 | 0.344799203 |
| chr7:15069377-15069615 | 0.833684412 | 0.116905648 | 0.344812708 |
| chr3:31919641-31919857 | 0.788419163 | 0.116913794 | 0.344823464 |
| chr2:240435152-240435766 | 0.542349805 | 0.116926688 | 0.344848217 |
| chr22:39242759-39243095 | 0.597918729 | 0.116946723 | 0.34489403 |
| chr3:180855409-180855786 | -0.64188106 | 0.116955487 | 0.344906602 |
| chrX:153708478-153708712 | 0.688710504 | 0.116963558 | 0.344913006 |
| chr14:23068901-23069174 | 0.587991348 | 0.116966662 | 0.344913006 |
| chr6:165704164-165704447 | -0.79029936 | 0.116999897 | 0.344997734 |
| chr3:171625104-171625519 | -0.649595019 | 0.117019221 | 0.345041435 |
| chr2:70935388-70935904 | -0.586134176 | 0.117028563 | 0.345046331 |
| chr7:93622454-93622753 | -0.620603923 | 0.117029888 | 0.345046331 |
| chr4:170193358-170193981 | -0.532264275 | 0.117050041 | 0.345086067 |
| chr5:134648331-134649134 | -0.524614217 | 0.117052372 | 0.345086067 |
| chr3:64476176-64476423 | 0.557875387 | 0.117066999 | 0.345115909 |
| chr5:168706630-168706903 | -0.658718862 | 0.117078704 | 0.345137136 |
| chr2:101518527-101519216 | -0.594658163 | 0.117089253 | 0.345154956 |
| chr17:63696610-63696871 | 0.593403234 | 0.117133412 | 0.345265847 |
| chr8:89735687-89735930 | 0.694705129 | 0.117135884 | 0.345265847 |
| chr6:41735499-41735855 | 0.562140523 | 0.117142979 | 0.345273477 |
| chr5:177930185-177930487 | -0.697090387 | 0.117148282 | 0.345275826 |
| chr16:527158-527774 | 0.528153363 | 0.11716702 | 0.345311171 |

| | | | |
|---|---|---|---|
| chr16:66751309-66752011 | 0.557110684 | 0.117169287 | 0.345311171 |
| chr2:120949394-120949733 | -0.614655749 | 0.117183501 | 0.345339779 |
| chr5:33903402-33903714 | 0.782737661 | 0.117228966 | 0.345460476 |
| chr12:102917104-102917888 | -0.540253414 | 0.117235645 | 0.345466872 |
| chr13:37058894-37060304 | -0.499660728 | 0.117260424 | 0.3455266 |
| chr9:86946882-86947381 | 0.61854373 | 0.117278202 | 0.345565696 |
| chr8:25614628-25615118 | -0.630768347 | 0.117300276 | 0.345597958 |
| chr2:24827506-24827932 | -0.566247906 | 0.117300888 | 0.345597958 |
| chr6:37354547-37354840 | 0.66736308 | 0.117302682 | 0.345597958 |
| chr7:21759686-21759926 | 0.714058876 | 0.117318603 | 0.345631575 |
| chr9:120936184-120936421 | 0.805051313 | 0.117331784 | 0.345657117 |
| chr11:60618692-60619014 | -0.669409733 | 0.117352969 | 0.3456999 |
| chr6:127956197-127956599 | -0.631967602 | 0.11735533 | 0.3456999 |
| chr11:134270366-134270615 | 0.713791398 | 0.117361585 | 0.345705034 |
| chr20:53608048-53608926 | 0.493913152 | 0.117379891 | 0.345732312 |
| chr4:72210200-72210639 | -0.620365774 | 0.117382307 | 0.345732312 |
| chr9:100437841-100438214 | -0.609155417 | 0.117384382 | 0.345732312 |
| chrX:107025641-107025874 | -0.743089005 | 0.117401009 | 0.345767993 |
| chr2:234633735-234634307 | 0.564617467 | 0.117413216 | 0.345786584 |
| chr12:85036123-85036700 | -0.53184164 | 0.117419917 | 0.345786584 |
| chr1:210711880-210712118 | -0.781394425 | 0.117422896 | 0.345786584 |
| chr7:116180282-116180707 | -0.629591457 | 0.117425373 | 0.345786584 |
| chr17:66025890-66026148 | -0.809695745 | 0.117432826 | 0.345795241 |
| chr2:62369519-62369814 | -0.71666162 | 0.117451023 | 0.345835536 |
| chr1:91220899-91221371 | -0.661927393 | 0.117457815 | 0.345842242 |
| chr17:62163895-62164242 | 0.586780456 | 0.117472479 | 0.345872129 |
| chr10:51699245-51700326 | -0.509189528 | 0.117483862 | 0.345892353 |
| chr7:37833626-37833860 | 0.710335338 | 0.117501578 | 0.34593122 |
| chr1:227793821-227794193 | 0.643122344 | 0.117510678 | 0.345944717 |
| chr6:53335212-53335497 | -0.704456194 | 0.117519177 | 0.345956447 |
| chr9:105656095-105656295 | -0.847065236 | 0.117531591 | 0.345976955 |
| chr2:42086557-42086845 | -0.672292225 | 0.117536158 | 0.345976955 |
| chr17:29384927-29385210 | -0.700405771 | 0.11753969 | 0.345976955 |
| chr6:144264286-144264524 | -0.776867541 | 0.117548769 | 0.345990389 |
| chr5:132301962-132302162 | -0.816671898 | 0.117562284 | 0.346016877 |
| chr10:732759-733167 | -0.658699556 | 0.117571382 | 0.34602302 |
| chr8:130015188-130017004 | 0.500274208 | 0.117574719 | 0.34602302 |
| chr5:141320205-141321358 | -0.512483706 | 0.11757792 | 0.34602302 |
| chr11:64439329-64439529 | 0.804455246 | 0.117591504 | 0.346049706 |
| chrX:54787037-54787245 | -0.826029442 | 0.117599198 | 0.346056348 |
| chr3:169041673-169042146 | -0.636362112 | 0.117602793 | 0.346056348 |
| chr14:67526607-67527146 | 0.550310547 | 0.117613774 | 0.346075369 |
| chr3:160498301-160498704 | -0.691057699 | 0.117633416 | 0.346110899 |
| chr2:10835041-10835644 | -0.532114155 | 0.117634883 | 0.346110899 |
| chr4:128572888-128573088 | 0.874451003 | 0.117667315 | 0.346193028 |
| chr13:38417619-38418159 | -0.555299561 | 0.117701847 | 0.346281328 |
| chr5:97425606-97425816 | -0.817153766 | 0.11771594 | 0.346309494 |
| chr3:19971521-19971723 | 0.804153049 | 0.117726228 | 0.346326462 |
| chr10:103847990-103848227 | -0.791938509 | 0.117753131 | 0.346392307 |
| chr16:68738098-68738729 | 0.555470146 | 0.117767578 | 0.346421504 |

| | | | |
|---|---|---|---|
| chr3:100006759-100007004 | -0.766749974 | 0.117801764 | 0.346508762 |
| chr20:53527289-53527489 | 0.73066242 | 0.117810873 | 0.346522253 |
| chr19:49150247-49150669 | 0.542832654 | 0.117839585 | 0.346593401 |
| chr20:40990523-40990977 | 0.545501568 | 0.117896097 | 0.346746305 |
| chr7:120502797-120503033 | 0.739672725 | 0.117901829 | 0.346749853 |
| chr2:215435574-215436382 | -0.535656318 | 0.117912491 | 0.346767901 |
| chr17:72287437-72287646 | 0.735383002 | 0.117920934 | 0.34677258 |
| chrX:149836133-149836354 | 0.78019854 | 0.117923134 | 0.34677258 |
| chr14:50848765-50849007 | 0.737729967 | 0.117929498 | 0.346777986 |
| chr4:172865641-172865923 | -0.738580316 | 0.11795944 | 0.346852722 |
| chr5:122258306-122258521 | -0.803736317 | 0.117967351 | 0.346862671 |
| chr5:55933880-55934636 | -0.538529334 | 0.117994989 | 0.346930624 |
| chrX:78065031-78065429 | -0.754779801 | 0.118005375 | 0.346933766 |
| chr10:107618303-107618848 | -0.560695592 | 0.118008551 | 0.346933766 |
| chr18:73774880-73775317 | -0.642931333 | 0.118009641 | 0.346933766 |
| chr5:41433051-41433256 | 0.875288085 | 0.118038957 | 0.347006636 |
| chr10:103890135-103890701 | 0.527808658 | 0.118047758 | 0.347019195 |
| chr7:46291072-46291464 | -0.642812386 | 0.11809116 | 0.347117829 |
| chr2:20113579-20114456 | -0.51026318 | 0.118093295 | 0.347117829 |
| chr19:3429622-3430069 | -0.665893151 | 0.118094902 | 0.347117829 |
| chr8:100680706-100681701 | 0.512066202 | 0.118122806 | 0.347179867 |
| chr14:52790919-52792189 | 0.4942943 | 0.118128681 | 0.347179867 |
| chr1:110433718-110434147 | -0.587495336 | 0.118129601 | 0.347179867 |
| chr22:31131751-31132125 | 0.585873977 | 0.118150335 | 0.347217029 |
| chr6:34657334-34657671 | -0.708185469 | 0.118151309 | 0.347217029 |
| chr5:15824324-15824538 | -0.745378087 | 0.118161138 | 0.347228241 |
| chr3:7407132-7407425 | -0.661310854 | 0.118164188 | 0.347228241 |
| chr6:97978702-97979628 | -0.508767799 | 0.118175678 | 0.347246427 |
| chr5:50962866-50963263 | -0.581036253 | 0.11817944 | 0.347246427 |
| chr18:3665880-3666166 | -0.682705865 | 0.118186442 | 0.347253684 |
| chr14:99969967-99970285 | 0.584929924 | 0.118206407 | 0.347289898 |
| chr10:131569967-131570170 | 0.782537897 | 0.118208412 | 0.347289898 |
| chr1:115713090-115713642 | -0.514297548 | 0.118212365 | 0.347289898 |
| chr7:38341215-38341760 | -0.540849652 | 0.118246203 | 0.347363022 |
| chr4:168200787-168201073 | -0.690335817 | 0.118246322 | 0.347363022 |
| chr9:84856261-84856529 | -0.750753487 | 0.118303204 | 0.347516796 |
| chr3:142931279-142931575 | -0.628963808 | 0.118320907 | 0.347555474 |
| chrX:106692741-106693752 | 0.525659298 | 0.118328332 | 0.347563959 |
| chr2:163349077-163349328 | 0.677728952 | 0.118354501 | 0.347610267 |
| chr8:97627075-97627340 | -0.743133569 | 0.118361366 | 0.347610267 |
| chr21:41878062-41878304 | -0.674869733 | 0.118361915 | 0.347610267 |
| chr15:92091189-92091427 | 0.691814134 | 0.118362244 | 0.347610267 |
| chr2:169573198-169574130 | -0.521308694 | 0.118370228 | 0.347620389 |
| chr5:145347061-145347399 | -0.663842713 | 0.118389736 | 0.347664356 |
| chr22:25400782-25401397 | -0.549873225 | 0.118419483 | 0.347738384 |
| chr13:47220423-47220863 | 0.602264795 | 0.118437915 | 0.347765421 |
| chr17:28634423-28634849 | 0.561120048 | 0.118441261 | 0.347765421 |
| chr2:25361919-25362389 | -0.532980358 | 0.118442307 | 0.347765421 |
| chr13:86838466-86838743 | 0.748165441 | 0.118453941 | 0.347786255 |
| chr2:109470237-109470955 | -0.548407359 | 0.11846319 | 0.347800082 |

| | | | |
|---|---|---|---|
| chr12:105053064-105053419 | -0.714691601 | 0.118494139 | 0.347877617 |
| chr8:128090989-128091554 | -0.526178269 | 0.118507007 | 0.347902066 |
| chr5:103119715-103120755 | -0.496188315 | 0.11851602 | 0.347909429 |
| chr1:9345981-9346274 | -0.718042041 | 0.118522009 | 0.347909429 |
| chr10:83225906-83226226 | 0.65441046 | 0.118523137 | 0.347909429 |
| chr14:40671901-40672101 | 0.835736672 | 0.118529712 | 0.3479154 |
| chr2:88671813-88672182 | -0.582223913 | 0.118537243 | 0.347916098 |
| chr13:72839821-72840220 | -0.594654882 | 0.118539031 | 0.347916098 |
| chr9:137254690-137254927 | 0.671483366 | 0.118547198 | 0.347921497 |
| chr8:97868787-97869856 | 0.512317962 | 0.118549952 | 0.347921497 |
| chr5:141557917-141558123 | -0.769540355 | 0.118559334 | 0.347935704 |
| chr17:8016707-8016933 | -0.74304513 | 0.118573324 | 0.347945021 |
| chr2:181058736-181058976 | -0.74304513 | 0.118573324 | 0.347945021 |
| chr2:189347568-189347768 | 0.834836786 | 0.118576132 | 0.347945021 |
| chr14:60222863-60223183 | 0.625622844 | 0.118585621 | 0.347959538 |
| chr11:60841833-60842253 | 0.532431331 | 0.118629743 | 0.348075675 |
| chr1:9234241-9235345 | 0.525289647 | 0.118643686 | 0.348097768 |
| chr10:50421330-50421594 | -0.658879712 | 0.118646359 | 0.348097768 |
| chr21:32990790-32991030 | 0.795174884 | 0.118659616 | 0.34811709 |
| chr10:119682074-119682443 | -0.619183788 | 0.118663948 | 0.34811709 |
| chr3:64219929-64220807 | 0.499460035 | 0.118666575 | 0.34811709 |
| chr3:100002804-100003454 | 0.532458164 | 0.118673182 | 0.348118347 |
| chr10:119101606-119101915 | 0.591449806 | 0.11867609 | 0.348118347 |
| chr17:30905910-30906737 | 0.554064151 | 0.118682788 | 0.348124668 |
| chr3:153209616-153209898 | 0.740251703 | 0.118693757 | 0.348143517 |
| chr3:135516301-135516595 | -0.668972985 | 0.118701477 | 0.348152831 |
| chr21:8213013-8213552 | 0.520784239 | 0.118739737 | 0.348233085 |
| chrX:64894546-64895009 | -0.64264471 | 0.118740257 | 0.348233085 |
| chr13:65612950-65613284 | -0.788494898 | 0.118742474 | 0.348233085 |
| chr1:79044970-79045405 | -0.576441365 | 0.11875047 | 0.348243208 |
| chr9:25716141-25716578 | -0.573648321 | 0.118756084 | 0.348246343 |
| chr14:80954768-80955577 | 0.529192812 | 0.118760812 | 0.348246878 |
| chr1:207629354-207629554 | 0.788392597 | 0.118767217 | 0.348252332 |
| chr3:129162231-129162453 | 0.649919054 | 0.118774199 | 0.348255824 |
| chr2:73981284-73981681 | -0.561676066 | 0.118777498 | 0.348255824 |
| chr6:135137113-135137428 | -0.785756856 | 0.118793634 | 0.348280415 |
| chr3:115092223-115092780 | -0.617397806 | 0.118794976 | 0.348280415 |
| chr4:15469701-15470016 | -0.688366451 | 0.118822169 | 0.34834681 |
| chr3:52385238-52385495 | 0.639977833 | 0.118845022 | 0.348400478 |
| chr6:156437269-156437909 | -0.597833128 | 0.118861446 | 0.348435294 |
| chr3:160032205-160032667 | -0.564559939 | 0.118876532 | 0.348463542 |
| chr2:178992761-178993046 | -0.84795179 | 0.118880178 | 0.348463542 |
| chr14:94173827-94174786 | 0.520125352 | 0.118905108 | 0.348523286 |
| chr2:195779561-195779793 | -0.778156719 | 0.118915425 | 0.348540191 |
| chr3:10085806-10086114 | 0.640523428 | 0.118932322 | 0.348576383 |
| chr5:88674071-88675446 | -0.497541619 | 0.118939036 | 0.348581685 |
| chr19:38543779-38544643 | 0.537068201 | 0.118947016 | 0.348581685 |
| chr14:95398128-95398395 | 0.706228742 | 0.118951698 | 0.348581685 |
| chr13:42883816-42884186 | -0.62932993 | 0.118962791 | 0.348581685 |
| chr14:93017443-93017656 | -0.72463361 | 0.118963744 | 0.348581685 |

| | | | |
|---|---|---|---|
| chr20:21048783-21049150 | 0.608957354 | 0.118964628 | 0.348581685 |
| chr11:85263426-85263801 | 0.655598295 | 0.118965977 | 0.348581685 |
| chr15:40903812-40904281 | 0.54630946 | 0.118977112 | 0.348591045 |
| chr10:71270307-71270555 | -0.708137837 | 0.11897827 | 0.348591045 |
| chr10:19592350-19592590 | -0.918875236 | 0.11901455 | 0.348676051 |
| chr2:101770505-101771001 | 0.639987133 | 0.119019989 | 0.348676051 |
| chr13:50365663-50366427 | -0.569722468 | 0.119023844 | 0.348676051 |
| chr6:126804244-126804503 | -0.713696959 | 0.119025486 | 0.348676051 |
| chr16:14301954-14302369 | 0.57907192 | 0.119036743 | 0.348695695 |
| chr1:28887763-28888090 | 0.636111774 | 0.119041977 | 0.348697697 |
| chr7:101216981-101217825 | -0.521817154 | 0.119047102 | 0.348699378 |
| chr2:38697190-38697408 | -0.813077854 | 0.119072518 | 0.348760492 |
| chr8:144810513-144810765 | 0.76925038 | 0.119088324 | 0.34878252 |
| chr11:47566150-47566386 | 0.627775883 | 0.119089143 | 0.34878252 |
| chr20:35277616-35278314 | 0.532961192 | 0.119118115 | 0.348849245 |
| chr1:183590360-183590624 | -0.793696695 | 0.119121031 | 0.348849245 |
| chr9:4759355-4759555 | -0.840726149 | 0.119129516 | 0.34886076 |
| chr2:60707703-60707954 | -0.734279124 | 0.119153241 | 0.3489169 |
| chr1:153922032-153922284 | 0.605780479 | 0.119160171 | 0.34892386 |
| chr5:105104953-105105215 | -0.831583819 | 0.119170394 | 0.34894046 |
| chr3:177777653-177778100 | -0.584125513 | 0.119180448 | 0.348956561 |
| chr2:186589636-186590690 | -0.496524724 | 0.119209787 | 0.349029128 |
| chr20:50267654-50267883 | 0.655575454 | 0.119215659 | 0.349032985 |
| chr6:29976479-29976809 | 0.608752218 | 0.119237478 | 0.349083525 |
| chr12:99455349-99455712 | -0.595968141 | 0.119249616 | 0.349105723 |
| chr8:78250759-78250977 | 0.793859264 | 0.119256656 | 0.349112993 |
| chrX:23901866-23902208 | -0.581116738 | 0.11926237 | 0.349116381 |
| chr11:25211878-25212214 | -0.680945805 | 0.119268029 | 0.34911961 |
| chr3:123258846-123259057 | -0.78999642 | 0.119283329 | 0.349151057 |
| chr7:88219270-88219584 | -0.602353679 | 0.119292353 | 0.349164133 |
| chr19:45076250-45076672 | 0.565040997 | 0.119310574 | 0.349204126 |
| chrX:53434237-53434733 | 0.551238637 | 0.119349711 | 0.349291748 |
| chr2:3125824-3126024 | 0.762595533 | 0.119354164 | 0.349291748 |
| chr15:61227462-61227912 | 0.690855402 | 0.119357055 | 0.349291748 |
| chr9:81522774-81523619 | -0.551017689 | 0.119362057 | 0.349291748 |
| chr16:86180562-86180828 | 0.736801687 | 0.119363305 | 0.349291748 |
| chr13:68850388-68850762 | -0.759037708 | 0.119374499 | 0.349311166 |
| chr17:12342679-12342894 | -0.814116973 | 0.119389385 | 0.349334888 |
| chr10:78047308-78047538 | 0.664991762 | 0.119391725 | 0.349334888 |
| chr18:13630160-13630413 | 0.736808228 | 0.119398789 | 0.349342219 |
| chr13:28819407-28819623 | -0.676589848 | 0.119411618 | 0.349343135 |
| chr15:48352521-48352835 | -0.686253449 | 0.119411646 | 0.349343135 |
| chr13:36428163-36428430 | -0.683352157 | 0.11941278 | 0.349343135 |
| chr21:45137204-45137404 | 0.739131209 | 0.119421133 | 0.349344866 |
| chr12:57861485-57861834 | 0.592293582 | 0.119422491 | 0.349344866 |
| chr2:67650767-67651135 | -0.657682143 | 0.119447533 | 0.349404782 |
| chr7:105429582-105430050 | -0.616282532 | 0.119484435 | 0.349484481 |
| chr9:87725688-87726657 | 0.504809701 | 0.119487643 | 0.349484481 |
| chr7:152833460-152833667 | -0.821862503 | 0.119488462 | 0.349484481 |
| chr15:80094182-80094449 | -0.775097008 | 0.119495132 | 0.34949065 |

| | | | |
|---|---|---|---|
| chr22:19632178-19632378 | 0.828135158 | 0.119502847 | 0.349499871 |
| chr14:56117832-56118496 | 0.570144872 | 0.119507975 | 0.34950153 |
| chr3:127823343-127823600 | 0.685890512 | 0.119542025 | 0.349574525 |
| chr20:37992754-37993069 | -0.682061505 | 0.11954206 | 0.349574525 |
| chr7:47979088-47979912 | -0.499267605 | 0.119553832 | 0.349595607 |
| chr17:71380318-71380805 | -0.640829264 | 0.119558512 | 0.349595952 |
| chr17:60214989-60215189 | 0.65139734 | 0.119579208 | 0.349643126 |
| chr11:6476004-6476255 | -0.619303759 | 0.119597624 | 0.349678283 |
| chr8:134720471-134720750 | 0.615117038 | 0.11960036 | 0.349678283 |
| chr3:48918419-48919029 | 0.561523783 | 0.119615437 | 0.34970902 |
| chr10:79150093-79150303 | -0.728509388 | 0.119631228 | 0.349741844 |
| chr20:57709846-57710743 | 0.495048385 | 0.119657683 | 0.349805837 |
| chr7:32427220-32427539 | -0.639878684 | 0.11968961 | 0.349885824 |
| chr10:29409400-29410040 | -0.542669001 | 0.119710543 | 0.349933664 |
| chr2:71331211-71332473 | -0.50587756 | 0.119731099 | 0.349980402 |
| chr6:40396496-40397073 | 0.553876887 | 0.119738489 | 0.349988652 |
| chr6:43060141-43060386 | 0.630348194 | 0.119752935 | 0.350010163 |
| chr12:40692050-40692564 | -0.571825286 | 0.119754985 | 0.350010163 |
| chr9:114405549-114405962 | 0.540868114 | 0.119777274 | 0.350061956 |
| chr1:173876818-173877108 | 0.630075759 | 0.119801051 | 0.350118093 |
| chr20:50715796-50716010 | 0.529749221 | 0.119809935 | 0.3501307 |
| chr7:101870310-101871124 | -0.540955846 | 0.119842607 | 0.350202042 |
| chr6:121042129-121042763 | 0.556209835 | 0.119843488 | 0.350202042 |
| chr2:42568497-42569447 | -0.537201006 | 0.119864753 | 0.350250823 |
| chr1:26695531-26695764 | 0.665581234 | 0.119876035 | 0.350270432 |
| chr3:133804981-133805193 | 0.680832193 | 0.119886725 | 0.350288311 |
| chr5:102754587-102756313 | -0.499237352 | 0.119912431 | 0.35035006 |
| chr8:55988955-55989355 | -0.562319805 | 0.11993701 | 0.350408511 |
| chr10:52682865-52683093 | 0.763666191 | 0.119952086 | 0.350415757 |
| chr14:96797047-96797945 | 0.503109145 | 0.119954341 | 0.350415757 |
| chr18:80210359-80210687 | -0.638790294 | 0.119958284 | 0.350415757 |
| chr11:85739917-85740123 | 0.767373709 | 0.119961457 | 0.350415757 |
| chr6:32154607-32154921 | 0.589007182 | 0.119962357 | 0.350415757 |
| chr17:58377269-58377683 | 0.537831697 | 0.119974701 | 0.350428003 |
| chr1:35268443-35269188 | 0.516314263 | 0.119975696 | 0.350428003 |
| chr3:45841785-45842427 | -0.548679917 | 0.119989782 | 0.350455785 |
| chr2:131683016-131683265 | -0.638029159 | 0.119999501 | 0.350470813 |
| chr5:140965959-140966956 | -0.532942499 | 0.120019309 | 0.350515303 |
| chr15:90902257-90903648 | 0.497407062 | 0.120025415 | 0.350519774 |
| chr9:129641991-129642622 | 0.523280069 | 0.120041667 | 0.350553877 |
| chr19:58499085-58499892 | 0.501801916 | 0.120070213 | 0.35061227 |
| chr5:34798631-34798849 | -0.879999432 | 0.12007539 | 0.35061227 |
| chr7:86905351-86905554 | -0.879999432 | 0.12007539 | 0.35061227 |
| chr12:93377460-93378828 | -0.495503163 | 0.120089675 | 0.350627773 |
| chr3:156589206-156589486 | -0.719327272 | 0.120094254 | 0.350627773 |
| chr2:138274402-138274719 | -0.688055055 | 0.120101112 | 0.350627773 |
| chrX:39012653-39013054 | -0.676242037 | 0.120102245 | 0.350627773 |
| chr17:30291967-30292411 | 0.635289112 | 0.120105776 | 0.350627773 |
| chr21:44765496-44765706 | 0.66369008 | 0.120108157 | 0.350627773 |
| chr8:108592214-108592489 | 0.720641091 | 0.120142195 | 0.350705433 |

| | | | |
|---|---|---|---|
| chr11:65181715-65181994 | 0.619474486 | 0.120148399 | 0.350705433 |
| chr3:130849000-130849605 | -0.597937576 | 0.120154328 | 0.350705433 |
| chr10:73887578-73888085 | -0.566533894 | 0.12015554 | 0.350705433 |
| chr1:1019383-1020096 | 0.526475862 | 0.120160882 | 0.350705433 |
| chr6:36667243-36667503 | 0.778778532 | 0.120165132 | 0.350705433 |
| chr11:122883511-122883723 | 0.76047174 | 0.120170443 | 0.350705433 |
| chr7:5406996-5407317 | 0.663202055 | 0.120171376 | 0.350705433 |
| chr1:8903997-8904210 | 0.755042126 | 0.12022445 | 0.350834876 |
| chr7:151875278-151875748 | -0.565894765 | 0.120224887 | 0.350834876 |
| chr14:49173779-49174098 | 0.618756279 | 0.120233671 | 0.350847145 |
| chr21:33347706-33348019 | 0.710103768 | 0.12027223 | 0.350946297 |
| chr3:197949654-197950674 | -0.508898543 | 0.12030056 | 0.351015595 |
| chr14:28330237-28330501 | -0.775068489 | 0.120306601 | 0.351019853 |
| chr22:19497246-19497500 | -0.735968904 | 0.120331003 | 0.351064359 |
| chr9:5839300-5839539 | -0.724016987 | 0.12033358 | 0.351064359 |
| chr22:29245288-29245831 | 0.592259671 | 0.1203356 | 0.351064359 |
| chrX:74383675-74383875 | 0.795086568 | 0.120352128 | 0.35109921 |
| chr2:20642404-20642800 | -0.542011038 | 0.120391339 | 0.351200229 |
| chr14:90760708-90761036 | 0.58496597 | 0.120421852 | 0.351275865 |
| chr21:38460711-38460911 | -0.819271141 | 0.12043861 | 0.351297859 |
| chr21:42097899-42098130 | 0.688430526 | 0.120442749 | 0.351297859 |
| chr17:8118754-8119318 | -0.52203334 | 0.120443146 | 0.351297859 |
| chr15:41507841-41508198 | 0.586576264 | 0.120450116 | 0.351304814 |
| chr16:66326943-66327390 | 0.647594428 | 0.120456476 | 0.351309992 |
| chr17:80426960-80427192 | 0.633781875 | 0.120463143 | 0.351311002 |
| chr5:39716397-39717103 | 0.515414542 | 0.120465992 | 0.351311002 |
| chr6:50815915-50816359 | -0.660462062 | 0.120476693 | 0.351328835 |
| chr16:175457-175918 | 0.582577258 | 0.12052688 | 0.351461812 |
| chr4:143355595-143355825 | -0.773152558 | 0.120552092 | 0.351521953 |
| chr10:18723301-18723735 | -0.691220421 | 0.120562141 | 0.351533083 |
| chr17:56939962-56940248 | 0.586828127 | 0.120565084 | 0.351533083 |
| chr1:217198759-217198959 | -0.861629082 | 0.120578725 | 0.351540478 |
| chr7:75866519-75866817 | -0.652582791 | 0.120578783 | 0.351540478 |
| chr1:215242140-215242373 | 0.76217517 | 0.120581384 | 0.351540478 |
| chr10:86967052-86967267 | -0.598637682 | 0.120591373 | 0.351556223 |
| chr5:172765080-172765764 | -0.504090304 | 0.12060689 | 0.351588081 |
| chr8:104477133-104477419 | 0.6485914 | 0.120623848 | 0.351624138 |
| chr13:46893913-46894617 | -0.521064908 | 0.120639888 | 0.351647654 |
| chr13:46665587-46665881 | 0.644511886 | 0.120648122 | 0.351647654 |
| chr13:49652615-49653066 | -0.59714909 | 0.120652459 | 0.351647654 |
| chr1:66332024-66332673 | -0.54304004 | 0.120654659 | 0.351647654 |
| chr18:54260634-54261221 | -0.576886747 | 0.120654862 | 0.351647654 |
| chr17:57068715-57069015 | -0.61728392 | 0.120660453 | 0.351650574 |
| chr11:72786295-72786820 | -0.516853851 | 0.120668692 | 0.351661208 |
| chr21:34888558-34888985 | 0.543944797 | 0.12069056 | 0.351711563 |
| chr7:149668961-149669385 | -0.609827964 | 0.120704107 | 0.351737663 |
| chr11:29020404-29020649 | -0.774236252 | 0.120710981 | 0.351744317 |
| chr9:19457442-19457802 | -0.605743788 | 0.120717372 | 0.351749563 |
| chr5:174982266-174982599 | 0.790882511 | 0.120756995 | 0.351826946 |
| chr6:143513448-143513661 | -0.70359457 | 0.120758788 | 0.351826946 |

| | | | |
|---|---|---|---|
| chr11:11931509-11931757 | -0.812993054 | 0.120766348 | 0.351826946 |
| chr11:2390809-2391094 | -0.673939532 | 0.120766706 | 0.351826946 |
| chr19:45439533-45439764 | 0.676735512 | 0.120766888 | 0.351826946 |
| chr10:130136026-130137438 | 0.507211142 | 0.120780623 | 0.35185358 |
| chr5:90439765-90440160 | -0.686960671 | 0.120814285 | 0.351938263 |
| chr7:148905395-148905614 | -0.782005983 | 0.12082165 | 0.351941346 |
| chr9:385869-386107 | 0.680578985 | 0.12082453 | 0.351941346 |
| chr10:80196504-80196834 | -0.694137599 | 0.120833874 | 0.351944948 |
| chr7:954208-954558 | 0.628170742 | 0.120839226 | 0.351944948 |
| chr1:109162384-109162620 | -0.666131001 | 0.120839546 | 0.351944948 |
| chr3:51384920-51385376 | 0.548292523 | 0.120864276 | 0.352003594 |
| chr2:88765724-88766084 | -0.6755398 | 0.120880693 | 0.352038027 |
| chr7:92245294-92247007 | -0.494744073 | 0.120916162 | 0.352108933 |
| chr11:119389959-119390246 | -0.694360672 | 0.12091714 | 0.352108933 |
| chr22:39634636-39635230 | 0.566151416 | 0.120921099 | 0.352108933 |
| chr1:183454350-183454850 | -0.559800434 | 0.120923422 | 0.352108933 |
| chr3:40476944-40477484 | 0.578800197 | 0.120961439 | 0.352206247 |
| chr2:43880594-43881395 | -0.530210897 | 0.120970011 | 0.352217821 |
| chr4:8410550-8411118 | 0.522828428 | 0.120995385 | 0.352268817 |
| chr5:130141772-130142063 | -0.654979291 | 0.120996721 | 0.352268817 |
| chr9:129278338-129279013 | -0.541619131 | 0.121001473 | 0.352269267 |
| chrX:47751680-47751988 | 0.694838812 | 0.121009632 | 0.352279636 |
| chr15:74128165-74128365 | -0.812357685 | 0.121061201 | 0.352416373 |
| chr7:148983343-148984121 | -0.50958007 | 0.121069643 | 0.35242257 |
| chr7:41711195-41712038 | -0.552749442 | 0.121072529 | 0.35242257 |
| chr9:137028793-137029095 | 0.582647643 | 0.121077383 | 0.352423311 |
| chr19:29389596-29390043 | 0.549460438 | 0.121090242 | 0.352447218 |
| chr1:200106991-200107558 | -0.538159524 | 0.12109651 | 0.352447218 |
| chr1:113617018-113617274 | 0.742819784 | 0.121101851 | 0.352447218 |
| chr3:12967037-12967481 | 0.537832652 | 0.121103995 | 0.352447218 |
| chr3:45761187-45761387 | 0.804593675 | 0.121114306 | 0.352462577 |
| chr4:28443805-28444129 | 0.696387463 | 0.121124558 | 0.352462577 |
| chr11:76838248-76838510 | 0.713851262 | 0.121125356 | 0.352462577 |
| chr8:106762869-106763447 | -0.520671743 | 0.121133752 | 0.352462577 |
| chr8:122467086-122467581 | -0.538745592 | 0.121136268 | 0.352462577 |
| chr11:66531947-66532185 | -0.753890139 | 0.121136873 | 0.352462577 |
| chr12:102740758-102741000 | -0.74761977 | 0.121147543 | 0.352480236 |
| chrX:49627007-49627396 | -0.717248612 | 0.121160046 | 0.352503228 |
| chr14:33696950-33697440 | -0.591067375 | 0.121173965 | 0.352530338 |
| chr10:45155843-45156092 | -0.760737216 | 0.121181506 | 0.352538893 |
| chr12:83584680-83584988 | -0.71732132 | 0.121198893 | 0.352573622 |
| chr1:202086919-202087431 | -0.519930324 | 0.121202647 | 0.352573622 |
| chr2:168961694-168961977 | -0.622993884 | 0.121220132 | 0.352600494 |
| chr10:71303242-71303536 | 0.619274012 | 0.121223839 | 0.352600494 |
| chr1:224565831-224566031 | -0.848434942 | 0.12122569 | 0.352600494 |
| chr11:74662563-74662791 | -0.832960202 | 0.121234146 | 0.352611703 |
| chr17:39402447-39402745 | 0.650826689 | 0.121239767 | 0.352614667 |
| chr10:29386423-29386625 | -0.868347917 | 0.121246188 | 0.352619959 |
| chr11:26583317-26583916 | -0.549117478 | 0.121258392 | 0.352642065 |
| chr19:4284563-4284813 | 0.742478602 | 0.121292328 | 0.35271862 |

| | | | |
|---|---|---|---|
| chr5:180934775-180935082 | -0.784299206 | 0.121293922 | 0.35271862 |
| chr11:76823640-76823892 | -0.681471972 | 0.121341105 | 0.352842435 |
| chr2:219001714-219002739 | -0.500996669 | 0.12136491 | 0.352898264 |
| chr2:21027469-21027761 | -0.69866583 | 0.121374047 | 0.352902035 |
| chr12:88499475-88499675 | -0.868699182 | 0.121375419 | 0.352902035 |
| chr3:61764370-61764765 | 0.54499405 | 0.121383116 | 0.352911023 |
| chr6:102554182-102554447 | -0.698804266 | 0.121401281 | 0.352942923 |
| chr15:81183015-81183236 | 0.722314659 | 0.121413754 | 0.352942923 |
| chr17:19897104-19897748 | -0.582153105 | 0.121414747 | 0.352942923 |
| chr3:168088429-168088688 | -0.622180314 | 0.121415535 | 0.352942923 |
| chr1:232329211-232329441 | 0.721976461 | 0.121417119 | 0.352942923 |
| chr7:145036686-145036911 | -0.858401451 | 0.121424054 | 0.352949691 |
| chr13:30848818-30849023 | -0.805265615 | 0.121454689 | 0.352997173 |
| chr9:112012066-112012771 | -0.546995834 | 0.121455201 | 0.352997173 |
| chr2:160043578-160043865 | -0.877888174 | 0.121458588 | 0.352997173 |
| chr17:50519321-50519527 | 0.712448306 | 0.121458817 | 0.352997173 |
| chr16:75174856-75175119 | 0.724028718 | 0.121479972 | 0.353045265 |
| chr1:200866813-200867177 | 0.593697515 | 0.121601761 | 0.353385804 |
| chr16:68264007-68264683 | 0.522752885 | 0.121655998 | 0.353527155 |
| chr19:18451010-18451289 | -0.7291984 | 0.121659628 | 0.353527155 |
| chr8:39618331-39618807 | -0.616457755 | 0.121671057 | 0.353537128 |
| chr1:9142257-9142512 | 0.795539283 | 0.121672288 | 0.353537128 |
| chr2:133886730-133886988 | -0.639892092 | 0.121697922 | 0.353562971 |
| chr1:175840168-175840622 | -0.573957863 | 0.121698319 | 0.353562971 |
| chr7:142485672-142485909 | -0.720892771 | 0.121699153 | 0.353562971 |
| chr11:22163429-22163937 | -0.634628135 | 0.12169964 | 0.353562971 |
| chr20:49969874-49970282 | -0.57112907 | 0.121705817 | 0.353567512 |
| chr11:20132420-20132737 | 0.644721289 | 0.121728162 | 0.353596134 |
| chr6:152727603-152727822 | 0.751787804 | 0.121728221 | 0.353596134 |
| chr2:24434602-24434831 | -0.841681863 | 0.12173053 | 0.353596134 |
| chr10:17273389-17273623 | -0.799585573 | 0.121734129 | 0.353596134 |
| chr10:59227195-59227485 | -0.773848913 | 0.121750579 | 0.353630511 |
| chr5:132776121-132776340 | -0.699291647 | 0.121764925 | 0.353652779 |
| chr5:120212096-120212339 | -0.804817694 | 0.121772715 | 0.353652779 |
| chr5:174752160-174752416 | 0.638472179 | 0.121784218 | 0.353652779 |
| chr12:53180329-53181446 | 0.500964912 | 0.121792455 | 0.353652779 |
| chr8:120345859-120346068 | 0.754861114 | 0.12179309 | 0.353652779 |
| chr19:53103248-53103895 | 0.511348148 | 0.121793781 | 0.353652779 |
| chr19:37898708-37898925 | 0.751667008 | 0.121794534 | 0.353652779 |
| chr15:44683273-44683606 | -0.624972388 | 0.121798239 | 0.353652779 |
| chrX:48494023-48494434 | 0.589000907 | 0.121803015 | 0.353652779 |
| chr11:119420185-119420438 | -0.774553234 | 0.121804401 | 0.353652779 |
| chr6:24761496-24761839 | -0.694768591 | 0.121813229 | 0.353663267 |
| chr5:53035810-53036058 | -0.72900598 | 0.121825908 | 0.353663267 |
| chr3:33482198-33482628 | 0.659812503 | 0.121827493 | 0.353663267 |
| chr4:169124736-169124936 | -0.846181312 | 0.121827792 | 0.353663267 |
| chr12:52143566-52143969 | 0.537845645 | 0.121831092 | 0.353663267 |
| chr1:218553079-218553617 | -0.540202287 | 0.12185264 | 0.353711443 |
| chr5:132495791-132497349 | -0.487870638 | 0.121856921 | 0.353711443 |
| chr21:37257354-37257587 | 0.748768098 | 0.121868307 | 0.353721188 |

| | | | |
|---|---|---|---|
| chr5:32710452-32710729 | -0.568065341 | 0.121869511 | 0.353721188 |
| chr6:170153128-170153565 | 0.584674467 | 0.121878829 | 0.353734833 |
| chr15:89017377-89017924 | 0.538957255 | 0.121889503 | 0.35374606 |
| chr13:30802944-30803251 | 0.604342 | 0.12189193 | 0.35374606 |
| chr16:85395513-85395914 | -0.555900794 | 0.121898159 | 0.353750737 |
| chr4:140884465-140884896 | -0.587746716 | 0.121905408 | 0.353758378 |
| chr20:53905954-53906703 | 0.49084874 | 0.121924508 | 0.353800402 |
| chr4:76504975-76505225 | 0.698132335 | 0.121936179 | 0.35382087 |
| chr17:76262907-76263176 | 0.627115434 | 0.121968988 | 0.353902668 |
| chr10:100878285-100878808 | 0.564830314 | 0.121973706 | 0.353902958 |
| chr17:50056483-50056740 | 0.59257619 | 0.121999409 | 0.353931663 |
| chr17:7836738-7837091 | -0.554434191 | 0.122002901 | 0.353931663 |
| chr16:50066215-50066686 | 0.545746421 | 0.122003456 | 0.353931663 |
| chr8:66688971-66689542 | -0.590086337 | 0.122003774 | 0.353931663 |
| chr14:99389251-99389574 | 0.607771314 | 0.122006996 | 0.353931663 |
| chr12:116878956-116879156 | 0.733938565 | 0.122011315 | 0.353931663 |
| chr2:164737657-164738040 | -0.619979506 | 0.122055484 | 0.354046386 |
| chrX:149527776-149528068 | 0.662266007 | 0.122087731 | 0.35412652 |
| chr2:11682020-11682567 | -0.531439436 | 0.122100122 | 0.354148541 |
| chr14:38971305-38971571 | -0.681370435 | 0.122104567 | 0.354148541 |
| chr12:107402483-107402746 | -0.670371024 | 0.122138791 | 0.354220089 |
| chr3:152961074-152961830 | -0.546327447 | 0.122142531 | 0.354220089 |
| chr16:10828797-10829309 | -0.616772405 | 0.12214478 | 0.354220089 |
| chr2:60794327-60794562 | -0.854608246 | 0.122153055 | 0.354220089 |
| chr14:35403580-35404295 | 0.50245483 | 0.122155806 | 0.354220089 |
| chr14:69152475-69153569 | 0.50458261 | 0.122156974 | 0.354220089 |
| chr3:42130470-42130818 | 0.629349586 | 0.122195604 | 0.354316249 |
| chr15:89888335-89888535 | 0.790900836 | 0.122199776 | 0.354316249 |
| chr19:43465528-43465856 | 0.562885623 | 0.122204008 | 0.354316249 |
| chr1:32197286-32197532 | -0.736245061 | 0.122216754 | 0.354339795 |
| chr15:29687537-29687835 | -0.702110109 | 0.122221691 | 0.354340701 |
| chr4:148056908-148057266 | -0.584727103 | 0.122229392 | 0.354349621 |
| chr9:101388555-101388837 | -0.712040707 | 0.122239339 | 0.354361711 |
| chr13:34906341-34906541 | 0.814121024 | 0.122242812 | 0.354361711 |
| chr3:34260698-34260907 | 0.817839477 | 0.122266437 | 0.354416788 |
| chr12:119667432-119668488 | 0.498116975 | 0.122272769 | 0.354421733 |
| chr1:246753788-246754449 | -0.534955298 | 0.122285245 | 0.354439922 |
| chr12:4142842-4143224 | -0.711125278 | 0.122296084 | 0.354439922 |
| chr5:132427369-132427942 | -0.551770733 | 0.122296867 | 0.354439922 |
| chr19:4278196-4278451 | 0.805543071 | 0.122297547 | 0.354439922 |
| chr1:103389988-103390254 | 0.706539182 | 0.122310543 | 0.354464178 |
| chr10:25113128-25113701 | -0.564564076 | 0.122346105 | 0.354553828 |
| chr1:201517711-201517994 | 0.576187485 | 0.122367661 | 0.354597616 |
| chr10:109542294-109542536 | -0.783631182 | 0.12237047 | 0.354597616 |
| chr2:178072438-178073374 | 0.514435222 | 0.122387479 | 0.354633492 |
| chr14:99430534-99430790 | 0.645635199 | 0.122417931 | 0.354672318 |
| chr17:34961085-34961665 | 0.562526431 | 0.122418855 | 0.354672318 |
| chr8:628147-628354 | -0.783446754 | 0.122421373 | 0.354672318 |
| chr5:172848408-172848871 | -0.576671641 | 0.122421608 | 0.354672318 |
| chr16:67465827-67466127 | 0.596729008 | 0.122424023 | 0.354672318 |

| | | | |
|---|---|---|---|
| chr1:197485455-197486399 | -0.502733239 | 0.122446911 | 0.354725215 |
| chr6:43509242-43509488 | 0.651647041 | 0.12246924 | 0.354771098 |
| chr17:61738237-61738461 | 0.538358611 | 0.122472009 | 0.354771098 |
| chr15:45863962-45864216 | 0.70580481 | 0.122477877 | 0.354772944 |
| chrX:101596035-101596686 | 0.544401143 | 0.122481907 | 0.354772944 |
| chr11:66996894-66997421 | -0.563013243 | 0.122490692 | 0.354784978 |
| chr7:42745420-42745620 | -0.855945588 | 0.122508652 | 0.354814525 |
| chr12:89016949-89017446 | -0.552550831 | 0.122510155 | 0.354814525 |
| chr2:3369171-3370103 | -0.523120027 | 0.122530605 | 0.354854431 |
| chr12:77049796-77050126 | 0.577087512 | 0.122533196 | 0.354854431 |
| chr3:100561447-100561877 | 0.622426639 | 0.122558723 | 0.354914944 |
| chr6:92942548-92942996 | 0.619286659 | 0.12257687 | 0.35495408 |
| chr3:17431004-17431345 | -0.621341453 | 0.122625692 | 0.355082037 |
| chr1:202193225-202193447 | 0.637576028 | 0.122650134 | 0.355133953 |
| chr20:59850067-59850339 | 0.630946743 | 0.122652891 | 0.355133953 |
| chr5:6372258-6372585 | 0.723535605 | 0.122674898 | 0.355184252 |
| chr3:171972968-171973259 | -0.724181009 | 0.122699237 | 0.355241296 |
| chrX:100685711-100686053 | -0.609157751 | 0.122710098 | 0.355259319 |
| chr1:197179082-197179397 | -0.652667368 | 0.122731284 | 0.355307229 |
| chr1:171188516-171189037 | -0.571352826 | 0.122742461 | 0.355326162 |
| chr3:64763382-64763686 | 0.535730202 | 0.12275402 | 0.355344691 |
| chr2:190695912-190696217 | -0.800590726 | 0.122758137 | 0.355344691 |
| chr20:64112243-64112672 | -0.567729732 | 0.122776152 | 0.355370926 |
| chr7:33191334-33191715 | -0.560143065 | 0.122776476 | 0.355370926 |
| chr1:206833764-206834066 | -0.606596774 | 0.12279479 | 0.355410508 |
| chr18:57748776-57749255 | -0.619466169 | 0.122835787 | 0.355515738 |
| chr19:55027759-55028092 | 0.654516962 | 0.12284515 | 0.355527409 |
| chr2:48900281-48900972 | -0.564848182 | 0.122849099 | 0.355527409 |
| chr22:37658220-37658906 | -0.511526619 | 0.122885273 | 0.355612605 |
| chr16:2022731-2023455 | -0.521276559 | 0.122888299 | 0.355612605 |
| chr7:98879360-98880035 | -0.525015864 | 0.122892461 | 0.355612605 |
| chr2:42513284-42513836 | -0.561680374 | 0.122903975 | 0.355632492 |
| chr21:8434052-8434311 | 0.528160424 | 0.122915307 | 0.355641974 |
| chr6:125749120-125750056 | 0.505080281 | 0.122916535 | 0.355641974 |
| chr20:40862188-40862388 | 0.779281942 | 0.122935794 | 0.355684264 |
| chr14:34403535-34403817 | 0.717131222 | 0.122962426 | 0.355718495 |
| chr7:90603373-90603603 | -0.791210961 | 0.122964074 | 0.355718495 |
| chr2:113278716-113278916 | 0.808812051 | 0.122967114 | 0.355718495 |
| chr7:149064817-149066159 | -0.494280049 | 0.122968233 | 0.355718495 |
| chr14:68511058-68511367 | 0.669202907 | 0.122970837 | 0.355718495 |
| chr14:44771385-44771901 | 0.53845049 | 0.122987564 | 0.355753449 |
| chr13:48232596-48233864 | -0.506801243 | 0.1230176 | 0.355817911 |
| chr12:120116367-120117575 | 0.499302176 | 0.123019136 | 0.355817911 |
| chr4:88242199-88242480 | -0.679056977 | 0.123037227 | 0.355856802 |
| chr6:131682483-131682789 | -0.714023538 | 0.123051605 | 0.355878595 |
| chr5:15935081-15935288 | -0.807953923 | 0.123057512 | 0.355878595 |
| chr7:6150889-6151278 | -0.664894742 | 0.123058696 | 0.355878595 |
| chr2:54498110-54498369 | -0.685667669 | 0.12307679 | 0.355917491 |
| chr1:111592919-111593223 | -0.722276728 | 0.123086874 | 0.355933216 |
| chr3:113051225-113051554 | -0.592828204 | 0.123098705 | 0.355953995 |

| | | | |
|---|---|---|---|
| chr5:170903204-170903417 | -0.802180438 | 0.123106448 | 0.355962953 |
| chr10:3892815-3893110 | -0.718849598 | 0.123114573 | 0.355973011 |
| chr19:43432921-43433459 | -0.618212485 | 0.123121757 | 0.355980352 |
| chr20:53639630-53639854 | 0.520205288 | 0.123130328 | 0.3559917 |
| chr10:115573079-115573407 | -0.626956999 | 0.123163035 | 0.356072825 |
| chr2:40238010-40239032 | -0.547940765 | 0.123192471 | 0.35613948 |
| chr19:55634802-55635212 | 0.568593739 | 0.123195387 | 0.35613948 |
| chr12:89225424-89226242 | -0.517740443 | 0.123203699 | 0.356150073 |
| chr4:10115853-10117389 | 0.489939257 | 0.123216546 | 0.356173774 |
| chr7:33841707-33841996 | -0.647922664 | 0.123223587 | 0.356180688 |
| chr8:57587180-57587646 | 0.594438479 | 0.123282741 | 0.356338232 |
| chr12:130192976-130193284 | -0.701764357 | 0.12333672 | 0.356480807 |
| chr6:128487122-128487600 | -0.597415669 | 0.123347893 | 0.356499652 |
| chr10:60141243-60141537 | -0.616180873 | 0.123364096 | 0.3565256 |
| chr4:147515530-147515836 | -0.721298343 | 0.123366177 | 0.3565256 |
| chr5:91314265-91314732 | -0.590020861 | 0.12337198 | 0.356528922 |
| chr3:25428084-25428317 | -0.755702502 | 0.12338354 | 0.356548882 |
| chr5:170100448-170100648 | 0.68901734 | 0.123413797 | 0.356622866 |
| chr1:197868935-197869299 | -0.55528507 | 0.12345594 | 0.356731191 |
| chr1:195173807-195174109 | 0.653842085 | 0.123461005 | 0.356732374 |
| chr20:844397-844720 | 0.569384127 | 0.123481739 | 0.356778831 |
| chr7:5569299-5570792 | -0.489296694 | 0.123486729 | 0.356779794 |
| chr21:41029520-41029852 | -0.667983499 | 0.123495696 | 0.356792247 |
| chr1:167056966-167057628 | -0.525405886 | 0.123504991 | 0.356799122 |
| chr5:154250766-154251021 | -0.838733117 | 0.123508324 | 0.356799122 |
| chr14:24944548-24944748 | 0.86609329 | 0.123512045 | 0.356799122 |
| chr9:20683566-20683789 | -0.655112183 | 0.123525243 | 0.356813556 |
| chr7:122637957-122638183 | -0.896487949 | 0.123526355 | 0.356813556 |
| chr8:27592288-27592569 | 0.663439592 | 0.123539498 | 0.356838067 |
| chr17:73099494-73099694 | 0.81441139 | 0.123551721 | 0.356854571 |
| chr3:70871983-70872242 | -0.750966097 | 0.123554526 | 0.356854571 |
| chr15:42374119-42374388 | 0.788857454 | 0.12356958 | 0.356884597 |
| chr11:61488535-61489293 | -0.515640013 | 0.123588044 | 0.356915981 |
| chr22:43110382-43110689 | 0.688691263 | 0.123589763 | 0.356915981 |
| chr2:217106715-217106953 | -0.833932957 | 0.123612794 | 0.356969039 |
| chr3:138434293-138434747 | 0.55882105 | 0.123620997 | 0.356972524 |
| chr19:17932483-17933125 | 0.545080946 | 0.123623319 | 0.356972524 |
| chr22:46742637-46743310 | 0.543327316 | 0.123652792 | 0.357041567 |
| chr5:1385760-1386640 | 0.512884537 | 0.123656549 | 0.357041567 |
| chrX:135596462-135596858 | 0.592333553 | 0.123669393 | 0.357065198 |
| chr1:64499966-64500353 | -0.598826903 | 0.123686801 | 0.357077852 |
| chr12:55728757-55730293 | 0.488061071 | 0.1236877 | 0.357077852 |
| chr1:182181209-182181581 | 0.597669393 | 0.123689912 | 0.357077852 |
| chr1:54121522-54122079 | -0.548579399 | 0.123692417 | 0.357077852 |
| chr6:10099693-10100042 | -0.646221836 | 0.12370177 | 0.357085571 |
| chr4:78566911-78567189 | -0.76547312 | 0.123706324 | 0.357085571 |
| chr17:66296329-66296623 | 0.638791323 | 0.123711007 | 0.357085571 |
| chr16:11915720-11916466 | 0.498250998 | 0.123722411 | 0.357085571 |
| chr1:215773580-215773780 | 0.762371905 | 0.123722741 | 0.357085571 |
| chr1:6427508-6428206 | -0.54516195 | 0.123723053 | 0.357085571 |

| | | | |
|---|---|---|---|
| chr16:79601423-79601700 | -0.643950629 | 0.123738866 | 0.35711776 |
| chr17:74380228-74380526 | 0.62296409 | 0.123764732 | 0.357178957 |
| chr12:80477439-80478263 | -0.508082217 | 0.123773708 | 0.357191408 |
| chr14:20630376-20631319 | -0.505553702 | 0.12379543 | 0.357230279 |
| chr7:31251541-31251779 | -0.739950507 | 0.123796502 | 0.357230279 |
| chr3:175223967-175224287 | 0.689943002 | 0.123811149 | 0.35725909 |
| chr16:28846071-28846827 | 0.502049011 | 0.123820051 | 0.357271323 |
| chr1:220521823-220522111 | -0.657946751 | 0.123842194 | 0.357321758 |
| chr13:112943585-112943897 | 0.662083949 | 0.123855498 | 0.357343413 |
| chr20:57142474-57143023 | 0.491531608 | 0.123860779 | 0.357343413 |
| chr7:93591128-93592046 | -0.527513825 | 0.123863691 | 0.357343413 |
| chr12:51421359-51421559 | 0.730200682 | 0.123873803 | 0.357347223 |
| chr14:22556987-22557697 | 0.532960759 | 0.123874339 | 0.357347223 |
| chr20:3159772-3160123 | 0.606481077 | 0.123880751 | 0.357349121 |
| chrX:74181896-74182341 | 0.562044357 | 0.123884324 | 0.357349121 |
| chr2:227466236-227466595 | -0.585772581 | 0.123892184 | 0.357358338 |
| chr3:189720817-189721097 | 0.739625462 | 0.123900133 | 0.357367816 |
| chr5:170604020-170604332 | -0.612594001 | 0.123926665 | 0.357430887 |
| chr14:28681701-28682044 | 0.591440281 | 0.123942598 | 0.357463384 |
| chr4:103046729-103047102 | -0.622732367 | 0.123952676 | 0.357478994 |
| chr10:23082026-23082287 | -0.745500217 | 0.123969026 | 0.357512691 |
| chr1:207020168-207020368 | -0.67119753 | 0.124011757 | 0.357622464 |
| chr18:9862808-9863008 | -0.915322171 | 0.124030139 | 0.35766201 |
| chr7:149310012-149310751 | -0.505846437 | 0.124051303 | 0.357709579 |
| chr7:91941507-91941745 | -0.634302525 | 0.124063465 | 0.357731187 |
| chr1:194342896-194343189 | -0.700097705 | 0.124074812 | 0.357742828 |
| chr22:49918015-49918759 | 0.517139771 | 0.12407684 | 0.357742828 |
| chr3:189270322-189270582 | -0.623524233 | 0.124089848 | 0.35776687 |
| chr1:114594126-114594440 | 0.558921681 | 0.124108573 | 0.357807394 |
| chr8:138713911-138714259 | 0.661060069 | 0.124119763 | 0.357815727 |
| chr14:20499205-20499686 | 0.634773611 | 0.124120804 | 0.357815727 |
| chr4:4575310-4575748 | 0.607179404 | 0.124172621 | 0.357947615 |
| chr10:133114063-133114395 | 0.653420783 | 0.124175897 | 0.357947615 |
| chr1:244740087-244740319 | 0.777553033 | 0.124203089 | 0.35801253 |
| chr17:714246-715259 | -0.531130127 | 0.124267259 | 0.358182113 |
| chr1:155978472-155978689 | 0.685750187 | 0.12427127 | 0.358182113 |
| chr15:33915577-33916165 | 0.533477612 | 0.124287379 | 0.358203594 |
| chr2:7378122-7378516 | 0.597013097 | 0.124288073 | 0.358203594 |
| chr4:110211346-110211593 | -0.698530031 | 0.124301168 | 0.358227861 |
| chr12:122952338-122952556 | 0.661194195 | 0.124334619 | 0.358278437 |
| chr7:35173124-35173363 | 0.787089676 | 0.124335578 | 0.358278437 |
| chr8:102528460-102529203 | -0.508166073 | 0.124338896 | 0.358278437 |
| chr13:26952266-26952478 | -0.74044435 | 0.124343225 | 0.358278437 |
| chr16:29811180-29811641 | 0.518930683 | 0.124349797 | 0.358278437 |
| chr1:164518815-164519046 | 0.751706088 | 0.124350848 | 0.358278437 |
| chr12:51823999-51824201 | 0.693994929 | 0.124351449 | 0.358278437 |
| chr1:230315184-230315384 | 0.805296463 | 0.124374954 | 0.358323868 |
| chr2:48471007-48471476 | -0.605301875 | 0.124376571 | 0.358323868 |
| chr17:64961881-64962146 | -0.607426865 | 0.124385841 | 0.358336483 |
| chr3:64368694-64368967 | 0.537278412 | 0.124390303 | 0.358336483 |

| | | | |
|---|---|---|---|
| chr1:165432489-165432700 | -0.862401334 | 0.124404149 | 0.358362899 |
| chr6:23803937-23804178 | -0.801876076 | 0.124412491 | 0.358373455 |
| chr13:40456998-40457339 | -0.676977234 | 0.124423296 | 0.358391105 |
| chr15:31364492-31364856 | -0.602831263 | 0.124428989 | 0.358394031 |
| chr11:119168580-119169630 | 0.511579214 | 0.124448016 | 0.358435361 |
| chr4:153318720-153319052 | -0.638561538 | 0.124453578 | 0.358437905 |
| chr5:177981743-177982432 | 0.552446783 | 0.124458314 | 0.358438074 |
| chr11:116736830-116737083 | 0.655830837 | 0.124512193 | 0.358579765 |
| chr6:17393207-17394408 | -0.499480427 | 0.124537252 | 0.358638454 |
| chr5:129410578-129410815 | -0.787919576 | 0.124549077 | 0.358659027 |
| chr10:80436217-80437030 | -0.510408595 | 0.124554086 | 0.358659973 |
| chr4:88378586-88379625 | -0.499874261 | 0.124588806 | 0.358746468 |
| chr2:46925579-46926019 | 0.635994739 | 0.124594276 | 0.358748735 |
| chr20:53946438-53947028 | 0.490388389 | 0.124621171 | 0.358812691 |
| chr1:147021797-147022173 | 0.629785444 | 0.124635833 | 0.358818933 |
| chr6:154978215-154978415 | 0.81770532 | 0.124638063 | 0.358818933 |
| chr2:117958063-117958263 | 0.757395893 | 0.124638866 | 0.358818933 |
| chr7:38196921-38197133 | 0.745336593 | 0.12464207 | 0.358818933 |
| chr19:15456830-15457030 | -0.748533321 | 0.124659516 | 0.358849459 |
| chr8:100309227-100310756 | 0.483535822 | 0.124662041 | 0.358849459 |
| chr3:185552534-185552816 | 0.733592742 | 0.124676678 | 0.358874169 |
| chrX:8364765-8365191 | -0.612478577 | 0.124679993 | 0.358874169 |
| chr5:149990668-149991086 | -0.598529596 | 0.124693689 | 0.358886203 |
| chr20:64044743-64045032 | 0.581872903 | 0.124696185 | 0.358886203 |
| chr2:18855095-18855337 | -0.706322175 | 0.124701621 | 0.358886203 |
| chr11:7575845-7576102 | -0.697439149 | 0.124702909 | 0.358886203 |
| chr17:61792510-61792881 | 0.508335636 | 0.124717008 | 0.3589133 |
| chr2:188978312-188978542 | 0.743326814 | 0.124739611 | 0.358964864 |
| chr11:59756867-59757212 | -0.565250738 | 0.12474443 | 0.358965251 |
| chr1:224356648-224357682 | 0.498755349 | 0.124752926 | 0.358976216 |
| chr8:127923782-127924062 | 0.610670867 | 0.124759254 | 0.358980944 |
| chr5:144170119-144171214 | -0.490592603 | 0.124766005 | 0.358981267 |
| chr2:85145531-85145774 | -0.778515325 | 0.12477661 | 0.358981267 |
| chr2:148254497-148254731 | 0.754035133 | 0.124778306 | 0.358981267 |
| chr5:113682648-113683123 | -0.623596997 | 0.124788687 | 0.358981267 |
| chr11:63767446-63767759 | 0.592924492 | 0.124793372 | 0.358981267 |
| chr4:1798330-1798610 | 0.747417106 | 0.124795349 | 0.358981267 |
| chr1:92485099-92485679 | -0.580359855 | 0.124796923 | 0.358981267 |
| chr2:14535172-14535635 | -0.53311008 | 0.124799261 | 0.358981267 |
| chr18:831021-831275 | -0.710585055 | 0.124801532 | 0.358981267 |
| chr2:103388076-103388320 | -0.718618883 | 0.124819635 | 0.359019862 |
| chr15:68559637-68559905 | 0.638647038 | 0.124825281 | 0.359022624 |
| chr5:132446760-132447004 | -0.665639807 | 0.124839731 | 0.359050705 |
| chr2:154482199-154482530 | -0.691948616 | 0.124860861 | 0.359097999 |
| chr4:134789600-134789902 | -0.729115795 | 0.124935113 | 0.35929806 |
| chr3:19889039-19889787 | 0.536211135 | 0.124963031 | 0.359364861 |
| chr3:136224108-136224862 | -0.569184411 | 0.12496991 | 0.35936663 |
| chr16:31073622-31073948 | 0.544124314 | 0.124973026 | 0.35936663 |
| chr11:64235056-64235351 | 0.653119437 | 0.12501265 | 0.359463853 |
| chr2:33670548-33670941 | -0.564272911 | 0.125016219 | 0.359463853 |

| | | | |
|---|---|---|---|
| chr17:66595725-66596086 | 0.651833923 | 0.125030486 | 0.359491385 |
| chr8:95701881-95702116 | 0.778171679 | 0.125048451 | 0.359529548 |
| chr4:93158063-93158555 | -0.697730327 | 0.125085221 | 0.35961325 |
| chr14:65065764-65066067 | 0.655789216 | 0.125088905 | 0.35961325 |
| chr17:76585726-76586222 | 0.581148179 | 0.125100499 | 0.35961325 |
| chr12:51238561-51239401 | 0.500545897 | 0.125102102 | 0.35961325 |
| chr3:134505491-134505837 | -0.616735534 | 0.125102681 | 0.35961325 |
| chr19:34973209-34973636 | 0.618101922 | 0.125108754 | 0.35961325 |
| chr3:50291812-50292029 | 0.677092216 | 0.125113206 | 0.35961325 |
| chr11:18698429-18698922 | 0.520066393 | 0.125115111 | 0.35961325 |
| chr16:2215568-2215768 | 0.653003453 | 0.125121227 | 0.35961734 |
| chr14:35533116-35534387 | 0.495186088 | 0.125133391 | 0.359638809 |
| chr5:160560314-160560570 | -0.801139947 | 0.125144748 | 0.35965796 |
| chr22:31207291-31207597 | 0.618158598 | 0.1251633 | 0.359684713 |
| chr14:31678714-31678947 | 0.765322951 | 0.125163445 | 0.359684713 |
| chr16:71611531-71611766 | -0.700667078 | 0.125173361 | 0.359699717 |
| chr7:127067040-127067257 | -0.816423705 | 0.12519729 | 0.359754988 |
| chr5:64768135-64769271 | -0.495035613 | 0.125208965 | 0.359775044 |
| chr3:192818867-192819385 | -0.560136816 | 0.125216733 | 0.359783872 |
| chr7:35025355-35026448 | 0.495371492 | 0.125242233 | 0.359841496 |
| chr10:71334037-71334474 | -0.657893287 | 0.125249676 | 0.359841496 |
| chr16:19696207-19696609 | 0.685266926 | 0.125250877 | 0.359841496 |
| chr11:87352697-87353132 | 0.55887555 | 0.125257449 | 0.359846883 |
| chr12:77925457-77925953 | -0.538407636 | 0.125266535 | 0.359859496 |
| chr20:45470755-45470970 | 0.667016636 | 0.125276485 | 0.359874587 |
| chr17:38718666-38718906 | 0.700810643 | 0.125335314 | 0.360030082 |
| chr1:81305995-81306436 | -0.638179286 | 0.125342471 | 0.360037144 |
| chr1:206586789-206587261 | 0.525518167 | 0.125377417 | 0.360124024 |
| chr14:91573904-91574891 | 0.499320685 | 0.125389106 | 0.360144096 |
| chr1:23742958-23743644 | 0.556642068 | 0.125393996 | 0.360144643 |
| chr22:46069762-46071015 | 0.51050182 | 0.125401212 | 0.360145047 |
| chr5:27241685-27242410 | -0.565238586 | 0.125403538 | 0.360145047 |
| chr4:2764133-2764453 | 0.627235931 | 0.125409511 | 0.360148702 |
| chr8:96766486-96766995 | -0.603249105 | 0.125425202 | 0.360177892 |
| chr4:143389580-143389818 | 0.680772784 | 0.12543212 | 0.360177892 |
| chr3:33096462-33097791 | 0.494996913 | 0.12543506 | 0.360177892 |
| chr21:39061837-39062096 | -0.679879221 | 0.12544818 | 0.360177892 |
| chr5:174419154-174419912 | -0.530888046 | 0.125455781 | 0.360177892 |
| chr9:84317558-84317765 | 0.764442481 | 0.125457609 | 0.360177892 |
| chrX:91752739-91753445 | -0.534852521 | 0.125458697 | 0.360177892 |
| chr20:15654135-15654505 | -0.587991646 | 0.125459138 | 0.360177892 |
| chr3:159829596-159829937 | -0.650329156 | 0.125467214 | 0.360177892 |
| chr18:52423060-52423305 | -0.793604487 | 0.125473369 | 0.360177892 |
| chr7:50615266-50615466 | -0.800609294 | 0.1254754 | 0.360177892 |
| chr5:55232408-55234084 | -0.491583942 | 0.125476083 | 0.360177892 |
| chr4:171641301-171641632 | -0.841743344 | 0.125508563 | 0.360257628 |
| chr10:62513033-62513265 | -0.811713593 | 0.125530157 | 0.360306114 |
| chrX:131802861-131803390 | 0.568762555 | 0.125557013 | 0.360369699 |
| chr5:90136580-90136804 | 0.718017128 | 0.125576653 | 0.360397269 |
| chr4:143000053-143000280 | 0.791531794 | 0.125583725 | 0.360397269 |

| | | | |
|---|---|---|---|
| chr1:23691581-23692146 | -0.566386507 | 0.125585905 | 0.360397269 |
| chr15:62832249-62832767 | -0.546011956 | 0.12559112 | 0.360397269 |
| chr1:200258467-200258775 | -0.570533416 | 0.12559147 | 0.360397269 |
| chr10:80896195-80896395 | 0.697055397 | 0.125597739 | 0.360397269 |
| chr18:48827911-48828144 | -0.706429037 | 0.125603488 | 0.360397269 |
| chr5:173917444-173917644 | -0.833309923 | 0.125606834 | 0.360397269 |
| chr3:133644807-133645173 | -0.614203069 | 0.125608951 | 0.360397269 |
| chr17:16935079-16935356 | -0.684846921 | 0.125630714 | 0.360446214 |
| chr20:12594676-12595057 | -0.665399741 | 0.125654341 | 0.360500504 |
| chr15:56245862-56246397 | 0.512029839 | 0.125662767 | 0.360511178 |
| chr2:142811474-142812423 | -0.554944688 | 0.125683924 | 0.360558376 |
| chr2:5195423-5195958 | -0.58593047 | 0.125695499 | 0.360578081 |
| chr19:4769531-4769878 | 0.631023954 | 0.125702387 | 0.360584341 |
| chr11:63016765-63016965 | 0.743926157 | 0.125735449 | 0.360630862 |
| chr8:94592370-94592872 | 0.547133554 | 0.12574499 | 0.360630862 |
| chr7:20458497-20458728 | 0.73313848 | 0.125745468 | 0.360630862 |
| chr7:36036705-36036972 | -0.618454513 | 0.125746668 | 0.360630862 |
| chr7:139284166-139284622 | -0.624360007 | 0.125746818 | 0.360630862 |
| chr21:40512470-40512930 | -0.607271815 | 0.125746845 | 0.360630862 |
| chr15:98890331-98890531 | 0.798373686 | 0.125766398 | 0.360662289 |
| chr2:165204046-165204269 | -0.81882926 | 0.125767217 | 0.360662289 |
| chr16:11297708-11297908 | 0.694697374 | 0.125793302 | 0.360723592 |
| chr16:71358469-71358804 | 0.572968362 | 0.125816698 | 0.36077718 |
| chr19:5680375-5681163 | 0.510269019 | 0.12582498 | 0.360787425 |
| chr11:67640092-67640298 | 0.713345382 | 0.12583064 | 0.360790154 |
| chr8:125334114-125334413 | 0.647088477 | 0.125844417 | 0.360816155 |
| chr3:39141141-39141724 | 0.530270631 | 0.125892732 | 0.360931256 |
| chr1:223155251-223155485 | -0.805400561 | 0.125893983 | 0.360931256 |
| chr17:10586796-10587226 | -0.746892604 | 0.125900239 | 0.360935686 |
| chr16:54193475-54194663 | -0.509626371 | 0.125907619 | 0.360936691 |
| chr16:57810353-57810576 | 0.766854746 | 0.12591001 | 0.360936691 |
| chr8:100526974-100527251 | 0.640870281 | 0.125925774 | 0.360968376 |
| chr11:121025045-121025245 | -0.882912909 | 0.125944132 | 0.361007493 |
| chr14:104985488-104986460 | 0.497722063 | 0.125961221 | 0.3610217 |
| chr3:64700537-64700993 | 0.509503056 | 0.125969866 | 0.3610217 |
| chr9:5230412-5230902 | 0.596211779 | 0.125971842 | 0.3610217 |
| chr1:162069488-162070695 | 0.501523725 | 0.125972463 | 0.3610217 |
| chr13:44872491-44872753 | 0.693451969 | 0.125975298 | 0.3610217 |
| chr15:65054374-65054630 | 0.586381875 | 0.125977359 | 0.3610217 |
| chr20:40673428-40673868 | 0.592935821 | 0.125982706 | 0.361023521 |
| chr1:168735002-168735202 | 0.760356509 | 0.126006735 | 0.361078876 |
| chr19:45670809-45671305 | 0.524345947 | 0.126013716 | 0.361085376 |
| chr20:10093306-10093564 | 0.641472801 | 0.126029352 | 0.361116674 |
| chr7:47968891-47969161 | -0.66669464 | 0.1260467 | 0.361152878 |
| chr11:575801-576629 | 0.51221341 | 0.126057683 | 0.361170842 |
| chr14:81252390-81252590 | -0.766131462 | 0.126085663 | 0.361237498 |
| chr6:18051755-18052033 | -0.711121062 | 0.126131455 | 0.361355181 |
| chr11:63869063-63869511 | 0.587462055 | 0.126147105 | 0.361386507 |
| chr11:22655080-22655341 | -0.754242275 | 0.126171542 | 0.361442998 |
| chr1:85599235-85599515 | -0.668477756 | 0.12618606 | 0.361471074 |

| | | | |
|---|---|---|---|
| chr9:41643536-41644115 | -0.559044133 | 0.12620041 | 0.361498667 |
| chr19:1275325-1275581 | 0.617230323 | 0.126209576 | 0.361504133 |
| chr6:104607404-104607852 | -0.638794279 | 0.126211754 | 0.361504133 |
| chr16:31148132-31148624 | 0.558443728 | 0.126224798 | 0.361527979 |
| chrX:154515958-154516929 | 0.50246019 | 0.126247597 | 0.361566364 |
| chr9:77455326-77455669 | -0.587563778 | 0.126250811 | 0.361566364 |
| chr17:61363891-61364198 | 0.506557549 | 0.126252357 | 0.361566364 |
| chr3:29937247-29937643 | 0.590661736 | 0.126259531 | 0.361573395 |
| chr15:61195251-61195495 | 0.69872017 | 0.126265671 | 0.361577464 |
| chr22:31249084-31249293 | 0.716685103 | 0.126285315 | 0.361620203 |
| chr20:50717860-50718326 | 0.490224397 | 0.126291449 | 0.361624253 |
| chr4:7972780-7973101 | 0.615511629 | 0.126329514 | 0.361719732 |
| chr7:26328948-26329229 | 0.658414918 | 0.126339057 | 0.361733538 |
| chr1:61711204-61711577 | 0.730308866 | 0.126349844 | 0.361750906 |
| chr2:5365895-5366454 | -0.567770466 | 0.126379046 | 0.361820993 |
| chr5:62094030-62094317 | -0.678178037 | 0.126388176 | 0.361833614 |
| chr20:38658927-38659237 | 0.596208071 | 0.126395242 | 0.361840321 |
| chr2:58341072-58341387 | -0.701499577 | 0.126411614 | 0.361864138 |
| chr9:15276627-15276982 | -0.638230314 | 0.126413006 | 0.361864138 |
| chr15:85381167-85381373 | 0.667086139 | 0.126417907 | 0.361864646 |
| chr13:44450435-44450874 | -0.578836545 | 0.12643015 | 0.361886173 |
| chr12:69518497-69518697 | -0.927890727 | 0.126478321 | 0.361992645 |
| chr7:3065820-3066020 | 0.723402666 | 0.126481178 | 0.361992645 |
| chr5:134758231-134759427 | -0.497889359 | 0.126481521 | 0.361992645 |
| chr2:238060429-238061920 | -0.498854582 | 0.126513414 | 0.3620704 |
| chr3:98894135-98894343 | -0.837569278 | 0.126525733 | 0.362079506 |
| chr10:118850497-118850843 | -0.643883863 | 0.126526047 | 0.362079506 |
| chr7:85695662-85695945 | -0.707482083 | 0.126553671 | 0.362145033 |
| chr12:114683783-114685058 | 0.487166601 | 0.126565234 | 0.362164594 |
| chr3:32052346-32052630 | 0.666099457 | 0.126583336 | 0.362202867 |
| chr8:56447973-56448226 | -0.736724941 | 0.126603382 | 0.362246699 |
| chrX:135343910-135344352 | 0.513839848 | 0.126625582 | 0.36228482 |
| chr11:113474860-113475073 | -0.803867621 | 0.126626162 | 0.36228482 |
| chr1:170563877-170564110 | 0.809308553 | 0.126644047 | 0.362322463 |
| chr7:103347475-103347846 | -0.534576773 | 0.126652277 | 0.362332479 |
| chr2:181954099-181954577 | -0.572645438 | 0.126675378 | 0.362383425 |
| chr17:72560251-72560711 | -0.543211415 | 0.126681739 | 0.362383425 |
| chr16:54425646-54425846 | -0.835061351 | 0.126687943 | 0.362383425 |
| chr21:42064077-42064447 | -0.630818914 | 0.126689003 | 0.362383425 |
| chr2:127909028-127909271 | -0.736850592 | 0.126698754 | 0.362397786 |
| chr19:44164499-44165213 | 0.516914804 | 0.126727733 | 0.362467145 |
| chr5:149646576-149646936 | -0.615590504 | 0.126733761 | 0.362470855 |
| chr3:15797762-15798303 | -0.595464403 | 0.126764157 | 0.362544258 |
| chr6:27375776-27376066 | -0.680292394 | 0.126786122 | 0.362583035 |
| chr14:22598126-22598564 | 0.555795698 | 0.126791915 | 0.362583035 |
| chr11:62981747-62982019 | 0.724266992 | 0.126792972 | 0.362583035 |
| chr12:67085650-67086147 | -0.595665187 | 0.126804842 | 0.362583035 |
| chr16:8880228-8880668 | 0.570858072 | 0.126805428 | 0.362583035 |
| chr10:75431589-75431991 | 0.631252538 | 0.126810302 | 0.362583035 |
| chr13:49691025-49691835 | -0.508950142 | 0.12681084 | 0.362583035 |

| | | | |
|---|---|---|---|
| chr1:45804070-45804270 | 0.750016041 | 0.126827553 | 0.362617291 |
| chr7:50782153-50783176 | -0.503122806 | 0.126851775 | 0.362673012 |
| chr6:137873358-137873698 | -0.708382515 | 0.126869219 | 0.36269643 |
| chr13:20721958-20722285 | 0.664910721 | 0.126869433 | 0.36269643 |
| chr1:212515259-212515644 | -0.702175327 | 0.126878137 | 0.362707779 |
| chr5:85225916-85226183 | -0.775563285 | 0.126906548 | 0.362775462 |
| chr2:47767906-47768246 | -0.591671615 | 0.126935889 | 0.362845801 |
| chr6:37818593-37819835 | 0.484744991 | 0.126959464 | 0.362866469 |
| chr6:10430112-10430351 | -0.726315105 | 0.126959686 | 0.362866469 |
| chr12:101839625-101840149 | -0.58280927 | 0.126961167 | 0.362866469 |
| chr1:216965550-216965754 | -0.777317919 | 0.126962376 | 0.362866469 |
| chr3:193225692-193226185 | -0.63929023 | 0.126966798 | 0.362866469 |
| chr2:53786615-53788123 | -0.489239267 | 0.126978538 | 0.362886485 |
| chrX:39895171-39895490 | 0.675248643 | 0.126997492 | 0.362927116 |
| chr15:74373634-74373976 | 0.550324381 | 0.127006893 | 0.362940446 |
| chr1:51235132-51235349 | 0.662285135 | 0.127021509 | 0.362957358 |
| chr16:89995826-89996050 | -0.691016517 | 0.127022285 | 0.362957358 |
| chr20:57254987-57255228 | 0.510180044 | 0.127029159 | 0.362959448 |
| chr11:46845485-46846504 | 0.498677748 | 0.127032491 | 0.362959448 |
| chr3:18330641-18331329 | -0.548496202 | 0.127082172 | 0.363084552 |
| chr11:63871902-63872404 | 0.541572246 | 0.127085754 | 0.363084552 |
| chr6:144195735-144196084 | 0.577109663 | 0.127091279 | 0.363084557 |
| chr12:93146416-93146702 | -0.742206167 | 0.127095233 | 0.363084557 |
| chr3:64112946-64113195 | 0.527431817 | 0.127116856 | 0.363132792 |
| chr18:50162553-50162981 | 0.621002687 | 0.127138849 | 0.363180331 |
| chr1:155063322-155064134 | 0.499617143 | 0.1271473 | 0.363180331 |
| chr8:138851455-138851777 | -0.598246677 | 0.127147717 | 0.363180331 |
| chr1:32107847-32108746 | 0.512250746 | 0.127159297 | 0.363199868 |
| chr3:99637658-99637911 | -0.754333779 | 0.127171537 | 0.363221288 |
| chr5:113859862-113860078 | -0.79933044 | 0.127177133 | 0.363223733 |
| chr19:37594216-37595339 | -0.506297157 | 0.12719474 | 0.363239861 |
| chr8:133768713-133769439 | -0.498402532 | 0.12719993 | 0.363239861 |
| chr2:117803526-117803998 | -0.637874484 | 0.127207923 | 0.363239861 |
| chr17:42180162-42180362 | 0.761038968 | 0.127208604 | 0.363239861 |
| chr1:53196596-53197672 | 0.510632134 | 0.127211054 | 0.363239861 |
| chr10:125494237-125494488 | -0.754700765 | 0.127219964 | 0.363239861 |
| chr1:106306502-106307132 | 0.508530449 | 0.127222947 | 0.363239861 |
| chr19:1604987-1605190 | -0.772233529 | 0.127224556 | 0.363239861 |
| chr2:149451708-149452292 | -0.56823803 | 0.127225446 | 0.363239861 |
| chr7:37181974-37182379 | -0.565610282 | 0.127244298 | 0.363280146 |
| chr11:83828003-83828297 | -0.646958735 | 0.127249917 | 0.363282652 |
| chr9:76634367-76634658 | -0.694292047 | 0.127260405 | 0.363297137 |
| chr17:61680778-61681138 | 0.514430926 | 0.127266062 | 0.363297137 |
| chr17:49604350-49604649 | 0.614204865 | 0.127269215 | 0.363297137 |
| chr10:93991887-93992258 | 0.645900137 | 0.127287898 | 0.363336933 |
| chr9:101601990-101602505 | -0.610438276 | 0.127305574 | 0.363365376 |
| chr10:8725692-8725892 | -0.787481573 | 0.127307347 | 0.363365376 |
| chr4:41130366-41130890 | 0.590683065 | 0.127337874 | 0.363426639 |
| chr11:19311397-19311623 | -0.789086307 | 0.127350706 | 0.363426639 |
| chr22:29388128-29388843 | 0.519219889 | 0.127350895 | 0.363426639 |

| | | | |
|---|---|---|---|
| chr3:79336555-79336755 | 0.850669949 | 0.127359175 | 0.363426639 |
| chr2:215216691-215217130 | -0.551118208 | 0.127360341 | 0.363426639 |
| chr1:89615923-89616329 | -0.611795771 | 0.127363607 | 0.363426639 |
| chr7:28926915-28927313 | 0.628520028 | 0.127364679 | 0.363426639 |
| chr14:104864291-104864770 | 0.523256687 | 0.127366756 | 0.363426639 |
| chr7:45324897-45325235 | -0.729481622 | 0.127385101 | 0.363465451 |
| chr5:19206976-19207462 | 0.569162054 | 0.127408176 | 0.363503251 |
| chr1:217964842-217965149 | -0.61769859 | 0.127410156 | 0.363503251 |
| chr5:163503107-163503456 | -0.556215782 | 0.127412581 | 0.363503251 |
| chr7:111953665-111953946 | -0.812418922 | 0.127422728 | 0.363517846 |
| chr1:109910530-109911017 | 0.558631696 | 0.127428661 | 0.363517846 |
| chr16:77374289-77374800 | 0.550364349 | 0.127432196 | 0.363517846 |
| chr11:46113042-46113264 | -0.734062142 | 0.127436675 | 0.363517846 |
| chr7:158802480-158802718 | -0.789556661 | 0.127454459 | 0.363545113 |
| chr5:127934794-127935286 | -0.519367444 | 0.127455722 | 0.363545113 |
| chr8:109116686-109117078 | 0.561265277 | 0.127513885 | 0.363697472 |
| chr11:122655506-122656693 | 0.516309673 | 0.12752221 | 0.363707678 |
| chr15:67049367-67049988 | -0.541019221 | 0.127554653 | 0.363786668 |
| chr3:192789392-192789767 | 0.666177913 | 0.127562781 | 0.363796309 |
| chr22:17941885-17942085 | -0.856046019 | 0.127568033 | 0.363797745 |
| chr3:105997828-105998363 | -0.623312074 | 0.127610461 | 0.363905197 |
| chr3:63938039-63938260 | 0.552332332 | 0.127648018 | 0.36399875 |
| chr21:41783379-41784250 | -0.507007703 | 0.127660471 | 0.3640037 |
| chr16:8963840-8964424 | 0.545581605 | 0.127662287 | 0.3640037 |
| chr11:45771258-45771818 | 0.583753548 | 0.127664006 | 0.3640037 |
| chr1:34781374-34781589 | 0.692863576 | 0.12768472 | 0.364049215 |
| chr8:8942632-8942866 | -0.767810791 | 0.127692208 | 0.364057018 |
| chr17:62416557-62417255 | -0.528023595 | 0.127711236 | 0.364097719 |
| chr3:115744623-115744928 | -0.654669734 | 0.127768076 | 0.364246214 |
| chr3:187064272-187064559 | -0.635438995 | 0.127791551 | 0.364299585 |
| chr11:77253092-77253321 | -0.780259162 | 0.127802359 | 0.36431684 |
| chr7:91940121-91941357 | -0.491055158 | 0.127809929 | 0.364324864 |
| chr5:72314895-72315130 | -0.734148937 | 0.127850873 | 0.36441686 |
| chr14:54348045-54348300 | -0.665373655 | 0.127852983 | 0.36441686 |
| chr6:5866871-5867101 | 0.722891935 | 0.127859185 | 0.36441686 |
| chr3:196660519-196660920 | 0.608876831 | 0.127862205 | 0.36441686 |
| chr13:39600849-39601051 | -0.736771116 | 0.127865982 | 0.36441686 |
| chr16:1572736-1572962 | -0.726575639 | 0.127874922 | 0.364428782 |
| chr11:64269147-64269360 | 0.613596924 | 0.127898689 | 0.364482959 |
| chr4:4673005-4673447 | 0.519013665 | 0.127928196 | 0.364553491 |
| chr12:7155516-7156047 | -0.553159857 | 0.127933996 | 0.364556461 |
| chr4:81260389-81260732 | -0.74208302 | 0.12794385 | 0.364569577 |
| chr12:19917015-19917286 | -0.733080252 | 0.127951614 | 0.364569577 |
| chr3:23804441-23804783 | -0.618388016 | 0.127952873 | 0.364569577 |
| chr3:105368231-105368456 | -0.570494725 | 0.127962562 | 0.364576747 |
| chr2:96064383-96064624 | 0.711452044 | 0.127964906 | 0.364576747 |
| chr11:117231913-117232806 | 0.512699681 | 0.128007069 | 0.36468331 |
| chr4:7939646-7940129 | 0.540385414 | 0.128016251 | 0.364695909 |
| chr11:125423276-125423524 | -0.761048923 | 0.128044382 | 0.364762264 |
| chr7:98418802-98419241 | -0.566332628 | 0.128049064 | 0.364762264 |

| | | | |
|---|---|---|---|
| chr19:2427518-2428016 | 0.570148781 | 0.128056578 | 0.364770109 |
| chr7:99374499-99375704 | -0.496050703 | 0.12813653 | 0.364967115 |
| chr3:106281118-106281551 | -0.588743571 | 0.128138708 | 0.364967115 |
| chr17:72602893-72603198 | -0.644931117 | 0.128143054 | 0.364967115 |
| chr16:2296654-2296878 | -0.733862442 | 0.128144792 | 0.364967115 |
| chr9:134325323-134325592 | 0.625336594 | 0.128152538 | 0.364975609 |
| chr3:187862889-187863140 | -0.779736857 | 0.128161796 | 0.364988269 |
| chr9:136800940-136801430 | 0.5197963 | 0.128175378 | 0.364988269 |
| chr1:203906991-203907273 | -0.612877203 | 0.128178917 | 0.364988269 |
| chr19:7507949-7508174 | 0.644255347 | 0.128179149 | 0.364988269 |
| chr7:106166801-106167026 | -0.804002138 | 0.128180801 | 0.364988269 |
| chr5:180085186-180085501 | 0.58885354 | 0.128186353 | 0.364990515 |
| chr10:116168445-116168645 | -0.803826865 | 0.12820476 | 0.365015796 |
| chr12:30588383-30588583 | -0.803826865 | 0.12820476 | 0.365015796 |
| chr13:45370992-45371321 | -0.563210489 | 0.128229936 | 0.365073911 |
| chr8:125859836-125860036 | 0.782081049 | 0.128241406 | 0.365082371 |
| chrX:106863032-106863591 | -0.547279206 | 0.128246445 | 0.365082371 |
| chr17:79810132-79810379 | 0.573921645 | 0.128247202 | 0.365082371 |
| chr7:16753317-16754556 | 0.479158576 | 0.128259768 | 0.365091709 |
| chr7:45984708-45984999 | 0.644315376 | 0.12826347 | 0.365091709 |
| chr7:135801142-135801358 | -0.702377191 | 0.128264777 | 0.365091709 |
| chr2:218216554-218216790 | -0.717413004 | 0.128271378 | 0.365096935 |
| chr1:52686248-52686576 | 0.620616626 | 0.128297445 | 0.365157565 |
| chr4:65998519-65999695 | -0.500792855 | 0.128315304 | 0.365194831 |
| chr16:87885360-87885731 | -0.667733134 | 0.12836308 | 0.365312636 |
| chr14:45252905-45253966 | 0.48144648 | 0.128366232 | 0.365312636 |
| chr18:3397039-3397713 | -0.5353741 | 0.128372696 | 0.365317463 |
| chr6:28674257-28674486 | -0.754310983 | 0.128382383 | 0.365331463 |
| chr6:155313971-155315235 | -0.483751159 | 0.128425986 | 0.36542029 |
| chr9:136727983-136728367 | 0.559665903 | 0.128427615 | 0.36542029 |
| chr20:41086564-41087502 | 0.505534898 | 0.128427906 | 0.36542029 |
| chr2:54456887-54457098 | 0.650608308 | 0.128447026 | 0.365461123 |
| chr4:179626761-179626982 | -0.798546841 | 0.128458359 | 0.3654728 |
| chr3:64777411-64777733 | 0.523486814 | 0.12846067 | 0.3654728 |
| chr6:33840110-33840440 | -0.608645655 | 0.128466103 | 0.365474686 |
| chr7:46544275-46544570 | -0.766761231 | 0.128494288 | 0.365541298 |
| chr19:50510898-50511673 | 0.50139922 | 0.128517007 | 0.365592357 |
| chr7:48110020-48110564 | -0.528469139 | 0.128533171 | 0.365622373 |
| chr3:172515559-172515879 | -0.689022794 | 0.128543825 | 0.365622373 |
| chr3:87733687-87734439 | -0.511746636 | 0.128545749 | 0.365622373 |
| chr18:54950293-54950549 | -0.685585016 | 0.128546646 | 0.365622373 |
| chr5:84113188-84113589 | -0.726467647 | 0.128582362 | 0.36570195 |
| chr1:52141302-52141539 | 0.710995845 | 0.12858417 | 0.36570195 |
| chr11:79174464-79174937 | -0.574688943 | 0.128602103 | 0.365730433 |
| chr15:90873952-90874376 | 0.552960408 | 0.128603731 | 0.365730433 |
| chr15:92082392-92082605 | 0.653711367 | 0.128613773 | 0.365739714 |
| chr1:195763074-195764149 | -0.494956256 | 0.1286177 | 0.365739714 |
| chr5:158382528-158382759 | 0.764548694 | 0.128621936 | 0.365739714 |
| chr1:157137461-157137661 | 0.647390574 | 0.128626088 | 0.365739714 |
| chr9:134572255-134572842 | -0.596295021 | 0.128645629 | 0.365781705 |

| | | | |
|---|---|---|---|
| chr5:56201173-56201724 | -0.59151224 | 0.128679958 | 0.365858105 |
| chr3:62241174-62241639 | -0.538714415 | 0.128682049 | 0.365858105 |
| chr10:56231438-56231788 | -0.626151115 | 0.128706989 | 0.365915437 |
| chr6:113970176-113971615 | -0.484411809 | 0.128724365 | 0.365939438 |
| chr4:68361121-68361601 | -0.53371748 | 0.128724983 | 0.365939438 |
| chr5:138316498-138316698 | -0.831007736 | 0.128741194 | 0.365971943 |
| chrX:10101246-10101481 | 0.749078613 | 0.128759222 | 0.366009613 |
| chr5:129043597-129043832 | -0.799608398 | 0.128776106 | 0.366044026 |
| chr11:49207952-49208855 | -0.525525287 | 0.128822518 | 0.366155044 |
| chr3:188975962-188976220 | -0.684897746 | 0.12882472 | 0.366155044 |
| chr8:82443541-82443821 | -0.621240421 | 0.1288329 | 0.366164711 |
| chr2:437751-438356 | -0.536155863 | 0.128857743 | 0.366221735 |
| chr3:141056485-141056931 | -0.631320912 | 0.128870852 | 0.366245406 |
| chr16:82033070-82033360 | 0.751794639 | 0.128887063 | 0.366277893 |
| chr3:126357744-126358081 | 0.580374857 | 0.128906646 | 0.366319957 |
| chr21:45820591-45820991 | 0.562507858 | 0.128926772 | 0.366345046 |
| chr14:94080838-94081789 | 0.49180004 | 0.128930535 | 0.366345046 |
| chr8:133851707-133852019 | 0.627236985 | 0.128934553 | 0.366345046 |
| chr7:98841794-98842072 | -0.693234661 | 0.128934599 | 0.366345046 |
| chr11:24389655-24389882 | -0.811889089 | 0.128943883 | 0.366357838 |
| chr9:119986340-119986548 | -0.775901154 | 0.128950583 | 0.36636329 |
| chr19:1438114-1438678 | 0.561792284 | 0.128981086 | 0.366429994 |
| chr11:133956715-133957581 | 0.508169678 | 0.128983626 | 0.366429994 |
| chr11:15707985-15708339 | -0.578104231 | 0.129033142 | 0.366550315 |
| chr12:29317728-29318123 | -0.61545375 | 0.129035547 | 0.366550315 |
| chr13:46896768-46897166 | -0.540497866 | 0.129061788 | 0.366604728 |
| chr4:76733520-76733783 | -0.704248614 | 0.129064271 | 0.366604728 |
| chr4:129136194-129136426 | 0.769902485 | 0.129090621 | 0.366660119 |
| chr15:69603795-69604318 | 0.54308122 | 0.129093342 | 0.366660119 |
| chr20:41339960-41341548 | 0.476590497 | 0.129107771 | 0.366686035 |
| chr6:154624110-154624434 | 0.655112448 | 0.129115601 | 0.366686035 |
| chr8:88438356-88438556 | 0.756136135 | 0.129117437 | 0.366686035 |
| chr11:49995618-49995828 | 0.760472269 | 0.129121609 | 0.366686035 |
| chr10:29106216-29106558 | 0.593475246 | 0.129142682 | 0.366732288 |
| chr8:142632163-142632374 | 0.642270334 | 0.129150703 | 0.366741473 |
| chr14:65816987-65817498 | 0.540144567 | 0.129165387 | 0.366769576 |
| chr20:44310334-44311574 | 0.490879118 | 0.12917848 | 0.366793164 |
| chr3:6125585-6125943 | -0.726141557 | 0.129218114 | 0.366848436 |
| chr11:44911374-44911591 | 0.72256346 | 0.129222176 | 0.366848436 |
| chr1:59718151-59718351 | -0.874512196 | 0.129223953 | 0.366848436 |
| chr20:46575575-46575775 | 0.73214787 | 0.129225218 | 0.366848436 |
| chr10:25903635-25903847 | -0.829565838 | 0.12922651 | 0.366848436 |
| chr7:36441728-36441928 | 0.766423581 | 0.129226673 | 0.366848436 |
| chr19:16542101-16542709 | 0.545592498 | 0.129238983 | 0.366869788 |
| chr2:239731575-239731853 | -0.75530561 | 0.129246789 | 0.366877677 |
| chr4:7167767-7168086 | 0.649149243 | 0.129251338 | 0.366877677 |
| chr9:129110724-129111702 | 0.494970424 | 0.129258839 | 0.366883554 |
| chr1:173405817-173406177 | -0.629424694 | 0.129262985 | 0.366883554 |
| chr4:104451659-104451967 | -0.710504428 | 0.129283858 | 0.366929204 |
| chrX:54447601-54447835 | -0.8405004 | 0.129291133 | 0.366930566 |

| | | | |
|---|---|---|---|
| chr7:18038466-18038887 | 0.584537504 | 0.129294796 | 0.366930566 |
| chr16:54423931-54424259 | -0.71694731 | 0.129298704 | 0.366930566 |
| chr3:150737171-150737784 | -0.529282338 | 0.129309056 | 0.366946354 |
| chr5:74241800-74242016 | 0.719845337 | 0.129314273 | 0.366947569 |
| chr13:81640079-81640529 | -0.67171574 | 0.129329594 | 0.366977453 |
| chr11:60193663-60193980 | -0.627954084 | 0.129344929 | 0.367007374 |
| chr11:36000874-36001372 | -0.568132388 | 0.129386536 | 0.367111836 |
| chr10:26046113-26046639 | -0.57710194 | 0.129432938 | 0.367221637 |
| chr19:5823648-5823891 | 0.656037753 | 0.129435048 | 0.367221637 |
| chr1:43366882-43367211 | 0.573724764 | 0.129440186 | 0.367221637 |
| chr1:6602529-6603157 | 0.531453052 | 0.129444405 | 0.367221637 |
| chr7:110413782-110414059 | -0.788835528 | 0.129476799 | 0.367282351 |
| chr19:55536465-55536911 | -0.622195657 | 0.12948006 | 0.367282351 |
| chr5:136082527-136082785 | -0.698915039 | 0.129480187 | 0.367282351 |
| chr1:248862991-248863473 | -0.516602633 | 0.129491114 | 0.367297257 |
| chr12:130872391-130872599 | 0.641381558 | 0.129495029 | 0.367297257 |
| chr7:87219297-87219982 | -0.509238951 | 0.129504122 | 0.367309453 |
| chr6:138007023-138007223 | -0.886635845 | 0.129511538 | 0.367316889 |
| chr11:2930054-2930577 | 0.53285057 | 0.129517043 | 0.367318904 |
| chr14:81589484-81589707 | -0.735272475 | 0.129524119 | 0.367325377 |
| chr4:50555998-50556288 | 0.551052004 | 0.129550654 | 0.367387033 |
| chr18:10520154-10520442 | -0.682343152 | 0.129559766 | 0.367399274 |
| chr2:113156395-113156608 | 0.680877941 | 0.129568807 | 0.367411315 |
| chr10:102634781-102635019 | 0.803184085 | 0.12957557 | 0.367414709 |
| chr2:189836793-189836993 | -0.851846766 | 0.129579594 | 0.367414709 |
| chr20:23637285-23638226 | 0.500170027 | 0.129599301 | 0.367456987 |
| chr5:81591794-81592050 | -0.777738541 | 0.129646529 | 0.367577293 |
| chr21:45724687-45724887 | 0.775587262 | 0.129672422 | 0.367637102 |
| chr4:164012629-164013047 | -0.584875805 | 0.129679592 | 0.367643827 |
| chr17:40201874-40202078 | 0.75922193 | 0.12970404 | 0.367699532 |
| chr8:11746352-11746928 | -0.58584352 | 0.129729613 | 0.367755419 |
| chr20:37345956-37346188 | 0.659176564 | 0.129733353 | 0.367755419 |
| chr15:88246807-88247156 | 0.556845868 | 0.129762567 | 0.367824624 |
| chr2:149329638-149331336 | -0.479558414 | 0.129773917 | 0.367843189 |
| chr11:68903572-68904011 | 0.55194664 | 0.129800679 | 0.367905436 |
| chr2:11365648-11365874 | 0.759482214 | 0.129811995 | 0.367923899 |
| chr2:54579507-54580213 | -0.510744722 | 0.129840359 | 0.367990677 |
| chr3:174002640-174003057 | -0.740250571 | 0.129850688 | 0.368006341 |
| chr19:52139765-52140240 | 0.526555339 | 0.129872091 | 0.368048682 |
| chr3:147838732-147838992 | -0.665421131 | 0.129875235 | 0.368048682 |
| chr10:102461740-102462019 | 0.631205733 | 0.129945781 | 0.368234981 |
| chr1:179866532-179867027 | -0.536436058 | 0.129952263 | 0.368239729 |
| chr20:49323687-49324086 | 0.570366286 | 0.129982212 | 0.368310975 |
| chr12:28190238-28191396 | -0.494727824 | 0.1299982 | 0.368334958 |
| chr8:144003432-144003890 | 0.545965426 | 0.130000291 | 0.368334958 |
| chr7:128892995-128893289 | -0.663314148 | 0.130010987 | 0.368341615 |
| chr17:64765330-64765733 | 0.592481953 | 0.13001454 | 0.368341615 |
| chr1:106233847-106234409 | 0.512631671 | 0.130017062 | 0.368341615 |
| chr17:56231808-56232008 | -0.756181335 | 0.130031195 | 0.368367552 |
| chr1:153608008-153609548 | 0.477207668 | 0.130035832 | 0.368367552 |

| | | | |
|---|---|---|---|
| chr9:82939562-82939813 | -0.722622793 | 0.13006465 | 0.368426838 |
| chr8:86108614-86108975 | -0.575063229 | 0.130066377 | 0.368426838 |
| chr19:43619042-43620227 | 0.493537636 | 0.13008032 | 0.368445864 |
| chr15:78565192-78566412 | 0.497520002 | 0.130082711 | 0.368445864 |
| chr1:199251092-199251352 | 0.751615655 | 0.130102734 | 0.36848837 |
| chr4:118942561-118942770 | -0.793184899 | 0.130107337 | 0.36848837 |
| chr12:131224138-131224838 | 0.503819029 | 0.130144003 | 0.368578592 |
| chr7:129984518-129984788 | 0.73832194 | 0.130168648 | 0.368634763 |
| chr3:5568535-5568746 | -0.793520061 | 0.130174098 | 0.368636572 |
| chr20:54605312-54605672 | 0.50615431 | 0.130184204 | 0.36865072 |
| chr19:1568337-1568573 | 0.761512256 | 0.13019193 | 0.36865072 |
| chr14:36773184-36773384 | 0.801937806 | 0.130193528 | 0.36865072 |
| chr7:157355593-157355794 | -0.799743997 | 0.130242111 | 0.368774659 |
| chr14:90646110-90646335 | 0.670274927 | 0.13025895 | 0.368808709 |
| chr1:73675082-73675288 | -0.815791872 | 0.130295916 | 0.368899741 |
| chr10:86229750-86229988 | 0.670530127 | 0.13031636 | 0.368934196 |
| chr12:50251917-50252284 | 0.58967012 | 0.13031835 | 0.368934196 |
| chr4:173608408-173609424 | -0.504000803 | 0.130322531 | 0.368934196 |
| chr11:123096355-123096717 | -0.665257116 | 0.130354369 | 0.369004122 |
| chr19:17209727-17210124 | 0.614998911 | 0.130356863 | 0.369004122 |
| chr11:937823-938090 | -0.735665751 | 0.130365319 | 0.369014425 |
| chr15:69861729-69862205 | 0.594409805 | 0.130381442 | 0.36904643 |
| chr22:41401905-41402506 | 0.548943904 | 0.130406551 | 0.369083041 |
| chr12:122031693-122031901 | 0.691454837 | 0.130409865 | 0.369083041 |
| chr3:127304059-127304812 | -0.49393288 | 0.130411364 | 0.369083041 |
| chr1:161749516-161750480 | 0.485388537 | 0.130421955 | 0.369083041 |
| chr19:35519478-35519704 | 0.697069932 | 0.13042323 | 0.369083041 |
| chr7:101411238-101411584 | -0.618171045 | 0.130427717 | 0.369083041 |
| chr8:37551707-37551920 | -0.805083166 | 0.130428095 | 0.369083041 |
| chr2:42930003-42930254 | -0.648945144 | 0.130449125 | 0.369128917 |
| chr13:42298502-42298724 | 0.748749796 | 0.130496037 | 0.369242569 |
| chr16:3106218-3107095 | 0.509037232 | 0.13050344 | 0.369242569 |
| chr8:119641396-119641668 | 0.691346899 | 0.130506186 | 0.369242569 |
| chrX:119672151-119672813 | 0.575536462 | 0.130508565 | 0.369242569 |
| chr16:23509797-23510943 | 0.497949521 | 0.130514634 | 0.369246106 |
| chr7:76303469-76303790 | -0.562083416 | 0.13052687 | 0.369263399 |
| chr21:45747807-45748149 | 0.604864322 | 0.130530385 | 0.369263399 |
| chr7:123285442-123285642 | -0.786686549 | 0.130537556 | 0.36927005 |
| chr3:84965192-84965394 | 0.742740822 | 0.130550004 | 0.36929163 |
| chr13:102870650-102870937 | -0.733307083 | 0.130563173 | 0.369315248 |
| chr11:67053904-67054173 | -0.556407064 | 0.130581253 | 0.369352756 |
| chr5:79025295-79025498 | -0.790093809 | 0.130599794 | 0.369390217 |
| chr17:17676258-17676467 | -0.727069541 | 0.130604139 | 0.369390217 |
| chr13:43851516-43851808 | 0.745827299 | 0.130614838 | 0.369398421 |
| chr2:240143846-240144098 | -0.75015091 | 0.130616682 | 0.369398421 |
| chr5:118775888-118776328 | 0.627411746 | 0.130625211 | 0.369398512 |
| chr12:102119623-102120733 | -0.484168714 | 0.130638777 | 0.369398512 |
| chr19:45075455-45075813 | 0.557888843 | 0.130642505 | 0.369398512 |
| chr10:49293463-49293735 | -0.796385093 | 0.130643665 | 0.369398512 |
| chr13:93202259-93202489 | -0.796385093 | 0.130643665 | 0.369398512 |

| | | | |
|---|---|---|---|
| chr2:39568180-39568401 | -0.731501012 | 0.130645641 | 0.369398512 |
| chr2:79494882-79495104 | -0.726664949 | 0.130680867 | 0.369484478 |
| chr11:37712323-37712528 | -0.780489036 | 0.130692027 | 0.369502397 |
| chr4:171903543-171903800 | -0.741386096 | 0.130701707 | 0.369516131 |
| chr1:185170184-185170773 | -0.520915811 | 0.130715654 | 0.369541927 |
| chr3:23576621-23576973 | 0.647340958 | 0.13073432 | 0.369574203 |
| chr5:76690961-76691335 | -0.714542027 | 0.130736717 | 0.369574203 |
| chr13:39810467-39810813 | -0.574249893 | 0.130745335 | 0.369574529 |
| chr12:122449350-122449744 | 0.577415722 | 0.130746479 | 0.369574529 |
| chr11:61816063-61817210 | 0.495360933 | 0.130799374 | 0.369704622 |
| chr1:40668915-40669182 | 0.632594047 | 0.130803276 | 0.369704622 |
| chr6:44128223-44128449 | 0.62445654 | 0.130807368 | 0.369704622 |
| chr6:122465521-122465721 | 0.79967779 | 0.130811803 | 0.369704622 |
| chr3:177394723-177394959 | -0.780041615 | 0.130852693 | 0.369793644 |
| chr20:36677200-36677492 | -0.741704394 | 0.130853807 | 0.369793644 |
| chr7:36312549-36313091 | -0.526267703 | 0.13085778 | 0.369793644 |
| chr1:96872817-96873060 | -0.781030212 | 0.130869461 | 0.369810525 |
| chr11:62974744-62975018 | -0.767834611 | 0.130873407 | 0.369810525 |
| chr12:31235324-31235599 | -0.67282396 | 0.130889876 | 0.369842873 |
| chr10:24251820-24252054 | 0.719308073 | 0.130898722 | 0.369842873 |
| chr3:127652184-127652469 | 0.594820039 | 0.130902656 | 0.369842873 |
| chr16:2339721-2340464 | 0.526103666 | 0.130904162 | 0.369842873 |
| chr10:58152801-58153032 | -0.680975051 | 0.130932868 | 0.369910336 |
| chr9:16192211-16192510 | -0.670315715 | 0.13094414 | 0.369927317 |
| chr19:49813484-49813791 | -0.566078674 | 0.130948535 | 0.369927317 |
| chr6:154995658-154996135 | 0.576747769 | 0.130954877 | 0.369931595 |
| chr14:49676407-49677145 | -0.537429345 | 0.130964384 | 0.369944813 |
| chr8:85521985-85522257 | 0.63317366 | 0.130974107 | 0.369958638 |
| chr3:8501391-8501994 | 0.48992697 | 0.131000859 | 0.370020346 |
| chr8:130463367-130463648 | 0.741991563 | 0.13101768 | 0.370020346 |
| chr1:201538390-201538635 | 0.616610176 | 0.131018608 | 0.370020346 |
| chr8:141164976-141165176 | 0.731217629 | 0.131019101 | 0.370020346 |
| chr12:79727811-79728054 | 0.796670177 | 0.131029023 | 0.370020346 |
| chr17:60465095-60465300 | 0.549404673 | 0.131031447 | 0.370020346 |
| chr14:91405791-91406078 | 0.637446712 | 0.131033361 | 0.370020346 |
| chr21:43691156-43691422 | -0.664303159 | 0.131034586 | 0.370020346 |
| chr5:23611528-23611728 | 0.795550333 | 0.131048934 | 0.370045917 |
| chr19:50330777-50330977 | 0.609499483 | 0.131053301 | 0.370045917 |
| chr6:85579598-85580009 | 0.581033597 | 0.131072204 | 0.37008469 |
| chr16:73048556-73048768 | 0.612019955 | 0.131076692 | 0.37008469 |
| chr3:113685225-113685468 | -0.68934604 | 0.131103202 | 0.3701459 |
| chr14:75654181-75654488 | 0.572257424 | 0.131119689 | 0.370167262 |
| chr17:2029883-2030246 | -0.554282783 | 0.131120431 | 0.370167262 |
| chr3:22595105-22595305 | 0.739234719 | 0.131126529 | 0.370170839 |
| chr7:28933359-28933679 | -0.68610468 | 0.131144419 | 0.370207703 |
| chr17:60305124-60305375 | 0.544150345 | 0.131160708 | 0.370225776 |
| chr2:108182572-108182823 | -0.724548937 | 0.13116319 | 0.370225776 |
| chr6:24431145-24431352 | -0.774940706 | 0.131165317 | 0.370225776 |
| chr1:61082789-61083166 | -0.589168103 | 0.131183281 | 0.370243464 |
| chr16:78576426-78576626 | 0.870150906 | 0.131191648 | 0.370243464 |

| | | | |
|---|---|---|---|
| chr6:34553332-34553578 | -0.653484142 | 0.131198674 | 0.370243464 |
| chr5:160065110-160065573 | 0.578971539 | 0.131200433 | 0.370243464 |
| chr10:36725588-36725823 | -0.767623972 | 0.131201853 | 0.370243464 |
| chr3:78659528-78659779 | -0.777556081 | 0.131210906 | 0.370243464 |
| chr14:21631163-21631605 | 0.616838006 | 0.131211951 | 0.370243464 |
| chr8:22561800-22562095 | 0.589938012 | 0.131212007 | 0.370243464 |
| chr7:101507761-101508119 | -0.597259815 | 0.131215072 | 0.370243464 |
| chr17:44774868-44775487 | 0.512970422 | 0.131229574 | 0.370270747 |
| chr14:91568688-91568931 | 0.667236845 | 0.131237801 | 0.370280326 |
| chr5:119267886-119268383 | -0.4926128 | 0.131243531 | 0.370282858 |
| chr4:40927664-40927891 | 0.705324136 | 0.131251628 | 0.370291732 |
| chr5:106122004-106122331 | -0.618100189 | 0.131256342 | 0.370291732 |
| chr7:150370079-150370328 | -0.620483647 | 0.131266496 | 0.370295595 |
| chr1:19799186-19799439 | -0.811624932 | 0.131273302 | 0.370295595 |
| chr15:67175111-67175311 | -0.744367459 | 0.131273807 | 0.370295595 |
| chr18:3187313-3187587 | -0.676854799 | 0.131277042 | 0.370295595 |
| chr4:121656004-121656533 | 0.578986577 | 0.131283064 | 0.370295789 |
| chr3:133208298-133208545 | -0.702864988 | 0.131286777 | 0.370295789 |
| chr7:51079502-51079773 | -0.740081566 | 0.131310935 | 0.370350295 |
| chr4:2408549-2408770 | 0.709469849 | 0.131318832 | 0.370358934 |
| chr6:108689196-108689868 | -0.527810935 | 0.131331449 | 0.370372654 |
| chr1:32063385-32063663 | -0.634164456 | 0.131341235 | 0.370372654 |
| chr16:2520752-2521191 | 0.560905394 | 0.13134126 | 0.370372654 |
| chr11:124970984-124971215 | 0.738914497 | 0.131343031 | 0.370372654 |
| chr2:153870362-153870648 | -0.637291315 | 0.131373316 | 0.370434538 |
| chr2:14903759-14904116 | -0.533004292 | 0.131374646 | 0.370434538 |
| chr17:75674738-75675208 | -0.597927152 | 0.131422763 | 0.370556575 |
| chr20:56712309-56712509 | 0.661527541 | 0.131451566 | 0.370624149 |
| chr17:31487574-31488177 | 0.513619638 | 0.131483893 | 0.370701655 |
| chr2:19138434-19138714 | 0.613484975 | 0.13149809 | 0.370727376 |
| chr3:87097670-87098461 | -0.508553967 | 0.131505154 | 0.370727376 |
| chr6:134109586-134109872 | -0.629534648 | 0.131507532 | 0.370727376 |
| chr4:51844545-51844974 | 0.512863843 | 0.131549857 | 0.37082628 |
| chr12:26884671-26885506 | -0.525190158 | 0.131552295 | 0.37082628 |
| chr10:11204690-11204974 | -0.711207279 | 0.131565924 | 0.370851056 |
| chr2:17257454-17257654 | 0.834895153 | 0.131578406 | 0.370872596 |
| chr3:52534956-52535167 | 0.63994451 | 0.13159832 | 0.370890801 |
| chr9:30774403-30774732 | -0.690789532 | 0.131602737 | 0.370890801 |
| chr6:140585897-140586158 | -0.770444391 | 0.131603652 | 0.370890801 |
| chr12:124730506-124730748 | 0.665341667 | 0.131604227 | 0.370890801 |
| chr1:218339320-218339563 | -0.63848055 | 0.131617991 | 0.370915949 |
| chr10:119509900-119510522 | -0.533363684 | 0.131630817 | 0.37093845 |
| chr8:141446828-141447296 | -0.590698095 | 0.131637673 | 0.370944128 |
| chr8:19267646-19267862 | 0.755554309 | 0.131671333 | 0.371025333 |
| chr2:187009742-187010072 | -0.777368954 | 0.131681846 | 0.371041313 |
| chr4:111602951-111603196 | -0.725342485 | 0.131701914 | 0.371084213 |
| chr15:74054084-74054350 | -0.646368262 | 0.131713997 | 0.37110114 |
| chr2:160181215-160181502 | -0.688257155 | 0.131717608 | 0.37110114 |
| chr16:71552773-71553272 | -0.58552072 | 0.131736592 | 0.371132814 |
| chr19:47436696-47437012 | 0.579988178 | 0.131746712 | 0.371132814 |

| | | | |
|---|---|---|---|
| chr5:5025792-5026338 | -0.546964164 | 0.131748637 | 0.371132814 |
| chr2:189440745-189441987 | -0.476289314 | 0.131750169 | 0.371132814 |
| chr4:78911737-78911937 | 0.815076558 | 0.131753069 | 0.371132814 |
| chr1:29453533-29453871 | 0.620886506 | 0.131784824 | 0.371179421 |
| chr7:141762093-141762387 | -0.852903786 | 0.13178902 | 0.371179421 |
| chr8:61913508-61913713 | -0.852903786 | 0.13178902 | 0.371179421 |
| chr4:105835250-105835534 | 0.727528679 | 0.131792314 | 0.371179421 |
| chr6:473848-474647 | -0.552869834 | 0.13179615 | 0.371179421 |
| chr7:132656913-132657370 | -0.616496652 | 0.131802338 | 0.371179421 |
| chr2:227658242-227658538 | 0.643444309 | 0.131803525 | 0.371179421 |
| chr7:94509363-94510964 | -0.487118159 | 0.131821312 | 0.371215868 |
| chr11:47407810-47408657 | 0.487895752 | 0.131828746 | 0.371217325 |
| chr7:123242546-123243207 | -0.508804039 | 0.131831519 | 0.371217325 |
| chr2:33134343-33134543 | -0.869450899 | 0.131846537 | 0.37124597 |
| chr12:19198385-19198703 | -0.719524918 | 0.131862603 | 0.371277566 |
| chr13:48769124-48769503 | -0.553457073 | 0.131870599 | 0.371281217 |
| chr12:949070-950127 | -0.494143769 | 0.131873591 | 0.371281217 |
| chrX:139891499-139891718 | 0.724670199 | 0.131891 | 0.371309836 |
| chr4:111641757-111642476 | -0.508548739 | 0.131897965 | 0.371309836 |
| chr8:27614302-27615021 | 0.515646295 | 0.131898294 | 0.371309836 |
| chr10:34985842-34986176 | -0.603361207 | 0.131915164 | 0.371343683 |
| chr8:27772437-27772832 | 0.557576997 | 0.131921232 | 0.371347122 |
| chrX:21973409-21974414 | -0.50011628 | 0.13193353 | 0.371368095 |
| chr20:59192961-59193324 | 0.571470028 | 0.131946737 | 0.371382782 |
| chr15:80309859-80310093 | -0.74308294 | 0.131948441 | 0.371382782 |
| chr3:14579908-14580117 | 0.706845652 | 0.13195891 | 0.371398606 |
| chr10:79825594-79825861 | -0.747946687 | 0.131986406 | 0.371462348 |
| chr13:99916505-99916898 | -0.596286565 | 0.131999837 | 0.371481805 |
| chr10:21516306-21517152 | -0.523757247 | 0.132003016 | 0.371481805 |
| chr2:15913052-15913411 | -0.641882256 | 0.132008984 | 0.371484955 |
| chr14:53180890-53181293 | -0.588786874 | 0.132016283 | 0.371491501 |
| chr5:133108581-133108816 | -0.741400083 | 0.132021007 | 0.371491501 |
| chr2:198901976-198902240 | -0.713811739 | 0.132027933 | 0.371497349 |
| chr10:124450204-124450445 | 0.742894037 | 0.132062419 | 0.371580737 |
| chr10:127012727-127013054 | -0.655973705 | 0.132071562 | 0.371583233 |
| chr3:120795081-120795419 | -0.572479638 | 0.132073005 | 0.371583233 |
| chr17:2336340-2336711 | -0.55335041 | 0.132119783 | 0.371700383 |
| chr18:63422061-63423130 | -0.479316399 | 0.132124346 | 0.371700383 |
| chr3:103992476-103992792 | -0.65833663 | 0.132136952 | 0.3717222 |
| chr10:25768315-25768528 | -0.828131628 | 0.13216034 | 0.371774344 |
| chrX:74458794-74459190 | -0.682781899 | 0.132187617 | 0.371837424 |
| chr17:18118393-18118728 | -0.630428988 | 0.132220368 | 0.371915898 |
| chr13:71247556-71247793 | -0.806899431 | 0.132231277 | 0.371931124 |
| chr6:115896581-115896827 | -0.727783708 | 0.132235489 | 0.371931124 |
| chr13:98840176-98840473 | 0.623371532 | 0.132246719 | 0.371949055 |
| chr6:33513322-33513576 | -0.734269264 | 0.132281103 | 0.372016587 |
| chr6:99219739-99220209 | -0.540712424 | 0.132284166 | 0.372016587 |
| chr3:62517211-62517411 | 0.614298359 | 0.132287942 | 0.372016587 |
| chr7:136115282-136115596 | -0.59551474 | 0.132290151 | 0.372016587 |
| chr14:45756297-45757077 | 0.519492896 | 0.132296982 | 0.372022144 |

| | | | |
|---|---|---|---|
| chr3:129565013-129565576 | 0.532884995 | 0.13231842 | 0.372068772 |
| chr4:104930828-104931498 | -0.536516471 | 0.132330526 | 0.372089158 |
| chr5:127564152-127564507 | -0.603443229 | 0.132370405 | 0.37217449 |
| chr16:70524604-70524896 | 0.61011823 | 0.132370588 | 0.37217449 |
| chr4:55184658-55185066 | -0.619396596 | 0.132389865 | 0.372215031 |
| chr1:106372534-106373103 | 0.511280916 | 0.132416073 | 0.372275055 |
| chr13:36345866-36347353 | -0.478214718 | 0.132431224 | 0.372303991 |
| chr2:73214022-73214746 | -0.500430237 | 0.132488235 | 0.372442538 |
| chr22:19718026-19718226 | 0.690062562 | 0.132498034 | 0.372442538 |
| chr7:99415606-99415881 | -0.709005301 | 0.132499833 | 0.372442538 |
| chr14:105020755-105021621 | 0.484583521 | 0.13249995 | 0.372442538 |
| chr7:101100154-101100487 | -0.618822155 | 0.132539012 | 0.372529859 |
| chr7:108454646-108455441 | -0.518673878 | 0.132541542 | 0.372529859 |
| chr15:55318800-55319514 | 0.498002792 | 0.132546268 | 0.372529859 |
| chr8:10334913-10335143 | -0.798017626 | 0.132550462 | 0.372529859 |
| chr20:57080838-57081164 | 0.552205304 | 0.132577776 | 0.372592955 |
| chr6:94941481-94941825 | -0.735303554 | 0.132583979 | 0.372596723 |
| chr7:87797888-87798185 | -0.692289502 | 0.132590513 | 0.372601418 |
| chr21:38158938-38159607 | -0.525835385 | 0.13260891 | 0.37263945 |
| chr19:18000967-18001167 | 0.753504211 | 0.132623544 | 0.372666906 |
| chr12:111578271-111578822 | 0.56666128 | 0.132652163 | 0.372733655 |
| chr4:72209785-72210011 | -0.737153976 | 0.132663823 | 0.372739751 |
| chr1:55916512-55917139 | -0.592388427 | 0.132667344 | 0.372739751 |
| chr2:25991487-25991863 | -0.606767535 | 0.132668927 | 0.372739751 |
| chr14:52950339-52951931 | 0.482613879 | 0.132686894 | 0.372766567 |
| chr21:43829499-43829824 | 0.603237372 | 0.132688201 | 0.372766567 |
| chr6:33289163-33289941 | 0.511240996 | 0.132709212 | 0.372811926 |
| chr5:43944440-43944696 | 0.722568572 | 0.132714809 | 0.372813979 |
| chr2:163735841-163737137 | -0.488404045 | 0.132735715 | 0.372859038 |
| chr11:10294556-10294877 | -0.712947795 | 0.132744606 | 0.372861828 |
| chr14:105710047-105710247 | 0.669236891 | 0.132746441 | 0.372861828 |
| chr3:148121932-148122238 | -0.747281649 | 0.132789583 | 0.372944703 |
| chr6:46042429-46042632 | -0.804534093 | 0.132793481 | 0.372944703 |
| chr6:53746919-53747155 | -0.804534093 | 0.132793481 | 0.372944703 |
| chr10:124389188-124389745 | 0.530811874 | 0.132795415 | 0.372944703 |
| chr12:102061182-102062324 | -0.485029926 | 0.132819322 | 0.372981253 |
| chr11:120169140-120169459 | 0.616481953 | 0.132827298 | 0.372981253 |
| chr9:128881804-128882680 | 0.496473886 | 0.13282751 | 0.372981253 |
| chr9:123467785-123467985 | 0.715960468 | 0.132828697 | 0.372981253 |
| chr1:204001287-204001808 | -0.543531801 | 0.132832769 | 0.372981253 |
| chr7:13989703-13989903 | 0.583566113 | 0.132841096 | 0.372990966 |
| chr1:182084925-182085289 | 0.548086013 | 0.132851371 | 0.373006146 |
| chr1:211469920-211470426 | -0.550875578 | 0.132891101 | 0.373092411 |
| chr5:79173615-79174056 | -0.561507666 | 0.132891834 | 0.373092411 |
| chr13:99656204-99656551 | 0.629964272 | 0.132899279 | 0.373099644 |
| chr11:64358890-64359577 | 0.52781022 | 0.13290971 | 0.373115256 |
| chr2:20496000-20496757 | -0.513945042 | 0.132915917 | 0.37311901 |
| chr8:108366641-108366848 | 0.769287632 | 0.132927167 | 0.373136921 |
| chr12:96091924-96092211 | -0.645754142 | 0.132938235 | 0.373154318 |
| chr5:178145927-178146810 | -0.485406036 | 0.133028162 | 0.373393062 |

| | | | |
|---|---|---|---|
| chr14:96762707-96763049 | 0.566430017 | 0.133043529 | 0.373409295 |
| chr12:98572092-98572351 | -0.657937144 | 0.133043692 | 0.373409295 |
| chr2:56108921-56109294 | 0.640739198 | 0.133053877 | 0.373424203 |
| chr7:100122151-100122447 | 0.560129779 | 0.133068898 | 0.37345268 |
| chr3:63705195-63705434 | 0.563617633 | 0.133086265 | 0.373487741 |
| chr10:104614288-104614602 | -0.648661306 | 0.133113014 | 0.373540786 |
| chr1:216830973-216831203 | -0.67224906 | 0.133114917 | 0.373540786 |
| chr12:67792092-67792538 | -0.611978235 | 0.133131841 | 0.373574597 |
| chr1:147877798-147878064 | 0.652815234 | 0.133141511 | 0.373579961 |
| chr10:69722027-69722266 | -0.729964106 | 0.133146948 | 0.373579961 |
| chr19:10654665-10655048 | 0.569437528 | 0.13314838 | 0.373579961 |
| chr17:31579385-31579617 | 0.74718753 | 0.133166692 | 0.37361574 |
| chr1:181033255-181034166 | -0.48723559 | 0.133170884 | 0.37361574 |
| chr10:585600-585800 | 0.70602386 | 0.133185056 | 0.373621621 |
| chr10:2599862-2600154 | 0.70602386 | 0.133185056 | 0.373621621 |
| chr14:24429769-24430356 | 0.502873585 | 0.133187609 | 0.373621621 |
| chr3:110439982-110440182 | -0.787676435 | 0.133199452 | 0.373641164 |
| chr6:106444990-106445236 | -0.781189354 | 0.133204666 | 0.373642113 |
| chr11:108497284-108497984 | -0.52686429 | 0.133223073 | 0.373672904 |
| chr3:22960411-22960687 | -0.640456984 | 0.133229755 | 0.373672904 |
| chr2:30239239-30239537 | 0.617100313 | 0.133230274 | 0.373672904 |
| chr14:40111031-40111276 | 0.737361225 | 0.133241368 | 0.37369034 |
| chr3:42014140-42014401 | -0.690797882 | 0.133262617 | 0.373724575 |
| chr2:84807176-84807885 | 0.563243631 | 0.133263329 | 0.373724575 |
| chr22:19121564-19122118 | 0.534419879 | 0.133278111 | 0.373752349 |
| chr3:137033032-137033232 | -0.671994772 | 0.133290086 | 0.373758002 |
| chr16:89947993-89948392 | -0.582083587 | 0.133290179 | 0.373758002 |
| chr15:77696389-77696600 | 0.669563139 | 0.133294761 | 0.373758002 |
| chr16:2513490-2514065 | 0.507073157 | 0.133319944 | 0.373810464 |
| chr11:123083065-123083299 | 0.686257669 | 0.133323228 | 0.373810464 |
| chr6:76861461-76861759 | -0.62135272 | 0.133333595 | 0.373825852 |
| chr15:72475021-72475295 | 0.587624882 | 0.13335104 | 0.373857031 |
| chr11:85253408-85253718 | -0.765721572 | 0.133354474 | 0.373857031 |
| chr15:100926392-100926601 | -0.775093102 | 0.13339528 | 0.373950237 |
| chr8:122575791-122576053 | 0.648837234 | 0.133404209 | 0.373950237 |
| chr2:121136456-121136685 | -0.696403474 | 0.133405919 | 0.373950237 |
| chr12:121888400-121889208 | 0.489345656 | 0.133410166 | 0.373950237 |
| chr15:99925832-99926492 | 0.537212426 | 0.133412123 | 0.373950237 |
| chr21:39094990-39095558 | 0.52499855 | 0.133427146 | 0.373978666 |
| chr12:14187673-14187873 | 0.852955653 | 0.133445005 | 0.37401504 |
| chr9:137255006-137255826 | 0.505843521 | 0.133453205 | 0.374024341 |
| chrX:48917155-48917538 | 0.534681061 | 0.133483499 | 0.374095561 |
| chr2:166437574-166437915 | -0.758542 | 0.133498725 | 0.374124549 |
| chr11:13114695-13115254 | -0.527732776 | 0.133546836 | 0.374226433 |
| chr7:150378239-150378597 | -0.533396533 | 0.133553275 | 0.374226433 |
| chr21:40841541-40842303 | -0.501311649 | 0.133554085 | 0.374226433 |
| chr16:65429500-65429774 | -0.763885631 | 0.133554616 | 0.374226433 |
| chr9:83231098-83231465 | -0.582472907 | 0.133565571 | 0.374243444 |
| chr17:59297284-59297484 | 0.657852765 | 0.133595093 | 0.374299227 |
| chr3:166969632-166969842 | 0.775766628 | 0.13359525 | 0.374299227 |

| | | | |
|---|---|---|---|
| chr8:107706051-107706282 | -0.673896743 | 0.133603287 | 0.374308058 |
| chr6:94903004-94903535 | -0.597035805 | 0.133632528 | 0.37437629 |
| chr6:75244242-75244585 | 0.580878767 | 0.133657516 | 0.374408382 |
| chr1:189416180-189416425 | 0.736808388 | 0.133659078 | 0.374408382 |
| chr20:25853901-25854250 | 0.569441252 | 0.133661289 | 0.374408382 |
| chr4:119039874-119040107 | -0.701410448 | 0.133664482 | 0.374408382 |
| chr15:28096029-28096411 | 0.553807779 | 0.133668415 | 0.374408382 |
| chr20:36436009-36436596 | 0.511614816 | 0.133700337 | 0.374468376 |
| chr16:740356-740683 | 0.613223074 | 0.133706213 | 0.374468376 |
| chr20:43728156-43728415 | 0.672660738 | 0.133706801 | 0.374468376 |
| chr2:110853602-110853802 | -0.789578909 | 0.133709382 | 0.374468376 |
| chr5:105587980-105588490 | -0.654393392 | 0.133720344 | 0.374485388 |
| chr5:181149332-181149532 | -0.774339992 | 0.133732769 | 0.374493865 |
| chr6:17281187-17281452 | 0.707187549 | 0.133733146 | 0.374493865 |
| chr14:59139438-59139664 | 0.768127615 | 0.133744637 | 0.374503227 |
| chr1:196059172-196059591 | -0.615553234 | 0.133748219 | 0.374503227 |
| chr7:107201901-107202220 | -0.6443797 | 0.133751153 | 0.374503227 |
| chr17:19866807-19868196 | 0.499269908 | 0.133760204 | 0.374514885 |
| chr15:92820209-92821191 | 0.47974735 | 0.133771751 | 0.37453353 |
| chr9:134365860-134366542 | 0.502795204 | 0.133791002 | 0.374573741 |
| chr22:34414600-34415171 | -0.572720438 | 0.133818284 | 0.374628528 |
| chr2:218322462-218324319 | -0.47731865 | 0.133820349 | 0.374628528 |
| chr2:214395043-214395377 | -0.649374339 | 0.133825659 | 0.374629705 |
| chr2:138455345-138456129 | -0.523134736 | 0.13383193 | 0.374633571 |
| chr10:34994656-34995059 | -0.553175765 | 0.133844133 | 0.374639798 |
| chr1:116672642-116672843 | 0.697168298 | 0.133848662 | 0.374639798 |
| chr2:81943408-81943655 | -0.821990126 | 0.13385299 | 0.374639798 |
| chr15:101252632-101252956 | 0.590921934 | 0.133853712 | 0.374639798 |
| chr3:184155159-184155878 | 0.526153489 | 0.133875289 | 0.374686503 |
| chr6:37473228-37473664 | 0.553121072 | 0.133887945 | 0.374708238 |
| chr6:73884614-73884911 | -0.755427956 | 0.133896507 | 0.374718512 |
| chr1:37946782-37947250 | 0.549071439 | 0.133905359 | 0.3747296 |
| chr1:112152330-112152654 | -0.754870121 | 0.133949699 | 0.374839992 |
| chr12:47408348-47408548 | 0.781318082 | 0.134000611 | 0.374968769 |
| chr1:208461683-208461978 | -0.700342989 | 0.134018581 | 0.375005357 |
| chr5:140988123-140988337 | 0.822428641 | 0.13404138 | 0.375046326 |
| chr4:168995663-168996044 | -0.69977746 | 0.134046571 | 0.375046326 |
| chr2:84741448-84741894 | -0.567521798 | 0.134047906 | 0.375046326 |
| chr15:59751895-59752135 | 0.753535223 | 0.13406718 | 0.375086554 |
| chr10:1100932-1101323 | -0.58927954 | 0.134093243 | 0.375145775 |
| chr2:61423758-61424011 | 0.696079318 | 0.134108885 | 0.375175838 |
| chr20:55416188-55416451 | 0.656526274 | 0.134115751 | 0.375181346 |
| chr6:30883910-30884602 | 0.498354978 | 0.134129318 | 0.3752056 |
| chr20:45336714-45336996 | 0.613624941 | 0.134141365 | 0.375225601 |
| chr14:43540747-43541602 | -0.51323684 | 0.134162096 | 0.375269217 |
| chr22:23913779-23914343 | 0.500189528 | 0.134166753 | 0.375269217 |
| chr17:57833150-57833638 | 0.522187671 | 0.134184204 | 0.375281436 |
| chr17:72676492-72676741 | 0.670004453 | 0.134185332 | 0.375281436 |
| chrX:54695910-54696333 | 0.649294126 | 0.134185815 | 0.375281436 |
| chr3:107431038-107432077 | 0.492525352 | 0.134223501 | 0.375373133 |

| | | | |
|---|---|---|---|
| chr7:113774291-113774543 | -0.722331577 | 0.134243189 | 0.37541449 |
| chr8:118488752-118488967 | 0.760054431 | 0.134252612 | 0.375427139 |
| chr22:48169259-48169597 | -0.686064982 | 0.134278174 | 0.375484917 |
| chr5:53481892-53482180 | -0.607678178 | 0.134298546 | 0.37552298 |
| chr17:52418917-52419674 | 0.511102148 | 0.134303318 | 0.37552298 |
| chr12:97552473-97552923 | -0.68026291 | 0.134311226 | 0.37552298 |
| chr12:46372170-46372513 | 0.548121708 | 0.134311389 | 0.37552298 |
| chr4:45663025-45663365 | -0.640168662 | 0.134327099 | 0.375553199 |
| chr1:116570296-116571295 | 0.4778642 | 0.1343475 | 0.375596532 |
| chr8:80145975-80146292 | -0.584459575 | 0.134364159 | 0.375629399 |
| chr1:240455276-240455558 | -0.65686169 | 0.134397403 | 0.375707661 |
| chr6:47073084-47073348 | -0.693040652 | 0.13440196 | 0.375707661 |
| chr6:121703196-121703521 | -0.63764462 | 0.134424443 | 0.3757568 |
| chr1:12618416-12618663 | -0.769892273 | 0.134437083 | 0.375762611 |
| chr20:44388408-44388861 | 0.530572162 | 0.134441196 | 0.375762611 |
| chr1:49297832-49298065 | -0.736152937 | 0.134448474 | 0.375762611 |
| chr7:150367957-150369462 | -0.479466402 | 0.134449759 | 0.375762611 |
| chr22:26461295-26461539 | 0.669278372 | 0.134451042 | 0.375762611 |
| chr2:43219176-43220279 | -0.49149858 | 0.134457392 | 0.375766652 |
| chr5:142699123-142699429 | 0.587712272 | 0.134464822 | 0.375773712 |
| chr1:119968699-119968899 | 0.735131078 | 0.134487171 | 0.375822459 |
| chr5:170184039-170184391 | -0.557583331 | 0.134504735 | 0.375857835 |
| chr6:116615933-116616626 | -0.491695565 | 0.134521419 | 0.375878763 |
| chr3:114418721-114419227 | -0.547250981 | 0.134522036 | 0.375878763 |
| chr4:18417924-18418573 | -0.517592815 | 0.134529411 | 0.375885663 |
| chr1:107683115-107683589 | -0.53377621 | 0.134544576 | 0.37591152 |
| chr10:53664855-53665079 | -0.719691514 | 0.134548477 | 0.37591152 |
| chr14:50397453-50397745 | -0.641604653 | 0.134560208 | 0.375930586 |
| chr17:41573698-41573981 | 0.719614355 | 0.134576253 | 0.375961704 |
| chr16:57486372-57486667 | 0.619996193 | 0.134584583 | 0.375971268 |
| chr22:37592333-37592892 | -0.58601581 | 0.134594334 | 0.375983247 |
| chrX:54495693-54496373 | -0.553773538 | 0.134598685 | 0.375983247 |
| chr3:176948061-176948597 | -0.534096572 | 0.134644713 | 0.376084676 |
| chr9:125199403-125199726 | -0.592894923 | 0.134644813 | 0.376084676 |
| chr2:102845894-102846096 | 0.784325455 | 0.134656341 | 0.376103167 |
| chr3:169563075-169563374 | -0.582796972 | 0.13467262 | 0.376134923 |
| chr4:136329115-136329632 | -0.572771153 | 0.134678417 | 0.376137403 |
| chr12:63918370-63918732 | -0.666451231 | 0.134700998 | 0.376186757 |
| chr10:32005085-32005374 | -0.654911971 | 0.134721357 | 0.376229901 |
| chr9:127715442-127716192 | 0.49445634 | 0.134733533 | 0.376243194 |
| chr17:63452414-63452626 | 0.589559763 | 0.134735937 | 0.376243194 |
| chrX:65582048-65582257 | 0.746362409 | 0.13475706 | 0.376288465 |
| chr10:105063880-105064107 | -0.749076032 | 0.134768361 | 0.376306306 |
| chr19:44856617-44856915 | -0.678437078 | 0.134785745 | 0.376341132 |
| chr3:28286647-28286954 | -0.617060362 | 0.134797436 | 0.376360061 |
| chr1:35807582-35808056 | 0.564105498 | 0.134808863 | 0.376378251 |
| chr4:152935633-152937278 | 0.476253871 | 0.134851695 | 0.376470429 |
| chr7:22872120-22872396 | 0.626850885 | 0.134852278 | 0.376470429 |
| chr13:19783327-19783570 | -0.645813708 | 0.134856619 | 0.376470429 |
| chr2:5713011-5713575 | -0.559561041 | 0.134881984 | 0.376518749 |

| | | | |
|---|---|---|---|
| chr7:77714910-77715180 | 0.772153677 | 0.134883756 | 0.376518749 |
| chr6:151668929-151669254 | 0.578097869 | 0.134890424 | 0.376523645 |
| chr3:145094432-145094934 | -0.564764428 | 0.134901969 | 0.376542155 |
| chr20:48785270-48785601 | 0.519936926 | 0.134925874 | 0.37659516 |
| chr10:14498985-14499445 | 0.588553401 | 0.134947762 | 0.3766199 |
| chr6:44087263-44087463 | 0.718871566 | 0.134952495 | 0.3766199 |
| chr4:139514516-139514726 | 0.696729191 | 0.134955543 | 0.3766199 |
| chr7:21507678-21508192 | 0.540304088 | 0.134959067 | 0.3766199 |
| chr17:4795953-4796296 | -0.596776466 | 0.134959315 | 0.3766199 |
| chr3:62993139-62993339 | 0.646476015 | 0.134978906 | 0.376638789 |
| chr2:121151807-121152114 | -0.661397022 | 0.134979219 | 0.376638789 |
| chr5:11372990-11373216 | -0.786852238 | 0.134997875 | 0.376638789 |
| chr3:133742187-133742962 | -0.557482666 | 0.134998576 | 0.376638789 |
| chr12:131454807-131455036 | 0.68321532 | 0.135002167 | 0.376638789 |
| chr20:15867917-15868217 | -0.667719522 | 0.135009084 | 0.376638789 |
| chr4:177159647-177159923 | -0.775400566 | 0.13501077 | 0.376638789 |
| chr7:155798032-155798267 | -0.775400566 | 0.13501077 | 0.376638789 |
| chr17:50318115-50318315 | 0.711632099 | 0.135015709 | 0.376638789 |
| chr16:74826223-74826441 | -0.744184851 | 0.135019165 | 0.376638789 |
| chr8:33545761-33545961 | 0.813691806 | 0.135020154 | 0.376638789 |
| chr12:2885238-2885579 | -0.604647968 | 0.135038143 | 0.376675257 |
| chr3:133663627-133664062 | -0.56333972 | 0.135046721 | 0.376685471 |
| chr17:78314086-78314629 | 0.52550776 | 0.13510202 | 0.376825997 |
| chr2:42788411-42788706 | -0.592081347 | 0.135116308 | 0.376852132 |
| chr13:50801371-50801671 | -0.629816356 | 0.135121493 | 0.376852875 |
| chr17:55446756-55447130 | -0.601335912 | 0.135129846 | 0.37685837 |
| chr4:89700846-89701051 | -0.850493817 | 0.1351333 | 0.37685837 |
| chr2:53634532-53634923 | -0.658575515 | 0.135157715 | 0.376906403 |
| chr19:44747305-44748783 | 0.477231433 | 0.135165242 | 0.376906403 |
| chr2:47198912-47199232 | -0.722065134 | 0.135165281 | 0.376906403 |
| chr17:60102725-60103039 | 0.521667303 | 0.135193859 | 0.376972373 |
| chr2:227324686-227324903 | -0.673874527 | 0.135199676 | 0.376974875 |
| chr6:146785052-146785406 | -0.599055082 | 0.135212367 | 0.376996544 |
| chr2:64685444-64685726 | -0.673498669 | 0.135230477 | 0.377033317 |
| chr12:56047168-56047415 | 0.637580316 | 0.135270174 | 0.377109238 |
| chr8:70534477-70534792 | -0.562661275 | 0.135271235 | 0.377109238 |
| chr4:103786147-103786392 | 0.682894663 | 0.135274964 | 0.377109238 |
| chr16:79291233-79291521 | 0.76815956 | 0.135277585 | 0.377109238 |
| chr11:110174255-110174462 | -0.737942906 | 0.135288278 | 0.377109238 |
| chr12:67220192-67220397 | -0.773030311 | 0.135290864 | 0.377109238 |
| chr4:38872742-38873056 | 0.617372947 | 0.135292159 | 0.377109238 |
| chr9:25519116-25519730 | -0.542041783 | 0.135309003 | 0.377142467 |
| chr4:124718781-124719175 | -0.56221022 | 0.135324826 | 0.377172849 |
| chr20:57155499-57155699 | 0.508621833 | 0.13533203 | 0.377179209 |
| chr2:117537053-117537368 | -0.680205593 | 0.13534206 | 0.377193442 |
| chr15:31325750-31325967 | 0.609284654 | 0.135353702 | 0.377212169 |
| chr12:99270270-99270476 | -0.800499572 | 0.135372836 | 0.377225886 |
| chr3:65320865-65321314 | -0.604401135 | 0.135374614 | 0.377225886 |
| chr10:91671043-91671328 | -0.602043439 | 0.135376488 | 0.377225886 |
| chr15:86232197-86232408 | 0.700136762 | 0.135378317 | 0.377225886 |

| | | | |
|---|---|---|---|
| chr3:123027489-123028032 | 0.56011821 | 0.135386188 | 0.377226137 |
| chr7:158973353-158973830 | -0.524550509 | 0.135391693 | 0.377226137 |
| chr2:122988732-122988943 | -0.769627928 | 0.135395703 | 0.377226137 |
| chr11:65569935-65570549 | 0.491955565 | 0.135398922 | 0.377226137 |
| chr17:28613435-28613762 | -0.661283961 | 0.135404267 | 0.377226137 |
| chr16:80540454-80541324 | -0.486384365 | 0.135409685 | 0.377226137 |
| chr11:95152070-95152350 | -0.567211714 | 0.135413269 | 0.377226137 |
| chr18:54983206-54983580 | -0.71227755 | 0.135417793 | 0.377226137 |
| chr22:28819243-28819802 | 0.521652939 | 0.135427725 | 0.377240088 |
| chr19:14081236-14081683 | 0.530912631 | 0.135446991 | 0.377273966 |
| chr1:165144840-165145115 | 0.663442664 | 0.135457241 | 0.377273966 |
| chr11:35275474-35275743 | 0.663442664 | 0.135457241 | 0.377273966 |
| chr3:42526614-42526882 | -0.731896902 | 0.135459582 | 0.377273966 |
| chr10:69743314-69743614 | 0.665824458 | 0.135466025 | 0.377278199 |
| chr7:39910045-39910352 | -0.678415004 | 0.135475689 | 0.377291399 |
| chr9:126489064-126489323 | 0.723431018 | 0.135481314 | 0.377293349 |
| chr3:105387115-105387502 | -0.678128961 | 0.135491683 | 0.377308514 |
| chr18:34726273-34727165 | -0.500636738 | 0.135509787 | 0.377345214 |
| chr9:118117087-118117449 | -0.634700281 | 0.135521658 | 0.377364555 |
| chr4:153677073-153677482 | -0.620677034 | 0.135538347 | 0.377386871 |
| chr4:132937830-132938091 | 0.687787408 | 0.135539523 | 0.377386871 |
| chr12:48995389-48995835 | 0.528154769 | 0.135566164 | 0.377447335 |
| chr14:101477059-101477365 | 0.571882636 | 0.135610424 | 0.377556843 |
| chr5:903103-903396 | -0.669465355 | 0.135620489 | 0.377560089 |
| chr7:140079104-140079331 | -0.779348709 | 0.135621445 | 0.377560089 |
| chr2:52358437-52358760 | -0.782409406 | 0.135636517 | 0.377577573 |
| chr17:66426437-66426656 | 0.631851484 | 0.135637581 | 0.377577573 |
| chr12:87151033-87151472 | -0.620477525 | 0.135653837 | 0.377604851 |
| chr3:129829646-129829917 | 0.69005912 | 0.13566255 | 0.377604851 |
| chr2:183179127-183179435 | -0.703833615 | 0.135664748 | 0.377604851 |
| chr12:132360675-132361000 | 0.706445839 | 0.135667092 | 0.377604851 |
| chr1:218683310-218683536 | -0.682579331 | 0.135706656 | 0.37769759 |
| chr3:23103743-23104103 | 0.59622643 | 0.13571027 | 0.37769759 |
| chr2:197613425-197613949 | -0.511567626 | 0.135728457 | 0.377729786 |
| chr16:28291952-28292152 | 0.65866434 | 0.135731699 | 0.377729786 |
| chr10:79444869-79445109 | -0.613593711 | 0.135739259 | 0.377737108 |
| chr1:109636411-109637015 | -0.532118823 | 0.135745448 | 0.37774061 |
| chr17:62882782-62883167 | -0.591278732 | 0.135766904 | 0.377786597 |
| chr13:106917602-106918357 | -0.524031355 | 0.135784924 | 0.377823017 |
| chr2:97360247-97360447 | 0.753641153 | 0.135798146 | 0.377846086 |
| chr2:231200841-231201214 | -0.582673388 | 0.135813058 | 0.377862813 |
| chr17:4161223-4162019 | -0.507045286 | 0.135816181 | 0.377862813 |
| chr1:27979695-27979942 | 0.756368867 | 0.135824032 | 0.377862813 |
| chr3:160407214-160407453 | -0.797822609 | 0.135825615 | 0.377862813 |
| chr20:43775876-43776141 | -0.747241201 | 0.135828815 | 0.377862813 |
| chr7:76306835-76307299 | -0.523425009 | 0.135836942 | 0.377871701 |
| chr3:122258830-122259065 | 0.689159161 | 0.13585054 | 0.377887869 |
| chr2:222088390-222088726 | -0.69049442 | 0.135853212 | 0.377887869 |
| chr20:41352207-41352443 | 0.594675464 | 0.135857549 | 0.377887869 |
| chr17:82021187-82021387 | 0.582642595 | 0.135865999 | 0.377895721 |

| | | | |
|---|---|---|---|
| chr2:73202207-73202432 | -0.71362488 | 0.135870236 | 0.377895721 |
| chr20:54930587-54930953 | 0.523782636 | 0.135885804 | 0.377925301 |
| chr3:57597061-57597941 | 0.47722496 | 0.135895827 | 0.377929627 |
| chr5:55694251-55694491 | -0.712042963 | 0.135898718 | 0.377929627 |
| chr8:135325582-135325793 | -0.894219354 | 0.135903554 | 0.377929627 |
| chr8:103111444-103111869 | 0.524764799 | 0.135908364 | 0.377929627 |
| chr15:49229532-49229765 | -0.708380849 | 0.135915445 | 0.377929627 |
| chr2:217684820-217685580 | 0.516936424 | 0.135919154 | 0.377929627 |
| chr20:53748882-53749136 | 0.493939466 | 0.135921886 | 0.377929627 |
| chr4:145216634-145216852 | -0.801715922 | 0.135943793 | 0.377964058 |
| chr17:19724135-19724398 | 0.644925537 | 0.135944135 | 0.377964058 |
| chr4:108714909-108715167 | 0.687090663 | 0.135965108 | 0.378008653 |
| chr9:116820031-116820231 | 0.847356273 | 0.135977991 | 0.378030755 |
| chr1:31584203-31584433 | 0.64351502 | 0.135989555 | 0.378049187 |
| chr10:52883886-52884578 | -0.485535231 | 0.136012563 | 0.378099431 |
| chr15:72971383-72971728 | -0.63260486 | 0.136020675 | 0.378105468 |
| chr16:86682835-86683035 | 0.785349959 | 0.136024604 | 0.378105468 |
| chr1:209433291-209433601 | -0.571996429 | 0.136041167 | 0.378125574 |
| chr12:131441047-131441261 | 0.670357429 | 0.13605025 | 0.378125574 |
| chr1:203256509-203257100 | -0.538596166 | 0.13605973 | 0.378125574 |
| chr1:192445894-192446179 | -0.671927268 | 0.136061802 | 0.378125574 |
| chr1:167601279-167601748 | -0.597792587 | 0.13606183 | 0.378125574 |
| chr2:161805728-161805967 | -0.768316791 | 0.136071925 | 0.378125574 |
| chr3:128794952-128795361 | -0.608145993 | 0.136073136 | 0.378125574 |
| chr1:165601683-165601933 | -0.595762483 | 0.136073389 | 0.378125574 |
| chr11:67645101-67645301 | 0.703148647 | 0.136076252 | 0.378125574 |
| chr6:20211318-20211649 | -0.628538595 | 0.136106071 | 0.378194719 |
| chr5:38371793-38372092 | -0.7758157 | 0.136123228 | 0.378228675 |
| chr15:88331211-88331477 | 0.622480158 | 0.136134935 | 0.378246098 |
| chr7:69775309-69775563 | -0.764372577 | 0.136149131 | 0.378246098 |
| chr1:106759062-106759268 | 0.549411653 | 0.136149227 | 0.378246098 |
| chr5:69166726-69167543 | -0.483241118 | 0.136151581 | 0.378246098 |
| chr17:18682092-18682534 | -0.573782787 | 0.136156969 | 0.378246098 |
| chr20:35447069-35447555 | -0.56688049 | 0.136159117 | 0.378246098 |
| chr2:108583924-108584265 | 0.572388054 | 0.136180823 | 0.378289632 |
| chr9:136799953-136800624 | 0.497899068 | 0.136185195 | 0.378289632 |
| chr20:53606441-53606777 | 0.492054801 | 0.13619118 | 0.378289632 |
| chr18:46324708-46324908 | -0.800839865 | 0.136194537 | 0.378289632 |
| chr8:110735217-110735637 | 0.612285858 | 0.136208559 | 0.378314867 |
| chr12:27383988-27384258 | -0.708904384 | 0.136228406 | 0.378356275 |
| chr1:205947001-205947418 | -0.513319633 | 0.136240436 | 0.378375972 |
| chr14:104137615-104137894 | 0.560688656 | 0.136252994 | 0.378397134 |
| chr14:104978317-104978614 | 0.553381316 | 0.136262527 | 0.378401195 |
| chr5:138752386-138752640 | -0.618946423 | 0.13626567 | 0.378401195 |
| chrX:19335092-19335318 | -0.790618097 | 0.136269272 | 0.378401195 |
| chr15:74048226-74048462 | 0.621687332 | 0.136276371 | 0.378407194 |
| chr3:63508340-63508719 | 0.532850818 | 0.136284601 | 0.378416332 |
| chr8:65912386-65912827 | -0.650060612 | 0.136311204 | 0.378476484 |
| chr1:201337694-201337976 | 0.614737799 | 0.13633899 | 0.378539917 |
| chr9:136438765-136439084 | 0.609666873 | 0.136400073 | 0.378695788 |

| | | | |
|---|---|---|---|
| chr2:88350595-88351530 | -0.499408015 | 0.136413029 | 0.378718038 |
| chr7:95527471-95527753 | -0.691182582 | 0.136436596 | 0.378769742 |
| chr11:36622843-36623043 | 0.762057988 | 0.13644367 | 0.378775656 |
| chr10:60810079-60810763 | -0.535243871 | 0.136461285 | 0.378810831 |
| chr10:930646-932419 | 0.467716167 | 0.136514226 | 0.378944064 |
| chr11:73185160-73186038 | -0.510016108 | 0.136530423 | 0.378975296 |
| chr19:48584219-48584591 | 0.559207534 | 0.136566589 | 0.379061953 |
| chr6:26026810-26027477 | -0.504265748 | 0.136581804 | 0.379090451 |
| chr13:52011305-52012542 | -0.477444165 | 0.136587347 | 0.379092104 |
| chr5:56270579-56271234 | -0.534206144 | 0.13659938 | 0.379111769 |
| chr13:75863692-75863892 | -0.857312512 | 0.136619962 | 0.379136562 |
| chr7:33091854-33092141 | 0.575548497 | 0.136621853 | 0.379136562 |
| chr1:248857810-248859583 | -0.4666598 | 0.136623158 | 0.379136562 |
| chr1:83819742-83819942 | -0.806769853 | 0.136631947 | 0.379143649 |
| chr3:72702560-72702760 | -0.810186663 | 0.136635608 | 0.379143649 |
| chr15:71079426-71079737 | -0.657317333 | 0.136641863 | 0.379147273 |
| chr2:128040321-128040527 | 0.702136176 | 0.136649842 | 0.379150006 |
| chr11:119018061-119019254 | 0.476636887 | 0.136652744 | 0.379150006 |
| chr18:2104810-2105190 | 0.599109329 | 0.136691217 | 0.379243018 |
| chr8:26053297-26053543 | -0.804100246 | 0.136721021 | 0.379311974 |
| chr11:9546114-9546323 | 0.745585817 | 0.136737935 | 0.379345163 |
| chr7:100691591-100691984 | -0.58255832 | 0.136749398 | 0.379359852 |
| chr7:83155431-83155713 | -0.788184856 | 0.13675888 | 0.379359852 |
| chr4:26161492-26161692 | 0.739669504 | 0.136763929 | 0.379359852 |
| chr11:61187335-61187855 | 0.56950954 | 0.136765756 | 0.379359852 |
| chr2:197785880-197786831 | -0.523431046 | 0.136767985 | 0.379359852 |
| chr2:237475270-237476092 | -0.515172689 | 0.136785492 | 0.379388449 |
| chr4:152050168-152050568 | 0.562474126 | 0.136789443 | 0.379388449 |
| chr22:22171444-22171901 | -0.616853496 | 0.136797063 | 0.379388449 |
| chr11:18775660-18776003 | -0.587922529 | 0.136798101 | 0.379388449 |
| chr1:169586195-169586896 | -0.517115342 | 0.136804246 | 0.379391759 |
| chr3:132642343-132642550 | 0.754592746 | 0.13681454 | 0.379392543 |
| chr5:53109394-53110245 | -0.479848987 | 0.136816675 | 0.379392543 |
| chr16:3112194-3112701 | 0.537372552 | 0.136819383 | 0.379392543 |
| chr17:7717098-7717972 | -0.503831289 | 0.13682835 | 0.379403679 |
| chr6:7618645-7618887 | -0.711845264 | 0.136845374 | 0.379437151 |
| chr9:107243799-107244190 | -0.535007233 | 0.136855813 | 0.379452365 |
| chr1:214658742-214659237 | -0.514002347 | 0.13687037 | 0.379469929 |
| chr12:50387039-50387343 | 0.679858422 | 0.136872053 | 0.379469929 |
| chr17:60515609-60515869 | 0.526623313 | 0.136893087 | 0.379514512 |
| chr3:57060444-57060722 | 0.668073407 | 0.13691277 | 0.379555347 |
| chr15:85296587-85296979 | 0.56710546 | 0.136932904 | 0.379597431 |
| chr16:85198478-85198690 | -0.678698529 | 0.13695176 | 0.379633622 |
| chr21:44455136-44455599 | 0.505949139 | 0.136955869 | 0.379633622 |
| chr10:91389681-91389935 | -0.696216857 | 0.136965756 | 0.379647296 |
| chr8:137492862-137493171 | -0.605308754 | 0.136974737 | 0.379658454 |
| chr8:38721331-38721686 | -0.646383134 | 0.136981073 | 0.379660258 |
| chrX:78157890-78158187 | -0.629194558 | 0.136985298 | 0.379660258 |
| chr17:64469738-64469960 | 0.645677743 | 0.137001782 | 0.379680145 |
| chr2:104965999-104966262 | 0.754325328 | 0.137002384 | 0.379680145 |

| | | | |
|---|---|---|---|
| chr13:20248104-20248483 | -0.684965762 | 0.137010911 | 0.379685829 |
| chr8:143596125-143596679 | 0.493317164 | 0.137014346 | 0.379685829 |
| chr2:167451493-167451818 | -0.661289973 | 0.137021647 | 0.37969233 |
| chr8:116703084-116703477 | -0.534969544 | 0.137058162 | 0.379770144 |
| chr9:93183895-93184310 | 0.553417232 | 0.137064313 | 0.379770144 |
| chrX:25144094-25144364 | -0.686686114 | 0.137067775 | 0.379770144 |
| chr17:3683066-3683302 | -0.747325856 | 0.137069554 | 0.379770144 |
| chr21:42495794-42496863 | -0.470488574 | 0.137082394 | 0.379791987 |
| chr21:31545059-31545345 | 0.647596892 | 0.137101367 | 0.379825775 |
| chr8:127809498-127810649 | 0.479812542 | 0.137104504 | 0.379825775 |
| chr6:25070730-25071044 | 0.645387645 | 0.137112107 | 0.379827323 |
| chr3:66499942-66500176 | 0.608499073 | 0.137114977 | 0.379827323 |
| chr12:71710614-71710921 | -0.738372129 | 0.13712165 | 0.379832076 |
| chr19:49581140-49581357 | 0.656034997 | 0.137131905 | 0.37984675 |
| chr2:30090878-30091176 | -0.655277268 | 0.137139435 | 0.379853877 |
| chr4:104904099-104904615 | -0.548185504 | 0.137152906 | 0.379877455 |
| chr13:42478453-42478846 | -0.60187745 | 0.137183829 | 0.379949372 |
| chr2:219003117-219003354 | -0.599877818 | 0.137194548 | 0.379957602 |
| chr5:163269724-163269944 | -0.798811495 | 0.137196719 | 0.379957602 |
| chr7:71736025-71736387 | -0.656379482 | 0.137211087 | 0.379983659 |
| chr7:137963023-137963536 | 0.545428936 | 0.137233909 | 0.380027167 |
| chr19:45927646-45928081 | 0.579323733 | 0.137238103 | 0.380027167 |
| chr6:139374178-139375716 | -0.469688253 | 0.137241676 | 0.380027167 |
| chr2:106461964-106462206 | -0.757002467 | 0.137281884 | 0.380119986 |
| chr7:49324671-49324893 | -0.788600129 | 0.137285119 | 0.380119986 |
| chr4:3037368-3037620 | 0.649522582 | 0.137298447 | 0.380143151 |
| chrX:27398937-27399254 | -0.6134106 | 0.137331105 | 0.380215222 |
| chr5:172666395-172666648 | -0.695729925 | 0.137334401 | 0.380215222 |
| chr5:52799343-52801081 | -0.471395039 | 0.137348487 | 0.380240481 |
| chr22:30605579-30605797 | 0.69469947 | 0.137360043 | 0.380242604 |
| chr11:117907161-117907612 | 0.508272996 | 0.137362744 | 0.380242604 |
| chr15:69973250-69973459 | 0.721790979 | 0.137377228 | 0.380242604 |
| chr4:138495022-138495326 | -0.632702507 | 0.137377523 | 0.380242604 |
| chr1:15834310-15834630 | 0.582836214 | 0.137378255 | 0.380242604 |
| chr9:22244219-22244447 | -0.773388938 | 0.137379029 | 0.380242604 |
| chr2:176821454-176821706 | -0.72215891 | 0.137399983 | 0.380266379 |
| chr12:6678943-6679181 | 0.810591428 | 0.137401376 | 0.380266379 |
| chr11:94237550-94238960 | 0.495258847 | 0.137402508 | 0.380266379 |
| chr9:75238202-75238889 | -0.530982209 | 0.137409108 | 0.38027091 |
| chr8:29087562-29087810 | -0.703795561 | 0.137418307 | 0.380282632 |
| chr10:26550818-26551245 | -0.592855497 | 0.137427456 | 0.380294216 |
| chr14:37211789-37212019 | 0.757062226 | 0.137480176 | 0.380414827 |
| chr12:107248730-107248930 | -0.792619008 | 0.137480971 | 0.380414827 |
| chr12:124321022-124321316 | 0.574211229 | 0.137489517 | 0.380424735 |
| chr1:229412105-229412421 | -0.588216858 | 0.137496712 | 0.380425279 |
| chr2:206559060-206560039 | 0.524437639 | 0.137499643 | 0.380425279 |
| chr17:82281754-82282169 | -0.56436039 | 0.137507876 | 0.380430038 |
| chr21:22399804-22400450 | 0.551477994 | 0.137511293 | 0.380430038 |
| chr5:172902690-172903220 | 0.503654474 | 0.137518457 | 0.380436121 |
| chr7:2860732-2860962 | 0.655629831 | 0.137530805 | 0.380456542 |

| | | | |
|---|---|---|---|
| chr10:5812729-5813971 | 0.472265068 | 0.137542375 | 0.380474814 |
| chr19:54209461-54209812 | 0.578226288 | 0.137561022 | 0.380480552 |
| chr4:164138469-164138669 | -0.829331874 | 0.137563524 | 0.380480552 |
| chr13:85828192-85828402 | -0.868548994 | 0.137570235 | 0.380480552 |
| chr14:48331249-48331739 | -0.597430233 | 0.137571822 | 0.380480552 |
| chr1:154335124-154335334 | 0.655635811 | 0.137572534 | 0.380480552 |
| chr14:39102454-39103718 | 0.470966798 | 0.137577876 | 0.380480552 |
| chr3:195784182-195784403 | 0.707264854 | 0.137579209 | 0.380480552 |
| chr12:79647910-79648203 | 0.648169939 | 0.137600074 | 0.380524238 |
| chr4:144174467-144174714 | -0.828768903 | 0.137604938 | 0.380524238 |
| chr14:74922761-74923448 | 0.505554141 | 0.137634077 | 0.38059108 |
| chr17:59876886-59877369 | 0.48935827 | 0.137649532 | 0.380616536 |
| chr7:154928234-154929461 | -0.483070768 | 0.137653217 | 0.380616536 |
| chr10:80599446-80599750 | -0.576379834 | 0.137670769 | 0.380644782 |
| chr11:49283275-49283650 | 0.577409805 | 0.137678958 | 0.380644782 |
| chr20:53959601-53959886 | 0.4962232 | 0.137679822 | 0.380644782 |
| chr9:35814904-35815656 | -0.483133558 | 0.137683304 | 0.380644782 |
| chr20:56818494-56818905 | -0.564524807 | 0.137728073 | 0.380754813 |
| chr14:104370271-104370520 | 0.600345289 | 0.137735621 | 0.380756499 |
| chr17:61536512-61536749 | 0.513829822 | 0.13774415 | 0.380756499 |
| chr3:45167506-45167850 | 0.59372166 | 0.137748833 | 0.380756499 |
| chr4:84531003-84531346 | -0.719905528 | 0.137749622 | 0.380756499 |
| chr18:46946554-46947396 | -0.504396517 | 0.137758549 | 0.380756499 |
| chr7:76358620-76359949 | -0.470691201 | 0.137762225 | 0.380756499 |
| chr2:240731519-240731723 | -0.810216489 | 0.137767984 | 0.380756499 |
| chr9:128655853-128656203 | 0.622724835 | 0.137768437 | 0.380756499 |
| chr15:70517229-70517562 | 0.554742625 | 0.137828783 | 0.38090954 |
| chr19:42360867-42361193 | -0.66919336 | 0.137840046 | 0.380913625 |
| chr8:109735238-109735570 | 0.546017247 | 0.137840204 | 0.380913625 |
| chr2:147843848-147844895 | -0.48884885 | 0.137847675 | 0.380920535 |
| chr3:71873135-71873457 | -0.71369248 | 0.137872579 | 0.380958623 |
| chr13:30395196-30395861 | -0.500909686 | 0.137873239 | 0.380958623 |
| chr3:64539582-64540044 | 0.49872725 | 0.137884976 | 0.380958623 |
| chr3:4338795-4339235 | -0.580129288 | 0.137885278 | 0.380958623 |
| chr1:222814872-222815614 | 0.505231136 | 0.137886318 | 0.380958623 |
| chr4:3942508-3942779 | -0.715685808 | 0.137908022 | 0.381004848 |
| chr1:220733010-220733468 | -0.533723158 | 0.137918899 | 0.381021159 |
| chr5:132184666-132184936 | -0.702887814 | 0.137953597 | 0.381103277 |
| chr19:10565725-10566210 | 0.5039875 | 0.137970804 | 0.38112369 |
| chr2:195600781-195601180 | -0.552577267 | 0.137970934 | 0.38112369 |
| chr12:104733694-104733948 | -0.659201218 | 0.138013248 | 0.381226833 |
| chr9:105557834-105558542 | -0.503174258 | 0.138040704 | 0.381277553 |
| chr16:50368311-50368552 | 0.682283198 | 0.138041562 | 0.381277553 |
| chr11:12122145-12122470 | -0.61200353 | 0.138049116 | 0.381284671 |
| chr10:86599739-86600008 | 0.62690812 | 0.138077426 | 0.381345035 |
| chr6:146598746-146599349 | -0.521782201 | 0.138080925 | 0.381345035 |
| chr3:88113524-88113926 | -0.574237365 | 0.138105748 | 0.381399842 |
| chr4:84503926-84504160 | 0.697111386 | 0.138113832 | 0.381408421 |
| chr3:25534046-25534339 | 0.701664502 | 0.138138582 | 0.38146302 |
| chr1:156691691-156692056 | 0.54649091 | 0.138163748 | 0.381518766 |

| | | | |
|---|---|---|---|
| chr7:2687480-2688438 | -0.500745933 | 0.138174063 | 0.381525651 |
| chr5:138032278-138033688 | -0.475910778 | 0.1381762 | 0.381525651 |
| chr2:215214221-215214771 | -0.515889868 | 0.138193966 | 0.381560956 |
| chr2:174486434-174487554 | -0.471188089 | 0.138228054 | 0.381641321 |
| chr15:58329875-58330837 | 0.476918765 | 0.13825354 | 0.381697934 |
| chr17:42566806-42567718 | 0.489657488 | 0.138281953 | 0.381755508 |
| chr10:57925026-57925308 | -0.645136182 | 0.138284359 | 0.381755508 |
| chr6:106085962-106086557 | -0.536273693 | 0.138300995 | 0.381787681 |
| chr3:64319484-64319833 | 0.502435928 | 0.138332488 | 0.381829449 |
| chr3:67790103-67790378 | -0.632204226 | 0.138333538 | 0.381829449 |
| chr15:80173356-80173661 | 0.577670557 | 0.138344246 | 0.381829449 |
| chr1:241627574-241627778 | -0.797121597 | 0.138345412 | 0.381829449 |
| chr11:103491851-103492087 | -0.797121597 | 0.138345412 | 0.381829449 |
| chr8:112801149-112801498 | 0.591005841 | 0.138346025 | 0.381829449 |
| chr1:220281055-220281586 | -0.564087317 | 0.138354781 | 0.38183986 |
| chr19:45862876-45863524 | 0.496340477 | 0.138381781 | 0.38187985 |
| chr3:113551787-113552192 | 0.588751298 | 0.138382524 | 0.38187985 |
| chr16:89972384-89973172 | -0.486968986 | 0.138384223 | 0.38187985 |
| chrX:115227019-115227255 | 0.712556262 | 0.13841176 | 0.381911828 |
| chr5:95727997-95728213 | -0.647712909 | 0.138413792 | 0.381911828 |
| chr1:235083604-235084062 | 0.548960055 | 0.138420643 | 0.381911828 |
| chr2:178891310-178891725 | -0.613369197 | 0.138422023 | 0.381911828 |
| chr4:160849442-160849950 | -0.548310708 | 0.138425118 | 0.381911828 |
| chrX:102899819-102900977 | -0.496427976 | 0.138425717 | 0.381911828 |
| chr2:1591949-1592305 | -0.555804331 | 0.138437728 | 0.381931213 |
| chr11:60701321-60701538 | 0.690501465 | 0.138474699 | 0.382019458 |
| chr8:22383515-22383821 | -0.665289205 | 0.138495955 | 0.38206434 |
| chr20:18405690-18405965 | -0.752577695 | 0.138503555 | 0.382065796 |
| chr14:103613308-103613811 | 0.584514544 | 0.138506455 | 0.382065796 |
| chr2:238239666-238240282 | 0.527117371 | 0.138527225 | 0.382109333 |
| chr3:113847645-113847963 | -0.648812708 | 0.138534095 | 0.382114526 |
| chr14:26960232-26960812 | 0.498115027 | 0.138545396 | 0.38213194 |
| chr14:23171501-23171943 | -0.627271312 | 0.138560956 | 0.382148622 |
| chr15:98705971-98706171 | 0.698716382 | 0.138564571 | 0.382148622 |
| chr5:141475639-141476861 | -0.480100303 | 0.138566406 | 0.382148622 |
| chr1:39576752-39577082 | 0.572264422 | 0.138578188 | 0.38216736 |
| chr1:170369425-170369631 | 0.712375025 | 0.138588744 | 0.382182715 |
| chr1:205504240-205505393 | -0.469568663 | 0.138612787 | 0.38223526 |
| chr5:119268437-119269117 | -0.477188643 | 0.138629338 | 0.382267144 |
| chr2:176129897-176130974 | 0.505176839 | 0.138668731 | 0.382362009 |
| chr3:98013752-98014057 | -0.642274501 | 0.138673979 | 0.38236272 |
| chr17:62456135-62456439 | 0.619931325 | 0.138694644 | 0.382379174 |
| chr1:16513423-16514638 | 0.488686698 | 0.138694903 | 0.382379174 |
| chr14:42192704-42193673 | 0.491352501 | 0.138694918 | 0.382379174 |
| chr3:16316877-16317099 | 0.680963393 | 0.138720715 | 0.382406396 |
| chr1:3650024-3650226 | 0.679217573 | 0.138721406 | 0.382406396 |
| chr9:89648650-89648850 | 0.768930776 | 0.13872448 | 0.382406396 |
| chr10:79239263-79239507 | -0.59897241 | 0.138724755 | 0.382406396 |
| chr5:154940900-154941463 | -0.493711767 | 0.138730611 | 0.382408782 |
| chr20:6940033-6940360 | -0.628101099 | 0.138759031 | 0.38247336 |

| | | | |
|---|---|---|---|
| chr14:36655892-36656538 | 0.504474851 | 0.138775809 | 0.382505847 |
| chr10:127407715-127408036 | 0.738368172 | 0.138793374 | 0.382528168 |
| chr3:113288117-113288341 | 0.617696569 | 0.138797544 | 0.382528168 |
| chr6:125268267-125268488 | -0.691177559 | 0.138798884 | 0.382528168 |
| chr2:110854376-110854658 | 0.614856548 | 0.138805086 | 0.382531501 |
| chr12:56224019-56224495 | 0.501639126 | 0.138831708 | 0.382577736 |
| chr17:40518731-40518972 | 0.655129716 | 0.138831849 | 0.382577736 |
| chr1:184408469-184408879 | -0.655251188 | 0.138847858 | 0.382588219 |
| chr9:120579473-120580460 | 0.484110196 | 0.138854691 | 0.382588219 |
| chr20:57210543-57210748 | 0.497875626 | 0.138855643 | 0.382588219 |
| chr12:459484-459999 | -0.604266919 | 0.13885995 | 0.382588219 |
| chr1:188292636-188293361 | -0.538917437 | 0.138860619 | 0.382588219 |
| chr5:139765497-139766216 | -0.513243139 | 0.138877299 | 0.382610033 |
| chr7:92447187-92448595 | -0.47670763 | 0.138878524 | 0.382610033 |
| chr2:179410936-179411258 | -0.611837418 | 0.138909699 | 0.382682162 |
| chr17:44014178-44014541 | 0.622062424 | 0.138920777 | 0.382698921 |
| chr6:52663718-52663933 | 0.613156043 | 0.138931526 | 0.382714773 |
| chr6:104683160-104683410 | -0.664599783 | 0.138937353 | 0.382717065 |
| chr16:28211058-28211565 | 0.529554629 | 0.138948794 | 0.382734821 |
| chr11:83724649-83725288 | -0.525173947 | 0.138963634 | 0.382761938 |
| chr7:131689162-131689457 | -0.588359631 | 0.138971966 | 0.382771126 |
| chr3:64006707-64007066 | 0.496903634 | 0.138993149 | 0.382815709 |
| chr1:178055157-178055463 | -0.587793088 | 0.13900098 | 0.382823519 |
| chr10:91745059-91745754 | 0.511076861 | 0.139022916 | 0.38287017 |
| chr9:74952177-74953328 | -0.482924302 | 0.139033993 | 0.382886914 |
| chr2:49026729-49026931 | 0.744477739 | 0.139047018 | 0.382909021 |
| chr17:80236987-80237229 | 0.832112945 | 0.139061733 | 0.382935782 |
| chr1:39739435-39739642 | 0.775032663 | 0.139073432 | 0.382941542 |
| chr16:1430894-1431129 | 0.640033227 | 0.139073821 | 0.382941542 |
| chr2:204031172-204031483 | 0.729100376 | 0.139115651 | 0.383042958 |
| chr6:132279553-132280074 | 0.559266213 | 0.139137298 | 0.383088795 |
| chr1:159584346-159584677 | -0.789033701 | 0.139145632 | 0.383097974 |
| chr2:210476204-210476986 | -0.487285269 | 0.139200421 | 0.383226664 |
| chr16:362133-362707 | 0.519013835 | 0.139202377 | 0.383226664 |
| chr6:137218572-137219711 | -0.477512416 | 0.139208874 | 0.383230781 |
| chr20:15699413-15699820 | -0.620303851 | 0.139222966 | 0.383254318 |
| chr6:36680400-36680727 | 0.579258512 | 0.139227427 | 0.383254318 |
| chr5:169089471-169089746 | -0.657671014 | 0.139253423 | 0.383312106 |
| chr17:8746077-8746577 | -0.594026905 | 0.139268069 | 0.38333865 |
| chr20:32743850-32744109 | 0.597100157 | 0.139314726 | 0.3834533 |
| chr4:10410805-10411196 | -0.622953677 | 0.139336451 | 0.38348226 |
| chr7:129612412-129612999 | -0.543541751 | 0.139339919 | 0.38348226 |
| chr5:34244285-34245078 | -0.507318471 | 0.139340263 | 0.38348226 |
| chr8:101157703-101157903 | 0.646999493 | 0.139346782 | 0.383486429 |
| chr6:17101757-17102261 | -0.565959057 | 0.139368016 | 0.38353109 |
| chr3:9986393-9987229 | 0.489130411 | 0.139391249 | 0.38357543 |
| chr14:68978725-68978964 | 0.571970858 | 0.139394141 | 0.38357543 |
| chr1:188706717-188707128 | 0.560037748 | 0.13941499 | 0.383619025 |
| chr6:152485737-152486175 | -0.674433346 | 0.139430672 | 0.383648398 |
| chr10:28496369-28497090 | 0.499339877 | 0.139444355 | 0.383672269 |

| | | | |
|---|---|---|---|
| chr8:128269299-128269853 | -0.511202161 | 0.139450352 | 0.383674992 |
| chr2:210181527-210182222 | -0.555486051 | 0.139458142 | 0.38367658 |
| chr10:77832271-77832524 | -0.71946988 | 0.139468982 | 0.38367658 |
| chr11:102109897-102112080 | -0.470686282 | 0.139470779 | 0.38367658 |
| chr13:67230023-67231334 | -0.469484429 | 0.139470958 | 0.38367658 |
| chr2:221205187-221205387 | 0.786426807 | 0.139478846 | 0.383684502 |
| chr7:152597397-152597897 | -0.618412123 | 0.139499914 | 0.383728682 |
| chr1:204257718-204257929 | 0.596846331 | 0.139521518 | 0.38377433 |
| chr14:105605647-105606224 | 0.562607221 | 0.139569519 | 0.383891876 |
| chr16:30064956-30065871 | 0.475609637 | 0.139574272 | 0.383891876 |
| chr1:107219532-107220272 | 0.481212152 | 0.139581716 | 0.383898571 |
| chr16:30096106-30096441 | 0.56690394 | 0.139605257 | 0.383949535 |
| chr1:171353195-171353444 | 0.732321668 | 0.139649165 | 0.384043319 |
| chr5:111757163-111758398 | -0.470156756 | 0.139649382 | 0.384043319 |
| chr5:91280370-91281012 | -0.494822738 | 0.139662434 | 0.384065429 |
| chr1:158348508-158348708 | 0.774570193 | 0.139671364 | 0.3840762 |
| chr21:36319802-36320859 | 0.475077524 | 0.139698849 | 0.384116211 |
| chr3:48609003-48609909 | 0.509659294 | 0.139700959 | 0.384116211 |
| chr2:121257361-121257561 | 0.767156574 | 0.139706591 | 0.384116211 |
| chr13:47385894-47386294 | 0.580824481 | 0.13970817 | 0.384116211 |
| chr15:78241015-78241246 | 0.682218064 | 0.13971098 | 0.384116211 |
| chr5:149962083-149962313 | -0.627044006 | 0.139723705 | 0.384137413 |
| chr3:33131667-33131990 | 0.619533803 | 0.139729478 | 0.38413931 |
| chr3:87186463-87187292 | -0.535845992 | 0.139734421 | 0.38413931 |
| chr10:68223671-68223922 | -0.706586994 | 0.139747922 | 0.38416264 |
| chr2:88187132-88187398 | -0.587056943 | 0.139753997 | 0.384165558 |
| chr3:142077465-142077665 | 0.80813409 | 0.139766893 | 0.384168623 |
| chr10:102644914-102645245 | 0.523775412 | 0.139769336 | 0.384168623 |
| chr9:127980327-127981530 | 0.48262588 | 0.139770153 | 0.384168623 |
| chr20:33666609-33667689 | 0.484505053 | 0.139786889 | 0.384190523 |
| chr1:99728820-99729144 | -0.589144612 | 0.13978815 | 0.384190523 |
| chr11:69515149-69515661 | -0.525370265 | 0.139803726 | 0.38421955 |
| chr5:135570159-135570514 | -0.565441468 | 0.139820295 | 0.384243217 |
| chr5:96112314-96112531 | -0.746681807 | 0.139822367 | 0.384243217 |
| chr14:24391899-24392224 | 0.572846776 | 0.139839791 | 0.38427554 |
| chr1:211260626-211261107 | -0.526451188 | 0.139844159 | 0.38427554 |
| chr6:48097770-48098115 | -0.759267933 | 0.139861127 | 0.384308382 |
| chr14:64515121-64515364 | 0.636838314 | 0.139881501 | 0.384350583 |
| chr1:24502152-24503152 | 0.504753461 | 0.139892195 | 0.384353737 |
| chr7:16794508-16795274 | 0.502342396 | 0.139897666 | 0.384353737 |
| chr10:94963818-94964201 | 0.565018808 | 0.139897697 | 0.384353737 |
| chr14:90757463-90757708 | -0.595056358 | 0.139913003 | 0.384382005 |
| chr3:117399369-117399686 | -0.743997442 | 0.139923013 | 0.384395724 |
| chr13:32911530-32911891 | -0.543524339 | 0.139983539 | 0.38454774 |
| chr14:43789453-43789728 | 0.616887134 | 0.139988386 | 0.38454774 |
| chr13:62250531-62250813 | -0.759201768 | 0.140001916 | 0.38457112 |
| chr6:152301520-152302562 | -0.486506295 | 0.140018912 | 0.384587992 |
| chr2:237572001-237572293 | -0.597825121 | 0.140020312 | 0.384587992 |
| chrX:43505833-43506139 | -0.715931262 | 0.140023116 | 0.384587992 |
| chr4:54683695-54684037 | -0.637464649 | 0.140034345 | 0.384605047 |

| | | | |
|---|---|---|---|
| chr17:69776763-69776977 | 0.755009053 | 0.140047714 | 0.384624243 |
| chr2:5842540-5842859 | 0.761540351 | 0.140051374 | 0.384624243 |
| chr11:86743854-86744056 | -0.794057556 | 0.140084658 | 0.384701863 |
| chr7:128425872-128426157 | -0.587734923 | 0.140099612 | 0.384715649 |
| chr2:27582806-27583197 | 0.569558717 | 0.14009972 | 0.384715649 |
| chr7:48528827-48529050 | -0.796554304 | 0.140107519 | 0.384723278 |
| chr4:139227766-139228093 | -0.604367815 | 0.140118469 | 0.384739558 |
| chr1:152001785-152002035 | 0.545878152 | 0.140134874 | 0.384770813 |
| chr17:44775517-44776171 | 0.482559293 | 0.140146245 | 0.384788247 |
| chr1:106914776-106915042 | 0.549582416 | 0.140155868 | 0.384800879 |
| chr19:12774975-12775803 | 0.49029026 | 0.140189109 | 0.384873948 |
| chr7:105313975-105314293 | -0.621061744 | 0.140192528 | 0.384873948 |
| chr1:180105077-180105497 | -0.58636441 | 0.140203702 | 0.384890833 |
| chr2:153166638-153166909 | -0.759562167 | 0.140222073 | 0.384927475 |
| chr9:133478642-133479684 | 0.486179542 | 0.140237445 | 0.384955881 |
| chr9:71782936-71783646 | -0.508374978 | 0.140243233 | 0.384957979 |
| chr18:76274685-76275048 | -0.621686439 | 0.14026313 | 0.384998802 |
| chr20:52312170-52312391 | 0.649362088 | 0.140278322 | 0.385026708 |
| chr2:175203589-175203988 | -0.588934161 | 0.140288499 | 0.385040849 |
| chr15:96052925-96053747 | 0.511224349 | 0.140299177 | 0.385056362 |
| chr3:107078616-107078851 | -0.794061039 | 0.140310117 | 0.385072595 |
| chr12:87848382-87848789 | 0.615307378 | 0.140335088 | 0.385127333 |
| chr3:128504990-128505295 | -0.638493006 | 0.140351582 | 0.385158802 |
| chr7:95199887-95200116 | -0.723011743 | 0.140371942 | 0.385200879 |
| chr17:7483794-7485012 | -0.470954429 | 0.140378987 | 0.385206415 |
| chr8:143013190-143013485 | -0.612388057 | 0.140396877 | 0.385232651 |
| chr8:118029794-118030008 | 0.729977643 | 0.140398603 | 0.385232651 |
| chr1:106210633-106210981 | 0.549216445 | 0.140407401 | 0.385242996 |
| chr14:103091422-103091854 | -0.538575153 | 0.140429509 | 0.385289857 |
| chr7:8217331-8217782 | 0.562271739 | 0.140434564 | 0.385289931 |
| chr6:143449693-143450249 | -0.506093904 | 0.140450263 | 0.385319204 |
| chr19:5261218-5261542 | 0.590947404 | 0.140463623 | 0.385342059 |
| chr14:92951747-92952257 | 0.524056163 | 0.140481182 | 0.385376432 |
| chr13:40536677-40537122 | 0.570718998 | 0.140497369 | 0.385403149 |
| chr7:10720135-10720335 | 0.713678864 | 0.140500981 | 0.385403149 |
| chr22:27474849-27475396 | 0.546198708 | 0.140507957 | 0.385408486 |
| chr8:107497852-107498131 | 0.572757054 | 0.140536464 | 0.385472796 |
| chr19:11056357-11056573 | -0.808087698 | 0.140541464 | 0.385472796 |
| chr8:126331266-126331475 | 0.663752245 | 0.14056503 | 0.385523633 |
| chr17:1574610-1574822 | -0.761211373 | 0.140580309 | 0.385551738 |
| chr19:2049666-2050203 | 0.533346742 | 0.140597986 | 0.38557449 |
| chr4:143531435-143531687 | -0.649763388 | 0.140601144 | 0.38557449 |
| chr19:18279550-18279788 | 0.5822039 | 0.140603799 | 0.38557449 |
| chr1:78004006-78005050 | -0.481300981 | 0.140608734 | 0.38557449 |
| chr21:41584503-41584703 | 0.708548323 | 0.14066178 | 0.38570615 |
| chr2:241637422-241638129 | 0.500460737 | 0.140684171 | 0.38575374 |
| chr14:89363179-89363379 | 0.737965635 | 0.140690799 | 0.38575811 |
| chr1:174799557-174799980 | -0.556744496 | 0.140727112 | 0.385843868 |
| chr18:48139321-48139704 | -0.595850234 | 0.14076401 | 0.385931225 |
| chr1:107104283-107104496 | 0.569136766 | 0.140796223 | 0.386005732 |

| | | | |
|---|---|---|---|
| chr2:179125019-179125280 | -0.745515733 | 0.140820251 | 0.386054135 |
| chr17:28416760-28417161 | -0.589078795 | 0.140823955 | 0.386054135 |
| chr8:94719356-94720485 | 0.467668285 | 0.140832062 | 0.386062547 |
| chr7:2321082-2321424 | 0.590339461 | 0.140880468 | 0.386179587 |
| chr5:101292258-101292739 | -0.607291104 | 0.140884837 | 0.386179587 |
| chr1:56569439-56570203 | -0.4797099 | 0.140892813 | 0.386187634 |
| chr6:154370621-154370876 | -0.687793576 | 0.140924581 | 0.386260893 |
| chr6:51934201-51934424 | -0.747138868 | 0.140969806 | 0.386368252 |
| chr2:133780193-133780413 | -0.703187758 | 0.140976 | 0.386368252 |
| chr3:148943552-148943922 | -0.524922181 | 0.140978885 | 0.386368252 |
| chr10:73592014-73592214 | -0.792275086 | 0.140988963 | 0.386368252 |
| chr13:91045560-91045807 | -0.792275086 | 0.140988963 | 0.386368252 |
| chr1:35715521-35715809 | 0.576232057 | 0.141025062 | 0.386453355 |
| chrX:96182424-96182745 | -0.584057561 | 0.141039342 | 0.386478665 |
| chr13:72781323-72782396 | -0.471278457 | 0.141049458 | 0.386492564 |
| chr3:120796024-120796370 | 0.626454247 | 0.141057457 | 0.386500661 |
| chr20:9884678-9884962 | -0.670559184 | 0.141062648 | 0.386501063 |
| chr1:214123205-214123580 | -0.663579147 | 0.14108677 | 0.386553333 |
| chr5:176837440-176837955 | -0.546262286 | 0.141123689 | 0.386568107 |
| chr6:136207784-136208132 | -0.729379185 | 0.141124362 | 0.386568107 |
| chr3:49018042-49018260 | 0.61738535 | 0.141125698 | 0.386568107 |
| chr10:72222945-72223145 | -0.847603033 | 0.141129042 | 0.386568107 |
| chr16:83606745-83607007 | -0.648785703 | 0.141135816 | 0.386568107 |
| chr3:159191524-159191790 | -0.711571616 | 0.141137349 | 0.386568107 |
| chr2:142130142-142131799 | -0.475464943 | 0.141139272 | 0.386568107 |
| chr5:141960718-141961082 | -0.557281938 | 0.141141461 | 0.386568107 |
| chr4:88290972-88291355 | -0.599782771 | 0.141142581 | 0.386568107 |
| chr18:77262881-77263438 | -0.615590988 | 0.141142614 | 0.386568107 |
| chr2:115194642-115194939 | -0.668689954 | 0.141156342 | 0.386591886 |
| chr1:223081162-223081709 | 0.494185641 | 0.141172433 | 0.386622137 |
| chr4:77240473-77240805 | -0.591654798 | 0.14118145 | 0.386633013 |
| chr10:88042550-88042827 | -0.659852583 | 0.141210101 | 0.386667405 |
| chr3:161883927-161884189 | 0.762933914 | 0.141211648 | 0.386667405 |
| chr3:173287400-173287739 | -0.63191038 | 0.141214066 | 0.386667405 |
| chr8:126488146-126488532 | 0.561322579 | 0.141214619 | 0.386667405 |
| chr4:110586059-110586529 | -0.595277843 | 0.141219241 | 0.386667405 |
| chrX:110401232-110401432 | 0.746633738 | 0.1412298 | 0.3866825 |
| chr10:100759527-100759783 | 0.633036009 | 0.141243219 | 0.38669513 |
| chr17:81097409-81097890 | -0.542810906 | 0.141244507 | 0.38669513 |
| chr14:39989061-39989267 | 0.749918448 | 0.141272866 | 0.386758951 |
| chr8:61789433-61789702 | -0.643171267 | 0.141285785 | 0.386780501 |
| chr19:45642070-45642404 | 0.536511799 | 0.141301811 | 0.386809417 |
| chr16:4288390-4288736 | 0.566445151 | 0.141307857 | 0.386809417 |
| chr7:51234993-51235199 | -0.732482 | 0.141311493 | 0.386809417 |
| chrX:130109687-130110068 | 0.546735003 | 0.141319734 | 0.386818157 |
| chr2:85531058-85531617 | -0.588548487 | 0.141330667 | 0.386834264 |
| chr8:58701797-58702110 | -0.58533441 | 0.141342424 | 0.386852625 |
| chr1:151060209-151060433 | 0.569754964 | 0.141352803 | 0.386854861 |
| chr20:57654332-57654607 | 0.52948097 | 0.141357528 | 0.386854861 |
| chr3:63152053-63152288 | 0.595761097 | 0.141362868 | 0.386854861 |

| | | | |
|---|---|---|---|
| chr5:130526700-130527088 | -0.557891854 | 0.141363437 | 0.386854861 |
| chr2:206353802-206354137 | -0.716410503 | 0.141370885 | 0.386861427 |
| chr20:57109110-57109643 | 0.469529184 | 0.141382647 | 0.386879796 |
| chr2:201158397-201158693 | -0.616863353 | 0.141395701 | 0.386880602 |
| chr21:44845743-44846171 | 0.540479777 | 0.141397084 | 0.386880602 |
| chr3:107470510-107470914 | -0.588314292 | 0.141398089 | 0.386880602 |
| chrX:80156822-80157437 | -0.56466626 | 0.141430704 | 0.386938688 |
| chr20:56787603-56787958 | 0.533228604 | 0.141431497 | 0.386938688 |
| chr4:137819876-137820275 | 0.562958483 | 0.141434468 | 0.386938688 |
| chr6:73876077-73876519 | -0.579441957 | 0.141454636 | 0.386980046 |
| chr3:135945224-135945587 | 0.684043515 | 0.14146275 | 0.386988425 |
| chr3:98812869-98813331 | -0.576253604 | 0.141481778 | 0.387026661 |
| chr17:69776392-69776592 | 0.786920187 | 0.141503859 | 0.387073246 |
| chr1:173076421-173076624 | 0.78290413 | 0.14151709 | 0.38709562 |
| chr1:86504990-86505208 | 0.707002266 | 0.141526547 | 0.387097988 |
| chr15:43511589-43511877 | 0.548760374 | 0.14152806 | 0.387097988 |
| chr16:10256839-10257133 | 0.632280259 | 0.141558238 | 0.387146411 |
| chr1:232334780-232335306 | -0.553717478 | 0.141560438 | 0.387146411 |
| chr15:90291649-90291951 | 0.565588693 | 0.141560922 | 0.387146411 |
| chr2:143705069-143705269 | 0.790755902 | 0.141568484 | 0.387153272 |
| chrX:86969327-86969901 | 0.569844271 | 0.141580129 | 0.387171301 |
| chr14:36520562-36520901 | 0.549500169 | 0.141612442 | 0.387238114 |
| chr1:109257442-109257676 | -0.609711807 | 0.141614669 | 0.387238114 |
| chr21:31798948-31799180 | -0.790345726 | 0.141648163 | 0.387281242 |
| chr14:85248923-85249316 | -0.604724534 | 0.141649966 | 0.387281242 |
| chr2:88919324-88919673 | -0.665616439 | 0.141650119 | 0.387281242 |
| chr21:33011491-33011730 | 0.730297184 | 0.141650659 | 0.387281242 |
| chr10:116026381-116026620 | -0.704099124 | 0.141664485 | 0.387305222 |
| chr10:105797304-105797830 | -0.574679553 | 0.141704625 | 0.387392169 |
| chr5:158013975-158014415 | -0.574487289 | 0.141709765 | 0.387392169 |
| chr7:141221086-141221324 | -0.729346255 | 0.141711455 | 0.387392169 |
| chr10:19231171-19231616 | -0.621256316 | 0.14172072 | 0.387403674 |
| chr13:33228604-33228871 | -0.60639672 | 0.141742627 | 0.387449212 |
| chr19:38401171-38401429 | -0.621494913 | 0.141747492 | 0.387449212 |
| chr1:192805890-192806153 | -0.570312327 | 0.141767462 | 0.38748073 |
| chr3:118213965-118214189 | -0.725473265 | 0.141769137 | 0.38748073 |
| chr20:47319001-47319209 | 0.49740379 | 0.141784559 | 0.387509061 |
| chr1:207822720-207823042 | 0.527582376 | 0.141793544 | 0.387519793 |
| chr7:74254047-74254930 | -0.493997466 | 0.141804131 | 0.387524023 |
| chr20:53967248-53967496 | 0.499422412 | 0.141805207 | 0.387524023 |
| chrX:136726992-136727248 | -0.714465894 | 0.141844153 | 0.387616629 |
| chr5:78222175-78222464 | -0.708587858 | 0.1418518 | 0.387623703 |
| chr17:28471727-28471935 | -0.77310935 | 0.14186213 | 0.387638106 |
| chr2:2899918-2900118 | -0.846628582 | 0.141882472 | 0.387679866 |
| chrX:153744377-153744593 | 0.663392646 | 0.141888914 | 0.387683643 |
| chr5:56996345-56996602 | -0.68597445 | 0.14190905 | 0.387711833 |
| chr7:149489429-149489844 | -0.568241906 | 0.141909352 | 0.387711833 |
| chr14:77070521-77070972 | 0.522255728 | 0.141936307 | 0.387771651 |
| chr2:67846162-67846791 | -0.5275634 | 0.14198991 | 0.387904264 |
| chr2:113806552-113806932 | 0.577090961 | 0.142006268 | 0.387935121 |

| | | | |
|---|---|---|---|
| chr2:28958695-28958986 | -0.577088548 | 0.142022339 | 0.387955917 |
| chr21:9366080-9366399 | 0.604942369 | 0.142024007 | 0.387955917 |
| chr7:76047667-76048273 | -0.490533285 | 0.142066378 | 0.388049305 |
| chr1:179251791-179251991 | 0.76915606 | 0.142068324 | 0.388049305 |
| chr5:30693787-30694217 | 0.580143221 | 0.142081526 | 0.388058218 |
| chr1:53811593-53811965 | -0.651393515 | 0.142081847 | 0.388058218 |
| chr11:88418537-88418948 | -0.580073003 | 0.14209259 | 0.388058218 |
| chr6:34527106-34527644 | -0.506585072 | 0.142094626 | 0.388058218 |
| chr10:52999343-52999624 | -0.789091889 | 0.142103652 | 0.388058218 |
| chrX:10633786-10634061 | 0.642376086 | 0.14210585 | 0.388058218 |
| chr3:99532188-99532625 | -0.533357471 | 0.142107039 | 0.388058218 |
| chr10:125900971-125901396 | 0.534625153 | 0.142113966 | 0.388063303 |
| chr18:76821674-76822598 | -0.499432514 | 0.142126371 | 0.388083348 |
| chrX:107725785-107726214 | 0.514233649 | 0.142131744 | 0.388084189 |
| chr10:134605-135089 | 0.506214396 | 0.142154157 | 0.388113848 |
| chr9:136644459-136644659 | 0.692557474 | 0.142155021 | 0.388113848 |
| chr20:53639943-53640261 | 0.48306165 | 0.142157802 | 0.388113848 |
| chrX:118345329-118346832 | 0.46642651 | 0.142174466 | 0.388145512 |
| chr7:5366073-5366321 | -0.606982188 | 0.142180025 | 0.388146858 |
| chrX:96575277-96575477 | -0.83509412 | 0.142185413 | 0.388147739 |
| chr21:39867188-39867621 | -0.527547149 | 0.142200065 | 0.388161785 |
| chr5:67493746-67494011 | -0.646818015 | 0.14220069 | 0.388161785 |
| chr2:177106094-177106580 | -0.569329003 | 0.142215022 | 0.388187076 |
| chr10:103482023-103482382 | 0.580063482 | 0.142229249 | 0.38821208 |
| chr14:50025842-50026069 | -0.718084527 | 0.142235311 | 0.388214796 |
| chr1:85027780-85027995 | -0.718052394 | 0.142265478 | 0.388283303 |
| chr6:7106863-7107984 | -0.490309464 | 0.142282244 | 0.388289834 |
| chr5:7015769-7016157 | 0.642034562 | 0.142282316 | 0.388289834 |
| chr2:118865157-118865440 | 0.624072155 | 0.142283287 | 0.388289834 |
| chr8:109690933-109692257 | -0.467594228 | 0.142288141 | 0.388289834 |
| chr8:39763610-39763959 | -0.668283621 | 0.142319844 | 0.388362517 |
| chr2:131413160-131413446 | -0.65687026 | 0.14233368 | 0.388381663 |
| chr14:90503641-90503925 | -0.632706308 | 0.142336998 | 0.388381663 |
| chr8:19757793-19758141 | 0.568417276 | 0.142346227 | 0.388382361 |
| chr11:71745658-71745998 | 0.560113634 | 0.142352425 | 0.388382361 |
| chr5:784554-785513 | 0.492206856 | 0.142352461 | 0.388382361 |
| chr18:5960859-5961059 | -0.873448934 | 0.142361 | 0.38839183 |
| chr7:129343106-129343345 | -0.749627097 | 0.142376089 | 0.388394679 |
| chrX:69035039-69035420 | -0.726685819 | 0.142377282 | 0.388394679 |
| chr2:161308052-161309005 | -0.479482633 | 0.142381645 | 0.388394679 |
| chr22:20064868-20065115 | 0.705485493 | 0.14238232 | 0.388394679 |
| chr10:6014252-6014452 | 0.692016576 | 0.142392363 | 0.388408245 |
| chr17:64037330-64037594 | 0.595008753 | 0.142406424 | 0.3884204 |
| chrX:119693064-119693501 | 0.564932668 | 0.142406957 | 0.3884204 |
| chr10:77005008-77005285 | -0.651096511 | 0.142417458 | 0.388425248 |
| chr5:75167418-75167699 | -0.678956467 | 0.142418873 | 0.388425248 |
| chr1:106924387-106924661 | 0.548115244 | 0.14242858 | 0.388437895 |
| chr7:3628486-3629032 | -0.601891035 | 0.142437042 | 0.388445903 |
| chr6:3053717-3054033 | 0.59550675 | 0.142441656 | 0.388445903 |
| chr5:141702134-141702613 | -0.56543176 | 0.14249434 | 0.38856282 |

| | | | |
|---|---|---|---|
| chr1:146911147-146911347 | 0.758047262 | 0.142494671 | 0.38856282 |
| chr1:223204630-223204918 | 0.747456603 | 0.142514434 | 0.388602881 |
| chr11:84687935-84688152 | -0.694862736 | 0.142566234 | 0.388707951 |
| chr13:96822938-96823188 | -0.702518452 | 0.142569174 | 0.388707951 |
| chr11:67410140-67410386 | 0.583157484 | 0.142578408 | 0.388707951 |
| chr2:219429336-219429567 | 0.687964416 | 0.14258173 | 0.388707951 |
| chr20:36575477-36575802 | 0.573056545 | 0.142582016 | 0.388707951 |
| chr2:19305478-19305965 | -0.578879133 | 0.142584317 | 0.388707951 |
| chr2:191631714-191632322 | -0.536585798 | 0.142588478 | 0.388707951 |
| chr6:160072320-160072632 | -0.667796189 | 0.142612696 | 0.38876014 |
| chr22:20567388-20567827 | 0.524724128 | 0.142618884 | 0.388763178 |
| chr13:19580010-19580262 | -0.750043578 | 0.142634489 | 0.388790328 |
| chr13:52651970-52653278 | -0.465759649 | 0.142642988 | 0.388790328 |
| chr13:33328778-33329058 | -0.740103234 | 0.142647939 | 0.388790328 |
| chr11:448010-448655 | 0.498006363 | 0.142649141 | 0.388790328 |
| chr11:45340079-45340415 | 0.569875038 | 0.14267236 | 0.388839779 |
| chrX:91414792-91414999 | -0.777441308 | 0.142681816 | 0.38885172 |
| chr10:35089883-35090994 | -0.485928903 | 0.142700307 | 0.388888281 |
| chr8:41529724-41529936 | 0.6573484 | 0.142726698 | 0.388943596 |
| chr7:1922370-1922740 | 0.553788902 | 0.14274065 | 0.388943596 |
| chr2:183951475-183952005 | -0.588238714 | 0.142740728 | 0.388943596 |
| chr13:99496244-99496444 | -0.627064764 | 0.142740909 | 0.388943596 |
| chr10:124864369-124864609 | 0.681759891 | 0.142749427 | 0.388950106 |
| chr17:59921420-59921793 | 0.49648354 | 0.14275345 | 0.388950106 |
| chr2:25000731-25001148 | -0.640273749 | 0.1427589 | 0.388951122 |
| chr6:41223769-41224131 | -0.691835399 | 0.142766269 | 0.388957371 |
| chr21:38139539-38140489 | 0.473545628 | 0.14277196 | 0.388959044 |
| chr8:119958026-119958255 | -0.798687911 | 0.142778529 | 0.38896311 |
| chr1:108830229-108830937 | 0.50006424 | 0.142818869 | 0.389059174 |
| chr13:95704563-95704767 | -0.821303458 | 0.142841931 | 0.389108163 |
| chr5:53799461-53800025 | -0.518271003 | 0.142851371 | 0.389120045 |
| chr16:23905077-23905436 | 0.650333551 | 0.142861264 | 0.389133158 |
| chr2:3756215-3756469 | 0.711341492 | 0.142875089 | 0.389156983 |
| chr5:173075569-173075833 | 0.661597105 | 0.142912951 | 0.389223 |
| chr14:59333772-59334010 | -0.806428101 | 0.142913643 | 0.389223 |
| chr1:220391377-220391834 | -0.62677016 | 0.142914566 | 0.389223 |
| chr9:66269767-66270742 | -0.499212881 | 0.142930883 | 0.389243919 |
| chr3:32240143-32240502 | 0.551978359 | 0.142932407 | 0.389243919 |
| chr18:48975652-48976352 | -0.519223226 | 0.142945565 | 0.389265915 |
| chr15:40358501-40358774 | 0.663397251 | 0.142953654 | 0.389274109 |
| chr11:20158685-20158953 | -0.696518298 | 0.142999197 | 0.389384288 |
| chr1:107162799-107163038 | 0.539820843 | 0.143012356 | 0.38940628 |
| chr8:113546561-113546888 | 0.635768479 | 0.143024192 | 0.38942467 |
| chr1:201909820-201910126 | -0.722399095 | 0.143043225 | 0.389425055 |
| chr2:16039085-16039285 | -0.793832702 | 0.143045522 | 0.389425055 |
| chr9:33290003-33291171 | -0.469193288 | 0.143049141 | 0.389425055 |
| chr10:71956132-71956438 | -0.629048867 | 0.143049552 | 0.389425055 |
| chr20:57229513-57229884 | 0.473541225 | 0.143049745 | 0.389425055 |
| chr2:70329107-70329510 | -0.648112385 | 0.143059403 | 0.38943751 |
| chr19:42280344-42280641 | 0.539980218 | 0.143076606 | 0.389470503 |

| | | | |
|---|---|---|---|
| chr9:129619976-129620204 | 0.661350557 | 0.143092118 | 0.38949889 |
| chr13:51222178-51222642 | -0.570571781 | 0.143097481 | 0.389499653 |
| chr6:45423213-45423462 | 0.618284979 | 0.143105935 | 0.389508824 |
| chr1:216093113-216093612 | -0.601675085 | 0.143122337 | 0.389539633 |
| chr16:89833590-89834351 | -0.489496289 | 0.143142343 | 0.389570925 |
| chr4:133069512-133069890 | 0.653698033 | 0.143146944 | 0.389570925 |
| chr4:146663728-146664363 | -0.545780423 | 0.143151152 | 0.389570925 |
| chr3:44861513-44862035 | -0.523981184 | 0.143154172 | 0.389570925 |
| chr3:118981571-118981978 | -0.634502493 | 0.143161926 | 0.389578189 |
| chr12:95534432-95534908 | -0.612470728 | 0.143175086 | 0.389600166 |
| chr21:46347739-46348071 | 0.589634026 | 0.143196971 | 0.389641458 |
| chr17:31095320-31095653 | 0.615195056 | 0.143200431 | 0.389641458 |
| chr10:10313509-10313823 | -0.645484084 | 0.143262651 | 0.389796913 |
| chr11:65662769-65663431 | 0.494742238 | 0.143272004 | 0.389799662 |
| chr3:197510183-197510383 | 0.671415316 | 0.143276782 | 0.389799662 |
| chr3:151856131-151856656 | 0.549040824 | 0.143278923 | 0.389799662 |
| chr1:6108915-6109352 | 0.582912241 | 0.143304909 | 0.389856514 |
| chr1:114032900-114033671 | 0.469140467 | 0.143317003 | 0.389875575 |
| chr9:136728775-136728976 | 0.643541078 | 0.14332495 | 0.389882268 |
| chr2:36618396-36618726 | -0.540262708 | 0.143329641 | 0.389882268 |
| chr1:42127791-42127991 | 0.727238354 | 0.143361545 | 0.389945726 |
| chr5:163461338-163461844 | -0.496540602 | 0.143363148 | 0.389945726 |
| chr17:64649168-64649470 | -0.589886713 | 0.143393162 | 0.389999198 |
| chr20:35284399-35285230 | 0.493016446 | 0.143397709 | 0.389999198 |
| chr1:204294384-204294735 | -0.569303917 | 0.143398076 | 0.389999198 |
| chr10:77756449-77756802 | -0.626020286 | 0.143411734 | 0.390002087 |
| chr1:37556789-37557068 | -0.640887928 | 0.143415321 | 0.390002087 |
| chr1:3530472-3530701 | 0.64113114 | 0.143415467 | 0.390002087 |
| chr13:100722632-100722885 | -0.800304232 | 0.143422033 | 0.390002087 |
| chr1:198130333-198130669 | -0.689651093 | 0.143425574 | 0.390002087 |
| chr18:46425015-46425302 | -0.620830005 | 0.143429678 | 0.390002087 |
| chr5:134227314-134227546 | -0.578174679 | 0.14344896 | 0.390040675 |
| chr11:47426102-47427337 | 0.466252878 | 0.14348391 | 0.39012186 |
| chr2:17365869-17366196 | -0.684758775 | 0.143493779 | 0.390134849 |
| chr7:82729501-82729711 | -0.764068438 | 0.143517998 | 0.390186853 |
| chr12:67468595-67468795 | 0.791121825 | 0.143526259 | 0.390195467 |
| chr8:125504585-125504785 | 0.635545136 | 0.143552983 | 0.390254272 |
| chr15:69576043-69576253 | 0.672879731 | 0.14356826 | 0.390281955 |
| chr21:41816419-41816877 | 0.570326959 | 0.14357637 | 0.390290155 |
| chr20:45300958-45301355 | 0.541253138 | 0.143591899 | 0.390318521 |
| chr1:26417009-26417248 | 0.756766668 | 0.143619383 | 0.390379382 |
| chr12:40224446-40225215 | -0.511362169 | 0.143637305 | 0.390414246 |
| chr10:93086880-93087080 | 0.78461196 | 0.143669233 | 0.390487175 |
| chr19:39539848-39540425 | 0.501765498 | 0.143676448 | 0.390492867 |
| chr5:116084291-116085566 | -0.473393807 | 0.14368152 | 0.390492867 |
| chr3:149367017-149367428 | -0.564755829 | 0.14369832 | 0.390522886 |
| chr18:53380134-53380400 | -0.739951765 | 0.143704166 | 0.390522886 |
| chr10:99358158-99358358 | 0.762579855 | 0.143707855 | 0.390522886 |
| chr14:79884910-79885338 | -0.684432327 | 0.14371398 | 0.390525679 |
| chr1:48677348-48677647 | -0.603408389 | 0.143740964 | 0.390585152 |

| | | | |
|---|---|---|---|
| chr1:217791790-217792002 | 0.69385715 | 0.143748151 | 0.390590831 |
| chr4:112012995-112013346 | -0.698071946 | 0.143754619 | 0.390594553 |
| chr16:27779424-27779624 | 0.772047644 | 0.143777176 | 0.390641991 |
| chr13:20182350-20182561 | 0.67567026 | 0.14379788 | 0.39068439 |
| chr8:116175209-116175472 | -0.650527604 | 0.143833939 | 0.390768502 |
| chr17:16873344-16873544 | 0.696138341 | 0.143871694 | 0.390854573 |
| chr6:41683546-41683913 | 0.573811824 | 0.143875822 | 0.390854573 |
| chr2:183123989-183125088 | -0.462951197 | 0.143882283 | 0.390858266 |
| chr20:34111150-34111417 | 0.667504753 | 0.143925462 | 0.390959088 |
| chr1:204131160-204131707 | -0.485272431 | 0.143929602 | 0.390959088 |
| chr1:155323715-155324848 | 0.463121577 | 0.143937037 | 0.390965424 |
| chr12:121856409-121856666 | 0.720142741 | 0.143975733 | 0.391046261 |
| chr12:93524445-93524772 | 0.605324143 | 0.143977005 | 0.391046261 |
| chr12:25249936-25250552 | -0.527843899 | 0.14400985 | 0.391106715 |
| chr7:107044068-107045088 | -0.499945426 | 0.144010878 | 0.391106715 |
| chr2:130988103-130988304 | -0.747251006 | 0.144014576 | 0.391106715 |
| chr20:57125552-57125795 | 0.489682757 | 0.144031876 | 0.391139833 |
| chr14:90785559-90785778 | 0.630572982 | 0.144042023 | 0.391153527 |
| chr6:132798190-132799520 | -0.474115701 | 0.144059035 | 0.391172216 |
| chr7:1513231-1513825 | -0.545207883 | 0.144063274 | 0.391172216 |
| chr10:113820562-113820813 | -0.803457817 | 0.144066941 | 0.391172216 |
| chr11:37110447-37110753 | -0.702605452 | 0.14407367 | 0.391172216 |
| chr6:217207-217416 | -0.727164227 | 0.144074432 | 0.391172216 |
| chrX:107484404-107484701 | -0.77543277 | 0.144099007 | 0.391212308 |
| chr16:81977346-81977680 | -0.693933343 | 0.14409941 | 0.391212308 |
| chr19:29811416-29812255 | 0.480533785 | 0.144111661 | 0.391218099 |
| chr4:76127933-76128144 | -0.769007569 | 0.1441139 | 0.391218099 |
| chr8:26670616-26670816 | -0.876977299 | 0.14411686 | 0.391218099 |
| chr5:64595283-64595486 | -0.725579078 | 0.144140522 | 0.39126847 |
| chr8:22165128-22165383 | 0.662630308 | 0.144150708 | 0.391282256 |
| chr10:778945-779215 | 0.616213257 | 0.144161928 | 0.391298035 |
| chr8:65834636-65834937 | -0.660623327 | 0.144166734 | 0.391298035 |
| chr12:99907043-99907285 | -0.71338923 | 0.144174778 | 0.391301967 |
| chr16:55712354-55712613 | -0.69106938 | 0.144181839 | 0.391301967 |
| chr20:43513894-43514640 | 0.474563066 | 0.144184423 | 0.391301967 |
| chr1:59926375-59927129 | -0.507327797 | 0.144188611 | 0.391301967 |
| chr13:73399287-73399653 | -0.601873595 | 0.14419768 | 0.391305861 |
| chr9:128127366-128127598 | 0.618522409 | 0.144200259 | 0.391305861 |
| chr1:246822237-246822474 | -0.65780177 | 0.144206651 | 0.391309347 |
| chr4:87622363-87622669 | -0.688505543 | 0.144214755 | 0.391317479 |
| chr12:12334437-12334655 | -0.714427597 | 0.144223102 | 0.391326269 |
| chr19:4153453-4153659 | 0.683067842 | 0.144253331 | 0.39139443 |
| chr18:11849809-11850362 | 0.53059243 | 0.144259051 | 0.391396092 |
| chr14:67667280-67667503 | 0.674840685 | 0.144292226 | 0.391462323 |
| chr2:9408892-9409092 | -0.770198712 | 0.144305784 | 0.391462323 |
| chr10:62608537-62608849 | -0.685214923 | 0.144307061 | 0.391462323 |
| chr14:35045986-35047099 | 0.461761848 | 0.144309025 | 0.391462323 |
| chr2:30847358-30847632 | 0.672644696 | 0.144309649 | 0.391462323 |
| chr3:32385080-32385280 | -0.87710213 | 0.144315556 | 0.391462323 |
| chr6:35146475-35146681 | -0.830742507 | 0.144328928 | 0.391462323 |

| | | | |
|---|---|---|---|
| chr6:152238469-152238729 | -0.830742507 | 0.144328928 | 0.391462323 |
| chr15:81280791-81281118 | -0.588339603 | 0.144329444 | 0.391462323 |
| chr14:89672765-89673139 | 0.587201952 | 0.144336649 | 0.391468009 |
| chr19:11346224-11346834 | 0.489057151 | 0.144355891 | 0.391504296 |
| chr5:35201406-35201606 | 0.669614535 | 0.144360247 | 0.391504296 |
| chr20:62288768-62289124 | -0.624481976 | 0.144414117 | 0.391636529 |
| chr20:48038173-48038586 | 0.486831115 | 0.144424484 | 0.391650782 |
| chr3:170483488-170483860 | -0.559770583 | 0.144430069 | 0.391652064 |
| chr9:29211620-29211912 | -0.588793321 | 0.144444171 | 0.39167281 |
| chr10:86801911-86802153 | 0.707601307 | 0.144447943 | 0.39167281 |
| chr1:10867147-10867513 | 0.576182153 | 0.144464473 | 0.391698916 |
| chr15:41184911-41185165 | 0.678851215 | 0.1444725 | 0.391698916 |
| chr10:100267344-100268046 | 0.506370481 | 0.144472907 | 0.391698916 |
| chr10:51035408-51035627 | -0.732642373 | 0.144483274 | 0.391712269 |
| chr16:28553771-28554602 | 0.483839834 | 0.144488057 | 0.391712269 |
| chr5:66697967-66698314 | -0.631420923 | 0.144546386 | 0.391856539 |
| chr7:139231761-139232112 | -0.549245942 | 0.144586238 | 0.391950707 |
| chr21:8464641-8464841 | 0.728327881 | 0.144609461 | 0.391999792 |
| chr7:149799846-149800110 | -0.621264836 | 0.144619123 | 0.392012114 |
| chr10:50501522-50501807 | -0.598145237 | 0.1446498 | 0.392081397 |
| chr4:45946788-45947050 | -0.765304898 | 0.144655779 | 0.392083735 |
| chr3:63523156-63523399 | 0.532697452 | 0.144692395 | 0.392158503 |
| chr10:5976671-5978341 | 0.467502163 | 0.1446936 | 0.392158503 |
| chr17:61870335-61870792 | 0.47834798 | 0.144704163 | 0.392173259 |
| chr2:156781589-156782263 | -0.518628213 | 0.144714384 | 0.392187088 |
| chr13:72726846-72728265 | -0.461275535 | 0.144749361 | 0.392268005 |
| chr7:142909604-142909804 | -0.80906839 | 0.144758415 | 0.392278665 |
| chr4:174469647-174470051 | -0.590172497 | 0.144771725 | 0.392300861 |
| chr2:47006914-47007130 | -0.64271978 | 0.144778923 | 0.392306491 |
| chr8:143401150-143401599 | 0.522622178 | 0.144791568 | 0.39232046 |
| chr22:37686171-37686974 | 0.523175431 | 0.144794319 | 0.39232046 |
| chr15:101349072-101349321 | -0.676833391 | 0.144805135 | 0.392335894 |
| chr1:219530771-219531606 | -0.47488113 | 0.144825484 | 0.392377152 |
| chr2:237438176-237438432 | -0.682781192 | 0.144845592 | 0.392406572 |
| chr16:83989287-83989712 | -0.538572717 | 0.144846585 | 0.392406572 |
| chr1:167540421-167540621 | 0.753294101 | 0.144868764 | 0.392452779 |
| chr3:39189701-39190075 | 0.586370114 | 0.144886808 | 0.392466042 |
| chr5:149970884-149971292 | -0.624069239 | 0.144887024 | 0.392466042 |
| chr10:97518305-97518580 | -0.613255101 | 0.144889026 | 0.392466042 |
| chr3:14369696-14369940 | 0.572989989 | 0.144896825 | 0.392473294 |
| chr1:211515284-211515805 | -0.537501314 | 0.144931405 | 0.392553081 |
| chr12:100539322-100539522 | -0.817020272 | 0.144946763 | 0.392580803 |
| chr20:53763843-53764130 | 0.482178895 | 0.144969464 | 0.392628409 |
| chr1:231475966-231476166 | -0.874220294 | 0.144984663 | 0.392647532 |
| chr8:42195199-42195437 | -0.717269054 | 0.144986774 | 0.392647532 |
| chr5:7746022-7746397 | -0.649158647 | 0.144992722 | 0.39264976 |
| chr7:6930112-6930476 | -0.632257955 | 0.145012546 | 0.392667835 |
| chr22:19431611-19432079 | 0.538270052 | 0.145014438 | 0.392667835 |
| chr10:101301525-101301786 | 0.630160179 | 0.145016607 | 0.392667835 |
| chr17:71909326-71909526 | 0.780176201 | 0.145026424 | 0.392667835 |

| | | | |
|---|---|---|---|
| chr10:65570082-65570978 | -0.482285614 | 0.145028884 | 0.392667835 |
| chr5:57220971-57221282 | -0.603872056 | 0.145034777 | 0.392667835 |
| chr9:109520520-109520805 | 0.568645629 | 0.145035269 | 0.392667835 |
| chr16:87764949-87766175 | 0.479703151 | 0.145045846 | 0.392679376 |
| chr5:117591045-117591269 | -0.737896122 | 0.145049782 | 0.392679376 |
| chr12:62724925-62725484 | -0.49992376 | 0.145074044 | 0.392731184 |
| chr11:118919677-118920086 | 0.49866597 | 0.14508027 | 0.392734163 |
| chr4:100096051-100096325 | -0.637130239 | 0.145094217 | 0.39275804 |
| chr22:21055676-21055876 | 0.718859821 | 0.145099401 | 0.392758199 |
| chr7:15906537-15906737 | 0.750038544 | 0.145133062 | 0.392835435 |
| chrX:39492465-39492804 | -0.665836357 | 0.145152011 | 0.392863399 |
| chr3:158139175-158139600 | -0.669947404 | 0.145158343 | 0.392863399 |
| chr18:61650288-61650539 | -0.650827539 | 0.145158776 | 0.392863399 |
| chr14:55753009-55753654 | 0.50204962 | 0.145179815 | 0.39289537 |
| chr7:34066768-34067376 | -0.503589925 | 0.145180844 | 0.39289537 |
| chr2:61004402-61004718 | -0.580793376 | 0.145206611 | 0.392951224 |
| chr10:77925976-77926539 | 0.520695063 | 0.145217593 | 0.392967065 |
| chr9:111099322-111099674 | -0.656913525 | 0.145228807 | 0.392976803 |
| chr13:83831812-83832108 | -0.602656278 | 0.145236933 | 0.392976803 |
| chrX:132954375-132954909 | 0.538693955 | 0.145254842 | 0.392976803 |
| chr12:59403327-59403603 | 0.677172908 | 0.145257783 | 0.392976803 |
| chr4:113398568-113398863 | 0.607820064 | 0.145259391 | 0.392976803 |
| chr17:40953904-40954139 | 0.718265421 | 0.145263433 | 0.392976803 |
| chr4:33482830-33483149 | 0.618674525 | 0.14527007 | 0.392976803 |
| chr1:197553338-197553635 | -0.640131136 | 0.14527063 | 0.392976803 |
| chr3:123817614-123817849 | -0.689963157 | 0.145271637 | 0.392976803 |
| chr4:80467017-80467255 | -0.807657423 | 0.14527248 | 0.392976803 |
| chr15:99749386-99749740 | 0.588641887 | 0.145285722 | 0.39299875 |
| chr2:100594947-100595339 | -0.60739883 | 0.145347495 | 0.393151968 |
| chr3:122405554-122405986 | 0.665268584 | 0.145365339 | 0.393157578 |
| chr4:48269018-48269226 | 0.632039924 | 0.14536725 | 0.393157578 |
| chr11:61503375-61503654 | 0.68831985 | 0.145367749 | 0.393157578 |
| chr14:87805237-87805484 | 0.677329373 | 0.145371162 | 0.393157578 |
| chr12:107918187-107918570 | -0.544304059 | 0.145375225 | 0.393157578 |
| chr21:30216000-30216333 | 0.619199171 | 0.14539718 | 0.393201783 |
| chr17:14300633-14301428 | -0.469642092 | 0.145401834 | 0.393201783 |
| chr1:58816306-58816586 | 0.539759662 | 0.145410188 | 0.393210497 |
| chr9:32384237-32385167 | -0.486559485 | 0.145427938 | 0.39322426 |
| chr17:79722459-79722728 | 0.609682127 | 0.14542995 | 0.39322426 |
| chr3:151943606-151943965 | -0.591074357 | 0.145430674 | 0.39322426 |
| chr2:157628381-157629540 | -0.466939667 | 0.145440049 | 0.393235719 |
| chrX:12309066-12309340 | 0.63121893 | 0.145445176 | 0.393235719 |
| chr5:106454675-106455037 | -0.577464166 | 0.145462932 | 0.393269847 |
| chr12:53159262-53159716 | 0.526538986 | 0.145546435 | 0.39348172 |
| chr2:31354877-31355147 | -0.741543043 | 0.145585717 | 0.393574033 |
| chr6:29977193-29977408 | 0.699751461 | 0.145628201 | 0.393670488 |
| chr1:104940450-104940823 | -0.599449534 | 0.145631672 | 0.393670488 |
| chr3:98359990-98360377 | 0.57726954 | 0.145640275 | 0.393679854 |
| chr1:218059987-218060445 | 0.64066341 | 0.14565864 | 0.393710503 |
| chr16:11586741-11587448 | 0.509970022 | 0.145664383 | 0.393710503 |

| | | | |
|---|---|---|---|
| chr9:97876865-97877624 | -0.506486599 | 0.145667028 | 0.393710503 |
| chr19:917758-917973 | 0.678340731 | 0.145698507 | 0.393768067 |
| chr3:188282775-188283211 | -0.592910544 | 0.145698605 | 0.393768067 |
| chr4:183796983-183797845 | -0.524141152 | 0.14576397 | 0.39393083 |
| chr3:63940502-63940823 | 0.504514968 | 0.145776059 | 0.393944439 |
| chr6:159450852-159451052 | -0.827793659 | 0.145779288 | 0.393944439 |
| chr18:24487715-24488034 | -0.595312885 | 0.145800842 | 0.39398008 |
| chr7:148976653-148976908 | -0.77554489 | 0.145802761 | 0.39398008 |
| chr6:152532510-152532945 | 0.555544231 | 0.145837003 | 0.394011706 |
| chr20:47391828-47392629 | 0.457662986 | 0.145841576 | 0.394011706 |
| chr5:32759988-32760305 | -0.736460425 | 0.145842749 | 0.394011706 |
| chr15:41283640-41284344 | 0.482946895 | 0.145850416 | 0.394011706 |
| chr16:11976310-11976877 | 0.518281196 | 0.14585119 | 0.394011706 |
| chr2:30862108-30862390 | -0.649716088 | 0.145853054 | 0.394011706 |
| chr10:24319494-24319798 | -0.635783322 | 0.145854274 | 0.394011706 |
| chr13:32205909-32206534 | -0.573403729 | 0.145855603 | 0.394011706 |
| chr14:63116041-63116241 | 0.822589445 | 0.145862202 | 0.394015642 |
| chr11:20477274-20477713 | 0.609480155 | 0.145911044 | 0.394120436 |
| chr7:12158482-12158956 | 0.529353901 | 0.145911284 | 0.394120436 |
| chr7:43892848-43893087 | -0.745847315 | 0.145917018 | 0.394122031 |
| chr14:56849921-56850121 | 0.766479506 | 0.145929855 | 0.39413615 |
| chr3:15207097-15207319 | 0.559999439 | 0.145937985 | 0.39413615 |
| chr11:93359457-93359766 | -0.626621621 | 0.145940748 | 0.39413615 |
| chr1:151594151-151594492 | 0.530817205 | 0.145942821 | 0.39413615 |
| chr17:59114305-59114582 | 0.501608353 | 0.145949542 | 0.394140409 |
| chr1:65174615-65174815 | -0.728516158 | 0.145963577 | 0.394164418 |
| chr5:135857743-135858419 | -0.51909938 | 0.14600078 | 0.394240886 |
| chr13:80473718-80474111 | -0.591131159 | 0.14601247 | 0.394240886 |
| chr3:107846239-107846630 | -0.524701063 | 0.14601254 | 0.394240886 |
| chr6:69835805-69836036 | -0.688829989 | 0.146016251 | 0.394240886 |
| chr6:4985784-4986033 | 0.791000624 | 0.146018063 | 0.394240886 |
| chr10:80290518-80290718 | -0.769100383 | 0.146022766 | 0.394240886 |
| chr5:154575854-154576133 | -0.663623548 | 0.146037562 | 0.394266942 |
| chrX:24859426-24859702 | -0.737818019 | 0.146044339 | 0.394271346 |
| chr5:35938531-35938914 | -0.516543513 | 0.146068401 | 0.394321535 |
| chr3:180134875-180135093 | -0.767632239 | 0.146073461 | 0.394321535 |
| chr3:63947231-63947508 | 0.532964445 | 0.146079057 | 0.394321535 |
| chr16:24539792-24540234 | 0.529838811 | 0.146083515 | 0.394321535 |
| chr20:54223730-54224051 | 0.497377229 | 0.146094686 | 0.394337796 |
| chr5:1345072-1345327 | 0.569346813 | 0.146107987 | 0.394359807 |
| chr3:24471551-24471804 | -0.80632999 | 0.146124195 | 0.394389659 |
| chr3:154781694-154781949 | -0.66946864 | 0.146160693 | 0.394470008 |
| chr6:170279928-170281039 | -0.471531122 | 0.146164261 | 0.394470008 |
| chr19:49546684-49547330 | -0.518082845 | 0.146204289 | 0.394564138 |
| chr19:17292401-17292684 | 0.575781597 | 0.146221212 | 0.394590234 |
| chrX:84501715-84502711 | -0.473821215 | 0.146227245 | 0.394590234 |
| chr14:78959795-78960009 | -0.719086795 | 0.146229408 | 0.394590234 |
| chr12:53500535-53501668 | 0.4591319 | 0.146246829 | 0.394623347 |
| chr10:61750890-61751394 | -0.520218663 | 0.146259127 | 0.394642631 |
| chr3:49018393-49018602 | 0.586398934 | 0.146289408 | 0.394710438 |

| | | | |
|---|---|---|---|
| chr7:55409929-55410136 | -0.732630993 | 0.146316079 | 0.394768499 |
| chr14:79342870-79343410 | -0.498771588 | 0.146344387 | 0.394830973 |
| chr3:56801632-56802464 | -0.519904378 | 0.146370382 | 0.394887201 |
| chr20:57108205-57109049 | 0.457022343 | 0.146380156 | 0.394899666 |
| chr10:72248877-72249084 | -0.733180975 | 0.146385898 | 0.394901251 |
| chr16:20806070-20807027 | 0.482258667 | 0.146401268 | 0.394928812 |
| chr6:96419572-96419772 | -0.831003212 | 0.146431714 | 0.39498313 |
| chr7:124058050-124058308 | -0.831003212 | 0.146431714 | 0.39498313 |
| chr4:120921796-120923198 | -0.471248805 | 0.146461634 | 0.395040696 |
| chr19:49543110-49543342 | 0.727346901 | 0.146463367 | 0.395040696 |
| chr21:42091831-42092031 | -0.758286045 | 0.146483917 | 0.395082217 |
| chr3:98023220-98023583 | -0.567299072 | 0.14650478 | 0.395124579 |
| chr11:20610152-20610550 | -0.528454738 | 0.146512711 | 0.395132059 |
| chr19:41754059-41754259 | 0.724471104 | 0.146523053 | 0.395134253 |
| chr4:113547532-113547982 | -0.503037926 | 0.146523838 | 0.395134253 |
| chr17:8397674-8398285 | -0.558198997 | 0.146533224 | 0.395145656 |
| chr20:46357070-46357334 | -0.616887376 | 0.146540838 | 0.395152281 |
| chr3:165185742-165186246 | -0.528179344 | 0.146568693 | 0.395213486 |
| chr10:107731925-107732141 | -0.786351211 | 0.146577173 | 0.395222443 |
| chr14:100240307-100240766 | 0.501557291 | 0.14658873 | 0.395233732 |
| chr10:132001360-132001964 | 0.49551511 | 0.146592904 | 0.395233732 |
| chr14:73508107-73508347 | 0.684183027 | 0.146605145 | 0.395233732 |
| chr19:12495222-12495643 | 0.557972417 | 0.146610998 | 0.395233732 |
| chr4:68447283-68447501 | 0.644884812 | 0.146612294 | 0.395233732 |
| chr19:17784468-17784690 | 0.706190632 | 0.146612309 | 0.395233732 |
| chr16:13866223-13866454 | 0.71333594 | 0.146623218 | 0.395249234 |
| chr1:188335986-188336419 | -0.606878379 | 0.146629885 | 0.395253301 |
| chr3:189167197-189167397 | -0.764724276 | 0.146639784 | 0.395253867 |
| chr16:9138411-9138616 | 0.727830217 | 0.146640412 | 0.395253867 |
| chr3:124313274-124313506 | 0.697898367 | 0.146650869 | 0.395268146 |
| chr6:116462028-116462228 | 0.764674038 | 0.146658192 | 0.395273979 |
| chr20:49484004-49484531 | 0.512137974 | 0.146667458 | 0.395285051 |
| chr4:152359873-152360377 | -0.592742135 | 0.146694356 | 0.395332147 |
| chr18:12304966-12305217 | 0.613446914 | 0.146695252 | 0.395332147 |
| chr7:131556026-131556516 | -0.566965547 | 0.146730834 | 0.395399778 |
| chr1:172687636-172687863 | 0.730403482 | 0.14673204 | 0.395399778 |
| chr16:20741035-20742063 | 0.475757338 | 0.146735829 | 0.395399778 |
| chr5:148306428-148306813 | -0.602546994 | 0.146754999 | 0.395429456 |
| chr9:21396195-21396405 | 0.715733746 | 0.146757334 | 0.395429456 |
| chr20:45993377-45993954 | 0.481264126 | 0.146762325 | 0.395429456 |
| chr1:47508681-47509014 | 0.607329624 | 0.14677619 | 0.395452909 |
| chr10:32650202-32650512 | -0.616476804 | 0.146841954 | 0.395616182 |
| chr3:50528306-50528506 | -0.814991296 | 0.146858861 | 0.395638227 |
| chr15:75115621-75115867 | 0.75339197 | 0.146860464 | 0.395638227 |
| chr12:52023546-52023996 | -0.519415055 | 0.146910531 | 0.395759191 |
| chr5:136028767-136029009 | 0.670637519 | 0.146934405 | 0.395803365 |
| chr8:102961874-102962133 | -0.711500694 | 0.146941863 | 0.395803365 |
| chr6:2634111-2634833 | -0.504910935 | 0.146942425 | 0.395803365 |
| chr15:22895435-22895860 | 0.529470442 | 0.146970752 | 0.395850418 |
| chr5:97162026-97162334 | -0.615216489 | 0.14697102 | 0.395850418 |

| | | | |
|---|---|---|---|
| chr6:20291688-20292017 | -0.559328858 | 0.146980347 | 0.395850418 |
| chr8:66541739-66542834 | -0.478165481 | 0.146983203 | 0.395850418 |
| chr7:101308671-101309465 | -0.522564122 | 0.146989175 | 0.395850418 |
| chr2:219571292-219572212 | 0.483232253 | 0.146994673 | 0.395850418 |
| chr1:176480033-176480233 | 0.790206504 | 0.146996058 | 0.395850418 |
| chr10:25725911-25726258 | -0.628496867 | 0.147005056 | 0.395860738 |
| chr19:53554128-53554880 | -0.531638347 | 0.147019214 | 0.395881242 |
| chr17:62995366-62995658 | -0.57719858 | 0.147023004 | 0.395881242 |
| chr1:27830159-27831380 | 0.476646982 | 0.147030224 | 0.395886771 |
| chr15:40367927-40368354 | 0.530673331 | 0.147035566 | 0.395887241 |
| chr1:151871455-151871715 | 0.616975624 | 0.147061256 | 0.395942497 |
| chr8:47987549-47987830 | 0.626436617 | 0.147105711 | 0.396048272 |
| chrX:38627696-38627979 | -0.708784735 | 0.147126228 | 0.39608959 |
| chr5:173808073-173808273 | 0.756313926 | 0.147149532 | 0.396138411 |
| chr14:24013962-24014363 | 0.542257626 | 0.147161107 | 0.396155652 |
| chr5:25766613-25766917 | -0.634955284 | 0.147173938 | 0.396176276 |
| chr5:77282147-77282929 | -0.492914298 | 0.147188771 | 0.396188846 |
| chr3:169012102-169012319 | 0.715359669 | 0.14718895 | 0.396188846 |
| chr19:45038864-45039653 | 0.498269314 | 0.147210617 | 0.396229849 |
| chr14:72776639-72776962 | 0.59796323 | 0.147214525 | 0.396229849 |
| chr1:191752435-191752635 | -0.78059872 | 0.147230642 | 0.396247178 |
| chr8:109366316-109366516 | -0.721840071 | 0.147231306 | 0.396247178 |
| chr16:19135992-19136443 | -0.572819351 | 0.14724229 | 0.39626282 |
| chr20:46748031-46748231 | 0.654358325 | 0.14727765 | 0.39634406 |
| chr22:28935865-28936129 | 0.662401271 | 0.147302781 | 0.39639777 |
| chr5:1183573-1183803 | 0.681348437 | 0.147311862 | 0.396408285 |
| chr2:5915116-5915360 | 0.780232445 | 0.147336671 | 0.396461119 |
| chr20:52083841-52084044 | 0.657391548 | 0.147342883 | 0.396463912 |
| chr3:172990241-172990441 | 0.824181499 | 0.147366247 | 0.396512855 |
| chr1:116092597-116093003 | 0.588410826 | 0.147408632 | 0.396612972 |
| chrX:104636335-104636622 | -0.642962556 | 0.1474201 | 0.3966299 |
| chr14:73759920-73760514 | 0.472449225 | 0.147426341 | 0.396632764 |
| chr19:38441713-38441948 | 0.671679258 | 0.147437437 | 0.39664869 |
| chr5:114018159-114018523 | -0.601760857 | 0.14744458 | 0.396651466 |
| chrX:140507785-140508481 | -0.479527642 | 0.14746125 | 0.396651466 |
| chr17:28661620-28662608 | -0.474866039 | 0.147467073 | 0.396651466 |
| chr7:7512785-7513186 | 0.595324378 | 0.147468917 | 0.396651466 |
| chr14:93789053-93789443 | 0.567009092 | 0.147472087 | 0.396651466 |
| chr20:10131703-10132672 | -0.487030317 | 0.147473704 | 0.396651466 |
| chr10:11823176-11823645 | 0.524314333 | 0.147483065 | 0.396651466 |
| chr7:112205718-112205989 | -0.56277925 | 0.147483722 | 0.396651466 |
| chr3:180679144-180680391 | -0.468025625 | 0.14748506 | 0.396651466 |
| chr5:73792820-73793096 | -0.728342358 | 0.147531825 | 0.396752894 |
| chr12:120086838-120087615 | 0.469356134 | 0.14753313 | 0.396752894 |
| chr14:70907190-70908143 | 0.478366448 | 0.147558289 | 0.396806628 |
| chr4:157514725-157515115 | -0.622253553 | 0.147569433 | 0.396808749 |
| chr5:174903372-174903712 | -0.639509478 | 0.147569436 | 0.396808749 |
| chr16:77169720-77169956 | 0.61268502 | 0.147582286 | 0.396812694 |
| chr7:12496906-12497796 | 0.494335718 | 0.14758393 | 0.396812694 |
| chr12:115473218-115473534 | 0.643302929 | 0.147586439 | 0.396812694 |

| | | | |
|---|---|---|---|
| chr3:73255223-73255423 | 0.801881728 | 0.147607202 | 0.396854591 |
| chr17:79933867-79934128 | 0.584672374 | 0.147635421 | 0.396916533 |
| chr5:95843270-95843470 | -0.719173035 | 0.147662031 | 0.396974147 |
| chr14:34461657-34463020 | 0.459625816 | 0.147681413 | 0.396990969 |
| chr11:65793260-65793494 | 0.586382251 | 0.147683038 | 0.396990969 |
| chr20:53848547-53848747 | 0.496059941 | 0.147686669 | 0.396990969 |
| chr7:44986104-44987098 | -0.465076789 | 0.147691039 | 0.396990969 |
| chr9:27247804-27248004 | 0.775003143 | 0.147694194 | 0.396990969 |
| chr7:19708421-19709539 | 0.461648426 | 0.147702259 | 0.396998718 |
| chr14:86035926-86036220 | -0.605978409 | 0.147722041 | 0.397037961 |
| chr1:47990360-47990560 | -0.738241701 | 0.147732704 | 0.397052692 |
| chr8:128343590-128344387 | -0.487907608 | 0.14775924 | 0.39709386 |
| chr13:31655911-31656132 | -0.73744845 | 0.147761965 | 0.39709386 |
| chr8:94638944-94639658 | 0.46376336 | 0.147763568 | 0.39709386 |
| chr2:41840658-41841590 | -0.463545004 | 0.147797833 | 0.397160632 |
| chr5:17096411-17096611 | -0.701975341 | 0.147798782 | 0.397160632 |
| chr19:55258765-55259662 | 0.464993621 | 0.147807408 | 0.397169883 |
| chr2:100893991-100894191 | -0.834990613 | 0.147825886 | 0.397177741 |
| chr3:158539519-158539775 | -0.679361207 | 0.147826295 | 0.397177741 |
| chr18:77207519-77207758 | -0.700649481 | 0.147827748 | 0.397177741 |
| chr3:139916236-139917142 | -0.488180422 | 0.147831067 | 0.397177741 |
| chr15:98865347-98865699 | 0.506050403 | 0.147849163 | 0.397212432 |
| chr2:60998979-60999311 | -0.590889811 | 0.147857197 | 0.397220088 |
| chr20:59525530-59526138 | 0.495813748 | 0.14786565 | 0.397228871 |
| chr12:91562261-91562474 | -0.748668099 | 0.147904572 | 0.397314641 |
| chr12:15527117-15527822 | -0.540190669 | 0.147907948 | 0.397314641 |
| chr20:9978404-9978910 | 0.550001506 | 0.147921552 | 0.397337253 |
| chrX:132368062-132368315 | 0.671003398 | 0.147927672 | 0.397339407 |
| chr5:54769078-54769342 | -0.712013115 | 0.147934993 | 0.397339407 |
| chr5:4012171-4012378 | 0.727679884 | 0.147941338 | 0.397339407 |
| chr9:98791270-98791488 | 0.621071833 | 0.147943144 | 0.397339407 |
| chr14:101964839-101965071 | 0.58900822 | 0.147948282 | 0.397339407 |
| chr1:223698262-223698571 | 0.568596068 | 0.147961316 | 0.397357039 |
| chr21:42174182-42174382 | -0.755983673 | 0.147965219 | 0.397357039 |
| chr6:139331233-139331612 | 0.583465512 | 0.147976257 | 0.397364609 |
| chr20:47334019-47334472 | 0.467419614 | 0.14797841 | 0.397364609 |
| chr14:105090509-105090808 | 0.526917661 | 0.147983746 | 0.39736501 |
| chr11:43515180-43515433 | 0.69765074 | 0.148028551 | 0.397471393 |
| chr21:42209902-42210261 | -0.560517339 | 0.148037301 | 0.397473933 |
| chr13:111437150-111437387 | -0.750514253 | 0.148039873 | 0.397473933 |
| chr7:37352919-37353214 | 0.570178661 | 0.14805465 | 0.397499681 |
| chr2:232624256-232624807 | -0.541582292 | 0.148086249 | 0.397570588 |
| chr3:145288916-145289175 | -0.722395423 | 0.148131344 | 0.397667826 |
| chr8:10547507-10547763 | -0.709349498 | 0.148132848 | 0.397667826 |
| chr3:17994428-17994628 | 0.776051962 | 0.148145212 | 0.397680462 |
| chr17:51667965-51668165 | 0.71920326 | 0.148147935 | 0.397680462 |
| chr10:125284878-125285506 | -0.53188456 | 0.148160164 | 0.397699355 |
| chr16:13957422-13958132 | -0.527941667 | 0.148169648 | 0.397710881 |
| chr1:45010507-45010724 | 0.619901629 | 0.148182709 | 0.397732004 |
| chr2:127068014-127068358 | -0.557000523 | 0.148190002 | 0.397737646 |

| | | | |
|---|---|---|---|
| chr15:75627772-75627977 | -0.736668131 | 0.148196597 | 0.397741415 |
| chr7:98826535-98827128 | -0.550005335 | 0.148202472 | 0.397743251 |
| chr3:141370073-141370534 | -0.508818826 | 0.148210223 | 0.397750121 |
| chr2:47905681-47907204 | -0.457654666 | 0.148219336 | 0.397751346 |
| chr9:128191551-128191940 | 0.554209162 | 0.148221061 | 0.397751346 |
| chr2:123514871-123515165 | -0.68852052 | 0.148235439 | 0.397775995 |
| chr1:200816901-200817101 | -0.765199094 | 0.148278863 | 0.397865622 |
| chr16:12289831-12290089 | -0.748295882 | 0.148279224 | 0.397865622 |
| chr20:58650750-58651798 | 0.459446524 | 0.148306735 | 0.397906684 |
| chr8:70403755-70403988 | 0.565924704 | 0.148308374 | 0.397906684 |
| chr10:58280441-58280685 | -0.750467351 | 0.148310107 | 0.397906684 |
| chr14:77139579-77139931 | 0.532567928 | 0.148342209 | 0.397978877 |
| chr10:90821305-90821677 | -0.688042602 | 0.148355491 | 0.398000576 |
| chr20:57389234-57389754 | 0.487816076 | 0.148369547 | 0.398024347 |
| chr3:173538491-173538754 | -0.69388854 | 0.148393404 | 0.39807441 |
| chr6:134379028-134379422 | -0.51284854 | 0.148401545 | 0.398082313 |
| chr2:70124044-70125417 | -0.469444321 | 0.148414865 | 0.398090311 |
| chrX:120734794-120735143 | -0.591710408 | 0.148414918 | 0.398090311 |
| chr16:85144405-85144629 | 0.640272842 | 0.14842382 | 0.398100252 |
| chr7:105691680-105692013 | 0.555184474 | 0.148434738 | 0.398115601 |
| chr4:96483377-96483701 | -0.669248981 | 0.148463237 | 0.398154163 |
| chr11:6618754-6619745 | -0.47150677 | 0.14846414 | 0.398154163 |
| chr1:204443200-204443683 | 0.487231746 | 0.148464705 | 0.398154163 |
| chr20:53669064-53669385 | 0.477410486 | 0.148483893 | 0.39818193 |
| chr7:47951413-47952098 | -0.497389187 | 0.148487275 | 0.39818193 |
| chr1:145823561-145824567 | 0.465547507 | 0.148493854 | 0.39818193 |
| chr8:37903886-37904436 | 0.515103395 | 0.148495845 | 0.39818193 |
| chr1:26279666-26280099 | 0.551549607 | 0.148516302 | 0.398222849 |
| chr6:152669986-152670249 | -0.742181946 | 0.148565757 | 0.398341514 |
| chr5:23220261-23220749 | 0.514536308 | 0.148630652 | 0.398501569 |
| chr9:38417036-38417298 | -0.773015875 | 0.14864117 | 0.398515825 |
| chr9:98611942-98612350 | -0.662868936 | 0.148661782 | 0.39855714 |
| chr4:158094815-158095015 | -0.76049234 | 0.1486675 | 0.398558524 |
| chr10:78293491-78293691 | -0.787740305 | 0.148680356 | 0.398579045 |
| chr20:63825352-63825593 | -0.677970556 | 0.148686855 | 0.398582521 |
| chr2:96824098-96824442 | 0.550863113 | 0.148704401 | 0.39861561 |
| chr2:65309463-65310337 | -0.466295964 | 0.148722355 | 0.398647597 |
| chr3:91891127-91891524 | 0.570243858 | 0.148726739 | 0.398647597 |
| chr1:215445141-215445341 | -0.749665943 | 0.148749394 | 0.398694373 |
| chr6:131915005-131915503 | -0.527730387 | 0.148771792 | 0.39874046 |
| chr20:58850733-58851116 | 0.497171801 | 0.148778236 | 0.398743781 |
| chr14:92726624-92726855 | 0.652079339 | 0.148789859 | 0.398760985 |
| chr10:66410453-66410747 | -0.765829346 | 0.148802346 | 0.398780504 |
| chr2:108215753-108216315 | -0.529180156 | 0.148831169 | 0.398843797 |
| chr12:66678995-66679195 | -0.764128859 | 0.148854556 | 0.398876983 |
| chr8:120605510-120606071 | -0.495811112 | 0.148857704 | 0.398876983 |
| chrX:154458171-154458437 | 0.594237371 | 0.14885917 | 0.398876983 |
| chr12:76405784-76406015 | -0.78046894 | 0.148869121 | 0.398889697 |
| chr2:188580860-188581128 | -0.702360913 | 0.148881343 | 0.398908495 |
| chr8:81890515-81890820 | 0.571328996 | 0.148898953 | 0.398934649 |

| | | | |
|---|---|---|---|
| chr2:112543175-112543397 | 0.63311822 | 0.148901517 | 0.398934649 |
| chr19:49301345-49301578 | -0.692957509 | 0.148944044 | 0.399034635 |
| chr20:57211021-57211590 | 0.459391519 | 0.14897248 | 0.399096862 |
| chr17:61284073-61284302 | 0.494228496 | 0.14898662 | 0.399120788 |
| chr5:153832054-153832332 | -0.588602324 | 0.149003605 | 0.399152334 |
| chr14:60551201-60551401 | 0.691681397 | 0.149030173 | 0.399209548 |
| chr1:106549745-106549945 | 0.532978647 | 0.149100707 | 0.399368642 |
| chr11:61173597-61173806 | 0.674504455 | 0.149103086 | 0.399368642 |
| chr20:48189276-48189516 | 0.507045849 | 0.149105202 | 0.399368642 |
| chr19:11355144-11355581 | 0.520333797 | 0.149116008 | 0.399383625 |
| chr12:6730270-6730654 | -0.680487427 | 0.149130839 | 0.399409387 |
| chr6:160088422-160088697 | 0.651274999 | 0.149170936 | 0.399502811 |
| chr19:33302396-33303274 | 0.474738292 | 0.149181656 | 0.399516949 |
| chrX:137418183-137418383 | 0.722858464 | 0.149191522 | 0.399516949 |
| chr2:26478021-26478223 | -0.673001869 | 0.149195873 | 0.399516949 |
| chr4:23368107-23368591 | -0.582838783 | 0.149199697 | 0.399516949 |
| chr3:155952414-155952662 | 0.645908954 | 0.149202285 | 0.399516949 |
| chr12:128259468-128259768 | -0.695737674 | 0.149213042 | 0.399531791 |
| chr14:106830636-106831016 | 0.553775561 | 0.149240064 | 0.399582653 |
| chr2:48496595-48496876 | -0.743219786 | 0.149242468 | 0.399582653 |
| chr10:74127141-74127344 | -0.794383488 | 0.149253411 | 0.39959799 |
| chr21:46334289-46334489 | -0.77705931 | 0.149269968 | 0.39961002 |
| chr4:121774676-121774876 | -0.77705931 | 0.149269968 | 0.39961002 |
| chr13:114143818-114144018 | -0.748260481 | 0.149277056 | 0.39961002 |
| chr3:11271811-11272878 | 0.464928952 | 0.149278766 | 0.39961002 |
| chr10:88460731-88460988 | 0.706347127 | 0.149284629 | 0.399611753 |
| chr3:147399116-147399456 | -0.529293978 | 0.149329467 | 0.399695642 |
| chr10:6667767-6667967 | 0.751578353 | 0.149331458 | 0.399695642 |
| chr12:121799621-121800423 | 0.469473342 | 0.14933777 | 0.399695642 |
| chr17:21020923-21021192 | -0.7854403 | 0.149337911 | 0.399695642 |
| chr19:10879184-10879472 | -0.635407703 | 0.14934205 | 0.399695642 |
| chr7:23722374-23723306 | 0.482139476 | 0.149359641 | 0.399728761 |
| chr6:42122646-42123074 | 0.515209497 | 0.149379673 | 0.399764885 |
| chr5:159133139-159133501 | -0.642483803 | 0.149388534 | 0.399764885 |
| chr10:88932490-88933006 | -0.506622438 | 0.149400986 | 0.399764885 |
| chr5:116864328-116864610 | 0.731470499 | 0.149405515 | 0.399764885 |
| chr7:115634662-115634879 | -0.781789698 | 0.149407175 | 0.399764885 |
| chr7:129780847-129781069 | -0.591288856 | 0.149411162 | 0.399764885 |
| chr1:32725149-32725474 | 0.593168088 | 0.149416731 | 0.399764885 |
| chr2:85266509-85266746 | 0.659213908 | 0.149418272 | 0.399764885 |
| chr21:38191427-38191790 | 0.598881391 | 0.149420095 | 0.399764885 |
| chr6:73610511-73610758 | -0.707004062 | 0.149434391 | 0.399789172 |
| chrX:67674225-67674534 | -0.764411655 | 0.149458688 | 0.399840214 |
| chr15:87047456-87047718 | 0.685053198 | 0.149465496 | 0.399844466 |
| chr18:23101702-23101902 | -0.747819505 | 0.149483814 | 0.399879509 |
| chr9:69054207-69054513 | -0.613112749 | 0.14949349 | 0.399891432 |
| chr3:19118491-19118695 | 0.738060259 | 0.149545749 | 0.400017258 |
| chr2:231613674-231614208 | 0.527377027 | 0.149551883 | 0.400019703 |
| chr10:55500982-55501307 | -0.586214094 | 0.149591451 | 0.400111131 |
| chr7:20329614-20331769 | 0.451888166 | 0.149598684 | 0.400111131 |

| | | | |
|---|---|---|---|
| chr1:179229879-179230159 | -0.598786115 | 0.14960173 | 0.400111131 |
| chr10:112804500-112804745 | 0.665850954 | 0.149609208 | 0.400117164 |
| chr11:58592915-58593136 | 0.733930822 | 0.149629167 | 0.400135756 |
| chr16:19691173-19691373 | -0.758830463 | 0.149630204 | 0.400135756 |
| chr20:53580748-53581108 | 0.47094756 | 0.149631826 | 0.400135756 |
| chr10:102133523-102133820 | 0.564002018 | 0.149639457 | 0.400142196 |
| chr15:38072287-38073398 | -0.475609594 | 0.149645461 | 0.400144285 |
| chr2:29033318-29033661 | -0.618816881 | 0.149664121 | 0.400180217 |
| chr1:107039899-107040177 | 0.527734245 | 0.149671899 | 0.400187048 |
| chr3:142078493-142078732 | 0.768370256 | 0.149682824 | 0.400188582 |
| chr4:89056986-89057496 | -0.511087456 | 0.149682919 | 0.400188582 |
| chr5:143183204-143183750 | -0.537856449 | 0.149694438 | 0.400205416 |
| chr5:170136138-170136596 | -0.567367268 | 0.149732443 | 0.400285819 |
| chr3:64255407-64255698 | 0.497407227 | 0.149739365 | 0.400285819 |
| chr12:75527009-75527263 | -0.650964098 | 0.149740185 | 0.400285819 |
| chr14:95427076-95427309 | 0.597595476 | 0.149756412 | 0.400302122 |
| chr8:93822368-93822697 | 0.516844048 | 0.149756732 | 0.400302122 |
| chr2:120695393-120695978 | -0.533721186 | 0.149762752 | 0.400304247 |
| chr1:172582830-172583222 | -0.627824988 | 0.149771147 | 0.400312723 |
| chr18:54269243-54270602 | -0.460426499 | 0.149777721 | 0.400313654 |
| chr12:49828111-49829050 | 0.467566351 | 0.149782469 | 0.400313654 |
| chr16:9194121-9194321 | 0.759020869 | 0.149787169 | 0.400313654 |
| chr1:206739056-206739350 | -0.537994277 | 0.149811836 | 0.400356092 |
| chr13:41942677-41943052 | -0.570843976 | 0.149813499 | 0.400356092 |
| chr22:23627479-23627771 | -0.625730461 | 0.149828664 | 0.40037317 |
| chr20:38304917-38305185 | 0.658751627 | 0.14983034 | 0.40037317 |
| chr2:11398821-11399078 | -0.652419517 | 0.149837768 | 0.400379058 |
| chr17:47220771-47221047 | -0.650950013 | 0.149844904 | 0.400384163 |
| chr6:31702893-31703631 | 0.483713452 | 0.149871312 | 0.400438393 |
| chr14:80202990-80203304 | -0.633525473 | 0.149875652 | 0.400438393 |
| chr1:153976786-153978260 | 0.462526344 | 0.149889384 | 0.400461118 |
| chr6:70242043-70242529 | -0.574751824 | 0.149896122 | 0.400465155 |
| chr12:131755372-131755587 | 0.685642684 | 0.149913362 | 0.400497249 |
| chr3:13885444-13885766 | 0.541411185 | 0.14993907 | 0.400535577 |
| chr12:15221073-15221639 | -0.508336015 | 0.149940367 | 0.400535577 |
| chr17:64056389-64056730 | 0.546493595 | 0.14994379 | 0.400535577 |
| chr10:32139351-32139630 | -0.62406695 | 0.149953761 | 0.400535577 |
| chr5:103475726-103476202 | -0.578797719 | 0.149958597 | 0.400535577 |
| chr3:61249939-61250389 | 0.496266562 | 0.149959073 | 0.400535577 |
| chrX:132132943-132133480 | -0.584992721 | 0.149981026 | 0.400567009 |
| chr17:19011113-19011408 | -0.600277835 | 0.149981297 | 0.400567009 |
| chr7:80918907-80919482 | -0.4747797 | 0.149991148 | 0.400571272 |
| chr7:139359373-139360884 | -0.452971075 | 0.14999594 | 0.400571272 |
| chr13:35879270-35879513 | -0.732006962 | 0.149998577 | 0.400571272 |
| chr6:167849865-167850178 | 0.641066049 | 0.150024321 | 0.400620943 |
| chr19:2527719-2527919 | -0.77186206 | 0.150029428 | 0.400620943 |
| chrX:134806963-134807821 | 0.504240823 | 0.150032862 | 0.400620943 |
| chr8:134478475-134478687 | 0.653145423 | 0.150048509 | 0.40064876 |
| chr4:114326349-114326699 | -0.705870146 | 0.150062014 | 0.400670858 |
| chr12:108563583-108563792 | -0.58825016 | 0.150070092 | 0.400678465 |

| | | | |
|---|---|---|---|
| chr3:146641320-146641630 | -0.595718291 | 0.150085171 | 0.400691728 |
| chr9:120702093-120702293 | 0.73364479 | 0.15008809 | 0.400691728 |
| chr5:142432999-142433272 | -0.667291433 | 0.150090748 | 0.400691728 |
| chr2:161872686-161872957 | -0.742337685 | 0.150142532 | 0.40081601 |
| chr5:16749651-16750068 | -0.568589789 | 0.15018133 | 0.400882769 |
| chr1:197769327-197769662 | -0.617929296 | 0.150182707 | 0.400882769 |
| chr10:77410022-77410525 | 0.504175897 | 0.150183236 | 0.400882769 |
| chr10:1055893-1056141 | 0.564519469 | 0.150193197 | 0.400895392 |
| chr9:81758261-81758521 | -0.72834869 | 0.150208542 | 0.400907429 |
| chr17:40140020-40140584 | 0.495929127 | 0.150217278 | 0.400907429 |
| chr18:76411006-76411416 | -0.629989707 | 0.150217652 | 0.400907429 |
| chr11:8603954-8604314 | -0.613634089 | 0.150218635 | 0.400907429 |
| chr17:38606453-38606680 | 0.632725882 | 0.150227223 | 0.400916382 |
| chr8:66406103-66406423 | -0.722956711 | 0.150236582 | 0.400927396 |
| chr4:124887736-124888155 | -0.556610243 | 0.150257874 | 0.400961967 |
| chr3:64546466-64546788 | 0.504971628 | 0.150260003 | 0.400961967 |
| chr15:101069063-101069604 | -0.511987972 | 0.150273155 | 0.400983098 |
| chr17:54806599-54806911 | 0.703015355 | 0.150298751 | 0.401037433 |
| chr5:88660532-88661648 | -0.456839357 | 0.150332156 | 0.401112596 |
| chr8:133411769-133412095 | -0.712635224 | 0.150372273 | 0.401205666 |
| chr1:188958468-188958792 | -0.576384759 | 0.150385316 | 0.401226495 |
| chr7:6272538-6273080 | -0.49096286 | 0.150421643 | 0.40130944 |
| chr4:115332097-115332429 | -0.617100965 | 0.15044122 | 0.401347695 |
| chr6:169789895-169790993 | -0.468828477 | 0.150454097 | 0.401368075 |
| chr2:37993214-37993414 | 0.749849859 | 0.150473182 | 0.401400998 |
| chr2:58441700-58441996 | -0.66555732 | 0.150476916 | 0.401400998 |
| chr5:115246409-115246698 | -0.607321624 | 0.150513747 | 0.401485267 |
| chr17:57968719-57968953 | 0.657116207 | 0.150521128 | 0.401490978 |
| chr12:49741114-49742350 | 0.457013058 | 0.150542177 | 0.401533146 |
| chr1:114554619-114554855 | 0.517373964 | 0.150558655 | 0.401563118 |
| chr6:95091880-95092165 | -0.646559603 | 0.150570308 | 0.40157255 |
| chr10:118581273-118581779 | -0.584803081 | 0.150572673 | 0.40157255 |
| chr7:3809910-3810130 | -0.783026378 | 0.15058763 | 0.40159138 |
| chrX:9688718-9689031 | -0.711459571 | 0.150595093 | 0.40159138 |
| chr5:1392268-1392468 | 0.696026035 | 0.150599399 | 0.40159138 |
| chr18:10560126-10560456 | -0.62453137 | 0.150600698 | 0.40159138 |
| chr13:46764099-46764368 | -0.537226187 | 0.150665195 | 0.401749385 |
| chr17:68195955-68196316 | 0.524880283 | 0.150679114 | 0.401756253 |
| chrX:17788485-17788753 | 0.672124721 | 0.150687224 | 0.401756253 |
| chr10:98331974-98332191 | -0.691830283 | 0.150690083 | 0.401756253 |
| chr9:84261172-84261480 | 0.577666215 | 0.150690119 | 0.401756253 |
| chrX:110853710-110853920 | 0.67203323 | 0.150693988 | 0.401756253 |
| chr9:127937125-127938019 | 0.482984603 | 0.150714157 | 0.40179293 |
| chr10:29163819-29164132 | -0.625479707 | 0.150723057 | 0.40179293 |
| chr12:96906846-96907575 | -0.475627441 | 0.15072424 | 0.40179293 |
| chr20:58855700-58856456 | 0.463339049 | 0.150729221 | 0.40179293 |
| chr7:91636177-91636423 | -0.727173814 | 0.150733964 | 0.40179293 |
| chr2:205903727-205904110 | -0.51669856 | 0.150742024 | 0.401800436 |
| chr2:180124381-180124725 | -0.621857648 | 0.150763558 | 0.401843856 |
| chr14:65070431-65070996 | 0.494652986 | 0.150793281 | 0.4019091 |

| | | | |
|---|---|---|---|
| chr15:33066736-33066948 | -0.672657884 | 0.15083863 | 0.40201583 |
| chr7:107924959-107925255 | -0.732345049 | 0.150860884 | 0.40201583 |
| chr11:65860233-65860712 | 0.478230249 | 0.150861505 | 0.40201583 |
| chr11:120335663-120337112 | 0.467638445 | 0.150861651 | 0.40201583 |
| chrX:107736891-107737344 | 0.524399764 | 0.150864183 | 0.40201583 |
| chr3:150409768-150410488 | 0.474078127 | 0.150864806 | 0.40201583 |
| chr2:71952868-71953424 | 0.520199604 | 0.150870709 | 0.402017579 |
| chr22:29306695-29307394 | 0.492128327 | 0.150888459 | 0.402050894 |
| chr2:14693963-14694221 | -0.676207664 | 0.150899969 | 0.40206758 |
| chr14:86038371-86038731 | -0.558831596 | 0.15092124 | 0.402110275 |
| chr2:183078302-183079273 | -0.461895102 | 0.150935437 | 0.402115543 |
| chr14:76696800-76697195 | 0.555254665 | 0.150937867 | 0.402115543 |
| chr3:101561455-101562206 | -0.467668909 | 0.150938962 | 0.402115543 |
| chrX:131841218-131842546 | -0.476509842 | 0.150957079 | 0.402149826 |
| chr14:60313241-60313443 | 0.714010792 | 0.150992379 | 0.402229881 |
| chr1:169509452-169509672 | -0.651736828 | 0.151017633 | 0.40228317 |
| chr10:84362078-84362484 | -0.617695677 | 0.151042316 | 0.402325505 |
| chr6:112087094-112088010 | -0.470414327 | 0.151044028 | 0.402325505 |
| chr1:217718864-217719272 | -0.527873026 | 0.151055973 | 0.402343338 |
| chrX:100979938-100981197 | -0.480902793 | 0.151078406 | 0.402389099 |
| chr10:121927265-121928545 | 0.465063412 | 0.151159876 | 0.402592096 |
| chr6:106782270-106782889 | 0.52274007 | 0.15117128 | 0.402608476 |
| chr16:66508322-66508528 | -0.745924731 | 0.151178748 | 0.402614369 |
| chr9:112751326-112751870 | 0.507600985 | 0.151201155 | 0.402647503 |
| chr5:119418777-119419067 | -0.581914484 | 0.151201699 | 0.402647503 |
| chr9:135086056-135086289 | 0.641977335 | 0.151222914 | 0.402690002 |
| chr13:51935840-51936240 | -0.576102156 | 0.151238273 | 0.402701917 |
| chr12:90909110-90909436 | -0.612235413 | 0.151244516 | 0.402701917 |
| chr19:1082003-1082931 | 0.460392045 | 0.151247603 | 0.402701917 |
| chr1:205983847-205984475 | 0.475384631 | 0.151248411 | 0.402701917 |
| chr6:37257445-37258438 | 0.467898814 | 0.151255894 | 0.402707848 |
| chr3:64711763-64711965 | 0.528160605 | 0.151270735 | 0.402733365 |
| chr4:168694531-168695142 | 0.516342923 | 0.15127715 | 0.40273645 |
| chr8:89849297-89849517 | 0.595558149 | 0.151287415 | 0.402749785 |
| chr1:63088079-63088279 | -0.790467307 | 0.151297052 | 0.402761447 |
| chr6:45922868-45923068 | -0.76475695 | 0.15132267 | 0.402803087 |
| chr7:55254538-55255368 | -0.476215931 | 0.151323208 | 0.402803087 |
| chr8:118878326-118879030 | -0.476112869 | 0.151355924 | 0.40286997 |
| chr7:100100452-100100987 | -0.480994372 | 0.151361582 | 0.40286997 |
| chr6:26165883-26166174 | 0.642433784 | 0.151364108 | 0.40286997 |
| chr19:11293790-11294208 | 0.543940658 | 0.151376321 | 0.40288848 |
| chr7:41704917-41705895 | -0.469397636 | 0.151393837 | 0.402921104 |
| chr8:131094196-131094447 | -0.741802752 | 0.151421919 | 0.402978853 |
| chr5:96175177-96175403 | -0.683507737 | 0.151426054 | 0.402978853 |
| chr7:30866904-30867104 | -0.816637624 | 0.151439144 | 0.402999691 |
| chr11:4493119-4493539 | 0.559756391 | 0.151450135 | 0.403001751 |
| chr4:74974495-74974820 | -0.766084835 | 0.151452093 | 0.403001751 |
| chr18:36066906-36068002 | -0.466678078 | 0.151455697 | 0.403001751 |
| chr22:43341646-43341864 | 0.75584291 | 0.151462702 | 0.403006394 |
| chr4:169104069-169104453 | -0.549549456 | 0.151470268 | 0.403012532 |

| | | | |
|---|---|---|---|
| chr8:134039283-134039565 | 0.60341437 | 0.151479821 | 0.403017799 |
| chr15:25850413-25850801 | -0.601985437 | 0.151482767 | 0.403017799 |
| chr7:66902137-66902355 | -0.768270997 | 0.151533817 | 0.403135398 |
| chr12:42012198-42012846 | -0.544113269 | 0.151539281 | 0.403135398 |
| chr10:97914757-97915140 | 0.512260708 | 0.151543918 | 0.403135398 |
| chr16:75029722-75030225 | -0.534015633 | 0.151551837 | 0.403135398 |
| chr8:129985515-129985990 | 0.510130652 | 0.151557277 | 0.403135398 |
| chr11:78645270-78645578 | -0.593766153 | 0.151558538 | 0.403135398 |
| chr4:74214861-74215111 | -0.690537404 | 0.151567493 | 0.403145222 |
| chr15:89411468-89411842 | 0.502496056 | 0.151584216 | 0.403164703 |
| chr8:130651045-130651462 | -0.576915488 | 0.15158534 | 0.403164703 |
| chr8:144045957-144046157 | 0.65566467 | 0.151592047 | 0.403168548 |
| chrX:128106722-128107004 | 0.663503303 | 0.151606925 | 0.403192472 |
| chr4:115234506-115234941 | -0.620448004 | 0.151611567 | 0.403192472 |
| chr1:188345073-188345288 | -0.659592285 | 0.151637129 | 0.403246456 |
| chr1:201955139-201955981 | 0.46908171 | 0.151646476 | 0.403257315 |
| chr5:53733953-53734218 | -0.673195954 | 0.151679313 | 0.403330638 |
| chr15:83352739-83353202 | -0.533773013 | 0.151689688 | 0.403338004 |
| chr13:55245150-55245622 | -0.599191065 | 0.151692611 | 0.403338004 |
| chr3:164380224-164380784 | -0.542445804 | 0.151709247 | 0.40336824 |
| chr1:3796388-3797047 | 0.495735791 | 0.151719286 | 0.403380934 |
| chr5:67235664-67236121 | -0.592147599 | 0.151742504 | 0.403425839 |
| chr15:41811480-41811680 | 0.69183651 | 0.151746706 | 0.403425839 |
| chr1:2240206-2240533 | 0.594636759 | 0.151768367 | 0.403469427 |
| chr12:48815238-48815644 | 0.505669114 | 0.151782526 | 0.403493069 |
| chr2:132430802-132431058 | 0.662291404 | 0.151792473 | 0.403505511 |
| chr8:132674956-132676082 | -0.473650544 | 0.151824949 | 0.403550798 |
| chr2:45807925-45808125 | 0.749077385 | 0.151826618 | 0.403550798 |
| chr6:156028382-156028661 | -0.767560893 | 0.151833605 | 0.403550798 |
| chr10:96340391-96340616 | 0.691551925 | 0.151834316 | 0.403550798 |
| chr2:172427278-172427900 | 0.537097937 | 0.151836147 | 0.403550798 |
| chr3:135736875-135737075 | 0.730659764 | 0.151844199 | 0.403550798 |
| chr3:30078783-30079039 | 0.70407394 | 0.151846376 | 0.403550798 |
| chr16:66934278-66934767 | 0.542594141 | 0.151859749 | 0.403556732 |
| chr15:76904707-76904968 | 0.590694799 | 0.15186372 | 0.403556732 |
| chr9:83622732-83623864 | -0.460413992 | 0.15186441 | 0.403556732 |
| chr17:61843222-61843736 | 0.465613208 | 0.151876551 | 0.403574999 |
| chr8:26307936-26308337 | 0.586405 | 0.151884164 | 0.403581232 |
| chr14:69439585-69439785 | 0.79409726 | 0.151901289 | 0.403604527 |
| chr15:68560003-68560220 | 0.660407033 | 0.151903466 | 0.403604527 |
| chr19:49559045-49559261 | -0.774580706 | 0.151985085 | 0.403797201 |
| chr7:69211802-69212061 | -0.735049328 | 0.151991391 | 0.403797201 |
| chr7:17710907-17711124 | 0.682045094 | 0.151991792 | 0.403797201 |
| chr2:128608909-128609149 | -0.70706901 | 0.152016841 | 0.403849746 |
| chr3:165254857-165255165 | -0.561645221 | 0.152039972 | 0.403882829 |
| chr5:24228840-24229040 | 0.736651705 | 0.152044724 | 0.403882829 |
| chr8:71819949-71820297 | 0.570052428 | 0.152045107 | 0.403882829 |
| chr7:74405319-74405621 | -0.61196514 | 0.152053638 | 0.403891487 |
| chr7:34675806-34676115 | 0.546580473 | 0.152066511 | 0.403901774 |
| chrX:135707902-135708330 | -0.524481287 | 0.152068053 | 0.403901774 |

| | | | |
|---|---|---|---|
| chr6:28922106-28922411 | 0.533347746 | 0.152089531 | 0.403944817 |
| chrX:68732965-68733207 | 0.693833531 | 0.152099933 | 0.403958443 |
| chr2:83827719-83827925 | -0.809876154 | 0.152108318 | 0.40396671 |
| chr11:104968971-104969405 | -0.58611947 | 0.152125553 | 0.40399848 |
| chr17:61444709-61445197 | 0.461021848 | 0.152181884 | 0.40413407 |
| chr6:80192385-80192666 | -0.708835161 | 0.152189416 | 0.404140065 |
| chr7:80876921-80877289 | -0.608534122 | 0.152207919 | 0.404173686 |
| chr14:91286481-91286801 | 0.554656595 | 0.152212627 | 0.404173686 |
| chr12:55940028-55940628 | 0.491836225 | 0.152225747 | 0.404194216 |
| chr3:144067345-144067577 | -0.713016149 | 0.152230909 | 0.404194216 |
| chr4:34314681-34315178 | -0.542921874 | 0.15225649 | 0.40424813 |
| chr10:52435396-52435686 | -0.607502274 | 0.152267496 | 0.404263344 |
| chr10:68559906-68560546 | -0.528155088 | 0.152283325 | 0.404291359 |
| chr9:101530169-101530400 | 0.593088373 | 0.152340333 | 0.404422083 |
| chr17:60198133-60198485 | 0.501316653 | 0.15234312 | 0.404422083 |
| chr2:190550186-190550462 | -0.658447758 | 0.15236219 | 0.404445559 |
| chr14:64841184-64841384 | 0.669696434 | 0.152363628 | 0.404445559 |
| chr14:101235212-101235551 | -0.591125584 | 0.152367799 | 0.404445559 |
| chr11:118004066-118004375 | -0.697546679 | 0.1523779 | 0.404458359 |
| chr7:148122515-148122724 | 0.765926167 | 0.152389277 | 0.404474545 |
| chr7:6238472-6238672 | 0.714319891 | 0.152404981 | 0.404502215 |
| chr12:67772654-67772949 | 0.649459922 | 0.152412596 | 0.404508415 |
| chr8:131999242-131999736 | 0.518474737 | 0.152431122 | 0.404533611 |
| chr16:3716794-3717913 | 0.475714681 | 0.152433337 | 0.404533611 |
| chr4:148302428-148302684 | 0.666968181 | 0.152437928 | 0.404533611 |
| chr2:227884497-227884773 | -0.633272628 | 0.152443553 | 0.404534526 |
| chr3:73024827-73025188 | -0.588196999 | 0.152471061 | 0.404593512 |
| chr16:81405744-81406296 | -0.508955061 | 0.152478914 | 0.404600338 |
| chr4:137532167-137532751 | -0.528181234 | 0.152539746 | 0.404747737 |
| chr21:43975539-43975930 | -0.583578114 | 0.152545956 | 0.404750198 |
| chr16:57244639-57245811 | 0.466157447 | 0.152559444 | 0.404765627 |
| chr14:66117589-66117789 | 0.695235128 | 0.152562484 | 0.404765627 |
| chr3:157572587-157572934 | -0.563079937 | 0.152567619 | 0.404765627 |
| chr3:113844682-113844992 | -0.59185883 | 0.152575787 | 0.404773282 |
| chr3:106923412-106923915 | -0.524841039 | 0.152581221 | 0.404773684 |
| chr14:38662828-38663192 | 0.568931924 | 0.152600859 | 0.404811766 |
| chr15:89356819-89357054 | 0.556920822 | 0.152609601 | 0.404820939 |
| chr12:115917177-115917544 | 0.541449409 | 0.15264053 | 0.404868206 |
| chr7:77834615-77834964 | -0.58199158 | 0.152642063 | 0.404868206 |
| chr2:191155528-191155978 | -0.550311577 | 0.15264459 | 0.404868206 |
| chr1:247119656-247120184 | -0.505757112 | 0.152648555 | 0.404868206 |
| chr6:145200011-145200273 | 0.64595207 | 0.152690858 | 0.404960603 |
| chr17:75326158-75326359 | -0.71663196 | 0.152693962 | 0.404960603 |
| chrX:153743641-153744158 | 0.519271225 | 0.152709133 | 0.404986821 |
| chr16:74700212-74701392 | -0.462359715 | 0.152719475 | 0.404992681 |
| chr20:54232431-54232687 | 0.504158599 | 0.152721915 | 0.404992681 |
| chr1:114239533-114239762 | 0.524912863 | 0.152734228 | 0.405007423 |
| chr20:3407009-3407283 | 0.597706654 | 0.152738866 | 0.405007423 |
| chr9:109410191-109410451 | 0.659628988 | 0.152743331 | 0.405007423 |
| chr2:17671074-17671274 | -0.743403048 | 0.152756571 | 0.405016244 |

| | | | |
|---|---|---|---|
| chr1:113510831-113511031 | 0.76677699 | 0.152757229 | 0.405016244 |
| chr9:15357096-15357305 | -0.832900055 | 0.152791706 | 0.405093636 |
| chr8:102152420-102152681 | 0.6310599 | 0.152840537 | 0.405201055 |
| chr20:17438682-17439039 | -0.609694923 | 0.152842798 | 0.405201055 |
| chr17:54544337-54544714 | -0.647438104 | 0.152889912 | 0.405311936 |
| chr9:82205402-82205669 | -0.813143929 | 0.152903456 | 0.405318567 |
| chr15:99468543-99468792 | 0.692667863 | 0.152905461 | 0.405318567 |
| chr3:34668977-34670024 | -0.481836451 | 0.152908284 | 0.405318567 |
| chr10:86688345-86688648 | 0.517574433 | 0.152915952 | 0.405324871 |
| chr3:49719390-49719889 | 0.504258347 | 0.152930836 | 0.405339682 |
| chr5:169980338-169980995 | -0.480718091 | 0.15293212 | 0.405339682 |
| chr1:68401509-68401935 | -0.514612029 | 0.152954519 | 0.405380185 |
| chr5:20972189-20972970 | -0.476887879 | 0.152960172 | 0.405380185 |
| chr13:112213343-112213870 | 0.493871487 | 0.152963273 | 0.405380185 |
| chr2:15560577-15561564 | -0.471061339 | 0.152973847 | 0.405385531 |
| chr1:182789258-182789923 | 0.489438694 | 0.152975872 | 0.405385531 |
| chr4:107183212-107183539 | -0.566468705 | 0.152986478 | 0.405393403 |
| chr2:5442783-5443051 | 0.619489691 | 0.152991913 | 0.405393403 |
| chr12:120430513-120430770 | 0.606465677 | 0.152999196 | 0.405393403 |
| chr11:13347962-13348167 | -0.69588787 | 0.153005126 | 0.405393403 |
| chr15:63047998-63049381 | 0.455750771 | 0.153005296 | 0.405393403 |
| chr17:38628489-38628689 | 0.764526567 | 0.153034661 | 0.405457185 |
| chr5:161007960-161008195 | -0.678944214 | 0.153041142 | 0.405460335 |
| chr9:706657-706919 | 0.591439974 | 0.153068126 | 0.405517804 |
| chrX:111447231-111447541 | 0.699141057 | 0.153074632 | 0.40552102 |
| chr8:61480689-61480893 | 0.721963748 | 0.153105967 | 0.405590009 |
| chr2:72862156-72863094 | -0.479303097 | 0.153138206 | 0.405661388 |
| chr7:23938943-23939314 | 0.508187273 | 0.153144235 | 0.405663333 |
| chr2:231701661-231701898 | -0.67674695 | 0.153152353 | 0.405670812 |
| chr12:122127250-122127450 | 0.67637286 | 0.153183643 | 0.405732346 |
| chr3:101753167-101754131 | -0.465202984 | 0.153186482 | 0.405732346 |
| chr2:101327743-101328379 | -0.494522574 | 0.15319756 | 0.405732346 |
| chr9:79955425-79955625 | -0.792761641 | 0.153199274 | 0.405732346 |
| chr14:37114031-37114263 | 0.654539482 | 0.15320206 | 0.405732346 |
| chr6:44110480-44110712 | -0.697520574 | 0.153219935 | 0.405765661 |
| chr1:182068981-182069353 | -0.536187834 | 0.153234931 | 0.405791348 |
| chr8:23421265-23421531 | 0.66565401 | 0.153253483 | 0.405826451 |
| chr13:33070800-33071000 | 0.731202254 | 0.153270538 | 0.405857589 |
| chr11:8234409-8234874 | -0.549842249 | 0.153285116 | 0.405876685 |
| chr4:68288-68488 | 0.66559877 | 0.153288344 | 0.405876685 |
| chr4:13262579-13263213 | 0.491443071 | 0.153295326 | 0.405881146 |
| chr17:57215334-57215553 | -0.627813357 | 0.153302002 | 0.405884795 |
| chr13:28796067-28796432 | -0.612038717 | 0.153335472 | 0.405959385 |
| chr17:44110884-44111504 | 0.500926931 | 0.153359972 | 0.406001268 |
| chrX:107676229-107677210 | -0.472615573 | 0.153371165 | 0.406001268 |
| chr3:64892540-64893120 | 0.467813007 | 0.153371633 | 0.406001268 |
| chr14:56856207-56856407 | -0.806178593 | 0.153372487 | 0.406001268 |
| chr22:49935893-49936574 | 0.492939704 | 0.153407374 | 0.40607959 |
| chr3:107317743-107317969 | 0.752394802 | 0.153415124 | 0.406086076 |
| chr12:67483727-67484457 | -0.475295864 | 0.153445423 | 0.406141599 |

| | | | |
|---|---|---|---|
| chr6:118875733-118876103 | -0.608964111 | 0.153450131 | 0.406141599 |
| chr1:224050666-224051129 | 0.548101784 | 0.153454932 | 0.406141599 |
| chr8:95225539-95225888 | 0.530597218 | 0.153457302 | 0.406141599 |
| chr20:46022482-46023169 | 0.486681968 | 0.153471631 | 0.406165491 |
| chr19:9934008-9934509 | 0.520143481 | 0.153477381 | 0.406166681 |
| chr5:115665563-115665763 | 0.734162952 | 0.15350025 | 0.40621317 |
| chr13:44141664-44142524 | -0.463206342 | 0.153534497 | 0.406289768 |
| chr14:93332140-93333447 | 0.454989675 | 0.153542376 | 0.406296587 |
| chr1:172744953-172745261 | -0.572737357 | 0.153548749 | 0.406299418 |
| chr8:39470995-39471896 | -0.506197529 | 0.153573613 | 0.406351176 |
| chr14:53002563-53002792 | 0.736869956 | 0.153597129 | 0.406386399 |
| chr16:70018304-70018676 | -0.567515708 | 0.153597532 | 0.406386399 |
| chr4:337157-338335 | 0.46012329 | 0.15362993 | 0.406445116 |
| chr13:34907257-34907476 | -0.709021763 | 0.153630334 | 0.406445116 |
| chr15:82709399-82709657 | 0.565438129 | 0.153641808 | 0.406461437 |
| chr2:69307015-69307736 | -0.48336729 | 0.153663592 | 0.406489518 |
| chr3:114178888-114179088 | -0.748883028 | 0.153667879 | 0.406489518 |
| chr20:47428195-47428571 | 0.463230661 | 0.153668338 | 0.406489518 |
| chr10:80925048-80925351 | 0.653193827 | 0.153687911 | 0.40652726 |
| chr14:67151108-67151614 | 0.565625229 | 0.153702396 | 0.406551538 |
| chr16:53434302-53434641 | 0.548499376 | 0.153711794 | 0.406562364 |
| chr1:155017178-155017839 | 0.474424481 | 0.153752556 | 0.406615245 |
| chr4:106409596-106410253 | 0.524928752 | 0.153756651 | 0.406615245 |
| chr5:172716890-172717107 | 0.575300856 | 0.153758382 | 0.406615245 |
| chr2:174748899-174749264 | -0.536183524 | 0.153759276 | 0.406615245 |
| chr15:61227981-61228217 | 0.714532938 | 0.153761055 | 0.406615245 |
| chr15:31366450-31366857 | -0.580627997 | 0.153763628 | 0.406615245 |
| chr7:71733046-71733678 | -0.540876109 | 0.15378151 | 0.406635197 |
| chr18:70337175-70337475 | -0.568616318 | 0.153781787 | 0.406635197 |
| chr6:111605468-111606806 | -0.459938524 | 0.153802314 | 0.40667544 |
| chr6:70520700-70521074 | -0.75970903 | 0.153808978 | 0.406679026 |
| chr1:77018786-77019076 | 0.618907379 | 0.153828755 | 0.406713738 |
| chr21:28517018-28517371 | -0.577984299 | 0.153832722 | 0.406713738 |
| chr1:215323567-215323888 | -0.626938218 | 0.15384384 | 0.406729098 |
| chr8:86494526-86494758 | 0.670699547 | 0.153874352 | 0.40679573 |
| chr14:23551548-23551940 | 0.484949367 | 0.153886515 | 0.406813846 |
| chr9:85512080-85512311 | -0.770696267 | 0.15389946 | 0.406834034 |
| chr2:120399156-120399525 | 0.553144211 | 0.153915388 | 0.406861373 |
| chr20:53819719-53820623 | 0.449133162 | 0.153920423 | 0.406861373 |
| chr4:170788437-170789108 | -0.531622677 | 0.153941696 | 0.406879486 |
| chr20:16760517-16760756 | 0.621729033 | 0.153949938 | 0.406879486 |
| chr3:62667774-62667988 | -0.550676105 | 0.153952345 | 0.406879486 |
| chr10:15902622-15903091 | -0.555972158 | 0.153957421 | 0.406879486 |
| chr3:99648035-99648380 | -0.548826691 | 0.153966163 | 0.406879486 |
| chr1:32351038-32352182 | 0.457981523 | 0.153970051 | 0.406879486 |
| chr15:30791219-30791537 | -0.643419587 | 0.153973328 | 0.406879486 |
| chr2:71460884-71461245 | -0.567584688 | 0.153973556 | 0.406879486 |
| chr2:232651695-232652186 | -0.54652159 | 0.153978558 | 0.406879486 |
| chr3:103991856-103992056 | -0.762901849 | 0.153980377 | 0.406879486 |
| chr14:85131498-85131846 | 0.554217652 | 0.15398972 | 0.40689014 |

| | | | |
|---|---|---|---|
| chr9:114738878-114739148 | 0.612474929 | 0.154010272 | 0.406930412 |
| chr1:188944449-188944716 | -0.681869595 | 0.154023512 | 0.406937891 |
| chr20:53849631-53849847 | 0.483951651 | 0.15402374 | 0.406937891 |
| chr2:204691933-204692272 | -0.62447464 | 0.154029035 | 0.406937891 |
| chr20:12207949-12208190 | -0.70576059 | 0.154045117 | 0.406950853 |
| chr2:44361066-44362784 | -0.451270456 | 0.15404533 | 0.406950853 |
| chr7:4849372-4849793 | -0.548032536 | 0.154049875 | 0.406950853 |
| chr5:138283191-138283611 | -0.507685857 | 0.154071225 | 0.406960534 |
| chr2:68487776-68488005 | -0.771312313 | 0.154074347 | 0.406960534 |
| chr2:109546169-109546370 | -0.771312313 | 0.154074347 | 0.406960534 |
| chr1:80392434-80393051 | -0.514499404 | 0.154078391 | 0.406960534 |
| chr1:206714522-206715054 | 0.48587576 | 0.154080096 | 0.406960534 |
| chr5:109141865-109142094 | -0.691450772 | 0.154087552 | 0.406966201 |
| chr2:215060844-215061127 | -0.669125204 | 0.154110781 | 0.40701352 |
| chr3:136142698-136142963 | -0.712431533 | 0.154127456 | 0.407043531 |
| chr14:44846132-44846516 | 0.584641082 | 0.154162803 | 0.407122849 |
| chr7:1555180-1556630 | 0.449972348 | 0.154188705 | 0.40716642 |
| chr12:56630174-56630495 | 0.518970657 | 0.15419109 | 0.40716642 |
| chr1:184663753-184664491 | -0.486103043 | 0.154195244 | 0.40716642 |
| chrX:129982201-129983155 | 0.460307774 | 0.154224669 | 0.407223611 |
| chrX:150898271-150899754 | 0.455455163 | 0.154227532 | 0.407223611 |
| chr3:196793227-196793615 | 0.54246857 | 0.154241 | 0.407245138 |
| chr3:105792327-105792774 | -0.549719388 | 0.15425743 | 0.407249992 |
| chr13:41060098-41062186 | -0.443719243 | 0.154257438 | 0.407249992 |
| chr7:96321215-96322488 | -0.458072379 | 0.15426261 | 0.407249992 |
| chrX:102858941-102859451 | -0.576086628 | 0.154271128 | 0.407249992 |
| chr1:29803594-29803850 | -0.709130996 | 0.154271752 | 0.407249992 |
| chr3:17833559-17833759 | 0.73253469 | 0.154274729 | 0.407249992 |
| chr2:19020422-19020770 | -0.69091062 | 0.154292722 | 0.407271377 |
| chr1:188782195-188782787 | -0.50840043 | 0.154306355 | 0.407271377 |
| chr4:87521872-87522438 | 0.533112836 | 0.154312954 | 0.407271377 |
| chr4:72295960-72296278 | -0.607108571 | 0.154313008 | 0.407271377 |
| chr19:47765687-47766131 | 0.535447953 | 0.154316063 | 0.407271377 |
| chr14:60534727-60534968 | 0.732708513 | 0.154322255 | 0.407271377 |
| chr14:75578135-75579423 | 0.461565351 | 0.154322377 | 0.407271377 |
| chr3:168094730-168096710 | -0.444003195 | 0.154325353 | 0.407271377 |
| chr11:44143925-44144427 | 0.531426945 | 0.154339202 | 0.407283295 |
| chr16:53435174-53435689 | 0.510674801 | 0.1543405 | 0.407283295 |
| chr20:34704078-34704794 | 0.484153768 | 0.154347528 | 0.407287815 |
| chr2:43605182-43605403 | -0.697146403 | 0.154359518 | 0.407305427 |
| chr14:95156639-95158072 | 0.451600573 | 0.154372786 | 0.407320103 |
| chr3:31980518-31982080 | 0.448965712 | 0.154376925 | 0.407320103 |
| chr9:124482468-124482733 | 0.600847812 | 0.154383968 | 0.407320103 |
| chrX:46325496-46325698 | -0.723333099 | 0.154386344 | 0.407320103 |
| chr20:9215649-9215856 | -0.728887633 | 0.154392746 | 0.407322968 |
| chr3:71416846-71417176 | -0.648325556 | 0.154411894 | 0.40735946 |
| chr3:195897332-195897566 | -0.602409243 | 0.154455393 | 0.407460185 |
| chr1:84944702-84945528 | -0.480170566 | 0.154464251 | 0.407469527 |
| chr12:25866924-25867231 | -0.68339767 | 0.154473815 | 0.407480727 |
| chr5:118638639-118639149 | 0.560899306 | 0.154550174 | 0.407668115 |

| | | | |
|---|---|---|---|
| chr16:9139990-9140221 | -0.635316117 | 0.15456887 | 0.407700281 |
| chr7:17640807-17641059 | 0.608936613 | 0.154574211 | 0.407700281 |
| chr19:53520492-53521667 | 0.461077708 | 0.154583346 | 0.407700281 |
| chr10:114654873-114655094 | 0.729319942 | 0.154583881 | 0.407700281 |
| chr1:204684706-204685106 | 0.522672993 | 0.154588973 | 0.407700281 |
| chr2:96931990-96932308 | -0.587678819 | 0.154603202 | 0.407723776 |
| chr13:34357569-34357982 | 0.589579706 | 0.154622415 | 0.407760409 |
| chr1:55337855-55338418 | -0.53760794 | 0.154636856 | 0.407784457 |
| chr16:1494244-1494646 | 0.541175886 | 0.154705512 | 0.407951465 |
| chr3:64699260-64699650 | 0.480975372 | 0.154719268 | 0.4079737 |
| chr11:819245-819851 | 0.497332247 | 0.154729886 | 0.407983811 |
| chr5:176559778-176560015 | -0.681096434 | 0.154733752 | 0.407983811 |
| chr10:83606209-83606719 | -0.553280722 | 0.154785236 | 0.408105515 |
| chr6:33585663-33586038 | 0.537341306 | 0.154807931 | 0.408136561 |
| chr8:8736940-8737232 | -0.62875793 | 0.154816708 | 0.408136561 |
| chr8:21116575-21117032 | -0.697671695 | 0.154817337 | 0.408136561 |
| chr16:68658357-68659013 | 0.507981153 | 0.154818318 | 0.408136561 |
| chr11:7612653-7613072 | -0.609663655 | 0.154831477 | 0.408157208 |
| chr15:44288463-44289261 | 0.474673112 | 0.154840461 | 0.40816685 |
| chr5:35617354-35618416 | -0.461656175 | 0.154858569 | 0.408192004 |
| chr8:64613021-64613221 | 0.726262524 | 0.154860658 | 0.408192004 |
| chr17:53032754-53033388 | -0.49231956 | 0.154892514 | 0.408261926 |
| chr1:22025727-22026177 | 0.53934812 | 0.154903289 | 0.408267506 |
| chr18:59918483-59918717 | -0.684687497 | 0.154905672 | 0.408267506 |
| chr20:51326303-51326516 | 0.604157645 | 0.154910616 | 0.408267506 |
| chr20:53572764-53573006 | 0.477703807 | 0.154923883 | 0.408276833 |
| chr5:21152640-21152961 | -0.603547286 | 0.154930943 | 0.408276833 |
| chr7:55415217-55415499 | -0.57463429 | 0.154934852 | 0.408276833 |
| chr2:19347568-19348571 | -0.453088019 | 0.154935468 | 0.408276833 |
| chr9:110248329-110248750 | -0.558033363 | 0.154949929 | 0.408289438 |
| chr10:52780105-52780428 | -0.522013749 | 0.154950909 | 0.408289438 |
| chr20:57159407-57159727 | 0.46383921 | 0.154972167 | 0.408331409 |
| chr1:185645661-185646017 | -0.578568521 | 0.154981331 | 0.408341513 |
| chr7:151076875-151077179 | -0.566551061 | 0.155007602 | 0.408375922 |
| chr2:1601110-1601397 | 0.572659438 | 0.155008722 | 0.408375922 |
| chr5:95822586-95823104 | -0.500682639 | 0.155011613 | 0.408375922 |
| chr5:171162644-171162882 | -0.63944246 | 0.15501571 | 0.408375922 |
| chr18:34153737-34153940 | -0.750278559 | 0.155027423 | 0.408382528 |
| chr7:101005134-101005352 | -0.77927097 | 0.155028877 | 0.408382528 |
| chr17:78865771-78866056 | 0.545023855 | 0.155039018 | 0.408390791 |
| chr4:118352589-118353337 | 0.476675602 | 0.155042673 | 0.408390791 |
| chr5:166633781-166634074 | -0.68437798 | 0.155048974 | 0.408393348 |
| chr12:93741717-93742174 | 0.56135347 | 0.155103951 | 0.408524111 |
| chr1:67500380-67500868 | -0.504972082 | 0.155131845 | 0.408551978 |
| chr17:80474554-80474879 | 0.550953989 | 0.155133553 | 0.408551978 |
| chr5:133967758-133968003 | -0.619971963 | 0.155135087 | 0.408551978 |
| chr3:62558590-62559082 | 0.472403183 | 0.155135859 | 0.408551978 |
| chr16:57377127-57378228 | -0.477183122 | 0.155158975 | 0.408593417 |
| chr1:227762439-227762747 | 0.553168697 | 0.15516226 | 0.408593417 |
| chr16:2980126-2980900 | 0.478707505 | 0.155174703 | 0.408601303 |

| | | | |
|---|---|---|---|
| chr4:154549871-154550840 | -0.458401234 | 0.15517592 | 0.408601303 |
| chr7:147412730-147413087 | 0.593646014 | 0.155189301 | 0.408622495 |
| chr18:76820298-76820637 | -0.573254282 | 0.155203403 | 0.408623505 |
| chr7:54468648-54468848 | -0.782267426 | 0.155203502 | 0.408623505 |
| chr2:134158227-134158458 | -0.760211493 | 0.155205683 | 0.408623505 |
| chr11:34537692-34538106 | 0.546805266 | 0.155237321 | 0.408692756 |
| chr6:10347568-10348097 | 0.539159788 | 0.155248027 | 0.408695055 |
| chr13:26859648-26859860 | -0.798025892 | 0.155251686 | 0.408695055 |
| chr11:128932224-128932438 | 0.773225538 | 0.155259179 | 0.408695055 |
| chr5:178042674-178043047 | -0.652725835 | 0.155259529 | 0.408695055 |
| chr1:231883508-231883718 | 0.706776519 | 0.155265078 | 0.408695621 |
| chr10:100508784-100509083 | 0.587300022 | 0.155274465 | 0.408698243 |
| chr3:22495259-22495459 | 0.756364195 | 0.155279228 | 0.408698243 |
| chr11:117946869-117947434 | 0.506415571 | 0.15528861 | 0.408698243 |
| chr15:65365247-65365568 | 0.548319619 | 0.155288712 | 0.408698243 |
| chr17:59073746-59073961 | 0.517513749 | 0.155292744 | 0.408698243 |
| chr4:56849372-56849606 | 0.705968916 | 0.155302992 | 0.408705578 |
| chr6:46990778-46991229 | -0.586507358 | 0.15531199 | 0.408705578 |
| chr22:27331720-27332012 | -0.731954498 | 0.155320079 | 0.408705578 |
| chr1:6470882-6471152 | 0.582056988 | 0.15532129 | 0.408705578 |
| chr2:44955659-44955962 | -0.558056144 | 0.155322202 | 0.408705578 |
| chr11:83256604-83256975 | -0.623416809 | 0.155369566 | 0.40881617 |
| chr12:105279462-105279769 | -0.626334426 | 0.155377909 | 0.408824083 |
| chr17:57033336-57033562 | 0.632507689 | 0.155392055 | 0.408847264 |
| chr2:85818034-85818256 | 0.655533073 | 0.155403076 | 0.408850745 |
| chr15:92188141-92188518 | -0.596081099 | 0.15540405 | 0.408850745 |
| chr8:47100677-47101025 | -0.673456266 | 0.155421772 | 0.408882552 |
| chr5:101942315-101942810 | -0.537547183 | 0.155429549 | 0.408882552 |
| chr1:107347397-107347687 | 0.502560596 | 0.155432149 | 0.408882552 |
| chr1:16545095-16545501 | -0.692859736 | 0.155466724 | 0.408959466 |
| chr17:2049590-2050580 | -0.474434056 | 0.15548153 | 0.408984373 |
| chr5:126171973-126172312 | -0.687849213 | 0.155512752 | 0.409033761 |
| chr6:82385179-82385379 | 0.76015035 | 0.155514561 | 0.409033761 |
| chr5:142145366-142145708 | -0.582955363 | 0.155516321 | 0.409033761 |
| chr1:2049879-2050518 | 0.485671544 | 0.155542341 | 0.409082136 |
| chrX:132217169-132218387 | 0.457155499 | 0.155545391 | 0.409082136 |
| chr8:133569694-133570160 | 0.482898446 | 0.1555593 | 0.409093436 |
| chr2:135744835-135745093 | -0.549327226 | 0.155561692 | 0.409093436 |
| chr5:139100937-139101152 | -0.721728404 | 0.155566035 | 0.409093436 |
| chr2:10057212-10058082 | -0.466081965 | 0.155571044 | 0.409093436 |
| chr7:123872263-123872627 | -0.540753187 | 0.155583637 | 0.409112511 |
| chr20:16701318-16701564 | 0.656432207 | 0.155601194 | 0.409130918 |
| chr1:211655210-211655560 | -0.54114606 | 0.155605462 | 0.409130918 |
| chr2:14572413-14573033 | 0.54523543 | 0.155612058 | 0.409130918 |
| chr7:107130505-107130791 | -0.681673272 | 0.155615945 | 0.409130918 |
| chr11:16935911-16936208 | -0.592747609 | 0.155620663 | 0.409130918 |
| chr2:119407734-119407968 | 0.617152286 | 0.155628053 | 0.409130918 |
| chr12:109762710-109762927 | 0.654363579 | 0.155636543 | 0.409130918 |
| chr17:59329631-59329831 | 0.542536807 | 0.155640953 | 0.409130918 |
| chr6:133817543-133817923 | -0.575938228 | 0.155642904 | 0.409130918 |

| | | | |
|---|---|---|---|
| chr12:112409101-112409877 | 0.482085446 | 0.155644034 | 0.409130918 |
| chr5:133993522-133993995 | 0.506557697 | 0.155666529 | 0.409167158 |
| chr11:77800557-77800776 | 0.714519198 | 0.155673121 | 0.409167158 |
| chr19:40403484-40403899 | 0.501076237 | 0.15567384 | 0.409167158 |
| chr20:20087867-20088105 | -0.802257143 | 0.155692114 | 0.409201152 |
| chr2:46565691-46566126 | -0.569612816 | 0.15570293 | 0.409215541 |
| chr2:22355863-22356073 | -0.751039309 | 0.155708415 | 0.409215922 |
| chr14:63179565-63179775 | 0.700236551 | 0.155723639 | 0.409241894 |
| chr20:54076068-54076766 | 0.450980946 | 0.15573367 | 0.409254218 |
| chr17:78345253-78345453 | -0.75628422 | 0.155762115 | 0.409314933 |
| chr1:228108946-228109961 | 0.463140222 | 0.155804825 | 0.409413125 |
| chr1:151898147-151898546 | 0.550530794 | 0.155822712 | 0.409446084 |
| chr19:41928335-41928535 | 0.687570351 | 0.155829363 | 0.40944952 |
| chr8:55881464-55881664 | -0.812314518 | 0.155868608 | 0.409530625 |
| chr8:132265708-132266040 | -0.672601804 | 0.15587092 | 0.409530625 |
| chr5:150538282-150538662 | -0.596239463 | 0.155919801 | 0.409645007 |
| chr22:31646652-31646927 | 0.680131728 | 0.155926151 | 0.409647645 |
| chr17:55264597-55265086 | 0.495064679 | 0.155941595 | 0.409674172 |
| chr5:38673676-38673876 | 0.73290027 | 0.155975229 | 0.409748482 |
| chr5:149128110-149128310 | -0.674181173 | 0.155992082 | 0.409778706 |
| chr17:39621996-39622264 | 0.596904018 | 0.156083996 | 0.4100061 |
| chr1:114002874-114003193 | 0.506760828 | 0.156099455 | 0.410017626 |
| chr9:135797036-135797236 | 0.717201473 | 0.156103216 | 0.410017626 |
| chr5:168288372-168288653 | 0.589078861 | 0.156107198 | 0.410017626 |
| chr6:116244282-116244653 | 0.541619371 | 0.156109788 | 0.410017626 |
| chr1:243941432-243941720 | -0.730545449 | 0.156118383 | 0.410019756 |
| chr5:61162885-61163207 | -0.542689049 | 0.156121302 | 0.410019756 |
| chr18:12423872-12424207 | 0.556749568 | 0.156139042 | 0.410035318 |
| chr12:88393427-88393774 | -0.590489716 | 0.156140647 | 0.410035318 |
| chr15:88450305-88450505 | 0.63597313 | 0.156148691 | 0.410035318 |
| chr8:126539569-126539842 | 0.598412235 | 0.156151209 | 0.410035318 |
| chr1:242019588-242020126 | -0.554313494 | 0.156153984 | 0.410035318 |
| chr8:143874250-143874770 | 0.482508361 | 0.156166828 | 0.41005499 |
| chr8:25184216-25185777 | 0.461766958 | 0.156179716 | 0.410074778 |
| chr2:217331495-217331968 | 0.508968135 | 0.156186771 | 0.41007925 |
| chr1:198686166-198686569 | 0.581976119 | 0.156227216 | 0.410167599 |
| chr4:105694859-105695061 | -0.781944912 | 0.15623336 | 0.410167599 |
| chr1:206798138-206798417 | -0.556437931 | 0.156237551 | 0.410167599 |
| chr12:31749150-31749710 | -0.476589172 | 0.156243868 | 0.410167599 |
| chr16:67279276-67279659 | 0.52536219 | 0.156249055 | 0.410167599 |
| chr2:108396903-108397142 | 0.661309572 | 0.156255 | 0.410167599 |
| chr15:87527911-87528136 | 0.722624065 | 0.156257892 | 0.410167599 |
| chr15:101613633-101613894 | 0.559397131 | 0.156277686 | 0.410205503 |
| chr2:98793258-98793530 | 0.627049104 | 0.156293002 | 0.410227498 |
| chr3:52471957-52472188 | -0.678467913 | 0.156296773 | 0.410227498 |
| chr5:10655227-10655577 | 0.569657705 | 0.15631315 | 0.410245287 |
| chr3:9809577-9810661 | 0.460748604 | 0.156316525 | 0.410245287 |
| chr1:106594509-106594762 | 0.497902431 | 0.156319613 | 0.410245287 |
| chr17:46894158-46894363 | -0.708216928 | 0.156337762 | 0.410278865 |
| chr14:39203419-39204373 | -0.481774155 | 0.15635705 | 0.410315428 |

| | | | |
|---|---|---|---|
| chr8:63427779-63428099 | -0.638099593 | 0.15637342 | 0.410344334 |
| chr15:29821796-29823125 | 0.470855172 | 0.156378813 | 0.410344432 |
| chr1:246573653-246573886 | -0.689433583 | 0.156403848 | 0.410396069 |
| chr5:74766782-74767808 | -0.463888624 | 0.156430766 | 0.410444499 |
| chr10:131982475-131982675 | 0.696887669 | 0.156433018 | 0.410444499 |
| chr11:77191369-77192450 | -0.456399108 | 0.156438815 | 0.410445653 |
| chr1:153352553-153352753 | 0.688721661 | 0.156450368 | 0.410461909 |
| chr5:139707763-139707984 | -0.628715053 | 0.156473189 | 0.410507727 |
| chrX:126770271-126770581 | 0.629470091 | 0.15651484 | 0.41060294 |
| chr2:30989796-30990552 | -0.497824076 | 0.156531173 | 0.410631729 |
| chr18:45722954-45723154 | -0.775502672 | 0.156536868 | 0.41063261 |
| chr20:62514309-62514548 | 0.594842645 | 0.156552903 | 0.410660615 |
| chr5:38420714-38421004 | -0.570209026 | 0.156573068 | 0.410699449 |
| chr16:11532187-11532464 | 0.590556042 | 0.156600991 | 0.410740738 |
| chr14:76894344-76894594 | 0.614775855 | 0.156601386 | 0.410740738 |
| chr20:48574458-48574758 | 0.482020883 | 0.15660489 | 0.410740738 |
| chr1:114482690-114483046 | 0.484248221 | 0.15662075 | 0.410768272 |
| chr17:78341499-78342016 | 0.53942256 | 0.156660283 | 0.410857892 |
| chr12:107519997-107520309 | -0.635544462 | 0.156667856 | 0.410863691 |
| chr7:143490801-143491050 | -0.64520125 | 0.156681668 | 0.41088585 |
| chr1:217372080-217372645 | -0.505290252 | 0.156693913 | 0.410903441 |
| chr1:247238437-247238911 | -0.593846278 | 0.156699101 | 0.410903441 |
| chr17:72725527-72725765 | 0.603449396 | 0.156724689 | 0.410946424 |
| chr8:66919396-66919651 | 0.636047636 | 0.15672622 | 0.410946424 |
| chr1:155241197-155241397 | 0.614131964 | 0.156732632 | 0.410949174 |
| chr9:81410098-81410569 | -0.550811492 | 0.156754461 | 0.410992345 |
| chr17:64699959-64700159 | -0.76361879 | 0.156785998 | 0.411057517 |
| chr12:11498661-11498949 | -0.672839456 | 0.156790047 | 0.411057517 |
| chr1:148679419-148679920 | 0.559871663 | 0.156796554 | 0.411060511 |
| chr8:89944396-89944706 | 0.681546998 | 0.156808474 | 0.411077695 |
| chr11:8813632-8813995 | -0.557823308 | 0.156834855 | 0.411132788 |
| chr12:33817718-33817958 | -0.769613784 | 0.15684079 | 0.411134281 |
| chr2:14907023-14907338 | -0.564140696 | 0.156860451 | 0.411171751 |
| chr19:9584375-9584780 | 0.558573915 | 0.156897765 | 0.411249474 |
| chr3:97197062-97197329 | -0.647857985 | 0.156901834 | 0.411249474 |
| chrX:50211527-50211857 | -0.590225628 | 0.156906203 | 0.411249474 |
| chr1:171226783-171227127 | -0.543066297 | 0.156929117 | 0.411286906 |
| chr11:4187540-4187934 | 0.517190595 | 0.156935779 | 0.411286906 |
| chr9:93201678-93202001 | -0.583515256 | 0.156939175 | 0.411286906 |
| chr17:49763496-49763772 | -0.555453735 | 0.156942336 | 0.411286906 |
| chr16:24685988-24686387 | -0.646271219 | 0.156947324 | 0.411286906 |
| chr10:107623320-107623576 | 0.648831628 | 0.156958312 | 0.411301634 |
| chr1:186482567-186483015 | -0.539338122 | 0.156969352 | 0.411316497 |
| chr2:200457535-200457767 | 0.745055043 | 0.156989921 | 0.411356328 |
| chr22:46773847-46774351 | -0.536417908 | 0.157011456 | 0.411384513 |
| chr11:34356647-34356992 | 0.542472977 | 0.157012192 | 0.411384513 |
| chr6:27903563-27903773 | -0.644333972 | 0.157016785 | 0.411384513 |
| chr3:125212409-125212842 | 0.537004081 | 0.157029387 | 0.411403462 |
| chr15:85794004-85795312 | 0.457633352 | 0.157044875 | 0.411416491 |
| chr9:83242074-83242491 | -0.575556897 | 0.157048045 | 0.411416491 |

| | | | |
|---|---|---|---|
| chr1:196317982-196318248 | -0.602278133 | 0.157051627 | 0.411416491 |
| chr20:10047513-10047943 | -0.599481662 | 0.157067036 | 0.411416491 |
| chr10:61943945-61944395 | -0.558897324 | 0.157067148 | 0.411416491 |
| chr2:191013509-191014571 | -0.452597666 | 0.157069138 | 0.411416491 |
| chr5:66306225-66306698 | -0.527018741 | 0.157076416 | 0.411416491 |
| chr3:102104223-102104423 | -0.780370553 | 0.157077315 | 0.411416491 |
| chr17:62704484-62704801 | -0.505326534 | 0.157086262 | 0.411425861 |
| chr12:53441085-53441901 | 0.457931896 | 0.157096872 | 0.411439587 |
| chr17:60096598-60097193 | 0.464309872 | 0.157104938 | 0.411446648 |
| chr8:87046294-87046664 | 0.578286758 | 0.157113177 | 0.411454161 |
| chr8:58608584-58608919 | -0.557536356 | 0.157123026 | 0.411456102 |
| chr13:36761901-36762559 | 0.496629018 | 0.157124658 | 0.411456102 |
| chr2:96761357-96761566 | -0.611238368 | 0.157166598 | 0.411536373 |
| chr22:17258061-17258454 | -0.611238368 | 0.157166598 | 0.411536373 |
| chr7:17741037-17741240 | 0.661418614 | 0.157173604 | 0.411536373 |
| chr2:63530729-63531302 | -0.524146389 | 0.1571823 | 0.411536373 |
| chr7:12595105-12595639 | 0.500634453 | 0.157182974 | 0.411536373 |
| chr20:53678064-53678273 | 0.481792469 | 0.157196785 | 0.411536373 |
| chr16:85558336-85558592 | 0.550786711 | 0.157198229 | 0.411536373 |
| chr10:128150029-128150526 | 0.510992575 | 0.15719828 | 0.411536373 |
| chr3:64179279-64179559 | 0.486961184 | 0.157230839 | 0.411597665 |
| chr15:41493077-41494120 | -0.475745429 | 0.157232435 | 0.411597665 |
| chr9:124852804-124853760 | 0.462712116 | 0.157253808 | 0.411639551 |
| chr7:137626443-137626812 | -0.598456697 | 0.15729193 | 0.411711765 |
| chr4:125977936-125978284 | -0.589746441 | 0.157292142 | 0.411711765 |
| chr18:58670924-58672252 | -0.454371525 | 0.157326806 | 0.411788431 |
| chr3:140015165-140015365 | 0.6435042 | 0.157333006 | 0.411790592 |
| chr15:101463201-101463625 | -0.549046037 | 0.157350615 | 0.411822613 |
| chr5:43389692-43390085 | 0.504613222 | 0.157356484 | 0.411823907 |
| chr7:156905864-156906064 | -0.686069679 | 0.157407606 | 0.411943629 |
| chr5:123511607-123512894 | -0.450437238 | 0.157426907 | 0.411980069 |
| chr6:133568938-133569856 | -0.495495599 | 0.157472902 | 0.412072703 |
| chr6:151325408-151325673 | -0.680580732 | 0.15747306 | 0.412072703 |
| chr1:226711255-226711543 | 0.580984588 | 0.157494006 | 0.412096654 |
| chr9:76825352-76825623 | -0.785233032 | 0.157505643 | 0.412096654 |
| chr5:34651917-34652144 | 0.689477512 | 0.157509445 | 0.412096654 |
| chr5:107466342-107466828 | -0.542348849 | 0.157513291 | 0.412096654 |
| chr9:111810484-111810974 | -0.57154984 | 0.157517364 | 0.412096654 |
| chr6:148558-148929 | -0.599144917 | 0.157523532 | 0.412096654 |
| chr11:84958267-84958686 | -0.556271811 | 0.157524616 | 0.412096654 |
| chr6:128696718-128697035 | 0.606572855 | 0.15752524 | 0.412096654 |
| chr11:82945718-82945997 | -0.670672453 | 0.15755374 | 0.412157141 |
| chr22:37457444-37457840 | -0.620152487 | 0.1575764 | 0.412202346 |
| chr2:74054098-74054340 | -0.716490013 | 0.157600115 | 0.412242383 |
| chr11:30322793-30323479 | -0.458448597 | 0.157604913 | 0.412242383 |
| chr1:151340911-151341212 | 0.519797154 | 0.157609951 | 0.412242383 |
| chr3:21751056-21751300 | 0.699775715 | 0.157613226 | 0.412242383 |
| chr3:492982-493219 | -0.700380178 | 0.157623218 | 0.412254444 |
| chr7:97532612-97533131 | -0.613342834 | 0.157651568 | 0.412300719 |
| chr3:135192653-135192854 | 0.779996415 | 0.157653081 | 0.412300719 |

| | | | |
|---|---|---|---|
| chr22:20031989-20032274 | -0.695491398 | 0.157666055 | 0.412300719 |
| chr17:61511605-61512010 | 0.460776197 | 0.157667579 | 0.412300719 |
| chr3:139312940-139313242 | -0.616248504 | 0.157667815 | 0.412300719 |
| chr1:120069351-120069798 | 0.552144569 | 0.157694749 | 0.412354441 |
| chr3:97278515-97278900 | 0.533779639 | 0.157699123 | 0.412354441 |
| chr20:54793407-54795328 | 0.441469604 | 0.157740439 | 0.412448401 |
| chr2:46873711-46873925 | -0.638525075 | 0.157780071 | 0.41252932 |
| chr15:82952185-82952901 | 0.498233911 | 0.157782155 | 0.41252932 |
| chr19:49640089-49641006 | 0.449604824 | 0.157795975 | 0.412544884 |
| chr22:18110614-18110981 | -0.589793495 | 0.157798876 | 0.412544884 |
| chr14:67420118-67420484 | 0.518081228 | 0.157840796 | 0.412640401 |
| chr2:24164212-24164751 | -0.504957162 | 0.157862343 | 0.412681764 |
| chr15:58227796-58227996 | -0.745601728 | 0.15786739 | 0.412681764 |
| chr5:72328153-72328430 | -0.700738249 | 0.1578771 | 0.412689115 |
| chr11:122203013-122203254 | 0.64739439 | 0.157881733 | 0.412689115 |
| chr3:22569659-22570060 | -0.553130471 | 0.157886361 | 0.412689115 |
| chr2:215071660-215071986 | -0.730366522 | 0.157926692 | 0.412775776 |
| chr8:103210190-103210581 | -0.556720702 | 0.157934433 | 0.412775776 |
| chr1:206909053-206910105 | -0.448291783 | 0.157937178 | 0.412775776 |
| chr11:64894327-64894605 | 0.571940436 | 0.157941064 | 0.412775776 |
| chr12:11171028-11171816 | -0.480780508 | 0.157978103 | 0.412858495 |
| chr18:48161186-48162292 | -0.462290473 | 0.157992656 | 0.412879555 |
| chr13:25771227-25771911 | 0.503229894 | 0.157996939 | 0.412879555 |
| chr7:31192953-31193282 | -0.585246688 | 0.158003681 | 0.412883092 |
| chr7:140506900-140507219 | -0.62931877 | 0.158026724 | 0.412925292 |
| chr18:76803847-76804123 | -0.569753856 | 0.158030608 | 0.412925292 |
| chr1:229143941-229144148 | 0.733203747 | 0.158038671 | 0.412932277 |
| chr7:83271280-83271515 | -0.69447781 | 0.158073903 | 0.41301025 |
| chr7:100629386-100629656 | -0.588583092 | 0.158084406 | 0.413017287 |
| chr17:2054440-2054787 | 0.54434893 | 0.158087377 | 0.413017287 |
| chr10:74433242-74433574 | -0.594183894 | 0.158105814 | 0.413051372 |
| chr20:57509732-57510001 | 0.527262195 | 0.15811243 | 0.413054573 |
| chr12:67275390-67275906 | -0.576818953 | 0.158126483 | 0.413077201 |
| chr5:178123557-178124753 | -0.443203171 | 0.158166449 | 0.413167517 |
| chr10:4771243-4771448 | 0.747516711 | 0.158184332 | 0.413200147 |
| chr17:57357306-57358480 | 0.451007177 | 0.158218561 | 0.413275469 |
| chr20:15297151-15297351 | 0.695822709 | 0.158225621 | 0.413279821 |
| chr1:183042443-183042643 | 0.732562965 | 0.158245567 | 0.41331783 |
| chr10:101067621-101068162 | 0.496660707 | 0.158296767 | 0.413436137 |
| chr2:219722345-219722633 | -0.623037901 | 0.158301655 | 0.413436137 |
| chr3:113231956-113232259 | 0.615003035 | 0.158321735 | 0.413474486 |
| chr12:101697249-101698648 | -0.454700915 | 0.158336199 | 0.413498169 |
| chr19:50759207-50759500 | -0.727873183 | 0.158366812 | 0.413564018 |
| chr1:197274733-197274972 | -0.7296096 | 0.158372365 | 0.413564424 |
| chr7:91034564-91034770 | -0.794087474 | 0.158377897 | 0.413564775 |
| chr22:41446751-41447174 | 0.484174294 | 0.158389161 | 0.413569631 |
| chr1:198423979-198424179 | 0.722536589 | 0.158390551 | 0.413569631 |
| chr20:51649342-51649582 | 0.585906865 | 0.158404987 | 0.41359323 |
| chr18:43984324-43984621 | -0.717262966 | 0.158410689 | 0.413594023 |
| chr12:54300828-54301093 | 0.575007598 | 0.158418408 | 0.41359435 |

| | | | |
|---|---|---|---|
| chr16:30091845-30092250 | 0.493847698 | 0.15842161 | 0.41359435 |
| chr7:152699238-152700088 | -0.495152785 | 0.158481083 | 0.413689649 |
| chr20:57104285-57104497 | 0.477109924 | 0.158482453 | 0.413689649 |
| chr18:74152963-74153168 | -0.666334504 | 0.158484535 | 0.413689649 |
| chr1:26038880-26039174 | -0.713109935 | 0.158485202 | 0.413689649 |
| chr9:97468657-97468857 | 0.674164574 | 0.158486245 | 0.413689649 |
| chr4:79820476-79820742 | -0.708722353 | 0.158490507 | 0.413689649 |
| chr2:52967024-52967466 | -0.602422059 | 0.1585119 | 0.413731395 |
| chr15:45314621-45314836 | -0.733747685 | 0.158522312 | 0.413737979 |
| chr16:73057652-73057890 | 0.581197451 | 0.158525223 | 0.413737979 |
| chr5:157172501-157172701 | -0.790883959 | 0.158531804 | 0.413741063 |
| chr4:2934713-2934913 | 0.59719171 | 0.158537281 | 0.413741264 |
| chr2:71226794-71227737 | -0.448636521 | 0.15855122 | 0.41376355 |
| chr5:117486901-117487496 | -0.506783533 | 0.158560482 | 0.413773627 |
| chr7:53100991-53101576 | -0.502898028 | 0.15857075 | 0.413786331 |
| chr20:62411667-62411907 | -0.534963756 | 0.158594164 | 0.413820323 |
| chr7:141072480-141073104 | -0.493843686 | 0.158606087 | 0.413820323 |
| chr3:189188485-189188758 | -0.663453909 | 0.1586068 | 0.413820323 |
| chr14:106011707-106011982 | 0.5653993 | 0.158607707 | 0.413820323 |
| chr17:69268572-69268820 | -0.667098488 | 0.158610781 | 0.413820323 |
| chr16:69150201-69150560 | 0.590755754 | 0.158672355 | 0.413966876 |
| chr20:53067903-53068103 | 0.738855171 | 0.158699404 | 0.414023347 |
| chr14:99381497-99381823 | -0.625007432 | 0.158724212 | 0.414073971 |
| chr8:125528472-125529118 | 0.480091771 | 0.158754284 | 0.414137244 |
| chr3:122356474-122356776 | -0.603058726 | 0.158759276 | 0.414137244 |
| chr16:3058772-3059751 | 0.483451363 | 0.158772722 | 0.414158219 |
| chr7:87927059-87927373 | -0.706202319 | 0.158801905 | 0.414206702 |
| chr2:163780296-163780550 | -0.649144912 | 0.158807052 | 0.414206702 |
| chr11:89557308-89557819 | -0.526931064 | 0.158807526 | 0.414206702 |
| chr7:16078109-16078557 | 0.556586507 | 0.15882452 | 0.414230466 |
| chr21:44455709-44456189 | 0.512475159 | 0.15882745 | 0.414230466 |
| chr6:13677851-13678278 | 0.518000862 | 0.158843401 | 0.414245422 |
| chrX:9429812-9430244 | 0.558322169 | 0.158843997 | 0.414245422 |
| chr6:87473665-87473920 | 0.576671102 | 0.158861316 | 0.414273833 |
| chr14:101009250-101009510 | -0.743875735 | 0.158865705 | 0.414273833 |
| chr1:213050888-213051840 | -0.462468946 | 0.158896317 | 0.414339559 |
| chr16:46908673-46908873 | 0.754195281 | 0.158909268 | 0.414359229 |
| chr5:172420313-172421044 | -0.467714819 | 0.158950618 | 0.414444089 |
| chr16:80804641-80805055 | 0.54090145 | 0.15895263 | 0.414444089 |
| chr13:76885115-76885975 | -0.474566759 | 0.158968438 | 0.414457131 |
| chr2:20457662-20458204 | 0.493571979 | 0.158970597 | 0.414457131 |
| chr19:39890143-39890642 | 0.566837969 | 0.158977579 | 0.414457131 |
| chr16:3059847-3060149 | 0.620974352 | 0.158984504 | 0.414457131 |
| chr10:30605733-30606071 | -0.675743225 | 0.158984678 | 0.414457131 |
| chr19:917080-917400 | -0.609640281 | 0.158991759 | 0.41446149 |
| chr1:235971565-235972133 | -0.501373731 | 0.159009751 | 0.414492503 |
| chr2:189150029-189150436 | -0.586559155 | 0.159014475 | 0.414492503 |
| chr1:223550419-223550619 | 0.69457877 | 0.159027277 | 0.41451177 |
| chr1:202614253-202614681 | -0.563869104 | 0.159058487 | 0.414579019 |
| chr17:61444127-61444544 | 0.455987736 | 0.159081771 | 0.414612783 |

| | | | |
|---|---|---|---|
| chr15:88701714-88701916 | 0.660395348 | 0.159082264 | 0.414612783 |
| chr5:134968076-134968639 | -0.529715709 | 0.159099395 | 0.414615379 |
| chr8:85828701-85829077 | 0.601359579 | 0.159100283 | 0.414615379 |
| chr20:53826045-53826245 | 0.483698808 | 0.159101332 | 0.414615379 |
| chr1:24705227-24705644 | 0.511960125 | 0.159109489 | 0.414615379 |
| chr4:170089168-170090477 | -0.453084016 | 0.159110316 | 0.414615379 |
| chr20:56346133-56346430 | 0.593125324 | 0.159125144 | 0.414639917 |
| chr8:129864176-129864470 | 0.708671183 | 0.159158224 | 0.414687566 |
| chr7:10941049-10941263 | 0.536793332 | 0.159159471 | 0.414687566 |
| chr14:50116165-50117019 | 0.465115412 | 0.159159925 | 0.414687566 |
| chr20:60078408-60078869 | -0.516837082 | 0.159165078 | 0.414687566 |
| chr15:89654902-89655102 | 0.672814043 | 0.159204842 | 0.414766073 |
| chr12:93742303-93742868 | -0.517472949 | 0.159209054 | 0.414766073 |
| chr17:55432867-55434034 | -0.452882598 | 0.15921145 | 0.414766073 |
| chr6:53945672-53946155 | -0.566185127 | 0.159243215 | 0.414822701 |
| chr16:84648181-84649043 | 0.468126026 | 0.159249498 | 0.414822701 |
| chr4:176319373-176320771 | -0.447385025 | 0.159257609 | 0.414822701 |
| chr17:44920081-44920396 | 0.562967353 | 0.159258087 | 0.414822701 |
| chr15:84840830-84841243 | -0.532503572 | 0.159264173 | 0.414822701 |
| chr11:75372259-75372530 | 0.573861026 | 0.159265671 | 0.414822701 |
| chr1:81549500-81549842 | -0.61285861 | 0.15927936 | 0.414842585 |
| chr6:49849637-49849872 | 0.652522883 | 0.159284133 | 0.414842585 |
| chr1:151024571-151024931 | 0.496273896 | 0.159317284 | 0.41491482 |
| chr13:32134328-32134594 | 0.68183485 | 0.15934524 | 0.414964843 |
| chr7:16982727-16982927 | 0.679281071 | 0.159352038 | 0.414964843 |
| chr8:86537992-86538192 | 0.689185058 | 0.15936225 | 0.414964843 |
| chr17:28750129-28750380 | -0.571944429 | 0.159366871 | 0.414964843 |
| chr8:37736211-37736801 | 0.482210676 | 0.159367944 | 0.414964843 |
| chr5:103487326-103487535 | -0.753892549 | 0.159368986 | 0.414964843 |
| chr12:101945377-101945708 | 0.555586031 | 0.159386996 | 0.414991841 |
| chr7:40483698-40484007 | -0.628997648 | 0.159394071 | 0.414991841 |
| chr8:130527861-130528091 | -0.65502393 | 0.159395603 | 0.414991841 |
| chr20:51386035-51386289 | 0.555534134 | 0.159430478 | 0.415068535 |
| chr1:44807260-44807674 | -0.519945618 | 0.159440958 | 0.415075871 |
| chr2:184138425-184138761 | -0.673231039 | 0.15944413 | 0.415075871 |
| chr7:87416748-87417024 | -0.651511234 | 0.159466588 | 0.415112868 |
| chr22:46071140-46071488 | 0.563838853 | 0.159469735 | 0.415112868 |
| chr2:191740228-191740487 | -0.670023311 | 0.159474594 | 0.415112868 |
| chr12:13962777-13963027 | -0.649178273 | 0.159487624 | 0.415132681 |
| chr12:89117393-89118068 | -0.484705553 | 0.159496583 | 0.415136958 |
| chr6:150053837-150054076 | -0.74150214 | 0.159500103 | 0.415136958 |
| chr13:33335516-33335716 | 0.780027024 | 0.159532473 | 0.415207103 |
| chr5:10307327-10308043 | -0.466530928 | 0.15954002 | 0.415211911 |
| chr19:18636122-18637080 | 0.453875681 | 0.159545158 | 0.415211911 |
| chr3:108022537-108022768 | 0.688834507 | 0.159553855 | 0.415220442 |
| chr17:71084161-71084553 | -0.564967479 | 0.159582404 | 0.415280633 |
| chr16:1379507-1379913 | 0.509962819 | 0.159618029 | 0.415359233 |
| chr11:63986146-63986919 | 0.465832451 | 0.159637259 | 0.415395167 |
| chr17:61480217-61480596 | 0.456146012 | 0.159643024 | 0.41539606 |
| chr14:96117081-96117800 | 0.472091994 | 0.159671366 | 0.415443574 |

| | | | |
|---|---|---|---|
| chr7:51430503-51430789 | -0.645595238 | 0.159672128 | 0.415443574 |
| chr3:120177018-120177335 | -0.621218051 | 0.159679788 | 0.415449395 |
| chr2:65413200-65413429 | 0.644605792 | 0.159702334 | 0.415493947 |
| chr20:26009666-26010305 | 0.536302389 | 0.159729917 | 0.415549915 |
| chr9:19404743-19405005 | 0.546576395 | 0.159734693 | 0.415549915 |
| chr17:14768950-14769235 | -0.619756557 | 0.159741464 | 0.415553421 |
| chr12:96516769-96517032 | -0.702074813 | 0.159754239 | 0.415572545 |
| chr6:139714119-139714589 | -0.564045909 | 0.159765897 | 0.415588761 |
| chr1:237864298-237865008 | -0.500998712 | 0.159817846 | 0.415695813 |
| chr12:127355594-127355838 | -0.755882773 | 0.159817902 | 0.415695813 |
| chr12:4604756-4605157 | -0.575350301 | 0.159831623 | 0.415710479 |
| chr2:196808323-196808606 | -0.660051465 | 0.159838044 | 0.415710479 |
| chrX:128518580-128518805 | -0.691993258 | 0.159839817 | 0.415710479 |
| chr10:31602699-31602949 | -0.604189794 | 0.159846183 | 0.415712926 |
| chr3:11236692-11236894 | 0.644460252 | 0.159864486 | 0.415746417 |
| chr11:85627762-85628966 | -0.444961352 | 0.159881741 | 0.415777178 |
| chr4:180057843-180058174 | -0.540583348 | 0.159907528 | 0.415830124 |
| chr3:171421410-171421610 | 0.740449281 | 0.159916322 | 0.415835454 |
| chr14:26841871-26842519 | -0.508331515 | 0.159920432 | 0.415835454 |
| chrX:141814173-141814792 | -0.487783018 | 0.159931589 | 0.415850353 |
| chr2:15940642-15940941 | 0.569530866 | 0.159942582 | 0.415859085 |
| chr9:120981718-120981918 | 0.68581633 | 0.159951378 | 0.415859085 |
| chr15:95958499-95958828 | 0.624076342 | 0.159952498 | 0.415859085 |
| chr3:45665179-45665501 | -0.62346927 | 0.159956951 | 0.415859085 |
| chr1:196071048-196071276 | -0.659658546 | 0.159966584 | 0.415859085 |
| chr11:103682280-103682844 | -0.520569065 | 0.159967512 | 0.415859085 |
| chr9:127568152-127568481 | 0.52616045 | 0.159991707 | 0.415897958 |
| chr7:145902247-145902447 | -0.780143266 | 0.159993321 | 0.415897958 |
| chr5:157354849-157355049 | -0.769968997 | 0.160004271 | 0.415912313 |
| chr2:164638305-164638505 | -0.793807925 | 0.160049107 | 0.416014746 |
| chr1:56414736-56415169 | 0.531493001 | 0.16010351 | 0.416142038 |
| chr15:90157950-90158503 | 0.468412992 | 0.160128927 | 0.416192877 |
| chr12:93979224-93979501 | -0.678802131 | 0.160133933 | 0.416192877 |
| chr3:154939380-154939645 | 0.667213546 | 0.160144974 | 0.416207454 |
| chr13:23714856-23716715 | -0.452137924 | 0.160171448 | 0.41626214 |
| chr8:137800606-137800865 | 0.719293969 | 0.160210664 | 0.41634469 |
| chr16:2267763-2268620 | 0.46679414 | 0.160214079 | 0.41634469 |
| chr1:12477825-12478603 | -0.505689666 | 0.160230206 | 0.41635901 |
| chr20:47220064-47220567 | 0.450659753 | 0.160232104 | 0.41635901 |
| chr7:43454901-43455356 | -0.567895482 | 0.160240092 | 0.41635901 |
| chr22:20289921-20290159 | 0.68363876 | 0.160241325 | 0.41635901 |
| chr1:35573180-35573380 | 0.668481609 | 0.160250111 | 0.416367718 |
| chr17:30832245-30832550 | 0.650800913 | 0.160256212 | 0.416369451 |
| chr8:142452799-142453064 | 0.577898391 | 0.16027357 | 0.41640043 |
| chr20:43449060-43449271 | 0.685811956 | 0.160309697 | 0.416480168 |
| chr12:96399285-96401556 | -0.443505389 | 0.16035168 | 0.416575114 |
| chr3:49120807-49121021 | 0.651283909 | 0.160375527 | 0.41661047 |
| chr17:1399403-1400498 | -0.471375871 | 0.160376164 | 0.41661047 |
| chr11:121023888-121024503 | 0.509947118 | 0.160402581 | 0.416664967 |
| chr5:94968574-94968885 | -0.584601204 | 0.160428855 | 0.41670768 |

| | | | |
|---|---|---|---|
| chr7:107642903-107643327 | -0.655127539 | 0.160432191 | 0.41670768 |
| chr1:28259183-28260502 | 0.452852619 | 0.160435339 | 0.41670768 |
| chr17:64808505-64808952 | -0.524271848 | 0.160467306 | 0.416776581 |
| chr6:44043664-44044016 | -0.580468469 | 0.160490016 | 0.416808552 |
| chr3:187066670-187066991 | -0.555508556 | 0.160490495 | 0.416808552 |
| chr21:45578475-45578858 | 0.513003831 | 0.160517678 | 0.41686502 |
| chr4:98506813-98507031 | -0.685360222 | 0.160523221 | 0.416865286 |
| chr12:29782997-29783197 | -0.772035354 | 0.160530751 | 0.416870712 |
| chr12:95003096-95003957 | -0.456389378 | 0.160561442 | 0.41693628 |
| chr16:88747965-88748208 | 0.549884753 | 0.160568064 | 0.416939344 |
| chr2:214683692-214684395 | -0.555683714 | 0.160577596 | 0.416949966 |
| chr7:88291809-88292248 | 0.562468659 | 0.160591526 | 0.416972006 |
| chr7:116525972-116526680 | -0.480181427 | 0.160608136 | 0.417001004 |
| chr5:134586783-134587544 | -0.469857323 | 0.160615988 | 0.417007259 |
| chr20:42614232-42614538 | 0.605457758 | 0.160646357 | 0.417071975 |
| chr19:35092836-35093088 | -0.718399304 | 0.160675113 | 0.417132497 |
| chr12:9412620-9412906 | -0.716167262 | 0.160695314 | 0.417170806 |
| chrX:111098842-111099106 | -0.607024581 | 0.160710768 | 0.417196791 |
| chr10:80468861-80469311 | 0.534226608 | 0.160763022 | 0.417318301 |
| chr8:13059535-13060110 | -0.537655375 | 0.160779782 | 0.41731947 |
| chr1:244864121-244864868 | 0.454985332 | 0.160794438 | 0.41731947 |
| chr2:130302388-130302672 | 0.615298227 | 0.160795265 | 0.41731947 |
| chr9:77794088-77794331 | -0.764060479 | 0.160797783 | 0.41731947 |
| chr8:113617891-113618285 | 0.580600448 | 0.160805898 | 0.41731947 |
| chrX:81195711-81196066 | -0.591256682 | 0.160813037 | 0.41731947 |
| chr1:46302184-46302384 | 0.605822391 | 0.160813159 | 0.41731947 |
| chr11:129324547-129324939 | 0.511948487 | 0.160816222 | 0.41731947 |
| chr6:7275717-7276252 | 0.538054556 | 0.160817323 | 0.41731947 |
| chr2:62411928-62412357 | -0.566114562 | 0.160817937 | 0.41731947 |
| chr2:11764188-11764766 | -0.479083664 | 0.160837697 | 0.417356611 |
| chr1:204036861-204037497 | -0.489908376 | 0.160844195 | 0.417359338 |
| chr6:41342053-41342344 | -0.660018531 | 0.160857935 | 0.417380856 |
| chr2:94759226-94759558 | -0.542754801 | 0.160898278 | 0.417463137 |
| chr13:113747832-113748164 | -0.694721721 | 0.160904405 | 0.417463137 |
| chr2:23824801-23825001 | -0.760145559 | 0.160916914 | 0.417463137 |
| chr15:68271833-68272408 | -0.482748674 | 0.160921374 | 0.417463137 |
| chr17:43878645-43878883 | 0.581766556 | 0.160922478 | 0.417463137 |
| chr17:39674821-39675307 | 0.504676012 | 0.160922515 | 0.417463137 |
| chr3:13896595-13896868 | 0.507951545 | 0.160927784 | 0.417463137 |
| chr14:94363469-94363669 | 0.684290253 | 0.16094188 | 0.417485569 |
| chr6:66985801-66986424 | -0.49561164 | 0.160954149 | 0.417503261 |
| chrX:153400161-153400372 | 0.679320896 | 0.16102469 | 0.417672097 |
| chr11:8218119-8218373 | 0.708041823 | 0.161034007 | 0.417682126 |
| chr1:85283539-85283885 | -0.519038232 | 0.161069266 | 0.417759438 |
| chr8:88944229-88944799 | -0.542931446 | 0.161075304 | 0.417760958 |
| chr8:141845015-141845401 | -0.65643135 | 0.161087728 | 0.417779037 |
| chr19:7482983-7483236 | -0.646526484 | 0.161188158 | 0.418024796 |
| chr18:48952544-48952992 | -0.493303034 | 0.161193933 | 0.418024796 |
| chr15:72155875-72156402 | 0.520159371 | 0.161198855 | 0.418024796 |
| chr3:165139742-165140421 | -0.503312303 | 0.16122183 | 0.41806775 |

| | | | |
|---|---|---|---|
| chr14:82611555-82611982 | -0.584229947 | 0.161226331 | 0.41806775 |
| chr18:74255444-74255646 | -0.737818639 | 0.161233041 | 0.418071001 |
| chr22:26483287-26484114 | 0.470071926 | 0.161250626 | 0.418102449 |
| chr14:53247696-53247932 | -0.780212214 | 0.161258997 | 0.418110006 |
| chr17:61658679-61659004 | 0.474462977 | 0.161270041 | 0.418124492 |
| chr5:139222209-139222428 | -0.726800917 | 0.161284773 | 0.418146637 |
| chr13:27620065-27620265 | 0.559242424 | 0.161294567 | 0.418146637 |
| chr1:218050381-218050707 | -0.586247632 | 0.161294954 | 0.418146637 |
| chr21:43733500-43733755 | -0.659322804 | 0.16130214 | 0.418151118 |
| chr10:112761228-112761437 | 0.671906436 | 0.161313332 | 0.418165983 |
| chr14:100740633-100740852 | -0.683747847 | 0.161337365 | 0.418214133 |
| chr14:69812912-69813145 | 0.628073633 | 0.161353536 | 0.418241904 |
| chr9:89022192-89022534 | 0.571011952 | 0.161372118 | 0.4182684 |
| chr5:121829094-121829294 | 0.727128943 | 0.161374676 | 0.4182684 |
| chr17:18203353-18203553 | 0.717200048 | 0.161445674 | 0.418438266 |
| chr9:5450345-5450732 | -0.54169119 | 0.161453104 | 0.418443368 |
| chr8:47541691-47542083 | 0.583471417 | 0.161520286 | 0.418600922 |
| chr11:73252924-73253164 | -0.757311796 | 0.161524821 | 0.418600922 |
| chr5:29005339-29005658 | -0.590952394 | 0.161534717 | 0.418605644 |
| chr5:91651545-91652246 | -0.521479605 | 0.161541007 | 0.418605644 |
| chr1:46183072-46183884 | 0.476870639 | 0.161548711 | 0.418605644 |
| chr9:19471795-19472253 | 0.534445094 | 0.161548831 | 0.418605644 |
| chr6:50030599-50031497 | -0.4698605 | 0.16155396 | 0.418605644 |
| chr16:76639931-76640540 | 0.478283594 | 0.161564738 | 0.418619415 |
| chr10:7687252-7687617 | 0.535609512 | 0.161572111 | 0.418624362 |
| chr15:69482684-69483155 | 0.483625755 | 0.161593628 | 0.41865257 |
| chr5:61745203-61745760 | -0.505117543 | 0.161593925 | 0.41865257 |
| chr10:104274727-104275689 | -0.467490396 | 0.161603796 | 0.418663987 |
| chr9:137520619-137520934 | 0.566462855 | 0.161626393 | 0.418700055 |
| chr1:106781841-106782041 | 0.525045181 | 0.16163392 | 0.418700055 |
| chr12:49580467-49580856 | 0.506569334 | 0.161634112 | 0.418700055 |
| chr8:100075348-100075548 | 0.726980704 | 0.161647374 | 0.418715065 |
| chr16:78080858-78081095 | 0.71694536 | 0.161650836 | 0.418715065 |
| chr11:77332649-77332988 | -0.667142335 | 0.16167581 | 0.418761243 |
| chr20:37543844-37544110 | 0.713389397 | 0.161679594 | 0.418761243 |
| chr6:10801323-10801602 | 0.6263315 | 0.161709111 | 0.418823536 |
| chr15:66765910-66766219 | 0.556849803 | 0.161726759 | 0.418855086 |
| chr1:43171822-43172804 | -0.467182527 | 0.16173944 | 0.418873771 |
| chr10:60282856-60283251 | -0.5530524 | 0.161762292 | 0.418901709 |
| chr5:149828246-149828837 | -0.490142808 | 0.161763785 | 0.418901709 |
| chr8:22130872-22132012 | -0.468724278 | 0.161766629 | 0.418901709 |
| chr15:77864803-77865105 | -0.564528847 | 0.161780503 | 0.418915814 |
| chr5:150296204-150296413 | -0.619344164 | 0.161783011 | 0.418915814 |
| chr2:72815876-72816232 | -0.550275589 | 0.161802318 | 0.41895165 |
| chr17:39739321-39739575 | 0.600943512 | 0.161824344 | 0.418994523 |
| chr3:147836376-147836592 | -0.729117447 | 0.161865795 | 0.419066679 |
| chr7:10939568-10940748 | 0.440711039 | 0.161866623 | 0.419066679 |
| chr12:65963100-65963374 | -0.627576292 | 0.161869061 | 0.419066679 |
| chr7:8035602-8036288 | 0.47159386 | 0.161877077 | 0.419066679 |
| chr1:6625057-6625642 | 0.509541861 | 0.161879559 | 0.419066679 |

| | | | |
|---|---|---|---|
| chr20:54471174-54471474 | 0.471393451 | 0.161926402 | 0.419173782 |
| chr3:176167263-176167765 | 0.488732028 | 0.161935369 | 0.419182832 |
| chr8:109720306-109720520 | 0.63347979 | 0.161946797 | 0.419198253 |
| chr20:61212471-61212889 | 0.609037177 | 0.161992831 | 0.419303245 |
| chr2:23033367-23033622 | 0.622866096 | 0.162011787 | 0.419324038 |
| chr8:116113532-116113786 | -0.669398554 | 0.162015118 | 0.419324038 |
| chr11:17979859-17980059 | 0.695421989 | 0.162021402 | 0.419324038 |
| chr2:109319104-109319311 | -0.727181456 | 0.162022754 | 0.419324038 |
| chr4:114807475-114808169 | -0.491157572 | 0.162036475 | 0.419332769 |
| chr3:166498377-166498950 | -0.542892992 | 0.162037073 | 0.419332769 |
| chr3:170414154-170414451 | -0.572892258 | 0.162067668 | 0.419393004 |
| chr9:137077339-137077555 | 0.677595901 | 0.162080176 | 0.419393004 |
| chr14:74362394-74362670 | 0.582170002 | 0.162084638 | 0.419393004 |
| chr19:7894889-7895202 | 0.573129043 | 0.162086751 | 0.419393004 |
| chr1:157074105-157074586 | 0.527964486 | 0.162090399 | 0.419393004 |
| chr11:47961362-47961663 | 0.578924503 | 0.16209319 | 0.419393004 |
| chr17:61285763-61286200 | 0.456397977 | 0.162101021 | 0.419397536 |
| chr10:86960412-86960858 | -0.480563403 | 0.162105889 | 0.419397536 |
| chr12:47797491-47797788 | -0.596935835 | 0.16212482 | 0.419422134 |
| chr1:195495997-195496197 | -0.740444068 | 0.16213462 | 0.419422134 |
| chr11:75748545-75748745 | -0.740444068 | 0.16213462 | 0.419422134 |
| chr8:84604734-84604934 | 0.648055735 | 0.162137292 | 0.419422134 |
| chr10:31770356-31770633 | -0.607617903 | 0.162150501 | 0.419442143 |
| chr2:126239535-126239844 | 0.64774768 | 0.162180149 | 0.419455536 |
| chr8:22579251-22579673 | 0.539615783 | 0.162181275 | 0.419455536 |
| chr7:4123779-4124137 | -0.601598229 | 0.162182552 | 0.419455536 |
| chr12:19482828-19483074 | -0.691959738 | 0.162183978 | 0.419455536 |
| chr4:168631378-168632196 | -0.495936273 | 0.16218401 | 0.419455536 |
| chr15:92751592-92751901 | 0.578464541 | 0.162188525 | 0.419455536 |
| chr14:105398121-105398781 | 0.466581204 | 0.162214068 | 0.419507435 |
| chr8:8288461-8288805 | -0.620256945 | 0.162249437 | 0.419570911 |
| chr17:71432934-71433193 | 0.716019931 | 0.162249564 | 0.419570911 |
| chr20:53779579-53779908 | 0.459518427 | 0.162256241 | 0.419572655 |
| chr3:142578286-142579231 | -0.448618214 | 0.16226119 | 0.419572655 |
| chr3:22983098-22983586 | -0.543857797 | 0.162271317 | 0.419584681 |
| chr8:73327751-73327951 | -0.776060348 | 0.162293612 | 0.419607965 |
| chr9:124258212-124259094 | 0.476212271 | 0.162294268 | 0.419607965 |
| chr17:68785890-68786141 | -0.603518342 | 0.162296751 | 0.419607965 |
| chr1:190754826-190755142 | -0.570305956 | 0.162315192 | 0.419624281 |
| chr12:64392280-64392488 | 0.62239421 | 0.162318194 | 0.419624281 |
| chr13:25574473-25574825 | 0.534085079 | 0.162319491 | 0.419624281 |
| chr1:150162145-150162373 | 0.574958774 | 0.162325014 | 0.419624401 |
| chr8:28891007-28891281 | 0.550715497 | 0.162331943 | 0.419628156 |
| chr11:118295330-118295530 | 0.684090214 | 0.162372522 | 0.419718891 |
| chr17:9924784-9925232 | -0.55635334 | 0.162380859 | 0.419721531 |
| chr1:179564396-179564687 | 0.634562269 | 0.162384499 | 0.419721531 |
| chr2:149006990-149007400 | 0.539055926 | 0.162396399 | 0.41973813 |
| chr1:153568555-153568869 | 0.503725147 | 0.162423575 | 0.419791185 |
| chr1:109161897-109162164 | -0.587736961 | 0.162427883 | 0.419791185 |
| chr7:44565556-44565809 | 0.565254539 | 0.162507967 | 0.419983994 |

| | | | |
|---|---|---|---|
| chr22:39176536-39176804 | -0.7670438 | 0.162529709 | 0.420026018 |
| chr9:121055671-121056069 | -0.536243844 | 0.162548347 | 0.420046942 |
| chr3:194271258-194271525 | -0.590436174 | 0.16254877 | 0.420046942 |
| chr12:75267731-75268124 | -0.652385413 | 0.162559332 | 0.420060067 |
| chr8:50378046-50378416 | -0.544521885 | 0.162570504 | 0.42007477 |
| chr16:15643511-15643777 | 0.568200284 | 0.162588885 | 0.420098631 |
| chr1:9129044-9129802 | 0.476038311 | 0.162592569 | 0.420098631 |
| chr3:172476991-172477690 | -0.500344056 | 0.162596186 | 0.420098631 |
| chr14:64194403-64194622 | 0.688140249 | 0.162637237 | 0.420190523 |
| chr14:100025107-100025558 | 0.488369896 | 0.162643001 | 0.420191246 |
| chr16:27897891-27898359 | -0.558887886 | 0.162655814 | 0.420210183 |
| chr14:95774421-95774621 | 0.686418995 | 0.162687779 | 0.42027859 |
| chr5:14180532-14180765 | 0.571889653 | 0.162695484 | 0.420284325 |
| chr1:200535492-200535947 | -0.492261178 | 0.162717113 | 0.420326028 |
| chr5:72319716-72320794 | -0.450653773 | 0.162729316 | 0.420332036 |
| chr4:88033319-88033575 | 0.658512092 | 0.162733839 | 0.420332036 |
| chr1:70966102-70966393 | -0.656677535 | 0.162735896 | 0.420332036 |
| chrX:71103546-71103955 | -0.555600479 | 0.162749936 | 0.42035413 |
| chr1:48228178-48228516 | -0.588868826 | 0.16276775 | 0.420385968 |
| chr13:29050336-29050630 | -0.674253098 | 0.162783466 | 0.420412389 |
| chr20:48354931-48355134 | 0.512596395 | 0.162789214 | 0.420413062 |
| chr4:148067563-148067834 | -0.62478706 | 0.162808165 | 0.420447834 |
| chr3:150227812-150228068 | 0.586501497 | 0.162823968 | 0.420460448 |
| chr1:9476762-9477135 | -0.619035172 | 0.162832024 | 0.420460448 |
| chr10:71696575-71697016 | -0.543196431 | 0.162844038 | 0.420460448 |
| chr16:4802057-4803152 | 0.447641266 | 0.162846037 | 0.420460448 |
| chr4:169045968-169046263 | -0.589340952 | 0.162849561 | 0.420460448 |
| chr17:67942823-67943197 | -0.504211616 | 0.162849768 | 0.420460448 |
| chr1:205329141-205329342 | -0.606010079 | 0.162851462 | 0.420460448 |
| chrX:100636028-100637171 | -0.452051464 | 0.162901247 | 0.420567468 |
| chr13:81987282-81987483 | -0.689106883 | 0.16290389 | 0.420567468 |
| chr1:43423580-43423823 | -0.681619922 | 0.162964482 | 0.42070972 |
| chr22:28800027-28801045 | 0.452714279 | 0.163001528 | 0.420764883 |
| chr15:66701894-66702366 | 0.474778375 | 0.163003259 | 0.420764883 |
| chr18:58269927-58270178 | 0.615537111 | 0.163003913 | 0.420764883 |
| chr19:40404035-40404558 | 0.488634439 | 0.163010533 | 0.420764883 |
| chr1:169781060-169781294 | -0.594626638 | 0.163013306 | 0.420764883 |
| chr3:34620099-34620362 | 0.619614233 | 0.163026604 | 0.420785031 |
| chr2:64312278-64312937 | -0.481019708 | 0.163035629 | 0.420786381 |
| chr5:127385908-127386288 | -0.544692296 | 0.16303811 | 0.420786381 |
| chr2:218648547-218648882 | 0.555119944 | 0.163047937 | 0.420797568 |
| chr5:16542074-16542852 | 0.47181135 | 0.163053612 | 0.420798041 |
| chr1:8703180-8703696 | 0.520756084 | 0.163084432 | 0.420860109 |
| chr1:26899907-26900607 | 0.491208198 | 0.163094992 | 0.420860109 |
| chr1:204751899-204752714 | -0.466247729 | 0.163097587 | 0.420860109 |
| chr7:6826155-6826453 | -0.559479116 | 0.163108261 | 0.420860109 |
| chr8:115978187-115978627 | -0.545386191 | 0.163108851 | 0.420860109 |
| chr2:37890969-37891298 | -0.552965995 | 0.163110618 | 0.420860109 |
| chr12:127898508-127898915 | -0.592592167 | 0.163116751 | 0.420861761 |
| chr18:75208905-75209789 | -0.461309002 | 0.163145901 | 0.420922796 |

| | | | |
|---|---|---|---|
| chr11:101852284-101853126 | -0.485356008 | 0.163165654 | 0.420959586 |
| chr2:10557524-10557748 | -0.624862268 | 0.163171906 | 0.420961542 |
| chr17:40027015-40027270 | 0.587492446 | 0.163194383 | 0.421005355 |
| chr7:41573043-41573315 | -0.656583502 | 0.163205735 | 0.421008846 |
| chr2:193584625-193584980 | -0.534238212 | 0.163206726 | 0.421008846 |
| chr11:83193317-83194804 | -0.440158965 | 0.163218161 | 0.421023037 |
| chr2:3002087-3002483 | -0.600053868 | 0.163223217 | 0.421023037 |
| chr8:24218511-24219098 | -0.510253869 | 0.163271616 | 0.421133703 |
| chr16:85894330-85894642 | 0.551795926 | 0.163286547 | 0.421150241 |
| chr4:53929991-53930313 | -0.581314658 | 0.163289021 | 0.421150241 |
| chr2:15955387-15956039 | 0.476972816 | 0.16329462 | 0.421150505 |
| chr8:94713488-94713688 | 0.652041769 | 0.16331063 | 0.421177621 |
| chr16:89289467-89289772 | -0.549265068 | 0.16333255 | 0.421219975 |
| chr1:180923580-180923798 | 0.58119411 | 0.163343354 | 0.421233659 |
| chr2:9830262-9831148 | -0.457532758 | 0.163361711 | 0.421266821 |
| chr7:155531673-155532136 | 0.517290963 | 0.163379569 | 0.421298692 |
| chr6:40431782-40432079 | -0.708376623 | 0.163399296 | 0.421335381 |
| chr2:19901174-19902535 | -0.444082019 | 0.16342438 | 0.421385881 |
| chr18:23515978-23516285 | 0.547960713 | 0.163467086 | 0.421481813 |
| chr14:21066987-21067189 | 0.638842467 | 0.163495237 | 0.421527329 |
| chr18:48445561-48445812 | -0.665690424 | 0.163495741 | 0.421527329 |
| chr17:17533975-17534242 | 0.638581713 | 0.163513011 | 0.421544454 |
| chr1:31669720-31670419 | 0.482288559 | 0.163513387 | 0.421544454 |
| chr6:28769857-28770327 | -0.548021253 | 0.163530062 | 0.42157326 |
| chr7:5822624-5823613 | -0.456143823 | 0.163544839 | 0.421597168 |
| chr18:51029554-51030954 | -0.450248751 | 0.163551511 | 0.421600185 |
| chr14:23631948-23632302 | 0.522021941 | 0.163560371 | 0.421608838 |
| chr2:1839207-1839503 | 0.599510739 | 0.163571731 | 0.421623938 |
| chr5:170331273-170331515 | -0.668581576 | 0.163586339 | 0.421647407 |
| chr5:172303661-172304015 | -0.620858243 | 0.163604219 | 0.421679309 |
| chr6:158644175-158645252 | -0.450358532 | 0.163610628 | 0.421681641 |
| chr2:96138557-96138929 | 0.520686467 | 0.163619821 | 0.42169115 |
| chr3:62148819-62149073 | 0.557649729 | 0.163640518 | 0.421725793 |
| chr17:61785959-61786292 | 0.466742105 | 0.16364427 | 0.421725793 |
| chr3:85081599-85081857 | -0.679043894 | 0.16365678 | 0.421734753 |
| chr18:7830428-7830774 | -0.582383739 | 0.163658755 | 0.421734753 |
| chr20:56929699-56929988 | -0.568308734 | 0.163666943 | 0.421741667 |
| chr5:1523848-1524407 | 0.47973197 | 0.163678214 | 0.421756528 |
| chr15:69762779-69763019 | 0.61235539 | 0.163689588 | 0.421771651 |
| chr9:93154301-93154510 | -0.695973423 | 0.163717659 | 0.421829795 |
| chr10:28997711-28997986 | -0.722717565 | 0.163726229 | 0.421837693 |
| chr15:88897208-88897702 | 0.492068636 | 0.163736078 | 0.421848884 |
| chr4:125283755-125284020 | -0.610079216 | 0.163746769 | 0.42184975 |
| chr2:170816233-170816433 | 0.635510741 | 0.163747426 | 0.42184975 |
| chr20:41451963-41452186 | -0.654886329 | 0.163756506 | 0.421858959 |
| chr9:138044345-138044545 | 0.738033571 | 0.163793997 | 0.421933554 |
| chr3:166453518-166453872 | -0.611093649 | 0.163801949 | 0.421933554 |
| chr1:106675575-106675916 | 0.485518679 | 0.163801982 | 0.421933554 |
| chr3:120559714-120559914 | -0.620316972 | 0.16381346 | 0.421939988 |
| chr3:154132881-154133294 | -0.595874625 | 0.163815494 | 0.421939988 |

| | | | |
|---|---|---|---|
| chr7:98051186-98051421 | -0.685210754 | 0.163836035 | 0.421956033 |
| chr4:37441320-37442048 | -0.47639241 | 0.16383898 | 0.421956033 |
| chr19:18341240-18341474 | -0.733624773 | 0.163846453 | 0.421956033 |
| chrX:121823496-121823740 | 0.633453451 | 0.163848116 | 0.421956033 |
| chr6:1436660-1436991 | 0.58617284 | 0.163854376 | 0.421956033 |
| chr14:61714044-61714330 | 0.703827379 | 0.163854765 | 0.421956033 |
| chr8:60397359-60397559 | 0.651177551 | 0.163873097 | 0.421989057 |
| chr9:129383605-129383832 | 0.640165177 | 0.163899541 | 0.42204297 |
| chr19:19170160-19170464 | -0.633966884 | 0.163920274 | 0.422070782 |
| chr17:61490659-61490911 | 0.463472663 | 0.163921359 | 0.422070782 |
| chr3:148288154-148288354 | -0.752493014 | 0.163927416 | 0.422072197 |
| chr4:82739157-82739698 | -0.488311208 | 0.163934324 | 0.422075798 |
| chr1:205396999-205397438 | -0.500988015 | 0.163968254 | 0.422148972 |
| chr10:98467107-98468401 | -0.457591818 | 0.164008536 | 0.42222633 |
| chr3:24914123-24914432 | 0.64945845 | 0.164012099 | 0.42222633 |
| chr9:136283449-136284070 | -0.488348789 | 0.164015946 | 0.42222633 |
| chr17:52556272-52556533 | -0.682610918 | 0.164020343 | 0.42222633 |
| chr1:43619205-43619632 | 0.579113994 | 0.164030081 | 0.422229222 |
| chr2:130738295-130738621 | -0.66151643 | 0.164032488 | 0.422229222 |
| chr5:67209202-67209592 | -0.515100496 | 0.164038505 | 0.422230528 |
| chr8:121641421-121642055 | -0.47634618 | 0.164047237 | 0.422238819 |
| chr1:147109743-147109943 | 0.610382146 | 0.164058064 | 0.422252501 |
| chr2:160114385-160115030 | -0.581998826 | 0.164079253 | 0.422292853 |
| chr8:22598575-22598864 | 0.574969122 | 0.164103673 | 0.42232012 |
| chr17:76125187-76125465 | 0.585122266 | 0.164105307 | 0.42232012 |
| chr2:71023599-71024143 | -0.490811107 | 0.164106678 | 0.42232012 |
| chr3:114179695-114179897 | -0.689297064 | 0.164111894 | 0.42232012 |
| chr6:12287741-12288676 | 0.467807602 | 0.164137179 | 0.422371 |
| chr1:240732284-240732621 | 0.574454341 | 0.164204677 | 0.422530501 |
| chr3:64012434-64012664 | 0.491830169 | 0.164221995 | 0.422548041 |
| chr10:28405221-28405569 | 0.521861555 | 0.164233699 | 0.422548041 |
| chr5:15472235-15472650 | -0.656571407 | 0.164234734 | 0.422548041 |
| chrX:38220573-38221002 | 0.573539025 | 0.164237243 | 0.422548041 |
| chr16:67948092-67948504 | -0.513658426 | 0.164239067 | 0.422548041 |
| chr6:135181005-135181538 | -0.46938782 | 0.164362186 | 0.4228506 |
| chr1:211416031-211417302 | -0.462748253 | 0.164374781 | 0.422868804 |
| chr11:112588248-112588465 | 0.668678492 | 0.16438954 | 0.422892574 |
| chr9:93058390-93058779 | -0.527612893 | 0.164406078 | 0.422919981 |
| chr13:78700193-78700393 | -0.807616449 | 0.164411233 | 0.422919981 |
| chr8:65643862-65644124 | -0.521042025 | 0.164422837 | 0.422935632 |
| chr1:76901233-76901561 | 0.578975183 | 0.164444854 | 0.422965491 |
| chrX:48597278-48598406 | 0.44603925 | 0.164446848 | 0.422965491 |
| chr12:89762669-89763205 | 0.482534411 | 0.164451005 | 0.422965491 |
| chr4:163122620-163122867 | -0.791529528 | 0.164467643 | 0.422994083 |
| chr9:1558371-1559288 | -0.490004682 | 0.164492825 | 0.423031107 |
| chr7:32747404-32747681 | 0.665376647 | 0.164498581 | 0.423031107 |
| chr10:5999363-5999969 | 0.519191283 | 0.164498601 | 0.423031107 |
| chr2:196967213-196967440 | -0.749564584 | 0.164533357 | 0.423104882 |
| chr20:45822858-45823215 | 0.513943784 | 0.164538333 | 0.423104882 |
| chr16:27169545-27169918 | 0.537542014 | 0.164547138 | 0.423113323 |

| | | | |
|---|---|---|---|
| chr8:53786528-53787055 | 0.510582165 | 0.164562527 | 0.423134515 |
| chr20:10179667-10180088 | -0.58869358 | 0.164566424 | 0.423134515 |
| chr8:93700093-93701210 | 0.450212936 | 0.164603407 | 0.423203444 |
| chr22:29772337-29772632 | 0.572427999 | 0.164604278 | 0.423203444 |
| chr3:151335743-151336001 | 0.713302812 | 0.164644641 | 0.423293014 |
| chr1:27934818-27935385 | 0.486400109 | 0.164674199 | 0.423354799 |
| chr7:152081235-152081735 | -0.592701611 | 0.164700723 | 0.423408783 |
| chr5:66633281-66633540 | -0.592732566 | 0.16470764 | 0.423412359 |
| chr8:111228949-111229167 | 0.738665918 | 0.164717064 | 0.42342238 |
| chr12:75623772-75624198 | -0.488946987 | 0.16472837 | 0.423437237 |
| chr10:69644605-69644957 | -0.573488079 | 0.164760471 | 0.423505002 |
| chr10:98425717-98426053 | -0.521717402 | 0.164765787 | 0.423505002 |
| chr6:89950925-89951135 | 0.704548343 | 0.164773826 | 0.423511459 |
| chr8:54169363-54169631 | 0.657798168 | 0.16478784 | 0.423533271 |
| chr7:1928577-1929196 | -0.500969346 | 0.164818793 | 0.423572533 |
| chr4:54363145-54363345 | 0.735599139 | 0.164825468 | 0.423572533 |
| chr2:224909941-224910601 | -0.511705915 | 0.164830596 | 0.423572533 |
| chr8:88859993-88860888 | -0.477758392 | 0.164832432 | 0.423572533 |
| chr14:106166267-106166601 | -0.684881588 | 0.16483284 | 0.423572533 |
| chr16:89043580-89043858 | 0.582326375 | 0.164836284 | 0.423572533 |
| chr20:34362935-34363634 | 0.479628447 | 0.164846609 | 0.423584858 |
| chr12:49366452-49367690 | 0.448491496 | 0.164854974 | 0.423592147 |
| chr1:155124531-155124776 | 0.536058226 | 0.164869927 | 0.423616361 |
| chr1:229507914-229508642 | 0.457003267 | 0.164878382 | 0.423623881 |
| chr9:89992304-89992515 | -0.760257234 | 0.164884168 | 0.423624542 |
| chr1:218090458-218090658 | -0.727481875 | 0.164893848 | 0.423635208 |
| chr8:71554142-71554595 | -0.594367124 | 0.16492393 | 0.423698286 |
| chr17:44324187-44324607 | 0.503574955 | 0.164940503 | 0.423726655 |
| chr19:29831053-29831253 | -0.721782216 | 0.16496156 | 0.423760111 |
| chr1:16677965-16678443 | -0.500756993 | 0.164964587 | 0.423760111 |
| chr5:35963330-35963776 | 0.55764075 | 0.164972795 | 0.42376699 |
| chr10:22805419-22805640 | 0.68464614 | 0.165003751 | 0.423821988 |
| chrX:84187040-84188749 | -0.450483557 | 0.165005269 | 0.423821988 |
| chr19:58304845-58305930 | 0.453235271 | 0.165030497 | 0.423862185 |
| chr2:68466917-68467901 | -0.442527029 | 0.165031983 | 0.423862185 |
| chr17:47673786-47673986 | 0.659627618 | 0.165057871 | 0.423914466 |
| chr15:89361754-89362027 | 0.522530867 | 0.165069695 | 0.423930624 |
| chr20:62367136-62367439 | 0.503160532 | 0.165078459 | 0.423938923 |
| chr1:79290911-79291371 | 0.531064229 | 0.165087459 | 0.423947827 |
| chr2:102540954-102541202 | -0.640422812 | 0.165098289 | 0.423955478 |
| chr2:215296397-215296617 | -0.647746127 | 0.165105147 | 0.423955478 |
| chr8:22066616-22067267 | 0.510645292 | 0.165107361 | 0.423955478 |
| chr18:28176999-28177247 | -0.687351642 | 0.165112571 | 0.423955478 |
| chr7:54759041-54759990 | -0.447269335 | 0.165141525 | 0.424009957 |
| chr13:30107803-30108009 | -0.621562881 | 0.165144856 | 0.424009957 |
| chr18:399291-399849 | -0.536863801 | 0.165160432 | 0.42403574 |
| chr10:95445894-95446260 | -0.532084818 | 0.165178519 | 0.424067966 |
| chr7:149610682-149610944 | 0.604436204 | 0.165184821 | 0.424069938 |
| chr19:56403803-56404516 | 0.502705398 | 0.165212631 | 0.424127122 |
| chr12:80598058-80598401 | -0.561799102 | 0.165218816 | 0.424128789 |

| | | | |
|---|---|---|---|
| chrX:81120668-81121320 | 0.493992495 | 0.165236949 | 0.424161127 |
| chr12:100573179-100574585 | -0.442185826 | 0.165249861 | 0.424180061 |
| chr12:35614848-35615131 | 0.539149929 | 0.165270227 | 0.424212502 |
| chr16:88195816-88196105 | 0.621937522 | 0.165273572 | 0.424212502 |
| chr2:173589673-173589905 | -0.70786262 | 0.165294787 | 0.424239332 |
| chr17:54536363-54536715 | 0.589453202 | 0.165295098 | 0.424239332 |
| chr14:24392342-24392679 | 0.493353257 | 0.165323202 | 0.424286081 |
| chr7:128154472-128154672 | -0.784171763 | 0.165324388 | 0.424286081 |
| chr7:111570054-111570336 | -0.670063451 | 0.165333728 | 0.424294442 |
| chr14:89270851-89271186 | -0.659425576 | 0.165345996 | 0.424294442 |
| chrX:110146605-110146985 | 0.539907021 | 0.16535467 | 0.424294442 |
| chr1:107324281-107324574 | 0.496212669 | 0.165357518 | 0.424294442 |
| chr5:168947303-168947559 | -0.605098431 | 0.165358527 | 0.424294442 |
| chr17:67653384-67653625 | -0.641779237 | 0.16536087 | 0.424294442 |
| chr6:170124851-170125182 | 0.530739342 | 0.165376524 | 0.424320396 |
| chr8:8893291-8893781 | 0.547886963 | 0.165438615 | 0.424450511 |
| chr5:69048319-69048598 | -0.628499197 | 0.165440609 | 0.424450511 |
| chr5:19733587-19733826 | -0.657345012 | 0.165444638 | 0.424450511 |
| chr7:8785105-8785406 | -0.577520745 | 0.165449393 | 0.424450511 |
| chr20:53938722-53939064 | 0.456265686 | 0.165469274 | 0.4244873 |
| chr8:101126140-101127481 | 0.441744674 | 0.165501911 | 0.424541413 |
| chr5:145658380-145658883 | -0.587653196 | 0.165506568 | 0.424541413 |
| chr12:27523645-27524633 | -0.447548161 | 0.165512275 | 0.424541413 |
| chr7:50709960-50710221 | 0.642843562 | 0.16551253 | 0.424541413 |
| chr20:18496717-18497440 | -0.473106383 | 0.165523704 | 0.424555862 |
| chrX:110002197-110003145 | 0.449791134 | 0.165563962 | 0.424644906 |
| chr2:102006290-102006520 | 0.654414572 | 0.165591333 | 0.424687738 |
| chr20:54782394-54783242 | -0.4418879 | 0.165591747 | 0.424687738 |
| chr20:9632941-9633291 | -0.602199465 | 0.165608944 | 0.424709288 |
| chr1:95072497-95073383 | -0.459282878 | 0.165611948 | 0.424709288 |
| chrX:48801250-48801467 | 0.561462012 | 0.165616779 | 0.424709288 |
| chr6:131124381-131124645 | 0.590067098 | 0.165624561 | 0.424715029 |
| chr22:47460870-47461498 | 0.501687651 | 0.165662088 | 0.424797045 |
| chr20:48620929-48621672 | 0.437454915 | 0.165704873 | 0.424880959 |
| chr20:57212567-57212773 | 0.466702851 | 0.165711115 | 0.424880959 |
| chr6:26188438-26189368 | -0.458462239 | 0.165711448 | 0.424880959 |
| chr2:25035691-25036213 | -0.515964252 | 0.165734704 | 0.424923027 |
| chr7:98085733-98085957 | 0.669018615 | 0.165738947 | 0.424923027 |
| chr22:20290648-20291060 | 0.586371173 | 0.165758665 | 0.424949618 |
| chr9:100426484-100427419 | -0.469871022 | 0.165760411 | 0.424949618 |
| chrX:77046162-77047033 | -0.474768965 | 0.165775767 | 0.424974767 |
| chr14:79960384-79960640 | 0.668351903 | 0.165786629 | 0.424988394 |
| chr2:18047880-18048271 | -0.515780617 | 0.165804535 | 0.425007905 |
| chr12:14588982-14589284 | -0.678879593 | 0.165805334 | 0.425007905 |
| chr14:83638769-83639071 | -0.575275058 | 0.16582205 | 0.425036533 |
| chr16:12087918-12088440 | -0.542696461 | 0.165832863 | 0.425037579 |
| chr3:64354976-64355205 | 0.499473068 | 0.165833552 | 0.425037579 |
| chr8:86289942-86291000 | 0.455868124 | 0.16586558 | 0.425097267 |
| chr6:29632400-29632600 | 0.614655201 | 0.165867936 | 0.425097267 |
| chr2:179936766-179937007 | -0.644737718 | 0.165878703 | 0.425110643 |

| | | | |
|---|---|---|---|
| chr8:108760437-108760929 | 0.51305936 | 0.165888716 | 0.425122084 |
| chr16:31002327-31002725 | 0.504917059 | 0.165894858 | 0.425123605 |
| chr14:66661900-66662207 | 0.607020937 | 0.165926764 | 0.425171648 |
| chr14:100639019-100639315 | 0.653965348 | 0.165926797 | 0.425171648 |
| chr10:98242974-98243174 | -0.766986679 | 0.1659327 | 0.425171648 |
| chr3:115871295-115871634 | -0.705852733 | 0.165935801 | 0.425171648 |
| chr13:36297404-36298280 | -0.469356844 | 0.16596448 | 0.42523091 |
| chr17:4263558-4264180 | -0.496444765 | 0.165975667 | 0.425245354 |
| chr4:85196012-85196681 | -0.498026654 | 0.165984898 | 0.425254786 |
| chr10:66925857-66926202 | -0.530501713 | 0.166037936 | 0.425376446 |
| chr5:112968788-112969112 | 0.542847613 | 0.16605276 | 0.4254002 |
| chr3:143674363-143674751 | -0.517570765 | 0.166100036 | 0.425507088 |
| chr16:30585273-30586142 | 0.455998263 | 0.166129295 | 0.425567812 |
| chr11:70760415-70761033 | -0.498136959 | 0.166152426 | 0.425594435 |
| chr6:6712564-6713151 | -0.52146036 | 0.166159385 | 0.425594435 |
| chr17:50956823-50957117 | -0.519124196 | 0.166160701 | 0.425594435 |
| chr3:189195415-189195724 | -0.637353602 | 0.166166984 | 0.425594435 |
| chr7:5677981-5678306 | 0.547699976 | 0.166169123 | 0.425594435 |
| chr22:38236785-38237111 | 0.589217665 | 0.166173014 | 0.425594435 |
| chr9:83132816-83133059 | 0.627132638 | 0.166187047 | 0.425616147 |
| chr7:135645348-135645610 | -0.594970194 | 0.166197286 | 0.425628143 |
| chr21:34367368-34367642 | 0.586193142 | 0.166287787 | 0.425845682 |
| chr9:14692860-14693847 | -0.475649494 | 0.166334355 | 0.425950702 |
| chr11:63945903-63946132 | 0.597939577 | 0.166362842 | 0.426009413 |
| chr11:113806486-113806734 | -0.696077511 | 0.166377967 | 0.426033906 |
| chr20:40415145-40415535 | 0.537405454 | 0.166393151 | 0.426046284 |
| chr17:28251630-28251952 | -0.555057698 | 0.166397661 | 0.426046284 |
| chr5:141758693-141758960 | -0.6454556 | 0.166399482 | 0.426046284 |
| chr7:142854590-142854872 | -0.562846825 | 0.166409952 | 0.426054746 |
| chr19:39498402-39498682 | 0.608641195 | 0.166416655 | 0.426054746 |
| chr16:27549518-27550473 | 0.443800585 | 0.166420501 | 0.426054746 |
| chr9:135075094-135075594 | 0.528970473 | 0.166427413 | 0.426054746 |
| chr19:49347814-49348192 | 0.525063903 | 0.166430589 | 0.426054746 |
| chr19:47483735-47484439 | 0.471316586 | 0.166437686 | 0.426058679 |
| chr3:166864952-166865717 | -0.488847578 | 0.166480983 | 0.426155276 |
| chr12:3333696-3334136 | -0.617505002 | 0.166488768 | 0.426160966 |
| chr2:55923328-55924161 | 0.467517235 | 0.166521186 | 0.426229709 |
| chr9:110058314-110058711 | -0.524887806 | 0.166555327 | 0.426292985 |
| chr1:151739773-151740317 | 0.468584901 | 0.166557034 | 0.426292985 |
| chr5:176793370-176794270 | -0.472126068 | 0.166564094 | 0.426296815 |
| chr9:122366324-122366531 | 0.705257495 | 0.166573392 | 0.426306371 |
| chr1:100037838-100038973 | -0.445248251 | 0.166579745 | 0.42630839 |
| chr20:41567228-41567464 | 0.589426941 | 0.166600017 | 0.42634603 |
| chr19:40716625-40717197 | 0.45596437 | 0.166609666 | 0.426356484 |
| chr13:21458432-21459842 | 0.441878159 | 0.166630502 | 0.426395562 |
| chr5:18698198-18698960 | -0.461419843 | 0.166648182 | 0.426426562 |
| chr10:98376486-98376882 | -0.527655911 | 0.166696005 | 0.426520817 |
| chr4:152655859-152656059 | 0.732392614 | 0.16669615 | 0.426520817 |
| chr13:70232399-70232618 | -0.778373194 | 0.166710851 | 0.426526513 |
| chr2:153584226-153584474 | -0.685340908 | 0.16671092 | 0.426526513 |

| | | | |
|---|---|---|---|
| chr14:88857748-88858177 | 0.5634222 | 0.166715076 | 0.426526513 |
| chr3:52084975-52085217 | 0.714633511 | 0.166721882 | 0.426529684 |
| chr10:1369199-1369479 | 0.608916254 | 0.166770884 | 0.426614292 |
| chr1:112638141-112638343 | 0.70759301 | 0.166771527 | 0.426614292 |
| chr21:43718761-43719583 | 0.4587972 | 0.166771657 | 0.426614292 |
| chr1:191153184-191153976 | 0.518640641 | 0.166780714 | 0.426618184 |
| chr5:155028048-155028363 | -0.617530735 | 0.166784314 | 0.426618184 |
| chr11:8033263-8033465 | 0.662979834 | 0.166839983 | 0.426737415 |
| chr4:132935012-132935335 | 0.615828515 | 0.166842066 | 0.426737415 |
| chr17:30116615-30117017 | 0.535986714 | 0.166857475 | 0.426752237 |
| chr5:141734885-141735161 | -0.554750729 | 0.166859 | 0.426752237 |
| chr5:133968043-133968948 | -0.4535708 | 0.166893892 | 0.42682723 |
| chr21:43342983-43343799 | 0.453716595 | 0.166913112 | 0.426862136 |
| chr6:110944654-110944863 | -0.75739447 | 0.166924327 | 0.426876569 |
| chr15:76059272-76059915 | 0.521241066 | 0.166962689 | 0.426960423 |
| chr2:197599397-197599996 | -0.491810119 | 0.166979779 | 0.426989875 |
| chr20:2872557-2873564 | 0.465664858 | 0.166994014 | 0.427004735 |
| chr15:98452271-98452471 | -0.728250391 | 0.166998261 | 0.427004735 |
| chr15:98017316-98017562 | 0.692279056 | 0.167002309 | 0.427004735 |
| chr7:100569383-100569630 | -0.628018317 | 0.167039432 | 0.427081819 |
| chr18:41745531-41745777 | -0.688430012 | 0.167046208 | 0.427081819 |
| chr15:65295684-65296790 | 0.442613528 | 0.167049178 | 0.427081819 |
| chr16:80744756-80745220 | -0.568354192 | 0.167070257 | 0.427121458 |
| chr1:100265717-100266860 | -0.454265424 | 0.167079032 | 0.427126112 |
| chr2:62467768-62468113 | -0.591381473 | 0.167083226 | 0.427126112 |
| chr18:23259638-23260874 | -0.445682711 | 0.167105646 | 0.427159392 |
| chr17:7233989-7234729 | -0.465400432 | 0.167107394 | 0.427159392 |
| chr16:75110664-75111741 | -0.43967477 | 0.167133051 | 0.427210723 |
| chr12:100199390-100199597 | -0.54917719 | 0.167147009 | 0.427221973 |
| chr2:142944757-142945161 | 0.566720525 | 0.167148604 | 0.427221973 |
| chr15:76265873-76266075 | 0.695296042 | 0.167175859 | 0.427277384 |
| chr8:15539695-15540857 | -0.455964974 | 0.167183202 | 0.427281897 |
| chrX:131277895-131278126 | 0.689273989 | 0.167190925 | 0.427287384 |
| chr5:42889391-42889698 | 0.53019672 | 0.167197274 | 0.427289359 |
| chr15:98751799-98752347 | 0.501705932 | 0.167211142 | 0.427299583 |
| chr7:17769814-17770201 | 0.499337341 | 0.167216445 | 0.427299583 |
| chr7:151315107-151315374 | -0.605343383 | 0.167218005 | 0.427299583 |
| chr10:87735243-87735581 | -0.563758128 | 0.167226377 | 0.427306725 |
| chr11:68386430-68386773 | -0.581089933 | 0.167285375 | 0.427443227 |
| chr2:138374075-138374300 | -0.718133514 | 0.167307251 | 0.427484866 |
| chr1:27772703-27773770 | 0.44242154 | 0.167313455 | 0.427486462 |
| chr14:21014725-21015234 | 0.51155429 | 0.167341808 | 0.427544648 |
| chr2:54557561-54559009 | -0.441226697 | 0.167354095 | 0.427561784 |
| chr13:42776360-42776901 | -0.508611558 | 0.167369197 | 0.42758611 |
| chr16:33767001-33767261 | 0.683443274 | 0.167375151 | 0.427587065 |
| chr5:99320547-99320893 | -0.565895541 | 0.167384309 | 0.427596204 |
| chr10:44864586-44865284 | -0.459744519 | 0.167410947 | 0.427649994 |
| chr3:48847760-48848082 | 0.533991482 | 0.167432975 | 0.427692005 |
| chr1:219133977-219134217 | 0.680092542 | 0.167438892 | 0.427692863 |
| chr15:68930215-68930866 | 0.470580163 | 0.167471918 | 0.427746155 |

| | | | |
|---|---|---|---|
| chr11:107017352-107019472 | -0.44408604 | 0.167475341 | 0.427746155 |
| chr12:119835193-119835400 | -0.770817521 | 0.167476503 | 0.427746155 |
| chr2:99205639-99206000 | 0.695235235 | 0.16749795 | 0.427777616 |
| chr19:35986984-35987349 | 0.559350655 | 0.167505535 | 0.427777616 |
| chr12:17929019-17929272 | -0.705060755 | 0.167505571 | 0.427777616 |
| chr14:41362217-41362440 | -0.673729294 | 0.167550956 | 0.427879261 |
| chr10:86078853-86079082 | 0.64415699 | 0.167564921 | 0.427900664 |
| chrX:104359336-104360125 | -0.536754739 | 0.167604071 | 0.427978077 |
| chr11:130434952-130435263 | 0.564359213 | 0.167606407 | 0.427978077 |
| chr2:214378426-214379180 | -0.458180205 | 0.167622419 | 0.427992038 |
| chr20:53929025-53929293 | 0.465534915 | 0.167626591 | 0.427992038 |
| chr14:75002403-75003150 | 0.44749428 | 0.167628632 | 0.427992038 |
| chr21:38821351-38821652 | -0.628265798 | 0.167652992 | 0.428025335 |
| chr10:924529-924794 | 0.581642825 | 0.16765745 | 0.428025335 |
| chrX:99481340-99481636 | 0.634481633 | 0.167658432 | 0.428025335 |
| chr2:129922857-129923807 | -0.471676752 | 0.167678891 | 0.428053665 |
| chr8:83040812-83041012 | 0.764350462 | 0.167680702 | 0.428053665 |
| chr1:17551053-17551499 | 0.607316511 | 0.167723761 | 0.428137974 |
| chr20:18632294-18632531 | 0.596073997 | 0.167724904 | 0.428137974 |
| chr8:133076262-133076538 | -0.592625189 | 0.167769175 | 0.428236714 |
| chr2:221254383-221254689 | -0.620473703 | 0.167782869 | 0.428257404 |
| chr2:73204630-73205095 | -0.495277821 | 0.167805874 | 0.428301854 |
| chr11:65574102-65574368 | 0.511322567 | 0.167822751 | 0.428319001 |
| chr5:13815419-13815656 | -0.708766541 | 0.167823772 | 0.428319001 |
| chr9:134455599-134455883 | 0.612715031 | 0.167842784 | 0.428353257 |
| chr4:8073183-8073570 | 0.613117833 | 0.167853975 | 0.428367547 |
| chr1:109980817-109981144 | -0.532471234 | 0.167878042 | 0.428404325 |
| chr8:109858626-109858852 | -0.7073679 | 0.167879568 | 0.428404325 |
| chr19:3275481-3275792 | 0.543791963 | 0.167904217 | 0.428452957 |
| chr1:153400412-153400612 | 0.694257241 | 0.167919446 | 0.428457088 |
| chr6:63440148-63440414 | -0.653439204 | 0.167926734 | 0.428457088 |
| chr7:42890794-42890994 | -0.785596179 | 0.167931529 | 0.428457088 |
| chrX:121239693-121239927 | -0.785596179 | 0.167931529 | 0.428457088 |
| chr10:49539442-49539936 | -0.47608295 | 0.167935006 | 0.428457088 |
| chr2:154375672-154376007 | -0.583052656 | 0.167939387 | 0.428457088 |
| chr17:67108440-67108651 | -0.727630614 | 0.167956151 | 0.428472385 |
| chr1:173148702-173149019 | -0.634020956 | 0.167956567 | 0.428472385 |
| chr4:2755947-2756688 | 0.466540314 | 0.167965525 | 0.428480971 |
| chr5:18305396-18305656 | 0.58227831 | 0.168000015 | 0.428554688 |
| chr16:11640123-11640861 | 0.468406985 | 0.168024277 | 0.428592899 |
| chr18:50266174-50266775 | -0.47473026 | 0.168026448 | 0.428592899 |
| chr14:102691870-102692152 | 0.608223338 | 0.168031775 | 0.428592899 |
| chr6:3348590-3349212 | -0.53116222 | 0.168051348 | 0.428608695 |
| chr2:191690932-191691338 | -0.522694025 | 0.168051553 | 0.428608695 |
| chr1:114853900-114854100 | 0.597447132 | 0.168054769 | 0.428608695 |
| chr7:144027171-144027399 | -0.735072567 | 0.168065249 | 0.428608695 |
| chr2:39228018-39228356 | 0.561634033 | 0.168065937 | 0.428608695 |
| chr12:14384872-14385661 | -0.488333944 | 0.168084926 | 0.428642856 |
| chr20:24949682-24950233 | -0.543947294 | 0.168118038 | 0.428713026 |
| chr7:122183305-122183637 | 0.636819189 | 0.168129446 | 0.428727849 |

| | | | |
|---|---|---|---|
| chr16:75565749-75567157 | -0.438849804 | 0.168140892 | 0.428730028 |
| chr10:24952735-24953360 | -0.519669671 | 0.168141491 | 0.428730028 |
| chr19:39045864-39046673 | 0.485584515 | 0.168178801 | 0.428810892 |
| chr8:126744503-126744845 | -0.589984009 | 0.168194884 | 0.428831431 |
| chr11:96272992-96273278 | -0.682300148 | 0.16820362 | 0.428831431 |
| chr19:42884946-42885191 | 0.681085888 | 0.168203669 | 0.428831431 |
| chr1:50286427-50286861 | 0.551195003 | 0.168209243 | 0.428831431 |
| chr19:38962210-38962496 | 0.605821574 | 0.16822198 | 0.428847987 |
| chr2:95402073-95402360 | 0.552999464 | 0.168226932 | 0.428847987 |
| chr21:41878451-41879941 | -0.448361915 | 0.168241486 | 0.428857494 |
| chr4:124262271-124262485 | 0.702651583 | 0.168244739 | 0.428857494 |
| chr2:159399851-159400233 | -0.582013145 | 0.168250723 | 0.428857494 |
| chr1:174466651-174466918 | -0.616417802 | 0.168253049 | 0.428857494 |
| chr1:246565741-246567106 | -0.431302742 | 0.168260008 | 0.428860964 |
| chr2:173872849-173873049 | 0.679857403 | 0.168266766 | 0.428863923 |
| chr17:61358916-61359155 | 0.456808164 | 0.168280261 | 0.428884054 |
| chr2:60429763-60430084 | -0.609856542 | 0.168296498 | 0.428911167 |
| chr17:54993134-54993433 | 0.534226816 | 0.168310752 | 0.428933227 |
| chr12:46133817-46134137 | 0.57176063 | 0.168332411 | 0.428974157 |
| chr17:60620113-60620321 | 0.51066922 | 0.168354246 | 0.429015532 |
| chr7:54785304-54785536 | -0.604658333 | 0.168366128 | 0.429031544 |
| chr9:136152698-136152928 | 0.628630524 | 0.168380711 | 0.42904755 |
| chr10:69696084-69696502 | -0.512957746 | 0.168383609 | 0.42904755 |
| chr6:30490029-30490349 | 0.524256266 | 0.168411635 | 0.429104693 |
| chr1:184805574-184805945 | -0.494941645 | 0.16843639 | 0.429153496 |
| chr4:183073580-183073874 | -0.596201641 | 0.168468248 | 0.429220393 |
| chr11:19905288-19905593 | -0.609271486 | 0.168486947 | 0.429253762 |
| chr13:32310158-32310609 | -0.515627594 | 0.16851469 | 0.429310166 |
| chr12:88763584-88763947 | -0.540159724 | 0.168534427 | 0.429346175 |
| chr5:90409178-90410325 | -0.433051365 | 0.168551059 | 0.429374269 |
| chr19:4124102-4124485 | 0.513104238 | 0.168588076 | 0.429451407 |
| chr10:74778494-74778694 | 0.74108938 | 0.168592549 | 0.429451407 |
| chr1:16079034-16079289 | 0.626246584 | 0.168599844 | 0.429455711 |
| chr19:8513566-8514406 | 0.452036148 | 0.168625595 | 0.429507025 |
| chr8:96132799-96133422 | -0.479676309 | 0.168638536 | 0.429515674 |
| chr13:28316180-28316381 | 0.685167467 | 0.168645769 | 0.429515674 |
| chr10:107264827-107265115 | -0.681600251 | 0.168645808 | 0.429515674 |
| chr14:96072662-96072920 | 0.638137621 | 0.168664922 | 0.429550077 |
| chrX:121047444-121048543 | 0.485255214 | 0.168685041 | 0.429587037 |
| chr1:65309266-65309621 | 0.507033671 | 0.168700073 | 0.429611042 |
| chr5:91162419-91163408 | -0.452102656 | 0.168716584 | 0.429630151 |
| chr1:156052910-156053376 | 0.482028519 | 0.168718792 | 0.429630151 |
| chr7:2266961-2267161 | -0.725541382 | 0.168778557 | 0.429758658 |
| chr5:31844704-31844916 | -0.757385885 | 0.168780475 | 0.429758658 |
| chr8:144466756-144467018 | 0.504333154 | 0.168799475 | 0.42979239 |
| chr12:59113802-59114277 | 0.546159267 | 0.168807057 | 0.42979239 |
| chr12:50070477-50070782 | 0.541031876 | 0.168812523 | 0.42979239 |
| chr7:109479457-109479748 | -0.585172577 | 0.168816159 | 0.42979239 |
| chr1:42209542-42210216 | -0.516423256 | 0.168846119 | 0.429854381 |
| chr18:11860005-11860292 | 0.627945033 | 0.168855804 | 0.429859857 |

| | | | |
|---|---|---|---|
| chr9:129028177-129028702 | 0.465582174 | 0.168859984 | 0.429859857 |
| chr13:49570218-49570434 | -0.731997246 | 0.16886737 | 0.429859857 |
| chr3:185559008-185559388 | 0.514594206 | 0.16887071 | 0.429859857 |
| chr16:57092248-57092526 | 0.639776606 | 0.168895946 | 0.429909812 |
| chr5:97405727-97405929 | -0.702114414 | 0.168927635 | 0.429976189 |
| chr19:54157000-54157275 | 0.521981044 | 0.168938062 | 0.429988446 |
| chr1:232508321-232508641 | -0.575463612 | 0.168974059 | 0.430041682 |
| chr7:123846093-123846293 | -0.783696102 | 0.168976414 | 0.430041682 |
| chr3:133661401-133662183 | -0.457461296 | 0.168980529 | 0.430041682 |
| chr17:38967321-38967927 | 0.49967393 | 0.168981428 | 0.430041682 |
| chr15:40421786-40422197 | 0.522148707 | 0.169002072 | 0.430079935 |
| chr18:21192069-21192357 | 0.616474203 | 0.169010959 | 0.430088265 |
| chr12:130916289-130916489 | 0.707098093 | 0.169029764 | 0.430121834 |
| chr18:48388946-48389193 | -0.841818142 | 0.169047395 | 0.430145218 |
| chr7:155214013-155215058 | -0.463075897 | 0.169055672 | 0.430145218 |
| chr6:123393343-123393632 | -0.636949768 | 0.169055794 | 0.430145218 |
| chr17:9050821-9051105 | -0.63458919 | 0.169068734 | 0.430163857 |
| chr4:17753619-17754293 | -0.528498066 | 0.16908843 | 0.430199685 |
| chr3:171569625-171569926 | -0.631199246 | 0.169106602 | 0.43023041 |
| chr19:33274367-33274653 | 0.581240703 | 0.169111737 | 0.43023041 |
| chr3:77159259-77159533 | -0.646492806 | 0.169128442 | 0.430243805 |
| chr14:68458912-68459503 | -0.500406181 | 0.169128784 | 0.430243805 |
| chr11:76494159-76494388 | 0.714067931 | 0.169133847 | 0.430243805 |
| chr5:14775785-14776078 | 0.58916362 | 0.169140444 | 0.430246301 |
| chr12:100229217-100230419 | -0.442679766 | 0.169155478 | 0.430270259 |
| chr5:37043744-37043962 | 0.677859619 | 0.169173432 | 0.430301643 |
| chr14:24436743-24437051 | 0.501486943 | 0.169180127 | 0.430304387 |
| chr2:43634669-43634917 | 0.589335354 | 0.169192866 | 0.430322505 |
| chr5:58582732-58583537 | -0.448667637 | 0.169199949 | 0.430323679 |
| chr4:134202595-134203101 | 0.532588569 | 0.16920456 | 0.430323679 |
| chr10:93745503-93745741 | -0.613636124 | 0.169216215 | 0.430339037 |
| chr3:25916334-25916825 | 0.509064566 | 0.169255129 | 0.430420488 |
| chr1:167043756-167044472 | -0.478562678 | 0.169259478 | 0.430420488 |
| chr6:166949992-166950223 | -0.638072751 | 0.169265813 | 0.430422312 |
| chr5:173868181-173868476 | 0.526658892 | 0.169271754 | 0.430423135 |
| chr18:62384226-62385258 | -0.46009195 | 0.16929718 | 0.430473501 |
| chr1:106643627-106643827 | 0.540037455 | 0.169314092 | 0.430500819 |
| chr4:144562473-144562684 | -0.72485393 | 0.16931916 | 0.430500819 |
| chr2:107687242-107687449 | -0.729135937 | 0.16932489 | 0.430501102 |
| chr15:96200575-96200964 | -0.583258337 | 0.169340169 | 0.430525663 |
| chr2:131434016-131434594 | -0.561967306 | 0.169350901 | 0.430538661 |
| chr7:47100903-47101195 | -0.550003077 | 0.169375202 | 0.430586155 |
| chr3:106610451-106610946 | -0.518441581 | 0.169404625 | 0.430620193 |
| chr18:14979097-14979297 | 0.752123597 | 0.169412056 | 0.430620193 |
| chr2:3780629-3781111 | -0.497459036 | 0.169413211 | 0.430620193 |
| chr2:48530598-48530887 | 0.538501152 | 0.169415853 | 0.430620193 |
| chr17:82212712-82212952 | 0.537579398 | 0.169416691 | 0.430620193 |
| chr9:3467696-3467898 | 0.716687849 | 0.169438352 | 0.430660963 |
| chr17:68961282-68961482 | 0.706759235 | 0.169476 | 0.430728372 |
| chr11:102733269-102733600 | -0.598024233 | 0.169476116 | 0.430728372 |

| | | | |
|---|---|---|---|
| chr9:38541708-38542005 | -0.721044273 | 0.169488724 | 0.430746127 |
| chr13:39616133-39616665 | -0.49929203 | 0.169508245 | 0.43078145 |
| chr11:65027217-65027984 | 0.456088368 | 0.169536329 | 0.430838533 |
| chr6:142518025-142518596 | -0.49576616 | 0.169545777 | 0.430842752 |
| chr17:62774000-62774407 | 0.519919788 | 0.169549235 | 0.430842752 |
| chr19:6501230-6501435 | 0.735007789 | 0.169562381 | 0.430855949 |
| chr8:141391833-141392396 | 0.464386425 | 0.169566201 | 0.430855949 |
| chr17:58647898-58648682 | 0.439382777 | 0.169571298 | 0.430855949 |
| chrX:31590669-31590966 | -0.632259844 | 0.169590727 | 0.430883029 |
| chr17:72640499-72640753 | 0.619183866 | 0.169593203 | 0.430883029 |
| chr9:76419801-76420130 | -0.564580155 | 0.16960194 | 0.430890939 |
| chr4:89226659-89227339 | -0.47223319 | 0.169613638 | 0.430906371 |
| chr1:21338748-21339035 | -0.56687586 | 0.16963039 | 0.430934642 |
| chr6:56714383-56714644 | 0.652518913 | 0.169681916 | 0.43105125 |
| chr10:43265942-43266370 | 0.502434839 | 0.169705233 | 0.431080994 |
| chr10:118423835-118424148 | -0.536143255 | 0.169709968 | 0.431080994 |
| chr17:40560171-40560598 | 0.509884527 | 0.169710503 | 0.431080994 |
| chr18:54191456-54191677 | -0.686371563 | 0.16972922 | 0.431114244 |
| chr5:137078387-137078685 | -0.648451425 | 0.169735438 | 0.431115747 |
| chr1:77281850-77282349 | 0.538311707 | 0.169752105 | 0.431143788 |
| chr4:139526979-139527399 | -0.510395121 | 0.169789728 | 0.431225049 |
| chr1:178238753-178239044 | 0.579612827 | 0.169798516 | 0.431229122 |
| chr4:39539253-39539736 | -0.50601609 | 0.169802587 | 0.431229122 |
| chr1:241749530-241750345 | 0.476517286 | 0.16981373 | 0.431243126 |
| chr3:62876161-62876361 | 0.510931231 | 0.169833789 | 0.431279772 |
| chr12:32947292-32947551 | 0.609148597 | 0.169869161 | 0.4313553 |
| chr9:86946307-86946724 | 0.532705108 | 0.169875197 | 0.431356334 |
| chr2:74833741-74834189 | 0.462662287 | 0.169884405 | 0.431365419 |
| chr21:30072522-30072801 | 0.624302092 | 0.169898073 | 0.431385829 |
| chr15:88638250-88639571 | 0.437376178 | 0.169930025 | 0.431417168 |
| chr2:30174934-30175164 | -0.731738581 | 0.16993206 | 0.431417168 |
| chr6:14758445-14758964 | -0.537881192 | 0.169940926 | 0.431417168 |
| chr16:62225734-62226059 | -0.534285738 | 0.169942613 | 0.431417168 |
| chr10:127649739-127650007 | -0.702266541 | 0.16994813 | 0.431417168 |
| chr12:116066694-116066924 | -0.702266541 | 0.16994813 | 0.431417168 |
| chr2:206556441-206556659 | -0.724578604 | 0.169949829 | 0.431417168 |
| chr1:207654322-207654666 | -0.545937652 | 0.169963342 | 0.431434473 |
| chr2:139031615-139032146 | -0.539845754 | 0.169967907 | 0.431434473 |
| chr5:19653600-19653802 | -0.781764959 | 0.169983751 | 0.431460395 |
| chr12:111813161-111813480 | 0.547799856 | 0.169999564 | 0.431486239 |
| chr10:27154130-27154743 | -0.461770506 | 0.170008418 | 0.431494418 |
| chr9:122092531-122093065 | 0.510158268 | 0.170021833 | 0.431514174 |
| chr1:154559517-154559906 | 0.487275777 | 0.17004782 | 0.43155512 |
| chr5:108557648-108557953 | -0.703040163 | 0.170054636 | 0.43155512 |
| chr21:32569345-32570128 | 0.469680944 | 0.170054863 | 0.43155512 |
| chr3:121096692-121097300 | 0.500711259 | 0.170064446 | 0.431565144 |
| chr20:24102297-24102599 | -0.608774265 | 0.170083709 | 0.431591407 |
| chr22:23376353-23376598 | 0.644665439 | 0.170094135 | 0.431591407 |
| chr1:16032885-16033148 | 0.57538946 | 0.17009586 | 0.431591407 |
| chr22:43385728-43385972 | 0.648133124 | 0.170097326 | 0.431591407 |

| | | | |
|---|---|---|---|
| chr11:74492736-74492936 | -0.621614574 | 0.170113976 | 0.43161217 |
| chr8:17802782-17803119 | 0.511916498 | 0.170116775 | 0.43161217 |
| chr8:100559161-100560023 | 0.444296269 | 0.170210089 | 0.431822466 |
| chr12:117968836-117969617 | 0.444916695 | 0.170216048 | 0.431822466 |
| chr17:55448841-55449071 | -0.634484952 | 0.170216569 | 0.431822466 |
| chr7:54278613-54279227 | -0.561539003 | 0.170224776 | 0.43182899 |
| chr1:106329671-106329941 | 0.515482588 | 0.170232658 | 0.431834687 |
| chr1:15584571-15584996 | -0.606071792 | 0.170275512 | 0.431929096 |
| chr7:50567742-50567983 | -0.669820309 | 0.17032096 | 0.432015823 |
| chr2:241665516-241665761 | 0.599236124 | 0.170320978 | 0.432015823 |
| chr20:1325701-1326033 | 0.533201513 | 0.170386273 | 0.432167135 |
| chrX:152830447-152831557 | 0.440937186 | 0.170407702 | 0.432207181 |
| chr8:75408611-75408919 | -0.622988924 | 0.170416221 | 0.432213521 |
| chr5:28789024-28789287 | -0.651547913 | 0.170421483 | 0.432213521 |
| chr5:86617674-86618720 | -0.441107024 | 0.170433056 | 0.432220232 |
| chr22:42867794-42868089 | 0.558918354 | 0.170435412 | 0.432220232 |
| chr21:45073235-45073677 | 0.505734061 | 0.170463635 | 0.4322775 |
| chr13:41055242-41055493 | -0.570077339 | 0.170471773 | 0.432283829 |
| chr19:49125325-49125811 | 0.48341429 | 0.170495311 | 0.432329209 |
| chr13:110527304-110528038 | 0.47764971 | 0.170507915 | 0.43234585 |
| chr11:119485289-119485549 | -0.61815557 | 0.170523653 | 0.43234585 |
| chr4:106213300-106213987 | -0.507859547 | 0.170524232 | 0.43234585 |
| chr15:81410317-81410629 | -0.558619637 | 0.170526068 | 0.43234585 |
| chr3:153257790-153258003 | 0.698890184 | 0.170530086 | 0.43234585 |
| chr12:131390220-131390886 | 0.478921194 | 0.170556413 | 0.432398288 |
| chr19:55549680-55549966 | 0.715546498 | 0.170569407 | 0.432409705 |
| chr4:95142973-95143185 | -0.734981816 | 0.170572203 | 0.432409705 |
| chr12:132754647-132754936 | 0.624217307 | 0.170655725 | 0.432607125 |
| chr2:63587997-63589366 | -0.430678919 | 0.170663359 | 0.432612165 |
| chr9:77456951-77457152 | -0.649610168 | 0.170674349 | 0.432625711 |
| chr17:50817255-50817547 | -0.583138395 | 0.170701424 | 0.432679387 |
| chr7:36937772-36937986 | 0.698496999 | 0.170706819 | 0.432679387 |
| chr14:75521358-75521952 | 0.475814757 | 0.170722596 | 0.432705061 |
| chr19:11814015-11814393 | 0.511731696 | 0.170751638 | 0.432764356 |
| chr14:74018585-74019735 | 0.443253988 | 0.17075805 | 0.432766293 |
| chr10:132662642-132662856 | -0.717560423 | 0.170772304 | 0.432788101 |
| chr1:106345104-106345552 | 0.48829175 | 0.17079849 | 0.432826802 |
| chr15:63389939-63390191 | 0.603837909 | 0.170798872 | 0.432826802 |
| chr2:178413295-178414217 | -0.455789439 | 0.170806636 | 0.432832162 |
| chr17:20843856-20844167 | 0.626533145 | 0.17082357 | 0.432860757 |
| chr16:81723116-81723488 | 0.494457735 | 0.170830289 | 0.432863468 |
| chr2:60550369-60550679 | -0.512923344 | 0.170843433 | 0.432882457 |
| chr2:216590861-216591084 | 0.697982282 | 0.170851218 | 0.432887868 |
| chr3:155745765-155746163 | -0.487880557 | 0.170869546 | 0.432907617 |
| chr4:127870995-127871366 | 0.605082625 | 0.170884053 | 0.432907617 |
| chr9:133615804-133616004 | 0.6811337 | 0.170884602 | 0.432907617 |
| chr5:43040318-43040628 | -0.491815632 | 0.170884721 | 0.432907617 |
| chr12:30807428-30807677 | 0.620196499 | 0.170887262 | 0.432907617 |
| chr11:86799752-86800005 | -0.612816028 | 0.17096146 | 0.433080527 |
| chr2:109417071-109417331 | 0.620292425 | 0.170971787 | 0.433080527 |

| | | | |
|---|---|---|---|
| chrX:76427485-76428442 | 0.476240729 | 0.170972473 | 0.433080527 |
| chr6:54874338-54874627 | -0.591482372 | 0.171014206 | 0.433161458 |
| chr11:70568011-70568831 | -0.471158624 | 0.171019781 | 0.433161458 |
| chr2:8809963-8810264 | -0.68728288 | 0.17102619 | 0.433161458 |
| chr7:144011508-144012000 | -0.614379528 | 0.171028558 | 0.433161458 |
| chr19:35495402-35495722 | 0.524460296 | 0.17103269 | 0.433161458 |
| chr17:55514035-55514301 | 0.580342894 | 0.171050505 | 0.433188475 |
| chr3:84825896-84826121 | -0.737448357 | 0.171065525 | 0.433188475 |
| chr11:61917483-61917977 | 0.529654957 | 0.171065746 | 0.433188475 |
| chr15:40952737-40953712 | 0.449135964 | 0.171065972 | 0.433188475 |
| chr11:123511096-123511561 | -0.49169378 | 0.171072222 | 0.433189986 |
| chr8:87695010-87695518 | 0.480344027 | 0.171078477 | 0.433191508 |
| chr17:57256888-57257236 | -0.497682859 | 0.17108752 | 0.433193857 |
| chr5:140841020-140841303 | -0.582849189 | 0.171090711 | 0.433193857 |
| chr11:10302815-10303396 | 0.493384329 | 0.171110371 | 0.433229317 |
| chr9:14030325-14030580 | -0.692874087 | 0.171123218 | 0.433240185 |
| chr18:39799581-39799802 | -0.650481661 | 0.171131223 | 0.433240185 |
| chr17:60018384-60020146 | 0.43209558 | 0.171131626 | 0.433240185 |
| chr21:31509144-31509973 | -0.454595938 | 0.1711443 | 0.433244409 |
| chr2:9703128-9703570 | -0.518900838 | 0.171144603 | 0.433244409 |
| chr5:176658113-176658545 | 0.499008545 | 0.171153468 | 0.433252536 |
| chr12:21680215-21680736 | -0.542957304 | 0.171184227 | 0.433316084 |
| chr20:57230208-57230543 | 0.446439349 | 0.171214495 | 0.433365067 |
| chr2:97590266-97590466 | -0.777992281 | 0.171215763 | 0.433365067 |
| chr20:37823364-37823606 | 0.680454226 | 0.171220546 | 0.433365067 |
| chr5:45521080-45521398 | -0.637013881 | 0.17123172 | 0.433368671 |
| chr8:100907755-100909027 | 0.437279697 | 0.171233282 | 0.433368671 |
| chr7:45506757-45506958 | -0.701223645 | 0.171249132 | 0.43339447 |
| chr3:104460759-104461013 | -0.652010691 | 0.171265128 | 0.433420639 |
| chr4:165131272-165131574 | -0.54851031 | 0.171302621 | 0.433487406 |
| chr10:30401670-30401870 | 0.717304777 | 0.171305084 | 0.433487406 |
| chr6:10429671-10429994 | -0.571379013 | 0.171308484 | 0.433487406 |
| chr2:237673144-237673358 | 0.651520709 | 0.171320905 | 0.433494472 |
| chr13:26551118-26551417 | 0.652955342 | 0.171324825 | 0.433494472 |
| chr17:61869787-61870044 | 0.472312466 | 0.171332689 | 0.433494472 |
| chr11:6353373-6353799 | -0.535692268 | 0.171333907 | 0.433494472 |
| chr4:118669260-118669525 | -0.663724487 | 0.171354735 | 0.433524174 |
| chrX:74118589-74119055 | -0.551813988 | 0.171359098 | 0.433524174 |
| chr5:34007603-34008587 | -0.44496979 | 0.17136262 | 0.433524174 |
| chr20:53700898-53701663 | 0.43292547 | 0.171374652 | 0.433540299 |
| chr6:157597349-157598033 | 0.476654465 | 0.171397422 | 0.433574654 |
| chr3:169914385-169914622 | 0.611525031 | 0.171400068 | 0.433574654 |
| chr15:70635440-70635660 | -0.795206084 | 0.171405208 | 0.433574654 |
| chr15:84599102-84599327 | 0.578937575 | 0.171426724 | 0.433582999 |
| chr1:100543714-100544039 | -0.539405679 | 0.171434426 | 0.433582999 |
| chr1:101820164-101820399 | -0.688814296 | 0.171436244 | 0.433582999 |
| chr6:156780408-156780672 | -0.560465608 | 0.171438594 | 0.433582999 |
| chr6:52881941-52882712 | -0.463128086 | 0.171443484 | 0.433582999 |
| chr2:69386606-69387629 | -0.446226384 | 0.171445815 | 0.433582999 |
| chr7:519093-519378 | 0.512727023 | 0.171449106 | 0.433582999 |

| | | | |
|---|---|---|---|
| chr12:31012088-31012594 | -0.562254031 | 0.171453777 | 0.433582999 |
| chr2:131527691-131528146 | -0.53053608 | 0.171475097 | 0.43360506 |
| chr1:94321835-94322108 | 0.617418553 | 0.171480909 | 0.43360506 |
| chr20:62703490-62703871 | 0.485451775 | 0.171483457 | 0.43360506 |
| chr3:193879530-193879755 | -0.636436392 | 0.171485137 | 0.43360506 |
| chr17:66502909-66503656 | 0.460928955 | 0.171524272 | 0.433689702 |
| chr1:37807552-37808649 | 0.4440375 | 0.171537576 | 0.433703707 |
| chr14:85443140-85443400 | -0.667558844 | 0.171541131 | 0.433703707 |
| chr13:65959301-65959682 | 0.581838963 | 0.171550731 | 0.433713667 |
| chr20:35607499-35607716 | 0.643638735 | 0.171575645 | 0.433762342 |
| chr20:54028316-54028543 | 0.480290562 | 0.171587226 | 0.433777309 |
| chr3:140276800-140277019 | 0.632404846 | 0.171596667 | 0.433786862 |
| chr8:142039299-142039546 | 0.561758118 | 0.1716283 | 0.433852516 |
| chr10:87846009-87846209 | -0.652868465 | 0.171640812 | 0.43386983 |
| chr6:28489831-28490197 | 0.519832997 | 0.171666157 | 0.433894004 |
| chr20:13153620-13153825 | -0.767673686 | 0.171668235 | 0.433894004 |
| chr17:18218687-18218913 | -0.694624236 | 0.171672429 | 0.433894004 |
| chr17:28644599-28645580 | -0.446438255 | 0.171673026 | 0.433894004 |
| chr19:43555212-43555515 | 0.595130759 | 0.171689338 | 0.433920918 |
| chr10:50788973-50789323 | -0.547664463 | 0.171709956 | 0.433958713 |
| chr16:76823133-76823333 | 0.641123614 | 0.171716695 | 0.433961431 |
| chr5:160345053-160345758 | -0.467625491 | 0.171735825 | 0.433995461 |
| chr1:80155468-80155668 | -0.738825533 | 0.171779555 | 0.434091654 |
| chr17:56499897-56500115 | -0.645043201 | 0.171800077 | 0.434128228 |
| chr11:72821597-72822967 | -0.437779145 | 0.17180796 | 0.434128228 |
| chr20:31924339-31924616 | -0.660640901 | 0.171811025 | 0.434128228 |
| chr3:37996850-37997153 | 0.493149249 | 0.171826906 | 0.434154038 |
| chr17:74923545-74923792 | 0.570449895 | 0.171864377 | 0.434234397 |
| chr3:156151819-156152375 | -0.482960363 | 0.1718922 | 0.434290373 |
| chrX:133923648-133923880 | -0.662882321 | 0.17192238 | 0.434352303 |
| chr3:10008611-10009321 | 0.450073869 | 0.171933951 | 0.434355745 |
| chr2:151927582-151927842 | -0.625230534 | 0.171941646 | 0.434355745 |
| chr8:126293788-126293995 | 0.631482109 | 0.171948452 | 0.434355745 |
| chr19:10219672-10219890 | 0.668888309 | 0.17195367 | 0.434355745 |
| chr1:182633458-182634411 | -0.440907434 | 0.171955727 | 0.434355745 |
| chr16:25256985-25257185 | 0.612204426 | 0.171966967 | 0.434355745 |
| chrX:131262788-131263071 | 0.570975993 | 0.17196924 | 0.434355745 |
| chr2:138507410-138507666 | -0.680496849 | 0.17197541 | 0.434355745 |
| chr8:124262163-124262394 | -0.587987683 | 0.171976945 | 0.434355745 |
| chr20:56249404-56249682 | -0.531906374 | 0.171980431 | 0.434355745 |
| chr1:197737212-197737584 | 0.576525575 | 0.172004499 | 0.434371659 |
| chrX:106998913-106999339 | -0.492718217 | 0.172004994 | 0.434371659 |
| chr7:12556788-12557046 | 0.690806356 | 0.17200932 | 0.434371659 |
| chr6:56663358-56663759 | -0.539305347 | 0.172014359 | 0.434371659 |
| chr8:118005020-118005266 | 0.655171459 | 0.172015077 | 0.434371659 |
| chr18:2851295-2851495 | -0.756547386 | 0.172028576 | 0.434382207 |
| chr8:143943675-143944183 | 0.466348593 | 0.172033277 | 0.434382207 |
| chr1:18912959-18913319 | 0.579779818 | 0.172046203 | 0.434382207 |
| chr1:235636354-235636586 | 0.633364532 | 0.172048357 | 0.434382207 |
| chr1:32413980-32414255 | 0.646265944 | 0.172051943 | 0.434382207 |

| | | | |
|---|---|---|---|
| chr11:57493474-57493717 | 0.604196932 | 0.172057521 | 0.434382207 |
| chr7:5174469-5174864 | -0.525613646 | 0.172058939 | 0.434382207 |
| chr8:83331068-83331389 | 0.574407958 | 0.172085742 | 0.434431057 |
| chr9:7146627-7146976 | -0.57315952 | 0.172092239 | 0.434431057 |
| chr20:39112109-39112417 | -0.566233781 | 0.172095297 | 0.434431057 |
| chr5:35030593-35030793 | -0.713763929 | 0.172113733 | 0.434463283 |
| chr3:112437047-112437485 | -0.597771055 | 0.172156866 | 0.434557845 |
| chr11:75340600-75340933 | -0.504395613 | 0.172171003 | 0.434568054 |
| chr1:206447784-206448073 | -0.538618229 | 0.172172254 | 0.434568054 |
| chr21:23452442-23453117 | -0.475035645 | 0.172188929 | 0.434590152 |
| chr1:244796665-244797317 | 0.500195531 | 0.172197789 | 0.434590152 |
| chr12:51005281-51006061 | 0.48392392 | 0.172208992 | 0.434590152 |
| chr5:142403453-142403970 | -0.483795319 | 0.172211248 | 0.434590152 |
| chr13:72985675-72985954 | -0.708663692 | 0.172213225 | 0.434590152 |
| chr14:46259637-46259837 | 0.736626958 | 0.17221504 | 0.434590152 |
| chr2:118321576-118322313 | -0.489644142 | 0.172241562 | 0.434637181 |
| chr2:33515367-33515613 | -0.762592158 | 0.172254496 | 0.434637181 |
| chr8:103194630-103194865 | -0.673895309 | 0.17225495 | 0.434637181 |
| chr5:75049903-75050120 | -0.604031009 | 0.172256366 | 0.434637181 |
| chr2:74147764-74148342 | -0.479270336 | 0.172267603 | 0.434645243 |
| chr19:19732970-19733392 | -0.503789434 | 0.172282012 | 0.434645243 |
| chr3:131331406-131332210 | -0.463997457 | 0.172282022 | 0.434645243 |
| chr1:116737354-116737593 | 0.564903225 | 0.172290833 | 0.434645243 |
| chr2:138814161-138814425 | -0.664073962 | 0.172296073 | 0.434645243 |
| chr2:110204740-110205278 | -0.504577379 | 0.172296746 | 0.434645243 |
| chr20:32744979-32745360 | 0.494375442 | 0.172299269 | 0.434645243 |
| chr4:159446518-159446891 | -0.497577327 | 0.172317972 | 0.434666584 |
| chr5:54742337-54742968 | -0.506982796 | 0.172319075 | 0.434666584 |
| chr17:75309235-75309844 | -0.480831803 | 0.172332331 | 0.434685713 |
| chr14:73777423-73777845 | 0.479561131 | 0.172347924 | 0.434704128 |
| chrX:147853805-147854032 | 0.683833712 | 0.172352674 | 0.434704128 |
| chr13:51523231-51523485 | -0.596440761 | 0.172356652 | 0.434704128 |
| chr12:120710062-120711256 | 0.456669292 | 0.172387427 | 0.434752542 |
| chr3:116053381-116053581 | 0.792100898 | 0.172391587 | 0.434752542 |
| chr3:61634693-61634955 | 0.496114559 | 0.172395814 | 0.434752542 |
| chr18:63238047-63238386 | -0.558470596 | 0.172406054 | 0.434752542 |
| chr11:77075455-77075660 | 0.619038313 | 0.1724062 | 0.434752542 |
| chr2:71545548-71545994 | 0.513582475 | 0.17242551 | 0.434752542 |
| chr20:48214721-48215015 | 0.472315687 | 0.17242746 | 0.434752542 |
| chr2:226797278-226797543 | -0.543617497 | 0.172429327 | 0.434752542 |
| chr3:15405567-15405966 | 0.580176673 | 0.172435889 | 0.434752542 |
| chr20:53921449-53921945 | 0.438257713 | 0.17244605 | 0.434752542 |
| chr5:91189952-91190183 | -0.696306029 | 0.172451548 | 0.434752542 |
| chr14:78158115-78158435 | 0.569642495 | 0.172452957 | 0.434752542 |
| chr5:103942644-103942940 | -0.657190314 | 0.172453576 | 0.434752542 |
| chr3:151166235-151166435 | -0.723802679 | 0.172455284 | 0.434752542 |
| chr1:51296826-51297371 | -0.481211542 | 0.172476788 | 0.434792447 |
| chr16:12561682-12561928 | 0.685975986 | 0.172487075 | 0.434804074 |
| chr12:113185344-113186056 | 0.449989776 | 0.172541473 | 0.434926892 |
| chr6:122563199-122563490 | -0.665768162 | 0.17259643 | 0.43505111 |

| | | | |
|---|---|---|---|
| chr1:113835720-113836020 | 0.48575478 | 0.172620308 | 0.435077243 |
| chr14:58184357-58184603 | 0.649425256 | 0.172624146 | 0.435077243 |
| chr6:37208020-37208362 | 0.509014235 | 0.17262705 | 0.435077243 |
| chr2:76748288-76748491 | -0.747860383 | 0.17262951 | 0.435077243 |
| chr3:195954365-195954737 | 0.505401767 | 0.172638076 | 0.435084521 |
| chr16:8441016-8441279 | -0.715499581 | 0.1726687 | 0.435147387 |
| chr17:73697823-73698152 | -0.551571741 | 0.172722997 | 0.435269905 |
| chr5:99032488-99032852 | -0.51487102 | 0.172730603 | 0.435274756 |
| chr17:79020774-79021013 | -0.721391423 | 0.172746143 | 0.4352996 |
| chr16:2514667-2514890 | 0.620947825 | 0.172757794 | 0.435303214 |
| chr15:77118881-77119283 | -0.644076526 | 0.172758939 | 0.435303214 |
| chr1:178548171-178548446 | 0.570824452 | 0.172766643 | 0.43530831 |
| chr3:75640985-75641750 | -0.459570249 | 0.172786275 | 0.435329531 |
| chr10:37417808-37418013 | -0.737963639 | 0.172786428 | 0.435329531 |
| chr5:123089195-123090461 | 0.446872435 | 0.172794528 | 0.435335623 |
| chr4:140740508-140740714 | -0.63255693 | 0.17289803 | 0.435582063 |
| chr11:69316997-69317263 | -0.628730084 | 0.172912283 | 0.435601611 |
| chr7:111855248-111855556 | -0.517820183 | 0.172917159 | 0.435601611 |
| chr8:87793584-87793913 | 0.608165021 | 0.172933187 | 0.435627664 |
| chr21:34912211-34912488 | 0.650463433 | 0.172939809 | 0.435630022 |
| chr16:80852339-80852594 | -0.594402179 | 0.172951725 | 0.435645718 |
| chr17:4565571-4565876 | -0.57099291 | 0.172962949 | 0.435659668 |
| chr9:34191564-34191826 | -0.690094701 | 0.172968935 | 0.435660424 |
| chr5:148998266-148998558 | -0.583403073 | 0.172979532 | 0.435672792 |
| chr1:60425608-60425867 | -0.620030293 | 0.173015507 | 0.435749078 |
| chr14:61368304-61368699 | 0.476961774 | 0.173026389 | 0.435759532 |
| chr19:48574724-48574996 | -0.614314765 | 0.173031032 | 0.435759532 |
| chr13:28636640-28637023 | 0.529046709 | 0.173038698 | 0.435764514 |
| chr10:122054147-122054371 | 0.606724834 | 0.173056055 | 0.435789974 |
| chr4:15963012-15963439 | 0.554586803 | 0.173060183 | 0.435789974 |
| chr8:123918839-123919124 | 0.575807017 | 0.173098858 | 0.435848994 |
| chr6:140494678-140495083 | -0.571555721 | 0.173106187 | 0.435848994 |
| chr3:142717263-142717607 | 0.554146498 | 0.173109958 | 0.435848994 |
| chr1:179826067-179826482 | 0.534398602 | 0.173117425 | 0.435848994 |
| chr18:59005346-59005663 | -0.608335729 | 0.173119985 | 0.435848994 |
| chr3:12759130-12760254 | 0.440963365 | 0.173124683 | 0.435848994 |
| chr21:44001415-44001690 | -0.675950405 | 0.173129483 | 0.435848994 |
| chr7:127233542-127233816 | -0.712305139 | 0.173133965 | 0.435848994 |
| chr3:11770127-11771469 | 0.433518618 | 0.173134815 | 0.435848994 |
| chr7:19681643-19681844 | 0.749550352 | 0.173145175 | 0.435860754 |
| chr11:121459573-121459839 | 0.616504445 | 0.173200213 | 0.435984976 |
| chr6:72787522-72787933 | -0.571522808 | 0.173211635 | 0.435999406 |
| chr22:29552875-29553146 | 0.604851331 | 0.173223057 | 0.436013832 |
| chr4:101169248-101169717 | -0.505211945 | 0.173253487 | 0.436063641 |
| chr3:174355202-174355877 | -0.504662366 | 0.173254228 | 0.436063641 |
| chr1:235415339-235415583 | 0.583310345 | 0.173261956 | 0.436068768 |
| chr2:193845522-193845940 | -0.590512536 | 0.173276639 | 0.436091398 |
| chr17:2786253-2786614 | -0.553728364 | 0.173303123 | 0.436143726 |
| chr20:58198491-58198693 | 0.55816422 | 0.173310146 | 0.436147077 |
| chr8:134905509-134906009 | -0.483424198 | 0.17333333 | 0.436173972 |

| | | | |
|---|---|---|---|
| chr10:8752117-8752780 | -0.536913038 | 0.173336932 | 0.436173972 |
| chr1:2044856-2045139 | -0.610260249 | 0.173337911 | 0.436173972 |
| chr7:112414901-112415208 | -0.583938631 | 0.173355224 | 0.436203212 |
| chr14:50560292-50561244 | 0.462860502 | 0.173377236 | 0.436222769 |
| chr17:75153467-75153972 | 0.481542342 | 0.1733793 | 0.436222769 |
| chr9:36166551-36166920 | 0.545371801 | 0.173380076 | 0.436222769 |
| chr9:137605019-137606099 | 0.439058748 | 0.173407387 | 0.436277157 |
| chr11:70713798-70714008 | 0.687984397 | 0.173416625 | 0.436277868 |
| chr8:11867892-11868300 | 0.498827304 | 0.173430747 | 0.436277868 |
| chr17:48823601-48823811 | 0.682116318 | 0.173434039 | 0.436277868 |
| chr4:77698383-77698721 | -0.593495075 | 0.173434054 | 0.436277868 |
| chr1:91131541-91131761 | -0.736862862 | 0.173436139 | 0.436277868 |
| chr5:138704940-138705430 | 0.531623118 | 0.173449309 | 0.436296674 |
| chr19:40613716-40614264 | 0.459668657 | 0.17345605 | 0.436299305 |
| chr17:63362287-63362534 | -0.605000352 | 0.173467397 | 0.436310955 |
| chr6:17721805-17722005 | 0.697397329 | 0.173481845 | 0.436310955 |
| chr19:5790491-5791420 | 0.450763143 | 0.173485861 | 0.436310955 |
| chr2:156341500-156343019 | -0.428057927 | 0.173487754 | 0.436310955 |
| chr3:140933215-140933518 | -0.634965551 | 0.173489153 | 0.436310955 |
| chr20:33373695-33374281 | 0.486103295 | 0.173509885 | 0.436328418 |
| chr12:121227235-121227701 | 0.488741907 | 0.173519185 | 0.436328418 |
| chr16:10369089-10369307 | 0.652999811 | 0.173528542 | 0.436328418 |
| chr8:104793356-104793778 | -0.498862774 | 0.173529885 | 0.436328418 |
| chr12:50084950-50085803 | 0.44437663 | 0.17353345 | 0.436328418 |
| chr6:145888074-145888430 | -0.565660349 | 0.173534856 | 0.436328418 |
| chr1:3901216-3901597 | 0.467105701 | 0.173535959 | 0.436328418 |
| chr1:239923785-239924026 | 0.737370936 | 0.173552964 | 0.436355847 |
| chr7:91953752-91953978 | -0.742104716 | 0.17356158 | 0.436355847 |
| chr1:247965851-247966280 | -0.583436824 | 0.173570859 | 0.436355847 |
| chrX:132119751-132120145 | 0.533324598 | 0.173574128 | 0.436355847 |
| chr7:116525400-116525864 | -0.527341391 | 0.173575342 | 0.436355847 |
| chr11:118679409-118679699 | 0.622607474 | 0.173596563 | 0.436394877 |
| chr4:72119845-72120613 | -0.497666092 | 0.173642692 | 0.436496517 |
| chr8:20231893-20232093 | -0.741629028 | 0.173650782 | 0.436502534 |
| chr17:62986394-62986625 | 0.575018692 | 0.173659427 | 0.436508534 |
| chr12:123571311-123571593 | -0.603415792 | 0.173664563 | 0.436508534 |
| chr10:79248814-79249014 | 0.62288066 | 0.173670443 | 0.436508993 |
| chr11:787979-788247 | -0.647863262 | 0.173698468 | 0.436565113 |
| chr8:86581105-86581369 | 0.603042814 | 0.173718218 | 0.436600429 |
| chr7:44307306-44307689 | -0.525230058 | 0.173742869 | 0.436648062 |
| chr7:132842912-132843150 | -0.714167691 | 0.17375184 | 0.436656286 |
| chr12:64088422-64088726 | 0.538261327 | 0.173778685 | 0.436709425 |
| chr4:180666258-180666526 | -0.654849773 | 0.173796411 | 0.436739648 |
| chr10:16436504-16437320 | -0.43881815 | 0.173812235 | 0.436765088 |
| chr16:30406901-30408179 | 0.441238865 | 0.173839678 | 0.436804939 |
| chr13:65202938-65203198 | -0.661950983 | 0.173840006 | 0.436804939 |
| chr1:186244800-186245309 | -0.499740215 | 0.173846183 | 0.436804939 |
| chr1:197615019-197615496 | -0.486325836 | 0.173861665 | 0.436804939 |
| chr20:53346255-53346739 | 0.496240628 | 0.173862237 | 0.436804939 |
| chr17:39573848-39574307 | 0.535597557 | 0.173862298 | 0.436804939 |

| | | | |
|---|---|---|---|
| chr2:231590602-231591009 | 0.50247707 | 0.173879926 | 0.436834902 |
| chr9:113767654-113767984 | -0.669407588 | 0.173908875 | 0.436893306 |
| chr7:3428057-3428257 | -0.767668794 | 0.173916994 | 0.436899379 |
| chr1:12549778-12549999 | 0.702523399 | 0.173933192 | 0.436925744 |
| chr7:10973533-10974393 | 0.430656878 | 0.173957439 | 0.436972326 |
| chr1:30807550-30807804 | -0.604343135 | 0.173970605 | 0.436991072 |
| chr5:119315431-119315957 | -0.491307364 | 0.17398646 | 0.437007049 |
| chrX:117376467-117376825 | -0.54732623 | 0.173988372 | 0.437007049 |
| chr3:197410313-197410552 | -0.716813063 | 0.17399439 | 0.43700784 |
| chr13:112606238-112606765 | -0.514435172 | 0.174004442 | 0.437018762 |
| chr10:11831701-11831967 | -0.568363845 | 0.174010961 | 0.437020808 |
| chrX:18620424-18620706 | 0.594032185 | 0.17402029 | 0.437021542 |
| chr7:4798180-4798679 | -0.481620854 | 0.174029219 | 0.437021542 |
| chr20:24469012-24469217 | 0.670621779 | 0.174032462 | 0.437021542 |
| chr8:95071382-95072124 | -0.444231712 | 0.174035798 | 0.437021542 |
| chr6:3239521-3239743 | 0.647788441 | 0.174039771 | 0.437021542 |
| chr4:169619923-169620781 | -0.433003154 | 0.17404941 | 0.437031425 |
| chr7:33716677-33716922 | 0.553395301 | 0.17406535 | 0.43705588 |
| chr3:102235768-102236004 | 0.61536485 | 0.174070558 | 0.43705588 |
| chr8:141716027-141716399 | 0.528048713 | 0.174091083 | 0.437080289 |
| chr5:42423225-42423856 | 0.448313412 | 0.174091688 | 0.437080289 |
| chr7:50119807-50120132 | -0.574943474 | 0.174125064 | 0.437149759 |
| chr7:30028039-30029142 | 0.450722959 | 0.17413384 | 0.437157469 |
| chr10:57049119-57049553 | -0.585743166 | 0.174158794 | 0.43720579 |
| chr6:143842755-143843797 | -0.435740453 | 0.174205539 | 0.437300909 |
| chr5:151697822-151698034 | -0.705411648 | 0.174213098 | 0.437300909 |
| chr5:43461503-43461776 | 0.631894668 | 0.174213806 | 0.437300909 |
| chr2:161135305-161135505 | 0.708772941 | 0.174271763 | 0.437432059 |
| chr22:20843682-20843892 | -0.69315754 | 0.174279948 | 0.437438273 |
| chr20:25685755-25686462 | -0.505043971 | 0.174319848 | 0.437524089 |
| chr14:90114646-90114911 | 0.565588379 | 0.174339472 | 0.437538745 |
| chr3:189041535-189041812 | -0.586879553 | 0.174340821 | 0.437538745 |
| chr12:88141535-88143059 | -0.427674228 | 0.174342818 | 0.437538745 |
| chr20:44738868-44739175 | 0.497400094 | 0.174360161 | 0.437567936 |
| chr7:138074676-138074916 | -0.710287355 | 0.174369056 | 0.437575928 |
| chr19:35309559-35309828 | -0.639491155 | 0.174385119 | 0.437601906 |
| chr17:60566279-60566683 | 0.455206929 | 0.174408321 | 0.437615149 |
| chr2:50529655-50529924 | -0.626615642 | 0.174409098 | 0.437615149 |
| chr15:55513885-55514204 | 0.51699389 | 0.174414612 | 0.437615149 |
| chr17:43832618-43832859 | 0.604576972 | 0.174416371 | 0.437615149 |
| chr15:68042655-68042892 | -0.67079004 | 0.174418953 | 0.437615149 |
| chr5:166595362-166595605 | -0.610684724 | 0.174458259 | 0.437699435 |
| chr4:55379026-55379512 | 0.468652675 | 0.174523129 | 0.437847849 |
| chr7:122093013-122093307 | -0.581104521 | 0.174537066 | 0.43786848 |
| chr1:51058756-51059095 | 0.541646385 | 0.174557816 | 0.437906196 |
| chr15:85168579-85169234 | 0.452271055 | 0.17456436 | 0.437908276 |
| chr2:49973898-49974573 | -0.462258673 | 0.174576437 | 0.437924234 |
| chr1:229420760-229421042 | -0.631196985 | 0.17459084 | 0.437937883 |
| chr13:91398341-91399000 | -0.468252527 | 0.174595379 | 0.437937883 |
| chr7:47581519-47582354 | 0.46649635 | 0.174599024 | 0.437937883 |

| | | | |
|---|---|---|---|
| chr12:113980629-113980838 | 0.644800289 | 0.174606143 | 0.437941402 |
| chr16:1048524-1048779 | 0.62729549 | 0.174613273 | 0.437944949 |
| chr20:41028316-41029300 | 0.428173963 | 0.174647238 | 0.438009461 |
| chr3:196097974-196098241 | 0.695482758 | 0.174650428 | 0.438009461 |
| chr20:50587485-50587685 | 0.554274795 | 0.174661671 | 0.438023322 |
| chr20:48078305-48078635 | 0.458369998 | 0.174678146 | 0.438038038 |
| chr15:45641348-45641743 | -0.530457967 | 0.174678973 | 0.438038038 |
| chr12:123085861-123086161 | -0.629505328 | 0.174691167 | 0.438054282 |
| chr2:191170208-191170606 | -0.508464193 | 0.174720272 | 0.438103596 |
| chr5:132807395-132807702 | -0.605039732 | 0.174722269 | 0.438103596 |
| chr7:28956004-28956332 | 0.540676441 | 0.174832647 | 0.438366016 |
| chr4:105551945-105552989 | -0.451088912 | 0.17487071 | 0.438429486 |
| chr11:16806728-16807171 | 0.505807315 | 0.17487131 | 0.438429486 |
| chr3:196596270-196596647 | -0.600597053 | 0.174875316 | 0.438429486 |
| chr6:165309123-165309771 | -0.515193001 | 0.17488449 | 0.438429486 |
| chr3:50272975-50273184 | 0.640037434 | 0.17489129 | 0.438429486 |
| chr1:179399442-179400060 | -0.494184179 | 0.174892293 | 0.438429486 |
| chrX:143648753-143649101 | -0.56659773 | 0.174919888 | 0.438484318 |
| chr19:19591087-19591364 | -0.675323059 | 0.17493355 | 0.43850422 |
| chr12:121046591-121046900 | -0.609142984 | 0.174994416 | 0.43864244 |
| chr3:123479112-123479319 | -0.735242833 | 0.175026992 | 0.438709744 |
| chr2:203648373-203648654 | -0.582584859 | 0.175037123 | 0.438720787 |
| chr18:65091873-65092075 | -0.753602515 | 0.175066217 | 0.438779357 |
| chr3:124761279-124761873 | 0.477359125 | 0.175084767 | 0.438811496 |
| chr7:50714949-50715212 | -0.620788698 | 0.175105644 | 0.438825112 |
| chr1:192095941-192096141 | 0.742235982 | 0.175109537 | 0.438825112 |
| chr3:61685613-61685926 | 0.487771701 | 0.175111558 | 0.438825112 |
| chr16:66552876-66553120 | -0.625602417 | 0.175113109 | 0.438825112 |
| chr9:23089185-23089385 | -0.739226281 | 0.175135061 | 0.43886577 |
| chr1:40952504-40952744 | 0.612093339 | 0.175153863 | 0.438877436 |
| chr7:134444158-134444440 | -0.579632267 | 0.175161339 | 0.438877436 |
| chr15:44243079-44243336 | 0.725642054 | 0.175162588 | 0.438877436 |
| chr7:114414700-114415094 | -0.57060508 | 0.175165746 | 0.438877436 |
| chr5:65664182-65664628 | -0.580499413 | 0.175172069 | 0.438877436 |
| chr5:146182268-146183295 | -0.428591632 | 0.175174083 | 0.438877436 |
| chr20:47093111-47093318 | -0.69841331 | 0.175195865 | 0.438904703 |
| chr1:75614649-75616121 | -0.440465223 | 0.175196423 | 0.438904703 |
| chr16:74391489-74391864 | 0.570759775 | 0.175209485 | 0.438923075 |
| chr5:77129514-77129760 | -0.614205739 | 0.175217921 | 0.438929858 |
| chr7:138477325-138477576 | -0.677566989 | 0.175227978 | 0.438939335 |
| chr18:65439247-65439590 | -0.611199251 | 0.175235503 | 0.438939335 |
| chr6:79777956-79778559 | -0.458570117 | 0.17523889 | 0.438939335 |
| chr1:149843461-149843832 | 0.5072054 | 0.175259147 | 0.438975723 |
| chr13:112984882-112985255 | -0.606655407 | 0.175312047 | 0.439093869 |
| chr20:49678405-49678720 | 0.597484936 | 0.175319333 | 0.439097765 |
| chr7:73841726-73841963 | -0.630700362 | 0.175325834 | 0.439099694 |
| chr14:37669194-37669579 | 0.478674331 | 0.175335741 | 0.439110153 |
| chr19:1592281-1592492 | 0.623084394 | 0.175348111 | 0.43912678 |
| chr13:51906974-51907248 | -0.63977648 | 0.175361841 | 0.439146812 |
| chr3:184354309-184354735 | 0.495991577 | 0.175369428 | 0.439151457 |

| | | | |
|---|---|---|---|
| chr1:39788783-39789551 | 0.46420963 | 0.175422498 | 0.439269997 |
| chr6:123802403-123803375 | -0.456821976 | 0.175455987 | 0.439332209 |
| chr2:163348007-163348597 | 0.460972326 | 0.17545881 | 0.439332209 |
| chr22:27918539-27920309 | 0.433658235 | 0.175466644 | 0.439337467 |
| chr17:29930133-29930600 | 0.521428392 | 0.175496571 | 0.439394832 |
| chrX:129637312-129637744 | 0.545246568 | 0.175501024 | 0.439394832 |
| chr17:12667352-12667756 | -0.554459494 | 0.175510199 | 0.439403446 |
| chr10:987811-988910 | 0.43213466 | 0.175534794 | 0.439450664 |
| chr16:451694-451894 | -0.704161935 | 0.175541093 | 0.439452073 |
| chr1:153990517-153991101 | 0.448637556 | 0.175554374 | 0.43945915 |
| chr15:31633059-31633340 | 0.685074087 | 0.17555539 | 0.43945915 |
| chr1:30700942-30701305 | 0.537306025 | 0.175574097 | 0.439483804 |
| chr11:123407436-123407646 | 0.66631374 | 0.175576711 | 0.439483804 |
| chr12:69416357-69416611 | -0.655679318 | 0.175595952 | 0.439517608 |
| chr1:54016037-54016811 | -0.45151185 | 0.175619859 | 0.439557499 |
| chr19:2168933-2169146 | 0.65629804 | 0.175623362 | 0.439557499 |
| chr10:97571612-97571948 | 0.509561043 | 0.175655371 | 0.439623252 |
| chr16:788349-788782 | 0.473564137 | 0.175677593 | 0.439664506 |
| chr1:183739025-183739378 | -0.610273351 | 0.175686899 | 0.43966565 |
| chr5:179519477-179519904 | 0.543892415 | 0.175689792 | 0.43966565 |
| chr1:106326491-106326691 | 0.543832057 | 0.175695264 | 0.43966565 |
| chr5:159510292-159510734 | -0.471741501 | 0.175754138 | 0.439798614 |
| chr10:14723392-14723909 | -0.57149137 | 0.175775524 | 0.439837764 |
| chr14:68816469-68816965 | 0.474885305 | 0.175832857 | 0.43996686 |
| chr1:194657229-194658403 | -0.44713762 | 0.175856525 | 0.440011712 |
| chr21:5154674-5155025 | 0.492111557 | 0.175867925 | 0.440021785 |
| chr15:97996088-97996640 | -0.523991235 | 0.175876044 | 0.440021785 |
| chr5:168422455-168422749 | -0.561963769 | 0.17588031 | 0.440021785 |
| chr16:31105686-31106191 | 0.516029805 | 0.175883522 | 0.440021785 |
| chr3:155184636-155184836 | -0.718080873 | 0.175905623 | 0.44005787 |
| chr11:16859610-16859925 | 0.554775691 | 0.175909432 | 0.44005787 |
| chr6:143634585-143634810 | -0.716343823 | 0.175926428 | 0.440069922 |
| chr15:40906197-40906743 | 0.460673511 | 0.175926441 | 0.440069922 |
| chr5:149904931-149905297 | -0.558557074 | 0.17593148 | 0.440069922 |
| chr18:51006522-51006740 | -0.707588555 | 0.175943883 | 0.440078428 |
| chr8:22401204-22401418 | 0.678529971 | 0.175946368 | 0.440078428 |
| chr6:56954310-56955791 | 0.432982738 | 0.175967127 | 0.44011572 |
| chr17:64036278-64036887 | -0.470051772 | 0.175972765 | 0.44011572 |
| chr20:43442582-43442803 | 0.670313163 | 0.175995603 | 0.440158472 |
| chr2:58046266-58047509 | -0.428993482 | 0.17600962 | 0.440169319 |
| chr7:34174762-34175629 | -0.437306185 | 0.17601143 | 0.440169319 |
| chr5:3310980-3311251 | -0.61749844 | 0.176037068 | 0.440209923 |
| chr14:98700741-98700963 | -0.703360482 | 0.176039157 | 0.440209923 |
| chr21:38833237-38833458 | -0.569289722 | 0.1760612 | 0.440239971 |
| chr1:179129811-179130203 | -0.564885501 | 0.176062664 | 0.440239971 |
| chr10:95617767-95618007 | 0.655718308 | 0.176077005 | 0.440261462 |
| chr5:62138581-62138801 | -0.653687961 | 0.176111315 | 0.440332881 |
| chr8:114636497-114637628 | -0.441060582 | 0.176129362 | 0.440358771 |
| chr6:24520987-24521379 | -0.527539987 | 0.176133164 | 0.440358771 |
| chr17:81495067-81495276 | 0.610769438 | 0.176212499 | 0.440542747 |

| | | | |
|---|---|---|---|
| chr2:156435462-156437008 | -0.428436678 | 0.176228699 | 0.440568873 |
| chr12:50025252-50025839 | 0.452775691 | 0.176235894 | 0.440572486 |
| chr3:130821015-130821311 | -0.591135706 | 0.176247491 | 0.440580164 |
| chr19:43934953-43935664 | 0.488326056 | 0.176250466 | 0.440580164 |
| chr15:101615892-101616360 | 0.481829524 | 0.176285833 | 0.440650959 |
| chr6:51490014-51490517 | 0.484283444 | 0.176292215 | 0.440650959 |
| chr5:170752200-170752471 | -0.625733034 | 0.17629604 | 0.440650959 |
| chr12:91577855-91578181 | -0.608035513 | 0.176368573 | 0.440817875 |
| chr13:36134050-36134272 | -0.599661715 | 0.176376119 | 0.440822357 |
| chr6:42142058-42142322 | 0.532565215 | 0.176382742 | 0.440824531 |
| chr4:151459364-151459656 | 0.585836474 | 0.176401518 | 0.440857076 |
| chr4:1693462-1693679 | 0.6169251 | 0.176421177 | 0.440879824 |
| chr15:84023738-84023960 | 0.664792737 | 0.176425388 | 0.440879824 |
| chr6:125267893-125268135 | -0.592397395 | 0.176427882 | 0.440879824 |
| chr4:184250860-184251128 | -0.688823078 | 0.176439519 | 0.440894525 |
| chr2:125680112-125680312 | 0.708698228 | 0.176453134 | 0.44091046 |
| chr3:196208150-196208474 | 0.503860061 | 0.176457405 | 0.44091046 |
| chr2:17540444-17540692 | 0.603471414 | 0.176479678 | 0.440933762 |
| chr3:156043899-156044183 | -0.543315225 | 0.176488153 | 0.440933762 |
| chr4:76305864-76307572 | 0.429605903 | 0.176488821 | 0.440933762 |
| chr7:55546949-55547230 | -0.657996751 | 0.176489749 | 0.440933762 |
| chr2:43443393-43443670 | -0.63626489 | 0.176516499 | 0.440982011 |
| chr1:1941187-1941475 | -0.691714188 | 0.176520572 | 0.440982011 |
| chr9:133417745-133418461 | 0.45770795 | 0.176552576 | 0.441039956 |
| chr6:144482165-144482415 | -0.644204074 | 0.176563024 | 0.441039956 |
| chr16:30187159-30187722 | 0.495985701 | 0.176563589 | 0.441039956 |
| chr10:89414119-89415647 | -0.435765609 | 0.176566791 | 0.441039956 |
| chr11:22456975-22457706 | -0.462055108 | 0.176585909 | 0.441058168 |
| chr4:32024467-32025099 | 0.483576584 | 0.176586298 | 0.441058168 |
| chr13:27341656-27341980 | -0.543240055 | 0.176591351 | 0.441058168 |
| chr7:30283962-30285159 | 0.430219816 | 0.176602392 | 0.44106704 |
| chr12:97447010-97447210 | -0.73062876 | 0.176611369 | 0.44106704 |
| chr10:58299530-58299922 | -0.528054235 | 0.176612172 | 0.44106704 |
| chr13:100721965-100722165 | 0.701760645 | 0.176627497 | 0.441083733 |
| chr2:74267137-74267377 | -0.61195021 | 0.176630369 | 0.441083733 |
| chr11:75366876-75367204 | 0.53000987 | 0.176639226 | 0.441084467 |
| chr11:11157295-11157542 | -0.715052757 | 0.176647865 | 0.441084467 |
| chr6:55578810-55579609 | -0.463946651 | 0.176651758 | 0.441084467 |
| chr5:135887388-135887723 | -0.49526864 | 0.17665369 | 0.441084467 |
| chr2:217943844-217944460 | -0.478949502 | 0.176692271 | 0.441166423 |
| chr10:82934514-82934773 | 0.621253622 | 0.176711237 | 0.441199401 |
| chr6:104887421-104887714 | 0.61993595 | 0.176752599 | 0.441284729 |
| chr8:81090527-81090900 | -0.520793443 | 0.176757726 | 0.441284729 |
| chr20:50724350-50725488 | -0.420787809 | 0.176766337 | 0.441284729 |
| chr5:44952358-44952765 | 0.524876791 | 0.17676845 | 0.441284729 |
| chr7:17618524-17619077 | 0.475361313 | 0.176777879 | 0.441284853 |
| chr8:107744816-107745016 | 0.680105203 | 0.176780018 | 0.441284853 |
| chr7:87564276-87564584 | -0.574103667 | 0.176787514 | 0.441289187 |
| chr20:25000280-25000601 | 0.535144028 | 0.176802485 | 0.441312181 |
| chr9:78030238-78031061 | -0.4542222 | 0.176816418 | 0.441332581 |

| | | | |
|---|---|---|---|
| chr13:68067069-68067476 | -0.599653174 | 0.176842175 | 0.441382492 |
| chr2:62775086-62775343 | -0.607192903 | 0.176862076 | 0.441417786 |
| chr16:23324728-23325086 | 0.542742307 | 0.176870803 | 0.441425188 |
| chr15:71926651-71926878 | 0.695342221 | 0.17687993 | 0.441429661 |
| chr18:59358598-59359838 | -0.43439323 | 0.176884118 | 0.441429661 |
| chrX:30935991-30936232 | -0.708559911 | 0.176903507 | 0.441463669 |
| chr5:95641794-95642102 | 0.5227605 | 0.176950776 | 0.441539939 |
| chr2:79131718-79131918 | 0.715580478 | 0.176952054 | 0.441539939 |
| chr2:28594296-28594813 | 0.485551623 | 0.176956065 | 0.441539939 |
| chr15:101250912-101251245 | 0.51194499 | 0.17695712 | 0.441539939 |
| chr2:154712479-154712989 | -0.574147739 | 0.176996271 | 0.441623247 |
| chr3:16178309-16178676 | 0.491749835 | 0.177037573 | 0.441706913 |
| chr12:76688717-76689019 | -0.611633763 | 0.177050415 | 0.441706913 |
| chr4:74531913-74532223 | -0.561524157 | 0.177051181 | 0.441706913 |
| chr9:76460387-76460642 | -0.559741828 | 0.177052862 | 0.441706913 |
| chr12:93025513-93025999 | 0.518333685 | 0.177058966 | 0.441707758 |
| chr1:230867994-230869316 | 0.440642884 | 0.177070478 | 0.441722096 |
| chr3:58111877-58112289 | 0.50940253 | 0.177092646 | 0.441763012 |
| chr2:139946823-139947026 | -0.716064629 | 0.177134039 | 0.441851884 |
| chr4:56410020-56410868 | -0.476928465 | 0.17714042 | 0.441853416 |
| chr6:43369229-43369955 | 0.445467501 | 0.177167271 | 0.441906007 |
| chr11:3840296-3840568 | 0.567399644 | 0.177225353 | 0.44203649 |
| chr10:71014274-71014474 | 0.653872807 | 0.177237877 | 0.442053338 |
| chr14:48553392-48553592 | 0.71108957 | 0.177257596 | 0.442088129 |
| chr5:108727458-108728769 | -0.434402001 | 0.17727219 | 0.442098481 |
| chr5:55307325-55308254 | -0.445380352 | 0.177273286 | 0.442098481 |
| chr19:13764462-13764670 | 0.67395222 | 0.177281962 | 0.442105727 |
| chr13:114043483-114043817 | -0.523563613 | 0.177293765 | 0.442120773 |
| chr3:61858103-61858368 | 0.501960907 | 0.177312797 | 0.442153843 |
| chr2:75821565-75821866 | 0.655101837 | 0.177320043 | 0.442157523 |
| chr14:38187551-38187980 | -0.519133707 | 0.177338208 | 0.442188241 |
| chrX:102155061-102156249 | -0.438179389 | 0.177343905 | 0.442188241 |
| chr14:68979036-68979714 | 0.438394483 | 0.177357514 | 0.442207784 |
| chr6:164881801-164882113 | -0.66601548 | 0.177377487 | 0.442243194 |
| chr6:134178265-134178800 | -0.464057928 | 0.177388445 | 0.44225349 |
| chr6:43129890-43130177 | 0.551236794 | 0.177393161 | 0.44225349 |
| chr5:178642366-178642839 | -0.543824425 | 0.177408378 | 0.442277037 |
| chr5:34898303-34898618 | 0.528843891 | 0.177456083 | 0.442358355 |
| chr16:30957067-30957366 | 0.49526276 | 0.177456221 | 0.442358355 |
| chr9:69877884-69878201 | -0.652533436 | 0.177458316 | 0.442358355 |
| chr4:9123591-9123933 | 0.526111237 | 0.177491807 | 0.442417987 |
| chr16:72502158-72502414 | 0.627004566 | 0.177493786 | 0.442417987 |
| chr3:183179378-183179799 | 0.52832598 | 0.177589247 | 0.442641529 |
| chr8:130621060-130621260 | 0.711908775 | 0.177600478 | 0.442655124 |
| chr3:194104447-194104710 | -0.629130112 | 0.177626145 | 0.442701311 |
| chr13:26253702-26254964 | 0.433249767 | 0.177630565 | 0.442701311 |
| chr2:138463570-138463822 | -0.771748416 | 0.17765159 | 0.442739312 |
| chr2:10460379-10460709 | -0.520585594 | 0.177666477 | 0.442743139 |
| chr1:170402621-170403051 | -0.640682507 | 0.177667223 | 0.442743139 |
| chr13:105816996-105817883 | -0.462760274 | 0.177672306 | 0.442743139 |

| | | | |
|---|---|---|---|
| chr17:61615364-61616299 | 0.426404734 | 0.177676239 | 0.442743139 |
| chr10:69921432-69921649 | 0.664276064 | 0.177691154 | 0.442765906 |
| chr8:141318138-141318374 | -0.599582741 | 0.177703057 | 0.442781167 |
| chr6:97440616-97440889 | -0.586869865 | 0.177714496 | 0.44279527 |
| chr1:73802741-73803241 | -0.627068063 | 0.177729334 | 0.442817839 |
| chr4:176054993-176055305 | -0.57378759 | 0.177738148 | 0.4428254 |
| chr10:69728368-69728570 | -0.6423487 | 0.177762539 | 0.442867753 |
| chr15:92205497-92205697 | 0.678606705 | 0.177766707 | 0.442867753 |
| chr4:52665680-52666040 | 0.523033042 | 0.1777828 | 0.442874946 |
| chr4:105193021-105193221 | 0.74813542 | 0.177783013 | 0.442874946 |
| chr10:124917551-124917889 | 0.52733174 | 0.177790556 | 0.442874946 |
| chr13:47463895-47464133 | 0.606823065 | 0.177792714 | 0.442874946 |
| chr4:25862106-25862480 | 0.589357101 | 0.177809434 | 0.442899679 |
| chr17:64718098-64718355 | 0.548120001 | 0.177814204 | 0.442899679 |
| chr12:56078960-56080389 | 0.429618707 | 0.177820788 | 0.442901681 |
| chr9:38227018-38227633 | -0.445227702 | 0.177832591 | 0.442901709 |
| chr20:32778447-32778823 | -0.514988916 | 0.177834456 | 0.442901709 |
| chr3:163780045-163780312 | -0.676326977 | 0.17783814 | 0.442901709 |
| chr5:108521883-108522428 | -0.499852335 | 0.177847616 | 0.442910912 |
| chr4:156677286-156677592 | -0.537720332 | 0.17790361 | 0.443025581 |
| chr5:78984455-78985547 | -0.442945401 | 0.177908406 | 0.443025581 |
| chr15:40754457-40754758 | 0.545232163 | 0.177911006 | 0.443025581 |
| chr10:33315906-33316106 | 0.788396681 | 0.177928967 | 0.443045633 |
| chr4:140012677-140013027 | 0.56460106 | 0.177930623 | 0.443045633 |
| chr14:58171146-58171482 | -0.481228487 | 0.177948424 | 0.443075557 |
| chr6:138107032-138107995 | 0.454472004 | 0.177957872 | 0.443084684 |
| chr8:93361150-93361429 | 0.548280276 | 0.177976048 | 0.443105238 |
| chr17:48908648-48908901 | -0.553838338 | 0.177977693 | 0.443105238 |
| chr13:39365101-39365532 | -0.483848906 | 0.178031866 | 0.443200018 |
| chr3:15186592-15187324 | -0.49823401 | 0.178032611 | 0.443200018 |
| chr16:2015276-2015815 | 0.500791558 | 0.178033115 | 0.443200018 |
| chr18:68790565-68790871 | -0.657730311 | 0.178053196 | 0.443235607 |
| chrX:56107224-56107937 | 0.474953816 | 0.178082431 | 0.443293981 |
| chr4:139575741-139576009 | 0.585059587 | 0.178088399 | 0.443294435 |
| chr11:27201946-27202308 | 0.527671103 | 0.17809983 | 0.443308487 |
| chr18:32261096-32261313 | -0.675472099 | 0.178107194 | 0.443312417 |
| chr8:115693614-115693860 | 0.607309124 | 0.178159288 | 0.443427674 |
| chr9:33466642-33466924 | -0.591066953 | 0.178174596 | 0.443439516 |
| chrX:39384255-39384555 | -0.648483839 | 0.17817562 | 0.443439516 |
| chr5:151406347-151406614 | -0.601246687 | 0.178183495 | 0.44344471 |
| chr9:131580374-131580617 | -0.672897597 | 0.1781949 | 0.44345869 |
| chr13:32683313-32683661 | -0.666396073 | 0.178227415 | 0.443525202 |
| chr7:74289209-74289527 | -0.542144567 | 0.178237081 | 0.443533694 |
| chr2:118882567-118883173 | -0.477429459 | 0.178242404 | 0.443533694 |
| chr6:144063621-144064791 | -0.440308518 | 0.178280411 | 0.443613861 |
| chr16:66357541-66357741 | -0.745144945 | 0.178290028 | 0.443623384 |
| chr19:45974016-45974270 | 0.60000606 | 0.178299322 | 0.443632105 |
| chr1:43558021-43558221 | -0.703445488 | 0.178327321 | 0.443687361 |
| chr3:64796845-64797441 | 0.438570169 | 0.17833314 | 0.443687433 |
| chr6:37222105-37222956 | 0.439281431 | 0.178362818 | 0.443746862 |

| | | | |
|---|---|---|---|
| chr3:187702543-187702804 | 0.688941597 | 0.178383897 | 0.443784895 |
| chr5:104645942-104646149 | -0.710030957 | 0.178404334 | 0.443821328 |
| chr6:5422418-5422637 | 0.609285493 | 0.178439685 | 0.443894859 |
| chr2:95031747-95032080 | 0.517851629 | 0.178452122 | 0.443905395 |
| chr8:94234254-94234780 | 0.474490348 | 0.178455508 | 0.443905395 |
| chr13:105139323-105140493 | -0.454509902 | 0.178485817 | 0.443953338 |
| chr2:3728149-3728427 | -0.600718794 | 0.178486369 | 0.443953338 |
| chr2:30621528-30621830 | -0.688579837 | 0.178512493 | 0.444001102 |
| chr2:111698902-111699166 | 0.556630915 | 0.178517161 | 0.444001102 |
| chr7:7829748-7830020 | -0.562572151 | 0.178523376 | 0.444002146 |
| chr16:88581479-88581679 | -0.674797076 | 0.178546461 | 0.444045147 |
| chrX:11144019-11144602 | -0.504441345 | 0.17856014 | 0.444058311 |
| chr1:89843194-89844143 | -0.462158783 | 0.178563345 | 0.444058311 |
| chr6:18269279-18269507 | 0.622550565 | 0.178585412 | 0.444098775 |
| chr8:144792855-144793101 | 0.52206039 | 0.178591821 | 0.444100299 |
| chr1:223992988-223993254 | -0.609206317 | 0.178601498 | 0.444109948 |
| chr3:110888521-110889502 | -0.44770399 | 0.178634512 | 0.444165319 |
| chr2:98058328-98058862 | -0.546361867 | 0.178635359 | 0.444165319 |
| chr1:38587020-38587295 | 0.584572391 | 0.178656048 | 0.444183946 |
| chr7:149595115-149595482 | -0.552896871 | 0.178663261 | 0.444183946 |
| chr20:45470048-45470644 | 0.48469644 | 0.178678112 | 0.444183946 |
| chr13:35772011-35772246 | 0.670751074 | 0.178683227 | 0.444183946 |
| chr11:78619698-78619935 | -0.668322364 | 0.178683657 | 0.444183946 |
| chr2:2935408-2935679 | 0.654230794 | 0.178689673 | 0.444183946 |
| chr1:94481476-94481706 | -0.701656007 | 0.178692899 | 0.444183946 |
| chr11:44899408-44899640 | -0.699382604 | 0.178700505 | 0.444183946 |
| chr12:113098215-113098851 | 0.455213722 | 0.178702505 | 0.444183946 |
| chr20:52972271-52972485 | -0.654464195 | 0.178706284 | 0.444183946 |
| chr5:32887318-32887827 | 0.51563401 | 0.178706618 | 0.444183946 |
| chr5:51860525-51861010 | 0.561989899 | 0.178753289 | 0.444285536 |
| chr14:90697650-90697915 | 0.557232347 | 0.178765414 | 0.444300956 |
| chr14:69139010-69139210 | 0.701782872 | 0.178775323 | 0.444300956 |
| chr11:8870414-8871531 | -0.466288946 | 0.178776889 | 0.444300956 |
| chr11:7548883-7549215 | -0.537070582 | 0.178798125 | 0.444339321 |
| chr2:154452952-154453299 | -0.597847789 | 0.178813321 | 0.444362674 |
| chr15:40874102-40874407 | 0.499035014 | 0.178822515 | 0.444371107 |
| chr5:117417518-117417901 | -0.654506521 | 0.178838306 | 0.444390629 |
| chr3:10263832-10264346 | 0.503231257 | 0.17884197 | 0.444390629 |
| chr4:111937430-111937749 | -0.558292232 | 0.17884955 | 0.444395052 |
| chr3:182165277-182165637 | 0.499386413 | 0.178861908 | 0.444411348 |
| chr5:18532352-18532703 | -0.646709314 | 0.178883785 | 0.444434526 |
| chr20:22836363-22836563 | -0.775227843 | 0.178888337 | 0.444434526 |
| chr7:76297366-76297736 | -0.507156386 | 0.178890849 | 0.444434526 |
| chr18:36634605-36634854 | -0.70599281 | 0.178898472 | 0.444434526 |
| chr14:89701158-89702196 | -0.441529848 | 0.178900509 | 0.444434526 |
| chr12:49894530-49894772 | 0.577954014 | 0.178906039 | 0.444434526 |
| chr17:21303018-21303335 | 0.524669763 | 0.178939388 | 0.444460545 |
| chr18:45642776-45642976 | -0.759172672 | 0.178940032 | 0.444460545 |
| chr18:78407127-78407327 | -0.759172672 | 0.178940032 | 0.444460545 |
| chr4:57303006-57303722 | -0.484285358 | 0.17894374 | 0.444460545 |

| | | | |
|---|---|---|---|
| chr10:91776397-91776631 | -0.620031576 | 0.178945516 | 0.444460545 |
| chr12:12092342-12092836 | -0.530005428 | 0.178976514 | 0.444511944 |
| chr14:38589646-38589862 | 0.64452279 | 0.178977813 | 0.444511944 |
| chr3:150786721-150786943 | 0.608342172 | 0.179004656 | 0.444554013 |
| chr4:70992482-70992700 | -0.585115701 | 0.17900926 | 0.444554013 |
| chr19:47391437-47391778 | 0.547557322 | 0.179012157 | 0.444554013 |
| chr19:1275675-1276109 | 0.469210052 | 0.17903109 | 0.444561239 |
| chr12:16359982-16360182 | 0.694349157 | 0.179036406 | 0.444561239 |
| chr12:122203266-122203872 | 0.457943143 | 0.179036898 | 0.444561239 |
| chr5:64784510-64784710 | 0.712015044 | 0.179038275 | 0.444561239 |
| chr2:88060396-88060720 | -0.54307311 | 0.179060806 | 0.444602777 |
| chr18:21611950-21612661 | 0.456042583 | 0.179094615 | 0.444672313 |
| chr17:43897894-43898302 | -0.507304934 | 0.179102704 | 0.444677988 |
| chr2:15492564-15492860 | -0.674094129 | 0.179114372 | 0.44468961 |
| chr12:56995-57340 | 0.527038685 | 0.179118992 | 0.44468961 |
| chr4:35908629-35909173 | -0.477391957 | 0.179126767 | 0.444694503 |
| chr4:80709465-80709754 | 0.641898921 | 0.179135991 | 0.444702993 |
| chr19:3093969-3094456 | 0.509316095 | 0.179158657 | 0.444744852 |
| chr11:64803191-64803483 | 0.515529395 | 0.179183742 | 0.444792713 |
| chr20:23920073-23920444 | 0.491363362 | 0.179202233 | 0.444803438 |
| chr19:7681328-7681601 | -0.582487884 | 0.17920507 | 0.444803438 |
| chr15:77993975-77994329 | 0.518195888 | 0.179210884 | 0.444803438 |
| chr2:106562843-106563219 | -0.541541842 | 0.179214689 | 0.444803438 |
| chr12:8827995-8828419 | -0.565993318 | 0.179217088 | 0.444803438 |
| chr8:58689080-58689566 | -0.534112612 | 0.179224128 | 0.444806503 |
| chr7:145005490-145006155 | -0.495003327 | 0.179254422 | 0.444867279 |
| chr6:44264893-44265821 | 0.442973988 | 0.179279189 | 0.444914333 |
| chr2:186780761-186781338 | -0.485682813 | 0.179286053 | 0.444914872 |
| chr1:177913141-177914033 | -0.457270732 | 0.17929102 | 0.444914872 |
| chr8:107599972-107600240 | 0.530557279 | 0.179299285 | 0.444920974 |
| chr14:95291154-95291481 | -0.553719483 | 0.1793083 | 0.444928934 |
| chr11:24833366-24833828 | -0.552439665 | 0.179321506 | 0.444947294 |
| chr7:146187386-146187586 | 0.804918835 | 0.179341971 | 0.444970062 |
| chr10:118543876-118544449 | 0.484257966 | 0.179342297 | 0.444970062 |
| chr3:134326845-134327304 | 0.493874673 | 0.179389914 | 0.44506348 |
| chr3:71859234-71859471 | 0.633460282 | 0.179393153 | 0.44506348 |
| chr6:167502087-167502307 | 0.641552444 | 0.179397374 | 0.44506348 |
| chr6:158474931-158475215 | 0.607912543 | 0.179425609 | 0.445119117 |
| chr11:46889557-46889921 | 0.505106882 | 0.179445276 | 0.445151301 |
| chr3:108863857-108864210 | -0.682451096 | 0.179456628 | 0.445151301 |
| chr12:1518719-1519010 | -0.614185596 | 0.179459977 | 0.445151301 |
| chr17:15171485-15171773 | -0.536184917 | 0.179467527 | 0.445151301 |
| chrX:135021634-135022808 | 0.433454317 | 0.179467631 | 0.445151301 |
| chr1:43453736-43454155 | 0.491350061 | 0.179484417 | 0.445164945 |
| chr6:90296707-90297076 | 0.506524773 | 0.179484752 | 0.445164945 |
| chr6:138086573-138086845 | 0.559999126 | 0.17950195 | 0.44519319 |
| chr1:114636320-114636630 | 0.494344143 | 0.179527324 | 0.44524171 |
| chr6:160668008-160668208 | 0.657482175 | 0.179541087 | 0.445253025 |
| chr6:7238845-7239209 | 0.49106278 | 0.179543509 | 0.445253025 |
| chr18:48547957-48548503 | -0.466838649 | 0.179579744 | 0.445328473 |

| | | | |
|---|---|---|---|
| chr2:28736142-28736391 | -0.649714979 | 0.17959432 | 0.445350206 |
| chr18:37253951-37254859 | -0.461534214 | 0.17961235 | 0.445380501 |
| chr5:172688755-172689224 | 0.495591894 | 0.179621393 | 0.445384567 |
| chr12:57941075-57942243 | -0.427895701 | 0.179625615 | 0.445384567 |
| chr1:206658421-206658811 | -0.477867194 | 0.179637831 | 0.445400444 |
| chr5:127326413-127326722 | -0.617647386 | 0.179654351 | 0.445426989 |
| chr8:115333760-115334450 | -0.467446338 | 0.179699854 | 0.44552539 |
| chr9:21789497-21789986 | -0.523976291 | 0.17974462 | 0.445621959 |
| chr13:21326146-21326561 | 0.534491922 | 0.179755715 | 0.445625982 |
| chr16:76650605-76650954 | 0.507575641 | 0.179762414 | 0.445625982 |
| chr11:16918378-16918905 | -0.497568896 | 0.17976369 | 0.445625982 |
| chr10:8725912-8726506 | -0.502730979 | 0.179790185 | 0.445677242 |
| chr3:58332543-58333601 | 0.461120203 | 0.179796509 | 0.445678499 |
| chr11:13238842-13239183 | -0.601866222 | 0.179813751 | 0.445706821 |
| chr22:31343362-31343587 | 0.514287888 | 0.179827306 | 0.445726001 |
| chr2:164914129-164914398 | -0.670717213 | 0.179843059 | 0.445727908 |
| chr13:50185724-50186490 | -0.48632728 | 0.179843941 | 0.445727908 |
| chr5:132445696-132445908 | -0.662221851 | 0.17984828 | 0.445727908 |
| chr2:154402179-154402474 | -0.541591811 | 0.179851345 | 0.445727908 |
| chr1:1307864-1308866 | 0.449295063 | 0.179899043 | 0.445828379 |
| chr17:64157640-64158016 | -0.516044965 | 0.179903522 | 0.445828379 |
| chr7:151249111-151249458 | -0.519127348 | 0.179909406 | 0.445828542 |
| chr10:133261864-133262260 | 0.478995494 | 0.1799238 | 0.445849792 |
| chr6:8462174-8462566 | -0.577208032 | 0.179964531 | 0.445922811 |
| chr4:135863537-135864777 | -0.428777622 | 0.179964907 | 0.445922811 |
| chr12:114625251-114625496 | -0.625632443 | 0.17998085 | 0.445944834 |
| chr7:156156832-156157159 | -0.535760007 | 0.179985435 | 0.445944834 |
| chr11:65507169-65508535 | -0.430542915 | 0.180008443 | 0.445987418 |
| chr19:48469565-48469862 | 0.497914941 | 0.180065902 | 0.446115354 |
| chr14:73058131-73059076 | 0.430858537 | 0.180085264 | 0.446148898 |
| chr2:185979706-185980179 | -0.541367908 | 0.180094081 | 0.446156315 |
| chr10:23345920-23346177 | -0.654172297 | 0.18010031 | 0.446157322 |
| chr17:26603599-26604452 | 0.426473044 | 0.180116983 | 0.446184199 |
| chr11:46808816-46809062 | 0.598681413 | 0.180130056 | 0.446202159 |
| chr4:168931172-168931472 | -0.631164825 | 0.180167341 | 0.446280089 |
| chr2:9797682-9798177 | -0.474237221 | 0.180183418 | 0.446288881 |
| chr6:6556899-6557376 | -0.552192359 | 0.18018376 | 0.446288881 |
| chr16:88236338-88236545 | 0.613324923 | 0.180188364 | 0.446288881 |
| chr2:1140305-1140724 | -0.540407954 | 0.180194797 | 0.446290388 |
| chr7:55571574-55572213 | 0.483998022 | 0.180222709 | 0.446345092 |
| chr18:36296536-36296893 | -0.5890873 | 0.180231106 | 0.44635146 |
| chr1:180554618-180554889 | 0.660909786 | 0.180286071 | 0.446473152 |
| chr16:58638297-58638546 | -0.638776751 | 0.180298831 | 0.44649032 |
| chr6:163392621-163392865 | -0.706697559 | 0.180328778 | 0.446550049 |
| chr1:209569538-209569765 | -0.603850951 | 0.180362864 | 0.44660559 |
| chr10:21402714-21402960 | -0.603850951 | 0.180362864 | 0.44660559 |
| chr6:13678540-13678863 | 0.582553922 | 0.180370728 | 0.44661063 |
| chr20:12358062-12358408 | -0.598033937 | 0.180402518 | 0.446665655 |
| chr6:116668226-116669020 | -0.439935698 | 0.180406913 | 0.446665655 |
| chr11:60559650-60559850 | 0.69982374 | 0.180411209 | 0.446665655 |

| | | | |
|---|---|---|---|
| chr1:2491969-2492351 | -0.60528889 | 0.180416269 | 0.446665655 |
| chr17:19111018-19111397 | 0.529349271 | 0.180433285 | 0.446677867 |
| chr1:217869218-217869481 | -0.596623689 | 0.180435258 | 0.446677867 |
| chr8:29737241-29737812 | -0.473850666 | 0.180438691 | 0.446677867 |
| chr10:71339890-71340422 | -0.489176525 | 0.180446529 | 0.44668284 |
| chr13:66666065-66666265 | -0.724193117 | 0.180490993 | 0.446766391 |
| chr6:127266412-127267843 | -0.426194047 | 0.180496141 | 0.446766391 |
| chr17:60490134-60490375 | 0.480327526 | 0.180501981 | 0.446766391 |
| chr14:68628228-68629326 | -0.441320281 | 0.180503604 | 0.446766391 |
| chr3:38459286-38459666 | 0.520323819 | 0.1805533 | 0.44686657 |
| chr6:94636855-94637388 | -0.493728877 | 0.180560459 | 0.44686657 |
| chr11:78417732-78418110 | -0.483268171 | 0.18056242 | 0.44686657 |
| chr2:20396019-20396535 | -0.471201005 | 0.180567407 | 0.44686657 |
| chr10:12993699-12993966 | -0.587601576 | 0.180573621 | 0.446867514 |
| chr20:49901407-49901607 | 0.623182499 | 0.180583593 | 0.446877758 |
| chr1:153222927-153223446 | 0.49898332 | 0.180615623 | 0.446935698 |
| chr15:32730722-32731465 | 0.476387185 | 0.180618672 | 0.446935698 |
| chr17:29838930-29839260 | -0.65459116 | 0.180665626 | 0.447037448 |
| chr7:30551051-30551437 | 0.487079148 | 0.180690284 | 0.447080298 |
| chr16:71214979-71215339 | -0.57701773 | 0.180694614 | 0.447080298 |
| chr7:2558749-2559176 | -0.461735162 | 0.180710776 | 0.44710585 |
| chr14:73490593-73492037 | 0.430340363 | 0.180720728 | 0.44711482 |
| chr6:35895662-35895913 | 0.614543965 | 0.180726072 | 0.44711482 |
| chr10:119442405-119442834 | -0.538663858 | 0.180749946 | 0.447135712 |
| chr8:130692207-130692640 | -0.507108904 | 0.180752338 | 0.447135712 |
| chr10:1110122-1110403 | 0.517850594 | 0.180755356 | 0.447135712 |
| chr5:129094102-129095449 | -0.420757485 | 0.180763446 | 0.447135712 |
| chr2:96843213-96843449 | 0.61251806 | 0.180763695 | 0.447135712 |
| chr11:128886480-128886704 | -0.633126075 | 0.180779578 | 0.447160565 |
| chr14:35341488-35342082 | 0.444159539 | 0.18079274 | 0.447178686 |
| chr1:147612313-147613030 | -0.436685486 | 0.180804745 | 0.447193944 |
| chr10:35568873-35569088 | -0.704747164 | 0.180822059 | 0.44722233 |
| chr6:12205573-12205842 | -0.751264869 | 0.180839938 | 0.447252113 |
| chr5:149645274-149645548 | 0.637122535 | 0.180851296 | 0.447265766 |
| chr6:27592467-27593059 | -0.44398869 | 0.180881411 | 0.447325806 |
| chr9:134175280-134175665 | 0.510153928 | 0.180894746 | 0.447342559 |
| chrX:102863818-102864082 | 0.55033158 | 0.180899862 | 0.447342559 |
| chr19:54231522-54232073 | 0.493723354 | 0.18090919 | 0.44735119 |
| chr8:134692642-134692845 | -0.610936531 | 0.180931565 | 0.447392079 |
| chr2:33598558-33598787 | -0.508238644 | 0.180964051 | 0.447457969 |
| chr21:45950719-45950950 | 0.59908481 | 0.180980961 | 0.44746839 |
| chr7:42618718-42619195 | -0.534900802 | 0.180982876 | 0.44746839 |
| chr3:37990645-37990845 | -0.639668997 | 0.180985786 | 0.44746839 |
| chr11:44786230-44786643 | -0.659365743 | 0.18099951 | 0.447487882 |
| chr1:38046520-38047056 | -0.505039696 | 0.181046169 | 0.447588796 |
| chr5:106843828-106844149 | -0.552729419 | 0.181065415 | 0.447615381 |
| chr20:52128536-52129048 | 0.489467586 | 0.181068606 | 0.447615381 |
| chr9:119368278-119368587 | -0.630778333 | 0.181084568 | 0.447640397 |
| chr2:143026437-143026762 | 0.675162138 | 0.181124621 | 0.447718033 |
| chr6:108115817-108116179 | 0.548704357 | 0.18112766 | 0.447718033 |

| | | | |
|---|---|---|---|
| chr1:95233328-95234805 | -0.433344369 | 0.18115916 | 0.447781449 |
| chr20:47438719-47439442 | 0.42200112 | 0.181179713 | 0.447805682 |
| chr14:23988799-23988999 | 0.676379606 | 0.181180652 | 0.447805682 |
| chr4:40283482-40283737 | -0.667057509 | 0.181204492 | 0.447850157 |
| chr2:143259483-143259754 | -0.629344964 | 0.181237664 | 0.447917694 |
| chr7:140005914-140006122 | -0.624095888 | 0.181249927 | 0.447933554 |
| chr22:21867536-21868234 | 0.465474457 | 0.181280819 | 0.447995449 |
| chr1:179838424-179838727 | -0.560391028 | 0.181340548 | 0.448128604 |
| chr19:4042689-4043114 | 0.532580691 | 0.181360583 | 0.448163659 |
| chr14:100221567-10022191C | -0.541188632 | 0.181370421 | 0.448173516 |
| chr4:57119180-57119628 | -0.543740791 | 0.181384336 | 0.448193446 |
| chr3:71414283-71414498 | -0.68410208 | 0.18141392 | 0.448252093 |
| chr7:53315855-53316341 | -0.478516409 | 0.181437887 | 0.448296856 |
| chr1:167401396-167401626 | 0.614430655 | 0.181453674 | 0.448321404 |
| chr14:94423464-94423756 | 0.552689676 | 0.181473967 | 0.448341617 |
| chr7:88140933-88141305 | 0.574672474 | 0.181476525 | 0.448341617 |
| chrX:138710487-138710807 | 0.571763154 | 0.181479409 | 0.448341617 |
| chr1:226308434-226309617 | 0.436662415 | 0.181526181 | 0.448442707 |
| chr7:103753085-103753302 | -0.72770817 | 0.181556072 | 0.448502091 |
| chr11:74921777-74921982 | -0.69768751 | 0.181578115 | 0.448523958 |
| chr20:49812298-49813379 | 0.432537828 | 0.1815801 | 0.448523958 |
| chr6:159978788-159979026 | -0.690754965 | 0.181582485 | 0.448523958 |
| chr2:227325029-227325777 | -0.444099583 | 0.18160317 | 0.448544599 |
| chr11:17021534-17022052 | 0.481798674 | 0.181603692 | 0.448544599 |
| chr15:30991122-30992319 | 0.441263736 | 0.181611591 | 0.448544599 |
| chr11:18412113-18412325 | -0.717055749 | 0.181614257 | 0.448544599 |
| chr22:29907751-29908146 | 0.515582045 | 0.181626235 | 0.448559722 |
| chr10:75245387-75246123 | -0.452581763 | 0.181634599 | 0.448564884 |
| chr1:204924223-204924925 | 0.449601473 | 0.181640033 | 0.448564884 |
| chr1:107327079-107327323 | 0.494472698 | 0.181647601 | 0.448569114 |
| chr2:112483141-112483368 | 0.59090924 | 0.181662298 | 0.448589319 |
| chr16:4425315-4426058 | 0.438283503 | 0.181667492 | 0.448589319 |
| chr6:143510946-143512047 | -0.438851415 | 0.181682083 | 0.448597314 |
| chrX:10156642-10157022 | 0.527384221 | 0.181682439 | 0.448597314 |
| chr5:180605761-180606244 | 0.49461362 | 0.181705921 | 0.448640837 |
| chr6:112296163-112296460 | -0.552176955 | 0.18171834 | 0.448656455 |
| chr5:168309123-168309564 | -0.525391676 | 0.181723958 | 0.448656455 |
| chr2:21042251-21042497 | -0.640980191 | 0.18175332 | 0.448714489 |
| chr22:18906904-18907214 | 0.534161305 | 0.181773227 | 0.448749176 |
| chr6:105606944-105607144 | 0.668782801 | 0.181785525 | 0.448765078 |
| chr7:16658029-16658250 | 0.586944575 | 0.181801571 | 0.448767246 |
| chr13:109786434-109787184 | 0.445441394 | 0.18180861 | 0.448767246 |
| chr3:150787830-150788153 | -0.521844106 | 0.181810596 | 0.448767246 |
| chr16:29691976-29692185 | 0.609200726 | 0.181811751 | 0.448767246 |
| chr6:11207085-11207611 | 0.496138671 | 0.181815688 | 0.448767246 |
| chr19:1507733-1508486 | 0.449274303 | 0.181880484 | 0.448907965 |
| chr18:8811216-8811496 | -0.649980026 | 0.181884416 | 0.448907965 |
| chr17:73196595-73196858 | -0.513408797 | 0.181920514 | 0.448982595 |
| chr11:88484836-88485105 | -0.576864233 | 0.181930976 | 0.448993952 |
| chr15:84929792-84930378 | -0.447386041 | 0.181942867 | 0.449008836 |

| | | | |
|---|---|---|---|
| chr16:31458979-31460246 | 0.422964872 | 0.181967259 | 0.44905457 |
| chr11:65366329-65366657 | 0.553894727 | 0.182006023 | 0.449135766 |
| chr6:130518235-130518456 | 0.718325665 | 0.182017531 | 0.449149697 |
| chr11:16820352-16820856 | 0.518698783 | 0.182025842 | 0.44915574 |
| chr4:154503698-154503972 | -0.622659204 | 0.182035583 | 0.449165312 |
| chr19:14517808-14519001 | 0.434996686 | 0.182041585 | 0.449165657 |
| chr20:31634232-31634505 | 0.556733656 | 0.182054496 | 0.449177721 |
| chr17:60558593-60559601 | 0.424153061 | 0.182063404 | 0.449177721 |
| chr16:89216558-89216759 | -0.624855482 | 0.182064061 | 0.449177721 |
| chr15:96428738-96429084 | 0.569556103 | 0.182097965 | 0.44922624 |
| chr7:32347337-32347593 | -0.554575795 | 0.182098751 | 0.44922624 |
| chr14:25371036-25371425 | 0.489050027 | 0.182101882 | 0.44922624 |
| chr6:75486932-75487428 | -0.551443555 | 0.18211125 | 0.44922624 |
| chr14:53285819-53286074 | -0.630046284 | 0.182113935 | 0.44922624 |
| chr11:58927443-58927753 | 0.52735965 | 0.182118904 | 0.44922624 |
| chr1:211135403-211135635 | -0.696216714 | 0.18212509 | 0.449227036 |
| chr4:82275908-82276247 | 0.562354401 | 0.182132877 | 0.449231781 |
| chr8:128450542-128450742 | 0.659128009 | 0.1821434 | 0.449232309 |
| chr19:7920529-7921275 | 0.447694887 | 0.182144817 | 0.449232309 |
| chr1:50605514-50605805 | -0.672002486 | 0.1821749 | 0.449279312 |
| chr6:99822781-99823313 | -0.489672398 | 0.182175602 | 0.449279312 |
| chr7:1247007-1247424 | -0.518714844 | 0.182183582 | 0.449284532 |
| chr11:95150890-95151965 | -0.43537724 | 0.18219297 | 0.449293223 |
| chr1:41574110-41574393 | 0.565809905 | 0.182215333 | 0.44933103 |
| chr17:28494833-28495051 | -0.59045342 | 0.182220029 | 0.44933103 |
| chr17:18946749-18946996 | 0.660540226 | 0.182231432 | 0.449334387 |
| chr16:3642647-3643191 | 0.471743206 | 0.182233119 | 0.449334387 |
| chr5:79706144-79706792 | -0.492097019 | 0.182243514 | 0.449337869 |
| chr14:36662554-36663021 | 0.457141197 | 0.18224626 | 0.449337869 |
| chr11:83025301-83025583 | -0.552049154 | 0.182284981 | 0.449418876 |
| chr6:30730797-30731189 | 0.491922497 | 0.182312932 | 0.449450123 |
| chr20:62335910-62336160 | 0.569315288 | 0.182319076 | 0.449450123 |
| chr11:65945626-65945885 | 0.611539422 | 0.18231945 | 0.449450123 |
| chr17:7014511-7015164 | 0.451160851 | 0.182321118 | 0.449450123 |
| chr6:4957555-4957917 | -0.577445965 | 0.182332123 | 0.449462791 |
| chr20:53660748-53661179 | 0.431450706 | 0.182344066 | 0.44947777 |
| chr2:227848684-227848884 | -0.734319282 | 0.182363714 | 0.449511741 |
| chr16:425192-425757 | 0.476765671 | 0.18237722 | 0.449523834 |
| chr17:17180107-17180454 | 0.543144803 | 0.182380353 | 0.449523834 |
| chr11:85809856-85810293 | -0.477772771 | 0.182397279 | 0.449539347 |
| chr4:38867463-38868119 | 0.465320337 | 0.182398381 | 0.449539347 |
| chr22:36028004-36029400 | 0.430668484 | 0.182427524 | 0.449584626 |
| chr21:29218874-29219380 | 0.513432367 | 0.182429444 | 0.449584626 |
| chr16:4846507-4847878 | 0.429069356 | 0.182434355 | 0.449584626 |
| chrX:5480136-5480656 | -0.557756135 | 0.182445396 | 0.44958762 |
| chrX:133921895-133922179 | -0.580489135 | 0.182447306 | 0.44958762 |
| chr4:137735702-137735953 | -0.695446662 | 0.18246049 | 0.449605649 |
| chr1:212595888-212596663 | -0.464725773 | 0.182475789 | 0.449628888 |
| chr20:9087694-9088201 | 0.53685821 | 0.182486695 | 0.449641301 |
| chr17:71230741-71231057 | 0.609166301 | 0.182495439 | 0.449648387 |

| | | | |
|---|---|---|---|
| chr19:44003759-44004043 | 0.654482474 | 0.182522891 | 0.449701565 |
| chr14:86091945-86092271 | -0.606946215 | 0.182538552 | 0.449725689 |
| chr10:80110178-80110534 | -0.569777532 | 0.182552347 | 0.449745214 |
| chr5:97044127-97044405 | -0.544974321 | 0.182574802 | 0.449786074 |
| chr10:80158627-80158928 | -0.549208633 | 0.182599867 | 0.449796862 |
| chr2:954424-954686 | -0.620222173 | 0.182601277 | 0.449796862 |
| chr1:36420848-36421150 | 0.546233703 | 0.182605923 | 0.449796862 |
| chr22:19554792-19555095 | -0.563697096 | 0.182611324 | 0.449796862 |
| chr2:121074768-121075564 | -0.438560326 | 0.1826138 | 0.449796862 |
| chr9:76460007-76460270 | -0.526191104 | 0.182614403 | 0.449796862 |
| chr3:12782424-12782857 | -0.509024077 | 0.182625013 | 0.449808536 |
| chr12:46372635-46373057 | 0.444239601 | 0.182677584 | 0.449923557 |
| chr3:53269922-53270540 | 0.455726937 | 0.182701172 | 0.449953313 |
| chr20:53723150-53723383 | 0.447269239 | 0.18270141 | 0.449953313 |
| chr2:54330202-54331281 | -0.433228959 | 0.182738075 | 0.450015872 |
| chr14:23295109-23295330 | 0.571439172 | 0.182739149 | 0.450015872 |
| chr6:17815371-17815665 | -0.534843955 | 0.182745806 | 0.450015872 |
| chr2:201116789-201117205 | -0.490453393 | 0.182750305 | 0.450015872 |
| chr19:45117716-45118321 | 0.470303665 | 0.182766601 | 0.450041536 |
| chr5:71601208-71601984 | -0.467975856 | 0.18278044 | 0.450061149 |
| chr16:8675806-8676041 | -0.594985618 | 0.182791194 | 0.450073168 |
| chr10:70532392-70532637 | -0.632633509 | 0.18279968 | 0.450079598 |
| chr3:179768078-179768431 | -0.514820332 | 0.182822703 | 0.450121193 |
| chr1:53326880-53327087 | 0.544342185 | 0.182828323 | 0.450121193 |
| chr2:119838173-119838464 | -0.593582011 | 0.182845526 | 0.450138948 |
| chr3:184727413-184727797 | -0.524093419 | 0.182847284 | 0.450138948 |
| chr12:52022863-52023112 | -0.562334628 | 0.182858609 | 0.450152364 |
| chr3:170651260-170651583 | 0.545574973 | 0.182922853 | 0.450293958 |
| chrX:53281103-53281837 | 0.454521802 | 0.18293352 | 0.450293958 |
| chr12:115885102-11588541[illegible] | 0.571890969 | 0.182936333 | 0.450293958 |
| chr6:129318431-129318720 | -0.658131228 | 0.182939634 | 0.450293958 |
| chr12:69284203-69284451 | 0.581142627 | 0.182948251 | 0.450295497 |
| chr11:28529664-28530078 | 0.588311176 | 0.182952013 | 0.450295497 |
| chr5:111772542-111772757 | -0.729539912 | 0.183007617 | 0.450410096 |
| chr12:112285811-112286022 | 0.695281154 | 0.18301033 | 0.450410096 |
| chr11:12687410-12688126 | -0.482353506 | 0.183020063 | 0.450417336 |
| chr3:49689445-49689881 | 0.516040249 | 0.183034605 | 0.450417336 |
| chr11:118359152-118359917 | 0.454756839 | 0.183035024 | 0.450417336 |
| chr1:244010262-244010550 | -0.606709577 | 0.183036786 | 0.450417336 |
| chr1:186917144-186917444 | -0.55175312 | 0.183047998 | 0.450430461 |
| chr13:85709789-85710129 | -0.619580686 | 0.1830611 | 0.450448236 |
| chr2:16079431-16079994 | -0.490571608 | 0.183089845 | 0.4505045 |
| chrX:36957310-36957700 | -0.523434558 | 0.183097965 | 0.450510012 |
| chr1:106619336-106620389 | 0.423572981 | 0.183116164 | 0.450540322 |
| chr4:78810549-78810814 | -0.667649482 | 0.183125599 | 0.450549068 |
| chrX:123450347-123450547 | 0.7169904 | 0.183132933 | 0.450552645 |
| chr13:27295509-27295744 | -0.662859077 | 0.183147836 | 0.450574843 |
| chr14:50600033-50600236 | 0.645125521 | 0.183162122 | 0.450595521 |
| chr2:136666089-136666564 | -0.59176666 | 0.183168102 | 0.450595766 |
| chrX:103674463-103674838 | -0.643898225 | 0.183178692 | 0.45060735 |

| | | | |
|---|---|---|---|
| chr12:92144146-92144405 | -0.472369918 | 0.183207952 | 0.450664859 |
| chr16:81141184-81141653 | -0.501230246 | 0.183214602 | 0.450666749 |
| chr7:101440477-101440695 | -0.639153177 | 0.18322555 | 0.450672061 |
| chr9:26271609-26271901 | -0.601554416 | 0.183228525 | 0.450672061 |
| chr17:61654711-61654911 | 0.471684133 | 0.183249492 | 0.450698247 |
| chr2:85334037-85334390 | -0.63067262 | 0.183250936 | 0.450698247 |
| chr4:87103908-87104192 | -0.617384082 | 0.183258777 | 0.450703065 |
| chr2:44956112-44956377 | -0.575921056 | 0.183279357 | 0.450739212 |
| chr6:93418149-93418349 | -0.572911666 | 0.183289225 | 0.450749012 |
| chr9:86187156-86187455 | 0.588719341 | 0.183315855 | 0.450786372 |
| chr8:48075140-48075340 | 0.659357803 | 0.183316183 | 0.450786372 |
| chr3:64482111-64482408 | 0.452379102 | 0.18333039 | 0.45080684 |
| chr6:76416847-76417062 | -0.716683319 | 0.183342754 | 0.450822775 |
| chr12:16349223-16349430 | 0.629943462 | 0.183352636 | 0.450832605 |
| chr10:62375057-62375402 | -0.511342102 | 0.183379492 | 0.45088187 |
| chr17:45468477-45468751 | 0.635816205 | 0.18338444 | 0.45088187 |
| chr9:134295341-134295545 | -0.661444589 | 0.18342163 | 0.450958836 |
| chr19:57632785-57633758 | 0.451580578 | 0.183473125 | 0.451056579 |
| chr11:102317074-102317800 | -0.452689615 | 0.183476594 | 0.451056579 |
| chr3:171168198-171168530 | -0.495634576 | 0.183482457 | 0.451056579 |
| chr2:216012147-216012495 | 0.49604163 | 0.183490411 | 0.451056579 |
| chr21:34073823-34074085 | 0.552942076 | 0.183496465 | 0.451056579 |
| chr4:118431996-118432229 | -0.704151655 | 0.183496706 | 0.451056579 |
| chrX:102494692-102494921 | -0.707663078 | 0.183515383 | 0.451088017 |
| chr20:16553822-16554056 | -0.695252869 | 0.183537234 | 0.451111185 |
| chr1:75092104-75092544 | -0.518508446 | 0.183544473 | 0.451111185 |
| chr4:158861896-158862111 | -0.663388193 | 0.183545317 | 0.451111185 |
| chr4:41015-41215 | 0.595555697 | 0.183552471 | 0.451111185 |
| chr1:32539309-32539532 | 0.598009544 | 0.183556628 | 0.451111185 |
| chr14:55051221-55052310 | 0.421485769 | 0.183560133 | 0.451111185 |
| chr15:96264879-96265217 | 0.560108901 | 0.183570523 | 0.451122251 |
| chr1:231970786-231970986 | 0.74795602 | 0.183598489 | 0.451176506 |
| chr21:28641770-28642073 | -0.6023372 | 0.183610723 | 0.451192098 |
| chr8:90758414-90758819 | 0.533697248 | 0.183631818 | 0.451201586 |
| chr17:66468783-66469053 | -0.587400254 | 0.183636655 | 0.451201586 |
| chr1:156689500-156689805 | 0.469318576 | 0.183641971 | 0.451201586 |
| chr16:56657444-56657991 | 0.465478889 | 0.183642638 | 0.451201586 |
| chr5:149450421-149450625 | 0.674833423 | 0.183644027 | 0.451201586 |
| chr12:103337196-103337450 | -0.731763178 | 0.183675705 | 0.451234448 |
| chr11:72162951-72163408 | -0.52684818 | 0.183675966 | 0.451234448 |
| chr4:24753447-24753962 | 0.466620737 | 0.18367611 | 0.451234448 |
| chr20:55066988-55067201 | 0.663070542 | 0.183680959 | 0.451234448 |
| chr5:81675892-81676368 | -0.55750094 | 0.183708163 | 0.451286808 |
| chr12:109855355-109855576 | 0.627344353 | 0.183714812 | 0.451288675 |
| chr3:170884336-170884646 | -0.672871417 | 0.18372508 | 0.45129943 |
| chr17:74864731-74865216 | 0.475224892 | 0.18374168 | 0.451315103 |
| chr17:42964111-42965114 | 0.445861067 | 0.183748381 | 0.451315103 |
| chr8:74088163-74088448 | -0.537469004 | 0.183758615 | 0.451315103 |
| chr20:16518359-16518617 | 0.559051403 | 0.18375944 | 0.451315103 |
| chr13:70391653-70392286 | -0.489065953 | 0.183763192 | 0.451315103 |

| | | | |
|---|---|---|---|
| chr7:156038631-156038881 | -0.627584582 | 0.183766802 | 0.451315103 |
| chr7:32110470-32110670 | 0.681740846 | 0.18377846 | 0.451327854 |
| chr11:27416844-27417096 | 0.609426481 | 0.183786915 | 0.451327854 |
| chr1:148402103-148403164 | 0.432812475 | 0.183793203 | 0.451327854 |
| chr14:63727885-63728569 | 0.441689958 | 0.183795555 | 0.451327854 |
| chr15:62125811-62126099 | 0.590728968 | 0.183839385 | 0.451421015 |
| chr1:246601359-246601581 | -0.580685276 | 0.183852852 | 0.451439617 |
| chr16:67246185-67247115 | 0.429764811 | 0.183884693 | 0.451492703 |
| chr6:86897746-86898404 | -0.530018871 | 0.183886257 | 0.451492703 |
| chr3:187619588-187619792 | 0.658152628 | 0.183897291 | 0.451505326 |
| chr13:38697967-38698284 | -0.516418932 | 0.183936505 | 0.451587136 |
| chr19:37719144-37719583 | 0.505642668 | 0.183943047 | 0.451588725 |
| chr1:48931638-48932177 | -0.549569227 | 0.183955749 | 0.45159293 |
| chr5:91499927-91500246 | -0.58433545 | 0.183956547 | 0.45159293 |
| chr1:115911370-115911617 | 0.5746496 | 0.183967721 | 0.451605891 |
| chr14:96038326-96039492 | 0.43153648 | 0.183976145 | 0.451609049 |
| chr14:23071913-23072232 | 0.470985518 | 0.183980795 | 0.451609049 |
| chr12:2624576-2625212 | -0.47800006 | 0.183991464 | 0.45162077 |
| chr9:128455755-128457098 | 0.428582039 | 0.184043306 | 0.451733548 |
| chr14:22334052-22334292 | 0.590184217 | 0.18407428 | 0.451783463 |
| chr20:61268543-61268743 | 0.606962813 | 0.184075434 | 0.451783463 |
| chr11:20363340-20364300 | 0.440919631 | 0.18411231 | 0.451847169 |
| chr10:93974872-93975246 | -0.614307554 | 0.184113185 | 0.451847169 |
| chr14:71139002-71139329 | 0.50353204 | 0.184136004 | 0.451861551 |
| chr20:54161117-54161445 | 0.446345627 | 0.184137676 | 0.451861551 |
| chr4:1720805-1721616 | 0.43654273 | 0.18414303 | 0.451861551 |
| chr8:65926163-65926363 | 0.762304556 | 0.184144306 | 0.451861551 |
| chr1:26500164-26501520 | -0.432711123 | 0.184148532 | 0.451861551 |
| chr14:99807367-99807591 | 0.620901987 | 0.184196739 | 0.451965369 |
| chr1:200509971-200510485 | -0.45735458 | 0.184213595 | 0.451992252 |
| chr16:23423424-23423664 | 0.647751456 | 0.184221991 | 0.45199838 |
| chr19:3976668-3976915 | 0.650360693 | 0.184257138 | 0.452070138 |
| chr5:126776466-126777238 | -0.423318038 | 0.184290116 | 0.452136572 |
| chr11:65743849-65744824 | -0.446120527 | 0.184335326 | 0.45223301 |
| chr1:150624157-150624494 | 0.47841334 | 0.184365228 | 0.452291887 |
| chr5:101442333-101442760 | -0.500545774 | 0.184385054 | 0.452314979 |
| chr12:26274437-26275072 | -0.465132844 | 0.184386447 | 0.452314979 |
| chr4:169612003-169612902 | -0.425248956 | 0.184418299 | 0.452366528 |
| chr1:226547791-226548020 | 0.579923619 | 0.184419269 | 0.452366528 |
| chr1:106832205-106832405 | 0.495467046 | 0.184441434 | 0.452406414 |
| chr16:67404596-67405351 | 0.436599266 | 0.184461233 | 0.452440494 |
| chr17:39687666-39688473 | 0.455937289 | 0.184493765 | 0.452505803 |
| chr11:77047640-77047970 | -0.477228862 | 0.184506911 | 0.452522869 |
| chr11:60970375-60970575 | 0.665104109 | 0.18451331 | 0.452522869 |
| chr20:23937167-23937667 | 0.501923743 | 0.184523338 | 0.452522869 |
| chr19:15836769-15837318 | -0.471140721 | 0.184524347 | 0.452522869 |
| chr3:96794434-96794819 | -0.624608053 | 0.1845488 | 0.452563586 |
| chr6:19812611-19812849 | -0.657135561 | 0.184552763 | 0.452563586 |
| chr20:354469-354841 | -0.535717653 | 0.184563343 | 0.452569459 |
| chr3:150314567-150314915 | 0.498500209 | 0.184566971 | 0.452569459 |

| | | | |
|---|---|---|---|
| chr16:31201922-31202560 | 0.464571889 | 0.184587908 | 0.452606314 |
| chr16:15093956-15094724 | 0.45690163 | 0.184610362 | 0.452646823 |
| chr8:110822082-110822473 | -0.485312492 | 0.184616384 | 0.452646823 |
| chr2:5852831-5853031 | -0.803041378 | 0.184622152 | 0.452646823 |
| chr11:76169035-76169264 | 0.599986442 | 0.184632128 | 0.452656799 |
| chr20:57013841-57014129 | 0.494492849 | 0.184640703 | 0.452663337 |
| chr20:54688440-54688864 | 0.437686695 | 0.184649654 | 0.452670798 |
| chr1:106569304-106569655 | 0.458270331 | 0.184692809 | 0.452744776 |
| chr8:125069754-125070407 | 0.451861411 | 0.184697072 | 0.452744776 |
| chr1:36835112-36835968 | -0.471520272 | 0.184697557 | 0.452744776 |
| chr1:157090114-157090495 | -0.490603732 | 0.184730633 | 0.452806838 |
| chr7:115146149-115146543 | -0.578804176 | 0.184734694 | 0.452806838 |
| chr10:26726193-26726395 | -0.65717445 | 0.184741197 | 0.452808292 |
| chr1:61057576-61058271 | -0.457154589 | 0.184748231 | 0.452811048 |
| chr3:48919430-48919678 | 0.638631281 | 0.184765727 | 0.452834709 |
| chr12:93410921-93411298 | -0.645589976 | 0.184769705 | 0.452834709 |
| chr5:91219885-91220085 | -0.644312452 | 0.1847843 | 0.452855995 |
| chr18:31610965-31611249 | -0.56162586 | 0.1847936 | 0.4528643 |
| chr3:127881899-127882154 | -0.684969816 | 0.184803201 | 0.45286859 |
| chr19:57915982-57916326 | 0.564381312 | 0.18480798 | 0.45286859 |
| chrX:103658676-103659565 | -0.449869021 | 0.184813082 | 0.45286859 |
| chr11:86978859-86979061 | 0.659509778 | 0.18482194 | 0.452875815 |
| chr1:10825049-10825249 | -0.748349479 | 0.184879267 | 0.452995545 |
| chr8:94948369-94948989 | 0.448841848 | 0.184882628 | 0.452995545 |
| chr10:17445225-17446019 | -0.454205178 | 0.184912251 | 0.45305364 |
| chr2:95991505-95991739 | 0.586821456 | 0.184930184 | 0.453073194 |
| chr8:142864914-142865326 | 0.52659438 | 0.184932058 | 0.453073194 |
| chr10:123995841-123996211 | 0.48469611 | 0.184954257 | 0.453100424 |
| chr8:80078366-80078661 | -0.560619889 | 0.184954999 | 0.453100424 |
| chr16:85674030-85674530 | -0.515494061 | 0.184971004 | 0.453125145 |
| chr20:58611063-58611626 | 0.433692518 | 0.184988687 | 0.453153975 |
| chr3:48717106-48717580 | 0.469377388 | 0.184998485 | 0.453163488 |
| chr3:160210793-160211236 | -0.539831695 | 0.185011912 | 0.453181891 |
| chr2:164514313-164515363 | -0.42371999 | 0.18502978 | 0.45321117 |
| chr8:70114758-70114958 | -0.740038157 | 0.185043804 | 0.453231032 |
| chr6:41278976-41279256 | -0.600325127 | 0.185063785 | 0.453249547 |
| chr3:45929103-45929446 | -0.622279527 | 0.185065481 | 0.453249547 |
| chr10:10677523-10677936 | 0.52970242 | 0.185069393 | 0.453249547 |
| chr5:38660208-38660707 | -0.446220626 | 0.185079318 | 0.453249547 |
| chr20:12925179-12925663 | 0.483730017 | 0.185085798 | 0.453249547 |
| chr11:2902024-2902631 | 0.456858197 | 0.185086856 | 0.453249547 |
| chr5:168591568-168591791 | 0.625789695 | 0.185104931 | 0.453279324 |
| chr10:93408979-93409179 | -0.705168601 | 0.185120856 | 0.45330212 |
| chr3:14203488-14203796 | 0.492758369 | 0.185126073 | 0.45330212 |
| chr5:16906248-16906545 | 0.5564402 | 0.185155757 | 0.453357819 |
| chr2:171346496-171346734 | -0.623244373 | 0.185160653 | 0.453357819 |
| chr2:230067426-230067634 | 0.658939951 | 0.185181985 | 0.453381112 |
| chr16:56401498-56401793 | -0.660769018 | 0.185182001 | 0.453381112 |
| chr20:5610708-5611362 | 0.447745751 | 0.185188675 | 0.453382964 |
| chr4:138865092-138865443 | -0.557380085 | 0.185203152 | 0.453403921 |

| | | | |
|---|---|---|---|
| chr22:29202443-29202766 | 0.564862444 | 0.185209609 | 0.453405242 |
| chr15:99487055-99487255 | -0.666997173 | 0.185229504 | 0.453427587 |
| chr20:40938084-40938723 | 0.462759657 | 0.185230572 | 0.453427587 |
| chr11:76096608-76096913 | -0.600106005 | 0.185253015 | 0.453468038 |
| chr21:39527746-39528313 | -0.459495466 | 0.185283615 | 0.453528453 |
| chr10:24724089-24725143 | -0.429992046 | 0.185294148 | 0.453539746 |
| chr2:138177625-138178066 | 0.567923981 | 0.185320187 | 0.453588992 |
| chr8:87166232-87166797 | -0.454852711 | 0.185345898 | 0.453622352 |
| chr2:6922449-6922749 | -0.520012401 | 0.185351143 | 0.453622352 |
| chr11:101345922-101346211 | -0.639790145 | 0.185352301 | 0.453622352 |
| chr22:37483484-37483684 | -0.823156903 | 0.185362449 | 0.453622352 |
| chr5:132426823-132427126 | -0.542111752 | 0.185363418 | 0.453622352 |
| chr20:58252021-58252379 | 0.500901768 | 0.185380107 | 0.453648704 |
| chr3:40505586-40506514 | 0.439706203 | 0.185397208 | 0.453675465 |
| chr3:145999897-146000122 | -0.688832067 | 0.185402885 | 0.453675465 |
| chr11:48011408-48011626 | 0.584365529 | 0.185412179 | 0.453683719 |
| chr14:22777452-22777705 | 0.589479063 | 0.185431321 | 0.453716069 |
| chr12:108911791-108912313 | 0.469826848 | 0.185470866 | 0.453798335 |
| chr15:99409038-99409391 | 0.486529371 | 0.185498499 | 0.453851455 |
| chr1:239898109-239898485 | -0.495112256 | 0.185509505 | 0.45386389 |
| chr16:4416287-4416938 | 0.448048293 | 0.185516478 | 0.453866457 |
| chr8:123456056-123456263 | 0.551849675 | 0.185535711 | 0.453894112 |
| chr10:7092732-7092932 | 0.66110511 | 0.185544343 | 0.453894112 |
| chr5:157142280-157143363 | -0.422862593 | 0.185548202 | 0.453894112 |
| chr11:30585327-30585620 | -0.55622601 | 0.185560139 | 0.453894112 |
| chr5:108748316-108749799 | -0.415980078 | 0.185561962 | 0.453894112 |
| chr11:66347656-66348551 | 0.430438589 | 0.185563325 | 0.453894112 |
| chr1:53889508-53890028 | -0.475636148 | 0.185584962 | 0.453932546 |
| chr3:59049563-59050433 | -0.439081335 | 0.185614728 | 0.453990861 |
| chr2:179860598-179861682 | -0.457569681 | 0.18562914 | 0.453997826 |
| chr17:16997540-16997853 | 0.538743098 | 0.185629426 | 0.453997826 |
| chr12:118034876-118035180 | 0.5380418 | 0.18563978 | 0.454008657 |
| chr3:14921149-14921560 | 0.456595594 | 0.185649968 | 0.454019081 |
| chr14:88488802-88489046 | 0.578141867 | 0.185655965 | 0.454019258 |
| chr11:24155839-24156164 | -0.55172655 | 0.185662746 | 0.454021348 |
| chr20:47297008-47297307 | 0.436784708 | 0.185671175 | 0.454027472 |
| chr3:141623464-141623764 | -0.663770514 | 0.185678665 | 0.454031297 |
| chr8:75648022-75648463 | -0.564520435 | 0.185702995 | 0.454072808 |
| chr12:100200395-100201231 | -0.428315566 | 0.185707984 | 0.454072808 |
| chr16:31442643-31443394 | 0.445375485 | 0.18571342 | 0.454072808 |
| chr3:132803884-132804157 | -0.631613271 | 0.185745823 | 0.454131996 |
| chr1:198309138-198309440 | 0.563117981 | 0.185749481 | 0.454131996 |
| chr7:30445990-30446190 | 0.67798291 | 0.185785216 | 0.454204869 |
| chr2:20962940-20963269 | -0.503026834 | 0.185797813 | 0.454221173 |
| chr5:88268377-88269704 | -0.420898025 | 0.185819659 | 0.454260087 |
| chr2:38431280-38431504 | -0.678750541 | 0.185829564 | 0.454269807 |
| chrX:24464706-24465782 | -0.428740591 | 0.185850518 | 0.454292664 |
| chr5:126679568-126679950 | -0.467263052 | 0.185850772 | 0.454292664 |
| chr5:113847757-113848195 | -0.552204127 | 0.185873571 | 0.454333901 |
| chr1:106680278-106680484 | 0.487525615 | 0.185884405 | 0.454345887 |

| | | | |
|---|---|---|---|
| chr3:181316579-181316779 | -0.742376495 | 0.185924364 | 0.454425625 |
| chr13:28459844-28460226 | -0.584333709 | 0.185928889 | 0.454425625 |
| chr4:128046382-128046582 | 0.680438699 | 0.185955036 | 0.454461654 |
| chrX:55640975-55641189 | -0.711736827 | 0.185955493 | 0.454461654 |
| chr2:105337111-105337917 | -0.440348221 | 0.185971809 | 0.454475361 |
| chr1:25905405-25905899 | 0.461168188 | 0.185972964 | 0.454475361 |
| chr15:43775426-43775767 | 0.472261774 | 0.186019713 | 0.454575106 |
| chr8:77053049-77053341 | -0.67203362 | 0.186043727 | 0.454613708 |
| chr5:9932889-9933276 | 0.571175005 | 0.186047376 | 0.454613708 |
| chr4:5615446-5615646 | 0.710120246 | 0.186076844 | 0.454671214 |
| chr11:86183247-86183569 | -0.587709352 | 0.186096422 | 0.454704553 |
| chr2:105398446-105399284 | 0.448130573 | 0.186117202 | 0.454740824 |
| chr2:181393460-181393863 | -0.537589688 | 0.186129885 | 0.454757311 |
| chr8:10838913-10839209 | 0.539370821 | 0.186153316 | 0.454800058 |
| chr8:75259486-75260206 | -0.480924127 | 0.186172915 | 0.454833437 |
| chr12:111368591-111369295 | 0.472999145 | 0.186198157 | 0.454873214 |
| chr10:33193400-33193673 | -0.59131141 | 0.18620255 | 0.454873214 |
| chr12:95623434-95623982 | -0.482108816 | 0.186207006 | 0.454873214 |
| chr22:37639421-37639683 | -0.579086644 | 0.186216296 | 0.454876344 |
| chr5:66143619-66145032 | -0.419827681 | 0.186220161 | 0.454876344 |
| chr10:30600016-30600231 | -0.774158414 | 0.186229841 | 0.454885489 |
| chr16:87087649-87088041 | -0.616310858 | 0.186246861 | 0.454912559 |
| chr19:51594290-51594901 | 0.462752273 | 0.186283013 | 0.454986359 |
| chr6:43263494-43263748 | 0.556424327 | 0.186314885 | 0.455028042 |
| chr2:108960074-108960335 | 0.59396948 | 0.186318326 | 0.455028042 |
| chr7:47435860-47436125 | -0.550284035 | 0.186323759 | 0.455028042 |
| chr2:191625056-191625338 | -0.636271545 | 0.186324087 | 0.455028042 |
| chr5:111530228-111530747 | -0.49103596 | 0.186329773 | 0.455028042 |
| chr17:40549436-40549683 | 0.560636397 | 0.186337996 | 0.45503362 |
| chr1:51170353-51170633 | 0.582584972 | 0.186348739 | 0.455035588 |
| chr21:31659226-31660428 | 0.423060298 | 0.186350679 | 0.455035588 |
| chr3:119795231-119795513 | -0.596944938 | 0.186358669 | 0.455040597 |
| chr17:29710898-29711308 | -0.539565679 | 0.186371579 | 0.455057619 |
| chr5:62256826-62257042 | -0.692237917 | 0.186385512 | 0.455077136 |
| chr5:145834786-145836020 | -0.428736502 | 0.186413155 | 0.455105082 |
| chr7:99408498-99409562 | -0.4325069 | 0.186416983 | 0.455105082 |
| chr8:92136121-92136611 | 0.524042461 | 0.186417036 | 0.455105082 |
| chr15:39172087-39172392 | 0.53167859 | 0.186420716 | 0.455105082 |
| chr8:98700778-98701136 | 0.515544396 | 0.186427011 | 0.45510595 |
| chr5:55994468-55995589 | -0.425269675 | 0.186443616 | 0.455131985 |
| chr21:42279661-42279861 | 0.620095543 | 0.186464129 | 0.455156976 |
| chr10:16699878-16700078 | -0.752077015 | 0.186465734 | 0.455156976 |
| chrX:30972228-30972562 | -0.512859722 | 0.186480905 | 0.455179507 |
| chr5:13218574-13218889 | -0.526667345 | 0.186494696 | 0.455198668 |
| chr10:11602526-11602726 | 0.75089344 | 0.186505291 | 0.455205504 |
| chr2:64873415-64873880 | -0.472558964 | 0.18651183 | 0.455205504 |
| chr17:44168414-44168837 | 0.485243348 | 0.186523488 | 0.455205504 |
| chr20:50272964-50273164 | -0.58078126 | 0.186530619 | 0.455205504 |
| chrX:100413279-100413573 | -0.506495172 | 0.186530642 | 0.455205504 |
| chr10:59906277-59907217 | -0.421404514 | 0.18653976 | 0.455205504 |

| | | | |
|---|---|---|---|
| chr11:450061-450417 | 0.480561249 | 0.186540467 | 0.455205504 |
| chr2:219104156-219104440 | -0.621940202 | 0.186545024 | 0.455205504 |
| chr3:184229336-184229949 | 0.478123325 | 0.186551505 | 0.455206822 |
| chr17:9576496-9576765 | 0.526847465 | 0.186561596 | 0.455216948 |
| chr4:105924420-105924667 | 0.625582242 | 0.186573832 | 0.455232307 |
| chr17:32341771-32342379 | 0.482931553 | 0.186606516 | 0.455276318 |
| chr6:27503576-27503867 | -0.519958391 | 0.186608996 | 0.455276318 |
| chr2:662628-663084 | -0.513833307 | 0.186609695 | 0.455276318 |
| chr7:98106624-98107378 | -0.440037958 | 0.186617178 | 0.455280077 |
| chr13:50131084-50131292 | -0.726959726 | 0.186631914 | 0.455292215 |
| chr2:186792839-186793066 | -0.700737871 | 0.186634037 | 0.455292215 |
| chr6:15254002-15254268 | -0.58997697 | 0.18664352 | 0.455300853 |
| chr20:13302750-13303170 | 0.543849087 | 0.186664972 | 0.455338688 |
| chr20:49320451-49320829 | 0.480386145 | 0.186676411 | 0.455352092 |
| chr1:108876572-108877410 | -0.429525617 | 0.186716636 | 0.455434132 |
| chr1:58035595-58035795 | -0.691763154 | 0.186725434 | 0.455434132 |
| chr13:76965802-76966124 | -0.563347383 | 0.186727875 | 0.455434132 |
| chr13:32910909-32911153 | -0.695268201 | 0.186804708 | 0.45560263 |
| chr4:26319011-26320205 | 0.420213506 | 0.186808852 | 0.45560263 |
| chr1:119713147-119713362 | 0.624873138 | 0.18681771 | 0.455609734 |
| chr20:6005709-6006378 | 0.443955416 | 0.186833997 | 0.455634951 |
| chr8:123866749-123866988 | 0.649406522 | 0.186864414 | 0.45567691 |
| chr6:138588926-138589329 | -0.551884946 | 0.18686715 | 0.45567691 |
| chr11:68764227-68764551 | 0.5115 | 0.186870867 | 0.45567691 |
| chr12:65160389-65160671 | 0.661295183 | 0.186874991 | 0.45567691 |
| chr1:218209742-218209985 | 0.589630761 | 0.186897543 | 0.455712606 |
| chr15:89306010-89306397 | 0.523616658 | 0.186902764 | 0.455712606 |
| chr14:89040231-89041602 | -0.425477323 | 0.186907473 | 0.455712606 |
| chr3:148372442-148372832 | -0.596348024 | 0.186918754 | 0.45572561 |
| chr4:113616500-113616720 | 0.642054783 | 0.18694724 | 0.455780559 |
| chr7:30026513-30027165 | 0.442180844 | 0.186965148 | 0.455806491 |
| chr1:155354863-155355130 | 0.66550842 | 0.186969774 | 0.455806491 |
| chrX:100587932-100588280 | -0.637654855 | 0.186986899 | 0.455833738 |
| chr2:48456390-48456672 | -0.717714599 | 0.187000102 | 0.455847964 |
| chr9:95103365-95103618 | 0.629158005 | 0.18700509 | 0.455847964 |
| chr18:40810477-40810719 | -0.651407484 | 0.187010583 | 0.455847964 |
| chr1:90723834-90724079 | -0.701843039 | 0.1870392 | 0.455896039 |
| chr1:164598505-164599070 | 0.463198183 | 0.187047603 | 0.455896039 |
| chr21:41366668-41367100 | -0.49907364 | 0.18705545 | 0.455896039 |
| chr7:73624443-73624770 | 0.483855315 | 0.187056906 | 0.455896039 |
| chr22:24010927-24011960 | 0.435363714 | 0.187060056 | 0.455896039 |
| chr15:78751028-78751298 | 0.574285288 | 0.187068513 | 0.455902151 |
| chr7:42389232-42389967 | -0.504609956 | 0.187078122 | 0.455911067 |
| chr3:47837717-47838010 | 0.590795534 | 0.187124549 | 0.456009707 |
| chr8:110138979-110139179 | 0.661273763 | 0.187141774 | 0.456037178 |
| chr2:68837287-68837578 | -0.537895632 | 0.187175018 | 0.456103684 |
| chr15:80743058-80743735 | 0.454549569 | 0.187191533 | 0.456129422 |
| chr13:77143498-77144182 | -0.447364634 | 0.187199925 | 0.456135363 |
| chr10:86644537-86644737 | 0.606295186 | 0.187248772 | 0.456239876 |
| chr1:60639957-60640677 | -0.447550726 | 0.187268805 | 0.456274179 |

| | | | |
|---|---|---|---|
| chr12:27710199-27711608 | -0.427750791 | 0.187281238 | 0.456275547 |
| chr7:155713508-155713745 | -0.692451754 | 0.187282575 | 0.456275547 |
| chr5:125050014-125050280 | -0.643850189 | 0.187292167 | 0.456275547 |
| chr2:127270680-127271139 | 0.510197349 | 0.187293186 | 0.456275547 |
| chr1:165403966-165404305 | -0.523062288 | 0.187308761 | 0.456298982 |
| chr1:152543948-152544221 | -0.611771595 | 0.187331419 | 0.45633967 |
| chr1:188877595-188877953 | -0.645689113 | 0.187337788 | 0.456340677 |
| chr12:97148965-97149352 | -0.490551799 | 0.187361023 | 0.456382768 |
| chr3:28726902-28727210 | -0.544654475 | 0.187375521 | 0.456403573 |
| chr16:16232391-16232647 | 0.570877943 | 0.187390189 | 0.456415476 |
| chr20:56815938-56816147 | 0.540309132 | 0.1873939 | 0.456415476 |
| chr3:127853744-127853979 | -0.601693996 | 0.187400244 | 0.456415476 |
| chr20:48025061-48025399 | 0.450838948 | 0.187409536 | 0.456415476 |
| chr15:39580222-39580590 | 0.46775787 | 0.187410192 | 0.456415476 |
| chr5:157887476-157887717 | -0.759583063 | 0.187423098 | 0.4564324 |
| chr14:31286748-31287123 | -0.502626499 | 0.187442634 | 0.456465469 |
| chr8:49796276-49796477 | -0.684151164 | 0.187457335 | 0.456486761 |
| chr14:104297648-104297848 | -0.666524372 | 0.18746906 | 0.456500804 |
| chr8:133836894-133837410 | -0.54031942 | 0.187486315 | 0.456528314 |
| chr2:11129837-11130419 | -0.448261165 | 0.187515738 | 0.456580386 |
| chr12:66352298-66352554 | 0.582917334 | 0.187519618 | 0.456580386 |
| chrX:65595415-65595831 | 0.489588892 | 0.187525859 | 0.456581074 |
| chr5:83394283-83394565 | -0.653136092 | 0.187557128 | 0.456634137 |
| chr10:65891065-65891830 | 0.455517545 | 0.187565308 | 0.456634137 |
| chr17:39636937-39637137 | 0.652589053 | 0.187565532 | 0.456634137 |
| chr5:60458774-60458974 | -0.6721902 | 0.187592677 | 0.456685713 |
| chr21:38350635-38350871 | -0.638834563 | 0.187607259 | 0.4567067 |
| chr14:24481580-24481982 | -0.478184502 | 0.187648557 | 0.456792722 |
| chr4:77862436-77863250 | -0.429361876 | 0.187655491 | 0.456795089 |
| chr1:196535786-196535986 | -0.654851382 | 0.18770591 | 0.456880127 |
| chr20:47314679-47314967 | 0.434301091 | 0.187707671 | 0.456880127 |
| chr12:122836000-122836417 | 0.497982595 | 0.187711049 | 0.456880127 |
| chr10:1435399-1435703 | 0.551992457 | 0.187714276 | 0.456880127 |
| chr17:31094577-31095062 | 0.466069166 | 0.187731143 | 0.456881682 |
| chr2:68251805-68252207 | -0.49036215 | 0.187731852 | 0.456881682 |
| chr6:39347893-39348126 | -0.684573836 | 0.187732804 | 0.456881682 |
| chr6:25006531-25006762 | -0.683681823 | 0.187763884 | 0.456942289 |
| chr2:208867291-208867892 | -0.502268975 | 0.187774575 | 0.456942289 |
| chr8:61810057-61810423 | 0.517943424 | 0.187775598 | 0.456942289 |
| chrX:40104770-40105000 | 0.558330656 | 0.187785064 | 0.456944818 |
| chr5:2038427-2038956 | 0.483432086 | 0.187794159 | 0.456944818 |
| chr6:141195270-141195810 | -0.519470271 | 0.187794528 | 0.456944818 |
| chr7:15015629-15016448 | 0.438146816 | 0.187804528 | 0.456954639 |
| chr3:24363302-24363547 | -0.636258679 | 0.187844931 | 0.457035079 |
| chr8:104639196-104639850 | 0.469048571 | 0.187849518 | 0.457035079 |
| chr2:120175054-120175336 | 0.590392941 | 0.187880785 | 0.457082123 |
| chr3:47522157-47522406 | 0.590392941 | 0.187880785 | 0.457082123 |
| chr5:11240445-11240669 | 0.635696638 | 0.187913187 | 0.457094366 |
| chr16:31025953-31026187 | 0.621380005 | 0.187916335 | 0.457094366 |
| chr3:53130280-53130659 | 0.477835385 | 0.187916625 | 0.457094366 |

| | | | |
|---|---|---|---|
| chr10:33273177-33273969 | -0.447537532 | 0.187920693 | 0.457094366 |
| chr1:42380685-42381091 | 0.531367261 | 0.187923536 | 0.457094366 |
| chr18:14074345-14074546 | 0.674554369 | 0.187924965 | 0.457094366 |
| chr3:130747377-130748342 | -0.431800018 | 0.187927576 | 0.457094366 |
| chr16:21158470-21159117 | 0.466559827 | 0.187940485 | 0.457111255 |
| chrX:101390454-101391511 | -0.42555588 | 0.187955106 | 0.457132304 |
| chr2:157572759-157573040 | -0.582870241 | 0.187974283 | 0.457163263 |
| chr8:31308818-31309090 | -0.607919474 | 0.187983378 | 0.457163263 |
| chr6:142845603-142846477 | -0.431957593 | 0.187985734 | 0.457163263 |
| chr16:69310887-69312181 | 0.422078619 | 0.188000575 | 0.457184843 |
| chr8:48007742-48008097 | -0.522767933 | 0.188057445 | 0.457308628 |
| chr5:160421642-160422219 | -0.435217222 | 0.18807213 | 0.457329825 |
| chr20:57228477-57228757 | 0.433153977 | 0.188104023 | 0.457386027 |
| chr11:16761454-16761712 | -0.623678532 | 0.188112874 | 0.457386027 |
| chr6:142300942-142302818 | -0.416348029 | 0.188113151 | 0.457386027 |
| chr4:107913326-107913632 | -0.575044704 | 0.188185073 | 0.457540328 |
| chr20:25336595-25336855 | 0.557221327 | 0.188188555 | 0.457540328 |
| chr10:11539233-11539708 | -0.536371447 | 0.188195443 | 0.457542557 |
| chr10:5890652-5890893 | 0.494715497 | 0.188202552 | 0.457545322 |
| chr10:18651292-18652064 | -0.432865616 | 0.18822003 | 0.457568289 |
| chr17:58492555-58492848 | 0.549290316 | 0.188223942 | 0.457568289 |
| chr3:12483854-12484912 | 0.419515527 | 0.188233099 | 0.457576032 |
| chr2:173914540-173914914 | -0.583763412 | 0.188240578 | 0.457579386 |
| chr1:28648119-28649508 | 0.420940074 | 0.188246423 | 0.457579386 |
| chr20:45934186-45934386 | 0.520677422 | 0.188282881 | 0.457653488 |
| chr3:32238492-32239171 | 0.45290275 | 0.188300438 | 0.457681644 |
| chr20:53927190-53927604 | 0.427601246 | 0.188308294 | 0.457686219 |
| chr8:72913124-72913447 | 0.622851678 | 0.188324144 | 0.457703317 |
| chr5:10140302-10140837 | 0.472737971 | 0.18832922 | 0.457703317 |
| chr3:63961876-63962151 | 0.510290462 | 0.188333374 | 0.457703317 |
| chr3:188947361-188948399 | -0.426590177 | 0.188339222 | 0.457703317 |
| chr8:15562572-15562862 | -0.580973238 | 0.188352145 | 0.457720205 |
| chr2:162118381-162118753 | -0.592214988 | 0.188377498 | 0.457767297 |
| chr10:73849096-73849296 | 0.589021646 | 0.188394634 | 0.45779442 |
| chr10:3142321-3142602 | 0.565543393 | 0.188423015 | 0.457847754 |
| chr3:50268809-50269224 | -0.475735928 | 0.188428533 | 0.457847754 |
| chr21:36835515-36835774 | 0.638626384 | 0.188489115 | 0.457977189 |
| chr2:183071138-183071582 | -0.464411071 | 0.188497629 | 0.457977189 |
| chr19:18995895-18996140 | -0.623380699 | 0.188504635 | 0.457977189 |
| chr15:31975758-31975966 | 0.633430507 | 0.188506493 | 0.457977189 |
| chr10:17673483-17674023 | 0.511934821 | 0.188514138 | 0.457977189 |
| chr19:32674777-32676021 | 0.417213501 | 0.188517665 | 0.457977189 |
| chrX:49291924-49292234 | -0.568340054 | 0.188529732 | 0.457991983 |
| chr15:66803213-66803413 | 0.643948224 | 0.188551978 | 0.458029157 |
| chr7:50479713-50479983 | -0.696541131 | 0.188558629 | 0.458029157 |
| chr15:56243103-56244077 | 0.424744196 | 0.188568711 | 0.458029157 |
| chr20:50662360-50662566 | 0.556791863 | 0.188568946 | 0.458029157 |
| chr6:110642682-110642882 | 0.697497163 | 0.188583023 | 0.458045421 |
| chr6:168390684-168391105 | 0.487066728 | 0.188587598 | 0.458045421 |
| chr1:166153645-166153912 | 0.679416868 | 0.18859987 | 0.45805113 |

| | | | |
|---|---|---|---|
| chr14:74712593-74713322 | 0.45041218 | 0.188601904 | 0.45805113 |
| chr2:45765812-45766246 | -0.521853918 | 0.188624565 | 0.458091646 |
| chr15:36579063-36580096 | -0.423048924 | 0.188644023 | 0.45812438 |
| chr2:20899980-20900180 | 0.719818719 | 0.188668699 | 0.458169786 |
| chr21:42117490-42117715 | 0.524046033 | 0.188686371 | 0.458198179 |
| chr1:108570195-108570471 | -0.527452393 | 0.188697283 | 0.458198657 |
| chr4:92387533-92387788 | -0.609618759 | 0.188698528 | 0.458198657 |
| chr2:10013019-10013227 | 0.639550976 | 0.188725598 | 0.458249866 |
| chr14:61537434-61538136 | 0.436454576 | 0.188740598 | 0.458271766 |
| chr12:109573053-109573946 | -0.419942351 | 0.188759408 | 0.458291303 |
| chr12:114760329-114760543 | 0.650928337 | 0.188760607 | 0.458291303 |
| chr17:46822207-46822669 | 0.540005832 | 0.188783138 | 0.458331483 |
| chr14:68671511-68671711 | -0.700495981 | 0.188792939 | 0.458340756 |
| chr5:146542291-146542572 | 0.539538823 | 0.188807158 | 0.458360751 |
| chr5:110726355-110727271 | -0.429175841 | 0.188869668 | 0.458463948 |
| chr5:146878072-146879114 | -0.446797888 | 0.188876821 | 0.458463948 |
| chr7:1458822-1459036 | 0.493646493 | 0.188888402 | 0.458463948 |
| chr6:148166447-148166647 | 0.666952494 | 0.188893151 | 0.458463948 |
| chr4:95124930-95125281 | -0.54066796 | 0.188898378 | 0.458463948 |
| chr5:2111026-2111237 | 0.698151068 | 0.188903121 | 0.458463948 |
| chr5:693251-693950 | 0.443284051 | 0.188903677 | 0.458463948 |
| chr1:106293436-106293680 | 0.510956298 | 0.188909835 | 0.458463948 |
| chr14:68590252-68590589 | 0.508068269 | 0.188916218 | 0.458463948 |
| chrX:39969058-39969326 | -0.626329845 | 0.188918596 | 0.458463948 |
| chrX:119475843-119476301 | -0.631044288 | 0.188920086 | 0.458463948 |
| chr6:30325747-30326139 | -0.48146068 | 0.188926805 | 0.458463948 |
| chr14:59464380-59465927 | 0.423925656 | 0.188929837 | 0.458463948 |
| chrX:103498380-103498680 | -0.639322503 | 0.188933435 | 0.458463948 |
| chr14:105181004-105181697 | 0.439317882 | 0.188955004 | 0.458496658 |
| chr19:55624094-55625030 | 0.428270865 | 0.188959274 | 0.458496658 |
| chr1:43707070-43707821 | -0.447069156 | 0.188964866 | 0.458496658 |
| chr9:71806853-71807306 | -0.498101119 | 0.188980588 | 0.458520284 |
| chr21:39713021-39713289 | 0.551992664 | 0.18899286 | 0.458535541 |
| chr1:226386656-226386856 | 0.675951003 | 0.189017702 | 0.458579307 |
| chr12:51047711-51048468 | 0.431315431 | 0.189022869 | 0.458579307 |
| chr17:72618008-72618593 | -0.465797343 | 0.189029272 | 0.458580322 |
| chr21:26023041-26023607 | -0.553490188 | 0.189053629 | 0.458611769 |
| chr4:105966091-105966401 | -0.589516365 | 0.189054206 | 0.458611769 |
| chr7:500528-500760 | -0.653058501 | 0.189092786 | 0.458677141 |
| chr6:32947838-32948116 | -0.659266831 | 0.189095355 | 0.458677141 |
| chr2:64562055-64562381 | -0.625967694 | 0.189101956 | 0.458677141 |
| chr12:130357866-130358199 | -0.600975967 | 0.189105099 | 0.458677141 |
| chr17:73156276-73156565 | 0.524180949 | 0.189126165 | 0.458713716 |
| chr17:10845165-10845441 | 0.519069848 | 0.189154725 | 0.458750227 |
| chr10:110852286-110852664 | 0.524169865 | 0.189157925 | 0.458750227 |
| chr1:114311129-114311834 | 0.433035119 | 0.18915918 | 0.458750227 |
| chr3:186931902-186932172 | -0.62166126 | 0.189190849 | 0.458812509 |
| chr2:46297573-46297859 | 0.528295793 | 0.18919839 | 0.458816275 |
| chr3:81581823-81582527 | -0.469527038 | 0.189231905 | 0.458883028 |
| chr5:171419939-171420149 | 0.538835569 | 0.189284029 | 0.458994901 |

| | | | |
|---|---|---|---|
| chr21:14864276-14864927 | -0.488257808 | 0.189310575 | 0.459044744 |
| chr9:113339823-113340555 | 0.439057774 | 0.189318707 | 0.459049936 |
| chr22:30391191-30391405 | 0.584432885 | 0.189358597 | 0.459065599 |
| chr4:99816622-99817067 | -0.49935549 | 0.189358849 | 0.459065599 |
| chr10:77037400-77037603 | -0.684410922 | 0.189365557 | 0.459065599 |
| chr1:200693430-200693651 | 0.594228044 | 0.189371711 | 0.459065599 |
| chr2:143333297-143333607 | 0.581973074 | 0.189375535 | 0.459065599 |
| chr7:153381635-153381835 | -0.703362661 | 0.18937711 | 0.459065599 |
| chr3:100992211-100992735 | -0.548870949 | 0.189378033 | 0.459065599 |
| chr3:183767083-183767393 | -0.548870949 | 0.189378033 | 0.459065599 |
| chr19:16097885-16098257 | 0.549984996 | 0.189379088 | 0.459065599 |
| chr1:40140428-40140628 | 0.669031644 | 0.189390043 | 0.459075369 |
| chr9:93133148-93133513 | 0.475140065 | 0.189395102 | 0.459075369 |
| chr11:117098311-117099159 | 0.44095509 | 0.18940648 | 0.459088425 |
| chr3:119690158-119690399 | -0.602195552 | 0.189421554 | 0.459110439 |
| chr3:67531331-67531926 | -0.475436549 | 0.189447959 | 0.459159914 |
| chr2:62705016-62705955 | -0.415512366 | 0.189466668 | 0.459179659 |
| chr10:24642714-24643393 | -0.496790449 | 0.189470863 | 0.459179659 |
| chr3:136862672-136862978 | 0.480496374 | 0.189474084 | 0.459179659 |
| chr10:88318827-88319088 | -0.629031321 | 0.189494897 | 0.459215572 |
| chr12:6835722-6835967 | -0.550173337 | 0.189534862 | 0.459297896 |
| chr9:127451266-127452576 | 0.422507545 | 0.189549072 | 0.459302439 |
| chr5:118009872-118010320 | -0.542458587 | 0.189552264 | 0.459302439 |
| chr5:79990728-79991443 | -0.460204425 | 0.18955472 | 0.459302439 |
| chr10:62015927-62016168 | -0.645625696 | 0.189566621 | 0.459316752 |
| chr9:128371007-128371679 | 0.448257257 | 0.189586238 | 0.459348551 |
| chr14:99580071-99580388 | 0.510706387 | 0.189594763 | 0.459348551 |
| chrX:101905065-101905407 | 0.546665134 | 0.18959773 | 0.459348551 |
| chr1:151990108-151990308 | 0.500741453 | 0.189612817 | 0.459357965 |
| chr12:3303873-3304078 | -0.696977507 | 0.189613606 | 0.459357965 |
| chr6:166788817-166789584 | -0.456272107 | 0.18962285 | 0.459365836 |
| chr14:46344225-46344701 | -0.484805231 | 0.189633447 | 0.459376983 |
| chr2:95345534-95346047 | 0.461421439 | 0.189655592 | 0.459416104 |
| chr15:73051486-73052814 | 0.424420967 | 0.18966961 | 0.459435536 |
| chr4:25792331-25792531 | 0.615522011 | 0.189677195 | 0.459439383 |
| chr7:134520140-134520369 | -0.63841058 | 0.189725859 | 0.459542731 |
| chr9:91775579-91775786 | 0.604875792 | 0.189732977 | 0.459545445 |
| chr9:131178332-131178586 | 0.623184791 | 0.189741701 | 0.459551031 |
| chr2:169827925-169828127 | 0.570817597 | 0.189747279 | 0.459551031 |
| chr6:72968778-72968984 | 0.635418263 | 0.189782893 | 0.459609723 |
| chr6:28537537-28537759 | -0.692148127 | 0.189783509 | 0.459609723 |
| chr2:235833967-235834280 | -0.590989861 | 0.189795328 | 0.459623819 |
| chr6:75813573-75813816 | -0.647512055 | 0.189835438 | 0.459651942 |
| chr5:107630944-107631244 | -0.694113953 | 0.189837033 | 0.459651942 |
| chr12:43938053-43938311 | -0.617354466 | 0.189844056 | 0.459651942 |
| chr6:31546596-31547007 | 0.451272442 | 0.189845697 | 0.459651942 |
| chr17:21456938-21457375 | 0.455502806 | 0.189848515 | 0.459651942 |
| chr12:47106420-47106654 | -0.657711915 | 0.189852696 | 0.459651942 |
| chr19:47241067-47241351 | 0.535663334 | 0.189854651 | 0.459651942 |
| chr4:189780429-189780687 | -0.639925536 | 0.189854933 | 0.459651942 |

| | | | |
|---|---|---|---|
| chr14:24377684-24377953 | 0.531246774 | 0.189862758 | 0.459656363 |
| chr13:42229138-42229517 | -0.593911826 | 0.189890221 | 0.459704598 |
| chr8:57578147-57578422 | 0.617205704 | 0.189901828 | 0.459704598 |
| chr2:176111283-176111653 | -0.518748788 | 0.18990253 | 0.459704598 |
| chr12:70808029-70808384 | -0.563068506 | 0.189908838 | 0.459704598 |
| chr2:127398264-127398628 | -0.473515799 | 0.18991268 | 0.459704598 |
| chr1:177440848-177441204 | -0.514633959 | 0.189928324 | 0.459727943 |
| chr3:113614509-113614790 | -0.589334525 | 0.189978605 | 0.459835123 |
| chr20:53834701-53835059 | 0.429759045 | 0.18998513 | 0.459836391 |
| chr16:76465315-76465655 | -0.490960324 | 0.190008119 | 0.459877506 |
| chr20:4227014-4227367 | 0.561429241 | 0.190028637 | 0.459912638 |
| chr9:132079655-132079997 | 0.54721922 | 0.19004865 | 0.459939128 |
| chr4:165440356-165441061 | -0.466235214 | 0.190051587 | 0.459939128 |
| chr6:154277180-154277675 | -0.50472064 | 0.190080415 | 0.459994364 |
| chr14:104689133-104689730 | 0.442293313 | 0.190088464 | 0.459999315 |
| chr20:44773429-44773629 | 0.640098244 | 0.190112532 | 0.460043029 |
| chr2:37083767-37084860 | -0.415215547 | 0.19014109 | 0.460097602 |
| chr1:147237248-147237906 | 0.440750811 | 0.19015003 | 0.460104706 |
| chr20:57212081-57212482 | 0.422030419 | 0.190163754 | 0.460123384 |
| chr21:45309027-45309360 | -0.51740483 | 0.190200146 | 0.460193172 |
| chrX:5236084-5236485 | -0.541411804 | 0.190204609 | 0.460193172 |
| chr1:234610127-234610452 | 0.465264579 | 0.19022384 | 0.460225168 |
| chr20:54252461-54252703 | 0.460834859 | 0.190248701 | 0.460270783 |
| chr12:25591244-25591475 | -0.611031415 | 0.190271965 | 0.460312531 |
| chr3:156630458-156630663 | -0.61858655 | 0.190302227 | 0.460364756 |
| chr3:119993856-119994089 | -0.641754854 | 0.190305568 | 0.460364756 |
| chr4:53365659-53366318 | -0.43381453 | 0.190321427 | 0.460386453 |
| chr7:155069990-155070320 | -0.506220039 | 0.190326555 | 0.460386453 |
| chr18:63317974-63318432 | -0.453392292 | 0.190367541 | 0.460471058 |
| chr5:52988604-52990059 | -0.420859718 | 0.190408471 | 0.460543167 |
| chr12:123062847-123063228 | 0.481588762 | 0.190413423 | 0.460543167 |
| chr1:153945322-153946927 | 0.417468768 | 0.190418568 | 0.460543167 |
| chr5:7370868-7371152 | 0.677631072 | 0.190422911 | 0.460543167 |
| chr3:64004370-64004776 | 0.438583761 | 0.190427405 | 0.460543167 |
| chr13:98468448-98468683 | -0.754636951 | 0.190433744 | 0.460543961 |
| chr12:92928528-92929947 | -0.412297015 | 0.190494409 | 0.460661613 |
| chr12:153107-153435 | -0.528024196 | 0.190494417 | 0.460661613 |
| chr8:12659485-12659751 | 0.597280107 | 0.190508588 | 0.460681343 |
| chr6:36678563-36678833 | 0.515896294 | 0.190527053 | 0.460688635 |
| chr17:81057103-81057686 | 0.447190196 | 0.190538234 | 0.460688635 |
| chr12:67592734-67593750 | -0.432015911 | 0.190539392 | 0.460688635 |
| chr12:50132543-50132743 | 0.652067563 | 0.190541432 | 0.460688635 |
| chr4:132557732-132557970 | -0.632330961 | 0.190541666 | 0.460688635 |
| chr9:93312581-93312937 | 0.502873636 | 0.190568713 | 0.460731416 |
| chr15:91894259-91894568 | -0.618062333 | 0.190571386 | 0.460731416 |
| chr1:209354162-209354649 | -0.496535951 | 0.1905826 | 0.460735206 |
| chr20:53852623-53852864 | 0.442737935 | 0.19058498 | 0.460735206 |
| chr11:82701952-82702152 | -0.671210657 | 0.190607312 | 0.460764014 |
| chr2:23414974-23415255 | -0.560115473 | 0.190608924 | 0.460764014 |
| chr15:69118249-69118449 | 0.683390842 | 0.19063491 | 0.460800227 |

| | | | |
|---|---|---|---|
| chr9:105447499-105448684 | -0.421822301 | 0.190635932 | 0.460800227 |
| chr14:55336280-55336575 | -0.559091703 | 0.1906508 | 0.460812039 |
| chr8:72098091-72098370 | -0.622109287 | 0.190652847 | 0.460812039 |
| chr2:111897810-111898613 | 0.454393849 | 0.19067669 | 0.46084779 |
| chr2:87662280-87662765 | -0.461401393 | 0.190683426 | 0.46084779 |
| chr10:100364778-100365212 | 0.486372028 | 0.190685682 | 0.46084779 |
| chr5:173793948-173794564 | -0.474935521 | 0.190708369 | 0.460888082 |
| chr4:8460101-8460458 | 0.508101387 | 0.190716755 | 0.460893813 |
| chr3:152539239-152539720 | -0.486363283 | 0.190724849 | 0.460898835 |
| chr10:117588578-117588781 | 0.725876767 | 0.190745314 | 0.460933753 |
| chr13:27379980-27380285 | -0.498467284 | 0.190768336 | 0.460960535 |
| chr8:137258908-137259191 | -0.6161126 | 0.190778255 | 0.460960535 |
| chr2:110115713-110116675 | 0.448371654 | 0.190778282 | 0.460960535 |
| chr22:50029969-50030578 | 0.474341627 | 0.190780461 | 0.460960535 |
| chrX:123569825-123570834 | 0.423755725 | 0.190795666 | 0.460966941 |
| chr1:185733800-185734688 | -0.438506321 | 0.190797337 | 0.460966941 |
| chr18:59823679-59824095 | -0.580280242 | 0.190801161 | 0.460966941 |
| chr8:143334434-143335482 | 0.427076322 | 0.190815719 | 0.460987578 |
| chr9:76894164-76894544 | -0.57063298 | 0.190825883 | 0.460997597 |
| chr10:102432475-102432894 | 0.444907735 | 0.190836173 | 0.461004777 |
| chr16:667785-668196 | 0.469169237 | 0.190842241 | 0.461004777 |
| chr20:34516152-34516801 | 0.449819268 | 0.190846905 | 0.461004777 |
| chr14:61091059-61091264 | -0.682134171 | 0.190859869 | 0.461021559 |
| chr1:106479821-106480068 | 0.50761029 | 0.190874575 | 0.461033037 |
| chr16:28616061-28616261 | 0.627386289 | 0.190876655 | 0.461033037 |
| chr1:233520591-233520931 | -0.551867799 | 0.190914168 | 0.461066568 |
| chr8:143369212-143369751 | 0.44168914 | 0.190926191 | 0.461066568 |
| chr15:73742796-73743473 | -0.460816067 | 0.190930537 | 0.461066568 |
| chr10:13456101-13456421 | -0.661462569 | 0.19093077 | 0.461066568 |
| chr2:113965734-113965974 | -0.605122162 | 0.190931875 | 0.461066568 |
| chr5:56244779-56245023 | -0.72648322 | 0.190933787 | 0.461066568 |
| chr3:84968420-84968635 | -0.681939621 | 0.190937647 | 0.461066568 |
| chr18:3449894-3450340 | 0.442111447 | 0.190938677 | 0.461066568 |
| chr14:36069525-36069950 | 0.469829986 | 0.190946442 | 0.461070788 |
| chr2:19178335-19178784 | -0.549263228 | 0.19097298 | 0.461082943 |
| chr18:9614529-9614856 | 0.527044948 | 0.190973722 | 0.461082943 |
| chr18:110534-110861 | 0.542650301 | 0.190976818 | 0.461082943 |
| chr14:105889799-105890497 | -0.468043644 | 0.190985013 | 0.461082943 |
| chr8:119223130-119223443 | -0.496211571 | 0.190986154 | 0.461082943 |
| chr10:1594882-1595082 | 0.627349883 | 0.190990911 | 0.461082943 |
| chr20:32136065-32136376 | -0.538877258 | 0.190997718 | 0.461082943 |
| chr8:22999198-22999620 | 0.513893036 | 0.190999617 | 0.461082943 |
| chr15:43745946-43746563 | 0.44916837 | 0.191011035 | 0.461095981 |
| chr4:143558765-143560115 | -0.423323303 | 0.191052871 | 0.461182441 |
| chr3:14423194-14423562 | 0.481929059 | 0.191062642 | 0.461191499 |
| chr14:75295159-75295401 | -0.545013547 | 0.191075567 | 0.461208169 |
| chr9:38226484-38226912 | -0.524932744 | 0.191101438 | 0.461255305 |
| chr7:102490605-102490872 | -0.552259736 | 0.191113167 | 0.461255305 |
| chr21:45555432-45556406 | 0.428677241 | 0.191116699 | 0.461255305 |
| chr15:89665896-89666111 | 0.569788368 | 0.191120007 | 0.461255305 |

| | | | |
|---|---|---|---|
| chr11:125141835-125142047 | -0.650947263 | 0.191125195 | 0.461255305 |
| chr3:75442550-75442774 | -0.712738697 | 0.191186537 | 0.461374306 |
| chr2:44291701-44292034 | -0.525995944 | 0.191186547 | 0.461374306 |
| chr13:60765962-60766365 | -0.536181959 | 0.191222988 | 0.461447714 |
| chr7:4361930-4362176 | 0.721474267 | 0.191229873 | 0.461449794 |
| chr7:31899011-31899820 | -0.446164849 | 0.191244134 | 0.461469673 |
| chr8:88008055-88008318 | 0.621029205 | 0.191292219 | 0.461569739 |
| chr11:17704058-17704305 | 0.613081346 | 0.191299951 | 0.461569739 |
| chr12:50224431-50224631 | 0.682012915 | 0.191309509 | 0.461569739 |
| chr17:61455192-61456380 | 0.408146636 | 0.191318698 | 0.461569739 |
| chr5:168774286-168774540 | -0.638434489 | 0.191324918 | 0.461569739 |
| chr6:104789945-104790162 | -0.638434489 | 0.191324918 | 0.461569739 |
| chr1:197994883-197995499 | -0.471962298 | 0.191335174 | 0.461569739 |
| chr3:8768908-8769691 | 0.435665839 | 0.191336174 | 0.461569739 |
| chr12:62123690-62124012 | 0.528327799 | 0.191339819 | 0.461569739 |
| chr9:120767387-120767842 | 0.506895479 | 0.191354268 | 0.461590063 |
| chr14:58414875-58415189 | 0.519905424 | 0.191361127 | 0.461592076 |
| chr14:81565940-81566238 | 0.509446187 | 0.191376114 | 0.461613694 |
| chr12:46371299-46372126 | 0.426635688 | 0.191416291 | 0.46169607 |
| chr7:42831648-42832043 | -0.485642328 | 0.191445947 | 0.461753065 |
| chr8:120810605-120811407 | -0.433204803 | 0.191458422 | 0.461761367 |
| chr2:187580741-187580941 | 0.622981773 | 0.191461442 | 0.461761367 |
| chr17:64313361-64314266 | 0.416546681 | 0.191485699 | 0.461805334 |
| chr16:76882195-76882424 | 0.567285194 | 0.191503725 | 0.461834269 |
| chr1:202034434-202035371 | 0.415573609 | 0.191527551 | 0.461877193 |
| chr11:96242956-96243592 | -0.467379813 | 0.191538911 | 0.461890049 |
| chr3:159018668-159018963 | 0.641348777 | 0.191562322 | 0.461931965 |
| chr13:62216526-62216751 | -0.651528568 | 0.191613936 | 0.462041888 |
| chr1:170689372-170689853 | -0.505710002 | 0.191624 | 0.462051614 |
| chr16:27920282-27920895 | -0.483189577 | 0.191639502 | 0.462074452 |
| chr19:35648059-35648408 | 0.465940491 | 0.191649878 | 0.46208493 |
| chr20:25195400-25197050 | 0.413764527 | 0.191662005 | 0.462099626 |
| chr4:65007873-65008151 | -0.585609549 | 0.191730887 | 0.462251158 |
| chr1:156584917-156585122 | 0.582701216 | 0.191807602 | 0.462419249 |
| chr5:59813723-59813978 | -0.561780188 | 0.191812678 | 0.462419249 |
| chr20:41628576-41628798 | 0.622243124 | 0.191833742 | 0.462455479 |
| chr10:3539268-3539468 | 0.598320741 | 0.19184498 | 0.462468022 |
| chr15:89129469-89129837 | -0.544839804 | 0.191858962 | 0.462487177 |
| chr4:82268398-82268925 | -0.48161833 | 0.191913189 | 0.46260334 |
| chr17:75267644-75267918 | 0.485826419 | 0.191936183 | 0.462644211 |
| chr17:82283253-82283522 | 0.546800966 | 0.19194514 | 0.462651247 |
| chr7:6978241-6978469 | 0.653735228 | 0.191965376 | 0.462675381 |
| chr19:35510367-35510736 | 0.480445037 | 0.191967229 | 0.462675381 |
| chr12:101407238-101408850 | -0.416503311 | 0.191984363 | 0.462702121 |
| chr5:149345153-149346165 | -0.419900798 | 0.191996568 | 0.462714593 |
| chr10:75111166-75112293 | -0.421710209 | 0.192007024 | 0.462714593 |
| chr1:228371285-228371631 | 0.494152384 | 0.19201192 | 0.462714593 |
| chr7:33831838-33832103 | -0.549124921 | 0.192013693 | 0.462714593 |
| chr6:28225091-28225661 | 0.464024653 | 0.192048832 | 0.462767289 |
| chr2:151409562-151410819 | -0.415848502 | 0.192053072 | 0.462767289 |

| | | | |
|---|---|---|---|
| chr14:102540394-102540594 | -0.610393736 | 0.192053679 | 0.462767289 |
| chr14:44449738-44450054 | 0.511109616 | 0.192111196 | 0.462887676 |
| chr5:43432399-43432925 | -0.449640512 | 0.192118724 | 0.462887676 |
| chr5:133421449-133421679 | -0.601720459 | 0.192121765 | 0.462887676 |
| chr1:10731145-10731474 | 0.608461711 | 0.192170267 | 0.462989977 |
| chr9:126612275-126612758 | -0.490152029 | 0.192185696 | 0.46301259 |
| chr1:50308945-50309189 | 0.600414163 | 0.192254289 | 0.463147502 |
| chr10:123133064-123133433 | 0.527330225 | 0.192262943 | 0.463147502 |
| chr6:52590085-52590328 | -0.587786363 | 0.1922638 | 0.463147502 |
| chr21:14863732-14864058 | -0.611886003 | 0.192265873 | 0.463147502 |
| chr10:4174147-4174399 | 0.627335697 | 0.192290713 | 0.463192776 |
| chr8:130324065-130324371 | 0.548883406 | 0.192338224 | 0.463292657 |
| chr17:38729999-38730557 | 0.469704844 | 0.192346153 | 0.463297192 |
| chr15:82953002-82953202 | 0.613848457 | 0.192373737 | 0.463334533 |
| chr3:161852003-161852292 | -0.481131119 | 0.192377648 | 0.463334533 |
| chr19:53889649-53890269 | 0.43274732 | 0.192382792 | 0.463334533 |
| chr8:141883304-141883717 | 0.546123252 | 0.192386897 | 0.463334533 |
| chr5:337734-338107 | 0.49536653 | 0.192397643 | 0.463334533 |
| chr10:69222359-69222906 | -0.49699854 | 0.192398214 | 0.463334533 |
| chr8:60576891-60577124 | -0.739677727 | 0.192403985 | 0.463334533 |
| chr9:129362266-129362619 | -0.504506024 | 0.192420645 | 0.46334745 |
| chr8:123763257-123763683 | 0.459942084 | 0.192421444 | 0.46334745 |
| chr1:214952695-214953091 | -0.522738733 | 0.192444022 | 0.463387255 |
| chr5:155822257-155822502 | -0.65374675 | 0.19246048 | 0.463412321 |
| chr7:34708487-34708687 | 0.706419166 | 0.192495665 | 0.463482476 |
| chr1:31626550-31626865 | -0.571843057 | 0.192502063 | 0.463483316 |
| chr14:60616154-60616998 | -0.449130821 | 0.192528357 | 0.46353206 |
| chr3:58025874-58026081 | 0.63817164 | 0.192547304 | 0.463558829 |
| chr5:162520683-162521227 | -0.500759935 | 0.192551576 | 0.463558829 |
| chr7:42856558-42857577 | -0.437913438 | 0.19259265 | 0.463628978 |
| chr7:94584194-94584429 | -0.680357885 | 0.192592816 | 0.463628978 |
| chr13:52128486-52129272 | -0.443179751 | 0.192601151 | 0.463634476 |
| chr2:210224642-210225329 | -0.477889315 | 0.192619491 | 0.463661881 |
| chr18:12420266-12420592 | 0.505140784 | 0.192624638 | 0.463661881 |
| chr3:66983276-66983500 | -0.690084858 | 0.192675113 | 0.463768809 |
| chr6:46491037-46492390 | -0.433725498 | 0.192697782 | 0.463807469 |
| chr1:38009132-38009488 | -0.524176063 | 0.192708581 | 0.463807469 |
| chr17:56698889-56699903 | -0.421851716 | 0.192709334 | 0.463807469 |
| chr8:94894512-94894822 | 0.457951161 | 0.192734468 | 0.463853392 |
| chr5:154118742-154119245 | -0.481996771 | 0.192746798 | 0.463868494 |
| chr11:26994169-26994879 | -0.519515833 | 0.192772934 | 0.463910082 |
| chr11:21511861-21512172 | 0.532070546 | 0.192777931 | 0.463910082 |
| chr9:95090523-95090723 | 0.647687232 | 0.192782242 | 0.463910082 |
| chr14:47278292-47278580 | -0.537224988 | 0.192802997 | 0.463945456 |
| chr15:74374115-74374910 | 0.429503511 | 0.192813918 | 0.463957165 |
| chr13:39783091-39783291 | -0.740702027 | 0.192820112 | 0.4639575 |
| chr10:8594721-8595096 | 0.54112852 | 0.192867238 | 0.46405632 |
| chr15:79812034-79812342 | -0.542959351 | 0.192891678 | 0.464100551 |
| chr3:10286602-10287222 | 0.465478378 | 0.192899019 | 0.464103639 |
| chr17:75012335-75013126 | 0.429549597 | 0.192923706 | 0.464148461 |

| | | | |
|---|---|---|---|
| chr14:86352471-86352728 | 0.609938258 | 0.192946078 | 0.46418771 |
| chr8:94187599-94188003 | 0.49023086 | 0.192956401 | 0.464189906 |
| chr17:78127951-78128434 | 0.452517166 | 0.192959107 | 0.464189906 |
| chr19:34365460-34365926 | 0.470708517 | 0.192993657 | 0.464253773 |
| chr1:175646515-175646756 | -0.592527249 | 0.192998165 | 0.464253773 |
| chr4:46648055-46648471 | 0.481015174 | 0.193003833 | 0.464253773 |
| chr9:36174718-36175064 | -0.577416639 | 0.193031636 | 0.464306074 |
| chr5:112228711-112228953 | -0.605361109 | 0.193051716 | 0.464317038 |
| chr14:74914026-74914226 | 0.578928371 | 0.193053392 | 0.464317038 |
| chr2:65639583-65639896 | -0.547631795 | 0.193056604 | 0.464317038 |
| chr11:43504596-43504873 | 0.545925779 | 0.193060433 | 0.464317038 |
| chr7:54484576-54484811 | -0.595895215 | 0.193074296 | 0.464335805 |
| chr13:46168407-46168761 | -0.479042246 | 0.193106225 | 0.464398016 |
| chr1:1778160-1779259 | 0.434787186 | 0.193125119 | 0.464428877 |
| chr15:78338617-78338846 | 0.632816207 | 0.193154509 | 0.464470696 |
| chr3:10248139-10249000 | 0.421406735 | 0.193157909 | 0.464470696 |
| chr16:84994502-84994708 | 0.605203553 | 0.193160694 | 0.464470696 |
| chr14:71541716-71541981 | 0.665817375 | 0.193169842 | 0.464478115 |
| chr21:7926259-7926530 | 0.461765348 | 0.193199011 | 0.464533675 |
| chr19:43827102-43828056 | 0.449681333 | 0.193283212 | 0.464702438 |
| chr1:11934229-11934999 | -0.454809249 | 0.193289634 | 0.464702438 |
| chr3:124564978-124565259 | 0.541842531 | 0.193292831 | 0.464702438 |
| chr4:142462955-142463643 | -0.518798711 | 0.193293459 | 0.464702438 |
| chr12:70264449-70264682 | -0.644690016 | 0.193328503 | 0.464747731 |
| chrX:7120002-7120484 | -0.536955738 | 0.193329936 | 0.464747731 |
| chr1:215526535-215526832 | -0.5138658 | 0.193330495 | 0.464747731 |
| chr3:153785997-153786243 | -0.559747147 | 0.193346302 | 0.464771147 |
| chr4:139454375-139454627 | 0.493981988 | 0.193365003 | 0.464801521 |
| chr2:230341883-230342574 | 0.464237587 | 0.193373536 | 0.464805566 |
| chr13:49247479-49248632 | -0.424792317 | 0.193378819 | 0.464805566 |
| chr12:100170474-100170705 | -0.576168677 | 0.193392686 | 0.464824317 |
| chr16:29228140-29228390 | -0.604986074 | 0.193418146 | 0.464870929 |
| chr1:221284106-221284416 | -0.61232003 | 0.193440592 | 0.464910292 |
| chr1:154324652-154324940 | 0.476815215 | 0.193458755 | 0.464915694 |
| chr17:64055976-64056317 | 0.505442918 | 0.193463618 | 0.464915694 |
| chr1:215878714-215879006 | -0.625777799 | 0.193465136 | 0.464915694 |
| chr2:39155067-39155330 | -0.658257752 | 0.19346711 | 0.464915694 |
| chr18:59402313-59403045 | -0.438359789 | 0.193482451 | 0.464937978 |
| chr5:157328385-157328969 | -0.45501842 | 0.193515786 | 0.464993225 |
| chr1:46940358-46940626 | -0.564858681 | 0.193517579 | 0.464993225 |
| chr14:93135477-93135882 | 0.484862684 | 0.193540292 | 0.465019832 |
| chr1:46548120-46548320 | 0.576769896 | 0.19354467 | 0.465019832 |
| chr17:68656741-68656954 | -0.678029654 | 0.193556988 | 0.465019832 |
| chr15:89791105-89791305 | 0.636829583 | 0.1935585 | 0.465019832 |
| chr12:42364729-42365047 | -0.585393734 | 0.193568243 | 0.465019832 |
| chr20:57224399-57224657 | 0.431253193 | 0.193570143 | 0.465019832 |
| chr20:58890434-58891688 | 0.413211156 | 0.193571778 | 0.465019832 |
| chr1:229514753-229515013 | 0.611976764 | 0.193577204 | 0.465019832 |
| chr18:76657403-76657699 | -0.717566094 | 0.193593767 | 0.465045039 |
| chr2:33598861-33599959 | -0.410607476 | 0.1936009 | 0.465047594 |

| | | | |
|---|---|---|---|
| chr17:57440556-57441471 | -0.432002392 | 0.193616782 | 0.465066867 |
| chr3:35908414-35908625 | 0.699958888 | 0.193628641 | 0.465066867 |
| chr9:113349360-113349913 | 0.461994903 | 0.193632311 | 0.465066867 |
| chr9:35072500-35073094 | -0.430439993 | 0.193633202 | 0.465066867 |
| chr17:62275559-62275915 | -0.577856846 | 0.193640256 | 0.465069232 |
| chr6:13797490-13797863 | 0.498230521 | 0.193648667 | 0.465074854 |
| chr22:46006386-46006680 | 0.531550144 | 0.193669311 | 0.465091399 |
| chrX:94961787-94962053 | -0.610644105 | 0.193671029 | 0.465091399 |
| chr15:31157709-31158084 | 0.543281186 | 0.193673766 | 0.465091399 |
| chr5:151312816-151313041 | -0.634802514 | 0.193684253 | 0.465093414 |
| chr3:2166595-2166870 | -0.568102125 | 0.193686745 | 0.465093414 |
| chr8:132215539-132216021 | -0.482390778 | 0.193696138 | 0.465100045 |
| chr17:61263084-61263471 | 0.429411013 | 0.193701646 | 0.465100045 |
| chr5:144848499-144848825 | -0.5134936 | 0.193708842 | 0.465102749 |
| chr13:39037536-39038882 | -0.409535452 | 0.193733153 | 0.4651247 |
| chr6:153494686-153495006 | -0.619603851 | 0.19373383 | 0.4651247 |
| chr6:85592896-85593096 | 0.55401276 | 0.193740081 | 0.4651247 |
| chr20:63552754-63553747 | -0.423352184 | 0.193742266 | 0.4651247 |
| chr3:150504592-150504820 | 0.645624078 | 0.193755426 | 0.46514172 |
| chr17:50757168-50757368 | -0.691380415 | 0.193764923 | 0.465149944 |
| chr1:214727001-214727626 | -0.464672995 | 0.193815055 | 0.465242053 |
| chr10:50552828-50553097 | -0.569941129 | 0.193815907 | 0.465242053 |
| chr8:33484579-33485539 | -0.423264005 | 0.193823207 | 0.465242053 |
| chr19:52786241-52787000 | 0.461276047 | 0.19382758 | 0.465242053 |
| chr14:93074174-93074446 | 0.594658572 | 0.193835745 | 0.465242892 |
| chr10:5715773-5716198 | 0.462051162 | 0.193840073 | 0.465242892 |
| chr12:112418533-112419604 | 0.422339463 | 0.193853297 | 0.465260058 |
| chr13:47511710-47511919 | 0.600909509 | 0.193869974 | 0.465285508 |
| chr2:85401867-85402104 | -0.596224566 | 0.193880413 | 0.465295165 |
| chr11:77088637-77088999 | 0.462482771 | 0.193886143 | 0.465295165 |
| chr2:106192970-106193209 | -0.539089078 | 0.193907885 | 0.465332769 |
| chr5:138694114-138694510 | 0.507405693 | 0.193919386 | 0.465345794 |
| chr13:51583699-51585064 | -0.407311588 | 0.193939785 | 0.46538017 |
| chr14:44961210-44961410 | 0.496508242 | 0.193946898 | 0.465382663 |
| chr3:124550535-124550748 | 0.592566919 | 0.193962845 | 0.465406355 |
| chr6:96499898-96500159 | -0.640489613 | 0.193997625 | 0.465470051 |
| chr2:79936355-79936685 | -0.510928545 | 0.194013985 | 0.465470051 |
| chr11:59565780-59566627 | -0.424384354 | 0.194019108 | 0.465470051 |
| chr11:3745203-3745436 | 0.648841515 | 0.194019371 | 0.465470051 |
| chr12:13739715-13739952 | -0.617404618 | 0.194019766 | 0.465470051 |
| chrX:98720424-98720641 | -0.616729244 | 0.194032069 | 0.465484992 |
| chr6:57274124-57274366 | -0.605847461 | 0.194045771 | 0.465503288 |
| chr5:16787358-16787562 | 0.594501318 | 0.194058071 | 0.465509633 |
| chr19:10700961-10701931 | 0.443938972 | 0.194060566 | 0.465509633 |
| chr9:134379008-134379254 | 0.52705161 | 0.194118371 | 0.465633716 |
| chr17:80575382-80575672 | -0.523544869 | 0.194138496 | 0.465667413 |
| chr8:126453913-126454320 | -0.527579681 | 0.19417021 | 0.465728904 |
| chr17:52677912-52678775 | 0.431654294 | 0.194179995 | 0.465734366 |
| chr3:169229519-169229719 | 0.695869306 | 0.194184644 | 0.465734366 |
| chr17:36703865-36704065 | 0.695306983 | 0.194197109 | 0.465749683 |

| | | | |
|---|---|---|---|
| chr4:42152251-42152511 | 0.640118925 | 0.194223139 | 0.465785005 |
| chr5:10030251-10030865 | -0.454995443 | 0.194223994 | 0.465785005 |
| chr11:66345005-66345356 | 0.453340975 | 0.194237246 | 0.465802206 |
| chr2:55860030-55860230 | -0.678011967 | 0.194252663 | 0.465824597 |
| chr15:66293196-66294317 | 0.430600626 | 0.194287644 | 0.465887583 |
| chr22:17257368-17257896 | -0.526686573 | 0.194296187 | 0.465887583 |
| chr15:45684404-45685041 | 0.441234539 | 0.194297169 | 0.465887583 |
| chr16:82358647-82359148 | 0.480961396 | 0.194311604 | 0.465907615 |
| chrX:93419651-93419997 | -0.615350201 | 0.19432527 | 0.465917907 |
| chr20:58754260-58754467 | 0.558334099 | 0.194328058 | 0.465917907 |
| chr8:100017936-100018278 | -0.526194462 | 0.19433851 | 0.465928388 |
| chr18:58942577-58942857 | -0.650303702 | 0.194359682 | 0.465964566 |
| chr17:51707261-51707525 | -0.520100273 | 0.194409077 | 0.466068405 |
| chr3:38165806-38166147 | 0.490123153 | 0.194421792 | 0.466084305 |
| chr17:75238738-75239443 | 0.45488265 | 0.194438287 | 0.466095508 |
| chr3:14546856-14547056 | -0.648154425 | 0.194438632 | 0.466095508 |
| chr9:129197794-129198041 | 0.622397807 | 0.194453952 | 0.46611765 |
| chr9:136992590-136992824 | 0.507907934 | 0.194472134 | 0.466146651 |
| chr4:100779760-100780018 | -0.5960848 | 0.194502567 | 0.466191061 |
| chr9:87766669-87767221 | 0.44221117 | 0.19450283 | 0.466191061 |
| chr22:34055911-34056111 | -0.688141499 | 0.194523636 | 0.466226346 |
| chr2:54657615-54657938 | -0.494534469 | 0.194538346 | 0.466247018 |
| chr20:51456389-51457112 | -0.421417003 | 0.194552542 | 0.466256331 |
| chr9:71758855-71759185 | -0.532635778 | 0.194554402 | 0.466256331 |
| chr11:119533972-119534363 | 0.521545918 | 0.194573058 | 0.466277051 |
| chr4:13544626-13545311 | 0.445881989 | 0.194575219 | 0.466277051 |
| chr10:35546871-35547238 | -0.542130851 | 0.19459293 | 0.466304909 |
| chr3:27943184-27943502 | -0.532376594 | 0.194608826 | 0.466328418 |
| chr14:100375554-100375781 | 0.533632432 | 0.194621064 | 0.466343158 |
| chr13:30551253-30551539 | -0.545619161 | 0.194641412 | 0.46637733 |
| chrX:86370253-86370596 | -0.614167893 | 0.194652838 | 0.466390124 |
| chr14:44149582-44149979 | -0.470821376 | 0.194668133 | 0.466412186 |
| chr12:69388429-69388676 | 0.563161381 | 0.194680581 | 0.466427032 |
| chr3:6262075-6262361 | -0.605286522 | 0.194686504 | 0.466427032 |
| chr10:90771465-90771732 | -0.679588531 | 0.194700125 | 0.46644508 |
| chr17:39235744-39236179 | -0.515426082 | 0.194750446 | 0.466551047 |
| chr8:141368907-141369274 | -0.50508409 | 0.19476228 | 0.466564809 |
| chr1:61149339-61149712 | 0.491354109 | 0.194770145 | 0.466569062 |
| chr3:50274303-50274555 | 0.56503482 | 0.194801306 | 0.466629121 |
| chr5:114789851-114790074 | -0.639314037 | 0.194823474 | 0.466667632 |
| chr17:65035286-65035564 | -0.603496157 | 0.194833279 | 0.4666697 |
| chr7:137656357-137656585 | -0.600175328 | 0.194836518 | 0.4666697 |
| chr5:91279701-91280143 | -0.482688022 | 0.194871821 | 0.466700314 |
| chr2:67476863-67477106 | -0.604727741 | 0.194877287 | 0.466700314 |
| chr17:76868238-76868498 | 0.546682817 | 0.194879485 | 0.466700314 |
| chr11:75490412-75490732 | 0.535153876 | 0.194881013 | 0.466700314 |
| chr2:42168947-42169943 | -0.41843948 | 0.194883554 | 0.466700314 |
| chr4:88673053-88673296 | 0.621990937 | 0.194889309 | 0.466700314 |
| chr11:14907304-14908119 | -0.432778979 | 0.194892517 | 0.466700314 |
| chr6:154509371-154511211 | -0.409042754 | 0.194898027 | 0.466700314 |

| | | | |
|---|---|---|---|
| chr2:106330694-106330921 | 0.602976558 | 0.194914407 | 0.466724951 |
| chr22:20606096-20606668 | -0.443398574 | 0.194925878 | 0.466733942 |
| chr21:42364547-42364747 | -0.539615895 | 0.194932515 | 0.466733942 |
| chr16:801158-801388 | 0.561640044 | 0.194936436 | 0.466733942 |
| chr14:68911641-68911974 | 0.489546789 | 0.194966083 | 0.466790338 |
| chr16:75250800-75251051 | 0.473463271 | 0.194976916 | 0.466801688 |
| chr17:36601217-36602073 | 0.43919565 | 0.194992925 | 0.466825013 |
| chr17:67942478-67942698 | -0.582940447 | 0.194998843 | 0.466825013 |
| chr11:9264706-9264906 | -0.675850956 | 0.195009434 | 0.466835781 |
| chr16:2459634-2460384 | 0.436995365 | 0.195094269 | 0.467024278 |
| chr6:33788252-33788684 | 0.499357825 | 0.195118765 | 0.467052087 |
| chr10:43044598-43045008 | 0.488004733 | 0.19512274 | 0.467052087 |
| chr3:157182951-157183201 | -0.579898093 | 0.195124173 | 0.467052087 |
| chr14:77111095-77111449 | -0.500541537 | 0.195167071 | 0.467140177 |
| chr6:142026293-142026874 | -0.479396926 | 0.195186441 | 0.467171945 |
| chr2:73156668-73157179 | -0.489409267 | 0.195207716 | 0.467208272 |
| chr17:60509074-60509278 | 0.4673017 | 0.195293727 | 0.467399529 |
| chr6:30238524-30238733 | 0.630216373 | 0.195308182 | 0.467419525 |
| chr5:64384824-64385083 | -0.572022914 | 0.195318127 | 0.467423086 |
| chr2:42196733-42197096 | -0.491926559 | 0.195324717 | 0.467423086 |
| chr3:196807692-196808150 | -0.545071127 | 0.195327971 | 0.467423086 |
| chr10:76500-76737 | 0.569052711 | 0.195356947 | 0.467477826 |
| chr1:158117851-158118232 | -0.475932484 | 0.195373181 | 0.467502072 |
| chr3:99824035-99824267 | -0.682747295 | 0.195389156 | 0.467525698 |
| chr6:19892100-19892765 | 0.456397839 | 0.195418076 | 0.467580295 |
| chr12:49644613-49644818 | 0.581791407 | 0.195432039 | 0.467595571 |
| chr7:101165425-101165781 | -0.479953712 | 0.195436665 | 0.467595571 |
| chr15:78056814-78057292 | -0.520645818 | 0.195459597 | 0.467635835 |
| chr2:8074018-8074269 | -0.675299163 | 0.19547077 | 0.467645231 |
| chr1:180923133-180923333 | 0.544324716 | 0.195478529 | 0.467645231 |
| chr12:111624640-111624840 | 0.666055185 | 0.195481834 | 0.467645231 |
| chr8:125729335-125729644 | -0.493669727 | 0.19550367 | 0.46768001 |
| chr5:65623923-65625440 | -0.41041874 | 0.19550858 | 0.46768001 |
| chr9:129496655-129496965 | 0.480005931 | 0.195522622 | 0.467687895 |
| chr5:7897037-7897376 | -0.505897169 | 0.195529822 | 0.467687895 |
| chr3:8417449-8417667 | 0.545059968 | 0.195532283 | 0.467687895 |
| chr22:26428682-26428928 | 0.580235745 | 0.195536291 | 0.467687895 |
| chr1:225845606-225846055 | 0.474877597 | 0.195572609 | 0.467760159 |
| chr21:43609695-43610112 | 0.489621658 | 0.195588947 | 0.467784632 |
| chr2:239927433-239927824 | 0.575810892 | 0.19560358 | 0.467804794 |
| chr14:24095392-24095592 | 0.53002233 | 0.195609587 | 0.467804794 |
| chr11:16959960-16960664 | -0.476565971 | 0.195661843 | 0.467904452 |
| chr1:83860403-83861314 | -0.420561517 | 0.195663472 | 0.467904452 |
| chr20:50603010-50603296 | 0.509298898 | 0.195775868 | 0.468158622 |
| chr1:87331253-87331678 | 0.47052539 | 0.195796238 | 0.46819272 |
| chr10:4982155-4982425 | 0.533171048 | 0.195806851 | 0.468200686 |
| chr1:99004988-99005222 | 0.657486052 | 0.19581179 | 0.468200686 |
| chr19:9075564-9075938 | -0.544333383 | 0.195817939 | 0.468200777 |
| chr8:19291512-19291803 | 0.556272047 | 0.195885102 | 0.46834592 |
| chr4:83010728-83010937 | 0.499719693 | 0.195892294 | 0.46834592 |

| | | | |
|---|---|---|---|
| chr12:77023697-77023978 | 0.618073591 | 0.195901851 | 0.46834592 |
| chr16:23435486-23435752 | 0.592164774 | 0.195903092 | 0.46834592 |
| chr10:75853170-75853401 | -0.67550405 | 0.195915075 | 0.468351239 |
| chr3:153052092-153052480 | 0.495683364 | 0.195917542 | 0.468351239 |
| chr7:64892482-64892682 | -0.726287719 | 0.195943172 | 0.468397894 |
| chr4:56362658-56363493 | -0.441863974 | 0.1959698 | 0.468446934 |
| chr4:54233169-54233643 | 0.471492456 | 0.195981533 | 0.468459286 |
| chr17:76279147-76279377 | 0.537008895 | 0.195987195 | 0.468459286 |
| chr15:57527516-57527870 | -0.53081821 | 0.196006824 | 0.468491588 |
| chr17:27609667-27610087 | -0.521544517 | 0.19602759 | 0.468526608 |
| chr13:43772541-43772756 | -0.708846067 | 0.196083078 | 0.468644606 |
| chr10:123796805-123797005 | 0.721261479 | 0.196093205 | 0.468644606 |
| chr14:66662441-66662675 | 0.601174199 | 0.196097428 | 0.468644606 |
| chr19:8113043-8113409 | -0.541316712 | 0.196102209 | 0.468644606 |
| chr12:62934252-62935487 | 0.410966392 | 0.196107541 | 0.468644606 |
| chr7:80139289-80139495 | -0.689587626 | 0.196121894 | 0.468664288 |
| chr1:106590492-106590716 | 0.468053239 | 0.196160346 | 0.468735796 |
| chr14:24052386-24052710 | 0.477446404 | 0.196164053 | 0.468735796 |
| chr17:39634456-39634656 | 0.653707724 | 0.196199793 | 0.468806578 |
| chr16:68529783-68530681 | 0.432102263 | 0.196210801 | 0.468818261 |
| chrX:108641659-108641922 | -0.656894015 | 0.196237434 | 0.468867275 |
| chr3:63047288-63047593 | 0.457199365 | 0.196250931 | 0.468875262 |
| chr5:123022636-123022978 | -0.635276419 | 0.196253015 | 0.468875262 |
| chr18:36809601-36809905 | -0.554386239 | 0.196303985 | 0.468982412 |
| chr1:203776747-203776966 | -0.574212622 | 0.196334896 | 0.468994847 |
| chr10:84762748-84762987 | -0.673872638 | 0.196335398 | 0.468994847 |
| chr16:68064720-68065003 | -0.564763181 | 0.196335757 | 0.468994847 |
| chr20:52438330-52438823 | -0.508744372 | 0.196341472 | 0.468994847 |
| chr1:201730821-201731214 | 0.499235731 | 0.196345179 | 0.468994847 |
| chr6:107238070-107238300 | 0.69578564 | 0.196356327 | 0.468994847 |
| chr1:27508249-27508511 | -0.640251964 | 0.196359613 | 0.468994847 |
| chr16:18801187-18802120 | 0.414800175 | 0.196371058 | 0.468994847 |
| chr7:30504713-30505386 | 0.424122237 | 0.196378255 | 0.468994847 |
| chr1:186541032-186541310 | -0.595465452 | 0.196380092 | 0.468994847 |
| chr10:15444260-15444529 | 0.617219103 | 0.196401076 | 0.468994847 |
| chr4:15964201-15964512 | 0.514336417 | 0.196407919 | 0.468994847 |
| chr11:6297903-6298134 | -0.63007327 | 0.196408054 | 0.468994847 |
| chr5:151298107-151298399 | -0.529084411 | 0.196412253 | 0.468994847 |
| chr5:21225290-21225947 | -0.47118175 | 0.196412667 | 0.468994847 |
| chr19:10510636-10510836 | 0.670934313 | 0.196416892 | 0.468994847 |
| chr8:22756083-22756340 | -0.594737194 | 0.196417747 | 0.468994847 |
| chr8:6779662-6779924 | -0.582819326 | 0.196424052 | 0.468994847 |
| chr17:81037333-81037717 | 0.45118198 | 0.196434316 | 0.468994847 |
| chr2:121077014-121077275 | 0.495152311 | 0.196442736 | 0.468994847 |
| chr10:1464992-1465428 | -0.487061725 | 0.196449002 | 0.468994847 |
| chr1:151575040-151575240 | 0.565741936 | 0.19645282 | 0.468994847 |
| chr1:112577912-112578185 | -0.642930978 | 0.196455369 | 0.468994847 |
| chr17:50482461-50482768 | 0.474883708 | 0.196459406 | 0.468994847 |
| chr8:106877791-106878045 | 0.670968344 | 0.196462212 | 0.468994847 |
| chr20:31517370-31517689 | 0.486361816 | 0.19647434 | 0.469009187 |

| | | | |
|---|---|---|---|
| chr17:47884066-47884717 | -0.438888408 | 0.196485684 | 0.469021654 |
| chr18:61435139-61435376 | -0.621994385 | 0.196507521 | 0.469059166 |
| chr5:61920022-61920445 | -0.571811271 | 0.196518724 | 0.469071295 |
| chr1:176878607-176878936 | -0.545377231 | 0.196531019 | 0.469086031 |
| chr7:43869187-43869890 | -0.420774816 | 0.196546011 | 0.4691072 |
| chr16:11255705-11256881 | 0.414149843 | 0.196566485 | 0.469141453 |
| chr13:31055729-31056029 | 0.553029966 | 0.19666565 | 0.469363509 |
| chr5:32852616-32853007 | -0.516572641 | 0.196680886 | 0.469373147 |
| chr15:57340745-57340954 | 0.674378979 | 0.19668194 | 0.469373147 |
| chr11:70382016-70382247 | -0.632565405 | 0.196697307 | 0.4693952 |
| chr20:23050100-23050560 | 0.47560161 | 0.196703625 | 0.469395657 |
| chr11:31509123-31510343 | -0.411758686 | 0.19671293 | 0.469401825 |
| chr11:29334287-29334545 | -0.615478337 | 0.196723086 | 0.469401825 |
| chr14:88463170-88463690 | 0.47085627 | 0.196724588 | 0.469401825 |
| chr1:47630695-47630895 | -0.64941192 | 0.196735205 | 0.469412539 |
| chr10:79116186-79116570 | -0.54509984 | 0.196754414 | 0.469429083 |
| chr2:196068176-196069416 | -0.410845621 | 0.19675628 | 0.469429083 |
| chr11:62680804-62681267 | 0.464602733 | 0.196760599 | 0.469429083 |
| chr14:80364351-80364630 | -0.568747596 | 0.196766645 | 0.469429083 |
| chr3:45226249-45226539 | -0.663731506 | 0.196782292 | 0.469451797 |
| chr3:39378149-39378440 | 0.512377783 | 0.196802149 | 0.469473012 |
| chr18:58541215-58541504 | -0.609510388 | 0.19680344 | 0.469473012 |
| chr14:52694839-52696161 | 0.414977512 | 0.196827089 | 0.46951481 |
| chr7:80542395-80542826 | -0.523751661 | 0.196841005 | 0.469533389 |
| chr9:86948496-86948764 | 0.537101717 | 0.196856885 | 0.46954239 |
| chr10:88389515-88389751 | -0.666511911 | 0.196857035 | 0.46954239 |
| chr13:26529489-26529711 | -0.622831083 | 0.196875647 | 0.469572165 |
| chr14:59244453-59244952 | 0.482404708 | 0.196918511 | 0.469636207 |
| chr11:125942044-125942313 | 0.543701824 | 0.196923211 | 0.469636207 |
| chr12:74316957-74317321 | 0.500975662 | 0.196931043 | 0.469636207 |
| chr17:10226904-10227191 | 0.581863374 | 0.196935173 | 0.469636207 |
| chr2:34012755-34013056 | -0.551236018 | 0.196940217 | 0.469636207 |
| chr6:89644928-89645209 | -0.596792696 | 0.196941092 | 0.469636207 |
| chr3:13895097-13895743 | 0.420415533 | 0.196946299 | 0.469636207 |
| chr16:21599011-21600183 | 0.418904946 | 0.196951532 | 0.469636207 |
| chr6:141562220-141562719 | -0.512932956 | 0.196967353 | 0.469643867 |
| chr13:46321644-46322347 | -0.472965564 | 0.196972369 | 0.469643867 |
| chr5:179167153-179167614 | -0.48558436 | 0.196976217 | 0.469643867 |
| chr1:223363031-223363801 | -0.439695599 | 0.196979261 | 0.469643867 |
| chr3:72628029-72628534 | -0.432263394 | 0.196992768 | 0.469645921 |
| chr4:44629285-44629553 | -0.656150977 | 0.197006573 | 0.469645921 |
| chrX:110822240-110822503 | -0.68327817 | 0.197018685 | 0.469645921 |
| chr6:46381534-46381868 | -0.652133596 | 0.197024979 | 0.469645921 |
| chr1:81800776-81801551 | -0.430348085 | 0.197025309 | 0.469645921 |
| chr6:6851028-6851338 | 0.60303438 | 0.1970359 | 0.469645921 |
| chr8:101266145-101266345 | 0.66122624 | 0.197038593 | 0.469645921 |
| chr4:89112272-89112525 | 0.494498801 | 0.197045704 | 0.469645921 |
| chr3:86389770-86390218 | 0.492004909 | 0.197047169 | 0.469645921 |
| chrX:47200457-47200657 | 0.711153044 | 0.19705164 | 0.469645921 |
| chr3:20265239-20265503 | -0.630017208 | 0.197052464 | 0.469645921 |

| | | | |
|---|---|---|---|
| chr1:117733608-117733840 | 0.636091061 | 0.197059649 | 0.469645921 |
| chr15:98805792-98806034 | 0.540000287 | 0.197059805 | 0.469645921 |
| chr2:172704184-172704702 | -0.522897631 | 0.197066093 | 0.4696463 |
| chr2:66553484-66553980 | -0.53459783 | 0.197104579 | 0.469723409 |
| chr6:32195072-32195272 | 0.525451413 | 0.197125413 | 0.469739833 |
| chr1:27044631-27044944 | 0.577679136 | 0.197127945 | 0.469739833 |
| chr5:151779772-151780162 | -0.59885134 | 0.197129863 | 0.469739833 |
| chr17:57348274-57348580 | -0.560178921 | 0.197142184 | 0.469743463 |
| chr13:34364702-34364988 | -0.587077721 | 0.197143647 | 0.469743463 |
| chr17:82838441-82838667 | -0.531524928 | 0.19717213 | 0.46979672 |
| chr6:116444234-116444449 | 0.624853486 | 0.197181627 | 0.469804739 |
| chr22:22564794-22565017 | -0.702510319 | 0.197201388 | 0.469828736 |
| chr4:128803331-128803708 | -0.568462612 | 0.197203962 | 0.469828736 |
| chr17:82528621-82529091 | 0.482670772 | 0.197227658 | 0.469870581 |
| chr8:135335311-135335757 | -0.50921219 | 0.197250005 | 0.46990921 |
| chr1:30684981-30685896 | 0.42612026 | 0.197257149 | 0.469911617 |
| chrX:120302591-120302791 | 0.660848595 | 0.197268745 | 0.469917639 |
| chr2:9261739-9261949 | 0.669590486 | 0.197271943 | 0.469917639 |
| chr1:184492709-184493051 | 0.61803858 | 0.197296143 | 0.469960677 |
| chr20:54557102-54557368 | 0.44957106 | 0.197304579 | 0.469966161 |
| chr4:139509756-139510013 | 0.547666774 | 0.197321955 | 0.469992939 |
| chr6:46283225-46283457 | -0.714565904 | 0.197339116 | 0.47000872 |
| chr16:2964889-2965249 | 0.454683597 | 0.197342231 | 0.47000872 |
| chr1:54153858-54154108 | 0.615918453 | 0.197349186 | 0.47000872 |
| chr10:73879713-73879959 | 0.522433817 | 0.197353117 | 0.47000872 |
| chr5:115262067-115263276 | -0.422839305 | 0.197382307 | 0.470063628 |
| chr8:96163601-96163968 | 0.518398774 | 0.197415226 | 0.470127412 |
| chr20:47676329-47676615 | 0.42421726 | 0.197459276 | 0.470193974 |
| chr1:107318964-107319317 | 0.434323685 | 0.197461622 | 0.470193974 |
| chr12:9708222-9708508 | 0.614675046 | 0.197462543 | 0.470193974 |
| chr8:67866690-67866937 | -0.627475923 | 0.197467723 | 0.470193974 |
| chr17:65074793-65075033 | -0.524576735 | 0.197485592 | 0.470217709 |
| chr7:70631358-70631643 | -0.675357381 | 0.197489965 | 0.470217709 |
| chr12:101855367-101855821 | -0.526993375 | 0.197499011 | 0.470224636 |
| chr3:120416803-120417908 | -0.41229182 | 0.197507871 | 0.470231118 |
| chrX:15738756-15738956 | 0.556534073 | 0.197519613 | 0.470244463 |
| chrX:148862608-148862853 | 0.639318223 | 0.197527146 | 0.470247787 |
| chr2:191358132-191358956 | 0.439174948 | 0.197550364 | 0.470281444 |
| chr14:33985129-33985394 | 0.553420752 | 0.197557249 | 0.470281444 |
| chr20:47740262-47740597 | 0.421879264 | 0.197559697 | 0.470281444 |
| chr9:137217185-137217563 | 0.460114188 | 0.197624166 | 0.470420102 |
| chr4:103935198-103935491 | -0.540515 | 0.197630225 | 0.470420102 |
| chr6:130760053-130760412 | -0.526683572 | 0.197636375 | 0.470420127 |
| chr8:120956692-120957226 | -0.436789555 | 0.197656987 | 0.470435776 |
| chr4:4859477-4859760 | 0.531857965 | 0.197659806 | 0.470435776 |
| chr13:21613835-21614035 | -0.713328192 | 0.197665732 | 0.470435776 |
| chr9:4933324-4933524 | 0.669205232 | 0.197672177 | 0.470435776 |
| chr5:157718560-157718849 | -0.52196666 | 0.197673648 | 0.470435776 |
| chr20:48359719-48360042 | 0.439694092 | 0.19769434 | 0.470470408 |
| chr14:23984981-23985351 | 0.527346945 | 0.197710562 | 0.470494399 |

| | | | |
|---|---|---|---|
| chr5:88690804-88691323 | -0.459658567 | 0.197736298 | 0.47054103 |
| chr1:182514933-182515235 | 0.578793822 | 0.197749849 | 0.470550555 |
| chr4:113851907-113852249 | -0.608232287 | 0.197752583 | 0.470550555 |
| chr20:11205224-11205473 | -0.576359683 | 0.197763726 | 0.470562456 |
| chr4:1793143-1793343 | 0.594018571 | 0.197795922 | 0.470623644 |
| chr5:172697231-172697837 | -0.439952296 | 0.197801726 | 0.470623644 |
| chr1:5481993-5482255 | 0.643367862 | 0.197820158 | 0.470649457 |
| chr12:65911348-65911561 | -0.700892104 | 0.19782486 | 0.470649457 |
| chr2:113437477-113438034 | 0.494688846 | 0.197837275 | 0.470664381 |
| chr4:55366787-55367057 | -0.686374467 | 0.197863412 | 0.470711946 |
| chr1:53541809-53542037 | 0.570677652 | 0.197893881 | 0.470769813 |
| chr15:99414907-99415182 | -0.586130309 | 0.197909724 | 0.470792886 |
| chr2:160221651-160222230 | 0.456896361 | 0.197918123 | 0.47079825 |
| chr1:183543649-183543954 | 0.496430472 | 0.197931422 | 0.470812887 |
| chr2:68447820-68448473 | -0.514242158 | 0.197936566 | 0.470812887 |
| chr7:101251810-101252635 | -0.420646386 | 0.197948369 | 0.470813308 |
| chr5:181151298-181151542 | 0.535182551 | 0.197949032 | 0.470813308 |
| chr1:111507634-111508527 | -0.435287957 | 0.197969238 | 0.470846752 |
| chr1:205774691-205775841 | -0.414557808 | 0.197979741 | 0.470857116 |
| chr1:33036040-33037240 | 0.416432249 | 0.198001146 | 0.470893407 |
| chr4:120843848-120844195 | -0.511530573 | 0.198022131 | 0.470928698 |
| chr5:136212336-136212536 | 0.600886145 | 0.198031601 | 0.470936602 |
| chr17:77477466-77477783 | -0.494943245 | 0.198042646 | 0.470948253 |
| chr3:44510450-44510806 | 0.529248561 | 0.198059117 | 0.470962669 |
| chr11:4175706-4176112 | -0.552775788 | 0.198061002 | 0.470962669 |
| chr5:113513396-113514379 | -0.419013317 | 0.198084114 | 0.47099142 |
| chr7:6871491-6871927 | 0.479715445 | 0.198085626 | 0.47099142 |
| chr2:171999612-172000333 | -0.431085318 | 0.198091534 | 0.47099142 |
| chr1:150876117-150877163 | 0.406977627 | 0.198107922 | 0.471015769 |
| chr2:232292718-232292929 | -0.668953061 | 0.198188747 | 0.471193316 |
| chr3:8223945-8224297 | 0.574751809 | 0.198213525 | 0.471237603 |
| chr14:72894007-72894518 | 0.429314105 | 0.198243756 | 0.471294852 |
| chr11:62337318-62337841 | -0.50408233 | 0.198297642 | 0.471399698 |
| chr7:75878731-75879514 | -0.437310829 | 0.198300163 | 0.471399698 |
| chr5:157026984-157027368 | 0.49831434 | 0.198315393 | 0.471421278 |
| chr22:30252370-30252965 | -0.468793402 | 0.198330879 | 0.471438603 |
| chr3:127696175-127696375 | 0.600001957 | 0.198342239 | 0.471438603 |
| chr8:143015876-143016253 | -0.459049027 | 0.198346449 | 0.471438603 |
| chr16:46554400-46554679 | -0.571807646 | 0.198353374 | 0.471438603 |
| chr17:81927617-81928168 | 0.430417459 | 0.198353445 | 0.471438603 |
| chr11:18589099-18589310 | -0.614939712 | 0.198363694 | 0.471448339 |
| chr5:150045627-150045875 | -0.605570132 | 0.198375791 | 0.471462465 |
| chr3:196684993-196685381 | -0.584196088 | 0.198383016 | 0.471465013 |
| chr5:136365247-136365449 | -0.681204996 | 0.198394811 | 0.471478375 |
| chr20:62668517-62668758 | 0.519089232 | 0.198400945 | 0.471478375 |
| chr6:878035-878483 | -0.571869612 | 0.198416141 | 0.471499748 |
| chr3:123979805-123980381 | -0.480359291 | 0.198422246 | 0.471499748 |
| chr3:150286408-150286921 | 0.463533598 | 0.198478962 | 0.471619891 |
| chr3:161080790-161080990 | 0.694961156 | 0.198500684 | 0.471656841 |
| chr6:66482608-66482960 | -0.567516861 | 0.19851775 | 0.471656841 |

| | | | |
|---|---|---|---|
| chr5:120817101-120817355 | -0.575498056 | 0.198524051 | 0.471656841 |
| chr16:28894140-28894871 | 0.4832144 | 0.198525422 | 0.471656841 |
| chr21:43374975-43375175 | 0.616968512 | 0.198536838 | 0.471656841 |
| chr2:176140286-176141332 | -0.419655684 | 0.198537656 | 0.471656841 |
| chr7:105934651-105934917 | -0.553349634 | 0.198546608 | 0.471656841 |
| chr1:247090373-247090885 | -0.47225086 | 0.198549359 | 0.471656841 |
| chr12:18430505-18430740 | -0.596263435 | 0.198549913 | 0.471656841 |
| chr4:112075176-112075542 | -0.502980091 | 0.198559464 | 0.471660327 |
| chr11:61158645-61158951 | 0.475633051 | 0.198564669 | 0.471660327 |
| chr2:241610036-241610661 | 0.44776658 | 0.198571407 | 0.471660327 |
| chr6:154934666-154934915 | 0.642365492 | 0.198576003 | 0.471660327 |
| chr16:80014257-80014655 | -0.543137721 | 0.198600983 | 0.471687576 |
| chr1:246798026-246798244 | -0.543351615 | 0.198604448 | 0.471687576 |
| chr8:66026425-66027132 | -0.426300082 | 0.198605943 | 0.471687576 |
| chr3:27228816-27229126 | -0.555161337 | 0.198640501 | 0.471748857 |
| chr10:32095087-32095487 | -0.537583831 | 0.198645899 | 0.471748857 |
| chr18:48538819-48539286 | -0.445812048 | 0.198650216 | 0.471748857 |
| chr9:128438445-128439298 | 0.448362465 | 0.198656753 | 0.471749759 |
| chr19:35269508-35269833 | 0.484765839 | 0.198669374 | 0.471765109 |
| chr1:35930321-35930521 | 0.603889988 | 0.198677493 | 0.471769768 |
| chr2:161586605-161586830 | -0.670215361 | 0.198692285 | 0.471790272 |
| chr2:59176831-59177031 | -0.695082507 | 0.198699394 | 0.47179253 |
| chr10:91043319-91043519 | -0.650368813 | 0.19873646 | 0.471865917 |
| chr20:57132307-57132553 | 0.43210433 | 0.198749678 | 0.471877552 |
| chr4:172892767-172893051 | -0.610960216 | 0.198754632 | 0.471877552 |
| chr8:121189209-121189409 | 0.650105029 | 0.198759836 | 0.471877552 |
| chr12:100996708-100996932 | -0.676854045 | 0.198781683 | 0.471914798 |
| chr2:214465811-214466090 | -0.603003921 | 0.19879374 | 0.471928801 |
| chr3:162285708-162286050 | 0.559461709 | 0.198864861 | 0.472083012 |
| chr7:36120118-36120797 | -0.465458921 | 0.198882228 | 0.472094022 |
| chr2:216682500-216682788 | 0.520645852 | 0.198887843 | 0.472094022 |
| chr1:201545925-201546514 | 0.438543216 | 0.198901908 | 0.472094022 |
| chr8:89718264-89718594 | 0.469917376 | 0.198906225 | 0.472094022 |
| chr5:78046061-78046278 | -0.716121639 | 0.198913693 | 0.472094022 |
| chr2:215294913-215295597 | -0.469002735 | 0.198916828 | 0.472094022 |
| chr8:125126021-125126221 | 0.624967485 | 0.198917891 | 0.472094022 |
| chr18:50610241-50610576 | -0.574797036 | 0.19891879 | 0.472094022 |
| chr2:218979845-218980354 | -0.459731431 | 0.19892803 | 0.47210133 |
| chr4:153083465-153083744 | 0.513171013 | 0.198951189 | 0.472141666 |
| chrX:115560901-115561510 | -0.443958705 | 0.198970718 | 0.472172743 |
| chr1:84852470-84852818 | -0.528495666 | 0.198976608 | 0.472172743 |
| chr19:12965374-12965712 | 0.519399938 | 0.198989474 | 0.472184745 |
| chr8:121631757-121632053 | 0.58100462 | 0.198993991 | 0.472184745 |
| chr22:18000869-18001988 | 0.422348623 | 0.199015175 | 0.472220386 |
| chr17:59885046-59885517 | 0.422053109 | 0.19902581 | 0.472230998 |
| chr10:32181737-32182038 | -0.59901778 | 0.199044341 | 0.472260344 |
| chr19:43438533-43438783 | 0.638545377 | 0.199056436 | 0.472274416 |
| chr7:157138238-157139412 | -0.41132317 | 0.19915406 | 0.472468445 |
| chr5:11008082-11008336 | -0.631472185 | 0.19917184 | 0.472468445 |
| chr1:156510656-156510864 | 0.598793415 | 0.199172534 | 0.472468445 |

| | | | |
|---|---|---|---|
| chr1:34681516-34681771 | 0.577123871 | 0.199172688 | 0.472468445 |
| chr10:82463318-82463940 | -0.477718306 | 0.199175304 | 0.472468445 |
| chr10:66257837-66258405 | 0.50805038 | 0.199184027 | 0.472468445 |
| chr10:50579770-50580566 | -0.452363986 | 0.199185878 | 0.472468445 |
| chr2:176118960-176119325 | 0.513203526 | 0.199187547 | 0.472468445 |
| chr11:59556347-59556636 | -0.477214791 | 0.199195084 | 0.472471697 |
| chrX:134372808-134373678 | 0.417348607 | 0.199204542 | 0.472473013 |
| chr1:151399200-151400228 | 0.413658854 | 0.199207971 | 0.472473013 |
| chr3:13648244-13648782 | 0.44156044 | 0.199214477 | 0.472473818 |
| chr17:63124648-63124913 | -0.537645398 | 0.199285399 | 0.472612725 |
| chr5:171907774-171908033 | -0.577232033 | 0.199289616 | 0.472612725 |
| chr16:3880661-3881201 | 0.45181777 | 0.199293426 | 0.472612725 |
| chr14:65321419-65322020 | 0.473358526 | 0.199297718 | 0.472612725 |
| chr4:124631739-124632193 | -0.453543582 | 0.19932301 | 0.472649023 |
| chr1:156584095-156584382 | 0.49717306 | 0.199325362 | 0.472649023 |
| chr3:173371593-173372176 | -0.483902777 | 0.19935887 | 0.47271385 |
| chr10:113701851-113702051 | -0.716906038 | 0.199375157 | 0.472737841 |
| chr20:56843909-56844124 | 0.514373384 | 0.19939054 | 0.472759524 |
| chr1:67559418-67559761 | -0.548190659 | 0.199396642 | 0.472759524 |
| chr8:108295762-108295991 | -0.664139386 | 0.199421441 | 0.472803691 |
| chr19:49487118-49487731 | 0.429607059 | 0.199442652 | 0.472839349 |
| chr14:77668023-77668293 | 0.57621267 | 0.199466369 | 0.472866852 |
| chr8:85290071-85290478 | 0.475604045 | 0.199466596 | 0.472866852 |
| chr9:127826240-127826440 | 0.636514856 | 0.199509988 | 0.472939426 |
| chrX:30895287-30895652 | -0.542603795 | 0.199519085 | 0.472939426 |
| chr17:81101696-81102053 | 0.468918158 | 0.199524379 | 0.472939426 |
| chr10:91681660-91682308 | -0.473937025 | 0.199526892 | 0.472939426 |
| chr9:34628558-34628758 | -0.687841029 | 0.199533379 | 0.472939426 |
| chr4:138182938-138183157 | -0.63589101 | 0.199537406 | 0.472939426 |
| chr2:189483959-189484330 | -0.475508935 | 0.199540416 | 0.472939426 |
| chr8:101474058-101474582 | -0.43282117 | 0.199549157 | 0.472944012 |
| chr6:116561428-116561704 | 0.527425857 | 0.199568569 | 0.472944012 |
| chr7:43261289-43261692 | -0.500003999 | 0.199568911 | 0.472944012 |
| chr2:197813132-197813505 | -0.556911946 | 0.199570256 | 0.472944012 |
| chr12:54232763-54233150 | -0.525208962 | 0.199573213 | 0.472944012 |
| chr17:57407549-57408192 | 0.428236167 | 0.199586853 | 0.472961707 |
| chr1:236141350-236141818 | -0.457501459 | 0.199599697 | 0.472977516 |
| chr21:36927730-36928007 | 0.578218482 | 0.199615434 | 0.473000179 |
| chr10:18215029-18215266 | -0.599551528 | 0.199631046 | 0.473011274 |
| chr10:124530176-124530747 | 0.449702443 | 0.199632463 | 0.473011274 |
| chrX:16807602-16807891 | -0.523626419 | 0.1996519 | 0.473042701 |
| chr19:34920503-34920722 | 0.690416908 | 0.199671484 | 0.473074473 |
| chr13:24671423-24671681 | -0.499555851 | 0.199688015 | 0.473082453 |
| chr6:31795154-31795357 | 0.593706717 | 0.199693474 | 0.473082453 |
| chr20:62914985-62915283 | -0.533928856 | 0.199700261 | 0.473082453 |
| chr5:103804729-103805038 | -0.626622347 | 0.199702625 | 0.473082453 |
| chr17:77591360-77591685 | 0.504797412 | 0.199710994 | 0.473082453 |
| chr1:197663145-197663950 | -0.435285943 | 0.199716015 | 0.473082453 |
| chr15:72998244-72998489 | -0.621555201 | 0.199718072 | 0.473082453 |
| chr14:34539140-34540186 | 0.409996908 | 0.199747485 | 0.47312193 |

| | | | |
|---|---|---|---|
| chr20:45856862-45857858 | 0.413906054 | 0.199749835 | 0.47312193 |
| chr1:149064621-149064882 | 0.665873309 | 0.199753262 | 0.47312193 |
| chr9:72579207-72579470 | -0.56063058 | 0.199782938 | 0.473168434 |
| chr19:38321712-38321928 | -0.593314853 | 0.199786488 | 0.473168434 |
| chr13:30916121-30916464 | -0.575351783 | 0.199791422 | 0.473168434 |
| chr19:4457746-4458222 | -0.456744313 | 0.199801832 | 0.473178463 |
| chr16:14974556-14975425 | 0.422715058 | 0.199820665 | 0.473208438 |
| chr8:108965871-108966421 | 0.493285418 | 0.199838024 | 0.47323492 |
| chr3:22862199-22862511 | 0.510765545 | 0.199879912 | 0.473302185 |
| chr4:116232098-116232298 | -0.707062008 | 0.199881881 | 0.473302185 |
| chr12:9407479-9407751 | -0.660799823 | 0.19988496 | 0.473302185 |
| chr7:34645015-34645341 | 0.556383241 | 0.199924906 | 0.473369915 |
| chr18:22169214-22169915 | 0.448092499 | 0.19992592 | 0.473369915 |
| chr3:132818216-132818447 | 0.63018636 | 0.199935483 | 0.47337793 |
| chr7:77679685-77680100 | -0.506930207 | 0.199955294 | 0.47339984 |
| chr6:149648171-149649348 | 0.410267807 | 0.199957094 | 0.47339984 |
| chr3:169772655-169773846 | -0.402498035 | 0.199966442 | 0.473401086 |
| chr9:77663725-77664269 | 0.47539083 | 0.199974448 | 0.473401086 |
| chr10:13299489-13299799 | 0.504681651 | 0.199976155 | 0.473401086 |
| chr15:63381631-63382461 | -0.425701192 | 0.200000929 | 0.473441071 |
| chr1:226333471-226333712 | 0.598263411 | 0.200006094 | 0.473441071 |
| chr5:27403402-27403644 | -0.697276675 | 0.200011583 | 0.473441071 |
| chr1:178532341-178532637 | 0.515232598 | 0.200023941 | 0.473455699 |
| chr6:109349544-109349833 | 0.538423311 | 0.20003878 | 0.473467998 |
| chr17:1147693-1148067 | -0.533422939 | 0.200041496 | 0.473467998 |
| chr3:20577286-20577486 | 0.670246316 | 0.200055689 | 0.473486964 |
| chr14:75982242-75982872 | 0.432579993 | 0.200066979 | 0.473496223 |
| chr2:241810996-241811264 | -0.562198738 | 0.20007196 | 0.473496223 |
| chr4:137124299-137124894 | -0.454596255 | 0.200082476 | 0.473506486 |
| chr8:58552715-58553974 | -0.412726096 | 0.200112692 | 0.473545194 |
| chr10:58781927-58782137 | -0.634110453 | 0.200123359 | 0.473545194 |
| chr10:65335452-65335690 | -0.634110453 | 0.200123359 | 0.473545194 |
| chr17:45102152-45102967 | -0.431299451 | 0.200123553 | 0.473545194 |
| chr15:100224628-100224964 | 0.564213094 | 0.200130877 | 0.4735479 |
| chr2:20635843-20636216 | 0.458427304 | 0.200160643 | 0.473603705 |
| chr22:37643339-37643619 | -0.528607395 | 0.200180566 | 0.47362963 |
| chr8:92053253-92053453 | 0.694262342 | 0.200183963 | 0.47362963 |
| chr2:177195212-177195484 | 0.561084255 | 0.200214185 | 0.473686509 |
| chr17:63405693-63405999 | 0.470602068 | 0.20026214 | 0.473785336 |
| chr5:74341349-74341574 | -0.678921448 | 0.200297 | 0.473838549 |
| chr6:89368186-89368386 | -0.678921448 | 0.200297 | 0.473838549 |
| chr21:43836762-43837045 | -0.660705859 | 0.200318661 | 0.473875161 |
| chr1:2412475-2412868 | 0.449784887 | 0.200355057 | 0.473946626 |
| chr2:893081-893838 | -0.425248737 | 0.200403083 | 0.474045598 |
| chr2:28716339-28716925 | -0.457002113 | 0.200430074 | 0.474090746 |
| chr19:54417662-54417862 | 0.650544603 | 0.200437981 | 0.474090746 |
| chr7:39888152-39888751 | -0.44269702 | 0.200446344 | 0.474090746 |
| chr6:165890090-165890375 | 0.565663692 | 0.200446919 | 0.474090746 |
| chr14:68306809-68307279 | 0.482266848 | 0.200459625 | 0.474091988 |
| chr1:71080512-71081520 | -0.409075086 | 0.200466651 | 0.474091988 |

| | | | |
|---|---|---|---|
| chrX:111869620-111869877 | 0.549876363 | 0.200470403 | 0.474091988 |
| chr6:87301161-87301586 | -0.520641623 | 0.200472194 | 0.474091988 |
| chr4:82852729-82853036 | -0.587473099 | 0.200489132 | 0.474117412 |
| chr20:46307145-46307361 | 0.539367578 | 0.200507439 | 0.474138264 |
| chr10:75926800-75927017 | -0.659889553 | 0.200511612 | 0.474138264 |
| chr15:45403169-45403787 | 0.450632851 | 0.200516514 | 0.474138264 |
| chr2:147812460-147812798 | -0.547887146 | 0.200524982 | 0.474143656 |
| chr6:68877253-68877457 | -0.705022795 | 0.200541682 | 0.474150917 |
| chr19:12885025-12885735 | -0.435363439 | 0.200545244 | 0.474150917 |
| chr6:44661341-44661575 | 0.575484772 | 0.200546617 | 0.474150917 |
| chr1:200139648-200140019 | -0.45101098 | 0.200553309 | 0.474152107 |
| chr14:102509720-102510153 | 0.435092677 | 0.200559859 | 0.474152962 |
| chr2:174614287-174614580 | 0.58843526 | 0.200574904 | 0.474161988 |
| chr2:82023573-82024004 | -0.546573532 | 0.200579712 | 0.474161988 |
| chr3:168558749-168558949 | 0.700749152 | 0.200582242 | 0.474161988 |
| chr3:89944445-89945169 | -0.440435324 | 0.200588714 | 0.474162659 |
| chr8:145051351-145051560 | -0.662406727 | 0.200607492 | 0.474192418 |
| chr18:49561861-49562138 | -0.598721018 | 0.200620895 | 0.474206701 |
| chr15:101773771-101774143 | 0.487052227 | 0.200635846 | 0.474206701 |
| chr14:26393985-26394577 | 0.487403885 | 0.200636094 | 0.474206701 |
| chr3:129908419-129908697 | 0.524206633 | 0.20063829 | 0.474206701 |
| chr14:20600982-20601242 | 0.607011899 | 0.200656625 | 0.474235407 |
| chr19:5686469-5687427 | 0.416189105 | 0.200695538 | 0.474312744 |
| chr7:38105924-38106235 | -0.513535624 | 0.200717422 | 0.474349834 |
| chr5:115313157-115313390 | 0.649755774 | 0.200740459 | 0.47437686 |
| chr8:101079201-101079486 | 0.509333641 | 0.200742034 | 0.47437686 |
| chr14:102774693-102775158 | -0.486387149 | 0.200748084 | 0.47437686 |
| chr6:135089308-135089517 | -0.620226019 | 0.200753623 | 0.47437686 |
| chr13:69324164-69324411 | -0.650459419 | 0.200773155 | 0.474408384 |
| chr11:118634765-118635311 | 0.456239185 | 0.200792301 | 0.474423425 |
| chr17:27226641-27226904 | -0.677694423 | 0.200794195 | 0.474423425 |
| chr5:130720405-130720802 | -0.500044262 | 0.200799333 | 0.474423425 |
| chr16:70654256-70654556 | 0.532195481 | 0.200804287 | 0.474423425 |
| chr7:111959052-111959318 | -0.618089644 | 0.200823116 | 0.474438379 |
| chr19:34241289-34241905 | 0.462491048 | 0.200825882 | 0.474438379 |
| chr3:136847947-136848178 | 0.667530715 | 0.200835876 | 0.474438379 |
| chr19:38315741-38316393 | 0.42524049 | 0.200840508 | 0.474438379 |
| chr12:62543576-62544068 | 0.500622198 | 0.200844537 | 0.474438379 |
| chr13:44364586-44365091 | -0.496621097 | 0.200847768 | 0.474438379 |
| chr2:190486076-190486538 | -0.519642859 | 0.200860903 | 0.474445052 |
| chr3:57755746-57756564 | 0.420164791 | 0.200862977 | 0.474445052 |
| chr3:90016541-90016809 | -0.557899354 | 0.200877043 | 0.474463609 |
| chr1:237163350-237163641 | -0.583951808 | 0.200883218 | 0.474463609 |
| chr8:105718025-105718374 | 0.460027956 | 0.20089479 | 0.474476315 |
| chr9:89605733-89606472 | 0.441630134 | 0.200915654 | 0.474510966 |
| chr5:12935763-12936059 | 0.572108472 | 0.200922412 | 0.47451219 |
| chr1:84996771-84997945 | -0.421366682 | 0.200928558 | 0.47451219 |
| chr1:151512853-151513411 | 0.420682368 | 0.200951518 | 0.474529136 |
| chr17:80124359-80124882 | 0.466980704 | 0.200953285 | 0.474529136 |
| chr12:55738229-55738646 | 0.467866472 | 0.200964739 | 0.474529136 |

| | | | |
|---|---|---|---|
| chr11:129771544-129771769 | -0.630600951 | 0.200970904 | 0.474529136 |
| chr19:40613233-40613561 | 0.463568502 | 0.200973633 | 0.474529136 |
| chr12:56521655-56521957 | -0.551312166 | 0.200976206 | 0.474529136 |
| chr5:95609874-95610157 | -0.667972735 | 0.200982156 | 0.474529136 |
| chr2:16782416-16782623 | 0.646548221 | 0.200985279 | 0.474529136 |
| chr20:45095835-45096137 | 0.516336691 | 0.200994653 | 0.474536645 |
| chr19:32340321-32340585 | 0.517828574 | 0.201003764 | 0.474541853 |
| chr15:81797884-81798482 | -0.513230737 | 0.201009339 | 0.474541853 |
| chr1:201724504-201724736 | 0.527157455 | 0.201015438 | 0.474541853 |
| chr6:45419546-45419988 | 0.4599275 | 0.201049235 | 0.474607015 |
| chr14:81710308-81710594 | 0.525710336 | 0.201065697 | 0.474631253 |
| chr2:199462903-199463766 | -0.422589851 | 0.201092021 | 0.474676515 |
| chr7:120272220-120272504 | -0.531744325 | 0.201097261 | 0.474676515 |
| chr14:106506981-106507226 | 0.6462984 | 0.201123211 | 0.47470174 |
| chr2:57906206-57906442 | 0.566964185 | 0.201124391 | 0.47470174 |
| chr3:165548206-165548554 | 0.532804427 | 0.201126534 | 0.47470174 |
| chr8:103635950-103636167 | 0.587942797 | 0.201149862 | 0.474736915 |
| chr10:122298834-122299216 | 0.501724265 | 0.201153829 | 0.474736915 |
| chr16:71501059-71501266 | -0.661430896 | 0.201166117 | 0.474751292 |
| chr3:185335847-185336114 | 0.558444656 | 0.20118018 | 0.474769859 |
| chr5:118887740-118887955 | -0.721909794 | 0.201196212 | 0.47479307 |
| chr12:94134129-94134576 | -0.507667816 | 0.201205623 | 0.474800655 |
| chr11:47259527-47259962 | 0.466904883 | 0.201218485 | 0.474816383 |
| chr11:66834282-66834576 | 0.501102698 | 0.201226969 | 0.47482178 |
| chr1:212034938-212036061 | -0.405561894 | 0.201235594 | 0.474827509 |
| chr20:53735612-53736054 | 0.411448476 | 0.201277065 | 0.474910737 |
| chr11:68209240-68209759 | -0.462717004 | 0.201296189 | 0.474941235 |
| chr16:2340597-2341165 | 0.429651268 | 0.201309697 | 0.474945819 |
| chr18:51135829-51136032 | -0.666706321 | 0.201310529 | 0.474945819 |
| chr5:850065-850359 | 0.538487669 | 0.201343883 | 0.475009884 |
| chr3:134364050-134364913 | 0.419826699 | 0.201359107 | 0.475031174 |
| chr21:34582887-34583087 | 0.623586334 | 0.2013964 | 0.475097549 |
| chr10:24238497-24238772 | -0.509650188 | 0.201404213 | 0.475097549 |
| chr2:186771377-186772627 | -0.404890874 | 0.201405844 | 0.475097549 |
| chr5:56429055-56429630 | -0.463345743 | 0.201413427 | 0.475100811 |
| chr10:124479188-124479401 | -0.736560808 | 0.20145034 | 0.475173254 |
| chr7:53114465-53114665 | -0.704226356 | 0.201457742 | 0.475176085 |
| chr5:2739810-2740041 | 0.589138809 | 0.201489729 | 0.475231798 |
| chr20:47989448-47990516 | -0.400777859 | 0.201493767 | 0.475231798 |
| chr5:93620511-93621774 | -0.410600277 | 0.201519601 | 0.475278098 |
| chr2:21281438-21281670 | -0.551676573 | 0.201543568 | 0.475282393 |
| chr11:57130908-57131351 | -0.54767923 | 0.201544585 | 0.475282393 |
| chr1:99327432-99327632 | -0.695182575 | 0.201546765 | 0.475282393 |
| chr13:33676064-33676484 | -0.487560677 | 0.201552918 | 0.475282393 |
| chr2:137944578-137945039 | -0.520016368 | 0.201557961 | 0.475282393 |
| chr6:143592384-143592636 | -0.601565013 | 0.201569391 | 0.475282393 |
| chr6:26047012-26047322 | 0.593829991 | 0.201582812 | 0.475282393 |
| chr9:133509403-133510056 | -0.45423143 | 0.201583282 | 0.475282393 |
| chr1:115051909-115052854 | -0.413687419 | 0.201586916 | 0.475282393 |
| chr19:44845529-44846435 | 0.411113885 | 0.201589495 | 0.475282393 |

| | | | |
|---|---|---|---|
| chr12:83389154-83389863 | -0.436415131 | 0.201589654 | 0.475282393 |
| chr6:43628580-43629025 | 0.455659284 | 0.201612 | 0.475295703 |
| chr8:66190896-66191267 | -0.506040523 | 0.201612493 | 0.475295703 |
| chr4:109859436-109859738 | 0.54164231 | 0.201613909 | 0.475295703 |
| chr17:43367661-43368539 | 0.424078756 | 0.201625568 | 0.475308564 |
| chr2:20822075-20823341 | -0.407614544 | 0.201642634 | 0.475334172 |
| chr7:90595831-90596672 | -0.452171323 | 0.201651923 | 0.475341443 |
| chr12:89949949-89950154 | -0.665978914 | 0.201681127 | 0.47539566 |
| chr10:42517923-42518170 | -0.670139908 | 0.20169754 | 0.475419721 |
| chr3:40309285-40310384 | 0.416372821 | 0.201729727 | 0.475472335 |
| chr8:53742151-53742438 | 0.53650039 | 0.201732272 | 0.475472335 |
| chr8:67339531-67339817 | 0.492259224 | 0.201755719 | 0.475493663 |
| chr3:14633881-14634140 | 0.505465879 | 0.201767505 | 0.475493663 |
| chr2:127535444-127535808 | -0.529212253 | 0.201769223 | 0.475493663 |
| chr2:70789603-70789888 | -0.536177314 | 0.201769975 | 0.475493663 |
| chr2:165778392-165778649 | -0.602848457 | 0.201772349 | 0.475493663 |
| chr1:6438085-6438321 | 0.552543121 | 0.201803292 | 0.475551957 |
| chr2:27683392-27683919 | -0.457908046 | 0.201834507 | 0.47560463 |
| chr21:42157101-42157301 | -0.642309511 | 0.201838059 | 0.47560463 |
| chr1:16512960-16513162 | 0.667186677 | 0.201845255 | 0.475606959 |
| chr19:46753861-46754151 | -0.521684233 | 0.20186666 | 0.47564277 |
| chr19:24054098-24054332 | 0.668537835 | 0.201894338 | 0.475683432 |
| chr3:82496922-82497468 | -0.494380174 | 0.201902331 | 0.475683432 |
| chrX:95321332-95321898 | -0.564816132 | 0.201909229 | 0.475683432 |
| chr7:100874743-100875786 | -0.416505802 | 0.201912108 | 0.475683432 |
| chr1:81174726-81175404 | -0.455839511 | 0.201914959 | 0.475683432 |
| chr12:101145980-10114618[illegible] | -0.654449143 | 0.201933779 | 0.475713144 |
| chr11:70460897-70461097 | -0.660274717 | 0.201953601 | 0.475745213 |
| chr6:141610798-141611095 | -0.581092282 | 0.201993286 | 0.475824072 |
| chr17:979684-980068 | -0.486132612 | 0.202013608 | 0.475857313 |
| chr13:107941443-10794177[illegible] | -0.569666802 | 0.202038386 | 0.475901049 |
| chr15:78000094-78000442 | -0.485703055 | 0.202046243 | 0.475904925 |
| chr1:196856185-196856390 | 0.642260377 | 0.202069512 | 0.475945104 |
| chr5:42604212-42604603 | 0.547625534 | 0.202107185 | 0.476019203 |
| chr5:78182682-78183198 | -0.502290246 | 0.202118368 | 0.476024428 |
| chr6:128570162-128571086 | -0.430635787 | 0.202123433 | 0.476024428 |
| chr5:157855436-157855729 | -0.516232807 | 0.202128041 | 0.476024428 |
| chr5:176898091-176898291 | -0.612017155 | 0.202154066 | 0.476046737 |
| chr19:4310677-4311193 | 0.481369854 | 0.202154841 | 0.476046737 |
| chr2:208501243-208501866 | -0.474398514 | 0.202156154 | 0.476046737 |
| chr11:27173697-27174519 | -0.440178032 | 0.202162366 | 0.476046737 |
| chr12:8914130-8915352 | -0.426731837 | 0.202218788 | 0.476164965 |
| chr1:111192816-111193033 | -0.656572445 | 0.202239029 | 0.476191179 |
| chr12:4307658-4308354 | -0.43120661 | 0.20224235 | 0.476191179 |
| chr2:212892240-212892592 | -0.496282402 | 0.202251207 | 0.4761974 |
| chr6:26272288-26272493 | 0.550486188 | 0.202264478 | 0.476214013 |
| chr2:26346651-26346870 | 0.522183698 | 0.202279662 | 0.476221882 |
| chr16:87581915-87582154 | 0.578865307 | 0.202285805 | 0.476221882 |
| chr6:35731446-35731833 | -0.490274022 | 0.202286466 | 0.476221882 |
| chr12:121538668-121538955 | 0.497172161 | 0.202308654 | 0.476259484 |

| | | | |
|---|---|---|---|
| chr1:27518130-27518401 | -0.61358667 | 0.202328546 | 0.476289888 |
| chr5:147857582-147857826 | -0.614051784 | 0.202337434 | 0.476289888 |
| chr3:134197757-134198074 | 0.528551327 | 0.202344896 | 0.476289888 |
| chr1:202112547-202113527 | -0.414397863 | 0.202346433 | 0.476289888 |
| chr5:114757264-114757464 | -0.736027469 | 0.202353893 | 0.476292815 |
| chr17:21119693-21119990 | -0.565613912 | 0.202377219 | 0.476333085 |
| chr20:20017064-20017715 | 0.426575693 | 0.202385216 | 0.476337276 |
| chr5:321603-322012 | 0.480035819 | 0.202396109 | 0.476348281 |
| chr9:33107308-33107578 | -0.585035705 | 0.202412349 | 0.476371872 |
| chr20:12838022-12838386 | -0.63359649 | 0.202433877 | 0.476407905 |
| chr17:5500995-5501387 | -0.559225163 | 0.202449292 | 0.476429549 |
| chr21:17689022-17689475 | 0.49266511 | 0.202463863 | 0.476449205 |
| chr7:129969769-129970423 | -0.464617904 | 0.202483259 | 0.476469704 |
| chr9:107819822-107820043 | 0.690640167 | 0.202492199 | 0.476469704 |
| chrX:136955006-136955206 | 0.690640167 | 0.202492199 | 0.476469704 |
| chr21:34544091-34544301 | 0.620119777 | 0.202497448 | 0.476469704 |
| chr4:139849202-139849751 | 0.45567267 | 0.202533269 | 0.476539356 |
| chr2:214154859-214155096 | -0.656760513 | 0.202542322 | 0.476546024 |
| chr14:103394168-103394486 | 0.532897809 | 0.202556642 | 0.476565081 |
| chr7:93715327-93715538 | -0.686332301 | 0.202571563 | 0.476585553 |
| chr8:47302879-47303642 | -0.444482585 | 0.202588679 | 0.476600248 |
| chr21:16942829-16943077 | 0.561564547 | 0.202592981 | 0.476600248 |
| chr3:157081611-157082249 | -0.420206335 | 0.202600428 | 0.476600248 |
| chr13:106570388-106570857 | -0.43871867 | 0.20260269 | 0.476600248 |
| chr20:32718835-32719257 | 0.516763691 | 0.202613883 | 0.476611947 |
| chr6:27693620-27694806 | -0.429986677 | 0.202627895 | 0.476626534 |
| chr9:36136515-36136988 | 0.489655231 | 0.202632525 | 0.476626534 |
| chr5:40659076-40659276 | 0.641501784 | 0.202649258 | 0.476632037 |
| chr12:12564092-12564345 | -0.51574575 | 0.202652686 | 0.476632037 |
| chr11:63035730-63035961 | -0.619215406 | 0.202656028 | 0.476632037 |
| chr16:75789060-75789520 | -0.495527866 | 0.202659747 | 0.476632037 |
| chr5:7904260-7904536 | -0.60340208 | 0.202681049 | 0.476667505 |
| chr1:21956671-21957351 | -0.465675642 | 0.20270336 | 0.476696903 |
| chr8:66228423-66228643 | -0.665139408 | 0.202705992 | 0.476696903 |
| chr9:1453730-1454054 | 0.578511217 | 0.202719188 | 0.476713305 |
| chr1:112808959-112809629 | -0.478072465 | 0.202766791 | 0.476810614 |
| chr9:109144036-109144347 | -0.55032332 | 0.202782381 | 0.476819133 |
| chr11:110478476-110478820 | -0.567889307 | 0.20278286 | 0.476819133 |
| chr1:223223059-223223487 | -0.561072312 | 0.202793774 | 0.476821171 |
| chr7:100852057-100853108 | -0.412599887 | 0.202800152 | 0.476821171 |
| chr4:128349919-128350169 | 0.564892556 | 0.20280885 | 0.476821171 |
| chr12:3204312-3204549 | 0.566636334 | 0.2028089 | 0.476821171 |
| chr11:126405868-126406296 | 0.461117625 | 0.202821476 | 0.476821171 |
| chr14:101591535-101591766 | -0.611074175 | 0.202822065 | 0.476821171 |
| chr6:137641538-137641789 | -0.60550798 | 0.202827287 | 0.476821171 |
| chr12:96035047-96036409 | -0.400831202 | 0.202846524 | 0.476833981 |
| chr2:145593788-145594199 | -0.523553147 | 0.202847951 | 0.476833981 |
| chr10:65571278-65571548 | -0.560515036 | 0.202851406 | 0.476833981 |
| chr4:160364746-160365785 | -0.422486349 | 0.202868317 | 0.476859102 |
| chr11:74417368-74417568 | 0.655737634 | 0.202879651 | 0.476871116 |

| | | | |
|---|---|---|---|
| chr1:106244528-106244935 | 0.461105826 | 0.202914355 | 0.476938057 |
| chr2:157403152-157403365 | 0.633148283 | 0.202922505 | 0.476942581 |
| chr11:14974770-14975473 | -0.452727719 | 0.202949136 | 0.476986643 |
| chr6:113879559-113879822 | 0.523507974 | 0.202953702 | 0.476986643 |
| chr1:155763467-155763715 | 0.592229585 | 0.202978859 | 0.477031135 |
| chr1:187696098-187696415 | -0.584115253 | 0.202989871 | 0.477042385 |
| chr10:100009557-100010381 | -0.44027033 | 0.203006146 | 0.477053477 |
| chr20:47312081-47312327 | 0.427959299 | 0.203007043 | 0.477053477 |
| chr6:617234-617845 | -0.48791439 | 0.203029044 | 0.477090546 |
| chr4:40622764-40623255 | -0.493587798 | 0.203050044 | 0.47712526 |
| chr1:214764467-214764954 | -0.484812061 | 0.20306901 | 0.477155192 |
| chr18:49492002-49492695 | -0.440667147 | 0.203083014 | 0.477162982 |
| chr5:51882574-51882939 | -0.681937649 | 0.20308963 | 0.477162982 |
| chr7:6842613-6843022 | 0.464769063 | 0.203091008 | 0.477162982 |
| chr6:151652442-151652662 | 0.579949837 | 0.203100054 | 0.477169606 |
| chr5:139403650-139404481 | -0.41222893 | 0.203146142 | 0.477263251 |
| chr1:116595961-116596546 | 0.459471269 | 0.203168076 | 0.477300148 |
| chr8:11448717-11448917 | -0.65438104 | 0.20319633 | 0.477351888 |
| chr19:47756370-47756570 | -0.661368141 | 0.203221123 | 0.47737731 |
| chr7:155994216-155994476 | -0.638497369 | 0.203222175 | 0.47737731 |
| chr6:42782666-42783208 | 0.460503247 | 0.203231528 | 0.47737731 |
| chr3:186752994-186753488 | -0.495926801 | 0.203232072 | 0.47737731 |
| chr3:99377053-99377282 | -0.715608045 | 0.203286789 | 0.477481854 |
| chr5:45454195-45454443 | 0.566515089 | 0.203289043 | 0.477481854 |
| chr7:54485224-54485613 | -0.505232041 | 0.203307789 | 0.477511247 |
| chr1:204560692-204560900 | 0.555562394 | 0.20331712 | 0.477518526 |
| chr1:47507871-47508653 | -0.458357596 | 0.203336731 | 0.477522853 |
| chr6:22148162-22148382 | -0.660779936 | 0.203342748 | 0.477522853 |
| chr12:96658190-96658454 | -0.582268329 | 0.203342843 | 0.477522853 |
| chr1:197024475-197024767 | 0.554182357 | 0.203343892 | 0.477522853 |
| chr12:48903964-48904189 | 0.643275644 | 0.203357443 | 0.477537368 |
| chr5:140725833-140726560 | 0.414442487 | 0.203362537 | 0.477537368 |
| chr8:81280174-81281039 | -0.42366627 | 0.20337334 | 0.477547401 |
| chr1:19251375-19252130 | 0.455874091 | 0.203379275 | 0.477547401 |
| chr22:36433329-36433571 | -0.633641448 | 0.203391592 | 0.477558843 |
| chr20:8759552-8759908 | -0.53769174 | 0.203396613 | 0.477558843 |
| chr5:149367187-149367857 | -0.431865062 | 0.203408731 | 0.477572661 |
| chr16:1773382-1773721 | 0.47891546 | 0.20342903 | 0.477605684 |
| chr20:57251016-57251338 | 0.41385495 | 0.203455827 | 0.477634188 |
| chr19:38662021-38662363 | 0.485638998 | 0.203458372 | 0.477634188 |
| chr1:161092575-161092851 | 0.494789231 | 0.203459871 | 0.477634188 |
| chr9:137453903-137454103 | 0.62567598 | 0.203480423 | 0.477650662 |
| chr20:54070255-54071005 | 0.400380333 | 0.203483044 | 0.477650662 |
| chr5:149391629-149391829 | 0.65838311 | 0.203485591 | 0.477650662 |
| chr9:38024129-38024641 | -0.475773097 | 0.203512522 | 0.477659457 |
| chr5:10663473-10663824 | -0.499838395 | 0.203515155 | 0.477659457 |
| chr3:55485150-55485361 | 0.636844548 | 0.203520097 | 0.477659457 |
| chr7:117949320-117949590 | -0.532535291 | 0.203521957 | 0.477659457 |
| chr14:73949729-73950783 | 0.404910708 | 0.203525844 | 0.477659457 |
| chr12:110888017-110888492 | 0.451312368 | 0.203526741 | 0.477659457 |

| | | | |
|---|---|---|---|
| chr1:45881632-45881934 | -0.569987981 | 0.203543392 | 0.477683904 |
| chr7:56004851-56005132 | 0.553562337 | 0.203558131 | 0.477703862 |
| chr1:149942340-149942571 | -0.544166138 | 0.203600375 | 0.477788367 |
| chr3:49029151-49029679 | 0.446885567 | 0.203635637 | 0.477842539 |
| chr19:11482585-11482829 | -0.569149961 | 0.20363637 | 0.477842539 |
| chr11:66183782-66184050 | -0.61410378 | 0.203649277 | 0.477842539 |
| chr19:12881329-12881675 | -0.48125776 | 0.20365387 | 0.477842539 |
| chr2:161237389-161238225 | -0.445398729 | 0.203654642 | 0.477842539 |
| chr4:114070811-114071351 | -0.49507273 | 0.203662769 | 0.477846976 |
| chr14:34922220-34922480 | 0.632750973 | 0.203691875 | 0.477900632 |
| chr9:131513527-131513727 | 0.622748243 | 0.203716298 | 0.477943299 |
| chr2:214410891-214411582 | -0.444767066 | 0.203723125 | 0.47794468 |
| chr16:3264092-3264853 | 0.422259207 | 0.203754387 | 0.478003387 |
| chr3:67210148-67210348 | -0.767822859 | 0.203774357 | 0.4780356 |
| chr19:40257123-40257445 | -0.536481102 | 0.203793662 | 0.478059067 |
| chr13:40340499-40340706 | -0.589437337 | 0.203796838 | 0.478059067 |
| chr17:73402160-73402559 | 0.517837979 | 0.203805539 | 0.47806484 |
| chr14:40417706-40418144 | 0.468069782 | 0.203823276 | 0.47809181 |
| chr15:39621292-39621538 | -0.638958311 | 0.203834551 | 0.478096837 |
| chr1:204198584-204199070 | -0.440630712 | 0.203837898 | 0.478096837 |
| chr1:25819785-25820200 | 0.450033885 | 0.203856312 | 0.478125389 |
| chr7:26151914-26152389 | 0.452729449 | 0.203869762 | 0.478142299 |
| chr4:25376744-25377580 | 0.413538021 | 0.203886596 | 0.478151096 |
| chr1:28249532-28249746 | -0.622379958 | 0.203889114 | 0.478151096 |
| chr1:185257170-185258047 | -0.464451809 | 0.203892613 | 0.478151096 |
| chr3:169762948-169763337 | -0.440589018 | 0.203898474 | 0.478151096 |
| chr14:105315194-105315751 | 0.443245317 | 0.203908473 | 0.478159908 |
| chr2:32427781-32428286 | -0.546725856 | 0.203925106 | 0.478184279 |
| chr12:95790001-95790767 | -0.464165223 | 0.203941638 | 0.47820841 |
| chr1:113675381-113675754 | -0.502578995 | 0.203951476 | 0.478211228 |
| chr2:75646206-75647434 | -0.412229221 | 0.20396041 | 0.478211228 |
| chr1:207868766-207869561 | -0.42882784 | 0.203963103 | 0.478211228 |
| chr6:55734384-55734584 | -0.684232558 | 0.203971525 | 0.478211228 |
| chr8:23084151-23084503 | 0.501221438 | 0.203978124 | 0.478211228 |
| chr3:181046335-181046583 | -0.651769915 | 0.203983351 | 0.478211228 |
| chr17:79993423-79993846 | 0.452373879 | 0.203986528 | 0.478211228 |
| chr19:36141141-36141535 | 0.44413679 | 0.204019922 | 0.478274881 |
| chr16:84176918-84177851 | -0.414183843 | 0.204036232 | 0.478296291 |
| chr17:56679817-56680092 | -0.511545446 | 0.20404154 | 0.478296291 |
| chr7:33807822-33808164 | -0.484640602 | 0.204055611 | 0.478302715 |
| chr5:131164386-131165728 | -0.402308589 | 0.204056765 | 0.478302715 |
| chr2:88212604-88212853 | -0.57253818 | 0.20412491 | 0.478447808 |
| chr10:125633645-125633845 | 0.609160448 | 0.204152643 | 0.478498174 |
| chr3:66573455-66573723 | -0.618126884 | 0.204179362 | 0.47854616 |
| chr4:128810907-128812590 | 0.402224142 | 0.2042038 | 0.478566504 |
| chr12:85076638-85077032 | 0.487431173 | 0.204206318 | 0.478566504 |
| chr18:9914929-9915179 | 0.511616992 | 0.204214518 | 0.478566504 |
| chr16:88856635-88857588 | -0.406765654 | 0.204214727 | 0.478566504 |
| chr6:79842093-79842415 | -0.526207668 | 0.204219271 | 0.478566504 |
| chr6:31834419-31835124 | 0.436460982 | 0.204253583 | 0.478632273 |

| | | | |
|---|---|---|---|
| chr12:113103760-113103973 | 0.576365929 | 0.204266534 | 0.478647981 |
| chr4:77366875-77367075 | -0.727863103 | 0.204279242 | 0.478663122 |
| chr6:122840035-122840235 | 0.662324741 | 0.204287411 | 0.478667625 |
| chr20:32445644-32445844 | 0.624312292 | 0.204345989 | 0.478790237 |
| chr12:7130771-7131399 | -0.460811271 | 0.204353482 | 0.478793153 |
| chr10:129466624-129467476 | 0.410948077 | 0.204378691 | 0.478837576 |
| chr11:129696047-129696388 | -0.486025147 | 0.204391677 | 0.47884413 |
| chr4:8427788-8429102 | 0.414239741 | 0.204393988 | 0.47884413 |
| chr1:113949157-113949432 | 0.46280398 | 0.204435707 | 0.47888007 |
| chr5:112995900-112996362 | -0.478126499 | 0.204439325 | 0.47888007 |
| chr7:45939011-45939234 | -0.629300006 | 0.204443467 | 0.47888007 |
| chr5:125846247-125846511 | -0.654279137 | 0.204444294 | 0.47888007 |
| chr3:164950455-164950809 | -0.582288882 | 0.204444314 | 0.47888007 |
| chr6:42217646-42218306 | 0.428619273 | 0.204447783 | 0.47888007 |
| chr1:61293331-61293548 | -0.716461433 | 0.204453078 | 0.47888007 |
| chr17:3012987-3013277 | 0.571835023 | 0.20449588 | 0.47894127 |
| chr19:49664199-49664932 | 0.41969202 | 0.204502298 | 0.47894127 |
| chr19:43639845-43640128 | 0.572249051 | 0.204503662 | 0.47894127 |
| chr13:85586414-85586908 | -0.488529431 | 0.204504209 | 0.47894127 |
| chr14:78164247-78164803 | -0.473841052 | 0.204530525 | 0.478967441 |
| chr4:58406814-58407195 | 0.583933868 | 0.204530911 | 0.478967441 |
| chr5:136006145-136006373 | -0.652450876 | 0.204542893 | 0.478967441 |
| chr7:52842125-52842741 | 0.444535488 | 0.204546745 | 0.478967441 |
| chr7:152491395-152491850 | -0.569330336 | 0.204551839 | 0.478967441 |
| chr19:12141227-12141429 | 0.653477443 | 0.20456962 | 0.478967441 |
| chr4:148090087-148090463 | 0.559822812 | 0.204569954 | 0.478967441 |
| chr12:48753452-48753732 | 0.580638663 | 0.204584325 | 0.478967441 |
| chr16:31700348-31700795 | 0.452082028 | 0.204586433 | 0.478967441 |
| chr14:53103963-53104272 | 0.53641561 | 0.204596844 | 0.478967441 |
| chr11:111253253-111253463 | -0.664663382 | 0.204601499 | 0.478967441 |
| chr17:42517574-42517829 | 0.514387643 | 0.204604633 | 0.478967441 |
| chr14:74881399-74882538 | 0.409772039 | 0.204606349 | 0.478967441 |
| chr10:24238958-24239299 | -0.489307518 | 0.204607946 | 0.478967441 |
| chr5:160165577-160166018 | -0.506003059 | 0.20460915 | 0.478967441 |
| chr17:48215672-48215989 | -0.560167559 | 0.204623252 | 0.47898582 |
| chr16:12388857-12389182 | -0.610524553 | 0.204646645 | 0.479025943 |
| chr4:116516035-116516437 | -0.49841159 | 0.204712692 | 0.479165904 |
| chr17:79006084-79006284 | 0.586893418 | 0.204756183 | 0.479253062 |
| chr12:68158532-68158827 | -0.556381381 | 0.204798293 | 0.479336983 |
| chr15:34588418-34588646 | -0.560726789 | 0.204812979 | 0.479356713 |
| chr3:3998275-3998517 | 0.600251964 | 0.204829232 | 0.47938011 |
| chr1:60979009-60979277 | -0.516162537 | 0.204839 | 0.479388329 |
| chr3:159789056-159789256 | 0.621074822 | 0.204881201 | 0.479463752 |
| chr12:72293807-72294007 | 0.694174396 | 0.204883743 | 0.479463752 |
| chr8:142824328-142824528 | 0.638242126 | 0.20491374 | 0.479519306 |
| chr10:122112546-122114039 | 0.406640723 | 0.204929048 | 0.479531525 |
| chr2:48946514-48946819 | -0.495752278 | 0.20493878 | 0.479531525 |
| chr9:107489433-107490457 | -0.409062916 | 0.204949906 | 0.479531525 |
| chr1:107464838-107465100 | -0.539557951 | 0.20495373 | 0.479531525 |
| chr10:108204490-108204736 | 0.652708717 | 0.204968795 | 0.479531525 |

| | | | |
|---|---|---|---|
| chr16:67150922-67151288 | 0.514358625 | 0.2049751 | 0.479531525 |
| chr7:35055436-35055640 | -0.604325194 | 0.204976725 | 0.479531525 |
| chr20:50085293-50086899 | 0.408136863 | 0.20498117 | 0.479531525 |
| chr1:225999040-225999853 | 0.413031732 | 0.204983295 | 0.479531525 |
| chr2:215931317-215931602 | 0.496545659 | 0.204985762 | 0.479531525 |
| chr10:44259141-44259687 | -0.476898011 | 0.204987805 | 0.479531525 |
| chr2:30821708-30821908 | 0.712201411 | 0.204996633 | 0.479537537 |
| chr6:167827756-167827988 | -0.649205783 | 0.205005545 | 0.479539786 |
| chr4:95166041-95166241 | -0.659285443 | 0.205010946 | 0.479539786 |
| chr8:86275113-86275487 | 0.484652277 | 0.20501637 | 0.479539786 |
| chr10:120948928-120949240 | 0.507818158 | 0.205032321 | 0.479562457 |
| chr5:7927819-7928078 | 0.534954339 | 0.205050465 | 0.479590253 |
| chr19:17266947-17267739 | 0.424667757 | 0.205079405 | 0.4796433 |
| chr17:75625946-75626249 | 0.454698932 | 0.205103757 | 0.479653478 |
| chr4:118034159-118034536 | 0.45101214 | 0.205104787 | 0.479653478 |
| chr6:125780137-125781300 | -0.405417817 | 0.205106874 | 0.479653478 |
| chr17:63710437-63710679 | -0.523505406 | 0.205113435 | 0.479653478 |
| chr7:115497540-115497879 | -0.656119624 | 0.205115057 | 0.479653478 |
| chr18:34322385-34322700 | -0.633385103 | 0.205133478 | 0.479681915 |
| chr4:173417018-173417314 | -0.503014892 | 0.205145898 | 0.479695786 |
| chr18:9722715-9723022 | 0.512408615 | 0.205163753 | 0.479695786 |
| chr17:61352228-61352596 | 0.411163144 | 0.205164711 | 0.479695786 |
| chr15:30894515-30894715 | 0.646156253 | 0.20516934 | 0.479695786 |
| chr19:17294033-17294339 | 0.506422043 | 0.205170712 | 0.479695786 |
| chr13:42327817-42328058 | -0.581308697 | 0.205186892 | 0.479717465 |
| chr11:118057351-118057760 | -0.498852736 | 0.205192506 | 0.479717465 |
| chr2:157751396-157751895 | -0.457777677 | 0.205203047 | 0.479723954 |
| chr21:32601010-32601210 | 0.6897178 | 0.205207803 | 0.479723954 |
| chr2:179562346-179562744 | -0.610349756 | 0.205232675 | 0.479742442 |
| chr1:113874390-113874873 | 0.429725527 | 0.205235683 | 0.479742442 |
| chr18:32714628-32714845 | -0.642721359 | 0.20524254 | 0.479742442 |
| chr6:169810342-169810643 | 0.500185407 | 0.20524769 | 0.479742442 |
| chr6:73898697-73898904 | 0.596364265 | 0.205248411 | 0.479742442 |
| chr19:7629469-7629927 | 0.451459764 | 0.205253279 | 0.479742442 |
| chr13:32390532-32390801 | 0.583873329 | 0.205297382 | 0.479798221 |
| chr19:46745637-46746074 | -0.474429251 | 0.205297728 | 0.479798221 |
| chr3:66037389-66037589 | 0.522142288 | 0.205301401 | 0.479798221 |
| chr16:11345016-11345741 | 0.428872246 | 0.205302191 | 0.479798221 |
| chr4:94127780-94128081 | -0.542248901 | 0.205309007 | 0.479799516 |
| chr19:58519022-58520460 | 0.404870823 | 0.205321397 | 0.479813836 |
| chr7:87918674-87919086 | -0.505040876 | 0.205354104 | 0.479875634 |
| chr7:66994763-66995017 | 0.485860548 | 0.205370329 | 0.479898912 |
| chr1:229270767-229271690 | 0.424084036 | 0.205398368 | 0.479949795 |
| chr3:94028490-94028753 | -0.58741131 | 0.205409504 | 0.479961181 |
| chr19:10695683-10696256 | 0.428996775 | 0.205435433 | 0.480007129 |
| chr6:165884852-165885132 | 0.554599464 | 0.20544284 | 0.480009797 |
| chr13:46707936-46708145 | 0.549631515 | 0.205459792 | 0.480034767 |
| chr18:57435050-57435693 | -0.427766571 | 0.205473482 | 0.480052115 |
| chr2:97112905-97113965 | -0.415742961 | 0.205482968 | 0.480059639 |
| chr1:192491947-192492147 | -0.651898161 | 0.205504724 | 0.480084626 |

| | | | |
|---|---|---|---|
| chr15:28021620-28021948 | -0.579374668 | 0.205506195 | 0.480084626 |
| chr4:67288937-67289244 | -0.623269427 | 0.205526409 | 0.480090387 |
| chr3:172691089-172691850 | 0.457720772 | 0.205528917 | 0.480090387 |
| chr3:64801646-64801856 | 0.453415074 | 0.205532294 | 0.480090387 |
| chr17:72727696-72728122 | 0.467700757 | 0.205535691 | 0.480090387 |
| chr12:34163424-34163686 | -0.562492198 | 0.205539989 | 0.480090387 |
| chr14:89973289-89973608 | 0.510077177 | 0.20562902 | 0.480267327 |
| chr9:34592994-34593345 | 0.507993232 | 0.205634113 | 0.480267327 |
| chr12:112012667-112014416 | 0.398747879 | 0.205634546 | 0.480267327 |
| chr10:8045119-8045411 | 0.466052446 | 0.20564385 | 0.480270534 |
| chr6:35392126-35392576 | 0.50176575 | 0.205648456 | 0.480270534 |
| chr2:27087014-27087396 | -0.45022776 | 0.205672357 | 0.480311714 |
| chr4:168831805-168832797 | -0.41829632 | 0.205707248 | 0.480378555 |
| chr3:196108158-196108583 | -0.527586117 | 0.205713818 | 0.480379255 |
| chr12:53974522-53974932 | 0.462587022 | 0.205797662 | 0.480560402 |
| chr7:118094067-118094362 | -0.53347086 | 0.205807372 | 0.48056843 |
| chr1:178575882-178576199 | -0.562315029 | 0.205868662 | 0.480696895 |
| chr4:153038966-153039342 | 0.526845922 | 0.205883088 | 0.48070623 |
| chr21:41167386-41168267 | -0.421907573 | 0.205887369 | 0.48070623 |
| chr1:216210025-216210430 | -0.51125022 | 0.205894091 | 0.48070623 |
| chr3:101466694-101467270 | -0.470470447 | 0.205897755 | 0.48070623 |
| chr11:67356882-67357147 | 0.480571522 | 0.205912742 | 0.480726572 |
| chr13:47337909-47338309 | 0.498107965 | 0.205929148 | 0.480750078 |
| chr8:63394699-63394920 | -0.603125635 | 0.205935359 | 0.480750078 |
| chr6:33589713-33589937 | 0.58528066 | 0.205952336 | 0.480775064 |
| chr1:200245321-200245527 | 0.592299715 | 0.20596157 | 0.48078197 |
| chr16:30922160-30923623 | 0.398576945 | 0.205968389 | 0.480783241 |
| chr2:40760098-40760421 | -0.509986029 | 0.205997056 | 0.480835508 |
| chr5:62427464-62427664 | -0.684819864 | 0.206008358 | 0.480840112 |
| chr16:69324320-69324608 | -0.544110474 | 0.206011702 | 0.480840112 |
| chr14:52960764-52960964 | 0.631832548 | 0.206020135 | 0.480840112 |
| chr5:149747522-149747863 | -0.508155673 | 0.20602413 | 0.480840112 |
| chr20:20780494-20780819 | -0.531162816 | 0.20606761 | 0.480918125 |
| chr13:46058682-46058933 | -0.62258776 | 0.206070109 | 0.480918125 |
| chr8:120720051-120720357 | 0.522263878 | 0.206082939 | 0.480933419 |
| chr12:124898523-124898825 | 0.50417665 | 0.206092238 | 0.480940471 |
| chr19:3505722-3506420 | 0.420913282 | 0.20610242 | 0.480949585 |
| chr19:45668920-45669313 | 0.470411593 | 0.206114164 | 0.480962342 |
| chr4:42048316-42048785 | -0.524576079 | 0.20613353 | 0.480992885 |
| chr20:53570572-53570996 | 0.410032634 | 0.206140092 | 0.480993548 |
| chr2:85989078-85989343 | 0.57862988 | 0.206176804 | 0.481064559 |
| chr15:75647453-75648432 | 0.422456145 | 0.206194249 | 0.481090613 |
| chr20:54694208-54694413 | 0.457386484 | 0.20620669 | 0.48110499 |
| chr19:41563238-41563793 | -0.45881876 | 0.206232172 | 0.481149776 |
| chr20:19463591-19463913 | 0.580345755 | 0.206238445 | 0.481149776 |
| chr4:77819939-77820724 | -0.41244816 | 0.206249635 | 0.481161233 |
| chr2:164458895-164459103 | -0.70209876 | 0.206265688 | 0.481169383 |
| chr20:16184378-16184638 | -0.70209876 | 0.206265688 | 0.481169383 |
| chr12:44972852-44973160 | -0.554447464 | 0.206272523 | 0.481170678 |
| chr7:45923090-45923480 | -0.532823933 | 0.206282659 | 0.481179673 |

| | | | |
|---|---|---|---|
| chr7:26278851-26279133 | 0.541819981 | 0.206302654 | 0.481211664 |
| chr4:95547697-95548458 | -0.459825592 | 0.206326249 | 0.481252051 |
| chr4:69060352-69060561 | -0.672467491 | 0.206336344 | 0.481260946 |
| chr19:49495881-49496572 | 0.412081209 | 0.206402422 | 0.481391357 |
| chr1:203282094-203282642 | 0.436012752 | 0.206404822 | 0.481391357 |
| chr19:44207308-44207811 | 0.482518038 | 0.206426063 | 0.481401559 |
| chr11:5620722-5621531 | 0.462110494 | 0.206427391 | 0.481401559 |
| chr10:79163023-79163431 | -0.4992088 | 0.206428044 | 0.481401559 |
| chr5:168334725-168335251 | -0.454728804 | 0.206440832 | 0.481404258 |
| chr17:45160953-45162000 | 0.409781817 | 0.206446284 | 0.481404258 |
| chr5:19519593-19519793 | -0.683604313 | 0.20644805 | 0.481404258 |
| chr16:87017431-87017728 | -0.565635774 | 0.206457452 | 0.48141153 |
| chr14:67172512-67172955 | 0.510187481 | 0.206471256 | 0.481429067 |
| chr18:13267341-13267639 | 0.539541428 | 0.206486677 | 0.481450373 |
| chr8:64567646-64567935 | -0.535075447 | 0.20649303 | 0.481450536 |
| chr3:108369668-108370049 | 0.52828446 | 0.206509984 | 0.481464184 |
| chr17:50372799-50373432 | -0.419411061 | 0.206517642 | 0.481464184 |
| chr3:101983859-101984182 | -0.623539349 | 0.206520577 | 0.481464184 |
| chr3:197458360-197458691 | -0.5389865 | 0.206524018 | 0.481464184 |
| chr12:52203094-52203754 | 0.426460803 | 0.206564159 | 0.481530304 |
| chr7:43700024-43700247 | 0.612300434 | 0.20656495 | 0.481530304 |
| chr13:33388241-33388732 | -0.470843875 | 0.206590509 | 0.481575236 |
| chr3:115919064-115919465 | -0.490016167 | 0.206598033 | 0.481578123 |
| chr17:58006484-58008096 | -0.394890335 | 0.206613373 | 0.48159923 |
| chr10:125231515-125231831 | 0.528012018 | 0.206654479 | 0.481635311 |
| chr20:956512-956807 | 0.571760137 | 0.206667282 | 0.481635311 |
| chr20:61154410-61154652 | 0.557443065 | 0.206667892 | 0.481635311 |
| chr18:74597297-74598755 | -0.405751687 | 0.206672761 | 0.481635311 |
| chr10:88114419-88114785 | -0.481698676 | 0.206677806 | 0.481635311 |
| chr1:107643812-107644202 | -0.513698574 | 0.206680643 | 0.481635311 |
| chr5:151613068-151613461 | -0.502881638 | 0.206688054 | 0.481635311 |
| chr12:100399733-100399990 | -0.633100835 | 0.206693636 | 0.481635311 |
| chr15:84845623-84845861 | 0.556297474 | 0.206693754 | 0.481635311 |
| chr8:33512536-33512913 | -0.507369447 | 0.206700057 | 0.481635311 |
| chr9:134689607-134689836 | 0.545475369 | 0.206708352 | 0.481635311 |
| chr6:76230692-76231025 | -0.504338943 | 0.206713626 | 0.481635311 |
| chr1:2545886-2546683 | 0.421207164 | 0.206720676 | 0.481635311 |
| chr3:164997618-164997941 | -0.582777831 | 0.206722856 | 0.481635311 |
| chr10:70169929-70170882 | -0.409611242 | 0.20672314 | 0.481635311 |
| chr3:197590531-197590749 | 0.616851713 | 0.206754389 | 0.481654607 |
| chr14:101707368-101707623 | 0.522169067 | 0.206758549 | 0.481654607 |
| chr10:61835143-61835408 | -0.542137505 | 0.206759725 | 0.481654607 |
| chr18:58496946-58497232 | -0.623551975 | 0.206760532 | 0.481654607 |
| chr8:97644908-97645357 | 0.439198432 | 0.206762853 | 0.481654607 |
| chr1:23528176-23528509 | -0.504819098 | 0.206824765 | 0.481756206 |
| chr9:102335023-102335223 | -0.724215847 | 0.206824943 | 0.481756206 |
| chr7:47371762-47372343 | -0.465637368 | 0.20682533 | 0.481756206 |
| chr19:51091981-51092213 | -0.568991315 | 0.206844737 | 0.481777952 |
| chr6:54699041-54699320 | 0.54649647 | 0.206847241 | 0.481777952 |
| chr17:67974591-67974815 | -0.576063161 | 0.206860516 | 0.481781337 |

| | | | |
|---|---|---|---|
| chr12:74803248-74803507 | -0.566269825 | 0.206865547 | 0.481781337 |
| chr12:106189667-106189867 | -0.692632694 | 0.206871131 | 0.481781337 |
| chr14:20682352-20683101 | 0.405037463 | 0.206873845 | 0.481781337 |
| chr12:66130014-66131339 | -0.404548662 | 0.206883331 | 0.481788785 |
| chr4:107708236-107708585 | 0.503528485 | 0.206900927 | 0.481815118 |
| chr7:147413263-147413627 | 0.48714774 | 0.206934137 | 0.481865416 |
| chr7:55018697-55019472 | 0.435228255 | 0.206935765 | 0.481865416 |
| chr16:31033148-31034432 | 0.406215515 | 0.206941392 | 0.481865416 |
| chr10:101817511-101817914 | 0.455672014 | 0.206951337 | 0.481873927 |
| chr2:39778695-39779517 | -0.417713772 | 0.206969244 | 0.481900979 |
| chr13:90838980-90839691 | -0.428085051 | 0.206978302 | 0.481907424 |
| chr12:79174759-79175031 | -0.550453555 | 0.206985045 | 0.481908481 |
| chr6:33092010-33092249 | -0.691970641 | 0.207011111 | 0.481928484 |
| chr16:72124899-72125210 | -0.481792205 | 0.207012407 | 0.481928484 |
| chr1:3856638-3857812 | 0.404478729 | 0.207012506 | 0.481928484 |
| chr7:7968313-7970008 | 0.395099948 | 0.207019443 | 0.481929992 |
| chr9:115591553-115591850 | 0.499634047 | 0.207040771 | 0.481965 |
| chr9:98117987-98118262 | -0.536288972 | 0.207049723 | 0.481971196 |
| chr5:24073037-24073523 | 0.483562968 | 0.207078525 | 0.482018696 |
| chr11:78574282-78575553 | -0.401183216 | 0.207086229 | 0.482018696 |
| chr20:21280599-21280991 | -0.607635373 | 0.207091973 | 0.482018696 |
| chr9:87716326-87716537 | 0.618987452 | 0.207097185 | 0.482018696 |
| chr21:33229669-33230755 | 0.425991876 | 0.207106554 | 0.482018696 |
| chr1:205142270-205142540 | -0.528375208 | 0.207110542 | 0.482018696 |
| chr17:64588848-64589093 | -0.665533832 | 0.207120233 | 0.482018696 |
| chr5:175460553-175460763 | 0.593919413 | 0.207123844 | 0.482018696 |
| chrX:37685407-37686281 | -0.450238577 | 0.207130013 | 0.482018696 |
| chr17:79839376-79839790 | 0.432607418 | 0.207137012 | 0.482018696 |
| chr5:129852801-129853095 | -0.495234585 | 0.207139328 | 0.482018696 |
| chr7:95596112-95596333 | -0.625326074 | 0.207163948 | 0.482058315 |
| chr20:63475339-63475565 | 0.519210496 | 0.207168936 | 0.482058315 |
| chr18:74896485-74896731 | -0.592034253 | 0.207191378 | 0.482095893 |
| chr10:51073653-51075008 | -0.409113375 | 0.207204858 | 0.482107781 |
| chr14:61526237-61526823 | 0.428309883 | 0.207209071 | 0.482107781 |
| chr16:83878933-83879154 | 0.572595799 | 0.207226032 | 0.482108163 |
| chrX:87141242-87141442 | -0.723406375 | 0.207228524 | 0.482108163 |
| chr3:72655781-72656124 | 0.516478545 | 0.2072314 | 0.482108163 |
| chr3:40457146-40457654 | 0.436120526 | 0.207234403 | 0.482108163 |
| chr2:177170203-177170500 | -0.551309153 | 0.20726137 | 0.482156258 |
| chr5:79112114-79112326 | -0.671793726 | 0.207340552 | 0.48230841 |
| chr6:56904698-56904941 | -0.593702369 | 0.207347664 | 0.48230841 |
| chr11:31240396-31240618 | -0.682333518 | 0.207348625 | 0.48230841 |
| chr16:76676117-76676861 | 0.411656223 | 0.207356817 | 0.48230841 |
| chr12:42111084-42111382 | 0.534739529 | 0.207358248 | 0.48230841 |
| chr12:19439281-19440812 | -0.411058904 | 0.207378915 | 0.482318075 |
| chr4:54626148-54626354 | -0.672860494 | 0.207380389 | 0.482318075 |
| chr2:10465205-10465460 | -0.540665012 | 0.20738226 | 0.482318075 |
| chr1:222546966-222547341 | -0.479214659 | 0.207387582 | 0.482318075 |
| chr22:38084161-38084394 | 0.593418573 | 0.207394085 | 0.482318561 |
| chr1:186204948-186205148 | -0.711850057 | 0.20740121 | 0.48232049 |

| | | | |
|---|---|---|---|
| chr20:53631782-53632420 | 0.401148306 | 0.207421373 | 0.482352739 |
| chr3:62917238-62917810 | 0.425595124 | 0.207444579 | 0.482392064 |
| chr6:129751736-129752228 | -0.47317425 | 0.207460739 | 0.482409458 |
| chr16:80619811-80620441 | 0.415229665 | 0.20746465 | 0.482409458 |
| chr3:27266727-27266927 | -0.717713402 | 0.207510349 | 0.482501075 |
| chr1:52365222-52365438 | 0.507146127 | 0.207527412 | 0.482526109 |
| chr2:25251257-25252745 | -0.408965839 | 0.207536149 | 0.48253178 |
| chr8:122638843-122639439 | -0.418731064 | 0.207558441 | 0.482568966 |
| chr9:70205690-70205998 | 0.508624833 | 0.207607939 | 0.482658248 |
| chr1:151993181-151994397 | 0.400785349 | 0.20760944 | 0.482658248 |
| chr1:155261389-155262646 | 0.403292654 | 0.207622508 | 0.482662304 |
| chr5:58069667-58070369 | -0.424829553 | 0.207623783 | 0.482662304 |
| chr16:76132293-76132735 | -0.499655424 | 0.207631406 | 0.482665379 |
| chr7:45035882-45036209 | 0.490565398 | 0.207643542 | 0.482678947 |
| chr6:116139974-116140273 | -0.52445273 | 0.20768842 | 0.482768622 |
| chr5:57298251-57298518 | -0.53438339 | 0.207705277 | 0.482777341 |
| chr2:88545250-88545561 | -0.489154032 | 0.207709767 | 0.482777341 |
| chr2:112750308-112750525 | 0.671399053 | 0.207711073 | 0.482777341 |
| chr13:42295147-42295418 | 0.643866341 | 0.207742259 | 0.48283518 |
| chr7:151754128-151754451 | 0.485048283 | 0.20780279 | 0.482961217 |
| chr15:34038180-34039418 | 0.415516474 | 0.207815794 | 0.482973194 |
| chr20:46997407-46997713 | 0.525488574 | 0.207822524 | 0.482973194 |
| chr5:104412821-104413282 | -0.510507095 | 0.207832449 | 0.482973194 |
| chr12:121626093-121627545 | 0.404741206 | 0.207836625 | 0.482973194 |
| chr6:135438960-135439431 | -0.539153504 | 0.20783946 | 0.482973194 |
| chr1:172399432-172399827 | -0.509905547 | 0.207856013 | 0.48299701 |
| chr13:52809323-52809680 | -0.464182159 | 0.207905218 | 0.483096697 |
| chr19:7068990-7070147 | 0.415579531 | 0.207940301 | 0.483134522 |
| chr1:218380124-218380899 | -0.41143373 | 0.207942905 | 0.483134522 |
| chr17:70630001-70630201 | -0.685771983 | 0.207946718 | 0.483134522 |
| chr19:6066701-6066901 | -0.685771983 | 0.207946718 | 0.483134522 |
| chr2:42457123-42457323 | -0.710828785 | 0.207974628 | 0.483152196 |
| chr2:96412896-96413190 | -0.547001787 | 0.207977558 | 0.483152196 |
| chr17:62718030-62718926 | -0.412518449 | 0.207978987 | 0.483152196 |
| chr5:96627827-96628054 | -0.614906114 | 0.207979547 | 0.483152196 |
| chr18:48423193-48423393 | -0.685765441 | 0.207986786 | 0.483154363 |
| chr5:100822359-100823364 | -0.42268425 | 0.207998812 | 0.483159709 |
| chr2:73212546-73212824 | -0.483656057 | 0.208006809 | 0.483159709 |
| chr10:68208440-68209000 | -0.445084837 | 0.208008004 | 0.483159709 |
| chr11:12264110-12264352 | 0.643125225 | 0.208028056 | 0.483189125 |
| chr4:141133044-141133799 | 0.421939688 | 0.2080383 | 0.483189125 |
| chr14:24333718-24333932 | 0.509802169 | 0.208051368 | 0.483189125 |
| chr1:151830691-151831102 | 0.433506077 | 0.208051862 | 0.483189125 |
| chr8:60280636-60281809 | -0.407763815 | 0.208062408 | 0.483189125 |
| chr11:69891361-69891857 | 0.467413006 | 0.208073068 | 0.483189125 |
| chr6:41470137-41470357 | -0.674930212 | 0.20807324 | 0.483189125 |
| chr6:35496634-35497962 | 0.408546499 | 0.208073548 | 0.483189125 |
| chr20:381431-381826 | 0.491937618 | 0.208078616 | 0.483189125 |
| chr22:38218609-38218880 | -0.579284927 | 0.20808373 | 0.483189125 |
| chr6:125201938-125202584 | -0.422362369 | 0.208097107 | 0.483198057 |

| | | | |
|---|---|---|---|
| chr19:19033381-19033982 | -0.458625194 | 0.208104651 | 0.483198057 |
| chr7:926998-927550 | 0.46633649 | 0.208112999 | 0.483198057 |
| chr8:98286223-98286452 | 0.5857826 | 0.208113662 | 0.483198057 |
| chr5:112920926-112922555 | -0.396082279 | 0.208124961 | 0.483198057 |
| chr1:111208952-111209376 | -0.487582448 | 0.208125414 | 0.483198057 |
| chr21:8441092-8441310 | 0.449757596 | 0.208134324 | 0.483201506 |
| chr17:61384484-61385298 | 0.396368303 | 0.208139512 | 0.483201506 |
| chr1:169595808-169596128 | -0.490547112 | 0.208150558 | 0.483212509 |
| chr1:233613657-233615774 | 0.401359171 | 0.208161862 | 0.48322354 |
| chr20:56690532-56691042 | -0.441487433 | 0.208192452 | 0.48322354 |
| chr5:170590364-170590564 | -0.670922832 | 0.208192912 | 0.48322354 |
| chr1:32650246-32650768 | 0.427012609 | 0.20819706 | 0.48322354 |
| chr1:159923692-159924025 | -0.449197172 | 0.208197376 | 0.48322354 |
| chr6:89242649-89242945 | -0.56449394 | 0.208197604 | 0.48322354 |
| chr12:11869529-11870092 | -0.543536345 | 0.208199456 | 0.48322354 |
| chr2:226791550-226792129 | -0.437971616 | 0.208223611 | 0.483264964 |
| chr3:11155513-11155777 | 0.510367855 | 0.208238161 | 0.483278118 |
| chr5:112757674-112757984 | -0.521540969 | 0.208241893 | 0.483278118 |
| chr11:86193562-86193860 | 0.476295605 | 0.208248361 | 0.483278491 |
| chr2:203883955-203884161 | 0.68899518 | 0.208259247 | 0.483289116 |
| chr20:13898091-13898344 | 0.549388223 | 0.208279416 | 0.483321281 |
| chr4:81032837-81033076 | -0.681412438 | 0.208295707 | 0.483344448 |
| chr3:104273082-104273413 | -0.526988653 | 0.20832337 | 0.483394 |
| chr20:53799051-53799435 | 0.40690642 | 0.208330275 | 0.483395382 |
| chr21:39925071-39925301 | 0.553091286 | 0.208343909 | 0.483412379 |
| chr8:132046674-132047234 | -0.424888252 | 0.208363456 | 0.483432279 |
| chr8:17755073-17755798 | -0.473742519 | 0.208365104 | 0.483432279 |
| chr3:50590899-50591118 | -0.603608013 | 0.208371588 | 0.483432683 |
| chr6:53130643-53131124 | -0.513303626 | 0.20840543 | 0.48349656 |
| chr20:19348572-19348991 | -0.586853606 | 0.208415833 | 0.483506055 |
| chr10:122276908-122277191 | -0.572778815 | 0.208445 | 0.483553659 |
| chr9:122056566-122056766 | 0.670768869 | 0.208448975 | 0.483553659 |
| chr8:126149064-126149264 | -0.680606354 | 0.208465572 | 0.48357752 |
| chr3:133423603-133423901 | -0.491083601 | 0.208474111 | 0.483582687 |
| chr1:230758160-230758454 | 0.53432964 | 0.20852237 | 0.483679987 |
| chr8:143070946-143071262 | 0.514322005 | 0.208539932 | 0.483706081 |
| chr12:98418732-98419029 | -0.521157563 | 0.208561385 | 0.483741198 |
| chr3:101693583-101693940 | -0.4535824 | 0.208572417 | 0.483752141 |
| chr15:47374391-47374610 | -0.658846748 | 0.208625434 | 0.483858354 |
| chr2:69436613-69437991 | -0.401078895 | 0.208631746 | 0.483858354 |
| chr3:187764363-187764615 | -0.670668825 | 0.208637156 | 0.483858354 |
| chr5:101347517-101348018 | -0.480919358 | 0.208664588 | 0.483895593 |
| chr21:25283805-25284109 | -0.578456822 | 0.208665844 | 0.483895593 |
| chr10:13707445-13707743 | -0.488134836 | 0.208703904 | 0.48396108 |
| chr5:114449897-114450465 | -0.447218584 | 0.208710519 | 0.48396108 |
| chr5:41758985-41759234 | -0.647863884 | 0.208713032 | 0.48396108 |
| chr17:51653967-51654274 | -0.496502128 | 0.208720772 | 0.483964381 |
| chr7:128387757-128388041 | -0.472121395 | 0.208763038 | 0.484047736 |
| chr19:3382314-3382587 | 0.624790832 | 0.208778033 | 0.484067856 |
| chr11:95107184-95108116 | -0.433116172 | 0.20879257 | 0.484086912 |

| | | | |
|---|---|---|---|
| chr6:83065437-83066024 | 0.423104855 | 0.20881118 | 0.484115412 |
| chr5:51810546-51810995 | 0.487522331 | 0.208824781 | 0.484132296 |
| chr7:105770351-105770617 | -0.623178459 | 0.208838636 | 0.484149769 |
| chr2:84958118-84958748 | 0.434018997 | 0.208865614 | 0.4841908 |
| chr3:115658348-115658680 | -0.505354892 | 0.208871831 | 0.4841908 |
| chr8:143989854-143990192 | 0.461130982 | 0.208875293 | 0.4841908 |
| chr20:37651898-37652170 | -0.570821253 | 0.208887268 | 0.484203909 |
| chr2:71727649-71728010 | 0.562331156 | 0.208917685 | 0.484259768 |
| chr7:148262749-148263208 | -0.477325269 | 0.208925679 | 0.484263646 |
| chr2:32932075-32932280 | -0.602846624 | 0.208949742 | 0.484304627 |
| chr2:82139946-82140478 | 0.448198277 | 0.208956 | 0.484304627 |
| chr4:56753087-56753287 | 0.635661106 | 0.208988636 | 0.484365555 |
| chr10:5488312-5489144 | -0.4088455 | 0.208994931 | 0.484365555 |
| chr7:10023006-10023482 | -0.468417594 | 0.209016338 | 0.48439712 |
| chr3:155253372-155253572 | -0.697018258 | 0.209021195 | 0.48439712 |
| chr21:28699686-28699886 | 0.680703127 | 0.209036037 | 0.484403411 |
| chr21:46215730-46215965 | 0.564901723 | 0.209036553 | 0.484403411 |
| chr14:64892200-64892519 | 0.634419336 | 0.20905634 | 0.48442886 |
| chr7:92076554-92076857 | -0.597324312 | 0.20906018 | 0.48442886 |
| chr4:86685333-86685752 | -0.508642342 | 0.209081076 | 0.484462629 |
| chr12:57243683-57243960 | 0.477772522 | 0.209104694 | 0.484502703 |
| chr12:42213048-42213400 | -0.529328159 | 0.209120493 | 0.484512702 |
| chr1:23497150-23497365 | 0.605161798 | 0.209125919 | 0.484512702 |
| chr17:14926516-14926735 | -0.632809365 | 0.209137932 | 0.484512702 |
| chr15:87348908-87349139 | 0.608110435 | 0.209142555 | 0.484512702 |
| chr13:38888267-38888729 | -0.463975687 | 0.209144905 | 0.484512702 |
| chr19:13880125-13880502 | 0.463732994 | 0.20914695 | 0.484512702 |
| chr7:73738708-73739298 | -0.412312655 | 0.209169562 | 0.484550436 |
| chr15:73927560-73928179 | 0.438656151 | 0.209191398 | 0.484586368 |
| chr6:43477346-43477941 | 0.434487389 | 0.209205872 | 0.484599639 |
| chr20:59788445-59788645 | 0.572111084 | 0.209209776 | 0.484599639 |
| chr12:129911546-129911746 | -0.679572337 | 0.209217 | 0.484601722 |
| chr20:46739822-46740174 | 0.485846352 | 0.209248801 | 0.484657999 |
| chr16:46629206-46629788 | 0.464826926 | 0.209253947 | 0.484657999 |
| chr9:31045950-31046166 | 0.640450414 | 0.20926879 | 0.484669712 |
| chr6:138106343-138106633 | 0.525772909 | 0.209272171 | 0.484669712 |
| chrX:140507259-140507617 | -0.456252681 | 0.20927798 | 0.484669712 |
| chr16:14309422-14309860 | 0.435667526 | 0.209344746 | 0.484809681 |
| chr20:55027358-55027594 | 0.574853545 | 0.209359748 | 0.484826412 |
| chr8:127565095-127565367 | 0.51302041 | 0.209364625 | 0.484826412 |
| chr1:217379437-217379794 | -0.498019764 | 0.209406669 | 0.484909117 |
| chr6:13014600-13014906 | -0.584383911 | 0.209426358 | 0.484933477 |
| chr11:98330226-98330521 | -0.612191944 | 0.209432297 | 0.484933477 |
| chr19:18557545-18558142 | -0.461980992 | 0.209436175 | 0.484933477 |
| chrY:11295508-11295720 | 0.66323361 | 0.209484133 | 0.485029862 |
| chr12:99387016-99387229 | -0.660967182 | 0.209501214 | 0.485053711 |
| chr12:122094058-122094289 | 0.597964925 | 0.20951249 | 0.485053711 |
| chr5:177133195-177134442 | -0.408430619 | 0.209513424 | 0.485053711 |
| chr3:99967723-99968075 | -0.615003882 | 0.209531747 | 0.485080226 |
| chr15:74382388-74382702 | 0.480053881 | 0.209537539 | 0.485080226 |

| | | | |
|---|---|---|---|
| chrX:13937535-13938704 | 0.430673007 | 0.209550089 | 0.485094624 |
| chr11:101003540-101003773 | -0.645205993 | 0.209587112 | 0.485159161 |
| chr7:47047026-47047257 | -0.594634131 | 0.209590632 | 0.485159161 |
| chr9:97598765-97599075 | 0.521150559 | 0.209598591 | 0.485162927 |
| chr1:218742970-218743184 | -0.581466013 | 0.209612381 | 0.485166593 |
| chr11:124745895-124746095 | -0.652476812 | 0.209612838 | 0.485166593 |
| chr1:165768443-165769557 | -0.404779345 | 0.209636837 | 0.485196035 |
| chr2:5916061-5916466 | -0.495852589 | 0.209638223 | 0.485196035 |
| chr5:135560846-135561144 | 0.504803572 | 0.20965023 | 0.485209168 |
| chr1:68819438-68819748 | -0.505142204 | 0.209667642 | 0.485231814 |
| chr12:78181600-78181927 | -0.509001996 | 0.209672681 | 0.485231814 |
| chr5:159209422-159210119 | -0.412435226 | 0.209723438 | 0.485334621 |
| chr5:36870499-36870782 | -0.520002563 | 0.20973485 | 0.485346371 |
| chr8:123677767-123677999 | 0.541788475 | 0.209763531 | 0.485398082 |
| chr19:5577751-5578509 | -0.454276759 | 0.209769998 | 0.485398387 |
| chr10:443405-443739 | 0.527985611 | 0.209778966 | 0.485404479 |
| chr7:111633670-111633929 | -0.603914213 | 0.209815816 | 0.485467492 |
| chr10:46216314-46216601 | 0.567219392 | 0.209832511 | 0.485467492 |
| chr20:43457124-43458568 | 0.400590791 | 0.20983692 | 0.485467492 |
| chr17:48261285-48261628 | -0.513826342 | 0.209837059 | 0.485467492 |
| chr14:59311583-59311932 | -0.517603198 | 0.209837878 | 0.485467492 |
| chr12:75329762-75330541 | -0.416699284 | 0.209915863 | 0.485622118 |
| chr3:192726999-192728105 | -0.405901072 | 0.209917389 | 0.485622118 |
| chr10:9051636-9051973 | 0.570412578 | 0.209948737 | 0.485679975 |
| chr1:214280434-214282178 | -0.405189086 | 0.209976533 | 0.485729612 |
| chr17:75877800-75878061 | 0.492673341 | 0.21000773 | 0.485750124 |
| chr10:45235200-45235665 | -0.473570729 | 0.210010083 | 0.485750124 |
| chr20:36093955-36094155 | 0.636363692 | 0.210011978 | 0.485750124 |
| chr1:189250226-189250477 | -0.653868699 | 0.210012099 | 0.485750124 |
| chr15:57376688-57377212 | 0.462563922 | 0.210017098 | 0.485750124 |
| chr2:84495621-84495945 | -0.538291986 | 0.210046494 | 0.485803449 |
| chr11:71215609-71216036 | -0.546556669 | 0.210058642 | 0.485816881 |
| chr2:164704358-164704586 | -0.638582955 | 0.210078533 | 0.485848218 |
| chr9:114587128-114588260 | 0.396679781 | 0.210098569 | 0.485879891 |
| chrX:74250007-74250341 | 0.507508933 | 0.210122145 | 0.485919749 |
| chr5:170056720-170057219 | 0.4834859 | 0.210138086 | 0.485922687 |
| chr7:54788586-54789065 | -0.442926011 | 0.210141495 | 0.485922687 |
| chr7:46493259-46493486 | -0.595985145 | 0.210147609 | 0.485922687 |
| chr14:31025525-31027159 | 0.393039646 | 0.210150045 | 0.485922687 |
| chr3:122680627-122681389 | -0.435328429 | 0.210160645 | 0.485922687 |
| chr17:76839761-76840018 | -0.547664407 | 0.210161467 | 0.485922687 |
| chr15:58077546-58077835 | -0.529704269 | 0.210173137 | 0.485935006 |
| chr12:71567628-71567957 | -0.588045791 | 0.210218374 | 0.486010267 |
| chr14:31127758-31128030 | -0.588045791 | 0.210218374 | 0.486010267 |
| chr20:58447780-58448176 | 0.442008078 | 0.210232491 | 0.486028238 |
| chr17:8998158-8998610 | -0.540165877 | 0.210238978 | 0.486028571 |
| chr15:58056113-58056313 | -0.72933412 | 0.210297855 | 0.486150016 |
| chr9:42900247-42900494 | 0.584559359 | 0.210315389 | 0.48617588 |
| chr3:13135039-13135448 | 0.50095543 | 0.210328607 | 0.486191768 |
| chr14:42224138-42224682 | 0.45232015 | 0.210338612 | 0.486200228 |

| | | | |
|---|---|---|---|
| chr10:61903176-61903979 | -0.424196044 | 0.210357644 | 0.486229551 |
| chr19:36139475-36140195 | 0.412957398 | 0.21036902 | 0.486241179 |
| chr13:110309670-110309873 | 0.729642627 | 0.210396174 | 0.486289271 |
| chr19:47244102-47244674 | 0.436268327 | 0.210412412 | 0.486312134 |
| chr1:108561764-108562059 | -0.46074883 | 0.210432484 | 0.486343853 |
| chr11:75231046-75231529 | -0.510691735 | 0.210447038 | 0.486362819 |
| chr2:132146849-132147083 | -0.644116956 | 0.210464258 | 0.486387947 |
| chr5:139880099-139880392 | -0.634192464 | 0.210486141 | 0.486396862 |
| chr3:156825831-156827080 | -0.394564762 | 0.210487651 | 0.486396862 |
| chr9:133335496-133335696 | 0.585362026 | 0.210489166 | 0.486396862 |
| chr2:217289896-217290208 | -0.579463232 | 0.210493508 | 0.486396862 |
| chr2:81324110-81324424 | 0.496652968 | 0.210549758 | 0.48649769 |
| chr1:196732617-196732817 | 0.630143938 | 0.210555498 | 0.48649769 |
| chr1:227734682-227735824 | 0.404455178 | 0.21055619 | 0.48649769 |
| chr12:219892-220237 | 0.482051879 | 0.210599353 | 0.486582746 |
| chr6:24494611-24495762 | -0.414109612 | 0.210669114 | 0.486729251 |
| chr11:71294087-71294297 | -0.645195057 | 0.210681348 | 0.486731578 |
| chr21:10751669-10751929 | 0.655511119 | 0.210686702 | 0.486731578 |
| chr8:136716800-136717079 | -0.644366899 | 0.210689179 | 0.486731578 |
| chr19:51421920-51422300 | 0.49073101 | 0.21069554 | 0.486731599 |
| chr3:4245559-4246431 | -0.424621659 | 0.210712336 | 0.486755724 |
| chr4:107606742-107606942 | 0.638421911 | 0.210727204 | 0.486775395 |
| chr2:156892414-156893247 | -0.427101933 | 0.210745874 | 0.486803846 |
| chr5:36079183-36079810 | -0.432332424 | 0.210760739 | 0.486806642 |
| chr9:129026544-129027123 | 0.423204139 | 0.2107685 | 0.486806642 |
| chr2:195612930-195613159 | 0.640549608 | 0.210773616 | 0.486806642 |
| chr1:8789636-8789928 | 0.540045117 | 0.210775929 | 0.486806642 |
| chr6:28922915-28924424 | 0.404203019 | 0.210778851 | 0.486806642 |
| chr12:62898709-62899032 | 0.516124778 | 0.210800735 | 0.486840699 |
| chr6:28855321-28855546 | 0.600554477 | 0.210806305 | 0.486840699 |
| chr2:74473045-74473321 | 0.547569528 | 0.210873288 | 0.486980714 |
| chr17:45147683-45147992 | 0.465613788 | 0.210921689 | 0.487077808 |
| chr1:71796099-71796346 | -0.596608927 | 0.210934836 | 0.487093486 |
| chr6:122839516-122839859 | -0.492277003 | 0.210950551 | 0.487115095 |
| chr17:75844290-75845590 | 0.399120545 | 0.210961617 | 0.487125968 |
| chr5:54701293-54701549 | 0.594237493 | 0.210984361 | 0.487143791 |
| chr18:11872560-11872828 | 0.549778852 | 0.210985528 | 0.487143791 |
| chr3:24853751-24854416 | 0.465523304 | 0.210988409 | 0.487143791 |
| chr2:20122968-20123193 | 0.528101589 | 0.211001893 | 0.487148549 |
| chr12:66861216-66861506 | 0.546436901 | 0.211003185 | 0.487148549 |
| chr17:58947811-58948033 | 0.466639671 | 0.211033013 | 0.487202732 |
| chr1:214823277-214823505 | -0.580103737 | 0.211044164 | 0.487213798 |
| chr5:13986060-13986448 | 0.49295714 | 0.211060602 | 0.487237065 |
| chr13:100035920-100036321 | 0.498819127 | 0.211077439 | 0.487261253 |
| chr19:6730289-6731036 | 0.429074352 | 0.211084187 | 0.487262152 |
| chr5:141084462-141084925 | -0.471270254 | 0.211107016 | 0.487292483 |
| chr20:53750835-53751387 | 0.398189709 | 0.211110047 | 0.487292483 |
| chr3:123605158-123605468 | 0.498265583 | 0.211142713 | 0.487342965 |
| chr19:56142238-56142740 | 0.433585295 | 0.211150579 | 0.487342965 |
| chr14:76848936-76849192 | 0.516158301 | 0.211155472 | 0.487342965 |

| | | | |
|---|---|---|---|
| chr8:127737594-127737956 | 0.422593239 | 0.211157358 | 0.487342965 |
| chr2:70293233-70294203 | -0.403493296 | 0.211197186 | 0.48740711 |
| chr5:94525393-94525797 | -0.619483072 | 0.211197874 | 0.48740711 |
| chr8:128744929-128745333 | 0.472071549 | 0.211253502 | 0.487520807 |
| chr2:138564717-138565421 | -0.439090979 | 0.211274107 | 0.487527082 |
| chr17:78128534-78128965 | 0.437251689 | 0.211275234 | 0.487527082 |
| chr4:174156289-174156563 | -0.583154013 | 0.211278028 | 0.487527082 |
| chr19:45455708-45456179 | 0.442247132 | 0.211285653 | 0.487527082 |
| chr2:76256599-76257107 | 0.447595789 | 0.211288035 | 0.487527082 |
| chrX:39688844-39689218 | 0.54419881 | 0.211306463 | 0.48755492 |
| chr18:63850069-63850366 | -0.513126684 | 0.211335852 | 0.487589685 |
| chrX:105763793-105764033 | -0.688240394 | 0.21133706 | 0.487589685 |
| chr8:60052188-60052482 | 0.590345867 | 0.211340621 | 0.487589685 |
| chr7:149690488-149691512 | -0.397135192 | 0.211381385 | 0.487669049 |
| chr8:98169940-98170583 | 0.43018055 | 0.211392637 | 0.487680325 |
| chr19:13169687-13170236 | 0.477862292 | 0.211399583 | 0.487681665 |
| chr5:32224006-32224398 | 0.50363621 | 0.211425029 | 0.487725683 |
| chr1:205515815-205516335 | -0.429912072 | 0.211436825 | 0.48773821 |
| chr13:108915753-108915968 | 0.530662244 | 0.211450783 | 0.487750822 |
| chr4:163111645-163111904 | -0.542859048 | 0.211459438 | 0.487750822 |
| chr18:12702436-12703304 | 0.424137615 | 0.211461389 | 0.487750822 |
| chr3:163786501-163786833 | 0.589419798 | 0.211488741 | 0.487799226 |
| chr1:226626471-226626745 | 0.525012699 | 0.211497789 | 0.487805412 |
| chr1:98991209-98991410 | -0.680428679 | 0.21152756 | 0.487854551 |
| chr11:75420386-75420720 | -0.513995416 | 0.211531828 | 0.487854551 |
| chr3:64099866-64100129 | 0.428842836 | 0.211545385 | 0.487871131 |
| chr2:157447711-157447914 | -0.692189553 | 0.211563053 | 0.487897193 |
| chr3:186425996-186426208 | 0.62143669 | 0.211603884 | 0.487976668 |
| chr15:69074901-69075239 | -0.472563741 | 0.211640982 | 0.488046277 |
| chr10:26879748-26880040 | -0.540502422 | 0.211646808 | 0.488046277 |
| chr22:30968440-30968750 | 0.483752748 | 0.211676068 | 0.48809906 |
| chr1:86474188-86474431 | -0.623186722 | 0.21168646 | 0.488108334 |
| chr15:96331469-96331755 | 0.450573883 | 0.211695068 | 0.488113494 |
| chr8:71827691-71828286 | -0.50198563 | 0.211774181 | 0.488272521 |
| chr2:190996327-190996713 | -0.462262869 | 0.211776783 | 0.488272521 |
| chr9:90895281-90895545 | -0.528178198 | 0.211827172 | 0.488374001 |
| chr19:35055746-35056069 | -0.513698925 | 0.211841028 | 0.488382075 |
| chr2:26401708-26402353 | -0.433038225 | 0.211846986 | 0.488382075 |
| chr3:72661078-72661409 | -0.498400196 | 0.211849795 | 0.488382075 |
| chr1:1723909-1724808 | 0.417861525 | 0.211858662 | 0.488387821 |
| chr3:163939936-163940405 | -0.460764325 | 0.211892058 | 0.488450111 |
| chr14:104665665-104666046 | 0.470972801 | 0.211908273 | 0.488472795 |
| chr17:79808735-79809179 | 0.445877063 | 0.211960032 | 0.488577406 |
| chr5:141878394-141878682 | 0.492752811 | 0.21198011 | 0.488608989 |
| chr16:14632615-14633164 | 0.413209007 | 0.212012833 | 0.488645759 |
| chr7:36115526-36115850 | -0.522165342 | 0.212014535 | 0.488645759 |
| chr21:41583328-41583657 | 0.495181925 | 0.21202883 | 0.488645759 |
| chr10:49464781-49465067 | -0.56661948 | 0.212031403 | 0.488645759 |
| chr11:110763498-110763873 | -0.56661948 | 0.212031403 | 0.488645759 |
| chr3:85270982-85271281 | -0.514361778 | 0.212034327 | 0.488645759 |

| | | | |
|---|---|---|---|
| chr10:79826374-79826910 | 0.449977543 | 0.212043444 | 0.488652073 |
| chr1:230642167-230642970 | 0.412738302 | 0.212065948 | 0.488689235 |
| chr1:207589584-207589878 | 0.519224605 | 0.212082906 | 0.488713615 |
| chrX:12946796-12947557 | -0.420214184 | 0.212102404 | 0.488743847 |
| chr20:51750145-51750766 | -0.403377534 | 0.212110023 | 0.488746706 |
| chr1:40143992-40144269 | -0.533781047 | 0.212126546 | 0.488770078 |
| chrX:100379219-100379688 | -0.477534154 | 0.212147106 | 0.488778794 |
| chr16:11405618-11405862 | -0.563739546 | 0.212147205 | 0.488778794 |
| chr1:156797367-156797568 | 0.587238656 | 0.212155735 | 0.488778794 |
| chr5:98772953-98774060 | -0.394184591 | 0.212155845 | 0.488778794 |
| chr13:99610637-99611001 | 0.517526488 | 0.212176332 | 0.488811296 |
| chr1:201946345-201946736 | -0.499164317 | 0.21218487 | 0.488816268 |
| chr5:103373786-103374198 | -0.506105256 | 0.212193848 | 0.488822255 |
| chr13:88000279-88000591 | -0.550958795 | 0.212209035 | 0.488837674 |
| chr8:22604431-22605154 | 0.422086955 | 0.212213301 | 0.488837674 |
| chr12:12990726-12991153 | 0.465574911 | 0.2122257 | 0.488851538 |
| chr14:97331172-97331422 | 0.626185186 | 0.212238335 | 0.488854701 |
| chr5:135064107-135064365 | 0.59685768 | 0.212239833 | 0.488854701 |
| chr11:63938044-63939058 | 0.403900315 | 0.212264756 | 0.48889741 |
| chr10:47208378-47208625 | -0.64293966 | 0.212297477 | 0.488958075 |
| chr1:61724566-61725640 | -0.407370393 | 0.212310243 | 0.488972781 |
| chr2:160384818-160385080 | -0.626376098 | 0.212322431 | 0.488977236 |
| chr3:23713038-23713276 | 0.566456543 | 0.212324941 | 0.488977236 |
| chr2:65238043-65238308 | -0.651809151 | 0.212333239 | 0.48898165 |
| chr17:61799912-61800214 | 0.414852491 | 0.212347108 | 0.488998892 |
| chr2:188629357-188629663 | -0.537637513 | 0.212360452 | 0.489002964 |
| chr5:59543603-59543817 | -0.636330021 | 0.21236164 | 0.489002964 |
| chr10:68406159-68407604 | -0.399081366 | 0.212371202 | 0.489010286 |
| chr3:62041787-62042059 | 0.468935954 | 0.212378244 | 0.489011804 |
| chr2:95374836-95375040 | 0.651247926 | 0.212391999 | 0.489028779 |
| chr13:50644153-50644999 | 0.425425441 | 0.212486926 | 0.489231999 |
| chr17:40648009-40648695 | 0.468684526 | 0.21249303 | 0.489231999 |
| chr3:167057570-167057957 | -0.507027259 | 0.212502976 | 0.489240198 |
| chr4:151766748-151767085 | -0.513979092 | 0.212511518 | 0.489245164 |
| chr12:119423348-119423595 | -0.603655656 | 0.212521771 | 0.489254068 |
| chr8:30911832-30912502 | -0.475341722 | 0.212546111 | 0.4892954 |
| chr2:10435016-10435232 | -0.512227889 | 0.212583443 | 0.489365754 |
| chr9:70113547-70113752 | -0.658889427 | 0.212589445 | 0.489365754 |
| chr9:34681718-34681989 | -0.621510312 | 0.212640018 | 0.489467464 |
| chr21:22477300-22477500 | -0.690620609 | 0.212650296 | 0.489476418 |
| chr9:98604057-98604337 | -0.559071647 | 0.212693247 | 0.489552348 |
| chr1:156751555-156752133 | 0.408064956 | 0.212696062 | 0.489552348 |
| chr6:34392357-34393104 | 0.425129717 | 0.212710575 | 0.489563575 |
| chr2:168948754-168949001 | -0.663216875 | 0.212717661 | 0.489563575 |
| chr20:53345469-53346059 | -0.475721297 | 0.212720108 | 0.489563575 |
| chrX:121013979-121014749 | 0.444722711 | 0.212739421 | 0.489593318 |
| chr1:1330839-1331363 | -0.47067064 | 0.21275557 | 0.489615776 |
| chr1:172090054-172090539 | -0.493189492 | 0.212765197 | 0.489623226 |
| chr2:43848278-43848531 | 0.552479277 | 0.21277948 | 0.489632466 |
| chr7:2523950-2524432 | -0.423807269 | 0.212781993 | 0.489632466 |

| | | | |
|---|---|---|---|
| chr6:99349265-99350157 | 0.412116501 | 0.212790422 | 0.489637158 |
| chr1:15128556-15128839 | 0.547144109 | 0.212799931 | 0.489644334 |
| chr3:3133521-3133868 | -0.522017741 | 0.212811037 | 0.489655184 |
| chr15:100713565-100713799 | 0.517401371 | 0.212821651 | 0.489664902 |
| chr8:28701943-28702191 | 0.535922093 | 0.212860173 | 0.489710698 |
| chr1:24642782-24643732 | 0.410594618 | 0.212860396 | 0.489710698 |
| chr1:27392034-27392778 | 0.421801229 | 0.212860729 | 0.489710698 |
| chr1:165258019-165258351 | 0.502717339 | 0.212867394 | 0.489711326 |
| chr9:112097711-112098108 | -0.50366253 | 0.2128797 | 0.489724934 |
| chr2:102479203-102479437 | 0.681763852 | 0.212897154 | 0.489749345 |
| chr8:120971013-120971213 | 0.603708098 | 0.212903095 | 0.489749345 |
| chr11:62612165-62613526 | 0.398275853 | 0.212915306 | 0.489762732 |
| chr10:130109800-130110024 | 0.489486827 | 0.212927041 | 0.489775022 |
| chr2:24805764-24806599 | -0.418821104 | 0.212947854 | 0.489804602 |
| chr20:15850410-15850684 | 0.616786479 | 0.212952686 | 0.489804602 |
| chr7:75638481-75639243 | -0.45234443 | 0.212992351 | 0.489881129 |
| chr9:130355833-130356033 | 0.61938219 | 0.213015124 | 0.489918799 |
| chr12:29171998-29172238 | -0.719606027 | 0.213024086 | 0.489922279 |
| chr1:234458461-234459000 | 0.480099218 | 0.213034193 | 0.489922279 |
| chr5:56460704-56460913 | -0.595613728 | 0.213035819 | 0.489922279 |
| chr13:40069259-40069522 | -0.517639549 | 0.213089486 | 0.49003099 |
| chr2:3483784-3484045 | 0.515329303 | 0.213106788 | 0.490042381 |
| chr20:59087871-59088176 | 0.479272273 | 0.21310723 | 0.490042381 |
| chr16:19095088-19095377 | -0.57272477 | 0.213120196 | 0.490057489 |
| chr1:29236314-29237012 | 0.438344317 | 0.213131403 | 0.490062549 |
| chr1:213015119-213016215 | -0.400257111 | 0.213135188 | 0.490062549 |
| chr7:8436882-8437278 | 0.425543896 | 0.21318649 | 0.490151272 |
| chr13:24580322-24580541 | -0.619380863 | 0.213186569 | 0.490151272 |
| chr15:89971526-89971915 | 0.455382773 | 0.21323558 | 0.490236853 |
| chr15:40865796-40866430 | -0.418250139 | 0.213238586 | 0.490236853 |
| chr20:57173748-57174217 | 0.397832223 | 0.213242986 | 0.490236853 |
| chr15:75347930-75349034 | 0.414342328 | 0.21325111 | 0.490236911 |
| chr22:26360309-26360796 | 0.491990121 | 0.213261337 | 0.490236911 |
| chr22:20027997-20028197 | 0.61240985 | 0.213268239 | 0.490236911 |
| chr16:67159258-67159458 | 0.515668646 | 0.21327563 | 0.490236911 |
| chr19:29941816-29942813 | 0.418549962 | 0.213277679 | 0.490236911 |
| chr7:95871220-95871623 | -0.584209188 | 0.21328295 | 0.490236911 |
| chr15:92655105-92655994 | 0.419615333 | 0.213299518 | 0.490236911 |
| chr8:133674682-133675089 | 0.482273274 | 0.213305946 | 0.490236911 |
| chr12:65983657-65983981 | -0.536575066 | 0.213306216 | 0.490236911 |
| chr6:53309167-53309377 | 0.649185576 | 0.213307582 | 0.490236911 |
| chr3:161313114-161313324 | 0.595653856 | 0.213313391 | 0.490236911 |
| chr17:65987791-65988310 | -0.449657609 | 0.213328533 | 0.490257006 |
| chr1:21638916-21639271 | 0.529948152 | 0.213376067 | 0.490351538 |
| chrX:118116214-118117270 | 0.437235983 | 0.213398002 | 0.490387239 |
| chr1:168879943-168880283 | 0.561455176 | 0.213409438 | 0.490397835 |
| chr3:141201840-141202086 | -0.652320025 | 0.213415413 | 0.490397835 |
| chr1:64898093-64898820 | -0.43491097 | 0.213429052 | 0.490414468 |
| chrX:40109011-40109262 | 0.511643864 | 0.213449966 | 0.49043717 |
| chr13:28138030-28138585 | -0.41852491 | 0.213451734 | 0.49043717 |

| | | | |
|---|---|---|---|
| chr1:36415683-36415958 | 0.5123686 | 0.213483125 | 0.490494588 |
| chr7:51666613-51666819 | -0.640519575 | 0.213497758 | 0.4905135 |
| chr5:114803772-114804100 | -0.525349391 | 0.213536918 | 0.490587995 |
| chr6:28837788-28839213 | 0.398652779 | 0.213542988 | 0.490587995 |
| chr9:71910666-71912050 | -0.392265878 | 0.213593834 | 0.490656527 |
| chr2:23199430-23199630 | 0.634059236 | 0.213595016 | 0.490656527 |
| chr1:63007129-63007375 | -0.676588853 | 0.213595127 | 0.490656527 |
| chr16:24883817-24884017 | 0.643172907 | 0.213598433 | 0.490656527 |
| chr11:72781027-72781472 | 0.434883622 | 0.213625029 | 0.49070291 |
| chr6:34696062-34697294 | -0.405153877 | 0.213632435 | 0.490705211 |
| chr5:56534712-56534998 | -0.616869867 | 0.213662909 | 0.490754717 |
| chr17:75105850-75106072 | 0.522288762 | 0.213668799 | 0.490754717 |
| chr9:96417105-96417608 | 0.434923403 | 0.213673202 | 0.490754717 |
| chr1:215942963-215943163 | -0.666614219 | 0.213680594 | 0.490756983 |
| chr3:135855915-135856197 | 0.549305464 | 0.213693633 | 0.490765286 |
| chrX:54029600-54029980 | 0.51968159 | 0.213702045 | 0.490765286 |
| chr8:18954780-18955118 | 0.522470345 | 0.213703424 | 0.490765286 |
| chr10:133134696-133135070 | 0.513127513 | 0.21378167 | 0.490930263 |
| chrX:107324185-107325347 | -0.401933117 | 0.213788385 | 0.490930969 |
| chr9:98071794-98072350 | -0.457136755 | 0.213850867 | 0.491054264 |
| chr19:46863901-46864239 | -0.452279483 | 0.213855122 | 0.491054264 |
| chr14:104681667-104682095 | 0.46660935 | 0.213861304 | 0.491054264 |
| chr3:155949282-155949839 | -0.451086372 | 0.213880334 | 0.491083245 |
| chr2:104434427-104434771 | -0.531968319 | 0.213901287 | 0.491116637 |
| chr7:149296295-149296555 | -0.572172089 | 0.213913233 | 0.491129349 |
| chr15:92044527-92044728 | 0.617119817 | 0.213928357 | 0.491135752 |
| chr9:124617497-124617806 | 0.52475187 | 0.213928842 | 0.491135752 |
| chr10:124743336-124743774 | 0.473977017 | 0.213955154 | 0.491169718 |
| chr10:99387255-99387497 | 0.596077233 | 0.213956457 | 0.491169718 |
| chr13:102916081-102916340 | -0.551138817 | 0.213967264 | 0.49117981 |
| chr16:68351841-68352302 | 0.4413398 | 0.213983598 | 0.491188541 |
| chr11:38461687-38461987 | 0.636912325 | 0.213983888 | 0.491188541 |
| chr1:206925268-206925715 | -0.432379188 | 0.214001942 | 0.491203317 |
| chr6:67778081-67778281 | -0.648374586 | 0.214003147 | 0.491203317 |
| chr10:73876169-73876524 | -0.470370666 | 0.214015467 | 0.4912136 |
| chr14:75978408-75978814 | -0.447633943 | 0.214020449 | 0.4912136 |
| chr19:51690748-51691071 | 0.455489791 | 0.214034689 | 0.491214064 |
| chr19:17403310-17403593 | -0.594359043 | 0.214042592 | 0.491214064 |
| chr6:113493921-113494490 | 0.42208464 | 0.214046544 | 0.491214064 |
| chr12:89525272-89526373 | -0.399621564 | 0.214056568 | 0.491214064 |
| chr11:83092703-83093371 | -0.409824271 | 0.214059566 | 0.491214064 |
| chr20:17681396-17681679 | 0.46319803 | 0.214065463 | 0.491214064 |
| chr5:7887062-7887390 | -0.588509737 | 0.214065527 | 0.491214064 |
| chr6:73798579-73798921 | -0.516572373 | 0.214092418 | 0.491261058 |
| chr19:4038489-4038689 | -0.652736654 | 0.214114741 | 0.491285915 |
| chr5:108701759-108702003 | -0.706945609 | 0.214116074 | 0.491285915 |
| chr1:201307751-201308448 | -0.402648451 | 0.214125617 | 0.491290309 |
| chr15:55897591-55898052 | 0.459575151 | 0.214130813 | 0.491290309 |
| chr9:83199346-83199817 | -0.507892105 | 0.214167419 | 0.491354728 |
| chr3:64263827-64265121 | 0.389672648 | 0.214171716 | 0.491354728 |

| | | | |
|---|---|---|---|
| chr14:74978622-74979021 | 0.4871254 | 0.214272952 | 0.49154897 |
| chr3:17039630-17039885 | 0.606686607 | 0.214273389 | 0.49154897 |
| chr14:94486176-94486837 | 0.430504862 | 0.214275628 | 0.49154897 |
| chr5:90783738-90784058 | 0.485009301 | 0.21428604 | 0.491558138 |
| chr3:136250979-136251236 | -0.472519062 | 0.214292456 | 0.491558141 |
| chr19:55080158-55080492 | 0.444849343 | 0.214309154 | 0.491581726 |
| chr3:157249035-157249258 | 0.568353592 | 0.214326376 | 0.491606513 |
| chr1:204763702-204764115 | -0.442330935 | 0.214363325 | 0.491676546 |
| chr13:110644882-110645233 | -0.505065058 | 0.214376279 | 0.491687084 |
| chr17:57121456-57121941 | -0.457489702 | 0.214380754 | 0.491687084 |
| chr4:16125876-16126458 | -0.452378639 | 0.214391165 | 0.491696244 |
| chr18:36284772-36285007 | -0.61852368 | 0.214411616 | 0.491702729 |
| chr4:54841109-54841644 | -0.511824485 | 0.214411866 | 0.491702729 |
| chr10:30522489-30523097 | -0.427272076 | 0.214413244 | 0.491702729 |
| chr8:22747802-22748023 | -0.563797367 | 0.214446266 | 0.491763738 |
| chr20:21096314-21096554 | 0.654070629 | 0.214493682 | 0.491843031 |
| chr3:108942480-108942704 | 0.654070629 | 0.214493682 | 0.491843031 |
| chr1:106799034-106799454 | 0.41328223 | 0.214512465 | 0.491871382 |
| chrX:94524887-94525232 | -0.628778837 | 0.214526222 | 0.491888206 |
| chr3:132399302-132399612 | -0.494174552 | 0.214569498 | 0.49196199 |
| chr4:17472475-17472720 | 0.617322715 | 0.214572488 | 0.49196199 |
| chr5:127176836-127177163 | -0.60834029 | 0.214577663 | 0.49196199 |
| chr3:57228441-57228671 | -0.582432316 | 0.214592531 | 0.491976795 |
| chr17:60212127-60212403 | 0.497699579 | 0.214596962 | 0.491976795 |
| chr15:63121601-63122734 | 0.400428022 | 0.214612302 | 0.491997241 |
| chr15:40814959-40815667 | 0.436743482 | 0.214621728 | 0.492004131 |
| chr1:46037078-46037361 | -0.523964037 | 0.214639799 | 0.492017754 |
| chr18:78774629-78774961 | -0.550500368 | 0.214640514 | 0.492017754 |
| chr11:66388861-66389466 | -0.450032177 | 0.214673714 | 0.492079138 |
| chr20:54938476-54940525 | 0.388307925 | 0.214682085 | 0.492083604 |
| chr15:42736705-42737488 | 0.416549856 | 0.214703005 | 0.492102181 |
| chr7:121421004-121421401 | -0.500960688 | 0.21470338 | 0.492102181 |
| chr8:95237191-95237498 | -0.474946457 | 0.214714289 | 0.492102181 |
| chr10:93858080-93858280 | 0.674307932 | 0.214715879 | 0.492102181 |
| chr8:80230814-80231309 | 0.437086654 | 0.214736923 | 0.492132836 |
| chr13:38788115-38788351 | -0.754876671 | 0.214742101 | 0.492132836 |
| chr6:149082300-149082523 | 0.550210193 | 0.214753624 | 0.492144524 |
| chr1:239308789-239309089 | -0.547521179 | 0.214796257 | 0.492227503 |
| chr1:204445889-204446178 | -0.452543224 | 0.214837375 | 0.492307006 |
| chr2:16012505-16013408 | -0.418493687 | 0.214850404 | 0.492322139 |
| chr5:52668771-52669204 | -0.525060189 | 0.214908353 | 0.492440198 |
| chr8:27325195-27325813 | 0.426675569 | 0.214928208 | 0.492470967 |
| chr1:228173566-228174200 | 0.416981465 | 0.214940202 | 0.492483721 |
| chr17:39952695-39953580 | 0.409134228 | 0.214957791 | 0.49250066 |
| chr2:177224551-177224943 | -0.494601352 | 0.21496045 | 0.49250066 |
| chr12:49842067-49843349 | 0.401552336 | 0.21497668 | 0.492523117 |
| chr1:218330113-218330361 | -0.517708646 | 0.214984388 | 0.492526051 |
| chr10:104291396-104291811 | 0.440444322 | 0.215003728 | 0.492555632 |
| chr2:186485704-186486836 | -0.39138096 | 0.215055171 | 0.492658754 |
| chr19:2950405-2951288 | -0.411960552 | 0.21506298 | 0.492661911 |

| | | | |
|---|---|---|---|
| chr9:132901929-132902176 | 0.536737156 | 0.21510636 | 0.492743714 |
| chr10:123621705-123622076 | 0.581590013 | 0.215111551 | 0.492743714 |
| chr7:152894416-152894616 | -0.622271726 | 0.215148825 | 0.492814363 |
| chr10:34826248-34826480 | -0.581599555 | 0.215184732 | 0.492876592 |
| chr8:101301397-101301732 | 0.46817455 | 0.215188858 | 0.492876592 |
| chr4:76611552-76611884 | -0.552291447 | 0.215198548 | 0.492881423 |
| chr8:101243729-101243991 | 0.53242873 | 0.215203832 | 0.492881423 |
| chr22:33325412-33325646 | 0.597061952 | 0.215218313 | 0.492899856 |
| chr2:159146970-159147170 | -0.676143625 | 0.215244591 | 0.492936084 |
| chr1:243786653-243786853 | 0.640423463 | 0.215246998 | 0.492936084 |
| chr19:29056145-29056463 | 0.537575128 | 0.215283617 | 0.49300521 |
| chr3:51995655-51996132 | 0.450367574 | 0.215293729 | 0.493013631 |
| chr21:45541576-45541985 | 0.45292853 | 0.21532835 | 0.493078175 |
| chr5:74394943-74395214 | -0.562707978 | 0.215338973 | 0.493087765 |
| chr9:73515812-73516012 | -0.682296277 | 0.215350368 | 0.493099122 |
| chr3:49469485-49470794 | 0.400706493 | 0.215367249 | 0.493123037 |
| chr2:19073614-19073864 | -0.568479684 | 0.215389012 | 0.493157459 |
| chr17:7280557-7281780 | -0.404239566 | 0.215395155 | 0.493157459 |
| chr10:34206132-34206332 | -0.675513108 | 0.215419418 | 0.493198274 |
| chr16:30090040-30090255 | 0.507369353 | 0.215429431 | 0.493206462 |
| chr13:44249557-44249832 | -0.539126691 | 0.21546605 | 0.493275558 |
| chr5:35230167-35230910 | -0.423275061 | 0.215478714 | 0.493285367 |
| chr15:43746615-43747343 | 0.444442556 | 0.215489082 | 0.493285367 |
| chr1:248807178-248807488 | -0.44663494 | 0.21549727 | 0.493285367 |
| chr17:59620517-59620780 | 0.432244483 | 0.215498418 | 0.493285367 |
| chr11:23990439-23990832 | -0.626591699 | 0.215502524 | 0.493285367 |
| chr7:47149437-47149663 | 0.683807473 | 0.215535706 | 0.493346583 |
| chr11:126340154-126340386 | -0.555377576 | 0.215580203 | 0.493433694 |
| chr16:81730087-81730815 | -0.426570374 | 0.215605032 | 0.493475783 |
| chr12:113358117-113359249 | 0.402675219 | 0.215613458 | 0.493480327 |
| chr4:126779457-126780153 | 0.447084568 | 0.215627826 | 0.493484901 |
| chr12:27076978-27077310 | -0.541050247 | 0.215628338 | 0.493484901 |
| chr14:59127198-59127480 | 0.518768724 | 0.215686056 | 0.493593037 |
| chr3:193712010-193712741 | 0.406822048 | 0.215688472 | 0.493593037 |
| chr2:63840327-63841938 | -0.391544987 | 0.215696617 | 0.493596935 |
| chr18:28189538-28189738 | -0.715160274 | 0.21572252 | 0.493634463 |
| chr2:10844058-10844468 | -0.486122342 | 0.215725901 | 0.493634463 |
| chrX:11191249-11191480 | -0.621186961 | 0.215739551 | 0.493641152 |
| chr11:119067398-119068170 | 0.408769701 | 0.215741709 | 0.493641152 |
| chr18:23053737-23054104 | -0.552408861 | 0.215767198 | 0.493684731 |
| chr3:146184734-146184946 | -0.589451975 | 0.21578344 | 0.49370715 |
| chr5:62739841-62740237 | -0.563511376 | 0.215802451 | 0.493718237 |
| chr20:49541824-49542223 | 0.457484499 | 0.215808778 | 0.493718237 |
| chr3:134794891-134795638 | -0.411375445 | 0.215819593 | 0.493718237 |
| chr3:69200382-69201075 | -0.406372607 | 0.215825247 | 0.493718237 |
| chr2:102443316-102443523 | 0.629148757 | 0.215827441 | 0.493718237 |
| chr2:2057181-2057451 | -0.584375303 | 0.215828129 | 0.493718237 |
| chr2:184067560-184067801 | -0.58397282 | 0.215833391 | 0.493718237 |
| chr3:148794325-148794641 | -0.498579022 | 0.215862145 | 0.493769272 |
| chr3:174862157-174862585 | -0.42012437 | 0.215879234 | 0.49379362 |

| | | | |
|---|---|---|---|
| chr8:51888202-51888427 | -0.604609398 | 0.2158923 | 0.493803759 |
| chr10:133307806-133308086 | 0.449168734 | 0.215896556 | 0.493803759 |
| chr14:92880834-92881704 | 0.424070433 | 0.215925023 | 0.493853592 |
| chr9:89593778-89594226 | -0.44122977 | 0.215931235 | 0.493853592 |
| chr19:33868849-33869049 | 0.674954814 | 0.215984787 | 0.493958181 |
| chr5:149581161-149581836 | 0.438462309 | 0.215989858 | 0.493958181 |
| chr8:30338490-30338811 | -0.538772253 | 0.216010314 | 0.493975886 |
| chr19:51688931-51689366 | 0.439655714 | 0.216010493 | 0.493975886 |
| chr20:13286605-13287012 | 0.480594098 | 0.216053219 | 0.494036088 |
| chr16:46972784-46974119 | 0.40902539 | 0.216056828 | 0.494036088 |
| chr5:34523160-34523461 | -0.524794723 | 0.216067091 | 0.494036088 |
| chr2:96769717-96770138 | -0.438133517 | 0.216070161 | 0.494036088 |
| chr22:20084715-20085047 | -0.520515802 | 0.216080508 | 0.494036088 |
| chr17:74862273-74862487 | 0.536855147 | 0.216085059 | 0.494036088 |
| chr6:3227436-3227676 | -0.694554258 | 0.216086664 | 0.494036088 |
| chr19:7522347-7522758 | 0.443616946 | 0.216092215 | 0.494036088 |
| chr22:21009681-21009900 | 0.575271747 | 0.216094849 | 0.494036088 |
| chr2:216283528-216283862 | -0.532329226 | 0.216110932 | 0.494058117 |
| chrX:132631729-132631937 | 0.604521251 | 0.216123224 | 0.494071476 |
| chr1:153986164-153986558 | 0.421561629 | 0.216160075 | 0.494140976 |
| chr3:177211958-177212513 | 0.449406383 | 0.216178354 | 0.494144227 |
| chr17:18363053-18363818 | -0.408914457 | 0.216178837 | 0.494144227 |
| chr12:79367173-79367622 | -0.492982248 | 0.216192179 | 0.494144227 |
| chrX:97189709-97190054 | -0.494224948 | 0.216197798 | 0.494144227 |
| chr12:12070775-12071245 | -0.513223231 | 0.216206274 | 0.494144227 |
| chr17:68771888-68772313 | -0.461251707 | 0.216206521 | 0.494144227 |
| chr1:242467606-242467891 | -0.499278842 | 0.216206641 | 0.494144227 |
| chr12:118161048-118161270 | 0.687136167 | 0.216228158 | 0.494178665 |
| chr9:88378315-88378515 | 0.636956675 | 0.216245131 | 0.494199112 |
| chr21:40993787-40994096 | -0.56893593 | 0.216250614 | 0.494199112 |
| chr17:21051717-21052114 | -0.561614371 | 0.216256454 | 0.494199112 |
| chr21:41832313-41832513 | 0.580016154 | 0.216269427 | 0.494212475 |
| chr10:123743474-123743674 | 0.574070328 | 0.216279614 | 0.494212475 |
| chr16:76081820-76082372 | -0.506871386 | 0.216281651 | 0.494212475 |
| chr17:77375946-77376191 | -0.567030072 | 0.216310823 | 0.494264392 |
| chr12:132686599-132687993 | 0.399796776 | 0.216319473 | 0.494269419 |
| chr12:109864346-109864666 | -0.495791227 | 0.21635495 | 0.49433463 |
| chr11:11966909-11967192 | 0.498552855 | 0.216360917 | 0.49433463 |
| chr9:86093757-86094252 | 0.436844477 | 0.216385991 | 0.494377177 |
| chr7:34906207-34906844 | -0.448934007 | 0.21640249 | 0.494384744 |
| chr1:160830602-160830814 | -0.654621997 | 0.216409652 | 0.494384744 |
| chr3:104604404-104604604 | -0.648109744 | 0.216410616 | 0.494384744 |
| chr16:56311268-56311506 | -0.588649616 | 0.216415112 | 0.494384744 |
| chr15:67848942-67849209 | -0.609603833 | 0.21642295 | 0.49438791 |
| chr2:156400199-156401574 | -0.388119139 | 0.216454212 | 0.494444582 |
| chr10:112183254-112184180 | -0.411391906 | 0.216461368 | 0.494446188 |
| chr6:132298399-132298645 | 0.620533151 | 0.216480929 | 0.494476128 |
| chr17:78241290-78241762 | 0.471333839 | 0.216493623 | 0.494490382 |
| chr4:75561770-75562397 | 0.41111072 | 0.216513472 | 0.494520978 |
| chr9:109969201-109969401 | 0.595272394 | 0.216532264 | 0.494540388 |

| | | | |
|---|---|---|---|
| chr2:201233264-201233741 | -0.421777081 | 0.216534879 | 0.494540388 |
| chr3:192467298-192467597 | -0.500614614 | 0.216551945 | 0.494546111 |
| chr10:19586648-19587374 | -0.464871714 | 0.216554938 | 0.494546111 |
| chr11:21323737-21324012 | -0.636330679 | 0.216556748 | 0.494546111 |
| chr1:212630035-212630563 | -0.449191967 | 0.216574185 | 0.494571192 |
| chr10:29583708-29583975 | -0.601720188 | 0.216584116 | 0.494575776 |
| chr14:66746370-66746645 | 0.52647633 | 0.216593443 | 0.494575776 |
| chr12:112387411-112387673 | 0.521154167 | 0.21659933 | 0.494575776 |
| chr9:90226840-90227338 | 0.469231443 | 0.216602012 | 0.494575776 |
| chr9:126662386-126662891 | -0.428589103 | 0.216610723 | 0.494580929 |
| chr1:3531295-3531779 | 0.436784141 | 0.216621888 | 0.494591684 |
| chr1:240947005-240947246 | 0.53324085 | 0.216628843 | 0.494592826 |
| chr5:67003750-67004809 | -0.405191977 | 0.216640253 | 0.494596559 |
| chr16:3100057-3100466 | 0.434917248 | 0.2166472 | 0.494596559 |
| chr2:23940181-23941047 | -0.406540314 | 0.216654314 | 0.494596559 |
| chr12:49567398-49567632 | 0.517675793 | 0.216656299 | 0.494596559 |
| chr17:52751181-52751723 | -0.501919569 | 0.216678948 | 0.494633528 |
| chr15:101641145-101641363 | 0.571554309 | 0.216701314 | 0.494669848 |
| chr10:46523506-46523937 | -0.458626629 | 0.21671497 | 0.494674031 |
| chr2:73244090-73244373 | -0.523185607 | 0.216716059 | 0.494674031 |
| chr19:2475888-2476242 | 0.452810016 | 0.216739656 | 0.494713156 |
| chr14:93178079-93178416 | 0.462038302 | 0.216753834 | 0.49473078 |
| chr3:58034823-58035107 | -0.619719287 | 0.216862078 | 0.494963097 |
| chr19:15837409-15837624 | -0.576137181 | 0.216897355 | 0.495014124 |
| chr6:56757681-56757951 | -0.576137181 | 0.216897355 | 0.495014124 |
| chr1:177969775-177970139 | -0.559129767 | 0.216908458 | 0.495024718 |
| chr3:155012661-155012905 | -0.527379375 | 0.216936513 | 0.495073999 |
| chr1:247111250-247112763 | -0.391565039 | 0.21696299 | 0.495117198 |
| chr17:66087709-66088464 | 0.443885321 | 0.216973385 | 0.495117198 |
| chr1:186082319-186082637 | -0.527294115 | 0.216974828 | 0.495117198 |
| chr8:133209276-133209596 | 0.46367356 | 0.216984763 | 0.495125123 |
| chr14:86373511-86374455 | 0.42778527 | 0.217002348 | 0.495150504 |
| chr4:76767902-76768732 | 0.420195138 | 0.217026809 | 0.495191571 |
| chr19:5352207-5352885 | 0.470786501 | 0.217052714 | 0.495235931 |
| chr4:78032462-78032800 | -0.549965472 | 0.217082311 | 0.495278947 |
| chr16:4360659-4360933 | 0.556368923 | 0.217087269 | 0.495278947 |
| chr15:100934891-100935192 | -0.51245942 | 0.217090959 | 0.495278947 |
| chr20:51542034-51542374 | 0.445320257 | 0.217098299 | 0.495279593 |
| chr7:87934723-87935180 | -0.474079087 | 0.21710417 | 0.495279593 |
| chr17:30378674-30379011 | 0.471392512 | 0.217119924 | 0.495300787 |
| chr3:13092157-13092497 | 0.443097492 | 0.217143539 | 0.495326349 |
| chr17:61554352-61554845 | 0.398010141 | 0.217148518 | 0.495326349 |
| chr14:53956515-53957082 | 0.416607007 | 0.217150523 | 0.495326349 |
| chr12:67193819-67194057 | 0.550967359 | 0.217168679 | 0.495353017 |
| chr6:120826095-120826557 | -0.533123778 | 0.217194356 | 0.495396838 |
| chr1:247210153-247211163 | -0.391387611 | 0.217205992 | 0.49540557 |
| chr10:97400127-97400585 | -0.435701591 | 0.217211116 | 0.49540557 |
| chr16:30400169-30400445 | 0.51659977 | 0.217232923 | 0.495425812 |
| chr17:79844821-79845040 | 0.51659977 | 0.217232923 | 0.495425812 |
| chr16:24340796-24340996 | 0.681687403 | 0.217240386 | 0.495428086 |

| | | | |
|---|---|---|---|
| chr1:206847537-206847909 | -0.468367122 | 0.217263579 | 0.495466233 |
| chr19:41421536-41421759 | -0.604529642 | 0.217271825 | 0.495470291 |
| chr17:64965766-64966262 | -0.440532035 | 0.217304593 | 0.495519555 |
| chr15:89004168-89004379 | 0.541727649 | 0.217308699 | 0.495519555 |
| chr1:179586195-179586891 | 0.411108559 | 0.217312829 | 0.495519555 |
| chr14:72561425-72561634 | 0.626961871 | 0.217327523 | 0.495538312 |
| chr3:64419958-64420203 | 0.44818752 | 0.21734261 | 0.495557965 |
| chr10:5678664-5678872 | -0.599869066 | 0.217388606 | 0.495648092 |
| chr4:177339840-177340160 | -0.50563381 | 0.2174214 | 0.495708111 |
| chr7:264794-265004 | 0.504012012 | 0.217439335 | 0.495732717 |
| chr2:60642398-60642693 | -0.520707833 | 0.217450248 | 0.495732717 |
| chr20:4221221-4221730 | 0.465859475 | 0.217454184 | 0.495732717 |
| chr21:44911296-44912266 | -0.414012608 | 0.217463765 | 0.495732717 |
| chr5:79548054-79548340 | -0.529873791 | 0.217469186 | 0.495732717 |
| chr5:98796243-98796668 | -0.458156252 | 0.217471011 | 0.495732717 |
| chr4:150078433-150079625 | -0.416591772 | 0.217479666 | 0.495737698 |
| chr20:53642804-53643083 | 0.406635451 | 0.217517221 | 0.495808552 |
| chr21:25708506-25708754 | 0.582673562 | 0.217539786 | 0.495837903 |
| chr14:50047287-50047554 | 0.553460332 | 0.21754304 | 0.495837903 |
| chr1:207393675-207393974 | 0.506839533 | 0.217549517 | 0.495837918 |
| chr17:59120601-59120814 | 0.469988868 | 0.217556156 | 0.495838299 |
| chr12:52879444-52880249 | 0.401321025 | 0.217623719 | 0.495977531 |
| chr2:177489787-177490071 | -0.58409292 | 0.217651709 | 0.496018394 |
| chr17:58809366-58809638 | 0.457780107 | 0.217659361 | 0.496018394 |
| chr16:81914591-81914820 | 0.584895727 | 0.217661069 | 0.496018394 |
| chr1:177212388-177212725 | -0.601287483 | 0.217668932 | 0.496021559 |
| chr9:20292022-20292272 | 0.615967925 | 0.217682736 | 0.496031376 |
| chr3:192862513-192862721 | 0.613238491 | 0.217689779 | 0.496031376 |
| chr6:44667990-44668452 | 0.486421396 | 0.217692661 | 0.496031376 |
| chr14:32366326-32366526 | 0.610763587 | 0.217704759 | 0.496044191 |
| chr17:63881376-63881669 | -0.466576362 | 0.217722083 | 0.496055095 |
| chr2:95414049-95414299 | -0.550446417 | 0.217722493 | 0.496055095 |
| chr8:60763320-60763521 | 0.599672926 | 0.217750033 | 0.496103091 |
| chr20:46255434-46255796 | 0.452719682 | 0.217759065 | 0.496108915 |
| chr10:128211916-128212249 | 0.475508501 | 0.217772589 | 0.496119835 |
| chr21:42219802-42220643 | -0.404868185 | 0.217776807 | 0.496119835 |
| chr9:109193412-109193655 | 0.567709735 | 0.217791538 | 0.496127895 |
| chr3:174863296-174863947 | -0.406914976 | 0.217793295 | 0.496127895 |
| chr9:121735055-121735321 | 0.519832903 | 0.217812852 | 0.496157695 |
| chr16:58495014-58495503 | -0.43256065 | 0.217834899 | 0.496193163 |
| chr1:108553918-108554449 | 0.415680352 | 0.217850936 | 0.496198323 |
| chr5:10689459-10690087 | -0.427156818 | 0.217855528 | 0.496198323 |
| chr1:155084649-155085650 | 0.392699347 | 0.217856592 | 0.496198323 |
| chr17:37643329-37643611 | -0.560987978 | 0.217881138 | 0.496239479 |
| chr2:9269382-9269590 | -0.642616364 | 0.21790166 | 0.496271468 |
| chr2:141699825-141700025 | -0.655891239 | 0.217916288 | 0.496271614 |
| chr2:190648233-190649628 | -0.405021382 | 0.2179164 | 0.496271614 |
| chr6:152948142-152948438 | -0.54636033 | 0.217921155 | 0.496271614 |
| chr1:207250630-207251063 | -0.497797908 | 0.217947255 | 0.496316301 |
| chr13:105441410-105441620 | -0.68237522 | 0.217962466 | 0.496336189 |

| | | | |
|---|---|---|---|
| chr12:100199813-100200373 | -0.411973348 | 0.218006869 | 0.496396954 |
| chr10:43255893-43256139 | -0.507559392 | 0.218006869 | 0.496396954 |
| chr4:161201000-161201427 | 0.468779356 | 0.218021064 | 0.496396954 |
| chr2:194254203-194254488 | -0.657565866 | 0.218022756 | 0.496396954 |
| chrX:19364058-19364421 | -0.578102441 | 0.218027869 | 0.496396954 |
| chr12:120469243-120470322 | 0.403046741 | 0.218031261 | 0.496396954 |
| chr11:117328549-117328749 | 0.542867462 | 0.2180345 | 0.496396954 |
| chr10:49941233-49942410 | -0.414916823 | 0.218073219 | 0.496461588 |
| chr6:97130948-97131272 | -0.593308782 | 0.218075848 | 0.496461588 |
| chr18:37444772-37445112 | -0.469580273 | 0.218089321 | 0.496477508 |
| chr2:58240618-58241639 | -0.39772137 | 0.218101186 | 0.496489768 |
| chr9:110329820-110330039 | 0.628194854 | 0.218113494 | 0.496503035 |
| chr14:74763041-74763591 | 0.417237227 | 0.218129062 | 0.496511546 |
| chr2:126042225-126042546 | 0.538060003 | 0.218130193 | 0.496511546 |
| chr5:2503805-2504031 | 0.594665505 | 0.21815965 | 0.496536114 |
| chr5:93039912-93040275 | 0.460072047 | 0.218161588 | 0.496536114 |
| chr5:143185553-143185804 | -0.539291348 | 0.218169493 | 0.496536114 |
| chr11:65422116-65422507 | 0.42266446 | 0.218172113 | 0.496536114 |
| chr12:26043621-26044175 | 0.453821008 | 0.218177771 | 0.496536114 |
| chr2:196315446-196315826 | -0.545219744 | 0.218179868 | 0.496536114 |
| chr16:50741409-50742378 | 0.398895533 | 0.218194031 | 0.496553595 |
| chr12:88566476-88566684 | -0.615420625 | 0.218201564 | 0.496555991 |
| chr3:104596666-104596930 | -0.540355031 | 0.218236814 | 0.496621459 |
| chr9:126644224-126644424 | 0.540697462 | 0.218254261 | 0.496646411 |
| chr14:94036555-94036876 | 0.452884932 | 0.218274827 | 0.496655088 |
| chr3:64792305-64792670 | 0.420332215 | 0.218275362 | 0.496655088 |
| chr15:59437592-59438614 | 0.434153046 | 0.218277519 | 0.496655088 |
| chr1:38078952-38079272 | -0.615223089 | 0.218300569 | 0.496675857 |
| chr3:183480383-183480667 | -0.518905532 | 0.218304821 | 0.496675857 |
| chr3:102000701-102000912 | -0.583125615 | 0.218306094 | 0.496675857 |
| chr4:143346296-143346562 | 0.528869193 | 0.2183473 | 0.496745084 |
| chr2:73753988-73754215 | -0.57779026 | 0.218349487 | 0.496745084 |
| chr3:146995172-146995594 | -0.576033053 | 0.218356972 | 0.496747361 |
| chr8:49519024-49519233 | -0.700646874 | 0.218369764 | 0.496761714 |
| chr1:183580426-183580889 | -0.438834463 | 0.218377177 | 0.496763829 |
| chr19:35702173-35703180 | 0.403289116 | 0.218417835 | 0.496841566 |
| chr7:146699314-146699717 | -0.571213962 | 0.218430148 | 0.496854825 |
| chr11:46593504-46594438 | -0.414186978 | 0.218448627 | 0.496882109 |
| chr11:1997871-1998182 | 0.519321403 | 0.218463339 | 0.496900822 |
| chr14:20999009-20999290 | 0.505946521 | 0.218490148 | 0.49693407 |
| chr3:89697272-89697738 | -0.443952681 | 0.218490928 | 0.49693407 |
| chr3:25647906-25648433 | -0.418405007 | 0.218519601 | 0.496975429 |
| chr8:8487628-8487865 | -0.61027735 | 0.218523242 | 0.496975429 |
| chr5:138559872-138560258 | 0.47960027 | 0.21852874 | 0.496975429 |
| chr5:38071769-38072033 | -0.662394942 | 0.218535057 | 0.496975429 |
| chr14:90533817-90534026 | -0.62016691 | 0.218576826 | 0.497039634 |
| chr6:6803528-6803960 | -0.494444928 | 0.218579812 | 0.497039634 |
| chr2:109332571-109332859 | -0.515488396 | 0.21858275 | 0.497039634 |
| chr20:52297338-52297693 | 0.464505808 | 0.218590585 | 0.497042698 |
| chr2:20466023-20466758 | -0.409853643 | 0.218604584 | 0.497047981 |

| | | | |
|---|---|---|---|
| chr1:240693946-240694253 | -0.590512798 | 0.218606916 | 0.497047981 |
| chr12:131218360-131218897 | -0.460258313 | 0.218612369 | 0.497047981 |
| chr2:171687209-171688244 | -0.404583329 | 0.218622309 | 0.497053925 |
| chr17:23016165-23016494 | 0.452217748 | 0.218634701 | 0.497053925 |
| chr15:41971939-41973080 | 0.399205104 | 0.218644474 | 0.497053925 |
| chr5:136174105-136174338 | -0.630175581 | 0.218645996 | 0.497053925 |
| chr10:89951457-89951701 | -0.658533076 | 0.218647419 | 0.497053925 |
| chr1:110783369-110783643 | 0.541893724 | 0.218684944 | 0.497124482 |
| chr1:56818919-56819796 | -0.415759485 | 0.218731585 | 0.497215757 |
| chr20:50152361-50152656 | 0.438153545 | 0.218756152 | 0.497256849 |
| chr3:193909544-193909749 | 0.635130464 | 0.218763229 | 0.497258185 |
| chr1:199046424-199047048 | -0.435949997 | 0.21876987 | 0.497258527 |
| chr1:151195299-151195603 | -0.422950427 | 0.218777616 | 0.497261383 |
| chr10:133258225-133258437 | 0.507526105 | 0.218791637 | 0.497277761 |
| chr7:30239013-30239217 | 0.60941206 | 0.218797801 | 0.497277761 |
| chr10:25465830-25466326 | -0.483671792 | 0.218810337 | 0.497291501 |
| chr5:163505334-163506407 | -0.392886217 | 0.218873851 | 0.497417431 |
| chr3:149507194-149507490 | 0.512900672 | 0.218882382 | 0.497417431 |
| chr2:205925824-205926252 | -0.441924737 | 0.218885222 | 0.497417431 |
| chr3:31562341-31562884 | -0.461737989 | 0.218893385 | 0.497421228 |
| chr6:97497755-97498269 | -0.457046258 | 0.218904974 | 0.497432809 |
| chr5:115543915-115544192 | -0.478322496 | 0.218917271 | 0.497446001 |
| chr14:105659177-105659555 | -0.501508239 | 0.218932278 | 0.497460254 |
| chr19:49056165-49056556 | 0.491400927 | 0.218936528 | 0.497460254 |
| chr10:34723774-34724122 | -0.525040381 | 0.218958636 | 0.497471524 |
| chr15:96312647-96312884 | -0.654423278 | 0.218960485 | 0.497471524 |
| chr15:85251005-85251463 | 0.461130071 | 0.218966593 | 0.497471524 |
| chr16:28864052-28864299 | 0.51843626 | 0.218973131 | 0.497471524 |
| chr8:85278964-85279396 | 0.445617664 | 0.218985678 | 0.497471524 |
| chr17:1684379-1685161 | -0.414687875 | 0.218986671 | 0.497471524 |
| chr18:9092520-9092737 | -0.726407518 | 0.218986936 | 0.497471524 |
| chr2:69616871-69617076 | -0.616112442 | 0.21901512 | 0.497508842 |
| chr4:1248294-1248919 | 0.431364146 | 0.21901635 | 0.497508842 |
| chr6:134621928-134622215 | -0.602678342 | 0.219040481 | 0.497534922 |
| chr9:132252509-132252927 | -0.526082455 | 0.219040818 | 0.497534922 |
| chr6:40894948-40895321 | 0.498695444 | 0.21905273 | 0.49754723 |
| chr5:14011040-14012018 | -0.393297934 | 0.219067272 | 0.49755891 |
| chr3:17841221-17841583 | -0.449718105 | 0.219070859 | 0.49755891 |
| chr9:121650283-121650525 | 0.523130978 | 0.21912553 | 0.497666567 |
| chr8:90948343-90948561 | 0.554078962 | 0.21913125 | 0.497666567 |
| chr3:124584493-124585393 | 0.412995036 | 0.219166462 | 0.497731441 |
| chr7:110363297-110363671 | -0.567028301 | 0.219172807 | 0.497731441 |
| chr15:71104601-71104973 | -0.548822872 | 0.219199252 | 0.49776239 |
| chr7:148987127-148987835 | -0.401213951 | 0.219203292 | 0.49776239 |
| chr8:95352319-95352519 | 0.578315643 | 0.219211575 | 0.49776239 |
| chr5:128121690-128122116 | -0.442518922 | 0.21921242 | 0.49776239 |
| chr11:73404134-73404359 | -0.551080554 | 0.219222311 | 0.497770096 |
| chr15:74901665-74901918 | 0.497517662 | 0.219232038 | 0.497777431 |
| chr3:83500515-83500842 | -0.554548713 | 0.219266898 | 0.497833041 |
| chr12:89019597-89019957 | -0.444470657 | 0.219269676 | 0.497833041 |

| | | | |
|---|---|---|---|
| chr3:148333092-148333478 | 0.480646888 | 0.219279602 | 0.497833041 |
| chr2:65227097-65228471 | -0.391113487 | 0.219282518 | 0.497833041 |
| chr14:25308086-25308626 | -0.456734657 | 0.219289295 | 0.497833675 |
| chr1:106651320-106651558 | 0.456854284 | 0.219307976 | 0.497843704 |
| chr14:25007002-25007451 | -0.519561165 | 0.219309941 | 0.497843704 |
| chr1:156281280-156281480 | 0.507088841 | 0.219325623 | 0.497843704 |
| chr16:28491885-28492433 | 0.419169666 | 0.219326002 | 0.497843704 |
| chr17:7242361-7243241 | -0.404971237 | 0.219332089 | 0.497843704 |
| chr15:86622685-86622932 | 0.633811822 | 0.219332697 | 0.497843704 |
| chr1:197146015-197147187 | -0.386512322 | 0.219345025 | 0.497850113 |
| chr3:9172790-9173108 | 0.47299576 | 0.219350445 | 0.497850113 |
| chr2:113242064-113242402 | -0.569842785 | 0.219355013 | 0.497850113 |
| chr19:49665551-49666221 | 0.413062537 | 0.219363542 | 0.497854724 |
| chr9:122213817-122214306 | 0.427768813 | 0.219379663 | 0.497872183 |
| chr2:52416457-52416662 | -0.657314355 | 0.219385256 | 0.497872183 |
| chr6:5891541-5892455 | -0.428611703 | 0.219390728 | 0.497872183 |
| chr17:5111715-5112275 | -0.440933659 | 0.219410969 | 0.49790337 |
| chr6:18209531-18209738 | 0.650294484 | 0.219418431 | 0.497905556 |
| chr5:44638392-44638914 | -0.421211364 | 0.219456546 | 0.497977299 |
| chrX:123648171-123649202 | 0.415509313 | 0.219469284 | 0.497991455 |
| chr12:105574289-105574574 | -0.566492444 | 0.21947946 | 0.497999799 |
| chr4:108620075-108621051 | -0.395946286 | 0.21950822 | 0.498048969 |
| chr10:106605839-106606096 | -0.535478544 | 0.219514131 | 0.498048969 |
| chrX:108000836-108001958 | -0.408326572 | 0.219549421 | 0.498114289 |
| chr12:97686237-97686482 | -0.680001573 | 0.219588851 | 0.498183339 |
| chr3:119380724-119381018 | 0.49307618 | 0.21959286 | 0.498183339 |
| chr2:94266743-94267073 | 0.455065359 | 0.219606649 | 0.498199871 |
| chr14:103510757-103510957 | 0.580134429 | 0.219631838 | 0.498242264 |
| chr9:86034922-86035172 | -0.576427368 | 0.219654128 | 0.498270151 |
| chr11:64777688-64777903 | 0.497824584 | 0.219672808 | 0.498270151 |
| chr3:122443750-122444144 | -0.450969465 | 0.21967485 | 0.498270151 |
| chr2:160142035-160142344 | 0.527163212 | 0.219675161 | 0.498270151 |
| chr2:11073164-11073582 | -0.461482422 | 0.219676646 | 0.498270151 |
| chr17:64342470-64344168 | 0.382989151 | 0.21973213 | 0.498381246 |
| chr15:95247050-95247844 | -0.410979682 | 0.219742122 | 0.498389157 |
| chr10:88152701-88153316 | -0.424198665 | 0.219835875 | 0.498587036 |
| chr22:21938030-21938787 | 0.417563983 | 0.219878343 | 0.498657566 |
| chr12:80607839-80608047 | -0.622520095 | 0.219879989 | 0.498657566 |
| chr2:37231195-37232058 | -0.397237991 | 0.21990099 | 0.498690433 |
| chr9:62838061-62838794 | -0.403472988 | 0.219926771 | 0.498734137 |
| chr19:58395164-58396009 | 0.409525362 | 0.219955353 | 0.498784192 |
| chr1:239407955-239408952 | 0.403311685 | 0.220008968 | 0.498876252 |
| chr17:75996799-75997107 | -0.508508038 | 0.220008972 | 0.498876252 |
| chr2:158464720-158464930 | -0.586752186 | 0.220034264 | 0.498918837 |
| chr20:50169402-50169611 | 0.488563651 | 0.220048421 | 0.498936174 |
| chr10:25059784-25060387 | -0.466717327 | 0.220085044 | 0.498990934 |
| chr5:115674869-115675224 | -0.466026849 | 0.220085597 | 0.498990934 |
| chr8:114454542-114454789 | 0.548989936 | 0.220146978 | 0.499115332 |
| chr8:144636392-144636592 | 0.660432364 | 0.22016315 | 0.499134089 |
| chr6:154289879-154290079 | -0.628262223 | 0.22016828 | 0.499134089 |

| | | | |
|---|---|---|---|
| chr2:84880985-84881412 | -0.497202366 | 0.220178132 | 0.499141656 |
| chr7:73719606-73719880 | 0.473562383 | 0.220186212 | 0.499145206 |
| chr1:61453113-61453721 | -0.436128772 | 0.22019913 | 0.499159721 |
| chr8:140337335-140337920 | -0.423397756 | 0.220210576 | 0.499164001 |
| chr10:70325452-70325738 | -0.56876211 | 0.220214047 | 0.499164001 |
| chr4:129015378-129015642 | -0.642442452 | 0.220262334 | 0.499238176 |
| chr3:195035277-195035518 | -0.582576564 | 0.220265301 | 0.499238176 |
| chr10:44310799-44311369 | -0.457175373 | 0.220266317 | 0.499238176 |
| chr1:42045694-42046123 | -0.562211277 | 0.220272861 | 0.499238239 |
| chr2:2472156-2472357 | 0.695599256 | 0.220290369 | 0.499263153 |
| chr3:25455534-25455773 | -0.607541597 | 0.22032599 | 0.499299222 |
| chr2:222775058-222775519 | -0.507927349 | 0.220330738 | 0.499299222 |
| chr17:17692784-17692991 | -0.520001463 | 0.220331969 | 0.499299222 |
| chr4:54440017-54440275 | -0.562679238 | 0.22033235 | 0.499299222 |
| chr12:57896521-57897392 | -0.416836249 | 0.220378857 | 0.49937764 |
| chr18:499546-499910 | -0.518215156 | 0.220388476 | 0.49937764 |
| chr3:172673174-172673528 | -0.448631321 | 0.220406771 | 0.49937764 |
| chr21:38853550-38853750 | -0.602922137 | 0.220409658 | 0.49937764 |
| chr10:83425368-83425568 | 0.664279639 | 0.220413051 | 0.49937764 |
| chr14:24984363-24984563 | 0.664279639 | 0.220413051 | 0.49937764 |
| chr16:58500534-58501031 | 0.449848341 | 0.220420425 | 0.49937764 |
| chr3:83467187-83467512 | -0.648927919 | 0.220423272 | 0.49937764 |
| chr8:82996333-82997288 | 0.415098818 | 0.220425611 | 0.49937764 |
| chr14:103075516-103076050 | -0.422673236 | 0.22044404 | 0.499404626 |
| chr4:184430274-184430523 | -0.642863146 | 0.220467802 | 0.49944369 |
| chr8:140192341-140192787 | 0.435981898 | 0.220479219 | 0.499449382 |
| chr15:69594085-69595051 | 0.393865054 | 0.220492505 | 0.499449382 |
| chr14:69605753-69606103 | -0.512987331 | 0.220494641 | 0.499449382 |
| chr3:8442133-8442357 | 0.495436952 | 0.220502466 | 0.499449382 |
| chr5:102506675-102507144 | -0.492340437 | 0.220504991 | 0.499449382 |
| chr8:120785749-120786378 | 0.428020961 | 0.220514474 | 0.499449382 |
| chr13:38349358-38350423 | -0.388216023 | 0.220519539 | 0.499449382 |
| chr10:66322268-66322537 | -0.613763764 | 0.220522462 | 0.499449382 |
| chr13:108961624-108961872 | -0.580329666 | 0.220540478 | 0.499474668 |
| chr14:103222170-103222370 | 0.659161557 | 0.220546663 | 0.499474668 |
| chr2:1631978-1632388 | -0.471814509 | 0.220566284 | 0.49950418 |
| chr2:1550816-1551209 | 0.476878742 | 0.220573692 | 0.49950418 |
| chr21:38229161-38229533 | -0.527127586 | 0.220579252 | 0.49950418 |
| chr20:49859834-49860208 | 0.434101731 | 0.220600686 | 0.499520861 |
| chr22:35154954-35155154 | -0.667862961 | 0.220602957 | 0.499520861 |
| chr16:19716174-19716514 | 0.478177835 | 0.220607843 | 0.499520861 |
| chr5:98865332-98865532 | 0.68806782 | 0.220612695 | 0.499520861 |
| chr8:143953240-143953590 | 0.43949322 | 0.220647992 | 0.499586019 |
| chr10:121696939-121697218 | -0.552124405 | 0.220690268 | 0.499650137 |
| chr16:31712930-31713717 | 0.411752928 | 0.220692783 | 0.499650137 |
| chr4:33104378-33104685 | -0.601943584 | 0.220695874 | 0.499650137 |
| chr11:4626238-4626550 | -0.557543036 | 0.220726428 | 0.499704547 |
| chr15:98852223-98852888 | 0.399324323 | 0.220763763 | 0.499774302 |
| chr8:128428149-128428572 | 0.431731004 | 0.220783115 | 0.499794677 |
| chr5:40755033-40755320 | 0.429580733 | 0.220785809 | 0.499794677 |

| | | | |
|---|---|---|---|
| chr4:71059626-71059925 | -0.539580425 | 0.220806673 | 0.499827141 |
| chr12:86342651-86342872 | 0.624349463 | 0.220875832 | 0.499957396 |
| chr5:61658591-61658949 | -0.490721019 | 0.220877265 | 0.499957396 |
| chr1:107017665-107017975 | 0.449546312 | 0.220894014 | 0.499974906 |
| chr3:77039082-77039479 | -0.472578487 | 0.220905005 | 0.499974906 |
| chr11:89498912-89499245 | -0.525838157 | 0.22090892 | 0.499974906 |
| chr10:30365307-30365625 | -0.526595234 | 0.22091349 | 0.499974906 |
| chr16:10594412-10595017 | -0.39949226 | 0.220917627 | 0.499974906 |
| chr5:86880713-86881210 | -0.408949931 | 0.220925974 | 0.499979028 |
| chr11:123261561-123261825 | 0.534583221 | 0.221002401 | 0.500137218 |
| chr4:41213840-41214088 | 0.480135909 | 0.221019313 | 0.500151196 |
| chr1:119675366-119675931 | -0.446478902 | 0.221021632 | 0.500151196 |
| chr1:46769155-46769369 | 0.595585387 | 0.221049334 | 0.500199111 |
| chr5:151742840-151743181 | -0.45727812 | 0.221106365 | 0.500298189 |
| chr13:44404189-44404587 | -0.450659437 | 0.221111241 | 0.500298189 |
| chr12:27332776-27333205 | -0.517583939 | 0.22111824 | 0.500298189 |
| chr9:84172748-84173427 | -0.446222949 | 0.221119237 | 0.500298189 |
| chr3:82132985-82133344 | -0.469597427 | 0.221142547 | 0.500336156 |
| chr17:78724519-78724818 | -0.539360322 | 0.221157348 | 0.500354868 |
| chr4:101958701-101958942 | 0.630228489 | 0.221174028 | 0.50037783 |
| chr22:24805885-24806316 | 0.492865501 | 0.221191002 | 0.500401458 |
| chr7:44639167-44639431 | 0.523842695 | 0.221255225 | 0.500531971 |
| chr10:101694056-101695319 | 0.404875999 | 0.221266427 | 0.500542534 |
| chr10:573322-573647 | -0.514405035 | 0.221292086 | 0.5005858 |
| chr4:74134101-74134545 | -0.474065657 | 0.221316564 | 0.500626393 |
| chr22:28663372-28663580 | -0.665831827 | 0.221325128 | 0.50062917 |
| chr5:15500917-15501750 | -0.423795774 | 0.22133086 | 0.50062917 |
| chr11:112280399-112280861 | 0.487517783 | 0.22139837 | 0.500767088 |
| chr3:174776213-174776415 | -0.665482935 | 0.221410795 | 0.50078041 |
| chr6:98287640-98287985 | 0.557318821 | 0.221421755 | 0.500790416 |
| chr3:5026075-5027246 | -0.390039312 | 0.221433622 | 0.500802473 |
| chr2:9778008-9778293 | -0.483469264 | 0.221446312 | 0.500805243 |
| chr1:169794381-169795464 | -0.390860464 | 0.221447919 | 0.500805243 |
| chr2:37004958-37005705 | -0.456256448 | 0.221489963 | 0.500873541 |
| chr19:34310469-34310953 | -0.447722264 | 0.221493966 | 0.500873541 |
| chr15:55799834-55800034 | 0.603658587 | 0.221506924 | 0.500873541 |
| chr8:28072787-28073030 | -0.582240287 | 0.221514886 | 0.500873541 |
| chr5:52787689-52788595 | -0.413828433 | 0.221519681 | 0.500873541 |
| chr20:9196281-9196481 | -0.681774243 | 0.221521968 | 0.500873541 |
| chr8:113466524-113466927 | 0.469448077 | 0.221531705 | 0.500873541 |
| chr12:102938478-102938678 | -0.716558283 | 0.221541795 | 0.500873541 |
| chr8:141308224-141308673 | 0.429877605 | 0.221542325 | 0.500873541 |
| chr5:168647069-168647474 | -0.532337177 | 0.221543488 | 0.500873541 |
| chr19:8444606-8445309 | 0.417890769 | 0.221558861 | 0.500882149 |
| chr9:89752075-89752404 | -0.523397731 | 0.22156037 | 0.500882149 |
| chr12:13061549-13062286 | -0.423202636 | 0.221578159 | 0.500907585 |
| chr3:125967233-125967609 | 0.488548451 | 0.221596618 | 0.500924239 |
| chr22:38681725-38682306 | -0.450598456 | 0.221598601 | 0.500924239 |
| chr12:111156310-111156634 | 0.531920796 | 0.221616066 | 0.500939238 |
| chr11:107799339-107799539 | 0.618322337 | 0.221631157 | 0.500939238 |

| | | | |
|---|---|---|---|
| chr15:77108120-77108320 | 0.60790851 | 0.2216358 | 0.500939238 |
| chr11:84642596-84642820 | 0.60790851 | 0.2216358 | 0.500939238 |
| chr5:132274950-132275213 | 0.568633506 | 0.221637925 | 0.500939238 |
| chr1:110875569-110875909 | -0.58674188 | 0.221656711 | 0.50096692 |
| chr3:113148133-113148797 | -0.404874464 | 0.221669089 | 0.500980117 |
| chr20:62986925-62987159 | 0.544866737 | 0.221681794 | 0.500994055 |
| chr13:21637404-21637625 | 0.582702243 | 0.22168876 | 0.50099502 |
| chr7:35985203-35985490 | 0.453403971 | 0.221736245 | 0.501087553 |
| chr1:116342874-116343079 | 0.5470308 | 0.221746525 | 0.501096005 |
| chr6:26249719-26250785 | -0.40564612 | 0.221759321 | 0.501110142 |
| chr19:52139219-52139618 | 0.454697366 | 0.221815392 | 0.501222063 |
| chr4:79274078-79274414 | 0.640076982 | 0.221865346 | 0.501303578 |
| chr3:180595531-180595889 | -0.575024158 | 0.221868449 | 0.501303578 |
| chr20:543429-544233 | 0.409825411 | 0.221871094 | 0.501303578 |
| chr2:37526395-37526633 | 0.616196151 | 0.221881388 | 0.501312054 |
| chr12:42372034-42372457 | -0.505745723 | 0.221900846 | 0.501341234 |
| chr5:118698877-118699209 | -0.562159179 | 0.221912487 | 0.501345958 |
| chr1:182889531-182889771 | -0.629043127 | 0.221922566 | 0.501345958 |
| chr1:198315501-198315716 | -0.629043127 | 0.221922566 | 0.501345958 |
| chr6:166420391-166420699 | -0.520828254 | 0.221952526 | 0.501398856 |
| chr1:31587795-31587995 | 0.584602927 | 0.221965669 | 0.501413766 |
| chr17:78168790-78169147 | 0.449416123 | 0.221977923 | 0.501422913 |
| chr20:57061387-57061637 | 0.485065573 | 0.221986962 | 0.501422913 |
| chr10:53045379-53045611 | -0.676521643 | 0.221989351 | 0.501422913 |
| chr10:15084325-15085394 | -0.394410396 | 0.222002181 | 0.501426234 |
| chr16:77450517-77451072 | 0.427294757 | 0.22200391 | 0.501426234 |
| chr13:23158798-23159067 | 0.466958309 | 0.222056093 | 0.501529313 |
| chr3:152466418-152467207 | -0.434884748 | 0.222079933 | 0.501568373 |
| chr5:159965629-159965885 | -0.513054265 | 0.222091583 | 0.501574542 |
| chr3:64379967-64380484 | 0.409158012 | 0.222095757 | 0.501574542 |
| chr17:53025776-53026265 | 0.466207573 | 0.222115769 | 0.501604953 |
| chr17:45681158-45681587 | -0.542872429 | 0.222134115 | 0.501622128 |
| chr3:70805266-70805889 | 0.470401779 | 0.222137844 | 0.501622128 |
| chr8:118897335-118897535 | 0.63434681 | 0.222143014 | 0.501622128 |
| chr1:38071909-38072115 | -0.585796646 | 0.222169413 | 0.501638923 |
| chr18:65753630-65753855 | -0.591812813 | 0.222172946 | 0.501638923 |
| chr17:72120765-72121114 | -0.420421485 | 0.222176012 | 0.501638923 |
| chr13:22418257-22418495 | -0.651238157 | 0.222179368 | 0.501638923 |
| chr3:166125270-166125497 | 0.603135025 | 0.22218896 | 0.501638923 |
| chr15:64958846-64959188 | 0.590186363 | 0.222193698 | 0.501638923 |
| chr3:138689166-138689406 | -0.625529495 | 0.222198335 | 0.501638923 |
| chr14:99604128-99604759 | 0.396515835 | 0.222207746 | 0.501638923 |
| chr17:39676261-39676523 | 0.504327317 | 0.222220571 | 0.501638923 |
| chr2:163132915-163133224 | -0.549091669 | 0.222220674 | 0.501638923 |
| chr10:121465599-121465820 | 0.640871474 | 0.222222468 | 0.501638923 |
| chr3:122768213-122768413 | -0.665482836 | 0.222231399 | 0.501644303 |
| chr19:19663211-19663487 | -0.558791005 | 0.22226058 | 0.501695394 |
| chr5:143209857-143210153 | -0.535599769 | 0.222279049 | 0.501708132 |
| chr4:155742421-155742621 | -0.64972239 | 0.222285867 | 0.501708132 |
| chr8:115203504-115203714 | -0.64972239 | 0.222285867 | 0.501708132 |

| | | | |
|---|---|---|---|
| chr12:64802242-64802533 | 0.476420212 | 0.222294208 | 0.501709628 |
| chr1:32753712-32754713 | 0.397991653 | 0.222304221 | 0.501709628 |
| chr3:63136159-63136895 | -0.396560235 | 0.222319778 | 0.501709628 |
| chr8:101137204-101137655 | 0.428574285 | 0.222322414 | 0.501709628 |
| chr3:72655050-72655250 | 0.593973106 | 0.222335019 | 0.501709628 |
| chr1:39378354-39378680 | -0.53007102 | 0.222335106 | 0.501709628 |
| chr1:221839807-221840217 | -0.485329987 | 0.222335827 | 0.501709628 |
| chr13:81540879-81541494 | -0.473623815 | 0.222338912 | 0.501709628 |
| chr13:112894141-112894470 | 0.513373531 | 0.22238089 | 0.501781258 |
| chr20:8018868-8020077 | -0.390179485 | 0.222383753 | 0.501781258 |
| chr5:135398140-135398811 | -0.425049397 | 0.222397031 | 0.501782393 |
| chr6:110681629-110681829 | 0.672964216 | 0.222397354 | 0.501782393 |
| chr2:20868439-20868659 | -0.654858416 | 0.22240491 | 0.501784664 |
| chr7:42931916-42932702 | -0.393450294 | 0.222456583 | 0.501871466 |
| chr18:9913225-9914870 | 0.38662876 | 0.222458555 | 0.501871466 |
| chr19:4976397-4976745 | -0.547392464 | 0.22247234 | 0.501871466 |
| chr11:47466951-47467377 | 0.471765276 | 0.222474715 | 0.501871466 |
| chr17:81711793-81712903 | 0.395189635 | 0.222476132 | 0.501871466 |
| chr11:46529268-46529661 | -0.447841979 | 0.222522015 | 0.501960026 |
| chr6:14144683-14144899 | 0.593649945 | 0.222528797 | 0.501960026 |
| chr7:2559424-2559712 | -0.45448419 | 0.222542219 | 0.501960026 |
| chr3:27768782-27769160 | 0.496548101 | 0.222543581 | 0.501960026 |
| chr11:75499829-75500090 | -0.567457451 | 0.222551105 | 0.501960026 |
| chr10:123642823-123643047 | 0.661003189 | 0.222554697 | 0.501960026 |
| chr12:48189505-48190296 | -0.449390627 | 0.222591708 | 0.502021448 |
| chr1:231651771-231652053 | 0.532495876 | 0.222598446 | 0.502021448 |
| chr5:134446788-134447258 | -0.467171092 | 0.222601586 | 0.502021448 |
| chr20:47500773-47501011 | 0.403380367 | 0.222611259 | 0.502026751 |
| chr3:159026422-159027119 | -0.417919251 | 0.222617041 | 0.502026751 |
| chr3:197546782-197546998 | -0.595873384 | 0.22266823 | 0.502108002 |
| chr8:128807527-128807831 | 0.558415905 | 0.222682576 | 0.502108002 |
| chr1:51558353-51558670 | 0.51108137 | 0.222691048 | 0.502108002 |
| chr10:75188034-75188283 | -0.539569859 | 0.222698163 | 0.502108002 |
| chrX:55160739-55161771 | -0.399583272 | 0.222704916 | 0.502108002 |
| chrX:2897629-2898241 | 0.432306077 | 0.222705985 | 0.502108002 |
| chr8:54090572-54090797 | -0.572433766 | 0.222711069 | 0.502108002 |
| chr3:166844833-166845087 | 0.591252549 | 0.222712017 | 0.502108002 |
| chr19:52095195-52095454 | 0.546030864 | 0.222714205 | 0.502108002 |
| chr3:24494041-24495942 | 0.38403233 | 0.222724618 | 0.502108002 |
| chr4:173307845-173308045 | -0.642311522 | 0.222727259 | 0.502108002 |
| chr4:112817860-112818639 | -0.415896317 | 0.222731707 | 0.502108002 |
| chr11:120511090-120511415 | 0.520334862 | 0.222740944 | 0.502114052 |
| chr10:71684200-71684606 | -0.504407231 | 0.222805439 | 0.502232903 |
| chr4:2262042-2262710 | 0.399328168 | 0.222806777 | 0.502232903 |
| chr1:203720340-203720606 | 0.522404351 | 0.222822047 | 0.502252549 |
| chr20:973748-974075 | 0.561197587 | 0.222858771 | 0.502302896 |
| chr12:89051101-89051410 | 0.533370245 | 0.222868001 | 0.502302896 |
| chr2:86622500-86622766 | -0.502893327 | 0.222870093 | 0.502302896 |
| chr2:120675535-120675811 | -0.563759391 | 0.222870606 | 0.502302896 |
| chr6:28020769-28021371 | -0.485522854 | 0.222896949 | 0.502347492 |

| | | | |
|---|---|---|---|
| chr17:68740351-68740551 | -0.544385523 | 0.222936713 | 0.502422331 |
| chr13:76991504-76992979 | -0.386125192 | 0.222975338 | 0.50246803 |
| chr7:7565921-7567519 | 0.381961898 | 0.222979812 | 0.50246803 |
| chr10:2137564-2137923 | -0.51128646 | 0.222981889 | 0.50246803 |
| chr6:80143521-80143843 | -0.5830544 | 0.222983222 | 0.50246803 |
| chr6:91031688-91031949 | 0.527426424 | 0.2229936 | 0.502476639 |
| chr21:17296550-17296753 | -0.652221454 | 0.223008301 | 0.502486214 |
| chr9:104763585-104764747 | -0.407394913 | 0.223010965 | 0.502486214 |
| chr18:70288472-70289743 | -0.405060041 | 0.223027545 | 0.502503372 |
| chr19:55173820-55174398 | -0.424202936 | 0.223038393 | 0.502503372 |
| chr5:112860940-112861653 | -0.397310793 | 0.223051339 | 0.502503372 |
| chr12:107520388-107520878 | -0.490777725 | 0.22305423 | 0.502503372 |
| chr2:169453187-169453574 | -0.488983452 | 0.223054743 | 0.502503372 |
| chr11:1947568-1947812 | 0.556911393 | 0.22305793 | 0.502503372 |
| chr9:12955668-12956181 | -0.441197959 | 0.223069265 | 0.502514132 |
| chr15:40486828-40487161 | -0.632743425 | 0.223086757 | 0.502538763 |
| chr5:35187831-35188610 | 0.409620912 | 0.223104757 | 0.502564536 |
| chr7:128010497-128010722 | -0.55059971 | 0.223161983 | 0.502678665 |
| chr1:239814252-239814636 | -0.46294624 | 0.223183346 | 0.502712007 |
| chr7:44072378-44072669 | -0.459571394 | 0.223213254 | 0.502750058 |
| chr2:178132341-178132583 | -0.563574319 | 0.223213362 | 0.502750058 |
| chr12:104671702-104672297 | -0.435142394 | 0.223260601 | 0.502841674 |
| chr7:38345416-38345944 | -0.421374052 | 0.223308876 | 0.502853861 |
| chr10:25142219-25142427 | -0.655444904 | 0.223313397 | 0.502853861 |
| chrX:50448618-50448818 | 0.634209012 | 0.223316096 | 0.502853861 |
| chr18:31101077-31102221 | 0.408648033 | 0.22331833 | 0.502853861 |
| chr14:31082048-31082329 | 0.48356967 | 0.223320178 | 0.502853861 |
| chr13:44489834-44490051 | 0.594164245 | 0.22333468 | 0.502853861 |
| chr6:111709736-111710207 | 0.459837427 | 0.223335195 | 0.502853861 |
| chr16:72664331-72665446 | -0.390195298 | 0.223339064 | 0.502853861 |
| chr3:84772846-84773164 | -0.534258682 | 0.223356398 | 0.502853861 |
| chr20:22453330-22453572 | -0.670962813 | 0.223365315 | 0.502853861 |
| chr2:26577817-26578094 | -0.467698323 | 0.22336744 | 0.502853861 |
| chr17:68759382-68760220 | -0.393208815 | 0.223369291 | 0.502853861 |
| chr5:173973248-173973918 | 0.427575622 | 0.223370536 | 0.502853861 |
| chr14:99563198-99563398 | 0.511128202 | 0.223372992 | 0.502853861 |
| chr16:27235365-27235674 | 0.494142448 | 0.223379783 | 0.502853861 |
| chr4:140050209-140050688 | -0.475409011 | 0.223380409 | 0.502853861 |
| chr2:166631975-166632242 | -0.56672259 | 0.223387474 | 0.502853861 |
| chr4:145618838-145619903 | -0.391335827 | 0.223389498 | 0.502853861 |
| chr2:26758404-26758604 | -0.586798993 | 0.223390705 | 0.502853861 |
| chr7:52726962-52727268 | 0.505426627 | 0.223400235 | 0.502860541 |
| chr17:4143521-4144281 | -0.414919996 | 0.223413189 | 0.502874926 |
| chr5:58459613-58461048 | -0.385392166 | 0.22342224 | 0.502878181 |
| chr14:49861671-49861952 | 0.446043283 | 0.223427761 | 0.502878181 |
| chr20:57117234-57117553 | 0.397073881 | 0.223434503 | 0.502878584 |
| chr2:188456579-188456860 | -0.522104905 | 0.223444524 | 0.502886365 |
| chr3:123702096-123702397 | 0.478625958 | 0.223481134 | 0.502944513 |
| chr3:111768566-111768769 | -0.629051354 | 0.223483488 | 0.502944513 |
| chr19:2478699-2479151 | -0.497623963 | 0.223500478 | 0.502955862 |

| | | | |
|---|---|---|---|
| chr3:154194932-154195178 | -0.630334034 | 0.223501659 | 0.502955862 |
| chr13:49308195-49308432 | 0.596324334 | 0.223515517 | 0.502972274 |
| chr2:68799122-68799415 | -0.566251803 | 0.223543378 | 0.503020196 |
| chr10:98446182-98447134 | -0.404185078 | 0.223561164 | 0.503045445 |
| chr7:112616646-112616890 | -0.688602311 | 0.223582607 | 0.503078923 |
| chr1:230426292-230426969 | 0.420542802 | 0.223590276 | 0.503081404 |
| chr7:152664615-152665055 | -0.464730218 | 0.223606258 | 0.503102002 |
| chr16:90046756-90047105 | -0.450553981 | 0.223634314 | 0.503102002 |
| chr1:115210389-115210760 | 0.492043656 | 0.223635189 | 0.503102002 |
| chr8:84182118-84183294 | 0.401617116 | 0.223635413 | 0.503102002 |
| chr19:50147499-50148458 | 0.402428666 | 0.223636739 | 0.503102002 |
| chr19:48491213-48491419 | 0.466673187 | 0.223638827 | 0.503102002 |
| chr14:77027743-77029016 | 0.383475535 | 0.223663806 | 0.50312863 |
| chr14:104827398-104827954 | 0.412728166 | 0.223668654 | 0.50312863 |
| chr1:114560468-114560866 | 0.424325145 | 0.223679002 | 0.50312863 |
| chr2:74441547-74442541 | -0.395893966 | 0.223686618 | 0.50312863 |
| chr10:28628165-28628484 | -0.476760798 | 0.223687944 | 0.50312863 |
| chr15:64877637-64877853 | 0.612453279 | 0.223690062 | 0.50312863 |
| chr2:100397896-100398174 | 0.570203128 | 0.223707166 | 0.503152332 |
| chr3:134677270-134678018 | -0.432090289 | 0.223730166 | 0.503189291 |
| chr3:96355710-96355958 | 0.639004121 | 0.223739891 | 0.503196394 |
| chr1:201740083-201740493 | -0.431109986 | 0.22379431 | 0.503304009 |
| chr1:217119569-217120081 | -0.446590034 | 0.223815153 | 0.503329407 |
| chr9:83583255-83583455 | -0.658142796 | 0.223821344 | 0.503329407 |
| chr16:82604254-82604475 | -0.559744033 | 0.22382531 | 0.503329407 |
| chrX:3393684-3393954 | -0.626787983 | 0.223842685 | 0.503353706 |
| chr1:125180018-125180473 | 0.401561579 | 0.223892216 | 0.503450311 |
| chr19:36190601-36190801 | 0.648327385 | 0.223905987 | 0.503454436 |
| chr9:32552174-32552775 | -0.408598479 | 0.223907191 | 0.503454436 |
| chr8:124333885-124334117 | -0.596112274 | 0.22396775 | 0.503575825 |
| chr6:44013021-44013273 | -0.644674717 | 0.224036797 | 0.503694411 |
| chr20:57223397-57224196 | 0.382472358 | 0.224042092 | 0.503694411 |
| chr12:68792966-68793325 | -0.432885152 | 0.224043839 | 0.503694411 |
| chr17:35930657-35930969 | 0.489003172 | 0.224056162 | 0.503694411 |
| chr2:85394441-85394650 | -0.5149994 | 0.224056249 | 0.503694411 |
| chr10:72625573-72626407 | -0.396279805 | 0.224059934 | 0.503694411 |
| chr3:51670812-51671752 | -0.404158137 | 0.224073614 | 0.503710385 |
| chr19:55437416-55437678 | 0.517715599 | 0.224147073 | 0.503851633 |
| chr7:55250294-55250740 | -0.435626949 | 0.224149826 | 0.503851633 |
| chr1:153024758-153025039 | 0.60743147 | 0.224156175 | 0.503851633 |
| chr14:27608431-27608744 | 0.497757227 | 0.224178619 | 0.503887298 |
| chr19:35757802-35758699 | 0.392037532 | 0.22418547 | 0.503887916 |
| chr1:92029658-92030519 | 0.44340528 | 0.224195092 | 0.503894762 |
| chr18:59517860-59518060 | -0.693527001 | 0.224215575 | 0.503926017 |
| chr1:48572637-48572997 | 0.463704609 | 0.2242401 | 0.503966355 |
| chr14:43312854-43313085 | 0.58531954 | 0.224276794 | 0.504019255 |
| chr15:70140265-70140465 | 0.58531954 | 0.224276794 | 0.504019255 |
| chr15:45790430-45790644 | -0.616633479 | 0.224307597 | 0.504073694 |
| chr1:216750075-216750377 | -0.468899385 | 0.2243378 | 0.504125902 |
| chr8:104688426-104688776 | 0.480797842 | 0.224347129 | 0.504125902 |

| | | | |
|---|---|---|---|
| chr20:46758131-46758347 | 0.576226179 | 0.224350567 | 0.504125902 |
| chr15:73273447-73273672 | -0.648204048 | 0.224360664 | 0.504133806 |
| chr8:103707316-103707872 | 0.418758791 | 0.22437193 | 0.504144337 |
| chr20:57213880-57214690 | 0.382368363 | 0.224391056 | 0.504172526 |
| chr2:27265022-27265242 | 0.526535111 | 0.22441838 | 0.504219134 |
| chr7:80428337-80428781 | -0.480336979 | 0.224440673 | 0.504233125 |
| chr12:34110054-34110961 | -0.424342622 | 0.224457728 | 0.504233125 |
| chr4:103091619-103091926 | -0.502591362 | 0.224461639 | 0.504233125 |
| chr11:47946980-47947582 | 0.429079411 | 0.224462092 | 0.504233125 |
| chr1:56821064-56821327 | -0.49928535 | 0.224462892 | 0.504233125 |
| chr1:51569776-51570228 | 0.439159907 | 0.224466687 | 0.504233125 |
| chr12:12349747-12350668 | -0.410172358 | 0.224470673 | 0.504233125 |
| chr1:32716428-32717042 | 0.406838741 | 0.224485352 | 0.504237167 |
| chr13:22270588-22270859 | -0.653739079 | 0.224489668 | 0.504237167 |
| chr1:153032975-153033193 | 0.547580959 | 0.224492215 | 0.504237167 |
| chr6:97474013-97474281 | -0.612403027 | 0.22449985 | 0.504238702 |
| chr10:48707465-48707665 | -0.628432287 | 0.22450606 | 0.504238702 |
| chr12:72395340-72395599 | -0.565189511 | 0.224519825 | 0.504248974 |
| chr19:18192696-18193332 | 0.406887986 | 0.224523795 | 0.504248974 |
| chr11:102811047-102811289 | -0.595272708 | 0.224530528 | 0.504249315 |
| chr1:109509173-109510055 | 0.407938379 | 0.224552044 | 0.504282855 |
| chrX:11770317-11770622 | 0.522779579 | 0.224566527 | 0.504300598 |
| chr18:51197388-51198483 | -0.383537102 | 0.224577579 | 0.504310637 |
| chr10:85547498-85547806 | -0.525358798 | 0.224592983 | 0.504330449 |
| chr17:82914372-82914690 | -0.483211526 | 0.224662325 | 0.504471373 |
| chr6:137799962-137800208 | -0.640819727 | 0.224676608 | 0.504480028 |
| chr16:2995713-2995937 | -0.522364672 | 0.224679347 | 0.504480028 |
| chr1:152460256-152460604 | 0.44116428 | 0.224694177 | 0.504484434 |
| chr1:196941385-196942168 | -0.435704203 | 0.224694478 | 0.504484434 |
| chr16:234336-235802 | 0.395326152 | 0.224702193 | 0.504486974 |
| chr6:132767962-132768564 | -0.405603734 | 0.224719629 | 0.50449793 |
| chr10:101986806-101987080 | 0.551268236 | 0.22473192 | 0.50449793 |
| chr12:71594294-71594632 | 0.486489716 | 0.224751562 | 0.50449793 |
| chr1:114386766-114387108 | 0.433183398 | 0.224757937 | 0.50449793 |
| chr8:103764370-103764570 | 0.593309325 | 0.224758779 | 0.50449793 |
| chr22:31159999-31160489 | 0.460269767 | 0.224758921 | 0.50449793 |
| chr10:8053377-8054176 | 0.389451174 | 0.224761247 | 0.50449793 |
| chr15:78702775-78702975 | 0.670827743 | 0.224763048 | 0.50449793 |
| chr1:241611056-241611408 | 0.468388227 | 0.224770149 | 0.50449793 |
| chr2:39120713-39121468 | -0.398920475 | 0.224772915 | 0.50449793 |
| chr9:137169253-137169786 | 0.429845521 | 0.224804479 | 0.504553993 |
| chr18:8425380-8425709 | -0.540924924 | 0.224832952 | 0.50459947 |
| chr5:13553962-13554166 | 0.599701922 | 0.224837912 | 0.50459947 |
| chr12:53625781-53626736 | 0.38663448 | 0.224861444 | 0.504637501 |
| chr6:80074903-80075156 | 0.628423835 | 0.224884225 | 0.504673846 |
| chr12:103859108-103859308 | 0.630151164 | 0.224896836 | 0.504687365 |
| chr3:168139919-168140269 | 0.489056715 | 0.224938264 | 0.504745387 |
| chr2:47339738-47340557 | -0.397472609 | 0.224938677 | 0.504745387 |
| chr16:434261-434517 | -0.686436481 | 0.224942454 | 0.504745387 |
| chr2:33373063-33373275 | -0.585779478 | 0.224976258 | 0.50480556 |

| | | | |
|---|---|---|---|
| chr3:120203315-120203534 | -0.606030243 | 0.224982447 | 0.50480556 |
| chr4:92270977-92271233 | -0.579825306 | 0.224996194 | 0.50480875 |
| chr12:123388739-123388940 | 0.513318428 | 0.224997045 | 0.50480875 |
| chr8:98020214-98020431 | 0.569621986 | 0.225021751 | 0.504844901 |
| chr7:32164385-32164976 | -0.409853926 | 0.225026336 | 0.504844901 |
| chr4:121072281-121072531 | 0.613743761 | 0.225042918 | 0.504852538 |
| chr5:42170586-42170786 | 0.613743761 | 0.225042918 | 0.504852538 |
| chr11:69462279-69462479 | 0.613639925 | 0.225055152 | 0.504865202 |
| chr20:57166866-57167174 | 0.396010079 | 0.225074811 | 0.504894522 |
| chr15:74799757-74800504 | 0.406370349 | 0.225097247 | 0.504930068 |
| chr1:106648720-106648992 | 0.437342224 | 0.225105819 | 0.504934515 |
| chr12:57093913-57094346 | -0.453289158 | 0.225128738 | 0.504971141 |
| chr7:1065780-1066106 | 0.464871426 | 0.225135666 | 0.504971898 |
| chr20:54382483-54382979 | 0.395852449 | 0.225180711 | 0.505050368 |
| chr11:64146167-64146397 | 0.522363079 | 0.225183834 | 0.505050368 |
| chr6:33571189-33571561 | -0.492731364 | 0.225192132 | 0.505054197 |
| chr1:180962228-180962511 | -0.475153484 | 0.225229164 | 0.505122465 |
| chr14:105489703-105490165 | 0.404971033 | 0.225238459 | 0.505128526 |
| chr18:9825601-9825905 | 0.524833403 | 0.225258759 | 0.505159266 |
| chr5:64229286-64229512 | 0.60882252 | 0.225273281 | 0.505177048 |
| chr2:102240834-102241203 | 0.472102826 | 0.225280418 | 0.505178267 |
| chr8:41146447-41146705 | -0.596017624 | 0.225304469 | 0.505215567 |
| chr16:73232758-73232984 | -0.581888549 | 0.225310239 | 0.505215567 |
| chr2:75896799-75897165 | -0.459509915 | 0.225356879 | 0.505288868 |
| chr22:38201682-38202183 | 0.425430135 | 0.225362377 | 0.505288868 |
| chr4:37722049-37722343 | -0.529882288 | 0.225362713 | 0.505288868 |
| chr13:113362979-113363772 | 0.419101341 | 0.22538445 | 0.505305018 |
| chr1:114282484-114283155 | 0.389080499 | 0.225384874 | 0.505305018 |
| chr9:121201026-121202461 | 0.386072891 | 0.2253897 | 0.505305018 |
| chr3:36082681-36083574 | -0.430408297 | 0.225422664 | 0.505364135 |
| chr12:124392868-124393528 | -0.416697735 | 0.225451258 | 0.505395188 |
| chr21:43727497-43728045 | 0.445512603 | 0.225453161 | 0.505395188 |
| chrX:102950973-102951911 | -0.4075238 | 0.225456984 | 0.505395188 |
| chr1:171817707-171818108 | -0.484759402 | 0.22547755 | 0.505395188 |
| chr12:10613268-10614105 | -0.418259916 | 0.225480447 | 0.505395188 |
| chr18:29002725-29003619 | -0.403037702 | 0.225481035 | 0.505395188 |
| chr11:3126613-3126813 | -0.623535233 | 0.225482688 | 0.505395188 |
| chr3:162998301-162998579 | -0.543896806 | 0.225523672 | 0.505449342 |
| chr1:109201204-109201708 | -0.433623095 | 0.225529306 | 0.505449342 |
| chr16:68902407-68902659 | -0.621330667 | 0.225530968 | 0.505449342 |
| chr20:54816723-54817063 | 0.431759771 | 0.225533235 | 0.505449342 |
| chr1:41123236-41123459 | -0.600147067 | 0.225547422 | 0.505466353 |
| chr17:67063187-67063458 | -0.501597116 | 0.225559076 | 0.505477684 |
| chr16:13919730-13920503 | 0.396357845 | 0.225576017 | 0.505492532 |
| chr6:52263837-52264047 | -0.587490955 | 0.225578896 | 0.505492532 |
| chr20:53647021-53647221 | 0.41733705 | 0.225592723 | 0.505508734 |
| chr20:44736273-44736579 | 0.45589999 | 0.225666963 | 0.505638069 |
| chr1:47437146-47438201 | 0.392436264 | 0.225667502 | 0.505638069 |
| chr13:18280028-18280357 | -0.495161696 | 0.225673881 | 0.505638069 |
| chr8:95297425-95297722 | -0.504405417 | 0.225676838 | 0.505638069 |

| | | | |
|---|---|---|---|
| chr19:43983802-43984305 | 0.447243444 | 0.225690613 | 0.505654147 |
| chr4:16916747-16917166 | -0.459234603 | 0.225706973 | 0.505676015 |
| chr10:122142692-122143504 | 0.409907525 | 0.225713702 | 0.505676304 |
| chr17:38861558-38861758 | 0.632579941 | 0.225724472 | 0.505685647 |
| chr17:75809057-75809277 | -0.510597288 | 0.225778912 | 0.505792819 |
| chr14:61485269-61485519 | 0.488071071 | 0.225791265 | 0.505805703 |
| chr5:17012391-17012710 | 0.500084948 | 0.225812789 | 0.505815837 |
| chr3:111860075-111860856 | -0.408166043 | 0.22581358 | 0.505815837 |
| chr3:10495111-10495369 | 0.520243272 | 0.225820699 | 0.505815837 |
| chr3:166923702-166923902 | -0.685452997 | 0.225822194 | 0.505815837 |
| chr20:48035980-48036267 | 0.410815091 | 0.225874579 | 0.505903034 |
| chr3:20274895-20275248 | 0.470851473 | 0.225878553 | 0.505903034 |
| chr6:119333754-119334070 | -0.499852821 | 0.225880931 | 0.505903034 |
| chr1:51551880-51552247 | 0.541958378 | 0.225905542 | 0.505943367 |
| chr9:36036755-36037311 | -0.475420421 | 0.225956506 | 0.506042715 |
| chr3:161031087-161031736 | -0.430332055 | 0.225968929 | 0.506055746 |
| chr1:185733169-185733369 | -0.64876644 | 0.225988718 | 0.506085272 |
| chr19:49987106-49987592 | -0.427780426 | 0.226012632 | 0.506124032 |
| chr3:23019725-23020821 | 0.397888752 | 0.226022635 | 0.50613164 |
| chr4:147019065-147019417 | -0.489454718 | 0.226035032 | 0.506144608 |
| chr11:96145186-96145545 | 0.507001529 | 0.226077862 | 0.50622054 |
| chr7:43943087-43944094 | -0.412073042 | 0.226082155 | 0.50622054 |
| chr3:196988193-196988537 | -0.502648447 | 0.226094313 | 0.506232969 |
| chr20:47712149-47713051 | 0.380470872 | 0.226116836 | 0.506268605 |
| chr7:157228932-157229190 | -0.526090861 | 0.226128048 | 0.506278914 |
| chr12:126157284-126157633 | -0.518205959 | 0.226149588 | 0.50629801 |
| chr3:71114396-71114632 | -0.655952918 | 0.226149792 | 0.50629801 |
| chr14:103582044-103582276 | 0.566884138 | 0.22616338 | 0.506313635 |
| chr6:31927822-31928094 | 0.618522265 | 0.226180726 | 0.506337674 |
| chr17:71424058-71424395 | -0.517068363 | 0.226204473 | 0.50637604 |
| chr1:246821445-246821645 | 0.595720476 | 0.226216349 | 0.506387833 |
| chr2:59961669-59961999 | -0.557905291 | 0.226240624 | 0.506427376 |
| chr13:44435081-44435850 | -0.387576394 | 0.226253579 | 0.506433984 |
| chr5:64632361-64632638 | -0.542783765 | 0.226256795 | 0.506433984 |
| chr4:22508545-22508745 | -0.674758798 | 0.226269451 | 0.50643901 |
| chr1:197915684-197916653 | -0.389747463 | 0.226272259 | 0.50643901 |
| chr15:75274416-75275111 | 0.410900716 | 0.226285854 | 0.506454642 |
| chr15:74073046-74073277 | 0.622961656 | 0.226315153 | 0.506492042 |
| chr1:28591008-28591251 | 0.619651199 | 0.226315785 | 0.506492042 |
| chr15:94526078-94526354 | -0.53540474 | 0.22632733 | 0.506503087 |
| chr20:8368704-8369276 | -0.452351547 | 0.226351577 | 0.506542555 |
| chr2:5625573-5625857 | -0.541675194 | 0.226366473 | 0.506552868 |
| chr4:184722998-184723239 | -0.604593278 | 0.226369408 | 0.506552868 |
| chr2:43636833-43637564 | -0.405615762 | 0.226383641 | 0.506569923 |
| chr17:17819762-17819962 | 0.499502461 | 0.226406508 | 0.506587791 |
| chr15:31977682-31978057 | 0.493398133 | 0.22641028 | 0.506587791 |
| chr14:45526814-45527637 | 0.405751715 | 0.226419963 | 0.506587791 |
| chr2:3651662-3652424 | -0.404528668 | 0.226423533 | 0.506587791 |
| chr15:63474864-63475070 | -0.539017213 | 0.226424683 | 0.506587791 |
| chr7:144738835-144739040 | -0.651763843 | 0.226451652 | 0.506618543 |

| | | | |
|---|---|---|---|
| chr8:138477356-138477592 | -0.651763843 | 0.226451652 | 0.506618543 |
| chr1:218402114-218402420 | -0.511275806 | 0.226464433 | 0.506632344 |
| chr22:21037919-21038119 | 0.611726118 | 0.226510484 | 0.506709186 |
| chr10:5684420-5685787 | 0.383798161 | 0.226512056 | 0.506709186 |
| chr21:26116397-26116745 | -0.478482102 | 0.226518621 | 0.506709186 |
| chr2:32893406-32893647 | -0.64907855 | 0.226544818 | 0.506730367 |
| chr17:76251989-76252530 | -0.447980152 | 0.226547443 | 0.506730367 |
| chr14:81408602-81408873 | 0.594749201 | 0.22655413 | 0.506730367 |
| chr10:97318840-97319972 | 0.390812601 | 0.226558914 | 0.506730367 |
| chrX:11111001-11111977 | -0.393025544 | 0.226561156 | 0.506730367 |
| chr14:92613114-92613377 | 0.504146518 | 0.226573498 | 0.506743179 |
| chr3:125936986-125937216 | -0.584249339 | 0.226623465 | 0.506840139 |
| chr1:70324793-70325058 | -0.61932503 | 0.226631318 | 0.506842908 |
| chr17:8139340-8139765 | -0.423510449 | 0.226655523 | 0.506864502 |
| chr22:17773126-17773568 | 0.43567079 | 0.226658758 | 0.506864502 |
| chr17:64918731-64920412 | -0.383668958 | 0.226660819 | 0.506864502 |
| chr16:2430362-2430629 | 0.480154712 | 0.226678917 | 0.50687594 |
| chr4:85909915-85910175 | -0.570859206 | 0.226679165 | 0.50687594 |
| chr4:175128887-175129124 | -0.608927924 | 0.226685998 | 0.506876427 |
| chr1:54323305-54323810 | -0.441208885 | 0.226732901 | 0.50696651 |
| chr4:9103226-9103468 | 0.577876971 | 0.226760171 | 0.507005731 |
| chr17:59972841-59973119 | 0.431988772 | 0.226763676 | 0.507005731 |
| chr3:20054491-20055185 | -0.407427942 | 0.226822796 | 0.507123116 |
| chr7:154841788-154842001 | -0.637883587 | 0.226850661 | 0.507170615 |
| chr9:71001872-71002177 | -0.592198816 | 0.226877409 | 0.507204579 |
| chr5:139388174-139388719 | 0.411549596 | 0.226883593 | 0.507204579 |
| chr5:149960334-149961243 | -0.391859655 | 0.226891377 | 0.507204579 |
| chr2:165406154-165406615 | -0.442062864 | 0.226892331 | 0.507204579 |
| chr4:674153-674975 | 0.396360389 | 0.226913594 | 0.507237314 |
| chr8:38769771-38769971 | -0.628105563 | 0.226928578 | 0.507256009 |
| chr8:64819831-64820195 | -0.5460991 | 0.226941934 | 0.507271065 |
| chr2:169528271-169528542 | -0.543411028 | 0.226953101 | 0.507273845 |
| chr3:106861822-106862107 | -0.614333013 | 0.226964086 | 0.507273845 |
| chr8:36773708-36773920 | 0.62574188 | 0.226967267 | 0.507273845 |
| chr2:196263440-196263640 | -0.674324354 | 0.226975522 | 0.507273845 |
| chr6:135498274-135498635 | -0.422597208 | 0.226982055 | 0.507273845 |
| chr1:146478116-146478578 | 0.465659929 | 0.2269829 | 0.507273845 |
| chr16:49739396-49739624 | -0.603128039 | 0.226997294 | 0.507291217 |
| chr19:19516034-19516329 | -0.533625215 | 0.227008554 | 0.507301586 |
| chr7:9227608-9227914 | -0.596496594 | 0.227037364 | 0.507351171 |
| chr21:45455370-45455584 | 0.511888906 | 0.227046492 | 0.50735677 |
| chr12:132908147-132909086 | 0.386879658 | 0.227062993 | 0.507360728 |
| chr1:204103915-204104950 | -0.391655148 | 0.227063385 | 0.507360728 |
| chr15:90374598-90374802 | 0.634353477 | 0.227068128 | 0.507360728 |
| chr3:177483181-177483455 | -0.53315459 | 0.227078071 | 0.50736815 |
| chr9:137584006-137584271 | 0.575115352 | 0.227091454 | 0.507383257 |
| chr4:87375711-87375983 | -0.562113892 | 0.227114681 | 0.507420354 |
| chr2:47402714-47404019 | -0.384629179 | 0.227129486 | 0.507420561 |
| chr16:2874891-2875182 | 0.513357925 | 0.227131051 | 0.507420561 |
| chr2:106193410-106193716 | 0.459598982 | 0.22713464 | 0.507420561 |

| | | | |
|---|---|---|---|
| chr17:58667879-58668542 | 0.388650369 | 0.227169882 | 0.507474004 |
| chr6:55543984-55544224 | -0.602860752 | 0.227172764 | 0.507474004 |
| chr3:152000148-152000348 | -0.686954791 | 0.227182179 | 0.507474004 |
| chr18:8882901-8883198 | -0.581443983 | 0.227185055 | 0.507474004 |
| chr5:34410751-34411063 | 0.500251294 | 0.227216275 | 0.507528946 |
| chrX:72131867-72132342 | 0.439186499 | 0.227248435 | 0.507577311 |
| chr10:25279647-25279963 | -0.632837707 | 0.227251177 | 0.507577311 |
| chr11:76097082-76097384 | 0.595660278 | 0.227273614 | 0.507612629 |
| chr21:39317349-39317684 | 0.418306223 | 0.227296706 | 0.507649407 |
| chr17:35808903-35809852 | 0.399507358 | 0.227324851 | 0.507697469 |
| chr5:149732050-149732316 | 0.49520096 | 0.227332518 | 0.507699794 |
| chr6:70412916-70414149 | -0.38923775 | 0.227339842 | 0.507701352 |
| chr7:158673385-158673589 | -0.627613204 | 0.227348681 | 0.507706035 |
| chr18:47149542-47151061 | -0.391100655 | 0.227361302 | 0.507706035 |
| chr16:23742857-23743057 | -0.657578104 | 0.227361818 | 0.507706035 |
| chr17:41505432-41505733 | -0.57185999 | 0.227376254 | 0.507720329 |
| chr2:194424911-194425159 | -0.569607402 | 0.227381471 | 0.507720329 |
| chr20:22806986-22807456 | 0.47412062 | 0.227393427 | 0.507732229 |
| chr9:98263816-98264075 | 0.564529429 | 0.227404511 | 0.507733771 |
| chr3:12967556-12968236 | 0.389163886 | 0.227407371 | 0.507733771 |
| chr6:122459121-122459579 | -0.433538323 | 0.227425499 | 0.507759452 |
| chr2:35871778-35871985 | -0.583197478 | 0.227468527 | 0.507833007 |
| chr7:81256992-81257247 | -0.591970885 | 0.2274717 | 0.507833007 |
| chr14:38922041-38922323 | 0.522221753 | 0.227481721 | 0.507840582 |
| chr11:21988535-21988799 | -0.556484529 | 0.227624341 | 0.50812092 |
| chr17:60004640-60005983 | 0.382582237 | 0.227629605 | 0.50812092 |
| chr3:129003805-129004108 | 0.468984611 | 0.227638255 | 0.50812092 |
| chr2:203327709-203329039 | -0.388804139 | 0.227639356 | 0.50812092 |
| chr2:196677441-196677697 | -0.624775926 | 0.227640454 | 0.50812092 |
| chr4:85919653-85919853 | -0.653470242 | 0.227667058 | 0.508151756 |
| chr3:23698922-23699214 | 0.545410214 | 0.227673399 | 0.508151756 |
| chr14:60327482-60327715 | -0.526229197 | 0.227677124 | 0.508151756 |
| chr15:90693789-90693989 | 0.575331182 | 0.227680796 | 0.508151756 |
| chr5:150129560-150130308 | -0.410779492 | 0.227721163 | 0.508227046 |
| chr5:97140344-97140563 | -0.564452819 | 0.227730901 | 0.50822815 |
| chr1:54810495-54810972 | 0.440893065 | 0.227734923 | 0.50822815 |
| chr10:29304564-29305029 | -0.490137945 | 0.227757808 | 0.508264418 |
| chr2:231666325-231666755 | 0.427736978 | 0.227764526 | 0.508264606 |
| chr19:35479366-35479566 | 0.591792403 | 0.227788349 | 0.508284257 |
| chr1:37691598-37691821 | 0.486628169 | 0.227788487 | 0.508284257 |
| chr7:3722185-3722889 | -0.438362213 | 0.227793233 | 0.508284257 |
| chr8:11456577-11457092 | 0.434616753 | 0.227813413 | 0.508297474 |
| chr8:101554339-101554547 | 0.585574666 | 0.227820948 | 0.508297474 |
| chr13:73372750-73373011 | -0.582839884 | 0.227822116 | 0.508297474 |
| chr12:19130897-19131380 | -0.462827855 | 0.227830274 | 0.508297474 |
| chr1:167474456-167474675 | -0.612722572 | 0.227832569 | 0.508297474 |
| chr4:73038914-73039118 | -0.632125421 | 0.227838959 | 0.508297474 |
| chr10:26696884-26698211 | -0.39500506 | 0.227863559 | 0.508337555 |
| chr17:65332972-65333246 | 0.503664438 | 0.227874647 | 0.508347488 |
| chr3:30996519-30996952 | 0.478650288 | 0.227887319 | 0.508360957 |

| | | | |
|---|---|---|---|
| chr10:122342695-122342895 | 0.64085424 | 0.227894142 | 0.508361377 |
| chr3:167913235-167913531 | -0.515040882 | 0.227945329 | 0.508460756 |
| chr2:167292361-167293293 | -0.393805833 | 0.227962206 | 0.508483601 |
| chr13:99501014-99501794 | -0.394612692 | 0.227977974 | 0.508503969 |
| chr1:115040604-115040959 | 0.467469084 | 0.227992948 | 0.508510313 |
| chr2:38692844-38693223 | 0.468618599 | 0.227994092 | 0.508510313 |
| chr21:38906922-38907578 | -0.3902043 | 0.228006173 | 0.508522456 |
| chr2:21633592-21633811 | -0.610411801 | 0.228013529 | 0.50852406 |
| chr2:188396825-188397025 | -0.643099413 | 0.228048263 | 0.508576631 |
| chr6:15949275-15949793 | 0.427507221 | 0.228051211 | 0.508576631 |
| chr15:75922530-75922763 | 0.660147766 | 0.228057013 | 0.508576631 |
| chr9:96383036-96383871 | 0.426368774 | 0.228075042 | 0.508602034 |
| chr12:54720-54926 | -0.586350592 | 0.228141384 | 0.508735169 |
| chr4:22897607-22897844 | 0.562215536 | 0.228172341 | 0.508770191 |
| chr11:17631788-17632046 | -0.565650589 | 0.228173926 | 0.508770191 |
| chr21:44927244-44928466 | -0.405568854 | 0.22817701 | 0.508770191 |
| chr11:107859639-10785983[illegible] | -0.540540003 | 0.228185532 | 0.508774387 |
| chr2:74227707-74228227 | -0.443036971 | 0.228193377 | 0.508777074 |
| chr2:10721514-10722165 | 0.439510961 | 0.228205315 | 0.508788886 |
| chr15:73239860-73240126 | -0.529792209 | 0.228219938 | 0.508806682 |
| chr20:57047334-57047841 | -0.398795582 | 0.228238085 | 0.508832335 |
| chr2:27134296-27134953 | -0.401631362 | 0.228257464 | 0.508860734 |
| chr1:230795328-230795558 | 0.56140372 | 0.228346334 | 0.509044043 |
| chr3:122315745-122315945 | -0.681407445 | 0.228364775 | 0.509070343 |
| chr1:155173083-155173551 | 0.401681553 | 0.228394369 | 0.509121501 |
| chr2:18520385-18520883 | 0.447798014 | 0.228423767 | 0.509156152 |
| chr14:66359449-66359665 | -0.599518251 | 0.228423946 | 0.509156152 |
| chr16:57694709-57694909 | -0.642610886 | 0.228429849 | 0.509156152 |
| chr1:202553738-202553987 | -0.534111243 | 0.228446077 | 0.509177511 |
| chr1:150213453-150214335 | -0.391934973 | 0.228486695 | 0.509220309 |
| chr4:17529671-17529923 | 0.561808175 | 0.228496566 | 0.509220309 |
| chr5:73729258-73729609 | -0.530898083 | 0.228498465 | 0.509220309 |
| chr6:170372469-170373286 | -0.435878575 | 0.228502378 | 0.509220309 |
| chr6:37303991-37304278 | 0.49373205 | 0.228510709 | 0.509220309 |
| chr6:105896795-105897004 | 0.596344747 | 0.228512716 | 0.509220309 |
| chr2:166516795-166517018 | -0.596308028 | 0.228521105 | 0.509220309 |
| chr6:151399535-151400047 | -0.434408529 | 0.228525003 | 0.509220309 |
| chr20:53876638-53878006 | 0.376390331 | 0.228525091 | 0.509220309 |
| chr12:89504671-89504871 | -0.641136643 | 0.228556905 | 0.509276388 |
| chr22:38114508-38114745 | 0.552921891 | 0.228571741 | 0.509294635 |
| chr14:50321054-50321644 | 0.412787774 | 0.228585187 | 0.509300916 |
| chr19:40216725-40217667 | 0.404039859 | 0.228587853 | 0.509300916 |
| chr16:88941863-88942142 | 0.467997561 | 0.228607034 | 0.509328842 |
| chr1:65235969-65236210 | -0.611694246 | 0.228621914 | 0.509347182 |
| chr1:44005746-44005968 | -0.532774879 | 0.228631024 | 0.509352668 |
| chr11:61532218-61532418 | 0.546642167 | 0.228644087 | 0.50936696 |
| chr20:58054662-58054976 | 0.465261761 | 0.228718186 | 0.509514975 |
| chr1:247529265-247529596 | -0.505678354 | 0.228723827 | 0.509514975 |
| chr10:71074216-71074759 | -0.442980181 | 0.228746319 | 0.509537437 |
| chr15:90293266-90293807 | 0.410093056 | 0.228747211 | 0.509537437 |

| | | | |
|---|---|---|---|
| chr1:200971978-200972755 | -0.39981731 | 0.228783256 | 0.509593417 |
| chr12:102824511-102824773 | -0.556885424 | 0.22878797 | 0.509593417 |
| chr15:62572963-62573169 | 0.634222639 | 0.228792294 | 0.509593417 |
| chr9:113913413-113913800 | 0.512321001 | 0.228802476 | 0.509601283 |
| chr7:1530481-1530902 | 0.4187315 | 0.228811445 | 0.509606445 |
| chr7:51428343-51428707 | -0.516926143 | 0.228833321 | 0.509627797 |
| chr8:124295323-124295523 | 0.568390276 | 0.228834334 | 0.509627797 |
| chr7:39720548-39720937 | -0.415319898 | 0.228845416 | 0.509632061 |
| chr12:72570668-72570927 | -0.566472442 | 0.228854943 | 0.509632061 |
| chrX:103845115-103845349 | 0.529677242 | 0.228860699 | 0.509632061 |
| chr17:7262331-7263503 | -0.389474028 | 0.228862854 | 0.509632061 |
| chr7:33278176-33278397 | 0.522159677 | 0.228883153 | 0.509662452 |
| chr3:32106300-32106752 | 0.405484328 | 0.228895349 | 0.50966497 |
| chr1:33219699-33220336 | 0.42676167 | 0.22890059 | 0.50966497 |
| chr10:100969518-100969727 | -0.584881458 | 0.228906608 | 0.50966497 |
| chr8:113980864-113981064 | -0.6483597 | 0.22891197 | 0.50966497 |
| chr2:208034145-208034549 | -0.447433512 | 0.228917543 | 0.50966497 |
| chr2:238285856-238286056 | -0.513978068 | 0.228937929 | 0.50969555 |
| chr9:68584467-68584854 | -0.468410957 | 0.228948942 | 0.509696395 |
| chr19:16111223-16111952 | 0.395728044 | 0.228962602 | 0.509696395 |
| chr17:61729772-61730219 | 0.390786454 | 0.228963635 | 0.509696395 |
| chr10:51331122-51331394 | -0.634002697 | 0.228964918 | 0.509696395 |
| chr3:164574273-164574547 | -0.617005368 | 0.229017647 | 0.509798963 |
| chr7:24721508-24722112 | 0.408488334 | 0.229031549 | 0.50980524 |
| chr6:106381962-106382162 | -0.648142503 | 0.229033773 | 0.50980524 |
| chr5:50588449-50588686 | 0.566382762 | 0.229062045 | 0.509846904 |
| chr11:120818980-120819199 | -0.602776931 | 0.229068866 | 0.509846904 |
| chr7:73811682-73811999 | -0.522528786 | 0.229072453 | 0.509846904 |
| chr20:3407741-3408318 | 0.419691803 | 0.229105896 | 0.509906525 |
| chr20:57130151-57130726 | 0.383160015 | 0.229153956 | 0.509998676 |
| chr10:63464879-63466442 | -0.391801539 | 0.229183339 | 0.510049254 |
| chr13:95478515-95478887 | 0.459736139 | 0.229233798 | 0.510146735 |
| chr2:26011674-26012900 | 0.38505602 | 0.229280476 | 0.510235795 |
| chr2:119167143-119167369 | 0.527975665 | 0.229294107 | 0.510251308 |
| chr3:159322408-159322643 | -0.586315876 | 0.229321042 | 0.510296427 |
| chr1:10474762-10475425 | 0.427457802 | 0.229359233 | 0.510360751 |
| chr20:45446011-45446295 | 0.482669755 | 0.22936327 | 0.510360751 |
| chr13:21124847-21125175 | 0.472279258 | 0.229377134 | 0.51037021 |
| chrX:134542401-134542821 | 0.494190098 | 0.22938795 | 0.51037021 |
| chr17:6651170-6651952 | -0.404400165 | 0.229397888 | 0.51037021 |
| chr10:5442108-5442346 | 0.552836995 | 0.229404113 | 0.51037021 |
| chr6:107075962-107076429 | -0.454953572 | 0.229409576 | 0.51037021 |
| chr3:172461602-172461909 | -0.478704818 | 0.229409637 | 0.51037021 |
| chr10:22615767-22615986 | -0.559052022 | 0.229414147 | 0.51037021 |
| chr1:215710337-215710697 | -0.451295353 | 0.229443755 | 0.510420335 |
| chr6:30679004-30679336 | 0.467172051 | 0.229452719 | 0.510420335 |
| chr15:70892094-70893194 | 0.388386125 | 0.229461133 | 0.510420335 |
| chr3:151426067-151426577 | -0.433529922 | 0.229463325 | 0.510420335 |
| chr18:37367250-37367450 | 0.661778347 | 0.229489492 | 0.510463723 |
| chr9:133459487-133460753 | 0.38249546 | 0.229503511 | 0.510472759 |

| | | | |
|---|---|---|---|
| chr21:39022868-39023075 | 0.539783955 | 0.229507534 | 0.510472759 |
| chr12:132700998-132701348 | 0.53751976 | 0.229513541 | 0.510472759 |
| chr11:21460221-21460426 | 0.61223774 | 0.229535692 | 0.510507208 |
| chr16:50152412-50153405 | 0.398165938 | 0.229597985 | 0.510630931 |
| chr7:142535076-142535915 | -0.422974843 | 0.22964393 | 0.510718288 |
| chr19:9818768-9819904 | 0.384558554 | 0.229733968 | 0.510862243 |
| chr10:33534561-33534778 | 0.576211231 | 0.229735448 | 0.510862243 |
| chr1:107593833-107594241 | 0.459858477 | 0.229736507 | 0.510862243 |
| chr8:103501184-103502012 | 0.389366455 | 0.229742417 | 0.510862243 |
| chr9:136687341-136687693 | 0.443315787 | 0.229744732 | 0.510862243 |
| chr9:134617185-134617535 | -0.582407702 | 0.229748663 | 0.510862243 |
| chr4:114217578-114217842 | -0.643721705 | 0.22978403 | 0.510926058 |
| chr8:30157666-30157869 | 0.551218233 | 0.229811496 | 0.510952004 |
| chrX:130120512-130121247 | 0.413915689 | 0.22981177 | 0.510952004 |
| chr8:800380-800699 | 0.651411299 | 0.229815705 | 0.510952004 |
| chr1:19800035-19800271 | -0.597429564 | 0.229825882 | 0.510959806 |
| chr7:120988585-120988792 | -0.580387426 | 0.229867679 | 0.511024617 |
| chr14:60722382-60723055 | 0.40182079 | 0.229868373 | 0.511024617 |
| chr19:3557421-3558113 | 0.413255488 | 0.229896139 | 0.511071516 |
| chr12:87439897-87440768 | -0.386680003 | 0.229909368 | 0.511086096 |
| chr1:44775288-44776224 | 0.404006455 | 0.229932085 | 0.511105092 |
| chr2:138501140-138502661 | -0.382101909 | 0.22993857 | 0.511105092 |
| chr5:31934246-31934483 | 0.522639689 | 0.229942986 | 0.511105092 |
| chr16:70523383-70524323 | 0.398594589 | 0.229944595 | 0.511105092 |
| chr19:3700451-3700903 | 0.472844343 | 0.229957307 | 0.51111852 |
| chr14:74508406-74508680 | 0.45433083 | 0.229966322 | 0.511123732 |
| chr1:203117867-203118106 | -0.496792894 | 0.22998257 | 0.511145018 |
| chr16:2501142-2501710 | 0.438388084 | 0.230020291 | 0.511199683 |
| chr6:144287115-144287315 | 0.530349873 | 0.230020509 | 0.511199683 |
| chr11:33241446-33241646 | 0.599768192 | 0.230069728 | 0.511294237 |
| chr7:82727007-82727477 | -0.50271909 | 0.230092768 | 0.511317894 |
| chr5:55355462-55355997 | -0.474499937 | 0.23009372 | 0.511317894 |
| chr14:34693103-34693566 | 0.43601385 | 0.230107419 | 0.511324669 |
| chr14:67694724-67695959 | 0.387359054 | 0.230110162 | 0.511324669 |
| chr19:32363085-32363607 | 0.432741762 | 0.230117797 | 0.511324669 |
| chr1:36223862-36225101 | 0.381223538 | 0.230129744 | 0.511324669 |
| chr1:166489658-166489858 | 0.604028872 | 0.230132312 | 0.511324669 |
| chr4:42296232-42296449 | -0.626886705 | 0.230143482 | 0.511324669 |
| chr7:30897092-30897323 | -0.626886705 | 0.230143482 | 0.511324669 |
| chr17:72477610-72477896 | 0.485458733 | 0.230173896 | 0.511362655 |
| chr20:54200998-54201681 | 0.381283669 | 0.230173927 | 0.511362655 |
| chr1:39410985-39411202 | 0.507094466 | 0.230208529 | 0.51140166 |
| chr3:125357104-125357664 | 0.442466908 | 0.230209916 | 0.51140166 |
| chr18:10525735-10526746 | 0.39263697 | 0.230211507 | 0.51140166 |
| chr13:45115024-45115224 | -0.664905566 | 0.230234865 | 0.511438721 |
| chr12:6371057-6371464 | -0.435672179 | 0.230265543 | 0.51149204 |
| chr5:124599612-124599956 | -0.539593183 | 0.230274895 | 0.511492552 |
| chr1:183551903-183552219 | -0.535659314 | 0.230279125 | 0.511492552 |
| chr8:14264114-14264960 | -0.439127282 | 0.230291134 | 0.511504398 |
| chr1:182484501-182485134 | -0.430305977 | 0.230328168 | 0.511571828 |

| | | | |
|---|---|---|---|
| chr4:55853382-55854071 | -0.395875927 | 0.230388364 | 0.511660308 |
| chr4:147837457-147837703 | 0.52167733 | 0.230403465 | 0.511660308 |
| chr20:50957840-50958109 | 0.459036443 | 0.230406239 | 0.511660308 |
| chr15:89243623-89244776 | 0.383375286 | 0.230406893 | 0.511660308 |
| chr7:76201733-76202197 | -0.461900786 | 0.230413612 | 0.511660308 |
| chr18:8052406-8052606 | -0.652815227 | 0.230414749 | 0.511660308 |
| chr18:73406670-73406881 | -0.652815227 | 0.230414749 | 0.511660308 |
| chr4:25913825-25914767 | 0.385992289 | 0.230427767 | 0.511674385 |
| chr2:229450628-229450828 | -0.670484324 | 0.230471334 | 0.511753901 |
| chr2:101244293-101245229 | -0.41244557 | 0.230479623 | 0.511753901 |
| chr8:121803145-121803374 | -0.578747445 | 0.230486785 | 0.511753901 |
| chr2:156383071-156383673 | -0.434174063 | 0.230493422 | 0.511753901 |
| chr10:62011575-62011927 | -0.460451901 | 0.230496971 | 0.511753901 |
| chr21:38838721-38839017 | -0.529706479 | 0.230509166 | 0.511761656 |
| chr20:54277034-54277420 | 0.403284082 | 0.230513822 | 0.511761656 |
| chr11:16605438-16605741 | -0.512273561 | 0.230525449 | 0.511772641 |
| chr17:74769861-74770282 | -0.436223284 | 0.230542787 | 0.511796303 |
| chr1:221908552-221908989 | -0.459569696 | 0.230568508 | 0.511838574 |
| chr14:35374405-35374670 | 0.529698187 | 0.230575291 | 0.511838803 |
| chr3:62464259-62464537 | 0.446214801 | 0.230611541 | 0.511904441 |
| chr22:19432176-19432735 | 0.40546957 | 0.230646864 | 0.51195224 |
| chr2:86562584-86563686 | -0.385575104 | 0.230650337 | 0.51195224 |
| chr14:55532927-55533238 | 0.490413277 | 0.230653119 | 0.51195224 |
| chr14:23010071-23010315 | 0.455336732 | 0.230690342 | 0.512020027 |
| chr1:223393648-223394077 | 0.490729178 | 0.230720553 | 0.512072247 |
| chr2:176830490-176830747 | -0.532913554 | 0.230730392 | 0.512079251 |
| chr6:136583126-136583369 | 0.558403715 | 0.23080486 | 0.512223425 |
| chr5:177599345-177599561 | -0.531636637 | 0.230808723 | 0.512223425 |
| chr2:42787934-42788343 | -0.449966999 | 0.230838282 | 0.512274187 |
| chr19:3721544-3721769 | 0.534243535 | 0.23086487 | 0.512318351 |
| chr4:17701076-17701653 | 0.479983658 | 0.23088058 | 0.512338376 |
| chr11:10750800-10751471 | 0.409350158 | 0.230904498 | 0.512355045 |
| chr1:215055396-215055635 | -0.533769755 | 0.230907584 | 0.512355045 |
| chr7:12210512-12211834 | 0.375661366 | 0.230908152 | 0.512355045 |
| chr13:92716826-92717026 | 0.661570944 | 0.230929537 | 0.512387657 |
| chr13:52159178-52159848 | -0.435246059 | 0.230955722 | 0.51242066 |
| chr5:175729805-175730187 | 0.425209931 | 0.23096502 | 0.51242066 |
| chr22:29217083-29217509 | -0.484086329 | 0.230965037 | 0.51242066 |
| chr12:22543629-22544907 | -0.381649423 | 0.230974362 | 0.51242066 |
| chr7:50746760-50746964 | -0.627915026 | 0.230977849 | 0.51242066 |
| chr8:86033227-86033732 | 0.433098337 | 0.231048313 | 0.512562141 |
| chr8:93369086-93369381 | 0.517258034 | 0.231063819 | 0.512581701 |
| chr20:49246629-49246838 | 0.516788297 | 0.231071589 | 0.512584096 |
| chr20:62616780-62617166 | 0.452715303 | 0.231099811 | 0.512616153 |
| chr8:141075603-141076185 | 0.415640563 | 0.231104102 | 0.512616153 |
| chr10:87817662-87819065 | -0.382109878 | 0.231106111 | 0.512616153 |
| chr3:193829795-193830107 | 0.480838872 | 0.231119957 | 0.512628443 |
| chr22:17847092-17847703 | 0.444342997 | 0.231134103 | 0.512628443 |
| chr11:48649181-48649457 | -0.618895667 | 0.231140631 | 0.512628443 |
| chr1:107926697-107926993 | 0.515242045 | 0.23114468 | 0.512628443 |

| | | | |
|---|---|---|---|
| chr11:9663706-9664751 | -0.404659181 | 0.231145103 | 0.512628443 |
| chr15:78366178-78366378 | -0.656438317 | 0.23117624 | 0.512681296 |
| chr8:124440219-124440705 | -0.408103676 | 0.231182317 | 0.512681296 |
| chr11:69674780-69675826 | 0.387907841 | 0.231215202 | 0.512739385 |
| chr8:137843339-137843544 | -0.572684331 | 0.231242708 | 0.512772237 |
| chr17:82153818-82154286 | 0.421452084 | 0.231249521 | 0.512772237 |
| chr7:73866227-73866524 | -0.560511949 | 0.231252086 | 0.512772237 |
| chr3:100392070-100392309 | 0.60287996 | 0.231256785 | 0.512772237 |
| chr17:77772420-77772963 | 0.433633299 | 0.231269104 | 0.512784712 |
| chr14:106872576-10687347 | 0.386075622 | 0.231276371 | 0.512785986 |
| chr6:21381655-21381970 | -0.545363244 | 0.23128432 | 0.512788772 |
| chr20:408534-408960 | 0.470179796 | 0.231305343 | 0.512820543 |
| chr9:33264079-33265226 | -0.381671998 | 0.231321707 | 0.512841985 |
| chr4:165206972-165208119 | -0.395910432 | 0.23133362 | 0.512853557 |
| chr4:22357252-22357470 | 0.588607964 | 0.231371938 | 0.512912737 |
| chr5:69517791-69518062 | -0.526359647 | 0.231373703 | 0.512912737 |
| chr6:83884494-83884781 | -0.565455635 | 0.2313875 | 0.512928484 |
| chr2:13414072-13414336 | -0.572492075 | 0.231400881 | 0.512943305 |
| chr18:12419793-12420184 | 0.455202053 | 0.231417051 | 0.512955235 |
| chr4:169619496-169619835 | -0.40874638 | 0.231419652 | 0.512955235 |
| chr2:11588269-11588652 | -0.462311548 | 0.23145338 | 0.513015155 |
| chr11:2256927-2257747 | 0.408693093 | 0.231463194 | 0.513022067 |
| chr1:117312504-117312763 | 0.51224365 | 0.231475408 | 0.513034299 |
| chr10:68836900-68837253 | -0.541886646 | 0.231490501 | 0.513039374 |
| chr6:36123823-36124023 | 0.549192748 | 0.23149109 | 0.513039374 |
| chr20:52379833-52380052 | -0.566277273 | 0.231512101 | 0.513060825 |
| chr7:138029200-138029450 | -0.610867797 | 0.231514161 | 0.513060825 |
| chr17:61554989-61555540 | 0.38585357 | 0.23155346 | 0.513128458 |
| chr4:119227570-119227800 | 0.659710537 | 0.231558073 | 0.513128458 |
| chr7:128444779-128445206 | -0.440605672 | 0.231564812 | 0.513128551 |
| chr6:127302826-127303033 | -0.612930309 | 0.231579635 | 0.513146557 |
| chr2:42835968-42836214 | -0.602057995 | 0.231608174 | 0.513194954 |
| chr3:155805351-155806580 | -0.381833961 | 0.231643812 | 0.51325229 |
| chr3:62399049-62399295 | 0.47161701 | 0.231648067 | 0.51325229 |
| chr20:20864469-20864674 | 0.568410901 | 0.231654146 | 0.51325229 |
| chrX:129115090-129115500 | 0.449215549 | 0.23166191 | 0.513254651 |
| chr6:130708133-130708746 | -0.422197085 | 0.231669731 | 0.513257138 |
| chr11:96463874-96464475 | -0.433920409 | 0.231694534 | 0.513284001 |
| chr11:44037349-44037666 | -0.505669974 | 0.231695254 | 0.513284001 |
| chr20:64038972-64039283 | 0.431990682 | 0.231718348 | 0.513318201 |
| chr2:120262828-120263084 | 0.543042926 | 0.23172409 | 0.513318201 |
| chr2:119294954-119295154 | -0.631287811 | 0.231786433 | 0.513441459 |
| chr3:162482289-162482529 | 0.540740684 | 0.231799803 | 0.513455746 |
| chr15:92521563-92521938 | 0.444408348 | 0.231806285 | 0.513455746 |
| chr2:74000087-74000460 | -0.464729959 | 0.231827191 | 0.513487208 |
| chr1:232600940-232601173 | -0.581980771 | 0.231837195 | 0.513494524 |
| chr1:236515172-236515707 | -0.471753201 | 0.231844841 | 0.513496616 |
| chr6:1396985-1397185 | 0.640952739 | 0.231853889 | 0.5134983 |
| chr14:76151616-76152712 | 0.382173414 | 0.231859005 | 0.5134983 |
| chr1:203040204-203040427 | 0.468509856 | 0.231874932 | 0.513518731 |

| | | | |
|---|---|---|---|
| chr7:158266975-158267401 | -0.484506694 | 0.23191578 | 0.513594349 |
| chr1:41449542-41449947 | -0.437704074 | 0.231948106 | 0.513651093 |
| chr14:93139202-93139571 | 0.428363328 | 0.231977685 | 0.513701748 |
| chr20:46688902-46689960 | 0.381418726 | 0.231993026 | 0.513712499 |
| chr1:151006764-151007064 | 0.409720647 | 0.231995949 | 0.513712499 |
| chr3:38003466-38003712 | 0.463600784 | 0.232008428 | 0.513724385 |
| chr10:43885950-43886150 | -0.666752992 | 0.232024769 | 0.513724385 |
| chr2:19480775-19480975 | -0.666752992 | 0.232024769 | 0.513724385 |
| chr1:9627091-9627317 | 0.587792599 | 0.232029107 | 0.513724385 |
| chr11:45103085-45103363 | -0.585858634 | 0.23203484 | 0.513724385 |
| chr17:74748943-74749146 | 0.479702825 | 0.23204908 | 0.513741067 |
| chr14:97169097-97169313 | 0.615289762 | 0.232100419 | 0.51380697 |
| chr17:67475414-67475914 | -0.416112161 | 0.232101763 | 0.51380697 |
| chr6:138800882-138801403 | -0.491359921 | 0.2321022 | 0.51380697 |
| chr11:92136872-92137113 | -0.644217197 | 0.23210567 | 0.51380697 |
| chr8:38552992-38553192 | 0.637800572 | 0.232113539 | 0.513809544 |
| chr5:175871588-175872026 | 0.436521973 | 0.232128436 | 0.513827677 |
| chrX:4844335-4844798 | 0.452229645 | 0.232160183 | 0.513883105 |
| chr5:168486191-168487447 | -0.381056618 | 0.23217973 | 0.513911526 |
| chr1:164256507-164256707 | 0.672122303 | 0.232208356 | 0.51396004 |
| chr18:73843656-73843952 | -0.556472103 | 0.232220027 | 0.513971025 |
| chr10:69438622-69439568 | -0.404557171 | 0.232230914 | 0.513980274 |
| chr1:37794308-37794508 | 0.569730986 | 0.232246983 | 0.513993862 |
| chr1:219432384-219432584 | -0.617437693 | 0.23225047 | 0.513993862 |
| chr8:128549273-128549683 | -0.473332152 | 0.232258162 | 0.513996041 |
| chr22:44804546-44804747 | 0.67146471 | 0.23229173 | 0.51405548 |
| chr16:12068022-12068222 | 0.583550483 | 0.232305188 | 0.514059693 |
| chr1:219157403-219157603 | 0.6044115 | 0.232310937 | 0.514059693 |
| chr8:69113084-69113332 | -0.559248343 | 0.232313761 | 0.514059693 |
| chr20:53949482-53949954 | 0.385163768 | 0.232373596 | 0.514177246 |
| chr14:100108426-100108626 | 0.516925371 | 0.232385271 | 0.514188231 |
| chr9:124312770-124312989 | -0.578778104 | 0.232399158 | 0.514204109 |
| chr1:171393468-171393684 | -0.587212355 | 0.232433396 | 0.514256602 |
| chr5:31532739-31533306 | -0.413271182 | 0.232442083 | 0.514256602 |
| chr2:21391759-21392013 | -0.538371921 | 0.232444921 | 0.514256602 |
| chr4:17597565-17597907 | 0.462469629 | 0.232454822 | 0.514256602 |
| chr8:109737615-109737986 | -0.440456021 | 0.232456441 | 0.514256602 |
| chr9:86126925-86127745 | -0.394937689 | 0.232479334 | 0.514292398 |
| chr6:75888260-75888515 | 0.647236906 | 0.232490283 | 0.514296277 |
| chr6:53795917-53796210 | 0.460165602 | 0.232494511 | 0.514296277 |
| chrX:87727672-87727877 | -0.565806691 | 0.232512682 | 0.514321623 |
| chr20:51220247-51220468 | 0.508472687 | 0.232522128 | 0.514327671 |
| chr2:217650807-217651085 | -0.51670093 | 0.232557596 | 0.514376647 |
| chr14:44316922-44317276 | 0.434095143 | 0.232557696 | 0.514376647 |
| chr7:149707481-149707931 | 0.418834447 | 0.232564521 | 0.514376893 |
| chr3:18126929-18127402 | -0.477398835 | 0.232586306 | 0.514410228 |
| chr12:100678399-100678672 | 0.561750069 | 0.232599427 | 0.514414713 |
| chr6:163413319-163414773 | -0.388233898 | 0.232601761 | 0.514414713 |
| chr22:23166578-23166907 | -0.527047023 | 0.232620385 | 0.514432284 |
| chr10:125061209-125061440 | 0.502055027 | 0.232623134 | 0.514432284 |

| | | | |
|---|---|---|---|
| chr4:6974477-6974686 | 0.583261709 | 0.232634447 | 0.514439014 |
| chr15:84715998-84717053 | 0.380213151 | 0.232642319 | 0.514439014 |
| chr8:86703324-86703733 | 0.428761847 | 0.232651334 | 0.514439014 |
| chr16:87100117-87100448 | 0.569133196 | 0.232653033 | 0.514439014 |
| chr3:37242804-37244360 | 0.382331082 | 0.232675967 | 0.514474878 |
| chr7:106069133-106069352 | -0.522837843 | 0.232701355 | 0.514491115 |
| chr9:94638694-94639048 | 0.479043442 | 0.232703104 | 0.514491115 |
| chr4:102731211-102731481 | -0.516537464 | 0.232703454 | 0.514491115 |
| chr14:39426688-39426924 | 0.482476475 | 0.232751136 | 0.514574728 |
| chr8:98085172-98085936 | 0.395435623 | 0.232754704 | 0.514574728 |
| chr3:165322075-165322293 | 0.597910826 | 0.232768768 | 0.514590974 |
| chr2:188291438-188293220 | -0.378762802 | 0.232781353 | 0.514603947 |
| chr16:67028639-67029290 | 0.413473274 | 0.232811945 | 0.514656729 |
| chr16:19907356-19907787 | 0.45612987 | 0.232818714 | 0.514656844 |
| chr1:197399988-197400519 | -0.424075999 | 0.232827211 | 0.514660778 |
| chr9:104991046-104992055 | -0.3992894 | 0.232848384 | 0.514679229 |
| chr14:39358850-39359342 | -0.442921128 | 0.232849638 | 0.514679229 |
| chr5:174214725-174215114 | 0.518891701 | 0.232867347 | 0.514679229 |
| chr6:110509711-110510054 | -0.500550714 | 0.232869179 | 0.514679229 |
| chr12:111685403-111686509 | 0.391180749 | 0.232875459 | 0.514679229 |
| chr7:5352413-5352740 | -0.5460317 | 0.232880419 | 0.514679229 |
| chr17:29406713-29406930 | -0.662588196 | 0.232884325 | 0.514679229 |
| chr19:42255059-42255360 | 0.457180518 | 0.232889295 | 0.514679229 |
| chr2:175282382-175282669 | -0.54901571 | 0.232962186 | 0.514821588 |
| chr22:20111142-20111400 | -0.505952378 | 0.23296715 | 0.514821588 |
| chr14:64372557-64372766 | -0.628489285 | 0.232986419 | 0.514848836 |
| chr18:34788281-34788488 | 0.606211695 | 0.232994169 | 0.514848836 |
| chr10:97861414-97861627 | 0.534058552 | 0.232999637 | 0.514848836 |
| chr9:136791627-136791916 | 0.439059641 | 0.23300856 | 0.514853705 |
| chr12:113736184-113736398 | -0.639720409 | 0.23302791 | 0.514876443 |
| chr6:98947188-98948560 | -0.3767863 | 0.23303229 | 0.514876443 |
| chrX:49155325-49156584 | -0.385431179 | 0.233060193 | 0.514923244 |
| chr6:111276521-111277167 | 0.41322815 | 0.23309214 | 0.514978978 |
| chrX:154534825-154535193 | 0.456352702 | 0.233129662 | 0.515047026 |
| chr8:121214312-121214512 | 0.640337383 | 0.233157897 | 0.515079217 |
| chr20:57187362-57187673 | 0.390995601 | 0.233163441 | 0.515079217 |
| chr1:231162387-231162587 | 0.570826974 | 0.2331644 | 0.515079217 |
| chr7:76220735-76221015 | -0.598420944 | 0.233186258 | 0.515112652 |
| chr2:15800946-15801154 | -0.569043014 | 0.233209672 | 0.515149523 |
| chr16:66549784-66550106 | 0.481674517 | 0.233253618 | 0.515231744 |
| chr20:62274402-62274828 | -0.468944949 | 0.233263248 | 0.515238161 |
| chr2:53202132-53202416 | -0.543818569 | 0.233279759 | 0.515255741 |
| chr5:96875781-96876768 | -0.409338538 | 0.233289348 | 0.515255741 |
| chr8:80611703-80612057 | -0.523241727 | 0.233291909 | 0.515255741 |
| chr7:151448363-151448941 | 0.447825593 | 0.233298105 | 0.515255741 |
| chr10:62610630-62611157 | -0.437344468 | 0.233322687 | 0.515295178 |
| chr20:40945806-40946356 | 0.418745145 | 0.233397471 | 0.515438053 |
| chr16:30377610-30377993 | 0.429821405 | 0.233402358 | 0.515438053 |
| chr8:42541098-42541391 | 0.523157999 | 0.233407561 | 0.515438053 |
| chr11:73660022-73660349 | 0.446505645 | 0.233426792 | 0.515465666 |

| | | | |
|---|---|---|---|
| chr1:25616157-25616762 | 0.435621262 | 0.233460662 | 0.515517221 |
| chr20:63957066-63958143 | -0.381184754 | 0.233465325 | 0.515517221 |
| chr1:145795346-145795586 | -0.594777941 | 0.233470323 | 0.515517221 |
| chr17:70089361-70090218 | -0.430885715 | 0.233486234 | 0.515527058 |
| chr19:18251874-18252096 | 0.591249402 | 0.233494025 | 0.515527058 |
| chr2:53969379-53969591 | -0.497418071 | 0.233494962 | 0.515527058 |
| chr10:93438611-93439275 | -0.397586417 | 0.233521798 | 0.515566249 |
| chr15:68673943-68674273 | 0.459206618 | 0.23352617 | 0.515566249 |
| chr2:8591035-8591329 | 0.49172287 | 0.233543881 | 0.515579675 |
| chr2:176269050-176270237 | -0.385350594 | 0.233549806 | 0.515579675 |
| chr15:82058053-82058253 | -0.660247401 | 0.233558704 | 0.515579675 |
| chr10:89251214-89252247 | -0.380695758 | 0.233559168 | 0.515579675 |
| chr2:27271674-27271890 | 0.592394233 | 0.233574243 | 0.515598101 |
| chr6:43427357-43428241 | 0.386259538 | 0.23359062 | 0.515604171 |
| chr10:70052437-70053057 | -0.438187675 | 0.233590718 | 0.515604171 |
| chr16:88838397-88838695 | -0.505421351 | 0.233597181 | 0.515604171 |
| chr15:41603806-41604180 | -0.4568453 | 0.233607051 | 0.515611105 |
| chr8:11467205-11467908 | 0.420578439 | 0.233624008 | 0.515616443 |
| chr9:25076373-25076573 | 0.628125305 | 0.233627819 | 0.515616443 |
| chr11:62599166-62599520 | 0.411260577 | 0.233632717 | 0.515616443 |
| chr14:33849161-33849703 | 0.419002547 | 0.233641802 | 0.515616443 |
| chr6:24359714-24360514 | -0.386940184 | 0.233643117 | 0.515616443 |
| chr1:161117449-161118393 | 0.390592413 | 0.233663846 | 0.515636741 |
| chr10:95841734-95842246 | -0.470156133 | 0.233665774 | 0.515636741 |
| chr20:47424913-47425197 | 0.394950953 | 0.233692837 | 0.515677548 |
| chr6:26312507-26313424 | -0.384671264 | 0.233697726 | 0.515677548 |
| chr3:99686978-99687339 | -0.492088279 | 0.233710727 | 0.515691385 |
| chr6:106927845-106928104 | 0.564785502 | 0.233735562 | 0.515724355 |
| chr1:56803592-56803901 | -0.477397605 | 0.233739131 | 0.515724355 |
| chr1:203089242-203089476 | -0.471377492 | 0.233751163 | 0.515736052 |
| chr10:6737277-6737613 | -0.507844257 | 0.233780771 | 0.515767152 |
| chr11:75815525-75815759 | 0.448317324 | 0.23378122 | 0.515767152 |
| chr8:143976625-143977380 | 0.400222788 | 0.233785453 | 0.515767152 |
| chr4:159723606-159723941 | -0.542026671 | 0.233793275 | 0.51576956 |
| chr2:2297177-2297448 | 0.573390331 | 0.233802651 | 0.515775394 |
| chr7:129218562-129218891 | -0.458959501 | 0.233810313 | 0.515777446 |
| chr7:37732247-37732545 | -0.475716227 | 0.233837936 | 0.515823531 |
| chr14:78272320-78272542 | 0.594103006 | 0.233862621 | 0.515863132 |
| chr11:33869442-33869987 | 0.476481253 | 0.233871864 | 0.515868669 |
| chr2:189640833-189641464 | -0.430680589 | 0.233885638 | 0.5158842 |
| chr15:92773555-92773823 | -0.587263831 | 0.233896877 | 0.51589414 |
| chr5:77630003-77631203 | -0.389444447 | 0.233916459 | 0.51592248 |
| chr20:53976000-53976200 | 0.411019192 | 0.233972987 | 0.516032302 |
| chr2:207311353-207311711 | 0.562129285 | 0.234010257 | 0.516099645 |
| chr11:134579470-134579698 | -0.580609598 | 0.234025815 | 0.516119103 |
| chr2:208395274-208395611 | -0.46388873 | 0.234069921 | 0.516201517 |
| chr4:143025005-143025710 | 0.435756078 | 0.234098449 | 0.516249572 |
| chr22:50474954-50475263 | -0.470784941 | 0.234108295 | 0.516255457 |
| chr3:123028128-123028375 | 0.520905955 | 0.234121293 | 0.516255457 |
| chr7:124152885-124153213 | -0.508506741 | 0.234124849 | 0.516255457 |

| | | | |
|---|---|---|---|
| chr17:4949186-4949533 | 0.459476096 | 0.234128069 | 0.516255457 |
| chr6:42548072-42548461 | 0.458941394 | 0.234140123 | 0.516257703 |
| chr1:62906520-62906903 | -0.535061184 | 0.234142563 | 0.516257703 |
| chr3:87167814-87168205 | -0.512977652 | 0.234150171 | 0.516259622 |
| chr3:62580592-62580792 | -0.525228017 | 0.234175874 | 0.516294105 |
| chr16:48609892-48610523 | 0.419804487 | 0.234181166 | 0.516294105 |
| chr21:39291408-39291732 | 0.49199948 | 0.234191894 | 0.516294105 |
| chr14:93184343-93185455 | 0.380232678 | 0.234192763 | 0.516294105 |
| chr11:75351602-75352018 | 0.416189226 | 0.234233641 | 0.516351267 |
| chrX:108439422-108440505 | -0.402792561 | 0.23424346 | 0.516351267 |
| chr12:40325398-40326713 | -0.399608456 | 0.234247889 | 0.516351267 |
| chr1:111920275-111920547 | 0.442371613 | 0.234253045 | 0.516351267 |
| chr13:45718175-45718385 | -0.583140299 | 0.23425536 | 0.516351267 |
| chr3:113494170-113494710 | -0.434987431 | 0.234270436 | 0.516351267 |
| chr1:47477705-47478914 | -0.395166478 | 0.234276141 | 0.516351267 |
| chr5:129438960-129439277 | -0.514841297 | 0.234276581 | 0.516351267 |
| chr5:158946635-158947074 | -0.509083923 | 0.234279343 | 0.516351267 |
| chr3:64559235-64559535 | 0.409282725 | 0.234344031 | 0.516478984 |
| chr10:126598078-126598395 | -0.478827351 | 0.234356336 | 0.516491246 |
| chr8:8892087-8893063 | 0.411238153 | 0.234394269 | 0.516550945 |
| chr7:151233106-151233468 | 0.438694228 | 0.234396907 | 0.516550945 |
| chr11:61481092-61481330 | -0.515567148 | 0.234411683 | 0.516568651 |
| chr12:46260575-46260796 | 0.550821307 | 0.234433758 | 0.51659524 |
| chr4:122167452-122167692 | -0.588763999 | 0.234443976 | 0.51659524 |
| chr4:147925189-147925432 | -0.588763999 | 0.234443976 | 0.51659524 |
| chr7:116524576-116524956 | -0.468755744 | 0.234453972 | 0.516599221 |
| chr13:46992419-46992807 | -0.427832512 | 0.234459267 | 0.516599221 |
| chr2:48420511-48421068 | -0.411817884 | 0.234480528 | 0.516631211 |
| chr3:192571293-192571593 | -0.502735629 | 0.234531201 | 0.516728002 |
| chr1:247087368-247087568 | 0.535671128 | 0.234610294 | 0.516887399 |
| chr19:57280173-57280769 | 0.397817962 | 0.234656569 | 0.516974485 |
| chr6:97501978-97502271 | -0.537214106 | 0.234684103 | 0.517020279 |
| chr9:88387346-88387555 | 0.511622386 | 0.234706923 | 0.517055686 |
| chr13:32537853-32539278 | -0.380229776 | 0.234736713 | 0.517063421 |
| chr17:54486748-54487285 | -0.438298873 | 0.234737585 | 0.517063421 |
| chr2:61004908-61005188 | -0.49783087 | 0.234737643 | 0.517063421 |
| chr2:174336727-174337414 | -0.433015632 | 0.234744627 | 0.517063421 |
| chr15:79333126-79333454 | 0.488265981 | 0.234751088 | 0.517063421 |
| chr5:31638709-31639482 | 0.39077144 | 0.23476397 | 0.517063421 |
| chr3:175121312-175121660 | -0.458689421 | 0.234767774 | 0.517063421 |
| chr13:22796073-22796282 | 0.56880814 | 0.234767966 | 0.517063421 |
| chr7:143837328-143837557 | -0.626770671 | 0.234777105 | 0.517063421 |
| chr10:87659418-87660112 | 0.434045699 | 0.234777916 | 0.517063421 |
| chr7:20648259-20648519 | 0.495483598 | 0.234815446 | 0.517131211 |
| chr2:10285539-10285961 | -0.439062353 | 0.234831455 | 0.517151605 |
| chr2:38690632-38691194 | 0.423534595 | 0.234874473 | 0.517231473 |
| chr22:17058183-17058674 | 0.463136324 | 0.234886146 | 0.517242314 |
| chrX:140764357-140765389 | 0.401272381 | 0.234913163 | 0.517286941 |
| chr15:81697157-81697496 | 0.555762844 | 0.234930931 | 0.517295038 |
| chr4:52659114-52660127 | 0.39483989 | 0.234932424 | 0.517295038 |

| | | | |
|---|---|---|---|
| chr5:148841566-148841883 | -0.550903097 | 0.234937094 | 0.517295038 |
| chr8:139705714-139706101 | 0.420176513 | 0.234973438 | 0.517346033 |
| chr14:55344835-55345035 | 0.610626194 | 0.234973758 | 0.517346033 |
| chr12:48197506-48197741 | -0.546433757 | 0.235009422 | 0.517393417 |
| chr1:119647671-119647947 | -0.515884452 | 0.235013608 | 0.517393417 |
| chr11:30323630-30323837 | -0.528738077 | 0.235016358 | 0.517393417 |
| chr4:166324340-166324739 | -0.450764816 | 0.23502229 | 0.517393417 |
| chr22:25235835-25236219 | -0.460385263 | 0.235044009 | 0.517426366 |
| chr9:94571822-94572022 | 0.671931473 | 0.235089016 | 0.517447121 |
| chr17:14205987-14206464 | -0.427690328 | 0.235094119 | 0.517447121 |
| chr20:57252325-57253191 | 0.372643753 | 0.235095695 | 0.517447121 |
| chr17:76072426-76072913 | 0.41501616 | 0.235101149 | 0.517447121 |
| chr6:140672000-140672200 | -0.662205503 | 0.235101853 | 0.517447121 |
| chr2:26942664-26942907 | -0.522839095 | 0.235101992 | 0.517447121 |
| chr15:49225602-49226126 | -0.444509457 | 0.235106009 | 0.517447121 |
| chr17:43907428-43907793 | -0.471004692 | 0.235107464 | 0.517447121 |
| chr15:56042558-56042911 | 0.474740988 | 0.235132108 | 0.517472784 |
| chr12:6062975-6063446 | 0.446183716 | 0.235132631 | 0.517472784 |
| chr15:40511604-40512144 | 0.42897654 | 0.235143404 | 0.51748163 |
| chr6:147201978-147202223 | -0.485553826 | 0.235155747 | 0.517493929 |
| chr17:66558335-66558549 | 0.52632509 | 0.235190952 | 0.517556539 |
| chr6:37018166-37018488 | 0.43463638 | 0.235209971 | 0.517583528 |
| chr21:35249721-35249924 | 0.639333075 | 0.235220032 | 0.517590801 |
| chr5:75566728-75567087 | -0.569658158 | 0.235231861 | 0.517601966 |
| chr2:231514191-231514834 | 0.388048978 | 0.235241091 | 0.517606593 |
| chr1:113674715-113674915 | -0.569322224 | 0.235249517 | 0.517606593 |
| chrX:7252168-7252586 | -0.423710485 | 0.235259212 | 0.517606593 |
| chrX:80808000-80808299 | -0.440289999 | 0.235260985 | 0.517606593 |
| chr10:33334060-33334260 | 0.549182182 | 0.235285409 | 0.517645466 |
| chr6:379254-379454 | 0.51919347 | 0.235298221 | 0.517658789 |
| chr11:58654066-58654339 | 0.505508263 | 0.23530923 | 0.517668145 |
| chr1:58699452-58700340 | -0.387055948 | 0.235332492 | 0.517704455 |
| chr4:128529228-128529431 | 0.553915382 | 0.2353549 | 0.517717616 |
| chr12:53746453-53747008 | 0.415243486 | 0.235358733 | 0.517717616 |
| chr7:50141979-50142179 | -0.643194549 | 0.235358745 | 0.517717616 |
| chr6:30101787-30102283 | -0.431373523 | 0.235375837 | 0.51774035 |
| chr14:50714543-50714973 | 0.464644094 | 0.235389931 | 0.517756488 |
| chr7:155815290-155815504 | -0.520899945 | 0.235402128 | 0.517768453 |
| chr18:62425309-62425973 | -0.432701151 | 0.235410605 | 0.517772237 |
| chr4:26372519-26372998 | -0.455476483 | 0.235430548 | 0.517801236 |
| chr8:11346852-11348226 | 0.389223238 | 0.235438474 | 0.517803804 |
| chr7:100483853-100484079 | -0.515677811 | 0.235449719 | 0.517813674 |
| chr12:73514720-73514983 | -0.526269482 | 0.235459236 | 0.51781974 |
| chr4:186782893-186783108 | -0.623459972 | 0.235466618 | 0.517821113 |
| chr2:158033066-158033361 | -0.588600757 | 0.235474475 | 0.51782353 |
| chr8:77606227-77606549 | -0.468194122 | 0.235489784 | 0.517827723 |
| chr4:99562925-99564223 | -0.377274221 | 0.235493839 | 0.517827723 |
| chr8:33857355-33857888 | -0.473543905 | 0.235503337 | 0.517827723 |
| chr2:174546675-174547507 | -0.406758737 | 0.235503415 | 0.517827723 |
| chr2:30871562-30871780 | -0.604686648 | 0.235541489 | 0.517888152 |

| | | | |
|---|---|---|---|
| chr3:134358404-134358782 | 0.443490086 | 0.235544416 | 0.517888152 |
| chr20:63434946-63435160 | -0.518479953 | 0.235569882 | 0.517929282 |
| chr1:44154081-44154304 | -0.689246982 | 0.235586605 | 0.517951188 |
| chr1:2655815-2656028 | 0.63158993 | 0.235593463 | 0.517951403 |
| chr12:124240565-124240791 | 0.580644452 | 0.235600488 | 0.517951986 |
| chr11:85269132-85269465 | -0.532079838 | 0.235621413 | 0.517983127 |
| chr6:40661854-40662055 | -0.645439112 | 0.235635236 | 0.517996959 |
| chr17:19897993-19898316 | 0.514409 | 0.235641226 | 0.517996959 |
| chr16:87552047-87552421 | 0.472728809 | 0.235662807 | 0.51801498 |
| chr7:81716052-81716319 | -0.538947459 | 0.235662945 | 0.51801498 |
| chr16:85151685-85151977 | -0.489380083 | 0.235671705 | 0.518017522 |
| chr5:150560976-150561288 | -0.456082596 | 0.23568267 | 0.518017522 |
| chr1:215258370-215258583 | -0.527134225 | 0.235684383 | 0.518017522 |
| chr6:55953482-55953720 | -0.581363506 | 0.23572493 | 0.518090883 |
| chr11:60942215-60942650 | 0.420285909 | 0.235736288 | 0.518090883 |
| chr12:57754733-57755060 | 0.404105538 | 0.235742458 | 0.518090883 |
| chr12:56464363-56464563 | 0.525473276 | 0.235744807 | 0.518090883 |
| chr3:63593766-63594285 | 0.432002534 | 0.235752462 | 0.518092845 |
| chr1:43763516-43763770 | 0.557996946 | 0.235774998 | 0.518127511 |
| chr6:33298584-33298900 | -0.448485773 | 0.235805987 | 0.518177288 |
| chr10:84927598-84927943 | -0.460683728 | 0.235811175 | 0.518177288 |
| chr6:43702519-43703301 | 0.400063201 | 0.235847945 | 0.518217004 |
| chr16:58392763-58392963 | 0.56524862 | 0.235854509 | 0.518217004 |
| chr3:58640595-58640956 | 0.479801468 | 0.235855489 | 0.518217004 |
| chr3:120595880-120596830 | -0.378457895 | 0.235856302 | 0.518217004 |
| chr3:90008270-90008470 | -0.693516025 | 0.235865034 | 0.518221329 |
| chr15:71220676-71220945 | 0.634081661 | 0.235886851 | 0.518243964 |
| chr6:31058394-31058598 | -0.618717267 | 0.235888863 | 0.518243964 |
| chr2:139685634-139685856 | -0.570677232 | 0.235905744 | 0.518249744 |
| chr10:62554694-62554926 | -0.538214878 | 0.235911275 | 0.518249744 |
| chr8:12755086-12755286 | 0.543981907 | 0.235911786 | 0.518249744 |
| chr7:81499766-81500156 | -0.53561638 | 0.2359236 | 0.518260838 |
| chr13:24247659-24247893 | -0.606916121 | 0.235933599 | 0.51826533 |
| chr8:122038327-122038550 | -0.604953449 | 0.235939172 | 0.51826533 |
| chr3:64140523-64140734 | 0.421369396 | 0.235948911 | 0.518268266 |
| chr20:30500493-30500896 | 0.458070653 | 0.235954037 | 0.518268266 |
| chr8:115660594-115661133 | -0.420971932 | 0.236005098 | 0.518365561 |
| chr13:49858222-49858474 | 0.543663674 | 0.236039406 | 0.518426056 |
| chr8:109346917-109347302 | -0.434832593 | 0.236046525 | 0.518426831 |
| chr11:69950072-69950372 | 0.512816794 | 0.236076704 | 0.518478251 |
| chr2:46360957-46361207 | 0.571557915 | 0.236092084 | 0.518488277 |
| chr3:94285708-94286230 | -0.472960285 | 0.236094803 | 0.518488277 |
| chr3:71583562-71584603 | -0.385676946 | 0.236107796 | 0.51850195 |
| chr14:44759221-44759718 | 0.455263415 | 0.236119118 | 0.518505781 |
| chr5:9099746-9099979 | 0.539594455 | 0.236127099 | 0.518505781 |
| chr7:12788008-12788208 | 0.607269367 | 0.236129842 | 0.518505781 |
| chr1:174583666-174584481 | -0.429073876 | 0.236180978 | 0.518603206 |
| chrX:49199997-49200479 | 0.412338652 | 0.236238683 | 0.51871505 |
| chr16:76959222-76959457 | 0.529776484 | 0.236270206 | 0.518766767 |
| chr15:101152865-101153228 | -0.453362143 | 0.236275778 | 0.518766767 |

| | | | |
|---|---|---|---|
| chr20:57006810-57007010 | 0.489812448 | 0.236284484 | 0.518771018 |
| chr1:157878142-157878370 | -0.593477731 | 0.236292317 | 0.518773351 |
| chr6:11299097-11299381 | -0.625086522 | 0.236323046 | 0.518825949 |
| chr10:43648340-43649274 | -0.392473432 | 0.236333338 | 0.518833678 |
| chr19:51338861-51339342 | 0.419315994 | 0.236344683 | 0.518843719 |
| chrX:36898341-36898858 | -0.439173716 | 0.236367318 | 0.518878543 |
| chr12:1690517-1691590 | -0.394784939 | 0.236398117 | 0.518931286 |
| chr4:21765855-21766107 | 0.541644946 | 0.236435577 | 0.51897948 |
| chr13:27372880-27373259 | 0.446840581 | 0.236445932 | 0.51897948 |
| chr8:22693439-22693653 | -0.480899049 | 0.236446218 | 0.51897948 |
| chr2:190578580-190579097 | -0.489448011 | 0.236451818 | 0.51897948 |
| chr18:44130384-44130584 | -0.657113684 | 0.236457338 | 0.51897948 |
| chr13:46759944-46760302 | -0.440794779 | 0.236460712 | 0.51897948 |
| chr6:116370492-116371847 | -0.383185051 | 0.236509558 | 0.519071819 |
| chr5:1846558-1846934 | 0.522580375 | 0.236518605 | 0.519076806 |
| chr1:152777601-152777842 | -0.571344258 | 0.236556701 | 0.5191391 |
| chr19:44141197-44141757 | 0.44170555 | 0.236562304 | 0.5191391 |
| chr5:170387884-170388311 | 0.434786005 | 0.236577304 | 0.5191391 |
| chrX:41380525-41380745 | -0.627104963 | 0.236580421 | 0.5191391 |
| chr5:88680230-88680786 | -0.405920495 | 0.236580866 | 0.5191391 |
| chr5:52809988-52810377 | 0.514009439 | 0.236608909 | 0.519171032 |
| chr20:53669608-53669825 | 0.404558971 | 0.236608969 | 0.519171032 |
| chr3:39416129-39416329 | -0.59400783 | 0.236625431 | 0.519187124 |
| chr19:1253006-1253254 | 0.456609434 | 0.236631212 | 0.519187124 |
| chr4:84167030-84167320 | -0.587204034 | 0.236636631 | 0.519187124 |
| chr11:86706091-86706778 | -0.419591691 | 0.236660279 | 0.51921591 |
| chr5:7856927-7857158 | -0.555422454 | 0.236665923 | 0.51921591 |
| chr5:60041950-60042510 | -0.464612116 | 0.23667008 | 0.51921591 |
| chr15:99928360-99928925 | 0.41696568 | 0.236691371 | 0.519247751 |
| chr13:110885883-110886294 | -0.508294763 | 0.236700897 | 0.519253784 |
| chr9:131046576-131047080 | 0.458216245 | 0.236763141 | 0.519375458 |
| chr10:27383959-27384159 | -0.633715386 | 0.236792404 | 0.51942478 |
| chr21:31580788-31581205 | 0.441667219 | 0.23681069 | 0.519450021 |
| chr5:145237996-145238242 | -0.534567973 | 0.236871537 | 0.519568617 |
| chr3:185365343-185365567 | -0.522357498 | 0.236891483 | 0.519597493 |
| chr3:5057771-5058272 | 0.418178275 | 0.236906296 | 0.519604064 |
| chr11:59557283-59557815 | -0.394462905 | 0.23691483 | 0.519604064 |
| chr3:126895712-126895972 | -0.519034972 | 0.236916399 | 0.519604064 |
| chr9:134411645-134411993 | -0.464939046 | 0.236921605 | 0.519604064 |
| chr6:29887880-29888464 | -0.520157231 | 0.236987931 | 0.519709617 |
| chr6:28943586-28943869 | 0.435295575 | 0.236990903 | 0.519709617 |
| chr13:60267467-60268211 | -0.411416935 | 0.236996096 | 0.519709617 |
| chr1:192382816-192383066 | -0.560011101 | 0.236996864 | 0.519709617 |
| chr4:109076753-109077060 | 0.534515335 | 0.237070644 | 0.51985653 |
| chr7:5586338-5586591 | -0.515820266 | 0.237078763 | 0.519859456 |
| chr6:102732926-102733244 | -0.481434816 | 0.237101395 | 0.519894204 |
| chr16:67198252-67198600 | -0.438435742 | 0.237124943 | 0.519905204 |
| chr5:81851467-81852149 | -0.400520308 | 0.237125422 | 0.519905204 |
| chr4:164949527-164949727 | -0.569201848 | 0.237126767 | 0.519905204 |
| chr1:25568341-25568723 | -0.493482346 | 0.237146737 | 0.519908861 |

| | | | |
|---|---|---|---|
| chr14:36661597-36662356 | 0.378066893 | 0.237148386 | 0.519908861 |
| chr7:3897531-3897741 | -0.565529983 | 0.237148791 | 0.519908861 |
| chr18:8962441-8962672 | -0.626440642 | 0.237175275 | 0.519952045 |
| chr3:18757924-18758441 | 0.410126108 | 0.237196308 | 0.519961079 |
| chr11:118930117-118930611 | 0.389913179 | 0.237198735 | 0.519961079 |
| chr1:112673664-112673891 | -0.516567398 | 0.23721093 | 0.519961079 |
| chr16:22205970-22206858 | 0.384677079 | 0.237212278 | 0.519961079 |
| chr6:136762348-136762828 | -0.507827146 | 0.237213326 | 0.519961079 |
| chr8:129631261-129631650 | -0.527951481 | 0.237223051 | 0.51996752 |
| chr22:36481344-36481776 | 0.4292927 | 0.237245162 | 0.51997749 |
| chr22:20429192-20429468 | 0.482842801 | 0.23725627 | 0.51997749 |
| chr11:67990896-67991096 | 0.581156138 | 0.237267343 | 0.51997749 |
| chr6:166627987-166628187 | 0.512753198 | 0.237269069 | 0.51997749 |
| chr12:117580449-117581093 | 0.422192767 | 0.237270087 | 0.51997749 |
| chr1:150168980-150169297 | 0.428727949 | 0.237273795 | 0.51997749 |
| chr17:8846803-8847023 | -0.629444394 | 0.237277818 | 0.51997749 |
| chr7:123367537-123367937 | -0.435689343 | 0.237291987 | 0.51997749 |
| chr4:112705558-112705982 | 0.462387044 | 0.23729657 | 0.51997749 |
| chr10:96765675-96765964 | 0.49065469 | 0.237297901 | 0.51997749 |
| chr11:116193733-116194035 | 0.526700357 | 0.237302248 | 0.51997749 |
| chr12:109567331-109567784 | -0.429913413 | 0.237349934 | 0.520059561 |
| chr5:132736856-132738046 | -0.379411242 | 0.237360372 | 0.520059561 |
| chr17:66310832-66311096 | 0.515369544 | 0.237370435 | 0.520059561 |
| chr8:143117415-143117913 | 0.419509348 | 0.237370483 | 0.520059561 |
| chr7:29106481-29106727 | 0.535129047 | 0.23737364 | 0.520059561 |
| chr4:6327274-6327515 | 0.502011146 | 0.237384349 | 0.520068154 |
| chr12:106955322-106956940 | -0.379147014 | 0.237400213 | 0.520086746 |
| chrX:153495388-153495651 | 0.495135006 | 0.2374103 | 0.520086746 |
| chr17:1268006-1268604 | 0.441132472 | 0.237413199 | 0.520086746 |
| chr12:1519373-1519597 | -0.611969943 | 0.23745202 | 0.520135863 |
| chr17:66603099-66603316 | 0.620628751 | 0.237453204 | 0.520135863 |
| chr8:98364672-98365046 | 0.489451981 | 0.237455985 | 0.520135863 |
| chr3:166901068-166901613 | -0.453776903 | 0.237473723 | 0.520159848 |
| chr12:121467036-121468079 | 0.386744343 | 0.237498056 | 0.520189391 |
| chr2:64898015-64898225 | -0.733687229 | 0.237500789 | 0.520189391 |
| chr3:73520797-73521567 | -0.406896795 | 0.23752708 | 0.520232105 |
| chr21:17549880-17550108 | 0.568756031 | 0.237536838 | 0.520238606 |
| chr11:88431847-88432190 | -0.608578001 | 0.237565056 | 0.520285534 |
| chr2:37880670-37880917 | -0.519379661 | 0.23758328 | 0.520310575 |
| chr5:170691680-170691899 | -0.570800837 | 0.237595267 | 0.520321954 |
| chr3:166864715-166864915 | 0.577090175 | 0.237613328 | 0.520335667 |
| chr10:34906071-34906331 | 0.500739857 | 0.23761511 | 0.520335667 |
| chr11:46898697-46899052 | 0.436045002 | 0.237623639 | 0.520339472 |
| chr1:45730534-45730764 | 0.560104005 | 0.237657909 | 0.520359892 |
| chr7:99325656-99326158 | -0.406153257 | 0.237664216 | 0.520359892 |
| chr12:66824271-66824483 | 0.557894485 | 0.237665135 | 0.520359892 |
| chr1:201828912-201829741 | 0.407643764 | 0.237665736 | 0.520359892 |
| chr8:72542544-72542819 | -0.523826467 | 0.237684146 | 0.520359892 |
| chr2:15591434-15592458 | -0.378199288 | 0.237688135 | 0.520359892 |
| chr3:62888140-62888559 | -0.393475878 | 0.237692969 | 0.520359892 |

| | | | |
|---|---|---|---|
| chr7:155796991-155797646 | -0.427887472 | 0.237699484 | 0.520359892 |
| chr17:70239144-70239351 | -0.64386579 | 0.237700448 | 0.520359892 |
| chr10:73886131-73886407 | -0.463865065 | 0.237700877 | 0.520359892 |
| chr12:127381093-127381381 | 0.515312459 | 0.237710794 | 0.520366735 |
| chr9:129320568-129321093 | 0.423895691 | 0.237727044 | 0.52038744 |
| chr22:36856311-36856613 | 0.502920424 | 0.237750425 | 0.520414315 |
| chr13:42728345-42728983 | -0.465629593 | 0.237752905 | 0.520414315 |
| chr10:81414103-81414399 | -0.529530661 | 0.237762973 | 0.520421485 |
| chr1:16904695-16905495 | 0.397930812 | 0.237784568 | 0.520436834 |
| chr5:6462429-6462637 | -0.609936891 | 0.237791901 | 0.520436834 |
| chr14:77133775-77134086 | 0.432211586 | 0.237794807 | 0.520436834 |
| chr16:22099047-22099253 | 0.579113443 | 0.237797154 | 0.520436834 |
| chr13:31161580-31162660 | -0.372261751 | 0.23781789 | 0.52046735 |
| chr7:44039008-44039575 | -0.387348616 | 0.237863208 | 0.520551661 |
| chr3:15205805-15206863 | 0.374238079 | 0.237883658 | 0.520570418 |
| chr9:90920917-90921271 | 0.560596411 | 0.237885367 | 0.520570418 |
| chr2:18482832-18483046 | 0.646612434 | 0.237909513 | 0.520581275 |
| chr8:126564005-126564889 | 0.391554375 | 0.237910603 | 0.520581275 |
| chr13:80239697-80239902 | 0.591335017 | 0.237910711 | 0.520581275 |
| chr7:135147629-135148724 | -0.393367638 | 0.237935173 | 0.520613428 |
| chr4:182806328-182806883 | -0.434478773 | 0.237946163 | 0.520613428 |
| chr9:116686778-116687450 | 0.408364844 | 0.237961656 | 0.520613428 |
| chr2:63692059-63692285 | -0.607043429 | 0.237970368 | 0.520613428 |
| chr2:222767262-222767526 | -0.607043429 | 0.237970368 | 0.520613428 |
| chr12:48156965-48157286 | -0.46755968 | 0.237978025 | 0.520613428 |
| chr16:4624233-4625310 | 0.382777672 | 0.237979093 | 0.520613428 |
| chr8:117303998-117304450 | -0.435405484 | 0.237979762 | 0.520613428 |
| chr11:116312980-116313222 | -0.607941761 | 0.238060963 | 0.520775895 |
| chr1:106902113-106902453 | 0.440464465 | 0.238069883 | 0.520775895 |
| chr2:102619006-102620084 | 0.389204065 | 0.238074418 | 0.520775895 |
| chr15:71764479-71764781 | 0.554051583 | 0.238138755 | 0.520887685 |
| chr6:8085092-8085444 | -0.481662448 | 0.238140622 | 0.520887685 |
| chr1:202118883-202119288 | 0.396918755 | 0.238146167 | 0.520887685 |
| chr5:179559339-179560399 | -0.376875787 | 0.238152716 | 0.520887685 |
| chr7:131106750-131106997 | -0.486196765 | 0.238159706 | 0.520887925 |
| chr16:89217326-89217868 | -0.408972091 | 0.238166422 | 0.520887925 |
| chr17:79836025-79836606 | 0.397475794 | 0.238181288 | 0.520898902 |
| chr6:33983101-33983444 | 0.469680274 | 0.238185037 | 0.520898902 |
| chr10:79127156-79127478 | -0.464089726 | 0.238201212 | 0.520919408 |
| chr22:19168213-19168629 | -0.439177846 | 0.238237308 | 0.520983476 |
| chr9:124834806-124835389 | 0.455437467 | 0.238264543 | 0.521028164 |
| chr21:46294780-46296452 | -0.383358815 | 0.238274356 | 0.521034751 |
| chr13:28321191-28321434 | 0.609022097 | 0.238287379 | 0.521048358 |
| chr9:82105815-82106031 | 0.5827569 | 0.238301366 | 0.521064073 |
| chr5:124651361-124651948 | -0.425361552 | 0.238343594 | 0.521141537 |
| chr14:32407786-32407986 | 0.619398672 | 0.238362559 | 0.521168132 |
| chr17:61289784-61290058 | 0.399685271 | 0.238409539 | 0.521255976 |
| chr2:42896864-42897073 | -0.589213285 | 0.23841997 | 0.521263908 |
| chr21:34888058-34888323 | 0.488630538 | 0.238428546 | 0.521267551 |
| chr10:8543976-8544827 | 0.401309191 | 0.238435242 | 0.521267551 |

| | | | |
|---|---|---|---|
| chr2:96753651-96753885 | -0.565498901 | 0.238495702 | 0.521378497 |
| chr10:91901834-91902039 | -0.641438981 | 0.2384996 | 0.521378497 |
| chr5:15132817-15133109 | -0.593816446 | 0.238508977 | 0.521384122 |
| chr20:46363601-46364631 | 0.379208768 | 0.238559984 | 0.521470497 |
| chr12:131871982-131872737 | 0.392163791 | 0.238562101 | 0.521470497 |
| chr8:4643902-4644155 | -0.570908168 | 0.238606184 | 0.521551977 |
| chr19:13861839-13862508 | 0.426554418 | 0.238624598 | 0.521577349 |
| chr9:119969134-119969434 | -0.474102907 | 0.238636511 | 0.521588509 |
| chr5:91266378-91266627 | -0.558387398 | 0.238644426 | 0.521590928 |
| chr12:19130464-19130820 | -0.44540246 | 0.238673901 | 0.521640472 |
| chr11:101327984-101328277 | -0.554149202 | 0.238687402 | 0.5216551 |
| chr6:137157583-137157912 | 0.472238428 | 0.238719211 | 0.521686722 |
| chr16:1943165-1943578 | 0.410060185 | 0.238721059 | 0.521686722 |
| chr17:40177318-40177786 | 0.440525357 | 0.238727236 | 0.521686722 |
| chr17:32267926-32268192 | -0.589335315 | 0.238729106 | 0.521686722 |
| chr1:109483187-109483520 | 0.438733729 | 0.238745426 | 0.521697949 |
| chr2:18883938-18884440 | -0.49205991 | 0.238754324 | 0.521697949 |
| chrX:109536444-109537678 | 0.37589959 | 0.238754669 | 0.521697949 |
| chr9:126606460-126606717 | 0.539118597 | 0.238801767 | 0.521780064 |
| chr11:4192859-4193130 | 0.561139992 | 0.238805869 | 0.521780064 |
| chr8:67462764-67463019 | 0.485994752 | 0.238828433 | 0.521814485 |
| chr1:50332988-50333429 | 0.436722617 | 0.238839622 | 0.521824053 |
| chr1:116569883-116570132 | 0.445070609 | 0.23884752 | 0.521826428 |
| chr12:56006864-56007135 | 0.464742971 | 0.238867113 | 0.521854356 |
| chr10:77332344-77332606 | 0.591338442 | 0.238880129 | 0.521867911 |
| chr8:89848269-89848799 | 0.405034589 | 0.23891106 | 0.521920603 |
| chr10:60686943-60687336 | -0.454847082 | 0.238925871 | 0.521937286 |
| chr20:57148973-57149251 | 0.391057446 | 0.238940097 | 0.521937286 |
| chr15:89632345-89632800 | 0.466469666 | 0.238943587 | 0.521937286 |
| chr3:16934424-16934683 | 0.580621149 | 0.238947155 | 0.521937286 |
| chr10:116671415-116671828 | 0.457682875 | 0.238952756 | 0.521937286 |
| chr15:78930906-78931338 | -0.418013777 | 0.238964019 | 0.521947008 |
| chr2:218896620-218897438 | -0.405813417 | 0.23898502 | 0.521978 |
| chr3:63364950-63365368 | -0.406938135 | 0.239010607 | 0.522000229 |
| chr21:33854764-33854972 | -0.588474352 | 0.239013413 | 0.522000229 |
| chr9:37913643-37914106 | -0.429997148 | 0.239015636 | 0.522000229 |
| chr17:55454886-55455303 | -0.481351242 | 0.23902362 | 0.522002787 |
| chr9:101791868-101792246 | -0.503962936 | 0.239049876 | 0.522038756 |
| chr10:31603193-31604098 | -0.38070417 | 0.239053716 | 0.522038756 |
| chr4:3781161-3781814 | 0.390254479 | 0.239066807 | 0.522052464 |
| chr1:155126495-155127526 | 0.382555044 | 0.239102626 | 0.522080358 |
| chr4:48723062-48723285 | 0.568163524 | 0.239105103 | 0.522080358 |
| chr7:135707587-135707872 | -0.51368422 | 0.239117456 | 0.522080358 |
| chr7:74464636-74464911 | -0.563490245 | 0.239122022 | 0.522080358 |
| chr18:57621287-57622404 | -0.380583457 | 0.239123634 | 0.522080358 |
| chr8:106208761-106209252 | 0.43438071 | 0.239131917 | 0.522080358 |
| chr19:2702755-2702968 | 0.542163124 | 0.239134351 | 0.522080358 |
| chr2:163375115-163375322 | -0.608353173 | 0.239140904 | 0.522080358 |
| chr18:31294441-31294697 | -0.608353173 | 0.239140904 | 0.522080358 |
| chr3:8327965-8328312 | -0.518669951 | 0.239153306 | 0.522092365 |

| | | | |
|---|---|---|---|
| chr6:158049470-158050024 | -0.429449674 | 0.239160031 | 0.522092365 |
| chr2:119126562-119126862 | -0.502853924 | 0.239191936 | 0.522147137 |
| chr5:62135753-62135990 | -0.614401805 | 0.239217045 | 0.522187072 |
| chr11:73217975-73218869 | 0.386278027 | 0.239227781 | 0.522193716 |
| chr2:48734633-48734947 | 0.571786009 | 0.23923571 | 0.522193716 |
| chr6:29737917-29738363 | -0.470731596 | 0.239244596 | 0.522193716 |
| chr4:159175536-159175940 | -0.448387693 | 0.239248437 | 0.522193716 |
| chr1:157014693-157014972 | -0.557951651 | 0.239262086 | 0.522193716 |
| chr17:43778854-43779151 | 0.473928403 | 0.239269277 | 0.522193716 |
| chr10:132904292-132904518 | 0.523399153 | 0.239270212 | 0.522193716 |
| chr1:237736679-237736916 | -0.58277645 | 0.239281333 | 0.522193716 |
| chr17:60593407-60593749 | 0.404346808 | 0.239283992 | 0.522193716 |
| chr2:3462264-3462557 | -0.46816976 | 0.23928824 | 0.522193716 |
| chr13:52455060-52456102 | -0.379963489 | 0.23932079 | 0.522249874 |
| chr1:72282704-72284280 | -0.377512527 | 0.239346862 | 0.522259908 |
| chr2:222561948-222562165 | -0.564718592 | 0.239348716 | 0.522259908 |
| chr12:49567686-49568554 | 0.386744602 | 0.239365729 | 0.522259908 |
| chrX:135003822-135004717 | 0.38027496 | 0.239368256 | 0.522259908 |
| chr18:70423993-70424200 | 0.566610109 | 0.239372829 | 0.522259908 |
| chr17:75653048-75653326 | 0.473832913 | 0.239380799 | 0.522259908 |
| chr11:81570749-81570964 | -0.585482812 | 0.239384746 | 0.522259908 |
| chr17:68229175-68229511 | 0.446782627 | 0.239396231 | 0.522259908 |
| chr5:68214899-68216294 | -0.376665169 | 0.239396884 | 0.522259908 |
| chr11:66126701-66126959 | -0.590705463 | 0.239402615 | 0.522259908 |
| chr3:185575545-185575761 | 0.533764789 | 0.239406681 | 0.522259908 |
| chr2:132416001-132417275 | -0.379083867 | 0.239407181 | 0.522259908 |
| chr19:45817707-45818060 | -0.485478351 | 0.239469486 | 0.522370069 |
| chr1:206684668-206685193 | -0.391944048 | 0.239471314 | 0.522370069 |
| chr4:109967747-109968166 | 0.451992885 | 0.239505259 | 0.522429241 |
| chr2:177416216-177416513 | -0.502439531 | 0.23953387 | 0.522476777 |
| chr6:106798256-106798681 | 0.459774306 | 0.239544217 | 0.52248447 |
| chr16:12455325-12455580 | 0.50337131 | 0.239551717 | 0.522485957 |
| chr2:161123677-161124217 | 0.426287983 | 0.239599743 | 0.52257583 |
| chr10:89514316-89514609 | -0.577992119 | 0.239617219 | 0.522596208 |
| chr1:84759286-84759497 | -0.619483733 | 0.239622727 | 0.522596208 |
| chr1:116666784-116668138 | 0.377645428 | 0.239650444 | 0.522641781 |
| chr4:87363402-87364014 | -0.41800594 | 0.239674499 | 0.522679365 |
| chr16:54930464-54931156 | 0.386639602 | 0.239710961 | 0.522744002 |
| chr19:37941938-37942292 | -0.513971618 | 0.239733967 | 0.522779293 |
| chr14:82218424-82218747 | 0.487148155 | 0.239767641 | 0.52279878 |
| chr1:98761370-98761588 | -0.610360349 | 0.239769038 | 0.52279878 |
| chr9:90419627-90419884 | -0.602489957 | 0.239769916 | 0.52279878 |
| chr19:36378612-36379432 | 0.397802365 | 0.239770196 | 0.52279878 |
| chr3:82707077-82707378 | -0.505733083 | 0.239781177 | 0.522807846 |
| chr4:57037996-57038576 | -0.415673373 | 0.239806161 | 0.522847442 |
| chr2:34461893-34462108 | -0.684162238 | 0.239830924 | 0.52288441 |
| chr5:132386631-132386831 | -0.533857573 | 0.239836765 | 0.52288441 |
| chr13:39777481-39778152 | -0.406969257 | 0.23984732 | 0.522892544 |
| chr1:64861751-64861957 | 0.63277214 | 0.239908233 | 0.522992956 |
| chr7:132225292-132225599 | -0.485603537 | 0.23990863 | 0.522992956 |

| | | | |
|---|---|---|---|
| chr5:160213555-160213847 | -0.466058096 | 0.239927455 | 0.522992956 |
| chr2:70524288-70524695 | -0.473301783 | 0.239928 | 0.522992956 |
| chr16:48164934-48165206 | 0.507914156 | 0.239931896 | 0.522992956 |
| chr19:30696557-30696820 | -0.605767105 | 0.239934332 | 0.522992956 |
| chr8:85841353-85841585 | -0.492337844 | 0.239990236 | 0.523099931 |
| chr5:149439875-149440270 | -0.410076218 | 0.24000034 | 0.523107072 |
| chr20:11274108-11274457 | 0.46786518 | 0.240007404 | 0.523107589 |
| chr2:158580770-158581153 | -0.437211369 | 0.240025242 | 0.523131588 |
| chr14:103736315-103736638 | 0.448271673 | 0.240053102 | 0.523177426 |
| chr19:32405168-32406368 | 0.378028057 | 0.240064197 | 0.523184517 |
| chr19:30682950-30683235 | 0.507297663 | 0.240070011 | 0.523184517 |
| chr6:8888522-8888781 | 0.559040247 | 0.240082194 | 0.523188765 |
| chr10:97869380-97870005 | -0.404567706 | 0.240093436 | 0.523188765 |
| chr9:119284198-119284405 | -0.610967793 | 0.240098105 | 0.523188765 |
| chr14:54797735-54797958 | 0.566719608 | 0.240099273 | 0.523188765 |
| chr2:25977156-25977390 | -0.565414968 | 0.240138789 | 0.52325999 |
| chr17:74354052-74354271 | 0.519926344 | 0.240146844 | 0.523262661 |
| chr9:112348303-112348687 | 0.431844255 | 0.240163967 | 0.52327251 |
| chr11:86511178-86511378 | -0.619010861 | 0.240165022 | 0.52327251 |
| chr4:100189778-100190676 | -0.412452548 | 0.240212123 | 0.52336025 |
| chr11:2269533-2269775 | 0.531779532 | 0.240264964 | 0.523452696 |
| chr2:160225924-160226195 | -0.512159833 | 0.240268217 | 0.523452696 |
| chr13:106521254-106521681 | 0.46145597 | 0.240282141 | 0.523454001 |
| chr7:149782445-149782697 | -0.486769289 | 0.240282479 | 0.523454001 |
| chr10:32650596-32651049 | -0.457722879 | 0.240302602 | 0.523482955 |
| chr8:116354472-116354716 | 0.496266602 | 0.24031275 | 0.523490177 |
| chr13:25678415-25678679 | 0.543228991 | 0.240328553 | 0.523504175 |
| chr12:42589643-42589979 | 0.447759667 | 0.24033284 | 0.523504175 |
| chr1:168089966-168090175 | -0.631911588 | 0.24037345 | 0.523556339 |
| chr8:53344300-53344566 | -0.534858603 | 0.240380497 | 0.523556339 |
| chr1:195005301-195005812 | 0.433033614 | 0.240383587 | 0.523556339 |
| chr8:125563480-125563848 | 0.460119354 | 0.24038412 | 0.523556339 |
| chr16:11396223-11396699 | 0.44750782 | 0.24039335 | 0.523561561 |
| chr13:33487886-33488230 | -0.461237572 | 0.240454442 | 0.523667629 |
| chr2:219253709-219254413 | 0.394211376 | 0.24045572 | 0.523667629 |
| chr6:72216257-72216629 | 0.43594457 | 0.24049087 | 0.523729293 |
| chr9:115883637-115884030 | -0.491783644 | 0.240520294 | 0.523778485 |
| chr2:11863602-11863984 | -0.419662552 | 0.240581675 | 0.523897263 |
| chr12:100157140-100157499 | -0.491430589 | 0.240597802 | 0.523917492 |
| chr17:70855405-70855783 | 0.482697844 | 0.240614999 | 0.523938856 |
| chr12:125065002-125065856 | 0.386876816 | 0.240621289 | 0.523938856 |
| chr11:60924209-60924849 | 0.403578148 | 0.24063774 | 0.523949589 |
| chr17:17682339-17682585 | 0.459809259 | 0.240649914 | 0.523949589 |
| chr1:17011700-17012213 | 0.426545948 | 0.240653981 | 0.523949589 |
| chr5:85647801-85648028 | 0.588000501 | 0.240674083 | 0.523949589 |
| chr1:184110308-184110612 | -0.527482119 | 0.240677052 | 0.523949589 |
| chr12:104456487-104457724 | 0.383396211 | 0.240689394 | 0.523949589 |
| chr1:196991602-196992730 | -0.374691336 | 0.240691312 | 0.523949589 |
| chr6:35496276-35496476 | 0.496861563 | 0.24069381 | 0.523949589 |
| chr22:12174962-12175212 | 0.464520973 | 0.240699871 | 0.523949589 |

| | | | |
|---|---|---|---|
| chr8:122160877-122161118 | -0.507959208 | 0.240700562 | 0.523949589 |
| chr12:120677808-120678008 | -0.678975008 | 0.240701437 | 0.523949589 |
| chr2:39696960-39697180 | -0.6061553 | 0.24073726 | 0.524000446 |
| chr12:120234558-120235216 | 0.38711773 | 0.240742633 | 0.524000446 |
| chr9:107412576-107412910 | -0.539881324 | 0.240749639 | 0.524000446 |
| chr5:151157490-151158579 | -0.388752779 | 0.240752156 | 0.524000446 |
| chr20:11912176-11912670 | -0.456728249 | 0.240784096 | 0.524055077 |
| chr15:75170759-75171155 | 0.510192354 | 0.240794431 | 0.524062685 |
| chr2:105294922-105295153 | -0.59399681 | 0.240802134 | 0.524064564 |
| chr15:89740101-89740411 | 0.46538576 | 0.240809216 | 0.524065091 |
| chr3:135106457-135106718 | -0.548900843 | 0.240822784 | 0.524075815 |
| chr3:9362741-9363699 | 0.376705754 | 0.240827823 | 0.524075815 |
| chrX:36662025-36662613 | -0.42608738 | 0.240866071 | 0.524133345 |
| chr7:25892863-25893349 | 0.46762304 | 0.240867972 | 0.524133345 |
| chr7:106736592-106736816 | -0.567031957 | 0.240874781 | 0.524133345 |
| chr2:174462653-174463089 | -0.492809418 | 0.240899387 | 0.52415934 |
| chr2:216151714-216151914 | -0.639522614 | 0.24090725 | 0.52415934 |
| chr3:186551417-186551649 | -0.639522614 | 0.24090725 | 0.52415934 |
| chr7:40134043-40135292 | -0.373536121 | 0.240953182 | 0.524244391 |
| chrX:73562930-73563909 | 0.393169285 | 0.240960141 | 0.524244646 |
| chr17:81277321-81277803 | 0.415866656 | 0.240991127 | 0.524297172 |
| chr3:143328514-143328788 | -0.538083867 | 0.241038119 | 0.52435495 |
| chr4:168695331-168695627 | -0.501826289 | 0.241038995 | 0.52435495 |
| chr1:218996481-218996795 | -0.491621716 | 0.241051637 | 0.52435495 |
| chr4:161642725-161643051 | -0.516626133 | 0.241055094 | 0.52435495 |
| chr7:100080973-100081199 | 0.53402488 | 0.241056786 | 0.52435495 |
| chr1:149124023-149124223 | -0.671945429 | 0.241058745 | 0.52435495 |
| chr3:52851339-52851786 | 0.443724819 | 0.241072401 | 0.52436977 |
| chr10:29998342-29998644 | -0.462863236 | 0.241115749 | 0.524443063 |
| chr10:14761905-14762183 | -0.605710475 | 0.241119786 | 0.524443063 |
| chr1:161227453-161227721 | 0.441334098 | 0.241126884 | 0.524443614 |
| chr11:35419291-35419523 | 0.578977933 | 0.241151708 | 0.524482718 |
| chr1:203396819-203397061 | -0.560268731 | 0.2411784 | 0.524517257 |
| chr1:218546114-218546638 | -0.434360212 | 0.241181279 | 0.524517257 |
| chrX:152941392-152942271 | 0.389388176 | 0.241200662 | 0.524530095 |
| chr4:73170400-73171214 | -0.384567315 | 0.241200874 | 0.524530095 |
| chrX:123881359-123881633 | 0.495292563 | 0.241220574 | 0.524532359 |
| chr6:112328795-112329178 | -0.524501212 | 0.241226733 | 0.524532359 |
| chr17:60759811-60760036 | 0.4103177 | 0.241227387 | 0.524532359 |
| chr11:67081522-67081870 | -0.429137924 | 0.241229298 | 0.524532359 |
| chr8:79655955-79656237 | -0.603421613 | 0.241247296 | 0.524556608 |
| chr8:66064955-66065243 | -0.517626813 | 0.241259442 | 0.524568132 |
| chrX:43693699-43694431 | 0.419694295 | 0.241282998 | 0.524604464 |
| chr20:48461111-48461910 | 0.377187121 | 0.241290284 | 0.524605417 |
| chr6:551363-551860 | -0.464751142 | 0.24132784 | 0.524659192 |
| chr4:153204947-153205348 | 0.429920196 | 0.241328712 | 0.524659192 |
| chr16:3646663-3646883 | 0.499208398 | 0.241341173 | 0.524667036 |
| chr11:94493470-94494597 | -0.380789879 | 0.241350735 | 0.524667036 |
| chr14:42378398-42378630 | 0.540528 | 0.241360608 | 0.524667036 |
| chr10:90695976-90696272 | -0.49304873 | 0.241364479 | 0.524667036 |

| | | | |
|---|---|---|---|
| chr6:42166474-42167094 | 0.398533379 | 0.241373804 | 0.524667036 |
| chr7:99142864-99143654 | -0.37635767 | 0.241379175 | 0.524667036 |
| chr7:118109658-118110119 | 0.457348862 | 0.241380252 | 0.524667036 |
| chr2:215238448-215238850 | -0.476577537 | 0.241416382 | 0.524730682 |
| chr15:81018315-81018533 | -0.68293219 | 0.241424811 | 0.524734117 |
| chr3:197297452-197299497 | -0.370448896 | 0.241434267 | 0.524739787 |
| chr9:33249464-33249668 | -0.572238796 | 0.241445953 | 0.524746458 |
| chr21:33302770-33303047 | -0.534487519 | 0.241461828 | 0.524746458 |
| chr4:89284284-89284594 | -0.597959545 | 0.241463853 | 0.524746458 |
| chrX:85930216-85930417 | -0.64542159 | 0.241464731 | 0.524746458 |
| chr8:103979883-103980234 | 0.453211986 | 0.241491879 | 0.524790571 |
| chr17:52762276-52763291 | -0.400356199 | 0.241502718 | 0.524799241 |
| chr5:107673407-107673977 | -0.396037778 | 0.241524028 | 0.524822084 |
| chr1:71094045-71094256 | -0.680335033 | 0.241533896 | 0.524822084 |
| chr1:6701631-6702267 | 0.414232611 | 0.241540353 | 0.524822084 |
| chr3:81352852-81353391 | -0.410574555 | 0.241542552 | 0.524822084 |
| chr14:51738827-51739060 | 0.53753074 | 0.241547477 | 0.524822084 |
| chr14:91369984-91370610 | -0.405194006 | 0.24155986 | 0.524834107 |
| chr9:88941590-88941902 | 0.511030828 | 0.241580568 | 0.524864215 |
| chr13:25756970-25757198 | -0.546367872 | 0.241608594 | 0.52491022 |
| chr12:58026455-58026728 | 0.533428651 | 0.241645251 | 0.524949322 |
| chr3:124809338-124809687 | 0.475403195 | 0.241651621 | 0.524949322 |
| chr17:27577414-27577628 | 0.580091495 | 0.241672178 | 0.524949322 |
| chr5:94660736-94661025 | -0.551460547 | 0.241676291 | 0.524949322 |
| chr12:6363259-6364363 | -0.383602006 | 0.241678843 | 0.524949322 |
| chr20:43590424-43591342 | 0.377910952 | 0.24167946 | 0.524949322 |
| chr15:78714892-78715115 | 0.535070484 | 0.241684047 | 0.524949322 |
| chr10:79264878-79265374 | -0.393121518 | 0.241685038 | 0.524949322 |
| chr2:177692860-177693067 | -0.613724048 | 0.241688252 | 0.524949322 |
| chr4:87559095-87559849 | -0.404174651 | 0.241730799 | 0.525026851 |
| chr3:134880451-134880741 | 0.548487365 | 0.241739201 | 0.525030219 |
| chr3:14034717-14035068 | 0.527631108 | 0.24175679 | 0.525044886 |
| chr3:38961050-38961250 | 0.611351825 | 0.24175966 | 0.525044886 |
| chr9:107172234-107172490 | -0.509745025 | 0.24180035 | 0.525118372 |
| chr15:55764457-55764701 | 0.518152115 | 0.241845482 | 0.525192643 |
| chr14:103880021-103880235 | 0.515239304 | 0.241848258 | 0.525192643 |
| chr15:91021718-91022797 | 0.386210391 | 0.241864772 | 0.525212478 |
| chr7:118538-118738 | -0.567476164 | 0.241871101 | 0.525212478 |
| chr22:18078413-18078654 | 0.486862962 | 0.241892783 | 0.525224782 |
| chr16:54928865-54929522 | 0.379743206 | 0.241907639 | 0.525224782 |
| chr5:60699481-60700736 | -0.370430648 | 0.241910827 | 0.525224782 |
| chr1:225923599-225924692 | 0.376934857 | 0.241912264 | 0.525224782 |
| chr7:76397357-76397732 | -0.415157924 | 0.241915591 | 0.525224782 |
| chr5:87267382-87269043 | -0.36760682 | 0.241921758 | 0.525224782 |
| chr14:103292890-103293090 | 0.534991398 | 0.241928439 | 0.525224782 |
| chr3:42159710-42160405 | -0.394539898 | 0.241931605 | 0.525224782 |
| chr10:100325260-100325460 | 0.614698218 | 0.241955969 | 0.525253818 |
| chr2:129397303-129397577 | -0.534320187 | 0.24195869 | 0.525253818 |
| chr12:124539462-124539662 | -0.616405405 | 0.241976248 | 0.525271579 |
| chr6:9851004-9851420 | 0.5020168 | 0.24198437 | 0.525271579 |

| | | | |
|---|---|---|---|
| chr10:44517603-44517931 | -0.518816644 | 0.241993191 | 0.525271579 |
| chr14:55138201-55138422 | -0.567313605 | 0.241994293 | 0.525271579 |
| chr1:211133346-211133755 | -0.531378819 | 0.242005255 | 0.525280492 |
| chr6:157763324-157763761 | -0.411523617 | 0.242014776 | 0.525286277 |
| chr15:79221668-79222001 | 0.455463969 | 0.242029833 | 0.525304079 |
| chr18:76348420-76348806 | -0.437931905 | 0.242045734 | 0.525323709 |
| chr16:1342836-1343464 | 0.383042393 | 0.242065333 | 0.525351365 |
| chr3:102536571-102536875 | -0.507175649 | 0.24209191 | 0.525394163 |
| chr16:75463736-75465121 | -0.367329081 | 0.242111112 | 0.525420953 |
| chr6:139134831-139135286 | -0.404980855 | 0.242119488 | 0.52542425 |
| chr15:56244369-56244569 | 0.475262652 | 0.242131388 | 0.525435192 |
| chr10:86253761-86254306 | -0.444631915 | 0.242145608 | 0.525451168 |
| chr12:80353300-80353560 | -0.60434359 | 0.242194055 | 0.525505191 |
| chr13:28805277-28805490 | -0.60434359 | 0.242194055 | 0.525505191 |
| chr4:85827431-85828499 | -0.402518347 | 0.242195046 | 0.525505191 |
| chr3:4218845-4219199 | 0.496981043 | 0.242197937 | 0.525505191 |
| chr8:97736153-97736886 | 0.407465505 | 0.242238701 | 0.525578756 |
| chrX:21940386-21941478 | -0.389317839 | 0.242259719 | 0.525609474 |
| chr6:63487272-63487476 | 0.644696578 | 0.242273096 | 0.525616528 |
| chr2:47617951-47618151 | 0.633812015 | 0.242278527 | 0.525616528 |
| chr5:154649604-154649810 | -0.619932682 | 0.24228851 | 0.525616528 |
| chr5:31291489-31291797 | -0.482456744 | 0.24229041 | 0.525616528 |
| chr5:178635461-178635755 | 0.562147814 | 0.242314797 | 0.52565455 |
| chr17:40533307-40533633 | 0.547341111 | 0.242322586 | 0.525656565 |
| chr5:34585535-34585849 | -0.520932508 | 0.24238193 | 0.52577041 |
| chr8:66036591-66036826 | 0.489477528 | 0.242395545 | 0.525785059 |
| chr9:13278339-13279012 | -0.431626648 | 0.242405262 | 0.525791252 |
| chr15:64731321-64731662 | 0.436828341 | 0.242434903 | 0.52584066 |
| chr18:23994480-23994680 | 0.571139911 | 0.242491167 | 0.525932953 |
| chrX:47658514-47659540 | 0.375695479 | 0.242491182 | 0.525932953 |
| chr12:2480176-2480376 | 0.626462074 | 0.242506692 | 0.525951705 |
| chr5:58999919-59000182 | -0.577112718 | 0.242525925 | 0.525964926 |
| chr14:31090537-31090756 | 0.532106531 | 0.242528712 | 0.525964926 |
| chr13:38842273-38842828 | 0.457036189 | 0.242533381 | 0.525964926 |
| chr6:29589523-29589758 | -0.603869546 | 0.242544184 | 0.525973467 |
| chr12:49276018-49276225 | 0.526796459 | 0.242563176 | 0.525999764 |
| chr15:25437926-25438521 | 0.429209032 | 0.242570926 | 0.526001684 |
| chr1:65002372-65003326 | -0.390004576 | 0.242621504 | 0.526087224 |
| chr11:118922371-118922650 | 0.418193399 | 0.242624172 | 0.526087224 |
| chr15:67937173-67937508 | -0.469065658 | 0.242632209 | 0.526087224 |
| chr14:53151625-53153859 | 0.369409122 | 0.242637838 | 0.526087224 |
| chr2:113117582-113118007 | -0.423458428 | 0.242663754 | 0.526128528 |
| chr19:33406924-33407261 | 0.46852962 | 0.242676815 | 0.526133064 |
| chr8:121132924-121133193 | -0.487846786 | 0.242679579 | 0.526133064 |
| chr1:26035804-26036295 | 0.434322155 | 0.242696373 | 0.526143433 |
| chr16:22374306-22375181 | 0.384803786 | 0.242698096 | 0.526143433 |
| chr1:228081732-228083043 | 0.375724837 | 0.242733022 | 0.52620426 |
| chr17:63484793-63485048 | 0.433399299 | 0.242744087 | 0.52621336 |
| chr10:45250084-45250313 | -0.577193954 | 0.242757217 | 0.526226934 |
| chr7:117312647-117312910 | -0.632782595 | 0.242779948 | 0.526256426 |

| | | | |
|---|---|---|---|
| chr7:1127087-1127303 | -0.502770581 | 0.242792813 | 0.526256426 |
| chr3:72310412-72311354 | -0.402119617 | 0.242794171 | 0.526256426 |
| chr1:15409156-15410040 | -0.409967124 | 0.242800107 | 0.526256426 |
| chr12:21313470-21313886 | -0.505518882 | 0.242812754 | 0.526256426 |
| chr4:55122383-55122586 | 0.618489113 | 0.242817423 | 0.526256426 |
| chr4:73446388-73446651 | -0.513643327 | 0.242818899 | 0.526256426 |
| chr6:17103782-17104100 | -0.522687191 | 0.242846222 | 0.526300755 |
| chr15:90165689-90166158 | 0.418711282 | 0.242877985 | 0.526354705 |
| chr10:43263618-43263861 | -0.510232995 | 0.242887531 | 0.526360505 |
| chr19:40165643-40165843 | -0.639311321 | 0.242926499 | 0.526425373 |
| chr3:159787795-159788071 | -0.525179826 | 0.242931205 | 0.526425373 |
| chr21:25385995-25386218 | -0.543307957 | 0.242950662 | 0.526452647 |
| chr5:154180119-154180646 | -0.429752318 | 0.242958471 | 0.52645468 |
| chr5:105105420-105105681 | -0.595220734 | 0.242990591 | 0.526490432 |
| chr3:12809992-12810709 | 0.377560236 | 0.242992531 | 0.526490432 |
| chr5:149474342-149474575 | -0.600895385 | 0.242999378 | 0.526490432 |
| chr16:83795371-83795580 | 0.611841539 | 0.243002455 | 0.526490432 |
| chr22:26672407-26673155 | 0.386201434 | 0.243033383 | 0.526536038 |
| chr8:74048899-74049364 | -0.454674181 | 0.243044063 | 0.526536038 |
| chr3:159881883-159882213 | -0.511149345 | 0.243046997 | 0.526536038 |
| chr6:152206497-152206745 | -0.503313629 | 0.243050992 | 0.526536038 |
| chr1:156500474-156501581 | 0.369804069 | 0.243068162 | 0.526558346 |
| chr3:194761408-194761629 | -0.580832141 | 0.24309451 | 0.526600536 |
| chr15:92848342-92848552 | -0.629367118 | 0.243106698 | 0.526609876 |
| chr6:83592944-83593144 | 0.603575693 | 0.243119047 | 0.526609876 |
| chr5:143570997-143571639 | -0.43311905 | 0.243127882 | 0.526609876 |
| chr16:15777932-15778216 | 0.527504269 | 0.243131512 | 0.526609876 |
| chr13:32285235-32285777 | -0.432117876 | 0.243133186 | 0.526609876 |
| chr1:110029571-110030080 | -0.434267511 | 0.2431456 | 0.526615189 |
| chr6:132813713-132814895 | -0.369413792 | 0.243149385 | 0.526615189 |
| chr3:42654097-42654345 | -0.560095821 | 0.243171899 | 0.526649063 |
| chr14:64504990-64505456 | 0.391195137 | 0.243194786 | 0.526662914 |
| chr12:124993507-124993914 | 0.429414528 | 0.243196397 | 0.526662914 |
| chr13:51832480-51832683 | -0.592455354 | 0.243198914 | 0.526662914 |
| chr6:125292217-125292417 | -0.678439071 | 0.243216847 | 0.526663196 |
| chr9:99855847-99856089 | 0.58168011 | 0.24322774 | 0.526663196 |
| chr12:124608898-124609518 | 0.404748356 | 0.243238136 | 0.526663196 |
| chr10:100997456-100998013 | 0.393204334 | 0.243244275 | 0.526663196 |
| chr2:47264425-47264635 | 0.573716341 | 0.243248462 | 0.526663196 |
| chr6:105136946-105137273 | 0.523897548 | 0.243249388 | 0.526663196 |
| chr11:67464008-67464227 | 0.467640849 | 0.243249546 | 0.526663196 |
| chr8:101790868-101791266 | -0.432792737 | 0.243254033 | 0.526663196 |
| chr15:65902449-65902779 | 0.457232665 | 0.243262224 | 0.526666049 |
| chr7:143974499-143974815 | 0.500727303 | 0.243269963 | 0.526667923 |
| chr17:79647975-79648175 | 0.58610561 | 0.243289501 | 0.52669534 |
| chr4:137139232-137139732 | -0.437081806 | 0.243321977 | 0.526750763 |
| chr14:34120257-34120497 | 0.535731395 | 0.243329151 | 0.526751412 |
| chr12:13278196-13278489 | -0.518113756 | 0.243339887 | 0.526753354 |
| chr8:30648495-30648775 | -0.53863684 | 0.243350119 | 0.526753354 |
| chr22:38283362-38284027 | 0.424306814 | 0.243361677 | 0.526753354 |

| | | | |
|---|---|---|---|
| chr6:153231067-153231461 | -0.482727142 | 0.243366738 | 0.526753354 |
| chr1:40315391-40315591 | 0.548948281 | 0.243367124 | 0.526753354 |
| chr3:16247336-16248119 | 0.395870573 | 0.243371297 | 0.526753354 |
| chr19:42216990-42218034 | 0.377231749 | 0.243391336 | 0.526781847 |
| chr1:154479860-154480336 | 0.441429141 | 0.243402389 | 0.526790888 |
| chr2:26316124-26316441 | 0.497780958 | 0.243431353 | 0.526838694 |
| chrX:46758186-46759620 | 0.385042175 | 0.243459374 | 0.52687098 |
| chr2:200578146-200578346 | 0.66889662 | 0.243470941 | 0.52687098 |
| chr8:134856541-134856788 | -0.571878185 | 0.243482362 | 0.52687098 |
| chr1:215388709-215388948 | 0.565080154 | 0.243491423 | 0.52687098 |
| chr6:16184163-16184363 | 0.589993745 | 0.243494038 | 0.52687098 |
| chr20:47984364-47984671 | 0.391501853 | 0.243500304 | 0.52687098 |
| chr16:31859457-31859759 | 0.495600536 | 0.243500828 | 0.52687098 |
| chr12:94546157-94546590 | 0.427755605 | 0.243501281 | 0.52687098 |
| chr19:50777750-50778361 | -0.442624606 | 0.24354038 | 0.526926034 |
| chr6:148559830-148560434 | -0.404336283 | 0.243540479 | 0.526926034 |
| chr10:35448572-35448772 | 0.605785325 | 0.243555695 | 0.526935647 |
| chr20:53738439-53738822 | 0.377952953 | 0.243558676 | 0.526935647 |
| chr8:37891827-37892122 | -0.580368947 | 0.243571383 | 0.52694826 |
| chr22:50516280-50516480 | 0.599041985 | 0.243595601 | 0.526985773 |
| chr11:126080085-126080555 | -0.438348188 | 0.243612681 | 0.526987615 |
| chr11:96272598-96272869 | -0.547258736 | 0.243620499 | 0.526987615 |
| chr20:62598691-62599060 | 0.420553407 | 0.243621724 | 0.526987615 |
| chr4:70948706-70949166 | -0.475486381 | 0.243630734 | 0.526987615 |
| chr5:9538652-9538913 | -0.542571867 | 0.243630841 | 0.526987615 |
| chr2:79855482-79855723 | -0.523145185 | 0.243667067 | 0.527016369 |
| chr12:62900742-62901302 | 0.410047778 | 0.243668326 | 0.527016369 |
| chr9:5339377-5340015 | -0.440394768 | 0.243671537 | 0.527016369 |
| chr5:168448987-168449555 | -0.415014548 | 0.243672435 | 0.527016369 |
| chr7:130363411-130363686 | 0.583974984 | 0.243678525 | 0.527016369 |
| chr11:111446540-111446902 | -0.457638195 | 0.243719052 | 0.527089141 |
| chr3:160811997-160812207 | -0.628778223 | 0.243779344 | 0.527194086 |
| chr11:50241695-50241932 | 0.63324847 | 0.243786399 | 0.527194086 |
| chr20:10479003-10479238 | 0.504155516 | 0.243788218 | 0.527194086 |
| chr10:79562192-79562474 | -0.485844297 | 0.243811825 | 0.527230256 |
| chr2:231985797-231986191 | -0.468324042 | 0.243824565 | 0.527242926 |
| chr12:49059914-49060139 | 0.494857178 | 0.243838614 | 0.527258426 |
| chr6:128598423-128598969 | 0.415391723 | 0.243854847 | 0.527278645 |
| chr15:94666584-94666876 | 0.48780547 | 0.243866637 | 0.527279195 |
| chr7:4261203-4261485 | -0.518099443 | 0.243876589 | 0.527279195 |
| chr7:39863901-39864224 | 0.505585231 | 0.2438793 | 0.527279195 |
| chr6:75817668-75818027 | -0.498956192 | 0.243883659 | 0.527279195 |
| chr6:89281828-89282154 | -0.456810855 | 0.243889509 | 0.527279195 |
| chr10:119658380-119658592 | 0.519128222 | 0.243905572 | 0.527299045 |
| chr10:67520906-67521265 | -0.476817994 | 0.243923849 | 0.527317427 |
| chr1:91016304-91016504 | -0.613887506 | 0.243939242 | 0.527317427 |
| chr20:57464921-57465289 | 0.394311677 | 0.243941236 | 0.527317427 |
| chr10:27899331-27899799 | -0.483399143 | 0.243944402 | 0.527317427 |
| chr12:67595393-67595695 | -0.435677357 | 0.243948485 | 0.527317427 |
| chr3:182947924-182948309 | -0.45577151 | 0.243961326 | 0.527320999 |

| | | | |
|---|---|---|---|
| chr15:88546156-88546897 | 0.376961799 | 0.243968916 | 0.527320999 |
| chr13:96076049-96076302 | -0.599339431 | 0.243970784 | 0.527320999 |
| chr10:71948704-71949005 | 0.526928328 | 0.243977727 | 0.527321131 |
| chr10:100045556-100045847 | -0.634107246 | 0.243996903 | 0.527347701 |
| chr7:144720166-144720482 | -0.478253082 | 0.244024244 | 0.527391916 |
| chr3:160618426-160618755 | 0.475962787 | 0.244049847 | 0.527418205 |
| chr7:100335537-100336520 | -0.380374567 | 0.244050174 | 0.527418205 |
| chr14:21191925-21193268 | 0.381644211 | 0.244074933 | 0.527429252 |
| chr12:9448106-9448554 | -0.428282953 | 0.244076785 | 0.527429252 |
| chr12:52495386-52495689 | 0.527866165 | 0.244086249 | 0.527429252 |
| chr5:10780267-10780562 | -0.467579626 | 0.244090786 | 0.527429252 |
| chr7:100802193-100802624 | -0.42330619 | 0.24409104 | 0.527429252 |
| chr4:144611922-144612205 | -0.535897169 | 0.244096587 | 0.527429252 |
| chrX:49029775-49030212 | 0.424621396 | 0.244183233 | 0.527601593 |
| chr17:61470641-61470919 | 0.382422417 | 0.244228316 | 0.527684122 |
| chr12:93340825-93341026 | -0.664258425 | 0.244248344 | 0.527710869 |
| chr8:138114192-138114661 | 0.473289818 | 0.24425548 | 0.527710869 |
| chr8:23160859-23161516 | -0.4214505 | 0.244261357 | 0.527710869 |
| chr11:63671186-63671985 | 0.388063294 | 0.244312768 | 0.527807057 |
| chr20:41173163-41173440 | 0.430554553 | 0.244327645 | 0.527824316 |
| chr1:31815731-31816236 | -0.419218195 | 0.244411888 | 0.527977942 |
| chr1:196743320-196743620 | -0.45662832 | 0.24441254 | 0.527977942 |
| chr7:33787601-33787812 | 0.559516908 | 0.244451075 | 0.5280463 |
| chr17:60028805-60029093 | 0.507918997 | 0.244466522 | 0.52806478 |
| chr7:22660824-22661094 | 0.478211694 | 0.244474591 | 0.528067321 |
| chr1:29230645-29231220 | 0.400265589 | 0.244489259 | 0.528084119 |
| chr5:134845687-134846450 | -0.383511787 | 0.244508744 | 0.528111316 |
| chr13:29658321-29658713 | -0.520964788 | 0.244529517 | 0.528125839 |
| chr5:12153694-12154032 | -0.500740831 | 0.244531968 | 0.528125839 |
| chr6:30326294-30327473 | 0.379964088 | 0.244536145 | 0.528125839 |
| chr7:47635134-47635758 | 0.405109406 | 0.244566061 | 0.528175561 |
| chr2:208595501-208596132 | -0.431717687 | 0.244602724 | 0.528239458 |
| chr19:47112133-47112546 | 0.410186999 | 0.244611517 | 0.528239458 |
| chr14:45976473-45976917 | -0.460835086 | 0.244616897 | 0.528239458 |
| chr8:9933645-9933874 | -0.588500941 | 0.244623224 | 0.528239458 |
| chr6:49890952-49891409 | 0.479970988 | 0.244654967 | 0.528277771 |
| chr15:92077359-92077752 | 0.410511374 | 0.244658993 | 0.528277771 |
| chr12:1927244-1927461 | -0.642661534 | 0.244669744 | 0.528277771 |
| chr10:12720731-12720982 | 0.535900259 | 0.244687824 | 0.528277771 |
| chr2:20208593-20208869 | -0.461391373 | 0.244689267 | 0.528277771 |
| chr17:10073190-10073438 | 0.552793627 | 0.244689954 | 0.528277771 |
| chr7:23546880-23547328 | 0.4009311 | 0.244693953 | 0.528277771 |
| chr11:30585818-30586933 | -0.385660154 | 0.244696123 | 0.528277771 |
| chr2:235506720-235507083 | 0.483103724 | 0.24473955 | 0.528356638 |
| chr10:125048033-125048375 | 0.449740697 | 0.244756011 | 0.528377289 |
| chr2:546152-546494 | 0.507255887 | 0.244779369 | 0.528412824 |
| chr2:66212481-66212681 | -0.623670276 | 0.24479139 | 0.528419259 |
| chr10:82190650-82191189 | -0.434331368 | 0.244796142 | 0.528419259 |
| chr5:122201332-122201532 | -0.641294452 | 0.244831906 | 0.52847506 |
| chr7:7628642-7628842 | 0.578684831 | 0.244835787 | 0.52847506 |

| | | | |
|---|---|---|---|
| chr5:136053597-136053979 | -0.426299052 | 0.24487276 | 0.528539975 |
| chr1:3553455-3553748 | -0.529329177 | 0.244906953 | 0.528589636 |
| chr5:24093086-24093324 | -0.627270429 | 0.244909565 | 0.528589636 |
| chr7:21427020-21428569 | 0.362022601 | 0.244924032 | 0.528591451 |
| chr9:121213243-121213554 | 0.460188128 | 0.244926305 | 0.528591451 |
| chr17:61800341-61800743 | 0.383253924 | 0.244932386 | 0.528591451 |
| chr20:53230356-53230610 | -0.520526744 | 0.244938037 | 0.528591451 |
| chr5:172451030-172451518 | 0.431319889 | 0.244944899 | 0.528591451 |
| chr12:128823758-128824804 | 0.380544924 | 0.244958952 | 0.528606889 |
| chr18:66872360-66872638 | -0.588139566 | 0.244970481 | 0.528616881 |
| chr19:43950914-43951269 | 0.43984324 | 0.244990485 | 0.528645159 |
| chr18:63476614-63477214 | -0.413262756 | 0.245010954 | 0.528674439 |
| chr1:35192004-35192313 | 0.439307131 | 0.24502821 | 0.528696784 |
| chr3:120507230-120507490 | -0.483135999 | 0.245063571 | 0.528758193 |
| chr12:11554083-11554418 | -0.550697959 | 0.245075661 | 0.528769388 |
| chr6:106858667-106858971 | 0.500318633 | 0.245087203 | 0.528779401 |
| chr12:108515933-108516133 | 0.482341894 | 0.245100196 | 0.528792544 |
| chr1:232950177-232950932 | 0.386981247 | 0.245109178 | 0.528797033 |
| chr13:50081599-50083064 | -0.369322403 | 0.245131238 | 0.528822136 |
| chr12:11668676-11668917 | -0.562011422 | 0.245135887 | 0.528822136 |
| chr10:22251750-22252144 | -0.490672126 | 0.245161329 | 0.528822136 |
| chr17:50527145-50527396 | -0.503887275 | 0.245162822 | 0.528822136 |
| chr16:80654253-80655189 | 0.4137872 | 0.245164373 | 0.528822136 |
| chr7:76278648-76278864 | -0.578827053 | 0.24516848 | 0.528822136 |
| chr1:161153412-161154351 | -0.38966723 | 0.245169126 | 0.528822136 |
| chr3:101636239-101636518 | 0.456472248 | 0.245188801 | 0.528824001 |
| chr7:74220912-74221207 | -0.525319186 | 0.245188976 | 0.528824001 |
| chr9:129140665-129140870 | 0.503859292 | 0.245190695 | 0.528824001 |
| chr20:54646954-54647164 | 0.398427541 | 0.245215768 | 0.528835719 |
| chr13:106700643-106700877 | 0.583316581 | 0.245218584 | 0.528835719 |
| chr3:63490133-63490429 | 0.430042902 | 0.245219957 | 0.528835719 |
| chr8:141127800-141129119 | 0.370146802 | 0.245229226 | 0.528835719 |
| chr2:68357983-68358385 | -0.426627155 | 0.245230638 | 0.528835719 |
| chr13:102799825-102800071 | -0.478809608 | 0.245270639 | 0.528907094 |
| chr3:63937709-63937984 | 0.407290066 | 0.245334415 | 0.529029734 |
| chr10:71840144-71840485 | -0.451991609 | 0.245350799 | 0.529031674 |
| chr2:106575909-106576386 | -0.399630885 | 0.245354397 | 0.529031674 |
| chr17:57096661-57096942 | 0.482316984 | 0.245356028 | 0.529031674 |
| chr20:16551668-16551916 | 0.527022326 | 0.245419871 | 0.529154441 |
| chr12:102883268-102883520 | -0.486465343 | 0.245427817 | 0.529156683 |
| chr2:31413932-31414201 | -0.534435897 | 0.245437416 | 0.529162488 |
| chr11:60899332-60899612 | 0.497773144 | 0.245452951 | 0.529180939 |
| chr11:76807253-76807745 | -0.431903664 | 0.245459787 | 0.529180939 |
| chr13:72960608-72960885 | -0.524725653 | 0.245469429 | 0.529182659 |
| chr1:202807434-202807829 | -0.400196756 | 0.245474397 | 0.529182659 |
| chr13:110914596-110914849 | 0.474492659 | 0.245488452 | 0.52919807 |
| chr2:235542779-235543550 | -0.411713482 | 0.245521774 | 0.529241938 |
| chr10:35822354-35822590 | -0.587446375 | 0.245532987 | 0.529241938 |
| chr14:19168354-19168615 | -0.595659846 | 0.245534229 | 0.529241938 |
| chr7:17370074-17370316 | -0.557314272 | 0.245540919 | 0.529241938 |

| | | | |
|---|---|---|---|
| chr7:113937378-113937958 | -0.490519068 | 0.245543338 | 0.529241938 |
| chr3:12545247-12545807 | -0.390663094 | 0.24555702 | 0.52925654 |
| chr3:129576840-129577058 | 0.504826369 | 0.245637074 | 0.529414191 |
| chr1:9910054-9911091 | 0.375658228 | 0.245695799 | 0.529525863 |
| chr6:108848024-108848996 | -0.379188218 | 0.245727834 | 0.529564416 |
| chr20:62795301-62795614 | -0.416414809 | 0.245733214 | 0.529564416 |
| chr16:5033427-5034087 | 0.399277886 | 0.245740203 | 0.529564416 |
| chr14:104820690-104821046 | 0.429362278 | 0.245741332 | 0.529564416 |
| chr14:30656214-30656711 | 0.429129238 | 0.245761575 | 0.529584008 |
| chr7:30172935-30173371 | 0.429401561 | 0.245764247 | 0.529584008 |
| chr14:49497782-49498055 | 0.45538226 | 0.245799386 | 0.529603383 |
| chr2:189751121-189751618 | -0.464683134 | 0.245808735 | 0.529603383 |
| chr4:69617130-69617386 | -0.55711493 | 0.245814936 | 0.529603383 |
| chr1:220776768-220777010 | -0.521695277 | 0.245823955 | 0.529603383 |
| chr8:22393142-22393408 | -0.595020698 | 0.245825339 | 0.529603383 |
| chr10:1110780-1110986 | 0.524507324 | 0.24582536 | 0.529603383 |
| chr9:91758465-91758665 | -0.599712105 | 0.245828127 | 0.529603383 |
| chr6:113414544-113414835 | 0.490457118 | 0.245828534 | 0.529603383 |
| chr13:32497658-32497871 | -0.614884168 | 0.245848125 | 0.529619694 |
| chr5:160008865-160009870 | -0.374202417 | 0.245849929 | 0.529619694 |
| chr20:25116127-25116361 | 0.553309605 | 0.245869352 | 0.529639764 |
| chr10:50623860-50625483 | -0.364478865 | 0.24587307 | 0.529639764 |
| chr4:138035786-138036299 | 0.453341678 | 0.245897553 | 0.529677612 |
| chr8:142624004-142624295 | 0.439906949 | 0.245914413 | 0.529699037 |
| chr18:54767097-54767510 | 0.421230991 | 0.245944807 | 0.52972895 |
| chr14:21940902-21941166 | -0.551342114 | 0.245949602 | 0.52972895 |
| chr13:49996692-49997778 | -0.379577321 | 0.245961474 | 0.52972895 |
| chr16:24538968-24539710 | 0.377993725 | 0.245961544 | 0.52972895 |
| chrX:16436606-16436927 | -0.53530547 | 0.245966089 | 0.52972895 |
| chr8:135394883-135395190 | -0.575570965 | 0.245969782 | 0.52972895 |
| chr16:19389234-19389578 | 0.463382318 | 0.246008748 | 0.529781952 |
| chr11:62694806-62695096 | 0.503048024 | 0.246012863 | 0.529781952 |
| chr21:37367791-37368042 | 0.490067018 | 0.246015134 | 0.529781952 |
| chr20:54804692-54804938 | 0.406555073 | 0.24605605 | 0.52985517 |
| chr16:71611021-71611395 | -0.415665801 | 0.246079198 | 0.52985858 |
| chr6:3751860-3752082 | 0.520772428 | 0.246083709 | 0.52985858 |
| chr6:69180390-69180655 | 0.514300679 | 0.246095478 | 0.52985858 |
| chr2:61404483-61404742 | -0.577830575 | 0.246095767 | 0.52985858 |
| chr13:44140206-44140550 | 0.421865586 | 0.246102551 | 0.52985858 |
| chr14:78566598-78567134 | 0.41423661 | 0.246103906 | 0.52985858 |
| chr8:63605523-63605797 | -0.541468239 | 0.246111626 | 0.52985858 |
| chr1:205899771-205899971 | -0.592013688 | 0.246112955 | 0.52985858 |
| chr22:21102824-21103178 | 0.440973313 | 0.246129376 | 0.529879043 |
| chr3:127822316-127822540 | 0.507336023 | 0.246147109 | 0.529899547 |
| chr7:8389699-8390566 | 0.375642341 | 0.24615551 | 0.529899547 |
| chr5:137779076-137779314 | -0.590035086 | 0.246159647 | 0.529899547 |
| chr11:113384686-113384923 | -0.58666554 | 0.246167727 | 0.529902053 |
| chr1:205668766-205669073 | -0.453219013 | 0.246190873 | 0.529936376 |
| chr6:27078705-27079231 | -0.42009193 | 0.246203098 | 0.529936376 |
| chr3:13619920-13620120 | 0.531666712 | 0.24620442 | 0.529936376 |

| | | | |
|---|---|---|---|
| chr3:4980362-4980572 | 0.424120446 | 0.246225279 | 0.529964694 |
| chr1:81179232-81179549 | -0.475791909 | 0.24623141 | 0.529964694 |
| chr16:2047647-2048257 | 0.398982132 | 0.246240132 | 0.529968579 |
| chr10:4267458-4267661 | -0.614353741 | 0.246249767 | 0.529974429 |
| chr22:35583095-35583307 | 0.582141677 | 0.246263756 | 0.52998965 |
| chr5:58886655-58887179 | -0.431669892 | 0.246280657 | 0.530003715 |
| chr18:48616015-48616373 | -0.425629438 | 0.246286235 | 0.530003715 |
| chr17:67980067-67980419 | -0.486325339 | 0.246302963 | 0.530003715 |
| chr10:1379735-1379956 | 0.513586916 | 0.246303814 | 0.530003715 |
| chr9:126082834-126083034 | 0.554967375 | 0.246304877 | 0.530003715 |
| chr9:37464816-37466247 | -0.364853025 | 0.246332421 | 0.530048099 |
| chr1:17502382-17502620 | 0.563995234 | 0.246348541 | 0.5300679 |
| chr12:57591461-57591931 | 0.410128884 | 0.246402441 | 0.530143055 |
| chrX:140040319-140040955 | -0.400195465 | 0.246412139 | 0.530143055 |
| chr16:75622757-75623686 | -0.37257307 | 0.246429816 | 0.530143055 |
| chr3:64745401-64745666 | 0.405421562 | 0.246433275 | 0.530143055 |
| chr6:163356441-163356670 | -0.647944827 | 0.246433919 | 0.530143055 |
| chr21:39215810-39216010 | 0.566290978 | 0.246434417 | 0.530143055 |
| chr14:84036844-84037081 | -0.566501413 | 0.24644201 | 0.530143055 |
| chr6:28982601-28982826 | -0.471964143 | 0.246444294 | 0.530143055 |
| chr3:159404668-159404938 | 0.522412519 | 0.24644574 | 0.530143055 |
| chr7:134637983-134638184 | -0.659145947 | 0.246455036 | 0.530148168 |
| chr1:215780251-215780451 | 0.545664866 | 0.246483652 | 0.530174944 |
| chr7:46153881-46154580 | -0.410395702 | 0.246484556 | 0.530174944 |
| chr5:44468260-44468546 | -0.61439244 | 0.246495312 | 0.530174944 |
| chr20:41136690-41138308 | 0.362968892 | 0.246496727 | 0.530174944 |
| chr4:176341040-176341307 | -0.498878441 | 0.24650208 | 0.530174944 |
| chr7:141701963-141702816 | -0.395853582 | 0.246513993 | 0.530179911 |
| chr15:66603640-66603998 | 0.476736085 | 0.246518228 | 0.530179911 |
| chr3:64729117-64729465 | 0.392949461 | 0.246532921 | 0.530196629 |
| chr17:39232087-39232287 | -0.576635329 | 0.246547501 | 0.530213101 |
| chr10:119750628-119750951 | 0.457097316 | 0.246603757 | 0.530319199 |
| chrX:81208456-81208793 | -0.50543641 | 0.24662205 | 0.530343654 |
| chr19:1180462-1180775 | -0.483051505 | 0.246641799 | 0.530353025 |
| chr14:73457854-73458981 | 0.370149555 | 0.246650266 | 0.530353025 |
| chr6:138115925-138116203 | 0.481790601 | 0.246651998 | 0.530353025 |
| chr17:50266708-50266944 | 0.476777591 | 0.246658764 | 0.530353025 |
| chr17:58740846-58741696 | -0.371052179 | 0.246661017 | 0.530353025 |
| chr9:36486999-36488152 | -0.385756501 | 0.246686275 | 0.53039245 |
| chr2:62529791-62530204 | -0.542240104 | 0.246725546 | 0.530462002 |
| chr16:1677938-1678543 | 0.396703404 | 0.246739076 | 0.530476205 |
| chr3:40188106-40188354 | -0.548569812 | 0.2467699 | 0.530527589 |
| chr9:71681726-71682257 | -0.448682186 | 0.246781232 | 0.530537066 |
| chr17:34361631-34361841 | 0.599791733 | 0.246789754 | 0.530540501 |
| chr1:70751644-70753132 | -0.375164788 | 0.246803614 | 0.530555411 |
| chr10:80454845-80455354 | 0.410688817 | 0.246827955 | 0.530592851 |
| chr3:57164566-57164766 | -0.645803959 | 0.246857968 | 0.530627593 |
| chr7:46722470-46722670 | -0.645803959 | 0.246857968 | 0.530627593 |
| chr1:84442257-84442573 | -0.498242024 | 0.246886535 | 0.530674113 |
| chr7:102274019-102274327 | -0.488913475 | 0.246896327 | 0.530680273 |

| | | | |
|---|---|---|---|
| chr17:21280086-21280362 | 0.43016915 | 0.246911643 | 0.530693296 |
| chr13:24995295-24995566 | 0.509727228 | 0.246916238 | 0.530693296 |
| chr3:170584575-170584886 | -0.484042748 | 0.246942089 | 0.530733969 |
| chr3:53134798-53134998 | 0.517877126 | 0.246969817 | 0.530778676 |
| chr20:42486641-42486981 | 0.458142496 | 0.246997936 | 0.530824219 |
| chr1:216765372-216765748 | 0.442625784 | 0.247020271 | 0.530857328 |
| chr8:126876608-126876994 | -0.437081927 | 0.247030391 | 0.530864189 |
| chr2:227107214-227107414 | -0.6456043 | 0.247060042 | 0.530910399 |
| chr12:130818278-130819407 | 0.377379283 | 0.247065753 | 0.530910399 |
| chr5:15469738-15469976 | 0.621326746 | 0.247073916 | 0.530913052 |
| chr12:124568455-124568776 | -0.449025753 | 0.24713851 | 0.531036959 |
| chr2:55432073-55432273 | -0.595428899 | 0.247155745 | 0.531059099 |
| chr13:95314445-95314823 | 0.446275676 | 0.247170871 | 0.531067738 |
| chr9:72024933-72025325 | -0.473737476 | 0.247173627 | 0.531067738 |
| chr8:76254840-76255208 | -0.461512447 | 0.247193213 | 0.531094927 |
| chr1:232047658-232047858 | 0.641515216 | 0.247202433 | 0.531099844 |
| chr8:61139011-61139331 | 0.46693519 | 0.247215806 | 0.531113682 |
| chrX:154596485-154596939 | 0.468561597 | 0.247249768 | 0.531171753 |
| chr5:133247951-133248170 | -0.524019709 | 0.2472687 | 0.531197531 |
| chr20:47400611-47401083 | 0.371141975 | 0.247298309 | 0.53121773 |
| chr21:33542001-33543394 | 0.367956221 | 0.24730037 | 0.53121773 |
| chr9:111399855-111400463 | -0.413411704 | 0.247303206 | 0.53121773 |
| chr20:53736226-53736621 | 0.376006889 | 0.247311743 | 0.53121773 |
| chr3:111717341-111717897 | -0.400828196 | 0.247312767 | 0.53121773 |
| chr10:91006335-91006624 | 0.444204722 | 0.247321045 | 0.531220618 |
| chr3:163878548-163878757 | -0.620858773 | 0.247357232 | 0.531283452 |
| chr17:7213990-7215332 | -0.370077253 | 0.247372045 | 0.531300374 |
| chrX:82230000-82230345 | -0.487673177 | 0.247388468 | 0.531311387 |
| chr8:143876145-143876425 | 0.447567792 | 0.247391041 | 0.531311387 |
| chr1:202401923-202402167 | 0.49277851 | 0.247400555 | 0.531313603 |
| chr9:4073464-4073783 | -0.499780625 | 0.247405941 | 0.531313603 |
| chr13:33310726-33311222 | -0.440845128 | 0.247434443 | 0.531359918 |
| chr18:37282375-37282790 | -0.474127338 | 0.247490951 | 0.531466372 |
| chr3:169871347-169871663 | -0.507528293 | 0.247511795 | 0.531496237 |
| chr15:83107274-83107533 | -0.541239702 | 0.247577658 | 0.531621661 |
| chr6:35452046-35452446 | 0.411338182 | 0.24758408 | 0.531621661 |
| chr12:73153319-73153690 | 0.501553427 | 0.247605083 | 0.531646246 |
| chr3:16855091-16855321 | -0.582098318 | 0.247609406 | 0.531646246 |
| chr4:45869261-45869557 | 0.513940101 | 0.247624379 | 0.531663496 |
| chr15:72230290-72230592 | 0.416832099 | 0.247634609 | 0.531670561 |
| chr11:66579067-66579496 | 0.439011548 | 0.24767135 | 0.531729439 |
| chr4:37977791-37978075 | 0.483931533 | 0.247685187 | 0.531729439 |
| chr6:13328038-13328945 | -0.376613049 | 0.247689674 | 0.531729439 |
| chr9:68779584-68780472 | -0.385944607 | 0.247697073 | 0.531729439 |
| chr5:16489315-16489727 | -0.459341927 | 0.247706387 | 0.531729439 |
| chr17:5418990-5419199 | 0.50871914 | 0.247707059 | 0.531729439 |
| chr2:101823316-101823580 | -0.498278993 | 0.247712233 | 0.531729439 |
| chr16:25225281-25225497 | 0.52820613 | 0.247717549 | 0.531729439 |
| chr3:149378618-149379095 | 0.448830273 | 0.247740407 | 0.531763607 |
| chr17:60054370-60055015 | 0.37638364 | 0.247749369 | 0.531767946 |

| | | | |
|---|---|---|---|
| chr10:102642001-102642668 | 0.380791089 | 0.247766067 | 0.53176926 |
| chr1:185344809-185345251 | -0.466396155 | 0.247779677 | 0.53176926 |
| chr18:52299471-52299673 | -0.625830698 | 0.247781876 | 0.53176926 |
| chr12:14257607-14258240 | -0.391704733 | 0.247785123 | 0.53176926 |
| chr16:57418822-57419143 | 0.455423501 | 0.247786508 | 0.53176926 |
| chr14:88168596-88168896 | 0.487430322 | 0.247791622 | 0.53176926 |
| chr8:1973452-1973925 | -0.487195448 | 0.247799181 | 0.531770589 |
| chr8:140213723-140214131 | 0.454059013 | 0.247845214 | 0.531854478 |
| chr10:42903436-42903656 | -0.619623786 | 0.247857342 | 0.531860397 |
| chr3:125165328-125165576 | 0.517233783 | 0.247864395 | 0.531860397 |
| chrX:84123456-84123708 | -0.570518912 | 0.247868796 | 0.531860397 |
| chr12:122265814-122267115 | 0.37542461 | 0.247890377 | 0.531891807 |
| chr16:85659825-85660154 | 0.428290386 | 0.247900039 | 0.531897644 |
| chr1:155206911-155207339 | 0.405087987 | 0.247913972 | 0.531912645 |
| chr20:57772398-57772928 | 0.387635206 | 0.247985882 | 0.532047777 |
| chr8:141279118-141279423 | -0.495336309 | 0.247990842 | 0.532047777 |
| chr12:49776203-49776798 | 0.419712204 | 0.248071873 | 0.532206721 |
| chr4:124581469-124581698 | -0.591986527 | 0.248091068 | 0.532211081 |
| chr4:152662770-152663099 | 0.563160031 | 0.248093281 | 0.532211081 |
| chr11:62705434-62706929 | 0.369708661 | 0.248094743 | 0.532211081 |
| chr20:54038207-54038581 | 0.38918722 | 0.248121664 | 0.532230591 |
| chr13:98650323-98650838 | -0.419942152 | 0.248129233 | 0.532230591 |
| chr19:42777118-42777352 | -0.594620999 | 0.248129424 | 0.532230591 |
| chr19:35744650-35745180 | 0.38479621 | 0.248133502 | 0.532230591 |
| chr2:173752206-173752463 | -0.587296849 | 0.248138569 | 0.532230591 |
| chr5:176180840-176181040 | 0.574161664 | 0.248174451 | 0.532258648 |
| chr6:55276821-55277044 | -0.591877825 | 0.248184815 | 0.532258648 |
| chr3:167734407-167735894 | -0.359286492 | 0.248184919 | 0.532258648 |
| chr12:119988422-119989449 | 0.368414135 | 0.248190321 | 0.532258648 |
| chr20:56890005-56890517 | -0.400036571 | 0.248190859 | 0.532258648 |
| chr2:158998847-158999277 | -0.46975276 | 0.248196853 | 0.532258648 |
| chr17:67689598-67689864 | -0.490795442 | 0.248200409 | 0.532258648 |
| chrX:63754468-63755559 | -0.373740887 | 0.248211109 | 0.532258648 |
| chr16:10981879-10982079 | 0.564198285 | 0.248224045 | 0.532258648 |
| chr12:70365555-70366535 | 0.376460193 | 0.248226897 | 0.532258648 |
| chr2:105144386-105144586 | 0.526002129 | 0.248228061 | 0.532258648 |
| chr17:81345605-81346536 | 0.374711832 | 0.248237887 | 0.532261663 |
| chr16:82329541-82330316 | 0.387646663 | 0.248243361 | 0.532261663 |
| chr9:96087042-96087242 | -0.636740487 | 0.248256827 | 0.532275641 |
| chr12:103799853-103800116 | -0.512122883 | 0.248272709 | 0.532294798 |
| chrX:142391025-142391735 | -0.415378357 | 0.248295766 | 0.53232887 |
| chr5:78699552-78700232 | -0.409604392 | 0.248302495 | 0.53232887 |
| chr17:8173039-8173985 | -0.388368544 | 0.248330933 | 0.532353116 |
| chr11:19742691-19742991 | 0.483536497 | 0.248331269 | 0.532353116 |
| chr3:59320076-59320529 | 0.457151922 | 0.24835117 | 0.532353116 |
| chr6:136967976-136968481 | -0.437317355 | 0.248352623 | 0.532353116 |
| chr16:52068543-52068829 | -0.586572777 | 0.248357295 | 0.532353116 |
| chr1:3982197-3982459 | -0.584349116 | 0.248364788 | 0.532353116 |
| chr10:60964202-60964447 | -0.522916553 | 0.248372473 | 0.532353116 |
| chr8:94242028-94242836 | 0.395081062 | 0.248405109 | 0.532353116 |

| | | | |
|---|---|---|---|
| chr18:45782740-45782959 | -0.559569947 | 0.248405876 | 0.532353116 |
| chr12:1907585-1908163 | 0.421987109 | 0.248410551 | 0.532353116 |
| chr11:17328968-17329209 | -0.589788434 | 0.248412235 | 0.532353116 |
| chr5:114377943-114378374 | -0.520529261 | 0.248413886 | 0.532353116 |
| chr12:51291726-51292355 | 0.448112956 | 0.248424249 | 0.532353116 |
| chr8:29752382-29752735 | 0.49790629 | 0.248431694 | 0.532353116 |
| chr16:19114359-19114623 | 0.500609562 | 0.248432727 | 0.532353116 |
| chr8:70393044-70393339 | -0.580943703 | 0.248445323 | 0.532353116 |
| chr10:103884724-103885017 | 0.490039013 | 0.248446309 | 0.532353116 |
| chr8:100509860-100510617 | 0.38577551 | 0.248459438 | 0.532353116 |
| chr21:33361285-33361586 | 0.460376713 | 0.248462572 | 0.532353116 |
| chr3:24773898-24774328 | 0.486547636 | 0.248469092 | 0.532353116 |
| chr20:56995182-56995479 | 0.439905702 | 0.24846987 | 0.532353116 |
| chr1:231806694-231807240 | 0.412704992 | 0.248470193 | 0.532353116 |
| chrX:30489426-30489656 | 0.531509148 | 0.248473604 | 0.532353116 |
| chr8:102067874-102068254 | 0.461640374 | 0.248507429 | 0.532410699 |
| chr11:121655330-121656539 | -0.3890705 | 0.248554972 | 0.532491998 |
| chr8:22081919-22082252 | -0.562890567 | 0.248559276 | 0.532491998 |
| chr2:96116474-96116750 | 0.452076979 | 0.248566737 | 0.532493095 |
| chr8:64399280-64399611 | -0.492228751 | 0.248577432 | 0.532501118 |
| chr10:70405508-70405797 | -0.454145456 | 0.24860894 | 0.532549814 |
| chr9:127965450-127965872 | 0.430374178 | 0.248614064 | 0.532549814 |
| chr8:78805332-78805577 | 0.480947038 | 0.248625412 | 0.532559232 |
| chr9:115395782-115395995 | -0.59999683 | 0.248648138 | 0.532593024 |
| chr8:50153818-50154265 | -0.502122942 | 0.24865758 | 0.53259836 |
| chr14:26820211-26820543 | 0.492649817 | 0.248675171 | 0.532613003 |
| chr1:11012152-11013284 | 0.378294317 | 0.248678319 | 0.532613003 |
| chr22:11916339-11916539 | 0.584260496 | 0.24870783 | 0.532638543 |
| chr11:70593848-70594067 | -0.591904806 | 0.24871049 | 0.532638543 |
| chr21:8230612-8230834 | 0.44854244 | 0.248717583 | 0.532638543 |
| chr16:25031279-25032318 | 0.375294659 | 0.248725236 | 0.532638543 |
| chr6:31197001-31198122 | 0.380141141 | 0.248730308 | 0.532638543 |
| chr17:3913755-3914141 | 0.447866776 | 0.248731953 | 0.532638543 |
| chr3:155753086-155753558 | -0.439773064 | 0.24874663 | 0.532655086 |
| chr4:61202168-61203475 | -0.386517483 | 0.248753587 | 0.532655097 |
| chr5:145903203-145903419 | 0.555327097 | 0.248767102 | 0.532669151 |
| chr7:322098-322451 | 0.418791254 | 0.248833496 | 0.532795984 |
| chr3:192566798-192567231 | 0.461265336 | 0.248840243 | 0.532795984 |
| chr17:70753085-70753285 | -0.643747008 | 0.248860965 | 0.532815976 |
| chr10:59854266-59854999 | 0.399703629 | 0.248866166 | 0.532815976 |
| chr3:102225435-102225725 | -0.463348438 | 0.248885594 | 0.532815976 |
| chr12:119208593-119208945 | -0.482905497 | 0.248887035 | 0.532815976 |
| chr1:109959773-109960269 | 0.415685458 | 0.248893814 | 0.532815976 |
| chr1:91959093-91959305 | -0.596441609 | 0.248898257 | 0.532815976 |
| chr1:161285565-161285809 | -0.596441609 | 0.248898257 | 0.532815976 |
| chr11:113401690-113402073 | -0.467748693 | 0.248917552 | 0.532828226 |
| chrX:109635306-109635617 | -0.489089603 | 0.248919272 | 0.532828226 |
| chr11:67996733-67997328 | 0.389656563 | 0.248932212 | 0.532828226 |
| chr6:38755174-38755558 | 0.445346225 | 0.248939648 | 0.532828226 |
| chr8:83163209-83163448 | 0.52457555 | 0.248940101 | 0.532828226 |

| | | | |
|---|---|---|---|
| chr15:70368081-70368427 | 0.522924841 | 0.248951316 | 0.532828226 |
| chr9:109199045-109199331 | -0.501085834 | 0.248952657 | 0.532828226 |
| chr1:16621048-16621380 | 0.415816692 | 0.248985923 | 0.532868856 |
| chr4:98113123-98113362 | -0.613768046 | 0.248996939 | 0.532868856 |
| chr6:68778095-68778324 | -0.613768046 | 0.248996939 | 0.532868856 |
| chr3:152574523-152574770 | -0.509224772 | 0.248999459 | 0.532868856 |
| chr6:67059626-67059953 | -0.534896895 | 0.249014135 | 0.532885382 |
| chrX:143664340-143664606 | 0.59720264 | 0.249028381 | 0.532896816 |
| chr20:31604073-31604526 | 0.383917273 | 0.249033388 | 0.532896816 |
| chr17:58755375-58757063 | -0.35986211 | 0.249045086 | 0.532906964 |
| chr15:35293454-35293683 | -0.595688571 | 0.249060207 | 0.532924437 |
| chr17:78716824-78717191 | -0.494742914 | 0.249106217 | 0.533008003 |
| chr1:32741808-32742008 | 0.582596911 | 0.249118079 | 0.533018498 |
| chr2:222899561-222899802 | 0.596852569 | 0.249140689 | 0.533024445 |
| chr16:66867826-66868086 | 0.534997576 | 0.249141572 | 0.533024445 |
| chr17:42423151-42423435 | 0.513667009 | 0.249143028 | 0.533024445 |
| chr8:125686376-125686576 | 0.512519794 | 0.249148685 | 0.533024445 |
| chr11:65170047-65170278 | -0.570555088 | 0.249252406 | 0.533231458 |
| chr6:106335106-106335382 | -0.471600652 | 0.249320966 | 0.533363237 |
| chr11:15208626-15209032 | -0.433398911 | 0.249329812 | 0.53336727 |
| chr14:77654732-77655493 | 0.391560978 | 0.24935187 | 0.533369622 |
| chr2:44086964-44087580 | -0.397208077 | 0.249355016 | 0.533369622 |
| chr3:113825232-113825521 | -0.459751871 | 0.249357356 | 0.533369622 |
| chr14:22570282-22570718 | 0.432773399 | 0.249358756 | 0.533369622 |
| chr12:67483402-67483640 | 0.497980359 | 0.24938362 | 0.533407916 |
| chr9:116693116-116693523 | 0.446102239 | 0.249427401 | 0.533486666 |
| chr5:65562835-65563728 | -0.37046698 | 0.249454627 | 0.533530005 |
| chr4:108004021-108004318 | 0.537318867 | 0.249482194 | 0.533561419 |
| chr5:171342462-171343053 | 0.429052 | 0.249483242 | 0.533561419 |
| chr6:31068919-31069119 | -0.627358511 | 0.249535686 | 0.533658684 |
| chr3:108489755-108490055 | -0.456218937 | 0.24955917 | 0.53366248 |
| chrX:42423041-42423272 | -0.635045946 | 0.249564832 | 0.53366248 |
| chr3:177596967-177597271 | -0.490155335 | 0.249566993 | 0.53366248 |
| chr8:143716120-143716493 | -0.413396008 | 0.249571744 | 0.53366248 |
| chr7:66095076-66095320 | 0.586977527 | 0.249572285 | 0.53366248 |
| chr3:171809592-171810810 | -0.366685441 | 0.24960057 | 0.533708068 |
| chr8:133954474-133954853 | 0.443116767 | 0.249612085 | 0.533710337 |
| chr3:106842153-106842437 | -0.561647845 | 0.249615562 | 0.533710337 |
| chr6:163821516-163821796 | -0.580113561 | 0.249627281 | 0.533720499 |
| chr1:51061055-51061259 | -0.606614161 | 0.249703148 | 0.53386781 |
| chr2:210663611-210663836 | -0.560078844 | 0.249721694 | 0.533892566 |
| chr16:30591582-30591785 | 0.652557347 | 0.249733629 | 0.533903184 |
| chr16:75265862-75266337 | 0.385379949 | 0.249769652 | 0.533965299 |
| chr14:77204368-77204736 | 0.416834262 | 0.249779958 | 0.533966603 |
| chr16:82661702-82661902 | -0.642818438 | 0.2497842 | 0.533966603 |
| chr1:153613017-153613243 | 0.439858962 | 0.249803719 | 0.533993432 |
| chr7:137636157-137636362 | -0.610432566 | 0.249824416 | 0.534022775 |
| chr11:23865648-23866039 | 0.437869834 | 0.249833645 | 0.534027605 |
| chr19:47471966-47472217 | 0.50959491 | 0.249850374 | 0.534048467 |
| chr7:155344717-155345313 | -0.39745933 | 0.249911452 | 0.534164117 |

| | | | |
|---|---|---|---|
| chr3:64113348-64113949 | 0.371704197 | 0.249923559 | 0.53416706 |
| chr2:19711605-19711852 | -0.548494932 | 0.249930512 | 0.53416706 |
| chr6:100001968-100002266 | -0.478211159 | 0.249940715 | 0.53416706 |
| chr9:71285150-71285508 | -0.478211159 | 0.249940715 | 0.53416706 |
| chr1:110630785-110631017 | -0.538205929 | 0.24995105 | 0.534174249 |
| chr10:120455314-120455517 | -0.60402887 | 0.250027735 | 0.53431005 |
| chr5:173825274-173825565 | 0.438107024 | 0.25002854 | 0.53431005 |
| chr5:36150861-36151348 | 0.404761261 | 0.250043249 | 0.534326579 |
| chr5:162227119-162227505 | -0.481700612 | 0.250054701 | 0.534336149 |
| chr6:104232061-104232335 | -0.474164055 | 0.250068577 | 0.534349751 |
| chr2:67234502-67234738 | -0.559773887 | 0.250075014 | 0.534349751 |
| chr2:88469348-88469662 | 0.521685755 | 0.250144452 | 0.534473426 |
| chr8:66228712-66229090 | -0.429454919 | 0.250146845 | 0.534473426 |
| chr12:100138422-100138637 | -0.469529114 | 0.250218931 | 0.53460811 |
| chr8:132480958-132481425 | -0.449412324 | 0.250228498 | 0.53460811 |
| chr8:95159637-95160414 | 0.392232218 | 0.250230812 | 0.53460811 |
| chr2:156536692-156537019 | -0.526383797 | 0.250283664 | 0.534706118 |
| chr14:54219974-54220392 | -0.551614306 | 0.250292877 | 0.534710891 |
| chr1:120175989-120176778 | 0.382702711 | 0.250306098 | 0.534721347 |
| chr13:49791798-49792092 | -0.448666177 | 0.250311729 | 0.534721347 |
| chr8:128766814-128767014 | 0.548411844 | 0.250330538 | 0.534746619 |
| chr3:11718754-11720871 | 0.359294894 | 0.250344779 | 0.534762132 |
| chr21:39096789-39097049 | -0.495612178 | 0.250357525 | 0.534769576 |
| chr8:115778258-115778544 | 0.491454841 | 0.250362223 | 0.534769576 |
| chr1:76331049-76331393 | 0.492190558 | 0.25040709 | 0.534850501 |
| chr7:47405674-47406427 | -0.390417778 | 0.250417601 | 0.534858043 |
| chr3:193885761-193886383 | -0.454867773 | 0.250429941 | 0.534869489 |
| chr12:60936022-60936267 | 0.523501109 | 0.250442936 | 0.53487247 |
| chr15:57561030-57561308 | 0.45762014 | 0.250445298 | 0.53487247 |
| chr19:55081143-55081740 | 0.392309955 | 0.250459117 | 0.534887075 |
| chrX:57929438-57929662 | 0.563822643 | 0.250482052 | 0.534918557 |
| chr10:75234107-75236268 | -0.360169479 | 0.250487821 | 0.534918557 |
| chr19:12801694-12802075 | 0.446189685 | 0.25050048 | 0.534930682 |
| chr1:52176400-52177123 | -0.412968111 | 0.250552951 | 0.53502782 |
| chr13:52845405-52845733 | -0.445827386 | 0.25057156 | 0.535042838 |
| chr5:79235590-79236654 | -0.3735806 | 0.25057395 | 0.535042838 |
| chr16:77722019-77723074 | 0.362179536 | 0.250592506 | 0.535048719 |
| chr11:16986494-16987585 | 0.373651063 | 0.250597306 | 0.535048719 |
| chr18:37565407-37565607 | -0.5409137 | 0.250597653 | 0.535048719 |
| chr3:126535605-126535833 | -0.571690374 | 0.250613003 | 0.535059727 |
| chr22:20038449-20038761 | 0.508087405 | 0.250616775 | 0.535059727 |
| chr18:48002977-48003336 | 0.502935846 | 0.250631825 | 0.535076949 |
| chr6:18062817-18063072 | 0.547733758 | 0.250659728 | 0.535117901 |
| chr17:60801404-60801727 | 0.413186575 | 0.250665879 | 0.535117901 |
| chr3:63146970-63147179 | 0.456788969 | 0.250671959 | 0.535117901 |
| chr16:23059555-23059880 | 0.585392598 | 0.250683889 | 0.535128461 |
| chr2:216082454-216082913 | -0.438200169 | 0.250718621 | 0.535187691 |
| chr8:86147714-86148106 | 0.454747599 | 0.250734879 | 0.535202244 |
| chr17:70779162-70779747 | -0.459825948 | 0.250745494 | 0.535202244 |
| chr11:129703219-129703657 | -0.492292264 | 0.250746394 | 0.535202244 |

| | | | |
|---|---|---|---|
| chr4:119041942-119042256 | 0.465100735 | 0.250791641 | 0.535283912 |
| chr3:184037884-184038136 | -0.553089141 | 0.250811242 | 0.535292111 |
| chr3:132316958-132317987 | -0.374898433 | 0.250812842 | 0.535292111 |
| chr20:53899573-53900012 | 0.371941059 | 0.250816441 | 0.535292111 |
| chr11:70196110-70196422 | 0.466487053 | 0.250853362 | 0.535351812 |
| chr1:107047533-107047733 | 0.449768083 | 0.250858389 | 0.535351812 |
| chr1:67552305-67552707 | -0.478673273 | 0.250876211 | 0.535366848 |
| chrX:130170850-130171443 | 0.398951112 | 0.250879409 | 0.535366848 |
| chr1:30782383-30782659 | 0.44187683 | 0.250895954 | 0.535386504 |
| chr2:110949075-110949305 | 0.607384072 | 0.250902594 | 0.535386504 |
| chr5:126368453-126368884 | -0.418825648 | 0.250988411 | 0.535543676 |
| chr11:3824707-3825044 | -0.466028987 | 0.250992516 | 0.535543676 |
| chr1:32180366-32180681 | 0.444555905 | 0.250997219 | 0.535543676 |
| chr8:133341903-133342139 | -0.591114557 | 0.251009303 | 0.535554544 |
| chr5:111461245-111461724 | -0.44305022 | 0.251044659 | 0.535615065 |
| chr2:9206182-9206913 | 0.391130468 | 0.251064447 | 0.535627842 |
| chr1:111864896-111865170 | 0.42298131 | 0.251070812 | 0.535627842 |
| chr17:60756075-60756585 | 0.371436583 | 0.251075412 | 0.535627842 |
| chr1:215583201-215583872 | -0.38680264 | 0.251078609 | 0.535627842 |
| chr4:108513206-108513457 | -0.547157252 | 0.251094303 | 0.535646408 |
| chr3:151808676-151809450 | -0.400144815 | 0.251107321 | 0.535653503 |
| chrX:71129155-71129495 | 0.494855166 | 0.251111611 | 0.535653503 |
| chr5:93801069-93801796 | -0.425994299 | 0.25112446 | 0.535666 |
| chr20:7980701-7980901 | 0.563074194 | 0.251187303 | 0.535785133 |
| chr14:90399570-90399828 | 0.430624612 | 0.251201373 | 0.535800229 |
| chr20:63545194-63545604 | -0.42526356 | 0.251228686 | 0.535843569 |
| chr17:13306503-13306905 | -0.488481624 | 0.251246986 | 0.535861629 |
| chr2:100427654-100428020 | -0.475719891 | 0.25125114 | 0.535861629 |
| chr6:72260624-72260824 | -0.632943003 | 0.251260362 | 0.53586638 |
| chr6:14996963-14997352 | -0.521042429 | 0.25126737 | 0.535866411 |
| chr6:72819640-72819965 | -0.590938922 | 0.251277932 | 0.535874021 |
| chr14:43005910-43006131 | 0.569102933 | 0.25128626 | 0.535876867 |
| chr6:154603761-154603969 | 0.575916002 | 0.251304893 | 0.535901687 |
| chr2:61846298-61846628 | -0.560244965 | 0.251327612 | 0.535920336 |
| chr12:102095737-102095993 | -0.561587323 | 0.251327627 | 0.535920336 |
| chr11:118791639-118791977 | 0.422725368 | 0.251362488 | 0.535979755 |
| chr7:149381286-149381643 | 0.437244322 | 0.251372349 | 0.535985867 |
| chr1:100835120-100835330 | -0.587378121 | 0.251422408 | 0.536077687 |
| chr8:136665509-136665990 | -0.468885285 | 0.251447864 | 0.536117044 |
| chr13:112689400-112690233 | -0.39739153 | 0.251458917 | 0.536125693 |
| chr1:46611738-46612096 | -0.44484832 | 0.251490521 | 0.536177758 |
| chr13:99354644-99354847 | 0.580084156 | 0.251497333 | 0.536177758 |
| chr21:23725669-23725869 | -0.64131957 | 0.251505505 | 0.536180262 |
| chr2:200305872-200306101 | -0.536593647 | 0.251567071 | 0.536296593 |
| chr5:78995032-78995300 | -0.608498026 | 0.251617114 | 0.536388351 |
| chr14:88916278-88916478 | 0.636492277 | 0.251643633 | 0.536429959 |
| chr2:42528891-42529744 | -0.41452901 | 0.251657728 | 0.53644508 |
| chr3:64254889-64255120 | 0.408168603 | 0.251669392 | 0.536455021 |
| chr16:87868908-87869874 | -0.377477658 | 0.25174939 | 0.536597508 |
| chr9:84789236-84789983 | 0.414436025 | 0.251750244 | 0.536597508 |

| | | | |
|---|---|---|---|
| chr18:34330972-34331848 | -0.390607988 | 0.251808752 | 0.536707286 |
| chr3:194168315-194168575 | -0.563812587 | 0.251816135 | 0.536708091 |
| chr16:82016359-82016569 | -0.537938473 | 0.251831224 | 0.536708635 |
| chr7:90070295-90070504 | 0.599275092 | 0.251837531 | 0.536708635 |
| chr18:21167739-21168027 | -0.536468447 | 0.251840584 | 0.536708635 |
| chr6:98040230-98040458 | -0.610700213 | 0.251846395 | 0.536708635 |
| chr1:163361219-163361491 | -0.48619951 | 0.251851413 | 0.536708635 |
| chr19:16076110-16076650 | 0.420807875 | 0.251859586 | 0.536711126 |
| chr6:86936867-86938052 | -0.374419077 | 0.251880195 | 0.536728963 |
| chr18:6781162-6781727 | -0.46090312 | 0.251881967 | 0.536728963 |
| chr3:158727823-158728094 | -0.465290801 | 0.251901637 | 0.53675595 |
| chr1:41381202-41381469 | -0.561049101 | 0.251917226 | 0.536769201 |
| chr3:154089470-154090306 | 0.383150571 | 0.25193292 | 0.536769201 |
| chr3:129507475-129507823 | -0.441608388 | 0.251935059 | 0.536769201 |
| chr1:169215132-169215392 | -0.643178039 | 0.251935877 | 0.536769201 |
| chr2:60560718-60561084 | -0.441198734 | 0.251971017 | 0.536814008 |
| chr16:85422364-85423120 | 0.388351225 | 0.25197355 | 0.536814008 |
| chr1:21937309-21937792 | 0.433025907 | 0.251977926 | 0.536814008 |
| chr16:2857851-2859093 | 0.368492355 | 0.251989635 | 0.536824029 |
| chr11:65859396-65859785 | 0.39208455 | 0.252003545 | 0.536838735 |
| chr19:55346267-55346467 | 0.56128175 | 0.252013819 | 0.536845695 |
| chr11:24509232-24509432 | 0.606860661 | 0.252033846 | 0.536873432 |
| chr8:144126423-144126798 | 0.425861012 | 0.252044617 | 0.536881451 |
| chr19:45091525-45091935 | 0.408126263 | 0.252066078 | 0.536912238 |
| chr21:41506865-41507704 | -0.387001662 | 0.252075535 | 0.536917455 |
| chr21:10330783-10331033 | 0.567025339 | 0.25208941 | 0.536932084 |
| chr9:39809825-39810218 | -0.456158057 | 0.252099716 | 0.536939108 |
| chr2:226791185-226791467 | 0.463428519 | 0.252114956 | 0.536956643 |
| chr5:10601247-10602215 | -0.371154898 | 0.252159388 | 0.537036347 |
| chr2:20446738-20447802 | -0.364288257 | 0.252189042 | 0.537073503 |
| chr1:75233214-75233653 | -0.466104802 | 0.25219226 | 0.537073503 |
| chr8:90870629-90870829 | 0.661777077 | 0.252197863 | 0.537073503 |
| chr14:90981805-90982005 | 0.609652882 | 0.252218955 | 0.537103492 |
| chr2:117835950-117836253 | 0.465672892 | 0.252236774 | 0.537111964 |
| chr5:146203065-146204063 | -0.365555902 | 0.252236953 | 0.537111964 |
| chr3:25441236-25441578 | -0.496486403 | 0.25225411 | 0.537113264 |
| chr11:8910826-8911664 | 0.388728985 | 0.25225986 | 0.537113264 |
| chr2:3956905-3957105 | 0.591878017 | 0.252261019 | 0.537113264 |
| chr11:77073487-77073748 | 0.488598867 | 0.25226682 | 0.537113264 |
| chr18:74760426-74760821 | 0.487391764 | 0.252272613 | 0.537113264 |
| chr4:144399953-144400322 | -0.477415756 | 0.252289978 | 0.53713531 |
| chr2:20001416-20001616 | -0.652006691 | 0.25230722 | 0.537141904 |
| chr3:120561308-120561716 | -0.418127763 | 0.252310613 | 0.537141904 |
| chr6:54267830-54268103 | -0.540910875 | 0.252314106 | 0.537141904 |
| chr13:41919957-41920298 | 0.510319202 | 0.2523449 | 0.537192536 |
| chr20:48034874-48035277 | 0.378768611 | 0.252352957 | 0.537194761 |
| chr7:72879780-72880231 | 0.457894571 | 0.252365363 | 0.537206246 |
| chr20:44683641-44683841 | 0.531502854 | 0.252413414 | 0.537293605 |
| chr1:36258421-36258985 | 0.422164761 | 0.252431208 | 0.537311417 |
| chr1:40316555-40316966 | -0.468507915 | 0.252435807 | 0.537311417 |

| | | | |
|---|---|---|---|
| chr8:90691920-90692584 | -0.431192413 | 0.25245395 | 0.537335107 |
| chr1:204320447-204320701 | 0.470013743 | 0.252461923 | 0.537337152 |
| chr7:21407811-21408013 | 0.559299933 | 0.252469265 | 0.537337852 |
| chr3:123629708-123629977 | -0.48135376 | 0.252519011 | 0.537428799 |
| chr12:53658965-53659311 | 0.535056187 | 0.252546032 | 0.537471378 |
| chrX:135344491-135345067 | 0.373609479 | 0.252556661 | 0.537472499 |
| chr3:153903526-153903754 | 0.538485561 | 0.252561562 | 0.537472499 |
| chr14:38721837-38722127 | 0.451381839 | 0.252567602 | 0.537472499 |
| chr22:36977409-36977664 | -0.534104105 | 0.252607052 | 0.537527755 |
| chr14:22940973-22941250 | 0.487255309 | 0.252607599 | 0.537527755 |
| chr20:19073601-19073829 | -0.574578273 | 0.252625243 | 0.537537319 |
| chr14:56491307-56491675 | 0.426055679 | 0.252635294 | 0.537537319 |
| chr4:147866797-147867171 | 0.419586179 | 0.252639819 | 0.537537319 |
| chr15:90451475-90451799 | 0.474037244 | 0.252640155 | 0.537537319 |
| chr7:98291806-98292051 | 0.525037311 | 0.252669756 | 0.537584198 |
| chr10:47377533-47377791 | -0.529075913 | 0.25267622 | 0.537584198 |
| chr9:113381954-113382392 | 0.417121864 | 0.252701397 | 0.537622837 |
| chr1:218367081-218367416 | 0.45835425 | 0.252709717 | 0.537625609 |
| chr8:143899874-143900088 | 0.478966143 | 0.252720499 | 0.53763362 |
| chr7:108660508-108660740 | -0.578943924 | 0.252739809 | 0.537654702 |
| chr13:52846160-52846509 | -0.421367083 | 0.252744443 | 0.537654702 |
| chr2:85537814-85538035 | -0.458872807 | 0.252765502 | 0.537684572 |
| chr17:64773690-64774892 | -0.360632351 | 0.25277657 | 0.537693189 |
| chr2:154762703-154762966 | -0.506189631 | 0.252793658 | 0.537702218 |
| chr9:128852053-128852324 | 0.529135778 | 0.25279598 | 0.537702218 |
| chr13:30818803-30819179 | -0.495884281 | 0.252801868 | 0.537702218 |
| chr22:31346556-31347094 | 0.402673516 | 0.252820149 | 0.537726174 |
| chr16:70346660-70347246 | 0.395158548 | 0.252840459 | 0.537754445 |
| chr5:134738980-134739433 | -0.398215837 | 0.252847909 | 0.537755364 |
| chr5:129119826-129120033 | -0.524643776 | 0.252861942 | 0.537770282 |
| chr10:24911625-24911909 | -0.585543593 | 0.252889354 | 0.537813652 |
| chr10:1526391-1526785 | -0.443100896 | 0.252900604 | 0.537815302 |
| chr6:99868182-99868441 | -0.505281979 | 0.252904168 | 0.537815302 |
| chr3:100521495-100521878 | -0.415033591 | 0.252926021 | 0.537846847 |
| chr7:55365356-55366730 | -0.360529518 | 0.252943911 | 0.537869962 |
| chr1:247672587-247672796 | -0.575755811 | 0.252959981 | 0.537889207 |
| chr15:67064995-67066095 | 0.372566877 | 0.25298879 | 0.537925445 |
| chr3:112529150-112529413 | 0.504955344 | 0.252997868 | 0.537925445 |
| chr12:78097160-78097404 | -0.583691875 | 0.252998085 | 0.537925445 |
| chr6:125046017-125046557 | -0.429237685 | 0.253033457 | 0.537985725 |
| chr3:151145793-151146389 | -0.399439572 | 0.253042657 | 0.537990357 |
| chr5:11903348-11904996 | 0.36409973 | 0.253075572 | 0.538045407 |
| chr5:4206162-4206487 | -0.5174999 | 0.253110793 | 0.538105357 |
| chr13:24182264-24182986 | 0.429299016 | 0.253155674 | 0.538185841 |
| chr1:244801290-244801497 | -0.546229211 | 0.253171497 | 0.538193562 |
| chr5:131170560-131171378 | -0.363125244 | 0.253184029 | 0.538193562 |
| chr6:12113210-12113441 | -0.591259874 | 0.253187402 | 0.538193562 |
| chr7:35248781-35249000 | -0.591259874 | 0.253187402 | 0.538193562 |
| chr6:53171855-53172250 | -0.415744564 | 0.25319728 | 0.538199628 |
| chr6:1424347-1424683 | 0.466810317 | 0.253228835 | 0.538245528 |

| | | | |
|---|---|---|---|
| chr3:196978664-196979194 | 0.416392526 | 0.25323662 | 0.538245528 |
| chr9:111516067-111516406 | -0.52735177 | 0.253239948 | 0.538245528 |
| chr6:14117567-14118149 | 0.429784933 | 0.253257388 | 0.538257778 |
| chr22:28882768-28883644 | 0.369129337 | 0.253259761 | 0.538257778 |
| chr2:111050920-111051363 | -0.470902186 | 0.253274035 | 0.538273183 |
| chr15:56647641-56647909 | 0.595061084 | 0.253293439 | 0.538299492 |
| chr16:27640236-27640507 | -0.479789159 | 0.253342937 | 0.538383501 |
| chr15:89906039-89906258 | 0.509378033 | 0.253347022 | 0.538383501 |
| chr2:64607832-64608168 | -0.411113738 | 0.25341963 | 0.538522864 |
| chr5:98084378-98084614 | 0.61889111 | 0.253427038 | 0.53852367 |
| chr13:75687034-75687464 | -0.458724322 | 0.253448664 | 0.538547861 |
| chr5:163173014-163173360 | -0.518665354 | 0.253452479 | 0.538547861 |
| chr1:194117622-194118399 | -0.382809367 | 0.253489406 | 0.538607297 |
| chr1:201649995-201650318 | 0.418310145 | 0.25349451 | 0.538607297 |
| chr1:170513194-170513483 | -0.496844341 | 0.253523878 | 0.538654759 |
| chr6:52399806-52400006 | -0.559837528 | 0.253537763 | 0.538659522 |
| chr1:53363364-53363640 | -0.508749059 | 0.25354018 | 0.538659522 |
| chr7:1872071-1872488 | 0.44732624 | 0.253549925 | 0.53866529 |
| chr7:30419883-30420093 | 0.593380144 | 0.253562144 | 0.538667056 |
| chr4:8918026-8918332 | 0.454647227 | 0.253564817 | 0.538667056 |
| chr7:54739791-54740161 | -0.55977402 | 0.253578298 | 0.538680761 |
| chr10:102241542-102241869 | 0.424761466 | 0.253592915 | 0.538696877 |
| chr19:893006-893626 | -0.414987204 | 0.253611985 | 0.53870895 |
| chr2:62706021-62706364 | -0.405557852 | 0.253614397 | 0.53870895 |
| chr5:135020282-135020557 | -0.51090922 | 0.253619691 | 0.53870895 |
| chr11:73275364-73275691 | 0.440348754 | 0.25365389 | 0.538766656 |
| chr22:29310937-29311352 | -0.411371213 | 0.253666913 | 0.538779381 |
| chr1:74732612-74733543 | -0.368362782 | 0.253712882 | 0.538862082 |
| chr1:34786510-34786814 | 0.443663919 | 0.253733356 | 0.538875562 |
| chr1:195791420-195792160 | -0.426865413 | 0.253735283 | 0.538875562 |
| chr7:159008157-159008357 | -0.6144046 | 0.253740328 | 0.538875562 |
| chr11:107457431-107458345 | -0.38738425 | 0.253758667 | 0.538899573 |
| chrX:102154347-102154574 | -0.498733087 | 0.253778468 | 0.538913666 |
| chr6:33580015-33580491 | 0.3927501 | 0.25377937 | 0.538913666 |
| chr14:26444801-26445064 | -0.542335249 | 0.253801342 | 0.538945387 |
| chr6:132209404-132209673 | -0.506088442 | 0.253822194 | 0.538968589 |
| chr7:45917643-45917930 | -0.471756906 | 0.253826336 | 0.538968589 |
| chr1:35202146-35202418 | 0.47891745 | 0.253841112 | 0.538985026 |
| chr4:82984480-82984821 | -0.47104498 | 0.253868309 | 0.539027838 |
| chr3:127544283-127544577 | -0.432185308 | 0.253887605 | 0.539053871 |
| chr20:34390973-34391173 | -0.624860818 | 0.253949177 | 0.539162018 |
| chr9:122005667-122006005 | -0.525115148 | 0.253952615 | 0.539162018 |
| chr11:70630314-70630659 | 0.481356904 | 0.253986096 | 0.539209952 |
| chr11:10933787-10934117 | 0.434231521 | 0.253999271 | 0.539209952 |
| chr15:101288981-101289231 | 0.490459097 | 0.254000552 | 0.539209952 |
| chr8:135730174-135730529 | -0.437527588 | 0.254003341 | 0.539209952 |
| chr15:83207966-83208249 | 0.45491923 | 0.254053486 | 0.539301459 |
| chr1:46516919-46517622 | -0.385358215 | 0.25406983 | 0.539314082 |
| chr11:10792439-10792826 | 0.451807139 | 0.254074152 | 0.539314082 |
| chr4:176277648-176277930 | -0.464879208 | 0.254084533 | 0.539314082 |

| | | | |
|---|---|---|---|
| chr19:7829435-7830290 | 0.36727432 | 0.254087586 | 0.539314082 |
| chr12:81381888-81382531 | 0.39425134 | 0.254106441 | 0.539339161 |
| chr9:137695210-137695490 | 0.480904859 | 0.254120976 | 0.539342086 |
| chr12:8231794-8231994 | -0.589598974 | 0.254121897 | 0.539342086 |
| chr20:37527692-37528376 | 0.391736315 | 0.254131585 | 0.539347708 |
| chr11:105609327-105609579 | -0.517978393 | 0.254159573 | 0.539392167 |
| chr16:76265461-76265770 | 0.440422135 | 0.254166834 | 0.539392637 |
| chr13:32660216-32660556 | 0.52250051 | 0.254179369 | 0.539404298 |
| chr7:151689622-151689954 | -0.462602964 | 0.254191799 | 0.539415737 |
| chr4:104394457-104394815 | -0.508096826 | 0.254205819 | 0.539430548 |
| chr14:43041040-43041240 | 0.676411005 | 0.254214316 | 0.539430551 |
| chr4:164938036-164938236 | 0.543918729 | 0.254223163 | 0.539430551 |
| chr14:22813403-22813649 | -0.539626198 | 0.25422694 | 0.539430551 |
| chr21:33428671-33429099 | -0.45922064 | 0.254260276 | 0.539486344 |
| chr2:68156949-68158384 | -0.361063189 | 0.254311819 | 0.539580765 |
| chr1:184956921-184957186 | -0.574357242 | 0.25433576 | 0.539616619 |
| chr10:114043558-114044098 | 0.397777171 | 0.254363876 | 0.539645814 |
| chr3:36943995-36945228 | 0.377169996 | 0.254370196 | 0.539645814 |
| chr2:168494525-168494745 | -0.527317346 | 0.254370649 | 0.539645814 |
| chr10:72325225-72325664 | -0.420037005 | 0.254430499 | 0.539757842 |
| chr3:16724042-16724242 | 0.535643305 | 0.254444516 | 0.539769858 |
| chr3:148366797-148367059 | 0.479362334 | 0.254461542 | 0.539769858 |
| chr3:36124169-36124427 | -0.516164621 | 0.254466973 | 0.539769858 |
| chr8:17496756-17497169 | -0.430509201 | 0.254470337 | 0.539769858 |
| chr5:73135444-73135736 | -0.496178991 | 0.254471386 | 0.539769858 |
| chr4:149795264-149795526 | 0.553247909 | 0.254482234 | 0.539777924 |
| chrX:129523291-129523555 | 0.526486533 | 0.254527785 | 0.539859597 |
| chr3:52705467-52706464 | 0.383105787 | 0.25456002 | 0.539913022 |
| chr4:118835639-118836344 | 0.370633242 | 0.254586897 | 0.539955081 |
| chrX:96372465-96373101 | 0.416573517 | 0.254596556 | 0.53996062 |
| chr1:150425936-150426136 | 0.524039524 | 0.254608325 | 0.539967633 |
| chr19:47863530-47863730 | 0.59017049 | 0.254613957 | 0.539967633 |
| chr20:47759023-47759232 | 0.405732643 | 0.254644334 | 0.540017109 |
| chr18:57706282-57706482 | 0.586610112 | 0.254661609 | 0.540038796 |
| chr2:208112675-208113359 | -0.410070582 | 0.25468432 | 0.540059875 |
| chr8:52340236-52340528 | -0.500923381 | 0.25468918 | 0.540059875 |
| chr2:79860626-79860876 | 0.4826625 | 0.254692694 | 0.540059875 |
| chr8:121639179-121639632 | -0.411983028 | 0.254700253 | 0.540060957 |
| chr17:17496854-17497193 | 0.419504501 | 0.254728092 | 0.54010504 |
| chr2:203720927-203721129 | -0.60967054 | 0.254741264 | 0.540116003 |
| chr21:41681319-41681701 | -0.399050875 | 0.254752065 | 0.540116003 |
| chr17:73783128-73783360 | -0.536212529 | 0.254754409 | 0.540116003 |
| chr13:98445323-98445694 | 0.423822436 | 0.254769493 | 0.540133036 |
| chr5:107608065-107608379 | -0.560534167 | 0.254783281 | 0.540147322 |
| chr17:45019336-45019679 | 0.453249466 | 0.254806069 | 0.540180688 |
| chr8:96180134-96180540 | 0.409187142 | 0.254844175 | 0.540246524 |
| chr13:85806614-85806850 | -0.556456856 | 0.254875338 | 0.540276917 |
| chr14:93023162-93023445 | 0.456976797 | 0.254876092 | 0.540276917 |
| chr13:38615162-38615381 | -0.647545008 | 0.254881782 | 0.540276917 |
| chr10:5867622-5868073 | 0.454262406 | 0.254892678 | 0.540276917 |

| | | | |
|---|---|---|---|
| chr6:108384068-108384349 | 0.552904298 | 0.254893768 | 0.540276917 |
| chr12:76037155-76037743 | -0.398555097 | 0.25491356 | 0.540303921 |
| chr11:1998409-1998739 | 0.466161062 | 0.254922812 | 0.540305909 |
| chr21:32258406-32258683 | 0.505879314 | 0.2549286 | 0.540305909 |
| chr5:126778173-126778657 | -0.376027679 | 0.254959205 | 0.540355827 |
| chr6:44221583-44221859 | 0.436586295 | 0.254969815 | 0.540363366 |
| chrX:128020684-128021202 | 0.388085724 | 0.254993651 | 0.540398935 |
| chr2:45610446-45610993 | -0.388816881 | 0.2550117 | 0.540422239 |
| chr9:33816637-33818041 | -0.36619624 | 0.255023041 | 0.540431325 |
| chr2:71001072-71001766 | -0.394464889 | 0.255072671 | 0.540521549 |
| chr12:79681282-79681534 | 0.520048099 | 0.255088012 | 0.540539109 |
| chr5:128082765-128084102 | -0.362677169 | 0.255102886 | 0.540555678 |
| chr7:32691675-32692264 | -0.402604507 | 0.255124466 | 0.540582499 |
| chr15:95196573-95196803 | -0.585823921 | 0.255129654 | 0.540582499 |
| chr16:87465389-87465636 | -0.477350597 | 0.25515667 | 0.540624793 |
| chr6:130308012-130308212 | 0.553763242 | 0.255167451 | 0.540632686 |
| chr3:15636309-15636566 | -0.534222273 | 0.255188093 | 0.540661471 |
| chr11:68708307-68708518 | 0.589839749 | 0.255220063 | 0.540714252 |
| chr11:70240468-70240741 | -0.539068225 | 0.255249282 | 0.540737626 |
| chr17:7797861-7798195 | -0.560232485 | 0.255251662 | 0.540737626 |
| chr13:36724226-36724498 | -0.520710167 | 0.255252267 | 0.540737626 |
| chr14:99962224-99962600 | -0.458642694 | 0.25526873 | 0.540757552 |
| chr12:119179304-119179907 | 0.400718467 | 0.255277894 | 0.540762015 |
| chrX:107766593-107767044 | -0.427337249 | 0.255343782 | 0.540886632 |
| chr5:155616824-155617088 | -0.495168998 | 0.255379249 | 0.540945018 |
| chr12:49622783-49623867 | 0.370123579 | 0.255385465 | 0.540945018 |
| chr20:63956275-63956858 | -0.380767297 | 0.255406204 | 0.540973992 |
| chr3:179929164-179929619 | -0.526156712 | 0.255418385 | 0.540978761 |
| chr12:104054815-104055074 | 0.563536402 | 0.255422576 | 0.540978761 |
| chr8:21915414-21915614 | -0.557282012 | 0.255479089 | 0.541083498 |
| chr16:30378092-30379118 | 0.36585377 | 0.255499811 | 0.541084474 |
| chr15:59372334-59373153 | 0.375095631 | 0.255503003 | 0.541084474 |
| chr8:95357663-95357872 | 0.525399013 | 0.255515564 | 0.541084474 |
| chr5:94693074-94693355 | -0.540371996 | 0.255518016 | 0.541084474 |
| chr15:93245270-93245533 | 0.578148958 | 0.255526845 | 0.541084474 |
| chr8:37841487-37841731 | 0.497515625 | 0.255528577 | 0.541084474 |
| chr2:12580052-12580268 | -0.61200558 | 0.255536044 | 0.541084474 |
| chr17:70551640-70551840 | -0.61200558 | 0.255536044 | 0.541084474 |
| chr5:114055276-114055654 | -0.452743505 | 0.255544251 | 0.541086898 |
| chr17:71161149-71161648 | -0.468526452 | 0.255554617 | 0.541093895 |
| chr3:148953161-148953393 | -0.518863633 | 0.255620209 | 0.541217819 |
| chrX:115560434-115560733 | -0.471873564 | 0.25565607 | 0.54127879 |
| chr10:68720630-68721768 | -0.365600627 | 0.255663668 | 0.541278879 |
| chr3:155744078-155745474 | -0.360401957 | 0.255670241 | 0.541278879 |
| chr2:29921519-29921897 | 0.418908952 | 0.255715126 | 0.541358948 |
| chr3:15148585-15148990 | -0.414599468 | 0.255727492 | 0.54137017 |
| chr22:17158506-17160174 | 0.362688486 | 0.255745902 | 0.541394184 |
| chr6:33200540-33200947 | 0.392483384 | 0.255758003 | 0.541404843 |
| chr1:169539047-169539383 | -0.454912406 | 0.255783807 | 0.541440585 |
| chr6:154234229-154234480 | 0.596927897 | 0.25578902 | 0.541440585 |

| | | | |
|---|---|---|---|
| chr15:74433858-74434073 | 0.498741032 | 0.255806613 | 0.541462867 |
| chr14:90758144-90758928 | 0.369654632 | 0.255832556 | 0.541500769 |
| chr1:200638722-200638967 | 0.497082547 | 0.255842749 | 0.541500769 |
| chr22:30396790-30397304 | 0.416773566 | 0.255846037 | 0.541500769 |
| chr3:155533697-155533897 | 0.620047216 | 0.255852788 | 0.541500769 |
| chr7:143321720-143322059 | -0.42583881 | 0.255893057 | 0.541571037 |
| chr14:34648797-34649119 | 0.497415252 | 0.255904131 | 0.541574637 |
| chr12:112524695-112524895 | 0.580487005 | 0.255913544 | 0.541574637 |
| chr20:51899762-51899976 | 0.46761536 | 0.25592361 | 0.541574637 |
| chr20:56629014-56629434 | 0.36976844 | 0.25592654 | 0.541574637 |
| chr11:86261390-86261826 | -0.437448639 | 0.255930099 | 0.541574637 |
| chr2:118227818-118228120 | -0.519987033 | 0.255966195 | 0.541636061 |
| chr11:117535496-117535763 | 0.479166688 | 0.255987138 | 0.541665419 |
| chr9:121074336-121075594 | 0.367118734 | 0.256037369 | 0.541755444 |
| chr6:148764772-148764972 | -0.642260174 | 0.256043824 | 0.541755444 |
| chr1:183318585-183318785 | 0.508370685 | 0.256052273 | 0.54175836 |
| chr5:170273872-170274284 | -0.405098577 | 0.256062189 | 0.54176438 |
| chr4:8306766-8307071 | -0.497454587 | 0.25607522 | 0.541776991 |
| chr14:51651153-51652685 | 0.359168435 | 0.256087813 | 0.541787953 |
| chr3:69079788-69080591 | 0.376561384 | 0.256094543 | 0.541787953 |
| chr2:38095839-38096676 | -0.365686672 | 0.256102956 | 0.541790792 |
| chr8:143407775-143408400 | 0.38219914 | 0.256122058 | 0.541812572 |
| chr17:50957135-50957490 | -0.445000077 | 0.256127394 | 0.541812572 |
| chr4:180059672-180059874 | -0.504911611 | 0.256134812 | 0.541813306 |
| chr12:48296646-48296880 | 0.552495721 | 0.256150361 | 0.541831238 |
| chr5:128682322-128682785 | 0.450524297 | 0.256203173 | 0.54192799 |
| chr16:10918765-10919065 | 0.443646488 | 0.256225101 | 0.541959412 |
| chr2:42567920-42568343 | -0.425009604 | 0.25624019 | 0.541976367 |
| chr17:38349178-38349446 | -0.493127313 | 0.256253424 | 0.541989395 |
| chr13:101785919-101786211 | -0.534400347 | 0.25626293 | 0.541994541 |
| chr2:74867932-74868470 | 0.38494472 | 0.256272469 | 0.541999754 |
| chr1:93846216-93846857 | -0.388832705 | 0.256293338 | 0.54202893 |
| chr2:27303498-27303702 | 0.489785023 | 0.256300828 | 0.542029809 |
| chrX:123833344-123833760 | 0.409649036 | 0.256326318 | 0.542068755 |
| chr12:19273870-19274281 | -0.448376366 | 0.256333965 | 0.542069965 |
| chr2:207624570-207626302 | -0.357442649 | 0.256341186 | 0.542070275 |
| chr5:122101731-122101986 | -0.515640565 | 0.256390138 | 0.542145505 |
| chr16:50397186-50397534 | -0.458174027 | 0.256390913 | 0.542145505 |
| chr8:53880531-53881068 | 0.420969387 | 0.256418582 | 0.542189049 |
| chr3:8663259-8663823 | 0.400504771 | 0.256440275 | 0.542217492 |
| chr12:94725823-94726100 | -0.523602437 | 0.256446186 | 0.542217492 |
| chr6:13378885-13379252 | -0.43263856 | 0.256460798 | 0.542233424 |
| chr12:6215935-6216526 | -0.400782251 | 0.256522531 | 0.54234898 |
| chr5:52657306-52657585 | -0.604679896 | 0.25653442 | 0.542359149 |
| chr7:50696809-50697019 | -0.564800187 | 0.256543002 | 0.54236233 |
| chr1:196516400-196516873 | -0.437953951 | 0.256577042 | 0.542419327 |
| chr20:43749761-43750111 | 0.444829792 | 0.256598772 | 0.542450299 |
| chr2:21363400-21363627 | -0.566564684 | 0.256632381 | 0.54250638 |
| chr14:20351024-20351224 | 0.554296081 | 0.256646361 | 0.542520966 |
| chr17:78351512-78352085 | -0.418344433 | 0.256654573 | 0.542523359 |

| | | | |
|---|---|---|---|
| chr5:4643261-4643461 | -0.607208608 | 0.256680923 | 0.54256409 |
| chr6:73585290-73585877 | -0.392415331 | 0.256705982 | 0.54260209 |
| chr19:37317189-37318114 | 0.375275099 | 0.256714492 | 0.542605109 |
| chr5:108642231-108642630 | -0.423505612 | 0.25674031 | 0.54264471 |
| chr1:224503114-224503946 | -0.396174545 | 0.256757788 | 0.542661707 |
| chr19:2977176-2977938 | 0.393204189 | 0.256771543 | 0.542661707 |
| chr12:74508856-74509640 | -0.390889851 | 0.256776512 | 0.542661707 |
| chr19:35643394-35643995 | 0.379961194 | 0.256776681 | 0.542661707 |
| chr2:45259153-45259403 | 0.48546067 | 0.256791131 | 0.54266735 |
| chr5:138610401-138610646 | 0.462387141 | 0.256794709 | 0.54266735 |
| chr11:12964032-12964428 | 0.438497366 | 0.256800598 | 0.54266735 |
| chr1:1574340-1575243 | 0.393970839 | 0.256811152 | 0.542671047 |
| chr1:117967487-117967780 | -0.545908037 | 0.256816513 | 0.542671047 |
| chr8:14860250-14860473 | -0.578066932 | 0.256834016 | 0.542693067 |
| chr13:30690349-30691085 | -0.400402019 | 0.256869297 | 0.542745529 |
| chr15:81265860-81266137 | -0.446461786 | 0.256875633 | 0.542745529 |
| chr1:154607430-154608401 | -0.381575103 | 0.256880095 | 0.542745529 |
| chr21:42258591-42258887 | -0.442762724 | 0.25690403 | 0.542781133 |
| chr2:42089358-42089662 | -0.467169594 | 0.256934588 | 0.54282082 |
| chr1:31684048-31684306 | 0.47445628 | 0.256936983 | 0.54282082 |
| chr5:172833692-172833953 | 0.463198642 | 0.256992418 | 0.542922967 |
| chr15:47196078-47196314 | -0.545687618 | 0.257032936 | 0.542993592 |
| chr2:165953257-165954371 | -0.359414468 | 0.257058036 | 0.543020512 |
| chr21:26843460-26843858 | -0.426975287 | 0.257072139 | 0.543020512 |
| chr17:82297884-82298578 | 0.380326331 | 0.25707321 | 0.543020512 |
| chr10:97320162-97320418 | 0.460279058 | 0.257078937 | 0.543020512 |
| chr9:7094132-7094332 | 0.599248054 | 0.257086345 | 0.543020512 |
| chr11:128802690-128802916 | -0.568129578 | 0.257095287 | 0.543020512 |
| chr12:80966083-80966328 | -0.568129578 | 0.257095287 | 0.543020512 |
| chr22:19217979-19218218 | -0.551485387 | 0.257112306 | 0.543041489 |
| chr5:134026541-134026908 | -0.421932282 | 0.257131864 | 0.543067827 |
| chr12:2958785-2959411 | 0.397133389 | 0.257156483 | 0.543103096 |
| chr3:160444480-160444820 | -0.38841559 | 0.257162739 | 0.543103096 |
| chr13:45011876-45012144 | 0.578950862 | 0.25717843 | 0.543106631 |
| chrX:52574672-52575002 | 0.455048189 | 0.25717859 | 0.543106631 |
| chr6:85865809-85866295 | -0.491349306 | 0.257195299 | 0.543126948 |
| chr1:154608620-154608907 | 0.476906212 | 0.257208542 | 0.543139945 |
| chr1:54481595-54481850 | -0.626404458 | 0.257242158 | 0.543195962 |
| chr10:44386572-44386876 | -0.494970551 | 0.257253142 | 0.543202871 |
| chr1:117210659-117211144 | 0.39689829 | 0.257263935 | 0.543202871 |
| chr14:100068490-100068825 | -0.489314291 | 0.257266699 | 0.543202871 |
| chr4:13544337-13544608 | 0.489727329 | 0.257279552 | 0.543215042 |
| chr2:216404083-216404289 | -0.567882645 | 0.257322037 | 0.543289772 |
| chr20:41125235-41125462 | 0.499386661 | 0.257331088 | 0.543293912 |
| chr10:43355599-43356442 | -0.379303114 | 0.257347223 | 0.543313007 |
| chr13:114339786-114340046 | -0.536644724 | 0.257361627 | 0.543327191 |
| chr16:66092449-66092659 | 0.54763849 | 0.257372372 | 0.543327191 |
| chr8:108887664-108888190 | -0.405846169 | 0.257375215 | 0.543327191 |
| chr6:151493405-151494945 | -0.361875445 | 0.25739028 | 0.543344023 |
| chr3:126679260-126679729 | -0.41059668 | 0.257402572 | 0.543355002 |

| | | | |
|---|---|---|---|
| chr2:5742070-5742513 | 0.450863772 | 0.257424058 | 0.543385387 |
| chr15:82750832-82751114 | 0.443590876 | 0.257440127 | 0.543404336 |
| chr2:86298661-86298933 | 0.519731148 | 0.257466958 | 0.543446 |
| chr14:92917021-92917747 | -0.393914259 | 0.257482883 | 0.543446693 |
| chrX:30888797-30889849 | -0.359324183 | 0.257488719 | 0.543446693 |
| chr6:107952577-107952902 | 0.453982897 | 0.257491482 | 0.543446693 |
| chr3:4821394-4821816 | 0.442164701 | 0.257495657 | 0.543446693 |
| chr4:48963126-48963446 | 0.438869135 | 0.257512709 | 0.543467714 |
| chr1:247343909-247344152 | -0.520851319 | 0.257534265 | 0.543498236 |
| chr10:124152276-124153767 | -0.366825693 | 0.257566746 | 0.543550432 |
| chr6:21587105-21588726 | -0.365260749 | 0.257585297 | 0.543550432 |
| chr1:191702305-191702516 | 0.559191867 | 0.257603979 | 0.543550432 |
| chr1:111570760-111570998 | -0.597038388 | 0.257604848 | 0.543550432 |
| chr14:65356281-65356854 | 0.382570627 | 0.257604953 | 0.543550432 |
| chr5:157965088-157965351 | 0.534737371 | 0.257605258 | 0.543550432 |
| chr17:60792271-60792790 | 0.372826324 | 0.257608655 | 0.543550432 |
| chr5:62403373-62404611 | -0.359338095 | 0.257636589 | 0.543570334 |
| chr7:54546464-54547155 | -0.39084869 | 0.257637413 | 0.543570334 |
| chr13:67438615-67438815 | -0.60930996 | 0.257645283 | 0.543570334 |
| chr6:122947372-122947745 | 0.4541262 | 0.257646464 | 0.543570334 |
| chr20:347566-348361 | 0.382790274 | 0.257682447 | 0.543629283 |
| chr7:67378237-67378508 | 0.478080038 | 0.257688595 | 0.543629283 |
| chrX:18104712-18104959 | 0.521201942 | 0.257710755 | 0.543634666 |
| chr16:78040630-78040869 | 0.527964925 | 0.257724884 | 0.543634666 |
| chr22:24146368-24146583 | -0.516630264 | 0.257728547 | 0.543634666 |
| chr3:11582282-11582533 | -0.516630264 | 0.257728547 | 0.543634666 |
| chr2:31333587-31333942 | -0.433752177 | 0.257730281 | 0.543634666 |
| chr8:127715326-127715576 | 0.489518413 | 0.257733717 | 0.543634666 |
| chr1:211260216-211260513 | 0.428453602 | 0.257763833 | 0.54368113 |
| chr10:86366242-86366674 | 0.417732458 | 0.257769937 | 0.54368113 |
| chr5:55823074-55823526 | -0.487125757 | 0.257791819 | 0.543699146 |
| chr2:118763768-118763976 | -0.548424291 | 0.257799248 | 0.543699146 |
| chr19:55207145-55207598 | 0.444611953 | 0.257799766 | 0.543699146 |
| chr14:50061062-50061339 | 0.495016803 | 0.257826133 | 0.543732943 |
| chr15:100879151-100879896 | 0.396457891 | 0.257829984 | 0.543732943 |
| chr6:27668527-27668797 | -0.46082226 | 0.257876417 | 0.543815898 |
| chr17:8158774-8159005 | -0.46696418 | 0.257908431 | 0.543868441 |
| chr1:161389660-161390736 | 0.361017644 | 0.257936444 | 0.543907592 |
| chr9:130578838-130579702 | 0.380287096 | 0.257943571 | 0.543907592 |
| chr20:1118339-1118779 | -0.415715224 | 0.257948293 | 0.543907592 |
| chr19:7926010-7926234 | 0.498003905 | 0.257955408 | 0.543907628 |
| chr17:62964458-62966545 | 0.352067265 | 0.258018337 | 0.544025033 |
| chr14:104876178-104876676 | 0.390249297 | 0.25802529 | 0.544025033 |
| chr1:106682078-106682405 | 0.394233726 | 0.258045089 | 0.544046279 |
| chr19:39792975-39793450 | 0.427334422 | 0.258049567 | 0.544046279 |
| chr7:146115596-146117026 | -0.367535513 | 0.25807265 | 0.544066925 |
| chr6:14280786-14281080 | -0.464811501 | 0.258073561 | 0.544066925 |
| chr12:116955714-116956019 | 0.453896557 | 0.258097822 | 0.544103102 |
| chr14:67674097-67675290 | 0.362950111 | 0.2581173 | 0.544120952 |
| chr14:97499652-97500529 | 0.389771548 | 0.258120492 | 0.544120952 |

| | | | |
|---|---|---|---|
| chr9:112366630-112367031 | -0.457726758 | 0.258157881 | 0.544184796 |
| chr9:113591638-113591953 | 0.428161471 | 0.25817879 | 0.5442139 |
| chr20:53712738-53712990 | 0.376406448 | 0.258199081 | 0.544236915 |
| chr5:36751939-36752169 | -0.509907435 | 0.258203914 | 0.544236915 |
| chr6:89021746-89022071 | 0.468105801 | 0.258211311 | 0.544237534 |
| chr11:45146963-45147682 | 0.39790229 | 0.258245705 | 0.544283996 |
| chr1:234111417-234111763 | 0.458401382 | 0.258252134 | 0.544283996 |
| chr12:119876700-119877799 | 0.367226128 | 0.258273624 | 0.544283996 |
| chr2:5291164-5291461 | -0.511986101 | 0.258276719 | 0.544283996 |
| chr10:116783321-116783530 | -0.567604384 | 0.258277066 | 0.544283996 |
| chr16:46688548-46689862 | 0.359160389 | 0.258289741 | 0.544283996 |
| chr2:90069348-90069643 | -0.463997111 | 0.258292695 | 0.544283996 |
| chr9:121663569-121663944 | 0.507486996 | 0.258295198 | 0.544283996 |
| chr3:21098796-21098996 | -0.677602351 | 0.258297286 | 0.544283996 |
| chr16:82344081-82344495 | -0.43750696 | 0.258305166 | 0.544285632 |
| chr1:219215447-219215858 | -0.448369786 | 0.25832384 | 0.544298372 |
| chrX:103250967-103251330 | 0.458467701 | 0.25832542 | 0.544298372 |
| chr2:173210568-173210866 | 0.477762156 | 0.258335005 | 0.544300343 |
| chr3:48989689-48990268 | 0.413840192 | 0.258340562 | 0.544300343 |
| chr15:89087857-89088631 | 0.362227706 | 0.258361878 | 0.544330285 |
| chr1:159798831-159799288 | -0.398184969 | 0.258386762 | 0.544367743 |
| chr2:212940852-212941091 | -0.548205964 | 0.258405396 | 0.544392032 |
| chr8:142763242-142763510 | 0.476344934 | 0.258435861 | 0.544441245 |
| chr10:72297571-72298462 | -0.372238349 | 0.258449837 | 0.544455719 |
| chr17:25788501-25788757 | 0.534139641 | 0.258458181 | 0.544456239 |
| chr8:127760791-127761067 | 0.45663488 | 0.258464295 | 0.544456239 |
| chr7:5592629-5593203 | -0.393686418 | 0.258507465 | 0.544532205 |
| chr19:43475206-43475496 | 0.482300933 | 0.25855121 | 0.544585691 |
| chr2:45263074-45263525 | 0.420069047 | 0.258561452 | 0.544585691 |
| chr4:100504578-100504914 | -0.517327967 | 0.258562643 | 0.544585691 |
| chr11:112225988-112226967 | 0.374948463 | 0.258562782 | 0.544585691 |
| chr16:47144328-47144658 | 0.459820522 | 0.258568394 | 0.544585691 |
| chr19:44259707-44260143 | 0.406397278 | 0.258587462 | 0.544610676 |
| chr10:34753079-34753299 | -0.553277513 | 0.258594472 | 0.544610676 |
| chr1:41241685-41242303 | -0.428463148 | 0.258620002 | 0.544613078 |
| chr11:82627797-82628143 | -0.494479674 | 0.258636516 | 0.544613078 |
| chr2:79111400-79111838 | -0.47538712 | 0.258636617 | 0.544613078 |
| chr20:31260365-31260666 | 0.49764857 | 0.258637316 | 0.544613078 |
| chr17:17681836-17682265 | 0.41588151 | 0.258637356 | 0.544613078 |
| chr5:34001442-34001642 | 0.558035884 | 0.258638259 | 0.544613078 |
| chr9:128818161-128818707 | 0.378049553 | 0.258673876 | 0.544667655 |
| chr1:7953903-7954612 | 0.397209496 | 0.258678395 | 0.544667655 |
| chr6:41941347-41942304 | 0.374810371 | 0.258689935 | 0.544676986 |
| chr12:116406435-116406956 | -0.393258979 | 0.258701595 | 0.54468106 |
| chr2:181742845-181743144 | -0.503266247 | 0.258718859 | 0.54468106 |
| chr11:124837544-124837783 | -0.486887806 | 0.258725833 | 0.54468106 |
| chr13:46160567-46160810 | -0.486887806 | 0.258725833 | 0.54468106 |
| chr6:135932597-135933139 | -0.414801239 | 0.258737543 | 0.54468106 |
| chr8:94079571-94080130 | 0.378080525 | 0.258740586 | 0.54468106 |
| chr15:34215876-34216076 | 0.571310746 | 0.258745228 | 0.54468106 |

| | | | |
|---|---|---|---|
| chr21:21238971-21239419 | 0.42804065 | 0.258748739 | 0.54468106 |
| chr14:56482341-56482756 | -0.47067094 | 0.258794285 | 0.54475395 |
| chr3:97204746-97204983 | -0.589553809 | 0.258799024 | 0.54475395 |
| chr2:136116847-136117054 | 0.479444101 | 0.258810744 | 0.54475395 |
| chr10:30751890-30752174 | -0.449283161 | 0.258812647 | 0.54475395 |
| chrX:36860798-36861485 | -0.418214679 | 0.258818914 | 0.54475395 |
| chr3:48598350-48598884 | -0.433831244 | 0.258860289 | 0.544826069 |
| chr6:119129912-119130501 | -0.398239442 | 0.258873635 | 0.544839193 |
| chr7:150628146-150628532 | -0.536668122 | 0.258883888 | 0.544842821 |
| chr1:197693026-197693296 | -0.472841741 | 0.258890952 | 0.544842821 |
| chr7:122847954-122848171 | -0.622315302 | 0.258906909 | 0.544842821 |
| chr4:169270163-169271787 | -0.358583325 | 0.25892231 | 0.544842821 |
| chr5:35089785-35090031 | -0.486697921 | 0.258930223 | 0.544842821 |
| chr1:150619891-150620204 | 0.418174351 | 0.258933127 | 0.544842821 |
| chr3:72318557-72318878 | 0.456913798 | 0.258941986 | 0.544842821 |
| chr6:106913622-106914139 | 0.416617219 | 0.258949542 | 0.544842821 |
| chr3:108402693-108402912 | 0.592944098 | 0.258958125 | 0.544842821 |
| chr10:92240525-92240835 | 0.447888208 | 0.258958387 | 0.544842821 |
| chr12:47307628-47308027 | -0.480546286 | 0.258964675 | 0.544842821 |
| chr8:140633986-140634540 | 0.389306734 | 0.258967197 | 0.544842821 |
| chr5:141100353-141100576 | 0.556444708 | 0.258967799 | 0.544842821 |
| chr13:46086359-46087062 | 0.408387545 | 0.259019584 | 0.544936807 |
| chr6:7008459-7009007 | 0.401839239 | 0.259046176 | 0.544964434 |
| chr12:110449751-110450618 | -0.381281474 | 0.25904694 | 0.544964434 |
| chr10:121887086-121887357 | -0.478002986 | 0.259057211 | 0.544971079 |
| chr3:104611323-104611553 | 0.510343378 | 0.259067289 | 0.544975323 |
| chr10:13536563-13537311 | -0.398857615 | 0.259073454 | 0.544975323 |
| chr16:29961569-29962362 | 0.374440681 | 0.259089941 | 0.544995044 |
| chr7:31663027-31663456 | -0.4134692 | 0.259142128 | 0.545089854 |
| chr5:115178468-115178727 | -0.568651234 | 0.259163745 | 0.545117358 |
| chr11:65857824-65858456 | 0.372540679 | 0.259169432 | 0.545117358 |
| chr7:33808257-33808497 | -0.523568076 | 0.25925443 | 0.545274186 |
| chr11:17384021-17384237 | 0.456039086 | 0.259260511 | 0.545274186 |
| chr10:96129657-96130878 | -0.365282916 | 0.259265343 | 0.545274186 |
| chr9:38008247-38008695 | -0.437408273 | 0.259276162 | 0.545281586 |
| chr12:108302420-108302831 | -0.438479349 | 0.259283095 | 0.545281586 |
| chr7:11298264-11298469 | 0.576557299 | 0.259295003 | 0.545285312 |
| chr5:126106180-126106380 | -0.595948435 | 0.2592991 | 0.545285312 |
| chr10:83677905-83678125 | -0.603932276 | 0.259320258 | 0.545314839 |
| chr17:28540714-28541027 | -0.442878895 | 0.259344621 | 0.545351104 |
| chr3:170307967-170308302 | 0.431812237 | 0.259383576 | 0.545418051 |
| chr8:91612596-91612939 | 0.428500187 | 0.259409687 | 0.545457987 |
| chr19:44844725-44845188 | 0.388827114 | 0.259428697 | 0.54546599 |
| chr3:32004213-32004509 | 0.483732856 | 0.259434786 | 0.54546599 |
| chr19:49225975-49226289 | -0.466685366 | 0.259441505 | 0.54546599 |
| chr14:39467883-39468497 | 0.393655691 | 0.259441969 | 0.54546599 |
| chr11:59550258-59551506 | -0.355061068 | 0.259452718 | 0.545473623 |
| chr12:32755096-32755644 | -0.38909329 | 0.259484895 | 0.545526303 |
| chr14:54901837-54903495 | 0.355495717 | 0.259502035 | 0.545547369 |
| chr5:111550316-111551062 | -0.404964689 | 0.259555188 | 0.545644139 |

| | | | |
|---|---|---|---|
| chr5:112289546-112289750 | -0.574787457 | 0.259562337 | 0.545644198 |
| chr16:88534104-88534457 | 0.425661497 | 0.259573886 | 0.545653506 |
| chr17:42773246-42773979 | 0.403890843 | 0.259594454 | 0.545681771 |
| chr6:36196126-36196816 | 0.395553691 | 0.259611826 | 0.545695226 |
| chr1:151196377-151196618 | 0.400939176 | 0.259615099 | 0.545695226 |
| chr13:49821465-49821764 | -0.475111399 | 0.259628093 | 0.545695666 |
| chr8:143018802-143019352 | 0.387919595 | 0.259633498 | 0.545695666 |
| chr5:128897622-128897903 | -0.613645475 | 0.259636674 | 0.545695666 |
| chr4:102705950-102706150 | -0.617269129 | 0.259653601 | 0.545716274 |
| chr12:78788839-78789127 | -0.615210155 | 0.259667999 | 0.545731565 |
| chr2:227163897-227164691 | 0.395172505 | 0.259681914 | 0.545741102 |
| chr2:31234457-31234657 | 0.501303652 | 0.259692102 | 0.545741102 |
| chr13:95300591-95301107 | 0.423315102 | 0.259693904 | 0.545741102 |
| chr8:63395155-63395461 | 0.477349104 | 0.259710727 | 0.545761486 |
| chr8:19312854-19314076 | 0.378748496 | 0.259720423 | 0.545766894 |
| chr7:92880255-92880536 | -0.542716035 | 0.259732942 | 0.545778233 |
| chr8:11413639-11414057 | -0.443864216 | 0.259743438 | 0.545780376 |
| chr20:60137732-60137999 | 0.475200473 | 0.259748208 | 0.545780376 |
| chr6:43951097-43951297 | 0.563416102 | 0.259783243 | 0.545839023 |
| chr8:98116721-98117864 | 0.356171108 | 0.259799516 | 0.545858246 |
| chr8:30082151-30083469 | -0.36236256 | 0.259843017 | 0.545908136 |
| chr5:73215651-73216665 | -0.377268311 | 0.25984449 | 0.545908136 |
| chr18:26185389-26185643 | -0.489580538 | 0.259844635 | 0.545908136 |
| chr6:121663538-121663795 | -0.591421916 | 0.259885177 | 0.545978341 |
| chr17:64472635-64473055 | -0.464968215 | 0.259903724 | 0.545994446 |
| chr19:41427213-41428381 | 0.369569385 | 0.259907095 | 0.545994446 |
| chr10:58268597-58269611 | -0.356289056 | 0.259919372 | 0.546005267 |
| chr3:97719162-97719501 | -0.442692475 | 0.259952097 | 0.546059041 |
| chr19:55080565-55080891 | 0.417981062 | 0.259977384 | 0.546097188 |
| chr19:4567702-4568074 | 0.494388641 | 0.259995239 | 0.546119721 |
| chr12:61546325-61546658 | 0.446445643 | 0.260002942 | 0.546120931 |
| chr11:14375025-14375511 | -0.459275308 | 0.26001632 | 0.546134059 |
| chr2:231961288-231962182 | 0.395570408 | 0.260042174 | 0.546167947 |
| chr11:61967191-61968074 | 0.370855475 | 0.260046711 | 0.546167947 |
| chr4:147429752-147429970 | -0.575593469 | 0.260060995 | 0.546182978 |
| chr4:169976483-169976731 | -0.569556227 | 0.260076589 | 0.546199102 |
| chr1:58754094-58754294 | -0.571932768 | 0.260088952 | 0.546199102 |
| chr4:6595795-6596301 | 0.401284678 | 0.260090058 | 0.546199102 |
| chr2:219206139-219207256 | -0.361936133 | 0.260152419 | 0.546315089 |
| chr1:2391453-2391935 | 0.381490893 | 0.260187495 | 0.546355578 |
| chr4:160026765-160027067 | -0.485614661 | 0.260196895 | 0.546355578 |
| chr1:32208879-32209336 | -0.399862035 | 0.260198151 | 0.546355578 |
| chr20:38073495-38073736 | 0.539085793 | 0.260200222 | 0.546355578 |
| chr10:8975557-8976095 | -0.403803283 | 0.260208356 | 0.546357685 |
| chrX:48475790-48476572 | 0.383453693 | 0.260230294 | 0.546374818 |
| chr9:76987613-76988107 | -0.461488176 | 0.260230777 | 0.546374818 |
| chr17:49899172-49899379 | 0.485935591 | 0.260240353 | 0.546379951 |
| chr6:105424201-105424497 | 0.463615694 | 0.260292098 | 0.546473616 |
| chr3:165572216-165572490 | -0.488151349 | 0.260323472 | 0.546499052 |
| chr2:78677442-78677642 | -0.579717683 | 0.260323831 | 0.546499052 |

| | | | |
|---|---|---|---|
| chr8:50115450-50116068 | 0.430726026 | 0.26032561 | 0.546499052 |
| chr18:23779787-23780031 | -0.531472287 | 0.260357697 | 0.546551437 |
| chr16:81165554-81166050 | -0.416840578 | 0.260403738 | 0.546633111 |
| chr1:175338535-175338947 | -0.408510953 | 0.260411244 | 0.546633892 |
| chr2:141666269-141666470 | -0.6178333 | 0.260437746 | 0.546674546 |
| chr12:67714347-67715030 | -0.413699663 | 0.260464364 | 0.546715442 |
| chr1:216153244-216153460 | -0.543566679 | 0.260496778 | 0.5467685 |
| chr11:18578918-18579655 | 0.391600521 | 0.260515698 | 0.546790113 |
| chr4:119000033-119000261 | 0.572095232 | 0.260521347 | 0.546790113 |
| chr3:10945794-10946003 | 0.468136059 | 0.260532236 | 0.54679799 |
| chr3:14178046-14179017 | 0.357419841 | 0.260548774 | 0.546806436 |
| chr6:76351349-76351618 | 0.473245075 | 0.260550534 | 0.546806436 |
| chr10:68302269-68302723 | -0.452455823 | 0.260601693 | 0.546898823 |
| chr22:19448165-19448456 | 0.43940036 | 0.260635186 | 0.546948062 |
| chr16:21952460-21953703 | 0.361517413 | 0.260639432 | 0.546948062 |
| chr5:946233-946557 | 0.498228656 | 0.260668517 | 0.546994115 |
| chr2:130371830-130372917 | 0.369575784 | 0.260730131 | 0.547108423 |
| chr3:14876901-14877338 | 0.402598433 | 0.260746364 | 0.547127503 |
| chr11:94183884-94184085 | -0.573334964 | 0.260793993 | 0.547212458 |
| chr6:31902008-31902638 | 0.384007092 | 0.260840708 | 0.54729549 |
| chr2:112665101-112665301 | -0.645161138 | 0.260867975 | 0.547323464 |
| chr9:114972680-114972890 | -0.632304317 | 0.260875678 | 0.547323464 |
| chr4:83362545-83363028 | -0.430231146 | 0.260878758 | 0.547323464 |
| chr9:14465516-14465816 | -0.462178241 | 0.260882613 | 0.547323464 |
| chr16:19396484-19396949 | -0.476247139 | 0.260895524 | 0.547335566 |
| chr22:22342889-22343204 | 0.455003229 | 0.260903222 | 0.547336728 |
| chr2:101238031-101238316 | 0.474901309 | 0.26093077 | 0.547379534 |
| chr8:109118658-109118964 | -0.477953329 | 0.260944807 | 0.547393994 |
| chr22:21643039-21643348 | 0.425576023 | 0.260994477 | 0.547483199 |
| chr13:46387154-46387470 | 0.432448295 | 0.261013001 | 0.547507068 |
| chr11:8247203-8247689 | -0.447050148 | 0.261048218 | 0.54755344 |
| chr7:17942331-17942541 | 0.442191422 | 0.261049401 | 0.54755344 |
| chr6:137341723-137342193 | -0.467850006 | 0.261058845 | 0.547558073 |
| chr19:49813026-49813384 | -0.440849954 | 0.261072798 | 0.547558073 |
| chr14:77151335-77151551 | 0.521655113 | 0.261076566 | 0.547558073 |
| chr3:149473638-149474095 | -0.38983455 | 0.261081553 | 0.547558073 |
| chr8:124561867-124562177 | 0.413787394 | 0.261089461 | 0.547558073 |
| chr10:125982361-125982561 | 0.610231687 | 0.261094487 | 0.547558073 |
| chr15:71251793-71252223 | -0.460526118 | 0.261124217 | 0.547597315 |
| chrX:133414573-133414780 | 0.557735607 | 0.261127492 | 0.547597315 |
| chr19:3985136-3985718 | -0.386849212 | 0.2611419 | 0.547611339 |
| chr8:139795467-139795710 | 0.469703537 | 0.261155135 | 0.547611339 |
| chr19:40487908-40488257 | 0.47925796 | 0.26115562 | 0.547611339 |
| chr6:27518435-27518655 | -0.529172241 | 0.261166895 | 0.547618513 |
| chr16:69108133-69108400 | 0.518547915 | 0.261173336 | 0.547618513 |
| chr10:4825789-4826799 | 0.365478224 | 0.261219372 | 0.547693295 |
| chr4:55125321-55125792 | 0.433485187 | 0.261223297 | 0.547693295 |
| chr12:3299942-3300220 | 0.447098936 | 0.261274366 | 0.547785379 |
| chr14:73602204-73602646 | 0.444339847 | 0.261288218 | 0.547799433 |
| chr6:44256516-44257881 | 0.361108107 | 0.261301931 | 0.547807035 |

| | | | |
|---|---|---|---|
| chr20:4109673-4109873 | -0.590140873 | 0.261306143 | 0.547807035 |
| chr3:167520760-167520962 | -0.626657514 | 0.261331732 | 0.547842263 |
| chr18:1663932-1664466 | -0.433261115 | 0.261343245 | 0.547842263 |
| chr20:57075641-57076641 | -0.363617704 | 0.261344397 | 0.547842263 |
| chr20:58429361-58429705 | -0.433553486 | 0.261385724 | 0.547905495 |
| chr2:178914324-178914569 | -0.527167826 | 0.261388863 | 0.547905495 |
| chr12:101012407-101012771 | -0.490732958 | 0.26148052 | 0.548082627 |
| chr12:54622911-54623187 | -0.491683125 | 0.26151962 | 0.548149587 |
| chr11:78284106-78284372 | 0.543832938 | 0.261528304 | 0.548152795 |
| chr8:65660482-65660782 | -0.583977226 | 0.261567335 | 0.548219605 |
| chr1:111959484-111959807 | 0.393971699 | 0.261586288 | 0.548244334 |
| chr2:32010606-32011438 | -0.368368093 | 0.261598283 | 0.548248324 |
| chr3:3291684-3292102 | 0.44677807 | 0.261602503 | 0.548248324 |
| chr11:74272272-74272667 | -0.515042925 | 0.261613239 | 0.548255829 |
| chr1:46735830-46736117 | 0.502108074 | 0.261621974 | 0.548259139 |
| chr17:50534262-50534619 | 0.390296525 | 0.261654889 | 0.548292149 |
| chr17:9885939-9886205 | -0.615798642 | 0.261659714 | 0.548292149 |
| chr9:77176860-77178373 | -0.354062007 | 0.261670858 | 0.548292149 |
| chr7:123131187-123131562 | -0.501310436 | 0.261672835 | 0.548292149 |
| chr8:110060301-110060883 | -0.371633957 | 0.261676696 | 0.548292149 |
| chr22:37745735-37746202 | 0.461801165 | 0.261680661 | 0.548292149 |
| chr1:154906732-154906943 | 0.565007327 | 0.261700588 | 0.548318908 |
| chr19:53792957-53793337 | 0.422134261 | 0.261716354 | 0.548328633 |
| chr2:43038879-43039181 | -0.497984983 | 0.261719542 | 0.548328633 |
| chr1:74652496-74653197 | -0.39502573 | 0.261735004 | 0.548346034 |
| chr5:4879730-4880078 | 0.416856857 | 0.261760853 | 0.548385193 |
| chr3:160169759-160169975 | -0.635632791 | 0.261771545 | 0.5483926 |
| chr2:208254116-208255628 | -0.353522552 | 0.261791778 | 0.548419993 |
| chr6:27334788-27335170 | -0.420628701 | 0.261817122 | 0.548458089 |
| chr2:61629915-61630115 | -0.630218909 | 0.261849797 | 0.548511542 |
| chr17:60408694-60409003 | 0.382020431 | 0.261863343 | 0.548524921 |
| chr15:81001248-81001670 | 0.40875835 | 0.261871025 | 0.548526017 |
| chr17:13602199-13602539 | -0.508107625 | 0.261902457 | 0.548559009 |
| chr10:32055486-32056634 | -0.366815828 | 0.261904429 | 0.548559009 |
| chr18:45966866-45967799 | -0.369525668 | 0.261908254 | 0.548559009 |
| chr1:215636625-215636978 | -0.440132255 | 0.261927058 | 0.548576701 |
| chr2:106085443-106085845 | 0.488138268 | 0.26193102 | 0.548576701 |
| chr6:31547339-31547709 | 0.400828996 | 0.261970008 | 0.548643359 |
| chr17:751712-752665 | -0.37448013 | 0.26199287 | 0.548676242 |
| chr4:82462481-82462783 | -0.459288591 | 0.26201539 | 0.548708407 |
| chr9:62801838-62802849 | -0.376054384 | 0.262038502 | 0.548741811 |
| chr10:103859249-103859529 | -0.474176608 | 0.262064458 | 0.548777913 |
| chr6:133921362-133921604 | -0.501187568 | 0.262076652 | 0.548777913 |
| chr8:135515484-135515721 | -0.538283098 | 0.262077228 | 0.548777913 |
| chr20:19427541-19427880 | 0.428211023 | 0.262139925 | 0.548894197 |
| chr3:87196880-87197255 | 0.437514291 | 0.262155001 | 0.548910567 |
| chr4:105196845-105197074 | -0.588618321 | 0.262169235 | 0.548910567 |
| chr4:172514345-172514578 | -0.588618321 | 0.262169235 | 0.548910567 |
| chr1:179484670-179484920 | 0.505509013 | 0.262184768 | 0.548928089 |
| chr9:126704690-126704982 | 0.450634562 | 0.262204936 | 0.548955314 |

| | | | |
|---|---|---|---|
| chr18:43277235-43277692 | -0.444142007 | 0.26221816 | 0.548968001 |
| chr1:160205162-160205580 | 0.414489939 | 0.262277489 | 0.549077207 |
| chr1:40097106-40097838 | 0.389720845 | 0.262295358 | 0.549099612 |
| chr5:138212862-138213893 | -0.358107314 | 0.262339995 | 0.549178054 |
| chrX:149962903-149963304 | 0.44474591 | 0.262406683 | 0.549302649 |
| chr15:60819852-60820202 | -0.541064584 | 0.262424677 | 0.549325309 |
| chr2:168639063-168639373 | -0.43906423 | 0.262457201 | 0.549378382 |
| chr12:52116275-52116822 | 0.391556018 | 0.262480341 | 0.549411809 |
| chr15:58269975-58270266 | 0.460597172 | 0.262494886 | 0.549427245 |
| chr9:69645448-69645979 | -0.386476983 | 0.262560953 | 0.549542427 |
| chr11:18700329-18700655 | 0.42988661 | 0.26256426 | 0.549542427 |
| chr16:88737433-88737644 | 0.446049757 | 0.26262685 | 0.549649507 |
| chr6:75800714-75801035 | -0.557434393 | 0.262630509 | 0.549649507 |
| chr5:133025998-133026822 | -0.365055718 | 0.262638398 | 0.549649507 |
| chr10:116788103-116788329 | 0.499451985 | 0.262645545 | 0.549649507 |
| chr5:108278873-108279164 | -0.470638712 | 0.262651766 | 0.549649507 |
| chr3:85801984-85802240 | -0.532450481 | 0.262664512 | 0.549649507 |
| chr2:20650349-20651762 | -0.355170931 | 0.262669005 | 0.549649507 |
| chr1:11478540-11479517 | 0.385129276 | 0.262672809 | 0.549649507 |
| chr5:38676546-38676803 | 0.631975674 | 0.262688078 | 0.549654983 |
| chrX:85808092-85808534 | -0.49079886 | 0.262689773 | 0.549654983 |
| chrX:120560455-120561787 | 0.35640505 | 0.262706676 | 0.549657308 |
| chr11:2099806-2100006 | 0.538565418 | 0.2627089 | 0.549657308 |
| chr12:132710171-132711420 | 0.363149456 | 0.262713941 | 0.549657308 |
| chr2:158045495-158045919 | -0.504885206 | 0.262719579 | 0.549657308 |
| chr2:121777167-121777494 | 0.494636888 | 0.262740401 | 0.549679823 |
| chr20:407624-408515 | 0.383696927 | 0.262749553 | 0.549679823 |
| chr4:98142776-98143862 | -0.356813705 | 0.262756478 | 0.549679823 |
| chr7:121337522-121337741 | -0.574255924 | 0.262759036 | 0.549679823 |
| chr10:125895671-125896225 | 0.398541484 | 0.262797921 | 0.549746159 |
| chr6:28807739-28808118 | 0.407176965 | 0.262821989 | 0.549781496 |
| chr14:69251282-69251482 | 0.555592437 | 0.262833167 | 0.549785992 |
| chr6:122494704-122495618 | -0.372981157 | 0.262838489 | 0.549785992 |
| chr17:81552294-81553041 | 0.375112199 | 0.262893239 | 0.549866694 |
| chr2:10136282-10136524 | -0.521204915 | 0.262893722 | 0.549866694 |
| chr9:108163664-108163988 | -0.520995128 | 0.262904267 | 0.549866694 |
| chr14:81553694-81554070 | -0.544886964 | 0.262905775 | 0.549866694 |
| chr1:73813802-73814034 | -0.56176166 | 0.262918574 | 0.549878453 |
| chr18:57820056-57820342 | -0.461262519 | 0.262935521 | 0.549898885 |
| chr3:21082471-21082836 | -0.472608152 | 0.262955252 | 0.549925141 |
| chr17:47698834-47699148 | -0.414306505 | 0.262986942 | 0.549976405 |
| chr3:124586634-124587373 | 0.374708844 | 0.263043813 | 0.550080323 |
| chr4:173370634-173371783 | -0.354989032 | 0.263069605 | 0.550119245 |
| chr18:26745427-26745627 | -0.596049518 | 0.263086753 | 0.55014009 |
| chr21:41069049-41069382 | -0.419448497 | 0.263123611 | 0.550182742 |
| chr8:23528420-23529741 | -0.365131441 | 0.263124711 | 0.550182742 |
| chr19:39038099-39038494 | -0.462584522 | 0.263128691 | 0.550182742 |
| chr6:44172724-44173019 | 0.428704354 | 0.263158392 | 0.550229829 |
| chr11:93850761-93850962 | -0.580946972 | 0.263230958 | 0.5503626 |
| chr5:149442535-149442871 | -0.407780051 | 0.263240092 | 0.5503626 |

| | | | |
|---|---|---|---|
| chr1:224454305-224454998 | 0.375297766 | 0.263245762 | 0.5503626 |
| chr20:3752038-3752778 | -0.387297859 | 0.263250624 | 0.5503626 |
| chr3:84447241-84447454 | -0.602207395 | 0.263274315 | 0.550377703 |
| chr9:114098160-114099465 | -0.357787591 | 0.263277967 | 0.550377703 |
| chr22:38793802-38794413 | -0.393224634 | 0.263279397 | 0.550377703 |
| chr15:75532545-75532749 | 0.535479711 | 0.263296205 | 0.550397823 |
| chr9:112112970-112113339 | 0.428807615 | 0.263314114 | 0.550420243 |
| chr11:78138998-78140095 | -0.362144074 | 0.263339317 | 0.550443943 |
| chr3:17692960-17693970 | -0.359649449 | 0.263341068 | 0.550443943 |
| chr6:27672017-27672647 | -0.379275968 | 0.263347003 | 0.550443943 |
| chr4:68411393-68411674 | 0.574834146 | 0.263360176 | 0.550456461 |
| chr14:68255903-68256177 | 0.511105196 | 0.263374272 | 0.550470907 |
| chrX:76172583-76173793 | -0.361001369 | 0.263405198 | 0.550520527 |
| chr1:39205085-39205318 | -0.642096733 | 0.263422213 | 0.550541072 |
| chr17:62300737-62301318 | 0.416389616 | 0.263454943 | 0.550594459 |
| chr17:22337614-22337821 | -0.567721764 | 0.263490653 | 0.550654069 |
| chr2:227703240-227703714 | 0.422490168 | 0.263555752 | 0.550766971 |
| chr19:18683383-18684072 | 0.396704299 | 0.263559053 | 0.550766971 |
| chr2:213221081-213221281 | -0.598792697 | 0.263566831 | 0.550768204 |
| chr10:84008836-84009160 | -0.467823112 | 0.263579446 | 0.550769536 |
| chr21:9162200-9162400 | -0.609817557 | 0.263583147 | 0.550769536 |
| chr14:26163728-26164580 | 0.376233943 | 0.263594296 | 0.550769536 |
| chr1:182993766-182994010 | -0.602116336 | 0.263605677 | 0.550769536 |
| chr10:125977254-125977464 | -0.602116336 | 0.263605677 | 0.550769536 |
| chr17:43839611-43840164 | 0.411148787 | 0.26361607 | 0.550769536 |
| chr11:33716318-33716828 | -0.462566512 | 0.263617786 | 0.550769536 |
| chr7:100673230-100673864 | -0.365783084 | 0.263668117 | 0.550851407 |
| chr17:6436018-6436370 | -0.448064396 | 0.26367692 | 0.550851407 |
| chr11:83521544-83521979 | -0.448714695 | 0.263678539 | 0.550851407 |
| chr20:50372803-50373003 | -0.468130407 | 0.263687176 | 0.55085443 |
| chr4:35541011-35541622 | -0.432228896 | 0.263711718 | 0.55089068 |
| chr13:51222651-51222961 | -0.524659984 | 0.263722603 | 0.550895853 |
| chr6:15690426-15690899 | -0.531602525 | 0.263728574 | 0.550895853 |
| chr12:71392561-71392913 | -0.421014076 | 0.263750075 | 0.550925746 |
| chr19:29841727-29841992 | 0.421204473 | 0.263770019 | 0.550938592 |
| chr5:176630404-176630722 | 0.444205571 | 0.263770605 | 0.550938592 |
| chr2:31675652-31675927 | 0.478112137 | 0.263802504 | 0.550975872 |
| chr12:28815287-28815614 | -0.606455215 | 0.263802835 | 0.550975872 |
| chr19:41426901-41427191 | 0.466662919 | 0.26384001 | 0.551038494 |
| chr9:123268537-123269174 | 0.364814861 | 0.263860024 | 0.551065274 |
| chr12:116773489-116773784 | 0.462850892 | 0.263873934 | 0.551079303 |
| chr5:168600314-168600757 | -0.39737621 | 0.263886574 | 0.551082999 |
| chr14:55411458-55412323 | 0.366709518 | 0.263890088 | 0.551082999 |
| chr13:36420403-36420937 | -0.391185046 | 0.26392317 | 0.551087407 |
| chr17:60240588-60240788 | 0.418212649 | 0.263923294 | 0.551087407 |
| chr20:50038481-50039061 | 0.3820718 | 0.263936043 | 0.551087407 |
| chr3:32467952-32468152 | 0.516829354 | 0.263938919 | 0.551087407 |
| chr22:38222835-38223175 | -0.496655557 | 0.263941984 | 0.551087407 |
| chr15:85164789-85165165 | 0.414319652 | 0.263944738 | 0.551087407 |
| chr16:70686026-70686264 | 0.435832639 | 0.2639517 | 0.551087407 |

| | | | |
|---|---|---|---|
| chr3:61731499-61731768 | 0.428207783 | 0.263953088 | 0.551087407 |
| chr1:63522862-63523973 | -0.360343859 | 0.263956929 | 0.551087407 |
| chr17:39979886-39980182 | -0.504852249 | 0.263972308 | 0.551104498 |
| chr6:53104080-53104530 | -0.459084942 | 0.263986305 | 0.551108884 |
| chr10:77826115-77826315 | -0.541412809 | 0.263988794 | 0.551108884 |
| chr2:174603416-174603629 | 0.570834414 | 0.263999685 | 0.551116605 |
| chr4:33060952-33061289 | 0.427180933 | 0.264023062 | 0.551150391 |
| chr20:33673703-33674516 | 0.383484162 | 0.264033682 | 0.551157544 |
| chr6:96394791-96395077 | 0.47575786 | 0.264065447 | 0.551208835 |
| chr1:151251787-151251996 | 0.442954354 | 0.264183947 | 0.55141679 |
| chr4:64314528-64314757 | -0.578070229 | 0.264193373 | 0.55141679 |
| chr10:80896544-80897343 | 0.367236493 | 0.264193622 | 0.55141679 |
| chr17:3737732-3738104 | -0.435544037 | 0.264193858 | 0.55141679 |
| chr1:168439560-168440168 | -0.377293916 | 0.264221938 | 0.551460376 |
| chr9:109903599-109903844 | 0.502648428 | 0.264254736 | 0.551479895 |
| chr10:23068185-23068484 | -0.494097286 | 0.26426008 | 0.551479895 |
| chr13:113208173-113208383 | -0.494097286 | 0.26426008 | 0.551479895 |
| chr3:16796810-16797150 | -0.494097286 | 0.26426008 | 0.551479895 |
| chr2:211867945-211868289 | -0.594581485 | 0.26427821 | 0.551482338 |
| chr6:131182959-131183392 | -0.458205193 | 0.264280542 | 0.551482338 |
| chr1:144560097-144560394 | -0.509535827 | 0.264288432 | 0.551482338 |
| chr13:31927370-31927855 | 0.457643924 | 0.26429004 | 0.551482338 |
| chr1:219304286-219304499 | -0.525304794 | 0.264311411 | 0.551503173 |
| chr1:156616066-156616402 | 0.440958474 | 0.26431442 | 0.551503173 |
| chr6:7200949-7201711 | 0.387966698 | 0.264336385 | 0.551531746 |
| chr8:25195501-25195701 | 0.578176413 | 0.264342511 | 0.551531746 |
| chrX:15619431-15619744 | -0.46257354 | 0.264377003 | 0.551588693 |
| chr4:183492949-183493149 | -0.631371332 | 0.26440631 | 0.551599129 |
| chr6:128295778-128295978 | -0.631371332 | 0.26440631 | 0.551599129 |
| chr2:42784646-42784846 | -0.50294525 | 0.264410004 | 0.551599129 |
| chr2:25184194-25184776 | 0.394142906 | 0.264413243 | 0.551599129 |
| chr7:47575895-47576776 | 0.360976085 | 0.264430018 | 0.551599129 |
| chr8:76648024-76648267 | -0.615726717 | 0.264431504 | 0.551599129 |
| chr9:95875320-95876242 | 0.358982144 | 0.264432398 | 0.551599129 |
| chr14:87927718-87927918 | 0.581163252 | 0.264455434 | 0.551617848 |
| chr17:54754527-54754826 | -0.486633887 | 0.264459349 | 0.551617848 |
| chr20:49957788-49958021 | 0.481297294 | 0.26446297 | 0.551617848 |
| chr8:66223800-66224026 | -0.50428243 | 0.264473399 | 0.551624585 |
| chr14:57317294-57317569 | -0.499516652 | 0.26450391 | 0.551654544 |
| chr16:85639519-85639882 | -0.415000445 | 0.26450536 | 0.551654544 |
| chr10:1006452-1006702 | -0.544952108 | 0.26451553 | 0.551654544 |
| chr6:33822594-33823000 | -0.436421478 | 0.264516561 | 0.551654544 |
| chr9:98192563-98193545 | 0.365977213 | 0.264536179 | 0.551680442 |
| chr3:185070563-185070775 | 0.500872129 | 0.264546463 | 0.551686872 |
| chr6:3063621-3064157 | 0.371789835 | 0.264559125 | 0.551698263 |
| chr1:55035952-55036161 | -0.566635795 | 0.264574281 | 0.551714853 |
| chr3:1988041-1988304 | -0.49064629 | 0.264586162 | 0.551724613 |
| chr14:75985627-75985935 | 0.40206703 | 0.264598219 | 0.55173474 |
| chr9:33228724-33229076 | -0.456119947 | 0.264638899 | 0.551804549 |
| chr1:149558108-149559043 | 0.362833484 | 0.264686389 | 0.551872181 |

| | | | |
|---|---|---|---|
| chr19:49198857-49199125 | -0.577705474 | 0.264691568 | 0.551872181 |
| chr10:79070798-79070998 | 0.456478934 | 0.264692943 | 0.551872181 |
| chr14:36184570-36185280 | -0.41382363 | 0.264707492 | 0.551887499 |
| chr6:108027649-108027917 | -0.557048857 | 0.264751933 | 0.551950314 |
| chr9:132878701-132879019 | -0.447592417 | 0.264760956 | 0.551950314 |
| chr9:99821584-99822105 | 0.364220751 | 0.264768446 | 0.551950314 |
| chr2:55574630-55574904 | -0.668601994 | 0.264772568 | 0.551950314 |
| chr1:217516320-217516680 | -0.407771078 | 0.264781929 | 0.551950314 |
| chr8:101070127-101070384 | -0.546816138 | 0.264787941 | 0.551950314 |
| chr20:9067574-9068146 | -0.372713707 | 0.264788887 | 0.551950314 |
| chr12:76442755-76442955 | 0.608509333 | 0.264795893 | 0.551950314 |
| chr15:92277803-92278127 | -0.43619399 | 0.264811237 | 0.551950314 |
| chr2:233354302-233354533 | -0.526315915 | 0.264816446 | 0.551950314 |
| chr6:36964498-36964747 | -0.495618852 | 0.26481686 | 0.551950314 |
| chr3:32502020-32503128 | 0.368835894 | 0.264864884 | 0.55203369 |
| chr2:88832646-88832875 | -0.582508911 | 0.264871272 | 0.55203369 |
| chr16:31215298-31216315 | 0.375874861 | 0.264898245 | 0.552066108 |
| chr3:38136861-38137816 | 0.356058325 | 0.264910514 | 0.552066108 |
| chr5:37370765-37371954 | -0.353789004 | 0.264916255 | 0.552066108 |
| chr3:113947334-113948881 | 0.354411307 | 0.264917698 | 0.552066108 |
| chr13:99176410-99176987 | 0.404584134 | 0.264922851 | 0.552066108 |
| chr2:62505521-62506650 | -0.359612869 | 0.264961735 | 0.55212558 |
| chr17:43677932-43678132 | 0.536792214 | 0.264966864 | 0.55212558 |
| chr14:101972389-101972711 | 0.439838997 | 0.264973008 | 0.55212558 |
| chrX:147911315-147912094 | 0.359584789 | 0.264992388 | 0.552150945 |
| chr12:79058455-79058751 | -0.472947662 | 0.265002465 | 0.552156928 |
| chr2:70425800-70426398 | -0.437213055 | 0.265018392 | 0.552175099 |
| chr14:37623528-37624139 | 0.375650854 | 0.265032332 | 0.552178713 |
| chr1:215023913-215024483 | -0.406431471 | 0.265037121 | 0.552178713 |
| chr15:92673274-92673539 | -0.538601119 | 0.265050608 | 0.552178713 |
| chr2:41410095-41410299 | -0.566855392 | 0.26505245 | 0.552178713 |
| chr16:70608544-70608807 | -0.511583028 | 0.265056159 | 0.552178713 |
| chr12:115433112-115433389 | -0.561990792 | 0.265067458 | 0.552187238 |
| chr16:30985500-30985787 | 0.425964074 | 0.265103882 | 0.552248101 |
| chr4:154349925-154350133 | -0.554223662 | 0.265133508 | 0.552293996 |
| chr7:30983636-30983836 | 0.522011015 | 0.265149266 | 0.552293996 |
| chr12:57845509-57845770 | -0.416132617 | 0.265149502 | 0.552293996 |
| chr5:148398332-148399153 | -0.384587953 | 0.265154746 | 0.552293996 |
| chr1:778315-779015 | 0.382726483 | 0.265177494 | 0.552296859 |
| chr17:19244822-19245154 | -0.456274503 | 0.2651838 | 0.552296859 |
| chr20:20593569-20593985 | -0.435381088 | 0.265184273 | 0.552296859 |
| chr18:28658160-28658360 | -0.610017176 | 0.26518547 | 0.552296859 |
| chr12:63968489-63968858 | -0.520017887 | 0.265195627 | 0.552296859 |
| chr14:76812276-76813194 | 0.359100561 | 0.265205004 | 0.552296859 |
| chr8:8333635-8334144 | -0.451445547 | 0.265206577 | 0.552296859 |
| chr5:79021090-79021509 | -0.451422135 | 0.265292272 | 0.552460305 |
| chr16:85000664-85001316 | 0.372560743 | 0.26532547 | 0.552487311 |
| chr2:180125340-180126103 | -0.387004178 | 0.265328451 | 0.552487311 |
| chr14:35353895-35354218 | 0.422575421 | 0.265329509 | 0.552487311 |
| chr4:68865234-68865442 | 0.513240685 | 0.265334082 | 0.552487311 |

| | | | |
|---|---|---|---|
| chr15:90265164-90266421 | 0.358402371 | 0.26535954 | 0.552519232 |
| chr11:106076690-106078119 | -0.351369161 | 0.265363835 | 0.552519232 |
| chr1:34549286-34549526 | 0.57650431 | 0.26538537 | 0.552549056 |
| chr7:36994481-36994696 | -0.540255378 | 0.265411337 | 0.552581331 |
| chr9:34500664-34500971 | -0.470255235 | 0.26541787 | 0.552581331 |
| chr6:145963714-145964906 | -0.357520105 | 0.265429406 | 0.552581331 |
| chr1:47432493-47432766 | -0.445796366 | 0.265429719 | 0.552581331 |
| chr16:33913662-33913908 | 0.54737543 | 0.265450704 | 0.552610005 |
| chr3:64474921-64476064 | 0.351837313 | 0.265464157 | 0.552621337 |
| chr3:9932610-9934511 | 0.348628234 | 0.265470572 | 0.552621337 |
| chr4:89001059-89001368 | -0.573030969 | 0.265502155 | 0.552672067 |
| chr12:88431362-88431562 | 0.561979375 | 0.265524972 | 0.552704547 |
| chr6:117399925-117400256 | 0.445489555 | 0.265545276 | 0.552731796 |
| chr7:134606386-134606849 | -0.430164764 | 0.265569292 | 0.552743084 |
| chr8:11065701-11065907 | -0.606354277 | 0.2655697 | 0.552743084 |
| chr1:36149015-36149914 | 0.358923289 | 0.265572341 | 0.552743084 |
| chr2:128322422-128322755 | 0.431472514 | 0.265590308 | 0.552765465 |
| chr1:17439674-17439885 | 0.561732615 | 0.265604942 | 0.552780907 |
| chrX:105324041-105324328 | -0.447767433 | 0.26569783 | 0.552959208 |
| chr3:138329161-138329513 | 0.405619676 | 0.265714078 | 0.552978003 |
| chr2:55048982-55049273 | -0.398035108 | 0.265728672 | 0.552993355 |
| chr8:53712455-53712789 | 0.491042904 | 0.265756516 | 0.553017932 |
| chr5:38845068-38845278 | -0.522659856 | 0.265757854 | 0.553017932 |
| chr7:157493007-157493298 | -0.475586558 | 0.265762135 | 0.553017932 |
| chr3:4645633-4645888 | -0.522240225 | 0.265781807 | 0.553043847 |
| chr3:124901424-124901624 | 0.555934761 | 0.265809982 | 0.553087454 |
| chr6:30884773-30884973 | 0.447219345 | 0.265823332 | 0.553100213 |
| chr3:127812805-127813064 | -0.551415537 | 0.265835933 | 0.553111413 |
| chr17:61635795-61636311 | 0.357365602 | 0.265845838 | 0.553117001 |
| chr1:160780157-160780368 | 0.607156726 | 0.26587537 | 0.553149781 |
| chr14:61456692-61457041 | 0.421172941 | 0.26588033 | 0.553149781 |
| chr16:84637024-84637294 | -0.479712018 | 0.265883251 | 0.553149781 |
| chr1:229342048-229343148 | 0.366308514 | 0.265912471 | 0.553195552 |
| chr10:12333472-12334354 | -0.371838571 | 0.26596053 | 0.553280509 |
| chr10:46284329-46285053 | 0.383365079 | 0.26599817 | 0.553343788 |
| chr5:23105337-23105578 | 0.592369694 | 0.266047816 | 0.553432041 |
| chr8:128056866-128057737 | -0.373161874 | 0.266065779 | 0.553454382 |
| chr15:90930050-90930262 | 0.41976679 | 0.266080602 | 0.55347019 |
| chr11:62546730-62547136 | -0.404552404 | 0.266115222 | 0.553511707 |
| chr16:76946996-76947233 | 0.488080314 | 0.266125573 | 0.553511707 |
| chr1:245856838-245857045 | -0.578311031 | 0.266129657 | 0.553511707 |
| chr15:80404306-80405121 | 0.383871801 | 0.266130056 | 0.553511707 |
| chr1:218528538-218528738 | -0.569640897 | 0.266139908 | 0.553511707 |
| chr5:115601940-115602330 | -0.452150768 | 0.266143905 | 0.553511707 |
| chr9:15513524-15513742 | -0.52154093 | 0.266151715 | 0.553512925 |
| chr18:80108705-80109921 | -0.35534266 | 0.266186951 | 0.553559951 |
| chr10:44857802-44858019 | -0.494278731 | 0.266188776 | 0.553559951 |
| chr20:41391728-41392139 | -0.396259029 | 0.266201102 | 0.55357056 |
| chr3:63588819-63589172 | 0.424966011 | 0.266212346 | 0.553578918 |
| chr8:11651605-11651866 | -0.539173384 | 0.266220629 | 0.553581117 |

| | | | |
|---|---|---|---|
| chr7:147833447-147833953 | 0.422727157 | 0.26624218 | 0.553596945 |
| chr17:60307579-60307980 | 0.372547485 | 0.266247575 | 0.553596945 |
| chr1:207691881-207692248 | -0.394267927 | 0.266249916 | 0.553596945 |
| chr8:142095555-142095795 | -0.53198257 | 0.266292478 | 0.553643471 |
| chr12:71972803-71973046 | -0.511199904 | 0.266306714 | 0.553643471 |
| chr22:19997050-19997343 | -0.539033551 | 0.266307079 | 0.553643471 |
| chr17:57113769-57114382 | -0.391705863 | 0.266307255 | 0.553643471 |
| chr10:13265230-13265430 | 0.6243224 | 0.266309341 | 0.553643471 |
| chr2:10036043-10036337 | -0.44569534 | 0.266315646 | 0.553643471 |
| chr5:95730824-95732374 | -0.354891567 | 0.266362975 | 0.553718507 |
| chr17:69235449-69235809 | -0.399181351 | 0.266367311 | 0.553718507 |
| chr20:297224-297888 | 0.400702737 | 0.266379866 | 0.553718507 |
| chr3:189961513-189961879 | -0.432417481 | 0.266380646 | 0.553718507 |
| chr3:130329645-130330089 | -0.474883468 | 0.266417317 | 0.553758393 |
| chr4:151113710-151113910 | 0.530037186 | 0.266422371 | 0.553758393 |
| chr2:61905331-61906035 | -0.372764518 | 0.266424621 | 0.553758393 |
| chr2:152717131-152719226 | -0.34598727 | 0.266428743 | 0.553758393 |
| chr17:60193576-60193915 | 0.385609895 | 0.266467632 | 0.553803662 |
| chr8:94357091-94357410 | 0.408560253 | 0.266468929 | 0.553803662 |
| chr3:20064929-20065281 | 0.455688977 | 0.266472206 | 0.553803662 |
| chr16:89822238-89822926 | -0.403145768 | 0.266493393 | 0.553811796 |
| chr17:5062478-5062715 | 0.550090989 | 0.266500864 | 0.553811796 |
| chr22:29882878-29883945 | 0.373048695 | 0.266512064 | 0.553811796 |
| chr7:120876990-120877234 | -0.540594762 | 0.266512165 | 0.553811796 |
| chr20:57178490-57179066 | 0.352112397 | 0.26651226 | 0.553811796 |
| chr16:84622856-84623056 | 0.475132177 | 0.266543421 | 0.553851109 |
| chr3:64688904-64689148 | -0.381654506 | 0.266545635 | 0.553851109 |
| chr3:186931188-186931729 | -0.420568324 | 0.266555668 | 0.553856937 |
| chr6:130885416-130885664 | 0.520126277 | 0.266565411 | 0.553862161 |
| chr4:187826820-187827106 | 0.526802455 | 0.266575815 | 0.553868758 |
| chr16:80726726-80727825 | -0.366307102 | 0.266590279 | 0.553883793 |
| chr14:35422459-35422790 | 0.449343851 | 0.266600792 | 0.553886845 |
| chr17:77494981-77495214 | 0.458126748 | 0.266606206 | 0.553886845 |
| chr10:27154904-27156220 | -0.353465049 | 0.266620394 | 0.553901303 |
| chr4:9531946-9532445 | 0.417912082 | 0.2666645 | 0.553954083 |
| chr4:87150626-87151129 | -0.437864575 | 0.266666296 | 0.553954083 |
| chr13:38670642-38670905 | -0.459699322 | 0.266667489 | 0.553954083 |
| chr14:91109289-91109543 | 0.47289344 | 0.26668669 | 0.553978951 |
| chr10:79128763-79129073 | 0.465008277 | 0.266706705 | 0.553998697 |
| chr9:108551363-108551919 | -0.423164909 | 0.266710656 | 0.553998697 |
| chr11:24635762-24636562 | 0.384773756 | 0.266745718 | 0.554056504 |
| chr15:90660569-90660798 | 0.471603815 | 0.266793161 | 0.554140026 |
| chr5:179811198-179811457 | -0.463750274 | 0.266827964 | 0.554197292 |
| chr12:132828717-132829488 | 0.364574923 | 0.266855573 | 0.554239611 |
| chr1:204628500-204628705 | 0.451906313 | 0.266885468 | 0.554282964 |
| chr19:56066258-56066741 | 0.422381164 | 0.266890915 | 0.554282964 |
| chr1:75618478-75618734 | 0.466062958 | 0.266925329 | 0.55433941 |
| chr2:172226432-172226641 | -0.543626987 | 0.26694412 | 0.554363409 |
| chr20:48065005-48065220 | 0.398738199 | 0.26701402 | 0.554493543 |
| chr2:188369389-188369661 | -0.469410701 | 0.267045514 | 0.554543914 |

| | | | |
|---|---|---|---|
| chr14:69574446-69574789 | 0.429667763 | 0.267071953 | 0.554583787 |
| chr2:23002333-23002686 | -0.489962985 | 0.267117377 | 0.55466308 |
| chr8:73449101-73449465 | -0.433284262 | 0.267137615 | 0.554671701 |
| chr5:72107236-72107516 | 0.461368285 | 0.267143163 | 0.554671701 |
| chr1:16913633-16914594 | 0.382888164 | 0.267143246 | 0.554671701 |
| chr17:83103623-83104196 | 0.372484896 | 0.267159557 | 0.5546777 |
| chr16:89721863-89722063 | -0.465433485 | 0.267161947 | 0.5546777 |
| chr4:149371075-149371466 | 0.442634731 | 0.267172642 | 0.5546777 |
| chr16:23735267-23735508 | -0.605281859 | 0.267175092 | 0.5546777 |
| chr5:154650235-154650450 | -0.552303921 | 0.267186379 | 0.554682159 |
| chr22:31292330-31292659 | 0.433885231 | 0.267191718 | 0.554682159 |
| chr9:114504400-114505029 | -0.382831649 | 0.267226563 | 0.554709801 |
| chr2:26292737-26292977 | -0.529018311 | 0.267229132 | 0.554709801 |
| chr14:52015593-52015971 | 0.405101995 | 0.267230287 | 0.554709801 |
| chr1:62963071-62963477 | -0.461966602 | 0.267233992 | 0.554709801 |
| chr4:39591341-39591638 | -0.552109132 | 0.267251263 | 0.554730625 |
| chr12:56591272-56591472 | 0.553046569 | 0.267266824 | 0.554747897 |
| chr12:26434428-26434885 | -0.457231461 | 0.267282763 | 0.554764585 |
| chr3:189841904-189842235 | -0.441253422 | 0.267289345 | 0.554764585 |
| chr10:84114521-84114834 | -0.547794772 | 0.267309884 | 0.554792187 |
| chr10:34684582-34684975 | -0.4898708 | 0.267337586 | 0.554834652 |
| chr4:67327392-67327592 | 0.547940244 | 0.267345982 | 0.554837048 |
| chrX:109393861-109394152 | 0.47825064 | 0.267380727 | 0.554894126 |
| chr1:107299518-107299764 | 0.408071812 | 0.267396914 | 0.554912691 |
| chr2:176814359-176814616 | 0.478069915 | 0.267410036 | 0.554924891 |
| chr17:51746011-51746335 | -0.433510347 | 0.267437402 | 0.554962293 |
| chr3:46163122-46164109 | 0.37509844 | 0.267442545 | 0.554962293 |
| chr8:110830323-110830774 | -0.402469118 | 0.267460576 | 0.55498468 |
| chr10:92405586-92405793 | 0.539786401 | 0.267481619 | 0.55501173 |
| chr21:45596770-45596979 | 0.510398361 | 0.267497611 | 0.55501173 |
| chr4:38663269-38664920 | 0.354232487 | 0.267501841 | 0.55501173 |
| chr4:168803239-168803541 | -0.573453284 | 0.267502586 | 0.55501173 |
| chr16:28524810-28525092 | 0.517393881 | 0.267531082 | 0.555047634 |
| chr1:242454739-242455104 | 0.439868536 | 0.267534379 | 0.555047634 |
| chr13:25375797-25376004 | 0.504964004 | 0.267561079 | 0.555086969 |
| chr8:103660361-103660583 | 0.504500289 | 0.267567828 | 0.555086969 |
| chr1:1079381-1080507 | -0.360226555 | 0.267614108 | 0.555154431 |
| chr6:36455551-36455819 | 0.468501886 | 0.267619929 | 0.555154431 |
| chr1:156599930-156600130 | 0.485235093 | 0.267622083 | 0.555154431 |
| chr9:111036499-111036775 | -0.517356299 | 0.267631893 | 0.555159752 |
| chr19:11243313-11243674 | 0.418872351 | 0.267646952 | 0.555174276 |
| chr6:28643334-28643705 | 0.400425284 | 0.267653523 | 0.555174276 |
| chr11:68410943-68411143 | -0.559070854 | 0.267660632 | 0.555174276 |
| chr18:50307670-50308020 | -0.404716697 | 0.267681657 | 0.555202857 |
| chr10:69883271-69883741 | -0.408530463 | 0.267702732 | 0.555231539 |
| chr17:60054030-60054298 | 0.383816952 | 0.267721128 | 0.555241977 |
| chr2:5022884-5023105 | 0.544407391 | 0.267722258 | 0.555241977 |
| chr6:12844782-12845036 | 0.537292309 | 0.267733014 | 0.555249256 |
| chr4:41216378-41216709 | 0.395569175 | 0.267746065 | 0.555261293 |
| chr5:139147341-139147662 | -0.467218577 | 0.267769529 | 0.555288221 |

| | | | |
|---|---|---|---|
| chr14:74294367-74294617 | 0.456854255 | 0.267773544 | 0.555288221 |
| chr6:16409794-16410232 | -0.497152908 | 0.267807039 | 0.555327861 |
| chr5:165057166-165057555 | -0.438532971 | 0.267810993 | 0.555327861 |
| chr4:103075716-103077007 | -0.360309867 | 0.267820075 | 0.555327861 |
| chr18:24176790-24177002 | -0.593350335 | 0.267836557 | 0.555327861 |
| chr4:86453538-86454232 | 0.376887507 | 0.267838639 | 0.555327861 |
| chr14:31492738-31492938 | 0.539452297 | 0.267838651 | 0.555327861 |
| chr9:129131695-129131982 | 0.459139234 | 0.267843393 | 0.555327861 |
| chr3:63371284-63371484 | 0.461034128 | 0.267897435 | 0.555417227 |
| chr6:161439110-161439362 | -0.511354419 | 0.267900993 | 0.555417227 |
| chr7:45937382-45937605 | -0.583878983 | 0.267930674 | 0.555455595 |
| chr2:149611133-149611333 | -0.594159587 | 0.267933998 | 0.555455595 |
| chr5:21481683-21482264 | 0.367180319 | 0.267946747 | 0.555461886 |
| chr2:196453695-196453895 | 0.537839404 | 0.267951531 | 0.555461886 |
| chr22:24768906-24769107 | 0.566003317 | 0.267964644 | 0.55547404 |
| chr10:18582300-18582556 | -0.554022251 | 0.267995785 | 0.555523565 |
| chr5:125835585-125835859 | -0.498424382 | 0.268015314 | 0.555549016 |
| chr10:46524722-46525101 | -0.417218886 | 0.268031327 | 0.555553559 |
| chr2:73926625-73927232 | -0.372493863 | 0.268033616 | 0.555553559 |
| chr10:21174899-21175172 | 0.436760206 | 0.268052276 | 0.555553559 |
| chr11:8328794-8329300 | -0.396722647 | 0.268052397 | 0.555553559 |
| chr11:67615437-67615696 | 0.438295666 | 0.268053758 | 0.555553559 |
| chr8:115154790-115154991 | -0.545143419 | 0.268063215 | 0.555558131 |
| chr3:119206400-119206600 | 0.585319374 | 0.268100336 | 0.555620035 |
| chr20:2751914-2752388 | -0.442023095 | 0.268113806 | 0.555629055 |
| chr17:1742154-1742388 | -0.587681766 | 0.268119191 | 0.555629055 |
| chr5:103561723-103561975 | -0.465205809 | 0.268136006 | 0.555637141 |
| chr17:12545944-12546166 | -0.533288519 | 0.268139564 | 0.555637141 |
| chr9:104743510-104743746 | -0.458930274 | 0.268144848 | 0.555637141 |
| chr5:88140743-88141815 | -0.351273195 | 0.268218869 | 0.555772575 |
| chr1:235511552-235512267 | 0.381016061 | 0.268251096 | 0.555772575 |
| chr16:81568405-81568639 | -0.471483609 | 0.268257741 | 0.555772575 |
| chr3:171187730-171188035 | -0.443924742 | 0.268259155 | 0.555772575 |
| chr3:10744056-10744351 | 0.464549396 | 0.268260015 | 0.555772575 |
| chr5:6812230-6812555 | -0.553565734 | 0.268262645 | 0.555772575 |
| chr4:158176794-158177172 | 0.433807829 | 0.268263672 | 0.555772575 |
| chr20:4686230-4686958 | 0.389464189 | 0.268268235 | 0.555772575 |
| chr14:47934831-47935349 | -0.441769158 | 0.268282039 | 0.555786145 |
| chr2:36785202-36785534 | -0.4233452 | 0.268298774 | 0.555805788 |
| chr7:149127581-149127979 | -0.42633881 | 0.268329891 | 0.555855222 |
| chr17:53686627-53686920 | 0.488716144 | 0.268359789 | 0.555902126 |
| chr1:203316329-203317102 | -0.35872878 | 0.268373672 | 0.555915855 |
| chr9:37747228-37747428 | -0.599787056 | 0.268390422 | 0.555935304 |
| chr4:35152063-35152269 | 0.553246285 | 0.268397572 | 0.555935304 |
| chr5:138068762-138069350 | 0.40113683 | 0.268452812 | 0.556002761 |
| chr11:69071638-69071892 | -0.522622396 | 0.268453788 | 0.556002761 |
| chr13:32587590-32587790 | -0.474551837 | 0.268455646 | 0.556002761 |
| chr14:40393237-40393470 | 0.549462507 | 0.268462528 | 0.556002761 |
| chr2:185187310-185187903 | -0.393716134 | 0.268472973 | 0.556002761 |
| chr3:154181406-154181739 | 0.416990089 | 0.268473678 | 0.556002761 |

| | | | |
|---|---|---|---|
| chr14:21771792-21772017 | 0.500426544 | 0.268502914 | 0.556048278 |
| chr22:49305348-49305690 | -0.507773306 | 0.268514632 | 0.556053922 |
| chr3:157248562-157248926 | -0.462862897 | 0.268520153 | 0.556053922 |
| chr1:226955029-226955229 | 0.580728806 | 0.268540612 | 0.55606741 |
| chr12:96219380-96220113 | -0.383270541 | 0.268541181 | 0.55606741 |
| chr2:43898402-43898801 | -0.445159853 | 0.268563015 | 0.556090164 |
| chr15:82566787-82567024 | 0.492999441 | 0.268566685 | 0.556090164 |
| chr3:97559456-97559778 | 0.547898744 | 0.268583341 | 0.556096489 |
| chr2:105790416-105790748 | -0.517813165 | 0.268584255 | 0.556096489 |
| chr1:65041481-65041799 | -0.439124141 | 0.268594286 | 0.556102232 |
| chr9:96617777-96617977 | 0.584536268 | 0.268611592 | 0.556123034 |
| chr11:110762782-110763065 | -0.59238818 | 0.268621659 | 0.55612885 |
| chr2:158745247-158745447 | -0.571836278 | 0.268642763 | 0.556157516 |
| chr16:70331763-70332098 | 0.531933507 | 0.268650057 | 0.556157589 |
| chr17:30731482-30732022 | -0.448025922 | 0.268676782 | 0.556197886 |
| chr15:31065888-31066148 | -0.516137484 | 0.268684831 | 0.556199522 |
| chr12:68661608-68661841 | -0.627525789 | 0.268701953 | 0.556219938 |
| chr5:61304503-61304870 | -0.421325736 | 0.268712122 | 0.556225962 |
| chr14:73591639-73592620 | 0.374094377 | 0.268738808 | 0.556252546 |
| chr15:42156281-42156962 | -0.403159845 | 0.268739484 | 0.556252546 |
| chr8:126824460-126825068 | -0.414825871 | 0.26874956 | 0.556258376 |
| chr4:89254806-89255094 | 0.46273832 | 0.268757135 | 0.556259028 |
| chr7:152445994-152446228 | -0.577702937 | 0.268798099 | 0.556328785 |
| chr2:113324805-113325084 | 0.493165169 | 0.268818126 | 0.556351414 |
| chr1:233705027-233705532 | 0.395848096 | 0.268823555 | 0.556351414 |
| chr15:48002675-48002914 | -0.5160802 | 0.268847393 | 0.556385721 |
| chr5:172622613-172622813 | -0.545942736 | 0.268859584 | 0.556388841 |
| chr8:98875938-98876138 | 0.586541449 | 0.268869488 | 0.556388841 |
| chr20:10671927-10672320 | -0.413838563 | 0.268872915 | 0.556388841 |
| chrX:135051398-135052553 | 0.352860387 | 0.268893438 | 0.556388841 |
| chr1:99849766-99850625 | -0.362767479 | 0.268897446 | 0.556388841 |
| chr7:32043049-32043273 | 0.488502282 | 0.268899776 | 0.556388841 |
| chr2:113876466-113877188 | 0.377049331 | 0.268905438 | 0.556388841 |
| chr14:44759770-44760038 | 0.524191829 | 0.268906992 | 0.556388841 |
| chr14:105999348-105999558 | 0.52725307 | 0.268917228 | 0.556394996 |
| chr4:13627379-13628410 | 0.362869525 | 0.268955293 | 0.556458726 |
| chr13:21176078-21177183 | 0.3511529 | 0.268974798 | 0.556483766 |
| chr11:12175689-12175898 | -0.571519124 | 0.268985716 | 0.556483766 |
| chrX:13028866-13029686 | -0.367181607 | 0.268989184 | 0.556483766 |
| chr3:154018123-154018396 | -0.47145283 | 0.268998475 | 0.556486993 |
| chr11:62013688-62013900 | -0.548375761 | 0.269005269 | 0.556486993 |
| chr20:45562568-45562791 | 0.552559177 | 0.269020122 | 0.556502695 |
| chr5:34967111-34967361 | -0.573085541 | 0.269035793 | 0.556520087 |
| chr11:17473392-17473766 | -0.403527942 | 0.269093106 | 0.556623616 |
| chr9:3827218-3827607 | -0.463571197 | 0.269128964 | 0.556682761 |
| chr4:99231654-99232223 | -0.449491168 | 0.269141986 | 0.556694667 |
| chr19:50050375-50050695 | 0.405876691 | 0.269164849 | 0.556726928 |
| chr5:92518337-92518821 | -0.419749942 | 0.269197879 | 0.556780216 |
| chr1:39893203-39893433 | -0.582954782 | 0.26921795 | 0.556801646 |
| chr9:32431994-32432241 | -0.56667951 | 0.269222774 | 0.556801646 |

| | | | |
|---|---|---|---|
| chr18:9102206-9103216 | 0.365572654 | 0.269269187 | 0.556821753 |
| chr13:51905868-51906089 | -0.543612101 | 0.269272299 | 0.556821753 |
| chr21:43883992-43884305 | 0.45575127 | 0.269274241 | 0.556821753 |
| chr6:161342213-161342437 | 0.591215701 | 0.269274919 | 0.556821753 |
| chr16:12819459-12819664 | 0.565922543 | 0.269288983 | 0.556821753 |
| chr3:36902740-36903121 | -0.447050752 | 0.269296145 | 0.556821753 |
| chr17:4498530-4498934 | 0.430487409 | 0.269299611 | 0.556821753 |
| chr7:75527884-75528589 | -0.371219045 | 0.269300439 | 0.556821753 |
| chr1:39129068-39129268 | 0.528929043 | 0.269300566 | 0.556821753 |
| chr11:61435595-61435795 | 0.534019279 | 0.269313489 | 0.556821753 |
| chr4:103415783-103416167 | -0.481792742 | 0.269318478 | 0.556821753 |
| chr13:39257756-39257956 | -0.558458194 | 0.269323999 | 0.556821753 |
| chr2:223836829-223838146 | -0.365540621 | 0.269326969 | 0.556821753 |
| chr16:77236038-77236783 | 0.36282317 | 0.269377782 | 0.556908207 |
| chr5:1799693-1800090 | 0.38112198 | 0.269387975 | 0.556908207 |
| chr21:41535887-41536925 | -0.367476477 | 0.269396795 | 0.556908207 |
| chr11:80998354-80998669 | -0.543367038 | 0.269397858 | 0.556908207 |
| chr3:100761261-100761561 | -0.44321841 | 0.269440166 | 0.556980639 |
| chr19:1249471-1249919 | 0.381605979 | 0.269480394 | 0.557038625 |
| chr22:31085280-31085480 | 0.498399855 | 0.26948625 | 0.557038625 |
| chr1:11749260-11749501 | 0.499184147 | 0.269505799 | 0.557038625 |
| chr6:10520719-10520932 | -0.528437243 | 0.269515578 | 0.557038625 |
| chr12:53379346-53380855 | 0.345457505 | 0.269520338 | 0.557038625 |
| chr16:31996367-31996646 | 0.500528291 | 0.269521421 | 0.557038625 |
| chr11:111429159-111429359 | -0.591286603 | 0.269522515 | 0.557038625 |
| chr10:98488667-98488867 | -0.595879238 | 0.269526376 | 0.557038625 |
| chr5:92051088-92051349 | -0.531735609 | 0.269540426 | 0.557052638 |
| chr21:29378797-29379213 | 0.393411555 | 0.269548768 | 0.557054853 |
| chr16:20763830-20764088 | 0.48742409 | 0.269559899 | 0.55706283 |
| chr20:50788059-50788648 | 0.350635567 | 0.26957613 | 0.557081349 |
| chr11:68213564-68214092 | 0.387678147 | 0.269588971 | 0.557092859 |
| chr11:14249398-14249598 | -0.554129134 | 0.269632103 | 0.557166964 |
| chr6:138227451-138227997 | 0.391331857 | 0.269650936 | 0.557186284 |
| chr6:143673701-143673995 | 0.474673827 | 0.269655997 | 0.557186284 |
| chr18:57959350-57959612 | -0.485471868 | 0.269678501 | 0.557217757 |
| chr13:114065037-114065757 | -0.372081211 | 0.269714688 | 0.557277501 |
| chr7:100538492-100539393 | -0.356999859 | 0.269733931 | 0.557302232 |
| chr13:110507699-110508170 | 0.437366606 | 0.269741766 | 0.557303392 |
| chr11:94767978-94768586 | -0.374889532 | 0.269752354 | 0.557310241 |
| chr2:181077165-181077403 | -0.516768521 | 0.269769874 | 0.557331408 |
| chr15:40694394-40695623 | -0.356549437 | 0.269789916 | 0.557357787 |
| chr6:152552658-152552899 | -0.513386801 | 0.269819502 | 0.55740388 |
| chr3:129026998-129027553 | 0.371006667 | 0.269856346 | 0.557464963 |
| chr17:68328510-68329618 | 0.348970976 | 0.269875255 | 0.557488995 |
| chr18:50374684-50375401 | -0.377936815 | 0.269922602 | 0.557571768 |
| chr12:3350378-3350639 | -0.549987243 | 0.269935238 | 0.557582837 |
| chr11:61699763-61700122 | -0.448642491 | 0.269943319 | 0.557584498 |
| chr19:33080765-33081286 | 0.396558458 | 0.269967657 | 0.557591786 |
| chr17:58932987-58933258 | 0.407540582 | 0.269973215 | 0.557591786 |
| chr10:68174534-68174734 | -0.620566699 | 0.269977212 | 0.557591786 |

| | | | |
|---|---|---|---|
| chr19:12802191-12802419 | 0.518583271 | 0.269982049 | 0.557591786 |
| chr3:150925558-150926157 | -0.383102701 | 0.269985744 | 0.557591786 |
| chr2:111068186-111068817 | -0.398035341 | 0.27000235 | 0.557591786 |
| chr1:42932169-42932373 | -0.502209342 | 0.270009097 | 0.557591786 |
| chr1:172401962-172402185 | -0.502209342 | 0.270009097 | 0.557591786 |
| chr14:60721672-60721990 | 0.395971757 | 0.270012341 | 0.557591786 |
| chr2:16525299-16525766 | -0.421077338 | 0.27004109 | 0.557636124 |
| chr10:80945147-80945584 | 0.417431546 | 0.270105035 | 0.557753139 |
| chr3:154252825-154253425 | -0.376878167 | 0.270114633 | 0.557757927 |
| chr1:193443259-193443506 | 0.550558427 | 0.270141092 | 0.557797531 |
| chr4:166463743-166463974 | -0.52417651 | 0.27018692 | 0.557871241 |
| chr1:229781631-229781859 | 0.518351039 | 0.270199288 | 0.557871241 |
| chr9:119963414-119963614 | -0.53668518 | 0.270199784 | 0.557871241 |
| chr4:173505898-173506424 | -0.388713814 | 0.270205913 | 0.557871241 |
| chr11:134293147-134293359 | 0.527762031 | 0.270248833 | 0.557944819 |
| chr12:101764497-101764726 | 0.544505724 | 0.270270217 | 0.557973934 |
| chr2:208024989-208025787 | -0.385078549 | 0.27034958 | 0.558122742 |
| chr10:75280866-75281348 | -0.494617373 | 0.270386727 | 0.55818439 |
| chr6:125986205-125987245 | -0.35138809 | 0.270414868 | 0.558227443 |
| chr9:137578479-137579105 | 0.373934268 | 0.270428715 | 0.558240989 |
| chr18:36933335-36933728 | -0.504477373 | 0.270466438 | 0.558303817 |
| chr1:228471880-228472087 | 0.532719169 | 0.270484818 | 0.558326718 |
| chr14:49380698-49380898 | 0.563124872 | 0.270519276 | 0.55834506 |
| chr13:47255831-47256068 | -0.489861339 | 0.27052078 | 0.55834506 |
| chr7:57417785-57418002 | -0.578592324 | 0.270521532 | 0.55834506 |
| chr10:78406993-78407605 | -0.385255406 | 0.270522852 | 0.55834506 |
| chr2:120917366-120917717 | 0.514578512 | 0.270548769 | 0.558382084 |
| chr11:33735887-33736691 | -0.361700661 | 0.270555366 | 0.558382084 |
| chr17:63752701-63753028 | 0.434368855 | 0.270583296 | 0.558424685 |
| chr8:6919421-6919735 | -0.571042676 | 0.270601238 | 0.558446672 |
| chr7:37208458-37208823 | 0.432040674 | 0.270611554 | 0.558452922 |
| chr3:184241609-184242191 | -0.415811038 | 0.270645714 | 0.558502622 |
| chr3:12706398-12706836 | 0.383503327 | 0.270659838 | 0.558502622 |
| chr3:130746263-130747084 | -0.359469965 | 0.270663049 | 0.558502622 |
| chr16:14405719-14405919 | 0.552694183 | 0.270671983 | 0.558502622 |
| chr6:99282543-99282797 | -0.50166527 | 0.270672083 | 0.558502622 |
| chr17:57459213-57459764 | -0.373632786 | 0.27067966 | 0.558503216 |
| chr1:202671503-202671753 | -0.442008518 | 0.270688421 | 0.558506252 |
| chr1:224304769-224305005 | -0.581169798 | 0.27070126 | 0.558517704 |
| chr3:15241441-15241672 | -0.58051279 | 0.270720184 | 0.558541707 |
| chr3:134291939-134292235 | -0.512547716 | 0.270753988 | 0.558580877 |
| chr12:92649948-92650330 | -0.413519957 | 0.270757856 | 0.558580877 |
| chr19:42296112-42296512 | 0.422445645 | 0.270765699 | 0.558580877 |
| chr17:17387949-17388502 | -0.377350677 | 0.270768329 | 0.558580877 |
| chr6:52552011-52552592 | 0.399264228 | 0.270780452 | 0.558590846 |
| chr6:81753742-81754212 | -0.37985547 | 0.270787849 | 0.558591066 |
| chr17:38887765-38888024 | 0.517577302 | 0.270816089 | 0.558634282 |
| chr1:114330352-114330687 | 0.385909241 | 0.270868047 | 0.558726418 |
| chr7:151552267-151553022 | -0.374676593 | 0.270881417 | 0.558738955 |
| chr14:54503441-54503894 | -0.388138205 | 0.270896103 | 0.558754205 |

| | | | |
|---|---|---|---|
| chr7:27150447-27151210 | 0.372757183 | 0.270906082 | 0.558759746 |
| chr19:5827781-5828396 | 0.3921672 | 0.270956688 | 0.55883555 |
| chr8:18179650-18179909 | -0.496601644 | 0.270957421 | 0.55883555 |
| chr14:96501722-96502896 | 0.349561333 | 0.270969328 | 0.558843297 |
| chr3:107927273-107928567 | -0.355649065 | 0.270982467 | 0.558843297 |
| chr8:125451865-125452597 | -0.359127795 | 0.270983058 | 0.558843297 |
| chr9:37591906-37592882 | -0.368756514 | 0.271005261 | 0.558860526 |
| chr19:33235368-33235651 | 0.442396398 | 0.271006 | 0.558860526 |
| chr20:44080803-44081276 | 0.403000865 | 0.271034814 | 0.558876644 |
| chr13:27833631-27833836 | -0.586371419 | 0.271035025 | 0.558876644 |
| chr2:10097308-10097743 | 0.40408041 | 0.271035698 | 0.558876644 |
| chr21:38614199-38614399 | -0.54113464 | 0.27104552 | 0.558881858 |
| chr20:53957844-53958521 | 0.350025945 | 0.271068137 | 0.558913453 |
| chr4:179290334-179290590 | -0.543534892 | 0.271115552 | 0.558971058 |
| chr7:50467854-50468375 | -0.388604176 | 0.271122057 | 0.558971058 |
| chr1:233721301-233721623 | -0.446299735 | 0.271122784 | 0.558971058 |
| chr5:43232044-43232336 | 0.473585044 | 0.271125256 | 0.558971058 |
| chr6:79813940-79814163 | 0.524782597 | 0.271177549 | 0.559063827 |
| chr21:38804634-38806371 | -0.344896066 | 0.271204256 | 0.559103842 |
| chr2:240586199-240586860 | 0.37711157 | 0.271230625 | 0.559143159 |
| chr8:38509788-38510774 | -0.378952079 | 0.271238648 | 0.559144655 |
| chr14:99622014-99622320 | -0.42257085 | 0.271250471 | 0.559153984 |
| chrX:49079756-49080312 | 0.38136872 | 0.271259446 | 0.559157443 |
| chr6:152982838-152984053 | -0.345546592 | 0.271296881 | 0.559219565 |
| chr15:63752256-63752478 | -0.561995908 | 0.271332517 | 0.559277974 |
| chr10:4663122-4663465 | 0.442070938 | 0.271413806 | 0.55943048 |
| chr1:58466660-58466889 | 0.514910401 | 0.271421435 | 0.559431156 |
| chrX:113929050-113929318 | 0.47302006 | 0.271430908 | 0.559435632 |
| chr6:1596278-1596517 | 0.478188996 | 0.27145118 | 0.559451139 |
| chr11:78639856-78640056 | 0.599927638 | 0.271453035 | 0.559451139 |
| chr7:12570135-12571175 | 0.356774174 | 0.271494499 | 0.559519365 |
| chr1:23483870-23484070 | 0.531594766 | 0.271500744 | 0.559519365 |
| chr1:211017156-211017681 | -0.45613451 | 0.271527605 | 0.559559672 |
| chr1:36203786-36203986 | -0.542869431 | 0.271535083 | 0.559560032 |
| chr8:31185333-31185533 | 0.537452809 | 0.271555789 | 0.559587652 |
| chr11:68214108-68214484 | 0.421563413 | 0.271573208 | 0.559608496 |
| chr10:31808456-31808914 | -0.456918393 | 0.271595613 | 0.559639615 |
| chr8:121617419-121617660 | -0.540946442 | 0.271630903 | 0.559693944 |
| chr1:26530138-26530338 | -0.568397918 | 0.271636589 | 0.559693944 |
| chr5:177585297-177585598 | -0.522258597 | 0.271701917 | 0.559813495 |
| chr6:6694326-6694684 | -0.403886096 | 0.27171102 | 0.559817198 |
| chr2:24083740-24085289 | -0.346906221 | 0.271772173 | 0.559915014 |
| chr19:2069788-2069988 | 0.590254823 | 0.271773111 | 0.559915014 |
| chr4:76950388-76950781 | 0.404132301 | 0.271811098 | 0.559978219 |
| chr2:111702155-111702420 | 0.458831394 | 0.271825598 | 0.559993036 |
| chr5:146947248-146947495 | -0.517470552 | 0.271851086 | 0.560016966 |
| chr13:60431154-60431372 | -0.579192718 | 0.27185452 | 0.560016966 |
| chr2:11822667-11823307 | -0.388836629 | 0.271861207 | 0.560016966 |
| chr20:38139210-38139514 | 0.464098223 | 0.271871129 | 0.560016966 |
| chr19:3306743-3306943 | 0.531357238 | 0.271877036 | 0.560016966 |

| | | | |
|---|---|---|---|
| chr10:133278429-133279246 | 0.367267188 | 0.271881067 | 0.560016966 |
| chr7:145463028-145463258 | -0.56586628 | 0.271914912 | 0.560071623 |
| chr17:41456303-41456552 | 0.528069241 | 0.271939606 | 0.560099683 |
| chr3:102741207-102741539 | -0.431784667 | 0.271943205 | 0.560099683 |
| chr14:59688144-59688757 | -0.409315515 | 0.271951891 | 0.560099683 |
| chr10:101031938-101032725 | -0.368021335 | 0.271957775 | 0.560099683 |
| chr10:113126540-113126813 | -0.60344818 | 0.271966104 | 0.560101783 |
| chr2:18794195-18794449 | 0.56487923 | 0.271987908 | 0.560127452 |
| chr5:20336063-20336379 | -0.502963175 | 0.271993189 | 0.560127452 |
| chr3:187709216-187709533 | -0.475431419 | 0.272011593 | 0.560150299 |
| chr17:61106220-61106619 | 0.418505276 | 0.272039499 | 0.560192709 |
| chr7:151612469-151612716 | -0.558251799 | 0.272049463 | 0.560198171 |
| chr2:73101677-73101917 | 0.558772713 | 0.272056974 | 0.560198583 |
| chr13:59603607-59603871 | -0.56023765 | 0.272066525 | 0.560199015 |
| chr20:20237335-20237625 | 0.495276164 | 0.272077745 | 0.560199015 |
| chr10:42999574-42999774 | 0.554376299 | 0.272079118 | 0.560199015 |
| chr10:44284279-44284602 | -0.477150018 | 0.272091725 | 0.560209919 |
| chr11:100841815-100842015 | 0.551494627 | 0.272105875 | 0.560212468 |
| chr20:54090157-54090502 | 0.364163842 | 0.272107585 | 0.560212468 |
| chr1:156284169-156284379 | 0.443467409 | 0.27212014 | 0.560221224 |
| chr20:9617107-9617378 | -0.465963701 | 0.272126461 | 0.560221224 |
| chr3:139628910-139629354 | 0.414734215 | 0.272136218 | 0.560226257 |
| chr9:124885624-124885824 | -0.585512003 | 0.272146559 | 0.560232494 |
| chr14:51552348-51552623 | -0.468380594 | 0.272156349 | 0.560237595 |
| chr10:80855464-80856014 | -0.425364588 | 0.272168044 | 0.560246617 |
| chr12:12032869-12033352 | -0.453828098 | 0.27218944 | 0.560275608 |
| chr1:101653460-101653682 | -0.521557988 | 0.272213528 | 0.560310139 |
| chr2:96509412-96509844 | 0.408206879 | 0.27222562 | 0.560316315 |
| chr13:34469263-34469904 | 0.381722883 | 0.272233197 | 0.560316315 |
| chrX:120629703-120630619 | 0.35294037 | 0.272239711 | 0.560316315 |
| chr6:143232905-143233182 | -0.495242915 | 0.272250266 | 0.560316315 |
| chr7:138936983-138937418 | -0.506752929 | 0.272253092 | 0.560316315 |
| chr3:62833511-62833832 | -0.40222563 | 0.272269907 | 0.56033587 |
| chr6:11044156-11044754 | -0.421409121 | 0.272283218 | 0.560348214 |
| chr21:43344466-43344729 | 0.442481496 | 0.272298124 | 0.56036384 |
| chr16:74286158-74286358 | 0.586708097 | 0.272309283 | 0.560368781 |
| chr1:67211708-67211928 | -0.560593827 | 0.272315152 | 0.560368781 |
| chr2:185441586-185441786 | -0.62472153 | 0.272354781 | 0.56040383 |
| chr2:221969626-221969834 | -0.62472153 | 0.272354781 | 0.56040383 |
| chr14:54488375-54489329 | 0.356198505 | 0.272358501 | 0.56040383 |
| chr6:85449775-85450609 | 0.379450337 | 0.27236144 | 0.56040383 |
| chrX:20527547-20527747 | -0.582382078 | 0.272374758 | 0.560408438 |
| chr12:26937789-26939001 | -0.347859182 | 0.272379496 | 0.560408438 |
| chr14:32745444-32746039 | 0.372861016 | 0.272389089 | 0.560408438 |
| chr14:50007350-50007833 | -0.396305689 | 0.272392935 | 0.560408438 |
| chr13:98419750-98420401 | -0.393314983 | 0.272409623 | 0.560427724 |
| chr17:45819771-45820011 | 0.561109924 | 0.272425661 | 0.560445671 |
| chr3:134250126-134251220 | 0.35986654 | 0.272457488 | 0.560484538 |
| chr8:94949182-94949851 | 0.360478082 | 0.272461769 | 0.560484538 |
| chr12:64659350-64659562 | 0.578741192 | 0.272466499 | 0.560484538 |

| | | | |
|---|---|---|---|
| chr7:135877882-135878200 | -0.453050945 | 0.272479857 | 0.560496968 |
| chr1:215576373-215576599 | -0.518353214 | 0.272579426 | 0.560686732 |
| chr9:112925121-112925321 | -0.562380657 | 0.272617613 | 0.560745363 |
| chr9:96031514-96031825 | 0.462336317 | 0.272622566 | 0.560745363 |
| chr2:73290245-73290537 | -0.416998289 | 0.272647245 | 0.560768955 |
| chr1:83641243-83641482 | -0.578827221 | 0.272656641 | 0.560768955 |
| chr13:112968378-112968811 | 0.430873788 | 0.272661459 | 0.560768955 |
| chr19:58347179-58347924 | 0.367388853 | 0.272663311 | 0.560768955 |
| chr14:56660030-56660257 | -0.523025604 | 0.272691396 | 0.560811664 |
| chr11:85606342-85606579 | -0.559350376 | 0.272702202 | 0.560812304 |
| chr7:47423408-47423767 | -0.454095514 | 0.272709478 | 0.560812304 |
| chr6:28725927-28726250 | -0.435404853 | 0.272713665 | 0.560812304 |
| chr18:76625473-76625673 | -0.60067254 | 0.272739683 | 0.560836323 |
| chr16:19395730-19396229 | -0.438190091 | 0.272742339 | 0.560836323 |
| chr4:120256109-120256842 | -0.388184296 | 0.272747303 | 0.560836323 |
| chr19:48619037-48619884 | 0.357219236 | 0.272765467 | 0.56085862 |
| chr4:145180193-145181230 | -0.348890433 | 0.272841013 | 0.560998902 |
| chr11:103108974-103110179 | -0.355509759 | 0.272857005 | 0.56101673 |
| chrX:53088619-53088926 | 0.405908501 | 0.272881956 | 0.561052975 |
| chr5:126847761-126848035 | -0.527975725 | 0.272919496 | 0.561115102 |
| chr1:24909535-24910049 | -0.430259876 | 0.272948547 | 0.561159773 |
| chr2:240983630-240983835 | 0.535595334 | 0.2730145 | 0.561260747 |
| chr6:11810066-11810279 | -0.545725069 | 0.273035213 | 0.561260747 |
| chr3:23900816-23901125 | -0.476233742 | 0.273045683 | 0.561260747 |
| chr14:105767531-105767926 | 0.454995925 | 0.273045692 | 0.561260747 |
| chr20:40962645-40963333 | 0.359393153 | 0.273046653 | 0.561260747 |
| chr11:84780770-84780990 | -0.579212244 | 0.273051728 | 0.561260747 |
| chr16:28070739-28071041 | 0.447058977 | 0.273052212 | 0.561260747 |
| chr2:197728466-197729810 | -0.357398935 | 0.273056261 | 0.561260747 |
| chr3:4720768-4720994 | -0.465117422 | 0.273071561 | 0.561277138 |
| chr3:87503956-87504428 | -0.410306609 | 0.273103601 | 0.561327935 |
| chr11:67792387-67792719 | -0.502799553 | 0.273162397 | 0.561423299 |
| chr5:43010652-43010924 | 0.416447403 | 0.273164653 | 0.561423299 |
| chr6:148342211-148343275 | -0.370568077 | 0.273173605 | 0.561426639 |
| chr17:74602079-74602382 | 0.460542118 | 0.273181993 | 0.56142882 |
| chr11:61985970-61986324 | 0.437212282 | 0.273199801 | 0.561450358 |
| chr6:11390006-11390382 | 0.439457321 | 0.273238515 | 0.561513944 |
| chr10:14296124-14296954 | -0.408625025 | 0.273245398 | 0.561513944 |
| chr15:36347627-36347840 | 0.553738044 | 0.273273427 | 0.561556481 |
| chr10:92239635-92240159 | 0.401334063 | 0.27329793 | 0.561591773 |
| chr12:6567956-6568715 | -0.379545064 | 0.27332789 | 0.561638274 |
| chr1:42992302-42992764 | -0.412886736 | 0.273362319 | 0.561693955 |
| chr19:29780013-29780221 | 0.594141053 | 0.273385213 | 0.561725933 |
| chr1:205879669-205879878 | 0.485192966 | 0.273419044 | 0.561780382 |
| chr14:103698852-103699202 | 0.413657845 | 0.273510426 | 0.561933238 |
| chr17:39754181-39754744 | 0.392925999 | 0.273514528 | 0.561933238 |
| chr14:23665957-23666186 | 0.462703579 | 0.273515441 | 0.561933238 |
| chr20:62209414-62209710 | 0.420829745 | 0.273545764 | 0.561980468 |
| chr2:217387706-217388434 | -0.379363946 | 0.273562949 | 0.562000704 |
| chr13:30971109-30971335 | -0.619137929 | 0.273588095 | 0.562002289 |

| | | | |
|---|---|---|---|
| chr14:21834771-21834971 | 0.592784402 | 0.273593382 | 0.562002289 |
| chr1:219221104-219221304 | -0.560706368 | 0.273607473 | 0.562002289 |
| chr11:63558640-63559069 | -0.434539184 | 0.273609238 | 0.562002289 |
| chr1:32626224-32626424 | 0.543234268 | 0.273624799 | 0.562002289 |
| chr9:92140348-92140626 | 0.439783776 | 0.273625686 | 0.562002289 |
| chr4:163333214-163333440 | -0.445877077 | 0.273628509 | 0.562002289 |
| chr5:127562347-127562573 | -0.556034709 | 0.273636224 | 0.562002289 |
| chr7:115475329-115475653 | -0.569351934 | 0.273636838 | 0.562002289 |
| chr4:72567788-72568571 | -0.361651901 | 0.273637068 | 0.562002289 |
| chr1:60455258-60455646 | -0.520343674 | 0.27365844 | 0.562022963 |
| chr1:41387675-41387927 | 0.478787063 | 0.273664336 | 0.562022963 |
| chr10:29958110-29958310 | -0.591315519 | 0.273670868 | 0.562022963 |
| chr1:196030324-196030558 | 0.538320948 | 0.273682837 | 0.562022963 |
| chr12:122974076-122975404 | 0.348862532 | 0.273683809 | 0.562022963 |
| chr22:36180282-36180620 | -0.465931199 | 0.273693425 | 0.562027648 |
| chr4:107740368-107740734 | -0.446921365 | 0.27370793 | 0.562042371 |
| chr15:90852377-90852631 | 0.485859351 | 0.273728754 | 0.562061244 |
| chr11:65589442-65589642 | 0.503385192 | 0.273747626 | 0.562061244 |
| chr11:48109931-48110441 | -0.39444228 | 0.273751674 | 0.562061244 |
| chr12:70645853-70646053 | 0.54961029 | 0.273753173 | 0.562061244 |
| chr17:4807757-4808002 | 0.494467664 | 0.273753799 | 0.562061244 |
| chr14:75535808-75536427 | 0.394231884 | 0.273799544 | 0.562140104 |
| chr8:108080469-108080789 | -0.409650975 | 0.273832859 | 0.562164482 |
| chr6:154478287-154478487 | -0.632067691 | 0.273833322 | 0.562164482 |
| chr1:117201963-117202168 | 0.523836805 | 0.273833954 | 0.562164482 |
| chr1:9110787-9111047 | 0.514760595 | 0.273840765 | 0.562164482 |
| chrX:136162903-136163184 | 0.474288812 | 0.273892347 | 0.562255309 |
| chr11:43644058-43644361 | -0.480074858 | 0.273928361 | 0.562302034 |
| chr3:10524865-10525211 | 0.424610313 | 0.273933494 | 0.562302034 |
| chr15:98779494-98780124 | -0.366820345 | 0.273944986 | 0.562302034 |
| chr2:212538532-212538959 | 0.411668337 | 0.273956589 | 0.562302034 |
| chr11:66957820-66958901 | 0.349814872 | 0.273957537 | 0.562302034 |
| chr20:53734247-53734638 | 0.354144475 | 0.273962385 | 0.562302034 |
| chr10:102229822-102230348 | 0.391049244 | 0.273966479 | 0.562302034 |
| chr11:30422895-30423394 | 0.431740837 | 0.274003003 | 0.562355009 |
| chr9:75083230-75083525 | -0.435339967 | 0.274010216 | 0.562355009 |
| chr17:43900377-43900837 | 0.40495223 | 0.274020744 | 0.562355009 |
| chr2:64018826-64019810 | -0.366414155 | 0.274021647 | 0.562355009 |
| chr1:93638570-93638787 | -0.568165637 | 0.274041115 | 0.562364838 |
| chr1:199533506-199533706 | -0.568165637 | 0.274041115 | 0.562364838 |
| chr18:79290472-79290721 | -0.550719831 | 0.274054618 | 0.562377486 |
| chr16:74998515-74999510 | -0.35410319 | 0.274108463 | 0.562458908 |
| chr12:32918979-32919408 | 0.508776092 | 0.274109595 | 0.562458908 |
| chr14:96194873-96195313 | -0.423648753 | 0.27412057 | 0.562458908 |
| chr7:92589692-92590644 | -0.350547447 | 0.2741273 | 0.562458908 |
| chr3:32177900-32178194 | 0.431486499 | 0.274131 | 0.562458908 |
| chr19:55180085-55180302 | 0.477013817 | 0.27415001 | 0.562482851 |
| chr1:217161288-217161500 | -0.560423664 | 0.274158904 | 0.562486038 |
| chr5:15580720-15580966 | 0.546253272 | 0.274180551 | 0.562515387 |
| chr14:100159538-100159878 | -0.394883314 | 0.274196022 | 0.562532065 |

| | | | |
|---|---|---|---|
| chr1:193381744-193382029 | -0.479896406 | 0.274236105 | 0.562599236 |
| chr12:39442379-39443699 | 0.346738037 | 0.274250063 | 0.562612807 |
| chr3:139535043-139535355 | -0.44465387 | 0.274278259 | 0.562654697 |
| chr6:32796887-32797261 | 0.487377633 | 0.274287423 | 0.562654697 |
| chr13:46962641-46962841 | -0.497826702 | 0.274303161 | 0.562654697 |
| chr12:68223025-68223500 | -0.377657637 | 0.274304825 | 0.562654697 |
| chr6:43246522-43247748 | 0.354425045 | 0.274307199 | 0.562654697 |
| chr3:172140472-172141263 | 0.362621481 | 0.274334755 | 0.562696155 |
| chr22:17837353-17837740 | 0.439372382 | 0.274356837 | 0.562719102 |
| chr5:150449221-150450236 | -0.352256323 | 0.27436063 | 0.562719102 |
| chr14:75200348-75200904 | 0.36628143 | 0.274382423 | 0.562748736 |
| chr17:60217275-60217542 | 0.411016907 | 0.274397123 | 0.56275857 |
| chr1:222617527-222618203 | 0.363230518 | 0.274403171 | 0.56275857 |
| chr6:133923996-133924270 | -0.466110871 | 0.274409252 | 0.56275857 |
| chr6:4203818-4204248 | 0.43867196 | 0.274423078 | 0.562771862 |
| chr4:168487058-168487294 | -0.552961581 | 0.274446879 | 0.562805609 |
| chr5:140575428-140575731 | 0.44165517 | 0.274471545 | 0.562841127 |
| chr7:106891781-106892099 | -0.495238241 | 0.274483336 | 0.562850243 |
| chr14:49767502-49769134 | 0.345748694 | 0.274530578 | 0.562932051 |
| chr1:234323031-234323313 | -0.481147856 | 0.274556248 | 0.562969622 |
| chr7:36949836-36950104 | 0.455350452 | 0.274574686 | 0.562992361 |
| chr2:75464648-75464868 | -0.511375223 | 0.274595856 | 0.563020703 |
| chr7:158977094-158977399 | -0.418761235 | 0.274616408 | 0.563047776 |
| chr17:60367160-60367401 | 0.38441224 | 0.274635172 | 0.563059113 |
| chr11:84317123-84317542 | -0.415762479 | 0.274637578 | 0.563059113 |
| chr7:52294793-52295040 | 0.573742946 | 0.274643983 | 0.563059113 |
| chr5:131634678-131636107 | -0.34379979 | 0.274662723 | 0.563082466 |
| chr14:98070639-98070844 | -0.598853804 | 0.274674829 | 0.563092217 |
| chr19:51703816-51704429 | 0.370109028 | 0.274716349 | 0.563162269 |
| chr16:14486595-14486795 | 0.544519024 | 0.274731367 | 0.563177987 |
| chr4:140743216-140743811 | -0.392984965 | 0.274753486 | 0.563208261 |
| chr19:2151490-2152033 | 0.367934293 | 0.274768757 | 0.563224497 |
| chr18:76492685-76493272 | -0.367629909 | 0.274826641 | 0.563328078 |
| chr20:63636976-63637300 | -0.446484613 | 0.274866738 | 0.56338235 |
| chr6:144013468-144013863 | -0.42755634 | 0.274869849 | 0.56338235 |
| chr20:3053969-3054189 | 0.553679534 | 0.274875176 | 0.56338235 |
| chr7:1160706-1160965 | 0.418681968 | 0.274891182 | 0.563396241 |
| chr14:93206638-93207812 | 0.348314761 | 0.274903827 | 0.563396241 |
| chr10:123994335-123994980 | 0.370543435 | 0.274904755 | 0.563396241 |
| chr3:126982320-126983121 | 0.36151735 | 0.274914681 | 0.563396241 |
| chr5:35273754-35273954 | -0.575485411 | 0.274924483 | 0.563396241 |
| chr1:225772293-225772687 | 0.37690728 | 0.274926072 | 0.563396241 |
| chr6:133945181-133945527 | -0.436552219 | 0.27494311 | 0.563416089 |
| chr17:32998293-32998663 | 0.487988528 | 0.274976239 | 0.563458584 |
| chr1:1918949-1919846 | 0.356622585 | 0.274981164 | 0.563458584 |
| chr20:47357256-47357980 | 0.342610239 | 0.274985908 | 0.563458584 |
| chr1:106194271-106194574 | 0.401849632 | 0.274993697 | 0.563459475 |
| chr2:122451135-122451388 | -0.564160334 | 0.275005847 | 0.563459911 |
| chr17:40803619-40803867 | 0.524711862 | 0.275008618 | 0.563459911 |
| chr1:194836889-194837089 | 0.593601159 | 0.275041325 | 0.563508033 |

| | | | |
|---|---|---|---|
| chr8:65568246-65568522 | -0.552439048 | 0.275046813 | 0.563508033 |
| chr19:46746796-46747003 | 0.465508702 | 0.275071132 | 0.563542788 |
| chr16:30571535-30572536 | 0.355908448 | 0.275079382 | 0.563544622 |
| chr7:1569820-1570778 | 0.357003158 | 0.275100733 | 0.563573294 |
| chr3:171825269-171825732 | -0.419576313 | 0.275162684 | 0.563685136 |
| chr2:44286762-44287188 | -0.453519794 | 0.275180609 | 0.563706787 |
| chr15:37135287-37135551 | -0.490596305 | 0.275192229 | 0.563715519 |
| chr1:106201257-106201812 | 0.372269757 | 0.275239878 | 0.563798053 |
| chr1:76154127-76155021 | -0.367863075 | 0.27530356 | 0.563913422 |
| chr8:115665872-115666072 | 0.516579891 | 0.275311504 | 0.563914619 |
| chr13:30328148-30328348 | -0.576486919 | 0.275335981 | 0.563949679 |
| chr4:25723641-25723854 | 0.566890335 | 0.275352845 | 0.563965392 |
| chr9:102223241-102224049 | -0.363427242 | 0.275365468 | 0.563965392 |
| chr1:111441289-111441867 | -0.43480904 | 0.275365734 | 0.563965392 |
| chr13:42047595-42049771 | -0.342587042 | 0.275381246 | 0.563982087 |
| chr2:29207119-29207400 | 0.498413173 | 0.275460286 | 0.564106707 |
| chr2:183193814-183194190 | 0.470207462 | 0.275461746 | 0.564106707 |
| chr4:16443622-16443869 | 0.510811329 | 0.275470942 | 0.564106707 |
| chr2:85729011-85729267 | 0.466720932 | 0.275471544 | 0.564106707 |
| chr15:40937930-40938399 | 0.386812512 | 0.275482037 | 0.564113117 |
| chr2:196925957-196927404 | -0.347092848 | 0.275491718 | 0.564117866 |
| chr17:69524977-69525177 | -0.521573744 | 0.275501832 | 0.5641235 |
| chr13:36918990-36920677 | -0.355656551 | 0.275513141 | 0.564131581 |
| chr1:114809412-114809645 | 0.424413264 | 0.27554935 | 0.564184176 |
| chr7:111853647-111853847 | -0.513372092 | 0.275553554 | 0.564184176 |
| chr14:90705834-90706034 | 0.585365483 | 0.275566918 | 0.564196462 |
| chr5:146427237-146427483 | -0.500422752 | 0.275585073 | 0.564218556 |
| chr17:76612945-76613777 | -0.368438788 | 0.275603363 | 0.564240925 |
| chr9:83538082-83539096 | -0.353228819 | 0.275624136 | 0.564268378 |
| chr11:79578508-79578708 | 0.557033521 | 0.275649521 | 0.564301241 |
| chrX:104309209-104309490 | -0.462573835 | 0.275660302 | 0.564301241 |
| chr20:44909800-44910467 | 0.364509284 | 0.275662283 | 0.564301241 |
| chr2:105859351-105859567 | -0.536367195 | 0.27567279 | 0.564307674 |
| chr11:43119663-43119863 | 0.585258029 | 0.275685634 | 0.564315562 |
| chr10:124532185-124532477 | 0.412748171 | 0.275691374 | 0.564315562 |
| chr11:102451784-102452098 | 0.451211209 | 0.275713215 | 0.564345192 |
| chr2:174067551-174067793 | -0.553362444 | 0.275748804 | 0.564395612 |
| chr3:132670901-132671287 | -0.483621045 | 0.275753804 | 0.564395612 |
| chr1:109152148-109152468 | 0.410869011 | 0.275769269 | 0.564395612 |
| chr1:205880061-205880265 | 0.526156917 | 0.275773941 | 0.564395612 |
| chr18:36297152-36298028 | -0.393640635 | 0.275774677 | 0.564395612 |
| chr1:58803025-58803332 | -0.505579397 | 0.27579087 | 0.564413676 |
| chr3:130098061-130098298 | -0.518086866 | 0.275802755 | 0.564422923 |
| chr14:88233910-88234251 | 0.442565941 | 0.275849519 | 0.564503548 |
| chr7:1124937-1125340 | -0.417011299 | 0.275898998 | 0.564589724 |
| chr19:39914115-39914354 | 0.549678155 | 0.275910668 | 0.564591505 |
| chr8:85393934-85394256 | 0.461792092 | 0.275920779 | 0.564591505 |
| chr12:4538162-4539402 | -0.362937282 | 0.275926701 | 0.564591505 |
| chr10:87622289-87622526 | -0.519516045 | 0.275929342 | 0.564591505 |
| chr14:79708885-79709111 | 0.516548204 | 0.27596267 | 0.564639195 |

| | | | |
|---|---|---|---|
| chr3:197002740-197003814 | 0.359756225 | 0.275970416 | 0.564639195 |
| chr5:132820098-132820442 | -0.418909385 | 0.275977689 | 0.564639195 |
| chr10:103908304-103908651 | -0.431575498 | 0.275982546 | 0.564639195 |
| chr9:133873950-133874697 | 0.366723718 | 0.275989496 | 0.564639195 |
| chr4:102826843-102828586 | -0.339371802 | 0.276005628 | 0.564657124 |
| chr16:70380520-70382065 | 0.345217495 | 0.276028767 | 0.564689385 |
| chr2:207692239-207692525 | -0.492386918 | 0.276063935 | 0.564746251 |
| chr1:115041183-115042005 | 0.362537348 | 0.276083202 | 0.56476912 |
| chr17:37358443-37358643 | 0.546814126 | 0.276089856 | 0.56476912 |
| chr20:58460749-58461011 | 0.433055016 | 0.276102817 | 0.564780113 |
| chr14:58362834-58363135 | 0.448179171 | 0.276109972 | 0.564780113 |
| chr3:37987590-37987790 | 0.541881803 | 0.27612347 | 0.564792645 |
| chr15:61835531-61835795 | 0.451925322 | 0.276137503 | 0.564806272 |
| chr4:153085206-153085529 | -0.421853777 | 0.276145169 | 0.564806874 |
| chr10:28291831-28292287 | -0.406623419 | 0.276155353 | 0.564812628 |
| chr3:79521270-79521595 | -0.450217884 | 0.276173146 | 0.564833941 |
| chr5:60677727-60678388 | -0.382614206 | 0.276184019 | 0.564841102 |
| chr17:57408452-57408652 | 0.445659635 | 0.276221649 | 0.564886268 |
| chr3:165322402-165322841 | -0.473070403 | 0.276222486 | 0.564886268 |
| chr6:29015514-29016064 | -0.411849014 | 0.276229364 | 0.564886268 |
| chr3:94973665-94974179 | -0.482391561 | 0.276241117 | 0.564886268 |
| chr13:28753557-28753883 | -0.403049347 | 0.276254847 | 0.564886268 |
| chr14:37546774-37547327 | 0.38402196 | 0.27626937 | 0.564886268 |
| chr6:82624275-82624564 | -0.45267435 | 0.276270366 | 0.564886268 |
| chr3:120153425-120153674 | -0.539677385 | 0.276272051 | 0.564886268 |
| chr4:183906145-183906421 | -0.546452162 | 0.276284029 | 0.564886268 |
| chr18:77131811-77132550 | -0.381721715 | 0.276288067 | 0.564886268 |
| chr4:82422023-82422301 | -0.501182868 | 0.276295204 | 0.564886268 |
| chr15:40245233-40245577 | 0.416885633 | 0.276306171 | 0.564886268 |
| chr11:12914603-12914846 | -0.529613266 | 0.276311194 | 0.564886268 |
| chr8:102411591-102413150 | 0.341904965 | 0.276319483 | 0.564886268 |
| chr3:155270933-155271565 | 0.395529976 | 0.27633035 | 0.564886268 |
| chr1:187624428-187624726 | -0.573842075 | 0.276333317 | 0.564886268 |
| chr1:247103813-247104680 | -0.344550973 | 0.2763499 | 0.564886268 |
| chr1:112507907-112508206 | 0.481962844 | 0.276352954 | 0.564886268 |
| chr6:147971043-147971243 | 0.52661746 | 0.276355763 | 0.564886268 |
| chr1:160244291-160244561 | -0.496827076 | 0.276358926 | 0.564886268 |
| chr12:95505716-95505916 | -0.6176112 | 0.276360923 | 0.564886268 |
| chrX:103614916-103615270 | -0.536584064 | 0.276422675 | 0.564997418 |
| chr8:134866827-134867173 | -0.489330142 | 0.276509189 | 0.565153696 |
| chr15:66567086-66567514 | 0.419698557 | 0.276513885 | 0.565153696 |
| chr6:159042627-159042921 | -0.489557692 | 0.276538013 | 0.565187934 |
| chr5:61409596-61409916 | -0.438744572 | 0.276551083 | 0.56519957 |
| chr12:132593747-132594360 | 0.361907276 | 0.276567697 | 0.565199659 |
| chr5:160787284-160787581 | -0.457208516 | 0.276582419 | 0.565199659 |
| chr10:84407471-84407689 | -0.547706505 | 0.276590173 | 0.565199659 |
| chr20:53884403-53885369 | 0.342204039 | 0.276592116 | 0.565199659 |
| chr7:36987116-36987461 | 0.418311586 | 0.276600869 | 0.565199659 |
| chr17:60649529-60649746 | 0.397435434 | 0.276601873 | 0.565199659 |
| chr2:113235193-113235914 | 0.37216756 | 0.276602762 | 0.565199659 |

| | | | |
|---|---|---|---|
| chr11:4182138-4182435 | 0.431422052 | 0.276687421 | 0.565355881 |
| chr19:1905040-1905845 | 0.365817609 | 0.276701069 | 0.565355881 |
| chr2:12433702-12433944 | -0.515540952 | 0.276704027 | 0.565355881 |
| chr4:167622774-167623016 | -0.536969821 | 0.276708729 | 0.565355881 |
| chrX:100423700-100423976 | 0.515056275 | 0.276737781 | 0.565389151 |
| chr2:33786657-33786857 | -0.63691337 | 0.276746319 | 0.565389151 |
| chr14:92692755-92692955 | -0.516819887 | 0.27674715 | 0.565389151 |
| chr19:17292771-17293103 | 0.403160324 | 0.276775345 | 0.565431677 |
| chr3:163167373-163168210 | -0.387264903 | 0.276808704 | 0.565484751 |
| chr17:58786379-58786618 | 0.407006679 | 0.27683893 | 0.56553142 |
| chr1:65459434-65459706 | -0.480240995 | 0.276848576 | 0.565533258 |
| chrX:154525797-154526037 | 0.514943442 | 0.276864121 | 0.565533258 |
| chr1:153112370-153112570 | -0.585416686 | 0.276866818 | 0.565533258 |
| chr17:15509485-15509700 | 0.572517781 | 0.276869353 | 0.565533258 |
| chr1:28886780-28887379 | 0.373917093 | 0.276884523 | 0.565549167 |
| chr1:200629783-200630081 | 0.462637631 | 0.276903512 | 0.565572878 |
| chr5:119350635-119350867 | -0.5504469 | 0.276948562 | 0.565649812 |
| chr9:88310446-88310686 | -0.527220488 | 0.2770022 | 0.565744285 |
| chr18:45384423-45384731 | -0.476035686 | 0.277034499 | 0.56579517 |
| chr16:11668032-11668481 | 0.400541114 | 0.277090586 | 0.565894635 |
| chr5:161944982-161945418 | -0.387936494 | 0.277099243 | 0.565897231 |
| chr6:113433260-113434033 | -0.368773601 | 0.277141604 | 0.565968656 |
| chr3:62319774-62320199 | 0.357610935 | 0.277150678 | 0.565972102 |
| chr6:23547898-23548127 | 0.515358364 | 0.27722009 | 0.566092027 |
| chr13:41194133-41194984 | -0.350501385 | 0.27722418 | 0.566092027 |
| chr22:18149615-18150362 | -0.378903954 | 0.27725764 | 0.566145263 |
| chr15:75809998-75810270 | -0.473890174 | 0.277270982 | 0.566147973 |
| chr17:79789748-79790080 | -0.471930403 | 0.277273744 | 0.566147973 |
| chr10:14153817-14154064 | -0.544863851 | 0.277284641 | 0.566155135 |
| chr2:235493564-235494087 | 0.401075781 | 0.277309641 | 0.566191092 |
| chrX:151402777-151403037 | 0.482421636 | 0.277336389 | 0.566230617 |
| chr2:28508328-28508681 | -0.447664334 | 0.277344308 | 0.566231697 |
| chr5:133993224-133993433 | 0.479330484 | 0.277352371 | 0.566233071 |
| chr9:91727376-91727576 | 0.524245527 | 0.277424399 | 0.566324295 |
| chr19:3572549-3573064 | 0.372657553 | 0.277425605 | 0.566324295 |
| chr11:12816629-12816833 | 0.574553302 | 0.277430301 | 0.566324295 |
| chr13:41458301-41458780 | -0.393531118 | 0.277433954 | 0.566324295 |
| chr15:98865768-98866374 | 0.372982217 | 0.27743401 | 0.566324295 |
| chr2:191030606-191030894 | -0.426098898 | 0.277455107 | 0.566347881 |
| chr12:125983058-125984103 | 0.347856175 | 0.277460348 | 0.566347881 |
| chr1:184075476-184075708 | -0.530166513 | 0.27747068 | 0.566353883 |
| chr2:112480616-112480816 | 0.494556339 | 0.277488172 | 0.566374499 |
| chr19:35645009-35645309 | -0.409145141 | 0.277508628 | 0.566401163 |
| chr12:53743708-53743915 | 0.462220868 | 0.277524459 | 0.566412699 |
| chr12:119974192-119974745 | 0.394932186 | 0.277534201 | 0.566412699 |
| chr8:122884965-122885281 | 0.402722745 | 0.277536457 | 0.566412699 |
| chrX:106959934-106960409 | -0.397753054 | 0.277552202 | 0.566429747 |
| chr1:90035957-90036216 | -0.596596228 | 0.27759499 | 0.566474012 |
| chr1:59043755-59044201 | -0.430093382 | 0.2775967 | 0.566474012 |
| chr2:18991645-18992151 | -0.397630984 | 0.27760967 | 0.566474012 |

| | | | |
|---|---|---|---|
| chr4:184343444-184344163 | -0.400901251 | 0.277609756 | 0.566474012 |
| chr20:17996177-17996420 | 0.494368236 | 0.277624977 | 0.566474012 |
| chr7:72326431-72326755 | -0.46734936 | 0.277625079 | 0.566474012 |
| chr20:47912208-47912453 | 0.392949612 | 0.277625644 | 0.566474012 |
| chr19:52270028-52270255 | 0.504086209 | 0.277647147 | 0.566502801 |
| chr7:91263443-91263663 | -0.468129182 | 0.277655051 | 0.566503843 |
| chr9:92325041-92326344 | 0.3499815 | 0.277693415 | 0.56656703 |
| chr10:75402176-75402561 | 0.385302895 | 0.27773337 | 0.566623393 |
| chr10:132291952-132292401 | 0.467177055 | 0.27773583 | 0.566623393 |
| chr19:4343405-4343667 | 0.414716703 | 0.277744809 | 0.566626625 |
| chr15:69523426-69523641 | 0.544929813 | 0.277753715 | 0.566627707 |
| chr5:181160479-181160780 | 0.405168343 | 0.277768251 | 0.566627707 |
| chr14:52551687-52551890 | 0.429952142 | 0.277774219 | 0.566627707 |
| chr8:102585827-102586129 | 0.395651858 | 0.27777492 | 0.566627707 |
| chr1:247951751-247952032 | -0.570307657 | 0.277785629 | 0.566634466 |
| chr11:117844016-117844381 | 0.443082472 | 0.277807979 | 0.566655601 |
| chr4:123555412-123556040 | 0.382881548 | 0.277810781 | 0.566655601 |
| chr2:15041465-15041665 | -0.594515157 | 0.277841462 | 0.566703095 |
| chr3:10549920-10550216 | -0.435007835 | 0.277868269 | 0.566742687 |
| chr19:17245960-17246214 | 0.457354551 | 0.277889949 | 0.566750099 |
| chr15:69056485-69056685 | -0.517300933 | 0.277890104 | 0.566750099 |
| chr14:88562545-88563675 | 0.345297779 | 0.277894093 | 0.566750099 |
| chr21:34128220-34128420 | -0.585470886 | 0.277946239 | 0.566818529 |
| chr17:60222119-60222449 | 0.380097528 | 0.277954516 | 0.566818529 |
| chr7:27160393-27160593 | 0.5668796 | 0.277963179 | 0.566818529 |
| chr16:23516444-23516691 | -0.517368404 | 0.277963848 | 0.566818529 |
| chr11:36056414-36056644 | -0.500593569 | 0.277971643 | 0.566818529 |
| chr1:56547765-56548284 | 0.395455929 | 0.277972032 | 0.566818529 |
| chr3:170741328-170741678 | -0.420921033 | 0.277986572 | 0.566833092 |
| chr1:202022990-202023401 | -0.359686535 | 0.278023258 | 0.566892812 |
| chr9:89171705-89171907 | 0.563159911 | 0.27803825 | 0.566908294 |
| chr1:162230024-162230510 | -0.393554792 | 0.278060745 | 0.566935832 |
| chr1:32760865-32761454 | -0.37326748 | 0.278066554 | 0.566935832 |
| chr2:27085881-27086252 | -0.398740368 | 0.278091576 | 0.566958905 |
| chr5:43192159-43193121 | 0.366920306 | 0.278092669 | 0.566958905 |
| chr6:5303092-5303646 | 0.410776917 | 0.278109951 | 0.566979051 |
| chr8:97587999-97588199 | 0.521240512 | 0.278155707 | 0.567057245 |
| chr1:87335216-87335690 | 0.429652573 | 0.27819409 | 0.567112561 |
| chr20:46438191-46438391 | 0.524347911 | 0.278199705 | 0.567112561 |
| chr1:153511019-153511221 | 0.472910989 | 0.278205045 | 0.567112561 |
| chr11:118233360-118233655 | -0.558350328 | 0.278215423 | 0.567118629 |
| chr9:112392205-112392603 | -0.438286822 | 0.278237561 | 0.567148668 |
| chr2:224039527-224039774 | 0.500991276 | 0.278250425 | 0.567159801 |
| chr16:87818312-87818512 | -0.549280598 | 0.278332519 | 0.567286274 |
| chr1:84620612-84621310 | -0.359932839 | 0.278340672 | 0.567286274 |
| chr16:30821545-30821745 | -0.623180928 | 0.278349995 | 0.567286274 |
| chrX:121407697-121407953 | -0.496459675 | 0.278350319 | 0.567286274 |
| chr3:71195482-71195786 | -0.456274156 | 0.278352146 | 0.567286274 |
| chrX:100895507-100895882 | -0.477003446 | 0.278356896 | 0.567286274 |
| chr9:120477472-120477977 | -0.397997707 | 0.278372438 | 0.567302861 |

| | | | |
|---|---|---|---|
| chr11:62560584-62560942 | 0.405715336 | 0.278391667 | 0.567326957 |
| chr8:22366928-22367568 | 0.385597036 | 0.278418938 | 0.567354267 |
| chr2:99337058-99337697 | 0.360944176 | 0.278420122 | 0.567354267 |
| chr19:47210757-47211007 | 0.472420875 | 0.278427281 | 0.567354267 |
| chr2:26076045-26076542 | -0.449178094 | 0.278439615 | 0.567357589 |
| chr14:71917025-71917225 | 0.553827209 | 0.278443721 | 0.567357589 |
| chr19:37655299-37656461 | 0.358475031 | 0.27845289 | 0.567361184 |
| chr7:148045546-148045821 | -0.515246834 | 0.278474879 | 0.567372738 |
| chr10:97542227-97542647 | 0.405234743 | 0.278485721 | 0.567372738 |
| chr11:62787114-62787654 | 0.361812716 | 0.278488003 | 0.567372738 |
| chr17:29347810-29348099 | -0.492310494 | 0.27848818 | 0.567372738 |
| chr17:61574746-61575542 | 0.342676839 | 0.278519999 | 0.567422478 |
| chr16:27080978-27081230 | 0.492780859 | 0.278544123 | 0.56745626 |
| chr20:38173516-38173716 | -0.555202539 | 0.278552853 | 0.56745626 |
| chr20:23799585-23799785 | 0.558376074 | 0.278560017 | 0.56745626 |
| chr11:44950735-44951054 | 0.439958072 | 0.278572377 | 0.56745626 |
| chr19:20052083-20052650 | 0.397156937 | 0.278573611 | 0.56745626 |
| chr13:69537356-69537813 | -0.449047722 | 0.278602434 | 0.5674848 |
| chr14:63689127-63689506 | -0.449047722 | 0.278602434 | 0.5674848 |
| chr6:43687642-43688256 | 0.372444214 | 0.278616984 | 0.56749935 |
| chr3:58306916-58307302 | 0.407363799 | 0.27865244 | 0.567545416 |
| chr6:129499074-129499343 | 0.467085312 | 0.278654414 | 0.567545416 |
| chr3:157785144-157785373 | -0.545346067 | 0.278675961 | 0.567574214 |
| chrX:41442025-41442800 | 0.363041128 | 0.278683451 | 0.567574381 |
| chr11:119315621-119315957 | -0.454407948 | 0.27870253 | 0.567598152 |
| chr15:87582234-87582668 | -0.418748792 | 0.278795834 | 0.567773081 |
| chr1:31412964-31413287 | 0.397005181 | 0.2788768 | 0.567922875 |
| chr15:69951548-69951865 | 0.456286351 | 0.278887332 | 0.567925663 |
| chr1:81912307-81912691 | -0.455073036 | 0.278892993 | 0.567925663 |
| chr7:76191844-76192185 | -0.423937245 | 0.278915704 | 0.567939704 |
| chr9:129802815-129803517 | 0.373942664 | 0.278917726 | 0.567939704 |
| chr6:52063566-52063839 | -0.495461288 | 0.278922125 | 0.567939704 |
| chr2:119641562-119641927 | 0.416225705 | 0.278967508 | 0.568017018 |
| chr6:21593359-21594376 | -0.35300408 | 0.278975192 | 0.56801757 |
| chr14:95915470-95915670 | 0.572424296 | 0.279010411 | 0.56806568 |
| chr6:81450209-81450471 | -0.55742653 | 0.279013649 | 0.56806568 |
| chr10:110610338-110610588 | -0.481343638 | 0.279043643 | 0.56807399 |
| chr9:93003622-93004428 | -0.371260708 | 0.279047644 | 0.56807399 |
| chr20:10434943-10435554 | -0.352970352 | 0.279048553 | 0.56807399 |
| chr4:143488631-143489381 | -0.395482794 | 0.279056236 | 0.56807399 |
| chr17:2011392-2012095 | -0.371383208 | 0.279056783 | 0.56807399 |
| chr3:21216685-21216989 | -0.516117083 | 0.279064978 | 0.56807399 |
| chr3:50321100-50321567 | -0.395958116 | 0.279069628 | 0.56807399 |
| chr2:75493316-75493754 | 0.418100277 | 0.279091837 | 0.568091942 |
| chr9:99307931-99308241 | 0.451374257 | 0.279096069 | 0.568091942 |
| chr20:39264549-39264863 | 0.452995677 | 0.279104766 | 0.568091942 |
| chr14:78758355-78758694 | -0.434319189 | 0.279114109 | 0.568091942 |
| chr5:59188576-59188777 | -0.571841676 | 0.279115518 | 0.568091942 |
| chr20:51712524-51713043 | -0.378998311 | 0.27913289 | 0.568112209 |
| chr20:31612876-31613211 | 0.383617508 | 0.279147586 | 0.568127028 |

| | | | |
|---|---|---|---|
| chr2:71241040-71241272 | -0.589424428 | 0.279167274 | 0.568152005 |
| chr12:131234842-131235164 | -0.455357307 | 0.279195183 | 0.568193713 |
| chr10:116742399-116743332 | 0.357317442 | 0.279219966 | 0.568229058 |
| chr10:113982685-113983541 | -0.363006885 | 0.279234656 | 0.568233493 |
| chr15:88647797-88648158 | 0.428338483 | 0.279241932 | 0.568233493 |
| chr14:91140793-91141051 | 0.447784508 | 0.279248301 | 0.568233493 |
| chr10:84375056-84375256 | 0.54335957 | 0.27925181 | 0.568233493 |
| chr1:186919425-186919625 | 0.514315989 | 0.279273569 | 0.568262677 |
| chr2:62787401-62787669 | -0.497925954 | 0.279295987 | 0.568293202 |
| chr2:190088943-190089184 | -0.515798096 | 0.279324454 | 0.568309408 |
| chr12:6383807-6384269 | 0.388100153 | 0.279327357 | 0.568309408 |
| chr2:70655247-70655511 | 0.528454167 | 0.279331422 | 0.568309408 |
| chr2:19369415-19369654 | -0.509121042 | 0.27933362 | 0.568309408 |
| chr7:134779368-134779738 | -0.441733299 | 0.279341825 | 0.568311012 |
| chr10:69179434-69180686 | -0.351730413 | 0.279440841 | 0.568494014 |
| chr8:80551213-80551475 | -0.542632405 | 0.279451801 | 0.568494014 |
| chr10:74293835-74294035 | -0.582287866 | 0.279454035 | 0.568494014 |
| chr17:78863974-78864227 | -0.491296345 | 0.279486265 | 0.568522643 |
| chr1:28517550-28517771 | -0.448331934 | 0.279487064 | 0.568522643 |
| chr16:85520872-85521378 | 0.373303478 | 0.279490367 | 0.568522643 |
| chr8:137847122-137847395 | -0.450192295 | 0.279501337 | 0.568529864 |
| chr17:72399222-72399468 | 0.473284811 | 0.279555597 | 0.568625137 |
| chr1:235073339-235074405 | 0.353562763 | 0.279570262 | 0.568630671 |
| chr9:129884454-129884732 | 0.423074637 | 0.279573989 | 0.568630671 |
| chr9:82923109-82923846 | -0.392360536 | 0.279580581 | 0.568630671 |
| chr7:132072832-132073073 | -0.55236714 | 0.27959383 | 0.568640636 |
| chr3:73614383-73614746 | -0.422621294 | 0.279606594 | 0.568640636 |
| chr9:69476725-69477007 | 0.539111094 | 0.27961438 | 0.568640636 |
| chr1:183633614-183634027 | -0.398916851 | 0.279615166 | 0.568640636 |
| chr4:6309333-6309552 | 0.481702556 | 0.279628522 | 0.568652704 |
| chr10:126902243-126902561 | -0.461010748 | 0.279659685 | 0.568694654 |
| chr6:147033285-147033589 | -0.505977489 | 0.279670263 | 0.568694654 |
| chr17:60425783-60426085 | 0.364219341 | 0.279678446 | 0.568694654 |
| chr20:59323671-59323871 | 0.506066585 | 0.279678838 | 0.568694654 |
| chr15:40764004-40764528 | 0.380837464 | 0.279692311 | 0.568701267 |
| chr16:85746389-85746656 | 0.421274199 | 0.279701355 | 0.568701267 |
| chr2:120456983-120457215 | 0.516728382 | 0.279705074 | 0.568701267 |
| chr16:22375316-22375516 | 0.497691142 | 0.279711779 | 0.568701267 |
| chr2:27169828-27170154 | -0.458557593 | 0.279724874 | 0.568706823 |
| chr5:74916740-74917289 | -0.404651723 | 0.279735052 | 0.568706823 |
| chrX:41971408-41971678 | -0.522482954 | 0.279736779 | 0.568706823 |
| chr1:107080419-107080619 | 0.430762893 | 0.279750737 | 0.568713338 |
| chr8:122742827-122743312 | -0.406095824 | 0.279754828 | 0.568713338 |
| chr10:30392509-30392848 | -0.416493785 | 0.279771426 | 0.568722123 |
| chr8:11787810-11788297 | -0.424817269 | 0.279785144 | 0.568722123 |
| chr17:13751257-13751719 | 0.435884286 | 0.279786344 | 0.568722123 |
| chr4:148829191-148829470 | 0.524793464 | 0.279788839 | 0.568722123 |
| chr1:168153389-168153703 | -0.603799485 | 0.27980298 | 0.568733968 |
| chr20:60107718-60107972 | 0.472700194 | 0.279816934 | 0.568733968 |
| chr5:553988-554211 | 0.472700194 | 0.279816934 | 0.568733968 |

| | | | |
|---|---|---|---|
| chr12:49347334-49348107 | 0.367637367 | 0.279834537 | 0.568754658 |
| chr7:64729174-64729543 | 0.446863575 | 0.279890572 | 0.568853459 |
| chr2:209208788-209209699 | 0.398745172 | 0.279908855 | 0.568875527 |
| chr22:32991777-32992045 | -0.525757108 | 0.279922415 | 0.568887998 |
| chr4:12852382-12852646 | -0.542047798 | 0.279944131 | 0.568902232 |
| chr2:159111586-159111862 | -0.467649377 | 0.279944269 | 0.568902232 |
| chr2:88903854-88904114 | -0.470301868 | 0.279967383 | 0.568915647 |
| chr8:124155935-124156165 | 0.496419601 | 0.279969828 | 0.568915647 |
| chr1:145990165-145990432 | -0.442160346 | 0.279973145 | 0.568915647 |
| chr13:33676664-33677016 | -0.376587433 | 0.27998988 | 0.56891749 |
| chr7:74477866-74478557 | -0.369241005 | 0.279993061 | 0.56891749 |
| chrX:133726123-133726416 | -0.523981163 | 0.279996327 | 0.56891749 |
| chr6:142926434-142927349 | -0.374785613 | 0.280033451 | 0.568928204 |
| chr1:217333962-217334345 | -0.391425774 | 0.280033846 | 0.568928204 |
| chr12:132284127-132284347 | -0.495708573 | 0.280034524 | 0.568928204 |
| chr21:15531952-15532252 | -0.575393222 | 0.280037239 | 0.568928204 |
| chr20:18879188-18879508 | -0.47025358 | 0.280045667 | 0.568928204 |
| chr8:93612994-93613194 | 0.562594686 | 0.280048624 | 0.568928204 |
| chr3:184448807-184449228 | -0.461568843 | 0.280056024 | 0.568928204 |
| chr6:28251649-28252476 | -0.376441073 | 0.280066164 | 0.568928204 |
| chr2:65416966-65417296 | 0.441163371 | 0.280073829 | 0.568928204 |
| chr15:39933569-39934520 | -0.354859818 | 0.280075851 | 0.568928204 |
| chr6:138468616-138469377 | 0.382459752 | 0.280084797 | 0.568931293 |
| chr5:90473314-90475224 | -0.339658285 | 0.2800942 | 0.56893531 |
| chr11:17544300-17544637 | 0.403547486 | 0.280108385 | 0.568942791 |
| chr4:156169000-156169430 | -0.453801204 | 0.280112733 | 0.568942791 |
| chrX:129843350-129844345 | 0.35324965 | 0.280138078 | 0.568979187 |
| chr1:117121358-117122674 | 0.340978817 | 0.280147688 | 0.568983623 |
| chr8:62248870-62249638 | -0.405051222 | 0.280160773 | 0.568995116 |
| chr1:156619444-156619665 | 0.482933155 | 0.280254443 | 0.569170271 |
| chr1:212656066-212656346 | -0.527525242 | 0.280309306 | 0.569252553 |
| chr17:45619926-45620485 | -0.400537219 | 0.28031556 | 0.569252553 |
| chr2:100730406-100730681 | 0.510040592 | 0.280317247 | 0.569252553 |
| chr1:67430098-67431240 | -0.34083305 | 0.280326319 | 0.569255888 |
| chr11:24100749-24101241 | -0.435255872 | 0.280379559 | 0.56930535 |
| chr1:110312625-110312929 | 0.425090149 | 0.280389683 | 0.56930535 |
| chr5:9920832-9921235 | 0.423481925 | 0.280394884 | 0.56930535 |
| chr12:54439608-54439940 | 0.417385674 | 0.280400433 | 0.56930535 |
| chr12:2812395-2813321 | -0.350787215 | 0.280400541 | 0.56930535 |
| chr20:63538791-63539081 | 0.383542377 | 0.28040204 | 0.56930535 |
| chr3:195708563-195708877 | 0.444261333 | 0.28040764 | 0.56930535 |
| chr6:72705043-72705314 | -0.45564958 | 0.280411775 | 0.56930535 |
| chr16:14489529-14489951 | -0.448156647 | 0.280418226 | 0.56930535 |
| chr16:50024895-50025584 | 0.363540848 | 0.280424976 | 0.56930535 |
| chr10:7301349-7301842 | -0.407334404 | 0.28047471 | 0.56939123 |
| chr13:39073846-39074319 | -0.417303733 | 0.280510953 | 0.569435642 |
| chr5:137220443-137220856 | -0.389275585 | 0.280513495 | 0.569435642 |
| chr9:75027486-75028740 | -0.341595236 | 0.280518882 | 0.569435642 |
| chr10:69357952-69358152 | -0.603335988 | 0.280531696 | 0.569441169 |
| chr6:54817340-54817659 | -0.539171184 | 0.280537054 | 0.569441169 |

| | | | |
|---|---|---|---|
| chr15:69686059-69686259 | 0.53958982 | 0.2805439 | 0.569441169 |
| chr1:197057184-197057393 | 0.574606869 | 0.280587108 | 0.569513785 |
| chr15:81061387-81061587 | -0.606564458 | 0.280617995 | 0.569561389 |
| chr14:88385236-88386224 | -0.34780269 | 0.280627487 | 0.569565568 |
| chr20:42179833-42180096 | -0.466968678 | 0.280653973 | 0.569584617 |
| chr6:85267670-85267953 | -0.568742049 | 0.280660335 | 0.569584617 |
| chr4:89836927-89838111 | -0.362527161 | 0.280662264 | 0.569584617 |
| chr2:76251385-76251650 | 0.484411405 | 0.280666608 | 0.569584617 |
| chr1:3530860-3531065 | 0.476285274 | 0.280703717 | 0.569633552 |
| chr8:101470414-101470797 | 0.389306297 | 0.280705589 | 0.569633552 |
| chr5:173056049-173056667 | -0.352269759 | 0.280718248 | 0.569644153 |
| chr16:22373833-22374116 | 0.451737975 | 0.28072823 | 0.569649323 |
| chr7:6279411-6279790 | -0.439193137 | 0.280786962 | 0.569745033 |
| chr9:2016513-2017056 | 0.377102277 | 0.280792427 | 0.569745033 |
| chr8:128767450-128767650 | 0.511628504 | 0.280797704 | 0.569745033 |
| chr10:125123697-125124039 | -0.468448051 | 0.280853393 | 0.569842938 |
| chr1:107944137-107944337 | 0.609237559 | 0.280865964 | 0.569846582 |
| chr10:61943678-61943878 | 0.518153446 | 0.28087511 | 0.569846582 |
| chr3:140898860-140899060 | 0.532768788 | 0.280878371 | 0.569846582 |
| chr9:33161545-33162037 | 0.379359784 | 0.280887127 | 0.569846582 |
| chr6:149962923-149964130 | -0.350804965 | 0.280892375 | 0.569846582 |
| chr1:156728335-156729007 | 0.350491629 | 0.280899913 | 0.569846787 |
| chr2:80073149-80073440 | -0.474369783 | 0.280920191 | 0.569872835 |
| chr1:244290094-244290294 | 0.541475242 | 0.280928557 | 0.569874719 |
| chr17:65198588-65199056 | -0.379964773 | 0.28094173 | 0.569886353 |
| chr6:81589982-81590249 | 0.523196189 | 0.280950473 | 0.569889001 |
| chr20:53763269-53763469 | 0.365861674 | 0.280961626 | 0.569896538 |
| chr5:146446774-146447860 | -0.340958771 | 0.280969702 | 0.569897832 |
| chr19:17555879-17556179 | 0.464898369 | 0.280986762 | 0.569910663 |
| chr14:37340512-37340721 | 0.506530937 | 0.280999535 | 0.569910663 |
| chr3:170869637-170870966 | -0.34125237 | 0.281002417 | 0.569910663 |
| chr2:49843453-49843686 | -0.493461741 | 0.28100578 | 0.569910663 |
| chr4:71161157-71161463 | -0.554681626 | 0.281022617 | 0.569929725 |
| chr2:174138230-174138482 | -0.514012305 | 0.28104495 | 0.569938862 |
| chr6:136110035-136110334 | -0.555096044 | 0.28104514 | 0.569938862 |
| chr16:181235-181934 | 0.384819656 | 0.281049437 | 0.569938862 |
| chr12:63862194-63862683 | -0.379030336 | 0.281066856 | 0.569959102 |
| chr20:3744981-3745422 | 0.379704211 | 0.281115517 | 0.570033357 |
| chr6:72570460-72570660 | -0.565548629 | 0.281124246 | 0.570033357 |
| chr20:15902570-15902835 | -0.517157016 | 0.281125793 | 0.570033357 |
| chr10:85876985-85877274 | -0.462982578 | 0.281146275 | 0.570059803 |
| chr14:57544948-57545209 | -0.554856286 | 0.281154919 | 0.570062243 |
| chr5:160080860-160081326 | -0.409425304 | 0.281174647 | 0.570080431 |
| chr1:212284627-212285651 | -0.340840257 | 0.281191292 | 0.570080431 |
| chr7:152504448-152504802 | 0.427218502 | 0.28119309 | 0.570080431 |
| chr10:73730170-73730933 | -0.364917345 | 0.28119365 | 0.570080431 |
| chr9:136885930-136886822 | 0.358666704 | 0.281221567 | 0.570121944 |
| chr2:75199537-75200895 | -0.35625225 | 0.281233368 | 0.570126364 |
| chr4:105708088-105709370 | -0.344022077 | 0.281238629 | 0.570126364 |
| chr7:149375707-149375955 | -0.481497085 | 0.281257867 | 0.57015028 |

| | | | |
|---|---|---|---|
| chr2:163155320-163155837 | -0.413955644 | 0.281271434 | 0.570162698 |
| chr11:62508296-62508848 | 0.390422624 | 0.281330041 | 0.570239018 |
| chr1:175221695-175222043 | -0.39668857 | 0.281331454 | 0.570239018 |
| chr21:15729457-15730573 | 0.349718538 | 0.28133536 | 0.570239018 |
| chr15:92831463-92832185 | 0.355749299 | 0.281338853 | 0.570239018 |
| chr22:46563336-46564055 | 0.382630526 | 0.281355373 | 0.57024514 |
| chr5:69414839-69415925 | -0.349093801 | 0.281362213 | 0.57024514 |
| chr1:39639393-39640240 | 0.370671781 | 0.2813642 | 0.57024514 |
| chr13:29923731-29924494 | 0.351494105 | 0.281374849 | 0.5702515 |
| chr1:35031556-35032286 | 0.354413586 | 0.281382224 | 0.5702515 |
| chr5:136062590-136062827 | -0.460299663 | 0.281389836 | 0.570251846 |
| chr5:126422947-126423898 | -0.350552299 | 0.281429976 | 0.57028005 |
| chr6:150147387-150147788 | -0.439745057 | 0.281435232 | 0.57028005 |
| chr5:88108632-88108906 | 0.527242251 | 0.281439218 | 0.57028005 |
| chr13:87672956-87673156 | -0.444392595 | 0.281440053 | 0.57028005 |
| chr7:140002371-140002798 | -0.433323862 | 0.281444254 | 0.57028005 |
| chr3:73809009-73809239 | 0.579847448 | 0.281457029 | 0.57028005 |
| chr16:70289056-70289850 | 0.348959848 | 0.281462495 | 0.57028005 |
| chr8:98383146-98383423 | 0.427542845 | 0.281463296 | 0.57028005 |
| chr3:151008798-151009090 | 0.437684248 | 0.281473819 | 0.570282889 |
| chrX:40146511-40146772 | 0.530938885 | 0.281482711 | 0.570282889 |
| chr1:202589225-202589637 | -0.386911327 | 0.281487025 | 0.570282889 |
| chr10:9302194-9302401 | 0.517592691 | 0.281495371 | 0.570284718 |
| chr19:56120332-56120675 | 0.401130256 | 0.281529967 | 0.570337063 |
| chr13:40789140-40790274 | -0.344548373 | 0.281543131 | 0.570337063 |
| chr11:79520170-79520416 | 0.496781271 | 0.281543539 | 0.570337063 |
| chr8:88896958-88897428 | 0.384959301 | 0.281574127 | 0.570383946 |
| chr7:124856299-124856676 | -0.439062569 | 0.281586751 | 0.570394438 |
| chr1:94025497-94025747 | -0.518470916 | 0.281626178 | 0.570440796 |
| chr10:30818670-30818870 | -0.601534443 | 0.281630314 | 0.570440796 |
| chr1:203929673-203930080 | -0.390108923 | 0.281634665 | 0.570440796 |
| chr1:59139500-59139747 | -0.510047319 | 0.281639416 | 0.570440796 |
| chr13:70310744-70311073 | -0.422228064 | 0.281695906 | 0.57054013 |
| chr19:48363084-48363438 | 0.375167833 | 0.281715457 | 0.570555428 |
| chr5:74684868-74685944 | -0.353839178 | 0.281718351 | 0.570555428 |
| chr1:207005031-207005315 | -0.388756938 | 0.281727729 | 0.57055934 |
| chr8:42896302-42897123 | 0.360030611 | 0.28174674 | 0.570582137 |
| chr7:102389690-102389957 | -0.552726376 | 0.28176062 | 0.570582137 |
| chr14:39266438-39267828 | 0.339978386 | 0.281761326 | 0.570582137 |
| chr10:61744602-61744823 | -0.589452916 | 0.28178012 | 0.570590415 |
| chr2:131516030-131516230 | 0.545813208 | 0.281782964 | 0.570590415 |
| chrX:129931207-129932764 | 0.338732511 | 0.281787754 | 0.570590415 |
| chr15:66191528-66191744 | 0.486248654 | 0.281812215 | 0.570624866 |
| chr7:38451930-38452144 | -0.500817369 | 0.281855786 | 0.570698009 |
| chr19:1690795-1691032 | 0.534015102 | 0.281885414 | 0.570742916 |
| chr8:98817853-98818195 | -0.449989011 | 0.281909605 | 0.570776814 |
| chr17:65294034-65294739 | -0.365575927 | 0.281930569 | 0.570804176 |
| chr22:40646329-40646556 | 0.502899139 | 0.281975747 | 0.570873534 |
| chr1:62334576-62334781 | 0.554303512 | 0.281997906 | 0.570873534 |
| chr2:161063687-161064232 | -0.389392583 | 0.282006838 | 0.570873534 |

| | | | |
|---|---|---|---|
| chr14:76514669-76514986 | 0.453265921 | 0.282008516 | 0.570873534 |
| chr6:166309218-166309517 | -0.43863409 | 0.28202213 | 0.570873534 |
| chr16:80084668-80084954 | -0.442002939 | 0.282023022 | 0.570873534 |
| chr2:100242852-100243259 | 0.416345843 | 0.282023831 | 0.570873534 |
| chr3:79517907-79518205 | 0.449499322 | 0.282045459 | 0.570873534 |
| chr5:141946372-141946617 | -0.537667629 | 0.282052308 | 0.570873534 |
| chr3:63142714-63143326 | 0.366755314 | 0.282053315 | 0.570873534 |
| chr16:83357276-83357602 | 0.527234679 | 0.282059025 | 0.570873534 |
| chr2:42562451-42563048 | -0.358390641 | 0.282062356 | 0.570873534 |
| chr1:64564421-64564623 | -0.579676634 | 0.282078641 | 0.570873534 |
| chr12:59780850-59781063 | -0.579676634 | 0.282078641 | 0.570873534 |
| chr5:38831767-38832123 | -0.432679694 | 0.282081151 | 0.570873534 |
| chr11:83285609-83286808 | -0.339133665 | 0.282084034 | 0.570873534 |
| chr2:235544448-235544817 | 0.447529468 | 0.282097389 | 0.570883927 |
| chr14:53260348-53260568 | 0.510009936 | 0.282104071 | 0.570883927 |
| chr10:24034613-24034813 | -0.60469195 | 0.28212097 | 0.570889403 |
| chr11:73271956-73272313 | 0.390998595 | 0.28212635 | 0.570889403 |
| chr16:74859543-74859791 | 0.534742731 | 0.282129129 | 0.570889403 |
| chr2:55231618-55232862 | -0.34028396 | 0.282141296 | 0.570893178 |
| chr2:175439462-175439780 | -0.501901584 | 0.282145896 | 0.570893178 |
| chr2:15031834-15032047 | -0.531040989 | 0.282158672 | 0.570893974 |
| chr6:140083404-140083882 | -0.422780314 | 0.282161191 | 0.570893974 |
| chr1:106418733-106418977 | 0.401561172 | 0.282196293 | 0.570949918 |
| chr8:93916109-93917919 | 0.335775826 | 0.282216113 | 0.570974943 |
| chr17:29744773-29745151 | -0.459433914 | 0.282234772 | 0.570997615 |
| chr18:74270153-74270420 | -0.539784676 | 0.282265682 | 0.571045073 |
| chr1:28184466-28184760 | -0.554591777 | 0.282277298 | 0.571053495 |
| chr17:80951938-80952138 | 0.512795605 | 0.282305231 | 0.571071925 |
| chr4:155026125-155026440 | 0.427941284 | 0.282307829 | 0.571071925 |
| chr16:81522095-81522399 | -0.41199232 | 0.282319528 | 0.571071925 |
| chr14:78448435-78448735 | 0.451023546 | 0.282323283 | 0.571071925 |
| chr3:128942443-128942719 | -0.498887123 | 0.28232856 | 0.571071925 |
| chr10:84118723-84118923 | -0.541249093 | 0.282336142 | 0.571071925 |
| chr17:5249618-5249833 | 0.5523693 | 0.282344282 | 0.571071925 |
| chr2:40554772-40555036 | -0.517750882 | 0.282349623 | 0.571071925 |
| chr17:16352758-16353059 | -0.521114174 | 0.282353486 | 0.571071925 |
| chr8:106419176-106419682 | 0.39710455 | 0.282364291 | 0.571078704 |
| chr9:94213560-94213837 | 0.4621114 | 0.282374707 | 0.571084695 |
| chr11:37858438-37858727 | -0.54827639 | 0.282394257 | 0.57109736 |
| chr13:30431355-30431619 | 0.504951961 | 0.282395876 | 0.57109736 |
| chr20:41572149-41572349 | 0.482183752 | 0.282422294 | 0.571135713 |
| chr11:9758018-9758459 | 0.387086799 | 0.282453742 | 0.571180809 |
| chr7:53209259-53209485 | -0.546459697 | 0.282468579 | 0.571180809 |
| chr15:65133481-65134103 | 0.363666855 | 0.282473239 | 0.571180809 |
| chr19:48445302-48446519 | 0.35260922 | 0.282480406 | 0.571180809 |
| chr14:92714742-92715032 | 0.439621759 | 0.282481866 | 0.571180809 |
| chr2:214830818-214831127 | -0.445464238 | 0.282521293 | 0.571245455 |
| chr2:205682252-205682879 | -0.391616306 | 0.282560248 | 0.571309145 |
| chr20:49937485-49937747 | 0.402895128 | 0.28257049 | 0.571314777 |
| chr14:91164673-91165109 | 0.372903915 | 0.282608886 | 0.571371423 |

| | | | |
|---|---|---|---|
| chr9:75601324-75601524 | 0.521650921 | 0.282613421 | 0.571371423 |
| chr1:41347382-41347687 | 0.438668767 | 0.282627809 | 0.571385435 |
| chr14:48970395-48970595 | 0.613007943 | 0.282641791 | 0.571385753 |
| chr2:52144593-52144824 | -0.558168833 | 0.282647919 | 0.571385753 |
| chr11:17036742-17037394 | -0.398566002 | 0.282654987 | 0.571385753 |
| chr2:10452367-10452658 | -0.404227845 | 0.282657795 | 0.571385753 |
| chrX:99173054-99173300 | -0.545817095 | 0.282694347 | 0.571412395 |
| chr16:11619202-11619586 | -0.462318142 | 0.282700226 | 0.571412395 |
| chr7:50068116-50068384 | -0.437569504 | 0.282700528 | 0.571412395 |
| chr1:89780707-89781046 | -0.455791505 | 0.282700805 | 0.571412395 |
| chr14:77882185-77882428 | 0.453756893 | 0.282713526 | 0.571421821 |
| chr2:97082382-97082763 | 0.451233565 | 0.282720384 | 0.571421821 |
| chr8:140653399-140653870 | 0.39303398 | 0.282738383 | 0.571443126 |
| chr17:51540245-51540446 | -0.507424125 | 0.282758312 | 0.571467515 |
| chr14:23056148-23056673 | 0.37183982 | 0.282765366 | 0.571467515 |
| chr18:72691971-72692846 | -0.384644938 | 0.282774684 | 0.571467527 |
| chr20:49928711-49929109 | 0.382695615 | 0.282791032 | 0.571467527 |
| chrX:77895274-77895861 | -0.352319469 | 0.282794443 | 0.571467527 |
| chr1:188489242-188489843 | 0.366178855 | 0.282795205 | 0.571467527 |
| chrX:123881715-123881935 | 0.529251372 | 0.282811159 | 0.571484693 |
| chr22:20123569-20124013 | 0.432307055 | 0.282820131 | 0.571487752 |
| chr17:46876303-46876777 | -0.387769586 | 0.282884739 | 0.57160323 |
| chr13:113428167-113428367 | 0.519158873 | 0.282898037 | 0.571615026 |
| chr14:91897179-91897769 | -0.378395129 | 0.282922542 | 0.571624416 |
| chr11:65418285-65419362 | 0.344424871 | 0.282923551 | 0.571624416 |
| chr20:45088297-45088559 | -0.477486936 | 0.282928029 | 0.571624416 |
| chr20:54056818-54057285 | 0.350470544 | 0.282934553 | 0.571624416 |
| chr12:96388096-96388296 | 0.576327038 | 0.282940645 | 0.571624416 |
| chr3:138347996-138348922 | 0.363158076 | 0.282947446 | 0.571624416 |
| chr10:58399952-58400307 | -0.547639832 | 0.282958669 | 0.571632016 |
| chr3:149925922-149926368 | -0.426197665 | 0.282966988 | 0.571633752 |
| chr11:120210749-120211101 | -0.479512719 | 0.282981865 | 0.571648733 |
| chr7:50142418-50142825 | -0.460464217 | 0.282994718 | 0.571653717 |
| chr22:44752366-44752822 | 0.413488583 | 0.282999345 | 0.571653717 |
| chr8:81870924-81871164 | 0.521316855 | 0.283006714 | 0.571653717 |
| chr18:79964954-79965863 | -0.363058721 | 0.283026796 | 0.571679211 |
| chr15:44219673-44219938 | 0.521794712 | 0.283100105 | 0.571812212 |
| chr16:85524917-85525233 | -0.40175991 | 0.283117658 | 0.571832591 |
| chr20:49045679-49045889 | -0.448195993 | 0.283153611 | 0.571880707 |
| chr17:5111435-5111635 | -0.565962935 | 0.283163642 | 0.571880707 |
| chr3:64561502-64561864 | 0.362964856 | 0.283163871 | 0.571880707 |
| chr17:56030154-56030410 | -0.502362168 | 0.283239596 | 0.572018563 |
| chr5:27072685-27072928 | 0.530143526 | 0.283258738 | 0.572042145 |
| chr16:64051859-64052106 | -0.555632244 | 0.283276778 | 0.572063499 |
| chr3:48634849-48635093 | 0.448958181 | 0.283300291 | 0.572078985 |
| chr20:45934524-45935618 | 0.341049696 | 0.283307786 | 0.572078985 |
| chrX:16785691-16787274 | -0.337733478 | 0.283309319 | 0.572078985 |
| chr12:66107414-66107785 | -0.399468573 | 0.283324896 | 0.572078985 |
| chr2:28894269-28895500 | -0.344269423 | 0.283325427 | 0.572078985 |
| chr2:212595062-212595363 | -0.482539901 | 0.283329244 | 0.572078985 |

| | | | |
|---|---|---|---|
| chr16:88634743-88635157 | -0.375811391 | 0.283341447 | 0.57208855 |
| chr13:77521988-77522326 | -0.482167108 | 0.28335577 | 0.572102393 |
| chr5:31929268-31929610 | 0.397168731 | 0.283376159 | 0.572113945 |
| chr15:48645545-48646192 | -0.416489594 | 0.283376425 | 0.572113945 |
| chr8:109575403-109575725 | 0.429099922 | 0.283397057 | 0.572128127 |
| chr20:36556509-36556850 | 0.44688836 | 0.283399049 | 0.572128127 |
| chr17:78226741-78227036 | 0.454284286 | 0.28340585 | 0.572128127 |
| chr5:1210393-1210733 | 0.405239263 | 0.283424296 | 0.57215029 |
| chrX:53284660-53284890 | 0.46121055 | 0.283462548 | 0.572195422 |
| chr14:63752832-63753132 | -0.469747447 | 0.283473583 | 0.572195422 |
| chr2:150487541-150487954 | -0.374320194 | 0.283473971 | 0.572195422 |
| chr2:156083485-156084083 | 0.38828255 | 0.283480948 | 0.572195422 |
| chr2:95053694-95053940 | 0.467435018 | 0.283483992 | 0.572195422 |
| chr3:16048664-16049049 | 0.396754224 | 0.283527939 | 0.572269051 |
| chr8:114789281-114789910 | -0.422316285 | 0.283547617 | 0.572293695 |
| chr5:115150586-115150798 | -0.562683611 | 0.283589522 | 0.572363196 |
| chr3:63932445-63932706 | 0.378655203 | 0.283647154 | 0.572464434 |
| chr11:18844810-18845065 | 0.487539534 | 0.283682035 | 0.572519751 |
| chr10:72274754-72274976 | 0.445894235 | 0.283694506 | 0.572525452 |
| chr2:98704767-98705440 | -0.419819803 | 0.283699804 | 0.572525452 |
| chr4:99179784-99179984 | 0.571970211 | 0.283709484 | 0.572529908 |
| chr19:37466882-37467660 | -0.379187042 | 0.283721177 | 0.572538426 |
| chr6:18022821-18023133 | 0.517388215 | 0.28373564 | 0.572552531 |
| chr19:36008372-36009005 | 0.36523952 | 0.283765298 | 0.572585655 |
| chr7:46253658-46253869 | -0.538140772 | 0.283767 | 0.572585655 |
| chr20:56974423-56975236 | -0.347313381 | 0.283850476 | 0.57273901 |
| chr8:102381074-102381274 | 0.572599469 | 0.283889867 | 0.572803408 |
| chr5:34552864-34553093 | 0.547533647 | 0.283930557 | 0.572870421 |
| chr14:55533977-55534315 | -0.428654907 | 0.284003224 | 0.57300195 |
| chr4:138241674-138242693 | -0.344867681 | 0.284022465 | 0.573025681 |
| chr14:77409188-77409471 | 0.455702008 | 0.284042781 | 0.57305158 |
| chr10:84204229-84204429 | -0.475100658 | 0.284064294 | 0.573079652 |
| chr20:47800966-47801633 | 0.343061915 | 0.284071654 | 0.573079652 |
| chr9:69086360-69086612 | -0.594496635 | 0.284093962 | 0.57309856 |
| chr20:50992981-50993583 | -0.362376006 | 0.284095986 | 0.57309856 |
| chr5:62268211-62268487 | -0.452895998 | 0.284149483 | 0.573191388 |
| chr19:13731633-13731987 | 0.47036724 | 0.284205113 | 0.573288513 |
| chr12:88035114-88036237 | -0.345285509 | 0.284226454 | 0.573316467 |
| chr16:30395031-30395649 | 0.375848293 | 0.284288289 | 0.573421235 |
| chr1:117067730-117068112 | 0.400654216 | 0.284293361 | 0.573421235 |
| chr2:85742529-85742905 | 0.3897652 | 0.28430306 | 0.573422704 |
| chr11:86608225-86608654 | 0.416480523 | 0.28430949 | 0.573422704 |
| chr1:61042936-61043399 | -0.409360683 | 0.284323779 | 0.573422704 |
| chr1:70896283-70896554 | -0.483889336 | 0.284324024 | 0.573422704 |
| chr7:134554074-134554295 | -0.548633803 | 0.28435056 | 0.573458792 |
| chr1:185813445-185813689 | -0.49236211 | 0.284356886 | 0.573458792 |
| chr8:94327231-94327644 | 0.469075521 | 0.284368529 | 0.573463375 |
| chr1:110893976-110894230 | -0.499493558 | 0.284374127 | 0.573463375 |
| chr2:190343524-190344501 | 0.367109427 | 0.284383833 | 0.573467855 |
| chr15:65318865-65319163 | 0.436012867 | 0.284394387 | 0.573474044 |

| | | | |
|---|---|---|---|
| chr7:50068918-50069562 | -0.367993816 | 0.284417823 | 0.573494024 |
| chr20:59608874-59609197 | 0.428146857 | 0.284419265 | 0.573494024 |
| chr5:135156803-135157003 | -0.573810464 | 0.284439103 | 0.573516999 |
| chr14:43649416-43649801 | 0.424502355 | 0.284445629 | 0.573516999 |
| chr14:99272926-99273623 | -0.356178555 | 0.284460884 | 0.573532665 |
| chr1:62230701-62231068 | -0.423245243 | 0.284487624 | 0.573553389 |
| chr6:98942312-98942580 | -0.454283766 | 0.28448818 | 0.573553389 |
| chr13:28415988-28416311 | -0.489804218 | 0.284493619 | 0.573553389 |
| chr16:76137974-76138632 | -0.382377195 | 0.284510736 | 0.573572804 |
| chr17:63889553-63889854 | -0.416039326 | 0.28452745 | 0.573591409 |
| chr18:72544230-72544640 | -0.398637782 | 0.28455087 | 0.573623531 |
| chr14:40252863-40253348 | 0.407576118 | 0.284602059 | 0.573711627 |
| chr4:172337538-172337878 | -0.513565774 | 0.284613449 | 0.573719494 |
| chr4:110851121-110851460 | -0.46867243 | 0.284631604 | 0.573740996 |
| chr1:212967539-212968110 | -0.388388349 | 0.284658632 | 0.573776934 |
| chr2:112979077-112979277 | -0.591110036 | 0.284665994 | 0.573776934 |
| chr3:197539557-197540192 | 0.365733423 | 0.284672038 | 0.573776934 |
| chr2:69178250-69178704 | -0.370295226 | 0.284679386 | 0.573776934 |
| chr1:15200468-15200981 | 0.416858557 | 0.284704146 | 0.573778077 |
| chr5:177948073-177948291 | -0.573001834 | 0.28470502 | 0.573778077 |
| chr1:241944819-241945363 | -0.397859378 | 0.284705156 | 0.573778077 |
| chr6:15956446-15956723 | -0.554220753 | 0.284709907 | 0.573778077 |
| chr19:44890342-44891375 | 0.339033665 | 0.284732709 | 0.573808937 |
| chr1:198072303-198072541 | 0.479854071 | 0.284751853 | 0.573832382 |
| chr11:797944-798655 | 0.374064335 | 0.284765397 | 0.573832382 |
| chr16:88296507-88296859 | 0.422548722 | 0.28476681 | 0.573832382 |
| chr18:2846410-2846714 | -0.499271587 | 0.284800341 | 0.573883317 |
| chr2:189674015-189675153 | -0.343947531 | 0.284807066 | 0.573883317 |
| chr7:47738003-47738203 | 0.498509488 | 0.28482677 | 0.573898733 |
| chr14:94330600-94330942 | 0.487568723 | 0.284829697 | 0.573898733 |
| chr12:53267784-53268624 | 0.364285782 | 0.284876531 | 0.573946253 |
| chr1:152458423-152458803 | 0.381335052 | 0.28488308 | 0.573946253 |
| chr9:113541104-113541590 | -0.38017614 | 0.28488777 | 0.573946253 |
| chr3:97011039-97011582 | 0.414725291 | 0.284893531 | 0.573946253 |
| chr10:1056227-1057290 | 0.342512326 | 0.284896967 | 0.573946253 |
| chr21:10331206-10331637 | 0.442129285 | 0.284898225 | 0.573946253 |
| chr2:158456167-158457064 | -0.343147178 | 0.284941029 | 0.574017393 |
| chr1:153775014-153776382 | 0.341998903 | 0.284949038 | 0.574018435 |
| chr2:134245399-134245893 | -0.418613032 | 0.284958293 | 0.574021988 |
| chr6:31763525-31763809 | 0.460850329 | 0.284967665 | 0.574025775 |
| chr9:35798713-35798957 | -0.579228571 | 0.284976493 | 0.574028468 |
| chr17:42276274-42276949 | 0.400064886 | 0.284984718 | 0.574029944 |
| chr3:87151366-87151753 | -0.467308959 | 0.285041199 | 0.574113898 |
| chr7:45387452-45387652 | 0.553923996 | 0.285041384 | 0.574113898 |
| chr20:52579655-52579855 | 0.617464301 | 0.285051178 | 0.574118533 |
| chr2:187532743-187533417 | 0.36900197 | 0.285075941 | 0.574153315 |
| chr3:25427202-25427419 | -0.553189852 | 0.285106761 | 0.574172751 |
| chr15:78682301-78682552 | -0.53641548 | 0.285107086 | 0.574172751 |
| chr20:51707258-51707574 | 0.418669203 | 0.285121532 | 0.574172751 |
| chrX:109943873-109944261 | -0.452870937 | 0.285129928 | 0.574172751 |

| | | | |
|---|---|---|---|
| chr20:47954384-47954616 | 0.373105487 | 0.285130007 | 0.574172751 |
| chr7:12130813-12131043 | 0.513001749 | 0.285130552 | 0.574172751 |
| chr1:183306493-183306937 | 0.407867808 | 0.285149306 | 0.574195425 |
| chr2:46698428-46699721 | -0.338576268 | 0.285157905 | 0.57419765 |
| chr11:67351274-67351530 | -0.421690527 | 0.285176987 | 0.574220983 |
| chr2:174688127-174688346 | -0.507995803 | 0.285197299 | 0.574246792 |
| chr2:156333940-156334914 | -0.344514918 | 0.285262429 | 0.574362839 |
| chr3:165855501-165855787 | 0.500429991 | 0.285270367 | 0.574363729 |
| chr3:100114359-100114897 | -0.446894514 | 0.285298061 | 0.574401549 |
| chr5:109409262-109410848 | -0.334717186 | 0.285304333 | 0.574401549 |
| chr17:16453228-16453622 | 0.404049367 | 0.285311641 | 0.574401549 |
| chr3:64318934-64319151 | 0.379220409 | 0.285339234 | 0.574442008 |
| chr12:89489923-89490189 | -0.509522744 | 0.285361745 | 0.574461183 |
| chr17:78112666-78112984 | 0.419604211 | 0.285363754 | 0.574461183 |
| chr6:132512561-132513604 | -0.342519721 | 0.285408029 | 0.574535218 |
| chr4:40363374-40363900 | 0.400232734 | 0.285446466 | 0.574587715 |
| chrX:102599006-102600001 | -0.346287298 | 0.285449106 | 0.574587715 |
| chr1:1692754-1693283 | 0.411886619 | 0.28549536 | 0.574665724 |
| chr3:165895896-165896301 | -0.421803129 | 0.285512935 | 0.574679149 |
| chr3:179322432-179323891 | -0.337604536 | 0.285522606 | 0.574679149 |
| chr1:3062244-3062571 | -0.571855112 | 0.28552453 | 0.574679149 |
| chr6:135851453-135851816 | -0.407634654 | 0.285533199 | 0.574681501 |
| chr11:67353175-67353954 | 0.34859977 | 0.285568857 | 0.574738172 |
| chr2:168148735-168149047 | -0.582766217 | 0.285605302 | 0.574796423 |
| chr10:84527237-84527912 | -0.366175014 | 0.285624168 | 0.574817574 |
| chr9:87704498-87704760 | 0.521493703 | 0.285630815 | 0.574817574 |
| chr8:50136524-50136819 | -0.504670945 | 0.28566764 | 0.574855547 |
| chr3:83798918-83799191 | -0.514100702 | 0.285668606 | 0.574855547 |
| chr9:121169117-121169357 | -0.547419336 | 0.285675916 | 0.574855547 |
| chr7:50404483-50404689 | -0.519918186 | 0.285679694 | 0.574855547 |
| chr2:222693960-222694186 | -0.572255179 | 0.28575688 | 0.574980663 |
| chr5:117057658-117057867 | -0.572255179 | 0.28575688 | 0.574980663 |
| chr1:198156173-198156424 | -0.373442349 | 0.285782856 | 0.575017831 |
| chr14:34714380-34715077 | 0.372357741 | 0.28580117 | 0.57503958 |
| chr4:110218133-110218431 | 0.523542405 | 0.285816301 | 0.575054922 |
| chr1:206856638-206857179 | -0.370510389 | 0.28587158 | 0.57513999 |
| chr6:157989948-157990252 | -0.434632762 | 0.285873594 | 0.57513999 |
| chr8:101123334-101123534 | 0.432604275 | 0.2858845 | 0.575146831 |
| chr12:62694493-62694806 | 0.501010304 | 0.285953605 | 0.575258775 |
| chr3:159123989-159124189 | 0.545155547 | 0.285955159 | 0.575258775 |
| chr8:138964815-138965180 | -0.441653808 | 0.285999004 | 0.575318875 |
| chr3:177827486-177827770 | 0.463858069 | 0.28600756 | 0.575318875 |
| chr4:10173337-10173596 | 0.463858069 | 0.28600756 | 0.575318875 |
| chr2:26606773-26607182 | 0.419313132 | 0.286028742 | 0.575346378 |
| chr6:142716145-142716351 | -0.578382541 | 0.286039748 | 0.575353414 |
| chr14:75277696-75277900 | 0.434694408 | 0.286074385 | 0.575407978 |
| chr12:100266335-100267914 | -0.334412916 | 0.286118393 | 0.575457639 |
| chr4:99663657-99664026 | -0.529348154 | 0.286125176 | 0.575457639 |
| chr17:61443845-61444055 | 0.361379032 | 0.28612909 | 0.575457639 |
| chr6:137705368-137705626 | -0.49788281 | 0.286129116 | 0.575457639 |

| | | | |
|---|---|---|---|
| chr13:18252841-18253150 | -0.472250569 | 0.286176903 | 0.57553864 |
| chr2:34486991-34487278 | -0.47108133 | 0.286193487 | 0.575556886 |
| chr20:59514886-59515367 | -0.388200394 | 0.286210298 | 0.575568194 |
| chr17:47850874-47851623 | 0.354860726 | 0.286214133 | 0.575568194 |
| chr13:23074650-23074873 | 0.555391554 | 0.286227618 | 0.575580206 |
| chr14:49687414-49688414 | 0.353537073 | 0.286236625 | 0.575583212 |
| chr9:113409992-113410911 | 0.342711952 | 0.286249135 | 0.575586005 |
| chr21:39425661-39425905 | 0.472053639 | 0.286253038 | 0.575586005 |
| chr15:74892269-74892844 | -0.384975968 | 0.286264925 | 0.575594802 |
| chr2:171415919-171416396 | 0.402501224 | 0.286274294 | 0.575598535 |
| chr9:34379399-34379674 | -0.40248937 | 0.286287952 | 0.575610892 |
| chr16:85516034-85516333 | -0.39424037 | 0.28633581 | 0.575663243 |
| chr20:47454708-47456154 | 0.331560198 | 0.286344524 | 0.575663243 |
| chr8:30358302-30358858 | -0.443244239 | 0.286348288 | 0.575663243 |
| chr4:128454329-128454800 | 0.388729748 | 0.286350278 | 0.575663243 |
| chr15:71199521-71199721 | 0.528976179 | 0.286351555 | 0.575663243 |
| chr6:99458377-99458622 | -0.502006109 | 0.286392217 | 0.575729882 |
| chr19:47663995-47664208 | -0.532911139 | 0.286437048 | 0.575784569 |
| chr17:52410889-52411146 | -0.491966739 | 0.286442513 | 0.575784569 |
| chr1:203480804-203481125 | -0.401030087 | 0.286448592 | 0.575784569 |
| chr1:193937396-193937675 | 0.435388619 | 0.286449944 | 0.575784569 |
| chr16:30445683-30446119 | 0.391186883 | 0.286456993 | 0.575784569 |
| chr5:118693000-118693415 | -0.436296007 | 0.286527936 | 0.575912057 |
| chr18:76897692-76897930 | -0.554733587 | 0.286559408 | 0.575959382 |
| chr4:94077070-94077270 | 0.572227434 | 0.286566515 | 0.575959382 |
| chr1:152440723-152441204 | 0.364984048 | 0.286584276 | 0.57597997 |
| chr16:85416213-85416798 | 0.357199665 | 0.286600881 | 0.575986623 |
| chr3:75415828-75416028 | 0.532743819 | 0.286602621 | 0.575986623 |
| chr10:52308245-52308620 | -0.426782661 | 0.286643945 | 0.576054564 |
| chr7:35037175-35038395 | 0.335396805 | 0.286651718 | 0.576055076 |
| chr2:8574330-8574555 | -0.429143094 | 0.286690547 | 0.576106723 |
| chr20:46772966-46773187 | 0.481055349 | 0.286703275 | 0.576106723 |
| chr16:70738390-70738741 | 0.408650496 | 0.286706393 | 0.576106723 |
| chr6:39166484-39166754 | -0.46198964 | 0.286707493 | 0.576106723 |
| chr16:84460198-84460467 | 0.483396782 | 0.286726507 | 0.576129819 |
| chr18:62324878-62325632 | -0.369084725 | 0.286749176 | 0.576151452 |
| chr7:92933813-92934480 | -0.354000686 | 0.286755947 | 0.576151452 |
| chr7:267062-267330 | 0.408496701 | 0.286762219 | 0.576151452 |
| chr14:52873249-52873482 | 0.490619827 | 0.286767351 | 0.576151452 |
| chr10:21517252-21517851 | -0.390967928 | 0.286780073 | 0.576157176 |
| chr2:25342404-25342720 | 0.387649783 | 0.286796248 | 0.576157176 |
| chr5:24548111-24548357 | 0.537588999 | 0.286798709 | 0.576157176 |
| chr20:46089171-46090262 | 0.344797807 | 0.286806036 | 0.576157176 |
| chr5:17228842-17229075 | 0.546236672 | 0.286807797 | 0.576157176 |
| chr3:105518831-105519162 | -0.454288534 | 0.286819709 | 0.57616487 |
| chr10:13212592-13212837 | -0.602412234 | 0.286826666 | 0.57616487 |
| chr11:67401148-67402177 | 0.344673061 | 0.286848373 | 0.576182407 |
| chr5:6377849-6378803 | 0.345497372 | 0.286850436 | 0.576182407 |
| chr14:90518948-90519233 | 0.403234501 | 0.286876828 | 0.576220314 |
| chr13:28658558-28659670 | -0.346371983 | 0.28688713 | 0.576225902 |

| | | | |
|---|---|---|---|
| chr2:43452222-43452667 | -0.436543701 | 0.286918433 | 0.57626409 |
| chr8:136164211-136164411 | 0.583046845 | 0.286925637 | 0.57626409 |
| chr7:12090994-12091518 | 0.390699014 | 0.286928706 | 0.57626409 |
| chr12:42590283-42590944 | -0.387389086 | 0.286957083 | 0.576290358 |
| chr20:59783118-59783386 | 0.398854189 | 0.286961209 | 0.576290358 |
| chr12:103387109-103387427 | -0.526278491 | 0.286964348 | 0.576290358 |
| chr6:115838785-115839191 | -0.403618147 | 0.286987614 | 0.576321974 |
| chr7:102332023-102332267 | -0.524040926 | 0.287033117 | 0.576398246 |
| chr19:47377366-47378174 | -0.371839677 | 0.287052799 | 0.576422159 |
| chr20:63627835-63628181 | -0.408346334 | 0.287060071 | 0.576422159 |
| chr3:137509081-137510132 | -0.359771524 | 0.287082237 | 0.576451561 |
| chr2:172052345-172052604 | -0.523797539 | 0.287116725 | 0.576505704 |
| chr2:238305050-238305478 | -0.438768268 | 0.287165634 | 0.576588799 |
| chr1:153122803-153123520 | 0.354566312 | 0.287182753 | 0.576599367 |
| chr2:176454951-176455240 | -0.441787507 | 0.287189704 | 0.576599367 |
| chr6:158205337-158205613 | -0.50116713 | 0.287202195 | 0.576599367 |
| chr2:59025201-59025491 | -0.458825622 | 0.287206357 | 0.576599367 |
| chr11:236232-236655 | 0.392195139 | 0.287208523 | 0.576599367 |
| chr3:147191858-147192311 | -0.393288293 | 0.287234556 | 0.576636523 |
| chr6:17705970-17707183 | -0.343820442 | 0.287269817 | 0.576692201 |
| chr10:48509970-48510220 | 0.466657831 | 0.287322812 | 0.576783476 |
| chr5:139787418-139787852 | 0.375740545 | 0.287344898 | 0.576809136 |
| chr1:155001588-155002622 | 0.338824124 | 0.28735065 | 0.576809136 |
| chr12:98002948-98003304 | -0.423114365 | 0.287405714 | 0.576904554 |
| chr10:22229032-22229744 | -0.349963918 | 0.287421605 | 0.57690908 |
| chr16:4474350-4474851 | 0.366388872 | 0.287428975 | 0.57690908 |
| chr6:49796713-49797500 | -0.383559467 | 0.287430557 | 0.57690908 |
| chr9:21334944-21335597 | -0.362713274 | 0.287481642 | 0.576996501 |
| chr9:129834776-129835612 | 0.35362847 | 0.287548236 | 0.577115041 |
| chr20:52001716-52002438 | -0.365715467 | 0.287569827 | 0.577120855 |
| chr3:131033562-131033783 | -0.569529565 | 0.287573729 | 0.577120855 |
| chr2:192197399-192197599 | -0.569529565 | 0.287573729 | 0.577120855 |
| chr1:217938128-217938343 | 0.537764381 | 0.287590857 | 0.577140112 |
| chr5:150485672-150487096 | 0.342913771 | 0.287603275 | 0.577149916 |
| chr1:165392736-165393247 | -0.386522283 | 0.287639174 | 0.57719115 |
| chr7:150003154-150003794 | 0.402337538 | 0.287639895 | 0.57719115 |
| chr3:157606501-157606976 | -0.381046453 | 0.28765739 | 0.57719115 |
| chr15:96332294-96332514 | 0.413433037 | 0.287664297 | 0.57719115 |
| chr19:48660813-48661368 | -0.386281873 | 0.287666205 | 0.57719115 |
| chr4:90113952-90114163 | 0.504181607 | 0.287670218 | 0.57719115 |
| chr10:74825832-74827103 | -0.333374755 | 0.287682975 | 0.57719115 |
| chr4:3410612-3410812 | 0.540572217 | 0.287684086 | 0.57719115 |
| chr7:116273302-116273524 | -0.538173007 | 0.28774316 | 0.577294556 |
| chr4:81471389-81472642 | -0.340167603 | 0.287754376 | 0.577301942 |
| chrX:135931936-135932204 | 0.437296811 | 0.287780186 | 0.577318149 |
| chr3:194472359-194472571 | -0.544768051 | 0.287780853 | 0.577318149 |
| chr8:123528241-123528541 | 0.419938056 | 0.287785058 | 0.577318149 |
| chr2:60926065-60926330 | -0.499423242 | 0.287832803 | 0.57739815 |
| chr11:119474721-119474921 | 0.497721804 | 0.287846382 | 0.57739815 |
| chr5:71827270-71827470 | -0.627950862 | 0.287847544 | 0.57739815 |

| | | | |
|---|---|---|---|
| chr3:19543177-19543595 | 0.390837182 | 0.28786449 | 0.57740652 |
| chr8:17699639-17699886 | -0.484349635 | 0.287866789 | 0.57740652 |
| chr18:12307984-12309084 | 0.357948536 | 0.287940464 | 0.57753918 |
| chr9:34664337-34664611 | -0.416450189 | 0.28796742 | 0.577568851 |
| chr8:111797234-111797582 | -0.413435614 | 0.287970336 | 0.577568851 |
| chr8:88320627-88320917 | 0.403961831 | 0.287982852 | 0.577568851 |
| chr11:67799868-67800099 | 0.529628129 | 0.287985408 | 0.577568851 |
| chr9:109469674-109470014 | -0.406891211 | 0.28802248 | 0.577618965 |
| chr17:4987839-4988039 | -0.511602352 | 0.288025473 | 0.577618965 |
| chr14:52706413-52706613 | 0.445480684 | 0.288085961 | 0.577712683 |
| chr1:111576551-111576938 | -0.449936584 | 0.288087284 | 0.577712683 |
| chr1:51752094-51752407 | -0.491425497 | 0.288107627 | 0.577723972 |
| chr6:87451971-87452228 | 0.494905816 | 0.288108235 | 0.577723972 |
| chr17:59095714-59095930 | 0.376325271 | 0.288119553 | 0.577723972 |
| chr13:20566410-20567729 | 0.338816101 | 0.288123073 | 0.577723972 |
| chr21:45059280-45059522 | 0.550191507 | 0.288179546 | 0.577820429 |
| chr12:53844898-53845150 | 0.490750339 | 0.288186261 | 0.577820429 |
| chr2:73128875-73129178 | -0.524728515 | 0.288204153 | 0.577841183 |
| chr19:48400134-48400425 | 0.413712687 | 0.288211881 | 0.577841556 |
| chr11:44894215-44894545 | -0.439481275 | 0.288224954 | 0.577852645 |
| chrX:120603679-120604425 | 0.350271482 | 0.28827363 | 0.577935112 |
| chr10:91798158-91799256 | -0.344920989 | 0.28828825 | 0.577949302 |
| chr18:58231780-58232229 | -0.430709145 | 0.288312093 | 0.577970825 |
| chr13:20120440-20121048 | -0.354613577 | 0.288318556 | 0.577970825 |
| chr13:36848645-36848961 | -0.434777277 | 0.288321616 | 0.577970825 |
| chr6:35419643-35419843 | 0.541138421 | 0.288384656 | 0.578035231 |
| chr1:75203623-75204072 | -0.408765172 | 0.288390287 | 0.578035231 |
| chrX:149950733-149951053 | 0.44937759 | 0.288390575 | 0.578035231 |
| chr9:94030318-94031100 | 0.357684136 | 0.288393211 | 0.578035231 |
| chr14:35292056-35293093 | 0.337881187 | 0.288393676 | 0.578035231 |
| chr11:70048622-70049160 | 0.361631547 | 0.288403668 | 0.578035231 |
| chr12:122030058-122030261 | -0.54882111 | 0.288409568 | 0.578035231 |
| chr14:99566047-99566384 | 0.406252706 | 0.288417091 | 0.578035231 |
| chr20:33554978-33555305 | 0.451672931 | 0.288421641 | 0.578035231 |
| chr8:65840846-65841067 | -0.463084064 | 0.288449492 | 0.578075859 |
| chr7:74275382-74275592 | -0.530896513 | 0.288462204 | 0.578075859 |
| chr8:26323397-26323685 | -0.482088992 | 0.288464547 | 0.578075859 |
| chr14:103104971-103105432 | 0.374864895 | 0.288473348 | 0.578078379 |
| chr2:177176343-177176645 | 0.494402085 | 0.288498577 | 0.578104335 |
| chr8:103539398-103539598 | 0.567160733 | 0.288501391 | 0.578104335 |
| chr7:100933758-100933958 | 0.579517504 | 0.288526593 | 0.578139717 |
| chr7:149674728-149675104 | 0.42354897 | 0.288550501 | 0.57816136 |
| chr1:119948686-119948959 | 0.446436259 | 0.288552486 | 0.57816136 |
| chr9:1083666-1084013 | 0.44338099 | 0.288589152 | 0.578219707 |
| chr5:19531319-19531789 | -0.400667053 | 0.288601345 | 0.578228663 |
| chr12:99680693-99681021 | -0.519818857 | 0.288624853 | 0.578228663 |
| chr16:67883774-67884056 | -0.438043286 | 0.288627613 | 0.578228663 |
| chr3:15310493-15311457 | 0.352090297 | 0.288631041 | 0.578228663 |
| chr5:50730457-50730676 | -0.543814564 | 0.288631355 | 0.578228663 |
| chr3:177295141-177295491 | -0.448774881 | 0.288639025 | 0.57822891 |

| | | | |
|---|---|---|---|
| chr2:104033509-104033718 | 0.510154842 | 0.288647455 | 0.578230681 |
| chr20:35698890-35699905 | 0.350064886 | 0.288660625 | 0.578241945 |
| chr12:131428037-131428404 | -0.432185701 | 0.28867057 | 0.578246749 |
| chr1:188701566-188701806 | -0.523741399 | 0.288688212 | 0.578256362 |
| chr20:35303915-35304164 | 0.440466601 | 0.288690463 | 0.578256362 |
| chr7:73324001-73324516 | -0.379244292 | 0.288719167 | 0.578283977 |
| chr12:6430320-6430775 | -0.45650022 | 0.288719344 | 0.578283977 |
| chr10:81018848-81019483 | -0.358349448 | 0.288779878 | 0.578359209 |
| chr1:82269463-82269787 | 0.477048951 | 0.288785918 | 0.578359209 |
| chr1:229229652-229229946 | -0.445688645 | 0.288794474 | 0.578359209 |
| chr7:6446985-6448686 | -0.334594358 | 0.288795083 | 0.578359209 |
| chr3:120146964-120147751 | 0.357521515 | 0.28879908 | 0.578359209 |
| chr15:87556511-87557315 | -0.38642331 | 0.288802501 | 0.578359209 |
| chrX:104921929-104922633 | -0.36272953 | 0.288809742 | 0.578359209 |
| chr8:96234654-96234912 | 0.410085203 | 0.288837073 | 0.578398825 |
| chr7:45111354-45112289 | -0.340345717 | 0.288875553 | 0.578460763 |
| chr1:147084040-147085175 | 0.341667762 | 0.288907128 | 0.578475039 |
| chr5:134592504-134592831 | -0.444202728 | 0.288917749 | 0.578475039 |
| chr12:71551623-71551827 | 0.552448677 | 0.288921159 | 0.578475039 |
| chr4:98929881-98930088 | 0.420201626 | 0.288922684 | 0.578475039 |
| chr10:116849251-116850001 | -0.346251064 | 0.288928124 | 0.578475039 |
| chr16:75484785-75485052 | -0.42754792 | 0.288934375 | 0.578475039 |
| chr19:665013-665271 | 0.47071775 | 0.288951374 | 0.578475039 |
| chr16:85029488-85029764 | 0.405577101 | 0.288954633 | 0.578475039 |
| chr9:137025460-137026121 | -0.358959748 | 0.288959105 | 0.578475039 |
| chr15:39404807-39405017 | 0.526226705 | 0.288969068 | 0.578475039 |
| chr1:174965497-174966477 | 0.356213123 | 0.288970998 | 0.578475039 |
| chr3:25022044-25022308 | -0.560019373 | 0.288973279 | 0.578475039 |
| chr8:124880180-124880380 | 0.518122756 | 0.288980891 | 0.578475162 |
| chr2:226835417-226836567 | -0.343306176 | 0.289030364 | 0.578559081 |
| chr12:53696257-53696672 | 0.368136284 | 0.289045614 | 0.578574493 |
| chr6:27777801-27778022 | -0.546251911 | 0.289065681 | 0.578599545 |
| chr10:113563451-113563839 | -0.391193498 | 0.289085508 | 0.578624116 |
| chr11:89147308-89147517 | -0.541424948 | 0.289097781 | 0.578631141 |
| chr11:82533354-82533554 | -0.581754945 | 0.289116656 | 0.578631141 |
| chr9:130771027-130771232 | 0.54463114 | 0.289117388 | 0.578631141 |
| chr3:38124786-38125072 | -0.422210057 | 0.289124884 | 0.578631141 |
| chr11:102664330-102664677 | -0.450658531 | 0.289127711 | 0.578631141 |
| chr12:87932976-87933286 | -0.517728868 | 0.289134328 | 0.578631141 |
| chr18:63225568-63225901 | -0.447544727 | 0.289146146 | 0.578638745 |
| chr12:45215403-45216292 | -0.347596375 | 0.289153232 | 0.578638745 |
| chr17:60095643-60096494 | 0.339259956 | 0.289170307 | 0.57864699 |
| chr13:106144074-106144426 | 0.435364629 | 0.289172456 | 0.57864699 |
| chr6:39251447-39251815 | -0.43160319 | 0.289203957 | 0.578694913 |
| chr17:29761053-29761680 | 0.3964653 | 0.289222238 | 0.57871638 |
| chr4:20251523-20253018 | 0.34289501 | 0.289238605 | 0.578734016 |
| chr10:102395458-102395789 | 0.381284031 | 0.289250615 | 0.578742932 |
| chrX:39983825-39984074 | 0.50636142 | 0.289330856 | 0.578888365 |
| chr1:224149831-224150046 | -0.533934229 | 0.289389051 | 0.578989683 |
| chr2:7870121-7870440 | -0.522943957 | 0.289406363 | 0.579009201 |

| | | | |
|---|---|---|---|
| chr1:35079164-35079938 | 0.369274764 | 0.289423183 | 0.5790138 |
| chr13:105814582-105814960 | -0.452712976 | 0.289429251 | 0.5790138 |
| chr3:195863126-195863345 | -0.521372652 | 0.289431333 | 0.5790138 |
| chr19:32748613-32748868 | 0.493663686 | 0.289460726 | 0.579057485 |
| chr1:175076271-175076487 | 0.478642154 | 0.28947044 | 0.579061798 |
| chr7:102279499-102279885 | -0.420894647 | 0.289495287 | 0.579096383 |
| chr21:38798538-38798749 | -0.501618925 | 0.289514563 | 0.579119824 |
| chr10:102120014-102120940 | 0.351259348 | 0.289554151 | 0.579169873 |
| chr13:29857139-29857417 | -0.438957428 | 0.289554701 | 0.579169873 |
| chr4:29757060-29757269 | -0.581219697 | 0.28957171 | 0.579179912 |
| chr3:197031407-197031638 | -0.477243801 | 0.289574838 | 0.579179912 |
| chr1:229229150-229229495 | 0.413485442 | 0.28959214 | 0.579199398 |
| chr2:24919062-24919329 | -0.430042184 | 0.289623637 | 0.579237746 |
| chr8:82841079-82841717 | 0.358854492 | 0.289626433 | 0.579237746 |
| chr8:80687213-80687560 | 0.405641762 | 0.289684563 | 0.579338883 |
| chrX:74449434-74449645 | 0.525479515 | 0.289709111 | 0.579370451 |
| chr9:68678496-68678768 | -0.513638252 | 0.289721438 | 0.579370451 |
| chr14:104752532-104753316 | 0.335292194 | 0.289723033 | 0.579370451 |
| chr12:62581791-62582081 | 0.429981351 | 0.289733685 | 0.579376631 |
| chr20:18152573-18153012 | -0.463746333 | 0.28975033 | 0.579386736 |
| chr22:32518813-32519052 | -0.58127195 | 0.289753861 | 0.579386736 |
| chr3:64189133-64189486 | 0.354651485 | 0.289776851 | 0.5793982 |
| chrX:110442694-110443065 | 0.400685734 | 0.289782543 | 0.5793982 |
| chr2:65435689-65436871 | 0.337993779 | 0.289787403 | 0.5793982 |
| chr16:1964731-1965395 | 0.357376395 | 0.289795495 | 0.5793982 |
| chr1:176119622-176119994 | 0.424867158 | 0.289797403 | 0.5793982 |
| chr5:74251261-74251486 | -0.537791859 | 0.289831475 | 0.579451201 |
| chr16:30893955-30894409 | 0.373026764 | 0.289848348 | 0.579469814 |
| chr4:53592766-53593016 | 0.423151487 | 0.289870169 | 0.579498319 |
| chr3:98733508-98733777 | -0.479578862 | 0.289904671 | 0.579542092 |
| chr7:101814119-101814542 | -0.389344626 | 0.28990916 | 0.579542092 |
| chr8:43140689-43140926 | 0.481387358 | 0.289915428 | 0.579542092 |
| chr13:64446457-64446662 | -0.549769319 | 0.289922319 | 0.579542092 |
| chr3:58584726-58585050 | -0.48408455 | 0.289967209 | 0.579596599 |
| chr14:51094832-51095434 | 0.343880825 | 0.289975249 | 0.579596599 |
| chr16:9139075-9139358 | 0.416795715 | 0.289978327 | 0.579596599 |
| chrX:107978812-107979019 | 0.53169267 | 0.289984417 | 0.579596599 |
| chr3:132034489-132035345 | -0.354256802 | 0.289987408 | 0.579596599 |
| chr17:74380635-74381160 | 0.383836583 | 0.290019904 | 0.579646428 |
| chr19:1837714-1837914 | 0.511596579 | 0.290069281 | 0.579716229 |
| chr6:111997093-111997467 | 0.403623023 | 0.290076195 | 0.579716229 |
| chr5:65721655-65722757 | -0.34181937 | 0.290082438 | 0.579716229 |
| chr6:28812017-28812226 | 0.46686342 | 0.290085502 | 0.579716229 |
| chr6:82337704-82338001 | -0.472635172 | 0.290092658 | 0.579716229 |
| chr7:149741655-149742706 | -0.356039354 | 0.290109893 | 0.57973555 |
| chr3:105956378-105956632 | -0.520183507 | 0.290119764 | 0.579740156 |
| chr10:111756428-111756884 | -0.404415865 | 0.290130409 | 0.579741956 |
| chr5:82165917-82166203 | -0.448810611 | 0.290138628 | 0.579741956 |
| chr5:150439245-150439517 | -0.447421783 | 0.290143363 | 0.579741956 |
| chr3:195914581-195915506 | -0.34386317 | 0.290156041 | 0.579752169 |

| | | | |
|---|---|---|---|
| chr11:40029242-40029648 | -0.461446185 | 0.29023116 | 0.579860811 |
| chr17:6853721-6854426 | -0.384133269 | 0.290237119 | 0.579860811 |
| chr15:34209526-34210441 | 0.339846215 | 0.290247455 | 0.579860811 |
| chr9:103789516-103790052 | -0.399756891 | 0.29025436 | 0.579860811 |
| chr1:203860876-203862189 | -0.333638763 | 0.290255094 | 0.579860811 |
| chr16:30650456-30651690 | 0.342791654 | 0.290262511 | 0.579860811 |
| chr10:62949476-62950472 | -0.362503412 | 0.290269131 | 0.579860811 |
| chr17:50660064-50661176 | -0.341130991 | 0.290270957 | 0.579860811 |
| chr18:59070422-59070773 | -0.434090387 | 0.29029005 | 0.579883833 |
| chr2:160110066-160110266 | 0.515537293 | 0.290306403 | 0.579901382 |
| chr19:31572495-31572695 | -0.594083702 | 0.290337979 | 0.579949336 |
| chr15:89647066-89647603 | 0.36932993 | 0.29037324 | 0.58000465 |
| chr22:31211877-31212724 | 0.362509466 | 0.290392462 | 0.580027925 |
| chr8:119832452-119833424 | -0.341603624 | 0.290400666 | 0.58002919 |
| chr4:163681299-163681553 | -0.466156763 | 0.290431592 | 0.580071704 |
| chr11:84437141-84438157 | -0.348490699 | 0.290437092 | 0.580071704 |
| chr1:216793391-216793664 | -0.444101239 | 0.290538369 | 0.580236358 |
| chr1:156941378-156941581 | 0.501853286 | 0.290548659 | 0.580236358 |
| chr4:39697691-39698881 | 0.342153418 | 0.290549376 | 0.580236358 |
| chrX:103503559-103503856 | 0.558847845 | 0.290549823 | 0.580236358 |
| chr11:129455028-129455401 | -0.426654346 | 0.290561678 | 0.580244909 |
| chr16:77660275-77660751 | 0.389367097 | 0.290569292 | 0.580244991 |
| chr17:32444134-32445413 | 0.34915518 | 0.290597122 | 0.580285442 |
| chr5:8247491-8247784 | -0.491715463 | 0.290645814 | 0.580364436 |
| chr14:91934613-91934923 | -0.554403448 | 0.29065183 | 0.580364436 |
| chr10:99540904-99541212 | -0.425456473 | 0.290663053 | 0.580371721 |
| chr11:44949631-44949854 | 0.498354301 | 0.290692786 | 0.58040122 |
| chr21:39321454-39322239 | 0.345048794 | 0.290697207 | 0.58040122 |
| chr16:86016443-86016683 | 0.433150689 | 0.290705448 | 0.58040122 |
| chrX:97296039-97296339 | -0.476535893 | 0.290708126 | 0.58040122 |
| chr12:89395753-89395953 | -0.541351848 | 0.290744233 | 0.580458184 |
| chr12:126784358-126784647 | -0.51384611 | 0.290778706 | 0.580511881 |
| chr15:35444704-35444972 | -0.548692071 | 0.290791382 | 0.580513696 |
| chr1:161439803-161440947 | 0.354005522 | 0.290794767 | 0.580513696 |
| chr2:169109821-169110099 | -0.456719709 | 0.290818786 | 0.580546519 |
| chr16:5043021-5043221 | -0.570081135 | 0.290866455 | 0.580626551 |
| chr6:13461652-13462025 | -0.460675632 | 0.290882266 | 0.580642985 |
| chr11:41201171-41201450 | -0.539590508 | 0.290907753 | 0.58066271 |
| chr4:46123968-46124360 | -0.438294103 | 0.290912048 | 0.58066271 |
| chr10:92892269-92892738 | 0.371662325 | 0.290914882 | 0.58066271 |
| chr6:17986762-17987271 | 0.38666982 | 0.290924518 | 0.580666817 |
| chr2:174465444-174465886 | -0.441746173 | 0.290943078 | 0.580685096 |
| chr19:17938275-17938475 | -0.578570853 | 0.290951323 | 0.580685096 |
| chr9:134419700-134420081 | 0.392773194 | 0.290957644 | 0.580685096 |
| chr12:52927236-52927739 | 0.376864531 | 0.29096399 | 0.580685096 |
| chr12:118271820-118272350 | 0.381060589 | 0.290993693 | 0.580729247 |
| chr7:102464738-102465317 | -0.355608103 | 0.291009554 | 0.580745775 |
| chr5:152402665-152403303 | -0.402522874 | 0.29104202 | 0.580795438 |
| chr1:941504-942277 | 0.362899422 | 0.291106418 | 0.58090882 |
| chr21:33552504-33552744 | -0.548701431 | 0.291136761 | 0.58095424 |

| | | | |
|---|---|---|---|
| chr9:136944510-136944953 | 0.373904409 | 0.291178781 | 0.581022958 |
| chr3:35924143-35924393 | 0.505178231 | 0.291232824 | 0.581115663 |
| chr4:85084507-85084721 | 0.566878249 | 0.291271138 | 0.581176978 |
| chr10:99296911-99297360 | 0.442749414 | 0.291285455 | 0.581183782 |
| chr1:221845935-221846193 | 0.495815918 | 0.291305302 | 0.581183782 |
| chr3:9827343-9827602 | 0.436082487 | 0.291310636 | 0.581183782 |
| chr1:150849844-150850044 | 0.49347173 | 0.291312929 | 0.581183782 |
| chr1:2642917-2643373 | 0.404297284 | 0.291321477 | 0.581183782 |
| chr6:27280066-27280439 | -0.395464997 | 0.291321653 | 0.581183782 |
| chr2:161238649-161239225 | -0.383333719 | 0.291330395 | 0.581183782 |
| chr3:138052270-138052580 | 0.424876159 | 0.291337704 | 0.581183782 |
| chr10:104305149-104306205 | -0.344696122 | 0.291342814 | 0.581183782 |
| chr10:109883971-109884706 | -0.364971028 | 0.291355099 | 0.581193158 |
| chr4:52552379-52552625 | 0.559006667 | 0.291373918 | 0.581208427 |
| chr1:156282158-156283134 | 0.341664299 | 0.291377924 | 0.581208427 |
| chr4:55335990-55336475 | -0.443384761 | 0.291389357 | 0.581216102 |
| chrX:22135923-22136217 | 0.464246716 | 0.291418103 | 0.581242172 |
| chr2:211371749-211372105 | 0.451093347 | 0.291425591 | 0.581242172 |
| chr8:75420816-75421016 | -0.588805018 | 0.291429188 | 0.581242172 |
| chr5:147949765-147949965 | 0.586055901 | 0.291434457 | 0.581242172 |
| chr8:78444846-78445514 | -0.373014106 | 0.291440356 | 0.581242172 |
| chr4:42865648-42865937 | -0.467862511 | 0.29146159 | 0.58126939 |
| chr6:121055551-121055901 | 0.39942524 | 0.291478261 | 0.581287506 |
| chr7:51776216-51776510 | -0.51931871 | 0.291489025 | 0.581293844 |
| chr16:28623401-28623777 | -0.445978894 | 0.291497423 | 0.581295462 |
| chr2:9570711-9570911 | -0.545428044 | 0.291507076 | 0.581299583 |
| chr5:132952848-132953156 | -0.435955437 | 0.2915161 | 0.581302449 |
| chr14:106627005-106627238 | 0.487162754 | 0.291549638 | 0.581344762 |
| chr1:26111337-26111950 | 0.373831308 | 0.291552494 | 0.581344762 |
| chr12:99253173-99253373 | 0.565957643 | 0.29156906 | 0.581362666 |
| chr6:26031925-26032390 | -0.358225486 | 0.291640725 | 0.581455094 |
| chr20:59939559-59941082 | 0.330740687 | 0.291641843 | 0.581455094 |
| chr8:123453570-123453780 | -0.439219786 | 0.291644116 | 0.581455094 |
| chr17:46286730-46287232 | 0.410786574 | 0.29164577 | 0.581455094 |
| chr2:241252015-241252215 | 0.572136443 | 0.291680342 | 0.58150889 |
| chr20:57114555-57115326 | 0.331702109 | 0.291712045 | 0.58152761 |
| chr7:128404997-128405197 | -0.451378561 | 0.291716723 | 0.58152761 |
| chr9:22107150-22107668 | -0.387119527 | 0.291717557 | 0.58152761 |
| chr5:26700829-26701123 | -0.420396725 | 0.29172009 | 0.58152761 |
| chr22:37538792-37539214 | -0.374834995 | 0.291740161 | 0.581531189 |
| chr2:240217350-240217990 | 0.354685274 | 0.291741377 | 0.581531189 |
| chr16:56424898-56425226 | 0.431374863 | 0.291744654 | 0.581531189 |
| chr22:42648960-42649160 | 0.526357149 | 0.291786681 | 0.581547984 |
| chr6:13614646-13616249 | 0.335837303 | 0.291791296 | 0.581547984 |
| chr6:27215074-27215276 | 0.479536201 | 0.291803776 | 0.581547984 |
| chr1:109739682-109740563 | -0.343398688 | 0.291804047 | 0.581547984 |
| chr3:197016109-197016569 | 0.415380731 | 0.29180724 | 0.581547984 |
| chr1:153958476-153958921 | 0.346352038 | 0.291816144 | 0.581547984 |
| chr12:131710520-131711412 | 0.362004853 | 0.291818767 | 0.581547984 |
| chrX:132022432-132022632 | 0.46810226 | 0.291821019 | 0.581547984 |

| | | | |
|---|---|---|---|
| chr19:39390608-39391704 | 0.338765283 | 0.291821389 | 0.581547984 |
| chr6:143940089-143940347 | -0.447338581 | 0.29184233 | 0.58157459 |
| chr5:157994961-157995385 | -0.396329548 | 0.291883567 | 0.58164164 |
| chr22:28663652-28663859 | -0.577618013 | 0.291897006 | 0.581653293 |
| chr10:63515486-63515748 | -0.509359731 | 0.291907767 | 0.581659608 |
| chr2:2291854-2292239 | -0.42497603 | 0.291922151 | 0.581661258 |
| chr3:107834565-107834783 | 0.56275334 | 0.291930545 | 0.581661258 |
| chr14:104342016-104342387 | 0.373738767 | 0.291936328 | 0.581661258 |
| chr2:16507689-16507947 | -0.453564099 | 0.29193896 | 0.581661258 |
| chr18:49655979-49656528 | -0.431746884 | 0.291948503 | 0.581663954 |
| chr6:131576196-131576396 | 0.554333223 | 0.291955496 | 0.581663954 |
| chr8:117947864-117948064 | -0.550763561 | 0.291981449 | 0.58169371 |
| chr1:214516360-214516646 | -0.483475448 | 0.291985615 | 0.58169371 |
| chr20:53926677-53927089 | 0.341516313 | 0.292019021 | 0.581745137 |
| chr6:98844921-98846113 | -0.345260271 | 0.292056486 | 0.581792521 |
| chr9:37753454-37754150 | -0.375923996 | 0.292059382 | 0.581792521 |
| chr6:91081505-91081807 | -0.471624345 | 0.292074981 | 0.581792521 |
| chr13:36226599-36226843 | 0.477775766 | 0.292077372 | 0.581792521 |
| chr15:44194569-44195862 | 0.339996878 | 0.292083226 | 0.581792521 |
| chr20:50749187-50749616 | 0.340602571 | 0.292088365 | 0.581792521 |
| chr2:3423850-3424230 | 0.402738412 | 0.292117713 | 0.581815261 |
| chr6:52419813-52421047 | -0.334280541 | 0.292121891 | 0.581815261 |
| chr13:32030992-32031883 | -0.350663186 | 0.292122562 | 0.581815261 |
| chr1:197675841-197676088 | -0.500370408 | 0.292131052 | 0.581817048 |
| chr11:62241952-62242446 | 0.379833744 | 0.292144777 | 0.581829259 |
| chr9:129139908-129140223 | 0.385807581 | 0.292165791 | 0.581850883 |
| chr7:47952292-47952493 | -0.494572831 | 0.292170822 | 0.581850883 |
| chr11:69301652-69302289 | -0.362035778 | 0.29220373 | 0.581901293 |
| chr10:30432989-30434765 | -0.335137492 | 0.292234167 | 0.581942527 |
| chr17:74190239-74190483 | 0.425374822 | 0.292240005 | 0.581942527 |
| chr12:122888363-122889169 | 0.359716775 | 0.29224722 | 0.581942527 |
| chr2:128316599-128317013 | -0.395452108 | 0.292266214 | 0.581965224 |
| chr6:42745969-42747272 | 0.335381552 | 0.292279615 | 0.581976785 |
| chr18:3449056-3449679 | 0.348681207 | 0.292318478 | 0.582039042 |
| chr6:9516227-9516603 | -0.39619768 | 0.292327936 | 0.58204275 |
| chrX:40003396-40003657 | -0.570907964 | 0.292367914 | 0.582097238 |
| chr20:33193264-33193572 | 0.475338332 | 0.292370497 | 0.582097238 |
| chr1:207032666-207033076 | -0.363979913 | 0.292405988 | 0.582152773 |
| chr20:51962311-51962681 | 0.38938943 | 0.292439768 | 0.582204897 |
| chr12:103758818-103759042 | -0.55512165 | 0.292467062 | 0.582234511 |
| chr2:119283686-119284347 | 0.359266702 | 0.29246984 | 0.582234511 |
| chr22:38335988-38336435 | 0.405104126 | 0.292493284 | 0.582266053 |
| chr3:50328060-50328630 | 0.369443654 | 0.292522977 | 0.582310035 |
| chr17:73310614-73311476 | 0.338539116 | 0.292544804 | 0.582338356 |
| chr1:236406635-236406835 | 0.539846723 | 0.292556007 | 0.582345527 |
| chr1:163171214-163171461 | -0.504958774 | 0.292598609 | 0.582415197 |
| chr14:89939449-89939990 | -0.370485604 | 0.292608507 | 0.58241977 |
| chr17:48023911-48024173 | 0.389502676 | 0.292626598 | 0.582440648 |
| chr13:34964860-34965155 | -0.48296978 | 0.292635518 | 0.582443273 |
| chr5:32093111-32093589 | -0.371460331 | 0.292657204 | 0.582471305 |

| | | | |
|---|---|---|---|
| chr18:62634825-62635176 | -0.475698074 | 0.292674831 | 0.582491256 |
| chr3:167748396-167748924 | 0.409417037 | 0.292715368 | 0.582556804 |
| chr1:78491264-78492067 | -0.363837459 | 0.29276493 | 0.582640306 |
| chr1:115012448-115012676 | 0.434926268 | 0.292774531 | 0.582644281 |
| chr21:45642921-45643804 | 0.336290207 | 0.292793678 | 0.582667251 |
| chr1:53070855-53071055 | 0.528182922 | 0.292802782 | 0.582670235 |
| chr3:143885872-143886106 | -0.463368299 | 0.292821505 | 0.582687427 |
| chr11:84714327-84714629 | -0.416366197 | 0.292833626 | 0.582687427 |
| chr9:90948194-90948487 | -0.515772883 | 0.292834235 | 0.582687427 |
| chr11:121292330-121293157 | 0.352739029 | 0.292856664 | 0.582716924 |
| chr6:36616912-36617448 | 0.380187318 | 0.292875007 | 0.58273829 |
| chr7:75957024-75957660 | -0.365069323 | 0.292888842 | 0.582750684 |
| chr17:82858182-82859020 | 0.359414591 | 0.292930659 | 0.582817151 |
| chr9:73199970-73200241 | -0.47890225 | 0.292941818 | 0.582817151 |
| chr1:147627512-147627824 | 0.401686336 | 0.292945067 | 0.582817151 |
| chr6:161373927-161374219 | -0.422108495 | 0.292971482 | 0.582841762 |
| chr3:122975366-122975575 | 0.468733479 | 0.292977944 | 0.582841762 |
| chr12:45991257-45991541 | 0.394633628 | 0.292980258 | 0.582841762 |
| chr3:27758136-27758387 | -0.50637792 | 0.293013511 | 0.58286786 |
| chr13:98484102-98485250 | -0.338457932 | 0.293016515 | 0.58286786 |
| chr1:70860860-70861110 | -0.546750794 | 0.293023016 | 0.58286786 |
| chr1:156193523-156194446 | 0.336255614 | 0.293023805 | 0.58286786 |
| chr1:228406570-228407355 | 0.345894342 | 0.293038609 | 0.582882176 |
| chr7:151017931-151018636 | -0.363037254 | 0.293069238 | 0.582927968 |
| chr5:140718554-140719466 | -0.334662447 | 0.293107146 | 0.582955596 |
| chr4:52545386-52545777 | 0.412898308 | 0.293108416 | 0.582955596 |
| chr12:51424943-51425319 | 0.391422233 | 0.293108786 | 0.582955596 |
| chr14:105467205-105467533 | 0.383216822 | 0.293113561 | 0.582955596 |
| chr1:3611576-3611880 | 0.42588059 | 0.293126732 | 0.582966659 |
| chr3:184134088-184134949 | 0.347479679 | 0.293141352 | 0.582980603 |
| chr14:56874461-56874716 | 0.462483273 | 0.293153591 | 0.582989812 |
| chr2:73984511-73984949 | -0.376571842 | 0.293169829 | 0.583006972 |
| chr8:101477244-101477583 | 0.409634815 | 0.2931943 | 0.583040504 |
| chr3:31157378-31157583 | -0.594925366 | 0.293221023 | 0.583067431 |
| chr4:93791650-93791865 | -0.544063853 | 0.29322306 | 0.583067431 |
| chr2:96117827-96118073 | 0.429930466 | 0.293231495 | 0.583069072 |
| chr14:63045316-63046146 | 0.364606549 | 0.293253288 | 0.583083862 |
| chr3:115783930-115785018 | -0.334149991 | 0.29325805 | 0.583083862 |
| chr10:28248524-28248815 | 0.519359236 | 0.293261763 | 0.583083862 |
| chr2:127197817-127198292 | 0.37657766 | 0.293326469 | 0.583184324 |
| chr3:39208514-39208796 | -0.469476531 | 0.293327513 | 0.583184324 |
| chr6:157273298-157273560 | 0.440381589 | 0.293397923 | 0.583309176 |
| chr1:146748763-146748985 | -0.540882916 | 0.293420528 | 0.583338981 |
| chr14:27739468-27740311 | 0.357880344 | 0.293444629 | 0.583371761 |
| chr2:12715981-12716280 | -0.379196721 | 0.29347197 | 0.583388705 |
| chr3:146589982-146590224 | -0.500913889 | 0.293473721 | 0.583388705 |
| chr5:38968722-38968958 | -0.500954351 | 0.293475994 | 0.583388705 |
| chr16:30104327-30104977 | -0.364101047 | 0.293501669 | 0.58341067 |
| chr18:21784639-21784885 | 0.46944417 | 0.29350735 | 0.58341067 |
| chr18:26202534-26202734 | -0.58989308 | 0.293514471 | 0.58341067 |

| | | | |
|---|---|---|---|
| chr20:45348287-45348617 | 0.382020673 | 0.293524592 | 0.58341067 |
| chr18:34978982-34979539 | -0.370791372 | 0.293525115 | 0.58341067 |
| chr2:71510317-71510563 | 0.422739304 | 0.293543121 | 0.583431326 |
| chr1:74197662-74198579 | -0.338316516 | 0.293567283 | 0.583452949 |
| chr6:43059204-43060068 | 0.341920627 | 0.293578173 | 0.583452949 |
| chr2:208929364-208929749 | -0.40216894 | 0.293581096 | 0.583452949 |
| chr13:85995869-85996308 | -0.446474721 | 0.293585819 | 0.583452949 |
| chr10:97427087-97427295 | 0.438545648 | 0.293592074 | 0.583452949 |
| chr7:37916916-37917708 | -0.364338356 | 0.29362161 | 0.583496512 |
| chr4:151117403-151117617 | -0.48988366 | 0.293665019 | 0.583504745 |
| chr17:37067916-37068174 | -0.522303245 | 0.29367251 | 0.583504745 |
| chr3:5122037-5123764 | 0.336493819 | 0.293677734 | 0.583504745 |
| chr17:39402107-39402338 | 0.507563844 | 0.293682165 | 0.583504745 |
| chr19:6361268-6361728 | 0.399957884 | 0.2936898 | 0.583504745 |
| chr10:18658900-18660106 | -0.337079677 | 0.293692823 | 0.583504745 |
| chr17:59296810-59297152 | 0.396092606 | 0.293707985 | 0.583504745 |
| chr12:72133584-72133878 | 0.460275095 | 0.29370989 | 0.583504745 |
| chr10:73246681-73247520 | -0.341986808 | 0.293714024 | 0.583504745 |
| chr4:7876154-7876354 | 0.543924752 | 0.293718645 | 0.583504745 |
| chr15:78756282-78756497 | -0.587225102 | 0.293728184 | 0.583504745 |
| chr9:129581441-129581687 | 0.48510227 | 0.29373292 | 0.583504745 |
| chr1:106534638-106534983 | 0.354862546 | 0.293733852 | 0.583504745 |
| chr21:34517611-34517825 | 0.505838788 | 0.293735138 | 0.583504745 |
| chr4:97498941-97499253 | -0.452680968 | 0.293746402 | 0.583504745 |
| chr14:60164436-60165625 | 0.338637323 | 0.293747726 | 0.583504745 |
| chr1:176249145-176249448 | -0.483132743 | 0.293755214 | 0.583504745 |
| chr11:129331869-129332132 | 0.479564445 | 0.293783283 | 0.583523976 |
| chr2:20093486-20093935 | -0.369344837 | 0.293784698 | 0.583523976 |
| chr19:11142507-11142985 | -0.376820545 | 0.293787743 | 0.583523976 |
| chr11:96486955-96487830 | -0.362026571 | 0.293811233 | 0.583555506 |
| chr5:132477551-132477772 | -0.468977404 | 0.293824145 | 0.583566023 |
| chr8:8701542-8702250 | 0.379219913 | 0.293854094 | 0.583610378 |
| chr11:31369427-31370277 | -0.342122658 | 0.293868245 | 0.583623355 |
| chr17:41647978-41648383 | 0.420916764 | 0.293891572 | 0.58363688 |
| chr18:31722417-31722741 | 0.420916764 | 0.293891572 | 0.58363688 |
| chr10:80407662-80409313 | -0.332855816 | 0.293907562 | 0.58363688 |
| chr1:209199822-209200119 | -0.461400733 | 0.29390898 | 0.58363688 |
| chr19:49057746-49058002 | 0.451259937 | 0.29391314 | 0.58363688 |
| chr1:218344863-218346365 | -0.334519804 | 0.293928752 | 0.58365152 |
| chr3:141659796-141660835 | 0.348797991 | 0.293935748 | 0.58365152 |
| chr14:67619399-67620369 | 0.337278087 | 0.293955941 | 0.583673842 |
| chr9:136758441-136758657 | 0.486635254 | 0.29396745 | 0.583673842 |
| chr2:218372648-218372908 | 0.507201799 | 0.293974275 | 0.583673842 |
| chr6:30768839-30769851 | -0.356465818 | 0.29397746 | 0.583673842 |
| chr17:49180383-49180608 | 0.465892455 | 0.293987947 | 0.583674634 |
| chr7:73851524-73851789 | -0.503814272 | 0.293993094 | 0.583674634 |
| chr17:50144986-50145186 | -0.464636366 | 0.294002537 | 0.583678258 |
| chr2:159285901-159287409 | -0.330307647 | 0.294017907 | 0.583683036 |
| chr4:90950999-90951448 | 0.398240296 | 0.294020179 | 0.583683036 |
| chr20:41956375-41956592 | 0.448799917 | 0.294030566 | 0.583688534 |

| | | | |
|---|---|---|---|
| chr3:176814322-176814644 | -0.39918496 | 0.294057395 | 0.583706078 |
| chr2:11778985-11779657 | -0.37496294 | 0.294063159 | 0.583706078 |
| chr1:223090107-223090374 | 0.437990627 | 0.294064289 | 0.583706078 |
| chr15:75625513-75625893 | 0.41501599 | 0.294069876 | 0.583706078 |
| chr3:126134176-126134429 | 0.434341111 | 0.294129194 | 0.583781075 |
| chr12:2034949-2035394 | -0.424579874 | 0.294135733 | 0.583781075 |
| chr22:25419887-25420087 | 0.563134887 | 0.294138528 | 0.583781075 |
| chr17:59904974-59905174 | 0.375759021 | 0.294138849 | 0.583781075 |
| chr14:97096525-97096812 | 0.491436946 | 0.294145754 | 0.583781075 |
| chr9:104747279-104748508 | -0.335900071 | 0.294179096 | 0.583809223 |
| chr2:2698658-2698858 | -0.529032877 | 0.29418016 | 0.583809223 |
| chr3:146421833-146422246 | 0.387098602 | 0.294183611 | 0.583809223 |
| chr2:201059028-201059248 | -0.572655962 | 0.294202219 | 0.583809223 |
| chr6:166381898-166383773 | -0.334541348 | 0.294203265 | 0.583809223 |
| chr10:114158027-114158251 | -0.526239568 | 0.294205653 | 0.583809223 |
| chr6:44133628-44133971 | 0.385888535 | 0.294217991 | 0.583818588 |
| chr6:3356134-3356795 | -0.373290627 | 0.294231386 | 0.583830047 |
| chr6:144330550-144331122 | -0.363472437 | 0.294247925 | 0.583847744 |
| chr8:108242900-108243205 | 0.419498776 | 0.294265016 | 0.583866538 |
| chr14:69818747-69819021 | 0.434173229 | 0.29429385 | 0.583894323 |
| chr3:9204357-9204557 | 0.482932627 | 0.294294261 | 0.583894323 |
| chr16:89458787-89458987 | 0.588681968 | 0.29431574 | 0.583921819 |
| chr22:33920898-33921388 | 0.357783657 | 0.294327751 | 0.583930529 |
| chr14:74160906-74161190 | 0.522814862 | 0.294361818 | 0.583982994 |
| chr12:32896414-32897561 | 0.338428454 | 0.294381897 | 0.584007708 |
| chr11:14499158-14500675 | -0.339917448 | 0.294423101 | 0.584074327 |
| chr3:139492103-139492337 | 0.468993055 | 0.294432729 | 0.584078307 |
| chr2:55838850-55839596 | -0.358109437 | 0.294440569 | 0.584078736 |
| chr5:141916062-141916476 | -0.40154388 | 0.294452373 | 0.584087031 |
| chr3:81496102-81496410 | -0.454631266 | 0.294513203 | 0.584192572 |
| chr7:133862010-133862211 | -0.561732419 | 0.294528515 | 0.584198568 |
| chr17:45043245-45043634 | -0.402047198 | 0.294531475 | 0.584198568 |
| chr1:161119628-161119927 | 0.396672154 | 0.294553737 | 0.584216851 |
| chr17:50497047-50497349 | -0.508780763 | 0.294562362 | 0.584216851 |
| chr2:30755951-30756508 | -0.350425696 | 0.294568339 | 0.584216851 |
| chr6:31529092-31529330 | -0.499922485 | 0.294571191 | 0.584216851 |
| chr16:28823612-28824721 | 0.337142117 | 0.294582366 | 0.584223892 |
| chr1:220841356-220841562 | -0.544990072 | 0.294594592 | 0.584233017 |
| chr1:56627808-56628008 | 0.558228533 | 0.294604955 | 0.584238446 |
| chr3:67263948-67264202 | 0.464217445 | 0.294631045 | 0.584270504 |
| chr11:75876207-75876461 | -0.574057497 | 0.294660367 | 0.584270504 |
| chr10:123109544-123109888 | -0.440218983 | 0.294662511 | 0.584270504 |
| chrX:125204881-125205869 | -0.337182422 | 0.294668806 | 0.584270504 |
| chr2:43463165-43463529 | 0.420237654 | 0.294670488 | 0.584270504 |
| chr2:45611101-45611510 | -0.359542181 | 0.294671274 | 0.584270504 |
| chr1:23838376-23838819 | -0.43482974 | 0.294677315 | 0.584270504 |
| chr11:1262520-1262902 | 0.39395011 | 0.294694892 | 0.584270504 |
| chr7:26452043-26452701 | 0.376260691 | 0.294701525 | 0.584270504 |
| chr16:2223086-2223745 | 0.344031257 | 0.294701558 | 0.584270504 |
| chr7:151175773-151176016 | -0.446209009 | 0.294704999 | 0.584270504 |

| | | | |
|---|---|---|---|
| chr7:149147009-149147603 | -0.361976108 | 0.294716746 | 0.584278675 |
| chr1:10835823-10836290 | 0.388568296 | 0.294733912 | 0.584297588 |
| chr13:21409646-21409935 | 0.456570048 | 0.294762882 | 0.584329539 |
| chr1:24847917-24848312 | 0.401567015 | 0.294786244 | 0.584329539 |
| chr1:214214039-214214261 | -0.488915322 | 0.294786406 | 0.584329539 |
| chr4:134920223-134920650 | -0.479621024 | 0.294786949 | 0.584329539 |
| chr3:30779713-30780110 | 0.427592412 | 0.294788159 | 0.584329539 |
| chr3:51637484-51637716 | -0.46081449 | 0.294802729 | 0.584343302 |
| chr14:80323379-80323596 | 0.496853165 | 0.294810958 | 0.584344496 |
| chr2:14287991-14288617 | -0.352456221 | 0.294819809 | 0.584346684 |
| chr1:154961299-154962158 | 0.347853254 | 0.294827314 | 0.584346684 |
| chr17:31566711-31566911 | 0.553461018 | 0.294865238 | 0.5844066 |
| chr6:170443487-170443779 | 0.420358584 | 0.294872798 | 0.5844066 |
| chrX:30127213-30127806 | -0.415102701 | 0.294920308 | 0.584476824 |
| chr14:24123566-24123861 | 0.42990177 | 0.29493096 | 0.584476824 |
| chr10:94617492-94617894 | -0.424334254 | 0.294936211 | 0.584476824 |
| chr14:93115187-93115705 | 0.348575106 | 0.294938744 | 0.584476824 |
| chr7:33287927-33288127 | -0.558375248 | 0.294949002 | 0.584482036 |
| chr16:89926093-89926605 | -0.360644655 | 0.294987098 | 0.584509395 |
| chr1:16707649-16708036 | 0.377198411 | 0.294987494 | 0.584509395 |
| chr22:49970683-49971192 | 0.361831443 | 0.294991485 | 0.584509395 |
| chr6:27312576-27313113 | 0.385529605 | 0.294998273 | 0.584509395 |
| chr14:65843446-65843772 | 0.48099518 | 0.29500095 | 0.584509395 |
| chr20:50741242-50741672 | 0.337725876 | 0.29502871 | 0.584546628 |
| chr17:12742168-12742552 | -0.434675061 | 0.295035 | 0.584546628 |
| chr8:81371139-81371366 | 0.487744264 | 0.29505112 | 0.584563451 |
| chrX:76148749-76149094 | 0.452490307 | 0.295067479 | 0.584580747 |
| chr1:36268860-36269256 | 0.435151265 | 0.295092435 | 0.584615072 |
| chr22:19051451-19051692 | -0.532639913 | 0.295108741 | 0.584618383 |
| chr13:103883128-103883489 | -0.396780453 | 0.295115913 | 0.584618383 |
| chr8:38268447-38269692 | -0.349504254 | 0.295116995 | 0.584618383 |
| chr1:233735504-233735811 | 0.462268482 | 0.295131549 | 0.584632097 |
| chr6:13873190-13874154 | -0.347744108 | 0.295168877 | 0.584690926 |
| chr4:41222332-41222532 | -0.577209144 | 0.295178759 | 0.584692707 |
| chr17:66900172-66900502 | 0.393558433 | 0.295185038 | 0.584692707 |
| chr17:70127274-70127581 | 0.424916131 | 0.295214036 | 0.584726504 |
| chr12:125518135-125518337 | 0.551609766 | 0.295217363 | 0.584726504 |
| chr2:166539330-166539701 | -0.414248592 | 0.295260753 | 0.584797328 |
| chr15:41989544-41989763 | 0.474451245 | 0.29527247 | 0.584805418 |
| chr2:219126534-219126920 | 0.402564376 | 0.295296889 | 0.584838664 |
| chr17:76501095-76501823 | 0.347536085 | 0.29537342 | 0.584975115 |
| chr3:138781128-138781508 | 0.4441406 | 0.2954568 | 0.58512436 |
| chr12:189117-189426 | -0.456227771 | 0.295471956 | 0.58512436 |
| chr2:14642540-14642821 | -0.509121515 | 0.295477126 | 0.58512436 |
| chr5:95960659-95962331 | -0.328556629 | 0.295479325 | 0.58512436 |
| chr18:11995475-11995784 | 0.409701803 | 0.295497892 | 0.585137317 |
| chr17:41930011-41930883 | 0.350396635 | 0.295516889 | 0.585137317 |
| chr6:28737790-28738045 | -0.49130264 | 0.295518226 | 0.585137317 |
| chr17:80650096-80650389 | 0.447086922 | 0.295519107 | 0.585137317 |
| chr15:85745101-85745301 | 0.571789821 | 0.295530876 | 0.585137317 |

| | | | |
|---|---|---|---|
| chr14:62179447-62179862 | 0.419672144 | 0.295531688 | 0.585137317 |
| chr1:204162417-204162876 | -0.345268391 | 0.295596808 | 0.585251128 |
| chr9:98118921-98119589 | -0.379686801 | 0.295616562 | 0.58525362 |
| chr4:156700229-156700436 | -0.573016413 | 0.295620982 | 0.58525362 |
| chr4:168183860-168184060 | -0.573016413 | 0.295620982 | 0.58525362 |
| chr19:18596796-18597027 | 0.494997674 | 0.295662273 | 0.58531454 |
| chr6:142462345-142462578 | -0.540901549 | 0.295667031 | 0.58531454 |
| chr2:44479479-44479763 | -0.46305257 | 0.29567937 | 0.585323843 |
| chr3:15427167-15427949 | 0.342482476 | 0.295689574 | 0.585328921 |
| chr20:57194587-57194978 | 0.337085496 | 0.295699916 | 0.585334272 |
| chr11:79762347-79762828 | -0.427000669 | 0.295715485 | 0.585347434 |
| chr8:127311484-127312270 | -0.342898492 | 0.295721845 | 0.585347434 |
| chr8:109213556-109213850 | 0.426942917 | 0.295783465 | 0.585454281 |
| chr3:53073339-53073555 | 0.458533213 | 0.295803935 | 0.585479674 |
| chr2:96888434-96889020 | -0.348115478 | 0.295830373 | 0.585509147 |
| chr11:82229419-82229707 | 0.456715506 | 0.295834109 | 0.585509147 |
| chr1:107150694-107150938 | 0.389233407 | 0.2958539 | 0.585533192 |
| chr14:73758337-73758623 | 0.377885308 | 0.295864317 | 0.585538685 |
| chr14:100204388-100204856 | -0.398482931 | 0.295874861 | 0.585544428 |
| chr15:59506023-59506225 | -0.558579787 | 0.295888164 | 0.58555563 |
| chr13:46211679-46212329 | -0.360143198 | 0.295929862 | 0.585605475 |
| chr12:50111580-50112529 | 0.331560391 | 0.295932434 | 0.585605475 |
| chr15:77376849-77377249 | 0.391093247 | 0.295936279 | 0.585605475 |
| chr14:81831542-81831747 | 0.518617686 | 0.295982292 | 0.585680336 |
| chr7:12322460-12322814 | 0.454438965 | 0.295989398 | 0.585680336 |
| chr12:67355664-67355903 | 0.505139438 | 0.296001359 | 0.585688879 |
| chr22:30325141-30325569 | 0.388200541 | 0.296024847 | 0.585700629 |
| chr12:70643800-70644268 | 0.402529379 | 0.296026472 | 0.585700629 |
| chr9:72364482-72366105 | -0.33020029 | 0.29603023 | 0.585700629 |
| chr17:62553185-62553497 | -0.420281758 | 0.296116548 | 0.585856284 |
| chr2:127701710-127702359 | -0.355511358 | 0.296156322 | 0.585919845 |
| chr9:121322304-121322515 | -0.489259116 | 0.296171999 | 0.585935731 |
| chr10:29318261-29318483 | 0.488316959 | 0.296249746 | 0.586074412 |
| chr8:72344375-72344755 | 0.422502117 | 0.296286684 | 0.586132354 |
| chr21:41898504-41898923 | 0.420288224 | 0.296319773 | 0.586182678 |
| chr4:8572162-8572565 | 0.448986143 | 0.296380597 | 0.586287864 |
| chr15:74937410-74938432 | 0.339984326 | 0.296390368 | 0.586292056 |
| chr12:52032877-52033321 | 0.352018382 | 0.296440098 | 0.586361099 |
| chr3:156751451-156752084 | -0.368218943 | 0.296448325 | 0.586361099 |
| chr20:57195487-57196041 | 0.332367876 | 0.296453031 | 0.586361099 |
| chr17:46835883-46836144 | 0.447832778 | 0.296455882 | 0.586361099 |
| chr12:127125950-127126248 | -0.483094972 | 0.296474981 | 0.586379686 |
| chr17:72030182-72030941 | -0.364635616 | 0.296480585 | 0.586379686 |
| chr6:70874570-70874903 | -0.52175775 | 0.296498441 | 0.586399865 |
| chr5:154857386-154858916 | -0.327467314 | 0.296529064 | 0.586444858 |
| chr22:22666605-22666829 | 0.490575937 | 0.296540011 | 0.586444858 |
| chr2:27966709-27966909 | -0.57558881 | 0.296546464 | 0.586444858 |
| chr12:6825524-6825882 | -0.400963387 | 0.296553508 | 0.586444858 |
| chr6:133952912-133953587 | -0.340474266 | 0.29656922 | 0.586444858 |
| chr1:76867467-76868021 | -0.37757832 | 0.296576283 | 0.586444858 |

| | | | |
|---|---|---|---|
| chr11:118522843-118523139 | 0.402815569 | 0.296579262 | 0.586444858 |
| chr9:35603857-35604509 | -0.340065904 | 0.296587627 | 0.586444858 |
| chr22:11027961-11028477 | 0.41876054 | 0.296590579 | 0.586444858 |
| chr16:70298957-70299848 | 0.351435704 | 0.296597728 | 0.586444858 |
| chr11:71284587-71284879 | -0.419833981 | 0.296608928 | 0.58645187 |
| chr17:59372519-59372953 | 0.368651276 | 0.296619608 | 0.586457852 |
| chr12:74664079-74664458 | 0.500784263 | 0.296655706 | 0.586500808 |
| chr10:7672594-7672824 | 0.543677114 | 0.296660878 | 0.586500808 |
| chr10:97714087-97715020 | 0.350020086 | 0.296669567 | 0.586500808 |
| chrX:86919332-86919996 | -0.39186745 | 0.296672599 | 0.586500808 |
| chr7:128947307-128947573 | -0.44591112 | 0.296679607 | 0.586500808 |
| chr3:106677153-106677526 | 0.44629861 | 0.29669128 | 0.586508752 |
| chr10:55174502-55174722 | -0.504149907 | 0.296714984 | 0.586527181 |
| chr16:67177974-67178717 | 0.342110056 | 0.296715912 | 0.586527181 |
| chr12:25820877-25821212 | -0.469113899 | 0.296743841 | 0.586567257 |
| chr12:57229980-57230369 | 0.397048405 | 0.296810207 | 0.586659829 |
| chr6:80457411-80457661 | -0.539629095 | 0.296813288 | 0.586659829 |
| chr1:52726263-52726899 | 0.359308213 | 0.296813643 | 0.586659829 |
| chrX:46446494-46446933 | -0.408562123 | 0.296824221 | 0.586661152 |
| chr22:21726476-21727002 | 0.394190629 | 0.296829625 | 0.586661152 |
| chr16:20774748-20775152 | 0.415124163 | 0.296839578 | 0.586665691 |
| chr4:135553069-135553530 | -0.424250591 | 0.296848449 | 0.58666809 |
| chr15:48811669-48811905 | -0.403299384 | 0.296886339 | 0.586727839 |
| chr2:75208769-75209074 | -0.461737043 | 0.29690513 | 0.586746463 |
| chr11:62804958-62805914 | 0.33635349 | 0.296915576 | 0.586746463 |
| chr12:85424727-85425199 | -0.419181638 | 0.296918735 | 0.586746463 |
| chr6:134641002-134641254 | 0.499145943 | 0.29692935 | 0.586752306 |
| chr12:86264390-86265575 | -0.336220163 | 0.296951022 | 0.586765001 |
| chr2:186498856-186499140 | 0.457744079 | 0.296953673 | 0.586765001 |
| chr11:86335129-86335626 | -0.435538429 | 0.296981406 | 0.586765001 |
| chr12:94092966-94093219 | -0.466880119 | 0.296983857 | 0.586765001 |
| chr1:174358365-174358679 | -0.454026609 | 0.296988284 | 0.586765001 |
| chr13:44128525-44128725 | -0.561781191 | 0.29699079 | 0.586765001 |
| chr14:44648994-44649194 | -0.561781191 | 0.29699079 | 0.586765001 |
| chr12:58148434-58148767 | -0.433532965 | 0.296997038 | 0.586765001 |
| chr10:97583981-97584632 | 0.356878403 | 0.297062818 | 0.586873264 |
| chr1:109090186-109091321 | -0.33473167 | 0.297067155 | 0.586873264 |
| chr3:111967684-111967990 | -0.417086612 | 0.297080725 | 0.58688494 |
| chr3:59575154-59575731 | 0.391889796 | 0.297091999 | 0.586892081 |
| chr5:2756244-2756508 | 0.399796116 | 0.297101373 | 0.586895467 |
| chr17:38606968-38607303 | 0.42233942 | 0.297171936 | 0.587019725 |
| chr7:3673613-3673813 | -0.547562564 | 0.297185747 | 0.587026393 |
| chr1:211258701-211259645 | -0.339261249 | 0.297190635 | 0.587026393 |
| chr9:33452694-33453404 | -0.371494451 | 0.297223642 | 0.587076457 |
| chr8:95223195-95223395 | 0.480121666 | 0.297264708 | 0.587110123 |
| chr3:42539892-42540138 | -0.446141368 | 0.297268468 | 0.587110123 |
| chr1:58823547-58823787 | -0.489715806 | 0.297268945 | 0.587110123 |
| chr15:38389985-38390315 | 0.425667419 | 0.2972775 | 0.587110123 |
| chr3:185733922-185734267 | -0.406941613 | 0.297278999 | 0.587110123 |
| chr20:51451114-51451360 | -0.417453968 | 0.297305043 | 0.587146425 |

| | | | |
|---|---|---|---|
| chr3:194135832-194136251 | -0.345280491 | 0.297347013 | 0.587214177 |
| chr6:143409403-143409691 | 0.453107313 | 0.297365817 | 0.587236177 |
| chr1:226101015-226101358 | -0.450104284 | 0.297391178 | 0.587257778 |
| chr10:58334721-58335991 | -0.337167777 | 0.297392084 | 0.587257778 |
| chr10:35427810-35428022 | -0.562089886 | 0.297417199 | 0.587289243 |
| chr12:89951242-89951926 | 0.355892053 | 0.297437893 | 0.587289243 |
| chr9:128192698-128193488 | 0.346250785 | 0.297438537 | 0.587289243 |
| chr1:190477768-190478034 | 0.420218751 | 0.297444442 | 0.587289243 |
| chr2:24104434-24104634 | -0.562084288 | 0.297450662 | 0.587289243 |
| chr3:32037654-32037959 | -0.442107634 | 0.297454007 | 0.587289243 |
| chr18:75208567-75208817 | -0.424409792 | 0.297479232 | 0.587323912 |
| chr3:129039513-129039812 | 0.429938211 | 0.297505793 | 0.587361217 |
| chr3:12841050-12842103 | 0.333251068 | 0.297528515 | 0.587390943 |
| chrX:146707736-146708433 | -0.377421745 | 0.297565208 | 0.587448246 |
| chr4:100581333-100581663 | -0.400218544 | 0.297579416 | 0.587461161 |
| chr1:231240519-231241553 | 0.343593074 | 0.297589613 | 0.587463009 |
| chr17:78522305-78522505 | 0.562727994 | 0.297595686 | 0.587463009 |
| chr1:50774810-50775014 | 0.558926313 | 0.297617725 | 0.587481896 |
| chr7:157707192-157707392 | -0.581001619 | 0.297623353 | 0.587481896 |
| chr15:87798812-87799159 | 0.486036632 | 0.297628256 | 0.587481896 |
| chr2:118855475-118855985 | 0.381128213 | 0.297642728 | 0.587482801 |
| chr7:5780875-5782127 | -0.334177435 | 0.297644049 | 0.587482801 |
| chr15:80455606-80455837 | -0.525438237 | 0.29768232 | 0.587536313 |
| chr2:223945033-223945580 | 0.379517133 | 0.297693039 | 0.587536313 |
| chr12:6778768-6779326 | -0.364656913 | 0.297694164 | 0.587536313 |
| chr3:98991499-98991893 | -0.447376724 | 0.297710565 | 0.587553546 |
| chr1:155308306-155309323 | 0.334888453 | 0.297731014 | 0.587577797 |
| chrX:131128622-131129334 | -0.376250296 | 0.29773819 | 0.587577797 |
| chr9:110089612-110090000 | 0.401086362 | 0.297748012 | 0.587582048 |
| chr3:17032478-17032761 | 0.451132292 | 0.297766355 | 0.587594426 |
| chr3:58049210-58049796 | -0.380925843 | 0.297769622 | 0.587594426 |
| chr2:162254612-162254863 | -0.547727193 | 0.297805523 | 0.587650136 |
| chr14:71221959-71222159 | 0.5887724 | 0.297828845 | 0.587681023 |
| chr17:60262999-60263247 | 0.370090916 | 0.297847507 | 0.587687897 |
| chr11:14357934-14359556 | 0.334542527 | 0.297852041 | 0.587687897 |
| chr5:11719870-11720197 | -0.414670956 | 0.297855339 | 0.587687897 |
| chr5:137760618-137761313 | 0.359421665 | 0.297887036 | 0.587724925 |
| chr10:126904896-126906307 | 0.343836831 | 0.297896765 | 0.587724925 |
| chr10:1089096-1089296 | -0.552303559 | 0.29791452 | 0.587724925 |
| chr1:234658227-234658447 | 0.468533369 | 0.297925752 | 0.587724925 |
| chr5:37489993-37490579 | -0.40254017 | 0.297926209 | 0.587724925 |
| chr2:131673947-131674256 | -0.38757471 | 0.297930124 | 0.587724925 |
| chr5:128905576-128905857 | 0.448946009 | 0.297939836 | 0.587724925 |
| chr8:10590164-10590501 | -0.42951136 | 0.297941331 | 0.587724925 |
| chr3:148204220-148204439 | -0.487343622 | 0.29794314 | 0.587724925 |
| chr10:117216588-117217125 | -0.373946235 | 0.297989242 | 0.587780219 |
| chr19:1490253-1491385 | 0.336507841 | 0.297990262 | 0.587780219 |
| chr7:99030201-99030401 | 0.559292363 | 0.297999941 | 0.587780219 |
| chr1:91851296-91851735 | -0.427605876 | 0.298001856 | 0.587780219 |
| chr5:10507629-10507911 | 0.471759816 | 0.298123753 | 0.587960117 |

| | | | |
|---|---|---|---|
| chr9:19102208-19103333 | -0.340564937 | 0.29812428 | 0.587960117 |
| chr18:23695322-23695539 | 0.531708777 | 0.298131175 | 0.587960117 |
| chr6:73929444-73929773 | -0.475402456 | 0.298141196 | 0.587960117 |
| chr15:100433375-100433604 | 0.466581314 | 0.298142152 | 0.587960117 |
| chr10:79984939-79985168 | -0.458457339 | 0.298142911 | 0.587960117 |
| chr5:141040780-141041475 | -0.34182977 | 0.298149784 | 0.587960117 |
| chr3:12562967-12563209 | 0.440602346 | 0.298154451 | 0.587960117 |
| chr13:32495280-32495721 | -0.42819296 | 0.298167522 | 0.587969199 |
| chr16:54286151-54287158 | 0.336787188 | 0.298174404 | 0.587969199 |
| chr15:65475246-65475446 | 0.582345401 | 0.298187661 | 0.587970589 |
| chr1:204941798-204942015 | -0.437786517 | 0.298190456 | 0.587970589 |
| chr6:117776565-117776792 | 0.491290623 | 0.298211058 | 0.58799608 |
| chr3:53551475-53551685 | -0.538505741 | 0.2982197 | 0.58799799 |
| chr2:30714055-30714325 | -0.418001575 | 0.298234205 | 0.588009117 |
| chr10:3043509-3043850 | 0.395933296 | 0.298250268 | 0.588009117 |
| chr3:187808485-187809400 | -0.344611558 | 0.298259426 | 0.588009117 |
| chr18:44731585-44732131 | -0.381504695 | 0.298264411 | 0.588009117 |
| chr14:72416751-72417204 | 0.404097588 | 0.298267986 | 0.588009117 |
| chr4:119314394-119314709 | 0.409671213 | 0.298272182 | 0.588009117 |
| chr2:149037774-149039173 | -0.336308875 | 0.298279063 | 0.588009117 |
| chr13:110108902-110109404 | 0.395662149 | 0.298295587 | 0.588022604 |
| chr1:231626471-231627640 | 0.345701585 | 0.298301253 | 0.588022604 |
| chr17:76631955-76632271 | 0.436510982 | 0.298311943 | 0.588028548 |
| chr3:177050848-177051048 | -0.59203698 | 0.29834551 | 0.588071417 |
| chr11:117222067-117222267 | 0.51032565 | 0.29834904 | 0.588071417 |
| chr1:221573702-221574036 | -0.476976527 | 0.298380608 | 0.588118148 |
| chr15:65304293-65305658 | 0.329093812 | 0.298389202 | 0.588118148 |
| chr1:94236931-94238017 | 0.351278889 | 0.298395776 | 0.588118148 |
| chr1:53953135-53953500 | 0.400884903 | 0.298419228 | 0.588149242 |
| chr3:94862762-94863030 | -0.446515429 | 0.298448935 | 0.588192662 |
| chr5:1409377-1409624 | 0.431457222 | 0.298470611 | 0.588220252 |
| chr7:137903640-137903905 | -0.42031879 | 0.298485496 | 0.588234456 |
| chr22:22164957-22165202 | -0.513490519 | 0.298496519 | 0.58824105 |
| chr1:147669441-147670977 | 0.328498659 | 0.298508916 | 0.588242589 |
| chr14:48561233-48561906 | -0.345655211 | 0.298512654 | 0.588242589 |
| chr20:11496362-11496585 | -0.566814755 | 0.298521938 | 0.588245754 |
| chr3:108221878-108223084 | -0.330822619 | 0.298585967 | 0.58835637 |
| chr1:183471035-183473147 | -0.323912993 | 0.29859343 | 0.58835637 |
| chr16:71422086-71422370 | 0.440334088 | 0.298609351 | 0.58837261 |
| chr8:95305413-95305699 | 0.406723345 | 0.298640816 | 0.588419476 |
| chr12:51026042-51026718 | 0.346177163 | 0.298650974 | 0.58842175 |
| chr21:45255600-45255871 | 0.463128784 | 0.298659413 | 0.58842175 |
| chr1:149970756-149970966 | 0.538193229 | 0.298665009 | 0.58842175 |
| chr14:44913388-44914024 | -0.348712482 | 0.298673807 | 0.588423955 |
| chr20:61966778-61967093 | 0.431406891 | 0.298697202 | 0.588454915 |
| chr10:83978977-83979222 | -0.528134363 | 0.298709937 | 0.58845746 |
| chr7:26061651-26062002 | 0.409615108 | 0.298713854 | 0.58845746 |
| chr1:248837742-248839373 | -0.324088182 | 0.298749061 | 0.588493442 |
| chr5:9313739-9313982 | -0.537913948 | 0.29875515 | 0.588493442 |
| chr4:165873280-165874070 | -0.358406806 | 0.29875516 | 0.588493442 |

| | | | |
|---|---|---|---|
| chr11:124836739-124836939 | -0.608498453 | 0.298780228 | 0.58852769 |
| chr1:66331631-66331901 | -0.442374252 | 0.298861958 | 0.588664138 |
| chr16:30805396-30806138 | 0.35815953 | 0.298864864 | 0.588664138 |
| chr8:39327444-39327644 | -0.576665044 | 0.298899177 | 0.588701209 |
| chr1:203145304-203145713 | -0.394274768 | 0.298902572 | 0.588701209 |
| chr3:196568130-196569383 | -0.338587654 | 0.298907336 | 0.588701209 |
| chr16:47461809-47462261 | 0.394357528 | 0.298914418 | 0.588701209 |
| chr2:27133894-27134111 | 0.450529929 | 0.298923086 | 0.588703148 |
| chr17:63139553-63139913 | -0.450310706 | 0.29896898 | 0.5887784 |
| chr1:86703920-86705570 | -0.331440181 | 0.298997206 | 0.588818852 |
| chr2:73736809-73737723 | -0.337065389 | 0.299018378 | 0.588845412 |
| chr2:179219047-179219476 | -0.432338423 | 0.299046256 | 0.588865278 |
| chr2:241508457-241509058 | 0.347880047 | 0.299046595 | 0.588865278 |
| chr14:40855686-40855886 | 0.562462031 | 0.299052489 | 0.588865278 |
| chr8:68490727-68491001 | 0.468899337 | 0.299063261 | 0.588865278 |
| chr4:95100993-95101213 | -0.584329313 | 0.299070213 | 0.588865278 |
| chr11:95066853-95067958 | -0.346263162 | 0.299078351 | 0.588865278 |
| chr17:74671285-74671485 | 0.606215129 | 0.299082263 | 0.588865278 |
| chr19:29665202-29665783 | 0.359234069 | 0.299092179 | 0.588869671 |
| chr14:21825215-21825496 | -0.423456603 | 0.299100194 | 0.588870318 |
| chr9:94925875-94926513 | 0.361060517 | 0.299128135 | 0.588910198 |
| chr12:120356613-120357057 | -0.374533331 | 0.299142905 | 0.588911611 |
| chr8:109362189-109362766 | -0.3619154 | 0.299144225 | 0.588911611 |
| chr11:114036108-114036467 | 0.409319643 | 0.299170091 | 0.588938654 |
| chr12:64549552-64549938 | 0.368814265 | 0.299173334 | 0.588938654 |
| chr12:61620906-61621273 | 0.491560241 | 0.299185624 | 0.588947716 |
| chr11:74381080-74381370 | 0.486086696 | 0.299193441 | 0.588947973 |
| chr16:1296854-1297220 | 0.417860872 | 0.299207455 | 0.588949866 |
| chr1:10724091-10724594 | 0.389922799 | 0.299209776 | 0.588949866 |
| chr17:28684411-28684655 | -0.501831528 | 0.299218028 | 0.588950868 |
| chr2:208983432-208983836 | -0.372660371 | 0.299225658 | 0.588950868 |
| chr17:81910874-81911785 | 0.337306894 | 0.299250051 | 0.58898375 |
| chr2:190468636-190469047 | 0.428791238 | 0.299264769 | 0.588991029 |
| chr3:1700763-1701010 | -0.532831076 | 0.29927681 | 0.588991029 |
| chr3:53375059-53375264 | -0.532831076 | 0.29927681 | 0.588991029 |
| chr2:37156170-37157450 | -0.336809798 | 0.299287209 | 0.588996367 |
| chr8:38770116-38770322 | -0.539274155 | 0.299304863 | 0.58901598 |
| chr1:208531466-208531767 | -0.489993901 | 0.299329145 | 0.589047328 |
| chr6:86856964-86857233 | -0.481882584 | 0.299336167 | 0.589047328 |
| chr5:118987820-118988890 | -0.328983595 | 0.299366838 | 0.589085115 |
| chr20:5126416-5127202 | 0.336540374 | 0.299378237 | 0.589085115 |
| chr8:83400436-83400709 | -0.4677781 | 0.299382426 | 0.589085115 |
| chr7:101437319-101437519 | -0.527325165 | 0.299392078 | 0.589085115 |
| chr13:21034576-21035006 | -0.469232063 | 0.299393811 | 0.589085115 |
| chr4:40645588-40645788 | -0.591568488 | 0.299401499 | 0.589085115 |
| chr1:44814972-44815217 | 0.43610125 | 0.299453533 | 0.589172366 |
| chr19:7534014-7534340 | 0.434071347 | 0.299475154 | 0.589199776 |
| chr7:86469742-86470241 | -0.400482372 | 0.299504139 | 0.58922527 |
| chr6:7261972-7262398 | -0.368923444 | 0.299507599 | 0.58922527 |
| chr2:45649943-45651372 | -0.334306113 | 0.299511182 | 0.58922527 |

| | | | |
|---|---|---|---|
| chr6:88165750-88166513 | 0.359764764 | 0.299530953 | 0.589241684 |
| chr6:113858086-113858349 | 0.426802374 | 0.299540153 | 0.589241684 |
| chr1:171801477-171801942 | 0.373459242 | 0.299546938 | 0.589241684 |
| chr13:113584242-113585010 | 0.350177971 | 0.299550287 | 0.589241684 |
| chr17:43546235-43546858 | 0.366449564 | 0.299568566 | 0.589262514 |
| chr17:59211001-59211216 | 0.421284482 | 0.299617315 | 0.589325649 |
| chr16:12610110-12610344 | -0.568351851 | 0.299625241 | 0.589325649 |
| chr1:46616664-46617183 | 0.370436996 | 0.29963192 | 0.589325649 |
| chr20:2524444-2524730 | -0.503780442 | 0.299634942 | 0.589325649 |
| chr1:54500106-54500306 | -0.550895055 | 0.29964213 | 0.589325649 |
| chr7:129376510-129376747 | -0.480032938 | 0.299646811 | 0.589325649 |
| chr16:85275598-85275942 | 0.380311995 | 0.299661786 | 0.589339973 |
| chrX:48897138-48897537 | 0.356535617 | 0.29970051 | 0.589393776 |
| chr8:123192642-123192882 | 0.437983882 | 0.299704527 | 0.589393776 |
| chr13:31094642-31094992 | 0.382073082 | 0.299720239 | 0.589409018 |
| chr4:52861833-52862607 | 0.34862107 | 0.299727663 | 0.589409018 |
| chr16:46789610-46790105 | 0.404379434 | 0.29973557 | 0.589409441 |
| chr4:95693711-95693926 | -0.547321596 | 0.29976074 | 0.589443809 |
| chr10:25157685-25157957 | -0.488277318 | 0.299787812 | 0.589481913 |
| chr1:236127161-236127399 | -0.504656973 | 0.299841407 | 0.589572169 |
| chr4:123096131-123096353 | 0.533570495 | 0.299862305 | 0.589590503 |
| chr1:114435148-114435449 | 0.366587079 | 0.299866121 | 0.589590503 |
| chr5:168348042-168348338 | -0.402204093 | 0.299961525 | 0.589762952 |
| chr17:66451584-66451823 | -0.468589073 | 0.29997039 | 0.589765247 |
| chr8:53692632-53693447 | 0.346356421 | 0.300019258 | 0.589836416 |
| chr2:207796645-207796865 | -0.500871546 | 0.300021984 | 0.589836416 |
| chr5:176945636-176946052 | 0.425460097 | 0.300040807 | 0.589847191 |
| chr1:39076626-39076877 | -0.533705461 | 0.300042862 | 0.589847191 |
| chr9:3181512-3181909 | -0.431494765 | 0.300061128 | 0.589867967 |
| chr2:26922793-26922993 | -0.517662849 | 0.300093673 | 0.589916811 |
| chr11:124837043-124837319 | -0.432642947 | 0.300116339 | 0.589944615 |
| chr13:98448436-98448654 | -0.546332823 | 0.300123836 | 0.589944615 |
| chr4:178224138-178224551 | -0.445991442 | 0.300130916 | 0.589944615 |
| chr1:11907768-11909875 | 0.32900144 | 0.300158855 | 0.589984397 |
| chr6:42428659-42428946 | -0.456702228 | 0.300174667 | 0.590000342 |
| chr17:19317357-19317606 | -0.491131888 | 0.300203289 | 0.590041463 |
| chr8:96644886-96645762 | 0.337805882 | 0.300234983 | 0.590088621 |
| chr19:32287858-32288130 | -0.476844517 | 0.300294261 | 0.590189987 |
| chr12:27243912-27244862 | -0.339045049 | 0.300313735 | 0.590213122 |
| chr17:67293767-67294110 | -0.383427185 | 0.30034065 | 0.590233708 |
| chr17:40100521-40101384 | -0.338194083 | 0.300342589 | 0.590233708 |
| chr1:16545815-16546140 | -0.453607548 | 0.300347319 | 0.590233708 |
| chr8:33805774-33806063 | -0.459716309 | 0.300362548 | 0.590248497 |
| chr2:173571139-173571857 | 0.365363295 | 0.300376824 | 0.590261243 |
| chr9:22455437-22455686 | -0.533476676 | 0.300384441 | 0.590261243 |
| chr3:193391818-193392164 | -0.474088671 | 0.300393798 | 0.590264492 |
| chr12:8081775-8082827 | -0.339961472 | 0.300403858 | 0.590268567 |
| chr8:103871265-103871648 | 0.40147285 | 0.300411279 | 0.590268567 |
| chr18:23689273-23689622 | 0.418835142 | 0.300419982 | 0.59027053 |
| chr4:123350283-123350484 | 0.490891382 | 0.300441563 | 0.590287243 |

| | | | |
|---|---|---|---|
| chr2:108228571-108228805 | -0.57177836 | 0.300448029 | 0.590287243 |
| chr3:37998893-37999366 | 0.35553385 | 0.300463266 | 0.590287243 |
| chr13:36490275-36490491 | 0.542622967 | 0.300463274 | 0.590287243 |
| chr16:88769654-88770252 | -0.344943338 | 0.300467007 | 0.590287243 |
| chrX:10051830-10052199 | -0.426809237 | 0.300490846 | 0.590292203 |
| chr16:63617934-63618237 | -0.428571789 | 0.300492884 | 0.590292203 |
| chr11:34468344-34468575 | -0.509194203 | 0.30049499 | 0.590292203 |
| chr4:30719463-30719746 | 0.385110702 | 0.300501219 | 0.590292203 |
| chr6:49531953-49532476 | -0.361596118 | 0.300513559 | 0.590292203 |
| chr12:131230222-131230422 | 0.502460884 | 0.300521268 | 0.590292203 |
| chr12:6229318-6229723 | -0.398333332 | 0.300528707 | 0.590292203 |
| chrX:140713529-140713847 | -0.50947803 | 0.300531164 | 0.590292203 |
| chr16:87871556-87871782 | 0.425147373 | 0.300542025 | 0.590298403 |
| chr6:89375430-89375871 | -0.369901061 | 0.300572601 | 0.590330445 |
| chr19:36837826-36838953 | 0.345979559 | 0.300573747 | 0.590330445 |
| chr5:19706320-19707363 | -0.351692902 | 0.300605809 | 0.59036328 |
| chr12:131949443-131950134 | 0.344034978 | 0.300609692 | 0.59036328 |
| chr8:95248739-95248985 | 0.455169037 | 0.30061358 | 0.59036328 |
| chr5:150523770-150524039 | 0.440214616 | 0.300653651 | 0.590426841 |
| chr15:55959808-55960035 | 0.509434032 | 0.300663775 | 0.590431589 |
| chr5:116766647-116766861 | 0.51260336 | 0.300674544 | 0.590437605 |
| chr2:176553695-176553992 | -0.415749539 | 0.300706991 | 0.590484442 |
| chr21:9324791-9325638 | 0.338570831 | 0.300719602 | 0.590484442 |
| chr13:83881887-83882259 | -0.423519154 | 0.300725239 | 0.590484442 |
| chr21:41045512-41045965 | -0.379732597 | 0.300733373 | 0.590484442 |
| chr14:23302708-23302980 | 0.377419097 | 0.300736927 | 0.590484442 |
| chr11:64844400-64844938 | 0.356766227 | 0.30074553 | 0.590486202 |
| chr13:27830033-27830279 | -0.434130792 | 0.300759946 | 0.590499375 |
| chr1:106998489-106998892 | 0.360210619 | 0.300787647 | 0.590526961 |
| chr1:19344478-19344678 | 0.531730582 | 0.300789411 | 0.590526961 |
| chr2:205776099-205776541 | -0.386877722 | 0.300803519 | 0.590539529 |
| chr11:76041259-76041575 | 0.444781635 | 0.300830955 | 0.590578259 |
| chr9:101398254-101399220 | -0.331417546 | 0.300841084 | 0.590583011 |
| chr1:184386339-184387823 | -0.331683387 | 0.300857194 | 0.590598716 |
| chr10:70348316-70348786 | -0.401808698 | 0.30086763 | 0.590598716 |
| chr17:49169474-49169678 | -0.503387478 | 0.300872255 | 0.590598716 |
| chr1:171985716-171986012 | -0.423417956 | 0.300879915 | 0.590598716 |
| chr1:160330213-160330456 | -0.516768064 | 0.30089225 | 0.590607797 |
| chr3:75287992-75288278 | 0.518432267 | 0.300905359 | 0.590618398 |
| chr2:65711053-65711397 | -0.412457511 | 0.300916355 | 0.590624851 |
| chr3:121995481-121995866 | 0.375939099 | 0.300944792 | 0.590641117 |
| chr7:914005-914536 | -0.370084999 | 0.300952577 | 0.590641117 |
| chr15:81000506-81001119 | 0.349665064 | 0.300959763 | 0.590641117 |
| chr17:77011425-77011673 | 0.522033146 | 0.300959848 | 0.590641117 |
| chr17:62755167-62755598 | -0.371398889 | 0.300963185 | 0.590641117 |
| chr15:98491273-98491473 | 0.532744354 | 0.301010218 | 0.59071829 |
| chr7:44471588-44472061 | -0.477378169 | 0.30102112 | 0.590724554 |
| chr1:235060542-235061368 | 0.337261254 | 0.301035929 | 0.590738487 |
| chr5:159986899-159987696 | -0.345417401 | 0.301052022 | 0.590754938 |
| chr3:23017771-23018145 | 0.379643379 | 0.30107894 | 0.590792628 |

| | | | |
|---|---|---|---|
| chr17:16363767-16364029 | -0.447527694 | 0.30113061 | 0.590878886 |
| chr11:120121719-120121919 | 0.523244 | 0.301176329 | 0.590953462 |
| chr20:58048239-58048441 | 0.438731831 | 0.301187893 | 0.590961018 |
| chr13:46796432-46797609 | -0.324469817 | 0.301210848 | 0.590985329 |
| chr12:84727953-84728434 | -0.42210879 | 0.301216166 | 0.590985329 |
| chr17:74417708-74418027 | -0.448349861 | 0.301224417 | 0.590985329 |
| chr6:53340522-53340751 | -0.534751745 | 0.301235319 | 0.590985329 |
| chr1:244549972-244550174 | 0.569829885 | 0.301239087 | 0.590985329 |
| chr1:25232952-25233391 | 0.393267202 | 0.301254239 | 0.590985329 |
| chr14:77013920-77014650 | -0.337998707 | 0.301254274 | 0.590985329 |
| chr14:105448636-105448931 | 0.405233812 | 0.301280046 | 0.591014854 |
| chr17:57268978-57269178 | 0.455900496 | 0.301284751 | 0.591014854 |
| chr6:15662150-15663264 | 0.338861039 | 0.301347003 | 0.591121837 |
| chr2:71378938-71379151 | -0.554392088 | 0.3013694 | 0.591150637 |
| chr7:6058595-6059732 | -0.330484043 | 0.301392449 | 0.591161702 |
| chr1:153384259-153384459 | 0.535174258 | 0.301393007 | 0.591161702 |
| chr8:99706464-99706734 | -0.445771957 | 0.301398307 | 0.591161702 |
| chr2:63888843-63889099 | -0.467207912 | 0.301405902 | 0.591161702 |
| chr17:59310993-59311477 | 0.347799235 | 0.301428856 | 0.59119159 |
| chr2:112869667-112869913 | -0.486434385 | 0.301461986 | 0.591207618 |
| chr5:134506082-134506868 | -0.346706087 | 0.301463634 | 0.591207618 |
| chr3:18982756-18982956 | 0.597308314 | 0.301466246 | 0.591207618 |
| chr6:29072149-29072487 | -0.408517815 | 0.301467892 | 0.591207618 |
| chr10:2898170-2898433 | 0.542964159 | 0.301495286 | 0.591225057 |
| chr7:98223273-98223925 | 0.37503888 | 0.301498403 | 0.591225057 |
| chr2:60961583-60961841 | -0.46578157 | 0.301503705 | 0.591225057 |
| chr3:152632711-152632941 | -0.522807516 | 0.301507649 | 0.591225057 |
| chr15:63870648-63870922 | 0.419509692 | 0.301606621 | 0.591391034 |
| chr8:122721211-122721548 | -0.425175011 | 0.301607729 | 0.591391034 |
| chr11:129895030-129896040 | 0.34197451 | 0.301638103 | 0.591435455 |
| chr7:100611868-100612694 | -0.337106526 | 0.301696454 | 0.59153473 |
| chr11:88543999-88544697 | -0.389565679 | 0.301704853 | 0.59153606 |
| chr3:122651386-122651990 | -0.384411844 | 0.301742736 | 0.591584758 |
| chr3:7171484-7171796 | 0.442898839 | 0.301752342 | 0.591584758 |
| chr16:11387059-11387336 | -0.441430358 | 0.301752853 | 0.591584758 |
| chr2:120762651-120762851 | 0.545399663 | 0.301766831 | 0.591597025 |
| chr6:53359100-53360054 | -0.333808439 | 0.301807843 | 0.591641816 |
| chr8:118415767-118415967 | -0.531919977 | 0.30180842 | 0.591641816 |
| chr6:92132537-92132767 | 0.48985904 | 0.301812843 | 0.591641816 |
| chr4:147616949-147618154 | -0.326920879 | 0.301845076 | 0.591689865 |
| chr18:9575536-9575800 | 0.445135745 | 0.301902942 | 0.591773612 |
| chr11:61567283-61567547 | 0.393228267 | 0.301904645 | 0.591773612 |
| chr7:100626515-100626887 | 0.370204873 | 0.301910969 | 0.591773612 |
| chr17:1770202-1770443 | -0.477669743 | 0.301938948 | 0.591804746 |
| chr6:149318061-149318426 | 0.394080174 | 0.3019423 | 0.591804746 |
| chr8:60718910-60719244 | -0.391453427 | 0.30195186 | 0.591808344 |
| chr17:80169736-80170258 | 0.359273438 | 0.302004846 | 0.591897053 |
| chr1:54973175-54973423 | -0.465134597 | 0.302055 | 0.591980207 |
| chrX:101260488-101260753 | 0.426452696 | 0.302080941 | 0.592015906 |
| chrX:137194227-137194495 | 0.458327683 | 0.302118753 | 0.592050696 |

| | | | |
|---|---|---|---|
| chr11:40186472-40187190 | 0.364366986 | 0.302123231 | 0.592050696 |
| chr2:239758290-239758852 | -0.399014389 | 0.302135213 | 0.592050696 |
| chr2:5121725-5122021 | -0.421338599 | 0.302137164 | 0.592050696 |
| chr5:76204784-76205141 | 0.405896322 | 0.302144038 | 0.592050696 |
| chr8:61734382-61734736 | -0.410495845 | 0.302149348 | 0.592050696 |
| chr18:26621211-26621786 | -0.364677845 | 0.302152782 | 0.592050696 |
| chr5:177456331-177456596 | 0.366283601 | 0.302230843 | 0.592167994 |
| chr9:109183190-109183481 | -0.40828766 | 0.302240092 | 0.592167994 |
| chr3:151839048-151839289 | 0.445736383 | 0.302241867 | 0.592167994 |
| chr11:73236923-73237250 | 0.396353708 | 0.302243558 | 0.592167994 |
| chr14:102795142-102795409 | -0.487289494 | 0.302270127 | 0.592168511 |
| chrX:123976770-123976970 | 0.527241326 | 0.302275343 | 0.592168511 |
| chr2:169840337-169840537 | 0.553777072 | 0.302277822 | 0.592168511 |
| chrX:155057380-155057876 | 0.368636962 | 0.302278966 | 0.592168511 |
| chr9:131238125-131238350 | 0.486701009 | 0.302282464 | 0.592168511 |
| chr18:62635467-62635667 | -0.552423762 | 0.302316604 | 0.592215396 |
| chr3:163701665-163702065 | -0.409144867 | 0.30232696 | 0.592215396 |
| chr21:42009166-42010654 | -0.324073216 | 0.302329585 | 0.592215396 |
| chr10:6088631-6089822 | 0.328993687 | 0.302347243 | 0.592234846 |
| chr17:57157987-57158207 | 0.469028547 | 0.302367658 | 0.592257675 |
| chr11:7651219-7651456 | -0.519394541 | 0.302383655 | 0.592257675 |
| chr17:39737787-39738800 | 0.347003202 | 0.302392675 | 0.592257675 |
| chr14:81512982-81513363 | -0.45754671 | 0.302404377 | 0.592257675 |
| chr4:92305385-92305818 | 0.392901865 | 0.302404876 | 0.592257675 |
| chr14:52724277-52724556 | 0.436682146 | 0.302405997 | 0.592257675 |
| chrX:130051081-130051414 | -0.502074169 | 0.302413005 | 0.592257675 |
| chr6:104525622-104526055 | -0.39653895 | 0.30243003 | 0.592266465 |
| chr3:17166455-17166655 | 0.511783762 | 0.302441908 | 0.592266465 |
| chr15:68817527-68817944 | 0.374796544 | 0.302443511 | 0.592266465 |
| chr2:739909-741210 | -0.331659574 | 0.302448412 | 0.592266465 |
| chr11:47469419-47469745 | 0.451994364 | 0.302457913 | 0.592269933 |
| chr17:82213366-82213834 | -0.361823112 | 0.302480153 | 0.592297194 |
| chr14:65961108-65961384 | 0.416492144 | 0.302487295 | 0.592297194 |
| chr16:84367621-84367869 | -0.408049812 | 0.302502214 | 0.592305549 |
| chr1:156659974-156660536 | 0.339909235 | 0.302507022 | 0.592305549 |
| chr8:86785245-86785647 | 0.376355371 | 0.302527492 | 0.592317645 |
| chr4:81919272-81919570 | -0.422621893 | 0.302528661 | 0.592317645 |
| chr19:42325177-42326227 | 0.331612509 | 0.302550803 | 0.592330845 |
| chr3:163874940-163875140 | 0.534066776 | 0.302550864 | 0.592330845 |
| chr2:231640637-231641040 | 0.403233368 | 0.302599069 | 0.592405328 |
| chr2:117623158-117623945 | -0.358324138 | 0.302610104 | 0.592405328 |
| chr8:106765242-106765572 | -0.366707827 | 0.302612103 | 0.592405328 |
| chr3:125990296-125990877 | 0.378953025 | 0.302631095 | 0.592412995 |
| chr3:19102634-19103458 | -0.347018985 | 0.302632157 | 0.592412995 |
| chr15:77908284-77908560 | 0.406632187 | 0.302639214 | 0.592412995 |
| chr11:37449214-37449449 | -0.506779483 | 0.302692963 | 0.592474094 |
| chr7:113710315-113710642 | -0.409260089 | 0.302695043 | 0.592474094 |
| chr4:140372841-140373098 | -0.384610198 | 0.302708347 | 0.592474094 |
| chr7:45217409-45217714 | -0.418691909 | 0.302708639 | 0.592474094 |
| chr17:83174141-83174997 | 0.336419645 | 0.302709089 | 0.592474094 |

| | | | |
|---|---|---|---|
| chr3:132536118-132536374 | 0.428447097 | 0.302723534 | 0.592487232 |
| chr2:61911576-61911808 | -0.498096384 | 0.302744509 | 0.592493168 |
| chrX:130330675-130330980 | 0.392462185 | 0.302744634 | 0.592493168 |
| chr15:68820846-68821193 | 0.376580717 | 0.302749765 | 0.592493168 |
| chr10:72331202-72332531 | -0.328310288 | 0.302777113 | 0.592531554 |
| chr2:159035924-159036443 | 0.37550321 | 0.302792193 | 0.592543815 |
| chr2:162343583-162344690 | -0.325547516 | 0.302798845 | 0.592543815 |
| chr15:44578482-44578776 | -0.435847739 | 0.302814127 | 0.592548561 |
| chr2:53422969-53423410 | -0.422340955 | 0.302816737 | 0.592548561 |
| chr7:6602222-6602792 | -0.344352039 | 0.302852503 | 0.592578748 |
| chr8:109642599-109642823 | -0.385026436 | 0.302870272 | 0.592578748 |
| chr18:48615613-48615850 | -0.452297344 | 0.302870358 | 0.592578748 |
| chr8:85480193-85480393 | -0.549453243 | 0.302871695 | 0.592578748 |
| chr7:112749870-112750129 | -0.496271546 | 0.302880239 | 0.592578748 |
| chr4:16225955-16226330 | 0.42219579 | 0.302884304 | 0.592578748 |
| chr2:147808474-147808899 | -0.409052942 | 0.302893463 | 0.592578748 |
| chr2:10454597-10454797 | 0.492973838 | 0.302894034 | 0.592578748 |
| chr1:185652292-185652610 | -0.462150457 | 0.302921847 | 0.592618031 |
| chr1:164166347-164166789 | -0.422311918 | 0.302942384 | 0.592643076 |
| chr9:34612877-34613121 | -0.44501171 | 0.302961905 | 0.59265292 |
| chr17:45149251-45149469 | 0.414474801 | 0.302962886 | 0.59265292 |
| chr2:223613005-223613472 | -0.397618514 | 0.302987481 | 0.592679187 |
| chr6:44223382-44224464 | 0.338641709 | 0.302995867 | 0.592679187 |
| chr8:107526345-107526595 | 0.445324599 | 0.303012862 | 0.592679187 |
| chr1:52002364-52002669 | 0.42570033 | 0.303016922 | 0.592679187 |
| chr6:138211133-138211370 | -0.532918786 | 0.303017933 | 0.592679187 |
| chr14:37594186-37596612 | 0.318960762 | 0.303022724 | 0.592679187 |
| chr14:64636713-64637018 | 0.392149626 | 0.303069539 | 0.592748095 |
| chr11:7251785-7252208 | 0.442706743 | 0.303087259 | 0.592748095 |
| chrX:25013003-25013708 | 0.369917475 | 0.303087387 | 0.592748095 |
| chr16:86975369-86975696 | -0.428848506 | 0.3030949 | 0.592748095 |
| chr10:101124402-101124622 | -0.537630056 | 0.303096635 | 0.592748095 |
| chr19:45443366-45444170 | -0.344666579 | 0.303118684 | 0.592776086 |
| chr1:64744249-64746141 | -0.330866183 | 0.30315261 | 0.5928273 |
| chr2:73234068-73234700 | -0.338860907 | 0.303172859 | 0.592851768 |
| chr12:3752919-3753177 | -0.472468267 | 0.303181083 | 0.59285272 |
| chr5:21495327-21495590 | 0.427100677 | 0.303224883 | 0.592923235 |
| chr14:30980970-30981270 | 0.477238927 | 0.303238099 | 0.592926236 |
| chr12:6235276-6235917 | 0.343050439 | 0.303241894 | 0.592926236 |
| chr2:88458701-88458901 | 0.548264474 | 0.30325849 | 0.592943134 |
| chr8:133449517-133449815 | 0.385243137 | 0.303276226 | 0.592943134 |
| chr4:23361560-23361805 | -0.497409945 | 0.303283042 | 0.592943134 |
| chr12:53199653-53200000 | 0.443973642 | 0.303283654 | 0.592943134 |
| chr18:23330492-23331850 | -0.336029738 | 0.303289229 | 0.592943134 |
| chr11:64835558-64835859 | 0.48735208 | 0.303312383 | 0.592970598 |
| chr1:96482263-96482601 | -0.508273502 | 0.303324405 | 0.592970598 |
| chr10:69465158-69465373 | -0.542757824 | 0.303331674 | 0.592970598 |
| chr15:43440881-43441138 | -0.518365715 | 0.303351079 | 0.592970598 |
| chr7:100960807-100961163 | -0.442264037 | 0.303351293 | 0.592970598 |
| chr9:128771685-128772607 | 0.339966909 | 0.303354341 | 0.592970598 |

| | | | |
|---|---|---|---|
| chr8:122816957-122817303 | 0.396932811 | 0.303373297 | 0.592970598 |
| chr8:47924945-47925190 | 0.516208806 | 0.303378687 | 0.592970598 |
| chr3:154954807-154955097 | 0.440123537 | 0.303381943 | 0.592970598 |
| chr2:88115676-88115907 | -0.540463634 | 0.303387038 | 0.592970598 |
| chr4:82677211-82677688 | 0.375965005 | 0.303390471 | 0.592970598 |
| chr17:72407432-72407836 | -0.402981169 | 0.303400929 | 0.592970598 |
| chr20:57748413-57749361 | 0.333073736 | 0.303403882 | 0.592970598 |
| chr16:89816116-89816349 | -0.423053361 | 0.303448802 | 0.593043261 |
| chr15:45162848-45163371 | -0.364938779 | 0.303476077 | 0.593081438 |
| chr10:73625239-73626270 | -0.338188753 | 0.303489032 | 0.593091629 |
| chr17:3892760-3893504 | -0.351495039 | 0.303507002 | 0.593096932 |
| chr10:6134187-6134484 | 0.486497387 | 0.303512936 | 0.593096932 |
| chr3:62964488-62964788 | 0.373378243 | 0.303514967 | 0.593096932 |
| chr21:41817304-41817539 | -0.429685417 | 0.303540823 | 0.593111475 |
| chr5:74567374-74567780 | -0.451580511 | 0.303551158 | 0.593111475 |
| chr11:17599008-17599249 | -0.44530114 | 0.303553372 | 0.593111475 |
| chr4:83504360-83504656 | -0.44530114 | 0.303553372 | 0.593111475 |
| chr12:114869556-114869960 | 0.469758082 | 0.303571329 | 0.593131435 |
| chr2:219638171-219638430 | 0.41048851 | 0.303603121 | 0.593165396 |
| chr5:106093918-106094649 | -0.357163791 | 0.303604194 | 0.593165396 |
| chr5:155839812-155840012 | -0.554040332 | 0.303621178 | 0.593183455 |
| chr11:130315449-130315662 | -0.495478017 | 0.30363331 | 0.593184657 |
| chr7:2364871-2365125 | 0.428477157 | 0.303644862 | 0.593184657 |
| chr16:85819241-85819557 | 0.392941748 | 0.303653969 | 0.593184657 |
| chr8:122794964-122795541 | -0.338331644 | 0.303655817 | 0.593184657 |
| chr4:143773959-143774449 | -0.362730568 | 0.303665187 | 0.593184657 |
| chrX:69192901-69193157 | 0.558369547 | 0.303668244 | 0.593184657 |
| chr1:23593595-23593855 | -0.495276972 | 0.303685664 | 0.593203563 |
| chr17:50273116-50273357 | 0.448252379 | 0.303699634 | 0.59321228 |
| chr10:31942821-31943162 | 0.456931416 | 0.303712262 | 0.59321228 |
| chr19:40598629-40598897 | 0.425715788 | 0.303723595 | 0.59321228 |
| chr13:86839489-86839689 | 0.560008439 | 0.303731072 | 0.59321228 |
| chr3:113557434-113557744 | -0.392639819 | 0.303734659 | 0.59321228 |
| chr3:128051494-128052781 | 0.330432543 | 0.303736579 | 0.59321228 |
| chr14:91036488-91036707 | -0.570761318 | 0.30375457 | 0.593232295 |
| chr15:75686271-75686471 | 0.447015302 | 0.303769826 | 0.593242094 |
| chr20:51725588-51725943 | -0.365700359 | 0.303781583 | 0.593242094 |
| chr7:154832559-154832879 | -0.433485972 | 0.303782814 | 0.593242094 |
| chr13:51801775-51802096 | -0.393124028 | 0.303825005 | 0.593309364 |
| chr1:32754826-32755035 | 0.427266144 | 0.303841433 | 0.593326323 |
| chr3:127697471-127697746 | 0.390386719 | 0.303870893 | 0.593342836 |
| chr3:175212905-175213640 | -0.379456711 | 0.303871943 | 0.593342836 |
| chr14:92348102-92348567 | -0.366931821 | 0.303880261 | 0.593342836 |
| chr16:11949608-11949922 | 0.418088452 | 0.303880864 | 0.593342836 |
| chr20:35776554-35776831 | 0.431820416 | 0.303925384 | 0.593410122 |
| chr1:43634755-43634955 | -0.623200114 | 0.303932419 | 0.593410122 |
| chr2:196468597-196468797 | -0.545915025 | 0.303944372 | 0.593410122 |
| chr15:66386472-66387625 | 0.329569251 | 0.30394677 | 0.593410122 |
| chr8:120397930-120398178 | -0.522408878 | 0.303954048 | 0.593410122 |
| chr5:157743170-157744233 | -0.333137905 | 0.303993791 | 0.593466111 |

| | | | |
|---|---|---|---|
| chr16:88770663-88771631 | -0.332316469 | 0.304002889 | 0.593466111 |
| chr4:139655125-139655423 | -0.40683678 | 0.304007498 | 0.593466111 |
| chr3:152707292-152707619 | -0.440859963 | 0.30401485 | 0.593466111 |
| chr1:167095354-167095554 | -0.5568239 | 0.304021453 | 0.593466111 |
| chr12:67647809-67649068 | -0.330451079 | 0.304081881 | 0.593563785 |
| chr12:91387738-91388121 | -0.430441321 | 0.30408789 | 0.593563785 |
| chr18:35289781-35290609 | -0.340086388 | 0.30409473 | 0.593563785 |
| chr1:217055175-217055473 | 0.450588764 | 0.304131526 | 0.593620485 |
| chr3:42135262-42135462 | 0.516069304 | 0.304149019 | 0.593629347 |
| chr3:73502767-73502998 | -0.47147526 | 0.304151561 | 0.593629347 |
| chr1:2611208-2611579 | 0.404662585 | 0.304164657 | 0.59363749 |
| chr18:36097080-36097346 | 0.504195203 | 0.304171229 | 0.59363749 |
| chr12:93419522-93419729 | -0.526470363 | 0.304182984 | 0.593645312 |
| chr14:54782784-54783042 | 0.44419423 | 0.304251059 | 0.593763045 |
| chr10:17387834-17388611 | -0.332229537 | 0.304287442 | 0.593818924 |
| chr9:15796489-15796886 | -0.526959887 | 0.304301209 | 0.593830665 |
| chr9:34328962-34329896 | -0.332228568 | 0.304339755 | 0.593880708 |
| chr6:4727642-4727932 | 0.527821137 | 0.304342354 | 0.593880708 |
| chr14:87945041-87945251 | 0.542637748 | 0.304410909 | 0.593985217 |
| chr11:26753909-26754263 | -0.41819305 | 0.304411416 | 0.593985217 |
| chr3:62140039-62140254 | 0.409889842 | 0.304442513 | 0.594030768 |
| chr5:74641859-74642292 | -0.38981882 | 0.304474036 | 0.594062684 |
| chr3:139288701-139289039 | 0.428050879 | 0.304474376 | 0.594062684 |
| chr10:89426223-89426550 | -0.422098073 | 0.30450815 | 0.594113451 |
| chr19:1205333-1205921 | -0.356107177 | 0.304517589 | 0.594116739 |
| chr12:85448609-85448868 | 0.447346867 | 0.304560234 | 0.594184811 |
| chr6:52664092-52664864 | 0.332989424 | 0.30459409 | 0.594235732 |
| chr2:61065497-61066691 | -0.330815447 | 0.304630693 | 0.594266261 |
| chrX:149371759-149371970 | -0.588430786 | 0.304635555 | 0.594266261 |
| chr11:130068923-130070049 | 0.328603717 | 0.304643807 | 0.594266261 |
| chrX:119391357-119391608 | 0.463279141 | 0.304647384 | 0.594266261 |
| chr3:64641753-64642221 | 0.341259241 | 0.304652433 | 0.594266261 |
| chr1:169781388-169781761 | -0.35838887 | 0.304663346 | 0.594266261 |
| chr12:102481774-102482121 | -0.424277282 | 0.304671042 | 0.594266261 |
| chr10:81839062-81839262 | 0.498328018 | 0.304671785 | 0.594266261 |
| chr11:58675628-58675851 | -0.535857703 | 0.304695776 | 0.594282799 |
| chr16:66512707-66513012 | -0.535857703 | 0.304695776 | 0.594282799 |
| chr20:47854467-47855125 | 0.330117072 | 0.304708728 | 0.594292933 |
| chr19:4717349-4717990 | -0.409329874 | 0.304731421 | 0.594322063 |
| chr12:103848231-103848480 | -0.495982709 | 0.304754538 | 0.594352021 |
| chr5:6745600-6745989 | 0.377529805 | 0.304772164 | 0.594371269 |
| chr5:39080604-39080893 | 0.423822176 | 0.304795026 | 0.594399919 |
| chr4:86922403-86922918 | -0.354262306 | 0.30480237 | 0.594399919 |
| chr5:139273378-139274465 | -0.333839457 | 0.304817174 | 0.594413659 |
| chr2:162732967-162733630 | -0.368292711 | 0.304847409 | 0.59445749 |
| chr3:58305982-58306695 | 0.341004442 | 0.304898268 | 0.594541536 |
| chr2:215839922-215840131 | 0.508473214 | 0.30492933 | 0.594567697 |
| chr17:60738218-60738498 | 0.353458464 | 0.304936683 | 0.594567697 |
| chr11:18171432-18171666 | -0.487468351 | 0.304939064 | 0.594567697 |
| chr1:186152578-186153336 | -0.346038576 | 0.304946038 | 0.594567697 |

| | | | |
|---|---|---|---|
| chr20:62838098-62838419 | -0.413126603 | 0.304965299 | 0.594567697 |
| chr5:127875552-127876103 | -0.353140115 | 0.304967926 | 0.594567697 |
| chr20:57190563-57191109 | 0.326938989 | 0.304972689 | 0.594567697 |
| chr4:73740471-73740707 | -0.48456668 | 0.304973762 | 0.594567697 |
| chr14:90726686-90726968 | 0.421389568 | 0.304981609 | 0.594567866 |
| chr1:10692414-10692693 | -0.486044037 | 0.304994227 | 0.594568511 |
| chr6:26054922-26056011 | -0.343337115 | 0.304997459 | 0.594568511 |
| chr13:32314959-32316249 | -0.327143974 | 0.305024858 | 0.594606794 |
| chr1:172325627-172325972 | 0.398554837 | 0.305039179 | 0.594619583 |
| chr2:25219717-25219957 | 0.45837872 | 0.305048106 | 0.594621857 |
| chr9:121612892-121613287 | 0.399375055 | 0.305068707 | 0.594646886 |
| chr5:89730121-89730375 | -0.495693512 | 0.30510033 | 0.594680183 |
| chr7:75994326-75995041 | -0.343420399 | 0.305107012 | 0.594680183 |
| chr15:59528859-59529248 | 0.378486855 | 0.305119577 | 0.594680183 |
| chr2:88831758-88831995 | -0.485921229 | 0.305121433 | 0.594680183 |
| chr8:694099-694671 | 0.378536581 | 0.305124595 | 0.594680183 |
| chr11:16788425-16788723 | -0.392015872 | 0.305145767 | 0.594706319 |
| chr8:119855197-119856582 | -0.327895185 | 0.305161606 | 0.594722062 |
| chr2:84682442-84682705 | -0.500802526 | 0.305192029 | 0.594766224 |
| chr12:79420182-79420553 | 0.372797495 | 0.305253578 | 0.594863657 |
| chr6:13908970-13909286 | -0.472960605 | 0.305257552 | 0.594863657 |
| chr20:64161991-64162191 | 0.446930235 | 0.305265447 | 0.594863913 |
| chr5:14648613-14648867 | 0.429004218 | 0.305277792 | 0.594872842 |
| chr3:125594914-125595996 | 0.328168044 | 0.305306221 | 0.594913109 |
| chr14:50829928-50831890 | 0.326595737 | 0.305314686 | 0.594914474 |
| chr8:134734549-134735115 | -0.364212415 | 0.305366821 | 0.59500093 |
| chr7:153244972-153245330 | -0.424138666 | 0.305378375 | 0.595008311 |
| chr11:10388096-10388474 | 0.420810252 | 0.305399239 | 0.595032673 |
| chr2:60565507-60565725 | -0.495465737 | 0.30540641 | 0.595032673 |
| chr5:96601784-96602043 | -0.43429665 | 0.305415444 | 0.595035144 |
| chr1:32215647-32215959 | 0.407566085 | 0.305465078 | 0.595116212 |
| chr8:120861315-120861879 | -0.364557418 | 0.305472588 | 0.595116212 |
| chr14:75283651-75284065 | -0.39361967 | 0.305485521 | 0.595126276 |
| chr14:102279603-102280207 | -0.376558052 | 0.305506888 | 0.595151817 |
| chr5:1593867-1595076 | 0.33144959 | 0.305514166 | 0.595151817 |
| chr14:54691617-54691877 | 0.41593176 | 0.305534748 | 0.595176294 |
| chr2:138544594-138544927 | -0.510228701 | 0.305542266 | 0.595176294 |
| chr1:154220089-154221135 | 0.327780616 | 0.305554042 | 0.595177437 |
| chr9:109627102-109627334 | 0.513502865 | 0.305563611 | 0.595177437 |
| chr8:113068312-113068666 | -0.399712993 | 0.305566308 | 0.595177437 |
| chr8:118209226-118209426 | 0.546266027 | 0.305580608 | 0.595177437 |
| chr1:177113368-177113791 | 0.420342306 | 0.305581692 | 0.595177437 |
| chr11:84390493-84390836 | -0.431087394 | 0.305666114 | 0.595326732 |
| chr4:54234883-54235157 | -0.427834852 | 0.305678199 | 0.595335137 |
| chr5:18256951-18257151 | 0.543260507 | 0.305713288 | 0.595388341 |
| chr17:81742171-81742429 | 0.486963165 | 0.305724771 | 0.595395572 |
| chr16:4548815-4549059 | 0.457877185 | 0.305739566 | 0.595409252 |
| chr1:147098423-147098704 | -0.413022053 | 0.305776027 | 0.595465122 |
| chr9:2477713-2478064 | -0.529860717 | 0.305786921 | 0.595471204 |
| chr7:36670810-36671160 | 0.397985896 | 0.30581257 | 0.595506017 |

| | | | |
|---|---|---|---|
| chr10:29812145-29812453 | 0.435711792 | 0.305839894 | 0.59554409 |
| chr16:57649010-57649367 | 0.397728927 | 0.305853902 | 0.59555623 |
| chr11:30673566-30674282 | -0.352250771 | 0.305882122 | 0.595581925 |
| chr12:27427907-27428205 | 0.436836534 | 0.305882643 | 0.595581925 |
| chr3:111674208-111675346 | -0.328588778 | 0.305911676 | 0.595623317 |
| chr5:112240067-112240281 | -0.520228808 | 0.305933651 | 0.595650969 |
| chr4:27176485-27176701 | -0.491428986 | 0.305945183 | 0.595658286 |
| chr1:206953935-206954245 | 0.398671115 | 0.305958457 | 0.595665484 |
| chr6:46015109-46015941 | 0.373764331 | 0.305969476 | 0.595665484 |
| chr5:172902182-172902578 | 0.365198085 | 0.30599599 | 0.595665484 |
| chr1:71506940-71507166 | 0.525865108 | 0.306001409 | 0.595665484 |
| chr17:82551879-82552079 | 0.47312524 | 0.306004584 | 0.595665484 |
| chr11:112961078-112961498 | -0.405043485 | 0.306006821 | 0.595665484 |
| chr4:128641697-128641914 | 0.494979225 | 0.306010743 | 0.595665484 |
| chr3:71894106-71894411 | -0.435913931 | 0.306015631 | 0.595665484 |
| chr1:160353226-160353426 | -0.524489781 | 0.306022313 | 0.595665484 |
| chr10:27865101-27865377 | -0.480873757 | 0.306030385 | 0.595665484 |
| chr17:68868069-68868269 | 0.562881128 | 0.306034396 | 0.595665484 |
| chr5:172383219-172383669 | 0.38503705 | 0.306057358 | 0.595695045 |
| chr4:55708014-55708247 | -0.51564597 | 0.306089913 | 0.595743277 |
| chr2:199910794-199912031 | -0.32438837 | 0.306100552 | 0.59574885 |
| chr6:144360532-144360732 | -0.55774464 | 0.306123176 | 0.595758981 |
| chr3:48223231-48223680 | -0.407563476 | 0.306127689 | 0.595758981 |
| chr5:116574688-116575153 | -0.343778331 | 0.306129083 | 0.595758981 |
| chrX:67544992-67545543 | 0.391565455 | 0.306137379 | 0.595759995 |
| chr16:71846461-71846783 | 0.399175314 | 0.306157277 | 0.595783586 |
| chr22:42947643-42947843 | 0.513764793 | 0.306202219 | 0.595855909 |
| chr10:28381828-28382248 | -0.361625188 | 0.306226614 | 0.595888248 |
| chr2:138893586-138894153 | -0.376496258 | 0.306257473 | 0.595933162 |
| chr12:50241306-50241541 | 0.454137285 | 0.306267672 | 0.595937872 |
| chr16:50696676-50696876 | -0.494491687 | 0.306283504 | 0.595953546 |
| chrX:23114016-23114583 | -0.38973565 | 0.306307779 | 0.595985643 |
| chr15:96345628-96345828 | -0.539415636 | 0.306325078 | 0.59600009 |
| chr13:51125605-51126121 | -0.390773365 | 0.306330761 | 0.59600009 |
| chr11:62456496-62456976 | -0.432604938 | 0.306346254 | 0.5960151 |
| chr5:32842843-32843081 | -0.476342574 | 0.306384954 | 0.596075258 |
| chr3:10154842-10155108 | 0.492764003 | 0.306395859 | 0.596081339 |
| chr12:34108609-34109014 | 0.436509315 | 0.306417575 | 0.596108451 |
| chr3:89049206-89049512 | -0.459914967 | 0.306452228 | 0.596160728 |
| chr11:13313084-13313601 | -0.358078923 | 0.306465409 | 0.596171235 |
| chr5:136132082-136132291 | -0.487782171 | 0.306480872 | 0.596186178 |
| chr19:33879927-33880354 | -0.456635473 | 0.306490022 | 0.596188842 |
| chr1:68759015-68759242 | -0.554075706 | 0.3065075 | 0.596207705 |
| chr13:32790716-32790964 | -0.440132229 | 0.306522053 | 0.596220876 |
| chr8:102647476-102648010 | 0.34341437 | 0.306529919 | 0.596221042 |
| chr8:86658809-86659030 | 0.478129902 | 0.306541455 | 0.596228344 |
| chr1:70874799-70875157 | 0.395014751 | 0.306549721 | 0.596229287 |
| chr15:70427066-70427511 | 0.37693682 | 0.306590611 | 0.596285046 |
| chr12:128387865-128388185 | -0.433889814 | 0.306593954 | 0.596285046 |
| chr3:45915786-45916312 | -0.401817468 | 0.306604404 | 0.596286615 |

| | | | |
|---|---|---|---|
| chr9:85610136-85610363 | -0.495868219 | 0.306611744 | 0.596286615 |
| chr1:173190412-173190922 | -0.38822663 | 0.306618107 | 0.596286615 |
| chr8:93740291-93741520 | 0.322110825 | 0.306649413 | 0.59633153 |
| chrX:21839380-21840189 | -0.351903976 | 0.30666386 | 0.59633153 |
| chr1:217250431-217250638 | 0.493701737 | 0.306671608 | 0.59633153 |
| chr1:150255406-150256200 | 0.337663121 | 0.306672334 | 0.59633153 |
| chr3:196969892-196970092 | 0.463456657 | 0.306696699 | 0.596363773 |
| chr1:245888844-245889241 | -0.431553416 | 0.306709744 | 0.596374004 |
| chr2:33475693-33476750 | -0.339526283 | 0.306736405 | 0.596410709 |
| chr8:61262684-61262884 | 0.543774367 | 0.306771149 | 0.59646313 |
| chr11:77637443-77638194 | -0.339930231 | 0.306780409 | 0.596465998 |
| chr1:119611651-119611932 | -0.445699864 | 0.306823593 | 0.596534822 |
| chr8:123563880-123564449 | -0.340554461 | 0.306872712 | 0.596615182 |
| chr7:73822281-73823072 | -0.338325064 | 0.306886166 | 0.596626201 |
| chr17:80260035-80260509 | 0.382937718 | 0.306919705 | 0.596676264 |
| chr21:39772254-39772492 | 0.53709441 | 0.306931882 | 0.596677382 |
| chr7:3399741-3400283 | -0.37931388 | 0.306939566 | 0.596677382 |
| chr7:102480022-102480443 | -0.424138433 | 0.306943641 | 0.596677382 |
| chr11:14148722-14148989 | -0.512628827 | 0.306995677 | 0.596763395 |
| chr5:21919479-21919756 | -0.451129292 | 0.307009535 | 0.596775195 |
| chr2:55687302-55687555 | -0.476840054 | 0.30703341 | 0.596806462 |
| chr5:21663082-21663338 | -0.467457904 | 0.307051624 | 0.596826725 |
| chr19:4025563-4025800 | 0.509272451 | 0.307095152 | 0.59689619 |
| chr22:46355453-46355715 | 0.436702451 | 0.307136052 | 0.596960545 |
| chr15:63019560-63020029 | 0.391843802 | 0.307145631 | 0.596964019 |
| chr8:54477951-54478414 | -0.429790575 | 0.307186672 | 0.597024422 |
| chr11:69683994-69684232 | -0.47651358 | 0.307192292 | 0.597024422 |
| chr3:90136823-90137062 | -0.518944275 | 0.307208823 | 0.597041405 |
| chr1:153049539-153049791 | -0.496918641 | 0.30721906 | 0.597046157 |
| chr14:75307796-75308052 | 0.423399157 | 0.307251315 | 0.597093695 |
| chr2:158745645-158745976 | -0.429025538 | 0.307264426 | 0.597104031 |
| chrX:119943110-119943976 | 0.335669283 | 0.307286272 | 0.597125589 |
| chr10:75209889-75211618 | -0.324701636 | 0.307291106 | 0.597125589 |
| chr9:125374156-125374550 | 0.408230868 | 0.307302891 | 0.597133347 |
| chr2:241950886-241951163 | 0.415131363 | 0.307343373 | 0.597196865 |
| chr9:132977774-132978021 | 0.467837296 | 0.307371467 | 0.597236307 |
| chr10:132164808-132165063 | 0.40280716 | 0.30738945 | 0.597256102 |
| chr3:127790893-127791936 | -0.341144692 | 0.307405951 | 0.597273018 |
| chr5:180596753-180597039 | 0.41814006 | 0.307428307 | 0.597301309 |
| chr4:78561602-78562036 | 0.402014207 | 0.307455605 | 0.5973392 |
| chr22:45911553-45912251 | -0.346284054 | 0.307466485 | 0.597345192 |
| chr8:30991426-30991678 | -0.482085576 | 0.307474412 | 0.597345446 |
| chr17:43760302-43760844 | 0.373832838 | 0.307483971 | 0.597348872 |
| chr7:11073913-11074113 | 0.506488963 | 0.307522807 | 0.597395145 |
| chr2:61176844-61178007 | -0.325535638 | 0.307532739 | 0.597395145 |
| chr3:74257389-74257589 | -0.541895101 | 0.30754225 | 0.597395145 |
| chr9:5846764-5847029 | 0.451824911 | 0.307542369 | 0.597395145 |
| chr5:103258310-103259781 | -0.320402253 | 0.307546774 | 0.597395145 |
| chr20:41664447-41664962 | 0.367059928 | 0.307584396 | 0.597453079 |
| chr11:108093525-108093744 | -0.516144888 | 0.307597775 | 0.597454065 |

| | | | |
|---|---|---|---|
| chr8:130596509-130597173 | -0.370170211 | 0.307600499 | 0.597454065 |
| chr15:91863262-91863515 | 0.486846656 | 0.30763904 | 0.597513777 |
| chr11:64240820-64242037 | 0.323420155 | 0.30769338 | 0.597604171 |
| chr12:11556140-11556439 | -0.518225869 | 0.307726946 | 0.597654214 |
| chr19:1455432-1456150 | 0.331458277 | 0.30778336 | 0.597748628 |
| chr5:173643270-173643474 | -0.486587903 | 0.30780441 | 0.597774358 |
| chr13:20043631-20043831 | -0.574075062 | 0.307817254 | 0.59778415 |
| chr6:83859381-83860427 | -0.327150549 | 0.307825162 | 0.597784357 |
| chr1:212950107-212950977 | -0.339745087 | 0.307845022 | 0.597784948 |
| chr1:165598007-165598669 | -0.337268952 | 0.307847243 | 0.597784948 |
| chr1:238252158-238252716 | -0.377255862 | 0.307861567 | 0.597784948 |
| chr7:17298141-17299494 | 0.322213183 | 0.307863127 | 0.597784948 |
| chrX:24054164-24055282 | -0.325027992 | 0.307864476 | 0.597784948 |
| chr5:25006750-25007278 | -0.369497725 | 0.307912795 | 0.59786362 |
| chr7:149261367-149261603 | -0.428528521 | 0.307929921 | 0.597881723 |
| chr16:84730484-84730771 | 0.390495602 | 0.307965452 | 0.597935558 |
| chr17:80064293-80064493 | 0.550791636 | 0.307987794 | 0.597963785 |
| chr2:48768598-48769040 | -0.378218116 | 0.308007242 | 0.59798639 |
| chr11:128831204-128831491 | -0.484520396 | 0.308038631 | 0.598010318 |
| chr8:97582481-97582873 | -0.380682855 | 0.308042076 | 0.598010318 |
| chr5:152593470-152593732 | -0.47994167 | 0.308045165 | 0.598010318 |
| chr11:124644393-124644952 | 0.361326531 | 0.308050785 | 0.598010318 |
| chr9:35114145-35114363 | -0.410545484 | 0.308073924 | 0.59801797 |
| chr10:115417601-115417975 | 0.40744337 | 0.308080357 | 0.59801797 |
| chr9:78239497-78239816 | -0.376192957 | 0.308085149 | 0.59801797 |
| chrX:101659111-101659493 | -0.401432717 | 0.308085946 | 0.59801797 |
| chr2:203534777-203535896 | 0.328781659 | 0.308096344 | 0.598023003 |
| chr17:43024453-43025206 | -0.342992086 | 0.308126393 | 0.598048961 |
| chr6:104623118-104623377 | -0.485897457 | 0.308128143 | 0.598048961 |
| chr17:60545875-60546231 | 0.346101485 | 0.308133132 | 0.598048961 |
| chr7:2802786-2803555 | 0.35588243 | 0.308156847 | 0.598079838 |
| chr1:52032825-52033089 | 0.42527886 | 0.308175521 | 0.59810093 |
| chr8:100123782-100124142 | 0.3729796 | 0.30819588 | 0.598118116 |
| chr14:103354510-103355143 | 0.346007059 | 0.308199987 | 0.598118116 |
| chrX:3707849-3708268 | -0.418859876 | 0.308215534 | 0.59812282 |
| chr5:179494125-179494337 | 0.491911568 | 0.308218024 | 0.59812282 |
| chr4:72505246-72505626 | -0.37848005 | 0.308233056 | 0.598136842 |
| chr1:153355541-153355839 | 0.381943028 | 0.30824925 | 0.598153118 |
| chr2:218282650-218283094 | 0.373151559 | 0.308267564 | 0.598173508 |
| chrX:87140652-87140871 | -0.497748747 | 0.308327352 | 0.598231668 |
| chr3:46693145-46693850 | 0.365951142 | 0.308333708 | 0.598231668 |
| chr3:19885525-19885725 | 0.463701419 | 0.308336045 | 0.598231668 |
| chr8:79982129-79982982 | -0.344886482 | 0.308339426 | 0.598231668 |
| chrX:125998453-125998704 | -0.503317348 | 0.308342682 | 0.598231668 |
| chr8:124751535-124752176 | -0.347901546 | 0.30834618 | 0.598231668 |
| chr15:77633371-77633841 | -0.353181571 | 0.30835219 | 0.598231668 |
| chr8:111252959-111253453 | 0.362902892 | 0.308369397 | 0.598249904 |
| chr8:57420546-57420746 | 0.523473885 | 0.308385062 | 0.598265146 |
| chr5:36693166-36693486 | -0.385280121 | 0.308422457 | 0.598322543 |
| chr1:207084361-207084561 | -0.515231761 | 0.308460146 | 0.598380508 |

| | | | |
|---|---|---|---|
| chr3:150721314-150722024 | 0.34853968 | 0.308472782 | 0.598388603 |
| chr6:2738451-2738659 | 0.514000744 | 0.308479938 | 0.598388603 |
| chr17:53150142-53150394 | -0.459203966 | 0.308488949 | 0.598390932 |
| chr2:28390148-28390797 | -0.344082457 | 0.308501044 | 0.598399246 |
| chr20:54378089-54378512 | 0.342173637 | 0.308516183 | 0.598400135 |
| chr1:215674276-215674494 | -0.473877037 | 0.30852196 | 0.598400135 |
| chr15:87349255-87349506 | -0.462644218 | 0.308524932 | 0.598400135 |
| chr2:160324261-160324469 | -0.580631707 | 0.308534485 | 0.598403517 |
| chr10:77787713-77787918 | 0.52881285 | 0.308545514 | 0.598409759 |
| chr1:169893276-169894479 | -0.319243882 | 0.308564916 | 0.598423084 |
| chr10:43703104-43703480 | 0.438671408 | 0.308570643 | 0.598423084 |
| chr18:5023465-5023665 | -0.537072487 | 0.308584009 | 0.598423084 |
| chr2:133390122-133390395 | -0.522677527 | 0.308586494 | 0.598423084 |
| chr18:50158872-50159308 | -0.359667551 | 0.308595597 | 0.598423084 |
| chr4:136025666-136025896 | 0.47860265 | 0.308599244 | 0.598423084 |
| chr19:55598864-55600431 | 0.320617097 | 0.308615005 | 0.598438501 |
| chr17:4638864-4639247 | -0.44785759 | 0.30866627 | 0.598522762 |
| chr16:84511222-84511615 | -0.371338794 | 0.308675188 | 0.598524908 |
| chr10:28903078-28903457 | -0.449506897 | 0.308685963 | 0.598530655 |
| chr20:38221252-38221452 | 0.47417264 | 0.30869826 | 0.598539351 |
| chr8:124215051-124215278 | 0.441571032 | 0.308712192 | 0.598551219 |
| chr8:124660372-124660582 | -0.443462265 | 0.308728489 | 0.598567669 |
| chr1:58687511-58687846 | -0.461020561 | 0.308738324 | 0.598571592 |
| chr10:17348441-17349137 | -0.339987493 | 0.308751368 | 0.598581735 |
| chr3:27483207-27484951 | 0.318965239 | 0.308761229 | 0.598585707 |
| chr16:27230761-27230971 | -0.461311259 | 0.308769955 | 0.598587478 |
| chr14:91127362-91127992 | 0.334777261 | 0.308818793 | 0.598667009 |
| chr7:121563288-121563576 | -0.459956439 | 0.308850722 | 0.598706616 |
| chr9:128829492-128830226 | 0.339263203 | 0.308854851 | 0.598706616 |
| chr8:82307615-82307913 | 0.408883831 | 0.308922419 | 0.598822445 |
| chr11:60913744-60915012 | 0.332569721 | 0.308933658 | 0.598829081 |
| chr10:33292033-33292252 | -0.494923028 | 0.308956976 | 0.598857721 |
| chr21:25039929-25040213 | 0.494228198 | 0.308964065 | 0.598857721 |
| chr3:162400784-162401173 | -0.411297108 | 0.308988254 | 0.598881748 |
| chr2:207556958-207557207 | -0.510447361 | 0.308998397 | 0.598881748 |
| chr20:56441779-56442116 | -0.408397231 | 0.309006718 | 0.598881748 |
| chr18:21997697-21998027 | -0.448272067 | 0.309007725 | 0.598881748 |
| chr21:40717316-40717534 | -0.510516635 | 0.309029975 | 0.598909721 |
| chrX:131289105-131289533 | 0.395227215 | 0.309065587 | 0.598957906 |
| chr22:26796785-26797052 | -0.53986586 | 0.309070472 | 0.598957906 |
| chr1:145918479-145920007 | 0.32079537 | 0.309097317 | 0.59899478 |
| chr21:21096880-21097374 | -0.37474436 | 0.309105509 | 0.598995506 |
| chr20:12910773-12911114 | -0.445612775 | 0.30911684 | 0.599002315 |
| chr4:78176570-78177291 | -0.383929777 | 0.309131502 | 0.599015576 |
| chr2:57720810-57721179 | -0.392553451 | 0.309146297 | 0.599027717 |
| chr7:102512684-102512884 | 0.537628635 | 0.309153403 | 0.599027717 |
| chr20:34190613-34190883 | 0.418481412 | 0.309183361 | 0.599070616 |
| chr12:50400605-50401483 | 0.33301832 | 0.309195404 | 0.599073554 |
| chr7:47088835-47089088 | -0.431625478 | 0.309217235 | 0.599073554 |
| chr1:100351182-100352636 | -0.320485216 | 0.309221047 | 0.599073554 |

| | | | |
|---|---|---|---|
| chr5:125412837-125413081 | -0.421564097 | 0.30922146 | 0.599073554 |
| chr5:62123992-62124329 | -0.464339287 | 0.30922397 | 0.599073554 |
| chr19:53334144-53334344 | 0.522330822 | 0.309262155 | 0.599116605 |
| chr14:61355745-61356346 | 0.338665781 | 0.309275291 | 0.599116605 |
| chr7:836532-836769 | 0.426566571 | 0.309275466 | 0.599116605 |
| chr11:38460903-38461315 | -0.3685221 | 0.309284792 | 0.599116605 |
| chr1:225480939-225481294 | 0.383627631 | 0.309285288 | 0.599116605 |
| chr17:60627203-60627425 | 0.366352545 | 0.309294477 | 0.599119259 |
| chr1:112614535-112614745 | 0.477728373 | 0.309302841 | 0.599120314 |
| chr19:35612431-35613228 | 0.342860072 | 0.309354868 | 0.59920453 |
| chr18:32549284-32549540 | -0.449153976 | 0.309363406 | 0.59920453 |
| chr20:49384562-49384768 | 0.421013474 | 0.309369779 | 0.59920453 |
| chr22:46036761-46036961 | 0.441650954 | 0.309410757 | 0.599268751 |
| chr6:118565361-118565645 | -0.431210893 | 0.309442331 | 0.599314754 |
| chr4:108649763-108651003 | -0.32505403 | 0.30949526 | 0.599402114 |
| chr5:108436672-108437024 | -0.461758139 | 0.309525522 | 0.59944557 |
| chr13:81487243-81487496 | -0.466296705 | 0.309581592 | 0.599539005 |
| chr1:146036023-146036284 | 0.349459081 | 0.309598879 | 0.599546992 |
| chr7:112160113-112160350 | -0.486157539 | 0.309607296 | 0.599546992 |
| chr6:42142377-42143021 | 0.351894268 | 0.30960919 | 0.599546992 |
| chr1:156766478-156767000 | 0.351916444 | 0.309638727 | 0.599574614 |
| chr5:86463415-86463808 | -0.426183375 | 0.309639813 | 0.599574614 |
| chr1:167082783-167083130 | -0.408833686 | 0.30964693 | 0.599574614 |
| chr9:109331373-109331843 | -0.351003706 | 0.30966807 | 0.599600396 |
| chr8:75275066-75275591 | -0.36644326 | 0.309735038 | 0.599706945 |
| chr14:82088987-82089245 | -0.454814471 | 0.309745347 | 0.599706945 |
| chr1:231252431-231252631 | -0.582097157 | 0.309746579 | 0.599706945 |
| chrX:120675708-120675973 | -0.501835579 | 0.309779423 | 0.599750009 |
| chr9:73684269-73684741 | -0.446891957 | 0.309785908 | 0.599750009 |
| chr7:539237-540328 | -0.323154911 | 0.309792303 | 0.599750009 |
| chr3:64073447-64073691 | 0.35444945 | 0.309824988 | 0.599790873 |
| chr17:62262959-62263209 | 0.46488222 | 0.309829067 | 0.599790873 |
| chr6:157281593-157281914 | -0.442154417 | 0.309839559 | 0.599795435 |
| chr17:72146101-72146341 | 0.421134799 | 0.309849586 | 0.599795435 |
| chr3:172637771-172638346 | -0.358542343 | 0.309854907 | 0.599795435 |
| chr20:32277409-32277782 | -0.397319818 | 0.309867487 | 0.599804632 |
| chr16:4116417-4116746 | 0.406552452 | 0.309891042 | 0.599805034 |
| chr3:184129014-184129270 | 0.448061283 | 0.309898947 | 0.599805034 |
| chr1:99808710-99809321 | -0.380027608 | 0.309902532 | 0.599805034 |
| chr7:127683397-127683616 | -0.541571818 | 0.309903436 | 0.599805034 |
| chr19:50724443-50725480 | 0.325883643 | 0.309906835 | 0.599805034 |
| chr17:57946723-57947024 | -0.400662023 | 0.30993162 | 0.599837852 |
| chr12:43882627-43882875 | -0.484097723 | 0.309970999 | 0.599898913 |
| chr8:8669803-8670044 | -0.513705347 | 0.309985054 | 0.599910962 |
| chr5:82641341-82641718 | -0.38077589 | 0.309993741 | 0.599912621 |
| chr19:53977571-53977797 | 0.409889334 | 0.310052627 | 0.600011425 |
| chr17:81254970-81255223 | 0.412936443 | 0.31010594 | 0.600092265 |
| chr12:55681091-55682008 | -0.333307857 | 0.310110064 | 0.600092265 |
| chr8:103020455-103021678 | 0.320993427 | 0.310146092 | 0.600141951 |
| chr12:47963307-47963946 | 0.349459659 | 0.310156793 | 0.600141951 |

| | | | |
|---|---|---|---|
| chr19:35154553-35154829 | 0.506059149 | 0.310161389 | 0.600141951 |
| chr4:51895839-51896103 | -0.419441959 | 0.31016707 | 0.600141951 |
| chr8:119086410-119086798 | -0.363676079 | 0.310188915 | 0.600169062 |
| chr16:30557882-30558733 | 0.333714141 | 0.310213062 | 0.600187405 |
| chr1:120558440-120559023 | 0.376698278 | 0.310214061 | 0.600187405 |
| chr7:32093423-32093785 | -0.402069851 | 0.310235372 | 0.600213481 |
| chr5:69527615-69527859 | -0.444479107 | 0.310255205 | 0.600233205 |
| chr14:103546779-103547092 | 0.372368005 | 0.310261234 | 0.600233205 |
| chr9:115013847-115014330 | 0.386872792 | 0.310273413 | 0.600237105 |
| chr15:69319893-69320265 | 0.427443286 | 0.310278918 | 0.600237105 |
| chr10:14991063-14991854 | -0.369884055 | 0.310294057 | 0.600251238 |
| chr9:110207960-110208295 | -0.423841325 | 0.310359725 | 0.60034072 |
| chr9:113463486-113463957 | 0.356581888 | 0.310370742 | 0.60034072 |
| chr3:3604174-3604554 | 0.435040202 | 0.310376891 | 0.60034072 |
| chr3:189146872-189147235 | -0.438199361 | 0.310386577 | 0.60034072 |
| chr10:21751511-21751841 | -0.474779565 | 0.310399692 | 0.60034072 |
| chr15:89206856-89207328 | 0.357191386 | 0.310408385 | 0.60034072 |
| chr6:26473785-26474407 | -0.345844823 | 0.310412244 | 0.60034072 |
| chr17:59492089-59492352 | 0.396741718 | 0.310417688 | 0.60034072 |
| chr17:57166739-57167192 | 0.379806318 | 0.310417935 | 0.60034072 |
| chr2:64865145-64865908 | -0.341809977 | 0.310418665 | 0.60034072 |
| chr11:1382872-1383505 | 0.367182764 | 0.310449581 | 0.600377222 |
| chr13:26037306-26037539 | -0.456097255 | 0.310469962 | 0.600377222 |
| chr14:73612419-73612773 | 0.428264036 | 0.310473451 | 0.600377222 |
| chr18:63016349-63016553 | -0.565856759 | 0.310483397 | 0.600377222 |
| chr2:135875511-135876761 | -0.321840634 | 0.310495585 | 0.600377222 |
| chr11:78798555-78798883 | -0.394140875 | 0.310501255 | 0.600377222 |
| chr9:88106488-88106726 | -0.476046474 | 0.310511871 | 0.600377222 |
| chr20:18276788-18277060 | -0.408470691 | 0.31051481 | 0.600377222 |
| chr9:27165673-27165992 | -0.45206977 | 0.310516516 | 0.600377222 |
| chr20:55752746-55752946 | 0.503309051 | 0.310517127 | 0.600377222 |
| chrX:41253438-41254212 | -0.383350546 | 0.310523731 | 0.600377222 |
| chr7:1096439-1096724 | -0.416387246 | 0.310539808 | 0.600393157 |
| chr12:38906449-38907035 | -0.36266253 | 0.310563863 | 0.600424512 |
| chr19:41957642-41958133 | 0.36387289 | 0.310610421 | 0.600499375 |
| chr16:88681038-88681383 | 0.375119523 | 0.310618752 | 0.600500328 |
| chr5:10551426-10551758 | 0.38512001 | 0.310645205 | 0.600536316 |
| chr4:151276411-151276611 | 0.547931913 | 0.310662969 | 0.600555506 |
| chr9:5186510-5186726 | -0.550016654 | 0.310704949 | 0.600618966 |
| chr10:70404419-70404619 | -0.458185955 | 0.310711474 | 0.600618966 |
| chr8:118415018-118415218 | 0.492252096 | 0.31073357 | 0.600646524 |
| chr2:45273580-45273788 | 0.483521147 | 0.310751843 | 0.600652766 |
| chr1:106717763-106718067 | 0.358528331 | 0.310752477 | 0.600652766 |
| chrX:110160559-110160797 | 0.443345978 | 0.310789474 | 0.600682415 |
| chr15:67453725-67453928 | 0.491963686 | 0.310794465 | 0.600682415 |
| chr1:187928424-187928637 | -0.502457542 | 0.310800826 | 0.600682415 |
| chr10:81108517-81109185 | 0.374791543 | 0.310810262 | 0.600682415 |
| chr9:32988978-32989211 | -0.524360492 | 0.310819973 | 0.600682415 |
| chr6:158781333-158781816 | 0.388119125 | 0.310824243 | 0.600682415 |
| chr4:7844346-7844581 | 0.447202306 | 0.310832079 | 0.600682415 |

| | | | |
|---|---|---|---|
| chr9:111891940-111892391 | -0.392633563 | 0.310833888 | 0.600682415 |
| chr10:80130056-80130574 | -0.38148542 | 0.310840034 | 0.600682415 |
| chr2:173038316-173038653 | 0.409128731 | 0.310863486 | 0.600682415 |
| chr8:133455476-133456036 | 0.354151769 | 0.310866651 | 0.600682415 |
| chr2:20661513-20661977 | -0.350140191 | 0.310869192 | 0.600682415 |
| chr19:35247592-35249145 | 0.322620178 | 0.310873346 | 0.600682415 |
| chr5:143418440-143418681 | -0.496987284 | 0.31087757 | 0.600682415 |
| chr1:32200518-32200811 | -0.418757026 | 0.310926315 | 0.600759525 |
| chr2:128386114-128386337 | -0.514875514 | 0.310952893 | 0.600759525 |
| chr13:42271282-42272751 | -0.317957951 | 0.310953762 | 0.600759525 |
| chr1:16145233-16145464 | -0.519746997 | 0.310955123 | 0.600759525 |
| chr4:137614597-137614797 | 0.529398478 | 0.310958461 | 0.600759525 |
| chr7:77121977-77122835 | -0.32626321 | 0.310964521 | 0.600759525 |
| chr11:34251090-34251539 | -0.405640739 | 0.310988821 | 0.600791323 |
| chr2:66434832-66435275 | -0.343369061 | 0.311006884 | 0.600798829 |
| chr21:43775792-43776904 | 0.330550428 | 0.311008389 | 0.600798829 |
| chr16:3400645-3401685 | 0.329981255 | 0.311028183 | 0.60080206 |
| chr2:59939296-59939626 | -0.510215957 | 0.311032399 | 0.60080206 |
| chr8:115992832-115993064 | -0.456946194 | 0.311033585 | 0.60080206 |
| chr8:134631621-134631892 | -0.473051964 | 0.311045865 | 0.600810636 |
| chr21:18759757-18760039 | 0.448069693 | 0.311082544 | 0.600856153 |
| chr16:81746184-81746561 | -0.387606176 | 0.311085114 | 0.600856153 |
| chr17:7887344-7887909 | -0.347529399 | 0.311119061 | 0.600906573 |
| chr5:145442355-145443033 | -0.362551671 | 0.3111873 | 0.601013063 |
| chr7:4815727-4815955 | -0.481540501 | 0.311189883 | 0.601013063 |
| chr3:166071282-166071686 | 0.4083944 | 0.311220812 | 0.601052637 |
| chr4:110130635-110130900 | 0.439162016 | 0.311226063 | 0.601052637 |
| chr20:57272915-57273424 | 0.321432542 | 0.311268909 | 0.601120232 |
| chr12:25250664-25251602 | -0.32853491 | 0.311294422 | 0.601154351 |
| chr3:50227316-50228154 | -0.339059278 | 0.311333031 | 0.601213759 |
| chr5:9575544-9575772 | -0.549656561 | 0.311353166 | 0.601237488 |
| chr17:74566231-74566431 | 0.467718281 | 0.311368118 | 0.601251208 |
| chr14:56579348-56580282 | 0.328812988 | 0.311401743 | 0.60128352 |
| chr18:28176286-28176584 | -0.458745664 | 0.311407568 | 0.60128352 |
| chr10:9964949-9965207 | -0.50170428 | 0.311428158 | 0.60128352 |
| chr1:175824280-175824480 | -0.557997581 | 0.311435394 | 0.60128352 |
| chr11:20364410-20364749 | 0.39462439 | 0.311442884 | 0.60128352 |
| chr15:101201506-101201722 | -0.486215912 | 0.31144311 | 0.60128352 |
| chr12:55926641-55928269 | 0.321358974 | 0.311444997 | 0.60128352 |
| chr17:59324285-59324507 | 0.423548979 | 0.311460114 | 0.60128352 |
| chr1:246735721-246735995 | 0.424326944 | 0.311462848 | 0.60128352 |
| chr1:215978694-215979038 | -0.463487603 | 0.311463325 | 0.60128352 |
| chr16:75130440-75130798 | -0.36944037 | 0.311480283 | 0.601301107 |
| chr8:132395882-132396169 | -0.418248004 | 0.311528839 | 0.601379691 |
| chr14:61559636-61560150 | -0.334571742 | 0.311580641 | 0.601449667 |
| chr17:19362303-19363183 | -0.328523607 | 0.311580787 | 0.601449667 |
| chr2:6883592-6883917 | -0.424466495 | 0.311605352 | 0.601481931 |
| chr7:14313153-14313998 | 0.333552019 | 0.311613255 | 0.601482035 |
| chr16:3509636-3509853 | 0.505802672 | 0.311621242 | 0.601482299 |
| chr15:83365528-83365732 | -0.526540836 | 0.311645767 | 0.601514484 |

| | | | |
|---|---|---|---|
| chr13:105904870-105905079 | -0.525766278 | 0.31167221 | 0.601550368 |
| chr7:123901183-123901435 | 0.449180598 | 0.311754601 | 0.601694232 |
| chr15:59688498-59689677 | 0.325280574 | 0.311780675 | 0.601725708 |
| chr1:247457385-247457659 | -0.4403095 | 0.311786616 | 0.601725708 |
| chr4:37440290-37440499 | 0.453818954 | 0.311810804 | 0.601753569 |
| chr21:43810816-43811295 | 0.358344618 | 0.311816759 | 0.601753569 |
| chr3:145915668-145916109 | -0.40682695 | 0.311934637 | 0.601935946 |
| chr2:217830535-217830735 | 0.526822026 | 0.311934864 | 0.601935946 |
| chr13:114035497-114036233 | 0.345328292 | 0.311939347 | 0.601935946 |
| chr2:203225488-203225715 | -0.501980305 | 0.311942687 | 0.601935946 |
| chr9:114502289-114502545 | 0.407634185 | 0.31203396 | 0.602096908 |
| chr4:129096039-129096457 | -0.364615837 | 0.312057179 | 0.602126547 |
| chr12:110704745-110705018 | -0.495992385 | 0.3120849 | 0.602164871 |
| chr12:116819254-116819815 | 0.347187668 | 0.31212809 | 0.602210617 |
| chr12:56634966-56635265 | 0.362568848 | 0.312130636 | 0.602210617 |
| chr4:128191359-128191666 | 0.420942798 | 0.312132187 | 0.602210617 |
| chr1:228735174-228735743 | 0.356567929 | 0.312147413 | 0.602224828 |
| chr1:152361306-152361506 | 0.519266851 | 0.31222855 | 0.602366199 |
| chr19:9834690-9835383 | 0.339603114 | 0.312279215 | 0.602448776 |
| chr10:49481916-49482185 | 0.46630414 | 0.31231407 | 0.602500847 |
| chr17:1115079-1115832 | 0.358008735 | 0.312324978 | 0.602506722 |
| chr8:123670413-123670668 | 0.428859338 | 0.312348894 | 0.602530024 |
| chr4:127997566-127998024 | -0.407961881 | 0.312369578 | 0.602530024 |
| chr8:22672991-22673300 | -0.395201975 | 0.312371116 | 0.602530024 |
| chr6:15953211-15953485 | -0.525187714 | 0.312372762 | 0.602530024 |
| chr1:181622130-181622561 | -0.434863631 | 0.312376376 | 0.602530024 |
| chr6:157221017-157221722 | -0.353077394 | 0.312387777 | 0.602532095 |
| chr21:42840634-42841054 | 0.392374394 | 0.312393177 | 0.602532095 |
| chr10:118641705-118642066 | -0.454543721 | 0.312406468 | 0.602542563 |
| chr13:24693961-24694430 | 0.369706406 | 0.312458767 | 0.602621708 |
| chr6:153701910-153702332 | 0.410185527 | 0.312465213 | 0.602621708 |
| chr1:39192157-39192402 | -0.461285316 | 0.312471098 | 0.602621708 |
| chr6:123965937-123966241 | -0.41818195 | 0.312495403 | 0.602653414 |
| chr14:98851593-98851908 | -0.502771016 | 0.312582654 | 0.602806506 |
| chr14:73662447-73662834 | 0.383524772 | 0.312613923 | 0.602851635 |
| chr3:67605581-67605900 | -0.489779612 | 0.312667166 | 0.602939136 |
| chr3:107753487-107753870 | -0.378810033 | 0.312699851 | 0.602986989 |
| chr2:96208141-96209222 | 0.325440771 | 0.312714928 | 0.603000886 |
| chr6:5379672-5379872 | 0.564774411 | 0.312735982 | 0.603026308 |
| chr2:74836544-74836753 | -0.428409017 | 0.312764723 | 0.603066551 |
| chr20:57200930-57201426 | 0.32238242 | 0.31277518 | 0.603071538 |
| chr5:20024218-20024418 | 0.509510944 | 0.31282883 | 0.603141488 |
| chr7:126589147-126589364 | 0.485110202 | 0.312831135 | 0.603141488 |
| chr3:165643088-165643411 | -0.406037418 | 0.312835074 | 0.603141488 |
| chr1:31230505-31230725 | -0.519723531 | 0.312858844 | 0.60317214 |
| chr11:62573616-62574647 | -0.325201696 | 0.312912997 | 0.603261364 |
| chr3:10950907-10951290 | 0.382036787 | 0.312936126 | 0.603290774 |
| chr6:33599397-33600009 | 0.351674314 | 0.313006683 | 0.603411615 |
| chr8:115467726-115467950 | -0.550098605 | 0.31302423 | 0.60343026 |
| chr4:174604209-174604535 | -0.466679129 | 0.313032316 | 0.603430666 |

| | | | |
|---|---|---|---|
| chr7:6556131-6556609 | -0.351047871 | 0.313052362 | 0.603454126 |
| chr6:20217549-20217818 | -0.463309552 | 0.313074041 | 0.603480734 |
| chr5:76175121-76175357 | -0.487540071 | 0.313093689 | 0.603503424 |
| chr1:156105999-156106685 | -0.332384211 | 0.313113323 | 0.603514637 |
| chr22:19806496-19807046 | 0.351860181 | 0.313117228 | 0.603514637 |
| chr3:107981426-107982054 | 0.350704195 | 0.313123135 | 0.603514637 |
| chrX:49073893-49074332 | 0.354476801 | 0.313145218 | 0.603532192 |
| chr5:64505858-64506699 | -0.349734851 | 0.313151046 | 0.603532192 |
| chr1:97920440-97921627 | -0.322492813 | 0.313155874 | 0.603532192 |
| chr11:61897725-61897941 | 0.453929314 | 0.313167623 | 0.603539656 |
| chr17:4948448-4949158 | -0.338861798 | 0.313196094 | 0.603579343 |
| chr1:202018145-202019431 | -0.316794035 | 0.313207888 | 0.603586892 |
| chr11:73227724-73228259 | -0.342874017 | 0.313243394 | 0.603640134 |
| chr18:79934389-79934917 | -0.410097673 | 0.313253711 | 0.603644834 |
| chr3:139187784-139188013 | 0.487464955 | 0.313285031 | 0.603690004 |
| chr5:138766455-138766690 | 0.527512662 | 0.313298376 | 0.603700537 |
| chr20:54139363-54139620 | 0.344024616 | 0.313315645 | 0.60371863 |
| chr2:43105900-43106145 | -0.451304315 | 0.313338024 | 0.60374657 |
| chr4:154928644-154928942 | -0.440594266 | 0.313362001 | 0.603777586 |
| chr1:150281350-150282973 | -0.314842066 | 0.313389154 | 0.603802126 |
| chr15:84859121-84859586 | 0.359052373 | 0.31339405 | 0.603802126 |
| chr17:50560436-50560795 | 0.356227638 | 0.313400678 | 0.603802126 |
| chr5:124655950-124656284 | -0.392191717 | 0.313406258 | 0.603802126 |
| chr2:161872098-161872476 | -0.419540244 | 0.313418473 | 0.603810477 |
| chr2:54503255-54503523 | -0.450066752 | 0.313505418 | 0.603962793 |
| chr2:28621158-28621365 | -0.476169429 | 0.313532027 | 0.603998868 |
| chr17:50308209-50308409 | -0.516505338 | 0.313568503 | 0.60405395 |
| chr1:29876937-29877225 | 0.459412073 | 0.313578588 | 0.60405819 |
| chr20:38925982-38926634 | 0.333871098 | 0.313614582 | 0.604085018 |
| chr5:157148923-157149200 | -0.433654832 | 0.313617206 | 0.604085018 |
| chrX:136767630-136768058 | -0.389308943 | 0.31361922 | 0.604085018 |
| chrX:56248545-56248808 | -0.493990522 | 0.313624051 | 0.604085018 |
| chr21:36183226-36183426 | 0.508586025 | 0.313635308 | 0.604091515 |
| chr12:102543444-102543802 | -0.417650102 | 0.313664269 | 0.604122725 |
| chr20:25407406-25408019 | -0.339744449 | 0.313667281 | 0.604122725 |
| chr12:30985865-30986148 | -0.432255161 | 0.313675678 | 0.604123712 |
| chr17:1488009-1488497 | -0.373119485 | 0.313721623 | 0.604197014 |
| chr17:50546911-50547599 | 0.334349153 | 0.313746284 | 0.604229321 |
| chr1:174262748-174263294 | 0.367135595 | 0.313795535 | 0.604308983 |
| chr5:120463927-120465091 | -0.349709275 | 0.313836433 | 0.604372555 |
| chr1:195867002-195867209 | 0.535525087 | 0.313869432 | 0.604419093 |
| chr13:31350160-31350366 | 0.51775254 | 0.313876377 | 0.604419093 |
| chr4:17636800-17637172 | 0.378214646 | 0.313913685 | 0.604452056 |
| chr11:111588904-111589490 | -0.419985309 | 0.313914893 | 0.604452056 |
| chr12:64403990-64405349 | 0.317243774 | 0.31391716 | 0.604452056 |
| chr17:75827636-75827876 | 0.414859238 | 0.313931175 | 0.604453613 |
| chr1:198254618-198254820 | -0.493630627 | 0.313933747 | 0.604453613 |
| chr1:199954730-199955048 | -0.37693894 | 0.313953057 | 0.604469396 |
| chr19:49339324-49339770 | 0.348226957 | 0.313957722 | 0.604469396 |
| chr6:143059657-143060032 | 0.3927291 | 0.313976024 | 0.604484821 |

| | | | |
|---|---|---|---|
| chr6:83709268-83709488 | 0.483964408 | 0.313981511 | 0.604484821 |
| chr7:11153747-11154014 | 0.415131876 | 0.313990898 | 0.604486335 |
| chr3:11847637-11847838 | 0.456126632 | 0.314000241 | 0.604486335 |
| chr3:159260135-159260384 | 0.430461052 | 0.314010977 | 0.604486335 |
| chr1:77979911-77980216 | -0.39307585 | 0.314013855 | 0.604486335 |
| chr11:65488168-65488430 | -0.405554126 | 0.314046708 | 0.604534389 |
| chr15:74842529-74843500 | 0.324657999 | 0.314065054 | 0.604539462 |
| chr12:52083472-52083692 | 0.437593718 | 0.314065123 | 0.604539462 |
| chrX:107759737-107760041 | -0.501195989 | 0.314076696 | 0.604546553 |
| chr2:207828325-207828918 | 0.36002215 | 0.314099164 | 0.604561638 |
| chr10:93813535-93813819 | 0.4004304 | 0.314101939 | 0.604561638 |
| chr3:17639793-17640095 | -0.448945418 | 0.314126766 | 0.604561638 |
| chr18:45386394-45386633 | -0.484797549 | 0.314138827 | 0.604561638 |
| chr17:18933112-18933329 | 0.503279954 | 0.314144444 | 0.604561638 |
| chr11:61021344-61022171 | -0.356312763 | 0.314148329 | 0.604561638 |
| chr12:85156647-85156934 | -0.4697002 | 0.314152865 | 0.604561638 |
| chr2:38665605-38666506 | 0.332179371 | 0.314155253 | 0.604561638 |
| chr5:160418732-160419571 | -0.327451745 | 0.314155545 | 0.604561638 |
| chr10:27099899-27100964 | -0.322725947 | 0.314185419 | 0.604603943 |
| chr5:162593925-162594253 | -0.418479206 | 0.31420119 | 0.604619107 |
| chr6:14253672-14253948 | 0.49845571 | 0.314216416 | 0.60463322 |
| chr1:152392718-152393283 | -0.351368793 | 0.314235026 | 0.604652067 |
| chr17:42028622-42028840 | 0.445451512 | 0.314244213 | 0.604652067 |
| chr4:3791989-3792205 | 0.488773442 | 0.314249884 | 0.604652067 |
| chr10:133520053-133520584 | -0.371640861 | 0.314296502 | 0.604718673 |
| chr9:5683456-5684352 | -0.358549912 | 0.314318108 | 0.604718673 |
| chr6:158518164-158518477 | -0.429235746 | 0.314323887 | 0.604718673 |
| chr1:31938215-31938816 | 0.335046024 | 0.314324902 | 0.604718673 |
| chr4:82852349-82852556 | 0.485735779 | 0.314328112 | 0.604718673 |
| chr21:41766828-41767335 | -0.343955111 | 0.314331854 | 0.604718673 |
| chr1:243792683-243792914 | 0.519218126 | 0.314356457 | 0.604750822 |
| chr2:201363824-201364273 | -0.385950363 | 0.314369646 | 0.60476101 |
| chr2:32332395-32332777 | -0.489370775 | 0.314449975 | 0.604900354 |
| chr2:212227853-212228053 | -0.54185705 | 0.314487922 | 0.604958163 |
| chr7:139074942-139075399 | -0.379907447 | 0.314528594 | 0.605008656 |
| chr5:28214258-28214592 | 0.420928679 | 0.314529963 | 0.605008656 |
| chr8:117841869-117842069 | -0.565130249 | 0.314563349 | 0.605057685 |
| chr8:125911981-125912253 | -0.418089791 | 0.31458439 | 0.605082969 |
| chr6:75205777-75206353 | 0.34530323 | 0.314598251 | 0.605083948 |
| chr4:53592138-53592348 | 0.443612167 | 0.314606354 | 0.605083948 |
| chr8:87580871-87581277 | 0.371067266 | 0.31460859 | 0.605083948 |
| chr12:31636733-31637032 | -0.453210221 | 0.31464293 | 0.605134804 |
| chr1:204410937-204412429 | -0.316520974 | 0.314673798 | 0.605151987 |
| chr22:23248558-23248768 | 0.501260593 | 0.314673906 | 0.605151987 |
| chr17:35747569-35747785 | 0.440865424 | 0.314675558 | 0.605151987 |
| chr15:95922485-95922757 | 0.468586149 | 0.31468357 | 0.605152207 |
| chr4:70038923-70039145 | -0.513449533 | 0.314700499 | 0.605169573 |
| chr12:51952399-51952599 | 0.466791168 | 0.314739211 | 0.605228826 |
| chr2:137202377-137202889 | -0.38649831 | 0.314777145 | 0.605284694 |
| chr2:70948514-70948967 | -0.373675192 | 0.314784063 | 0.605284694 |

| | | | |
|---|---|---|---|
| chr14:22982058-22982359 | 0.39301447 | 0.314851093 | 0.605398389 |
| chr14:104205892-104206111 | 0.453845192 | 0.314869435 | 0.605414448 |
| chr8:106193761-106194024 | 0.441739708 | 0.314875247 | 0.605414448 |
| chr20:14860376-14860852 | -0.430135321 | 0.314892959 | 0.605433311 |
| chr6:133340568-133340870 | -0.519962151 | 0.314918019 | 0.605466301 |
| chr5:10284035-10284337 | 0.362715735 | 0.31492837 | 0.605467541 |
| chr14:68363385-68363748 | 0.391382783 | 0.314938083 | 0.605467541 |
| chr1:183723925-183724294 | 0.423651281 | 0.31494237 | 0.605467541 |
| chr3:164983716-164983916 | -0.598748634 | 0.314960131 | 0.605482724 |
| chr12:46125227-46125455 | 0.47408016 | 0.314986484 | 0.605482724 |
| chrX:105190260-105190585 | -0.430989041 | 0.314994409 | 0.605482724 |
| chr12:76927317-76927526 | -0.522201851 | 0.314994641 | 0.605482724 |
| chr14:42201020-42201243 | -0.522201851 | 0.314994641 | 0.605482724 |
| chr7:30135450-30136173 | 0.338983106 | 0.314997681 | 0.605482724 |
| chr20:31604995-31605580 | 0.329096179 | 0.315028275 | 0.60552634 |
| chr6:7479883-7480269 | 0.412480935 | 0.315043724 | 0.605540845 |
| chr1:89018393-89018877 | -0.404500265 | 0.315061187 | 0.605559218 |
| chr17:42744268-42745222 | 0.332909146 | 0.315094168 | 0.605607418 |
| chr1:8489918-8490361 | 0.416532811 | 0.315138923 | 0.605678244 |
| chr14:24058333-24058716 | 0.373858781 | 0.315177714 | 0.605737604 |
| chr15:74905967-74907117 | 0.32912883 | 0.31524072 | 0.605831706 |
| chr1:156130050-156130628 | 0.34323962 | 0.31524249 | 0.605831706 |
| chr22:21702818-21703133 | 0.497335991 | 0.315280195 | 0.605873821 |
| chr19:41310034-41310746 | 0.348532303 | 0.31528022 | 0.605873821 |
| chrX:9462918-9463383 | 0.340244842 | 0.315299823 | 0.605896298 |
| chr7:6306547-6306857 | 0.436143427 | 0.31531051 | 0.605901637 |
| chr1:2801719-2801979 | -0.45387401 | 0.315329829 | 0.605923566 |
| chr3:53114003-53114229 | 0.469518286 | 0.31536294 | 0.605971992 |
| chr6:158245788-158246137 | -0.419350714 | 0.315380064 | 0.6059897 |
| chr18:22799188-22799479 | 0.478008683 | 0.315395598 | 0.606004351 |
| chr3:14848015-14848510 | 0.35820828 | 0.315447708 | 0.606083771 |
| chrX:119791236-119792246 | 0.32422074 | 0.315452753 | 0.606083771 |
| chr9:131304812-131305376 | -0.366243155 | 0.315480365 | 0.606110869 |
| chr8:124490887-124491722 | 0.326352661 | 0.315482677 | 0.606110869 |
| chr1:60281347-60281693 | -0.381177568 | 0.315504858 | 0.606138285 |
| chr12:54579734-54580041 | 0.376627883 | 0.315518097 | 0.606148521 |
| chr14:91654776-91655062 | 0.47151243 | 0.315549153 | 0.606192985 |
| chr8:126820884-126821210 | -0.41788736 | 0.315572153 | 0.606195877 |
| chr3:65575385-65575724 | -0.466888596 | 0.315576401 | 0.606195877 |
| chr12:110403100-110403870 | -0.343571191 | 0.315587675 | 0.606195877 |
| chr11:93043175-93043573 | -0.40172808 | 0.315592952 | 0.606195877 |
| chr17:44307775-44308723 | 0.326902426 | 0.315593474 | 0.606195877 |
| chr1:42146924-42147208 | 0.394590297 | 0.315610927 | 0.606195877 |
| chr17:2401033-2401362 | -0.356239748 | 0.315611544 | 0.606195877 |
| chr12:48474367-48474895 | -0.387433973 | 0.315624314 | 0.606195877 |
| chr19:5977734-5978466 | 0.340616031 | 0.315625942 | 0.606195877 |
| chr5:55188739-55188995 | -0.469837289 | 0.315629774 | 0.606195877 |
| chr16:22321281-22321513 | 0.444066163 | 0.315639695 | 0.606199736 |
| chr8:26374039-26374458 | 0.396200763 | 0.315674686 | 0.606251743 |
| chr3:160400141-160400472 | 0.361171432 | 0.3156826 | 0.606251746 |

| | | | |
|---|---|---|---|
| chrX:8987684-8987910 | -0.540515813 | 0.315696556 | 0.606263353 |
| chr7:151759435-151759831 | -0.38865852 | 0.315716126 | 0.606285739 |
| chr8:55990628-55991064 | 0.386848123 | 0.315752517 | 0.606340427 |
| chr1:54499552-54499937 | -0.421553841 | 0.315799484 | 0.606391675 |
| chr6:81816458-81816750 | 0.45240912 | 0.315800371 | 0.606391675 |
| chr3:148697680-148698068 | 0.408650826 | 0.315812854 | 0.606391675 |
| chr15:88132151-88132351 | 0.526664713 | 0.315821956 | 0.606391675 |
| chr22:38718524-38718770 | 0.451486263 | 0.315835807 | 0.606391675 |
| chr2:178478991-178479343 | -0.346009204 | 0.315837933 | 0.606391675 |
| chr7:43468378-43468578 | 0.538794578 | 0.315847914 | 0.606391675 |
| chr15:44862900-44863105 | 0.513734393 | 0.315848327 | 0.606391675 |
| chr8:98817318-98817767 | 0.358191636 | 0.315856225 | 0.606391675 |
| chr16:72924971-72925278 | 0.387721398 | 0.315858345 | 0.606391675 |
| chrX:135024056-135024301 | 0.427678194 | 0.315884503 | 0.606419021 |
| chr1:109953399-109953599 | -0.62892245 | 0.315888418 | 0.606419021 |
| chrX:7927052-7927990 | -0.329780096 | 0.315898556 | 0.606423289 |
| chr9:121550504-121550805 | 0.392622076 | 0.315923168 | 0.606448708 |
| chr17:50426566-50426828 | 0.406392401 | 0.315930079 | 0.606448708 |
| chr1:73635154-73635354 | -0.554014096 | 0.315935542 | 0.606448708 |
| chr8:103980411-103980969 | 0.349572885 | 0.315965049 | 0.606478083 |
| chr20:53974840-53975110 | 0.339187723 | 0.315966675 | 0.606478083 |
| chr3:64611561-64611913 | 0.340975825 | 0.316050365 | 0.606609102 |
| chrX:154473751-154474154 | 0.370711907 | 0.316050768 | 0.606609102 |
| chr5:83927178-83927423 | -0.440041512 | 0.316078819 | 0.606647746 |
| chrX:88886870-88887396 | -0.342526334 | 0.316091212 | 0.606656335 |
| chr21:45880442-45880769 | 0.388345076 | 0.316112652 | 0.606682287 |
| chr11:809244-810045 | 0.335343511 | 0.316135052 | 0.606708983 |
| chr12:58067470-58068149 | 0.35691869 | 0.316146609 | 0.606708983 |
| chr16:88118984-88119296 | 0.504850129 | 0.316150317 | 0.606708983 |
| chr10:48444974-48445360 | -0.370038909 | 0.316182734 | 0.606755 |
| chr15:75029332-75029628 | 0.372398737 | 0.316190133 | 0.606755 |
| chr1:158177655-158178074 | 0.381724246 | 0.316202027 | 0.6067588 |
| chr8:126327290-126327593 | -0.406130237 | 0.316207951 | 0.6067588 |
| chr9:137618182-137619079 | 0.331517451 | 0.31622406 | 0.606774514 |
| chrX:93673067-93673856 | -0.352295244 | 0.3162514 | 0.606807215 |
| chr9:109556159-109556468 | 0.405594949 | 0.316256941 | 0.606807215 |
| chr13:29080931-29081355 | -0.486594372 | 0.31627637 | 0.606829298 |
| chr6:100880859-100881881 | -0.323010272 | 0.31628484 | 0.606829782 |
| chr22:35452934-35453134 | 0.505643231 | 0.316294837 | 0.606829782 |
| chr6:153493556-153493941 | -0.417268061 | 0.316300381 | 0.606829782 |
| chr6:6057596-6057796 | -0.511363188 | 0.316318589 | 0.606837172 |
| chr6:163700938-163701399 | 0.413890052 | 0.316320073 | 0.606837172 |
| chr7:29442230-29442488 | -0.471676091 | 0.316339316 | 0.606858894 |
| chr16:2236324-2236530 | -0.467557347 | 0.316355456 | 0.606860037 |
| chr2:71530783-71531089 | -0.390524783 | 0.316355752 | 0.606860037 |
| chr1:207228423-207228730 | -0.445523301 | 0.316364053 | 0.606860766 |
| chr8:30554125-30554477 | -0.429764942 | 0.31637716 | 0.606870716 |
| chr12:56116196-56116915 | 0.326881019 | 0.316420454 | 0.606938061 |
| chr1:204506169-204507278 | -0.314506647 | 0.316437631 | 0.606938061 |
| chr1:233791130-233791393 | 0.439142181 | 0.316440118 | 0.606938061 |

| | | | |
|---|---|---|---|
| chr6:110618957-110619157 | -0.570614289 | 0.316443954 | 0.606938061 |
| chr14:105021664-105021988 | 0.353184561 | 0.316502178 | 0.607011422 |
| chr1:36187786-36188589 | 0.343053024 | 0.316504809 | 0.607011422 |
| chr11:119666414-119666688 | 0.417234084 | 0.316507734 | 0.607011422 |
| chr3:129086937-129087194 | 0.45085558 | 0.316515252 | 0.607011422 |
| chr4:139295309-139296061 | -0.331706814 | 0.316522206 | 0.607011422 |
| chr4:153634778-153635081 | -0.428694935 | 0.316530984 | 0.607011422 |
| chr22:21735428-21736207 | 0.335682781 | 0.316537657 | 0.607011422 |
| chr12:5002999-5003422 | -0.402874674 | 0.316553321 | 0.607026267 |
| chr11:71605254-71605454 | 0.483390458 | 0.316566194 | 0.607035761 |
| chr12:92869865-92870118 | -0.507676034 | 0.316618718 | 0.607121285 |
| chr5:2124495-2124706 | -0.54371672 | 0.316638749 | 0.6071445 |
| chr1:62849118-62849406 | 0.429125108 | 0.31666964 | 0.607178555 |
| chr15:64151294-64151764 | 0.355090332 | 0.316673986 | 0.607178555 |
| chr12:120495595-120496592 | 0.327060089 | 0.316683938 | 0.607178555 |
| chr7:83162103-83163263 | -0.318752193 | 0.31669693 | 0.607178555 |
| chr6:44475656-44476305 | 0.344873452 | 0.316700102 | 0.607178555 |
| chr21:10470381-10470581 | 0.594837537 | 0.316711284 | 0.607178555 |
| chr1:172484174-172484897 | -0.343807428 | 0.31671247 | 0.607178555 |
| chr6:110875149-110876515 | 0.321491642 | 0.316719904 | 0.607178555 |
| chr1:224329791-224330648 | 0.33005731 | 0.316728219 | 0.607179304 |
| chr7:77415844-77416793 | -0.338472774 | 0.316776693 | 0.607242167 |
| chr1:12019139-12019635 | -0.343460653 | 0.316776861 | 0.607242167 |
| chr10:99172169-99172446 | 0.45836842 | 0.316819545 | 0.607307175 |
| chr11:118247000-118247360 | 0.46525628 | 0.316828766 | 0.607307175 |
| chr9:95336424-95336624 | 0.532521815 | 0.316834551 | 0.607307175 |
| chr14:73059160-73059360 | 0.419804254 | 0.316851986 | 0.6073254 |
| chr10:69563448-69563679 | -0.518065787 | 0.316880954 | 0.607358414 |
| chr21:36209613-36209962 | 0.416400192 | 0.316885063 | 0.607358414 |
| chr5:2802575-2803114 | -0.379837129 | 0.316893581 | 0.607359547 |
| chr2:174247831-174249093 | -0.3193768 | 0.316935362 | 0.607399048 |
| chr6:38262112-38262312 | 0.499039364 | 0.316947724 | 0.607399048 |
| chr20:6828876-6829098 | -0.463113375 | 0.31695246 | 0.607399048 |
| chr8:2000726-2000926 | 0.499144064 | 0.316956694 | 0.607399048 |
| chr18:28177328-28177913 | -0.373033438 | 0.316956968 | 0.607399048 |
| chr1:13543949-13544470 | -0.42392397 | 0.316972378 | 0.607399048 |
| chr10:104472460-104472821 | -0.411761946 | 0.316981586 | 0.607399048 |
| chr10:30534631-30534916 | -0.429096592 | 0.316983979 | 0.607399048 |
| chr1:110163610-110163845 | -0.44910024 | 0.316985536 | 0.607399048 |
| chr5:90866357-90866627 | -0.468515686 | 0.317001088 | 0.607413658 |
| chr14:43503591-43503878 | 0.415992499 | 0.317033808 | 0.607460624 |
| chr20:56654746-56654946 | 0.445397265 | 0.317041455 | 0.607460624 |
| chr7:4681999-4682314 | 0.384419866 | 0.317108924 | 0.607574704 |
| chr11:19259752-19260053 | 0.418784935 | 0.317131807 | 0.607603354 |
| chr17:1998053-1998319 | -0.444635582 | 0.317168456 | 0.607658376 |
| chr5:170105753-170105953 | -0.486993019 | 0.317185273 | 0.607675401 |
| chr20:1900278-1900659 | -0.396248597 | 0.317200324 | 0.607675999 |
| chr17:74760756-74761096 | -0.363362291 | 0.317207623 | 0.607675999 |
| chr13:39352115-39352330 | -0.537562334 | 0.317209378 | 0.607675999 |
| chr22:30212578-30212914 | -0.436301276 | 0.317229335 | 0.607699038 |

| | | | |
|---|---|---|---|
| chr1:48061236-48061468 | -0.464120169 | 0.317250605 | 0.60772459 |
| chr17:49435102-49435410 | 0.380879819 | 0.317270383 | 0.607732215 |
| chr12:122078465-122078846 | 0.366177078 | 0.317270449 | 0.607732215 |
| chr1:152397573-152397831 | 0.386880603 | 0.317281614 | 0.607738409 |
| chr14:31751018-31751218 | 0.544964178 | 0.317293206 | 0.60774542 |
| chr3:11995065-11995439 | -0.399640995 | 0.31731013 | 0.607762644 |
| chr5:129460964-129461396 | -0.348624695 | 0.317331428 | 0.607775809 |
| chr10:11913409-11913627 | 0.51472372 | 0.317339979 | 0.607775809 |
| chr13:36163354-36163554 | -0.501905609 | 0.3173408 | 0.607775809 |
| chr13:60156036-60156297 | -0.45080613 | 0.317356373 | 0.607790443 |
| chr15:38218368-38218627 | -0.560394715 | 0.317369012 | 0.607799456 |
| chr3:156943416-156943782 | -0.379259512 | 0.317388012 | 0.607820651 |
| chr10:91298270-91298889 | -0.340855323 | 0.317418484 | 0.607863815 |
| chr8:129748344-129748607 | 0.424175377 | 0.317434334 | 0.607878975 |
| chr3:107523870-107524076 | 0.397808719 | 0.317466724 | 0.607925807 |
| chr2:33681430-33681859 | -0.430355045 | 0.317484434 | 0.607944528 |
| chr22:22348123-22348454 | -0.427162316 | 0.317494308 | 0.607948241 |
| chr16:69287092-69287292 | 0.550273306 | 0.317540172 | 0.607982472 |
| chr8:102628249-102628818 | -0.361112248 | 0.317543225 | 0.607982472 |
| chr15:68053881-68055004 | 0.324968399 | 0.317543655 | 0.607982472 |
| chr15:42471695-42471910 | 0.473347486 | 0.317543924 | 0.607982472 |
| chr5:134958355-134958556 | -0.554688846 | 0.317614568 | 0.608100601 |
| chr13:49110069-49110355 | -0.434187699 | 0.317631647 | 0.608100601 |
| chr2:65686845-65687179 | -0.463796849 | 0.317632187 | 0.608100601 |
| chr6:31739595-31740152 | 0.329062687 | 0.317644138 | 0.608100601 |
| chr11:72589859-72590133 | -0.427019678 | 0.317652705 | 0.608100601 |
| chr5:101174803-101175954 | -0.333196234 | 0.31765324 | 0.608100601 |
| chr7:144178423-144178727 | -0.415443088 | 0.3177395 | 0.60824414 |
| chr13:110222107-110222482 | -0.448286019 | 0.317744097 | 0.60824414 |
| chr5:113950744-113951002 | 0.467087094 | 0.317798427 | 0.60832629 |
| chr1:179002293-179003164 | -0.336397719 | 0.31780289 | 0.60832629 |
| chr3:154971945-154972267 | -0.481086262 | 0.317826137 | 0.608355591 |
| chr17:18225253-18225651 | -0.368584476 | 0.317874153 | 0.608432298 |
| chr19:29716618-29716829 | -0.459950493 | 0.317887453 | 0.608433707 |
| chr21:38261619-38261910 | -0.476180154 | 0.317896033 | 0.608433707 |
| chr5:168630740-168631420 | -0.361975574 | 0.317898711 | 0.608433707 |
| chr15:77501061-77501263 | 0.459696813 | 0.317927082 | 0.608472809 |
| chr16:79600668-79601035 | -0.359198644 | 0.317988207 | 0.608554259 |
| chr2:141864940-141865220 | -0.501389537 | 0.317992201 | 0.608554259 |
| chr12:104734383-104734799 | -0.438268979 | 0.317993467 | 0.608554259 |
| chr1:39762890-39763118 | -0.493248223 | 0.318009128 | 0.60856903 |
| chr16:89201566-89202255 | -0.327349227 | 0.318037165 | 0.608604523 |
| chr7:65731566-65731941 | -0.444021246 | 0.318043561 | 0.608604523 |
| chr3:139934518-139936163 | 0.320374659 | 0.318053728 | 0.608608779 |
| chr19:12757653-12757968 | 0.421411617 | 0.318067423 | 0.608619786 |
| chr9:90700281-90700511 | 0.470961671 | 0.318083219 | 0.608634811 |
| chr17:60204476-60204847 | 0.352886262 | 0.318097234 | 0.608644321 |
| chr6:136249687-136250934 | -0.317256539 | 0.318104763 | 0.608644321 |
| chr13:87259477-87259756 | -0.470174409 | 0.318112625 | 0.608644321 |
| chr7:86643708-86644288 | -0.382948018 | 0.318146997 | 0.608644321 |

| | | | |
|---|---|---|---|
| chr12:7994848-7995632 | -0.342643671 | 0.318152705 | 0.608644321 |
| chr14:56994873-56995431 | -0.399466922 | 0.318153424 | 0.608644321 |
| chr6:66558366-66558636 | -0.480873249 | 0.318164797 | 0.608644321 |
| chr20:37902188-37902411 | -0.446162795 | 0.318170491 | 0.608644321 |
| chr1:191027211-191027552 | 0.42815031 | 0.318171625 | 0.608644321 |
| chr12:12665236-12665677 | -0.402417298 | 0.318172367 | 0.608644321 |
| chr15:84914965-84915495 | 0.332118448 | 0.318177363 | 0.608644321 |
| chr7:35249925-35250125 | -0.518963124 | 0.31818351 | 0.608644321 |
| chr1:108813355-108813629 | -0.404429528 | 0.318216932 | 0.608679318 |
| chr15:99213282-99213499 | 0.473328274 | 0.318217694 | 0.608679318 |
| chr2:165573851-165574267 | -0.403958389 | 0.318226028 | 0.608680065 |
| chr17:31008428-31008628 | 0.504001889 | 0.318253466 | 0.608707196 |
| chr8:57994309-57995096 | -0.342801729 | 0.318256101 | 0.608707196 |
| chr1:161258543-161259007 | 0.382114984 | 0.318302403 | 0.608751164 |
| chr7:33888215-33888517 | 0.416528564 | 0.318304703 | 0.608751164 |
| chr2:179738763-179739126 | -0.489475013 | 0.318310858 | 0.608751164 |
| chr11:88016468-88016668 | -0.540976743 | 0.318318617 | 0.608751164 |
| chr1:53107627-53107891 | 0.412254488 | 0.318318813 | 0.608751164 |
| chr19:8002322-8002681 | -0.433574302 | 0.3183309 | 0.60875542 |
| chr20:54984140-54984348 | -0.509508161 | 0.318336929 | 0.60875542 |
| chr15:76330382-76330591 | -0.482212654 | 0.318371516 | 0.608796127 |
| chr16:23060720-23061013 | 0.436166325 | 0.318374107 | 0.608796127 |
| chr12:100926740-100927009 | -0.434506274 | 0.318384452 | 0.608800716 |
| chr1:164611772-164612368 | -0.338239361 | 0.318401002 | 0.608817168 |
| chr6:151564718-151565018 | 0.398853579 | 0.318437338 | 0.608871453 |
| chr5:76386438-76386724 | -0.431908803 | 0.318459026 | 0.608880709 |
| chr10:114520003-114520345 | 0.453257534 | 0.318464092 | 0.608880709 |
| chr9:38047632-38048087 | 0.378762382 | 0.318469439 | 0.608880709 |
| chr11:83086995-83087322 | -0.395133252 | 0.318473965 | 0.608880709 |
| chr8:1831773-1832088 | 0.415682768 | 0.318515905 | 0.608938244 |
| chr14:76564535-76564735 | -0.547559111 | 0.318519953 | 0.608938244 |
| chr14:73935794-73936007 | 0.470428339 | 0.318539153 | 0.608950303 |
| chr4:139497826-139498135 | -0.422624538 | 0.318542156 | 0.608950303 |
| chr3:106457274-106457690 | -0.395944666 | 0.318573584 | 0.608983802 |
| chr2:162582425-162582843 | -0.403091871 | 0.318575576 | 0.608983802 |
| chr16:87706307-87706809 | 0.338079531 | 0.318585602 | 0.608987776 |
| chr20:23799289-23799524 | 0.437250018 | 0.318603906 | 0.609007571 |
| chr4:172938972-172939193 | -0.567823678 | 0.318625739 | 0.609034111 |
| chr18:46158595-46158795 | -0.5381191 | 0.318646697 | 0.609055629 |
| chr11:3000238-3001130 | 0.336184752 | 0.318652894 | 0.609055629 |
| chr10:119527481-119527703 | -0.521628333 | 0.318691522 | 0.609092555 |
| chr13:96089742-96090174 | 0.384916519 | 0.318696046 | 0.609092555 |
| chr12:92739135-92739370 | -0.5029351 | 0.318699478 | 0.609092555 |
| chr7:27095714-27096777 | 0.332507563 | 0.318704011 | 0.609092555 |
| chr2:95389255-95389521 | -0.42919334 | 0.318719199 | 0.609106389 |
| chrX:34521635-34521928 | -0.437888957 | 0.318738468 | 0.609123802 |
| chr9:97983892-97985068 | 0.330456368 | 0.31874421 | 0.609123802 |
| chr4:39365861-39366625 | -0.336876367 | 0.318770139 | 0.609156377 |
| chr19:17778202-17778850 | 0.341417085 | 0.318777156 | 0.609156377 |
| chr5:83608183-83608487 | -0.412065606 | 0.318791306 | 0.609168225 |

| | | | |
|---|---|---|---|
| chr2:27916001-27916281 | 0.491434758 | 0.31880959 | 0.609187052 |
| chr7:32890536-32890756 | 0.370678579 | 0.31882304 | 0.609187052 |
| chr11:746808-747044 | 0.442603749 | 0.318831306 | 0.609187052 |
| chr5:23978086-23978286 | -0.51580431 | 0.318832961 | 0.609187052 |
| chr5:41636667-41636947 | 0.425049256 | 0.318844382 | 0.609191216 |
| chr6:118800452-118801081 | -0.355427235 | 0.318851042 | 0.609191216 |
| chr2:75637561-75637924 | -0.401430346 | 0.318878965 | 0.609229374 |
| chr7:51299885-51300085 | -0.506625415 | 0.318935516 | 0.609293341 |
| chr1:114435614-114435855 | 0.392493586 | 0.318941664 | 0.609293341 |
| chr2:134843185-134843611 | -0.406398301 | 0.318942531 | 0.609293341 |
| chr2:71191039-71191530 | -0.434681808 | 0.318944254 | 0.609293341 |
| chr8:134367732-134367932 | 0.478975647 | 0.318953002 | 0.609294861 |
| chr11:64878458-64879207 | 0.32439176 | 0.318970549 | 0.609302295 |
| chr3:146927028-146927324 | -0.504196783 | 0.318972797 | 0.609302295 |
| chr6:70686619-70686862 | -0.483436338 | 0.318986313 | 0.609306229 |
| chr12:12611490-12612416 | -0.33401437 | 0.318994718 | 0.609306229 |
| chr3:170553028-170553248 | -0.54693574 | 0.319008858 | 0.609306229 |
| chrX:12701729-12702014 | 0.446896877 | 0.319013892 | 0.609306229 |
| chr13:27406322-27406733 | 0.384241602 | 0.319014617 | 0.609306229 |
| chr11:69644676-69644885 | -0.420760819 | 0.31902613 | 0.609313031 |
| chr3:94062559-94063448 | -0.317504883 | 0.319123599 | 0.609468073 |
| chr4:2652920-2653148 | 0.483702275 | 0.319126919 | 0.609468073 |
| chr10:55263783-55264226 | -0.374742763 | 0.31913117 | 0.609468073 |
| chr20:49825836-49826073 | -0.481458115 | 0.319142605 | 0.609474718 |
| chr2:161160092-161161409 | -0.315041582 | 0.319155065 | 0.609483324 |
| chr8:134566519-134566901 | -0.387257708 | 0.319180246 | 0.609498754 |
| chr8:110976763-110977242 | 0.383865454 | 0.319181478 | 0.609498754 |
| chr7:27130334-27131461 | 0.3314811 | 0.319190512 | 0.609498754 |
| chr7:112239372-112239642 | -0.556101552 | 0.319194964 | 0.609498754 |
| chrX:71935875-71936201 | 0.486864722 | 0.319203673 | 0.609500196 |
| chr17:60165194-60165398 | 0.342211069 | 0.31921286 | 0.60950069 |
| chr7:1441532-1441776 | 0.416553051 | 0.319235043 | 0.60950069 |
| chr15:70093575-70093925 | 0.347515098 | 0.319235539 | 0.60950069 |
| chr19:10928285-10929116 | 0.324232696 | 0.319241351 | 0.60950069 |
| chr5:141923324-141924352 | -0.320654008 | 0.31925153 | 0.60950069 |
| chr14:55416901-55417150 | -0.432993479 | 0.31925166 | 0.60950069 |
| chr15:78720183-78720596 | -0.385131061 | 0.31927329 | 0.609515194 |
| chr16:68520580-68521089 | -0.35901256 | 0.31927971 | 0.609515194 |
| chr16:77171783-77172201 | 0.355536963 | 0.319283122 | 0.609515194 |
| chr21:44801662-44802695 | 0.333674045 | 0.319292121 | 0.609517188 |
| chr2:9178342-9178904 | 0.364885556 | 0.319321056 | 0.609557237 |
| chr6:56381895-56382121 | 0.496311528 | 0.319336865 | 0.60956517 |
| chr4:141635911-141636877 | -0.337086597 | 0.319342262 | 0.60956517 |
| chr2:39783181-39783701 | -0.364616669 | 0.319349078 | 0.60956517 |
| chr20:18406296-18406584 | 0.472039747 | 0.319396863 | 0.609641194 |
| chr22:18024261-18024942 | 0.362059437 | 0.319417789 | 0.609665948 |
| chr2:164495576-164495899 | 0.413266574 | 0.319449831 | 0.609711918 |
| chr3:83460099-83460351 | -0.486191939 | 0.319473369 | 0.60973009 |
| chr19:34359063-34359853 | 0.317882985 | 0.319475267 | 0.60973009 |
| chr1:171507020-171507359 | -0.431309389 | 0.319505354 | 0.609772323 |

| | | | |
|---|---|---|---|
| chr2:200888546-200890053 | -0.313319216 | 0.319530891 | 0.609805872 |
| chr1:178657281-178657584 | -0.433368582 | 0.319541622 | 0.609811162 |
| chr2:228425373-228426012 | -0.378324958 | 0.319574311 | 0.609852912 |
| chr5:53231374-53231714 | 0.498072239 | 0.319579417 | 0.609852912 |
| chr19:15479716-15479955 | -0.461890146 | 0.319598953 | 0.609875003 |
| chr14:73785330-73786303 | 0.324273528 | 0.319609595 | 0.609880121 |
| chr16:27374897-27375407 | 0.351220034 | 0.319627113 | 0.609894926 |
| chr3:172035134-172035393 | -0.383002409 | 0.319633273 | 0.609894926 |
| chr12:54304228-54304486 | 0.415397808 | 0.319725496 | 0.610019697 |
| chr3:23743299-23744153 | -0.338693612 | 0.319729522 | 0.610019697 |
| chr7:12674911-12675397 | 0.360606226 | 0.319729867 | 0.610019697 |
| chr13:59318414-59318705 | -0.465803679 | 0.319730508 | 0.610019697 |
| chr10:112140010-112140492 | 0.350093204 | 0.319763707 | 0.610031321 |
| chr13:42477847-42478065 | 0.51726809 | 0.319767196 | 0.610031321 |
| chr11:61595329-61595795 | 0.38312208 | 0.319774395 | 0.610031321 |
| chr6:132820661-132821183 | -0.368099972 | 0.319780677 | 0.610031321 |
| chr19:29605750-29606649 | 0.328587246 | 0.319781015 | 0.610031321 |
| chr19:46714098-46714740 | -0.341639519 | 0.319796854 | 0.610031321 |
| chr16:8673980-8674812 | 0.330568977 | 0.319798801 | 0.610031321 |
| chr22:18077719-18078104 | 0.355636158 | 0.319800294 | 0.610031321 |
| chr14:79437917-79438314 | -0.41730672 | 0.319863236 | 0.610095882 |
| chrX:143788328-143788550 | 0.481137259 | 0.319863242 | 0.610095882 |
| chr2:203352852-203353123 | 0.439966589 | 0.319863435 | 0.610095882 |
| chr3:139427589-139427789 | 0.467506078 | 0.319870909 | 0.610095882 |
| chr17:82030487-82030781 | 0.364245688 | 0.319878401 | 0.610095882 |
| chr2:142682569-142682812 | -0.501509174 | 0.319887648 | 0.610095882 |
| chr3:194583511-194584598 | -0.327595848 | 0.319889876 | 0.610095882 |
| chr10:25280478-25280718 | -0.527828601 | 0.319900377 | 0.610100724 |
| chr8:143430634-143431332 | 0.325276013 | 0.319911568 | 0.610106767 |
| chr6:170433164-170433399 | 0.491016628 | 0.319919471 | 0.610106767 |
| chr19:46014764-46015640 | 0.337138388 | 0.319947941 | 0.610145876 |
| chr1:159132362-159132563 | -0.527234042 | 0.319970121 | 0.610168328 |
| chr10:79275468-79275840 | -0.413457939 | 0.319976292 | 0.610168328 |
| chr6:2790671-2790871 | 0.521298456 | 0.319983605 | 0.610168328 |
| chr20:39049790-39050448 | 0.350307813 | 0.320050096 | 0.610278788 |
| chr17:77286174-77287421 | -0.313762767 | 0.320057461 | 0.610278788 |
| chr12:97395684-97395914 | -0.508161582 | 0.320072509 | 0.610292294 |
| chr2:393658-394271 | -0.382516748 | 0.320169285 | 0.610447723 |
| chr1:106957817-106958093 | 0.360341471 | 0.320175367 | 0.610447723 |
| chr6:160103453-160103678 | -0.474164974 | 0.320177927 | 0.610447723 |
| chr17:77086546-77086813 | 0.385049556 | 0.320198943 | 0.610472603 |
| chr9:12814394-12814767 | 0.387097437 | 0.320219336 | 0.610494321 |
| chr11:67659457-67659714 | 0.411980195 | 0.32022627 | 0.610494321 |
| chr19:50758701-50759026 | -0.461302652 | 0.32024241 | 0.610509901 |
| chr15:97239119-97239485 | -0.397524664 | 0.320260351 | 0.61052006 |
| chr5:152330854-152331124 | -0.482365997 | 0.320263675 | 0.61052006 |
| chr17:50338133-50338340 | -0.555557707 | 0.320290824 | 0.610538692 |
| chr5:7850567-7851385 | -0.328514897 | 0.320296106 | 0.610538692 |
| chr13:41458919-41459179 | -0.381207164 | 0.320306213 | 0.610538692 |
| chr5:33550536-33550736 | -0.539629264 | 0.32031121 | 0.610538692 |

| | | | |
|---|---|---|---|
| chr6:68778714-68778963 | -0.456472915 | 0.32031329 | 0.610538692 |
| chr9:93288450-93288705 | -0.472576608 | 0.320336987 | 0.61056315 |
| chr5:139968692-139969172 | -0.357288194 | 0.320342058 | 0.61056315 |
| chr12:121443244-121443575 | 0.376031717 | 0.320380224 | 0.610606903 |
| chr15:69131704-69131904 | -0.56649651 | 0.3203885 | 0.610606903 |
| chr3:161821370-161821570 | -0.514053486 | 0.320388921 | 0.610606903 |
| chr12:66775921-66776135 | 0.532081329 | 0.320401663 | 0.610608935 |
| chr1:202835864-202836094 | 0.404104699 | 0.320405926 | 0.610608935 |
| chr21:18384737-18385001 | -0.538740792 | 0.320418568 | 0.610617841 |
| chr9:83956038-83956294 | -0.434422207 | 0.320479377 | 0.610718535 |
| chr1:162284985-162285898 | 0.341591214 | 0.320494076 | 0.610731356 |
| chrX:80205635-80206051 | -0.424590769 | 0.320520147 | 0.610745391 |
| chr4:140825798-140826702 | -0.340561273 | 0.320525748 | 0.610745391 |
| chr2:111116941-111117324 | -0.350324659 | 0.32053259 | 0.610745391 |
| chr7:130380838-130381038 | 0.514932469 | 0.320533325 | 0.610745391 |
| chr2:130964566-130964839 | 0.401810132 | 0.32054238 | 0.610747457 |
| chr2:96714329-96714962 | -0.36450635 | 0.320566608 | 0.610759729 |
| chr16:11797203-11797875 | 0.334062606 | 0.320568318 | 0.610759729 |
| chrX:129540053-129541195 | 0.330722747 | 0.320578352 | 0.610759729 |
| chr10:119445436-119445727 | 0.395051249 | 0.320580705 | 0.610759729 |
| chr4:118629778-118630031 | -0.480200143 | 0.320603965 | 0.610773568 |
| chr17:77854632-77854852 | -0.475433848 | 0.320615054 | 0.610773568 |
| chr3:13496291-13496636 | 0.362891235 | 0.320615212 | 0.610773568 |
| chr1:215657619-215657880 | -0.448786428 | 0.320623846 | 0.610773568 |
| chr20:44027893-44028164 | 0.443797451 | 0.320627825 | 0.610773568 |
| chr14:57512164-57512364 | 0.54091761 | 0.32066826 | 0.610813463 |
| chr5:161230281-161230525 | -0.527165252 | 0.32068725 | 0.610813463 |
| chr15:63188963-63189838 | 0.321967424 | 0.320689311 | 0.610813463 |
| chr11:25766561-25767067 | 0.394424217 | 0.320697079 | 0.610813463 |
| chr2:155100290-155100592 | -0.439693555 | 0.320698823 | 0.610813463 |
| chr6:151716280-151716812 | 0.369985689 | 0.320700135 | 0.610813463 |
| chr3:15268005-15268255 | 0.401038152 | 0.320721073 | 0.610813463 |
| chr12:67451248-67451569 | -0.440392022 | 0.320721205 | 0.610813463 |
| chr1:50968900-50969492 | -0.34786617 | 0.320723334 | 0.610813463 |
| chr13:46931660-46931861 | -0.473568958 | 0.320728486 | 0.610813463 |
| chr12:81128960-81129486 | -0.365970597 | 0.3207802 | 0.610896767 |
| chr10:90293580-90294145 | -0.363776958 | 0.320792696 | 0.61090074 |
| chr3:107767803-107768003 | 0.510315873 | 0.320798233 | 0.61090074 |
| chr10:101570647-101570928 | 0.403660623 | 0.320819366 | 0.610925801 |
| chr8:11986576-11986960 | -0.441725137 | 0.320845036 | 0.6109595 |
| chr12:121015917-121016713 | 0.335414122 | 0.320866417 | 0.61097476 |
| chrX:82961192-82961421 | -0.49234121 | 0.320868998 | 0.61097476 |
| chr19:55516836-55517439 | 0.337571519 | 0.320884878 | 0.610979898 |
| chr3:34011357-34011562 | 0.458780257 | 0.320887729 | 0.610979898 |
| chr12:103929429-103930645 | -0.312132145 | 0.320895618 | 0.610979898 |
| chr13:45309352-45309774 | -0.345428381 | 0.320947677 | 0.611063833 |
| chr7:121062691-121062934 | -0.521145335 | 0.320968603 | 0.611088491 |
| chr15:67024404-67024689 | -0.389257959 | 0.321020405 | 0.611149371 |
| chr2:149366330-149366684 | -0.437497784 | 0.321024055 | 0.611149371 |
| chr19:1040898-1041269 | 0.353012267 | 0.321024509 | 0.611149371 |

| | | | |
|---|---|---|---|
| chr16:70114531-70114893 | 0.387469702 | 0.321054524 | 0.611191328 |
| chr3:147447871-147448116 | -0.493991859 | 0.321086683 | 0.611237362 |
| chr1:2212561-2212893 | 0.436309324 | 0.321099177 | 0.611245961 |
| chr3:120693454-120693780 | 0.417886723 | 0.321127694 | 0.611285058 |
| chr3:161171697-161172197 | -0.368929216 | 0.321145701 | 0.611298817 |
| chr1:157132567-157133094 | -0.34151694 | 0.321150878 | 0.611298817 |
| chr9:124869423-124869913 | 0.356358129 | 0.321171085 | 0.611322093 |
| chr14:59919804-59920180 | -0.423267174 | 0.321224074 | 0.611393485 |
| chr9:132582107-132582390 | 0.409720823 | 0.321231819 | 0.611393485 |
| chr16:8572944-8573204 | -0.477642024 | 0.321240854 | 0.611393485 |
| chr11:66720148-66720864 | -0.325072273 | 0.321252312 | 0.611393485 |
| chr2:196291314-196291947 | 0.351343525 | 0.321261212 | 0.611393485 |
| chr9:34646192-34647324 | -0.324303161 | 0.321265753 | 0.611393485 |
| chr9:130042529-130043593 | 0.330858523 | 0.321272792 | 0.611393485 |
| chr5:36073658-36074008 | 0.407484401 | 0.32127905 | 0.611393485 |
| chr20:42530949-42531157 | 0.401832248 | 0.321280406 | 0.611393485 |
| chr19:54463601-54465012 | 0.312772403 | 0.321295669 | 0.611407345 |
| chr22:11915750-11916046 | 0.452726296 | 0.321315895 | 0.611430648 |
| chr21:37533170-37533700 | -0.367347941 | 0.321336936 | 0.611444089 |
| chr10:102449601-102449970 | 0.367707584 | 0.321338918 | 0.611444089 |
| chr14:97053137-97053337 | 0.554931038 | 0.321385053 | 0.611516689 |
| chr3:114213918-114214510 | 0.365786807 | 0.321428287 | 0.611581769 |
| chr2:38777826-38778745 | -0.327640298 | 0.32143522 | 0.611581769 |
| chr22:24247668-24247943 | 0.419824247 | 0.321462405 | 0.611618306 |
| chr3:158672124-158673122 | -0.317282197 | 0.321497359 | 0.611666011 |
| chr1:206479551-206479940 | -0.349125507 | 0.321513117 | 0.611666011 |
| chr7:39949204-39950564 | -0.310287766 | 0.321516347 | 0.611666011 |
| chr2:155879909-155880212 | -0.489733538 | 0.321520314 | 0.611666011 |
| chr8:108786945-108787856 | -0.317127401 | 0.321527393 | 0.611666011 |
| chr2:36578922-36579252 | 0.432054352 | 0.321535701 | 0.611666628 |
| chr1:168136628-168137607 | 0.340771653 | 0.321567773 | 0.611688654 |
| chr20:57276923-57278185 | 0.309555825 | 0.321570664 | 0.611688654 |
| chr13:25233040-25233240 | 0.473901162 | 0.321571229 | 0.611688654 |
| chr12:42325800-42326500 | 0.331312278 | 0.321582129 | 0.611694203 |
| chr7:99472547-99473115 | -0.332839725 | 0.321606613 | 0.611713197 |
| chr14:60649951-60650312 | 0.376324481 | 0.321608081 | 0.611713197 |
| chr1:117562565-117562867 | 0.3854299 | 0.321618953 | 0.61171869 |
| chr1:36099599-36100315 | 0.339350637 | 0.321635707 | 0.611733996 |
| chr9:126146350-126146787 | 0.400317381 | 0.321642968 | 0.611733996 |
| chr18:12727046-12727451 | 0.423661191 | 0.321657133 | 0.611745752 |
| chr2:162221004-162221394 | -0.371175218 | 0.321686369 | 0.611771074 |
| chr7:143620838-143621102 | -0.446975746 | 0.321686416 | 0.611771074 |
| chr17:41548354-41548554 | -0.444671272 | 0.321708257 | 0.611797425 |
| chr12:24562388-24562676 | -0.429883099 | 0.3217245 | 0.611810121 |
| chr5:154642487-154642968 | -0.366203742 | 0.321730903 | 0.611810121 |
| chrX:10190417-10190635 | -0.47728614 | 0.321814077 | 0.611953101 |
| chr2:153593040-153593370 | -0.452466574 | 0.321823648 | 0.611956112 |
| chr1:186532390-186532629 | 0.536537122 | 0.321886746 | 0.612060905 |
| chr15:84570153-84570642 | 0.361531543 | 0.321905876 | 0.612082091 |
| chr2:182668748-182668997 | -0.429587077 | 0.321926047 | 0.612094581 |

| | | | |
|---|---|---|---|
| chr7:124099863-124100063 | -0.57280017 | 0.321936246 | 0.612094581 |
| chr3:52278046-52279061 | 0.326693923 | 0.321936752 | 0.612094581 |
| chr11:62245299-62245507 | 0.504920993 | 0.321946172 | 0.612094581 |
| chr15:76390506-76390863 | 0.403049716 | 0.321952387 | 0.612094581 |
| chr1:60858392-60858616 | -0.517731074 | 0.321974872 | 0.612122141 |
| chr1:234478131-234479348 | 0.318502783 | 0.322001799 | 0.612139291 |
| chr1:207730399-207730686 | -0.447703753 | 0.322002276 | 0.612139291 |
| chr14:105880795-105881198 | 0.398528039 | 0.32200786 | 0.612139291 |
| chrX:98831912-98832281 | -0.418244858 | 0.322031266 | 0.612147383 |
| chr9:125538008-125538396 | 0.38660193 | 0.322039175 | 0.612147383 |
| chr17:11644237-11644710 | -0.390380669 | 0.322040563 | 0.612147383 |
| chr18:49728660-49728860 | -0.542670832 | 0.32204765 | 0.612147383 |
| chr19:54172508-54173446 | 0.32472826 | 0.322052328 | 0.612147383 |
| chr14:95993710-95993997 | 0.469567996 | 0.322066631 | 0.612147383 |
| chr20:58965194-58965394 | 0.444045177 | 0.322073435 | 0.612147383 |
| chr2:203551731-203552168 | -0.415490092 | 0.32207603 | 0.612147383 |
| chr17:60577134-60577887 | -0.315264743 | 0.322094401 | 0.612153502 |
| chr20:50483424-50483855 | -0.337751038 | 0.322095228 | 0.612153502 |
| chr15:40843932-40844636 | 0.333437323 | 0.322114194 | 0.612174363 |
| chr8:141629228-141629945 | 0.348399958 | 0.322137337 | 0.612203162 |
| chr3:64040148-64040360 | 0.354526392 | 0.322183258 | 0.612275246 |
| chr1:228457418-228458770 | 0.314878878 | 0.322212557 | 0.612315739 |
| chr17:42608845-42610034 | 0.319647031 | 0.32222529 | 0.612324747 |
| chr5:172443543-172443747 | -0.492818944 | 0.322254014 | 0.612364146 |
| chr1:222589663-222590316 | -0.335463247 | 0.322274771 | 0.612368176 |
| chr3:164806152-164806575 | -0.369680411 | 0.32228275 | 0.612368176 |
| chr19:12722017-12722220 | 0.480643052 | 0.322287174 | 0.612368176 |
| chr1:155086093-155086317 | 0.371640174 | 0.322288103 | 0.612368176 |
| chr1:17377272-17377828 | 0.348675035 | 0.322320291 | 0.612414148 |
| chr5:175908562-175909270 | -0.347188703 | 0.322355556 | 0.612465965 |
| chr19:7903394-7904006 | 0.327330807 | 0.322399087 | 0.612511973 |
| chr6:41936333-41936588 | 0.39950302 | 0.322403122 | 0.612511973 |
| chr1:166603265-166603735 | -0.354359157 | 0.322403753 | 0.612511973 |
| chr13:75635782-75637118 | -0.323642456 | 0.322427632 | 0.612542151 |
| chr5:151652100-151652418 | -0.380412987 | 0.322479514 | 0.612625526 |
| chr2:26846335-26846701 | 0.384212385 | 0.322498082 | 0.612645611 |
| chr5:60649323-60649622 | -0.38768526 | 0.322530169 | 0.612691375 |
| chr8:124138156-124139033 | 0.327429124 | 0.32254681 | 0.612707796 |
| chr6:82363421-82364908 | -0.314874311 | 0.32255944 | 0.61271287 |
| chr10:121608657-121609000 | 0.388817266 | 0.322573444 | 0.61271287 |
| chr5:7146977-7147231 | 0.4802438 | 0.32257347 | 0.61271287 |
| chr12:54527122-54527355 | -0.467298702 | 0.322592841 | 0.612733733 |
| chr3:109369585-109369820 | 0.433917355 | 0.322601175 | 0.612733733 |
| chr2:113626988-113627665 | 0.336360937 | 0.322608445 | 0.612733733 |
| chr20:62232255-62233191 | -0.318952961 | 0.322651482 | 0.612786053 |
| chr8:144373618-144374351 | 0.32336382 | 0.322663316 | 0.612786053 |
| chr8:128754250-128754959 | 0.361935929 | 0.322663947 | 0.612786053 |
| chr8:43092989-43094078 | 0.326257473 | 0.322668549 | 0.612786053 |
| chr3:12663270-12664718 | 0.312138709 | 0.322675979 | 0.612786053 |
| chr12:93154456-93154656 | -0.509145678 | 0.322697795 | 0.612800834 |

| | | | |
|---|---|---|---|
| chr12:129187030-129187230 | 0.512499071 | 0.322699758 | 0.612800834 |
| chr17:82962962-82963536 | 0.333660161 | 0.322741182 | 0.612864308 |
| chr6:165987107-165987307 | 0.490902008 | 0.322784406 | 0.612931198 |
| chr20:52917804-52918087 | -0.496316805 | 0.322813794 | 0.612967932 |
| chr13:110555848-110556104 | 0.453131195 | 0.322824059 | 0.612967932 |
| chr7:131722069-131722299 | -0.585940971 | 0.322827751 | 0.612967932 |
| chr7:76235125-76235541 | -0.342730783 | 0.322855475 | 0.613005383 |
| chr2:208482131-208483099 | -0.334765622 | 0.322868354 | 0.613011535 |
| chr3:8416988-8417280 | 0.392641738 | 0.322874716 | 0.613011535 |
| chr3:14548290-14548652 | 0.358081143 | 0.322892683 | 0.613030457 |
| chr15:38683648-38683965 | -0.420982094 | 0.322906848 | 0.613035309 |
| chr2:210170367-210171873 | -0.310064635 | 0.322912288 | 0.613035309 |
| chr5:75037114-75037710 | -0.334972022 | 0.322926516 | 0.613035309 |
| chr8:67717723-67717959 | 0.564278182 | 0.322927242 | 0.613035309 |
| chr1:36088563-36089435 | 0.33109742 | 0.322947046 | 0.613057715 |
| chr4:76585927-76586424 | 0.36535325 | 0.3229835 | 0.613111726 |
| chr12:111396983-111397189 | 0.456503025 | 0.323028704 | 0.613169986 |
| chr5:132746878-132747340 | 0.366045777 | 0.323030196 | 0.613169986 |
| chr20:44651998-44652376 | 0.379706295 | 0.323047575 | 0.613187785 |
| chr5:44035427-44036077 | -0.352267461 | 0.323103452 | 0.613278654 |
| chr4:973459-973858 | 0.357373063 | 0.323137842 | 0.613328735 |
| chr8:96313480-96313837 | -0.38582172 | 0.323148476 | 0.613333727 |
| chr7:150745125-150745357 | 0.453756778 | 0.32316681 | 0.613341354 |
| chr20:63482870-63483183 | 0.381431503 | 0.323168505 | 0.613341354 |
| chr3:154059784-154060254 | -0.327813542 | 0.323180452 | 0.61334166 |
| chr9:122228638-122228841 | -0.488193094 | 0.323184675 | 0.61334166 |
| chr4:74306686-74306932 | -0.452350047 | 0.323220033 | 0.613393569 |
| chr5:25706940-25707292 | 0.422307765 | 0.323236325 | 0.613409294 |
| chr10:106795687-106795958 | 0.467954795 | 0.32328262 | 0.61347194 |
| chr20:56808867-56809135 | -0.370704796 | 0.323285349 | 0.61347194 |
| chr17:59802174-59802440 | 0.408017449 | 0.323295116 | 0.613475281 |
| chr17:29779288-29779584 | -0.488021705 | 0.323346038 | 0.613556714 |
| chr8:37594998-37595198 | 0.486982477 | 0.32338697 | 0.613596911 |
| chr12:121793863-121794575 | 0.322816759 | 0.323389677 | 0.613596911 |
| chr15:32609576-32609898 | -0.439076332 | 0.323391246 | 0.613596911 |
| chr4:91840347-91840841 | 0.351135972 | 0.323442841 | 0.61366741 |
| chr19:42386836-42387582 | 0.338043866 | 0.32344442 | 0.61366741 |
| chr5:68071894-68072143 | -0.563172052 | 0.323454182 | 0.613670736 |
| chr17:59121359-59121701 | 0.340717383 | 0.323475725 | 0.613696413 |
| chr13:30870346-30870877 | 0.330810704 | 0.323496321 | 0.613697514 |
| chr14:100176746-100177006 | -0.454473552 | 0.323498969 | 0.613697514 |
| chr2:72719406-72719792 | -0.416889653 | 0.323500334 | 0.613697514 |
| chr8:134906094-134906473 | -0.362268009 | 0.323567831 | 0.61380065 |
| chr16:35560467-35560754 | 0.440121612 | 0.323570722 | 0.61380065 |
| chr11:118929416-118929818 | 0.332155122 | 0.323597278 | 0.613832083 |
| chrX:66728758-66729308 | 0.360660035 | 0.323603314 | 0.613832083 |
| chr21:8442201-8442512 | 0.324015778 | 0.323659319 | 0.613923117 |
| chr13:30158840-30159084 | 0.423977111 | 0.323681813 | 0.613950586 |
| chr2:126658008-126658208 | -0.526358253 | 0.323696415 | 0.613955318 |
| chr12:49110545-49111148 | 0.325142609 | 0.323700333 | 0.613955318 |

| | | | |
|---|---|---|---|
| chr5:17305775-17306730 | 0.312700673 | 0.323745082 | 0.613998542 |
| chr1:34946222-34946649 | -0.384812732 | 0.323766888 | 0.613998542 |
| chr10:62082689-62082966 | -0.445898773 | 0.323770967 | 0.613998542 |
| chr16:81036653-81036938 | 0.396334501 | 0.32377809 | 0.613998542 |
| chr20:50246887-50247255 | 0.353487564 | 0.323780507 | 0.613998542 |
| chr18:9333912-9335400 | 0.310913825 | 0.323785497 | 0.613998542 |
| chr13:51347252-51347452 | -0.532371892 | 0.323787229 | 0.613998542 |
| chr13:109412444-109412644 | -0.532371892 | 0.323787229 | 0.613998542 |
| chr13:20703991-20704903 | 0.320007879 | 0.323836874 | 0.614077485 |
| chr12:6662592-6663495 | -0.322807295 | 0.323845334 | 0.61407833 |
| chr8:108816971-108817273 | 0.451136105 | 0.323868053 | 0.614106213 |
| chr20:54186191-54186525 | 0.330550915 | 0.32393657 | 0.614220931 |
| chr15:40845880-40846210 | 0.352400672 | 0.323953263 | 0.614234861 |
| chr21:39205383-39205617 | 0.479556318 | 0.323959949 | 0.614234861 |
| chr1:22451617-22452451 | 0.333108636 | 0.323977306 | 0.614235167 |
| chr1:156677242-156677597 | 0.364365274 | 0.323988424 | 0.614235167 |
| chr7:6290397-6290597 | 0.517866972 | 0.323989694 | 0.614235167 |
| chr14:95361480-95361735 | 0.425937458 | 0.323996981 | 0.614235167 |
| chr5:105506746-105507227 | -0.380684816 | 0.324000193 | 0.614235167 |
| chr9:21812943-21813521 | -0.352886956 | 0.324024394 | 0.614261305 |
| chr1:189774737-189775113 | -0.375670582 | 0.324031949 | 0.614261305 |
| chr13:44116669-44116904 | 0.492163898 | 0.32403803 | 0.614261305 |
| chrX:71870719-71871085 | 0.370032605 | 0.324068931 | 0.614291419 |
| chr5:115625484-115626683 | -0.313380217 | 0.324075513 | 0.614291419 |
| chr3:115890259-115890499 | 0.512467726 | 0.324077968 | 0.614291419 |
| chr1:181135425-181135667 | 0.36613242 | 0.324100953 | 0.614319792 |
| chr3:13494446-13494937 | 0.350288969 | 0.324170252 | 0.614435945 |
| chr4:102420634-102420834 | -0.522641086 | 0.32421752 | 0.614487727 |
| chr3:128875611-128875867 | -0.376378773 | 0.324220047 | 0.614487727 |
| chr20:50689729-50691396 | 0.306728421 | 0.324221631 | 0.614487727 |
| chr16:71808313-71809833 | -0.313203425 | 0.3242362 | 0.614500139 |
| chr16:2537465-2538284 | 0.329054737 | 0.324278168 | 0.614541835 |
| chr7:73855484-73855848 | -0.373587524 | 0.32428211 | 0.614541835 |
| chr11:105968041-105968277 | -0.561969882 | 0.324282262 | 0.614541835 |
| chr4:148054384-148054614 | -0.434498255 | 0.324312254 | 0.614583473 |
| chr16:15888128-15888945 | 0.318477187 | 0.324320351 | 0.614583615 |
| chr10:34816524-34816905 | -0.381412068 | 0.324335787 | 0.614591541 |
| chr1:180178840-180179146 | 0.368488329 | 0.324344775 | 0.614591541 |
| chr5:146007327-146007583 | 0.442086491 | 0.324348596 | 0.614591541 |
| chr5:149729975-149730484 | -0.345200276 | 0.324378779 | 0.614633534 |
| chr7:116786813-116787039 | -0.469977223 | 0.324425328 | 0.614692043 |
| chr5:181098831-181099251 | 0.421381268 | 0.324425703 | 0.614692043 |
| chr1:163535899-163536138 | 0.475588052 | 0.324458127 | 0.614729907 |
| chr1:145858650-145860151 | 0.309737556 | 0.324474985 | 0.614729907 |
| chr4:76179759-76180190 | 0.364912848 | 0.324482471 | 0.614729907 |
| chrX:30196375-30196660 | -0.414902794 | 0.324484667 | 0.614729907 |
| chr18:5979925-5980263 | -0.430933065 | 0.324485801 | 0.614729907 |
| chr13:31797094-31797492 | -0.397763373 | 0.324526991 | 0.614788546 |
| chr16:4253303-4254338 | 0.315064437 | 0.324536611 | 0.614788546 |
| chr2:74908740-74909541 | 0.34203464 | 0.324549382 | 0.614788546 |

| | | | |
|---|---|---|---|
| chr7:8724130-8724669 | -0.36165782 | 0.324552717 | 0.614788546 |
| chr2:168541989-168542231 | -0.50993263 | 0.324564896 | 0.614788546 |
| chr2:179495132-179495402 | -0.50993263 | 0.324564896 | 0.614788546 |
| chr2:4106954-4107154 | -0.56282083 | 0.324590959 | 0.614810729 |
| chr1:205841479-205841679 | 0.443248764 | 0.324592655 | 0.614810729 |
| chr6:20454301-20454745 | -0.433414075 | 0.324609444 | 0.614827331 |
| chr3:167184554-167184912 | -0.369936137 | 0.324628881 | 0.61484476 |
| chr20:33035063-33035319 | 0.45612587 | 0.324634695 | 0.61484476 |
| chr10:84552548-84552785 | -0.437931978 | 0.324662366 | 0.614876549 |
| chr13:46973719-46974341 | -0.339337725 | 0.324668925 | 0.614876549 |
| chr1:212791325-212792423 | -0.324908533 | 0.324675554 | 0.614876549 |
| chr15:32029465-32029685 | 0.439958594 | 0.324687518 | 0.61488401 |
| chr3:177810183-177810447 | -0.494240415 | 0.324730832 | 0.614920441 |
| chr4:70521788-70521998 | -0.494240415 | 0.324730832 | 0.614920441 |
| chr5:131589572-131589777 | -0.494240415 | 0.324730832 | 0.614920441 |
| chr21:33915375-33916095 | 0.326289354 | 0.324745276 | 0.614932595 |
| chr12:64758715-64759852 | 0.314635345 | 0.324792719 | 0.614964121 |
| chr20:60868621-60868958 | -0.420058815 | 0.324795265 | 0.614964121 |
| chr16:56370879-56371276 | -0.373221294 | 0.324795545 | 0.614964121 |
| chrX:116240110-116240310 | 0.503443334 | 0.324798601 | 0.614964121 |
| chr5:62752739-62753314 | -0.344721915 | 0.324809757 | 0.614964121 |
| chr11:12412322-12412602 | -0.453252059 | 0.324810081 | 0.614964121 |
| chr6:154675615-154675970 | -0.344592481 | 0.32482936 | 0.614985427 |
| chr6:3933080-3933394 | 0.441523293 | 0.324844173 | 0.614986921 |
| chr8:142912363-142912584 | 0.452827556 | 0.32485291 | 0.614986921 |
| chr8:110733984-110734187 | 0.464449034 | 0.324856183 | 0.614986921 |
| chr3:154616461-154616661 | 0.451381572 | 0.324862254 | 0.614986921 |
| chr7:143362165-143362700 | -0.385096043 | 0.324877448 | 0.614993577 |
| chr20:4760638-4760838 | 0.536059396 | 0.324895578 | 0.614993577 |
| chr7:34586658-34586858 | 0.478218737 | 0.324899231 | 0.614993577 |
| chr13:114132542-114132745 | 0.485276425 | 0.324901546 | 0.614993577 |
| chr3:37741699-37741907 | 0.466762567 | 0.324905902 | 0.614993577 |
| chr2:152743115-152743315 | -0.502622498 | 0.324942292 | 0.615047263 |
| chr5:90558373-90559244 | -0.311449836 | 0.324951753 | 0.615049978 |
| chr6:26103594-26104347 | 0.327467085 | 0.32497567 | 0.61506683 |
| chr4:129747810-129748010 | 0.481132103 | 0.324981249 | 0.61506683 |
| chr5:217733-218965 | 0.315125169 | 0.324984738 | 0.61506683 |
| chr18:267452-268866 | -0.316669705 | 0.325010089 | 0.615099615 |
| chr6:47595873-47596124 | -0.469206708 | 0.325034668 | 0.615130939 |
| chrX:149512301-149512719 | 0.346284406 | 0.325059864 | 0.615151248 |
| chr19:39846116-39846689 | 0.348280926 | 0.325061456 | 0.615151248 |
| chr16:27925751-27925960 | -0.487960827 | 0.325097837 | 0.615204902 |
| chr12:89814483-89814741 | -0.547511303 | 0.325114368 | 0.615215842 |
| chr5:131005991-131006394 | 0.381294131 | 0.325119677 | 0.615215842 |
| chr1:199551989-199552274 | -0.410936141 | 0.325149953 | 0.615257939 |
| chr5:61408545-61409076 | -0.321533459 | 0.325170557 | 0.61528173 |
| chr10:88990247-88991488 | -0.321651947 | 0.325188656 | 0.615287336 |
| chr10:37146439-37146787 | -0.421982361 | 0.32519197 | 0.615287336 |
| chr6:152100064-152100264 | 0.488361542 | 0.325198567 | 0.615287336 |
| chr3:149028521-149028721 | 0.547125904 | 0.325210464 | 0.615287336 |

| | | | |
|---|---|---|---|
| chr14:76761218-76761994 | 0.33663234 | 0.325216338 | 0.615287336 |
| chr7:140923571-140923941 | -0.392311687 | 0.325229058 | 0.615287336 |
| chr15:64811810-64812109 | -0.434525282 | 0.32522973 | 0.615287336 |
| chr4:57055859-57056141 | -0.5160988 | 0.325271281 | 0.61535075 |
| chr7:881494-881874 | 0.355416251 | 0.32528918 | 0.615357683 |
| chr16:89759936-89760228 | 0.427540519 | 0.325291008 | 0.615357683 |
| chr20:54881669-54882339 | 0.319195701 | 0.325345027 | 0.615444677 |
| chr14:92759369-92759569 | 0.537115001 | 0.325380295 | 0.615496197 |
| chr2:191020154-191020873 | -0.31702341 | 0.32539005 | 0.615499455 |
| chr10:125160835-125161721 | 0.324614926 | 0.32541594 | 0.615533231 |
| chr22:43923665-43923901 | 0.452037671 | 0.325434271 | 0.615543481 |
| chr17:16991480-16991695 | 0.447320935 | 0.325437425 | 0.615543481 |
| chr9:89603733-89604050 | 0.364675148 | 0.325488137 | 0.615615671 |
| chr11:33773711-33773981 | -0.443652694 | 0.325496408 | 0.615615671 |
| chr11:2700608-2700916 | 0.404202515 | 0.325505424 | 0.615615671 |
| chr3:112519373-112519762 | 0.371755878 | 0.32550773 | 0.615615671 |
| chr14:77087997-77088903 | 0.318638281 | 0.325552232 | 0.615684637 |
| chr1:223531828-223532068 | 0.447017326 | 0.325567723 | 0.615698738 |
| chr13:35255390-35255694 | -0.487463496 | 0.32559534 | 0.615724925 |
| chr4:7719428-7719813 | 0.436336394 | 0.325597802 | 0.615724925 |
| chr10:34049969-34050169 | -0.521886736 | 0.325609411 | 0.615724925 |
| chr8:113067834-113068059 | 0.486975939 | 0.325613714 | 0.615724925 |
| chr6:63078540-63078763 | -0.473452977 | 0.325668994 | 0.615814261 |
| chr22:45974498-45974732 | -0.444433333 | 0.325722499 | 0.615878553 |
| chr9:133207606-133208116 | -0.422878511 | 0.325732257 | 0.615878553 |
| chr12:2003941-2004760 | -0.335952333 | 0.325741835 | 0.615878553 |
| chr15:73944902-73945203 | -0.424765428 | 0.325742801 | 0.615878553 |
| chr6:31137533-31137953 | -0.397709413 | 0.32574686 | 0.615878553 |
| chr10:43984753-43984955 | 0.489075547 | 0.325751222 | 0.615878553 |
| chr10:26043149-26043653 | -0.366503318 | 0.325773466 | 0.615905411 |
| chr9:97983562-97983817 | 0.410214492 | 0.325783911 | 0.615909962 |
| chr4:115852507-115852763 | -0.411630923 | 0.32580798 | 0.615912846 |
| chr3:165321439-165321761 | 0.491180772 | 0.325808049 | 0.615912846 |
| chr13:111344781-111345154 | 0.372482569 | 0.32580975 | 0.615912846 |
| chr16:25066716-25067665 | 0.325784909 | 0.32581759 | 0.615912846 |
| chrX:112867530-112868066 | -0.365053445 | 0.325833054 | 0.615918163 |
| chr10:71974578-71974867 | -0.432953488 | 0.325846957 | 0.615918163 |
| chr15:66903706-66904000 | -0.435789011 | 0.325851559 | 0.615918163 |
| chr8:80579991-80580198 | -0.464093502 | 0.325852556 | 0.615918163 |
| chr6:83511844-83512230 | -0.365490743 | 0.325897233 | 0.615981629 |
| chr10:90579645-90580031 | 0.424306235 | 0.325902212 | 0.615981629 |
| chr9:130712430-130713331 | 0.317247305 | 0.325937951 | 0.616033984 |
| chr1:117562986-117563441 | -0.364625688 | 0.325958936 | 0.616058448 |
| chr14:66791609-66791809 | 0.595835609 | 0.326009972 | 0.616139709 |
| chr12:49060207-49061281 | 0.31915808 | 0.326031789 | 0.616165744 |
| chr15:77910380-77910976 | 0.337117159 | 0.326051528 | 0.616187849 |
| chr7:150778514-150779045 | -0.375171318 | 0.326069004 | 0.616200432 |
| chr10:1142651-1142981 | -0.436677614 | 0.326081975 | 0.616200432 |
| chr18:46468551-46468751 | -0.479645677 | 0.326082312 | 0.616200432 |
| chr5:126579541-126579782 | 0.514954761 | 0.32610245 | 0.616223288 |

| | | | |
|---|---|---|---|
| chr15:88878599-88879067 | 0.346407349 | 0.326117611 | 0.61623674 |
| chr1:66543519-66543781 | -0.438596879 | 0.326148575 | 0.616280052 |
| chr16:69424217-69425174 | -0.321246653 | 0.32615771 | 0.616282116 |
| chr20:43539838-43540038 | -0.551845905 | 0.326169441 | 0.616289083 |
| chr5:151264211-151264553 | -0.388994276 | 0.326183825 | 0.61629025 |
| chr2:94994820-94995020 | 0.49797601 | 0.326186145 | 0.61629025 |
| chr7:105024841-105025041 | 0.483883962 | 0.326281799 | 0.616441228 |
| chr6:136317041-136317339 | -0.395902614 | 0.326282144 | 0.616441228 |
| chr8:138951516-138951723 | -0.484950693 | 0.326299357 | 0.616458548 |
| chr4:124904002-124904930 | -0.328733493 | 0.326330818 | 0.616502785 |
| chr2:64997019-64997240 | -0.463550903 | 0.32640826 | 0.616633884 |
| chr15:45586938-45588010 | 0.317494249 | 0.326416692 | 0.61663461 |
| chr17:60299009-60299218 | 0.356432333 | 0.326443592 | 0.616670222 |
| chr10:12720323-12720675 | 0.37285318 | 0.326460674 | 0.616673351 |
| chr10:84783294-84783654 | -0.382400184 | 0.326461344 | 0.616673351 |
| chr12:89352367-89353095 | -0.319970696 | 0.326493663 | 0.616719196 |
| chr3:12556387-12557595 | 0.312823368 | 0.32655538 | 0.616820567 |
| chr17:60441059-60441637 | 0.318763844 | 0.326570306 | 0.616827598 |
| chr10:56360908-56361599 | -0.316040004 | 0.326575202 | 0.616827598 |
| chr10:48289721-48289958 | -0.440297497 | 0.326611922 | 0.616881746 |
| chr3:82049831-82050065 | -0.545585904 | 0.32662436 | 0.616890031 |
| chr5:139197899-139198640 | -0.327990762 | 0.326651233 | 0.61692558 |
| chr9:34139792-34139994 | -0.53476364 | 0.326678564 | 0.616951804 |
| chr8:22668799-22669311 | 0.351779748 | 0.326689663 | 0.616951804 |
| chr7:148988063-148988298 | -0.384776777 | 0.326691834 | 0.616951804 |
| chr17:35568571-35568795 | 0.449527225 | 0.326697326 | 0.616951804 |
| chr1:177872439-177873071 | 0.354918317 | 0.326757223 | 0.61704971 |
| chr6:78810039-78810475 | 0.362815292 | 0.326839074 | 0.617189065 |
| chr1:155085757-155085974 | 0.370636728 | 0.326858577 | 0.617210683 |
| chr12:6961170-6963330 | -0.308756608 | 0.326889161 | 0.617253223 |
| chr4:17511658-17512355 | 0.350001827 | 0.326903384 | 0.617264868 |
| chr17:45248277-45248494 | 0.399422055 | 0.326926265 | 0.617292861 |
| chr6:157823010-157823749 | 0.346626133 | 0.326938978 | 0.617301654 |
| chrX:104804085-104804298 | 0.497181549 | 0.327010715 | 0.617420259 |
| chr3:148955585-148955832 | 0.423534366 | 0.327041521 | 0.617420259 |
| chr3:79885403-79885892 | 0.34576623 | 0.327046565 | 0.617420259 |
| chr10:110729410-110729882 | -0.38864401 | 0.327056695 | 0.617420259 |
| chr8:96315990-96316220 | 0.396463771 | 0.327068111 | 0.617420259 |
| chr2:202237758-202238965 | -0.315357139 | 0.327071157 | 0.617420259 |
| chr16:73171345-73171778 | -0.379240237 | 0.32707474 | 0.617420259 |
| chr6:43476851-43477051 | 0.475990879 | 0.327076669 | 0.617420259 |
| chr5:19880545-19881332 | -0.335296085 | 0.327078002 | 0.617420259 |
| chrX:4379191-4379391 | 0.504838525 | 0.327082375 | 0.617420259 |
| chr21:9346289-9346531 | 0.460324561 | 0.327148773 | 0.617530382 |
| chr8:121485348-121485634 | 0.452579333 | 0.327189094 | 0.61759096 |
| chr19:58451123-58451987 | 0.319499372 | 0.327196985 | 0.61759096 |
| chr16:2205074-2205918 | 0.324708982 | 0.327214327 | 0.617608478 |
| chr4:101730544-101730836 | -0.399060097 | 0.32724785 | 0.617639183 |
| chr11:1763869-1764421 | -0.361880545 | 0.327259598 | 0.617639183 |
| chrX:85358710-85359549 | -0.359877222 | 0.327263359 | 0.617639183 |

| | | | |
|---|---|---|---|
| chr9:84947372-84947573 | 0.490847126 | 0.327270576 | 0.617639183 |
| chr6:100657808-100658099 | -0.4601308 | 0.327277826 | 0.617639183 |
| chr3:195105427-195105830 | 0.394971706 | 0.32727896 | 0.617639183 |
| chr17:7177452-7177672 | 0.454278417 | 0.327317638 | 0.617691019 |
| chr1:114368113-114368385 | 0.356871133 | 0.327324425 | 0.617691019 |
| chr2:30807263-30807678 | -0.337371476 | 0.327330612 | 0.617691019 |
| chr11:71928089-71929421 | -0.315828921 | 0.327342912 | 0.617699018 |
| chr17:60008427-60008690 | -0.357003659 | 0.327373827 | 0.617730859 |
| chr3:98817525-98817985 | -0.3765508 | 0.32737591 | 0.617730859 |
| chrX:107775560-107776252 | 0.330279324 | 0.327386118 | 0.617734909 |
| chr6:49543175-49543474 | -0.418575643 | 0.327405471 | 0.617756211 |
| chrX:102832687-102832933 | -0.473343697 | 0.327490929 | 0.61790224 |
| chr8:111236343-111236673 | -0.35005249 | 0.327517548 | 0.617919647 |
| chr11:95108217-95108495 | -0.423764088 | 0.32752041 | 0.617919647 |
| chr7:128035780-128035993 | -0.442933638 | 0.327524348 | 0.617919647 |
| chr15:75036216-75036742 | 0.344341233 | 0.327536663 | 0.617927665 |
| chr12:108403890-108404090 | 0.538228224 | 0.32756758 | 0.617940128 |
| chr12:1566511-1566753 | -0.487772536 | 0.327569778 | 0.617940128 |
| chr14:96362751-96363934 | 0.313770548 | 0.327569953 | 0.617940128 |
| chr3:74267617-74267910 | -0.412142696 | 0.327578917 | 0.617940128 |
| chr9:31848562-31849180 | -0.397843048 | 0.327583593 | 0.617940128 |
| chr3:194985070-194985469 | 0.390248868 | 0.327597429 | 0.617951014 |
| chr3:114173247-114173474 | -0.497565116 | 0.327631193 | 0.617999489 |
| chr4:24973496-24974251 | 0.347173156 | 0.327660062 | 0.61803873 |
| chr3:63228193-63228442 | 0.370722975 | 0.327681314 | 0.6180636 |
| chr3:129633897-129634471 | 0.337955649 | 0.327716693 | 0.618115115 |
| chr4:85147571-85147797 | -0.477981535 | 0.32773278 | 0.618122683 |
| chr10:119267376-119267599 | 0.477313753 | 0.32773684 | 0.618122683 |
| chr12:71663021-71664677 | -0.306040659 | 0.327766938 | 0.618158993 |
| chr6:149484693-149484932 | -0.416752775 | 0.327775745 | 0.618158993 |
| chr19:2739920-2740376 | 0.355677203 | 0.327783345 | 0.618158993 |
| chr5:122774579-122775716 | -0.313004628 | 0.327792212 | 0.618158993 |
| chr6:75601584-75603012 | -0.312587877 | 0.32779643 | 0.618158993 |
| chr3:149334500-149334700 | 0.496446503 | 0.327822934 | 0.61818078 |
| chr3:148911713-148911932 | -0.472983071 | 0.327824119 | 0.61818078 |
| chr12:31362885-31363140 | -0.470232441 | 0.327847046 | 0.6182088 |
| chr5:55928236-55928518 | -0.435139148 | 0.327862204 | 0.618214678 |
| chr5:157348133-157348333 | -0.527255118 | 0.3278663 | 0.618214678 |
| chr5:63923922-63924213 | 0.394357092 | 0.32787941 | 0.618224184 |
| chr7:101702492-101702721 | -0.441100314 | 0.327909652 | 0.618262319 |
| chr3:154120807-154122473 | -0.307611167 | 0.327915773 | 0.618262319 |
| chr3:85572501-85572701 | -0.503909719 | 0.327925634 | 0.618264925 |
| chr10:103241280-103242067 | 0.32321225 | 0.327933294 | 0.618264925 |
| chr14:60550566-60550858 | 0.431478211 | 0.327966091 | 0.618311546 |
| chr21:33324284-33325672 | 0.316031077 | 0.327996043 | 0.618352799 |
| chr8:25183596-25183908 | -0.383329024 | 0.328005183 | 0.618354816 |
| chr1:197763379-197763657 | 0.407306786 | 0.328031353 | 0.618374886 |
| chr17:74646746-74646946 | -0.541186792 | 0.32803197 | 0.618374886 |
| chr12:54159154-54159536 | 0.357073401 | 0.328058363 | 0.618409426 |
| chr15:44296708-44296928 | 0.45030435 | 0.328070577 | 0.618417234 |

| | | | |
|---|---|---|---|
| chr4:141220682-141221681 | -0.321787716 | 0.328078656 | 0.618417249 |
| chr11:71589264-71589509 | -0.455368234 | 0.328121646 | 0.618470719 |
| chr14:77457121-77458231 | 0.309161247 | 0.328123166 | 0.618470719 |
| chr10:80589609-80589840 | -0.483661348 | 0.328203753 | 0.618578106 |
| chr15:41804399-41804657 | 0.454764191 | 0.328204358 | 0.618578106 |
| chr18:12934552-12934804 | 0.454764191 | 0.328204358 | 0.618578106 |
| chr17:61580061-61580664 | 0.313237016 | 0.328231072 | 0.618613239 |
| chr4:24763396-24764593 | 0.320427367 | 0.328265491 | 0.61866289 |
| chr1:11541321-11541524 | -0.495328084 | 0.328290274 | 0.61869438 |
| chr19:3035591-3035933 | 0.349292692 | 0.328310264 | 0.618714054 |
| chr14:99574131-99574682 | 0.338286245 | 0.328330078 | 0.618714054 |
| chr3:134350741-134351345 | -0.323409187 | 0.328339746 | 0.618714054 |
| chr7:39519910-39520463 | -0.341091448 | 0.328339797 | 0.618714054 |
| chr2:210577513-210578034 | -0.388461873 | 0.328341088 | 0.618714054 |
| chr15:82530522-82531321 | 0.322912024 | 0.328361936 | 0.61872664 |
| chr3:163951283-163951691 | -0.402428316 | 0.32836451 | 0.61872664 |
| chr10:29918987-29919187 | -0.531278648 | 0.328371993 | 0.61872664 |
| chr7:34740211-34740411 | 0.493325431 | 0.328412334 | 0.618749042 |
| chr12:51269300-51270585 | 0.319118041 | 0.328415102 | 0.618749042 |
| chr7:100089177-100089464 | -0.433710358 | 0.328418062 | 0.618749042 |
| chr7:77331948-77332191 | -0.445224314 | 0.328426691 | 0.618749042 |
| chr5:160188149-160188971 | -0.337203491 | 0.328429637 | 0.618749042 |
| chr7:43990154-43990433 | 0.405394721 | 0.328432334 | 0.618749042 |
| chr15:65132989-65133421 | 0.352039594 | 0.328446492 | 0.6187605 |
| chr5:72525853-72526101 | -0.494869702 | 0.328463663 | 0.618777636 |
| chr5:174338332-174338560 | -0.477347134 | 0.328482152 | 0.618796287 |
| chr18:43034628-43035238 | -0.329792249 | 0.328489715 | 0.618796287 |
| chr1:36155712-36156285 | 0.327942302 | 0.328519866 | 0.6188077 |
| chr4:102868512-102869755 | -0.309972225 | 0.328523329 | 0.6188077 |
| chr3:19866590-19867098 | 0.36245064 | 0.328523452 | 0.6188077 |
| chr12:124914007-124914265 | 0.368621788 | 0.328528078 | 0.6188077 |
| chr4:110196306-110197430 | -0.317974883 | 0.328550349 | 0.61882345 |
| chr10:70902812-70903037 | 0.451396805 | 0.328564013 | 0.61882345 |
| chr2:1897202-1897476 | -0.465801411 | 0.328571986 | 0.61882345 |
| chr14:58455591-58456237 | 0.336106521 | 0.328572152 | 0.61882345 |
| chr4:103019368-103020240 | -0.317862925 | 0.32858227 | 0.61882345 |
| chr9:136226091-136226408 | 0.387476153 | 0.328584898 | 0.61882345 |
| chr8:81780514-81781073 | -0.347036418 | 0.328599971 | 0.618836627 |
| chr17:42981029-42981515 | 0.362789859 | 0.328647349 | 0.61891064 |
| chr16:2423167-2423417 | 0.419609916 | 0.328675271 | 0.618948009 |
| chr8:107068653-107068975 | 0.359806787 | 0.328687521 | 0.618955865 |
| chr11:68946652-68946983 | -0.407326532 | 0.328697119 | 0.618958727 |
| chr11:27506234-27507218 | -0.311413346 | 0.328714377 | 0.618976014 |
| chr20:59560621-59560873 | 0.386821495 | 0.328741201 | 0.619011313 |
| chr2:8692863-8693613 | -0.345488395 | 0.328754357 | 0.619020872 |
| chr7:5513297-5515000 | -0.307050529 | 0.328768268 | 0.619031853 |
| chr16:56519489-56520285 | 0.330512509 | 0.32881364 | 0.619093963 |
| chr6:130843515-130843898 | -0.370111478 | 0.328823349 | 0.619093963 |
| chr12:54370240-54370610 | 0.359859395 | 0.328826686 | 0.619093963 |
| chr2:55412156-55412489 | -0.471908483 | 0.328833574 | 0.619093963 |

| | | | |
|---|---|---|---|
| chr20:36717767-36717998 | -0.498670791 | 0.328862678 | 0.61911852 |
| chr14:89739971-89740708 | 0.342957687 | 0.328862778 | 0.61911852 |
| chr10:93511359-93511767 | -0.408319266 | 0.32890256 | 0.619175436 |
| chr18:47174739-47174974 | 0.437815058 | 0.328909172 | 0.619175436 |
| chr12:116697393-116698455 | 0.323739115 | 0.328940791 | 0.619206062 |
| chr6:104859228-104860184 | -0.317636288 | 0.328947472 | 0.619206062 |
| chr3:165526486-165526717 | -0.490117678 | 0.328952803 | 0.619206062 |
| chr15:72796843-72797405 | 0.359035056 | 0.328961221 | 0.619206062 |
| chr5:97296943-97297665 | -0.336299813 | 0.32897623 | 0.619206062 |
| chr1:200640081-200640297 | -0.436914389 | 0.328985183 | 0.619206062 |
| chr1:58620753-58621314 | 0.337590231 | 0.328987762 | 0.619206062 |
| chr5:96344794-96345074 | -0.393654978 | 0.328990092 | 0.619206062 |
| chrX:110162915-110163132 | 0.43042331 | 0.329003545 | 0.619216174 |
| chr20:54141679-54141961 | 0.333388957 | 0.329029285 | 0.619223712 |
| chr17:60210366-60210619 | 0.384257819 | 0.329030033 | 0.619223712 |
| chr10:110431786-110431986 | -0.48629707 | 0.329046667 | 0.619223712 |
| chr12:11666203-11666458 | -0.48629707 | 0.329046667 | 0.619223712 |
| chr10:26781214-26781422 | -0.489781793 | 0.329053078 | 0.619223712 |
| chr2:165793496-165794919 | -0.308941599 | 0.329066898 | 0.619223712 |
| chr13:43258805-43259077 | 0.443972896 | 0.329068037 | 0.619223712 |
| chr2:162615425-162615665 | -0.487670956 | 0.329079326 | 0.619223712 |
| chr13:19116498-19116787 | 0.455663576 | 0.329080285 | 0.619223712 |
| chr3:186783145-186784529 | -0.31588217 | 0.329101289 | 0.619244489 |
| chr3:49673503-49674715 | 0.312902823 | 0.32910811 | 0.619244489 |
| chr11:129923304-129923565 | -0.439193746 | 0.329119935 | 0.619244489 |
| chr3:187862493-187862693 | -0.525688216 | 0.329125713 | 0.619244489 |
| chr3:113897284-113897639 | -0.360149935 | 0.329131735 | 0.619244489 |
| chr16:1679904-1680109 | 0.43314304 | 0.329143647 | 0.619251695 |
| chr3:158261634-158261948 | -0.431006207 | 0.329187795 | 0.619319549 |
| chr17:7842139-7842665 | -0.337843327 | 0.329211323 | 0.61934387 |
| chr17:60628514-60628751 | 0.349180005 | 0.32922086 | 0.61934387 |
| chr12:72346319-72346697 | -0.364565533 | 0.329224972 | 0.61934387 |
| chr1:40625139-40625418 | 0.403569648 | 0.32923593 | 0.619349278 |
| chr8:22605188-22605499 | 0.428904633 | 0.32924827 | 0.619357286 |
| chr4:153684898-153685098 | 0.509596461 | 0.329267827 | 0.619372773 |
| chr19:1897603-1897963 | -0.402226949 | 0.32927267 | 0.619372773 |
| chr6:30907757-30908520 | -0.324010913 | 0.329307399 | 0.619422892 |
| chr1:199010827-199011085 | -0.459714789 | 0.329317373 | 0.619426447 |
| chr4:145172012-145172250 | -0.475560473 | 0.329326407 | 0.619428233 |
| chr19:47047800-47048275 | 0.344966382 | 0.329359442 | 0.619462522 |
| chr10:11881426-11881626 | -0.459520451 | 0.329367024 | 0.619462522 |
| chr8:141942176-141942432 | 0.393818036 | 0.329378091 | 0.619462522 |
| chr5:170017872-170018132 | 0.461933586 | 0.329380874 | 0.619462522 |
| chr14:49892295-49893774 | 0.308497932 | 0.329386434 | 0.619462522 |
| chr17:15650881-15652085 | 0.312976274 | 0.329397534 | 0.619462522 |
| chrX:133792525-133792997 | 0.387456205 | 0.329401229 | 0.619462522 |
| chrX:53081831-53082578 | 0.319138831 | 0.329431216 | 0.61950371 |
| chr16:53510528-53510728 | 0.480843295 | 0.329497001 | 0.619612213 |
| chr21:45449021-45449484 | -0.395108161 | 0.329516363 | 0.619633415 |
| chr5:114863143-114863457 | -0.567434806 | 0.329536314 | 0.619655725 |

| | | | |
|---|---|---|---|
| chr10:53210005-53210239 | -0.456392216 | 0.32957071 | 0.619705194 |
| chr7:155787259-155787508 | -0.450063768 | 0.329590769 | 0.619727704 |
| chr17:8435399-8436685 | -0.317127675 | 0.329611692 | 0.619751836 |
| chr4:145481550-145483080 | -0.31239241 | 0.329624571 | 0.619760394 |
| chr3:144085981-144086315 | -0.378605975 | 0.329632421 | 0.619760394 |
| chr6:77699888-77700816 | -0.324855839 | 0.329674836 | 0.619824933 |
| chr12:14937780-14938042 | -0.460408334 | 0.329689605 | 0.619837489 |
| chr14:68806698-68807845 | -0.32155984 | 0.32973042 | 0.619899015 |
| chr3:182259649-182259862 | -0.525786375 | 0.329748211 | 0.619917251 |
| chr1:153626941-153627986 | 0.31262581 | 0.329757065 | 0.619918687 |
| chr2:182745129-182745329 | 0.491884772 | 0.32976786 | 0.619923771 |
| chr13:27270540-27271480 | 0.320650607 | 0.329794968 | 0.619927193 |
| chr14:64060140-64060470 | -0.409563519 | 0.329801511 | 0.619927193 |
| chr6:166460945-166461208 | -0.489451675 | 0.32980294 | 0.619927193 |
| chr3:105962537-105962827 | 0.457372473 | 0.329804961 | 0.619927193 |
| chr1:43021665-43021865 | 0.494280615 | 0.329810134 | 0.619927193 |
| chr7:3200549-3200897 | -0.384879905 | 0.329827484 | 0.619944597 |
| chr22:41447502-41448112 | 0.327875569 | 0.329920386 | 0.620104003 |
| chr1:62294297-62294503 | -0.485409981 | 0.329936743 | 0.620107986 |
| chr4:141307200-141307717 | -0.372338004 | 0.329939523 | 0.620107986 |
| chr5:69234415-69235315 | -0.317149189 | 0.32995032 | 0.620107986 |
| chr1:198911374-198911669 | -0.463212943 | 0.329954877 | 0.620107986 |
| chr5:28344670-28345337 | -0.34895326 | 0.329981325 | 0.620136783 |
| chr3:64399240-64399722 | 0.315150273 | 0.329986387 | 0.620136783 |
| chr8:127620444-127620708 | 0.421935194 | 0.329998093 | 0.620143573 |
| chr9:127758857-127759122 | -0.445964113 | 0.330092494 | 0.62030576 |
| chr20:10090637-10090839 | -0.46603056 | 0.330106577 | 0.620317011 |
| chr3:122306259-122306633 | -0.404915165 | 0.330160718 | 0.620403534 |
| chr1:31229021-31229319 | -0.430494337 | 0.3301728 | 0.620411024 |
| chr5:149665472-149665740 | 0.463485149 | 0.330185709 | 0.62041561 |
| chr1:151443886-151444200 | 0.35408672 | 0.330191435 | 0.62041561 |
| chr2:68441740-68442063 | -0.384585024 | 0.330208982 | 0.620433366 |
| chr17:56009678-56010037 | 0.431129189 | 0.330259227 | 0.620512555 |
| chr3:61997158-61997417 | 0.365182651 | 0.330333308 | 0.620610995 |
| chr2:96800983-96801277 | -0.380763883 | 0.330339022 | 0.620610995 |
| chr7:30564270-30564839 | 0.351577376 | 0.330340179 | 0.620610995 |
| chr7:146928377-146929092 | -0.336254723 | 0.330349895 | 0.620610995 |
| chr3:149315783-149316148 | -0.465618065 | 0.330352119 | 0.620610995 |
| chr1:53485555-53485963 | 0.372709497 | 0.330385162 | 0.620629373 |
| chr10:31927892-31929651 | -0.309319168 | 0.330396552 | 0.620629373 |
| chr6:154578262-154578462 | -0.530693162 | 0.33040476 | 0.620629373 |
| chr17:58136763-58136983 | 0.406336356 | 0.330416729 | 0.620629373 |
| chr2:96857594-96858659 | -0.310643965 | 0.330417172 | 0.620629373 |
| chr2:152740793-152741057 | -0.427353244 | 0.330428617 | 0.620629373 |
| chr8:125737728-125738038 | 0.415146599 | 0.330446157 | 0.620629373 |
| chr1:211493370-211493580 | 0.511375873 | 0.330450142 | 0.620629373 |
| chr12:70724548-70724794 | -0.444350975 | 0.330479854 | 0.620629373 |
| chr8:21041096-21041305 | -0.494696877 | 0.330481948 | 0.620629373 |
| chr16:89917941-89918515 | -0.340570734 | 0.330486567 | 0.620629373 |
| chr1:190453603-190453927 | -0.397719145 | 0.330493265 | 0.620629373 |

| | | | |
|---|---|---|---|
| chr6:34791453-34792710 | 0.324212471 | 0.330496521 | 0.620629373 |
| chr17:63564551-63564858 | -0.38842327 | 0.330496644 | 0.620629373 |
| chr5:145825717-145825960 | 0.454545006 | 0.330496824 | 0.620629373 |
| chr19:3500496-3501077 | 0.350261395 | 0.330500998 | 0.620629373 |
| chr2:167551421-167551741 | 0.462608476 | 0.330502292 | 0.620629373 |
| chr3:63906678-63906896 | 0.358089333 | 0.330508486 | 0.620629373 |
| chrX:106953040-106953328 | -0.435816333 | 0.330515799 | 0.620629373 |
| chr7:139645972-139646267 | -0.419564404 | 0.330532908 | 0.62064629 |
| chr1:167267599-167268145 | -0.362189322 | 0.330620069 | 0.620792374 |
| chr11:126440827-126441222 | 0.362046572 | 0.330633583 | 0.620792374 |
| chr10:74266085-74266404 | -0.407569995 | 0.330646333 | 0.620792374 |
| chr17:80005820-80006046 | 0.40012245 | 0.330646425 | 0.620792374 |
| chr13:110561102-110562193 | 0.321965996 | 0.330651217 | 0.620792374 |
| chr22:21567587-21568047 | 0.378241127 | 0.330701524 | 0.620809532 |
| chr11:73760241-73760450 | 0.397780216 | 0.330705215 | 0.620809532 |
| chr7:80623955-80624155 | -0.533754137 | 0.330705503 | 0.620809532 |
| chr12:69633086-69633320 | 0.445512079 | 0.330719191 | 0.620809532 |
| chr15:37291955-37292261 | -0.454006092 | 0.330722894 | 0.620809532 |
| chr1:156337966-156339002 | 0.308321916 | 0.330731734 | 0.620809532 |
| chr2:16506512-16506742 | -0.447886601 | 0.330736067 | 0.620809532 |
| chr16:88803484-88804025 | -0.329146256 | 0.3307414 | 0.620809532 |
| chr9:120936557-120936831 | 0.411850204 | 0.330741859 | 0.620809532 |
| chr8:105933134-105933396 | 0.390163253 | 0.330745561 | 0.620809532 |
| chr4:40840990-40841369 | -0.35878843 | 0.330756818 | 0.620809532 |
| chr7:148535172-148535663 | -0.388187818 | 0.330770179 | 0.620809532 |
| chr14:95035671-95035871 | 0.474216066 | 0.330775479 | 0.620809532 |
| chr5:132190233-132190532 | -0.429497773 | 0.330792466 | 0.620809532 |
| chr14:94824005-94824227 | 0.428972255 | 0.33079303 | 0.620809532 |
| chr22:32202474-32202693 | 0.421212193 | 0.330800182 | 0.620809532 |
| chr1:151510975-151512126 | 0.307675558 | 0.330800965 | 0.620809532 |
| chr1:190633578-190634112 | 0.359388304 | 0.330806197 | 0.620809532 |
| chr13:37837054-37837369 | -0.438555992 | 0.330824976 | 0.620829569 |
| chr5:9401270-9401470 | -0.513041152 | 0.330852761 | 0.620857946 |
| chr9:111975716-111976000 | -0.448895357 | 0.330856303 | 0.620857946 |
| chr7:1046754-1046988 | 0.37360309 | 0.330867577 | 0.620863897 |
| chr9:76219857-76220064 | -0.500869278 | 0.330901837 | 0.620898127 |
| chrX:93434712-93435658 | -0.338252674 | 0.330902026 | 0.620898127 |
| chr4:122576914-122577181 | 0.446944401 | 0.330914386 | 0.620906114 |
| chr9:95073021-95073317 | 0.414684026 | 0.330934292 | 0.620928258 |
| chr11:57476466-57476812 | 0.383738391 | 0.330980139 | 0.620992589 |
| chr16:78099182-78100321 | 0.304740167 | 0.330984787 | 0.620992589 |
| chr14:49240504-49240704 | -0.515381525 | 0.331052354 | 0.62110415 |
| chr3:120159996-120160431 | -0.342609412 | 0.331075136 | 0.621123838 |
| chr6:30647634-30647940 | 0.358502872 | 0.331081992 | 0.621123838 |
| chr1:56578266-56578610 | 0.369640246 | 0.331093868 | 0.621123838 |
| chr4:87838281-87838574 | 0.431199111 | 0.331095273 | 0.621123838 |
| chr8:126820484-126820751 | -0.428370631 | 0.331126274 | 0.621164416 |
| chr2:208265782-208266641 | -0.311447362 | 0.331133118 | 0.621164416 |
| chr10:789028-789332 | 0.359051187 | 0.331157399 | 0.621194756 |
| chr4:110315311-110315780 | -0.424746621 | 0.331240617 | 0.621319478 |

| | | | |
|---|---|---|---|
| chr10:81696879-81697079 | 0.507711181 | 0.331252398 | 0.621319478 |
| chr16:58236397-58236781 | -0.390495781 | 0.331270517 | 0.621319478 |
| chr12:83106890-83107207 | -0.403341603 | 0.331275584 | 0.621319478 |
| chr17:83029985-83030185 | 0.469228485 | 0.331281873 | 0.621319478 |
| chr17:61673086-61673790 | 0.309586072 | 0.33129836 | 0.621319478 |
| chr12:121223732-121223976 | -0.423857602 | 0.331304007 | 0.621319478 |
| chrX:57918997-57919420 | 0.401234766 | 0.331313848 | 0.621319478 |
| chr2:86632744-86633021 | 0.406423592 | 0.331314732 | 0.621319478 |
| chr6:7243318-7243818 | -0.370236926 | 0.331315244 | 0.621319478 |
| chr9:69131424-69131762 | -0.48763639 | 0.331323557 | 0.621319478 |
| chr6:43181849-43182642 | 0.328787982 | 0.331337022 | 0.621319478 |
| chr2:11343616-11345374 | -0.308667425 | 0.331338313 | 0.621319478 |
| chr4:101970606-101970806 | 0.524752042 | 0.33133971 | 0.621319478 |
| chr12:53675703-53676512 | 0.310640193 | 0.331345521 | 0.621319478 |
| chr1:53470128-53470434 | -0.40559762 | 0.331368849 | 0.621348016 |
| chr3:156842926-156843181 | -0.433009502 | 0.331381076 | 0.621355737 |
| chr5:140639056-140640169 | -0.312514322 | 0.331399454 | 0.621359864 |
| chr21:25562093-25562531 | 0.414819515 | 0.331407003 | 0.621359864 |
| chr2:7879125-7879360 | -0.495857596 | 0.331414982 | 0.621359864 |
| chr1:63441631-63442088 | -0.356892261 | 0.331421519 | 0.621359864 |
| chr15:41558581-41558863 | 0.407498704 | 0.331425828 | 0.621359864 |
| chr3:53881625-53882457 | -0.332505521 | 0.331433045 | 0.621359864 |
| chr7:80090156-80090450 | 0.484226295 | 0.331448863 | 0.621359864 |
| chr14:101843742-101844056 | -0.481972995 | 0.331450664 | 0.621359864 |
| chrX:102835627-102836085 | -0.395063791 | 0.331456262 | 0.621359864 |
| chr22:41021521-41022401 | 0.336441273 | 0.331486521 | 0.621376242 |
| chr5:40904740-40905063 | -0.382818914 | 0.331488553 | 0.621376242 |
| chr9:137551432-137552396 | 0.31864888 | 0.331498824 | 0.621376242 |
| chr8:126384338-126384607 | -0.371922969 | 0.331500109 | 0.621376242 |
| chr7:149090177-149091842 | -0.306531363 | 0.331505547 | 0.621376242 |
| chr1:23743749-23743968 | 0.481527906 | 0.331516648 | 0.621381849 |
| chr8:101332748-101333298 | 0.339279725 | 0.331534185 | 0.621399519 |
| chr1:88892482-88893235 | -0.323018206 | 0.331548558 | 0.621411257 |
| chr12:69534470-69534827 | -0.398649907 | 0.331602539 | 0.621484414 |
| chr7:1455233-1455755 | 0.320667977 | 0.331603812 | 0.621484414 |
| chr20:48350684-48350952 | 0.341760526 | 0.331614418 | 0.62148909 |
| chr8:90791632-90792323 | 0.34058849 | 0.331700183 | 0.62163462 |
| chr1:107855969-107856197 | 0.517108789 | 0.331708299 | 0.621634624 |
| chr2:15058137-15058372 | -0.491149572 | 0.33172659 | 0.621653699 |
| chr2:3857027-3857227 | 0.534270457 | 0.331741495 | 0.621660057 |
| chr2:241636626-241637402 | 0.329665011 | 0.33174621 | 0.621660057 |
| chr7:73715534-73716105 | -0.330109412 | 0.331808474 | 0.621761528 |
| chr4:138522119-138522742 | 0.356386887 | 0.331826464 | 0.621774014 |
| chr8:8418646-8419009 | -0.457045219 | 0.331831367 | 0.621774014 |
| chr1:197356746-197357303 | -0.336076598 | 0.331847154 | 0.62178839 |
| chr1:148528706-148529674 | -0.314747746 | 0.331859076 | 0.621790466 |
| chr2:66432881-66433092 | -0.413333957 | 0.331876187 | 0.621790466 |
| chr4:118969341-118969645 | 0.374599876 | 0.331876825 | 0.621790466 |
| chr3:181087170-181087462 | -0.520834182 | 0.331880722 | 0.621790466 |
| chr1:192580489-192580811 | 0.388375517 | 0.33193454 | 0.621876091 |

| | | | |
|---|---|---|---|
| chr13:49792439-49793279 | -0.311983209 | 0.33195503 | 0.621899271 |
| chr18:46658184-46658531 | -0.384345092 | 0.331989266 | 0.621948204 |
| chr8:121144948-121145162 | -0.426075424 | 0.332008979 | 0.621969927 |
| chr7:20785364-20785871 | 0.362468341 | 0.332026664 | 0.62198785 |
| chr10:72082095-72082376 | -0.425760211 | 0.33203925 | 0.621996221 |
| chr5:65035243-65035994 | -0.322551347 | 0.332054964 | 0.621999997 |
| chr19:18161442-18161922 | -0.385513836 | 0.332057502 | 0.621999997 |
| chr1:106487192-106487438 | 0.37317325 | 0.332068305 | 0.622005029 |
| chr21:41680536-41680805 | -0.369466816 | 0.332083352 | 0.622018007 |
| chr17:82874472-82874857 | -0.356237167 | 0.332159366 | 0.622122769 |
| chr7:149460251-149461176 | -0.325963112 | 0.332159609 | 0.622122769 |
| chr1:212367803-212368330 | -0.360354694 | 0.332163641 | 0.622122769 |
| chr5:13831389-13831756 | -0.426631369 | 0.332205481 | 0.622174272 |
| chr20:61329254-61329493 | 0.410615454 | 0.332209968 | 0.622174272 |
| chr20:48070587-48070891 | 0.338645919 | 0.332223466 | 0.622174272 |
| chr17:7835383-7836010 | -0.321756309 | 0.332223619 | 0.622174272 |
| chr5:34438312-34438512 | 0.527400099 | 0.332258185 | 0.622202966 |
| chr6:91276141-91276652 | -0.366323284 | 0.33226086 | 0.622202966 |
| chr13:99416415-99416615 | -0.514981555 | 0.332263303 | 0.622202966 |
| chr7:104848511-104848765 | -0.423990468 | 0.33231754 | 0.622281511 |
| chr8:4837927-4838127 | -0.522457002 | 0.332321489 | 0.622281511 |
| chr7:32957098-32958378 | 0.306905087 | 0.332329651 | 0.622281587 |
| chr17:45316562-45317621 | 0.308606606 | 0.332351302 | 0.622297031 |
| chr6:34546821-34547030 | -0.395303298 | 0.332354142 | 0.622297031 |
| chr14:69101281-69101741 | 0.371606343 | 0.332373738 | 0.622318515 |
| chr6:2596328-2596693 | 0.447433221 | 0.332400584 | 0.62234037 |
| chr9:136866210-136866714 | 0.346689281 | 0.332425998 | 0.62234037 |
| chr3:4744448-4744833 | 0.394233686 | 0.332433137 | 0.62234037 |
| chr8:117663428-117663665 | 0.507943868 | 0.332433521 | 0.62234037 |
| chr1:83704250-83704628 | -0.421929288 | 0.332439224 | 0.62234037 |
| chr1:190124125-190124394 | -0.421929288 | 0.332439224 | 0.62234037 |
| chr21:43318062-43318276 | 0.445927121 | 0.332442266 | 0.62234037 |
| chr2:10752150-10752728 | -0.335246822 | 0.332476081 | 0.622388465 |
| chr2:117856573-117856828 | 0.520146987 | 0.33252745 | 0.622469419 |
| chr1:143184995-143185263 | 0.446224146 | 0.33255687 | 0.622509283 |
| chr4:88178094-88178929 | 0.335988153 | 0.332589847 | 0.622545631 |
| chr22:46335566-46336205 | 0.359302738 | 0.332592538 | 0.622545631 |
| chr1:198314823-198315123 | -0.418480838 | 0.33261123 | 0.622560243 |
| chr8:22440653-22441536 | 0.334195367 | 0.332616594 | 0.622560243 |
| chr16:81329704-81329906 | -0.468318088 | 0.332644345 | 0.622596976 |
| chr6:3126416-3126721 | 0.416434263 | 0.332737584 | 0.622729881 |
| chr16:54062384-54063117 | -0.376176958 | 0.332745035 | 0.622729881 |
| chr16:11213217-11213426 | 0.470175144 | 0.332747961 | 0.622729881 |
| chr3:33113739-33114856 | 0.312817182 | 0.332753507 | 0.622729881 |
| chr3:149042540-149042740 | -0.505844422 | 0.332760446 | 0.622729881 |
| chr15:82943202-82943552 | -0.351142417 | 0.332764118 | 0.622729881 |
| chr11:48042027-48042304 | 0.420992441 | 0.332822401 | 0.622821881 |
| chr20:23349702-23350926 | 0.306971356 | 0.332834018 | 0.622821881 |
| chrX:57910173-57910412 | -0.482018801 | 0.332837728 | 0.622821881 |
| chr4:128552850-128553655 | 0.333646974 | 0.332845793 | 0.622821881 |

| | | | |
|---|---|---|---|
| chr18:80247419-80247800 | 0.364597692 | 0.332864486 | 0.622841648 |
| chr11:12409913-12410158 | -0.469751129 | 0.332892158 | 0.622870674 |
| chr21:46143574-46144226 | 0.338579759 | 0.332896257 | 0.622870674 |
| chr13:80339947-80340581 | -0.34106761 | 0.33291331 | 0.622887371 |
| chr3:186806092-186807067 | -0.319218422 | 0.332941255 | 0.622909973 |
| chr22:23233191-23233526 | -0.461855881 | 0.332941649 | 0.622909973 |
| chr4:105682740-105683452 | 0.350239647 | 0.332957416 | 0.622924261 |
| chr6:154343722-154344335 | -0.312257879 | 0.333041968 | 0.623059205 |
| chr10:5889479-5890384 | 0.307795511 | 0.333045807 | 0.623059205 |
| chr17:78198845-78199349 | 0.328392691 | 0.333055383 | 0.623061575 |
| chr5:107808490-107808766 | -0.445421304 | 0.333064259 | 0.623061575 |
| chr4:78612438-78612725 | -0.418091716 | 0.333071469 | 0.623061575 |
| chr5:150758292-150759416 | -0.307786111 | 0.333097384 | 0.62309484 |
| chr11:60763705-60764031 | -0.392155782 | 0.333109109 | 0.62310156 |
| chr13:28056991-28057253 | 0.454545879 | 0.333117518 | 0.62310208 |
| chr11:83724016-83724410 | 0.379936983 | 0.333137799 | 0.623117966 |
| chr8:60473264-60473557 | -0.388079731 | 0.333142276 | 0.623117966 |
| chr4:155033604-155033820 | -0.518833029 | 0.333157949 | 0.62313207 |
| chr6:71855456-71855689 | -0.466266479 | 0.333183797 | 0.623158438 |
| chr19:47427337-47427564 | 0.394991266 | 0.333194828 | 0.623158438 |
| chr20:25294478-25294756 | -0.395120748 | 0.333196445 | 0.623158438 |
| chr15:100721722-100722301 | 0.335292221 | 0.333210376 | 0.62316928 |
| chr14:68795707-68796342 | 0.320897145 | 0.333225524 | 0.6231824 |
| chr1:96845541-96845818 | -0.407962518 | 0.333269281 | 0.623249019 |
| chr4:145651529-145651729 | 0.539259304 | 0.333286593 | 0.623266184 |
| chr2:9908749-9908964 | -0.510718795 | 0.333325138 | 0.623305989 |
| chr20:25178894-25179483 | 0.347695492 | 0.333332289 | 0.623305989 |
| chr11:119121174-119121937 | 0.310082402 | 0.333335387 | 0.623305989 |
| chr12:102769478-102769871 | -0.414109017 | 0.333340418 | 0.623305989 |
| chr9:22200303-22200723 | -0.429167939 | 0.333386638 | 0.623363964 |
| chr2:48315239-48315783 | -0.331969851 | 0.333387694 | 0.623363964 |
| chr15:85897717-85898030 | 0.439947124 | 0.33339917 | 0.62337021 |
| chr1:9681337-9681549 | -0.543345483 | 0.33345493 | 0.623444431 |
| chr16:72836983-72837183 | 0.464817614 | 0.333455139 | 0.623444431 |
| chr20:36573866-36574097 | 0.409251372 | 0.333494745 | 0.623503266 |
| chr8:80807862-80808184 | 0.366346425 | 0.333535496 | 0.623546165 |
| chr15:63166347-63166556 | -0.493706261 | 0.333536137 | 0.623546165 |
| chr14:60723317-60723884 | 0.328021096 | 0.333542104 | 0.623546165 |
| chr7:75073041-75074146 | -0.322487422 | 0.333569204 | 0.623581613 |
| chr17:79991660-79992653 | 0.31931499 | 0.333587514 | 0.623585771 |
| chr14:73644476-73645326 | 0.32242634 | 0.333597911 | 0.623585771 |
| chr10:32346082-32347479 | -0.307068284 | 0.333598264 | 0.623585771 |
| chr19:40609906-40610258 | 0.336860158 | 0.333603982 | 0.623585771 |
| chr3:61920030-61920266 | 0.372303884 | 0.333629066 | 0.623617445 |
| chr21:31873934-31874245 | 0.425396995 | 0.333678626 | 0.623694345 |
| chr2:75053555-75053802 | 0.425082088 | 0.333695114 | 0.623694345 |
| chr1:33295481-33295898 | 0.380567168 | 0.333716725 | 0.623694345 |
| chr8:143267012-143267642 | 0.327994253 | 0.33372016 | 0.623694345 |
| chr7:88306381-88306612 | -0.527138955 | 0.333731246 | 0.623694345 |
| chr20:47677707-47678227 | 0.306493764 | 0.333732398 | 0.623694345 |

| | | | |
|---|---|---|---|
| chr10:75202769-75203048 | 0.429182947 | 0.333740298 | 0.623694345 |
| chr11:77471627-77472088 | 0.363538493 | 0.333742203 | 0.623694345 |
| chr3:4712203-4712427 | -0.455169787 | 0.333743466 | 0.623694345 |
| chr11:20599798-20600050 | -0.432611857 | 0.33375792 | 0.623706145 |
| chr8:142114743-142114955 | 0.431797434 | 0.333794254 | 0.623746113 |
| chr3:140060465-140060667 | -0.501388811 | 0.333799101 | 0.623746113 |
| chr6:34656807-34657007 | -0.533281891 | 0.333810942 | 0.623746113 |
| chr11:14300811-14301121 | 0.418664751 | 0.33381187 | 0.623746113 |
| chr1:109773659-109774006 | 0.338554992 | 0.333835766 | 0.623756062 |
| chr18:75479124-75479365 | -0.493567504 | 0.333838585 | 0.623756062 |
| chr1:32759437-32759916 | 0.329960809 | 0.333841617 | 0.623756062 |
| chr17:59114624-59114922 | 0.333382604 | 0.333874454 | 0.623792296 |
| chr18:79712111-79712783 | -0.356979704 | 0.333877292 | 0.623792296 |
| chr5:148145921-148146308 | -0.345351656 | 0.333890446 | 0.623801662 |
| chr10:90863012-90863222 | -0.509640973 | 0.333941582 | 0.623880692 |
| chr17:60727663-60728280 | 0.317421431 | 0.333949032 | 0.623880692 |
| chr15:41581174-41581390 | 0.43457818 | 0.333989273 | 0.623940657 |
| chr21:21101888-21102219 | -0.383534625 | 0.334009322 | 0.623962899 |
| chr11:100671540-10067181б | 0.41754143 | 0.334031904 | 0.623989871 |
| chr5:10801952-10802437 | 0.376235214 | 0.334043089 | 0.623995553 |
| chr5:11822705-11822912 | -0.514966485 | 0.334088831 | 0.624065784 |
| chr3:9843120-9844289 | 0.307336304 | 0.334133051 | 0.62411535 |
| chr4:64954208-64954862 | 0.348678277 | 0.334139843 | 0.62411535 |
| chr7:2612444-2612873 | -0.349339175 | 0.33414406 | 0.62411535 |
| chrX:74902799-74903233 | 0.380609222 | 0.334152659 | 0.62411535 |
| chr3:23728833-23729164 | -0.397127856 | 0.334156093 | 0.62411535 |
| chr4:122224320-122224665 | 0.435985579 | 0.334179828 | 0.624144467 |
| chr2:119753340-119753765 | -0.367254624 | 0.33420961 | 0.624172981 |
| chrX:103220509-103220729 | 0.45150046 | 0.334211387 | 0.624172981 |
| chr12:57239130-57239440 | 0.354588832 | 0.334264712 | 0.624255678 |
| chr11:112290047-11229037ε | 0.387135155 | 0.334271994 | 0.624255678 |
| chr9:81070751-81070987 | -0.490756803 | 0.334280108 | 0.624255678 |
| chr8:39989802-39990310 | -0.360881929 | 0.334303258 | 0.624283693 |
| chr1:107011087-107011383 | 0.374068564 | 0.334315258 | 0.624290887 |
| chr11:19919287-19919626 | -0.410440857 | 0.334326037 | 0.624295801 |
| chr7:155297012-155298119 | -0.308358327 | 0.334341403 | 0.62430928 |
| chr1:112491258-112491535 | -0.520676372 | 0.334425061 | 0.624436316 |
| chr8:143281502-143281875 | 0.338269086 | 0.334425735 | 0.624436316 |
| chr12:121990372-12199063C | 0.499094654 | 0.334462603 | 0.624489938 |
| chr14:73251562-73251762 | 0.440957319 | 0.33447437 | 0.624496691 |
| chr1:25543454-25544027 | 0.366014329 | 0.334508466 | 0.624545132 |
| chr9:137236533-137236764 | 0.379857012 | 0.334530441 | 0.624556844 |
| chr16:48385274-48385678 | 0.361360354 | 0.334532418 | 0.624556844 |
| chr5:61055536-61055760 | -0.453252793 | 0.334539192 | 0.624556844 |
| chr3:53800123-53800351 | -0.501494893 | 0.3345512 | 0.624559739 |
| chr2:85418341-85419025 | -0.318983523 | 0.334560288 | 0.624559739 |
| chr1:43536266-43536821 | 0.33625141 | 0.334565196 | 0.624559739 |
| chr5:112339059-112339322 | 0.431367708 | 0.334574198 | 0.624561327 |
| chrX:48026605-48026987 | 0.374273076 | 0.334590963 | 0.624573677 |
| chr22:29267646-29268771 | 0.310367953 | 0.334597117 | 0.624573677 |

| | | | |
|---|---|---|---|
| chr15:75786332-75786569 | 0.450407987 | 0.334610114 | 0.624582722 |
| chr10:97400857-97401795 | -0.311859334 | 0.334625555 | 0.624596328 |
| chr12:64610056-64611276 | 0.312106653 | 0.334647244 | 0.624621595 |
| chr8:105908303-105908673 | -0.368917648 | 0.334692725 | 0.624676645 |
| chr9:68500451-68501369 | -0.323449704 | 0.334693043 | 0.624676645 |
| chr3:30596183-30596506 | -0.429008441 | 0.334716346 | 0.624704922 |
| chr5:163564398-163564610 | -0.451614803 | 0.334814266 | 0.624872456 |
| chrX:31423952-31424389 | -0.409071431 | 0.334900562 | 0.625018288 |
| chr14:55519495-55519806 | 0.370843943 | 0.334919665 | 0.625038716 |
| chr20:2663839-2664453 | -0.359692893 | 0.334958479 | 0.625095926 |
| chr11:93850174-93850683 | -0.38029357 | 0.334997961 | 0.625154382 |
| chr11:93570809-93571121 | -0.456817893 | 0.33501653 | 0.625167055 |
| chr3:25789773-25790583 | 0.311394531 | 0.33502107 | 0.625167055 |
| chr13:36571275-36571636 | -0.431442567 | 0.335074316 | 0.625251187 |
| chr16:21520118-21520724 | 0.324032125 | 0.335088963 | 0.625255037 |
| chr2:152912119-152912461 | -0.423955742 | 0.3350927 | 0.625255037 |
| chr1:89762999-89763359 | -0.385213262 | 0.335143153 | 0.62533288 |
| chr16:1959236-1959632 | 0.348022719 | 0.335150741 | 0.62533288 |
| chr8:95313265-95313525 | 0.421236585 | 0.335169649 | 0.625352932 |
| chr12:123808277-123808499 | -0.45667806 | 0.335184584 | 0.625365569 |
| chr15:31215426-31215774 | 0.354425476 | 0.335210273 | 0.625398271 |
| chr16:50530734-50530997 | -0.501706009 | 0.335279752 | 0.625484666 |
| chr2:86426340-86426577 | -0.497245973 | 0.335280545 | 0.625484666 |
| chr10:110471622-110471972 | -0.434467849 | 0.33528107 | 0.625484666 |
| chr4:73187466-73187721 | -0.46593834 | 0.335307799 | 0.625519299 |
| chr9:128275571-128276798 | 0.307349402 | 0.335317137 | 0.62552149 |
| chr21:31731146-31732734 | 0.305321037 | 0.335329054 | 0.625521719 |
| chr5:130109241-130109658 | 0.422201809 | 0.335333588 | 0.625521719 |
| chr14:61196828-61197195 | 0.386403404 | 0.335350089 | 0.625537271 |
| chrX:125826896-125827096 | 0.4923218 | 0.335383564 | 0.625575118 |
| chr19:46529567-46530383 | 0.335973523 | 0.335386707 | 0.625575118 |
| chrX:43949694-43949894 | -0.538033556 | 0.335410368 | 0.625604022 |
| chrX:130024119-130024560 | 0.347638752 | 0.335444807 | 0.625653027 |
| chr11:108467035-108467732 | -0.315197089 | 0.335461475 | 0.625655201 |
| chr18:12327734-12328296 | -0.349905294 | 0.335462304 | 0.625655201 |
| chr3:171736519-171736787 | -0.474044759 | 0.335482648 | 0.625677915 |
| chr4:54384623-54385034 | -0.391729588 | 0.335505349 | 0.625699768 |
| chr12:127108691-127108967 | -0.445627824 | 0.335510697 | 0.625699768 |
| chr5:36305770-36306091 | 0.415060509 | 0.335522779 | 0.625700571 |
| chr12:47234707-47235041 | -0.425380685 | 0.335530277 | 0.625700571 |
| chr2:134460488-134460846 | 0.355494773 | 0.335538806 | 0.625700571 |
| chr18:63025207-63025689 | -0.377393478 | 0.335543793 | 0.625700571 |
| chr11:133968423-133968651 | 0.507655064 | 0.335584033 | 0.625757579 |
| chr1:196723188-196724243 | -0.313422269 | 0.335593265 | 0.625757579 |
| chr1:37537371-37537736 | -0.427504553 | 0.335598865 | 0.625757579 |
| chr3:161104462-161105540 | -0.310226485 | 0.335637801 | 0.625813771 |
| chr6:17086274-17086829 | -0.364438559 | 0.335652166 | 0.625813771 |
| chr3:21894558-21894778 | 0.488349468 | 0.335653504 | 0.625813771 |
| chr18:35040976-35041777 | -0.318104577 | 0.335683515 | 0.625854497 |
| chr1:52843646-52844175 | -0.370380947 | 0.335697125 | 0.625864643 |

| | | | |
|---|---|---|---|
| chr8:143431405-143431700 | 0.363561723 | 0.335707311 | 0.625868404 |
| chr20:50587970-50588408 | 0.346939467 | 0.335746393 | 0.625926038 |
| chr16:87560483-87561285 | 0.313168563 | 0.335779452 | 0.625972438 |
| chr11:77473473-77474469 | 0.318810555 | 0.335821734 | 0.62603603 |
| chr5:151051673-151052015 | -0.364583243 | 0.335840305 | 0.626055418 |
| chr8:124726982-124728283 | 0.309706947 | 0.335858399 | 0.626067415 |
| chr11:320271-320901 | 0.36785958 | 0.335869425 | 0.626067415 |
| chr6:41456337-41457302 | -0.317946738 | 0.335871253 | 0.626067415 |
| chr15:48085097-48085383 | -0.42075469 | 0.335879629 | 0.626067797 |
| chr6:10694262-10695224 | -0.312559807 | 0.335895336 | 0.626069035 |
| chr1:204806909-204807749 | -0.312321077 | 0.335896635 | 0.626069035 |
| chr19:32691565-32691985 | 0.34855811 | 0.335935257 | 0.626125791 |
| chr3:73681762-73681982 | 0.472348498 | 0.335943852 | 0.626126581 |
| chr3:160003609-160003953 | 0.367983381 | 0.335952426 | 0.62612733 |
| chr11:65817959-65818627 | 0.337199399 | 0.33596194 | 0.626129831 |
| chr1:161473258-161473729 | -0.385215171 | 0.335972795 | 0.626129885 |
| chr10:91551911-91552131 | 0.428446691 | 0.335985666 | 0.626129885 |
| chr2:121897634-121897901 | 0.413473327 | 0.335986484 | 0.626129885 |
| chr17:57605562-57605887 | 0.400554415 | 0.335997161 | 0.626134554 |
| chr9:112793246-112793567 | 0.392727052 | 0.336007208 | 0.626138048 |
| chr15:76336360-76336838 | 0.320714814 | 0.336037185 | 0.626167849 |
| chrX:108091063-108092396 | -0.303253445 | 0.336039544 | 0.626167849 |
| chr8:142809425-142809625 | -0.50244339 | 0.33605783 | 0.62618428 |
| chr1:165717993-165718198 | 0.48537123 | 0.336064707 | 0.62618428 |
| chr10:60768911-60769171 | -0.446475413 | 0.336077313 | 0.626192542 |
| chr10:125024588-125024797 | 0.459720345 | 0.336086338 | 0.62619413 |
| chr17:62179571-62180186 | 0.328726911 | 0.336147497 | 0.626292852 |
| chr9:28213009-28213272 | 0.42546474 | 0.336225277 | 0.626410448 |
| chrX:105821481-105822428 | -0.325398026 | 0.336226965 | 0.626410448 |
| chr8:134261659-134261999 | 0.376062037 | 0.336237258 | 0.626414394 |
| chr13:28854843-28855078 | -0.469679182 | 0.336262556 | 0.626437742 |
| chr6:80338321-80338532 | -0.490960557 | 0.336266142 | 0.626437742 |
| chr13:44050535-44050750 | -0.508397835 | 0.33629089 | 0.626451411 |
| chr8:123066123-123066390 | -0.399683899 | 0.336297468 | 0.626451411 |
| chr12:46201465-46201745 | -0.405405998 | 0.336298007 | 0.626451411 |
| chr7:44596038-44596238 | 0.492040382 | 0.336327573 | 0.626484435 |
| chr10:1244077-1244345 | 0.436467848 | 0.336332088 | 0.626484435 |
| chr7:94657705-94658722 | -0.301055672 | 0.3363444 | 0.626492138 |
| chr3:162488223-162488423 | -0.527308833 | 0.336376024 | 0.626525865 |
| chr3:153396429-153396727 | -0.5031134 | 0.336381913 | 0.626525865 |
| chr1:205631031-205632526 | -0.301001273 | 0.336387037 | 0.626525865 |
| chr9:95426656-95427354 | 0.324604827 | 0.33640649 | 0.626546866 |
| chr17:63697795-63698260 | -0.321744096 | 0.336430535 | 0.626576419 |
| chr14:68669174-68669518 | 0.378396334 | 0.336447267 | 0.626592352 |
| chr5:73625663-73626842 | -0.321813261 | 0.336460968 | 0.626602638 |
| chr1:167533828-167534131 | 0.381205102 | 0.336485293 | 0.626632708 |
| chr6:125363203-125363784 | 0.348920693 | 0.336504002 | 0.626639562 |
| chr2:64203807-64204160 | -0.4104871 | 0.336509499 | 0.626639562 |
| chr1:43552817-43553033 | 0.394571959 | 0.336513508 | 0.626639562 |
| chr7:55887070-55887668 | 0.347800198 | 0.336540484 | 0.626667958 |

| | | | |
|---|---|---|---|
| chr20:63176737-63177004 | 0.389448872 | 0.336545115 | 0.626667958 |
| chr9:103245532-103245796 | -0.461582638 | 0.336575522 | 0.626709347 |
| chr9:117160299-117160530 | 0.468512853 | 0.336591384 | 0.626723652 |
| chr1:246152197-246152409 | -0.513814078 | 0.336606024 | 0.626735682 |
| chr17:10876006-10876262 | -0.431182866 | 0.336635011 | 0.626774423 |
| chr6:156399567-156399935 | -0.458289881 | 0.336661228 | 0.626787922 |
| chr2:88961431-88961714 | -0.440891756 | 0.336665643 | 0.626787922 |
| chr10:84267559-84267921 | -0.366019767 | 0.336666802 | 0.626787922 |
| chr3:153160521-153160999 | -0.318994536 | 0.336703326 | 0.626839415 |
| chr9:128370732-128370932 | 0.459206267 | 0.336710823 | 0.626839415 |
| chr5:10546376-10546576 | -0.490364997 | 0.336756395 | 0.626891655 |
| chr15:89760768-89761025 | 0.385811683 | 0.336759746 | 0.626891655 |
| chr3:63565390-63565590 | 0.420869401 | 0.33677535 | 0.626891655 |
| chr11:45180210-45180410 | 0.445298304 | 0.336777384 | 0.626891655 |
| chr22:42621532-42621828 | -0.431915456 | 0.336779791 | 0.626891655 |
| chr7:1530983-1531226 | 0.382664652 | 0.33681323 | 0.626938668 |
| chr2:218908754-218909518 | -0.324028059 | 0.336834337 | 0.626962725 |
| chr1:219519636-219519985 | -0.371983738 | 0.336844775 | 0.626966922 |
| chr8:138642052-138642305 | -0.488242682 | 0.336894092 | 0.627029111 |
| chr15:40873598-40874023 | 0.333469772 | 0.336894553 | 0.627029111 |
| chr2:131801785-131802427 | -0.329420374 | 0.336938259 | 0.62706084 |
| chr13:37934304-37934512 | -0.537120099 | 0.336946455 | 0.62706084 |
| chr9:99820994-99821237 | 0.362649634 | 0.336947113 | 0.62706084 |
| chr11:47934515-47934764 | 0.391745024 | 0.336949761 | 0.62706084 |
| chrX:49039854-49040220 | -0.344988395 | 0.33695252 | 0.62706084 |
| chr7:101688769-101688972 | -0.476324158 | 0.336985536 | 0.627098868 |
| chrX:71185219-71185512 | 0.398292019 | 0.336992548 | 0.627098868 |
| chr1:162790262-162790753 | 0.33407856 | 0.336997507 | 0.627098868 |
| chr15:34342831-34343422 | 0.323443076 | 0.337012353 | 0.627111264 |
| chr3:11846211-11847152 | 0.31249816 | 0.337035911 | 0.627139871 |
| chr3:146218921-146219170 | -0.493078721 | 0.33704794 | 0.627147023 |
| chr6:44100108-44100381 | -0.411543227 | 0.337189669 | 0.627395502 |
| chr4:18021093-18022521 | 0.302980275 | 0.337228275 | 0.627452098 |
| chr3:24203947-24204178 | 0.431835783 | 0.337236965 | 0.627453031 |
| chr1:92079983-92080936 | -0.317475151 | 0.337245927 | 0.62745447 |
| chr1:173451986-173452394 | -0.400776046 | 0.337258895 | 0.627457326 |
| chr5:137753880-137755220 | -0.304622451 | 0.33727437 | 0.627457326 |
| chr1:202571774-202572001 | -0.419496981 | 0.337279507 | 0.627457326 |
| chr5:164013126-164013374 | -0.420361914 | 0.337280218 | 0.627457326 |
| chr6:121932198-121932467 | -0.504566921 | 0.337300386 | 0.627479611 |
| chr19:39403450-39404168 | 0.319123045 | 0.33731388 | 0.627489477 |
| chr18:13812033-13812379 | 0.415683772 | 0.337367169 | 0.627573372 |
| chr1:113758678-113760078 | -0.302349729 | 0.337382592 | 0.627584071 |
| chr22:20020695-20021181 | 0.333183245 | 0.337389301 | 0.627584071 |
| chr1:212558232-212559449 | -0.311299007 | 0.33743818 | 0.627655657 |
| chr21:29476434-29476776 | -0.388683113 | 0.337444169 | 0.627655657 |
| chr1:176886657-176886965 | -0.453243707 | 0.337457131 | 0.627662127 |
| chr19:2494682-2495210 | -0.391139928 | 0.337464031 | 0.627662127 |
| chr7:76310689-76310915 | -0.39066564 | 0.337491053 | 0.627697149 |
| chr3:179132008-179132922 | -0.329246699 | 0.337515711 | 0.627719067 |

| | | | |
|---|---|---|---|
| chr12:76030898-76032433 | 0.303809329 | 0.337519222 | 0.627719067 |
| chr6:148509902-148510202 | 0.495630831 | 0.337557757 | 0.627775496 |
| chr7:74307013-74307213 | -0.503678406 | 0.337584946 | 0.627804733 |
| chr21:42073637-42074013 | 0.392831708 | 0.337589865 | 0.627804733 |
| chr3:14768822-14769289 | -0.375109112 | 0.337606871 | 0.627821121 |
| chr6:118650678-118652040 | 0.305492014 | 0.337662332 | 0.627909019 |
| chr10:14687198-14687801 | 0.369588847 | 0.337695265 | 0.627955019 |
| chr9:120714339-120714996 | 0.315037952 | 0.337762372 | 0.628025674 |
| chr17:60592053-60592361 | 0.331308313 | 0.337762703 | 0.628025674 |
| chr1:185720739-185720939 | -0.500683344 | 0.337766046 | 0.628025674 |
| chr12:105337583-105337826 | -0.500683344 | 0.337766046 | 0.628025674 |
| chr4:3484421-3485307 | -0.319627109 | 0.337799797 | 0.628073187 |
| chr2:134120393-134121160 | -0.322811137 | 0.337827428 | 0.628109321 |
| chr4:152708283-152709131 | 0.324128461 | 0.337857199 | 0.628144526 |
| chr16:73070488-73071057 | -0.316896402 | 0.337862759 | 0.628144526 |
| chr7:102875960-102876200 | -0.5061734 | 0.337889795 | 0.628179547 |
| chr3:119293793-119294773 | 0.31727933 | 0.337901701 | 0.628186439 |
| chr2:98648868-98649214 | 0.455900305 | 0.337915734 | 0.628187152 |
| chr14:73825657-73825858 | 0.444250473 | 0.337918481 | 0.628187152 |
| chr12:105383470-105383670 | -0.48303625 | 0.33793946 | 0.628200293 |
| chr3:113111280-113111689 | 0.35792189 | 0.337947306 | 0.628200293 |
| chr20:8106054-8106534 | -0.422430296 | 0.337950146 | 0.628200293 |
| chr19:38838314-38838901 | 0.33911078 | 0.337962766 | 0.628208512 |
| chr8:105285042-105285250 | -0.503332344 | 0.337991102 | 0.628245941 |
| chr11:17813732-17813970 | 0.434024402 | 0.338011113 | 0.628267896 |
| chr3:99945346-99945546 | -0.488968247 | 0.338044889 | 0.628282045 |
| chr15:33994043-33994368 | -0.428091095 | 0.338045762 | 0.628282045 |
| chr6:75350891-75351230 | 0.384033812 | 0.338045857 | 0.628282045 |
| chr11:24779809-24780054 | 0.509784388 | 0.338054076 | 0.628282045 |
| chr2:102806298-102806586 | -0.403240873 | 0.338066828 | 0.628282045 |
| chr3:41961389-41962671 | -0.305538875 | 0.338067923 | 0.628282045 |
| chr21:9819031-9819261 | 0.470337859 | 0.338089427 | 0.628306769 |
| chr5:129271180-129271642 | -0.358758191 | 0.338108894 | 0.628316515 |
| chr18:79394155-79394375 | 0.446256169 | 0.338123242 | 0.628316515 |
| chr2:72150311-72150805 | -0.334119306 | 0.338128969 | 0.628316515 |
| chr4:144572677-144572885 | -0.507798609 | 0.338129439 | 0.628316515 |
| chr19:39406532-39406953 | 0.335809806 | 0.338135673 | 0.628316515 |
| chr9:33445535-33445794 | 0.42228074 | 0.338144515 | 0.628317709 |
| chr22:42719199-42719438 | -0.495918175 | 0.338185483 | 0.628378593 |
| chr3:101229683-101230042 | 0.371951887 | 0.338209963 | 0.62840884 |
| chr8:144337023-144337288 | 0.39789627 | 0.338225517 | 0.628422502 |
| chr19:29844310-29845479 | 0.302013361 | 0.338245344 | 0.628437359 |
| chr1:9182244-9182520 | 0.40742742 | 0.338249917 | 0.628437359 |
| chr9:94630744-94631233 | 0.334168174 | 0.338259259 | 0.628439478 |
| chr4:83570003-83570346 | -0.401180919 | 0.338275279 | 0.628454002 |
| chr19:44718532-44718904 | 0.377958233 | 0.338289831 | 0.628465799 |
| chr5:73902650-73903005 | -0.42658876 | 0.338311126 | 0.628483049 |
| chr1:58460580-58460822 | -0.46278691 | 0.338315521 | 0.628483049 |
| chr8:118951457-118952394 | 0.315949039 | 0.338348553 | 0.628519749 |
| chr12:3748218-3748808 | 0.348741769 | 0.338356067 | 0.628519749 |

| | | | |
|---|---|---|---|
| chr2:130355786-130356738 | 0.319721709 | 0.338364135 | 0.628519749 |
| chr6:138987400-138988706 | -0.311278406 | 0.338368088 | 0.628519749 |
| chr11:1260097-1260369 | 0.396317076 | 0.338384518 | 0.628524069 |
| chr12:100141940-100143344 | -0.303974942 | 0.33838682 | 0.628524069 |
| chr9:137035226-137035744 | 0.34035168 | 0.338414262 | 0.628559804 |
| chr3:100533251-100533514 | -0.429699869 | 0.338424866 | 0.628564262 |
| chr8:58658635-58660018 | -0.305526742 | 0.338434384 | 0.628566704 |
| chr12:88299703-88300059 | -0.452208599 | 0.338460849 | 0.628600619 |
| chr15:71302339-71302549 | -0.500792072 | 0.338478726 | 0.628618584 |
| chr2:27628109-27629278 | 0.319078036 | 0.338488504 | 0.628621508 |
| chr9:6080078-6080757 | -0.315873164 | 0.338514793 | 0.628640078 |
| chr12:57462331-57462982 | -0.318111088 | 0.338516902 | 0.628640078 |
| chr14:55102187-55102470 | -0.434715862 | 0.338523117 | 0.628640078 |
| chr10:85105734-85105953 | -0.45350134 | 0.338579963 | 0.628698793 |
| chr12:117857775-117858032 | -0.42376867 | 0.338580948 | 0.628698793 |
| chr11:67993784-67994024 | 0.386934004 | 0.338583827 | 0.628698793 |
| chr3:133895444-133895977 | 0.33355552 | 0.338597227 | 0.628698793 |
| chr1:189693086-189693357 | 0.419468036 | 0.338599991 | 0.628698793 |
| chr8:126159035-126159306 | 0.416102241 | 0.338619234 | 0.628698793 |
| chr12:68365026-68365827 | -0.317093839 | 0.338619386 | 0.628698793 |
| chr13:41614867-41615089 | -0.474626665 | 0.338620376 | 0.628698793 |
| chr5:18772246-18772850 | -0.351158338 | 0.33862865 | 0.628698919 |
| chr17:4731031-4732009 | -0.309836913 | 0.338637779 | 0.628700635 |
| chrX:150848901-150849310 | 0.374071608 | 0.338650964 | 0.628709881 |
| chr5:113487322-113489037 | -0.30634608 | 0.338664167 | 0.628719158 |
| chr5:16666699-16667356 | -0.340917939 | 0.338685977 | 0.628738617 |
| chr14:92747537-92747901 | 0.374107352 | 0.338697237 | 0.628738617 |
| chr12:122980415-122981167 | 0.329417659 | 0.338699266 | 0.628738617 |
| chr5:6780874-6781081 | 0.443744397 | 0.338723911 | 0.628751014 |
| chr6:56247310-56247916 | -0.357966333 | 0.338727615 | 0.628751014 |
| chr18:24734114-24734355 | -0.430566074 | 0.338730562 | 0.628751014 |
| chr14:95307534-95307734 | 0.440154335 | 0.338752123 | 0.628775803 |
| chr12:49298107-49298434 | 0.348853095 | 0.33878507 | 0.628821725 |
| chrX:135973055-135974661 | 0.306329645 | 0.338807654 | 0.628848411 |
| chr17:68774263-68774949 | -0.32022071 | 0.338816723 | 0.62885001 |
| chr6:57722612-57722888 | -0.451674802 | 0.338872444 | 0.628938193 |
| chr7:10632800-10633069 | 0.430059699 | 0.338906339 | 0.628985866 |
| chr14:54642813-54643252 | 0.397391681 | 0.338926974 | 0.629008928 |
| chr3:165196478-165196729 | 0.419255188 | 0.338953267 | 0.629041306 |
| chr13:73849358-73849567 | -0.507081011 | 0.338960839 | 0.629041306 |
| chr17:68613878-68614285 | -0.34242452 | 0.338994146 | 0.62908788 |
| chr7:26376904-26377397 | 0.345333691 | 0.339021278 | 0.629119637 |
| chr6:128507570-128507870 | -0.417385666 | 0.33902768 | 0.629119637 |
| chrX:26586735-26586984 | -0.504737774 | 0.339041007 | 0.62912913 |
| chr5:123099199-123100233 | -0.320459022 | 0.339051746 | 0.629133821 |
| chr12:119545143-119545343 | 0.471287032 | 0.339079554 | 0.629155341 |
| chr12:4320475-4321722 | -0.31176221 | 0.339079766 | 0.629155341 |
| chr15:84980295-84981900 | 0.301000679 | 0.339120899 | 0.629216426 |
| chr15:35697568-35697869 | -0.476365716 | 0.339170364 | 0.629292965 |
| chr14:99717199-99717506 | 0.431851714 | 0.339186853 | 0.629297575 |

| | | | |
|---|---|---|---|
| chr1:175373864-175374133 | -0.390283337 | 0.339190481 | 0.629297575 |
| chr7:8998814-8999235 | -0.411366761 | 0.339197487 | 0.629297575 |
| chr3:173690494-173691046 | 0.363197284 | 0.339234775 | 0.629351515 |
| chr11:69479566-69480282 | -0.33740088 | 0.339245996 | 0.629357095 |
| chr9:89310311-89311768 | 0.301390607 | 0.339268509 | 0.629383621 |
| chr2:55268641-55269498 | -0.310897892 | 0.339294044 | 0.629400565 |
| chr12:56993356-56993556 | 0.525584687 | 0.339294071 | 0.629400565 |
| chr4:98778239-98778513 | -0.442180866 | 0.33930552 | 0.629406564 |
| chr9:35812047-35812359 | -0.364286484 | 0.339343606 | 0.629449089 |
| chr12:6927658-6927949 | 0.394393989 | 0.339344875 | 0.629449089 |
| chr12:12813063-12813667 | -0.332926077 | 0.339362136 | 0.629465869 |
| chr15:40439732-40440221 | -0.327142882 | 0.339413484 | 0.629545872 |
| chr9:31807670-31807905 | -0.45719253 | 0.339493244 | 0.629676809 |
| chrX:100732080-100732433 | 0.402399247 | 0.339500514 | 0.629676809 |
| chr17:19388002-19388229 | 0.40409979 | 0.339518554 | 0.629695026 |
| chr9:134655784-134656163 | -0.417662305 | 0.339551403 | 0.629740708 |
| chr2:19071802-19072084 | -0.412604606 | 0.33956338 | 0.629747678 |
| chr6:37499511-37500363 | 0.310031853 | 0.339575277 | 0.629748304 |
| chr5:23800320-23800712 | -0.363827471 | 0.339580156 | 0.629748304 |
| chr3:166685320-166685726 | 0.382874159 | 0.339606059 | 0.629766682 |
| chr8:143368442-143369132 | 0.310724315 | 0.339606504 | 0.629766682 |
| chr20:54629672-54630513 | 0.30434756 | 0.33962384 | 0.629774496 |
| chr12:116881013-116881363 | 0.371813408 | 0.339638477 | 0.629774496 |
| chr5:152271529-152271790 | -0.422567626 | 0.33964126 | 0.629774496 |
| chr20:47150636-47150976 | 0.339614804 | 0.339643595 | 0.629774496 |
| chr11:63916465-63916753 | 0.383950932 | 0.339692779 | 0.629850453 |
| chr5:23352447-23352752 | 0.378730363 | 0.339741562 | 0.629925662 |
| chr3:71390007-71390284 | -0.440782914 | 0.339807676 | 0.630033001 |
| chr2:108594943-108595217 | -0.479138523 | 0.339871547 | 0.630136174 |
| chr20:47983529-47983774 | 0.33198642 | 0.339893499 | 0.630161626 |
| chr5:35851273-35852066 | -0.361535735 | 0.339915386 | 0.630186955 |
| chr12:69439914-69440380 | 0.346140941 | 0.339931138 | 0.630191401 |
| chr2:197309926-197311101 | 0.315218667 | 0.339934233 | 0.630191401 |
| chr2:14751425-14752111 | 0.341499641 | 0.339959614 | 0.630223205 |
| chr1:231986523-231987401 | -0.328616595 | 0.339997975 | 0.630257437 |
| chr10:94165053-94165621 | -0.343206334 | 0.340005356 | 0.630257437 |
| chr18:5978532-5978925 | -0.446639392 | 0.340009564 | 0.630257437 |
| chr10:43606005-43607028 | 0.318777327 | 0.340019978 | 0.630257437 |
| chr2:19673263-19673857 | -0.352661343 | 0.340019993 | 0.630257437 |
| chr7:53350373-53350573 | -0.517161405 | 0.340027432 | 0.630257437 |
| chr7:99374091-99374432 | -0.361426333 | 0.340045965 | 0.630276541 |
| chr16:28863117-28864039 | 0.314082707 | 0.340082373 | 0.630328776 |
| chr11:57205063-57205445 | -0.365629808 | 0.340115619 | 0.630375148 |
| chr5:180455913-180456171 | 0.428961601 | 0.340128811 | 0.630375603 |
| chr6:33317622-33318214 | 0.316686159 | 0.340135185 | 0.630375603 |
| chr18:48963323-48963560 | -0.491698335 | 0.340153331 | 0.630375603 |
| chr4:152784834-152785136 | -0.372389125 | 0.340153776 | 0.630375603 |
| chr1:39547064-39547277 | 0.486914686 | 0.340157 | 0.630375603 |
| chr1:47424510-47424867 | -0.400966198 | 0.340168884 | 0.630382379 |
| chrX:74293380-74293731 | 0.350366568 | 0.340181201 | 0.630389959 |

| | | | |
|---|---|---|---|
| chr6:17585808-17586190 | -0.33702212 | 0.34021355 | 0.630434657 |
| chr7:12801085-12801738 | 0.328115911 | 0.340228617 | 0.630447329 |
| chr9:36190485-36191263 | -0.312799556 | 0.340241405 | 0.63045578 |
| chr2:100381231-100381439 | -0.472840047 | 0.340259462 | 0.630473991 |
| chr2:189555812-189556377 | -0.368076725 | 0.340292388 | 0.63050727 |
| chr20:54112111-54112464 | 0.317605115 | 0.34029388 | 0.63050727 |
| chr1:219544597-219544939 | -0.480511967 | 0.34033617 | 0.630557868 |
| chr10:128210226-128210619 | 0.375141905 | 0.340337647 | 0.630557868 |
| chr12:48183051-48183884 | 0.324684704 | 0.340355736 | 0.630576135 |
| chr20:4014934-4015406 | 0.362772503 | 0.340373951 | 0.630594635 |
| chr1:242475352-242475575 | 0.455619807 | 0.340408139 | 0.630627859 |
| chr1:163933597-163933920 | -0.425962648 | 0.340435495 | 0.630627859 |
| chr1:177959784-177960042 | -0.403962129 | 0.340436648 | 0.630627859 |
| chr1:63466834-63467044 | -0.535682798 | 0.340441267 | 0.630627859 |
| chr1:178311339-178311613 | -0.535682798 | 0.340441267 | 0.630627859 |
| chr10:118521998-118522198 | -0.535682798 | 0.340441267 | 0.630627859 |
| chr3:112109448-112109761 | -0.391856893 | 0.3404594 | 0.630634956 |
| chr4:109207203-109207484 | -0.450549217 | 0.340461559 | 0.630634956 |
| chr7:128455463-128456326 | -0.312863675 | 0.340480389 | 0.630652091 |
| chr11:125410131-125410333 | -0.49105888 | 0.340491446 | 0.630652091 |
| chr2:152189994-152190210 | -0.491087325 | 0.340495502 | 0.630652091 |
| chr15:42548189-42549339 | 0.317175908 | 0.340523272 | 0.630688281 |
| chr6:56842940-56844477 | 0.30473662 | 0.340552724 | 0.630727582 |
| chr3:36540987-36541187 | -0.513505082 | 0.340571597 | 0.630732045 |
| chr5:31967780-31968027 | -0.513505082 | 0.340571597 | 0.630732045 |
| chr22:38162511-38162900 | 0.434594095 | 0.340610033 | 0.630746857 |
| chr4:118890643-118890939 | 0.408488025 | 0.340610046 | 0.630746857 |
| chr8:102788865-102789324 | 0.328600406 | 0.340617321 | 0.630746857 |
| chr5:1633619-1634372 | 0.340771245 | 0.340620155 | 0.630746857 |
| chr10:125538863-125539501 | 0.35096167 | 0.340620754 | 0.630746857 |
| chr20:62142532-62143605 | 0.312216146 | 0.340636596 | 0.630760948 |
| chr2:69564995-69565217 | -0.482262021 | 0.340675955 | 0.630818585 |
| chr5:67206971-67207322 | -0.380311927 | 0.340689211 | 0.630825424 |
| chr2:133900921-133901173 | 0.43983074 | 0.340696114 | 0.630825424 |
| chr3:171771123-171771650 | -0.3384722 | 0.340718079 | 0.630850848 |
| chr14:23413910-23414122 | 0.440100817 | 0.340743888 | 0.63088339 |
| chr1:43285453-43285843 | 0.345740853 | 0.340761055 | 0.630897167 |
| chr5:132829909-132830445 | 0.333940692 | 0.340767797 | 0.630897167 |
| chr8:125219266-125219650 | 0.359817446 | 0.340780507 | 0.630905454 |
| chr1:24199359-24199559 | -0.526127807 | 0.340812902 | 0.630950184 |
| chrX:19372233-19373047 | -0.334067438 | 0.340838212 | 0.630950347 |
| chr6:108560147-108560370 | -0.369881582 | 0.34083978 | 0.630950347 |
| chr5:69369193-69370235 | -0.306390324 | 0.340840315 | 0.630950347 |
| chr22:43883114-43883334 | -0.455897056 | 0.340845929 | 0.630950347 |
| chr2:3651009-3651277 | -0.366459675 | 0.34086595 | 0.630968942 |
| chr16:86577305-86577505 | -0.496612887 | 0.340872444 | 0.630968942 |
| chr7:43810624-43810886 | -0.39265234 | 0.340909872 | 0.631014419 |
| chr6:56570830-56571267 | 0.381938815 | 0.340913483 | 0.631014419 |
| chr5:2992744-2993017 | 0.445101899 | 0.340955714 | 0.63106405 |
| chr9:83061972-83063485 | -0.299853944 | 0.340956769 | 0.63106405 |

| | | | |
|---|---|---|---|
| chrX:110012728-110013243 | 0.342127623 | 0.340989892 | 0.631110111 |
| chr8:119574207-119574457 | -0.499914002 | 0.341007187 | 0.631123466 |
| chr20:63088982-63089510 | -0.34173061 | 0.341013581 | 0.631123466 |
| chr10:3668420-3668907 | 0.340713401 | 0.341031582 | 0.631132475 |
| chr1:6701204-6701404 | 0.462259997 | 0.341044405 | 0.631132475 |
| chr19:38263976-38265048 | 0.306683355 | 0.341047257 | 0.631132475 |
| chr7:55307149-55307406 | -0.422362436 | 0.341054504 | 0.631132475 |
| chr15:96644976-96645304 | -0.458841611 | 0.341062306 | 0.631132475 |
| chr6:145075482-145075682 | 0.528714483 | 0.341067871 | 0.631132475 |
| chr17:74443016-74443281 | -0.403311828 | 0.341109953 | 0.631185479 |
| chr5:38922137-38922646 | -0.360112122 | 0.34111299 | 0.631185479 |
| chr16:71542323-71542670 | -0.349998052 | 0.34113088 | 0.63120334 |
| chr1:47894796-47894996 | -0.47972009 | 0.341139381 | 0.631203827 |
| chr15:82141000-82141228 | 0.471605203 | 0.341160039 | 0.631226807 |
| chr1:42913358-42913659 | 0.391443608 | 0.341172297 | 0.631234244 |
| chr9:128159746-128160412 | 0.31695212 | 0.341199541 | 0.631239577 |
| chr8:143309758-143310094 | 0.405874629 | 0.341203024 | 0.631239577 |
| chr6:79586616-79586866 | 0.437872251 | 0.341205108 | 0.631239577 |
| chr7:6403986-6404271 | 0.374238972 | 0.341211264 | 0.631239577 |
| chr15:77496824-77497596 | -0.327265923 | 0.341222236 | 0.631239577 |
| chr6:43037896-43038179 | -0.415271515 | 0.34122461 | 0.631239577 |
| chr2:244774-245066 | 0.461258638 | 0.341264693 | 0.631298487 |
| chr1:54564386-54564624 | 0.437885656 | 0.341279485 | 0.631303948 |
| chr12:96902145-96902387 | -0.416131478 | 0.341284124 | 0.631303948 |
| chr17:78445695-78445916 | 0.448876158 | 0.341294991 | 0.631308808 |
| chr3:179562537-179563832 | -0.298765451 | 0.341401806 | 0.631484476 |
| chr7:44112613-44114121 | -0.301974059 | 0.341406442 | 0.631484476 |
| chr19:15222958-15223751 | -0.338232398 | 0.341429528 | 0.631499516 |
| chr8:145052021-145052673 | 0.335252775 | 0.341431057 | 0.631499516 |
| chr22:41413438-41414295 | 0.321029987 | 0.341475981 | 0.631554138 |
| chr9:125449657-125449938 | 0.410952186 | 0.341479526 | 0.631554138 |
| chr6:160991249-160992621 | 0.312109311 | 0.341485317 | 0.631554138 |
| chr16:72104885-72105261 | -0.40894298 | 0.341503886 | 0.631573237 |
| chr4:87006282-87007571 | -0.304017446 | 0.341518615 | 0.631585232 |
| chr11:88337309-88338113 | -0.330731046 | 0.341531739 | 0.631594258 |
| chr7:155852141-155852341 | -0.518750749 | 0.34154324 | 0.63159998 |
| chr10:14953421-14954589 | -0.304870246 | 0.341553355 | 0.63159998 |
| chr4:178226462-178226662 | -0.471688681 | 0.341560331 | 0.63159998 |
| chr9:23598629-23599462 | 0.321291222 | 0.34158105 | 0.63159998 |
| chr5:134140504-134140916 | -0.341947617 | 0.341585122 | 0.63159998 |
| chr20:20465370-20465707 | 0.413176814 | 0.341604659 | 0.63159998 |
| chr8:112134779-112135001 | -0.490238644 | 0.341611279 | 0.63159998 |
| chr1:199493578-199494072 | 0.380677159 | 0.341613705 | 0.63159998 |
| chr8:117934463-117934663 | -0.496735293 | 0.341614249 | 0.63159998 |
| chr3:104628203-104628533 | -0.389327313 | 0.341617264 | 0.63159998 |
| chr2:198296739-198297064 | -0.411651146 | 0.341653437 | 0.631640365 |
| chr6:110543002-110543720 | -0.321807092 | 0.341658244 | 0.631640365 |
| chr10:61834514-61835101 | -0.345291926 | 0.341665296 | 0.631640365 |
| chr3:133662255-133662573 | -0.351882497 | 0.341692132 | 0.631640365 |
| chr6:35687846-35688256 | -0.357372233 | 0.341700816 | 0.631640365 |

| | | | |
|---|---|---|---|
| chr21:46524152-46524386 | 0.483703841 | 0.341709097 | 0.631640365 |
| chr3:162418205-162418479 | -0.466328039 | 0.341712432 | 0.631640365 |
| chr12:3236839-3237139 | -0.426802027 | 0.341712567 | 0.631640365 |
| chr5:11766121-11766321 | 0.452822895 | 0.341713299 | 0.631640365 |
| chr10:25174353-25175245 | -0.318368809 | 0.341752149 | 0.631696937 |
| chr4:70901763-70903079 | -0.305836753 | 0.34178393 | 0.631739263 |
| chr3:33440104-33441361 | 0.298799299 | 0.341791537 | 0.631739263 |
| chr4:127400808-127401109 | -0.393029621 | 0.341835353 | 0.631782351 |
| chr1:116764167-116764421 | 0.402436201 | 0.341844269 | 0.631782351 |
| chr1:1504769-1505741 | 0.322568702 | 0.34184767 | 0.631782351 |
| chr4:6916559-6917122 | 0.335318453 | 0.341847831 | 0.631782351 |
| chr6:129936610-129936863 | 0.427434832 | 0.341856761 | 0.631783617 |
| chr1:151859712-151859968 | 0.383671025 | 0.341880595 | 0.631796375 |
| chr2:171272029-171272325 | -0.406132119 | 0.341882428 | 0.631796375 |
| chr5:167723329-167723807 | 0.374092589 | 0.341888402 | 0.631796375 |
| chr2:166375406-166376680 | -0.311085906 | 0.341929606 | 0.63185728 |
| chr7:110766932-110767261 | 0.425957575 | 0.341951948 | 0.631883327 |
| chr2:218356540-218357076 | -0.345909825 | 0.342015414 | 0.631961385 |
| chr5:75236615-75237331 | -0.323069607 | 0.342018435 | 0.631961385 |
| chr16:3183252-3183738 | 0.332567521 | 0.342033615 | 0.631961385 |
| chr5:151214076-151214458 | -0.401777837 | 0.342034772 | 0.631961385 |
| chr2:166127899-166128103 | -0.515020383 | 0.342043677 | 0.631961385 |
| chr20:18026171-18026371 | -0.515020383 | 0.342043677 | 0.631961385 |
| chr13:36282854-36283226 | -0.384674129 | 0.342061885 | 0.631979787 |
| chr6:168370327-168370573 | -0.494255994 | 0.342142554 | 0.632055365 |
| chr11:107999871-108000083 | 0.4565268 | 0.342143724 | 0.632055365 |
| chr5:5886927-5887257 | -0.377138119 | 0.34215065 | 0.632055365 |
| chr2:179711439-179711730 | -0.404530087 | 0.342154119 | 0.632055365 |
| chr14:81509714-81510336 | 0.341488581 | 0.34216025 | 0.632055365 |
| chr6:170522695-170523390 | 0.352055773 | 0.342165899 | 0.632055365 |
| chr4:139144498-139145553 | -0.325656211 | 0.342166022 | 0.632055365 |
| chr1:14955519-14955830 | -0.445969743 | 0.342168784 | 0.632055365 |
| chr3:191499392-191500050 | -0.351408389 | 0.342187742 | 0.632057083 |
| chr14:66069780-66070229 | -0.358982228 | 0.342191853 | 0.632057083 |
| chr3:182018451-182018683 | 0.459028746 | 0.342200627 | 0.632057083 |
| chr11:83491935-83492151 | -0.499450542 | 0.342205011 | 0.632057083 |
| chr15:65980902-65981206 | -0.365699489 | 0.342212161 | 0.632057083 |
| chr10:102478900-102479266 | 0.360846537 | 0.342219208 | 0.632057083 |
| chr8:56970896-56971096 | 0.522156604 | 0.342251942 | 0.632091273 |
| chr2:39123897-39124709 | -0.306558001 | 0.342258842 | 0.632091273 |
| chr12:48887117-48887365 | 0.395489555 | 0.342262468 | 0.632091273 |
| chr12:401382-401970 | -0.320887721 | 0.34229164 | 0.632129912 |
| chr1:198136370-198136570 | 0.48645935 | 0.34234159 | 0.632205388 |
| chr4:48033887-48034150 | -0.447136735 | 0.342355224 | 0.632205388 |
| chr1:217242453-217242654 | -0.493462675 | 0.342357262 | 0.632205388 |
| chr9:92790834-92791074 | 0.434786997 | 0.342365676 | 0.632205688 |
| chr5:15013935-15014226 | 0.372706876 | 0.342399405 | 0.632252735 |
| chr12:66048562-66048762 | -0.537813908 | 0.342426031 | 0.632286662 |
| chr3:19940632-19940925 | 0.388040057 | 0.342455092 | 0.632325085 |
| chr8:19014455-19014679 | -0.457468796 | 0.342487048 | 0.63236885 |

| | | | |
|---|---|---|---|
| chr8:103877994-103878194 | 0.485308268 | 0.342498427 | 0.632374623 |
| chr7:35185590-35187072 | -0.305464922 | 0.342521878 | 0.632379327 |
| chr11:86632825-86633105 | 0.401062369 | 0.342522392 | 0.632379327 |
| chr2:10405435-10405701 | 0.368651837 | 0.342525734 | 0.632379327 |
| chr12:131327789-131328029 | -0.474195157 | 0.342611106 | 0.632516586 |
| chr14:77663328-77663665 | -0.386465595 | 0.34261659 | 0.632516586 |
| chr8:23287916-23288264 | 0.369981388 | 0.342632232 | 0.632526025 |
| chr9:35903108-35903357 | -0.451117632 | 0.342641864 | 0.632526025 |
| chr16:70653575-70654028 | 0.372562151 | 0.342646468 | 0.632526025 |
| chr16:85306124-85306677 | -0.326951737 | 0.342665879 | 0.632532361 |
| chr9:109489360-109489605 | 0.443618158 | 0.342666412 | 0.632532361 |
| chr2:101831324-101831562 | -0.455640164 | 0.342688966 | 0.632554462 |
| chr19:8321055-8321714 | 0.332430428 | 0.342694895 | 0.632554462 |
| chr8:132489616-132489821 | -0.454150413 | 0.342726375 | 0.632597328 |
| chrX:10003873-10004292 | 0.369906676 | 0.342759693 | 0.632606403 |
| chr17:57721078-57721322 | -0.433197759 | 0.342763811 | 0.632606403 |
| chr7:98106062-98106454 | -0.392315567 | 0.342769425 | 0.632606403 |
| chr2:113198659-113199523 | 0.310440779 | 0.342775468 | 0.632606403 |
| chr21:42218785-42219242 | 0.341419135 | 0.342776633 | 0.632606403 |
| chr4:148051560-148051848 | -0.409023543 | 0.342780829 | 0.632606403 |
| chr1:65247669-65248382 | 0.349491876 | 0.34279364 | 0.632614809 |
| chr2:131031636-131031860 | -0.468549897 | 0.342811201 | 0.63261761 |
| chr10:110311377-110311757 | 0.374933373 | 0.34281167 | 0.63261761 |
| chr7:81527747-81527947 | -0.50854426 | 0.342869839 | 0.632709716 |
| chr5:180790740-180791191 | -0.388869145 | 0.342933222 | 0.632773562 |
| chr4:98027805-98028080 | 0.4252783 | 0.342936438 | 0.632773562 |
| chr8:144526804-144527219 | 0.333466058 | 0.342937486 | 0.632773562 |
| chr2:167127197-167127469 | -0.429938267 | 0.34294573 | 0.632773562 |
| chr20:17721692-17721933 | -0.429938267 | 0.34294573 | 0.632773562 |
| chr7:150378792-150379834 | -0.305409057 | 0.342962267 | 0.632780917 |
| chr2:152465663-152465895 | -0.446575691 | 0.342966233 | 0.632780917 |
| chr17:64345198-64345444 | 0.327748515 | 0.342997006 | 0.632822455 |
| chr20:31553315-31553545 | 0.440268857 | 0.343031126 | 0.632858954 |
| chr3:155727767-155728175 | -0.337625611 | 0.34304967 | 0.632858954 |
| chr2:96930611-96930926 | -0.385119128 | 0.343049673 | 0.632858954 |
| chr16:30874749-30875635 | 0.317789986 | 0.343049826 | 0.632858954 |
| chr14:64726467-64726791 | 0.375977138 | 0.343074615 | 0.632877814 |
| chr15:85485984-85486278 | 0.413070854 | 0.343076569 | 0.632877814 |
| chr22:38052370-38052583 | -0.463065087 | 0.343101844 | 0.632900084 |
| chr10:75294657-75295502 | -0.321725402 | 0.343105162 | 0.632900084 |
| chr10:34703957-34704157 | -0.540481821 | 0.343199282 | 0.63304644 |
| chr1:165075783-165076100 | -0.380448949 | 0.343201027 | 0.63304644 |
| chr2:71276037-71277170 | -0.30621988 | 0.343214145 | 0.633047326 |
| chr3:155411904-155412163 | -0.425552825 | 0.343218032 | 0.633047326 |
| chr12:53287791-53288013 | 0.424943819 | 0.343238023 | 0.63306896 |
| chr12:107872688-107872933 | -0.466909803 | 0.343276324 | 0.633124362 |
| chr16:74709473-74709704 | 0.395608752 | 0.343307229 | 0.633166121 |
| chr16:4956213-4956421 | 0.391353573 | 0.343348244 | 0.633212117 |
| chr20:47319445-47320163 | 0.297633158 | 0.343348696 | 0.633212117 |
| chr2:178478504-178478780 | -0.338388461 | 0.343370942 | 0.633237901 |

| | | | |
|---|---|---|---|
| chr4:26857334-26858083 | 0.310330156 | 0.343404215 | 0.633246584 |
| chr10:230810-231048 | 0.432523252 | 0.343410044 | 0.633246584 |
| chr9:21154002-21154299 | -0.417605609 | 0.343411913 | 0.633246584 |
| chr12:116277525-116278095 | 0.321627641 | 0.343416118 | 0.633246584 |
| chr7:64948709-64948956 | -0.427436894 | 0.343416973 | 0.633246584 |
| chr2:3518416-3518616 | -0.375823214 | 0.343426019 | 0.633248024 |
| chr3:172685732-172685932 | 0.494947755 | 0.343468347 | 0.633310227 |
| chr20:51231433-51231748 | 0.395570606 | 0.343476283 | 0.633310227 |
| chr2:69160041-69160246 | -0.487973838 | 0.34348645 | 0.633313732 |
| chr12:6924312-6924611 | 0.363516044 | 0.343501597 | 0.633326419 |
| chr1:110407132-110408155 | 0.311042441 | 0.343559997 | 0.633400674 |
| chr5:42514071-42514271 | 0.54876001 | 0.343562915 | 0.633400674 |
| chr2:46164041-46164675 | 0.33017674 | 0.34356667 | 0.633400674 |
| chr12:39474843-39475873 | -0.320921144 | 0.343614597 | 0.633471781 |
| chr3:33452438-33452940 | 0.358182871 | 0.343626404 | 0.633471781 |
| chr12:62913458-62913702 | 0.40197843 | 0.343630042 | 0.633471781 |
| chr2:166384757-166384997 | -0.459514522 | 0.343644257 | 0.633482745 |
| chr9:88272935-88273213 | 0.425863551 | 0.343664826 | 0.633485772 |
| chr18:34439732-34440087 | -0.462528848 | 0.343665733 | 0.633485772 |
| chrX:103152189-103152554 | -0.349286909 | 0.343676754 | 0.633485772 |
| chr9:103202792-103203170 | -0.418327186 | 0.34367897 | 0.633485772 |
| chr12:45490258-45490693 | 0.345062239 | 0.343695077 | 0.633500221 |
| chr4:119067450-119068294 | -0.35192701 | 0.343717378 | 0.633524965 |
| chr12:80918852-80919127 | -0.42310739 | 0.343731468 | 0.633524965 |
| chr6:169654661-169655153 | 0.346420042 | 0.343734515 | 0.633524965 |
| chr17:73287076-73287276 | 0.39688791 | 0.343741574 | 0.633524965 |
| chr6:154040178-154040378 | 0.486712052 | 0.34375204 | 0.633529015 |
| chr5:73496015-73496238 | -0.438256837 | 0.343818979 | 0.633637141 |
| chr4:106315526-106316741 | -0.300405988 | 0.343842304 | 0.633664887 |
| chr8:142609084-142609463 | 0.336878293 | 0.343860994 | 0.633684089 |
| chr20:54007347-54007607 | 0.320241657 | 0.343871736 | 0.633688645 |
| chr9:122144998-122145216 | -0.494397743 | 0.343934737 | 0.6337895 |
| chr6:111052173-111052373 | 0.496067785 | 0.343950049 | 0.633802473 |
| chr9:78031120-78031386 | -0.382587411 | 0.343969407 | 0.633822901 |
| chr1:26857094-26857564 | 0.349159076 | 0.343989983 | 0.633837361 |
| chr12:89470821-89471094 | -0.432097483 | 0.343997574 | 0.633837361 |
| chr5:90113149-90113637 | -0.376098947 | 0.34400207 | 0.633837361 |
| chr2:70006263-70006523 | 0.427244249 | 0.34402588 | 0.633865987 |
| chr1:181070713-181070913 | 0.468666229 | 0.344042081 | 0.633880595 |
| chr15:74598083-74598793 | -0.320078416 | 0.344066274 | 0.63389123 |
| chr1:86833610-86833943 | -0.389942003 | 0.344071685 | 0.63389123 |
| chr12:389187-389770 | -0.334485453 | 0.344120409 | 0.63389123 |
| chr5:141814146-141814368 | 0.424045917 | 0.34412594 | 0.63389123 |
| chr18:34661586-34661816 | -0.444306023 | 0.344129213 | 0.63389123 |
| chr2:181581620-181581873 | -0.444306023 | 0.344129213 | 0.63389123 |
| chr14:42750741-42751176 | -0.458749127 | 0.344131154 | 0.63389123 |
| chr2:75480963-75481246 | 0.402522821 | 0.344131334 | 0.63389123 |
| chr1:61852224-61852424 | -0.494178311 | 0.344136326 | 0.63389123 |
| chr2:215966701-215967335 | 0.324806411 | 0.344137003 | 0.63389123 |
| chr9:98538742-98539013 | 0.419760744 | 0.344141026 | 0.63389123 |

| | | | |
|---|---|---|---|
| chr2:1657447-1657730 | -0.415293737 | 0.344147129 | 0.63389123 |
| chrX:98405335-98405766 | -0.398736147 | 0.344157421 | 0.633894951 |
| chr20:53870453-53870742 | 0.309166192 | 0.344168115 | 0.633899408 |
| chr20:62382775-62383018 | 0.360992969 | 0.344187089 | 0.633912873 |
| chr16:68237332-68238112 | 0.303954916 | 0.344192801 | 0.633912873 |
| chr19:6767114-6768191 | 0.310259144 | 0.344200245 | 0.633912873 |
| chr8:96601199-96601416 | 0.525241546 | 0.34421691 | 0.633928327 |
| chr19:12964985-12965251 | 0.404241723 | 0.344226294 | 0.633930373 |
| chr7:3474333-3474901 | -0.388119366 | 0.344264543 | 0.633985575 |
| chr10:18400349-18400630 | 0.403422776 | 0.344289935 | 0.634003882 |
| chr6:63272483-63272683 | -0.535343065 | 0.344310031 | 0.634003882 |
| chr7:46407093-46407384 | 0.434843261 | 0.34431387 | 0.634003882 |
| chr22:33302061-33302338 | 0.39758946 | 0.344321031 | 0.634003882 |
| chr20:53892936-53893529 | 0.302106201 | 0.344327298 | 0.634003882 |
| chr10:73529458-73529719 | 0.443644389 | 0.344340333 | 0.634003882 |
| chr17:74657507-74657799 | 0.3648844 | 0.344344637 | 0.634003882 |
| chr3:119160421-119160671 | 0.422213171 | 0.344353964 | 0.634003882 |
| chr9:136808043-136808477 | 0.352467868 | 0.344356683 | 0.634003882 |
| chr7:5529615-5531561 | -0.295815829 | 0.344357228 | 0.634003882 |
| chr2:172153-172407 | -0.425642198 | 0.34437658 | 0.634024275 |
| chr13:32228317-32228674 | -0.374939316 | 0.344401653 | 0.634055202 |
| chr1:228933757-228934003 | 0.472975621 | 0.344411805 | 0.634058657 |
| chr16:84884146-84884346 | 0.476560887 | 0.344440247 | 0.634069192 |
| chr9:93827469-93827808 | 0.37728312 | 0.344444677 | 0.634069192 |
| chr6:101115164-101115392 | -0.491647249 | 0.344447072 | 0.634069192 |
| chr17:66006724-66006949 | -0.437770072 | 0.344450628 | 0.634069192 |
| chr12:95918305-95918857 | -0.354753599 | 0.344460882 | 0.634072834 |
| chr5:141326061-141326757 | -0.344625208 | 0.344506695 | 0.63414193 |
| chr3:153072844-153073172 | 0.407169923 | 0.344518912 | 0.634149184 |
| chr12:49346628-49346901 | 0.387185828 | 0.34453676 | 0.634156834 |
| chrX:96406616-96407884 | -0.308399295 | 0.344539621 | 0.634156834 |
| chr20:25280202-25280540 | -0.377286156 | 0.344569102 | 0.634195863 |
| chr1:204791774-204792108 | -0.355130294 | 0.344579843 | 0.634200398 |
| chr5:71482220-71482489 | -0.436143526 | 0.344615784 | 0.634242651 |
| chr1:202789716-202790049 | 0.353827705 | 0.344621482 | 0.634242651 |
| chr2:97995398-97996616 | 0.304693532 | 0.344627633 | 0.634242651 |
| chr4:107265978-107266182 | 0.468066336 | 0.344646137 | 0.634261471 |
| chr2:214410274-214410736 | -0.374001228 | 0.344666768 | 0.634284204 |
| chr5:66891513-66891713 | 0.559494626 | 0.34467639 | 0.634284786 |
| chr3:138479407-138479675 | -0.44786509 | 0.344683641 | 0.634284786 |
| chr7:19851465-19851842 | 0.365208214 | 0.344707824 | 0.634309081 |
| chr8:99818286-99818844 | -0.379263298 | 0.3447134 | 0.634309081 |
| chr1:155688258-155689335 | 0.305026584 | 0.344723345 | 0.634312147 |
| chr13:21811953-21812215 | -0.480299323 | 0.344744927 | 0.634319175 |
| chr5:84089077-84089303 | -0.468682744 | 0.344750446 | 0.634319175 |
| chr10:110207417-110207731 | 0.342886488 | 0.344752 | 0.634319175 |
| chr12:16332042-16332446 | 0.367026857 | 0.344778451 | 0.634335064 |
| chr2:133171469-133171932 | -0.408163366 | 0.344782445 | 0.634335064 |
| chr2:11629692-11630025 | -0.396367276 | 0.344785472 | 0.634335064 |
| chr17:48830639-48831639 | -0.307629641 | 0.344807811 | 0.634360931 |

| | | | |
|---|---|---|---|
| chr13:89514563-89514877 | -0.446308552 | 0.34483255 | 0.634391213 |
| chr1:234987384-234987983 | 0.327431239 | 0.344849147 | 0.634393592 |
| chr3:140051395-140051688 | -0.404508325 | 0.344853849 | 0.634393592 |
| chr7:99616429-99617314 | -0.321437971 | 0.344858681 | 0.634393592 |
| chr17:60044789-60044989 | 0.336464136 | 0.344884642 | 0.634426116 |
| chrX:152904541-152904836 | 0.385384988 | 0.344902474 | 0.634432557 |
| chr3:153997430-153997670 | -0.467223808 | 0.344904703 | 0.634432557 |
| chr8:127919469-127919693 | 0.380701434 | 0.344916917 | 0.63443677 |
| chr3:63995976-63996182 | 0.331942639 | 0.344923554 | 0.63443677 |
| chr3:119353306-119353618 | 0.37813644 | 0.344936055 | 0.634444534 |
| chr19:46614270-46614512 | 0.420448462 | 0.344954705 | 0.634463607 |
| chr1:13699936-13700522 | -0.370415574 | 0.344979349 | 0.634493703 |
| chr3:4021168-4021420 | -0.471990125 | 0.345004598 | 0.63451052 |
| chr2:113438199-113438473 | 0.390836168 | 0.345005279 | 0.63451052 |
| chr7:6630330-6630692 | 0.34474693 | 0.345019917 | 0.63451052 |
| chr15:78177447-78177811 | -0.375648067 | 0.345023483 | 0.63451052 |
| chr5:178196786-178197210 | 0.370839104 | 0.345029898 | 0.63451052 |
| chr6:116680858-116681971 | -0.301313424 | 0.345087296 | 0.634600845 |
| chr1:85308563-85308763 | -0.49359256 | 0.345111309 | 0.634629772 |
| chr1:164735591-164736004 | -0.398117136 | 0.345134567 | 0.634657309 |
| chr2:4955984-4956275 | -0.409476999 | 0.345174518 | 0.634715541 |
| chr17:63425435-63425926 | -0.318221855 | 0.345227378 | 0.6347794 |
| chr3:159781732-159782166 | 0.369682038 | 0.345238859 | 0.6347794 |
| chr20:34515474-34515819 | -0.39611289 | 0.345242725 | 0.6347794 |
| chr19:15418459-15419456 | 0.312608083 | 0.345265839 | 0.6347794 |
| chr3:87151846-87152352 | -0.388017979 | 0.345276806 | 0.6347794 |
| chr7:73842117-73842711 | -0.333500767 | 0.34529082 | 0.6347794 |
| chr1:112267720-112268066 | -0.463024479 | 0.34529327 | 0.6347794 |
| chr2:85503783-85504064 | -0.435807657 | 0.345302084 | 0.6347794 |
| chr17:42107709-42108097 | -0.345867288 | 0.345304396 | 0.6347794 |
| chr12:6904589-6905221 | -0.315962385 | 0.345314355 | 0.6347794 |
| chr16:80767842-80768042 | 0.468625932 | 0.345326861 | 0.6347794 |
| chr3:175062371-175062586 | 0.472904918 | 0.345338502 | 0.6347794 |
| chr14:36320036-36321268 | 0.3026731 | 0.345361364 | 0.6347794 |
| chr20:42204377-42204802 | -0.325773266 | 0.34536283 | 0.6347794 |
| chr3:11642800-11643160 | 0.354008747 | 0.345366961 | 0.6347794 |
| chr17:7843898-7844187 | -0.358440021 | 0.345367731 | 0.6347794 |
| chr11:121132292-121132681 | -0.401074098 | 0.345368406 | 0.6347794 |
| chr1:98051215-98051586 | -0.422801564 | 0.345369161 | 0.6347794 |
| chr16:19555101-19556005 | 0.30851697 | 0.345370867 | 0.6347794 |
| chr10:69861437-69861784 | -0.373423195 | 0.345396959 | 0.6347794 |
| chr19:46078616-46078884 | -0.425379968 | 0.345397146 | 0.6347794 |
| chr10:14653384-14654199 | -0.31361076 | 0.345406611 | 0.6347794 |
| chr12:3242390-3242638 | -0.449399326 | 0.345406621 | 0.6347794 |
| chr17:7913028-7913228 | -0.492868736 | 0.345411559 | 0.6347794 |
| chr20:36050480-36051373 | 0.320324407 | 0.345416359 | 0.6347794 |
| chr3:162190886-162191104 | -0.475534007 | 0.345438019 | 0.634801116 |
| chr2:38835084-38835327 | -0.475621273 | 0.345451028 | 0.634801116 |
| chr2:75591963-75592361 | 0.351491697 | 0.345453031 | 0.634801116 |
| chr10:99553451-99553716 | 0.407544524 | 0.345500687 | 0.634852687 |

| | | | |
|---|---|---|---|
| chr8:53286746-53286977 | -0.447225863 | 0.345535975 | 0.634852687 |
| chr19:35708436-35708636 | -0.51253685 | 0.345545405 | 0.634852687 |
| chr2:139685351-139685551 | -0.51253685 | 0.345545405 | 0.634852687 |
| chr2:25649938-25650179 | -0.51253685 | 0.345545405 | 0.634852687 |
| chr17:31580958-31581380 | 0.355142193 | 0.34556148 | 0.634852687 |
| chr15:65286144-65287255 | 0.306788477 | 0.345563829 | 0.634852687 |
| chr1:224641076-224641397 | 0.363208732 | 0.345566799 | 0.634852687 |
| chr1:201648246-201648499 | 0.380810705 | 0.345568861 | 0.634852687 |
| chr6:131910008-131910296 | -0.41230286 | 0.345570778 | 0.634852687 |
| chr10:73167684-73168735 | -0.308046315 | 0.345578762 | 0.634852687 |
| chr2:75278460-75278711 | -0.486229177 | 0.345580522 | 0.634852687 |
| chr1:169367410-169368717 | -0.297736023 | 0.345603225 | 0.634860898 |
| chr16:3123700-3124995 | 0.300469922 | 0.345606809 | 0.634860898 |
| chr5:143891484-143891684 | 0.545217869 | 0.345609848 | 0.634860898 |
| chr2:18331580-18331936 | -0.48160663 | 0.345639068 | 0.634899351 |
| chr13:50909546-50910819 | -0.309008255 | 0.345690064 | 0.634977803 |
| chr5:37717371-37717838 | -0.36052333 | 0.34571826 | 0.635014371 |
| chr1:113488741-113489105 | 0.369082743 | 0.345744142 | 0.63503158 |
| chr19:17830041-17831232 | 0.309861017 | 0.345744204 | 0.63503158 |
| chr5:72307937-72308742 | -0.304949299 | 0.345756892 | 0.635039661 |
| chr7:7026694-7027058 | 0.423966022 | 0.345804739 | 0.635112316 |
| chr21:41865243-41865589 | 0.38760484 | 0.345847141 | 0.635174967 |
| chr3:45089042-45089354 | 0.445916503 | 0.345885392 | 0.635229993 |
| chr4:42673168-42673368 | -0.519893028 | 0.345898865 | 0.635238337 |
| chr20:25861384-25861799 | 0.343436374 | 0.345919292 | 0.635238337 |
| chr7:44884411-44884867 | -0.347154965 | 0.345921157 | 0.635238337 |
| chr12:107671106-107671342 | -0.445222905 | 0.345923098 | 0.635238337 |
| chr1:197201955-197202221 | -0.339582817 | 0.345943357 | 0.635260315 |
| chr10:107914469-107915667 | 0.315282582 | 0.345973324 | 0.635286418 |
| chr12:55817270-55817489 | 0.376714688 | 0.345974154 | 0.635286418 |
| chr15:90301035-90301755 | -0.324708714 | 0.345996053 | 0.6352985 |
| chr5:157253477-157253833 | 0.353283469 | 0.345997317 | 0.6352985 |
| chr7:39911267-39911467 | -0.487846373 | 0.346054691 | 0.635388621 |
| chr14:40414568-40414824 | 0.422312728 | 0.346073858 | 0.635408586 |
| chr10:80530273-80530709 | -0.329211758 | 0.34609206 | 0.635426781 |
| chr16:13517717-13518067 | -0.506129957 | 0.346102807 | 0.63542781 |
| chr8:107804774-107805032 | 0.44447621 | 0.346109207 | 0.63542781 |
| chr9:129175858-129176139 | 0.357164192 | 0.346118058 | 0.635428835 |
| chr16:30697952-30699577 | 0.298069763 | 0.346130501 | 0.635436453 |
| chr8:126744012-126744386 | 0.379032474 | 0.346139972 | 0.635438615 |
| chr6:54758414-54758623 | -0.489807283 | 0.346161435 | 0.635443018 |
| chr17:3557664-3558142 | -0.357378 | 0.346166775 | 0.635443018 |
| chr6:63260397-63260697 | 0.42319303 | 0.34616725 | 0.635443018 |
| chr12:80504580-80504943 | -0.444869529 | 0.34617675 | 0.635445233 |
| chr3:156256915-156257127 | 0.476106905 | 0.346219091 | 0.635507731 |
| chr11:68447204-68447560 | 0.35260828 | 0.346268488 | 0.635583177 |
| chr17:60012816-60013030 | -0.37989348 | 0.346279187 | 0.635587587 |
| chr15:40893903-40894591 | 0.310025283 | 0.346289931 | 0.63559044 |
| chr20:47114150-47114399 | 0.447336849 | 0.346297331 | 0.63559044 |
| chr20:32031407-32031786 | -0.343368047 | 0.346306287 | 0.635591652 |

| | | | |
|---|---|---|---|
| chr8:143974880-143975316 | 0.333314474 | 0.346346555 | 0.635624777 |
| chr10:71623465-71623671 | -0.438514594 | 0.346347898 | 0.635624777 |
| chr6:107490339-107490631 | -0.397070067 | 0.346352507 | 0.635624777 |
| chr2:192604231-192604694 | -0.362349744 | 0.346357517 | 0.635624777 |
| chr12:56189168-56190915 | 0.298088635 | 0.346384246 | 0.635634035 |
| chr1:93878418-93878747 | 0.373434192 | 0.346389893 | 0.635634035 |
| chr7:27171682-27172077 | -0.394525462 | 0.346397477 | 0.635634035 |
| chr21:17512425-17513188 | 0.31901 | 0.346401528 | 0.635634035 |
| chr3:37814689-37814901 | 0.489061727 | 0.346404041 | 0.635634035 |
| chr17:79708672-79708872 | 0.458469597 | 0.346423261 | 0.63564499 |
| chr6:85923033-85923358 | -0.387652512 | 0.34643006 | 0.63564499 |
| chr12:108226782-108227100 | -0.463570515 | 0.346434898 | 0.63564499 |
| chr8:82774928-82775184 | -0.487358349 | 0.346452287 | 0.635661674 |
| chr9:14314487-14315688 | -0.304569015 | 0.346520498 | 0.635771601 |
| chr10:24952315-24952623 | -0.412434492 | 0.346533829 | 0.635780836 |
| chr6:110039456-110040387 | -0.323600714 | 0.34655536 | 0.635805113 |
| chr17:44324622-44325085 | 0.325351471 | 0.346566259 | 0.635809885 |
| chrX:119656286-119656718 | 0.351479995 | 0.346623097 | 0.635873179 |
| chr2:144441108-144441357 | -0.499000803 | 0.346639236 | 0.635873179 |
| chr3:194285577-194285777 | -0.499000803 | 0.346639236 | 0.635873179 |
| chr2:58870801-58871001 | -0.499000803 | 0.346639236 | 0.635873179 |
| chr3:53155619-53156207 | 0.312559709 | 0.346642253 | 0.635873179 |
| chr14:30005233-30005499 | 0.39732745 | 0.346651356 | 0.635874655 |
| chr5:62412406-62413354 | -0.299001355 | 0.346674514 | 0.635885803 |
| chr15:97899011-97899343 | 0.367840337 | 0.34667506 | 0.635885803 |
| chr1:220511276-220511476 | 0.466842709 | 0.346692128 | 0.635885803 |
| chr15:75328746-75328953 | 0.450459764 | 0.346696044 | 0.635885803 |
| chr5:141636285-141637642 | -0.299493887 | 0.346698929 | 0.635885803 |
| chr9:31937666-31938042 | 0.360314522 | 0.346726286 | 0.635906599 |
| chr14:61525165-61526158 | 0.304965078 | 0.346733759 | 0.635906599 |
| chr13:35448737-35449151 | -0.41767026 | 0.346735165 | 0.635906599 |
| chr2:141077486-141077744 | 0.436160604 | 0.346758365 | 0.635910102 |
| chr7:102390712-102391083 | -0.353104505 | 0.34675948 | 0.635910102 |
| chr3:36958698-36959044 | 0.349880042 | 0.346761973 | 0.635910102 |
| chr8:123823407-123823846 | -0.346985919 | 0.346784403 | 0.635936016 |
| chr16:70288689-70288958 | 0.384676986 | 0.346795542 | 0.635940625 |
| chr3:9396135-9397867 | 0.292335547 | 0.346809645 | 0.635940625 |
| chr5:172714731-172715295 | 0.317298893 | 0.346817638 | 0.635940625 |
| chr20:3218499-3218851 | 0.375389851 | 0.346820116 | 0.635940625 |
| chr6:73695651-73696496 | -0.320612173 | 0.346861361 | 0.635990111 |
| chr3:196502923-196504172 | -0.305113638 | 0.346863704 | 0.635990111 |
| chr17:44968277-44968734 | 0.341399473 | 0.346897553 | 0.636036953 |
| chr2:127024743-127025998 | -0.307272562 | 0.346914253 | 0.636052353 |
| chr7:100450421-100451037 | -0.326460557 | 0.346961757 | 0.636091934 |
| chr4:79917254-79917489 | -0.444250883 | 0.346962583 | 0.636091934 |
| chr8:126301046-126301257 | -0.399831241 | 0.346966465 | 0.636091934 |
| chr5:56669569-56669825 | -0.470940111 | 0.346977017 | 0.636091934 |
| chr7:76510078-76510797 | 0.321254548 | 0.346977349 | 0.636091934 |
| chr13:23028460-23028666 | 0.492973997 | 0.347011305 | 0.636123997 |
| chr5:180494077-180495123 | 0.304775698 | 0.347011443 | 0.636123997 |

| | | | |
|---|---|---|---|
| chrX:130268114-130269123 | 0.306567919 | 0.347037581 | 0.636138683 |
| chr11:46871418-46871878 | -0.400595451 | 0.347038747 | 0.636138683 |
| chr3:74351182-74351413 | 0.4997187 | 0.347044362 | 0.636138683 |
| chr9:32551353-32551553 | 0.404982985 | 0.347108462 | 0.636240958 |
| chr1:22172392-22172640 | -0.518428269 | 0.34713842 | 0.63628065 |
| chr3:61648299-61648690 | 0.331677687 | 0.347175426 | 0.63631413 |
| chr13:44854606-44854922 | -0.416250988 | 0.347184606 | 0.63631413 |
| chr19:18571396-18572102 | 0.323928826 | 0.347187469 | 0.63631413 |
| chr9:37651236-37651775 | -0.39582567 | 0.347189904 | 0.63631413 |
| chr3:75792337-75792606 | 0.452709072 | 0.347205802 | 0.636328047 |
| chr13:22915375-22916132 | -0.325012203 | 0.347249668 | 0.636389029 |
| chr14:57268182-57269399 | 0.300739162 | 0.347255688 | 0.636389029 |
| chr2:112960386-112960692 | -0.421946883 | 0.347280153 | 0.636408848 |
| chr2:27442269-27442676 | -0.36864225 | 0.347293254 | 0.636408848 |
| chr4:143514494-143514711 | 0.368476968 | 0.347297417 | 0.636408848 |
| chr7:87058778-87060215 | -0.30312655 | 0.347299726 | 0.636408848 |
| chr15:70771750-70771950 | 0.521736052 | 0.347323144 | 0.636421816 |
| chr20:23245501-23245943 | -0.367280777 | 0.347331433 | 0.636421816 |
| chr11:82494982-82495463 | -0.346973941 | 0.34733172 | 0.636421816 |
| chr11:62652825-62653781 | 0.304609758 | 0.347344458 | 0.636429936 |
| chr16:80751180-80751522 | -0.359869529 | 0.347360509 | 0.636444126 |
| chr1:226703048-226703910 | 0.330402983 | 0.34740742 | 0.636514858 |
| chr15:91247636-91248019 | 0.3832415 | 0.347431599 | 0.636543936 |
| chr5:154446629-154447021 | 0.349600955 | 0.347448012 | 0.636558786 |
| chr6:31718295-31718744 | 0.333679956 | 0.347473091 | 0.636570095 |
| chr6:141549976-141550253 | -0.486579353 | 0.347473508 | 0.636570095 |
| chr2:118061451-118061705 | 0.407188353 | 0.347479108 | 0.636570095 |
| chr1:114472553-114472762 | 0.360825171 | 0.347552809 | 0.636659452 |
| chr6:26421212-26421784 | -0.328837232 | 0.347557744 | 0.636659452 |
| chr8:109973871-109975151 | -0.297278337 | 0.347560448 | 0.636659452 |
| chr5:171152430-171152748 | -0.404265528 | 0.347561121 | 0.636659452 |
| chr2:43773800-43774567 | -0.317587422 | 0.347573667 | 0.636667212 |
| chr9:69035576-69036352 | -0.31173757 | 0.347594765 | 0.636690639 |
| chr3:16550849-16551236 | 0.326696862 | 0.347610714 | 0.636702071 |
| chr2:94208793-94209358 | 0.331278477 | 0.347617626 | 0.636702071 |
| chr5:172834086-172834516 | 0.335734273 | 0.347626491 | 0.636703087 |
| chr8:125957574-125958159 | -0.331614249 | 0.347657747 | 0.636745115 |
| chr7:11237734-11238164 | -0.367360388 | 0.347671739 | 0.636755521 |
| chr19:5923561-5923811 | -0.536578031 | 0.347693355 | 0.636779889 |
| chr13:102798743-102799501 | -0.307602374 | 0.347714645 | 0.63680366 |
| chr9:2857629-2857855 | 0.459980364 | 0.347760777 | 0.636844912 |
| chr13:23065544-23065959 | -0.395839537 | 0.34776885 | 0.636844912 |
| chr6:150817847-150818093 | 0.414220449 | 0.347771791 | 0.636844912 |
| chr17:76210947-76211160 | -0.53904754 | 0.347791549 | 0.636844912 |
| chr5:32734216-32734776 | -0.3985022 | 0.347796539 | 0.636844912 |
| chr2:714086-714785 | -0.331923508 | 0.347807921 | 0.636844912 |
| chr1:17520040-17520435 | -0.350213574 | 0.347808299 | 0.636844912 |
| chr1:177209385-177209585 | 0.483437514 | 0.347811974 | 0.636844912 |
| chr14:68295399-68295599 | 0.483437514 | 0.347811974 | 0.636844912 |
| chr1:51960330-51960586 | -0.463685128 | 0.347867975 | 0.636932231 |

| | | | |
|---|---|---|---|
| chr10:34740493-34740747 | -0.46217675 | 0.347883248 | 0.636944975 |
| chr15:44853344-44853608 | 0.419179266 | 0.347981817 | 0.637110222 |
| chr16:83926628-83927358 | 0.307636639 | 0.348044832 | 0.637205452 |
| chrX:101460728-101461018 | -0.426127464 | 0.348050463 | 0.637205452 |
| chr1:30788063-30788285 | 0.454888393 | 0.348067596 | 0.637221593 |
| chr20:8039223-8039463 | -0.4210674 | 0.348084734 | 0.637235528 |
| chrX:9788974-9789235 | -0.388388757 | 0.348091841 | 0.637235528 |
| chr5:91941653-91941963 | 0.436773628 | 0.348130679 | 0.637275237 |
| chr11:71179920-71180232 | -0.447446451 | 0.348138183 | 0.637275237 |
| chr2:96065452-96065684 | -0.44206212 | 0.348143972 | 0.637275237 |
| chr7:118590191-118590436 | -0.503011121 | 0.34815446 | 0.637275237 |
| chr7:44123161-44124487 | -0.300517773 | 0.348155117 | 0.637275237 |
| chr10:82397903-82398175 | -0.422144329 | 0.348176291 | 0.637298431 |
| chr11:109946294-109946571 | 0.403241974 | 0.348184424 | 0.637298431 |
| chr17:41535904-41536612 | -0.313220549 | 0.348231794 | 0.637350289 |
| chr6:131844097-131844625 | -0.343453615 | 0.348237651 | 0.637350289 |
| chr12:117837344-117837794 | 0.36118202 | 0.348239932 | 0.637350289 |
| chr4:100635780-100636336 | -0.336893089 | 0.348246028 | 0.637350289 |
| chr2:6826507-6826784 | 0.408764276 | 0.348267952 | 0.637364738 |
| chr13:34913173-34913380 | 0.47575344 | 0.34827056 | 0.637364738 |
| chr11:130002114-130003740 | 0.29598734 | 0.348300972 | 0.637405171 |
| chr1:150578741-150580153 | 0.296943457 | 0.348318596 | 0.637422199 |
| chr3:127694401-127694705 | 0.363260545 | 0.348332383 | 0.637430604 |
| chr17:82717021-82717444 | 0.348096364 | 0.348339827 | 0.637430604 |
| chr3:103440591-103441030 | 0.369137983 | 0.348404953 | 0.637522959 |
| chr3:64360266-64360490 | 0.337235689 | 0.34841494 | 0.637522959 |
| chr7:112383696-112384102 | -0.39331571 | 0.348423522 | 0.637522959 |
| chr14:50955360-50956149 | 0.300382959 | 0.348423578 | 0.637522959 |
| chrX:77785741-77786670 | -0.320599498 | 0.348434327 | 0.637527402 |
| chr7:2757378-2757630 | -0.42400609 | 0.34847773 | 0.63759159 |
| chr20:36886903-36887563 | 0.332429026 | 0.3484904 | 0.637599548 |
| chr9:98072563-98072887 | 0.382628344 | 0.348510777 | 0.637610549 |
| chr2:8513829-8514047 | 0.452526934 | 0.348516306 | 0.637610549 |
| chr12:102749718-102749918 | -0.567279522 | 0.348521378 | 0.637610549 |
| chr11:64305907-64306620 | -0.31576818 | 0.348532737 | 0.637616107 |
| chr1:206448959-206449160 | -0.396524017 | 0.34855897 | 0.637648873 |
| chr16:75206300-75206553 | 0.396017434 | 0.348576363 | 0.637665467 |
| chr20:25103599-25104495 | 0.33590384 | 0.348586524 | 0.637668832 |
| chr1:193104510-193105847 | -0.294873026 | 0.348602641 | 0.637683088 |
| chr12:111758106-111758562 | 0.342402272 | 0.348625025 | 0.637687912 |
| chr8:104990214-104990443 | -0.408194588 | 0.348626261 | 0.637687912 |
| chr2:217822369-217822569 | -0.459767339 | 0.348630245 | 0.637687912 |
| chr3:142279993-142280337 | 0.350882118 | 0.348659813 | 0.637726772 |
| chr3:139202171-139202884 | -0.329078805 | 0.348698477 | 0.637782267 |
| chr11:36744586-36744881 | -0.378327369 | 0.348714006 | 0.637795445 |
| chr4:37527786-37528054 | -0.416757535 | 0.348755407 | 0.637855941 |
| chr6:117344551-117344934 | -0.407100799 | 0.348780167 | 0.637886 |
| chr1:54904076-54904344 | -0.474264989 | 0.348829013 | 0.637960108 |
| chr11:107858001-107859359 | -0.303849648 | 0.348855121 | 0.637981903 |
| chr17:7833803-7834279 | -0.32478864 | 0.348862087 | 0.637981903 |

| | | | |
|---|---|---|---|
| chr5:94524893-94525204 | -0.40476734 | 0.34886591 | 0.637981903 |
| chr1:85214775-85215447 | -0.350791674 | 0.348906904 | 0.638034775 |
| chr13:48975516-48976954 | -0.298265443 | 0.348919982 | 0.638034775 |
| chr5:176810165-176810583 | -0.360728101 | 0.348933092 | 0.638034775 |
| chr2:195304394-195304651 | -0.480714448 | 0.348934221 | 0.638034775 |
| chr7:111763260-111763460 | -0.484960912 | 0.348936457 | 0.638034775 |
| chr3:168456293-168456493 | 0.489433122 | 0.348961464 | 0.638046679 |
| chr1:161427616-161428332 | -0.318997267 | 0.34896463 | 0.638046679 |
| chr10:5503405-5503904 | 0.312408942 | 0.348970184 | 0.638046679 |
| chr2:219352535-219352783 | -0.420500126 | 0.348976276 | 0.638046679 |
| chr7:8290658-8291314 | 0.318737149 | 0.348999801 | 0.638068438 |
| chr1:246712419-246712619 | -0.517468817 | 0.349022085 | 0.638068438 |
| chr2:35973158-35973472 | -0.389439848 | 0.349027985 | 0.638068438 |
| chr3:158858713-158858913 | 0.476440786 | 0.349030054 | 0.638068438 |
| chr8:143739929-143740322 | 0.326881604 | 0.349030386 | 0.638068438 |
| chr10:69407853-69408136 | -0.371430825 | 0.349044486 | 0.638068438 |
| chr3:40386850-40387298 | 0.389921921 | 0.34904647 | 0.638068438 |
| chr13:20902025-20903325 | 0.301395941 | 0.34906926 | 0.638094876 |
| chr6:36593467-36594727 | 0.303605835 | 0.34909199 | 0.638121203 |
| chr6:33425626-33426398 | 0.315986325 | 0.34913824 | 0.63817346 |
| chr19:32373734-32374254 | 0.325851111 | 0.349138522 | 0.63817346 |
| chr2:194550789-194551079 | 0.430728123 | 0.349145565 | 0.63817346 |
| chr6:3013528-3013831 | -0.386970556 | 0.349208584 | 0.638273422 |
| chr6:50849938-50850150 | 0.439689535 | 0.349234659 | 0.638286723 |
| chr17:57401948-57402427 | 0.335584825 | 0.34925212 | 0.638286723 |
| chr6:4020971-4021981 | 0.304372339 | 0.349253379 | 0.638286723 |
| chr19:55160542-55160995 | 0.327149991 | 0.349258075 | 0.638286723 |
| chr3:109647751-109647978 | -0.438263182 | 0.349267426 | 0.638286723 |
| chr17:59941747-59942075 | 0.33401033 | 0.349281016 | 0.638286723 |
| chr2:43798354-43798595 | -0.474411434 | 0.349282705 | 0.638286723 |
| chr3:134656085-134656285 | -0.482529118 | 0.349284767 | 0.638286723 |
| chr3:155325379-155325765 | -0.359939764 | 0.349300767 | 0.638286723 |
| chr6:26156176-26156434 | -0.357362158 | 0.349306288 | 0.638286723 |
| chr1:6927871-6928094 | -0.462422505 | 0.349307495 | 0.638286723 |
| chr9:126965890-126966090 | 0.487583627 | 0.349316697 | 0.638288315 |
| chr8:86522353-86522839 | -0.330896739 | 0.349349064 | 0.638316587 |
| chr17:73308308-73308618 | 0.326996651 | 0.349352124 | 0.638316587 |
| chr16:46830627-46831728 | 0.304234481 | 0.349357161 | 0.638316587 |
| chr12:20950260-20950480 | -0.454110489 | 0.349369112 | 0.638323202 |
| chr12:76739114-76739314 | 0.480875237 | 0.349434293 | 0.638427068 |
| chr3:62522284-62522845 | 0.318185858 | 0.349461812 | 0.638462123 |
| chr3:101091522-101091957 | -0.377397073 | 0.34949348 | 0.638493901 |
| chr11:18203870-18204155 | -0.416859811 | 0.349495953 | 0.638493901 |
| chr11:133078638-133078838 | -0.488738853 | 0.349504205 | 0.638493901 |
| chr12:13135394-13136249 | -0.314566189 | 0.349545038 | 0.638553273 |
| chr3:63228659-63229007 | -0.316878452 | 0.349563501 | 0.638560323 |
| chr7:150047521-150047844 | -0.449219259 | 0.349565565 | 0.638560323 |
| chr6:61032662-61033102 | 0.39490072 | 0.349587655 | 0.638573635 |
| chr3:193791652-193791967 | 0.364643597 | 0.349589521 | 0.638573635 |
| chrX:109268354-109268708 | -0.393137512 | 0.349624948 | 0.638623123 |

| | | | |
|---|---|---|---|
| chr13:22926116-22927523 | 0.297857896 | 0.349647309 | 0.638645485 |
| chr13:114234684-114235491 | 0.327549351 | 0.349653861 | 0.638645485 |
| chr3:78848869-78849202 | -0.402553196 | 0.349675531 | 0.638669842 |
| chr10:70867019-70867284 | 0.446942379 | 0.349710687 | 0.638675883 |
| chr15:66127872-66128083 | -0.475634632 | 0.349715889 | 0.638675883 |
| chr5:10424173-10424464 | 0.377625527 | 0.349720519 | 0.638675883 |
| chr11:4393194-4394073 | -0.320908558 | 0.349727814 | 0.638675883 |
| chr8:18563073-18563508 | 0.342355957 | 0.349736491 | 0.638675883 |
| chr19:37370696-37371577 | 0.311034309 | 0.349738805 | 0.638675883 |
| chr7:44795828-44797652 | -0.294033073 | 0.349743819 | 0.638675883 |
| chr11:116641586-116641836 | -0.428293402 | 0.349746306 | 0.638675883 |
| chr17:44123568-44124234 | 0.319742055 | 0.349757533 | 0.638675883 |
| chr10:100535562-100536274 | 0.3162193 | 0.349762192 | 0.638675883 |
| chr7:106009411-106009900 | -0.325258538 | 0.349778763 | 0.63869092 |
| chr2:72144667-72145130 | 0.352993495 | 0.349816405 | 0.638744431 |
| chr1:178037478-178038189 | -0.318887778 | 0.349911127 | 0.638902163 |
| chr4:124846359-124846630 | -0.477187445 | 0.349925744 | 0.638913627 |
| chr19:5719998-5720846 | 0.305698322 | 0.349962996 | 0.638966418 |
| chr6:15441338-15441757 | -0.420302547 | 0.349990778 | 0.639000539 |
| chr5:143977951-143978216 | -0.465476361 | 0.349998364 | 0.639000539 |
| chr4:72929647-72930232 | 0.334721385 | 0.350030699 | 0.639027282 |
| chr17:80345812-80346012 | 0.496996973 | 0.350030883 | 0.639027282 |
| chr16:2927054-2927369 | 0.368332407 | 0.350038147 | 0.639027282 |
| chr16:379239-379712 | -0.354162947 | 0.350046372 | 0.639027282 |
| chr8:110047652-110048137 | -0.370296631 | 0.350056762 | 0.639031025 |
| chr3:63015589-63016087 | 0.335364159 | 0.350082772 | 0.639063282 |
| chr3:47421031-47421404 | -0.350861495 | 0.350095937 | 0.639072088 |
| chr1:194763525-194763829 | -0.415345328 | 0.350113984 | 0.639089805 |
| chr1:154767419-154767761 | -0.422846628 | 0.350156964 | 0.639153034 |
| chr19:40798580-40799319 | 0.302860703 | 0.350168997 | 0.639159772 |
| chr17:59165911-59166279 | 0.327047769 | 0.350196899 | 0.639195472 |
| chr1:151384260-151384460 | 0.417802702 | 0.350207244 | 0.639195472 |
| chr6:27684322-27684857 | -0.313969859 | 0.350213582 | 0.639195472 |
| chr1:116466065-116466578 | 0.329092788 | 0.350264751 | 0.639273635 |
| chr19:38277493-38277717 | -0.423456037 | 0.350293888 | 0.639311586 |
| chr1:181514538-181514822 | -0.372717999 | 0.350348342 | 0.639395738 |
| chr5:179414410-179414625 | -0.43863715 | 0.350361974 | 0.639405387 |
| chr15:72184236-72184840 | -0.342107237 | 0.350402463 | 0.639453484 |
| chr7:143215126-143215464 | -0.367789696 | 0.35040502 | 0.639453484 |
| chr18:111348-112007 | 0.333199794 | 0.350438627 | 0.639493907 |
| chr20:1108441-1108701 | 0.463257495 | 0.350450434 | 0.639493907 |
| chr5:17413323-17413620 | -0.405929197 | 0.350452209 | 0.639493907 |
| chr9:93592660-93592973 | 0.372569969 | 0.35048909 | 0.639541096 |
| chr2:20135609-20135967 | -0.344990181 | 0.350497972 | 0.639541096 |
| chr13:40420337-40420537 | -0.466662927 | 0.350503109 | 0.639541096 |
| chr12:55817570-55818379 | 0.297015947 | 0.350526601 | 0.639567882 |
| chr12:79738146-79738461 | 0.370678384 | 0.350536999 | 0.639567882 |
| chrX:24693363-24694295 | -0.298149228 | 0.350542831 | 0.639567882 |
| chr16:1983862-1984375 | 0.323035211 | 0.350575022 | 0.639597846 |
| chr13:27374731-27374986 | 0.372966525 | 0.350588345 | 0.639597846 |

| | | | |
|---|---|---|---|
| chr2:1538662-1539125 | -0.35004585 | 0.350592144 | 0.639597846 |
| chr22:24594661-24594904 | 0.408149568 | 0.350609137 | 0.639597846 |
| chr7:134686850-134687133 | -0.444002834 | 0.35063288 | 0.639597846 |
| chr5:43602377-43604191 | 0.299616419 | 0.350635009 | 0.639597846 |
| chr9:3741246-3741455 | -0.506052513 | 0.350635754 | 0.639597846 |
| chr18:56358678-56358961 | -0.417273099 | 0.350636431 | 0.639597846 |
| chr2:30385420-30385641 | 0.452491782 | 0.350641038 | 0.639597846 |
| chr3:42202237-42202437 | 0.543852555 | 0.350642728 | 0.639597846 |
| chr16:54242278-54242567 | -0.421771857 | 0.350702703 | 0.639692016 |
| chr5:149922764-149923060 | -0.451029535 | 0.350743296 | 0.639739419 |
| chr1:108660630-108661636 | -0.304801144 | 0.350753081 | 0.639739419 |
| chr11:75583617-75583897 | -0.42963747 | 0.350753739 | 0.639739419 |
| chr9:72444467-72445026 | -0.374567523 | 0.350808798 | 0.63980693 |
| chr15:55301377-55301787 | 0.413917743 | 0.350817534 | 0.63980693 |
| chr1:169031530-169031880 | -0.411424463 | 0.350831733 | 0.63980693 |
| chrX:112247737-112248011 | 0.409184057 | 0.350836028 | 0.63980693 |
| chr14:23187136-23187372 | 0.416552799 | 0.350839665 | 0.63980693 |
| chr7:41224465-41225096 | 0.335575961 | 0.350847459 | 0.63980693 |
| chr6:29792374-29793109 | 0.337646447 | 0.350849205 | 0.63980693 |
| chr20:53984698-53984934 | 0.317407426 | 0.350858578 | 0.639808795 |
| chr5:13606044-13606486 | -0.338845958 | 0.350870319 | 0.639810729 |
| chr1:247256759-247257051 | 0.451728451 | 0.350885702 | 0.639810729 |
| chr17:29180082-29180836 | 0.323345861 | 0.350886757 | 0.639810729 |
| chr9:96169479-96169793 | -0.4195963 | 0.350893039 | 0.639810729 |
| chr8:90237600-90237862 | 0.444326401 | 0.350915347 | 0.639822549 |
| chr4:103994185-103994414 | -0.468243154 | 0.350941531 | 0.639822549 |
| chr5:56072433-56072679 | -0.468243154 | 0.350941531 | 0.639822549 |
| chr7:816329-817155 | -0.306152974 | 0.350941857 | 0.639822549 |
| chr12:106205321-106205707 | -0.371097975 | 0.35094678 | 0.639822549 |
| chr2:237471488-237471797 | 0.365655894 | 0.35095134 | 0.639822549 |
| chr20:64285654-64286213 | 0.348992161 | 0.350965868 | 0.639822549 |
| chr9:114681541-114681806 | 0.372764143 | 0.350966325 | 0.639822549 |
| chr15:48975706-48976404 | -0.331060426 | 0.350990564 | 0.639847385 |
| chr6:165632312-165632596 | -0.393882162 | 0.351013014 | 0.639847385 |
| chr16:81515675-81515890 | -0.432884421 | 0.351015341 | 0.639847385 |
| chr3:162835272-162835689 | -0.397846315 | 0.351020945 | 0.639847385 |
| chr6:32087673-32087873 | -0.48532948 | 0.3510371 | 0.639847385 |
| chr2:104431767-104432030 | 0.416817731 | 0.351040429 | 0.639847385 |
| chr7:460177-460428 | -0.390666859 | 0.351044284 | 0.639847385 |
| chr21:31604637-31604919 | 0.380579979 | 0.351049227 | 0.639847385 |
| chr5:178205231-178205565 | 0.332604888 | 0.351057481 | 0.639847385 |
| chr12:22121310-22121566 | -0.454072182 | 0.351063455 | 0.639847385 |
| chr1:9539808-9540134 | 0.378512972 | 0.351103902 | 0.639905881 |
| chr17:2393739-2394128 | -0.329499878 | 0.351122074 | 0.639908871 |
| chr3:107794705-107794930 | -0.427116936 | 0.351130277 | 0.639908871 |
| chr3:139491378-139491634 | -0.453966264 | 0.351130597 | 0.639908871 |
| chr19:45467497-45468074 | -0.311477092 | 0.351146865 | 0.639923298 |
| chr8:116734450-116734753 | 0.399475789 | 0.351221356 | 0.640022682 |
| chr7:96709291-96710278 | -0.312423963 | 0.351225562 | 0.640022682 |
| chr6:119742254-119742732 | -0.364016102 | 0.351234357 | 0.640022682 |

| | | | |
|---|---|---|---|
| chr3:153291144-153291421 | -0.376862975 | 0.351234812 | 0.640022682 |
| chr12:94560890-94561107 | -0.487567802 | 0.35125246 | 0.64003962 |
| chr14:37985933-37986256 | 0.377627504 | 0.351273291 | 0.640062354 |
| chr22:42519641-42520022 | 0.38467616 | 0.351288709 | 0.640075228 |
| chr7:122305132-122305381 | -0.398341311 | 0.351318897 | 0.640104802 |
| chr7:55951664-55952261 | 0.325753919 | 0.351321648 | 0.640104802 |
| chr9:100028171-100028521 | 0.37177371 | 0.351342352 | 0.640123329 |
| chr3:63940101-63940320 | 0.340029912 | 0.351348526 | 0.640123329 |
| chr5:15008444-15008916 | -0.314269332 | 0.351391739 | 0.640163764 |
| chr8:52695529-52695754 | 0.467876718 | 0.351399347 | 0.640163764 |
| chr10:96496324-96496524 | 0.459289478 | 0.351403276 | 0.640163764 |
| chr16:81076523-81077639 | -0.294420384 | 0.351404139 | 0.640163764 |
| chr12:118052477-118052868 | 0.349864207 | 0.351428362 | 0.640192671 |
| chrX:34893792-34894012 | -0.480081583 | 0.351484094 | 0.640278976 |
| chr11:45180662-45181167 | 0.348899127 | 0.351504761 | 0.6403014 |
| chr14:80088370-80089225 | -0.320796985 | 0.35151559 | 0.640305904 |
| chr5:74076430-74076630 | -0.471661307 | 0.351544043 | 0.640342509 |
| chr20:324124-324876 | 0.308611141 | 0.351569266 | 0.64037323 |
| chr2:70070862-70071190 | -0.369063734 | 0.351614764 | 0.640434242 |
| chr2:148926696-148926896 | -0.472018083 | 0.351619478 | 0.640434242 |
| chr3:38455329-38455823 | 0.316496535 | 0.351643548 | 0.640459639 |
| chr5:91383206-91383814 | -0.300485204 | 0.351665595 | 0.640459639 |
| chr2:64728154-64728365 | -0.438842025 | 0.351665922 | 0.640459639 |
| chr1:166975715-166975994 | -0.390281366 | 0.351666857 | 0.640459639 |
| chr3:88149094-88150365 | -0.293648444 | 0.35175183 | 0.640598552 |
| chr10:96089624-96090509 | -0.302605274 | 0.351766253 | 0.640598552 |
| chr2:10934261-10934645 | -0.383297596 | 0.351769674 | 0.640598552 |
| chr11:118928030-118928996 | 0.296552966 | 0.351776574 | 0.640598552 |
| chr19:39112278-39112665 | 0.378863674 | 0.351788497 | 0.640600458 |
| chr3:111577830-111578131 | 0.353455926 | 0.351794342 | 0.640600458 |
| chr17:63232552-63232824 | 0.386149699 | 0.351820687 | 0.640633205 |
| chr5:180090959-180091578 | -0.332640162 | 0.35188151 | 0.640726631 |
| chr9:35657414-35658800 | -0.29532648 | 0.351893853 | 0.640726631 |
| chr5:139864135-139864337 | 0.431724467 | 0.35189708 | 0.640726631 |
| chr7:152341493-152341762 | -0.444005849 | 0.351946919 | 0.640783485 |
| chr11:62497137-62497430 | -0.427220705 | 0.351950229 | 0.640783485 |
| chrX:115968365-115969308 | 0.304968055 | 0.351963547 | 0.640783485 |
| chr8:69290627-69290833 | -0.539377998 | 0.351965597 | 0.640783485 |
| chr22:30547339-30547586 | 0.457057899 | 0.35197012 | 0.640783485 |
| chr19:750714-751446 | 0.320644404 | 0.352007989 | 0.640833863 |
| chr20:53075049-53075264 | -0.461501784 | 0.352032044 | 0.640833863 |
| chr19:16185086-16185569 | 0.33750863 | 0.352038174 | 0.640833863 |
| chr13:41773406-41773606 | 0.524776874 | 0.352039631 | 0.640833863 |
| chr11:117985871-117986748 | 0.311833488 | 0.352046207 | 0.640833863 |
| chr1:154936441-154937680 | 0.29706132 | 0.352047973 | 0.640833863 |
| chr7:91089413-91089887 | -0.393007071 | 0.352075713 | 0.640854937 |
| chr14:55089559-55089850 | 0.358648937 | 0.35209238 | 0.640854937 |
| chr15:34183005-34183455 | -0.333451927 | 0.352098101 | 0.640854937 |
| chr6:52821872-52822080 | 0.491897542 | 0.3521004 | 0.640854937 |
| chr11:9572845-9573908 | 0.298995009 | 0.352101369 | 0.640854937 |

| | | | |
|---|---|---|---|
| chr7:99489131-99489511 | -0.405830824 | 0.352186106 | 0.640984881 |
| chr6:30571142-30571978 | 0.303912003 | 0.352189495 | 0.640984881 |
| chr1:80593796-80594574 | 0.309680864 | 0.352206529 | 0.641000658 |
| chr19:48255614-48256112 | 0.352974626 | 0.352247573 | 0.641053315 |
| chr8:114090935-114091144 | -0.467126268 | 0.352254369 | 0.641053315 |
| chr10:100482254-100482900 | 0.327890166 | 0.352260562 | 0.641053315 |
| chr5:43007349-43008865 | 0.295715028 | 0.352275194 | 0.641057804 |
| chr6:123777601-123777858 | -0.428074478 | 0.352279762 | 0.641057804 |
| chr7:6081023-6081892 | -0.297181474 | 0.352297535 | 0.641074922 |
| chr5:2989683-2989899 | -0.468990296 | 0.352320625 | 0.641093272 |
| chrX:31350232-31350689 | -0.377506556 | 0.352324353 | 0.641093272 |
| chr18:13680672-13680905 | -0.494202017 | 0.352364799 | 0.641151642 |
| chr22:31496029-31497173 | 0.304146067 | 0.352387652 | 0.641177997 |
| chr13:52127637-52128084 | -0.347185825 | 0.352433121 | 0.641245503 |
| chr3:138468905-138469229 | 0.403955331 | 0.352457208 | 0.641268311 |
| chr1:119870903-119871326 | 0.366329086 | 0.352462395 | 0.641268311 |
| chr14:102319431-102319798 | 0.333284184 | 0.35248491 | 0.64129171 |
| chr7:70599085-70599338 | -0.461122755 | 0.352491995 | 0.64129171 |
| chr19:3061287-3061571 | 0.460542865 | 0.352514966 | 0.641302409 |
| chr2:47175561-47177129 | -0.294464114 | 0.35251861 | 0.641302409 |
| chr5:119334980-119335442 | -0.335470018 | 0.352522985 | 0.641302409 |
| chr12:112148162-112148362 | 0.450887644 | 0.352560576 | 0.641353595 |
| chr14:77053247-77053600 | 0.351056586 | 0.352567862 | 0.641353595 |
| chr6:26331421-26331633 | -0.399490633 | 0.352586033 | 0.641366723 |
| chr16:74297347-74297673 | 0.338017479 | 0.352611721 | 0.641366723 |
| chr10:122648723-122649042 | 0.406090058 | 0.352617418 | 0.641366723 |
| chr18:623977-624260 | 0.404817111 | 0.352623482 | 0.641366723 |
| chr11:64304944-64305650 | 0.301158245 | 0.352635988 | 0.641366723 |
| chr17:55479232-55479433 | -0.455861098 | 0.35264024 | 0.641366723 |
| chr17:41528142-41528620 | 0.319385704 | 0.352641669 | 0.641366723 |
| chrX:66638529-66639582 | -0.316758332 | 0.352645718 | 0.641366723 |
| chr5:53430764-53430964 | -0.516193946 | 0.352653499 | 0.641366723 |
| chr9:137590417-137591014 | 0.314758652 | 0.352662383 | 0.641366723 |
| chr13:110590573-110591008 | 0.341984082 | 0.352667433 | 0.641366723 |
| chr20:31661480-31662056 | 0.324017871 | 0.352677952 | 0.641366723 |
| chr5:170790638-170790861 | -0.450836331 | 0.352683896 | 0.641366723 |
| chr8:132140617-132141040 | -0.369176642 | 0.352694584 | 0.641370936 |
| chr20:48620497-48620842 | 0.301881947 | 0.352705984 | 0.641376445 |
| chr13:38601649-38601849 | 0.490109041 | 0.352714712 | 0.641377095 |
| chr1:8848861-8849119 | 0.436011153 | 0.352724021 | 0.641378802 |
| chr4:181701370-181701579 | 0.497706777 | 0.352736789 | 0.641386796 |
| chr7:129433647-129434789 | -0.305767041 | 0.352783315 | 0.64143594 |
| chr7:91264100-91265549 | -0.295621485 | 0.352784921 | 0.64143594 |
| chr7:120272862-120273205 | -0.383588101 | 0.352794938 | 0.64143594 |
| chr2:128306045-128306700 | 0.306508882 | 0.352797301 | 0.64143594 |
| chr19:55484950-55485388 | 0.328515947 | 0.35285164 | 0.641490389 |
| chr2:20098237-20098554 | -0.346688108 | 0.352857847 | 0.641490389 |
| chr5:94850333-94850629 | -0.36628934 | 0.352858197 | 0.641490389 |
| chr2:211033858-211034316 | -0.391586959 | 0.352867857 | 0.641490389 |
| chr6:136922024-136922224 | 0.465621489 | 0.35286911 | 0.641490389 |

| | | | |
|---|---|---|---|
| chr13:42216117-42216317 | 0.487165679 | 0.352912915 | 0.641554802 |
| chr7:52328321-52328666 | -0.373346965 | 0.352955339 | 0.641616453 |
| chr4:124694359-124694626 | 0.400271087 | 0.352963576 | 0.641616453 |
| chr18:49460070-49461115 | -0.308193665 | 0.352972805 | 0.641618007 |
| chr3:185768548-185768777 | -0.428778056 | 0.352987903 | 0.641630231 |
| chr17:4789675-4790474 | 0.323057487 | 0.353007217 | 0.641650116 |
| chr12:541618-541818 | -0.503482432 | 0.353050994 | 0.641714465 |
| chr21:29298639-29299716 | 0.30413303 | 0.353101002 | 0.641771543 |
| chr13:45340411-45342057 | -0.289523025 | 0.353104493 | 0.641771543 |
| chr12:121797070-121797275 | 0.357112368 | 0.353107524 | 0.641771543 |
| chr5:66596002-66597159 | -0.303313473 | 0.353162775 | 0.641856736 |
| chr8:23161811-23162106 | 0.396913009 | 0.353206172 | 0.641915674 |
| chr1:39408518-39409397 | 0.302039066 | 0.353218049 | 0.641915674 |
| chr19:28291127-28291397 | 0.432042434 | 0.353220337 | 0.641915674 |
| chr6:36623521-36623771 | 0.417742324 | 0.353234351 | 0.641925836 |
| chr6:151690406-151690683 | -0.3795767 | 0.353242684 | 0.641925836 |
| chrX:21985267-21985617 | 0.357432888 | 0.353255165 | 0.641933291 |
| chr11:71800392-71800857 | -0.398736413 | 0.353267437 | 0.641940368 |
| chr2:101027891-101028091 | 0.495269844 | 0.353298204 | 0.641972217 |
| chr7:43511131-43511358 | -0.416918562 | 0.353302649 | 0.641972217 |
| chr17:45903084-45903318 | 0.412132622 | 0.353319955 | 0.641972217 |
| chr19:45667694-45668224 | 0.333153757 | 0.353324674 | 0.641972217 |
| chrX:12894136-12894383 | -0.448711828 | 0.35333337 | 0.641972217 |
| chr2:210195564-210195764 | -0.48284731 | 0.353342702 | 0.641972217 |
| chr8:61120784-61121279 | 0.370353464 | 0.353352965 | 0.641972217 |
| chrX:70423119-70423450 | 0.384755759 | 0.353362687 | 0.641972217 |
| chr3:189509183-189509386 | -0.489728762 | 0.353364216 | 0.641972217 |
| chr1:164590181-164591079 | 0.309090426 | 0.353393774 | 0.641972217 |
| chr3:153640599-153640865 | -0.460410966 | 0.353397026 | 0.641972217 |
| chr16:73714238-73714438 | -0.467779473 | 0.353403269 | 0.641972217 |
| chr22:37674940-37675690 | -0.316320335 | 0.353409168 | 0.641972217 |
| chr2:2844227-2844442 | -0.427162855 | 0.353410257 | 0.641972217 |
| chr2:5777112-5777450 | 0.363264423 | 0.353410641 | 0.641972217 |
| chr17:44344888-44345441 | 0.305820942 | 0.353432118 | 0.641995134 |
| chr7:43758337-43759299 | -0.307229383 | 0.353440014 | 0.641995134 |
| chr15:75646958-75647158 | 0.39885679 | 0.353449761 | 0.64199762 |
| chr14:105773980-105774330 | 0.384182427 | 0.353479166 | 0.642035811 |
| chr6:116268526-116268935 | -0.331040537 | 0.353489186 | 0.642038791 |
| chr10:119872066-119873663 | 0.296933103 | 0.353503171 | 0.642048971 |
| chr22:17742457-17742737 | -0.392484188 | 0.353515441 | 0.642053952 |
| chr22:27382117-27382355 | 0.450603994 | 0.353522672 | 0.642053952 |
| chr12:95259491-95259750 | -0.465712529 | 0.353538015 | 0.642066599 |
| chr8:17676269-17676606 | 0.368939028 | 0.353554589 | 0.642081481 |
| chr19:12666654-12666865 | 0.406536909 | 0.353630998 | 0.642188104 |
| chr6:166951438-166951638 | 0.424833061 | 0.353640964 | 0.642188104 |
| chr2:95159781-95160323 | 0.33669154 | 0.353649353 | 0.642188104 |
| chr17:74863051-74863315 | -0.410628 | 0.353651761 | 0.642188104 |
| chr1:200279806-200280133 | 0.332765783 | 0.353655207 | 0.642188104 |
| chr8:123190856-123191162 | -0.342148401 | 0.353706301 | 0.642260186 |
| chr8:116800308-116800997 | 0.313189071 | 0.353715712 | 0.642260186 |

| | | | |
|---|---|---|---|
| chr12:124544382-124544697 | -0.380058127 | 0.353720049 | 0.642260186 |
| chr4:115154930-115155130 | 0.468021307 | 0.353740663 | 0.642274409 |
| chr17:31842614-31842814 | 0.482957322 | 0.353750303 | 0.642274409 |
| chr1:160078672-160078958 | -0.375792404 | 0.35376055 | 0.642274409 |
| chr3:188202493-188202761 | -0.399973344 | 0.353761411 | 0.642274409 |
| chrX:41923298-41923885 | 0.326422813 | 0.353771548 | 0.642277593 |
| chr18:36075915-36076406 | -0.356175313 | 0.353797439 | 0.64230938 |
| chr3:69762646-69762850 | -0.503585471 | 0.353819854 | 0.642320442 |
| chr15:71232770-71233053 | -0.45328096 | 0.353825826 | 0.642320442 |
| chr1:156601106-156602086 | 0.312812057 | 0.353828681 | 0.642320442 |
| chr17:81200897-81201097 | 0.407809488 | 0.353846462 | 0.642323445 |
| chr10:50561824-50562072 | -0.413217408 | 0.353859688 | 0.642323445 |
| chr11:46120799-46122015 | -0.297259837 | 0.353866603 | 0.642323445 |
| chr11:60762891-60763137 | 0.435139446 | 0.353874565 | 0.642323445 |
| chr1:111738789-111739982 | 0.299097586 | 0.353875893 | 0.642323445 |
| chr3:72446999-72447980 | 0.301141084 | 0.353886457 | 0.642323445 |
| chr13:19487659-19488012 | 0.402888788 | 0.353889017 | 0.642323445 |
| chr1:59010688-59010916 | -0.51939702 | 0.353913315 | 0.642333721 |
| chr9:89664881-89665275 | 0.335109309 | 0.353915095 | 0.642333721 |
| chr5:23492139-23492339 | 0.452247094 | 0.353919828 | 0.642333721 |
| chr15:70508728-70508980 | 0.387530898 | 0.353952863 | 0.64237253 |
| chr8:118621137-118622561 | -0.293997236 | 0.353957979 | 0.64237253 |
| chr10:4434281-4434534 | 0.486066017 | 0.353997056 | 0.642413246 |
| chr3:14402074-14403359 | 0.292952547 | 0.354000858 | 0.642413246 |
| chr12:31589827-31591420 | -0.308386382 | 0.354005566 | 0.642413246 |
| chr13:20495629-20495866 | 0.402531991 | 0.354043635 | 0.642463285 |
| chr19:10960386-10961352 | 0.300662834 | 0.354063925 | 0.642463285 |
| chr1:181556643-181556859 | 0.439894672 | 0.354063943 | 0.642463285 |
| chr8:144509760-144509977 | 0.357625948 | 0.354074933 | 0.642463285 |
| chr14:85471676-85472091 | 0.371517878 | 0.354075065 | 0.642463285 |
| chr8:61479877-61480462 | -0.359653068 | 0.354097821 | 0.642489361 |
| chr1:63615847-63616085 | -0.429956113 | 0.354145668 | 0.642560961 |
| chr17:74720750-74720950 | -0.503191402 | 0.354164815 | 0.642573098 |
| chr6:71420042-71421143 | 0.297097887 | 0.35416913 | 0.642573098 |
| chr1:162151943-162152977 | -0.302974828 | 0.354191756 | 0.642598933 |
| chr18:33802965-33803265 | -0.426535863 | 0.354217984 | 0.642631301 |
| chr9:137067859-137068098 | 0.35832956 | 0.354252586 | 0.64267886 |
| chr7:32381117-32381340 | 0.389139234 | 0.354288634 | 0.642729039 |
| chr8:128035034-128035644 | 0.302572836 | 0.354315559 | 0.642762668 |
| chr12:106682909-106683179 | -0.44315659 | 0.354351484 | 0.642812618 |
| chr3:72048569-72048769 | 0.487544091 | 0.354366881 | 0.642825331 |
| chr3:171640487-171640926 | -0.36162383 | 0.354388739 | 0.642849762 |
| chr7:103347952-103348199 | -0.376296747 | 0.354405557 | 0.642865051 |
| chr11:27184034-27184234 | 0.456428253 | 0.354421017 | 0.642877874 |
| chr7:100157158-100157402 | -0.432276952 | 0.354434561 | 0.642885787 |
| chr11:126355208-126356955 | -0.299536764 | 0.35444216 | 0.642885787 |
| chr10:13302063-13302883 | 0.301873212 | 0.354484312 | 0.642947022 |
| chr4:128427733-128428380 | 0.316314526 | 0.354504444 | 0.642955324 |
| chr16:31868657-31868936 | 0.419400284 | 0.354505672 | 0.642955324 |
| chr11:2975649-2975943 | -0.383388048 | 0.354531642 | 0.642972209 |

| | | | |
|---|---|---|---|
| chr8:129687465-129687760 | -0.371309373 | 0.354531765 | 0.642972209 |
| chr3:112074734-112075530 | 0.316036763 | 0.354543831 | 0.642978045 |
| chr1:194616208-194616668 | -0.399849605 | 0.354551766 | 0.642978045 |
| chr12:6250443-6250643 | -0.53615303 | 0.354584943 | 0.643022993 |
| chr12:116533234-116533558 | 0.331129444 | 0.354597529 | 0.643030599 |
| chr16:81035358-81036470 | 0.300130442 | 0.354617605 | 0.643044939 |
| chr3:150079079-150079280 | 0.412217201 | 0.354622222 | 0.643044939 |
| chr1:155078645-155078936 | 0.34377571 | 0.354651254 | 0.643052594 |
| chr1:5509932-5510132 | 0.513400063 | 0.354654295 | 0.643052594 |
| chr17:54889023-54889279 | 0.405474901 | 0.354662713 | 0.643052594 |
| chrY:56840429-56840766 | 0.39176435 | 0.354667187 | 0.643052594 |
| chr7:87291215-87291587 | -0.409746887 | 0.354668407 | 0.643052594 |
| chr3:151461567-151461813 | 0.449348834 | 0.354682789 | 0.643063455 |
| chr20:9323707-9323940 | 0.414967157 | 0.354697167 | 0.643070732 |
| chr1:219469809-219470011 | 0.468009106 | 0.354703589 | 0.643070732 |
| chr14:55123054-55123374 | 0.396652074 | 0.354722061 | 0.643089006 |
| chr17:58679179-58680011 | 0.296272179 | 0.354738873 | 0.643095903 |
| chr10:104630861-104631061 | 0.485141728 | 0.354748068 | 0.643095903 |
| chr20:18187047-18187285 | -0.462768946 | 0.354751044 | 0.643095903 |
| chr10:12195406-12196648 | -0.297523231 | 0.354784272 | 0.64312859 |
| chr2:15577123-15577468 | -0.335467461 | 0.354785862 | 0.64312859 |
| chr1:201009081-201009430 | 0.338812356 | 0.354813244 | 0.643163009 |
| chr17:76132604-76132965 | 0.38075434 | 0.354844145 | 0.643203806 |
| chr17:50055533-50056328 | 0.301545084 | 0.3548581 | 0.643213884 |
| chr3:94234198-94234710 | -0.344275668 | 0.354870355 | 0.643220881 |
| chr12:95858453-95859308 | -0.299524733 | 0.354880866 | 0.643223607 |
| chr2:90385845-90386079 | 0.48642403 | 0.354888648 | 0.643223607 |
| chr12:74922071-74922417 | 0.423396247 | 0.354921609 | 0.643268131 |
| chr7:13990992-13991862 | 0.29403106 | 0.354940314 | 0.643277605 |
| chr9:77053446-77053746 | -0.478902967 | 0.354943627 | 0.643277605 |
| chr5:73519464-73520121 | -0.320850533 | 0.35497007 | 0.643298712 |
| chr3:13283210-13283429 | 0.391341888 | 0.354972065 | 0.643298712 |
| chr4:122205775-122206020 | -0.455525982 | 0.354981835 | 0.643301204 |
| chr3:119468709-119469690 | 0.298130109 | 0.355023711 | 0.643361874 |
| chr12:122834607-122834853 | 0.389718605 | 0.355034909 | 0.643366951 |
| chr17:49101010-49101930 | 0.306149816 | 0.355043629 | 0.643367538 |
| chr7:36510088-36510310 | -0.448706751 | 0.355054518 | 0.643371783 |
| chr12:71134670-71134870 | 0.508990802 | 0.355064666 | 0.643371783 |
| chr8:93118525-93118855 | -0.402754963 | 0.355071162 | 0.643371783 |
| chr1:28144608-28144896 | -0.448424928 | 0.355119796 | 0.643444689 |
| chr2:423603-423810 | -0.481054828 | 0.355140811 | 0.643466998 |
| chr2:64488842-64489084 | -0.448523063 | 0.355148904 | 0.643466998 |
| chr3:172750160-172751494 | -0.291102829 | 0.35521141 | 0.64356503 |
| chr11:68271203-68272489 | -0.30012982 | 0.355222888 | 0.643570607 |
| chr1:224481580-224481981 | 0.343159047 | 0.355234629 | 0.643576662 |
| chr10:52951870-52952179 | -0.359978794 | 0.355264344 | 0.64360207 |
| chr2:26355429-26355917 | -0.339741024 | 0.355270593 | 0.64360207 |
| chr12:118372550-118373320 | 0.297925353 | 0.355278963 | 0.64360207 |
| chr16:10737122-10737367 | 0.387648296 | 0.355282252 | 0.64360207 |
| chr1:114165142-114165525 | 0.31365384 | 0.355303134 | 0.643622294 |

| | | | |
|---|---|---|---|
| chr14:95611495-95611700 | -0.453323029 | 0.355310216 | 0.643622294 |
| chr2:20438563-20438836 | -0.371433589 | 0.355318995 | 0.643622979 |
| chr14:93959390-93959825 | -0.329823436 | 0.355406221 | 0.643759397 |
| chr2:10465462-10466211 | -0.31539781 | 0.355411109 | 0.643759397 |
| chr6:161122328-161122562 | -0.501297879 | 0.355426697 | 0.64376547 |
| chr18:21111192-21112278 | 0.299307543 | 0.355439627 | 0.64376547 |
| chr5:158594696-158595555 | 0.306098314 | 0.355439668 | 0.64376547 |
| chr8:60640493-60640760 | -0.512939368 | 0.355473092 | 0.643810789 |
| chr18:11994309-11994509 | -0.498667132 | 0.355484153 | 0.643815604 |
| chr9:127423651-127425154 | 0.300239513 | 0.355511327 | 0.643837057 |
| chr3:86990419-86990772 | 0.356173024 | 0.355512804 | 0.643837057 |
| chr6:3382385-3382647 | -0.432603745 | 0.355532161 | 0.643845799 |
| chr2:131046488-131046783 | -0.440553463 | 0.35553776 | 0.643845799 |
| chr6:53725853-53726103 | -0.400054615 | 0.35554284 | 0.643845799 |
| chr7:87151951-87152997 | -0.293933281 | 0.355555169 | 0.643851929 |
| chr8:144019737-144020237 | 0.336019676 | 0.35556303 | 0.643851929 |
| chr7:118195912-118196171 | -0.435381097 | 0.355581605 | 0.643870348 |
| chr11:69218374-69218648 | -0.396906106 | 0.355609988 | 0.643898248 |
| chr7:121107985-121108242 | -0.424279576 | 0.355613821 | 0.643898248 |
| chr17:50199723-50200993 | 0.293490475 | 0.355623682 | 0.643900887 |
| chr3:86245435-86245659 | -0.445255565 | 0.355635928 | 0.643906882 |
| chr19:47113698-47113974 | -0.34549839 | 0.3556438 | 0.643906882 |
| chr15:80289583-80289789 | -0.534940557 | 0.355678057 | 0.643939988 |
| chr3:193782364-193782571 | 0.404667812 | 0.355678894 | 0.643939988 |
| chr12:110048019-110048777 | -0.314208886 | 0.355702423 | 0.643967371 |
| chr12:109005282-109005508 | 0.471176172 | 0.355715619 | 0.643976045 |
| chr1:153963201-153964391 | 0.292105771 | 0.355772765 | 0.644056212 |
| chr17:67068411-67068611 | -0.460033868 | 0.355776712 | 0.644056212 |
| chr6:149031751-149032775 | -0.31533479 | 0.355840174 | 0.644114664 |
| chr11:47853566-47853793 | 0.393662512 | 0.355841898 | 0.644114664 |
| chr6:165834558-165835076 | -0.3251877 | 0.355855483 | 0.644114664 |
| chr19:42301901-42302971 | 0.29843609 | 0.355859324 | 0.644114664 |
| chr3:190513444-190515126 | -0.293545069 | 0.355866526 | 0.644114664 |
| chr9:37403900-37404250 | 0.357622489 | 0.355868927 | 0.644114664 |
| chr3:152743691-152744002 | 0.411088228 | 0.355869827 | 0.644114664 |
| chr7:64562640-64563363 | 0.30932652 | 0.355880228 | 0.644114664 |
| chr1:164659669-164660585 | 0.31137273 | 0.355884659 | 0.644114664 |
| chr6:53493395-53493698 | -0.394602212 | 0.355896497 | 0.644120875 |
| chr20:53685233-53685603 | 0.300026016 | 0.355927007 | 0.64416088 |
| chr1:206135241-206135653 | -0.357877575 | 0.355991507 | 0.644262394 |
| chr1:117059398-117060975 | 0.289603397 | 0.35601199 | 0.644284247 |
| chr15:64702889-64703735 | 0.302143007 | 0.356027413 | 0.64429694 |
| chr7:4631442-4632254 | -0.303403017 | 0.356047629 | 0.644306304 |
| chr2:128799865-128800239 | -0.410452659 | 0.356049405 | 0.644306304 |
| chr1:195615650-195615913 | -0.456069403 | 0.356063364 | 0.644316347 |
| chr4:162499583-162499813 | -0.446071206 | 0.356100952 | 0.644363291 |
| chr12:107685505-107686298 | -0.303350982 | 0.356111009 | 0.644363291 |
| chr8:107935629-107935829 | -0.494854684 | 0.356118193 | 0.644363291 |
| chr14:106661707-106661938 | -0.449713381 | 0.356130963 | 0.644363291 |
| chr13:41131999-41133119 | -0.294714752 | 0.356131354 | 0.644363291 |

| | | | |
|---|---|---|---|
| chr19:4535077-4535521 | 0.335530343 | 0.356145771 | 0.644373846 |
| chr13:32427179-32428439 | -0.299532774 | 0.356161774 | 0.644373846 |
| chr2:26462573-26462847 | -0.420429174 | 0.356169946 | 0.644373846 |
| chr20:44609930-44610387 | 0.316479767 | 0.356170827 | 0.644373846 |
| chr1:207692418-207692771 | -0.354736634 | 0.356203078 | 0.644416977 |
| chr6:3226698-3227163 | -0.372066429 | 0.356252046 | 0.644478899 |
| chr19:41642022-41642239 | -0.447501739 | 0.356254128 | 0.644478899 |
| chr1:114833471-114833808 | 0.347653343 | 0.356288273 | 0.644514827 |
| chr14:102086480-102086680 | 0.379389573 | 0.356290811 | 0.644514827 |
| chr17:21287788-21288301 | 0.312238338 | 0.356313148 | 0.644540017 |
| chr14:76913201-76913928 | 0.305806669 | 0.35635359 | 0.644597954 |
| chr2:113273100-113273411 | 0.366319499 | 0.356383868 | 0.644637506 |
| chr4:152408569-152408877 | -0.431835405 | 0.356407199 | 0.644664488 |
| chrX:78104493-78105173 | -0.316595683 | 0.356420578 | 0.64467347 |
| chr5:127280530-127280736 | -0.489136486 | 0.356431328 | 0.644677695 |
| chr5:34545435-34545763 | -0.447422655 | 0.356442033 | 0.644681839 |
| chr3:125214129-125214370 | -0.466759095 | 0.356453005 | 0.644686467 |
| chr8:103749935-103750209 | 0.373445299 | 0.356490589 | 0.644712999 |
| chr2:113890683-113891099 | -0.332698843 | 0.356499579 | 0.644712999 |
| chr11:1652763-1653065 | 0.426982794 | 0.356500855 | 0.644712999 |
| chr12:120439717-120440052 | 0.340901649 | 0.356504229 | 0.644712999 |
| chr12:131835136-131835400 | -0.377508099 | 0.356514777 | 0.644712999 |
| chr5:174581381-174581698 | -0.427161546 | 0.356518607 | 0.644712999 |
| chr18:7272803-7273018 | -0.508441819 | 0.356529399 | 0.644712999 |
| chr14:103692975-103693386 | -0.357127188 | 0.356534989 | 0.644712999 |
| chr22:19171046-19171417 | 0.347102172 | 0.356545065 | 0.644716004 |
| chr2:26771036-26771354 | -0.364175708 | 0.356559838 | 0.644727485 |
| chr5:149959416-149959809 | -0.342374146 | 0.356568243 | 0.644727485 |
| chr2:153478461-153479311 | -0.304511962 | 0.356603335 | 0.644775721 |
| chr8:133296989-133298170 | 0.290974103 | 0.356613417 | 0.644778735 |
| chr6:131116852-131117137 | -0.444139685 | 0.35663099 | 0.64478537 |
| chr6:11414620-11414920 | -0.39189186 | 0.356648975 | 0.64478537 |
| chr7:66557589-66557906 | -0.367183904 | 0.356649831 | 0.64478537 |
| chr16:70778927-70779337 | -0.319121165 | 0.356650747 | 0.64478537 |
| chr16:5097705-5098804 | 0.290514124 | 0.356669677 | 0.644787913 |
| chr17:71571242-71571494 | -0.456211149 | 0.35667206 | 0.644787913 |
| chr1:235128362-235129166 | 0.298219668 | 0.3566774 | 0.644787913 |
| chr17:59038869-59039182 | 0.312567034 | 0.356700047 | 0.644794226 |
| chr1:33612963-33613178 | -0.424418334 | 0.356705221 | 0.644794226 |
| chr1:217953308-217953842 | -0.309661823 | 0.356708347 | 0.644794226 |
| chr17:50933483-50933878 | -0.335296033 | 0.356718806 | 0.644794226 |
| chr14:30317711-30318065 | -0.377295154 | 0.356731357 | 0.644794226 |
| chr7:157253941-157254201 | -0.430267184 | 0.356731383 | 0.644794226 |
| chr10:132962175-132962451 | -0.376318228 | 0.356753334 | 0.644818691 |
| chr10:17787129-17787480 | -0.377346008 | 0.356776343 | 0.644829434 |
| chr17:81538107-81538307 | -0.453991088 | 0.356776734 | 0.644829434 |
| chr5:113606276-113606878 | -0.321501023 | 0.356784525 | 0.644829434 |
| chr12:2752591-2752943 | -0.389690298 | 0.356815308 | 0.644869858 |
| chr6:150418245-150418566 | -0.402080942 | 0.356829988 | 0.644878289 |
| chr11:27362721-27363537 | -0.299041661 | 0.356836805 | 0.644878289 |

| | | | |
|---|---|---|---|
| chr10:71512829-71513199 | 0.351211604 | 0.356862887 | 0.644887415 |
| chr20:47663598-47664436 | 0.288577502 | 0.356869538 | 0.644887415 |
| chr2:231884732-231884975 | -0.451865615 | 0.356880085 | 0.644887415 |
| chr1:25859256-25859672 | 0.35064767 | 0.356886747 | 0.644887415 |
| chr8:109803204-109803404 | 0.457097731 | 0.356890262 | 0.644887415 |
| chr8:144291869-144292448 | 0.312913891 | 0.356896911 | 0.644887415 |
| chr14:46250149-46250385 | -0.446905136 | 0.35690077 | 0.644887415 |
| chr10:14074516-14074818 | 0.404960221 | 0.356911042 | 0.644890767 |
| chr18:35999096-35999296 | 0.476154182 | 0.356932887 | 0.644899766 |
| chr12:126156829-126157156 | -0.386513481 | 0.356935431 | 0.644899766 |
| chr16:24070942-24071203 | 0.412003665 | 0.356947041 | 0.644899766 |
| chr14:77760443-77761944 | 0.29199232 | 0.356952972 | 0.644899766 |
| chr6:150408866-150409066 | -0.514751505 | 0.356964264 | 0.644899766 |
| chr12:130335574-130335774 | 0.449203715 | 0.356966522 | 0.644899766 |
| chr4:22962505-22962733 | -0.458746029 | 0.357000681 | 0.644941596 |
| chr6:13795145-13795361 | 0.412729034 | 0.357006511 | 0.644941596 |
| chr17:21599134-21599396 | -0.457517628 | 0.357058381 | 0.645020094 |
| chr8:140358688-140359362 | -0.341300716 | 0.357072057 | 0.645029591 |
| chr6:1227881-1228160 | -0.462344131 | 0.357085484 | 0.64503864 |
| chr16:67160655-67161248 | -0.309290806 | 0.357133682 | 0.645103576 |
| chr11:34358476-34358735 | 0.373039617 | 0.357138271 | 0.645103576 |
| chr19:40445947-40446147 | -0.394877929 | 0.357150315 | 0.645108719 |
| chr4:2418195-2419109 | 0.305974931 | 0.357157957 | 0.645108719 |
| chr12:53017994-53018194 | 0.475788572 | 0.357171248 | 0.645117518 |
| chr18:2337661-2337874 | -0.488401138 | 0.357196211 | 0.645147011 |
| chr3:83440117-83440389 | -0.45877548 | 0.357204416 | 0.645147011 |
| chr10:24722733-24723215 | -0.337835615 | 0.35723648 | 0.645189713 |
| chr12:26832411-26833648 | -0.299137956 | 0.357263237 | 0.645208372 |
| chr3:188176435-188176878 | 0.336059475 | 0.357263652 | 0.645208372 |
| chr6:42563554-42564663 | 0.297236889 | 0.357298043 | 0.645225489 |
| chrX:103699228-103699642 | -0.331718657 | 0.357298744 | 0.645225489 |
| chr16:4693017-4694390 | 0.293911387 | 0.35730436 | 0.645225489 |
| chr11:64304331-64304678 | 0.328507943 | 0.357306814 | 0.645225489 |
| chr1:107430534-107431012 | 0.358844814 | 0.357318256 | 0.645225745 |
| chr22:20298957-20299949 | 0.301039723 | 0.357323798 | 0.645225745 |
| chr13:46603156-46603660 | -0.320259215 | 0.357377681 | 0.645307835 |
| chrX:70033381-70033704 | 0.391574964 | 0.357430944 | 0.645377632 |
| chr2:179876348-179876592 | -0.42953439 | 0.357433181 | 0.645377632 |
| chr11:32154410-32154618 | 0.449841021 | 0.357489541 | 0.645464184 |
| chr1:193242080-193242417 | -0.426150374 | 0.357498044 | 0.645464327 |
| chr4:163587946-163588215 | -0.491732062 | 0.357529327 | 0.645487795 |
| chr1:114971130-114971330 | 0.412441886 | 0.357548533 | 0.645487795 |
| chr10:50600297-50600566 | 0.412441886 | 0.357548533 | 0.645487795 |
| chr7:32905160-32905715 | -0.302283065 | 0.357549534 | 0.645487795 |
| chr17:48160607-48161047 | 0.349154681 | 0.357553163 | 0.645487795 |
| chr2:178443542-178443796 | -0.485350783 | 0.357570056 | 0.645495287 |
| chr9:129610605-129611291 | 0.320961736 | 0.357574162 | 0.645495287 |
| chr1:171566571-171566842 | -0.399810921 | 0.357591771 | 0.645496406 |
| chr11:8336176-8336941 | -0.31753456 | 0.357607871 | 0.645496406 |
| chr8:95685453-95685657 | 0.470891793 | 0.35762607 | 0.645496406 |

| | | | |
|---|---|---|---|
| chr10:80260695-80260966 | 0.359035731 | 0.357635301 | 0.645496406 |
| chr22:39399581-39400059 | 0.341697626 | 0.357639362 | 0.645496406 |
| chr5:9207905-9208109 | 0.426328162 | 0.35764774 | 0.645496406 |
| chr9:131393456-131394166 | -0.315530746 | 0.35766223 | 0.645496406 |
| chr5:109290833-109291071 | -0.469512478 | 0.357667171 | 0.645496406 |
| chr1:223143818-223144188 | -0.349819447 | 0.357671573 | 0.645496406 |
| chrX:78103745-78104389 | 0.300724696 | 0.357690535 | 0.645496406 |
| chr2:241355030-241355693 | -0.304813094 | 0.357695112 | 0.645496406 |
| chr15:71093103-71093641 | -0.352289779 | 0.357696799 | 0.645496406 |
| chr7:75046426-75046679 | -0.424799747 | 0.357697224 | 0.645496406 |
| chr16:83877381-83877581 | -0.488575883 | 0.357712127 | 0.645496406 |
| chr15:63833048-63834196 | 0.305701448 | 0.357716474 | 0.645496406 |
| chr12:104288669-104289029 | -0.327461885 | 0.357722475 | 0.645496406 |
| chr1:119367591-119367861 | -0.443127819 | 0.357726759 | 0.645496406 |
| chr1:184974617-184974824 | -0.443127819 | 0.357726759 | 0.645496406 |
| chr10:98950014-98950228 | 0.412214613 | 0.357734846 | 0.645496406 |
| chr1:31593962-31594313 | 0.362994342 | 0.357769638 | 0.645543982 |
| chr5:36503792-36503992 | 0.537525553 | 0.3578134 | 0.645607698 |
| chr11:6234467-6235575 | -0.290485341 | 0.357821802 | 0.645607698 |
| chr18:49041965-49042424 | -0.349341875 | 0.357852304 | 0.645633562 |
| chr16:20691747-20691991 | 0.435347719 | 0.357855434 | 0.645633562 |
| chr10:132248894-132249319 | 0.322054302 | 0.357861416 | 0.645633562 |
| chr2:60439577-60439777 | -0.471373202 | 0.357873523 | 0.64563738 |
| chr16:72467211-72467487 | -0.418948042 | 0.357880384 | 0.64563738 |
| chr22:17558441-17558761 | 0.358905327 | 0.357891115 | 0.645641538 |
| chr6:17230495-17230749 | -0.543667301 | 0.357906588 | 0.645648146 |
| chr18:11689009-11689475 | -0.388187887 | 0.357911631 | 0.645648146 |
| chr18:47930160-47931691 | -0.292291288 | 0.357929227 | 0.645664687 |
| chr11:66745026-66745752 | 0.306956519 | 0.357975782 | 0.645733465 |
| chr7:41578669-41578907 | 0.459030369 | 0.357994175 | 0.64575144 |
| chr17:74089906-74090113 | 0.427835176 | 0.358007109 | 0.645759568 |
| chr14:26276772-26277096 | 0.391712935 | 0.358020137 | 0.64576739 |
| chr13:60351415-60351680 | -0.44430866 | 0.358028301 | 0.64576739 |
| chr14:90599856-90600396 | -0.337904752 | 0.358052496 | 0.645795828 |
| chr13:94596023-94596453 | -0.317837936 | 0.358069839 | 0.645805506 |
| chr17:62968512-62968757 | -0.33215705 | 0.358074719 | 0.645805506 |
| chr5:11251244-11251626 | -0.358179886 | 0.358104865 | 0.645833767 |
| chr3:18420584-18420963 | -0.362614879 | 0.358107246 | 0.645833767 |
| chr20:38143384-38143682 | -0.388187194 | 0.358138484 | 0.645874901 |
| chr9:70925021-70925306 | -0.430365911 | 0.358148825 | 0.645878348 |
| chr5:115745985-115746202 | 0.475080021 | 0.358175362 | 0.645911003 |
| chrX:82335842-82336092 | 0.416095719 | 0.358196474 | 0.645933872 |
| chr4:128591642-128591842 | 0.483011984 | 0.358210345 | 0.645943683 |
| chr2:16900358-16900716 | -0.368063012 | 0.358246423 | 0.645979182 |
| chr11:77914261-77914876 | -0.343877917 | 0.358246892 | 0.645979182 |
| chr1:244047729-244048697 | 0.294368127 | 0.358268271 | 0.645996572 |
| chr13:112985654-112986020 | -0.405982778 | 0.358273398 | 0.645996572 |
| chr16:20147072-20147291 | 0.420629355 | 0.358295151 | 0.646020592 |
| chr2:217326298-217326728 | -0.360343187 | 0.358322089 | 0.646041962 |
| chr1:113557324-113557753 | 0.360609852 | 0.358337118 | 0.646041962 |

| | | | |
|---|---|---|---|
| chr1:27144457-27144726 | -0.37871098 | 0.358338471 | 0.646041962 |
| chr12:9448688-9449030 | -0.367497111 | 0.35834073 | 0.646041962 |
| chr8:126579871-126580618 | -0.309792338 | 0.358350791 | 0.6460449 |
| chr4:99808329-99808620 | -0.384892959 | 0.358412446 | 0.646129037 |
| chr9:129751894-129752106 | -0.43788371 | 0.358414325 | 0.646129037 |
| chr5:11734327-11734783 | 0.34239715 | 0.358435058 | 0.646146684 |
| chr1:46724848-46725544 | 0.332396122 | 0.358440981 | 0.646146684 |
| chr16:89053454-89053736 | -0.371644095 | 0.358451518 | 0.646150252 |
| chr1:218496602-218497052 | -0.312725583 | 0.358467754 | 0.646150252 |
| chr3:62135993-62136516 | 0.317520755 | 0.358468259 | 0.646150252 |
| chr11:114399957-114400984 | 0.298813375 | 0.358478778 | 0.646154012 |
| chr2:14831561-14831761 | -0.530565553 | 0.358489608 | 0.646158334 |
| chr3:150370081-150370602 | 0.333062255 | 0.358536693 | 0.646227999 |
| chr13:75691279-75691493 | 0.462357499 | 0.358549583 | 0.646236032 |
| chr9:132309165-132309419 | 0.427111555 | 0.35857245 | 0.646262043 |
| chr5:98786501-98787164 | 0.338306124 | 0.358597676 | 0.646278043 |
| chr6:113689678-113689962 | -0.418257012 | 0.358604228 | 0.646278043 |
| chr15:83210596-83211031 | 0.314462649 | 0.358606631 | 0.646278043 |
| chr1:156379654-156379939 | 0.384118143 | 0.35864181 | 0.64632624 |
| chr11:118522337-118522537 | 0.466788343 | 0.358651381 | 0.646326808 |
| chr12:130870800-130872133 | 0.291699345 | 0.358658996 | 0.646326808 |
| chr1:226082942-226084155 | 0.294912774 | 0.358698863 | 0.64638345 |
| chr12:73278058-73278258 | -0.510946194 | 0.358748777 | 0.646458192 |
| chr11:72126060-72126260 | 0.5004253 | 0.358783271 | 0.646505146 |
| chr8:71327559-71327879 | -0.349562634 | 0.358813777 | 0.64654491 |
| chr10:99658754-99659786 | -0.297349625 | 0.358866438 | 0.646624593 |
| chr4:87270246-87270448 | 0.476037147 | 0.358882831 | 0.646638924 |
| chr5:141155537-141155805 | 0.43618538 | 0.358929985 | 0.64670868 |
| chr17:59846273-59847118 | -0.287075351 | 0.358949356 | 0.646724141 |
| chr5:35229402-35229778 | 0.325700153 | 0.358963624 | 0.646724141 |
| chr17:49361378-49362703 | 0.290568656 | 0.358963888 | 0.646724141 |
| chr2:26806347-26806547 | 0.442958113 | 0.358983767 | 0.646744749 |
| chr11:560692-561011 | 0.337003493 | 0.358997816 | 0.646754852 |
| chr8:47188483-47188886 | 0.369039752 | 0.359018448 | 0.646766071 |
| chr5:138180300-138180577 | -0.360861979 | 0.359034429 | 0.646766071 |
| chr3:151283013-151283287 | 0.382958186 | 0.359041091 | 0.646766071 |
| chr5:17024801-17025039 | 0.436756106 | 0.359056191 | 0.646766071 |
| chr5:132180241-132180441 | 0.436756106 | 0.359056191 | 0.646766071 |
| chr3:164254793-164255020 | -0.453687142 | 0.359068375 | 0.646766071 |
| chr5:134511075-134511613 | -0.329394334 | 0.359075425 | 0.646766071 |
| chr3:9067434-9067680 | 0.386304771 | 0.35907745 | 0.646766071 |
| chr15:89334360-89335855 | 0.286745808 | 0.359080012 | 0.646766071 |
| chr1:16644313-16645073 | 0.311118268 | 0.359094331 | 0.646776659 |
| chr6:36442737-36443373 | 0.325522722 | 0.359104599 | 0.646779948 |
| chr15:74144337-74144692 | -0.359414603 | 0.359125576 | 0.64678716 |
| chr14:33631495-33631695 | 0.43820921 | 0.359126978 | 0.64678716 |
| chr4:117528191-117528752 | -0.345967342 | 0.359139697 | 0.64678716 |
| chr6:91744553-91744835 | 0.391822749 | 0.359148181 | 0.64678716 |
| chr17:71968274-71968474 | 0.517392785 | 0.359151213 | 0.64678716 |
| chr16:85610565-85611428 | -0.293252776 | 0.359164769 | 0.64678716 |

| | | | |
|---|---|---|---|
| chr19:4201222-4201471 | 0.453495836 | 0.359167692 | 0.64678716 |
| chr3:47513221-47513969 | 0.30432454 | 0.359214638 | 0.646849005 |
| chr13:70579987-70580252 | 0.412265545 | 0.359218919 | 0.646849005 |
| chr6:44074516-44074833 | 0.357483522 | 0.35923685 | 0.646853954 |
| chr11:469736-469980 | 0.465879897 | 0.359243909 | 0.646853954 |
| chr12:67726840-67727392 | -0.363201208 | 0.359250498 | 0.646853954 |
| chr10:58588749-58589142 | -0.397734381 | 0.359255436 | 0.646853954 |
| chr3:133409959-133410159 | -0.489453462 | 0.359269125 | 0.646854523 |
| chr14:67502994-67503194 | -0.496950523 | 0.359272636 | 0.646854523 |
| chr7:38363542-38363774 | -0.43879715 | 0.359306212 | 0.646881831 |
| chr19:54209921-54210294 | 0.337585116 | 0.359309012 | 0.646881831 |
| chr14:24135682-24136672 | 0.294088297 | 0.359314923 | 0.646881831 |
| chr16:56194342-56194792 | 0.341467819 | 0.359321573 | 0.646881831 |
| chr5:178232012-178232870 | 0.300054535 | 0.359335622 | 0.646891922 |
| chr4:4669240-4669440 | -0.467969834 | 0.359353668 | 0.646909211 |
| chr2:238203101-238204074 | -0.31102732 | 0.359392495 | 0.646948081 |
| chr2:63408522-63408867 | -0.434329623 | 0.359398613 | 0.646948081 |
| chr13:46134897-46135578 | -0.297981411 | 0.359413968 | 0.646948081 |
| chr7:115135752-115135966 | -0.525712382 | 0.359418803 | 0.646948081 |
| chr15:85789136-85789639 | 0.314874025 | 0.359431654 | 0.646948081 |
| chr1:170367600-170367852 | 0.451788751 | 0.359448634 | 0.646948081 |
| chr11:45918743-45918997 | -0.325264283 | 0.359454397 | 0.646948081 |
| chr20:61915683-61915957 | -0.443321919 | 0.359454899 | 0.646948081 |
| chr1:170531641-170532698 | -0.291559004 | 0.359455375 | 0.646948081 |
| chr17:58447080-58447882 | -0.305852136 | 0.359459694 | 0.646948081 |
| chrX:22134613-22135712 | -0.301786943 | 0.359468744 | 0.646949025 |
| chr2:216692139-216692605 | -0.331101009 | 0.359477105 | 0.646949025 |
| chr6:30626595-30627025 | 0.33246348 | 0.359518252 | 0.647003495 |
| chr4:17490535-17491086 | -0.312170453 | 0.35952426 | 0.647003495 |
| chrX:100571853-100572082 | 0.459121997 | 0.359597808 | 0.647120655 |
| chr15:60299804-60300034 | -0.438493112 | 0.359609431 | 0.647126373 |
| chr16:10601431-10601750 | 0.384079891 | 0.359632055 | 0.64714394 |
| chr4:145061292-145061659 | -0.401408965 | 0.359636085 | 0.64714394 |
| chr17:21409118-21409318 | 0.529069212 | 0.35966766 | 0.647185558 |
| chr3:27312074-27312419 | -0.34774421 | 0.35972748 | 0.647277998 |
| chr3:26118209-26118491 | -0.446037959 | 0.359746471 | 0.647296969 |
| chr16:56681852-56682584 | 0.311641325 | 0.359784902 | 0.647350916 |
| chr2:231795542-231795958 | 0.342871274 | 0.359819267 | 0.647392422 |
| chr3:184513606-184514261 | 0.329670184 | 0.359836446 | 0.647392422 |
| chr22:19756874-19757074 | 0.464500722 | 0.35983694 | 0.647392422 |
| chr13:34375892-34376092 | 0.495166513 | 0.359841767 | 0.647392422 |
| chr4:155442263-155442591 | -0.36594204 | 0.359857737 | 0.647405953 |
| chr1:149636262-149637188 | -0.292842372 | 0.359908631 | 0.647475175 |
| chr4:138938374-138938858 | -0.37682753 | 0.359913114 | 0.647475175 |
| chr3:11059564-11059829 | 0.360930336 | 0.359951711 | 0.647529406 |
| chr16:57994878-57995108 | 0.424316211 | 0.359989585 | 0.647582334 |
| chr20:41316940-41317414 | 0.306423558 | 0.360019191 | 0.647620389 |
| chr11:6363069-6363438 | -0.343101081 | 0.360033122 | 0.647630243 |
| chr10:46992951-46993585 | -0.334682978 | 0.360055508 | 0.647655308 |
| chr5:170633059-170633731 | -0.311466903 | 0.360072593 | 0.647670834 |

| | | | |
|---|---|---|---|
| chrX:25019892-25020402 | -0.342574255 | 0.360089628 | 0.647686272 |
| chr1:1433252-1433622 | 0.342922397 | 0.360099591 | 0.647688988 |
| chr1:231039824-231040024 | 0.431627487 | 0.360118681 | 0.647698534 |
| chr15:100627805-100628135 | -0.358486414 | 0.360146433 | 0.647698534 |
| chr17:66676364-66676882 | -0.305281395 | 0.360148924 | 0.647698534 |
| chr6:27650590-27651124 | -0.321060059 | 0.360152325 | 0.647698534 |
| chr18:33495377-33495637 | -0.429277355 | 0.360157071 | 0.647698534 |
| chr12:97487493-97487835 | -0.363270137 | 0.36016875 | 0.647698534 |
| chr10:49593058-49593374 | -0.356633735 | 0.360178929 | 0.647698534 |
| chr17:47790052-47790417 | -0.348352018 | 0.3601855 | 0.647698534 |
| chr14:96434686-96435119 | 0.365165341 | 0.360186424 | 0.647698534 |
| chr5:151241339-151241672 | 0.355295655 | 0.36018943 | 0.647698534 |
| chr4:3388063-3388319 | 0.415788141 | 0.360203281 | 0.647708239 |
| chr16:78308138-78308570 | -0.35043092 | 0.36021557 | 0.647715136 |
| chr19:45113700-45113985 | -0.362554694 | 0.360229334 | 0.64771947 |
| chr12:55829317-55830281 | 0.291649411 | 0.36024418 | 0.64771947 |
| chr7:32034020-32034382 | -0.334878868 | 0.360254621 | 0.64771947 |
| chr1:106472772-106473283 | 0.309331645 | 0.36026147 | 0.64771947 |
| chr18:1585946-1586146 | -0.504724457 | 0.360267866 | 0.64771947 |
| chr20:62921240-62921542 | 0.345131766 | 0.360268701 | 0.64771947 |
| chr4:40046358-40046679 | -0.404346869 | 0.360290022 | 0.647740076 |
| chr2:174573725-174573925 | -0.488718307 | 0.360302178 | 0.647740076 |
| chr17:60770095-60770295 | 0.317327749 | 0.360305523 | 0.647740076 |
| chr7:7011263-7011629 | 0.402195818 | 0.360318436 | 0.647741 |
| chr1:66924630-66925792 | -0.287471595 | 0.36033658 | 0.647741 |
| chr3:128617286-128618346 | -0.295782422 | 0.36036211 | 0.647741 |
| chr2:73431805-73432010 | -0.504502395 | 0.36036284 | 0.647741 |
| chr1:193058530-193060279 | -0.285222927 | 0.360363942 | 0.647741 |
| chr6:75213830-75214090 | -0.429428451 | 0.360366008 | 0.647741 |
| chr19:3006133-3006397 | -0.478727616 | 0.360373667 | 0.647741 |
| chr2:54150692-54150970 | -0.478727616 | 0.360373667 | 0.647741 |
| chr20:51768063-51768295 | 0.336848133 | 0.360399269 | 0.647766928 |
| chr15:88484666-88485413 | -0.312326583 | 0.360405 | 0.647766928 |
| chr1:215340197-215340436 | -0.42707491 | 0.360418044 | 0.647775178 |
| chr1:76595055-76595479 | -0.397600057 | 0.360428117 | 0.647778088 |
| chr1:198756520-198756781 | -0.399886516 | 0.360471863 | 0.647841513 |
| chr12:49321929-49322179 | 0.388100846 | 0.36050256 | 0.647881485 |
| chr6:91445739-91446010 | -0.44674132 | 0.360514984 | 0.647888618 |
| chr8:74259974-74260174 | 0.494117179 | 0.360528428 | 0.647897583 |
| chr12:93441730-93442688 | -0.291013435 | 0.360540374 | 0.647903854 |
| chr18:48446518-48446920 | -0.356608034 | 0.360593128 | 0.647983457 |
| chr3:8987932-8988453 | 0.338257584 | 0.360620655 | 0.648017726 |
| chr9:91037050-91037576 | 0.322553382 | 0.360651179 | 0.648057378 |
| chr1:172615000-172615200 | 0.463664436 | 0.360682937 | 0.648099245 |
| chr13:110415654-110416015 | -0.394924202 | 0.360720925 | 0.648152305 |
| chr4:140052188-140052390 | 0.463759601 | 0.360763199 | 0.648203158 |
| chr3:71498042-71498512 | -0.364485378 | 0.360766147 | 0.648203158 |
| chr1:99000606-99000807 | -0.495289742 | 0.360787389 | 0.648226125 |
| chr4:114659427-114659729 | -0.367284608 | 0.360809641 | 0.648250904 |
| chr7:38433966-38434260 | 0.3749736 | 0.36085502 | 0.648305904 |

| | | | |
|---|---|---|---|
| chr16:88633553-88634040 | -0.31137765 | 0.360857175 | 0.648305904 |
| chr15:79381012-79381252 | 0.4169258 | 0.360866643 | 0.648307713 |
| chr3:63900517-63900835 | 0.323634966 | 0.360880681 | 0.648317731 |
| chr21:43333003-43333226 | 0.409564225 | 0.360890736 | 0.648320593 |
| chr16:21935117-21935439 | 0.366345012 | 0.360908738 | 0.648337734 |
| chr15:45093217-45093469 | -0.461091274 | 0.360938629 | 0.648350197 |
| chr8:22140817-22141473 | 0.312475548 | 0.360940668 | 0.648350197 |
| chr2:202831360-202831909 | -0.341542963 | 0.360952735 | 0.648350197 |
| chr12:6195681-6196168 | -0.325754152 | 0.360953262 | 0.648350197 |
| chr6:125956530-125957500 | -0.293579826 | 0.360957985 | 0.648350197 |
| chr12:40107611-40107856 | -0.381644821 | 0.360982278 | 0.648378634 |
| chr8:87958945-87959226 | -0.39865863 | 0.360999056 | 0.64838398 |
| chr4:145609878-145610130 | -0.407387215 | 0.361002179 | 0.64838398 |
| chr9:129550292-129550669 | -0.430273815 | 0.361045661 | 0.648446877 |
| chr12:50103794-50104119 | 0.332561874 | 0.361107969 | 0.648543581 |
| chr20:58388816-58389968 | 0.287357727 | 0.361119169 | 0.648548494 |
| chr16:28951260-28951589 | 0.343481058 | 0.361139542 | 0.648569881 |
| chr8:106617751-106618340 | -0.321813371 | 0.361176474 | 0.648610811 |
| chr6:111631746-111632175 | -0.3724683 | 0.361179263 | 0.648610811 |
| chr10:8616281-8616576 | 0.361493296 | 0.361188484 | 0.648612168 |
| chr10:91643788-91644079 | -0.377940146 | 0.36120665 | 0.64861935 |
| chr1:115981491-115981725 | 0.385595682 | 0.361216788 | 0.64861935 |
| chr2:184981345-184981777 | 0.378344075 | 0.361217879 | 0.64861935 |
| chr13:28059273-28059517 | 0.405290017 | 0.361248434 | 0.648659016 |
| chr16:28975087-28975330 | 0.371744882 | 0.361268994 | 0.648680731 |
| chr5:143596347-143596626 | 0.38721869 | 0.361287324 | 0.648698443 |
| chr18:50287111-50288383 | -0.29173167 | 0.361331375 | 0.648755213 |
| chr1:155062780-155062994 | 0.35824062 | 0.361344058 | 0.648755213 |
| chr3:149568741-149569266 | -0.338633036 | 0.361344342 | 0.648755213 |
| chr8:141158832-141159061 | 0.432175399 | 0.361364747 | 0.648767811 |
| chr4:105471603-105471862 | -0.35019118 | 0.361375132 | 0.648767811 |
| chr1:14909581-14909781 | 0.525867148 | 0.361376761 | 0.648767811 |
| chr10:22683175-22683897 | -0.32185154 | 0.361402967 | 0.648799657 |
| chr11:71425724-71426244 | -0.323993452 | 0.361429046 | 0.648831272 |
| chr1:6235323-6235523 | -0.461996909 | 0.361444389 | 0.648843613 |
| chr4:170026115-170026973 | -0.297243772 | 0.361465574 | 0.648852492 |
| chr12:106138704-106139884 | 0.302378472 | 0.361476011 | 0.648852492 |
| chr6:52128018-52128225 | -0.492170061 | 0.361479196 | 0.648852492 |
| chr17:82339267-82339674 | -0.36428342 | 0.361483208 | 0.648852492 |
| chr9:12775762-12776679 | -0.304596522 | 0.361521041 | 0.648895521 |
| chr12:64600209-64600542 | 0.356179419 | 0.361524117 | 0.648895521 |
| chr3:72458386-72458770 | 0.357196692 | 0.361546311 | 0.648920156 |
| chr3:67325415-67325615 | -0.470099509 | 0.361569802 | 0.648947117 |
| chr2:163778058-163778762 | -0.326251133 | 0.361599857 | 0.648985858 |
| chr1:167013412-167013612 | 0.537192909 | 0.36163366 | 0.649031323 |
| chr1:106268031-106268439 | -0.30608291 | 0.361644499 | 0.649035574 |
| chr6:147192356-147193009 | -0.339532926 | 0.361664501 | 0.649056269 |
| chr9:131106031-131106231 | -0.526326553 | 0.361715539 | 0.649119989 |
| chrX:134827240-134827667 | -0.418912791 | 0.36171695 | 0.649119989 |
| chr7:32210721-32211086 | -0.34772688 | 0.361746939 | 0.649158602 |

| | | | |
|---|---|---|---|
| chr3:179604259-179605369 | -0.291491483 | 0.361759124 | 0.649165264 |
| chr6:116969299-116969499 | 0.45707373 | 0.361810394 | 0.649238461 |
| chr1:109783445-109783645 | -0.42216001 | 0.361816861 | 0.649238461 |
| chr20:11302464-11302664 | 0.50890193 | 0.361844887 | 0.649273545 |
| chr22:25564271-25564980 | 0.324492572 | 0.36187324 | 0.649303904 |
| chr4:99947038-99947241 | -0.361640785 | 0.361883548 | 0.649303904 |
| chr8:104512613-104512822 | -0.43665492 | 0.361892457 | 0.649303904 |
| chr6:151763395-151763698 | 0.335856992 | 0.361895702 | 0.649303904 |
| chr5:173615927-173616259 | 0.327342923 | 0.361917223 | 0.649312552 |
| chr11:107590786-107591538 | 0.329582881 | 0.361931032 | 0.649312552 |
| chr12:107093434-107094328 | -0.291410719 | 0.361932879 | 0.649312552 |
| chr19:6218508-6218796 | 0.403214369 | 0.361945702 | 0.649312552 |
| chr3:181850794-181851527 | -0.305989008 | 0.361948807 | 0.649312552 |
| chr5:132370897-132371787 | -0.299252843 | 0.361953418 | 0.649312552 |
| chr14:100031552-100032294 | 0.338366932 | 0.361959842 | 0.649312552 |
| chr10:94544880-94545134 | -0.372707604 | 0.361975963 | 0.649312869 |
| chr16:69396105-69396382 | -0.409118627 | 0.361976968 | 0.649312869 |
| chr2:122136154-122136354 | -0.48635808 | 0.361991766 | 0.649324212 |
| chr12:109518821-109519021 | 0.456633625 | 0.36208579 | 0.649471645 |
| chr1:230978583-230979456 | 0.292649017 | 0.362092241 | 0.649471645 |
| chr1:202045867-202047472 | -0.285923315 | 0.362115582 | 0.649471645 |
| chr12:18320039-18320343 | -0.384930761 | 0.362134204 | 0.649471645 |
| chr5:140699928-140700776 | -0.307912401 | 0.362139103 | 0.649471645 |
| chr3:4472839-4473039 | 0.438079 | 0.362139773 | 0.649471645 |
| chr3:39051718-39052190 | -0.316054363 | 0.36215156 | 0.649471645 |
| chr13:43786631-43787789 | -0.299576705 | 0.362156805 | 0.649471645 |
| chr22:45269081-45269524 | 0.383678365 | 0.362160788 | 0.649471645 |
| chr17:64985213-64985547 | -0.350562966 | 0.362173511 | 0.649471645 |
| chr22:44684703-44685178 | 0.323471036 | 0.362174349 | 0.649471645 |
| chr4:152352625-152353085 | -0.318814836 | 0.362175674 | 0.649471645 |
| chr2:67840495-67840829 | -0.349742075 | 0.362227003 | 0.649547662 |
| chr16:55465127-55465523 | 0.360053188 | 0.362237952 | 0.649547662 |
| chr3:49554007-49554543 | 0.339892281 | 0.362243496 | 0.649547662 |
| chr10:68826899-68827640 | -0.315339185 | 0.362269592 | 0.649579254 |
| chr1:180270287-180270553 | -0.34942665 | 0.362300699 | 0.649614301 |
| chr5:138542834-138543993 | -0.288406502 | 0.36230926 | 0.649614301 |
| chr10:76265418-76265618 | 0.50990501 | 0.362314572 | 0.649614301 |
| chr2:162710211-162710782 | -0.341647138 | 0.362330048 | 0.649626847 |
| chr19:1864854-1865054 | 0.412694071 | 0.362353649 | 0.649653961 |
| chr1:151539436-151540775 | 0.285414903 | 0.362407588 | 0.649735463 |
| chr8:88189184-88189448 | 0.442452718 | 0.362418414 | 0.649739669 |
| chr4:169902670-169903257 | -0.325314932 | 0.362432493 | 0.649749707 |
| chr13:110616268-110616558 | -0.37229383 | 0.362443347 | 0.649753963 |
| chr4:40149968-40150168 | -0.511833083 | 0.362485813 | 0.649807242 |
| chr5:75457659-75458044 | 0.359873419 | 0.362490028 | 0.649807242 |
| chrX:149631458-149632196 | 0.311860208 | 0.362501787 | 0.649813117 |
| chr6:135423686-135424044 | -0.380392492 | 0.362529069 | 0.649846821 |
| chr1:27601775-27601991 | -0.398103161 | 0.362588229 | 0.649922702 |
| chr9:6412879-6413797 | 0.298790117 | 0.362588365 | 0.649922702 |
| chr16:28011997-28012231 | -0.45516171 | 0.362603599 | 0.649934802 |

| | | | |
|---|---|---|---|
| chr17:72134730-72134939 | -0.43156791 | 0.362612188 | 0.649934993 |
| chr12:40105328-40106462 | -0.295284594 | 0.362650117 | 0.649966409 |
| chr11:83819986-83820265 | 0.413488349 | 0.362656182 | 0.649966409 |
| chr14:56174863-56175153 | 0.433568171 | 0.362663134 | 0.649966409 |
| chrX:134172163-134172432 | 0.431133685 | 0.362663646 | 0.649966409 |
| chr3:196609319-196609622 | -0.369870614 | 0.36267254 | 0.649967145 |
| chr9:133335994-133336523 | 0.320013441 | 0.362685588 | 0.649970868 |
| chr15:33701391-33701706 | -0.363408578 | 0.36269937 | 0.649970868 |
| chr1:204624079-204624327 | -0.343091646 | 0.362700065 | 0.649970868 |
| chr1:153452538-153452755 | 0.441497267 | 0.362709764 | 0.649973047 |
| chr11:31997755-31998169 | -0.420240796 | 0.362730232 | 0.649984459 |
| chr21:22575160-22575497 | 0.395356468 | 0.362733098 | 0.649984459 |
| chr4:127261418-127261877 | -0.327794188 | 0.362757157 | 0.650012368 |
| chr2:67322545-67322931 | -0.375181297 | 0.362824494 | 0.650084428 |
| chr8:13194271-13194593 | -0.431387632 | 0.362824891 | 0.650084428 |
| chr6:3456841-3457353 | 0.315030954 | 0.362825078 | 0.650084428 |
| chr19:12933507-12933834 | 0.356674858 | 0.36283525 | 0.650084428 |
| chr12:26097097-26097340 | -0.427381615 | 0.362848433 | 0.650084428 |
| chr3:105366255-105367013 | -0.291211656 | 0.362858544 | 0.650084428 |
| chr22:16955914-16956198 | 0.415722171 | 0.362863228 | 0.650084428 |
| chr20:18725934-18726134 | -0.482722764 | 0.362865246 | 0.650084428 |
| chr3:126836048-126836248 | -0.468883254 | 0.362877403 | 0.650091007 |
| chr5:97357984-97358308 | 0.415163828 | 0.362894774 | 0.650106926 |
| chr3:46913729-46913929 | 0.445667213 | 0.362911911 | 0.650122427 |
| chr8:75324169-75324841 | -0.318099419 | 0.362938603 | 0.650151424 |
| chr14:23320693-23322208 | 0.286420184 | 0.362945068 | 0.650151424 |
| chr4:122732273-122733486 | -0.287184307 | 0.362960448 | 0.650163773 |
| chr17:49016813-49017067 | -0.396036811 | 0.362979238 | 0.650182232 |
| chr6:151658851-151659302 | -0.330907962 | 0.363051694 | 0.650296816 |
| chr1:23168734-23169032 | 0.363273809 | 0.363075589 | 0.650324415 |
| chr9:75087652-75088883 | -0.289741112 | 0.363126978 | 0.650401256 |
| chr16:1913910-1914237 | -0.382240353 | 0.363160295 | 0.650445725 |
| chr5:73764686-73764936 | 0.49995479 | 0.363170552 | 0.650448893 |
| chr1:112804254-112804494 | 0.386917326 | 0.363188178 | 0.650460165 |
| chr1:143729391-143729715 | 0.355060437 | 0.36319838 | 0.650460165 |
| chr7:82572247-82572766 | -0.345425895 | 0.363202314 | 0.650460165 |
| chr16:2682149-2683196 | 0.300608429 | 0.36321899 | 0.650474826 |
| chr2:27081159-27081541 | -0.321052131 | 0.363243742 | 0.650503949 |
| chr2:74250238-74250602 | -0.414100075 | 0.36331715 | 0.650615961 |
| chr11:66068326-66068526 | 0.399605664 | 0.363333559 | 0.650615961 |
| chr5:132385403-132385621 | -0.394273238 | 0.363333967 | 0.650615961 |
| chr20:8722359-8722620 | -0.387839091 | 0.363340254 | 0.650615961 |
| chr12:121803907-121804107 | 0.380082934 | 0.363363007 | 0.650641498 |
| chr19:43463154-43464364 | 0.288768564 | 0.36339004 | 0.650673256 |
| chr16:2970038-2970456 | 0.321693078 | 0.363397727 | 0.650673256 |
| chr1:39025910-39027008 | 0.299968816 | 0.363407859 | 0.650676193 |
| chr5:116731459-116731826 | 0.383979647 | 0.363439141 | 0.650715998 |
| chr18:45807269-45807470 | -0.4364776 | 0.363447076 | 0.650715998 |
| chr13:45417598-45418690 | -0.290829751 | 0.363546636 | 0.650879042 |
| chr19:43766504-43766736 | -0.403353059 | 0.363564323 | 0.650895499 |

| | | | |
|---|---|---|---|
| chr17:15490663-15490863 | 0.484879665 | 0.363576498 | 0.650902087 |
| chr3:132210694-132211303 | 0.346322422 | 0.363592323 | 0.650915209 |
| chr13:97221694-97221920 | -0.465821759 | 0.363611867 | 0.650934988 |
| chr11:118659393-118659890 | 0.312064832 | 0.363643013 | 0.650975537 |
| chr7:80874049-80874257 | -0.498091786 | 0.363678924 | 0.651024612 |
| chr3:14461737-14462085 | 0.341689023 | 0.363710611 | 0.651066125 |
| chr4:140755871-140757269 | -0.287205621 | 0.363726209 | 0.651078837 |
| chr12:116737738-116738677 | 0.290830358 | 0.363803042 | 0.651201156 |
| chr8:111931114-111931500 | 0.324443706 | 0.363814909 | 0.651202272 |
| chr14:23034518-23035459 | 0.293155771 | 0.363820663 | 0.651202272 |
| chr3:174613113-174613557 | -0.375175008 | 0.363837853 | 0.651209254 |
| chr11:62324724-62325227 | 0.334577952 | 0.363855916 | 0.651209254 |
| chr2:173037990-173038234 | 0.422517243 | 0.36385666 | 0.651209254 |
| chr10:15088396-15089010 | -0.304540857 | 0.36385856 | 0.651209254 |
| chr12:131410594-131411308 | -0.324794084 | 0.363876523 | 0.651219734 |
| chr3:33883493-33883713 | -0.421521519 | 0.363886235 | 0.651219734 |
| chr8:101326969-101327310 | 0.338759348 | 0.363889913 | 0.651219734 |
| chr18:24362847-24363327 | -0.38217072 | 0.363921049 | 0.651260244 |
| chr17:193194-193397 | -0.490045579 | 0.363946087 | 0.65128984 |
| chr7:75485779-75486697 | -0.313476612 | 0.364037329 | 0.651432194 |
| chr7:151687684-151687920 | -0.413518444 | 0.364058349 | 0.651432194 |
| chr12:123900866-123901249 | -0.371982407 | 0.364062059 | 0.651432194 |
| chr3:42507296-42507659 | 0.342040321 | 0.364073843 | 0.651432194 |
| chr15:90387632-90388898 | 0.286328217 | 0.364074756 | 0.651432194 |
| chr15:74956609-74957490 | 0.302648281 | 0.364076647 | 0.651432194 |
| chr7:43985959-43986421 | -0.324544274 | 0.36408927 | 0.651439568 |
| chr10:117084854-117085127 | -0.481949515 | 0.364117402 | 0.651474689 |
| chr18:31710897-31711375 | 0.358857701 | 0.364129213 | 0.65148061 |
| chr1:153633474-153634603 | 0.288504369 | 0.364159944 | 0.651520377 |
| chr1:207052675-207053841 | -0.28489869 | 0.364202392 | 0.651581108 |
| chr5:39100889-39101310 | 0.339434306 | 0.364261474 | 0.651658595 |
| chr14:76744625-76744893 | 0.370839993 | 0.364262714 | 0.651658595 |
| chr1:206461852-206462157 | 0.326251229 | 0.36429754 | 0.651705683 |
| chr10:75408968-75409907 | -0.294419416 | 0.364307928 | 0.65170905 |
| chr12:68113757-68113960 | -0.467950023 | 0.364371044 | 0.651800687 |
| chr2:27421085-27421364 | 0.382674324 | 0.364376166 | 0.651800687 |
| chr8:116548290-116548612 | -0.372374293 | 0.364432412 | 0.651872229 |
| chr11:77722202-77722415 | -0.540388977 | 0.364447894 | 0.651872229 |
| chr7:8751527-8751873 | 0.374655336 | 0.364452589 | 0.651872229 |
| chr12:66149929-66150298 | -0.374314194 | 0.36446669 | 0.651872229 |
| chr19:18292364-18293100 | -0.331421316 | 0.364474801 | 0.651872229 |
| chr11:67371504-67371704 | 0.372155709 | 0.36448079 | 0.651872229 |
| chr7:149239393-149240062 | -0.306533392 | 0.364480825 | 0.651872229 |
| chr7:154744337-154744537 | -0.497543611 | 0.364484482 | 0.651872229 |
| chr19:48169547-48171027 | 0.288161196 | 0.364492729 | 0.651872229 |
| chr2:10027152-10027591 | -0.380647612 | 0.36450907 | 0.651886237 |
| chr13:96052733-96054041 | -0.291315524 | 0.364537261 | 0.651920907 |
| chr11:45252508-45252712 | -0.460986087 | 0.364545472 | 0.651920907 |
| chr21:42107822-42108415 | -0.314256123 | 0.364571963 | 0.651953064 |
| chr10:104231826-104232864 | -0.292989684 | 0.364580553 | 0.651953211 |

| | | | |
|---|---|---|---|
| chr5:141877097-141877531 | -0.328958863 | 0.364609398 | 0.651989576 |
| chr18:32769356-32769741 | 0.387507854 | 0.364672408 | 0.65208703 |
| chr1:154982976-154983665 | 0.296415553 | 0.36471645 | 0.652150565 |
| chr4:8158689-8159196 | 0.306031615 | 0.364730344 | 0.652155856 |
| chr5:149805870-149806231 | -0.359880383 | 0.364736432 | 0.652155856 |
| chr11:17000773-17001227 | -0.366746839 | 0.364756456 | 0.65217644 |
| chr2:202635619-202636201 | -0.315222018 | 0.364773509 | 0.652191711 |
| chr11:64803589-64804077 | 0.32322147 | 0.36480162 | 0.652217575 |
| chr1:109099782-109100801 | -0.29155868 | 0.364804999 | 0.652217575 |
| chr1:145720664-145721674 | -0.296820258 | 0.364816854 | 0.652223551 |
| chr5:7436460-7436885 | 0.349337102 | 0.36483032 | 0.652232408 |
| chr22:25048819-25049066 | 0.413294054 | 0.364866311 | 0.652281533 |
| chr19:40502648-40503025 | 0.361460722 | 0.364916906 | 0.652342981 |
| chr4:148441971-148443073 | -0.288171235 | 0.364922512 | 0.652342981 |
| chr3:123960838-123961820 | 0.297035397 | 0.364926225 | 0.652342981 |
| chr16:83946837-83947372 | -0.344360097 | 0.364943764 | 0.652359114 |
| chr10:131860817-131861051 | 0.433021817 | 0.364952742 | 0.652359944 |
| chr3:62004012-62004326 | 0.3307747 | 0.364985108 | 0.652380514 |
| chr16:2603143-2604331 | 0.299300218 | 0.364992582 | 0.652380514 |
| chr16:89490416-89491222 | -0.290312493 | 0.364993948 | 0.652380514 |
| chr16:89167453-89167700 | -0.34683786 | 0.365006895 | 0.652380514 |
| chr12:3236535-3236737 | -0.469050347 | 0.365017971 | 0.652380514 |
| chr19:10653557-10654501 | 0.289646886 | 0.36502119 | 0.652380514 |
| chr11:78046246-78046797 | -0.322798499 | 0.365038857 | 0.652380514 |
| chr6:158242656-158242927 | 0.377870506 | 0.365048288 | 0.652380514 |
| chr7:38318772-38319177 | 0.312379357 | 0.36505044 | 0.652380514 |
| chr7:50418057-50418610 | -0.313438429 | 0.365054881 | 0.652380514 |
| chr5:149870283-149870534 | -0.398638161 | 0.365057907 | 0.652380514 |
| chr16:24033253-24033487 | -0.463497217 | 0.365071412 | 0.652389434 |
| chr14:43894556-43894880 | 0.34703445 | 0.365093181 | 0.652413119 |
| chr10:86366760-86366977 | 0.368850872 | 0.365119596 | 0.652445105 |
| chr7:44325078-44325278 | -0.454600667 | 0.365151902 | 0.652473492 |
| chr3:12891942-12892277 | 0.342238533 | 0.365152512 | 0.652473492 |
| chr20:50575951-50576166 | 0.358500956 | 0.365190577 | 0.65252629 |
| chr3:27914058-27914456 | -0.426772366 | 0.365205538 | 0.652537805 |
| chr8:129826187-129826598 | 0.318530253 | 0.365250844 | 0.652544646 |
| chr6:2911761-2912006 | 0.385945191 | 0.365251537 | 0.652544646 |
| chr1:182315114-182315552 | -0.373730467 | 0.365252956 | 0.652544646 |
| chr12:6554142-6554638 | -0.338103801 | 0.36526249 | 0.652544646 |
| chr8:48597612-48597835 | 0.489292652 | 0.365266028 | 0.652544646 |
| chr9:111828294-111828546 | -0.496142141 | 0.365266277 | 0.652544646 |
| chr1:158246657-158247111 | -0.343144735 | 0.365268981 | 0.652544646 |
| chr17:14196745-14197038 | -0.42280063 | 0.36530832 | 0.652596867 |
| chr14:68404709-68404909 | -0.473854721 | 0.365321372 | 0.652596867 |
| chr12:120702803-120703003 | 0.50863395 | 0.365326852 | 0.652596867 |
| chr3:32296756-32297070 | 0.369399917 | 0.365338316 | 0.652596867 |
| chr8:74053420-74053636 | 0.446181649 | 0.365340798 | 0.652596867 |
| chr11:77636698-77636907 | 0.406947731 | 0.365361998 | 0.652606246 |
| chr15:95182266-95182588 | -0.445502272 | 0.365363083 | 0.652606246 |
| chr15:101723873-101724759 | 0.288686655 | 0.365372812 | 0.652608411 |

| | | | |
|---|---|---|---|
| chr3:193805469-193805907 | 0.332580669 | 0.365397033 | 0.652634763 |
| chr10:30687696-30687932 | -0.44228837 | 0.365404601 | 0.652634763 |
| chr17:73360051-73360328 | 0.368721787 | 0.365427699 | 0.652654204 |
| chr7:48424828-48425088 | -0.411188489 | 0.365432522 | 0.652654204 |
| chr17:8410091-8410673 | -0.351336384 | 0.365496056 | 0.65275246 |
| chr2:78992478-78993483 | -0.295974064 | 0.365530262 | 0.652787092 |
| chrX:69387617-69387971 | -0.378623933 | 0.365549564 | 0.652787092 |
| chr20:44475641-44476607 | 0.297269897 | 0.365550867 | 0.652787092 |
| chr1:158177029-158177452 | -0.359232711 | 0.365556714 | 0.652787092 |
| chr7:95772326-95772949 | 0.329019522 | 0.365558045 | 0.652787092 |
| chr8:74350003-74351005 | -0.291779236 | 0.365588008 | 0.652825382 |
| chr5:115841382-115842543 | -0.286686578 | 0.365602541 | 0.652836119 |
| chr3:9902294-9903475 | -0.299754525 | 0.365628507 | 0.652850198 |
| chr16:10944271-10944781 | 0.308977985 | 0.365633944 | 0.652850198 |
| chr1:98430164-98430428 | -0.472717615 | 0.365635986 | 0.652850198 |
| chr10:11830912-11831300 | -0.375006905 | 0.365654337 | 0.65286775 |
| chr15:48177583-48178593 | -0.302179981 | 0.365676246 | 0.652874365 |
| chr5:178094265-178094584 | 0.365887578 | 0.365679428 | 0.652874365 |
| chr3:97927750-97928168 | 0.344708085 | 0.365683604 | 0.652874365 |
| chr17:57918377-57918620 | 0.355565293 | 0.365693871 | 0.652877482 |
| chrX:39759745-39760374 | -0.342005871 | 0.365709198 | 0.652888163 |
| chr13:27284054-27284325 | -0.404249293 | 0.365716895 | 0.652888163 |
| chr5:116921430-116921630 | -0.496079252 | 0.36574783 | 0.652928176 |
| chr13:98145115-98145329 | 0.409607644 | 0.365763747 | 0.652941377 |
| chr6:29649707-29650389 | -0.299003016 | 0.365808995 | 0.653006938 |
| chr3:186609512-186609794 | -0.409982737 | 0.365824494 | 0.653019391 |
| chrX:132092515-132092853 | 0.366273224 | 0.365853235 | 0.653048013 |
| chr16:21559711-21559922 | 0.500767653 | 0.365868544 | 0.653048013 |
| chr5:32709757-32710352 | -0.304388359 | 0.365878982 | 0.653048013 |
| chr16:50065741-50066125 | 0.312169913 | 0.365880895 | 0.653048013 |
| chr5:96823051-96823359 | -0.39795829 | 0.365883143 | 0.653048013 |
| chr3:100197753-100197984 | -0.464000668 | 0.365927863 | 0.653097406 |
| chr3:141384181-141384383 | -0.464000668 | 0.365927863 | 0.653097406 |
| chr3:107946806-107947006 | 0.493280246 | 0.365948583 | 0.653119173 |
| chr19:40718984-40719205 | 0.360694771 | 0.365986931 | 0.653172399 |
| chr16:85330783-85331102 | 0.342050973 | 0.366024396 | 0.65320953 |
| chr12:68863895-68864299 | -0.341892669 | 0.366024786 | 0.65320953 |
| chr1:167637811-167638011 | -0.482397754 | 0.366051865 | 0.653240922 |
| chr3:33427804-33428063 | -0.441501505 | 0.366059427 | 0.653240922 |
| chr2:182077545-182078089 | -0.329426166 | 0.36608647 | 0.653273965 |
| chr19:18999044-18999316 | -0.511494191 | 0.366102302 | 0.653276698 |
| chr9:38687551-38687845 | 0.377342577 | 0.366111213 | 0.653276698 |
| chr5:11724829-11725317 | -0.310641445 | 0.36611358 | 0.653276698 |
| chr16:117787-118152 | -0.343033097 | 0.366146141 | 0.653318294 |
| chr12:56221243-56222435 | 0.287359518 | 0.366153944 | 0.653318294 |
| chr4:187499794-187500262 | 0.361071517 | 0.366166861 | 0.653326128 |
| chr2:203651358-203651665 | 0.362244826 | 0.366207885 | 0.653375899 |
| chr12:94101914-94102114 | 0.412294403 | 0.366218167 | 0.653375899 |
| chr6:136783704-136783992 | -0.451887304 | 0.366220338 | 0.653375899 |
| chr8:97595783-97596160 | 0.325306003 | 0.366257617 | 0.653427194 |

| | | | |
|---|---|---|---|
| chr1:28552511-28553682 | 0.292229878 | 0.366278671 | 0.653447159 |
| chr2:207720898-207721224 | -0.402771135 | 0.366285864 | 0.653447159 |
| chr2:170212572-170212820 | 0.504269414 | 0.366309408 | 0.653468426 |
| chr5:130555484-130555706 | -0.450103902 | 0.366314842 | 0.653468426 |
| chr2:86232451-86232784 | -0.437018651 | 0.366352175 | 0.653510064 |
| chr2:85561014-85561943 | -0.296422352 | 0.366358607 | 0.653510064 |
| chr1:50977663-50978512 | 0.325624666 | 0.366372197 | 0.653510064 |
| chr16:23557016-23558131 | 0.286978346 | 0.366372299 | 0.653510064 |
| chr3:28317486-28317786 | 0.358201584 | 0.36639243 | 0.653523113 |
| chr2:16321992-16322208 | 0.433517184 | 0.366396673 | 0.653523113 |
| chr14:85414921-85415693 | -0.308239256 | 0.366408156 | 0.653528381 |
| chr3:186514337-186514572 | -0.476550798 | 0.366435177 | 0.653561363 |
| chr8:118111652-118112287 | 0.300315374 | 0.366447695 | 0.653568475 |
| chr16:77615128-77615390 | 0.4111048 | 0.366465388 | 0.653584818 |
| chrX:130224864-130225163 | 0.367966376 | 0.366481396 | 0.653598155 |
| chr14:79356171-79356371 | 0.466764102 | 0.366520735 | 0.653648136 |
| chr5:176610058-176610353 | -0.343041014 | 0.366534213 | 0.653648136 |
| chr16:28924519-28925671 | 0.285480493 | 0.366535014 | 0.653648136 |
| chr17:47821410-47822019 | 0.305480683 | 0.36655949 | 0.653662272 |
| chr20:44248172-44248468 | 0.378501161 | 0.366562514 | 0.653662272 |
| chr7:6367612-6367890 | 0.408910463 | 0.366575456 | 0.653662272 |
| chr10:97355788-97356068 | 0.38481111 | 0.366577064 | 0.653662272 |
| chr12:79274876-79275091 | 0.435812259 | 0.366639384 | 0.653758182 |
| chr14:57389745-57391617 | 0.285657124 | 0.36668899 | 0.653831421 |
| chr1:225445756-225445994 | -0.439819963 | 0.36670731 | 0.653834935 |
| chr6:43635478-43636347 | 0.307105756 | 0.366708028 | 0.653834935 |
| chr2:129813362-129813793 | -0.367309718 | 0.366727579 | 0.653840809 |
| chr17:68081309-68081811 | -0.333929446 | 0.366734037 | 0.653840809 |
| chr21:39788285-39788803 | -0.306753569 | 0.366736922 | 0.653840809 |
| chr1:201538703-201539580 | 0.297325617 | 0.366788559 | 0.653917235 |
| chr9:5437507-5438111 | -0.308520761 | 0.366796858 | 0.653917235 |
| chr10:48605005-48605259 | -0.389744247 | 0.366813179 | 0.653930198 |
| chr1:217477514-217477957 | -0.359523183 | 0.366821198 | 0.653930198 |
| chr5:32329315-32329809 | 0.315486647 | 0.366876277 | 0.653999576 |
| chr5:11289607-11290010 | 0.38013441 | 0.366877186 | 0.653999576 |
| chr1:35808487-35808752 | 0.396648756 | 0.366888679 | 0.654004848 |
| chr2:190469239-190469597 | 0.3456736 | 0.366927017 | 0.654057971 |
| chr3:2921930-2922130 | 0.433730461 | 0.366950285 | 0.65408423 |
| chr13:21595274-21595784 | -0.343680616 | 0.366963693 | 0.654092914 |
| chr14:40682231-40682537 | -0.403113792 | 0.366980802 | 0.654108193 |
| chr7:149145011-149145396 | -0.307472672 | 0.367003844 | 0.654126604 |
| chrX:49058817-49059061 | 0.381714148 | 0.367008205 | 0.654126604 |
| chr1:161367593-161368298 | -0.322298842 | 0.367028889 | 0.654138637 |
| chr6:117829651-117830166 | -0.329471014 | 0.367042822 | 0.654138637 |
| chr11:125429023-125429498 | 0.3503531 | 0.367064526 | 0.654138637 |
| chr3:14371824-14372349 | 0.312522273 | 0.367072367 | 0.654138637 |
| chr1:83052935-83053145 | -0.450861233 | 0.367076985 | 0.654138637 |
| chr18:31648988-31649221 | -0.471556069 | 0.367101946 | 0.654138637 |
| chr2:17148478-17148678 | -0.471556069 | 0.367101946 | 0.654138637 |
| chr10:123043424-123043694 | 0.397437698 | 0.367115876 | 0.654138637 |

| | | | |
|---|---|---|---|
| chr6:2938641-2939006 | -0.332814841 | 0.367116235 | 0.654138637 |
| chr14:74084169-74085325 | 0.283082621 | 0.367121982 | 0.654138637 |
| chr2:169827212-169827755 | -0.30366989 | 0.367137664 | 0.654138637 |
| chr12:81077330-81078251 | -0.289805945 | 0.367140732 | 0.654138637 |
| chr15:74817318-74817574 | 0.351528923 | 0.367145561 | 0.654138637 |
| chr16:83995830-83996353 | 0.313227958 | 0.367155069 | 0.654138637 |
| chrX:7593213-7593783 | -0.395843369 | 0.367158055 | 0.654138637 |
| chr1:205256135-205256503 | 0.309181609 | 0.367173044 | 0.654138637 |
| chr5:135094263-135094553 | -0.423845493 | 0.367177898 | 0.654138637 |
| chr12:116658328-116659168 | -0.315926547 | 0.36718322 | 0.654138637 |
| chr7:149153828-149154417 | -0.312421221 | 0.36718368 | 0.654138637 |
| chr3:161316251-161316512 | 0.375556051 | 0.367185701 | 0.654138637 |
| chr4:94140717-94140977 | -0.450874627 | 0.367225379 | 0.654194114 |
| chr11:78228458-78228770 | -0.362362419 | 0.367290867 | 0.654268241 |
| chr6:153814156-153814429 | 0.44184306 | 0.367293702 | 0.654268241 |
| chr3:94249588-94249826 | 0.3963359 | 0.367300018 | 0.654268241 |
| chr2:138335160-138335360 | -0.532600464 | 0.367301146 | 0.654268241 |
| chr9:133715721-133716084 | -0.384640937 | 0.367376646 | 0.65436861 |
| chr15:63402890-63403154 | 0.372918951 | 0.367379444 | 0.65436861 |
| chr7:37201685-37202125 | 0.348785565 | 0.367383113 | 0.65436861 |
| chr19:36127616-36128316 | 0.302832961 | 0.367430326 | 0.654437491 |
| chr1:86502735-86502935 | -0.47093158 | 0.367465527 | 0.654469761 |
| chr13:36831991-36832217 | -0.47093158 | 0.367465527 | 0.654469761 |
| chr17:45270988-45271320 | -0.437261714 | 0.36749255 | 0.654497748 |
| chr12:66756287-66756554 | -0.434294039 | 0.367498325 | 0.654497748 |
| chr3:4650582-4650913 | -0.344949057 | 0.367538219 | 0.654553584 |
| chr6:20401193-20401977 | -0.287771984 | 0.367575237 | 0.654604295 |
| chr6:11344319-11344787 | -0.34251852 | 0.367601972 | 0.654631255 |
| chr11:129148342-129148797 | 0.371203139 | 0.367607462 | 0.654631255 |
| chr9:74773942-74774166 | -0.412286964 | 0.367641732 | 0.654677065 |
| chr2:222423931-222424632 | -0.328049744 | 0.367669147 | 0.654710669 |
| chr11:35894142-35894486 | -0.372871246 | 0.367703221 | 0.654756107 |
| chr11:68297698-68298266 | -0.323877604 | 0.367711754 | 0.654756107 |
| chr1:212857598-212858927 | -0.288435057 | 0.367742779 | 0.654781245 |
| chr21:14069997-14070610 | -0.317168225 | 0.367742963 | 0.654781245 |
| chr17:326300-326623 | -0.488129955 | 0.367819586 | 0.654902457 |
| chr5:133610945-133611225 | 0.371161953 | 0.367836467 | 0.654907269 |
| chr16:56861229-56861574 | -0.397252165 | 0.367839383 | 0.654907269 |
| chr20:50624382-50625021 | 0.293896645 | 0.3678621 | 0.654921189 |
| chr14:45978705-45978917 | 0.426448939 | 0.367864296 | 0.654921189 |
| chr5:16615805-16617319 | 0.291164863 | 0.367888134 | 0.65494841 |
| chr14:95431309-95431704 | 0.371619421 | 0.367929017 | 0.655005975 |
| chr20:50635717-50636247 | 0.302752738 | 0.367938603 | 0.655007822 |
| chr6:35764448-35764889 | -0.337640606 | 0.367970205 | 0.65503679 |
| chr20:50019443-50020233 | 0.292002944 | 0.367971974 | 0.65503679 |
| chr17:75896948-75897201 | 0.364644205 | 0.368003964 | 0.655078517 |
| chr3:176872716-176873114 | 0.327915046 | 0.36803632 | 0.655120894 |
| chr7:16801157-16801421 | 0.351467466 | 0.368061851 | 0.65515112 |
| chr1:165888273-165888549 | 0.443443102 | 0.368081388 | 0.655170677 |
| chr8:18208963-18209190 | 0.398033927 | 0.368091206 | 0.655172931 |

| | | | |
|---|---|---|---|
| chr1:240687876-240688324 | 0.318442155 | 0.368126442 | 0.655220429 |
| chr8:101583411-101583682 | 0.370945771 | 0.368150776 | 0.655248519 |
| chr6:119331773-119331973 | 0.454306926 | 0.368196986 | 0.655315544 |
| chr6:118674743-118675191 | -0.381913141 | 0.368247165 | 0.655389628 |
| chr6:25788589-25788888 | -0.375355189 | 0.368284249 | 0.655440404 |
| chr1:21869921-21870287 | 0.362534783 | 0.368339672 | 0.655523815 |
| chr20:57197863-57198378 | 0.285755513 | 0.368353096 | 0.65553248 |
| chr1:246450868-246451189 | 0.373868379 | 0.368362687 | 0.655534323 |
| chr5:180719102-180719302 | -0.448184892 | 0.368374297 | 0.655539758 |
| chr10:9402091-9402846 | -0.306182405 | 0.368386284 | 0.655545765 |
| chr5:176199100-176199692 | 0.346699454 | 0.368395477 | 0.655545765 |
| chr1:198665510-198665710 | -0.4363629 | 0.368403339 | 0.655545765 |
| chr15:70378159-70378430 | -0.386605407 | 0.36841463 | 0.655550632 |
| chr1:88180558-88180860 | -0.391876516 | 0.368437858 | 0.655576739 |
| chr6:34431859-34432059 | 0.479351589 | 0.368452094 | 0.655582047 |
| chr3:172304023-172304223 | -0.467759996 | 0.368457953 | 0.655582047 |
| chr11:16737909-16739140 | -0.287461977 | 0.368474257 | 0.655595832 |
| chr1:245157018-245157391 | -0.348336631 | 0.368483197 | 0.655596515 |
| chr22:50190762-50191091 | 0.421550295 | 0.368500526 | 0.655612123 |
| chr4:140594948-140595502 | -0.306245573 | 0.368519931 | 0.655631424 |
| chr8:101460745-101460945 | 0.408860933 | 0.368581192 | 0.655709146 |
| chr1:205748866-205750545 | -0.280126134 | 0.368587828 | 0.655709146 |
| chrX:84729722-84730260 | -0.32243888 | 0.36860763 | 0.655709146 |
| chr2:25376825-25377143 | -0.346472265 | 0.368611564 | 0.655709146 |
| chr15:78709359-78709755 | 0.334251935 | 0.368615395 | 0.655709146 |
| chr5:177312854-177313166 | 0.325042758 | 0.368624805 | 0.655709146 |
| chr17:55356562-55357186 | -0.317053756 | 0.368627026 | 0.655709146 |
| chr2:127479267-127479532 | -0.441045954 | 0.368633917 | 0.655709146 |
| chr10:21495359-21495559 | -0.442451169 | 0.368647844 | 0.655709146 |
| chr5:119070514-119071795 | -0.282550412 | 0.368649194 | 0.655709146 |
| chr10:54706915-54707393 | -0.370006269 | 0.368710419 | 0.655792433 |
| chr2:219736316-219736533 | 0.421863279 | 0.368713137 | 0.655792433 |
| chr8:93286149-93286438 | 0.401507823 | 0.368728399 | 0.655804354 |
| chr1:155026715-155026957 | 0.416723649 | 0.368743057 | 0.655808988 |
| chr4:120198305-120198505 | -0.513366506 | 0.368748123 | 0.655808988 |
| chr20:63442195-63442467 | 0.373421983 | 0.368767327 | 0.655827921 |
| chr16:30957490-30958038 | 0.304009342 | 0.368805866 | 0.655881236 |
| chr20:5844366-5844649 | -0.474594797 | 0.368856593 | 0.655950721 |
| chr19:41263357-41264452 | 0.285464731 | 0.368862059 | 0.655950721 |
| chr8:125329639-125329947 | 0.390462361 | 0.368886143 | 0.655978325 |
| chr1:207565568-207566007 | -0.354960725 | 0.368918426 | 0.655993798 |
| chr19:45496913-45497351 | -0.31022227 | 0.368927212 | 0.655993798 |
| chr1:248890595-248891057 | -0.376500485 | 0.368929009 | 0.655993798 |
| chr3:187799335-187799535 | -0.472854625 | 0.368929625 | 0.655993798 |
| chr22:27880930-27881131 | 0.470866742 | 0.368937652 | 0.655993798 |
| chr2:202279045-202279277 | -0.40608071 | 0.368954226 | 0.656008045 |
| chr17:7141903-7142108 | 0.427145338 | 0.36906296 | 0.656171246 |
| chr2:69932049-69932249 | -0.507362468 | 0.369063142 | 0.656171246 |
| chr1:6962341-6963405 | 0.285233762 | 0.369072404 | 0.656172489 |
| chr2:179991475-179991675 | -0.463611206 | 0.369115928 | 0.656234643 |

| | | | |
|---|---|---|---|
| chr9:82606783-82607043 | -0.42735042 | 0.369131533 | 0.656237104 |
| chr10:48649463-48649773 | -0.397638898 | 0.369140516 | 0.656237104 |
| chr2:185275840-185276658 | 0.304152958 | 0.369143006 | 0.656237104 |
| chr12:120437714-120438494 | 0.290530368 | 0.369197468 | 0.656311698 |
| chr13:91162252-91162467 | -0.452756705 | 0.369208533 | 0.656311698 |
| chr5:37495299-37495499 | 0.438002926 | 0.369214578 | 0.656311698 |
| chr8:15237298-15237691 | -0.403119057 | 0.369219229 | 0.656311698 |
| chr8:113202357-113202574 | -0.445454899 | 0.369245334 | 0.656328529 |
| chr18:395181-395381 | -0.48241526 | 0.369247138 | 0.656328529 |
| chr13:28138783-28139642 | -0.291400062 | 0.369254394 | 0.656328529 |
| chr20:55088415-55088645 | -0.429698327 | 0.369269142 | 0.656339515 |
| chr4:151031943-151032221 | -0.400257385 | 0.369279809 | 0.65634325 |
| chr7:102572473-102572877 | 0.333437375 | 0.369304754 | 0.656372361 |
| chr12:12356774-12357773 | -0.294291616 | 0.369325893 | 0.656394706 |
| chr1:17437669-17437961 | 0.406863327 | 0.36936272 | 0.656444932 |
| chr14:50859983-50860372 | 0.347651739 | 0.369393327 | 0.6564841 |
| chr13:29676648-29677214 | -0.328894485 | 0.369419362 | 0.656515142 |
| chr20:6058699-6058915 | -0.474279282 | 0.369429772 | 0.656518416 |
| chr2:47272169-47272925 | -0.309971126 | 0.369443485 | 0.656522113 |
| chr5:138231267-138231467 | 0.444623295 | 0.369448989 | 0.656522113 |
| chr3:137110460-137110702 | 0.385626771 | 0.369459136 | 0.656524917 |
| chr14:91177418-91177677 | 0.358654739 | 0.369479052 | 0.656545082 |
| chr4:166165897-166166165 | 0.385410717 | 0.36951161 | 0.656587708 |
| chr3:158734162-158734368 | 0.415827644 | 0.369524739 | 0.65659581 |
| chr4:6308632-6308977 | 0.345892287 | 0.369550297 | 0.656614233 |
| chr14:56847512-56847900 | -0.35131849 | 0.369552246 | 0.656614233 |
| chr6:92849714-92849971 | 0.372036637 | 0.369579105 | 0.656646728 |
| chr3:18988645-18988973 | 0.359764356 | 0.369598427 | 0.656665833 |
| chr1:59921230-59921985 | 0.314215611 | 0.369609394 | 0.65667009 |
| chr15:64728190-64728557 | 0.330893183 | 0.369652814 | 0.656732006 |
| chr10:68900643-68901779 | -0.286094119 | 0.369672367 | 0.656740575 |
| chr6:3963016-3963278 | -0.424238536 | 0.36967478 | 0.656740575 |
| chr10:124612751-124612951 | 0.421094914 | 0.369693898 | 0.656753879 |
| chr1:154326267-154326582 | 0.329289484 | 0.369699412 | 0.656753879 |
| chr17:63305879-63306097 | -0.440015439 | 0.369721889 | 0.656778581 |
| chr14:63727485-63727750 | 0.347370689 | 0.369749786 | 0.65681291 |
| chr16:3282289-3283740 | 0.287027621 | 0.369781229 | 0.656842131 |
| chr1:16636656-16636856 | -0.481836391 | 0.369783905 | 0.656842131 |
| chr19:12945579-12945981 | 0.329477386 | 0.369791953 | 0.656842131 |
| chr7:90211492-90212378 | -0.300733243 | 0.369810801 | 0.656844522 |
| chr16:30334624-30335640 | 0.296401417 | 0.369814353 | 0.656844522 |
| chr1:36285539-36285739 | 0.463559824 | 0.369829315 | 0.656844522 |
| chr10:80058819-80059118 | -0.38909373 | 0.369837059 | 0.656844522 |
| chr11:71481254-71481956 | -0.299335619 | 0.369838099 | 0.656844522 |
| chr11:86218255-86219016 | -0.304524122 | 0.369844734 | 0.656844522 |
| chr1:211944835-211945035 | -0.468587619 | 0.369875446 | 0.65688384 |
| chr7:99525730-99525949 | -0.494605877 | 0.36989323 | 0.656884973 |
| chr7:38264655-38264891 | -0.494605877 | 0.36989323 | 0.656884973 |
| chr7:43362978-43363326 | -0.394576979 | 0.369917061 | 0.656910285 |
| chr4:90031018-90031293 | 0.38857056 | 0.36992463 | 0.656910285 |

| | | | |
|---|---|---|---|
| chrX:154541825-154542750 | 0.298033497 | 0.369956826 | 0.656945117 |
| chr7:4103440-4103823 | -0.359123895 | 0.369961392 | 0.656945117 |
| chr2:131681697-131682932 | -0.290826469 | 0.369976088 | 0.656955331 |
| chr1:161792722-161792922 | 0.43278794 | 0.369984292 | 0.656955331 |
| chr12:114663157-114663357 | -0.501366432 | 0.370027458 | 0.657016751 |
| chr20:53331520-53331875 | 0.336536051 | 0.370055917 | 0.657050624 |
| chr11:46338936-46339189 | -0.361928638 | 0.370063685 | 0.657050624 |
| chr2:211958055-211958380 | 0.37497662 | 0.370075461 | 0.657056307 |
| chr2:71154826-71155250 | -0.316862137 | 0.370105942 | 0.657095199 |
| chr10:34638469-34638730 | -0.397694573 | 0.37017631 | 0.657204903 |
| chr15:46312183-46312468 | -0.386471442 | 0.370197625 | 0.65722571 |
| chr3:171811583-171811861 | -0.340476823 | 0.370208403 | 0.65722571 |
| chr7:158992246-158992539 | -0.387129061 | 0.370213762 | 0.65722571 |
| chr8:133571072-133572765 | 0.281773114 | 0.370260503 | 0.657293458 |
| chr2:202911339-202912580 | -0.288393486 | 0.370293229 | 0.657307316 |
| chr8:65706003-65706340 | -0.367382008 | 0.370294779 | 0.657307316 |
| chr5:107630210-107630629 | -0.314685257 | 0.370298 | 0.657307316 |
| chr8:125828775-125829077 | -0.338267458 | 0.370302624 | 0.657307316 |
| chr15:73972896-73973427 | 0.319552308 | 0.370322302 | 0.657327019 |
| chr1:201999644-202000006 | 0.34653251 | 0.370336463 | 0.657336928 |
| chr19:45391439-45391747 | 0.40012019 | 0.370366609 | 0.657363406 |
| chr18:35497389-35498182 | -0.292243891 | 0.37036854 | 0.657363406 |
| chr3:169553584-169553962 | 0.391608721 | 0.370393265 | 0.657377066 |
| chr21:41605446-41605908 | -0.327253623 | 0.370393395 | 0.657377066 |
| chr20:62426779-62427854 | 0.284903232 | 0.370430726 | 0.657428094 |
| chr21:34201646-34202308 | -0.323938564 | 0.370477107 | 0.65749518 |
| chr1:222711317-222711541 | 0.349533135 | 0.370501963 | 0.657524062 |
| chr8:28701121-28701925 | 0.30588885 | 0.370532114 | 0.657562342 |
| chr11:68905612-68906197 | -0.303558063 | 0.37054529 | 0.657570494 |
| chr12:95837660-95837901 | -0.455122026 | 0.370576177 | 0.657610075 |
| chr3:180989144-180990170 | -0.288943558 | 0.370626759 | 0.657684605 |
| chr15:74614616-74615961 | 0.286347308 | 0.370638762 | 0.657690608 |
| chr13:27425897-27426458 | 0.299389199 | 0.370656758 | 0.657690608 |
| chr7:138195356-138195627 | -0.389452692 | 0.370664351 | 0.657690608 |
| chr17:38608675-38608890 | -0.434617302 | 0.370664477 | 0.657690608 |
| chr3:112136895-112137404 | -0.347087875 | 0.370679777 | 0.657702526 |
| chr10:115093249-115094038 | -0.312907594 | 0.370689879 | 0.657705219 |
| chr21:25700523-25701195 | 0.309341823 | 0.370700274 | 0.657708432 |
| chr10:86261718-86262425 | -0.309567372 | 0.370710027 | 0.657710507 |
| chr8:30384360-30385275 | 0.309131789 | 0.370814739 | 0.657881054 |
| chr1:217393893-217394258 | -0.347894313 | 0.370831556 | 0.65788829 |
| chr6:16238552-16238825 | -0.427094003 | 0.370843685 | 0.65788829 |
| chr2:676070-677839 | -0.278776159 | 0.370852096 | 0.65788829 |
| chr6:90648958-90649158 | -0.496954545 | 0.370853163 | 0.65788829 |
| chrX:83818943-83819905 | -0.296744516 | 0.370869089 | 0.657901311 |
| chr1:24744886-24745699 | 0.293505807 | 0.370878306 | 0.65790243 |
| chr16:9225564-9225793 | -0.449484563 | 0.370901979 | 0.657914109 |
| chr4:74793701-74794863 | -0.286332148 | 0.370902063 | 0.657914109 |
| chr11:47565191-47566008 | 0.299771706 | 0.370922686 | 0.657935459 |
| chr2:113249264-113249496 | 0.431884153 | 0.370967699 | 0.658000069 |

| | | | |
|---|---|---|---|
| chr20:50818449-50818742 | -0.345988226 | 0.37100762 | 0.658043109 |
| chr11:44564740-44565774 | -0.287637951 | 0.37100914 | 0.658043109 |
| chr5:54889915-54890185 | 0.411279321 | 0.3710486 | 0.658088313 |
| chr15:65184449-65184695 | 0.353633914 | 0.371060396 | 0.658088313 |
| chr2:208072375-208072629 | -0.38728914 | 0.371065608 | 0.658088313 |
| chr2:60581610-60582151 | -0.320121788 | 0.371068981 | 0.658088313 |
| chr17:63998091-63998668 | 0.300997631 | 0.371086009 | 0.658101051 |
| chr9:129704246-129704630 | -0.388528842 | 0.371093341 | 0.658101051 |
| chr12:128823123-128823749 | 0.317181605 | 0.371112152 | 0.658119178 |
| chr22:41366859-41367619 | -0.303344627 | 0.371156617 | 0.658182797 |
| chr1:10209869-10210664 | 0.298150991 | 0.371185413 | 0.658218628 |
| chr1:208021905-208022126 | -0.412878699 | 0.371245209 | 0.658309429 |
| chr3:163043123-163043525 | -0.335198162 | 0.371264874 | 0.658322663 |
| chr12:68240056-68240442 | -0.344328802 | 0.371269856 | 0.658322663 |
| chr1:33255339-33255824 | 0.308653645 | 0.371335879 | 0.658424496 |
| chr3:101573491-101575075 | -0.280576459 | 0.371344667 | 0.658424841 |
| chr1:2208264-2208595 | 0.365593808 | 0.371359826 | 0.658436482 |
| chr12:112992903-112993103 | -0.46711779 | 0.371391763 | 0.658465115 |
| chr14:47949042-47949319 | -0.389404496 | 0.371393162 | 0.658465115 |
| chr17:61425360-61426410 | 0.279921219 | 0.371404313 | 0.658469648 |
| chr15:86915752-86916383 | -0.317722614 | 0.371422727 | 0.658487058 |
| chr12:46354757-46354995 | 0.417280608 | 0.371452043 | 0.658523795 |
| chr19:47729261-47729900 | 0.314659513 | 0.371476404 | 0.658551746 |
| chr7:50685451-50686195 | -0.304484947 | 0.371512383 | 0.658592061 |
| chr1:148161484-148161711 | 0.396682511 | 0.371516336 | 0.658592061 |
| chr5:1686414-1686615 | 0.444705345 | 0.371538982 | 0.658597741 |
| chr15:88338043-88338281 | 0.400326369 | 0.371553392 | 0.658597741 |
| chr21:17611564-17612379 | 0.313153684 | 0.371554268 | 0.658597741 |
| chr5:62925490-62925840 | -0.396496672 | 0.37157448 | 0.658597741 |
| chr2:96251744-96251966 | 0.456184647 | 0.371575455 | 0.658597741 |
| chr20:20017800-20018190 | 0.33842074 | 0.371582099 | 0.658597741 |
| chr14:51129242-51129544 | -0.481772587 | 0.371587916 | 0.658597741 |
| chr17:73710192-73710539 | 0.346969933 | 0.371588303 | 0.658597741 |
| chr22:42662092-42662600 | -0.368188139 | 0.371597137 | 0.658598164 |
| chr7:1919639-1919842 | 0.376120276 | 0.371615059 | 0.658603898 |
| chr2:29072779-29072979 | -0.470220087 | 0.371617563 | 0.658603898 |
| chr9:132669644-132670672 | 0.292597662 | 0.37167436 | 0.658665243 |
| chr19:1479154-1479656 | 0.328668719 | 0.371675981 | 0.658665243 |
| chr17:81602757-81602989 | -0.474761248 | 0.371677966 | 0.658665243 |
| chr8:10964923-10965123 | 0.490841493 | 0.371750776 | 0.658779037 |
| chr13:113208517-113209626 | -0.290469047 | 0.37176319 | 0.658785798 |
| chr1:202341420-202342632 | -0.282599555 | 0.371781216 | 0.65878977 |
| chr12:52859892-52860612 | -0.299652831 | 0.371789232 | 0.65878977 |
| chrX:144657994-144658259 | -0.456039954 | 0.371801305 | 0.65878977 |
| chr14:64430035-64430264 | 0.422610902 | 0.371822473 | 0.65878977 |
| chr10:101830233-101831168 | 0.288506977 | 0.371824048 | 0.65878977 |
| chr6:136822186-136823274 | -0.286856382 | 0.371830288 | 0.65878977 |
| chr7:35849306-35849591 | 0.387775748 | 0.371833331 | 0.65878977 |
| chr5:90606615-90606867 | -0.39550449 | 0.371834215 | 0.65878977 |
| chr1:94069650-94069850 | 0.468204366 | 0.371854971 | 0.658811311 |

| | | | |
|---|---|---|---|
| chr2:86079948-86080220 | -0.372986333 | 0.371870379 | 0.658823375 |
| chr14:35985594-35985809 | 0.42680508 | 0.371887347 | 0.658838204 |
| chr17:68315337-68315572 | 0.379694967 | 0.371938858 | 0.658906159 |
| chr2:176188110-176188839 | 0.313948125 | 0.371942904 | 0.658906159 |
| chr13:85266425-85266665 | 0.455651086 | 0.371955208 | 0.658912722 |
| chrX:53071557-53072074 | 0.319111904 | 0.371979714 | 0.65892716 |
| chr4:108834811-108835070 | -0.453868085 | 0.371985965 | 0.65892716 |
| chr14:91276332-91276791 | 0.31512399 | 0.371989157 | 0.65892716 |
| chr6:117482241-117483626 | 0.282764 | 0.372005301 | 0.658940523 |
| chr2:193885563-193885787 | -0.469049649 | 0.372039936 | 0.658979941 |
| chr17:15828748-15829171 | -0.338438828 | 0.372044756 | 0.658979941 |
| chr2:10911740-10912054 | -0.349178964 | 0.372065728 | 0.659001854 |
| chr12:9662578-9663115 | 0.357422775 | 0.372112612 | 0.65906966 |
| chr8:112465108-112465322 | 0.442202867 | 0.372124551 | 0.659069823 |
| chr1:21782543-21783869 | 0.291410704 | 0.372129907 | 0.659069823 |
| chr17:81386890-81387152 | 0.372378569 | 0.37215397 | 0.659078686 |
| chr15:85380096-85381047 | 0.285824091 | 0.372154793 | 0.659078686 |
| chr10:127888189-127888389 | -0.477284875 | 0.372160717 | 0.659078686 |
| chr10:48287776-48288616 | -0.305228777 | 0.372180317 | 0.659088191 |
| chr16:87653341-87653552 | 0.376807645 | 0.372183723 | 0.659088191 |
| chr6:87547505-87547896 | 0.447303406 | 0.372191889 | 0.659088191 |
| chr12:85394879-85395088 | -0.453585503 | 0.372206357 | 0.659098579 |
| chr17:54967671-54969416 | -0.281410193 | 0.37221599 | 0.659100404 |
| chr10:3184224-3184435 | -0.449953245 | 0.372255014 | 0.659154273 |
| chr5:140647110-140648273 | -0.28749219 | 0.372304439 | 0.659226555 |
| chr20:9039930-9040180 | -0.459979155 | 0.372316203 | 0.659231911 |
| chr2:9207079-9208303 | -0.293168241 | 0.372324671 | 0.659231911 |
| chrX:140762158-140762408 | 0.407022625 | 0.372381808 | 0.659306834 |
| chr20:62955713-62956322 | -0.311628321 | 0.372384196 | 0.659306834 |
| chr20:9458484-9458768 | 0.388712779 | 0.372448844 | 0.659406056 |
| chrX:55202240-55202612 | -0.349351331 | 0.372481441 | 0.659421251 |
| chr8:129431313-129432299 | 0.293948984 | 0.37248215 | 0.659421251 |
| chr8:69632044-69632263 | 0.418553765 | 0.372490937 | 0.659421251 |
| chr8:117727403-117727654 | -0.407585598 | 0.37249185 | 0.659421251 |
| chr11:8721204-8721565 | 0.338354249 | 0.37254313 | 0.659496793 |
| chr10:101168423-101168623 | 0.508434941 | 0.37258583 | 0.659554323 |
| chr21:43045180-43045380 | -0.535679403 | 0.3726014 | 0.659554323 |
| chr1:211513968-211514527 | -0.3117469 | 0.372601452 | 0.659554323 |
| chr13:49443480-49444857 | -0.281925913 | 0.372616933 | 0.65956649 |
| chr8:103010501-103010888 | -0.327150868 | 0.37263052 | 0.659575303 |
| chr9:128625915-128626139 | -0.422749296 | 0.372641115 | 0.659576547 |
| chr9:126777883-126778083 | 0.428677999 | 0.372655739 | 0.659576547 |
| chr9:88950520-88950745 | 0.449222981 | 0.372663713 | 0.659576547 |
| chr5:78295539-78295978 | -0.312548078 | 0.372665656 | 0.659576547 |
| chr18:11962565-11962919 | -0.442028124 | 0.372679853 | 0.659586438 |
| chr6:56315124-56315335 | -0.468277857 | 0.372709519 | 0.659615771 |
| chr5:32173290-32175151 | 0.282545306 | 0.372734949 | 0.659615771 |
| chr10:62187509-62187813 | -0.440880248 | 0.372744786 | 0.659615771 |
| chr19:17947371-17948350 | 0.299837418 | 0.372746194 | 0.659615771 |
| chr19:49149148-49149640 | 0.296065485 | 0.372747614 | 0.659615771 |

| | | | |
|---|---|---|---|
| chr16:30747866-30748922 | 0.289574184 | 0.372748078 | 0.659615771 |
| chr13:49935639-49937098 | -0.282720353 | 0.372761946 | 0.659624168 |
| chr6:47780737-47781057 | -0.363513317 | 0.372770041 | 0.659624168 |
| chr3:63820527-63821022 | -0.297574096 | 0.372796263 | 0.659628532 |
| chr9:137286980-137287331 | 0.351861246 | 0.372797782 | 0.659628532 |
| chr1:28872123-28872392 | 0.441479884 | 0.372798334 | 0.659628532 |
| chr21:41283215-41283548 | -0.345783802 | 0.37283708 | 0.65967034 |
| chr2:71157554-71157797 | -0.390860136 | 0.372839181 | 0.65967034 |
| chr19:56105176-56105411 | 0.403760637 | 0.37287397 | 0.659704782 |
| chr17:58605427-58605649 | 0.320179445 | 0.372875867 | 0.659704782 |
| chr8:122988775-122988975 | 0.400731297 | 0.372885105 | 0.659705893 |
| chr1:62082642-62083008 | -0.374559023 | 0.372897453 | 0.659712507 |
| chr21:42086236-42086579 | 0.337857522 | 0.372915191 | 0.659714186 |
| chr9:62863353-62863706 | -0.331332399 | 0.372915623 | 0.659714186 |
| chr2:30217630-30218288 | -0.3051295 | 0.37293454 | 0.659732421 |
| chr20:761049-761278 | -0.441847533 | 0.372950696 | 0.659745769 |
| chr5:154189872-154190868 | -0.291384391 | 0.372960382 | 0.659747045 |
| chr12:103784370-103784603 | -0.465847684 | 0.372968638 | 0.659747045 |
| chr2:206693504-206694019 | -0.358137556 | 0.372981157 | 0.659753958 |
| chr4:45741956-45742234 | -0.380210681 | 0.372994571 | 0.659762455 |
| chr5:38382845-38383225 | 0.335958859 | 0.373030063 | 0.659798747 |
| chr19:53592993-53593193 | 0.439115951 | 0.373032311 | 0.659798747 |
| chr2:34482805-34483026 | 0.420112023 | 0.373045329 | 0.659806541 |
| chr5:177468733-177468998 | -0.386360597 | 0.373120952 | 0.659910672 |
| chr20:58829262-58829462 | 0.396738338 | 0.373121428 | 0.659910672 |
| chr14:57712204-57712404 | -0.465375874 | 0.373193885 | 0.660023586 |
| chr1:43348485-43349681 | 0.285502321 | 0.373268734 | 0.660071799 |
| chr7:151725912-151726376 | -0.332175144 | 0.373269436 | 0.660071799 |
| chr2:14123343-14124104 | -0.308071815 | 0.37327466 | 0.660071799 |
| chr2:45890-47245 | 0.289116725 | 0.373276638 | 0.660071799 |
| chr2:67450386-67450587 | -0.482892545 | 0.373285767 | 0.660071799 |
| chr2:129957966-129958285 | -0.340448189 | 0.373287122 | 0.660071799 |
| chr14:50311701-50313075 | 0.279364035 | 0.373289844 | 0.660071799 |
| chr17:72118635-72118846 | 0.359301955 | 0.373299476 | 0.660071799 |
| chr1:162497642-162498524 | 0.297976703 | 0.373300772 | 0.660071799 |
| chr7:128434856-128435563 | -0.314727045 | 0.37330833 | 0.660071799 |
| chr15:59138164-59138510 | -0.343073618 | 0.373321023 | 0.660071799 |
| chr10:83917563-83917763 | -0.429291641 | 0.373325454 | 0.660071799 |
| chr9:2466125-2466444 | 0.369136022 | 0.373341518 | 0.660071799 |
| chr5:167295445-167295913 | -0.356565025 | 0.373341751 | 0.660071799 |
| chr2:43609408-43609641 | 0.365295623 | 0.373363083 | 0.660094284 |
| chr2:11407114-11407324 | -0.423252286 | 0.373389686 | 0.660126085 |
| chr17:60505854-60506593 | 0.286335176 | 0.373457811 | 0.660231291 |
| chr3:154323778-154324787 | -0.28645916 | 0.37347256 | 0.660242132 |
| chrX:104641301-104641503 | -0.488257136 | 0.373497624 | 0.660271207 |
| chr17:48816747-48817591 | 0.292280598 | 0.37351522 | 0.660279786 |
| chr22:19291309-19291509 | 0.453887142 | 0.373523156 | 0.660279786 |
| chr22:33883595-33884119 | 0.347742673 | 0.373528329 | 0.660279786 |
| chr16:684171-684664 | -0.311964331 | 0.373559761 | 0.660320115 |
| chr7:33987149-33987613 | 0.344645872 | 0.373586759 | 0.660352603 |

| | | | |
|---|---|---|---|
| chr5:154972539-154972811 | -0.381055951 | 0.373598703 | 0.660358481 |
| chr18:39841455-39842097 | -0.325480133 | 0.37362462 | 0.660389057 |
| chr5:59768375-59768970 | 0.333137866 | 0.373662658 | 0.66042315 |
| chr3:11096464-11097169 | -0.30693392 | 0.373667142 | 0.66042315 |
| chr16:22190379-22191534 | 0.278985506 | 0.373676037 | 0.66042315 |
| chr14:35121613-35123052 | 0.279619156 | 0.373680851 | 0.66042315 |
| chr1:199917400-199917600 | 0.50937833 | 0.373687005 | 0.66042315 |
| chr19:44741438-44742286 | 0.301018886 | 0.373714582 | 0.660456654 |
| chr21:34745583-34745819 | 0.402939814 | 0.373732446 | 0.660472992 |
| chr6:160105071-160105319 | -0.471994201 | 0.373765679 | 0.660516487 |
| chr10:99370969-99371202 | 0.385241967 | 0.373780938 | 0.660528219 |
| chr5:95646296-95647334 | -0.282092249 | 0.373791809 | 0.660532195 |
| chr20:57290871-57291397 | 0.30823903 | 0.373801227 | 0.660533604 |
| chr3:11539756-11540044 | 0.333855492 | 0.373833509 | 0.660575415 |
| chr1:70745200-70745523 | -0.353820405 | 0.373867597 | 0.660586956 |
| chr3:141366521-141366745 | 0.412074491 | 0.373880112 | 0.660586956 |
| chr1:50368282-50368788 | -0.328300229 | 0.37390175 | 0.660586956 |
| chr10:92434962-92435357 | -0.347512521 | 0.373904185 | 0.660586956 |
| chr3:9249256-9250093 | 0.28973802 | 0.373911103 | 0.660586956 |
| chr12:11177232-11177478 | 0.405708118 | 0.373919701 | 0.660586956 |
| chr5:41843492-41843816 | -0.378078956 | 0.373925634 | 0.660586956 |
| chrX:134883388-134883666 | 0.397078373 | 0.373926817 | 0.660586956 |
| chr20:24900475-24901294 | -0.313917503 | 0.373929623 | 0.660586956 |
| chr5:167911341-167911774 | -0.344179631 | 0.373937266 | 0.660586956 |
| chr12:52904733-52905358 | 0.288396907 | 0.3739398 | 0.660586956 |
| chr12:113184646-113185192 | 0.301376772 | 0.373950908 | 0.660586956 |
| chr2:104332622-104333217 | 0.313296766 | 0.373955956 | 0.660586956 |
| chr5:35038927-35039178 | -0.463148902 | 0.37396074 | 0.660586956 |
| chr20:64129106-64129394 | -0.419011877 | 0.373980153 | 0.660592298 |
| chr16:47141760-47142048 | 0.375202251 | 0.373981007 | 0.660592298 |
| chr9:89795103-89795303 | 0.459496846 | 0.374047346 | 0.660674468 |
| chr19:14558493-14558714 | 0.503012795 | 0.37405501 | 0.660674468 |
| chr5:142024747-142025284 | -0.317487497 | 0.374059371 | 0.660674468 |
| chr12:7806969-7807188 | -0.428584729 | 0.374062016 | 0.660674468 |
| chr3:132343095-132343302 | -0.428824581 | 0.374082807 | 0.660687157 |
| chr17:64448463-64448677 | -0.455671942 | 0.374090658 | 0.660687157 |
| chr20:25309292-25309521 | -0.395483586 | 0.374095068 | 0.660687157 |
| chr19:8389857-8390477 | 0.293446671 | 0.374114007 | 0.660705377 |
| chr9:86281861-86283218 | 0.282616054 | 0.374123223 | 0.660706423 |
| chr22:39532604-39533149 | 0.303938271 | 0.374164864 | 0.660764282 |
| chr7:1255154-1255354 | 0.381805657 | 0.374173233 | 0.660764282 |
| chr9:88969652-88969933 | -0.377129732 | 0.37420301 | 0.660801637 |
| chr1:38123419-38123878 | 0.359737103 | 0.374232318 | 0.660828426 |
| chr14:45919959-45920611 | 0.315154059 | 0.374238857 | 0.660828426 |
| chr22:36123890-36124314 | -0.339769281 | 0.374244054 | 0.660828426 |
| chr20:41617808-41619167 | 0.280479402 | 0.374266712 | 0.660841444 |
| chr6:3885674-3885888 | 0.42252695 | 0.374272649 | 0.660841444 |
| chr17:77760897-77761106 | -0.469462137 | 0.3742773 | 0.660841444 |
| chr22:37933897-37934344 | 0.321152757 | 0.374296077 | 0.660854063 |
| chr16:68310464-68311508 | 0.292362879 | 0.37430551 | 0.660854063 |

| | | | |
|---|---|---|---|
| chr17:42458562-42458990 | 0.326892635 | 0.374315558 | 0.660854063 |
| chr16:29925501-29926649 | 0.292098659 | 0.374318947 | 0.660854063 |
| chr2:43612500-43612809 | -0.351632824 | 0.374338463 | 0.660873292 |
| chr20:24469346-24469837 | -0.354250801 | 0.374351013 | 0.660880221 |
| chr13:31002456-31002740 | -0.432626132 | 0.374406752 | 0.66094926 |
| chr2:25977964-25978207 | -0.406895479 | 0.374420729 | 0.66094926 |
| chr8:28594070-28594287 | 0.476019492 | 0.374424429 | 0.66094926 |
| chr16:81212861-81213334 | 0.329085067 | 0.37442589 | 0.66094926 |
| chr5:155260902-155261204 | -0.375067348 | 0.374437311 | 0.66094926 |
| chr4:159487285-159487726 | -0.330938327 | 0.374441877 | 0.66094926 |
| chr17:59871559-59871784 | 0.311463547 | 0.374476042 | 0.660994339 |
| chr17:69084147-69084347 | 0.453585221 | 0.374501158 | 0.661006286 |
| chr1:181152994-181153245 | 0.382593321 | 0.374505192 | 0.661006286 |
| chr11:119381185-119382077 | -0.289318822 | 0.374510668 | 0.661006286 |
| chr22:28020062-28020318 | 0.400719915 | 0.374517318 | 0.661006286 |
| chr5:146109566-146109775 | 0.483463649 | 0.374530612 | 0.661014523 |
| chr22:30386920-30387136 | 0.397884083 | 0.374549726 | 0.661029928 |
| chr5:151108185-151108414 | -0.383074243 | 0.374557997 | 0.661029928 |
| chr19:11374496-11375360 | 0.290519791 | 0.374574184 | 0.661029928 |
| chr8:71975329-71975556 | -0.406802734 | 0.374576247 | 0.661029928 |
| chr17:41800923-41801920 | 0.301411659 | 0.374582476 | 0.661029928 |
| chr9:98158325-98158681 | 0.335254577 | 0.374598838 | 0.661037991 |
| chr16:85027345-85028237 | -0.289204976 | 0.374604299 | 0.661037991 |
| chr5:175444575-175444996 | 0.333317051 | 0.374615129 | 0.661041878 |
| chr3:70738346-70738712 | 0.367416143 | 0.374637777 | 0.661056825 |
| chr11:64285116-64285967 | 0.293029488 | 0.374651446 | 0.661056825 |
| chr4:142434916-142435206 | -0.404510047 | 0.374655455 | 0.661056825 |
| chr17:30970752-30971413 | 0.326355258 | 0.37466124 | 0.661056825 |
| chr5:180325039-180325245 | -0.426433983 | 0.374666737 | 0.661056825 |
| chr6:82331362-82331893 | -0.344199586 | 0.374682925 | 0.661070163 |
| chr4:77818121-77819181 | -0.287421199 | 0.374701478 | 0.661073492 |
| chr8:22010108-22010838 | 0.302824194 | 0.374702066 | 0.661073492 |
| chr20:37095566-37096452 | 0.300310067 | 0.374712267 | 0.661076266 |
| chr20:46406114-46407396 | 0.283288607 | 0.374770857 | 0.661161996 |
| chr12:7919652-7920096 | 0.37359057 | 0.374785291 | 0.661161996 |
| chr8:134886478-134886789 | -0.334434618 | 0.374786747 | 0.661161996 |
| chr3:113339706-113339933 | 0.4148988 | 0.374808547 | 0.661185231 |
| chr16:84333143-84333816 | 0.31445302 | 0.374822988 | 0.661195483 |
| chr15:40746526-40746784 | 0.388919541 | 0.374870769 | 0.661264545 |
| chr17:81926007-81926234 | 0.393905965 | 0.374946702 | 0.661383263 |
| chr3:172560093-172560306 | -0.423442618 | 0.374973138 | 0.661414669 |
| chr9:135804833-135805129 | 0.444476166 | 0.374988804 | 0.661427076 |
| chr5:177311450-177311691 | 0.33393767 | 0.375005146 | 0.661436599 |
| chr2:213151499-213152419 | -0.288991058 | 0.375017881 | 0.661436599 |
| chr9:14314012-14314216 | 0.391094543 | 0.375020101 | 0.661436599 |
| chr20:31496964-31497266 | 0.367805058 | 0.375044847 | 0.661453434 |
| chr22:36277763-36278008 | 0.40475171 | 0.375049127 | 0.661453434 |
| chr13:113162844-113163405 | -0.359157274 | 0.375057573 | 0.661453434 |
| chr4:133264202-133264677 | -0.322475326 | 0.375064177 | 0.661453434 |
| chr20:58744317-58744517 | 0.387382287 | 0.375082299 | 0.661463631 |

| | | | |
|---|---|---|---|
| chr12:55797182-55797388 | 0.458348891 | 0.375093516 | 0.661463631 |
| chr17:81636336-81637321 | -0.294617713 | 0.375103735 | 0.661463631 |
| chr1:161097631-161098751 | 0.287424322 | 0.37510449 | 0.661463631 |
| chr20:35171691-35172164 | -0.331205849 | 0.375125768 | 0.661471101 |
| chr1:30905814-30906139 | -0.399113859 | 0.375132863 | 0.661471101 |
| chr1:234599699-234600537 | 0.293451263 | 0.375134625 | 0.661471101 |
| chr1:151372197-151372942 | 0.289586167 | 0.375146114 | 0.661476137 |
| chr14:52706699-52707718 | 0.283623523 | 0.37516805 | 0.661499594 |
| chr2:25534727-25534990 | -0.457059521 | 0.375182664 | 0.661510138 |
| chr13:37487811-37488123 | -0.476471882 | 0.375200949 | 0.661527155 |
| chr9:123327598-123328214 | 0.29742266 | 0.37522121 | 0.661547655 |
| chr17:62470724-62471079 | 0.319994504 | 0.375233426 | 0.661552293 |
| chr14:70809002-70810231 | 0.281426602 | 0.375241108 | 0.661552293 |
| chr12:69070396-69070685 | 0.369965618 | 0.37528254 | 0.66160299 |
| chr16:15146934-15147243 | -0.371394676 | 0.375287134 | 0.66160299 |
| chr12:104109175-104109480 | 0.363790747 | 0.375317703 | 0.661641658 |
| chr3:134224396-134224883 | 0.324729121 | 0.375344255 | 0.661664781 |
| chrX:68396872-68397365 | -0.364081473 | 0.37534809 | 0.661664781 |
| chr6:156883586-156883893 | -0.455005677 | 0.375387846 | 0.661719639 |
| chr16:84725928-84726582 | 0.308080761 | 0.375414721 | 0.661730175 |
| chr7:25094244-25094542 | -0.339522437 | 0.375416792 | 0.661730175 |
| chr19:38434358-38434558 | -0.478642918 | 0.375424332 | 0.661730175 |
| chr2:218995997-218996464 | 0.308945797 | 0.375433628 | 0.661730175 |
| chr4:684024-684229 | 0.377588394 | 0.375437005 | 0.661730175 |
| chr5:139148245-139148742 | -0.304919167 | 0.375472913 | 0.661765487 |
| chr18:12396426-12396858 | 0.325428363 | 0.375474313 | 0.661765487 |
| chr14:47538935-47539558 | 0.304257985 | 0.375489926 | 0.661769225 |
| chr19:15150468-15150705 | 0.430104847 | 0.375496716 | 0.661769225 |
| chr1:119712409-119712907 | 0.341496494 | 0.375509673 | 0.661769225 |
| chr14:23727437-23727913 | 0.323951188 | 0.375510981 | 0.661769225 |
| chr17:51797511-51797711 | 0.479803277 | 0.375543832 | 0.661797463 |
| chr3:119559534-119559910 | 0.332366683 | 0.375544278 | 0.661797463 |
| chr9:135067450-135067761 | -0.439473006 | 0.375570361 | 0.661799761 |
| chr17:42659094-42659761 | 0.306117602 | 0.375571637 | 0.661799761 |
| chr2:204109651-204110051 | -0.317309042 | 0.375573759 | 0.661799761 |
| chr11:65422644-65423135 | 0.300035057 | 0.375580131 | 0.661799761 |
| chr19:12564082-12564327 | 0.417886698 | 0.375621415 | 0.661857286 |
| chr7:100148531-100149156 | -0.289348361 | 0.375667259 | 0.661884828 |
| chr5:157294804-157295065 | -0.425307696 | 0.375671479 | 0.661884828 |
| chr11:35617749-35618005 | -0.395917184 | 0.375709186 | 0.661884828 |
| chr9:88938211-88938479 | -0.425356535 | 0.375710177 | 0.661884828 |
| chr19:38453067-38453348 | -0.413705284 | 0.375712415 | 0.661884828 |
| chr17:59572413-59572816 | 0.352966413 | 0.375715729 | 0.661884828 |
| chr12:124970896-124971114 | -0.452780796 | 0.375723109 | 0.661884828 |
| chr18:25921529-25921923 | -0.351724331 | 0.375725595 | 0.661884828 |
| chr2:95121484-95122454 | 0.291524099 | 0.375726849 | 0.661884828 |
| chr16:83952736-83953996 | 0.280801211 | 0.375728487 | 0.661884828 |
| chr17:6666097-6666343 | 0.435979559 | 0.375743225 | 0.661884828 |
| chr1:118318346-118318582 | 0.446990975 | 0.375753341 | 0.661884828 |
| chr11:119729065-119730377 | -0.288970405 | 0.375757763 | 0.661884828 |

| | | | |
|---|---|---|---|
| chr17:56257159-56257442 | -0.379959243 | 0.375779263 | 0.661884828 |
| chr8:10547852-10548458 | -0.317808107 | 0.375784139 | 0.661884828 |
| chr7:101094857-101095105 | -0.390787043 | 0.375794429 | 0.661884828 |
| chr1:156767061-156768042 | 0.282327642 | 0.375799812 | 0.661884828 |
| chr12:123601360-123602510 | 0.289553995 | 0.375802674 | 0.661884828 |
| chr17:82668449-82668909 | 0.332515285 | 0.375810354 | 0.661884828 |
| chr11:18727626-18727918 | 0.355569965 | 0.375813119 | 0.661884828 |
| chr9:130682483-130682811 | 0.340619445 | 0.375818451 | 0.661884828 |
| chr16:90011556-90011756 | 0.477004213 | 0.375859079 | 0.661941167 |
| chr2:85222917-85223220 | -0.400350186 | 0.375880353 | 0.661963419 |
| chr9:84668203-84668785 | -0.306231427 | 0.375893959 | 0.661972164 |
| chr8:144509458-144509698 | 0.338967877 | 0.375922174 | 0.662006638 |
| chr9:115495330-115495773 | -0.369135943 | 0.375945135 | 0.662031857 |
| chr8:133350682-133350914 | 0.425106049 | 0.375969129 | 0.662038939 |
| chr2:44167713-44168959 | -0.276856522 | 0.375980816 | 0.662038939 |
| chr20:51917975-51918584 | 0.30963876 | 0.375986891 | 0.662038939 |
| chr3:185825143-185825400 | -0.395484118 | 0.375993733 | 0.662038939 |
| chr11:65013538-65014866 | 0.288806034 | 0.375998876 | 0.662038939 |
| chr5:61347224-61347491 | -0.393659752 | 0.376000999 | 0.662038939 |
| chr2:227471905-227473103 | 0.293031311 | 0.376051404 | 0.662083234 |
| chr2:122063759-122063977 | -0.488507444 | 0.376066676 | 0.662083234 |
| chr4:9152902-9153678 | 0.28656081 | 0.376070408 | 0.662083234 |
| chr10:110211110-110211537 | -0.295620308 | 0.376076071 | 0.662083234 |
| chr4:26583568-26584422 | 0.294930979 | 0.376082892 | 0.662083234 |
| chrX:153922848-153923066 | 0.402178575 | 0.376105225 | 0.662083234 |
| chr16:58000677-58001818 | 0.292724654 | 0.376110403 | 0.662083234 |
| chr19:53432220-53432420 | 0.417568504 | 0.376113282 | 0.662083234 |
| chr17:8190161-8190413 | -0.345215829 | 0.376113917 | 0.662083234 |
| chr1:154405013-154405701 | 0.300570163 | 0.376129076 | 0.662083234 |
| chr2:191052241-191052441 | -0.454133204 | 0.376136296 | 0.662083234 |
| chr12:66234031-66234464 | -0.296971716 | 0.376140168 | 0.662083234 |
| chr12:95998629-95998937 | -0.380590193 | 0.37614616 | 0.662083234 |
| chr9:99266429-99266786 | -0.392032645 | 0.376153629 | 0.662083234 |
| chr16:3681300-3681500 | 0.499164655 | 0.376155769 | 0.662083234 |
| chr17:60355957-60356586 | 0.28527768 | 0.376178298 | 0.662107679 |
| chr17:4534422-4534786 | -0.355540092 | 0.376198461 | 0.662120873 |
| chr11:70637182-70637475 | 0.406771511 | 0.376203077 | 0.662120873 |
| chr7:100049414-100050104 | -0.307958001 | 0.376218774 | 0.662132952 |
| chr3:172648011-172648269 | -0.390351117 | 0.376227224 | 0.662132952 |
| chr12:132657413-132657673 | -0.448138301 | 0.376246922 | 0.66215241 |
| chr17:77331394-77331825 | 0.328351499 | 0.376273817 | 0.662184533 |
| chr11:75140541-75140823 | -0.370147325 | 0.376301727 | 0.66220361 |
| chr7:148264970-148265247 | -0.365863087 | 0.376301942 | 0.66220361 |
| chr5:179858286-179859458 | -0.282509954 | 0.376318857 | 0.662218167 |
| chr6:159762742-159762976 | 0.352310244 | 0.376353102 | 0.662249718 |
| chr17:67778671-67779611 | -0.285600986 | 0.376354072 | 0.662249718 |
| chr8:142874375-142874612 | 0.404974349 | 0.376381227 | 0.66227689 |
| chr11:61429173-61430461 | 0.288317753 | 0.376386801 | 0.66227689 |
| chr8:48008175-48008939 | -0.293012526 | 0.376422502 | 0.662324498 |
| chr1:227481551-227481850 | 0.377826183 | 0.37647958 | 0.662367419 |

| | | | |
|---|---|---|---|
| chr17:77306314-77306669 | -0.365073419 | 0.376482666 | 0.662367419 |
| chr7:120806000-120806255 | -0.395497984 | 0.376483956 | 0.662367419 |
| chr5:37379028-37379663 | -0.298250688 | 0.376486449 | 0.662367419 |
| chr1:187626189-187626389 | -0.474473186 | 0.376490118 | 0.662367419 |
| chr4:158826976-158827209 | -0.405031373 | 0.37653931 | 0.662414862 |
| chr11:8963989-8965260 | 0.28501125 | 0.376547695 | 0.662414862 |
| chr5:27472087-27472346 | 0.433318604 | 0.376557664 | 0.662414862 |
| chr20:4015516-4015981 | 0.312133686 | 0.376565199 | 0.662414862 |
| chr2:225270898-225271126 | -0.457928116 | 0.376585424 | 0.662414862 |
| chr2:19059012-19059354 | -0.338518366 | 0.376619189 | 0.662414862 |
| chr2:186590866-186591070 | -0.356224166 | 0.376619401 | 0.662414862 |
| chr10:15480036-15480374 | 0.466519511 | 0.376628232 | 0.662414862 |
| chr5:44691860-44692171 | -0.394212912 | 0.376629583 | 0.662414862 |
| chr19:49200397-49200695 | 0.362117325 | 0.376632631 | 0.662414862 |
| chr3:148158305-148158555 | -0.384367479 | 0.376632655 | 0.662414862 |
| chr1:219660616-219661449 | -0.298479236 | 0.376633787 | 0.662414862 |
| chr11:9265029-9265728 | 0.308026688 | 0.376653245 | 0.662414862 |
| chr17:59316923-59317234 | 0.319877217 | 0.376659856 | 0.662414862 |
| chr8:94528632-94529150 | 0.303687039 | 0.376666217 | 0.662414862 |
| chr11:102780508-102780946 | -0.383572072 | 0.376672812 | 0.662414862 |
| chr4:169613152-169613449 | -0.329516168 | 0.376673003 | 0.662414862 |
| chr21:39851967-39852282 | -0.366513168 | 0.376681997 | 0.662414862 |
| chr8:124160719-124160919 | -0.494102008 | 0.376688912 | 0.662414862 |
| chr2:230411879-230412228 | 0.386128613 | 0.376691773 | 0.662414862 |
| chr17:42979881-42980311 | -0.337033211 | 0.376698635 | 0.662414862 |
| chr17:68263164-68263712 | -0.319629906 | 0.376764238 | 0.662502314 |
| chr2:26961549-26962153 | -0.311946191 | 0.376765659 | 0.662502314 |
| chr6:43334518-43334742 | 0.41843794 | 0.376817718 | 0.662578648 |
| chr8:131036996-131037196 | 0.46305987 | 0.376843948 | 0.662609565 |
| chr10:75022386-75022663 | -0.378975995 | 0.376858258 | 0.662619521 |
| chr14:88928671-88928903 | 0.432871283 | 0.376890079 | 0.662660264 |
| chr7:37713784-37714047 | -0.438244907 | 0.376906658 | 0.662674208 |
| chr2:36376312-36376524 | -0.404727747 | 0.376974938 | 0.662779049 |
| chr19:5436709-5436929 | 0.430818979 | 0.377000338 | 0.662808496 |
| chr12:132478736-132478936 | 0.4106435 | 0.377039732 | 0.662862546 |
| chr21:44300534-44300744 | 0.359748036 | 0.377076113 | 0.662881157 |
| chr2:217650318-217650548 | -0.43817607 | 0.377078875 | 0.662881157 |
| chr9:124209733-124209933 | 0.489217498 | 0.377084157 | 0.662881157 |
| chr16:10521085-10521363 | 0.416134638 | 0.377084923 | 0.662881157 |
| chrX:134914353-134915787 | 0.280809439 | 0.377111155 | 0.662901552 |
| chr5:86153096-86153296 | -0.491223075 | 0.377114649 | 0.662901552 |
| chr2:233288660-233289009 | -0.399263846 | 0.37712759 | 0.662901552 |
| chr17:781939-782710 | -0.309079243 | 0.377131132 | 0.662901552 |
| chr7:6658966-6659956 | -0.290947624 | 0.377144046 | 0.662909045 |
| chr10:80498644-80499150 | -0.318717116 | 0.377221897 | 0.663030674 |
| chrX:16882246-16882446 | -0.439733294 | 0.377242924 | 0.663045159 |
| chr16:85245706-85246161 | 0.32046986 | 0.377247445 | 0.663045159 |
| chr12:91109082-91109609 | -0.341741136 | 0.377287934 | 0.663101111 |
| chr10:15715910-15716165 | -0.421354999 | 0.377307723 | 0.663108749 |
| chr20:45872605-45873141 | 0.296336166 | 0.377309588 | 0.663108749 |

| | | | |
|---|---|---|---|
| chr10:43407699-43409545 | -0.276999662 | 0.377337077 | 0.663137043 |
| chr3:9792142-9793712 | 0.277100205 | 0.377342996 | 0.663137043 |
| chr7:139598639-139598892 | 0.355707242 | 0.377386755 | 0.663198732 |
| chr6:125254425-125254634 | -0.44530146 | 0.377402813 | 0.663209122 |
| chr2:61016853-61018313 | -0.279830624 | 0.377409978 | 0.663209122 |
| chr9:72577248-72577776 | -0.325632672 | 0.377450903 | 0.663259115 |
| chr1:6998312-6998614 | 0.365940104 | 0.377463571 | 0.663259115 |
| chr3:149283508-149283983 | 0.330658127 | 0.377464396 | 0.663259115 |
| chr1:106473771-106474439 | 0.291289119 | 0.377482102 | 0.663275017 |
| chr13:95301347-95302140 | 0.303791018 | 0.377494437 | 0.66328148 |
| chr9:78031664-78032726 | -0.2862178 | 0.377555443 | 0.663373459 |
| chr3:101512513-101513661 | -0.284794696 | 0.377580265 | 0.663400222 |
| chr17:17613712-17614017 | 0.365836484 | 0.377587992 | 0.663400222 |
| chr10:24956771-24957094 | -0.445190564 | 0.377622782 | 0.663445262 |
| chr8:138698653-138699204 | -0.305774662 | 0.377630944 | 0.663445262 |
| chr14:35883731-35883959 | 0.379045772 | 0.377655967 | 0.663459023 |
| chr8:116374885-116375140 | 0.475185281 | 0.377656095 | 0.663459023 |
| chr1:97138126-97138612 | -0.336791761 | 0.377681409 | 0.663482609 |
| chr1:999976-1000607 | 0.299688252 | 0.377693734 | 0.663482609 |
| chr1:29197495-29197830 | 0.371563335 | 0.377695498 | 0.663482609 |
| chr8:61139499-61139719 | 0.417659834 | 0.377730901 | 0.663529588 |
| chr17:79087645-79087905 | 0.346674255 | 0.377739745 | 0.663529911 |
| chr4:135306603-135306803 | 0.459396691 | 0.37774882 | 0.663530642 |
| chr15:88537298-88537498 | 0.493206344 | 0.377772898 | 0.663557723 |
| chr6:132653245-132653445 | -0.495929738 | 0.377782511 | 0.663559397 |
| chr1:228500947-228501474 | 0.322613907 | 0.377815289 | 0.663601758 |
| chr16:89320381-89320772 | -0.342135067 | 0.37783779 | 0.663625628 |
| chr14:60656662-60657753 | 0.285744958 | 0.377846202 | 0.663625628 |
| chr11:125111031-125111256 | -0.391467414 | 0.377873538 | 0.663658428 |
| chr1:26530395-26530745 | 0.361176059 | 0.377896291 | 0.663683176 |
| chr8:70532864-70533234 | -0.345355609 | 0.377922706 | 0.663685925 |
| chr14:77301026-77301314 | 0.34962025 | 0.377925693 | 0.663685925 |
| chr7:20744857-20745282 | 0.330493162 | 0.377928147 | 0.663685925 |
| chr21:42516174-42516440 | 0.34462389 | 0.377935185 | 0.663685925 |
| chr2:240021536-240021792 | -0.405946919 | 0.377945968 | 0.663685925 |
| chr7:104983580-104984396 | -0.297193221 | 0.377951604 | 0.663685925 |
| chr12:109597685-109598215 | -0.310208833 | 0.377962432 | 0.663685925 |
| chr14:81464388-81464951 | -0.337929531 | 0.377967151 | 0.663685925 |
| chr3:151439448-151440165 | -0.303328597 | 0.378013227 | 0.663751621 |
| chr1:220747852-220748974 | 0.285099566 | 0.378038192 | 0.663780246 |
| chr4:106245221-106245457 | 0.39931696 | 0.378081745 | 0.663788829 |
| chr3:119810150-119810482 | -0.340952452 | 0.378089676 | 0.663788829 |
| chr20:49187279-49188733 | 0.282662459 | 0.378096359 | 0.663788829 |
| chr7:2172637-2173258 | 0.30505794 | 0.378099144 | 0.663788829 |
| chr1:74767558-74767759 | -0.444500031 | 0.378105945 | 0.663788829 |
| chrX:120485627-120486564 | 0.292637945 | 0.378106077 | 0.663788829 |
| chr3:197029738-197030052 | 0.335854479 | 0.378138235 | 0.663788829 |
| chr4:153432984-153433482 | -0.314005046 | 0.378141764 | 0.663788829 |
| chr1:226479371-226479857 | -0.321374009 | 0.378149555 | 0.663788829 |
| chr7:73433611-73433818 | 0.400463996 | 0.378151471 | 0.663788829 |

| | | | |
|---|---|---|---|
| chr16:87324829-87325123 | 0.368648942 | 0.378154327 | 0.663788829 |
| chr1:241583154-241583376 | 0.475930434 | 0.378157973 | 0.663788829 |
| chr14:104155330-104155530 | -0.440582547 | 0.378166952 | 0.663788829 |
| chr9:26955903-26956765 | -0.294231893 | 0.378180865 | 0.663788829 |
| chr7:99990917-99991117 | 0.427281816 | 0.378181691 | 0.663788829 |
| chr9:101727740-101727940 | 0.427281816 | 0.378181691 | 0.663788829 |
| chr18:24767018-24767290 | -0.441112496 | 0.378216153 | 0.663834111 |
| chr1:153968147-153968347 | 0.336031311 | 0.378305293 | 0.663955561 |
| chr4:87985621-87985971 | 0.329754029 | 0.378310055 | 0.663955561 |
| chr1:201190597-201190937 | 0.317910136 | 0.378311345 | 0.663955561 |
| chr1:46643362-46643795 | 0.369318124 | 0.378328177 | 0.663969894 |
| chr5:170597383-170597791 | -0.329690339 | 0.378361517 | 0.664008946 |
| chr9:69549527-69549727 | 0.452900672 | 0.378383348 | 0.664008946 |
| chr15:75640075-75640763 | 0.287561732 | 0.378386143 | 0.664008946 |
| chr5:96934676-96936273 | -0.27430396 | 0.378389105 | 0.664008946 |
| chr3:117019112-117019429 | -0.374565941 | 0.378401184 | 0.664008946 |
| chr8:79684528-79684935 | 0.332258567 | 0.378402425 | 0.664008946 |
| chr2:231706469-231706728 | -0.337040357 | 0.378415855 | 0.664017305 |
| chr2:89585566-89585979 | 0.321794131 | 0.378454927 | 0.664070658 |
| chr7:17215545-17215900 | 0.329413054 | 0.378464244 | 0.664071798 |
| chr17:45050468-45051165 | 0.3084405 | 0.378489783 | 0.664101402 |
| chr3:141460147-141460473 | 0.323113333 | 0.378505186 | 0.664113221 |
| chr5:11940959-11941281 | -0.427472414 | 0.378520193 | 0.66411978 |
| chr12:65414952-65415152 | 0.485365 | 0.378526259 | 0.66411978 |
| chr3:129121452-129122445 | 0.279809774 | 0.378569208 | 0.664173421 |
| chr1:113388660-113389095 | -0.304619796 | 0.378574169 | 0.664173421 |
| chr1:197587887-197588115 | 0.426776723 | 0.378592968 | 0.664180943 |
| chr5:43020033-43020315 | -0.325608129 | 0.378598892 | 0.664180943 |
| chr8:29542235-29542647 | 0.336644321 | 0.378604462 | 0.664180943 |
| chr5:173217713-173217913 | -0.469671149 | 0.378641577 | 0.664230846 |
| chr2:68362032-68362355 | -0.334183933 | 0.378663935 | 0.664252069 |
| chr3:150515242-150515539 | 0.392001003 | 0.378675579 | 0.664252069 |
| chr1:160198665-160198865 | 0.420791184 | 0.378679683 | 0.664252069 |
| chr1:158284106-158284306 | -0.48348371 | 0.378710883 | 0.66429159 |
| chr3:160754618-160756330 | -0.276368891 | 0.378723668 | 0.66429881 |
| chr1:112508463-112508707 | 0.400024577 | 0.378758448 | 0.664344607 |
| chr17:68745346-68745546 | -0.407587458 | 0.378781959 | 0.664355572 |
| chr16:3165924-3166257 | 0.334282656 | 0.378782041 | 0.664355572 |
| chr10:69637006-69637238 | 0.380168438 | 0.378790975 | 0.664356035 |
| chr10:6779123-6779427 | -0.385784559 | 0.378800726 | 0.664357929 |
| chr12:52243941-52244412 | 0.308151758 | 0.378850065 | 0.664429254 |
| chr3:54122111-54122324 | -0.486176768 | 0.378866818 | 0.664443427 |
| chr17:64506133-64507695 | -0.27427783 | 0.37887801 | 0.664447847 |
| chr5:14665234-14665434 | 0.351620591 | 0.378925072 | 0.664515171 |
| chr2:142980594-142981027 | -0.350829626 | 0.378942547 | 0.664530607 |
| chr10:95560591-95561690 | -0.287059258 | 0.378975697 | 0.664561207 |
| chr20:10647501-10647790 | -0.342611896 | 0.378977343 | 0.664561207 |
| chr14:24093783-24095065 | 0.279395603 | 0.379011056 | 0.66460271 |
| chr1:37690398-37690835 | 0.296961848 | 0.379018358 | 0.66460271 |
| chr14:68569898-68570553 | 0.30255928 | 0.379084513 | 0.664694152 |

| | | | |
|---|---|---|---|
| chr12:79012813-79013065 | 0.410917826 | 0.379093872 | 0.664694152 |
| chr3:64207386-64207716 | -0.298069189 | 0.379096532 | 0.664694152 |
| chrX:140815849-140816181 | 0.356294052 | 0.379132538 | 0.664742073 |
| chr14:92686730-92687385 | 0.298989506 | 0.379141998 | 0.664743448 |
| chr4:3852418-3852644 | -0.476639613 | 0.379232045 | 0.664865672 |
| chr12:33520765-33521658 | -0.289373748 | 0.379236047 | 0.664865672 |
| chr6:137415677-137415942 | -0.401524546 | 0.379239023 | 0.664865672 |
| chr3:46565858-46566058 | -0.480643129 | 0.379246418 | 0.664865672 |
| chr6:12010906-12012129 | -0.279735009 | 0.379340382 | 0.664997446 |
| chrX:97481225-97481577 | -0.397639114 | 0.379342236 | 0.664997446 |
| chr17:43785701-43786034 | -0.421006296 | 0.379353172 | 0.664997446 |
| chr9:18208256-18208563 | -0.410686545 | 0.379362299 | 0.664997446 |
| chr1:30717964-30719076 | 0.277851801 | 0.379364978 | 0.664997446 |
| chr8:127833093-127833486 | 0.314917562 | 0.37946358 | 0.66515507 |
| chr4:114606373-114606656 | 0.390692084 | 0.379490767 | 0.665187508 |
| chr3:179666495-179667134 | -0.3311981 | 0.379510105 | 0.665205585 |
| chr8:56059988-56060463 | 0.334102018 | 0.379520627 | 0.665205585 |
| chr19:46771073-46771319 | -0.384864148 | 0.379540363 | 0.665205585 |
| chr8:23681934-23683339 | 0.290318872 | 0.37954169 | 0.665205585 |
| chr3:120285044-120285382 | 0.373559776 | 0.379555422 | 0.665205585 |
| chr17:2054834-2055170 | -0.329288498 | 0.379558192 | 0.665205585 |
| chr19:55674865-55675569 | 0.296983562 | 0.379561852 | 0.665205585 |
| chr11:130314303-130315278 | 0.287830087 | 0.379583147 | 0.665227691 |
| chr8:65348204-65348446 | 0.463609358 | 0.379601774 | 0.66524512 |
| chr3:150703187-150704369 | 0.27872985 | 0.379630198 | 0.665279716 |
| chr6:56851806-56852339 | -0.340032601 | 0.379652794 | 0.665284425 |
| chr1:113103090-113103402 | -0.368087652 | 0.379657757 | 0.665284425 |
| chr10:123008252-123009363 | 0.284875524 | 0.379658933 | 0.665284425 |
| chr7:100081396-100082103 | -0.293480736 | 0.379686163 | 0.665316301 |
| chr3:63007625-63007862 | 0.379613244 | 0.37969449 | 0.665316301 |
| chr20:50679789-50680315 | 0.28105375 | 0.379759819 | 0.665406456 |
| chr6:79762424-79762719 | -0.380390029 | 0.37976331 | 0.665406456 |
| chr18:13003418-13003850 | 0.329436545 | 0.379772069 | 0.665406587 |
| chr11:125428598-125428798 | -0.476173599 | 0.379854149 | 0.665522543 |
| chr17:45218965-45219256 | 0.354463445 | 0.379855621 | 0.665522543 |
| chr12:27781003-27781387 | -0.347211293 | 0.379865731 | 0.665525039 |
| chr11:10076940-10077187 | 0.453722734 | 0.379876726 | 0.66552865 |
| chr4:19739081-19739376 | 0.377024385 | 0.379885164 | 0.66552865 |
| chr16:82230069-82230348 | -0.376166957 | 0.379900748 | 0.665531853 |
| chr7:74026679-74027107 | -0.36050616 | 0.379904364 | 0.665531853 |
| chr6:9510967-9511685 | -0.309146582 | 0.379950562 | 0.665597566 |
| chr6:35185297-35185500 | -0.444443695 | 0.379960593 | 0.665599921 |
| chr12:76878232-76879486 | -0.280734392 | 0.380021245 | 0.66569095 |
| chr16:88936109-88936309 | 0.374236239 | 0.380097661 | 0.665759462 |
| chr20:56689988-56690426 | -0.319422605 | 0.380103636 | 0.665759462 |
| chr10:43436261-43437597 | -0.282155884 | 0.38010549 | 0.665759462 |
| chr20:47383810-47384602 | 0.275093399 | 0.380108709 | 0.665759462 |
| chr18:57802860-57803856 | -0.28699558 | 0.38011391 | 0.665759462 |
| chr2:235754608-235755100 | -0.352756754 | 0.38011543 | 0.665759462 |
| chr5:20497306-20497686 | 0.365447944 | 0.380121179 | 0.665759462 |

| | | | |
|---|---|---|---|
| chr12:1661675-1662280 | 0.324367543 | 0.380150191 | 0.665795056 |
| chr8:143202736-143203026 | 0.360195956 | 0.380166544 | 0.665808478 |
| chr6:170289988-170290937 | 0.297873732 | 0.380183155 | 0.665822352 |
| chr4:120299537-120299915 | 0.397137299 | 0.380222618 | 0.665869056 |
| chr5:612623-612823 | 0.382229695 | 0.380230674 | 0.665869056 |
| chr11:48025342-48025542 | -0.429289145 | 0.380235894 | 0.665869056 |
| chr18:3296713-3297762 | -0.286797489 | 0.380267558 | 0.665909287 |
| chr11:8263854-8264100 | 0.398722147 | 0.380316396 | 0.665969395 |
| chr11:73360543-73360793 | -0.357829172 | 0.380319266 | 0.665969395 |
| chr11:20042282-20042494 | 0.43431155 | 0.3803315 | 0.665975597 |
| chr9:7570220-7570421 | -0.470367714 | 0.380358865 | 0.666008294 |
| chr1:26862809-26863188 | 0.320672634 | 0.380373083 | 0.66601797 |
| chr18:72998575-72998775 | -0.506827879 | 0.380386313 | 0.666025915 |
| chr11:73647465-73647846 | 0.320256468 | 0.380399535 | 0.666033846 |
| chr21:46254536-46254760 | -0.418631836 | 0.38043328 | 0.666056543 |
| chr3:39174902-39175139 | -0.380467704 | 0.380440925 | 0.666056543 |
| chr7:149873263-149874404 | 0.280744537 | 0.380441753 | 0.666056543 |
| chr1:203136532-203136754 | 0.351588008 | 0.380454432 | 0.666056543 |
| chr2:25757966-25758239 | 0.422770148 | 0.380455962 | 0.666056543 |
| chr6:3157167-3157882 | 0.300721597 | 0.38049806 | 0.666115024 |
| chr8:88191719-88191999 | -0.423904313 | 0.380529848 | 0.666134908 |
| chr17:77121854-77122100 | -0.385757175 | 0.380544764 | 0.666134908 |
| chr1:197736628-197736900 | -0.374918872 | 0.380550026 | 0.666134908 |
| chr10:49594615-49594851 | -0.39742545 | 0.380550215 | 0.666134908 |
| chr12:125049289-125050193 | 0.282399811 | 0.380552887 | 0.666134908 |
| chr18:811767-813119 | -0.281780943 | 0.380568423 | 0.666146885 |
| chr11:2197239-2197452 | 0.393007413 | 0.380602747 | 0.666191746 |
| chr7:43302447-43302647 | 0.425084681 | 0.380619381 | 0.666196585 |
| chr13:24002192-24002552 | -0.413262457 | 0.380622901 | 0.666196585 |
| chr4:76251411-76252233 | -0.297459511 | 0.380632677 | 0.666198478 |
| chrX:27444630-27444879 | -0.419622816 | 0.380679916 | 0.666257251 |
| chr2:120343419-120343637 | 0.387543726 | 0.380692343 | 0.666257251 |
| chr2:157302596-157302840 | 0.387543726 | 0.380692343 | 0.666257251 |
| chr6:87639206-87639406 | 0.490406793 | 0.380712286 | 0.666272998 |
| chr5:115831245-115831648 | -0.388199402 | 0.380718732 | 0.666272998 |
| chr18:24426365-24427075 | -0.299084422 | 0.380742767 | 0.666299842 |
| chr12:87617911-87618120 | -0.482149621 | 0.380755956 | 0.666307704 |
| chr2:11901625-11901847 | -0.447526625 | 0.380785919 | 0.66634492 |
| chr18:62388870-62389224 | 0.349167695 | 0.380816948 | 0.666368625 |
| chr2:237662657-237663029 | -0.341852662 | 0.380820937 | 0.666368625 |
| chr2:202848603-202848980 | -0.425286968 | 0.380829594 | 0.666368625 |
| chr7:1412659-1413271 | -0.290562331 | 0.380834252 | 0.666368625 |
| chr15:31435384-31435650 | 0.361934858 | 0.380862505 | 0.666402843 |
| chr15:77161121-77161383 | 0.399756481 | 0.380882904 | 0.666416389 |
| chr7:45854301-45854642 | -0.35609717 | 0.380887642 | 0.666416389 |
| chr4:78644760-78645023 | -0.389675988 | 0.380904759 | 0.666431119 |
| chr10:68988484-68989584 | -0.28195717 | 0.380930382 | 0.666460731 |
| chr20:56400085-56400348 | 0.347869474 | 0.380949371 | 0.666478148 |
| chr4:118033191-118034080 | 0.290102554 | 0.380958807 | 0.666478148 |
| chr3:143648760-143649061 | 0.358171463 | 0.380966432 | 0.666478148 |

| | | | |
|---|---|---|---|
| chr21:26844118-26844379 | 0.366391702 | 0.381019227 | 0.666555291 |
| chr6:32844497-32845007 | -0.305406053 | 0.381059502 | 0.666610529 |
| chr6:73739211-73739425 | -0.43819963 | 0.381078676 | 0.666628851 |
| chr3:169771882-169772390 | -0.28483972 | 0.38111467 | 0.666653952 |
| chr11:120235037-120235338 | -0.346712698 | 0.381125421 | 0.666653952 |
| chr1:216482358-216482647 | -0.40344981 | 0.381139526 | 0.666653952 |
| chr3:189779304-189779889 | 0.329775201 | 0.381142059 | 0.666653952 |
| chr1:86395715-86396520 | -0.295108216 | 0.381142993 | 0.666653952 |
| chr5:157265740-157266810 | -0.286448986 | 0.38114827 | 0.666653952 |
| chr2:96960304-96960759 | -0.310527747 | 0.381153929 | 0.666653952 |
| chr7:141288779-141288979 | -0.502619444 | 0.381178553 | 0.66668138 |
| chr11:955173-955373 | -0.469956778 | 0.381187012 | 0.66668138 |
| chr14:85528612-85528928 | -0.352040164 | 0.381223544 | 0.666716535 |
| chr5:180429036-180429271 | 0.458398098 | 0.381224515 | 0.666716535 |
| chr5:177591946-177592831 | -0.291383732 | 0.38123618 | 0.666721717 |
| chrX:116749961-116750359 | 0.319024744 | 0.381263143 | 0.666747508 |
| chr5:15095891-15096252 | 0.36490904 | 0.381268331 | 0.666747508 |
| chr9:89498011-89498363 | 0.326614045 | 0.381305653 | 0.666797556 |
| chr3:128709655-128710348 | 0.299204995 | 0.381324737 | 0.66681571 |
| chr17:4902916-4903355 | -0.372194296 | 0.381352733 | 0.666830846 |
| chr4:73620138-73620785 | 0.311059116 | 0.381356867 | 0.666830846 |
| chr11:58923565-58923815 | -0.404293788 | 0.381359501 | 0.666830846 |
| chr10:100346452-100347289 | 0.28915364 | 0.381394926 | 0.66687757 |
| chr13:41310379-41310727 | -0.325040474 | 0.381414556 | 0.666892006 |
| chrX:44872353-44873494 | 0.284004682 | 0.381428266 | 0.666892006 |
| chr20:10654597-10654943 | 0.361474803 | 0.381429293 | 0.666892006 |
| chr11:62194095-62194297 | 0.455623083 | 0.381470308 | 0.666936949 |
| chr7:5558935-5559232 | 0.303370031 | 0.3814755 | 0.666936949 |
| chr17:50845273-50845892 | -0.300405411 | 0.381481111 | 0.666936949 |
| chr17:66702571-66702950 | -0.348261627 | 0.381493945 | 0.66694417 |
| chr5:110267771-110268016 | -0.42171001 | 0.381524666 | 0.666980689 |
| chr2:156483972-156484465 | 0.348428944 | 0.381534766 | 0.666980689 |
| chr1:70737271-70737506 | -0.424818596 | 0.381540949 | 0.666980689 |
| chr3:193828530-193828859 | -0.36277887 | 0.381584797 | 0.667037858 |
| chr8:86609149-86609432 | 0.332514267 | 0.381591063 | 0.667037858 |
| chr10:43026159-43027017 | -0.293241396 | 0.381683517 | 0.66718425 |
| chr14:89955496-89956459 | 0.279979631 | 0.381698945 | 0.667195998 |
| chr5:21469754-21470105 | -0.345902095 | 0.381714212 | 0.667207462 |
| chr8:101345899-101346152 | 0.404416581 | 0.38173627 | 0.667230798 |
| chr3:10249159-10249577 | 0.314386538 | 0.381770248 | 0.667264677 |
| chr16:71630948-71631261 | 0.341376636 | 0.381774458 | 0.667264677 |
| chr19:51689784-51690032 | 0.344712084 | 0.381781778 | 0.667264677 |
| chrY:56870743-56871084 | 0.36929668 | 0.381823762 | 0.667322833 |
| chr19:45308052-45308552 | 0.319608594 | 0.381837416 | 0.667331475 |
| chr1:144560732-144561223 | -0.311736818 | 0.3818504 | 0.667334956 |
| chr7:32905803-32906185 | -0.307360568 | 0.381865245 | 0.667334956 |
| chr9:130432948-130433148 | -0.405988137 | 0.381865536 | 0.667334956 |
| chr2:70553264-70554508 | -0.280864377 | 0.381907865 | 0.667369789 |
| chrX:12137953-12138197 | -0.459369752 | 0.381919864 | 0.667369789 |
| chr5:138331326-138332270 | -0.280432848 | 0.381932449 | 0.667369789 |

| | | | |
|---|---|---|---|
| chr7:138701539-138701753 | -0.520710419 | 0.381932683 | 0.667369789 |
| chr15:40871688-40872571 | -0.282555198 | 0.381936193 | 0.667369789 |
| chr12:54028562-54028767 | -0.418428363 | 0.381937728 | 0.667369789 |
| chr2:127407745-127408810 | 0.286476304 | 0.381981685 | 0.667420732 |
| chr17:82716041-82716843 | -0.284554602 | 0.381984304 | 0.667420732 |
| chr20:25399175-25399395 | 0.403407402 | 0.381999627 | 0.667431481 |
| chr8:127914366-127914631 | -0.334836858 | 0.382007877 | 0.667431481 |
| chr6:26457504-26458468 | -0.293101617 | 0.382024572 | 0.667445429 |
| chr13:24192566-24193045 | -0.372913325 | 0.382042134 | 0.667449018 |
| chr21:26715981-26716227 | 0.405189943 | 0.382053433 | 0.667449018 |
| chr2:241558492-241559182 | -0.309124274 | 0.382059077 | 0.667449018 |
| chr7:72827908-72828611 | -0.283028239 | 0.38206147 | 0.667449018 |
| chr12:2783932-2784356 | -0.341290072 | 0.382132764 | 0.667558346 |
| chr6:12873771-12873971 | 0.46481932 | 0.38214798 | 0.667565549 |
| chr8:42051933-42052897 | 0.289054619 | 0.382154312 | 0.667565549 |
| chr11:126147592-126147828 | -0.474723102 | 0.382183841 | 0.667588447 |
| chr1:109197510-109197710 | -0.401414607 | 0.382191212 | 0.667588447 |
| chr10:5294074-5294467 | -0.328399763 | 0.382198122 | 0.667588447 |
| chr3:156776615-156776942 | -0.417064832 | 0.382226525 | 0.667588447 |
| chr2:41806395-41806631 | -0.435266199 | 0.382228902 | 0.667588447 |
| chr1:107739399-107740090 | -0.309142788 | 0.382230493 | 0.667588447 |
| chr8:60680074-60680659 | 0.318385668 | 0.382232329 | 0.667588447 |
| chr6:131807554-131808731 | 0.287818274 | 0.382237122 | 0.667588447 |
| chr17:50719123-50720076 | 0.281687264 | 0.38227872 | 0.667645881 |
| chr12:91001419-91001619 | -0.468747132 | 0.382297419 | 0.667655067 |
| chr5:51399140-51399345 | 0.448712534 | 0.382324178 | 0.667655067 |
| chr12:64082877-64083086 | -0.387664981 | 0.382324636 | 0.667655067 |
| chr6:27478703-27479162 | -0.302335705 | 0.382325365 | 0.667655067 |
| chr8:80103163-80103470 | -0.343746105 | 0.382327829 | 0.667655067 |
| chr17:28398071-28398597 | -0.348878889 | 0.382336261 | 0.667655067 |
| chrX:49840762-49841097 | -0.370949025 | 0.382365168 | 0.667689798 |
| chr6:127554534-127554785 | -0.394273468 | 0.382374074 | 0.667689798 |
| chr2:191302521-191303078 | -0.311533659 | 0.382389911 | 0.667689798 |
| chr2:175004510-175005875 | 0.285414022 | 0.382391007 | 0.667689798 |
| chr16:75089468-75089985 | 0.308396594 | 0.382416332 | 0.667695787 |
| chr4:8117283-8117681 | 0.308860424 | 0.382420384 | 0.667695787 |
| chr9:129202719-129203349 | 0.304002477 | 0.382420579 | 0.667695787 |
| chr7:130242261-130242815 | -0.35438864 | 0.382440514 | 0.667706913 |
| chr12:6778149-6778492 | -0.392861329 | 0.382451969 | 0.667706913 |
| chr22:31586080-31586843 | -0.307713495 | 0.382458959 | 0.667706913 |
| chr10:101050327-101050686 | -0.341705676 | 0.382461809 | 0.667706913 |
| chr17:31858589-31858915 | -0.373360608 | 0.382480513 | 0.667712749 |
| chr17:59847657-59847896 | -0.290166386 | 0.38248258 | 0.667712749 |
| chr1:3668815-3669188 | 0.349434968 | 0.382495193 | 0.667719555 |
| chr4:142473472-142475076 | -0.276233533 | 0.382520373 | 0.667748298 |
| chr20:9421665-9421887 | -0.45666243 | 0.382568363 | 0.667816857 |
| chr2:36770559-36771047 | -0.326961563 | 0.382581348 | 0.667824309 |
| chr6:18302828-18303052 | 0.460628044 | 0.382630822 | 0.667895454 |
| chr1:39691328-39691643 | 0.322107611 | 0.382745916 | 0.668081135 |
| chr7:12686377-12687785 | 0.271823464 | 0.382772664 | 0.668097047 |

| | | | |
|---|---|---|---|
| chr12:54201270-54201702 | 0.32028022 | 0.382773213 | 0.668097047 |
| chr19:1045183-1045599 | 0.309998351 | 0.38278119 | 0.668097047 |
| chr12:123250682-123251436 | -0.293164886 | 0.382802307 | 0.668118684 |
| chr1:160261620-160262734 | 0.282554645 | 0.382820998 | 0.668136087 |
| chr4:41361414-41361712 | -0.426639529 | 0.382860714 | 0.668190184 |
| chr1:107626377-107626860 | 0.317916409 | 0.382875039 | 0.668199964 |
| chr15:98868580-98868845 | 0.33360123 | 0.382885105 | 0.668202312 |
| chr18:63585699-63586000 | -0.447660349 | 0.382896622 | 0.668207192 |
| chr1:147840953-147841476 | 0.311243778 | 0.382909271 | 0.668214047 |
| chr5:43066835-43067188 | 0.299399074 | 0.382933567 | 0.668241227 |
| chr8:124556516-124556803 | 0.323101419 | 0.382967934 | 0.668276633 |
| chr7:144410645-144410961 | -0.40104788 | 0.382988333 | 0.668276633 |
| chr8:15644155-15644392 | -0.436453292 | 0.382991253 | 0.668276633 |
| chr20:63272586-63272992 | 0.308846766 | 0.382997086 | 0.668276633 |
| chr3:37986858-37987314 | -0.313511503 | 0.383003132 | 0.668276633 |
| chr1:184929893-184930589 | -0.295366205 | 0.383006187 | 0.668276633 |
| chr2:106582125-106582411 | -0.334667972 | 0.383025015 | 0.668284776 |
| chr8:99168198-99168472 | 0.374545183 | 0.383028298 | 0.668284776 |
| chr3:5194729-5195100 | -0.335975658 | 0.383063179 | 0.668314477 |
| chr14:63308816-63309106 | -0.368478458 | 0.383069829 | 0.668314477 |
| chr1:114485439-114485661 | 0.320212789 | 0.383071487 | 0.668314477 |
| chr1:17412091-17412543 | 0.321455873 | 0.383109848 | 0.668366183 |
| chr1:235866575-235867670 | 0.28099207 | 0.383122762 | 0.668373494 |
| chr9:86353633-86354931 | 0.276658492 | 0.383141738 | 0.668389348 |
| chr1:206912575-206912781 | -0.329181632 | 0.383153184 | 0.668389348 |
| chr6:142853162-142853954 | -0.294538276 | 0.38316567 | 0.668389348 |
| chr4:88591820-88592081 | -0.354778903 | 0.383166742 | 0.668389348 |
| chr7:40570223-40570423 | -0.532572908 | 0.383211693 | 0.668452541 |
| chr17:1562728-1563003 | 0.370727442 | 0.383259597 | 0.668520884 |
| chr4:77052335-77052587 | -0.386125033 | 0.383280145 | 0.668541506 |
| chr1:54070111-54070398 | 0.363784236 | 0.383296777 | 0.668555297 |
| chr22:50600108-50600308 | 0.442225876 | 0.383353593 | 0.668617013 |
| chr7:130875886-130876131 | -0.460263247 | 0.38335567 | 0.668617013 |
| chr7:137846212-137846887 | -0.323177908 | 0.383358338 | 0.668617013 |
| chr8:69894876-69895162 | -0.445780141 | 0.383395471 | 0.668666556 |
| chr17:51435686-51436053 | -0.304083655 | 0.383435853 | 0.668704402 |
| chr18:41096593-41096844 | -0.436379571 | 0.383437929 | 0.668704402 |
| chr1:157311214-157311414 | 0.451355001 | 0.383443353 | 0.668704402 |
| chr2:97347896-97348187 | -0.431288865 | 0.383489566 | 0.668737399 |
| chr14:33165392-33165601 | 0.434073587 | 0.383499636 | 0.668737399 |
| chr14:68718335-68718854 | -0.320367326 | 0.383511409 | 0.668737399 |
| chr11:69442978-69443192 | -0.447080034 | 0.383518758 | 0.668737399 |
| chr1:219555269-219555623 | -0.349141729 | 0.38352329 | 0.668737399 |
| chr6:93419449-93420090 | -0.282653964 | 0.383538879 | 0.668737399 |
| chr1:240026776-240027209 | -0.375347121 | 0.383544216 | 0.668737399 |
| chr3:129535101-129535420 | -0.344170219 | 0.383546423 | 0.668737399 |
| chr7:65770433-65770999 | 0.329973521 | 0.383546995 | 0.668737399 |
| chr1:145445239-145445495 | -0.42419891 | 0.383549551 | 0.668737399 |
| chr9:92687025-92687249 | 0.395947525 | 0.383617847 | 0.668827654 |
| chr16:68377715-68377992 | 0.368598774 | 0.383618775 | 0.668827654 |

| | | | |
|---|---|---|---|
| chr11:65614151-65614447 | 0.310391668 | 0.383652347 | 0.668870966 |
| chr4:74228727-74229038 | -0.376553477 | 0.383666178 | 0.668879861 |
| chrX:140091199-140091768 | 0.338297361 | 0.383698741 | 0.66892141 |
| chr2:99335849-99336704 | 0.28218752 | 0.383718831 | 0.668941215 |
| chr10:69507720-69508151 | -0.318177094 | 0.383747342 | 0.668975698 |
| chr10:96765280-96765538 | 0.391853362 | 0.383757772 | 0.66897866 |
| chr6:70371579-70371793 | -0.435730869 | 0.38378422 | 0.669009544 |
| chr8:48501476-48502062 | -0.3014792 | 0.383794248 | 0.669011804 |
| chr2:97590832-97591146 | -0.364839809 | 0.383840328 | 0.66904691 |
| chr11:44456952-44457261 | -0.397675824 | 0.383855036 | 0.66904691 |
| chr13:53782454-53782795 | -0.354326883 | 0.383856066 | 0.66904691 |
| chr19:35066834-35067386 | 0.317816215 | 0.383867469 | 0.66904691 |
| chr2:224585264-224586148 | -0.278309231 | 0.383869519 | 0.66904691 |
| chr5:96893821-96894062 | -0.441056448 | 0.38387096 | 0.66904691 |
| chr8:117232085-117232360 | -0.419881808 | 0.38387551 | 0.66904691 |
| chr1:188883074-188883274 | 0.464381924 | 0.383888482 | 0.669054301 |
| chr3:8111181-8111575 | 0.309413816 | 0.383910406 | 0.669077291 |
| chr21:27759137-27759396 | -0.392987435 | 0.383935343 | 0.6690974 |
| chr3:134485026-134485228 | -0.369536776 | 0.383939409 | 0.6690974 |
| chr19:55376636-55376895 | -0.387329863 | 0.384013313 | 0.669210974 |
| chr9:13181660-13182017 | 0.350624193 | 0.384038273 | 0.669228302 |
| chr14:81927139-81927566 | 0.347806741 | 0.384040725 | 0.669228302 |
| chr1:199610907-199611278 | -0.362081059 | 0.38406787 | 0.669228895 |
| chr17:81192418-81192860 | -0.327236236 | 0.384076494 | 0.669228895 |
| chr1:5745815-5746170 | -0.357918991 | 0.384078689 | 0.669228895 |
| chr15:96336942-96337279 | 0.337608017 | 0.384081266 | 0.669228895 |
| chr1:17239432-17239687 | 0.370942845 | 0.384084736 | 0.669228895 |
| chr12:47758892-47759335 | 0.312948682 | 0.384097033 | 0.669235103 |
| chr3:12763348-12763612 | 0.328284002 | 0.384115875 | 0.669252713 |
| chr5:37248852-37249920 | -0.284414842 | 0.384168595 | 0.669329348 |
| chr15:90932499-90932731 | 0.322457304 | 0.384204377 | 0.669375715 |
| chr15:89546812-89547021 | 0.431424473 | 0.384212679 | 0.669375715 |
| chr6:153791123-153791389 | 0.391498296 | 0.384238976 | 0.669406308 |
| chr12:101830270-101831306 | -0.277276545 | 0.384263228 | 0.669408749 |
| chr12:110501328-110502372 | -0.288013695 | 0.384265616 | 0.669408749 |
| chrX:106966229-106967103 | -0.284355806 | 0.384266587 | 0.669408749 |
| chr17:68511542-68512954 | -0.271690767 | 0.384285958 | 0.669421784 |
| chr1:58903332-58904292 | 0.281384103 | 0.384293121 | 0.669421784 |
| chr2:206127747-206127977 | -0.416558115 | 0.38430772 | 0.669421784 |
| chrX:130575886-130576091 | -0.450384437 | 0.384309016 | 0.669421784 |
| chr6:51473005-51473532 | 0.298666868 | 0.384329303 | 0.669441903 |
| chrX:3512246-3512453 | -0.457167718 | 0.384349357 | 0.669449256 |
| chr2:219510214-219510433 | -0.403858992 | 0.384350998 | 0.669449256 |
| chr15:34101295-34102422 | 0.277941898 | 0.384382172 | 0.669478865 |
| chr11:94845610-94846148 | 0.317445885 | 0.384386589 | 0.669478865 |
| chr10:73998931-73999324 | -0.323551486 | 0.384394211 | 0.669478865 |
| chr5:95758883-95759310 | -0.338718065 | 0.384406267 | 0.669484646 |
| chr2:175414952-175415152 | -0.45438353 | 0.384437149 | 0.669523212 |
| chr4:41214361-41215141 | 0.281849447 | 0.38450465 | 0.6696205 |
| chr11:63681123-63682199 | 0.284130712 | 0.38451049 | 0.6696205 |

| | | | |
|---|---|---|---|
| chr3:141186822-141187022 | 0.433586776 | 0.384536028 | 0.669649754 |
| chr12:48865339-48866084 | 0.298089067 | 0.384589099 | 0.669726953 |
| chr6:26157379-26158502 | -0.278157387 | 0.384603848 | 0.669737415 |
| chr20:953932-954132 | 0.4556531 | 0.384640872 | 0.669749835 |
| chr13:101544512-101544911 | -0.332767976 | 0.384646518 | 0.669749835 |
| chr14:72100549-72100873 | -0.377086764 | 0.3846522 | 0.669749835 |
| chr1:203682662-203683003 | -0.337905973 | 0.384657317 | 0.669749835 |
| chr16:88264499-88264746 | 0.421379688 | 0.384671397 | 0.669749835 |
| chr3:168847378-168847578 | 0.430620974 | 0.384676155 | 0.669749835 |
| chr17:64282018-64282512 | 0.304331175 | 0.384681524 | 0.669749835 |
| chrX:141133066-141134062 | 0.305418793 | 0.384683 | 0.669749835 |
| chr6:37137443-37137815 | -0.436420196 | 0.384689648 | 0.669749835 |
| chr9:36555816-36556016 | -0.476917186 | 0.384701529 | 0.669750561 |
| chr16:67247379-67247768 | 0.293227844 | 0.384707547 | 0.669750561 |
| chr7:4042263-4042609 | 0.386807438 | 0.38474369 | 0.669796193 |
| chr10:27241770-27242410 | -0.285425433 | 0.384768056 | 0.669796193 |
| chr7:31129135-31129434 | -0.370102885 | 0.384769097 | 0.669796193 |
| chr8:108133650-108133874 | -0.419092694 | 0.384774566 | 0.669796193 |
| chr4:152679699-152680365 | 0.298580894 | 0.384779021 | 0.669796193 |
| chr5:73031045-73031338 | -0.387332279 | 0.384792159 | 0.669796193 |
| chr8:53588143-53588399 | -0.403493726 | 0.384813545 | 0.669796193 |
| chr12:121003805-121004017 | -0.454770499 | 0.384815495 | 0.669796193 |
| chr4:79241791-79242291 | 0.341535724 | 0.384816878 | 0.669796193 |
| chr1:219067079-219067352 | -0.35688534 | 0.384824354 | 0.669796193 |
| chr15:100266266-100266830 | 0.341461692 | 0.384829916 | 0.669796193 |
| chr6:36682889-36683100 | -0.385022732 | 0.384841305 | 0.669800801 |
| chr3:27568371-27568707 | -0.339068718 | 0.384859758 | 0.669803097 |
| chr17:82236363-82236663 | 0.315267205 | 0.384865729 | 0.669803097 |
| chr3:16060302-16060796 | 0.305054379 | 0.384868849 | 0.669803097 |
| chr14:23665436-23665756 | -0.367576405 | 0.384921074 | 0.669873791 |
| chr1:23019166-23019831 | 0.310761437 | 0.384929805 | 0.669873791 |
| chr1:234876680-234877079 | 0.373097528 | 0.384935697 | 0.669873791 |
| chr17:82031941-82032392 | -0.306136343 | 0.384956472 | 0.669894728 |
| chr10:16891823-16892074 | -0.445876143 | 0.384984622 | 0.669915517 |
| chr11:57598054-57598288 | -0.447626524 | 0.384992373 | 0.669915517 |
| chr14:90396543-90397277 | 0.281656826 | 0.384994647 | 0.669915517 |
| chr1:31964110-31964460 | 0.324271758 | 0.385022519 | 0.669945831 |
| chr6:79878302-79878931 | -0.327474177 | 0.385029556 | 0.669945831 |
| chr12:67943421-67943666 | -0.397891045 | 0.385051833 | 0.66996938 |
| chr4:174585776-174586181 | -0.443808596 | 0.385084204 | 0.670004491 |
| chr5:10502693-10502893 | 0.396181682 | 0.385090038 | 0.670004491 |
| chr17:67630061-67630261 | 0.379457257 | 0.385100671 | 0.670004491 |
| chr8:48821930-48822175 | 0.435364657 | 0.38510699 | 0.670004491 |
| chr14:106790499-106790800 | -0.376098923 | 0.385171148 | 0.670090585 |
| chr2:74715042-74715945 | -0.285881858 | 0.385173966 | 0.670090585 |
| chr6:106871988-106872423 | -0.31884272 | 0.385185257 | 0.670095014 |
| chr8:85463162-85464404 | 0.275419095 | 0.385196675 | 0.670095147 |
| chr20:56874991-56875616 | 0.294471807 | 0.385202825 | 0.670095147 |
| chr3:20127953-20128180 | 0.448703317 | 0.385230442 | 0.670127976 |
| chr14:90476903-90477137 | -0.469676684 | 0.385254135 | 0.670153975 |

| | | | |
|---|---|---|---|
| chr1:37859302-37860436 | 0.279139837 | 0.385268632 | 0.67016398 |
| chr14:102304720-102305741 | -0.281856044 | 0.385288314 | 0.670183 |
| chr11:78311730-78312104 | -0.34402244 | 0.385300381 | 0.670188777 |
| chr2:128291323-128291931 | 0.298065182 | 0.385317715 | 0.670203713 |
| chr1:154199059-154199291 | 0.36737302 | 0.385374478 | 0.670287227 |
| chr12:109995671-109996574 | -0.288751356 | 0.385389187 | 0.670297597 |
| chr7:30478055-30479083 | 0.290996334 | 0.385416623 | 0.670329971 |
| chr16:87383056-87383493 | 0.300638064 | 0.385425298 | 0.670329971 |
| chr2:190618051-190618344 | -0.391707416 | 0.385493517 | 0.670392697 |
| chr3:48151656-48151998 | 0.399087154 | 0.38549443 | 0.670392697 |
| chr6:28104575-28105191 | 0.304786342 | 0.385496276 | 0.670392697 |
| chr12:121611396-121611611 | 0.407191049 | 0.385503033 | 0.670392697 |
| chr3:128624943-128625143 | 0.429287587 | 0.385517197 | 0.670392697 |
| chr6:131519606-131519980 | -0.333463986 | 0.385520357 | 0.670392697 |
| chr14:24036155-24037119 | 0.28681814 | 0.3855291 | 0.670392697 |
| chr2:6865252-6866708 | -0.2818682 | 0.385535466 | 0.670392697 |
| chr19:2163622-2164201 | 0.318714751 | 0.385540108 | 0.670392697 |
| chr13:55517416-55517616 | -0.501356467 | 0.385561772 | 0.670415151 |
| chr10:77949185-77949499 | -0.371704067 | 0.385581098 | 0.670424149 |
| chr19:40716101-40716423 | -0.319222407 | 0.385584446 | 0.670424149 |
| chr11:10808613-10809642 | 0.282962039 | 0.385601234 | 0.670438125 |
| chrX:18886055-18886255 | -0.421057257 | 0.385616896 | 0.670450144 |
| chr8:143572672-143572928 | -0.351228187 | 0.385636639 | 0.670469255 |
| chr7:12731135-12731567 | 0.350771442 | 0.385677196 | 0.670524553 |
| chr18:57586030-57587014 | -0.289162852 | 0.385740633 | 0.670619627 |
| chr1:84505598-84506898 | -0.274438539 | 0.385755535 | 0.670630318 |
| chr17:50684827-50685093 | 0.334239851 | 0.385784014 | 0.670656314 |
| chr1:160951424-160951661 | 0.373496762 | 0.385805603 | 0.670656314 |
| chr19:2461906-2462449 | -0.33917467 | 0.385808649 | 0.670656314 |
| chr16:14070587-14071712 | 0.288277756 | 0.385812137 | 0.670656314 |
| chr12:106148977-106149396 | -0.367466759 | 0.385814253 | 0.670656314 |
| chr8:97775539-97776578 | 0.276654111 | 0.385846428 | 0.670697027 |
| chr4:68076255-68076471 | 0.436883911 | 0.385857699 | 0.670701405 |
| chr12:102983345-102983762 | -0.426664842 | 0.385901869 | 0.670762964 |
| chr11:17013056-17014564 | 0.280770263 | 0.385946353 | 0.670789012 |
| chr11:117391369-117391569 | 0.459665748 | 0.385946542 | 0.670789012 |
| chr3:193787957-193788422 | 0.328202339 | 0.385959245 | 0.670789012 |
| chr6:30344957-30345623 | 0.296858076 | 0.385968748 | 0.670789012 |
| chr1:148459216-148459538 | -0.416232322 | 0.38597047 | 0.670789012 |
| chr19:17469467-17470542 | 0.280622098 | 0.38598368 | 0.670789012 |
| chr11:67682027-67682367 | 0.368599224 | 0.385986205 | 0.670789012 |
| chr2:54722957-54723969 | -0.274952981 | 0.385986891 | 0.670789012 |
| chr4:75939823-75941297 | 0.277203867 | 0.385996776 | 0.670790977 |
| chr2:122051933-122052287 | -0.387353828 | 0.386027945 | 0.670814676 |
| chr7:105791079-105791294 | -0.420648343 | 0.386031147 | 0.670814676 |
| chr22:19854744-19855994 | 0.282556831 | 0.386044834 | 0.670814676 |
| chr16:4293944-4294316 | 0.324772325 | 0.386049744 | 0.670814676 |
| chr14:99272104-99272500 | -0.299334395 | 0.386054187 | 0.670814676 |
| chr10:77251293-77251621 | -0.381583247 | 0.386063691 | 0.670815976 |
| chr19:5031675-5031903 | 0.395487115 | 0.386100003 | 0.670863859 |

| | | | |
|---|---|---|---|
| chr8:99982289-99982768 | 0.319828877 | 0.386122381 | 0.67087782 |
| chr16:10877486-10877779 | 0.37864042 | 0.38612555 | 0.67087782 |
| chr19:7506189-7506958 | 0.285009138 | 0.386136326 | 0.670881331 |
| chr18:37504617-37504817 | -0.420643228 | 0.38618124 | 0.670944152 |
| chr10:125584927-125585338 | -0.335924795 | 0.386221768 | 0.670986612 |
| chr8:122858325-122858526 | 0.377383195 | 0.386227121 | 0.670986612 |
| chr10:71213375-71213586 | 0.369431093 | 0.386237465 | 0.670986612 |
| chr3:166977543-166978052 | -0.349878937 | 0.386240708 | 0.670986612 |
| chr19:10952767-10953168 | -0.329560024 | 0.386261039 | 0.671006718 |
| chr2:241541112-241541493 | 0.309307676 | 0.386308203 | 0.671048828 |
| chr14:88468341-88468620 | -0.386225298 | 0.386315356 | 0.671048828 |
| chrX:136880085-136881032 | 0.282158895 | 0.386325067 | 0.671048828 |
| chr13:99980464-99980758 | -0.345841564 | 0.386331193 | 0.671048828 |
| chr17:1253839-1254109 | 0.399737472 | 0.38633727 | 0.671048828 |
| chr7:8365401-8366058 | 0.298636507 | 0.386338256 | 0.671048828 |
| chrX:99722121-99722612 | 0.333949665 | 0.386349488 | 0.671048828 |
| chr21:43740771-43741138 | -0.337616885 | 0.386355342 | 0.671048828 |
| chr8:127900548-127901020 | -0.304751953 | 0.386365123 | 0.671050604 |
| chr4:151569064-151569270 | 0.460732076 | 0.386378586 | 0.671058776 |
| chr12:62466142-62467622 | 0.276063932 | 0.386393856 | 0.671061374 |
| chr11:130431304-130431553 | -0.396809126 | 0.386405862 | 0.671061374 |
| chr13:35953634-35953962 | -0.350589061 | 0.386406356 | 0.671061374 |
| chr6:133951920-133952120 | -0.367044865 | 0.386452366 | 0.671126067 |
| chr17:7403576-7404902 | -0.276695684 | 0.386502122 | 0.671184931 |
| chr8:96327956-96328352 | 0.302019497 | 0.386503781 | 0.671184931 |
| chr5:150458790-150459493 | -0.292310518 | 0.386546786 | 0.671244399 |
| chr14:36747651-36747927 | 0.359759231 | 0.38656349 | 0.671258193 |
| chr11:118435845-118436615 | 0.277932216 | 0.386578818 | 0.671260006 |
| chr12:119225593-119225793 | 0.450761079 | 0.386586689 | 0.671260006 |
| chr10:84264520-84265073 | -0.301240868 | 0.386605678 | 0.671260006 |
| chr20:45392814-45393066 | 0.376031723 | 0.386610099 | 0.671260006 |
| chr5:18465688-18465888 | -0.417419314 | 0.386622416 | 0.671260006 |
| chr10:120981619-120981985 | -0.358256917 | 0.386624234 | 0.671260006 |
| chr12:96015710-96016064 | -0.348440464 | 0.386625859 | 0.671260006 |
| chr5:157731211-157732123 | -0.276087861 | 0.386698693 | 0.671368578 |
| chr12:49284101-49284311 | -0.424653034 | 0.386705918 | 0.671368578 |
| chr1:201972624-201973018 | 0.332724219 | 0.3867397 | 0.671412016 |
| chr13:44446305-44446505 | 0.455186357 | 0.38675005 | 0.671414772 |
| chr2:30621917-30622737 | -0.314211613 | 0.386784137 | 0.671458735 |
| chr18:50559737-50560250 | -0.329702223 | 0.386803829 | 0.671477706 |
| chr3:39498510-39498935 | -0.330516846 | 0.38681509 | 0.671482042 |
| chr19:44513043-44513332 | -0.369250257 | 0.386829022 | 0.671484075 |
| chr2:113079188-113079700 | 0.302454221 | 0.386835339 | 0.671484075 |
| chr1:224615797-224616670 | -0.295463474 | 0.386842552 | 0.671484075 |
| chr2:200643642-200644414 | -0.334821289 | 0.386886605 | 0.671545328 |
| chr6:32952961-32953524 | -0.293632609 | 0.386912368 | 0.671564622 |
| chr6:52234151-52234542 | -0.357777587 | 0.38691525 | 0.671564622 |
| chr1:147120657-147121057 | 0.351572582 | 0.386944915 | 0.671590287 |
| chr5:151924621-151924832 | -0.41997937 | 0.386947566 | 0.671590287 |
| chr15:92631838-92632040 | -0.400771183 | 0.386958799 | 0.67159457 |

| | | | |
|---|---|---|---|
| chr12:93156326-93156535 | -0.43092693 | 0.386989668 | 0.671618496 |
| chr5:159571576-159572325 | -0.315828562 | 0.386990115 | 0.671618496 |
| chr3:94549005-94549247 | -0.411395954 | 0.387038606 | 0.671687438 |
| chr16:75233861-75234519 | 0.290992693 | 0.387060493 | 0.671710208 |
| chr5:137743065-137743351 | -0.381202771 | 0.387085683 | 0.671738708 |
| chr10:115689548-115689762 | -0.461906607 | 0.387108872 | 0.671763736 |
| chr6:128519071-128521100 | -0.272049209 | 0.387146265 | 0.671801198 |
| chr16:30729411-30729654 | -0.442151157 | 0.387150733 | 0.671801198 |
| chr13:19089060-19089629 | -0.317510271 | 0.387162623 | 0.671801198 |
| chr13:28718356-28719666 | 0.276252352 | 0.387165531 | 0.671801198 |
| chr11:85949011-85949295 | 0.391729531 | 0.387187572 | 0.671824229 |
| chr2:199861266-199861553 | 0.449575054 | 0.387203972 | 0.671837471 |
| chr16:2251389-2251927 | 0.29943381 | 0.38723966 | 0.671872379 |
| chr1:117113280-117113591 | -0.407401793 | 0.387244812 | 0.671872379 |
| chr20:23215655-23215949 | -0.430980035 | 0.387255889 | 0.671872379 |
| chr1:241974390-241974616 | 0.409280526 | 0.387263964 | 0.671872379 |
| chr1:40774265-40774500 | -0.40863006 | 0.387267934 | 0.671872379 |
| chr16:30011077-30011277 | 0.405648073 | 0.387311067 | 0.671931997 |
| chr10:122374910-122375375 | 0.316452457 | 0.387341195 | 0.671967513 |
| chr5:174356614-174356814 | -0.461150143 | 0.387349079 | 0.671967513 |
| chr8:94248057-94248283 | 0.411467227 | 0.387393678 | 0.672018717 |
| chr2:68709278-68709922 | -0.322750422 | 0.387399996 | 0.672018717 |
| chr5:163459562-163461044 | -0.270941791 | 0.387412699 | 0.672018717 |
| chr17:59914303-59914594 | 0.290011525 | 0.387422553 | 0.672018717 |
| chr1:51218636-51218836 | 0.437780724 | 0.387422757 | 0.672018717 |
| chr14:52729729-52730833 | 0.272727059 | 0.387431219 | 0.672018717 |
| chr3:12135035-12135408 | 0.317047298 | 0.387499056 | 0.672121169 |
| chr4:150109599-150109818 | -0.437741817 | 0.387513856 | 0.672131029 |
| chr3:62212080-62212382 | -0.302045892 | 0.38752994 | 0.672131029 |
| chr16:11293444-11293644 | 0.388595357 | 0.387531056 | 0.672131029 |
| chr2:141511077-141511277 | 0.470647357 | 0.387557342 | 0.672153877 |
| chr6:160519750-160520804 | -0.291477393 | 0.387561774 | 0.672153877 |
| chr14:23352565-23353042 | 0.307821039 | 0.387586202 | 0.672164004 |
| chr5:149719792-149720024 | -0.437508582 | 0.387590859 | 0.672164004 |
| chr6:132484215-132484445 | 0.387004824 | 0.387593931 | 0.672164004 |
| chr6:26311564-26311979 | -0.302586847 | 0.387606775 | 0.672171065 |
| chr1:46223477-46223792 | 0.361374508 | 0.387615865 | 0.672171616 |
| chr8:118349676-118349948 | -0.387289156 | 0.387643728 | 0.672191378 |
| chr3:170221889-170223535 | -0.270903412 | 0.387652974 | 0.672191378 |
| chr5:160039844-160040392 | 0.316069465 | 0.38765358 | 0.672191378 |
| chr5:133489722-133489955 | -0.407217345 | 0.387672985 | 0.672194653 |
| chr10:63520716-63522266 | -0.275472375 | 0.387676515 | 0.672194653 |
| chr4:151729994-151730425 | -0.363733057 | 0.387681787 | 0.672194653 |
| chrX:46903665-46904403 | 0.293208485 | 0.387691124 | 0.67219563 |
| chr17:73643216-73643704 | 0.316475839 | 0.387711375 | 0.67220214 |
| chr17:64220800-64221000 | 0.373070538 | 0.387712424 | 0.67220214 |
| chr1:151311475-151311893 | 0.29692444 | 0.387731661 | 0.672219248 |
| chr1:24070887-24071139 | 0.410690424 | 0.387745404 | 0.672219248 |
| chr2:64616163-64616422 | 0.40872371 | 0.387748611 | 0.672219248 |
| chr10:59847971-59848226 | 0.373040318 | 0.387763242 | 0.672229403 |

| | | | |
|---|---|---|---|
| chr2:1136128-1136350 | -0.425183452 | 0.387797925 | 0.672274318 |
| chr19:47647976-47648989 | 0.280892917 | 0.387816901 | 0.672288163 |
| chr1:17520558-17520800 | -0.395489266 | 0.387823459 | 0.672288163 |
| chr3:173395042-173396159 | -0.277480377 | 0.387837603 | 0.672297472 |
| chr12:13234084-13234351 | -0.392450988 | 0.387859599 | 0.672307046 |
| chr13:25632197-25632464 | -0.395602364 | 0.387860675 | 0.672307046 |
| chr1:26431981-26432682 | 0.301169539 | 0.387870898 | 0.672309556 |
| chr20:25584749-25585863 | 0.286225204 | 0.387892106 | 0.672327944 |
| chr3:137347221-137347448 | -0.437301713 | 0.387899056 | 0.672327944 |
| chr4:112199096-112199457 | 0.338369953 | 0.387907864 | 0.672328003 |
| chr14:38867637-38867876 | 0.390952656 | 0.387993313 | 0.672438834 |
| chr16:75647317-75648761 | -0.272191878 | 0.387999751 | 0.672438834 |
| chr1:205488132-205488516 | -0.301528875 | 0.388005532 | 0.672438834 |
| chr1:16566965-16567328 | 0.350380834 | 0.388006913 | 0.672438834 |
| chr18:50183637-50183938 | -0.34857151 | 0.3880224 | 0.672450463 |
| chr6:79600893-79601313 | 0.357172989 | 0.388036354 | 0.672459436 |
| chr2:127591472-127591787 | -0.349405281 | 0.388050229 | 0.672468273 |
| chr11:86669008-86669289 | -0.358450611 | 0.388086816 | 0.672516464 |
| chr11:67312595-67313006 | 0.298734377 | 0.388112374 | 0.672545544 |
| chr2:207408466-207408791 | 0.426947907 | 0.388159404 | 0.672611828 |
| chr14:68248494-68248758 | 0.401484926 | 0.388206396 | 0.672667655 |
| chr7:66099623-66100471 | -0.285089852 | 0.388209179 | 0.672667655 |
| chr1:19653630-19654102 | 0.33416074 | 0.388221342 | 0.672670281 |
| chr1:156356883-156357420 | 0.30343361 | 0.388228463 | 0.672670281 |
| chr10:124308703-124308903 | -0.470682398 | 0.388237355 | 0.672670281 |
| chr9:114047827-114048097 | 0.366033047 | 0.388254398 | 0.672670281 |
| chr8:126068818-126069411 | -0.306612929 | 0.38825459 | 0.672670281 |
| chr13:38363087-38363312 | -0.438654353 | 0.388265039 | 0.672673175 |
| chr3:47824738-47825364 | -0.307129686 | 0.388282541 | 0.672688287 |
| chr11:12044280-12044613 | 0.32944549 | 0.388335632 | 0.672765054 |
| chr13:32381511-32381849 | -0.355210798 | 0.388416601 | 0.672885617 |
| chr5:103450539-103450851 | 0.36219344 | 0.388422788 | 0.672885617 |
| chr1:201559613-201560101 | 0.288365135 | 0.388432226 | 0.672886754 |
| chr9:122453155-122453516 | -0.346269261 | 0.38847236 | 0.672927323 |
| chr8:26867219-26867498 | -0.378746638 | 0.38847321 | 0.672927323 |
| chr14:73752732-73753033 | 0.320314375 | 0.388485588 | 0.672933553 |
| chr8:58459460-58459877 | 0.362265606 | 0.388518149 | 0.67297474 |
| chr15:99426023-99426338 | 0.351741958 | 0.388539323 | 0.672987133 |
| chr2:192286201-192286410 | 0.439829491 | 0.38854287 | 0.672987133 |
| chr8:133552814-133553014 | 0.420647206 | 0.388552031 | 0.672987787 |
| chr3:97882940-97883219 | -0.401460244 | 0.388599706 | 0.673049645 |
| chr6:166106180-166106412 | 0.394710399 | 0.388605313 | 0.673049645 |
| chr17:74212891-74213669 | 0.287021737 | 0.388619681 | 0.673059316 |
| chr12:102563866-102564374 | -0.338870532 | 0.388666384 | 0.673124988 |
| chrX:45186963-45187277 | -0.35951966 | 0.388703082 | 0.673173327 |
| chr4:137070654-137070854 | -0.442185271 | 0.388716862 | 0.673174531 |
| chr4:121156818-121157506 | -0.323818308 | 0.388722524 | 0.673174531 |
| chr11:14891081-14891337 | -0.327909392 | 0.388744693 | 0.673174531 |
| chr2:8986661-8986987 | -0.34846271 | 0.38875366 | 0.673174531 |
| chr1:6414361-6414686 | 0.411718088 | 0.388762737 | 0.673174531 |

| | | | |
|---|---|---|---|
| chr15:96171026-96171481 | -0.342900984 | 0.388762861 | 0.673174531 |
| chr1:27725487-27726537 | 0.274985429 | 0.388771411 | 0.673174531 |
| chr17:59910991-59911244 | 0.31052478 | 0.388774062 | 0.673174531 |
| chr19:3948528-3948728 | -0.467851831 | 0.388793343 | 0.673192703 |
| chr7:76166229-76167082 | -0.285318408 | 0.38882013 | 0.673205706 |
| chr11:85787936-85788178 | -0.390539102 | 0.388828698 | 0.673205706 |
| chr17:1716118-1716910 | -0.279361811 | 0.388832553 | 0.673205706 |
| chr11:57781559-57781976 | 0.308511638 | 0.388835997 | 0.673205706 |
| chr19:796799-797527 | 0.282011479 | 0.388886573 | 0.673278057 |
| chr2:142989355-142989556 | -0.460353164 | 0.38890755 | 0.673299161 |
| chr13:34201431-34202020 | 0.321629942 | 0.388953032 | 0.673326723 |
| chr8:80895370-80895760 | 0.333310525 | 0.388954168 | 0.673326723 |
| chr3:171774460-171774743 | 0.386274249 | 0.388957286 | 0.673326723 |
| chr17:83180890-83181299 | 0.319228542 | 0.38896907 | 0.673326723 |
| chrX:107857764-107858330 | 0.326916325 | 0.388977454 | 0.673326723 |
| chr20:41694084-41694396 | 0.316487512 | 0.388982013 | 0.673326723 |
| chr3:147392050-147392950 | -0.275204511 | 0.38899353 | 0.673326723 |
| chr1:149646286-149646643 | -0.37457693 | 0.388993771 | 0.673326723 |
| chr3:161276142-161276395 | -0.359217267 | 0.38908186 | 0.673463986 |
| chr10:117004503-117006005 | -0.274250818 | 0.389096112 | 0.673473441 |
| chr13:94416483-94416806 | -0.350463097 | 0.389106552 | 0.673476298 |
| chr1:16155036-16155527 | 0.344445719 | 0.389160157 | 0.673541725 |
| chr1:107339804-107340085 | 0.313166781 | 0.389165487 | 0.673541725 |
| chr22:24894541-24894916 | 0.338838328 | 0.389170724 | 0.673541725 |
| chr2:191514845-191515045 | -0.478383273 | 0.389207515 | 0.673572046 |
| chr1:212749837-212750156 | -0.371344127 | 0.38921133 | 0.673572046 |
| chr20:58295135-58295564 | 0.311727526 | 0.389214617 | 0.673572046 |
| chr7:64733751-64733951 | -0.442068123 | 0.389234708 | 0.673582476 |
| chr4:154618732-154619078 | -0.342237072 | 0.389238225 | 0.673582476 |
| chr12:45762488-45762770 | -0.355995887 | 0.389248995 | 0.6735859 |
| chr16:228697-228897 | -0.428831288 | 0.38932108 | 0.673680659 |
| chr8:35488450-35488663 | -0.433995305 | 0.389323435 | 0.673680659 |
| chr13:43081833-43082110 | -0.458242146 | 0.389330131 | 0.673680659 |
| chr14:93474292-93474540 | -0.486039778 | 0.389380876 | 0.673753251 |
| chr19:14061722-14061940 | -0.452238548 | 0.389392358 | 0.673757904 |
| chr2:63615121-63615472 | 0.345675 | 0.389407395 | 0.673768708 |
| chr13:71879341-71879542 | -0.45982629 | 0.389446092 | 0.673820446 |
| chr2:201071305-201072212 | -0.279692536 | 0.389468758 | 0.673829259 |
| chr1:175152051-175152359 | -0.322448489 | 0.389468774 | 0.673829259 |
| chr1:16534987-16535376 | 0.346173673 | 0.38947833 | 0.673830578 |
| chr20:57055490-57055711 | 0.36962122 | 0.38949175 | 0.673838579 |
| chr14:41449054-41449506 | -0.316899898 | 0.389506676 | 0.673849188 |
| chr4:113959161-113959604 | 0.321836899 | 0.389515946 | 0.673850011 |
| chr15:100759738-100759954 | 0.382937514 | 0.389534523 | 0.67386474 |
| chr2:45541899-45542134 | -0.445708592 | 0.389558924 | 0.67386474 |
| chr4:89307506-89308409 | 0.295960148 | 0.389567469 | 0.67386474 |
| chr16:87417756-87417956 | -0.413237534 | 0.389570222 | 0.67386474 |
| chr1:225875273-225875904 | 0.305706703 | 0.389572644 | 0.67386474 |
| chr8:121150864-121151391 | -0.300331626 | 0.389577589 | 0.67386474 |
| chr16:84096595-84096987 | 0.303643311 | 0.389586023 | 0.67386474 |

| | | | |
|---|---|---|---|
| chrX:11239286-11239660 | 0.342670507 | 0.38962254 | 0.67390342 |
| chr14:90061053-90061283 | 0.373638449 | 0.389625976 | 0.67390342 |
| chr7:116418093-116418300 | -0.417447303 | 0.389640199 | 0.673912809 |
| chr5:178730375-178731337 | -0.280465638 | 0.389669481 | 0.673948241 |
| chr3:94004361-94004664 | -0.337786318 | 0.38970914 | 0.67400162 |
| chr8:74320339-74321884 | 0.274214715 | 0.389720353 | 0.674005799 |
| chr8:60516325-60517697 | 0.274068414 | 0.389745174 | 0.674026107 |
| chr20:62133883-62134155 | -0.373162644 | 0.389764425 | 0.674026107 |
| chr1:115507270-115507474 | 0.376865493 | 0.389770143 | 0.674026107 |
| chr12:3658031-3658231 | -0.494617355 | 0.389772985 | 0.674026107 |
| chr14:63677793-63678477 | 0.283889809 | 0.389781439 | 0.674026107 |
| chr12:58310819-58311151 | -0.375549796 | 0.389784876 | 0.674026107 |
| chr17:17440135-17440335 | 0.431157982 | 0.389799848 | 0.674036784 |
| chr2:169479197-169480063 | -0.282168329 | 0.389813096 | 0.674036856 |
| chr19:47152709-47152999 | 0.38819759 | 0.389827034 | 0.674036856 |
| chr17:1648767-1649945 | -0.282755592 | 0.389834317 | 0.674036856 |
| chr1:228165280-228166773 | 0.269594912 | 0.389836942 | 0.674036856 |
| chr17:61467387-61468073 | 0.271306537 | 0.389846977 | 0.674036856 |
| chr5:100706956-100707180 | -0.43358445 | 0.389854693 | 0.674036856 |
| chr6:157193777-157194642 | 0.296932759 | 0.389861468 | 0.674036856 |
| chr13:41263118-41264012 | -0.277035793 | 0.389877046 | 0.67404858 |
| chr14:68935763-68936116 | -0.35691573 | 0.38991842 | 0.674101124 |
| chr7:97871721-97872766 | -0.276115273 | 0.389925034 | 0.674101124 |
| chr19:38387426-38388554 | 0.274648375 | 0.389955887 | 0.674126558 |
| chr2:130157380-130157683 | -0.376688245 | 0.389957342 | 0.674126558 |
| chr7:88357384-88357628 | -0.439025634 | 0.389984192 | 0.674157765 |
| chr12:34133739-34134097 | -0.332700613 | 0.390008956 | 0.674181169 |
| chr18:41883166-41883478 | 0.37781919 | 0.390015329 | 0.674181169 |
| chr11:118545676-118546005 | -0.377681734 | 0.390032048 | 0.674194861 |
| chr3:169564935-169565424 | -0.309712642 | 0.390052277 | 0.674214618 |
| chr8:96134514-96134859 | 0.309816344 | 0.390077128 | 0.674242363 |
| chr4:139643622-139643895 | -0.47525456 | 0.390101372 | 0.674265415 |
| chr11:119094879-119095881 | 0.273314659 | 0.390108064 | 0.674265415 |
| chr21:45970735-45970935 | 0.406846156 | 0.390117709 | 0.674266874 |
| chr3:164524284-164524530 | -0.394444839 | 0.390149159 | 0.674303377 |
| chr11:368318-368591 | 0.41234422 | 0.390156429 | 0.674303377 |
| chr1:23979957-23980653 | 0.292074056 | 0.39017089 | 0.674313159 |
| chr11:129023995-129024376 | -0.320133974 | 0.390209654 | 0.674349949 |
| chr17:52106570-52106946 | -0.343141213 | 0.390209779 | 0.674349949 |
| chr20:9112485-9112687 | 0.489840787 | 0.390225091 | 0.674361202 |
| chr13:24053399-24053767 | -0.312021236 | 0.390236278 | 0.674365324 |
| chr16:84228432-84228646 | 0.373228299 | 0.39027178 | 0.674404703 |
| chrX:70433810-70434873 | 0.279366086 | 0.390285328 | 0.674404703 |
| chr3:177622511-177623179 | 0.331892811 | 0.390285471 | 0.674404703 |
| chr14:39169544-39170816 | 0.271401103 | 0.390301894 | 0.674416505 |
| chr1:2227597-2228072 | 0.296381204 | 0.390323597 | 0.674416505 |
| chr11:79667474-79667849 | 0.336137249 | 0.39032456 | 0.674416505 |
| chr8:142726415-142727887 | 0.272353207 | 0.390332242 | 0.674416505 |
| chr1:151934048-151934407 | 0.316274381 | 0.39033949 | 0.674416505 |
| chr17:27506691-27506946 | -0.368527392 | 0.390345346 | 0.674416505 |

| | | | |
|---|---|---|---|
| chr11:125886973-125888337 | -0.273566494 | 0.390360307 | 0.674416505 |
| chr5:33649919-33650123 | 0.460085289 | 0.390368297 | 0.674416505 |
| chr8:60233276-60233755 | -0.324912291 | 0.390371517 | 0.674416505 |
| chr3:129961130-129961512 | -0.357229578 | 0.390383094 | 0.674421299 |
| chr8:92698356-92698556 | 0.428415381 | 0.390417438 | 0.674461823 |
| chr18:79069618-79069830 | -0.380344859 | 0.390424156 | 0.674461823 |
| chr7:120792034-120792482 | -0.332436662 | 0.390440025 | 0.67447403 |
| chr4:140523752-140524750 | -0.273997463 | 0.390454589 | 0.674483983 |
| chr1:44721127-44722305 | 0.282573069 | 0.390499649 | 0.674546613 |
| chr7:46492493-46492853 | 0.360627765 | 0.3905207 | 0.674567769 |
| chr3:63878644-63879090 | 0.293229772 | 0.390540657 | 0.674587034 |
| chr17:42024777-42025268 | -0.32177756 | 0.390549561 | 0.674587205 |
| chr1:198848275-198848518 | 0.412772391 | 0.390629533 | 0.67471013 |
| chr1:201127278-201127509 | 0.326743719 | 0.390640202 | 0.674713348 |
| chr17:52828230-52828509 | -0.354392765 | 0.390666471 | 0.674731591 |
| chr11:7512975-7514249 | 0.277905309 | 0.390668376 | 0.674731591 |
| chr2:20617889-20618365 | 0.290354295 | 0.390681808 | 0.674736449 |
| chr11:75561947-75562921 | 0.278585533 | 0.390688801 | 0.674736449 |
| chr5:162248191-162248583 | -0.344982531 | 0.390697792 | 0.674736768 |
| chr1:25997655-25998704 | 0.281898754 | 0.390741561 | 0.674771492 |
| chr22:11341448-11341671 | -0.460842637 | 0.390746778 | 0.674771492 |
| chr8:81235306-81235622 | 0.352208138 | 0.390752112 | 0.674771492 |
| chr4:123399414-123399762 | -0.334672436 | 0.390758563 | 0.674771492 |
| chr12:119061221-119061488 | -0.434672237 | 0.390768385 | 0.674771492 |
| chr5:26807288-26807770 | 0.353812126 | 0.390776125 | 0.674771492 |
| chrX:25007166-25007456 | -0.403371777 | 0.390779544 | 0.674771492 |
| chr13:99045934-99046794 | -0.291820471 | 0.390791772 | 0.6747774 |
| chr12:43915010-43915269 | -0.393127149 | 0.390855868 | 0.674858251 |
| chr20:57160562-57160782 | 0.2868327 | 0.390866656 | 0.674858251 |
| chr7:133434285-133434485 | -0.470813032 | 0.390876239 | 0.674858251 |
| chr12:67849573-67850248 | -0.324075076 | 0.390893367 | 0.674858251 |
| chr10:93090868-93091143 | 0.4338804 | 0.390900988 | 0.674858251 |
| chr16:80527366-80527697 | -0.344435948 | 0.390911767 | 0.674858251 |
| chr19:45406299-45406785 | 0.299819424 | 0.390926643 | 0.674858251 |
| chr7:1144218-1144686 | -0.321510844 | 0.390936206 | 0.674858251 |
| chr7:43582464-43583865 | -0.270998254 | 0.390937139 | 0.674858251 |
| chr12:53321145-53322053 | 0.278360185 | 0.390943523 | 0.674858251 |
| chr5:122478830-122479484 | -0.291627988 | 0.390943533 | 0.674858251 |
| chr2:120313293-120313518 | 0.379316671 | 0.390949311 | 0.674858251 |
| chr2:160077968-160078256 | -0.465967566 | 0.390953095 | 0.674858251 |
| chr1:187491503-187491802 | 0.385782642 | 0.390974521 | 0.674880033 |
| chr10:80100895-80101273 | -0.307028205 | 0.390992884 | 0.674896526 |
| chr7:152025249-152026076 | -0.286149828 | 0.391006224 | 0.674904347 |
| chr12:57087077-57088373 | -0.271322293 | 0.391020261 | 0.674913372 |
| chr8:97277113-97277331 | 0.354217211 | 0.391049894 | 0.674939628 |
| chr16:76618927-76619127 | 0.450889978 | 0.39105309 | 0.674939628 |
| chr2:234881677-234881989 | -0.375527711 | 0.391081414 | 0.67497331 |
| chr7:3089546-3089786 | -0.397357327 | 0.391120015 | 0.675024728 |
| chr3:38495992-38496888 | 0.281924204 | 0.391145353 | 0.675033721 |
| chr5:134039506-134039879 | -0.322697031 | 0.391149402 | 0.675033721 |

| | | | |
|---|---|---|---|
| chr15:71115979-71116385 | 0.319571207 | 0.391151656 | 0.675033721 |
| chr6:7108242-7108809 | 0.303813111 | 0.391177873 | 0.675055024 |
| chr4:153232735-153233040 | -0.366512125 | 0.39118162 | 0.675055024 |
| chr8:143246360-143247721 | 0.270858047 | 0.391218383 | 0.675103259 |
| chr3:159732208-159732408 | 0.432691547 | 0.391278671 | 0.675192089 |
| chr3:48739910-48740198 | 0.39087898 | 0.391297613 | 0.675209569 |
| chr1:109282801-109283594 | -0.277126742 | 0.391387777 | 0.67530959 |
| chr15:93333995-93334242 | 0.395423444 | 0.391393883 | 0.67530959 |
| chr2:219278945-219279640 | 0.281599515 | 0.391395381 | 0.67530959 |
| chr2:27211381-27212724 | -0.272837992 | 0.391398335 | 0.67530959 |
| chr17:66201676-66202080 | -0.326434864 | 0.391404952 | 0.67530959 |
| chrX:109644549-109644907 | -0.366439076 | 0.391428741 | 0.67530959 |
| chr11:122559328-122560273 | -0.279401304 | 0.391429637 | 0.67530959 |
| chr5:82351900-82352113 | -0.391359393 | 0.39143491 | 0.67530959 |
| chr12:14365288-14366131 | -0.27533361 | 0.391449674 | 0.67530959 |
| chr2:52386457-52386680 | -0.439658021 | 0.391452274 | 0.67530959 |
| chr1:168365766-168366270 | 0.333578426 | 0.391452526 | 0.67530959 |
| chr5:81667042-81667292 | -0.411650866 | 0.391507254 | 0.675388797 |
| chr20:63042901-63043101 | -0.439777144 | 0.391553079 | 0.675448598 |
| chr15:91572151-91572554 | 0.329816091 | 0.39155955 | 0.675448598 |
| chr7:100157840-100158040 | -0.389610388 | 0.391570167 | 0.675451706 |
| chr16:55508851-55509804 | 0.29457236 | 0.39162106 | 0.675523891 |
| chr1:205406165-205406430 | -0.382833328 | 0.391630055 | 0.675523891 |
| chr20:44552801-44553079 | -0.408721378 | 0.391640628 | 0.675523891 |
| chr20:41525057-41525541 | -0.322323222 | 0.391647279 | 0.675523891 |
| chr4:78057258-78057476 | 0.355098633 | 0.39169268 | 0.675555053 |
| chr7:143380188-143381833 | -0.27069409 | 0.391699943 | 0.675555053 |
| chr14:22496577-22496777 | -0.448264967 | 0.391707351 | 0.675555053 |
| chr15:79896201-79897572 | 0.276273444 | 0.391707537 | 0.675555053 |
| chr3:14301663-14301884 | -0.437656234 | 0.391712962 | 0.675555053 |
| chr20:9359518-9359955 | -0.348034733 | 0.391718246 | 0.675555053 |
| chr10:71212421-71212743 | 0.332070418 | 0.391755868 | 0.675604729 |
| chr5:112160501-112161342 | -0.277213416 | 0.391781778 | 0.675630223 |
| chr2:26785527-26786231 | -0.27957854 | 0.391788286 | 0.675630223 |
| chr3:72746828-72747066 | 0.396950053 | 0.39181609 | 0.675662963 |
| chr8:108206613-108207175 | 0.307224218 | 0.391835268 | 0.675680828 |
| chr3:20185388-20185769 | 0.297406139 | 0.391844331 | 0.67568125 |
| chr6:136334961-136335161 | 0.439302887 | 0.391868947 | 0.67570849 |
| chr1:177599121-177599345 | -0.426615104 | 0.391881983 | 0.675715761 |
| chr17:45385047-45385269 | 0.401938853 | 0.391900327 | 0.675732186 |
| chr5:60843159-60844665 | -0.275493673 | 0.391916624 | 0.675745079 |
| chr9:91005069-91005279 | 0.470353742 | 0.391931546 | 0.675755601 |
| chr10:74206747-74207090 | 0.331117782 | 0.391998795 | 0.675856342 |
| chr11:64729852-64730091 | 0.394922287 | 0.392032463 | 0.675870642 |
| chr16:86782135-86782497 | -0.388402692 | 0.392032689 | 0.675870642 |
| chr13:49953684-49953947 | 0.377420666 | 0.392035827 | 0.675870642 |
| chr7:135274523-135274820 | -0.350951333 | 0.392042373 | 0.675870642 |
| chr11:94776389-94776615 | 0.410805873 | 0.392056442 | 0.675879691 |
| chr3:119660276-119660503 | -0.462670038 | 0.392098426 | 0.675935668 |
| chr2:20184164-20184872 | -0.294767079 | 0.392108989 | 0.675935668 |

| | | | |
|---|---|---|---|
| chr6:99424719-99426163 | -0.269757057 | 0.392115378 | 0.675935668 |
| chr22:37199148-37199742 | 0.292776479 | 0.392147446 | 0.675965059 |
| chr19:10471810-10472158 | -0.405563241 | 0.392160929 | 0.675965059 |
| chr17:76141172-76141568 | -0.30696451 | 0.392162311 | 0.675965059 |
| chr5:77144632-77144837 | -0.443275678 | 0.392167717 | 0.675965059 |
| chr12:96658663-96658863 | -0.463322052 | 0.39220697 | 0.676017511 |
| chr19:13152719-13153161 | 0.301443166 | 0.392217991 | 0.6760213 |
| chr12:131474337-131474678 | 0.341815876 | 0.392237215 | 0.676022483 |
| chr8:28363402-28363662 | 0.394777009 | 0.392242926 | 0.676022483 |
| chr7:36785172-36785444 | 0.386087492 | 0.392245146 | 0.676022483 |
| chr19:28242409-28242624 | -0.454584698 | 0.39225674 | 0.67602726 |
| chr3:151135588-151135788 | -0.437307299 | 0.392274129 | 0.676042022 |
| chr2:42360854-42361923 | -0.271725513 | 0.392297375 | 0.676047402 |
| chr11:62855645-62856383 | 0.279576373 | 0.392306018 | 0.676047402 |
| chr19:19405396-19405821 | -0.325811941 | 0.392312302 | 0.676047402 |
| chr6:52918125-52918742 | 0.301957595 | 0.392325805 | 0.676047402 |
| chr9:14331263-14331678 | -0.370276244 | 0.392328502 | 0.676047402 |
| chr6:170260768-170261014 | 0.366535597 | 0.39233019 | 0.676047402 |
| chr11:69344135-69344357 | 0.451057355 | 0.392339831 | 0.676048812 |
| chr17:62064777-62065242 | 0.288236074 | 0.392385525 | 0.676112344 |
| chr9:136137260-136137480 | 0.430757064 | 0.392403375 | 0.676127896 |
| chr8:116471839-116472067 | 0.348251042 | 0.392415841 | 0.676134035 |
| chr4:127632464-127633434 | -0.275891807 | 0.392429135 | 0.676134035 |
| chr2:20051275-20052276 | -0.27066963 | 0.392433411 | 0.676134035 |
| chr5:141743467-141743670 | -0.415157707 | 0.392451718 | 0.676149213 |
| chr7:42401003-42401320 | -0.417786043 | 0.392467146 | 0.676149213 |
| chr5:141877625-141877827 | -0.353072033 | 0.392474458 | 0.676149213 |
| chr9:19202294-19202494 | -0.469275234 | 0.392488761 | 0.676149213 |
| chr10:84548965-84549234 | -0.339865278 | 0.392492582 | 0.676149213 |
| chr2:172084348-172085987 | 0.272349647 | 0.392495167 | 0.676149213 |
| chr1:26826128-26826329 | -0.435110042 | 0.392509872 | 0.676159343 |
| chr7:99929076-99929904 | -0.302174045 | 0.392583337 | 0.676260828 |
| chr14:93857863-93858140 | 0.366822763 | 0.392586436 | 0.676260828 |
| chr11:121446793-121446993 | 0.439425151 | 0.392597552 | 0.676264773 |
| chr4:102620625-102620951 | 0.320935832 | 0.392634169 | 0.676290917 |
| chr1:46622521-46622806 | -0.369304695 | 0.392634699 | 0.676290917 |
| chr1:246938822-246939154 | 0.359203023 | 0.392645628 | 0.676290917 |
| chr1:117540914-117541363 | -0.310747263 | 0.392648035 | 0.676290917 |
| chr20:49964945-49965267 | 0.315706365 | 0.392667484 | 0.676302043 |
| chr10:55287479-55287685 | -0.449964052 | 0.392676275 | 0.676302043 |
| chr12:52969962-52970569 | -0.291178563 | 0.392680974 | 0.676302043 |
| chr17:59830074-59830363 | 0.294112512 | 0.392714316 | 0.676344264 |
| chr4:145179577-145180075 | -0.288788815 | 0.392738965 | 0.676362095 |
| chr6:158718564-158718957 | 0.350717048 | 0.3927454 | 0.676362095 |
| chrX:130902789-130903630 | 0.27352051 | 0.392764859 | 0.676362095 |
| chr11:119188417-119188812 | -0.360005521 | 0.392765299 | 0.676362095 |
| chr2:58762175-58762571 | -0.33829026 | 0.392768806 | 0.676362095 |
| chr12:47705536-47706276 | 0.284198222 | 0.392822489 | 0.676439337 |
| chr11:117809284-117809808 | 0.301972929 | 0.392832158 | 0.676440785 |
| chrX:40266818-40267541 | 0.316032244 | 0.39289101 | 0.676511811 |

| | | | |
|---|---|---|---|
| chr4:151675211-151675475 | 0.412425395 | 0.392893321 | 0.676511811 |
| chr2:221635994-221636305 | -0.381507731 | 0.392899893 | 0.676511811 |
| chr14:91427170-91427734 | -0.294614866 | 0.392921717 | 0.676534185 |
| chr1:156225365-156225750 | 0.323169029 | 0.392964825 | 0.676586586 |
| chr6:99736329-99736648 | -0.36851268 | 0.392971597 | 0.676586586 |
| chr12:96047990-96048317 | 0.376408464 | 0.392987425 | 0.676586586 |
| chr3:182714071-182714425 | -0.334160322 | 0.392996721 | 0.676586586 |
| chr10:111360223-111360709 | -0.310712108 | 0.393004897 | 0.676586586 |
| chr5:12292706-12292911 | 0.450455958 | 0.393005131 | 0.676586586 |
| chr13:46120042-46120460 | 0.316705671 | 0.393039946 | 0.676599879 |
| chr13:110412147-110412516 | 0.334669135 | 0.393046925 | 0.676599879 |
| chr6:131627824-131628559 | -0.275062335 | 0.393054847 | 0.676599879 |
| chr1:7783746-7784472 | 0.281723495 | 0.393068733 | 0.676599879 |
| chr13:30673710-30674522 | 0.2860165 | 0.39308722 | 0.676599879 |
| chr19:43784989-43785619 | 0.307504076 | 0.393088775 | 0.676599879 |
| chr15:101088789-101089159 | -0.315217993 | 0.393097168 | 0.676599879 |
| chr7:993093-993564 | 0.289606136 | 0.393098804 | 0.676599879 |
| chr6:31399675-31400136 | 0.324430508 | 0.393109831 | 0.676599879 |
| chr18:23134009-23134643 | -0.292499549 | 0.393115677 | 0.676599879 |
| chr7:73578108-73579310 | -0.274538292 | 0.393128498 | 0.676599879 |
| chr5:172939151-172939361 | 0.366760572 | 0.39316449 | 0.676599879 |
| chr6:75849135-75850040 | 0.294967423 | 0.393177556 | 0.676599879 |
| chr1:152182507-152182721 | -0.383127871 | 0.393180318 | 0.676599879 |
| chr12:83823458-83824039 | -0.299062611 | 0.393191638 | 0.676599879 |
| chr1:155061557-155061990 | 0.296816752 | 0.393199425 | 0.676599879 |
| chr8:141174820-141175207 | -0.313971101 | 0.393203691 | 0.676599879 |
| chr5:139011281-139011558 | -0.410102172 | 0.393205267 | 0.676599879 |
| chr17:82037234-82037941 | 0.283271272 | 0.393208491 | 0.676599879 |
| chr13:91134285-91134564 | -0.37994425 | 0.393210338 | 0.676599879 |
| chr5:36060215-36060415 | 0.427826139 | 0.393212152 | 0.676599879 |
| chr19:37779181-37779997 | 0.297692575 | 0.393213016 | 0.676599879 |
| chr9:127199344-127199552 | 0.372927795 | 0.393215951 | 0.676599879 |
| chr12:76879569-76879785 | -0.33860665 | 0.39323348 | 0.676614846 |
| chr5:64635151-64635385 | 0.398579935 | 0.393266305 | 0.676618522 |
| chr5:138633023-138633425 | -0.32926043 | 0.39326777 | 0.676618522 |
| chr8:138928455-138928754 | -0.421838689 | 0.393269941 | 0.676618522 |
| chr1:154154877-154155077 | 0.434199065 | 0.393270939 | 0.676618522 |
| chr5:14144150-14145480 | 0.272385374 | 0.393298553 | 0.676624284 |
| chr17:32937026-32937236 | 0.435425141 | 0.39330078 | 0.676624284 |
| chr2:177334994-177335194 | 0.435425141 | 0.39330078 | 0.676624284 |
| chr7:151860064-151860311 | -0.417070859 | 0.393319134 | 0.676640668 |
| chr14:105570584-105570977 | -0.351509176 | 0.39333644 | 0.676655248 |
| chr6:166005602-166005888 | 0.39408614 | 0.393352507 | 0.676667695 |
| chr1:172788246-172788475 | -0.444966032 | 0.393383092 | 0.676694681 |
| chr7:157355958-157356199 | -0.408582911 | 0.393398163 | 0.676694681 |
| chr6:56727169-56727812 | -0.302957712 | 0.393402413 | 0.676694681 |
| chr3:99817095-99818337 | -0.271056102 | 0.393403521 | 0.676694681 |
| chr17:65122923-65123637 | -0.285491317 | 0.393421762 | 0.676710458 |
| chr5:172904649-172905342 | -0.282491843 | 0.393441563 | 0.676710458 |
| chr12:117907333-117907630 | -0.378855594 | 0.393453111 | 0.676710458 |

| | | | |
|---|---|---|---|
| chr12:105235657-105236447 | -0.28078981 | 0.393456009 | 0.676710458 |
| chr16:84367961-84368701 | -0.277410552 | 0.393456852 | 0.676710458 |
| chr7:76314274-76314665 | -0.293988563 | 0.393488945 | 0.676750465 |
| chr1:101996746-101997218 | -0.345418736 | 0.393504981 | 0.676758065 |
| chr17:80147048-80148066 | 0.276355699 | 0.393511029 | 0.676758065 |
| chr12:54214628-54215376 | 0.289214945 | 0.393523012 | 0.676763483 |
| chr2:177242496-177242973 | -0.318526973 | 0.393571914 | 0.676832392 |
| chr9:121720511-121720711 | -0.428236876 | 0.39361788 | 0.676896247 |
| chr7:35216126-35216447 | -0.357941216 | 0.39364977 | 0.676911259 |
| chr4:112145065-112146299 | -0.272648997 | 0.393656263 | 0.676911259 |
| chr3:23866137-23866337 | 0.420589545 | 0.393661947 | 0.676911259 |
| chr4:102318557-102318757 | 0.420589545 | 0.393661947 | 0.676911259 |
| chr1:37448638-37449115 | -0.309229519 | 0.393688244 | 0.676940082 |
| chr2:30888990-30889260 | -0.417029274 | 0.393699612 | 0.676940082 |
| chr22:19178660-19179234 | 0.284051359 | 0.393705214 | 0.676940082 |
| chr19:29760109-29760451 | 0.322307759 | 0.393726228 | 0.67694202 |
| chr10:114103386-114103586 | -0.437978596 | 0.393728086 | 0.67694202 |
| chr1:67850744-67850971 | 0.413281354 | 0.393743191 | 0.67694202 |
| chr7:135714446-135715104 | -0.300766068 | 0.393752044 | 0.67694202 |
| chr2:205183574-205184033 | 0.319686128 | 0.393759456 | 0.67694202 |
| chr14:67599608-67600637 | 0.275871245 | 0.393769811 | 0.67694202 |
| chr7:156970223-156970502 | -0.378788172 | 0.393770757 | 0.67694202 |
| chr10:100705143-100705344 | -0.474822809 | 0.393784871 | 0.67694202 |
| chr8:140159655-140159942 | 0.345974688 | 0.393794152 | 0.67694202 |
| chr3:187699931-187700134 | 0.433863659 | 0.393802436 | 0.67694202 |
| chr6:16331382-16331617 | 0.403980912 | 0.393803524 | 0.67694202 |
| chr15:75119908-75120202 | -0.424690943 | 0.393819076 | 0.676946373 |
| chr7:100272152-100272504 | 0.337516264 | 0.393823726 | 0.676946373 |
| chr3:34196867-34197260 | 0.326055496 | 0.393838442 | 0.676956482 |
| chr11:125433494-125433800 | -0.35108594 | 0.39387672 | 0.67700709 |
| chr19:13151578-13152036 | 0.296316564 | 0.393898731 | 0.677029736 |
| chr14:88792076-88793509 | 0.274945782 | 0.393929218 | 0.677065365 |
| chr1:224203643-224204095 | 0.308332591 | 0.393938681 | 0.677065365 |
| chr3:137792708-137792991 | -0.373722336 | 0.39394597 | 0.677065365 |
| chr3:13134573-13134935 | 0.363482376 | 0.393971397 | 0.677071283 |
| chr2:46543215-46544494 | -0.274122276 | 0.393971485 | 0.677071283 |
| chr7:100813862-100814337 | -0.347685763 | 0.393975923 | 0.677071283 |
| chr10:19817317-19817829 | -0.305100368 | 0.393992383 | 0.677084385 |
| chr8:144094489-144095181 | 0.280460637 | 0.394003008 | 0.677087458 |
| chr1:232868781-232869441 | 0.303619263 | 0.394026858 | 0.677113257 |
| chr5:127289999-127290217 | 0.389016723 | 0.394077413 | 0.677169759 |
| chr6:31307570-31307794 | 0.389016723 | 0.394077413 | 0.677169759 |
| chr20:44217375-44217575 | 0.396607247 | 0.394086905 | 0.677170448 |
| chr6:3473088-3473375 | -0.415119241 | 0.39409549 | 0.677170448 |
| chr22:50342740-50343415 | 0.308033551 | 0.394113122 | 0.677185559 |
| chr6:36387409-36387643 | -0.431685282 | 0.394185502 | 0.677284458 |
| chr22:21665796-21666535 | 0.288494723 | 0.394188358 | 0.677284458 |
| chr8:97142139-97142652 | -0.33981442 | 0.394248807 | 0.677373129 |
| chr2:68062681-68063435 | -0.285369404 | 0.394273936 | 0.677401115 |
| chr6:111777444-111777667 | -0.416545572 | 0.394363096 | 0.677508868 |

| | | | |
|---|---|---|---|
| chr1:40944365-40944663 | -0.369497711 | 0.39436554 | 0.677508868 |
| chr4:143254562-143254775 | -0.462255656 | 0.394367975 | 0.677508868 |
| chr5:17145921-17146135 | 0.406982101 | 0.394372021 | 0.677508868 |
| chr14:89035055-89035321 | -0.419879369 | 0.394389106 | 0.677523028 |
| chr20:56775509-56775844 | 0.346282364 | 0.394401765 | 0.677529584 |
| chr1:243850172-243850920 | 0.299831029 | 0.394418594 | 0.677537969 |
| chr10:110499420-110499757 | 0.323142503 | 0.394432841 | 0.677537969 |
| chr13:76980122-76980480 | -0.339231771 | 0.394433174 | 0.677537969 |
| chr8:3596463-3596805 | -0.419255153 | 0.394467998 | 0.677577887 |
| chr11:65150713-65151375 | -0.305105577 | 0.394474099 | 0.677577887 |
| chr19:40392739-40393217 | 0.308364161 | 0.394504736 | 0.677601351 |
| chr17:78378441-78379168 | 0.297241669 | 0.394505446 | 0.677601351 |
| chr11:10451072-10451467 | 0.333620433 | 0.394532184 | 0.677621331 |
| chr16:68744753-68745219 | 0.297934319 | 0.394534766 | 0.677621331 |
| chr2:209936388-209936645 | -0.437048259 | 0.39454478 | 0.677623342 |
| chr3:148962546-148962907 | -0.319520621 | 0.39455371 | 0.67762349 |
| chr14:103539128-103539651 | 0.301495598 | 0.394573939 | 0.677643042 |
| chr8:66774584-66775758 | -0.270994142 | 0.39459594 | 0.677665638 |
| chr6:37463576-37463917 | -0.392177825 | 0.394650907 | 0.677744073 |
| chr2:76953773-76954103 | -0.351428745 | 0.394659302 | 0.677744073 |
| chr7:23090530-23090752 | 0.413354809 | 0.394677319 | 0.677759822 |
| chr22:19767120-19767331 | 0.374096474 | 0.394712564 | 0.677805157 |
| chr15:83436480-83436854 | 0.344685841 | 0.394779761 | 0.677833939 |
| chr12:89524472-89525030 | -0.280735003 | 0.394783463 | 0.677833939 |
| chr5:116576079-116576461 | -0.293810959 | 0.394783724 | 0.677833939 |
| chr9:130212536-130212780 | -0.407394709 | 0.394784089 | 0.677833939 |
| chr12:100794475-100795052 | -0.322596505 | 0.394786964 | 0.677833939 |
| chr15:65610439-65611707 | 0.273495927 | 0.394793265 | 0.677833939 |
| chr13:108740630-108740928 | -0.407380328 | 0.394803353 | 0.677833939 |
| chr15:62059778-62060956 | -0.272562665 | 0.394808567 | 0.677833939 |
| chr5:154835242-154835619 | -0.33052466 | 0.394808943 | 0.677833939 |
| chr14:100076255-100076546 | 0.344744393 | 0.394823616 | 0.677834828 |
| chr7:32207043-32207308 | -0.337591533 | 0.394834413 | 0.677834828 |
| chr3:22345554-22345755 | -0.503803158 | 0.394844847 | 0.677834828 |
| chr4:82881744-82881944 | -0.503803158 | 0.394844847 | 0.677834828 |
| chr1:185585589-185586316 | 0.309679545 | 0.394867128 | 0.677855705 |
| chr2:15856373-15856593 | -0.418833139 | 0.394874702 | 0.677855705 |
| chr5:86970968-86971434 | -0.343288729 | 0.394896128 | 0.6778773 |
| chr2:10497865-10498233 | -0.320637913 | 0.394909265 | 0.677884662 |
| chr16:2174277-2174569 | 0.395732386 | 0.394951184 | 0.677928232 |
| chr22:41517557-41517964 | 0.332911728 | 0.394952342 | 0.677928232 |
| chr2:96022654-96023084 | -0.312375647 | 0.394975404 | 0.677948915 |
| chr3:186616971-186617374 | -0.309363024 | 0.394982088 | 0.677948915 |
| chr10:110617445-110617645 | 0.435152381 | 0.394993349 | 0.677953058 |
| chr3:162586594-162586798 | 0.41879121 | 0.395019027 | 0.677981942 |
| chr5:64811096-64811348 | 0.421258799 | 0.395033632 | 0.677989625 |
| chr20:59042053-59043335 | 0.267963646 | 0.3950412 | 0.677989625 |
| chr3:171614591-171614867 | -0.346064911 | 0.395058339 | 0.678000356 |
| chr3:851163-851389 | -0.426976667 | 0.39506515 | 0.678000356 |
| chr18:66124503-66124703 | 0.508140331 | 0.395090524 | 0.678028716 |

| | | | |
|---|---|---|---|
| chr18:12059838-12060124 | 0.351799328 | 0.395139955 | 0.67809836 |
| chr15:63503193-63503555 | 0.301658497 | 0.395160405 | 0.678118265 |
| chr19:55986544-55986879 | -0.393750054 | 0.395180452 | 0.678137479 |
| chr5:93677409-93677619 | -0.430743589 | 0.395223846 | 0.678194137 |
| chr5:142713140-142713729 | -0.287795767 | 0.395232228 | 0.678194137 |
| chr7:72829184-72829619 | -0.290283314 | 0.395240757 | 0.678194137 |
| chrX:49163578-49164422 | 0.280394244 | 0.395264153 | 0.678194137 |
| chr10:97800491-97800691 | -0.449271468 | 0.395265734 | 0.678194137 |
| chr6:42901486-42901850 | -0.332607026 | 0.395266576 | 0.678194137 |
| chr7:65981702-65982493 | -0.288231729 | 0.395286359 | 0.678212894 |
| chr3:39107062-39108013 | 0.277920337 | 0.395334613 | 0.678280497 |
| chr13:45879535-45880005 | -0.330145922 | 0.395346447 | 0.678285613 |
| chr1:38991597-38992096 | 0.315184126 | 0.395367145 | 0.678305936 |
| chr10:104406743-104406993 | -0.408186207 | 0.395393447 | 0.678335871 |
| chr19:18152406-18152740 | 0.323432176 | 0.395455919 | 0.678427858 |
| chr2:172931420-172931650 | 0.419307617 | 0.395481789 | 0.678457049 |
| chr3:28667651-28668188 | -0.308959623 | 0.395498705 | 0.678470878 |
| chr8:68827950-68828179 | -0.460947965 | 0.395534848 | 0.67851769 |
| chr16:78325692-78325904 | 0.423809686 | 0.39557719 | 0.678575133 |
| chr20:53739037-53739246 | 0.290410509 | 0.395641262 | 0.678650602 |
| chr6:34391693-34392151 | 0.318012093 | 0.395642404 | 0.678650602 |
| chr5:135641707-135642354 | -0.302831289 | 0.395647756 | 0.678650602 |
| chrX:23667214-23668091 | -0.278737356 | 0.395657662 | 0.6786524 |
| chr6:159451160-159451366 | -0.441830963 | 0.395686202 | 0.678686161 |
| chr1:145891742-145892668 | 0.270142202 | 0.395746976 | 0.678775206 |
| chr12:131555110-131555562 | -0.33188949 | 0.395776651 | 0.67879975 |
| chr12:75195877-75196107 | -0.46716247 | 0.395779004 | 0.67879975 |
| chr12:120630587-120630907 | -0.361214118 | 0.395793724 | 0.678807961 |
| chr22:39319305-39319936 | 0.295985864 | 0.39580151 | 0.678807961 |
| chr1:166975328-166975528 | -0.445993287 | 0.395822778 | 0.678827024 |
| chr1:14149461-14149736 | -0.446720067 | 0.395830344 | 0.678827024 |
| chr18:9771536-9771791 | 0.362734772 | 0.395860798 | 0.678864057 |
| chr11:74546707-74547159 | -0.304520629 | 0.395898526 | 0.678875879 |
| chr8:68773623-68774144 | 0.32641012 | 0.395900989 | 0.678875879 |
| chr6:125802439-125802639 | 0.489154432 | 0.395907036 | 0.678875879 |
| chr10:5943444-5943889 | -0.333712342 | 0.395914877 | 0.678875879 |
| chr3:185660245-185660707 | -0.323366905 | 0.395919716 | 0.678875879 |
| chr6:28985430-28985782 | 0.303262737 | 0.395924258 | 0.678875879 |
| chr16:81572716-81572916 | -0.424684677 | 0.395932528 | 0.678875879 |
| chr3:138923456-138923834 | 0.312825749 | 0.395942946 | 0.678875879 |
| chr12:106773977-106775110 | -0.271272019 | 0.395947432 | 0.678875879 |
| chr2:206800451-206800879 | -0.329401338 | 0.395956358 | 0.678875991 |
| chr2:156006348-156006694 | 0.351936197 | 0.395986691 | 0.678912806 |
| chr12:123043507-123043845 | 0.341437581 | 0.396028069 | 0.678956901 |
| chr15:35019498-35019798 | -0.373470415 | 0.396030132 | 0.678956901 |
| chrX:115940814-115941224 | 0.306771859 | 0.396039267 | 0.678957185 |
| chr1:114473116-114473752 | -0.277049876 | 0.39604802 | 0.678957185 |
| chr11:116791253-116791626 | -0.336645832 | 0.396074294 | 0.678961131 |
| chr3:23671982-23672383 | -0.364062987 | 0.396074982 | 0.678961131 |
| chr7:101169821-101170084 | -0.335539118 | 0.396077837 | 0.678961131 |

| | | | |
|---|---|---|---|
| chr18:54192970-54193179 | -0.431630447 | 0.396085766 | 0.678961131 |
| chr5:143014080-143014379 | -0.375739641 | 0.396146789 | 0.679050543 |
| chr8:1973065-1973324 | -0.435329956 | 0.39616359 | 0.679064151 |
| chr1:63917963-63918163 | 0.458739423 | 0.396179744 | 0.679072021 |
| chr11:70068006-70068441 | 0.311665299 | 0.396185907 | 0.679072021 |
| chr6:107079480-107079741 | -0.377352925 | 0.396206265 | 0.679073943 |
| chr19:18117757-18118270 | 0.333353883 | 0.396218186 | 0.679073943 |
| chr7:148142744-148142992 | -0.371129288 | 0.396221403 | 0.679073943 |
| chr6:143745338-143746009 | 0.299540449 | 0.396222479 | 0.679073943 |
| chr21:18717407-18718030 | -0.339525292 | 0.396280704 | 0.679158543 |
| chr20:54729203-54729729 | 0.280543484 | 0.396296331 | 0.679170133 |
| chr10:63083966-63084288 | -0.353690489 | 0.396324188 | 0.679174532 |
| chr10:61179774-61180128 | 0.358788615 | 0.396329762 | 0.679174532 |
| chr3:173411745-173411945 | -0.460349161 | 0.396333636 | 0.679174532 |
| chr19:1027521-1027771 | 0.356259074 | 0.396340102 | 0.679174532 |
| chr12:71900132-71900365 | -0.413079336 | 0.396343217 | 0.679174532 |
| chr7:42161812-42162131 | -0.383429483 | 0.396361013 | 0.679189836 |
| chr19:1383224-1383992 | 0.285127392 | 0.396392297 | 0.679214824 |
| chr10:91880281-91880793 | -0.293996613 | 0.396393324 | 0.679214824 |
| chr14:76373380-76373842 | 0.316390117 | 0.396410719 | 0.67922944 |
| chr22:21641940-21642539 | 0.282409313 | 0.396437467 | 0.679258488 |
| chr20:63537242-63538386 | 0.269016258 | 0.396445402 | 0.679258488 |
| chr1:149936721-149937064 | 0.296136707 | 0.396463113 | 0.67926064 |
| chr8:108585821-108586186 | 0.315877803 | 0.396464388 | 0.67926064 |
| chr1:177255447-177255828 | -0.339174581 | 0.396505211 | 0.679311638 |
| chr7:101173915-101174267 | -0.303710868 | 0.396511886 | 0.679311638 |
| chr17:65136954-65137716 | -0.277180316 | 0.396564524 | 0.679386629 |
| chr18:62714686-62715690 | -0.270626834 | 0.396598879 | 0.679430294 |
| chr1:234523514-234523936 | 0.310550541 | 0.396648638 | 0.679500345 |
| chr12:52980837-52981282 | 0.294421197 | 0.396692662 | 0.67956057 |
| chr3:1771360-1771666 | 0.401957067 | 0.396724755 | 0.679600352 |
| chr15:41773996-41775020 | 0.275762454 | 0.396740713 | 0.679601098 |
| chr10:11165972-11166384 | -0.354630948 | 0.396742929 | 0.679601098 |
| chr1:240004210-240004410 | -0.479722758 | 0.396753968 | 0.679604814 |
| chrX:17522278-17522653 | -0.404226771 | 0.39677884 | 0.679632225 |
| chr4:93199634-93199928 | -0.368448043 | 0.3968096 | 0.67964739 |
| chr20:50663437-50663789 | 0.310853486 | 0.396811156 | 0.67964739 |
| chr1:220090099-220090299 | -0.36272565 | 0.396819783 | 0.67964739 |
| chr2:19020095-19020318 | -0.397849242 | 0.396823175 | 0.67964739 |
| chr10:117165819-117166100 | -0.346485015 | 0.396839601 | 0.679660331 |
| chr11:75022940-75023221 | -0.34390925 | 0.396855671 | 0.679664104 |
| chrX:127255850-127256085 | 0.374516244 | 0.396859544 | 0.679664104 |
| chr14:53568534-53568734 | 0.465274182 | 0.396874473 | 0.679674479 |
| chr6:134317771-134318036 | -0.442862388 | 0.396887424 | 0.679676217 |
| chr3:69143708-69143957 | 0.374627491 | 0.396893229 | 0.679676217 |
| chr16:229015-229635 | 0.29557521 | 0.396912925 | 0.679694755 |
| chrX:124127603-124127844 | 0.382117901 | 0.396958355 | 0.679747566 |
| chr16:53502552-53503952 | 0.268322151 | 0.396965053 | 0.679747566 |
| chr17:65531954-65532154 | 0.34835697 | 0.396970379 | 0.679747566 |
| chr1:217363544-217363822 | 0.388220917 | 0.397007108 | 0.679779331 |

| | | | |
|---|---|---|---|
| chr3:30683848-30684190 | -0.399002338 | 0.39701257 | 0.679779331 |
| chr5:93569849-93570150 | -0.440512865 | 0.397015544 | 0.679779331 |
| chr1:200016999-200017254 | -0.37029981 | 0.397037589 | 0.679801884 |
| chr16:1971469-1972098 | 0.293871338 | 0.397099752 | 0.679893127 |
| chr6:121334265-121335492 | -0.271912041 | 0.397118358 | 0.67990979 |
| chr20:53996636-53996952 | 0.282893502 | 0.397157843 | 0.679962199 |
| chr4:109302909-109303462 | -0.330186698 | 0.397181458 | 0.679973736 |
| chr1:55143735-55144077 | 0.323662238 | 0.39718233 | 0.679973736 |
| chr9:121744438-121744641 | 0.402532156 | 0.3971958 | 0.679981602 |
| chr20:5112505-5113572 | 0.275982286 | 0.397227887 | 0.680017665 |
| chr21:21997678-21997932 | -0.470332504 | 0.397234615 | 0.680017665 |
| chr4:152063244-152063526 | 0.37549798 | 0.397248138 | 0.680025621 |
| chr14:36028263-36028476 | 0.453554621 | 0.397326679 | 0.680127709 |
| chr18:56637931-56639365 | -0.268412678 | 0.397330564 | 0.680127709 |
| chr4:87973436-87973907 | 0.345294799 | 0.397334482 | 0.680127709 |
| chrX:96063906-96064133 | -0.440346696 | 0.397350198 | 0.680127709 |
| chr4:172893570-172893807 | -0.452141091 | 0.397363176 | 0.680127709 |
| chr17:7219774-7219996 | -0.354636697 | 0.397369771 | 0.680127709 |
| chr4:88144891-88145346 | 0.297636165 | 0.397369909 | 0.680127709 |
| chr4:15654620-15656008 | 0.272670039 | 0.397405161 | 0.680172852 |
| chr19:17841559-17842737 | 0.282863766 | 0.397418048 | 0.680179716 |
| chr6:145888513-145888747 | 0.428746809 | 0.397456156 | 0.680229743 |
| chr6:57046384-57047030 | 0.29050863 | 0.397476038 | 0.68024511 |
| chr3:30150694-30151515 | -0.304329725 | 0.397500914 | 0.68024511 |
| chr16:16021991-16022410 | 0.318564773 | 0.397515966 | 0.68024511 |
| chr1:55021945-55022454 | -0.288559266 | 0.397528662 | 0.68024511 |
| chr4:109427528-109427893 | -0.327468825 | 0.397535662 | 0.68024511 |
| chr6:10127325-10127532 | 0.42406441 | 0.397540296 | 0.68024511 |
| chr8:135456679-135457946 | -0.2731397 | 0.397544295 | 0.68024511 |
| chr5:75336155-75337381 | -0.266323677 | 0.39755319 | 0.68024511 |
| chr4:102308470-102308738 | 0.377794247 | 0.397557975 | 0.68024511 |
| chr13:99605948-99606269 | -0.338460181 | 0.397573912 | 0.68024511 |
| chr17:9196295-9196729 | -0.320457194 | 0.397582292 | 0.68024511 |
| chr8:138546595-138546861 | 0.424059471 | 0.397586018 | 0.68024511 |
| chr6:70955365-70955736 | -0.353434704 | 0.397586674 | 0.68024511 |
| chr7:12244115-12244526 | 0.309918053 | 0.397589426 | 0.68024511 |
| chr6:150418869-150419354 | 0.340834796 | 0.397630946 | 0.680300957 |
| chr5:176490356-176490556 | -0.428533341 | 0.397667645 | 0.680348553 |
| chr7:44326025-44326319 | 0.320230567 | 0.397730786 | 0.680431659 |
| chr1:112395902-112396553 | -0.27808107 | 0.397733982 | 0.680431659 |
| chr12:67025565-67025807 | -0.390506405 | 0.397793165 | 0.680517714 |
| chr12:57242812-57243026 | -0.34439568 | 0.397809276 | 0.680530081 |
| chr6:99568069-99569258 | -0.269190006 | 0.397862339 | 0.680590664 |
| chr14:78595890-78596090 | 0.396709394 | 0.397862455 | 0.680590664 |
| chr3:77825172-77825895 | 0.292129935 | 0.397898972 | 0.680621464 |
| chr8:126302482-126302930 | 0.297337941 | 0.397903655 | 0.680621464 |
| chr2:200905337-200905627 | -0.362004437 | 0.397907109 | 0.680621464 |
| chr19:12681022-12682042 | 0.278150841 | 0.397920448 | 0.680629087 |
| chr1:26284889-26285218 | -0.360688483 | 0.397975556 | 0.680708152 |
| chr8:115668477-115669765 | -0.27467558 | 0.39799034 | 0.680718242 |

| | | | |
|---|---|---|---|
| chr14:69650961-69651199 | 0.357731191 | 0.398040259 | 0.680788426 |
| chr5:133077595-133077816 | 0.435337302 | 0.398059822 | 0.68080669 |
| chr14:23301250-23302597 | 0.266576937 | 0.398069254 | 0.680807625 |
| chr1:196243765-196244111 | -0.37761085 | 0.39809812 | 0.680841796 |
| chr6:41786507-41787796 | 0.273291758 | 0.398131987 | 0.680878096 |
| chr19:40570052-40571137 | 0.269531008 | 0.398137118 | 0.680878096 |
| chr7:3043112-3043312 | -0.441735698 | 0.398167588 | 0.680911057 |
| chr1:46413669-46413916 | 0.354115856 | 0.398174165 | 0.680911057 |
| chr3:15646046-15646346 | -0.387009658 | 0.398225285 | 0.680959354 |
| chr19:49506622-49506910 | 0.342063577 | 0.398228111 | 0.680959354 |
| chrX:100671223-100671517 | 0.358169406 | 0.398229069 | 0.680959354 |
| chr2:210002240-210003256 | -0.270694497 | 0.398271278 | 0.681016332 |
| chr20:10673875-10674562 | 0.272362407 | 0.398335491 | 0.681110932 |
| chr3:32667063-32667505 | 0.337913768 | 0.398344831 | 0.681111703 |
| chr20:50987552-50988106 | 0.291176848 | 0.398358247 | 0.681114975 |
| chr1:202154797-202155740 | -0.27304578 | 0.398364523 | 0.681114975 |
| chr3:170310153-170310699 | -0.283899415 | 0.398394608 | 0.681150604 |
| chr6:169859428-169859762 | -0.377264603 | 0.398403141 | 0.681150604 |
| chr7:97928286-97928637 | -0.340993194 | 0.398427749 | 0.681172531 |
| chr14:104708679-104709000 | 0.302113131 | 0.398447402 | 0.681172531 |
| chr21:33199881-33200123 | 0.368935961 | 0.398449141 | 0.681172531 |
| chr10:92912584-92912896 | 0.388075663 | 0.398451526 | 0.681172531 |
| chr6:37210498-37210787 | 0.367700027 | 0.398485991 | 0.681188966 |
| chr19:39328173-39328393 | -0.400434049 | 0.398493494 | 0.681188966 |
| chr12:107760525-107761864 | -0.269368685 | 0.398513067 | 0.681188966 |
| chr5:69332606-69333005 | 0.294321447 | 0.398513187 | 0.681188966 |
| chr5:171307223-171307564 | -0.357848687 | 0.398517154 | 0.681188966 |
| chr20:52628553-52628901 | -0.439023024 | 0.398517258 | 0.681188966 |
| chr17:74737648-74737912 | -0.33890072 | 0.398523372 | 0.681188966 |
| chr9:133992203-133992538 | 0.31200135 | 0.398556914 | 0.681229397 |
| chr18:37578864-37579226 | -0.312323676 | 0.398564807 | 0.681229397 |
| chr20:48187019-48187319 | 0.290390418 | 0.398615938 | 0.681295195 |
| chr2:217434114-217434706 | -0.298382575 | 0.398621086 | 0.681295195 |
| chr16:68764944-68765144 | 0.42805571 | 0.39866162 | 0.681349274 |
| chr7:44162894-44163330 | -0.329540614 | 0.398682641 | 0.681370002 |
| chr3:155995596-155995922 | -0.317792003 | 0.398713359 | 0.681407302 |
| chr1:62989353-62989724 | 0.384910747 | 0.398733866 | 0.681422436 |
| chr12:107607544-107607933 | -0.311203502 | 0.398740001 | 0.681422436 |
| chr3:178859808-178860561 | -0.287565346 | 0.398784666 | 0.681483566 |
| chr11:108524476-108525084 | 0.302211536 | 0.398810819 | 0.68151306 |
| chr19:7098769-7099737 | 0.286850093 | 0.398822308 | 0.681517493 |
| chr1:181126492-181127556 | 0.273638343 | 0.398860947 | 0.68156832 |
| chr16:87635533-87636712 | 0.27472211 | 0.398880674 | 0.681586829 |
| chrX:101450146-101450441 | 0.349766235 | 0.398901624 | 0.681598824 |
| chr10:32957935-32958948 | -0.272391252 | 0.398905485 | 0.681598824 |
| chr6:27230208-27230967 | -0.289326359 | 0.398940521 | 0.681643488 |
| chr3:57078500-57078811 | 0.356753946 | 0.39897516 | 0.681687472 |
| chr19:38851879-38852081 | 0.323904579 | 0.398990781 | 0.681698961 |
| chr5:81233009-81233439 | -0.365871669 | 0.399007359 | 0.681712085 |
| chr10:29179202-29179402 | -0.494038313 | 0.399049318 | 0.681758675 |

| | | | |
|---|---|---|---|
| chr19:48399802-48400026 | 0.351891564 | 0.399052424 | 0.681758675 |
| chr8:143569187-143569618 | 0.28760897 | 0.399071684 | 0.68177133 |
| chr12:107646728-107646977 | 0.435575738 | 0.399077627 | 0.68177133 |
| chr21:45542339-45542619 | 0.331350359 | 0.399088448 | 0.681774616 |
| chr12:131104666-131105058 | 0.318965541 | 0.399116403 | 0.681784006 |
| chr8:17156364-17157272 | 0.289792027 | 0.399126133 | 0.681784006 |
| chr22:24231284-24231652 | -0.367122714 | 0.399126392 | 0.681784006 |
| chr6:28802487-28803067 | -0.284129503 | 0.399129537 | 0.681784006 |
| chr1:201704146-201704578 | 0.287750003 | 0.399162331 | 0.681824824 |
| chr4:84924398-84924795 | -0.310615691 | 0.399201312 | 0.681876207 |
| chr17:44503036-44503772 | 0.283041222 | 0.399216064 | 0.681886204 |
| chr1:168861487-168862230 | 0.289550064 | 0.399259754 | 0.681945628 |
| chr2:138323862-138324334 | -0.327197102 | 0.399304859 | 0.681989712 |
| chr8:37153202-37153519 | -0.355960553 | 0.399325482 | 0.681989712 |
| chr21:29002651-29003037 | -0.321866672 | 0.399326245 | 0.681989712 |
| chrX:129603915-129604285 | 0.295779732 | 0.399335917 | 0.681989712 |
| chr12:124480174-124481275 | -0.28374082 | 0.399337296 | 0.681989712 |
| chr6:31971651-31972709 | 0.271369207 | 0.399338968 | 0.681989712 |
| chr19:32652899-32653099 | 0.439468701 | 0.399393569 | 0.682062534 |
| chr6:80106127-80107289 | -0.267542273 | 0.399406183 | 0.682062534 |
| chr5:73437178-73437378 | 0.395583858 | 0.399408314 | 0.682062534 |
| chr1:248851025-248851235 | 0.364822416 | 0.399431793 | 0.682083088 |
| chr19:11089291-11089609 | 0.321070609 | 0.399438154 | 0.682083088 |
| chr12:52874067-52874620 | 0.280241739 | 0.399447712 | 0.682084208 |
| chr8:94612894-94613322 | 0.302148967 | 0.399466433 | 0.682100975 |
| chr7:76379647-76380665 | -0.272768947 | 0.399487303 | 0.682121409 |
| chr3:62109543-62109797 | 0.3287528 | 0.399498364 | 0.682125095 |
| chr4:142405670-142405892 | 0.411002063 | 0.39952008 | 0.682146259 |
| chr5:21496165-21496445 | -0.360127936 | 0.399549049 | 0.682146259 |
| chr4:128461263-128461812 | 0.323197205 | 0.399549446 | 0.682146259 |
| chr1:67347720-67347956 | 0.384253635 | 0.399553168 | 0.682146259 |
| chr9:128819223-128819451 | 0.343036557 | 0.399555273 | 0.682146259 |
| chr10:34319534-34319923 | -0.31881423 | 0.399635244 | 0.682267588 |
| chr3:76024725-76024982 | -0.410257739 | 0.399646185 | 0.682271066 |
| chr3:5285543-5285786 | 0.458582349 | 0.399655421 | 0.682271632 |
| chr1:168585220-168585462 | -0.448068983 | 0.399686105 | 0.682308812 |
| chr20:61949293-61949784 | 0.314953247 | 0.399752845 | 0.682407541 |
| chr7:75615057-75615286 | 0.419130162 | 0.399821788 | 0.682472021 |
| chr7:43636287-43636497 | 0.419130162 | 0.399821788 | 0.682472021 |
| chr1:189692820-189693075 | 0.387170414 | 0.399832353 | 0.682472021 |
| chr19:45450351-45450558 | -0.373676134 | 0.399833703 | 0.682472021 |
| chr4:16227066-16227365 | 0.343482874 | 0.399835152 | 0.682472021 |
| chr2:215086580-215086780 | 0.45988431 | 0.399867194 | 0.682511509 |
| chr4:124940413-124940622 | -0.402926504 | 0.399887638 | 0.6825312 |
| chr15:73800884-73801129 | -0.395336501 | 0.399897471 | 0.682532779 |
| chr1:21963956-21964245 | 0.396604528 | 0.399911218 | 0.682541038 |
| chr2:95992614-95993076 | -0.334929529 | 0.399934143 | 0.68256496 |
| chr15:84142863-84143357 | 0.30041443 | 0.400046382 | 0.682726353 |
| chr7:38369871-38370212 | 0.313986763 | 0.400052583 | 0.682726353 |
| chr9:33374890-33375222 | -0.331493496 | 0.40005675 | 0.682726353 |

| | | | |
|---|---|---|---|
| chr6:1147427-1147762 | -0.453645112 | 0.400064349 | 0.682726353 |
| chr15:42736203-42736552 | 0.339480544 | 0.400081607 | 0.682739066 |
| chr7:35709497-35709768 | -0.369577224 | 0.400104146 | 0.682739066 |
| chr2:127810548-127811690 | -0.272532122 | 0.400106757 | 0.682739066 |
| chr4:112050458-112050658 | -0.484280514 | 0.400123215 | 0.682739066 |
| chr6:75932068-75932352 | -0.338188099 | 0.400137008 | 0.682739066 |
| chr11:75547588-75547867 | 0.338911582 | 0.400139953 | 0.682739066 |
| chr1:32886598-32886909 | -0.342244371 | 0.400148232 | 0.682739066 |
| chr11:115990058-115990361 | -0.397849846 | 0.400149019 | 0.682739066 |
| chr1:181141505-181141877 | 0.313834716 | 0.400151993 | 0.682739066 |
| chr10:3873798-3874055 | -0.449402006 | 0.400166257 | 0.682748201 |
| chr1:234610591-234611528 | 0.270248677 | 0.40017859 | 0.682754039 |
| chr2:35040308-35040508 | -0.512558317 | 0.4001885 | 0.682755744 |
| chr3:20514296-20514505 | -0.48423135 | 0.400200211 | 0.682760521 |
| chrX:24053553-24053835 | -0.340092993 | 0.400228019 | 0.68279276 |
| chr13:44971779-44972219 | -0.32789769 | 0.40024862 | 0.68279446 |
| chr12:68486507-68486757 | 0.37958544 | 0.400253737 | 0.68279446 |
| chr17:80988866-80989511 | 0.291287244 | 0.400255749 | 0.68279446 |
| chr5:17890581-17890829 | -0.400116343 | 0.40026756 | 0.682799407 |
| chr20:41692566-41693921 | 0.265766683 | 0.400315156 | 0.682865396 |
| chr17:74943410-74943638 | -0.410457809 | 0.40034243 | 0.682881428 |
| chr2:46500122-46500580 | -0.294193827 | 0.400342736 | 0.682881428 |
| chr19:48061576-48061928 | 0.360171231 | 0.400359835 | 0.682881428 |
| chr3:4633125-4633700 | -0.305073523 | 0.400360203 | 0.682881428 |
| chr16:31179629-31180909 | 0.268678798 | 0.400424128 | 0.682960397 |
| chr14:36519880-36520348 | 0.316802854 | 0.400424328 | 0.682960397 |
| chr9:90642738-90643229 | 0.326435774 | 0.40047381 | 0.683006421 |
| chr6:51855038-51855549 | -0.318572734 | 0.400477669 | 0.683006421 |
| chr11:65254031-65254231 | 0.427149869 | 0.400478054 | 0.683006421 |
| chr2:149597573-149598259 | -0.295741128 | 0.400542006 | 0.683100285 |
| chr15:89998185-89998400 | 0.346233347 | 0.400571897 | 0.683136057 |
| chr5:73451318-73451747 | -0.319582764 | 0.400615947 | 0.683184219 |
| chr8:23472447-23472673 | -0.412423273 | 0.400617971 | 0.683184219 |
| chr17:70636628-70636828 | 0.396043112 | 0.400630967 | 0.683191177 |
| chrX:135086504-135086793 | 0.355489381 | 0.400705799 | 0.683271586 |
| chr11:14589461-14589868 | -0.343199196 | 0.400707596 | 0.683271586 |
| chr10:8075658-8075858 | -0.391381556 | 0.400710872 | 0.683271586 |
| chr3:180601885-180602817 | -0.272461747 | 0.40072168 | 0.683271586 |
| chr10:105585770-105586038 | 0.371584909 | 0.400722707 | 0.683271586 |
| chr11:119108204-119108510 | 0.302505432 | 0.400770367 | 0.683311569 |
| chr12:109818940-109819390 | -0.2989229 | 0.400770881 | 0.683311569 |
| chr2:85391310-85391516 | -0.363177658 | 0.400783054 | 0.683311569 |
| chr14:67218854-67219148 | 0.394511986 | 0.400784202 | 0.683311569 |
| chr10:3300413-3300613 | -0.473659593 | 0.400790746 | 0.683311569 |
| chr9:35116798-35117119 | -0.307720254 | 0.400821347 | 0.683348535 |
| chr5:159208064-159208321 | 0.35208488 | 0.400868296 | 0.683413372 |
| chr14:99518424-99518804 | 0.301246089 | 0.400904557 | 0.683459983 |
| chr11:34345256-34345644 | -0.344879726 | 0.400914371 | 0.683461507 |
| chr14:61252811-61253103 | 0.321460968 | 0.400934792 | 0.683467588 |
| chr3:182212400-182212796 | -0.337226216 | 0.400943501 | 0.683467588 |

| | | | |
|---|---|---|---|
| chr15:64949526-64949795 | 0.38026118 | 0.400944697 | 0.683467588 |
| chr1:184310511-184311008 | 0.318314175 | 0.400968251 | 0.683490347 |
| chr13:23741984-23742184 | 0.465180497 | 0.400976581 | 0.683490347 |
| chr17:60328601-60329041 | 0.279158037 | 0.40098481 | 0.683490347 |
| chr8:18347654-18347940 | 0.397392897 | 0.400995308 | 0.683493037 |
| chr7:74453701-74454036 | -0.346255487 | 0.401021826 | 0.68351842 |
| chr17:76523147-76523644 | 0.316733281 | 0.401028041 | 0.68351842 |
| chrX:39689542-39689973 | -0.372091833 | 0.401059901 | 0.683557517 |
| chr3:126008417-126008758 | 0.350246249 | 0.401082547 | 0.683580909 |
| chr2:241374523-241374884 | -0.357572237 | 0.401099133 | 0.683593972 |
| chr13:91155490-91155847 | -0.353667624 | 0.40111981 | 0.683614006 |
| chr5:87781155-87781670 | -0.306975883 | 0.401188155 | 0.683715276 |
| chr1:181404891-181405463 | -0.298783239 | 0.401217922 | 0.683737982 |
| chrX:149392946-149393236 | -0.357001855 | 0.401219376 | 0.683737982 |
| chr11:119505174-119505374 | 0.4555775 | 0.401228249 | 0.683737982 |
| chr1:107689855-107690061 | -0.352249167 | 0.401296 | 0.683827344 |
| chr5:106561858-106562174 | -0.45498188 | 0.401298537 | 0.683827344 |
| chr19:4065655-4066310 | 0.279852699 | 0.401314866 | 0.68383289 |
| chr20:18566999-18567970 | 0.274369517 | 0.401319641 | 0.68383289 |
| chr17:40111695-40111965 | 0.334690678 | 0.401361269 | 0.683883342 |
| chr1:204398376-204398907 | -0.281182902 | 0.401367706 | 0.683883342 |
| chr18:32439888-32440089 | 0.449657153 | 0.401376026 | 0.683883342 |
| chr19:58407772-58409005 | 0.271633184 | 0.401405672 | 0.683915668 |
| chr20:64062852-64063184 | 0.29064426 | 0.40141285 | 0.683915668 |
| chr8:97170656-97171148 | 0.318397481 | 0.401462347 | 0.68397265 |
| chr8:127206966-127207299 | -0.312962775 | 0.401464148 | 0.68397265 |
| chr2:85577365-85578189 | -0.27967392 | 0.401476134 | 0.683975174 |
| chr8:22237386-22237610 | -0.492307573 | 0.401496386 | 0.683975174 |
| chr20:62365796-62366695 | 0.268835981 | 0.40150498 | 0.683975174 |
| chr3:172710713-172711767 | -0.267795564 | 0.401513634 | 0.683975174 |
| chr1:27603932-27604313 | 0.307288348 | 0.401535449 | 0.683975174 |
| chr4:179307561-179307869 | -0.377642601 | 0.401538012 | 0.683975174 |
| chr1:155856905-155857596 | 0.282637622 | 0.401546941 | 0.683975174 |
| chr15:83010865-83012124 | 0.270100598 | 0.401547889 | 0.683975174 |
| chr6:106912988-106913453 | 0.292258067 | 0.401548612 | 0.683975174 |
| chr4:68349472-68350511 | -0.267927979 | 0.401563259 | 0.683975174 |
| chr17:73189628-73189871 | 0.324880501 | 0.401563823 | 0.683975174 |
| chr8:54758890-54759095 | -0.436430992 | 0.401588482 | 0.68399108 |
| chr10:110746323-110746747 | 0.322842653 | 0.401591014 | 0.68399108 |
| chr11:43021478-43021809 | -0.331042142 | 0.401634978 | 0.684041794 |
| chr1:16872614-16872984 | -0.391785991 | 0.401638645 | 0.684041794 |
| chr6:27807477-27808387 | -0.271449473 | 0.401680265 | 0.684097472 |
| chr17:64797785-64798047 | -0.340720102 | 0.401702243 | 0.684112399 |
| chr11:126510693-126510956 | 0.391595203 | 0.401706887 | 0.684112399 |
| chr12:132809299-132809671 | 0.359320635 | 0.401721115 | 0.684115018 |
| chr6:139164059-139164516 | -0.337020877 | 0.401726281 | 0.684115018 |
| chr15:73994466-73994905 | -0.282712645 | 0.401757723 | 0.684148411 |
| chr1:169245688-169245972 | -0.390293593 | 0.401763748 | 0.684148411 |
| chr3:108406675-108406948 | 0.34269811 | 0.401791398 | 0.684158037 |
| chr12:62505760-62506096 | 0.355429235 | 0.401791584 | 0.684158037 |

| | | | |
|---|---|---|---|
| chr7:434979-435232 | -0.335155902 | 0.401796188 | 0.684158037 |
| chr3:113136073-113136391 | -0.36738173 | 0.401808753 | 0.684164229 |
| chr8:121811882-121812091 | 0.383832582 | 0.401842757 | 0.684185853 |
| chr19:14136428-14137622 | 0.269589546 | 0.401843768 | 0.684185853 |
| chr17:15448823-15449074 | -0.412854681 | 0.401848241 | 0.684185853 |
| chr7:37666501-37666985 | 0.338381085 | 0.401869595 | 0.684207007 |
| chrX:142606731-142606958 | 0.402393348 | 0.401889003 | 0.684224846 |
| chr16:87279135-87279335 | -0.385164409 | 0.401905904 | 0.684238417 |
| chr6:27902331-27903369 | -0.268319254 | 0.401915865 | 0.684240171 |
| chr11:16945915-16946260 | 0.316731663 | 0.401937933 | 0.684262538 |
| chr3:34680584-34680925 | -0.338507971 | 0.401971445 | 0.684304384 |
| chr7:50359828-50360177 | -0.385862434 | 0.401991518 | 0.684316582 |
| chr6:25359418-25359814 | -0.409616241 | 0.402014561 | 0.684316582 |
| chr13:50357638-50358125 | 0.293945956 | 0.402024696 | 0.684316582 |
| chr18:58144717-58145061 | -0.366435193 | 0.40202852 | 0.684316582 |
| chr1:157035005-157035224 | -0.453402822 | 0.402040321 | 0.684316582 |
| chr3:4303140-4304118 | -0.267437213 | 0.402040543 | 0.684316582 |
| chr7:14695647-14696079 | 0.309606082 | 0.402041127 | 0.684316582 |
| chr13:36999865-37001435 | -0.263327154 | 0.402089384 | 0.684383516 |
| chr1:85550728-85551123 | -0.361442543 | 0.402112408 | 0.684398458 |
| chr10:121900622-121900911 | -0.371831776 | 0.402116027 | 0.684398458 |
| chr2:56749183-56749519 | -0.349274187 | 0.402134519 | 0.684407684 |
| chr8:11474082-11474937 | 0.283232488 | 0.402147149 | 0.684407684 |
| chr22:50209914-50210354 | -0.370165868 | 0.402152847 | 0.684407684 |
| chr10:70132293-70132943 | -0.289177227 | 0.402157176 | 0.684407684 |
| chr16:27924866-27925079 | -0.400018103 | 0.402180801 | 0.684417812 |
| chr21:8440468-8440725 | 0.29016878 | 0.402180992 | 0.684417812 |
| chr14:93011592-93012250 | -0.288442949 | 0.40219771 | 0.684431061 |
| chr20:50441742-50441957 | 0.370142695 | 0.402223856 | 0.684460352 |
| chr8:120033801-120034001 | 0.433778289 | 0.402234388 | 0.684463074 |
| chr4:82201907-82202107 | -0.449512706 | 0.402247447 | 0.684463735 |
| chr8:88715724-88715957 | 0.372905504 | 0.402252643 | 0.684463735 |
| chr8:101505582-101505840 | 0.350958631 | 0.402266701 | 0.684472455 |
| chr1:55245967-55246176 | -0.457345335 | 0.402282555 | 0.684484231 |
| chr11:48315638-48315838 | 0.414870927 | 0.402339717 | 0.684523563 |
| chr5:32613124-32613330 | -0.406322773 | 0.40235095 | 0.684523563 |
| chr5:69181993-69182203 | -0.406322773 | 0.40235095 | 0.684523563 |
| chr1:205604179-205604379 | -0.414199661 | 0.402357545 | 0.684523563 |
| chr22:26512154-26512826 | 0.292348196 | 0.402373292 | 0.684523563 |
| chr4:172034062-172034382 | -0.33182365 | 0.402373609 | 0.684523563 |
| chr12:103878941-103879167 | -0.43613243 | 0.402380474 | 0.684523563 |
| chr16:726896-727378 | 0.288647257 | 0.402390253 | 0.684523563 |
| chr15:60526314-60526551 | -0.425788845 | 0.402396093 | 0.684523563 |
| chr5:150779180-150779472 | -0.321206266 | 0.402396492 | 0.684523563 |
| chr13:54241745-54242026 | -0.429724872 | 0.402403942 | 0.684523563 |
| chr11:61129769-61130191 | 0.295403816 | 0.402418249 | 0.684532704 |
| chr2:30225053-30225418 | 0.338007361 | 0.402461581 | 0.684591215 |
| chr7:26397958-26399428 | -0.280014061 | 0.402487835 | 0.684620675 |
| chr17:77288766-77289038 | -0.30707433 | 0.402500196 | 0.684626502 |
| chr10:53029914-53030466 | 0.315434245 | 0.402570412 | 0.68470556 |

| | | | |
|---|---|---|---|
| chr2:70086235-70087773 | -0.265754992 | 0.402570598 | 0.68470556 |
| chr3:49339634-49340857 | -0.286268874 | 0.402573484 | 0.68470556 |
| chr16:71412686-71413010 | -0.324849606 | 0.402612584 | 0.684730017 |
| chr7:151027861-151029012 | -0.263843607 | 0.40261459 | 0.684730017 |
| chr14:99274363-99275066 | -0.274376472 | 0.402614673 | 0.684730017 |
| chr16:57185935-57186685 | 0.272612808 | 0.40262806 | 0.684737587 |
| chr2:239246565-239246765 | -0.460841176 | 0.402645635 | 0.684751669 |
| chr7:92178806-92179359 | -0.290652798 | 0.402654214 | 0.684751669 |
| chr13:98523230-98523489 | 0.456798833 | 0.402668545 | 0.68475804 |
| chr1:21539948-21540232 | 0.365548456 | 0.402675834 | 0.68475804 |
| chr11:123438159-123438603 | 0.344612468 | 0.402706178 | 0.684781279 |
| chr7:26078999-26079252 | 0.341324745 | 0.402709116 | 0.684781279 |
| chr5:10810653-10810905 | -0.413665593 | 0.402716311 | 0.684781279 |
| chr6:37353783-37354394 | 0.283890187 | 0.402729575 | 0.684788636 |
| chr9:133127105-133127336 | -0.423601028 | 0.402767423 | 0.68479196 |
| chr19:10581636-10581836 | -0.434178214 | 0.402775739 | 0.68479196 |
| chr2:152155298-152155498 | -0.434178214 | 0.402775739 | 0.68479196 |
| chr4:105932584-105932784 | -0.434178214 | 0.402775739 | 0.68479196 |
| chr12:105860534-105861273 | 0.28501989 | 0.40278073 | 0.68479196 |
| chr3:52287656-52288206 | 0.284561057 | 0.402798759 | 0.68479196 |
| chr21:39631008-39631361 | 0.372109003 | 0.402815599 | 0.68479196 |
| chr17:29155329-29155798 | -0.310864466 | 0.402820168 | 0.68479196 |
| chr17:49887093-49888080 | 0.26725731 | 0.402821178 | 0.68479196 |
| chr14:56855820-56856109 | -0.3738677 | 0.40282984 | 0.68479196 |
| chr15:32677260-32677625 | -0.3738677 | 0.40282984 | 0.68479196 |
| chr10:89260355-89260726 | 0.308041307 | 0.402848109 | 0.684807822 |
| chr20:55057893-55058093 | -0.437619446 | 0.402881091 | 0.684848696 |
| chr2:101124817-101125316 | -0.30916779 | 0.402907244 | 0.684877959 |
| chr4:98262066-98262278 | 0.343212805 | 0.402942712 | 0.684891701 |
| chr2:70498109-70498449 | -0.341142998 | 0.402944772 | 0.684891701 |
| chr6:30597376-30597662 | 0.340571197 | 0.402959677 | 0.684891701 |
| chr5:56904-57285 | -0.331511646 | 0.402960569 | 0.684891701 |
| chr15:32030148-32030771 | 0.291605072 | 0.402964579 | 0.684891701 |
| chr3:170690451-170690677 | -0.372226019 | 0.40296896 | 0.684891701 |
| chr8:144472254-144472649 | 0.290682157 | 0.403010027 | 0.684938902 |
| chr11:65095998-65096973 | 0.27627116 | 0.403021784 | 0.684938902 |
| chr5:71632082-71632600 | 0.299650597 | 0.403025429 | 0.684938902 |
| chr9:38069161-38070085 | -0.270523034 | 0.403032488 | 0.684938902 |
| chr2:116510155-116510570 | 0.325850492 | 0.403060878 | 0.684940887 |
| chr14:27528369-27528698 | 0.313278873 | 0.403064414 | 0.684940887 |
| chr1:182869187-182869653 | 0.315603343 | 0.403070563 | 0.684940887 |
| chr8:22109087-22109889 | 0.282169729 | 0.40307262 | 0.684940887 |
| chr2:170973307-170974625 | -0.268387527 | 0.403078668 | 0.684940887 |
| chr10:95295620-95295944 | -0.355485459 | 0.403087292 | 0.684940887 |
| chr3:148861303-148861635 | -0.325918008 | 0.403101439 | 0.684944876 |
| chr15:80831272-80831701 | -0.35829919 | 0.403107518 | 0.684944876 |
| chr16:48117571-48117815 | 0.38405522 | 0.403138678 | 0.684982632 |
| chr3:141232351-141232678 | 0.316539434 | 0.403168026 | 0.685017308 |
| chr17:60327437-60328586 | 0.264153601 | 0.403188849 | 0.685037497 |
| chr15:68306644-68306934 | -0.315165156 | 0.403232438 | 0.685096365 |

| | | | |
|---|---|---|---|
| chr6:26233603-26233875 | -0.340437776 | 0.40327342 | 0.685150801 |
| chr6:32853108-32853492 | 0.333414673 | 0.403295463 | 0.685173058 |
| chr6:110341985-110342278 | -0.373567747 | 0.403314083 | 0.685189501 |
| chr19:45069192-45069413 | -0.399145655 | 0.403341307 | 0.685220558 |
| chr8:100950360-100950907 | 0.271538456 | 0.403371282 | 0.685256288 |
| chr9:133463233-133463476 | 0.32087985 | 0.403418754 | 0.68532174 |
| chr14:60562177-60562508 | -0.323616936 | 0.403448604 | 0.685324243 |
| chr5:128674917-128675140 | -0.389074034 | 0.403453298 | 0.685324243 |
| chr14:50231184-50232449 | 0.267151504 | 0.403456447 | 0.685324243 |
| chr8:142393871-142394071 | -0.443474782 | 0.40345754 | 0.685324243 |
| chr17:7293840-7294104 | -0.329348189 | 0.403464948 | 0.685324243 |
| chr1:196934024-196934268 | -0.390186749 | 0.403492578 | 0.68534752 |
| chr3:112327317-112327723 | 0.321205096 | 0.403496541 | 0.68534752 |
| chr2:36507382-36507674 | -0.331343448 | 0.403565028 | 0.685448651 |
| chr14:24051846-24052325 | 0.304451188 | 0.40358406 | 0.685465783 |
| chr18:59084245-59084483 | -0.385557422 | 0.403620056 | 0.685511724 |
| chr14:99605278-99605600 | 0.294993854 | 0.403682404 | 0.685602419 |
| chr3:161272593-161272869 | 0.343203762 | 0.40377631 | 0.68573356 |
| chr10:108911751-108912483 | -0.284251017 | 0.403777518 | 0.68573356 |
| chr1:248849681-248849881 | 0.387686971 | 0.403806795 | 0.685759247 |
| chr2:117470359-117471058 | 0.310286496 | 0.403810543 | 0.685759247 |
| chr9:89009092-89009301 | 0.400862036 | 0.40382733 | 0.685772555 |
| chr17:1829250-1830295 | -0.280668726 | 0.403847704 | 0.685790534 |
| chr7:7386620-7386847 | 0.385212978 | 0.403855818 | 0.685790534 |
| chr2:215498475-215498678 | 0.425667986 | 0.40387825 | 0.685799896 |
| chr3:25782839-25784095 | -0.265263106 | 0.403891715 | 0.685799896 |
| chr13:36107763-36107963 | -0.455787456 | 0.403900338 | 0.685799896 |
| chr12:67415532-67415789 | 0.365591766 | 0.40390273 | 0.685799896 |
| chr2:234951615-234952157 | -0.306907404 | 0.403920037 | 0.685799896 |
| chr5:5934637-5934953 | -0.408927702 | 0.403924729 | 0.685799896 |
| chr17:50161298-50161847 | 0.282955604 | 0.403930273 | 0.685799896 |
| chr2:206260375-206260608 | -0.385397249 | 0.403943133 | 0.685799896 |
| chr13:42861564-42861771 | -0.467668644 | 0.403943271 | 0.685799896 |
| chr3:155232573-155233126 | 0.321649883 | 0.403985579 | 0.685799896 |
| chr1:172449843-172450835 | -0.272205661 | 0.403992698 | 0.685799896 |
| chr2:221572179-221572572 | -0.296787853 | 0.403995093 | 0.685799896 |
| chr3:133452660-133452920 | 0.34656821 | 0.404001696 | 0.685799896 |
| chr2:109510124-109510577 | 0.311838312 | 0.404002753 | 0.685799896 |
| chr2:190649862-190650141 | -0.349594654 | 0.404006035 | 0.685799896 |
| chr6:125314142-125314462 | -0.336636357 | 0.404011402 | 0.685799896 |
| chr14:60839239-60839814 | -0.344031914 | 0.404013488 | 0.685799896 |
| chr14:73852216-73853050 | 0.27559136 | 0.404027267 | 0.685808092 |
| chr14:105352290-105352755 | 0.302627323 | 0.404039773 | 0.685814126 |
| chr19:10898888-10899098 | -0.375398164 | 0.404077835 | 0.685863538 |
| chr11:66568539-66568785 | 0.36031903 | 0.404098979 | 0.685874831 |
| chr20:62508642-62509000 | -0.311066706 | 0.404107815 | 0.685874831 |
| chr3:32422258-32422474 | 0.398795981 | 0.404111342 | 0.685874831 |
| chr3:195908496-195908795 | 0.316490715 | 0.404143937 | 0.685883372 |
| chr7:55572751-55573027 | 0.36890585 | 0.404152846 | 0.685883372 |
| chr1:211113684-211113924 | -0.417168505 | 0.404163628 | 0.685883372 |

| | | | |
|---|---|---|---|
| chr15:98548143-98548993 | -0.284892612 | 0.404169262 | 0.685883372 |
| chr12:85053655-85054070 | -0.339770096 | 0.404171281 | 0.685883372 |
| chr10:31031478-31032622 | -0.263092454 | 0.404175395 | 0.685883372 |
| chr6:41424515-41424715 | 0.396583571 | 0.40418351 | 0.685883372 |
| chr16:11742154-11743274 | 0.265865634 | 0.404187986 | 0.685883372 |
| chr17:59154545-59155902 | -0.259404925 | 0.404226921 | 0.685904959 |
| chr6:105384077-105384449 | -0.386234097 | 0.404235768 | 0.685904959 |
| chr19:39506521-39506778 | -0.403394583 | 0.404239884 | 0.685904959 |
| chr17:52181777-52181977 | 0.457125907 | 0.404244982 | 0.685904959 |
| chr4:159202817-159203300 | -0.32001122 | 0.404252923 | 0.685904959 |
| chr9:77977291-77977491 | 0.417970361 | 0.404254418 | 0.685904959 |
| chr14:22956418-22957637 | 0.271864944 | 0.404308998 | 0.68595238 |
| chr6:63560826-63561330 | -0.308077258 | 0.404313422 | 0.68595238 |
| chr9:121621086-121621354 | 0.369610714 | 0.404317419 | 0.68595238 |
| chr8:72918664-72918939 | -0.380855126 | 0.404318177 | 0.68595238 |
| chr14:70832205-70832405 | -0.452188825 | 0.404365283 | 0.686017109 |
| chr2:148674110-148674354 | -0.386369726 | 0.404376822 | 0.686021496 |
| chr2:127719615-127719871 | 0.433592508 | 0.404439958 | 0.68610015 |
| chr15:34969124-34970097 | -0.269229959 | 0.404445663 | 0.68610015 |
| chr20:9442620-9443258 | -0.280021347 | 0.404450048 | 0.68610015 |
| chr2:148020122-148021641 | -0.261484075 | 0.404465584 | 0.686102473 |
| chr3:160133813-160134056 | -0.379929628 | 0.404469326 | 0.686102473 |
| chr12:12562514-12563128 | 0.280874644 | 0.404527487 | 0.686176406 |
| chr3:132731104-132731418 | 0.344796481 | 0.4045319 | 0.686176406 |
| chr16:69339186-69340125 | 0.273747336 | 0.40454377 | 0.686176406 |
| chr6:136226296-136226592 | -0.344109971 | 0.404551957 | 0.686176406 |
| chr3:75614372-75614631 | -0.394501031 | 0.404586559 | 0.686176406 |
| chr9:121545947-121546452 | 0.293162154 | 0.404588928 | 0.686176406 |
| chr12:69156941-69157395 | -0.29449201 | 0.404596585 | 0.686176406 |
| chr1:60874828-60875054 | -0.455839037 | 0.40460166 | 0.686176406 |
| chr11:36747094-36747628 | -0.288771319 | 0.404604855 | 0.686176406 |
| chr3:115913157-115913441 | -0.346611493 | 0.404608982 | 0.686176406 |
| chrX:20067206-20067485 | 0.346252618 | 0.404616291 | 0.686176406 |
| chr20:35947510-35947770 | 0.365067318 | 0.404620375 | 0.686176406 |
| chr2:68225192-68225581 | -0.336500552 | 0.404650884 | 0.686199924 |
| chr5:181223021-181223836 | 0.263180224 | 0.404652154 | 0.686199924 |
| chr9:5628423-5629789 | -0.266861419 | 0.404665454 | 0.68620729 |
| chr10:102420674-102421652 | 0.267598785 | 0.404680003 | 0.686209118 |
| chr3:63450619-63450830 | 0.366561314 | 0.404692409 | 0.686209118 |
| chr5:67746225-67746439 | 0.412811652 | 0.404693399 | 0.686209118 |
| chr21:44509570-44509917 | 0.308240904 | 0.404704738 | 0.686213158 |
| chr10:58414516-58414899 | -0.330159754 | 0.404720428 | 0.686224578 |
| chr3:107512435-107512747 | 0.363120806 | 0.404741084 | 0.686240076 |
| chr15:75058260-75058585 | -0.370520739 | 0.404752962 | 0.686240076 |
| chr11:12986440-12986640 | 0.470722121 | 0.404756437 | 0.686240076 |
| chr2:64323327-64323549 | 0.383156298 | 0.40477595 | 0.686257974 |
| chr19:55503697-55504324 | -0.284731954 | 0.404795063 | 0.686275192 |
| chr8:64239709-64240160 | -0.33529266 | 0.404805949 | 0.686278463 |
| chr1:167052489-167052781 | 0.311814651 | 0.404815392 | 0.686279287 |
| chr19:35153697-35153979 | 0.35701985 | 0.40486111 | 0.686323797 |

| | | | |
|---|---|---|---|
| chr3:150538648-150539330 | 0.288184587 | 0.404864031 | 0.686323797 |
| chr14:74492958-74494254 | 0.263073469 | 0.404868518 | 0.686323797 |
| chr5:143674331-143674591 | 0.427422649 | 0.404880965 | 0.686329711 |
| chr17:6939122-6939392 | 0.421932508 | 0.404931771 | 0.686400649 |
| chr3:120567472-120568068 | 0.295442547 | 0.404965825 | 0.686443189 |
| chr5:31551914-31552176 | -0.377368166 | 0.405016953 | 0.686514666 |
| chr2:200901481-200901784 | -0.304322436 | 0.40502918 | 0.686520204 |
| chr18:12657615-12658376 | 0.285853948 | 0.405042855 | 0.686528196 |
| chr6:80128541-80128858 | -0.351117407 | 0.405079979 | 0.6865613 |
| chr5:54330028-54330472 | 0.325406628 | 0.405088633 | 0.6865613 |
| chr10:69045972-69046206 | 0.42805816 | 0.405089267 | 0.6865613 |
| chr8:108850201-108850598 | 0.345168232 | 0.405129105 | 0.686613631 |
| chr10:101310604-101310963 | -0.371047478 | 0.405149481 | 0.686632978 |
| chr6:135538066-135538320 | -0.414610412 | 0.405196633 | 0.6866977 |
| chr10:116470814-116471337 | 0.320370141 | 0.405212366 | 0.686709175 |
| chr12:108671770-108671970 | -0.469211049 | 0.405239741 | 0.686740377 |
| chr11:119721123-119721385 | 0.36229187 | 0.405258669 | 0.686757265 |
| chr16:30598206-30598661 | 0.325253836 | 0.405274032 | 0.686768111 |
| chr9:75077225-75077452 | -0.417739527 | 0.405318278 | 0.686827899 |
| chr2:20191385-20191599 | 0.354662846 | 0.405362065 | 0.686874131 |
| chr1:110619409-110619673 | -0.344117529 | 0.40536904 | 0.686874131 |
| chr1:204975676-204975979 | -0.298873517 | 0.405372454 | 0.686874131 |
| chr8:79767572-79768747 | 0.266782442 | 0.405394824 | 0.686896845 |
| chr8:128277475-128278095 | 0.293523587 | 0.40541645 | 0.686904452 |
| chr1:17059664-17059968 | 0.362373282 | 0.405417244 | 0.686904452 |
| chr20:23357824-23358486 | 0.266726835 | 0.405435423 | 0.686920064 |
| chr1:169711567-169711922 | -0.297830449 | 0.405464387 | 0.68692731 |
| chr8:112217908-112218598 | -0.325172351 | 0.405469805 | 0.68692731 |
| chr13:34361677-34361958 | 0.38064302 | 0.405484406 | 0.68692731 |
| chr14:68211555-68211924 | -0.342954244 | 0.405496927 | 0.68692731 |
| chr1:206738315-206738530 | -0.345910401 | 0.405497816 | 0.68692731 |
| chr4:112515518-112516224 | 0.292785513 | 0.405508944 | 0.68692731 |
| chr8:103487332-103487587 | 0.38732168 | 0.405511074 | 0.68692731 |
| chr10:33005648-33005961 | -0.326962567 | 0.405511421 | 0.68692731 |
| chr1:53860897-53861387 | 0.309694814 | 0.405533079 | 0.686947915 |
| chr8:100847123-100848027 | 0.268870761 | 0.405541515 | 0.686947915 |
| chr9:130317766-130317966 | 0.453307573 | 0.405560052 | 0.686964128 |
| chr13:97257170-97257496 | -0.341928834 | 0.405575204 | 0.686974605 |
| chr20:32902550-32902844 | 0.361266548 | 0.405633124 | 0.687057524 |
| chr6:151648081-151648366 | 0.331194181 | 0.405665795 | 0.687079981 |
| chr6:117674875-117675607 | -0.268668284 | 0.405666955 | 0.687079981 |
| chr17:29148731-29149194 | 0.280341118 | 0.405678622 | 0.687079981 |
| chr8:56992935-56994502 | -0.264454526 | 0.405690455 | 0.687079981 |
| chr2:26679071-26679271 | 0.443355256 | 0.405694204 | 0.687079981 |
| chr6:26553393-26553661 | 0.297636564 | 0.405704984 | 0.687079981 |
| chr1:228436515-228436715 | 0.454971752 | 0.405709152 | 0.687079981 |
| chr6:43170670-43171564 | 0.266634602 | 0.405719885 | 0.687082971 |
| chrX:36640123-36640367 | -0.405036231 | 0.405751606 | 0.687106566 |
| chr8:98825210-98826578 | 0.263428022 | 0.405751753 | 0.687106566 |
| chr16:84065627-84065862 | -0.371635808 | 0.405763449 | 0.687111188 |

| | | | |
|---|---|---|---|
| chr5:138352290-138353558 | -0.268503525 | 0.405792784 | 0.687145676 |
| chrX:71911084-71911336 | 0.420945612 | 0.405812656 | 0.687160754 |
| chr4:102299718-102300007 | 0.364109736 | 0.405819624 | 0.687160754 |
| chr1:157505667-157506011 | -0.325141755 | 0.405866498 | 0.687208933 |
| chr18:35326931-35327131 | -0.457451953 | 0.40587526 | 0.687208933 |
| chr7:2886989-2887240 | 0.332566358 | 0.405880362 | 0.687208933 |
| chr6:7725852-7726230 | 0.303898842 | 0.405891808 | 0.687208933 |
| chr16:89246869-89247985 | -0.266802425 | 0.405892922 | 0.687208933 |
| chr13:51797730-51798058 | -0.322209371 | 0.405954815 | 0.687298536 |
| chr14:44342949-44343149 | 0.445502238 | 0.405965102 | 0.687300766 |
| chr12:75458028-75458228 | -0.462554319 | 0.405984396 | 0.687318243 |
| chr3:139319885-139320138 | -0.38689223 | 0.406015217 | 0.687355235 |
| chr21:45421326-45421777 | 0.308601628 | 0.406025395 | 0.68735728 |
| chr2:3581575-3581834 | -0.398947694 | 0.406050408 | 0.687374713 |
| chr9:99140757-99141023 | 0.400522961 | 0.406053635 | 0.687374713 |
| chr1:44029938-44031185 | 0.272266279 | 0.406093729 | 0.687427397 |
| chr1:150268324-150269420 | 0.265494724 | 0.406146175 | 0.687500988 |
| chr16:8723091-8723291 | 0.460248237 | 0.40622953 | 0.687623924 |
| chr19:49641763-49642317 | 0.273787128 | 0.406236749 | 0.687623924 |
| chr15:63466706-63467087 | -0.313491354 | 0.406256 | 0.687638773 |
| chr7:32810726-32810942 | 0.432079713 | 0.40626347 | 0.687638773 |
| chr15:89998437-89998727 | 0.340173029 | 0.406298416 | 0.687669365 |
| chr3:170320591-170320926 | 0.335186343 | 0.406304408 | 0.687669365 |
| chr3:187988163-187988368 | -0.4429736 | 0.406313636 | 0.687669365 |
| chr19:4007194-4007874 | 0.276129395 | 0.406322342 | 0.687669365 |
| chr5:155126597-155126932 | 0.384706988 | 0.406326419 | 0.687669365 |
| chr3:158801619-158802758 | -0.264874621 | 0.406352325 | 0.68769802 |
| chr1:246897133-246897333 | 0.455084916 | 0.406368861 | 0.687710815 |
| chr3:184561085-184562237 | 0.263762814 | 0.406388289 | 0.687728505 |
| chr12:109589805-109590102 | -0.372580053 | 0.4064261 | 0.687777301 |
| chr17:67994492-67994759 | -0.305449953 | 0.40643711 | 0.687780743 |
| chr14:67584053-67584325 | 0.345705354 | 0.406448801 | 0.687781393 |
| chr21:42262649-42263218 | 0.291606235 | 0.406455447 | 0.687781393 |
| chr11:66051730-66052561 | 0.270454181 | 0.406479155 | 0.687798857 |
| chr8:80211286-80211574 | -0.336483547 | 0.40648372 | 0.687798857 |
| chr16:17101804-17102011 | 0.423909715 | 0.40649857 | 0.687808795 |
| chr8:122443392-122443732 | -0.344167653 | 0.406509542 | 0.68781217 |
| chr11:85827509-85828971 | -0.262849154 | 0.406521796 | 0.687817716 |
| chr6:25265739-25266023 | -0.394784325 | 0.406532375 | 0.687820427 |
| chr1:156153356-156153696 | 0.310381718 | 0.406555367 | 0.687832135 |
| chr9:83167229-83167451 | -0.390053903 | 0.406557249 | 0.687832135 |
| chr19:48516522-48516734 | -0.410612923 | 0.406576756 | 0.687837331 |
| chr10:133158927-133159520 | -0.295522755 | 0.406578275 | 0.687837331 |
| chr18:58043524-58043836 | -0.323093901 | 0.406609739 | 0.687875374 |
| chr2:206158951-206160315 | -0.26212988 | 0.406645247 | 0.687920256 |
| chr19:44776433-44776932 | 0.303565954 | 0.406656459 | 0.687924034 |
| chr3:9769487-9770115 | 0.284013139 | 0.406667675 | 0.687927821 |
| chr6:105134972-105135385 | -0.335556485 | 0.406693279 | 0.687955944 |
| chr20:32206557-32206780 | -0.360611181 | 0.406722759 | 0.687976407 |
| chr6:127972031-127972383 | 0.337219822 | 0.406723334 | 0.687976407 |

| | | | |
|---|---|---|---|
| chr1:106232122-106232778 | 0.2727886 | 0.406753058 | 0.688011497 |
| chr5:9958355-9958577 | 0.417833173 | 0.406796543 | 0.688061616 |
| chr4:104678250-104678682 | -0.387630817 | 0.406800648 | 0.688061616 |
| chr9:107425265-107425563 | -0.352422071 | 0.406811424 | 0.688064653 |
| chr9:36257918-36258796 | -0.285371877 | 0.40684329 | 0.688098108 |
| chr11:86571514-86571896 | -0.297138258 | 0.406851226 | 0.688098108 |
| chr2:72892814-72893014 | 0.353537933 | 0.406858145 | 0.688098108 |
| chr3:12719949-12720612 | -0.281082625 | 0.406881173 | 0.688121866 |
| chr22:46035603-46036222 | -0.273337473 | 0.406910379 | 0.68815607 |
| chr10:9219973-9220256 | 0.368955492 | 0.406931744 | 0.688177012 |
| chr1:85230678-85230967 | -0.337388395 | 0.406986115 | 0.688253771 |
| chr13:108911797-108912172 | 0.323658172 | 0.407010585 | 0.688261148 |
| chr2:97112386-97112812 | 0.314609618 | 0.40701264 | 0.688261148 |
| chr7:75025605-75025814 | 0.40000104 | 0.407017425 | 0.688261148 |
| chr16:86065907-86066107 | 0.390674718 | 0.407053196 | 0.688281532 |
| chr2:121356105-121356347 | 0.390674718 | 0.407053196 | 0.688281532 |
| chr1:199287778-199287978 | -0.435963688 | 0.407056428 | 0.688281532 |
| chr5:17249261-17249583 | 0.315963411 | 0.407068745 | 0.688287169 |
| chr13:36931375-36931714 | -0.384903342 | 0.407083388 | 0.68829674 |
| chr2:171564572-171564951 | 0.379696445 | 0.407112884 | 0.688302167 |
| chr7:2313845-2315030 | -0.264982916 | 0.40711668 | 0.688302167 |
| chr9:33229361-33230007 | -0.27506253 | 0.407137297 | 0.688302167 |
| chr3:179046737-179047373 | -0.330530987 | 0.407140498 | 0.688302167 |
| chr2:84959713-84960077 | -0.316978316 | 0.407142172 | 0.688302167 |
| chr9:122414053-122414738 | -0.312806017 | 0.407142251 | 0.688302167 |
| chr3:57227488-57228163 | 0.288702172 | 0.407157299 | 0.688302167 |
| chr2:231687153-231687849 | 0.292264681 | 0.407167134 | 0.688302167 |
| chr20:53725013-53725542 | 0.263928513 | 0.407167446 | 0.688302167 |
| chr12:106345804-106346186 | -0.308588935 | 0.407189463 | 0.6883242 |
| chr22:29517550-29517861 | 0.367536313 | 0.407237392 | 0.688345718 |
| chr18:54224375-54225108 | -0.270359876 | 0.407248638 | 0.688345718 |
| chr6:117346223-117346474 | -0.380739851 | 0.40726656 | 0.688345718 |
| chr5:95486391-95486704 | 0.408653704 | 0.407273497 | 0.688345718 |
| chr17:38742480-38742701 | -0.422007463 | 0.40727835 | 0.688345718 |
| chr22:43143102-43143594 | -0.311465317 | 0.407284552 | 0.688345718 |
| chr3:160929749-160929955 | -0.414604436 | 0.407284602 | 0.688345718 |
| chr4:84202900-84203200 | -0.414604436 | 0.407284602 | 0.688345718 |
| chr8:73207867-73208449 | -0.287450024 | 0.407286244 | 0.688345718 |
| chr5:173535757-173536436 | 0.286391774 | 0.407296131 | 0.688345718 |
| chr1:46903644-46903989 | 0.342598799 | 0.407306809 | 0.688345718 |
| chr6:56764047-56764326 | 0.350480144 | 0.407315709 | 0.688345718 |
| chr17:60553060-60554130 | 0.263913883 | 0.407320545 | 0.688345718 |
| chr11:85540726-85540926 | -0.419646742 | 0.407327963 | 0.688345718 |
| chr12:113461769-113461977 | 0.399395251 | 0.407393012 | 0.688440461 |
| chr3:32335116-32335316 | 0.42824416 | 0.407412788 | 0.688456736 |
| chr7:74364196-74364466 | -0.394194436 | 0.407420614 | 0.688456736 |
| chr2:202932854-202933361 | 0.320370042 | 0.407445718 | 0.688483974 |
| chr9:69672705-69673249 | -0.295852058 | 0.407466271 | 0.688495793 |
| chr7:50962850-50963392 | -0.297600716 | 0.407478362 | 0.688495793 |
| chr6:145148289-145148791 | -0.290326801 | 0.407479669 | 0.688495793 |

| | | | |
|---|---|---|---|
| chr10:132265915-132266179 | 0.3806708 | 0.407524973 | 0.688547527 |
| chr2:169364197-169364888 | 0.292894231 | 0.40752826 | 0.688547527 |
| chr13:98576071-98576586 | 0.312011968 | 0.407566219 | 0.688595055 |
| chr7:151046924-151047556 | -0.282457366 | 0.407574364 | 0.688595055 |
| chr8:108821582-108821873 | 0.356896063 | 0.407589292 | 0.68859886 |
| chr1:109483619-109484553 | -0.266500871 | 0.40759459 | 0.68859886 |
| chrX:105191970-105192272 | -0.352823204 | 0.407620705 | 0.68862229 |
| chr3:81006027-81006317 | -0.363027002 | 0.407626433 | 0.68862229 |
| chrX:47556047-47556686 | 0.282224773 | 0.407656242 | 0.688657463 |
| chr10:4842588-4842855 | 0.39823416 | 0.407681559 | 0.688685049 |
| chr2:65187266-65187587 | -0.330200946 | 0.407695279 | 0.688693041 |
| chr1:164748255-164748603 | -0.349044616 | 0.407705999 | 0.688695966 |
| chr1:184745963-184746212 | 0.397160729 | 0.40773644 | 0.688724802 |
| chr4:112636592-112637675 | -0.261357404 | 0.407741046 | 0.688724802 |
| chr15:89441443-89441654 | 0.362622545 | 0.407769489 | 0.688757661 |
| chr5:132064702-132065010 | -0.306668577 | 0.407785536 | 0.68876077 |
| chr8:41701751-41702227 | 0.339664276 | 0.407790026 | 0.68876077 |
| chr17:60088291-60088561 | 0.280869958 | 0.407798296 | 0.68876077 |
| chr13:95698098-95698298 | 0.437197993 | 0.407837182 | 0.688811263 |
| chr14:66070613-66071050 | -0.308129096 | 0.407857981 | 0.688831208 |
| chr1:207397657-207397988 | -0.326368207 | 0.407884499 | 0.688849997 |
| chr20:47356680-47356981 | 0.26806685 | 0.407890278 | 0.688849997 |
| chr3:188962031-188962245 | -0.40915228 | 0.407896077 | 0.688849997 |
| chr8:123295903-123296164 | 0.356340414 | 0.407930131 | 0.68886773 |
| chr2:105179803-105180182 | -0.321720828 | 0.407934465 | 0.68886773 |
| chr4:108774878-108775136 | -0.400111729 | 0.407938925 | 0.68886773 |
| chr6:120378613-120379164 | -0.305414901 | 0.407942539 | 0.68886773 |
| chr5:11791717-11792199 | 0.290345421 | 0.407965838 | 0.688891892 |
| chr15:62112656-62113480 | 0.28528433 | 0.407980579 | 0.688901601 |
| chr11:71579603-71579873 | -0.38449037 | 0.408032071 | 0.688973365 |
| chr3:14114475-14114675 | 0.384999543 | 0.408048351 | 0.688985671 |
| chr7:38411481-38411762 | -0.34789124 | 0.408094476 | 0.689048369 |
| chr17:29252155-29252355 | 0.40976322 | 0.408150232 | 0.689120023 |
| chr20:21084700-21084900 | -0.469460885 | 0.408162062 | 0.689120023 |
| chr3:141275950-141276377 | -0.278819076 | 0.408168025 | 0.689120023 |
| chr21:31117378-31117585 | 0.393739745 | 0.40818043 | 0.689120023 |
| chr21:44858295-44859107 | -0.292948589 | 0.408181883 | 0.689120023 |
| chr9:91734438-91734718 | -0.364926174 | 0.408195416 | 0.689127686 |
| chr18:47772696-47773003 | -0.346601016 | 0.408227022 | 0.68916586 |
| chr5:126777510-126777903 | -0.285771079 | 0.408249153 | 0.689188037 |
| chr7:100691132-100691361 | 0.363494338 | 0.408273355 | 0.689213709 |
| chr14:58152540-58152813 | 0.34354036 | 0.40828913 | 0.6892243 |
| chr20:61120675-61120952 | 0.329842536 | 0.40829762 | 0.6892243 |
| chr8:130442905-130444367 | -0.265441239 | 0.408349655 | 0.689296953 |
| chr6:2932156-2932735 | -0.296086545 | 0.408386057 | 0.689324027 |
| chr11:66880890-66881156 | 0.315303145 | 0.408390943 | 0.689324027 |
| chr10:14592190-14592501 | -0.36786073 | 0.408392684 | 0.689324027 |
| chr5:6837544-6837834 | 0.40883937 | 0.408418366 | 0.689352191 |
| chr4:77599474-77599694 | 0.375672434 | 0.408435967 | 0.689366712 |
| chr9:68999686-68999960 | -0.377811136 | 0.408475794 | 0.689414139 |

| | | | |
|---|---|---|---|
| chr6:27867427-27867717 | -0.322534867 | 0.408494128 | 0.689414139 |
| chr2:178306437-178306711 | -0.357231945 | 0.408499497 | 0.689414139 |
| chr1:152167551-152167772 | 0.392692305 | 0.40850355 | 0.689414139 |
| chr1:16217231-16217769 | 0.279217379 | 0.408509054 | 0.689414139 |
| chr11:69148416-69148715 | -0.388474344 | 0.408532927 | 0.689427598 |
| chr15:101015315-101015670 | 0.329076812 | 0.408535025 | 0.689427598 |
| chr1:154627033-154628139 | 0.272386313 | 0.408577008 | 0.689475484 |
| chr2:61895659-61895887 | -0.408700754 | 0.408581397 | 0.689475484 |
| chr12:32672002-32672202 | 0.475133584 | 0.408596105 | 0.689484669 |
| chr1:202888361-202889435 | -0.260388175 | 0.408604837 | 0.689484669 |
| chr6:149546882-149547375 | -0.281007371 | 0.408691683 | 0.689615161 |
| chr1:156625734-156626123 | 0.322866503 | 0.408702485 | 0.689615161 |
| chr4:102344356-102345995 | -0.264126479 | 0.408709171 | 0.689615161 |
| chr14:51988977-51990265 | 0.265866173 | 0.408736756 | 0.689646519 |
| chr1:180188179-180188515 | 0.294459272 | 0.4088065 | 0.689736002 |
| chr19:45404967-45406193 | 0.265936372 | 0.408807794 | 0.689736002 |
| chr6:110646844-110647178 | -0.377155403 | 0.408829123 | 0.689748511 |
| chr5:180097433-180097743 | 0.337508552 | 0.408833212 | 0.689748511 |
| chr11:130894478-130894744 | -0.382486951 | 0.408927127 | 0.689877446 |
| chr2:230848002-230848272 | -0.406655991 | 0.408927643 | 0.689877446 |
| chr3:142856956-142857402 | -0.281592119 | 0.408966241 | 0.689927372 |
| chr6:161831568-161831956 | -0.416140752 | 0.408982684 | 0.689939921 |
| chr12:6735691-6735912 | 0.394465265 | 0.408993644 | 0.68994322 |
| chr2:11418056-11418307 | -0.432328482 | 0.409010605 | 0.689952396 |
| chr3:36707744-36707955 | -0.403764116 | 0.409017386 | 0.689952396 |
| chr2:169324647-169324894 | -0.423587663 | 0.409031813 | 0.689952396 |
| chr6:99119508-99119734 | -0.3666531 | 0.409035102 | 0.689952396 |
| chr9:12784414-12785785 | -0.263812381 | 0.409063631 | 0.689972995 |
| chr19:55631758-55631958 | 0.392475604 | 0.409065324 | 0.689972995 |
| chr15:77959391-77959790 | 0.303677476 | 0.409097293 | 0.690011728 |
| chr19:53898799-53899356 | 0.291970573 | 0.409120029 | 0.690018455 |
| chr3:164162781-164163202 | -0.368380704 | 0.409120084 | 0.690018455 |
| chr1:195180548-195180798 | -0.383239497 | 0.409128298 | 0.690018455 |
| chr20:33810964-33811514 | 0.280837662 | 0.409168006 | 0.690062662 |
| chr14:104784176-104784410 | -0.356073148 | 0.409175565 | 0.690062662 |
| chr16:33589221-33589475 | 0.383952612 | 0.409181527 | 0.690062662 |
| chr1:39720092-39720357 | -0.376839952 | 0.409216755 | 0.690103246 |
| chr5:18527076-18527349 | -0.369828998 | 0.409223605 | 0.690103246 |
| chr10:84026427-84026802 | -0.385429475 | 0.409254971 | 0.690137714 |
| chr2:236449183-236449705 | -0.31628974 | 0.409262059 | 0.690137714 |
| chr20:58246409-58246711 | 0.307016766 | 0.409281998 | 0.690156149 |
| chr20:62571827-62572051 | -0.361625401 | 0.409309634 | 0.690176812 |
| chr15:78113270-78113702 | 0.307759906 | 0.409321755 | 0.690176812 |
| chr1:247063955-247064155 | -0.385269548 | 0.409327647 | 0.690176812 |
| chr22:38056953-38057617 | 0.286642854 | 0.409330282 | 0.690176812 |
| chr1:55785052-55785348 | 0.370784959 | 0.409339297 | 0.690176824 |
| chr17:81702812-81703724 | 0.273963236 | 0.409357401 | 0.690192162 |
| chr6:33248323-33248974 | -0.286900261 | 0.409408915 | 0.690257871 |
| chr12:13098143-13098436 | -0.327930112 | 0.409414391 | 0.690257871 |
| chr21:15192286-15192602 | 0.372126307 | 0.409434814 | 0.690274502 |

| | | | |
|---|---|---|---|
| chr3:194685661-194686432 | 0.276936769 | 0.409455065 | 0.690274502 |
| chr2:195318602-195318802 | 0.428992809 | 0.409455398 | 0.690274502 |
| chr6:11172493-11172997 | -0.326536514 | 0.409462991 | 0.690274502 |
| chr20:63682313-63682935 | -0.270873265 | 0.409469299 | 0.690274502 |
| chr18:12271550-12272168 | 0.285588799 | 0.409528886 | 0.69035321 |
| chr2:99489487-99489761 | 0.320600897 | 0.409540479 | 0.69035321 |
| chr1:110176392-110176592 | 0.388769144 | 0.409543018 | 0.69035321 |
| chr1:205284429-205285119 | -0.277760884 | 0.409582284 | 0.690375821 |
| chr14:93234723-93235117 | -0.324037851 | 0.409587525 | 0.690375821 |
| chr1:62783528-62784314 | -0.267252782 | 0.409592472 | 0.690375821 |
| chr11:67667928-67668150 | 0.362793304 | 0.409592473 | 0.690375821 |
| chr6:94144782-94145190 | -0.31511938 | 0.409604626 | 0.690381119 |
| chr16:85689434-85689964 | 0.285426519 | 0.409615769 | 0.690384713 |
| chr8:109644501-109645642 | -0.261077408 | 0.409652883 | 0.690432079 |
| chr1:89523989-89525741 | -0.262925797 | 0.409701459 | 0.690498761 |
| chr15:95826850-95827206 | 0.329313205 | 0.40971065 | 0.690499064 |
| chr4:117084884-117085847 | -0.272803884 | 0.409743793 | 0.690534921 |
| chr6:29848058-29848315 | 0.355079845 | 0.409752572 | 0.690534921 |
| chr8:90986023-90986277 | 0.405884899 | 0.409763196 | 0.690534921 |
| chr12:64672359-64672790 | -0.288216625 | 0.409775918 | 0.690534921 |
| chr3:23320719-23321573 | 0.274089086 | 0.409780298 | 0.690534921 |
| chr15:82680084-82680924 | 0.269792921 | 0.409786 | 0.690534921 |
| chr3:139588443-139588805 | 0.333776301 | 0.409813103 | 0.690552845 |
| chr8:128088316-128089020 | -0.269137173 | 0.409814661 | 0.690552845 |
| chr1:54794757-54795296 | 0.303179346 | 0.409846032 | 0.690590518 |
| chr2:47675126-47675454 | 0.364733368 | 0.409858687 | 0.690596655 |
| chr11:73671535-73671848 | 0.347811659 | 0.409872188 | 0.690604217 |
| chr9:35870314-35870514 | -0.480201173 | 0.40990309 | 0.690639175 |
| chr12:106986698-106987391 | -0.269846106 | 0.409922601 | 0.690639175 |
| chr15:84654742-84655021 | 0.294303709 | 0.409928505 | 0.690639175 |
| chr2:27033250-27033499 | 0.347736086 | 0.40992899 | 0.690639175 |
| chr11:83250112-83250321 | -0.389720989 | 0.409945677 | 0.690652104 |
| chr8:91891262-91891991 | 0.289919874 | 0.409967956 | 0.690674451 |
| chr19:32693522-32693722 | 0.38193776 | 0.409977677 | 0.690675642 |
| chr22:19517046-19517246 | -0.421744795 | 0.409987187 | 0.690676478 |
| chr16:58661242-58661516 | 0.340694701 | 0.410001483 | 0.690685375 |
| chr1:94993843-94994557 | -0.300024402 | 0.410053067 | 0.690743113 |
| chr7:795045-795326 | -0.404171104 | 0.410061743 | 0.690743113 |
| chr1:169550294-169550643 | 0.309150487 | 0.41007086 | 0.690743113 |
| chr17:15944918-15945550 | -0.309566543 | 0.410071816 | 0.690743113 |
| chr1:224780880-224781193 | 0.37417836 | 0.410172054 | 0.69088404 |
| chr1:161972143-161972448 | 0.362505112 | 0.410173514 | 0.69088404 |
| chr8:47512368-47512773 | 0.304874592 | 0.410211724 | 0.690933212 |
| chr6:89917841-89918169 | -0.380434938 | 0.410267606 | 0.691012145 |
| chr11:129704572-129704990 | -0.38329591 | 0.410314094 | 0.691075254 |
| chr15:63945317-63946266 | -0.269555767 | 0.410353428 | 0.691124971 |
| chr17:62680492-62680788 | -0.318674194 | 0.410367798 | 0.691124971 |
| chr8:107671619-107672048 | 0.301592883 | 0.410370673 | 0.691124971 |
| chr13:50123853-50125081 | -0.261578413 | 0.410390143 | 0.69114257 |
| chr2:113407676-113408151 | -0.309567839 | 0.410417676 | 0.6911662 |

| | | | |
|---|---|---|---|
| chr17:68196501-68197590 | 0.266217856 | 0.410422215 | 0.6911662 |
| chr11:86041929-86042261 | 0.329344596 | 0.410473997 | 0.691233778 |
| chr4:116180403-116180766 | 0.368775414 | 0.410480386 | 0.691233778 |
| chr4:6280369-6280873 | -0.304289895 | 0.410498217 | 0.691248613 |
| chr8:47961186-47961606 | 0.286378821 | 0.410530547 | 0.691263561 |
| chr1:243472126-243472358 | -0.44632618 | 0.410542441 | 0.691263561 |
| chrX:14528669-14529010 | 0.349780253 | 0.410550577 | 0.691263561 |
| chr9:98172607-98173302 | -0.284127592 | 0.410551841 | 0.691263561 |
| chr20:31587596-31587977 | 0.347050037 | 0.410553179 | 0.691263561 |
| chr2:85396698-85397423 | -0.276125624 | 0.410561224 | 0.691263561 |
| chr1:95116690-95117993 | -0.26992958 | 0.410581755 | 0.691282465 |
| chr1:93680716-93681368 | 0.286982898 | 0.410593584 | 0.691282465 |
| chr13:75610525-75610807 | 0.358265193 | 0.410599518 | 0.691282465 |
| chr4:146520886-146521153 | -0.330358074 | 0.410617156 | 0.691283239 |
| chr2:10408341-10408648 | -0.295787057 | 0.410618022 | 0.691283239 |
| chr16:71484053-71484826 | -0.269817756 | 0.41064811 | 0.691311546 |
| chr3:63201057-63201276 | 0.332691841 | 0.41065288 | 0.691311546 |
| chr6:27791245-27791622 | -0.293630723 | 0.410700298 | 0.691367204 |
| chr1:40769833-40770151 | 0.352286732 | 0.410703989 | 0.691367204 |
| chr20:61874664-61874993 | 0.310881517 | 0.410717559 | 0.691374858 |
| chr5:107409166-107409429 | -0.421075362 | 0.410734152 | 0.6913876 |
| chr8:98259949-98260907 | -0.277893726 | 0.410749906 | 0.691398931 |
| chrX:35890773-35891065 | -0.355637813 | 0.410796284 | 0.691461807 |
| chr15:72199755-72200089 | -0.311432721 | 0.410814282 | 0.69147691 |
| chr20:53708294-53708565 | 0.271734304 | 0.41084438 | 0.691497382 |
| chr2:64523966-64525176 | -0.26080427 | 0.410844494 | 0.691497382 |
| chr19:4831523-4831769 | 0.395177013 | 0.410878379 | 0.691518239 |
| chr9:21111002-21111212 | -0.427860051 | 0.410888636 | 0.691518239 |
| chr13:52210927-52211810 | -0.273283195 | 0.410902499 | 0.691518239 |
| chr8:144671431-144671652 | -0.386311992 | 0.410908928 | 0.691518239 |
| chr5:8955555-8955824 | -0.40048553 | 0.410914174 | 0.691518239 |
| chr5:32916661-32916995 | -0.33541124 | 0.410917992 | 0.691518239 |
| chr5:53745974-53746202 | 0.430730726 | 0.410920061 | 0.691518239 |
| chr1:167538701-167538973 | -0.401973159 | 0.410938468 | 0.691534026 |
| chr12:116426142-116426782 | -0.284104242 | 0.410979005 | 0.691587053 |
| chr17:54608727-54608953 | 0.396648013 | 0.411010805 | 0.691625377 |
| chr3:187738886-187739157 | -0.314687891 | 0.411028484 | 0.691639937 |
| chr3:62318414-62319639 | -0.256861859 | 0.411054269 | 0.691654212 |
| chr19:33119185-33119407 | 0.398491643 | 0.411056604 | 0.691654212 |
| chr18:58263041-58263469 | 0.322557226 | 0.411064048 | 0.691654212 |
| chr4:127870274-127870579 | -0.429761794 | 0.411077368 | 0.691661191 |
| chr20:25024699-25024978 | -0.399337687 | 0.41108625 | 0.691661191 |
| chr8:73848520-73848854 | 0.357820922 | 0.411136396 | 0.691720674 |
| chr14:84716562-84716762 | 0.417822219 | 0.411139659 | 0.691720674 |
| chr1:195376186-195376603 | -0.310724175 | 0.411154789 | 0.691730941 |
| chr8:94657035-94657688 | 0.281673425 | 0.411164952 | 0.691732851 |
| chr21:10542349-10542777 | -0.360742789 | 0.411182224 | 0.69174672 |
| chr19:50476093-50476559 | 0.297580397 | 0.411259373 | 0.69185501 |
| chr10:61453330-61453530 | -0.438925653 | 0.411264652 | 0.69185501 |
| chr3:4314781-4314991 | -0.396096729 | 0.411333022 | 0.691944943 |

| | | | |
|---|---|---|---|
| chr12:92145565-92146532 | -0.258584679 | 0.411342189 | 0.691944943 |
| chr1:202926604-202927666 | -0.258873902 | 0.411350439 | 0.691944943 |
| chr8:95285080-95285405 | -0.36236093 | 0.411357672 | 0.691944943 |
| chr5:126525652-126526138 | -0.343710232 | 0.411365636 | 0.691944943 |
| chr17:48585222-48585649 | -0.311426422 | 0.411372295 | 0.691944943 |
| chr6:79310497-79310764 | -0.416265916 | 0.411427111 | 0.692021955 |
| chr1:204320122-204320332 | 0.343389039 | 0.411445231 | 0.69203045 |
| chr14:88189407-88189779 | 0.299854297 | 0.411460409 | 0.69203045 |
| chr20:64120479-64120758 | 0.322293948 | 0.411461044 | 0.69203045 |
| chr16:29738633-29738886 | 0.35018677 | 0.411468289 | 0.69203045 |
| chr1:243859934-243860194 | -0.451220121 | 0.411502662 | 0.69205179 |
| chr14:105005955-105006225 | 0.367886092 | 0.411506002 | 0.69205179 |
| chr7:46938130-46938403 | -0.43833503 | 0.411508074 | 0.69205179 |
| chr8:143733763-143734238 | -0.296979262 | 0.411528388 | 0.692070764 |
| chr3:162865548-162865827 | -0.374294132 | 0.411541544 | 0.692077699 |
| chr18:50967844-50968603 | 0.285272674 | 0.411569965 | 0.692110303 |
| chr13:26698520-26699029 | -0.295665097 | 0.411592557 | 0.692133104 |
| chr12:103626556-103626822 | 0.344848685 | 0.411627322 | 0.692163663 |
| chr3:38164681-38165678 | 0.260540973 | 0.411628797 | 0.692163663 |
| chr12:50581289-50581819 | 0.314104899 | 0.411651817 | 0.692187183 |
| chr17:27970138-27970421 | 0.401595751 | 0.411731996 | 0.692277696 |
| chr1:237851281-237851514 | -0.390430179 | 0.411733371 | 0.692277696 |
| chr3:23115234-23115531 | 0.342463975 | 0.411735372 | 0.692277696 |
| chrX:15674669-15675963 | -0.259407932 | 0.411741786 | 0.692277696 |
| chr14:55271022-55272263 | 0.259820575 | 0.411753622 | 0.692282404 |
| chr2:212148025-212148262 | -0.450366682 | 0.411763077 | 0.69228311 |
| chr4:38728945-38729145 | -0.433636299 | 0.41179722 | 0.69228608 |
| chr1:168569719-168570018 | -0.375911233 | 0.411798859 | 0.69228608 |
| chr18:35386628-35387444 | -0.28726437 | 0.411804033 | 0.69228608 |
| chr22:19019310-19019510 | 0.414332003 | 0.411804483 | 0.69228608 |
| chr1:204207101-204207559 | -0.281445426 | 0.411810018 | 0.69228608 |
| chr3:136410681-136410981 | -0.424217479 | 0.411830043 | 0.692304554 |
| chr3:3233309-3233509 | 0.451388213 | 0.411848423 | 0.692319573 |
| chr20:5119320-5120423 | 0.264889007 | 0.411857049 | 0.692319573 |
| chr3:110314590-110314961 | -0.323692953 | 0.41186678 | 0.692320743 |
| chr4:17810218-17811467 | 0.260750998 | 0.411902454 | 0.692365519 |
| chr16:57535911-57536665 | 0.281407053 | 0.411930003 | 0.692396637 |
| chr8:29286709-29287346 | 0.284986354 | 0.41194996 | 0.692414991 |
| chr14:105440288-105440758 | 0.305175806 | 0.411995401 | 0.69246911 |
| chr5:166594602-166594914 | 0.331056513 | 0.412000232 | 0.69246911 |
| chr16:89565563-89566363 | -0.268397616 | 0.412031286 | 0.692506114 |
| chr12:14567740-14568043 | -0.345477755 | 0.41204422 | 0.692507567 |
| chr1:226806991-226807309 | -0.334235976 | 0.412050227 | 0.692507567 |
| chr19:47289408-47290052 | -0.293921757 | 0.412060018 | 0.692508833 |
| chr14:70074010-70074245 | 0.375503147 | 0.412079301 | 0.692512295 |
| chr2:48314096-48315145 | -0.258811396 | 0.412080154 | 0.692512295 |
| chr4:766042-766380 | -0.372862739 | 0.412092172 | 0.692517302 |
| chr14:100150224-100150545 | 0.32914188 | 0.412171179 | 0.69262007 |
| chr1:46442173-46442933 | 0.277222047 | 0.412171404 | 0.69262007 |
| chr3:98732121-98733393 | 0.268157544 | 0.412215942 | 0.69267972 |

| | | | |
|---|---|---|---|
| chr16:30844781-30845278 | 0.316375172 | 0.4122259 | 0.692681262 |
| chr7:32204370-32204766 | -0.278288174 | 0.412238123 | 0.692686611 |
| chr11:104352011-104352353 | -0.348128191 | 0.41226968 | 0.692691869 |
| chr20:41213949-41214149 | 0.410855269 | 0.412275599 | 0.692691869 |
| chr4:76681000-76681200 | 0.410855269 | 0.412275599 | 0.692691869 |
| chr9:137436468-137436727 | 0.382611122 | 0.412277414 | 0.692691869 |
| chr18:50297740-50298179 | -0.313301604 | 0.412291917 | 0.692699749 |
| chr1:167454792-167455371 | -0.3043765 | 0.412304427 | 0.692699749 |
| chr1:116432707-116433598 | 0.279592557 | 0.412309225 | 0.692699749 |
| chr13:113759071-113759682 | 0.294932696 | 0.412330408 | 0.692720148 |
| chr6:18434460-18434774 | -0.372615784 | 0.412347427 | 0.692733551 |
| chr19:35114633-35115195 | -0.289994961 | 0.412358188 | 0.69273644 |
| chr12:80919853-80920062 | -0.424461844 | 0.412435962 | 0.692828405 |
| chr11:70203977-70204345 | 0.320294938 | 0.412438416 | 0.692828405 |
| chr2:3398905-3399158 | -0.376595567 | 0.412444051 | 0.692828405 |
| chr10:109773616-109773854 | -0.445109315 | 0.412453023 | 0.692828405 |
| chr14:66624333-66624686 | -0.315092697 | 0.412472372 | 0.692828405 |
| chr4:122652435-122652645 | 0.416983066 | 0.412479041 | 0.692828405 |
| chr12:131960144-131960358 | 0.446269139 | 0.412481694 | 0.692828405 |
| chr12:56119437-56119690 | 0.298962426 | 0.412485268 | 0.692828405 |
| chr6:84119362-84119576 | -0.428547615 | 0.412495689 | 0.692830722 |
| chr12:131736917-131737436 | 0.301261338 | 0.412514514 | 0.692839873 |
| chr16:138507-139077 | -0.285372785 | 0.412519222 | 0.692839873 |
| chr22:36066209-36066483 | 0.353792936 | 0.412536622 | 0.692853909 |
| chr5:111309027-111309230 | -0.4523839 | 0.412580081 | 0.692911711 |
| chr9:6703550-6704705 | -0.264267404 | 0.412599866 | 0.69292975 |
| chr14:24141112-24142036 | 0.260186448 | 0.412619982 | 0.692948346 |
| chr2:231601712-231601951 | 0.331768594 | 0.412633355 | 0.692955616 |
| chr9:137070270-137071000 | 0.273409483 | 0.412651778 | 0.692968014 |
| chr12:56443248-56443540 | -0.403336547 | 0.412683141 | 0.692968014 |
| chr12:103642600-103642845 | 0.408536797 | 0.412691093 | 0.692968014 |
| chr1:61143204-61143404 | 0.382563488 | 0.41270963 | 0.692968014 |
| chr12:81451044-81451657 | 0.29291042 | 0.412711203 | 0.692968014 |
| chr1:247525496-247526052 | -0.331814636 | 0.412714686 | 0.692968014 |
| chr10:5094279-5094956 | 0.292481947 | 0.412723682 | 0.692968014 |
| chr18:47485263-47485484 | 0.441281513 | 0.41272469 | 0.692968014 |
| chr7:46260161-46260506 | 0.363222388 | 0.412730621 | 0.692968014 |
| chr18:36017729-36017966 | -0.416571979 | 0.412736529 | 0.692968014 |
| chr1:106193006-106193766 | 0.275186803 | 0.412740221 | 0.692968014 |
| chr1:164601875-164602369 | 0.279100527 | 0.412756066 | 0.692979431 |
| chr15:75347174-75347864 | 0.284817019 | 0.412767485 | 0.692983418 |
| chr10:11742674-11742876 | -0.399075625 | 0.412785301 | 0.692998145 |
| chr12:79013071-79013410 | 0.359858821 | 0.412809972 | 0.693024378 |
| chr14:98906545-98906826 | -0.331515362 | 0.412853773 | 0.693082726 |
| chr14:47442231-47442565 | 0.314392093 | 0.412943895 | 0.693190411 |
| chr21:33901799-33902062 | -0.370527136 | 0.412944602 | 0.693190411 |
| chr3:161315975-161316221 | 0.332777049 | 0.412945059 | 0.693190411 |
| chr15:90351797-90352944 | 0.264108017 | 0.412955215 | 0.693192272 |
| chr5:176613869-176614230 | -0.300535541 | 0.413014311 | 0.693276285 |
| chr1:248750430-248750736 | -0.337156539 | 0.41305343 | 0.693326759 |

| | | | |
|---|---|---|---|
| chr1:189021816-189022172 | 0.377022745 | 0.41307508 | 0.693347911 |
| chr15:42385519-42385781 | 0.359533032 | 0.413094495 | 0.693362427 |
| chr5:16160794-16161103 | 0.318990186 | 0.413101827 | 0.693362427 |
| chr9:100505537-100505834 | -0.371980047 | 0.41311381 | 0.693367353 |
| chr3:53254857-53255213 | 0.299967671 | 0.413124041 | 0.693369335 |
| chr5:36777135-36777647 | -0.306478984 | 0.4131495 | 0.693396876 |
| chr14:40221430-40221831 | 0.290808551 | 0.413179475 | 0.693431995 |
| chr3:58217186-58217502 | -0.325918195 | 0.413202848 | 0.693456033 |
| chr1:43991267-43992282 | 0.266649689 | 0.413219378 | 0.693457261 |
| chr12:56957307-56957786 | 0.310554837 | 0.413221681 | 0.693457261 |
| chr3:61894872-61895356 | 0.277330629 | 0.413253265 | 0.69348143 |
| chr3:27720805-27722565 | 0.257258905 | 0.413254184 | 0.69348143 |
| chr20:58209628-58210300 | 0.271037274 | 0.413293503 | 0.693524096 |
| chr6:152091881-152092226 | -0.325869447 | 0.413298511 | 0.693524096 |
| chr19:48720556-48721308 | -0.269115888 | 0.413306763 | 0.693524096 |
| chr10:96422319-96422632 | -0.393056103 | 0.413316273 | 0.693524866 |
| chr2:138493290-138493710 | -0.293848945 | 0.413334342 | 0.693539997 |
| chr4:92091400-92091624 | 0.392649009 | 0.413362486 | 0.693547323 |
| chr3:156448373-156448634 | -0.343332533 | 0.413365215 | 0.693547323 |
| chr12:105330242-105330969 | -0.276203478 | 0.413365863 | 0.693547323 |
| chrX:7121402-7121778 | -0.402822525 | 0.413385358 | 0.693560543 |
| chr5:138501577-138501872 | 0.352314673 | 0.413391846 | 0.693560543 |
| chr14:103141041-103141485 | 0.311278506 | 0.413408757 | 0.693573729 |
| chr5:113824399-113824600 | 0.396632019 | 0.413451665 | 0.693622013 |
| chr14:99682819-99683243 | 0.28924585 | 0.413459806 | 0.693622013 |
| chr1:242651880-242652114 | 0.407158624 | 0.413464694 | 0.693622013 |
| chr6:111210316-111210644 | -0.337731864 | 0.413478766 | 0.693630433 |
| chr3:124886085-124887783 | 0.26159888 | 0.413545989 | 0.693728014 |
| chr5:113804384-113804588 | -0.423471974 | 0.413560246 | 0.693736743 |
| chr2:157336276-157336477 | -0.419487734 | 0.413584979 | 0.693750429 |
| chrX:24748102-24748777 | -0.298177917 | 0.413586513 | 0.693750429 |
| chr19:47012887-47013488 | 0.287521605 | 0.413626008 | 0.693801489 |
| chr11:59102175-59102498 | -0.323253646 | 0.413648884 | 0.693811596 |
| chr11:15101184-15101414 | -0.408860611 | 0.413656629 | 0.693811596 |
| chr5:179732155-179732364 | -0.348844718 | 0.413667967 | 0.693811596 |
| chr3:141474265-141474523 | -0.36203385 | 0.413677485 | 0.693811596 |
| chr6:144560090-144560445 | -0.363043833 | 0.413680289 | 0.693811596 |
| chr12:108856602-108858219 | 0.262262223 | 0.413686364 | 0.693811596 |
| chr13:21998395-21998644 | 0.385698606 | 0.413698934 | 0.693817492 |
| chr1:156383017-156383217 | 0.355997604 | 0.413771542 | 0.693924073 |
| chr14:91361270-91361735 | 0.310357977 | 0.41378609 | 0.693928074 |
| chr15:81106500-81106727 | 0.371293197 | 0.41379204 | 0.693928074 |
| chr2:212659573-212659864 | 0.388586784 | 0.413838082 | 0.693953929 |
| chr17:59857126-59857671 | 0.258952844 | 0.413861635 | 0.693953929 |
| chr5:125027211-125027708 | 0.303520252 | 0.413864473 | 0.693953929 |
| chr6:8083388-8083643 | -0.396575366 | 0.413867254 | 0.693953929 |
| chr1:201512760-201513312 | 0.269751197 | 0.413872728 | 0.693953929 |
| chr5:101291529-101291734 | -0.429358347 | 0.413873723 | 0.693953929 |
| chr9:20640433-20640957 | -0.327748779 | 0.413874995 | 0.693953929 |
| chr22:49951028-49951690 | 0.273561152 | 0.413879912 | 0.693953929 |

| | | | |
|---|---|---|---|
| chr17:13002948-13003428 | 0.319640978 | 0.413966536 | 0.694083983 |
| chr11:6418942-6419608 | 0.298819996 | 0.413997301 | 0.694120377 |
| chr20:40874812-40875098 | 0.353633878 | 0.414008383 | 0.694123767 |
| chr5:139787936-139788324 | -0.299838074 | 0.414036014 | 0.694147664 |
| chr17:74493426-74493636 | 0.379984806 | 0.414040754 | 0.694147664 |
| chr14:67067475-67067825 | 0.327441306 | 0.41405498 | 0.694156325 |
| chr19:50369139-50369535 | 0.301828934 | 0.414082908 | 0.694187957 |
| chr5:80082855-80083140 | -0.346103869 | 0.414106609 | 0.694198264 |
| chr8:133120410-133120855 | -0.318137224 | 0.414107176 | 0.694198264 |
| chr6:39377549-39377816 | 0.36837495 | 0.414126562 | 0.694199294 |
| chr10:94677704-94677933 | 0.404325701 | 0.414149443 | 0.694199294 |
| chr2:85243732-85244192 | -0.315537196 | 0.414155747 | 0.694199294 |
| chr1:54122202-54122402 | -0.37427733 | 0.414164634 | 0.694199294 |
| chr13:43016361-43016699 | 0.407314945 | 0.414165807 | 0.694199294 |
| chr11:43944676-43945083 | 0.328465034 | 0.414175558 | 0.694199294 |
| chr1:188689667-188689883 | -0.435643999 | 0.414178739 | 0.694199294 |
| chr7:46281135-46281463 | 0.32725975 | 0.414180271 | 0.694199294 |
| chr6:716274-716474 | -0.443370819 | 0.414194958 | 0.694208725 |
| chr1:201397115-201397315 | 0.324172224 | 0.414226336 | 0.694227853 |
| chr4:123060105-123060357 | -0.361076134 | 0.414226505 | 0.694227853 |
| chr3:101484827-101485205 | -0.380500932 | 0.414240653 | 0.694227853 |
| chr12:6200288-6200691 | 0.308823177 | 0.414242612 | 0.694227853 |
| chr10:101041266-101041484 | 0.329446431 | 0.414275658 | 0.694232599 |
| chr4:94406806-94407048 | -0.380119045 | 0.414286569 | 0.694232599 |
| chr19:42244732-42245011 | 0.302949825 | 0.414286691 | 0.694232599 |
| chr6:135054279-135055422 | -0.260180847 | 0.414295131 | 0.694232599 |
| chr6:159941813-159942013 | -0.427176844 | 0.41429558 | 0.694232599 |
| chr2:12326847-12327171 | -0.340387893 | 0.414306712 | 0.694232599 |
| chr3:189125112-189125332 | -0.412490435 | 0.414308868 | 0.694232599 |
| chr7:16791926-16792201 | 0.365893696 | 0.414348562 | 0.694283929 |
| chr6:98833802-98834972 | -0.260660226 | 0.414441189 | 0.69442395 |
| chr12:58877024-58877626 | -0.301727756 | 0.414474203 | 0.69446408 |
| chr17:79665020-79665680 | -0.287743419 | 0.414490245 | 0.694475772 |
| chr12:48814386-48814900 | 0.281515328 | 0.414502179 | 0.694480582 |
| chr6:100327143-100327352 | -0.424608596 | 0.414523583 | 0.694501257 |
| chr2:130188126-130188329 | 0.475260853 | 0.414539807 | 0.694513253 |
| chr6:37018626-37019025 | 0.297360303 | 0.414580505 | 0.69456625 |
| chr2:7745482-7745742 | -0.365371796 | 0.414621354 | 0.694609715 |
| chr8:88327155-88329098 | 0.254952647 | 0.41462458 | 0.694609715 |
| chr6:8434898-8436159 | -0.262726343 | 0.414639081 | 0.694618822 |
| chr19:5622654-5623225 | 0.273743351 | 0.41467112 | 0.694657307 |
| chr9:120531721-120532036 | 0.317374197 | 0.414752568 | 0.694778559 |
| chr2:11446779-11447005 | -0.357409799 | 0.414797587 | 0.694835557 |
| chr5:39624512-39624859 | 0.300745713 | 0.41480473 | 0.694835557 |
| chr20:59031852-59033076 | 0.256507885 | 0.414884665 | 0.694926821 |
| chr22:20702573-20703440 | 0.262813204 | 0.414886343 | 0.694926821 |
| chr20:47527107-47527874 | 0.259026817 | 0.414893456 | 0.694926821 |
| chr14:91509409-91510985 | 0.258030209 | 0.414895492 | 0.694926821 |
| chr17:17891046-17891314 | -0.358133394 | 0.414913775 | 0.694928609 |
| chr17:50884459-50884882 | 0.315664632 | 0.414914698 | 0.694928609 |

| | | | |
|---|---|---|---|
| chr21:16419193-16419705 | -0.319909922 | 0.414932822 | 0.694934175 |
| chr16:21501905-21502744 | 0.262941272 | 0.4149438 | 0.694934175 |
| chr10:110871492-11087269! | -0.258349455 | 0.41494523 | 0.694934175 |
| chr17:44533501-44533829 | 0.33576881 | 0.414981308 | 0.694979406 |
| chr22:29580740-29581724 | 0.269928737 | 0.414990791 | 0.694980096 |
| chrX:15269610-15270440 | -0.291336171 | 0.415000757 | 0.694981597 |
| chr11:74592215-74593308 | -0.268476543 | 0.415052727 | 0.695049507 |
| chr8:70745571-70745804 | -0.398604645 | 0.415066031 | 0.695049507 |
| chr1:205565878-205566142 | 0.31593576 | 0.415068522 | 0.695049507 |
| chr13:34942990-34943875 | 0.271138161 | 0.415090037 | 0.695070344 |
| chr5:14326325-14326604 | -0.336049583 | 0.415138339 | 0.695136035 |
| chr17:1491101-1491385 | 0.332290671 | 0.415171966 | 0.695177151 |
| chr1:77825347-77825623 | 0.354889281 | 0.415219699 | 0.695241882 |
| chr5:149618759-149619135 | -0.308108305 | 0.415263002 | 0.695299194 |
| chr2:134118318-134118525 | -0.335853053 | 0.415276777 | 0.695307064 |
| chr15:83084975-83085257 | 0.354912486 | 0.415356479 | 0.695425315 |
| chr15:90934927-90935697 | 0.260152664 | 0.415380191 | 0.695443157 |
| chr1:169648859-169649104 | -0.433349926 | 0.415388228 | 0.695443157 |
| chr5:52987812-52988015 | -0.354142654 | 0.415394364 | 0.695443157 |
| chr16:68737033-68737461 | 0.284090437 | 0.415404449 | 0.695444846 |
| chr19:38402816-38403443 | 0.273025314 | 0.415445874 | 0.695499 |
| chr7:100132705-100132969 | -0.358287659 | 0.415464135 | 0.695514375 |
| chr7:21986334-21986534 | 0.413342452 | 0.415504254 | 0.695547319 |
| chr19:48634489-48634800 | 0.288714055 | 0.415517493 | 0.695547319 |
| chr12:110826711-110826992 | 0.347107488 | 0.41552432 | 0.695547319 |
| chr20:56468010-56468829 | -0.260096418 | 0.415526013 | 0.695547319 |
| chr2:143745053-143745253 | 0.415234586 | 0.415529202 | 0.695547319 |
| chr20:62500256-62500494 | 0.320348581 | 0.415569677 | 0.695599873 |
| chr12:2876498-2877867 | 0.268573276 | 0.415584163 | 0.695608925 |
| chrX:110944340-110945516 | 0.269671605 | 0.415614001 | 0.695631278 |
| chr1:23531244-23531456 | 0.367238662 | 0.415621616 | 0.695631278 |
| chr3:155996001-155996378 | -0.290961898 | 0.415624754 | 0.695631278 |
| chr19:45779250-45780004 | 0.269329134 | 0.415645219 | 0.695650335 |
| chr15:67335285-67335498 | -0.443614629 | 0.415657473 | 0.695655649 |
| chr20:47740678-47741415 | 0.256038284 | 0.415709128 | 0.695726903 |
| chrX:48631414-48631768 | 0.319800621 | 0.415728861 | 0.695744732 |
| chr12:78000121-78000350 | -0.376847849 | 0.415768477 | 0.695795833 |
| chr12:97308692-97308892 | -0.419901698 | 0.41579056 | 0.695817593 |
| chr9:127825428-127825693 | 0.375282067 | 0.415861106 | 0.69592045 |
| chr3:131502507-131503210 | -0.267609422 | 0.415929672 | 0.696003758 |
| chr21:37551741-37552120 | -0.369909315 | 0.415944248 | 0.696003758 |
| chr10:31935356-31935564 | -0.441239507 | 0.415948166 | 0.696003758 |
| chr10:80924374-80924665 | -0.344352732 | 0.415955256 | 0.696003758 |
| chr11:94973228-94973990 | -0.279519968 | 0.415957301 | 0.696003758 |
| chr1:239879241-239879815 | -0.285007922 | 0.415973605 | 0.696003758 |
| chr14:37583239-37584519 | 0.255591902 | 0.415974473 | 0.696003758 |
| chr1:27914177-27915152 | 0.266784687 | 0.415999155 | 0.696029856 |
| chr11:117819148-117819383 | 0.330551691 | 0.416055141 | 0.696108329 |
| chr6:21589299-21589499 | -0.344634402 | 0.416085074 | 0.696136232 |
| chr17:51298731-51299081 | 0.31226487 | 0.416089989 | 0.696136232 |

| | | | |
|---|---|---|---|
| chr7:30800776-30800995 | 0.381350995 | 0.416160403 | 0.696238835 |
| chr5:178515010-178515369 | 0.351503368 | 0.416192296 | 0.696275614 |
| chr14:64760237-64760437 | 0.333555092 | 0.416200561 | 0.696275614 |
| chr11:95849165-95849365 | 0.4095177 | 0.416236942 | 0.696321273 |
| chr3:42819450-42819795 | -0.367452809 | 0.416246186 | 0.696321535 |
| chr6:90476852-90477052 | -0.419053427 | 0.416255495 | 0.696321905 |
| chr22:41001736-41002089 | 0.358125006 | 0.416353386 | 0.69647012 |
| chr17:72479219-72479488 | 0.340580974 | 0.416362276 | 0.69647012 |
| chr10:125522039-125522483 | -0.299241188 | 0.416386118 | 0.696494796 |
| chr3:71724512-71725174 | 0.286574446 | 0.416396609 | 0.69649714 |
| chr1:33634837-33635187 | 0.321106236 | 0.416425322 | 0.696509475 |
| chr20:51161555-51161830 | 0.339051998 | 0.416427579 | 0.696509475 |
| chr6:7173867-7174233 | -0.361296833 | 0.416434882 | 0.696509475 |
| chr3:160956845-160957139 | -0.345639703 | 0.416451443 | 0.696509475 |
| chr3:86695810-86696202 | 0.321173647 | 0.416457875 | 0.696509475 |
| chr10:103069648-103069848 | 0.419964755 | 0.416458525 | 0.696509475 |
| chr8:102537618-102537858 | 0.338310521 | 0.416503905 | 0.696570167 |
| chr13:102724890-102725138 | -0.397000336 | 0.416531902 | 0.696591042 |
| chr3:122754909-122755237 | -0.320362534 | 0.416544856 | 0.696591042 |
| chr5:142548731-142549126 | -0.310022461 | 0.416548747 | 0.696591042 |
| chrX:106220314-106220514 | -0.456840846 | 0.416552752 | 0.696591042 |
| chr17:57892308-57893273 | -0.26564422 | 0.416587994 | 0.696634773 |
| chr3:64225146-64225617 | 0.269056953 | 0.416604767 | 0.696645039 |
| chr12:54009121-54009546 | -0.316888906 | 0.416612811 | 0.696645039 |
| chr18:6683153-6683768 | -0.290987383 | 0.416622714 | 0.696645039 |
| chrX:129427337-129427577 | 0.363995836 | 0.416630501 | 0.696645039 |
| chr21:26169116-26169383 | -0.327685705 | 0.416671285 | 0.696685974 |
| chr4:113979286-113980050 | 0.280690886 | 0.416673168 | 0.696685974 |
| chr8:130802631-130802919 | -0.33450969 | 0.416692482 | 0.696703065 |
| chr9:111598838-111599921 | -0.271631784 | 0.416723805 | 0.696740234 |
| chr4:146175568-146176507 | -0.266973748 | 0.416797987 | 0.696836619 |
| chr3:109055513-109055818 | 0.379047447 | 0.416799643 | 0.696836619 |
| chr9:34048743-34049664 | -0.260237917 | 0.416846235 | 0.696897589 |
| chr2:108989072-108989436 | -0.327084268 | 0.416854301 | 0.696897589 |
| chr1:146560887-146561154 | -0.399347689 | 0.416893964 | 0.69694869 |
| chr19:18418840-18419483 | 0.284992331 | 0.41690798 | 0.696956915 |
| chr3:100657427-100657664 | -0.426955706 | 0.416917518 | 0.696957654 |
| chr1:3454260-3454621 | 0.311622302 | 0.416933451 | 0.696962679 |
| chr10:98423700-98424021 | -0.296942691 | 0.416938716 | 0.696962679 |
| chr2:73385442-73386700 | -0.257774465 | 0.416961922 | 0.696986265 |
| chr16:23219069-23219310 | -0.424353594 | 0.416976254 | 0.696995016 |
| chr14:35818430-35818630 | 0.357672782 | 0.416990061 | 0.697002889 |
| chr10:69757573-69758149 | -0.298831516 | 0.416999903 | 0.697004136 |
| chr20:63973674-63974144 | -0.291916169 | 0.417018978 | 0.697020813 |
| chr17:4909098-4909407 | 0.345075465 | 0.417031469 | 0.697023866 |
| chr9:124415126-124415769 | 0.269384793 | 0.417040655 | 0.697023866 |
| chr19:35869690-35870229 | 0.282893052 | 0.417048095 | 0.697023866 |
| chr16:15950368-15950691 | 0.323668534 | 0.417058131 | 0.697025436 |
| chr6:21360495-21360826 | 0.341628215 | 0.417096946 | 0.697075102 |
| chr10:50470629-50471514 | -0.269772967 | 0.417128866 | 0.697100105 |

| | | | |
|---|---|---|---|
| chr2:38532832-38533032 | 0.38419724 | 0.417130102 | 0.697100105 |
| chr17:77270020-77270337 | 0.322046276 | 0.417158604 | 0.697108632 |
| chr1:210239859-210240123 | -0.363471037 | 0.41716136 | 0.697108632 |
| chr8:103796725-103797055 | -0.316368201 | 0.417162499 | 0.697108632 |
| chr8:133000097-133000416 | 0.326856582 | 0.417175324 | 0.69711486 |
| chr1:156082338-156082696 | 0.298439838 | 0.41720119 | 0.69711489 |
| chr10:4725980-4726326 | 0.357530423 | 0.417216606 | 0.69711489 |
| chr10:80253732-80254249 | -0.298413116 | 0.417218034 | 0.69711489 |
| chr9:96853634-96854758 | 0.263973107 | 0.417220406 | 0.69711489 |
| chr17:18260250-18261052 | -0.268008276 | 0.417223273 | 0.69711489 |
| chr20:42376296-42376676 | 0.292859812 | 0.41722993 | 0.69711489 |
| chr2:206989242-206989703 | -0.304682473 | 0.417257513 | 0.697145774 |
| chr3:87350511-87350954 | 0.31607772 | 0.417277324 | 0.697148995 |
| chr3:4271919-4272186 | -0.346195986 | 0.417279083 | 0.697148995 |
| chr5:177111715-177112181 | -0.296375474 | 0.417286737 | 0.697148995 |
| chr14:97872972-97873780 | -0.285142736 | 0.417301326 | 0.697155096 |
| chr9:92293525-92294082 | 0.273940833 | 0.417323173 | 0.697155096 |
| chr14:23006619-23007433 | 0.261481234 | 0.41732439 | 0.697155096 |
| chr8:102653617-102654586 | 0.255675338 | 0.417326783 | 0.697155096 |
| chr10:100454452-100455052 | -0.297709439 | 0.417343104 | 0.69716716 |
| chr19:54102589-54102996 | 0.285998698 | 0.417363869 | 0.697174842 |
| chr15:63599600-63601273 | 0.257947444 | 0.417388798 | 0.697174842 |
| chr6:28488925-28489176 | -0.359304216 | 0.41740021 | 0.697174842 |
| chr17:20801658-20801875 | 0.369762962 | 0.417408793 | 0.697174842 |
| chr2:175167556-175168849 | -0.254741293 | 0.417413925 | 0.697174842 |
| chr17:82424728-82424928 | 0.378806344 | 0.417418935 | 0.697174842 |
| chr9:94668909-94669109 | 0.403750538 | 0.417420654 | 0.697174842 |
| chr15:101217512-101217961 | 0.305209071 | 0.417424738 | 0.697174842 |
| chr17:44065918-44066413 | 0.295617454 | 0.417429592 | 0.697174842 |
| chr8:109867616-109867816 | 0.407559627 | 0.417478548 | 0.697236602 |
| chr22:20858272-20859113 | 0.264138016 | 0.41748477 | 0.697236602 |
| chr12:132761847-132762581 | 0.274316593 | 0.417537105 | 0.697308806 |
| chr2:209977729-209977949 | -0.405918541 | 0.417581024 | 0.697366954 |
| chr7:73832141-73832487 | -0.289160077 | 0.417592679 | 0.697371219 |
| chr20:41650301-41650962 | 0.264569643 | 0.417631456 | 0.697420775 |
| chr12:121203569-121203811 | 0.393294113 | 0.417672606 | 0.697474293 |
| chr8:68331056-68331681 | -0.28029958 | 0.417721849 | 0.697503539 |
| chr4:54227890-54228334 | -0.320760919 | 0.417728105 | 0.697503539 |
| chr14:58151367-58151636 | 0.351919096 | 0.417745035 | 0.697503539 |
| chr10:35192854-35193091 | -0.375180515 | 0.417750776 | 0.697503539 |
| chr19:14405903-14406528 | 0.272806547 | 0.417754894 | 0.697503539 |
| chr20:62143626-62143979 | 0.296346155 | 0.417759938 | 0.697503539 |
| chr11:112175143-112175657 | -0.303269578 | 0.417760835 | 0.697503539 |
| chr19:48390589-48390983 | 0.305929181 | 0.417762946 | 0.697503539 |
| chr8:115667538-115667888 | -0.30375602 | 0.417776896 | 0.697511631 |
| chr5:149114580-149115016 | -0.303561467 | 0.417812502 | 0.69755588 |
| chr19:16496025-16496705 | 0.278619793 | 0.417861702 | 0.69762282 |
| chr2:197803034-197803273 | 0.341928913 | 0.417872223 | 0.697625185 |
| chr3:15695078-15695384 | -0.429830058 | 0.417891555 | 0.697632147 |
| chr19:48675391-48675673 | 0.335705911 | 0.417900103 | 0.697632147 |

| | | | |
|---|---|---|---|
| chr2:172693128-172693401 | -0.373431357 | 0.417903708 | 0.697632147 |
| chr16:89873425-89874681 | -0.255874409 | 0.417920739 | 0.697645378 |
| chr7:77811832-77812036 | 0.440817979 | 0.417937044 | 0.697657397 |
| chr14:64764668-64765076 | 0.286186155 | 0.41796083 | 0.697681903 |
| chr4:47945917-47946180 | -0.421352889 | 0.417990616 | 0.697701223 |
| chr6:49542655-49542855 | -0.421352889 | 0.417990616 | 0.697701223 |
| chr7:6092903-6093134 | -0.36996533 | 0.418002818 | 0.697706392 |
| chr10:101272085-101272505 | -0.340483242 | 0.41801572 | 0.697709649 |
| chr12:39619734-39620200 | -0.314611338 | 0.418022981 | 0.697709649 |
| chr17:70717852-70718073 | -0.402378764 | 0.418039341 | 0.697713264 |
| chr13:20924900-20925136 | 0.39901278 | 0.418046087 | 0.697713264 |
| chr19:54126154-54126363 | -0.420286185 | 0.418052464 | 0.697713264 |
| chr2:514046-514246 | -0.457292799 | 0.418062605 | 0.697714991 |
| chr11:72782555-72782802 | -0.308261772 | 0.418095679 | 0.69775499 |
| chr1:200033458-200033754 | -0.331945168 | 0.418108159 | 0.697758486 |
| chr14:87491915-87492230 | -0.322807719 | 0.418115987 | 0.697758486 |
| chr6:56314767-56315003 | -0.395848875 | 0.418169895 | 0.697783147 |
| chr19:51394018-51394723 | 0.271196781 | 0.418172424 | 0.697783147 |
| chr2:182715567-182716982 | -0.256002346 | 0.418180471 | 0.697783147 |
| chr2:188028696-188028940 | -0.399395518 | 0.418187175 | 0.697783147 |
| chr1:1300343-1300609 | -0.387910482 | 0.418193211 | 0.697783147 |
| chr12:116673677-116673888 | 0.402929994 | 0.418196663 | 0.697783147 |
| chr1:153703461-153703661 | 0.421061159 | 0.418201275 | 0.697783147 |
| chr4:119627598-119628191 | 0.278118698 | 0.418205438 | 0.697783147 |
| chr12:48818938-48819268 | 0.300609487 | 0.418214118 | 0.697783147 |
| chr2:19312995-19313407 | -0.356178517 | 0.418221832 | 0.697783147 |
| chr16:85569823-85571297 | -0.258424923 | 0.418254825 | 0.697816754 |
| chr12:14802935-14804045 | -0.260263937 | 0.418260189 | 0.697816754 |
| chr6:27687474-27687982 | -0.283272079 | 0.418293417 | 0.697856995 |
| chr20:60263044-60263331 | -0.314335947 | 0.418349259 | 0.697934961 |
| chr1:37487237-37487614 | 0.30485862 | 0.418404137 | 0.698011317 |
| chr17:60148352-60148755 | 0.263867094 | 0.418442364 | 0.698031277 |
| chr12:118979876-118980148 | -0.384782416 | 0.418452984 | 0.698031277 |
| chr9:121320090-121320423 | 0.306976739 | 0.41846116 | 0.698031277 |
| chr2:3635089-3635327 | 0.397786308 | 0.418463003 | 0.698031277 |
| chr2:168348111-168348381 | -0.347745509 | 0.418475508 | 0.698031277 |
| chr1:85582344-85582703 | 0.29057067 | 0.418475539 | 0.698031277 |
| chr17:69610209-69610409 | 0.414964515 | 0.418479872 | 0.698031277 |
| chr9:79489325-79490037 | -0.272614128 | 0.418513224 | 0.698058933 |
| chr1:43640354-43640554 | 0.44792819 | 0.418517723 | 0.698058933 |
| chrX:3788697-3789185 | -0.333233993 | 0.418523783 | 0.698058933 |
| chr10:41882595-41882795 | 0.414279654 | 0.418539822 | 0.698059482 |
| chr3:1501491-1502164 | -0.300690125 | 0.418547286 | 0.698059482 |
| chr2:172038658-172038962 | -0.342186812 | 0.418551444 | 0.698059482 |
| chr14:53187384-53187691 | -0.34343341 | 0.418573754 | 0.698081496 |
| chr20:25863147-25863437 | 0.300943946 | 0.418584428 | 0.698084103 |
| chr11:46239025-46239454 | 0.291780237 | 0.418610699 | 0.698111503 |
| chr15:38868343-38868640 | -0.326695184 | 0.418637039 | 0.698111503 |
| chr15:40787473-40787864 | -0.290617437 | 0.418642643 | 0.698111503 |
| chr2:112481924-112482180 | -0.334899106 | 0.418647459 | 0.698111503 |

| | | | |
|---|---|---|---|
| chr8:62102823-62103054 | -0.407973856 | 0.418651275 | 0.698111503 |
| chr2:121957826-121958074 | 0.422510213 | 0.418659982 | 0.698111503 |
| chr2:218847597-218848032 | 0.291118606 | 0.418664635 | 0.698111503 |
| chr12:57175894-57176156 | 0.338018005 | 0.418769761 | 0.698259714 |
| chr4:17561220-17561855 | -0.300813689 | 0.418776988 | 0.698259714 |
| chr4:159748199-159748519 | -0.388450613 | 0.418780858 | 0.698259714 |
| chr10:124092348-124093115 | 0.270472195 | 0.418796281 | 0.698270235 |
| chr19:46776873-46777077 | -0.398199837 | 0.418807121 | 0.698273114 |
| chr4:9152097-9152785 | 0.269370809 | 0.418831063 | 0.698283402 |
| chr3:105584383-105584760 | -0.352278902 | 0.418848517 | 0.698283402 |
| chr1:239893126-239893642 | -0.302933848 | 0.418848934 | 0.698283402 |
| chr10:26945991-26946374 | -0.32465344 | 0.418849745 | 0.698283402 |
| chr8:142002276-142003176 | -0.277969065 | 0.418860074 | 0.698285428 |
| chr15:89957960-89958439 | 0.285451381 | 0.418873416 | 0.698292478 |
| chr10:21533465-21535262 | -0.256273913 | 0.418883086 | 0.698293406 |
| chr6:28728539-28729519 | 0.264875561 | 0.418892449 | 0.698293822 |
| chr7:157097659-157097919 | -0.407556407 | 0.418941931 | 0.698349649 |
| chr4:987205-987738 | 0.295564923 | 0.418944167 | 0.698349649 |
| chr14:94129228-94129855 | 0.276932668 | 0.418978837 | 0.698392247 |
| chr17:51119952-51120179 | -0.31031487 | 0.419007067 | 0.698424111 |
| chr2:74119945-74120759 | -0.268220872 | 0.419030654 | 0.698448232 |
| chr9:38620389-38621447 | -0.260784293 | 0.419055639 | 0.698474683 |
| chr5:108381197-108382688 | -0.254005092 | 0.419139657 | 0.698593778 |
| chr10:69360667-69361236 | 0.284603328 | 0.419145326 | 0.698593778 |
| chr4:13654515-13654883 | 0.310971412 | 0.41917303 | 0.698624755 |
| chr3:148379781-148380149 | -0.299524605 | 0.419196453 | 0.698648597 |
| chr14:65353220-65353461 | -0.364522257 | 0.419245999 | 0.698691439 |
| chr1:189429613-189429924 | -0.32291199 | 0.419246499 | 0.698691439 |
| chr2:175270346-175270592 | 0.433133995 | 0.419249514 | 0.698691439 |
| chr18:59837475-59837855 | -0.318735379 | 0.419265374 | 0.698702672 |
| chr20:42102386-42102656 | 0.33556769 | 0.419280865 | 0.698712717 |
| chr10:112115802-112116348 | -0.305183713 | 0.419289639 | 0.698712717 |
| chr2:220687252-220687617 | -0.323178022 | 0.419299537 | 0.698714015 |
| chr2:148509041-148509846 | -0.271061589 | 0.419309734 | 0.698715811 |
| chr6:152993496-152993735 | -0.37597132 | 0.419332018 | 0.698723203 |
| chr12:47717510-47717860 | -0.299157371 | 0.419332408 | 0.698723203 |
| chr16:2714172-2714706 | 0.297569456 | 0.419358588 | 0.698751631 |
| chr1:150611552-150611783 | 0.346779469 | 0.419370573 | 0.698756405 |
| chr6:170552915-170553490 | -0.265763224 | 0.419384058 | 0.698763678 |
| chr17:8221824-8223196 | -0.256165352 | 0.419401713 | 0.698770949 |
| chr5:30401384-30401782 | 0.362809098 | 0.419408634 | 0.698770949 |
| chr17:28896879-28897156 | 0.34588233 | 0.41941578 | 0.698770949 |
| chr2:74958274-74959192 | -0.259600537 | 0.419433713 | 0.698785632 |
| chr19:43527059-43527569 | 0.295659689 | 0.419464708 | 0.69880755 |
| chr7:147626784-147627097 | -0.402290302 | 0.419467759 | 0.69880755 |
| chr9:33922983-33923220 | -0.384081778 | 0.41948551 | 0.69880755 |
| chr22:23145108-23145568 | 0.27820296 | 0.419491739 | 0.69880755 |
| chr20:3781133-3781671 | 0.277920994 | 0.41949247 | 0.69880755 |
| chr4:47926581-47927292 | -0.283129422 | 0.419526459 | 0.698836402 |
| chr22:50027690-50028117 | 0.320015099 | 0.419528031 | 0.698836402 |

| | | | |
|---|---|---|---|
| chr11:62791885-62792186 | 0.293019565 | 0.419570603 | 0.698878986 |
| chr1:102191829-102192118 | 0.347783832 | 0.419571838 | 0.698878986 |
| chr3:107850673-107851135 | -0.28642731 | 0.419583369 | 0.698883001 |
| chr3:42150171-42150402 | -0.407177241 | 0.419606887 | 0.698906979 |
| chr10:109852448-109852776 | -0.313235834 | 0.41962487 | 0.69892174 |
| chr2:65055745-65056947 | -0.256420493 | 0.419689021 | 0.699013393 |
| chr19:531547-531864 | 0.311281893 | 0.419709678 | 0.699028126 |
| chr13:108215393-108215692 | 0.309825896 | 0.419716113 | 0.699028126 |
| chr10:103772215-103772845 | 0.28839121 | 0.419742474 | 0.699056835 |
| chr11:111436702-111437088 | 0.308065698 | 0.419759914 | 0.699070687 |
| chrX:32408427-32408707 | -0.405037803 | 0.41978346 | 0.699086001 |
| chr17:50426022-50426493 | 0.286201469 | 0.419787358 | 0.699086001 |
| chr8:141058010-141058555 | 0.280114794 | 0.419810689 | 0.699094715 |
| chr16:75305289-75305917 | -0.266700808 | 0.419819362 | 0.699094715 |
| chr3:136070966-136071386 | -0.300881065 | 0.419819962 | 0.699094715 |
| chrX:104370171-104370539 | -0.294704219 | 0.419855711 | 0.699121339 |
| chr15:85879224-85879725 | 0.311262223 | 0.419863332 | 0.699121339 |
| chr6:139134058-139134258 | -0.381519747 | 0.419864334 | 0.699121339 |
| chr2:108928459-108928667 | 0.366493246 | 0.419872447 | 0.699121339 |
| chr2:237486897-237487264 | -0.31278912 | 0.419903345 | 0.699157593 |
| chr10:21194539-21195041 | -0.301046493 | 0.419945484 | 0.699207007 |
| chr4:37157561-37157761 | -0.448115172 | 0.419951273 | 0.699207007 |
| chr3:149618831-149619031 | 0.404269392 | 0.419987916 | 0.699239671 |
| chr1:94445759-94446047 | 0.36640762 | 0.419996495 | 0.699239671 |
| chr8:38792883-38793439 | -0.297106756 | 0.419998269 | 0.699239671 |
| chr4:88411470-88411850 | -0.346596693 | 0.420021663 | 0.699245424 |
| chr5:169227044-169227351 | -0.37035499 | 0.420025834 | 0.699245424 |
| chr14:77804732-77804985 | -0.331584236 | 0.420029103 | 0.699245424 |
| chr6:107866886-107867260 | -0.367600043 | 0.420039294 | 0.699247197 |
| chr10:31620300-31620587 | -0.38346899 | 0.420054578 | 0.69925745 |
| chr11:236789-237262 | 0.294011998 | 0.420093296 | 0.699283372 |
| chr9:86947637-86948009 | 0.338564798 | 0.420100858 | 0.699283372 |
| chr2:85595196-85596150 | -0.258918587 | 0.420101039 | 0.699283372 |
| chr13:23020287-23020602 | 0.302294028 | 0.420106656 | 0.699283372 |
| chr3:130002735-130003392 | 0.282557122 | 0.420155118 | 0.699348847 |
| chr17:60044424-60044672 | 0.280804965 | 0.420178523 | 0.699372611 |
| chr6:157667191-157667391 | 0.436210663 | 0.42020365 | 0.699399241 |
| chr1:62175645-62175863 | -0.376458117 | 0.420219783 | 0.699410901 |
| chr6:34196347-34196770 | 0.319011218 | 0.420261221 | 0.699464675 |
| chr14:103384919-103386284 | 0.256705007 | 0.420304508 | 0.699490201 |
| chr5:132464240-132464678 | -0.295991573 | 0.420324792 | 0.699490201 |
| chr17:51608320-51609143 | -0.278096621 | 0.420336167 | 0.699490201 |
| chr12:83133196-83133413 | -0.430621713 | 0.420349137 | 0.699490201 |
| chr12:104361069-104361269 | -0.430621713 | 0.420349137 | 0.699490201 |
| chr2:39107210-39107415 | -0.430621713 | 0.420349137 | 0.699490201 |
| chr6:160733098-160733440 | 0.326935244 | 0.420358416 | 0.699490201 |
| chr17:44557004-44557508 | 0.288655346 | 0.420358599 | 0.699490201 |
| chr1:205485341-205485813 | -0.271134657 | 0.420359246 | 0.699490201 |
| chr7:44573600-44574561 | -0.259669898 | 0.420367849 | 0.699490201 |
| chr14:102878797-102879032 | -0.370186043 | 0.420380231 | 0.69949517 |

| | | | |
|---|---|---|---|
| chr5:94737777-94738231 | -0.30857524 | 0.420389093 | 0.69949517 |
| chr1:26471969-26472591 | 0.283705088 | 0.420426327 | 0.699541933 |
| chr6:44387150-44388241 | -0.259616671 | 0.420456714 | 0.699577301 |
| chr4:176277232-176277452 | -0.358687362 | 0.420467407 | 0.699579902 |
| chr18:45839366-45839713 | -0.304294776 | 0.420488235 | 0.699599364 |
| chr10:47024746-47025068 | 0.297420953 | 0.420561887 | 0.699706712 |
| chr20:59750977-59751441 | 0.272444733 | 0.42057779 | 0.699717977 |
| chrX:70259777-70260145 | -0.392872697 | 0.42064766 | 0.699779968 |
| chr6:6788543-6788752 | -0.415135966 | 0.420653266 | 0.699779968 |
| chr2:105237944-105238247 | 0.314675712 | 0.420661937 | 0.699779968 |
| chr17:68827777-68827981 | -0.399930413 | 0.420664071 | 0.699779968 |
| chr10:73997421-73998271 | -0.257892931 | 0.420666247 | 0.699779968 |
| chr2:7611663-7611995 | -0.384614693 | 0.420669848 | 0.699779968 |
| chr8:103644028-103644454 | 0.339737402 | 0.420699133 | 0.699803482 |
| chr6:44111662-44111961 | -0.363991343 | 0.42070225 | 0.699803482 |
| chr17:16584212-16584746 | -0.279566292 | 0.420744441 | 0.699858471 |
| chr2:27706055-27706742 | -0.279818531 | 0.420773736 | 0.699892005 |
| chr19:10870754-10871538 | 0.259056175 | 0.42079096 | 0.699905461 |
| chr6:44350097-44350297 | -0.444456491 | 0.420804096 | 0.699912116 |
| chr3:25663455-25663748 | 0.309930163 | 0.420826444 | 0.699934092 |
| chr5:81299668-81299869 | 0.352687414 | 0.42085629 | 0.69996854 |
| chr4:152957907-152958107 | 0.42201366 | 0.420902342 | 0.70001688 |
| chr1:145964307-145965167 | 0.257218622 | 0.420903626 | 0.70001688 |
| chr10:44974969-44975247 | 0.299870978 | 0.420950683 | 0.700079945 |
| chr12:79344909-79345204 | 0.307267866 | 0.420972072 | 0.700100322 |
| chr7:74465072-74465349 | -0.358626208 | 0.420992546 | 0.700119175 |
| chr2:113203959-113204450 | -0.290727454 | 0.421020989 | 0.70015128 |
| chr12:91893567-91893809 | 0.371223655 | 0.421035806 | 0.700160724 |
| chr17:44309133-44309333 | 0.359435105 | 0.421067623 | 0.70019563 |
| chr9:114341803-114342003 | 0.40053385 | 0.421085058 | 0.70019563 |
| chr2:215269435-215269722 | 0.366888498 | 0.4210855 | 0.70019563 |
| chr1:118081176-118081766 | -0.310897125 | 0.421093349 | 0.70019563 |
| chr20:47071919-47072173 | -0.370825866 | 0.421171505 | 0.700301269 |
| chr3:149657512-149659073 | -0.251765314 | 0.42117516 | 0.700301269 |
| chr10:68600121-68600723 | -0.285827122 | 0.421186968 | 0.700305706 |
| chr6:34756502-34756827 | -0.313801049 | 0.421207218 | 0.700324179 |
| chr1:193185867-193186109 | 0.369576907 | 0.421228944 | 0.700345104 |
| chr6:170516977-170517336 | 0.312968897 | 0.421303664 | 0.700438297 |
| chr9:135412596-135412959 | 0.324528963 | 0.421308415 | 0.700438297 |
| chr1:2526116-2526402 | 0.316756874 | 0.421316695 | 0.700438297 |
| chr2:128543608-128544009 | -0.315805187 | 0.421348086 | 0.700438297 |
| chr2:63050051-63051098 | -0.256962613 | 0.421353922 | 0.700438297 |
| chr17:50858509-50858767 | 0.34612829 | 0.421354442 | 0.700438297 |
| chr1:208244156-208244692 | -0.283865565 | 0.421356027 | 0.700438297 |
| chr16:79171997-79172197 | -0.416460245 | 0.421358127 | 0.700438297 |
| chr5:18397435-18397771 | -0.312903332 | 0.421369842 | 0.700442575 |
| chr1:231603678-231604005 | -0.340096778 | 0.421382182 | 0.700447891 |
| chr8:95885697-95885943 | 0.393479251 | 0.421413673 | 0.700481611 |
| chr3:6272231-6273182 | -0.291319509 | 0.421427859 | 0.700481611 |
| chr8:52713147-52714038 | -0.272605854 | 0.421429972 | 0.700481611 |

| | | | |
|---|---|---|---|
| chr1:151624901-151625101 | 0.389834021 | 0.421439036 | 0.700481611 |
| chr1:226870314-226871150 | 0.269683991 | 0.421453938 | 0.700491186 |
| chr10:78033508-78034231 | -0.262803818 | 0.421463899 | 0.700492546 |
| chr4:83012170-83013404 | 0.253505708 | 0.421494562 | 0.700525254 |
| chr4:169949070-169949412 | 0.393181849 | 0.421501863 | 0.700525254 |
| chr5:158626188-158626604 | -0.328374444 | 0.421531794 | 0.700559803 |
| chrX:151505832-151506087 | 0.383918408 | 0.421545685 | 0.700567692 |
| chr12:87951553-87951812 | -0.368165081 | 0.421571683 | 0.700595703 |
| chr4:128288787-128289022 | 0.292322603 | 0.421604179 | 0.700634511 |
| chr20:50929376-50929692 | 0.282159153 | 0.421655472 | 0.700700008 |
| chr1:180486115-180486362 | -0.401750775 | 0.421662353 | 0.700700008 |
| chr2:34435104-34435304 | -0.431874871 | 0.421677344 | 0.700700008 |
| chr20:59744095-59744485 | -0.284522855 | 0.421680576 | 0.700700008 |
| chr6:53961987-53962287 | 0.347790634 | 0.421695131 | 0.700700008 |
| chr3:37214142-37214363 | 0.383990996 | 0.421698461 | 0.700700008 |
| chr1:81007078-81007476 | -0.337062049 | 0.421753235 | 0.700723117 |
| chr10:7408446-7409441 | -0.269434667 | 0.421771148 | 0.700723117 |
| chr2:240584647-240584975 | 0.318470689 | 0.42177605 | 0.700723117 |
| chr15:78750034-78750245 | -0.416402006 | 0.421782033 | 0.700723117 |
| chr16:85962643-85962843 | -0.416402006 | 0.421782033 | 0.700723117 |
| chr11:11621568-11621768 | 0.418172849 | 0.421785659 | 0.700723117 |
| chr4:82500692-82500938 | 0.366335923 | 0.421787079 | 0.700723117 |
| chr2:239824265-239824659 | -0.315160614 | 0.421804124 | 0.700723117 |
| chr17:46940812-46941107 | -0.307460649 | 0.421816567 | 0.700723117 |
| chr2:161570463-161570663 | -0.424579984 | 0.421819262 | 0.700723117 |
| chr5:3512179-3512379 | -0.424579984 | 0.421819262 | 0.700723117 |
| chr5:178070044-178070253 | 0.333654427 | 0.421825118 | 0.700723117 |
| chr2:85637385-85637847 | -0.293170479 | 0.421831256 | 0.700723117 |
| chr1:51087107-51087411 | -0.368893575 | 0.421845283 | 0.700731226 |
| chr17:79114592-79115111 | -0.290905259 | 0.421860602 | 0.700741481 |
| chr15:69414639-69415111 | 0.289242332 | 0.421876632 | 0.700752917 |
| chr15:93072833-93073404 | 0.278723795 | 0.421911703 | 0.700791414 |
| chr10:23826451-23826698 | 0.35726214 | 0.421918101 | 0.700791414 |
| chr16:70800436-70801445 | 0.260155633 | 0.421985052 | 0.700887424 |
| chrX:21479886-21480179 | -0.34922755 | 0.422004685 | 0.70090484 |
| chr2:215189401-215189682 | 0.402294655 | 0.422016272 | 0.700908892 |
| chr5:24735282-24735736 | 0.296544232 | 0.422045747 | 0.70092948 |
| chr1:27342357-27342580 | -0.396956917 | 0.422046964 | 0.70092948 |
| chr9:128863566-128863986 | -0.305204273 | 0.422087698 | 0.700981936 |
| chr17:4882114-4882322 | -0.395402644 | 0.42210577 | 0.700989686 |
| chr8:85841859-85842090 | 0.343877045 | 0.422111896 | 0.700989686 |
| chr19:41363691-41364437 | 0.261916829 | 0.42211981 | 0.700989686 |
| chr4:157528411-157528735 | -0.396893383 | 0.422133997 | 0.700998052 |
| chr12:42332338-42332560 | 0.394977727 | 0.422185855 | 0.701068973 |
| chr10:63977349-63977764 | -0.292809992 | 0.422217486 | 0.701096835 |
| chr12:111298810-111299163 | 0.321013797 | 0.422228753 | 0.701096835 |
| chr9:23821428-23822186 | -0.287156211 | 0.422236002 | 0.701096835 |
| chr14:95660934-95661173 | -0.377113509 | 0.422246615 | 0.701096835 |
| chr16:73047442-73048438 | 0.260426689 | 0.422248384 | 0.701096835 |
| chr6:124349183-124349514 | -0.437922127 | 0.422279013 | 0.701105638 |

| | | | |
|---|---|---|---|
| chr20:32665341-32665585 | 0.37649061 | 0.422281505 | 0.701105638 |
| chr22:37817669-37817942 | 0.3495012 | 0.422290282 | 0.701105638 |
| chr1:157193662-157193975 | 0.308833897 | 0.422290841 | 0.701105638 |
| chr6:28953094-28953440 | 0.29018077 | 0.422299436 | 0.701105638 |
| chr14:80977412-80977638 | 0.352572586 | 0.422331276 | 0.701136986 |
| chr14:104564014-104564263 | 0.330586724 | 0.422336619 | 0.701136986 |
| chr12:7162663-7163153 | 0.300527644 | 0.422430579 | 0.701268965 |
| chr9:120897178-120897511 | 0.362878825 | 0.422443403 | 0.701268965 |
| chr8:98164610-98165140 | 0.271676408 | 0.422443575 | 0.701268965 |
| chr8:130796824-130797043 | -0.41158617 | 0.422474229 | 0.701283628 |
| chr16:57462383-57463000 | 0.29647176 | 0.422474236 | 0.701283628 |
| chr3:164451469-164451740 | -0.342856955 | 0.42249636 | 0.701283628 |
| chr17:18129128-18129353 | 0.333158479 | 0.422496627 | 0.701283628 |
| chr10:127907526-127907728 | -0.374554548 | 0.42249817 | 0.701283628 |
| chr19:46717065-46717353 | -0.303785468 | 0.4225512 | 0.701347494 |
| chr10:397475-398001 | -0.298056837 | 0.422558409 | 0.701347494 |
| chr19:54325217-54325427 | -0.416060382 | 0.422564108 | 0.701347494 |
| chr17:1276208-1276474 | -0.381523121 | 0.422587058 | 0.701370392 |
| chr1:114505144-114505641 | 0.258940667 | 0.422608645 | 0.701379184 |
| chr5:40679043-40680249 | -0.253119164 | 0.422610663 | 0.701379184 |
| chr10:124627208-124627430 | -0.369517412 | 0.422643714 | 0.701418269 |
| chr10:81820501-81820765 | 0.356758616 | 0.422652521 | 0.701418269 |
| chr1:228102992-228103659 | 0.265439764 | 0.422669911 | 0.701422961 |
| chr3:38738662-38738929 | -0.385230293 | 0.422677765 | 0.701422961 |
| chr17:75114122-75114322 | 0.353280451 | 0.422682812 | 0.701422961 |
| chr17:76584562-76584762 | 0.406370974 | 0.422696886 | 0.701431126 |
| chr10:85612866-85613252 | -0.312259099 | 0.42272728 | 0.701443841 |
| chr4:98657602-98657813 | -0.363380969 | 0.422740504 | 0.701443841 |
| chr2:235170963-235171272 | -0.338177991 | 0.422752584 | 0.701443841 |
| chr16:66717108-66717521 | 0.3416052 | 0.422756369 | 0.701443841 |
| chr16:23148918-23149941 | 0.263848498 | 0.422760575 | 0.701443841 |
| chr13:39406828-39407028 | -0.374438361 | 0.422766726 | 0.701443841 |
| chr15:31080606-31080846 | -0.382662628 | 0.422768631 | 0.701443841 |
| chr1:37732462-37732755 | -0.411787238 | 0.422810837 | 0.70145114 |
| chr1:201493960-201494260 | -0.300566948 | 0.422821384 | 0.70145114 |
| chr1:1748798-1749008 | 0.408874424 | 0.422828619 | 0.70145114 |
| chr6:75350666-75350869 | -0.400929942 | 0.422829306 | 0.70145114 |
| chr1:15603266-15603801 | -0.298917252 | 0.422832823 | 0.70145114 |
| chr7:38368112-38368534 | -0.277280981 | 0.422834567 | 0.70145114 |
| chr12:11754301-11754560 | -0.434788499 | 0.422837113 | 0.70145114 |
| chr3:128329221-128329447 | 0.390268061 | 0.422868927 | 0.701488729 |
| chr5:117013792-117013992 | -0.45091785 | 0.422883781 | 0.701498183 |
| chr7:29988421-29988740 | 0.322846672 | 0.422907266 | 0.701521953 |
| chr10:12605480-12606407 | 0.266959934 | 0.422930936 | 0.701532521 |
| chr4:6250362-6251003 | 0.28463149 | 0.422931949 | 0.701532521 |
| chr12:14845403-14845614 | -0.423354111 | 0.42294691 | 0.701533613 |
| chr16:4271633-4272362 | 0.266251456 | 0.422950918 | 0.701533613 |
| chr22:20394442-20394693 | 0.34592559 | 0.422962458 | 0.701537566 |
| chr1:50969574-50970247 | 0.273462938 | 0.422984699 | 0.701546866 |
| chr11:66637935-66638983 | -0.256510184 | 0.422988646 | 0.701546866 |

| | | | |
|---|---|---|---|
| chr10:68087381-68087749 | -0.342572645 | 0.422995532 | 0.701546866 |
| chr5:96336275-96336660 | 0.337202651 | 0.423036537 | 0.701599686 |
| chr7:54525654-54525893 | -0.42092152 | 0.423055764 | 0.701605255 |
| chr8:141589611-141589868 | -0.356550572 | 0.423058208 | 0.701605255 |
| chr8:67064805-67065137 | -0.286539452 | 0.423072451 | 0.701613691 |
| chr1:67792062-67792339 | 0.358745961 | 0.423132052 | 0.701644962 |
| chr10:67779981-67780282 | -0.3449381 | 0.423143326 | 0.701644962 |
| chr1:182446122-182446414 | -0.319058794 | 0.423144751 | 0.701644962 |
| chr14:57211949-57212241 | -0.382468637 | 0.423162492 | 0.701644962 |
| chr3:122911852-122912087 | 0.390684113 | 0.42316409 | 0.701644962 |
| chr1:167556137-167556394 | 0.329068615 | 0.42316966 | 0.701644962 |
| chr3:117454328-117454531 | -0.421581179 | 0.423170078 | 0.701644962 |
| chr20:47502793-47503222 | 0.256384142 | 0.423172042 | 0.701644962 |
| chr4:75630155-75631095 | -0.262067327 | 0.42317966 | 0.701644962 |
| chr8:103298517-103299840 | 0.255580533 | 0.42318288 | 0.701644962 |
| chr8:27727301-27727600 | 0.337872697 | 0.423208403 | 0.701672096 |
| chr5:126973187-126973549 | -0.383688207 | 0.423229457 | 0.701690733 |
| chr7:74587502-74587738 | -0.410925175 | 0.423247854 | 0.701690733 |
| chr4:78775456-78777110 | -0.254769575 | 0.423253133 | 0.701690733 |
| chr12:71654251-71654554 | 0.32838461 | 0.423256275 | 0.701690733 |
| chr12:121124401-121124942 | -0.324618979 | 0.423278305 | 0.701712073 |
| chr9:89686187-89686467 | 0.372384973 | 0.423333711 | 0.701788741 |
| chr7:156491784-156492097 | -0.407290861 | 0.423362959 | 0.701809619 |
| chr3:45938817-45939021 | 0.422325471 | 0.423364624 | 0.701809619 |
| chr10:7054937-7055250 | 0.324554164 | 0.423383885 | 0.701826364 |
| chr1:153277871-153278240 | 0.306984558 | 0.423407326 | 0.701850038 |
| chr7:42273212-42273620 | -0.322572831 | 0.423438365 | 0.701886304 |
| chr4:154082792-154083056 | 0.339515532 | 0.423471421 | 0.701913861 |
| chr2:215397743-215398022 | -0.355341309 | 0.423473311 | 0.701913861 |
| chr19:10602250-10602524 | 0.344479194 | 0.423495958 | 0.701918727 |
| chr22:38181628-38182230 | 0.268287215 | 0.423501927 | 0.701918727 |
| chr9:1232490-1232750 | 0.387924455 | 0.42350373 | 0.701918727 |
| chr18:10490682-10491035 | 0.307325973 | 0.423521802 | 0.701933497 |
| chr15:99981288-99981566 | 0.341042821 | 0.42353151 | 0.701934403 |
| chr19:34733402-34734673 | 0.261997443 | 0.423548147 | 0.701946793 |
| chr10:17607527-17608017 | -0.296510341 | 0.423558118 | 0.701948135 |
| chr14:105419725-105419933 | 0.352603166 | 0.423574086 | 0.701959417 |
| chr10:112314916-112315178 | 0.356083351 | 0.423588525 | 0.701968162 |
| chr12:93087278-93087724 | -0.308920214 | 0.423623354 | 0.70200243 |
| chr7:149838068-149839059 | -0.257587209 | 0.423627527 | 0.70200243 |
| chr16:89122411-89122656 | -0.347056853 | 0.423636964 | 0.702002887 |
| chr19:18154141-18154341 | 0.363651038 | 0.423647748 | 0.702005574 |
| chr18:25503519-25503982 | -0.306889857 | 0.423672749 | 0.702021592 |
| chr10:70364514-70364936 | 0.296085845 | 0.423681804 | 0.702021592 |
| chr15:91055901-91056258 | 0.302598903 | 0.423684901 | 0.702021592 |
| chr6:12504427-12504631 | 0.398379849 | 0.423711582 | 0.702048279 |
| chr12:92785103-92785490 | -0.285329899 | 0.423719332 | 0.702048279 |
| chr5:54309750-54311101 | -0.255741868 | 0.423749893 | 0.702079699 |
| chr22:49930779-49931356 | 0.291358174 | 0.423756621 | 0.702079699 |
| chr1:182962551-182962866 | -0.294129612 | 0.423769249 | 0.702085439 |

| | | | |
|---|---|---|---|
| chr10:91796347-91796586 | -0.322617214 | 0.423782388 | 0.702092027 |
| chrX:40012626-40012890 | -0.337349704 | 0.423815701 | 0.70212028 |
| chr14:58272575-58272787 | 0.366171019 | 0.423817769 | 0.70212028 |
| chr19:47125269-47125481 | 0.395664463 | 0.423829478 | 0.702124498 |
| chr2:91780789-91781080 | -0.434492831 | 0.423891862 | 0.702212662 |
| chr5:81507023-81507249 | -0.384024214 | 0.4239131 | 0.702232662 |
| chr6:75934941-75935356 | -0.296226035 | 0.423925432 | 0.702237908 |
| chr3:152269873-152270147 | -0.294477335 | 0.423978717 | 0.702310992 |
| chr1:149844230-149844479 | 0.31734205 | 0.424006991 | 0.702329958 |
| chr1:176691640-176691974 | 0.306995402 | 0.424008499 | 0.702329958 |
| chr18:37309787-37310164 | -0.311917206 | 0.42403385 | 0.702342045 |
| chr3:155853784-155855254 | -0.250077188 | 0.424034129 | 0.702342045 |
| chr1:225004511-225004711 | -0.423230098 | 0.424073544 | 0.702371046 |
| chr5:69217542-69218566 | -0.255545824 | 0.424076146 | 0.702371046 |
| chr2:241010151-241010557 | -0.344814087 | 0.424088753 | 0.702371046 |
| chr1:230104691-230105008 | 0.339652937 | 0.424093928 | 0.702371046 |
| chr1:189825917-189826226 | -0.401768582 | 0.424097471 | 0.702371046 |
| chr2:74421238-74422471 | -0.255570734 | 0.424135134 | 0.702418239 |
| chr2:24359725-24361187 | -0.259162044 | 0.424157358 | 0.702439492 |
| chr9:4762547-4762796 | -0.372390525 | 0.424166303 | 0.702439492 |
| chr6:13486699-13488074 | 0.25726776 | 0.42420966 | 0.702490675 |
| chr15:78437683-78438812 | 0.258605284 | 0.424215546 | 0.702490675 |
| chr3:33231546-33231746 | 0.418869935 | 0.42423314 | 0.702496778 |
| chr22:42079440-42080217 | 0.264775645 | 0.42424006 | 0.702496778 |
| chr6:31717600-31718014 | 0.304452684 | 0.424246736 | 0.702496778 |
| chr1:201496022-201496239 | -0.325547235 | 0.424270862 | 0.702503812 |
| chr1:161201873-161202760 | 0.259100294 | 0.424289312 | 0.702503812 |
| chr14:67704540-67704740 | 0.393310602 | 0.424294076 | 0.702503812 |
| chr4:84498669-84498935 | -0.373382836 | 0.424296312 | 0.702503812 |
| chr10:57217857-57218799 | -0.267648649 | 0.424304211 | 0.702503812 |
| chr13:32036930-32037142 | -0.428858577 | 0.424340246 | 0.702503812 |
| chr4:124691897-124692177 | -0.356924327 | 0.424348556 | 0.702503812 |
| chr10:101018953-101019327 | -0.299355872 | 0.424348589 | 0.702503812 |
| chr7:100965836-100966784 | -0.265276014 | 0.424356789 | 0.702503812 |
| chr2:85001636-85001965 | -0.37116868 | 0.424362418 | 0.702503812 |
| chr2:48221855-48222337 | -0.308059648 | 0.424375161 | 0.702503812 |
| chr1:32925301-32925686 | 0.342203666 | 0.424375384 | 0.702503812 |
| chr5:113345306-113345506 | -0.426635947 | 0.424376727 | 0.702503812 |
| chr1:206521012-206521587 | 0.265474451 | 0.424392211 | 0.702503812 |
| chr15:32679491-32679691 | -0.417790378 | 0.424395828 | 0.702503812 |
| chr13:36619800-36620018 | -0.441326755 | 0.424397679 | 0.702503812 |
| chr13:34941594-34942835 | -0.256692417 | 0.424427505 | 0.702538005 |
| chr20:56758510-56759298 | -0.262813489 | 0.424449156 | 0.702558667 |
| chr6:30877706-30878043 | 0.305259526 | 0.424473809 | 0.702584296 |
| chr2:68694719-68694919 | 0.465955785 | 0.424501725 | 0.702615323 |
| chr7:138483573-138483773 | -0.424004613 | 0.424579481 | 0.702728841 |
| chr10:122102686-122102886 | -0.399432871 | 0.424600664 | 0.702748723 |
| chrX:49878876-49879812 | 0.262649541 | 0.424612251 | 0.70275272 |
| chr7:112550501-112550800 | -0.321999218 | 0.424627307 | 0.702762458 |
| chr5:78590-78838 | 0.329650876 | 0.424639354 | 0.702762717 |

| | | | |
|---|---|---|---|
| chr4:119550924-119551124 | 0.4251458 | 0.424665608 | 0.702762717 |
| chr7:33320982-33321183 | 0.4251458 | 0.424665608 | 0.702762717 |
| chr19:44654480-44654680 | 0.381547818 | 0.424672386 | 0.702762717 |
| chr12:93234752-93235005 | -0.34579845 | 0.424679888 | 0.702762717 |
| chr7:104362835-104363083 | -0.466228311 | 0.424688515 | 0.702762717 |
| chr12:121786347-121786831 | 0.276288022 | 0.424691666 | 0.702762717 |
| chr8:119863076-119863443 | 0.311050388 | 0.42471653 | 0.702770068 |
| chr13:38065318-38066055 | -0.285356437 | 0.424726581 | 0.702770068 |
| chr8:1960105-1960337 | 0.391075681 | 0.424748796 | 0.702770068 |
| chr3:185199747-185199985 | -0.397674632 | 0.424750204 | 0.702770068 |
| chr5:129255696-129255942 | -0.397674632 | 0.424750204 | 0.702770068 |
| chr17:8138501-8139250 | -0.263012487 | 0.42475114 | 0.702770068 |
| chr3:125972238-125972759 | 0.293000996 | 0.424769629 | 0.702785483 |
| chr17:28842550-28843105 | 0.27493615 | 0.424786807 | 0.702798729 |
| chr19:50704251-50704470 | 0.377897036 | 0.424812433 | 0.702825951 |
| chr15:67023656-67024219 | -0.274643393 | 0.424855481 | 0.702850656 |
| chr5:40437262-40437631 | -0.366050662 | 0.424860952 | 0.702850656 |
| chr4:133682532-133682844 | -0.399262977 | 0.424862021 | 0.702850656 |
| chr17:42548029-42548530 | 0.29006915 | 0.424864058 | 0.702850656 |
| chr3:128608179-128608727 | 0.27905732 | 0.424877504 | 0.702857724 |
| chr11:65111605-65112088 | 0.275319761 | 0.424901421 | 0.702879072 |
| chr11:64917095-64917651 | 0.261295147 | 0.424922509 | 0.702879072 |
| chr1:154855854-154856077 | -0.401055598 | 0.424925793 | 0.702879072 |
| chr13:46648098-46648514 | -0.268151113 | 0.424933197 | 0.702879072 |
| chr5:50601622-50601952 | -0.314272308 | 0.42494487 | 0.702879072 |
| chrX:16719158-16720074 | -0.255111397 | 0.424945448 | 0.702879072 |
| chr12:100154743-100155234 | -0.284941431 | 0.424956434 | 0.70288207 |
| chr3:125647336-125647689 | -0.373207861 | 0.424978143 | 0.702890404 |
| chr1:197200804-197201839 | -0.250631196 | 0.424984548 | 0.702890404 |
| chr13:98672878-98673473 | 0.296037804 | 0.424988993 | 0.702890404 |
| chr11:12831902-12832102 | -0.419969382 | 0.425030047 | 0.702933342 |
| chr5:8281285-8281521 | -0.440920009 | 0.425033303 | 0.702933342 |
| chr2:85367200-85367400 | -0.366488392 | 0.42505069 | 0.70293966 |
| chr6:25991702-25992283 | -0.286590694 | 0.425055471 | 0.70293966 |
| chr15:49169876-49170612 | -0.260386367 | 0.425069963 | 0.702948454 |
| chr8:17602663-17602911 | -0.358408936 | 0.425093186 | 0.702957691 |
| chr2:17753229-17754710 | -0.251620358 | 0.425093898 | 0.702957691 |
| chr3:23761439-23761852 | 0.28492767 | 0.425132783 | 0.702988647 |
| chr2:32039328-32040170 | -0.258453739 | 0.425140482 | 0.702988647 |
| chrX:64214017-64214368 | -0.336373365 | 0.425141073 | 0.702988647 |
| chr7:141937474-141937718 | 0.441774728 | 0.425158491 | 0.702988647 |
| chrX:73850490-73850690 | 0.441774728 | 0.425158491 | 0.702988647 |
| chr15:65400734-65401065 | 0.303477237 | 0.425175908 | 0.702989603 |
| chr20:17651904-17652179 | 0.313360086 | 0.425177419 | 0.702989603 |
| chr3:159059540-159059776 | 0.378903017 | 0.425191454 | 0.702997639 |
| chr4:89640784-89641527 | 0.285170688 | 0.425231893 | 0.703048264 |
| chr2:62297739-62297945 | -0.380538098 | 0.425240424 | 0.703048264 |
| chr8:105660536-105661058 | 0.297927598 | 0.42530254 | 0.703135788 |
| chr11:36266062-36266327 | 0.322877443 | 0.425373334 | 0.703237655 |
| chr1:155745182-155745903 | 0.268320379 | 0.425407215 | 0.703275508 |

| | | | |
|---|---|---|---|
| chr18:5488788-5489110 | 0.359561039 | 0.425414587 | 0.703275508 |
| chr6:33199939-33200154 | 0.317394772 | 0.425429863 | 0.703285587 |
| chr19:21505470-21505670 | -0.419514842 | 0.425454249 | 0.703289059 |
| chr16:88020391-88020591 | 0.370864992 | 0.425454753 | 0.703289059 |
| chr12:122137739-122138207 | 0.282895201 | 0.425462865 | 0.703289059 |
| chr2:211269456-211269707 | 0.377391067 | 0.425468678 | 0.703289059 |
| chr8:88525904-88526150 | 0.358289532 | 0.425558381 | 0.703422161 |
| chr9:81071145-81071345 | 0.426851815 | 0.425586989 | 0.703436614 |
| chrX:38985498-38985723 | 0.426851815 | 0.425586989 | 0.703436614 |
| chr7:129367544-129368588 | -0.269099557 | 0.425602402 | 0.703436614 |
| chr11:925711-925928 | 0.384312653 | 0.425609955 | 0.703436614 |
| chr19:38647264-38647875 | 0.266438487 | 0.425613028 | 0.703436614 |
| chr11:93614490-93614763 | -0.388279883 | 0.425633043 | 0.703444361 |
| chr20:44785870-44786101 | 0.360607578 | 0.425636077 | 0.703444361 |
| chr7:48875316-48875810 | -0.295428368 | 0.425654251 | 0.703447401 |
| chr5:43067245-43067580 | 0.283838622 | 0.425656278 | 0.703447401 |
| chr4:99205153-99205415 | -0.388197395 | 0.425682411 | 0.703474962 |
| chr6:73870907-73871264 | -0.303355187 | 0.425691594 | 0.703474962 |
| chr5:128494489-128494748 | -0.392276287 | 0.425700498 | 0.703474962 |
| chr2:37196153-37197336 | -0.255831143 | 0.425717498 | 0.703487882 |
| chr9:35095923-35096786 | -0.266744471 | 0.425727415 | 0.703489098 |
| chr1:45444906-45445241 | -0.349883643 | 0.425747452 | 0.703507036 |
| chr7:75738521-75739878 | -0.25504094 | 0.425788277 | 0.703559323 |
| chr3:354367-354897 | -0.293147957 | 0.425821856 | 0.703595362 |
| chr9:120875910-120877766 | 0.253167307 | 0.425828453 | 0.703595362 |
| chr2:189661041-189661784 | -0.253516234 | 0.425856287 | 0.703599234 |
| chr5:78360829-78361292 | -0.278637517 | 0.425857803 | 0.703599234 |
| chr9:39464149-39464807 | -0.283902031 | 0.425858345 | 0.703599234 |
| chr5:180291606-180291811 | 0.325336461 | 0.425979638 | 0.703755283 |
| chr7:28685685-28686208 | -0.308492533 | 0.42599644 | 0.703755283 |
| chr16:84005856-84006107 | -0.354155584 | 0.426003062 | 0.703755283 |
| chr8:66459772-66460362 | 0.299059837 | 0.426006183 | 0.703755283 |
| chr3:34374593-34374944 | 0.342459272 | 0.426012459 | 0.703755283 |
| chr1:174074864-174075064 | -0.412328181 | 0.426015386 | 0.703755283 |
| chr16:77974823-77975240 | 0.311849513 | 0.426020327 | 0.703755283 |
| chr3:17413001-17413309 | 0.327749127 | 0.426032067 | 0.703755283 |
| chr2:59182374-59182648 | -0.347559591 | 0.426035457 | 0.703755283 |
| chrX:137151495-137151695 | 0.400959826 | 0.426088692 | 0.703812794 |
| chr2:147806949-147807162 | -0.410592856 | 0.426092993 | 0.703812794 |
| chr2:191501637-191501885 | -0.39349262 | 0.42609783 | 0.703812794 |
| chr11:1458889-1459252 | -0.317124576 | 0.426155997 | 0.703882275 |
| chr9:110090016-110090431 | 0.322316567 | 0.426158267 | 0.703882275 |
| chr15:78740465-78740833 | 0.321646942 | 0.426190246 | 0.703915185 |
| chr2:89048369-89048687 | 0.353038858 | 0.426196566 | 0.703915185 |
| chr1:31701081-31701389 | -0.33613368 | 0.426243638 | 0.703977755 |
| chr17:59521300-59522017 | 0.265974862 | 0.426263159 | 0.703994821 |
| chr3:38651286-38651572 | 0.323300511 | 0.426287433 | 0.704001529 |
| chr8:120995274-120995514 | 0.403663297 | 0.426287881 | 0.704001529 |
| chr7:12170509-12170798 | 0.334169522 | 0.426294784 | 0.704001529 |
| chr12:116358890-116359348 | 0.299717589 | 0.426315226 | 0.704020114 |

| | | | |
|---|---|---|---|
| chr5:160508858-160509133 | 0.308048133 | 0.426336167 | 0.704039364 |
| chr8:41612869-41613075 | 0.391695394 | 0.426349774 | 0.704039364 |
| chr1:220798425-220798759 | -0.327584636 | 0.426354448 | 0.704039364 |
| chr14:56996833-56997063 | 0.412502443 | 0.426393776 | 0.704078956 |
| chr1:25247283-25247966 | 0.267683639 | 0.426396802 | 0.704078956 |
| chr17:15563287-15563849 | 0.290504173 | 0.426411725 | 0.704088424 |
| chr5:141757513-141757722 | 0.347272755 | 0.42646095 | 0.704145618 |
| chr17:74770568-74771026 | 0.278184836 | 0.426471119 | 0.704145618 |
| chr1:156456433-156457856 | 0.249250436 | 0.426481657 | 0.704145618 |
| chr2:217786091-217786300 | 0.385840206 | 0.426483361 | 0.704145618 |
| chr8:126571304-126572225 | -0.256709027 | 0.426492313 | 0.704145618 |
| chr17:56149623-56149973 | -0.32475829 | 0.426558248 | 0.704232318 |
| chr5:142356712-142357539 | 0.277863313 | 0.426563208 | 0.704232318 |
| chr10:86649126-86649388 | 0.312813101 | 0.426584954 | 0.704250294 |
| chr12:12403942-12404306 | -0.356298226 | 0.426592478 | 0.704250294 |
| chr7:50092544-50092843 | -0.34898524 | 0.426607668 | 0.704260196 |
| chr6:26594791-26595060 | -0.313732046 | 0.426621049 | 0.704267111 |
| chr17:28726484-28726741 | 0.302896194 | 0.426648476 | 0.704285648 |
| chr15:59622381-59622590 | 0.372749666 | 0.426658862 | 0.704285648 |
| chr18:75070785-75071256 | 0.309866756 | 0.426669483 | 0.704285648 |
| chr8:122490207-122490407 | -0.3577737 | 0.426671123 | 0.704285648 |
| chr9:89759516-89760050 | 0.275354716 | 0.426678236 | 0.704285648 |
| chr20:62409941-62410765 | 0.259014024 | 0.42669011 | 0.704289638 |
| chr15:77569527-77569877 | -0.3226157 | 0.426699036 | 0.704289638 |
| chr17:38748330-38748576 | 0.331440431 | 0.426716083 | 0.704296149 |
| chr1:205438768-205439204 | -0.279025611 | 0.426729353 | 0.704296149 |
| chrX:19887691-19888820 | 0.263148225 | 0.426730557 | 0.704296149 |
| chr10:27503923-27504840 | -0.257176333 | 0.426743973 | 0.704303121 |
| chr1:15152941-15153199 | 0.344914388 | 0.426837846 | 0.704426184 |
| chr15:98320178-98320459 | 0.379213118 | 0.426844218 | 0.704426184 |
| chr6:53547678-53548548 | -0.255074474 | 0.426856334 | 0.704426184 |
| chr11:66782255-66782570 | 0.353987035 | 0.426857271 | 0.704426184 |
| chr1:149813288-149813845 | 0.275994663 | 0.426864506 | 0.704426184 |
| chr17:57739985-57740471 | 0.281281339 | 0.426933853 | 0.70452545 |
| chr6:11496249-11496545 | -0.311945807 | 0.427000672 | 0.704620539 |
| chr17:16536846-16537202 | -0.29155512 | 0.427020588 | 0.704627833 |
| chr19:56120753-56121887 | 0.252834262 | 0.427023485 | 0.704627833 |
| chr1:147171911-147173393 | 0.250032844 | 0.427033314 | 0.704628879 |
| chr8:129891695-129891895 | -0.406100152 | 0.427045397 | 0.704633642 |
| chr6:124294654-124294931 | -0.370033598 | 0.427061673 | 0.704645323 |
| chr15:79310652-79311469 | 0.265686396 | 0.427093032 | 0.704678804 |
| chr10:27252217-27252796 | -0.273130757 | 0.427100358 | 0.704678804 |
| chr6:39229044-39229810 | -0.270257265 | 0.427112551 | 0.704683747 |
| chr3:158434492-158434739 | -0.358448748 | 0.427122809 | 0.704685498 |
| chr2:230864610-230865004 | 0.310701273 | 0.427160223 | 0.704711054 |
| chr8:133218607-133218856 | 0.388385792 | 0.427164655 | 0.704711054 |
| chr17:77746329-77746539 | -0.42371141 | 0.427169633 | 0.704711054 |
| chr12:85575731-85576044 | 0.322123858 | 0.427175088 | 0.704711054 |
| chr17:76182230-76182551 | -0.341336699 | 0.427184721 | 0.704711773 |
| chr1:22780811-22781022 | 0.394539441 | 0.427210561 | 0.704739226 |

| | | | |
|---|---|---|---|
| chr20:50982257-50982695 | -0.284629734 | 0.427252721 | 0.704787258 |
| chr6:11096250-11096492 | -0.323834638 | 0.427258074 | 0.704787258 |
| chr3:111989075-111989419 | -0.30063897 | 0.427286707 | 0.704810823 |
| chrX:107726675-107727159 | 0.286017881 | 0.427290756 | 0.704810823 |
| chr13:23530552-23530792 | -0.381543426 | 0.427328335 | 0.704857635 |
| chr14:36525554-36525836 | 0.300522004 | 0.427348007 | 0.704874909 |
| chr16:87391706-87392411 | 0.256021061 | 0.427375531 | 0.704905132 |
| chr16:88505442-88505694 | 0.31279296 | 0.427406307 | 0.704932555 |
| chr1:186985974-186986207 | 0.348783726 | 0.427410557 | 0.704932555 |
| chr8:133645905-133646245 | 0.291482584 | 0.427430136 | 0.704949672 |
| chr5:140400685-140402038 | -0.24938983 | 0.427474133 | 0.70500706 |
| chr8:102655577-102656223 | 0.253525741 | 0.427497144 | 0.705026899 |
| chr9:96875301-96876179 | -0.265589175 | 0.427504565 | 0.705026899 |
| chr5:65747685-65747994 | -0.355382768 | 0.427529703 | 0.705053182 |
| chr1:27306942-27307158 | 0.404225561 | 0.427555558 | 0.705080644 |
| chr2:136384696-136385141 | -0.297132115 | 0.427591878 | 0.705121474 |
| chr3:158943820-158944237 | 0.324823247 | 0.427602497 | 0.705121474 |
| chr1:207823716-207824410 | -0.258725287 | 0.42761578 | 0.705121474 |
| chr2:126777355-126777711 | 0.286373534 | 0.427617384 | 0.705121474 |
| chr5:75717007-75717819 | -0.255655039 | 0.42762633 | 0.705121474 |
| chr10:71590048-71590433 | -0.352921539 | 0.42765698 | 0.705156838 |
| chr1:144551134-144552475 | 0.248682293 | 0.427666479 | 0.705157327 |
| chr17:68019620-68020738 | -0.249455442 | 0.427686081 | 0.705160027 |
| chr20:18983927-18984133 | -0.371778889 | 0.427686523 | 0.705160027 |
| chr5:132445278-132445498 | -0.355381004 | 0.427699307 | 0.705165931 |
| chr2:212940123-212940452 | -0.296994275 | 0.427747768 | 0.705215617 |
| chr2:10757325-10758024 | -0.272642704 | 0.42776331 | 0.705215617 |
| chr2:120252493-120253842 | 0.251500313 | 0.427771287 | 0.705215617 |
| chr3:119676835-119677805 | 0.253541305 | 0.42777173 | 0.705215617 |
| chr5:150801035-150801891 | -0.271329685 | 0.42778371 | 0.705215617 |
| chr1:185696517-185697160 | -0.276846085 | 0.427784666 | 0.705215617 |
| chr7:55186294-55186505 | -0.370688754 | 0.427811236 | 0.705244246 |
| chr14:61894510-61894799 | -0.359375668 | 0.427825816 | 0.705253108 |
| chr11:12287560-12287877 | 0.310206804 | 0.42784674 | 0.705272427 |
| chr14:97106341-97106644 | -0.35781408 | 0.427874923 | 0.70530371 |
| chr5:170571363-170571712 | -0.340175063 | 0.427896857 | 0.705324692 |
| chr2:127226085-127227321 | 0.255410989 | 0.427918931 | 0.705345904 |
| chr3:42536377-42536709 | 0.316577143 | 0.427937258 | 0.705345906 |
| chr1:110122118-110122323 | -0.432564429 | 0.427937343 | 0.705345906 |
| chr9:33161074-33161360 | -0.31080663 | 0.427947952 | 0.705348218 |
| chr20:3753111-3753330 | -0.328291613 | 0.427965393 | 0.705361791 |
| chr3:64166677-64167056 | 0.261018044 | 0.427986826 | 0.705381943 |
| chr1:165610988-165611241 | -0.403150001 | 0.428029737 | 0.705416119 |
| chr21:45073740-45074012 | 0.310551824 | 0.428030775 | 0.705416119 |
| chr12:31959197-31959938 | -0.2729479 | 0.428035181 | 0.705416119 |
| chr20:57180982-57181290 | 0.257368946 | 0.428064409 | 0.705449114 |
| chr1:96075799-96075999 | 0.412887195 | 0.428097779 | 0.705488934 |
| chr2:64453866-64454948 | 0.260489399 | 0.428141224 | 0.705534925 |
| chr7:6008725-6009777 | -0.251635665 | 0.428144102 | 0.705534925 |
| chr19:266950-267720 | 0.272391819 | 0.428204051 | 0.705609944 |

| | | | |
|---|---|---|---|
| chr19:53867452-53867961 | 0.264753227 | 0.428223824 | 0.705609944 |
| chr14:93262432-93262726 | 0.334327646 | 0.42823197 | 0.705609944 |
| chr7:63900084-63900446 | 0.282720084 | 0.42825118 | 0.705609944 |
| chrX:53714482-53715105 | 0.285716723 | 0.428259744 | 0.705609944 |
| chr3:66997889-66998854 | -0.253009805 | 0.428268149 | 0.705609944 |
| chr8:130357406-130357899 | -0.282031535 | 0.428268486 | 0.705609944 |
| chr5:173416662-173416914 | 0.355920224 | 0.428269386 | 0.705609944 |
| chr3:9727030-9727257 | 0.293001902 | 0.428272507 | 0.705609944 |
| chr1:151058626-151059853 | 0.25047123 | 0.428368479 | 0.705752888 |
| chr6:87929821-87930191 | 0.300188039 | 0.428405226 | 0.705798254 |
| chr20:19275814-19276127 | -0.3191211 | 0.428419406 | 0.705806439 |
| chr14:92748353-92748909 | 0.263402242 | 0.428473516 | 0.705865962 |
| chr12:91388488-91389044 | 0.298924806 | 0.42847396 | 0.705865962 |
| chr7:12518939-12519839 | 0.259820437 | 0.428519704 | 0.705926144 |
| chr1:31998458-31998658 | -0.425187416 | 0.428565879 | 0.705975371 |
| chr21:42234157-42235162 | -0.259400424 | 0.428574599 | 0.705975371 |
| chrX:100424071-100424355 | -0.334535789 | 0.428577228 | 0.705975371 |
| chr4:83490953-83491176 | -0.385879098 | 0.428607878 | 0.706008692 |
| chr8:106705048-106705429 | -0.294117878 | 0.428616585 | 0.706008692 |
| chr9:87714800-87715074 | -0.384082769 | 0.428625099 | 0.706008692 |
| chr17:47001160-47001432 | -0.329293362 | 0.428644295 | 0.706009414 |
| chr9:94782979-94783609 | 0.264417834 | 0.428646127 | 0.706009414 |
| chr1:165631926-165632341 | 0.283975682 | 0.42865318 | 0.706009414 |
| chr2:135913831-135914050 | -0.40513504 | 0.428672681 | 0.706026357 |
| chr5:81785318-81785577 | -0.361941095 | 0.428731268 | 0.706107672 |
| chr10:79292502-79292702 | 0.376115991 | 0.428751768 | 0.706126257 |
| chr17:27354260-27354558 | 0.326626582 | 0.428775561 | 0.706143337 |
| chr1:7961128-7962043 | 0.256854265 | 0.428780571 | 0.706143337 |
| chr6:31515556-31515816 | -0.410299036 | 0.428803615 | 0.706160763 |
| chr18:80253948-80254249 | 0.314052163 | 0.428809584 | 0.706160763 |
| chr12:101476040-101476352 | 0.406095797 | 0.428835132 | 0.70616981 |
| chr13:70505863-70506161 | 0.338704446 | 0.428835812 | 0.70616981 |
| chr6:87968656-87968984 | 0.356849266 | 0.428842727 | 0.70616981 |
| chr20:59253819-59254110 | 0.327973771 | 0.428867545 | 0.706195501 |
| chr14:58298978-58299213 | 0.298008196 | 0.428881084 | 0.706202618 |
| chr3:64496724-64497002 | 0.269044598 | 0.428895768 | 0.70621162 |
| chr7:30778211-30778582 | -0.336258739 | 0.428950081 | 0.706271598 |
| chr20:56568480-56568904 | 0.289173039 | 0.428957151 | 0.706271598 |
| chr5:149095021-149095534 | -0.287747381 | 0.428969314 | 0.706271598 |
| chr11:9758613-9758970 | 0.309487421 | 0.428978505 | 0.706271598 |
| chr3:120622784-120623871 | -0.254772119 | 0.42898141 | 0.706271598 |
| chr8:132062821-132063039 | -0.374199561 | 0.428987499 | 0.706271598 |
| chr6:135252024-135252333 | 0.331374111 | 0.429015466 | 0.706281046 |
| chr10:71377171-71377692 | 0.27755384 | 0.42902046 | 0.706281046 |
| chr17:48618738-48618938 | 0.356827329 | 0.429020891 | 0.706281046 |
| chr1:205658924-205659328 | 0.270701807 | 0.429060808 | 0.706326675 |
| chr1:161089360-161089794 | 0.266695875 | 0.429093972 | 0.706326675 |
| chr12:49069493-49070317 | 0.258020153 | 0.429098325 | 0.706326675 |
| chr15:73809892-73810092 | -0.395559684 | 0.429106799 | 0.706326675 |
| chr3:196650857-196651252 | -0.329364559 | 0.429119611 | 0.706326675 |

| | | | |
|---|---|---|---|
| chr16:10971582-10971790 | -0.385160302 | 0.4291304 | 0.706326675 |
| chr7:35425009-35425215 | 0.372528406 | 0.429135067 | 0.706326675 |
| chr9:108130938-108131373 | 0.348320818 | 0.429137991 | 0.706326675 |
| chr12:131242998-131243225 | -0.374948006 | 0.429147786 | 0.706326675 |
| chr2:42025207-42025426 | -0.407606955 | 0.429149353 | 0.706326675 |
| chr11:129966375-129966602 | -0.380050714 | 0.429150009 | 0.706326675 |
| chr9:105244167-105245705 | -0.250766192 | 0.429232133 | 0.706436633 |
| chr18:1914514-1914787 | -0.337370443 | 0.429239995 | 0.706436633 |
| chr2:131106210-131106639 | -0.287938941 | 0.429244477 | 0.706436633 |
| chr5:168295583-168295838 | 0.300521663 | 0.429266396 | 0.706457532 |
| chr11:72683835-72684035 | 0.336009694 | 0.429297351 | 0.706492255 |
| chr5:32451056-32451307 | -0.352343064 | 0.429305936 | 0.706492255 |
| chr4:73729639-73729856 | 0.370886944 | 0.429317294 | 0.706495774 |
| chr17:63950986-63951284 | -0.279929362 | 0.429329353 | 0.706500445 |
| chr14:102246349-102246549 | 0.39376449 | 0.429359699 | 0.706535208 |
| chr19:57819486-57820102 | 0.272979477 | 0.429452847 | 0.70667331 |
| chr3:6799441-6799694 | -0.363111273 | 0.429484009 | 0.706695554 |
| chr16:23678448-23679017 | 0.268111799 | 0.429484811 | 0.706695554 |
| chr1:77779421-77780330 | -0.256895997 | 0.429506345 | 0.70671581 |
| chr15:70507850-70508377 | 0.273117366 | 0.429602645 | 0.706844899 |
| chr8:109755734-109756054 | 0.359225616 | 0.429606297 | 0.706844899 |
| chr8:115276380-115276760 | -0.386948144 | 0.429612473 | 0.706844899 |
| chr3:15202728-15202973 | 0.300617793 | 0.429636957 | 0.706870003 |
| chr9:121486040-121486371 | 0.355996523 | 0.42965734 | 0.70688836 |
| chr7:97104700-97104988 | -0.337766198 | 0.429687019 | 0.706922009 |
| chr10:86716872-86717094 | -0.351425828 | 0.429727852 | 0.706936573 |
| chr5:174317271-174317564 | 0.307727842 | 0.429734747 | 0.706936573 |
| chr16:50720007-50720337 | -0.328916856 | 0.429739403 | 0.706936573 |
| chr3:67940649-67940882 | -0.40932954 | 0.429747573 | 0.706936573 |
| chrX:47803090-47803547 | 0.317422421 | 0.429750241 | 0.706936573 |
| chr12:85836746-85837067 | 0.350309418 | 0.429761374 | 0.706936573 |
| chr6:170575320-170575525 | 0.37887113 | 0.429770951 | 0.706936573 |
| chr1:75842569-75843008 | 0.29834888 | 0.429773977 | 0.706936573 |
| chr11:63478876-63479255 | -0.325496998 | 0.429786187 | 0.706936573 |
| chr13:104389877-104390077 | -0.482560006 | 0.429788134 | 0.706936573 |
| chr22:31122389-31122597 | 0.398706938 | 0.429818395 | 0.70695631 |
| chr7:98727311-98727534 | -0.425898505 | 0.429818586 | 0.70695631 |
| chr11:74948599-74949538 | -0.256905793 | 0.429830368 | 0.706960513 |
| chr2:87076252-87076890 | 0.277354497 | 0.429870023 | 0.707010559 |
| chr7:44044337-44045399 | -0.256234221 | 0.429888621 | 0.707025736 |
| chr11:16606004-16606715 | -0.27475573 | 0.429898659 | 0.707025736 |
| chr6:26204382-26205036 | -0.264602625 | 0.429906933 | 0.707025736 |
| chr4:140135842-140136042 | 0.427785381 | 0.429926307 | 0.707042422 |
| chr8:132651466-132651792 | -0.327733283 | 0.429949715 | 0.707065743 |
| chr5:142459434-142459852 | 0.38816838 | 0.42998333 | 0.707105846 |
| chr20:45419502-45419871 | 0.292643462 | 0.430005676 | 0.707125639 |
| chr7:32862514-32862813 | 0.346706032 | 0.430013823 | 0.707125639 |
| chr6:63066189-63066437 | -0.380534751 | 0.430042221 | 0.707126784 |
| chr3:66740372-66740752 | -0.312655242 | 0.430043212 | 0.707126784 |
| chr3:153260598-153260892 | 0.346585295 | 0.43004767 | 0.707126784 |

| | | | |
|---|---|---|---|
| chr17:68204773-68205024 | 0.294811854 | 0.43005959 | 0.707126784 |
| chr3:105232727-105232927 | -0.409346598 | 0.430060663 | 0.707126784 |
| chr13:44026323-44026523 | 0.410134221 | 0.430076067 | 0.707136937 |
| chr1:173429936-173430201 | -0.318340616 | 0.430108152 | 0.707163497 |
| chr20:52235535-52235884 | -0.355071709 | 0.430116302 | 0.707163497 |
| chr10:79071180-79071447 | -0.289958289 | 0.430128909 | 0.707163497 |
| chr5:151692279-151692718 | 0.306245457 | 0.430129138 | 0.707163497 |
| chr20:428857-429084 | -0.401524158 | 0.430157182 | 0.707194431 |
| chr19:6737270-6738172 | 0.255991753 | 0.430172525 | 0.707204481 |
| chr1:175022916-175023965 | -0.250560149 | 0.430192227 | 0.707221695 |
| chr10:67884147-67885376 | -0.251726999 | 0.430205512 | 0.707228362 |
| chr1:65719179-65719391 | 0.426393788 | 0.430256925 | 0.707297707 |
| chr11:67614043-67614491 | -0.293202357 | 0.430289399 | 0.70733094 |
| chr14:50619602-50619894 | 0.32841374 | 0.430317155 | 0.70733094 |
| chr4:90158307-90159348 | 0.266786912 | 0.430320862 | 0.70733094 |
| chr12:122880792-122881215 | 0.322826829 | 0.430323994 | 0.70733094 |
| chr6:12602434-12602748 | 0.320502368 | 0.430325936 | 0.70733094 |
| chr10:120286465-120286691 | 0.403432595 | 0.43033253 | 0.70733094 |
| chr2:126528220-126528579 | -0.341559554 | 0.430343193 | 0.707333294 |
| chr4:2441590-2442169 | -0.283770968 | 0.430375732 | 0.707368918 |
| chr5:28576069-28576432 | -0.306379318 | 0.430383331 | 0.707368918 |
| chr15:23039056-23040050 | 0.258930662 | 0.430404529 | 0.707373997 |
| chr18:3666331-3666591 | -0.325149768 | 0.430404885 | 0.707373997 |
| chr13:28495113-28495400 | -0.341291687 | 0.430428788 | 0.707398108 |
| chr4:100558668-100558898 | 0.403017539 | 0.430439106 | 0.707399892 |
| chr22:38871912-38872592 | 0.285795911 | 0.43046481 | 0.707418072 |
| chr15:43882018-43882250 | -0.371484803 | 0.430468633 | 0.707418072 |
| chr15:40776719-40777256 | -0.285788034 | 0.430510326 | 0.707450384 |
| chr6:10616397-10616751 | -0.406716141 | 0.430511822 | 0.707450384 |
| chr7:6705934-6707398 | -0.249378275 | 0.430515994 | 0.707450384 |
| chr12:63843436-63844840 | 0.248856827 | 0.4305396 | 0.707474002 |
| chr16:30996921-30997521 | 0.267244138 | 0.430549885 | 0.70747573 |
| chr5:79474014-79474241 | -0.44001806 | 0.430566221 | 0.707481372 |
| chr12:121399599-121400592 | 0.255011534 | 0.430578849 | 0.707481372 |
| chr16:19930020-19930279 | -0.393102725 | 0.430581019 | 0.707481372 |
| chr9:88370351-88370651 | 0.396976427 | 0.430679218 | 0.7075972 |
| chr9:92969665-92969897 | 0.396976427 | 0.430679218 | 0.7075972 |
| chr6:82178181-82178390 | 0.396976427 | 0.430679218 | 0.7075972 |
| chr22:19179370-19179698 | 0.290523383 | 0.430693796 | 0.707605979 |
| chr1:85276065-85277482 | -0.246907933 | 0.430724407 | 0.707641097 |
| chr6:105094282-105094484 | -0.416090926 | 0.430773972 | 0.707688049 |
| chr3:71729891-71730231 | 0.303583219 | 0.430779664 | 0.707688049 |
| chr2:27889999-27891386 | -0.251704868 | 0.430780694 | 0.707688049 |
| chr12:108956527-108956849 | 0.361217326 | 0.430836329 | 0.707740798 |
| chrX:101550286-101551487 | -0.255846258 | 0.430837112 | 0.707740798 |
| chr6:33299092-33299989 | -0.255790943 | 0.430840513 | 0.707740798 |
| chr2:86631877-86632319 | -0.308170239 | 0.430878138 | 0.70778743 |
| chr20:34069825-34070098 | 0.330841982 | 0.430897866 | 0.707797677 |
| chr9:22084713-22085074 | -0.316670996 | 0.430902851 | 0.707797677 |
| chr2:161250925-161251258 | -0.3134277 | 0.430943974 | 0.707848037 |

| | | | |
|---|---|---|---|
| chr11:35916707-35917116 | -0.38224596 | 0.430951986 | 0.707848037 |
| chr17:56973634-56973879 | 0.314041053 | 0.431038022 | 0.707961276 |
| chrX:15687110-15687369 | -0.365431492 | 0.431039408 | 0.707961276 |
| chr17:49032201-49032497 | 0.298405258 | 0.431055957 | 0.707973281 |
| chr1:116508126-116508354 | 0.363348074 | 0.431077673 | 0.707993773 |
| chr14:23379872-23380280 | 0.284574697 | 0.431092788 | 0.708003421 |
| chr16:75754214-75754498 | -0.334288173 | 0.431122761 | 0.708014841 |
| chr8:135828612-135828892 | 0.336366904 | 0.431140146 | 0.708014841 |
| chr20:32819647-32820687 | 0.258865233 | 0.431143067 | 0.708014841 |
| chr16:28291700-28291935 | 0.356329681 | 0.43114445 | 0.708014841 |
| chr7:36279304-36279524 | 0.355068105 | 0.431145943 | 0.708014841 |
| chr5:154631934-154632161 | -0.37811981 | 0.431163196 | 0.708027999 |
| chr12:118120886-118121399 | 0.29653944 | 0.431178762 | 0.708028208 |
| chr19:6393028-6393764 | 0.271801759 | 0.431181805 | 0.708028208 |
| chr17:64355576-64355776 | -0.349059029 | 0.431198369 | 0.708040013 |
| chr13:20021513-20021782 | -0.338823022 | 0.431207475 | 0.708040013 |
| chr2:88497727-88498010 | -0.355209656 | 0.431334718 | 0.708233767 |
| chr5:14871774-14872669 | -0.265332457 | 0.431393466 | 0.70831505 |
| chr6:53179891-53180175 | -0.384727919 | 0.431428713 | 0.70833449 |
| chr2:199850599-199851590 | -0.256070175 | 0.431434444 | 0.70833449 |
| chrX:141879529-141879956 | 0.303906274 | 0.431441078 | 0.70833449 |
| chr11:1705432-1706055 | 0.281306233 | 0.431443195 | 0.70833449 |
| chr12:85867051-85867524 | -0.293805869 | 0.431454189 | 0.70833449 |
| chr11:32188829-32189098 | -0.37068178 | 0.431473008 | 0.70833449 |
| chr13:44350833-44351521 | -0.29308551 | 0.431473795 | 0.70833449 |
| chr1:22053207-22053457 | 0.350832207 | 0.431479261 | 0.70833449 |
| chr3:158733746-158734071 | 0.319714496 | 0.431502815 | 0.708337964 |
| chr15:57461538-57461773 | 0.364285248 | 0.431511623 | 0.708337964 |
| chr5:72816055-72817441 | -0.254397677 | 0.431517856 | 0.708337964 |
| chr6:77465938-77466299 | -0.303578687 | 0.43152585 | 0.708337964 |
| chr14:91858647-91858918 | 0.331891738 | 0.431556303 | 0.708337964 |
| chr14:90191059-90191290 | 0.371325394 | 0.431565861 | 0.708337964 |
| chr15:99474913-99475113 | 0.414764701 | 0.431569506 | 0.708337964 |
| chr1:190085078-190085278 | -0.392077784 | 0.431585539 | 0.708337964 |
| chr1:235759009-235759231 | -0.392077784 | 0.431585539 | 0.708337964 |
| chr19:54030006-54030211 | -0.403959344 | 0.431590604 | 0.708337964 |
| chr12:123706034-123706234 | -0.463355311 | 0.431597495 | 0.708337964 |
| chr19:10587167-10587443 | 0.34152272 | 0.431599122 | 0.708337964 |
| chr7:1182859-1183328 | 0.28546721 | 0.431601558 | 0.708337964 |
| chr11:75248633-75248982 | 0.296793412 | 0.431633743 | 0.708375613 |
| chr2:148604546-148604785 | -0.398211735 | 0.43170388 | 0.708475544 |
| chr2:74653755-74654835 | -0.250430266 | 0.431747936 | 0.708504697 |
| chr17:58903219-58903426 | 0.313080238 | 0.431750647 | 0.708504697 |
| chr1:34742687-34742984 | -0.336782084 | 0.431758631 | 0.708504697 |
| chr11:118321759-118322126 | -0.336782084 | 0.431758631 | 0.708504697 |
| chr1:179105382-179105738 | -0.280296635 | 0.431779493 | 0.708512138 |
| chr20:47799289-47799511 | 0.277201996 | 0.431781659 | 0.708512138 |
| chr9:114682035-114682409 | 0.283150347 | 0.431793636 | 0.708516618 |
| chr15:74950431-74950924 | -0.287658766 | 0.431853839 | 0.708600228 |
| chr1:190065057-190065262 | -0.428890989 | 0.431886412 | 0.708637181 |

| | | | |
|---|---|---|---|
| chr14:102762242-102762485 | -0.388638864 | 0.431909167 | 0.708637181 |
| chr5:172040202-172040721 | -0.29426409 | 0.43192391 | 0.708637181 |
| chr1:204637627-204637868 | 0.336550694 | 0.431943068 | 0.708637181 |
| chr10:12716157-12716409 | -0.397891909 | 0.431943444 | 0.708637181 |
| chr2:136468994-136469308 | 0.33664919 | 0.43194448 | 0.708637181 |
| chr13:73096242-73096618 | -0.347479724 | 0.431951918 | 0.708637181 |
| chr19:17993201-17993467 | 0.349867122 | 0.431960739 | 0.708637181 |
| chr14:102316855-102317694 | 0.252983782 | 0.431969114 | 0.708637181 |
| chr3:159266871-159267155 | 0.341872721 | 0.431972819 | 0.708637181 |
| chr17:31072246-31072503 | 0.366930845 | 0.431980323 | 0.708637181 |
| chr7:93551171-93551371 | -0.454707054 | 0.431987898 | 0.708637181 |
| chr3:139265329-139265529 | 0.414846856 | 0.43199659 | 0.708637181 |
| chr17:4986777-4987832 | 0.263477938 | 0.432028008 | 0.708673546 |
| chr16:8791980-8792224 | 0.347146724 | 0.43205118 | 0.708681815 |
| chr19:12866402-12866602 | 0.409506918 | 0.432051547 | 0.708681815 |
| chr1:146400179-146400380 | 0.378756282 | 0.432083354 | 0.708703644 |
| chr13:23493675-23493943 | 0.378756282 | 0.432083354 | 0.708703644 |
| chr10:133261259-133261664 | 0.285594356 | 0.43210391 | 0.708716356 |
| chr20:17536960-17537454 | 0.276908907 | 0.432109603 | 0.708716356 |
| chr1:153566250-153567101 | 0.257814907 | 0.432139905 | 0.708747414 |
| chr11:81237254-81237462 | 0.368600137 | 0.432147039 | 0.708747414 |
| chr2:42787272-42787756 | -0.273764751 | 0.432171978 | 0.708773145 |
| chr1:180680588-180680788 | 0.405937301 | 0.432184896 | 0.708775625 |
| chr3:88311829-88312213 | 0.312308561 | 0.432191991 | 0.708775625 |
| chr6:82101230-82101968 | -0.276061134 | 0.432208557 | 0.708777877 |
| chr11:126365365-126365903 | -0.294904815 | 0.432220954 | 0.708777877 |
| chr1:2534855-2535106 | 0.34610675 | 0.432221116 | 0.708777877 |
| chr2:98936100-98936667 | 0.303005127 | 0.43224077 | 0.708784988 |
| chr5:19795251-19795569 | -0.307889642 | 0.432247747 | 0.708784988 |
| chr3:8714004-8714238 | 0.319131458 | 0.43225996 | 0.708784988 |
| chr1:113608420-113608620 | 0.365175245 | 0.432270816 | 0.708784988 |
| chr1:16155712-16156198 | 0.316153725 | 0.432271704 | 0.708784988 |
| chr4:972621-973062 | 0.305522898 | 0.43233259 | 0.708859703 |
| chr7:150898557-150898855 | -0.321109328 | 0.432335773 | 0.708859703 |
| chrX:106726576-106726859 | -0.375760444 | 0.432350192 | 0.708861183 |
| chr11:126368868-126369152 | 0.326095063 | 0.432355179 | 0.708861183 |
| chr17:64496167-64497679 | -0.246694107 | 0.432371169 | 0.708864173 |
| chr3:88058333-88059888 | -0.246223046 | 0.432375506 | 0.708864173 |
| chr4:139734806-139735259 | 0.304350081 | 0.432452122 | 0.708960319 |
| chr3:152474674-152474874 | -0.371284485 | 0.432478121 | 0.708960319 |
| chr22:30224230-30224430 | 0.400272782 | 0.432484192 | 0.708960319 |
| chr20:49608900-49609218 | 0.287883395 | 0.43249732 | 0.708960319 |
| chr7:100642536-100643055 | -0.299522194 | 0.432497354 | 0.708960319 |
| chr9:93533824-93534112 | -0.363510294 | 0.432501704 | 0.708960319 |
| chr1:218048918-218049127 | -0.360552353 | 0.432504086 | 0.708960319 |
| chr1:144545846-144546950 | 0.249596402 | 0.432508172 | 0.708960319 |
| chr1:115807836-115808036 | 0.359326317 | 0.432529602 | 0.708971092 |
| chr5:74648850-74649114 | -0.361282049 | 0.432535477 | 0.708971092 |
| chr17:19648163-19649240 | -0.253163463 | 0.432542503 | 0.708971092 |
| chrX:154420465-154420665 | 0.402744754 | 0.432552016 | 0.708971519 |

| | | | |
|---|---|---|---|
| chr2:4354634-4354987 | -0.307597278 | 0.432568252 | 0.708982964 |
| chr1:119275473-119275733 | -0.397379064 | 0.432581883 | 0.708990139 |
| chr13:48291943-48292185 | -0.396021094 | 0.4326248 | 0.709027565 |
| chr14:60207310-60207524 | -0.405024942 | 0.432626639 | 0.709027565 |
| chr19:30669472-30669687 | -0.373134268 | 0.432638927 | 0.709027565 |
| chr8:140727380-140727610 | 0.403233615 | 0.432641732 | 0.709027565 |
| chr14:64914113-64914917 | 0.25353941 | 0.432654092 | 0.709032657 |
| chr5:144467760-144468098 | -0.315333547 | 0.432703322 | 0.709084273 |
| chr2:143513045-143513386 | -0.348238426 | 0.432704098 | 0.709084273 |
| chr8:108691311-108691511 | 0.363374229 | 0.432740331 | 0.70912555 |
| chr8:137334681-137334930 | -0.327760822 | 0.432747796 | 0.70912555 |
| chr1:231590687-231590892 | -0.397679417 | 0.432777152 | 0.709135262 |
| chr21:25734464-25735881 | 0.248169912 | 0.432777777 | 0.709135262 |
| chr3:128845208-128846515 | -0.252800453 | 0.432784161 | 0.709135262 |
| chr3:8324772-8325108 | -0.297825708 | 0.432792503 | 0.709135262 |
| chr4:121484626-121484885 | 0.390165075 | 0.432808552 | 0.709135262 |
| chr11:120952792-120953138 | 0.298472871 | 0.432809252 | 0.709135262 |
| chr11:9607125-9607366 | 0.371111621 | 0.432848968 | 0.709185169 |
| chr6:138692053-138693870 | -0.251704544 | 0.432870852 | 0.709202868 |
| chr12:92574391-92574718 | -0.32829249 | 0.432885612 | 0.709202868 |
| chr10:4815309-4815518 | 0.355047952 | 0.43290884 | 0.709202868 |
| chr9:128552991-128553212 | 0.358761955 | 0.432920504 | 0.709202868 |
| chr3:193592776-193593988 | -0.248748689 | 0.432934817 | 0.709202868 |
| chr8:123980718-123980935 | 0.318723259 | 0.432938934 | 0.709202868 |
| chr13:99926562-99926783 | 0.402221781 | 0.432940163 | 0.709202868 |
| chr6:36766215-36766457 | 0.33083136 | 0.432946158 | 0.709202868 |
| chrX:103282834-103283227 | -0.31340413 | 0.432970661 | 0.709202868 |
| chr2:230684865-230685082 | -0.378042823 | 0.432970726 | 0.709202868 |
| chr7:74511658-74512028 | -0.370879733 | 0.432974776 | 0.709202868 |
| chr4:17562245-17562526 | 0.335805894 | 0.432975706 | 0.709202868 |
| chr2:111029608-111029844 | -0.371663374 | 0.432989171 | 0.709202868 |
| chr2:105125366-105125776 | -0.313417378 | 0.432997644 | 0.709202868 |
| chr9:81656674-81657127 | 0.329182122 | 0.432998608 | 0.709202868 |
| chr6:53794055-53795495 | -0.250057303 | 0.433032209 | 0.709242741 |
| chr15:98597676-98597884 | 0.397223291 | 0.433047021 | 0.70925184 |
| chr10:119207919-119208131 | 0.342235003 | 0.433066786 | 0.709269052 |
| chr6:45175747-45176178 | -0.34292205 | 0.433086161 | 0.709285622 |
| chr14:49867871-49868661 | 0.252866395 | 0.43309785 | 0.709289605 |
| chr2:88160051-88160456 | -0.343058964 | 0.433126735 | 0.709313884 |
| chr8:144078399-144078946 | 0.263801819 | 0.433134786 | 0.709313884 |
| chr3:139906354-139906645 | -0.374734379 | 0.433140447 | 0.709313884 |
| chr5:173625020-173625452 | -0.316613422 | 0.433178956 | 0.709361786 |
| chr4:56183458-56183658 | -0.402180212 | 0.433220739 | 0.70940316 |
| chr7:12802021-12802598 | -0.272860076 | 0.433222739 | 0.70940316 |
| chr16:8621354-8622244 | 0.259784997 | 0.4332555 | 0.709420554 |
| chr11:68972768-68972982 | 0.376060448 | 0.433266109 | 0.709420554 |
| chr18:50281190-50282003 | -0.251887456 | 0.433281886 | 0.709420554 |
| chr14:94599162-94599646 | 0.289132431 | 0.433284654 | 0.709420554 |
| chr16:48623160-48623719 | 0.281911557 | 0.433285346 | 0.709420554 |
| chr4:37502961-37503276 | 0.354834008 | 0.433299706 | 0.709420554 |

| | | | |
|---|---|---|---|
| chr4:77618654-77619034 | -0.29727987 | 0.433305357 | 0.709420554 |
| chr16:67993105-67993670 | 0.28239294 | 0.43330743 | 0.709420554 |
| chr5:20522324-20522534 | -0.438185226 | 0.433322627 | 0.709427957 |
| chr16:88639319-88639520 | -0.324512779 | 0.433335549 | 0.709427957 |
| chr6:89792792-89793209 | -0.295691388 | 0.433339728 | 0.709427957 |
| chr3:13484999-13485569 | -0.270992589 | 0.433367805 | 0.709451406 |
| chr11:67248547-67248996 | 0.275867811 | 0.43337257 | 0.709451406 |
| chr10:103966601-103967498 | -0.264329921 | 0.433384383 | 0.709452329 |
| chr1:145943674-145943948 | -0.367170576 | 0.433400638 | 0.709452329 |
| chr11:16924633-16926233 | 0.253382383 | 0.433400911 | 0.709452329 |
| chr17:46758699-46758899 | 0.426741129 | 0.433443115 | 0.709506256 |
| chr17:41785797-41787228 | 0.247688715 | 0.433454838 | 0.709506377 |
| chr6:75208918-75209236 | -0.288039271 | 0.433481774 | 0.709506377 |
| chr10:12782522-12782722 | -0.411572648 | 0.433483587 | 0.709506377 |
| chr12:44000345-44001020 | 0.279796604 | 0.433487103 | 0.709506377 |
| chr20:50713754-50714068 | 0.256626029 | 0.433489488 | 0.709506377 |
| chr19:36041898-36042098 | -0.411693033 | 0.433504328 | 0.709514063 |
| chr5:177457627-177458015 | -0.263539145 | 0.433520319 | 0.709514063 |
| chr2:127453392-127453912 | 0.287240755 | 0.433525398 | 0.709514063 |
| chr2:81598496-81598776 | 0.317702193 | 0.433531223 | 0.709514063 |
| chr8:128127342-128127804 | -0.293118181 | 0.433542773 | 0.70951781 |
| chr11:844473-844993 | -0.293186805 | 0.433560545 | 0.709531741 |
| chr14:37153662-37154077 | -0.350007712 | 0.433577164 | 0.709543784 |
| chr19:54208868-54209078 | -0.365767877 | 0.433637264 | 0.709613256 |
| chr6:26022189-26022399 | 0.314812927 | 0.433639898 | 0.709613256 |
| chr16:8567643-8568099 | 0.275230887 | 0.433651813 | 0.709613256 |
| chr15:75204803-75205075 | 0.284990061 | 0.433656661 | 0.709613256 |
| chr1:157363639-157363965 | 0.315335586 | 0.43368272 | 0.709630861 |
| chr18:46374952-46375268 | -0.346968701 | 0.433690126 | 0.709630861 |
| chrX:135206501-135206813 | 0.317624594 | 0.433695205 | 0.709630861 |
| chr1:197980249-197980793 | -0.301262906 | 0.433714167 | 0.709646734 |
| chr12:83004673-83004926 | 0.353115726 | 0.433768352 | 0.709705084 |
| chr6:20403281-20403778 | -0.260065825 | 0.433768354 | 0.709705084 |
| chr18:46916987-46917864 | -0.269091518 | 0.433796612 | 0.709736163 |
| chr18:62201923-62202169 | -0.370988514 | 0.433831085 | 0.709777409 |
| chr2:225761537-225761737 | -0.443449419 | 0.433840846 | 0.709778223 |
| chr3:32420233-32420495 | 0.352886915 | 0.433852004 | 0.709781322 |
| chr21:31671404-31671758 | -0.357728292 | 0.433886669 | 0.709805762 |
| chr1:39883963-39884168 | 0.39962511 | 0.433890237 | 0.709805762 |
| chr1:15847218-15847914 | 0.274773976 | 0.433894733 | 0.709805762 |
| chr10:78947343-78947543 | -0.39982253 | 0.43395592 | 0.7098907 |
| chr1:179877662-179878135 | -0.263885343 | 0.43402419 | 0.709987222 |
| chr6:35317895-35318910 | 0.260024478 | 0.434061022 | 0.710022074 |
| chr3:186980000-186980252 | 0.35804699 | 0.434064029 | 0.710022074 |
| chr22:30250793-30251055 | -0.35776383 | 0.434097636 | 0.710048396 |
| chr4:40908679-40909181 | -0.283177785 | 0.434098654 | 0.710048396 |
| chr9:108761910-108762132 | 0.363664113 | 0.434118769 | 0.710063317 |
| chr12:101570116-101570365 | -0.377000347 | 0.43412631 | 0.710063317 |
| chr17:50317109-50317494 | 0.292212112 | 0.434138529 | 0.710065969 |
| chr6:31664852-31666240 | 0.250688282 | 0.434146466 | 0.710065969 |

| | | | |
|---|---|---|---|
| chr3:185282716-185283423 | 0.258633005 | 0.434201378 | 0.710130036 |
| chr3:192879285-192880093 | 0.260117285 | 0.434209721 | 0.710130036 |
| chr18:35343824-35344719 | -0.25279608 | 0.434213442 | 0.710130036 |
| chr2:112345197-112346175 | 0.267331784 | 0.434248803 | 0.71017271 |
| chr7:99013581-99013794 | -0.422787506 | 0.434266907 | 0.710174844 |
| chr17:60571611-60571846 | 0.276208236 | 0.434276514 | 0.710174844 |
| chr10:28302524-28303489 | -0.248898534 | 0.434277914 | 0.710174844 |
| chr11:71283904-71284450 | 0.271493011 | 0.434336843 | 0.710248472 |
| chr14:92105862-92106941 | 0.248746355 | 0.434356913 | 0.710248472 |
| chr7:46778548-46778860 | -0.391723362 | 0.434373918 | 0.710248472 |
| chr17:60676995-60678438 | -0.242270481 | 0.434378877 | 0.710248472 |
| chr14:59631372-59631572 | 0.366727122 | 0.434384852 | 0.710248472 |
| chr11:117226478-117226781 | 0.311730665 | 0.434385375 | 0.710248472 |
| chr4:139665628-139666424 | -0.262322049 | 0.434387824 | 0.710248472 |
| chr19:3275918-3276906 | 0.251585996 | 0.434432396 | 0.710294308 |
| chr5:174019779-174020734 | -0.264600856 | 0.434451417 | 0.710294308 |
| chr10:44959218-44960046 | -0.256269769 | 0.434451527 | 0.710294308 |
| chr12:57001466-57001745 | 0.333780936 | 0.434453139 | 0.710294308 |
| chr3:107660511-107660711 | 0.425806313 | 0.434462713 | 0.710294308 |
| chr5:54229279-54229739 | 0.309452742 | 0.434471479 | 0.710294308 |
| chr6:129575168-129575869 | -0.283776417 | 0.434499225 | 0.710324513 |
| chr20:46220858-46221136 | 0.331261146 | 0.434536414 | 0.710370154 |
| chr8:139651177-139651401 | 0.344272902 | 0.434556195 | 0.710374607 |
| chr14:66403338-66403555 | -0.356216852 | 0.434566527 | 0.710374607 |
| chr11:34052888-34053200 | -0.321873339 | 0.434566952 | 0.710374607 |
| chr16:58249728-58250354 | 0.277823134 | 0.434594975 | 0.710402705 |
| chr19:10251718-10252344 | 0.262368308 | 0.434602683 | 0.710402705 |
| chr16:67667890-67668351 | 0.27353239 | 0.434632493 | 0.710435205 |
| chr14:49830154-49830388 | 0.372881272 | 0.43464111 | 0.710435205 |
| chr10:125160176-125160735 | 0.276832882 | 0.434678918 | 0.710480122 |
| chr4:119268310-119268754 | -0.333317738 | 0.434687135 | 0.710480122 |
| chr1:66242741-66243032 | 0.322382565 | 0.434707516 | 0.710498279 |
| chr1:152210968-152211402 | 0.310085561 | 0.434724723 | 0.710510756 |
| chr1:108560971-108561324 | -0.274201974 | 0.434733696 | 0.710510756 |
| chr19:10460926-10461881 | -0.258315228 | 0.434751309 | 0.710515564 |
| chr7:134750151-134750442 | -0.361781727 | 0.434768269 | 0.710515564 |
| chr11:21567719-21568297 | -0.287708137 | 0.434768746 | 0.710515564 |
| chr5:151354319-151354591 | -0.352419017 | 0.434775618 | 0.710515564 |
| chr12:44876079-44877167 | -0.255873583 | 0.434791547 | 0.710515564 |
| chr1:187051647-187051902 | -0.342265224 | 0.434792276 | 0.710515564 |
| chr1:162800262-162800601 | 0.312411621 | 0.43480635 | 0.71052341 |
| chr12:118181524-118181867 | 0.313393015 | 0.434837063 | 0.710547819 |
| chr7:73792414-73793283 | -0.259625341 | 0.434839834 | 0.710547819 |
| chr6:24284084-24284313 | -0.369421837 | 0.434851156 | 0.710551166 |
| chr15:98421607-98421865 | -0.323807413 | 0.434914933 | 0.710640224 |
| chr16:75267863-75268240 | -0.270212664 | 0.43495565 | 0.710677735 |
| chr8:135567898-135568149 | -0.393802828 | 0.43496228 | 0.710677735 |
| chr2:112582316-112582555 | -0.321922953 | 0.434970123 | 0.710677735 |
| chr3:713822-714031 | 0.430477653 | 0.434974991 | 0.710677735 |
| chr3:163476245-163476731 | -0.304599457 | 0.435000051 | 0.710703525 |

| | | | |
|---|---|---|---|
| chr2:74391214-74392278 | -0.260359673 | 0.435023538 | 0.71071823 |
| chr6:143450346-143451372 | -0.251776652 | 0.435035973 | 0.71071823 |
| chr3:129352081-129352368 | -0.301890177 | 0.435041348 | 0.71071823 |
| chr6:79076892-79077291 | 0.273978613 | 0.435046154 | 0.71071823 |
| chr12:121536664-121537517 | 0.262064158 | 0.435056243 | 0.710719559 |
| chr4:87659289-87659579 | 0.330908827 | 0.435071559 | 0.710729425 |
| chr17:60768473-60769795 | 0.243945224 | 0.435093997 | 0.710750927 |
| chr3:157865268-157865740 | -0.286097807 | 0.435121789 | 0.710767098 |
| chr11:65918959-65920133 | 0.249786404 | 0.435122448 | 0.710767098 |
| chr5:170267919-170268172 | 0.326142903 | 0.435160694 | 0.710814418 |
| chr5:23411950-23412364 | 0.295478036 | 0.435193778 | 0.710853304 |
| chr15:73684208-73684460 | 0.330826294 | 0.435207194 | 0.710860064 |
| chr11:76169740-76169967 | -0.348147908 | 0.435220873 | 0.710867253 |
| chr12:116222451-116222770 | -0.388991093 | 0.435234435 | 0.71087285 |
| chr5:152689312-152689546 | -0.384209428 | 0.435251003 | 0.71087285 |
| chr10:93580795-93581031 | -0.380005656 | 0.435252133 | 0.71087285 |
| chr6:90664006-90664214 | -0.40235111 | 0.435263275 | 0.710875896 |
| chr17:57747411-57747877 | 0.277943024 | 0.435274804 | 0.710879572 |
| chr5:150436089-150436523 | 0.285552543 | 0.435311595 | 0.710924506 |
| chr17:82751350-82752858 | 0.249482327 | 0.435324198 | 0.710929934 |
| chr8:93037978-93038195 | -0.430704251 | 0.435349994 | 0.710956909 |
| chr11:69545704-69545904 | 0.388360761 | 0.435409242 | 0.711017105 |
| chr16:85212469-85212700 | 0.326180138 | 0.43541503 | 0.711017105 |
| chr7:43421962-43422162 | 0.429464689 | 0.435424216 | 0.711017105 |
| chr17:21098889-21099731 | -0.256928944 | 0.435446815 | 0.711017105 |
| chrX:13669206-13669685 | 0.270624995 | 0.435451583 | 0.711017105 |
| chr16:2882495-2883706 | 0.248600582 | 0.43545181 | 0.711017105 |
| chr20:54920434-54920889 | 0.258009849 | 0.435451811 | 0.711017105 |
| chr1:167617306-167617619 | -0.376375231 | 0.435479815 | 0.711047679 |
| chr5:44603485-44604280 | -0.268677674 | 0.435504358 | 0.711050373 |
| chr1:46220884-46221084 | 0.417604547 | 0.435504453 | 0.711050373 |
| chr17:82592850-82593459 | -0.271022249 | 0.435515784 | 0.711050373 |
| chr14:52961137-52961454 | 0.293186901 | 0.43552418 | 0.711050373 |
| chr4:67713241-67713455 | 0.359789157 | 0.435527866 | 0.711050373 |
| chr4:137909938-137910212 | 0.344614837 | 0.435553121 | 0.711076455 |
| chr2:182867020-182867343 | -0.302292934 | 0.4356325 | 0.711182654 |
| chr3:169764334-169765320 | -0.252482909 | 0.435637773 | 0.711182654 |
| chr8:144819968-144820245 | -0.351792893 | 0.435646016 | 0.711182654 |
| chr21:38299893-38300093 | -0.429100688 | 0.435657715 | 0.7111866 |
| chr6:100518608-100518808 | 0.400139505 | 0.435688947 | 0.711222432 |
| chr15:98442463-98442739 | -0.311274584 | 0.435727716 | 0.711261664 |
| chr1:31977708-31978105 | -0.312550703 | 0.435731546 | 0.711261664 |
| chr5:107542347-107542846 | -0.283419693 | 0.435766135 | 0.711287243 |
| chr2:158255374-158255591 | 0.411120534 | 0.43577124 | 0.711287243 |
| chr10:15859766-15860733 | -0.258735814 | 0.435783324 | 0.711287243 |
| chr3:13112317-13112565 | 0.310318744 | 0.435793602 | 0.711287243 |
| chr2:151857655-151857971 | -0.359720016 | 0.435802709 | 0.711287243 |
| chr4:87636641-87636841 | 0.4298515 | 0.435802914 | 0.711287243 |
| chr19:46991244-46991509 | -0.344495258 | 0.435831404 | 0.71131859 |
| chr2:196078273-196078610 | -0.297390098 | 0.435843047 | 0.711322441 |

| | | | |
|---|---|---|---|
| chr10:8103661-8103949 | -0.331952121 | 0.435883433 | 0.71136434 |
| chr7:73647886-73648095 | 0.399778403 | 0.435895018 | 0.71136434 |
| chr1:188188398-188188723 | -0.32224799 | 0.435896572 | 0.71136434 |
| chr5:84274927-84275212 | -0.340829975 | 0.435910921 | 0.711372606 |
| chr3:184468500-184468717 | 0.365843742 | 0.435952566 | 0.71141428 |
| chr10:101602767-101603045 | 0.326507205 | 0.435955027 | 0.71141428 |
| chr8:127772236-127772447 | -0.380870134 | 0.435972918 | 0.711428324 |
| chr2:11688123-11688767 | -0.257572336 | 0.435989365 | 0.71144001 |
| chr12:57229477-57229937 | 0.269088392 | 0.436034729 | 0.711498881 |
| chr6:47309397-47310209 | -0.255470731 | 0.436068118 | 0.711538211 |
| chr2:158007410-158007823 | -0.315719358 | 0.436082902 | 0.711547183 |
| chr17:70621698-70621960 | -0.341287718 | 0.436127647 | 0.711605038 |
| chr8:16371508-16371762 | -0.375740679 | 0.436167029 | 0.711638161 |
| chr2:162524253-162524498 | -0.371686077 | 0.436173953 | 0.711638161 |
| chr17:48907976-48908563 | -0.255984288 | 0.436200395 | 0.711638161 |
| chr16:68770643-68770843 | -0.430901384 | 0.43620661 | 0.711638161 |
| chr20:54829208-54829586 | 0.258087623 | 0.436212907 | 0.711638161 |
| chr2:69082440-69082979 | -0.296164466 | 0.436213069 | 0.711638161 |
| chr6:159461295-159461517 | -0.370286134 | 0.43621665 | 0.711638161 |
| chr1:180269903-180270161 | -0.313036662 | 0.436222249 | 0.711638161 |
| chr5:13781830-13782108 | 0.366970026 | 0.436238922 | 0.71165021 |
| chr9:80664912-80665128 | -0.383552983 | 0.436263969 | 0.711675917 |
| chr17:54694342-54694589 | -0.374253501 | 0.43627634 | 0.711680946 |
| chr4:22522078-22522318 | -0.371442981 | 0.436312179 | 0.711715295 |
| chr5:150783311-150783807 | -0.282832775 | 0.436327192 | 0.711715295 |
| chr4:175239172-175239408 | -0.37958645 | 0.43633328 | 0.711715295 |
| chrX:136496544-136498033 | 0.248416704 | 0.436358214 | 0.711715295 |
| chr6:143677799-143678748 | -0.263554846 | 0.436367114 | 0.711715295 |
| chr2:164992903-164993185 | -0.351325596 | 0.436375954 | 0.711715295 |
| chr5:92050818-92051023 | -0.409461784 | 0.436385794 | 0.711715295 |
| chr1:205827683-205827984 | 0.285909694 | 0.436387759 | 0.711715295 |
| chr12:14769918-14770522 | -0.263244493 | 0.436387975 | 0.711715295 |
| chr8:67490372-67490576 | -0.397763416 | 0.436394478 | 0.711715295 |
| chr10:125201960-125202248 | -0.367922135 | 0.436399572 | 0.711715295 |
| chr6:87393078-87393370 | 0.326095787 | 0.436416947 | 0.711728482 |
| chr3:13451295-13452493 | 0.248239558 | 0.436459565 | 0.711782836 |
| chr10:69801129-69801532 | -0.291836238 | 0.436523877 | 0.71186155 |
| chr1:198307321-198307803 | -0.280811057 | 0.436526413 | 0.71186155 |
| chr21:34614978-34615724 | 0.265145589 | 0.436546049 | 0.711871741 |
| chr15:88089151-88089364 | 0.36329131 | 0.436551243 | 0.711871741 |
| chr1:74689894-74690099 | 0.397391319 | 0.436571211 | 0.711874269 |
| chr8:94813210-94813742 | 0.273481397 | 0.436571375 | 0.711874269 |
| chr4:78059613-78059881 | 0.303969605 | 0.436584338 | 0.711880257 |
| chr12:2944657-2944991 | -0.330866114 | 0.436627098 | 0.71193483 |
| chr5:154444878-154445187 | -0.28946432 | 0.436667527 | 0.711981288 |
| chr14:100145181-100145402 | -0.33547753 | 0.436674175 | 0.711981288 |
| chr12:74021856-74022202 | -0.291535383 | 0.436694346 | 0.711990779 |
| chr11:65329993-65330208 | 0.375084058 | 0.436698581 | 0.711990779 |
| chr11:72693520-72693788 | -0.318965355 | 0.436723938 | 0.712016971 |
| chr10:132421145-132421345 | 0.337468494 | 0.43673665 | 0.712019383 |

| | | | |
|---|---|---|---|
| chr12:88993765-88993980 | -0.409146793 | 0.436744003 | 0.712019383 |
| chr17:67433394-67433663 | -0.341752847 | 0.436769636 | 0.712046023 |
| chr6:168284265-168284465 | -0.429386663 | 0.436806034 | 0.712079544 |
| chr16:2829950-2830265 | 0.336683808 | 0.436808785 | 0.712079544 |
| chr1:205703606-205703906 | 0.302616807 | 0.436828652 | 0.71209678 |
| chr18:48023672-48024079 | -0.297823834 | 0.436845896 | 0.712109219 |
| chr12:52921000-52921272 | -0.341578961 | 0.43685487 | 0.712109219 |
| chr6:4707040-4707472 | -0.305130114 | 0.436882041 | 0.712124346 |
| chr17:58841298-58841527 | 0.287994346 | 0.436882738 | 0.712124346 |
| chr12:48350683-48351398 | 0.26991886 | 0.436914458 | 0.712160901 |
| chr1:77744642-77744895 | -0.351622152 | 0.436959154 | 0.712218602 |
| chr1:54406247-54406644 | -0.288410006 | 0.437009011 | 0.712284715 |
| chr11:59615184-59615439 | 0.298078768 | 0.437018387 | 0.712284845 |
| chr1:83055829-83056123 | 0.312487097 | 0.437027836 | 0.712285094 |
| chr12:82383929-82384193 | -0.357890729 | 0.437046408 | 0.712297002 |
| chr2:216114482-216115565 | -0.248636001 | 0.437057742 | 0.712297002 |
| chr19:53544787-53544987 | -0.43998365 | 0.437063031 | 0.712297002 |
| chr10:7410904-7411104 | 0.341361681 | 0.437080879 | 0.712310939 |
| chr16:11576631-11577266 | 0.275662889 | 0.437126612 | 0.712370318 |
| chr6:137588512-137589773 | -0.25256751 | 0.437144111 | 0.712383684 |
| chr11:65477096-65478079 | 0.253214506 | 0.437210084 | 0.712474893 |
| chr1:217731084-217731314 | -0.413018008 | 0.437218677 | 0.712474893 |
| chr7:9231904-9232104 | -0.414963492 | 0.437258954 | 0.712525373 |
| chr8:23692841-23693045 | 0.386760035 | 0.437268979 | 0.712526555 |
| chr3:169966064-169967682 | -0.246673672 | 0.437318625 | 0.712592299 |
| chr17:9817162-9817559 | -0.298450379 | 0.437354961 | 0.712636352 |
| chr1:15975291-15976271 | 0.263561645 | 0.437397388 | 0.712690328 |
| chr2:200578878-200579147 | -0.341766571 | 0.437417167 | 0.7127074 |
| chr12:51383588-51384003 | 0.305974156 | 0.437432169 | 0.712716688 |
| chrX:103774143-103774393 | 0.332576289 | 0.437461815 | 0.712735007 |
| chr8:38830749-38831005 | 0.386513184 | 0.437471319 | 0.712735007 |
| chr9:4298412-4298612 | 0.386513184 | 0.437471319 | 0.712735007 |
| chr2:237774720-237775200 | -0.307363521 | 0.43750145 | 0.712768941 |
| chr7:100366565-100367012 | -0.286682102 | 0.437531986 | 0.712803534 |
| chr11:75035906-75036245 | -0.311436681 | 0.437572697 | 0.712839697 |
| chr16:85688887-85689347 | 0.258982195 | 0.43757806 | 0.712839697 |
| chr3:107429911-107430262 | 0.278069885 | 0.437582093 | 0.712839697 |
| chr8:120939021-120939591 | -0.262049992 | 0.437670413 | 0.712916373 |
| chr9:137445337-137445837 | 0.278895994 | 0.437670883 | 0.712916373 |
| chr17:2930091-2930387 | 0.329174042 | 0.437673903 | 0.712916373 |
| chr3:156675397-156675750 | -0.268951438 | 0.437678366 | 0.712916373 |
| chr7:98781090-98781290 | -0.431187915 | 0.437681595 | 0.712916373 |
| chr14:76704445-76704683 | -0.366944355 | 0.437684987 | 0.712916373 |
| chr6:13624854-13625054 | -0.423655372 | 0.43769639 | 0.712919791 |
| chr4:80336503-80336925 | -0.277921011 | 0.437733908 | 0.712929512 |
| chr13:26496692-26496924 | 0.331405731 | 0.437735827 | 0.712929512 |
| chr11:3417027-3417266 | -0.365656547 | 0.437739055 | 0.712929512 |
| chr11:67095023-67095280 | 0.320995597 | 0.437742221 | 0.712929512 |
| chr11:93089573-93089773 | -0.426496227 | 0.437748881 | 0.712929512 |
| chr15:78108346-78108630 | 0.288610017 | 0.437804861 | 0.712993627 |

| | | | |
|---|---|---|---|
| chr8:22244537-22245691 | 0.249561383 | 0.437812509 | 0.712993627 |
| chr2:235669181-235669636 | 0.319597631 | 0.437816164 | 0.712993627 |
| chr4:128663504-128663728 | -0.353869417 | 0.437892654 | 0.713102185 |
| chr13:54416179-54416458 | -0.390593098 | 0.437901438 | 0.713102185 |
| chr15:90192013-90192375 | 0.294020385 | 0.437916994 | 0.713105961 |
| chr1:115805753-115805959 | -0.38513651 | 0.437925369 | 0.713105961 |
| chr4:160848797-160849390 | -0.291507472 | 0.437931678 | 0.713105961 |
| chr6:4008455-4008703 | 0.368648526 | 0.43794117 | 0.713106263 |
| chr1:180507955-180508157 | -0.379469394 | 0.437962464 | 0.713113882 |
| chr19:46416318-46416862 | -0.301190701 | 0.437971575 | 0.713113882 |
| chr1:46302939-46303956 | 0.250798413 | 0.437979566 | 0.713113882 |
| chr8:138478791-138479134 | -0.351689773 | 0.437994912 | 0.713113882 |
| chrX:72298063-72298305 | -0.376444092 | 0.43801251 | 0.713113882 |
| chr1:203625617-203625940 | -0.284170019 | 0.438013864 | 0.713113882 |
| chr1:65309763-65310350 | -0.264475353 | 0.438019492 | 0.713113882 |
| chr6:99520845-99521990 | -0.2504039 | 0.438020304 | 0.713113882 |
| chr17:76279539-76279840 | 0.298430416 | 0.438038136 | 0.713126114 |
| chr3:172202836-172203040 | -0.396657742 | 0.438052203 | 0.713126114 |
| chr11:67506009-67506418 | -0.266738856 | 0.438055739 | 0.713126114 |
| chr1:201604978-201605417 | -0.291624809 | 0.438148787 | 0.713262437 |
| chr4:3955304-3956337 | 0.245405333 | 0.438182419 | 0.713302032 |
| chr1:185559908-185560207 | 0.31993399 | 0.438226136 | 0.713358041 |
| chr10:2978180-2978396 | 0.399174458 | 0.438264087 | 0.713403734 |
| chr16:87939509-87939780 | -0.364577857 | 0.438272827 | 0.713403734 |
| chr4:80471781-80472032 | -0.347205772 | 0.438291511 | 0.713417932 |
| chr12:122730748-122731049 | -0.338924043 | 0.438307093 | 0.713417932 |
| chr10:88553267-88553514 | -0.365391705 | 0.438322808 | 0.713417932 |
| chr7:90629373-90629621 | -0.365284785 | 0.438327716 | 0.713417932 |
| chr11:59809953-59811025 | -0.25107474 | 0.438328104 | 0.713417932 |
| chr6:42984000-42984918 | 0.251699292 | 0.438345278 | 0.71343073 |
| chr22:26331030-26331307 | 0.332943096 | 0.438371898 | 0.713457926 |
| chr17:50518647-50518884 | -0.372624682 | 0.438384879 | 0.713457926 |
| chr6:51559141-51559363 | -0.362185345 | 0.438392356 | 0.713457926 |
| chr20:47535057-47535655 | 0.245585351 | 0.43840824 | 0.713457926 |
| chr3:172536376-172536912 | 0.287259921 | 0.43842458 | 0.713457926 |
| chr1:211928319-211928667 | 0.349487271 | 0.438431311 | 0.713457926 |
| chr15:72358817-72359269 | -0.291399197 | 0.438433026 | 0.713457926 |
| chr9:14433806-14434190 | 0.295517704 | 0.438440675 | 0.713457926 |
| chr15:66456502-66456766 | -0.382315702 | 0.438455074 | 0.713457926 |
| chr8:109607755-109608259 | -0.273962636 | 0.43845706 | 0.713457926 |
| chr4:77698015-77698215 | -0.419338497 | 0.438464414 | 0.713457926 |
| chr5:109688528-109690886 | -0.242373801 | 0.438498846 | 0.713492327 |
| chr2:28810064-28811075 | -0.265068749 | 0.438504179 | 0.713492327 |
| chr19:47920178-47920465 | -0.301455905 | 0.438521535 | 0.713503261 |
| chr5:17469388-17469588 | 0.429131952 | 0.438529522 | 0.713503261 |
| chrX:46327491-46327928 | -0.310413895 | 0.438553579 | 0.71352725 |
| chr5:134511706-134512495 | -0.265160186 | 0.438581833 | 0.713558069 |
| chr5:36060498-36060853 | -0.323050229 | 0.438615262 | 0.713594583 |
| chr3:141305525-141305818 | 0.309383541 | 0.438622903 | 0.713594583 |
| chr14:62126883-62127170 | -0.337528475 | 0.438639237 | 0.713598583 |

| | | | |
|---|---|---|---|
| chr11:10878694-10878906 | 0.420115954 | 0.438643988 | 0.713598583 |
| chr2:14507042-14507897 | -0.27937273 | 0.438691928 | 0.713661421 |
| chr14:81625439-81625763 | 0.292581804 | 0.438726142 | 0.713701926 |
| chr11:129323105-129323313 | 0.382955352 | 0.438736399 | 0.71370346 |
| chr4:45442608-45442833 | -0.344703116 | 0.438755211 | 0.713718909 |
| chr13:23578725-23579767 | 0.254155506 | 0.438795327 | 0.713769012 |
| chr2:54085171-54085421 | -0.32931128 | 0.43881709 | 0.713777226 |
| chr8:144373319-144373528 | 0.312412989 | 0.438820489 | 0.713777226 |
| chr9:113876205-113876460 | -0.377145233 | 0.4388392 | 0.713777226 |
| chr8:94272364-94272668 | 0.293445752 | 0.438842197 | 0.713777226 |
| chr12:122730273-122730626 | -0.331657117 | 0.438846954 | 0.713777226 |
| chr9:2241799-2242225 | -0.290829674 | 0.438859594 | 0.713782633 |
| chr3:112814411-112814616 | -0.37463533 | 0.438871629 | 0.713787055 |
| chr5:61330634-61331301 | -0.255924925 | 0.438893418 | 0.713800664 |
| chr9:129080673-129081523 | 0.249559129 | 0.438899414 | 0.713800664 |
| chr1:114969138-114969348 | 0.339360796 | 0.438907944 | 0.713800664 |
| chr22:17368677-17368897 | 0.361335526 | 0.438944732 | 0.71383425 |
| chr2:140964222-140964653 | -0.298888219 | 0.438947228 | 0.71383425 |
| chr5:24050972-24051421 | 0.325344362 | 0.438978908 | 0.713870617 |
| chr3:157073825-157074117 | -0.339201462 | 0.43902002 | 0.713918142 |
| chr7:149125754-149127233 | -0.245443189 | 0.439028987 | 0.713918142 |
| chr10:71285127-71285351 | 0.333102004 | 0.439036084 | 0.713918142 |
| chr3:63795103-63795872 | -0.247392235 | 0.439047861 | 0.713922141 |
| chr14:51866111-51866387 | -0.388674154 | 0.43910547 | 0.714000665 |
| chr6:27651300-27651735 | -0.29755816 | 0.439124628 | 0.714016664 |
| chr13:36841457-36841680 | -0.369281382 | 0.439165115 | 0.714067342 |
| chr20:52313417-52313861 | -0.302444893 | 0.439208195 | 0.714112164 |
| chr11:82718169-82718520 | 0.305737884 | 0.439211321 | 0.714112164 |
| chr1:147504974-147505276 | 0.32321272 | 0.439234167 | 0.714134156 |
| chr14:104723604-104723854 | 0.295634876 | 0.439246093 | 0.714138392 |
| chr8:130042053-130042819 | -0.253341497 | 0.439259922 | 0.714145722 |
| chr1:215181925-215182334 | -0.303468214 | 0.439279466 | 0.714162344 |
| chr3:81734822-81735423 | -0.300093511 | 0.439334246 | 0.714236248 |
| chr10:3120154-3120354 | 0.345284824 | 0.439372489 | 0.714283265 |
| chr9:3962484-3962756 | -0.404481273 | 0.439393274 | 0.714301899 |
| chr1:195190418-195190742 | 0.354181749 | 0.439417136 | 0.714325535 |
| chr17:47941205-47941935 | -0.248909129 | 0.439447466 | 0.714350465 |
| chr1:111422608-111423391 | -0.268643993 | 0.439451117 | 0.714350465 |
| chr10:29445525-29445810 | -0.333406036 | 0.439482739 | 0.714386711 |
| chr16:89827600-89828775 | -0.255805048 | 0.439531347 | 0.714450567 |
| chr17:16386524-16387446 | -0.263859028 | 0.439549597 | 0.714465077 |
| chr20:63580686-63581159 | -0.273982501 | 0.439582545 | 0.714503474 |
| chr13:112444008-112444400 | -0.301840586 | 0.439613784 | 0.71451266 |
| chr1:204827039-204827317 | -0.296259091 | 0.439619456 | 0.71451266 |
| chr16:3134386-3135335 | 0.253486506 | 0.439626762 | 0.71451266 |
| chr2:96413298-96413544 | -0.341664152 | 0.439645356 | 0.71451266 |
| chr3:15108118-15108443 | 0.291738993 | 0.43965241 | 0.71451266 |
| chr6:105991869-105993020 | -0.254533595 | 0.439653457 | 0.71451266 |
| chr1:110196996-110197216 | 0.315078225 | 0.439658401 | 0.71451266 |
| chr4:76213269-76214420 | 0.249199532 | 0.439668412 | 0.71451266 |

| | | | |
|---|---|---|---|
| chr2:38274661-38275079 | -0.308165082 | 0.439679554 | 0.71451266 |
| chr17:28335269-28336511 | 0.245812942 | 0.439681448 | 0.71451266 |
| chr8:70147954-70148469 | -0.289205418 | 0.439706441 | 0.714527938 |
| chr14:21104516-21104815 | 0.275368171 | 0.439709501 | 0.714527938 |
| chr8:80085085-80085359 | 0.283798824 | 0.439720651 | 0.714530905 |
| chr10:119650876-119651898 | 0.251661973 | 0.439735878 | 0.714540494 |
| chr5:600743-601084 | 0.335477155 | 0.439753184 | 0.714553462 |
| chr1:156705506-156706491 | 0.251358855 | 0.439782063 | 0.714585232 |
| chr4:138576679-138577026 | 0.312828261 | 0.439799692 | 0.714598724 |
| chr8:22589134-22589346 | 0.357884813 | 0.43984745 | 0.714640416 |
| chr8:130245761-130245961 | 0.399577637 | 0.439855374 | 0.714640416 |
| chr8:140635635-140636615 | 0.250889207 | 0.439864219 | 0.714640416 |
| chr1:232805216-232806234 | 0.262761838 | 0.439864375 | 0.714640416 |
| chr3:110881120-110881440 | -0.311657389 | 0.439876274 | 0.714640416 |
| chr8:22095331-22095782 | -0.303718486 | 0.439889982 | 0.714640416 |
| chr11:63888480-63888972 | 0.304760554 | 0.439890639 | 0.714640416 |
| chr12:121513779-121514235 | -0.285310202 | 0.439926942 | 0.714676512 |
| chr4:82499823-82500127 | -0.339865515 | 0.439942897 | 0.714676512 |
| chr17:48625777-48626036 | 0.329944762 | 0.439945132 | 0.714676512 |
| chr21:29298310-29298522 | 0.351995541 | 0.439950167 | 0.714676512 |
| chr2:7133585-7133845 | -0.363949314 | 0.439970912 | 0.714678108 |
| chr21:41786685-41786885 | -0.363949314 | 0.439970912 | 0.714678108 |
| chr21:33551953-33552162 | -0.405716957 | 0.439979132 | 0.714678108 |
| chr1:182782191-182782675 | 0.278784615 | 0.439996424 | 0.714679087 |
| chr3:146543888-146545041 | -0.244330264 | 0.440002386 | 0.714679087 |
| chr12:47754870-47755073 | 0.322819242 | 0.440016183 | 0.714679087 |
| chr4:131370892-131371198 | 0.359079507 | 0.440017044 | 0.714679087 |
| chr6:110483564-110483882 | 0.322047474 | 0.440067171 | 0.714733027 |
| chr10:72364131-72364658 | -0.276137792 | 0.440072831 | 0.714733027 |
| chr4:72568951-72569402 | -0.286926299 | 0.44008493 | 0.714733027 |
| chr6:19838004-19839238 | 0.254616777 | 0.440095681 | 0.714733027 |
| chr1:201895924-201896223 | 0.291391143 | 0.440096894 | 0.714733027 |
| chr12:32756250-32756558 | -0.279146759 | 0.440131769 | 0.714774263 |
| chr5:36241220-36242564 | 0.245547636 | 0.440140942 | 0.714774263 |
| chr11:79400605-79401043 | -0.295921312 | 0.44019552 | 0.714847744 |
| chr5:151771299-151772779 | -0.241534135 | 0.440221108 | 0.714859697 |
| chr15:62241223-62241480 | 0.334132021 | 0.440225858 | 0.714859697 |
| chr17:28812078-28813029 | -0.253969286 | 0.440235115 | 0.714859697 |
| chr5:115151122-115151514 | -0.354071941 | 0.440253084 | 0.714859697 |
| chr7:55433185-55433590 | -0.278590143 | 0.44025505 | 0.714859697 |
| chr10:1380367-1380640 | -0.351723087 | 0.440263408 | 0.714859697 |
| chr5:80654038-80655294 | -0.246672498 | 0.440268188 | 0.714859697 |
| chr15:70241379-70241604 | -0.402088619 | 0.440285997 | 0.714868024 |
| chr1:115239160-115239391 | 0.391746656 | 0.440299629 | 0.714868024 |
| chr2:15360312-15360575 | -0.325967806 | 0.440301306 | 0.714868024 |
| chr4:52037748-52038854 | -0.25897573 | 0.440343309 | 0.714921071 |
| chr2:75209191-75209499 | -0.35847447 | 0.440366128 | 0.714942969 |
| chr8:81841498-81842940 | 0.245016175 | 0.440413474 | 0.715004687 |
| chr16:87236120-87236327 | 0.347951301 | 0.440428367 | 0.715013715 |
| chr17:19977354-19978461 | -0.255710262 | 0.440441291 | 0.715019547 |

| | | | |
|---|---|---|---|
| chr3:159941135-159941335 | -0.424149153 | 0.440483144 | 0.715072343 |
| chr3:113726799-113726999 | -0.440622119 | 0.440537277 | 0.715145069 |
| chr16:18789975-18790637 | 0.25560379 | 0.440593533 | 0.715221238 |
| chr1:99849441-99849717 | 0.321020434 | 0.440623205 | 0.715250717 |
| chr10:114279657-114280289 | 0.269062792 | 0.440630362 | 0.715250717 |
| chr5:92019523-92019726 | -0.411613606 | 0.44064044 | 0.715251923 |
| chr3:157543078-157543531 | 0.274525195 | 0.440673518 | 0.715290463 |
| chr20:6074156-6074495 | 0.322404818 | 0.440695646 | 0.715311227 |
| chr1:119903188-119903388 | -0.407749676 | 0.440773359 | 0.715405679 |
| chr1:174011971-174012171 | -0.407749676 | 0.440773359 | 0.715405679 |
| chr8:143933931-143934131 | -0.350519381 | 0.440792459 | 0.715405679 |
| chr17:80544670-80545254 | -0.276880832 | 0.44080151 | 0.715405679 |
| chr7:17250766-17250966 | 0.430811435 | 0.44080558 | 0.715405679 |
| chr8:58078980-58079253 | 0.359754474 | 0.440809857 | 0.715405679 |
| chr18:59140329-59140593 | -0.315604819 | 0.440824723 | 0.715414651 |
| chrX:2928628-2929689 | 0.248843933 | 0.440850279 | 0.715440973 |
| chr7:6955760-6955990 | -0.358993987 | 0.440873095 | 0.715457863 |
| chr7:95545014-95545223 | -0.403174469 | 0.440882506 | 0.715457863 |
| chr6:71012902-71013114 | 0.354232371 | 0.440888699 | 0.715457863 |
| chr1:206634367-206634577 | 0.310336709 | 0.440902454 | 0.715465031 |
| chr20:57435555-57436278 | 0.250872916 | 0.440914707 | 0.715469762 |
| chr4:142353208-142353624 | -0.312804462 | 0.4409704 | 0.715540888 |
| chr6:44245964-44247391 | 0.243360809 | 0.440977216 | 0.715540888 |
| chr1:26864572-26864800 | -0.346064549 | 0.44100243 | 0.715566647 |
| chr13:102800182-102800827 | -0.260045551 | 0.441035776 | 0.715600536 |
| chr8:6136270-6136610 | -0.306024384 | 0.441041995 | 0.715600536 |
| chr10:11875257-11875457 | 0.362825748 | 0.441053043 | 0.715603309 |
| chrX:130108649-130109251 | 0.273060512 | 0.441077497 | 0.715618604 |
| chr1:214637000-214637530 | 0.276520501 | 0.441081149 | 0.715618604 |
| chr12:133181239-133181976 | 0.26679887 | 0.441106649 | 0.715632456 |
| chr17:1724381-1724900 | -0.286939592 | 0.441108367 | 0.715632456 |
| chr15:36195441-36196011 | -0.288274588 | 0.441122575 | 0.715634554 |
| chr2:233251389-233252069 | -0.271081278 | 0.441131458 | 0.715634554 |
| chr20:41734429-41734754 | 0.27684367 | 0.441138571 | 0.715634554 |
| chr1:151889055-151889277 | 0.347405804 | 0.441147019 | 0.715634554 |
| chr8:38966612-38966812 | -0.41345473 | 0.441172982 | 0.71566152 |
| chr3:156735536-156735796 | -0.371396859 | 0.441189174 | 0.715664847 |
| chr17:47229654-47230284 | 0.276949392 | 0.441201744 | 0.715664847 |
| chr14:95534274-95535160 | 0.244845279 | 0.441208227 | 0.715664847 |
| chr8:120890997-120891559 | 0.287809743 | 0.441212394 | 0.715664847 |
| chr5:157143478-157143751 | -0.312524652 | 0.441239324 | 0.715693378 |
| chr16:68558992-68559192 | 0.43509707 | 0.441263083 | 0.715708476 |
| chr2:128063438-128063927 | -0.280742093 | 0.441267314 | 0.715708476 |
| chr5:16456060-16456616 | -0.29350035 | 0.441302055 | 0.7157452 |
| chr7:159191883-159192083 | 0.412385121 | 0.441317143 | 0.7157452 |
| chr21:18087040-18087240 | 0.445003143 | 0.441325739 | 0.7157452 |
| chr1:153922524-153923753 | 0.243360789 | 0.441327321 | 0.7157452 |
| chr10:33005347-33005570 | -0.340406429 | 0.441339512 | 0.715749823 |
| chr19:52048425-52049177 | 0.274554038 | 0.441410561 | 0.715849895 |
| chr20:61387534-61387968 | -0.293136039 | 0.441446015 | 0.715892241 |

| | | | |
|---|---|---|---|
| chr13:40672512-40672717 | -0.368712062 | 0.441465094 | 0.715908028 |
| chr6:69796081-69797341 | -0.242592223 | 0.441492992 | 0.715938118 |
| chr8:96373761-96373961 | 0.438124517 | 0.441530023 | 0.715983015 |
| chr22:43915573-43915884 | -0.392063477 | 0.441546145 | 0.715994004 |
| chr8:78720402-78720669 | 0.356314543 | 0.441561156 | 0.716002255 |
| chr18:12883493-12884521 | 0.249319533 | 0.441569922 | 0.716002255 |
| chr3:4564930-4565282 | 0.318276055 | 0.441610808 | 0.716053399 |
| chr3:15331711-15333005 | 0.247676286 | 0.441623305 | 0.716058509 |
| chr3:63739979-63740537 | 0.2532482 | 0.441645795 | 0.71606949 |
| chr5:51383027-51383891 | 0.25947329 | 0.441648769 | 0.71606949 |
| chr7:25216809-25217089 | -0.376287168 | 0.441686879 | 0.716116127 |
| chr10:30220074-30220393 | -0.341368966 | 0.441706528 | 0.716132831 |
| chr19:46638349-46639381 | 0.253927551 | 0.441722182 | 0.716141004 |
| chr2:122191383-122191844 | -0.296038222 | 0.441730262 | 0.716141004 |
| chr1:234960765-234960965 | 0.39471422 | 0.44174163 | 0.716144281 |
| chr6:27502754-27503196 | -0.278603478 | 0.44176936 | 0.716167729 |
| chr15:97295621-97296453 | -0.254271965 | 0.441774787 | 0.716167729 |
| chr3:175988578-175988851 | -0.409426067 | 0.441784304 | 0.716168005 |
| chr18:23703138-23703411 | 0.326910939 | 0.441821601 | 0.716213313 |
| chr11:65525328-65525689 | 0.288912126 | 0.441861256 | 0.716259032 |
| chr6:144215847-144216920 | -0.243380809 | 0.4418685 | 0.716259032 |
| chr18:42596266-42596566 | -0.335024247 | 0.441902412 | 0.716298849 |
| chr17:81832638-81833724 | 0.249817501 | 0.441930062 | 0.716320028 |
| chr18:45993392-45993701 | -0.365712396 | 0.441937658 | 0.716320028 |
| chr1:959199-959458 | 0.309188843 | 0.441943524 | 0.716320028 |
| chr6:100078364-100078902 | -0.283459583 | 0.441957297 | 0.716327199 |
| chr10:110681103-110681314 | -0.378916685 | 0.44197038 | 0.716333252 |
| chr1:247532036-247532268 | -0.358989792 | 0.442002034 | 0.716369402 |
| chr3:11643226-11643707 | 0.277224884 | 0.442016755 | 0.716378107 |
| chr6:99393660-99394535 | -0.251321298 | 0.442028806 | 0.716382486 |
| chr5:49599514-49599763 | -0.350501843 | 0.442047572 | 0.716392687 |
| chrX:56232196-56232551 | 0.304194872 | 0.4420538 | 0.716392687 |
| chr4:74157678-74158639 | -0.247297811 | 0.442086839 | 0.716409602 |
| chr11:66328359-66328653 | 0.286406122 | 0.442088577 | 0.716409602 |
| chr11:76788250-76789117 | -0.248462816 | 0.442092287 | 0.716409602 |
| chr15:101980678-101981023 | 0.321246411 | 0.442123594 | 0.716425607 |
| chr3:47281569-47281940 | 0.309531772 | 0.442123738 | 0.716425607 |
| chr6:30075628-30076319 | -0.273574054 | 0.442133068 | 0.716425607 |
| chr8:143716683-143717027 | -0.286136994 | 0.442144766 | 0.716425607 |
| chr1:206556943-206557731 | 0.253939149 | 0.442148914 | 0.716425607 |
| chrX:137565690-137566682 | 0.251554036 | 0.442164099 | 0.716435061 |
| chr1:115469598-115470035 | 0.273505956 | 0.442192795 | 0.716458139 |
| chr9:137302209-137302750 | -0.268802803 | 0.4422076 | 0.716458139 |
| chr9:90773943-90774143 | -0.424447545 | 0.442231393 | 0.716458139 |
| chr19:7920125-7920435 | 0.288392166 | 0.44223307 | 0.716458139 |
| chr1:155208487-155209380 | 0.253323105 | 0.442233871 | 0.716458139 |
| chr3:65953309-65953851 | 0.258289009 | 0.442240183 | 0.716458139 |
| chr1:197939243-197939451 | -0.388182438 | 0.442243796 | 0.716458139 |
| chr3:59052909-59053109 | -0.398025352 | 0.442303016 | 0.716538928 |
| chr17:3863772-3863999 | -0.35468805 | 0.442321674 | 0.716544101 |

| | | | |
|---|---|---|---|
| chr1:197068560-197069044 | 0.29209378 | 0.442332062 | 0.716544101 |
| chr2:27262945-27263253 | 0.292370405 | 0.442335374 | 0.716544101 |
| chr10:101031064-101031558 | 0.263823954 | 0.442343616 | 0.716544101 |
| chr8:61023526-61023869 | -0.295018742 | 0.442369787 | 0.716571346 |
| chr9:137577962-137578182 | 0.334367148 | 0.442382996 | 0.716577595 |
| chr6:117907979-117908211 | 0.372002023 | 0.442420709 | 0.71660182 |
| chr6:31180651-31181054 | 0.286754822 | 0.442422374 | 0.71660182 |
| chr5:14946800-14947379 | 0.276019377 | 0.44242601 | 0.71660182 |
| chr14:73771784-73772190 | -0.288872926 | 0.442442085 | 0.716608961 |
| chr1:243254882-243256430 | -0.239812521 | 0.442458642 | 0.716608961 |
| chrX:146224460-146224745 | -0.329602676 | 0.442465791 | 0.716608961 |
| chr1:87450713-87451450 | -0.298546149 | 0.442467828 | 0.716608961 |
| chr15:68577885-68578896 | 0.25988599 | 0.442494807 | 0.716635338 |
| chr2:155708168-155708368 | -0.422092753 | 0.442505366 | 0.716635338 |
| chr5:19865819-19866019 | 0.376269131 | 0.442512173 | 0.716635338 |
| chr3:17686641-17686945 | -0.33653639 | 0.442578263 | 0.71671089 |
| chr14:24953309-24953556 | -0.368976138 | 0.44258378 | 0.71671089 |
| chr5:129541635-129541984 | -0.278088781 | 0.442586887 | 0.71671089 |
| chr2:1617595-1618011 | 0.29122434 | 0.442614004 | 0.716739654 |
| chr17:79631540-79631740 | 0.399311728 | 0.442668282 | 0.716803649 |
| chr5:25874371-25874699 | -0.371583369 | 0.442674055 | 0.716803649 |
| chr2:27050293-27051806 | -0.239727078 | 0.442681589 | 0.716803649 |
| chr7:22782668-22782953 | 0.29576849 | 0.44272438 | 0.716857789 |
| chr13:27251134-27252039 | -0.25125518 | 0.442766631 | 0.716911052 |
| chr5:16335300-16335728 | 0.308580076 | 0.442810596 | 0.716915593 |
| chr3:129428076-129429107 | -0.250963681 | 0.442818544 | 0.716915593 |
| chr13:37305243-37305443 | -0.414639059 | 0.442826353 | 0.716915593 |
| chr2:197384361-197384561 | -0.396460856 | 0.442826495 | 0.716915593 |
| chr8:141026884-141027213 | 0.320856453 | 0.442839972 | 0.716915593 |
| chr5:120926560-120926768 | 0.3721117 | 0.442844428 | 0.716915593 |
| chr6:31940276-31940476 | 0.3721117 | 0.442844428 | 0.716915593 |
| chr15:68257033-68257453 | 0.29117264 | 0.442845725 | 0.716915593 |
| chr19:55632314-55632698 | 0.293660177 | 0.442854362 | 0.716915593 |
| chr7:80611759-80612168 | -0.309891162 | 0.442866755 | 0.716915593 |
| chr5:9544402-9545689 | -0.247400787 | 0.442872358 | 0.716915593 |
| chr20:31833425-31833696 | 0.306505438 | 0.44288584 | 0.716917253 |
| chr11:40722049-40722532 | 0.283333923 | 0.442905516 | 0.716917253 |
| chr2:55223334-55224041 | -0.258690577 | 0.442907252 | 0.716917253 |
| chr3:52244253-52244473 | -0.379536465 | 0.442910809 | 0.716917253 |
| chr4:78951937-78952177 | -0.354272238 | 0.442926094 | 0.716926849 |
| chr3:133927862-133928070 | 0.355243751 | 0.442948538 | 0.716948033 |
| chr17:66602814-66603014 | 0.381993413 | 0.442967917 | 0.716964252 |
| chr2:216081297-216081985 | -0.257715135 | 0.442981591 | 0.71697124 |
| chr1:153670714-153672145 | 0.24161117 | 0.44301106 | 0.716991959 |
| chrX:15745228-15745465 | 0.356467868 | 0.443013107 | 0.716991959 |
| chr22:23289836-23290236 | 0.307349214 | 0.443066507 | 0.717047674 |
| chr1:110011640-110011969 | 0.294171987 | 0.44307426 | 0.717047674 |
| chr5:42908511-42909391 | 0.249717658 | 0.443084237 | 0.717047674 |
| chr17:17001660-17002163 | 0.285105519 | 0.443084966 | 0.717047674 |
| chr11:13480223-13480791 | -0.268023576 | 0.443122377 | 0.717093072 |

| | | | |
|---|---|---|---|
| chr10:17257717-17257970 | 0.341059298 | 0.443136876 | 0.717101391 |
| chr2:46889053-46889595 | -0.276762794 | 0.443155864 | 0.717102165 |
| chr9:214588-215500 | 0.261227547 | 0.443156073 | 0.717102165 |
| chr12:121712168-121712927 | 0.261389903 | 0.443237692 | 0.717219092 |
| chr10:13371859-13372254 | -0.286898028 | 0.443282172 | 0.717275918 |
| chr7:43195316-43195586 | -0.346915765 | 0.443297088 | 0.717284906 |
| chr6:52652087-52652367 | -0.306756642 | 0.443346166 | 0.717347791 |
| chr4:56048600-56050006 | 0.25161129 | 0.443354676 | 0.717347791 |
| chr2:158795160-158795635 | -0.30462414 | 0.443383452 | 0.717379202 |
| chr8:110613936-110614136 | 0.38154057 | 0.443400442 | 0.717391543 |
| chr20:25397947-25398147 | -0.399276439 | 0.443415469 | 0.717400707 |
| chr6:40509249-40509505 | 0.393210859 | 0.443483261 | 0.717495237 |
| chr13:108919091-108919957 | -0.253055602 | 0.443584997 | 0.717634789 |
| chr2:186848808-186849611 | -0.260792567 | 0.44358825 | 0.717634789 |
| chr3:167873038-167873288 | -0.345606148 | 0.443617556 | 0.717667047 |
| chr11:64240417-64240712 | 0.276756194 | 0.443630124 | 0.717672227 |
| chr10:117684635-117685294 | 0.264827643 | 0.443648934 | 0.717687504 |
| chr4:104065049-104065306 | -0.343225899 | 0.443665056 | 0.717698432 |
| chr12:3872959-3873646 | -0.258248911 | 0.443702554 | 0.717743938 |
| chr10:46106733-46106991 | -0.368048359 | 0.44373973 | 0.717787466 |
| chrX:20029805-20030037 | 0.347853355 | 0.443748199 | 0.717787466 |
| chr20:50764020-50764615 | 0.241662767 | 0.443768753 | 0.71780556 |
| chr10:114349922-114350266 | 0.322994478 | 0.443796428 | 0.717835172 |
| chr12:119274673-119274922 | -0.373009151 | 0.443861883 | 0.717925889 |
| chr10:57339092-57339376 | -0.328099718 | 0.443872334 | 0.717927638 |
| chr10:31961309-31961671 | 0.296049453 | 0.443882839 | 0.717929474 |
| chr6:129501459-129501771 | -0.31538433 | 0.443900085 | 0.717942212 |
| chr8:86558202-86558495 | -0.297640024 | 0.443920146 | 0.717959504 |
| chr13:24627118-24627518 | -0.296662102 | 0.443995075 | 0.718065531 |
| chr3:149581888-149582088 | -0.370425036 | 0.444006781 | 0.718066497 |
| chr2:58554045-58554245 | -0.425974088 | 0.444014416 | 0.718066497 |
| chr17:61416624-61417206 | 0.240710687 | 0.444041409 | 0.718070581 |
| chr19:45885931-45886235 | 0.303743538 | 0.444044587 | 0.718070581 |
| chr9:116737062-116737396 | -0.299239059 | 0.444045056 | 0.718070581 |
| chr1:162250080-162250332 | -0.380294037 | 0.444070461 | 0.718093612 |
| chr18:9474757-9475802 | 0.242532751 | 0.444078042 | 0.718093612 |
| chr1:186374393-186374613 | -0.280904834 | 0.444100517 | 0.7181148 |
| chr3:11662473-11662739 | 0.319000087 | 0.444119248 | 0.718129934 |
| chr19:50357142-50357486 | 0.298181526 | 0.444131224 | 0.718134143 |
| chr2:223041584-223041796 | -0.42255856 | 0.444167981 | 0.718178421 |
| chr6:63935250-63935511 | 0.436395777 | 0.444184242 | 0.718180416 |
| chr10:102678775-102679300 | 0.281002422 | 0.444188294 | 0.718180416 |
| chr9:108272590-108272863 | -0.314361655 | 0.444202832 | 0.718180416 |
| chr3:21255103-21255848 | -0.252635553 | 0.444215749 | 0.718180416 |
| chr3:146625195-146625650 | -0.320203648 | 0.44421608 | 0.718180416 |
| chr2:158885193-158885839 | -0.30134769 | 0.44424522 | 0.718211486 |
| chr12:105379110-105379336 | -0.385950849 | 0.444254044 | 0.718211486 |
| chr19:48636965-48637174 | -0.303158701 | 0.444276519 | 0.718221823 |
| chr15:98901527-98901733 | -0.374083045 | 0.444293052 | 0.718221823 |
| chr1:187281162-187281362 | 0.405692135 | 0.444297533 | 0.718221823 |

| | | | |
|---|---|---|---|
| chr1:2283493-2283693 | 0.439967944 | 0.444297933 | 0.718221823 |
| chr1:156145977-156146264 | 0.291984421 | 0.444344461 | 0.71822812 |
| chr5:171953092-171953292 | -0.431706526 | 0.444348608 | 0.71822812 |
| chr6:144094980-144096039 | -0.243070066 | 0.444355994 | 0.71822812 |
| chrX:135066314-135066602 | 0.333074802 | 0.444364275 | 0.71822812 |
| chr1:159924425-159925902 | 0.24232766 | 0.444365812 | 0.71822812 |
| chr14:81144259-81144859 | -0.27824268 | 0.444367131 | 0.71822812 |
| chr12:96907693-96908136 | -0.285891839 | 0.444389013 | 0.71822812 |
| chr2:179263651-179265020 | -0.248449388 | 0.444396519 | 0.71822812 |
| chr20:31547008-31547539 | 0.267321346 | 0.44440061 | 0.71822812 |
| chr14:77046508-77046904 | 0.311842636 | 0.444401786 | 0.71822812 |
| chr1:205344256-205344695 | -0.27372276 | 0.444404938 | 0.71822812 |
| chr4:70687788-70689116 | -0.241909903 | 0.44443707 | 0.718250462 |
| chr6:102619620-102619847 | -0.364965037 | 0.44443751 | 0.718250462 |
| chr5:178895686-178896211 | -0.279724362 | 0.444453003 | 0.718251889 |
| chr8:108890418-108890661 | -0.302135292 | 0.444464398 | 0.718251889 |
| chr1:10961021-10961495 | 0.292921317 | 0.444466515 | 0.718251889 |
| chr15:33309698-33309923 | 0.397468146 | 0.444483647 | 0.718263517 |
| chr15:64961185-64961502 | 0.354045406 | 0.444492459 | 0.718263517 |
| chr2:45643640-45643958 | 0.307959441 | 0.444508819 | 0.718274806 |
| chr9:90958546-90958749 | -0.414900061 | 0.444539974 | 0.718299449 |
| chr10:103476385-103476642 | -0.316661541 | 0.444542819 | 0.718299449 |
| chrX:129905728-129906299 | 0.254626721 | 0.444597944 | 0.718373372 |
| chr8:92812274-92812622 | 0.302046045 | 0.444646902 | 0.718437328 |
| chr4:84206260-84206552 | 0.336847895 | 0.444672106 | 0.7184629 |
| chr3:64687850-64688760 | -0.240923141 | 0.444687747 | 0.718466247 |
| chr3:62665654-62666387 | -0.247701576 | 0.444692931 | 0.718466247 |
| chr5:122258946-122259210 | -0.399558018 | 0.44472324 | 0.718500066 |
| chr11:84671313-84671651 | 0.324744458 | 0.444800798 | 0.718591507 |
| chr11:94046845-94047148 | -0.327383975 | 0.444815663 | 0.718591507 |
| chr22:22586951-22587390 | -0.310660631 | 0.444816048 | 0.718591507 |
| chr10:2619505-2619789 | -0.323805141 | 0.444817352 | 0.718591507 |
| chr15:42581384-42581667 | -0.32919852 | 0.444847771 | 0.718625496 |
| chr19:5164938-5165138 | -0.433146739 | 0.444891338 | 0.718680724 |
| chr1:162627807-162628007 | 0.378443606 | 0.444960324 | 0.718777011 |
| chr19:7145786-7146167 | -0.391808998 | 0.44498647 | 0.718788486 |
| chr11:61254460-61254724 | -0.312057943 | 0.444990132 | 0.718788486 |
| chr2:236986592-236986853 | 0.348058735 | 0.445005901 | 0.718788486 |
| chrX:86047101-86048066 | -0.250900226 | 0.445012117 | 0.718788486 |
| chr5:77181593-77181793 | -0.413385555 | 0.445014333 | 0.718788486 |
| chr17:75749874-75750204 | 0.273324004 | 0.445056537 | 0.718816743 |
| chr2:102841273-102841541 | -0.324835579 | 0.445058976 | 0.718816743 |
| chrX:126813317-126813573 | 0.439430724 | 0.445081427 | 0.718816743 |
| chr17:67824697-67825057 | 0.25859179 | 0.445094839 | 0.718816743 |
| chr4:121822860-121824331 | -0.24261165 | 0.445100546 | 0.718816743 |
| chr15:92917881-92918249 | 0.285285483 | 0.445101864 | 0.718816743 |
| chr10:106606231-106606465 | -0.368127034 | 0.445106073 | 0.718816743 |
| chr14:35826020-35827080 | 0.23977818 | 0.445106878 | 0.718816743 |
| chr2:232295657-232296286 | -0.286522894 | 0.445142605 | 0.718835325 |
| chr5:173030622-173030876 | 0.450161069 | 0.445143657 | 0.718835325 |

| | | | |
|---|---|---|---|
| chr11:58524205-58524417 | -0.36441886 | 0.445146529 | 0.718835325 |
| chr18:70496993-70497193 | -0.424386955 | 0.445181036 | 0.718842195 |
| chr9:136118050-136119391 | 0.244761458 | 0.44518741 | 0.718842195 |
| chr18:46072154-46072645 | -0.284966392 | 0.445205406 | 0.718842195 |
| chr19:45730744-45731265 | 0.260869417 | 0.445208271 | 0.718842195 |
| chr13:21951750-21952011 | 0.372433487 | 0.445215609 | 0.718842195 |
| chr11:108593488-108594000 | -0.252106143 | 0.445224961 | 0.718842195 |
| chr18:305154-305414 | -0.422428923 | 0.445229259 | 0.718842195 |
| chr17:43751733-43751933 | -0.44176558 | 0.445231731 | 0.718842195 |
| chr5:1948943-1949198 | 0.33235472 | 0.445240914 | 0.718842195 |
| chr3:18946577-18946921 | -0.295169938 | 0.4452446 | 0.718842195 |
| chr8:101466753-101466953 | 0.322025702 | 0.445263638 | 0.71885233 |
| chr9:137513886-137514095 | 0.340791498 | 0.44527556 | 0.71885233 |
| chr1:217568464-217568664 | -0.376194016 | 0.445279023 | 0.71885233 |
| chr19:11339205-11339405 | 0.415499534 | 0.445289381 | 0.718853907 |
| chr3:42012868-42013272 | -0.311816145 | 0.445315358 | 0.718880696 |
| chr5:11791375-11791575 | 0.373949886 | 0.445334928 | 0.718890339 |
| chr7:149330931-149331304 | -0.289830495 | 0.445340096 | 0.718890339 |
| chr16:4746478-4746747 | -0.374748745 | 0.44537671 | 0.718923059 |
| chr8:42502482-42502682 | -0.386482172 | 0.445379131 | 0.718923059 |
| chr12:53999602-54000162 | -0.269889483 | 0.445392287 | 0.71892915 |
| chr6:27279584-27279852 | -0.35226161 | 0.445425055 | 0.718948088 |
| chr6:127641856-127642324 | -0.304343683 | 0.445426662 | 0.718948088 |
| chr2:19048455-19049459 | -0.249911729 | 0.445432169 | 0.718948088 |
| chr11:125536222-125536725 | -0.340169889 | 0.44549387 | 0.71903253 |
| chr21:31558659-31560125 | 0.23963006 | 0.44555979 | 0.719099366 |
| chr5:174751397-174752066 | -0.269593573 | 0.445560541 | 0.719099366 |
| chr7:1920114-1920435 | -0.290414193 | 0.445581587 | 0.719099366 |
| chr2:163667141-163667666 | 0.317372555 | 0.44559423 | 0.719099366 |
| chr12:71639338-71639869 | -0.295816901 | 0.445610559 | 0.719099366 |
| chr5:137016800-137017232 | 0.326051469 | 0.445622076 | 0.719099366 |
| chr10:30059476-30059744 | 0.330186526 | 0.445625927 | 0.719099366 |
| chr4:74768841-74769545 | -0.292625883 | 0.445626164 | 0.719099366 |
| chr3:165482693-165482967 | -0.365442216 | 0.445626817 | 0.719099366 |
| chr3:141644700-141645206 | 0.275733179 | 0.445636146 | 0.719099366 |
| chr20:17822032-17822279 | 0.324048 | 0.445638515 | 0.719099366 |
| chr2:68715306-68715777 | -0.275659805 | 0.445672101 | 0.719138418 |
| chr6:660098-660373 | -0.375637117 | 0.445686357 | 0.719138688 |
| chr6:131135144-131136200 | -0.25252651 | 0.44569104 | 0.719138688 |
| chr4:60464124-60464440 | -0.299551437 | 0.445782308 | 0.719246115 |
| chr3:128725844-128726988 | 0.244660364 | 0.445799691 | 0.719246115 |
| chr11:62601269-62602171 | 0.245009552 | 0.445808017 | 0.719246115 |
| chr11:64847131-64847465 | 0.275831273 | 0.445813275 | 0.719246115 |
| chr7:94643798-94644100 | 0.333801291 | 0.44581732 | 0.719246115 |
| chr20:50570518-50570825 | 0.287525592 | 0.445818966 | 0.719246115 |
| chr1:154558613-154559390 | 0.24565107 | 0.445823327 | 0.719246115 |
| chr19:41563879-41564175 | -0.346481735 | 0.445834267 | 0.719248621 |
| chrX:68498403-68499347 | 0.251721744 | 0.445859097 | 0.719273533 |
| chr2:207395067-207395267 | -0.409856201 | 0.445889854 | 0.719308007 |
| chr11:24496652-24497460 | -0.256072842 | 0.445971652 | 0.719420617 |

| | | | |
|---|---|---|---|
| chr1:177615321-177615593 | 0.320732455 | 0.445984643 | 0.719420617 |
| chr17:8221377-8221668 | -0.27084902 | 0.445990858 | 0.719420617 |
| chr20:49706411-49706630 | -0.402780533 | 0.446011774 | 0.719420617 |
| chrX:80809723-80810284 | -0.246802852 | 0.446014827 | 0.719420617 |
| chr17:78653346-78653546 | 0.400208257 | 0.446015995 | 0.719420617 |
| chr2:190979951-190980748 | -0.250626238 | 0.446034112 | 0.719434695 |
| chr14:77032885-77034405 | 0.239698754 | 0.446082308 | 0.719487378 |
| chr11:60705267-60705489 | 0.405319807 | 0.446088173 | 0.719487378 |
| chr17:63839243-63839508 | -0.262488559 | 0.446103495 | 0.719487378 |
| chr1:112783069-112783353 | -0.387024008 | 0.446104335 | 0.719487378 |
| chrX:97310363-97310679 | -0.315548881 | 0.446152131 | 0.719549319 |
| chrX:98300261-98300921 | -0.280818096 | 0.446180815 | 0.719580434 |
| chr12:79407860-79408368 | 0.260385329 | 0.446195 | 0.719588165 |
| chr3:126433612-126434036 | 0.304219105 | 0.446241456 | 0.71963753 |
| chr2:11028484-11028766 | 0.332475754 | 0.44626489 | 0.71963753 |
| chr15:34991438-34991672 | -0.322317167 | 0.446267376 | 0.71963753 |
| chr10:110354239-110354650 | -0.28329879 | 0.446269267 | 0.71963753 |
| chr6:91167890-91168133 | 0.337807164 | 0.446278905 | 0.71963753 |
| chr8:116851247-116851451 | 0.381265192 | 0.446286384 | 0.71963753 |
| chr14:37650111-37651285 | 0.241564436 | 0.446291354 | 0.71963753 |
| chr14:76914680-76915293 | 0.250769835 | 0.446314187 | 0.719659202 |
| chr6:10316273-10316775 | -0.264872686 | 0.446331216 | 0.719661836 |
| chr3:97988820-97989024 | -0.381028719 | 0.446335929 | 0.719661836 |
| chr6:170150433-170150641 | 0.332702754 | 0.446364555 | 0.719661836 |
| chr3:129211771-129212233 | 0.292016041 | 0.446375589 | 0.719661836 |
| chr22:18109785-18110463 | 0.281658633 | 0.446376099 | 0.719661836 |
| chr13:51803450-51804511 | -0.245508105 | 0.446378077 | 0.719661836 |
| chr11:85810503-85811578 | -0.246938366 | 0.446383691 | 0.719661836 |
| chr11:37320955-37321324 | 0.300870039 | 0.446390959 | 0.719661836 |
| chr6:32415471-32415673 | 0.381007425 | 0.446401044 | 0.719662952 |
| chr16:75064485-75065084 | -0.26279646 | 0.446428417 | 0.71969194 |
| chr7:149714371-149715362 | 0.248304135 | 0.446451195 | 0.719712243 |
| chr14:101350461-101350880 | -0.346468795 | 0.446466086 | 0.719712243 |
| chr17:64837559-64837802 | -0.317165038 | 0.446471207 | 0.719712243 |
| chr16:728164-728644 | 0.261688649 | 0.446478584 | 0.719712243 |
| chr19:1876039-1876389 | 0.302176599 | 0.44651183 | 0.719745884 |
| chr10:64376268-64376504 | -0.366993543 | 0.44651824 | 0.719745884 |
| chr16:74695850-74696238 | -0.25981254 | 0.446545992 | 0.719775476 |
| chr9:4489824-4490523 | -0.270655959 | 0.4465755 | 0.719805215 |
| chr5:176543151-176544267 | -0.248046285 | 0.446583231 | 0.719805215 |
| chr9:27573158-27574267 | 0.246984828 | 0.446594722 | 0.719808595 |
| chr10:68331351-68333053 | -0.239332757 | 0.446606024 | 0.71981167 |
| chr9:113602125-113602405 | -0.325837553 | 0.446666036 | 0.719875501 |
| chr7:128427338-128427538 | -0.337852652 | 0.446677838 | 0.719875501 |
| chr10:103367780-103368136 | 0.271318924 | 0.446680199 | 0.719875501 |
| chr10:89302069-89302457 | -0.325938362 | 0.446707325 | 0.719875501 |
| chr2:55168691-55168891 | 0.383419518 | 0.446708163 | 0.719875501 |
| chr2:143336562-143336762 | 0.383419518 | 0.446708163 | 0.719875501 |
| chr16:10628083-10628567 | 0.275294994 | 0.446711394 | 0.719875501 |
| chr1:200076171-200076525 | 0.296802077 | 0.446770726 | 0.719955973 |

| | | | |
|---|---|---|---|
| chr12:71780298-71780621 | 0.325035301 | 0.446816726 | 0.719994999 |
| chr1:63593173-63594623 | 0.250870388 | 0.446831384 | 0.719994999 |
| chr2:131434751-131434951 | -0.470577615 | 0.446832531 | 0.719994999 |
| chr2:136810429-136810629 | -0.470577615 | 0.446832531 | 0.719994999 |
| chr12:11486652-11486984 | -0.325915361 | 0.446860469 | 0.720024874 |
| chr1:51616841-51617825 | 0.24818109 | 0.446891163 | 0.720059189 |
| chr2:108533383-108534702 | -0.241581438 | 0.446935111 | 0.720109684 |
| chr4:80267010-80267210 | 0.376786849 | 0.446941298 | 0.720109684 |
| chr15:89916830-89917030 | 0.371548505 | 0.446963655 | 0.720130563 |
| chr16:2003890-2004165 | 0.28940586 | 0.447000379 | 0.720160668 |
| chr16:75376522-75376907 | -0.30330922 | 0.447001138 | 0.720160668 |
| chr8:33681532-33681755 | -0.393875331 | 0.447051484 | 0.720220614 |
| chr7:140230976-140231906 | -0.257300858 | 0.447061012 | 0.720220614 |
| chr14:102185322-102185699 | -0.285435533 | 0.447066545 | 0.720220614 |
| chr5:148163390-148163634 | -0.383759028 | 0.447107319 | 0.720271157 |
| chr16:76278997-76279385 | 0.267482562 | 0.447143534 | 0.72031041 |
| chr17:81513371-81513682 | 0.279194056 | 0.447150488 | 0.72031041 |
| chr17:56653937-56654279 | 0.27488573 | 0.447183899 | 0.720349088 |
| chr16:58666575-58666788 | -0.40673819 | 0.44721481 | 0.720380076 |
| chr8:96164345-96164665 | 0.272311984 | 0.44722194 | 0.720380076 |
| chr8:22574333-22574704 | 0.276844887 | 0.447252497 | 0.720408549 |
| chr11:71080310-71080593 | 0.298568748 | 0.44725842 | 0.720408549 |
| chr2:61620620-61620936 | -0.329998348 | 0.447280229 | 0.720428532 |
| chr14:28012702-28013351 | 0.279156574 | 0.447308156 | 0.720458368 |
| chr11:728058-728375 | 0.309392052 | 0.447330374 | 0.720474711 |
| chr2:183171707-183172140 | 0.323726097 | 0.447337108 | 0.720474711 |
| chr16:28822509-28823151 | 0.258196337 | 0.447366632 | 0.720507117 |
| chr2:74327311-74327573 | 0.36795324 | 0.447412207 | 0.720565371 |
| chr22:26763951-26764348 | 0.298096752 | 0.447443043 | 0.720584782 |
| chr17:67246538-67247011 | 0.254086373 | 0.447443068 | 0.720584782 |
| chr4:102129523-102129774 | 0.395109545 | 0.447473982 | 0.720619422 |
| chr10:8800332-8800616 | -0.393523994 | 0.447520034 | 0.720663441 |
| chr3:120661178-120662299 | -0.25550365 | 0.447520127 | 0.720663441 |
| chr2:105546622-105546879 | 0.341370212 | 0.447570841 | 0.720692985 |
| chr10:12041937-12043544 | -0.239119425 | 0.447587963 | 0.720692985 |
| chr3:125399018-125399297 | -0.351166957 | 0.447592443 | 0.720692985 |
| chr4:168679018-168679328 | 0.319087856 | 0.447592446 | 0.720692985 |
| chr9:114889961-114890272 | -0.349747531 | 0.447610419 | 0.720692985 |
| chr2:47335093-47335506 | -0.260712531 | 0.447615958 | 0.720692985 |
| chr7:4952048-4952482 | -0.315488569 | 0.447616769 | 0.720692985 |
| chr10:34649389-34649589 | -0.408422914 | 0.447656635 | 0.720692985 |
| chr21:39444814-39445014 | 0.315142678 | 0.447660464 | 0.720692985 |
| chr10:102460922-102461575 | 0.258978763 | 0.44766348 | 0.720692985 |
| chr15:58612177-58612540 | -0.361527536 | 0.447667078 | 0.720692985 |
| chr9:93577438-93578101 | 0.25883067 | 0.447671324 | 0.720692985 |
| chr14:79810329-79810588 | 0.360873798 | 0.447685189 | 0.720692985 |
| chr18:63367063-63368126 | -0.241673872 | 0.447688904 | 0.720692985 |
| chr3:119518074-119518310 | 0.391064059 | 0.447689059 | 0.720692985 |
| chr2:50144088-50144638 | -0.281137254 | 0.447689338 | 0.720692985 |
| chr1:157132103-157132461 | -0.298767781 | 0.447698372 | 0.720692985 |

| | | | |
|---|---|---|---|
| chr17:996507-997508 | -0.25214567 | 0.447708337 | 0.720693885 |
| chr7:46685111-46685311 | 0.411521352 | 0.447753172 | 0.720750915 |
| chr7:75565179-75565760 | -0.290571832 | 0.447797152 | 0.72079192 |
| chr3:11944091-11944390 | -0.308924017 | 0.447802131 | 0.72079192 |
| chr11:69400875-69401187 | -0.350195168 | 0.447806867 | 0.72079192 |
| chr18:43789026-43789493 | -0.304398207 | 0.447828079 | 0.720803861 |
| chr20:50434663-50434916 | -0.305145955 | 0.4478331 | 0.720803861 |
| chr10:28284066-28284556 | 0.268144329 | 0.447874164 | 0.720854813 |
| chr6:10163059-10163837 | 0.260920752 | 0.447896183 | 0.72087511 |
| chr10:29149903-29150252 | -0.339104287 | 0.44793242 | 0.720915927 |
| chr10:100681263-100681463 | -0.424997203 | 0.447940361 | 0.720915927 |
| chr17:47884760-47885136 | -0.294769033 | 0.447973994 | 0.720953061 |
| chr2:137975209-137976107 | -0.250769375 | 0.447982253 | 0.720953061 |
| chr15:56052414-56052692 | 0.320733492 | 0.448009898 | 0.720964621 |
| chr6:28589251-28589824 | -0.250696273 | 0.448018113 | 0.720964621 |
| chr16:88928370-88928740 | -0.268223308 | 0.448018349 | 0.720964621 |
| chr1:222455144-222455679 | -0.274561192 | 0.448027074 | 0.720964621 |
| chr3:169863414-169863937 | -0.267505856 | 0.448050864 | 0.720987763 |
| chr3:104610944-104611231 | -0.375062443 | 0.448075485 | 0.720997956 |
| chr3:154615563-154615979 | -0.29948425 | 0.448080415 | 0.720997956 |
| chr1:211578665-211579512 | -0.244534057 | 0.448085985 | 0.720997956 |
| chr3:107938071-107938402 | -0.340015305 | 0.448098171 | 0.720997956 |
| chr3:101196153-101196416 | -0.326136387 | 0.448104844 | 0.720997956 |
| chr1:148151906-148152456 | 0.256455277 | 0.448113657 | 0.720997956 |
| chr2:179692398-179692916 | -0.288344343 | 0.448149371 | 0.721024269 |
| chr1:151909124-151910086 | 0.239953427 | 0.448156066 | 0.721024269 |
| chr2:3268037-3268377 | 0.328495692 | 0.44815825 | 0.721024269 |
| chr3:63936552-63936881 | -0.258548851 | 0.448188569 | 0.721024269 |
| chr12:56315468-56316549 | 0.24385454 | 0.448192874 | 0.721024269 |
| chr1:206384432-206384676 | -0.385869448 | 0.448209674 | 0.721024269 |
| chr11:57499558-57499884 | 0.288608503 | 0.44820998 | 0.721024269 |
| chr20:3795628-3796629 | 0.245359045 | 0.448214873 | 0.721024269 |
| chr1:119549720-119549933 | 0.337411813 | 0.448229479 | 0.721024269 |
| chr1:226679889-226680391 | 0.274949316 | 0.448229702 | 0.721024269 |
| chr5:148011777-148012098 | 0.297539878 | 0.448242149 | 0.721024269 |
| chr6:113857168-113857451 | 0.286843624 | 0.448242933 | 0.721024269 |
| chr10:94545212-94546104 | -0.247460334 | 0.448268061 | 0.72103697 |
| chr1:212488667-212488900 | 0.369529177 | 0.44826965 | 0.72103697 |
| chr11:33722563-33722860 | 0.321439735 | 0.448279738 | 0.721038061 |
| chr1:112011370-112011773 | 0.278299986 | 0.448301317 | 0.721049892 |
| chr2:172465543-172466034 | -0.288264 | 0.448315845 | 0.721049892 |
| chr13:44778890-44779139 | -0.328080373 | 0.448329672 | 0.721049892 |
| chr1:154182750-154183607 | 0.243500966 | 0.448334247 | 0.721049892 |
| chr8:118811978-118812390 | 0.303363865 | 0.448346423 | 0.721049892 |
| chr4:76420932-76421575 | 0.283687488 | 0.448353734 | 0.721049892 |
| chr13:36453587-36454026 | -0.296727583 | 0.448362022 | 0.721049892 |
| chr1:192808250-192809326 | -0.243214637 | 0.448362378 | 0.721049892 |
| chr3:45418517-45418717 | 0.389646331 | 0.448378728 | 0.721061052 |
| chr9:20551815-20552085 | -0.336660502 | 0.448398213 | 0.721077253 |
| chr1:231777888-231778606 | 0.255799786 | 0.448449601 | 0.721117167 |

| | | | |
|---|---|---|---|
| chr19:246847-247473 | -0.283142261 | 0.448450267 | 0.721117167 |
| chr8:102634297-102634541 | 0.296565093 | 0.448455382 | 0.721117167 |
| chr3:159884845-159885071 | -0.346660022 | 0.448484173 | 0.721117167 |
| chr3:150410731-150411018 | 0.277192301 | 0.448487818 | 0.721117167 |
| chr20:56247968-56248363 | -0.260025285 | 0.448487872 | 0.721117167 |
| chr5:150817032-150817251 | -0.389411892 | 0.448488913 | 0.721117167 |
| chr10:59958556-59958876 | 0.351004616 | 0.448519883 | 0.721143849 |
| chr2:211954707-211954948 | 0.365305209 | 0.448524331 | 0.721143849 |
| chr9:4740644-4742029 | -0.246162981 | 0.448553148 | 0.721175049 |
| chr12:48197882-48198100 | -0.372242868 | 0.44857968 | 0.721187537 |
| chr1:152188646-152188911 | 0.28745823 | 0.448589393 | 0.721187537 |
| chr1:201085988-201086257 | 0.282829406 | 0.448595821 | 0.721187537 |
| chr5:155394294-155394618 | -0.319859819 | 0.448603021 | 0.721187537 |
| chr20:25865701-25865964 | 0.29161178 | 0.448608992 | 0.721187537 |
| chr6:122710724-122711094 | -0.400155949 | 0.448629509 | 0.721187537 |
| chr8:113171807-113172067 | -0.344612062 | 0.448634911 | 0.721187537 |
| chr1:10866787-10867054 | -0.3358829 | 0.448656041 | 0.721187537 |
| chr3:153353359-153353622 | 0.365861367 | 0.448658971 | 0.721187537 |
| chr8:7354876-7355519 | -0.270767909 | 0.448666875 | 0.721187537 |
| chr6:125099537-125099962 | -0.296360473 | 0.448667489 | 0.721187537 |
| chr9:128298561-128299362 | 0.252332392 | 0.448673863 | 0.721187537 |
| chr9:133356716-133356971 | -0.311201152 | 0.448684585 | 0.721189643 |
| chr12:104057821-104058024 | -0.404410315 | 0.448698219 | 0.721196429 |
| chr9:120928814-120929285 | 0.289871252 | 0.448747156 | 0.721203092 |
| chr9:70602499-70602719 | -0.336289351 | 0.448747248 | 0.721203092 |
| chrX:135477593-135477892 | 0.320296024 | 0.448764909 | 0.721203092 |
| chr6:142088903-142089250 | 0.296690694 | 0.448766385 | 0.721203092 |
| chr21:39456690-39456922 | -0.340807926 | 0.448770214 | 0.721203092 |
| chr6:63565545-63565753 | -0.390400553 | 0.448776006 | 0.721203092 |
| chr8:129692340-129692654 | -0.298881559 | 0.448776522 | 0.721203092 |
| chr12:82019593-82019829 | -0.392436337 | 0.448785673 | 0.721203092 |
| chr10:24519048-24519296 | -0.334318405 | 0.448787077 | 0.721203092 |
| chr1:195948325-195948555 | -0.356200223 | 0.448804465 | 0.721215908 |
| chr14:30562368-30562800 | 0.271653007 | 0.448831506 | 0.721239838 |
| chr11:15649852-15650776 | -0.248716152 | 0.448849733 | 0.721239838 |
| chr2:42693050-42693289 | -0.365510723 | 0.448853717 | 0.721239838 |
| chr3:52198310-52198630 | 0.339204297 | 0.44887082 | 0.721239838 |
| chr11:64448831-64449144 | 0.2954844 | 0.448880066 | 0.721239838 |
| chr21:39062201-39062598 | -0.288813925 | 0.448882978 | 0.721239838 |
| chr3:83217501-83219135 | -0.241249259 | 0.448885247 | 0.721239838 |
| chr2:169694047-169694878 | -0.245640718 | 0.448915701 | 0.721273644 |
| chr5:175365820-175366086 | 0.349602258 | 0.448963917 | 0.721308552 |
| chr19:39125534-39125921 | 0.272493327 | 0.448968381 | 0.721308552 |
| chr12:70258670-70258899 | -0.365002645 | 0.448973995 | 0.721308552 |
| chr11:776515-776715 | -0.385223658 | 0.448975082 | 0.721308552 |
| chr8:15040293-15040659 | -0.3549559 | 0.448988553 | 0.721315069 |
| chr1:201229891-201230091 | 0.344426073 | 0.449012948 | 0.721339137 |
| chr19:35168321-35168608 | 0.31708574 | 0.449114392 | 0.721480763 |
| chr5:11384919-11385233 | 0.302809156 | 0.449127744 | 0.721480763 |
| chr14:101938748-101939011 | 0.350442095 | 0.449136905 | 0.721480763 |

| | | | |
|---|---|---|---|
| chr17:49630030-49630361 | -0.345075474 | 0.44915255 | 0.721480763 |
| chr11:133835310-133835510 | -0.377367033 | 0.449170875 | 0.721480763 |
| chr1:75723953-75725085 | -0.242289513 | 0.449174657 | 0.721480763 |
| chr11:6480986-6481944 | -0.245095451 | 0.449180405 | 0.721480763 |
| chr19:32655326-32655526 | 0.414984595 | 0.449184173 | 0.721480763 |
| chr11:59786998-59787368 | -0.284034698 | 0.449185852 | 0.721480763 |
| chr14:23953436-23954461 | 0.245555523 | 0.449207519 | 0.72150044 |
| chr4:64713292-64713528 | -0.33728246 | 0.449237947 | 0.721534189 |
| chr12:24562068-24562268 | -0.372929062 | 0.449256743 | 0.721549251 |
| chr6:5063678-5063878 | -0.415744853 | 0.449309191 | 0.721594654 |
| chr17:48791050-48791353 | -0.294191697 | 0.449311617 | 0.721594654 |
| chr1:22976360-22976833 | 0.286093057 | 0.449313264 | 0.721594654 |
| chr6:28742792-28743115 | -0.283963727 | 0.44933224 | 0.721610003 |
| chr2:163266864-163267153 | -0.391822499 | 0.449406153 | 0.721687972 |
| chr6:96626832-96627067 | -0.344799401 | 0.449408327 | 0.721687972 |
| chr19:55146220-55146625 | 0.297123646 | 0.449409045 | 0.721687972 |
| chr10:82009781-82009981 | 0.419915753 | 0.449422266 | 0.721694076 |
| chr1:54488065-54488682 | 0.256731266 | 0.449454548 | 0.721730789 |
| chr14:45716578-45717063 | 0.277208572 | 0.449476653 | 0.72175116 |
| chr15:78299054-78299886 | 0.247535155 | 0.449487188 | 0.721752951 |
| chr8:69946172-69946442 | 0.329221287 | 0.449499799 | 0.721758075 |
| chr7:18610444-18610650 | 0.36661678 | 0.449522933 | 0.721780096 |
| chr3:12471114-12471688 | 0.275039235 | 0.449546122 | 0.721802204 |
| chr1:219991099-219991371 | -0.387292848 | 0.449606634 | 0.721877081 |
| chr17:29568215-29569448 | 0.245247481 | 0.449611599 | 0.721877081 |
| chr8:90571128-90571337 | 0.337413958 | 0.449623412 | 0.721880921 |
| chr3:142091774-142091974 | -0.431445091 | 0.449646736 | 0.721901995 |
| chr10:13708257-13708457 | 0.368405279 | 0.449655381 | 0.721901995 |
| chr11:78045832-78046045 | -0.322807257 | 0.449665073 | 0.72190243 |
| chr5:100430331-100430568 | -0.37228273 | 0.449689443 | 0.721926428 |
| chr12:46382978-46384297 | 0.239360114 | 0.449730614 | 0.721977396 |
| chr9:95666201-95666401 | 0.401500997 | 0.44981145 | 0.722090636 |
| chr9:73107551-73107887 | -0.324967397 | 0.449823762 | 0.722090636 |
| chr13:71867168-71867473 | 0.314435599 | 0.449829425 | 0.722090636 |
| chr15:63247132-63247379 | 0.376206708 | 0.449902205 | 0.722146468 |
| chr7:34839658-34840005 | -0.306000249 | 0.44990813 | 0.722146468 |
| chr8:13122053-13122340 | -0.351916662 | 0.449916379 | 0.722146468 |
| chr4:77863449-77863789 | -0.29948447 | 0.449922282 | 0.722146468 |
| chr15:73683688-73683927 | 0.332930996 | 0.44992744 | 0.722146468 |
| chr7:8444457-8444709 | 0.31180827 | 0.449933112 | 0.722146468 |
| chr6:118551171-118551393 | -0.369469608 | 0.44993527 | 0.722146468 |
| chr9:19208525-19209080 | 0.285469832 | 0.449940113 | 0.722146468 |
| chr5:112586852-112587102 | -0.384619693 | 0.449955731 | 0.722146468 |
| chr6:30672366-30673352 | 0.25372562 | 0.449986244 | 0.722146468 |
| chr3:24299787-24300485 | -0.273809538 | 0.449991486 | 0.722146468 |
| chr2:135741274-135742720 | -0.237882701 | 0.449992793 | 0.722146468 |
| chr11:86846179-86846379 | 0.38748735 | 0.450003351 | 0.722146468 |
| chr1:171405075-171405418 | -0.273698508 | 0.450004783 | 0.722146468 |
| chr6:151616097-151616356 | -0.334564296 | 0.450011555 | 0.722146468 |
| chr11:110076511-110076885 | 0.33594928 | 0.450015003 | 0.722146468 |

| | | | |
|---|---|---|---|
| chr3:57100171-57100371 | 0.374243076 | 0.450030411 | 0.722150319 |
| chr2:83343035-83343235 | -0.381733022 | 0.450048973 | 0.722150319 |
| chr14:59055518-59055730 | 0.349736029 | 0.450053437 | 0.722150319 |
| chr3:126567955-126568206 | 0.343237409 | 0.450055102 | 0.722150319 |
| chr15:40088568-40088848 | -0.329894873 | 0.450070568 | 0.722160013 |
| chr8:126083708-126084062 | -0.279051952 | 0.45009452 | 0.72218332 |
| chr8:127734970-127736483 | -0.236240528 | 0.450109531 | 0.722192283 |
| chr22:29603081-29604144 | 0.244011011 | 0.450137709 | 0.722222371 |
| chr11:66545659-66546456 | 0.25004183 | 0.450174621 | 0.722266471 |
| chr19:556043-556269 | -0.38432366 | 0.450201478 | 0.722294435 |
| chr5:11785781-11786162 | 0.316356007 | 0.450294705 | 0.72242888 |
| chr20:49916432-49916844 | -0.257171843 | 0.450316098 | 0.722448075 |
| chr6:133286819-133287059 | -0.344224321 | 0.45037149 | 0.722507169 |
| chr14:101628456-101628727 | -0.319451355 | 0.450371792 | 0.722507169 |
| chr8:56437867-56438152 | 0.334740258 | 0.450383461 | 0.722510762 |
| chr2:118202722-118203096 | -0.28855751 | 0.450407898 | 0.722534838 |
| chr14:77707290-77708381 | 0.245120778 | 0.450449119 | 0.722574169 |
| chr1:966894-967118 | -0.343966449 | 0.450451374 | 0.722574169 |
| chr22:23240951-23241214 | -0.34570177 | 0.450460708 | 0.722574169 |
| chr14:89000581-89000900 | 0.304371591 | 0.450491662 | 0.722608694 |
| chr12:95072803-95073908 | -0.244061942 | 0.450512441 | 0.722623061 |
| chr6:149450681-149450907 | 0.326851971 | 0.45051948 | 0.722623061 |
| chr2:230713371-230713756 | -0.311138057 | 0.450541688 | 0.722643554 |
| chr17:61821003-61821386 | 0.246749073 | 0.45056658 | 0.722659667 |
| chr7:55354757-55355022 | -0.394485007 | 0.450579782 | 0.722659667 |
| chr1:231091829-231092305 | 0.293179939 | 0.450580029 | 0.722659667 |
| chr9:127608162-127608694 | 0.265229618 | 0.450592784 | 0.722664998 |
| chr10:128088282-128088634 | -0.300554385 | 0.45062241 | 0.722692291 |
| chr10:9342944-9343144 | 0.398913628 | 0.450628665 | 0.722692291 |
| chr10:83871523-83871880 | 0.290597532 | 0.450693875 | 0.722781743 |
| chr6:73121806-73122006 | 0.380076309 | 0.450708723 | 0.722790426 |
| chr1:28173867-28174067 | 0.383325066 | 0.450739956 | 0.722825385 |
| chr6:112254426-112254626 | 0.410383394 | 0.450751573 | 0.722828886 |
| chr8:141228519-141228786 | -0.30764156 | 0.450790446 | 0.722848849 |
| chr4:3704319-3704528 | 0.406913545 | 0.450790706 | 0.722848849 |
| chr2:240136198-240136554 | 0.28244488 | 0.450792324 | 0.722848849 |
| chr1:201500729-201500978 | 0.28757352 | 0.450818034 | 0.722855885 |
| chr5:32768043-32768313 | 0.354867057 | 0.450836396 | 0.722855885 |
| chr9:129168090-129168320 | 0.343581506 | 0.450846017 | 0.722855885 |
| chr1:106358616-106358816 | 0.292251688 | 0.450859624 | 0.722855885 |
| chr20:7691695-7692146 | -0.303500643 | 0.450868698 | 0.722855885 |
| chr10:9149316-9149761 | 0.294942733 | 0.450876657 | 0.722855885 |
| chr8:102197146-102197767 | 0.260819828 | 0.450880286 | 0.722855885 |
| chr1:85580491-85581180 | 0.253272235 | 0.450896888 | 0.722855885 |
| chr10:46558626-46558948 | 0.298332143 | 0.45089834 | 0.722855885 |
| chr2:5070391-5070612 | -0.390669767 | 0.450905416 | 0.722855885 |
| chr10:132389594-132389811 | 0.317260319 | 0.450913351 | 0.722855885 |
| chr21:38219088-38219438 | -0.297909631 | 0.450915573 | 0.722855885 |
| chr9:97895758-97896362 | -0.273006005 | 0.450924163 | 0.722855885 |
| chr8:60347721-60348009 | -0.33572033 | 0.450929679 | 0.722855885 |

| | | | |
|---|---|---|---|
| chr7:5426528-5426867 | -0.270619277 | 0.450938222 | 0.722855885 |
| chr5:113939976-113940448 | -0.276943012 | 0.450961427 | 0.722877959 |
| chr16:72093268-72094435 | -0.238284035 | 0.450984018 | 0.722897384 |
| chr5:156037671-156037995 | -0.358601145 | 0.450992414 | 0.722897384 |
| chr17:30340184-30340457 | -0.340156299 | 0.451078825 | 0.723020768 |
| chr10:8061909-8062205 | 0.289644232 | 0.451113064 | 0.723029063 |
| chr5:135426124-135426674 | 0.276393376 | 0.451118561 | 0.723029063 |
| chr2:9104013-9104254 | -0.336202957 | 0.451121032 | 0.723029063 |
| chr1:8525900-8526225 | 0.31216106 | 0.451121746 | 0.723029063 |
| chr2:54457100-54457329 | 0.330939588 | 0.451141624 | 0.723041917 |
| chr5:82312380-82312843 | -0.283929168 | 0.451148639 | 0.723041917 |
| chr8:88594306-88594506 | -0.418784766 | 0.451163939 | 0.723051314 |
| chr13:26482608-26482925 | -0.292879084 | 0.451232927 | 0.723146752 |
| chr9:92634776-92635079 | 0.300419625 | 0.451260747 | 0.723151934 |
| chr2:156455398-156455598 | 0.367164667 | 0.451263519 | 0.723151934 |
| chr14:39431425-39431788 | 0.252941538 | 0.451264475 | 0.723151934 |
| chr3:64193035-64193513 | 0.250969278 | 0.451283095 | 0.723166649 |
| chr1:36486866-36487304 | 0.296190163 | 0.451328087 | 0.723212145 |
| chr1:27858078-27858605 | 0.270164334 | 0.451330364 | 0.723212145 |
| chr10:119014470-119014700 | -0.401015601 | 0.451352453 | 0.723213938 |
| chr17:2965587-2965873 | 0.338316737 | 0.45135758 | 0.723213938 |
| chr3:153296988-153297188 | 0.387365309 | 0.451359799 | 0.723213938 |
| chr3:95418564-95418764 | 0.384083217 | 0.45139023 | 0.723230373 |
| chr20:52001296-52001539 | -0.323741299 | 0.451391927 | 0.723230373 |
| chr19:49151619-49152195 | 0.245869063 | 0.451398373 | 0.723230373 |
| chr5:36090084-36090382 | -0.297289099 | 0.451408079 | 0.723230801 |
| chr20:19688406-19688711 | 0.306527714 | 0.451453381 | 0.723260127 |
| chr5:111699352-111699552 | 0.4099327 | 0.45145737 | 0.723260127 |
| chrX:78157385-78157616 | 0.343260163 | 0.451458038 | 0.723260127 |
| chr1:65066425-65068273 | -0.240928517 | 0.45146414 | 0.723260127 |
| chr5:39642716-39642964 | 0.394389589 | 0.45152166 | 0.723325021 |
| chr22:32378706-32378990 | 0.324944842 | 0.451523528 | 0.723325021 |
| chr8:126547203-126547734 | -0.259611373 | 0.451541074 | 0.723338007 |
| chr2:15637329-15637611 | 0.303658774 | 0.45159085 | 0.723402621 |
| chr6:158071808-158072029 | 0.372481489 | 0.45160189 | 0.723405181 |
| chr17:81722179-81722763 | 0.255729388 | 0.451625951 | 0.723417191 |
| chr8:81302534-81302815 | -0.334251199 | 0.45162827 | 0.723417191 |
| chr3:125374567-125374899 | 0.287408806 | 0.451667094 | 0.72344587 |
| chr3:184135065-184135736 | -0.252592468 | 0.451669192 | 0.72344587 |
| chr14:99363147-99363347 | -0.400302686 | 0.451678491 | 0.72344587 |
| chr16:89658271-89658563 | -0.286121236 | 0.451692004 | 0.72344587 |
| chr21:42781455-42781898 | -0.270402161 | 0.451693383 | 0.72344587 |
| chr6:49550357-49551599 | -0.244221181 | 0.451758509 | 0.723535054 |
| chrX:104098231-104098581 | -0.307715825 | 0.451786445 | 0.723549692 |
| chr2:168455800-168457186 | -0.2402422 | 0.451788902 | 0.723549692 |
| chr15:65791528-65792819 | 0.240547525 | 0.451795979 | 0.723549692 |
| chr11:66257183-66258749 | 0.240389978 | 0.451807026 | 0.723552262 |
| chr5:78648036-78649240 | -0.250000754 | 0.451860009 | 0.723621987 |
| chr8:48318588-48319288 | -0.259433467 | 0.451892872 | 0.723655383 |
| chr5:149564762-149565161 | 0.267364299 | 0.451899751 | 0.723655383 |

| | | | |
|---|---|---|---|
| chr1:98053903-98054510 | -0.261633173 | 0.451918631 | 0.723662372 |
| chr20:35568667-35568967 | -0.328506763 | 0.451940941 | 0.723662372 |
| chr8:98165903-98166105 | -0.339610139 | 0.451942171 | 0.723662372 |
| chr15:31326054-31327246 | 0.238982617 | 0.451956481 | 0.723662372 |
| chr2:215038415-215038684 | -0.37638167 | 0.451965623 | 0.723662372 |
| chr1:201021098-201021300 | 0.300856164 | 0.451969867 | 0.723662372 |
| chr13:21603445-21604417 | 0.239864569 | 0.451970228 | 0.723662372 |
| chr14:65411627-65413119 | 0.23628464 | 0.452018452 | 0.723707418 |
| chr1:206469866-206471029 | 0.240041852 | 0.452026044 | 0.723707418 |
| chr9:109887875-109888247 | -0.32897561 | 0.452026698 | 0.723707418 |
| chr8:19314178-19314575 | -0.296519675 | 0.452070561 | 0.723739929 |
| chr19:50205287-50205529 | 0.296618602 | 0.452074734 | 0.723739929 |
| chr6:53863171-53863437 | -0.330862653 | 0.452080746 | 0.723739929 |
| chr22:38272092-38272292 | 0.37883231 | 0.452087552 | 0.723739929 |
| chr18:13558185-13558512 | 0.333232631 | 0.452094232 | 0.723739929 |
| chr20:42205987-42206497 | -0.257540525 | 0.452113615 | 0.723755838 |
| chr5:25858062-25858370 | 0.29794362 | 0.452140345 | 0.723783506 |
| chr4:88539225-88539831 | -0.262736244 | 0.452155594 | 0.723792795 |
| chr8:11902256-11902728 | 0.275121499 | 0.452194457 | 0.723836073 |
| chr14:100246781-100247124 | 0.281866213 | 0.452201524 | 0.723836073 |
| chr9:101311782-101312048 | -0.334572446 | 0.452221803 | 0.723847449 |
| chr11:18410430-18410792 | -0.28883106 | 0.452242131 | 0.723847449 |
| chr12:106106922-106107176 | -0.330233101 | 0.452253375 | 0.723847449 |
| chr8:80023688-80023895 | 0.357447988 | 0.452257643 | 0.723847449 |
| chr3:168249225-168250416 | -0.240234782 | 0.45225973 | 0.723847449 |
| chr6:166342369-166343128 | -0.255277064 | 0.452281333 | 0.723847449 |
| chr11:72382038-72382295 | -0.370800274 | 0.452285037 | 0.723847449 |
| chr4:176377632-176378028 | -0.318138335 | 0.4522877 | 0.723847449 |
| chr15:78427892-78428104 | -0.389797378 | 0.452306196 | 0.723847449 |
| chr11:9460329-9461565 | 0.238203425 | 0.452306678 | 0.723847449 |
| chr5:38265575-38266158 | -0.276041049 | 0.452312547 | 0.723847449 |
| chr6:143962562-143962814 | -0.340217259 | 0.452353925 | 0.723884297 |
| chr9:121400164-121400424 | -0.358796744 | 0.452354552 | 0.723884297 |
| chr1:161539788-161540579 | -0.253485795 | 0.452373132 | 0.723884297 |
| chr9:95341625-95341852 | 0.349292934 | 0.452373363 | 0.723884297 |
| chr17:19237226-19238459 | -0.249453092 | 0.452402735 | 0.723887612 |
| chr10:87861586-87862775 | -0.237890227 | 0.452404572 | 0.723887612 |
| chr5:172502920-172503468 | 0.274330487 | 0.452406625 | 0.723887612 |
| chr2:207511954-207512189 | 0.367660229 | 0.452413225 | 0.723887612 |
| chr20:46903882-46904082 | -0.38302725 | 0.452424697 | 0.723890852 |
| chr14:23569602-23570208 | -0.253492897 | 0.452546449 | 0.724069143 |
| chr4:93828341-93828809 | 0.281400506 | 0.45256229 | 0.724069143 |
| chr3:8302080-8302287 | 0.3622213 | 0.452588535 | 0.724069143 |
| chr6:3848910-3849528 | 0.25903812 | 0.452589558 | 0.724069143 |
| chr11:68257413-68257884 | -0.273150926 | 0.452609134 | 0.724069143 |
| chr1:52127557-52127767 | 0.380594103 | 0.452613147 | 0.724069143 |
| chr15:88999207-88999407 | 0.380594103 | 0.452613147 | 0.724069143 |
| chr5:131796274-131797772 | -0.236492608 | 0.452617071 | 0.724069143 |
| chr5:149371387-149371702 | 0.282684358 | 0.452621176 | 0.724069143 |
| chr14:20613101-20614284 | -0.234863262 | 0.452640677 | 0.724072484 |

| | | | |
|---|---|---|---|
| chr3:62532939-62533139 | 0.306942765 | 0.452656699 | 0.724072484 |
| chr11:6443413-6443718 | -0.305286206 | 0.452660209 | 0.724072484 |
| chr4:13075199-13075454 | -0.362087077 | 0.452661064 | 0.724072484 |
| chr6:123091016-123091326 | -0.334371489 | 0.452674732 | 0.724079231 |
| chr17:58658524-58660015 | -0.233021219 | 0.452702007 | 0.724107742 |
| chr3:15398898-15399245 | 0.287986394 | 0.452724681 | 0.724128894 |
| chr10:24886166-24886651 | 0.282554214 | 0.452752014 | 0.724157496 |
| chr18:3012543-3013686 | 0.243881613 | 0.452778771 | 0.724161235 |
| chr4:88158247-88159591 | 0.245402646 | 0.452782727 | 0.724161235 |
| chr16:22297025-22298707 | 0.235856637 | 0.452786702 | 0.724161235 |
| chr12:64451681-64453154 | 0.237486514 | 0.452808902 | 0.724161235 |
| chr13:29925555-29925755 | 0.305629574 | 0.452809274 | 0.724161235 |
| chr1:153727461-153728612 | 0.239895519 | 0.452817336 | 0.724161235 |
| chr17:11987007-11987208 | -0.385549619 | 0.45282051 | 0.724161235 |
| chr1:92298530-92299371 | -0.24810853 | 0.452841964 | 0.72418043 |
| chr14:22871148-22873102 | 0.235972065 | 0.452855365 | 0.724186747 |
| chr5:90529012-90530352 | -0.23650389 | 0.452891504 | 0.724229423 |
| chr2:196639115-196640249 | -0.239403392 | 0.452917832 | 0.724256409 |
| chr5:144142400-144142633 | 0.329563147 | 0.452939029 | 0.72427519 |
| chr3:185796542-185797067 | -0.275174616 | 0.452959619 | 0.724280095 |
| chr3:75446920-75447120 | -0.427704095 | 0.452961002 | 0.724280095 |
| chr8:125301504-125301704 | 0.347579073 | 0.45300464 | 0.724325628 |
| chr14:99980498-99980850 | 0.304827186 | 0.453008675 | 0.724325628 |
| chr14:69708890-69709165 | 0.305492557 | 0.45303036 | 0.724325628 |
| chr19:53906839-53907076 | 0.343468475 | 0.453036311 | 0.724325628 |
| chr3:17021700-17021955 | -0.365503517 | 0.453040524 | 0.724325628 |
| chr6:28737261-28737492 | 0.356424368 | 0.453046197 | 0.724325628 |
| chr20:45651831-45652065 | 0.356750244 | 0.453080418 | 0.724351056 |
| chr19:2328500-2328753 | 0.33709929 | 0.453095546 | 0.724351056 |
| chr5:174205086-174205310 | 0.363973623 | 0.453098414 | 0.724351056 |
| chr8:142402795-142403625 | -0.247394845 | 0.453099916 | 0.724351056 |
| chr12:105083592-105084846 | -0.243586707 | 0.453119697 | 0.724366107 |
| chr19:1103697-1104193 | 0.257877402 | 0.453128238 | 0.724366107 |
| chr7:127759526-127759782 | -0.390513657 | 0.453152076 | 0.724380311 |
| chr8:142602913-142603246 | -0.317335158 | 0.453156032 | 0.724380311 |
| chr9:110005862-110006062 | -0.432180099 | 0.453167971 | 0.724384283 |
| chr14:104724084-104724284 | 0.308305038 | 0.45318231 | 0.724392093 |
| chr2:171530049-171530286 | -0.39250006 | 0.453192858 | 0.724393841 |
| chr16:71564558-71565808 | -0.238284392 | 0.453284325 | 0.724477176 |
| chr1:159177296-159177650 | -0.29102373 | 0.453285136 | 0.724477176 |
| chr2:20242698-20243384 | -0.263059909 | 0.453285492 | 0.724477176 |
| chrX:69286077-69286496 | 0.319449916 | 0.453290062 | 0.724477176 |
| chr1:42923926-42924758 | 0.250097327 | 0.45329227 | 0.724477176 |
| chr14:31934730-31934975 | 0.329527146 | 0.453310817 | 0.724491707 |
| chr3:179942193-179942828 | -0.252468949 | 0.453340881 | 0.724524643 |
| chr16:2467627-2467928 | 0.311412563 | 0.453374756 | 0.724561108 |
| chr14:20487455-20487655 | -0.409972015 | 0.45338261 | 0.724561108 |
| chr14:77862240-77862569 | 0.280117334 | 0.453399458 | 0.724565601 |
| chr13:78978456-78978692 | -0.348443422 | 0.453412328 | 0.724565601 |
| chr19:510676-510876 | 0.340523516 | 0.45341379 | 0.724565601 |

| | | | |
|---|---|---|---|
| chr4:140727886-140728879 | 0.246821281 | 0.453452437 | 0.724583486 |
| chr13:96020156-96020434 | 0.321674053 | 0.453464569 | 0.724583486 |
| chr4:103150723-103151016 | -0.343260806 | 0.453475553 | 0.724583486 |
| chr3:105611504-105612056 | -0.280157765 | 0.45347844 | 0.724583486 |
| chr12:68686517-68687496 | -0.238805973 | 0.453480371 | 0.724583486 |
| chr9:35111453-35111894 | 0.253409315 | 0.453487499 | 0.724583486 |
| chr7:2425386-2425684 | -0.314319741 | 0.453491178 | 0.724583486 |
| chr7:75442436-75442738 | -0.355845351 | 0.453508307 | 0.724595744 |
| chr11:134868594-134868844 | -0.344384036 | 0.453602409 | 0.724716141 |
| chr10:30026805-30027690 | -0.252447581 | 0.453602578 | 0.724716141 |
| chr11:47847826-47848983 | 0.24014765 | 0.45363972 | 0.724751955 |
| chr1:215682294-215682494 | -0.349976924 | 0.453651279 | 0.724751955 |
| chr3:132342283-132342516 | -0.361365817 | 0.453659846 | 0.724751955 |
| chr5:17951288-17952426 | 0.238892028 | 0.453662829 | 0.724751955 |
| chr7:51306824-51307099 | -0.361589006 | 0.453694196 | 0.724776746 |
| chr7:33836975-33837175 | -0.351243789 | 0.453699235 | 0.724776746 |
| chr10:110498120-110498336 | 0.310499495 | 0.453710019 | 0.724776746 |
| chr1:54801292-54801686 | -0.301655503 | 0.453716183 | 0.724776746 |
| chrX:68433028-68434190 | -0.242732263 | 0.453732034 | 0.724785818 |
| chr17:63985220-63985420 | -0.324602434 | 0.45374082 | 0.724785818 |
| chr8:19408429-19408835 | -0.314379657 | 0.45375024 | 0.724785818 |
| chr10:11533501-11533785 | -0.342644965 | 0.453785605 | 0.724814823 |
| chr1:215474038-215474320 | -0.320298462 | 0.453787318 | 0.724814823 |
| chr1:66726022-66726394 | -0.318032483 | 0.453801751 | 0.724822766 |
| chr8:79615465-79615745 | -0.343384875 | 0.453819665 | 0.72483627 |
| chr20:57140417-57140714 | 0.246225336 | 0.453841878 | 0.724856638 |
| chr2:62341068-62341403 | 0.334361364 | 0.45386484 | 0.724867812 |
| chr5:12662786-12663011 | -0.409108319 | 0.453867795 | 0.724867812 |
| chr1:30275206-30275452 | -0.386817588 | 0.453930719 | 0.724924394 |
| chr1:156892391-156892643 | 0.304204232 | 0.453932089 | 0.724924394 |
| chr5:73207528-73207804 | 0.359609093 | 0.453939787 | 0.724924394 |
| chr1:197521204-197521598 | -0.301042446 | 0.453941067 | 0.724924394 |
| chr19:8390553-8391054 | 0.254690895 | 0.453957019 | 0.72493476 |
| chr1:22191874-22192157 | -0.424381476 | 0.454014317 | 0.72501115 |
| chr3:89852063-89852263 | 0.366836051 | 0.454024662 | 0.725012559 |
| chr11:101583167-101584070 | -0.262479102 | 0.454047261 | 0.725033537 |
| chr5:171750268-171750469 | 0.360887069 | 0.45407272 | 0.72505908 |
| chr10:19835362-19835618 | -0.338593309 | 0.454095272 | 0.725067587 |
| chr3:109561122-109561322 | -0.424508655 | 0.454096973 | 0.725067587 |
| chr8:38808639-38809129 | 0.292875108 | 0.454122592 | 0.725093383 |
| chr17:42832971-42833703 | 0.254706091 | 0.454149313 | 0.725120938 |
| chr2:240557856-240558140 | -0.301191551 | 0.45417605 | 0.725148516 |
| chr6:47477336-47478596 | -0.237916758 | 0.454199414 | 0.725152052 |
| chr5:132419604-132419971 | -0.278717378 | 0.454200354 | 0.725152052 |
| chr3:150407868-150409712 | 0.235283556 | 0.454212812 | 0.725152052 |
| chr20:60365256-60365481 | -0.353907067 | 0.45421612 | 0.725152052 |
| chr3:20039804-20041068 | 0.239018878 | 0.454231758 | 0.725161908 |
| chr7:130142811-130143023 | 0.362789885 | 0.454276186 | 0.725217725 |
| chr1:109298706-109298964 | -0.306895564 | 0.454287457 | 0.725219373 |
| chr3:186529357-186529644 | -0.325343104 | 0.454305378 | 0.725219373 |

| | | | |
|---|---|---|---|
| chr3:69410247-69410683 | 0.305760402 | 0.454305613 | 0.725219373 |
| chr2:41895306-41895511 | -0.389984214 | 0.454350321 | 0.725258497 |
| chr2:143431512-143431737 | -0.389984214 | 0.454350321 | 0.725258497 |
| chr3:123678103-123678430 | 0.287608615 | 0.454358518 | 0.725258497 |
| chr11:66789579-66790120 | -0.272209873 | 0.454398062 | 0.725306508 |
| chr14:41575422-41575699 | 0.335088818 | 0.454457414 | 0.72537826 |
| chr7:1269011-1269211 | 0.324833845 | 0.454461948 | 0.72537826 |
| chr2:68252393-68253285 | -0.238709762 | 0.454488724 | 0.725405887 |
| chr14:24194843-24196069 | 0.241831603 | 0.454510818 | 0.72542604 |
| chr2:231364946-231365802 | 0.249510507 | 0.454520728 | 0.725426745 |
| chr17:50270172-50270477 | -0.289492856 | 0.454560923 | 0.725474052 |
| chr1:27341833-27342089 | -0.331376009 | 0.454571224 | 0.725474052 |
| chr1:226127797-226128251 | -0.257413736 | 0.454578773 | 0.725474052 |
| chr19:1848450-1848983 | 0.26516118 | 0.454614107 | 0.725515331 |
| chr8:90521162-90521538 | 0.290681648 | 0.454651369 | 0.725540099 |
| chr3:65751184-65751419 | 0.352061221 | 0.45466807 | 0.725540099 |
| chr5:154583656-154584549 | -0.250152719 | 0.454682491 | 0.725540099 |
| chr10:76311813-76312087 | 0.327477197 | 0.454687107 | 0.725540099 |
| chr22:20079731-20080678 | 0.247165943 | 0.454696831 | 0.725540099 |
| chr9:72592875-72593077 | -0.39424091 | 0.454701047 | 0.725540099 |
| chr21:30340990-30341190 | 0.389221942 | 0.454710773 | 0.725540099 |
| chr2:172954038-172954412 | 0.294099683 | 0.454723163 | 0.725540099 |
| chr12:112787701-112787945 | -0.403970645 | 0.45472382 | 0.725540099 |
| chr3:79575737-79575996 | -0.390968789 | 0.454736984 | 0.725540099 |
| chr3:1231220-1231576 | -0.318373011 | 0.454748211 | 0.725540099 |
| chr7:44606102-44607254 | -0.236377917 | 0.454756582 | 0.725540099 |
| chr12:111597388-111597644 | 0.303684954 | 0.454769418 | 0.725540099 |
| chr12:48977265-48977646 | 0.287977259 | 0.454774372 | 0.725540099 |
| chr1:159790362-159790685 | -0.310946887 | 0.454781132 | 0.725540099 |
| chr1:204692819-204693089 | -0.310946887 | 0.454781132 | 0.725540099 |
| chr6:27583308-27583619 | -0.305235061 | 0.454800198 | 0.725550975 |
| chr11:22177236-22177514 | -0.332159806 | 0.454813111 | 0.725550975 |
| chr3:12175945-12176200 | 0.308359291 | 0.454819713 | 0.725550975 |
| chr5:163571862-163572267 | -0.314579664 | 0.454825826 | 0.725550975 |
| chr3:126474967-126475313 | 0.267627059 | 0.45484309 | 0.725563409 |
| chr12:131894201-131894744 | 0.249908003 | 0.454873746 | 0.725597205 |
| chr4:124437877-124438169 | 0.326195226 | 0.454926318 | 0.725636326 |
| chr20:9617953-9618326 | 0.27004202 | 0.454936426 | 0.725636326 |
| chr4:185227363-185227601 | -0.356972878 | 0.454937669 | 0.725636326 |
| chr2:162422978-162423178 | -0.401909281 | 0.454947802 | 0.725636326 |
| chr7:58434461-58434661 | 0.381289572 | 0.454956419 | 0.725636326 |
| chr18:61892925-61894845 | -0.240294923 | 0.454960563 | 0.725636326 |
| chr20:45592625-45592889 | 0.340474262 | 0.454970392 | 0.725636326 |
| chr6:119349362-119350702 | -0.241910171 | 0.454987606 | 0.725636326 |
| chr3:127590266-127591584 | 0.237730048 | 0.454998862 | 0.725636326 |
| chr10:81873955-81875186 | 0.245927265 | 0.455000889 | 0.725636326 |
| chr1:54887085-54887635 | -0.254452141 | 0.45500912 | 0.725636326 |
| chr5:179895884-179896110 | -0.337092644 | 0.455011915 | 0.725636326 |
| chr3:48685495-48686404 | 0.251279404 | 0.455068965 | 0.725712204 |
| chr20:63978795-63979073 | -0.272312559 | 0.455084376 | 0.725721675 |

| | | | |
|---|---|---|---|
| chr11:72434055-72434857 | -0.241626875 | 0.455136605 | 0.72577266 |
| chr8:128251825-128252133 | -0.278807964 | 0.455142136 | 0.72577266 |
| chr17:40417594-40418295 | 0.247327918 | 0.455144764 | 0.72577266 |
| chrY:56837492-56837692 | 0.363181275 | 0.455154497 | 0.725773076 |
| chr2:136423465-136423962 | -0.261519423 | 0.455209661 | 0.725831678 |
| chr1:188128747-188129178 | 0.29172085 | 0.455212682 | 0.725831678 |
| chr20:54012067-54012360 | 0.250481397 | 0.455219667 | 0.725831678 |
| chr2:106104386-106104691 | -0.308408941 | 0.455243464 | 0.725854517 |
| chr2:30692462-30692662 | 0.350379668 | 0.455255317 | 0.725858313 |
| chr1:47781127-47781427 | 0.308801337 | 0.455284406 | 0.725862308 |
| chr20:34215978-34216412 | -0.323434889 | 0.455285926 | 0.725862308 |
| chr19:57410947-57411709 | 0.260213716 | 0.455286243 | 0.725862308 |
| chr11:1882587-1882787 | 0.378316167 | 0.455331045 | 0.725918632 |
| chr5:148383566-148384457 | -0.240738625 | 0.455349966 | 0.725933692 |
| chr14:41656795-41657046 | -0.445191277 | 0.455369718 | 0.725946342 |
| chr11:694950-695950 | -0.244039532 | 0.45537685 | 0.725946342 |
| chr18:63439339-63439598 | -0.348454166 | 0.455403623 | 0.725965254 |
| chr12:123763188-123763477 | -0.398868118 | 0.455413209 | 0.725965254 |
| chr10:119139516-119139813 | -0.317329753 | 0.455417137 | 0.725965254 |
| chr19:29694595-29694917 | 0.277148352 | 0.455449712 | 0.726002077 |
| chr19:49712380-49712728 | -0.274777736 | 0.455466415 | 0.726013598 |
| chr8:123156677-123157944 | -0.233871386 | 0.45549892 | 0.726028242 |
| chr2:205745739-205745970 | -0.357654695 | 0.455511827 | 0.726028242 |
| chr9:33024779-33026109 | -0.239323359 | 0.455515384 | 0.726028242 |
| chr20:11267307-11267577 | -0.35195178 | 0.45551985 | 0.726028242 |
| chr7:2355234-2355456 | 0.301586768 | 0.455530329 | 0.726028242 |
| chr1:189249710-189250066 | -0.328602338 | 0.455532455 | 0.726028242 |
| chr10:98045813-98046120 | -0.308196357 | 0.455585208 | 0.72607388 |
| chr11:10307938-10308252 | -0.308196357 | 0.455585208 | 0.72607388 |
| chr1:68231206-68231884 | 0.271181672 | 0.455596738 | 0.72607388 |
| chr8:53944837-53945890 | 0.250049844 | 0.455598994 | 0.72607388 |
| chr16:70136658-70137056 | 0.294377649 | 0.455620437 | 0.726083781 |
| chr18:34912039-34912467 | 0.278736934 | 0.455634866 | 0.726083781 |
| chr15:89321809-89322079 | 0.291460843 | 0.455637483 | 0.726083781 |
| chr18:78241940-78242200 | -0.352788013 | 0.455652431 | 0.726083781 |
| chr5:171207325-171207576 | -0.368867119 | 0.455652587 | 0.726083781 |
| chr11:70650019-70650630 | 0.266381884 | 0.455671183 | 0.726095867 |
| chr1:150974663-150974982 | 0.262245723 | 0.455679125 | 0.726095867 |
| chr6:139977595-139978394 | 0.249462018 | 0.455692845 | 0.72610263 |
| chr17:78923705-78924072 | 0.286032821 | 0.455728884 | 0.726144954 |
| chr17:41521424-41521663 | 0.303697837 | 0.455780612 | 0.726188472 |
| chr8:130297937-130298142 | -0.381593519 | 0.455786088 | 0.726188472 |
| chr5:13111215-13111485 | 0.36550229 | 0.455790525 | 0.726188472 |
| chr6:41682135-41682481 | 0.287331304 | 0.455795745 | 0.726188472 |
| chr5:149938990-149939263 | -0.319920497 | 0.455803583 | 0.726188472 |
| chr1:198385824-198386065 | 0.338949134 | 0.455819019 | 0.726197964 |
| chr2:241507813-241508454 | 0.257830384 | 0.455838798 | 0.726214375 |
| chr7:151736449-151736855 | 0.2960081 | 0.455855564 | 0.726225985 |
| chr20:56980795-56981205 | 0.267602198 | 0.455868036 | 0.726230755 |
| chr17:58768132-58768332 | 0.289476233 | 0.455894467 | 0.726253052 |

| | | | |
|---|---|---|---|
| chr19:13774242-13774805 | -0.297397104 | 0.45591068 | 0.726253052 |
| chr1:30670478-30670709 | -0.393005543 | 0.455915266 | 0.726253052 |
| chr8:73563130-73563367 | 0.384996003 | 0.455919945 | 0.726253052 |
| chr4:80461522-80461766 | -0.38677976 | 0.455934085 | 0.726260476 |
| chr5:160399457-160400322 | -0.241869251 | 0.455948068 | 0.726267652 |
| chr18:76394795-76395064 | -0.321852311 | 0.455966724 | 0.726281086 |
| chr8:53881582-53881813 | -0.356002096 | 0.455986633 | 0.726281086 |
| chr3:5986932-5987570 | -0.285500718 | 0.455991668 | 0.726281086 |
| chr5:179622759-179624295 | -0.235117398 | 0.45599819 | 0.726281086 |
| chr1:217332720-217332981 | 0.294134354 | 0.456003895 | 0.726281086 |
| chr12:14567252-14567633 | -0.30190527 | 0.456048665 | 0.726282254 |
| chr15:42045271-42045565 | -0.316988927 | 0.456052365 | 0.726282254 |
| chr13:35921108-35921364 | -0.360727322 | 0.456057445 | 0.726282254 |
| chr2:219542913-219544211 | 0.235414679 | 0.456070391 | 0.726282254 |
| chr7:51309804-51310032 | -0.368670316 | 0.456080708 | 0.726282254 |
| chr1:61588520-61589032 | -0.283574616 | 0.456081219 | 0.726282254 |
| chr9:76393690-76395229 | -0.237504427 | 0.456095241 | 0.726282254 |
| chr8:86124953-86125440 | 0.266315985 | 0.456111687 | 0.726282254 |
| chr2:148829674-148829911 | -0.342391809 | 0.45611761 | 0.726282254 |
| chr2:110747686-110747989 | 0.316869547 | 0.456118066 | 0.726282254 |
| chr5:17882708-17883520 | -0.25590561 | 0.456120349 | 0.726282254 |
| chr3:45388069-45388761 | 0.262074624 | 0.456121267 | 0.726282254 |
| chrX:110418245-110418771 | 0.259544695 | 0.456127853 | 0.726282254 |
| chr3:83928396-83928662 | 0.317437516 | 0.456168436 | 0.726317516 |
| chr14:102545669-102545925 | -0.339758068 | 0.456178188 | 0.726317516 |
| chr12:60266474-60266700 | 0.350938414 | 0.456178436 | 0.726317516 |
| chr10:133438882-133439083 | 0.352417123 | 0.456192763 | 0.726318564 |
| chr8:122864749-122865252 | -0.27550065 | 0.456198053 | 0.726318564 |
| chr9:111896310-111897377 | 0.244203346 | 0.456219816 | 0.72633812 |
| chr19:45130867-45131379 | 0.26382113 | 0.456240105 | 0.726353669 |
| chr2:179595418-179595698 | -0.381379209 | 0.456248542 | 0.726353669 |
| chr2:166361348-166361549 | -0.406649895 | 0.456267689 | 0.72636906 |
| chr14:23910538-23910928 | 0.289523008 | 0.456304714 | 0.72639843 |
| chr17:40218652-40219795 | 0.257302299 | 0.456305098 | 0.72639843 |
| chr15:75775340-75775615 | 0.321643665 | 0.456324218 | 0.726413775 |
| chr3:169575418-169575637 | 0.357498952 | 0.456352042 | 0.726442975 |
| chr4:81913441-81913677 | -0.340612261 | 0.456375864 | 0.726461007 |
| chr16:30657832-30659232 | 0.23425978 | 0.456394838 | 0.726461007 |
| chr16:4767086-4767583 | 0.249766245 | 0.456400285 | 0.726461007 |
| chr2:67213825-67214036 | 0.356029269 | 0.456407544 | 0.726461007 |
| chr3:176855899-176856561 | -0.254832333 | 0.456410775 | 0.726461007 |
| chr3:15362555-15362783 | -0.320169263 | 0.456434197 | 0.726475802 |
| chr16:15949394-15949666 | 0.306637868 | 0.456439033 | 0.726475802 |
| chr17:43798082-43798298 | 0.372630289 | 0.456453014 | 0.726482964 |
| chr2:24490668-24491321 | -0.257382035 | 0.456467191 | 0.726490438 |
| chr12:6925811-6926230 | 0.283314577 | 0.456490906 | 0.72651309 |
| chr2:148926983-148927413 | -0.336230599 | 0.45650985 | 0.726526779 |
| chr19:56076521-56076759 | 0.335781978 | 0.456518471 | 0.726526779 |
| chr5:171324442-171324648 | -0.389344824 | 0.456540862 | 0.726536963 |
| chr1:40783141-40783440 | 0.315527546 | 0.456543835 | 0.726536963 |

| | | | |
|---|---|---|---|
| chr1:10507491-10507806 | -0.324504575 | 0.456566123 | 0.726552554 |
| chr14:77615925-77617229 | 0.233765958 | 0.456575052 | 0.726552554 |
| chr14:104137965-104138722 | 0.245998742 | 0.456582079 | 0.726552554 |
| chr11:126366265-126366505 | -0.325028348 | 0.45661944 | 0.72655952 |
| chr14:99393087-99393410 | 0.291136521 | 0.456628047 | 0.72655952 |
| chr12:80321645-80322146 | -0.273477567 | 0.456631102 | 0.72655952 |
| chr2:39120337-39120635 | -0.278810136 | 0.456632035 | 0.72655952 |
| chr9:110623544-110623828 | 0.324068401 | 0.456633868 | 0.72655952 |
| chr1:114466354-114467035 | 0.243545335 | 0.456653298 | 0.726575348 |
| chr9:111037476-111037948 | -0.290223363 | 0.456704127 | 0.72662486 |
| chr3:75503393-75503719 | -0.284305991 | 0.456709002 | 0.72662486 |
| chr8:85742208-85742465 | 0.406000709 | 0.456712866 | 0.72662486 |
| chr4:139666724-139666985 | -0.295273464 | 0.456729199 | 0.726635757 |
| chr16:80317853-80318116 | 0.321260512 | 0.456750843 | 0.72665182 |
| chr2:71129652-71130812 | -0.236611523 | 0.45676487 | 0.72665182 |
| chr11:66045146-66045529 | 0.272557473 | 0.456767747 | 0.72665182 |
| chr22:45979036-45979313 | 0.308961363 | 0.456777277 | 0.726651894 |
| chr10:87342246-87343344 | -0.246038978 | 0.456805824 | 0.726674285 |
| chr1:64824068-64824540 | -0.285459614 | 0.456814426 | 0.726674285 |
| chr18:34352845-34353045 | -0.433649716 | 0.456819803 | 0.726674285 |
| chr9:126781828-126782078 | -0.350520158 | 0.456834484 | 0.726682551 |
| chr7:4883484-4884105 | 0.244786475 | 0.456846232 | 0.726685994 |
| chr2:123511906-123512166 | -0.375522346 | 0.456855616 | 0.726685994 |
| chr3:150238042-150239348 | 0.236254667 | 0.456889809 | 0.726725296 |
| chr10:97685653-97685895 | -0.296291464 | 0.456977126 | 0.726840379 |
| chr20:57106403-57106645 | 0.246433761 | 0.456988935 | 0.726840379 |
| chr19:51367451-51368060 | 0.243014752 | 0.457005019 | 0.726840379 |
| chr5:153489616-153489897 | -0.363849597 | 0.457011324 | 0.726840379 |
| chr3:134308414-134308738 | 0.265335121 | 0.457016621 | 0.726840379 |
| chr13:94614193-94614420 | -0.380189497 | 0.457019078 | 0.726840379 |
| chr7:73734741-73735246 | -0.256482799 | 0.457042407 | 0.726862394 |
| chr19:40777417-40777753 | -0.262538641 | 0.457062361 | 0.726866839 |
| chr12:120904045-120904808 | 0.253992113 | 0.457064174 | 0.726866839 |
| chr12:76611528-76611994 | -0.289369648 | 0.457090935 | 0.726880392 |
| chr9:35518179-35518497 | -0.288221327 | 0.45709167 | 0.726880392 |
| chr20:51318227-51318569 | -0.273940395 | 0.457198752 | 0.727020513 |
| chr12:89353338-89354463 | -0.237319878 | 0.457199248 | 0.727020513 |
| chr5:155663041-155663332 | -0.369511623 | 0.457208249 | 0.727020513 |
| chr1:203871124-203871430 | -0.286190654 | 0.457243054 | 0.727060769 |
| chr11:119496943-119497205 | -0.345357869 | 0.457264838 | 0.727066818 |
| chr4:142499819-142500210 | -0.323898449 | 0.457279214 | 0.727066818 |
| chr13:39311849-39312196 | -0.322074004 | 0.457281375 | 0.727066818 |
| chr10:94279957-94280435 | 0.281385413 | 0.457284814 | 0.727066818 |
| chr1:203839148-203839355 | -0.333354985 | 0.457305542 | 0.727084686 |
| chr6:47698301-47698859 | -0.258120874 | 0.457346265 | 0.727112815 |
| chr16:67872746-67873275 | 0.263093013 | 0.457369635 | 0.727112815 |
| chr6:43516580-43517465 | 0.243158933 | 0.457374659 | 0.727112815 |
| chr1:180333640-180334432 | -0.264805299 | 0.457379269 | 0.727112815 |
| chr9:35814451-35814657 | -0.309510701 | 0.457382666 | 0.727112815 |
| chr17:79005387-79005741 | 0.272342504 | 0.457396344 | 0.727112815 |

| | | | |
|---|---|---|---|
| chr2:97740577-97740789 | -0.331601902 | 0.457396393 | 0.727112815 |
| chr8:78665439-78666803 | 0.234935973 | 0.457404199 | 0.727112815 |
| chr8:128859224-128859651 | 0.267777739 | 0.45740864 | 0.727112815 |
| chr1:205321227-205322082 | -0.238511942 | 0.457424014 | 0.727122169 |
| chr8:125429180-125430509 | -0.231363579 | 0.457447523 | 0.727143469 |
| chr3:193872240-193872631 | -0.259934372 | 0.457457186 | 0.727143469 |
| chr7:44490309-44491090 | -0.245815481 | 0.457465883 | 0.727143469 |
| chr7:129977914-129978162 | 0.378473484 | 0.457487756 | 0.727163151 |
| chr2:24971310-24972325 | -0.236830039 | 0.457514651 | 0.727179569 |
| chr4:38673701-38673901 | -0.363503431 | 0.457517067 | 0.727179569 |
| chr12:122940244-122940502 | -0.363288846 | 0.457541675 | 0.727203598 |
| chr1:23544795-23545004 | 0.383664126 | 0.457559202 | 0.72721637 |
| chr5:139648650-139648984 | -0.272677301 | 0.457571073 | 0.727220152 |
| chr7:150421077-150421420 | -0.281058251 | 0.457596545 | 0.727245551 |
| chr2:171522802-171524016 | -0.251183532 | 0.457671034 | 0.727348847 |
| chr5:123611186-123611386 | 0.419546678 | 0.457699935 | 0.72737969 |
| chr10:457869-458078 | 0.391748192 | 0.457714601 | 0.727387911 |
| chr10:71836158-71836488 | -0.341606753 | 0.457762777 | 0.727449384 |
| chr17:55026783-55027247 | -0.301439034 | 0.457778496 | 0.727459277 |
| chr5:23506437-23507715 | -0.233430781 | 0.457801062 | 0.727480048 |
| chr7:101608705-101609258 | -0.257534099 | 0.457826761 | 0.727502834 |
| chr19:6110392-6110992 | 0.258785494 | 0.457834391 | 0.727502834 |
| chr8:122536239-122536439 | -0.365793616 | 0.457849808 | 0.727511932 |
| chr7:2554863-2556012 | -0.237649305 | 0.457859106 | 0.727511932 |
| chr11:41245771-41246086 | -0.33698774 | 0.457870249 | 0.727513119 |
| chr16:19072201-19072532 | 0.286704289 | 0.457885623 | 0.727513119 |
| chrX:78139121-78140113 | -0.240882394 | 0.457888337 | 0.727513119 |
| chr5:23749705-23749905 | 0.400524595 | 0.457947032 | 0.727591289 |
| chr17:43124490-43124882 | 0.302214664 | 0.457984428 | 0.727635616 |
| chr22:22409459-22409717 | 0.339685167 | 0.458038821 | 0.727706944 |
| chr14:70317829-70318151 | 0.317007956 | 0.458067216 | 0.727720913 |
| chr11:108027158-108027457 | -0.368269864 | 0.458076441 | 0.727720913 |
| chr5:43017856-43018513 | 0.246598427 | 0.458083419 | 0.727720913 |
| chr5:17419757-17419986 | 0.362455068 | 0.458085603 | 0.727720913 |
| chr1:109490723-109490935 | 0.298312802 | 0.458100437 | 0.72772939 |
| chr2:98909377-98909819 | 0.286568467 | 0.4581168 | 0.727733921 |
| chr21:33605838-33606038 | 0.361404529 | 0.458122284 | 0.727733921 |
| chr13:34748998-34749439 | -0.272549076 | 0.458165161 | 0.727786942 |
| chr1:158149570-158150051 | -0.263074137 | 0.458188823 | 0.72780944 |
| chr20:18413150-18413617 | -0.284891636 | 0.458232873 | 0.727864322 |
| chr7:151819033-151819594 | -0.253617438 | 0.458262509 | 0.727890127 |
| chr1:201982267-201983455 | 0.24502893 | 0.458268118 | 0.727890127 |
| chr3:88307689-88307990 | -0.317149404 | 0.458300081 | 0.727900136 |
| chr13:44810197-44810400 | -0.390035772 | 0.458302603 | 0.727900136 |
| chr4:37729348-37729624 | -0.403216796 | 0.458308193 | 0.727900136 |
| chr5:40688170-40688541 | 0.269202607 | 0.458312419 | 0.727900136 |
| chr10:84196140-84196493 | -0.28857384 | 0.458333916 | 0.727917961 |
| chr10:99711207-99711487 | -0.324255065 | 0.458342642 | 0.727917961 |
| chrX:26918643-26918945 | -0.347564273 | 0.458362894 | 0.727918804 |
| chr19:40576427-40577254 | 0.245592133 | 0.458373768 | 0.727918804 |

| | | | |
|---|---|---|---|
| chr7:31633726-31634286 | -0.261054072 | 0.458376694 | 0.727918804 |
| chr4:37648086-37648635 | -0.252288684 | 0.458381174 | 0.727918804 |
| chr22:41196900-41197956 | 0.249969157 | 0.458439639 | 0.7279912 |
| chr13:32019677-32019907 | -0.36954148 | 0.458457979 | 0.7279912 |
| chr2:29309175-29309414 | 0.346120345 | 0.458463495 | 0.7279912 |
| chr10:91941771-91942067 | 0.366686501 | 0.458466616 | 0.7279912 |
| chr3:97956148-97956348 | 0.403592716 | 0.458474268 | 0.7279912 |
| chr1:115771561-115772318 | -0.243520076 | 0.458491136 | 0.728002898 |
| chr3:17554499-17554814 | -0.382358705 | 0.458521302 | 0.728035709 |
| chr11:706956-707257 | 0.295073732 | 0.458585739 | 0.728109218 |
| chr1:206725623-206725874 | -0.280941376 | 0.458586604 | 0.728109218 |
| chr12:110060251-110060519 | -0.350479743 | 0.458623741 | 0.728153094 |
| chr17:82205750-82206610 | 0.251826028 | 0.458718679 | 0.728288734 |
| chr1:180881953-180882435 | 0.273352403 | 0.458763712 | 0.728338347 |
| chr1:2174302-2174623 | -0.326078608 | 0.458774327 | 0.728338347 |
| chr13:44576814-44578307 | -0.232358691 | 0.458778445 | 0.728338347 |
| chr7:2477430-2478045 | 0.259956684 | 0.45879704 | 0.728352778 |
| chr5:127073322-127073695 | -0.275410893 | 0.458830318 | 0.728380651 |
| chr16:84511693-84511893 | -0.337573869 | 0.458836078 | 0.728380651 |
| chr12:12266486-12267526 | -0.242009633 | 0.458864594 | 0.728380651 |
| chr3:48998536-48998816 | -0.3858297 | 0.458871635 | 0.728380651 |
| chr20:22490212-22490412 | -0.3858297 | 0.458871635 | 0.728380651 |
| chr3:100125961-100126161 | -0.3858297 | 0.458871635 | 0.728380651 |
| chr1:150886971-150887295 | -0.309643227 | 0.458887281 | 0.728390397 |
| chr1:246475459-246475679 | -0.384249668 | 0.45890548 | 0.728404195 |
| chr1:24413451-24413904 | -0.278724366 | 0.458936142 | 0.728437773 |
| chr6:17589379-17589780 | -0.263693763 | 0.458965354 | 0.728469049 |
| chr17:60167634-60167882 | 0.253353834 | 0.459060889 | 0.728593548 |
| chr2:19266902-19267162 | -0.372388908 | 0.459062811 | 0.728593548 |
| chr6:134111081-134111407 | 0.29926273 | 0.459076282 | 0.728599835 |
| chr2:219836766-219836966 | 0.385451917 | 0.45910243 | 0.728626243 |
| chr3:21752826-21753072 | -0.312737575 | 0.459118984 | 0.728637424 |
| chr17:44086619-44087305 | 0.251887566 | 0.459160941 | 0.728688918 |
| chr16:29973428-29974072 | 0.242795938 | 0.459201615 | 0.728722187 |
| chr22:40651672-40652612 | -0.246712711 | 0.459205561 | 0.728722187 |
| chr6:30721334-30721582 | 0.284182039 | 0.459211388 | 0.728722187 |
| chr15:101546051-101546348 | 0.332581072 | 0.459229082 | 0.728722187 |
| chr1:207084704-207085011 | -0.280718541 | 0.459229458 | 0.728722187 |
| chr9:89000987-89001323 | -0.300178139 | 0.459251987 | 0.728727542 |
| chr15:98313044-98313417 | -0.320111967 | 0.459258921 | 0.728727542 |
| chr11:17498201-17498412 | 0.333725377 | 0.459261364 | 0.728727542 |
| chr2:7097275-7097888 | -0.259381824 | 0.459317226 | 0.728741393 |
| chr4:1615048-1615705 | 0.251895995 | 0.459320582 | 0.728741393 |
| chr19:18104394-18104658 | -0.327102337 | 0.459320893 | 0.728741393 |
| chr17:18163707-18163907 | -0.370327679 | 0.459332043 | 0.728741393 |
| chr17:61551241-61551682 | 0.238763405 | 0.459336329 | 0.728741393 |
| chr13:33230823-33231094 | -0.365122047 | 0.459337954 | 0.728741393 |
| chr15:70974086-70974371 | -0.34983409 | 0.459357978 | 0.728741393 |
| chr2:75609088-75609346 | -0.315131789 | 0.459371853 | 0.728741393 |
| chr11:109865879-109866131 | -0.364703084 | 0.459385737 | 0.728741393 |

| | | | |
|---|---|---|---|
| chr15:32662666-32662875 | 0.370345739 | 0.459386863 | 0.728741393 |
| chr3:108090355-108091910 | -0.23407472 | 0.459390408 | 0.728741393 |
| chr14:106808654-106808854 | 0.388613942 | 0.459403158 | 0.728741393 |
| chr5:13957109-13957409 | -0.288952365 | 0.459409278 | 0.728741393 |
| chr11:65663552-65664005 | 0.273174181 | 0.459414584 | 0.728741393 |
| chr12:82685734-82686100 | -0.245073886 | 0.459416497 | 0.728741393 |
| chr6:27019663-27020720 | 0.240345148 | 0.459422267 | 0.728741393 |
| chr19:48993121-48994019 | 0.238171623 | 0.459454686 | 0.728774295 |
| chr7:44131136-44131489 | -0.286632843 | 0.459462033 | 0.728774295 |
| chr9:34707245-34707508 | -0.379516325 | 0.459493127 | 0.728808528 |
| chr5:1316136-1316664 | 0.252905667 | 0.459545137 | 0.728875934 |
| chr1:87131769-87132238 | -0.286282204 | 0.459600731 | 0.728937376 |
| chr10:119489612-119490245 | -0.268359934 | 0.459611471 | 0.728937376 |
| chr16:31074124-31074767 | 0.242781138 | 0.459616929 | 0.728937376 |
| chr15:59144466-59144748 | 0.345403661 | 0.459621929 | 0.728937376 |
| chr6:144150033-144150926 | -0.249800372 | 0.459646093 | 0.728960611 |
| chr17:61409535-61410000 | 0.234593202 | 0.459669746 | 0.728983035 |
| chr16:8173164-8173364 | -0.430906681 | 0.459698193 | 0.728993418 |
| chr3:61910329-61910562 | 0.293441685 | 0.459710281 | 0.728993418 |
| chr9:17794309-17794509 | -0.390769451 | 0.459720199 | 0.728993418 |
| chr11:62840266-62842240 | 0.231216634 | 0.459724035 | 0.728993418 |
| chr5:104533574-104533918 | -0.349652982 | 0.459730738 | 0.728993418 |
| chr12:56333594-56334475 | 0.242951723 | 0.459733379 | 0.728993418 |
| chr8:119178449-119178781 | 0.323663821 | 0.459769556 | 0.729035696 |
| chr19:46000920-46001243 | 0.288408884 | 0.459787029 | 0.729048316 |
| chr8:91048290-91048551 | 0.351595821 | 0.459810121 | 0.729057624 |
| chr1:37509273-37509523 | 0.354428019 | 0.459817013 | 0.729057624 |
| chr17:54749217-54749528 | 0.282525171 | 0.459821445 | 0.729057624 |
| chr2:96010363-96010667 | -0.305193378 | 0.459902614 | 0.729154692 |
| chr2:8476406-8476633 | -0.34835952 | 0.459912374 | 0.729154692 |
| chr17:32141986-32142479 | 0.28755589 | 0.459920664 | 0.729154692 |
| chr21:38352678-38352878 | 0.336947816 | 0.459920731 | 0.729154692 |
| chr8:144103610-144105097 | 0.23323627 | 0.459967811 | 0.729214244 |
| chr16:265743-265949 | -0.350705848 | 0.459981191 | 0.729220367 |
| chr19:45423270-45424071 | 0.241051305 | 0.459994023 | 0.729225623 |
| chr10:120996518-120996752 | 0.339344306 | 0.460014002 | 0.729242208 |
| chr2:176465047-176465554 | -0.2739934 | 0.460028615 | 0.729245555 |
| chr19:40750355-40751359 | 0.235358533 | 0.460035149 | 0.729245555 |
| chr21:44817375-44818355 | 0.241206473 | 0.460059007 | 0.729268288 |
| chr5:16736270-16736673 | 0.27057852 | 0.460084771 | 0.72929404 |
| chr15:75720918-75721402 | 0.275711891 | 0.460102846 | 0.729300557 |
| chr8:93666188-93666519 | 0.301245905 | 0.460111908 | 0.729300557 |
| chr3:171650877-171651150 | -0.334189176 | 0.46013212 | 0.729300557 |
| chr13:44899985-44900232 | -0.332733603 | 0.460145043 | 0.729300557 |
| chr14:44579032-44579447 | -0.302356957 | 0.460156071 | 0.729300557 |
| chr1:16007219-16007468 | 0.346302445 | 0.460156942 | 0.729300557 |
| chr3:105222764-105223051 | -0.323817474 | 0.46016001 | 0.729300557 |
| chr17:45404087-45404298 | -0.380971579 | 0.460165027 | 0.729300557 |
| chr6:27871973-27872533 | -0.25879571 | 0.460195435 | 0.729333663 |
| chr4:40587917-40588182 | 0.354027047 | 0.460226225 | 0.729367374 |

| | | | |
|---|---|---|---|
| chr19:38931215-38931428 | 0.318754899 | 0.460241883 | 0.729377102 |
| chr17:60371716-60372152 | 0.244679936 | 0.460275563 | 0.729410183 |
| chr5:172293257-172293798 | 0.256717037 | 0.460309447 | 0.729410183 |
| chr10:64645267-64645475 | -0.398984983 | 0.46032257 | 0.729410183 |
| chr1:93399813-93400266 | 0.288843859 | 0.460326455 | 0.729410183 |
| chr4:153765862-153766173 | -0.352542118 | 0.460331418 | 0.729410183 |
| chr4:92406407-92406714 | -0.411126736 | 0.460336694 | 0.729410183 |
| chr3:146833006-146833270 | 0.371500811 | 0.460338316 | 0.729410183 |
| chr10:12265708-12266051 | -0.296116593 | 0.460338913 | 0.729410183 |
| chr11:66002066-66003069 | 0.239371367 | 0.460403368 | 0.729497227 |
| chr17:44326774-44327182 | 0.262106843 | 0.460445945 | 0.729537903 |
| chr7:4774714-4776266 | -0.234268833 | 0.460448082 | 0.729537903 |
| chr11:66363997-66364244 | -0.303512079 | 0.460468127 | 0.729554575 |
| chr8:89995358-89995937 | -0.251965572 | 0.460486143 | 0.729555027 |
| chr17:54900458-54901764 | -0.233524932 | 0.460487455 | 0.729555027 |
| chr6:27241476-27241879 | -0.30197715 | 0.460498744 | 0.729557827 |
| chr9:136994260-136994511 | -0.298149929 | 0.460575007 | 0.729663562 |
| chr9:128127606-128128798 | 0.239687889 | 0.460602709 | 0.729692362 |
| chr2:160784117-160784341 | 0.386902846 | 0.460614061 | 0.729695258 |
| chr5:173616392-173617029 | -0.241191435 | 0.460637454 | 0.729717231 |
| chr3:46979476-46979879 | 0.263982581 | 0.460697453 | 0.729794915 |
| chr8:133249300-133250253 | -0.245260752 | 0.46071084 | 0.729794915 |
| chr15:55768914-55769151 | 0.324786734 | 0.460736218 | 0.729794915 |
| chr20:49333905-49334158 | 0.294635495 | 0.460737983 | 0.729794915 |
| chr3:59664128-59664734 | -0.271235376 | 0.460743383 | 0.729794915 |
| chr9:114465938-114466524 | -0.279584071 | 0.460770416 | 0.729794915 |
| chr6:79234559-79235011 | -0.244794232 | 0.460772393 | 0.729794915 |
| chr12:7189083-7190047 | -0.248044424 | 0.460778598 | 0.729794915 |
| chr11:8338115-8338353 | -0.364894244 | 0.460781212 | 0.729794915 |
| chr8:142815923-142816123 | 0.315105006 | 0.460781739 | 0.729794915 |
| chr14:66070316-66070526 | -0.344425954 | 0.460848399 | 0.72988426 |
| chr19:38729743-38729988 | -0.32944819 | 0.460857201 | 0.72988426 |
| chr5:52367137-52367337 | -0.390415504 | 0.460892799 | 0.729914126 |
| chr5:1800614-1800825 | 0.300228079 | 0.46090838 | 0.729914126 |
| chr1:174575746-174576559 | -0.256198693 | 0.460910915 | 0.729914126 |
| chr19:982551-982951 | 0.278598442 | 0.460914164 | 0.729914126 |
| chrX:16892833-16893109 | 0.327952752 | 0.460931379 | 0.729926303 |
| chr16:67653157-67653357 | 0.348716059 | 0.460950003 | 0.729938675 |
| chr19:7515735-7516005 | -0.36306957 | 0.460967718 | 0.729938675 |
| chr3:50154577-50155109 | -0.260306174 | 0.460969661 | 0.729938675 |
| chr8:144684873-144685457 | 0.249344405 | 0.460977298 | 0.729938675 |
| chr15:69035691-69035962 | -0.306645635 | 0.460999435 | 0.729958643 |
| chr13:33697064-33697426 | -0.295014625 | 0.461039739 | 0.729971344 |
| chr7:131945454-131945688 | -0.352042422 | 0.461040035 | 0.729971344 |
| chr8:144413805-144414070 | 0.293534858 | 0.461044731 | 0.729971344 |
| chr10:62244928-62245335 | -0.28608467 | 0.461045565 | 0.729971344 |
| chr1:204072743-204073819 | -0.234470734 | 0.461060982 | 0.729980671 |
| chr16:73010028-73010423 | -0.276171958 | 0.461083473 | 0.730001195 |
| chr20:45812156-45813095 | 0.236673911 | 0.461105793 | 0.730021449 |
| chr12:113104125-113104355 | 0.335982524 | 0.461122061 | 0.73003212 |

| | | | |
|---|---|---|---|
| chr16:30124407-30124663 | 0.309358265 | 0.461138965 | 0.730043797 |
| chr5:178461169-178461467 | -0.296299855 | 0.461168019 | 0.730053689 |
| chr10:73843778-73843978 | -0.372322101 | 0.461174233 | 0.730053689 |
| chr7:34430282-34430928 | -0.260825282 | 0.461177017 | 0.730053689 |
| chr5:158673324-158673625 | 0.349225901 | 0.461195447 | 0.730053689 |
| chr16:11851150-11852464 | 0.235638761 | 0.461203421 | 0.730053689 |
| chr1:114558635-114558893 | 0.26553702 | 0.461215674 | 0.730053689 |
| chr17:40647529-40647966 | 0.301502223 | 0.461220301 | 0.730053689 |
| chr10:96709745-96710002 | -0.40275176 | 0.461221437 | 0.730053689 |
| chr3:119600776-119601100 | 0.290726663 | 0.461236696 | 0.730062761 |
| chr5:2556467-2556713 | -0.364435612 | 0.46125166 | 0.730071364 |
| chr15:87038501-87038782 | -0.333380067 | 0.461269484 | 0.730083654 |
| chr1:204417123-204417966 | -0.238521606 | 0.46128503 | 0.730083654 |
| chr16:87550814-87551014 | -0.364465249 | 0.461291374 | 0.730083654 |
| chr2:169695425-169696041 | -0.252483472 | 0.461297538 | 0.730083654 |
| chr5:143066608-143067007 | -0.30688178 | 0.46132172 | 0.730093605 |
| chr5:6576950-6577188 | -0.326607464 | 0.461322882 | 0.730093605 |
| chr4:150867122-150867364 | -0.348396096 | 0.461353944 | 0.730117397 |
| chr7:25215705-25215918 | -0.381922493 | 0.461356973 | 0.730117397 |
| chr1:42923047-42923311 | 0.293323742 | 0.461376228 | 0.730132788 |
| chr12:46268404-46269838 | 0.234847736 | 0.46147151 | 0.730260663 |
| chr3:156509758-156510032 | -0.313211543 | 0.461476094 | 0.730260663 |
| chr17:72270626-72270827 | 0.317124216 | 0.461520839 | 0.730316386 |
| chr14:72753739-72753939 | 0.362587863 | 0.46153177 | 0.7303186 |
| chr11:112086231-112087203 | 0.24154137 | 0.461562224 | 0.730334771 |
| chr4:80365963-80366272 | -0.30413295 | 0.461572492 | 0.730334771 |
| chr1:84298394-84299592 | -0.246256037 | 0.461578192 | 0.730334771 |
| chr11:68622590-68622790 | -0.415805461 | 0.461580116 | 0.730334771 |
| chr3:146250441-146251356 | -0.244402531 | 0.461592707 | 0.730339612 |
| chr1:32393934-32394994 | 0.234434606 | 0.461615574 | 0.730360711 |
| chr12:62827167-62827387 | 0.349877457 | 0.461646441 | 0.730373776 |
| chr10:4733095-4733665 | 0.251158981 | 0.461651327 | 0.730373776 |
| chr22:37267061-37267326 | -0.362519114 | 0.461652428 | 0.730373776 |
| chr21:38234824-38235127 | 0.273101422 | 0.461684958 | 0.73041016 |
| chr1:31162646-31163050 | -0.288233906 | 0.461780578 | 0.730546352 |
| chr4:40629489-40631123 | 0.235355384 | 0.461801435 | 0.730564264 |
| chr3:93470262-93470885 | 0.230920299 | 0.461861491 | 0.730644186 |
| chr6:88982729-88982974 | -0.343797254 | 0.461881652 | 0.730660994 |
| chr5:13514904-13515104 | -0.401984088 | 0.461898865 | 0.730673139 |
| chr16:4476014-4477129 | 0.238747842 | 0.461911295 | 0.730677717 |
| chr9:98848515-98848871 | 0.277694207 | 0.461936209 | 0.730702042 |
| chr17:83079429-83079707 | 0.269406597 | 0.46196993 | 0.730737801 |
| chr10:118754100-118754305 | -0.308853625 | 0.461982902 | 0.730737801 |
| chr12:57280700-57280900 | -0.387098842 | 0.462001566 | 0.730737801 |
| chr5:31544599-31544903 | -0.399639774 | 0.462002999 | 0.730737801 |
| chr16:2905098-2906161 | 0.233101514 | 0.4620065 | 0.730737801 |
| chr3:99789364-99789707 | -0.270964012 | 0.462033229 | 0.730756506 |
| chr19:45632428-45632897 | -0.27011487 | 0.4620374 | 0.730756506 |
| chr2:70190140-70191333 | -0.233893706 | 0.462069007 | 0.730762083 |
| chr5:65101117-65101389 | -0.291752721 | 0.462074008 | 0.730762083 |

| | | | |
|---|---|---|---|
| chr16:83996675-83997155 | -0.272873154 | 0.462076771 | 0.730762083 |
| chr4:39446557-39447605 | 0.237400142 | 0.462079076 | 0.730762083 |
| chr14:102726374-102726654 | -0.309850296 | 0.462105528 | 0.730777936 |
| chr11:68003565-68004323 | 0.245704754 | 0.462108175 | 0.730777936 |
| chr22:17774484-17774734 | 0.310817174 | 0.46214893 | 0.730801804 |
| chr10:93767668-93767950 | -0.330295248 | 0.46216435 | 0.730801804 |
| chrX:68647308-68648069 | -0.259905783 | 0.462166843 | 0.730801804 |
| chr13:82878640-82878876 | -0.347811856 | 0.462183321 | 0.730801804 |
| chr5:31232306-31232642 | -0.301064092 | 0.462187421 | 0.730801804 |
| chr1:200488497-200489049 | -0.248465198 | 0.462187836 | 0.730801804 |
| chr12:109093297-109093749 | 0.247137793 | 0.462190032 | 0.730801804 |
| chr22:38706236-38706436 | -0.343436691 | 0.462207849 | 0.730814895 |
| chr21:46318170-46318474 | 0.296500384 | 0.462225431 | 0.730827613 |
| chr9:126558101-126558335 | -0.341779919 | 0.462247818 | 0.73084721 |
| chr8:94640235-94641401 | 0.230664921 | 0.462256902 | 0.73084721 |
| chr17:75689432-75689699 | -0.30263241 | 0.462268256 | 0.73085008 |
| chr6:33073636-33073836 | 0.387712508 | 0.462282085 | 0.730856864 |
| chr17:69326467-69327885 | -0.230682432 | 0.462296598 | 0.730864728 |
| chr17:39154116-39154732 | 0.247794973 | 0.462306807 | 0.730865789 |
| chr17:38752234-38752875 | 0.248963702 | 0.462382461 | 0.730970308 |
| chr1:246529373-246529600 | -0.301491087 | 0.462397169 | 0.730978479 |
| chr5:51275916-51276188 | -0.345924198 | 0.46241034 | 0.730984218 |
| chr3:15685902-15686248 | 0.278449467 | 0.462456276 | 0.731033743 |
| chr4:88369221-88369574 | -0.323448585 | 0.462466658 | 0.731033743 |
| chr10:16116487-16116696 | -0.388303815 | 0.462470291 | 0.731033743 |
| chr3:63740682-63740999 | 0.253257205 | 0.462482442 | 0.731035788 |
| chr20:14775900-14776194 | -0.318995281 | 0.462490666 | 0.731035788 |
| chr12:4857832-4858060 | -0.338465183 | 0.462556939 | 0.73112546 |
| chr12:120496726-120496935 | 0.331220101 | 0.462587547 | 0.731146355 |
| chr17:80741810-80742102 | 0.350469778 | 0.462589243 | 0.731146355 |
| chr3:86969818-86970067 | -0.328698825 | 0.462601047 | 0.73114993 |
| chr14:45133931-45134906 | 0.231791795 | 0.462645038 | 0.731204375 |
| chr18:12612946-12613207 | -0.391151995 | 0.462679896 | 0.731244384 |
| chr22:26589282-26589581 | 0.304968786 | 0.462720197 | 0.731292994 |
| chr18:23101950-23102287 | 0.29951241 | 0.462740474 | 0.731309956 |
| chr15:34588062-34588294 | -0.320637384 | 0.462774243 | 0.73134824 |
| chr22:42978467-42978857 | -0.339248251 | 0.462795017 | 0.731353094 |
| chr1:174770082-174770282 | -0.375301678 | 0.462803424 | 0.731353094 |
| chr1:147599226-147600495 | 0.231243697 | 0.46280595 | 0.731353094 |
| chr6:82086746-82087305 | 0.27503713 | 0.462827647 | 0.731372298 |
| chr15:48330914-48331133 | 0.317112252 | 0.462858056 | 0.73138443 |
| chr22:26503143-26503353 | 0.399363209 | 0.462859749 | 0.73138443 |
| chr9:122465093-122465339 | 0.324554223 | 0.462863961 | 0.73138443 |
| chr20:17939090-17939317 | 0.303552012 | 0.462894967 | 0.731418341 |
| chr1:228139754-228140720 | 0.234452323 | 0.462925167 | 0.731418514 |
| chr12:69620819-69621137 | -0.362959563 | 0.462935959 | 0.731418514 |
| chr16:31142098-31142787 | 0.238842121 | 0.462947219 | 0.731418514 |
| chr3:185498383-185499337 | 0.23864598 | 0.462948551 | 0.731418514 |
| chr7:124311624-124311847 | -0.411088206 | 0.462951138 | 0.731418514 |
| chr16:31694467-31694667 | 0.374329205 | 0.462952352 | 0.731418514 |

| | | | |
|---|---|---|---|
| chr19:55247614-55247974 | -0.290241546 | 0.463042646 | 0.731499519 |
| chrX:48890998-48892042 | 0.238285636 | 0.46304449 | 0.731499519 |
| chr12:97567725-97568246 | -0.250766758 | 0.463045498 | 0.731499519 |
| chr18:49486884-49487615 | -0.247218559 | 0.463048006 | 0.731499519 |
| chr10:128536247-128536560 | -0.373347543 | 0.463051358 | 0.731499519 |
| chr19:374086-374321 | -0.342769884 | 0.463083317 | 0.731520538 |
| chr3:10983339-10983614 | 0.303425401 | 0.463101725 | 0.731520538 |
| chr6:64389669-64389952 | -0.333504816 | 0.463105507 | 0.731520538 |
| chr5:170070832-170071096 | 0.30468485 | 0.463106276 | 0.731520538 |
| chr8:29559946-29560197 | -0.407665551 | 0.463112399 | 0.731520538 |
| chr5:12041432-12041632 | -0.367136092 | 0.463154059 | 0.731562958 |
| chr5:176959083-176959444 | 0.316691148 | 0.46315835 | 0.731562958 |
| chr19:57746586-57747218 | 0.255401543 | 0.463175057 | 0.731574266 |
| chr20:44105106-44105329 | 0.334230063 | 0.463206174 | 0.731601134 |
| chr8:89596112-89596341 | 0.344343405 | 0.463211164 | 0.731601134 |
| chr1:162874011-162874596 | -0.262134729 | 0.463227869 | 0.731612437 |
| chr6:139250560-139250882 | -0.28921635 | 0.463257023 | 0.731628937 |
| chrX:21942027-21942235 | -0.331176993 | 0.463257413 | 0.731628937 |
| chr3:142908067-142908328 | -0.357581801 | 0.463290836 | 0.731658458 |
| chr10:5015025-5015302 | 0.328255757 | 0.463295203 | 0.731658458 |
| chr2:10936186-10936599 | 0.274724636 | 0.463333754 | 0.731695018 |
| chr2:169983510-169983820 | -0.33188433 | 0.463347856 | 0.731695018 |
| chr5:43120692-43121926 | 0.230121626 | 0.463360633 | 0.731695018 |
| chr9:97889982-97890375 | 0.297425132 | 0.463364 | 0.731695018 |
| chr22:19447232-19447956 | 0.249291202 | 0.463372363 | 0.731695018 |
| chr11:64317389-64318287 | 0.236518251 | 0.46337565 | 0.731695018 |
| chr9:95874916-95875216 | 0.270954411 | 0.46339214 | 0.731705977 |
| chr3:39000018-39000374 | 0.30775213 | 0.463442543 | 0.731755811 |
| chr10:60680285-60680619 | -0.339801553 | 0.463449861 | 0.731755811 |
| chr1:189929636-189929889 | 0.351859489 | 0.46345235 | 0.731755811 |
| chr20:12970300-12970796 | -0.291297064 | 0.463490125 | 0.731800374 |
| chr3:46982208-46982408 | 0.323072738 | 0.463510867 | 0.73181595 |
| chr16:53054695-53055415 | 0.249783477 | 0.463521776 | 0.73181595 |
| chr8:128712708-128712963 | -0.389187949 | 0.463528643 | 0.73181595 |
| chr17:45176208-45176415 | -0.344812436 | 0.463572028 | 0.731862841 |
| chr11:87038325-87038533 | -0.337662759 | 0.463577447 | 0.731862841 |
| chr9:15143004-15143204 | 0.350069402 | 0.46360123 | 0.731885309 |
| chr2:65316104-65316451 | 0.279158633 | 0.463651477 | 0.731949553 |
| chr10:110447140-110447393 | -0.374568878 | 0.463683594 | 0.731961722 |
| chr6:39303207-39303487 | 0.296696573 | 0.463684869 | 0.731961722 |
| chr11:57761559-57762042 | -0.261279053 | 0.463693815 | 0.731961722 |
| chr20:56872617-56872935 | -0.275230368 | 0.463697397 | 0.731961722 |
| chr14:103528899-103529325 | 0.253005822 | 0.463760192 | 0.732012711 |
| chr2:119367306-119367822 | 0.25250217 | 0.463770062 | 0.732012711 |
| chr4:87419502-87419842 | -0.284897284 | 0.463783396 | 0.732012711 |
| chr10:88439999-88440331 | 0.33154464 | 0.46378639 | 0.732012711 |
| chr4:82401455-82401944 | -0.274048502 | 0.463795081 | 0.732012711 |
| chr14:42053899-42054303 | -0.300964512 | 0.46380039 | 0.732012711 |
| chr3:73622135-73622408 | -0.302579825 | 0.463801408 | 0.732012711 |
| chr15:40150208-40150518 | -0.318125108 | 0.463815536 | 0.732012711 |

| | | | |
|---|---|---|---|
| chr1:247078050-247079055 | -0.232128753 | 0.463815681 | 0.732012711 |
| chr9:136410491-136410960 | 0.27485298 | 0.463911966 | 0.732149592 |
| chr22:44701807-44702325 | 0.264650426 | 0.463928757 | 0.73216101 |
| chr5:138463848-138465660 | -0.22846481 | 0.464007564 | 0.732270299 |
| chr3:179272467-179272720 | 0.352648911 | 0.464017407 | 0.732270642 |
| chr19:16324374-16324927 | 0.276914641 | 0.464026895 | 0.732270642 |
| chr2:31047906-31048168 | -0.363737774 | 0.464042134 | 0.732273017 |
| chr21:31098516-31098901 | 0.266318045 | 0.464047514 | 0.732273017 |
| chr22:50775186-50775478 | 0.303352343 | 0.464091597 | 0.732315321 |
| chr6:164709716-164709988 | -0.401985837 | 0.464093437 | 0.732315321 |
| chr5:60287206-60287501 | -0.303678862 | 0.464114199 | 0.732325847 |
| chr11:87463591-87463815 | -0.362007324 | 0.464119223 | 0.732325847 |
| chr4:71229801-71230003 | -0.385131131 | 0.464133848 | 0.732329153 |
| chr1:167342874-167343179 | -0.303204568 | 0.464140434 | 0.732329153 |
| chrX:46447916-46448116 | -0.390757291 | 0.464182361 | 0.732380225 |
| chr3:29065489-29065689 | -0.367964422 | 0.464195957 | 0.732384108 |
| chr2:200471349-200471549 | 0.356446274 | 0.464203939 | 0.732384108 |
| chr1:54069764-54069964 | 0.38867209 | 0.464227429 | 0.732401439 |
| chr5:57693135-57693335 | -0.374342752 | 0.464234041 | 0.732401439 |
| chr17:60011522-60012061 | 0.247463131 | 0.464282235 | 0.73245615 |
| chr17:2095069-2095318 | -0.342892302 | 0.464287839 | 0.73245615 |
| chr17:66570471-66570678 | -0.341173224 | 0.46434388 | 0.732512378 |
| chr3:193780766-193781339 | -0.262493205 | 0.464350885 | 0.732512378 |
| chr14:74501251-74501474 | 0.314779975 | 0.46435216 | 0.732512378 |
| chr7:66204921-66206025 | -0.244752271 | 0.464379432 | 0.732540317 |
| chr3:33439660-33439992 | 0.265409048 | 0.464402189 | 0.732561134 |
| chr5:155906088-155906288 | -0.384634289 | 0.464448504 | 0.732619111 |
| chr1:247118886-247119313 | -0.244727844 | 0.464468249 | 0.732635173 |
| chr21:33403681-33404542 | 0.254112659 | 0.46449265 | 0.732642844 |
| chr1:147242191-147243301 | 0.231042542 | 0.464501429 | 0.732642844 |
| chr3:109398893-109399247 | -0.30717316 | 0.464507474 | 0.732642844 |
| chr6:47665170-47665430 | -0.337182105 | 0.464511359 | 0.732642844 |
| chr1:40039834-40041054 | 0.233361153 | 0.464525125 | 0.732649475 |
| chr9:121992606-121992981 | -0.29441274 | 0.464535114 | 0.732650148 |
| chr11:133153285-133153505 | -0.41318879 | 0.464608977 | 0.732718009 |
| chr5:10145937-10146278 | 0.277421521 | 0.46462528 | 0.732718009 |
| chrX:104310038-104310552 | 0.274991303 | 0.464626401 | 0.732718009 |
| chr15:69088328-69088548 | -0.361641157 | 0.46462757 | 0.732718009 |
| chr19:48110455-48110963 | 0.264752427 | 0.46463205 | 0.732718009 |
| chr6:2952569-2953239 | 0.247824678 | 0.464635518 | 0.732718009 |
| chr1:173867646-173869024 | -0.227912823 | 0.464673098 | 0.732762192 |
| chr17:80873174-80873392 | 0.394278597 | 0.464696302 | 0.732783701 |
| chr5:72808578-72808841 | 0.34746067 | 0.464718221 | 0.732788916 |
| chr6:1995899-1996106 | -0.37910374 | 0.464718736 | 0.732788916 |
| chr8:144798483-144799363 | 0.242082243 | 0.464772874 | 0.732849059 |
| chr2:219476285-219476524 | -0.30505307 | 0.464778784 | 0.732849059 |
| chrX:128975721-128976058 | 0.2884279 | 0.464785571 | 0.732849059 |
| chr10:27783661-27784056 | -0.29700905 | 0.464830841 | 0.73289285 |
| chr3:143971230-143972981 | -0.227994182 | 0.464842395 | 0.73289285 |
| chr5:174323876-174324223 | -0.285989543 | 0.464863135 | 0.73289285 |

| | | | |
|---|---|---|---|
| chr4:167523783-167524415 | 0.264346326 | 0.464875242 | 0.73289285 |
| chrX:134761920-134762120 | 0.382552865 | 0.46488903 | 0.73289285 |
| chr9:98678344-98678544 | 0.382552865 | 0.46488903 | 0.73289285 |
| chr2:11320799-11321058 | -0.330747214 | 0.464889404 | 0.73289285 |
| chr6:57021238-57021452 | 0.355834003 | 0.464889864 | 0.73289285 |
| chr1:177329974-177330201 | -0.355000336 | 0.464905 | 0.732901632 |
| chr13:44759574-44760331 | -0.249336092 | 0.464915317 | 0.732902817 |
| chr17:60492202-60492507 | 0.249796202 | 0.464942482 | 0.732930562 |
| chr14:77213377-77213732 | 0.259892123 | 0.464979217 | 0.732973389 |
| chr7:43764097-43764367 | -0.29041355 | 0.465006767 | 0.733001738 |
| chr15:37100285-37100894 | 0.24671087 | 0.465019516 | 0.733006755 |
| chr1:153364272-153364551 | 0.296848732 | 0.465031044 | 0.733009847 |
| chr3:29576738-29577029 | -0.386456734 | 0.465063686 | 0.733010343 |
| chr1:160291331-160291705 | 0.275431023 | 0.46507902 | 0.733010343 |
| chr10:63901272-63901518 | 0.350459905 | 0.465087377 | 0.733010343 |
| chr7:1758608-1758849 | -0.314755754 | 0.465091933 | 0.733010343 |
| chr5:90722893-90723093 | 0.407794425 | 0.465109355 | 0.733010343 |
| chr1:227894130-227894469 | 0.296897008 | 0.465110853 | 0.733010343 |
| chr2:42385190-42385489 | -0.276804003 | 0.465114287 | 0.733010343 |
| chr11:66015853-66016053 | 0.332905168 | 0.465120336 | 0.733010343 |
| chr2:191812153-191812515 | -0.28247248 | 0.465123062 | 0.733010343 |
| chr10:18443404-18443626 | -0.429480473 | 0.465127025 | 0.733010343 |
| chr4:136060793-136061248 | 0.301277987 | 0.465177947 | 0.733056829 |
| chr5:24397894-24398094 | -0.394596811 | 0.465185329 | 0.733056829 |
| chr10:119618649-119619023 | 0.296185713 | 0.465201696 | 0.733056829 |
| chr16:89860626-89861209 | 0.256480231 | 0.465207587 | 0.733056829 |
| chr8:70577033-70577233 | 0.404304817 | 0.465217808 | 0.733056829 |
| chr17:45468913-45469172 | -0.335549769 | 0.46521978 | 0.733056829 |
| chr1:45583229-45585155 | -0.230454793 | 0.465237899 | 0.733056829 |
| chr11:103091451-10309176[illegible] | -0.279067625 | 0.46524018 | 0.733056829 |
| chr12:103956580-103957865 | -0.23342569 | 0.465248636 | 0.733056829 |
| chr15:66462140-66462340 | -0.386049282 | 0.465252194 | 0.733056829 |
| chr5:168578980-168580055 | -0.230984644 | 0.465265915 | 0.733063374 |
| chr6:111258920-111259858 | -0.233335653 | 0.465289901 | 0.733082221 |
| chr9:135501145-135501433 | 0.276410109 | 0.465299357 | 0.733082221 |
| chr6:19804409-19805400 | -0.237651149 | 0.46530658 | 0.733082221 |
| chr19:10403423-10404041 | 0.247699479 | 0.465328404 | 0.733101531 |
| chrX:9995810-9996121 | -0.31857573 | 0.465347781 | 0.733116985 |
| chr9:21235736-21236011 | -0.36987057 | 0.46537125 | 0.733124081 |
| chr8:14211130-14211330 | -0.422542471 | 0.465371422 | 0.733124081 |
| chr3:155766189-155766512 | 0.340868095 | 0.465388983 | 0.733136674 |
| chr9:6845991-6846191 | 0.388727787 | 0.465401435 | 0.733141217 |
| chr18:42048221-42048473 | -0.339162862 | 0.465449957 | 0.733187116 |
| chr18:45850055-45850288 | -0.344049452 | 0.465459391 | 0.733187116 |
| chr14:91911646-91912005 | -0.282836847 | 0.46546413 | 0.733187116 |
| chr20:61463631-61463904 | -0.325629344 | 0.46547708 | 0.733187116 |
| chr13:43303098-43303538 | -0.285367312 | 0.465479929 | 0.733187116 |
| chr3:194520155-194520488 | -0.316889569 | 0.465487986 | 0.733187116 |
| chr7:33177905-33178207 | 0.313570439 | 0.465504347 | 0.733197815 |
| chr9:33076110-33076403 | 0.288778706 | 0.46553662 | 0.733206694 |

| | | | |
|---|---|---|---|
| chr1:237672824-237673038 | -0.374893493 | 0.465538569 | 0.733206694 |
| chrX:13646113-13646409 | -0.310623792 | 0.465538692 | 0.733206694 |
| chr1:112637387-112637714 | -0.319059287 | 0.46557734 | 0.733252491 |
| chr2:159615075-159616614 | -0.233529223 | 0.465620989 | 0.733306162 |
| chr22:22720120-22720432 | -0.313555642 | 0.465666265 | 0.733362394 |
| chr10:86520932-86521161 | 0.290430608 | 0.465688174 | 0.733368997 |
| chr6:163945850-163946188 | 0.306975432 | 0.4656896 | 0.733368997 |
| chr17:14302634-14302862 | -0.300360435 | 0.465718031 | 0.733377678 |
| chr5:7392463-7392788 | -0.311385842 | 0.465728785 | 0.733377678 |
| chr12:46492591-46492855 | 0.319406771 | 0.465730331 | 0.733377678 |
| chr1:32133347-32133929 | -0.268215646 | 0.465733398 | 0.733377678 |
| chr5:52964247-52964489 | -0.331019811 | 0.465916127 | 0.733649886 |
| chr15:55969105-55969945 | 0.248105751 | 0.465925415 | 0.733649886 |
| chr3:9649082-9649898 | 0.233562124 | 0.465940121 | 0.733657966 |
| chr2:175187325-175187763 | -0.278985838 | 0.465963895 | 0.733680323 |
| chr1:232261559-232261772 | 0.402111847 | 0.466000981 | 0.733685303 |
| chr14:95740964-95741615 | 0.244870794 | 0.466001243 | 0.733685303 |
| chr14:35944798-35945193 | 0.273072271 | 0.466003318 | 0.733685303 |
| chr1:116817614-116818025 | 0.25853124 | 0.466005359 | 0.733685303 |
| chr5:81108477-81108688 | -0.375087133 | 0.466040865 | 0.733725987 |
| chr10:24617774-24618293 | -0.271977401 | 0.466050352 | 0.733725987 |
| chr11:78420306-78420598 | 0.27917697 | 0.46607558 | 0.733750628 |
| chr11:33161077-33161843 | 0.236522466 | 0.466156696 | 0.733863252 |
| chr10:23870130-23870338 | 0.35488856 | 0.466172404 | 0.733863409 |
| chr5:10201547-10201747 | -0.39162936 | 0.466175951 | 0.733863409 |
| chr19:46674836-46675131 | -0.283627495 | 0.466199987 | 0.733868068 |
| chr7:36366605-36367709 | -0.231885051 | 0.466202991 | 0.733868068 |
| chr16:73170686-73170972 | 0.301990729 | 0.466225685 | 0.733868068 |
| chr7:78134349-78134563 | 0.336015558 | 0.466232787 | 0.733868068 |
| chr14:105600909-105601109 | -0.370827319 | 0.46623417 | 0.733868068 |
| chr17:20801947-20802147 | -0.375907402 | 0.466236377 | 0.733868068 |
| chr3:17494427-17494650 | 0.374647856 | 0.466298459 | 0.73395071 |
| chr11:72725290-72725557 | 0.297782036 | 0.466323787 | 0.733963982 |
| chr4:15238727-15238992 | 0.34938989 | 0.466326049 | 0.733963982 |
| chr22:17201327-17201560 | -0.330625316 | 0.466344081 | 0.733966219 |
| chr1:200038987-200040200 | 0.234988657 | 0.466346629 | 0.733966219 |
| chr3:69324403-69324679 | -0.303459901 | 0.466400843 | 0.734031142 |
| chr4:118888611-118889254 | -0.256029243 | 0.46640704 | 0.734031142 |
| chr1:93448581-93448984 | 0.285665986 | 0.466435328 | 0.734050344 |
| chr1:227542357-227542910 | 0.261223049 | 0.466443196 | 0.734050344 |
| chr3:130759768-130760025 | -0.33214034 | 0.466447981 | 0.734050344 |
| chr11:111978395-111978681 | 0.294838404 | 0.466460514 | 0.73405499 |
| chrX:130529653-130529869 | 0.369112348 | 0.466479088 | 0.734069144 |
| chr20:35284123-35284384 | -0.30108676 | 0.466509586 | 0.734090169 |
| chr8:142355334-142355544 | -0.323226868 | 0.466512512 | 0.734090169 |
| chr11:39732126-39732346 | -0.382774417 | 0.466523403 | 0.734090169 |
| chr6:83417917-83418215 | 0.309376024 | 0.466530772 | 0.734090169 |
| chr14:39113909-39114716 | 0.232533348 | 0.46656053 | 0.734104239 |
| chr1:148334123-148335419 | 0.228948236 | 0.466564798 | 0.734104239 |
| chr3:118761051-118761328 | -0.333979514 | 0.466568456 | 0.734104239 |

| | | | |
|---|---|---|---|
| chr19:34134239-34134900 | -0.25071125 | 0.466588069 | 0.734120023 |
| chr20:33489585-33490655 | 0.237051807 | 0.46662964 | 0.734170355 |
| chr7:121558407-121558662 | -0.342546838 | 0.466726169 | 0.734307149 |
| chr10:123154649-123155177 | 0.250899205 | 0.466754476 | 0.734322539 |
| chr3:22743507-22743858 | -0.286733238 | 0.466755117 | 0.734322539 |
| chr7:7653603-7653939 | -0.291510224 | 0.466776838 | 0.734332252 |
| chr1:202058143-202058868 | 0.237519524 | 0.466780459 | 0.734332252 |
| chr1:6259833-6260256 | 0.267858956 | 0.466808167 | 0.734346032 |
| chrX:119573788-119575280 | 0.232244908 | 0.466808386 | 0.734346032 |
| chr9:127574469-127575027 | 0.246757119 | 0.466832089 | 0.734368242 |
| chr2:172977484-172977693 | 0.335620266 | 0.466853273 | 0.734386489 |
| chr15:95911625-95912754 | -0.24560343 | 0.466878134 | 0.73440315 |
| chr1:108730675-108730894 | 0.349154416 | 0.466883034 | 0.73440315 |
| chr6:1641289-1641770 | -0.253318177 | 0.466922637 | 0.734450367 |
| chr10:103193047-103193535 | 0.265276809 | 0.466947082 | 0.734450659 |
| chr15:96356958-96357439 | 0.265124431 | 0.466955586 | 0.734450659 |
| chr22:38700760-38701282 | -0.251830044 | 0.466959084 | 0.734450659 |
| chr12:132414893-132415267 | 0.262291024 | 0.466961164 | 0.734450659 |
| chr7:1504158-1504806 | 0.240837877 | 0.466972487 | 0.734453393 |
| chr16:29899205-29900896 | 0.232581547 | 0.466995798 | 0.734460018 |
| chr16:28211814-28212452 | 0.244196842 | 0.466995871 | 0.734460018 |
| chr20:59385598-59385798 | 0.307682106 | 0.467054022 | 0.734529849 |
| chrX:23902496-23902748 | -0.307372471 | 0.467059444 | 0.734529849 |
| chr2:36650024-36650422 | -0.262558133 | 0.467081619 | 0.734545374 |
| chrX:91434612-91435133 | -0.279183216 | 0.467097481 | 0.734545374 |
| chr10:70265005-70265473 | 0.277001784 | 0.467098076 | 0.734545374 |
| chr14:89482141-89482375 | -0.334432332 | 0.467113591 | 0.734547789 |
| chr11:93197122-93198275 | -0.235944601 | 0.467118785 | 0.734547789 |
| chr13:39023781-39024208 | 0.256014563 | 0.467149682 | 0.734552552 |
| chr2:20633952-20634281 | -0.279551332 | 0.467162346 | 0.734552552 |
| chr2:30422528-30422737 | 0.386201984 | 0.467175958 | 0.734552552 |
| chr19:49808788-49809080 | 0.268876933 | 0.467186356 | 0.734552552 |
| chr6:28764107-28764517 | -0.312617515 | 0.467187595 | 0.734552552 |
| chr3:46981848-46982126 | 0.287296157 | 0.467188151 | 0.734552552 |
| chr2:172859805-172860251 | -0.314956781 | 0.467195242 | 0.734552552 |
| chr10:101042615-101043328 | -0.239611341 | 0.467198508 | 0.734552552 |
| chr19:6862788-6863050 | 0.328527303 | 0.467251835 | 0.734620137 |
| chr2:177344446-177344738 | -0.326754595 | 0.467260669 | 0.734620137 |
| chr14:61347410-61347662 | 0.367074904 | 0.467287251 | 0.734643482 |
| chr22:20116651-20117599 | 0.237305021 | 0.467295414 | 0.734643482 |
| chr17:63974176-63974451 | -0.276311975 | 0.467304282 | 0.734643482 |
| chr3:119125850-119126829 | -0.24931295 | 0.467362699 | 0.734720245 |
| chr12:89768922-89769755 | -0.236695739 | 0.467377493 | 0.734721012 |
| chr1:191897893-191898116 | -0.389265573 | 0.467382365 | 0.734721012 |
| chr17:8186584-8188089 | -0.229086369 | 0.467445699 | 0.734796767 |
| chr2:142944147-142944645 | 0.273993755 | 0.467459033 | 0.734796767 |
| chr3:148779049-148779249 | -0.433375728 | 0.467459325 | 0.734796767 |
| chr6:102816389-102817040 | -0.27303093 | 0.467486024 | 0.73482366 |
| chr5:112976185-112977292 | -0.23004511 | 0.467506229 | 0.734840344 |
| chr7:65080883-65081695 | -0.249564747 | 0.467537708 | 0.734840526 |

| | | | |
|---|---|---|---|
| chr2:43142971-43143251 | -0.313023821 | 0.467543465 | 0.734840526 |
| chr9:35646708-35647269 | -0.252330789 | 0.467552718 | 0.734840526 |
| chr3:69306440-69306673 | -0.323266074 | 0.467554006 | 0.734840526 |
| chr9:78121542-78121794 | -0.322612983 | 0.467554297 | 0.734840526 |
| chr10:44878663-44878996 | -0.286427149 | 0.467577838 | 0.734859441 |
| chr3:129314837-129315221 | 0.255145992 | 0.467585513 | 0.734859441 |
| chr3:53493756-53494675 | 0.253490802 | 0.467613015 | 0.734887589 |
| chr14:50242762-50243177 | -0.303606727 | 0.467657289 | 0.734942095 |
| chr11:64165884-64166775 | 0.235042861 | 0.467688113 | 0.734975462 |
| chr1:206041553-206041905 | 0.283359091 | 0.467721265 | 0.735000902 |
| chr3:118884708-118884966 | -0.298725404 | 0.46773017 | 0.735000902 |
| chr1:23368734-23369256 | 0.258242381 | 0.467734158 | 0.735000902 |
| chr4:64781802-64782002 | -0.387941089 | 0.467742672 | 0.735000902 |
| chr1:70220984-70222022 | -0.234478095 | 0.467759949 | 0.735012978 |
| chr20:33443694-33444242 | 0.260763351 | 0.467772397 | 0.735017464 |
| chr7:153062043-153062243 | -0.363228002 | 0.467784451 | 0.735021332 |
| chr6:28880912-28881619 | 0.252364084 | 0.467800352 | 0.735031243 |
| chr14:32933617-32933897 | 0.302965275 | 0.467828099 | 0.735059767 |
| chr13:33440878-33441373 | -0.274164306 | 0.467860981 | 0.735083958 |
| chr21:39333570-39333908 | 0.287722691 | 0.467862683 | 0.735083958 |
| chr4:145097742-145099023 | -0.229779827 | 0.467876699 | 0.735090907 |
| chrX:140733776-140734086 | -0.292632805 | 0.467902961 | 0.735117094 |
| chr12:22529365-22529613 | -0.337360656 | 0.467942689 | 0.735155416 |
| chr1:41672228-41672574 | -0.285096506 | 0.467946542 | 0.735155416 |
| chr1:218705938-218706265 | -0.283834298 | 0.467988748 | 0.735206649 |
| chr11:79371139-79371419 | -0.32423853 | 0.468029685 | 0.735255885 |
| chr2:105576270-105576742 | -0.279113547 | 0.468050204 | 0.735273044 |
| chr1:206631167-206631396 | -0.2818477 | 0.468094534 | 0.735318913 |
| chr3:138285006-138285253 | -0.311544566 | 0.468098596 | 0.735318913 |
| chr2:31737676-31737883 | -0.402275539 | 0.468122305 | 0.735341081 |
| chr11:12461542-12461906 | -0.325905718 | 0.468153271 | 0.735364683 |
| chr3:109081864-109082393 | -0.279694607 | 0.468156525 | 0.735364683 |
| chr12:125533494-125533694 | -0.38955501 | 0.468178364 | 0.735383912 |
| chr16:69754425-69755267 | 0.239173664 | 0.468208866 | 0.735416746 |
| chr5:65722826-65723150 | -0.275287328 | 0.468222391 | 0.735422914 |
| chr19:41176078-41176465 | -0.32905253 | 0.468253203 | 0.735456234 |
| chr12:21668-21935 | 0.354917004 | 0.468266204 | 0.735461578 |
| chr3:142586802-142587319 | -0.256825584 | 0.468296734 | 0.73547798 |
| chr4:140609997-140610380 | 0.276215544 | 0.468306374 | 0.73547798 |
| chr3:50299130-50299914 | 0.233654293 | 0.468307178 | 0.73547798 |
| chr3:8755576-8756128 | 0.268007048 | 0.468315042 | 0.73547798 |
| chr4:46390369-46390597 | -0.358603237 | 0.468325462 | 0.735479269 |
| chr7:155921939-155922447 | -0.266185296 | 0.468354041 | 0.735498662 |
| chr8:118909545-118909745 | -0.41256687 | 0.468357009 | 0.735498662 |
| chr4:59381058-59381366 | -0.324110596 | 0.468383036 | 0.735521761 |
| chr22:30088947-30089277 | 0.317336287 | 0.468397871 | 0.735521761 |
| chr3:183765443-183765780 | -0.288922783 | 0.468400516 | 0.735521761 |
| chr1:155227345-155227862 | 0.23634051 | 0.468442583 | 0.735572744 |
| chr9:34441415-34441615 | 0.397694142 | 0.468464044 | 0.735591367 |
| chr13:39602481-39603904 | -0.234899451 | 0.468479405 | 0.735597906 |

| | | | |
|---|---|---|---|
| chr15:89556647-89556996 | 0.275539914 | 0.468487408 | 0.735597906 |
| chr8:97822058-97822342 | -0.296608042 | 0.468513342 | 0.735623551 |
| chr3:179898049-179898339 | -0.330411528 | 0.468545422 | 0.735631497 |
| chr1:157489399-157489629 | -0.339739471 | 0.468545514 | 0.735631497 |
| chr4:75848310-75848612 | 0.295088488 | 0.468550301 | 0.735631497 |
| chr12:121918073-121918287 | -0.361992867 | 0.468556806 | 0.735631497 |
| chr7:4683697-4684195 | -0.261225397 | 0.468598744 | 0.735661538 |
| chrX:48945969-48946200 | 0.37200644 | 0.468601941 | 0.735661538 |
| chr8:133886125-133886438 | 0.287676643 | 0.468604744 | 0.735661538 |
| chr15:62066756-62067214 | 0.254515012 | 0.46861981 | 0.735670117 |
| chr5:19322788-19323081 | -0.317144916 | 0.468675312 | 0.735742174 |
| chr10:90592995-90593204 | -0.364052954 | 0.468749704 | 0.735817205 |
| chr6:121849558-121849853 | -0.303289585 | 0.468754697 | 0.735817205 |
| chr20:36745602-36746518 | 0.246434939 | 0.468758176 | 0.735817205 |
| chr4:157015196-157015407 | -0.416189302 | 0.468761521 | 0.735817205 |
| chr5:132410360-132411668 | -0.238648376 | 0.46877856 | 0.735828878 |
| chr1:193111652-193111972 | -0.277100986 | 0.468803167 | 0.735852428 |
| chr4:104860580-104860860 | 0.310986721 | 0.468828788 | 0.735877569 |
| chr5:133700567-133700767 | -0.373602084 | 0.468860546 | 0.735912341 |
| chr20:34709407-34710058 | 0.247948907 | 0.46887404 | 0.735918446 |
| chr3:108588975-108590153 | -0.227919808 | 0.468903537 | 0.735949668 |
| chr10:122374100-122374791 | 0.243144332 | 0.468986419 | 0.736064674 |
| chr3:62233108-62233454 | -0.262577594 | 0.469015462 | 0.736082033 |
| chr6:2970751-2972073 | 0.239419868 | 0.469016693 | 0.736082033 |
| chr17:40099033-40099462 | 0.251606641 | 0.469031429 | 0.736090084 |
| chr5:154834112-154834368 | 0.364005801 | 0.469048023 | 0.736101048 |
| chr18:26649753-26650078 | 0.295185829 | 0.469061017 | 0.736106365 |
| chr7:44200714-44201408 | -0.238307755 | 0.46907154 | 0.736107801 |
| chr9:127224162-127225505 | 0.229602798 | 0.469140853 | 0.736158008 |
| chr9:34376041-34376370 | 0.284395009 | 0.469146094 | 0.736158008 |
| chr9:69759404-69760471 | -0.238958907 | 0.469155046 | 0.736158008 |
| chr11:27718424-27719286 | -0.249733954 | 0.469173499 | 0.736158008 |
| chr8:82937333-82937985 | 0.276956067 | 0.469174237 | 0.736158008 |
| chr6:39309944-39310423 | 0.271915286 | 0.469178052 | 0.736158008 |
| chr14:69376020-69376319 | 0.32752411 | 0.469182466 | 0.736158008 |
| chr17:50689980-50690355 | -0.271135139 | 0.469205222 | 0.736158008 |
| chr12:111599043-111600248 | 0.23483061 | 0.469225469 | 0.736158008 |
| chr12:24991537-24991784 | -0.355452743 | 0.469231714 | 0.736158008 |
| chr17:15479358-15479558 | 0.390713801 | 0.469232799 | 0.736158008 |
| chr1:235367108-235367836 | 0.232795604 | 0.469233752 | 0.736158008 |
| chr20:52731176-52731502 | -0.356513185 | 0.469238778 | 0.736158008 |
| chr5:123429035-123429431 | -0.294141712 | 0.46923937 | 0.736158008 |
| chr1:62391455-62391691 | -0.350211595 | 0.469249801 | 0.736158008 |
| chr2:72174112-72174312 | 0.386926137 | 0.469257255 | 0.736158008 |
| chr5:159175526-159175950 | 0.272315252 | 0.469295616 | 0.736203114 |
| chr15:97903925-97904606 | 0.246567579 | 0.469330202 | 0.736242297 |
| chr1:184446544-184446744 | 0.356170954 | 0.469384902 | 0.736313031 |
| chrX:149546694-149547211 | 0.275042622 | 0.469432245 | 0.736360544 |
| chr16:82263567-82263804 | 0.306727901 | 0.469434411 | 0.736360544 |
| chr22:30387332-30387823 | 0.268564617 | 0.469461163 | 0.736387431 |

| | | | |
|---|---|---|---|
| chr10:103598845-103599116 | 0.297850834 | 0.469504881 | 0.736426797 |
| chr16:1272231-1272606 | -0.295938556 | 0.46950732 | 0.736426797 |
| chr7:71752152-71752518 | -0.312335038 | 0.469515093 | 0.736426797 |
| chr3:58040146-58040466 | -0.29946116 | 0.469538873 | 0.73644902 |
| chr20:41317513-41318416 | 0.230871532 | 0.469555046 | 0.736454016 |
| chr10:70145710-70147253 | -0.229247848 | 0.46956276 | 0.736454016 |
| chr12:63944704-63944997 | 0.317556457 | 0.469576536 | 0.736454016 |
| chr5:148730961-148731291 | -0.282446049 | 0.469589773 | 0.736454016 |
| chr12:132207169-132207369 | 0.395083172 | 0.469612612 | 0.736454016 |
| chr19:2462906-2463288 | 0.282890542 | 0.469616091 | 0.736454016 |
| chr12:103965557-103966359 | -0.230763896 | 0.469618426 | 0.736454016 |
| chr3:148727235-148727590 | 0.267239575 | 0.46961895 | 0.736454016 |
| chr5:107806775-107807072 | -0.315847278 | 0.469657045 | 0.736498682 |
| chr16:5092298-5092627 | 0.269850584 | 0.469691357 | 0.736537415 |
| chr2:74399903-74400393 | -0.285835297 | 0.469734552 | 0.736579104 |
| chr15:93063365-93063677 | -0.288085164 | 0.469737169 | 0.736579104 |
| chr19:10695254-10695468 | -0.334794394 | 0.469764951 | 0.736607593 |
| chr1:93847086-93848283 | -0.231591261 | 0.469800001 | 0.736647478 |
| chr10:86377137-86377575 | 0.272765513 | 0.469819375 | 0.736662781 |
| chr2:197434524-197435654 | -0.229680006 | 0.469840299 | 0.736664585 |
| chr6:52576547-52577614 | 0.234381999 | 0.469861199 | 0.736664585 |
| chr19:50143014-50143238 | 0.351961615 | 0.469867832 | 0.736664585 |
| chr2:161134553-161134803 | 0.351961615 | 0.469867832 | 0.736664585 |
| chr2:119265178-119265495 | 0.286872765 | 0.469868597 | 0.736664585 |
| chr22:37197660-37198031 | 0.269737943 | 0.46988836 | 0.736676945 |
| chr1:225809908-225810527 | 0.246005487 | 0.469895709 | 0.736676945 |
| chr1:230863479-230863833 | 0.274392312 | 0.469924222 | 0.736693953 |
| chr3:156816491-156817748 | -0.228628625 | 0.469925787 | 0.736693953 |
| chr2:20584217-20584864 | 0.249217334 | 0.469944504 | 0.736698119 |
| chr8:104492090-104492728 | 0.269911261 | 0.469947674 | 0.736698119 |
| chr12:13404213-13404413 | -0.376392769 | 0.470005292 | 0.736773368 |
| chr15:88454924-88455455 | 0.247012671 | 0.470014962 | 0.736773454 |
| chr12:45727420-45728082 | -0.237535025 | 0.47004768 | 0.736793183 |
| chr14:91264227-91264827 | 0.256203508 | 0.470085045 | 0.736793183 |
| chr20:54711680-54712238 | 0.23621018 | 0.470088219 | 0.736793183 |
| chr14:57987665-57987937 | 0.302779633 | 0.470089554 | 0.736793183 |
| chr2:63555975-63556265 | 0.319430921 | 0.470090369 | 0.736793183 |
| chr20:18793868-18794404 | 0.248502825 | 0.470107193 | 0.736793183 |
| chr8:108942399-108942839 | -0.264950895 | 0.470118173 | 0.736793183 |
| chr15:85039640-85039944 | -0.354831701 | 0.470121658 | 0.736793183 |
| chr18:36828249-36829846 | -0.228344059 | 0.470122006 | 0.736793183 |
| chr1:114173832-114174741 | 0.232202229 | 0.470142619 | 0.736793183 |
| chr12:55715898-55716779 | 0.233104746 | 0.470150566 | 0.736793183 |
| chr15:101276802-101277925 | -0.231348206 | 0.470151184 | 0.736793183 |
| chr4:146033024-146033229 | -0.404679105 | 0.470152556 | 0.736793183 |
| chr19:35006442-35006716 | 0.290846539 | 0.470220957 | 0.736880032 |
| chr2:173354504-173355665 | -0.245369893 | 0.470232483 | 0.736880032 |
| chr4:107824296-107825461 | -0.231285515 | 0.470236826 | 0.736880032 |
| chr7:26178715-26178949 | 0.326975297 | 0.470257056 | 0.736896664 |
| chr1:44168934-44169177 | 0.345274725 | 0.470281288 | 0.736919564 |

| | | | |
|---|---|---|---|
| chr14:77799896-77800772 | 0.233086449 | 0.470315361 | 0.736932967 |
| chr8:127788825-127789078 | 0.287440332 | 0.470323808 | 0.736932967 |
| chr6:128511494-128511737 | 0.334227906 | 0.470331707 | 0.736932967 |
| chr10:42782289-42783151 | 0.235923699 | 0.470336783 | 0.736932967 |
| chr10:67550881-67551210 | -0.319134202 | 0.470342623 | 0.736932967 |
| chr1:194535409-194535756 | 0.31690801 | 0.470347548 | 0.736932967 |
| chr1:167721054-167721466 | 0.268932225 | 0.470382665 | 0.736962961 |
| chr3:69052823-69053029 | 0.323243345 | 0.470385928 | 0.736962961 |
| chr2:232444459-232444689 | -0.341319762 | 0.470456959 | 0.737041415 |
| chr1:172443633-172444514 | -0.235548708 | 0.470464524 | 0.737041415 |
| chr10:74150249-74151647 | -0.229037588 | 0.470464861 | 0.737041415 |
| chr4:75289810-75290234 | -0.29723873 | 0.470553598 | 0.737165361 |
| chr20:12358669-12358918 | -0.404270102 | 0.470578517 | 0.737189326 |
| chr5:78467876-78468078 | 0.373569479 | 0.470598731 | 0.737205921 |
| chr20:53950038-53950409 | 0.234604753 | 0.470621484 | 0.737212626 |
| chr2:144162716-144162975 | -0.315783392 | 0.470624885 | 0.737212626 |
| chr9:97238675-97238935 | 0.315913582 | 0.470631876 | 0.737212626 |
| chr11:124875946-124876915 | 0.236154944 | 0.470682687 | 0.737274591 |
| chr3:197906073-197906384 | 0.302774443 | 0.470690678 | 0.737274591 |
| chr3:196914930-196915158 | -0.372109614 | 0.470730826 | 0.737322404 |
| chr11:1834191-1834391 | -0.369110244 | 0.470764549 | 0.737360152 |
| chr22:41576967-41577167 | 0.349742596 | 0.470789536 | 0.737364745 |
| chr15:73305890-73306168 | 0.298832035 | 0.470798849 | 0.737364745 |
| chr4:25668221-25668451 | -0.375422168 | 0.470805655 | 0.737364745 |
| chr12:51520339-51520746 | -0.307022668 | 0.47081952 | 0.737364745 |
| chr14:104100266-104101274 | -0.23970487 | 0.470820372 | 0.737364745 |
| chr5:173984058-173984526 | 0.282125203 | 0.470825222 | 0.737364745 |
| chr22:46367679-46367972 | 0.296950573 | 0.470844307 | 0.737368228 |
| chr3:64718287-64718519 | 0.267124088 | 0.470847761 | 0.737368228 |
| chr5:24445348-24445952 | -0.24892393 | 0.470863644 | 0.737368228 |
| chr1:178530319-178530627 | -0.332234099 | 0.47087951 | 0.737368228 |
| chr20:12287561-12287982 | 0.277474754 | 0.470881798 | 0.737368228 |
| chr11:11621959-11622385 | 0.273522514 | 0.470885186 | 0.737368228 |
| chr15:34366200-34367928 | 0.228650993 | 0.470930139 | 0.737423549 |
| chr9:88387673-88388956 | 0.229651641 | 0.470949083 | 0.737429647 |
| chr2:121999313-121999749 | -0.275782429 | 0.470953282 | 0.737429647 |
| chr8:124147719-124147946 | 0.297498171 | 0.470982266 | 0.737459961 |
| chr4:111164674-111164874 | 0.395241432 | 0.471051832 | 0.737553815 |
| chr7:7306335-7306591 | 0.317496198 | 0.471077974 | 0.737579674 |
| chr3:64444889-64445924 | 0.226111076 | 0.471099328 | 0.737598037 |
| chr16:86874231-86874438 | 0.323880452 | 0.47110921 | 0.737598436 |
| chr20:35616862-35617097 | 0.270905344 | 0.47114498 | 0.737621572 |
| chr3:49132439-49133891 | 0.235076303 | 0.471149446 | 0.737621572 |
| chr9:99299696-99299896 | 0.371977112 | 0.471161585 | 0.737621572 |
| chr14:65019989-65020339 | 0.272227138 | 0.471179902 | 0.737621572 |
| chr10:125905766-125906025 | 0.386562764 | 0.471180718 | 0.737621572 |
| chr13:24670978-24671278 | -0.282764194 | 0.471181747 | 0.737621572 |
| chr12:43758218-43759708 | -0.227839837 | 0.47123877 | 0.737695767 |
| chr3:45986323-45986712 | 0.292687442 | 0.471254292 | 0.737704995 |
| chr2:21317176-21317472 | -0.352324251 | 0.471364929 | 0.737859937 |

| | | | |
|---|---|---|---|
| chr8:100145683-100146022 | -0.261465716 | 0.471372531 | 0.737859937 |
| chr17:74453806-74454618 | -0.239451277 | 0.471451254 | 0.737936707 |
| chr9:37594458-37594883 | -0.276341335 | 0.471454158 | 0.737936707 |
| chr19:9917210-9917475 | -0.353769289 | 0.47145952 | 0.737936707 |
| chr22:50270386-50270586 | 0.37175778 | 0.471460098 | 0.737936707 |
| chr1:229625338-229626669 | 0.234233491 | 0.471473186 | 0.737942118 |
| chr19:53537232-53538746 | 0.231323142 | 0.471495904 | 0.737960402 |
| chr7:92474018-92474515 | -0.274128105 | 0.47150413 | 0.737960402 |
| chr17:43360445-43361853 | 0.229633798 | 0.471550414 | 0.738017767 |
| chr8:21784366-21784607 | -0.374959755 | 0.471618387 | 0.738094132 |
| chr19:30719460-30719733 | 0.365623658 | 0.471618473 | 0.738094132 |
| chr3:71444814-71445088 | -0.34887587 | 0.471630758 | 0.738098284 |
| chr17:61389946-61390252 | 0.233141518 | 0.47168771 | 0.738149493 |
| chr11:119144844-119145226 | -0.262670708 | 0.471691107 | 0.738149493 |
| chr10:34973745-34974216 | 0.257637544 | 0.471695233 | 0.738149493 |
| chr1:170881997-170882305 | 0.307424816 | 0.471711006 | 0.738149493 |
| chr19:36053951-36054989 | 0.229027307 | 0.471713839 | 0.738149493 |
| chr2:41814965-41815303 | -0.295389731 | 0.471724362 | 0.738149493 |
| chr19:35775745-35775963 | 0.299632795 | 0.471730915 | 0.738149493 |
| chr15:72226648-72227006 | 0.250929351 | 0.47174167 | 0.738151247 |
| chr9:117487468-117488117 | -0.244691698 | 0.47176123 | 0.738166778 |
| chr1:220113262-220113462 | -0.380381601 | 0.471774615 | 0.738172647 |
| chr5:132905887-132906317 | -0.274770985 | 0.471794848 | 0.738189231 |
| chr10:47438808-47439225 | -0.2662128 | 0.471850008 | 0.738226701 |
| chr3:2765064-2765528 | -0.27887345 | 0.47185069 | 0.738226701 |
| chr11:82314890-82315463 | -0.264535024 | 0.471862503 | 0.738226701 |
| chr1:27527860-27529264 | -0.234595509 | 0.471868844 | 0.738226701 |
| chrX:54786012-54786212 | -0.399966274 | 0.471873695 | 0.738226701 |
| chr19:57578806-57579228 | 0.259970602 | 0.471876603 | 0.738226701 |
| chr3:129934364-129934816 | 0.269967571 | 0.471892621 | 0.738226878 |
| chrX:16883310-16883527 | -0.349237087 | 0.471895986 | 0.738226878 |
| chr5:19401028-19401233 | -0.386342515 | 0.471928138 | 0.738258528 |
| chr12:69936246-69936671 | 0.255838745 | 0.471935488 | 0.738258528 |
| chr1:178501397-178501612 | 0.325644373 | 0.471967543 | 0.738280795 |
| chr10:84114112-84114449 | -0.271390653 | 0.471968993 | 0.738280795 |
| chr10:74991436-74991749 | -0.317964129 | 0.471979468 | 0.738282108 |
| chr17:67401202-67401440 | 0.319851839 | 0.472013942 | 0.738311008 |
| chr13:26895266-26895663 | -0.299847875 | 0.472019569 | 0.738311008 |
| chr15:65032996-65033410 | -0.253894467 | 0.472038317 | 0.738311008 |
| chr1:161045067-161046376 | 0.230465741 | 0.472039972 | 0.738311008 |
| chr11:86499662-86500415 | -0.246969455 | 0.472046122 | 0.738311008 |
| chr8:143995831-143996235 | 0.263140037 | 0.472071643 | 0.738335853 |
| chr20:47196152-47197104 | 0.224909222 | 0.472088726 | 0.7383475 |
| chr6:53723191-53723669 | -0.254544519 | 0.472106213 | 0.738359779 |
| chr11:106305384-106305686 | -0.302855729 | 0.472119367 | 0.73836528 |
| chr8:26410030-26410422 | 0.3156989 | 0.472150888 | 0.738399449 |
| chr16:88910452-88910875 | -0.27390132 | 0.472168438 | 0.738399449 |
| chr2:112626521-112626990 | 0.266729009 | 0.472170126 | 0.738399449 |
| chr8:121055839-121056100 | -0.360229305 | 0.47219242 | 0.738419242 |
| chr8:143291039-143291687 | 0.242857231 | 0.472230773 | 0.738461737 |

| | | | |
|---|---|---|---|
| chr7:5704490-5704790 | 0.287608805 | 0.472238869 | 0.738461737 |
| chr1:204377797-204378507 | 0.237928387 | 0.47226786 | 0.738492 |
| chr7:39464870-39465299 | -0.292089369 | 0.472288612 | 0.738503273 |
| chr1:41492330-41492544 | 0.320866184 | 0.472296132 | 0.738503273 |
| chr6:26311149-26311349 | 0.309355851 | 0.472303985 | 0.738503273 |
| chr19:42075291-42075491 | 0.338096089 | 0.472416249 | 0.738659223 |
| chr8:47997273-47997605 | -0.277272404 | 0.472423002 | 0.738659223 |
| chr9:6015357-6016254 | 0.245686891 | 0.472451066 | 0.738688029 |
| chr17:10815314-10815634 | -0.304702155 | 0.47246143 | 0.73868916 |
| chr3:90015930-90016149 | -0.359882891 | 0.472481821 | 0.738705968 |
| chr17:76088092-76088292 | 0.349854837 | 0.472513722 | 0.738740771 |
| chr1:156838979-156839179 | 0.311953789 | 0.472538257 | 0.738764055 |
| chr5:11733585-11734267 | 0.249289174 | 0.472558946 | 0.738781326 |
| chr3:171286563-171286891 | -0.317656338 | 0.472575152 | 0.738791588 |
| chr2:31233769-31234253 | -0.2449699 | 0.472600762 | 0.738802324 |
| chr2:203484323-203484645 | -0.304031214 | 0.47261504 | 0.738802324 |
| chr8:144396937-144397245 | 0.268102069 | 0.472615162 | 0.738802324 |
| chr14:74465528-74465784 | 0.322611928 | 0.472620588 | 0.738802324 |
| chr6:162343297-162343520 | -0.396663845 | 0.472643224 | 0.738822635 |
| chr6:27157493-27157719 | -0.320919183 | 0.472654502 | 0.738825192 |
| chr11:28140469-28140669 | -0.38848605 | 0.472690193 | 0.738865909 |
| chr8:139675782-139676098 | 0.261370517 | 0.472700848 | 0.738867491 |
| chr6:17600056-17601311 | -0.231142306 | 0.472782214 | 0.738971864 |
| chr12:45299065-45299707 | -0.246373929 | 0.472786911 | 0.738971864 |
| chr6:107704317-107704639 | -0.347522438 | 0.472813803 | 0.738998821 |
| chr20:54149592-54149792 | 0.251128223 | 0.472829642 | 0.739008502 |
| chr5:142734690-142734951 | -0.333620344 | 0.472855343 | 0.739009616 |
| chr5:3757445-3757645 | -0.431711721 | 0.472868335 | 0.739009616 |
| chr17:27323641-27323937 | 0.290568463 | 0.47287123 | 0.739009616 |
| chr1:223486437-223486748 | -0.286953496 | 0.472875101 | 0.739009616 |
| chr19:8311907-8312177 | 0.296879518 | 0.472878579 | 0.739009616 |
| chr19:44879800-44880017 | 0.338151828 | 0.472916946 | 0.73902402 |
| chr2:16665794-16665994 | 0.338151828 | 0.472916946 | 0.73902402 |
| chrX:49053332-49053619 | 0.286195997 | 0.472941696 | 0.73902402 |
| chr1:166258110-166258393 | 0.301277525 | 0.472941754 | 0.73902402 |
| chr3:62332815-62333033 | 0.309859563 | 0.472949212 | 0.73902402 |
| chr2:74454653-74455380 | -0.233136515 | 0.472976405 | 0.73902402 |
| chr11:3063077-3063343 | 0.337974755 | 0.472979002 | 0.73902402 |
| chr9:122454551-122454812 | 0.320482479 | 0.472979461 | 0.73902402 |
| chr7:100705305-100706646 | -0.231202247 | 0.472979499 | 0.73902402 |
| chr3:19065106-19065342 | -0.333557911 | 0.472991498 | 0.73902402 |
| chr6:72622571-72623475 | -0.241269871 | 0.473001875 | 0.73902402 |
| chr17:17641691-17642124 | 0.275191545 | 0.473003537 | 0.73902402 |
| chr16:71648992-71649234 | 0.305094173 | 0.473019245 | 0.739027228 |
| chr13:19618982-19619208 | -0.320172609 | 0.473040431 | 0.739027228 |
| chr9:2843541-2844491 | -0.232367986 | 0.47304292 | 0.739027228 |
| chr20:53711273-53712007 | 0.225901477 | 0.473047234 | 0.739027228 |
| chr16:53022480-53022693 | 0.339301021 | 0.473053815 | 0.739027228 |
| chr4:73573803-73574024 | -0.352243424 | 0.473139169 | 0.739139126 |
| chr7:128791086-128791528 | -0.269794505 | 0.473148962 | 0.739139126 |

| | | | |
|---|---|---|---|
| chr20:48921005-48922593 | 0.222611893 | 0.473155497 | 0.739139126 |
| chr22:19300077-19300292 | -0.348609376 | 0.473164028 | 0.739139126 |
| chr8:133406659-133406859 | 0.318032653 | 0.473194326 | 0.739171386 |
| chr11:34307093-34307552 | 0.25885234 | 0.473215393 | 0.739175489 |
| chr5:17189400-17189920 | 0.260688987 | 0.473222993 | 0.739175489 |
| chrX:45515252-45515496 | -0.345879105 | 0.473225894 | 0.739175489 |
| chr17:51407499-51407968 | 0.258725473 | 0.473284535 | 0.739248796 |
| chr8:141169059-141169451 | -0.257923225 | 0.473292122 | 0.739248796 |
| chr17:81484331-81484725 | 0.258976879 | 0.473302139 | 0.739249373 |
| chr1:67479148-67479591 | 0.285921569 | 0.473329738 | 0.739265017 |
| chr8:120651182-120651479 | 0.326800458 | 0.473341099 | 0.739265017 |
| chrX:110003581-110003781 | 0.326800458 | 0.473341099 | 0.739265017 |
| chr16:67660086-67661158 | 0.229009566 | 0.473357727 | 0.739270222 |
| chr15:34753428-34753628 | 0.390073311 | 0.473364649 | 0.739270222 |
| chr1:106439190-106439618 | 0.254818244 | 0.473373377 | 0.739270222 |
| chr1:184561215-184561542 | -0.346802515 | 0.473422157 | 0.739331332 |
| chr19:29685251-29685964 | -0.26752053 | 0.473433653 | 0.739334218 |
| chr6:138278305-138278676 | 0.265557555 | 0.473485309 | 0.739397261 |
| chr6:143575389-143575658 | -0.303424621 | 0.473493335 | 0.739397261 |
| chr5:167725852-167726084 | -0.373689501 | 0.473517323 | 0.739397261 |
| chr11:33065624-33065884 | 0.295687632 | 0.473524067 | 0.739397261 |
| chr1:68975633-68975854 | -0.368705979 | 0.473533085 | 0.739397261 |
| chr10:104028213-104028526 | -0.305879274 | 0.473540052 | 0.739397261 |
| chr14:24270797-24271885 | 0.228374465 | 0.473541573 | 0.739397261 |
| chr2:219327778-219328002 | 0.350673148 | 0.473566948 | 0.739421814 |
| chr6:163956675-163956926 | -0.337826181 | 0.473590276 | 0.739443171 |
| chr2:225002131-225002356 | -0.342045689 | 0.473768367 | 0.73969522 |
| chr3:113484968-113485821 | -0.247964287 | 0.473771014 | 0.73969522 |
| chr17:78214014-78214788 | -0.244968365 | 0.473803886 | 0.73973147 |
| chr2:237502494-237502800 | -0.282940439 | 0.473819889 | 0.739741381 |
| chr1:205577613-205577880 | -0.284052071 | 0.473867478 | 0.739800605 |
| chr2:160269435-160270564 | 0.236826711 | 0.473884211 | 0.739804527 |
| chr6:131062725-131062945 | 0.30658486 | 0.473902733 | 0.739804527 |
| chr1:41889753-41890088 | -0.31396775 | 0.473905305 | 0.739804527 |
| chr17:66965120-66966301 | 0.229573804 | 0.473908611 | 0.739804527 |
| chr1:66765029-66765305 | -0.321136569 | 0.473944241 | 0.739845075 |
| chr1:182410205-182410531 | -0.301611433 | 0.473957185 | 0.739850208 |
| chr2:28821372-28821687 | 0.297389337 | 0.47397419 | 0.739855383 |
| chr7:129582672-129582888 | -0.369401291 | 0.473979812 | 0.739855383 |
| chr9:95004117-95004367 | 0.335648712 | 0.473995013 | 0.739864037 |
| chr19:34428142-34428724 | 0.254266218 | 0.474016817 | 0.739883 |
| chr6:90586605-90587561 | -0.228863075 | 0.474040657 | 0.739895357 |
| chr12:62260986-62261192 | -0.291293672 | 0.474044047 | 0.739895357 |
| chr10:88865550-88865779 | 0.3342783 | 0.474055834 | 0.739898682 |
| chr20:32091961-32092283 | -0.303586572 | 0.474127669 | 0.739995727 |
| chr3:9915794-9916238 | 0.249560924 | 0.474139778 | 0.739998313 |
| chr5:31826188-31826398 | -0.367669664 | 0.474148641 | 0.739998313 |
| chr20:34824455-34824756 | 0.289765227 | 0.474207691 | 0.740075397 |
| chr14:102591420-102591670 | -0.291638598 | 0.474220639 | 0.74008053 |
| chr3:7144777-7145096 | 0.289220162 | 0.474255001 | 0.740119083 |

| | | | |
|---|---|---|---|
| chr3:140898379-140898682 | 0.279892889 | 0.474294771 | 0.740156336 |
| chr7:48102110-48102355 | -0.313629901 | 0.474298192 | 0.740156336 |
| chr11:57322213-57322928 | -0.234781334 | 0.47431684 | 0.740170361 |
| chr17:3935132-3935565 | -0.284125822 | 0.474372457 | 0.740226821 |
| chr17:4832774-4833876 | -0.23555961 | 0.474378007 | 0.740226821 |
| chr2:3645351-3645586 | -0.34895975 | 0.474382003 | 0.740226821 |
| chr19:38989188-38989451 | -0.34156659 | 0.474391965 | 0.740227292 |
| chr9:32572323-32573444 | -0.234372243 | 0.474430737 | 0.740268679 |
| chr4:150297905-150298105 | 0.364931376 | 0.474443806 | 0.740268679 |
| chr9:109640219-109640628 | 0.283477024 | 0.474452442 | 0.740268679 |
| chrX:124211412-124211782 | -0.298507247 | 0.47446065 | 0.740268679 |
| chr4:77465383-77465583 | 0.346749105 | 0.474466796 | 0.740268679 |
| chr8:90504312-90504571 | 0.312406269 | 0.474486606 | 0.740284513 |
| chr8:3818990-3819230 | -0.379811971 | 0.474526449 | 0.740297367 |
| chr4:65865600-65865800 | -0.36741369 | 0.474539217 | 0.740297367 |
| chr12:89005531-89005773 | -0.36741369 | 0.474539217 | 0.740297367 |
| chr2:118222005-118222320 | 0.303060781 | 0.474542158 | 0.740297367 |
| chr15:40249515-40249974 | 0.271539056 | 0.474555129 | 0.740297367 |
| chr5:151284513-151284805 | -0.353631043 | 0.474559505 | 0.740297367 |
| chr11:13052094-13052623 | 0.267606172 | 0.474562476 | 0.740297367 |
| chr14:78836825-78837240 | -0.257506264 | 0.474582135 | 0.740312961 |
| chr10:77392000-77392218 | -0.319901124 | 0.474601753 | 0.740328492 |
| chr12:110345178-110345653 | 0.254446639 | 0.474635715 | 0.740353908 |
| chr2:29920587-29920813 | -0.355500843 | 0.474637371 | 0.740353908 |
| chr16:67935201-67936195 | 0.230077567 | 0.474657699 | 0.740370543 |
| chr13:45506039-45506323 | 0.345243181 | 0.474681402 | 0.740372993 |
| chr3:129942254-129942680 | -0.256904834 | 0.474693477 | 0.740372993 |
| chr5:74903315-74903743 | -0.261847321 | 0.474697828 | 0.740372993 |
| chr20:48697655-48698806 | 0.223740434 | 0.474708741 | 0.740372993 |
| chr17:3723465-3724031 | 0.2482499 | 0.474713597 | 0.740372993 |
| chr3:9730893-9731843 | 0.226106163 | 0.474717245 | 0.740372993 |
| chr6:19024823-19025073 | -0.311703928 | 0.474742842 | 0.740397843 |
| chr3:184145044-184145541 | 0.26299795 | 0.474866497 | 0.740548689 |
| chr3:192803056-192803256 | 0.364464381 | 0.474866852 | 0.740548689 |
| chr17:28710680-28711652 | -0.233801631 | 0.474868559 | 0.740548689 |
| chr11:134393977-134394177 | -0.407762375 | 0.474905968 | 0.740591955 |
| chr17:8164151-8164518 | 0.284461297 | 0.47492888 | 0.740598017 |
| chr17:67200590-67201069 | -0.258951284 | 0.474929187 | 0.740598017 |
| chr1:150394249-150394467 | 0.329709272 | 0.474941772 | 0.740602569 |
| chr6:119682968-119683642 | -0.254778502 | 0.474962991 | 0.74061284 |
| chr11:33630905-33631128 | -0.342250964 | 0.474983588 | 0.74061284 |
| chr3:161268611-161268935 | -0.254836146 | 0.474994331 | 0.74061284 |
| chr11:68342314-68343173 | 0.234425518 | 0.474994716 | 0.74061284 |
| chr6:156390654-156390880 | -0.359170087 | 0.474996687 | 0.74061284 |
| chr8:107497013-107497585 | -0.248555017 | 0.475026458 | 0.740631769 |
| chr15:44550335-44550535 | 0.358187184 | 0.47502816 | 0.740631769 |
| chr8:114846695-114846933 | -0.33025322 | 0.475048577 | 0.740635383 |
| chr19:40465525-40466523 | 0.227860439 | 0.475064587 | 0.740635383 |
| chr19:18831763-18831984 | 0.341021949 | 0.475065747 | 0.740635383 |
| chr2:120283711-120283915 | 0.368618565 | 0.475069142 | 0.740635383 |

| | | | |
|---|---|---|---|
| chr10:131964840-131965146 | -0.375926138 | 0.475124026 | 0.740705877 |
| chr22:40856109-40857377 | -0.235231357 | 0.475136412 | 0.740710116 |
| chr12:42145234-42145459 | -0.311920418 | 0.475148691 | 0.740714186 |
| chr3:185176857-185177057 | 0.385306876 | 0.475223953 | 0.740816441 |
| chr15:99273042-99273420 | -0.280996308 | 0.475266924 | 0.740845044 |
| chr2:165016883-165017086 | -0.35319644 | 0.475267569 | 0.740845044 |
| chr14:67485873-67486073 | 0.344111596 | 0.475279054 | 0.740845044 |
| chr1:192722869-192723069 | 0.384088592 | 0.475290518 | 0.740845044 |
| chr11:36122664-36123036 | -0.321666075 | 0.475290645 | 0.740845044 |
| chr6:33431003-33431311 | 0.283295995 | 0.475310397 | 0.74086076 |
| chr12:56103947-56105095 | 0.226818237 | 0.475349977 | 0.740907381 |
| chr19:18293228-18293609 | 0.278087571 | 0.475377152 | 0.740934665 |
| chr5:172106692-172107154 | -0.262600532 | 0.475398691 | 0.740941782 |
| chr7:33886215-33886418 | -0.329099902 | 0.475401058 | 0.740941782 |
| chr19:14495995-14496587 | -0.246085918 | 0.475411558 | 0.740943076 |
| chr3:119145393-119145908 | 0.255572083 | 0.47543262 | 0.740960829 |
| chr7:102909805-102910070 | -0.391353764 | 0.475458245 | 0.740985694 |
| chr8:117414453-117414662 | 0.340781622 | 0.475488985 | 0.741013369 |
| chr3:166837362-166837950 | -0.250989975 | 0.475501464 | 0.741013369 |
| chr12:54378872-54380015 | 0.23258421 | 0.475505016 | 0.741013369 |
| chr17:81387346-81387608 | 0.296247196 | 0.475549845 | 0.741068157 |
| chr1:20696462-20697195 | 0.244389385 | 0.475565983 | 0.741078234 |
| chr14:61235141-61235409 | -0.327263995 | 0.475582889 | 0.741089507 |
| chr1:235368010-235368286 | -0.291780766 | 0.475607075 | 0.741108593 |
| chr3:50321681-50321911 | -0.310090276 | 0.475631301 | 0.741108593 |
| chrX:141643098-141643303 | -0.401549516 | 0.475653457 | 0.741108593 |
| chr2:20232230-20233171 | 0.232965359 | 0.475654106 | 0.741108593 |
| chr17:50026843-50027138 | -0.286850661 | 0.475655767 | 0.741108593 |
| chr7:115085479-115085797 | -0.302711546 | 0.47567177 | 0.741108593 |
| chr2:1823282-1823551 | 0.309553965 | 0.475674032 | 0.741108593 |
| chr1:152060261-152060461 | 0.325169368 | 0.475688919 | 0.741108593 |
| chr5:16935469-16937364 | 0.224588575 | 0.475691001 | 0.741108593 |
| chr2:84906091-84906489 | -0.26239326 | 0.475694253 | 0.741108593 |
| chr2:25851937-25852203 | -0.327678552 | 0.475701533 | 0.741108593 |
| chr16:88764991-88765347 | 0.249280253 | 0.475731848 | 0.741140753 |
| chr14:75246658-75246858 | -0.389364835 | 0.475747589 | 0.741150206 |
| chr8:110452032-110452247 | 0.34034332 | 0.475769085 | 0.741166365 |
| chr3:15617823-15618100 | -0.375146066 | 0.475777307 | 0.741166365 |
| chr3:169149153-169149807 | -0.237419196 | 0.475830146 | 0.741228008 |
| chr22:38278792-38279083 | -0.327567452 | 0.475839822 | 0.741228008 |
| chr1:201793024-201793472 | -0.280698985 | 0.475850658 | 0.741228008 |
| chr14:48536335-48536644 | -0.296732155 | 0.475855573 | 0.741228008 |
| chr1:64564965-64565217 | 0.325001165 | 0.475876088 | 0.741241296 |
| chr17:35804769-35805116 | 0.276733604 | 0.475883451 | 0.741241296 |
| chr1:57425005-57425270 | -0.341499 | 0.475902942 | 0.741252023 |
| chr5:1966006-1966206 | -0.385900923 | 0.475918552 | 0.741252023 |
| chr1:166813956-166814193 | 0.324970699 | 0.475919361 | 0.741252023 |
| chr14:30223347-30223853 | 0.247606308 | 0.475936616 | 0.74126383 |
| chr20:49405559-49405810 | 0.312051345 | 0.475948622 | 0.741267461 |
| chr1:225446092-225446525 | 0.269268808 | 0.475989492 | 0.741308694 |

| | | | |
|---|---|---|---|
| chr20:47991567-47992366 | -0.226381806 | 0.4759979 | 0.741308694 |
| chr13:90838446-90838909 | -0.286151126 | 0.476004121 | 0.741308694 |
| chr1:199431816-199432042 | 0.347834681 | 0.476032482 | 0.741337795 |
| chr6:26056927-26057127 | -0.337050465 | 0.47604497 | 0.741341383 |
| chr21:42202111-42202311 | 0.38908156 | 0.476054137 | 0.741341383 |
| chr16:74005642-74005980 | 0.304509387 | 0.476081083 | 0.741368278 |
| chr5:127909134-127909348 | 0.320192297 | 0.476095058 | 0.741371561 |
| chr17:42154717-42155327 | 0.238049629 | 0.476111869 | 0.741371561 |
| chr14:49852318-49854191 | 0.224509537 | 0.476112219 | 0.741371561 |
| chr15:84563652-84563852 | -0.391883738 | 0.476123522 | 0.741373917 |
| chr15:95259480-95259849 | 0.274501872 | 0.476133083 | 0.741373917 |
| chr20:63747821-63748021 | -0.312276118 | 0.476147276 | 0.74138095 |
| chr9:15306101-15307765 | -0.226762362 | 0.476159165 | 0.741384396 |
| chr17:61837120-61837567 | 0.22978465 | 0.476174873 | 0.741393789 |
| chr3:157352046-157352246 | -0.352387547 | 0.476189949 | 0.741399717 |
| chr1:203027363-203027622 | 0.271661972 | 0.476198033 | 0.741399717 |
| chr2:149017507-149017714 | 0.312042196 | 0.476210641 | 0.741404282 |
| chr14:58832052-58832348 | -0.306534511 | 0.476279879 | 0.74147202 |
| chr5:88883062-88883824 | -0.248541934 | 0.47628005 | 0.74147202 |
| chr1:37448149-37448468 | 0.277869557 | 0.476283181 | 0.74147202 |
| chr1:151839487-151840264 | 0.229434386 | 0.476301772 | 0.741485897 |
| chr9:130940915-130941618 | 0.243403051 | 0.476365199 | 0.741569571 |
| chr1:19882118-19882666 | 0.251059798 | 0.476376129 | 0.74157152 |
| chr10:92612279-92612479 | -0.394863988 | 0.476409349 | 0.741608167 |
| chr10:87203419-87204456 | -0.239607999 | 0.476430073 | 0.74162536 |
| chr2:88669515-88669919 | -0.286662301 | 0.476453331 | 0.74164337 |
| chr16:3500700-3501282 | -0.247367499 | 0.476461002 | 0.74164337 |
| chr5:134632309-134633110 | -0.236948503 | 0.476488704 | 0.741671423 |
| chr6:128124788-128125016 | -0.349617392 | 0.47651205 | 0.741688368 |
| chr2:197356186-197356499 | -0.289845801 | 0.47651895 | 0.741688368 |
| chr2:175180818-175181964 | -0.230763164 | 0.47654561 | 0.741704084 |
| chr17:5255253-5255536 | -0.328175284 | 0.476571824 | 0.741704084 |
| chrX:138287260-138287516 | -0.335594791 | 0.476575798 | 0.741704084 |
| chr3:134352276-134352663 | -0.252759055 | 0.476576202 | 0.741704084 |
| chr3:172623307-172623533 | -0.310819487 | 0.476598379 | 0.741704084 |
| chr2:201118514-201119169 | -0.242834498 | 0.476608547 | 0.741704084 |
| chr16:89700874-89701229 | 0.249353486 | 0.476630949 | 0.741704084 |
| chr2:180763637-180763866 | -0.343080821 | 0.476643265 | 0.741704084 |
| chr22:16917452-16917674 | -0.343080821 | 0.476643265 | 0.741704084 |
| chr13:30752307-30752538 | -0.345762734 | 0.47664362 | 0.741704084 |
| chr8:97266107-97266692 | -0.244369693 | 0.476651632 | 0.741704084 |
| chrX:115561596-115562552 | 0.239776434 | 0.476666041 | 0.741704084 |
| chr2:31011525-31011960 | 0.271092545 | 0.476684826 | 0.741704084 |
| chr20:58822271-58822561 | 0.265149639 | 0.476687573 | 0.741704084 |
| chr7:153379837-153380108 | -0.346759347 | 0.476696347 | 0.741704084 |
| chr17:59843136-59844459 | -0.22150669 | 0.476705543 | 0.741704084 |
| chr9:130320807-130321134 | 0.292488432 | 0.476718106 | 0.741704084 |
| chr22:21657252-21658370 | 0.235701637 | 0.476727682 | 0.741704084 |
| chr8:87633850-87634179 | 0.293020526 | 0.476730803 | 0.741704084 |
| chr6:99434968-99435236 | -0.30762691 | 0.476743583 | 0.741704084 |

| | | | |
|---|---|---|---|
| chr20:19628595-19628884 | -0.304914209 | 0.476766693 | 0.741704084 |
| chr6:22363210-22363501 | 0.294226844 | 0.476771182 | 0.741704084 |
| chr3:8962932-8963920 | 0.228530951 | 0.476777519 | 0.741704084 |
| chr20:45342323-45342643 | 0.257768494 | 0.476793916 | 0.741704084 |
| chr13:38866616-38866910 | -0.31222179 | 0.476794912 | 0.741704084 |
| chr12:115872854-115873397 | 0.239963144 | 0.47681219 | 0.741704084 |
| chr9:96566454-96567481 | -0.239881265 | 0.476825361 | 0.741704084 |
| chr6:151763797-151764592 | 0.232332478 | 0.476827529 | 0.741704084 |
| chr6:121048549-121048934 | -0.298646077 | 0.476834469 | 0.741704084 |
| chr8:38975393-38975630 | -0.353505868 | 0.47683572 | 0.741704084 |
| chr2:9644624-9645076 | 0.259460859 | 0.476837932 | 0.741704084 |
| chr2:20452328-20452880 | -0.246978659 | 0.47687284 | 0.741704084 |
| chr5:142507576-142507839 | -0.329386744 | 0.476881966 | 0.741704084 |
| chr4:107094353-107094820 | -0.277944194 | 0.476884665 | 0.741704084 |
| chr1:193651600-193651888 | 0.341594753 | 0.476899183 | 0.741704084 |
| chr2:220228624-220228934 | -0.292885596 | 0.476899868 | 0.741704084 |
| chr20:740228-740435 | 0.330734138 | 0.476910277 | 0.741704084 |
| chr11:62621347-62622495 | 0.225987458 | 0.476922637 | 0.741704084 |
| chr2:105919952-105920152 | -0.357596933 | 0.476929363 | 0.741704084 |
| chr9:4315639-4315946 | 0.299302521 | 0.476940208 | 0.741704084 |
| chr3:158917978-158918270 | -0.284234177 | 0.476946126 | 0.741704084 |
| chr20:9359189-9359389 | -0.411794806 | 0.476948163 | 0.741704084 |
| chr4:148705510-148705712 | 0.356603981 | 0.476952512 | 0.741704084 |
| chrX:103464142-103465108 | -0.233457994 | 0.476959016 | 0.741704084 |
| chr1:110209968-110210519 | -0.251243832 | 0.476964649 | 0.741704084 |
| chr1:110585536-110585736 | 0.388300658 | 0.47700236 | 0.741747673 |
| chr2:10120163-10120807 | -0.232820458 | 0.477016142 | 0.741751914 |
| chr2:207374927-207375138 | -0.379525294 | 0.477024449 | 0.741751914 |
| chr11:118916908-118917145 | 0.264908113 | 0.477061652 | 0.741783397 |
| chr1:147496655-147496924 | -0.296763026 | 0.47707 | 0.741783397 |
| chr1:169858619-169858819 | 0.363199536 | 0.477073739 | 0.741783397 |
| chr3:72738608-72739380 | 0.24488956 | 0.477085641 | 0.741785372 |
| chr12:13127486-13127763 | 0.315285106 | 0.477099708 | 0.741785372 |
| chr5:57184398-57184831 | -0.26560464 | 0.477104053 | 0.741785372 |
| chr10:98028256-98028524 | -0.298957922 | 0.4771277 | 0.741794737 |
| chr16:89615464-89615697 | -0.322757771 | 0.477129438 | 0.741794737 |
| chr17:62870206-62870517 | -0.273189431 | 0.477160079 | 0.741820805 |
| chr2:197452922-197454081 | -0.223128958 | 0.477165569 | 0.741820805 |
| chr1:221741772-221743387 | -0.223432321 | 0.477179195 | 0.741826938 |
| chr2:112600041-112600373 | -0.284716835 | 0.477265202 | 0.741945591 |
| chr2:31365600-31365800 | -0.388466619 | 0.477380811 | 0.74209716 |
| chr22:21629103-21629592 | 0.244761114 | 0.47738207 | 0.74209716 |
| chr6:138773091-138774141 | -0.226281579 | 0.477419772 | 0.742140711 |
| chr2:86221038-86221443 | -0.270782574 | 0.477429494 | 0.742140767 |
| chr19:40831913-40832320 | 0.283709367 | 0.477492277 | 0.742223302 |
| chr15:58467087-58467551 | -0.26483599 | 0.47754485 | 0.742286457 |
| chr12:69595720-69595982 | -0.289883741 | 0.477552281 | 0.742286457 |
| chr1:70205080-70206114 | -0.230286565 | 0.477601341 | 0.742347653 |
| chr1:235067464-235068182 | 0.235488712 | 0.477642561 | 0.742396663 |
| chr3:4985921-4986380 | -0.249125813 | 0.477660421 | 0.742400099 |

| | | | |
|---|---|---|---|
| chr12:61768966-61769332 | 0.323128363 | 0.477671663 | 0.742400099 |
| chr7:33050800-33051093 | 0.289592047 | 0.477673839 | 0.742400099 |
| chr5:25134260-25134470 | -0.373569321 | 0.477771763 | 0.74250282 |
| chr14:101008801-101009060 | 0.334229599 | 0.477776962 | 0.74250282 |
| chr6:54847455-54847739 | -0.275359821 | 0.477779637 | 0.74250282 |
| chr8:75619604-75619804 | 0.412772517 | 0.477798785 | 0.74250282 |
| chr6:151649079-151649324 | 0.357803094 | 0.477812593 | 0.74250282 |
| chr15:43133170-43134179 | 0.236204946 | 0.477813869 | 0.74250282 |
| chr2:68722818-68723063 | -0.31804115 | 0.477828302 | 0.74250282 |
| chr14:96391251-96392397 | 0.225894399 | 0.477830817 | 0.74250282 |
| chrX:110665733-110665993 | 0.319943191 | 0.477832623 | 0.74250282 |
| chr1:41432752-41433357 | -0.252132286 | 0.477844108 | 0.74250282 |
| chr19:5142335-5142590 | -0.347168591 | 0.477846527 | 0.74250282 |
| chr1:144567320-144568065 | -0.247139465 | 0.477884117 | 0.742546171 |
| chr12:122063192-122063775 | 0.249032135 | 0.477932567 | 0.742576515 |
| chrX:111680938-111681980 | 0.225959867 | 0.47794226 | 0.742576515 |
| chrX:107777013-107777638 | -0.241354758 | 0.477942365 | 0.742576515 |
| chr3:69120938-69121245 | -0.31799863 | 0.477942411 | 0.742576515 |
| chr1:40257560-40258545 | -0.230614796 | 0.477959167 | 0.742583288 |
| chr2:175028161-175028691 | -0.266925677 | 0.477966153 | 0.742583288 |
| chr16:15055893-15056643 | 0.235573047 | 0.477988805 | 0.742589173 |
| chr4:90357197-90357418 | 0.329953686 | 0.477989324 | 0.742589173 |
| chr3:196142280-196143265 | 0.246648719 | 0.478011155 | 0.742599154 |
| chr4:69983991-69984492 | -0.268918979 | 0.478015132 | 0.742599154 |
| chr13:82627100-82627601 | -0.270759402 | 0.47802619 | 0.742600492 |
| chr5:41519788-41520188 | -0.293809036 | 0.47803829 | 0.742600492 |
| chr2:46616457-46617517 | -0.227907004 | 0.478045069 | 0.742600492 |
| chr14:103921033-103921898 | 0.232570796 | 0.478097414 | 0.742666749 |
| chr20:48075844-48076432 | 0.231392108 | 0.478150238 | 0.742733747 |
| chr3:141303438-141303722 | -0.298820223 | 0.478178302 | 0.742747394 |
| chr1:39575702-39575939 | 0.30520001 | 0.478178411 | 0.742747394 |
| chr20:58087122-58087593 | 0.250073773 | 0.478196718 | 0.742758256 |
| chr5:43041801-43042313 | 0.241969445 | 0.478204791 | 0.742758256 |
| chr1:17053713-17054529 | 0.23307809 | 0.478221961 | 0.742769867 |
| chr13:36894701-36894901 | -0.400007115 | 0.478284003 | 0.742800598 |
| chr5:21802706-21802965 | 0.317397565 | 0.47830068 | 0.742800598 |
| chr1:237913437-237913667 | -0.345654372 | 0.478316026 | 0.742800598 |
| chr3:42099847-42100149 | 0.295791772 | 0.478316285 | 0.742800598 |
| chr4:27861392-27861709 | 0.286893439 | 0.478341193 | 0.742800598 |
| chr6:136326065-136326302 | -0.341663841 | 0.478342031 | 0.742800598 |
| chr10:95042860-95043091 | -0.38709661 | 0.478343185 | 0.742800598 |
| chr16:13874039-13874244 | -0.38709661 | 0.478343185 | 0.742800598 |
| chr16:46746991-46747191 | -0.38709661 | 0.478343185 | 0.742800598 |
| chr1:159893096-159893296 | -0.356292511 | 0.478347827 | 0.742800598 |
| chr4:78565978-78566292 | 0.278916998 | 0.478348384 | 0.742800598 |
| chr12:116863275-116863555 | 0.285539574 | 0.478414716 | 0.74287639 |
| chrX:149903994-149904455 | 0.291353114 | 0.478419557 | 0.74287639 |
| chr6:163199579-163199965 | 0.283076053 | 0.478446786 | 0.74287639 |
| chr12:73685278-73685478 | 0.385614363 | 0.478451834 | 0.74287639 |
| chr1:114418180-114418429 | 0.264274886 | 0.478463793 | 0.74287639 |

| | | | |
|---|---|---|---|
| chr14:26531995-26532449 | 0.274016028 | 0.478465044 | 0.74287639 |
| chr1:149848704-149849173 | -0.253159374 | 0.478465061 | 0.74287639 |
| chr18:36120511-36120711 | 0.354756223 | 0.478494678 | 0.742893263 |
| chr16:53780162-53780362 | 0.37662419 | 0.478495319 | 0.742893263 |
| chr15:66252409-66252903 | 0.265469427 | 0.47850532 | 0.742893737 |
| chr7:34064548-34064935 | 0.260828443 | 0.478553763 | 0.742948758 |
| chr10:99620317-99621716 | -0.222008111 | 0.478574124 | 0.742948758 |
| chr16:81403528-81403926 | 0.247755429 | 0.478577093 | 0.742948758 |
| chr22:29837951-29838571 | -0.239567557 | 0.478579544 | 0.742948758 |
| chr7:6857559-6858095 | -0.245789122 | 0.478607645 | 0.74297136 |
| chr2:27723329-27723632 | -0.28387781 | 0.478633799 | 0.74297136 |
| chr9:33722003-33722500 | -0.271111014 | 0.478634212 | 0.74297136 |
| chr12:106357485-106358640 | -0.22552907 | 0.478642022 | 0.74297136 |
| chr3:152417319-152417568 | -0.360612549 | 0.478642586 | 0.74297136 |
| chr2:188983277-188983675 | -0.380895978 | 0.478667941 | 0.74297719 |
| chr20:12928274-12928505 | 0.366473777 | 0.478671543 | 0.74297719 |
| chr15:58440907-58441172 | 0.326554591 | 0.478675433 | 0.74297719 |
| chr16:23640976-23641699 | 0.231924183 | 0.478708354 | 0.743013238 |
| chr22:46250069-46250842 | 0.252101535 | 0.478735086 | 0.743039677 |
| chr17:6640135-6641159 | -0.235989034 | 0.478759912 | 0.743063158 |
| chr16:27737965-27738348 | -0.287536958 | 0.478793589 | 0.743100374 |
| chr17:71854936-71855206 | 0.334979338 | 0.478808908 | 0.743109098 |
| chr1:188499783-188500035 | 0.306347869 | 0.478818738 | 0.743109301 |
| chr9:112819846-112820203 | 0.29168257 | 0.478834652 | 0.743118948 |
| chr11:129703800-129704141 | -0.309930624 | 0.4788823 | 0.743170057 |
| chr10:86252-86555 | 0.311808663 | 0.478893516 | 0.743170057 |
| chr1:45339532-45340788 | -0.230001116 | 0.478896682 | 0.743170057 |
| chr2:38076047-38077509 | -0.222836909 | 0.478939398 | 0.743221293 |
| chr10:96017260-96017644 | -0.278217088 | 0.478960942 | 0.743239672 |
| chr2:169716006-169716207 | -0.344862248 | 0.478988265 | 0.743251966 |
| chr3:122101173-122101395 | -0.344862248 | 0.478988265 | 0.743251966 |
| chr14:55129067-55130494 | 0.227583144 | 0.479036388 | 0.743311587 |
| chr2:158034324-158034575 | -0.301674759 | 0.479096824 | 0.743390309 |
| chr20:56995661-56996130 | 0.241669675 | 0.479118937 | 0.743409566 |
| chr1:41450792-41451163 | -0.274911771 | 0.479143912 | 0.743409654 |
| chr19:47348854-47349476 | 0.233714242 | 0.479146913 | 0.743409654 |
| chr3:120802475-120802675 | 0.356523863 | 0.479148101 | 0.743409654 |
| chr9:98445523-98445857 | -0.311319518 | 0.479223804 | 0.743512054 |
| chrX:109952137-109952381 | -0.334715429 | 0.479252429 | 0.743529585 |
| chr10:104045476-104045856 | -0.273514384 | 0.479254511 | 0.743529585 |
| chr1:184222603-184222924 | -0.269419169 | 0.479282205 | 0.743557495 |
| chr5:67617113-67617358 | 0.34612882 | 0.479325031 | 0.743593693 |
| chr1:41282737-41282968 | -0.318717797 | 0.479332687 | 0.743593693 |
| chr22:44735881-44736086 | 0.328768442 | 0.479334652 | 0.743593693 |
| chr3:100603507-100603858 | 0.267259666 | 0.479376467 | 0.743643505 |
| chrX:71118416-71119100 | -0.250445186 | 0.479391478 | 0.743651736 |
| chr11:113895447-113895654 | -0.368375766 | 0.479437828 | 0.743680426 |
| chr4:37988688-37988962 | 0.336182017 | 0.479438029 | 0.743680426 |
| chr1:94061490-94061760 | 0.29068116 | 0.479439091 | 0.743680426 |
| chr17:82235976-82236244 | 0.269414633 | 0.479461739 | 0.743685278 |

| | | | |
|---|---|---|---|
| chr17:59989162-59989362 | 0.326101464 | 0.479487321 | 0.743685278 |
| chrX:8653918-8654327 | -0.260394203 | 0.4794984 | 0.743685278 |
| chr17:42618571-42618878 | 0.292956365 | 0.479531625 | 0.743685278 |
| chr6:117712808-117713008 | 0.431109641 | 0.479537904 | 0.743685278 |
| chr20:19291148-19291528 | 0.304632411 | 0.479540785 | 0.743685278 |
| chr4:49032464-49032709 | -0.347213227 | 0.479543331 | 0.743685278 |
| chr8:56604323-56604620 | 0.296150733 | 0.479549092 | 0.743685278 |
| chr17:5014888-5015152 | 0.370339103 | 0.479551742 | 0.743685278 |
| chr20:19283565-19283918 | 0.307737631 | 0.479561472 | 0.743685278 |
| chr1:170074253-170075033 | -0.224854659 | 0.479567535 | 0.743685278 |
| chr1:86913974-86914961 | -0.226877575 | 0.479567923 | 0.743685278 |
| chr3:188152801-188153363 | 0.239828038 | 0.479568395 | 0.743685278 |
| chr10:100529590-10053019€ | -0.247614826 | 0.479584884 | 0.743694872 |
| chr4:70576775-70577094 | -0.35377136 | 0.479600683 | 0.743694872 |
| chr10:122063142-12206371 | 0.245288063 | 0.479614199 | 0.743694872 |
| chr20:58829547-58829775 | 0.296894922 | 0.479617245 | 0.743694872 |
| chr2:139018394-139018640 | -0.365392432 | 0.479623112 | 0.743694872 |
| chr9:129036036-129037021 | 0.229439622 | 0.479649004 | 0.743719968 |
| chr12:121095971-12109617 | 0.344123493 | 0.479677457 | 0.743736674 |
| chr4:143711609-143712101 | -0.303877145 | 0.479679191 | 0.743736674 |
| chr11:65888106-65888790 | 0.231329958 | 0.479690504 | 0.743739164 |
| chr14:92379626-92379826 | 0.333015114 | 0.479720081 | 0.743761922 |
| chr9:113620531-113620884 | 0.284642153 | 0.479732206 | 0.743761922 |
| chr12:122225529-12222667 | 0.223344728 | 0.479734302 | 0.743761922 |
| chr1:181424691-181425421 | -0.234719128 | 0.479779192 | 0.743816466 |
| chr14:60091229-60092512 | -0.224142067 | 0.479816557 | 0.743848164 |
| chr1:224182861-224184020 | 0.228328801 | 0.479827814 | 0.743848164 |
| chr22:41666680-41667156 | -0.270900175 | 0.479829918 | 0.743848164 |
| chr8:81025501-81025921 | -0.267612897 | 0.479838484 | 0.743848164 |
| chr6:34248396-34249338 | 0.233617207 | 0.479848177 | 0.743848164 |
| chr14:68979792-68980060 | 0.265089631 | 0.479869941 | 0.743853369 |
| chr3:136644823-136645034 | 0.367903588 | 0.479879073 | 0.743853369 |
| chr16:3184119-3184381 | 0.273515095 | 0.479894243 | 0.743853369 |
| chr12:25958158-25959444 | -0.236546815 | 0.479899555 | 0.743853369 |
| chr6:153002327-153003249 | -0.227584594 | 0.47990715 | 0.743853369 |
| chr3:48194489-48194824 | -0.319763455 | 0.479916804 | 0.743853369 |
| chr17:60453663-60453883 | 0.25227371 | 0.479921174 | 0.743853369 |
| chr2:131104559-131106104 | -0.226430225 | 0.479961406 | 0.743853369 |
| chr12:53421260-53421884 | 0.238582364 | 0.479962611 | 0.743853369 |
| chr12:49235263-49235464 | 0.349472538 | 0.479965573 | 0.743853369 |
| chr14:99793670-99793893 | 0.349472538 | 0.479965573 | 0.743853369 |
| chr2:187174720-187174924 | -0.37332155 | 0.479976774 | 0.743853369 |
| chr17:58621338-58621718 | 0.239250727 | 0.479977741 | 0.743853369 |
| chr3:4986470-4987395 | 0.227580041 | 0.480032251 | 0.743908359 |
| chr13:49399703-49400183 | 0.255721764 | 0.480035415 | 0.743908359 |
| chr8:11853685-11854365 | -0.242481457 | 0.480061803 | 0.743908359 |
| chr13:32587007-32587422 | -0.256659805 | 0.480065744 | 0.743908359 |
| chr20:50121537-50122148 | -0.242992872 | 0.480070016 | 0.743908359 |
| chrX:106611554-106612500 | -0.236257718 | 0.480071476 | 0.743908359 |
| chr6:107114888-107115276 | -0.272497634 | 0.480096059 | 0.743930545 |

| | | | |
|---|---|---|---|
| chr12:112109270-112109578 | 0.268693012 | 0.480105211 | 0.743930545 |
| chr15:31221981-31222181 | -0.398829212 | 0.480126796 | 0.743944991 |
| chr1:30380082-30380689 | 0.254877567 | 0.480133953 | 0.743944991 |
| chr5:9344792-9345065 | -0.328985875 | 0.480158517 | 0.743966004 |
| chr3:129053438-129053973 | -0.25493399 | 0.480166934 | 0.743966004 |
| chr1:161763531-161763783 | 0.286794066 | 0.480200186 | 0.744002481 |
| chr19:48259989-48260383 | 0.29248587 | 0.480216813 | 0.744013197 |
| chr2:217785505-217785990 | 0.257596765 | 0.480253389 | 0.74405482 |
| chr3:25593986-25594335 | 0.272603333 | 0.480278302 | 0.744067561 |
| chr19:6753722-6753995 | 0.32806248 | 0.480281034 | 0.744067561 |
| chr5:892492-893351 | 0.235912898 | 0.480313101 | 0.744102194 |
| chr18:3261399-3263089 | 0.222280562 | 0.480409733 | 0.74423685 |
| chr3:157411838-157412102 | 0.344647919 | 0.480424433 | 0.744244575 |
| chrX:86806829-86807039 | -0.369069294 | 0.480454704 | 0.744276421 |
| chr6:27575970-27576343 | -0.282547731 | 0.480472078 | 0.744288287 |
| chr12:89256023-89256344 | 0.296892261 | 0.480514582 | 0.744327103 |
| chr22:20144170-20144384 | 0.370909557 | 0.480516564 | 0.744327103 |
| chr10:6918367-6918567 | 0.326624902 | 0.48054733 | 0.744359713 |
| chr2:1564196-1564406 | -0.358250795 | 0.480561393 | 0.744366447 |
| chr6:36130431-36131036 | -0.263364182 | 0.480584379 | 0.744387004 |
| chr2:42132778-42133507 | 0.240789894 | 0.480614886 | 0.744407115 |
| chr17:60171083-60171698 | 0.226155259 | 0.480616794 | 0.744407115 |
| chr1:208245269-208245522 | -0.296965871 | 0.480649989 | 0.744430493 |
| chr8:121176632-121176908 | -0.286680787 | 0.480651537 | 0.744430493 |
| chrX:69503519-69504669 | 0.229715032 | 0.480661034 | 0.744430493 |
| chr1:904638-904847 | 0.336698967 | 0.480681209 | 0.744446691 |
| chr5:138222812-138223157 | 0.292808526 | 0.480714095 | 0.744467527 |
| chr6:6676866-6677148 | 0.292808526 | 0.480714095 | 0.744467527 |
| chr19:3708186-3708924 | -0.250410579 | 0.480758547 | 0.744511441 |
| chr13:21371579-21371779 | 0.366814298 | 0.480761884 | 0.744511441 |
| chr12:132083638-132084744 | 0.227861675 | 0.480778866 | 0.744522692 |
| chr2:95231606-95231944 | -0.287943249 | 0.480799392 | 0.744527751 |
| chr14:103572074-103572299 | 0.322646792 | 0.480801567 | 0.744527751 |
| chr1:27489738-27490740 | 0.228445702 | 0.480825617 | 0.744549946 |
| chr6:142692937-142693251 | -0.313719975 | 0.480843992 | 0.744563352 |
| chr6:12751777-12752046 | -0.333809836 | 0.480885053 | 0.744598083 |
| chr4:147157920-147158617 | 0.254593046 | 0.480885857 | 0.744598083 |
| chr12:100668935-100669253 | -0.263153193 | 0.480933635 | 0.744657014 |
| chr5:15999391-15999591 | 0.376359347 | 0.480986744 | 0.744724197 |
| chr2:223727570-223727795 | -0.368765688 | 0.481035674 | 0.744784905 |
| chr9:89201281-89201557 | 0.357010344 | 0.48109165 | 0.744856521 |
| chr11:21861390-21861738 | -0.336304606 | 0.481127508 | 0.744896987 |
| chr9:127897282-127897773 | 0.249252954 | 0.481175268 | 0.744919911 |
| chr7:14734783-14735488 | -0.2417204 | 0.481176453 | 0.744919911 |
| chr21:23887659-23887916 | -0.352964755 | 0.481177317 | 0.744919911 |
| chr4:17780844-17781194 | -0.321067097 | 0.481181202 | 0.744919911 |
| chr16:14285275-14286406 | 0.225971993 | 0.481199084 | 0.744927337 |
| chr14:20412881-20413681 | 0.22825316 | 0.481205443 | 0.744927337 |
| chr5:149192396-149192613 | -0.368409605 | 0.481229101 | 0.74494891 |
| chr6:126170971-126171452 | 0.294996865 | 0.481266356 | 0.744969159 |

| | | | |
|---|---|---|---|
| chrX:74234957-74235157 | 0.386766147 | 0.481269634 | 0.744969159 |
| chr20:31072654-31072866 | -0.359028073 | 0.481277679 | 0.744969159 |
| chr3:87082868-87083383 | -0.259530101 | 0.481297399 | 0.744969159 |
| chr3:119821025-119821273 | -0.311690055 | 0.481298576 | 0.744969159 |
| chr17:9577166-9577433 | 0.300777717 | 0.481300518 | 0.744969159 |
| chr14:58672636-58672836 | 0.384982587 | 0.48136692 | 0.745042872 |
| chr18:45621009-45621267 | -0.315432757 | 0.481377259 | 0.745042872 |
| chr6:79233532-79234169 | -0.230770087 | 0.48137977 | 0.745042872 |
| chr11:71266291-71266620 | -0.309279189 | 0.481389684 | 0.745042872 |
| chr20:32536305-32537159 | 0.244440441 | 0.481396759 | 0.745042872 |
| chrX:102977210-102977453 | -0.349667653 | 0.48142052 | 0.745057247 |
| chr12:79689837-79690052 | -0.264704059 | 0.481425495 | 0.745057247 |
| chr2:215982388-215983057 | -0.229952077 | 0.481449595 | 0.745079495 |
| chr3:14780898-14781277 | -0.263392278 | 0.48147699 | 0.745106841 |
| chr5:177370451-177371163 | -0.230850124 | 0.48148725 | 0.74510767 |
| chr15:92733923-92734236 | -0.35826715 | 0.481545261 | 0.745182392 |
| chr11:70212439-70212817 | -0.295538872 | 0.481581659 | 0.745220095 |
| chr2:47776065-47776265 | -0.365150375 | 0.481595591 | 0.745220095 |
| chr17:39401500-39401794 | 0.293586455 | 0.481598802 | 0.745220095 |
| chr1:202810499-202811146 | -0.227015613 | 0.481625686 | 0.745228854 |
| chr19:17424978-17425192 | -0.349564005 | 0.481631239 | 0.745228854 |
| chr22:21045480-21046066 | 0.249108799 | 0.481633641 | 0.745228854 |
| chr7:129214046-129214268 | -0.312299795 | 0.481683188 | 0.745290467 |
| chr16:29848497-29849142 | 0.264016132 | 0.481729645 | 0.74531728 |
| chr19:48391377-48391735 | 0.264016132 | 0.481729645 | 0.74531728 |
| chr2:232300999-232301552 | 0.2580064 | 0.481734884 | 0.74531728 |
| chrX:44253409-44253609 | -0.381790757 | 0.481740893 | 0.74531728 |
| chr3:144903913-144904224 | -0.330793683 | 0.481758781 | 0.74531728 |
| chr5:141592100-141592320 | -0.347884911 | 0.481758881 | 0.74531728 |
| chr12:16165516-16165731 | -0.355029326 | 0.481773868 | 0.745321935 |
| chr2:75640584-75640878 | -0.269410721 | 0.481804404 | 0.745321935 |
| chr20:3783107-3783433 | 0.257940277 | 0.481817434 | 0.745321935 |
| chr1:51729042-51730256 | 0.226995627 | 0.481817853 | 0.745321935 |
| chr1:198056270-198056641 | -0.258229074 | 0.481821437 | 0.745321935 |
| chr3:51538191-51538835 | 0.269694028 | 0.481832341 | 0.745321935 |
| chr1:76868099-76868448 | 0.263188655 | 0.481844458 | 0.745321935 |
| chr4:67809577-67809927 | -0.32989102 | 0.481847891 | 0.745321935 |
| chr3:185907376-185907576 | 0.358329146 | 0.481849435 | 0.745321935 |
| chr19:2841079-2841609 | 0.241508642 | 0.481885789 | 0.74536312 |
| chr9:113807459-113807895 | 0.280465969 | 0.481900338 | 0.745370578 |
| chr15:74872863-74873556 | 0.232511078 | 0.481928909 | 0.745386438 |
| chr13:37888681-37888949 | 0.343038524 | 0.481934007 | 0.745386438 |
| chr13:36905080-36905331 | -0.371058485 | 0.481939776 | 0.745386438 |
| chr13:25727385-25727601 | 0.318094969 | 0.481967198 | 0.745409287 |
| chr2:217707105-217707560 | -0.263798753 | 0.481974007 | 0.745409287 |
| chr3:62589299-62589690 | 0.246450998 | 0.482005108 | 0.745427583 |
| chr9:35115849-35116568 | -0.230646483 | 0.482005294 | 0.745427583 |
| chr1:153238528-153238915 | 0.266986391 | 0.482016029 | 0.74542914 |
| chr3:155964050-155964327 | -0.331786166 | 0.482055704 | 0.745459629 |
| chr9:81689463-81690148 | -0.223166449 | 0.48206229 | 0.745459629 |

| | | | |
|---|---|---|---|
| chr10:95601156-95601553 | 0.253615619 | 0.482064931 | 0.745459629 |
| chr13:51452336-51453887 | -0.220925655 | 0.482100432 | 0.745499481 |
| chr10:93289063-93289290 | 0.304317985 | 0.482111716 | 0.745501885 |
| chr5:112707138-112708270 | -0.223472315 | 0.482128902 | 0.745513414 |
| chr5:113971128-113971839 | -0.254265773 | 0.482157508 | 0.745542603 |
| chr3:119034161-119035283 | -0.225459023 | 0.48218436 | 0.745569077 |
| chr7:5425863-5426166 | -0.25011768 | 0.482200311 | 0.745569867 |
| chr12:5078563-5078763 | 0.409076633 | 0.482210665 | 0.745569867 |
| chr5:168291363-168292585 | -0.225289654 | 0.482214063 | 0.745569867 |
| chr12:112891907-112892107 | -0.396796982 | 0.482226944 | 0.745574739 |
| chrX:47802583-47802883 | 0.27171292 | 0.482247728 | 0.745591829 |
| chr2:3030900-3031246 | -0.291002264 | 0.482307503 | 0.745654254 |
| chr1:110116889-110117257 | -0.252485308 | 0.482307568 | 0.745654254 |
| chr1:107529481-107529860 | -0.268479072 | 0.482338382 | 0.74567813 |
| chr14:38277745-38277984 | 0.339317381 | 0.482342475 | 0.74567813 |
| chr2:25673325-25674088 | -0.239957927 | 0.482384513 | 0.745728072 |
| chr17:67958015-67958245 | 0.338994311 | 0.482397246 | 0.745732711 |
| chr5:176388470-176389565 | -0.221795984 | 0.482409875 | 0.745737188 |
| chr7:26081715-26082027 | 0.293649684 | 0.482446471 | 0.74576924 |
| chr4:9495497-9495794 | 0.313355689 | 0.482459736 | 0.74576924 |
| chr6:26032493-26032707 | -0.328122147 | 0.482459809 | 0.74576924 |
| chr16:3458356-3458774 | 0.26874952 | 0.4824734 | 0.745775203 |
| chr20:47725309-47725899 | 0.221470575 | 0.482511733 | 0.745819411 |
| chr6:109344196-109344571 | 0.26928816 | 0.482526592 | 0.745827333 |
| chr12:2284835-2285068 | 0.325019549 | 0.48260696 | 0.745936508 |
| chr5:175501922-175502296 | -0.271607908 | 0.48262952 | 0.745955302 |
| chr1:159854112-159855069 | 0.234253865 | 0.482638591 | 0.745955302 |
| chr10:60048176-60048469 | -0.319960481 | 0.48265123 | 0.74595979 |
| chr14:34301225-34301425 | 0.347333806 | 0.482674285 | 0.745968791 |
| chr12:100828013-100828384 | -0.323515119 | 0.482695312 | 0.745968791 |
| chr12:107903572-107904225 | -0.244002421 | 0.482704163 | 0.745968791 |
| chr3:7034758-7034981 | -0.352863062 | 0.482707884 | 0.745968791 |
| chr7:48180022-48180233 | -0.352863062 | 0.482707884 | 0.745968791 |
| chr2:42342445-42343162 | -0.234268774 | 0.482715468 | 0.745968791 |
| chr3:130699836-130700105 | -0.306926674 | 0.482729964 | 0.745976147 |
| chr17:48080244-48080500 | -0.294776671 | 0.482753734 | 0.745996197 |
| chr5:34421261-34421980 | -0.277572994 | 0.48276241 | 0.745996197 |
| chr16:89994976-89995247 | -0.314417757 | 0.482796052 | 0.74599622 |
| chr1:198297610-198297810 | 0.36582271 | 0.482796314 | 0.74599622 |
| chr2:85603305-85603533 | -0.277973411 | 0.482801265 | 0.74599622 |
| chr21:42309485-42309909 | -0.259674282 | 0.482811088 | 0.74599622 |
| chr3:126259485-126259735 | -0.304053487 | 0.482811106 | 0.74599622 |
| chr10:28250818-28251037 | 0.362341685 | 0.482821489 | 0.74599722 |
| chr2:106269928-106270136 | 0.333766108 | 0.482844206 | 0.746017277 |
| chr19:47273438-47274941 | 0.221944987 | 0.482875951 | 0.74605128 |
| chr14:80003164-80003478 | 0.299685599 | 0.482897516 | 0.746051827 |
| chr16:82138482-82139428 | -0.233627284 | 0.482899184 | 0.746051827 |
| chr9:109289977-109290377 | -0.260745885 | 0.482905516 | 0.746051827 |
| chr13:47385056-47385634 | 0.254175239 | 0.482936461 | 0.746084592 |
| chr5:57862320-57862980 | -0.229858219 | 0.482960333 | 0.746106429 |

| | | | |
|---|---|---|---|
| chr3:126407758-126407977 | 0.32209858 | 0.482998036 | 0.746149631 |
| chr6:115882232-115882516 | 0.339041515 | 0.48302463 | 0.746152312 |
| chr11:12288638-12288879 | 0.29462782 | 0.483026761 | 0.746152312 |
| chr14:21098613-21098844 | 0.291237834 | 0.483032735 | 0.746152312 |
| chr1:4323156-4323396 | -0.388396551 | 0.483038724 | 0.746152312 |
| chr13:100392818-100393018 | -0.378922762 | 0.483078729 | 0.746182243 |
| chr13:47512812-47513031 | 0.327488783 | 0.483085854 | 0.746182243 |
| chr20:33651081-33651364 | -0.282609814 | 0.483095203 | 0.746182243 |
| chr8:93769378-93769578 | 0.359888188 | 0.483105867 | 0.746182243 |
| chr7:84194954-84195154 | -0.36185984 | 0.483106793 | 0.746182243 |
| chr3:129360764-129360970 | 0.339451297 | 0.483169712 | 0.746264382 |
| chr19:14072002-14072392 | 0.269475953 | 0.483188961 | 0.746279068 |
| chr2:70248154-70249174 | -0.222293792 | 0.483229214 | 0.746313928 |
| chr15:55587707-55587907 | 0.341345536 | 0.483240061 | 0.746313928 |
| chr1:81670352-81670580 | 0.336564293 | 0.483240752 | 0.746313928 |
| chr3:8492146-8492346 | -0.356817897 | 0.483279923 | 0.746359379 |
| chr3:481746-482013 | 0.292314922 | 0.483316245 | 0.746397981 |
| chr4:100093441-100093824 | -0.250024198 | 0.483324401 | 0.746397981 |
| chr2:73779558-73780385 | -0.233840191 | 0.483336014 | 0.746400872 |
| chr3:119301064-119301264 | -0.379295299 | 0.483359334 | 0.746408578 |
| chr6:114580855-114581459 | 0.276923358 | 0.483360487 | 0.746408578 |
| chr6:153767070-153767488 | -0.30720996 | 0.483405959 | 0.746463752 |
| chr17:49695144-49695742 | 0.250027948 | 0.483457856 | 0.746497395 |
| chr11:75706290-75706665 | 0.254046643 | 0.483471765 | 0.746497395 |
| chr6:31308606-31308879 | 0.316955729 | 0.483475153 | 0.746497395 |
| chr2:120442854-120443309 | 0.241280757 | 0.483486355 | 0.746497395 |
| chr19:37906068-37907376 | 0.226956547 | 0.483486839 | 0.746497395 |
| chr2:113700183-113700738 | 0.261312208 | 0.483487309 | 0.746497395 |
| chr22:37198476-37198805 | -0.272882467 | 0.483495944 | 0.746497395 |
| chr10:110823504-110823841 | 0.266628398 | 0.483515656 | 0.746512787 |
| chr5:18010111-18010355 | -0.339515678 | 0.483539528 | 0.7465346 |
| chr14:104856193-104856396 | 0.329966891 | 0.483574572 | 0.746554754 |
| chr12:79635025-79635321 | 0.323141769 | 0.483581853 | 0.746554754 |
| chr6:124712180-124712923 | -0.242145269 | 0.483590213 | 0.746554754 |
| chr10:24516522-24516722 | 0.396746502 | 0.483591555 | 0.746554754 |
| chr6:42226849-42227209 | -0.275517129 | 0.48364373 | 0.746595715 |
| chr10:97999835-98000100 | -0.352988632 | 0.483666975 | 0.746595715 |
| chr11:70569025-70569401 | -0.299811459 | 0.483675438 | 0.746595715 |
| chr1:232260876-232261218 | -0.274570529 | 0.483678822 | 0.746595715 |
| chr5:102910375-102910625 | -0.358222289 | 0.483682471 | 0.746595715 |
| chr9:134325839-134326313 | 0.253102627 | 0.483706494 | 0.746595715 |
| chr17:72176923-72177207 | 0.279274999 | 0.483720395 | 0.746595715 |
| chr3:64578474-64578683 | 0.252796935 | 0.483720782 | 0.746595715 |
| chr14:77045621-77045821 | 0.331239334 | 0.483727441 | 0.746595715 |
| chr2:238320178-238320613 | 0.249430528 | 0.483753447 | 0.746595715 |
| chr10:78109284-78109552 | -0.37392068 | 0.483758871 | 0.746595715 |
| chr12:75334275-75334812 | -0.23321531 | 0.483763419 | 0.746595715 |
| chr2:132268068-132268906 | -0.230629011 | 0.483767218 | 0.746595715 |
| chr2:18154897-18155097 | -0.346761429 | 0.483768546 | 0.746595715 |
| chr4:125751205-125751474 | -0.309703225 | 0.483771279 | 0.746595715 |

| | | | |
|---|---|---|---|
| chr17:79844136-79844359 | 0.296923135 | 0.48377399 | 0.746595715 |
| chr17:7687247-7687653 | 0.266722774 | 0.483804982 | 0.746619946 |
| chr20:5073007-5073598 | 0.257396856 | 0.483809179 | 0.746619946 |
| chr6:54960846-54961334 | -0.330976397 | 0.483819637 | 0.746621048 |
| chr1:153460635-153460884 | -0.325353494 | 0.483830094 | 0.746622147 |
| chr11:7571691-7572006 | 0.271354513 | 0.483845411 | 0.746630746 |
| chr8:95024443-95025419 | 0.222958692 | 0.48386317 | 0.746643114 |
| chr11:64316963-64317172 | 0.288179789 | 0.483874864 | 0.746645059 |
| chr14:20430532-20430737 | 0.339513446 | 0.483891995 | 0.746645059 |
| chr13:19888809-19889163 | 0.307471165 | 0.483893664 | 0.746645059 |
| chr6:31775251-31775792 | 0.255475424 | 0.483908799 | 0.746653376 |
| chr3:46909679-46909879 | 0.321482214 | 0.483919622 | 0.746653518 |
| chr5:122858530-122858756 | 0.348170801 | 0.48392838 | 0.746653518 |
| chr1:26285935-26286156 | 0.334194651 | 0.483960157 | 0.746687511 |
| chr10:11898259-11899005 | -0.24101548 | 0.484013994 | 0.746755537 |
| chr9:133742995-133743351 | -0.254748097 | 0.484035828 | 0.746774187 |
| chr18:63237446-63237740 | -0.29882584 | 0.48407251 | 0.746801993 |
| chr22:50562598-50563403 | 0.239826982 | 0.484073344 | 0.746801993 |
| chr18:34729344-34729579 | -0.300450406 | 0.484086821 | 0.746807748 |
| chr20:54228841-54229129 | 0.242196301 | 0.484126192 | 0.746839536 |
| chr17:76725930-76727365 | 0.223190744 | 0.484131241 | 0.746839536 |
| chr6:10555495-10556285 | -0.265019571 | 0.484136668 | 0.746839536 |
| chr2:121038924-121039150 | -0.356050082 | 0.484172375 | 0.746879582 |
| chr5:146409337-146409712 | -0.308645988 | 0.484199669 | 0.746890298 |
| chr12:47759473-47759692 | 0.305429478 | 0.484206656 | 0.746890298 |
| chr6:114058540-114058877 | -0.3190073 | 0.484208565 | 0.746890298 |
| chrX:49059271-49060085 | 0.242643519 | 0.484247639 | 0.746934213 |
| chr11:31341021-31341272 | 0.321137937 | 0.484257609 | 0.746934213 |
| chr10:87214749-87214958 | 0.366287234 | 0.484275783 | 0.746934213 |
| chr13:95430686-95430886 | 0.333937144 | 0.484290172 | 0.746934213 |
| chr5:133111246-133111550 | 0.277617377 | 0.484304114 | 0.746934213 |
| chr1:200409540-200410915 | -0.221262826 | 0.484306118 | 0.746934213 |
| chr4:138090439-138090802 | 0.270755234 | 0.484312568 | 0.746934213 |
| chr5:12014622-12014832 | -0.390963111 | 0.484315021 | 0.746934213 |
| chr3:73624588-73625611 | -0.230764349 | 0.484328998 | 0.746940734 |
| chr2:38115289-38115600 | -0.28932959 | 0.484347058 | 0.746953552 |
| chr5:54985349-54985649 | 0.308974172 | 0.484369747 | 0.746973508 |
| chr4:24396066-24396346 | 0.31091884 | 0.484383787 | 0.746980126 |
| chr20:10427140-10427393 | 0.309094459 | 0.484454489 | 0.747058808 |
| chr1:86915109-86915512 | -0.259884657 | 0.484457693 | 0.747058808 |
| chr17:12928778-12929066 | -0.316387918 | 0.484464059 | 0.747058808 |
| chr3:47475463-47476269 | 0.232868233 | 0.48448353 | 0.747071553 |
| chr4:133668366-133668639 | 0.325036629 | 0.484491824 | 0.747071553 |
| chr1:205301231-205301483 | -0.27676977 | 0.48450956 | 0.747083867 |
| chr20:46880955-46881176 | 0.346481683 | 0.484522373 | 0.747088589 |
| chr14:99407642-99407887 | -0.366724668 | 0.484548324 | 0.747113569 |
| chr2:168293824-168294070 | -0.331810449 | 0.484593847 | 0.747151844 |
| chr3:79032791-79033043 | 0.311826169 | 0.484594317 | 0.747151844 |
| chr5:29367500-29367754 | 0.32099535 | 0.484602401 | 0.747151844 |
| chr1:44793570-44794092 | -0.250207998 | 0.484637193 | 0.747172958 |

| | | | |
|---|---|---|---|
| chr17:3048708-3049483 | -0.234785535 | 0.484637533 | 0.747172958 |
| chr3:113054520-113054720 | -0.364176502 | 0.48464535 | 0.747172958 |
| chr5:10481673-10481966 | 0.267952731 | 0.484701943 | 0.747237606 |
| chr10:511036-511650 | -0.235037561 | 0.484715008 | 0.747237606 |
| chr1:117599801-117600171 | 0.295501637 | 0.484721732 | 0.747237606 |
| chr3:88807617-88807946 | 0.324820312 | 0.484726292 | 0.747237606 |
| chr11:76868595-76868795 | -0.37756989 | 0.484739331 | 0.747242672 |
| chr3:33980345-33980735 | 0.272945373 | 0.484753833 | 0.747249994 |
| chr2:101307925-101309172 | 0.225724003 | 0.484796131 | 0.747289657 |
| chr2:131659064-131659312 | 0.370967608 | 0.484799069 | 0.747289657 |
| chr1:219312746-219312976 | -0.324627409 | 0.48481237 | 0.747295126 |
| chr1:93681462-93682086 | 0.259388036 | 0.484833031 | 0.747311939 |
| chr17:61356371-61357227 | 0.218534142 | 0.484867515 | 0.747349973 |
| chr5:136133526-136133917 | -0.267297337 | 0.48489466 | 0.747349973 |
| chr1:41868250-41868809 | -0.260574662 | 0.484895793 | 0.747349973 |
| chr3:192130503-192130880 | -0.27023216 | 0.484903141 | 0.747349973 |
| chr3:183886760-183886986 | 0.302953191 | 0.484910051 | 0.747349973 |
| chr9:5818085-5818285 | -0.370876882 | 0.484916862 | 0.747349973 |
| chr14:94026084-94026539 | 0.237620906 | 0.484925982 | 0.747349973 |
| chr1:152185502-152185920 | 0.241233169 | 0.484940955 | 0.747355615 |
| chr10:106624844-106625161 | 0.275041703 | 0.484962135 | 0.747355615 |
| chr7:74993875-74994296 | 0.261836039 | 0.484979491 | 0.747355615 |
| chr1:172609565-172609857 | 0.283214947 | 0.484982098 | 0.747355615 |
| chr16:31202696-31202906 | 0.312707155 | 0.484992416 | 0.747355615 |
| chr14:99320149-99320605 | -0.266258479 | 0.484998245 | 0.747355615 |
| chr10:35640925-35641395 | -0.256497026 | 0.485007068 | 0.747355615 |
| chr1:6437039-6437299 | 0.305643679 | 0.485009427 | 0.747355615 |
| chr3:153042848-153043605 | -0.238293934 | 0.485034037 | 0.747355615 |
| chr15:45457408-45457835 | -0.275594439 | 0.485036453 | 0.747355615 |
| chr19:9140302-9140502 | 0.35652698 | 0.485036935 | 0.747355615 |
| chr14:96146043-96146579 | 0.257197916 | 0.485047337 | 0.747356614 |
| chr1:154329365-154329565 | 0.306567716 | 0.485080018 | 0.747381293 |
| chr2:152175318-152176666 | -0.221573416 | 0.485087174 | 0.747381293 |
| chr7:66930239-66930697 | -0.274787558 | 0.485099318 | 0.747381293 |
| chr6:20126844-20127050 | 0.320856108 | 0.485102371 | 0.747381293 |
| chr2:170929710-170929982 | -0.258467048 | 0.485139258 | 0.747398425 |
| chr15:79922923-79924230 | 0.220674685 | 0.485148897 | 0.747398425 |
| chr18:64134407-64134681 | 0.341567867 | 0.48515617 | 0.747398425 |
| chr2:35273719-35274093 | -0.340146517 | 0.485166134 | 0.747398425 |
| chr9:90032406-90032655 | -0.353597135 | 0.485170043 | 0.747398425 |
| chr1:191282117-191282317 | -0.37509622 | 0.485179393 | 0.747398425 |
| chr21:32578150-32578509 | -0.289249958 | 0.485181771 | 0.747398425 |
| chr7:151897247-151897769 | -0.260606222 | 0.485229098 | 0.747456303 |
| chr3:143970873-143971130 | -0.248077744 | 0.485277351 | 0.747515604 |
| chr14:85885317-85885835 | 0.2633645 | 0.485290535 | 0.747520885 |
| chrX:74535369-74535580 | 0.319209849 | 0.485332132 | 0.74756993 |
| chr11:13210459-13210664 | -0.338109396 | 0.485361294 | 0.747596127 |
| chr16:65117896-65118316 | 0.267821228 | 0.485368653 | 0.747596127 |
| chr1:181134575-181135321 | 0.22714402 | 0.485378863 | 0.747596824 |
| chr6:66496797-66497014 | 0.365010136 | 0.485423992 | 0.747650825 |

| | | | |
|---|---|---|---|
| chr13:27826339-27826929 | -0.241500351 | 0.485433438 | 0.747650825 |
| chr2:183037212-183039100 | -0.219680704 | 0.485444929 | 0.747653495 |
| chr2:113583563-113584178 | 0.248783956 | 0.485467363 | 0.747671504 |
| chr3:132079779-132079979 | -0.41676199 | 0.485476138 | 0.747671504 |
| chr13:33542164-33543129 | -0.219832847 | 0.485503055 | 0.747697929 |
| chr13:98577120-98577719 | 0.267667729 | 0.485557931 | 0.747767411 |
| chr11:68839223-68839582 | 0.242484402 | 0.485594597 | 0.74777056 |
| chr3:166029669-166029869 | 0.355830313 | 0.485594895 | 0.74777056 |
| chr2:200509234-200510534 | 0.227326246 | 0.485596737 | 0.74777056 |
| chr19:35420290-35420775 | 0.26649154 | 0.485599012 | 0.74777056 |
| chr16:30046233-30046485 | 0.324154794 | 0.485609259 | 0.747771311 |
| chr17:49494737-49495427 | -0.245960351 | 0.485621815 | 0.747775617 |
| chr14:104116301-104117392 | 0.228125907 | 0.485642786 | 0.747778726 |
| chr17:58317431-58318051 | -0.231405215 | 0.485643353 | 0.747778726 |
| chr12:61993779-61994050 | 0.283366507 | 0.48567015 | 0.747799097 |
| chr6:106979490-106979698 | 0.313316798 | 0.485676102 | 0.747799097 |
| chr11:63523192-63523481 | 0.34479916 | 0.485698587 | 0.747808487 |
| chr4:151324827-151325866 | 0.237240317 | 0.48570172 | 0.747808487 |
| chr1:156725816-156726145 | 0.246191841 | 0.485754231 | 0.747846983 |
| chr14:61320792-61322228 | 0.222136868 | 0.485758408 | 0.747846983 |
| chr4:80072393-80073353 | -0.230901999 | 0.485764636 | 0.747846983 |
| chr16:4894845-4895146 | -0.298702404 | 0.485765763 | 0.747846983 |
| chr8:125654188-125654515 | 0.270532814 | 0.485794745 | 0.747876574 |
| chr17:63445491-63447520 | -0.216714419 | 0.485809172 | 0.74787975 |
| chr3:122679130-122679742 | -0.243252746 | 0.48581633 | 0.74787975 |
| chr6:33322746-33323574 | 0.226197879 | 0.485864912 | 0.747939513 |
| chr2:89763374-89763616 | 0.357763903 | 0.485892068 | 0.747966037 |
| chr1:201376808-201377168 | 0.240462687 | 0.48590478 | 0.747966037 |
| chr19:6516016-6516615 | 0.2350696 | 0.485911428 | 0.747966037 |
| chr3:171834280-171834480 | 0.344772728 | 0.48593863 | 0.747968193 |
| chr21:38404200-38404601 | -0.294405504 | 0.485940804 | 0.747968193 |
| chr18:46148830-46149050 | -0.344553706 | 0.485942114 | 0.747968193 |
| chr10:97649025-97649271 | -0.305773766 | 0.485962385 | 0.747984369 |
| chr2:202034444-202035064 | 0.244069645 | 0.48601815 | 0.748055175 |
| chr19:10544238-10544580 | 0.281521871 | 0.486029674 | 0.748057885 |
| chr17:31541637-31541887 | 0.340869007 | 0.486081001 | 0.74809068 |
| chrX:20266480-20266680 | -0.310147664 | 0.486094738 | 0.74809068 |
| chr17:61583099-61583657 | 0.22148562 | 0.486099075 | 0.74809068 |
| chr8:87936081-87936281 | 0.3452487 | 0.486099196 | 0.74809068 |
| chr13:70568658-70568862 | -0.406186931 | 0.486099799 | 0.74809068 |
| chr12:11943971-11944171 | -0.415863415 | 0.486170813 | 0.748184942 |
| chrX:103918106-103918373 | 0.269580124 | 0.486188924 | 0.74818776 |
| chr3:17798099-17798344 | 0.297502352 | 0.486192174 | 0.74818776 |
| chr2:54665465-54665742 | -0.317518792 | 0.486224403 | 0.74822233 |
| chr4:1413015-1413215 | 0.379991834 | 0.486277892 | 0.748289612 |
| chr14:32583693-32583936 | 0.338854488 | 0.486305203 | 0.748316609 |
| chr4:143399322-143399753 | 0.260296552 | 0.486343814 | 0.748358668 |
| chr3:188036385-188037050 | -0.244086738 | 0.486352069 | 0.748358668 |
| chr12:95343240-95343440 | 0.348335313 | 0.486369 | 0.748369691 |
| chr6:139587997-139588593 | 0.248211277 | 0.486406062 | 0.748411689 |

| | | | |
|---|---|---|---|
| chr10:60086232-60086808 | 0.271866989 | 0.486436145 | 0.748433527 |
| chr2:125472061-125472579 | 0.260935401 | 0.486446374 | 0.748433527 |
| chr2:48552272-48552473 | -0.405762505 | 0.486467164 | 0.748433527 |
| chr1:38872801-38873821 | 0.227602182 | 0.486468439 | 0.748433527 |
| chr8:124425292-124426475 | 0.222622276 | 0.486469094 | 0.748433527 |
| chr2:138491219-138491658 | 0.249641192 | 0.486484485 | 0.748442178 |
| chr10:90857636-90858147 | -0.271554302 | 0.486537585 | 0.748508841 |
| chrX:30149201-30149630 | -0.246377606 | 0.486559892 | 0.748528129 |
| chr1:176880966-176881352 | -0.256302487 | 0.486584501 | 0.748550959 |
| chr14:73500504-73500876 | 0.286567743 | 0.486650622 | 0.748615541 |
| chr17:64602497-64602899 | -0.277150765 | 0.486652259 | 0.748615541 |
| chr1:117535089-117535409 | -0.267299323 | 0.486655792 | 0.748615541 |
| chr17:61180410-61180786 | 0.265848371 | 0.486717108 | 0.748648082 |
| chr20:36861948-36862473 | 0.240132304 | 0.486720917 | 0.748648082 |
| chr12:82358080-82359076 | -0.219765418 | 0.486725808 | 0.748648082 |
| chr14:90510230-90510501 | 0.256391118 | 0.486728006 | 0.748648082 |
| chr1:162560710-162562167 | 0.21885393 | 0.486728122 | 0.748648082 |
| chr12:62602326-62603935 | 0.219045396 | 0.48673557 | 0.748648082 |
| chr3:4905366-4905576 | 0.326004311 | 0.486782307 | 0.748668656 |
| chr16:57735167-57736055 | 0.224208336 | 0.48681172 | 0.748668656 |
| chr1:226197105-226197316 | 0.319377952 | 0.486828544 | 0.748668656 |
| chr21:17818382-17819905 | 0.224016804 | 0.486829221 | 0.748668656 |
| chr10:28532167-28533239 | -0.2233896 | 0.486843103 | 0.748668656 |
| chr7:12458571-12458867 | 0.267910593 | 0.486845604 | 0.748668656 |
| chr10:62817817-62818269 | -0.242897587 | 0.486848886 | 0.748668656 |
| chr15:73633051-73633843 | 0.226779282 | 0.486849505 | 0.748668656 |
| chr3:139113449-139113649 | -0.336595225 | 0.486851318 | 0.748668656 |
| chr22:26665583-26665783 | 0.342317489 | 0.486859902 | 0.748668656 |
| chr3:123642339-123642731 | -0.244313278 | 0.486861663 | 0.748668656 |
| chr21:33269103-33269427 | -0.276126322 | 0.486880207 | 0.748668656 |
| chr17:47618532-47618746 | 0.34045194 | 0.48689344 | 0.748668656 |
| chr15:75307860-75308394 | 0.26289367 | 0.486895352 | 0.748668656 |
| chrX:16946223-16947487 | 0.227794279 | 0.48689551 | 0.748668656 |
| chr1:84668278-84668507 | -0.367869707 | 0.486932728 | 0.748710859 |
| chr15:62390143-62390723 | 0.241150977 | 0.486966771 | 0.748715207 |
| chr3:153088255-153088462 | -0.360356595 | 0.486971492 | 0.748715207 |
| chr5:172454060-172454260 | -0.316870923 | 0.486984816 | 0.748715207 |
| chr17:38558192-38558566 | 0.280477422 | 0.486989822 | 0.748715207 |
| chr6:36423109-36424118 | -0.230384059 | 0.4869928 | 0.748715207 |
| chr14:97777674-97777880 | 0.327305448 | 0.486994186 | 0.748715207 |
| chr17:50852090-50852390 | -0.274162879 | 0.487004477 | 0.748716006 |
| chr2:211068502-211068845 | -0.310158188 | 0.487019326 | 0.748720472 |
| chr1:200307549-200307788 | 0.284524642 | 0.487026925 | 0.748720472 |
| chr1:155277100-155277540 | 0.230881011 | 0.48704273 | 0.748729747 |
| chr6:87079056-87079393 | -0.267662922 | 0.487096449 | 0.748762663 |
| chr21:5128353-5128645 | 0.281298394 | 0.487108309 | 0.748762663 |
| chr2:235514594-235514919 | -0.320252799 | 0.48710856 | 0.748762663 |
| chr18:24271315-24272314 | 0.230525904 | 0.487108646 | 0.748762663 |
| chr1:9874886-9875219 | -0.292194038 | 0.487121475 | 0.748762663 |
| chr5:26846933-26847835 | -0.23031795 | 0.487122775 | 0.748762663 |

| | | | |
|---|---|---|---|
| chr5:142601054-142601705 | -0.266461601 | 0.487176869 | 0.748806673 |
| chr5:33890314-33890978 | 0.256601184 | 0.487181534 | 0.748806673 |
| chr20:842246-842598 | -0.270323663 | 0.487193912 | 0.748806673 |
| chr2:49616903-49617112 | -0.336294689 | 0.487202752 | 0.748806673 |
| chr17:13337874-13338085 | -0.37927483 | 0.487211683 | 0.748806673 |
| chr5:16791144-16791373 | 0.313139136 | 0.487232262 | 0.748806673 |
| chr4:129093052-129094019 | -0.22101228 | 0.487236574 | 0.748806673 |
| chr7:44981073-44981417 | -0.27782271 | 0.487240879 | 0.748806673 |
| chr13:28750792-28750992 | -0.323890018 | 0.487248649 | 0.748806673 |
| chr2:216626221-216626511 | 0.283818616 | 0.487249133 | 0.748806673 |
| chr8:106856335-106856844 | -0.247681568 | 0.487264032 | 0.748811593 |
| chr19:14380937-14381449 | -0.267167746 | 0.48727285 | 0.748811593 |
| chr11:95705346-95705879 | -0.273414724 | 0.48728606 | 0.748811593 |
| chr1:156591281-156592257 | 0.231329559 | 0.487306325 | 0.748811593 |
| chr1:202101259-202101459 | -0.282731657 | 0.48731808 | 0.748811593 |
| chr6:97394216-97394489 | 0.297262023 | 0.487322106 | 0.748811593 |
| chr2:88015743-88017266 | -0.221862762 | 0.487327748 | 0.748811593 |
| chr2:96580857-96581069 | 0.353870613 | 0.487330517 | 0.748811593 |
| chr5:173144307-173145308 | -0.222478771 | 0.487352561 | 0.748830448 |
| chr12:71186756-71186956 | -0.373416636 | 0.487403199 | 0.748888197 |
| chr15:98647305-98648700 | 0.220280147 | 0.487409693 | 0.748888197 |
| chr3:115146590-115148169 | -0.216635593 | 0.487429282 | 0.748903277 |
| chr17:7052090-7052324 | 0.328432994 | 0.487449451 | 0.748919249 |
| chr10:87504412-87505684 | -0.223666882 | 0.487482193 | 0.748954536 |
| chr5:136216340-136216540 | -0.347826011 | 0.487533957 | 0.749015209 |
| chr10:89275242-89275947 | -0.249739237 | 0.487541236 | 0.749015209 |
| chrX:46573311-46574338 | 0.231177901 | 0.487580068 | 0.749019986 |
| chrX:135520429-135522212 | 0.218888053 | 0.487595375 | 0.749019986 |
| chr1:243344077-243344284 | -0.358567171 | 0.487602387 | 0.749019986 |
| chr1:231748780-231749140 | -0.319851836 | 0.487606357 | 0.749019986 |
| chr20:48012431-48012955 | 0.222745346 | 0.487609667 | 0.749019986 |
| chr1:151189626-151190414 | 0.224286584 | 0.487617837 | 0.749019986 |
| chr6:26327495-26328349 | -0.226676737 | 0.487619577 | 0.749019986 |
| chr5:105582330-105582539 | -0.365026863 | 0.48762548 | 0.749019986 |
| chr22:31345656-31346111 | 0.241779422 | 0.487632325 | 0.749019986 |
| chr9:126721865-126722091 | -0.328262582 | 0.487645769 | 0.749025621 |
| chrX:137080207-137080490 | 0.317542166 | 0.487657927 | 0.749029281 |
| chr3:134527087-134527388 | 0.290599779 | 0.487680329 | 0.749048675 |
| chr1:225975792-225976320 | 0.242696139 | 0.487724947 | 0.749092709 |
| chr5:141682201-141682410 | 0.336124058 | 0.487728552 | 0.749092709 |
| chr4:155376408-155377360 | -0.219463391 | 0.487763534 | 0.749124677 |
| chrX:5480833-5481033 | -0.393482047 | 0.487769886 | 0.749124677 |
| chr5:16894289-16895046 | 0.225598736 | 0.487786947 | 0.749124677 |
| chr1:63327601-63327946 | -0.256287756 | 0.487788473 | 0.749124677 |
| chr2:170126132-170126349 | -0.372857051 | 0.487821434 | 0.749152801 |
| chr6:45721750-45721954 | 0.372343084 | 0.487835159 | 0.749152801 |
| chr19:4304355-4304716 | -0.286612264 | 0.487836118 | 0.749152801 |
| chr1:201283339-201283646 | 0.250985091 | 0.487862827 | 0.749160919 |
| chr15:91706479-91706761 | 0.283387217 | 0.487865117 | 0.749160919 |
| chr1:215656971-215657386 | 0.255143268 | 0.487870736 | 0.749160919 |

| | | | |
|---|---|---|---|
| chr3:111338062-111338575 | -0.250626652 | 0.487934812 | 0.749220617 |
| chr6:33197482-33197807 | 0.258196576 | 0.487936917 | 0.749220617 |
| chr11:34371547-34371747 | 0.371235624 | 0.487938947 | 0.749220617 |
| chr16:85612722-85613368 | -0.225313566 | 0.487949138 | 0.749221251 |
| chr9:113061577-113061781 | 0.343684291 | 0.48797569 | 0.749247005 |
| chr7:40213424-40213624 | -0.407671694 | 0.48798902 | 0.749252459 |
| chr5:151252419-151253631 | -0.219381916 | 0.488026982 | 0.74929573 |
| chrX:22117678-22117933 | -0.336478534 | 0.488054504 | 0.749322972 |
| chr4:150010579-150010895 | -0.287592056 | 0.488135822 | 0.749424615 |
| chr17:8192215-8192474 | -0.262869864 | 0.488140269 | 0.749424615 |
| chr3:143001000-143002394 | -0.218822776 | 0.488152955 | 0.749429075 |
| chr12:95473771-95474581 | -0.223500084 | 0.488165059 | 0.749432642 |
| chr1:200302080-200302973 | -0.229349223 | 0.488180637 | 0.749441541 |
| chr7:83684799-83685357 | -0.239027073 | 0.48819794 | 0.74944245 |
| chr12:119328206-119328442 | -0.364589535 | 0.488202105 | 0.74944245 |
| chr5:146706167-146706367 | 0.386843434 | 0.488210572 | 0.74944245 |
| chr3:7800423-7800667 | 0.337352577 | 0.488245823 | 0.749481548 |
| chr4:123465394-123465627 | 0.355754979 | 0.48827096 | 0.749503728 |
| chr6:106629180-106630134 | -0.226873979 | 0.488284842 | 0.749503728 |
| chr3:119667026-119667237 | -0.385241666 | 0.488289618 | 0.749503728 |
| chr19:40348102-40348837 | 0.227308934 | 0.488309888 | 0.749510336 |
| chr1:42816594-42817580 | 0.227783307 | 0.488313487 | 0.749510336 |
| chr5:157390689-157390963 | 0.304427247 | 0.488352744 | 0.749555576 |
| chr19:47284210-47284673 | -0.271640048 | 0.488375505 | 0.749569389 |
| chr11:95254269-95254655 | -0.265265758 | 0.488381309 | 0.749569389 |
| chr6:111573188-111573609 | 0.283584762 | 0.488444948 | 0.749652047 |
| chr19:52189529-52190617 | 0.220053107 | 0.488483302 | 0.749695895 |
| chr14:100156041-100156386 | 0.24984341 | 0.48853413 | 0.749736475 |
| chr2:237322645-237322845 | 0.348958732 | 0.488543057 | 0.749736475 |
| chr11:45922512-45922741 | -0.258051113 | 0.488550206 | 0.749736475 |
| chr3:162697975-162698188 | -0.335259634 | 0.488552911 | 0.749736475 |
| chr5:69528013-69528492 | -0.241304776 | 0.488558667 | 0.749736475 |
| chr2:9306874-9307144 | -0.306262171 | 0.488616143 | 0.749797589 |
| chr2:20781747-20781965 | -0.330866869 | 0.488629052 | 0.749797589 |
| chr16:2128286-2128844 | -0.252704505 | 0.488635047 | 0.749797589 |
| chr2:218367775-218367979 | -0.335222395 | 0.488640028 | 0.749797589 |
| chr1:181189857-181190426 | 0.237067316 | 0.48864742 | 0.749797589 |
| chr11:47552231-47553688 | 0.221141232 | 0.48866615 | 0.749811313 |
| chr7:94834146-94834777 | 0.235897079 | 0.488704661 | 0.749817287 |
| chr1:176073553-176074081 | 0.252992294 | 0.48870485 | 0.749817287 |
| chr2:10900239-10900557 | -0.283280687 | 0.48870679 | 0.749817287 |
| chr7:963274-963609 | 0.267397201 | 0.488716101 | 0.749817287 |
| chr2:70257330-70258895 | -0.219418345 | 0.488718973 | 0.749817287 |
| chr1:231337480-231338474 | 0.221794728 | 0.488742705 | 0.749838684 |
| chr4:67544918-67545858 | -0.222235904 | 0.488761604 | 0.749852665 |
| chr19:4471550-4472266 | 0.225298398 | 0.488788738 | 0.749872608 |
| chr6:158060031-158060765 | -0.246177778 | 0.488822767 | 0.749872608 |
| chr2:187581480-187581680 | -0.370093555 | 0.488834609 | 0.749872608 |
| chr10:112137445-112137724 | 0.275524611 | 0.488859647 | 0.749872608 |
| chr14:99628895-99629233 | 0.255543753 | 0.488861905 | 0.749872608 |

| | | | |
|---|---|---|---|
| chr18:34121192-34121529 | -0.294646531 | 0.488864754 | 0.749872608 |
| chr8:73746023-73747660 | 0.219888339 | 0.488878327 | 0.749872608 |
| chr6:86723681-86723881 | 0.346878808 | 0.488878595 | 0.749872608 |
| chr11:66694671-66695128 | 0.23707794 | 0.488890345 | 0.749872608 |
| chr15:88476664-88476929 | 0.291155605 | 0.488892093 | 0.749872608 |
| chr3:50235682-50236368 | 0.226512637 | 0.488893741 | 0.749872608 |
| chr2:63555393-63555593 | 0.37953462 | 0.48889523 | 0.749872608 |
| chr17:63741281-63742972 | -0.216635263 | 0.488907726 | 0.749872608 |
| chr3:83217028-83217231 | -0.311802198 | 0.488921403 | 0.749872608 |
| chr6:157764556-157764792 | -0.311802198 | 0.488921403 | 0.749872608 |
| chr3:169390116-169391222 | -0.230524575 | 0.488942585 | 0.749882665 |
| chr1:92831768-92832442 | -0.227599134 | 0.488947534 | 0.749882665 |
| chr16:50265285-50265538 | -0.32150213 | 0.488960494 | 0.749887531 |
| chr2:76314615-76314863 | 0.31420031 | 0.488971324 | 0.749889131 |
| chr12:124981236-124981520 | -0.318955832 | 0.488995378 | 0.749908531 |
| chr16:4272737-4273123 | 0.250032478 | 0.489003548 | 0.749908531 |
| chr10:104498251-104498818 | -0.258219525 | 0.489034197 | 0.749940524 |
| chr2:7109097-7109297 | -0.378040213 | 0.489069036 | 0.749967394 |
| chr5:79236916-79237868 | -0.228444872 | 0.489075651 | 0.749967394 |
| chr18:3385301-3385635 | -0.354118493 | 0.489081083 | 0.749967394 |
| chr4:5518313-5518786 | -0.313506522 | 0.489129402 | 0.750026478 |
| chr13:111460550-111460777 | 0.362125685 | 0.489165785 | 0.750060058 |
| chr15:88904792-88905041 | 0.267441767 | 0.489179012 | 0.750060058 |
| chr4:53601107-53601936 | -0.228005649 | 0.489202093 | 0.750060058 |
| chr3:97252336-97252921 | -0.255831386 | 0.489206699 | 0.750060058 |
| chr5:51691824-51692043 | -0.363702227 | 0.489215692 | 0.750060058 |
| chr21:44752824-44753180 | 0.269808323 | 0.489217884 | 0.750060058 |
| chr12:105107190-105108353 | -0.221073167 | 0.489219826 | 0.750060058 |
| chr5:128667222-128667443 | -0.339609898 | 0.489230297 | 0.750061105 |
| chr3:10392799-10392999 | 0.336757594 | 0.489261442 | 0.75008332 |
| chr5:148443961-148444281 | -0.292520023 | 0.489264366 | 0.75008332 |
| chr14:96816472-96816673 | 0.385799378 | 0.489286095 | 0.750101624 |
| chr9:131762232-131762696 | 0.269361324 | 0.48938061 | 0.750199659 |
| chr8:26088770-26089218 | 0.309475979 | 0.489386603 | 0.750199659 |
| chr12:2682518-2682792 | -0.309262145 | 0.489391398 | 0.750199659 |
| chr19:38224009-38224827 | 0.229053168 | 0.489394529 | 0.750199659 |
| chr10:86560010-86560282 | -0.305147484 | 0.489398997 | 0.750199659 |
| chr2:167264242-167264720 | -0.27285834 | 0.489418251 | 0.750214165 |
| chr11:17230104-17230368 | -0.337063369 | 0.489466574 | 0.750253257 |
| chr9:126805808-126806073 | 0.285203037 | 0.489488603 | 0.750253257 |
| chr10:116848950-116849179 | 0.286839641 | 0.489488605 | 0.750253257 |
| chr19:2475202-2475486 | 0.289504495 | 0.48949173 | 0.750253257 |
| chr10:74189981-74190469 | 0.25163524 | 0.489492712 | 0.750253257 |
| chr15:48877716-48878543 | -0.227715863 | 0.489507981 | 0.750255407 |
| chr10:120460885-120461107 | -0.386793456 | 0.489513698 | 0.750255407 |
| chr6:153131143-153131670 | 0.253250708 | 0.489528338 | 0.750257533 |
| chr10:63132782-63133967 | -0.218647518 | 0.489534668 | 0.750257533 |
| chr8:80864753-80865134 | -0.283696245 | 0.489561882 | 0.750263525 |
| chr19:49118611-49119417 | 0.225371815 | 0.489588399 | 0.750263525 |
| chr16:74367965-74368661 | -0.228869674 | 0.489594399 | 0.750263525 |

| | | | |
|---|---|---|---|
| chr5:73375653-73375869 | 0.375932484 | 0.48960668 | 0.750263525 |
| chr3:102224240-102224568 | -0.267793958 | 0.489611076 | 0.750263525 |
| chr1:64269138-64269461 | -0.266130471 | 0.489615336 | 0.750263525 |
| chr7:100428207-100430031 | -0.216999916 | 0.489624486 | 0.750263525 |
| chr16:352603-353379 | 0.221764735 | 0.489626599 | 0.750263525 |
| chr2:39436504-39438256 | -0.218263856 | 0.489626704 | 0.750263525 |
| chr2:11584213-11584623 | -0.250537867 | 0.48964936 | 0.750283237 |
| chr5:5120103-5120303 | -0.393393121 | 0.489665719 | 0.750293027 |
| chr12:108884272-108884714 | 0.267752668 | 0.489675333 | 0.750293027 |
| chr6:151324232-151324575 | -0.278109203 | 0.489701135 | 0.750317556 |
| chrX:64976130-64976886 | -0.23509333 | 0.489779123 | 0.750422043 |
| chr12:54000320-54000629 | 0.271252381 | 0.489801511 | 0.750425384 |
| chr17:5078078-5078945 | 0.241381028 | 0.489804228 | 0.750425384 |
| chr9:90953913-90954113 | 0.375435052 | 0.48981215 | 0.750425384 |
| chr17:64027415-64027668 | -0.281114862 | 0.489835488 | 0.750425384 |
| chr11:119681090-119681694 | -0.254038213 | 0.489847587 | 0.750425384 |
| chr2:72886927-72888177 | -0.22261412 | 0.489856577 | 0.750425384 |
| chr3:4711373-4711927 | -0.237810493 | 0.489862585 | 0.750425384 |
| chr10:77111738-77112179 | -0.28144873 | 0.48987642 | 0.750425384 |
| chr3:108031803-108032232 | 0.269361855 | 0.48988432 | 0.750425384 |
| chr15:98662968-98663423 | 0.252872994 | 0.48989182 | 0.750425384 |
| chr1:17406196-17406484 | 0.300579497 | 0.489898761 | 0.750425384 |
| chr7:27136466-27136737 | 0.286650208 | 0.48989883 | 0.750425384 |
| chr17:59059860-59060208 | 0.235862945 | 0.489933236 | 0.750447812 |
| chr9:81116324-81116547 | -0.399428849 | 0.489941307 | 0.750447812 |
| chr3:151837048-151837581 | 0.246928022 | 0.489953232 | 0.750447812 |
| chr6:34689866-34690073 | -0.374767314 | 0.489959588 | 0.750447812 |
| chr8:48619905-48620532 | -0.236090496 | 0.489962442 | 0.750447812 |
| chr1:110003862-110004469 | -0.235461035 | 0.489975445 | 0.750452727 |
| chr18:35006041-35006320 | -0.325255108 | 0.489995897 | 0.750466771 |
| chr5:173304001-173304207 | -0.295786475 | 0.490004203 | 0.750466771 |
| chr7:75914421-75915447 | -0.226725842 | 0.490036862 | 0.750501788 |
| chr10:43945027-43945232 | -0.350183263 | 0.490047656 | 0.750503318 |
| chr7:36945928-36946545 | 0.23511866 | 0.490069663 | 0.75052202 |
| chr2:97663426-97664880 | 0.219297707 | 0.490092757 | 0.75053617 |
| chr3:75190264-75190472 | -0.337476424 | 0.490098493 | 0.75053617 |
| chr15:20192986-20193281 | -0.329717988 | 0.490148732 | 0.750586349 |
| chr7:148109995-148110493 | -0.249761252 | 0.490157243 | 0.750586349 |
| chr12:31103771-31104017 | -0.318021527 | 0.490160647 | 0.750586349 |
| chr1:87200961-87201460 | -0.26910658 | 0.490174425 | 0.750592445 |
| chr3:112642924-112643226 | -0.248824438 | 0.490208918 | 0.750626419 |
| chr2:119677980-119678465 | 0.247351152 | 0.490216204 | 0.750626419 |
| chr15:92059169-92059446 | -0.333835287 | 0.490248055 | 0.750660187 |
| chr8:95549665-95549940 | -0.334011278 | 0.49026168 | 0.750661311 |
| chr10:1097311-1097691 | -0.292247762 | 0.490277435 | 0.750661311 |
| chr1:164126359-164127152 | -0.222450798 | 0.49027818 | 0.750661311 |
| chr12:84503003-84503506 | 0.247420878 | 0.490291187 | 0.750666227 |
| chr14:78969100-78969520 | -0.266131747 | 0.49030683 | 0.750675177 |
| chr2:241881168-241882549 | -0.217561859 | 0.490330715 | 0.750694488 |
| chr19:45093090-45093349 | 0.28759153 | 0.490348955 | 0.750694488 |

| | | | |
|---|---|---|---|
| chr3:102529538-102529816 | 0.28759153 | 0.490348955 | 0.750694488 |
| chr15:82261932-82263010 | -0.223427195 | 0.490361981 | 0.750694488 |
| chrX:130248807-130249171 | -0.329390768 | 0.490382934 | 0.750694488 |
| chr2:85369372-85369981 | -0.245849018 | 0.490383228 | 0.750694488 |
| chrX:135740484-135741080 | -0.264331933 | 0.490394202 | 0.750694488 |
| chr1:40708801-40709386 | -0.254160752 | 0.490397822 | 0.750694488 |
| chr10:74213132-74213526 | -0.309732984 | 0.490411644 | 0.750700649 |
| chr1:54199488-54200718 | 0.222994615 | 0.490425952 | 0.750707554 |
| chr6:40818867-40819067 | -0.379106808 | 0.490442989 | 0.750718634 |
| chr2:218444513-218445108 | -0.2304993 | 0.490481344 | 0.75074519 |
| chr8:47738719-47739198 | 0.242661169 | 0.490494548 | 0.75074519 |
| chr8:42337909-42338866 | 0.229025378 | 0.490496867 | 0.75074519 |
| chr1:162613516-162614034 | 0.254367973 | 0.490524611 | 0.75074519 |
| chr1:106516649-106517081 | 0.247673328 | 0.4905247 | 0.75074519 |
| chr7:42918711-42918911 | -0.428822347 | 0.490528183 | 0.75074519 |
| chr8:2001617-2001999 | -0.28411569 | 0.490542923 | 0.75074519 |
| chr2:160290688-160290935 | 0.373660558 | 0.490552939 | 0.75074519 |
| chr7:100897227-100897603 | -0.278909222 | 0.490554299 | 0.75074519 |
| chr13:40193434-40193662 | -0.363425507 | 0.490558318 | 0.75074519 |
| chr20:47181148-47181687 | 0.218401295 | 0.490593675 | 0.750780605 |
| chr12:122064049-122064362 | -0.266412807 | 0.4906279 | 0.750780605 |
| chr10:124791230-124792506 | 0.224462665 | 0.490639193 | 0.750780605 |
| chr17:66727991-66728307 | -0.250159862 | 0.490641058 | 0.750780605 |
| chr3:133488803-133489519 | -0.236701128 | 0.490648618 | 0.750780605 |
| chr1:180953282-180954634 | -0.219651417 | 0.490652305 | 0.750780605 |
| chr20:59985893-59986288 | 0.259120333 | 0.490654604 | 0.750780605 |
| chr4:85910392-85910875 | -0.300955476 | 0.490659847 | 0.750780605 |
| chr5:78569500-78569700 | 0.353672603 | 0.490685057 | 0.750804186 |
| chr1:15100466-15100689 | 0.365762179 | 0.490732078 | 0.750858399 |
| chr17:58083080-58084391 | 0.21735699 | 0.490740086 | 0.750858399 |
| chr11:72812882-72813388 | -0.236283044 | 0.490783643 | 0.750896507 |
| chr6:28950868-28951238 | 0.246598074 | 0.490784593 | 0.750896507 |
| chr12:95317989-95318632 | 0.238122085 | 0.49080108 | 0.750906737 |
| chr8:130129683-130129895 | 0.337263881 | 0.4908856 | 0.751021054 |
| chr6:8101907-8102902 | -0.220465094 | 0.490939768 | 0.751066487 |
| chr14:75003180-75003411 | 0.283576689 | 0.490944584 | 0.751066487 |
| chr3:160691236-160691437 | -0.346152751 | 0.490944703 | 0.751066487 |
| chr5:154844748-154845288 | -0.232960568 | 0.490962123 | 0.751078141 |
| chr3:88203573-88203942 | -0.265114576 | 0.490977163 | 0.751086154 |
| chr6:9503367-9503721 | -0.287005371 | 0.491017361 | 0.751125657 |
| chr10:84352274-84352651 | -0.254710206 | 0.491022592 | 0.751125657 |
| chr8:124345094-124345302 | 0.333552935 | 0.491057668 | 0.75114521 |
| chr12:104014561-104014761 | -0.362211177 | 0.491064783 | 0.75114521 |
| chr14:82999894-83000094 | -0.362211177 | 0.491064783 | 0.75114521 |
| chr11:75214368-75214672 | 0.260379489 | 0.491103152 | 0.751180278 |
| chr17:39056807-39057341 | -0.252033132 | 0.491107317 | 0.751180278 |
| chr8:102768147-102768433 | -0.284929168 | 0.491118216 | 0.751181953 |
| chr9:131372989-131374067 | -0.222450154 | 0.491137822 | 0.751196945 |
| chr12:121209769-121210771 | 0.23072522 | 0.491153558 | 0.751206019 |
| chr1:218163946-218164796 | -0.229291719 | 0.491208103 | 0.751274448 |

| | | | |
|---|---|---|---|
| chr12:101917466-101918063 | 0.25336638 | 0.491225371 | 0.751285861 |
| chr2:217756369-217756880 | -0.260893032 | 0.491239024 | 0.751291746 |
| chr14:105045806-105046248 | 0.247704179 | 0.491258182 | 0.751296121 |
| chrX:47523396-47523726 | 0.271160647 | 0.491261495 | 0.751296121 |
| chr17:66952698-66953049 | -0.240535338 | 0.491272808 | 0.751297509 |
| chr15:43791835-43792463 | 0.223357543 | 0.491282013 | 0.751297509 |
| chr17:68740610-68741356 | -0.230504389 | 0.491297719 | 0.751304031 |
| chr10:50417812-50418598 | -0.23556123 | 0.491333015 | 0.751304031 |
| chr17:48663231-48663450 | 0.311428682 | 0.491365364 | 0.751304031 |
| chr2:106542803-106543463 | -0.242345081 | 0.491370221 | 0.751304031 |
| chr2:95338151-95338361 | 0.381399914 | 0.49138256 | 0.751304031 |
| chr12:107782156-107782356 | 0.381399914 | 0.49138256 | 0.751304031 |
| chr11:74929813-74930056 | -0.301115192 | 0.491395764 | 0.751304031 |
| chr17:69019766-69020026 | -0.369857489 | 0.491400407 | 0.751304031 |
| chr3:194056743-194057136 | -0.266176607 | 0.491404468 | 0.751304031 |
| chrX:72004398-72004660 | -0.333082715 | 0.491406145 | 0.751304031 |
| chr1:32332314-32332751 | 0.250380157 | 0.491409729 | 0.751304031 |
| chr13:75469162-75469417 | -0.29700245 | 0.491411979 | 0.751304031 |
| chr1:149885748-149886019 | 0.241352212 | 0.491424631 | 0.751304031 |
| chr5:65924918-65926179 | -0.220082609 | 0.491429309 | 0.751304031 |
| chr19:7197746-7198599 | 0.23819824 | 0.491436703 | 0.751304031 |
| chr4:76198579-76198896 | 0.296937902 | 0.491443163 | 0.751304031 |
| chr20:64102122-64102388 | -0.287131228 | 0.491472458 | 0.751333825 |
| chr1:207620136-207620680 | -0.240706905 | 0.491493226 | 0.751347527 |
| chr17:57783125-57783651 | 0.235949908 | 0.491501032 | 0.751347527 |
| chr15:62995068-62995462 | 0.283221876 | 0.491536225 | 0.751372338 |
| chr20:33731899-33733113 | 0.222940722 | 0.491567081 | 0.751372338 |
| chr3:168889496-168890136 | -0.246708107 | 0.491572928 | 0.751372338 |
| chr2:108448412-108449865 | 0.218667905 | 0.491577198 | 0.751372338 |
| chr13:99436641-99437394 | -0.243762668 | 0.491580371 | 0.751372338 |
| chr10:677733-677966 | 0.331543912 | 0.491582354 | 0.751372338 |
| chr17:78250849-78251825 | 0.226614155 | 0.491585906 | 0.751372338 |
| chr13:23087935-23088222 | 0.293675984 | 0.491616801 | 0.751389932 |
| chr2:177394306-177394524 | 0.34004987 | 0.49161703 | 0.751389932 |
| chr2:129979652-129980002 | -0.27986131 | 0.491663896 | 0.751445434 |
| chr12:79566491-79566800 | 0.286703784 | 0.491672958 | 0.751445434 |
| chr13:26880995-26881595 | -0.260762698 | 0.491691452 | 0.75145871 |
| chr3:170046699-170047270 | -0.240488398 | 0.491702665 | 0.751460858 |
| chr19:10315448-10316137 | 0.230797494 | 0.491724546 | 0.751473485 |
| chr3:10772481-10773044 | 0.243350021 | 0.491730542 | 0.751473485 |
| chr5:132283414-132283885 | -0.272003763 | 0.491773301 | 0.751522451 |
| chr6:116287696-116287896 | -0.361569385 | 0.4917822 | 0.751522451 |
| chr1:117675032-117675283 | -0.276795092 | 0.491795275 | 0.751527444 |
| chr20:36447153-36447404 | 0.318288522 | 0.491838844 | 0.75156061 |
| chr3:16557185-16557428 | -0.351641358 | 0.491855395 | 0.75156061 |
| chr2:73283497-73284025 | -0.247536938 | 0.491859998 | 0.75156061 |
| chr14:51744889-51745196 | 0.301139475 | 0.491861667 | 0.75156061 |
| chr1:68232336-68232768 | -0.250601225 | 0.491866022 | 0.75156061 |
| chr1:19715576-19716021 | -0.256493848 | 0.49199258 | 0.751738996 |
| chr14:77400294-77400494 | 0.37100104 | 0.49201881 | 0.751764083 |

| | | | |
|---|---|---|---|
| chr4:43843622-43843822 | 0.360565505 | 0.49205692 | 0.75180732 |
| chr1:32650840-32651670 | 0.222702929 | 0.492091346 | 0.751836009 |
| chr19:38975254-38975629 | 0.27023316 | 0.492095321 | 0.751836009 |
| chr4:73964953-73965231 | 0.357128119 | 0.492140586 | 0.751851259 |
| chr11:125404900-125405155 | 0.295687425 | 0.492144478 | 0.751851259 |
| chr1:50023318-50024383 | -0.231265412 | 0.492145347 | 0.751851259 |
| chrX:138711674-138712347 | 0.238893578 | 0.492162686 | 0.751851259 |
| chr20:18057247-18057636 | 0.254002167 | 0.49216983 | 0.751851259 |
| chr16:67278675-67279126 | 0.252253369 | 0.492176372 | 0.751851259 |
| chr12:65300795-65301178 | -0.254722456 | 0.492176374 | 0.751851259 |
| chr4:33625863-33626063 | -0.351328608 | 0.492186634 | 0.751851259 |
| chr6:132823088-132823292 | -0.326752133 | 0.492193615 | 0.751851259 |
| chr3:112013398-112013863 | -0.270139614 | 0.492216248 | 0.751870842 |
| chr10:116080715-116081028 | 0.304345576 | 0.492227656 | 0.751873279 |
| chr5:149482787-149483065 | -0.270774162 | 0.492266419 | 0.751914654 |
| chr11:102288000-102288249 | 0.314194193 | 0.492274369 | 0.751914654 |
| chr17:29390873-29391073 | 0.315361797 | 0.492287381 | 0.751919539 |
| chr11:17439313-17439872 | 0.247489436 | 0.492366161 | 0.752024877 |
| chr1:179867680-179868110 | 0.246098903 | 0.492380603 | 0.752031944 |
| chr6:133916689-133916889 | 0.336065101 | 0.492401123 | 0.752048295 |
| chr14:69531203-69531648 | -0.26060887 | 0.49242266 | 0.752059088 |
| chr4:116016280-116016739 | 0.275053759 | 0.49242782 | 0.752059088 |
| chr1:203273035-203273611 | -0.226911547 | 0.49244137 | 0.752064792 |
| chr4:163192053-163192253 | 0.376846292 | 0.492484548 | 0.752115742 |
| chr20:51752639-51752887 | -0.257594981 | 0.492518654 | 0.752149171 |
| chr6:153024385-153024739 | -0.270571586 | 0.49252607 | 0.752149171 |
| chr16:84475477-84475956 | -0.254952057 | 0.492565215 | 0.752155671 |
| chr8:75952077-75952766 | -0.254066337 | 0.492572692 | 0.752155671 |
| chr11:119321653-119322128 | -0.243439114 | 0.492577636 | 0.752155671 |
| chr2:217510885-217511090 | 0.319048882 | 0.492580351 | 0.752155671 |
| chr6:27866118-27866453 | -0.265447335 | 0.492585457 | 0.752155671 |
| chr19:13847229-13847429 | 0.31761507 | 0.492593576 | 0.752155671 |
| chr18:75231150-75231354 | -0.351041006 | 0.492599041 | 0.752155671 |
| chr9:89942031-89942242 | -0.387315068 | 0.492612395 | 0.752161073 |
| chr5:94111137-94112055 | -0.219646745 | 0.492637874 | 0.752182163 |
| chr16:15998521-15998761 | -0.348190736 | 0.492652818 | 0.752182163 |
| chr1:116311820-116312678 | 0.220177264 | 0.492655814 | 0.752182163 |
| chr14:106586195-106586395 | 0.336204425 | 0.492672401 | 0.752182163 |
| chr16:81896009-81896245 | 0.327263787 | 0.492683873 | 0.752182163 |
| chr2:63078905-63079207 | -0.28631192 | 0.492685109 | 0.752182163 |
| chr2:101150606-101151743 | 0.222346502 | 0.492702519 | 0.752193755 |
| chr20:25719091-25719451 | 0.280287326 | 0.49271467 | 0.752197319 |
| chr6:137094422-137094662 | -0.333337684 | 0.492762272 | 0.752250807 |
| chr1:246907565-246907765 | -0.313648889 | 0.4927715 | 0.752250807 |
| chr15:99486335-99486855 | -0.24570748 | 0.492784742 | 0.752250807 |
| chr10:87019836-87020290 | 0.274320349 | 0.492788977 | 0.752250807 |
| chr20:62766768-62766994 | -0.277099367 | 0.492807342 | 0.752263854 |
| chr16:84266399-84266776 | 0.238826285 | 0.492825489 | 0.752274798 |
| chr10:28922793-28923039 | 0.348637208 | 0.492834148 | 0.752274798 |
| chr11:133947282-133947718 | 0.263250762 | 0.492862006 | 0.752296014 |

| | | | |
|---|---|---|---|
| chr4:677981-678194 | -0.283720589 | 0.492876379 | 0.752296014 |
| chr15:100517167-100517367 | -0.363068043 | 0.492877501 | 0.752296014 |
| chr16:27193429-27193629 | 0.321435032 | 0.492905738 | 0.75230914 |
| chr17:44087387-44087587 | 0.321435032 | 0.492905738 | 0.75230914 |
| chr5:122468409-122468652 | -0.342074011 | 0.492953028 | 0.75233639 |
| chr2:11465482-11466904 | -0.217800375 | 0.492958869 | 0.75233639 |
| chr12:6310649-6311080 | -0.275570453 | 0.492961696 | 0.75233639 |
| chr5:16776695-16776933 | 0.298399041 | 0.492962867 | 0.75233639 |
| chr10:44253833-44254044 | 0.347565876 | 0.492980585 | 0.752348445 |
| chr12:53231876-53232104 | 0.2700301 | 0.493016148 | 0.752387734 |
| chr16:81873899-81874208 | 0.276565379 | 0.493045073 | 0.752394875 |
| chr10:79195454-79195746 | 0.268846132 | 0.493047292 | 0.752394875 |
| chrX:35380873-35381367 | -0.264178233 | 0.493059491 | 0.752394875 |
| chr2:218395950-218396732 | 0.221608603 | 0.493060106 | 0.752394875 |
| chr10:8097650-8097989 | -0.297602541 | 0.493082063 | 0.752413395 |
| chr9:128098128-128098668 | 0.233922125 | 0.493117559 | 0.752452574 |
| chr15:69987711-69988321 | 0.241244576 | 0.493172167 | 0.752520916 |
| chr9:1470204-1470637 | -0.278262026 | 0.493215669 | 0.752556002 |
| chr11:17388711-17389801 | 0.2205749 | 0.493218547 | 0.752556002 |
| chr5:14405347-14405606 | -0.299253465 | 0.493224626 | 0.752556002 |
| chr17:30599923-30600965 | 0.229709832 | 0.493240083 | 0.752562343 |
| chr16:81401085-81401464 | -0.248079496 | 0.493248425 | 0.752562343 |
| chr20:52190754-52191026 | -0.267893174 | 0.493271858 | 0.752571913 |
| chr12:47779345-47779677 | 0.257095886 | 0.493274342 | 0.752571913 |
| chr4:89901919-89902120 | -0.362281897 | 0.493310961 | 0.752593952 |
| chr11:82634154-82634388 | -0.328248303 | 0.493315585 | 0.752593952 |
| chr14:103562077-103563359 | 0.221263835 | 0.493322158 | 0.752593952 |
| chr4:129350858-129351243 | 0.27068078 | 0.493334899 | 0.752593952 |
| chr17:42221920-42222120 | -0.357407853 | 0.493337898 | 0.752593952 |
| chr19:5339807-5340007 | 0.330321279 | 0.493407421 | 0.752685025 |
| chr9:96666664-96667090 | 0.279864244 | 0.493452269 | 0.752738454 |
| chr6:119261000-119261821 | -0.246759671 | 0.493478261 | 0.752763116 |
| chr15:100433722-100434082 | 0.283121424 | 0.493498118 | 0.752764668 |
| chr16:9091048-9091781 | 0.218024533 | 0.493501818 | 0.752764668 |
| chr1:116785168-116785581 | 0.291162389 | 0.493508752 | 0.752764668 |
| chr16:67159787-67160050 | -0.26528541 | 0.49353193 | 0.752785036 |
| chr20:38633769-38633986 | 0.316918505 | 0.493545349 | 0.75279052 |
| chr19:44915037-44915280 | -0.308485113 | 0.493576119 | 0.752822466 |
| chr7:11636449-11636759 | 0.331529321 | 0.493593059 | 0.752833317 |
| chr4:177555065-177555279 | -0.347454803 | 0.493634374 | 0.752853209 |
| chr8:80170995-80171854 | 0.220494284 | 0.493636839 | 0.752853209 |
| chr19:57599505-57600365 | 0.233035881 | 0.493640825 | 0.752853209 |
| chr20:29765059-29765338 | 0.30857423 | 0.493646037 | 0.752853209 |
| chr3:42443882-42444199 | 0.289352769 | 0.493657993 | 0.752853209 |
| chr1:169546300-169546745 | -0.245150819 | 0.493665054 | 0.752853209 |
| chr6:113858578-113859276 | 0.225829867 | 0.493690946 | 0.75287771 |
| chr1:106221902-106222110 | 0.279221832 | 0.493721733 | 0.752909676 |
| chr16:58684259-58684981 | 0.238120361 | 0.493767282 | 0.752950613 |
| chr4:90419153-90419596 | 0.260654531 | 0.493788223 | 0.752950613 |
| chr11:43312680-43312960 | -0.276408315 | 0.493791458 | 0.752950613 |

| | | | |
|---|---|---|---|
| chr14:64972098-64972778 | 0.235514252 | 0.493791757 | 0.752950613 |
| chr5:6765670-6766008 | 0.241670385 | 0.493800659 | 0.752950613 |
| chr3:170078557-170078820 | -0.290707106 | 0.493807539 | 0.752950613 |
| chr13:44917513-44918745 | -0.21419915 | 0.493826242 | 0.752956243 |
| chr8:99012443-99013981 | 0.2170315 | 0.493839564 | 0.752956243 |
| chr3:149936781-149937367 | 0.244311981 | 0.493852847 | 0.752956243 |
| chr10:87771970-87772223 | 0.331394835 | 0.493865277 | 0.752956243 |
| chr2:97890191-97890419 | -0.329034477 | 0.493865378 | 0.752956243 |
| chr12:112190604-112190947 | -0.283079497 | 0.493877001 | 0.752956243 |
| chr5:78615820-78616020 | -0.36776972 | 0.49388002 | 0.752956243 |
| chr3:41199051-41200091 | 0.226199471 | 0.49390639 | 0.752981465 |
| chr20:32064739-32065008 | 0.29951283 | 0.49399631 | 0.753088902 |
| chr6:31580766-31581194 | 0.256444934 | 0.493996519 | 0.753088902 |
| chr6:27127107-27127405 | -0.267138398 | 0.494020549 | 0.75311055 |
| chr3:122732907-122733479 | -0.272241898 | 0.494044855 | 0.753124973 |
| chr10:125316635-125316895 | -0.323659121 | 0.494062201 | 0.753124973 |
| chr5:160391135-160391382 | -0.329774457 | 0.4940626 | 0.753124973 |
| chr2:85538438-85540079 | -0.215604441 | 0.494069326 | 0.753124973 |
| chr20:59183813-59184087 | 0.270939954 | 0.494118529 | 0.75318499 |
| chr4:39527103-39528350 | 0.216370804 | 0.494136567 | 0.753197502 |
| chr8:33471718-33472087 | -0.279376895 | 0.494169573 | 0.753221986 |
| chr14:87899301-87899557 | -0.302262091 | 0.494172291 | 0.753221986 |
| chr18:23143900-23144153 | -0.296866604 | 0.494192859 | 0.753238352 |
| chr8:87903660-87903920 | 0.314301325 | 0.494209046 | 0.753241036 |
| chr5:112691705-112692084 | -0.24640442 | 0.494214281 | 0.753241036 |
| chr7:47587273-47587473 | 0.311661546 | 0.494243258 | 0.753270218 |
| chr21:10145261-10145518 | 0.32334121 | 0.49429268 | 0.753330556 |
| chr2:26764037-26764465 | -0.248529878 | 0.49432419 | 0.753361482 |
| chr13:26459274-26459658 | -0.316499976 | 0.494338981 | 0.753361482 |
| chr20:63519747-63521340 | 0.216816951 | 0.494350465 | 0.753361482 |
| chrX:132022737-132024306 | 0.216693049 | 0.4943523 | 0.753361482 |
| chr12:124994544-124994883 | -0.272944857 | 0.494426334 | 0.753459319 |
| chr19:17334778-17334988 | 0.288149367 | 0.494458901 | 0.753493962 |
| chr15:41492771-41492981 | -0.32138113 | 0.494485783 | 0.753504332 |
| chr1:87318923-87319123 | -0.363287391 | 0.494489171 | 0.753504332 |
| chr5:173283691-173284142 | 0.243846523 | 0.494500897 | 0.753504332 |
| chr8:100625673-100626045 | 0.259683607 | 0.494508454 | 0.753504332 |
| chr9:121700133-121700447 | 0.259964851 | 0.494514876 | 0.753504332 |
| chr12:48965506-48965762 | -0.300809627 | 0.494553315 | 0.75354359 |
| chr6:87944493-87945004 | 0.258593053 | 0.494575305 | 0.75354359 |
| chr1:202418038-202418252 | 0.332564788 | 0.494577973 | 0.75354359 |
| chr7:95686207-95686425 | 0.327588788 | 0.494583519 | 0.75354359 |
| chr21:45527451-45527839 | 0.269059456 | 0.494592653 | 0.75354359 |
| chr4:138577741-138578224 | -0.270983754 | 0.494599648 | 0.75354359 |
| chr18:12539943-12540188 | 0.323029865 | 0.494648588 | 0.753603168 |
| chr16:84925168-84925453 | 0.277518431 | 0.494662304 | 0.75360908 |
| chr2:138434663-138434863 | 0.339903919 | 0.494701265 | 0.753609403 |
| chr4:160246738-160247013 | -0.3363897 | 0.494706172 | 0.753609403 |
| chr11:2929313-2929710 | 0.242838816 | 0.494708034 | 0.753609403 |
| chr6:139028407-139029818 | -0.218683131 | 0.494711 | 0.753609403 |

| | | | |
|---|---|---|---|
| chr1:55064065-55064430 | -0.256233916 | 0.494718988 | 0.753609403 |
| chr1:79745971-79746297 | -0.280540614 | 0.494721529 | 0.753609403 |
| chr6:160041472-160041740 | -0.374559252 | 0.494732801 | 0.753611593 |
| chrX:55908988-55909250 | -0.3132764 | 0.494749162 | 0.753621532 |
| chr18:839362-839581 | 0.3135223 | 0.494801367 | 0.75368607 |
| chr4:137226975-137227319 | 0.284003533 | 0.494825097 | 0.753693385 |
| chr1:208045615-208045993 | -0.275423249 | 0.494825843 | 0.753693385 |
| chr12:79711250-79711692 | -0.276183585 | 0.494850486 | 0.753702441 |
| chr13:36818807-36819758 | -0.218393815 | 0.494851462 | 0.753702441 |
| chr15:40420753-40421197 | -0.24030364 | 0.494866254 | 0.753709989 |
| chr1:110088108-110088432 | 0.26885531 | 0.494885449 | 0.753724241 |
| chr3:27293917-27294265 | 0.280448709 | 0.494906983 | 0.753738235 |
| chr1:206448210-206448791 | 0.232610607 | 0.494914311 | 0.753738235 |
| chr1:114458203-114458524 | -0.239430488 | 0.494926989 | 0.753742562 |
| chr14:34467916-34468201 | 0.318634884 | 0.494959794 | 0.753777539 |
| chr5:154097065-154097354 | 0.291340058 | 0.494980724 | 0.753794431 |
| chr3:155782533-155783016 | -0.248434279 | 0.49505598 | 0.753879415 |
| chrX:41302248-41302733 | 0.247785353 | 0.495056207 | 0.753879415 |
| chr11:111299498-111299698 | -0.340190005 | 0.495081352 | 0.753902725 |
| chr17:41772184-41772543 | 0.245936785 | 0.495103284 | 0.753921138 |
| chr1:116334031-116334355 | 0.265769597 | 0.495145053 | 0.753955814 |
| chr1:225792301-225792501 | -0.381799128 | 0.495145736 | 0.753955814 |
| chr17:29617956-29618349 | 0.247756844 | 0.495165529 | 0.75395741 |
| chr5:150459740-150459959 | -0.30576457 | 0.495166464 | 0.75395741 |
| chr6:75546483-75546723 | -0.322611865 | 0.495179793 | 0.753962722 |
| chr11:59135887-59136342 | -0.263560756 | 0.495218957 | 0.754004796 |
| chr6:128806455-128806691 | -0.379176148 | 0.495227106 | 0.754004796 |
| chr9:138066121-138066450 | -0.302723905 | 0.495239686 | 0.754008966 |
| chr4:74805658-74805873 | 0.321283237 | 0.495261668 | 0.754021759 |
| chr11:32176297-32176561 | -0.32164699 | 0.49526777 | 0.754021759 |
| chr6:33590288-33590488 | -0.342827535 | 0.495334671 | 0.754108629 |
| chr6:8109218-8109603 | -0.289742151 | 0.495400682 | 0.754174262 |
| chr5:173302854-173303121 | 0.266341486 | 0.495406964 | 0.754174262 |
| chr3:171156658-171156871 | 0.335515867 | 0.495418034 | 0.754174262 |
| chr1:203084581-203084837 | -0.26604129 | 0.495429955 | 0.754174262 |
| chr6:27823657-27823857 | -0.338567436 | 0.49543403 | 0.754174262 |
| chr10:115355253-115355498 | -0.334900535 | 0.495436839 | 0.754174262 |
| chr10:8660989-8661189 | 0.387699674 | 0.495512608 | 0.754266002 |
| chr6:26271300-26271691 | -0.243057058 | 0.49552065 | 0.754266002 |
| chr13:26557152-26558086 | -0.22011264 | 0.49556115 | 0.754266002 |
| chr17:28952246-28952701 | 0.224970521 | 0.495567714 | 0.754266002 |
| chr5:317092-317466 | 0.249923588 | 0.495580853 | 0.754266002 |
| chr2:191677357-191678960 | -0.214264604 | 0.495596979 | 0.754266002 |
| chr2:140401565-140401868 | -0.346926473 | 0.495597608 | 0.754266002 |
| chr3:158717517-158717717 | -0.346926473 | 0.495597608 | 0.754266002 |
| chr12:124359548-124360061 | 0.230664767 | 0.495624329 | 0.754266002 |
| chr13:107166221-107166493 | -0.372143234 | 0.495624447 | 0.754266002 |
| chr10:93518429-93518659 | 0.350537843 | 0.495624722 | 0.754266002 |
| chr3:141454956-141456298 | 0.217875294 | 0.495627965 | 0.754266002 |
| chr5:87924271-87924810 | -0.24664767 | 0.495630428 | 0.754266002 |

| | | | |
|---|---|---|---|
| chr21:45407919-45408219 | 0.28541096 | 0.495634921 | 0.754266002 |
| chr19:24033288-24033834 | 0.250419639 | 0.495678066 | 0.754298515 |
| chr1:13828823-13829155 | 0.288766748 | 0.495683992 | 0.754298515 |
| chr2:139353971-139354171 | -0.374491999 | 0.495695693 | 0.754298515 |
| chr11:45899296-45900221 | -0.22054182 | 0.495699456 | 0.754298515 |
| chr14:96364085-96364333 | 0.267955146 | 0.495705508 | 0.754298515 |
| chr3:49929242-49930161 | -0.231436609 | 0.495717541 | 0.754298572 |
| chr1:109397072-109397278 | 0.295784177 | 0.495725234 | 0.754298572 |
| chr4:20699971-20700801 | 0.221658946 | 0.495758064 | 0.754314638 |
| chr12:63901023-63901245 | 0.305950325 | 0.495768189 | 0.754314638 |
| chr12:50825363-50825604 | -0.346958306 | 0.495778575 | 0.754314638 |
| chr16:1782422-1783162 | 0.221186431 | 0.495783268 | 0.754314638 |
| chr17:38605936-38606221 | 0.279470662 | 0.495793942 | 0.754314638 |
| chr1:185822067-185822285 | 0.349935029 | 0.495802361 | 0.754314638 |
| chr11:2428319-2428519 | 0.325020245 | 0.495805729 | 0.754314638 |
| chr1:155135045-155135395 | 0.238216059 | 0.495816441 | 0.754314638 |
| chr1:70528543-70529052 | 0.249934018 | 0.495824394 | 0.754314638 |
| chr14:24263129-24263329 | 0.28656068 | 0.495848468 | 0.754336285 |
| chr1:26959703-26959903 | 0.34712633 | 0.495863239 | 0.754340656 |
| chr3:25095157-25095468 | -0.352341461 | 0.495875459 | 0.754340656 |
| chr12:57307771-57308049 | 0.319323591 | 0.495880876 | 0.754340656 |
| chr9:136774729-136774929 | 0.333254686 | 0.495924722 | 0.754392377 |
| chr10:13383166-13383403 | -0.319001756 | 0.495969764 | 0.754445916 |
| chr6:135234107-135234325 | -0.357982103 | 0.496027917 | 0.754519397 |
| chr2:106143686-106144056 | 0.292928371 | 0.496101333 | 0.754599329 |
| chr12:66688918-66689166 | 0.297248438 | 0.496109002 | 0.754599329 |
| chr9:131733585-131733971 | 0.25335978 | 0.49611001 | 0.754599329 |
| chr15:92666819-92667873 | -0.226690623 | 0.496130144 | 0.754614973 |
| chr3:14981755-14982229 | 0.278115004 | 0.496151498 | 0.754628198 |
| chr1:2133070-2133773 | 0.224439032 | 0.496161794 | 0.754628198 |
| chr7:45225323-45225523 | -0.346489701 | 0.496180391 | 0.754628198 |
| chr11:130916329-130916780 | -0.242500164 | 0.496182526 | 0.754628198 |
| chr4:176794607-176794950 | 0.263778937 | 0.496189589 | 0.754628198 |
| chr4:27010438-27010638 | 0.344715958 | 0.496197931 | 0.754628198 |
| chr1:198465678-198465890 | 0.312456418 | 0.496218857 | 0.754645045 |
| chr1:233876069-233876304 | 0.342486264 | 0.496229684 | 0.75464594 |
| chr6:158231798-158233193 | -0.223838571 | 0.496239144 | 0.75464594 |
| chr1:146051379-146052751 | 0.214036202 | 0.496285396 | 0.754700114 |
| chr22:31943799-31945016 | 0.218599186 | 0.496302985 | 0.754700114 |
| chr13:30125584-30125982 | -0.248725592 | 0.496323625 | 0.754700114 |
| chr8:101204826-101206555 | 0.212695833 | 0.496328343 | 0.754700114 |
| chr11:120316817-120317477 | -0.230347673 | 0.49633828 | 0.754700114 |
| chr3:94071954-94072168 | 0.329854244 | 0.49634047 | 0.754700114 |
| chr3:17233391-17233687 | 0.277378088 | 0.496343715 | 0.754700114 |
| chr17:28743123-28744148 | 0.220003608 | 0.496379364 | 0.754704921 |
| chr9:129790388-129790753 | 0.264369806 | 0.496379427 | 0.754704921 |
| chr7:151083371-151083601 | 0.268733335 | 0.496382547 | 0.754704921 |
| chr21:39612569-39613175 | -0.240425974 | 0.496387363 | 0.754704921 |
| chr3:183817832-183818149 | 0.304355641 | 0.496398106 | 0.754704921 |
| chr5:161432952-161433153 | -0.393728013 | 0.496414396 | 0.754704921 |

| | | | |
|---|---|---|---|
| chr10:22436699-22437741 | -0.22843984 | 0.496425196 | 0.754704921 |
| chr17:12902764-12903078 | 0.288893421 | 0.496425674 | 0.754704921 |
| chr9:38432947-38433201 | -0.305631018 | 0.49647622 | 0.754766789 |
| chr1:26110869-26111223 | 0.273808466 | 0.496486623 | 0.754767628 |
| chr13:73598297-73598584 | -0.35277853 | 0.496497372 | 0.754768995 |
| chr5:138337405-138338582 | -0.218200539 | 0.496508601 | 0.754770791 |
| chr13:32001235-32001633 | -0.248924649 | 0.496535283 | 0.754770791 |
| chr17:60654340-60654670 | 0.239220739 | 0.496535386 | 0.754770791 |
| chr8:1835288-1835493 | -0.336212977 | 0.496538847 | 0.754770791 |
| chr17:18780959-18781433 | 0.237115541 | 0.496547807 | 0.754770791 |
| chr15:70854724-70854924 | 0.351189004 | 0.496562637 | 0.75477836 |
| chrX:106404535-106404853 | -0.276697558 | 0.496596164 | 0.754807822 |
| chr2:86675631-86676107 | -0.251527627 | 0.496601721 | 0.754807822 |
| chr19:14206827-14207201 | 0.277863246 | 0.496640699 | 0.754852092 |
| chr2:216540002-216540288 | -0.292437164 | 0.496692612 | 0.754898713 |
| chr14:81169953-81170926 | -0.218099286 | 0.496700844 | 0.754898713 |
| chr1:44405134-44405344 | -0.337524578 | 0.49670093 | 0.754898713 |
| chr8:98045196-98046468 | 0.214792381 | 0.496713832 | 0.754903349 |
| chr2:73071208-73071964 | -0.220371084 | 0.496800821 | 0.755004942 |
| chr9:79233621-79233899 | -0.299457359 | 0.496802628 | 0.755004942 |
| chr1:223129396-223129659 | -0.293032924 | 0.496810239 | 0.755004942 |
| chr17:42667507-42667932 | 0.251279328 | 0.496825639 | 0.75501337 |
| chr6:32037856-32038329 | -0.311270578 | 0.496847694 | 0.755031912 |
| chr19:14689538-14690310 | 0.231052449 | 0.496876198 | 0.755060252 |
| chr6:153056316-153056606 | -0.31731767 | 0.496893334 | 0.755071318 |
| chr12:51371679-51371879 | 0.358079365 | 0.496922699 | 0.755100964 |
| chrX:108331159-108331727 | -0.250412282 | 0.496939533 | 0.75511157 |
| chr8:6405678-6406764 | -0.229821782 | 0.496950551 | 0.755113336 |
| chr12:66301984-66302980 | -0.216967821 | 0.497012308 | 0.755177393 |
| chr6:57038399-57038660 | 0.355300162 | 0.497012419 | 0.755177393 |
| chr12:78727110-78727402 | -0.328675375 | 0.497032875 | 0.755183085 |
| chr15:89575034-89575668 | 0.219885589 | 0.497035877 | 0.755183085 |
| chr5:23494542-23494952 | 0.250757853 | 0.497054516 | 0.75519643 |
| chr5:149431029-149431455 | -0.249749884 | 0.49712939 | 0.755294662 |
| chr12:12745-12995 | -0.340194115 | 0.497147668 | 0.755294662 |
| chr12:64780220-64781932 | 0.21499438 | 0.497152669 | 0.755294662 |
| chr16:89837733-89838079 | -0.247300215 | 0.497158601 | 0.755294662 |
| chr2:29196849-29197155 | 0.271542632 | 0.497181502 | 0.755300768 |
| chr21:33479297-33480528 | 0.216219715 | 0.497183861 | 0.755300768 |
| chr20:53258727-53258994 | 0.298864657 | 0.497192192 | 0.755300768 |
| chr1:172826083-172826299 | 0.328821667 | 0.497233746 | 0.755307758 |
| chr3:121433908-121434198 | -0.291527387 | 0.497235052 | 0.755307758 |
| chr8:117519975-117521269 | -0.215299977 | 0.49724162 | 0.755307758 |
| chr8:144095947-144096277 | 0.24457203 | 0.497241645 | 0.755307758 |
| chr17:6551960-6552748 | 0.239201477 | 0.497246509 | 0.755307758 |
| chr7:117873365-117873654 | -0.295861721 | 0.497255939 | 0.755307758 |
| chr1:52306519-52306721 | 0.338230722 | 0.497266774 | 0.755309244 |
| chr7:139447154-139447545 | -0.251575318 | 0.497304814 | 0.755341162 |
| chr1:51878161-51879382 | 0.215348838 | 0.497307504 | 0.755341162 |
| chr22:32474252-32475485 | 0.221413811 | 0.497326072 | 0.75534886 |

| | | | |
|---|---|---|---|
| chr15:69462076-69463190 | 0.218670024 | 0.497340969 | 0.75534886 |
| chr6:71510161-71510405 | -0.332000763 | 0.497349523 | 0.75534886 |
| chr8:12194082-12194522 | 0.239575456 | 0.49736062 | 0.75534886 |
| chr1:116268182-116268555 | 0.247814868 | 0.497372846 | 0.75534886 |
| chr3:28019424-28019738 | -0.285901454 | 0.497376967 | 0.75534886 |
| chr14:88977894-88978142 | 0.277601679 | 0.497390748 | 0.75534886 |
| chr6:26341413-26342084 | 0.227492728 | 0.497391438 | 0.75534886 |
| chr4:8961398-8961684 | 0.292004149 | 0.49743202 | 0.755395518 |
| chr2:219541877-219542088 | 0.259002452 | 0.497462287 | 0.755410331 |
| chr19:459990-460352 | 0.310255506 | 0.497466063 | 0.755410331 |
| chr5:65699691-65700027 | 0.335738268 | 0.497471351 | 0.755410331 |
| chr3:64484122-64484939 | 0.2141538 | 0.497494577 | 0.755430627 |
| chr8:110060988-110061203 | -0.287705825 | 0.497506072 | 0.755432253 |
| chr2:27735166-27736091 | -0.23368186 | 0.497515366 | 0.755432253 |
| chr9:79501438-79501638 | -0.379859157 | 0.497528691 | 0.755437516 |
| chr19:290066-290785 | 0.233510598 | 0.497567983 | 0.755482206 |
| chr9:19048904-19049915 | -0.219378908 | 0.497630809 | 0.755562624 |
| chr4:107760551-107760867 | -0.284839463 | 0.497653943 | 0.755582777 |
| chr5:133040691-133040973 | 0.288111288 | 0.497684638 | 0.755614408 |
| chr10:83992400-83992745 | -0.26069259 | 0.497703245 | 0.755627686 |
| chr19:2611161-2611448 | -0.336923872 | 0.497720656 | 0.755639147 |
| chr2:201860608-201860942 | 0.272490573 | 0.497774529 | 0.755684153 |
| chr1:226100428-226100786 | -0.299376767 | 0.49777744 | 0.755684153 |
| chr3:172809281-172809490 | -0.335342921 | 0.497788746 | 0.755684153 |
| chr1:27349288-27350050 | 0.236153905 | 0.49780522 | 0.755684153 |
| chr2:158901296-158901945 | -0.241953933 | 0.497807784 | 0.755684153 |
| chr2:43426508-43426862 | 0.279631478 | 0.497809476 | 0.755684153 |
| chr2:218445518-218445841 | -0.262104468 | 0.497822714 | 0.755689277 |
| chr7:46628668-46628900 | -0.353383812 | 0.497858578 | 0.755728747 |
| chr10:82282783-82283362 | -0.260699711 | 0.497875385 | 0.75573284 |
| chr14:70821982-70822308 | 0.27194993 | 0.497896063 | 0.75573284 |
| chr7:39733978-39734289 | -0.244673095 | 0.49790406 | 0.75573284 |
| chr3:132855016-132855240 | 0.339137547 | 0.497916526 | 0.75573284 |
| chr2:240060997-240061224 | -0.320427259 | 0.497919797 | 0.75573284 |
| chr5:133423837-133424071 | -0.295285536 | 0.497920453 | 0.75573284 |
| chr4:25401138-25401596 | 0.277392182 | 0.497951317 | 0.755751272 |
| chr7:66247302-66247759 | 0.280215675 | 0.497952324 | 0.755751272 |
| chr1:44406205-44406414 | 0.343295124 | 0.497965212 | 0.755755862 |
| chr6:136565846-136566535 | -0.227103898 | 0.497982822 | 0.755765563 |
| chr18:22932926-22934058 | -0.22307682 | 0.498007849 | 0.755765563 |
| chr11:67606452-67607076 | 0.228910838 | 0.498009648 | 0.755765563 |
| chr6:129733410-129733626 | -0.340713363 | 0.498011058 | 0.755765563 |
| chr9:137044499-137044765 | -0.2691356 | 0.498042887 | 0.755777751 |
| chr5:112692162-112692389 | 0.309929718 | 0.498043678 | 0.755777751 |
| chr1:202160546-202160784 | -0.256897345 | 0.49804868 | 0.755777751 |
| chr9:121640067-121640348 | 0.295660331 | 0.498061998 | 0.755782993 |
| chr2:133568187-133568474 | 0.305515695 | 0.498101639 | 0.755828176 |
| chr1:33350290-33350666 | 0.250893233 | 0.498143919 | 0.755877364 |
| chr19:49929066-49930227 | 0.217673123 | 0.498195495 | 0.75591191 |
| chr11:46369470-46369780 | 0.248365098 | 0.49820034 | 0.75591191 |

| | | | |
|---|---|---|---|
| chr14:83079124-83079768 | -0.257979127 | 0.498203256 | 0.75591191 |
| chr6:71886172-71887370 | -0.227916274 | 0.498206148 | 0.75591191 |
| chrX:102712346-102713707 | -0.218602415 | 0.498224811 | 0.755923916 |
| chr5:159703053-159703292 | -0.341932573 | 0.498233792 | 0.755923916 |
| chrX:4263082-4263282 | 0.38577523 | 0.498244404 | 0.755925048 |
| chr1:226061759-226063139 | 0.214668301 | 0.498281537 | 0.75595498 |
| chr5:177311816-177312818 | -0.215835068 | 0.498283864 | 0.75595498 |
| chr22:25110624-25111339 | -0.228647416 | 0.498311846 | 0.755982463 |
| chr5:61293134-61293393 | 0.283681979 | 0.498343472 | 0.756015473 |
| chr2:27293226-27293674 | 0.261154185 | 0.498364239 | 0.756017744 |
| chr22:45240920-45241219 | -0.283391752 | 0.498364703 | 0.756017744 |
| chr14:92690375-92690649 | 0.251585068 | 0.498395561 | 0.756022842 |
| chr3:138594034-138595127 | -0.214055206 | 0.498397559 | 0.756022842 |
| chr11:8343112-8343408 | -0.307247858 | 0.498397664 | 0.756022842 |
| chr15:59283799-59284354 | -0.233777603 | 0.498457214 | 0.756080397 |
| chr1:144101703-144101956 | 0.297633093 | 0.498460412 | 0.756080397 |
| chr8:13133098-13133691 | -0.21984763 | 0.498468305 | 0.756080397 |
| chr7:129271639-129272120 | -0.267354569 | 0.498482376 | 0.756080397 |
| chr15:45430263-45430758 | -0.259675409 | 0.498484944 | 0.756080397 |
| chr8:132768579-132768854 | 0.28164523 | 0.498515451 | 0.756098096 |
| chr14:78352439-78352824 | 0.297355197 | 0.498521392 | 0.756098096 |
| chr20:3767605-3768251 | 0.218524093 | 0.498534635 | 0.756098096 |
| chr4:112031725-112032047 | -0.282003332 | 0.498536085 | 0.756098096 |
| chr3:23095033-23095431 | -0.24997282 | 0.498608294 | 0.756192644 |
| chr11:28110167-28110439 | -0.251751979 | 0.498624864 | 0.756201694 |
| chr6:34237934-34238359 | -0.263515692 | 0.498637997 | 0.756201694 |
| chr3:164826908-164827108 | 0.323038204 | 0.498645782 | 0.756201694 |
| chr5:146041251-146041759 | -0.249251978 | 0.498661578 | 0.756201694 |
| chr8:6830176-6830377 | 0.385238709 | 0.498663609 | 0.756201694 |
| chr4:38860965-38861180 | 0.315683248 | 0.498756394 | 0.756319901 |
| chr8:133735633-133735833 | -0.334487771 | 0.4987613 | 0.756319901 |
| chr4:138555626-138555912 | -0.327420844 | 0.498783403 | 0.75633845 |
| chr10:119663945-119664406 | 0.249343246 | 0.498807048 | 0.756354835 |
| chr16:89193127-89193385 | -0.258131851 | 0.498834236 | 0.756354835 |
| chr18:7174779-7174979 | 0.359129962 | 0.498836584 | 0.756354835 |
| chr19:54024038-54024238 | 0.359129962 | 0.498836584 | 0.756354835 |
| chr6:85015429-85015832 | -0.260640582 | 0.498843565 | 0.756354835 |
| chr10:104546869-104547069 | -0.347444772 | 0.49890009 | 0.756403742 |
| chr2:99180760-99181560 | -0.224456126 | 0.498900852 | 0.756403742 |
| chr3:14352093-14352354 | 0.281290929 | 0.498905436 | 0.756403742 |
| chr8:124147196-124147396 | 0.308440592 | 0.498917353 | 0.756406842 |
| chr1:186404542-186404769 | -0.325214485 | 0.498928219 | 0.75640835 |
| chr5:73515264-73515787 | 0.270342497 | 0.499005863 | 0.75649799 |
| chr2:218010137-218010853 | 0.240915048 | 0.499011196 | 0.75649799 |
| chr17:65106416-65106949 | -0.246405519 | 0.499016965 | 0.75649799 |
| chr12:122526555-122527569 | 0.216884635 | 0.499069866 | 0.756563218 |
| chr21:45612175-45612505 | 0.274122731 | 0.499139683 | 0.756653085 |
| chr8:142029999-142030199 | -0.344119157 | 0.499148897 | 0.756653085 |
| chrX:109862411-109862777 | 0.269450094 | 0.499172814 | 0.756670757 |
| chr10:100656634-100656918 | 0.295049135 | 0.499189855 | 0.756670757 |

| | | | |
|---|---|---|---|
| chr17:76240848-76241126 | 0.279932489 | 0.499192887 | 0.756670757 |
| chr5:119420250-119420612 | -0.242188986 | 0.499200057 | 0.756670757 |
| chr17:48492622-48492958 | 0.282416171 | 0.499212191 | 0.756674181 |
| chr18:79988038-79989127 | -0.223529751 | 0.499239322 | 0.756700335 |
| chr10:121862702-12186291€ | 0.334317897 | 0.499266547 | 0.756724735 |
| chr3:10958121-10958372 | 0.296976977 | 0.499285003 | 0.756724735 |
| chr3:71247639-71247955 | -0.276388936 | 0.499285049 | 0.756724735 |
| chr2:47250750-47251023 | -0.320637572 | 0.499320658 | 0.756754169 |
| chr1:17125801-17126001 | 0.342234958 | 0.499332613 | 0.756754169 |
| chr21:44012905-44013308 | 0.262189029 | 0.499334098 | 0.756754169 |
| chr20:10218435-10219120 | 0.226525204 | 0.499398477 | 0.756831897 |
| chr7:7354648-7355073 | 0.230579011 | 0.499405141 | 0.756831897 |
| chr1:216859303-216859512 | 0.332680649 | 0.499417883 | 0.756836239 |
| chr7:22575623-22575873 | 0.319330993 | 0.499434616 | 0.756843165 |
| chr2:28683188-28683724 | -0.238067855 | 0.49949129 | 0.756843165 |
| chr3:2303627-2303827 | 0.336276166 | 0.499496894 | 0.756843165 |
| chr19:38304004-38304315 | 0.252238557 | 0.49949834 | 0.756843165 |
| chr7:1193683-1194358 | 0.232733991 | 0.499501982 | 0.756843165 |
| chr1:116982756-116982990 | 0.311720249 | 0.499507068 | 0.756843165 |
| chr16:28950521-28951147 | 0.220803083 | 0.499517698 | 0.756843165 |
| chr3:100590221-100590853 | 0.245566351 | 0.499527139 | 0.756843165 |
| chr14:54509333-54510493 | 0.214627658 | 0.499528457 | 0.756843165 |
| chr17:46225290-46225721 | -0.249652437 | 0.499531197 | 0.756843165 |
| chr2:211093777-211094036 | -0.302499318 | 0.499551655 | 0.756843165 |
| chr12:92055912-92056201 | -0.291531197 | 0.499555975 | 0.756843165 |
| chr11:2165206-2165406 | 0.312517053 | 0.499568415 | 0.756843165 |
| chr12:50039244-50039473 | 0.312517053 | 0.499568415 | 0.756843165 |
| chr1:151534739-151534970 | 0.253810333 | 0.499570618 | 0.756843165 |
| chr3:19855454-19855654 | -0.35378816 | 0.499618479 | 0.756873354 |
| chr9:92738532-92739150 | -0.239155536 | 0.499619651 | 0.756873354 |
| chr3:56556799-56557482 | -0.237540921 | 0.49962088 | 0.756873354 |
| chr19:46678147-46678450 | -0.263733641 | 0.499630057 | 0.756873354 |
| chr10:84076604-84076834 | 0.305348575 | 0.499686802 | 0.756930252 |
| chr18:76572615-76572971 | -0.277002205 | 0.499687375 | 0.756930252 |
| chr2:143162260-143162584 | -0.271491486 | 0.499720669 | 0.756965721 |
| chr17:59979038-59979307 | -0.269301941 | 0.499766541 | 0.757020242 |
| chr11:12599486-12599750 | -0.314535735 | 0.499781768 | 0.75702172 |
| chr18:26656551-26657220 | 0.245444269 | 0.499790871 | 0.75702172 |
| chr16:68245048-68245984 | 0.222416898 | 0.499797157 | 0.75702172 |
| chr11:75723215-75723577 | -0.252939458 | 0.499825845 | 0.757050208 |
| chr17:45355206-45355506 | -0.291941004 | 0.499860961 | 0.757088429 |
| chr19:47697969-47698542 | 0.23932889 | 0.499876654 | 0.757097233 |
| chr1:174158905-174160472 | -0.211637311 | 0.499917982 | 0.757137707 |
| chr6:5078344-5078619 | -0.28417072 | 0.499924691 | 0.757137707 |
| chr3:180693633-180693887 | -0.327880657 | 0.499933022 | 0.757137707 |
| chr5:75050401-75050679 | -0.275569332 | 0.499970956 | 0.757156762 |
| chr6:10415047-10415330 | 0.247196458 | 0.499972029 | 0.757156762 |
| chr3:44648525-44649199 | 0.242342582 | 0.499975249 | 0.757156762 |
| chr2:205025834-205026195 | -0.290214993 | 0.500015941 | 0.75720342 |
| chrX:38801025-38801611 | -0.224102311 | 0.50005725 | 0.75725101 |

| | | | |
|---|---|---|---|
| chr2:230762427-230762644 | -0.333466379 | 0.500080939 | 0.757271917 |
| chr3:183193726-183194236 | 0.253253812 | 0.500113184 | 0.757292832 |
| chr6:73451662-73452487 | 0.231251201 | 0.50011786 | 0.757292832 |
| chr1:196660973-196661272 | -0.260656509 | 0.500124402 | 0.757292832 |
| chr8:102627486-102627758 | 0.270588318 | 0.500166356 | 0.757341393 |
| chr4:147984054-147984362 | 0.258261543 | 0.500185604 | 0.757355572 |
| chr2:113674324-113674529 | -0.355429968 | 0.500196359 | 0.75735689 |
| chr3:64549967-64550204 | 0.241263655 | 0.500220023 | 0.757377753 |
| chr1:117754157-117754364 | 0.294329912 | 0.500238234 | 0.757390361 |
| chr12:70123089-70123289 | 0.349760542 | 0.500254351 | 0.757399796 |
| chr6:7542040-7542496 | 0.246802558 | 0.500324292 | 0.75749008 |
| chr5:175797805-175798052 | -0.293785292 | 0.500340954 | 0.75749008 |
| chr9:134286670-134286893 | -0.343165055 | 0.500346414 | 0.75749008 |
| chr12:119479199-119479755 | 0.232254439 | 0.500353527 | 0.75749008 |
| chr6:16964919-16965254 | -0.262886281 | 0.500405347 | 0.757553563 |
| chr19:3626751-3627326 | 0.235659911 | 0.500451575 | 0.75759139 |
| chr2:218068874-218069132 | -0.291511235 | 0.500490018 | 0.75759139 |
| chr3:126522279-126522578 | 0.263550022 | 0.500497787 | 0.75759139 |
| chr1:67771360-67771560 | 0.345571809 | 0.500502235 | 0.75759139 |
| chr10:102450087-102451109 | 0.223772323 | 0.500507843 | 0.75759139 |
| chrX:72306959-72307317 | -0.318721635 | 0.500509504 | 0.75759139 |
| chr1:187265017-187265311 | -0.272583104 | 0.500513939 | 0.75759139 |
| chr17:9292197-9292406 | -0.334751529 | 0.500516075 | 0.75759139 |
| chr16:79581655-79581934 | -0.350985923 | 0.50051932 | 0.75759139 |
| chr2:212776928-212777459 | -0.251978993 | 0.50053203 | 0.757595662 |
| chr22:50305903-50306156 | 0.283508297 | 0.500576262 | 0.757647644 |
| chr8:100721951-100722981 | 0.211233428 | 0.500592794 | 0.757657701 |
| chr10:70672150-70672364 | -0.33453636 | 0.50060348 | 0.757658908 |
| chr2:111659458-111659830 | 0.28281619 | 0.500629097 | 0.757662902 |
| chr3:172213990-172214266 | -0.295924871 | 0.500635168 | 0.757662902 |
| chr13:49585044-49586315 | -0.220595047 | 0.50063784 | 0.757662902 |
| chr9:136807739-136807993 | 0.265706111 | 0.500645672 | 0.757662902 |
| chr16:68528581-68529031 | 0.246949616 | 0.500657543 | 0.757665901 |
| chr11:128905307-128906230 | 0.226759906 | 0.50071024 | 0.757730363 |
| chr13:38641632-38642007 | -0.281821938 | 0.500735631 | 0.757730363 |
| chr6:158849915-158850260 | -0.266902907 | 0.500740844 | 0.757730363 |
| chr4:143513005-143514255 | -0.211799552 | 0.500742977 | 0.757730363 |
| chr4:104395260-104395597 | -0.302597849 | 0.500756974 | 0.757730363 |
| chr5:103015058-103015370 | -0.282286213 | 0.500759474 | 0.757730363 |
| chr3:6994407-6994608 | 0.331345146 | 0.500784239 | 0.757740567 |
| chr9:123584897-123585097 | 0.352430455 | 0.500790787 | 0.757740567 |
| chr7:105309816-105310145 | -0.286989564 | 0.500795885 | 0.757740567 |
| chr12:51024925-51025130 | 0.288120945 | 0.500805779 | 0.757740574 |
| chr3:16536054-16536329 | 0.322632242 | 0.500827878 | 0.757748325 |
| chr11:16841586-16841786 | -0.363996338 | 0.50084057 | 0.757748325 |
| chr11:38791405-38791622 | -0.363996338 | 0.50084057 | 0.757748325 |
| chr1:226424541-226424811 | -0.29061904 | 0.500856719 | 0.757757795 |
| chr3:143644473-143644781 | -0.287546941 | 0.500870207 | 0.757763238 |
| chr2:50492537-50492766 | 0.291322029 | 0.500894497 | 0.757784473 |
| chr3:73101092-73101314 | 0.311660731 | 0.500904022 | 0.757784473 |

| | | | |
|---|---|---|---|
| chr12:56985752-56986124 | 0.258061091 | 0.500925555 | 0.757802087 |
| chr16:71845218-71846382 | -0.216533537 | 0.50094532 | 0.757817024 |
| chr7:73593761-73594000 | -0.310230477 | 0.500987316 | 0.757865592 |
| chr6:142945031-142947729 | -0.209913148 | 0.50101129 | 0.757886896 |
| chr14:83639215-83639947 | -0.225782001 | 0.501031752 | 0.757902886 |
| chrX:24676498-24676730 | -0.311249046 | 0.501066667 | 0.757910709 |
| chr18:10787376-10787632 | -0.315225819 | 0.501073111 | 0.757910709 |
| chr19:5903717-5905073 | 0.212354817 | 0.5010779 | 0.757910709 |
| chr2:10447937-10448832 | 0.219228214 | 0.501090495 | 0.757910709 |
| chr14:23306575-23307162 | 0.220239958 | 0.501092661 | 0.757910709 |
| chr14:55213075-55213521 | 0.246550985 | 0.501096273 | 0.757910709 |
| chr1:202379800-202380666 | 0.218703785 | 0.501148554 | 0.757958183 |
| chr1:94820148-94820751 | 0.250594474 | 0.501171182 | 0.757958183 |
| chr8:128152996-128153509 | 0.22190974 | 0.50117472 | 0.757958183 |
| chr12:93541685-93542303 | -0.240169251 | 0.501189256 | 0.757958183 |
| chr11:130178862-130179083 | -0.349017185 | 0.501206744 | 0.757958183 |
| chr7:48067669-48067971 | -0.347042849 | 0.501214785 | 0.757958183 |
| chr17:28380205-28380600 | -0.248929221 | 0.501216162 | 0.757958183 |
| chr5:43306062-43306305 | 0.296291957 | 0.501219772 | 0.757958183 |
| chr1:203026168-203026901 | -0.218020812 | 0.501230527 | 0.757958183 |
| chr14:67240554-67242047 | 0.211644746 | 0.501236231 | 0.757958183 |
| chr13:33931467-33931805 | -0.266616983 | 0.501236474 | 0.757958183 |
| chr12:47637109-47637599 | 0.249876229 | 0.501266425 | 0.757988515 |
| chr10:73495237-73496445 | -0.219033442 | 0.501295196 | 0.758011346 |
| chr1:71087119-71087320 | -0.345568916 | 0.501307217 | 0.758011346 |
| chr2:141554897-141555126 | -0.310260389 | 0.501311202 | 0.758011346 |
| chr17:63675583-63676284 | -0.22493535 | 0.501340138 | 0.758031613 |
| chr1:23916866-23917110 | 0.313949277 | 0.501347386 | 0.758031613 |
| chr1:156114138-156114858 | -0.219690115 | 0.501358075 | 0.758031613 |
| chr3:377150-377609 | -0.25571733 | 0.501364179 | 0.758031613 |
| chr11:20672709-20672981 | -0.333761746 | 0.50139527 | 0.758063663 |
| chr14:34981904-34983375 | 0.211088229 | 0.501443145 | 0.758121086 |
| chr14:96774463-96774663 | 0.368769033 | 0.501455597 | 0.758123212 |
| chr12:85821735-85822035 | -0.298670179 | 0.501464339 | 0.758123212 |
| chr6:30646845-30647555 | -0.22271074 | 0.501494072 | 0.758142868 |
| chr7:143041612-143041921 | -0.32080245 | 0.50149713 | 0.758142868 |
| chr11:8688715-8689322 | 0.222015649 | 0.501514708 | 0.758144916 |
| chr1:106467114-106467880 | 0.221858296 | 0.501518274 | 0.758144916 |
| chr8:67062002-67062726 | -0.221662605 | 0.501548515 | 0.758175672 |
| chr2:119431444-119432048 | 0.24455968 | 0.501586913 | 0.758218759 |
| chr10:70528778-70528978 | -0.325222674 | 0.50161264 | 0.75824269 |
| chrX:14927539-14927773 | 0.336250869 | 0.501628988 | 0.758252442 |
| chr4:185325813-185326018 | -0.330057367 | 0.501653253 | 0.758263156 |
| chr10:99589590-99589793 | -0.349664689 | 0.501657395 | 0.758263156 |
| chr14:103260302-103260502 | -0.345630007 | 0.501665764 | 0.758263156 |
| chr3:121834125-121835588 | -0.213867084 | 0.501678262 | 0.758267088 |
| chr1:151967989-151968380 | 0.231961285 | 0.501700262 | 0.758285383 |
| chr6:41039238-41039757 | 0.239408839 | 0.501745454 | 0.758338729 |
| chr15:101050659-101051078 | -0.267176086 | 0.501787411 | 0.758360757 |
| chr11:80727073-80727276 | -0.352358979 | 0.501801173 | 0.758360757 |

| | | | |
|---|---|---|---|
| chr6:69866792-69867880 | -0.224249368 | 0.501805715 | 0.758360757 |
| chr7:122198693-122198918 | -0.346504674 | 0.501810743 | 0.758360757 |
| chr8:119663677-119664061 | -0.311108216 | 0.501816342 | 0.758360757 |
| chr20:4686967-4687471 | -0.254493376 | 0.501825192 | 0.758360757 |
| chr1:69566942-69568502 | 0.222119773 | 0.501829311 | 0.758360757 |
| chr5:74389695-74390198 | -0.276096619 | 0.501840032 | 0.758362002 |
| chrX:72276938-72277571 | 0.230294017 | 0.501877739 | 0.758401779 |
| chr20:58481450-58481879 | 0.232293188 | 0.501886151 | 0.758401779 |
| chr20:57091355-57091555 | -0.266160142 | 0.501911772 | 0.758414731 |
| chr5:38258444-38259421 | -0.230677502 | 0.501914518 | 0.758414731 |
| chr10:79157410-79157908 | -0.237272052 | 0.501952428 | 0.758447357 |
| chr3:126414886-126415139 | 0.307628085 | 0.501955907 | 0.758447357 |
| chr6:89940355-89940768 | -0.267959329 | 0.501987215 | 0.758479707 |
| chr19:55179631-55179880 | -0.288177557 | 0.501998826 | 0.758482292 |
| chr15:68138709-68139318 | 0.25310099 | 0.502025513 | 0.758496202 |
| chr11:3049879-3050167 | -0.276600974 | 0.50202783 | 0.758496202 |
| chr17:12730954-12731536 | -0.231497231 | 0.502075523 | 0.758553303 |
| chr11:35295700-35296201 | 0.248608522 | 0.502085719 | 0.75855375 |
| chr2:222860698-222861664 | -0.223904171 | 0.502122339 | 0.758584746 |
| chr2:217325901-217326141 | -0.344842401 | 0.502133715 | 0.758584746 |
| chr3:127073951-127074435 | 0.238103262 | 0.502135937 | 0.758584746 |
| chr19:40715562-40716030 | 0.231248806 | 0.502162795 | 0.758609788 |
| chr7:50649711-50650054 | 0.294689083 | 0.502172314 | 0.758609788 |
| chr3:67292507-67293316 | -0.241316996 | 0.502207089 | 0.758647363 |
| chr2:237291543-237291745 | -0.349517012 | 0.502318478 | 0.758800671 |
| chr2:241768306-241768533 | -0.365474473 | 0.502344833 | 0.758825522 |
| chr1:109540635-109540835 | 0.319023075 | 0.502372606 | 0.758852514 |
| chr1:197327833-197328182 | 0.273037019 | 0.5024339 | 0.75893014 |
| chr10:103736835-103737134 | -0.276183685 | 0.502483369 | 0.758970206 |
| chr3:15051447-15051920 | -0.228595792 | 0.502490937 | 0.758970206 |
| chr4:53706511-53706715 | -0.33552544 | 0.502496335 | 0.758970206 |
| chr15:33735733-33735965 | -0.324546135 | 0.502508506 | 0.758970206 |
| chr13:78121575-78121795 | -0.360134891 | 0.502514232 | 0.758970206 |
| chr7:3302129-3302561 | -0.249880231 | 0.502527699 | 0.758970206 |
| chr14:61526913-61527113 | 0.277828581 | 0.502529763 | 0.758970206 |
| chr2:219509660-219510134 | 0.24380936 | 0.502559457 | 0.759000092 |
| chr2:70034912-70035486 | -0.253375964 | 0.502590804 | 0.759032475 |
| chr17:81871008-81871801 | 0.220315163 | 0.502603621 | 0.75903687 |
| chr11:68354330-68354746 | -0.258670621 | 0.502650185 | 0.759081127 |
| chr8:38030170-38031741 | 0.216667119 | 0.502661558 | 0.759081127 |
| chr14:105504343-105504567 | 0.308998088 | 0.502666275 | 0.759081127 |
| chr10:72272916-72273385 | -0.228231775 | 0.502679928 | 0.759081127 |
| chr11:79940105-79940546 | 0.244928037 | 0.502688735 | 0.759081127 |
| chr11:33257079-33257870 | 0.224773749 | 0.502698317 | 0.759081127 |
| chr3:169835497-169835761 | -0.316119165 | 0.502702274 | 0.759081127 |
| chr19:49866985-49867722 | 0.223035021 | 0.502756358 | 0.759144055 |
| chr14:65751641-65751841 | 0.350754551 | 0.502763763 | 0.759144055 |
| chr17:9696660-9696890 | -0.372744642 | 0.502790106 | 0.75915645 |
| chr2:240513943-240514163 | -0.366250856 | 0.502809916 | 0.75915645 |
| chr16:4936950-4937486 | 0.239391142 | 0.502819773 | 0.75915645 |

| | | | |
|---|---|---|---|
| chr4:35253180-35253955 | -0.239971592 | 0.502831959 | 0.75915645 |
| chr19:44002566-44003588 | 0.230046665 | 0.502832405 | 0.75915645 |
| chr2:73070599-73071064 | -0.236944467 | 0.502837642 | 0.75915645 |
| chr3:117996944-117997835 | -0.22140644 | 0.502841327 | 0.75915645 |
| chr15:57706351-57707238 | -0.221065609 | 0.502886061 | 0.759189881 |
| chr20:17569021-17570837 | 0.210520562 | 0.502894153 | 0.759189881 |
| chr4:112327896-112328179 | -0.281877159 | 0.502937288 | 0.759189881 |
| chrX:14095705-14096064 | -0.272504599 | 0.502938939 | 0.759189881 |
| chr19:33041765-33042106 | 0.282559685 | 0.502940439 | 0.759189881 |
| chr22:41403772-41404193 | 0.241906968 | 0.502940865 | 0.759189881 |
| chr17:81891400-81892133 | 0.218718916 | 0.502946739 | 0.759189881 |
| chr22:46013414-46014283 | 0.226195902 | 0.502953257 | 0.759189881 |
| chr14:78021984-78022319 | -0.262992679 | 0.502965396 | 0.759189881 |
| chr14:92544078-92544442 | -0.279080859 | 0.502968499 | 0.759189881 |
| chrX:127968319-127968520 | 0.341439646 | 0.502981611 | 0.759189881 |
| chr1:66271250-66272319 | 0.222027659 | 0.502989053 | 0.759189881 |
| chr3:196163757-196163957 | -0.36002962 | 0.502992278 | 0.759189881 |
| chr2:74198258-74199389 | -0.214208304 | 0.503015507 | 0.759209986 |
| chr11:118111285-118111528 | -0.321390847 | 0.503045059 | 0.759217598 |
| chr3:101548852-101549052 | 0.326151644 | 0.503049846 | 0.759217598 |
| chr13:28166473-28166700 | -0.332691603 | 0.503050276 | 0.759217598 |
| chr10:50991660-50992277 | -0.253189153 | 0.503063338 | 0.759222358 |
| chr10:103555331-103555801 | -0.260361577 | 0.503095861 | 0.759223982 |
| chr5:10441060-10442432 | -0.211663314 | 0.503097156 | 0.759223982 |
| chr16:48165311-48165783 | 0.253716221 | 0.503097236 | 0.759223982 |
| chr17:55238112-55239200 | -0.21291037 | 0.503104049 | 0.759223982 |
| chr15:98650529-98650739 | 0.268746932 | 0.503121398 | 0.759235209 |
| chr14:79131562-79132145 | 0.248223379 | 0.50316768 | 0.759259709 |
| chr16:71499828-71500032 | 0.311164485 | 0.503173271 | 0.759259709 |
| chr2:38750778-38752023 | -0.211097191 | 0.503176719 | 0.759259709 |
| chr4:88456379-88457236 | -0.225035707 | 0.503177269 | 0.759259709 |
| chr3:101381939-101382139 | -0.335375353 | 0.503199778 | 0.759278719 |
| chr10:84429955-84430399 | -0.249175691 | 0.503223504 | 0.759299567 |
| chr5:271049-272001 | 0.214069687 | 0.503255024 | 0.759332174 |
| chr1:203216291-203216535 | 0.257841982 | 0.503273725 | 0.75933765 |
| chr2:43200144-43200344 | -0.328768746 | 0.503285447 | 0.75933765 |
| chr3:148820950-148821150 | 0.343443131 | 0.503315991 | 0.75933765 |
| chr2:227325910-227326153 | -0.283140387 | 0.503316179 | 0.75933765 |
| chr16:88793639-88793999 | -0.2696705 | 0.503318204 | 0.75933765 |
| chr14:68747383-68747811 | -0.265806495 | 0.503320613 | 0.75933765 |
| chr3:120710423-120710707 | 0.272957082 | 0.503328025 | 0.75933765 |
| chr5:88884008-88884410 | -0.265778704 | 0.503346051 | 0.759349894 |
| chr16:3263523-3263962 | 0.240102428 | 0.503394855 | 0.759403161 |
| chr15:59022038-59022423 | -0.313491824 | 0.503401182 | 0.759403161 |
| chr20:53866680-53867054 | 0.214816173 | 0.50341504 | 0.759404566 |
| chr17:79042432-79042632 | 0.301695714 | 0.503421935 | 0.759404566 |
| chr3:94336124-94336677 | -0.247691365 | 0.503487055 | 0.759487846 |
| chr1:51422068-51422321 | -0.329933404 | 0.503502974 | 0.759489189 |
| chr5:15383495-15383696 | -0.327675478 | 0.503512633 | 0.759489189 |
| chr8:107667744-107667944 | 0.319155019 | 0.503517682 | 0.759489189 |

| | | | |
|---|---|---|---|
| chr1:65527690-65528107 | -0.278649347 | 0.503544774 | 0.759491023 |
| chr20:4289920-4290237 | 0.293731586 | 0.503550954 | 0.759491023 |
| chr3:172298222-172298620 | 0.260607242 | 0.5035575 | 0.759491023 |
| chr22:46762128-46762819 | 0.236745353 | 0.503568627 | 0.759491023 |
| chr2:74855969-74856174 | -0.336218346 | 0.503581742 | 0.759491023 |
| chr3:5893770-5894041 | -0.336218346 | 0.503581742 | 0.759491023 |
| chr17:80008440-80008720 | 0.26625421 | 0.503588615 | 0.759491023 |
| chrX:18353995-18354259 | -0.320542615 | 0.503598195 | 0.759491023 |
| chr14:44626742-44626942 | 0.337013665 | 0.503608409 | 0.759491478 |
| chr5:112896536-112896821 | -0.263730318 | 0.503668705 | 0.75956746 |
| chr3:45555185-45555570 | -0.281974749 | 0.503730983 | 0.759646428 |
| chr6:125533841-125534346 | -0.262379113 | 0.503755561 | 0.759666158 |
| chr11:60775396-60775916 | 0.231662428 | 0.503763895 | 0.759666158 |
| chr8:40106513-40106767 | -0.353486973 | 0.503785064 | 0.759683129 |
| chr6:169950645-169950903 | 0.285451668 | 0.503810311 | 0.75970625 |
| chr1:152941156-152941362 | 0.316195452 | 0.503845409 | 0.759732096 |
| chr6:157981499-157982068 | 0.240800607 | 0.503859638 | 0.759732096 |
| chr2:215971786-215972071 | -0.27566097 | 0.503865613 | 0.759732096 |
| chr18:41075008-41075669 | -0.255061544 | 0.503872701 | 0.759732096 |
| chr5:38713180-38713500 | -0.317849754 | 0.50387957 | 0.759732096 |
| chr2:232365670-232365944 | -0.30586354 | 0.50388985 | 0.759732096 |
| chr1:19596612-19597151 | -0.233293523 | 0.503896859 | 0.759732096 |
| chr3:138961317-138961568 | -0.322154191 | 0.503915741 | 0.759745614 |
| chr5:148179318-148179533 | -0.346480677 | 0.503927908 | 0.75974901 |
| chr12:52964628-52964881 | 0.276151097 | 0.503954047 | 0.759773468 |
| chr17:50487254-50487992 | -0.237095382 | 0.503977725 | 0.759784094 |
| chr17:77473972-77474198 | -0.317018882 | 0.503980927 | 0.759784094 |
| chr17:73215460-73215912 | 0.248750011 | 0.504020391 | 0.759828639 |
| chr17:56914684-56915056 | -0.241830724 | 0.504041947 | 0.759846185 |
| chr21:40772800-40773034 | -0.351512144 | 0.504058199 | 0.759855735 |
| chr4:119559072-119559307 | 0.328483692 | 0.504074137 | 0.759864811 |
| chr18:12947255-12948671 | 0.21126197 | 0.504094357 | 0.759880342 |
| chr18:27605328-27605584 | -0.307941462 | 0.504126526 | 0.759913884 |
| chr2:110212605-110212871 | 0.301316484 | 0.504137398 | 0.759915323 |
| chr20:42382716-42382916 | 0.262735569 | 0.504188143 | 0.759976863 |
| chr12:63961882-63962134 | 0.332089689 | 0.504205161 | 0.75997808 |
| chr9:15510248-15510566 | 0.263878512 | 0.504208787 | 0.75997808 |
| chr3:2097565-2099017 | -0.214440421 | 0.504281825 | 0.760043995 |
| chr11:57560239-57560531 | 0.271642991 | 0.504293133 | 0.760043995 |
| chr17:50603397-50603649 | 0.266189954 | 0.504296734 | 0.760043995 |
| chr17:64025758-64026170 | -0.230974873 | 0.504299996 | 0.760043995 |
| chr11:14969865-14970146 | -0.269499581 | 0.504307114 | 0.760043995 |
| chr22:42292474-42292674 | 0.332025856 | 0.504312035 | 0.760043995 |
| chr18:13933934-13934285 | 0.295370746 | 0.504339956 | 0.760058393 |
| chr8:119207939-119209959 | -0.210032789 | 0.504341428 | 0.760058393 |
| chr5:107422424-107422801 | -0.3317353 | 0.50437596 | 0.760085986 |
| chr19:57814487-57815276 | 0.22049332 | 0.504387951 | 0.760085986 |
| chr16:20084690-20084980 | 0.299929575 | 0.504389497 | 0.760085986 |
| chr16:12803404-12804121 | 0.22100678 | 0.504402769 | 0.760091037 |
| chr8:111358171-111358465 | -0.31455352 | 0.504433581 | 0.760119901 |

| | | | |
|---|---|---|---|
| chr16:67396277-67396663 | -0.260211176 | 0.504441764 | 0.760119901 |
| chr3:57079207-57079672 | 0.254526485 | 0.504465013 | 0.760139985 |
| chr5:58502097-58502407 | 0.31535891 | 0.504496639 | 0.760172691 |
| chr11:61081232-61081511 | -0.313847344 | 0.504565079 | 0.760248711 |
| chr3:138097453-138097835 | 0.246189121 | 0.504566935 | 0.760248711 |
| chr19:4402260-4402686 | 0.234989658 | 0.504600563 | 0.760271333 |
| chr2:106863416-106863688 | 0.306613303 | 0.504601794 | 0.760271333 |
| chr5:83718286-83718492 | -0.279711972 | 0.504620447 | 0.760284487 |
| chr7:34036378-34036789 | -0.262469262 | 0.50464829 | 0.760302882 |
| chr3:147361849-147362153 | 0.298736476 | 0.504652501 | 0.760302882 |
| chr14:23237683-23238090 | -0.237465632 | 0.504671467 | 0.760305853 |
| chrX:48648416-48648616 | 0.337803777 | 0.504674319 | 0.760305853 |
| chr2:54115278-54116157 | -0.214267853 | 0.504703417 | 0.76033474 |
| chr17:37488885-37489145 | 0.263849003 | 0.504720031 | 0.76034482 |
| chr3:128665462-128665721 | -0.326530391 | 0.504730446 | 0.76034556 |
| chr2:96305082-96305996 | 0.220241913 | 0.504772477 | 0.760389396 |
| chr17:74971896-74972904 | 0.217199358 | 0.50478921 | 0.760389396 |
| chr6:151014335-151014652 | 0.269553461 | 0.504790293 | 0.760389396 |
| chr9:124150093-124150293 | -0.348938936 | 0.50479924 | 0.760389396 |
| chr11:554856-555208 | -0.270036082 | 0.504843334 | 0.760429514 |
| chr7:2511439-2511639 | 0.282600713 | 0.504847473 | 0.760429514 |
| chr4:7431796-7432029 | -0.319529389 | 0.504855647 | 0.760429514 |
| chr1:30911404-30911631 | 0.27631963 | 0.504881811 | 0.760453974 |
| chr11:107928016-10792873[illegible] | 0.218082581 | 0.504898044 | 0.760463475 |
| chr5:44808578-44809991 | -0.211456411 | 0.504916702 | 0.760476628 |
| chr10:52313685-52315067 | -0.211014415 | 0.504939749 | 0.760496392 |
| chr2:165874469-165874760 | -0.280009467 | 0.50496351 | 0.760517229 |
| chr1:214648939-214649320 | -0.271890825 | 0.504974998 | 0.760519583 |
| chr20:63230898-63231138 | 0.266231635 | 0.50503876 | 0.760600062 |
| chr1:232776691-232776966 | -0.334365314 | 0.505048288 | 0.760600062 |
| chr19:41403267-41403530 | -0.336706299 | 0.505100172 | 0.760643929 |
| chr22:30289399-30289910 | 0.228816468 | 0.50510252 | 0.760643929 |
| chr19:43533110-43534386 | 0.211486797 | 0.505107198 | 0.760643929 |
| chr10:73644184-73644465 | -0.296447679 | 0.505129988 | 0.760663298 |
| chr16:88792359-88792760 | -0.254521892 | 0.505143298 | 0.760668392 |
| chr17:75940673-75941419 | 0.222211181 | 0.50515754 | 0.760670948 |
| chr6:136921711-136921932 | -0.350691401 | 0.505164851 | 0.760670948 |
| chr6:119978356-119978580 | 0.316823528 | 0.505200649 | 0.760709903 |
| chr3:71681330-71681530 | -0.336560875 | 0.505242192 | 0.760757506 |
| chr12:32755724-32756123 | -0.22523195 | 0.505272924 | 0.760777856 |
| chr7:149284956-149285668 | -0.216145531 | 0.505275565 | 0.760777856 |
| chrX:10769276-10769501 | -0.385590247 | 0.505299709 | 0.760799259 |
| chrX:39134024-39134304 | -0.310571951 | 0.505361734 | 0.760877696 |
| chr9:113634562-113634762 | 0.347562423 | 0.505372798 | 0.760878206 |
| chr2:32063039-32064496 | -0.21097152 | 0.50539949 | 0.760878206 |
| chr12:4648505-4649218 | -0.251690706 | 0.505403178 | 0.760878206 |
| chr2:71798406-71798698 | -0.261381406 | 0.505417441 | 0.760878206 |
| chr14:69261247-69261456 | -0.321452979 | 0.505420902 | 0.760878206 |
| chrX:134796155-134797433 | 0.215562976 | 0.505421893 | 0.760878206 |
| chr3:98986084-98986452 | 0.285807193 | 0.505431586 | 0.760878206 |

| | | | |
|---|---|---|---|
| chr12:98838466-98838743 | -0.310542448 | 0.505455653 | 0.760899489 |
| chr7:106709756-106709957 | -0.367957436 | 0.505487343 | 0.760902112 |
| chr11:47641980-47642919 | 0.212983467 | 0.50550357 | 0.760902112 |
| chr16:10876988-10877334 | 0.274202107 | 0.505516085 | 0.760902112 |
| chr4:138233883-138234278 | 0.300117363 | 0.505522862 | 0.760902112 |
| chr17:17205853-17207110 | -0.213057056 | 0.505547949 | 0.760902112 |
| chr8:106080859-106081350 | 0.241370623 | 0.505549439 | 0.760902112 |
| chr5:1522098-1522298 | 0.294215631 | 0.505555472 | 0.760902112 |
| chr7:95342937-95343178 | 0.354143392 | 0.505572979 | 0.760902112 |
| chr2:228568230-228568507 | -0.316340442 | 0.505582879 | 0.760902112 |
| chr12:76952942-76953585 | -0.23555058 | 0.505591126 | 0.760902112 |
| chr3:127628912-127629312 | 0.253011987 | 0.505598992 | 0.760902112 |
| chr4:48943486-48943688 | 0.320878003 | 0.50560872 | 0.760902112 |
| chr15:98519068-98519319 | 0.287973022 | 0.505610259 | 0.760902112 |
| chr10:125045701-125046059 | 0.258978871 | 0.505612048 | 0.760902112 |
| chr1:153783366-153784212 | 0.21890721 | 0.505615368 | 0.760902112 |
| chr10:26698300-26698558 | -0.285227788 | 0.505616286 | 0.760902112 |
| chr12:63806129-63806467 | 0.277421933 | 0.505702008 | 0.761011667 |
| chr3:5891601-5892235 | -0.256452357 | 0.505708949 | 0.761011667 |
| chr11:65040768-65041590 | 0.220225829 | 0.505738408 | 0.761041053 |
| chr12:57771671-57773023 | 0.210237817 | 0.50575275 | 0.761047687 |
| chr3:42979189-42979725 | 0.26209278 | 0.50576363 | 0.761049114 |
| chr20:13839912-13840116 | -0.344943556 | 0.505832469 | 0.761108114 |
| chr7:67371860-67372100 | -0.29885147 | 0.50583264 | 0.761108114 |
| chr8:139799970-139800225 | -0.29885147 | 0.50583264 | 0.761108114 |
| chr12:106226898-106227098 | 0.325331957 | 0.505871691 | 0.761136281 |
| chr5:54867304-54867616 | -0.338642366 | 0.505874502 | 0.761136281 |
| chr11:59058316-59058757 | 0.239982362 | 0.505881877 | 0.761136281 |
| chr2:168773732-168774088 | -0.255345368 | 0.505891094 | 0.761136281 |
| chr18:9135933-9137526 | 0.210284941 | 0.505910282 | 0.761150205 |
| chr1:230825040-230825544 | 0.237280179 | 0.505947649 | 0.761185448 |
| chr17:56715719-56716256 | -0.233217724 | 0.505953576 | 0.761185448 |
| chr1:173255300-173255518 | 0.311717587 | 0.505966105 | 0.761189352 |
| chr5:6733016-6733429 | -0.261793108 | 0.505999305 | 0.761224353 |
| chr2:45585314-45585570 | 0.300949629 | 0.506029751 | 0.761255209 |
| chr10:5446353-5446899 | 0.223495877 | 0.506059527 | 0.761285056 |
| chr16:28945548-28945970 | 0.233909218 | 0.506070198 | 0.761286163 |
| chr7:97971442-97972710 | -0.220603697 | 0.506090151 | 0.761292678 |
| chr2:219368788-219369262 | -0.25050522 | 0.506094401 | 0.761292678 |
| chr7:33757615-33757815 | -0.328087781 | 0.506126336 | 0.761325771 |
| chr14:59263137-59263402 | 0.278493144 | 0.506150991 | 0.761347911 |
| chr8:136020491-136020913 | 0.259712482 | 0.506163968 | 0.761352485 |
| chr5:40797500-40798934 | 0.209863983 | 0.506197105 | 0.761383274 |
| chr17:60067975-60068353 | 0.226068344 | 0.506209516 | 0.761383274 |
| chrX:52934499-52935718 | 0.213810677 | 0.506214248 | 0.761383274 |
| chr14:20690827-20691322 | 0.240033965 | 0.506232915 | 0.761389575 |
| chr2:191867131-191867407 | 0.281078099 | 0.506238311 | 0.761389575 |
| chr11:2191047-2191389 | -0.28951005 | 0.506261482 | 0.761409479 |
| chr16:50210607-50210826 | -0.349699977 | 0.506275226 | 0.761415203 |
| chr9:134703006-134703447 | 0.243868217 | 0.506306989 | 0.761419835 |

| | | | |
|---|---|---|---|
| chr10:132752385-132752589 | -0.351370123 | 0.506307759 | 0.761419835 |
| chr19:49817934-49818446 | 0.236253016 | 0.506320561 | 0.761419835 |
| chr4:177051542-177051791 | -0.30638831 | 0.506328179 | 0.761419835 |
| chr15:59689819-59690069 | -0.267507355 | 0.50633341 | 0.761419835 |
| chr8:103370968-103371868 | 0.216601337 | 0.506341769 | 0.761419835 |
| chr9:134641485-134641759 | 0.259612715 | 0.506347867 | 0.761419835 |
| chr20:53748143-53748756 | 0.209837225 | 0.506360576 | 0.761424003 |
| chr9:112486181-112487271 | -0.220510227 | 0.506384352 | 0.761444812 |
| chr17:2511487-2512101 | -0.229401831 | 0.506413244 | 0.761473312 |
| chr10:63819827-63820028 | 0.344015529 | 0.506426788 | 0.761478735 |
| chr13:44403415-44404135 | -0.217184222 | 0.506437105 | 0.761479304 |
| chr5:35815520-35815942 | -0.262863166 | 0.506458123 | 0.761495964 |
| chr6:150395486-150395763 | -0.297742775 | 0.506505858 | 0.761552793 |
| chr16:66406447-66407222 | 0.231711673 | 0.506543746 | 0.761568431 |
| chr17:3724169-3724588 | -0.25949618 | 0.506545789 | 0.761568431 |
| chr10:73872044-73872261 | -0.279593534 | 0.506550717 | 0.761568431 |
| chr18:50578829-50579047 | 0.31853034 | 0.506556016 | 0.761568431 |
| chr4:28314427-28314705 | -0.331044314 | 0.506597617 | 0.761616031 |
| chr17:78681178-78681552 | -0.28453801 | 0.506620453 | 0.761630518 |
| chr8:130518335-130519298 | -0.230685325 | 0.506627134 | 0.761630518 |
| chr1:84433835-84434069 | -0.322668037 | 0.506656438 | 0.761659628 |
| chr16:89153746-89154507 | 0.227556305 | 0.506682909 | 0.761684479 |
| chr3:138125653-138125926 | 0.284252975 | 0.506694367 | 0.761686758 |
| chr13:33445381-33445718 | -0.282254939 | 0.506742617 | 0.761744346 |
| chr18:9971373-9971577 | 0.324849418 | 0.506818832 | 0.761841962 |
| chr22:38549020-38549392 | -0.281795505 | 0.506832131 | 0.761841962 |
| chr5:138641672-138642141 | -0.246561303 | 0.506850486 | 0.761841962 |
| chr14:105411040-105411468 | 0.240619509 | 0.506850936 | 0.761841962 |
| chr3:53072679-53072995 | 0.265508033 | 0.50685727 | 0.761841962 |
| chr6:118736800-118737063 | 0.328848115 | 0.50686777 | 0.761842801 |
| chr6:150668432-150668712 | -0.299068671 | 0.506934883 | 0.761928727 |
| chr20:63912455-63912687 | -0.291574763 | 0.506947233 | 0.761928762 |
| chr21:27047761-27048041 | -0.345999283 | 0.506954794 | 0.761928762 |
| chr19:1725638-1725906 | -0.319855387 | 0.506984647 | 0.761946629 |
| chr12:40152873-40153082 | 0.352500527 | 0.50698657 | 0.761946629 |
| chr10:98510376-98510597 | 0.338037413 | 0.507054912 | 0.761950467 |
| chr1:45018611-45019212 | -0.238570654 | 0.507067193 | 0.761950467 |
| chr2:202634473-202635478 | -0.213895212 | 0.507069345 | 0.761950467 |
| chr5:127029558-127031084 | -0.209853432 | 0.507073785 | 0.761950467 |
| chr20:31268703-31268903 | -0.39989962 | 0.507074265 | 0.761950467 |
| chr1:185670398-185670640 | 0.308413627 | 0.507089236 | 0.761950467 |
| chr14:91138531-91138839 | -0.287731575 | 0.507089874 | 0.761950467 |
| chr17:42677952-42678581 | 0.219889849 | 0.507096895 | 0.761950467 |
| chr20:35757096-35757416 | 0.283203211 | 0.507098463 | 0.761950467 |
| chr9:98241832-98242183 | 0.287146418 | 0.507100454 | 0.761950467 |
| chr1:159683672-159683872 | -0.367634778 | 0.507101396 | 0.761950467 |
| chr6:143060225-143061340 | -0.215693067 | 0.50711467 | 0.761950467 |
| chr15:75802092-75802615 | 0.236255201 | 0.507129912 | 0.761950467 |
| chr17:63258363-63258584 | -0.307767498 | 0.507130976 | 0.761950467 |
| chr4:120870369-120870613 | -0.305547664 | 0.507158611 | 0.761950467 |

| | | | |
|---|---|---|---|
| chr1:243758588-243758808 | 0.360760657 | 0.507160303 | 0.761950467 |
| chr10:84148082-84148282 | 0.360760657 | 0.507160303 | 0.761950467 |
| chr5:43083920-43084439 | 0.265725377 | 0.507168121 | 0.761950467 |
| chr15:33201796-33202240 | -0.231389128 | 0.507187332 | 0.761964388 |
| chr2:219498351-219499088 | 0.2134217 | 0.507242403 | 0.762032182 |
| chr1:108049438-108049865 | 0.259977656 | 0.507260574 | 0.76204454 |
| chr4:82552997-82553508 | 0.23401289 | 0.507296182 | 0.762083091 |
| chr7:155645081-155645337 | -0.272376722 | 0.507308379 | 0.762084781 |
| chr10:116272688-116273052 | -0.255191036 | 0.50732645 | 0.762084781 |
| chr3:61919456-61919679 | 0.254201655 | 0.507333298 | 0.762084781 |
| chr5:33300142-33300481 | -0.292747372 | 0.507344431 | 0.762084781 |
| chr2:44938690-44939061 | 0.273485037 | 0.507355884 | 0.762084781 |
| chr6:102813470-102813857 | -0.238303816 | 0.507357514 | 0.762084781 |
| chr20:45880938-45881519 | 0.221620128 | 0.507366929 | 0.762084781 |
| chr5:172843933-172844160 | 0.320874102 | 0.507406965 | 0.762129976 |
| chr3:36965286-36965764 | 0.244960387 | 0.507424057 | 0.762140708 |
| chr17:57903497-57903697 | 0.272974093 | 0.507507387 | 0.762248761 |
| chr2:74440234-74441361 | -0.211917576 | 0.507524775 | 0.762248761 |
| chr5:132369429-132370268 | -0.217036942 | 0.507525842 | 0.762248761 |
| chr2:84926657-84927300 | -0.222862981 | 0.507542496 | 0.762255127 |
| chr1:183685178-183685451 | -0.288329768 | 0.507549977 | 0.762255127 |
| chr4:138876013-138876523 | 0.259869981 | 0.507569451 | 0.762269433 |
| chr14:58426552-58428120 | 0.208704082 | 0.507586431 | 0.762279993 |
| chr15:59632702-59633095 | -0.256578249 | 0.507598863 | 0.762283723 |
| chr8:43057359-43057590 | 0.295783548 | 0.507648778 | 0.762343741 |
| chr3:141940155-141940876 | -0.229236204 | 0.507678816 | 0.762373907 |
| chr17:59786284-59787071 | 0.216622945 | 0.507708973 | 0.762404252 |
| chr8:73976044-73976588 | 0.222646331 | 0.507729347 | 0.762410886 |
| chr2:200905655-200906038 | -0.243597266 | 0.507733292 | 0.762410886 |
| chr1:195055140-195055340 | -0.336252306 | 0.507772037 | 0.762453109 |
| chr2:38315790-38316087 | -0.282253008 | 0.507781312 | 0.762453109 |
| chr9:34256160-34256573 | -0.272898893 | 0.507817734 | 0.762492856 |
| chr17:31008647-31008847 | 0.333491412 | 0.507839896 | 0.76251119 |
| chr6:149941577-149942495 | 0.222555928 | 0.50787012 | 0.762524946 |
| chr12:80316413-80316725 | 0.277296616 | 0.507880411 | 0.762524946 |
| chr9:7052655-7053054 | -0.256751903 | 0.50789049 | 0.762524946 |
| chr7:5486721-5487394 | -0.220807029 | 0.507891498 | 0.762524946 |
| chr4:147518864-147519230 | 0.251143747 | 0.507898817 | 0.762524946 |
| chr4:63854288-63854509 | -0.336018397 | 0.507921863 | 0.762544605 |
| chr9:119004987-119005357 | -0.300895078 | 0.507941375 | 0.762550494 |
| chr20:58892104-58892733 | 0.219369347 | 0.50794569 | 0.762550494 |
| chr15:75738467-75739164 | 0.21967022 | 0.507967344 | 0.762568061 |
| chr3:64501074-64502041 | 0.210072811 | 0.507981825 | 0.762574859 |
| chr2:196799346-196799954 | -0.223080112 | 0.507998798 | 0.762584561 |
| chr9:79670604-79670916 | 0.346577475 | 0.508008193 | 0.762584561 |
| chr2:151827908-151828986 | -0.212430627 | 0.508041605 | 0.762619777 |
| chr13:42970820-42971130 | -0.261840507 | 0.508078734 | 0.762660569 |
| chr17:4939757-4940790 | -0.214846436 | 0.508108494 | 0.76269016 |
| chr13:47455513-47455820 | 0.266502462 | 0.508118355 | 0.76269016 |
| chr8:87017085-87017387 | 0.297241889 | 0.508147495 | 0.762718959 |

| | | | |
|---|---|---|---|
| chr4:41360431-41360951 | 0.246736891 | 0.508179073 | 0.76274411 |
| chr7:16891938-16892494 | 0.239280259 | 0.508206075 | 0.76274411 |
| chr2:150610681-150611025 | 0.30260368 | 0.508214006 | 0.76274411 |
| chr17:68413498-68414265 | -0.229134659 | 0.508217508 | 0.76274411 |
| chr1:241542208-241543032 | 0.229561363 | 0.508240866 | 0.76274411 |
| chr16:89948562-89948810 | 0.272967339 | 0.508241107 | 0.76274411 |
| chr5:174910276-174910602 | -0.260982013 | 0.508242283 | 0.76274411 |
| chr6:27532124-27532474 | -0.268001051 | 0.508243889 | 0.76274411 |
| chr2:70976953-70977675 | 0.231963566 | 0.508278947 | 0.762781782 |
| chr2:84302942-84303418 | -0.252193882 | 0.508317327 | 0.762783805 |
| chr1:160376607-160376840 | -0.332963205 | 0.508320563 | 0.762783805 |
| chr10:93542058-93542274 | 0.303717154 | 0.508322687 | 0.762783805 |
| chr19:45468969-45469999 | 0.215012098 | 0.508324378 | 0.762783805 |
| chr7:46132416-46132743 | 0.262947913 | 0.508330071 | 0.762783805 |
| chr3:14557350-14557580 | -0.308342224 | 0.508383367 | 0.762837726 |
| chr1:108338761-108338989 | 0.336957679 | 0.508411181 | 0.762837726 |
| chr14:87916135-87916335 | 0.366400282 | 0.508415791 | 0.762837726 |
| chr19:8580785-8581055 | -0.316785854 | 0.508430676 | 0.762837726 |
| chr1:206961115-206961357 | -0.275309395 | 0.508435765 | 0.762837726 |
| chr14:75542500-75543182 | -0.229006305 | 0.508437263 | 0.762837726 |
| chr10:73743906-73744913 | -0.213666521 | 0.508437959 | 0.762837726 |
| chr8:99212181-99212394 | -0.332219463 | 0.508445651 | 0.762837726 |
| chrX:78314663-78314982 | -0.2958672 | 0.508457916 | 0.76284119 |
| chr1:39919539-39919849 | 0.266282596 | 0.508484289 | 0.762855554 |
| chr11:35618616-35619431 | -0.225372976 | 0.508487402 | 0.762855554 |
| chr19:43951353-43951588 | 0.306069483 | 0.508527305 | 0.762887791 |
| chr3:141250394-141250645 | -0.301399695 | 0.508528803 | 0.762887791 |
| chr21:42975013-42975213 | 0.274348029 | 0.508607142 | 0.762990376 |
| chr8:88610210-88610909 | 0.236035896 | 0.508630605 | 0.763010635 |
| chr14:68965514-68965740 | 0.280896174 | 0.50869439 | 0.763063918 |
| chr11:65780814-65781435 | 0.230775192 | 0.508700804 | 0.763063918 |
| chr16:73180364-73180564 | -0.310689249 | 0.508701782 | 0.763063918 |
| chr12:81610869-81611606 | -0.224286701 | 0.50870823 | 0.763063918 |
| chrX:140572647-140573057 | 0.261109589 | 0.508715918 | 0.763063918 |
| chr9:129885460-129886091 | 0.226561172 | 0.508762792 | 0.763094659 |
| chr5:164544145-164545061 | -0.237747371 | 0.508770465 | 0.763094659 |
| chr2:202148708-202149296 | -0.257149581 | 0.508770914 | 0.763094659 |
| chr12:113334772-113335539 | 0.221173949 | 0.508776249 | 0.763094659 |
| chr17:80254770-80255034 | 0.263165929 | 0.508793625 | 0.763105783 |
| chr11:63974210-63975110 | 0.215327004 | 0.508806112 | 0.763109573 |
| chr6:54865312-54865512 | -0.342510362 | 0.508855289 | 0.763168392 |
| chr2:232605760-232606514 | 0.231103596 | 0.508886407 | 0.763200122 |
| chr10:97686898-97687730 | 0.211030813 | 0.508899192 | 0.763204359 |
| chr7:37577181-37577401 | 0.341113094 | 0.508932773 | 0.763239782 |
| chr16:1414652-1415013 | 0.272415172 | 0.508959451 | 0.76325253 |
| chr6:4774823-4776680 | 0.208890466 | 0.508966937 | 0.76325253 |
| chr6:51494402-51494661 | -0.292015815 | 0.508983657 | 0.76325253 |
| chr6:31070649-31071098 | 0.259027042 | 0.508985925 | 0.76325253 |
| chr2:40256676-40256957 | -0.272732618 | 0.50899108 | 0.76325253 |
| chr1:147273825-147274045 | 0.283479438 | 0.509004692 | 0.763258005 |

| | | | |
|---|---|---|---|
| chr10:13409592-13410318 | -0.24525207 | 0.509027003 | 0.763276522 |
| chr3:88005160-88005376 | -0.310595149 | 0.509042384 | 0.763284494 |
| chr1:202135197-202135546 | -0.222205543 | 0.509062713 | 0.763284494 |
| chr17:28598858-28600178 | -0.211255789 | 0.509074994 | 0.763284494 |
| chr1:33086473-33086695 | 0.280457437 | 0.509088611 | 0.763284494 |
| chr19:770299-770530 | 0.347818398 | 0.509088747 | 0.763284494 |
| chr5:136058098-136058891 | -0.226708178 | 0.509092089 | 0.763284494 |
| chr5:176593976-176594185 | -0.300667067 | 0.509104651 | 0.763288393 |
| chr6:49410030-49410262 | 0.301912591 | 0.509128511 | 0.76329854 |
| chr6:42048491-42048924 | -0.244515247 | 0.509131343 | 0.76329854 |
| chr10:86755639-86757526 | 0.206991623 | 0.509155257 | 0.763312057 |
| chr3:152183543-152183887 | -0.252195557 | 0.509160283 | 0.763312057 |
| chr2:119174501-119174757 | 0.26871486 | 0.509174707 | 0.763318746 |
| chr6:33248997-33249693 | -0.216764868 | 0.509196844 | 0.763320067 |
| chr16:74117882-74118083 | -0.320719996 | 0.509198531 | 0.763320067 |
| chr11:108498116-108498699 | 0.248147001 | 0.509205475 | 0.763320067 |
| chr12:101847415-101847711 | -0.334965409 | 0.509217365 | 0.763322957 |
| chr1:203006309-203007924 | -0.207167844 | 0.509233983 | 0.763323837 |
| chr8:119767735-119768101 | -0.24680497 | 0.50925763 | 0.763323837 |
| chr1:153365078-153365848 | -0.217049617 | 0.509261626 | 0.763323837 |
| chr14:67514705-67515781 | 0.221288214 | 0.509265924 | 0.763323837 |
| chr10:19945353-19945553 | -0.375381076 | 0.509267763 | 0.763323837 |
| chr22:39360734-39361447 | 0.24813 | 0.509284046 | 0.763333311 |
| chr11:42625123-42625360 | -0.308093303 | 0.509321682 | 0.763374788 |
| chr5:133052513-133053105 | 0.22526519 | 0.509359974 | 0.763393454 |
| chr20:44449326-44449587 | -0.32065909 | 0.509362501 | 0.763393454 |
| chr10:116273425-116274033 | 0.23288179 | 0.509369178 | 0.763393454 |
| chr11:63813034-63814133 | 0.222445492 | 0.509378719 | 0.763393454 |
| chr1:40477130-40477620 | 0.260257213 | 0.509392809 | 0.763393454 |
| chr16:72729367-72729904 | -0.262039284 | 0.509393915 | 0.763393454 |
| chr8:135997317-135997791 | -0.266605324 | 0.509404256 | 0.76339402 |
| chr1:31842920-31843312 | 0.272824564 | 0.509421251 | 0.763402909 |
| chr3:129825007-129825239 | 0.28332307 | 0.509430114 | 0.763402909 |
| chr9:97803357-97804068 | 0.224167788 | 0.509441318 | 0.76340477 |
| chr14:42927297-42927607 | 0.283790324 | 0.509466291 | 0.763427261 |
| chr17:28174665-28174942 | -0.292329199 | 0.509493478 | 0.76345307 |
| chr3:108006202-108006635 | -0.252947145 | 0.509521237 | 0.763467816 |
| chr9:129111715-129112087 | -0.257090383 | 0.509523247 | 0.763467816 |
| chr1:159000722-159000957 | -0.317960057 | 0.509555556 | 0.763501297 |
| chr16:25233640-25234138 | -0.241171071 | 0.509590528 | 0.763538767 |
| chr17:62888089-62888553 | 0.24037136 | 0.509751603 | 0.763765176 |
| chr4:174855171-174855446 | -0.36030611 | 0.509778897 | 0.763785201 |
| chr9:77488539-77488796 | 0.30922166 | 0.509785049 | 0.763785201 |
| chr11:114762528-114762728 | -0.362286614 | 0.509794873 | 0.763785201 |
| chr3:166291367-166291719 | 0.301597538 | 0.509821432 | 0.763786887 |
| chr17:59722763-59723124 | -0.253984691 | 0.509830851 | 0.763786887 |
| chr11:73376040-73376851 | 0.214505078 | 0.509834029 | 0.763786887 |
| chr3:160449197-160450178 | -0.207250422 | 0.509835871 | 0.763786887 |
| chr9:14322186-14323103 | -0.217286486 | 0.50985798 | 0.763798944 |
| chr14:64503317-64504884 | 0.206247973 | 0.509870669 | 0.763798944 |

| | | | |
|---|---|---|---|
| chr8:129761575-129762006 | 0.28673769 | 0.509873825 | 0.763798944 |
| chr4:118835077-118835495 | 0.233473643 | 0.509893754 | 0.763813865 |
| chr1:204287569-204287852 | -0.238670958 | 0.50994929 | 0.763877216 |
| chr10:93437739-93438335 | -0.230515974 | 0.509955983 | 0.763877216 |
| chr3:146382413-146382729 | -0.271936107 | 0.509972523 | 0.763887058 |
| chr8:127401942-127402211 | 0.257359849 | 0.509996759 | 0.763903292 |
| chr22:21457003-21457561 | 0.256727472 | 0.510008677 | 0.763903292 |
| chr21:34857931-34858178 | 0.296029777 | 0.510013271 | 0.763903292 |
| chr5:5492209-5492687 | -0.253730176 | 0.510033801 | 0.76390393 |
| chr10:80378129-80378547 | -0.241153774 | 0.510041568 | 0.76390393 |
| chr17:73311926-73312622 | 0.212678879 | 0.510056843 | 0.76390393 |
| chr3:170128955-170129460 | 0.258736922 | 0.510067156 | 0.76390393 |
| chr5:106755146-106755346 | 0.32963147 | 0.510070963 | 0.76390393 |
| chr1:27234162-27234848 | 0.220344588 | 0.510082945 | 0.76390393 |
| chr20:25255186-25255643 | 0.26914834 | 0.51008791 | 0.76390393 |
| chr6:57274490-57274931 | -0.257768377 | 0.510093454 | 0.76390393 |
| chr18:3058411-3058713 | -0.301047474 | 0.510113173 | 0.76391853 |
| chr8:125557371-125557587 | 0.311501993 | 0.51013285 | 0.763933065 |
| chr9:128341450-128341699 | -0.258765858 | 0.510160202 | 0.763945206 |
| chr15:75367747-75368785 | 0.209926474 | 0.510160898 | 0.763945206 |
| chr1:35557068-35557965 | 0.212669163 | 0.510182238 | 0.763953321 |
| chr20:1503692-1503969 | 0.282653507 | 0.510193166 | 0.763953321 |
| chr9:130998370-130998611 | 0.286611822 | 0.510196228 | 0.763953321 |
| chr17:67125896-67126096 | -0.309553293 | 0.510260456 | 0.764019462 |
| chr10:80453761-80454828 | 0.213263251 | 0.510284874 | 0.764019462 |
| chr2:157784407-157784797 | -0.28330009 | 0.510285447 | 0.764019462 |
| chr6:33417845-33418656 | 0.211463199 | 0.510288378 | 0.764019462 |
| chr19:41411543-41412068 | -0.25331923 | 0.510295998 | 0.764019462 |
| chr10:91635467-91635707 | -0.276966679 | 0.510300844 | 0.764019462 |
| chr3:52704736-52705357 | -0.248827225 | 0.510310199 | 0.764019462 |
| chr8:144444058-144444907 | 0.218368009 | 0.510364598 | 0.764075411 |
| chr2:24219942-24220218 | -0.318649579 | 0.510367513 | 0.764075411 |
| chr11:111601922-111603104 | 0.213339774 | 0.510391891 | 0.764096979 |
| chr14:102139025-102140785 | 0.207171012 | 0.510411214 | 0.764110977 |
| chr1:153390743-153391311 | 0.235920413 | 0.510452608 | 0.764143913 |
| chr1:179365221-179366370 | -0.211443307 | 0.510471347 | 0.764143913 |
| chr2:68282889-68283120 | -0.33696003 | 0.510477814 | 0.764143913 |
| chr11:36672950-36673156 | 0.339793133 | 0.510484894 | 0.764143913 |
| chr10:1004001-1004509 | 0.24905599 | 0.510488776 | 0.764143913 |
| chr1:45011229-45011505 | 0.250650612 | 0.510499558 | 0.764143913 |
| chr10:1360056-1360302 | 0.295634653 | 0.510503025 | 0.764143913 |
| chrX:3815985-3816651 | -0.220980696 | 0.51051813 | 0.764149608 |
| chr10:77926579-77927128 | -0.234592663 | 0.510526776 | 0.764149608 |
| chr19:1269000-1269861 | 0.215237711 | 0.510539235 | 0.764153329 |
| chr6:2999223-3000420 | 0.209822425 | 0.51055136 | 0.764156551 |
| chr14:71309891-71310232 | 0.274947646 | 0.510569766 | 0.764167578 |
| chr22:37677196-37677481 | -0.258752116 | 0.510578674 | 0.764167578 |
| chr4:156951006-156951246 | -0.313452754 | 0.510667103 | 0.764260006 |
| chr20:1005645-1005927 | 0.286185216 | 0.510677782 | 0.764260006 |
| chr2:29224739-29224958 | 0.318541949 | 0.510680209 | 0.764260006 |

| | | | |
|---|---|---|---|
| chr17:70004523-70004723 | 0.397894318 | 0.510680327 | 0.764260006 |
| chr2:54983558-54983758 | -0.364874787 | 0.510693312 | 0.76426451 |
| chr14:77320222-77321583 | 0.205696358 | 0.510712903 | 0.764278902 |
| chr11:82226006-82226217 | 0.305603233 | 0.510731162 | 0.764291299 |
| chr14:81353787-81353987 | 0.349581982 | 0.510741144 | 0.76429131 |
| chr3:140972960-140973166 | 0.337777076 | 0.510758997 | 0.764302175 |
| chr6:127515544-127517039 | -0.211745302 | 0.510792877 | 0.764302175 |
| chr11:66282328-66282965 | -0.223546821 | 0.510796708 | 0.764302175 |
| chr1:109277441-109277815 | -0.228585396 | 0.510797809 | 0.764302175 |
| chr10:24208664-24209028 | -0.25501859 | 0.51081584 | 0.764302175 |
| chr11:74071102-74071547 | 0.241484744 | 0.510840202 | 0.764302175 |
| chr3:149825340-149825832 | 0.256891417 | 0.510840778 | 0.764302175 |
| chr8:90280952-90281152 | 0.34193274 | 0.510845677 | 0.764302175 |
| chr14:89417059-89418279 | 0.209945361 | 0.510880504 | 0.764302175 |
| chr5:83563604-83564429 | -0.224809268 | 0.51088271 | 0.764302175 |
| chr10:81894539-81894748 | -0.358187535 | 0.510884901 | 0.764302175 |
| chr12:123383572-123384309 | 0.213338584 | 0.51088549 | 0.764302175 |
| chr3:79767936-79768162 | 0.30803713 | 0.51089346 | 0.764302175 |
| chr19:8005951-8006180 | -0.322110429 | 0.510900995 | 0.764302175 |
| chr14:61381537-61382006 | -0.251313557 | 0.51090983 | 0.764302175 |
| chr6:45377341-45378623 | 0.209556369 | 0.510916487 | 0.764302175 |
| chr8:140616094-140616294 | 0.317503903 | 0.510917979 | 0.764302175 |
| chr10:133308478-133309641 | 0.209011326 | 0.510940302 | 0.764320647 |
| chr9:136842279-136842583 | 0.246896123 | 0.510976371 | 0.764359679 |
| chr18:74217610-74217831 | -0.361346075 | 0.511001526 | 0.764367538 |
| chr6:152595632-152595889 | 0.29957568 | 0.511001576 | 0.764367538 |
| chr5:178793253-178793490 | 0.276545436 | 0.511055997 | 0.764434018 |
| chr9:81450960-81451173 | 0.360533666 | 0.511078635 | 0.764450263 |
| chr6:7153926-7154127 | -0.353267456 | 0.511089106 | 0.764450263 |
| chr1:111236920-111237125 | -0.319185278 | 0.511096788 | 0.764450263 |
| chr12:80472969-80473169 | 0.298335201 | 0.511110358 | 0.764455637 |
| chr14:48008649-48008853 | -0.373817973 | 0.511125819 | 0.764463839 |
| chr6:50265062-50265291 | -0.342389156 | 0.5111521 | 0.764477834 |
| chr12:87779072-87779391 | 0.276395253 | 0.511161227 | 0.764477834 |
| chr20:50790439-50791089 | 0.206236944 | 0.511165107 | 0.764477834 |
| chr17:82534425-82534625 | 0.310714385 | 0.511184661 | 0.764491647 |
| chr14:88492242-88492652 | 0.23318821 | 0.511194299 | 0.764491647 |
| chr7:151059720-151059927 | 0.250525275 | 0.511220985 | 0.764516636 |
| chr8:78705312-78705699 | 0.280577478 | 0.51124051 | 0.764530912 |
| chr1:109274509-109275170 | -0.214248987 | 0.511288489 | 0.764583999 |
| chr11:64449583-64449966 | 0.242503424 | 0.511295966 | 0.764583999 |
| chr2:31138359-31138901 | -0.238688649 | 0.511331092 | 0.764606827 |
| chr3:37855442-37855771 | 0.263595615 | 0.51133119 | 0.764606827 |
| chr20:57350886-57352027 | 0.209793425 | 0.51136256 | 0.764623923 |
| chr14:49527857-49528390 | 0.236058804 | 0.511379719 | 0.764623923 |
| chr4:48016240-48017597 | -0.207054442 | 0.511388467 | 0.764623923 |
| chr16:76048979-76049269 | -0.329245382 | 0.511391594 | 0.764623923 |
| chr8:17779991-17780252 | -0.319992352 | 0.511392519 | 0.764623923 |
| chr16:81366408-81366678 | 0.270331536 | 0.511437207 | 0.764668037 |
| chr14:45083774-45084751 | 0.207702584 | 0.511457021 | 0.764668037 |

| | | | |
|---|---|---|---|
| chr2:113041942-113042233 | 0.289089419 | 0.511457941 | 0.764668037 |
| chr14:39016901-39017855 | -0.214847499 | 0.511461942 | 0.764668037 |
| chr4:155353933-155354241 | 0.275152048 | 0.511500896 | 0.764700421 |
| chr1:201941550-201942334 | -0.229115387 | 0.511503562 | 0.764700421 |
| chr12:31271931-31272233 | -0.275219641 | 0.511525225 | 0.764717885 |
| chr3:114519401-114519761 | -0.270937896 | 0.511539688 | 0.764724588 |
| chr3:27357720-27357976 | 0.272523753 | 0.511563065 | 0.764734609 |
| chr14:24279402-24280059 | -0.228590479 | 0.511572302 | 0.764734609 |
| chr12:751139-751574 | -0.235777375 | 0.511576334 | 0.764734609 |
| chr22:37316919-37317137 | 0.304195434 | 0.511596477 | 0.764745345 |
| chr6:42452189-42452432 | 0.260448613 | 0.511603477 | 0.764745345 |
| chr5:142914110-142914310 | -0.367726637 | 0.511646576 | 0.764794849 |
| chr2:140808854-140809174 | -0.276116956 | 0.511698216 | 0.764851054 |
| chr10:21989606-21989903 | -0.283912241 | 0.511704142 | 0.764851054 |
| chr16:15489506-15489910 | -0.276897738 | 0.511715531 | 0.764853158 |
| chr15:96355122-96355330 | 0.284737176 | 0.511766266 | 0.76491407 |
| chr16:8936467-8936775 | 0.27072377 | 0.511785802 | 0.764928347 |
| chr2:86105431-86106559 | -0.210614835 | 0.511826985 | 0.764961257 |
| chr5:173011657-173011862 | -0.31024493 | 0.511827788 | 0.764961257 |
| chr9:112784369-112784569 | 0.33424153 | 0.511852794 | 0.764965334 |
| chr5:56097611-56098081 | -0.257150787 | 0.511860467 | 0.764965334 |
| chr7:157273608-157273866 | -0.257150787 | 0.511860467 | 0.764965334 |
| chr11:64778441-64779279 | 0.209095119 | 0.511890918 | 0.764995922 |
| chr11:32448317-32448553 | -0.323319185 | 0.511919036 | 0.765013981 |
| chr19:49937751-49938179 | 0.265484311 | 0.51192297 | 0.765013981 |
| chr15:92850206-92850533 | -0.297551422 | 0.511935899 | 0.765018381 |
| chr3:101707055-101707638 | -0.244494603 | 0.511965513 | 0.765046322 |
| chr3:114299358-114299955 | -0.223560345 | 0.511974566 | 0.765046322 |
| chr15:44298936-44299136 | 0.356626686 | 0.511994333 | 0.765046648 |
| chr17:66594700-66595000 | -0.30469499 | 0.511994754 | 0.765046648 |
| chr9:40916119-40916319 | -0.376763435 | 0.512034764 | 0.765079311 |
| chr5:177336178-177336485 | -0.259087624 | 0.512042631 | 0.765079311 |
| chr7:107040824-107041024 | -0.359764083 | 0.512049254 | 0.765079311 |
| chrX:17614224-17614502 | -0.317382572 | 0.512056553 | 0.765079311 |
| chr3:64355532-64355829 | 0.224970544 | 0.512116946 | 0.765150784 |
| chr16:84308138-84308487 | 0.24813008 | 0.512125574 | 0.765150784 |
| chr6:89145487-89146878 | -0.211551121 | 0.512143134 | 0.765150784 |
| chr2:176001445-176002553 | -0.210426605 | 0.512144333 | 0.765150784 |
| chrX:55211058-55211558 | -0.26337377 | 0.512164235 | 0.765165598 |
| chr6:106540996-106541259 | -0.353850054 | 0.512184337 | 0.765178416 |
| chr12:76820971-76821259 | -0.289737799 | 0.512201364 | 0.765178416 |
| chr11:79270072-79270326 | -0.291224681 | 0.512211705 | 0.765178416 |
| chr7:142188875-142189166 | -0.312912306 | 0.512212761 | 0.765178416 |
| chr14:92456705-92456993 | 0.261719361 | 0.512222806 | 0.765178504 |
| chr3:62273500-62273763 | 0.262576558 | 0.512241058 | 0.765190852 |
| chr7:129952319-129952551 | -0.272595087 | 0.512267366 | 0.765215232 |
| chr3:188749038-188749363 | -0.284303421 | 0.512290334 | 0.765218138 |
| chr5:32753796-32754095 | -0.323168577 | 0.512291468 | 0.765218138 |
| chr17:16014014-16014214 | 0.32460625 | 0.512303618 | 0.765218138 |
| chr3:70753666-70753976 | 0.275730151 | 0.512309408 | 0.765218138 |

| | | | |
|---|---|---|---|
| chr1:9950353-9950817 | 0.242114831 | 0.512325334 | 0.765218138 |
| chr13:77593055-77593400 | -0.263477803 | 0.512347547 | 0.765218138 |
| chr2:152098752-152099020 | -0.287955609 | 0.512352246 | 0.765218138 |
| chr8:126425114-126425345 | 0.279839739 | 0.512356634 | 0.765218138 |
| chr14:96204501-96205051 | -0.21935535 | 0.512359194 | 0.765218138 |
| chr3:113890161-113890669 | -0.236792767 | 0.512389968 | 0.765249184 |
| chr6:130019370-130019635 | -0.322174936 | 0.512415711 | 0.765272714 |
| chr12:26853856-26854157 | -0.289173541 | 0.512437712 | 0.765282622 |
| chr9:84669845-84670123 | -0.293953726 | 0.512450095 | 0.765282622 |
| chr11:30911687-30911887 | -0.356442148 | 0.512452308 | 0.765282622 |
| chr17:75548990-75550418 | -0.206622628 | 0.512463183 | 0.765283947 |
| chr2:111924820-111925020 | 0.356069531 | 0.512501427 | 0.765288405 |
| chr9:96654385-96655682 | -0.214746431 | 0.512510277 | 0.765288405 |
| chr14:22314260-22314460 | 0.317158276 | 0.512513078 | 0.765288405 |
| chr16:53373239-53374085 | 0.228466779 | 0.512531703 | 0.765288405 |
| chr2:109499656-109499913 | 0.264644992 | 0.512532793 | 0.765288405 |
| chr5:155546093-155546398 | -0.300933224 | 0.512533208 | 0.765288405 |
| chr1:119544252-119544457 | -0.359966839 | 0.512543439 | 0.765288405 |
| chr11:73760734-73761721 | 0.209005148 | 0.512550488 | 0.765288405 |
| chr7:55473275-55473500 | -0.336526285 | 0.512564406 | 0.765288405 |
| chr6:73512894-73513173 | 0.270283282 | 0.512566047 | 0.765288405 |
| chr1:180863232-180863482 | -0.316852233 | 0.512621542 | 0.765348938 |
| chr3:124468248-124468538 | 0.272937231 | 0.512626567 | 0.765348938 |
| chr8:75483379-75483641 | 0.27248089 | 0.512643188 | 0.765358839 |
| chr11:43130674-43131012 | -0.293823002 | 0.512687371 | 0.765397751 |
| chr1:203475722-203475924 | -0.281674939 | 0.51268923 | 0.765397751 |
| chr5:170273155-170273356 | -0.327606115 | 0.512742065 | 0.765400591 |
| chr11:65122607-65122872 | 0.270639792 | 0.512746005 | 0.765400591 |
| chr2:1972497-1972732 | -0.29924363 | 0.512750732 | 0.765400591 |
| chr9:117086842-117087042 | 0.337954706 | 0.512753814 | 0.765400591 |
| chr1:25875453-25875807 | -0.263783179 | 0.512758707 | 0.765400591 |
| chr5:173753441-173753782 | 0.249843987 | 0.512774001 | 0.765400591 |
| chr5:127918916-127919332 | 0.252144174 | 0.512778746 | 0.765400591 |
| chr17:61235514-61236630 | 0.204309042 | 0.512779659 | 0.765400591 |
| chr3:9782859-9783059 | 0.297058847 | 0.512781036 | 0.765400591 |
| chr17:76671073-76672067 | 0.211822748 | 0.512793045 | 0.765403606 |
| chr13:20773163-20774347 | 0.209750619 | 0.512811992 | 0.765416976 |
| chr13:44988977-44989901 | -0.21094565 | 0.512882291 | 0.765506991 |
| chr3:62238528-62238737 | 0.259261173 | 0.512896382 | 0.765513111 |
| chr10:79981460-79982586 | 0.213567385 | 0.512907539 | 0.765514851 |
| chr5:59380542-59380811 | -0.326497159 | 0.512920559 | 0.765519373 |
| chr2:95728932-95729261 | -0.267055893 | 0.512947277 | 0.765544337 |
| chr15:85858606-85859132 | 0.236403659 | 0.512959592 | 0.765547805 |
| chr14:75296132-75296343 | 0.272724281 | 0.5129921 | 0.765581409 |
| chr1:75672797-75673070 | 0.297998819 | 0.513012581 | 0.765597063 |
| chrX:103585322-103586006 | -0.225421457 | 0.513064112 | 0.765659052 |
| chr22:50531552-50531819 | -0.317768138 | 0.513077766 | 0.765664516 |
| chr17:7107477-7107679 | -0.356357375 | 0.513096041 | 0.765674993 |
| chr1:62131186-62131609 | -0.259688448 | 0.513104772 | 0.765674993 |
| chr5:148650817-148651045 | -0.358964904 | 0.513146831 | 0.765722842 |

| | | | |
|---|---|---|---|
| chr6:56292503-56292877 | -0.270590441 | 0.513165954 | 0.765736465 |
| chr14:35884053-35884387 | 0.280797044 | 0.513181805 | 0.765745205 |
| chr16:88194041-88194257 | 0.29697198 | 0.513195984 | 0.76575145 |
| chr11:80109920-80110201 | -0.330265649 | 0.513217019 | 0.765767924 |
| chr5:80570370-80570747 | -0.282922582 | 0.513234386 | 0.765778456 |
| chr14:75589544-75589744 | 0.333668963 | 0.513248997 | 0.765778456 |
| chr13:99578690-99578963 | 0.288674327 | 0.51325406 | 0.765778456 |
| chr10:47311997-47312540 | -0.232035124 | 0.513270583 | 0.765779809 |
| chr17:45234767-45234967 | 0.328845401 | 0.513277123 | 0.765779809 |
| chr19:49113717-49114719 | -0.216845076 | 0.51328495 | 0.765779809 |
| chr16:70478682-70478903 | -0.31264922 | 0.513311837 | 0.765805011 |
| chr1:226125274-226125508 | 0.257785943 | 0.513341777 | 0.765834767 |
| chr12:18546553-18546803 | -0.314442754 | 0.513355813 | 0.765840795 |
| chrX:67695033-67695343 | -0.307497123 | 0.513393051 | 0.765881437 |
| chr5:138178434-138179479 | -0.210334376 | 0.513405561 | 0.765885188 |
| chr2:199366500-199366778 | -0.283424841 | 0.513435752 | 0.765900934 |
| chr10:103395828-103396075 | 0.28112851 | 0.513436108 | 0.765900934 |
| chr7:63900559-63901246 | -0.215301275 | 0.513452023 | 0.765907467 |
| chr17:60798573-60798806 | 0.239291017 | 0.51346048 | 0.765907467 |
| chr14:94838998-94839289 | 0.313371808 | 0.513478294 | 0.765919129 |
| chr6:30214404-30214695 | -0.267696909 | 0.513511994 | 0.76594425 |
| chr11:77031120-77031320 | -0.323571772 | 0.513524525 | 0.76594425 |
| chr1:240967147-240967411 | 0.301176249 | 0.513525124 | 0.76594425 |
| chr3:119322503-119323783 | 0.211933338 | 0.513541439 | 0.765953673 |
| chr2:119389606-119389902 | 0.285111841 | 0.513564193 | 0.765958911 |
| chr10:42878249-42878506 | -0.315438306 | 0.513564943 | 0.765958911 |
| chr16:5062851-5063068 | 0.308872681 | 0.513606247 | 0.765990693 |
| chr18:34976848-34977067 | 0.308872681 | 0.513606247 | 0.765990693 |
| chr2:46224773-46225099 | -0.324529255 | 0.51361687 | 0.765991627 |
| chr22:37849185-37849713 | -0.236941623 | 0.513630473 | 0.765997005 |
| chr16:89816468-89817130 | -0.218330093 | 0.513665048 | 0.766020348 |
| chr2:238227078-238227295 | -0.335710744 | 0.51367365 | 0.766020348 |
| chr8:103479488-103479709 | -0.344693706 | 0.513691129 | 0.766020348 |
| chr10:65838850-65839193 | 0.276709501 | 0.513696061 | 0.766020348 |
| chr1:195785992-195786218 | -0.334008021 | 0.513696112 | 0.766020348 |
| chr3:33231856-33232060 | 0.355145743 | 0.513710393 | 0.766026076 |
| chr21:46184165-46184547 | -0.253526782 | 0.513729425 | 0.766026076 |
| chr6:113432868-113433193 | 0.273424867 | 0.513729946 | 0.766026076 |
| chr1:35769285-35770401 | 0.209595717 | 0.513744072 | 0.766032232 |
| chr14:63563548-63564078 | -0.228547907 | 0.513762169 | 0.766044308 |
| chr4:113394705-113395034 | 0.249523962 | 0.513785995 | 0.766064926 |
| chr5:71455372-71456424 | -0.211229188 | 0.513825532 | 0.766107954 |
| chr17:60456616-60456823 | 0.235612974 | 0.513840982 | 0.766107954 |
| chr15:62306895-62307235 | 0.262432386 | 0.513853088 | 0.766107954 |
| chr2:58842644-58842887 | 0.306032819 | 0.513854847 | 0.766107954 |
| chr9:78667679-78667915 | -0.361721363 | 0.51387115 | 0.766115276 |
| chr1:191636499-191636870 | 0.26248841 | 0.513879756 | 0.766115276 |
| chr5:180802207-180803207 | 0.212657593 | 0.513895915 | 0.766124461 |
| chr12:122775097-122775510 | -0.254735545 | 0.513906553 | 0.766125414 |
| chr2:109046900-109047107 | 0.346869278 | 0.513950692 | 0.766171159 |

| | | | |
|---|---|---|---|
| chr15:40857421-40858005 | 0.220163327 | 0.513968157 | 0.766171159 |
| chr8:100365885-100366181 | -0.270789951 | 0.513971013 | 0.766171159 |
| chr19:10748946-10749454 | 0.239298739 | 0.51398403 | 0.766171159 |
| chr2:231612107-231612461 | -0.252720804 | 0.513991427 | 0.766171159 |
| chr3:36792368-36792704 | 0.267613305 | 0.513997234 | 0.766171159 |
| chr8:132144936-132145229 | 0.263165331 | 0.514074311 | 0.766239194 |
| chr16:77943918-77944120 | 0.310277472 | 0.514081369 | 0.766239194 |
| chr15:65638631-65638910 | 0.264222658 | 0.514087835 | 0.766239194 |
| chr8:26571300-26571511 | -0.312601362 | 0.514091392 | 0.766239194 |
| chr11:119030223-119031350 | 0.20783036 | 0.514092878 | 0.766239194 |
| chr1:68197925-68198199 | 0.27853757 | 0.514105357 | 0.766242888 |
| chr20:61254400-61254659 | -0.296124387 | 0.514142087 | 0.766269787 |
| chr15:63420315-63420819 | -0.250973422 | 0.514143405 | 0.766269787 |
| chr10:68550811-68551121 | 0.254143541 | 0.514157615 | 0.76627606 |
| chr1:109263356-109264000 | 0.218420106 | 0.514206333 | 0.76633376 |
| chr5:13976891-13977426 | -0.228464219 | 0.514220409 | 0.766334547 |
| chr15:96893410-96893610 | 0.332758006 | 0.514228788 | 0.766334547 |
| chrX:3549478-3549930 | -0.258472577 | 0.514247768 | 0.766334547 |
| chr11:12696918-12697168 | 0.293958825 | 0.514254301 | 0.766334547 |
| chr18:13546065-13546334 | 0.336918651 | 0.514256868 | 0.766334547 |
| chr3:133452963-133453318 | -0.23563614 | 0.514267696 | 0.766335778 |
| chr9:114624565-114625078 | -0.237347799 | 0.514292779 | 0.76634582 |
| chr11:103091964-103092477 | -0.219209677 | 0.514301833 | 0.76634582 |
| chr2:121902189-121902619 | -0.256485434 | 0.514310371 | 0.76634582 |
| chr8:100125768-100125971 | -0.332084656 | 0.514314441 | 0.76634582 |
| chr10:79347277-79348309 | -0.213230457 | 0.514335727 | 0.766362634 |
| chr15:62075918-62076310 | 0.238916369 | 0.514349532 | 0.766366253 |
| chr15:74489641-74490094 | 0.247578407 | 0.51435816 | 0.766366253 |
| chr17:55641634-55642128 | -0.236952786 | 0.514371673 | 0.766366259 |
| chr1:205061623-205062010 | 0.228242038 | 0.514378254 | 0.766366259 |
| chr1:52702932-52703615 | -0.219529721 | 0.514397509 | 0.766366259 |
| chr8:125700674-125700949 | -0.237105568 | 0.514398256 | 0.766366259 |
| chr17:73192300-73193648 | 0.209517483 | 0.514408174 | 0.766366259 |
| chr3:27659664-27659941 | -0.280530035 | 0.514447168 | 0.766409452 |
| chrX:70454435-70455889 | -0.207640102 | 0.51446885 | 0.766418945 |
| chr8:139213394-139213823 | -0.260336984 | 0.514473546 | 0.766418945 |
| chr3:183159435-183159709 | 0.275518695 | 0.514513178 | 0.766442989 |
| chr17:67510048-67510721 | -0.233207965 | 0.514519579 | 0.766442989 |
| chr12:62799189-62799414 | 0.290820115 | 0.514521125 | 0.766442989 |
| chr2:29309637-29309886 | -0.313593181 | 0.514529697 | 0.766442989 |
| chr3:43592061-43592261 | -0.364853536 | 0.514573563 | 0.76649343 |
| chr14:73472103-73472467 | 0.245673756 | 0.514623105 | 0.766551708 |
| chr15:85865118-85865383 | 0.276207232 | 0.514640795 | 0.766551708 |
| chr14:20327309-20327569 | -0.277455133 | 0.51465507 | 0.766551708 |
| chr2:29261684-29261938 | 0.305378145 | 0.514666331 | 0.766551708 |
| chr10:5222853-5223053 | -0.355145568 | 0.514667125 | 0.766551708 |
| chr9:76632609-76632879 | -0.355910946 | 0.514672713 | 0.766551708 |
| chr22:37906099-37906367 | 0.301624444 | 0.514712394 | 0.766582068 |
| chr1:215290316-215290563 | -0.319997296 | 0.514719388 | 0.766582068 |
| chr8:41844834-41845206 | -0.282623905 | 0.514730844 | 0.766582068 |

| | | | |
|---|---|---|---|
| chr8:123285927-123286135 | -0.300168617 | 0.514733116 | 0.766582068 |
| chr10:5089865-5090204 | 0.243067535 | 0.514828896 | 0.766709808 |
| chr9:128067167-128068318 | 0.206484358 | 0.514887384 | 0.766782009 |
| chr5:16649237-16649590 | -0.282757917 | 0.514897941 | 0.766782827 |
| chr16:4341033-4341256 | 0.320177975 | 0.514911086 | 0.7667875 |
| chr2:3557912-3559012 | -0.20647765 | 0.51495485 | 0.766830017 |
| chr3:149489133-149489375 | -0.273861849 | 0.514959653 | 0.766830017 |
| chr22:35399874-35400588 | 0.228546075 | 0.51501324 | 0.76689491 |
| chr3:31768881-31769081 | -0.360442562 | 0.515045074 | 0.766919114 |
| chr14:62691980-62692568 | 0.225303839 | 0.515058112 | 0.766919114 |
| chr5:155101584-155101916 | -0.311928414 | 0.515059522 | 0.766919114 |
| chr12:57522042-57522787 | 0.212227437 | 0.515092884 | 0.766953886 |
| chr3:128487681-128488183 | 0.225002152 | 0.515137089 | 0.767004801 |
| chr16:84119293-84119807 | -0.220024251 | 0.515157028 | 0.767019584 |
| chr11:65420016-65420225 | 0.254209362 | 0.515182353 | 0.767042385 |
| chr3:25837133-25837412 | 0.285050626 | 0.515232763 | 0.767102533 |
| chr17:64337084-64337473 | 0.213166899 | 0.515244445 | 0.767105021 |
| chr7:25862113-25862948 | 0.206141062 | 0.515308758 | 0.767185863 |
| chr3:45995484-45996101 | 0.232523249 | 0.515361364 | 0.76724925 |
| chr3:9302378-9302911 | 0.219432229 | 0.515371361 | 0.76724925 |
| chr10:62108997-62109369 | -0.259636962 | 0.515423256 | 0.767311599 |
| chr11:70397729-70398756 | 0.210019757 | 0.51546286 | 0.767341741 |
| chr6:10180571-10180857 | -0.323017461 | 0.515463532 | 0.767341741 |
| chr1:58155396-58155663 | -0.286421475 | 0.515485935 | 0.767360182 |
| chr11:65487443-65488041 | -0.212739669 | 0.515520072 | 0.767383848 |
| chr14:25049745-25049973 | 0.299810434 | 0.515530329 | 0.767383848 |
| chr11:2880685-2880885 | -0.378139165 | 0.515538071 | 0.767383848 |
| chr16:23803600-23803805 | 0.310258795 | 0.515543118 | 0.767383848 |
| chr1:56516891-56517300 | 0.232527348 | 0.515561768 | 0.767383848 |
| chrX:150575445-150575832 | -0.262180826 | 0.515561924 | 0.767383848 |
| chr8:124990729-124990938 | 0.279977228 | 0.515593415 | 0.767400906 |
| chr8:128469711-128469952 | 0.279977228 | 0.515593415 | 0.767400906 |
| chrX:149539224-149541124 | 0.202704825 | 0.515636194 | 0.767449669 |
| chr20:34523248-34523494 | -0.315956419 | 0.515695391 | 0.767512853 |
| chr10:15013544-15013894 | -0.250225002 | 0.515702103 | 0.767512853 |
| chr11:13487682-13488017 | -0.243912802 | 0.515708696 | 0.767512853 |
| chr14:27614524-27614724 | 0.33575519 | 0.515744087 | 0.767550616 |
| chr6:144008005-144008442 | -0.262771953 | 0.515759132 | 0.767552178 |
| chr12:50387549-50387864 | -0.274544887 | 0.515765172 | 0.767552178 |
| chr8:1897507-1897707 | 0.329213441 | 0.515787822 | 0.767570978 |
| chr14:102437906-102438333 | 0.249118462 | 0.515802914 | 0.767578528 |
| chr18:23918518-23918799 | -0.314512777 | 0.515853303 | 0.767622381 |
| chr4:85369768-85370017 | -0.304113909 | 0.515857583 | 0.767622381 |
| chr12:69935616-69936114 | -0.237608314 | 0.515873236 | 0.767622381 |
| chr12:114858441-114858666 | -0.357970479 | 0.515883107 | 0.767622381 |
| chr22:20956783-20957025 | 0.288921918 | 0.515889674 | 0.767622381 |
| chr20:54117358-54117625 | 0.225149767 | 0.515892906 | 0.767622381 |
| chr6:105402182-105403755 | -0.207849384 | 0.51590251 | 0.767622381 |
| chr1:224852958-224853168 | -0.3322454 | 0.51593502 | 0.76764094 |
| chr11:22617947-22618162 | -0.3322454 | 0.51593502 | 0.76764094 |

| | | | |
|---|---|---|---|
| chr19:46212195-46212512 | -0.31348783 | 0.515952222 | 0.767643661 |
| chr14:91417269-91419092 | 0.205281292 | 0.515960653 | 0.767643661 |
| chr14:44701651-44701892 | 0.291821727 | 0.515966905 | 0.767643661 |
| chr19:45523633-45524057 | -0.23381233 | 0.515994156 | 0.767669299 |
| chr19:38374348-38375116 | 0.214411669 | 0.516027725 | 0.767704334 |
| chr20:59994698-59995142 | 0.230436192 | 0.516051002 | 0.767712484 |
| chr3:154872655-154872855 | -0.345411713 | 0.516053242 | 0.767712484 |
| chr12:24960269-24960767 | -0.262483785 | 0.516072058 | 0.767725571 |
| chr12:11878206-11878776 | -0.252583451 | 0.516085357 | 0.767726779 |
| chr1:43147546-43148026 | -0.229817588 | 0.51610176 | 0.767726779 |
| chr12:105420716-105421004 | -0.272009394 | 0.516121607 | 0.767726779 |
| chr8:109639652-109639905 | -0.252797767 | 0.516123058 | 0.767726779 |
| chr11:32297606-32297889 | -0.283212511 | 0.516123998 | 0.767726779 |
| chr12:98990715-98991035 | -0.270267482 | 0.516132988 | 0.767726779 |
| chr21:15064448-15065908 | 0.210372331 | 0.516178228 | 0.767753474 |
| chr8:88881967-88882271 | 0.279403954 | 0.516194321 | 0.767753474 |
| chr1:246773313-246773692 | 0.259155798 | 0.516196235 | 0.767753474 |
| chr3:51388593-51388993 | 0.231929288 | 0.516196754 | 0.767753474 |
| chr8:41827851-41829125 | 0.213709589 | 0.516215124 | 0.767753474 |
| chr7:101308133-101308425 | -0.255805484 | 0.516228266 | 0.767753474 |
| chr22:36506704-36507410 | 0.224625771 | 0.516235299 | 0.767753474 |
| chr12:48681640-48682705 | 0.209457803 | 0.516238593 | 0.767753474 |
| chr7:2709844-2710044 | -0.287804339 | 0.516253084 | 0.767753474 |
| chr14:55026680-55027660 | 0.207378323 | 0.516264816 | 0.767753474 |
| chr9:35828896-35829585 | -0.224740601 | 0.516273882 | 0.767753474 |
| chr1:171984982-171985201 | -0.299968349 | 0.516288513 | 0.767753474 |
| chr14:46581175-46581550 | 0.280014022 | 0.516296329 | 0.767753474 |
| chr16:84672367-84672645 | 0.247485446 | 0.516310043 | 0.767753474 |
| chrX:103214769-103215255 | 0.235921225 | 0.516312652 | 0.767753474 |
| chr4:112604638-112604932 | -0.270182977 | 0.51633844 | 0.767753474 |
| chr8:90000725-90002174 | 0.204095475 | 0.516342047 | 0.767753474 |
| chr6:3001901-3002181 | 0.24973864 | 0.516343692 | 0.767753474 |
| chr19:10086082-10086409 | 0.280181652 | 0.516352996 | 0.767753474 |
| chr11:72751477-72751769 | -0.246789834 | 0.516355435 | 0.767753474 |
| chr3:52770542-52771523 | 0.223395003 | 0.516361355 | 0.767753474 |
| chr1:111222882-111223229 | 0.26120904 | 0.516426661 | 0.767781595 |
| chr1:167803287-167803660 | 0.236284004 | 0.516429117 | 0.767781595 |
| chr15:64153555-64153943 | 0.225759018 | 0.516429349 | 0.767781595 |
| chr19:45507200-45507750 | -0.222073745 | 0.516429468 | 0.767781595 |
| chr7:154777191-154777487 | -0.306820343 | 0.51643037 | 0.767781595 |
| chr2:42788929-42789351 | -0.234645598 | 0.516454734 | 0.767795579 |
| chr15:101307043-101307243 | -0.309091023 | 0.516463024 | 0.767795579 |
| chr6:84626235-84626505 | -0.276529285 | 0.516484888 | 0.767795579 |
| chr11:72813871-72815160 | 0.206341843 | 0.516485624 | 0.767795579 |
| chr16:2777129-2777908 | 0.222241382 | 0.516503165 | 0.767795579 |
| chr11:5684417-5685525 | 0.219331863 | 0.516503436 | 0.767795579 |
| chr1:192488127-192488377 | -0.29435555 | 0.51650992 | 0.767795579 |
| chr2:198930290-198930540 | -0.317924328 | 0.516528583 | 0.767808425 |
| chr15:90716782-90717896 | 0.20684114 | 0.516623758 | 0.767915659 |
| chr8:50398218-50398524 | 0.26392649 | 0.516626899 | 0.767915659 |

| | | | |
|---|---|---|---|
| chr6:95293994-95294573 | -0.241763458 | 0.516630789 | 0.767915659 |
| chr7:82214-82434 | -0.278455187 | 0.516687991 | 0.767982702 |
| chr17:63919186-63919559 | 0.229596743 | 0.516695939 | 0.767982702 |
| chr2:172951608-172951865 | 0.276181517 | 0.516739725 | 0.768032883 |
| chr6:139230490-139230811 | 0.286660403 | 0.516758667 | 0.768046139 |
| chr17:62986833-62987482 | -0.223560226 | 0.5167866 | 0.768072757 |
| chr13:21849381-21849652 | 0.286247639 | 0.51680665 | 0.768087657 |
| chr1:119744065-119744396 | 0.238546523 | 0.516848935 | 0.768135602 |
| chr19:49158071-49158508 | 0.244853352 | 0.516877871 | 0.768154326 |
| chr1:56758173-56758496 | -0.273521687 | 0.516881584 | 0.768154326 |
| chr5:138411104-138411347 | -0.296323854 | 0.516906602 | 0.768176607 |
| chr21:26149144-26149346 | 0.328826688 | 0.516936492 | 0.768206127 |
| chr6:35485940-35486565 | 0.218150529 | 0.516946762 | 0.76820649 |
| chr6:78676057-78676988 | 0.214232273 | 0.517000281 | 0.768271121 |
| chr5:79829321-79829654 | -0.251937881 | 0.517025503 | 0.7682937 |
| chr6:39724831-39725898 | -0.209919725 | 0.517052289 | 0.768300066 |
| chr17:74210322-74210669 | -0.24743461 | 0.517063981 | 0.768300066 |
| chr12:100404804-100405197 | -0.240493482 | 0.517067542 | 0.768300066 |
| chr15:72375542-72376540 | 0.211191913 | 0.517074192 | 0.768300066 |
| chr2:54574186-54574434 | -0.284258347 | 0.517079922 | 0.768300066 |
| chr20:10864443-10864697 | 0.308865065 | 0.517136074 | 0.768368598 |
| chr19:17076539-17076746 | 0.303733915 | 0.517192921 | 0.768438162 |
| chr7:2468646-2469270 | 0.223976159 | 0.517211762 | 0.768451254 |
| chr8:65948389-65949008 | -0.21752933 | 0.517227062 | 0.768454073 |
| chr1:145465188-145465990 | 0.206641504 | 0.51723969 | 0.768454073 |
| chr19:38917726-38918494 | -0.232268241 | 0.517243747 | 0.768454073 |
| chr2:14137506-14137993 | -0.254843568 | 0.517259275 | 0.768457486 |
| chr6:90286924-90287215 | -0.265365278 | 0.517314346 | 0.768457486 |
| chr9:78178559-78178786 | 0.338551721 | 0.517330157 | 0.768457486 |
| chr7:66108580-66108831 | 0.368511976 | 0.517338983 | 0.768457486 |
| chr1:106917669-106918015 | -0.25310171 | 0.517339021 | 0.768457486 |
| chr17:67963316-67963531 | -0.269428561 | 0.51735208 | 0.768457486 |
| chr19:29121781-29122107 | 0.275017348 | 0.517352482 | 0.768457486 |
| chr18:74342535-74342768 | -0.353637856 | 0.517356175 | 0.768457486 |
| chr2:190440905-190441170 | -0.353637856 | 0.517356175 | 0.768457486 |
| chr1:45490678-45491731 | 0.205952425 | 0.517356615 | 0.768457486 |
| chr16:720399-721012 | 0.222416402 | 0.517364711 | 0.768457486 |
| chr5:174571750-174572020 | 0.261147225 | 0.517366394 | 0.768457486 |
| chr19:17305394-17305887 | 0.218904748 | 0.517382063 | 0.768465862 |
| chr19:4598289-4598608 | 0.308237707 | 0.517433162 | 0.768526862 |
| chr15:77836466-77836675 | 0.29173396 | 0.517462984 | 0.768556257 |
| chr16:57093097-57093297 | 0.333911917 | 0.517499499 | 0.768595592 |
| chr4:38753108-38753503 | -0.270624627 | 0.517521398 | 0.768613218 |
| chr19:44770586-44771047 | 0.228521033 | 0.517560028 | 0.768652944 |
| chr4:78363455-78363781 | -0.297178361 | 0.517568209 | 0.768652944 |
| chr15:97301262-97301549 | -0.270504763 | 0.517609756 | 0.768699747 |
| chr3:163698152-163698938 | -0.221552764 | 0.517658671 | 0.768738225 |
| chr1:32291637-32292410 | 0.212555099 | 0.517666622 | 0.768738225 |
| chr3:192514064-192515728 | -0.205667711 | 0.517671928 | 0.768738225 |
| chr22:17706579-17706885 | 0.27544681 | 0.517682702 | 0.768738225 |

| | | | |
|---|---|---|---|
| chr1:171484862-171486443 | -0.20334911 | 0.51768583 | 0.768738225 |
| chr20:56187863-56188174 | -0.244192544 | 0.517704043 | 0.768750372 |
| chr5:168454586-168454992 | -0.253419315 | 0.51771757 | 0.768752811 |
| chr2:65432231-65432970 | 0.208300361 | 0.517725752 | 0.768752811 |
| chr14:82009076-82009367 | -0.305320021 | 0.51775003 | 0.768771301 |
| chr12:118377937-118378137 | 0.286704212 | 0.517770407 | 0.768771301 |
| chr16:73478679-73478879 | 0.325344935 | 0.517780918 | 0.768771301 |
| chr8:63220428-63220628 | 0.327751183 | 0.517782043 | 0.768771301 |
| chr6:118755879-118756262 | 0.293675576 | 0.517788371 | 0.768771301 |
| chr6:92810931-92811418 | -0.253063426 | 0.517834164 | 0.768824394 |
| chr12:82359200-82359402 | -0.254298102 | 0.517875379 | 0.768870687 |
| chr9:129488416-129489503 | -0.209846005 | 0.517900399 | 0.768892935 |
| chr2:158953958-158954297 | -0.275650448 | 0.517911916 | 0.768895136 |
| chr8:48172013-48172441 | 0.269009319 | 0.517932408 | 0.768902831 |
| chr17:7573558-7574013 | 0.23003884 | 0.517947125 | 0.768902831 |
| chr20:49914813-49915226 | 0.219260977 | 0.517947205 | 0.768902831 |
| chr11:102452511-102453179 | -0.237669679 | 0.517965999 | 0.768915834 |
| chr2:172995922-172996188 | 0.276845525 | 0.51799309 | 0.768941153 |
| chr11:131038688-131038888 | -0.336850028 | 0.518007611 | 0.768947811 |
| chr17:48804865-48805422 | -0.231348577 | 0.51808006 | 0.768994157 |
| chr21:41632052-41632477 | -0.257593476 | 0.518083637 | 0.768994157 |
| chr2:120011443-120011813 | 0.254576733 | 0.518087727 | 0.768994157 |
| chr3:52197763-52198269 | 0.24779232 | 0.518088898 | 0.768994157 |
| chr22:31248072-31248706 | 0.225737345 | 0.518089013 | 0.768994157 |
| chr2:204483570-204484108 | -0.242692237 | 0.518112675 | 0.76901438 |
| chr16:84952256-84952523 | 0.248078083 | 0.518136301 | 0.769034551 |
| chr2:197263055-197263255 | -0.361908616 | 0.518155851 | 0.769048671 |
| chr3:145316380-145316678 | -0.287558591 | 0.518214231 | 0.769101485 |
| chr17:10790577-10791348 | -0.215926001 | 0.518225766 | 0.769101485 |
| chr20:45340462-45340720 | -0.262008737 | 0.518231064 | 0.769101485 |
| chrX:133985494-133986172 | 0.223417929 | 0.518231586 | 0.769101485 |
| chr7:44712244-44712448 | -0.305466927 | 0.518290788 | 0.769163765 |
| chr6:139364475-139364745 | -0.305531858 | 0.518293628 | 0.769163765 |
| chr12:97716412-97716641 | -0.309585014 | 0.518361886 | 0.769241065 |
| chr2:637014-637338 | 0.261066143 | 0.518365794 | 0.769241065 |
| chr2:171811217-171811480 | -0.339807082 | 0.518387649 | 0.769242236 |
| chr7:73626947-73627195 | -0.278226311 | 0.518395776 | 0.769242236 |
| chr3:167709992-167710601 | -0.243712347 | 0.518397322 | 0.769242236 |
| chr9:4792213-4793451 | -0.206067435 | 0.518411535 | 0.769242236 |
| chr7:92691599-92691868 | -0.294336971 | 0.518429856 | 0.769242236 |
| chr1:32760077-32760566 | 0.217092749 | 0.518431808 | 0.769242236 |
| chr2:226796786-226797084 | -0.250808682 | 0.51843686 | 0.769242236 |
| chr17:47810233-47810858 | 0.217179275 | 0.518467969 | 0.769273498 |
| chr13:32669095-32669326 | -0.296656289 | 0.518479503 | 0.769275715 |
| chr9:109244556-109244890 | 0.250242018 | 0.518534341 | 0.769342182 |
| chr11:73931303-73931606 | 0.265202462 | 0.518571278 | 0.769382086 |
| chr2:84904983-84905971 | -0.208559632 | 0.518600624 | 0.769401217 |
| chr10:59709397-59710078 | 0.217556134 | 0.518614449 | 0.769401217 |
| chr11:118918011-118918583 | -0.212588602 | 0.518620759 | 0.769401217 |
| chr14:52568175-52568380 | -0.333889891 | 0.518624338 | 0.769401217 |

| | | | |
|---|---|---|---|
| chr8:11958962-11959186 | -0.324568601 | 0.518647877 | 0.76942124 |
| chr8:144767423-144767676 | -0.29435137 | 0.518669398 | 0.76943827 |
| chr9:135962126-135962368 | -0.28362082 | 0.51869289 | 0.769443382 |
| chr6:93036900-93037100 | -0.346731239 | 0.518693742 | 0.769443382 |
| chr12:75511066-75512015 | -0.207295933 | 0.518711281 | 0.769443382 |
| chr12:118874818-118875182 | 0.265055908 | 0.518713013 | 0.769443382 |
| chr6:8063727-8064757 | -0.207397997 | 0.518739026 | 0.769461049 |
| chr8:97594911-97595759 | -0.214911008 | 0.518750943 | 0.769461049 |
| chr1:239878739-239878956 | 0.329614117 | 0.518755049 | 0.769461049 |
| chr9:130689937-130690573 | -0.215895181 | 0.518784449 | 0.769489761 |
| chr4:24470267-24470467 | 0.311055964 | 0.51883798 | 0.769554264 |
| chr16:67480408-67480736 | 0.240757097 | 0.518850692 | 0.769558222 |
| chr17:82022762-82023477 | 0.205968434 | 0.518888125 | 0.769598844 |
| chr1:32062474-32062873 | 0.23351011 | 0.518900649 | 0.769602523 |
| chr14:81247109-81248139 | -0.216982253 | 0.518928141 | 0.769613014 |
| chr1:92441753-92442333 | -0.247124774 | 0.518937196 | 0.769613014 |
| chr8:127738561-127738811 | 0.228509715 | 0.518937855 | 0.769613014 |
| chr8:130764602-130764865 | 0.272024978 | 0.518958566 | 0.769615174 |
| chr20:54091191-54091420 | 0.22108555 | 0.5189594 | 0.769615174 |
| chr2:135709937-135710213 | -0.273001263 | 0.519008743 | 0.769656964 |
| chr2:53969848-53971567 | -0.205693901 | 0.519020507 | 0.769656964 |
| chr1:20440883-20441083 | -0.348510629 | 0.519028059 | 0.769656964 |
| chr1:11272773-11273861 | -0.213280579 | 0.519039206 | 0.769656964 |
| chr22:29733826-29734177 | 0.267703869 | 0.519048288 | 0.769656964 |
| chr16:86270755-86270955 | -0.367340213 | 0.519051403 | 0.769656964 |
| chr17:45135290-45135775 | 0.221350686 | 0.519071434 | 0.769656964 |
| chr12:125005706-125006078 | 0.277475045 | 0.519077355 | 0.769656964 |
| chr10:7787551-7787775 | 0.254670597 | 0.519077984 | 0.769656964 |
| chr10:79487443-79487948 | -0.231242697 | 0.519115217 | 0.769697276 |
| chr3:156762562-156762886 | 0.273237147 | 0.519144631 | 0.769724492 |
| chrX:134626507-134626707 | 0.342446019 | 0.519173168 | 0.769724492 |
| chr1:196817382-196817668 | -0.304167416 | 0.519173448 | 0.769724492 |
| chrX:12804733-12804965 | -0.309049656 | 0.519173755 | 0.769724492 |
| chr6:73774355-73774843 | 0.25424684 | 0.519222234 | 0.769781472 |
| chr10:131997710-131998099 | 0.236329595 | 0.519239326 | 0.769788963 |
| chr8:143986098-143986779 | -0.221054042 | 0.51924738 | 0.769788963 |
| chr4:184163867-184164079 | -0.321588666 | 0.519258303 | 0.769790261 |
| chr12:6970236-6971075 | 0.211735685 | 0.519271218 | 0.769794515 |
| chr5:171946155-171946355 | -0.352265836 | 0.519300446 | 0.769818737 |
| chr12:48795590-48796199 | 0.225199402 | 0.519307652 | 0.769818737 |
| chr19:17778972-17779873 | -0.221261332 | 0.519341764 | 0.76985441 |
| chr10:128035599-128035819 | -0.309867784 | 0.519387533 | 0.769907363 |
| chr1:184590464-184590889 | 0.265322777 | 0.51939953 | 0.769910252 |
| chr10:60816615-60816875 | -0.296848407 | 0.519440523 | 0.76995612 |
| chr1:16244471-16244671 | 0.348592913 | 0.519507109 | 0.769996049 |
| chr3:138025565-138025821 | -0.331067768 | 0.519511084 | 0.769996049 |
| chr15:64461124-64461656 | -0.228573514 | 0.519511579 | 0.769996049 |
| chr1:201494459-201494687 | -0.260259857 | 0.519516557 | 0.769996049 |
| chr1:150221656-150221878 | 0.288341411 | 0.519517707 | 0.769996049 |
| chr3:46930788-46931266 | 0.240352673 | 0.519528268 | 0.769996808 |

| | | | |
|---|---|---|---|
| chr2:101252255-101253052 | 0.208021806 | 0.519574951 | 0.770051101 |
| chr11:110296116-110297139 | -0.217280584 | 0.519600469 | 0.770064107 |
| chr9:84667636-84668064 | 0.247213515 | 0.519609064 | 0.770064107 |
| chr6:27890688-27890991 | -0.244779561 | 0.519613877 | 0.770064107 |
| chr15:40103206-40103853 | 0.212101369 | 0.519633205 | 0.770066535 |
| chr3:5994670-5994870 | -0.331085021 | 0.519635616 | 0.770066535 |
| chr10:65555293-65555550 | -0.286943487 | 0.519680432 | 0.770087947 |
| chr10:92122806-92123536 | 0.223877927 | 0.519681144 | 0.770087947 |
| chr7:35315383-35315583 | 0.337097144 | 0.519681946 | 0.770087947 |
| chr8:13080693-13080974 | -0.295765765 | 0.519690266 | 0.770087947 |
| chr6:31795413-31796379 | 0.214388557 | 0.519706313 | 0.770096833 |
| chr4:153095457-153095944 | -0.232775665 | 0.51972131 | 0.770104162 |
| chr5:27073190-27073404 | -0.325153752 | 0.519737534 | 0.77011331 |
| chr8:51979912-51980191 | -0.297658161 | 0.519765083 | 0.770139237 |
| chr17:69569061-69569653 | -0.226898785 | 0.519817999 | 0.770189711 |
| chr2:47782630-47784313 | -0.202543126 | 0.519835204 | 0.770189711 |
| chr5:62305532-62307064 | -0.203087458 | 0.519846092 | 0.770189711 |
| chr17:66695021-66695221 | -0.340708332 | 0.519855227 | 0.770189711 |
| chr21:44297712-44297927 | 0.265007934 | 0.519859171 | 0.770189711 |
| chr5:26941744-26942046 | 0.303832458 | 0.519859459 | 0.770189711 |
| chr17:31437362-31437662 | 0.258680953 | 0.519911177 | 0.77025144 |
| chr1:226121495-226122266 | 0.212587372 | 0.519946814 | 0.770280131 |
| chr6:17156192-17156634 | -0.301724498 | 0.519950649 | 0.770280131 |
| chr3:113912970-113913336 | -0.241009959 | 0.519966168 | 0.770288228 |
| chr15:63083618-63083849 | -0.307490418 | 0.519991861 | 0.770301206 |
| chr4:110631606-110632450 | -0.210477367 | 0.519995035 | 0.770301206 |
| chr12:47858906-47859119 | 0.323841357 | 0.520026608 | 0.770333084 |
| chr2:137847462-137847938 | -0.256796603 | 0.520082413 | 0.770400855 |
| chr1:247007504-247008409 | -0.203130619 | 0.520100321 | 0.770406416 |
| chr14:32076053-32077455 | 0.201988502 | 0.520106276 | 0.770406416 |
| chr17:16286059-16286259 | -0.332209117 | 0.520131546 | 0.770428953 |
| chr12:49018109-49019046 | 0.209243074 | 0.520159621 | 0.770446437 |
| chr13:62156975-62157236 | 0.288906938 | 0.520172849 | 0.770446437 |
| chr16:4615996-4616824 | 0.205997729 | 0.520173515 | 0.770446437 |
| chr19:40634384-40634714 | -0.281550909 | 0.520196297 | 0.770465287 |
| chr2:156934039-156934265 | -0.305608999 | 0.520240877 | 0.770502113 |
| chr3:128650097-128651243 | 0.208587779 | 0.520241273 | 0.770502113 |
| chr8:143942556-143942913 | 0.220472104 | 0.520252203 | 0.770503407 |
| chr12:1634595-1634911 | -0.287582893 | 0.520262317 | 0.770503494 |
| chr20:55415650-55415882 | -0.288817588 | 0.52040085 | 0.770671777 |
| chr15:69475819-69476314 | 0.236351332 | 0.520403887 | 0.770671777 |
| chr17:78618308-78618508 | -0.326246871 | 0.520420037 | 0.770671777 |
| chr6:35727969-35728227 | 0.254682258 | 0.52042206 | 0.770671777 |
| chrX:123432873-123433117 | 0.295112162 | 0.520432401 | 0.770671777 |
| chr6:150717489-150717771 | -0.286012147 | 0.520436294 | 0.770671777 |
| chr17:41972856-41973354 | 0.233838515 | 0.520505812 | 0.770752027 |
| chr6:86175206-86175491 | 0.284924764 | 0.520510606 | 0.770752027 |
| chr2:128736641-128736961 | -0.257797587 | 0.520525077 | 0.770758559 |
| chr9:109119200-109120376 | -0.206362881 | 0.520546202 | 0.770774945 |
| chr3:144162060-144162384 | -0.2617496 | 0.520603364 | 0.770834902 |

| | | | |
|---|---|---|---|
| chr12:12612527-12612727 | -0.315559914 | 0.520606815 | 0.770834902 |
| chr8:23457127-23458269 | 0.212686599 | 0.520703536 | 0.770963214 |
| chr17:10737172-10737435 | 0.270228882 | 0.520732179 | 0.770990725 |
| chr1:42003492-42003809 | -0.278768803 | 0.520777225 | 0.771017736 |
| chr9:105157160-105157605 | -0.257517122 | 0.520778946 | 0.771017736 |
| chr1:110963720-110964346 | -0.207909814 | 0.520785735 | 0.771017736 |
| chr3:63068089-63068416 | 0.223739665 | 0.520793366 | 0.771017736 |
| chr14:69397948-69399027 | 0.206202036 | 0.520800735 | 0.771017736 |
| chr20:50359779-50359980 | 0.274916416 | 0.520885833 | 0.771112761 |
| chr17:3667955-3669059 | 0.206944345 | 0.520895238 | 0.771112761 |
| chrX:134764155-134764594 | 0.222126754 | 0.52090098 | 0.771112761 |
| chr1:36384933-36385275 | 0.230865602 | 0.520908291 | 0.771112761 |
| chr11:67288334-67289749 | 0.206937789 | 0.520915241 | 0.771112761 |
| chr1:202056695-202057712 | -0.20305291 | 0.520927856 | 0.771116537 |
| chr14:53055005-53055243 | 0.295940052 | 0.52096898 | 0.771162514 |
| chr7:26864133-26865222 | 0.204768581 | 0.520987729 | 0.771165778 |
| chrX:40105833-40106398 | -0.231040829 | 0.52100126 | 0.771165778 |
| chr13:24892578-24892778 | -0.3436443 | 0.521010547 | 0.771165778 |
| chr8:80084146-80084818 | 0.207671156 | 0.521011444 | 0.771165778 |
| chr1:185917706-185917960 | 0.296930929 | 0.521025808 | 0.771172142 |
| chr2:215619100-215619577 | -0.25769083 | 0.52104431 | 0.77118463 |
| chr1:207698875-207699227 | -0.230049784 | 0.521076256 | 0.771217016 |
| chr3:151543480-151543833 | -0.271902656 | 0.52108827 | 0.7712199 |
| chr14:37197437-37198401 | 0.206957393 | 0.52110858 | 0.771235062 |
| chr10:86611573-86611780 | 0.283377941 | 0.521171287 | 0.771301767 |
| chr10:102169468-10217002( | 0.225635911 | 0.521173784 | 0.771301767 |
| chr19:4811668-4812002 | 0.275440824 | 0.521205099 | 0.77132292 |
| chr9:105290454-105290669 | 0.299292052 | 0.521208211 | 0.77132292 |
| chr8:99893052-99894045 | 0.20843964 | 0.521251958 | 0.771366056 |
| chr2:136214758-136214967 | 0.321117352 | 0.521257494 | 0.771366056 |
| chr1:152035863-152036073 | 0.245909949 | 0.521281704 | 0.77137411 |
| chr12:101823960-101824385 | -0.229125399 | 0.521283071 | 0.77137411 |
| chr7:45260045-45260403 | 0.249573287 | 0.521295531 | 0.771377652 |
| chr3:88032201-88032401 | -0.352469825 | 0.521367644 | 0.771445291 |
| chr3:137171863-137172063 | -0.352469825 | 0.521367644 | 0.771445291 |
| chr15:71819513-71819766 | -0.343419865 | 0.521371446 | 0.771445291 |
| chr3:141105864-141106174 | 0.259050207 | 0.521436868 | 0.771519102 |
| chr5:112166203-112166803 | -0.238394243 | 0.521442094 | 0.771519102 |
| chr7:149178979-149179228 | 0.320845579 | 0.521451538 | 0.771519102 |
| chr13:27578566-27578809 | -0.298890723 | 0.521484835 | 0.771553468 |
| chr3:184371550-184371873 | 0.240307327 | 0.521526825 | 0.771569906 |
| chr17:48878867-48879312 | 0.227925618 | 0.521559704 | 0.771569906 |
| chr14:20662784-20663610 | -0.209181912 | 0.521564435 | 0.771569906 |
| chr8:120084612-120084815 | -0.324670696 | 0.521567313 | 0.771569906 |
| chr9:108919606-108919846 | -0.324670696 | 0.521567313 | 0.771569906 |
| chr1:201506563-201507376 | 0.208005543 | 0.521590747 | 0.771569906 |
| chr3:75422103-75422465 | 0.24599549 | 0.521594243 | 0.771569906 |
| chr6:33075668-33076026 | -0.25151421 | 0.521619475 | 0.771569906 |
| chr11:86301958-86302784 | -0.207609967 | 0.521623615 | 0.771569906 |
| chr9:18442709-18443045 | -0.263655738 | 0.521642739 | 0.771569906 |

| | | | |
|---|---|---|---|
| chr1:26529536-26530096 | -0.237379881 | 0.521651163 | 0.771569906 |
| chr3:89246403-89246627 | -0.337049736 | 0.521659639 | 0.771569906 |
| chr3:165770858-165771076 | -0.337049736 | 0.521659639 | 0.771569906 |
| chr2:19140552-19141308 | 0.207715336 | 0.521677041 | 0.771569906 |
| chr7:30469296-30469790 | 0.231742397 | 0.521687529 | 0.771569906 |
| chr11:6315827-6316119 | -0.2815513 | 0.521697948 | 0.771569906 |
| chr7:5494577-5494939 | 0.231264244 | 0.521703363 | 0.771569906 |
| chr1:36576120-36576384 | 0.270871329 | 0.521704782 | 0.771569906 |
| chr19:14529137-14529971 | 0.210503681 | 0.52170535 | 0.771569906 |
| chr10:124743959-124744683 | 0.219494454 | 0.521717872 | 0.771569906 |
| chr1:40508468-40508884 | 0.243994943 | 0.521721158 | 0.771569906 |
| chr20:47983021-47983289 | 0.218695852 | 0.521721855 | 0.771569906 |
| chr3:62187005-62187317 | -0.219452862 | 0.521730488 | 0.771569906 |
| chr13:22735511-22735711 | 0.326556797 | 0.521747116 | 0.771569906 |
| chr19:50137056-50137338 | 0.302861205 | 0.52174769 | 0.771569906 |
| chr20:63234750-63235088 | -0.232786734 | 0.521777618 | 0.771585888 |
| chr5:2806325-2806546 | -0.31463406 | 0.521782507 | 0.771585888 |
| chr17:82098108-82099072 | 0.204997415 | 0.521806034 | 0.771585888 |
| chr2:240832195-240832575 | -0.271544655 | 0.521816029 | 0.771585888 |
| chr15:98311603-98312149 | -0.24600101 | 0.521824072 | 0.771585888 |
| chr14:93936338-93936615 | 0.253818891 | 0.521824426 | 0.771585888 |
| chr2:206230417-206230779 | 0.285745057 | 0.521851839 | 0.771585888 |
| chr3:185066775-185067021 | -0.319731392 | 0.521865909 | 0.771585888 |
| chr2:45936321-45936688 | -0.232745747 | 0.521868575 | 0.771585888 |
| chr8:107187213-107187664 | 0.240364124 | 0.521869559 | 0.771585888 |
| chr5:42320-42587 | 0.302639069 | 0.521870513 | 0.771585888 |
| chr1:200107657-200107887 | -0.283881685 | 0.521879338 | 0.771585888 |
| chr9:129139278-129139572 | 0.246135649 | 0.521910417 | 0.771616949 |
| chr9:90433006-90433925 | 0.213169302 | 0.521937503 | 0.771633274 |
| chr1:9823909-9824414 | 0.234455283 | 0.52195734 | 0.771633274 |
| chrX:102517322-102517669 | -0.242031441 | 0.521960944 | 0.771633274 |
| chr17:60415091-60415351 | 0.222936306 | 0.521963003 | 0.771633274 |
| chr7:138245392-138245592 | -0.336738985 | 0.521971813 | 0.771633274 |
| chr2:187500033-187500280 | -0.257119183 | 0.521994795 | 0.771645944 |
| chr7:31736317-31736739 | 0.225518316 | 0.522004101 | 0.771645944 |
| chr3:27632426-27633508 | 0.213963488 | 0.522010595 | 0.771645944 |
| chr12:38138607-38139271 | 0.227252938 | 0.522029023 | 0.77165694 |
| chr6:151900634-151901931 | -0.204282813 | 0.522068221 | 0.77165694 |
| chr6:106728328-106728607 | 0.297508629 | 0.522074194 | 0.77165694 |
| chr4:188372976-188373299 | -0.292587119 | 0.522087055 | 0.77165694 |
| chr18:21391157-21391408 | -0.303890259 | 0.522087252 | 0.77165694 |
| chr18:31950471-31950760 | -0.334021436 | 0.522092224 | 0.77165694 |
| chr10:132904767-132905090 | -0.266023854 | 0.52210045 | 0.77165694 |
| chr20:63707229-63707477 | 0.245731196 | 0.522102745 | 0.77165694 |
| chr14:94809722-94809922 | 0.324238665 | 0.522133526 | 0.77165694 |
| chr6:36743661-36743861 | -0.333802027 | 0.522149257 | 0.77165694 |
| chr2:111122338-111122730 | 0.228672076 | 0.52215381 | 0.77165694 |
| chr2:170235985-170236282 | 0.280801554 | 0.522169967 | 0.77165694 |
| chr6:87718947-87719461 | 0.232340477 | 0.522184013 | 0.77165694 |
| chr4:39514912-39515338 | 0.254294524 | 0.522191565 | 0.77165694 |

| | | | |
|---|---|---|---|
| chr13:36335301-36335594 | 0.315994059 | 0.52220907 | 0.77165694 |
| chr3:61664212-61664412 | 0.250009168 | 0.52221939 | 0.77165694 |
| chr22:23433237-23433462 | -0.299489546 | 0.522231631 | 0.77165694 |
| chr12:54444311-54444571 | -0.306587151 | 0.522234971 | 0.77165694 |
| chr16:1413765-1414203 | -0.252872201 | 0.522237093 | 0.77165694 |
| chr6:129621316-129621584 | -0.297516611 | 0.522240456 | 0.77165694 |
| chr5:137497379-137497647 | -0.286319896 | 0.522252962 | 0.77165694 |
| chr4:137490574-137490791 | -0.381800244 | 0.522266801 | 0.77165694 |
| chr8:13100550-13100801 | -0.381800244 | 0.522266801 | 0.77165694 |
| chr12:54394354-54394558 | 0.30116428 | 0.522274395 | 0.77165694 |
| chr15:82108569-82108824 | 0.25274506 | 0.522279702 | 0.77165694 |
| chr12:108234270-108234496 | -0.333711586 | 0.522291697 | 0.77165694 |
| chr21:16070180-16070596 | -0.308440294 | 0.522291862 | 0.77165694 |
| chr16:50245498-50245949 | 0.252352505 | 0.522300021 | 0.77165694 |
| chr3:179713824-179714194 | -0.271708911 | 0.522329098 | 0.771678852 |
| chr10:82938314-82938751 | 0.243945969 | 0.522334995 | 0.771678852 |
| chr1:108125489-108125923 | -0.222358352 | 0.522367851 | 0.771701256 |
| chr1:35568935-35569225 | 0.257971091 | 0.522371159 | 0.771701256 |
| chr8:66033791-66034008 | 0.280103637 | 0.522380375 | 0.771701256 |
| chr20:53963859-53964305 | 0.208912217 | 0.522399176 | 0.77171107 |
| chr4:88320716-88321327 | 0.225372324 | 0.522407161 | 0.77171107 |
| chr1:211520628-211520859 | -0.28613466 | 0.522455868 | 0.771758261 |
| chr11:64355078-64355319 | 0.270445497 | 0.522459251 | 0.771758261 |
| chr6:53180301-53180501 | -0.35564578 | 0.522481764 | 0.771765742 |
| chr1:117606891-117607242 | -0.23265958 | 0.52248446 | 0.771765742 |
| chr1:28812965-28813165 | -0.34233894 | 0.522523874 | 0.771795088 |
| chr18:50161912-50162113 | 0.297193985 | 0.522524473 | 0.771795088 |
| chr5:132866173-132867060 | -0.20594296 | 0.522581809 | 0.771859048 |
| chr2:15665710-15666096 | 0.265384607 | 0.522618585 | 0.771859048 |
| chr3:143071494-143071694 | 0.309585798 | 0.522620149 | 0.771859048 |
| chr7:87312319-87312540 | -0.281677594 | 0.522624308 | 0.771859048 |
| chr7:47228585-47229057 | -0.218300308 | 0.52262973 | 0.771859048 |
| chr18:48662216-48662475 | -0.285954691 | 0.522635534 | 0.771859048 |
| chr12:7109180-7109380 | 0.318429549 | 0.522638291 | 0.771859048 |
| chr2:190477933-190478133 | -0.347798607 | 0.52266661 | 0.771885994 |
| chr20:19130467-19130667 | 0.356846766 | 0.522686555 | 0.771900571 |
| chr20:6006447-6006816 | -0.26153729 | 0.52273598 | 0.771958682 |
| chr6:88983545-88983840 | -0.258535964 | 0.522763485 | 0.771984423 |
| chr4:148528015-148528259 | 0.341949116 | 0.522775888 | 0.77198786 |
| chr19:46016799-46017147 | 0.238486232 | 0.522798776 | 0.771996627 |
| chr11:83156234-83157512 | -0.199723188 | 0.522804103 | 0.771996627 |
| chr11:118156411-118156733 | 0.245854616 | 0.522812051 | 0.771996627 |
| chr16:50806320-50806786 | 0.253136831 | 0.522875977 | 0.772050281 |
| chr14:20632479-20633458 | -0.206323763 | 0.522882566 | 0.772050281 |
| chr14:70257572-70257901 | -0.254929017 | 0.522891391 | 0.772050281 |
| chr8:124592771-124593235 | -0.225776346 | 0.522897882 | 0.772050281 |
| chr2:175464023-175464224 | -0.326794238 | 0.522898767 | 0.772050281 |
| chr19:50383711-50384747 | 0.205062148 | 0.522952346 | 0.772114511 |
| chr6:52612668-52613037 | 0.295537994 | 0.522997342 | 0.772166066 |
| chr2:57759863-57760218 | -0.23916876 | 0.523026993 | 0.772179285 |

| | | | |
|---|---|---|---|
| chr5:78649477-78649677 | -0.329618461 | 0.523031812 | 0.772179285 |
| chr10:68074565-68075578 | -0.205006277 | 0.523036529 | 0.772179285 |
| chr18:3771317-3771668 | -0.244757193 | 0.523094096 | 0.772249395 |
| chr17:75514981-75515278 | -0.227931931 | 0.523127871 | 0.772284377 |
| chr16:23013620-23013820 | -0.345663209 | 0.523145811 | 0.772287301 |
| chr12:24463437-24463637 | -0.345104082 | 0.52315001 | 0.772287301 |
| chr5:150410662-150410867 | -0.270787516 | 0.523202142 | 0.77230522 |
| chr8:140459664-140459893 | 0.231124277 | 0.523210243 | 0.77230522 |
| chr13:31538168-31538372 | -0.317148245 | 0.523228118 | 0.77230522 |
| chr21:6371683-6372377 | 0.200485077 | 0.52323435 | 0.77230522 |
| chr7:149144646-149144914 | -0.245914897 | 0.523238604 | 0.77230522 |
| chr14:41636542-41637305 | 0.21547805 | 0.523242432 | 0.77230522 |
| chr16:84893654-84893870 | -0.31278868 | 0.523242641 | 0.77230522 |
| chr1:75991553-75991889 | -0.247812829 | 0.523248643 | 0.77230522 |
| chr6:127420522-127420914 | 0.256210952 | 0.523252863 | 0.77230522 |
| chr22:23858315-23858526 | 0.275377108 | 0.523303198 | 0.772358759 |
| chr4:125314682-125315339 | 0.207003362 | 0.523309297 | 0.772358759 |
| chr10:101141714-101141997 | -0.27318151 | 0.523336251 | 0.772383663 |
| chr20:13262077-13262421 | -0.260417263 | 0.523394981 | 0.772427721 |
| chr2:241561888-241562183 | 0.248261473 | 0.523404392 | 0.772427721 |
| chr22:19844363-19844589 | 0.31572304 | 0.523418316 | 0.772427721 |
| chr22:23750664-23751863 | 0.209064633 | 0.523422285 | 0.772427721 |
| chr3:2323716-2323924 | 0.344178468 | 0.523438108 | 0.772427721 |
| chr2:241549783-241550000 | -0.294032976 | 0.523450092 | 0.772427721 |
| chr5:135551895-135552211 | -0.260239612 | 0.523450516 | 0.772427721 |
| chr2:163368932-163369473 | 0.220073954 | 0.523454733 | 0.772427721 |
| chr1:151611734-151612982 | 0.2034733 | 0.523470148 | 0.772427721 |
| chr9:35604647-35605328 | -0.216124924 | 0.52347611 | 0.772427721 |
| chr1:218200917-218201210 | -0.275102031 | 0.523478961 | 0.772427721 |
| chr10:84537457-84537685 | -0.257735315 | 0.523494321 | 0.772427721 |
| chr10:94362539-94363394 | -0.212617684 | 0.523509677 | 0.772427721 |
| chr2:102032327-102032719 | -0.265050976 | 0.523524122 | 0.772427721 |
| chr13:44468355-44468599 | -0.312430877 | 0.523526986 | 0.772427721 |
| chr5:150010118-150010338 | -0.330129083 | 0.5235274 | 0.772427721 |
| chr11:61871147-61871596 | 0.228625306 | 0.523562106 | 0.772464054 |
| chrX:147920446-147920656 | -0.329062774 | 0.52357911 | 0.772474267 |
| chr17:54426319-54426575 | 0.316982237 | 0.523594887 | 0.772482669 |
| chr5:172770570-172771540 | 0.20304554 | 0.523624647 | 0.7725117 |
| chr13:29707737-29708445 | -0.22101744 | 0.523653035 | 0.772517829 |
| chr10:71850630-71851637 | -0.206110685 | 0.523658223 | 0.772517829 |
| chr10:117374942-117375928 | -0.210877379 | 0.523679004 | 0.772517829 |
| chr15:90954640-90955548 | 0.204140435 | 0.523680031 | 0.772517829 |
| chr11:11620174-11620685 | -0.235234625 | 0.523681907 | 0.772517829 |
| chrX:140665035-140665235 | 0.323065406 | 0.523693125 | 0.772517829 |
| chr17:81666245-81667064 | -0.210266269 | 0.523699376 | 0.772517829 |
| chr12:57487774-57488262 | 0.222187007 | 0.523737321 | 0.772558928 |
| chr3:153595248-153595546 | -0.253625562 | 0.523783211 | 0.772611747 |
| chr8:94163276-94163476 | 0.317395916 | 0.523828408 | 0.77266354 |
| chr3:50260005-50260458 | 0.236481011 | 0.523847586 | 0.772676954 |
| chr1:226186092-226187139 | 0.210138826 | 0.523904453 | 0.772745958 |

| | | | |
|---|---|---|---|
| chr10:100986712-100987719 | 0.213825277 | 0.523930721 | 0.7727607 |
| chr1:93620926-93621126 | -0.344725864 | 0.523934619 | 0.7727607 |
| chr2:47388146-47388617 | -0.227249089 | 0.523952995 | 0.772772929 |
| chr12:106247283-106247769 | 0.22768178 | 0.523991159 | 0.77281434 |
| chr11:47176657-47177877 | 0.202157582 | 0.524084493 | 0.772937116 |
| chr17:48131251-48131532 | -0.252265968 | 0.524154923 | 0.773023275 |
| chr1:203128107-203128419 | 0.231343312 | 0.524175067 | 0.773023275 |
| chr12:57766119-57767234 | -0.209114917 | 0.524182002 | 0.773023275 |
| chr8:103445554-103445880 | -0.235333528 | 0.524183267 | 0.773023275 |
| chr15:65049330-65049841 | -0.218237396 | 0.52419353 | 0.773023532 |
| chr12:124273700-124274062 | 0.248774426 | 0.524205682 | 0.773026574 |
| chr16:17211796-17212065 | -0.293590786 | 0.524216012 | 0.773026929 |
| chr12:56164307-56164730 | -0.250569051 | 0.524275512 | 0.773099792 |
| chr7:88546781-88547123 | -0.251023652 | 0.524287061 | 0.773101944 |
| chr11:47376641-47376857 | -0.281085776 | 0.52433151 | 0.773141483 |
| chr2:135530425-135531801 | -0.199795455 | 0.524334056 | 0.773141483 |
| chr5:151447319-151448105 | -0.209508613 | 0.524349174 | 0.773147378 |
| chr14:69766811-69768044 | 0.201360382 | 0.524358234 | 0.773147378 |
| chr16:85057322-85057540 | -0.282168398 | 0.524373429 | 0.773154903 |
| chr8:86182322-86182807 | 0.224592317 | 0.524421689 | 0.773204375 |
| chr14:55534507-55534769 | 0.313747356 | 0.524427164 | 0.773204375 |
| chr3:63172164-63172466 | -0.239472875 | 0.5244528 | 0.773227293 |
| chr6:116779670-116779948 | -0.300866863 | 0.524472258 | 0.773241102 |
| chr15:100914855-100915135 | 0.249612525 | 0.524504515 | 0.773266785 |
| chr4:109798930-109799173 | -0.304456017 | 0.524516115 | 0.773266785 |
| chr12:63206987-63207208 | -0.312538606 | 0.524519954 | 0.773266785 |
| chr15:101006976-101007230 | -0.241465015 | 0.524561059 | 0.773309692 |
| chr12:109154449-109155350 | 0.205184573 | 0.524569243 | 0.773309692 |
| chr17:64165595-64165826 | -0.299376747 | 0.524580206 | 0.773310975 |
| chr11:31800486-31800686 | 0.341450573 | 0.524592937 | 0.773314865 |
| chr5:173629776-173630041 | -0.255285557 | 0.524645963 | 0.773341271 |
| chr10:1448392-1449049 | -0.231634502 | 0.524655245 | 0.773341271 |
| chr17:60237585-60237820 | 0.231340162 | 0.524661109 | 0.773341271 |
| chr11:129881667-129881868 | 0.324047243 | 0.524661315 | 0.773341271 |
| chr13:109610622-109610847 | 0.324047243 | 0.524661315 | 0.773341271 |
| chr11:19077034-19077249 | -0.293079003 | 0.524696722 | 0.773378583 |
| chr19:1795887-1796087 | -0.350102039 | 0.524725639 | 0.773379038 |
| chr14:23141859-23142634 | -0.220165793 | 0.524736058 | 0.773379038 |
| chr12:57621839-57622060 | 0.286620417 | 0.524737794 | 0.773379038 |
| chr6:43884262-43884462 | -0.318811037 | 0.524745502 | 0.773379038 |
| chr17:50345268-50345651 | 0.231291737 | 0.524747498 | 0.773379038 |
| chr12:83690545-83690926 | 0.290919784 | 0.524784342 | 0.773390854 |
| chr11:66718704-66719040 | 0.225190097 | 0.524790854 | 0.773390854 |
| chr17:43388244-43389009 | 0.210554416 | 0.524795646 | 0.773390854 |
| chr2:9323637-9323837 | -0.331762153 | 0.524802097 | 0.773390854 |
| chr13:36981244-36981492 | -0.295439501 | 0.524814273 | 0.773390854 |
| chr11:67374058-67374357 | 0.252439075 | 0.524822478 | 0.773390854 |
| chr11:113796773-113796973 | 0.332725846 | 0.524833117 | 0.773390854 |
| chr17:55472916-55473259 | -0.26360647 | 0.524836264 | 0.773390854 |
| chr22:37519189-37519598 | -0.270481017 | 0.524884978 | 0.773447762 |

| | | | |
|---|---|---|---|
| chr5:119565117-119565332 | 0.340151876 | 0.524926336 | 0.773472859 |
| chr9:127734679-127735609 | 0.208246736 | 0.524946688 | 0.773472859 |
| chr15:57559867-57560146 | 0.266254404 | 0.524958435 | 0.773472859 |
| chr12:80589137-80589383 | 0.297913401 | 0.524972288 | 0.773472859 |
| chr20:59193562-59193762 | -0.281432911 | 0.524979397 | 0.773472859 |
| chr5:57173134-57174644 | -0.199331689 | 0.524980239 | 0.773472859 |
| chr20:23668411-23668898 | 0.241608953 | 0.524983596 | 0.773472859 |
| chr12:95963052-95963443 | -0.253178018 | 0.524985736 | 0.773472859 |
| chr20:44496199-44496433 | 0.318092233 | 0.525007988 | 0.773472859 |
| chr5:49658563-49658807 | 0.296970163 | 0.525011485 | 0.773472859 |
| chr1:183418096-183418585 | 0.235602302 | 0.52501305 | 0.773472859 |
| chr11:66855854-66857093 | 0.202954565 | 0.525032721 | 0.773483597 |
| chr1:183915759-183916420 | -0.219803908 | 0.525047536 | 0.773483597 |
| chr2:170324641-170324900 | -0.282527404 | 0.525050623 | 0.773483597 |
| chr11:506265-507553 | 0.203939044 | 0.525070529 | 0.773495674 |
| chr10:37774086-37774291 | -0.343437406 | 0.525082157 | 0.773495674 |
| chr12:121499943-121500288 | 0.275134576 | 0.525095771 | 0.773495674 |
| chr13:40770685-40771561 | -0.2028746 | 0.525099201 | 0.773495674 |
| chr13:43878941-43880439 | -0.202514032 | 0.525127822 | 0.773522963 |
| chr3:37175490-37176924 | 0.203993707 | 0.525174446 | 0.773576769 |
| chr5:149381601-149382235 | -0.226525977 | 0.525184966 | 0.773577394 |
| chr12:69397607-69397856 | -0.330668587 | 0.525213379 | 0.773604374 |
| chr16:569545-569948 | 0.247205676 | 0.525224849 | 0.773606398 |
| chr4:152505682-152505959 | -0.289229353 | 0.525235206 | 0.773606781 |
| chr10:28393900-28394475 | -0.246617299 | 0.525260605 | 0.773619023 |
| chr3:3200739-3200974 | 0.285141308 | 0.525263711 | 0.773619023 |
| chr20:23351414-23351624 | 0.256635751 | 0.525304014 | 0.773651073 |
| chr14:26382001-26382201 | 0.373527539 | 0.525305666 | 0.773651073 |
| chr2:30446703-30446903 | -0.298671152 | 0.52534116 | 0.773677399 |
| chr11:81497484-81497684 | -0.337909553 | 0.525358752 | 0.773677399 |
| chr4:154348728-154348928 | -0.337909553 | 0.525358752 | 0.773677399 |
| chr16:85289263-85289540 | -0.262561656 | 0.525363931 | 0.773677399 |
| chr2:61793219-61794050 | -0.216061956 | 0.52539188 | 0.773703689 |
| chr1:113072575-113073802 | -0.201928716 | 0.525465816 | 0.773797697 |
| chr14:56686598-56686802 | 0.29270904 | 0.525543083 | 0.773896607 |
| chr4:15882674-15882942 | 0.289563255 | 0.525594391 | 0.773954068 |
| chr12:14111249-14111791 | -0.253558592 | 0.525602306 | 0.773954068 |
| chr3:134211943-134212159 | -0.293515082 | 0.525624711 | 0.773963566 |
| chr21:34073124-34073758 | 0.225274964 | 0.525628959 | 0.773963566 |
| chr5:180208881-180209263 | -0.2610695 | 0.525654238 | 0.773985915 |
| chr19:47112657-47113518 | 0.20598613 | 0.525699996 | 0.774038415 |
| chr2:26562336-26562894 | 0.230749671 | 0.525720117 | 0.774044128 |
| chr11:69615116-69615350 | 0.332793164 | 0.525736324 | 0.774044128 |
| chr2:215920183-215920685 | 0.2365993 | 0.525738603 | 0.774044128 |
| chr14:73886368-73887377 | 0.203889379 | 0.52576108 | 0.774044128 |
| chr16:71607604-71608082 | 0.233001129 | 0.52576288 | 0.774044128 |
| chr2:95274222-95274954 | -0.20887451 | 0.525766058 | 0.774044128 |
| chr15:72722149-72722482 | 0.270840486 | 0.52577459 | 0.774044128 |
| chr8:31185768-31186025 | 0.271675658 | 0.525789016 | 0.774050492 |
| chr19:48965077-48965472 | 0.222756077 | 0.525806834 | 0.774059527 |

| | | | |
|---|---|---|---|
| chr20:50179276-50179569 | -0.241777124 | 0.525815357 | 0.774059527 |
| chr8:133695770-133696062 | 0.26753085 | 0.525844144 | 0.77407614 |
| chr10:35641540-35642730 | -0.206365062 | 0.525846848 | 0.77407614 |
| chr1:234612573-234613002 | 0.219039652 | 0.525908187 | 0.774149597 |
| chr3:169083358-169083591 | -0.311443881 | 0.525916956 | 0.774149597 |
| chr8:133382066-133382340 | 0.26393719 | 0.525930714 | 0.774154978 |
| chr16:58652784-58653040 | -0.33052392 | 0.525946149 | 0.774162678 |
| chr6:35467997-35468843 | 0.208713141 | 0.525966847 | 0.774162678 |
| chr5:80184764-80185022 | -0.293639773 | 0.525972322 | 0.774162678 |
| chr9:16133790-16134178 | 0.243988224 | 0.525981643 | 0.774162678 |
| chr2:168912968-168913217 | -0.36233977 | 0.525986464 | 0.774162678 |
| chr2:166628480-166628970 | 0.232163386 | 0.526033578 | 0.77421715 |
| chr9:108710520-108710804 | -0.305563417 | 0.526083574 | 0.774262787 |
| chr11:119101547-119102748 | 0.200953334 | 0.526084796 | 0.774262787 |
| chr6:43874301-43874611 | 0.255257478 | 0.526120785 | 0.77429422 |
| chr5:132134432-132134703 | -0.297035962 | 0.526132805 | 0.77429422 |
| chr10:101095394-101095931 | -0.210986957 | 0.526136469 | 0.77429422 |
| chr5:43609808-43610124 | -0.260752915 | 0.526172597 | 0.774308412 |
| chr10:102729293-102729610 | 0.255619317 | 0.526176774 | 0.774308412 |
| chr20:51595021-51595372 | 0.221251383 | 0.526178375 | 0.774308412 |
| chr10:93072276-93073042 | 0.215866251 | 0.526186535 | 0.774308412 |
| chr12:63319433-63319893 | -0.241606094 | 0.526226431 | 0.774352249 |
| chr8:106858221-106858550 | -0.237694827 | 0.526257274 | 0.774382763 |
| chr8:99651912-99652164 | -0.3478475 | 0.526283488 | 0.774406465 |
| chr11:14418889-14419141 | 0.289809076 | 0.526307491 | 0.774424588 |
| chr1:1013259-1014107 | 0.222577082 | 0.526316019 | 0.774424588 |
| chr14:106879061-106879825 | 0.224665268 | 0.526365297 | 0.77448056 |
| chr11:33802190-33803081 | -0.221679481 | 0.526374274 | 0.77448056 |
| chr2:20012249-20012514 | -0.267371352 | 0.526416361 | 0.774527611 |
| chr20:5949828-5950028 | -0.327386958 | 0.526446041 | 0.774556407 |
| chr8:127400667-127401001 | -0.237400918 | 0.526460115 | 0.774562241 |
| chr2:75528240-75528560 | 0.246733422 | 0.526487408 | 0.774578573 |
| chr20:50376870-50377221 | -0.234690896 | 0.526492156 | 0.774578573 |
| chr6:32169246-32169515 | -0.290430123 | 0.526501543 | 0.774578573 |
| chr2:86314474-86314713 | 0.316515276 | 0.526547361 | 0.774631106 |
| chr5:128201060-128201708 | -0.213240012 | 0.526596266 | 0.774688179 |
| chr15:98085120-98085336 | -0.348519972 | 0.526612912 | 0.774697792 |
| chr5:58259706-58260223 | -0.246571402 | 0.526663558 | 0.77473971 |
| chr2:65576893-65577196 | -0.261121647 | 0.526680904 | 0.77473971 |
| chr10:84328211-84329244 | 0.200912815 | 0.526685371 | 0.77473971 |
| chr8:52495860-52496421 | -0.235096223 | 0.526687594 | 0.77473971 |
| chr18:58121989-58122220 | 0.300772185 | 0.526691962 | 0.77473971 |
| chr10:9188055-9189086 | 0.202159171 | 0.526736228 | 0.77478995 |
| chr6:3363342-3363728 | -0.233554946 | 0.526755783 | 0.774799376 |
| chrX:11100196-11100481 | 0.263909427 | 0.52676286 | 0.774799376 |
| chr8:106845740-106846332 | -0.220174082 | 0.526838578 | 0.774887905 |
| chr16:87854418-87854939 | -0.218759653 | 0.526843275 | 0.774887905 |
| chr9:108854106-108854423 | -0.262499315 | 0.526883042 | 0.774931519 |
| chr21:39715895-39716124 | -0.292899044 | 0.527024691 | 0.775124976 |
| chr1:188769087-188770468 | 0.209018577 | 0.527040604 | 0.775133502 |

| | | | |
|---|---|---|---|
| chr3:195732883-195733244 | 0.301934323 | 0.527072187 | 0.775165073 |
| chr14:81236877-81237157 | -0.302991141 | 0.527095037 | 0.775183798 |
| chr1:151716749-151717604 | 0.210198704 | 0.527125261 | 0.775193631 |
| chr6:136791775-136792020 | -0.26356301 | 0.527142534 | 0.775193631 |
| chr15:67079499-67079771 | 0.268097373 | 0.527150879 | 0.775193631 |
| chr17:45889435-45889811 | 0.269271202 | 0.527156066 | 0.775193631 |
| chr6:56995400-56995690 | -0.274612108 | 0.527179612 | 0.775193631 |
| chr13:32246421-32246820 | -0.224070607 | 0.527184379 | 0.775193631 |
| chr12:70756212-70756526 | -0.273652896 | 0.527185224 | 0.775193631 |
| chr20:43666652-43668116 | 0.203156398 | 0.527187108 | 0.775193631 |
| chr10:44374054-44374445 | -0.283234488 | 0.527192776 | 0.775193631 |
| chr3:14673615-14674097 | -0.238932071 | 0.527241945 | 0.775251052 |
| chr19:10835450-10836713 | 0.200116975 | 0.527281077 | 0.775280567 |
| chr22:33210214-33210791 | -0.248874819 | 0.527286973 | 0.775280567 |
| chr8:98835073-98835273 | 0.309356803 | 0.527292533 | 0.775280567 |
| chr9:89747802-89748431 | 0.225922759 | 0.527302491 | 0.775280567 |
| chr6:34462467-34462780 | -0.245215784 | 0.527330172 | 0.775306389 |
| chr9:89021308-89021589 | 0.281119785 | 0.527356067 | 0.775322884 |
| chrX:107741840-107742040 | 0.311286942 | 0.527374352 | 0.775322884 |
| chr9:127417009-127417824 | -0.216433001 | 0.527383471 | 0.775322884 |
| chr12:69879027-69879695 | -0.216646687 | 0.527395129 | 0.775322884 |
| chr8:89886213-89886617 | -0.22993859 | 0.527400478 | 0.775322884 |
| chr12:6608046-6608671 | -0.214672452 | 0.527407327 | 0.775322884 |
| chr16:68272665-68272880 | 0.343204659 | 0.527412223 | 0.775322884 |
| chr16:67367130-67367374 | -0.326703707 | 0.527434798 | 0.775341195 |
| chrX:15816184-15816393 | -0.332378554 | 0.527499356 | 0.77542122 |
| chr5:84255391-84255627 | -0.297038984 | 0.527522565 | 0.77543596 |
| chr12:45604382-45604853 | -0.270403851 | 0.527529624 | 0.77543596 |
| chr3:134199989-134200209 | -0.329269899 | 0.527585716 | 0.775503534 |
| chrX:42119110-42119501 | -0.24487163 | 0.527604207 | 0.7755046 |
| chr7:9194405-9194658 | -0.339366081 | 0.527611746 | 0.7755046 |
| chr20:32109312-32110259 | 0.206008919 | 0.527616804 | 0.7755046 |
| chr4:51842601-51843584 | 0.200499017 | 0.527657188 | 0.775537176 |
| chr12:68406344-68406593 | 0.268299659 | 0.527659211 | 0.775537176 |
| chr11:57345588-57345994 | -0.226849076 | 0.527677978 | 0.775549883 |
| chr17:32007002-32007763 | 0.226238121 | 0.527700819 | 0.775568013 |
| chr17:7455402-7455890 | -0.23317172 | 0.527710558 | 0.775568013 |
| chr10:96720420-96720653 | 0.282632213 | 0.527727632 | 0.775578231 |
| chr19:50090782-50091057 | 0.28698709 | 0.527750077 | 0.77559634 |
| chr10:73940421-73941117 | 0.225112635 | 0.527784443 | 0.775631968 |
| chr3:165854612-165855095 | -0.253008146 | 0.527797924 | 0.77563545 |
| chr13:51896325-51896652 | -0.257304189 | 0.527814883 | 0.77563545 |
| chr16:2474926-2475215 | 0.228564547 | 0.52781718 | 0.77563545 |
| chr7:94580647-94580847 | -0.379115281 | 0.52784915 | 0.775643405 |
| chr1:24283913-24284113 | 0.358973513 | 0.527852963 | 0.775643405 |
| chr12:101062370-101062570 | 0.358973513 | 0.527852963 | 0.775643405 |
| chr1:203340104-203340423 | -0.224695814 | 0.527863311 | 0.775643736 |
| chr5:6759469-6759768 | 0.250482643 | 0.52791973 | 0.775711762 |
| chr11:20648015-20648319 | 0.265616369 | 0.527969183 | 0.775754129 |
| chr11:134050524-134050808 | 0.258766031 | 0.527981236 | 0.775754129 |

| | | | |
|---|---|---|---|
| chr6:83067424-83068303 | -0.204126309 | 0.527983945 | 0.775754129 |
| chr17:42535883-42536790 | 0.210824104 | 0.528001033 | 0.775754129 |
| chr10:29141901-29142101 | 0.32838885 | 0.528008178 | 0.775754129 |
| chr8:134528080-134528280 | 0.32371333 | 0.52800931 | 0.775754129 |
| chr2:86337430-86337630 | 0.284577416 | 0.528027464 | 0.775765927 |
| chr10:29630141-29630958 | -0.220349997 | 0.528073189 | 0.775804668 |
| chr6:122800368-122800609 | 0.323571107 | 0.528074084 | 0.775804668 |
| chr1:176499597-176499854 | -0.321089074 | 0.528121179 | 0.775858981 |
| chr20:55655772-55656211 | -0.257458899 | 0.52816847 | 0.775900943 |
| chr16:75516561-75517737 | -0.199389667 | 0.528169995 | 0.775900943 |
| chr1:39980425-39980986 | -0.227069106 | 0.528184875 | 0.775907925 |
| chr3:154678267-154678467 | -0.336028929 | 0.528204572 | 0.775921984 |
| chr10:34815694-34815896 | -0.274475441 | 0.528244818 | 0.775966228 |
| chr20:34676418-34677448 | 0.205754887 | 0.528290647 | 0.776012756 |
| chr3:95329692-95329902 | -0.322576676 | 0.528306496 | 0.776012756 |
| chr3:49093574-49093837 | 0.25346151 | 0.528306875 | 0.776012756 |
| chr1:51535073-51535355 | 0.287405757 | 0.528339654 | 0.776037106 |
| chr3:101026477-101026764 | -0.309684463 | 0.528346919 | 0.776037106 |
| chr8:127629847-127630047 | 0.29428813 | 0.528359586 | 0.776037106 |
| chr3:155978408-155978623 | -0.316888734 | 0.528374331 | 0.776037106 |
| chr7:54774147-54774520 | -0.223292925 | 0.528378778 | 0.776037106 |
| chr12:12803203-12803587 | -0.245643465 | 0.528397864 | 0.776037106 |
| chr3:14651307-14652269 | 0.201129969 | 0.528399348 | 0.776037106 |
| chr6:20212016-20212992 | -0.204398438 | 0.528404478 | 0.776037106 |
| chr2:160406938-160407425 | -0.232133945 | 0.528426303 | 0.776054285 |
| chr10:95693569-95694415 | -0.201778712 | 0.528478366 | 0.776102458 |
| chr11:106021727-10602275 | -0.201608374 | 0.528479363 | 0.776102458 |
| chr1:16624953-16625224 | 0.24080999 | 0.528523988 | 0.776153117 |
| chr16:86563544-86564155 | 0.222242312 | 0.528548564 | 0.776174332 |
| chr3:4491948-4493703 | 0.196966879 | 0.528601128 | 0.776235654 |
| chr1:70487709-70488239 | 0.227684437 | 0.528610584 | 0.776235654 |
| chr17:75904575-75905375 | 0.204230185 | 0.528621652 | 0.77623703 |
| chr7:6664649-6664849 | -0.31435866 | 0.528668069 | 0.776289563 |
| chr1:152047599-152048769 | -0.197200115 | 0.52867769 | 0.776289563 |
| chr3:94587126-94587753 | -0.21765235 | 0.528708121 | 0.77631937 |
| chr14:98887015-98887349 | -0.253351833 | 0.528725872 | 0.776330557 |
| chr1:115112823-115113163 | 0.239668798 | 0.528753468 | 0.776349854 |
| chr10:12068502-12069433 | -0.201502219 | 0.528766385 | 0.776349854 |
| chr1:112830669-112830869 | 0.340502886 | 0.528789018 | 0.776349854 |
| chr10:29130152-29130531 | -0.23929936 | 0.528792188 | 0.776349854 |
| chr8:67749394-67749690 | -0.267446161 | 0.528798829 | 0.776349854 |
| chr2:15939653-15939945 | -0.266565676 | 0.528799808 | 0.776349854 |
| chr3:52153471-52155066 | 0.203028456 | 0.528837528 | 0.776360417 |
| chr8:102906236-102906597 | -0.228508626 | 0.528841283 | 0.776360417 |
| chr15:31296726-31297044 | 0.259852472 | 0.528841897 | 0.776360417 |
| chr1:145977541-145977801 | 0.227270042 | 0.528859086 | 0.776360417 |
| chr10:82377514-82378040 | 0.246983975 | 0.528866003 | 0.776360417 |
| chr11:64250993-64251610 | 0.220163216 | 0.528867797 | 0.776360417 |
| chr19:58504486-58505076 | 0.227408855 | 0.528900017 | 0.77639284 |
| chr2:207829074-207829600 | -0.244750311 | 0.5289681 | 0.776465519 |

| | | | |
|---|---|---|---|
| chr3:58009998-58010459 | -0.245469801 | 0.528969795 | 0.776465519 |
| chr15:90249225-90249856 | 0.216869484 | 0.529006284 | 0.77649846 |
| chr16:84116307-84117417 | 0.201442002 | 0.529012504 | 0.77649846 |
| chr4:173288394-173288640 | 0.300305263 | 0.529038433 | 0.776521643 |
| chr20:63627020-63627577 | -0.211792441 | 0.529079776 | 0.77656745 |
| chr1:53063668-53064020 | 0.2764926 | 0.529132064 | 0.77662932 |
| chr10:102352149-102352358 | -0.329795586 | 0.529163803 | 0.77665334 |
| chr8:138945193-138945414 | 0.30950515 | 0.529168702 | 0.77665334 |
| chr11:65726797-65726997 | -0.332734615 | 0.529259901 | 0.776738664 |
| chr7:936630-936897 | -0.273822748 | 0.529281196 | 0.776738664 |
| chr1:147495008-147495334 | 0.228928118 | 0.529287706 | 0.776738664 |
| chr6:154868154-154868420 | -0.277419637 | 0.529289352 | 0.776738664 |
| chr3:100368630-100368865 | -0.323727152 | 0.529290548 | 0.776738664 |
| chr22:46296215-46297032 | 0.213505358 | 0.529292446 | 0.776738664 |
| chr7:36201403-36201678 | -0.280656312 | 0.529306858 | 0.776738664 |
| chr9:123107940-123108308 | -0.273780668 | 0.529307935 | 0.776738664 |
| chr8:273811-274165 | 0.282510268 | 0.529340393 | 0.776769614 |
| chr17:49014319-49014779 | 0.232429234 | 0.529349301 | 0.776769614 |
| chr17:26881171-26881598 | 0.230846607 | 0.529372718 | 0.776789099 |
| chr6:17622223-17622507 | -0.352183925 | 0.529392126 | 0.776802702 |
| chr11:125062963-125063361 | 0.24255663 | 0.529412529 | 0.776816885 |
| chr20:50874237-50874492 | 0.263519365 | 0.529428223 | 0.776816885 |
| chr14:104085182-104085631 | 0.234076086 | 0.529433684 | 0.776816885 |
| chr20:58839463-58839791 | 0.231472311 | 0.529450083 | 0.776816885 |
| chr19:45770264-45771028 | 0.202970814 | 0.529452483 | 0.776816885 |
| chr12:94532832-94533138 | -0.244690558 | 0.5294793 | 0.776824959 |
| chr21:29372440-29372648 | 0.289973165 | 0.529497536 | 0.776824959 |
| chr4:155403946-155404146 | -0.330008242 | 0.529501853 | 0.776824959 |
| chr8:30227720-30227920 | -0.356407574 | 0.529506438 | 0.776824959 |
| chr11:59536979-59538140 | -0.221967898 | 0.529508679 | 0.776824959 |
| chr8:47044191-47044472 | 0.284353573 | 0.529521798 | 0.776829332 |
| chr20:19964477-19964832 | 0.272804178 | 0.529536599 | 0.776836172 |
| chr17:41918783-41919839 | 0.203367298 | 0.529555155 | 0.77684852 |
| chrX:55908654-55908945 | 0.273605388 | 0.529568734 | 0.776853567 |
| chr14:41448616-41448816 | -0.330727082 | 0.529593955 | 0.776864412 |
| chr8:95849096-95849414 | 0.2389215 | 0.529600883 | 0.776864412 |
| chr8:94553136-94553930 | 0.200504202 | 0.529606543 | 0.776864412 |
| chr1:93079727-93079927 | -0.308677733 | 0.529636336 | 0.776893241 |
| chr14:49879688-49880134 | -0.22301599 | 0.529656463 | 0.776907891 |
| chr18:12645284-12646397 | 0.21572083 | 0.529693296 | 0.776947045 |
| chr1:108316434-108316811 | 0.254795748 | 0.529721137 | 0.776973009 |
| chr1:211249738-211249971 | -0.324683433 | 0.529737901 | 0.776976944 |
| chr1:15849365-15849696 | 0.257174576 | 0.529745017 | 0.776976944 |
| chr1:161013480-161014123 | 0.212085501 | 0.529754242 | 0.776976944 |
| chr3:38842254-38842546 | -0.308691556 | 0.529765871 | 0.776979128 |
| chr10:23417278-23417478 | -0.334729636 | 0.529784807 | 0.776992027 |
| chr10:132162989-132164028 | 0.200052786 | 0.529848005 | 0.777058185 |
| chr17:3695685-3695943 | -0.29193263 | 0.529853046 | 0.777058185 |
| chr1:195269203-195269403 | -0.322935776 | 0.52986034 | 0.777058185 |
| chr17:65278183-65278895 | -0.214135528 | 0.529904543 | 0.777108136 |

| | | | |
|---|---|---|---|
| chr20:35619314-35619733 | -0.220404482 | 0.529972882 | 0.777175441 |
| chr10:119667604-119668059 | 0.229950905 | 0.529977365 | 0.777175441 |
| chr2:223989989-223990214 | 0.314429218 | 0.529980867 | 0.777175441 |
| chr1:230066614-230068160 | 0.201632715 | 0.529993428 | 0.777178988 |
| chr2:216808916-216809165 | -0.320565414 | 0.530021912 | 0.777205881 |
| chr3:22685624-22685946 | 0.23944041 | 0.530053014 | 0.777220934 |
| chr15:73704831-73705122 | 0.242133778 | 0.530056025 | 0.777220934 |
| chr2:27298654-27299186 | 0.236253573 | 0.530062607 | 0.777220934 |
| chr5:52424570-52424779 | -0.308951221 | 0.530090928 | 0.777247586 |
| chr7:151100847-151101119 | -0.307166607 | 0.530106223 | 0.777255138 |
| chr20:33638803-33639130 | -0.303942593 | 0.530163342 | 0.77730556 |
| chr17:7356058-7356318 | -0.263690083 | 0.53016428 | 0.77730556 |
| chr5:104011885-104012085 | -0.339580604 | 0.530177628 | 0.77730556 |
| chr16:88953323-88953675 | 0.270254682 | 0.530181191 | 0.77730556 |
| chr6:34448070-34448283 | -0.286238949 | 0.530241235 | 0.777372847 |
| chr8:17912406-17912862 | -0.245375306 | 0.530251462 | 0.777372847 |
| chr20:56716876-56717207 | 0.225063351 | 0.530263573 | 0.777372847 |
| chr1:200621313-200621646 | -0.222889182 | 0.530267667 | 0.777372847 |
| chr9:127931198-127931398 | 0.256862512 | 0.530310321 | 0.777409602 |
| chr1:87328161-87328979 | -0.206963529 | 0.530317984 | 0.777409602 |
| chr5:95743342-95743542 | -0.33191945 | 0.530328555 | 0.777409602 |
| chr16:4870164-4870430 | 0.308959998 | 0.53034804 | 0.777409602 |
| chr16:50468392-50468597 | 0.308959998 | 0.53034804 | 0.777409602 |
| chr3:84958501-84959434 | -0.200859881 | 0.530362468 | 0.777409602 |
| chr1:28296766-28297142 | 0.245086382 | 0.530363761 | 0.777409602 |
| chr12:106987877-106988178 | -0.2457807 | 0.530381691 | 0.777421011 |
| chr5:169511818-169512242 | -0.260287315 | 0.530453459 | 0.777511332 |
| chr12:110102311-110102635 | -0.277296151 | 0.530485276 | 0.777525985 |
| chr1:35641211-35642199 | 0.206662175 | 0.530505362 | 0.777525985 |
| chr18:34322810-34323010 | -0.339043176 | 0.530505739 | 0.777525985 |
| chr10:73018706-73018906 | 0.317740424 | 0.530513062 | 0.777525985 |
| chr19:49452981-49453822 | 0.203445038 | 0.530514193 | 0.777525985 |
| chr5:151448177-151448476 | -0.251300921 | 0.530556309 | 0.77757131 |
| chr11:103609501-103609725 | -0.307983277 | 0.530568241 | 0.77757131 |
| chr13:29594317-29596161 | -0.197416392 | 0.530584415 | 0.77757131 |
| chr20:56358628-56359755 | 0.197633815 | 0.530590364 | 0.77757131 |
| chr20:35208468-35208684 | -0.30791735 | 0.53059586 | 0.77757131 |
| chr10:18610565-18610919 | 0.259898511 | 0.530613902 | 0.777582492 |
| chr3:136179280-136179511 | -0.301719356 | 0.530626857 | 0.777582492 |
| chr2:110735436-110735718 | -0.291016338 | 0.530633935 | 0.777582492 |
| chr3:69012935-69014150 | 0.200490503 | 0.53066887 | 0.777618814 |
| chr2:113137479-113137727 | -0.30409638 | 0.530744184 | 0.777703087 |
| chr1:205229451-205229823 | -0.220623018 | 0.530756532 | 0.777703087 |
| chr22:37608301-37609117 | -0.206557269 | 0.53076705 | 0.777703087 |
| chr7:47751044-47751350 | 0.287867322 | 0.530769237 | 0.777703087 |
| chr12:94764278-94764538 | -0.264870301 | 0.530778359 | 0.777703087 |
| chr1:64965859-64967035 | -0.202421804 | 0.530787279 | 0.777703087 |
| chr4:146582974-146583194 | 0.305280948 | 0.530806967 | 0.777717061 |
| chr6:27518879-27519832 | -0.212949023 | 0.530830153 | 0.777722126 |
| chr11:96464779-96465057 | -0.276457098 | 0.530830724 | 0.777722126 |

| | | | |
|---|---|---|---|
| chr1:244918663-244919561 | 0.220135879 | 0.530848638 | 0.777731292 |
| chr8:33334552-33334970 | -0.274450669 | 0.530857281 | 0.777731292 |
| chr17:45395116-45395401 | 0.27356964 | 0.530909677 | 0.777793183 |
| chr5:59923125-59923458 | -0.294181342 | 0.530937959 | 0.777809734 |
| chr1:16895957-16897385 | 0.204693035 | 0.53095136 | 0.777809734 |
| chr21:40660934-40661169 | -0.316428946 | 0.530968271 | 0.777809734 |
| chr6:88238172-88238429 | 0.276850519 | 0.530970922 | 0.777809734 |
| chr12:57865024-57865778 | -0.214745697 | 0.530981521 | 0.777809734 |
| chr10:102399314-102399820 | 0.219985616 | 0.530981882 | 0.777809734 |
| chr15:38251204-38252838 | -0.199128393 | 0.5310032 | 0.777826091 |
| chr1:203410518-203411044 | -0.228792769 | 0.531035025 | 0.777848307 |
| chr3:38805828-38806030 | 0.313478407 | 0.531052381 | 0.777848307 |
| chr12:115998163-115998544 | -0.279482634 | 0.53106855 | 0.777848307 |
| chr19:29881203-29881403 | 0.333521282 | 0.531070677 | 0.777848307 |
| chr15:84600288-84601290 | 0.201152224 | 0.531125215 | 0.777848307 |
| chr20:1392855-1393520 | 0.216338725 | 0.531127823 | 0.777848307 |
| chr16:31095692-31096062 | 0.244327575 | 0.531130672 | 0.777848307 |
| chr6:96521364-96522688 | -0.199731143 | 0.531135934 | 0.777848307 |
| chr8:33590872-33591268 | -0.242780445 | 0.531146293 | 0.777848307 |
| chr6:90701455-90701724 | -0.27833378 | 0.531154302 | 0.777848307 |
| chr16:19375686-19375908 | -0.33681072 | 0.531157707 | 0.777848307 |
| chr9:125189260-125190288 | 0.205379763 | 0.531163968 | 0.777848307 |
| chr10:38402330-38403429 | -0.207124022 | 0.531169161 | 0.777848307 |
| chr17:82772639-82772850 | 0.28989825 | 0.531171804 | 0.777848307 |
| chr7:6200497-6200727 | -0.324575303 | 0.531180069 | 0.777848307 |
| chr1:205121620-205122606 | -0.198608695 | 0.531188198 | 0.777848307 |
| chr3:114362847-114363090 | 0.27491524 | 0.531190946 | 0.777848307 |
| chr8:6648858-6649272 | -0.263586126 | 0.531210206 | 0.777861644 |
| chr7:3109348-3109905 | -0.227434767 | 0.531235888 | 0.777884384 |
| chr18:23586073-23587169 | 0.201283304 | 0.531303112 | 0.777965564 |
| chr1:112619553-112620152 | -0.211579383 | 0.531315668 | 0.777965564 |
| chr3:150422144-150422428 | -0.261670603 | 0.531321787 | 0.777965564 |
| chr12:54368654-54369364 | -0.213039151 | 0.531359224 | 0.778005513 |
| chr3:148076198-148076473 | 0.258155 | 0.531417879 | 0.778076526 |
| chr6:135497307-135498137 | -0.201356908 | 0.531450877 | 0.778108633 |
| chr5:173302136-173302774 | -0.212158706 | 0.531460118 | 0.778108633 |
| chr2:171433739-171434922 | -0.198089008 | 0.531481941 | 0.778111759 |
| chr6:74343653-74344056 | -0.293582646 | 0.531484986 | 0.778111759 |
| chr16:85253166-85253492 | -0.235796427 | 0.531502417 | 0.778111759 |
| chr12:20798661-20798861 | -0.338612972 | 0.531502874 | 0.778111759 |
| chr8:131262042-131262838 | -0.222015035 | 0.531521164 | 0.778119555 |
| chr20:62791501-62791720 | -0.235775082 | 0.531545797 | 0.778119555 |
| chr1:117093899-117094200 | 0.276387998 | 0.531559097 | 0.778119555 |
| chr18:7170540-7170743 | -0.332449158 | 0.531563951 | 0.778119555 |
| chr2:100883360-100883564 | -0.309896373 | 0.531572085 | 0.778119555 |
| chr11:124763176-124763393 | 0.295815854 | 0.531577715 | 0.778119555 |
| chr17:29389879-29390722 | -0.208109684 | 0.531609363 | 0.778119555 |
| chr3:150668166-150668371 | -0.314476714 | 0.531617167 | 0.778119555 |
| chr14:80927678-80927891 | -0.307201017 | 0.531618995 | 0.778119555 |
| chr7:150974425-150975351 | -0.206942828 | 0.531641191 | 0.778119555 |

| | | | |
|---|---|---|---|
| chr3:15305304-15305630 | -0.235313112 | 0.531644565 | 0.778119555 |
| chr1:201946911-201947136 | -0.313698805 | 0.531648817 | 0.778119555 |
| chr1:174258113-174258592 | -0.231415361 | 0.531650081 | 0.778119555 |
| chr8:49797838-49798174 | -0.261879357 | 0.53165696 | 0.778119555 |
| chr16:29469735-29470006 | -0.308618266 | 0.531660529 | 0.778119555 |
| chr22:50200766-50201116 | 0.250560198 | 0.531678215 | 0.778130577 |
| chr3:61917157-61917496 | 0.231460302 | 0.531690665 | 0.778133934 |
| chr19:10388928-10389311 | -0.258823811 | 0.531702856 | 0.778136913 |
| chr14:67819392-67819951 | -0.213986127 | 0.531737083 | 0.77817214 |
| chr15:77617524-77617894 | 0.236459205 | 0.53175196 | 0.778178056 |
| chr20:36217337-36217585 | -0.279729857 | 0.531761437 | 0.778178056 |
| chr6:154408897-154409147 | 0.265022364 | 0.531786063 | 0.778199231 |
| chr1:8878544-8879445 | 0.206159767 | 0.53186198 | 0.778275995 |
| chr3:38024334-38025434 | 0.196603599 | 0.531867538 | 0.778275995 |
| chr2:102064277-102064578 | -0.261822616 | 0.531890456 | 0.778275995 |
| chr19:28989207-28989407 | -0.332989012 | 0.531897397 | 0.778275995 |
| chr20:14878779-14878979 | -0.332989012 | 0.531897397 | 0.778275995 |
| chr7:34596955-34597234 | -0.250335518 | 0.531914424 | 0.778275995 |
| chr2:36646281-36646564 | -0.264078413 | 0.531924228 | 0.778275995 |
| chr15:73915026-73915354 | 0.252915493 | 0.531936922 | 0.778275995 |
| chr11:47963901-47964101 | -0.306237376 | 0.531947795 | 0.778275995 |
| chr8:103207389-103207614 | 0.264847584 | 0.531954355 | 0.778275995 |
| chr1:180230979-180231920 | 0.203626043 | 0.531977239 | 0.778275995 |
| chr20:15709060-15709340 | 0.274277239 | 0.531989565 | 0.778275995 |
| chr3:137264439-137264694 | 0.314472146 | 0.531993879 | 0.778275995 |
| chr16:714863-715163 | 0.240164998 | 0.531998301 | 0.778275995 |
| chr6:89372052-89372395 | 0.239624234 | 0.53199987 | 0.778275995 |
| chr16:56451146-56451975 | 0.206405384 | 0.532005023 | 0.778275995 |
| chr10:32253301-32254154 | 0.212217873 | 0.532021838 | 0.778275995 |
| chr7:133275038-133275305 | -0.320902499 | 0.532029093 | 0.778275995 |
| chr13:39526056-39526337 | -0.321476404 | 0.53203151 | 0.778275995 |
| chr17:42017282-42018087 | 0.208810221 | 0.532051417 | 0.778279367 |
| chr14:69532697-69532952 | 0.275055878 | 0.53205413 | 0.778279367 |
| chr1:16895550-16895755 | -0.30354432 | 0.532091309 | 0.77831419 |
| chrX:54357997-54358293 | -0.274655931 | 0.532098252 | 0.77831419 |
| chr15:59174846-59175239 | -0.232318288 | 0.532132533 | 0.77831795 |
| chr10:80664101-80664440 | 0.228596797 | 0.532140773 | 0.77831795 |
| chr10:71811624-71812194 | 0.231157495 | 0.532147006 | 0.77831795 |
| chr4:81844804-81845010 | 0.30252531 | 0.532149656 | 0.77831795 |
| chr9:122060023-122060250 | 0.294072409 | 0.532152261 | 0.77831795 |
| chr6:118080697-118080897 | -0.356849111 | 0.532161769 | 0.77831795 |
| chr1:246767848-246768082 | 0.271763992 | 0.5321847 | 0.778332256 |
| chr1:200217818-200218018 | 0.284750982 | 0.53219936 | 0.778332256 |
| chr19:12946120-12946390 | -0.256598881 | 0.532202025 | 0.778332256 |
| chr8:41577534-41578756 | -0.204461986 | 0.532235806 | 0.778366804 |
| chr1:160193114-160193455 | 0.229341914 | 0.532261676 | 0.778389781 |
| chr1:231077614-231077814 | 0.349037788 | 0.53227351 | 0.77839223 |
| chr16:70454278-70455072 | 0.218596035 | 0.532296814 | 0.778411453 |
| chr3:67655389-67655616 | -0.285505006 | 0.532322308 | 0.778422957 |
| chr16:85172533-85172857 | -0.240410749 | 0.5323395 | 0.778422957 |

| | | | |
|---|---|---|---|
| chr3:154220290-154220511 | -0.323085194 | 0.532343695 | 0.778422957 |
| chr2:176187689-176187894 | 0.286743968 | 0.532345318 | 0.778422957 |
| chrX:130975991-130976220 | 0.280393091 | 0.532360128 | 0.778429758 |
| chr7:45542283-45542483 | -0.339984837 | 0.532418646 | 0.778500467 |
| chr1:33654346-33654546 | 0.308901361 | 0.532444651 | 0.778515634 |
| chr20:41135251-41135658 | 0.212004585 | 0.532454964 | 0.778515634 |
| chr2:67397081-67398113 | -0.203680228 | 0.5324595 | 0.778515634 |
| chr8:64266917-64267211 | -0.271982051 | 0.532485143 | 0.778538271 |
| chr2:68643410-68643964 | -0.2281101 | 0.532509062 | 0.778557984 |
| chr12:108731261-108732413 | -0.196440942 | 0.532518948 | 0.778557984 |
| chr8:72957218-72957418 | -0.34589466 | 0.532581126 | 0.778608366 |
| chr2:15334707-15335066 | -0.247173323 | 0.53258115 | 0.778608366 |
| chr10:43204876-43205528 | -0.211247048 | 0.532583893 | 0.778608366 |
| chr18:75183501-75183892 | -0.25860999 | 0.532646626 | 0.77867906 |
| chr16:85799039-85800455 | 0.200191433 | 0.532661318 | 0.77867906 |
| chr3:31754605-31754819 | -0.303327927 | 0.532674867 | 0.77867906 |
| chr1:201713440-201713701 | 0.245687443 | 0.532680742 | 0.77867906 |
| chrX:54529696-54530821 | -0.20504189 | 0.532685917 | 0.77867906 |
| chr3:147532311-147532620 | -0.269334362 | 0.532694125 | 0.77867906 |
| chr15:40530693-40531033 | -0.291030756 | 0.532719124 | 0.77867906 |
| chr5:139638312-139638624 | -0.229288948 | 0.532721037 | 0.77867906 |
| chr5:179907674-179908202 | -0.219740787 | 0.532728619 | 0.77867906 |
| chr17:69348611-69348832 | -0.303131297 | 0.532733876 | 0.77867906 |
| chr15:66700796-66701686 | 0.201336798 | 0.532750647 | 0.77868872 |
| chr11:43358687-43359861 | -0.202061226 | 0.532772966 | 0.778696918 |
| chr10:22894799-22895097 | -0.274720827 | 0.532780485 | 0.778696918 |
| chr1:32179298-32180213 | 0.204699116 | 0.532786744 | 0.778696918 |
| chr11:45815792-45816143 | 0.247033869 | 0.532825973 | 0.778736872 |
| chr14:24231695-24233234 | 0.198207867 | 0.532834408 | 0.778736872 |
| chr7:36091661-36091861 | -0.343053564 | 0.532846752 | 0.77874006 |
| chr1:106473384-106473600 | 0.235751561 | 0.532913099 | 0.778801501 |
| chr2:222318014-222318710 | 0.221100735 | 0.532927086 | 0.778801501 |
| chr11:78155941-78156830 | 0.212556053 | 0.532929324 | 0.778801501 |
| chr10:69375253-69376120 | -0.213943524 | 0.532935903 | 0.778801501 |
| chr11:79512321-79512521 | -0.32046638 | 0.532944202 | 0.778801501 |
| chr5:178305959-178306159 | 0.278326264 | 0.532957131 | 0.778801501 |
| chr4:151035302-151035991 | -0.207770216 | 0.532959942 | 0.778801501 |
| chr6:26122876-26123482 | -0.205845742 | 0.532988216 | 0.778827963 |
| chr1:28369426-28370606 | 0.203958209 | 0.533010352 | 0.778838682 |
| chr16:75054144-75054544 | 0.233468269 | 0.533015881 | 0.778838682 |
| chr5:10366724-10366936 | 0.297619653 | 0.533044956 | 0.778845298 |
| chr1:2256576-2256776 | -0.342995148 | 0.533049239 | 0.778845298 |
| chr13:33634768-33634968 | 0.308448759 | 0.533050903 | 0.778845298 |
| chr1:205273473-205273889 | -0.211027818 | 0.533061704 | 0.778846228 |
| chr7:44348441-44348689 | -0.293261366 | 0.533077251 | 0.778854091 |
| chr1:184433105-184433305 | 0.299270722 | 0.533130583 | 0.778900985 |
| chr14:69603650-69603891 | 0.261538469 | 0.533136422 | 0.778900985 |
| chr11:39365583-39366159 | -0.2573908 | 0.533139843 | 0.778900985 |
| chr11:2933401-2933957 | 0.21627674 | 0.533165406 | 0.77892348 |
| chr2:96745727-96745986 | 0.295912627 | 0.533199907 | 0.778959031 |

| | | | |
|---|---|---|---|
| chr8:73227351-73227578 | 0.315759865 | 0.533248252 | 0.779014806 |
| chr22:35589288-35589488 | 0.295929164 | 0.533266186 | 0.779018563 |
| chr15:97245867-97246307 | 0.259663112 | 0.533295289 | 0.779018563 |
| chr9:98215885-98216119 | 0.276962058 | 0.53330765 | 0.779018563 |
| chr4:143321312-143321544 | -0.300915344 | 0.533310579 | 0.779018563 |
| chr3:169661465-169663420 | -0.195338042 | 0.533314032 | 0.779018563 |
| chr7:99014471-99014911 | 0.26667923 | 0.533327259 | 0.779018563 |
| chr19:2154505-2154909 | 0.236235084 | 0.533329019 | 0.779018563 |
| chr8:109042164-109042436 | 0.260381739 | 0.533340766 | 0.779018563 |
| chr2:41853683-41854014 | -0.258595937 | 0.533342327 | 0.779018563 |
| chr1:184051179-184052274 | -0.201485542 | 0.53336842 | 0.779041825 |
| chr2:112941729-112942132 | 0.238519176 | 0.533415565 | 0.779082968 |
| chr4:1374189-1374389 | 0.354496761 | 0.533416924 | 0.779082968 |
| chr3:187714412-187714758 | -0.23273677 | 0.533460501 | 0.779131763 |
| chr10:102485109-102485309 | -0.366840432 | 0.533481731 | 0.779147918 |
| chr21:33127385-33127648 | 0.27704951 | 0.53350913 | 0.779173081 |
| chr16:89105786-89106002 | -0.281738203 | 0.533528401 | 0.779184391 |
| chr16:30740319-30740795 | 0.21623775 | 0.533538771 | 0.779184391 |
| chr7:113394214-113394440 | -0.338235043 | 0.533547381 | 0.779184391 |
| chr2:131802500-131802942 | -0.216496902 | 0.533562844 | 0.779192122 |
| chr7:95367657-95368006 | -0.259633933 | 0.533592605 | 0.77921144 |
| chr6:154344504-154344739 | -0.250115486 | 0.533596412 | 0.77921144 |
| chr10:49164650-49164863 | 0.278604256 | 0.533629049 | 0.779220161 |
| chr1:22275837-22276149 | -0.255367744 | 0.533637066 | 0.779220161 |
| chr5:127072843-127073203 | -0.25612364 | 0.533643986 | 0.779220161 |
| chr12:56028679-56029339 | 0.222262392 | 0.533655509 | 0.779220161 |
| chr11:65333534-65334370 | 0.207763369 | 0.533658002 | 0.779220161 |
| chr1:215894731-215894971 | -0.282460242 | 0.533663402 | 0.779220161 |
| chr2:149169466-149170215 | -0.217611736 | 0.533680274 | 0.77922143 |
| chr19:41597622-41597881 | 0.281665516 | 0.53368461 | 0.77922143 |
| chr2:173815906-173816165 | -0.319983776 | 0.533708349 | 0.779241242 |
| chr19:42420119-42420448 | 0.245115479 | 0.533732472 | 0.779247211 |
| chr8:56320007-56320691 | 0.213502539 | 0.533732777 | 0.779247211 |
| chr16:71961315-71961794 | -0.22693492 | 0.53376115 | 0.779267093 |
| chr6:50850267-50850754 | 0.221324799 | 0.533766736 | 0.779267093 |
| chr1:21439741-21440278 | -0.22644924 | 0.533802608 | 0.779304615 |
| chr14:101073240-101073683 | -0.219428799 | 0.533819916 | 0.779315035 |
| chr2:89004407-89004607 | -0.344969629 | 0.533836706 | 0.779324698 |
| chr4:85166507-85166761 | 0.299882056 | 0.533894656 | 0.779383983 |
| chr2:10487068-10487591 | 0.226486109 | 0.533910183 | 0.779383983 |
| chr3:166844099-166844346 | 0.28143871 | 0.533921697 | 0.779383983 |
| chr1:28511667-28512041 | -0.248039377 | 0.533921751 | 0.779383983 |
| chr17:65737251-65737494 | -0.300405009 | 0.533928175 | 0.779383983 |
| chr17:59963917-59965526 | -0.193151956 | 0.533986855 | 0.779438963 |
| chr15:96524821-96525030 | -0.306771806 | 0.533994687 | 0.779438963 |
| chr2:238426136-238427074 | 0.210587386 | 0.534002875 | 0.779438963 |
| chr1:200131540-200131775 | -0.257032113 | 0.534016165 | 0.779438963 |
| chr12:54358874-54359669 | 0.202991963 | 0.534016703 | 0.779438963 |
| chr17:79993148-79993348 | 0.258720305 | 0.534091722 | 0.77953361 |
| chr8:110559592-110559934 | 0.235104708 | 0.534113402 | 0.779537705 |

| | | | |
|---|---|---|---|
| chr8:102810414-102811122 | 0.197926435 | 0.534114876 | 0.779537705 |
| chr8:139997177-139997436 | -0.289924472 | 0.534189746 | 0.779604532 |
| chr5:177446306-177447658 | -0.198909323 | 0.534195486 | 0.779604532 |
| chr7:44349389-44349708 | -0.300925918 | 0.53420375 | 0.779604532 |
| chr7:44018882-44019555 | -0.204850264 | 0.534214802 | 0.779604532 |
| chr1:21655821-21656042 | 0.283153954 | 0.534227365 | 0.779604532 |
| chr22:42614458-42615806 | -0.19948547 | 0.534229202 | 0.779604532 |
| chr6:151706324-151706806 | -0.221328479 | 0.534231886 | 0.779604532 |
| chr10:110225678-110227602 | -0.196654894 | 0.534249068 | 0.779614758 |
| chr6:90859687-90859887 | -0.322141958 | 0.534278318 | 0.779639635 |
| chr19:20039348-20039669 | 0.260363461 | 0.534286466 | 0.779639635 |
| chr15:89472655-89473148 | 0.226240893 | 0.534330749 | 0.779670048 |
| chr14:88053619-88053832 | 0.327990068 | 0.534332857 | 0.779670048 |
| chr4:38893690-38894008 | -0.300823564 | 0.534337835 | 0.779670048 |
| chr20:53600870-53601458 | 0.198139655 | 0.534373671 | 0.77970749 |
| chr2:24972576-24972781 | 0.261954921 | 0.534391412 | 0.779718528 |
| chr2:24471706-24471980 | -0.282039675 | 0.534417416 | 0.779730232 |
| chr9:9931597-9931817 | -0.308701747 | 0.534419786 | 0.779730232 |
| chr11:44979609-44979925 | 0.243272253 | 0.534456742 | 0.77974844 |
| chr5:69501972-69502253 | -0.285351543 | 0.534464164 | 0.77974844 |
| chr19:16771555-16771802 | 0.314326242 | 0.534464819 | 0.77974844 |
| chr20:20549286-20549648 | 0.251888525 | 0.534472972 | 0.77974844 |
| chr3:111859007-111859912 | -0.20710595 | 0.534507835 | 0.779784454 |
| chr3:127597984-127598800 | 0.202263493 | 0.534537261 | 0.779800269 |
| chr15:65359897-65360125 | 0.296358716 | 0.53453903 | 0.779800269 |
| chr19:14564213-14564882 | -0.222957253 | 0.534579052 | 0.779843808 |
| chr17:71633768-71633971 | -0.340465564 | 0.534621528 | 0.779881938 |
| chr1:233025138-233025523 | -0.250638724 | 0.53462565 | 0.779881938 |
| chr9:80787479-80787940 | -0.255079587 | 0.534635725 | 0.779881938 |
| chr16:4097857-4098126 | -0.279784037 | 0.534657992 | 0.77988406 |
| chr8:81720499-81721726 | 0.19678709 | 0.534665947 | 0.77988406 |
| chr14:29434281-29434481 | -0.348094488 | 0.534667716 | 0.77988406 |
| chr2:171433121-171433573 | -0.219736711 | 0.53479173 | 0.780006466 |
| chr5:109438903-109439110 | 0.320472607 | 0.534800731 | 0.780006466 |
| chr14:80356532-80356751 | -0.287975613 | 0.53480132 | 0.780006466 |
| chr8:40174779-40175065 | -0.265206623 | 0.534801953 | 0.780006466 |
| chr9:120742940-120743309 | -0.240985274 | 0.534809263 | 0.780006466 |
| chr6:122789040-122790237 | -0.198072372 | 0.534820493 | 0.780006466 |
| chr22:41544322-41544986 | 0.20666671 | 0.534827093 | 0.780006466 |
| chr1:22751256-22751493 | -0.301495326 | 0.534833073 | 0.780006466 |
| chr2:43881665-43882014 | -0.228009971 | 0.534895684 | 0.78008293 |
| chr1:93179628-93180973 | 0.19870117 | 0.534911966 | 0.780091828 |
| chr3:98039249-98039922 | -0.205131348 | 0.534978453 | 0.780162751 |
| chr22:37843712-37843912 | 0.308990259 | 0.534980962 | 0.780162751 |
| chr12:108692165-108692366 | -0.277553572 | 0.534994307 | 0.78016325 |
| chr21:13877069-13877269 | -0.325617165 | 0.535013749 | 0.78016325 |
| chr20:46332952-46333288 | -0.257083094 | 0.535019341 | 0.78016325 |
| chrX:48883773-48884146 | -0.280743034 | 0.535026435 | 0.78016325 |
| chr5:10595758-10596292 | -0.222529184 | 0.53503938 | 0.78016325 |
| chr11:73420243-73420562 | -0.289963652 | 0.535042396 | 0.78016325 |

| | | | |
|---|---|---|---|
| chr1:202438853-202439291 | -0.239662487 | 0.53508634 | 0.780201072 |
| chr15:43800404-43800805 | 0.220240885 | 0.535088699 | 0.780201072 |
| chr1:203263424-203263863 | -0.23467444 | 0.535118222 | 0.780229271 |
| chr10:59708813-59709207 | -0.238025477 | 0.535138583 | 0.78024411 |
| chr20:49277964-49278816 | 0.196893264 | 0.53518705 | 0.780295484 |
| chr8:91393489-91393908 | -0.227099175 | 0.535195983 | 0.780295484 |
| chr17:79521421-79521640 | 0.312569104 | 0.535204369 | 0.780295484 |
| chr6:4829436-4829684 | 0.313828687 | 0.535239706 | 0.780332155 |
| chr13:27177960-27178160 | -0.305926835 | 0.535263829 | 0.780352476 |
| chr5:177071830-177072154 | 0.240869108 | 0.535351449 | 0.780465367 |
| chr4:106993936-106994188 | -0.327841513 | 0.535421108 | 0.780552067 |
| chr13:110870055-110870702 | -0.209375852 | 0.535457271 | 0.780560823 |
| chr11:73258595-73258795 | 0.304797663 | 0.535476581 | 0.780560823 |
| chr17:62693230-62693496 | -0.226246826 | 0.535479114 | 0.780560823 |
| chr17:47776631-47777085 | -0.240553372 | 0.535482708 | 0.780560823 |
| chr5:141177622-141177861 | 0.295792064 | 0.53548327 | 0.780560823 |
| chr5:78633692-78633900 | -0.313048559 | 0.535488237 | 0.780560823 |
| chr14:72981491-72981731 | -0.317673133 | 0.53550793 | 0.780567176 |
| chr1:180229899-180230457 | 0.213826177 | 0.535541007 | 0.780567176 |
| chr16:72092910-72093140 | -0.237601571 | 0.535542635 | 0.780567176 |
| chr19:47130365-47131185 | 0.205771637 | 0.535549216 | 0.780567176 |
| chr13:74134850-74135092 | -0.243211008 | 0.535551487 | 0.780567176 |
| chr2:11676558-11676931 | -0.22307377 | 0.535553719 | 0.780567176 |
| chr20:42215625-42215896 | 0.275894737 | 0.535571178 | 0.780577775 |
| chr19:14888875-14889086 | -0.335425841 | 0.535593348 | 0.780583083 |
| chr7:55279993-55280307 | -0.242334708 | 0.535595195 | 0.780583083 |
| chr22:19070500-19070709 | -0.329790756 | 0.535606267 | 0.780584373 |
| chr4:85370391-85370782 | 0.233968781 | 0.53565729 | 0.780643885 |
| chr13:31409759-31410038 | -0.265684313 | 0.535708191 | 0.780703217 |
| chr5:919386-919706 | 0.247314364 | 0.535734604 | 0.78071846 |
| chr22:24952128-24953325 | 0.200075983 | 0.535748298 | 0.78071846 |
| chr8:6909745-6910158 | -0.262147259 | 0.535750588 | 0.78071846 |
| chr11:64269711-64270565 | 0.198959909 | 0.535766562 | 0.78071846 |
| chr3:196787458-196788110 | -0.227812237 | 0.535769635 | 0.78071846 |
| chr17:40418394-40418594 | -0.283058563 | 0.535783257 | 0.78071846 |
| chr19:40785621-40785821 | 0.331233689 | 0.535789976 | 0.78071846 |
| chr20:64079483-64080203 | 0.210253397 | 0.535822805 | 0.78075145 |
| chr20:2470444-2471178 | 0.209238757 | 0.535846323 | 0.780766175 |
| chr11:30882540-30882820 | -0.327820617 | 0.535863481 | 0.780766175 |
| chr2:196719101-196719314 | -0.327820617 | 0.535863481 | 0.780766175 |
| chr7:4831742-4832569 | -0.204122209 | 0.53589786 | 0.780768058 |
| chr7:10874518-10874718 | 0.310093926 | 0.535899201 | 0.780768058 |
| chr2:63594706-63594906 | -0.318554175 | 0.535903511 | 0.780768058 |
| chr15:98800488-98800905 | -0.240046418 | 0.535910871 | 0.780768058 |
| chr1:204359827-204360210 | 0.228166018 | 0.535915722 | 0.780768058 |
| chr16:11675634-11675844 | -0.2886192 | 0.53595804 | 0.780814863 |
| chr1:114480665-114481084 | 0.212425209 | 0.535987146 | 0.78084242 |
| chr17:50001290-50001613 | -0.249437135 | 0.536006786 | 0.780853447 |
| chrX:136783170-136783524 | 0.238995413 | 0.536015097 | 0.780853447 |
| chr13:44990106-44990411 | -0.240353024 | 0.536034135 | 0.780866335 |

| | | | |
|---|---|---|---|
| chr8:1701191-1701586 | 0.248293867 | 0.536055477 | 0.780882578 |
| chr15:79853545-79853858 | 0.25274327 | 0.536085549 | 0.780899596 |
| chr3:129249199-129249924 | 0.203460193 | 0.536087542 | 0.780899596 |
| chr16:29943895-29944437 | 0.215863804 | 0.536129202 | 0.780916936 |
| chr7:99471734-99472168 | -0.229653699 | 0.536138802 | 0.780916936 |
| chrX:107838874-107839143 | -0.309556715 | 0.536147241 | 0.780916936 |
| chr8:33566218-33567595 | -0.199272481 | 0.536147776 | 0.780916936 |
| chr16:83807478-83808662 | -0.19784448 | 0.536150405 | 0.780916936 |
| chr1:205354355-205354595 | 0.240971987 | 0.536180018 | 0.780945222 |
| chr20:63707546-63708154 | 0.208993934 | 0.536203268 | 0.780964241 |
| chr1:57824501-57824886 | -0.244072096 | 0.536250507 | 0.781010231 |
| chr14:101980641-101980943 | 0.26547722 | 0.53625523 | 0.781010231 |
| chrX:68197053-68197261 | 0.293764357 | 0.536279618 | 0.781027952 |
| chr10:90787773-90788235 | -0.243300041 | 0.536297259 | 0.781027952 |
| chr2:240989774-240990145 | 0.228136242 | 0.536297978 | 0.781027952 |
| chr1:1615111-1615952 | 0.203786252 | 0.536311285 | 0.781031666 |
| chr19:3785479-3785843 | 0.260891196 | 0.536322258 | 0.781031666 |
| chr8:85855097-85855662 | -0.227418782 | 0.536331107 | 0.781031666 |
| chr2:71672377-71672663 | 0.278594044 | 0.536365929 | 0.78106753 |
| chr9:469757-470661 | 0.210352013 | 0.536408644 | 0.781100061 |
| chr4:37890414-37891559 | 0.201711513 | 0.536408657 | 0.781100061 |
| chr14:38627300-38627529 | 0.303029792 | 0.536420105 | 0.781101886 |
| chr11:102807754-102808296 | 0.222693193 | 0.536445621 | 0.781124196 |
| chr9:85804880-85806001 | 0.212467997 | 0.536513494 | 0.781208181 |
| chr10:110609750-110610017 | -0.270504437 | 0.536529539 | 0.781216698 |
| chr16:30526710-30527705 | 0.197578796 | 0.536586436 | 0.781266821 |
| chr4:77385456-77385686 | -0.272838936 | 0.536608804 | 0.781266821 |
| chr2:241929906-241930217 | -0.280321278 | 0.53661769 | 0.781266821 |
| chr12:117874921-117875186 | 0.28053011 | 0.536626477 | 0.781266821 |
| chr1:1906305-1906505 | 0.284812436 | 0.536654558 | 0.781266821 |
| chr8:124037601-124038099 | -0.213586264 | 0.536656093 | 0.781266821 |
| chr6:100107518-100107794 | -0.275796578 | 0.536674279 | 0.781266821 |
| chr5:170791355-170791645 | -0.252573502 | 0.536684724 | 0.781266821 |
| chr17:7931703-7932336 | -0.2046068 | 0.536695519 | 0.781266821 |
| chr3:5187233-5188974 | -0.193237641 | 0.536705282 | 0.781266821 |
| chr2:74412816-74413409 | -0.221892011 | 0.536705419 | 0.781266821 |
| chr19:10744187-10744569 | -0.229479358 | 0.536708843 | 0.781266821 |
| chr3:153161085-153162513 | -0.192755175 | 0.536718647 | 0.781266821 |
| chr12:131222287-131222564 | -0.265787629 | 0.536721719 | 0.781266821 |
| chr18:58043982-58045334 | -0.203764195 | 0.536724112 | 0.781266821 |
| chr7:51141502-51141773 | -0.275685818 | 0.536730944 | 0.781266821 |
| chr11:46315094-46315495 | -0.215709124 | 0.536737301 | 0.781266821 |
| chr7:100119220-100119982 | -0.206867097 | 0.536771507 | 0.781301769 |
| chr14:77376588-77377946 | 0.198871912 | 0.536807981 | 0.781328517 |
| chr3:110916638-110917052 | -0.246768602 | 0.536820703 | 0.781328517 |
| chr11:20355295-20355559 | 0.26374525 | 0.536829845 | 0.781328517 |
| chr19:49580140-49580804 | 0.212133673 | 0.536830673 | 0.781328517 |
| chr10:84665942-84666312 | -0.242608957 | 0.536853669 | 0.781347146 |
| chr20:35771765-35772455 | 0.203186189 | 0.536883712 | 0.781376029 |
| chr7:73436061-73436721 | -0.21837537 | 0.536895936 | 0.781378978 |

| | | | |
|---|---|---|---|
| chr15:81039420-81039659 | -0.326893512 | 0.536914984 | 0.781391857 |
| chr3:101495510-101495958 | -0.227748245 | 0.536925764 | 0.781392704 |
| chr8:92062413-92062792 | -0.260582743 | 0.536973813 | 0.781447788 |
| chr8:86791757-86792418 | -0.22373761 | 0.537006009 | 0.7814798 |
| chr3:134449320-134449590 | -0.251956025 | 0.537035908 | 0.781481277 |
| chr1:156281792-156282035 | -0.248357203 | 0.537044594 | 0.781481277 |
| chr2:181656462-181657265 | -0.204369351 | 0.537050786 | 0.781481277 |
| chr8:94404771-94405127 | 0.260166814 | 0.537054083 | 0.781481277 |
| chr1:111203596-111204718 | -0.199562519 | 0.53705802 | 0.781481277 |
| chr12:112938160-112938636 | -0.228394453 | 0.537109576 | 0.781541456 |
| chr20:37315651-37315893 | -0.301867774 | 0.537135561 | 0.78155882 |
| chr6:154156030-154156505 | -0.219062704 | 0.537146651 | 0.78155882 |
| chr6:95577162-95578283 | -0.199108611 | 0.537153745 | 0.78155882 |
| chr3:66763150-66763929 | 0.20732401 | 0.537168436 | 0.78155882 |
| chr1:206746526-206746768 | -0.271700252 | 0.537174932 | 0.78155882 |
| chr14:28679001-28679299 | 0.270055342 | 0.53718807 | 0.78155882 |
| chr2:172735348-172735583 | -0.321531182 | 0.537203111 | 0.78155882 |
| chr2:184611612-184611812 | -0.321531182 | 0.537203111 | 0.78155882 |
| chr3:20225870-20226084 | 0.290408735 | 0.53722337 | 0.78157198 |
| chr21:33374728-33375145 | -0.289332008 | 0.537232558 | 0.78157198 |
| chr11:71448158-71448955 | 0.200775216 | 0.537249155 | 0.781581285 |
| chr2:28566752-28567121 | 0.228695327 | 0.537263023 | 0.781586622 |
| chr7:75157047-75157252 | -0.326508607 | 0.537316234 | 0.78164919 |
| chr3:151276240-151276487 | -0.309996124 | 0.537407273 | 0.781766785 |
| chrX:39984328-39984802 | 0.247856758 | 0.537442274 | 0.781802857 |
| chr1:203166307-203166845 | 0.206475442 | 0.537489212 | 0.781805455 |
| chr16:22316603-22316900 | 0.285744122 | 0.53749275 | 0.781805455 |
| chr2:169828476-169828701 | -0.248237506 | 0.537496333 | 0.781805455 |
| chr17:8346210-8346569 | -0.278756959 | 0.537501155 | 0.781805455 |
| chr2:231198251-231198797 | 0.219493522 | 0.537501795 | 0.781805455 |
| chr2:74845192-74846022 | -0.203951743 | 0.537510623 | 0.781805455 |
| chr1:171403346-171403803 | 0.221494701 | 0.537520418 | 0.781805455 |
| chr18:11908122-11909100 | 0.203752259 | 0.537525687 | 0.781805455 |
| chrX:37993899-37994226 | -0.29178331 | 0.537595366 | 0.78189165 |
| chr8:85973425-85973704 | 0.261384689 | 0.537614317 | 0.78189165 |
| chr11:57324681-57325585 | -0.199722124 | 0.537617504 | 0.78189165 |
| chr15:100928867-100929311 | 0.220701493 | 0.537625768 | 0.78189165 |
| chr3:63576798-63577716 | -0.197485931 | 0.537672603 | 0.781944922 |
| chr3:123652421-123652707 | 0.230000951 | 0.537693991 | 0.781961186 |
| chr11:57476925-57477267 | -0.230144179 | 0.537720813 | 0.78198535 |
| chr3:62529305-62529602 | 0.21842918 | 0.537732244 | 0.781986184 |
| chr7:41059308-41059771 | -0.227118834 | 0.537741798 | 0.781986184 |
| chr6:38702436-38703386 | 0.204261829 | 0.537819041 | 0.782083668 |
| chr1:172719703-172719969 | -0.253587906 | 0.537844809 | 0.782105157 |
| chr17:81553049-81553784 | 0.205220704 | 0.537854233 | 0.782105157 |
| chr5:180291903-180292821 | 0.199782488 | 0.537911022 | 0.782172891 |
| chr8:126433818-126434024 | -0.281727701 | 0.53794016 | 0.782186745 |
| chr15:92654191-92654469 | 0.225342302 | 0.537940967 | 0.782186745 |
| chr16:21501608-21501816 | 0.271255029 | 0.53795506 | 0.782192395 |
| chr12:6312456-6313186 | -0.210243744 | 0.537967426 | 0.782195532 |

| | | | |
|---|---|---|---|
| chr14:21498648-21498849 | 0.280698757 | 0.537995445 | 0.782221427 |
| chr3:108620082-108620720 | 0.221768948 | 0.538006716 | 0.782222972 |
| chr10:132684618-132684818 | 0.338967594 | 0.538029455 | 0.782238922 |
| chr8:107813448-107813820 | 0.221215352 | 0.538038105 | 0.782238922 |
| chr2:85615918-85616481 | -0.205962085 | 0.538054176 | 0.7822442 |
| chr3:4722171-4722409 | 0.247477949 | 0.538062153 | 0.7822442 |
| chr5:102222881-102223087 | -0.298094779 | 0.538077986 | 0.782252375 |
| chr4:4985922-4986122 | -0.332726494 | 0.538120523 | 0.782299372 |
| chr2:72993124-72993408 | -0.252810732 | 0.538172125 | 0.782350283 |
| chr8:119264012-119264244 | -0.273959578 | 0.538183808 | 0.782350283 |
| chr6:10147053-10147253 | -0.327491332 | 0.538211033 | 0.782350283 |
| chrX:54808033-54809265 | 0.196968527 | 0.538216212 | 0.782350283 |
| chr6:122609889-122611467 | -0.195220666 | 0.53821999 | 0.782350283 |
| chr6:121510663-121511117 | -0.252831966 | 0.538223943 | 0.782350283 |
| chr3:18086154-18086391 | -0.300498928 | 0.538227017 | 0.782350283 |
| chr3:103991536-103991750 | -0.302307522 | 0.538286071 | 0.782388435 |
| chr12:524029-524413 | -0.294883715 | 0.538286562 | 0.782388435 |
| chr3:158626876-158627237 | -0.298380561 | 0.538294599 | 0.782388435 |
| chrX:10082760-10083677 | -0.205689176 | 0.538296534 | 0.782388435 |
| chr11:61730381-61730612 | 0.279378219 | 0.538304319 | 0.782388435 |
| chr4:145936171-145937760 | 0.197989553 | 0.538352871 | 0.782431568 |
| chr1:67053059-67054638 | -0.196622163 | 0.538354419 | 0.782431568 |
| chr3:157159396-157161057 | -0.192783626 | 0.53837013 | 0.782439561 |
| chr16:57775572-57775893 | -0.245224476 | 0.538382363 | 0.782442499 |
| chr20:4108740-4108942 | 0.299536933 | 0.538405081 | 0.782455992 |
| chr4:113181296-113181591 | -0.257268126 | 0.538412071 | 0.782455992 |
| chr5:141349905-141350411 | -0.228970066 | 0.538437579 | 0.782478221 |
| chr16:75789646-75789846 | 0.304524537 | 0.538461131 | 0.782497606 |
| chr14:45698487-45698827 | -0.251729237 | 0.538477429 | 0.782506449 |
| chr4:50662224-50662507 | 0.290908471 | 0.538529057 | 0.782566632 |
| chr1:165631500-165631855 | -0.229116952 | 0.538545294 | 0.782575386 |
| chr19:47310919-47311119 | -0.354554069 | 0.538563576 | 0.78258711 |
| chr1:40373247-40373874 | -0.207092181 | 0.538595491 | 0.782618643 |
| chr2:96637820-96638888 | -0.200628946 | 0.538640987 | 0.78266991 |
| chrX:18983762-18985180 | 0.196541509 | 0.538661218 | 0.782677988 |
| chr16:52896-53974 | 0.203344629 | 0.538675387 | 0.782677988 |
| chr17:4807449-4807672 | -0.291778097 | 0.538678552 | 0.782677988 |
| chr17:42527244-42527622 | 0.239472703 | 0.538699852 | 0.782677988 |
| chr13:113741942-113742283 | -0.262569657 | 0.538707245 | 0.782677988 |
| chr5:31581582-31581782 | 0.335556471 | 0.538707835 | 0.782677988 |
| chr8:144147458-144148410 | 0.196406985 | 0.538741813 | 0.782712512 |
| chr1:184212998-184213339 | -0.239832968 | 0.538785481 | 0.782760952 |
| chr12:47904583-47905525 | 0.201089867 | 0.538795586 | 0.782760952 |
| chr2:21042901-21043414 | -0.226241504 | 0.538839886 | 0.782806193 |
| chr7:123309604-123309833 | -0.309053812 | 0.5388487 | 0.782806193 |
| chr2:46780102-46780346 | 0.293326157 | 0.538858015 | 0.782806193 |
| chr19:49876564-49876972 | 0.207185602 | 0.538867593 | 0.782806193 |
| chr6:116633755-116633972 | -0.305295354 | 0.538895432 | 0.782831055 |
| chr18:69179913-69180177 | -0.305300922 | 0.53890514 | 0.782831055 |
| chrX:30135087-30135338 | -0.321486329 | 0.538927733 | 0.782849032 |

| | | | |
|---|---|---|---|
| chr5:158175000-158175201 | -0.310304312 | 0.538938984 | 0.782850535 |
| chr3:170180594-170182124 | -0.193600495 | 0.538969869 | 0.782880555 |
| chr14:58668041-58668241 | 0.307434798 | 0.538988612 | 0.782892939 |
| chr17:18951973-18952173 | -0.308034045 | 0.539002589 | 0.782898399 |
| chr15:45522642-45523246 | 0.220554669 | 0.539019156 | 0.782907621 |
| chr4:73653661-73653980 | 0.279013113 | 0.539067446 | 0.782962919 |
| chr14:81006189-81006484 | 0.290586838 | 0.539098456 | 0.782993117 |
| chr11:62671510-62672118 | 0.207196047 | 0.53912023 | 0.782997099 |
| chr5:10331356-10331807 | -0.227196846 | 0.539121635 | 0.782997099 |
| chr17:45132496-45133122 | 0.20715139 | 0.53915589 | 0.783032007 |
| chr20:31822370-31823120 | 0.206866719 | 0.539177447 | 0.783048473 |
| chr18:57210302-57210657 | -0.256525513 | 0.539195727 | 0.783060178 |
| chr6:41027766-41028760 | -0.206220885 | 0.539245021 | 0.783085793 |
| chr17:54814020-54814826 | 0.212581181 | 0.539249374 | 0.783085793 |
| chr19:45340576-45340858 | -0.252254892 | 0.539262725 | 0.783085793 |
| chrX:76279092-76279292 | -0.341364263 | 0.539270965 | 0.783085793 |
| chr15:91049803-91050136 | 0.237040954 | 0.53927163 | 0.783085793 |
| chr11:19116856-19117933 | -0.197609647 | 0.539274686 | 0.783085793 |
| chr14:90331562-90332319 | 0.197316772 | 0.539308922 | 0.783120667 |
| chr5:180071660-180072303 | 0.210010636 | 0.539358454 | 0.783177749 |
| chr7:1813369-1813721 | 0.253215158 | 0.53938929 | 0.783180157 |
| chr20:24918192-24918619 | 0.23937722 | 0.539391186 | 0.783180157 |
| chr15:77107687-77107954 | 0.259415831 | 0.539399184 | 0.783180157 |
| chr17:79658891-79659161 | -0.27350872 | 0.539400998 | 0.783180157 |
| chr13:38372810-38373274 | -0.245064998 | 0.539425852 | 0.783201403 |
| chr10:102245044-102245820 | -0.200062336 | 0.539442183 | 0.783210273 |
| chr12:66409374-66409634 | -0.264939655 | 0.539467134 | 0.783231658 |
| chr6:159692618-159693929 | 0.200193849 | 0.539477628 | 0.783232052 |
| chr11:76217679-76217879 | -0.311490177 | 0.539532986 | 0.783297581 |
| chr14:55599203-55599879 | 0.216492467 | 0.539573285 | 0.783341245 |
| chr4:18602527-18603332 | -0.221133876 | 0.539603921 | 0.783358585 |
| chr21:8438364-8439022 | 0.193870629 | 0.539615002 | 0.783358585 |
| chr6:43769549-43770342 | 0.200686357 | 0.5396159 | 0.783358585 |
| chr19:29730534-29730779 | 0.275327033 | 0.539629189 | 0.783363035 |
| chr1:21270212-21270438 | 0.326184547 | 0.539687226 | 0.783432443 |
| chr10:84291486-84291727 | 0.276324579 | 0.539718817 | 0.783455777 |
| chr17:48636539-48636967 | 0.241930433 | 0.53972375 | 0.783455777 |
| chr9:112379079-112380576 | 0.19456817 | 0.539734077 | 0.783455925 |
| chr3:27467588-27467865 | -0.263856662 | 0.539767148 | 0.783489086 |
| chr16:75481569-75482032 | -0.235573814 | 0.539784783 | 0.783499842 |
| chr4:139603710-139603910 | 0.30691947 | 0.539824372 | 0.783542462 |
| chr7:36128421-36128969 | -0.218261274 | 0.539847835 | 0.783561675 |
| chr17:17303693-17303896 | -0.288342778 | 0.539860158 | 0.783564718 |
| chr19:54136918-54137912 | 0.193564929 | 0.539928266 | 0.783648727 |
| chr19:43755569-43755784 | 0.248946737 | 0.539976128 | 0.783703349 |
| chr5:10576441-10576684 | 0.282508099 | 0.540017127 | 0.783748007 |
| chr4:17576617-17577951 | 0.193770882 | 0.540067201 | 0.783795617 |
| chr2:3741470-3741768 | -0.294703562 | 0.54007039 | 0.783795617 |
| chr5:14983773-14984026 | 0.275028956 | 0.540134171 | 0.783822333 |
| chr20:40715601-40715801 | -0.337094744 | 0.540149453 | 0.783822333 |

| | | | |
|---|---|---|---|
| chr7:48088275-48089821 | 0.194657559 | 0.540149829 | 0.783822333 |
| chr8:110718621-110718922 | -0.319446511 | 0.540166449 | 0.783822333 |
| chr8:135701310-135701535 | -0.30292421 | 0.540172371 | 0.783822333 |
| chr11:11297655-11298017 | -0.270495677 | 0.540188505 | 0.783822333 |
| chr10:87961065-87961265 | 0.316047456 | 0.540191561 | 0.783822333 |
| chr5:151786565-151786773 | 0.342007091 | 0.540198217 | 0.783822333 |
| chr14:22852894-22853772 | 0.200735729 | 0.540211052 | 0.783822333 |
| chr8:106863980-106864242 | 0.309080186 | 0.540213109 | 0.783822333 |
| chr5:147782400-147782989 | -0.217020664 | 0.540225008 | 0.783822333 |
| chr5:177086442-177087466 | -0.198348741 | 0.540236174 | 0.783822333 |
| chr7:34425247-34425615 | -0.253164072 | 0.540250842 | 0.783822333 |
| chr21:30146641-30146912 | 0.270508832 | 0.540252346 | 0.783822333 |
| chr6:141562880-141563133 | -0.281354923 | 0.540255786 | 0.783822333 |
| chr1:161523569-161524772 | -0.201270915 | 0.540264022 | 0.783822333 |
| chr15:77070652-77071390 | 0.207058601 | 0.540269275 | 0.783822333 |
| chr3:158109662-158110968 | -0.193435798 | 0.540283562 | 0.783822333 |
| chr1:43454311-43454762 | 0.228122888 | 0.540291078 | 0.783822333 |
| chr1:231931886-231932167 | -0.324286251 | 0.540296365 | 0.783822333 |
| chr8:78393475-78393725 | 0.274962845 | 0.540313852 | 0.783822333 |
| chr8:134885877-134886122 | 0.274962845 | 0.540313852 | 0.783822333 |
| chr2:229714225-229714897 | 0.217425948 | 0.540327489 | 0.783827275 |
| chrX:56995046-56995998 | -0.206021859 | 0.540342932 | 0.783834838 |
| chr16:31056263-31056527 | -0.328977093 | 0.54037405 | 0.783865137 |
| chr10:126749102-126749383 | 0.262106071 | 0.540410231 | 0.78389528 |
| chr4:13298605-13299210 | 0.216282328 | 0.540415291 | 0.78389528 |
| chr19:45506448-45507020 | 0.20647357 | 0.540490688 | 0.783960573 |
| chr16:67928573-67928816 | -0.25435797 | 0.540503778 | 0.783960573 |
| chr1:19487832-19488032 | -0.309091812 | 0.540505577 | 0.783960573 |
| chr8:119645091-119645340 | -0.273219629 | 0.54050682 | 0.783960573 |
| chr2:129764425-129764718 | 0.260076678 | 0.540511461 | 0.783960573 |
| chr3:137692866-137693205 | -0.238087765 | 0.540527255 | 0.783968641 |
| chr12:46218548-46219217 | 0.226423906 | 0.540595642 | 0.784052987 |
| chr7:25456910-25457208 | 0.274723583 | 0.540630169 | 0.784082452 |
| chr1:241362244-241362638 | -0.265964758 | 0.540642662 | 0.784082452 |
| chrX:149504926-149505584 | 0.200756834 | 0.540646657 | 0.784082452 |
| chr7:6160278-6160498 | -0.279988639 | 0.540668396 | 0.784086393 |
| chr2:62195495-62196742 | -0.196688822 | 0.540669841 | 0.784086393 |
| chr16:83968112-83968312 | 0.265368833 | 0.540697254 | 0.784087987 |
| chr4:98479374-98479909 | -0.223757607 | 0.540700178 | 0.784087987 |
| chr10:38010373-38011512 | -0.1940208 | 0.540703106 | 0.784087987 |
| chr3:5644707-5644975 | 0.283047938 | 0.5407149 | 0.784087987 |
| chr8:42270811-42271957 | -0.20300468 | 0.540730686 | 0.784087987 |
| chr4:151801678-151801884 | 0.299550552 | 0.54073234 | 0.784087987 |
| chr2:125694896-125695103 | 0.27572507 | 0.540773365 | 0.784121115 |
| chr4:103099716-103100173 | -0.212523477 | 0.540776428 | 0.784121115 |
| chr19:50679396-50679697 | 0.244240826 | 0.540785887 | 0.784121115 |
| chr10:125745053-125745357 | -0.297143002 | 0.540837541 | 0.784181173 |
| chr3:4687127-4687608 | 0.204982202 | 0.540867934 | 0.78418129 |
| chr11:18383659-18384090 | -0.236785066 | 0.540869081 | 0.78418129 |
| chr20:47798506-47798714 | 0.220044611 | 0.540889742 | 0.78418129 |

| | | | |
|---|---|---|---|
| chr17:77542774-77543079 | 0.238854894 | 0.540913852 | 0.78418129 |
| chr3:62753600-62753811 | -0.264226472 | 0.540918015 | 0.78418129 |
| chr3:194686736-194686943 | 0.280658404 | 0.540924463 | 0.78418129 |
| chr1:45206306-45206950 | 0.226615239 | 0.540927121 | 0.78418129 |
| chr2:104872742-104873829 | -0.200818374 | 0.540927161 | 0.78418129 |
| chr5:2541543-2541851 | -0.279792258 | 0.540933609 | 0.78418129 |
| chr8:15702371-15702632 | -0.303686192 | 0.540939966 | 0.78418129 |
| chr15:93018587-93018887 | -0.28501455 | 0.541005446 | 0.784261376 |
| chr3:69051339-69052755 | -0.194462696 | 0.541063122 | 0.784330145 |
| chr16:81625744-81626038 | -0.247054998 | 0.541080243 | 0.784332076 |
| chr3:36868939-36869139 | -0.320361243 | 0.541084926 | 0.784332076 |
| chr3:3179145-3179991 | -0.199221313 | 0.541115769 | 0.784358807 |
| chr4:87093399-87093880 | 0.228466839 | 0.541123841 | 0.784358807 |
| chr7:5178385-5178652 | -0.284134253 | 0.541138638 | 0.784365417 |
| chr7:34091401-34091601 | -0.28603289 | 0.54117623 | 0.784397353 |
| chr18:711621-712062 | 0.222070463 | 0.541181145 | 0.784397353 |
| chr21:23968652-23969284 | 0.214086024 | 0.541223496 | 0.784429883 |
| chr11:111765693-111766926 | -0.199474546 | 0.54123023 | 0.784429883 |
| chr17:81394971-81395542 | 0.199682589 | 0.541242932 | 0.784429883 |
| chr6:34746079-34746365 | -0.288460857 | 0.54124541 | 0.784429883 |
| chr3:144303180-144303424 | -0.311013334 | 0.541258237 | 0.784429883 |
| chr2:120012781-120013748 | 0.199512252 | 0.541275686 | 0.784429883 |
| chr19:46855932-46856414 | -0.213979024 | 0.541279706 | 0.784429883 |
| chr3:117853838-117854366 | -0.232008388 | 0.54128549 | 0.784429883 |
| chr3:97018811-97019011 | 0.324374612 | 0.541299015 | 0.784434647 |
| chr2:234165418-234165691 | 0.29868339 | 0.541354983 | 0.784477017 |
| chr5:152286325-152286530 | -0.320311663 | 0.541359754 | 0.784477017 |
| chr10:94804031-94804292 | -0.296827094 | 0.541371778 | 0.784477017 |
| chr6:83051794-83052060 | -0.284832926 | 0.54138732 | 0.784477017 |
| chr14:90816369-90816576 | 0.275187352 | 0.541392076 | 0.784477017 |
| chr9:90102303-90102678 | 0.232590209 | 0.541398578 | 0.784477017 |
| chr7:100585566-100586776 | -0.193479202 | 0.54139992 | 0.784477017 |
| chr17:5191525-5192192 | 0.206214295 | 0.541437493 | 0.784514768 |
| chr12:26125714-26126322 | -0.218278867 | 0.541455033 | 0.784514768 |
| chr19:45016959-45017285 | 0.258099851 | 0.541463981 | 0.784514768 |
| chr15:89690190-89691200 | 0.198025698 | 0.541466929 | 0.784514768 |
| chr15:77010360-77010635 | 0.264837236 | 0.541505435 | 0.784555724 |
| chr9:120338506-120338706 | 0.311908505 | 0.541559019 | 0.784618522 |
| chr19:49589933-49590874 | 0.197506113 | 0.541597681 | 0.784649604 |
| chr9:133114528-133114872 | 0.242988515 | 0.541600953 | 0.784649604 |
| chr5:151047896-151048150 | 0.241472803 | 0.54163488 | 0.784663618 |
| chr8:79890607-79891320 | 0.217515355 | 0.541646225 | 0.784663618 |
| chr2:105143405-105144082 | 0.205845789 | 0.541649321 | 0.784663618 |
| chr19:1930237-1930513 | -0.295328849 | 0.541658444 | 0.784663618 |
| chr12:117890238-117890453 | 0.326593007 | 0.54166183 | 0.784663618 |
| chr4:85969021-85969262 | -0.309637023 | 0.541691801 | 0.784667574 |
| chr5:38207404-38207668 | -0.289752398 | 0.541696238 | 0.784667574 |
| chr9:22446596-22447718 | 0.197949018 | 0.541701898 | 0.784667574 |
| chr9:138211922-138212122 | 0.311045123 | 0.541705524 | 0.784667574 |
| chr19:5703776-5704010 | 0.304784738 | 0.54172587 | 0.784682212 |

| | | | |
|---|---|---|---|
| chr3:149752070-149753056 | -0.198258466 | 0.54177436 | 0.784737614 |
| chr8:10332807-10333028 | -0.295295196 | 0.541815773 | 0.784762801 |
| chr9:92618683-92619011 | -0.260973527 | 0.541818921 | 0.784762801 |
| chr6:113596854-113597227 | 0.231168471 | 0.541822476 | 0.784762801 |
| chr3:63791908-63792278 | -0.205546416 | 0.541838568 | 0.784771275 |
| chr14:20610037-20610316 | 0.210707731 | 0.541851804 | 0.78477561 |
| chr8:106608537-106608934 | -0.219407846 | 0.541891702 | 0.78480691 |
| chr9:4662062-4662522 | -0.208454264 | 0.541914259 | 0.78480691 |
| chr17:45197683-45198208 | 0.22461926 | 0.541920787 | 0.78480691 |
| chr2:10709349-10709893 | 0.216872742 | 0.541928411 | 0.78480691 |
| chr1:27383044-27383564 | 0.217091587 | 0.541928862 | 0.78480691 |
| chr5:43396832-43397306 | 0.228825277 | 0.541939916 | 0.78480691 |
| chr14:54175445-54175647 | 0.298465785 | 0.541945113 | 0.78480691 |
| chr1:32992449-32992766 | 0.249047505 | 0.541965531 | 0.784821645 |
| chr14:65274239-65274688 | 0.231294244 | 0.541980317 | 0.784828225 |
| chr9:100352204-100353214 | -0.197029975 | 0.542020628 | 0.784871765 |
| chr10:30002465-30002760 | -0.261786968 | 0.542038735 | 0.784877038 |
| chr12:116916804-116917043 | 0.280970426 | 0.542051888 | 0.784877038 |
| chr3:115975577-115975862 | 0.285741706 | 0.542070969 | 0.784877038 |
| chr1:35718488-35719298 | 0.206646937 | 0.542088997 | 0.784877038 |
| chr17:48457101-48457608 | -0.217370077 | 0.542092917 | 0.784877038 |
| chr19:35947411-35947707 | 0.240860021 | 0.542093208 | 0.784877038 |
| chr20:25389812-25391214 | 0.194173249 | 0.542116283 | 0.784877038 |
| chr2:119366597-119367284 | 0.19740625 | 0.542130059 | 0.784877038 |
| chr6:107727556-107727858 | 0.273711983 | 0.542163695 | 0.784877038 |
| chr19:51339664-51340107 | 0.20611263 | 0.542177398 | 0.784877038 |
| chr5:13601146-13601505 | -0.273274172 | 0.542187115 | 0.784877038 |
| chr5:174675817-174676038 | 0.313166805 | 0.542187352 | 0.784877038 |
| chr5:57779468-57779720 | 0.269183259 | 0.542188177 | 0.784877038 |
| chr20:36574742-36575048 | 0.247979676 | 0.542191021 | 0.784877038 |
| chr1:206664235-206664511 | 0.228240632 | 0.542191631 | 0.784877038 |
| chr3:56494700-56494977 | 0.251439906 | 0.542194043 | 0.784877038 |
| chr8:66022193-66022526 | 0.218002895 | 0.542198409 | 0.784877038 |
| chr1:169928512-169929219 | 0.226278783 | 0.542209403 | 0.784878124 |
| chr10:45465013-45465352 | 0.247555248 | 0.542225483 | 0.784886573 |
| chr5:174724206-174725347 | -0.200860493 | 0.542286564 | 0.78496016 |
| chr1:184143398-184143856 | 0.232565949 | 0.542314665 | 0.784974378 |
| chr2:61763715-61764831 | -0.193055154 | 0.542325075 | 0.784974378 |
| chr1:108643610-108643830 | -0.284950436 | 0.542336675 | 0.784974378 |
| chr13:46289066-46289324 | 0.243688271 | 0.542337365 | 0.784974378 |
| chr19:19320336-19320919 | -0.216596446 | 0.542407965 | 0.785061733 |
| chr1:180154285-180155037 | -0.19883888 | 0.542425049 | 0.78506798 |
| chr16:2194948-2195148 | -0.295979525 | 0.542432773 | 0.78506798 |
| chr17:39547710-39548213 | 0.243198469 | 0.542490857 | 0.785125934 |
| chr17:50312400-50312603 | -0.295274223 | 0.542493309 | 0.785125934 |
| chr14:60151678-60152084 | -0.26886551 | 0.542517504 | 0.785127985 |
| chr10:97590234-97590438 | 0.301370215 | 0.542517692 | 0.785127985 |
| chr12:110039456-110039656 | -0.299129418 | 0.542525467 | 0.785127985 |
| chr12:92080401-92080912 | 0.221464837 | 0.542542647 | 0.785138018 |
| chr7:145082575-145082880 | -0.284180381 | 0.542591089 | 0.785193292 |

| | | | |
|---|---|---|---|
| chr8:102872291-102872688 | 0.229764304 | 0.542634052 | 0.785228159 |
| chr7:2844027-2844830 | -0.196557292 | 0.542635679 | 0.785228159 |
| chr16:56989410-56989814 | 0.245977443 | 0.542674901 | 0.785270085 |
| chr12:54596574-54596809 | -0.28015198 | 0.542715446 | 0.785313924 |
| chr8:144517043-144517243 | 0.252169687 | 0.542736063 | 0.785328926 |
| chr10:86103512-86103737 | -0.262358116 | 0.542815207 | 0.785428613 |
| chr10:101893823-101894159 | -0.288765698 | 0.542829856 | 0.785434978 |
| chr11:73974001-73974226 | -0.264132194 | 0.542840405 | 0.78543541 |
| chr7:5665459-5665682 | -0.267034508 | 0.542890761 | 0.785493436 |
| chr8:18083444-18085105 | -0.194242954 | 0.542929055 | 0.785534009 |
| chr3:132416862-132418403 | -0.191003509 | 0.542965951 | 0.785568357 |
| chr17:76641611-76641983 | 0.262461197 | 0.542978042 | 0.785568357 |
| chr9:118170093-118170337 | -0.288943018 | 0.542983553 | 0.785568357 |
| chr9:89435417-89435885 | -0.21492527 | 0.543005239 | 0.785584899 |
| chr19:35828757-35829006 | 0.293212187 | 0.543023605 | 0.785596637 |
| chr1:46171303-46172304 | -0.199762359 | 0.543049624 | 0.785619446 |
| chr2:64767771-64768495 | -0.208198592 | 0.543076618 | 0.785643664 |
| chr5:132300769-132301393 | -0.219545206 | 0.543091716 | 0.785646116 |
| chr22:29359362-29359864 | 0.241290582 | 0.54309882 | 0.785646116 |
| chr18:2547579-2547815 | 0.286186923 | 0.543172713 | 0.785738174 |
| chr2:5042749-5043020 | -0.315132818 | 0.543210488 | 0.785777983 |
| chr9:33720061-33720345 | -0.268757401 | 0.543233092 | 0.785795847 |
| chr7:102244645-102244924 | -0.253979898 | 0.543244421 | 0.785797399 |
| chr14:20961261-20961462 | -0.32332944 | 0.543265492 | 0.785807286 |
| chr15:84955371-84955913 | 0.216103926 | 0.543271767 | 0.785807286 |
| chr15:64067525-64067914 | 0.228262746 | 0.543296038 | 0.785815239 |
| chr4:112112699-112113718 | -0.201296694 | 0.543297777 | 0.785815239 |
| chr10:123635831-123636083 | -0.298508765 | 0.54336827 | 0.7858942 |
| chr17:43368748-43369001 | 0.242672028 | 0.543372883 | 0.7858942 |
| chr19:2507192-2507392 | 0.302250649 | 0.543443765 | 0.785975082 |
| chr1:156851784-156851984 | 0.297358549 | 0.543449321 | 0.785975082 |
| chr9:114138601-114138808 | -0.305398751 | 0.543498755 | 0.78603174 |
| chr19:17390950-17391447 | -0.227307517 | 0.543510663 | 0.786034125 |
| chr15:41904367-41905022 | -0.22568312 | 0.54352658 | 0.78604231 |
| chr1:206635178-206636264 | -0.193845418 | 0.54355783 | 0.786061618 |
| chr17:19004610-19005155 | 0.214969032 | 0.54356045 | 0.786061618 |
| chr1:1231936-1232361 | 0.217560406 | 0.543573592 | 0.786064109 |
| chr1:185156399-185157625 | -0.190504853 | 0.54358269 | 0.786064109 |
| chr7:117582235-117582605 | -0.280793442 | 0.543603901 | 0.786079946 |
| chr16:85321947-85322319 | -0.228691705 | 0.543638502 | 0.786100675 |
| chr1:156104085-156104422 | -0.219856991 | 0.543660533 | 0.786100675 |
| chr3:172203968-172204222 | 0.254848548 | 0.543669069 | 0.786100675 |
| chr4:110622597-110623480 | -0.194914309 | 0.54368622 | 0.786100675 |
| chr8:85911513-85911801 | -0.266485654 | 0.543689912 | 0.786100675 |
| chr1:156692599-156693559 | 0.196463161 | 0.54371021 | 0.786100675 |
| chr2:69828997-69830665 | -0.191864834 | 0.543710806 | 0.786100675 |
| chr10:63722605-63722869 | 0.268137553 | 0.543714871 | 0.786100675 |
| chr9:127244836-127245614 | -0.202893415 | 0.543719089 | 0.786100675 |
| chr3:72938705-72939133 | 0.231522226 | 0.54372083 | 0.786100675 |
| chr19:53853007-53853241 | -0.281675298 | 0.543749159 | 0.786120443 |

| | | | |
|---|---|---|---|
| chr1:119139924-119141386 | -0.193235453 | 0.543756319 | 0.786120443 |
| chr3:146326685-146326885 | 0.289903768 | 0.543765282 | 0.786120443 |
| chr7:36440964-36441206 | 0.264944586 | 0.543802581 | 0.78614617 |
| chr12:7092530-7092751 | -0.288073165 | 0.543803627 | 0.78614617 |
| chr5:93306060-93306350 | -0.268066762 | 0.543813859 | 0.78614617 |
| chr10:62724631-62724831 | -0.308046971 | 0.543868894 | 0.786198895 |
| chr7:101602460-101602783 | -0.237302625 | 0.543881151 | 0.786198895 |
| chr1:108691661-108693624 | -0.190884721 | 0.543899987 | 0.786198895 |
| chr20:51513336-51513700 | -0.25301556 | 0.543909043 | 0.786198895 |
| chr1:46477486-46477766 | 0.240141032 | 0.54391166 | 0.786198895 |
| chr20:60300031-60300292 | -0.237761807 | 0.543912452 | 0.786198895 |
| chr19:10416569-10417111 | -0.210215494 | 0.543922156 | 0.786198895 |
| chr1:62784414-62784614 | -0.271571106 | 0.543937923 | 0.786206854 |
| chrX:52261431-52262284 | -0.201029151 | 0.543968151 | 0.786221957 |
| chr6:112618889-112619089 | -0.310272607 | 0.543968894 | 0.786221957 |
| chr4:150960166-150960395 | -0.271631355 | 0.543986089 | 0.786227331 |
| chr6:41936004-41936252 | 0.267159721 | 0.543993134 | 0.786227331 |
| chr4:122577874-122578114 | -0.27350933 | 0.544039339 | 0.786279278 |
| chrX:125436941-125437224 | 0.274388256 | 0.544056046 | 0.786285215 |
| chr4:64617828-64618141 | -0.266181262 | 0.544078633 | 0.786285215 |
| chr7:42691025-42691337 | -0.244884416 | 0.544098039 | 0.786285215 |
| chr3:108732188-108732392 | 0.312972741 | 0.544105203 | 0.786285215 |
| chr2:191218347-191218628 | 0.237216835 | 0.544106282 | 0.786285215 |
| chr4:82428935-82431243 | -0.189230424 | 0.544111715 | 0.786285215 |
| chr17:51056868-51057318 | 0.239907757 | 0.544133183 | 0.786285215 |
| chr20:54219898-54220133 | -0.216490082 | 0.544136295 | 0.786285215 |
| chr16:11250242-11250520 | 0.28066334 | 0.544138508 | 0.786285215 |
| chr11:21811575-21812055 | -0.230932529 | 0.544146065 | 0.786285215 |
| chr11:18384200-18384536 | 0.244227557 | 0.544174719 | 0.786299983 |
| chr1:53220130-53220849 | 0.19876179 | 0.544182336 | 0.786299983 |
| chr8:86176159-86176402 | -0.253171976 | 0.544187071 | 0.786299983 |
| chr16:56318408-56318679 | 0.25270384 | 0.544228468 | 0.786328862 |
| chr1:201482072-201482427 | 0.203037927 | 0.54424652 | 0.786328862 |
| chr6:28218589-28219147 | -0.222940996 | 0.544248634 | 0.786328862 |
| chr2:101342184-101342532 | -0.230981306 | 0.544250084 | 0.786328862 |
| chr6:46170494-46171358 | -0.195383112 | 0.544258371 | 0.786328862 |
| chr8:128744493-128744712 | -0.290637829 | 0.544275609 | 0.786332562 |
| chr5:6766383-6767106 | -0.198009713 | 0.544281456 | 0.786332562 |
| chr3:16101562-16101762 | 0.292176874 | 0.544310889 | 0.786351252 |
| chr2:135984768-135986201 | -0.191943841 | 0.544314919 | 0.786351252 |
| chr10:80504380-80504735 | -0.2413588 | 0.544351601 | 0.786389419 |
| chr4:121272585-121272820 | -0.294651695 | 0.544382529 | 0.786419271 |
| chr16:27268485-27269250 | -0.202146569 | 0.544402874 | 0.786433834 |
| chr5:170429094-170429346 | -0.255997625 | 0.544431854 | 0.786451015 |
| chr10:37857038-37858435 | 0.19194558 | 0.544435295 | 0.786451015 |
| chr9:131349971-131350556 | -0.220753905 | 0.544455269 | 0.786459958 |
| chr8:37826513-37826861 | -0.247938131 | 0.544462015 | 0.786459958 |
| chr9:87678705-87679030 | -0.237430483 | 0.544498678 | 0.786483921 |
| chr12:8890606-8891116 | -0.253157591 | 0.544504944 | 0.786483921 |
| chr10:34728939-34729267 | -0.270300968 | 0.544514109 | 0.786483921 |

| | | | |
|---|---|---|---|
| chr5:151719134-151719564 | -0.251816754 | 0.544522288 | 0.786483921 |
| chr4:88593021-88593552 | 0.202827765 | 0.544529926 | 0.786483921 |
| chr9:109401335-109401851 | 0.223170705 | 0.544543695 | 0.786488982 |
| chrX:120857175-120857712 | -0.225408635 | 0.544572536 | 0.786511147 |
| chr1:205459000-205459269 | -0.226466375 | 0.544579924 | 0.786511147 |
| chr19:10896671-10897169 | 0.219319896 | 0.544596711 | 0.786511147 |
| chr9:76303031-76303254 | -0.317028176 | 0.544600101 | 0.786511147 |
| chr5:34656088-34657060 | 0.203008257 | 0.544640923 | 0.786544029 |
| chr1:37692119-37692776 | 0.198780641 | 0.544645401 | 0.786544029 |
| chr8:87175663-87175959 | 0.26781781 | 0.544653665 | 0.786544029 |
| chr3:47282382-47283067 | -0.206304763 | 0.544673329 | 0.786544953 |
| chr1:91500493-91501629 | -0.196617269 | 0.544674835 | 0.786544953 |
| chr16:88264946-88265270 | 0.250901939 | 0.544688052 | 0.786549215 |
| chr7:99365607-99365893 | -0.251646786 | 0.544707015 | 0.786550384 |
| chr3:151316673-151317044 | -0.232094359 | 0.544715622 | 0.786550384 |
| chr6:32838117-32838990 | 0.205941042 | 0.544753469 | 0.786550384 |
| chr5:34673831-34674036 | 0.299691697 | 0.544761677 | 0.786550384 |
| chr10:90723720-90724081 | -0.269861119 | 0.544762413 | 0.786550384 |
| chr6:105087683-105087883 | 0.312701847 | 0.544766171 | 0.786550384 |
| chr2:118881430-118881918 | 0.209238645 | 0.544769192 | 0.786550384 |
| chr12:47097337-47097554 | -0.33973787 | 0.544770984 | 0.786550384 |
| chr7:43042765-43043030 | -0.315827941 | 0.54478935 | 0.786562079 |
| chr5:42424355-42425076 | 0.205493155 | 0.544813671 | 0.786582373 |
| chr13:68927436-68927739 | -0.255605415 | 0.544825003 | 0.786583912 |
| chr8:21856266-21856471 | -0.333099026 | 0.544862075 | 0.786622612 |
| chr1:113042713-113043016 | -0.257250173 | 0.544875933 | 0.786627797 |
| chr1:209435683-209436040 | -0.225012005 | 0.544930579 | 0.786670177 |
| chr6:75349640-75349849 | -0.334609816 | 0.544941815 | 0.786670177 |
| chr6:155177428-155177650 | -0.334609816 | 0.544941815 | 0.786670177 |
| chr2:240560621-240561519 | 0.195401207 | 0.544966291 | 0.786670177 |
| chr22:22110961-22111240 | 0.256004005 | 0.544972437 | 0.786670177 |
| chr14:75230248-75230791 | 0.222595145 | 0.54498448 | 0.786670177 |
| chr4:84167644-84167897 | 0.296145283 | 0.544997196 | 0.786670177 |
| chr11:38250761-38250961 | 0.322310659 | 0.545001288 | 0.786670177 |
| chr12:74537272-74538426 | -0.192221388 | 0.545019916 | 0.786670177 |
| chr10:11260261-11260537 | -0.263084151 | 0.545026454 | 0.786670177 |
| chr6:154995185-154995470 | 0.259021497 | 0.545027706 | 0.786670177 |
| chr10:73785125-73785775 | -0.200392956 | 0.545029043 | 0.786670177 |
| chr6:35544725-35545353 | -0.23493758 | 0.54504122 | 0.786670177 |
| chr6:73733122-73733447 | -0.226555059 | 0.545049025 | 0.786670177 |
| chr1:23554798-23555234 | 0.213775635 | 0.545066987 | 0.786679293 |
| chr19:32343669-32343878 | -0.26268675 | 0.545075874 | 0.786679293 |
| chrX:150772204-150772514 | 0.2396943 | 0.545109257 | 0.786696801 |
| chr6:30720053-30720427 | 0.212797757 | 0.545112051 | 0.786696801 |
| chr9:125745929-125746900 | 0.194819546 | 0.545138723 | 0.786696801 |
| chr1:157170850-157171060 | -0.272172377 | 0.545142423 | 0.786696801 |
| chr9:122827990-122828998 | 0.197788141 | 0.545155309 | 0.786696801 |
| chr19:49478766-49479206 | 0.214527007 | 0.545176251 | 0.786696801 |
| chr3:130106030-130106351 | -0.227996289 | 0.545176848 | 0.786696801 |
| chr19:40798173-40798373 | 0.249575344 | 0.545183783 | 0.786696801 |

| | | | |
|---|---|---|---|
| chr10:16817077-16818004 | -0.191746424 | 0.545188799 | 0.786696801 |
| chr5:140151792-140151992 | -0.278532596 | 0.545190678 | 0.786696801 |
| chr15:82124745-82124952 | 0.31007566 | 0.545240989 | 0.786754582 |
| chr22:37260903-37261201 | 0.295924539 | 0.545269581 | 0.786770069 |
| chrX:147804030-147804230 | 0.323700384 | 0.545307605 | 0.786770069 |
| chr8:87342804-87343079 | -0.263896583 | 0.545310217 | 0.786770069 |
| chr1:43934833-43935085 | -0.242659152 | 0.545320076 | 0.786770069 |
| chr13:98430914-98431471 | 0.216169391 | 0.54532106 | 0.786770069 |
| chr8:127478822-127479046 | 0.284806141 | 0.5453224 | 0.786770069 |
| chr2:62423267-62423553 | -0.250263138 | 0.545323599 | 0.786770069 |
| chr3:129490249-129490524 | 0.225099871 | 0.545337904 | 0.786771365 |
| chr21:45387042-45387242 | 0.275711728 | 0.545345034 | 0.786771365 |
| chr2:170816755-170817206 | 0.225264107 | 0.545358152 | 0.786775476 |
| chr6:31307891-31308356 | 0.211827703 | 0.545373751 | 0.786783167 |
| chr17:68221774-68222152 | 0.228379649 | 0.545418404 | 0.786832771 |
| chr5:150409808-150410309 | -0.209928494 | 0.545470838 | 0.786893597 |
| chr15:57524223-57524497 | -0.304544473 | 0.545497186 | 0.786894737 |
| chr1:202213462-202213662 | -0.252750246 | 0.545499366 | 0.786894737 |
| chr5:25329557-25329787 | -0.301779855 | 0.545502437 | 0.786894737 |
| chr8:70906079-70906323 | 0.28579414 | 0.545531644 | 0.786922053 |
| chr19:7619271-7619756 | 0.224589973 | 0.545550734 | 0.786933332 |
| chr20:18058692-18059349 | -0.203107133 | 0.54557509 | 0.786933332 |
| chr11:120247749-120247979 | -0.335233604 | 0.54558293 | 0.786933332 |
| chr1:148353373-148354042 | -0.195693109 | 0.545589898 | 0.786933332 |
| chr1:235417163-235417452 | 0.249414618 | 0.545605872 | 0.786933332 |
| chr1:48743906-48744332 | -0.258082953 | 0.54561447 | 0.786933332 |
| chr12:110100257-110100726 | -0.263891159 | 0.545622902 | 0.786933332 |
| chr2:25041244-25042496 | -0.192834133 | 0.54562833 | 0.786933332 |
| chr2:187054565-187054765 | 0.314551312 | 0.545637404 | 0.786933332 |
| chr7:45073679-45074131 | -0.215154114 | 0.545642166 | 0.786933332 |
| chr17:56809776-56810002 | 0.256821504 | 0.545684006 | 0.786978861 |
| chr13:38915473-38916046 | -0.214013191 | 0.545697269 | 0.786982684 |
| chr17:60126764-60129087 | 0.1880917 | 0.545707199 | 0.786982684 |
| chr13:103140115-103140503 | -0.294850221 | 0.545720275 | 0.786986729 |
| chr3:157088551-157089672 | -0.191071611 | 0.54575425 | 0.787020912 |
| chr14:20625204-20625782 | -0.196539898 | 0.545829973 | 0.787109952 |
| chr2:3785325-3785639 | -0.249923224 | 0.545836539 | 0.787109952 |
| chr17:35578811-35579045 | 0.318419134 | 0.545863902 | 0.787120278 |
| chr13:47164455-47164684 | 0.274131056 | 0.545864245 | 0.787120278 |
| chr6:89369057-89369290 | -0.267848688 | 0.545891511 | 0.787144781 |
| chr14:93984982-93985304 | 0.23493895 | 0.545915355 | 0.787164349 |
| chr13:98556929-98557540 | 0.23074614 | 0.545932171 | 0.787173782 |
| chr14:20989589-20990578 | 0.195074637 | 0.545946086 | 0.787179032 |
| chr13:43565420-43565646 | 0.317392021 | 0.545957882 | 0.787181228 |
| chr14:41280798-41281087 | 0.254516194 | 0.54598696 | 0.787208341 |
| chr4:158176175-158176603 | 0.226945177 | 0.54602192 | 0.787243082 |
| chr3:185280821-185281021 | 0.305941533 | 0.546031604 | 0.787243082 |
| chr7:17021214-17021535 | 0.2422738 | 0.546056067 | 0.787263537 |
| chr4:150215386-150216058 | -0.201561928 | 0.546076954 | 0.787278838 |
| chr12:46984027-46984283 | -0.276732892 | 0.546092625 | 0.787286617 |

| | | | |
|---|---|---|---|
| chr2:128321821-128322169 | 0.217082653 | 0.546118677 | 0.787309362 |
| chr17:28233927-28234271 | 0.227976432 | 0.546140816 | 0.787326464 |
| chr8:91578096-91578381 | 0.268490439 | 0.546157473 | 0.787334327 |
| chr3:5974903-5975112 | 0.323447311 | 0.546166821 | 0.787334327 |
| chr1:204213615-204214642 | -0.192013653 | 0.546203206 | 0.787335775 |
| chr6:100257968-100258187 | -0.307820843 | 0.546205483 | 0.787335775 |
| chr10:71486510-71486843 | 0.231074713 | 0.546207433 | 0.787335775 |
| chr3:137409550-137409788 | -0.301252114 | 0.546208928 | 0.787335775 |
| chr17:57955164-57955935 | -0.199516766 | 0.546245025 | 0.78735518 |
| chr5:156552336-156552653 | -0.29266266 | 0.546261526 | 0.78735518 |
| chr10:110210708-110210931 | 0.246836351 | 0.546278932 | 0.78735518 |
| chr1:166841065-166841332 | -0.23521783 | 0.546279089 | 0.78735518 |
| chr11:75867833-75868064 | -0.304319087 | 0.546291576 | 0.78735518 |
| chr17:73970167-73970367 | 0.296746544 | 0.546292659 | 0.78735518 |
| chr4:25107843-25108326 | -0.236717807 | 0.54629432 | 0.78735518 |
| chr1:30929977-30930205 | 0.295708536 | 0.546336283 | 0.787399205 |
| chr4:101089298-101089542 | 0.307283789 | 0.546354202 | 0.787399205 |
| chr3:18100543-18100754 | 0.340237027 | 0.546355696 | 0.787399205 |
| chr5:21145565-21145765 | 0.298454506 | 0.546386006 | 0.787428076 |
| chr8:103073887-103074338 | 0.217939774 | 0.546401561 | 0.787432462 |
| chr9:121121484-121122012 | -0.205780269 | 0.546409603 | 0.787432462 |
| chr12:46261131-46261441 | 0.221483857 | 0.546421619 | 0.787434969 |
| chr4:1166974-1168062 | 0.202074719 | 0.546446532 | 0.78745606 |
| chr1:925594-925810 | 0.269783552 | 0.546483489 | 0.787494506 |
| chr1:201740938-201741139 | -0.240366491 | 0.546496845 | 0.787498395 |
| chr20:431438-431771 | 0.238601675 | 0.546506743 | 0.787498395 |
| chr16:69132093-69133084 | 0.200163833 | 0.546536903 | 0.787527044 |
| chr20:47389758-47390471 | 0.190134879 | 0.546547238 | 0.787527126 |
| chr10:8856831-8857227 | 0.235498927 | 0.546568617 | 0.787535694 |
| chr14:67618673-67618925 | -0.231739878 | 0.546573741 | 0.787535694 |
| chr2:190615543-190615774 | 0.283884647 | 0.546604289 | 0.787555321 |
| chr3:111745923-111746262 | 0.277866723 | 0.546607919 | 0.787555321 |
| chr1:39738628-39739356 | -0.207581304 | 0.546621461 | 0.787560023 |
| chr10:26860502-26861424 | -0.191310077 | 0.546633387 | 0.787562397 |
| chr17:7329219-7329862 | -0.202737845 | 0.546646274 | 0.787566155 |
| chrX:50204290-50204932 | -0.212391744 | 0.546714842 | 0.787650133 |
| chr16:46883801-46884432 | 0.200675706 | 0.546762498 | 0.78765392 |
| chr10:92547731-92547931 | 0.301453821 | 0.546766911 | 0.78765392 |
| chr14:33357893-33358094 | 0.301453821 | 0.546766911 | 0.78765392 |
| chr18:74526560-74526760 | 0.301453821 | 0.546766911 | 0.78765392 |
| chr1:193452476-193452676 | -0.320803309 | 0.546773645 | 0.78765392 |
| chr4:108762553-108763276 | 0.207817288 | 0.546785958 | 0.78765392 |
| chr6:116877066-116877539 | -0.233012274 | 0.546789429 | 0.78765392 |
| chr15:78308510-78308710 | 0.257850323 | 0.546813124 | 0.787673243 |
| chr4:130808583-130808845 | -0.257938856 | 0.546824204 | 0.787674396 |
| chr13:113489984-113490252 | -0.2490514 | 0.546851534 | 0.787680442 |
| chr17:58113027-58113357 | 0.216422189 | 0.546856599 | 0.787680442 |
| chr5:151757310-151757855 | 0.205168748 | 0.546859241 | 0.787680442 |
| chr9:27529352-27530040 | 0.215843675 | 0.546894709 | 0.787704105 |
| chr3:138417529-138417956 | 0.212601865 | 0.546900499 | 0.787704105 |

| | | | |
|---|---|---|---|
| chr7:43536616-43536973 | -0.263764854 | 0.546906511 | 0.787704105 |
| chr2:129526901-129527151 | -0.302752694 | 0.546950033 | 0.787751982 |
| chr11:95231054-95231728 | -0.193510025 | 0.546963952 | 0.787757222 |
| chr8:57343414-57343688 | 0.249509566 | 0.54697843 | 0.787763266 |
| chr4:99655355-99655576 | -0.313957834 | 0.546995126 | 0.787772504 |
| chr8:106909004-106909209 | 0.291524706 | 0.547070477 | 0.787866214 |
| chr14:105490613-105491432 | 0.190437556 | 0.547089632 | 0.787878993 |
| chr6:131949907-131950227 | -0.221029957 | 0.547115095 | 0.787896938 |
| chr15:91480525-91480725 | 0.297704175 | 0.547123907 | 0.787896938 |
| chr6:53746515-53746736 | 0.286033271 | 0.547132942 | 0.787896938 |
| chr12:47870546-47870848 | -0.255087067 | 0.54714351 | 0.787897048 |
| chr6:52861234-52861434 | -0.331060836 | 0.54715742 | 0.787897048 |
| chr1:47192999-47193199 | 0.275399179 | 0.547163867 | 0.787897048 |
| chr12:62813187-62813552 | 0.23298329 | 0.547206135 | 0.787943105 |
| chr1:40899106-40899904 | 0.20096233 | 0.547229033 | 0.787961268 |
| chr5:22467093-22467293 | 0.323282471 | 0.547252847 | 0.78798075 |
| chr19:45922426-45922847 | -0.250434869 | 0.547270688 | 0.787991631 |
| chr1:211830492-211831118 | -0.21952697 | 0.547296108 | 0.787996706 |
| chr10:125624889-12562515€ | -0.285591204 | 0.547300804 | 0.787996706 |
| chr12:58855672-58855872 | 0.288521454 | 0.547305065 | 0.787996706 |
| chr8:38467848-38468333 | 0.234117116 | 0.547358921 | 0.788059438 |
| chr17:2303009-2304311 | -0.195751731 | 0.547382239 | 0.788078202 |
| chr8:124888768-124889111 | 0.206165295 | 0.547440607 | 0.788147426 |
| chr8:17800833-17802009 | 0.198478032 | 0.547503352 | 0.78822295 |
| chr14:94636665-94636965 | -0.256110562 | 0.547516252 | 0.788226712 |
| chr15:98003544-98003882 | 0.254311291 | 0.547555832 | 0.788259116 |
| chr10:27558545-27558828 | 0.264150926 | 0.547559336 | 0.788259116 |
| chr10:3289889-3290156 | 0.258777191 | 0.547591152 | 0.788290107 |
| chr19:40823308-40823578 | -0.305096375 | 0.547678984 | 0.788401734 |
| chr16:30761959-30762615 | 0.196832316 | 0.547737294 | 0.78847086 |
| chr11:33997005-33997330 | 0.22727069 | 0.547782404 | 0.788509323 |
| chr1:204380425-204380793 | -0.208669172 | 0.547784596 | 0.788509323 |
| chr7:129713538-129713756 | -0.311707498 | 0.547799882 | 0.788513566 |
| chr14:29636240-29636512 | 0.262425665 | 0.547812323 | 0.788513566 |
| chr14:42308770-42309006 | -0.296542477 | 0.547818416 | 0.788513566 |
| chr5:78269225-78269474 | -0.311728435 | 0.54784937 | 0.788543308 |
| chr1:23562364-23562683 | 0.237342226 | 0.547879266 | 0.788571524 |
| chr2:135904504-135904829 | 0.270706104 | 0.547939643 | 0.788643232 |
| chr2:218876683-218876982 | -0.239321446 | 0.547954768 | 0.788643232 |
| chr10:97072888-97073092 | -0.266994832 | 0.547959964 | 0.788643232 |
| chr17:7204900-7205265 | 0.242018019 | 0.547979854 | 0.788657044 |
| chr17:43389187-43389398 | 0.248860826 | 0.548008639 | 0.788683657 |
| chr3:72875349-72875812 | -0.237213655 | 0.548027221 | 0.788691756 |
| chr1:106217871-106218154 | -0.215975962 | 0.548041739 | 0.788691756 |
| chr8:143951732-143953184 | -0.190990769 | 0.548045146 | 0.788691756 |
| chr1:179312717-179312966 | 0.291361655 | 0.548187078 | 0.788857145 |
| chr3:176403503-176403911 | -0.225697841 | 0.548188412 | 0.788857145 |
| chr21:33361689-33361987 | -0.276767201 | 0.548194961 | 0.788857145 |
| chr3:191136396-191136674 | -0.320957457 | 0.548205587 | 0.788857145 |
| chr3:103794849-103795098 | -0.299572667 | 0.548223362 | 0.788857145 |

| | | | |
|---|---|---|---|
| chr2:66201314-66201576 | -0.297082982 | 0.54823435 | 0.788857145 |
| chr12:52895322-52896475 | -0.190992977 | 0.54823455 | 0.788857145 |
| chr16:88510354-88510855 | 0.208696449 | 0.548242434 | 0.788857145 |
| chr17:8288437-8288792 | -0.230329999 | 0.548253584 | 0.788858373 |
| chr1:161020589-161021635 | 0.190006459 | 0.548273563 | 0.788872307 |
| chr12:120290763-120293318 | -0.187737532 | 0.548301597 | 0.788890497 |
| chr12:111753247-111753502 | 0.250219508 | 0.548316994 | 0.788890497 |
| chr21:42833500-42833700 | -0.250965641 | 0.548317093 | 0.788890497 |
| chr1:165871458-165871780 | -0.262655584 | 0.548346567 | 0.788908278 |
| chr20:58980882-58981759 | 0.19006774 | 0.548359087 | 0.788908278 |
| chr14:97584569-97584831 | -0.280858096 | 0.54836034 | 0.788908278 |
| chr14:71215439-71215651 | -0.296112325 | 0.548402123 | 0.788953576 |
| chr10:89224430-89224699 | -0.297460222 | 0.548414138 | 0.788956048 |
| chr4:74675773-74676002 | -0.264350025 | 0.548483191 | 0.789018046 |
| chr3:13533830-13534261 | -0.229898181 | 0.548483373 | 0.789018046 |
| chr8:22587758-22588006 | 0.277642466 | 0.548488126 | 0.789018046 |
| chr21:39451547-39452067 | -0.209151059 | 0.548511086 | 0.789029208 |
| chr3:169668986-169669529 | 0.200579691 | 0.548517626 | 0.789029208 |
| chr6:138403317-138405148 | -0.190126348 | 0.54853512 | 0.789029208 |
| chrX:149570663-149571072 | 0.222391863 | 0.548537076 | 0.789029208 |
| chr11:77127181-77127611 | 0.219496625 | 0.548547754 | 0.789029755 |
| chr14:75640915-75641153 | -0.280216041 | 0.548576369 | 0.789056102 |
| chr2:104758396-104758693 | -0.264903388 | 0.548687296 | 0.789197186 |
| chr8:109848353-109848553 | 0.301897526 | 0.548695055 | 0.789197186 |
| chr16:20300284-20300774 | 0.221206652 | 0.548715884 | 0.789212331 |
| chr22:24532049-24532365 | 0.232398764 | 0.548740115 | 0.789228576 |
| chr16:18926219-18927161 | 0.194280559 | 0.54874778 | 0.789228576 |
| chr1:197902311-197903349 | -0.189637739 | 0.548765998 | 0.789231623 |
| chr1:115608220-115608489 | -0.23898661 | 0.548775528 | 0.789231623 |
| chr5:149035386-149035586 | 0.305517983 | 0.548780799 | 0.789231623 |
| chr17:55501605-55502037 | 0.236006566 | 0.548850739 | 0.789292186 |
| chr8:65794634-65795056 | -0.287497839 | 0.548851827 | 0.789292186 |
| chr12:113547418-113547660 | -0.289482715 | 0.548853814 | 0.789292186 |
| chr17:59845546-59846149 | -0.188596561 | 0.548869972 | 0.789300608 |
| chr1:115906702-115907067 | 0.239453696 | 0.548887973 | 0.789311681 |
| chr15:90002318-90002643 | -0.233887895 | 0.548905599 | 0.789316266 |
| chr9:105413611-105413879 | 0.275035876 | 0.548911765 | 0.789316266 |
| chr7:33018638-33018838 | 0.323949413 | 0.548953914 | 0.789347248 |
| chr7:37516776-37516976 | 0.323949413 | 0.548953914 | 0.789347248 |
| chr2:30165046-30165360 | -0.237796454 | 0.549001107 | 0.789400293 |
| chr3:186760385-186760645 | 0.266352385 | 0.549032955 | 0.789424788 |
| chr12:64256264-64256501 | 0.287276975 | 0.549038905 | 0.789424788 |
| chr8:8227600-8228684 | -0.19170572 | 0.549049051 | 0.789424788 |
| chr4:129362745-129362989 | -0.283297827 | 0.549083406 | 0.78945937 |
| chr2:38875291-38876658 | -0.191861857 | 0.549130884 | 0.789507867 |
| chrX:54182805-54183790 | -0.19844976 | 0.549160449 | 0.789507867 |
| chr2:74507208-74508489 | -0.189726742 | 0.549173028 | 0.789507867 |
| chr17:73231879-73233445 | -0.189808466 | 0.549200106 | 0.789507867 |
| chr4:28728776-28729080 | -0.277051863 | 0.549205438 | 0.789507867 |
| chr14:60302920-60303120 | -0.31229314 | 0.549208333 | 0.789507867 |

| | | | |
|---|---|---|---|
| chr11:134033963-134034353 | 0.215524127 | 0.549217841 | 0.789507867 |
| chr20:32337000-32337215 | 0.325605803 | 0.549220069 | 0.789507867 |
| chr5:51033202-51033402 | 0.325605803 | 0.549220069 | 0.789507867 |
| chr9:105426110-105426375 | -0.294271839 | 0.549220176 | 0.789507867 |
| chr9:126752442-126752743 | 0.256322349 | 0.549231332 | 0.789509092 |
| chr15:48963865-48964070 | -0.282689953 | 0.54924952 | 0.789520425 |
| chr5:1479614-1479942 | 0.227595185 | 0.549265122 | 0.78952804 |
| chr16:273575-273935 | -0.232945073 | 0.549291975 | 0.789551827 |
| chr8:96405013-96405282 | -0.270874683 | 0.549376369 | 0.78964963 |
| chr22:22959205-22959442 | 0.274893069 | 0.54938465 | 0.78964963 |
| chr14:52823527-52823847 | 0.286879561 | 0.549390934 | 0.78964963 |
| chr2:215002624-215002824 | -0.329219674 | 0.549450449 | 0.789714405 |
| chr10:137070-137407 | -0.220191485 | 0.549456614 | 0.789714405 |
| chr5:77852514-77852864 | -0.253840657 | 0.549476522 | 0.789728205 |
| chrX:71532746-71533395 | 0.204894416 | 0.549488037 | 0.789729942 |
| chr9:104863296-104863655 | -0.253292229 | 0.549531965 | 0.789776336 |
| chr5:173315527-173315935 | -0.220579556 | 0.549540933 | 0.789776336 |
| chr3:155869916-155871903 | -0.186547816 | 0.549552356 | 0.789777939 |
| chr3:42263331-42263710 | 0.252319787 | 0.549565858 | 0.789778744 |
| chr20:57391119-57391508 | 0.201005723 | 0.549573531 | 0.789778744 |
| chr4:144597288-144597516 | -0.321511064 | 0.549594988 | 0.789794767 |
| chr10:79929443-79929722 | 0.250453635 | 0.549630126 | 0.78980463 |
| chr17:72986108-72986705 | 0.20935088 | 0.54963173 | 0.78980463 |
| chr14:69625449-69625788 | -0.266649388 | 0.549632775 | 0.78980463 |
| chr1:155562549-155563419 | 0.194528554 | 0.549644973 | 0.789807346 |
| chr3:9864419-9864625 | 0.288321344 | 0.549678331 | 0.789831526 |
| chr5:128349314-128349514 | -0.295088933 | 0.549682417 | 0.789831526 |
| chr7:42152506-42152955 | -0.22775723 | 0.549703654 | 0.78984723 |
| chr13:46753313-46753574 | -0.245895211 | 0.549722424 | 0.789859388 |
| chr17:18261235-18261599 | -0.209443339 | 0.549750835 | 0.789884233 |
| chr15:47471567-47471801 | 0.263322917 | 0.549768224 | 0.789884233 |
| chr15:64521732-64522137 | -0.255471878 | 0.549770642 | 0.789884233 |
| chr6:109251212-109251544 | 0.248287632 | 0.549805729 | 0.789919832 |
| chr1:223992416-223992887 | -0.20895529 | 0.549835285 | 0.789945836 |
| chr5:140710507-140711575 | -0.191139196 | 0.549844448 | 0.789945836 |
| chr19:30801244-30801444 | -0.341595034 | 0.54989041 | 0.789997056 |
| chr2:42829147-42829371 | 0.268686532 | 0.549933507 | 0.790035284 |
| chr3:128174480-128174763 | -0.291577553 | 0.549937641 | 0.790035284 |
| chr6:149545210-149546809 | -0.188784952 | 0.549952954 | 0.790037818 |
| chr6:3542054-3542474 | -0.247329994 | 0.549960027 | 0.790037818 |
| chr9:136893469-136893770 | -0.291529387 | 0.549999675 | 0.790079143 |
| chr11:110405642-110405971 | 0.241006132 | 0.550011405 | 0.790079143 |
| chr9:125143242-125144098 | 0.195128159 | 0.550019728 | 0.790079143 |
| chr10:118241283-118241585 | 0.250368025 | 0.550045875 | 0.79010189 |
| chr8:142427846-142428054 | 0.299047831 | 0.550075549 | 0.790113609 |
| chr8:30803404-30803615 | 0.310392505 | 0.550076417 | 0.790113609 |
| chr6:31203303-31203687 | -0.243013486 | 0.55010175 | 0.790113609 |
| chr10:101835978-101836299 | -0.229818345 | 0.550104741 | 0.790113609 |
| chr2:227406079-227406829 | 0.209865813 | 0.550105592 | 0.790113609 |
| chr15:65184859-65185949 | 0.191662625 | 0.550192061 | 0.790196488 |

| | | | |
|---|---|---|---|
| chr7:99504362-99505368 | -0.192238513 | 0.550194155 | 0.790196488 |
| chr6:49097326-49098115 | -0.201146356 | 0.550194235 | 0.790196488 |
| chr5:44925631-44926054 | -0.265501964 | 0.550234803 | 0.79023994 |
| chr5:151164666-151165108 | -0.231680945 | 0.5502522 | 0.790250114 |
| chr19:680364-681208 | 0.193234329 | 0.550335891 | 0.790343934 |
| chr17:51167084-51167375 | 0.21097012 | 0.550341659 | 0.790343934 |
| chr7:115038125-115038325 | -0.31603338 | 0.550360263 | 0.790343934 |
| chr2:162123673-162124144 | -0.242793821 | 0.550362768 | 0.790343934 |
| chr19:4861106-4861617 | 0.226033699 | 0.550376319 | 0.790343934 |
| chr2:207071852-207072052 | 0.317158864 | 0.550379416 | 0.790343934 |
| chr22:50675081-50675303 | 0.298774514 | 0.550461845 | 0.790444891 |
| chr8:85865478-85866013 | -0.217019009 | 0.550491463 | 0.790444891 |
| chr14:100201740-100202027 | -0.253422144 | 0.550494264 | 0.790444891 |
| chr3:63351200-63351510 | -0.205993455 | 0.550500307 | 0.790444891 |
| chr1:53572433-53573210 | 0.207260949 | 0.550501301 | 0.790444891 |
| chr19:13116589-13116935 | 0.247191979 | 0.550537469 | 0.790467709 |
| chr17:78254482-78254752 | 0.23221625 | 0.550543904 | 0.790467709 |
| chr1:109249358-109249596 | -0.240606413 | 0.550554644 | 0.790467709 |
| chr5:170211900-170212114 | -0.293127666 | 0.550565216 | 0.790467709 |
| chr1:200669273-200670629 | -0.186056878 | 0.550568775 | 0.790467709 |
| chr21:42892720-42893826 | -0.191738598 | 0.55059345 | 0.790481498 |
| chr8:123181924-123182874 | -0.191444454 | 0.550601911 | 0.790481498 |
| chr8:48723607-48723807 | -0.320800154 | 0.550614301 | 0.790481498 |
| chr21:38172082-38172376 | 0.23991477 | 0.550619646 | 0.790481498 |
| chr17:1259217-1259417 | -0.301494221 | 0.550640132 | 0.790496098 |
| chrX:65528795-65529031 | 0.27188178 | 0.550657374 | 0.790506039 |
| chr1:229093629-229093917 | 0.244262244 | 0.550722295 | 0.790569599 |
| chr12:94710991-94711208 | -0.305292587 | 0.55073106 | 0.790569599 |
| chrX:74924392-74925620 | -0.198997309 | 0.550741088 | 0.790569599 |
| chr2:207529001-207530368 | -0.187886201 | 0.550742921 | 0.790569599 |
| chr5:73470558-73470804 | -0.328564495 | 0.550776618 | 0.790603159 |
| chr7:56079317-56079793 | 0.21844573 | 0.550798957 | 0.790620414 |
| chr9:35756562-35757689 | 0.197925562 | 0.55085796 | 0.790682191 |
| chr3:10025936-10026933 | 0.197609307 | 0.550862634 | 0.790682191 |
| chr1:149712504-149712946 | -0.208412774 | 0.550876461 | 0.790687226 |
| chr5:684714-685335 | -0.24149706 | 0.550918779 | 0.790718518 |
| chr6:170295815-170296122 | 0.239259409 | 0.550918902 | 0.790718518 |
| chr19:588646-588865 | 0.261845178 | 0.550959952 | 0.790735467 |
| chr10:130190107-130190720 | 0.201232035 | 0.550997513 | 0.790735467 |
| chr8:105781833-105782062 | 0.254702446 | 0.550998826 | 0.790735467 |
| chr13:59636367-59636670 | 0.314594176 | 0.551016284 | 0.790735467 |
| chrX:16445081-16445493 | -0.258827532 | 0.551025087 | 0.790735467 |
| chr12:71754352-71755693 | -0.187837961 | 0.551027602 | 0.790735467 |
| chr17:58595257-58595830 | -0.192185403 | 0.551039809 | 0.790735467 |
| chr12:47079466-47079897 | -0.227298146 | 0.55104977 | 0.790735467 |
| chr18:80033240-80034796 | -0.187945036 | 0.55105195 | 0.790735467 |
| chr2:208466363-208466714 | -0.252352336 | 0.551054221 | 0.790735467 |
| chr19:36213995-36215385 | 0.18976593 | 0.551055629 | 0.790735467 |
| chr1:109689025-109689232 | 0.263354292 | 0.551074386 | 0.790735467 |
| chr7:37901342-37901759 | -0.235729188 | 0.551074726 | 0.790735467 |

| | | | |
|---|---|---|---|
| chr1:109075659-109076341 | 0.196375801 | 0.55107519 | 0.790735467 |
| chr4:99950139-99950970 | -0.189328329 | 0.551087209 | 0.790737905 |
| chr2:88452380-88452872 | 0.223843673 | 0.551132906 | 0.790788666 |
| chr1:54887648-54887848 | -0.249629354 | 0.55114337 | 0.790788871 |
| chr3:34035827-34036402 | -0.223025239 | 0.5512043 | 0.790861485 |
| chr16:66961075-66961437 | -0.242183847 | 0.551228421 | 0.790881284 |
| chr17:7046217-7046683 | 0.220157916 | 0.551262865 | 0.790915892 |
| chr15:87958978-87959442 | 0.218629097 | 0.551283596 | 0.790930826 |
| chr12:6890780-6891562 | 0.197235154 | 0.551324784 | 0.790941639 |
| chr2:177161754-177161965 | 0.287002219 | 0.551331869 | 0.790941639 |
| chr4:65223669-65223885 | 0.287012993 | 0.551335254 | 0.790941639 |
| chr16:58392218-58392623 | 0.224580406 | 0.551341919 | 0.790941639 |
| chr11:70450830-70451096 | 0.277200678 | 0.551363232 | 0.790941639 |
| chr8:141258253-141258511 | -0.266534218 | 0.551368396 | 0.790941639 |
| chr6:27188105-27189188 | -0.205091207 | 0.551369172 | 0.790941639 |
| chr1:2526514-2526800 | 0.244340023 | 0.551391911 | 0.790941639 |
| chr11:125569101-125570016 | 0.198424121 | 0.551393896 | 0.790941639 |
| chr5:76755018-76755249 | -0.296405058 | 0.55140566 | 0.790941639 |
| chr16:84738318-84738990 | -0.202704328 | 0.551427183 | 0.790941639 |
| chr2:70509575-70509775 | -0.297295028 | 0.551429547 | 0.790941639 |
| chr17:52827656-52828084 | -0.235783231 | 0.551439623 | 0.790941639 |
| chr14:61837619-61838015 | 0.218866801 | 0.551441973 | 0.790941639 |
| chr9:37785491-37785794 | -0.259239957 | 0.551445972 | 0.790941639 |
| chr10:49762076-49762530 | -0.215770169 | 0.551462281 | 0.790947323 |
| chr10:116597655-116597879 | 0.288308106 | 0.551476644 | 0.790947323 |
| chr6:63635374-63636738 | 0.189901401 | 0.551480904 | 0.790947323 |
| chr11:130159559-130160394 | 0.201225777 | 0.551500514 | 0.790948696 |
| chr11:70399541-70400058 | 0.208947651 | 0.551511434 | 0.790948696 |
| chr4:169079874-169080634 | -0.208856005 | 0.551522737 | 0.790948696 |
| chr7:142297999-142298272 | -0.256241334 | 0.551523151 | 0.790948696 |
| chr11:46557508-46557708 | 0.307480871 | 0.551542794 | 0.790956514 |
| chr8:110419635-110419856 | 0.335399723 | 0.551549249 | 0.790956514 |
| chr7:76262993-76263275 | -0.232635381 | 0.5515766 | 0.7909586 |
| chr1:178724761-178726598 | 0.189566224 | 0.551580527 | 0.7909586 |
| chr1:224922374-224922941 | 0.213367511 | 0.551594666 | 0.7909586 |
| chr1:147640750-147641197 | -0.201435771 | 0.55159589 | 0.7909586 |
| chr2:135926656-135926941 | 0.252827304 | 0.551618627 | 0.7909586 |
| chr11:80720285-80720656 | 0.219424334 | 0.551621927 | 0.7909586 |
| chr6:30570797-30570997 | 0.260753532 | 0.551624716 | 0.7909586 |
| chr7:147196997-147197346 | -0.230893019 | 0.551634843 | 0.7909586 |
| chr1:88061592-88061808 | -0.311971165 | 0.551643609 | 0.7909586 |
| chr2:217899555-217899903 | 0.214601695 | 0.55166069 | 0.790968289 |
| chr12:98725901-98726178 | 0.258305481 | 0.551716207 | 0.791033087 |
| chr19:35124384-35124873 | 0.212394006 | 0.55173125 | 0.791036697 |
| chr1:186330876-186331128 | -0.259834462 | 0.551749838 | 0.791036697 |
| chr7:87153088-87153343 | -0.22911347 | 0.551751903 | 0.791036697 |
| chr19:53394915-53395818 | 0.203064655 | 0.551762953 | 0.791036697 |
| chr18:10695122-10695455 | -0.246071055 | 0.551770344 | 0.791036697 |
| chr11:564153-564514 | 0.22322587 | 0.551825675 | 0.791101219 |
| chr2:219530165-219530515 | 0.218663122 | 0.551853302 | 0.791126023 |

| | | | |
|---|---|---|---|
| chr18:48222159-48222472 | -0.254651537 | 0.551886267 | 0.791151678 |
| chr15:33700713-33701208 | -0.214886322 | 0.551891848 | 0.791151678 |
| chr6:109978201-109978857 | 0.218458265 | 0.551948004 | 0.791217375 |
| chr6:89432853-89433465 | -0.20508266 | 0.552014745 | 0.791298244 |
| chr8:140321470-140321936 | 0.234216831 | 0.55204562 | 0.791327699 |
| chr18:12990914-12991863 | 0.193339753 | 0.552137316 | 0.791402342 |
| chr16:8797255-8798323 | 0.191845329 | 0.55214683 | 0.791402342 |
| chr16:81581191-81581624 | 0.206209187 | 0.552147386 | 0.791402342 |
| chr3:183447223-183448035 | -0.206661824 | 0.552153294 | 0.791402342 |
| chr10:5583555-5583893 | 0.241988566 | 0.552156629 | 0.791402342 |
| chr21:40291066-40291402 | -0.278538278 | 0.552159664 | 0.791402342 |
| chr11:15276709-15276909 | 0.348439238 | 0.552195617 | 0.791424264 |
| chr13:37410095-37410295 | 0.348439238 | 0.552195617 | 0.791424264 |
| chr4:37390026-37390337 | -0.246413066 | 0.552213035 | 0.791425886 |
| chr18:46097856-46098723 | -0.20125794 | 0.552220273 | 0.791425886 |
| chr1:85109424-85109727 | -0.25523011 | 0.552227736 | 0.791425886 |
| chr4:164753947-164754147 | 0.294023271 | 0.552243915 | 0.791429981 |
| chr10:77533585-77534273 | -0.217324292 | 0.552251251 | 0.791429981 |
| chr19:2944701-2945599 | 0.195307866 | 0.552263717 | 0.791433043 |
| chr13:99200213-99201003 | -0.194271437 | 0.552283576 | 0.791441853 |
| chr9:91761425-91761681 | -0.306561392 | 0.552290523 | 0.791441853 |
| chr5:166746090-166746290 | -0.338494372 | 0.552306907 | 0.79145053 |
| chr20:41746686-41747008 | 0.226335404 | 0.552361877 | 0.791514499 |
| chrX:103176220-103176651 | 0.232701827 | 0.55241143 | 0.791570703 |
| chr11:20027227-20027575 | 0.240029431 | 0.55243894 | 0.791585927 |
| chrX:19334113-19334443 | -0.249995056 | 0.552442717 | 0.791585927 |
| chr2:129469492-129469692 | 0.316221247 | 0.552472414 | 0.791599436 |
| chr12:63668007-63668693 | -0.219884502 | 0.552472807 | 0.791599436 |
| chr1:193197623-193197923 | -0.293282149 | 0.552488305 | 0.791606838 |
| chr5:181092105-181092319 | -0.308819513 | 0.552501182 | 0.791610485 |
| chr11:48061325-48061705 | 0.217962582 | 0.552524681 | 0.791614651 |
| chr1:197924599-197924879 | -0.224767777 | 0.552524752 | 0.791614651 |
| chr7:42479892-42480358 | 0.22962393 | 0.552541623 | 0.791624021 |
| chr15:79028455-79028655 | 0.262366863 | 0.552571816 | 0.791648402 |
| chr9:77751527-77751792 | -0.257680806 | 0.552589466 | 0.791648402 |
| chr7:64902953-64903958 | 0.196594833 | 0.552599867 | 0.791648402 |
| chr2:109516389-109516709 | 0.230670462 | 0.552600492 | 0.791648402 |
| chr4:108572626-108572851 | -0.285744205 | 0.5526103 | 0.791648402 |
| chr12:122123253-122123515 | -0.258419635 | 0.552642707 | 0.791666111 |
| chr6:4489490-4490408 | 0.195185201 | 0.552643326 | 0.791666111 |
| chr20:54250751-54251697 | -0.186914922 | 0.552685917 | 0.791711068 |
| chr5:111223273-111223928 | -0.233451129 | 0.552695375 | 0.791711068 |
| chr9:123349886-123350280 | 0.215758172 | 0.552713692 | 0.791722505 |
| chr20:53592583-53592824 | 0.195584459 | 0.552751256 | 0.791761511 |
| chrX:136200556-136200825 | 0.262241612 | 0.552782257 | 0.791779185 |
| chr9:13279166-13279836 | -0.217891598 | 0.55279223 | 0.791779185 |
| chr16:28974534-28974982 | 0.20590534 | 0.552794595 | 0.791779185 |
| chr18:9366144-9366662 | -0.207920723 | 0.552816443 | 0.791790853 |
| chr13:33119928-33120541 | 0.199615004 | 0.552836901 | 0.791790853 |
| chr3:11379197-11379555 | 0.230001214 | 0.552878265 | 0.791790853 |

| | | | |
|---|---|---|---|
| chr5:171877612-171878032 | -0.27468935 | 0.552891234 | 0.791790853 |
| chr19:1513873-1514176 | 0.241897287 | 0.552893757 | 0.791790853 |
| chr2:219713174-219713409 | -0.282863578 | 0.552894306 | 0.791790853 |
| chr11:74726555-74727109 | -0.207301784 | 0.552894331 | 0.791790853 |
| chr2:106596929-106597197 | 0.257695202 | 0.552903701 | 0.791790853 |
| chr4:140428165-140428365 | 0.298610813 | 0.552904507 | 0.791790853 |
| chr5:172282875-172284284 | 0.189877171 | 0.552926474 | 0.791790853 |
| chr15:92709118-92709860 | 0.212683637 | 0.552931156 | 0.791790853 |
| chr13:25286943-25287754 | 0.196512799 | 0.552936984 | 0.791790853 |
| chr19:1194449-1194853 | 0.233391355 | 0.55293708 | 0.791790853 |
| chr13:113393743-113394319 | 0.223876304 | 0.552999841 | 0.791865925 |
| chr5:135662788-135663042 | -0.27894589 | 0.553014094 | 0.791871537 |
| chr8:107181477-107181739 | -0.293348573 | 0.553062307 | 0.791925774 |
| chr12:85441878-85442080 | -0.31085136 | 0.553081635 | 0.79193865 |
| chr12:57182975-57183373 | -0.209466799 | 0.553095875 | 0.79194424 |
| chr7:102236707-102237029 | -0.245039289 | 0.553125522 | 0.791960093 |
| chr4:154559389-154559798 | 0.216272685 | 0.553127618 | 0.791960093 |
| chr3:151227563-151228217 | -0.206905581 | 0.553146549 | 0.791969904 |
| chr14:80288526-80288726 | 0.341399317 | 0.553155143 | 0.791969904 |
| chr17:46922732-46923685 | 0.196894089 | 0.553176983 | 0.791986375 |
| chr12:68610310-68611256 | 0.192093357 | 0.553254441 | 0.792082471 |
| chr14:55581092-55581322 | 0.241236351 | 0.553290641 | 0.792119498 |
| chr3:100103845-100104245 | -0.247849254 | 0.553313097 | 0.792136846 |
| chr20:33645036-33645236 | 0.288536325 | 0.553332618 | 0.792149992 |
| chr2:75241494-75241801 | -0.268695647 | 0.553398258 | 0.79222916 |
| chr5:180401286-180401486 | -0.310175887 | 0.553443365 | 0.792274295 |
| chr17:63390642-63390927 | -0.257031585 | 0.553450466 | 0.792274295 |
| chrX:80982712-80982912 | 0.301938574 | 0.553476765 | 0.792297139 |
| chrX:10833339-10833868 | -0.220809434 | 0.55350468 | 0.792322297 |
| chr12:113093117-113093461 | 0.21855124 | 0.55352652 | 0.792336267 |
| chr17:41548823-41549549 | -0.20130251 | 0.553553504 | 0.792336267 |
| chr22:45671480-45672136 | 0.21693939 | 0.553556048 | 0.792336267 |
| chr16:66873124-66873421 | 0.250380748 | 0.553560128 | 0.792336267 |
| chr20:25059128-25059420 | -0.234644148 | 0.553566144 | 0.792336267 |
| chr7:100149238-100149647 | -0.220268834 | 0.553589675 | 0.792340589 |
| chr9:135343866-135344117 | 0.276462371 | 0.553589845 | 0.792340589 |
| chr3:132180267-132180845 | 0.210872907 | 0.553603401 | 0.792345192 |
| chr2:178428365-178428863 | 0.223904955 | 0.553639515 | 0.792351883 |
| chr8:112463260-112463598 | -0.237895799 | 0.553639523 | 0.792351883 |
| chr17:50865962-50868077 | -0.184129467 | 0.553666527 | 0.792351883 |
| chr14:31787452-31787703 | -0.29016146 | 0.553668324 | 0.792351883 |
| chr17:58304088-58304684 | -0.198795399 | 0.553680573 | 0.792351883 |
| chr4:68497952-68498152 | -0.31502827 | 0.553692181 | 0.792351883 |
| chr5:90421152-90421561 | -0.218212268 | 0.553707589 | 0.792351883 |
| chr1:40979272-40980516 | 0.191286242 | 0.553709768 | 0.792351883 |
| chr11:126494663-126494966 | 0.300118129 | 0.553718099 | 0.792351883 |
| chr1:146228759-146229103 | 0.249669087 | 0.553720725 | 0.792351883 |
| chr7:37284753-37285108 | -0.23036925 | 0.553721828 | 0.792351883 |
| chr5:19790894-19791141 | -0.270994821 | 0.553752859 | 0.792381489 |
| chr4:158902593-158902970 | -0.223067647 | 0.553768447 | 0.792388995 |

| | | | |
|---|---|---|---|
| chrX:95076598-95076822 | -0.264897697 | 0.553778874 | 0.792389119 |
| chr2:19933396-19933699 | -0.274750762 | 0.553790603 | 0.792391104 |
| chr5:176447522-176448736 | -0.18799731 | 0.553836673 | 0.792442224 |
| chr3:172982816-172983171 | 0.24967135 | 0.553857137 | 0.792445164 |
| chr21:41727240-41727689 | -0.232969144 | 0.553859412 | 0.792445164 |
| chr6:26383721-26383967 | 0.271681151 | 0.553876108 | 0.792450448 |
| chr11:10931048-10931981 | 0.207271751 | 0.55388379 | 0.792450448 |
| chr3:69231877-69232165 | 0.254008952 | 0.553922836 | 0.792491515 |
| chr2:240232942-240234020 | 0.190322931 | 0.553972539 | 0.792547826 |
| chr2:128595163-128595429 | 0.268597808 | 0.554010088 | 0.792571949 |
| chr21:46189589-46189811 | 0.268597808 | 0.554010088 | 0.792571949 |
| chr16:81143861-81144486 | 0.201959578 | 0.554060793 | 0.792615389 |
| chr7:15889556-15890084 | 0.213535756 | 0.554061142 | 0.792615389 |
| chr4:122921977-122923526 | -0.188386396 | 0.554095585 | 0.792620218 |
| chr17:73420900-73421314 | -0.210708632 | 0.554105568 | 0.792620218 |
| chr14:49620146-49621381 | 0.187188406 | 0.554117166 | 0.792620218 |
| chr15:100002964-100003178 | 0.30159916 | 0.554122391 | 0.792620218 |
| chr11:85680872-85681100 | -0.270446682 | 0.554126585 | 0.792620218 |
| chr21:44874696-44874896 | -0.270446682 | 0.554126585 | 0.792620218 |
| chr4:42041370-42041644 | -0.310309612 | 0.554150674 | 0.792628575 |
| chr3:160225017-160226248 | 0.193073419 | 0.554153117 | 0.792628575 |
| chr5:11619487-11620006 | 0.226019691 | 0.554172715 | 0.79264181 |
| chr19:1174221-1174421 | 0.272601957 | 0.554234602 | 0.79271553 |
| chr18:32019204-32019555 | 0.229788808 | 0.554252102 | 0.792721213 |
| chr5:61253299-61253556 | -0.312872886 | 0.554259267 | 0.792721213 |
| chr2:38947123-38947971 | -0.200269678 | 0.554310136 | 0.792773434 |
| chr4:141375850-141376250 | -0.228605348 | 0.554316472 | 0.792773434 |
| chr17:28571366-28571868 | -0.200072052 | 0.554342241 | 0.792785253 |
| chr4:158729762-158730312 | -0.222691325 | 0.554353247 | 0.792785253 |
| chr20:34044514-34044963 | -0.258114004 | 0.554355776 | 0.792785253 |
| chr15:70313468-70313819 | 0.273424373 | 0.554381365 | 0.79280705 |
| chr7:115356225-115356515 | -0.270318384 | 0.554408261 | 0.792830716 |
| chrX:102659764-102660517 | 0.197296358 | 0.554444283 | 0.792867431 |
| chr5:11428097-11428297 | 0.316272594 | 0.554503508 | 0.792931598 |
| chr6:19211250-19211574 | -0.262317219 | 0.554509851 | 0.792931598 |
| chr4:56976789-56977264 | 0.197299961 | 0.554521168 | 0.792932983 |
| chr19:51366827-51367063 | -0.233522686 | 0.554548735 | 0.792957539 |
| chr3:128993543-128994515 | 0.19600227 | 0.554559039 | 0.792957539 |
| chr16:67428128-67428488 | 0.219188465 | 0.554586955 | 0.792959991 |
| chr11:21552474-21552705 | 0.309660019 | 0.554588443 | 0.792959991 |
| chr2:191434151-191434475 | -0.245265441 | 0.554591801 | 0.792959991 |
| chr17:53302711-53302948 | -0.285308461 | 0.55461004 | 0.792971273 |
| chr5:110360350-110360670 | -0.315862071 | 0.554624565 | 0.792977243 |
| chr1:56578707-56579611 | 0.196485932 | 0.554651594 | 0.792982381 |
| chr21:42923881-42924184 | -0.233470085 | 0.554659077 | 0.792982381 |
| chr7:44534646-44534917 | -0.237370314 | 0.554659206 | 0.792982381 |
| chr17:71237876-71238122 | -0.256163663 | 0.554701869 | 0.793028578 |
| chr12:93677019-93678129 | -0.189684757 | 0.554728133 | 0.79304327 |
| chr7:49791114-49791424 | -0.236243572 | 0.554732846 | 0.79304327 |
| chr17:3635538-3636971 | -0.190373756 | 0.554763611 | 0.793072454 |

| | | | |
|---|---|---|---|
| chr9:37784706-37785442 | -0.201321867 | 0.55483237 | 0.793155951 |
| chr1:152142018-152142361 | 0.218820946 | 0.554889412 | 0.79320124 |
| chr5:94044661-94044861 | 0.328063328 | 0.554894332 | 0.79320124 |
| chr7:42014679-42014974 | -0.274344202 | 0.554907751 | 0.79320124 |
| chr17:35063565-35063983 | -0.234614598 | 0.554913465 | 0.79320124 |
| chr2:65861681-65861881 | -0.316513335 | 0.554915811 | 0.79320124 |
| chr2:20231404-20232029 | -0.195631779 | 0.554968519 | 0.793237204 |
| chr3:135907834-135908515 | -0.213788897 | 0.554973445 | 0.793237204 |
| chr3:21256995-21257195 | -0.2874216 | 0.554978272 | 0.793237204 |
| chr4:109977999-109978250 | -0.290484871 | 0.554982381 | 0.793237204 |
| chr15:42650408-42650790 | 0.229076962 | 0.555005103 | 0.793254882 |
| chr7:100247220-100247720 | -0.222371595 | 0.555070143 | 0.793284524 |
| chr1:13965860-13966128 | 0.280457584 | 0.555080443 | 0.793284524 |
| chr6:30103110-30103616 | -0.217137776 | 0.555083141 | 0.793284524 |
| chrX:117366175-117366534 | -0.249078021 | 0.555084403 | 0.793284524 |
| chr3:197627229-197627429 | 0.288798924 | 0.555087961 | 0.793284524 |
| chr16:62965984-62966184 | 0.288798924 | 0.555087961 | 0.793284524 |
| chr19:50414849-50415101 | 0.250734897 | 0.555118941 | 0.793292195 |
| chr1:167553123-167554023 | -0.190649839 | 0.55512335 | 0.793292195 |
| chr10:124088056-124088363 | -0.231152276 | 0.555124588 | 0.793292195 |
| chr2:209961025-209961382 | 0.22401761 | 0.555134742 | 0.793292195 |
| chr11:83836822-83837298 | -0.210933228 | 0.555219557 | 0.793397216 |
| chr3:9175516-9175789 | 0.239163873 | 0.555228943 | 0.793397216 |
| chr12:80916338-80916538 | -0.297977219 | 0.555265239 | 0.793434284 |
| chr7:151116012-151116437 | 0.208468311 | 0.555303092 | 0.793466011 |
| chr8:144517436-144518308 | 0.193031327 | 0.555334723 | 0.793466011 |
| chr15:39835632-39835832 | -0.33999833 | 0.55534339 | 0.793466011 |
| chr12:121060395-121060894 | -0.23112674 | 0.555345429 | 0.793466011 |
| chr19:40385162-40385448 | -0.305894188 | 0.555349479 | 0.793466011 |
| chr17:62109454-62109831 | -0.195994443 | 0.555349576 | 0.793466011 |
| chr12:97637964-97638164 | 0.307310886 | 0.55541028 | 0.793530602 |
| chr3:122793490-122794331 | -0.193863789 | 0.555424096 | 0.793530602 |
| chr12:65169176-65170703 | 0.186267866 | 0.555425853 | 0.793530602 |
| chr2:172102567-172103284 | 0.195827456 | 0.555445657 | 0.793544099 |
| chrX:136250560-136250922 | 0.243275576 | 0.555457896 | 0.793546788 |
| chr16:67260676-67260876 | 0.300765727 | 0.555474819 | 0.793556169 |
| chr2:97734235-97734438 | 0.278736119 | 0.555508358 | 0.793589287 |
| chr1:20493845-20494097 | 0.279941969 | 0.555523844 | 0.793596614 |
| chr14:24201337-24201640 | 0.25037672 | 0.555548509 | 0.793606468 |
| chr5:160118583-160118783 | -0.225621494 | 0.555576656 | 0.793606468 |
| chr12:75390664-75391651 | -0.186743681 | 0.555577628 | 0.793606468 |
| chr17:64369406-64369701 | -0.256436962 | 0.555578047 | 0.793606468 |
| chr14:69090268-69090575 | 0.242760751 | 0.555582529 | 0.793606468 |
| chr19:33684200-33684544 | 0.237614875 | 0.555597443 | 0.793612976 |
| chr1:109493828-109494417 | -0.195595096 | 0.555609268 | 0.793615073 |
| chr12:104381712-104382167 | 0.218055836 | 0.555672769 | 0.793690979 |
| chr16:84037022-84037284 | 0.266768962 | 0.555707056 | 0.79371787 |
| chr1:167721568-167722687 | 0.189408139 | 0.555712313 | 0.79371787 |
| chr9:21183117-21183386 | -0.281438904 | 0.555764933 | 0.793737006 |
| chr14:21476309-21477636 | 0.185706869 | 0.555775336 | 0.793737006 |

| | | | |
|---|---|---|---|
| chr14:92794736-92794966 | -0.226992886 | 0.555784737 | 0.793737006 |
| chr2:75677290-75677705 | -0.209199935 | 0.55579459 | 0.793737006 |
| chr22:38656090-38657414 | -0.197045927 | 0.55579817 | 0.793737006 |
| chr4:83484647-83485522 | -0.190693161 | 0.555798313 | 0.793737006 |
| chrX:78029734-78029934 | 0.30409102 | 0.55580182 | 0.793737006 |
| chr11:43278644-43278992 | 0.223623211 | 0.555808584 | 0.793737006 |
| chr3:66195277-66195701 | -0.232344319 | 0.555867623 | 0.793749595 |
| chr2:42008168-42008368 | 0.320044212 | 0.555875036 | 0.793749595 |
| chr19:10231234-10231635 | 0.222016557 | 0.555877103 | 0.793749595 |
| chr2:854498-854747 | 0.25923196 | 0.555886314 | 0.793749595 |
| chr8:121154436-121154942 | -0.203898674 | 0.555887854 | 0.793749595 |
| chr2:170770087-170771544 | -0.189361048 | 0.5558933 | 0.793749595 |
| chr7:100675530-100676025 | 0.205402977 | 0.555900713 | 0.793749595 |
| chr11:129364143-129364534 | -0.22082824 | 0.555916111 | 0.793749595 |
| chr11:2699871-2700528 | 0.19996761 | 0.55592036 | 0.793749595 |
| chr8:94822420-94823881 | 0.184932096 | 0.555920992 | 0.793749595 |
| chr20:47764438-47764670 | 0.205018028 | 0.555941883 | 0.793764631 |
| chr14:100238448-100239216 | 0.190779127 | 0.555970187 | 0.793767508 |
| chr5:141004140-141004547 | 0.226557458 | 0.555981196 | 0.793767508 |
| chr12:97003783-97004033 | -0.308651842 | 0.555981886 | 0.793767508 |
| chr2:241271907-241272116 | -0.258686979 | 0.555986817 | 0.793767508 |
| chr2:38376275-38377913 | -0.18547779 | 0.555995695 | 0.793767508 |
| chr3:170864594-170865011 | -0.225559874 | 0.556008243 | 0.793770632 |
| chr12:96194146-96194644 | -0.21794727 | 0.556020194 | 0.793772904 |
| chr3:152834490-152835597 | 0.189420763 | 0.556045205 | 0.79379382 |
| chr3:32611390-32611590 | 0.325468546 | 0.556078566 | 0.793826655 |
| chr8:88289736-88290121 | -0.206303009 | 0.5561087 | 0.793854883 |
| chr21:8208547-8209197 | 0.188979283 | 0.556129327 | 0.793869537 |
| chr3:36904762-36904980 | 0.269112946 | 0.556179493 | 0.793925851 |
| chr1:60848550-60848867 | -0.272970281 | 0.5561895 | 0.793925851 |
| chr12:104063766-104064914 | -0.189488436 | 0.556253953 | 0.793989342 |
| chr5:1772703-1772982 | 0.236913708 | 0.556254704 | 0.793989342 |
| chr3:99748077-99748358 | 0.253214655 | 0.556288064 | 0.794022169 |
| chr3:166560365-166560746 | 0.246396088 | 0.556306393 | 0.794033539 |
| chr12:98653970-98654209 | -0.284227915 | 0.556330801 | 0.794038871 |
| chr17:4366510-4367114 | -0.198050916 | 0.556330854 | 0.794038871 |
| chr2:26345527-26346472 | -0.188000568 | 0.556344556 | 0.794043635 |
| chr17:29591725-29592259 | 0.201776206 | 0.556373807 | 0.794056727 |
| chr4:52051209-52052231 | 0.194108585 | 0.556374455 | 0.794056727 |
| chr12:103434074-103434608 | -0.216168032 | 0.556390729 | 0.794062636 |
| chr8:143635781-143636714 | 0.194327887 | 0.556402012 | 0.794062636 |
| chr17:17476789-17477147 | 0.211901303 | 0.556409686 | 0.794062636 |
| chr10:85547935-85548375 | -0.221177001 | 0.556476034 | 0.7941243 |
| chrX:110605773-110606486 | -0.208426483 | 0.556483139 | 0.7941243 |
| chr19:15332523-15332723 | 0.309261386 | 0.556483987 | 0.7941243 |
| chr15:69394627-69394897 | 0.239866651 | 0.556500278 | 0.794128843 |
| chr8:115838102-115838843 | 0.192585274 | 0.556516158 | 0.794128843 |
| chr15:58182631-58182865 | -0.282844527 | 0.556518263 | 0.794128843 |
| chr12:6278109-6278434 | -0.225161576 | 0.556529115 | 0.794129036 |
| chr3:45037477-45038022 | -0.224871949 | 0.556539127 | 0.794129036 |

| | | | |
|---|---|---|---|
| chr3:525467-525675 | 0.318052229 | 0.556584199 | 0.794178561 |
| chr1:41484155-41484891 | -0.200518782 | 0.556600068 | 0.794186414 |
| chr21:8434793-8435344 | 0.187908281 | 0.556631511 | 0.794210987 |
| chr3:151201260-151201613 | -0.277971104 | 0.556646585 | 0.794210987 |
| chr1:50081027-50081510 | -0.231037664 | 0.556648386 | 0.794210987 |
| chr17:48074409-48074624 | 0.270936025 | 0.556664749 | 0.794219544 |
| chr14:81267938-81268509 | 0.201474745 | 0.556687916 | 0.794237808 |
| chr17:67468917-67469732 | -0.203426627 | 0.55674478 | 0.794304147 |
| chr12:43676308-43676512 | -0.274984832 | 0.556770896 | 0.794326616 |
| chr9:15552205-15552444 | 0.244959801 | 0.556876032 | 0.794461818 |
| chr1:116485262-116485689 | 0.203383351 | 0.556935555 | 0.794505232 |
| chr3:168417063-168417377 | 0.242295634 | 0.556940865 | 0.794505232 |
| chr17:9802300-9802992 | -0.219719572 | 0.556946346 | 0.794505232 |
| chr6:56955836-56956197 | -0.21592539 | 0.55695711 | 0.794505232 |
| chr1:245106939-245107300 | 0.233763237 | 0.556958308 | 0.794505232 |
| chr3:75434792-75436175 | -0.182702617 | 0.55700181 | 0.794552494 |
| chr5:152505434-152505841 | 0.227618608 | 0.557021059 | 0.794565161 |
| chr15:76190884-76191120 | 0.245403043 | 0.557055031 | 0.794598827 |
| chr1:201882817-201883020 | -0.299530995 | 0.557068084 | 0.794602654 |
| chr15:63857174-63857505 | 0.250579634 | 0.55708895 | 0.794617624 |
| chr3:146883173-146883496 | -0.256695186 | 0.557142226 | 0.794678821 |
| chr16:11371808-11372013 | -0.306061734 | 0.557154232 | 0.794681153 |
| chr5:43042781-43043071 | -0.220155268 | 0.557202717 | 0.794725404 |
| chr5:141564140-141564582 | -0.209177801 | 0.557218076 | 0.794725404 |
| chr9:136051822-136052572 | 0.197899821 | 0.557223154 | 0.794725404 |
| chr20:32207466-32208029 | 0.20142336 | 0.557226744 | 0.794725404 |
| chr4:98939859-98940095 | -0.289609683 | 0.557243766 | 0.794734888 |
| chr3:54433866-54434144 | -0.269166307 | 0.55727293 | 0.794753434 |
| chr14:34030922-34031122 | 0.261876197 | 0.557283262 | 0.794753434 |
| chr1:8702188-8702481 | -0.268218961 | 0.557293187 | 0.794753434 |
| chr2:177332428-177332657 | -0.270436783 | 0.55729826 | 0.794753434 |
| chr13:112942701-112943260 | 0.203981433 | 0.55731963 | 0.794758605 |
| chr14:40368803-40369003 | -0.301924455 | 0.557322631 | 0.794758605 |
| chr6:21883869-21884236 | -0.247849856 | 0.557358123 | 0.794762741 |
| chr19:39154412-39155396 | -0.19339346 | 0.557361914 | 0.794762741 |
| chr8:132666414-132666656 | -0.301818782 | 0.557366087 | 0.794762741 |
| chr20:54067505-54068190 | 0.184271079 | 0.557367986 | 0.794762741 |
| chr20:61966224-61966468 | -0.252493606 | 0.557377394 | 0.794762741 |
| chr2:146927195-146927613 | -0.235830083 | 0.557389504 | 0.794765219 |
| chr9:137492354-137492956 | 0.212239893 | 0.55740639 | 0.794774506 |
| chr7:149771600-149772305 | 0.191372426 | 0.557482502 | 0.794850806 |
| chr7:83947346-83947601 | -0.270354513 | 0.557488211 | 0.794850806 |
| chr4:180449416-180449773 | -0.222148186 | 0.557491023 | 0.794850806 |
| chr6:36759196-36759496 | 0.231085785 | 0.557503912 | 0.794854393 |
| chr19:14090737-14091747 | 0.201026576 | 0.557522714 | 0.794855278 |
| chr3:170046328-170046615 | 0.216866217 | 0.557534143 | 0.794855278 |
| chr14:21526123-21526612 | -0.209512154 | 0.557535655 | 0.794855278 |
| chr11:124673316-124674307 | -0.193833955 | 0.557583812 | 0.794891187 |
| chr14:59965071-59965430 | -0.220316323 | 0.557590746 | 0.794891187 |
| chrX:23504042-23504710 | 0.207022049 | 0.557592333 | 0.794891187 |

| | | | |
|---|---|---|---|
| chr18:64208522-64208764 | 0.32140144 | 0.557602554 | 0.794891187 |
| chr1:153312591-153313075 | 0.225278591 | 0.5576269 | 0.794891187 |
| chr11:10540609-10541558 | 0.198857469 | 0.557631407 | 0.794891187 |
| chr11:120280715-120281111 | 0.237017928 | 0.55763635 | 0.794891187 |
| chr3:196632052-196633222 | 0.191830565 | 0.557643835 | 0.794891187 |
| chr11:59770119-59770401 | 0.242796257 | 0.557677689 | 0.794924655 |
| chrX:109839965-109840232 | -0.300019838 | 0.557711606 | 0.79492754 |
| chr13:43785611-43786492 | -0.192717428 | 0.557723202 | 0.79492754 |
| chr2:36965933-36967142 | -0.186015467 | 0.557729382 | 0.79492754 |
| chr7:1231492-1231692 | 0.278502387 | 0.557743883 | 0.79492754 |
| chr10:72273441-72274390 | -0.187206036 | 0.557747622 | 0.79492754 |
| chr3:71544895-71545230 | -0.31495895 | 0.557748869 | 0.79492754 |
| chr17:64231004-64231209 | -0.278144857 | 0.557752336 | 0.79492754 |
| chr5:15103309-15103514 | 0.275968747 | 0.557767383 | 0.794934199 |
| chr2:85988642-85988978 | -0.256549572 | 0.557789232 | 0.794950553 |
| chr3:26736125-26736521 | 0.253466638 | 0.557816573 | 0.794974731 |
| chr3:130755631-130756006 | 0.231153316 | 0.557849099 | 0.7950063 |
| chr2:11669812-11670254 | 0.21353471 | 0.557873241 | 0.795025917 |
| chr3:164169970-164170291 | -0.258810348 | 0.557903259 | 0.795053908 |
| chr12:79766701-79766947 | -0.299810007 | 0.557955971 | 0.795113652 |
| chr1:225567878-225568094 | -0.305355884 | 0.557965936 | 0.795113652 |
| chr1:39132877-39133520 | -0.213541441 | 0.557986218 | 0.795127767 |
| chr14:35814416-35814735 | -0.222717186 | 0.558055152 | 0.795211208 |
| chr11:65250418-65250620 | -0.285845855 | 0.558083109 | 0.795215243 |
| chr4:95102519-95102729 | -0.298676923 | 0.558085412 | 0.795215243 |
| chr4:123772748-123773217 | 0.214418727 | 0.5581141 | 0.795215243 |
| chr13:40993500-40993847 | -0.240606298 | 0.558114718 | 0.795215243 |
| chr3:90182350-90182815 | -0.226081299 | 0.558115967 | 0.795215243 |
| chr18:23540069-23540419 | -0.225982098 | 0.558133363 | 0.795215243 |
| chr1:215380446-215380822 | -0.227239672 | 0.558137545 | 0.795215243 |
| chr22:29306172-29306581 | 0.211480658 | 0.558141011 | 0.795215243 |
| chr11:9748484-9748825 | 0.236163154 | 0.558175191 | 0.795226609 |
| chr8:123979887-123980214 | -0.230700614 | 0.558182985 | 0.795226609 |
| chr6:161550325-161550605 | 0.262865693 | 0.558194653 | 0.795226609 |
| chr17:76856579-76856844 | 0.259508174 | 0.558196077 | 0.795226609 |
| chr2:8678711-8679408 | 0.191730338 | 0.558206085 | 0.795226609 |
| chr5:56013439-56013740 | 0.246261529 | 0.55821126 | 0.795226609 |
| chr18:44678957-44679216 | -0.239549047 | 0.558226135 | 0.795233015 |
| chr3:156010818-156011196 | 0.235326552 | 0.55823716 | 0.795233935 |
| chr11:110201441-110201641 | 0.304100979 | 0.558251657 | 0.795239802 |
| chr10:80535655-80536505 | -0.190152077 | 0.558273582 | 0.795247579 |
| chr7:100065749-100065949 | -0.321956118 | 0.558277874 | 0.795247579 |
| chr9:109499741-109500117 | -0.211144092 | 0.55834607 | 0.795329936 |
| chr11:68021381-68021602 | -0.246757535 | 0.558368956 | 0.795334388 |
| chr5:9361541-9362117 | -0.213049891 | 0.558384245 | 0.795334388 |
| chr5:69007844-69008090 | 0.257679497 | 0.558385794 | 0.795334388 |
| chr1:88678480-88678704 | 0.283421014 | 0.558398716 | 0.795334388 |
| chr1:236168802-236169061 | -0.282004857 | 0.558401096 | 0.795334388 |
| chr4:87659631-87660073 | 0.215216948 | 0.558431236 | 0.795362532 |
| chr2:3379488-3380175 | -0.189593776 | 0.558474251 | 0.795395888 |

| | | | |
|---|---|---|---|
| chr16:56931795-56932648 | 0.19622033 | 0.558475962 | 0.795395888 |
| chr21:44338955-44339717 | 0.190554637 | 0.558493368 | 0.795395888 |
| chr14:77059579-77060070 | -0.211226235 | 0.558496729 | 0.795395888 |
| chr9:117970696-117970973 | -0.296415113 | 0.558508315 | 0.795395888 |
| chr14:51780208-51780415 | 0.253766786 | 0.558518511 | 0.795395888 |
| chr6:30541749-30541958 | 0.281976205 | 0.558527321 | 0.795395888 |
| chr9:128688553-128689633 | 0.190844544 | 0.558542322 | 0.795402468 |
| chr21:25790186-25790805 | -0.218358677 | 0.558557679 | 0.795409554 |
| chr14:55864912-55865137 | 0.303857426 | 0.558570267 | 0.795412697 |
| chr8:96227231-96227589 | 0.223259275 | 0.558645804 | 0.795505478 |
| chr11:15202609-15203020 | -0.239042139 | 0.558685179 | 0.795530371 |
| chr8:122782260-122782613 | -0.200454396 | 0.558687444 | 0.795530371 |
| chr12:31403190-31403427 | -0.269162757 | 0.558719641 | 0.795530371 |
| chr17:64847874-64848303 | -0.21946088 | 0.558725298 | 0.795530371 |
| chr4:128863298-128863815 | 0.211133692 | 0.558727285 | 0.795530371 |
| chr8:96493439-96495524 | 0.183552048 | 0.558734455 | 0.795530371 |
| chr1:247180502-247180770 | -0.265219817 | 0.558735962 | 0.795530371 |
| chr17:79874412-79874678 | 0.261720997 | 0.558767401 | 0.795546685 |
| chr17:51585561-51585991 | -0.215808825 | 0.558768186 | 0.795546685 |
| chr5:142611138-142611424 | 0.24065976 | 0.558780318 | 0.795549176 |
| chr6:109304048-109304498 | -0.24107653 | 0.55880709 | 0.79556434 |
| chr1:206583736-206584003 | 0.217224391 | 0.558811735 | 0.79556434 |
| chr5:86624215-86624471 | -0.304776064 | 0.558881684 | 0.795649141 |
| chr2:149070849-149071441 | -0.220416822 | 0.558907975 | 0.795671786 |
| chr21:39236010-39236210 | -0.300097706 | 0.558921973 | 0.79567693 |
| chr20:53757566-53758008 | 0.187175011 | 0.55895727 | 0.795687246 |
| chr14:89416052-89416404 | -0.199688319 | 0.558971675 | 0.795687246 |
| chr3:143134353-143134684 | -0.256756617 | 0.558972741 | 0.795687246 |
| chr1:155000669-155001508 | 0.186310516 | 0.558979959 | 0.795687246 |
| chr9:2017294-2017630 | 0.22007935 | 0.558981142 | 0.795687246 |
| chr12:46176554-46176768 | 0.310879856 | 0.559005897 | 0.79570628 |
| chr4:146945579-146946295 | -0.196423525 | 0.559015283 | 0.79570628 |
| chr7:34497178-34497467 | 0.243063243 | 0.559057859 | 0.795726557 |
| chr3:142211093-142211301 | 0.303628115 | 0.559060684 | 0.795726557 |
| chr3:162786126-162786326 | 0.303628115 | 0.559060684 | 0.795726557 |
| chr8:78470688-78471067 | 0.23148367 | 0.559077926 | 0.795736316 |
| chr8:20542866-20543238 | -0.270282992 | 0.559101864 | 0.795741806 |
| chr5:93915117-93915406 | -0.2431525 | 0.559108684 | 0.795741806 |
| chr7:75951500-75951965 | -0.220709681 | 0.559142399 | 0.795741806 |
| chr19:35485570-35485770 | 0.329109208 | 0.559143691 | 0.795741806 |
| chr4:82044110-82044743 | 0.207427513 | 0.559148545 | 0.795741806 |
| chr5:178113145-178114274 | -0.186735443 | 0.559159721 | 0.795741806 |
| chrX:103420474-103420674 | 0.319391834 | 0.559170308 | 0.795741806 |
| chr7:3334524-3334766 | -0.307203284 | 0.559170865 | 0.795741806 |
| chrX:7147329-7148878 | 0.186758416 | 0.559187553 | 0.795741806 |
| chr6:146857278-146857817 | 0.212682834 | 0.559189555 | 0.795741806 |
| chr11:67302793-67302993 | -0.24171698 | 0.559199359 | 0.795741806 |
| chr7:34652741-34653039 | -0.257792796 | 0.559209255 | 0.795741806 |
| chr19:39480528-39481327 | 0.193377052 | 0.559216792 | 0.795741806 |
| chr4:932123-932666 | 0.204321024 | 0.559236352 | 0.795747476 |

| | | | |
|---|---|---|---|
| chr4:48269294-48270377 | 0.186019021 | 0.559248714 | 0.795747476 |
| chr21:21921679-21921927 | -0.265873776 | 0.559257243 | 0.795747476 |
| chr19:639455-639903 | -0.226079176 | 0.559264865 | 0.795747476 |
| chr2:218186348-218186643 | 0.249770841 | 0.559272704 | 0.795747476 |
| chr14:105480682-105482490 | 0.183324092 | 0.559317599 | 0.795796578 |
| chr12:94615913-94616489 | -0.220110986 | 0.559360474 | 0.795811046 |
| chr14:40206729-40207143 | -0.266113094 | 0.559367203 | 0.795811046 |
| chr17:27455696-27456047 | -0.238080505 | 0.559369609 | 0.795811046 |
| chr3:181712317-181712710 | -0.228524009 | 0.559377636 | 0.795811046 |
| chr8:69140422-69140822 | -0.249604018 | 0.5593797 | 0.795811046 |
| chr16:76653469-76653840 | 0.210895947 | 0.559487598 | 0.795949772 |
| chr1:154820166-154820444 | 0.270090238 | 0.559515917 | 0.795964173 |
| chr8:95268507-95270213 | -0.181876295 | 0.559518498 | 0.795964173 |
| chr19:633479-633871 | 0.212878035 | 0.559551089 | 0.795995758 |
| chr15:77083748-77084357 | -0.200054052 | 0.559592137 | 0.796039372 |
| chr1:45521613-45522312 | 0.193497978 | 0.559648922 | 0.796105371 |
| chr15:40421332-40421579 | -0.249921431 | 0.559727257 | 0.796202022 |
| chr6:35568973-35569173 | 0.283854799 | 0.559753252 | 0.796224218 |
| chr2:112906570-112906823 | -0.286484049 | 0.559776102 | 0.796241938 |
| chr7:42911591-42912538 | -0.188274773 | 0.559794304 | 0.796242718 |
| chr15:94119946-94120285 | 0.218360058 | 0.559797434 | 0.796242718 |
| chr3:101562308-101562716 | 0.206099648 | 0.55983474 | 0.796281 |
| chr10:52451890-52452293 | -0.235705667 | 0.559849307 | 0.796286937 |
| chr6:170116574-170116863 | -0.256800948 | 0.559910933 | 0.796358487 |
| chr11:49526237-49526558 | 0.245181664 | 0.559927181 | 0.796358487 |
| chr13:52449880-52450963 | -0.188924984 | 0.559937521 | 0.796358487 |
| chr16:4183112-4184376 | -0.188357385 | 0.559950673 | 0.796358487 |
| chr16:84175306-84175664 | 0.205117291 | 0.559951578 | 0.796358487 |
| chr10:83562462-83562667 | -0.279767635 | 0.559981918 | 0.796363936 |
| chr19:44671537-44671899 | 0.233457034 | 0.559985438 | 0.796363936 |
| chr3:113838581-113839291 | -0.194102136 | 0.55998659 | 0.796363936 |
| chr22:36409980-36410624 | 0.202629119 | 0.560001227 | 0.79636997 |
| chr11:124170874-124171510 | -0.201907886 | 0.560037111 | 0.79640622 |
| chr17:45719842-45720091 | 0.275510051 | 0.560092805 | 0.796470638 |
| chr9:100598580-100599098 | -0.194498865 | 0.560106142 | 0.796474822 |
| chr6:26607421-26607646 | -0.299539107 | 0.56017476 | 0.796544105 |
| chr14:23285962-23286865 | 0.185591214 | 0.560175656 | 0.796544105 |
| chr9:82532541-82532798 | 0.311643864 | 0.560189769 | 0.796549391 |
| chr11:65261483-65262345 | 0.192096455 | 0.560251263 | 0.796622047 |
| chr3:69084717-69085396 | -0.194945355 | 0.560313415 | 0.796695637 |
| chr14:77009036-77009387 | -0.209646142 | 0.560385064 | 0.796777566 |
| chr4:118077278-118077591 | -0.238990939 | 0.560391833 | 0.796777566 |
| chr3:163849285-163849510 | -0.315235969 | 0.560426766 | 0.796812449 |
| chr1:78617436-78618032 | -0.213004124 | 0.560455937 | 0.796830771 |
| chr8:9151522-9151757 | 0.251877744 | 0.560463208 | 0.796830771 |
| chr1:155167336-155168201 | 0.192097014 | 0.560470851 | 0.796830771 |
| chr8:90709095-90709349 | 0.275259428 | 0.560504456 | 0.796833877 |
| chr8:118012480-118012680 | -0.318014324 | 0.560523953 | 0.796833877 |
| chr22:18119923-18120148 | -0.285854716 | 0.560547028 | 0.796833877 |
| chr3:114370894-114371110 | -0.299345744 | 0.560555906 | 0.796833877 |

| | | | |
|---|---|---|---|
| chr12:113220694-113221933 | 0.183888557 | 0.560559433 | 0.796833877 |
| chr17:65055556-65057236 | -0.182756094 | 0.560562433 | 0.796833877 |
| chr10:96043124-96044555 | -0.187795601 | 0.560579822 | 0.796833877 |
| chr6:138161732-138162729 | 0.192618727 | 0.560586488 | 0.796833877 |
| chr7:74083497-74083863 | -0.217473428 | 0.560587741 | 0.796833877 |
| chr3:185270556-185270834 | -0.234638396 | 0.560588819 | 0.796833877 |
| chr17:73216288-73216501 | -0.300904047 | 0.560596878 | 0.796833877 |
| chr15:67044418-67044658 | 0.271625925 | 0.560601357 | 0.796833877 |
| chr12:116709480-116709680 | -0.279929414 | 0.560614318 | 0.796833877 |
| chr3:8569932-8570179 | 0.298475318 | 0.56061863 | 0.796833877 |
| chr8:99859892-99860131 | 0.244274593 | 0.560646058 | 0.796857611 |
| chr7:46363734-46364244 | -0.239187716 | 0.560669507 | 0.796857611 |
| chr6:158518584-158518849 | 0.235513339 | 0.56067037 | 0.796857611 |
| chr7:151705494-151705865 | -0.231787972 | 0.560676927 | 0.796857611 |
| chr17:48068661-48068881 | 0.257882289 | 0.560721343 | 0.79686509 |
| chr11:777377-777687 | 0.230466828 | 0.560729014 | 0.79686509 |
| chr16:85267637-85267857 | 0.270364154 | 0.560729367 | 0.79686509 |
| chr9:81690596-81690876 | -0.20986039 | 0.560732167 | 0.79686509 |
| chr14:102516492-102517134 | 0.200499774 | 0.560745553 | 0.79686509 |
| chr6:132963058-132963333 | 0.248938014 | 0.56074594 | 0.79686509 |
| chr1:152109765-152111174 | 0.185629697 | 0.560754989 | 0.79686509 |
| chr7:136566994-136567249 | -0.301756491 | 0.560785421 | 0.796893556 |
| chr9:34048403-34048737 | -0.219584226 | 0.560810132 | 0.796913891 |
| chr1:16236179-16237416 | -0.191108757 | 0.560826011 | 0.796921676 |
| chr20:50670126-50671090 | 0.181916175 | 0.560910671 | 0.797027195 |
| chr11:75525739-75526303 | 0.195497219 | 0.560966561 | 0.797080898 |
| chr9:127418060-127418421 | -0.234086465 | 0.56096927 | 0.797080898 |
| chr6:140921511-140921894 | -0.250394553 | 0.561001811 | 0.797112353 |
| chr8:119873445-119874884 | -0.182988206 | 0.56107531 | 0.797202002 |
| chr17:58952122-58952697 | 0.194261978 | 0.56110056 | 0.797212772 |
| chr6:109705223-109705516 | 0.231367768 | 0.561103698 | 0.797212772 |
| chr9:95510689-95511395 | 0.193737977 | 0.561125633 | 0.797229153 |
| chr6:153553504-153553981 | -0.220050221 | 0.561137438 | 0.797231144 |
| chr11:77259956-77260156 | -0.324668337 | 0.561195527 | 0.797298888 |
| chr2:48572731-48573268 | -0.206441765 | 0.56123544 | 0.797317506 |
| chr8:29314808-29315052 | 0.264845471 | 0.561239766 | 0.797317506 |
| chr19:29524095-29524295 | 0.287391059 | 0.561239849 | 0.797317506 |
| chr20:17854361-17854590 | 0.247973834 | 0.561300877 | 0.79738282 |
| chr20:56796418-56796668 | 0.234697417 | 0.561314482 | 0.79738282 |
| chr16:1351744-1351986 | 0.249166309 | 0.561317044 | 0.79738282 |
| chr4:98994990-98996408 | 0.183553899 | 0.561349967 | 0.797414806 |
| chr9:5832329-5833364 | -0.192996108 | 0.561406256 | 0.797479981 |
| chr16:2214762-2215537 | 0.188612776 | 0.561418071 | 0.797481979 |
| chr1:201120380-201120580 | 0.262756678 | 0.56143001 | 0.797484153 |
| chr5:181187531-181188685 | -0.18239777 | 0.561441345 | 0.797485471 |
| chr2:73925084-73925284 | 0.27581235 | 0.561470046 | 0.797488485 |
| chr3:52001494-52001697 | 0.27581235 | 0.561470046 | 0.797488485 |
| chrX:130401385-130402408 | 0.192581037 | 0.561474692 | 0.797488485 |
| chrX:121581987-121582239 | 0.289792432 | 0.561507218 | 0.797497833 |
| chr3:180912112-180913009 | -0.187327401 | 0.561508073 | 0.797497833 |

| | | | |
|---|---|---|---|
| chr17:75565072-75565515 | 0.200055744 | 0.561516413 | 0.797497833 |
| chr19:41785685-41785916 | -0.285701939 | 0.561522906 | 0.797497833 |
| chr2:69742664-69742906 | -0.250399187 | 0.561534804 | 0.797499949 |
| chr17:54976462-54976678 | 0.296244556 | 0.561559201 | 0.797505381 |
| chrX:47560861-47561757 | 0.188303513 | 0.561565185 | 0.797505381 |
| chr6:7541144-7542007 | 0.192150034 | 0.561569853 | 0.797505381 |
| chr6:41318668-41319167 | -0.211277114 | 0.561595273 | 0.797526698 |
| chrX:49043064-49044076 | 0.184580455 | 0.561610614 | 0.797533703 |
| chr3:153970850-153971050 | -0.291036143 | 0.561651134 | 0.797571591 |
| chr14:89500607-89500807 | -0.298627446 | 0.561658645 | 0.797571591 |
| chr6:16239167-16239918 | 0.204399844 | 0.561668522 | 0.797571591 |
| chr1:79568875-79569075 | 0.336759007 | 0.561730488 | 0.79763812 |
| chr1:244299486-244300142 | -0.206449296 | 0.561736193 | 0.79763812 |
| chr14:20454465-20455791 | 0.182261121 | 0.561767913 | 0.797668379 |
| chr14:75452933-75453709 | -0.192509003 | 0.56179254 | 0.797688564 |
| chr1:178501745-178502052 | -0.229638542 | 0.561822595 | 0.797703544 |
| chr1:200638190-200638412 | 0.245933179 | 0.561834594 | 0.797703544 |
| chr13:52194094-52195082 | -0.194388375 | 0.561840575 | 0.797703544 |
| chr2:206135025-206135225 | 0.330514141 | 0.561855145 | 0.797703544 |
| chr3:163101231-163101431 | 0.330514141 | 0.561855145 | 0.797703544 |
| chr17:18182909-18183975 | -0.185234281 | 0.561883984 | 0.797729708 |
| chr3:181003283-181003500 | 0.303701763 | 0.561916704 | 0.797750878 |
| chr11:116732790-116733010 | 0.257957038 | 0.561919718 | 0.797750878 |
| chr8:138914442-138914896 | 0.21187556 | 0.561941401 | 0.797763785 |
| chr4:153204241-153204556 | 0.215484445 | 0.561949633 | 0.797763785 |
| chr5:23812729-23813052 | 0.237365277 | 0.561998054 | 0.797817744 |
| chr6:31826857-31827953 | 0.184706359 | 0.562021104 | 0.797835683 |
| chr8:123766258-123766516 | -0.205761924 | 0.562075232 | 0.797897741 |
| chr7:42337658-42337863 | -0.322438416 | 0.562097957 | 0.797915217 |
| chr12:109875607-109875827 | 0.255925467 | 0.562153383 | 0.797973955 |
| chr21:6362106-6362616 | 0.211025648 | 0.562164171 | 0.797973955 |
| chr8:123502256-123502473 | 0.262210305 | 0.562170579 | 0.797973955 |
| chr7:50835177-50835469 | -0.262284193 | 0.562211211 | 0.798016847 |
| chr10:97640222-97640993 | 0.184963424 | 0.562298846 | 0.798126452 |
| chr8:95758699-95758989 | 0.229598546 | 0.562322409 | 0.798145113 |
| chr19:39133369-39133790 | -0.233017759 | 0.562438886 | 0.798285263 |
| chr3:135439111-135439938 | -0.197988654 | 0.562445655 | 0.798285263 |
| chr11:111918611-111919356 | 0.197715443 | 0.562452405 | 0.798285263 |
| chr7:18894239-18894440 | 0.270168953 | 0.562492985 | 0.798318688 |
| chr17:34980262-34981409 | 0.192159405 | 0.562496793 | 0.798318688 |
| chr12:109020793-109021290 | 0.211011471 | 0.56251328 | 0.798327299 |
| chr12:119938031-119938316 | 0.266016499 | 0.562577411 | 0.798403526 |
| chr1:181190477-181190865 | 0.20870037 | 0.562604338 | 0.798416248 |
| chr10:3502646-3503033 | -0.234519767 | 0.562607215 | 0.798416248 |
| chr1:240475804-240476232 | 0.232567851 | 0.56266215 | 0.798425622 |
| chr1:41322831-41323181 | 0.224605109 | 0.562667272 | 0.798425622 |
| chr17:79578864-79579112 | 0.282269802 | 0.562668229 | 0.798425622 |
| chr2:44044213-44044704 | 0.201032963 | 0.562669046 | 0.798425622 |
| chr8:95133144-95134396 | 0.181446534 | 0.562676707 | 0.798425622 |
| chr12:123744842-123745152 | 0.252692381 | 0.562686383 | 0.798425622 |

| | | | |
|---|---|---|---|
| chr3:22373205-22373405 | 0.305079452 | 0.56268952 | 0.798425622 |
| chr7:74417486-74417743 | -0.260855573 | 0.562707008 | 0.798425622 |
| chr4:70704232-70706138 | -0.183451635 | 0.562707603 | 0.798425622 |
| chr8:20436786-20437171 | -0.247965764 | 0.562773422 | 0.798504225 |
| chr6:160956170-160956475 | -0.296045987 | 0.562839013 | 0.798580233 |
| chr4:27164041-27164291 | -0.310867976 | 0.562847836 | 0.798580233 |
| chr6:72182539-72182755 | 0.307540265 | 0.562884935 | 0.798618081 |
| chr11:134244763-134244973 | -0.293658714 | 0.562899391 | 0.798623804 |
| chr17:50055029-50055379 | 0.20068488 | 0.562929089 | 0.7986411 |
| chr12:108950972-108951421 | -0.230306548 | 0.562936261 | 0.7986411 |
| chr4:86791252-86791452 | -0.324228697 | 0.562946903 | 0.7986411 |
| chr2:167689502-167689882 | -0.240601146 | 0.562953275 | 0.7986411 |
| chr2:161319180-161319522 | 0.302794776 | 0.562992042 | 0.79868131 |
| chr17:58640535-58640742 | 0.202511292 | 0.563040458 | 0.798701194 |
| chr1:236517328-236517559 | -0.263236008 | 0.563053674 | 0.798701194 |
| chr3:33787379-33787707 | 0.234035223 | 0.563068199 | 0.798701194 |
| chr16:67846410-67847329 | 0.187610818 | 0.563072804 | 0.798701194 |
| chr2:1990873-1991092 | -0.288547418 | 0.563102575 | 0.798701194 |
| chr2:39008101-39008338 | -0.288547418 | 0.563102575 | 0.798701194 |
| chr13:106129350-106129626 | 0.274507711 | 0.563109547 | 0.798701194 |
| chr5:32852323-32852523 | -0.301616507 | 0.563120627 | 0.798701194 |
| chr8:55589405-55589632 | -0.301568329 | 0.563127806 | 0.798701194 |
| chr1:20476386-20476741 | -0.243246302 | 0.563129334 | 0.798701194 |
| chr1:102097482-102097793 | -0.243246302 | 0.563129334 | 0.798701194 |
| chr17:5468563-5469248 | -0.198471183 | 0.563131145 | 0.798701194 |
| chr5:85048647-85049466 | 0.200737306 | 0.563142358 | 0.798702313 |
| chr3:50112578-50112808 | -0.287136565 | 0.563211426 | 0.798774285 |
| chr2:100984926-100985230 | -0.263174155 | 0.563213953 | 0.798774285 |
| chr8:69511536-69511752 | 0.290823519 | 0.5632802 | 0.798853453 |
| chr10:109903123-109903516 | -0.238537993 | 0.563298371 | 0.798860562 |
| chr1:204301262-204301904 | -0.195322024 | 0.563325376 | 0.798860562 |
| chr12:1630344-1630598 | -0.267254185 | 0.563330227 | 0.798860562 |
| chr22:38982225-38982499 | 0.248634412 | 0.563338637 | 0.798860562 |
| chr13:37560519-37560719 | -0.348200556 | 0.563344757 | 0.798860562 |
| chr8:117847624-117848061 | 0.214727234 | 0.563347769 | 0.798860562 |
| chr18:11518984-11519279 | 0.284240858 | 0.56336006 | 0.798863207 |
| chr3:196941788-196943338 | -0.1835898 | 0.563386473 | 0.798885876 |
| chr22:31313373-31313719 | 0.237066297 | 0.563407737 | 0.798901244 |
| chr20:54062719-54063293 | 0.183760629 | 0.563419097 | 0.798902568 |
| chr8:49913785-49914020 | -0.276994384 | 0.563459276 | 0.798944754 |
| chr19:55095495-55095853 | 0.250837376 | 0.563474127 | 0.798951027 |
| chr19:17496292-17496629 | 0.23421512 | 0.56351043 | 0.798979362 |
| chr20:46151889-46152216 | 0.235085212 | 0.563520095 | 0.798979362 |
| chr1:100430400-100430666 | 0.298639451 | 0.563531662 | 0.798979362 |
| chr17:66076762-66076962 | 0.284010388 | 0.563535821 | 0.798979362 |
| chr3:49856226-49857049 | 0.198448227 | 0.563554986 | 0.798982128 |
| chr14:82721110-82721337 | -0.298592492 | 0.563565842 | 0.798982128 |
| chr7:123682720-123682960 | 0.294238206 | 0.563569055 | 0.798982128 |
| chr1:52296320-52296624 | 0.242219485 | 0.563594404 | 0.799003282 |
| chr2:110854790-110855090 | -0.257439278 | 0.563619393 | 0.799015395 |

| | | | |
|---|---|---|---|
| chr14:21103556-21104370 | 0.185834729 | 0.563651257 | 0.799015395 |
| chr5:177581573-177581807 | -0.273658299 | 0.563652598 | 0.799015395 |
| chr1:202237090-202237417 | -0.199446989 | 0.563660351 | 0.799015395 |
| chr6:14664696-14664906 | -0.312178847 | 0.56367804 | 0.799015395 |
| chr9:42951298-42951528 | -0.312178847 | 0.56367804 | 0.799015395 |
| chr14:54396422-54397674 | 0.182816655 | 0.563686758 | 0.799015395 |
| chr3:149564221-149564421 | 0.259354819 | 0.563696323 | 0.799015395 |
| chr10:112446643-112447693 | -0.189078027 | 0.563701137 | 0.799015395 |
| chr16:30370326-30371347 | 0.182260758 | 0.563707228 | 0.799015395 |
| chr7:4700699-4700956 | 0.282506092 | 0.563729718 | 0.799032493 |
| chr17:8073635-8073869 | 0.249883114 | 0.563786401 | 0.799097388 |
| chr2:181104602-181104839 | 0.269616139 | 0.563796361 | 0.799097388 |
| chr16:68328607-68328925 | 0.215555095 | 0.563856082 | 0.79916725 |
| chr14:70416581-70417260 | 0.200477472 | 0.563878784 | 0.799180576 |
| chr2:105930796-105930996 | 0.318967126 | 0.563886345 | 0.799180576 |
| chr4:65293694-65294575 | -0.212483998 | 0.563914502 | 0.7992057 |
| chr14:63276840-63277062 | -0.268900363 | 0.563983142 | 0.799288196 |
| chr12:132283696-132283929 | -0.281563021 | 0.564004749 | 0.799304033 |
| chr12:121009793-121010078 | 0.250273517 | 0.564027198 | 0.799307426 |
| chr3:182162397-182162665 | -0.249737208 | 0.564028006 | 0.799307426 |
| chr3:155526760-155527000 | 0.272914234 | 0.5640824 | 0.799357623 |
| chr11:30143164-30143441 | -0.247375788 | 0.564084293 | 0.799357623 |
| chr8:95127008-95127374 | -0.239669668 | 0.564108637 | 0.799377337 |
| chr19:10404118-10404491 | -0.216961138 | 0.564213767 | 0.799487153 |
| chr17:16353515-16353737 | 0.313423934 | 0.56421559 | 0.799487153 |
| chr1:40868299-40868644 | 0.218449843 | 0.564225801 | 0.799487153 |
| chr14:84587126-84587611 | -0.227242318 | 0.564227869 | 0.799487153 |
| chr4:154982289-154982841 | -0.213474039 | 0.564258426 | 0.799501548 |
| chr17:59059138-59059778 | 0.18763639 | 0.564258897 | 0.799501548 |
| chr8:94579245-94579445 | 0.301876827 | 0.564292558 | 0.799513767 |
| chr12:51262768-51262968 | 0.293915179 | 0.564298824 | 0.799513767 |
| chr14:53412984-53413184 | 0.293915179 | 0.564298824 | 0.799513767 |
| chr17:3844019-3844230 | -0.272173089 | 0.564310147 | 0.799515026 |
| chr7:1176167-1176671 | -0.202498477 | 0.564327996 | 0.79952553 |
| chr1:70410617-70411705 | -0.182892882 | 0.564376523 | 0.7995444 |
| chr13:45473711-45474025 | -0.247740303 | 0.564378188 | 0.7995444 |
| chr3:48429397-48430280 | 0.193810894 | 0.564378726 | 0.7995444 |
| chr7:74288778-74289021 | -0.259733609 | 0.564394895 | 0.7995444 |
| chr1:33165235-33166100 | -0.205694969 | 0.564401847 | 0.7995444 |
| chr2:148876583-148877242 | -0.192093226 | 0.564403925 | 0.7995444 |
| chr17:44941852-44942084 | -0.291168822 | 0.564416906 | 0.799548007 |
| chr1:116389512-116389712 | 0.264713575 | 0.564437217 | 0.799561997 |
| chr11:12556769-12557192 | -0.224653604 | 0.564498226 | 0.799632527 |
| chr15:90184029-90185537 | 0.183456061 | 0.564509158 | 0.799632527 |
| chr5:138752919-138754449 | -0.183800606 | 0.564518314 | 0.799632527 |
| chr12:102793265-102793465 | -0.34768536 | 0.564550312 | 0.799663068 |
| chr1:46348095-46348474 | 0.211382051 | 0.564572377 | 0.799679539 |
| chr7:74635647-74636496 | 0.198702938 | 0.564583105 | 0.799679952 |
| chr3:152079599-152079799 | 0.30125249 | 0.564608425 | 0.799687239 |
| chr2:70923463-70923758 | 0.250468048 | 0.564609123 | 0.799687239 |

| | | | |
|---|---|---|---|
| chr19:39445346-39445746 | 0.202743576 | 0.564633022 | 0.799706305 |
| chr6:15642361-15642602 | -0.287416713 | 0.564649857 | 0.799715367 |
| chr11:75031988-75032397 | -0.21498344 | 0.564702189 | 0.799768967 |
| chr18:44679309-44679566 | -0.259371821 | 0.564717064 | 0.799768967 |
| chr6:125205060-125205491 | -0.225950612 | 0.564719016 | 0.799768967 |
| chr3:177084603-177084987 | -0.230090938 | 0.564736954 | 0.79977959 |
| chr20:50916348-50916548 | 0.263327351 | 0.564767685 | 0.799800543 |
| chr2:176177806-176178890 | -0.194253856 | 0.564772627 | 0.799800543 |
| chr10:11687832-11688124 | 0.217432051 | 0.564796743 | 0.799819914 |
| chr7:74174002-74174895 | 0.194489044 | 0.564809013 | 0.799822506 |
| chr1:34755360-34755586 | -0.276864888 | 0.564840729 | 0.7998439 |
| chr11:67345733-67346441 | -0.200169468 | 0.564844998 | 0.7998439 |
| chr4:37977191-37977631 | 0.219946834 | 0.564869707 | 0.799862704 |
| chr3:45210542-45210861 | -0.23497511 | 0.564879155 | 0.799862704 |
| chr2:100562444-100563539 | 0.181757459 | 0.564890743 | 0.799864331 |
| chr9:92877532-92878388 | 0.197026212 | 0.564905324 | 0.799870195 |
| chr1:205367852-205368183 | -0.230783672 | 0.564923764 | 0.799876466 |
| chr10:119165153-119166170 | -0.187830534 | 0.564936461 | 0.799876466 |
| chr10:26606357-26606887 | -0.223588969 | 0.56494107 | 0.799876466 |
| chr3:112304687-112304887 | -0.297782511 | 0.564952505 | 0.799877875 |
| chr17:67995989-67996192 | -0.226593477 | 0.5649712 | 0.799889564 |
| chr13:50319410-50320199 | -0.202327629 | 0.565018997 | 0.799942453 |
| chr3:138834155-138835370 | 0.184675902 | 0.565060174 | 0.79998597 |
| chr1:149962103-149962339 | 0.236320311 | 0.565081991 | 0.799993693 |
| chr10:5125459-5125762 | -0.277340533 | 0.565096952 | 0.799993693 |
| chr10:123427091-123427385 | -0.277340533 | 0.565096952 | 0.799993693 |
| chr16:89171215-89171840 | -0.186063398 | 0.565118338 | 0.800009188 |
| chr20:46014200-46014621 | 0.215016826 | 0.56516741 | 0.8000607 |
| chr22:32797007-32797292 | -0.270540413 | 0.565175609 | 0.8000607 |
| chr11:70191623-70191823 | 0.283511304 | 0.565207185 | 0.800090617 |
| chr8:145026625-145027213 | 0.191139701 | 0.565232381 | 0.800111502 |
| chrX:100768764-100769102 | 0.222595999 | 0.565283008 | 0.800162674 |
| chr1:145280580-145281851 | 0.180720836 | 0.565291929 | 0.800162674 |
| chr3:179811668-179811947 | -0.253233393 | 0.56529986 | 0.800162674 |
| chr14:32576150-32576454 | 0.227049647 | 0.565357125 | 0.800228947 |
| chr10:119006682-119007330 | 0.195418229 | 0.565424082 | 0.800308937 |
| chr6:150072292-150072505 | -0.256448059 | 0.565473714 | 0.800352001 |
| chr18:22559932-22560226 | -0.253740245 | 0.565475398 | 0.800352001 |
| chr5:132557429-132557801 | -0.22187497 | 0.565506389 | 0.800381081 |
| chr17:28854591-28855376 | 0.195291626 | 0.565535148 | 0.800405814 |
| chr1:172532153-172533653 | -0.17915369 | 0.565544757 | 0.800405814 |
| chr5:38193137-38193408 | 0.242916555 | 0.565573016 | 0.800431023 |
| chr16:89701463-89702367 | -0.182521666 | 0.565623401 | 0.800480373 |
| chr20:36439490-36439765 | 0.245884381 | 0.56562878 | 0.800480373 |
| chr5:76083911-76084181 | -0.282125254 | 0.565697723 | 0.800563155 |
| chr13:46659611-46660105 | 0.204498326 | 0.565782293 | 0.800627914 |
| chr19:34814944-34815144 | 0.305481188 | 0.565793558 | 0.800627914 |
| chr15:91302710-91303154 | 0.220121391 | 0.565794439 | 0.800627914 |
| chr6:46652482-46653642 | -0.188328458 | 0.565794506 | 0.800627914 |
| chr1:55012690-55012921 | -0.283230768 | 0.565795728 | 0.800627914 |

| | | | |
|---|---|---|---|
| chr1:147699984-147700430 | -0.20723628 | 0.565850866 | 0.80069115 |
| chr5:141778641-141778876 | 0.277511328 | 0.565920848 | 0.800775387 |
| chr13:32277984-32278197 | -0.300843191 | 0.565959678 | 0.800806524 |
| chr15:98241125-98241537 | -0.240356971 | 0.565963756 | 0.800806524 |
| chr17:63773261-63774688 | -0.179248415 | 0.565976767 | 0.800810147 |
| chr3:137065331-137065629 | -0.261611997 | 0.566015165 | 0.800849687 |
| chr12:26918747-26919181 | -0.231364729 | 0.566073232 | 0.800917056 |
| chr2:236167682-236167968 | 0.257610421 | 0.566112026 | 0.800948398 |
| chr15:89875060-89875299 | 0.253498305 | 0.56611629 | 0.800948398 |
| chr2:18824237-18824791 | -0.200711034 | 0.566136551 | 0.800962273 |
| chr17:50044272-50044472 | -0.292767432 | 0.566148224 | 0.800963999 |
| chr10:48523823-48524023 | -0.296456948 | 0.566166453 | 0.800975 |
| chr2:200713022-200713473 | -0.205946635 | 0.56618922 | 0.800992419 |
| chr8:95101133-95101595 | 0.20665163 | 0.566207291 | 0.801003195 |
| chr8:27774176-27774848 | 0.192167818 | 0.566219789 | 0.801006087 |
| chr20:53863977-53865641 | 0.177657203 | 0.56623969 | 0.801019451 |
| chr2:9630298-9631484 | -0.181250457 | 0.566272814 | 0.801043903 |
| chr5:16813785-16814038 | 0.270060206 | 0.566281013 | 0.801043903 |
| chr13:41204554-41204787 | -0.298366831 | 0.566298208 | 0.801043903 |
| chr7:2631517-2631989 | -0.186730882 | 0.566316677 | 0.801043903 |
| chr10:93596637-93596837 | 0.280464608 | 0.566322811 | 0.801043903 |
| chr1:16206205-16207196 | 0.189123117 | 0.566349695 | 0.801043903 |
| chr4:121997974-121998345 | 0.235077427 | 0.566354753 | 0.801043903 |
| chr7:89887358-89887605 | -0.305340117 | 0.566357469 | 0.801043903 |
| chr10:14501924-14502158 | 0.287131332 | 0.566360676 | 0.801043903 |
| chr3:130111329-130112692 | -0.17908317 | 0.56636152 | 0.801043903 |
| chr5:1271265-1271465 | -0.312520975 | 0.566404853 | 0.801078667 |
| chr8:42893778-42893988 | 0.240111205 | 0.56640701 | 0.801078667 |
| chr1:182391130-182392413 | 0.183084416 | 0.56643492 | 0.801085792 |
| chr10:107814879-10781532C | -0.237182721 | 0.566438997 | 0.801085792 |
| chr2:169789770-169790328 | -0.199935291 | 0.566443412 | 0.801085792 |
| chr10:95532931-95533131 | 0.323928914 | 0.566463105 | 0.801098856 |
| chr21:45334676-45334908 | -0.286705479 | 0.566489035 | 0.80112074 |
| chr5:39547633-39547845 | -0.286117273 | 0.566558078 | 0.801194927 |
| chr14:68140344-68140544 | -0.273026051 | 0.566582605 | 0.801194927 |
| chr14:81435460-81437040 | -0.181741537 | 0.566590949 | 0.801194927 |
| chr19:48190357-48190911 | 0.199089058 | 0.566600839 | 0.801194927 |
| chr6:143176028-143176334 | 0.264185778 | 0.566625578 | 0.801194927 |
| chr3:12149097-12149312 | -0.282720815 | 0.566626616 | 0.801194927 |
| chr15:84516824-84517069 | 0.272038317 | 0.566634692 | 0.801194927 |
| chr1:32448351-32448856 | 0.214360767 | 0.566644498 | 0.801194927 |
| chr3:157437323-157437665 | 0.237882344 | 0.56665065 | 0.801194927 |
| chr8:140790757-140791312 | -0.208405826 | 0.566656279 | 0.801194927 |
| chr18:37537861-37538237 | -0.219523966 | 0.566660904 | 0.801194927 |
| chr6:121187774-121188346 | -0.207520511 | 0.566666971 | 0.801194927 |
| chr1:37096541-37096782 | -0.277746375 | 0.566685732 | 0.801206668 |
| chr4:7116223-7116474 | 0.28649504 | 0.566730578 | 0.801255288 |
| chr5:42949328-42949586 | -0.251457519 | 0.56674111 | 0.801255393 |
| chr16:88452608-88453359 | 0.184327265 | 0.566755352 | 0.801260745 |
| chr10:128324183-128324403 | -0.285820038 | 0.56682048 | 0.801338034 |

| | | | |
|---|---|---|---|
| chr6:106973762-106975480 | 0.182577002 | 0.566861123 | 0.801366386 |
| chr8:119200710-119201099 | -0.209519171 | 0.566871505 | 0.801366386 |
| chr15:34360337-34360672 | 0.206170204 | 0.566874807 | 0.801366386 |
| chr1:177181342-177182285 | -0.18980737 | 0.566882369 | 0.801366386 |
| chr22:41970070-41970270 | -0.311610234 | 0.566904969 | 0.8013712 |
| chr16:3150385-3151053 | 0.184538391 | 0.566906691 | 0.8013712 |
| chr5:5136064-5136276 | 0.264876838 | 0.566943375 | 0.801407285 |
| chr2:85772388-85772645 | 0.240864777 | 0.566978428 | 0.801407285 |
| chr5:73437548-73437821 | 0.240864777 | 0.566978428 | 0.801407285 |
| chr7:66654128-66655320 | 0.183113999 | 0.566979692 | 0.801407285 |
| chr11:134223819-134225209 | 0.182593244 | 0.566994288 | 0.801407285 |
| chr3:14507293-14507554 | 0.246313863 | 0.566994974 | 0.801407285 |
| chr7:45124033-45124393 | -0.245548343 | 0.567027541 | 0.801432211 |
| chr17:42121203-42121489 | -0.230589841 | 0.567033529 | 0.801432211 |
| chr11:75507947-75508521 | -0.211736863 | 0.567059627 | 0.801454314 |
| chr6:39892627-39892836 | 0.269549741 | 0.56707646 | 0.801463322 |
| chr20:31593216-31594105 | -0.185723106 | 0.567122943 | 0.801514233 |
| chr11:7281146-7281354 | 0.282981325 | 0.567149444 | 0.801534511 |
| chr5:112367004-112367308 | -0.228643187 | 0.567158213 | 0.801534511 |
| chr17:8237147-8237490 | -0.241933781 | 0.567187357 | 0.801560915 |
| chr10:60114784-60115080 | -0.258483217 | 0.567221224 | 0.801568933 |
| chr15:91032929-91033510 | 0.194047673 | 0.567224368 | 0.801568933 |
| chr7:44082195-44083102 | -0.181678331 | 0.567224415 | 0.801568933 |
| chr12:68745558-68747041 | -0.181402708 | 0.567241842 | 0.801578093 |
| chr12:67268403-67270121 | -0.178911813 | 0.567255306 | 0.801578093 |
| chr3:63667280-63667521 | -0.225137532 | 0.567262281 | 0.801578093 |
| chr11:130636821-130637461 | -0.213947456 | 0.567328479 | 0.801651924 |
| chr9:92502011-92502362 | 0.223445811 | 0.567335455 | 0.801651924 |
| chr11:7600293-7600695 | -0.203624559 | 0.567357442 | 0.801664137 |
| chr10:96375057-96375778 | -0.18898152 | 0.567365024 | 0.801664137 |
| chr1:205919013-205919281 | 0.210776217 | 0.567412987 | 0.801710613 |
| chr1:81800319-81800519 | -0.279057573 | 0.567419143 | 0.801710613 |
| chr12:119444513-119444831 | -0.236817184 | 0.567445488 | 0.801710613 |
| chr5:139637469-139638115 | -0.192646273 | 0.567450061 | 0.801710613 |
| chr14:77321701-77321947 | 0.20622744 | 0.567466687 | 0.801710613 |
| chr12:121672978-121673236 | 0.241258299 | 0.567475315 | 0.801710613 |
| chr2:212905502-212906443 | -0.186135862 | 0.567486315 | 0.801710613 |
| chr8:102569130-102569465 | 0.223621256 | 0.567491295 | 0.801710613 |
| chr3:195542764-195544018 | -0.184140082 | 0.567512511 | 0.801710613 |
| chr1:175844935-175845145 | -0.263044504 | 0.567523608 | 0.801710613 |
| chr8:142614377-142615189 | -0.189154288 | 0.567526432 | 0.801710613 |
| chr13:24359210-24359548 | -0.240163367 | 0.567532114 | 0.801710613 |
| chr3:86990921-86991703 | -0.190083003 | 0.567536917 | 0.801710613 |
| chr7:35219482-35219728 | 0.272743535 | 0.5675444 | 0.801710613 |
| chr20:47272789-47273164 | 0.183782257 | 0.567557169 | 0.80171387 |
| chr5:148860845-148861080 | -0.272358487 | 0.567572518 | 0.80172077 |
| chr12:49887996-49888259 | 0.260417034 | 0.567590827 | 0.801731852 |
| chr8:23105278-23106214 | 0.204871786 | 0.567680777 | 0.801840967 |
| chr7:5085422-5085622 | 0.243972885 | 0.567697503 | 0.801840967 |
| chr18:7022254-7022613 | -0.228504065 | 0.56769947 | 0.801840967 |

| | | | |
|---|---|---|---|
| chr4:83583955-83584388 | 0.229649508 | 0.567721273 | 0.801845373 |
| chr3:11620344-11620623 | -0.255528017 | 0.567723519 | 0.801845373 |
| chr3:107206231-107206431 | -0.276014913 | 0.567745788 | 0.801862044 |
| chr5:74640321-74641639 | 0.182848842 | 0.567759475 | 0.801866594 |
| chr13:49301239-49301553 | -0.231498801 | 0.567770572 | 0.801867487 |
| chr16:88087529-88087931 | -0.246572308 | 0.567793738 | 0.801883256 |
| chr3:105858271-105858534 | -0.280064801 | 0.567813005 | 0.801883256 |
| chr7:25855000-25855371 | -0.201366867 | 0.567813134 | 0.801883256 |
| chr15:85840537-85840934 | 0.205052391 | 0.567865406 | 0.801942295 |
| chrX:74437556-74437818 | 0.27377115 | 0.567898382 | 0.801966804 |
| chr3:144585299-144585690 | 0.249801139 | 0.567915596 | 0.801966804 |
| chr19:8525418-8526165 | 0.184430815 | 0.567927401 | 0.801966804 |
| chr2:150485955-150486485 | 0.196042865 | 0.567930248 | 0.801966804 |
| chr12:752203-753529 | -0.184778995 | 0.567935093 | 0.801966804 |
| chr18:54959237-54959607 | -0.242693654 | 0.567957611 | 0.801973478 |
| chr15:70362938-70363249 | 0.231240273 | 0.567960753 | 0.801973478 |
| chr3:120244239-120244627 | 0.211131284 | 0.568006657 | 0.802008172 |
| chr16:67185453-67185653 | 0.241293016 | 0.568008557 | 0.802008172 |
| chr2:69997424-69997624 | -0.293687755 | 0.568016725 | 0.802008172 |
| chr6:97291982-97292182 | -0.292126326 | 0.56804668 | 0.802032379 |
| chr14:101533780-101533980 | -0.287171245 | 0.568073228 | 0.802032379 |
| chr8:127800315-127801753 | -0.180959178 | 0.568075586 | 0.802032379 |
| chr1:163528759-163529037 | -0.274542407 | 0.568075739 | 0.802032379 |
| chr5:57650240-57650639 | -0.218357822 | 0.568128592 | 0.802077488 |
| chr1:163561580-163561851 | 0.269764913 | 0.5681351 | 0.802077488 |
| chr3:174614126-174614326 | -0.301966532 | 0.568150565 | 0.802077488 |
| chr2:139992014-139992214 | 0.321221798 | 0.568152615 | 0.802077488 |
| chr4:120156195-120156470 | 0.256267691 | 0.568167327 | 0.802077488 |
| chr8:89474324-89474524 | -0.290517252 | 0.568174094 | 0.802077488 |
| chr1:167442103-167442504 | 0.203163075 | 0.568180965 | 0.802077488 |
| chr9:101449091-101449714 | -0.192490527 | 0.568242506 | 0.802149585 |
| chr1:115546073-115546493 | 0.199261755 | 0.568271826 | 0.802174149 |
| chr1:226814725-226815686 | 0.192546934 | 0.568293069 | 0.802174149 |
| chr17:80175536-80175736 | -0.299366314 | 0.56832254 | 0.802174149 |
| chr14:71166465-71166666 | 0.259924121 | 0.5683261 | 0.802174149 |
| chr11:32829552-32830671 | -0.188423692 | 0.568327444 | 0.802174149 |
| chr1:47726122-47726602 | 0.209622517 | 0.5683304 | 0.802174149 |
| chr2:105287800-105288084 | -0.252547554 | 0.568339714 | 0.802174149 |
| chr3:169677973-169678440 | 0.240542204 | 0.568346563 | 0.802174149 |
| chr13:27171286-27172455 | 0.184489356 | 0.568354131 | 0.802174149 |
| chr1:204254188-204254476 | -0.21508544 | 0.568364626 | 0.802174185 |
| chr1:160251938-160252291 | -0.214473337 | 0.568397896 | 0.802180536 |
| chr12:110742706-110743302 | -0.194757984 | 0.568401118 | 0.802180536 |
| chr8:123041527-123042910 | -0.178682964 | 0.56841539 | 0.802180536 |
| chr12:119229869-119230124 | -0.242396647 | 0.568444776 | 0.802180536 |
| chr8:43139145-43139434 | -0.232286817 | 0.568449041 | 0.802180536 |
| chr8:88898849-88899332 | 0.207006449 | 0.568452811 | 0.802180536 |
| chr3:75361991-75362724 | -0.18880237 | 0.568459322 | 0.802180536 |
| chr7:152823338-152823680 | -0.241204098 | 0.568462766 | 0.802180536 |
| chr15:59724989-59725246 | 0.280650791 | 0.568463349 | 0.802180536 |

| | | | |
|---|---|---|---|
| chr6:162442715-162442966 | 0.277567421 | 0.568480399 | 0.802189822 |
| chr3:23129658-23130270 | -0.1910185 | 0.568491898 | 0.802191274 |
| chr15:83446706-83448273 | 0.182358598 | 0.56851452 | 0.802208422 |
| chr1:215318750-215318979 | -0.297461142 | 0.568551235 | 0.802234406 |
| chr1:109923481-109923815 | -0.227801874 | 0.568553874 | 0.802234406 |
| chr15:100247156-100247671 | 0.200946101 | 0.568579241 | 0.802255426 |
| chr18:34493100-34494490 | -0.182274261 | 0.568595068 | 0.802262983 |
| chr2:96547340-96547601 | -0.242696642 | 0.568652065 | 0.802282214 |
| chr3:163056948-163057263 | -0.227811022 | 0.568652602 | 0.802282214 |
| chr7:150963004-150963235 | -0.241736202 | 0.568654606 | 0.802282214 |
| chr17:28676047-28676356 | -0.244509456 | 0.568665481 | 0.802282214 |
| chr1:182614777-182615280 | 0.200905442 | 0.568680488 | 0.802282214 |
| chr1:156657552-156657752 | 0.236203108 | 0.568695165 | 0.802282214 |
| chr1:65075414-65075931 | -0.212793144 | 0.568696997 | 0.802282214 |
| chr17:67221130-67221370 | 0.236235442 | 0.568698918 | 0.802282214 |
| chr2:134657794-134658001 | 0.300696185 | 0.568702934 | 0.802282214 |
| chr3:98592738-98592938 | 0.267072695 | 0.568751753 | 0.802325392 |
| chr8:100149980-100150308 | -0.206165514 | 0.568754483 | 0.802325392 |
| chr7:101378891-101379091 | -0.297185768 | 0.568785108 | 0.802353822 |
| chr11:35485692-35486141 | -0.213859842 | 0.568840566 | 0.802400167 |
| chr11:43880546-43880972 | -0.203572162 | 0.568847134 | 0.802400167 |
| chr6:30489262-30489975 | 0.191544094 | 0.568849379 | 0.802400167 |
| chr3:175378339-175378539 | 0.271951899 | 0.568892009 | 0.802442888 |
| chr11:86800268-86800839 | -0.197068725 | 0.56890061 | 0.802442888 |
| chr3:93973157-93974248 | -0.182517376 | 0.568920734 | 0.802450519 |
| chr19:4803871-4804071 | 0.284969168 | 0.568926966 | 0.802450519 |
| chr15:82263288-82263613 | 0.216672268 | 0.568962119 | 0.802471565 |
| chr15:76217078-76217309 | -0.284281442 | 0.568962833 | 0.802471565 |
| chr4:721374-721909 | -0.21744971 | 0.56898987 | 0.802484019 |
| chr4:80215503-80215738 | -0.301217122 | 0.568993319 | 0.802484019 |
| chr20:23626084-23626707 | -0.210103451 | 0.569028502 | 0.802484019 |
| chr4:56301564-56301832 | 0.243710993 | 0.569030915 | 0.802484019 |
| chr17:82974499-82974741 | 0.261536543 | 0.569033892 | 0.802484019 |
| chr5:19820372-19820572 | 0.307574636 | 0.569034504 | 0.802484019 |
| chr12:123094756-123095210 | -0.222291449 | 0.569089466 | 0.80254676 |
| chr10:46030126-46030872 | -0.185759249 | 0.569137065 | 0.802588752 |
| chr4:164700440-164700685 | -0.27374997 | 0.569140193 | 0.802588752 |
| chr5:146815095-146815348 | -0.298544157 | 0.569184497 | 0.802636458 |
| chr8:100951863-100953226 | 0.177454919 | 0.569270196 | 0.802735928 |
| chr15:93372695-93373028 | 0.249975675 | 0.56928208 | 0.802735928 |
| chr20:35146831-35147653 | 0.193425283 | 0.569286466 | 0.802735928 |
| chr10:70222469-70222922 | 0.228028028 | 0.569303067 | 0.802744564 |
| chr7:151519206-151519554 | -0.212479423 | 0.569323931 | 0.80275921 |
| chr6:87139327-87139537 | -0.291159479 | 0.569334886 | 0.802759885 |
| chr15:88274603-88274870 | 0.228300949 | 0.569346294 | 0.802761197 |
| chrX:48835486-48836008 | -0.22102207 | 0.569357013 | 0.802761539 |
| chrX:66110255-66110474 | -0.313018838 | 0.569380217 | 0.802779483 |
| chr19:6372412-6372890 | 0.197404965 | 0.569428716 | 0.802833089 |
| chr21:46184648-46185178 | 0.203634365 | 0.569445073 | 0.802834774 |
| chr10:69801717-69802040 | 0.219905399 | 0.569461449 | 0.802834774 |

| | | | |
|---|---|---|---|
| chr6:53795684-53795902 | -0.229600874 | 0.569469514 | 0.802834774 |
| chr19:38115397-38115708 | 0.250688698 | 0.569471822 | 0.802834774 |
| chr9:38088065-38088265 | -0.291054145 | 0.569605768 | 0.803005583 |
| chr17:13182956-13183156 | -0.290993175 | 0.569613942 | 0.803005583 |
| chr2:18861427-18861627 | 0.31521687 | 0.569653008 | 0.803045881 |
| chr1:147541352-147541563 | 0.267535248 | 0.569673093 | 0.803046807 |
| chr5:82364561-82364795 | -0.287115033 | 0.569687124 | 0.803046807 |
| chr20:18057811-18058097 | 0.203635946 | 0.569691771 | 0.803046807 |
| chr3:170054273-170054592 | 0.222942295 | 0.569697672 | 0.803046807 |
| chr12:42111477-42111841 | 0.22414313 | 0.569706068 | 0.803046807 |
| chr2:112054835-112055637 | 0.182049421 | 0.56972157 | 0.803053885 |
| chr11:9538628-9538828 | -0.327190042 | 0.56981553 | 0.803150729 |
| chr6:49570825-49571055 | -0.276238724 | 0.569821721 | 0.803150729 |
| chr7:89623107-89623333 | -0.276238724 | 0.569821721 | 0.803150729 |
| chr7:104848076-104848321 | -0.273374069 | 0.569861064 | 0.803191406 |
| chr6:117601875-117603046 | -0.179988386 | 0.569878424 | 0.8032011 |
| chr3:4736557-4736852 | -0.246445531 | 0.569958328 | 0.803298943 |
| chr17:18968095-18968415 | -0.221193425 | 0.569982846 | 0.803318722 |
| chr10:32155728-32156406 | -0.193684046 | 0.570005416 | 0.803322799 |
| chr1:202719638-202719974 | -0.200071211 | 0.570006707 | 0.803322799 |
| chr14:104888267-104888554 | 0.229461268 | 0.570026799 | 0.803336339 |
| chr19:46346731-46347640 | 0.187981718 | 0.57003961 | 0.803339618 |
| chr21:45981243-45981706 | -0.21857589 | 0.570081187 | 0.803372832 |
| chr16:2904213-2904450 | 0.218747851 | 0.570085081 | 0.803372832 |
| chr9:133030105-133031103 | -0.1858435 | 0.570100185 | 0.803372832 |
| chr15:64835651-64836394 | 0.18909505 | 0.570105784 | 0.803372832 |
| chr17:62830592-62830792 | 0.262765944 | 0.570115603 | 0.803372832 |
| chr3:166367974-166368443 | -0.225203063 | 0.570135039 | 0.803381622 |
| chr2:128098298-128098498 | -0.326039621 | 0.570142811 | 0.803381622 |
| chr8:142383389-142383593 | 0.264926217 | 0.570174819 | 0.803411949 |
| chr7:138324981-138325762 | -0.195710725 | 0.570205966 | 0.803441062 |
| chr9:136807367-136807622 | 0.2270575 | 0.570236983 | 0.803458056 |
| chr4:139761407-139761787 | -0.220938258 | 0.570238999 | 0.803458056 |
| chr6:27479246-27479823 | -0.196913761 | 0.570303034 | 0.803533506 |
| chr17:76103235-76104030 | 0.189649344 | 0.570340358 | 0.803558012 |
| chr3:141789349-141789613 | -0.244267276 | 0.570341402 | 0.803558012 |
| chr15:82809169-82810389 | 0.181770369 | 0.570397177 | 0.80360893 |
| chr15:70636900-70637100 | 0.285568506 | 0.570421081 | 0.80360893 |
| chr12:81992553-81992763 | -0.31209807 | 0.570422879 | 0.80360893 |
| chr1:23451989-23452389 | 0.208365107 | 0.570423322 | 0.80360893 |
| chr9:131336240-131336532 | 0.234574598 | 0.570439003 | 0.80360893 |
| chr7:156892485-156893766 | -0.179981072 | 0.57044047 | 0.80360893 |
| chr3:5290953-5291153 | -0.293529559 | 0.570494027 | 0.803669602 |
| chr3:25904374-25904574 | -0.309134851 | 0.570555862 | 0.803741934 |
| chr18:57772248-57772730 | -0.217101121 | 0.570582282 | 0.803760565 |
| chr17:5534069-5534338 | -0.306224731 | 0.570592401 | 0.803760565 |
| chr8:106577611-106577980 | -0.208889028 | 0.570600558 | 0.803760565 |
| chr1:150568181-150568830 | 0.184287068 | 0.570627915 | 0.803784325 |
| chr21:38787431-38788249 | -0.187118501 | 0.570649079 | 0.803790412 |
| chr2:127107012-127107929 | 0.192528664 | 0.570661981 | 0.803790412 |

| | | | |
|---|---|---|---|
| chr12:121589925-121590515 | 0.202344393 | 0.570663708 | 0.803790412 |
| chr17:81023789-81024056 | 0.231175015 | 0.570714845 | 0.803847663 |
| chr4:87235811-87236532 | -0.205706729 | 0.57073229 | 0.803854816 |
| chr16:2127780-2127996 | -0.272763191 | 0.570743213 | 0.803854816 |
| chr1:181557374-181557722 | 0.231642494 | 0.570751397 | 0.803854816 |
| chr1:109655607-109656493 | 0.188909502 | 0.570802293 | 0.803903349 |
| chr3:4670409-4670609 | 0.252900108 | 0.570813335 | 0.803903349 |
| chr11:117776691-117776891 | -0.311153197 | 0.570821843 | 0.803903349 |
| chr20:57031396-57031608 | 0.223607272 | 0.570830277 | 0.803903349 |
| chr5:66633797-66634127 | -0.220005673 | 0.570838316 | 0.803903349 |
| chr7:44020241-44020655 | -0.204209117 | 0.570884546 | 0.803953679 |
| chr8:88811535-88811959 | -0.219921429 | 0.570907349 | 0.803962222 |
| chr5:139147826-139148046 | -0.241525526 | 0.570911598 | 0.803962222 |
| chr17:63838546-63838920 | -0.186287947 | 0.571023002 | 0.804104324 |
| chr2:207746067-207746433 | -0.218900319 | 0.571048483 | 0.804119775 |
| chr21:26842975-26843269 | -0.214275752 | 0.571054964 | 0.804119775 |
| chr1:35192796-35193580 | 0.183919757 | 0.571083502 | 0.804145183 |
| chr11:66950108-66950623 | 0.202240071 | 0.571120166 | 0.804182031 |
| chr17:19507514-19508692 | -0.182426755 | 0.57119773 | 0.804276467 |
| chr1:40438506-40438741 | 0.292829805 | 0.571238213 | 0.80431211 |
| chr9:84397104-84397635 | -0.216233684 | 0.571260691 | 0.80431211 |
| chr4:84965560-84967320 | -0.179513168 | 0.57126746 | 0.80431211 |
| chr4:160568531-160568893 | -0.292712111 | 0.571271363 | 0.80431211 |
| chr11:32071888-32072123 | 0.245759506 | 0.57127553 | 0.80431211 |
| chr17:62870693-62870964 | -0.235041964 | 0.571335034 | 0.804370848 |
| chr8:120593173-120593476 | -0.234237695 | 0.571338245 | 0.804370848 |
| chr9:114653364-114653829 | 0.204286789 | 0.571369779 | 0.80438352 |
| chr17:75978717-75980018 | 0.18116858 | 0.571370363 | 0.80438352 |
| chr20:23340562-23340958 | -0.222592914 | 0.571378741 | 0.80438352 |
| chr14:22887321-22887608 | 0.211963343 | 0.571392654 | 0.804384592 |
| chr4:83639946-83640187 | -0.276251057 | 0.571400498 | 0.804384592 |
| chr9:137294685-137295062 | -0.227808092 | 0.571449561 | 0.804438881 |
| chr19:41353629-41354414 | -0.193495837 | 0.571526162 | 0.804531932 |
| chr19:55476130-55477190 | 0.184811196 | 0.571544952 | 0.804535146 |
| chr10:49512204-49512562 | 0.227818943 | 0.571550384 | 0.804535146 |
| chr10:93376418-93376715 | -0.247037022 | 0.571568933 | 0.804535146 |
| chr20:45933719-45934014 | 0.205576172 | 0.571583362 | 0.804535146 |
| chr3:49093923-49094768 | 0.184736685 | 0.571588007 | 0.804535146 |
| chr4:106238854-106239449 | -0.207663553 | 0.571598202 | 0.804535146 |
| chr16:87550281-87550678 | 0.214522687 | 0.571612278 | 0.804535146 |
| chr20:7047293-7047493 | -0.325627444 | 0.571612445 | 0.804535146 |
| chr11:83292918-83293250 | 0.217846708 | 0.571648758 | 0.804571475 |
| chr2:72638832-72639103 | -0.271645764 | 0.571671024 | 0.804575833 |
| chr3:149480359-149480753 | 0.197635123 | 0.571672855 | 0.804575833 |
| chr15:81827890-81828108 | -0.312854543 | 0.571724809 | 0.804602769 |
| chr5:179550241-179551402 | -0.183932708 | 0.571727537 | 0.804602769 |
| chr16:89489166-89489532 | -0.202477987 | 0.571735464 | 0.804602769 |
| chr20:31313187-31313762 | -0.199816381 | 0.571738921 | 0.804602769 |
| chr3:129485689-129486287 | -0.189028556 | 0.571748737 | 0.804602769 |
| chr3:175956890-175957385 | 0.201364685 | 0.571764208 | 0.804602769 |

| | | | |
|---|---|---|---|
| chr13:41222294-41222593 | -0.212236776 | 0.571765501 | 0.804602769 |
| chr13:106535367-106536757 | 0.180484422 | 0.57180578 | 0.804644673 |
| chr18:48975333-48975533 | -0.293198064 | 0.571821875 | 0.804652544 |
| chr7:18666190-18666497 | 0.225401695 | 0.571892774 | 0.80471943 |
| chr12:3365990-3366218 | 0.27636791 | 0.571899009 | 0.80471943 |
| chr6:108133560-108134257 | 0.192884142 | 0.571901276 | 0.80471943 |
| chr17:56798965-56799238 | -0.226303293 | 0.571911416 | 0.80471943 |
| chr8:109189658-109189968 | -0.248762182 | 0.57194371 | 0.804735972 |
| chr10:50605342-50605736 | -0.209751029 | 0.571963064 | 0.804735972 |
| chr8:70327799-70328231 | -0.200609694 | 0.571980122 | 0.804735972 |
| chrX:107372539-107372739 | 0.325379175 | 0.57198084 | 0.804735972 |
| chr3:29280692-29281124 | -0.226552457 | 0.571985957 | 0.804735972 |
| chr16:66947240-66947491 | 0.246721408 | 0.571997712 | 0.804735972 |
| chr1:205688222-205688436 | -0.229814313 | 0.571998383 | 0.804735972 |
| chr11:83410726-83411261 | -0.220054276 | 0.572016601 | 0.804735972 |
| chr15:82540113-82541457 | 0.179854476 | 0.572017697 | 0.804735972 |
| chr8:94474502-94475668 | 0.178796598 | 0.572047165 | 0.804751261 |
| chr4:107734208-107734446 | -0.258309172 | 0.572049571 | 0.804751261 |
| chr13:30335134-30335416 | -0.225226883 | 0.572064224 | 0.8047571 |
| chr10:22325522-22325920 | -0.224193219 | 0.572079345 | 0.804763596 |
| chr1:156726630-156726846 | 0.231796941 | 0.572121963 | 0.804797422 |
| chr10:90718290-90718490 | -0.313238992 | 0.572124583 | 0.804797422 |
| chr1:205672136-205672469 | -0.197869029 | 0.572134901 | 0.804797422 |
| chr2:118186250-118186775 | 0.212117456 | 0.5721497 | 0.804803465 |
| chr5:168314756-168315014 | -0.223067254 | 0.572181046 | 0.804815734 |
| chr12:110582348-110583595 | -0.183593406 | 0.572181515 | 0.804815734 |
| chr1:85329054-85329315 | -0.278609322 | 0.572189934 | 0.804815734 |
| chr1:25608569-25608818 | 0.284250855 | 0.572206253 | 0.804823914 |
| chr7:4437380-4437659 | -0.263211543 | 0.572233547 | 0.804847529 |
| chr17:50472654-50473092 | -0.213634209 | 0.572251209 | 0.804857596 |
| chr2:79887823-79888319 | -0.21235821 | 0.572279259 | 0.804882273 |
| chrX:152642008-152642278 | 0.254498921 | 0.572292554 | 0.804886198 |
| chr4:126396827-126397027 | -0.286699417 | 0.572339225 | 0.804932681 |
| chr14:26293973-26294224 | 0.253488848 | 0.572353438 | 0.804932681 |
| chr19:11558850-11559707 | 0.194442761 | 0.5723635 | 0.804932681 |
| chr20:46715217-46715722 | 0.197363177 | 0.572367626 | 0.804932681 |
| chr13:36992408-36992652 | -0.26902 | 0.572408332 | 0.804950595 |
| chr1:147553307-147553515 | 0.282419881 | 0.57241446 | 0.804950595 |
| chr1:10793806-10794285 | -0.209811945 | 0.572416361 | 0.804950595 |
| chr16:81413520-81413923 | -0.234998624 | 0.572422385 | 0.804950595 |
| chr16:85350988-85351301 | -0.206652031 | 0.572444918 | 0.804967507 |
| chr1:17119370-17119615 | 0.265605719 | 0.572467157 | 0.804984007 |
| chr3:47802173-47803599 | 0.18275223 | 0.572494056 | 0.805007058 |
| chr13:19601700-19602809 | 0.183203744 | 0.572523938 | 0.805026008 |
| chrX:69222168-69222458 | 0.233719652 | 0.572528546 | 0.805026008 |
| chr20:56462427-56462643 | 0.252750579 | 0.572546135 | 0.805035966 |
| chr11:59754002-59754266 | -0.218376242 | 0.57256976 | 0.805054412 |
| chr17:44608546-44608861 | 0.257123372 | 0.572593253 | 0.80507267 |
| chr3:174821442-174821672 | -0.29169741 | 0.572616278 | 0.80509027 |
| chrX:70133086-70134065 | -0.182550649 | 0.572650055 | 0.805096579 |

| | | | |
|---|---|---|---|
| chr16:18969419-18969619 | 0.287386948 | 0.572651853 | 0.805096579 |
| chr19:46009708-46010038 | -0.231892522 | 0.572659612 | 0.805096579 |
| chr17:28405867-28406342 | -0.227360643 | 0.572663814 | 0.805096579 |
| chr3:185823706-185823906 | 0.279127362 | 0.572673302 | 0.805096579 |
| chr6:170389737-170390272 | 0.223433789 | 0.572713588 | 0.805136134 |
| chr5:175796843-175797579 | 0.191738247 | 0.572737668 | 0.805136134 |
| chr15:43185085-43185734 | 0.195130182 | 0.572741769 | 0.805136134 |
| chr1:228174454-228174955 | -0.200575314 | 0.57274347 | 0.805136134 |
| chr15:38474172-38474640 | 0.214993113 | 0.572816378 | 0.805223853 |
| chr7:49635784-49636111 | -0.251390791 | 0.572834172 | 0.805234093 |
| chr12:3655585-3655950 | -0.230634325 | 0.572881175 | 0.805285252 |
| chr2:101113388-101113665 | 0.225138539 | 0.572891586 | 0.805285252 |
| chr2:10380089-10380347 | -0.243395886 | 0.57291347 | 0.80530124 |
| chr4:166655232-166655991 | -0.196110315 | 0.573031544 | 0.805435848 |
| chrX:143633444-143634767 | -0.185920071 | 0.57303703 | 0.805435848 |
| chr10:21403656-21403943 | 0.236394518 | 0.573040769 | 0.805435848 |
| chr19:4759861-4760061 | -0.293697998 | 0.573071821 | 0.805456539 |
| chr1:54222015-54222387 | 0.237302421 | 0.573079282 | 0.805456539 |
| chr5:123000018-123000280 | 0.237634127 | 0.573087545 | 0.805456539 |
| chr9:113657836-113658381 | 0.198295869 | 0.573100243 | 0.805456539 |
| chr20:25623138-25624630 | 0.178228992 | 0.57310805 | 0.805456539 |
| chr19:3366327-3366527 | 0.288779685 | 0.5731253 | 0.805466007 |
| chr21:38132317-38132523 | -0.261440376 | 0.573136567 | 0.805467068 |
| chr1:229070143-229070561 | 0.208116504 | 0.573165419 | 0.805490121 |
| chr2:75374400-75374677 | 0.255563437 | 0.573186731 | 0.805490121 |
| chrX:155235924-155236215 | 0.249694588 | 0.573200836 | 0.805490121 |
| chr2:64727862-64728067 | 0.285139403 | 0.573203047 | 0.805490121 |
| chr7:28956553-28957060 | 0.21298711 | 0.573205533 | 0.805490121 |
| chr8:126275865-126276065 | -0.302723399 | 0.573224095 | 0.805497529 |
| chr19:3036068-3036535 | 0.199896408 | 0.573245569 | 0.805497529 |
| chr6:53728663-53728945 | -0.243191345 | 0.573260299 | 0.805497529 |
| chr20:54133424-54133700 | 0.190238291 | 0.573263697 | 0.805497529 |
| chr9:87974274-87975096 | 0.182771963 | 0.573268174 | 0.805497529 |
| chr1:156212743-156213707 | 0.183024654 | 0.57327388 | 0.805497529 |
| chr11:72183458-72183739 | -0.232221816 | 0.573288576 | 0.805503406 |
| chr11:126211110-126212247 | 0.180893205 | 0.573337231 | 0.805545193 |
| chr10:62701339-62701606 | -0.286725431 | 0.573339343 | 0.805545193 |
| chr1:196244315-196244516 | 0.267330857 | 0.573364197 | 0.805548586 |
| chr6:51603773-51604125 | -0.252705722 | 0.573371457 | 0.805548586 |
| chr14:37150607-37150858 | 0.249510675 | 0.573373297 | 0.805548586 |
| chr2:145752300-145752780 | -0.222323633 | 0.573465172 | 0.805662891 |
| chr8:72402688-72402958 | -0.242185685 | 0.573539286 | 0.80575224 |
| chr3:161560085-161560609 | -0.197071957 | 0.573565443 | 0.805774213 |
| chr3:32780872-32781158 | 0.218950425 | 0.573581264 | 0.80578083 |
| chr15:45022439-45023669 | -0.182494366 | 0.573591185 | 0.80578083 |
| chr4:38188538-38189056 | -0.211068402 | 0.57363961 | 0.805834083 |
| chr13:30422073-30422466 | -0.200935115 | 0.573659154 | 0.805846764 |
| chr8:54101368-54102703 | -0.179772692 | 0.573687441 | 0.80587018 |
| chr6:119116975-119117197 | -0.279537997 | 0.573697958 | 0.80587018 |
| chr6:42783634-42783846 | -0.245446821 | 0.573707376 | 0.80587018 |

| | | | |
|---|---|---|---|
| chr19:39122091-39122358 | -0.240519861 | 0.573719576 | 0.805872544 |
| chr1:160284839-160285337 | 0.192771113 | 0.573734327 | 0.80587849 |
| chr1:161088784-161089006 | 0.232025233 | 0.573748749 | 0.805883974 |
| chr15:42575043-42575911 | -0.183507193 | 0.573768013 | 0.805884807 |
| chr6:37576544-37576756 | -0.288594903 | 0.573770378 | 0.805884807 |
| chr16:87945297-87946105 | 0.179526215 | 0.57379944 | 0.805910853 |
| chr1:246837805-246838153 | -0.220382427 | 0.573859273 | 0.805930254 |
| chr17:67272567-67273095 | -0.197878562 | 0.573878041 | 0.805930254 |
| chr19:47729014-47729240 | 0.256274811 | 0.573888373 | 0.805930254 |
| chr11:82900156-82901923 | -0.176057513 | 0.573893781 | 0.805930254 |
| chr2:45440238-45440441 | -0.323943937 | 0.573894289 | 0.805930254 |
| chr8:82557677-82557956 | -0.257072739 | 0.573897968 | 0.805930254 |
| chr10:109757712-109757929 | 0.284456117 | 0.573905808 | 0.805930254 |
| chr6:56595576-56595868 | 0.243936727 | 0.573906456 | 0.805930254 |
| chr7:39623036-39624338 | -0.178913623 | 0.573914367 | 0.805930254 |
| chr2:55961330-55961580 | -0.268937744 | 0.573918436 | 0.805930254 |
| chr10:451075-451275 | 0.307910936 | 0.573931768 | 0.805934205 |
| chr8:103511851-103512577 | 0.188304349 | 0.573976815 | 0.805942162 |
| chr3:106317236-106317507 | -0.291645094 | 0.573983395 | 0.805942162 |
| chrX:75747094-75747482 | 0.232426293 | 0.573984936 | 0.805942162 |
| chr4:121800943-121801825 | -0.18636243 | 0.573997019 | 0.805942162 |
| chr1:204363284-204364002 | -0.182964229 | 0.574008725 | 0.805942162 |
| chr20:16242029-16242334 | -0.237692797 | 0.574026482 | 0.805942162 |
| chr3:161854166-161854724 | -0.199342007 | 0.574035819 | 0.805942162 |
| chr7:44847604-44848796 | -0.178054184 | 0.57404294 | 0.805942162 |
| chr16:15649955-15650351 | 0.202487889 | 0.574068123 | 0.805942162 |
| chr5:40288857-40289142 | 0.267032353 | 0.574069948 | 0.805942162 |
| chr4:142985266-142985489 | 0.273489994 | 0.57408558 | 0.805942162 |
| chrX:101942975-101943249 | -0.243448976 | 0.574090419 | 0.805942162 |
| chr1:31373318-31373827 | -0.216617968 | 0.574099842 | 0.805942162 |
| chr1:112698916-112699287 | -0.203516526 | 0.574100069 | 0.805942162 |
| chr9:125261466-125262360 | -0.189273512 | 0.574104055 | 0.805942162 |
| chr15:34721966-34722382 | 0.220085625 | 0.574105729 | 0.805942162 |
| chr1:108312557-108312864 | 0.269512488 | 0.574139068 | 0.805974198 |
| chr10:33155825-33156209 | -0.209531008 | 0.57415098 | 0.805976154 |
| chr17:40615612-40615944 | 0.224366183 | 0.574164942 | 0.805980988 |
| chr15:74889805-74890206 | 0.202309423 | 0.574200009 | 0.806015445 |
| chr10:103493475-103493675 | -0.261896266 | 0.574234563 | 0.806049182 |
| chr3:122021746-122022474 | -0.190927807 | 0.574248793 | 0.80605439 |
| chr14:52552127-52553037 | 0.177488399 | 0.57428105 | 0.806080723 |
| chr9:93951092-93951418 | 0.234706344 | 0.574293356 | 0.806080723 |
| chr6:142944543-142944745 | -0.218881379 | 0.574299114 | 0.806080723 |
| chr11:129732725-129733304 | 0.213882824 | 0.574329966 | 0.806109261 |
| chr2:219177732-219178404 | 0.183844974 | 0.57436648 | 0.806145744 |
| chr1:99146121-99146810 | -0.199382984 | 0.57438703 | 0.80615982 |
| chr5:129254984-129255184 | -0.296930937 | 0.574462729 | 0.806251296 |
| chr6:7145757-7146769 | -0.188781524 | 0.574473541 | 0.806251701 |
| chr8:61467601-61467810 | 0.260827425 | 0.574501588 | 0.806266587 |
| chr3:108477523-108477863 | -0.218403083 | 0.574505192 | 0.806266587 |
| chr21:28745718-28746039 | -0.231653455 | 0.574565228 | 0.806336073 |

| | | | |
|---|---|---|---|
| chr17:42561937-42562278 | 0.215434144 | 0.574591373 | 0.806340377 |
| chr10:128285037-128285277 | -0.303714449 | 0.574597989 | 0.806340377 |
| chr1:110612210-110612433 | 0.231544181 | 0.574606706 | 0.806340377 |
| chr18:59139296-59140232 | -0.180507807 | 0.574610389 | 0.806340377 |
| chr19:1207470-1207912 | -0.217761799 | 0.574647458 | 0.806377626 |
| chr2:127293459-127294379 | 0.185845922 | 0.574671099 | 0.806396032 |
| chr20:33181974-33182174 | -0.288651895 | 0.574702235 | 0.806403926 |
| chr4:72303193-72303393 | -0.288651895 | 0.574702235 | 0.806403926 |
| chr3:136277279-136278037 | -0.196318504 | 0.574747495 | 0.806403926 |
| chr4:97648039-97648239 | 0.298094636 | 0.574747828 | 0.806403926 |
| chr13:94444841-94445145 | 0.238261443 | 0.574765772 | 0.806403926 |
| chr2:38725982-38726361 | -0.209775645 | 0.574768264 | 0.806403926 |
| chr9:22447829-22448223 | 0.216613094 | 0.57477645 | 0.806403926 |
| chr1:67489749-67489949 | -0.305683131 | 0.57478832 | 0.806403926 |
| chr17:31859101-31859472 | 0.222322035 | 0.57478945 | 0.806403926 |
| chr11:34274613-34275392 | 0.180999657 | 0.574793199 | 0.806403926 |
| chr1:10704649-10704890 | -0.286995699 | 0.574797292 | 0.806403926 |
| chr5:112929038-112929518 | -0.211179244 | 0.574803018 | 0.806403926 |
| chr17:44222320-44222520 | 0.239407875 | 0.574822755 | 0.806416851 |
| chr1:43577576-43577912 | -0.240073249 | 0.574846361 | 0.806431989 |
| chr1:66929945-66930931 | -0.179763484 | 0.574854595 | 0.806431989 |
| chr3:127547791-127548235 | 0.198687106 | 0.574892032 | 0.806469742 |
| chr17:1639681-1640037 | 0.220917146 | 0.574916301 | 0.806479824 |
| chr8:83729581-83729781 | 0.289017802 | 0.574963968 | 0.806479824 |
| chr2:110677758-110678373 | -0.190968005 | 0.574964659 | 0.806479824 |
| chr1:217847735-217847935 | -0.303177974 | 0.574967251 | 0.806479824 |
| chr19:12610517-12611115 | 0.198433258 | 0.57500312 | 0.806479824 |
| chr5:23139679-23139992 | 0.224615101 | 0.575017004 | 0.806479824 |
| chr11:613548-613851 | -0.22051136 | 0.5750204 | 0.806479824 |
| chr1:48767120-48767415 | -0.27903461 | 0.575032525 | 0.806479824 |
| chr10:90484623-90484876 | 0.277472087 | 0.575033743 | 0.806479824 |
| chr1:184986381-184986641 | 0.230814624 | 0.575037088 | 0.806479824 |
| chr10:24513270-24513506 | 0.274380419 | 0.575042916 | 0.806479824 |
| chr8:123447768-123448092 | -0.207897516 | 0.575053762 | 0.806479824 |
| chr1:111905122-111905339 | 0.217243706 | 0.575065957 | 0.806479824 |
| chr11:12496159-12496448 | 0.237557379 | 0.575066324 | 0.806479824 |
| chr6:89050230-89050454 | -0.275440704 | 0.575074629 | 0.806479824 |
| chr7:107862-108071 | -0.26924622 | 0.575075932 | 0.806479824 |
| chr5:30439225-30439561 | -0.261346772 | 0.575087539 | 0.806479824 |
| chr5:14663823-14665138 | 0.17711666 | 0.575089126 | 0.806479824 |
| chr22:17628301-17629121 | 0.186903891 | 0.575108997 | 0.806479824 |
| chr4:167021596-167022052 | -0.21753 | 0.575116014 | 0.806479824 |
| chr1:218514209-218514410 | -0.265451287 | 0.575120253 | 0.806479824 |
| chr6:2764587-2766034 | 0.177654523 | 0.575141865 | 0.80649537 |
| chr10:71134951-71135151 | 0.278847717 | 0.575162426 | 0.806509442 |
| chr5:77016001-77016413 | 0.236790051 | 0.575186375 | 0.806528264 |
| chr20:56829343-56830169 | 0.18490506 | 0.575212712 | 0.806542429 |
| chr8:116100135-116100482 | 0.213433705 | 0.575223097 | 0.806542429 |
| chr19:42242531-42243481 | 0.178804222 | 0.575228056 | 0.806542429 |
| chr6:97925396-97925825 | -0.236721595 | 0.575261592 | 0.806553037 |

| | | | |
|---|---|---|---|
| chr11:72752329-72753040 | -0.183435929 | 0.575266277 | 0.806553037 |
| chr5:42971291-42971567 | 0.27559304 | 0.575267201 | 0.806553037 |
| chr4:140359537-140360459 | 0.193040547 | 0.575304339 | 0.806569279 |
| chr12:112198172-112198586 | 0.211200143 | 0.575307998 | 0.806569279 |
| chr12:131682236-131682524 | 0.267318215 | 0.575312161 | 0.806569279 |
| chr7:33243351-33243646 | -0.225573075 | 0.575320892 | 0.806569279 |
| chr1:235450496-235450743 | 0.291151838 | 0.575355121 | 0.806580782 |
| chr14:56886514-56886781 | 0.263558056 | 0.57535791 | 0.806580782 |
| chr3:38025750-38026033 | -0.207337562 | 0.575370578 | 0.806580782 |
| chr2:128262711-128262946 | -0.270621845 | 0.575371204 | 0.806580782 |
| chr2:219521391-219521619 | 0.236848646 | 0.57545346 | 0.806676127 |
| chr12:131883295-131883885 | 0.206574403 | 0.57546174 | 0.806676127 |
| chr6:134746186-134746394 | -0.317598634 | 0.575470802 | 0.806676127 |
| chr1:244460737-244461973 | -0.178711591 | 0.575502274 | 0.806693667 |
| chr2:3518694-3519771 | -0.176224669 | 0.575504371 | 0.806693667 |
| chr22:33921809-33922050 | 0.237686921 | 0.575564573 | 0.806744519 |
| chr11:1827175-1827555 | -0.214392195 | 0.575581576 | 0.806744519 |
| chr19:35628420-35629193 | 0.182723862 | 0.575582319 | 0.806744519 |
| chr16:782720-783333 | -0.193721372 | 0.575582765 | 0.806744519 |
| chr8:109316232-109316536 | -0.219188024 | 0.575629552 | 0.806747941 |
| chr2:24075958-24077164 | -0.174800762 | 0.575638218 | 0.806747941 |
| chr3:52831625-52831879 | 0.242805718 | 0.575647551 | 0.806747941 |
| chr11:1947105-1947476 | 0.217468824 | 0.57565611 | 0.806747941 |
| chr21:39182923-39184213 | -0.175966178 | 0.575658063 | 0.806747941 |
| chr1:47709567-47709797 | 0.240839422 | 0.575659888 | 0.806747941 |
| chr8:69025732-69026175 | -0.193824241 | 0.575666045 | 0.806747941 |
| chr1:110113560-110113864 | -0.225595382 | 0.575669438 | 0.806747941 |
| chr4:118865017-118865504 | 0.19445986 | 0.575750464 | 0.806846735 |
| chr2:214674442-214674699 | -0.274297549 | 0.575762452 | 0.806848777 |
| chr2:137675910-137676301 | -0.205526955 | 0.575779553 | 0.806857985 |
| chr14:72199512-72199712 | -0.285621371 | 0.575859222 | 0.80691656 |
| chr17:45893979-45894677 | 0.191061569 | 0.575862799 | 0.80691656 |
| chr10:29119508-29119714 | 0.252993449 | 0.575863019 | 0.80691656 |
| chr14:91956763-91956986 | 0.282141939 | 0.575863477 | 0.80691656 |
| chr21:46058691-46059089 | -0.232079846 | 0.575894083 | 0.80691941 |
| chr10:120997323-120997645 | 0.221922711 | 0.575897276 | 0.80691941 |
| chr4:125484458-125484750 | -0.292780731 | 0.575904409 | 0.80691941 |
| chr10:110007270-110008617 | -0.181632384 | 0.575907635 | 0.80691941 |
| chr19:46829227-46829449 | 0.261902251 | 0.575918921 | 0.8069196 |
| chr8:73673072-73673308 | -0.267974992 | 0.575933994 | 0.8069196 |
| chr18:75809963-75810163 | 0.306503485 | 0.575939365 | 0.8069196 |
| chr14:44613151-44613689 | -0.212790658 | 0.57600732 | 0.806966422 |
| chr8:113437716-113438617 | -0.18418598 | 0.576009976 | 0.806966422 |
| chr13:113105280-113106065 | 0.204882173 | 0.576023791 | 0.806966422 |
| chr2:178522583-178523922 | -0.182989513 | 0.576024005 | 0.806966422 |
| chr20:53654623-53655578 | -0.175310234 | 0.576033689 | 0.806966422 |
| chr19:58513779-58514483 | 0.184718218 | 0.576035975 | 0.806966422 |
| chr11:57293092-57293292 | 0.27805877 | 0.576062027 | 0.806984152 |
| chr2:206154321-206154535 | -0.253836022 | 0.576069695 | 0.806984152 |
| chr14:100486559-100486759 | 0.281270412 | 0.57609212 | 0.807000812 |

| | | | |
|---|---|---|---|
| chr12:81268113-81268922 | -0.185583779 | 0.576106253 | 0.807005856 |
| chr16:88760674-88761029 | -0.204684551 | 0.576121229 | 0.807008942 |
| chr18:13862991-13863523 | 0.222577413 | 0.576134889 | 0.807008942 |
| chr8:84626135-84626512 | -0.201361214 | 0.57614431 | 0.807008942 |
| chr13:51924090-51924290 | 0.275110293 | 0.57615528 | 0.807008942 |
| chr1:30719267-30719545 | 0.205254156 | 0.576161118 | 0.807008942 |
| chr11:64148709-64148913 | -0.268771955 | 0.576202984 | 0.807035705 |
| chr18:8696766-8697087 | 0.230610145 | 0.576218613 | 0.807035705 |
| chr17:28879114-28879418 | 0.233997819 | 0.57621956 | 0.807035705 |
| chr14:49972430-49973122 | -0.205636474 | 0.576227317 | 0.807035705 |
| chr11:47118907-47119546 | 0.190831787 | 0.576232889 | 0.807035705 |
| chr1:114955849-114956083 | 0.234035874 | 0.576247936 | 0.807042028 |
| chr12:124838086-124838686 | -0.193930165 | 0.576262011 | 0.807046989 |
| chr5:63125821-63126172 | -0.223765531 | 0.576277478 | 0.807047351 |
| chr17:37609307-37609774 | 0.198219504 | 0.576283335 | 0.807047351 |
| chr8:123548269-123548487 | -0.259450155 | 0.576297701 | 0.807052719 |
| chr11:45846990-45847661 | 0.179353998 | 0.576332859 | 0.807076208 |
| chr9:19464645-19465435 | -0.18978803 | 0.576338179 | 0.807076208 |
| chr11:33736764-33737228 | -0.206233613 | 0.576346074 | 0.807076208 |
| chr7:44196483-44196688 | -0.236222225 | 0.576379611 | 0.807091042 |
| chr20:20408007-20408209 | -0.272740396 | 0.576386189 | 0.807091042 |
| chr16:67226844-67227488 | 0.188528318 | 0.576388268 | 0.807091042 |
| chr3:63644456-63645980 | -0.175050696 | 0.576406882 | 0.807102357 |
| chr7:44618727-44619310 | 0.196059466 | 0.576465283 | 0.807169382 |
| chr10:29124786-29125459 | 0.194719024 | 0.576503968 | 0.807206688 |
| chr20:62796253-62796605 | -0.192002703 | 0.576516293 | 0.807206688 |
| chr1:146386979-146388158 | 0.177133743 | 0.576523531 | 0.807206688 |
| chr8:73174045-73174720 | -0.190342836 | 0.576549048 | 0.807216587 |
| chr15:101558366-101558632 | -0.226906665 | 0.576551672 | 0.807216587 |
| chr1:23949224-23949503 | 0.265108533 | 0.576614272 | 0.807271802 |
| chr8:140200217-140200553 | 0.238975312 | 0.576617266 | 0.807271802 |
| chr2:3377369-3378450 | -0.178676853 | 0.576627253 | 0.807271802 |
| chr14:71721090-71721414 | 0.226353495 | 0.576637004 | 0.807271802 |
| chr2:16104347-16104588 | 0.279158492 | 0.576643787 | 0.807271802 |
| chr4:147719126-147719326 | 0.301982173 | 0.576690809 | 0.807285722 |
| chr1:205524108-205524601 | 0.192682753 | 0.576699745 | 0.807285722 |
| chr1:27568352-27568798 | 0.189061019 | 0.576712131 | 0.807285722 |
| chr8:93974912-93975497 | 0.195616497 | 0.576714892 | 0.807285722 |
| chr16:78770549-78771026 | -0.204069545 | 0.576715496 | 0.807285722 |
| chr3:23888599-23888886 | 0.235934799 | 0.576716946 | 0.807285722 |
| chr8:52008473-52008829 | -0.206588693 | 0.576817012 | 0.807411043 |
| chr3:87298135-87298590 | -0.218232372 | 0.576852478 | 0.807445936 |
| chr3:65342144-65342603 | 0.218832772 | 0.576896516 | 0.807492827 |
| chr4:77095381-77095796 | -0.212260764 | 0.57691061 | 0.807497803 |
| chr8:115998302-115998533 | 0.250537984 | 0.576937263 | 0.807520358 |
| chr7:150570646-150570900 | -0.283165107 | 0.576951969 | 0.807526191 |
| chr4:151276821-151277080 | -0.254954408 | 0.576996137 | 0.807573258 |
| chr19:48492780-48493739 | 0.175756309 | 0.577058405 | 0.807645656 |
| chr5:2715134-2715419 | -0.234649038 | 0.577100738 | 0.807671447 |
| chr10:114821374-114822955 | -0.181400873 | 0.577106604 | 0.807671447 |

| | | | |
|---|---|---|---|
| chr20:45044099-45044323 | 0.282512524 | 0.577108455 | 0.807671447 |
| chr14:20413802-20414094 | 0.216337679 | 0.577126521 | 0.807681978 |
| chrX:117969390-117969629 | -0.285186819 | 0.577144151 | 0.807686324 |
| chr1:246689384-246690114 | -0.181387491 | 0.577151108 | 0.807686324 |
| chr3:102229104-102229370 | -0.225102192 | 0.57716125 | 0.807686324 |
| chr5:174402292-174402515 | 0.282526236 | 0.577172204 | 0.807686901 |
| chr15:72271955-72273086 | 0.177468026 | 0.577195849 | 0.807705239 |
| chr12:64760280-64760618 | -0.211676437 | 0.577226858 | 0.80773388 |
| chr4:87849077-87849317 | 0.248384396 | 0.577264532 | 0.807771846 |
| chr11:65454777-65455403 | 0.188601971 | 0.577288737 | 0.807790963 |
| chr15:92881965-92883199 | 0.175086771 | 0.577301571 | 0.80779417 |
| chr7:47475075-47476254 | -0.17638534 | 0.577316428 | 0.807794473 |
| chr1:199684684-199685079 | -0.203291665 | 0.577322873 | 0.807794473 |
| chr7:52918078-52918313 | 0.26339079 | 0.577406129 | 0.807894538 |
| chr11:83048582-83048782 | -0.288188738 | 0.577426846 | 0.807894538 |
| chr17:44249864-44250105 | -0.266460937 | 0.577428559 | 0.807894538 |
| chr18:57970465-57970665 | -0.296552751 | 0.577436564 | 0.807894538 |
| chr4:7082040-7082335 | 0.24453998 | 0.577471508 | 0.807928676 |
| chr11:49780830-49781191 | 0.216658554 | 0.577512334 | 0.807962629 |
| chr15:37891796-37892136 | -0.224658396 | 0.577524551 | 0.807962629 |
| chr12:13990-14190 | 0.278733556 | 0.577536606 | 0.807962629 |
| chr19:10289192-10289655 | 0.202394518 | 0.577537955 | 0.807962629 |
| chr20:54132603-54133099 | -0.180853228 | 0.577564958 | 0.807982065 |
| chr2:161280725-161281445 | 0.187017846 | 0.577576658 | 0.807982065 |
| chr10:14877826-14879392 | -0.17610231 | 0.577595505 | 0.807982065 |
| chr8:55970871-55971341 | 0.210128495 | 0.577602279 | 0.807982065 |
| chr1:25239603-25240327 | -0.183977023 | 0.577604573 | 0.807982065 |
| chr4:44996245-44996699 | 0.20969387 | 0.5776485 | 0.808002474 |
| chr10:85537601-85537801 | 0.31066749 | 0.577660647 | 0.808002474 |
| chr7:32039414-32039670 | -0.224359212 | 0.577662113 | 0.808002474 |
| chr11:12366294-12366531 | -0.260914453 | 0.57767284 | 0.808002474 |
| chr7:49099610-49099870 | -0.251741118 | 0.577674745 | 0.808002474 |
| chr4:142846194-142847602 | 0.178068883 | 0.577682435 | 0.808002474 |
| chr4:40797471-40797695 | -0.305824365 | 0.577717127 | 0.808036247 |
| chr6:168877886-168878126 | -0.294672111 | 0.577766308 | 0.808068275 |
| chr6:81076098-81076314 | -0.294672111 | 0.577766308 | 0.808068275 |
| chr1:197787742-197788003 | 0.217722531 | 0.577774797 | 0.808068275 |
| chr8:94982210-94982480 | 0.246602408 | 0.577792695 | 0.808068275 |
| chr22:38923809-38924165 | -0.210087583 | 0.577792757 | 0.808068275 |
| chr6:160300196-160300589 | -0.250712338 | 0.577828523 | 0.808103546 |
| chr5:15798093-15798378 | 0.238848186 | 0.577846538 | 0.80811399 |
| chr15:72286026-72286244 | 0.248173125 | 0.577861398 | 0.808120022 |
| chrX:134254032-134254232 | 0.279963747 | 0.577882654 | 0.808134999 |
| chr3:67594022-67594247 | -0.271655036 | 0.577912067 | 0.808161381 |
| chr16:12340458-12340672 | 0.270781205 | 0.577926828 | 0.808167264 |
| chr21:44011971-44012410 | 0.200078674 | 0.577947843 | 0.808167264 |
| chr14:64960708-64961289 | 0.193568778 | 0.577947916 | 0.808167264 |
| chr1:84234135-84234828 | 0.206195602 | 0.57797027 | 0.808183773 |
| chr2:113084348-113084564 | 0.248164515 | 0.578009169 | 0.808187446 |
| chr14:72723379-72723579 | 0.297871947 | 0.578018718 | 0.808187446 |

| | | | |
|---|---|---|---|
| chr15:41838353-41839018 | -0.186200021 | 0.578028466 | 0.808187446 |
| chr1:153967230-153967939 | 0.179673976 | 0.578028698 | 0.808187446 |
| chr9:128786829-128787463 | -0.194996133 | 0.578028957 | 0.808187446 |
| chr5:152218547-152219093 | 0.201388265 | 0.578036183 | 0.808187446 |
| chr10:17616643-17617993 | 0.178092749 | 0.578101979 | 0.808264691 |
| chr1:230702964-230703164 | -0.313246928 | 0.578146065 | 0.80831158 |
| chr17:7394105-7395025 | -0.187543706 | 0.578186778 | 0.808340601 |
| chr3:163721829-163722029 | 0.296359512 | 0.578200182 | 0.808340601 |
| chr5:41290842-41291123 | -0.264588487 | 0.578200725 | 0.808340601 |
| chr17:79301257-79301653 | 0.227128862 | 0.578209022 | 0.808340601 |
| chr17:51371710-51371968 | -0.220221318 | 0.578234888 | 0.808356726 |
| chr13:44410401-44410845 | -0.211420586 | 0.57825579 | 0.808356726 |
| chr4:88574093-88574330 | -0.277827839 | 0.578265511 | 0.808356726 |
| chr2:165015089-165015380 | 0.237217123 | 0.578281588 | 0.808356726 |
| chr6:4794447-4794772 | 0.257404793 | 0.578291044 | 0.808356726 |
| chr1:208108554-208108754 | -0.296154137 | 0.578293913 | 0.808356726 |
| chrX:149470894-149471293 | -0.206591745 | 0.578294406 | 0.808356726 |
| chr2:161747060-161747260 | 0.316517103 | 0.578341647 | 0.808392241 |
| chr12:99811324-99811532 | -0.27530024 | 0.578352022 | 0.808392241 |
| chr11:47980548-47981071 | 0.204835456 | 0.578360007 | 0.808392241 |
| chr20:2476068-2476418 | -0.232401046 | 0.578368617 | 0.808392241 |
| chr1:56831959-56832225 | -0.239626579 | 0.578372565 | 0.808392241 |
| chr18:13286884-13287096 | -0.320624639 | 0.578387412 | 0.808398247 |
| chr1:108200825-108201078 | 0.202860388 | 0.578443463 | 0.808454139 |
| chr1:207434152-207434352 | -0.290579933 | 0.578461217 | 0.808454139 |
| chr3:177520882-177521667 | -0.18582069 | 0.578464846 | 0.808454139 |
| chr3:151402308-151402637 | 0.252885911 | 0.578469607 | 0.808454139 |
| chrX:65534299-65535117 | -0.182804053 | 0.578509202 | 0.808487585 |
| chr9:109316260-109316646 | -0.201575927 | 0.578527149 | 0.808487585 |
| chr1:99405287-99405537 | -0.284848655 | 0.57853053 | 0.808487585 |
| chr19:34345904-34346201 | 0.22734331 | 0.578535745 | 0.808487585 |
| chr7:18357737-18358015 | 0.233211594 | 0.578558352 | 0.808504433 |
| chr2:190407813-190409054 | 0.181999366 | 0.578577582 | 0.808504494 |
| chr4:25045814-25046043 | -0.275317019 | 0.578579499 | 0.808504494 |
| chr1:230120949-230121228 | 0.238429659 | 0.578599135 | 0.808517187 |
| chr2:10914328-10914588 | -0.220009646 | 0.57861288 | 0.808521649 |
| chr8:130528385-130528666 | 0.218764635 | 0.57862468 | 0.808523393 |
| chr1:162191947-162192615 | 0.20247613 | 0.57864179 | 0.808526623 |
| chr11:116454187-116454397 | 0.282964344 | 0.578648096 | 0.808526623 |
| chr2:10016023-10016292 | 0.259038329 | 0.57867006 | 0.80853838 |
| chr20:5127430-5127659 | 0.237045394 | 0.578677615 | 0.80853838 |
| chr3:31567064-31567339 | 0.232395589 | 0.57871524 | 0.80856555 |
| chr17:19766481-19766912 | 0.214639155 | 0.578723503 | 0.80856555 |
| chr5:14264621-14264975 | -0.192356595 | 0.57874576 | 0.80856555 |
| chr17:75405023-75406281 | 0.175877514 | 0.578748628 | 0.80856555 |
| chr1:157369927-157370237 | 0.227090897 | 0.578749824 | 0.80856555 |
| chr14:72741327-72741527 | 0.288557648 | 0.578769734 | 0.808578102 |
| chr6:150906237-150906443 | 0.28076889 | 0.578779914 | 0.808578102 |
| chr1:236774971-236775275 | 0.239329626 | 0.578794963 | 0.808584383 |
| chr3:75258870-75259122 | -0.269634611 | 0.578849914 | 0.808646406 |

| | | | |
|---|---|---|---|
| chr2:205568707-205568907 | 0.288356639 | 0.578904127 | 0.808707397 |
| chr14:30621190-30622739 | 0.175058761 | 0.578916281 | 0.808709631 |
| chr3:101792055-101792255 | -0.300435127 | 0.578930109 | 0.808714204 |
| chr12:121624723-121625051 | 0.222286576 | 0.57895713 | 0.808727787 |
| chr20:54387833-54388126 | 0.18860125 | 0.578960942 | 0.808727787 |
| chr6:138555665-138555912 | 0.284433667 | 0.578986581 | 0.808748857 |
| chr12:94603481-94603763 | -0.238506166 | 0.57899785 | 0.808749024 |
| chr1:109138340-109138775 | -0.229769511 | 0.579007811 | 0.808749024 |
| chr5:111764033-111764306 | 0.25899928 | 0.579035498 | 0.808772953 |
| chr9:120023909-120024173 | -0.295199502 | 0.579085277 | 0.8088203 |
| chr2:203578854-203579646 | -0.182477124 | 0.579090507 | 0.8088203 |
| chr13:41440272-41440610 | -0.254859023 | 0.579160335 | 0.808903085 |
| chr19:42191242-42191613 | 0.252491284 | 0.579187943 | 0.808926898 |
| chr14:24160884-24161764 | -0.180144674 | 0.579213806 | 0.808948275 |
| chr8:106623929-106624385 | -0.22090695 | 0.579239989 | 0.808970097 |
| chr1:93446857-93447482 | -0.205494187 | 0.579272173 | 0.809000299 |
| chr16:68959137-68959489 | -0.243262732 | 0.579282873 | 0.809000497 |
| chr16:75695824-75696052 | 0.269898202 | 0.579322599 | 0.809013212 |
| chr10:91091211-91091418 | 0.28253358 | 0.579343028 | 0.809013212 |
| chr19:14117643-14118220 | 0.201032044 | 0.579346445 | 0.809013212 |
| chr17:11880741-11881094 | -0.219804743 | 0.579347969 | 0.809013212 |
| chr10:847871-848302 | 0.228144641 | 0.579349125 | 0.809013212 |
| chr22:24707347-24707635 | 0.263193099 | 0.579355328 | 0.809013212 |
| chr7:155803938-155804248 | -0.224516136 | 0.579392189 | 0.80904994 |
| chr1:172933903-172934423 | 0.200641264 | 0.579432242 | 0.809075944 |
| chr2:203623669-203623874 | 0.262110915 | 0.579433 | 0.809075944 |
| chr6:33815885-33816192 | -0.232137055 | 0.579443368 | 0.809075944 |
| chr11:62380806-62381095 | -0.231102347 | 0.579470061 | 0.809075944 |
| chr2:27321011-27321431 | 0.186931272 | 0.579473629 | 0.809075944 |
| chr12:89911183-89911463 | -0.281437387 | 0.579474167 | 0.809075944 |
| chr19:14550283-14550776 | 0.216129618 | 0.579509446 | 0.809102623 |
| chr22:20956334-20956713 | 0.206026501 | 0.579532497 | 0.809102623 |
| chr14:65861978-65862254 | -0.261016754 | 0.57953304 | 0.809102623 |
| chr1:27154717-27155307 | 0.202422442 | 0.579535514 | 0.809102623 |
| chr18:46273747-46274260 | 0.200419176 | 0.57960137 | 0.809153709 |
| chr13:44373201-44373437 | -0.273960201 | 0.579609618 | 0.809153709 |
| chr15:31280120-31280412 | -0.273286505 | 0.579610575 | 0.809153709 |
| chrX:102720529-102721381 | -0.189622026 | 0.579614346 | 0.809153709 |
| chr7:104896170-104896456 | 0.296698987 | 0.57963885 | 0.809163838 |
| chr3:62229942-62230216 | -0.204695293 | 0.579649892 | 0.809163838 |
| chr10:95530138-95530352 | 0.308908387 | 0.579653283 | 0.809163838 |
| chr20:9838385-9839449 | -0.179638618 | 0.579673402 | 0.809171189 |
| chr5:14542876-14543180 | -0.240768299 | 0.579679671 | 0.809171189 |
| chr11:124740601-124740879 | -0.225499759 | 0.579697494 | 0.809181327 |
| chr16:690207-690554 | 0.220522683 | 0.579712679 | 0.809187781 |
| chr9:122159163-122160031 | -0.181656618 | 0.579794761 | 0.809273752 |
| chr7:36342482-36342784 | -0.252010925 | 0.579807661 | 0.809273752 |
| chrX:46912838-46913346 | 0.203778991 | 0.579807773 | 0.809273752 |
| chr2:70871469-70872075 | -0.19045941 | 0.579820197 | 0.809273752 |
| chr20:54549809-54550084 | 0.194793812 | 0.579827079 | 0.809273752 |

| | | | |
|---|---|---|---|
| chr17:53765677-53765910 | -0.30928956 | 0.579855099 | 0.809288675 |
| chr5:35246414-35246751 | -0.214267492 | 0.579871746 | 0.809288675 |
| chr15:46918782-46919376 | -0.197532722 | 0.579873633 | 0.809288675 |
| chr11:6298638-6298964 | -0.238920607 | 0.579880019 | 0.809288675 |
| chr8:42842641-42843702 | 0.176806538 | 0.579927096 | 0.809314428 |
| chr1:155273059-155273757 | 0.18349986 | 0.579939601 | 0.809314428 |
| chr3:159616348-159616548 | 0.282033 | 0.579946427 | 0.809314428 |
| chr11:118291033-118291297 | 0.240794054 | 0.579951366 | 0.809314428 |
| chr11:47161095-47161450 | 0.203071813 | 0.579954959 | 0.809314428 |
| chr2:39838478-39838692 | -0.276264351 | 0.579961847 | 0.809314428 |
| chr15:64681486-64681796 | 0.213379666 | 0.580039786 | 0.809400208 |
| chr12:57227666-57227979 | -0.230695155 | 0.580044886 | 0.809400208 |
| chrX:40176527-40176884 | 0.21857304 | 0.580055008 | 0.809400208 |
| chr11:77819372-77819663 | -0.197934529 | 0.580103926 | 0.809447657 |
| chr15:38440292-38441034 | -0.193814828 | 0.580117363 | 0.809447657 |
| chr9:2728051-2728338 | -0.265180233 | 0.580120705 | 0.809447657 |
| chr7:155560181-155560481 | -0.262621131 | 0.58013579 | 0.809447963 |
| chr12:55072144-55072384 | 0.259050469 | 0.580142052 | 0.809447963 |
| chr15:39581690-39582001 | 0.204849122 | 0.580174037 | 0.809470767 |
| chr9:129413313-129414065 | -0.183917493 | 0.580179804 | 0.809470767 |
| chr10:100019857-100020137 | 0.242126927 | 0.58019009 | 0.809470767 |
| chr1:155135511-155136226 | 0.180450253 | 0.580270882 | 0.809568746 |
| chr10:1508456-1508656 | 0.303639722 | 0.580283528 | 0.809571648 |
| chr2:84631140-84631406 | 0.229980914 | 0.58030027 | 0.809573151 |
| chr15:60478602-60479646 | 0.179118691 | 0.580305737 | 0.809573151 |
| chr3:30768742-30768959 | -0.278505866 | 0.580368479 | 0.80964261 |
| chr8:27583620-27583957 | 0.216216496 | 0.580402209 | 0.80964261 |
| chr1:170984062-170984312 | -0.283674519 | 0.580404208 | 0.80964261 |
| chr3:58568682-58568900 | -0.298842388 | 0.580426966 | 0.80964261 |
| chr17:28548214-28548456 | -0.220594505 | 0.580429803 | 0.80964261 |
| chr16:21083857-21084057 | 0.266249779 | 0.580432231 | 0.80964261 |
| chr16:85297066-85297387 | -0.21224823 | 0.580436181 | 0.80964261 |
| chr4:121726720-121727032 | -0.216534502 | 0.580440059 | 0.80964261 |
| chr3:19485087-19485361 | 0.252038172 | 0.580491948 | 0.809684118 |
| chr14:91582846-91583050 | 0.269058349 | 0.580507464 | 0.809684118 |
| chr1:50280011-50280221 | 0.271804831 | 0.580519543 | 0.809684118 |
| chr13:42011775-42012428 | -0.188797073 | 0.580523992 | 0.809684118 |
| chr7:54773579-54774070 | -0.206157841 | 0.580530834 | 0.809684118 |
| chr10:84715886-84716202 | -0.221376352 | 0.580545221 | 0.809684118 |
| chr1:51344263-51344472 | 0.256592482 | 0.58056213 | 0.809684118 |
| chr8:90510028-90510228 | 0.257712467 | 0.580583852 | 0.809684118 |
| chr6:43013308-43014460 | 0.176588708 | 0.580589227 | 0.809684118 |
| chr14:44979996-44980196 | 0.274907391 | 0.58059272 | 0.809684118 |
| chr19:12611204-12611412 | 0.267480438 | 0.580593473 | 0.809684118 |
| chr6:170529821-170530137 | 0.222813119 | 0.580596623 | 0.809684118 |
| chr9:93119889-93120190 | 0.224437207 | 0.58061066 | 0.809688956 |
| chr13:43179663-43179863 | -0.308681531 | 0.580660306 | 0.809743452 |
| chr21:17454253-17455274 | -0.176075893 | 0.580675257 | 0.809749564 |
| chr10:29858446-29858793 | -0.228913152 | 0.580698492 | 0.809753721 |
| chr3:10344065-10344439 | 0.216036905 | 0.580699374 | 0.809753721 |

| | | | |
|---|---|---|---|
| chr1:21634540-21634990 | 0.210804195 | 0.58074554 | 0.809803359 |
| chr2:58155263-58155463 | -0.289501318 | 0.580780808 | 0.809837799 |
| chr5:158560256-158560600 | -0.220294492 | 0.580808569 | 0.809850134 |
| chr5:1360801-1361031 | -0.265174233 | 0.580810793 | 0.809850134 |
| chr3:66345148-66345358 | 0.285024564 | 0.580833932 | 0.80986766 |
| chr12:121296575-121297581 | 0.180584034 | 0.580868015 | 0.809886504 |
| chr3:5731181-5731418 | -0.263016604 | 0.580868586 | 0.809886504 |
| chr10:99930807-99931320 | -0.20481004 | 0.580928379 | 0.809946569 |
| chr2:41118600-41118869 | -0.281681166 | 0.580932808 | 0.809946569 |
| chr10:110600556-110600797 | -0.271603912 | 0.580965232 | 0.809975626 |
| chr6:45535774-45536210 | 0.220020694 | 0.580978115 | 0.809975626 |
| chr10:12348732-12349593 | -0.194883659 | 0.580985362 | 0.809975626 |
| chr8:54010522-54010724 | 0.305396241 | 0.58105062 | 0.810010375 |
| chr15:31870336-31871414 | 0.17969378 | 0.58107152 | 0.810010375 |
| chr5:95710374-95710740 | 0.21691954 | 0.581072657 | 0.810010375 |
| chr17:82216800-82217045 | -0.226233417 | 0.58108241 | 0.810010375 |
| chr20:59207810-59208067 | 0.216412913 | 0.581083842 | 0.810010375 |
| chr12:116354714-116354914 | -0.302532504 | 0.581086295 | 0.810010375 |
| chr11:57457387-57457679 | 0.217928914 | 0.581092664 | 0.810010375 |
| chr17:71447685-71447998 | 0.230943763 | 0.581094859 | 0.810010375 |
| chr12:33248487-33248884 | 0.229872614 | 0.581119073 | 0.810029391 |
| chr11:76152190-76152472 | -0.214873912 | 0.581151197 | 0.810059433 |
| chr14:66315425-66315846 | 0.224170628 | 0.581161867 | 0.810059569 |
| chr10:81082583-81082793 | 0.277650381 | 0.581182591 | 0.810073719 |
| chr15:76963647-76963914 | -0.248224227 | 0.581207948 | 0.810094326 |
| chr10:88935398-88935705 | -0.20974672 | 0.581286954 | 0.81018854 |
| chr10:72198398-72198598 | -0.311181033 | 0.581296986 | 0.81018854 |
| chr4:11635519-11635719 | 0.267054414 | 0.581314609 | 0.81018854 |
| chr17:64147314-64147536 | -0.214868542 | 0.58132133 | 0.81018854 |
| chr17:58854979-58855479 | -0.191775858 | 0.581337256 | 0.81018854 |
| chr2:131016786-131017520 | -0.200213352 | 0.581338986 | 0.81018854 |
| chrX:49123400-49124101 | 0.190823817 | 0.581360673 | 0.810204029 |
| chr12:62914046-62914285 | 0.261709755 | 0.581387293 | 0.810211932 |
| chr6:27251048-27252124 | -0.179530947 | 0.581387492 | 0.810211932 |
| chr6:33828391-33828638 | -0.265153989 | 0.581417495 | 0.810236924 |
| chr15:83380419-83380946 | 0.198349432 | 0.581426575 | 0.810236924 |
| chr6:33297399-33298284 | -0.178113476 | 0.581469517 | 0.810255443 |
| chr6:167307210-167307416 | -0.32014369 | 0.581471037 | 0.810255443 |
| chr3:141737913-141738847 | 0.175480731 | 0.581477095 | 0.810255443 |
| chr21:42330801-42331196 | 0.232430986 | 0.581489157 | 0.810255443 |
| chr22:45863972-45864219 | 0.252826436 | 0.581494907 | 0.810255443 |
| chr12:101490336-101490622 | -0.225630346 | 0.581503313 | 0.810255443 |
| chr17:46193339-46194291 | -0.181579363 | 0.581514988 | 0.810256977 |
| chr11:134252796-134254016 | 0.178500143 | 0.581535085 | 0.810264694 |
| chr17:7381376-7381699 | -0.238226122 | 0.581550155 | 0.810264694 |
| chr21:44012543-44012833 | 0.218657812 | 0.581556554 | 0.810264694 |
| chr18:12376747-12377973 | 0.173840842 | 0.581573703 | 0.810264694 |
| chr5:115544287-115545369 | -0.175261445 | 0.58157656 | 0.810264694 |
| chr19:50205830-50206184 | -0.208531605 | 0.581585163 | 0.810264694 |
| chr12:22304446-22304713 | 0.261658674 | 0.581594551 | 0.810264694 |

| | | | |
|---|---|---|---|
| chr15:77419921-77421272 | -0.176085868 | 0.581605738 | 0.810265548 |
| chr8:15493972-15494206 | 0.302052786 | 0.581639578 | 0.810297959 |
| chr1:98661276-98662602 | -0.178287104 | 0.581665734 | 0.810310783 |
| chr15:66862147-66862371 | 0.241962394 | 0.58167115 | 0.810310783 |
| chr14:69001421-69001704 | 0.232776555 | 0.581687494 | 0.810310783 |
| chr2:221845061-221845261 | -0.298111359 | 0.581691085 | 0.810310783 |
| chr2:39513644-39514006 | 0.221040803 | 0.581709684 | 0.810315982 |
| chr1:11926226-11926702 | -0.205318609 | 0.581715968 | 0.810315982 |
| chr5:103864091-103864321 | 0.263986935 | 0.581741139 | 0.810336217 |
| chr16:931873-932105 | 0.269643727 | 0.581751646 | 0.810336217 |
| chr21:31548722-31548957 | -0.274913584 | 0.581827182 | 0.810375999 |
| chr3:149300105-149300527 | -0.204590712 | 0.581833525 | 0.810375999 |
| chr6:137708043-137708255 | 0.264193257 | 0.581835805 | 0.810375999 |
| chr3:104904162-104904362 | -0.287208728 | 0.581836492 | 0.810375999 |
| chr6:3064228-3064488 | 0.208420096 | 0.581842283 | 0.810375999 |
| chr5:150357388-150358288 | -0.17882097 | 0.581843664 | 0.810375999 |
| chr8:80732431-80732761 | 0.243384703 | 0.58185964 | 0.81038352 |
| chr13:21110817-21111101 | 0.224786482 | 0.581883139 | 0.810401517 |
| chrX:20680981-20681232 | -0.279607657 | 0.581899021 | 0.810408906 |
| chr4:99047812-99048012 | -0.287080662 | 0.581954421 | 0.810470465 |
| chr7:4307532-4307742 | -0.279461481 | 0.581966989 | 0.810470465 |
| chr19:984578-984832 | -0.230550337 | 0.581983094 | 0.810470465 |
| chr1:174021534-174022967 | -0.173299849 | 0.581985532 | 0.810470465 |
| chr3:151460732-151461353 | -0.190238746 | 0.582034215 | 0.810504384 |
| chr2:200306283-200307527 | -0.178913112 | 0.582034744 | 0.810504384 |
| chr17:68911804-68912014 | -0.248722038 | 0.582041623 | 0.810504384 |
| chr15:75178467-75178734 | -0.256519736 | 0.582093905 | 0.810562456 |
| chr20:43914476-43915060 | 0.205921331 | 0.582115757 | 0.810578155 |
| chr20:31895174-31895669 | -0.205279011 | 0.582134921 | 0.810590109 |
| chr16:24010555-24010755 | -0.272720199 | 0.582175758 | 0.81063224 |
| chr5:179527541-179527814 | -0.259198149 | 0.582275217 | 0.810755995 |
| chr19:16463595-16464102 | 0.214048321 | 0.582330625 | 0.810810522 |
| chr8:125357418-125357754 | -0.231019314 | 0.582335542 | 0.810810522 |
| chr2:229715000-229715340 | 0.204158106 | 0.582361347 | 0.810815372 |
| chr7:208249-208491 | 0.240035343 | 0.582362494 | 0.810815372 |
| chr1:85307991-85308479 | -0.197180782 | 0.582370771 | 0.810815372 |
| chr6:41545864-41546966 | 0.179685084 | 0.582418405 | 0.810866958 |
| chr4:103197809-103198794 | -0.178006926 | 0.582431772 | 0.810867438 |
| chr1:40862187-40862966 | -0.17900675 | 0.582440619 | 0.810867438 |
| chr10:125162307-125162507 | -0.244290923 | 0.582450498 | 0.810867438 |
| chr3:150237413-150237787 | -0.201930795 | 0.582489378 | 0.810906832 |
| chr17:7350133-7350474 | 0.215857617 | 0.582533215 | 0.810947693 |
| chr2:95076623-95077373 | -0.182081772 | 0.582539897 | 0.810947693 |
| chr2:121476261-121476487 | -0.267019458 | 0.582585025 | 0.810986053 |
| chr3:58019893-58020155 | -0.284126604 | 0.582588621 | 0.810986053 |
| chr3:48088623-48089464 | 0.182047708 | 0.582605062 | 0.810993211 |
| chr17:58863683-58863966 | 0.195057447 | 0.582625463 | 0.810993211 |
| chr5:91182800-91183000 | -0.307739854 | 0.582625516 | 0.810993211 |
| chr9:128551967-128552848 | 0.181686018 | 0.582646266 | 0.81100736 |
| chrX:138756246-138756530 | -0.241201969 | 0.582706829 | 0.811076926 |

| | | | |
|---|---|---|---|
| chr10:102132475-102133352 | 0.178882073 | 0.5827599 | 0.811116295 |
| chr12:59150008-59150361 | -0.214913634 | 0.582763745 | 0.811116295 |
| chr10:103716265-103716677 | -0.199319375 | 0.58276687 | 0.811116295 |
| chr1:161159035-161160005 | 0.179398347 | 0.582792204 | 0.811126649 |
| chr7:150550678-150550878 | -0.300166193 | 0.582795482 | 0.811126649 |
| chr16:77043238-77043438 | 0.270149029 | 0.582823058 | 0.811133112 |
| chr7:100088735-100089159 | 0.210134487 | 0.582831407 | 0.811133112 |
| chr4:106935246-106935447 | 0.298871018 | 0.582839152 | 0.811133112 |
| chr1:226697759-226698408 | 0.190193531 | 0.582845917 | 0.811133112 |
| chr13:28591076-28591301 | -0.29643902 | 0.582853155 | 0.811133112 |
| chr20:32017888-32018921 | 0.178089241 | 0.582863642 | 0.811133112 |
| chr1:164558673-164559426 | 0.175173562 | 0.582926042 | 0.811193151 |
| chrX:108735631-108737092 | 0.178552479 | 0.582927959 | 0.811193151 |
| chr4:139030697-139030971 | 0.225163752 | 0.5829396 | 0.811194618 |
| chr18:65750452-65751682 | 0.179757393 | 0.582967992 | 0.811219393 |
| chr11:71059642-71059842 | 0.276332735 | 0.582983682 | 0.811222257 |
| chr12:10723026-10723683 | -0.190385037 | 0.582991224 | 0.811222257 |
| chr20:44540902-44541165 | -0.288019236 | 0.583002439 | 0.811223131 |
| chr4:69823426-69823972 | 0.19502708 | 0.583055475 | 0.811282194 |
| chr3:10173529-10174208 | 0.206555169 | 0.583071202 | 0.811289345 |
| chr12:56634239-56634552 | 0.194654828 | 0.583159673 | 0.81139771 |
| chr1:97415291-97415577 | -0.268919797 | 0.583176935 | 0.811406993 |
| chr20:25574499-25574724 | -0.258960979 | 0.583206415 | 0.811414591 |
| chr2:149005790-149006357 | -0.204290963 | 0.58321685 | 0.811414591 |
| chr17:49968955-49969282 | 0.195199647 | 0.583223468 | 0.811414591 |
| chr4:93828904-93829174 | -0.236129543 | 0.583231027 | 0.811414591 |
| chr1:113140629-113140860 | 0.240087883 | 0.583238363 | 0.811414591 |
| chr8:128242135-128242621 | -0.188753743 | 0.58324594 | 0.811414591 |
| chr6:159969509-159970207 | -0.201519745 | 0.583256525 | 0.811414591 |
| chr12:57127684-57128582 | -0.181625844 | 0.583316608 | 0.811458132 |
| chr12:3034183-3034563 | 0.213994064 | 0.583318605 | 0.811458132 |
| chr14:63361775-63362094 | 0.218550371 | 0.583335916 | 0.811458132 |
| chr10:86399788-86400824 | 0.174469054 | 0.583337638 | 0.811458132 |
| chr8:64240276-64240708 | -0.229386294 | 0.583340774 | 0.811458132 |
| chr2:19256262-19256658 | -0.209104947 | 0.583376802 | 0.811479789 |
| chr17:43334160-43334486 | 0.219348659 | 0.583377525 | 0.811479789 |
| chr14:103333327-103334526 | 0.173491611 | 0.583411205 | 0.811511907 |
| chr19:53425044-53425307 | 0.257904054 | 0.583442002 | 0.811540012 |
| chr5:81195049-81195399 | -0.217728637 | 0.583476923 | 0.811572911 |
| chr2:73232673-73233828 | -0.173716224 | 0.583493078 | 0.811572911 |
| chr5:90929512-90929712 | -0.282044304 | 0.58349743 | 0.811572911 |
| chr2:164955071-164955743 | -0.194003062 | 0.583510701 | 0.811573411 |
| chr15:44728509-44729611 | 0.176796905 | 0.583528624 | 0.811573411 |
| chr5:90873874-90874288 | 0.249814889 | 0.583536866 | 0.811573411 |
| chr2:121095283-121095656 | 0.205124519 | 0.583540753 | 0.811573411 |
| chrX:47955924-47956340 | 0.241490124 | 0.583550749 | 0.811573411 |
| chr11:106604170-106604370 | -0.272770259 | 0.583689363 | 0.811749657 |
| chr15:77875190-77875458 | 0.233282556 | 0.583698665 | 0.811749657 |
| chr1:204808093-204808333 | -0.216396994 | 0.583711563 | 0.811752861 |
| chr6:3457685-3458075 | 0.198215455 | 0.583748349 | 0.811789284 |

| | | | |
|---|---|---|---|
| chr3:32816657-32817277 | 0.205552741 | 0.583782714 | 0.811804887 |
| chr10:35501971-35502227 | -0.281698043 | 0.583790928 | 0.811804887 |
| chr2:191555099-191555355 | 0.23847838 | 0.583791353 | 0.811804887 |
| chr1:171741330-171742884 | -0.173020227 | 0.583813977 | 0.811820408 |
| chr19:7636950-7637531 | 0.200419249 | 0.583823705 | 0.811820408 |
| chr16:1812848-1813066 | 0.247700492 | 0.583857022 | 0.811852003 |
| chr11:73729840-73730091 | -0.256276571 | 0.58393056 | 0.811939522 |
| chr9:99105365-99105981 | -0.197667644 | 0.58396854 | 0.811977597 |
| chr20:63949007-63949457 | -0.186793446 | 0.58404235 | 0.812065489 |
| chr9:34546863-34547220 | -0.213672757 | 0.584053361 | 0.812066064 |
| chr8:69833737-69835214 | -0.177603042 | 0.584064572 | 0.812066915 |
| chr1:217721754-217721970 | -0.241510527 | 0.584087701 | 0.812084338 |
| chr3:24478844-24479054 | -0.257322429 | 0.58411395 | 0.812085526 |
| chr7:147789535-147789735 | -0.304563401 | 0.584128926 | 0.812085526 |
| chr11:33038963-33040118 | 0.175958661 | 0.584130253 | 0.812085526 |
| chr7:65006212-65006992 | -0.180969324 | 0.58413095 | 0.812085526 |
| chr3:165572595-165572980 | -0.210100926 | 0.584165384 | 0.812089887 |
| chr6:33788718-33789373 | -0.189383195 | 0.584173253 | 0.812089887 |
| chr1:235104222-235104675 | 0.198874572 | 0.584174838 | 0.812089887 |
| chrX:101485144-101485629 | 0.203533963 | 0.584176482 | 0.812089887 |
| chr16:86093782-86094025 | 0.253089011 | 0.584217094 | 0.81213161 |
| chr17:67374682-67374970 | -0.18901608 | 0.58423526 | 0.812139611 |
| chr11:100753239-100753459 | 0.262356554 | 0.584244049 | 0.812139611 |
| chr5:179818894-179819299 | 0.189893326 | 0.584271786 | 0.812163433 |
| chr5:107571432-107572529 | 0.180515788 | 0.584288578 | 0.812172042 |
| chr6:134093261-134093493 | -0.255146092 | 0.584334616 | 0.812221301 |
| chr1:153431972-153432321 | -0.221321668 | 0.584365309 | 0.812236613 |
| chr8:687940-688508 | 0.196168767 | 0.584371835 | 0.812236613 |
| chr15:69133646-69133847 | 0.286016754 | 0.58439135 | 0.812236613 |
| chr3:107643063-107643378 | -0.233375531 | 0.584393269 | 0.812236613 |
| chr15:68229071-68230312 | 0.176742563 | 0.584398636 | 0.812236613 |
| chr11:60363488-60363843 | -0.228917344 | 0.584416938 | 0.812242533 |
| chr11:119374176-119374469 | 0.206990203 | 0.584424096 | 0.812242533 |
| chr10:23461594-23461823 | -0.314952443 | 0.584446565 | 0.812259028 |
| chr13:74575732-74576001 | -0.256155779 | 0.584461652 | 0.812265263 |
| chr5:134370875-134371801 | -0.175057417 | 0.584492958 | 0.812294037 |
| chr12:79374629-79374881 | 0.237868268 | 0.584508632 | 0.812301086 |
| chr2:21125177-21125415 | 0.276337311 | 0.584524483 | 0.812308381 |
| chr15:69957249-69957512 | 0.236461094 | 0.584536483 | 0.812310325 |
| chr15:94582372-94582605 | 0.2529608 | 0.584551752 | 0.812316811 |
| chr5:180484495-180484745 | -0.250678836 | 0.584567213 | 0.812323564 |
| chr11:133927642-133928916 | -0.176712876 | 0.584602785 | 0.812339938 |
| chr1:228459218-228459514 | 0.202938372 | 0.584606411 | 0.812339938 |
| chr15:77607239-77607737 | 0.203448463 | 0.584610802 | 0.812339938 |
| chr12:60616002-60616312 | -0.259963642 | 0.584626431 | 0.812346923 |
| chr17:61568232-61569226 | 0.170997696 | 0.58465039 | 0.812351012 |
| chr10:43881391-43881591 | 0.280266324 | 0.584650577 | 0.812351012 |
| chr17:1516573-1517107 | 0.185824957 | 0.584696335 | 0.81237712 |
| chr10:104121480-104122337 | -0.174594155 | 0.584727442 | 0.81237712 |
| chr8:98377595-98378119 | 0.196525633 | 0.584741162 | 0.81237712 |

| | | | |
|---|---|---|---|
| chr2:46499805-46500005 | 0.25721996 | 0.584741256 | 0.81237712 |
| chr22:29209086-29209319 | 0.25721996 | 0.584741256 | 0.81237712 |
| chr6:27688100-27688554 | -0.183631542 | 0.584744624 | 0.81237712 |
| chr8:125854874-125855074 | -0.271403873 | 0.584751103 | 0.81237712 |
| chr6:112029255-112029553 | 0.235964438 | 0.584754187 | 0.81237712 |
| chr17:58746649-58747037 | 0.192584697 | 0.584774052 | 0.812387963 |
| chr10:102230458-102230741 | 0.217181786 | 0.584783197 | 0.812387963 |
| chr1:10398379-10399700 | 0.17676096 | 0.584796724 | 0.812392026 |
| chr1:226939374-226940717 | 0.173327446 | 0.584816262 | 0.812404439 |
| chr4:139099435-139099769 | 0.214256341 | 0.584861295 | 0.812452267 |
| chr7:2688532-2689364 | 0.18307517 | 0.584898548 | 0.812489286 |
| chr10:43596151-43596361 | -0.280860642 | 0.584916412 | 0.812499371 |
| chr1:174204597-174204918 | -0.251338412 | 0.584936003 | 0.812511855 |
| chr3:197627606-197628293 | -0.185212429 | 0.584955659 | 0.812524428 |
| chr8:108261495-108261704 | -0.254428116 | 0.584971456 | 0.812531641 |
| chr3:94875086-94875472 | -0.252473002 | 0.585062097 | 0.81264281 |
| chr1:28914151-28915425 | -0.178482445 | 0.585083067 | 0.812650836 |
| chr12:13211959-13212211 | 0.250584536 | 0.585097511 | 0.812650836 |
| chr10:49418705-49419463 | -0.180235522 | 0.585099693 | 0.812650836 |
| chr13:37069144-37069399 | -0.24491512 | 0.585121159 | 0.812651587 |
| chr6:28105398-28105598 | -0.278574701 | 0.585121446 | 0.812651587 |
| chr17:64688644-64688915 | -0.230159889 | 0.585134265 | 0.812654662 |
| chr11:75343487-75343897 | 0.191243025 | 0.585218155 | 0.812756438 |
| chr5:38809769-38810124 | -0.210285453 | 0.585257522 | 0.812792354 |
| chr1:42455910-42457100 | -0.180673487 | 0.585265231 | 0.812792354 |
| chr2:53786257-53786552 | 0.211719647 | 0.585294063 | 0.812817662 |
| chr1:193519526-193519751 | 0.280425326 | 0.585323556 | 0.812836802 |
| chr7:37052753-37052975 | 0.25685279 | 0.585329062 | 0.812836802 |
| chr8:102862859-102864854 | 0.170068113 | 0.585399475 | 0.812887194 |
| chr15:29403860-29404062 | -0.284459546 | 0.585422945 | 0.812887194 |
| chr10:73810804-73812123 | -0.174638312 | 0.585426982 | 0.812887194 |
| chr5:138643059-138643312 | -0.233994544 | 0.585440444 | 0.812887194 |
| chr3:120418966-120419219 | 0.206024576 | 0.585443537 | 0.812887194 |
| chr8:33512989-33513989 | -0.174888952 | 0.585444778 | 0.812887194 |
| chr11:22829196-22830197 | -0.172639048 | 0.585459 | 0.812887194 |
| chr1:237013160-237013399 | -0.286110217 | 0.585459966 | 0.812887194 |
| chr2:96832752-96833473 | -0.183750628 | 0.585460962 | 0.812887194 |
| chr20:56735033-56735782 | 0.178600377 | 0.58547144 | 0.812887194 |
| chr5:36017673-36017893 | 0.256707129 | 0.585482059 | 0.812887208 |
| chr7:122352345-122352612 | -0.276763133 | 0.585516799 | 0.81292071 |
| chr3:142987113-142987380 | -0.250981617 | 0.585539006 | 0.812936813 |
| chr8:118862108-118862532 | -0.209132763 | 0.585559681 | 0.812950786 |
| chr8:12755402-12755840 | -0.191151302 | 0.585577158 | 0.812959834 |
| chr16:75197711-75198047 | 0.212098455 | 0.585587418 | 0.812959834 |
| chr15:40859237-40859491 | 0.217653762 | 0.585652251 | 0.813035109 |
| chrX:47541143-47541523 | -0.201794863 | 0.58569477 | 0.813075747 |
| chr8:78232338-78232648 | 0.209694598 | 0.585702747 | 0.813075747 |
| chr2:54704761-54704963 | -0.281777947 | 0.585723483 | 0.813089803 |
| chr9:98837967-98838219 | 0.270508882 | 0.585737123 | 0.813094006 |
| chr16:89700048-89700602 | -0.178575539 | 0.585750082 | 0.813097264 |

| | | | |
|---|---|---|---|
| chr1:196661418-196661821 | -0.230354508 | 0.585798167 | 0.81314928 |
| chr14:64303394-64303806 | 0.202103352 | 0.585832537 | 0.81316583 |
| chr9:80885951-80886166 | -0.263631505 | 0.585833919 | 0.81316583 |
| chr15:73375952-73376284 | -0.219655899 | 0.585841927 | 0.81316583 |
| chr3:62554278-62554659 | 0.182534852 | 0.585901849 | 0.813229945 |
| chr10:16843303-16843527 | -0.276662197 | 0.585909345 | 0.813229945 |
| chr19:12919145-12919526 | 0.20271077 | 0.58596953 | 0.813296682 |
| chr10:112296307-112296748 | -0.218492099 | 0.585978657 | 0.813296682 |
| chr19:58150669-58150950 | 0.221982474 | 0.586076 | 0.813417053 |
| chr10:97709607-97709835 | 0.243697435 | 0.586145408 | 0.813491806 |
| chr17:60039259-60040056 | -0.172670399 | 0.586151094 | 0.813491806 |
| chr8:118941243-118941480 | 0.259843078 | 0.586188511 | 0.813529 |
| chr16:22948816-22949039 | 0.260955584 | 0.586234145 | 0.813569635 |
| chr2:200751554-200751942 | 0.239531713 | 0.58624471 | 0.813569635 |
| chr5:86461272-86461740 | 0.207060898 | 0.586249644 | 0.813569635 |
| chr17:63490335-63490606 | 0.207280264 | 0.58626518 | 0.813576459 |
| chr13:21152999-21153199 | 0.28555275 | 0.586303098 | 0.813614343 |
| chr15:64989273-64990381 | 0.175029267 | 0.586320604 | 0.813623901 |
| chr6:125584377-125585071 | 0.185026765 | 0.586373429 | 0.813682468 |
| chr11:65381929-65382149 | 0.218314255 | 0.58638438 | 0.813682929 |
| chr12:121365828-121366159 | -0.252861612 | 0.586412621 | 0.813707381 |
| chr10:23742788-23743090 | 0.246094365 | 0.586473729 | 0.813777437 |
| chr8:32549097-32549341 | 0.258040369 | 0.586614567 | 0.813903193 |
| chr6:27293839-27294064 | -0.237433808 | 0.586616721 | 0.813903193 |
| chr1:61704521-61704721 | 0.303394782 | 0.586625184 | 0.813903193 |
| chr16:3045358-3046046 | 0.189261359 | 0.586632819 | 0.813903193 |
| chr17:40054271-40054838 | 0.193154267 | 0.586634545 | 0.813903193 |
| chrX:15854215-15855668 | -0.173779493 | 0.586635495 | 0.813903193 |
| chr2:20106756-20107112 | -0.205859511 | 0.586638715 | 0.813903193 |
| chr17:81238802-81239641 | 0.175510933 | 0.586660687 | 0.8139052 |
| chr4:78669499-78669746 | -0.250364678 | 0.586661406 | 0.8139052 |
| chr20:59222298-59222826 | 0.189536794 | 0.586706756 | 0.813950756 |
| chr1:108559775-108560149 | -0.192544766 | 0.586719968 | 0.813950756 |
| chr10:88457668-88457935 | -0.249800216 | 0.586738589 | 0.813950756 |
| chr8:106427321-106427647 | -0.208696372 | 0.586743076 | 0.813950756 |
| chr11:86676557-86676939 | -0.218385743 | 0.586747358 | 0.813950756 |
| chr6:153512191-153512462 | 0.229685069 | 0.586812144 | 0.813959818 |
| chrX:19343536-19344464 | 0.178040416 | 0.586818286 | 0.813959818 |
| chr9:35477222-35477574 | -0.223348094 | 0.586828135 | 0.813959818 |
| chr2:218399597-218400034 | -0.188529012 | 0.586832489 | 0.813959818 |
| chr17:30336502-30336937 | -0.242425692 | 0.586846038 | 0.813959818 |
| chr5:38314640-38314899 | 0.276419799 | 0.586847584 | 0.813959818 |
| chr7:149735174-149735374 | 0.244213757 | 0.586847736 | 0.813959818 |
| chr3:13308410-13308917 | 0.19444081 | 0.586850872 | 0.813959818 |
| chr3:172039326-172041024 | -0.171047188 | 0.586854865 | 0.813959818 |
| chr1:179128544-179128858 | 0.210591901 | 0.586871722 | 0.813959818 |
| chr5:16803868-16804200 | -0.229807793 | 0.586876101 | 0.813959818 |
| chr6:30742222-30743196 | 0.17732853 | 0.586888576 | 0.813959818 |
| chr16:84186797-84187306 | 0.182885968 | 0.586896046 | 0.813959818 |
| chr16:72400323-72400608 | -0.264006828 | 0.586902613 | 0.813959818 |

| | | | |
|---|---|---|---|
| chr5:133754641-133754862 | -0.277958403 | 0.586937058 | 0.813992855 |
| chr19:13163423-13164001 | 0.194401685 | 0.586948914 | 0.813994564 |
| chr5:64147969-64148176 | 0.258036683 | 0.586987698 | 0.814021249 |
| chr6:18122378-18123280 | -0.183096716 | 0.586989403 | 0.814021249 |
| chr3:189458458-189458919 | -0.199887615 | 0.587025205 | 0.814054565 |
| chr17:60560130-60561060 | 0.172630108 | 0.587044924 | 0.814054565 |
| chr17:68198077-68198463 | -0.185945055 | 0.5870453 | 0.814054565 |
| chr2:136117387-136118742 | 0.174222912 | 0.587064839 | 0.814066926 |
| chr19:53868607-53869792 | 0.175266651 | 0.58711278 | 0.814118672 |
| chr14:74302620-74303464 | 0.1778102 | 0.587130109 | 0.814127967 |
| chr3:94036936-94037269 | -0.206165786 | 0.587153961 | 0.814132799 |
| chr7:51366896-51367390 | -0.192948431 | 0.587154844 | 0.814132799 |
| chr12:13000178-13000974 | -0.177331974 | 0.587168562 | 0.814137087 |
| chr15:67254139-67255220 | 0.174298998 | 0.58720558 | 0.814173121 |
| chr19:39111601-39111815 | 0.246457566 | 0.587215802 | 0.814173121 |
| chr3:64233133-64233458 | 0.182487606 | 0.587229794 | 0.814176649 |
| chr16:1610626-1610932 | 0.202560768 | 0.587239598 | 0.814176649 |
| chr1:224513751-224514045 | -0.226708483 | 0.587268561 | 0.814202071 |
| chr20:50354733-50355045 | -0.212537759 | 0.587294942 | 0.814211347 |
| chr14:102063082-102063665 | 0.188018612 | 0.587303876 | 0.814211347 |
| chr15:99104659-99105368 | 0.179819921 | 0.587313177 | 0.814211347 |
| chr16:83533201-83533568 | -0.215890635 | 0.587317757 | 0.814211347 |
| chr13:30114737-30115343 | -0.182718036 | 0.58733383 | 0.814218899 |
| chr17:67359930-67360182 | -0.222653618 | 0.587361201 | 0.814228917 |
| chr17:64216551-64216832 | -0.224344616 | 0.58736231 | 0.814228917 |
| chr12:111313614-111314047 | 0.218082296 | 0.587380897 | 0.814231104 |
| chr13:38755887-38756087 | 0.306049227 | 0.587385141 | 0.814231104 |
| chr4:140373201-140374235 | -0.171798177 | 0.587396573 | 0.814232221 |
| chr2:9002957-9003253 | -0.215447677 | 0.587445466 | 0.814260935 |
| chr1:192506714-192506969 | -0.258375277 | 0.587478965 | 0.814260935 |
| chr14:32203101-32203714 | 0.175405032 | 0.587494447 | 0.814260935 |
| chr6:31541643-31542676 | -0.174404447 | 0.587495793 | 0.814260935 |
| chr11:88231953-88232153 | 0.300076387 | 0.587497467 | 0.814260935 |
| chr8:142634722-142634988 | -0.223371322 | 0.587536068 | 0.814260935 |
| chr9:120619797-120620017 | 0.264029109 | 0.58754007 | 0.814260935 |
| chr1:178493202-178493587 | -0.242854373 | 0.587546433 | 0.814260935 |
| chr10:58267874-58268398 | -0.178716558 | 0.58755991 | 0.814260935 |
| chr2:20066112-20066623 | 0.198429228 | 0.587560356 | 0.814260935 |
| chr12:124917129-124917811 | 0.175806519 | 0.587572669 | 0.814260935 |
| chr1:31769797-31770175 | -0.2183966 | 0.587573248 | 0.814260935 |
| chr2:240998179-240998903 | 0.189213747 | 0.587574025 | 0.814260935 |
| chr5:160887729-160888157 | -0.223096686 | 0.587578465 | 0.814260935 |
| chr14:91835779-91836928 | 0.173761871 | 0.58758128 | 0.814260935 |
| chr7:76201431-76201647 | 0.247945266 | 0.58758732 | 0.814260935 |
| chr6:30885572-30885960 | -0.195911557 | 0.587601088 | 0.814263357 |
| chr19:10250153-10250386 | 0.240301206 | 0.587610321 | 0.814263357 |
| chr16:31108072-31108503 | 0.210208722 | 0.587635135 | 0.814270248 |
| chr1:147723727-147724273 | 0.190416656 | 0.587639433 | 0.814270248 |
| chr2:211833352-211833577 | 0.290121537 | 0.587647176 | 0.814270248 |
| chr21:46457976-46459279 | 0.17274187 | 0.587677254 | 0.814296554 |

| | | | |
|---|---|---|---|
| chr2:85231277-85231638 | 0.22637437 | 0.587697592 | 0.814296554 |
| chr8:59223055-59223343 | -0.253747136 | 0.587698042 | 0.814296554 |
| chr14:34356322-34356522 | 0.277989667 | 0.587713881 | 0.81429727 |
| chr11:101914781-101915542 | 0.179869135 | 0.587719814 | 0.81429727 |
| chr15:82139871-82140099 | -0.255974064 | 0.587731325 | 0.814298494 |
| chr17:60391566-60392952 | -0.167967751 | 0.587749187 | 0.814301008 |
| chr8:80131485-80131685 | 0.279700534 | 0.587756231 | 0.814301008 |
| chr11:75328674-75329063 | 0.189856429 | 0.587768573 | 0.814301008 |
| chr4:101347284-101348241 | -0.173003907 | 0.58777565 | 0.814301008 |
| chr2:105744662-105745810 | -0.185555236 | 0.587800053 | 0.814306386 |
| chr1:191690973-191691322 | 0.203312839 | 0.587811127 | 0.814306386 |
| chr2:220179311-220179660 | -0.233441829 | 0.587811415 | 0.814306386 |
| chr11:19071213-19071600 | 0.195246153 | 0.587825755 | 0.814311529 |
| chr22:40177773-40178003 | 0.245121562 | 0.587861049 | 0.814345699 |
| chr5:174735787-174736537 | -0.185082718 | 0.58791059 | 0.814353741 |
| chr17:60421854-60422676 | 0.17216261 | 0.58793147 | 0.814353741 |
| chr20:41476245-41476469 | -0.243861442 | 0.587933876 | 0.814353741 |
| chr12:65631573-65632079 | -0.195817462 | 0.587943808 | 0.814353741 |
| chr10:129808261-129808709 | -0.234381447 | 0.587949134 | 0.814353741 |
| chr3:11060750-11061252 | 0.192577056 | 0.587957568 | 0.814353741 |
| chr12:56763188-56763388 | -0.276353433 | 0.587957915 | 0.814353741 |
| chr4:107737016-107737259 | -0.244036478 | 0.587961226 | 0.814353741 |
| chr15:65376839-65377783 | 0.180820972 | 0.587966062 | 0.814353741 |
| chr2:25091167-25091376 | 0.274124377 | 0.587973136 | 0.814353741 |
| chr5:135457509-135457712 | -0.238360328 | 0.587992441 | 0.814365758 |
| chr6:27617693-27618513 | -0.187566142 | 0.588006433 | 0.814370417 |
| chr5:86807025-86807316 | -0.278566822 | 0.588043123 | 0.814385155 |
| chr9:136267122-136267715 | 0.18374157 | 0.588058108 | 0.814385155 |
| chr19:44954643-44955640 | 0.178259519 | 0.5880648 | 0.814385155 |
| chr20:62407240-62407561 | -0.194886917 | 0.588100501 | 0.814385155 |
| chr9:3525713-3527050 | 0.171148081 | 0.588102292 | 0.814385155 |
| chr19:50133775-50134040 | 0.270660212 | 0.588111677 | 0.814385155 |
| chr6:42929029-42929926 | 0.175408093 | 0.588113171 | 0.814385155 |
| chrX:47644763-47644973 | 0.267812173 | 0.588117242 | 0.814385155 |
| chr5:176415722-176416901 | -0.175523134 | 0.588125989 | 0.814385155 |
| chr3:184335758-184336638 | -0.175369933 | 0.588132152 | 0.814385155 |
| chr13:100588319-100589209 | -0.182544345 | 0.588141836 | 0.814385155 |
| chr10:123995398-123995716 | 0.217290394 | 0.588144618 | 0.814385155 |
| chr16:589149-590162 | 0.174823171 | 0.588215573 | 0.814468685 |
| chr1:89987285-89987546 | 0.246590207 | 0.588245901 | 0.814495052 |
| chr18:57715605-57716022 | -0.238024845 | 0.588255876 | 0.814495052 |
| chr19:38657917-38658141 | -0.259310157 | 0.588285497 | 0.814519043 |
| chr13:21327435-21327805 | -0.247209531 | 0.588306024 | 0.814519043 |
| chr10:53214541-53214752 | -0.292954082 | 0.588312609 | 0.814519043 |
| chr11:72244665-72244869 | -0.252053919 | 0.588315724 | 0.814519043 |
| chr3:155283706-155283961 | 0.248472049 | 0.588332051 | 0.81452693 |
| chr3:42080443-42081027 | -0.200102928 | 0.588343282 | 0.814527762 |
| chr19:55117563-55118201 | -0.180727169 | 0.588412848 | 0.814596489 |
| chr3:181137799-181138069 | -0.28834253 | 0.588414188 | 0.814596489 |
| chr9:135170993-135171193 | 0.284722289 | 0.588439848 | 0.814609965 |

| | | | |
|---|---|---|---|
| chr11:3378775-3379483 | 0.179736285 | 0.588445185 | 0.814609965 |
| chr7:2063653-2064097 | -0.207470822 | 0.588484847 | 0.814650152 |
| chr20:25696168-25697179 | 0.174616404 | 0.588505628 | 0.814658238 |
| chr12:89542988-89543329 | -0.206198597 | 0.588544479 | 0.814658238 |
| chr3:190120740-190121205 | -0.191515035 | 0.588548963 | 0.814658238 |
| chr2:69455518-69455729 | 0.270499135 | 0.588550445 | 0.814658238 |
| chr21:44892880-44893092 | -0.245104524 | 0.588554307 | 0.814658238 |
| chr10:94027216-94027434 | 0.273658329 | 0.588566777 | 0.814658238 |
| chr15:85103185-85103423 | -0.266832137 | 0.588569363 | 0.814658238 |
| chr7:636565-636770 | 0.254654695 | 0.588575745 | 0.814658238 |
| chr17:79777758-79778351 | -0.190490214 | 0.588622921 | 0.814697101 |
| chr3:159805119-159805660 | -0.191133997 | 0.588663257 | 0.814697101 |
| chr17:75601244-75601735 | -0.185752285 | 0.588672253 | 0.814697101 |
| chr17:78140298-78140833 | 0.17495918 | 0.588674606 | 0.814697101 |
| chr6:54856056-54856300 | 0.294578849 | 0.588679839 | 0.814697101 |
| chr12:57240519-57241000 | 0.187797634 | 0.588681226 | 0.814697101 |
| chr17:75504330-75504609 | 0.19946374 | 0.588682087 | 0.814697101 |
| chr8:37762314-37762929 | 0.182603367 | 0.588688884 | 0.814697101 |
| chr14:75660733-75661754 | 0.170968908 | 0.588703385 | 0.814702454 |
| chr6:157020943-157022375 | -0.173471225 | 0.588726544 | 0.814719789 |
| chr20:62910263-62910472 | -0.266097776 | 0.588776776 | 0.814774587 |
| chr1:232211239-232211483 | 0.248952526 | 0.588841501 | 0.814849439 |
| chr3:171703869-171704366 | -0.194296727 | 0.588898435 | 0.814893153 |
| chr17:78661400-78661686 | -0.274474935 | 0.588900328 | 0.814893153 |
| chr5:145697327-145697539 | -0.283351289 | 0.58890956 | 0.814893153 |
| chr5:17138875-17139112 | -0.29726251 | 0.588917125 | 0.814893153 |
| chr21:45554864-45555133 | -0.212384743 | 0.588943381 | 0.814893153 |
| chr8:142943682-142943882 | 0.255488795 | 0.588950579 | 0.814893153 |
| chr2:55281817-55282919 | -0.17635888 | 0.588963688 | 0.814893153 |
| chr11:66267163-66267871 | 0.177357263 | 0.58896466 | 0.814893153 |
| chr1:8313957-8314159 | -0.280892726 | 0.588968807 | 0.814893153 |
| chr21:45308707-45308918 | 0.244234018 | 0.588993624 | 0.814912775 |
| chr1:44355007-44355685 | 0.185965856 | 0.589034371 | 0.81494108 |
| chr1:22642088-22642364 | -0.247038407 | 0.589051552 | 0.81494108 |
| chr2:175309821-175310155 | -0.230698293 | 0.58905361 | 0.81494108 |
| chr6:109482542-109483781 | 0.175069922 | 0.589056626 | 0.81494108 |
| chr6:159968519-159969247 | -0.181089771 | 0.589074503 | 0.814951098 |
| chr13:42671108-42671420 | -0.291221315 | 0.589159704 | 0.815037199 |
| chr9:135967451-135968057 | 0.194668646 | 0.589166371 | 0.815037199 |
| chr12:120686236-120687251 | 0.171070977 | 0.589168651 | 0.815037199 |
| chr9:137150238-137151554 | -0.173297908 | 0.589187633 | 0.815048743 |
| chr12:78102280-78102548 | -0.228579416 | 0.589202976 | 0.815055252 |
| chr8:125931555-125931895 | -0.218346122 | 0.589231592 | 0.81507769 |
| chr8:121164541-121164776 | 0.233622567 | 0.589249404 | 0.81507769 |
| chr5:163489505-163489840 | -0.234267891 | 0.589260993 | 0.81507769 |
| chr9:136991832-136992503 | 0.178552779 | 0.589278611 | 0.81507769 |
| chr5:134465755-134467059 | -0.171959472 | 0.589281887 | 0.81507769 |
| chr9:97696704-97697702 | 0.175919375 | 0.589286006 | 0.81507769 |
| chr17:74748262-74748807 | 0.177866722 | 0.58929366 | 0.81507769 |
| chr6:11428873-11429134 | 0.255170267 | 0.589329148 | 0.815097482 |

| | | | |
|---|---|---|---|
| chr1:71047209-71048387 | -0.177045462 | 0.589366226 | 0.815097482 |
| chr8:66020661-66021578 | 0.178944191 | 0.589368475 | 0.815097482 |
| chr16:3171955-3172551 | -0.185616892 | 0.58939025 | 0.815097482 |
| chr2:127526035-127527598 | 0.171632369 | 0.58939674 | 0.815097482 |
| chr12:115398306-115398754 | 0.206009051 | 0.58940652 | 0.815097482 |
| chr14:49598197-49599441 | 0.171286108 | 0.589435227 | 0.815097482 |
| chr19:10717360-10718157 | 0.174186058 | 0.589435741 | 0.815097482 |
| chr2:32326240-32326826 | -0.183249335 | 0.589459761 | 0.815097482 |
| chr1:48498141-48498418 | -0.238259931 | 0.589461037 | 0.815097482 |
| chr4:92047546-92047779 | 0.252901548 | 0.589465366 | 0.815097482 |
| chr16:85448993-85449199 | 0.225571094 | 0.589467399 | 0.815097482 |
| chr15:82750034-82750745 | -0.177960986 | 0.58947334 | 0.815097482 |
| chr1:9428566-9429413 | -0.1802493 | 0.589477923 | 0.815097482 |
| chr17:64081851-64082497 | 0.184533935 | 0.589509392 | 0.815097482 |
| chr2:168246392-168247098 | -0.185779928 | 0.589517448 | 0.815097482 |
| chr7:99270807-99271065 | 0.257623475 | 0.58952508 | 0.815097482 |
| chr4:109702759-109703844 | 0.173661492 | 0.589535294 | 0.815097482 |
| chr11:86291888-86292151 | 0.226779485 | 0.589550391 | 0.815097482 |
| chr15:43510255-43511432 | 0.172848532 | 0.589556297 | 0.815097482 |
| chr6:107848235-107848795 | -0.192345633 | 0.589560365 | 0.815097482 |
| chr3:19467619-19468047 | -0.214497633 | 0.589561089 | 0.815097482 |
| chr14:74608943-74609684 | 0.180918842 | 0.589561265 | 0.815097482 |
| chr16:72874961-72875161 | 0.293287303 | 0.589563279 | 0.815097482 |
| chr3:143775420-143775686 | -0.240458962 | 0.589592147 | 0.815108896 |
| chr18:76421047-76421663 | 0.20075742 | 0.589598051 | 0.815108896 |
| chr2:46986815-46987482 | -0.188250951 | 0.589604243 | 0.815108896 |
| chr8:101635542-101636122 | 0.182289923 | 0.589614087 | 0.815108896 |
| chr5:132295261-132295462 | 0.212438051 | 0.589648771 | 0.815131144 |
| chr20:54648482-54649793 | -0.167760732 | 0.589671719 | 0.815131144 |
| chrX:103878754-103879024 | 0.229702621 | 0.589691418 | 0.815131144 |
| chr19:18346227-18346427 | 0.28258151 | 0.589691543 | 0.815131144 |
| chr8:133799344-133799544 | 0.280275114 | 0.58969828 | 0.815131144 |
| chr2:178804806-178805042 | 0.262412703 | 0.589701139 | 0.815131144 |
| chrX:50332689-50332944 | -0.264941571 | 0.589704648 | 0.815131144 |
| chrX:95764931-95765204 | -0.230357151 | 0.589727539 | 0.815148079 |
| chr8:122673315-122674001 | -0.181243182 | 0.589764831 | 0.815184921 |
| chr14:106373418-106373919 | -0.201261306 | 0.589781327 | 0.815193017 |
| chr20:31562899-31563114 | 0.217733309 | 0.589810944 | 0.815209324 |
| chr19:1407122-1407629 | -0.190728865 | 0.589820083 | 0.815209324 |
| chr1:156740591-156741565 | 0.17214993 | 0.589825044 | 0.815209324 |
| chr10:119030417-119030643 | 0.222784729 | 0.589850541 | 0.815229859 |
| chr16:19410407-19410862 | 0.210521954 | 0.58986359 | 0.815233189 |
| chr8:100143971-100144302 | -0.207207162 | 0.58989214 | 0.815244595 |
| chrX:7132295-7133007 | -0.180820158 | 0.589901218 | 0.815244595 |
| chr2:9951046-9952324 | 0.172384052 | 0.589903762 | 0.815244595 |
| chr1:93078982-93079632 | -0.183930804 | 0.589933885 | 0.81527152 |
| chr1:217403271-217403549 | -0.227201839 | 0.589949764 | 0.815278759 |
| chr2:66433972-66434707 | -0.183207695 | 0.589975555 | 0.815299696 |
| chr15:92900079-92900607 | -0.179049844 | 0.589989239 | 0.815303902 |
| chr1:50873375-50874130 | 0.18998792 | 0.590013482 | 0.815306095 |

| | | | |
|---|---|---|---|
| chr10:48804835-48805151 | -0.243408526 | 0.590020497 | 0.815306095 |
| chr3:27659195-27659559 | 0.197538373 | 0.590022747 | 0.815306095 |
| chr2:143284138-143284378 | -0.259225573 | 0.590064133 | 0.815348578 |
| chr1:34924846-34925134 | 0.217604121 | 0.590098713 | 0.815372532 |
| chr11:27471926-27473073 | -0.171940661 | 0.590102751 | 0.815372532 |
| chr16:83793234-83793720 | -0.195538132 | 0.590168998 | 0.815428973 |
| chr8:143038765-143039342 | 0.202061832 | 0.590182151 | 0.815428973 |
| chr10:92592370-92593504 | -0.173094701 | 0.590188095 | 0.815428973 |
| chr11:67482833-67483458 | -0.179342219 | 0.590196133 | 0.815428973 |
| chr9:101119260-101119512 | 0.23437732 | 0.59019681 | 0.815428973 |
| chr14:45268375-45269071 | 0.185844684 | 0.590223796 | 0.815451553 |
| chr1:151970672-151970872 | -0.237445395 | 0.59026314 | 0.815491206 |
| chr11:123020922-123021221 | -0.242239398 | 0.590278545 | 0.815497786 |
| chr11:14623080-14623284 | -0.300950592 | 0.590298567 | 0.815507305 |
| chr3:170189103-170189453 | 0.196180427 | 0.590306722 | 0.815507305 |
| chr19:41951651-41951941 | -0.231361732 | 0.59035262 | 0.815556009 |
| chr2:169798616-169799650 | -0.17599278 | 0.590370432 | 0.81556591 |
| chr10:114937988-114938735 | -0.178898312 | 0.590382509 | 0.815567891 |
| chr2:220224750-220225633 | -0.175703239 | 0.590438957 | 0.815619513 |
| chr7:43925859-43927008 | -0.173483626 | 0.590441168 | 0.815619513 |
| chr1:71050079-71050303 | -0.227987819 | 0.590523613 | 0.815708944 |
| chr13:24878474-24878674 | 0.282350466 | 0.5905272 | 0.815708944 |
| chr6:110981324-110982547 | -0.174263005 | 0.590551923 | 0.815728388 |
| chr2:133641432-133641940 | -0.177328937 | 0.590576662 | 0.815747855 |
| chr17:63698641-63699368 | -0.178868479 | 0.590606007 | 0.815773682 |
| chr17:75096192-75096482 | 0.20622886 | 0.590622881 | 0.815782283 |
| chr19:4968566-4969143 | 0.180917307 | 0.59064299 | 0.815795352 |
| chr12:63826056-63826408 | -0.208355903 | 0.590677983 | 0.815828979 |
| chr8:142816317-142816658 | 0.195892491 | 0.590817405 | 0.816006835 |
| chr9:94554105-94555343 | 0.172367117 | 0.590839137 | 0.816022141 |
| chr7:5189901-5190863 | -0.174306873 | 0.590877439 | 0.816060332 |
| chr6:161439517-161439737 | 0.268826807 | 0.590904944 | 0.816083609 |
| chr17:59043543-59043752 | 0.193165843 | 0.590923361 | 0.816094334 |
| chr20:58919280-58919529 | -0.238124829 | 0.590996592 | 0.81618076 |
| chr2:74178092-74179470 | -0.172216097 | 0.591016108 | 0.816183279 |
| chrX:8748945-8749164 | -0.282730595 | 0.591019721 | 0.816183279 |
| chr1:153394085-153394293 | -0.258009103 | 0.59103162 | 0.816185001 |
| chr4:108350366-108350713 | -0.238129341 | 0.591096107 | 0.816243366 |
| chr16:67884720-67885054 | -0.202440153 | 0.591102591 | 0.816243366 |
| chr6:24357836-24358217 | -0.189143852 | 0.591113326 | 0.816243366 |
| chr2:201450965-201452252 | 0.170764557 | 0.591116496 | 0.816243366 |
| chr22:46798624-46798833 | -0.256934489 | 0.591136727 | 0.816244486 |
| chr17:30824566-30825193 | -0.193415641 | 0.591161115 | 0.816244486 |
| chr20:55122380-55122609 | -0.268040915 | 0.591167642 | 0.816244486 |
| chr14:32957868-32958068 | 0.280886219 | 0.591168123 | 0.816244486 |
| chr15:45278991-45279794 | 0.172454595 | 0.5911857 | 0.816244486 |
| chr6:35731922-35732182 | 0.213160027 | 0.591190691 | 0.816244486 |
| chr17:56761114-56761421 | 0.222036274 | 0.591191877 | 0.816244486 |
| chr20:45629546-45629851 | 0.217749764 | 0.591205794 | 0.816248993 |
| chr7:1137913-1138893 | -0.173245415 | 0.591268357 | 0.816297244 |

| | | | |
|---|---|---|---|
| chr6:12331316-12331788 | 0.197624116 | 0.591277726 | 0.816297244 |
| chr3:112086008-112086914 | -0.174781169 | 0.591293357 | 0.816297244 |
| chr5:134145882-134146348 | -0.197538612 | 0.591293654 | 0.816297244 |
| chr2:30872135-30872612 | -0.209384855 | 0.59129401 | 0.816297244 |
| chr19:2060983-2061870 | 0.177929112 | 0.591334219 | 0.816338046 |
| chr16:73010449-73010772 | 0.201790648 | 0.591353011 | 0.81634928 |
| chr1:47436543-47436975 | 0.195196895 | 0.591368631 | 0.816356134 |
| chr12:64080417-64081030 | -0.179554118 | 0.591392532 | 0.81637442 |
| chr1:3453170-3453677 | 0.196661302 | 0.591437654 | 0.816422 |
| chr7:155598830-155599097 | -0.234318536 | 0.591455875 | 0.816430572 |
| chr1:114485989-114486192 | 0.20796149 | 0.591469047 | 0.816430572 |
| chr17:75514350-75514656 | -0.186284533 | 0.591481667 | 0.816430572 |
| chr5:10850336-10850599 | 0.257079766 | 0.591487867 | 0.816430572 |
| chr4:173038180-173038411 | 0.289525855 | 0.591497141 | 0.816430572 |
| chr5:57499463-57499693 | -0.244450521 | 0.59151299 | 0.816437742 |
| chr1:58814164-58815949 | -0.16892625 | 0.59159276 | 0.816533135 |
| chr10:12711138-12711368 | -0.265893139 | 0.591606377 | 0.816537221 |
| chr8:80991754-80992249 | 0.204089778 | 0.591634678 | 0.816561574 |
| chr10:133257486-13325805€ | 0.185428299 | 0.591651481 | 0.816562272 |
| chr5:18170253-18170453 | 0.283563533 | 0.591656498 | 0.816562272 |
| chr21:39742568-39743040 | 0.193129665 | 0.591667373 | 0.816562572 |
| chr11:34640281-34640649 | 0.196356702 | 0.591716865 | 0.816616167 |
| chr3:165041550-165041777 | -0.268994024 | 0.591773718 | 0.816679919 |
| chr6:133405722-133406074 | -0.278055859 | 0.591808619 | 0.816702293 |
| chr3:149085580-149086963 | -0.169057437 | 0.59181146 | 0.816702293 |
| chr9:134068022-134068405 | -0.207295429 | 0.591828113 | 0.816702293 |
| chrX:106898666-106898875 | 0.243352289 | 0.591832566 | 0.816702293 |
| chr15:90047267-90047467 | 0.238331809 | 0.591859634 | 0.816722131 |
| chr14:44961584-44962791 | 0.168074881 | 0.591873226 | 0.816722131 |
| chr1:28235848-28236528 | 0.181022853 | 0.591878918 | 0.816722131 |
| chr12:70730060-70730565 | -0.213288943 | 0.591890742 | 0.816723737 |
| chr14:93681884-93682130 | 0.286708481 | 0.591954857 | 0.816797498 |
| chr9:130445606-130445836 | -0.215645272 | 0.591981475 | 0.816819517 |
| chr2:130728061-130728265 | -0.236668336 | 0.592013612 | 0.816841261 |
| chr5:141968641-141969682 | -0.173229994 | 0.592034309 | 0.816841261 |
| chr16:27363920-27364126 | -0.242919916 | 0.592051225 | 0.816841261 |
| chr4:77464112-77464366 | 0.232413033 | 0.59205637 | 0.816841261 |
| chr1:173103364-173103652 | -0.271433759 | 0.592067142 | 0.816841261 |
| chr2:241686286-241687504 | 0.174015106 | 0.592068061 | 0.816841261 |
| chr3:16703383-16703656 | 0.254502677 | 0.592084521 | 0.816841261 |
| chr17:64975149-64976302 | -0.167912415 | 0.592090089 | 0.816841261 |
| chr3:142929151-142929447 | -0.215855374 | 0.59209318 | 0.816841261 |
| chr4:77433430-77433685 | 0.238878893 | 0.59213118 | 0.816878977 |
| chr1:85642239-85642733 | -0.213758734 | 0.592146367 | 0.816881296 |
| chr2:221084733-221084938 | 0.284276811 | 0.592160191 | 0.816881296 |
| chr8:98926778-98927074 | -0.21518595 | 0.592164844 | 0.816881296 |
| chr13:54185753-54186037 | -0.252829471 | 0.592197023 | 0.816910979 |
| chr13:94154421-94154628 | -0.285015443 | 0.59223925 | 0.816941188 |
| chr6:27609368-27609568 | 0.27856718 | 0.592256535 | 0.816941188 |
| chr1:171393816-171394193 | 0.203278924 | 0.592272498 | 0.816941188 |

| | | | |
|---|---|---|---|
| chr17:7023863-7024127 | -0.241187407 | 0.592281388 | 0.816941188 |
| chr4:147979574-147980140 | -0.205914486 | 0.592284962 | 0.816941188 |
| chr8:102108008-102108356 | -0.208661923 | 0.592293411 | 0.816941188 |
| chr7:100126692-100127715 | -0.175973854 | 0.592311563 | 0.816941188 |
| chr3:79603148-79603532 | 0.208299894 | 0.592322164 | 0.816941188 |
| chr4:87644383-87644829 | 0.230369882 | 0.592325198 | 0.816941188 |
| chr8:101603583-101603956 | 0.180478709 | 0.592333858 | 0.816941188 |
| chr13:48896353-48896553 | -0.298951625 | 0.592344652 | 0.816941188 |
| chr6:28863381-28864451 | 0.173793381 | 0.592346866 | 0.816941188 |
| chr22:29821338-29821709 | 0.234381969 | 0.592371255 | 0.816960119 |
| chr8:70668813-70669655 | -0.171528294 | 0.5923908 | 0.816964655 |
| chr4:53377331-53378212 | -0.177097667 | 0.592395868 | 0.816964655 |
| chr6:42779135-42779342 | 0.242275073 | 0.592414718 | 0.816975946 |
| chr11:14519647-14520833 | -0.172555522 | 0.592484969 | 0.81705812 |
| chr17:29111859-29112169 | 0.21019117 | 0.592503627 | 0.817065579 |
| chr5:54715293-54715551 | -0.286701724 | 0.592517334 | 0.817065579 |
| chr14:95693388-95693696 | 0.224171368 | 0.592528473 | 0.817065579 |
| chr8:121776256-121776754 | 0.185792321 | 0.592533032 | 0.817065579 |
| chrX:107718179-107718402 | 0.218237908 | 0.592580655 | 0.817116543 |
| chr1:242075135-242075335 | -0.293596664 | 0.592608552 | 0.817135233 |
| chr3:141011573-141011860 | -0.219356551 | 0.592615538 | 0.817135233 |
| chr11:69353692-69353892 | -0.266991469 | 0.592659495 | 0.817181137 |
| chr4:173333733-173333957 | -0.197038114 | 0.59270519 | 0.817229438 |
| chr14:30965966-30966166 | -0.280415218 | 0.59273297 | 0.817253035 |
| chr11:62542756-62543073 | -0.191069145 | 0.592744233 | 0.817253858 |
| chr9:112921491-112921753 | 0.234800782 | 0.592786377 | 0.817291577 |
| chr12:118089788-118089988 | 0.290929629 | 0.592802234 | 0.817291577 |
| chr17:64262328-64263918 | -0.169665741 | 0.592811854 | 0.817291577 |
| chr5:84634643-84634885 | -0.282158578 | 0.592814256 | 0.817291577 |
| chr4:56959101-56959301 | 0.262931755 | 0.592835855 | 0.817293114 |
| chr7:26667495-26667848 | 0.213806085 | 0.592843884 | 0.817293114 |
| chr13:107203189-107203545 | -0.222007466 | 0.592847371 | 0.817293114 |
| chr4:88592178-88592908 | 0.17530721 | 0.592894175 | 0.817335144 |
| chr8:126698962-126699499 | 0.181523526 | 0.592917819 | 0.817335144 |
| chr3:36729493-36729752 | 0.239177125 | 0.592922363 | 0.817335144 |
| chr7:44477412-44477764 | -0.202004754 | 0.592931757 | 0.817335144 |
| chr14:106343137-106343337 | -0.282197386 | 0.592940784 | 0.817335144 |
| chr3:164949069-164949305 | 0.283293388 | 0.592941862 | 0.817335144 |
| chr16:89744167-89744434 | 0.238408547 | 0.592968722 | 0.817357465 |
| chr2:7044357-7044613 | 0.226013522 | 0.593047816 | 0.817443653 |
| chr1:183655564-183655809 | 0.224916849 | 0.593054183 | 0.817443653 |
| chr14:99415765-99415995 | -0.254594845 | 0.593063254 | 0.817443653 |
| chr10:85642232-85643049 | 0.180652239 | 0.593107327 | 0.817489695 |
| chr10:78029412-78030020 | -0.17991484 | 0.593118368 | 0.817490208 |
| chr11:83706687-83706954 | -0.294243767 | 0.593145392 | 0.817510864 |
| chr6:26553752-26554394 | -0.175375619 | 0.593154694 | 0.817510864 |
| chr18:10062311-10062729 | -0.22272806 | 0.593171831 | 0.817519778 |
| chr5:126676145-126676354 | -0.228613224 | 0.593193805 | 0.817524428 |
| chr20:32457500-32457743 | 0.228515123 | 0.593196544 | 0.817524428 |
| chr19:7944306-7944568 | 0.221747181 | 0.593245579 | 0.8175773 |

| | | | |
|---|---|---|---|
| chrX:69109686-69110316 | 0.18375882 | 0.59325938 | 0.817581614 |
| chr5:141182243-141182671 | -0.21904004 | 0.593273095 | 0.817584194 |
| chr14:77580597-77581227 | 0.206458483 | 0.593287196 | 0.817584194 |
| chr9:122264404-122265524 | 0.172427112 | 0.593293263 | 0.817584194 |
| chr20:51306764-51307193 | 0.184622277 | 0.593358365 | 0.817659202 |
| chr17:56679275-56679475 | 0.235982277 | 0.59340576 | 0.817696842 |
| chr10:108598900-108599100 | 0.285747054 | 0.593415804 | 0.817696842 |
| chr8:17412938-17414103 | -0.176229768 | 0.593417695 | 0.817696842 |
| chr10:5434053-5434532 | 0.209600509 | 0.593439061 | 0.817711578 |
| chr1:151047578-151048978 | 0.168404269 | 0.593452435 | 0.817715301 |
| chr1:36464080-36464858 | 0.17878648 | 0.593466175 | 0.817719254 |
| chr19:16659729-16660522 | 0.176417786 | 0.593490428 | 0.817719254 |
| chr17:41965761-41966301 | -0.180098367 | 0.593493023 | 0.817719254 |
| chr6:57960911-57961670 | 0.184795829 | 0.593502523 | 0.817719254 |
| chr5:107977768-107977968 | 0.272008981 | 0.593508665 | 0.817719254 |
| chr4:180060227-180060470 | -0.215771967 | 0.593536914 | 0.817726045 |
| chr20:57198996-57199213 | 0.177903782 | 0.593538329 | 0.817726045 |
| chr1:244383655-244383873 | 0.275651633 | 0.59354561 | 0.817726045 |
| chr16:84141234-84141529 | -0.200435976 | 0.593557821 | 0.817728166 |
| chr19:51002558-51003035 | -0.193084192 | 0.593614683 | 0.817790524 |
| chr22:34099258-34099516 | -0.23780159 | 0.593624431 | 0.817790524 |
| chr14:34305770-34305970 | 0.274115101 | 0.593635707 | 0.817791355 |
| chr1:200558908-200559233 | -0.204959166 | 0.593656303 | 0.817805024 |
| chr2:27489144-27490135 | -0.17034818 | 0.593720166 | 0.817878295 |
| chr1:176515400-176515680 | 0.227123598 | 0.593739537 | 0.817890276 |
| chr8:55972256-55972456 | 0.262437166 | 0.593758264 | 0.817901368 |
| chr9:12884832-12885242 | -0.226431785 | 0.593774468 | 0.817902969 |
| chr17:65587998-65588466 | 0.190839772 | 0.593787596 | 0.817902969 |
| chr8:81590911-81591153 | 0.239446694 | 0.59379145 | 0.817902969 |
| chr11:82695533-82696445 | -0.174362689 | 0.593817256 | 0.817923812 |
| chr8:142403812-142404079 | -0.21672634 | 0.593830209 | 0.817926949 |
| chr4:46329148-46329506 | -0.212417175 | 0.593851093 | 0.817941011 |
| chr10:939173-939431 | 0.258579149 | 0.593893975 | 0.817985371 |
| chr16:10578510-10579188 | -0.177848361 | 0.593920716 | 0.818007498 |
| chr10:33330575-33330775 | -0.255943515 | 0.594004813 | 0.818102682 |
| chr8:98690853-98691096 | 0.239334901 | 0.59402194 | 0.818102682 |
| chr3:32865151-32865599 | 0.232427527 | 0.594024015 | 0.818102682 |
| chr8:70493168-70493989 | -0.180257943 | 0.594033757 | 0.818102682 |
| chr1:54945198-54945436 | -0.224996539 | 0.594043211 | 0.818102682 |
| chr8:90772176-90772380 | 0.287477749 | 0.594065722 | 0.818118979 |
| chr10:43390783-43390998 | -0.232453275 | 0.594078984 | 0.818122539 |
| chr17:9644500-9645684 | -0.171215081 | 0.594092799 | 0.81812686 |
| chr10:103275987-103276187 | -0.265947379 | 0.594115183 | 0.81814298 |
| chr8:38104665-38105965 | 0.173198128 | 0.594136304 | 0.818143047 |
| chr3:132379245-132379459 | -0.289801497 | 0.594145705 | 0.818143047 |
| chr12:59595759-59597205 | 0.169622391 | 0.594147264 | 0.818143047 |
| chr6:16752454-16752776 | 0.249601958 | 0.594185773 | 0.818181369 |
| chr1:149939020-149939718 | -0.173574302 | 0.594267716 | 0.818279497 |
| chr8:26451227-26451567 | -0.198547 | 0.594279609 | 0.818281169 |
| chr14:61831816-61832113 | 0.266473145 | 0.594292026 | 0.818283561 |

| | | | |
|---|---|---|---|
| chr4:39638231-39639698 | 0.169648007 | 0.594416268 | 0.818439923 |
| chr15:78374533-78375478 | 0.186395449 | 0.59444391 | 0.818451466 |
| chr16:83989939-83990274 | -0.22149125 | 0.594453809 | 0.818451466 |
| chr1:151146372-151147320 | 0.170413047 | 0.594467293 | 0.818451466 |
| chr1:6392622-6394144 | 0.17317131 | 0.594468271 | 0.818451466 |
| chr17:58363849-58364082 | -0.214722923 | 0.59447806 | 0.818451466 |
| chr19:10332948-10333817 | -0.175214144 | 0.594498917 | 0.818465475 |
| chr16:70165016-70165467 | -0.197689382 | 0.594519111 | 0.818478571 |
| chr8:118961941-118962299 | -0.213695907 | 0.594537579 | 0.818489289 |
| chr10:114472284-114472484 | 0.280374479 | 0.594571718 | 0.818515632 |
| chr21:39341434-39341823 | -0.218943064 | 0.594583016 | 0.818515632 |
| chr10:117496527-117496748 | 0.258910974 | 0.59460509 | 0.818515632 |
| chr10:28722061-28722841 | -0.189925676 | 0.594609777 | 0.818515632 |
| chr15:61051285-61051737 | -0.214972886 | 0.594610126 | 0.818515632 |
| chr12:4030941-4031673 | 0.184794204 | 0.594627908 | 0.818516551 |
| chr8:139023129-139023445 | -0.210412942 | 0.594632159 | 0.818516551 |
| chr5:43042454-43042758 | 0.192483761 | 0.594685365 | 0.818568679 |
| chr10:37976129-37976982 | -0.169591102 | 0.59469526 | 0.818568679 |
| chrX:24647190-24647641 | -0.188742128 | 0.594707788 | 0.818568679 |
| chr17:74736219-74736713 | 0.193707069 | 0.59471555 | 0.818568679 |
| chr1:116878080-116878475 | 0.210141979 | 0.594723445 | 0.818568679 |
| chr17:81736155-81736405 | 0.209210329 | 0.594754384 | 0.818596558 |
| chr1:147213595-147213795 | 0.257576898 | 0.594785168 | 0.818599266 |
| chr22:32137668-32138160 | 0.21136332 | 0.594794365 | 0.818599266 |
| chr17:59561063-59561341 | -0.193708776 | 0.594800694 | 0.818599266 |
| chr3:177360445-177360753 | 0.20149135 | 0.594809622 | 0.818599266 |
| chr6:27602339-27602763 | -0.187465506 | 0.594809769 | 0.818599266 |
| chr5:149357497-149358631 | -0.170827259 | 0.594837247 | 0.818622379 |
| chr14:49721773-49722016 | 0.242277176 | 0.594855992 | 0.818633472 |
| chr4:13483588-13485273 | 0.168886936 | 0.594868834 | 0.818636442 |
| chr16:89918624-89918854 | -0.211258936 | 0.594950942 | 0.818734731 |
| chr20:57022314-57022677 | 0.185770451 | 0.594964188 | 0.818738255 |
| chr1:19596164-19596364 | -0.263463427 | 0.595014566 | 0.818787081 |
| chr1:19209612-19210678 | 0.178994555 | 0.595021041 | 0.818787081 |
| chr15:77543385-77543984 | 0.181890474 | 0.595093897 | 0.818872629 |
| chr2:102355549-102356101 | 0.191098151 | 0.595129236 | 0.818906549 |
| chr5:43401904-43402147 | -0.227386868 | 0.595185309 | 0.818957181 |
| chr15:64905392-64905783 | -0.215461267 | 0.595187408 | 0.818957181 |
| chr20:3811452-3812207 | 0.171320319 | 0.595222782 | 0.818985493 |
| chrX:16739595-16739816 | -0.250031696 | 0.595239279 | 0.818985493 |
| chr17:67139860-67140140 | -0.219977932 | 0.59524005 | 0.818985493 |
| chr19:6828219-6829508 | 0.171284322 | 0.595291391 | 0.819041425 |
| chr1:94417880-94418781 | -0.174304757 | 0.59532174 | 0.819068473 |
| chr4:1365508-1366117 | 0.195022291 | 0.595338477 | 0.819076793 |
| chr3:63364565-63364833 | -0.200089062 | 0.595364544 | 0.81909795 |
| chr8:115139239-115139527 | 0.239263203 | 0.595379369 | 0.819103638 |
| chr15:95799860-95800144 | 0.229003662 | 0.595441306 | 0.819152196 |
| chr17:64776826-64777101 | -0.187392974 | 0.595455512 | 0.819152196 |
| chr10:45971538-45972052 | 0.205681315 | 0.595456485 | 0.819152196 |
| chr19:45640824-45641605 | -0.174643274 | 0.595461307 | 0.819152196 |

| | | | |
|---|---|---|---|
| chr2:64831288-64832136 | -0.18095765 | 0.59548173 | 0.819152196 |
| chr4:184472618-184473335 | -0.18375886 | 0.595488158 | 0.819152196 |
| chr17:58387818-58388221 | 0.186710608 | 0.5954895 | 0.819152196 |
| chr3:58842548-58842848 | -0.226033673 | 0.595534023 | 0.819187919 |
| chr3:160398934-160400045 | -0.16792571 | 0.595542406 | 0.819187919 |
| chr16:53781369-53781569 | 0.288594679 | 0.595547543 | 0.819187919 |
| chr2:58614369-58614666 | 0.270328272 | 0.59557688 | 0.819189788 |
| chr6:107980337-107980570 | 0.270328272 | 0.59557688 | 0.819189788 |
| chr1:231421231-231422873 | 0.170099512 | 0.595580976 | 0.819189788 |
| chr19:23273899-23274427 | 0.208422041 | 0.595601137 | 0.819202813 |
| chr17:28925395-28925639 | -0.27849104 | 0.595667359 | 0.819275578 |
| chr9:15135167-15135526 | 0.210537657 | 0.595675425 | 0.819275578 |
| chr3:59809810-59810068 | -0.256292322 | 0.595691818 | 0.819283417 |
| chr10:19815628-19817118 | -0.171909932 | 0.595703456 | 0.819284718 |
| chr3:57803573-57803940 | 0.237330739 | 0.595729361 | 0.819305639 |
| chr1:72845421-72845869 | 0.213191772 | 0.595779397 | 0.819331057 |
| chr6:79869605-79870114 | -0.191623495 | 0.595787433 | 0.819331057 |
| chr1:106831202-106831932 | 0.170114177 | 0.595795347 | 0.819331057 |
| chr3:10738707-10739443 | 0.183777879 | 0.595800519 | 0.819331057 |
| chr1:156096654-156096966 | 0.195960106 | 0.595801308 | 0.819331057 |
| chr5:74240893-74241401 | 0.199395788 | 0.595813171 | 0.819332665 |
| chr6:97437068-97437290 | -0.280608163 | 0.595839287 | 0.819353874 |
| chr3:52445002-52445245 | 0.212821041 | 0.595896308 | 0.819417579 |
| chr10:102072945-102073145 | 0.248509542 | 0.595908656 | 0.819419852 |
| chr12:57614246-57614506 | 0.213114055 | 0.595925268 | 0.81942799 |
| chr17:28591774-28591988 | -0.269228528 | 0.595960284 | 0.819457895 |
| chr15:30625593-30626560 | -0.176070732 | 0.595968406 | 0.819457895 |
| chr17:17840130-17840549 | -0.183773192 | 0.595986629 | 0.819468246 |
| chr16:81625440-81625640 | -0.248589522 | 0.596017086 | 0.819495257 |
| chr8:28493835-28494537 | 0.182012522 | 0.596027664 | 0.819495257 |
| chr20:44747113-44747955 | 0.176611941 | 0.596041418 | 0.819499462 |
| chr9:128339918-128341065 | 0.171512488 | 0.596068647 | 0.819522194 |
| chrX:48021011-48021932 | -0.179412628 | 0.596112771 | 0.819540505 |
| chr1:114979329-114979605 | -0.204876785 | 0.596114633 | 0.819540505 |
| chr4:81153587-81154420 | -0.181583821 | 0.596120203 | 0.819540505 |
| chrX:52990038-52990280 | -0.244996508 | 0.596124749 | 0.819540505 |
| chr19:33521572-33522199 | 0.183435688 | 0.596145943 | 0.819547356 |
| chr5:170623600-170624086 | -0.196967665 | 0.596151124 | 0.819547356 |
| chr4:110119162-110119706 | 0.188728745 | 0.596197896 | 0.819596949 |
| chr12:82627549-82628236 | -0.188927902 | 0.596269924 | 0.819668115 |
| chr19:55453000-55453260 | 0.237866483 | 0.596271058 | 0.819668115 |
| chr1:202693015-202693290 | 0.199281024 | 0.596309274 | 0.819697209 |
| chr6:93721419-93721689 | 0.227213683 | 0.596313619 | 0.819697209 |
| chr14:92937592-92937851 | -0.216859576 | 0.596338664 | 0.819713189 |
| chr17:51640095-51640377 | -0.222756938 | 0.596346641 | 0.819713189 |
| chr22:25202431-25202632 | 0.264306916 | 0.596368429 | 0.819728432 |
| chr9:122796896-122797244 | -0.267585412 | 0.596460374 | 0.819840107 |
| chr1:196997579-196997779 | -0.262418987 | 0.596482436 | 0.819855725 |
| chr3:138961923-138962195 | 0.2514073 | 0.596494436 | 0.819857511 |
| chr1:47179060-47179915 | 0.172868996 | 0.596542839 | 0.819908786 |

| | | | |
|---|---|---|---|
| chr8:80067147-80067406 | 0.242844034 | 0.596622315 | 0.819908786 |
| chr19:52049281-52049481 | 0.280007781 | 0.596623818 | 0.819908786 |
| chr11:74069104-74069474 | 0.196138918 | 0.596646762 | 0.819908786 |
| chr10:100481504-100482077 | 0.183958806 | 0.596650744 | 0.819908786 |
| chr4:76234009-76234992 | -0.175167623 | 0.596656956 | 0.819908786 |
| chr17:60160919-60161198 | 0.176710952 | 0.596658695 | 0.819908786 |
| chr3:24197977-24198197 | -0.261720496 | 0.596659027 | 0.819908786 |
| chr10:13787856-13788074 | -0.264039603 | 0.596661992 | 0.819908786 |
| chr9:87662997-87663248 | 0.244969772 | 0.596664525 | 0.819908786 |
| chr21:39737502-39738114 | 0.17958942 | 0.596672175 | 0.819908786 |
| chr19:49877211-49878279 | 0.166970635 | 0.596673251 | 0.819908786 |
| chr13:98751417-98751621 | 0.275080947 | 0.596681117 | 0.819908786 |
| chr1:156505319-156505604 | 0.192548682 | 0.59668667 | 0.819908786 |
| chr11:18659175-18659462 | -0.218221194 | 0.596692252 | 0.819908786 |
| chr15:41928913-41929526 | 0.185946571 | 0.59670609 | 0.819913098 |
| chr14:29016255-29016592 | 0.209585583 | 0.596727038 | 0.819927177 |
| chr4:140861413-140861640 | 0.24286256 | 0.596747516 | 0.819940611 |
| chr5:9560099-9560299 | 0.253719538 | 0.596792603 | 0.819961605 |
| chr11:66744391-66744865 | 0.179749723 | 0.596792674 | 0.819961605 |
| chr2:197515498-197516803 | -0.16797415 | 0.596814675 | 0.819961605 |
| chr17:8156250-8156905 | -0.171236178 | 0.596828624 | 0.819961605 |
| chr4:153643490-153643765 | -0.213246735 | 0.596832386 | 0.819961605 |
| chr10:110918423-110919706 | -0.166027802 | 0.596840978 | 0.819961605 |
| chr2:112645215-112646223 | 0.168884013 | 0.596848868 | 0.819961605 |
| chr16:57628368-57628713 | -0.205221598 | 0.596859066 | 0.819961605 |
| chr4:128310414-128310645 | 0.271048508 | 0.596859108 | 0.819961605 |
| chr12:46164244-46164485 | 0.26142714 | 0.596892628 | 0.819982595 |
| chr2:150860-151179 | 0.277941865 | 0.596915886 | 0.819982595 |
| chr2:25521582-25521782 | -0.318859449 | 0.596916727 | 0.819982595 |
| chr8:93754621-93755436 | -0.17006076 | 0.596917193 | 0.819982595 |
| chr22:47129919-47130449 | -0.194248267 | 0.596932639 | 0.819984503 |
| chr9:133115150-133115350 | 0.252394522 | 0.596946543 | 0.819984503 |
| chr15:31129028-31129562 | 0.201397432 | 0.596950687 | 0.819984503 |
| chr6:156776505-156776748 | -0.230647686 | 0.596973924 | 0.819986254 |
| chr9:33001091-33002020 | -0.173008596 | 0.596982993 | 0.819986254 |
| chr11:121112570-121112907 | 0.221457621 | 0.596984067 | 0.819986254 |
| chr8:27515943-27516143 | 0.298241311 | 0.597037187 | 0.820044516 |
| chr18:79245610-79246074 | 0.186307901 | 0.597073286 | 0.820079398 |
| chr10:100680824-100681024 | -0.277307433 | 0.597106719 | 0.82010513 |
| chr17:70271735-70271956 | 0.282991705 | 0.597113427 | 0.82010513 |
| chr3:51499199-51500453 | 0.169833528 | 0.597163354 | 0.820159001 |
| chr17:51764379-51764826 | -0.18783727 | 0.597201623 | 0.820183304 |
| chr6:16947610-16947810 | -0.271780784 | 0.597212721 | 0.820183304 |
| chr8:37546835-37547243 | -0.224593656 | 0.597213638 | 0.820183304 |
| chr1:230742603-230742871 | 0.226659269 | 0.597223866 | 0.820183304 |
| chr8:102656350-102656597 | 0.187141213 | 0.597269932 | 0.820223139 |
| chr17:50532340-50532835 | 0.181245349 | 0.597274282 | 0.820223139 |
| chr1:51896938-51897139 | 0.270883965 | 0.597316611 | 0.820234058 |
| chr1:206776319-206776814 | 0.187196524 | 0.597323503 | 0.820234058 |
| chr16:57091570-57091770 | 0.274298304 | 0.597344947 | 0.820234058 |

| | | | |
|---|---|---|---|
| chr4:42656801-42657842 | -0.170147335 | 0.597360299 | 0.820234058 |
| chr8:87021335-87021683 | -0.231888812 | 0.597364048 | 0.820234058 |
| chr11:83071390-83072453 | -0.167054942 | 0.597364287 | 0.820234058 |
| chr2:202588607-202589153 | -0.186905824 | 0.59736533 | 0.820234058 |
| chr21:38228093-38228359 | -0.223841044 | 0.597384882 | 0.820234058 |
| chr2:84314848-84315316 | -0.189839949 | 0.59739735 | 0.820234058 |
| chr1:181469552-181470067 | -0.194663426 | 0.597398494 | 0.820234058 |
| chr8:135465920-135466259 | 0.23626837 | 0.597399987 | 0.820234058 |
| chr14:89954375-89955287 | 0.167757703 | 0.597427444 | 0.820257057 |
| chr8:124148010-124148231 | 0.223248504 | 0.59745031 | 0.820270767 |
| chr14:44896892-44898059 | 0.16571353 | 0.59745884 | 0.820270767 |
| chr5:151757889-151759009 | -0.172369456 | 0.597482525 | 0.820279359 |
| chr8:127589699-127589908 | 0.237865429 | 0.597491178 | 0.820279359 |
| chr20:25237793-25238159 | 0.193726054 | 0.597513244 | 0.820279359 |
| chr1:24216408-24216871 | -0.219363803 | 0.59753126 | 0.820279359 |
| chr3:152268428-152269747 | -0.166792814 | 0.59756112 | 0.820279359 |
| chr6:54874718-54874946 | 0.234605519 | 0.597565264 | 0.820279359 |
| chr10:36900071-36900432 | -0.200235711 | 0.597569831 | 0.820279359 |
| chr1:84753561-84753919 | -0.212653377 | 0.597569953 | 0.820279359 |
| chr5:138354983-138355280 | 0.206220692 | 0.597572082 | 0.820279359 |
| chr8:122785770-122786502 | -0.167663602 | 0.597572153 | 0.820279359 |
| chr3:15378017-15378587 | -0.177301819 | 0.597608132 | 0.820283646 |
| chr20:38646186-38646766 | 0.197128741 | 0.597615845 | 0.820283646 |
| chr20:52061065-52061401 | 0.204044772 | 0.597625408 | 0.820283646 |
| chr18:48818445-48818858 | -0.27275501 | 0.597625847 | 0.820283646 |
| chr10:127563958-127564253 | -0.234488175 | 0.597628805 | 0.820283646 |
| chrX:10189651-10189941 | -0.21876783 | 0.597655589 | 0.820291621 |
| chr1:45750198-45751131 | 0.16974856 | 0.597656026 | 0.820291621 |
| chr5:72653603-72653900 | 0.256817482 | 0.59766704 | 0.820292044 |
| chr17:1717008-1717335 | -0.187003624 | 0.597726575 | 0.820359061 |
| chr22:36481878-36482340 | 0.182835155 | 0.597749942 | 0.820376436 |
| chr8:122133263-122133485 | 0.257120458 | 0.597777735 | 0.820399885 |
| chr6:41673778-41674056 | 0.268047302 | 0.597793067 | 0.820406233 |
| chr11:67986314-67986543 | 0.281495923 | 0.59781452 | 0.82042098 |
| chr8:132898374-132898712 | 0.206032621 | 0.597863326 | 0.820468703 |
| chr6:4018246-4019031 | -0.171570287 | 0.59787071 | 0.820468703 |
| chr10:24557203-24557515 | 0.210302507 | 0.597917262 | 0.820513376 |
| chr3:172717553-172717852 | -0.200298851 | 0.597927168 | 0.820513376 |
| chr17:67390644-67390942 | 0.201483894 | 0.597935389 | 0.820513376 |
| chr15:89767040-89767731 | 0.186900494 | 0.597947071 | 0.820514712 |
| chr16:66603831-66605068 | 0.175495336 | 0.597973695 | 0.820524705 |
| chr9:21381989-21382395 | 0.225095584 | 0.597975771 | 0.820524705 |
| chr8:97643468-97644810 | 0.166628504 | 0.59801186 | 0.82055953 |
| chr20:10078496-10078764 | 0.248045215 | 0.598042212 | 0.820571541 |
| chr10:28432256-28432496 | -0.234881924 | 0.598048174 | 0.820571541 |
| chr2:201882569-201882950 | -0.236553939 | 0.598052741 | 0.820571541 |
| chr18:51531517-51531734 | 0.270123947 | 0.598068437 | 0.820578382 |
| chr18:45802527-45803338 | -0.174141572 | 0.598098995 | 0.820605616 |
| chr8:37851482-37851700 | 0.234309675 | 0.598114267 | 0.820611874 |
| chr16:57824933-57825203 | 0.220701762 | 0.598128237 | 0.820616348 |

| | | | |
|---|---|---|---|
| chr11:33543049-33543249 | 0.259213104 | 0.598189295 | 0.820666081 |
| chr21:8443152-8443432 | 0.180180885 | 0.598203796 | 0.820666081 |
| chr15:40477926-40478126 | -0.257249197 | 0.598205667 | 0.820666081 |
| chr8:117330126-117330547 | -0.201307493 | 0.598209039 | 0.820666081 |
| chr13:106567179-106567439 | -0.213535756 | 0.59821804 | 0.820666081 |
| chr17:78084994-78085194 | 0.265174851 | 0.598239336 | 0.820675128 |
| chr19:45882781-45883073 | -0.207203484 | 0.598254127 | 0.820675128 |
| chr2:189783455-189785039 | -0.165933093 | 0.598264892 | 0.820675128 |
| chr17:81142560-81143113 | 0.186974957 | 0.598269148 | 0.820675128 |
| chr14:80226392-80226641 | -0.279711596 | 0.598278188 | 0.820675128 |
| chr4:147845064-147845264 | -0.276391848 | 0.598327207 | 0.820698718 |
| chr2:64913261-64913684 | -0.194334874 | 0.598327832 | 0.820698718 |
| chr3:195106320-195107037 | -0.189209298 | 0.598334835 | 0.820698718 |
| chr1:54581683-54582007 | -0.207680797 | 0.598355745 | 0.820698718 |
| chr16:148808-149008 | 0.257765325 | 0.598376222 | 0.820698718 |
| chr1:151718110-151718444 | 0.190877669 | 0.598399961 | 0.820698718 |
| chr11:17730602-17731383 | 0.175647995 | 0.598404978 | 0.820698718 |
| chr20:18144501-18144733 | -0.26220846 | 0.598409586 | 0.820698718 |
| chr19:16628124-16628627 | 0.181739078 | 0.598414864 | 0.820698718 |
| chr21:34351790-34352036 | -0.236238102 | 0.598418736 | 0.820698718 |
| chr11:117928896-117929841 | 0.173942494 | 0.598423777 | 0.820698718 |
| chr9:96449638-96450812 | 0.17042511 | 0.598423917 | 0.820698718 |
| chr6:27592167-27592382 | -0.239613951 | 0.598439391 | 0.82070525 |
| chr20:7113759-7114181 | 0.197411437 | 0.598470437 | 0.82072197 |
| chr7:20991570-20991770 | 0.258943421 | 0.598473006 | 0.82072197 |
| chr1:214753076-214753282 | -0.266292755 | 0.598508873 | 0.820742992 |
| chr9:27436271-27436671 | 0.209855189 | 0.598513817 | 0.820742992 |
| chr4:123397944-123399151 | 0.170268155 | 0.598520469 | 0.820742992 |
| chr2:234544968-234545233 | 0.241599167 | 0.598541143 | 0.820756652 |
| chr19:45493158-45493475 | 0.200268035 | 0.598564315 | 0.820773738 |
| chr16:67480797-67481527 | 0.173371586 | 0.598585299 | 0.820787824 |
| chr5:35926308-35926790 | 0.190793699 | 0.598600318 | 0.820793729 |
| chr2:1750854-1751369 | 0.189497519 | 0.598643277 | 0.820797051 |
| chr12:98438729-98438969 | 0.252421943 | 0.598646722 | 0.820797051 |
| chr17:60078004-60079804 | -0.163115027 | 0.598676825 | 0.820797051 |
| chr3:101951415-101951657 | -0.246202188 | 0.598678694 | 0.820797051 |
| chr3:48504142-48504360 | -0.242873921 | 0.598678761 | 0.820797051 |
| chr17:82839719-82840710 | -0.174089264 | 0.598684027 | 0.820797051 |
| chr2:164560063-164560492 | -0.231565426 | 0.598705097 | 0.820797051 |
| chr11:7657357-7657718 | 0.205542856 | 0.598706819 | 0.820797051 |
| chr22:20130287-20130587 | 0.202685652 | 0.59870871 | 0.820797051 |
| chr10:35635543-35635842 | -0.214950422 | 0.598709863 | 0.820797051 |
| chr20:3118001-3118299 | -0.219416338 | 0.598720588 | 0.820797068 |
| chr13:113001428-113001859 | 0.195507881 | 0.598770354 | 0.820841785 |
| chr15:62254450-62254662 | -0.259534553 | 0.598797114 | 0.820841785 |
| chr14:73851551-73852160 | 0.171395657 | 0.598803259 | 0.820841785 |
| chr3:118817543-118818044 | 0.195497999 | 0.598803408 | 0.820841785 |
| chr16:8863838-8864038 | -0.227371707 | 0.598810893 | 0.820841785 |
| chr1:43270435-43271138 | -0.179213266 | 0.598817484 | 0.820841785 |
| chr3:62992720-62993061 | 0.204905246 | 0.598891036 | 0.820905421 |

| | | | |
|---|---|---|---|
| chr5:138077096-138077391 | -0.213523848 | 0.598908681 | 0.820905421 |
| chr6:19873117-19873318 | -0.30292565 | 0.598909876 | 0.820905421 |
| chr6:21937638-21937838 | -0.30292565 | 0.598909876 | 0.820905421 |
| chr3:108053056-108053683 | 0.185796393 | 0.598917476 | 0.820905421 |
| chr8:81641521-81641808 | 0.229490501 | 0.598947048 | 0.820931269 |
| chr1:202075431-202075639 | 0.217842757 | 0.598959645 | 0.820933849 |
| chr17:1710329-1710529 | 0.236927346 | 0.598979617 | 0.82094243 |
| chr16:85353792-85354161 | -0.187940976 | 0.598987334 | 0.82094243 |
| chr1:30418361-30418618 | -0.236851992 | 0.599027056 | 0.820969692 |
| chr3:159988538-159989428 | -0.173756926 | 0.599028654 | 0.820969692 |
| chr3:189355411-189355875 | -0.204032911 | 0.599044617 | 0.820976884 |
| chr2:101001859-101002961 | -0.171449095 | 0.599060417 | 0.820977742 |
| chr12:124556249-124556486 | -0.254415423 | 0.599066672 | 0.820977742 |
| chr5:43312812-43313959 | 0.165244293 | 0.59909049 | 0.820982676 |
| chr5:173719693-173720038 | 0.210303668 | 0.599091701 | 0.820982676 |
| chr6:157036282-157036484 | -0.260585642 | 0.599134298 | 0.821019548 |
| chr16:3028201-3028566 | 0.186089775 | 0.599140039 | 0.821019548 |
| chrX:10189393-10189650 | -0.234245215 | 0.599157513 | 0.821024459 |
| chr8:66195714-66195961 | 0.245092475 | 0.599165053 | 0.821024459 |
| chr20:48102109-48102398 | 0.18092607 | 0.599183087 | 0.821033854 |
| chr16:74107807-74108191 | 0.193128078 | 0.599202193 | 0.821033854 |
| chr17:61862835-61864192 | -0.16294547 | 0.599204055 | 0.821033854 |
| chr6:47631321-47631955 | 0.198912979 | 0.599221678 | 0.821034828 |
| chr3:119883825-119884131 | 0.210839614 | 0.599226197 | 0.821034828 |
| chr22:25154332-25154673 | 0.216841003 | 0.599243842 | 0.821044323 |
| chr7:8952620-8952820 | 0.285812564 | 0.59926003 | 0.821051821 |
| chr6:158559729-158560216 | -0.197943283 | 0.599274602 | 0.821052226 |
| chr1:52918304-52918504 | -0.28145527 | 0.599287895 | 0.821052226 |
| chr7:45154647-45154973 | 0.195712222 | 0.599292473 | 0.821052226 |
| chr5:158980106-158980538 | -0.23921658 | 0.599344739 | 0.821088142 |
| chr2:102472982-102473319 | 0.215824548 | 0.599346647 | 0.821088142 |
| chr17:29344532-29344769 | -0.274019913 | 0.599359857 | 0.821088142 |
| chr2:142877471-142877707 | -0.219452815 | 0.599371221 | 0.821088142 |
| chr9:112890372-112890645 | 0.230472616 | 0.599372269 | 0.821088142 |
| chr3:60770671-60770871 | -0.270159174 | 0.599413829 | 0.821130396 |
| chr9:129459205-129459686 | -0.191848212 | 0.599455136 | 0.821172301 |
| chr14:88013777-88014088 | 0.206222455 | 0.59947593 | 0.821184074 |
| chr12:48788660-48788985 | 0.216176513 | 0.599489358 | 0.821184074 |
| chr2:173965200-173965974 | -0.171020394 | 0.599505055 | 0.821184074 |
| chr11:125091274-125091674 | 0.222321417 | 0.599511097 | 0.821184074 |
| chr2:101696583-101697840 | 0.174759447 | 0.599526279 | 0.821184074 |
| chr17:78168351-78168671 | 0.190548776 | 0.599528035 | 0.821184074 |
| chr16:10580006-10580209 | 0.216551915 | 0.599565529 | 0.821215806 |
| chr6:35804955-35805434 | -0.200354832 | 0.599572637 | 0.821215806 |
| chr5:160119209-160120204 | -0.165075503 | 0.599597119 | 0.821234657 |
| chr8:77940540-77940814 | -0.255078022 | 0.5996736 | 0.821324728 |
| chr7:30226914-30227138 | -0.250155081 | 0.59971284 | 0.821363789 |
| chr11:94152387-94152607 | -0.262422552 | 0.599758559 | 0.821394805 |
| chr4:76086987-76087490 | 0.195811203 | 0.599770569 | 0.821394805 |
| chr9:135500462-135500943 | 0.182353011 | 0.599778314 | 0.821394805 |

| | | | |
|---|---|---|---|
| chr2:207710870-207712379 | -0.164909489 | 0.599778366 | 0.821394805 |
| chr17:50094886-50095460 | 0.173163114 | 0.599790015 | 0.821396077 |
| chr8:144630122-144630508 | 0.227332378 | 0.599814795 | 0.821415332 |
| chr4:87058670-87059073 | -0.208240289 | 0.599826673 | 0.821416917 |
| chr1:71593536-71593736 | -0.262983652 | 0.599863355 | 0.821452469 |
| chr10:48332606-48332859 | -0.236004319 | 0.599933749 | 0.821534184 |
| chrX:149825173-149825748 | -0.206221413 | 0.599952047 | 0.821541117 |
| chr3:50611668-50612204 | -0.18207022 | 0.599964602 | 0.821541117 |
| chr4:112766507-112766707 | -0.267506074 | 0.599970978 | 0.821541117 |
| chr20:56391651-56392969 | 0.163915901 | 0.599995866 | 0.821547328 |
| chr12:95217022-95218387 | -0.165202482 | 0.599998683 | 0.821547328 |
| chr22:16717619-16718381 | -0.187514182 | 0.60000768 | 0.821547328 |
| chr3:42502225-42502719 | 0.185753222 | 0.600052198 | 0.821578181 |
| chr17:81845232-81845432 | 0.236233041 | 0.600053059 | 0.821578181 |
| chr13:31184518-31184718 | -0.283012485 | 0.600062381 | 0.821578181 |
| chr17:74757498-74757783 | 0.195136968 | 0.600094805 | 0.821607894 |
| chr11:85854594-85855480 | -0.168281935 | 0.600140216 | 0.821655385 |
| chr1:204287987-204288677 | -0.169381166 | 0.600158416 | 0.821665622 |
| chr17:8991642-8991842 | -0.272418549 | 0.600172915 | 0.82167079 |
| chr1:173202431-173202658 | 0.228638798 | 0.600204775 | 0.821691064 |
| chr2:215138565-215139081 | -0.199995348 | 0.600209172 | 0.821691064 |
| chr1:112004998-112005243 | 0.20827941 | 0.600232411 | 0.821708197 |
| chr14:38528075-38528470 | -0.216311734 | 0.60024425 | 0.821709723 |
| chr15:58467719-58467974 | -0.2319807 | 0.600263631 | 0.821721066 |
| chr10:85995783-85996170 | -0.203393539 | 0.600291589 | 0.821721066 |
| chr11:62012945-62013145 | 0.265751339 | 0.600292902 | 0.821721066 |
| chr1:1162747-1162958 | 0.229431077 | 0.600300525 | 0.821721066 |
| chr12:121537554-121537799 | 0.225464309 | 0.600316608 | 0.821721066 |
| chr5:50521914-50522215 | -0.260974761 | 0.600322321 | 0.821721066 |
| chr1:203593664-203594203 | -0.181896847 | 0.600335392 | 0.821721066 |
| chr6:87701787-87702844 | -0.166875787 | 0.600338331 | 0.821721066 |
| chr8:9555581-9556223 | 0.174448691 | 0.600357557 | 0.821722589 |
| chr2:50357884-50358084 | 0.304384891 | 0.600360892 | 0.821722589 |
| chr3:37945982-37946408 | -0.215654495 | 0.600376202 | 0.821728866 |
| chr17:72518729-72519586 | -0.175369869 | 0.600397748 | 0.821741123 |
| chr1:114083146-114083638 | 0.174136188 | 0.600406607 | 0.821741123 |
| chr3:123586199-123586491 | -0.193603299 | 0.600420176 | 0.821745016 |
| chr12:115871791-115872006 | 0.216006826 | 0.600467841 | 0.821786891 |
| chr14:23328081-23328366 | 0.211336737 | 0.600472223 | 0.821786891 |
| chr11:79611105-79611405 | -0.240475448 | 0.600512705 | 0.821817743 |
| chr5:65076350-65076681 | -0.200304684 | 0.600528319 | 0.821817743 |
| chr19:30453845-30454072 | 0.295883872 | 0.600531746 | 0.821817743 |
| chr10:93554819-93555127 | -0.20262374 | 0.600543145 | 0.821817743 |
| chr1:167132620-167132855 | 0.306757332 | 0.60055912 | 0.821817743 |
| chr12:108580973-108581173 | 0.306757332 | 0.60055912 | 0.821817743 |
| chr9:84615255-84615534 | -0.258944951 | 0.600587995 | 0.821830316 |
| chr14:101947833-101948708 | 0.166281934 | 0.600589759 | 0.821830316 |
| chr11:64810283-64811183 | 0.172027887 | 0.600612489 | 0.821846742 |
| chrX:116462567-116463251 | 0.178295234 | 0.600647115 | 0.82186759 |
| chr10:5665874-5666954 | 0.172551978 | 0.600649178 | 0.82186759 |

| | | | |
|---|---|---|---|
| chr6:85958945-85959223 | 0.260242106 | 0.600694218 | 0.82191454 |
| chr1:96966590-96966996 | -0.2095004 | 0.600718544 | 0.821924252 |
| chr7:141695817-141696133 | 0.207598346 | 0.60072277 | 0.821924252 |
| chr20:36951154-36952042 | -0.172650866 | 0.600787946 | 0.821949944 |
| chr4:187354877-187355077 | -0.295321521 | 0.600788861 | 0.821949944 |
| chr6:116218183-116218411 | -0.295321521 | 0.600788861 | 0.821949944 |
| chr19:45769106-45769727 | -0.173850484 | 0.600793506 | 0.821949944 |
| chr1:160291817-160292035 | -0.249177596 | 0.600795183 | 0.821949944 |
| chr3:23389108-23389502 | -0.200627547 | 0.600817175 | 0.821965354 |
| chr20:22248121-22248358 | -0.265588246 | 0.600830848 | 0.821969384 |
| chr8:60045964-60046164 | -0.289194644 | 0.600876869 | 0.821990792 |
| chr6:170561657-170561900 | 0.234063054 | 0.600878681 | 0.821990792 |
| chr9:85900566-85900838 | 0.234063054 | 0.600878681 | 0.821990792 |
| chr2:43038515-43038715 | -0.255358994 | 0.600912841 | 0.822001053 |
| chr2:134917879-134919591 | -0.163965113 | 0.600915935 | 0.822001053 |
| chr6:89918321-89918521 | -0.272784221 | 0.600918365 | 0.822001053 |
| chr15:97623994-97624196 | -0.29437578 | 0.600968056 | 0.822050536 |
| chr13:30453867-30454210 | -0.203723572 | 0.600975997 | 0.822050536 |
| chr2:75153906-75154625 | 0.174654181 | 0.601028615 | 0.822107834 |
| chr2:241802079-241802354 | -0.233049064 | 0.601054488 | 0.822115385 |
| chr18:12597772-12597991 | -0.268930535 | 0.601061033 | 0.822115385 |
| chr11:118565644-118566446 | 0.177007786 | 0.601066324 | 0.822115385 |
| chr2:105037401-105038632 | -0.165793319 | 0.601089142 | 0.822131919 |
| chr6:89610064-89610401 | 0.207778973 | 0.601131251 | 0.822168648 |
| chr5:25461910-25462166 | 0.241014408 | 0.601141087 | 0.822168648 |
| chr3:142723776-142725078 | -0.165192271 | 0.601148187 | 0.822168648 |
| chr3:182962370-182962713 | -0.24057306 | 0.601170088 | 0.822183927 |
| chr20:17768326-17768537 | 0.249494875 | 0.601208103 | 0.822200098 |
| chr3:127679717-127679943 | 0.23381501 | 0.60121893 | 0.822200098 |
| chr16:87779037-87779463 | 0.180286676 | 0.601220937 | 0.822200098 |
| chr3:185560170-185560462 | 0.224759479 | 0.601224835 | 0.822200098 |
| chr13:109998668-109999238 | 0.183471275 | 0.601243135 | 0.82221045 |
| chr4:39976418-39978250 | 0.166851436 | 0.601254956 | 0.822211941 |
| chr7:47610875-47611179 | -0.223655369 | 0.601286218 | 0.822227593 |
| chr1:152121684-152121889 | 0.254323058 | 0.601287864 | 0.822227593 |
| chr2:98935107-98935307 | 0.239960696 | 0.601303212 | 0.822228319 |
| chr17:47978328-47978685 | 0.196031995 | 0.601310294 | 0.822228319 |
| chr8:104442410-104442620 | 0.250535727 | 0.601331258 | 0.822228319 |
| chr5:133746925-133747125 | -0.289634592 | 0.601331319 | 0.822228319 |
| chr5:41856676-41856992 | 0.20119653 | 0.60134804 | 0.82223651 |
| chr3:75218659-75218885 | 0.243757517 | 0.601372253 | 0.822254943 |
| chr5:139453985-139454185 | 0.265251992 | 0.601396163 | 0.822272962 |
| chr14:37230654-37230917 | 0.235419789 | 0.601461396 | 0.822337548 |
| chr15:84671971-84672239 | -0.229033931 | 0.601464865 | 0.822337548 |
| chr17:29929350-29930034 | -0.192975971 | 0.601490497 | 0.822357919 |
| chr11:71972555-71972871 | -0.242206935 | 0.601509196 | 0.822368811 |
| chr4:51868051-51868389 | 0.234648883 | 0.601533208 | 0.822372395 |
| chr6:137309210-137309662 | -0.207103461 | 0.601533283 | 0.822372395 |
| chr2:120737924-120738143 | 0.265974983 | 0.601570893 | 0.822389045 |
| chr10:83078046-83078266 | -0.293870098 | 0.60157479 | 0.822389045 |

| | | | |
|---|---|---|---|
| chr14:24605472-24605739 | 0.215510047 | 0.601577662 | 0.822389045 |
| chr19:29212995-29213470 | -0.178403877 | 0.601602308 | 0.822395768 |
| chr10:122879066-122880274 | 0.167415619 | 0.601604046 | 0.822395768 |
| chr7:47444796-47444997 | -0.260548901 | 0.60163798 | 0.822427484 |
| chr2:133981491-133981843 | 0.262170718 | 0.601678705 | 0.822455412 |
| chr20:58293479-58293762 | 0.197345893 | 0.601679879 | 0.822455412 |
| chr3:4524390-4525002 | 0.175564249 | 0.601705082 | 0.82247519 |
| chr12:64083175-64083523 | 0.206351717 | 0.601735898 | 0.822502639 |
| chr2:38508353-38508691 | 0.198519883 | 0.601784612 | 0.822554553 |
| chr11:62545545-62546257 | -0.173275044 | 0.601805732 | 0.822568746 |
| chr4:17447773-17447973 | 0.27923959 | 0.601822347 | 0.822576783 |
| chr5:175478015-175478705 | -0.178359685 | 0.601865234 | 0.822612414 |
| chr8:86430131-86430410 | -0.238511307 | 0.601869888 | 0.822612414 |
| chr3:151739698-151739915 | 0.262510415 | 0.601916195 | 0.822661031 |
| chr6:31862506-31863858 | 0.166489704 | 0.601975818 | 0.822727844 |
| chr20:9393338-9393626 | -0.21352429 | 0.601990311 | 0.822732977 |
| chr11:78921682-78921955 | -0.238050285 | 0.602003547 | 0.822733238 |
| chr21:6369259-6369511 | 0.173398444 | 0.602045141 | 0.822733238 |
| chr9:115593604-115593845 | 0.234922385 | 0.602047237 | 0.822733238 |
| chr17:20095364-20095594 | -0.244439361 | 0.602048701 | 0.822733238 |
| chr1:228646644-228648482 | 0.168197485 | 0.602053745 | 0.822733238 |
| chr16:88737793-88738062 | 0.21076967 | 0.602054928 | 0.822733238 |
| chr18:28961754-28962043 | -0.247054399 | 0.602107497 | 0.822790403 |
| chr5:10563559-10563887 | 0.201717603 | 0.602174777 | 0.822825502 |
| chr2:171940994-171941257 | -0.239940037 | 0.602186426 | 0.822825502 |
| chr7:66493230-66494211 | -0.169604511 | 0.602188678 | 0.822825502 |
| chr13:36115116-36115650 | 0.189289399 | 0.602194189 | 0.822825502 |
| chr12:22101060-22101312 | -0.245963499 | 0.602196103 | 0.822825502 |
| chr17:70493419-70493619 | -0.28871008 | 0.602197616 | 0.822825502 |
| chr3:13229459-13229932 | 0.178953619 | 0.602236627 | 0.822864133 |
| chr5:146533325-146533625 | -0.20710299 | 0.602248 | 0.822864999 |
| chr10:101249542-101249812 | 0.257245355 | 0.602304361 | 0.822909749 |
| chr2:182586487-182586869 | 0.21761594 | 0.602308336 | 0.822909749 |
| chr8:101504624-101504964 | 0.190392515 | 0.602312972 | 0.822909749 |
| chr2:12168378-12168734 | -0.197490975 | 0.602332 | 0.822916652 |
| chr12:9049658-9049898 | -0.267980003 | 0.602355599 | 0.822916652 |
| chr14:93113405-93113605 | 0.221234267 | 0.602365897 | 0.822916652 |
| chr10:47894668-47894868 | 0.280795895 | 0.602370956 | 0.822916652 |
| chr1:54018057-54018287 | 0.239199544 | 0.602371724 | 0.822916652 |
| chr14:64703221-64703546 | 0.197337259 | 0.602397577 | 0.822937298 |
| chr18:21956160-21956361 | 0.257522939 | 0.602420909 | 0.822954499 |
| chr7:41065787-41065987 | -0.271643071 | 0.602449713 | 0.822979175 |
| chr5:90135839-90136317 | -0.185061621 | 0.60254446 | 0.823084498 |
| chr10:76547479-76547878 | -0.227687272 | 0.602548298 | 0.823084498 |
| chr19:32787428-32788008 | -0.197499325 | 0.602590332 | 0.823127243 |
| chr20:53291618-53291867 | -0.241436954 | 0.60265282 | 0.823197925 |
| chr8:99705838-99706103 | -0.224741648 | 0.602670327 | 0.823207163 |
| chr5:150217961-150218488 | 0.187637411 | 0.602685296 | 0.82320793 |
| chr12:93570020-93571364 | 0.168117031 | 0.602700367 | 0.82320793 |
| chr20:53412979-53413513 | -0.183611595 | 0.60270312 | 0.82320793 |

| | | | |
|---|---|---|---|
| chr13:24389662-24389926 | -0.246254027 | 0.602772203 | 0.823284305 |
| chr18:46293894-46294220 | 0.218520535 | 0.602786568 | 0.823284305 |
| chr17:69213451-69213808 | -0.200667564 | 0.602791271 | 0.823284305 |
| chr17:66906614-66906974 | 0.188542584 | 0.602883884 | 0.823396117 |
| chr10:88744044-88744335 | 0.218246349 | 0.602921438 | 0.823423231 |
| chr8:89856564-89857160 | -0.173123201 | 0.60292523 | 0.823423231 |
| chr12:121295822-121296488 | 0.17783725 | 0.602956756 | 0.823433344 |
| chr13:27388329-27388553 | -0.240824979 | 0.602961887 | 0.823433344 |
| chr11:13439931-13440477 | -0.187533327 | 0.602964874 | 0.823433344 |
| chr5:149471516-149471833 | 0.224554893 | 0.60298184 | 0.823435868 |
| chr17:60640519-60640836 | -0.170962905 | 0.602998417 | 0.823435868 |
| chr20:50115855-50116196 | 0.177220459 | 0.603004879 | 0.823435868 |
| chr11:134346003-134346222 | 0.278299736 | 0.603013118 | 0.823435868 |
| chr3:62201458-62201766 | 0.183779426 | 0.603020457 | 0.823435868 |
| chr12:39765057-39765259 | 0.266512077 | 0.603079901 | 0.823502364 |
| chr7:149625460-149625890 | 0.185343813 | 0.603097224 | 0.823511344 |
| chr2:203238676-203240135 | -0.164117351 | 0.603112001 | 0.823516845 |
| chr17:20944333-20944636 | -0.210141578 | 0.603175166 | 0.823588417 |
| chr1:236727098-236727492 | 0.237703091 | 0.60319788 | 0.823596296 |
| chr1:36385492-36386331 | 0.166081652 | 0.603202434 | 0.823596296 |
| chr21:40143836-40144090 | 0.217080156 | 0.603239109 | 0.823631693 |
| chr9:88970036-88970259 | -0.250088637 | 0.603257107 | 0.823641591 |
| chr14:25090990-25091374 | 0.195893993 | 0.603278666 | 0.823656349 |
| chr7:4641378-4642507 | -0.164869644 | 0.603294666 | 0.823663517 |
| chr13:41256979-41257179 | 0.285126569 | 0.60331277 | 0.823673558 |
| chr12:123458056-123459052 | 0.173283366 | 0.603327982 | 0.823679649 |
| chr3:184803244-184803553 | -0.213951744 | 0.603343562 | 0.82368436 |
| chr14:96194336-96194549 | 0.250350274 | 0.603356993 | 0.82368436 |
| chr18:34492712-34492986 | 0.204331483 | 0.603363682 | 0.82368436 |
| chr7:37031875-37032345 | 0.179690835 | 0.60338044 | 0.823690409 |
| chr20:38408605-38408927 | -0.209747721 | 0.603389614 | 0.823690409 |
| chr13:26051144-26051684 | -0.195127893 | 0.603401216 | 0.823691573 |
| chr3:85821210-85821532 | 0.205490624 | 0.603421106 | 0.823695092 |
| chr8:73893024-73893224 | 0.252795447 | 0.603425294 | 0.823695092 |
| chr10:6108999-6109209 | 0.259724186 | 0.603513638 | 0.823786378 |
| chr12:38316419-38317139 | -0.176091836 | 0.603513672 | 0.823786378 |
| chr7:43705613-43705813 | -0.2807689 | 0.603526837 | 0.823789673 |
| chr15:90336722-90336922 | 0.258218434 | 0.603562972 | 0.82382432 |
| chr15:73899418-73899624 | -0.239777661 | 0.603593287 | 0.823851023 |
| chr2:236569645-236569974 | -0.209291672 | 0.603626547 | 0.823860249 |
| chrX:102768490-102769610 | -0.16716471 | 0.603627866 | 0.823860249 |
| chr11:77411849-77412234 | 0.195127357 | 0.603642124 | 0.823860249 |
| chr20:35604431-35604671 | 0.24332344 | 0.603643056 | 0.823860249 |
| chr2:128232902-128233898 | -0.173498789 | 0.603657945 | 0.823865895 |
| chr12:55738876-55739092 | 0.208863943 | 0.603721611 | 0.823938109 |
| chr1:199057448-199057713 | -0.253177884 | 0.603796248 | 0.824025294 |
| chr3:162569223-162569694 | 0.198259055 | 0.603836207 | 0.82406515 |
| chr20:31722745-31723078 | 0.18024713 | 0.603863427 | 0.824087619 |
| chr13:46146916-46147194 | -0.20672953 | 0.60390472 | 0.824108929 |
| chr5:36265491-36265748 | 0.253676921 | 0.603908378 | 0.824108929 |

| | | | |
|---|---|---|---|
| chr22:44697438-44697709 | -0.232771442 | 0.603911308 | 0.824108929 |
| chrX:125204181-125204624 | 0.181991498 | 0.60392387 | 0.824111394 |
| chrX:112840467-112841907 | 0.164621761 | 0.604001332 | 0.824187026 |
| chr2:187853540-187853788 | -0.258955507 | 0.604003598 | 0.824187026 |
| chr8:132192004-132192311 | -0.22501243 | 0.604011564 | 0.824187026 |
| chr6:77620334-77620617 | 0.204807055 | 0.604074359 | 0.824258033 |
| chr3:73508385-73508682 | -0.214900746 | 0.60410777 | 0.824261016 |
| chr1:33227963-33228182 | -0.279588342 | 0.604108201 | 0.824261016 |
| chr15:71975252-71975926 | -0.177986658 | 0.604116148 | 0.824261016 |
| chr18:41324312-41324710 | -0.219040309 | 0.604119576 | 0.824261016 |
| chr17:31901447-31902244 | -0.172613963 | 0.604139641 | 0.824273716 |
| chr10:133127836-133128036 | 0.233574362 | 0.604162008 | 0.824277631 |
| chr4:67988398-67988753 | 0.199382127 | 0.604164026 | 0.824277631 |
| chr2:28955586-28955800 | -0.243833221 | 0.604239942 | 0.824366527 |
| chr6:51844195-51844585 | 0.219640462 | 0.604263689 | 0.824384246 |
| chr8:135762301-135762509 | -0.261922808 | 0.604292866 | 0.824409373 |
| chr16:73196692-73197114 | -0.193833097 | 0.604380213 | 0.824513855 |
| chr3:194381405-194381907 | 0.19367899 | 0.604407308 | 0.824529184 |
| chr1:204692425-204692646 | 0.212886407 | 0.604424528 | 0.824529184 |
| chr6:37103311-37103534 | 0.228733997 | 0.60444731 | 0.824529184 |
| chr1:241605232-241605486 | 0.216559998 | 0.604451813 | 0.824529184 |
| chr1:153533172-153533500 | 0.20513838 | 0.604457408 | 0.824529184 |
| chr12:56645854-56646466 | 0.169566508 | 0.604459993 | 0.824529184 |
| chr5:78537902-78538113 | 0.237699696 | 0.604466776 | 0.824529184 |
| chr2:46657069-46657285 | 0.242725205 | 0.604506805 | 0.824555513 |
| chr5:147892746-147893049 | -0.198351633 | 0.604507601 | 0.824555513 |
| chr1:55694180-55694685 | 0.186329077 | 0.604526182 | 0.824566179 |
| chr6:73986087-73986287 | 0.272946704 | 0.604556621 | 0.824593019 |
| chr15:67264630-67264830 | 0.29266715 | 0.604593871 | 0.824629147 |
| chr11:115503921-115505130 | -0.166972861 | 0.604632719 | 0.824667453 |
| chr19:28308569-28308769 | 0.277824538 | 0.604695359 | 0.824708164 |
| chr10:33126212-33126452 | -0.26077343 | 0.604703705 | 0.824708164 |
| chr1:11979855-11980850 | 0.168809136 | 0.604717269 | 0.824708164 |
| chr17:81328881-81330489 | -0.163577365 | 0.604727983 | 0.824708164 |
| chr1:202657319-202657597 | -0.1976045 | 0.604728726 | 0.824708164 |
| chr6:149847436-149847872 | 0.191380919 | 0.604729855 | 0.824708164 |
| chr9:124861381-124862585 | 0.166383167 | 0.604737911 | 0.824708164 |
| chr5:16690317-16690606 | -0.197307197 | 0.604754527 | 0.824716146 |
| chr10:60450906-60451207 | -0.259395207 | 0.604798082 | 0.82475193 |
| chr19:18521932-18522914 | 0.166715511 | 0.604802295 | 0.82475193 |
| chr5:27124699-27124899 | -0.275615903 | 0.604813713 | 0.824752822 |
| chr2:70203574-70203774 | -0.262043226 | 0.604850909 | 0.824757012 |
| chr1:27395507-27395707 | -0.229860366 | 0.604855769 | 0.824757012 |
| chr20:47801771-47802085 | 0.174567167 | 0.60486853 | 0.824757012 |
| chr8:66099912-66100133 | -0.258183431 | 0.60487864 | 0.824757012 |
| chr20:50071294-50071615 | 0.183624103 | 0.604891978 | 0.824757012 |
| chr1:185358335-185358648 | -0.235235585 | 0.604897636 | 0.824757012 |
| chr9:103397870-103398289 | -0.190388264 | 0.604923092 | 0.824757012 |
| chr1:15741433-15741720 | 0.212076785 | 0.604923915 | 0.824757012 |
| chr2:60880749-60882150 | -0.164109833 | 0.604930813 | 0.824757012 |

| | | | |
|---|---|---|---|
| chr3:157879474-157879947 | -0.206073425 | 0.604936042 | 0.824757012 |
| chr2:119476225-119476696 | 0.182161028 | 0.604939395 | 0.824757012 |
| chr3:52536328-52536732 | 0.178516863 | 0.604945953 | 0.824757012 |
| chr6:107565269-107565591 | 0.223948349 | 0.604977633 | 0.82477326 |
| chr7:18846472-18847237 | 0.181186397 | 0.604989854 | 0.82477326 |
| chr6:148177634-148177963 | 0.211734313 | 0.604990163 | 0.82477326 |
| chr5:171273281-171273557 | 0.221976626 | 0.60502515 | 0.824806282 |
| chr17:5281859-5282402 | -0.180795701 | 0.605097334 | 0.82489001 |
| chr21:42228054-42228413 | 0.189420484 | 0.605161422 | 0.824943631 |
| chr6:143267764-143267979 | -0.282714751 | 0.6051638 | 0.824943631 |
| chr6:83512258-83512722 | -0.184678825 | 0.605168966 | 0.824943631 |
| chr21:39313262-39314041 | 0.165878539 | 0.605189419 | 0.824949486 |
| chr3:126309494-126309737 | -0.244900541 | 0.605194795 | 0.824949486 |
| chr6:102525332-102525620 | 0.256956824 | 0.60520901 | 0.824954187 |
| chr12:71731658-71731858 | 0.24712178 | 0.60523116 | 0.824964769 |
| chr1:203784481-203784681 | -0.27537275 | 0.605238307 | 0.824964769 |
| chr8:91033803-91034286 | 0.19141207 | 0.605258661 | 0.824977838 |
| chr2:26274242-26274587 | -0.252841218 | 0.605278429 | 0.824990105 |
| chr10:133393903-133394696 | 0.169445834 | 0.605301965 | 0.825004254 |
| chr1:35708748-35708956 | -0.275073502 | 0.605329099 | 0.825004254 |
| chr1:215220322-215220522 | -0.259838639 | 0.605335289 | 0.825004254 |
| chr3:142180855-142181060 | 0.25995095 | 0.605335664 | 0.825004254 |
| chr4:131644671-131644988 | -0.209633827 | 0.605351171 | 0.825004254 |
| chr9:127943706-127944074 | -0.191310007 | 0.605361646 | 0.825004254 |
| chr12:71938630-71939246 | -0.180732059 | 0.60536418 | 0.825004254 |
| chr22:30823064-30823309 | -0.240775521 | 0.605416556 | 0.825046466 |
| chr1:17494202-17494497 | 0.222338167 | 0.605416689 | 0.825046466 |
| chr17:76452400-76453561 | 0.169273337 | 0.60547312 | 0.825108694 |
| chr11:125410681-125410940 | 0.261350837 | 0.60550556 | 0.825130579 |
| chr14:68824501-68825162 | 0.17762131 | 0.605510717 | 0.825130579 |
| chr11:110435334-110435546 | -0.270763455 | 0.605529202 | 0.825138026 |
| chr16:74623256-74623570 | 0.199150037 | 0.605537721 | 0.825138026 |
| chr1:186304985-186305271 | 0.203635081 | 0.60554866 | 0.825138259 |
| chr4:109406382-109406746 | 0.209404944 | 0.605564494 | 0.82514516 |
| chr2:130229064-130229540 | 0.185074526 | 0.605588844 | 0.825162945 |
| chr5:38891690-38891948 | -0.255803734 | 0.605599084 | 0.825162945 |
| chr7:122304381-122305057 | -0.173747906 | 0.605632383 | 0.825183816 |
| chr17:60139880-60140154 | 0.170744659 | 0.605641423 | 0.825183816 |
| chr3:188294781-188295078 | -0.240365652 | 0.605660211 | 0.825183816 |
| chr1:151156504-151156743 | 0.196174781 | 0.605700804 | 0.825183816 |
| chr4:113673505-113673841 | -0.209604921 | 0.605702615 | 0.825183816 |
| chr10:62817022-62817431 | -0.186437551 | 0.605725402 | 0.825183816 |
| chr8:104738797-104739005 | 0.263056632 | 0.605734345 | 0.825183816 |
| chr5:5422032-5423022 | 0.163163772 | 0.605744749 | 0.825183816 |
| chr9:130444615-130445067 | -0.180707911 | 0.605748654 | 0.825183816 |
| chrX:83416354-83416583 | -0.26698928 | 0.605752679 | 0.825183816 |
| chr1:35345352-35345606 | -0.232244362 | 0.605755563 | 0.825183816 |
| chr6:152750759-152750989 | -0.262021074 | 0.605755635 | 0.825183816 |
| chr3:168047219-168047597 | -0.189117257 | 0.605761858 | 0.825183816 |
| chr1:56390368-56390568 | -0.280911938 | 0.605773373 | 0.825183816 |

| | | | |
|---|---|---|---|
| chr14:88824172-88825062 | -0.165862646 | 0.605779121 | 0.825183816 |
| chr7:35231352-35231744 | -0.203278192 | 0.605790815 | 0.825183816 |
| chr14:100192735-100193590 | 0.166226635 | 0.605801731 | 0.825183816 |
| chr20:58066016-58066271 | 0.219607854 | 0.605808648 | 0.825183816 |
| chr1:203268963-203269400 | -0.178271299 | 0.605827726 | 0.825183816 |
| chr6:31462793-31463269 | -0.181169266 | 0.605830851 | 0.825183816 |
| chr19:6424791-6425634 | 0.167346702 | 0.605847495 | 0.825183816 |
| chr15:40765388-40765825 | -0.18784729 | 0.605851332 | 0.825183816 |
| chr11:66408455-66408735 | 0.205304383 | 0.60587013 | 0.825187875 |
| chr10:120851018-120851739 | 0.171294974 | 0.605881091 | 0.825187875 |
| chr17:64418585-64419053 | -0.189910209 | 0.605886621 | 0.825187875 |
| chr19:21020347-21020819 | -0.192219111 | 0.605917853 | 0.825214689 |
| chr19:40090338-40091133 | 0.176775022 | 0.605927849 | 0.825214689 |
| chr17:81295963-81296348 | 0.186606136 | 0.605977326 | 0.825253659 |
| chr16:54422661-54423034 | 0.21423387 | 0.605981448 | 0.825253659 |
| chr1:32449193-32449430 | -0.2456818 | 0.605999528 | 0.825253659 |
| chr6:28816965-28817499 | -0.177237096 | 0.605999545 | 0.825253659 |
| chr4:72169225-72169986 | -0.183677701 | 0.606039098 | 0.825292855 |
| chr2:105736063-105736445 | 0.209911509 | 0.606072234 | 0.825301995 |
| chr16:66521537-66521856 | -0.265207534 | 0.606076109 | 0.825301995 |
| chr2:237085490-237086495 | 0.176741839 | 0.606078123 | 0.825301995 |
| chr19:47414469-47414771 | 0.206909787 | 0.606128424 | 0.825346656 |
| chr10:92843135-92843373 | -0.215944062 | 0.606138457 | 0.825346656 |
| chr1:185954793-185955049 | 0.270410867 | 0.606150134 | 0.825346656 |
| chr15:81155621-81155831 | 0.245999612 | 0.606154008 | 0.825346656 |
| chr10:75401037-75402058 | -0.169520614 | 0.606277148 | 0.825489219 |
| chr17:59406797-59407465 | 0.171377484 | 0.606282893 | 0.825489219 |
| chr2:219176262-219177360 | -0.168100765 | 0.60629103 | 0.825489219 |
| chr1:196807345-196807550 | -0.303544014 | 0.606321756 | 0.825502077 |
| chr8:141726921-141727203 | 0.21631894 | 0.606323254 | 0.825502077 |
| chr15:101269957-101270157 | -0.235186671 | 0.606340663 | 0.825502077 |
| chr4:72496263-72496513 | -0.251534838 | 0.606346015 | 0.825502077 |
| chr15:98493916-98494294 | 0.200209112 | 0.606369578 | 0.825502077 |
| chr3:160039075-160039308 | 0.244692949 | 0.606372825 | 0.825502077 |
| chr3:82410336-82410777 | -0.198212117 | 0.606393792 | 0.825502077 |
| chr5:111427838-111428082 | -0.258458996 | 0.606394443 | 0.825502077 |
| chr7:66628149-66629369 | -0.164706709 | 0.606397437 | 0.825502077 |
| chr21:14114307-14114628 | -0.217702284 | 0.606423948 | 0.825507773 |
| chr15:64740439-64740686 | -0.220295018 | 0.606430244 | 0.825507773 |
| chr1:160990332-160990749 | 0.207010554 | 0.606433942 | 0.825507773 |
| chr5:134512566-134512801 | 0.222828423 | 0.606446559 | 0.825510281 |
| chr7:40977623-40977950 | 0.234490223 | 0.606471562 | 0.825518159 |
| chr8:98650315-98650985 | 0.184291227 | 0.606473894 | 0.825518159 |
| chr9:15510614-15511597 | -0.171093968 | 0.606504026 | 0.825544508 |
| chr12:102563067-102563405 | -0.21974066 | 0.606527831 | 0.825551503 |
| chr17:47099790-47100699 | -0.170784266 | 0.606530713 | 0.825551503 |
| chr5:176732863-176733142 | -0.235388908 | 0.606556571 | 0.825572033 |
| chr2:115164601-115164827 | 0.285415829 | 0.606585663 | 0.825588121 |
| chr4:99949051-99949348 | 0.181905236 | 0.606589941 | 0.825588121 |
| chr22:28741637-28742751 | 0.163185781 | 0.606653791 | 0.825660357 |

| | | | |
|---|---|---|---|
| chr5:57137900-57138449 | -0.180971945 | 0.606670274 | 0.825668124 |
| chr12:74897508-74897772 | -0.272557388 | 0.606714395 | 0.825689416 |
| chr1:109368569-109368783 | -0.258025637 | 0.606716737 | 0.825689416 |
| chr7:98400489-98401598 | -0.166507407 | 0.606718247 | 0.825689416 |
| chr17:17356132-17356363 | -0.210732434 | 0.606733744 | 0.825693915 |
| chrX:110167894-110168424 | -0.191068112 | 0.606748016 | 0.825693915 |
| chr2:175366833-175367171 | -0.210851664 | 0.606780473 | 0.825693915 |
| chr14:82251596-82251797 | -0.266223717 | 0.60678425 | 0.825693915 |
| chr22:20916808-20917603 | 0.168044442 | 0.606786922 | 0.825693915 |
| chr15:45634457-45635858 | 0.164460267 | 0.606792426 | 0.825693915 |
| chr5:39073356-39075267 | -0.162347033 | 0.606799709 | 0.825693915 |
| chr18:24397116-24398279 | 0.164587603 | 0.606807762 | 0.825693915 |
| chr11:22245012-22245351 | -0.202526534 | 0.606911267 | 0.82582009 |
| chr8:82862731-82863341 | -0.180769945 | 0.606940878 | 0.825822323 |
| chr4:55948548-55949579 | 0.165645756 | 0.606946891 | 0.825822323 |
| chr3:134313287-134313873 | 0.179341404 | 0.606960633 | 0.825822323 |
| chr12:66279200-66279659 | -0.203378587 | 0.606966812 | 0.825822323 |
| chr2:234288975-234289213 | 0.27122676 | 0.606983443 | 0.825822323 |
| chr7:64794151-64794827 | -0.171125269 | 0.606999206 | 0.825822323 |
| chr20:23446695-23446895 | 0.244252908 | 0.607007254 | 0.825822323 |
| chr17:4815335-4815535 | 0.258715188 | 0.607009161 | 0.825822323 |
| chr9:100472673-100472953 | 0.223780193 | 0.607009909 | 0.825822323 |
| chr20:59223145-59223417 | 0.202521374 | 0.607049225 | 0.82585298 |
| chr2:17878293-17880085 | -0.162563953 | 0.607053999 | 0.82585298 |
| chr1:206544630-206545146 | 0.178334478 | 0.607066206 | 0.825854923 |
| chr8:95202974-95203208 | 0.209350445 | 0.60708029 | 0.82585942 |
| chr7:38177750-38178879 | -0.162495076 | 0.6071388 | 0.825924353 |
| chr1:223202470-223202670 | -0.277380896 | 0.607184104 | 0.825968469 |
| chr6:99993309-99993509 | 0.316901374 | 0.607207624 | 0.825968469 |
| chr19:55385278-55386028 | 0.168653796 | 0.607209667 | 0.825968469 |
| chr5:56669084-56669464 | -0.212679528 | 0.60721435 | 0.825968469 |
| chr18:10560497-10560847 | -0.201918304 | 0.607272407 | 0.826017928 |
| chr3:128323594-128323941 | -0.203859726 | 0.607273191 | 0.826017928 |
| chr1:198939269-198939640 | 0.1737074 | 0.607287598 | 0.826017928 |
| chr21:42145090-42145307 | -0.244870509 | 0.607293831 | 0.826017928 |
| chr15:43329752-43330956 | 0.167454711 | 0.607319431 | 0.826032685 |
| chr1:38012344-38013279 | 0.166943012 | 0.607326242 | 0.826032685 |
| chr2:177263397-177265539 | -0.161176633 | 0.607360756 | 0.826064965 |
| chr18:26813052-26813415 | -0.212367127 | 0.607376596 | 0.826071589 |
| chr7:106603334-106603683 | -0.219041166 | 0.607388388 | 0.826071589 |
| chr17:50681371-50681616 | 0.202622136 | 0.607399717 | 0.826071589 |
| chrX:138763938-138764138 | -0.259497429 | 0.607408751 | 0.826071589 |
| chr17:59837053-59838084 | -0.159828843 | 0.607450817 | 0.826106908 |
| chrX:86969981-86970306 | 0.218130304 | 0.607483364 | 0.826106908 |
| chrX:151176219-151176600 | 0.213802713 | 0.607485769 | 0.826106908 |
| chr1:54052651-54054134 | -0.16208109 | 0.607497881 | 0.826106908 |
| chr10:124878387-124878673 | 0.213571784 | 0.607512643 | 0.826106908 |
| chr1:197786336-197786536 | 0.248595775 | 0.607523901 | 0.826106908 |
| chr4:83252832-83253087 | -0.221544859 | 0.607563412 | 0.826106908 |
| chr8:63964251-63964451 | -0.275061657 | 0.607563466 | 0.826106908 |

| | | | |
|---|---|---|---|
| chr8:106908755-106908955 | -0.275061657 | 0.607563466 | 0.826106908 |
| chr1:40949000-40949266 | -0.216902516 | 0.607574544 | 0.826106908 |
| chr4:121462740-121462953 | 0.243699275 | 0.607581103 | 0.826106908 |
| chr19:12696447-12696760 | 0.220296102 | 0.607585428 | 0.826106908 |
| chr17:19992807-19993328 | -0.180967096 | 0.607592649 | 0.826106908 |
| chr3:99875913-99876473 | 0.180152601 | 0.607596116 | 0.826106908 |
| chr20:53529284-53529500 | 0.229086214 | 0.607596445 | 0.826106908 |
| chr3:8954983-8955362 | 0.199701806 | 0.607615153 | 0.826117685 |
| chr16:58629511-58630709 | 0.163362238 | 0.607641756 | 0.826139195 |
| chr6:72271640-72271938 | 0.222300217 | 0.607658882 | 0.82614782 |
| chr10:124010942-124011200 | 0.231298158 | 0.607698232 | 0.826176649 |
| chr8:51526098-51526399 | -0.206682512 | 0.607708432 | 0.826176649 |
| chr10:107409599-107409847 | -0.236207535 | 0.607715216 | 0.826176649 |
| chr5:181191377-181192160 | 0.163587161 | 0.607724335 | 0.826176649 |
| chr14:102568105-102568678 | -0.188249186 | 0.607733999 | 0.826176649 |
| chr10:95865190-95865517 | -0.206269808 | 0.607804131 | 0.82624497 |
| chr11:59560687-59561153 | -0.175748234 | 0.607805822 | 0.82624497 |
| chr1:84689838-84691199 | -0.162404214 | 0.607829995 | 0.826263171 |
| chr15:31605399-31605940 | 0.185508092 | 0.607849832 | 0.826267048 |
| chr3:95329377-95329624 | 0.249618442 | 0.60786426 | 0.826267048 |
| chr16:15576297-15576764 | 0.201781586 | 0.607865198 | 0.826267048 |
| chr3:183115712-183116354 | 0.177875342 | 0.607904624 | 0.82630598 |
| chr7:42024615-42024815 | 0.268699867 | 0.607948507 | 0.82635097 |
| chr5:103364240-103364567 | -0.234206087 | 0.607968919 | 0.826364055 |
| chr19:54114884-54115950 | 0.167922105 | 0.607996135 | 0.826386388 |
| chr8:104588075-104590204 | 0.160031008 | 0.608035847 | 0.826408011 |
| chr6:45454395-45454595 | 0.30121158 | 0.608049069 | 0.826408011 |
| chr17:78421408-78421995 | 0.172444051 | 0.608054754 | 0.826408011 |
| chr11:73597814-73598865 | 0.163428848 | 0.608055186 | 0.826408011 |
| chr4:47122735-47123045 | -0.22632162 | 0.608092124 | 0.826417435 |
| chr6:110958125-110959138 | 0.168872968 | 0.608092787 | 0.826417435 |
| chr17:20320750-20321182 | -0.21442412 | 0.608094477 | 0.826417435 |
| chrX:149871960-149872353 | 0.189581708 | 0.60814395 | 0.826470012 |
| chr1:215175886-215176086 | -0.27057077 | 0.608160826 | 0.826478288 |
| chr8:20376366-20376593 | -0.264539891 | 0.608202773 | 0.826520633 |
| chr8:143362952-143363204 | 0.232140816 | 0.608247214 | 0.82656379 |
| chr21:46286109-46286805 | 0.169791384 | 0.608263667 | 0.82656379 |
| chr20:47768460-47768873 | 0.170552907 | 0.608269866 | 0.82656379 |
| chr4:74283728-74283928 | -0.284295073 | 0.608293789 | 0.82656379 |
| chr3:142595654-142596951 | -0.162998813 | 0.60829833 | 0.82656379 |
| chr3:139718472-139718691 | 0.235274437 | 0.608328188 | 0.82656379 |
| chr6:34425851-34426366 | 0.175905127 | 0.608329221 | 0.82656379 |
| chr6:166675756-166676540 | 0.180630394 | 0.608330626 | 0.82656379 |
| chr17:68133557-68133781 | 0.216096937 | 0.608350252 | 0.82656379 |
| chr17:42050858-42051168 | 0.221767688 | 0.608353929 | 0.82656379 |
| chr9:34620373-34620716 | 0.189153259 | 0.608356543 | 0.82656379 |
| chr4:153536174-153536404 | -0.252795796 | 0.60836738 | 0.82656379 |
| chr6:28349778-28350431 | -0.18583872 | 0.608376395 | 0.82656379 |
| chr4:87304523-87304973 | 0.216144502 | 0.608389076 | 0.82656379 |
| chr19:18281489-18282151 | 0.172333109 | 0.608406127 | 0.82656379 |

| | | | |
|---|---|---|---|
| chr3:156554592-156555759 | -0.161519649 | 0.608413724 | 0.82656379 |
| chr18:46333391-46334116 | 0.176468967 | 0.608417919 | 0.82656379 |
| chr13:95213473-95213839 | 0.224725382 | 0.608440131 | 0.826579311 |
| chr1:243433001-243433443 | -0.194292682 | 0.608453192 | 0.826582399 |
| chr14:43736982-43737182 | 0.276101913 | 0.608468704 | 0.826586543 |
| chr11:118997590-118998811 | 0.164487176 | 0.608478777 | 0.826586543 |
| chr14:100304891-100305260 | -0.183370171 | 0.608488606 | 0.826586543 |
| chr15:44243928-44244139 | -0.272649181 | 0.608500183 | 0.826587615 |
| chr3:75595037-75595237 | 0.285710734 | 0.608544579 | 0.826622074 |
| chr16:87777575-87778322 | -0.170008016 | 0.60858187 | 0.826622074 |
| chr14:101242892-101243285 | -0.211582114 | 0.608587385 | 0.826622074 |
| chr3:155760267-155760467 | 0.285674608 | 0.608599884 | 0.826622074 |
| chr19:8526277-8526539 | -0.197667602 | 0.608609226 | 0.826622074 |
| chr16:25048547-25048783 | -0.244882143 | 0.608612852 | 0.826622074 |
| chr9:126499220-126499679 | -0.196166241 | 0.608627541 | 0.826622074 |
| chr9:36873345-36873562 | -0.250089008 | 0.608631804 | 0.826622074 |
| chr2:143263323-143263595 | -0.225841442 | 0.608633506 | 0.826622074 |
| chr11:65354229-65354919 | 0.17112765 | 0.608644768 | 0.826622074 |
| chr16:81445279-81445698 | 0.182614442 | 0.608645849 | 0.826622074 |
| chr2:56183598-56186083 | -0.160669019 | 0.60865501 | 0.826622074 |
| chr21:23401393-23401790 | -0.216648101 | 0.608778369 | 0.826774955 |
| chr2:6103920-6104291 | 0.208743226 | 0.608839515 | 0.826841785 |
| chr2:47632079-47632344 | -0.205039085 | 0.60884916 | 0.826841785 |
| chr16:30021209-30021892 | 0.170102132 | 0.608884154 | 0.826864433 |
| chr8:24387802-24388032 | -0.255330006 | 0.60889319 | 0.826864433 |
| chr12:27013699-27014633 | -0.170913646 | 0.608898212 | 0.826864433 |
| chr8:17922459-17923782 | -0.163340535 | 0.608941173 | 0.826904598 |
| chr1:241998416-241998868 | -0.189646073 | 0.60894983 | 0.826904598 |
| chr17:49975655-49975901 | 0.215606145 | 0.608970513 | 0.826904598 |
| chr6:79570528-79571293 | -0.184661952 | 0.608970957 | 0.826904598 |
| chr4:112099701-112100236 | 0.182630031 | 0.608997498 | 0.826925983 |
| chr8:43139786-43140601 | 0.17116999 | 0.609032825 | 0.826945121 |
| chr5:78920527-78920877 | 0.222114276 | 0.609046669 | 0.826945121 |
| chr1:174709102-174709460 | -0.217549804 | 0.609048216 | 0.826945121 |
| chr10:78386666-78386970 | -0.239773714 | 0.609062922 | 0.826945121 |
| chr17:8391761-8392103 | 0.218989651 | 0.609065555 | 0.826945121 |
| chr17:17041911-17042521 | 0.171593372 | 0.609083828 | 0.826955278 |
| chr22:46542374-46542626 | 0.217119171 | 0.609096268 | 0.826957514 |
| chr3:136642743-136643141 | 0.202878654 | 0.609108777 | 0.826959843 |
| chr15:65862650-65863144 | -0.188771444 | 0.609243261 | 0.827127772 |
| chr4:116702977-116703467 | 0.185966099 | 0.609275695 | 0.827157149 |
| chr7:100435746-100436804 | -0.167787015 | 0.609307779 | 0.827182713 |
| chr18:27921959-27922229 | -0.236006222 | 0.609339946 | 0.827182713 |
| chr1:11642466-11642687 | -0.24634138 | 0.6093631 | 0.827182713 |
| chr3:134357454-134358321 | 0.168395383 | 0.609364586 | 0.827182713 |
| chr10:71729397-71729708 | -0.208403702 | 0.609372024 | 0.827182713 |
| chr1:203148144-203148430 | 0.186143926 | 0.609379298 | 0.827182713 |
| chr14:95265465-95265778 | -0.20361662 | 0.609395648 | 0.827182713 |
| chr8:1763605-1764827 | 0.164415085 | 0.609396947 | 0.827182713 |
| chr15:77024351-77024565 | -0.22304756 | 0.609414659 | 0.827182713 |

| | | | |
|---|---|---|---|
| chr1:39105588-39106178 | 0.190458384 | 0.609423158 | 0.827182713 |
| chr10:87093986-87095866 | 0.160516735 | 0.609441327 | 0.827182713 |
| chr11:110092474-110093796 | 0.170035763 | 0.609451704 | 0.827182713 |
| chr8:105424705-105425393 | 0.170129466 | 0.609455336 | 0.827182713 |
| chr17:7836313-7836546 | -0.21046834 | 0.609459746 | 0.827182713 |
| chr19:15836383-15836689 | 0.225712913 | 0.60946613 | 0.827182713 |
| chr3:101676146-101677526 | -0.159577968 | 0.609467256 | 0.827182713 |
| chr4:118270337-118270758 | -0.2062085 | 0.609480455 | 0.827185975 |
| chr5:115602347-115602547 | -0.2453378 | 0.609503863 | 0.827203092 |
| chr5:61138964-61139439 | -0.201002323 | 0.609521208 | 0.82721198 |
| chr3:63853194-63853479 | -0.178854439 | 0.609557153 | 0.827217816 |
| chr3:138608469-138609435 | 0.165548765 | 0.609565756 | 0.827217816 |
| chr6:2846348-2846581 | -0.213902352 | 0.609572356 | 0.827217816 |
| chr5:43066305-43066710 | 0.178040184 | 0.60958198 | 0.827217816 |
| chr8:22549582-22549832 | -0.229300782 | 0.609587422 | 0.827217816 |
| chr15:31293840-31294050 | 0.237198996 | 0.609590285 | 0.827217816 |
| chr15:85800470-85800831 | 0.184717141 | 0.609626015 | 0.827238661 |
| chr10:91220367-91221484 | -0.164213056 | 0.609627238 | 0.827238661 |
| chr6:151807069-151808589 | -0.163217023 | 0.60965083 | 0.827240554 |
| chr15:69001975-69002262 | -0.253805037 | 0.609654794 | 0.827240554 |
| chr2:26982371-26982665 | -0.210050023 | 0.609661022 | 0.827240554 |
| chr8:127848814-127849034 | 0.195252916 | 0.609738475 | 0.827324791 |
| chr19:1592663-1593209 | 0.186864465 | 0.609744698 | 0.827324791 |
| chr15:78396947-78397259 | -0.20590698 | 0.609770262 | 0.827342681 |
| chr8:128024785-128025942 | -0.162896357 | 0.609779479 | 0.827342681 |
| chr1:148150855-148151264 | -0.195557444 | 0.609805654 | 0.8273546 |
| chr1:203217288-203217544 | 0.195660658 | 0.609809859 | 0.8273546 |
| chrX:115650963-115651219 | 0.22757255 | 0.609834923 | 0.827360904 |
| chr5:55534166-55535504 | -0.164377724 | 0.609836102 | 0.827360904 |
| chr11:20443719-20443919 | -0.263792244 | 0.609851923 | 0.827367719 |
| chr8:85031994-85032275 | -0.216186074 | 0.60993729 | 0.827437006 |
| chr17:58686964-58687529 | 0.164202311 | 0.609950812 | 0.827437006 |
| chr14:70707266-70707604 | -0.20481371 | 0.609955267 | 0.827437006 |
| chr11:58567987-58568187 | -0.262418353 | 0.60995583 | 0.827437006 |
| chr1:36374352-36375001 | 0.170874911 | 0.609956989 | 0.827437006 |
| chr17:58631142-58632502 | -0.158330601 | 0.609977462 | 0.827450129 |
| chr4:119455753-119456017 | -0.191208717 | 0.61002013 | 0.827457419 |
| chr15:99755817-99756104 | 0.202379688 | 0.610024061 | 0.827457419 |
| chr14:70113157-70113399 | 0.22889143 | 0.610036158 | 0.827457419 |
| chr12:87667310-87667964 | -0.185490424 | 0.610041659 | 0.827457419 |
| chr1:53326012-53326807 | 0.164097635 | 0.610047534 | 0.827457419 |
| chr11:66677509-66678830 | 0.162297233 | 0.610047631 | 0.827457419 |
| chr3:148631400-148631831 | 0.197422092 | 0.610078805 | 0.827475701 |
| chr11:69640984-69641382 | 0.177438228 | 0.610095376 | 0.827475701 |
| chr8:80249573-80249934 | -0.199317643 | 0.610100668 | 0.827475701 |
| chr5:143403419-143404637 | 0.163011462 | 0.610104307 | 0.827475701 |
| chr1:112298614-112298925 | 0.223711418 | 0.610132148 | 0.827498813 |
| chr5:120684723-120685309 | -0.19321102 | 0.610192983 | 0.827516429 |
| chr5:23526636-23526938 | 0.180751831 | 0.610208823 | 0.827516429 |
| chr11:62753569-62754607 | 0.161951416 | 0.610220017 | 0.827516429 |

| | | | |
|---|---|---|---|
| chr6:110528130-110528330 | 0.25651738 | 0.61022837 | 0.827516429 |
| chr6:7425047-7425312 | 0.222539397 | 0.610228993 | 0.827516429 |
| chr4:79576410-79576737 | 0.206658296 | 0.610238419 | 0.827516429 |
| chr1:179755658-179755869 | 0.258057406 | 0.610239732 | 0.827516429 |
| chr15:64841159-64841939 | 0.165127891 | 0.610283383 | 0.827516429 |
| chr10:102114754-102115182 | 0.178010037 | 0.61028697 | 0.827516429 |
| chr9:110194330-110194530 | 0.247960835 | 0.610289851 | 0.827516429 |
| chr8:93664777-93664977 | -0.265754671 | 0.610306082 | 0.827516429 |
| chr7:127186093-127186310 | 0.261207441 | 0.610323841 | 0.827516429 |
| chr8:68528864-68529087 | 0.261207441 | 0.610323841 | 0.827516429 |
| chr3:170496472-170496690 | -0.215467447 | 0.610325748 | 0.827516429 |
| chr2:70143532-70143772 | -0.186512604 | 0.610327996 | 0.827516429 |
| chr19:43662585-43662787 | 0.264719706 | 0.610331528 | 0.827516429 |
| chr17:75204662-75206006 | 0.162514863 | 0.610337824 | 0.827516429 |
| chr10:110857690-110857890 | -0.264917894 | 0.610339536 | 0.827516429 |
| chr16:66977660-66978008 | 0.20059805 | 0.610365449 | 0.827520232 |
| chr11:117643879-117644126 | -0.264963847 | 0.610375226 | 0.827520232 |
| chr10:38528162-38529464 | 0.169775584 | 0.610387346 | 0.827520232 |
| chr1:47710535-47710751 | 0.228771102 | 0.610391239 | 0.827520232 |
| chr1:162207183-162207651 | 0.188222599 | 0.610396401 | 0.827520232 |
| chr1:171245486-171245836 | 0.191889947 | 0.610407142 | 0.827520232 |
| chrX:101415524-101415850 | 0.204123893 | 0.610420442 | 0.827523622 |
| chr3:190718169-190718369 | -0.255568253 | 0.610445663 | 0.827531802 |
| chr5:138536717-138537047 | -0.211151467 | 0.610448077 | 0.827531802 |
| chr9:93566387-93567127 | -0.175000953 | 0.610473226 | 0.827551254 |
| chr20:11553602-11553852 | 0.246687701 | 0.610484094 | 0.827551346 |
| chr4:77881971-77882202 | -0.257331347 | 0.610507189 | 0.827568011 |
| chrX:17736766-17737708 | -0.163131755 | 0.610526906 | 0.827568451 |
| chr14:40074318-40074570 | 0.235895353 | 0.610529115 | 0.827568451 |
| chr5:87286927-87287227 | 0.242826299 | 0.610560538 | 0.827578208 |
| chr10:58065317-58065806 | -0.196495665 | 0.610565214 | 0.827578208 |
| chr4:68129722-68130151 | 0.182438166 | 0.610568716 | 0.827578208 |
| chr15:74995270-74995736 | 0.181254075 | 0.610596721 | 0.827601527 |
| chr1:22420547-22420760 | 0.247876862 | 0.6106428 | 0.827631482 |
| chr17:60393084-60393303 | -0.169825216 | 0.610666458 | 0.827631482 |
| chr1:28669103-28669701 | 0.187641362 | 0.61068624 | 0.827631482 |
| chr8:127911136-127911466 | -0.195264362 | 0.610692592 | 0.827631482 |
| chr1:115595643-115596189 | 0.176001918 | 0.610693759 | 0.827631482 |
| chr1:149792874-149793553 | -0.175817915 | 0.610695792 | 0.827631482 |
| chr20:32040573-32041003 | 0.187786924 | 0.610714764 | 0.827631482 |
| chr6:7910428-7911629 | -0.167669847 | 0.610718284 | 0.827631482 |
| chr7:3922262-3922565 | 0.202913506 | 0.610728852 | 0.827631482 |
| chr1:201739476-201740010 | -0.170062264 | 0.610736249 | 0.827631482 |
| chr18:59150311-59150657 | -0.246390568 | 0.610737638 | 0.827631482 |
| chr8:76308040-76308875 | -0.167720298 | 0.610749961 | 0.827633545 |
| chr12:123519287-123519620 | 0.206997861 | 0.610809318 | 0.827699342 |
| chr17:10616514-10616932 | -0.186182135 | 0.610891198 | 0.82777154 |
| chr2:108636190-108636907 | -0.17171096 | 0.610894854 | 0.82777154 |
| chr1:161530638-161531455 | 0.163851029 | 0.610903771 | 0.82777154 |
| chr19:12736727-12737618 | 0.167475457 | 0.610905811 | 0.82777154 |

| | | | |
|---|---|---|---|
| chr17:66715785-66716112 | 0.188405313 | 0.61091732 | 0.827772497 |
| chr1:146007136-146007558 | -0.185672932 | 0.610934501 | 0.827775868 |
| chr9:114487645-114487960 | -0.190238081 | 0.610941415 | 0.827775868 |
| chr10:59873015-59873532 | 0.18079556 | 0.610990901 | 0.827828279 |
| chr7:1269688-1269889 | -0.229570772 | 0.611015216 | 0.827846585 |
| chr10:93702357-93703052 | -0.169695601 | 0.611056679 | 0.827888122 |
| chr17:57340459-57341144 | -0.16952351 | 0.611069632 | 0.827891033 |
| chr16:27605620-27605820 | 0.289146163 | 0.611105508 | 0.827924998 |
| chr14:54773996-54774319 | 0.17912954 | 0.611124931 | 0.827936674 |
| chr1:174875345-174875631 | 0.212522508 | 0.611137082 | 0.827938496 |
| chr12:44948590-44949252 | -0.167815403 | 0.611151507 | 0.8279434 |
| chr3:195804172-195804771 | -0.190446632 | 0.611169258 | 0.82795281 |
| chr4:157045580-157045864 | -0.204125549 | 0.611182881 | 0.827955378 |
| chr5:76004845-76005075 | -0.288922224 | 0.611192766 | 0.827955378 |
| chr14:75063531-75064451 | 0.167467824 | 0.611208806 | 0.827962469 |
| chr4:47704710-47705082 | -0.204656782 | 0.611221988 | 0.827965687 |
| chr5:32724591-32724886 | -0.251567001 | 0.611257583 | 0.827990701 |
| chr1:118635656-118636478 | -0.16961559 | 0.611262066 | 0.827990701 |
| chr22:40818850-40819881 | -0.172869442 | 0.611280947 | 0.827995501 |
| chr10:4141612-4141872 | -0.251423378 | 0.611287221 | 0.827995501 |
| chr20:1894441-1894818 | 0.219196707 | 0.611357459 | 0.828072788 |
| chr8:132581753-132582043 | -0.248053795 | 0.611371862 | 0.828072788 |
| chr8:138700383-138700687 | -0.191051479 | 0.611376703 | 0.828072788 |
| chr10:3783793-3784258 | 0.172876007 | 0.61140031 | 0.828080055 |
| chr16:85370003-85370327 | 0.189765127 | 0.611403682 | 0.828080055 |
| chr3:183436721-183436971 | -0.276244811 | 0.611455678 | 0.828135839 |
| chr15:44408785-44409018 | -0.29299465 | 0.61148453 | 0.828152919 |
| chr11:117071374-117071589 | 0.263794251 | 0.611489906 | 0.828152919 |
| chr10:123153935-123154553 | 0.165183592 | 0.61151298 | 0.828159071 |
| chr5:128964951-128965983 | -0.167063239 | 0.611516065 | 0.828159071 |
| chr6:9467782-9468280 | -0.183463609 | 0.611568685 | 0.828197979 |
| chr7:83643891-83644123 | 0.229756445 | 0.611575541 | 0.828197979 |
| chr10:103451877-103452663 | 0.172455385 | 0.611577636 | 0.828197979 |
| chr11:17063467-17063751 | -0.199612704 | 0.61158803 | 0.828197979 |
| chr12:56724999-56725740 | -0.166231434 | 0.611674217 | 0.828280193 |
| chr2:197499647-197500749 | -0.159326093 | 0.611678489 | 0.828280193 |
| chr3:124786719-124787035 | -0.208699693 | 0.611681171 | 0.828280193 |
| chr4:76430850-76431087 | -0.252631463 | 0.611703048 | 0.828295178 |
| chr10:132185618-132185915 | 0.20079613 | 0.611714741 | 0.828296373 |
| chr9:2621310-2621654 | 0.196926658 | 0.611764033 | 0.828340717 |
| chr22:20418327-20418527 | 0.237908196 | 0.611769111 | 0.828340717 |
| chr14:96574872-96575223 | -0.215175414 | 0.611799884 | 0.828358132 |
| chr11:119352875-119353324 | -0.19554034 | 0.611803595 | 0.828358132 |
| chr11:67629896-67631065 | 0.161587353 | 0.611836077 | 0.828387474 |
| chr7:455655-455998 | 0.194565682 | 0.611847978 | 0.828388949 |
| chr12:13001150-13001365 | 0.216691334 | 0.6118943 | 0.828431645 |
| chr9:137114354-137115351 | 0.173752711 | 0.611903283 | 0.828431645 |
| chr7:36473417-36474237 | -0.174185683 | 0.611914547 | 0.828431645 |
| chr6:95087295-95087739 | -0.209240021 | 0.611922761 | 0.828431645 |
| chr13:27449903-27450837 | 0.160590176 | 0.61196545 | 0.828436648 |

| | | | |
|---|---|---|---|
| chr13:34894803-34895003 | 0.270727646 | 0.61196783 | 0.828436648 |
| chr3:157174565-157175572 | -0.159879483 | 0.611968518 | 0.828436648 |
| chr11:66172166-66172413 | 0.247148939 | 0.611969705 | 0.828436648 |
| chr2:128812502-128812731 | -0.271811878 | 0.611998841 | 0.828442911 |
| chr1:226407523-226408674 | 0.164614583 | 0.612005771 | 0.828442911 |
| chr1:106620438-106620984 | 0.165506947 | 0.612006768 | 0.828442911 |
| chr11:34915771-34916971 | 0.161641243 | 0.612062265 | 0.828503398 |
| chr11:76721814-76722226 | 0.184992508 | 0.612084729 | 0.828519169 |
| chr4:89438198-89438722 | -0.194303505 | 0.612103019 | 0.82852929 |
| chr1:109397729-109398547 | -0.166960385 | 0.612131901 | 0.828553748 |
| chr2:224650934-224651167 | -0.279020927 | 0.612152195 | 0.828564516 |
| chr2:64721501-64721724 | 0.249285694 | 0.612161484 | 0.828564516 |
| chr6:28212224-28213285 | -0.166697052 | 0.612211793 | 0.828617972 |
| chr10:62076952-62077175 | -0.229655445 | 0.612289216 | 0.828708124 |
| chr11:71792726-71793004 | 0.205821858 | 0.612305406 | 0.828715398 |
| chr3:167114956-167115156 | -0.265116202 | 0.612332406 | 0.828737302 |
| chr3:112527150-112527425 | -0.255914352 | 0.612366112 | 0.828755259 |
| chr1:44354718-44354980 | 0.21697094 | 0.612367306 | 0.828755259 |
| chr11:66421620-66421968 | 0.187328438 | 0.612422089 | 0.828806188 |
| chr3:53891287-53892300 | 0.165225854 | 0.612432357 | 0.828806188 |
| chr9:98255148-98256407 | 0.161469652 | 0.612455838 | 0.828806188 |
| chr5:149857582-149857833 | -0.226627797 | 0.612458277 | 0.828806188 |
| chr10:103549174-103549450 | -0.239982677 | 0.612459106 | 0.828806188 |
| chr19:46829926-46830353 | -0.198200684 | 0.612486484 | 0.828806188 |
| chr19:49474086-49474359 | 0.208128881 | 0.612490206 | 0.828806188 |
| chr6:138372880-138373080 | -0.283720197 | 0.612491473 | 0.828806188 |
| chr1:180754691-180754986 | 0.230914195 | 0.61252922 | 0.828842629 |
| chrX:73127483-73128313 | 0.171683728 | 0.612541388 | 0.828844458 |
| chr16:86888222-86888422 | -0.256668218 | 0.612562953 | 0.828854962 |
| chr9:99411173-99411402 | -0.272195814 | 0.612570786 | 0.828854962 |
| chr10:13099638-13100763 | 0.166579757 | 0.612584612 | 0.828859032 |
| chr15:40430310-40430732 | -0.206642054 | 0.612646724 | 0.828928436 |
| chr2:202141065-202141479 | -0.224866648 | 0.612675148 | 0.828952255 |
| chr13:53250353-53250608 | -0.234910727 | 0.612698423 | 0.828969108 |
| chr3:7546264-7546617 | 0.209629286 | 0.612776205 | 0.829059707 |
| chr4:126405389-126405704 | -0.214642694 | 0.612788919 | 0.829062269 |
| chr15:87121788-87122058 | 0.218222335 | 0.612821032 | 0.829091077 |
| chr10:15167768-15169232 | -0.159384865 | 0.612842614 | 0.829105636 |
| chr10:96586874-96587451 | -0.173663176 | 0.612854024 | 0.829106433 |
| chr1:201214497-201214755 | -0.193062385 | 0.612872735 | 0.829117107 |
| chr14:28116733-28117052 | 0.209623745 | 0.612900657 | 0.829138551 |
| chr5:90604603-90604877 | -0.232838995 | 0.612910719 | 0.829138551 |
| chr12:52025696-52026151 | 0.174738037 | 0.61292105 | 0.829138551 |
| chr3:109224303-109224698 | 0.193065171 | 0.612932597 | 0.829139534 |
| chr7:26376142-26376843 | 0.182025874 | 0.612954687 | 0.829154776 |
| chrX:78124371-78124571 | 0.253105383 | 0.612971372 | 0.829162708 |
| chr19:45186844-45187136 | -0.238692179 | 0.613033797 | 0.829226463 |
| chr15:66851195-66851538 | 0.183769511 | 0.613040158 | 0.829226463 |
| chr1:25267834-25268119 | 0.235955227 | 0.613050971 | 0.829226463 |
| chr1:212995485-212995818 | 0.211652599 | 0.613093026 | 0.829242434 |

| | | | |
|---|---|---|---|
| chr5:134047257-134047489 | 0.234066526 | 0.613111063 | 0.829242434 |
| chr5:166845185-166845414 | 0.252942631 | 0.613113879 | 0.829242434 |
| chr1:117841895-117842671 | 0.181006141 | 0.613120704 | 0.829242434 |
| chr12:88180244-88180525 | -0.224499751 | 0.613125985 | 0.829242434 |
| chr14:80157604-80157869 | -0.269159658 | 0.613127713 | 0.829242434 |
| chrX:54440102-54440895 | -0.171516619 | 0.61315143 | 0.829245607 |
| chr3:124521084-124521433 | 0.2084912 | 0.613151704 | 0.829245607 |
| chr4:39182060-39182877 | -0.162127813 | 0.613200866 | 0.82928419 |
| chr12:67555046-67555753 | -0.176415674 | 0.613201879 | 0.82928419 |
| chr1:197731452-197731745 | -0.19876332 | 0.613248122 | 0.829332091 |
| chr6:3023479-3024345 | -0.171506593 | 0.613326767 | 0.829423808 |
| chr5:161846831-161847127 | -0.216769667 | 0.613393121 | 0.829495973 |
| chr4:48578353-48578689 | -0.209764626 | 0.613401782 | 0.829495973 |
| chr2:98601666-98601866 | 0.248297158 | 0.6134258 | 0.829503401 |
| chr6:96724725-96724926 | 0.254552528 | 0.61344752 | 0.829503401 |
| chr2:12716362-12717127 | -0.166244153 | 0.613465593 | 0.829503401 |
| chr12:120528963-120529525 | -0.178655125 | 0.613477811 | 0.829503401 |
| chr18:21600137-21601359 | 0.16045656 | 0.613478902 | 0.829503401 |
| chr15:70762561-70764016 | -0.161208934 | 0.613479986 | 0.829503401 |
| chr22:20981765-20982033 | 0.210711245 | 0.613492555 | 0.829503401 |
| chr2:96536440-96536781 | 0.190029054 | 0.613493882 | 0.829503401 |
| chr3:113205327-113205836 | -0.195471701 | 0.613546736 | 0.829560226 |
| chr19:58558539-58559851 | 0.159862482 | 0.613589362 | 0.829588816 |
| chr1:26921336-26922104 | 0.174521006 | 0.613589536 | 0.829588816 |
| chr3:196837897-196838180 | -0.236359658 | 0.613627682 | 0.829615095 |
| chr7:1666327-1666582 | 0.208753525 | 0.613669147 | 0.829615095 |
| chr7:76502674-76502945 | -0.290482659 | 0.613677537 | 0.829615095 |
| chr1:37795281-37795634 | 0.201320848 | 0.61367871 | 0.829615095 |
| chr6:142518680-142518880 | -0.259472341 | 0.61368878 | 0.829615095 |
| chr12:98515117-98515843 | -0.163866274 | 0.613694115 | 0.829615095 |
| chr5:32116163-32116363 | -0.262574135 | 0.61369676 | 0.829615095 |
| chr3:123068201-123068466 | 0.202600386 | 0.613696987 | 0.829615095 |
| chr1:228622039-228622332 | -0.229640047 | 0.613706419 | 0.829615095 |
| chr14:61388204-61388679 | 0.184926831 | 0.613723421 | 0.829623442 |
| chr2:114261499-114261699 | -0.290586781 | 0.613737504 | 0.829627685 |
| chr1:154272184-154272814 | 0.169774139 | 0.613755584 | 0.829627685 |
| chr17:17972157-17973111 | -0.160024463 | 0.613759042 | 0.829627685 |
| chr1:203589514-203589714 | -0.22942734 | 0.613783535 | 0.829639549 |
| chr10:9733873-9734131 | -0.227928621 | 0.613818913 | 0.829639549 |
| chr17:62703015-62704329 | -0.160364719 | 0.613835144 | 0.829639549 |
| chrX:114344808-114345120 | -0.211098718 | 0.613838743 | 0.829639549 |
| chr3:72915917-72916117 | -0.284152191 | 0.613845948 | 0.829639549 |
| chr8:111152566-111152913 | -0.212332496 | 0.613849569 | 0.829639549 |
| chr1:215557303-215557584 | -0.219886257 | 0.613850612 | 0.829639549 |
| chr11:61281197-61281878 | 0.16968703 | 0.613860545 | 0.829639549 |
| chr2:42095941-42096265 | -0.200435451 | 0.613865268 | 0.829639549 |
| chr8:24415005-24415435 | -0.193345605 | 0.613889565 | 0.829657752 |
| chr1:53739148-53739649 | 0.183297261 | 0.613957211 | 0.829726348 |
| chr19:13162659-13163300 | 0.167034222 | 0.613961979 | 0.829726348 |
| chr1:152004051-152004608 | 0.172950433 | 0.614012871 | 0.829774404 |

| | | | |
|---|---|---|---|
| chr1:214678403-214678655 | -0.217987451 | 0.614019197 | 0.829774404 |
| chr14:106009144-106009528 | 0.198911601 | 0.614065443 | 0.829822264 |
| chr19:41997191-41997391 | 0.253652925 | 0.614079601 | 0.829826762 |
| chr7:6435460-6435961 | 0.178089888 | 0.614098478 | 0.829837635 |
| chr17:78169409-78169968 | 0.17119273 | 0.614122928 | 0.829848549 |
| chr5:10857058-10857261 | 0.230664803 | 0.614130891 | 0.829848549 |
| chr1:226520094-226520425 | 0.246497076 | 0.614159182 | 0.829848549 |
| chr10:103748481-103748824 | 0.224240268 | 0.614184053 | 0.829848549 |
| chr17:59173048-59173282 | -0.210221876 | 0.614187855 | 0.829848549 |
| chr15:100339048-100339304 | 0.190382088 | 0.614195007 | 0.829848549 |
| chr19:10424350-10424601 | 0.209217704 | 0.61420083 | 0.829848549 |
| chr8:58534889-58535231 | -0.206745343 | 0.614204007 | 0.829848549 |
| chr2:231674983-231675256 | 0.216826452 | 0.614204029 | 0.829848549 |
| chr6:2209317-2209589 | -0.225509647 | 0.614220032 | 0.829855539 |
| chr3:107471084-107471292 | -0.252753181 | 0.61424003 | 0.829858981 |
| chr4:152779389-152780760 | 0.158859955 | 0.614248412 | 0.829858981 |
| chr10:70537516-70537885 | -0.195641016 | 0.614264857 | 0.829858981 |
| chr6:45607866-45608139 | -0.244429096 | 0.614265902 | 0.829858981 |
| chr2:195562547-195563018 | -0.19172337 | 0.614303835 | 0.829895595 |
| chr14:61750921-61751463 | 0.172760426 | 0.614317512 | 0.829899439 |
| chr15:48520599-48521037 | -0.203576312 | 0.614342114 | 0.829918043 |
| chr20:20369260-20369774 | 0.188506906 | 0.614391677 | 0.829946716 |
| chr16:73058014-73058233 | 0.223265764 | 0.614393244 | 0.829946716 |
| chr19:4342427-4343277 | 0.163032345 | 0.61441048 | 0.829946716 |
| chr16:73176641-73176841 | 0.23509404 | 0.614423965 | 0.829946716 |
| chr13:32585978-32586740 | -0.164490378 | 0.614424592 | 0.829946716 |
| chr17:4365853-4366348 | -0.183713012 | 0.614438628 | 0.829946716 |
| chr1:39098572-39099003 | -0.186115814 | 0.614456693 | 0.829946716 |
| chr18:62584380-62584641 | -0.204652586 | 0.61446279 | 0.829946716 |
| chr19:48738615-48739231 | 0.170702415 | 0.614465178 | 0.829946716 |
| chr3:157137427-157138204 | -0.16839964 | 0.614479519 | 0.829946716 |
| chr17:56787020-56787260 | -0.207787211 | 0.614485345 | 0.829946716 |
| chr2:24435877-24436206 | 0.223798061 | 0.614504067 | 0.829946716 |
| chr13:51070161-51070436 | -0.220756395 | 0.614509338 | 0.829946716 |
| chr15:83383682-83383987 | 0.207692701 | 0.614514983 | 0.829946716 |
| chr1:27575115-27575562 | 0.190034125 | 0.614542466 | 0.829963112 |
| chr1:219567341-219567701 | 0.187250985 | 0.614557408 | 0.829963112 |
| chr11:68038590-68039108 | 0.17810844 | 0.614559619 | 0.829963112 |
| chr19:4186504-4186723 | 0.226218606 | 0.614597827 | 0.830000083 |
| chr21:15210027-15210634 | -0.173376931 | 0.614608703 | 0.830000141 |
| chr17:60591557-60591900 | 0.169564077 | 0.614623102 | 0.830004958 |
| chr17:75154202-75154924 | -0.163507694 | 0.614666478 | 0.8300316 |
| chr17:60185002-60185232 | 0.182556363 | 0.614679429 | 0.8300316 |
| chr11:71955771-71955971 | -0.279180648 | 0.614683285 | 0.8300316 |
| chr3:15953779-15954150 | -0.211796677 | 0.614686161 | 0.8300316 |
| chr8:68330430-68330902 | 0.179439996 | 0.614736119 | 0.830070547 |
| chr10:97037342-97038362 | 0.165205039 | 0.614736671 | 0.830070547 |
| chr2:223641133-223641607 | 0.18507838 | 0.614757677 | 0.830084284 |
| chr16:1325662-1325862 | -0.276191176 | 0.614790965 | 0.830104687 |
| chr1:156709821-156710156 | -0.190614147 | 0.614818105 | 0.830104687 |

| | | | |
|---|---|---|---|
| chr19:40420685-40421468 | -0.167899036 | 0.614844232 | 0.830104687 |
| chr6:33607812-33608039 | -0.270441608 | 0.614849175 | 0.830104687 |
| chr7:150101467-150101671 | -0.270441608 | 0.614849175 | 0.830104687 |
| chr9:127460370-127460591 | -0.270441608 | 0.614849175 | 0.830104687 |
| chr1:228664054-228664254 | -0.274122887 | 0.614854599 | 0.830104687 |
| chr12:98456651-98457312 | 0.179268962 | 0.614876874 | 0.830104687 |
| chr3:134008471-134008813 | 0.198638829 | 0.614884678 | 0.830104687 |
| chr8:86140910-86141310 | -0.180276401 | 0.614885191 | 0.830104687 |
| chr10:133360125-133360433 | -0.207603194 | 0.614898007 | 0.830104687 |
| chr13:42968921-42969437 | 0.185117767 | 0.614924769 | 0.830104687 |
| chr12:88958170-88958422 | 0.262911144 | 0.614941428 | 0.830104687 |
| chr11:71431270-71431470 | -0.240025234 | 0.614952224 | 0.830104687 |
| chr1:215476131-215476802 | 0.174570162 | 0.614966074 | 0.830104687 |
| chr20:54172536-54173297 | 0.159256837 | 0.614966516 | 0.830104687 |
| chr15:42920702-42921407 | -0.166123635 | 0.614970969 | 0.830104687 |
| chr6:131597526-131597726 | -0.276717369 | 0.614987941 | 0.830104687 |
| chr16:56576977-56577345 | 0.195673303 | 0.614989422 | 0.830104687 |
| chr9:27572694-27572956 | 0.213085006 | 0.614989463 | 0.830104687 |
| chr1:200500075-200500290 | -0.238231541 | 0.615070127 | 0.83019894 |
| chr1:110053916-110054155 | 0.20876357 | 0.615092168 | 0.830207747 |
| chr11:31810126-31810606 | -0.198540028 | 0.615098321 | 0.830207747 |
| chr9:94198538-94198844 | 0.243396011 | 0.615124436 | 0.830228369 |
| chr5:82277810-82278827 | -0.161368065 | 0.61514568 | 0.830242418 |
| chrX:137052060-137052802 | -0.172672496 | 0.615188803 | 0.830274232 |
| chr8:52714188-52715062 | -0.164969255 | 0.615190924 | 0.830274232 |
| chr17:61103689-61103889 | 0.233802339 | 0.615230752 | 0.830313258 |
| chr4:99024918-99025217 | 0.206008448 | 0.615241722 | 0.830313258 |
| chr3:105904716-105904925 | 0.278033567 | 0.61525235 | 0.830313258 |
| chr5:39217773-39218116 | -0.196222999 | 0.615273169 | 0.83032673 |
| chr16:84340268-84340821 | -0.173060473 | 0.615298966 | 0.830331138 |
| chr1:111015333-111015533 | -0.27359308 | 0.615306437 | 0.830331138 |
| chr13:55199228-55199500 | -0.224774071 | 0.615308945 | 0.830331138 |
| chr3:156763684-156763884 | 0.279728797 | 0.615368467 | 0.830396834 |
| chr9:132409932-132410298 | 0.202605269 | 0.615413892 | 0.830437243 |
| chr5:126339269-126339478 | -0.264960274 | 0.615423398 | 0.830437243 |
| chr16:87492729-87493347 | 0.168225528 | 0.615453112 | 0.830437243 |
| chr5:2751439-2752482 | 0.160210599 | 0.615456209 | 0.830437243 |
| chr12:84128727-84128990 | 0.234397716 | 0.615476086 | 0.830437243 |
| chr12:39912466-39913060 | 0.175299446 | 0.615478684 | 0.830437243 |
| chr16:807537-808055 | 0.181034729 | 0.615484558 | 0.830437243 |
| chr6:98769560-98769842 | 0.208949668 | 0.615485116 | 0.830437243 |
| chr19:47555592-47555805 | 0.25980817 | 0.615502961 | 0.830446697 |
| chr18:26863112-26863805 | 0.177864591 | 0.615569535 | 0.830507372 |
| chr16:30813841-30814998 | -0.165173781 | 0.61556961 | 0.830507372 |
| chr15:70099462-70099895 | 0.165724099 | 0.615631303 | 0.830575981 |
| chr4:129390900-129391205 | 0.264454943 | 0.615645906 | 0.830578071 |
| chr20:18587621-18588606 | 0.162179125 | 0.615654532 | 0.830578071 |
| chr17:78186492-78187675 | 0.159616111 | 0.6157132 | 0.830627907 |
| chr8:123022244-123022772 | -0.174203938 | 0.615715558 | 0.830627907 |
| chr19:29259174-29259508 | -0.209656066 | 0.615729713 | 0.830627907 |

| | | | |
|---|---|---|---|
| chr17:27332373-27333430 | -0.160454003 | 0.615734835 | 0.830627907 |
| chr8:86173759-86174291 | -0.175038702 | 0.615830689 | 0.830727884 |
| chr1:112390831-112391262 | -0.179869338 | 0.615832359 | 0.830727884 |
| chr17:77846512-77846824 | 0.213073012 | 0.615841472 | 0.830727884 |
| chr5:177553623-177554297 | 0.172358124 | 0.615867137 | 0.830745143 |
| chr8:105092438-105092696 | 0.209632301 | 0.615875951 | 0.830745143 |
| chr3:18963192-18964175 | -0.167423959 | 0.615923544 | 0.830794715 |
| chr2:96550033-96550492 | 0.178832517 | 0.615935086 | 0.830795658 |
| chr2:85611479-85612270 | -0.164290167 | 0.615981723 | 0.830843938 |
| chr1:37479721-37480066 | -0.179293613 | 0.616021314 | 0.830872422 |
| chr2:208738651-208738920 | 0.248224545 | 0.616024528 | 0.830872422 |
| chr17:35088618-35090006 | 0.163918257 | 0.616086509 | 0.830924445 |
| chr7:27142686-27144016 | 0.162117153 | 0.616098741 | 0.830924445 |
| chr6:170025287-170025623 | -0.191593834 | 0.616106199 | 0.830924445 |
| chr7:48041218-48041418 | -0.277586966 | 0.616122355 | 0.830924445 |
| chr1:246506444-246507680 | -0.159917385 | 0.616131716 | 0.830924445 |
| chr7:99438316-99439507 | -0.162839224 | 0.616132257 | 0.830924445 |
| chr7:51549539-51549786 | 0.247920512 | 0.616142972 | 0.830924445 |
| chr1:115088896-115089121 | -0.207962469 | 0.616149855 | 0.830924445 |
| chr4:10063769-10064003 | -0.239750401 | 0.61617483 | 0.830936864 |
| chr16:3157063-3157627 | -0.165072019 | 0.616180753 | 0.830936864 |
| chr10:63158212-63158471 | -0.217822275 | 0.616200701 | 0.830949139 |
| chr12:124559399-124559754 | 0.209543539 | 0.616235958 | 0.830959163 |
| chr2:190372415-190372647 | 0.234687312 | 0.616240063 | 0.830959163 |
| chr17:55502051-55502360 | -0.227957227 | 0.616248186 | 0.830959163 |
| chr7:64849186-64849470 | -0.266403136 | 0.616251514 | 0.830959163 |
| chr10:35306858-35307334 | 0.173438607 | 0.616272317 | 0.830962736 |
| chr3:62632861-62633533 | -0.16266432 | 0.616277661 | 0.830962736 |
| chr12:113497708-113498130 | -0.213809024 | 0.616286699 | 0.830962736 |
| chrX:69493241-69493449 | 0.256750165 | 0.616313064 | 0.830968797 |
| chr13:39145004-39145260 | 0.213450828 | 0.616316678 | 0.830968797 |
| chr11:47243601-47243811 | -0.214963213 | 0.616323729 | 0.830968797 |
| chr17:70622415-70622852 | 0.189810095 | 0.616355792 | 0.830976618 |
| chr16:75860714-75861002 | -0.218028251 | 0.616357569 | 0.830976618 |
| chr7:7762516-7762718 | 0.220591688 | 0.616362065 | 0.830976618 |
| chr12:116533628-116534155 | 0.180148087 | 0.616394374 | 0.83100046 |
| chr11:123740963-123741853 | 0.167190084 | 0.61640144 | 0.83100046 |
| chr20:19027171-19027616 | 0.194363383 | 0.616459224 | 0.831037321 |
| chr7:72924907-72925585 | 0.175290646 | 0.616461629 | 0.831037321 |
| chr3:134769763-134770019 | 0.234784707 | 0.616470541 | 0.831037321 |
| chr12:7970258-7970474 | -0.2543512 | 0.616487621 | 0.831037321 |
| chr11:95789353-95790694 | -0.157691038 | 0.616490967 | 0.831037321 |
| chr2:206781640-206781871 | -0.234462055 | 0.616493858 | 0.831037321 |
| chr2:74697885-74698132 | 0.248124258 | 0.616505753 | 0.831038736 |
| chr1:198399460-198399681 | 0.243578101 | 0.616551926 | 0.831071357 |
| chr7:116442149-116442737 | -0.178840894 | 0.616562186 | 0.831071357 |
| chr12:12890519-12891765 | 0.158515424 | 0.616562492 | 0.831071357 |
| chr3:194935075-194935294 | -0.282656962 | 0.616596362 | 0.83108777 |
| chr4:104015748-104015948 | -0.282656962 | 0.616596362 | 0.83108777 |
| chr20:47985095-47985308 | 0.178013157 | 0.616633236 | 0.831122851 |

| | | | |
|---|---|---|---|
| chr13:19960066-19960400 | 0.174784735 | 0.616646586 | 0.831126225 |
| chr6:28940364-28940564 | -0.199411254 | 0.616662968 | 0.831133684 |
| chr14:69503835-69504035 | 0.2306829 | 0.616687467 | 0.831149203 |
| chr9:98974698-98975329 | -0.180414849 | 0.616696177 | 0.831149203 |
| chr12:65823889-65825303 | 0.166511453 | 0.616713846 | 0.831158396 |
| chr12:6945991-6946762 | 0.161822623 | 0.616757007 | 0.831187209 |
| chr14:25938507-25938707 | 0.273413587 | 0.616757376 | 0.831187209 |
| chr3:169029433-169029656 | 0.259842782 | 0.616767769 | 0.831187209 |
| chr12:98515907-98516681 | -0.164477546 | 0.616795064 | 0.831200143 |
| chr1:227563343-227564244 | 0.161683697 | 0.616808786 | 0.831200143 |
| chr2:153229021-153229567 | -0.177554772 | 0.61680991 | 0.831200143 |
| chr2:119218456-119218730 | -0.209991867 | 0.616834462 | 0.831218609 |
| chr16:24853868-24854218 | -0.214584197 | 0.616854911 | 0.831226284 |
| chr4:133320615-133320868 | 0.225499184 | 0.616880635 | 0.831226284 |
| chr2:53460393-53460677 | -0.223640338 | 0.616886542 | 0.831226284 |
| chr18:23282026-23282633 | -0.178416657 | 0.616890264 | 0.831226284 |
| chr10:128124963-128125184 | 0.207133937 | 0.616918858 | 0.831226284 |
| chr21:38669251-38669484 | -0.222035791 | 0.616922397 | 0.831226284 |
| chr12:57547058-57547445 | 0.175062076 | 0.616925992 | 0.831226284 |
| chr17:49227486-49227735 | 0.19009617 | 0.616926944 | 0.831226284 |
| chrX:103375997-103377373 | -0.163045893 | 0.616956568 | 0.831251581 |
| chr1:21334734-21335040 | -0.201872004 | 0.617009804 | 0.831293505 |
| chr5:174693824-174694266 | 0.190858673 | 0.617037274 | 0.831293505 |
| chr6:139996474-139996730 | -0.224495822 | 0.617039253 | 0.831293505 |
| chr6:124823781-124823988 | -0.246748445 | 0.617058216 | 0.831293505 |
| chr8:95628053-95628282 | -0.246748445 | 0.617058216 | 0.831293505 |
| chr11:75445368-75445743 | 0.18487539 | 0.617058483 | 0.831293505 |
| chr18:37564846-37565106 | -0.220560375 | 0.617069451 | 0.831293505 |
| chr15:74540367-74541451 | 0.16261745 | 0.617083508 | 0.831293505 |
| chr13:20966404-20966859 | -0.204271913 | 0.617099395 | 0.831293505 |
| chr2:177198221-177198421 | -0.245135493 | 0.617105255 | 0.831293505 |
| chr17:51891762-51891962 | 0.249075242 | 0.617107027 | 0.831293505 |
| chr17:62278361-62278635 | -0.241332123 | 0.617137282 | 0.831319646 |
| chr5:168679450-168679739 | -0.200426993 | 0.617149342 | 0.831321277 |
| chr8:141392492-141392890 | 0.178276378 | 0.617178524 | 0.831334814 |
| chr6:46129349-46130506 | -0.16045628 | 0.617182002 | 0.831334814 |
| chr5:75460581-75460787 | -0.276899072 | 0.617191941 | 0.831334814 |
| chr1:64137173-64137428 | -0.217259104 | 0.617219552 | 0.831357391 |
| chr2:60554621-60554923 | -0.194305873 | 0.617251853 | 0.831386283 |
| chr2:70135939-70136143 | 0.243104104 | 0.617264274 | 0.831387763 |
| chr3:38756898-38757171 | -0.250512267 | 0.617280953 | 0.831387763 |
| chr15:94324098-94324520 | 0.193835867 | 0.617285503 | 0.831387763 |
| chr1:198158788-198159429 | -0.161015336 | 0.617352726 | 0.831442172 |
| chr8:123416095-123417313 | -0.155828372 | 0.617363393 | 0.831442172 |
| chr8:14802762-14803158 | -0.21733955 | 0.617380187 | 0.831442172 |
| chr2:197704858-197706112 | -0.158701185 | 0.617380263 | 0.831442172 |
| chr14:66038446-66038724 | -0.243333502 | 0.617382616 | 0.831442172 |
| chr6:116100383-116101750 | -0.158464173 | 0.617391008 | 0.831442172 |
| chr6:16759965-16760533 | -0.174456785 | 0.617438772 | 0.83146576 |
| chr5:10855739-10855953 | 0.226883194 | 0.617439101 | 0.83146576 |

| | | | |
|---|---|---|---|
| chr1:160790303-160790545 | -0.26127134 | 0.617443668 | 0.83146576 |
| chrX:136954004-136954670 | -0.187460231 | 0.61745193 | 0.83146576 |
| chr11:73660607-73661648 | 0.162324221 | 0.617533389 | 0.831552169 |
| chr10:89702121-89702376 | 0.1895432 | 0.617537803 | 0.831552169 |
| chr3:123881059-123881531 | -0.192694021 | 0.617560785 | 0.831556693 |
| chr2:182666962-182667266 | -0.206404192 | 0.617568154 | 0.831556693 |
| chr1:27366828-27367118 | -0.190805554 | 0.61758445 | 0.831556693 |
| chr1:197180129-197180627 | -0.176983258 | 0.617584573 | 0.831556693 |
| chr13:30865359-30865664 | 0.187790026 | 0.617657354 | 0.831640075 |
| chrX:116961995-116962274 | 0.221090771 | 0.617717834 | 0.831692822 |
| chr11:78000124-78000528 | -0.186459695 | 0.617726022 | 0.831692822 |
| chr17:79045911-79046179 | 0.204656296 | 0.617738767 | 0.831692822 |
| chr4:38328184-38328590 | -0.189883246 | 0.617739947 | 0.831692822 |
| chr2:60907930-60908610 | -0.193876854 | 0.617759485 | 0.831704514 |
| chr6:43489734-43490114 | 0.198809459 | 0.617822421 | 0.83176183 |
| chr2:62287956-62288171 | -0.221520396 | 0.617823768 | 0.83176183 |
| chr9:32550465-32551250 | -0.163236909 | 0.617876619 | 0.831810091 |
| chr1:149390087-149390888 | -0.167090472 | 0.617881329 | 0.831810091 |
| chr3:197668190-197668424 | 0.235725411 | 0.617908465 | 0.831819921 |
| chr14:101961170-101961440 | -0.194253996 | 0.617911186 | 0.831819921 |
| chr11:74532379-74532839 | 0.19555802 | 0.617923466 | 0.831819921 |
| chr19:18543482-18544238 | 0.181245282 | 0.617932055 | 0.831819921 |
| chr3:86180782-86180982 | 0.262888541 | 0.617948255 | 0.831823883 |
| chr1:201435629-201436226 | -0.164464254 | 0.61795671 | 0.831823883 |
| chr3:68467537-68467810 | 0.218080284 | 0.617988265 | 0.83184607 |
| chr18:12271206-12271420 | -0.23527809 | 0.617994906 | 0.83184607 |
| chr1:149928113-149928481 | -0.17021275 | 0.618015309 | 0.831853691 |
| chr7:34504555-34504928 | -0.211294503 | 0.618022281 | 0.831853691 |
| chr1:27402397-27402614 | -0.233287753 | 0.618043867 | 0.831857341 |
| chr5:66741422-66741833 | 0.189548715 | 0.618046706 | 0.831857341 |
| chr2:9813861-9814170 | -0.18658663 | 0.618083352 | 0.831876937 |
| chr21:14215894-14216766 | -0.171703822 | 0.618100532 | 0.831876937 |
| chr17:75034802-75035545 | 0.159064831 | 0.618108198 | 0.831876937 |
| chr10:73965884-73966383 | 0.193392913 | 0.618111429 | 0.831876937 |
| chr17:65561448-65562155 | -0.165518967 | 0.61811555 | 0.831876937 |
| chr6:29927091-29927489 | -0.202166881 | 0.618146884 | 0.831904495 |
| chr10:43395763-43397541 | -0.157637022 | 0.618184486 | 0.831931077 |
| chr10:32105853-32106113 | -0.239318119 | 0.618188351 | 0.831931077 |
| chr16:80171224-80171544 | -0.245345399 | 0.618230946 | 0.831973788 |
| chr22:37953252-37954012 | -0.171802211 | 0.618251849 | 0.831987305 |
| chr6:50608425-50608680 | 0.225441364 | 0.618281034 | 0.832011967 |
| chr8:109164033-109164292 | 0.228851207 | 0.61830924 | 0.832024224 |
| chr19:2503828-2504103 | 0.228677518 | 0.618332126 | 0.832024224 |
| chr13:45655088-45655289 | -0.252938925 | 0.618332194 | 0.832024224 |
| chr9:97710672-97711176 | 0.191917248 | 0.618349836 | 0.832024224 |
| chr10:71888735-71889297 | 0.188696218 | 0.618354927 | 0.832024224 |
| chr21:33801657-33802112 | 0.188359855 | 0.618360336 | 0.832024224 |
| chr8:73240407-73240681 | 0.217500307 | 0.618366154 | 0.832024224 |
| chr4:34953382-34953739 | 0.184457531 | 0.618386373 | 0.832036817 |
| chr12:66066690-66066890 | 0.263394418 | 0.61845905 | 0.832119993 |

| | | | |
|---|---|---|---|
| chr6:74932660-74932868 | -0.268603445 | 0.618502142 | 0.832163358 |
| chr16:58339520-58339720 | 0.255316693 | 0.618530011 | 0.832175697 |
| chr22:21020901-21021126 | -0.235089377 | 0.618533034 | 0.832175697 |
| chr1:150532606-150532930 | 0.181540478 | 0.618544795 | 0.832176908 |
| chr9:29691852-29692307 | -0.200499261 | 0.61861219 | 0.832252967 |
| chr14:55103021-55103616 | 0.16966011 | 0.618637269 | 0.832272094 |
| chr17:82519019-82519758 | -0.170447966 | 0.618717254 | 0.832360807 |
| chr12:102863466-102863718 | -0.264871043 | 0.61872815 | 0.832360807 |
| chr2:116696265-116696465 | -0.266731393 | 0.6187358 | 0.832360807 |
| chr21:40663096-40663380 | -0.229013395 | 0.618760987 | 0.83236533 |
| chr8:9002600-9003472 | -0.164571642 | 0.618793357 | 0.83236533 |
| chr12:57430343-57431317 | -0.16324237 | 0.618800958 | 0.83236533 |
| chr22:36239222-36239824 | 0.184706598 | 0.618814627 | 0.83236533 |
| chr1:155859238-155859787 | 0.165843322 | 0.618816532 | 0.83236533 |
| chr7:151277077-151277365 | 0.195692561 | 0.618823009 | 0.83236533 |
| chr6:31749778-31750016 | 0.237351209 | 0.618832943 | 0.83236533 |
| chr10:14975757-14976001 | -0.237402013 | 0.61884235 | 0.83236533 |
| chr17:68247321-68248554 | -0.155922867 | 0.618842972 | 0.83236533 |
| chr9:32230408-32230608 | -0.278749324 | 0.618847795 | 0.83236533 |
| chr10:25227266-25227519 | 0.246168329 | 0.618868886 | 0.832379086 |
| chr11:69638984-69639386 | 0.17020027 | 0.618883339 | 0.832383913 |
| chr7:7477701-7478279 | 0.190691503 | 0.6188977 | 0.832388618 |
| chr17:17186135-17186335 | -0.236897247 | 0.618920439 | 0.83240459 |
| chr10:50614224-50614857 | -0.177271896 | 0.618985339 | 0.832476414 |
| chr10:101781134-101781339 | -0.20868043 | 0.619004123 | 0.832476414 |
| chr6:42260582-42261140 | -0.183430399 | 0.619006437 | 0.832476414 |
| chr8:133952105-133952352 | -0.213281642 | 0.619037742 | 0.832498135 |
| chr14:99683629-99684614 | 0.159098853 | 0.619044318 | 0.832498135 |
| chr13:104571643-104571954 | 0.205893208 | 0.619065132 | 0.83250443 |
| chr1:112501770-112502321 | 0.182802198 | 0.619081254 | 0.83250443 |
| chr6:9248137-9248429 | -0.245312082 | 0.619081595 | 0.83250443 |
| chr3:149120231-149120680 | 0.180292782 | 0.61909315 | 0.832505359 |
| chr3:62455040-62455240 | 0.196057759 | 0.619125889 | 0.832534773 |
| chr12:67594929-67595212 | -0.198188703 | 0.619153969 | 0.83255792 |
| chr17:47188897-47189699 | -0.161287748 | 0.619175893 | 0.83257279 |
| chr9:78235659-78236845 | -0.159200931 | 0.619208516 | 0.83258844 |
| chr12:22335097-22336164 | -0.166027785 | 0.619223386 | 0.83258844 |
| chr15:100349670-100350549 | -0.161227653 | 0.6192255 | 0.83258844 |
| chr7:45325438-45325666 | 0.257015626 | 0.619230997 | 0.83258844 |
| chr6:83199756-83200204 | -0.181480763 | 0.619256125 | 0.832607616 |
| chr19:48745266-48745482 | 0.227199323 | 0.619267716 | 0.83260859 |
| chr4:75555460-75555662 | -0.251008144 | 0.619328595 | 0.832661755 |
| chr6:31123013-31123357 | 0.200999182 | 0.619328993 | 0.832661755 |
| chr17:57084661-57086631 | 0.155345878 | 0.619368115 | 0.832696153 |
| chr2:47063677-47063886 | -0.228203764 | 0.619384059 | 0.832696153 |
| chr13:21297810-21298918 | 0.159096163 | 0.619397635 | 0.832696153 |
| chr10:60209940-60210182 | 0.228093457 | 0.619398048 | 0.832696153 |
| chr3:79868022-79868401 | 0.187892679 | 0.619481601 | 0.832793867 |
| chr19:1402728-1403092 | -0.198673368 | 0.619508219 | 0.832798938 |
| chr12:129914674-129914988 | -0.210974129 | 0.619509741 | 0.832798938 |

| | | | |
|---|---|---|---|
| chr13:27770861-27771093 | 0.227036519 | 0.61951798 | 0.832798938 |
| chr13:41239859-41240190 | -0.221481973 | 0.619534535 | 0.832806582 |
| chr5:174972364-174972606 | -0.245000531 | 0.619566086 | 0.832828101 |
| chr20:21027612-21027920 | 0.195717373 | 0.619572282 | 0.832828101 |
| chr8:84181734-84182001 | -0.213635647 | 0.61959009 | 0.832832466 |
| chr17:27380232-27380448 | 0.241684614 | 0.619597268 | 0.832832466 |
| chr2:36812248-36812530 | -0.213582554 | 0.619610008 | 0.83283498 |
| chr2:202178406-202178653 | -0.229233629 | 0.619645105 | 0.832867544 |
| chr7:92057351-92057718 | 0.184551002 | 0.61971327 | 0.832944553 |
| chr11:68798331-68798711 | -0.212794339 | 0.619745792 | 0.832972397 |
| chr19:43595949-43596704 | 0.160462438 | 0.619761865 | 0.832972397 |
| chr2:47344961-47345770 | -0.163346468 | 0.619774077 | 0.832972397 |
| chr3:12984224-12984568 | 0.176270261 | 0.619777471 | 0.832972397 |
| chr18:50410125-50410336 | -0.275269143 | 0.619800168 | 0.832988291 |
| chr15:76931279-76932244 | 0.160562019 | 0.619878648 | 0.833058765 |
| chr11:69122677-69123443 | 0.16147604 | 0.619884275 | 0.833058765 |
| chr2:45786412-45786652 | -0.248809884 | 0.619885222 | 0.833058765 |
| chr13:30979391-30979650 | 0.208088925 | 0.619914055 | 0.833082902 |
| chr8:144147020-144147364 | 0.176883319 | 0.619958013 | 0.833124239 |
| chr1:226348988-226349188 | -0.267579191 | 0.619971378 | 0.833124239 |
| chr1:67713473-67713849 | -0.21049636 | 0.619983121 | 0.833124239 |
| chr9:79134749-79134961 | -0.250439354 | 0.619988308 | 0.833124239 |
| chr17:76272330-76272530 | 0.230324185 | 0.6200299 | 0.833141317 |
| chr9:95509231-95509472 | 0.200973069 | 0.620031493 | 0.833141317 |
| chr14:96583860-96584585 | 0.166572155 | 0.620033636 | 0.833141317 |
| chr10:71860280-71860830 | 0.178373104 | 0.620046751 | 0.833144329 |
| chr12:49609672-49610164 | 0.178184317 | 0.620060879 | 0.833148701 |
| chr14:81324149-81324364 | -0.240528601 | 0.620116697 | 0.83320909 |
| chr17:41666813-41667324 | -0.187848551 | 0.620146076 | 0.833213326 |
| chr12:78562723-78562982 | 0.216973183 | 0.620146186 | 0.833213326 |
| chr7:33895308-33895714 | 0.171580354 | 0.620152473 | 0.833213326 |
| chr15:95024547-95025040 | 0.211632118 | 0.620169344 | 0.833221383 |
| chr16:80614939-80615221 | 0.179725234 | 0.620192301 | 0.833233181 |
| chr10:86824240-86824598 | -0.205376208 | 0.620199874 | 0.833233181 |
| chr3:63309693-63309913 | 0.219007959 | 0.620224021 | 0.833251012 |
| chr1:29490361-29490610 | -0.249759687 | 0.620246282 | 0.833266309 |
| chr3:135503688-135504212 | -0.183077546 | 0.620269265 | 0.833282574 |
| chr10:26598954-26599188 | 0.264902448 | 0.620286745 | 0.833291447 |
| chr21:40718489-40718730 | -0.257015678 | 0.620312854 | 0.833311912 |
| chr3:15098556-15099836 | 0.157091977 | 0.620340013 | 0.833322565 |
| chr14:66932587-66932880 | 0.202259972 | 0.620349186 | 0.833322565 |
| chr8:144316062-144316381 | 0.192938061 | 0.620353412 | 0.833322565 |
| chr17:67162682-67163164 | -0.197706509 | 0.620388335 | 0.833354867 |
| chr20:17968526-17969645 | 0.15750727 | 0.62047859 | 0.833442366 |
| chr17:45425004-45426121 | 0.16255715 | 0.620480055 | 0.833442366 |
| chr14:64966732-64967424 | -0.16032432 | 0.620486105 | 0.833442366 |
| chr10:92243442-92243642 | 0.231064735 | 0.620515444 | 0.833455511 |
| chr2:120825337-120825537 | -0.255421302 | 0.620517646 | 0.833455511 |
| chr11:36564421-36564721 | 0.209213939 | 0.620621673 | 0.833580623 |
| chr11:118622996-118623584 | -0.172321638 | 0.620650708 | 0.833604253 |

| | | | |
|---|---|---|---|
| chr1:88608202-88608421 | -0.259513517 | 0.620661025 | 0.833604253 |
| chr3:158644244-158644959 | -0.159330584 | 0.620710746 | 0.83365642 |
| chr1:201364590-201364838 | 0.202790523 | 0.620726295 | 0.833662691 |
| chr7:1377148-1377348 | 0.247434961 | 0.620747511 | 0.833676572 |
| chr3:128068127-128068350 | 0.219136982 | 0.620769268 | 0.833682769 |
| chr8:143345935-143346165 | 0.258804348 | 0.620773886 | 0.833682769 |
| chr13:28531706-28531906 | 0.234964007 | 0.620873183 | 0.83379887 |
| chr19:49476178-49476572 | 0.188506918 | 0.620882101 | 0.83379887 |
| chr4:39526783-39527000 | -0.208364054 | 0.620906023 | 0.833816381 |
| chr19:36687144-36688069 | 0.163790461 | 0.620961115 | 0.833867656 |
| chr1:196954348-196954548 | 0.255541365 | 0.62096597 | 0.833867656 |
| chr1:56968271-56968549 | -0.234220264 | 0.62100602 | 0.833906822 |
| chrX:131068935-131069402 | 0.19577356 | 0.621037301 | 0.833931393 |
| chr2:38245150-38245359 | -0.231217886 | 0.621056864 | 0.833931393 |
| chr16:636158-637319 | 0.158029756 | 0.621070381 | 0.833931393 |
| chr2:187520292-187520583 | 0.219816071 | 0.621095648 | 0.833931393 |
| chr6:84728617-84728889 | -0.235199568 | 0.621102253 | 0.833931393 |
| chr1:169485181-169486524 | -0.155477982 | 0.621120645 | 0.833931393 |
| chrX:98850522-98851001 | -0.1862109 | 0.621122409 | 0.833931393 |
| chr8:121669811-121670011 | 0.22963588 | 0.621136783 | 0.833931393 |
| chr18:57778512-57778859 | -0.195671019 | 0.621140812 | 0.833931393 |
| chr9:114080844-114081134 | 0.202295829 | 0.621145432 | 0.833931393 |
| chr13:46052179-46053202 | -0.158044455 | 0.621162692 | 0.833931393 |
| chr1:37454890-37455190 | -0.192940991 | 0.621165147 | 0.833931393 |
| chr4:156475280-156475617 | -0.209001129 | 0.621165806 | 0.833931393 |
| chr20:62141748-62142034 | -0.198861213 | 0.621206018 | 0.833970766 |
| chr6:151451879-151452763 | -0.156728283 | 0.621233007 | 0.833992387 |
| chr4:110410866-110411958 | -0.167478798 | 0.621290397 | 0.834003732 |
| chr14:67427166-67427483 | 0.184189406 | 0.621296577 | 0.834003732 |
| chr14:76605091-76605300 | 0.244636911 | 0.621305908 | 0.834003732 |
| chr2:11788800-11789437 | 0.170407168 | 0.621306454 | 0.834003732 |
| chr1:182595735-182595980 | -0.200845739 | 0.62131446 | 0.834003732 |
| chr8:42541497-42542424 | -0.169331048 | 0.621317641 | 0.834003732 |
| chr5:6496046-6496289 | 0.220424 | 0.621317651 | 0.834003732 |
| chr5:10627706-10628197 | -0.165912387 | 0.621346007 | 0.834011835 |
| chr11:128884598-128884990 | 0.192748078 | 0.621354933 | 0.834011835 |
| chr20:18137378-18138396 | 0.161698783 | 0.621362639 | 0.834011835 |
| chr19:38142314-38142711 | 0.194540162 | 0.621372461 | 0.834011835 |
| chr7:10265935-10266214 | -0.230257989 | 0.621385746 | 0.834011835 |
| chr10:100446474-100446760 | -0.262992281 | 0.621388996 | 0.834011835 |
| chr14:90274149-90274448 | 0.189338841 | 0.621416095 | 0.834033597 |
| chr3:97764139-97765189 | -0.157479846 | 0.62147257 | 0.834094785 |
| chr16:593593-593801 | 0.20282129 | 0.62149183 | 0.834106023 |
| chr5:157431919-157432156 | -0.226214388 | 0.621516801 | 0.834124927 |
| chr3:86774175-86774378 | -0.246360182 | 0.621537378 | 0.834137933 |
| chr10:113678907-113680603 | -0.156489303 | 0.621553312 | 0.834144707 |
| chr12:122400476-122400780 | 0.186784499 | 0.621564319 | 0.834144868 |
| chr6:28895481-28896550 | 0.157700392 | 0.621598605 | 0.834167595 |
| chr11:67241080-67241347 | -0.191694562 | 0.621611465 | 0.834167595 |
| chr19:50794303-50794743 | 0.181083873 | 0.621634784 | 0.834167595 |

| | | | |
|---|---|---|---|
| chrX:52920294-52921729 | -0.158168589 | 0.621640209 | 0.834167595 |
| chr2:111883612-111884587 | 0.160789897 | 0.621658593 | 0.834167595 |
| chr9:13088837-13089083 | 0.232265415 | 0.621694196 | 0.834167595 |
| chr8:98546999-98547251 | 0.22018173 | 0.621696273 | 0.834167595 |
| chr8:3726632-3726839 | 0.242628103 | 0.621726403 | 0.834167595 |
| chr7:128937699-128938360 | -0.172147506 | 0.621733862 | 0.834167595 |
| chr22:31884462-31884718 | -0.251331878 | 0.621735199 | 0.834167595 |
| chr9:5304575-5304784 | -0.2361508 | 0.62173639 | 0.834167595 |
| chr11:6288123-6288499 | -0.207862697 | 0.621738365 | 0.834167595 |
| chr1:168888962-168889270 | 0.222698284 | 0.62173865 | 0.834167595 |
| chr8:40201831-40202031 | 0.237575444 | 0.621743928 | 0.834167595 |
| chr16:78925078-78925650 | -0.189757788 | 0.621744556 | 0.834167595 |
| chr15:65463216-65463416 | -0.270600493 | 0.621791333 | 0.834215747 |
| chr3:31724726-31725116 | 0.18782691 | 0.621860961 | 0.834294553 |
| chr1:116233058-116233258 | 0.237540396 | 0.621898337 | 0.834323631 |
| chr6:98539872-98540135 | -0.237291613 | 0.621911722 | 0.834323631 |
| chr2:216059612-216059851 | -0.267028923 | 0.62192552 | 0.834323631 |
| chr19:32692136-32692526 | 0.19416739 | 0.621936025 | 0.834323631 |
| chr15:90868394-90868751 | 0.171165044 | 0.621937078 | 0.834323631 |
| chr17:63990137-63990337 | -0.203846874 | 0.621964615 | 0.834329637 |
| chr17:51259662-51261149 | -0.154020821 | 0.621979416 | 0.834329637 |
| chr14:81990238-81990611 | -0.203644126 | 0.621986155 | 0.834329637 |
| chr6:131950340-131950912 | -0.16932563 | 0.621988933 | 0.834329637 |
| chr3:194313841-194314069 | 0.225311678 | 0.622002548 | 0.834329637 |
| chr10:69317650-69318946 | -0.158366401 | 0.622006889 | 0.834329637 |
| chr14:25833410-25833690 | -0.235786058 | 0.622061832 | 0.834388728 |
| chr17:68456263-68456463 | 0.185265952 | 0.622076864 | 0.8343936 |
| chr1:201385327-201385729 | 0.16455011 | 0.622087244 | 0.8343936 |
| chr17:58711418-58711689 | -0.173991968 | 0.62221476 | 0.834537519 |
| chr14:21455679-21456662 | 0.156080539 | 0.622221987 | 0.834537519 |
| chr17:81263040-81263323 | 0.228794912 | 0.622231513 | 0.834537519 |
| chr13:35826722-35826954 | -0.231408444 | 0.62223811 | 0.834537519 |
| chr10:50067531-50068095 | -0.174382608 | 0.622265881 | 0.834550358 |
| chr8:80103912-80104248 | -0.216777088 | 0.622269467 | 0.834550358 |
| chr19:47231176-47231583 | 0.183023226 | 0.622309132 | 0.834588947 |
| chr1:167066441-167066993 | -0.170203372 | 0.622330422 | 0.834599314 |
| chr17:28719428-28720275 | -0.159928286 | 0.622339126 | 0.834599314 |
| chr5:154038232-154039325 | -0.159611711 | 0.62234954 | 0.834599314 |
| chr1:113978819-113979731 | 0.156019298 | 0.622376833 | 0.834611635 |
| chr6:26537281-26538587 | -0.155431172 | 0.622380512 | 0.834611635 |
| chr12:27333351-27333682 | 0.21176756 | 0.622424601 | 0.834649563 |
| chrX:47619808-47620165 | 0.194806704 | 0.622452635 | 0.834649563 |
| chr16:80336992-80337240 | 0.242336182 | 0.622465148 | 0.834649563 |
| chr3:52006050-52006344 | 0.235046893 | 0.622467294 | 0.834649563 |
| chr8:125476954-125477385 | 0.178130412 | 0.622471087 | 0.834649563 |
| chr1:167714229-167715746 | -0.155066393 | 0.622474154 | 0.834649563 |
| chr20:50509864-50511142 | -0.156007332 | 0.622500882 | 0.834650823 |
| chr8:73312402-73312700 | -0.202399712 | 0.622502172 | 0.834650823 |
| chrX:81209131-81209369 | -0.245590172 | 0.622507773 | 0.834650823 |
| chr11:90222374-90223546 | -0.161088176 | 0.622578959 | 0.834728513 |

| | | | |
|---|---|---|---|
| chr16:20869033-20869328 | -0.244711558 | 0.622591022 | 0.834728513 |
| chr17:8375802-8376927 | -0.164898221 | 0.622598399 | 0.834728513 |
| chr10:117442941-117443280 | -0.191358267 | 0.622684681 | 0.834791457 |
| chr3:186841728-186841994 | 0.214570308 | 0.622685242 | 0.834791457 |
| chr1:151941517-151941721 | -0.20742348 | 0.622698441 | 0.834791457 |
| chr3:87792734-87793584 | 0.172466529 | 0.622709027 | 0.834791457 |
| chr9:92176112-92176361 | -0.214104115 | 0.622735742 | 0.834791457 |
| chr12:3300476-3300848 | -0.184549075 | 0.622739381 | 0.834791457 |
| chr1:207077852-207078151 | -0.178267141 | 0.622739563 | 0.834791457 |
| chr8:70088400-70088669 | -0.235325272 | 0.622750042 | 0.834791457 |
| chr12:57069487-57069797 | -0.208814872 | 0.622753563 | 0.834791457 |
| chr3:167882135-167882412 | -0.208416652 | 0.622759779 | 0.834791457 |
| chr3:195909129-195909984 | 0.163235489 | 0.622776072 | 0.834791457 |
| chr2:74632314-74632624 | -0.228225765 | 0.622776086 | 0.834791457 |
| chr8:105674751-105675141 | -0.205955512 | 0.622798792 | 0.834807289 |
| chr2:9843103-9844384 | -0.156405525 | 0.622826957 | 0.83481705 |
| chr7:30740485-30741196 | 0.162846986 | 0.622827865 | 0.83481705 |
| chr1:24610073-24610273 | 0.240451421 | 0.622907356 | 0.834904888 |
| chr4:118278210-118279192 | -0.155699042 | 0.622921281 | 0.834904888 |
| chr20:266995-267275 | -0.233040403 | 0.622926087 | 0.834904888 |
| chr2:213153687-213154003 | -0.183493327 | 0.622966979 | 0.834940261 |
| chr12:15532469-15532712 | -0.219780008 | 0.622974273 | 0.834940261 |
| chr12:110552383-110552583 | -0.263304456 | 0.623002327 | 0.834958175 |
| chr12:33407181-33407432 | 0.242950053 | 0.623014178 | 0.834958175 |
| chr3:113863630-113863843 | 0.241794476 | 0.62302033 | 0.834958175 |
| chr11:94128263-94129630 | -0.159386942 | 0.623051636 | 0.834971964 |
| chr12:83827833-83828121 | -0.219289835 | 0.623052414 | 0.834971964 |
| chr6:109094321-109095675 | 0.154853437 | 0.623064818 | 0.834973984 |
| chr1:32500204-32501027 | 0.172042602 | 0.623137585 | 0.835056894 |
| chr9:88886770-88887062 | 0.210804631 | 0.623151369 | 0.835060761 |
| chr9:122991495-122991746 | -0.261677258 | 0.623164702 | 0.835064024 |
| chr2:119691661-119692030 | -0.216929349 | 0.623193757 | 0.835077795 |
| chr19:55353930-55354265 | 0.194026943 | 0.623196776 | 0.835077795 |
| chr17:81393758-81394298 | -0.166155992 | 0.623215212 | 0.835087894 |
| chrX:107439598-107439820 | -0.225264227 | 0.623237943 | 0.835103697 |
| chr18:22683495-22683695 | 0.251899645 | 0.623248803 | 0.835103697 |
| chr12:71685496-71686675 | -0.156643464 | 0.623264944 | 0.83511072 |
| chr6:167815382-167816227 | -0.166505053 | 0.623306467 | 0.835140852 |
| chr11:59106594-59107372 | 0.159158713 | 0.623309231 | 0.835140852 |
| chr8:103795317-103795782 | 0.183436909 | 0.623350097 | 0.835163737 |
| chr2:164268054-164268397 | 0.217430919 | 0.623364656 | 0.835163737 |
| chr17:68301316-68301931 | -0.162148815 | 0.62337074 | 0.835163737 |
| chr12:56127806-56128622 | 0.156650024 | 0.623381421 | 0.835163737 |
| chr1:166692213-166692502 | 0.232051492 | 0.623381972 | 0.835163737 |
| chr1:106360350-106360609 | -0.176144319 | 0.62339171 | 0.835163737 |
| chr16:87129943-87130169 | -0.242301256 | 0.623434201 | 0.835206059 |
| chr20:31991097-31991358 | 0.231044412 | 0.62349354 | 0.835253507 |
| chr12:12724170-12724370 | -0.214601366 | 0.623498658 | 0.835253507 |
| chr4:16196535-16196765 | 0.233154109 | 0.623515048 | 0.835253507 |
| chr14:53217090-53218189 | -0.156318927 | 0.623519049 | 0.835253507 |

| | | | |
|---|---|---|---|
| chr3:67311378-67311909 | 0.189989206 | 0.623533511 | 0.835253507 |
| chrX:47836142-47837028 | 0.159667097 | 0.623535024 | 0.835253507 |
| chr13:26051768-26052258 | -0.172651136 | 0.623558621 | 0.835270514 |
| chr4:56978057-56978361 | -0.182980876 | 0.623611142 | 0.835326264 |
| chr6:107318216-107318443 | -0.233717674 | 0.623682423 | 0.83540714 |
| chr18:65876275-65876681 | 0.182114449 | 0.623709814 | 0.835429225 |
| chr3:96813123-96815176 | -0.154546572 | 0.623722414 | 0.835431497 |
| chrX:83845453-83845752 | -0.198267725 | 0.623748669 | 0.835438023 |
| chr2:219642459-219642881 | 0.166577292 | 0.623764585 | 0.835438023 |
| chr19:41570265-41570472 | 0.262468188 | 0.623782632 | 0.835438023 |
| chr1:171017672-171018041 | 0.214146428 | 0.623795307 | 0.835438023 |
| chr8:29340264-29340535 | -0.222337679 | 0.623799152 | 0.835438023 |
| chr9:122099228-122099626 | 0.192074221 | 0.623802832 | 0.835438023 |
| chr7:32489903-32490616 | 0.158426455 | 0.62380361 | 0.835438023 |
| chr1:212413985-212415109 | -0.16174499 | 0.623818732 | 0.835443673 |
| chr16:68840925-68841125 | -0.244181476 | 0.623859891 | 0.835467622 |
| chr11:33808302-33808502 | -0.250155818 | 0.623867873 | 0.835467622 |
| chr11:615601-616157 | 0.175630527 | 0.623872812 | 0.835467622 |
| chr7:36278807-36279242 | 0.183287463 | 0.623880229 | 0.835467622 |
| chr1:220271748-220272909 | -0.156645914 | 0.62390057 | 0.835480259 |
| chr6:169831666-169831961 | -0.197121863 | 0.623913338 | 0.835482756 |
| chr17:50169096-50169662 | -0.166979467 | 0.623952063 | 0.83552001 |
| chr7:150444458-150444658 | 0.244801393 | 0.624007186 | 0.835578271 |
| chr17:79401194-79401406 | 0.233816777 | 0.624018081 | 0.835578271 |
| chr2:72859780-72860055 | -0.199122582 | 0.62403326 | 0.835578271 |
| chr1:49802028-49802866 | -0.17398774 | 0.624039192 | 0.835578271 |
| chr1:171007477-171007700 | 0.268475703 | 0.62405052 | 0.835578837 |
| chr1:218361344-218361673 | -0.183528774 | 0.624069098 | 0.835589111 |
| chr17:19447146-19447423 | -0.211336428 | 0.624085246 | 0.835594183 |
| chr1:15526382-15526964 | 0.16107728 | 0.624096476 | 0.835594183 |
| chr3:132253262-132253514 | -0.215364348 | 0.624116391 | 0.835594183 |
| chr5:174800103-174800357 | -0.234280633 | 0.624116509 | 0.835594183 |
| chr16:85086732-85087068 | 0.183095806 | 0.624249361 | 0.835740067 |
| chr5:21191809-21192102 | -0.205693742 | 0.624260983 | 0.835740067 |
| chr2:74587675-74587915 | -0.216044995 | 0.624271055 | 0.835740067 |
| chr3:64551448-64552381 | -0.156235328 | 0.624276689 | 0.835740067 |
| chr12:55996823-55997645 | -0.161203777 | 0.624280007 | 0.835740067 |
| chr7:34781776-34782055 | 0.214350671 | 0.624314651 | 0.835771842 |
| chr5:152181344-152181781 | 0.185478544 | 0.62436272 | 0.835818257 |
| chr11:73964325-73964761 | 0.17083534 | 0.624373009 | 0.835818257 |
| chr1:189775354-189775712 | 0.199840815 | 0.624387064 | 0.835818257 |
| chr10:67700069-67700269 | -0.249277379 | 0.624397348 | 0.835818257 |
| chr6:4344578-4344778 | -0.256782208 | 0.624421443 | 0.835818257 |
| chr6:22471069-22471269 | -0.256782208 | 0.624421443 | 0.835818257 |
| chr13:45654653-45654927 | -0.226161315 | 0.624435186 | 0.835818257 |
| chr1:58163010-58163359 | -0.196640553 | 0.624436589 | 0.835818257 |
| chr13:46672508-46672748 | 0.20118193 | 0.62445154 | 0.835820068 |
| chr10:59622335-59622671 | -0.201000993 | 0.624459758 | 0.835820068 |
| chr14:50882975-50883472 | 0.178550931 | 0.624555054 | 0.835886434 |
| chr1:27500062-27500416 | 0.227339626 | 0.624556043 | 0.835886434 |

| | | | |
|---|---|---|---|
| chr14:24215483-24216229 | 0.159745705 | 0.62456389 | 0.835886434 |
| chr5:157858293-157859369 | -0.161516365 | 0.624586832 | 0.835886434 |
| chr14:70665417-70666319 | 0.16135923 | 0.62458686 | 0.835886434 |
| chr10:29672715-29672953 | -0.256743272 | 0.624601974 | 0.835886434 |
| chr7:332569-333335 | 0.166853244 | 0.62460488 | 0.835886434 |
| chr7:45628276-45628481 | -0.261438374 | 0.624613432 | 0.835886434 |
| chr4:40924579-40924839 | -0.207342879 | 0.624642326 | 0.835886434 |
| chr16:27639809-27640050 | -0.21147376 | 0.624645023 | 0.835886434 |
| chr2:30955900-30956156 | 0.243223782 | 0.624645768 | 0.835886434 |
| chr1:173476634-173478039 | -0.153827755 | 0.624650763 | 0.835886434 |
| chr11:10366734-10367036 | 0.217445607 | 0.624651162 | 0.835886434 |
| chr6:29017169-29017405 | 0.214083599 | 0.624710942 | 0.83595183 |
| chr17:68465943-68466436 | -0.169241967 | 0.624744607 | 0.835954234 |
| chr1:1157431-1158202 | 0.171363154 | 0.624751861 | 0.835954234 |
| chr10:5009219-5009439 | 0.231136978 | 0.624754816 | 0.835954234 |
| chr8:96775758-96775985 | 0.236862781 | 0.624758793 | 0.835954234 |
| chr20:20664858-20665058 | 0.24177858 | 0.624767289 | 0.835954234 |
| chr7:26200243-26200483 | -0.175650257 | 0.62478492 | 0.835963227 |
| chr1:168342708-168343015 | -0.198945575 | 0.624801223 | 0.835966905 |
| chr3:10703043-10703366 | -0.182959043 | 0.62480949 | 0.835966905 |
| chr11:64743574-64744108 | 0.172892099 | 0.624824075 | 0.835971823 |
| chr1:247424737-247424937 | 0.2195528 | 0.624838485 | 0.835976505 |
| chr5:175867697-175868639 | -0.159210628 | 0.624850256 | 0.835977656 |
| chr10:99845774-99846088 | -0.196828869 | 0.624872528 | 0.835980978 |
| chr7:12583984-12584289 | 0.221980602 | 0.62487456 | 0.835980978 |
| chr4:139028427-139028708 | 0.206826273 | 0.624898031 | 0.835997782 |
| chr3:42476629-42477184 | 0.170619504 | 0.624927672 | 0.836022839 |
| chr6:118776583-118776918 | -0.194627287 | 0.624941112 | 0.836026222 |
| chr1:243850986-243851268 | 0.213245103 | 0.624989068 | 0.836075779 |
| chr14:45426667-45427033 | 0.179841708 | 0.625021869 | 0.83610506 |
| chr1:33430391-33431523 | 0.156457128 | 0.625039045 | 0.836113439 |
| chr10:97081686-97082146 | 0.192439842 | 0.625051754 | 0.836115843 |
| chr9:84255095-84255488 | 0.200393559 | 0.625092692 | 0.836150371 |
| chr16:33336791-33337016 | 0.250562735 | 0.625099391 | 0.836150371 |
| chr10:90986348-90986599 | -0.2303407 | 0.625113519 | 0.836154672 |
| chr1:223123763-223123984 | -0.231420057 | 0.62513765 | 0.83617163 |
| chr18:51621237-51621437 | -0.256270374 | 0.625148023 | 0.83617163 |
| chr6:24645681-24646752 | -0.162761386 | 0.625174994 | 0.836180774 |
| chr1:107676172-107676392 | -0.236902633 | 0.62518452 | 0.836180774 |
| chr14:72615435-72615755 | -0.199004019 | 0.625187598 | 0.836180774 |
| chr16:1420413-1421208 | 0.162050535 | 0.62521778 | 0.836201566 |
| chr12:56710185-56710452 | -0.237868923 | 0.625224971 | 0.836201566 |
| chr3:115814809-115815129 | -0.193825685 | 0.625243339 | 0.836211536 |
| chr2:41782420-41782897 | 0.181162973 | 0.625291271 | 0.836260093 |
| chr19:54448374-54449981 | 0.155771991 | 0.625317847 | 0.836260093 |
| chr19:14536912-14537587 | 0.171450554 | 0.625318036 | 0.836260093 |
| chr19:35268045-35268485 | 0.183203395 | 0.625342879 | 0.836260093 |
| chr6:15036900-15037182 | -0.235940633 | 0.62534544 | 0.836260093 |
| chr2:101401958-101402220 | -0.235905404 | 0.625353178 | 0.836260093 |
| chr5:150635031-150635299 | -0.212870636 | 0.625359638 | 0.836260093 |

| | | | |
|---|---|---|---|
| chr6:121686293-121686564 | 0.245918112 | 0.625379802 | 0.836260093 |
| chr17:9744920-9745120 | -0.251229843 | 0.625383291 | 0.836260093 |
| chr3:164686327-164686536 | -0.252475964 | 0.625399642 | 0.836260093 |
| chr1:155186882-155187123 | 0.206454824 | 0.625399701 | 0.836260093 |
| chr8:139405252-139406097 | -0.166582595 | 0.625415375 | 0.836264688 |
| chr2:85290746-85290946 | 0.261815435 | 0.625430593 | 0.836264688 |
| chr16:88559080-88559335 | -0.196611271 | 0.625435879 | 0.836264688 |
| chr17:57448397-57448622 | -0.207239535 | 0.625471724 | 0.836280124 |
| chr11:95923747-95924620 | -0.169525202 | 0.625477163 | 0.836280124 |
| chr1:44739374-44740400 | -0.160252152 | 0.625480167 | 0.836280124 |
| chr17:74203355-74204090 | -0.159493355 | 0.625497242 | 0.836288361 |
| chr18:79396182-79396664 | 0.183594888 | 0.625516147 | 0.836299044 |
| chr2:155356282-155356702 | -0.199191292 | 0.625541929 | 0.836318921 |
| chr19:3710371-3710751 | 0.19356769 | 0.625582943 | 0.83635127 |
| chr2:120228579-120228838 | -0.223596119 | 0.625587956 | 0.83635127 |
| chr20:63183341-63183620 | -0.18796417 | 0.625665862 | 0.836440828 |
| chr1:202643634-202643874 | 0.203027962 | 0.625701805 | 0.836474286 |
| chr8:114230127-114230327 | -0.259768475 | 0.625714925 | 0.836477231 |
| chr5:98975288-98975563 | -0.232258742 | 0.625762542 | 0.836493451 |
| chr1:224851533-224851792 | 0.210895617 | 0.625769424 | 0.836493451 |
| chrX:74535779-74536069 | 0.188069742 | 0.625772552 | 0.836493451 |
| chr20:8142152-8142468 | -0.226952338 | 0.625798868 | 0.836493451 |
| chrX:108072077-108072297 | 0.221290903 | 0.625800748 | 0.836493451 |
| chr3:137521000-137521255 | 0.255158367 | 0.625811545 | 0.836493451 |
| chr9:89526077-89526323 | -0.247656778 | 0.625812498 | 0.836493451 |
| chr22:23173881-23174243 | -0.186056295 | 0.625823515 | 0.836493451 |
| chr2:200124653-200124853 | 0.24533116 | 0.625856788 | 0.836493451 |
| chr2:224331759-224331959 | 0.24533116 | 0.625856788 | 0.836493451 |
| chr14:24806240-24806483 | -0.218709938 | 0.625857422 | 0.836493451 |
| chr1:207320959-207322188 | -0.15264612 | 0.625858064 | 0.836493451 |
| chr5:16195462-16195662 | -0.241583112 | 0.625888099 | 0.836493468 |
| chr5:95239093-95239295 | 0.232634915 | 0.625893359 | 0.836493468 |
| chr8:103124878-103125170 | -0.188003237 | 0.625899377 | 0.836493468 |
| chr17:59830772-59831450 | 0.156368226 | 0.625909716 | 0.836493468 |
| chr6:157274310-157275397 | -0.159496637 | 0.625912662 | 0.836493468 |
| chr14:53513362-53513703 | 0.241969201 | 0.625927752 | 0.836499044 |
| chr18:28995348-28995587 | 0.22039332 | 0.625974881 | 0.836547438 |
| chr1:161420776-161421150 | 0.18499458 | 0.625987135 | 0.836549223 |
| chr10:22340175-22341466 | 0.162752944 | 0.626020596 | 0.836561695 |
| chrX:150503905-150504116 | 0.229287207 | 0.626023935 | 0.836561695 |
| chr22:41800383-41801166 | 0.172852089 | 0.626033419 | 0.836561695 |
| chr15:75579144-75579822 | -0.169466627 | 0.62604014 | 0.836561695 |
| chr6:104896994-104897194 | -0.277226959 | 0.626133292 | 0.836671581 |
| chr4:7049766-7049978 | -0.244186726 | 0.626207551 | 0.8367428 |
| chr3:146817708-146817997 | -0.208970812 | 0.626208431 | 0.8367428 |
| chr8:120622763-120623178 | 0.190998041 | 0.626223842 | 0.836748182 |
| chr15:37143414-37143689 | 0.207970959 | 0.6262343 | 0.836748182 |
| chr16:68645106-68645951 | 0.165818086 | 0.626254902 | 0.836751632 |
| chr3:14947105-14948726 | 0.152418867 | 0.626258722 | 0.836751632 |
| chr9:98998903-98999215 | 0.190761496 | 0.626322023 | 0.836799895 |

| | | | |
|---|---|---|---|
| chr5:153959730-153959930 | -0.225782713 | 0.626322216 | 0.836799895 |
| chr5:74327117-74327317 | -0.286382129 | 0.626327608 | 0.836799895 |
| chr12:57604632-57605450 | 0.166236637 | 0.626365247 | 0.836821446 |
| chr8:106521825-106522152 | -0.180793826 | 0.626365581 | 0.836821446 |
| chr14:77181215-77182436 | 0.154934932 | 0.626422903 | 0.836867073 |
| chr2:20213093-20213515 | 0.168418709 | 0.626427102 | 0.836867073 |
| chr19:10194537-10195245 | 0.161183214 | 0.626434605 | 0.836867073 |
| chr6:142329590-142329950 | 0.220207693 | 0.626448417 | 0.836867073 |
| chr8:94896507-94896742 | 0.179817175 | 0.626454343 | 0.836867073 |
| chr6:149061050-149061729 | -0.170564047 | 0.626478921 | 0.83687192 |
| chr12:6264657-6265186 | 0.180685423 | 0.626488157 | 0.83687192 |
| chr15:80140751-80140998 | 0.262435232 | 0.626499202 | 0.83687192 |
| chr11:65497310-65497941 | -0.160885314 | 0.62650166 | 0.83687192 |
| chr8:120811987-120812465 | -0.16898272 | 0.626515257 | 0.836875493 |
| chr8:95386643-95386861 | -0.220550254 | 0.626529253 | 0.836879598 |
| chr2:189555333-189555638 | -0.205332219 | 0.626600329 | 0.836948338 |
| chr10:119973215-119973660 | 0.170871893 | 0.626602561 | 0.836948338 |
| chr3:7543373-7543578 | -0.244710103 | 0.626664943 | 0.837017071 |
| chr14:19801125-19801325 | -0.26666765 | 0.626705008 | 0.837055992 |
| chr3:142976617-142976901 | -0.203897812 | 0.626770701 | 0.837128862 |
| chr2:28452482-28452806 | 0.186200652 | 0.626788334 | 0.837128862 |
| chr17:63013174-63013470 | -0.221121092 | 0.626792342 | 0.837128862 |
| chr1:185316150-185316946 | -0.157806031 | 0.626820033 | 0.83714421 |
| chr1:204383633-204383937 | 0.188686662 | 0.626825685 | 0.83714421 |
| chr18:22913686-22914248 | 0.167485426 | 0.626864607 | 0.837161784 |
| chr2:152126720-152127009 | -0.228359461 | 0.626880598 | 0.837161784 |
| chr16:19284233-19284502 | 0.218261599 | 0.626880933 | 0.837161784 |
| chr15:93997633-93997833 | 0.27364983 | 0.626882547 | 0.837161784 |
| chr10:29318624-29318829 | 0.25660294 | 0.626901488 | 0.837168049 |
| chr11:7604441-7604938 | -0.174652971 | 0.62690909 | 0.837168049 |
| chr10:80143732-80144669 | -0.160675865 | 0.626947362 | 0.837194515 |
| chr1:10794441-10794641 | 0.245115289 | 0.626956935 | 0.837194515 |
| chr1:181038401-181038785 | -0.16652454 | 0.626961688 | 0.837194515 |
| chr10:22714383-22715024 | -0.178904074 | 0.626983131 | 0.837195048 |
| chr2:144331764-144333188 | -0.15341843 | 0.62698394 | 0.837195048 |
| chr2:11477962-11478232 | -0.198612016 | 0.627031161 | 0.837221144 |
| chr20:32790144-32790344 | 0.238990887 | 0.627033835 | 0.837221144 |
| chr15:81813302-81813509 | 0.227574997 | 0.627036263 | 0.837221144 |
| chr14:51239929-51240732 | 0.155844942 | 0.627051427 | 0.837226801 |
| chr10:69914676-69914940 | -0.209944826 | 0.627063994 | 0.837228991 |
| chr22:44975380-44975817 | 0.199954862 | 0.627097219 | 0.837258763 |
| chr17:44345541-44346056 | 0.171735375 | 0.627128035 | 0.837282081 |
| chr1:115782865-115783295 | 0.197996666 | 0.627137693 | 0.837282081 |
| chr5:178232924-178233124 | -0.203869558 | 0.627147467 | 0.837282081 |
| chr4:153142999-153143495 | -0.178930496 | 0.627192231 | 0.837313453 |
| chr16:48244115-48244787 | 0.172033999 | 0.627192821 | 0.837313453 |
| chr12:53231025-53231225 | 0.202960892 | 0.627222242 | 0.837335648 |
| chr15:67998749-67999135 | 0.211662891 | 0.627231303 | 0.837335648 |
| chr4:37999427-37999710 | -0.215795578 | 0.627256909 | 0.837337748 |
| chr4:140094913-140095227 | -0.215795578 | 0.627256909 | 0.837337748 |

| | | | |
|---|---|---|---|
| chr10:114507488-114507688 | -0.251082771 | 0.62726566 | 0.837337748 |
| chr1:110830009-110830351 | -0.190419664 | 0.62729475 | 0.837351269 |
| chr7:44138863-44139182 | -0.195302873 | 0.627297646 | 0.837351269 |
| chr14:61475334-61475599 | 0.209502319 | 0.627338436 | 0.837386536 |
| chr9:32581601-32581934 | -0.206712293 | 0.627351082 | 0.837386536 |
| chr3:157208603-157208880 | -0.245264019 | 0.627356852 | 0.837386536 |
| chr3:122247605-122247828 | 0.24301995 | 0.627381497 | 0.83739066 |
| chr6:79077760-79078984 | -0.155510725 | 0.627392248 | 0.83739066 |
| chr7:151172615-151173128 | 0.175366362 | 0.62740087 | 0.83739066 |
| chr3:171829931-171830334 | -0.168545963 | 0.627403657 | 0.83739066 |
| chr3:133041271-133041535 | -0.197923054 | 0.627430999 | 0.837412567 |
| chr2:67003025-67003225 | 0.284450463 | 0.627455278 | 0.837422493 |
| chr18:39523222-39523543 | -0.232111693 | 0.627460295 | 0.837422493 |
| chr11:83956500-83956700 | -0.261845971 | 0.627514106 | 0.837463129 |
| chr17:41835358-41835721 | -0.225928994 | 0.627517086 | 0.837463129 |
| chr14:46048159-46048677 | -0.1703258 | 0.627529724 | 0.837463129 |
| chr6:100493250-100493847 | 0.181864568 | 0.627534462 | 0.837463129 |
| chr11:2924441-2924740 | 0.190729789 | 0.627554977 | 0.837473356 |
| chr14:33676066-33676527 | 0.174163907 | 0.62756674 | 0.837473356 |
| chr20:56629534-56631169 | 0.151590302 | 0.627574916 | 0.837473356 |
| chr14:24678887-24679243 | 0.185942275 | 0.62766127 | 0.837574006 |
| chr2:96554485-96554691 | -0.216795668 | 0.627686256 | 0.837592761 |
| chr19:51702157-51702662 | -0.166789629 | 0.627781423 | 0.837692935 |
| chrX:3026194-3026537 | -0.211568113 | 0.627783192 | 0.837692935 |
| chr2:39180102-39180682 | -0.17128107 | 0.627810012 | 0.8377103 |
| chr14:79997963-79998255 | 0.212639638 | 0.627818071 | 0.8377103 |
| chr12:79105002-79105228 | -0.257317093 | 0.627907122 | 0.837799172 |
| chr10:93465709-93466723 | -0.155454513 | 0.627917207 | 0.837799172 |
| chr5:14489716-14490153 | 0.175314559 | 0.627917479 | 0.837799172 |
| chr10:43455042-43456058 | 0.156679477 | 0.627944994 | 0.837809832 |
| chr6:147202390-147204279 | -0.153305997 | 0.627959896 | 0.837809832 |
| chr4:56907501-56909638 | 0.153190778 | 0.627968337 | 0.837809832 |
| chr3:9749340-9750450 | 0.152599515 | 0.627969205 | 0.837809832 |
| chr3:13479454-13480683 | 0.153424367 | 0.62799204 | 0.837822585 |
| chr20:34312567-34313041 | -0.181781009 | 0.628000634 | 0.837822585 |
| chr20:31861496-31862296 | 0.162052504 | 0.628055052 | 0.837880597 |
| chr2:113831206-113831547 | 0.213757307 | 0.628097244 | 0.837900389 |
| chr5:4868172-4868425 | 0.211966416 | 0.628102304 | 0.837900389 |
| chr16:72066778-72066999 | -0.221107068 | 0.628112506 | 0.837900389 |
| chr18:596628-597456 | -0.161417958 | 0.628119818 | 0.837900389 |
| chr1:47175420-47175624 | -0.216445038 | 0.628133822 | 0.837900389 |
| chr15:73897440-73897640 | 0.219796683 | 0.628146166 | 0.837900389 |
| chr2:214809002-214810340 | -0.151949244 | 0.628149188 | 0.837900389 |
| chr4:108168557-108169325 | 0.162920137 | 0.628161488 | 0.837900389 |
| chr10:55450100-55450300 | -0.242841677 | 0.628168307 | 0.837900389 |
| chr22:26445500-26445710 | -0.249143394 | 0.628181767 | 0.837903756 |
| chr9:21813885-21814145 | -0.200295652 | 0.628197998 | 0.837910819 |
| chrX:109997898-109998136 | -0.206800565 | 0.628220884 | 0.837925778 |
| chr1:173420193-173420667 | -0.170391129 | 0.628251866 | 0.837925778 |
| chr5:67963992-67964500 | 0.185594424 | 0.628261461 | 0.837925778 |

| | | | |
|---|---|---|---|
| chr21:22855684-22855949 | -0.222781141 | 0.628263718 | 0.837925778 |
| chr8:100116710-100116955 | 0.253301199 | 0.628263893 | 0.837925778 |
| chr14:21308178-21309152 | -0.153801273 | 0.62828317 | 0.837936903 |
| chr9:131336910-131337120 | 0.234403877 | 0.628297485 | 0.83794141 |
| chr3:66971868-66972150 | -0.198059406 | 0.628318895 | 0.837955379 |
| chr2:7037790-7038089 | -0.194438187 | 0.628346822 | 0.837978038 |
| chr2:207768945-207770518 | -0.152596376 | 0.628360874 | 0.837982193 |
| chr8:72944020-72944340 | -0.192648217 | 0.628442963 | 0.838073477 |
| chr1:64915766-64916265 | 0.163065939 | 0.628451199 | 0.838073477 |
| chr9:27333213-27333533 | 0.197936511 | 0.628530289 | 0.838136265 |
| chr2:42809783-42811374 | -0.151223562 | 0.628538463 | 0.838136265 |
| chr5:130995447-130995647 | -0.262828823 | 0.628541952 | 0.838136265 |
| chr17:57106575-57106848 | 0.187257188 | 0.628542037 | 0.838136265 |
| chr2:24047063-24047931 | -0.156178891 | 0.628556325 | 0.838140517 |
| chr20:45548110-45548357 | 0.228859006 | 0.628567841 | 0.838140517 |
| chr7:2519546-2519860 | 0.170873356 | 0.628578041 | 0.838140517 |
| chr18:58183290-58183694 | -0.182908599 | 0.628594941 | 0.838148465 |
| chr15:74616105-74616305 | 0.199044624 | 0.628626293 | 0.838175683 |
| chr14:100703667-100704391 | 0.165086993 | 0.628658256 | 0.838198368 |
| chr7:151123930-151124274 | 0.174620531 | 0.628665605 | 0.838198368 |
| chr17:63780474-63780879 | 0.178648954 | 0.628676124 | 0.838198368 |
| chr19:45485113-45486088 | 0.15835497 | 0.628708494 | 0.838221681 |
| chr15:88465904-88466157 | 0.210412596 | 0.628722643 | 0.838221681 |
| chrX:74251554-74251902 | 0.190122336 | 0.628726429 | 0.838221681 |
| chr12:10009934-10010320 | -0.205681174 | 0.628770202 | 0.838253352 |
| chr18:37322402-37322646 | 0.207512997 | 0.628772065 | 0.838253352 |
| chr20:54195455-54195958 | 0.154820273 | 0.628807476 | 0.838285975 |
| chr6:170129743-170130132 | 0.181682966 | 0.62884933 | 0.838327015 |
| chr11:62921823-62922424 | 0.169890072 | 0.628870454 | 0.838327015 |
| chr20:58965892-58966221 | -0.193001346 | 0.628873685 | 0.838327015 |
| chr17:57518722-57518922 | 0.21948926 | 0.628886829 | 0.838327015 |
| chr8:122721639-122722142 | 0.171701973 | 0.628892966 | 0.838327015 |
| chr2:113308741-113309099 | 0.21334342 | 0.628925964 | 0.838352406 |
| chr4:73069042-73070295 | -0.152328613 | 0.628941065 | 0.838352406 |
| chr19:49851391-49851591 | -0.225820816 | 0.628960456 | 0.838352406 |
| chr17:27805465-27805918 | 0.179351124 | 0.628975996 | 0.838352406 |
| chr12:109692567-109692805 | -0.224862561 | 0.628992109 | 0.838352406 |
| chr5:148957714-148957981 | -0.215474707 | 0.628995415 | 0.838352406 |
| chr19:42253840-42254761 | -0.163298661 | 0.629001403 | 0.838352406 |
| chr8:126688069-126688278 | -0.259660432 | 0.629003035 | 0.838352406 |
| chr5:149834299-149834547 | -0.199255968 | 0.629011575 | 0.838352406 |
| chr8:94894900-94896432 | 0.151250299 | 0.629021427 | 0.838352406 |
| chr19:37719624-37720061 | 0.173195101 | 0.629059745 | 0.838388892 |
| chr11:74749160-74749633 | -0.167786399 | 0.629086666 | 0.838410188 |
| chr15:41661794-41662046 | 0.218453671 | 0.62915478 | 0.838478599 |
| chr6:79630688-79631745 | 0.158665454 | 0.629159883 | 0.838478599 |
| chr16:11303328-11303947 | 0.171284812 | 0.629171128 | 0.838479001 |
| chr7:73769702-73769916 | -0.201726401 | 0.629242799 | 0.83855993 |
| chr4:72111000-72111200 | 0.23736467 | 0.629285504 | 0.838590987 |
| chr15:66046965-66047218 | 0.227650911 | 0.629287993 | 0.838590987 |

| | | | |
|---|---|---|---|
| chr3:143698043-143698270 | 0.234876718 | 0.629306497 | 0.838601061 |
| chr8:87502024-87502224 | 0.260288892 | 0.629372812 | 0.838661977 |
| chr17:78585804-78586057 | 0.23604641 | 0.629391012 | 0.838661977 |
| chr5:171405622-171405826 | 0.23604641 | 0.629391012 | 0.838661977 |
| chr2:180077391-180077618 | 0.244405236 | 0.629404024 | 0.838661977 |
| chr8:91343369-91343587 | 0.248940181 | 0.629411728 | 0.838661977 |
| chr8:106060558-106060779 | -0.235583162 | 0.629417882 | 0.838661977 |
| chr14:99480733-99481795 | 0.152070073 | 0.629467503 | 0.838713509 |
| chr11:58788396-58788746 | -0.182454335 | 0.629532706 | 0.8387858 |
| chr6:42961041-42961348 | -0.203747192 | 0.629552887 | 0.838798102 |
| chr20:62489480-62489859 | 0.179648682 | 0.629565033 | 0.8387997 |
| chr17:39199904-39200512 | -0.171765219 | 0.629607772 | 0.83883184 |
| chr3:16478611-16478917 | 0.19284711 | 0.629611051 | 0.83883184 |
| chr1:202808131-202808428 | 0.172004137 | 0.629623765 | 0.838834193 |
| chr5:140114295-140114943 | -0.171456438 | 0.629645847 | 0.838845965 |
| chr2:84881735-84881941 | 0.235619731 | 0.629654496 | 0.838845965 |
| chr7:25951100-25951806 | 0.16423229 | 0.629732236 | 0.838921114 |
| chr1:205703177-205703493 | 0.185448202 | 0.629753638 | 0.838921114 |
| chr1:97192812-97193486 | 0.169610986 | 0.629755402 | 0.838921114 |
| chr11:104254447-104254737 | -0.206355583 | 0.629759773 | 0.838921114 |
| chr2:84882161-84882503 | -0.190501803 | 0.629767487 | 0.838921114 |
| chr16:70065460-70066435 | -0.161589599 | 0.629776597 | 0.838921114 |
| chr13:41273697-41273969 | -0.246896694 | 0.629792428 | 0.838927616 |
| chr5:135834488-135835046 | -0.171795437 | 0.629825538 | 0.838957135 |
| chr1:176897415-176897637 | -0.221758527 | 0.629849186 | 0.838974051 |
| chr11:75399290-75400055 | 0.158447594 | 0.629873448 | 0.838991783 |
| chr1:233635924-233636469 | -0.174814062 | 0.629887389 | 0.838995768 |
| chr1:180416097-180416360 | -0.215590573 | 0.62990712 | 0.83899762 |
| chr1:147243434-147243652 | 0.18387454 | 0.629920687 | 0.83899762 |
| chrX:12790865-12792219 | -0.15851919 | 0.629926349 | 0.83899762 |
| chr15:66853454-66853812 | -0.203127706 | 0.629947555 | 0.83899762 |
| chr6:85580120-85580344 | -0.227212677 | 0.629953372 | 0.83899762 |
| chr4:110344217-110344425 | -0.244207411 | 0.629974001 | 0.83899762 |
| chr20:46971166-46971652 | 0.160676502 | 0.629977188 | 0.83899762 |
| chr12:82740724-82740984 | 0.227749415 | 0.629982964 | 0.83899762 |
| chr11:3130711-3130977 | 0.202054712 | 0.629987329 | 0.83899762 |
| chr10:86344718-86344954 | -0.197031361 | 0.630026196 | 0.839019367 |
| chr15:43798708-43799277 | -0.162110867 | 0.630042178 | 0.839019367 |
| chr1:47001811-47002040 | -0.241467526 | 0.630045339 | 0.839019367 |
| chr3:150521583-150521838 | -0.222439051 | 0.630054864 | 0.839019367 |
| chr6:156752620-156753324 | -0.177979998 | 0.630058408 | 0.839019367 |
| chr9:101533335-101534401 | -0.153653445 | 0.630102808 | 0.839044213 |
| chr2:202376581-202377001 | 0.1695713 | 0.630112305 | 0.839044213 |
| chr14:25381619-25381819 | -0.245680741 | 0.630124219 | 0.839044213 |
| chr12:7018109-7019115 | -0.154978554 | 0.630126075 | 0.839044213 |
| chr20:42589912-42590215 | 0.187864114 | 0.630131818 | 0.839044213 |
| chr5:174532525-174532942 | -0.167609627 | 0.630143682 | 0.839045428 |
| chr5:134019694-134020003 | -0.197077705 | 0.630239176 | 0.839143295 |
| chr8:108603261-108603461 | -0.243168854 | 0.63026234 | 0.839143295 |
| chr11:73907295-73907523 | -0.256416446 | 0.630268287 | 0.839143295 |

| | | | |
|---|---|---|---|
| chr12:52870336-52870836 | 0.165732964 | 0.630271201 | 0.839143295 |
| chr12:55737116-55737542 | -0.165769907 | 0.63028772 | 0.839143295 |
| chr4:170071932-170072209 | -0.24562776 | 0.630289537 | 0.839143295 |
| chr16:63617139-63617544 | 0.195833339 | 0.630293844 | 0.839143295 |
| chr17:75271049-75271402 | 0.174525233 | 0.630313875 | 0.839155383 |
| chr17:68435983-68436763 | -0.155070772 | 0.630373751 | 0.839220515 |
| chr3:158105676-158106043 | 0.174952062 | 0.630405653 | 0.839248405 |
| chr16:80615740-80616121 | -0.169419997 | 0.630434587 | 0.839272343 |
| chr1:188093942-188094180 | -0.245367309 | 0.630500002 | 0.839344843 |
| chr6:5143676-5143920 | 0.221753334 | 0.630538774 | 0.839381874 |
| chr1:21345410-21346065 | -0.168980139 | 0.630562399 | 0.839385502 |
| chr15:92803726-92804500 | -0.155417839 | 0.630564197 | 0.839385502 |
| chr5:23332746-23332946 | 0.235352043 | 0.630574363 | 0.839385502 |
| chr13:41646836-41647036 | 0.258684241 | 0.630640393 | 0.839437124 |
| chr16:74730025-74730689 | -0.164534435 | 0.630652865 | 0.839437124 |
| chr11:68843797-68844195 | 0.166945343 | 0.630654816 | 0.839437124 |
| chr22:20157007-20157359 | 0.181606197 | 0.630656966 | 0.839437124 |
| chr11:58923258-58923515 | -0.220233827 | 0.630683071 | 0.839448843 |
| chr9:122261527-122261843 | 0.17863399 | 0.630687682 | 0.839448843 |
| chr1:91021208-91022507 | 0.154631889 | 0.630700032 | 0.839450699 |
| chr10:95132992-95133241 | 0.214347036 | 0.630738966 | 0.839487938 |
| chr1:52032520-52032789 | -0.204644907 | 0.630785207 | 0.839525659 |
| chr3:87900178-87900450 | 0.224167906 | 0.630789222 | 0.839525659 |
| chr5:121378732-121379105 | -0.199012405 | 0.630812073 | 0.839540755 |
| chr1:179156064-179156272 | 0.244584498 | 0.63082342 | 0.839540755 |
| chr20:9149208-9149431 | -0.251421029 | 0.630833435 | 0.839540755 |
| chr17:66204193-66204588 | 0.187698901 | 0.630847878 | 0.839545395 |
| chr1:241531276-241531846 | 0.171543857 | 0.63088061 | 0.839572972 |
| chr12:23055119-23055321 | -0.245243494 | 0.630890515 | 0.839572972 |
| chr17:20008770-20009808 | 0.160926242 | 0.630914115 | 0.839582981 |
| chr10:101587820-101588471 | -0.163982017 | 0.630926753 | 0.839582981 |
| chr3:194672047-194672736 | 0.160686999 | 0.630945284 | 0.839582981 |
| chr2:138476936-138477442 | 0.190050988 | 0.630952206 | 0.839582981 |
| chr13:76039001-76039260 | -0.22368603 | 0.630952823 | 0.839582981 |
| chr14:56181664-56182144 | 0.183518156 | 0.630967014 | 0.839587283 |
| chr14:50039740-50040701 | -0.159397548 | 0.631031416 | 0.839658397 |
| chr13:19818181-19818616 | -0.166582301 | 0.631063275 | 0.839686207 |
| chr15:82045817-82047627 | 0.151060594 | 0.631078689 | 0.839687951 |
| chr4:151532457-151532695 | -0.240215781 | 0.631086503 | 0.839687951 |
| chr11:134360256-134360604 | -0.190052387 | 0.631128738 | 0.839706049 |
| chr12:95398155-95398439 | -0.250443082 | 0.631139695 | 0.839706049 |
| chr13:107211112-107211325 | -0.250443082 | 0.631139695 | 0.839706049 |
| chr19:14005676-14006061 | -0.182032989 | 0.6311482 | 0.839706049 |
| chr8:99865909-99866214 | -0.179271133 | 0.631182866 | 0.839706049 |
| chr11:65652454-65653154 | 0.160516561 | 0.631193209 | 0.839706049 |
| chrX:63930168-63930408 | 0.226904667 | 0.631193332 | 0.839706049 |
| chr10:13152354-13152626 | -0.227001572 | 0.631195796 | 0.839706049 |
| chr1:62760463-62760773 | -0.188132395 | 0.631212244 | 0.839706049 |
| chr17:49214674-49214907 | 0.207174679 | 0.631220133 | 0.839706049 |
| chr10:99485124-99485484 | -0.20636535 | 0.631231035 | 0.839706049 |

| | | | |
|---|---|---|---|
| chr2:20617206-20617471 | -0.185755109 | 0.631231614 | 0.839706049 |
| chrX:24491776-24492096 | -0.182158416 | 0.631272272 | 0.839745556 |
| chr15:78149039-78150319 | 0.154970272 | 0.631304395 | 0.839748772 |
| chr1:220874271-220874472 | 0.233555216 | 0.631306249 | 0.839748772 |
| chr17:4365195-4365485 | -0.183746132 | 0.631307568 | 0.839748772 |
| chr15:34048568-34048768 | 0.261035309 | 0.631332782 | 0.839757143 |
| chr22:17171178-17172295 | 0.155815185 | 0.631346394 | 0.839757143 |
| chr21:32392519-32393767 | 0.153787441 | 0.631346741 | 0.839757143 |
| chr7:95608769-95609124 | -0.20863815 | 0.631370563 | 0.839774251 |
| chr8:88554477-88554978 | -0.160236364 | 0.631391462 | 0.83978747 |
| chr17:51850309-51850509 | -0.227282455 | 0.631453708 | 0.839855683 |
| chr17:75027912-75028122 | 0.227836583 | 0.631469982 | 0.839862748 |
| chr14:100851159-100851592 | 0.17371148 | 0.63148699 | 0.839870791 |
| chr21:40372527-40372727 | -0.282681719 | 0.631520972 | 0.839880611 |
| chr4:142720237-142720519 | -0.2173563 | 0.631524023 | 0.839880611 |
| chr11:128849243-128849584 | 0.228672844 | 0.631529461 | 0.839880611 |
| chr13:40666030-40666883 | -0.156953252 | 0.631538219 | 0.839880611 |
| chr3:106680905-106681185 | 0.199411433 | 0.631557666 | 0.839891896 |
| chr7:7099427-7099963 | -0.163817189 | 0.631573612 | 0.839898525 |
| chr7:37433208-37433480 | 0.199023592 | 0.631588478 | 0.839903717 |
| chr6:122471106-122472240 | -0.153459494 | 0.631607639 | 0.839914621 |
| chr4:122003312-122003688 | -0.190115614 | 0.63163398 | 0.839934929 |
| chr11:111983121-111983407 | 0.199765395 | 0.631644835 | 0.839934929 |
| chrX:48508783-48509267 | 0.177826355 | 0.631673301 | 0.839945756 |
| chr3:121855602-121855851 | -0.221294554 | 0.631674901 | 0.839945756 |
| chr2:231681100-231681333 | -0.216412562 | 0.63170249 | 0.839967864 |
| chr5:170902907-170903107 | 0.267815742 | 0.631725431 | 0.839983792 |
| chr6:18990445-18990744 | -0.196548551 | 0.631754991 | 0.839998802 |
| chr10:48976257-48976532 | 0.199034426 | 0.631758646 | 0.839998802 |
| chr14:39187982-39188222 | 0.195446802 | 0.63182314 | 0.840056234 |
| chr12:116366432-116366736 | -0.218748358 | 0.631823768 | 0.840056234 |
| chr19:3284929-3286069 | 0.152734781 | 0.631854838 | 0.840082967 |
| chr17:75018819-75019024 | 0.258124317 | 0.631958593 | 0.840206335 |
| chr11:45917557-45918340 | -0.152273426 | 0.632013201 | 0.840258839 |
| chr15:43776630-43777787 | 0.154949941 | 0.632020016 | 0.840258839 |
| chr15:57732590-57732790 | -0.256511938 | 0.632038925 | 0.840269398 |
| chr1:150979367-150979759 | 0.163359092 | 0.632063011 | 0.840273991 |
| chr10:49001407-49001692 | 0.201518923 | 0.632064313 | 0.840273991 |
| chr4:41334252-41334520 | 0.211942515 | 0.632081869 | 0.840282752 |
| chr11:69654379-69655318 | 0.154340462 | 0.632103965 | 0.840297547 |
| chrX:49001961-49002686 | -0.159608277 | 0.632181685 | 0.840377294 |
| chr2:85707317-85707726 | 0.179733824 | 0.63218589 | 0.840377294 |
| chr5:16178796-16179160 | 0.175988191 | 0.632234962 | 0.840427946 |
| chr11:78365478-78365678 | 0.256461744 | 0.632257196 | 0.840442921 |
| chr20:37178167-37178772 | 0.165067907 | 0.632301223 | 0.84048604 |
| chr5:56588590-56588851 | 0.243672516 | 0.632311572 | 0.84048604 |
| chr6:36699490-36699690 | -0.241173128 | 0.632346531 | 0.840503562 |
| chr4:121869573-121870803 | -0.151924292 | 0.632346694 | 0.840503562 |
| chr8:127215024-127215259 | 0.204718611 | 0.632405326 | 0.840550928 |
| chr14:59905830-59906260 | 0.204393788 | 0.632410434 | 0.840550928 |

| | | | |
|---|---|---|---|
| chr19:47226086-47226413 | -0.175692354 | 0.632415239 | 0.840550928 |
| chr1:48292887-48293180 | -0.235504501 | 0.632432096 | 0.840558239 |
| chr20:13638290-13639474 | -0.151018753 | 0.63244268 | 0.840558239 |
| chr4:56467286-56468337 | 0.153856957 | 0.632515722 | 0.840630446 |
| chr3:141765395-141765710 | -0.212884436 | 0.632518952 | 0.840630446 |
| chr17:81860408-81861472 | 0.150821226 | 0.632540935 | 0.840645081 |
| chr11:17276202-17277480 | 0.152925168 | 0.632569844 | 0.84066892 |
| chr11:65539803-65541284 | 0.151704862 | 0.632587829 | 0.840669669 |
| chr15:63891046-63891522 | 0.175667813 | 0.632592351 | 0.840669669 |
| chr20:38860421-38860802 | 0.201504011 | 0.632604893 | 0.840671757 |
| chr7:34437320-34437520 | -0.254613041 | 0.63262594 | 0.840685145 |
| chr5:34486261-34486501 | -0.211694946 | 0.632644991 | 0.840689066 |
| chr1:107572627-107573237 | 0.16948526 | 0.632654292 | 0.840689066 |
| chr11:2089544-2089823 | -0.193339144 | 0.632663584 | 0.840689066 |
| chr11:59142565-59143472 | -0.156927006 | 0.632683474 | 0.840689066 |
| chr11:30488507-30488955 | 0.181084009 | 0.63268375 | 0.840689066 |
| chr5:65092858-65093117 | 0.24048791 | 0.632706954 | 0.840701543 |
| chr14:101832957-101833161 | -0.266970313 | 0.632715084 | 0.840701543 |
| chr1:196002242-196002829 | -0.161503249 | 0.632749115 | 0.840719473 |
| chr17:77280952-77281723 | 0.157761736 | 0.632758787 | 0.840719473 |
| chrX:108304343-108304543 | 0.248703926 | 0.632787284 | 0.840719473 |
| chr8:11769434-11770195 | -0.159270131 | 0.632790681 | 0.840719473 |
| chr5:157948787-157949354 | -0.168531121 | 0.632793624 | 0.840719473 |
| chr4:78551461-78552726 | -0.154892261 | 0.632794412 | 0.840719473 |
| chr1:236109388-236109618 | 0.198410222 | 0.632821143 | 0.840740409 |
| chr9:112717605-112718545 | 0.152997454 | 0.632841288 | 0.840750511 |
| chr9:133148879-133149836 | 0.154916371 | 0.632857152 | 0.840750511 |
| chr15:101132635-101132835 | 0.223718108 | 0.632861664 | 0.840750511 |
| chr17:16535547-16536270 | -0.156783625 | 0.632877663 | 0.840757187 |
| chr5:14035947-14036418 | 0.171382505 | 0.632905274 | 0.84077929 |
| chr14:104171457-104171693 | -0.205510369 | 0.632933616 | 0.840785576 |
| chr4:40316932-40317624 | 0.172449844 | 0.632938425 | 0.840785576 |
| chr1:174393503-174393703 | -0.257549331 | 0.632957436 | 0.840785576 |
| chr10:51032123-51032323 | -0.257549331 | 0.632957436 | 0.840785576 |
| chr2:201129236-201129436 | -0.247028724 | 0.632964872 | 0.840785576 |
| chr14:23469151-23470050 | 0.153291254 | 0.632996466 | 0.840812639 |
| chr14:57066934-57067253 | -0.203714952 | 0.633011453 | 0.840812639 |
| chr11:70270011-70270847 | 0.153775104 | 0.633018166 | 0.840812639 |
| chr11:120807501-120807917 | -0.188212468 | 0.633069862 | 0.840833479 |
| chr2:70994322-70994580 | 0.189635711 | 0.633096765 | 0.840833479 |
| chr19:35716832-35718045 | 0.154000147 | 0.633104112 | 0.840833479 |
| chr13:32259151-32259449 | -0.214797601 | 0.633107322 | 0.840833479 |
| chr8:26661844-26662074 | 0.226637469 | 0.633112061 | 0.840833479 |
| chr22:50507552-50508361 | -0.160202121 | 0.633123585 | 0.840833479 |
| chr15:41521401-41521602 | 0.249795398 | 0.633127816 | 0.840833479 |
| chr10:30695977-30696418 | -0.193801256 | 0.633147123 | 0.840833479 |
| chr2:43595493-43596707 | -0.151966187 | 0.633168116 | 0.840833479 |
| chr2:74778218-74778740 | -0.187890588 | 0.633174224 | 0.840833479 |
| chr7:873371-873628 | 0.178892071 | 0.633177401 | 0.840833479 |
| chr11:43802606-43802806 | 0.269735658 | 0.633187942 | 0.840833479 |

| | | | |
|---|---|---|---|
| chr9:133486231-133486635 | 0.172746095 | 0.633192048 | 0.840833479 |
| chr7:66306076-66306372 | -0.240491829 | 0.633194148 | 0.840833479 |
| chr2:241701232-241703231 | 0.149880481 | 0.633228227 | 0.840833479 |
| chr20:8082123-8082368 | 0.241578638 | 0.633228547 | 0.840833479 |
| chr8:128377704-128377970 | 0.216583461 | 0.633231217 | 0.840833479 |
| chr2:59301825-59302180 | -0.184493644 | 0.633231384 | 0.840833479 |
| chr14:96204209-96204421 | 0.208236687 | 0.633274289 | 0.84086378 |
| chr13:31199556-31200960 | 0.152684227 | 0.633284221 | 0.84086378 |
| chr17:34998305-34998510 | -0.233873779 | 0.633287126 | 0.84086378 |
| chrX:68462780-68462994 | -0.244050274 | 0.633316303 | 0.840881282 |
| chr1:9825388-9825905 | 0.168650996 | 0.633325661 | 0.840881282 |
| chr2:241150090-241150807 | 0.157380807 | 0.633335767 | 0.840881282 |
| chr9:37120029-37121092 | -0.154630852 | 0.633357909 | 0.840881282 |
| chr11:70270982-70271648 | 0.163955838 | 0.633358622 | 0.840881282 |
| chr14:24365702-24365973 | 0.176763996 | 0.633379425 | 0.840881282 |
| chrX:48737045-48737989 | -0.154738677 | 0.633386006 | 0.840881282 |
| chrX:103586151-103586627 | 0.180157909 | 0.633391155 | 0.840881282 |
| chr8:38382254-38383044 | 0.159563549 | 0.633399077 | 0.840881282 |
| chr6:32853506-32854283 | 0.160245237 | 0.633423101 | 0.840898606 |
| chr14:100553955-100554557 | 0.171880675 | 0.633443483 | 0.840911095 |
| chr10:48658366-48658608 | -0.19707346 | 0.633546591 | 0.841033402 |
| chr2:8803808-8804466 | 0.166124556 | 0.633588866 | 0.841066232 |
| chr11:45885370-45885901 | -0.165932476 | 0.633604515 | 0.841066232 |
| chr19:47868951-47869221 | 0.203960236 | 0.633608074 | 0.841066232 |
| chr4:188143818-188144027 | 0.241267179 | 0.633633949 | 0.841066232 |
| chr9:6023995-6024422 | 0.202491398 | 0.633636419 | 0.841066232 |
| chr10:72171147-72171648 | 0.183140303 | 0.633637183 | 0.841066232 |
| chr21:41507958-41508404 | 0.173418291 | 0.633659412 | 0.841067005 |
| chr14:40280732-40281167 | 0.194275445 | 0.633659719 | 0.841067005 |
| chr11:14643397-14644777 | -0.153307614 | 0.633680998 | 0.841080678 |
| chr1:31564148-31564348 | 0.242733297 | 0.633709955 | 0.841104543 |
| chr3:13650452-13650822 | 0.168599733 | 0.633768114 | 0.841167165 |
| chr4:100311193-100311556 | -0.203321411 | 0.633790395 | 0.841178721 |
| chr8:15657609-15657928 | -0.207874283 | 0.63382312 | 0.841178721 |
| chr10:69093344-69093579 | -0.23604617 | 0.633826379 | 0.841178721 |
| chr17:44708949-44709293 | 0.214603092 | 0.63382864 | 0.841178721 |
| chr11:74493023-74494000 | -0.15202497 | 0.633837369 | 0.841178721 |
| chr6:108080462-108080667 | 0.245936439 | 0.633860867 | 0.841178721 |
| chr2:109260924-109261139 | 0.230157236 | 0.633863006 | 0.841178721 |
| chr4:154329348-154329783 | -0.199087592 | 0.6338805 | 0.841178721 |
| chr15:74725435-74726323 | 0.161116334 | 0.633880601 | 0.841178721 |
| chr2:3303642-3304157 | -0.168676441 | 0.633886604 | 0.841178721 |
| chr1:206106625-206106852 | -0.195286475 | 0.633898102 | 0.841179411 |
| chr1:58546011-58547096 | -0.151895649 | 0.633931515 | 0.841205806 |
| chr18:2655046-2656661 | -0.15173079 | 0.63393995 | 0.841205806 |
| chr2:85648724-85648924 | 0.231706318 | 0.633967857 | 0.841228269 |
| chr14:61694849-61696379 | 0.149634542 | 0.633988499 | 0.841241091 |
| chr3:49939819-49940561 | 0.163389694 | 0.634049031 | 0.841276858 |
| chr14:32826179-32826469 | -0.228526669 | 0.634057144 | 0.841276858 |
| chr20:49277472-49277700 | 0.181650263 | 0.634057853 | 0.841276858 |

| | | | |
|---|---|---|---|
| chr15:78348051-78348644 | -0.162085051 | 0.634059373 | 0.841276858 |
| chr1:205227700-205228016 | -0.169055963 | 0.634097729 | 0.841308468 |
| chr3:122382876-122383640 | -0.155293472 | 0.63410729 | 0.841308468 |
| chr4:152997772-152998161 | 0.180446778 | 0.634116137 | 0.841308468 |
| chr21:42056417-42057721 | -0.152543418 | 0.634159029 | 0.841335405 |
| chr2:219298731-219299048 | -0.181774057 | 0.634168665 | 0.841335405 |
| chr1:37534482-37534834 | 0.187057762 | 0.63417795 | 0.841335405 |
| chr1:119674824-119675130 | -0.178222049 | 0.634180361 | 0.841335405 |
| chrX:136251129-136251924 | 0.160983167 | 0.634192321 | 0.841336704 |
| chr15:77821533-77821733 | 0.212870453 | 0.634212924 | 0.84134947 |
| chr9:137217708-137217908 | -0.218203813 | 0.634235803 | 0.841365254 |
| chr6:52503788-52504792 | -0.151633486 | 0.634255105 | 0.841375479 |
| chr17:75645402-75645976 | 0.160956659 | 0.634265473 | 0.841375479 |
| chr12:79886956-79887218 | 0.201630872 | 0.634287348 | 0.841389931 |
| chr1:198650506-198650817 | 0.221948226 | 0.634326111 | 0.841417991 |
| chr7:149027853-149029277 | -0.149775684 | 0.634330464 | 0.841417991 |
| chr17:75261108-75262337 | 0.153963232 | 0.634372802 | 0.841446112 |
| chr14:100396095-100396359 | 0.258306301 | 0.634373628 | 0.841446112 |
| chr18:48308085-48308386 | 0.19818223 | 0.634385033 | 0.841446673 |
| chr2:529376-529696 | -0.2008968 | 0.634430856 | 0.841492886 |
| chr6:17116225-17116425 | -0.253145627 | 0.634463408 | 0.841519472 |
| chr15:76840928-76841180 | -0.237319778 | 0.634472865 | 0.841519472 |
| chr17:60525325-60526653 | -0.147614244 | 0.634499791 | 0.841540617 |
| chr10:97553553-97553846 | 0.18803934 | 0.634516459 | 0.841548158 |
| chr10:133474770-133474985 | 0.256571694 | 0.634561969 | 0.841593948 |
| chr1:2194711-2195139 | 0.177028645 | 0.634583589 | 0.841604039 |
| chr6:31821357-31821909 | 0.174931138 | 0.634591545 | 0.841604039 |
| chr17:58896328-58896656 | 0.16905396 | 0.63460949 | 0.841612584 |
| chr14:79631237-79631572 | 0.198482743 | 0.634624471 | 0.841612584 |
| chr2:151392375-151392736 | -0.201341346 | 0.634637675 | 0.841612584 |
| chr9:135478840-135479131 | -0.195365368 | 0.634641924 | 0.841612584 |
| chr13:97975716-97977063 | -0.152200863 | 0.634661077 | 0.841623417 |
| chr6:137944988-137945422 | -0.187889528 | 0.634674968 | 0.841627272 |
| chr16:20901422-20902018 | 0.159482748 | 0.634702438 | 0.841649133 |
| chr1:176949021-176949433 | -0.170870347 | 0.634736634 | 0.841679912 |
| chr7:6637131-6637402 | 0.184574693 | 0.634783035 | 0.841716427 |
| chr5:93581226-93581893 | -0.166676396 | 0.634793133 | 0.841716427 |
| chr6:26233962-26234162 | 0.228800922 | 0.634798986 | 0.841716427 |
| chr7:79056094-79056384 | -0.195918178 | 0.634815279 | 0.841716427 |
| chr3:146327125-146327485 | 0.184745373 | 0.634819098 | 0.841716427 |
| chr3:130026181-130026680 | -0.177842698 | 0.634838239 | 0.841727242 |
| chr15:32042336-32042536 | 0.246876037 | 0.634853947 | 0.841729351 |
| chr5:171622136-171622401 | 0.210447938 | 0.634868608 | 0.841729351 |
| chr5:134927124-134927346 | -0.235398074 | 0.634873318 | 0.841729351 |
| chr5:157476907-157477239 | -0.189143009 | 0.634891329 | 0.841729351 |
| chr14:92926307-92926532 | -0.214331217 | 0.634894758 | 0.841729351 |
| chr1:58584576-58584843 | -0.210811245 | 0.634920904 | 0.841733243 |
| chr19:42081648-42081848 | 0.243376645 | 0.63492682 | 0.841733243 |
| chr5:12856152-12856491 | -0.25428006 | 0.63493065 | 0.841733243 |
| chr5:127517146-127518487 | -0.150724801 | 0.634985282 | 0.841786315 |

| | | | |
|---|---|---|---|
| chr6:38795565-38795765 | 0.248788927 | 0.634999063 | 0.841786315 |
| chr3:164533376-164533576 | 0.275628006 | 0.635012041 | 0.841786315 |
| chr16:16049323-16049603 | 0.235001781 | 0.63501704 | 0.841786315 |
| chr14:49464972-49465244 | -0.199609701 | 0.635028219 | 0.841786315 |
| chr10:5199274-5200120 | 0.161662704 | 0.6350366 | 0.841786315 |
| chr17:21099779-21099998 | -0.207007992 | 0.635056218 | 0.841797757 |
| chr17:52804864-52805158 | 0.224250121 | 0.63507656 | 0.841810159 |
| chr3:23052626-23052972 | 0.196484329 | 0.635120696 | 0.841838925 |
| chr4:73662724-73662942 | 0.233327843 | 0.635149825 | 0.841838925 |
| chr3:188839227-188839607 | 0.198258592 | 0.635151558 | 0.841838925 |
| chr5:134524317-134524575 | 0.174749962 | 0.635152769 | 0.841838925 |
| chr9:133356410-133356694 | 0.180286572 | 0.635153197 | 0.841838925 |
| chr1:163069241-163069549 | 0.196115827 | 0.635190969 | 0.841874426 |
| chr20:1118836-1119061 | -0.229152355 | 0.635254556 | 0.841930265 |
| chr4:75099395-75099665 | 0.237230996 | 0.635255075 | 0.841930265 |
| chr1:20575247-20575489 | 0.239672444 | 0.635268584 | 0.841933605 |
| chr10:6920066-6920347 | 0.193234609 | 0.635299729 | 0.841954733 |
| chrX:6302470-6302776 | 0.210744564 | 0.635313673 | 0.841954733 |
| chr8:123388130-123388388 | -0.204587745 | 0.635317491 | 0.841954733 |
| chr12:76490742-76490993 | -0.241974121 | 0.635389132 | 0.842035112 |
| chr17:79728667-79728967 | -0.191630837 | 0.63545804 | 0.842097584 |
| chr1:106344441-106345027 | 0.160543498 | 0.635462965 | 0.842097584 |
| chr20:38299991-38300344 | 0.181546839 | 0.635469244 | 0.842097584 |
| chr4:165378697-165379772 | -0.153984442 | 0.635519118 | 0.842136788 |
| chr6:28981989-28982362 | 0.162165745 | 0.63552081 | 0.842136788 |
| chr20:35664383-35665215 | 0.158842921 | 0.635583936 | 0.842205872 |
| chr1:159913774-159914181 | -0.175734514 | 0.635619019 | 0.842237795 |
| chr5:174157182-174157453 | -0.220260268 | 0.635652844 | 0.842268049 |
| chr16:16078517-16078717 | 0.24972186 | 0.635669117 | 0.842275046 |
| chr20:50380743-50380943 | -0.207689475 | 0.635683512 | 0.842276225 |
| chr1:168225497-168226737 | 0.154617124 | 0.635691992 | 0.842276225 |
| chr6:28790671-28790872 | -0.22064192 | 0.635734602 | 0.842291727 |
| chrX:112679493-112680328 | 0.164143959 | 0.635736921 | 0.842291727 |
| chr4:151760665-151761581 | -0.152492279 | 0.635743157 | 0.842291727 |
| chrX:7218837-7219390 | 0.18972908 | 0.635749641 | 0.842291727 |
| chr1:150567786-150568021 | 0.180383692 | 0.63577236 | 0.842291727 |
| chr11:60830852-60831580 | 0.164392303 | 0.635787441 | 0.842291727 |
| chr1:204286807-204287423 | -0.153797864 | 0.635802299 | 0.842291727 |
| chr14:21997354-21997676 | -0.201060256 | 0.635802627 | 0.842291727 |
| chr3:65439669-65439966 | -0.201060256 | 0.635802627 | 0.842291727 |
| chr7:33494500-33494898 | 0.189965739 | 0.635853942 | 0.842337116 |
| chr1:41900692-41900903 | -0.248437155 | 0.635861345 | 0.842337116 |
| chr3:184289392-184289853 | -0.174405634 | 0.635875621 | 0.842337116 |
| chr22:17181335-17181591 | -0.253104217 | 0.635887317 | 0.842337116 |
| chr11:122781393-122781777 | 0.186640427 | 0.63590276 | 0.842337116 |
| chr8:131859591-131859843 | 0.210495495 | 0.635911322 | 0.842337116 |
| chr2:31729718-31730091 | 0.183705192 | 0.635923945 | 0.842337116 |
| chr1:228236825-228237142 | -0.198839733 | 0.635924836 | 0.842337116 |
| chr4:153127903-153128161 | -0.203527233 | 0.635942906 | 0.842346489 |
| chr3:19174050-19174267 | 0.2410816 | 0.635966392 | 0.842363036 |

| | | | |
|---|---|---|---|
| chr12:133129859-133130761 | 0.15280954 | 0.635990221 | 0.842374679 |
| chr14:72847806-72848006 | 0.236696108 | 0.63599717 | 0.842374679 |
| chr1:181391380-181391763 | -0.182353996 | 0.63605927 | 0.842442368 |
| chrX:101622812-101623829 | -0.155594358 | 0.636086202 | 0.842463476 |
| chr17:80082771-80083035 | 0.205788066 | 0.636108289 | 0.842478165 |
| chr15:70097801-70098134 | 0.163723161 | 0.63612741 | 0.842488928 |
| chr17:79811447-79811698 | -0.179704873 | 0.636172809 | 0.842534491 |
| chr6:27237143-27237718 | 0.169996726 | 0.636192864 | 0.842546488 |
| chr11:111944314-111944514 | 0.250762762 | 0.63627476 | 0.842640383 |
| chr1:54819150-54819586 | -0.178604556 | 0.636297453 | 0.842655872 |
| chr1:202348333-202349598 | -0.149543064 | 0.636343454 | 0.84268331 |
| chr14:24656965-24657172 | -0.251164768 | 0.636346095 | 0.84268331 |
| chr15:40440624-40441215 | 0.159156527 | 0.636351165 | 0.84268331 |
| chr16:14656463-14656701 | 0.242931868 | 0.636382403 | 0.842710112 |
| chr13:33447146-33447465 | -0.177333396 | 0.636412032 | 0.842724687 |
| chr2:231056436-231057212 | 0.157780302 | 0.636415406 | 0.842724687 |
| chr11:30196734-30196934 | 0.234960603 | 0.63644661 | 0.842748147 |
| chr17:60638742-60639114 | -0.153468126 | 0.63646181 | 0.842748147 |
| chr16:72832608-72832889 | 0.187553147 | 0.636469828 | 0.842748147 |
| chr3:141646809-141647139 | -0.180586303 | 0.636487571 | 0.842748147 |
| chr3:39152059-39153120 | 0.151373832 | 0.63649477 | 0.842748147 |
| chr2:136214409-136214684 | 0.200784336 | 0.636499116 | 0.842748147 |
| chr10:13614626-13615076 | 0.184002069 | 0.636520499 | 0.842761896 |
| chr9:34522914-34523470 | 0.165954495 | 0.636566445 | 0.842783296 |
| chr1:79123352-79123552 | -0.274833831 | 0.63656651 | 0.842783296 |
| chr12:118135855-118136838 | 0.153747549 | 0.63656966 | 0.842783296 |
| chr6:42050329-42050906 | 0.163776404 | 0.636604813 | 0.842815275 |
| chr1:94448650-94449196 | 0.183075026 | 0.636642383 | 0.842844524 |
| chr12:45991754-45992188 | 0.166044713 | 0.636648907 | 0.842844524 |
| chr16:4461172-4461405 | -0.222930024 | 0.636670411 | 0.842853566 |
| chr2:26605431-26605631 | -0.227764961 | 0.636677736 | 0.842853566 |
| chr14:58297471-58298770 | 0.147809552 | 0.636711143 | 0.842867192 |
| chr5:95066889-95067177 | -0.228181809 | 0.636714604 | 0.842867192 |
| chr15:58066323-58066638 | -0.195613767 | 0.636721031 | 0.842867192 |
| chr2:184598292-184598960 | -0.163298327 | 0.636742683 | 0.842872053 |
| chr6:114108237-114108736 | 0.172296984 | 0.636746704 | 0.842872053 |
| chr14:74459675-74459899 | 0.198701437 | 0.636768765 | 0.842886695 |
| chr9:13034275-13034524 | -0.207362848 | 0.63678748 | 0.842889316 |
| chr20:40706998-40707271 | -0.207056823 | 0.636792747 | 0.842889316 |
| chr19:55454660-55454933 | 0.209208673 | 0.636804503 | 0.842890316 |
| chr17:68089897-68090215 | -0.178830502 | 0.636830306 | 0.842909909 |
| chr1:205812703-205813908 | -0.149044068 | 0.636862314 | 0.842925865 |
| chr16:78122626-78123130 | 0.165604805 | 0.636864364 | 0.842925865 |
| chr17:77140215-77141248 | 0.151002403 | 0.636899261 | 0.842953774 |
| chr1:188668160-188668421 | -0.195489333 | 0.636907453 | 0.842953774 |
| chr3:155608071-155608582 | -0.192517758 | 0.636927529 | 0.842965785 |
| chr11:84564679-84564904 | -0.225704911 | 0.636962105 | 0.842994939 |
| chr10:76984954-76985188 | -0.26014982 | 0.636985403 | 0.842994939 |
| chr10:17643455-17644740 | -0.151413453 | 0.636995272 | 0.842994939 |
| chr22:17830033-17830330 | 0.18361306 | 0.636995575 | 0.842994939 |

| | | | |
|---|---|---|---|
| chr15:41895274-41895481 | -0.225566372 | 0.637017111 | 0.842994939 |
| chr2:223989274-223989730 | -0.188065817 | 0.637023601 | 0.842994939 |
| chr17:17811789-17811992 | -0.187080129 | 0.637033098 | 0.842994939 |
| chr6:13426054-13426275 | 0.235946709 | 0.637037574 | 0.842994939 |
| chr1:148254712-148255306 | 0.172250442 | 0.637050666 | 0.842997705 |
| chr14:100376060-100376705 | 0.155630915 | 0.637075772 | 0.843016369 |
| chr1:202188875-202189342 | -0.158343749 | 0.637187876 | 0.843146379 |
| chr1:25895464-25895664 | 0.220981002 | 0.63719603 | 0.843146379 |
| chr8:47167246-47167680 | -0.17171768 | 0.637228751 | 0.843175115 |
| chr3:22563895-22564644 | -0.158481919 | 0.637266118 | 0.843209997 |
| chr2:194867481-194867694 | -0.232617849 | 0.637290415 | 0.843227584 |
| chr6:81451923-81452123 | 0.238649512 | 0.637331173 | 0.843266951 |
| chrX:101407560-101408758 | -0.148386178 | 0.637362217 | 0.843293465 |
| chr12:51678180-51678420 | 0.206144151 | 0.637375324 | 0.843296244 |
| chr4:127586999-127587292 | -0.174990999 | 0.637408432 | 0.843325486 |
| chr1:113021774-113021996 | 0.244787467 | 0.637443246 | 0.843347677 |
| chr3:70737701-70738338 | -0.165565656 | 0.637447218 | 0.843347677 |
| chr22:50485819-50486052 | -0.221385383 | 0.637475907 | 0.843371071 |
| chr3:30073626-30073868 | 0.223519348 | 0.637497904 | 0.843377386 |
| chr10:68987758-68987975 | 0.201562295 | 0.637502694 | 0.843377386 |
| chr20:7893383-7893583 | 0.238461026 | 0.637522234 | 0.843388674 |
| chr6:140053384-140053584 | 0.270548859 | 0.637558816 | 0.843397461 |
| chr2:61258241-61258512 | -0.248618583 | 0.637563718 | 0.843397461 |
| chr10:101060571-101061788 | 0.151378675 | 0.63756513 | 0.843397461 |
| chr5:32144698-32144898 | 0.234387016 | 0.637572905 | 0.843397461 |
| chr12:27703740-27703985 | 0.213103294 | 0.63759036 | 0.843402441 |
| chr14:64387309-64388758 | 0.147888881 | 0.637607902 | 0.843402441 |
| chr11:47969350-47969640 | 0.216456696 | 0.637609692 | 0.843402441 |
| chr7:5450829-5451267 | -0.175440987 | 0.637626188 | 0.843404403 |
| chr9:83707214-83708841 | 0.149491087 | 0.63763319 | 0.843404403 |
| chr15:74789352-74789657 | 0.173180242 | 0.637666054 | 0.843419351 |
| chr3:100260568-100261509 | -0.149514043 | 0.637666506 | 0.843419351 |
| chr13:82291958-82292869 | -0.160632428 | 0.637714284 | 0.843430165 |
| chr3:108493921-108494245 | -0.176916596 | 0.637715423 | 0.843430165 |
| chr17:68911079-68911419 | -0.189839726 | 0.637717057 | 0.843430165 |
| chr5:88964616-88965350 | 0.159311007 | 0.63772353 | 0.843430165 |
| chr3:128188839-128189092 | -0.228467605 | 0.63772972 | 0.843430165 |
| chr1:193905450-193905650 | 0.224476373 | 0.637800523 | 0.843503608 |
| chr8:6407065-6407502 | -0.184320276 | 0.637807269 | 0.843503608 |
| chr19:37506277-37507127 | 0.1666616 | 0.637833268 | 0.843523433 |
| chr3:111539630-111540063 | -0.184042634 | 0.637850093 | 0.843526035 |
| chr2:15940205-15940513 | 0.187992103 | 0.637857253 | 0.843526035 |
| chr4:15906047-15906449 | -0.186333037 | 0.637879633 | 0.843541072 |
| chr8:111140306-111141832 | -0.155120257 | 0.637906125 | 0.843552484 |
| chr10:5360297-5360614 | -0.221703849 | 0.637910282 | 0.843552484 |
| chr3:50121604-50121882 | -0.193141465 | 0.637934513 | 0.843569968 |
| chr17:75004951-75005157 | 0.222638345 | 0.637951642 | 0.84357806 |
| chr6:14953241-14953488 | -0.250556218 | 0.637968378 | 0.843585633 |
| chr19:4867534-4868124 | 0.164718645 | 0.637989044 | 0.843598401 |
| chr19:54256324-54256570 | -0.237959303 | 0.63801068 | 0.843612452 |

| | | | |
|---|---|---|---|
| chr9:21392281-21392643 | -0.171688291 | 0.638035314 | 0.843614374 |
| chr1:153535980-153536344 | -0.174940986 | 0.638039628 | 0.843614374 |
| chr14:102097392-102098085 | -0.156723711 | 0.638045164 | 0.843614374 |
| chr9:107488499-107488915 | 0.175085543 | 0.638065338 | 0.843626489 |
| chr9:99659161-99659419 | -0.226149492 | 0.6381188 | 0.843653719 |
| chr19:11504904-11506038 | 0.153250189 | 0.638120837 | 0.843653719 |
| chr1:85494895-85495174 | -0.203960847 | 0.638129595 | 0.843653719 |
| chr14:51979130-51979412 | 0.193203368 | 0.638129975 | 0.843653719 |
| chr8:21918414-21918677 | 0.192443006 | 0.638155538 | 0.843672958 |
| chr12:122097910-122098278 | -0.17455955 | 0.638187103 | 0.843695598 |
| chr1:188343952-188344186 | -0.223593216 | 0.638206652 | 0.843695598 |
| chr5:181243607-181244549 | 0.151870805 | 0.638219494 | 0.843695598 |
| chr5:127757583-127757851 | 0.185930086 | 0.638223765 | 0.843695598 |
| chr12:101759546-101759754 | 0.229974354 | 0.638241356 | 0.843695598 |
| chr20:53578590-53579315 | 0.147881342 | 0.638258149 | 0.843695598 |
| chr11:65888973-65889173 | 0.194279695 | 0.638266829 | 0.843695598 |
| chr6:10413878-10414358 | -0.165838561 | 0.638268341 | 0.843695598 |
| chr10:69515596-69515974 | 0.181507685 | 0.638271763 | 0.843695598 |
| chr1:227794495-227795000 | -0.177194383 | 0.638325686 | 0.843752319 |
| chr2:171921993-171923077 | -0.151660475 | 0.638357083 | 0.843766969 |
| chr20:54152253-54152901 | 0.149988188 | 0.638358793 | 0.843766969 |
| chr5:54895794-54896044 | -0.227128239 | 0.63837525 | 0.843772426 |
| chr2:85585000-85585272 | 0.199997391 | 0.638384946 | 0.843772426 |
| chr14:100306223-100306777 | 0.158094685 | 0.638403421 | 0.843781341 |
| chr20:19396765-19397168 | 0.194384907 | 0.638413715 | 0.843781341 |
| chr2:184646302-184646612 | 0.199496419 | 0.63845476 | 0.843821034 |
| chr3:135040294-135040626 | -0.232643213 | 0.638474087 | 0.843832022 |
| chr16:89637360-89637651 | 0.19683413 | 0.638497349 | 0.843833091 |
| chr10:3467351-3467892 | 0.167905114 | 0.638500817 | 0.843833091 |
| chr16:19067282-19068285 | 0.151286548 | 0.638507934 | 0.843833091 |
| chr11:19557614-19557989 | 0.177394995 | 0.63857524 | 0.843896481 |
| chr8:101491947-101492914 | 0.147634447 | 0.638577928 | 0.843896481 |
| chrX:123803317-123803917 | 0.159242154 | 0.638637926 | 0.843961214 |
| chr15:69388196-69388698 | 0.171301825 | 0.638676181 | 0.84399424 |
| chr15:98292486-98292815 | -0.179971904 | 0.63870038 | 0.84399424 |
| chr5:134873865-134875191 | -0.154215196 | 0.638708694 | 0.84399424 |
| chr3:144244771-144244985 | -0.238320735 | 0.638726056 | 0.84399424 |
| chr21:38712138-38712403 | -0.19577531 | 0.638727036 | 0.84399424 |
| chr7:21011139-21011549 | -0.181612179 | 0.638729007 | 0.84399424 |
| chr12:92408167-92408397 | -0.222407503 | 0.638748552 | 0.844005511 |
| chr8:88416318-88416518 | -0.231653535 | 0.638786585 | 0.844041209 |
| chr8:119416579-119417311 | -0.162811287 | 0.638810952 | 0.844051112 |
| chr10:97701412-97701988 | 0.16527211 | 0.63882483 | 0.844051112 |
| chr3:195259475-195260572 | -0.153511549 | 0.638836141 | 0.844051112 |
| chr8:43055963-43056999 | 0.155929794 | 0.638858483 | 0.844051112 |
| chr1:26408631-26408832 | -0.260235329 | 0.638861807 | 0.844051112 |
| chr7:106058446-106058856 | -0.1661046 | 0.638863135 | 0.844051112 |
| chr7:103446035-103446278 | 0.221891787 | 0.638871189 | 0.844051112 |
| chr20:35269323-35269558 | -0.207635486 | 0.638907742 | 0.844069754 |
| chr20:2101611-2102976 | 0.154282233 | 0.638909392 | 0.844069754 |

| | | | |
|---|---|---|---|
| chr17:58517491-58518951 | -0.145833883 | 0.638918348 | 0.844069754 |
| chr4:152937834-152938074 | -0.192883131 | 0.638934025 | 0.844072295 |
| chr10:93680485-93680731 | 0.207960062 | 0.638951863 | 0.844072295 |
| chr12:101019387-101019602 | -0.24748166 | 0.63895332 | 0.844072295 |
| chr5:80407548-80408710 | -0.148857718 | 0.6389715 | 0.844081759 |
| chr4:94731908-94732196 | 0.20911046 | 0.63899894 | 0.844094152 |
| chr18:72965491-72966006 | 0.171983278 | 0.639002914 | 0.844094152 |
| chr2:241734387-241735699 | 0.151480955 | 0.639026809 | 0.844111163 |
| chr2:132791703-132791903 | 0.239627483 | 0.639077796 | 0.844112655 |
| chr16:1920919-1921182 | 0.202417992 | 0.6390824 | 0.844112655 |
| chr6:42783348-42783620 | -0.188666359 | 0.639084937 | 0.844112655 |
| chr8:10928396-10929132 | -0.170026536 | 0.639100617 | 0.844112655 |
| chr2:47368881-47370177 | -0.149570857 | 0.639101228 | 0.844112655 |
| chr4:70577194-70577756 | -0.204595673 | 0.63910642 | 0.844112655 |
| chr3:196789776-196790122 | 0.189637575 | 0.639111711 | 0.844112655 |
| chr6:111872866-111873920 | 0.152942022 | 0.639126711 | 0.844112655 |
| chr9:110340436-110340636 | -0.244549334 | 0.639129469 | 0.844112655 |
| chr17:64779996-64780949 | 0.148913892 | 0.639138104 | 0.844112655 |
| chr14:59281774-59282031 | 0.226957566 | 0.639152061 | 0.84411542 |
| chr4:173038607-173038817 | -0.240652883 | 0.639162231 | 0.84411542 |
| chr12:77065112-77066228 | -0.148335095 | 0.639223933 | 0.844182357 |
| chr11:83956283-83956483 | 0.231808978 | 0.639272969 | 0.844232565 |
| chr3:41994864-41995158 | -0.215703657 | 0.639291674 | 0.844242716 |
| chr1:148345450-148345961 | 0.172989194 | 0.639326193 | 0.844244001 |
| chr7:34235675-34236575 | -0.155145181 | 0.639330279 | 0.844244001 |
| chr1:109864390-109864631 | 0.227980654 | 0.639334373 | 0.844244001 |
| chr1:199462796-199463045 | 0.198205029 | 0.63933672 | 0.844244001 |
| chr6:3732575-3732822 | -0.226308436 | 0.639350761 | 0.8442456 |
| chr10:103078796-103079052 | 0.214337955 | 0.639379123 | 0.8442456 |
| chr8:22548850-22549496 | 0.168790373 | 0.639379486 | 0.8442456 |
| chrX:149487380-149487655 | 0.192126348 | 0.639382005 | 0.8442456 |
| chr20:11919069-11919269 | -0.248772864 | 0.639412619 | 0.844271475 |
| chr9:109345360-109346106 | -0.156838 | 0.639434783 | 0.844286191 |
| chr17:67979832-67980032 | -0.217255857 | 0.639458897 | 0.844286195 |
| chr11:73779130-73780006 | -0.150009795 | 0.639463567 | 0.844286195 |
| chr6:166626869-166627770 | 0.149802498 | 0.639467843 | 0.844286195 |
| chr6:159252928-159253167 | -0.219090721 | 0.63948215 | 0.844287497 |
| chr11:12544068-12544286 | 0.219480343 | 0.639490866 | 0.844287497 |
| chr6:11232004-11232977 | -0.153258947 | 0.639505188 | 0.844291857 |
| chr1:94854370-94854976 | 0.161524876 | 0.639527715 | 0.844294073 |
| chr9:93095362-93096465 | 0.148759362 | 0.639528905 | 0.844294073 |
| chr5:54217952-54218193 | -0.23327011 | 0.639553738 | 0.844306541 |
| chr11:66430039-66430305 | 0.197102349 | 0.639560387 | 0.844306541 |
| chr19:40690522-40691050 | 0.15985079 | 0.639637412 | 0.844393677 |
| chr17:68558382-68558582 | -0.237665505 | 0.639653402 | 0.844400238 |
| chr8:93694147-93694414 | 0.195220604 | 0.639682715 | 0.844424385 |
| chr17:63522865-63523469 | -0.154227938 | 0.639718381 | 0.84445666 |
| chr17:20706999-20707289 | 0.217405558 | 0.639729207 | 0.84445666 |
| chr14:68792876-68793630 | -0.152718112 | 0.639742394 | 0.84445952 |
| chr12:90691172-90691417 | -0.248180498 | 0.639761843 | 0.844470644 |

| | | | |
|---|---|---|---|
| chr8:107895587-107896040 | -0.193152478 | 0.639828354 | 0.844543662 |
| chr17:64146398-64146707 | 0.169372711 | 0.639839205 | 0.844543662 |
| chr15:93087845-93088608 | 0.15980242 | 0.639852133 | 0.844546177 |
| chr2:7083759-7084179 | -0.196461145 | 0.6398799 | 0.844568279 |
| chr5:611659-612494 | 0.152606818 | 0.639916911 | 0.84459571 |
| chr20:47979297-47979538 | 0.169318176 | 0.639946554 | 0.84459571 |
| chr6:131951032-131951568 | 0.160546206 | 0.639947016 | 0.84459571 |
| chr12:62933243-62933566 | 0.167444089 | 0.639949384 | 0.84459571 |
| chr15:78859398-78859787 | 0.173088523 | 0.639955798 | 0.84459571 |
| chr1:206175551-206175794 | -0.234449296 | 0.640007915 | 0.844649945 |
| chr6:30886053-30886610 | 0.161401087 | 0.640044148 | 0.844663727 |
| chrX:74180124-74180343 | 0.219862489 | 0.640046279 | 0.844663727 |
| chr1:63701236-63701444 | 0.239609718 | 0.640058129 | 0.844663727 |
| chr6:35343130-35343377 | -0.20775287 | 0.640072372 | 0.844663727 |
| chr1:52368464-52368670 | -0.19769525 | 0.640099922 | 0.844663727 |
| chr6:28356155-28356506 | -0.185082307 | 0.640105439 | 0.844663727 |
| chr3:69740087-69740399 | 0.173709569 | 0.640107459 | 0.844663727 |
| chr16:67865435-67865670 | -0.220658902 | 0.640125893 | 0.844663727 |
| chr4:102449886-102450086 | -0.240497714 | 0.640131223 | 0.844663727 |
| chr5:122030292-122030492 | -0.240497714 | 0.640131223 | 0.844663727 |
| chr3:156201045-156201275 | -0.235497653 | 0.64013962 | 0.844663727 |
| chr9:93020930-93021412 | -0.187769741 | 0.640174769 | 0.844695559 |
| chr14:61568073-61569010 | -0.155934686 | 0.640205002 | 0.844716186 |
| chr13:26947060-26947272 | 0.20695399 | 0.640236562 | 0.844716186 |
| chr1:223610145-223610472 | -0.211806442 | 0.640255263 | 0.844716186 |
| chr10:103917598-103918759 | 0.148368641 | 0.640269008 | 0.844716186 |
| chr14:53942966-53943215 | -0.21856985 | 0.640271434 | 0.844716186 |
| chr5:62460561-62460761 | -0.241707606 | 0.640279316 | 0.844716186 |
| chr10:95183355-95184186 | -0.156689282 | 0.640281988 | 0.844716186 |
| chr10:45008544-45008988 | 0.170015921 | 0.640289488 | 0.844716186 |
| chr3:112467668-112467953 | -0.234332546 | 0.640289622 | 0.844716186 |
| chr19:33750171-33750371 | -0.26595353 | 0.640319284 | 0.844733208 |
| chr5:180625356-180625579 | 0.198635023 | 0.640324573 | 0.844733208 |
| chrX:146887056-146887406 | 0.180374324 | 0.640345546 | 0.844744818 |
| chr9:132800149-132800493 | 0.191169918 | 0.640355423 | 0.844744818 |
| chr7:100022700-100023023 | -0.188524119 | 0.640487398 | 0.844893912 |
| chr8:50020193-50020393 | -0.243556432 | 0.640497215 | 0.844893912 |
| chr6:68343961-68344261 | 0.1819097 | 0.640501524 | 0.844893912 |
| chr3:40457-40727 | 0.220738773 | 0.640606757 | 0.845018178 |
| chr10:100285925-100287021 | 0.152909753 | 0.640621068 | 0.845022508 |
| chr17:41835807-41836063 | 0.206859637 | 0.640642775 | 0.845036594 |
| chr3:187890251-187890859 | -0.173348502 | 0.640658897 | 0.845043312 |
| chr15:99669280-99669532 | -0.236932308 | 0.640691779 | 0.845065857 |
| chr4:75652197-75652525 | 0.202420889 | 0.640698047 | 0.845065857 |
| chr3:119579343-119580091 | 0.156228469 | 0.640737428 | 0.845074214 |
| chr6:120230063-120230396 | -0.195865635 | 0.640745818 | 0.845074214 |
| chr17:57021451-57022017 | 0.165931433 | 0.640746195 | 0.845074214 |
| chr7:76294760-76294960 | -0.191653897 | 0.640772892 | 0.845074214 |
| chr5:134102030-134102265 | 0.195828887 | 0.640787681 | 0.845074214 |
| chr12:79934301-79935653 | -0.146997279 | 0.64079257 | 0.845074214 |

| | | | |
|---|---|---|---|
| chr8:140456945-140458604 | 0.146156148 | 0.640830714 | 0.845074214 |
| chr9:25602436-25602710 | 0.196613416 | 0.640833311 | 0.845074214 |
| chr3:197487224-197487424 | 0.255736647 | 0.640836494 | 0.845074214 |
| chr10:109224020-109224267 | -0.216765676 | 0.640839854 | 0.845074214 |
| chr7:138396344-138396629 | -0.22057215 | 0.640851067 | 0.845074214 |
| chr11:94741364-94741641 | 0.203604143 | 0.640855361 | 0.845074214 |
| chr14:103500245-103500520 | 0.203604143 | 0.640855361 | 0.845074214 |
| chr22:45309511-45310321 | 0.155406117 | 0.640860114 | 0.845074214 |
| chr2:43995464-43996669 | -0.148445598 | 0.64086982 | 0.845074214 |
| chr14:64097446-64097675 | 0.216836488 | 0.640908233 | 0.84507669 |
| chr6:118264076-118264276 | 0.249814458 | 0.640917381 | 0.84507669 |
| chr7:101386904-101387158 | 0.21336528 | 0.640935955 | 0.84507669 |
| chr7:80334177-80334450 | -0.253622477 | 0.640945731 | 0.84507669 |
| chr3:186108504-186108741 | 0.197574266 | 0.640947341 | 0.84507669 |
| chr17:60161695-60162071 | 0.152337634 | 0.640953737 | 0.84507669 |
| chr4:82798618-82799094 | 0.165995262 | 0.640959881 | 0.84507669 |
| chr13:29472373-29472669 | 0.197231105 | 0.640959931 | 0.84507669 |
| chr1:234691692-234691892 | 0.249373769 | 0.640980464 | 0.84508922 |
| chr4:160836489-160836832 | -0.178227452 | 0.641047864 | 0.84516354 |
| chr1:101288480-101288919 | 0.175546602 | 0.641081937 | 0.845193919 |
| chr8:80578106-80578877 | 0.160177899 | 0.641098578 | 0.845201316 |
| chr6:12234047-12234294 | -0.211369555 | 0.641126963 | 0.845224194 |
| chr10:344567-344907 | 0.201456502 | 0.64114566 | 0.845234301 |
| chr14:61586063-61586282 | 0.230401953 | 0.6411697 | 0.845238907 |
| chr3:126703699-126704840 | -0.150314387 | 0.641171217 | 0.845238907 |
| chr17:60421287-60421507 | 0.163822395 | 0.641182934 | 0.845239811 |
| chr5:148979026-148979366 | -0.196904761 | 0.641214505 | 0.845266887 |
| chr1:179846375-179846575 | 0.262332787 | 0.641252626 | 0.845302596 |
| chr2:242088453-242089004 | 0.165732458 | 0.641302728 | 0.845343165 |
| chr1:175154707-175155189 | 0.173497108 | 0.641305467 | 0.845343165 |
| chr8:105881515-105881746 | 0.225671467 | 0.641327846 | 0.845352139 |
| chr4:82218412-82218612 | -0.244438558 | 0.641336675 | 0.845352139 |
| chr6:32940929-32941232 | 0.189173661 | 0.641345373 | 0.845352139 |
| chr8:143616890-143618321 | 0.148137117 | 0.641365908 | 0.845364663 |
| chr6:156776932-156777615 | 0.16051529 | 0.641391454 | 0.845371978 |
| chr1:223504717-223505003 | -0.208043971 | 0.641400949 | 0.845371978 |
| chr1:42835886-42836261 | -0.229953238 | 0.641405143 | 0.845371978 |
| chr20:25139130-25139468 | -0.182847503 | 0.64141559 | 0.845371978 |
| chr3:128170951-128171221 | 0.193701061 | 0.64145739 | 0.845412528 |
| chr2:112958901-112959150 | 0.196416715 | 0.641475335 | 0.845421637 |
| chrX:102516458-102517042 | -0.161644175 | 0.641499944 | 0.845439528 |
| chr11:18526294-18527375 | 0.147301168 | 0.641517589 | 0.84544824 |
| chr20:1185748-1186024 | 0.220060314 | 0.641538638 | 0.845461438 |
| chr3:149624081-149624393 | 0.203692288 | 0.641560332 | 0.845475487 |
| chr2:177799612-177799869 | -0.205364495 | 0.641610457 | 0.84549823 |
| chr1:48249876-48250139 | 0.205488667 | 0.641632353 | 0.84549823 |
| chr14:53110060-53110297 | 0.231904976 | 0.641634272 | 0.84549823 |
| chr14:35215127-35215628 | 0.180803427 | 0.641634912 | 0.84549823 |
| chr17:7997290-7997531 | 0.232756816 | 0.641642906 | 0.84549823 |
| chr19:33819974-33820576 | 0.16531931 | 0.641650217 | 0.84549823 |

| | | | |
|---|---|---|---|
| chr4:6546649-6546889 | -0.228225357 | 0.641660822 | 0.84549823 |
| chr16:81350305-81350515 | 0.220970948 | 0.641674221 | 0.84549823 |
| chr1:151365051-151365251 | 0.198130271 | 0.64171164 | 0.84549823 |
| chr12:77720381-77720654 | -0.234490975 | 0.641726403 | 0.84549823 |
| chr2:75549441-75550111 | -0.15431304 | 0.641733079 | 0.84549823 |
| chr3:39291852-39292165 | 0.183789908 | 0.64173712 | 0.84549823 |
| chr20:36122181-36122852 | -0.164208509 | 0.641751884 | 0.84549823 |
| chr3:45593754-45594137 | 0.181727425 | 0.641764903 | 0.84549823 |
| chr16:69859726-69859978 | 0.212641962 | 0.641765019 | 0.84549823 |
| chr1:44652958-44653218 | 0.197527925 | 0.64177192 | 0.84549823 |
| chr2:110678551-110678783 | -0.211498482 | 0.641773974 | 0.84549823 |
| chr5:66757275-66757671 | -0.192235684 | 0.641776214 | 0.84549823 |
| chr1:146472197-146473038 | -0.159068359 | 0.641791548 | 0.845503894 |
| chr6:31169323-31169699 | 0.196980776 | 0.641838994 | 0.845551861 |
| chr20:35541772-35542582 | 0.153210664 | 0.641871799 | 0.84558054 |
| chr4:145322782-145323223 | -0.211265472 | 0.641922398 | 0.845623713 |
| chr8:130821355-130821555 | -0.265732177 | 0.641926643 | 0.845623713 |
| chr16:48229693-48230004 | 0.208149158 | 0.641987294 | 0.845679505 |
| chr1:156489095-156489358 | 0.196138293 | 0.641993616 | 0.845679505 |
| chr2:8602863-8603190 | 0.185576708 | 0.642002506 | 0.845679505 |
| chr3:149258946-149259232 | 0.211825939 | 0.642013144 | 0.845679505 |
| chr18:50784768-50785012 | -0.209153538 | 0.64204411 | 0.845705755 |
| chr4:40675237-40675494 | 0.231235345 | 0.642080424 | 0.845739049 |
| chr20:51562354-51563043 | 0.156245524 | 0.642104701 | 0.845756487 |
| chr13:41960344-41961458 | -0.150760265 | 0.642147068 | 0.845797753 |
| chr14:22712933-22713486 | -0.173220214 | 0.642212698 | 0.845860122 |
| chr20:5100806-5101030 | 0.226076768 | 0.642216499 | 0.845860122 |
| chr8:98293429-98294536 | 0.150480956 | 0.642278347 | 0.845908546 |
| chr3:155347853-155348076 | 0.229639125 | 0.642283607 | 0.845908546 |
| chr8:50198400-50198622 | -0.242362427 | 0.642286385 | 0.845908546 |
| chr5:32489718-32490046 | 0.182625277 | 0.642328994 | 0.845950123 |
| chr16:4733907-4735024 | 0.149038508 | 0.64235065 | 0.845964103 |
| chr12:57888036-57888276 | -0.200127436 | 0.642376444 | 0.845983532 |
| chr13:113458331-113458531 | 0.237989667 | 0.642427461 | 0.846036179 |
| chr12:51217533-51218420 | 0.159887853 | 0.642492422 | 0.846107185 |
| chr16:80526535-80526948 | 0.187965043 | 0.64250607 | 0.846110616 |
| chr3:63108004-63108218 | 0.180788773 | 0.642550684 | 0.846118126 |
| chr4:87985080-87985280 | -0.232853343 | 0.642555964 | 0.846118126 |
| chr16:16127659-16127959 | -0.198455566 | 0.642557851 | 0.846118126 |
| chr3:108437458-108437705 | 0.225877252 | 0.642560497 | 0.846118126 |
| chr20:53217122-53217389 | 0.203058463 | 0.642566986 | 0.846118126 |
| chr8:105797629-105797853 | 0.220429271 | 0.642591265 | 0.846135554 |
| chr8:90644878-90646629 | 0.145378984 | 0.642618002 | 0.846142782 |
| chr17:44947300-44948122 | 0.156483896 | 0.64261884 | 0.846142782 |
| chr6:108782322-108782660 | -0.190632251 | 0.642661496 | 0.846184406 |
| chr6:30716999-30717826 | 0.149437533 | 0.642681922 | 0.846187905 |
| chr15:60916639-60916960 | 0.216459364 | 0.642686241 | 0.846187905 |
| chrX:71160526-71160825 | -0.205576219 | 0.642742638 | 0.846247618 |
| chr1:227727999-227728990 | 0.152212604 | 0.642782728 | 0.84627616 |
| chr17:71473272-71473501 | -0.199355633 | 0.642786406 | 0.84627616 |

| | | | |
|---|---|---|---|
| chr2:204260174-204260434 | -0.205263872 | 0.642842164 | 0.846323531 |
| chr8:99605031-99605288 | -0.21193005 | 0.642844477 | 0.846323531 |
| chr14:100252090-100252387 | -0.232442131 | 0.642886124 | 0.846361337 |
| chr12:39103588-39103934 | -0.184735506 | 0.642903187 | 0.846361337 |
| chr2:218959263-218959770 | 0.162437029 | 0.642906331 | 0.846361337 |
| chr5:27815836-27816748 | -0.153717788 | 0.642940915 | 0.846378014 |
| chr4:139705132-139705416 | -0.228678271 | 0.642969797 | 0.846378014 |
| chr10:126979253-126979592 | 0.208651744 | 0.642971898 | 0.846378014 |
| chr6:24402643-24403605 | -0.150293824 | 0.642972041 | 0.846378014 |
| chr14:63799109-63799325 | 0.247601497 | 0.64297423 | 0.846378014 |
| chrX:129957300-129957645 | -0.180319289 | 0.643004623 | 0.846394947 |
| chr11:72414147-72414491 | -0.182171284 | 0.643009186 | 0.846394947 |
| chr20:62642014-62643096 | 0.15100228 | 0.64302096 | 0.846395904 |
| chr6:39362079-39362349 | 0.195813142 | 0.643051147 | 0.846411542 |
| chr6:154882107-154882353 | -0.246832358 | 0.643068142 | 0.846411542 |
| chr4:75513817-75515104 | -0.146952218 | 0.643068214 | 0.846411542 |
| chr11:85647416-85648418 | -0.148552716 | 0.64309304 | 0.846411542 |
| chr3:165953552-165953868 | -0.217489966 | 0.643098014 | 0.846411542 |
| chr19:34926708-34927334 | 0.171145827 | 0.64309912 | 0.846411542 |
| chr3:51974910-51975220 | 0.192750828 | 0.643124109 | 0.846417959 |
| chr1:106465580-106465851 | 0.178000058 | 0.643126088 | 0.846417959 |
| chr12:120317211-120317959 | -0.149259161 | 0.643164801 | 0.846446917 |
| chr17:4998230-4998467 | -0.194921856 | 0.643182045 | 0.846446917 |
| chr11:46389406-46389711 | 0.169118713 | 0.643188381 | 0.846446917 |
| chr5:9156362-9156648 | -0.197685907 | 0.643192279 | 0.846446917 |
| chr3:138941355-138941643 | -0.182877913 | 0.643218436 | 0.846464139 |
| chr1:192756885-192757167 | 0.240459889 | 0.643227461 | 0.846464139 |
| chr16:3457429-3458227 | 0.155643961 | 0.64324655 | 0.846474721 |
| chr3:167628937-167629269 | -0.219266423 | 0.643257789 | 0.846474974 |
| chrX:100654358-100654581 | 0.211656747 | 0.643356623 | 0.846590492 |
| chr1:158121736-158121945 | -0.196565162 | 0.643368346 | 0.846591379 |
| chr10:48215505-48215796 | -0.189818603 | 0.643395576 | 0.846607359 |
| chr2:121736447-121737352 | 0.1475281 | 0.643402588 | 0.846607359 |
| chr12:76083815-76085223 | -0.145657762 | 0.643437394 | 0.846625792 |
| chr1:79568498-79568770 | 0.196110116 | 0.643454805 | 0.846625792 |
| chr7:15370218-15370418 | 0.248630933 | 0.643460409 | 0.846625792 |
| chr3:139343830-139344690 | -0.151095958 | 0.643460795 | 0.846625792 |
| chr7:43954565-43954845 | 0.183630801 | 0.643502864 | 0.846666606 |
| chr6:17012047-17013174 | -0.155889812 | 0.643517683 | 0.846671565 |
| chr2:24016331-24016777 | -0.169770409 | 0.643550819 | 0.846700623 |
| chrX:12950945-12951173 | -0.203131787 | 0.643564172 | 0.846703652 |
| chr1:114378376-114379123 | 0.150273441 | 0.643603269 | 0.846714974 |
| chr12:52148507-52149006 | 0.156726797 | 0.643609891 | 0.846714974 |
| chr3:13251901-13252101 | 0.221721953 | 0.64361049 | 0.846714974 |
| chr10:102834804-102835283 | -0.182591222 | 0.643620587 | 0.846714974 |
| chr7:2116201-2116616 | 0.177377956 | 0.64362803 | 0.846714974 |
| chr6:96923933-96925330 | -0.146606298 | 0.643709622 | 0.846801559 |
| chr13:38685994-38687462 | -0.146619154 | 0.643728053 | 0.846801559 |
| chr14:64625470-64626009 | 0.166593599 | 0.643730928 | 0.846801559 |
| chr3:149992757-149993037 | 0.208963141 | 0.643738054 | 0.846801559 |

| | | | |
|---|---|---|---|
| chr19:17022237-17022631 | -0.19037644 | 0.643755142 | 0.846809499 |
| chr8:93057714-93058134 | 0.174975718 | 0.643794421 | 0.846846628 |
| chr6:27726578-27727054 | -0.161024215 | 0.643869621 | 0.846931007 |
| chr1:170136100-170136572 | 0.170340754 | 0.643907725 | 0.846964622 |
| chr3:170809340-170809834 | -0.163443409 | 0.643919354 | 0.846964622 |
| chr1:10472224-10472765 | -0.165309282 | 0.643930825 | 0.846964622 |
| chr12:6493142-6494317 | -0.149244672 | 0.643950224 | 0.846964622 |
| chr14:94136699-94137111 | 0.168140811 | 0.643953555 | 0.846964622 |
| chr2:64649695-64649938 | -0.180935591 | 0.643961499 | 0.846964622 |
| chr1:183719375-183719623 | 0.20743032 | 0.64400567 | 0.847008178 |
| chr12:116599672-116599878 | 0.22853878 | 0.644021971 | 0.847015078 |
| chr22:29156611-29156924 | -0.189144356 | 0.644058322 | 0.847048347 |
| chr6:43804385-43804741 | -0.197113722 | 0.644083696 | 0.84706718 |
| chr2:227407632-227408008 | -0.186351387 | 0.644098566 | 0.847072197 |
| chr22:32206772-32207269 | 0.169868651 | 0.644130138 | 0.847078203 |
| chr1:151790048-151791613 | 0.144715069 | 0.644130588 | 0.847078203 |
| chr6:11538361-11538588 | -0.194354331 | 0.64413866 | 0.847078203 |
| chr9:126228878-126229216 | -0.176783413 | 0.644148582 | 0.847078203 |
| chr8:106674085-106674285 | 0.229796602 | 0.64415841 | 0.847078203 |
| chr2:200756651-200757379 | -0.172447113 | 0.644174221 | 0.847084457 |
| chr10:126568831-126569031 | 0.255244476 | 0.644200478 | 0.847104126 |
| chr4:83034327-83035532 | -0.145936573 | 0.644216737 | 0.847104126 |
| chr19:1260862-1261225 | -0.171033771 | 0.644222345 | 0.847104126 |
| chr14:91464809-91465009 | 0.238730257 | 0.644240963 | 0.84711407 |
| chr2:8684696-8686130 | -0.147245767 | 0.644280577 | 0.847151621 |
| chr16:11076459-11076674 | 0.236652786 | 0.644302412 | 0.847165793 |
| chr8:47260436-47261493 | 0.153192862 | 0.644350358 | 0.847205674 |
| chr17:7315102-7315514 | 0.174355542 | 0.644366735 | 0.847205674 |
| chr2:70861885-70862190 | -0.189297862 | 0.644376904 | 0.847205674 |
| chr12:71415676-71416126 | 0.183156869 | 0.644388939 | 0.847205674 |
| chr20:37615875-37616075 | 0.222321059 | 0.644400555 | 0.847205674 |
| chr16:81198782-81199045 | 0.188514066 | 0.644401 | 0.847205674 |
| chr22:22753555-22753818 | -0.201136277 | 0.644410142 | 0.847205674 |
| chr19:55653681-55655175 | 0.144845553 | 0.644427763 | 0.847214303 |
| chr5:132813553-132813801 | -0.210810019 | 0.644470465 | 0.847255906 |
| chr7:42763331-42763566 | -0.214866363 | 0.644507438 | 0.847289976 |
| chr12:98955362-98955605 | -0.20108229 | 0.644539312 | 0.847306482 |
| chr16:28093495-28093702 | -0.221516874 | 0.644551557 | 0.847306482 |
| chr8:27426207-27426520 | -0.190579997 | 0.644553169 | 0.847306482 |
| chr6:44630393-44630752 | 0.181314027 | 0.644594035 | 0.847345666 |
| chr13:112969270-112969576 | 0.202371101 | 0.644635357 | 0.847385447 |
| chr4:54539481-54539848 | 0.185881926 | 0.644685418 | 0.84742301 |
| chr10:21195235-21195742 | -0.166439631 | 0.644686052 | 0.84742301 |
| chr1:51192282-51192482 | 0.237968723 | 0.644714983 | 0.847446501 |
| chr3:20185853-20186667 | 0.146348167 | 0.644753195 | 0.84748219 |
| chr17:55601817-55602118 | -0.186227285 | 0.644771669 | 0.847491934 |
| chr9:128702267-128702874 | 0.159470832 | 0.644808562 | 0.847516453 |
| chr2:69914216-69915307 | -0.145125208 | 0.644812445 | 0.847516453 |
| chr14:106421581-106422023 | 0.182304345 | 0.64485623 | 0.847559463 |
| chr7:41979838-41980038 | 0.223137234 | 0.644896503 | 0.847571464 |

| | | | |
|---|---|---|---|
| chr20:44467423-44467623 | -0.243315487 | 0.644903611 | 0.847571464 |
| chr22:18122260-18122460 | -0.243315487 | 0.644903611 | 0.847571464 |
| chr7:56033865-56034710 | 0.153241772 | 0.644930819 | 0.847571464 |
| chr1:174999198-175000605 | -0.143844547 | 0.644931512 | 0.847571464 |
| chr3:128680414-128681398 | -0.147388396 | 0.64493173 | 0.847571464 |
| chr8:138371400-138371644 | 0.220995958 | 0.644943222 | 0.847572029 |
| chr8:92835865-92836065 | 0.247422863 | 0.64497784 | 0.847602985 |
| chr3:10754044-10754252 | 0.215933542 | 0.645017426 | 0.84764047 |
| chr3:52455313-52455666 | -0.17251619 | 0.645053141 | 0.847664663 |
| chr1:20633721-20634145 | 0.172666922 | 0.645057962 | 0.847664663 |
| chr5:132759126-132759415 | 0.189147061 | 0.645101973 | 0.847699438 |
| chr3:64422219-64422419 | 0.174274866 | 0.645124887 | 0.847699438 |
| chr2:46780955-46781155 | -0.233198057 | 0.645152721 | 0.847699438 |
| chr6:16183023-16183390 | -0.204723822 | 0.645216672 | 0.847699438 |
| chr12:70577081-70577322 | -0.213496358 | 0.645217817 | 0.847699438 |
| chr8:119152942-119153142 | -0.265163242 | 0.645221779 | 0.847699438 |
| chrX:97921475-97921685 | -0.265163242 | 0.645221779 | 0.847699438 |
| chr14:35336629-35337039 | 0.162843381 | 0.645222803 | 0.847699438 |
| chr3:167290110-167290310 | 0.250163216 | 0.645226102 | 0.847699438 |
| chr2:229587197-229587397 | -0.245259035 | 0.645227172 | 0.847699438 |
| chr1:51214956-51215364 | -0.194389311 | 0.645227278 | 0.847699438 |
| chr11:15074663-15075334 | -0.157535402 | 0.645228175 | 0.847699438 |
| chr3:24088797-24089085 | -0.203867794 | 0.645230579 | 0.847699438 |
| chr3:195960870-195961523 | 0.164899514 | 0.645239313 | 0.847699438 |
| chrX:70197331-70197531 | -0.241553592 | 0.645266607 | 0.847710904 |
| chr4:83906043-83906280 | 0.22403802 | 0.645281979 | 0.847710904 |
| chr2:216693161-216693432 | 0.182941512 | 0.645295681 | 0.847710904 |
| chr8:6707831-6709162 | -0.146472626 | 0.645310773 | 0.847710904 |
| chr13:46126591-46126941 | 0.161531709 | 0.645311337 | 0.847710904 |
| chr2:217366970-217367618 | 0.174191796 | 0.645314422 | 0.847710904 |
| chr17:48908903-48909301 | 0.16015002 | 0.645345741 | 0.847726502 |
| chr20:44151666-44151866 | -0.238015826 | 0.645348423 | 0.847726502 |
| chr2:198374458-198374700 | 0.221579348 | 0.645374651 | 0.847746421 |
| chr15:92904043-92904710 | 0.146524375 | 0.645405563 | 0.84776818 |
| chr3:152645900-152646252 | -0.196306339 | 0.645413344 | 0.84776818 |
| chr12:56359882-56360888 | 0.147231444 | 0.645469535 | 0.847827454 |
| chr8:11066917-11067291 | -0.192690585 | 0.645491524 | 0.847833538 |
| chr9:129823374-129824447 | -0.154945004 | 0.645496297 | 0.847833538 |
| chr19:35644116-35644744 | 0.151854911 | 0.645552493 | 0.847862782 |
| chr2:84089982-84090197 | -0.223083222 | 0.645557402 | 0.847862782 |
| chr8:67711821-67712687 | -0.159643902 | 0.645564526 | 0.847862782 |
| chr15:40823605-40823960 | 0.174211469 | 0.64559159 | 0.847862782 |
| chr3:170356653-170358529 | -0.143235368 | 0.64559571 | 0.847862782 |
| chr19:4909838-4910186 | -0.181579935 | 0.645595969 | 0.847862782 |
| chr13:19075178-19075449 | -0.21553721 | 0.645605115 | 0.847862782 |
| chr6:135850923-135851144 | -0.226892588 | 0.645607086 | 0.847862782 |
| chrX:48911447-48912390 | -0.152901107 | 0.645635934 | 0.847886134 |
| chr2:149989529-149990108 | -0.173477994 | 0.645653216 | 0.847894298 |
| chr12:54853983-54854478 | 0.166954414 | 0.645679473 | 0.847914248 |
| chr5:26812906-26813357 | -0.200348805 | 0.645714719 | 0.847934351 |

| | | | |
|---|---|---|---|
| chr1:150560948-150562401 | 0.14355827 | 0.645738939 | 0.847934351 |
| chr1:184754153-184755489 | -0.143705994 | 0.645743523 | 0.847934351 |
| chr17:77957883-77958943 | 0.14677436 | 0.645750109 | 0.847934351 |
| chr1:67367781-67368071 | -0.196127783 | 0.645750114 | 0.847934351 |
| chr7:141073203-141073595 | -0.171540495 | 0.645765434 | 0.847939935 |
| chr3:188206019-188206354 | 0.191118392 | 0.645808993 | 0.84795395 |
| chr9:14846821-14847102 | -0.202955066 | 0.64580936 | 0.84795395 |
| chr4:145710036-145710333 | 0.212633618 | 0.645816184 | 0.84795395 |
| chr11:74398292-74399227 | -0.148788318 | 0.645820374 | 0.84795395 |
| chr1:61446530-61447056 | -0.184100095 | 0.645865835 | 0.847999108 |
| chr16:30123113-30123506 | 0.166405595 | 0.645914219 | 0.848048104 |
| chr13:113013115-113013777 | -0.156810725 | 0.64594632 | 0.848070421 |
| chr1:89994674-89995666 | -0.148792782 | 0.645953354 | 0.848070421 |
| chr5:10987926-10988252 | 0.17081038 | 0.645992535 | 0.84810733 |
| chr16:58733673-58734598 | -0.152692846 | 0.646005416 | 0.84810971 |
| chr14:23071367-23071801 | -0.15800065 | 0.646037151 | 0.84812434 |
| chr3:138979754-138980079 | -0.185656167 | 0.646038698 | 0.84812434 |
| chr5:1008688-1009048 | 0.179375898 | 0.646108941 | 0.848202023 |
| chr10:95289667-95289917 | -0.192797669 | 0.646143824 | 0.848233284 |
| chr2:20474244-20474824 | -0.15725132 | 0.646169349 | 0.848239041 |
| chr3:131217480-131217812 | 0.196825416 | 0.64617035 | 0.848239041 |
| chr19:19628110-19628384 | -0.200089169 | 0.646201236 | 0.848265052 |
| chr14:77097674-77098792 | 0.144749943 | 0.646215057 | 0.848268663 |
| chr6:38933531-38933741 | -0.245858398 | 0.646245585 | 0.848294204 |
| chr7:54543279-54543479 | -0.226426599 | 0.646257378 | 0.848295151 |
| chr15:33074995-33075271 | -0.194826133 | 0.646287571 | 0.84832025 |
| chr2:212537144-212538142 | -0.158457731 | 0.646311177 | 0.84833039 |
| chr11:44065848-44066663 | -0.155014979 | 0.646328457 | 0.84833039 |
| chr10:91787745-91788150 | -0.179865077 | 0.646329177 | 0.84833039 |
| chr16:27154958-27155516 | -0.175276488 | 0.646339583 | 0.84833039 |
| chr5:14108106-14108364 | -0.194360906 | 0.646390882 | 0.848380765 |
| chr14:49153132-49153384 | 0.222066805 | 0.646400108 | 0.848380765 |
| chr3:63076494-63076727 | 0.172920986 | 0.646423302 | 0.848396674 |
| chr10:82928685-82928898 | 0.229533833 | 0.646436426 | 0.848399367 |
| chr22:39437211-39437503 | 0.203316327 | 0.646456225 | 0.848410819 |
| chr1:201399143-201399899 | 0.14942785 | 0.646482946 | 0.848431356 |
| chr17:7314720-7314974 | -0.195519574 | 0.646515711 | 0.848447511 |
| chr20:62182614-62183411 | 0.151757668 | 0.646524796 | 0.848447511 |
| chr5:135399040-135399587 | 0.160876193 | 0.646528475 | 0.848447511 |
| chr11:46247540-46247865 | -0.178539691 | 0.646561976 | 0.848449535 |
| chr5:173802840-173803160 | -0.20352383 | 0.646573009 | 0.848449535 |
| chr3:163632760-163633026 | -0.203493451 | 0.646581242 | 0.848449535 |
| chr5:149320790-149321244 | -0.169400153 | 0.646592883 | 0.848449535 |
| chr16:84088799-84089020 | -0.211080816 | 0.646596304 | 0.848449535 |
| chr5:6706024-6706293 | -0.202215205 | 0.646607209 | 0.848449535 |
| chr20:59955208-59955595 | 0.156493212 | 0.64660753 | 0.848449535 |
| chr16:85970781-85971016 | 0.210471702 | 0.646637836 | 0.848462896 |
| chr1:236523022-236524737 | 0.145194307 | 0.646658152 | 0.848462896 |
| chrX:119605503-119606690 | 0.148303304 | 0.646663376 | 0.848462896 |
| chr9:35719257-35719457 | -0.2582336 | 0.646672168 | 0.848462896 |

| | | | |
|---|---|---|---|
| chr5:62223429-62223690 | -0.216310149 | 0.646673079 | 0.848462896 |
| chr11:59754861-59756160 | 0.147609035 | 0.646723942 | 0.848481658 |
| chr8:75406941-75408218 | -0.148348129 | 0.646725269 | 0.848481658 |
| chr6:107675079-107675878 | -0.155528688 | 0.646727599 | 0.848481658 |
| chr15:86213274-86213721 | -0.20056355 | 0.646738581 | 0.848481658 |
| chr17:51535860-51536168 | -0.173306072 | 0.646742747 | 0.848481658 |
| chr10:128125903-128127117 | 0.149188367 | 0.646785934 | 0.848523788 |
| chr3:184565783-184566086 | 0.165715127 | 0.646901948 | 0.84864206 |
| chr6:51398868-51399114 | 0.204227245 | 0.646912097 | 0.84864206 |
| chr15:75903604-75904645 | -0.148359184 | 0.646914014 | 0.84864206 |
| chr3:173037957-173038421 | -0.187479796 | 0.646926267 | 0.84864206 |
| chr6:116253281-116254492 | -0.146880271 | 0.646931465 | 0.84864206 |
| chr14:26137208-26137455 | 0.219717493 | 0.646968326 | 0.848663547 |
| chr8:128032839-128033356 | 0.15229599 | 0.646973751 | 0.848663547 |
| chr20:62516401-62516701 | -0.182901414 | 0.646982413 | 0.848663547 |
| chr17:74744621-74744920 | 0.167056971 | 0.646995456 | 0.848663547 |
| chr1:235160741-235161610 | 0.150189927 | 0.647003225 | 0.848663547 |
| chr4:80335486-80336322 | -0.149764415 | 0.647044011 | 0.848697314 |
| chr3:136391607-136391943 | 0.178586807 | 0.647058771 | 0.848697314 |
| chr5:1004166-1004383 | -0.192108602 | 0.647062197 | 0.848697314 |
| chr2:69643512-69644580 | -0.148263912 | 0.647101725 | 0.848734631 |
| chr3:33372695-33373072 | -0.204024899 | 0.647127128 | 0.848746425 |
| chr17:57597839-57598141 | 0.165883724 | 0.647139321 | 0.848746425 |
| chr3:124730201-124730902 | -0.154872626 | 0.647143949 | 0.848746425 |
| chr1:245156422-245156811 | 0.160153389 | 0.64721469 | 0.84881904 |
| chr6:153401839-153402141 | -0.209799357 | 0.647221471 | 0.84881904 |
| chr4:86498622-86498834 | 0.210853265 | 0.647232984 | 0.84881961 |
| chr3:132352618-132352881 | 0.18727398 | 0.647247436 | 0.848824035 |
| chr13:25494199-25494468 | -0.19841295 | 0.64726607 | 0.848833944 |
| chr6:34756936-34757696 | -0.152579241 | 0.647291006 | 0.848852117 |
| chr4:42397321-42397590 | 0.232894587 | 0.647319362 | 0.848859807 |
| chr21:37914298-37914534 | -0.265945657 | 0.647355654 | 0.848859807 |
| chr19:2204093-2204298 | -0.265945657 | 0.647355654 | 0.848859807 |
| chr14:72933877-72934077 | 0.248476078 | 0.647358844 | 0.848859807 |
| chr11:108222368-108223425 | -0.144796385 | 0.647380149 | 0.848859807 |
| chr10:108544326-108544822 | -0.17668395 | 0.64738409 | 0.848859807 |
| chr12:12724698-12725954 | -0.146863562 | 0.647395774 | 0.848859807 |
| chr2:24015626-24016063 | -0.160822414 | 0.647396259 | 0.848859807 |
| chr14:58401631-58401871 | 0.199453322 | 0.647399148 | 0.848859807 |
| chr15:66803784-66804035 | -0.235301853 | 0.647407655 | 0.848859807 |
| chr3:195442396-195443519 | -0.145738174 | 0.647422448 | 0.848864676 |
| chr8:70113728-70113941 | -0.232279231 | 0.647453414 | 0.848890751 |
| chr11:129309071-129309271 | 0.253349237 | 0.647495306 | 0.848920086 |
| chr17:61201189-61201536 | 0.169032517 | 0.647502231 | 0.848920086 |
| chr22:25447656-25448244 | 0.157963345 | 0.647509026 | 0.848920086 |
| chr15:58957290-58957898 | 0.161175014 | 0.64758727 | 0.848991565 |
| chr4:26330241-26330451 | 0.209059221 | 0.647587792 | 0.848991565 |
| chr3:149277561-149277925 | 0.175241149 | 0.647596787 | 0.848991565 |
| chr12:120056832-120057032 | 0.231406538 | 0.647616577 | 0.849002983 |
| chr1:62530602-62530847 | 0.196115928 | 0.64763978 | 0.849013372 |

| | | | |
|---|---|---|---|
| chr6:132598478-132598801 | -0.186340929 | 0.647646663 | 0.849013372 |
| chr11:117619072-117619347 | 0.198836174 | 0.647717072 | 0.849091146 |
| chr5:157971701-157971901 | 0.23749666 | 0.64773863 | 0.849100263 |
| chr2:156181877-156182576 | -0.150164933 | 0.647762658 | 0.849100263 |
| chr1:64948045-64948540 | -0.179269883 | 0.64776398 | 0.849100263 |
| chr17:78013654-78014116 | -0.18153048 | 0.647774668 | 0.849100263 |
| chr19:57850609-57850839 | 0.208669904 | 0.647779435 | 0.849100263 |
| chr2:125654248-125654573 | -0.184645088 | 0.647841364 | 0.849164635 |
| chr17:55774118-55774447 | -0.199693613 | 0.64785396 | 0.849164635 |
| chr1:27391618-27391848 | 0.18563131 | 0.647861792 | 0.849164635 |
| chr7:157544652-157544860 | -0.233609862 | 0.647902486 | 0.849203447 |
| chr3:39434929-39435129 | -0.241063491 | 0.64793708 | 0.84920369 |
| chr5:115349381-115349664 | 0.196434298 | 0.647939323 | 0.84920369 |
| chr20:45322960-45323193 | 0.187897245 | 0.647955461 | 0.84920369 |
| chr17:49132697-49133020 | 0.179789316 | 0.647956701 | 0.84920369 |
| chr9:125374768-125375033 | 0.182645036 | 0.647958087 | 0.84920369 |
| chr10:110671864-110672784 | -0.16274926 | 0.647977918 | 0.849215155 |
| chr17:21223906-21224111 | -0.218248339 | 0.648020605 | 0.84924174 |
| chr1:145215832-145216296 | -0.183853003 | 0.648020802 | 0.84924174 |
| chr2:58427582-58428084 | -0.17221997 | 0.648044287 | 0.84924174 |
| chr18:11947628-11947831 | -0.240795253 | 0.648046586 | 0.84924174 |
| chr7:12839421-12840458 | 0.147037225 | 0.648053621 | 0.84924174 |
| chr7:75092604-75093186 | 0.164611509 | 0.648091023 | 0.849276228 |
| chr17:78736363-78737097 | 0.155821732 | 0.648115469 | 0.849293738 |
| chr15:98177977-98178316 | -0.173514535 | 0.64814967 | 0.84932403 |
| chr15:72866710-72866961 | -0.23548174 | 0.648210286 | 0.849364585 |
| chr12:116390284-116391060 | -0.150211865 | 0.648221605 | 0.849364585 |
| chr6:126054179-126054695 | 0.164538812 | 0.64822252 | 0.849364585 |
| chr16:1175821-1176063 | 0.223015592 | 0.64824943 | 0.849364585 |
| chr10:70382347-70382978 | 0.158177717 | 0.648261531 | 0.849364585 |
| chr20:33291673-33292026 | -0.195134024 | 0.648264791 | 0.849364585 |
| chr4:41281095-41281371 | 0.193602273 | 0.648272457 | 0.849364585 |
| chr5:28620289-28620790 | 0.196630401 | 0.648303981 | 0.849364585 |
| chr7:123208837-123209152 | -0.180927403 | 0.64831017 | 0.849364585 |
| chr4:111283163-111283365 | 0.224388737 | 0.648310839 | 0.849364585 |
| chr12:76423252-76423452 | -0.234054261 | 0.648332884 | 0.849364585 |
| chr12:80270070-80270301 | -0.214159593 | 0.648348214 | 0.849364585 |
| chr17:60382107-60382349 | 0.16168096 | 0.648357755 | 0.849364585 |
| chr9:6007724-6008738 | -0.146051307 | 0.648359146 | 0.849364585 |
| chr16:68448318-68448962 | 0.148703871 | 0.648363992 | 0.849364585 |
| chr1:58432911-58433177 | -0.205120891 | 0.648368467 | 0.849364585 |
| chr2:88880394-88880783 | -0.177771239 | 0.648369066 | 0.849364585 |
| chr1:33055189-33055999 | 0.155467599 | 0.648459654 | 0.849468731 |
| chr3:157201732-157202035 | -0.193511551 | 0.648540031 | 0.8495595 |
| chr3:166734521-166735222 | -0.156027815 | 0.648559287 | 0.8495702 |
| chr6:166956470-166957626 | -0.148188342 | 0.648596638 | 0.849604603 |
| chr2:27771316-27772093 | 0.152352273 | 0.648660607 | 0.84967387 |
| chrX:148049539-148049747 | 0.254171342 | 0.648678899 | 0.849683305 |
| chr12:56751816-56752647 | -0.154073965 | 0.648703046 | 0.849697559 |
| chr9:90985062-90985578 | 0.184364101 | 0.648726358 | 0.849697559 |

| | | | |
|---|---|---|---|
| chr12:33296046-33296248 | -0.271036551 | 0.648727468 | 0.849697559 |
| chr17:29292730-29293026 | -0.185097699 | 0.648734139 | 0.849697559 |
| chr5:53560569-53561163 | -0.166059412 | 0.648755785 | 0.849704825 |
| chr6:127342998-127343739 | -0.150023662 | 0.648761866 | 0.849704825 |
| chr19:17303320-17303615 | -0.181390537 | 0.648779298 | 0.849713133 |
| chr12:108628413-108628794 | -0.181628455 | 0.648794145 | 0.849714214 |
| chr11:65961258-65961873 | -0.153551526 | 0.648802304 | 0.849714214 |
| chr17:45794975-45795175 | -0.248200484 | 0.648820406 | 0.849716858 |
| chr1:28346103-28346497 | 0.200975525 | 0.648838084 | 0.849716858 |
| chr18:58781570-58781946 | 0.180109615 | 0.648848757 | 0.849716858 |
| chr10:97534827-97535087 | -0.183154946 | 0.648868986 | 0.849716858 |
| chr17:79905885-79906085 | 0.215157811 | 0.648870307 | 0.849716858 |
| chr7:98251374-98252682 | -0.145680792 | 0.64887086 | 0.849716858 |
| chr3:188929139-188929382 | -0.213719995 | 0.648887179 | 0.849723705 |
| chr9:34457721-34458031 | -0.17756544 | 0.648917023 | 0.849729639 |
| chr4:140252544-140253509 | -0.151887766 | 0.648931167 | 0.849729639 |
| chr1:226109550-226110848 | 0.146973885 | 0.648949674 | 0.849729639 |
| chr1:156651926-156652362 | -0.170332245 | 0.648954326 | 0.849729639 |
| chr1:41516124-41516415 | 0.175958023 | 0.648961354 | 0.849729639 |
| chr19:40702594-40702813 | 0.203085465 | 0.648975105 | 0.849729639 |
| chr15:92683335-92683800 | 0.166866625 | 0.648978027 | 0.849729639 |
| chr3:140552569-140552812 | 0.221404097 | 0.64898043 | 0.849729639 |
| chr9:128231987-128232197 | -0.23524445 | 0.649046201 | 0.849801234 |
| chr14:24188176-24189095 | 0.147038082 | 0.649082531 | 0.849813668 |
| chr11:122882677-122883491 | 0.160081234 | 0.649090599 | 0.849813668 |
| chr6:89484906-89485190 | -0.234881332 | 0.649111106 | 0.849813668 |
| chr1:107056433-107058065 | -0.141267209 | 0.649148529 | 0.849813668 |
| chr8:115563872-115564159 | 0.20883052 | 0.649149531 | 0.849813668 |
| chr11:126194195-126194407 | 0.237818266 | 0.649159812 | 0.849813668 |
| chr10:127853961-127854245 | -0.18867659 | 0.649159962 | 0.849813668 |
| chr12:108337436-108337636 | 0.216027702 | 0.649163179 | 0.849813668 |
| chr10:93816220-93816446 | -0.213585088 | 0.649168895 | 0.849813668 |
| chr4:68724520-68724736 | -0.241579478 | 0.649169892 | 0.849813668 |
| chr9:74940602-74940978 | -0.165852448 | 0.649208726 | 0.849813668 |
| chr2:130391165-130391946 | -0.154793723 | 0.649215743 | 0.849813668 |
| chr2:96265450-96266630 | 0.145512425 | 0.649215959 | 0.849813668 |
| chr5:97534124-97534487 | 0.176019447 | 0.64922416 | 0.849813668 |
| chr8:119755445-119755722 | -0.218605276 | 0.64922611 | 0.849813668 |
| chr5:169581693-169581939 | 0.177584612 | 0.649233153 | 0.849813668 |
| chr5:139438994-139439984 | -0.148351126 | 0.649246323 | 0.849816389 |
| chr10:119177824-119179129 | 0.145588294 | 0.649294373 | 0.849839256 |
| chr7:5614000-5614261 | -0.206053445 | 0.64930744 | 0.849839256 |
| chr17:50944008-50945005 | 0.145800686 | 0.649320266 | 0.849839256 |
| chr5:147511023-147511224 | 0.22123157 | 0.64933962 | 0.849839256 |
| chr14:75572215-75572441 | 0.19894782 | 0.649340084 | 0.849839256 |
| chr9:127771030-127771778 | 0.147240893 | 0.649348746 | 0.849839256 |
| chr9:129460270-129460617 | 0.162508822 | 0.6493541 | 0.849839256 |
| chr10:100968911-100969447 | 0.161416298 | 0.649356637 | 0.849839256 |
| chr2:20350209-20351873 | -0.142538197 | 0.649363615 | 0.849839256 |
| chr11:85663910-85665523 | -0.143726462 | 0.649512472 | 0.850002371 |

| | | | |
|---|---|---|---|
| chr20:62873693-62874048 | 0.168989201 | 0.64952419 | 0.850002371 |
| chr16:81778842-81780012 | -0.146548608 | 0.649528899 | 0.850002371 |
| chr2:241875883-241876097 | -0.210007569 | 0.649532625 | 0.850002371 |
| chr8:54073829-54074033 | 0.250435791 | 0.649549879 | 0.850010432 |
| chr22:21982696-21982950 | -0.198681233 | 0.649569719 | 0.850019331 |
| chr3:195558607-195558934 | -0.184203783 | 0.649580474 | 0.850019331 |
| chr1:176405753-176405978 | 0.224060485 | 0.64958996 | 0.850019331 |
| chr12:103877961-103878268 | -0.177767124 | 0.649602353 | 0.850020808 |
| chr8:135135825-135136027 | -0.230761037 | 0.649613276 | 0.850020808 |
| chr3:146503057-146504231 | 0.146525012 | 0.649665402 | 0.850074498 |
| chr4:110193808-110194134 | -0.158312003 | 0.649684381 | 0.850078048 |
| chr1:165887932-165888192 | 0.193364005 | 0.649691344 | 0.850078048 |
| chr14:91882360-91882561 | -0.238330822 | 0.649701399 | 0.850078048 |
| chr8:17576458-17577849 | -0.148948397 | 0.649747748 | 0.850106421 |
| chr6:25992426-25993094 | -0.150862502 | 0.649754033 | 0.850106421 |
| chr22:45553617-45554342 | 0.168409442 | 0.649762203 | 0.850106421 |
| chr20:60300537-60300789 | 0.185125475 | 0.649767463 | 0.850106421 |
| chr4:154743621-154744435 | -0.14870185 | 0.649782274 | 0.850111283 |
| chr19:45994976-45995647 | -0.147296336 | 0.649811713 | 0.850135283 |
| chr4:105633199-105633616 | -0.15944099 | 0.649845227 | 0.850164611 |
| chr17:81982342-81982612 | 0.174435182 | 0.649869685 | 0.850182093 |
| chr7:51067772-51067986 | 0.220659649 | 0.649926253 | 0.850240612 |
| chr17:79607928-79608360 | 0.168063037 | 0.649936609 | 0.850240612 |
| chr11:47443112-47443462 | -0.172507796 | 0.64999095 | 0.850297182 |
| chr3:29440359-29440869 | -0.180490818 | 0.650023508 | 0.850325256 |
| chr3:155884352-155884870 | 0.163050356 | 0.650065004 | 0.850336422 |
| chr4:88659232-88659711 | 0.164742135 | 0.650068559 | 0.850336422 |
| chr5:169826709-169826914 | -0.242572881 | 0.650069114 | 0.850336422 |
| chr14:105733911-105734146 | 0.205873353 | 0.650076435 | 0.850336422 |
| chr1:151090263-151090516 | 0.206650389 | 0.650115451 | 0.850347767 |
| chr3:127607653-127607894 | 0.207446823 | 0.650117256 | 0.850347767 |
| chr9:15018391-15018618 | 0.225524621 | 0.65014075 | 0.850347767 |
| chr8:117863599-117863846 | -0.198382146 | 0.65014129 | 0.850347767 |
| chr10:130110198-130111753 | 0.142124493 | 0.65014197 | 0.850347767 |
| chr17:48960894-48961162 | -0.196647479 | 0.650151696 | 0.850347767 |
| chr12:75171204-75171693 | -0.184848844 | 0.650188902 | 0.850381914 |
| chr17:9982227-9982556 | -0.195953277 | 0.650203523 | 0.850386522 |
| chr5:177303517-177303885 | -0.160070107 | 0.650233613 | 0.850399913 |
| chr3:139120274-139120482 | -0.215898718 | 0.650254 | 0.850399913 |
| chr13:28796629-28796940 | 0.167882361 | 0.65026754 | 0.850399913 |
| chr10:82887595-82887887 | -0.22035271 | 0.650272404 | 0.850399913 |
| chr2:61694382-61695191 | 0.148753956 | 0.650275103 | 0.850399913 |
| chr20:17611217-17611689 | 0.163717141 | 0.650280354 | 0.850399913 |
| chr17:48100542-48101878 | -0.144823956 | 0.650293463 | 0.850401852 |
| chr17:17710783-17711613 | -0.150314795 | 0.650304034 | 0.850401852 |
| chr17:57834096-57834647 | 0.151842661 | 0.650318724 | 0.850406548 |
| chr9:94300255-94300455 | -0.228933158 | 0.650381198 | 0.850424388 |
| chr2:85935820-85936086 | -0.234347193 | 0.650388317 | 0.850424388 |
| chr4:39135248-39135766 | 0.165575538 | 0.650395136 | 0.850424388 |
| chr3:150469673-150470000 | -0.17501126 | 0.650399854 | 0.850424388 |

| | | | |
|---|---|---|---|
| chr14:96263962-96264198 | -0.196884328 | 0.650400337 | 0.850424388 |
| chr20:23361538-23362261 | 0.144898903 | 0.650400711 | 0.850424388 |
| chr13:21477017-21477377 | 0.167004257 | 0.650442599 | 0.850424388 |
| chr9:102356070-102356441 | -0.191562287 | 0.650458724 | 0.850424388 |
| chr16:74862754-74863047 | -0.182793533 | 0.650485199 | 0.850424388 |
| chr6:55684924-55685290 | -0.194319798 | 0.650489478 | 0.850424388 |
| chr2:234952687-234952887 | -0.216838987 | 0.650493848 | 0.850424388 |
| chr13:23406930-23407389 | 0.172351266 | 0.650501946 | 0.850424388 |
| chr3:169813722-169814038 | -0.168045919 | 0.650502329 | 0.850424388 |
| chr14:101953615-101954024 | -0.184796475 | 0.65050495 | 0.850424388 |
| chr11:62553177-62553774 | -0.154765797 | 0.650523944 | 0.850424388 |
| chr20:5205438-5205705 | -0.192733048 | 0.6505275 | 0.850424388 |
| chr17:64457574-64457922 | 0.182421003 | 0.6505386 | 0.850424388 |
| chr15:40906969-40907535 | -0.15834864 | 0.650550668 | 0.850424388 |
| chr4:39458406-39459977 | 0.143504392 | 0.65055287 | 0.850424388 |
| chr10:17453333-17453986 | 0.152794743 | 0.650554346 | 0.850424388 |
| chr5:69093397-69094230 | -0.145350535 | 0.650592606 | 0.850430786 |
| chr4:83328556-83328770 | -0.207678164 | 0.650602941 | 0.850430786 |
| chr5:62354099-62354374 | -0.201936023 | 0.650604285 | 0.850430786 |
| chr3:125482042-125482272 | 0.222825905 | 0.650607026 | 0.850430786 |
| chr5:38208855-38209106 | -0.20390687 | 0.650614735 | 0.850430786 |
| chr1:119711705-119712240 | 0.160867432 | 0.650631135 | 0.850437528 |
| chr3:169218388-169218716 | -0.176366736 | 0.650642091 | 0.850437528 |
| chr6:142139014-142139245 | -0.245023857 | 0.650692515 | 0.850488927 |
| chr2:137659647-137659961 | -0.188670833 | 0.650716982 | 0.850503244 |
| chr2:131543068-131544018 | 0.149782792 | 0.650734749 | 0.850503244 |
| chr12:62794014-62794284 | -0.18870766 | 0.650750866 | 0.850503244 |
| chr17:76540569-76541374 | -0.147724268 | 0.650762953 | 0.850503244 |
| chr9:76458907-76459935 | -0.146153886 | 0.650766796 | 0.850503244 |
| chr3:135271958-135272165 | -0.231096715 | 0.650770069 | 0.850503244 |
| chr16:80608292-80608583 | -0.194935545 | 0.650802352 | 0.850503489 |
| chr1:151346805-151347775 | -0.143868347 | 0.650807211 | 0.850503489 |
| chr10:73781777-73782208 | 0.160079736 | 0.650824008 | 0.850503489 |
| chr3:30693590-30693813 | 0.219333754 | 0.650832562 | 0.850503489 |
| chr11:48172894-48173191 | -0.178852728 | 0.650833101 | 0.850503489 |
| chr1:107016609-107016959 | 0.160535683 | 0.650836856 | 0.850503489 |
| chr9:89919125-89919677 | -0.166450802 | 0.6508709 | 0.850533472 |
| chr3:10432219-10432580 | -0.182620014 | 0.650956808 | 0.850619219 |
| chr15:99076453-99077062 | -0.160360436 | 0.650965787 | 0.850619219 |
| chr2:169221888-169222097 | 0.229255868 | 0.650974221 | 0.850619219 |
| chr8:69026321-69026579 | -0.214616033 | 0.650983236 | 0.850619219 |
| chr17:75747891-75748173 | -0.171656138 | 0.650992025 | 0.850619219 |
| chr18:24014063-24015263 | 0.149946097 | 0.651014068 | 0.850633514 |
| chr2:64888911-64889263 | -0.181327134 | 0.651061874 | 0.850681472 |
| chr17:49223470-49223754 | 0.170290238 | 0.651095146 | 0.850710439 |
| chr13:29496151-29496602 | -0.158562863 | 0.651171147 | 0.850794569 |
| chr1:171780831-171782181 | -0.141880277 | 0.65118262 | 0.850794569 |
| chr2:140014025-140014225 | 0.240334391 | 0.651192847 | 0.850794569 |
| chr19:4723659-4724798 | 0.145833927 | 0.651206938 | 0.850798472 |
| chr2:102736098-102737296 | 0.147986727 | 0.651263757 | 0.85084143 |

| | | | |
|---|---|---|---|
| chr2:5938071-5938932 | -0.153907875 | 0.651268393 | 0.85084143 |
| chr3:56467070-56467401 | 0.17512466 | 0.651273132 | 0.85084143 |
| chr2:201780384-201781728 | -0.143473707 | 0.651298782 | 0.850848088 |
| chr5:177317986-177318426 | -0.16737205 | 0.651300437 | 0.850848088 |
| chr20:53828035-53829127 | 0.141532551 | 0.651353907 | 0.850903434 |
| chr13:52236259-52236540 | 0.203540629 | 0.651392285 | 0.850938831 |
| chr10:17910203-17910403 | -0.247789988 | 0.651420966 | 0.850938831 |
| chr9:134443194-134443836 | -0.154733287 | 0.65142381 | 0.850938831 |
| chr3:4438269-4438709 | 0.159293439 | 0.651428937 | 0.850938831 |
| chr8:113378693-113378910 | 0.210789168 | 0.651436532 | 0.850938831 |
| chr15:91737330-91737642 | -0.204792935 | 0.651470753 | 0.85096744 |
| chr10:49407660-49408114 | 0.165368624 | 0.651483569 | 0.85096744 |
| chr1:202236348-202236708 | -0.151031103 | 0.651491752 | 0.85096744 |
| chr2:219073297-219073616 | -0.214834925 | 0.65151645 | 0.850985193 |
| chr15:81274883-81275388 | 0.161847857 | 0.651543811 | 0.851005906 |
| chr1:150995927-150996179 | -0.189341906 | 0.651554923 | 0.851005906 |
| chr22:19962948-19963382 | 0.151785024 | 0.651565627 | 0.851005906 |
| chr3:32121153-32121463 | -0.191083644 | 0.651587525 | 0.851007087 |
| chr10:80552280-80552542 | -0.219021016 | 0.651600411 | 0.851007087 |
| chr17:60254286-60254672 | -0.156048131 | 0.651601588 | 0.851007087 |
| chr3:31958735-31958963 | -0.216845473 | 0.651620991 | 0.851007087 |
| chr1:205543209-205544320 | 0.14408949 | 0.651622064 | 0.851007087 |
| chr7:38222481-38222862 | 0.172695409 | 0.651635744 | 0.851010448 |
| chr8:74632088-74632303 | 0.215388132 | 0.651669634 | 0.851025978 |
| chr9:121281774-121282230 | -0.169142083 | 0.651669849 | 0.851025978 |
| chr10:42866751-42866956 | 0.228548603 | 0.651692191 | 0.851026392 |
| chr15:40755143-40755585 | 0.154695542 | 0.65169238 | 0.851026392 |
| chr2:42794456-42794771 | -0.183441524 | 0.651708282 | 0.851032653 |
| chr5:175640401-175640601 | 0.234520278 | 0.651740205 | 0.851059836 |
| chr2:73268514-73270021 | -0.145466186 | 0.651774577 | 0.851090215 |
| chr1:77605753-77606245 | 0.175603124 | 0.651798765 | 0.851090353 |
| chr20:60057393-60057693 | 0.162262705 | 0.651801933 | 0.851090353 |
| chr1:3319996-3320220 | -0.214621852 | 0.651817766 | 0.851090353 |
| chr4:82579020-82579227 | 0.2152502 | 0.651819114 | 0.851090353 |
| chrX:75188332-75188680 | -0.182939212 | 0.651910139 | 0.851192322 |
| chr10:132023110-132023379 | 0.186482814 | 0.651919426 | 0.851192322 |
| chr20:38826118-38826370 | -0.210844141 | 0.651945009 | 0.851211221 |
| chr17:50904269-50905665 | 0.142784257 | 0.651966566 | 0.851224861 |
| chr14:91060501-91061235 | 0.142594865 | 0.6519895 | 0.851231666 |
| chr4:47462856-47464005 | -0.143973771 | 0.651993997 | 0.851231666 |
| chr1:151254291-151255151 | 0.143621076 | 0.652010225 | 0.85123484 |
| chr4:87219587-87220858 | -0.14340503 | 0.652057946 | 0.85123484 |
| chr6:41181928-41182367 | -0.19670808 | 0.652062706 | 0.85123484 |
| chr17:46002338-46002853 | 0.164588899 | 0.652074345 | 0.85123484 |
| chr11:129767167-129767433 | -0.198278509 | 0.652077541 | 0.85123484 |
| chr3:14508923-14509221 | 0.16855152 | 0.65207775 | 0.85123484 |
| chr4:145154688-145154893 | 0.219386175 | 0.652080508 | 0.85123484 |
| chr13:36999136-36999572 | 0.150126603 | 0.652089586 | 0.85123484 |
| chr19:41230665-41230940 | 0.222807907 | 0.652105366 | 0.85123484 |
| chr2:32356446-32358020 | -0.142043948 | 0.652107523 | 0.85123484 |

| | | | |
|---|---|---|---|
| chr8:30733155-30733406 | 0.217981305 | 0.652153385 | 0.851274518 |
| chr8:116912670-116913084 | -0.173802683 | 0.652161973 | 0.851274518 |
| chr6:84779432-84779713 | 0.219426341 | 0.652185614 | 0.851274518 |
| chr10:71896432-71896697 | 0.199671602 | 0.652190328 | 0.851274518 |
| chr6:31681375-31681575 | 0.212149913 | 0.652193469 | 0.851274518 |
| chrX:10173354-10174111 | 0.153660824 | 0.652249171 | 0.851331146 |
| chr11:44632818-44633056 | -0.223375537 | 0.652259076 | 0.851331146 |
| chr13:113314916-113315301 | -0.178332566 | 0.652282481 | 0.851347192 |
| chr17:67276116-67276479 | -0.171061501 | 0.652300963 | 0.851348105 |
| chr5:27652584-27652904 | -0.191275707 | 0.652305403 | 0.851348105 |
| chr20:53162321-53162699 | -0.189453933 | 0.652346699 | 0.851375974 |
| chr9:81687073-81687294 | 0.166970231 | 0.652357497 | 0.851375974 |
| chr7:149872773-149873062 | -0.16654529 | 0.652386848 | 0.851375974 |
| chr10:69716796-69717276 | -0.171981028 | 0.652391827 | 0.851375974 |
| chr2:8571835-8572245 | 0.158205657 | 0.652393193 | 0.851375974 |
| chr22:31561281-31561682 | 0.17615294 | 0.652398185 | 0.851375974 |
| chr14:55446183-55446403 | -0.201262416 | 0.652415084 | 0.851375974 |
| chr17:9645843-9646185 | -0.170303789 | 0.652415647 | 0.851375974 |
| chr12:109775280-109775647 | 0.169627527 | 0.652443222 | 0.851379803 |
| chr4:47913831-47915137 | -0.142843812 | 0.652447968 | 0.851379803 |
| chr6:18154921-18156317 | 0.14196059 | 0.65245506 | 0.851379803 |
| chr9:12774849-12775077 | 0.201475653 | 0.652463027 | 0.851379803 |
| chr19:11155670-11156201 | -0.151570012 | 0.652512295 | 0.851429592 |
| chr9:124151041-124151251 | -0.244630497 | 0.652559219 | 0.851463198 |
| chr2:97700198-97700493 | 0.178962673 | 0.652560275 | 0.851463198 |
| chrX:154409243-154409487 | 0.19077347 | 0.652593692 | 0.851492301 |
| chrX:101357129-101357708 | 0.164428925 | 0.652649409 | 0.851541783 |
| chr22:19122328-19122839 | 0.148886973 | 0.652653843 | 0.851541783 |
| chr5:120139069-120139341 | -0.229653073 | 0.652682467 | 0.851554346 |
| chr15:76533436-76533643 | 0.222489582 | 0.652685699 | 0.851554346 |
| chr12:33093634-33094132 | 0.164835736 | 0.652739029 | 0.851609424 |
| chr6:52403629-52403986 | -0.200573703 | 0.652800512 | 0.851675137 |
| chr11:102495463-102495830 | 0.191386589 | 0.65284562 | 0.851708895 |
| chr16:10566699-10566977 | 0.193905362 | 0.652848618 | 0.851708895 |
| chr2:36701274-36701850 | -0.161402658 | 0.652890222 | 0.85171752 |
| chr17:82273334-82273909 | -0.155395607 | 0.652910022 | 0.85171752 |
| chr15:48285707-48286301 | 0.168244719 | 0.652920233 | 0.85171752 |
| chr10:526829-527114 | 0.181015753 | 0.652933129 | 0.85171752 |
| chr1:181043064-181043401 | 0.172945837 | 0.652940085 | 0.85171752 |
| chr4:105145773-105147174 | 0.14453892 | 0.652947183 | 0.85171752 |
| chr12:57803365-57803777 | -0.164875304 | 0.652951283 | 0.85171752 |
| chr18:63119646-63119857 | -0.241138254 | 0.652954068 | 0.85171752 |
| chr13:102772927-102774236 | 0.144867009 | 0.652960877 | 0.85171752 |
| chr15:77503341-77503584 | -0.19134105 | 0.652966387 | 0.85171752 |
| chr21:40693863-40694063 | -0.235757769 | 0.652988174 | 0.851726035 |
| chr5:58056438-58056787 | -0.238105504 | 0.652995147 | 0.851726035 |
| chr6:110682373-110682677 | -0.188319486 | 0.653013163 | 0.851735035 |
| chrX:44542628-44543327 | 0.149401789 | 0.653055083 | 0.851775211 |
| chr18:26226070-26227121 | 0.145021242 | 0.653083808 | 0.851798177 |
| chr1:32963870-32964249 | -0.174403162 | 0.653096361 | 0.851800051 |

| | | | |
|---|---|---|---|
| chr11:6612110-6613051 | -0.148249102 | 0.653126908 | 0.851823722 |
| chr11:18115310-18115823 | -0.168094314 | 0.653136745 | 0.851823722 |
| chr3:67930336-67930730 | 0.189349713 | 0.653152809 | 0.851830174 |
| chr19:703618-704552 | 0.146739082 | 0.653228701 | 0.851898327 |
| chr1:10032435-10033641 | 0.144002056 | 0.653237848 | 0.851898327 |
| chr16:74205972-74206181 | 0.23211009 | 0.653238421 | 0.851898327 |
| chr9:104926247-104926879 | 0.159644051 | 0.653302453 | 0.851967332 |
| chr1:225962403-225962637 | -0.232337143 | 0.653336724 | 0.851990828 |
| chr7:628069-628543 | -0.171052523 | 0.65334271 | 0.851990828 |
| chr17:1561812-1562017 | -0.225307396 | 0.653355117 | 0.851992507 |
| chr1:203674142-203674551 | -0.163489429 | 0.653385126 | 0.852013762 |
| chr8:21919406-21919851 | 0.159915802 | 0.653393655 | 0.852013762 |
| chr16:1794289-1794656 | -0.186239877 | 0.653430316 | 0.852046031 |
| chr7:32891063-32893031 | 0.140309535 | 0.653462383 | 0.852046031 |
| chr1:28437868-28438509 | -0.160498803 | 0.653464799 | 0.852046031 |
| chr1:201540117-201540547 | 0.166676391 | 0.653467616 | 0.852046031 |
| chr12:94102213-94102625 | -0.160627896 | 0.653482247 | 0.852046031 |
| chr1:202916672-202916967 | -0.199810952 | 0.653485123 | 0.852046031 |
| chr2:35655-36336 | -0.147236373 | 0.653496635 | 0.852046543 |
| chr2:69948300-69948574 | -0.234259178 | 0.653509171 | 0.852048389 |
| chr3:136343456-136344287 | 0.148442768 | 0.653526631 | 0.852056655 |
| chr20:53679726-53680523 | 0.14112194 | 0.6535412 | 0.852061151 |
| chr6:144581873-144582079 | -0.219111564 | 0.653556219 | 0.852066233 |
| chr1:93995953-93996153 | -0.233362672 | 0.653577281 | 0.852070275 |
| chrX:4788092-4788292 | 0.252470555 | 0.653582454 | 0.852070275 |
| chr1:46667867-46668789 | 0.147698849 | 0.65359268 | 0.852070275 |
| chr17:45099251-45099451 | 0.197639937 | 0.653619409 | 0.852090131 |
| chr13:106567630-106568450 | -0.150320496 | 0.653630153 | 0.852090131 |
| chr7:74067874-74068198 | -0.172580078 | 0.653671326 | 0.852129309 |
| chr2:86396506-86396763 | 0.19410021 | 0.6536832 | 0.85213029 |
| chr20:8963276-8963631 | 0.210470005 | 0.653707476 | 0.852136066 |
| chr12:124883558-124883834 | -0.197935347 | 0.653709874 | 0.852136066 |
| chr8:104077558-104077779 | -0.20058997 | 0.653727197 | 0.85214415 |
| chr2:117917564-117917792 | -0.205128688 | 0.653775048 | 0.852192026 |
| chr3:189430625-189430885 | -0.21760916 | 0.653789815 | 0.852196778 |
| chr3:23646660-23646860 | 0.220410491 | 0.653807805 | 0.852205729 |
| chr2:48440153-48441418 | -0.141950277 | 0.653826318 | 0.852215364 |
| chr10:80841898-80842143 | 0.226858312 | 0.65384286 | 0.852222428 |
| chr1:39539224-39539538 | -0.184781945 | 0.653872264 | 0.852226409 |
| chr7:32727869-32729119 | -0.1429293 | 0.653875827 | 0.852226409 |
| chr1:168178568-168179555 | 0.144066229 | 0.653879282 | 0.852226409 |
| chr2:48530055-48530499 | 0.166861441 | 0.653899963 | 0.852238867 |
| chrX:140141526-140141799 | -0.202656884 | 0.653926471 | 0.852258918 |
| chr1:159211629-159211829 | 0.247778884 | 0.653944069 | 0.852267356 |
| chr3:170978368-170978734 | 0.203551605 | 0.653957606 | 0.852270504 |
| chr6:73262712-73263676 | -0.153900067 | 0.653970086 | 0.852272272 |
| chr2:153064143-153064343 | -0.272008699 | 0.654026879 | 0.852322565 |
| chr10:38094042-38094980 | -0.147478294 | 0.654030925 | 0.852322565 |
| chr8:120984784-120985170 | -0.176221886 | 0.654098609 | 0.852396273 |
| chr12:63151515-63152422 | 0.153220444 | 0.654120308 | 0.852410052 |

| | | | |
|---|---|---|---|
| chr15:82672625-82672886 | -0.233838878 | 0.654151018 | 0.852430144 |
| chr3:121760973-121761184 | -0.222553296 | 0.654165283 | 0.852430144 |
| chr10:114551201-114551653 | -0.16895231 | 0.654180467 | 0.852430144 |
| chr7:108613703-108614209 | -0.165129907 | 0.65419807 | 0.852430144 |
| chr5:178153572-178154408 | -0.142458635 | 0.654199314 | 0.852430144 |
| chrX:101931401-101932336 | -0.146940659 | 0.65421158 | 0.852430144 |
| chr17:4142862-4143405 | -0.156040263 | 0.654213602 | 0.852430144 |
| chr2:231786182-231787023 | -0.152356966 | 0.654266128 | 0.852472439 |
| chr10:22002864-22004070 | -0.142910388 | 0.654268313 | 0.852472439 |
| chr15:41230647-41231739 | 0.145159525 | 0.654297612 | 0.85248452 |
| chr17:58541289-58541517 | 0.17787723 | 0.654299837 | 0.85248452 |
| chr3:169687423-169687762 | 0.201665275 | 0.65431415 | 0.852487566 |
| chr7:48734769-48735090 | 0.194932783 | 0.654334203 | 0.852487566 |
| chr11:2420738-2421009 | -0.186536587 | 0.654347943 | 0.852487566 |
| chr3:18847608-18847859 | -0.199451143 | 0.654357321 | 0.852487566 |
| chr18:74459810-74460179 | -0.192931959 | 0.654357805 | 0.852487566 |
| chr1:205711145-205711474 | 0.165257484 | 0.654387739 | 0.852512069 |
| chr11:57530481-57531183 | 0.146760626 | 0.654424886 | 0.852545967 |
| chr6:114342272-114343809 | -0.145607068 | 0.654448469 | 0.852552112 |
| chr3:46660870-46661195 | -0.18524068 | 0.654451856 | 0.852552112 |
| chr22:45105253-45105493 | -0.216235604 | 0.654466345 | 0.852556491 |
| chr15:40593745-40594810 | -0.143086565 | 0.654491129 | 0.852569365 |
| chr6:40587306-40588167 | -0.153921393 | 0.654498481 | 0.852569365 |
| chr2:10072606-10072954 | -0.168331474 | 0.65452287 | 0.852570955 |
| chr16:90022536-90022889 | -0.167372404 | 0.654544 | 0.852570955 |
| chr10:15720165-15720428 | -0.207517942 | 0.654545419 | 0.852570955 |
| chr20:58308722-58310390 | 0.140252754 | 0.654553167 | 0.852570955 |
| chr16:10385800-10386475 | 0.149160404 | 0.654556123 | 0.852570955 |
| chr17:17812263-17812494 | -0.175345831 | 0.654566463 | 0.852570955 |
| chr9:92716618-92716818 | -0.215064832 | 0.654631075 | 0.852640617 |
| chr2:66433188-66433777 | -0.150930643 | 0.654670367 | 0.852676675 |
| chr8:2071944-2072244 | -0.227704196 | 0.654681016 | 0.852676675 |
| chr14:93130368-93130608 | 0.186556518 | 0.654697632 | 0.852683822 |
| chr10:6239225-6239613 | 0.160803458 | 0.654717729 | 0.852695503 |
| chr8:102054896-102055116 | -0.223100157 | 0.654788584 | 0.852773289 |
| chr3:161231628-161232208 | -0.169856106 | 0.654854123 | 0.852844148 |
| chr14:52272431-52272901 | 0.171756506 | 0.654876603 | 0.852844278 |
| chr11:45771983-45772271 | -0.188661409 | 0.654880982 | 0.852844278 |
| chr5:181169537-181170212 | -0.150200293 | 0.654887614 | 0.852844278 |
| chr6:106566567-106566767 | -0.234453277 | 0.654925783 | 0.852871089 |
| chr6:102849910-102850410 | -0.160833797 | 0.654930464 | 0.852871089 |
| chrX:135186104-135186420 | -0.183711735 | 0.654952131 | 0.852884809 |
| chr21:46323670-46324997 | -0.140915858 | 0.654975232 | 0.852900396 |
| chr2:153784785-153785023 | 0.237096822 | 0.655036639 | 0.852965864 |
| chr5:147690648-147690948 | -0.179019078 | 0.65504821 | 0.852966435 |
| chr11:1869970-1870207 | 0.234051288 | 0.655078914 | 0.85299192 |
| chr2:68829269-68829677 | 0.176928243 | 0.655109253 | 0.853005571 |
| chr13:29925990-29926190 | -0.198815097 | 0.655111663 | 0.853005571 |
| chr17:41662751-41663008 | 0.180793974 | 0.65519624 | 0.85306017 |
| chr14:61860368-61860912 | 0.166598297 | 0.655196692 | 0.85306017 |

| | | | |
|---|---|---|---|
| chr20:62482158-62482651 | -0.157894442 | 0.655201038 | 0.85306017 |
| chr4:17549136-17549434 | 0.186282623 | 0.655211008 | 0.85306017 |
| chr12:14928943-14929358 | 0.165885466 | 0.655218784 | 0.85306017 |
| chr8:56028880-56029134 | 0.213991854 | 0.655220396 | 0.85306017 |
| chr8:138686551-138686760 | -0.194133663 | 0.655261162 | 0.853066619 |
| chr10:117166154-117166386 | -0.195689692 | 0.655263243 | 0.853066619 |
| chr16:57185545-57185759 | -0.190056869 | 0.655272869 | 0.853066619 |
| chr10:72215634-72216512 | -0.144622973 | 0.65527868 | 0.853066619 |
| chr17:79770188-79770402 | -0.215684574 | 0.655294064 | 0.853066619 |
| chr15:90903858-90904158 | 0.166285042 | 0.655310013 | 0.853066619 |
| chr12:13102142-13102342 | 0.207282358 | 0.655316166 | 0.853066619 |
| chr17:57010003-57010367 | -0.16178062 | 0.655331777 | 0.853066619 |
| chr22:23539565-23539915 | 0.172378301 | 0.655332954 | 0.853066619 |
| chr19:571113-571373 | 0.19642573 | 0.655336683 | 0.853066619 |
| chr2:182767608-182767880 | -0.18481632 | 0.655365427 | 0.853089543 |
| chr12:22588906-22589160 | -0.237829552 | 0.655422255 | 0.853148077 |
| chr17:13182536-13182802 | -0.211484821 | 0.655432664 | 0.853148077 |
| chr9:109497530-109497978 | 0.166790086 | 0.655467045 | 0.853165977 |
| chr2:154623864-154624139 | -0.205352935 | 0.655468685 | 0.853165977 |
| chr10:92960-93171 | 0.206242792 | 0.655482587 | 0.853169579 |
| chr2:68169333-68169571 | -0.213376721 | 0.655495293 | 0.853171624 |
| chr4:48726553-48726920 | -0.177731906 | 0.655554044 | 0.853233599 |
| chr2:197996980-197997239 | -0.241417184 | 0.655640804 | 0.853332026 |
| chr15:74227088-74227298 | 0.213779356 | 0.65569446 | 0.853387364 |
| chr4:82561490-82562411 | 0.152390434 | 0.655727715 | 0.853409825 |
| chr11:79809554-79809923 | -0.186941485 | 0.655733994 | 0.853409825 |
| chr3:134374074-134375468 | 0.141091997 | 0.655785495 | 0.853449496 |
| chr10:97676728-97676967 | 0.183084686 | 0.65579003 | 0.853449496 |
| chr5:172187822-172188963 | 0.142760118 | 0.655797891 | 0.853449496 |
| chr3:63096294-63097493 | 0.140842403 | 0.655834504 | 0.853471865 |
| chr5:29811974-29812204 | -0.211086595 | 0.655837357 | 0.853471865 |
| chr8:138165685-138165985 | 0.182326522 | 0.655946676 | 0.85358155 |
| chr8:66428443-66429910 | 0.141293206 | 0.655961519 | 0.85358155 |
| chr8:43141149-43141365 | 0.201331216 | 0.655962516 | 0.85358155 |
| chr4:123564762-123565147 | 0.160751131 | 0.655974558 | 0.85358155 |
| chr17:60308268-60308513 | 0.157759864 | 0.655998966 | 0.85358155 |
| chr8:66613918-66614181 | -0.172763984 | 0.656002296 | 0.85358155 |
| chr8:96856988-96857188 | 0.230072826 | 0.656009047 | 0.85358155 |
| chr13:35025304-35025767 | -0.154148435 | 0.656015088 | 0.85358155 |
| chr5:472061-472747 | 0.150494036 | 0.656026438 | 0.85358155 |
| chr12:26126440-26126735 | -0.178774558 | 0.656033044 | 0.85358155 |
| chr1:119316741-119317388 | -0.168249197 | 0.656063449 | 0.853602092 |
| chr12:47715370-47715620 | 0.188248427 | 0.656071113 | 0.853602092 |
| chr15:74770928-74771375 | 0.156921799 | 0.656106552 | 0.853633707 |
| chr4:27971-28189 | 0.208521214 | 0.65613293 | 0.85364421 |
| chr3:163957279-163957498 | -0.208955386 | 0.656145575 | 0.85364421 |
| chr20:62386529-62387266 | 0.146678221 | 0.656163671 | 0.85364421 |
| chr10:1420635-1421376 | 0.151435061 | 0.656172196 | 0.85364421 |
| chr2:218466862-218467152 | 0.22392841 | 0.656178735 | 0.85364421 |
| chr11:42969002-42969202 | -0.261404179 | 0.656189364 | 0.85364421 |

| | | | |
|---|---|---|---|
| chr14:61280478-61281763 | 0.139999677 | 0.656192612 | 0.85364421 |
| chr3:10991936-10992164 | -0.217206616 | 0.656203899 | 0.8536444 |
| chrX:100614960-100615263 | -0.190413713 | 0.656255402 | 0.853693177 |
| chr3:160928267-160928618 | -0.177530197 | 0.656263678 | 0.853693177 |
| chr17:77254895-77255416 | -0.160707952 | 0.656278859 | 0.853698431 |
| chr8:119535219-119535616 | -0.182215062 | 0.656290274 | 0.853698788 |
| chr4:83455296-83456306 | -0.141120874 | 0.656320295 | 0.853723345 |
| chr10:5412136-5413046 | 0.14465237 | 0.656334945 | 0.853727908 |
| chr6:52839542-52839963 | -0.163129565 | 0.656407207 | 0.85376241 |
| chrX:75273751-75274688 | 0.147659235 | 0.656407688 | 0.85376241 |
| chr3:119390838-119391038 | 0.240081338 | 0.656413214 | 0.85376241 |
| chr1:183087039-183087311 | -0.202081989 | 0.656424149 | 0.85376241 |
| chr11:72831509-72831759 | -0.186522829 | 0.656435647 | 0.85376241 |
| chr11:47193887-47194525 | 0.14978966 | 0.65644689 | 0.85376241 |
| chr1:40802144-40802660 | 0.158643259 | 0.656451972 | 0.85376241 |
| chr11:62054593-62054885 | 0.187587046 | 0.656453123 | 0.85376241 |
| chr20:3818408-3818632 | 0.177909321 | 0.656461752 | 0.85376241 |
| chr6:52284151-52285176 | -0.141495847 | 0.65647458 | 0.853764602 |
| chr12:18261091-18261627 | -0.174342129 | 0.656521235 | 0.853799526 |
| chr6:43884010-43884210 | 0.204836146 | 0.65652372 | 0.853799526 |
| chr17:2960516-2960768 | 0.204109898 | 0.656573417 | 0.853849665 |
| chr17:64345694-64346251 | 0.143948918 | 0.656586072 | 0.853851629 |
| chr4:109676932-109677176 | -0.218593868 | 0.656632818 | 0.853875779 |
| chr22:23768309-23768568 | 0.171443918 | 0.656646012 | 0.853875779 |
| chr1:166838770-166840304 | -0.1405802 | 0.656652813 | 0.853875779 |
| chr9:92115022-92115825 | 0.145752326 | 0.656654522 | 0.853875779 |
| chr12:16329636-16329964 | 0.185336618 | 0.656660362 | 0.853875779 |
| chr1:31644496-31645425 | 0.142564081 | 0.656726243 | 0.853946954 |
| chr19:43575357-43576103 | 0.147729424 | 0.656804962 | 0.85403482 |
| chr7:47604844-47605528 | 0.154472887 | 0.656859704 | 0.85406068 |
| chr13:64675303-64675710 | -0.172864468 | 0.656869466 | 0.85406068 |
| chr11:72482228-72482610 | 0.16372931 | 0.656872505 | 0.85406068 |
| chr11:21871641-21871908 | 0.240991028 | 0.656884913 | 0.85406068 |
| chr2:74865496-74865752 | 0.1881495 | 0.656890832 | 0.85406068 |
| chr2:27152066-27152393 | -0.175687261 | 0.656891729 | 0.85406068 |
| chr2:208373522-208373765 | 0.202201168 | 0.656924342 | 0.854066704 |
| chr10:132320632-132320909 | 0.2037475 | 0.656957095 | 0.854066704 |
| chr2:143635466-143635666 | 0.239700907 | 0.656957192 | 0.854066704 |
| chr9:34611954-34612301 | 0.163182674 | 0.656958114 | 0.854066704 |
| chr3:138186876-138188154 | -0.142008135 | 0.656962191 | 0.854066704 |
| chr3:138299910-138300180 | 0.195627434 | 0.656963241 | 0.854066704 |
| chr16:83954082-83954331 | -0.164958403 | 0.656990128 | 0.854087167 |
| chr11:61947657-61947988 | 0.18042182 | 0.65700752 | 0.854095286 |
| chr2:227766379-227766637 | -0.198506023 | 0.657072783 | 0.854151548 |
| chr10:48306171-48307396 | -0.141837671 | 0.657073095 | 0.854151548 |
| chr17:48024631-48024831 | 0.175131973 | 0.657102057 | 0.854171949 |
| chr2:3443785-3444058 | -0.20852358 | 0.657128984 | 0.854171949 |
| chr12:111286028-111286228 | 0.225093989 | 0.657131944 | 0.854171949 |
| chr2:177604514-177604747 | -0.217618484 | 0.657133381 | 0.854171949 |
| chr5:17204044-17204336 | 0.176092726 | 0.657148551 | 0.854177178 |

| | | | |
|---|---|---|---|
| chr2:84894021-84894703 | -0.153611894 | 0.657180537 | 0.854191795 |
| chr12:62434733-62436350 | 0.141018985 | 0.657182093 | 0.854191795 |
| chr9:119562147-119562432 | -0.214810606 | 0.657230221 | 0.854223808 |
| chr2:65640151-65640802 | -0.161050387 | 0.657234177 | 0.854223808 |
| chr14:106551351-106551582 | -0.212227666 | 0.657240168 | 0.854223808 |
| chr2:95022297-95022640 | 0.160101167 | 0.657260309 | 0.854235496 |
| chr9:117400073-117400642 | 0.153060667 | 0.657369857 | 0.854320394 |
| chr16:88743807-88744130 | 0.161589548 | 0.657378073 | 0.854320394 |
| chr5:18837570-18837877 | -0.216437463 | 0.657382334 | 0.854320394 |
| chr11:78416399-78417562 | -0.143844642 | 0.657390938 | 0.854320394 |
| chr8:125196413-125196623 | 0.218967528 | 0.657397525 | 0.854320394 |
| chr3:93979767-93980693 | -0.139996862 | 0.65740346 | 0.854320394 |
| chr17:48790709-48790925 | -0.200939862 | 0.6574105 | 0.854320394 |
| chr1:156751066-156751453 | -0.155033877 | 0.657416463 | 0.854320394 |
| chr17:7990057-7990510 | -0.168115395 | 0.65742598 | 0.854320394 |
| chr14:76784760-76785179 | 0.149241129 | 0.657451568 | 0.854339157 |
| chr11:102010934-102011437 | -0.164169032 | 0.657526956 | 0.854408141 |
| chr17:58270516-58270798 | -0.164169032 | 0.657526956 | 0.854408141 |
| chr21:28884938-28885564 | -0.147902085 | 0.657587125 | 0.854438422 |
| chr7:148883423-148883863 | -0.155847824 | 0.657594171 | 0.854438422 |
| chr3:173711572-173712231 | 0.15574947 | 0.657603006 | 0.854438422 |
| chr1:42165565-42166330 | 0.147672897 | 0.657616292 | 0.854438422 |
| chr10:97463676-97464005 | -0.163961407 | 0.657616692 | 0.854438422 |
| chr1:201346426-201346942 | 0.151648388 | 0.657617168 | 0.854438422 |
| chr8:125589954-125590396 | -0.156020963 | 0.657630538 | 0.854441305 |
| chr19:40442912-40443495 | 0.145640455 | 0.65766292 | 0.854468889 |
| chr1:39852217-39853173 | 0.153039918 | 0.657680722 | 0.854477529 |
| chr3:128343706-128343926 | 0.189738928 | 0.65772214 | 0.85450501 |
| chr11:71580118-71580333 | -0.197279327 | 0.657735051 | 0.85450501 |
| chr7:55192498-55192698 | 0.220376892 | 0.657735331 | 0.85450501 |
| chr5:174528927-174529181 | 0.185855247 | 0.657766393 | 0.854517741 |
| chr6:42452870-42453435 | 0.153659211 | 0.657767434 | 0.854517741 |
| chr15:40239061-40239591 | 0.157462011 | 0.657860746 | 0.854624474 |
| chr17:43021145-43021540 | -0.163293274 | 0.657898096 | 0.854640532 |
| chr2:87660283-87660502 | -0.223283612 | 0.657898528 | 0.854640532 |
| chr4:110970678-110971042 | 0.169153536 | 0.657924593 | 0.854640532 |
| chr1:217881551-217882282 | -0.150788966 | 0.657926101 | 0.854640532 |
| chr19:45164691-45164998 | 0.166876798 | 0.657964719 | 0.854640532 |
| chr3:127045042-127045344 | 0.170083906 | 0.657965388 | 0.854640532 |
| chr14:99380596-99380883 | -0.197103441 | 0.657968826 | 0.854640532 |
| chr15:78195728-78196034 | -0.167321876 | 0.657990652 | 0.854640532 |
| chr10:99392500-99392876 | 0.169492942 | 0.657997655 | 0.854640532 |
| chr3:58187687-58188008 | 0.173602614 | 0.658008096 | 0.854640532 |
| chr15:78415502-78415733 | 0.192471707 | 0.658010163 | 0.854640532 |
| chr16:64983952-64984301 | -0.1804304 | 0.658015309 | 0.854640532 |
| chr11:35364054-35364367 | -0.220086902 | 0.658018171 | 0.854640532 |
| chr16:9134820-9135081 | 0.182888674 | 0.658041486 | 0.854640532 |
| chr4:77146343-77146782 | 0.181081583 | 0.658050963 | 0.854640532 |
| chr3:19738717-19738988 | -0.201890176 | 0.658062166 | 0.854640532 |
| chr17:59869468-59869808 | 0.145513718 | 0.658072222 | 0.854640532 |

| | | | |
|---|---|---|---|
| chr14:21092507-21092913 | -0.15779242 | 0.65809355 | 0.854640532 |
| chr5:159341810-159342313 | -0.156496497 | 0.658095272 | 0.854640532 |
| chr20:50360150-50360761 | 0.154168247 | 0.658096187 | 0.854640532 |
| chr2:9808768-9809344 | -0.149623338 | 0.658116965 | 0.85465303 |
| chr3:190307253-190307516 | 0.226703316 | 0.658161203 | 0.8546742 |
| chr9:93159998-93160246 | 0.19379162 | 0.658162649 | 0.8546742 |
| chr19:40771349-40771742 | -0.204875746 | 0.658213931 | 0.8546742 |
| chr5:174045474-174045863 | -0.160674289 | 0.658216094 | 0.8546742 |
| chr5:180809716-180811119 | -0.138725566 | 0.658219384 | 0.8546742 |
| chr5:171263683-171264166 | -0.183545588 | 0.658228322 | 0.8546742 |
| chr3:58020241-58020487 | 0.221542889 | 0.658229773 | 0.8546742 |
| chr17:75109686-75110124 | 0.153592731 | 0.658238942 | 0.8546742 |
| chr7:97951618-97951952 | -0.173362375 | 0.658259675 | 0.8546742 |
| chr2:8681723-8682159 | -0.147260068 | 0.658265676 | 0.8546742 |
| chr1:26177216-26177632 | -0.164200016 | 0.658269372 | 0.8546742 |
| chr8:60412539-60412909 | -0.166128278 | 0.658269599 | 0.8546742 |
| chr9:129103032-129103361 | 0.171198259 | 0.658288613 | 0.8546742 |
| chr6:67679564-67679982 | 0.176583401 | 0.658301346 | 0.8546742 |
| chr16:28292954-28293447 | 0.165625765 | 0.658309177 | 0.8546742 |
| chr10:29804248-29804613 | -0.183118131 | 0.658319662 | 0.8546742 |
| chr14:48853077-48853458 | 0.199945113 | 0.658322892 | 0.8546742 |
| chr11:66883267-66883542 | -0.172726787 | 0.658371859 | 0.85471956 |
| chr7:1203840-1204058 | 0.214046602 | 0.658380141 | 0.85471956 |
| chr19:18879581-18880009 | -0.181262083 | 0.658499519 | 0.854860055 |
| chr16:14629917-14630595 | -0.145946444 | 0.658519418 | 0.854861707 |
| chr6:72218160-72218488 | 0.172313414 | 0.658523105 | 0.854861707 |
| chr1:67700640-67701265 | -0.160900511 | 0.658568787 | 0.854906525 |
| chr5:154754534-154755577 | -0.140658016 | 0.658597167 | 0.854910221 |
| chr1:117654489-117655376 | -0.152156127 | 0.658604354 | 0.854910221 |
| chr7:154913734-154914076 | -0.17132025 | 0.658610782 | 0.854910221 |
| chr13:45766837-45767238 | -0.197393595 | 0.65863756 | 0.854910221 |
| chr20:2750795-2751197 | -0.174131813 | 0.658655144 | 0.854910221 |
| chr1:46401998-46402583 | -0.149276535 | 0.658662576 | 0.854910221 |
| chr2:232440645-232440884 | 0.207838039 | 0.658669456 | 0.854910221 |
| chr6:48068333-48069051 | 0.161957587 | 0.658671704 | 0.854910221 |
| chr14:64465152-64466151 | 0.148777391 | 0.658675481 | 0.854910221 |
| chr10:43409842-43410065 | -0.168652284 | 0.658697796 | 0.854910221 |
| chr1:96756882-96757082 | 0.244539415 | 0.658704021 | 0.854910221 |
| chr8:89882573-89883054 | 0.147288449 | 0.658706714 | 0.854910221 |
| chr20:62121342-62123300 | 0.138606257 | 0.658716682 | 0.854910221 |
| chrX:40580405-40581115 | 0.146742599 | 0.658734958 | 0.854919459 |
| chr17:29097266-29097466 | 0.213890049 | 0.658782262 | 0.854966371 |
| chr15:31272475-31272729 | -0.201622739 | 0.658799548 | 0.854974322 |
| chr3:47164637-47164880 | -0.202964445 | 0.658838748 | 0.855010714 |
| chr5:36876518-36877541 | 0.142030294 | 0.658876388 | 0.855045079 |
| chr2:170476069-170476334 | -0.224158654 | 0.658912519 | 0.855068073 |
| chr6:156719613-156720460 | -0.146426628 | 0.65892026 | 0.855068073 |
| chr12:6321886-6322165 | 0.178216963 | 0.658927585 | 0.855068073 |
| chr6:52551242-52551749 | 0.159461709 | 0.658953867 | 0.85508165 |
| chr17:60586349-60586969 | 0.141664515 | 0.658960367 | 0.85508165 |

| | | | |
|---|---|---|---|
| chr14:38522863-38523069 | -0.244608798 | 0.658975905 | 0.85508733 |
| chr17:39980334-39981039 | 0.148364645 | 0.659009842 | 0.855116887 |
| chr1:221579512-221579726 | -0.226322294 | 0.659024265 | 0.85512112 |
| chr5:142421391-142421597 | 0.234633452 | 0.659057915 | 0.8551503 |
| chr5:135034132-135034553 | 0.163229121 | 0.659075197 | 0.855158243 |
| chr1:24567900-24568445 | -0.159787752 | 0.659094603 | 0.855168942 |
| chr8:73286423-73286718 | -0.210051656 | 0.659128841 | 0.855198883 |
| chr5:20024597-20024908 | 0.182244615 | 0.659152721 | 0.855215385 |
| chr2:177148711-177149251 | 0.156448054 | 0.659182789 | 0.855228796 |
| chr17:39244765-39245027 | 0.197989889 | 0.659185381 | 0.855228796 |
| chr22:31629351-31629711 | -0.171223947 | 0.659234111 | 0.855277536 |
| chr15:87368397-87368674 | -0.209423161 | 0.659257131 | 0.855292921 |
| chr11:18588281-18589024 | 0.147310904 | 0.659280503 | 0.85530876 |
| chr16:77190333-77191469 | 0.137806588 | 0.659332772 | 0.855309491 |
| chr7:10973018-10973458 | 0.14304073 | 0.659335573 | 0.855309491 |
| chr10:103712960-103713222 | 0.179123039 | 0.659342001 | 0.855309491 |
| chr20:47576580-47577187 | 0.139653029 | 0.659343489 | 0.855309491 |
| chr15:39801899-39802248 | -0.165062821 | 0.659352723 | 0.855309491 |
| chr6:142323366-142323982 | -0.142693672 | 0.659353875 | 0.855309491 |
| chr4:99462868-99463187 | 0.181790207 | 0.659359205 | 0.855309491 |
| chr18:24023374-24023801 | 0.16375109 | 0.659387772 | 0.855332067 |
| chr3:194861158-194861358 | -0.231187117 | 0.659423174 | 0.85535719 |
| chr1:244833863-244834261 | -0.155689178 | 0.659429466 | 0.85535719 |
| chr19:45584160-45585167 | 0.14001285 | 0.659459594 | 0.855371644 |
| chr2:74261152-74261402 | 0.192858245 | 0.65946477 | 0.855371644 |
| chrX:104136243-104136794 | -0.1640586 | 0.659481396 | 0.855371644 |
| chr7:108413901-108414144 | 0.220336075 | 0.659485264 | 0.855371644 |
| chr15:41806416-41806879 | 0.161513304 | 0.659518156 | 0.855399826 |
| chr6:105484407-105485049 | -0.170316031 | 0.659572671 | 0.855456053 |
| chr6:166955695-166955990 | -0.18395499 | 0.659644596 | 0.855534686 |
| chr8:109519805-109520238 | -0.166805364 | 0.659663782 | 0.855534686 |
| chr6:155174764-155175106 | 0.183588055 | 0.659666796 | 0.855534686 |
| chr2:217886700-217887053 | 0.173204325 | 0.659685394 | 0.855539258 |
| chr2:196314722-196314994 | 0.194461076 | 0.659692653 | 0.855539258 |
| chr12:42922157-42922383 | 0.198288932 | 0.659722034 | 0.855552401 |
| chr11:60906068-60906973 | 0.144049198 | 0.659731138 | 0.855552401 |
| chr10:102055623-102056467 | 0.145394456 | 0.659736285 | 0.855552401 |
| chr17:62626695-62627864 | 0.140932702 | 0.659748927 | 0.855554316 |
| chr1:226723660-226723873 | 0.222907293 | 0.659765829 | 0.855561754 |
| chrX:13488181-13488532 | 0.183869456 | 0.659811023 | 0.855605879 |
| chr10:74889435-74889635 | -0.218197779 | 0.659840516 | 0.855616041 |
| chr12:11228380-11228779 | -0.191286172 | 0.659841192 | 0.855616041 |
| chr3:186097860-186098104 | -0.205351152 | 0.659882859 | 0.855648882 |
| chr9:71758319-71758668 | -0.175027198 | 0.659909447 | 0.855648882 |
| chr11:13971034-13971296 | -0.199236515 | 0.659912626 | 0.855648882 |
| chr15:60507812-60508093 | -0.18293877 | 0.659918411 | 0.855648882 |
| chr1:186375001-186376197 | -0.137401473 | 0.659922355 | 0.855648882 |
| chr12:120124011-120124211 | 0.224180942 | 0.659952332 | 0.855670621 |
| chr3:186108874-186109556 | 0.142374109 | 0.659961456 | 0.855670621 |
| chr11:1309560-1310183 | 0.151467847 | 0.659985745 | 0.855687634 |

| | | | |
|---|---|---|---|
| chr1:183022943-183024280 | 0.140066552 | 0.660052869 | 0.855760181 |
| chr12:113965813-11396670[illegible] | -0.143037967 | 0.660079427 | 0.855780134 |
| chr16:73956696-73956949 | 0.207863829 | 0.660096513 | 0.855787804 |
| chr1:42957920-42959302 | -0.141385107 | 0.660205199 | 0.855898318 |
| chr5:124812824-124813060 | -0.228562598 | 0.660207946 | 0.855898318 |
| chr12:54680423-54680762 | -0.171582248 | 0.660218978 | 0.855898318 |
| chr2:88482109-88482346 | -0.230623714 | 0.660231922 | 0.855898318 |
| chr9:69421976-69422527 | -0.17223958 | 0.660242333 | 0.855898318 |
| chr6:27303693-27303924 | -0.216918228 | 0.660259685 | 0.855898318 |
| chr6:156686552-156687969 | -0.143421848 | 0.660259949 | 0.855898318 |
| chr2:91583393-91583594 | -0.234993369 | 0.660277626 | 0.855906753 |
| chr10:104622620-104622932 | -0.173447187 | 0.660300271 | 0.855921627 |
| chr9:89593026-89593377 | -0.161975059 | 0.660313296 | 0.855924031 |
| chr1:174232875-174233075 | -0.216401618 | 0.660344971 | 0.855943523 |
| chr1:61742078-61743283 | -0.140407853 | 0.660350676 | 0.855943523 |
| chr17:43315158-43316307 | 0.142836776 | 0.66040117 | 0.855994492 |
| chr20:22684087-22684351 | -0.183559016 | 0.660421471 | 0.856003888 |
| chr14:92121367-92122379 | 0.143582078 | 0.660430762 | 0.856003888 |
| chr14:37171815-37172819 | 0.140958395 | 0.660539729 | 0.856096866 |
| chr3:129183072-129184519 | 0.140193436 | 0.66054089 | 0.856096866 |
| chr16:89923033-89923442 | -0.153074128 | 0.660549128 | 0.856096866 |
| chr10:15096745-15097840 | -0.139251812 | 0.66055549 | 0.856096866 |
| chr14:55297893-55298432 | 0.159230996 | 0.660571981 | 0.856096866 |
| chr5:81544675-81545108 | 0.171237895 | 0.66057896 | 0.856096866 |
| chr1:121020700-121021007 | 0.18508995 | 0.66060461 | 0.856096866 |
| chr12:105319830-10532010[illegible] | -0.194338614 | 0.660608491 | 0.856096866 |
| chr14:64344242-64344471 | -0.19180239 | 0.660624551 | 0.856096866 |
| chr4:121057417-121057699 | -0.214259709 | 0.6606254 | 0.856096866 |
| chr7:22435853-22436105 | -0.214259709 | 0.6606254 | 0.856096866 |
| chr4:83109809-83110825 | 0.144084786 | 0.660674193 | 0.856116037 |
| chr10:85280086-85280321 | -0.221251953 | 0.660674911 | 0.856116037 |
| chr2:202374987-202376522 | -0.138795484 | 0.660695015 | 0.856116037 |
| chr9:36872939-36873217 | -0.212495892 | 0.660705487 | 0.856116037 |
| chr21:14064315-14064553 | 0.198491835 | 0.660706174 | 0.856116037 |
| chr8:94797574-94797827 | 0.199937011 | 0.660709898 | 0.856116037 |
| chr4:47716093-47716331 | -0.21507064 | 0.660718406 | 0.856116037 |
| chr1:152461548-152461907 | 0.156522689 | 0.66073564 | 0.85612389 |
| chr7:148242190-148242727 | -0.153292962 | 0.660765442 | 0.856148026 |
| chr13:46545957-46546759 | 0.144285681 | 0.660778934 | 0.856150323 |
| chr3:69451387-69451765 | 0.19776825 | 0.660789562 | 0.856150323 |
| chr20:62227918-62228426 | -0.15089738 | 0.660811534 | 0.856164314 |
| chr15:67520522-67521804 | -0.140773985 | 0.660833854 | 0.856178754 |
| chr12:109052252-109053375 | -0.144058698 | 0.660861005 | 0.856199454 |
| chr5:66597515-66597795 | -0.173860486 | 0.660898989 | 0.856221514 |
| chr3:63724064-63724276 | 0.157203886 | 0.660900381 | 0.856221514 |
| chr7:76069159-76069611 | -0.15591359 | 0.660932974 | 0.856249262 |
| chr12:111405188-11140602[illegible] | 0.143496787 | 0.660945014 | 0.856250382 |
| chr1:161382208-161382452 | 0.211225363 | 0.660980783 | 0.85627762 |
| chr7:1028164-1029209 | 0.140248609 | 0.660989226 | 0.85627762 |
| chr11:111878586-111879677 | 0.140261818 | 0.660999565 | 0.85627762 |

| | | | |
|---|---|---|---|
| chrX:69094038-69094318 | 0.195501259 | 0.661042162 | 0.856318324 |
| chr21:43938991-43939420 | -0.156001481 | 0.661076252 | 0.856348007 |
| chr15:74388061-74388294 | -0.192847655 | 0.66112072 | 0.856391131 |
| chr21:31866068-31866483 | -0.161397588 | 0.661172561 | 0.856443805 |
| chr1:91165237-91165587 | 0.17207117 | 0.661184115 | 0.856444294 |
| chr1:65254046-65255444 | 0.140004275 | 0.66121446 | 0.85646784 |
| chr17:36755580-36755874 | -0.187408866 | 0.661224649 | 0.85646784 |
| chr11:3454952-3455152 | 0.227577634 | 0.661248361 | 0.856484075 |
| chr12:76443344-76443566 | 0.21650383 | 0.661269048 | 0.856488277 |
| chr15:59371987-59372194 | -0.19099531 | 0.661273962 | 0.856488277 |
| chr14:83999666-83999929 | 0.21019143 | 0.66129049 | 0.856495206 |
| chr18:50819911-50820236 | -0.174779794 | 0.66132356 | 0.85652356 |
| chr1:6554426-6554843 | 0.172981361 | 0.661353274 | 0.856547566 |
| chr5:160370309-160370747 | -0.175592704 | 0.661412081 | 0.856598135 |
| chr6:99905560-99905857 | -0.180202757 | 0.661414678 | 0.856598135 |
| chr6:142207276-142207728 | 0.16398277 | 0.661456766 | 0.856638164 |
| chr19:36137059-36137310 | 0.176878065 | 0.661486593 | 0.8566561 |
| chr5:173822496-173822851 | -0.164119077 | 0.661492976 | 0.8566561 |
| chr9:77150845-77151086 | 0.202559477 | 0.661524231 | 0.856682097 |
| chr1:150515220-150516461 | 0.138341021 | 0.66155367 | 0.856705742 |
| chr11:109939285-109939559 | -0.203788974 | 0.661590492 | 0.856738945 |
| chr1:26306275-26306889 | -0.152099369 | 0.661611418 | 0.856751565 |
| chr20:46592709-46592909 | 0.216430634 | 0.661634994 | 0.856767614 |
| chr10:71319023-71319856 | 0.146658288 | 0.661665703 | 0.856789744 |
| chr16:71494136-71494360 | 0.211847139 | 0.661675262 | 0.856789744 |
| chr16:30941920-30942283 | 0.157899577 | 0.661685629 | 0.856789744 |
| chr3:3126289-3127894 | -0.137787187 | 0.66170548 | 0.856800968 |
| chrX:130958794-130959505 | 0.158487516 | 0.66172677 | 0.856807906 |
| chr17:20121481-20121872 | -0.184424161 | 0.661738067 | 0.856807906 |
| chr5:4511958-4512546 | -0.16189889 | 0.661744385 | 0.856807906 |
| chr16:81261119-81261353 | 0.196414836 | 0.661876526 | 0.856964518 |
| chr7:73342475-73342904 | 0.176597355 | 0.661908179 | 0.856991019 |
| chr5:69359383-69359605 | 0.238569663 | 0.661972363 | 0.857043268 |
| chr8:138935989-138936625 | -0.154310848 | 0.661978787 | 0.857043268 |
| chr8:10839410-10840229 | -0.14416788 | 0.661982091 | 0.857043268 |
| chr11:12673808-12674983 | -0.140340586 | 0.66200427 | 0.857046707 |
| chr3:194369763-194370021 | -0.193347366 | 0.662007118 | 0.857046707 |
| chr22:33294573-33294773 | 0.250845639 | 0.662035963 | 0.857068413 |
| chr14:58395552-58396600 | 0.141035802 | 0.662067692 | 0.857068413 |
| chr12:88684317-88684555 | -0.211138221 | 0.662081312 | 0.857068413 |
| chr7:30403502-30404013 | -0.158711885 | 0.662082839 | 0.857068413 |
| chr6:134822934-134823259 | -0.171783959 | 0.662091675 | 0.857068413 |
| chr2:130727268-130727940 | 0.145104879 | 0.662101553 | 0.857068413 |
| chr12:65193306-65193542 | -0.205123639 | 0.662102183 | 0.857068413 |
| chrX:9786220-9786737 | 0.16711455 | 0.662130769 | 0.857090936 |
| chr12:107685024-107685327 | -0.167763429 | 0.662183853 | 0.857136686 |
| chr19:18476796-18477009 | -0.230070826 | 0.662190463 | 0.857136686 |
| chr1:161427092-161427406 | 0.201142495 | 0.662199672 | 0.857136686 |
| chr6:157184309-157184534 | -0.196012577 | 0.662210943 | 0.857136795 |
| chr4:2302802-2303002 | 0.24061662 | 0.662291478 | 0.857226556 |

| | | | |
|---|---|---|---|
| chr16:46757390-46757685 | -0.20224844 | 0.662304684 | 0.857229168 |
| chr13:33254016-33254267 | 0.200642926 | 0.662318397 | 0.857232436 |
| chr3:127405006-127405418 | -0.169917197 | 0.662334971 | 0.857239407 |
| chr1:113785936-113786466 | -0.15756595 | 0.662348801 | 0.857242827 |
| chr5:132777142-132777544 | -0.171827778 | 0.662362006 | 0.857245438 |
| chr3:97971628-97972674 | 0.140101392 | 0.662390215 | 0.857267466 |
| chr9:7402174-7402374 | 0.219622999 | 0.662429845 | 0.857293216 |
| chr19:53980924-53981158 | 0.17935712 | 0.662432489 | 0.857293216 |
| chr6:91475856-91476317 | -0.161878146 | 0.662449699 | 0.857301009 |
| chr11:109914148-109914385 | -0.203986203 | 0.662476603 | 0.857309832 |
| chr19:50657948-50658764 | 0.139668839 | 0.662478894 | 0.857309832 |
| chr19:36772338-36773269 | 0.14981852 | 0.662516907 | 0.857335307 |
| chr4:87353577-87353777 | 0.213441971 | 0.662551592 | 0.857335307 |
| chr14:78917549-78917749 | 0.232114507 | 0.662562993 | 0.857335307 |
| chr12:87151702-87152136 | -0.180278859 | 0.662574537 | 0.857335307 |
| chr2:165293760-165294089 | -0.170729824 | 0.662576618 | 0.857335307 |
| chr11:19547158-19547358 | -0.261970133 | 0.662586288 | 0.857335307 |
| chr11:18395711-18396594 | 0.138963415 | 0.662586862 | 0.857335307 |
| chr20:9408124-9408390 | -0.199902285 | 0.662588093 | 0.857335307 |
| chr5:22431579-22431788 | -0.216293192 | 0.662620541 | 0.857348336 |
| chr6:79984361-79984579 | -0.216293192 | 0.662620541 | 0.857348336 |
| chr2:12126430-12126630 | -0.237803341 | 0.66266211 | 0.857387643 |
| chr13:113545537-113545880 | -0.198285793 | 0.662694746 | 0.857405269 |
| chr12:100826421-100826650 | -0.208597885 | 0.662710241 | 0.857405269 |
| chr3:163071713-163072081 | -0.169785325 | 0.662719911 | 0.857405269 |
| chr6:26322465-26323370 | -0.141155193 | 0.662720493 | 0.857405269 |
| chr8:117044729-117045061 | 0.166234069 | 0.662745516 | 0.857423165 |
| chr1:228013391-228013631 | 0.186638214 | 0.662760867 | 0.857428548 |
| chr2:221572947-221573843 | -0.143376221 | 0.662774744 | 0.857432023 |
| chr12:81644613-81644813 | -0.243688304 | 0.662795438 | 0.857444318 |
| chr2:128090902-128091531 | 0.144915367 | 0.662807282 | 0.857445163 |
| chr3:19506944-19507286 | -0.196611423 | 0.662831831 | 0.857452432 |
| chr19:35740271-35741059 | 0.142154651 | 0.662835282 | 0.857452432 |
| chr2:37168217-37168677 | 0.167298819 | 0.6628603 | 0.857470319 |
| chr1:27012719-27013540 | -0.145549165 | 0.662952353 | 0.85757492 |
| chr6:109440225-109441761 | -0.138834552 | 0.662972687 | 0.857584494 |
| chr1:219459696-219460184 | -0.160743961 | 0.662982139 | 0.857584494 |
| chr4:52721901-52722838 | -0.141094489 | 0.663008771 | 0.857604465 |
| chr14:22973603-22973944 | 0.159752215 | 0.663038383 | 0.857614486 |
| chr16:2911559-2912137 | -0.14887461 | 0.663038904 | 0.857614486 |
| chr14:65135336-65135536 | -0.231330372 | 0.663087173 | 0.857647963 |
| chr15:33737770-33737970 | -0.231330372 | 0.663087173 | 0.857647963 |
| chr5:112551498-112551949 | -0.179059927 | 0.66310328 | 0.857654319 |
| chr3:127473302-127473534 | -0.19669299 | 0.663121332 | 0.85766319 |
| chr15:63276850-63277644 | 0.148239994 | 0.663189726 | 0.857737171 |
| chr2:98758214-98758442 | -0.241503344 | 0.663216632 | 0.857745047 |
| chr7:124155454-124155661 | -0.232696228 | 0.663218205 | 0.857745047 |
| chr19:1826330-1826538 | 0.219029575 | 0.663230885 | 0.857746969 |
| chr14:92609341-92609651 | 0.181516275 | 0.663248359 | 0.857749503 |
| chr2:104804205-104804521 | -0.201034777 | 0.663255234 | 0.857749503 |

| | | | |
|---|---|---|---|
| chr10:132791082-132791293 | 0.216409148 | 0.663299426 | 0.857777698 |
| chr11:35362727-35362954 | 0.216409148 | 0.663299426 | 0.857777698 |
| chr2:3605144-3605344 | 0.207476481 | 0.663342024 | 0.857803831 |
| chr2:129748750-129748990 | 0.207476481 | 0.663342024 | 0.857803831 |
| chr1:43366266-43366829 | 0.150323323 | 0.663386802 | 0.857825529 |
| chr10:59604565-59604772 | -0.217814132 | 0.66338871 | 0.857825529 |
| chr3:142439416-142440198 | 0.150129699 | 0.663398629 | 0.857825529 |
| chr4:104352341-104352554 | 0.23845147 | 0.663414781 | 0.857825529 |
| chr5:175052596-175052796 | 0.23845147 | 0.663414781 | 0.857825529 |
| chr6:72784002-72784230 | -0.225081134 | 0.663438016 | 0.857841096 |
| chr5:158000730-158000960 | -0.242643778 | 0.663473392 | 0.857853621 |
| chr5:10202175-10202394 | -0.225802919 | 0.663489426 | 0.857853621 |
| chr10:77165798-77166051 | -0.190121188 | 0.663489648 | 0.857853621 |
| chr2:198127710-198127922 | 0.230913044 | 0.663492486 | 0.857853621 |
| chr1:241591366-241591665 | 0.172858678 | 0.663507563 | 0.857858638 |
| chr4:109998863-109999078 | -0.216688436 | 0.663532743 | 0.857868825 |
| chr11:64653125-64653365 | -0.185044278 | 0.663541943 | 0.857868825 |
| chr22:20016723-20017096 | 0.157664741 | 0.663549825 | 0.857868825 |
| chr7:5012940-5013218 | 0.205613845 | 0.663560226 | 0.857868825 |
| chr12:81261910-81262110 | -0.236997513 | 0.663587641 | 0.857876573 |
| chr2:211652310-211652557 | -0.224467774 | 0.663598502 | 0.857876573 |
| chr11:9137732-9138104 | -0.17203711 | 0.66360197 | 0.857876573 |
| chr14:77040119-77040836 | 0.145464545 | 0.663636188 | 0.857876573 |
| chr11:73876427-73877365 | 0.139650584 | 0.663642032 | 0.857876573 |
| chr9:109320955-109321332 | -0.153595061 | 0.663645136 | 0.857876573 |
| chr1:156471435-156472409 | 0.14345221 | 0.663660813 | 0.857876573 |
| chr16:71501680-71501931 | 0.19348781 | 0.663688039 | 0.857876573 |
| chr15:43824292-43824927 | -0.150211523 | 0.663705139 | 0.857876573 |
| chr10:61472202-61472402 | -0.221974376 | 0.663718845 | 0.857876573 |
| chr4:95637060-95637309 | 0.210436152 | 0.66372246 | 0.857876573 |
| chr1:84020271-84020544 | 0.190719406 | 0.663724796 | 0.857876573 |
| chr10:133026831-133027031 | 0.236195791 | 0.663730026 | 0.857876573 |
| chr5:23208907-23209339 | 0.156974774 | 0.663732924 | 0.857876573 |
| chr16:74108320-74108557 | 0.200234546 | 0.663734163 | 0.857876573 |
| chr9:83041576-83042121 | 0.159863975 | 0.663783887 | 0.857925408 |
| chr6:3432040-3432376 | -0.184605354 | 0.66379434 | 0.857925408 |
| chr14:39174622-39175881 | 0.136505228 | 0.663827078 | 0.857951221 |
| chr15:89898670-89899000 | -0.155362019 | 0.663836706 | 0.857951221 |
| chr1:207737758-207738009 | 0.220049371 | 0.663872462 | 0.857972904 |
| chr11:61333005-61333546 | 0.148933961 | 0.663875878 | 0.857972904 |
| chr10:107610842-107611502 | -0.14884713 | 0.663916461 | 0.857998971 |
| chr10:121573383-121573832 | -0.189334737 | 0.663918444 | 0.857998971 |
| chr5:56264162-56264785 | -0.160366902 | 0.663938869 | 0.858010896 |
| chr1:114620458-114620751 | 0.150788225 | 0.663970843 | 0.858024902 |
| chr13:23778868-23779152 | -0.217917921 | 0.663972104 | 0.858024902 |
| chr3:17877435-17877768 | -0.188155606 | 0.664038591 | 0.858085623 |
| chr17:61450057-61450971 | 0.135797277 | 0.66405802 | 0.858085623 |
| chr16:80447210-80447410 | -0.220553508 | 0.664068433 | 0.858085623 |
| chr10:117168156-117168485 | 0.162620502 | 0.664096188 | 0.858085623 |
| chr2:201653033-201653432 | -0.195080597 | 0.664097142 | 0.858085623 |

| | | | |
|---|---|---|---|
| chr8:97831335-97832076 | 0.144655223 | 0.664097356 | 0.858085623 |
| chr8:109858866-109859256 | -0.176747489 | 0.664100508 | 0.858085623 |
| chr15:82091016-82091268 | 0.203301113 | 0.664109598 | 0.858085623 |
| chr17:7561679-7562023 | 0.160046568 | 0.664141188 | 0.858085623 |
| chr2:64988326-64989540 | -0.14082681 | 0.664142259 | 0.858085623 |
| chr12:116427158-116427818 | 0.150594356 | 0.66414228 | 0.858085623 |
| chr3:134212872-134213275 | 0.158830618 | 0.664160386 | 0.858094547 |
| chr7:121547556-121547899 | -0.181872866 | 0.664176503 | 0.8581009 |
| chr15:82197989-82198354 | 0.167834133 | 0.664192103 | 0.858103677 |
| chr9:98205765-98205978 | -0.197505382 | 0.664204858 | 0.858103677 |
| chr7:33044063-33044636 | 0.151836057 | 0.664233406 | 0.858103677 |
| chr6:37029369-37029581 | 0.215124904 | 0.664233476 | 0.858103677 |
| chr1:35198950-35199195 | 0.204220908 | 0.664234648 | 0.858103677 |
| chr5:167244651-167244965 | -0.176724021 | 0.664247773 | 0.858106165 |
| chr10:24618397-24618804 | -0.16566164 | 0.664279161 | 0.858132246 |
| chr2:108982291-108982680 | -0.165802324 | 0.664345677 | 0.858196021 |
| chr18:35580738-35581295 | -0.163394452 | 0.664354955 | 0.858196021 |
| chr4:156378211-156378411 | -0.225585852 | 0.664362131 | 0.858196021 |
| chr4:87422337-87423088 | -0.144570244 | 0.664411815 | 0.858245732 |
| chr18:10189856-10190134 | 0.2198701 | 0.66447034 | 0.858306621 |
| chr4:8777397-8777610 | 0.201505216 | 0.664481356 | 0.858306621 |
| chr3:186566949-186567508 | -0.144341861 | 0.664530463 | 0.858355582 |
| chr3:149080357-149080559 | -0.217015061 | 0.664550762 | 0.858363898 |
| chr22:24102005-24102277 | -0.193800126 | 0.664565266 | 0.858363898 |
| chr3:196075234-196075462 | 0.228820157 | 0.664570509 | 0.858363898 |
| chr1:9867508-9867797 | -0.190172432 | 0.664588854 | 0.858373123 |
| chr20:3107693-3107893 | -0.22074082 | 0.664614998 | 0.858392422 |
| chr16:1705618-1706318 | 0.147880627 | 0.664652251 | 0.858412221 |
| chr12:54207651-54208089 | 0.161459866 | 0.664652734 | 0.858412221 |
| chr10:89267054-89267356 | -0.201708442 | 0.664687445 | 0.858442581 |
| chr5:77158963-77159163 | 0.244392382 | 0.66472693 | 0.858479105 |
| chr5:167049119-167049331 | -0.193729924 | 0.664744111 | 0.858486825 |
| chr9:88686073-88686445 | -0.216334963 | 0.664759146 | 0.858491771 |
| chr17:7583539-7584226 | -0.144682892 | 0.664790475 | 0.858517761 |
| chr20:8275634-8275834 | 0.219312852 | 0.664831803 | 0.858556662 |
| chr20:18304639-18304920 | 0.203935793 | 0.664845998 | 0.858560522 |
| chrX:35568498-35568811 | -0.185368434 | 0.664960529 | 0.858693952 |
| chr1:52662711-52662980 | 0.200388588 | 0.664998376 | 0.858726293 |
| chr11:65078180-65078432 | 0.185317011 | 0.665007988 | 0.858726293 |
| chr12:101670094-101670328 | -0.194219076 | 0.665034402 | 0.858745929 |
| chr3:24520909-24522162 | 0.140243019 | 0.665061823 | 0.858750217 |
| chr17:44409293-44409571 | -0.227231536 | 0.665071294 | 0.858750217 |
| chr12:6891660-6891860 | -0.200586351 | 0.66507458 | 0.858750217 |
| chr3:63862795-63865033 | -0.134236376 | 0.665094082 | 0.858750217 |
| chr7:1506884-1507265 | -0.152119274 | 0.665104731 | 0.858750217 |
| chr5:539611-539891 | -0.193398739 | 0.665104968 | 0.858750217 |
| chr8:111964239-111965141 | 0.145598647 | 0.665119213 | 0.858754138 |
| chr19:8208895-8209553 | 0.160014988 | 0.665145773 | 0.85877396 |
| chr5:175092904-175093313 | 0.185185479 | 0.665167233 | 0.858780179 |
| chr8:99358597-99358853 | 0.17920927 | 0.665173006 | 0.858780179 |

| | | | |
|---|---|---|---|
| chr8:16077504-16077729 | -0.229034181 | 0.665192742 | 0.858791189 |
| chr3:140093219-140093538 | -0.187494833 | 0.665235548 | 0.858818731 |
| chr7:73769993-73770923 | -0.140514671 | 0.665236492 | 0.858818731 |
| chr14:69535328-69535583 | -0.179924403 | 0.665253079 | 0.858825675 |
| chr15:94028898-94029166 | 0.191102915 | 0.665292324 | 0.858861868 |
| chr17:16414995-16415295 | 0.172378103 | 0.665346468 | 0.858917293 |
| chr17:13030552-13030786 | 0.189832834 | 0.6654175 | 0.858956685 |
| chr14:75176040-75177179 | 0.137674416 | 0.665431921 | 0.858956685 |
| chr20:50931247-50931949 | -0.144188241 | 0.665445602 | 0.858956685 |
| chr10:44236466-44237221 | 0.158855864 | 0.665449864 | 0.858956685 |
| chr4:68375922-68376540 | -0.147469772 | 0.66545331 | 0.858956685 |
| chr18:76256697-76257010 | -0.203803977 | 0.665458897 | 0.858956685 |
| chr1:114669560-114670885 | -0.134309949 | 0.665460792 | 0.858956685 |
| chr2:47712490-47712690 | 0.229177176 | 0.665472551 | 0.858956685 |
| chr1:152089304-152089652 | -0.170639737 | 0.665477874 | 0.858956685 |
| chr17:82736748-82736948 | 0.212797072 | 0.665518267 | 0.858976676 |
| chr2:160205336-160205955 | -0.150016801 | 0.665521138 | 0.858976676 |
| chr17:17591130-17592237 | -0.139669698 | 0.665526994 | 0.858976676 |
| chr4:190021009-190021523 | -0.174998219 | 0.665569208 | 0.858995906 |
| chr1:114219402-114219810 | 0.145354519 | 0.665573778 | 0.858995906 |
| chr15:66497143-66498366 | 0.140425681 | 0.665586408 | 0.858995906 |
| chr10:99406265-99406553 | -0.197938203 | 0.665590682 | 0.858995906 |
| chr1:204080189-204080808 | -0.139784867 | 0.665598128 | 0.858995906 |
| chr15:68306239-68306461 | 0.179754194 | 0.665609158 | 0.858995906 |
| chr7:2353884-2354951 | -0.137797847 | 0.66568124 | 0.859047101 |
| chr3:140927812-140928012 | -0.239448882 | 0.665681765 | 0.859047101 |
| chr1:237562339-237562558 | 0.205965368 | 0.665688928 | 0.859047101 |
| chr6:7140738-7141525 | -0.146289906 | 0.665694752 | 0.859047101 |
| chr3:23805623-23806196 | 0.147022005 | 0.665716446 | 0.859047101 |
| chr3:190139386-190139690 | 0.235173567 | 0.665721028 | 0.859047101 |
| chr20:64131372-64131776 | 0.154999336 | 0.665727308 | 0.859047101 |
| chr22:22848428-22848764 | -0.178144524 | 0.66574305 | 0.859052946 |
| chr5:151075707-151076108 | 0.159315512 | 0.665761868 | 0.859058436 |
| chr3:180595041-180595301 | -0.202742001 | 0.665769727 | 0.859058436 |
| chr19:11597197-11597819 | 0.147095228 | 0.665791245 | 0.859070737 |
| chr1:192877694-192877920 | 0.210202428 | 0.665804442 | 0.859070737 |
| chr3:57943084-57943306 | 0.216087913 | 0.66582382 | 0.859070737 |
| chr19:47035257-47036290 | -0.141758453 | 0.665824108 | 0.859070737 |
| chrX:101206524-101206796 | -0.186265215 | 0.665843796 | 0.859081673 |
| chr13:52195333-52195818 | -0.157447649 | 0.66587652 | 0.859092791 |
| chr7:5592299-5592519 | -0.178932542 | 0.665879921 | 0.859092791 |
| chr14:86646715-86646952 | 0.212337288 | 0.665886049 | 0.859092791 |
| chr1:147642064-147642299 | -0.169893893 | 0.66590222 | 0.859092872 |
| chr15:98428313-98429006 | -0.145564599 | 0.665909432 | 0.859092872 |
| chr1:85464482-85465599 | -0.13913374 | 0.665919748 | 0.859092872 |
| chr1:176717126-176717425 | -0.177570364 | 0.665936104 | 0.859099508 |
| chr16:85384582-85385440 | -0.141553821 | 0.665952515 | 0.859106215 |
| chr10:6647955-6648279 | -0.168706516 | 0.665973441 | 0.859112008 |
| chr10:84574192-84574478 | -0.178292016 | 0.666004098 | 0.859112008 |
| chr2:169824408-169825200 | 0.142402344 | 0.666027015 | 0.859112008 |

| | | | |
|---|---|---|---|
| chr9:137351942-137352327 | 0.184780928 | 0.666032841 | 0.859112008 |
| chr3:153513273-153513554 | -0.197867709 | 0.666034725 | 0.859112008 |
| chr14:25400716-25400983 | 0.200605729 | 0.666035459 | 0.859112008 |
| chr14:67729087-67729502 | -0.17010141 | 0.666035492 | 0.859112008 |
| chr6:49676643-49676857 | -0.223222292 | 0.666060557 | 0.859115291 |
| chr1:20633277-20633572 | 0.175027145 | 0.666088826 | 0.859115291 |
| chr14:40368319-40368695 | -0.159017412 | 0.666096885 | 0.859115291 |
| chr2:168119565-168119874 | -0.183176309 | 0.666110525 | 0.859115291 |
| chr16:75432873-75434227 | -0.135959915 | 0.666114268 | 0.859115291 |
| chr8:130065333-130065586 | 0.177532158 | 0.666115345 | 0.859115291 |
| chr18:741969-742169 | 0.21446861 | 0.666116524 | 0.859115291 |
| chr1:201126668-201127184 | 0.145733786 | 0.6661382 | 0.859121855 |
| chr6:26026295-26026643 | -0.154660077 | 0.66614511 | 0.859121855 |
| chr15:83707793-83708027 | -0.233405784 | 0.666155251 | 0.859121855 |
| chr6:153073526-153073794 | -0.220099498 | 0.666255315 | 0.859236442 |
| chr2:186256593-186256830 | 0.204766588 | 0.666273501 | 0.859245433 |
| chr6:137866517-137867843 | -0.138481163 | 0.666293814 | 0.859257167 |
| chr13:40498370-40498737 | 0.195584823 | 0.666352115 | 0.859317891 |
| chr15:90934347-90934652 | 0.146541215 | 0.666388668 | 0.859350566 |
| chr10:3652224-3652435 | -0.219570347 | 0.666404429 | 0.859356426 |
| chr3:124837738-124838095 | -0.157896218 | 0.666465285 | 0.859418216 |
| chr2:29923231-29923440 | -0.199922137 | 0.6664767 | 0.859418216 |
| chr1:119582844-119583134 | 0.189875156 | 0.666490974 | 0.859418216 |
| chr10:30510582-30510884 | -0.181582508 | 0.66649721 | 0.859418216 |
| chr1:204507449-204508020 | -0.1411818 | 0.666508485 | 0.859418291 |
| chr8:21909021-21909383 | -0.180752286 | 0.666522184 | 0.859421492 |
| chr22:37681648-37681878 | 0.182923816 | 0.666631927 | 0.859542823 |
| chr20:17505173-17505524 | 0.171118785 | 0.666638718 | 0.859542823 |
| chr15:90086887-90087153 | 0.16284052 | 0.666658354 | 0.859553678 |
| chr2:62857162-62857362 | -0.234708862 | 0.666705523 | 0.859593647 |
| chr4:153683992-153684778 | -0.150149466 | 0.666716029 | 0.859593647 |
| chr1:172933069-172933276 | -0.215961351 | 0.666732115 | 0.859593647 |
| chr12:80925246-80925679 | -0.160784739 | 0.666737447 | 0.859593647 |
| chr17:75784377-75785354 | 0.136330899 | 0.666745447 | 0.859593647 |
| chr3:188370226-188370436 | 0.24771627 | 0.666775191 | 0.859595538 |
| chr1:75785621-75786490 | -0.13826096 | 0.666781434 | 0.859595538 |
| chr1:77900701-77901070 | 0.176126083 | 0.666790114 | 0.859595538 |
| chr1:119872595-119872893 | -0.194321824 | 0.666798319 | 0.859595538 |
| chr11:76380701-76381820 | -0.138075211 | 0.666803007 | 0.859595538 |
| chr7:220510-220737 | -0.213438812 | 0.66682183 | 0.859605341 |
| chr4:134132047-134132828 | -0.152650328 | 0.666836375 | 0.859606742 |
| chr17:17823567-17824208 | 0.143041834 | 0.666851075 | 0.859606742 |
| chr3:59247507-59247848 | -0.175419796 | 0.666856573 | 0.859606742 |
| chr14:23555755-23556540 | 0.138944002 | 0.666876649 | 0.859618159 |
| chr10:28663773-28664171 | -0.16664338 | 0.666893676 | 0.859625645 |
| chr11:76210604-76210830 | 0.177558164 | 0.666916459 | 0.859639468 |
| chr7:120502511-120502772 | -0.198047116 | 0.666926838 | 0.859639468 |
| chr15:74695884-74696626 | 0.149800375 | 0.666943192 | 0.859646087 |
| chr8:86260916-86261116 | 0.202597882 | 0.666955964 | 0.859648088 |
| chr3:41268062-41268366 | 0.181964057 | 0.666979888 | 0.859664463 |

| | | | |
|---|---|---|---|
| chr1:201539641-201539903 | 0.1643564 | 0.666991896 | 0.859665478 |
| chr11:36405453-36405653 | 0.246791132 | 0.667003493 | 0.859665965 |
| chr3:113403816-113404016 | -0.23783093 | 0.667034999 | 0.859671963 |
| chr1:229217924-229218138 | 0.206072137 | 0.66703813 | 0.859671963 |
| chr8:41490225-41490785 | 0.147634618 | 0.667041805 | 0.859671963 |
| chr5:163566037-163566274 | 0.191772404 | 0.667065376 | 0.859687593 |
| chr5:139168439-139168647 | 0.222151102 | 0.667095424 | 0.859687593 |
| chrX:73944068-73944620 | -0.153066678 | 0.667104313 | 0.859687593 |
| chr18:79678847-79679963 | -0.136447298 | 0.667107752 | 0.859687593 |
| chr3:122514033-122515311 | 0.137525397 | 0.667120724 | 0.859687593 |
| chr1:159072050-159072302 | 0.192315688 | 0.667123255 | 0.859687593 |
| chr2:70995394-70995739 | 0.155575725 | 0.667132472 | 0.859687593 |
| chr4:153531072-153531281 | -0.214077393 | 0.667150034 | 0.859693558 |
| chr22:17329030-17329538 | 0.156355465 | 0.667166038 | 0.859693558 |
| chr5:58496592-58496959 | -0.201979388 | 0.667179323 | 0.859693558 |
| chr9:128361886-128362420 | 0.150796554 | 0.667189241 | 0.859693558 |
| chr3:56447082-56447387 | 0.18267658 | 0.667201922 | 0.859693558 |
| chr2:241844081-241844341 | -0.19636032 | 0.667214062 | 0.859693558 |
| chr17:59930361-59930661 | 0.151748759 | 0.667217363 | 0.859693558 |
| chr5:159515717-159516162 | -0.168444993 | 0.66722686 | 0.859693558 |
| chr19:11535945-11536316 | 0.181557344 | 0.667253095 | 0.859710114 |
| chr10:117174700-117175647 | 0.144244058 | 0.66726215 | 0.859710114 |
| chr1:154973654-154975149 | 0.134723932 | 0.667316725 | 0.859725092 |
| chr1:45303438-45304241 | 0.138877949 | 0.667327201 | 0.859725092 |
| chr8:143609488-143610129 | 0.139742223 | 0.667330859 | 0.859725092 |
| chr6:40912386-40912754 | -0.166854725 | 0.667332448 | 0.859725092 |
| chr13:68064352-68064787 | -0.190340984 | 0.667334535 | 0.859725092 |
| chr8:120951213-120951518 | -0.156605429 | 0.6673618 | 0.859725092 |
| chr17:21322993-21323322 | -0.156462468 | 0.667386286 | 0.859725092 |
| chr16:54080488-54080993 | -0.160434073 | 0.667389754 | 0.859725092 |
| chr17:28475426-28475646 | -0.190725973 | 0.667390432 | 0.859725092 |
| chr5:38636637-38637123 | -0.152308677 | 0.667392314 | 0.859725092 |
| chr10:83657675-83657875 | 0.233440899 | 0.667425919 | 0.859725092 |
| chr8:38468382-38468870 | 0.157529512 | 0.667429669 | 0.859725092 |
| chr3:193041214-193041438 | 0.212398849 | 0.66743086 | 0.859725092 |
| chr18:27185071-27185329 | 0.212398849 | 0.66743086 | 0.859725092 |
| chr6:119484485-119484928 | -0.164344567 | 0.667486943 | 0.859780194 |
| chr12:6851827-6852798 | -0.136433415 | 0.667499757 | 0.859780194 |
| chr2:61889681-61889881 | -0.192579936 | 0.667507301 | 0.859780194 |
| chr17:73429425-73429683 | -0.170555903 | 0.667525699 | 0.859789438 |
| chr1:189708562-189709249 | -0.150142154 | 0.667549244 | 0.859803678 |
| chr6:44469787-44470378 | -0.147651615 | 0.667559197 | 0.859803678 |
| chr12:72232103-72232303 | -0.231822093 | 0.667578635 | 0.859814261 |
| chr22:49781158-49781358 | 0.243589484 | 0.667612532 | 0.859843465 |
| chr1:155278257-155278657 | 0.151996215 | 0.667673577 | 0.859888489 |
| chr1:172476308-172476548 | -0.193730752 | 0.667674561 | 0.859888489 |
| chr11:3091823-3092152 | 0.165727249 | 0.667690669 | 0.859888489 |
| chr13:42415610-42415853 | 0.224578351 | 0.667697738 | 0.859888489 |
| chr1:68321145-68321399 | -0.201343683 | 0.667704548 | 0.859888489 |
| chr3:14474646-14474924 | -0.207853666 | 0.667715486 | 0.859888489 |

| | | | |
|---|---|---|---|
| chr14:31378123-31378380 | 0.196511397 | 0.667726048 | 0.859888489 |
| chr3:162868233-162868514 | -0.208932024 | 0.667746276 | 0.859900086 |
| chr1:157496093-157496424 | 0.173530541 | 0.667827552 | 0.859987617 |
| chr17:17303131-17303482 | 0.166976034 | 0.667844314 | 0.859987617 |
| chr1:161480673-161480913 | 0.214468782 | 0.667847918 | 0.859987617 |
| chr8:122780076-122780967 | 0.136607255 | 0.66790797 | 0.860026794 |
| chr22:31036606-31036889 | 0.196871376 | 0.667912745 | 0.860026794 |
| chr1:150293602-150294582 | -0.138437976 | 0.667923063 | 0.860026794 |
| chr2:64144159-64144828 | -0.137981964 | 0.667923239 | 0.860026794 |
| chr5:147577038-147577253 | -0.213716235 | 0.667949812 | 0.860046556 |
| chr7:149662559-149663149 | -0.148507188 | 0.667963828 | 0.860050151 |
| chr15:95894712-95894922 | -0.210333302 | 0.667985102 | 0.860054611 |
| chr3:160921023-160921344 | 0.17800469 | 0.667989742 | 0.860054611 |
| chr5:90740454-90740711 | 0.210848316 | 0.668004295 | 0.860058898 |
| chr3:134453712-134453934 | 0.199324307 | 0.668086328 | 0.860150061 |
| chr15:88834352-88834710 | 0.163080388 | 0.668131991 | 0.860176459 |
| chr2:241515565-241515765 | -0.254780619 | 0.668133002 | 0.860176459 |
| chr21:45630210-45630470 | 0.207581124 | 0.668146105 | 0.860176459 |
| chr15:40891925-40892125 | -0.181479225 | 0.668151736 | 0.860176459 |
| chr20:41373333-41373621 | -0.155838597 | 0.668225872 | 0.860257448 |
| chr18:74310446-74310836 | -0.159756411 | 0.668238369 | 0.860259082 |
| chr14:73136197-73137248 | 0.13831027 | 0.668253058 | 0.8602619 |
| chr4:60612163-60612371 | -0.22166352 | 0.668263012 | 0.8602619 |
| chr1:228155267-228155604 | 0.172064097 | 0.668279303 | 0.860268418 |
| chr4:100493783-100494194 | 0.161234946 | 0.66830595 | 0.860288267 |
| chr20:23889743-23890422 | -0.15035674 | 0.668329459 | 0.860301442 |
| chr19:45677262-45677817 | 0.143700727 | 0.668338641 | 0.860301442 |
| chr6:32894053-32895129 | -0.140101215 | 0.668361843 | 0.860316856 |
| chr1:117268593-117269543 | 0.138907412 | 0.668380176 | 0.860320178 |
| chr3:172722561-172722947 | -0.158603671 | 0.668386881 | 0.860320178 |
| chr3:129992768-129993234 | -0.159571203 | 0.668400353 | 0.860323067 |
| chr3:77563130-77563330 | -0.214336593 | 0.668452911 | 0.860370208 |
| chr7:65269804-65270111 | 0.182804858 | 0.668459435 | 0.860370208 |
| chr19:55165962-55166385 | 0.150160256 | 0.668473352 | 0.860373668 |
| chr2:183248932-183249250 | -0.178543538 | 0.668495622 | 0.860387879 |
| chr1:44047803-44048003 | 0.209495857 | 0.66852015 | 0.860404995 |
| chr16:75116570-75116990 | -0.144898329 | 0.668535313 | 0.860410057 |
| chr11:86068230-86069688 | -0.134616388 | 0.668613936 | 0.860465214 |
| chr6:154450345-154450815 | 0.158107437 | 0.668626371 | 0.860465214 |
| chr10:126787334-126787534 | -0.229475708 | 0.66864046 | 0.860465214 |
| chr15:96327319-96327529 | 0.17204056 | 0.668640567 | 0.860465214 |
| chr5:36725137-36725340 | -0.214677663 | 0.668641875 | 0.860465214 |
| chr6:143549487-143549689 | -0.223235646 | 0.66864555 | 0.860465214 |
| chr3:24011526-24011736 | 0.237074719 | 0.668678882 | 0.860493656 |
| chr10:20271518-20271727 | 0.215271473 | 0.668693259 | 0.860497705 |
| chr5:33843311-33843524 | -0.233289559 | 0.668708417 | 0.86050276 |
| chr15:43826804-43827093 | 0.178056654 | 0.668742963 | 0.860531927 |
| chrX:39908221-39908665 | -0.173790031 | 0.668753545 | 0.860531927 |
| chr7:6558921-6559240 | -0.155176896 | 0.66880999 | 0.860575908 |
| chr12:69639304-69639702 | -0.161229974 | 0.668878158 | 0.860575908 |

| | | | |
|---|---|---|---|
| chr2:180761359-180761572 | -0.219542196 | 0.668893718 | 0.860575908 |
| chr2:241492468-241492702 | -0.198199382 | 0.668908171 | 0.860575908 |
| chrX:74421228-74421623 | 0.157110416 | 0.66892373 | 0.860575908 |
| chr6:166556952-166557262 | -0.181103096 | 0.668937659 | 0.860575908 |
| chr13:110615253-110616187 | -0.142673067 | 0.668943308 | 0.860575908 |
| chr4:142436045-142436286 | 0.198614198 | 0.668945661 | 0.860575908 |
| chr6:124795957-124796189 | 0.198614198 | 0.668945661 | 0.860575908 |
| chr1:145451997-145452213 | 0.217543783 | 0.668965216 | 0.860575908 |
| chr4:124711936-124713615 | -0.134271698 | 0.668967697 | 0.860575908 |
| chr1:226946678-226946928 | 0.204932492 | 0.668972863 | 0.860575908 |
| chr17:31525403-31525825 | -0.164785031 | 0.668981896 | 0.860575908 |
| chr7:36049141-36049341 | 0.226092995 | 0.668986203 | 0.860575908 |
| chr11:86099874-86100098 | -0.212911662 | 0.669004676 | 0.860575908 |
| chr14:39231619-39231905 | -0.19299051 | 0.669011763 | 0.860575908 |
| chr5:180619282-180619484 | -0.204368141 | 0.669014023 | 0.860575908 |
| chr11:67261331-67261682 | -0.162925506 | 0.669030238 | 0.860575908 |
| chr11:6603309-6604637 | 0.13874231 | 0.669040912 | 0.860575908 |
| chr1:200620454-200621066 | -0.137403045 | 0.669050936 | 0.860575908 |
| chr1:89149284-89149524 | -0.216531693 | 0.669058114 | 0.860575908 |
| chr1:201309434-201309830 | 0.142057188 | 0.669060761 | 0.860575908 |
| chr6:93418538-93419239 | 0.143511453 | 0.669063597 | 0.860575908 |
| chr2:222572374-222572657 | -0.192271717 | 0.669073962 | 0.860575908 |
| chr18:37565965-37566452 | 0.166889122 | 0.66909574 | 0.860575908 |
| chr17:49191767-49193294 | -0.133481678 | 0.669122671 | 0.860575908 |
| chr3:135800294-135800576 | -0.197832763 | 0.669123058 | 0.860575908 |
| chr14:75966950-75967261 | -0.184657058 | 0.669128038 | 0.860575908 |
| chr2:202104306-202104592 | 0.180225399 | 0.669139159 | 0.860575908 |
| chr20:34311788-34311994 | 0.209032195 | 0.669140794 | 0.860575908 |
| chr4:118749518-118749842 | 0.187122454 | 0.66914293 | 0.860575908 |
| chr10:95656225-95657168 | 0.138259138 | 0.66914713 | 0.860575908 |
| chr16:83356144-83356491 | 0.180370876 | 0.669174482 | 0.86059664 |
| chrX:103918608-103919700 | -0.138178983 | 0.669195092 | 0.860604234 |
| chr17:47946956-47947456 | 0.143991937 | 0.669202851 | 0.860604234 |
| chr13:28819717-28820882 | -0.140786418 | 0.669234264 | 0.860630187 |
| chr6:27297833-27298211 | -0.161826864 | 0.669266141 | 0.860642369 |
| chr7:15007554-15007833 | -0.175785248 | 0.669266202 | 0.860642369 |
| chr13:31629264-31629464 | -0.225009966 | 0.669285039 | 0.860652148 |
| chr12:34350569-34350769 | -0.188365041 | 0.669325903 | 0.860678042 |
| chr20:50742029-50742251 | 0.14330147 | 0.669327641 | 0.860678042 |
| chr2:68646405-68646628 | -0.182982461 | 0.669347668 | 0.86068935 |
| chr21:43646569-43647001 | -0.157272535 | 0.669360741 | 0.860691188 |
| chr3:48635379-48636113 | -0.145803159 | 0.669383524 | 0.860691188 |
| chr15:78860040-78860905 | 0.142218318 | 0.669385016 | 0.860691188 |
| chr15:72117034-72118729 | 0.134507902 | 0.669394029 | 0.860691188 |
| chr3:151067099-151067575 | -0.159269155 | 0.669411772 | 0.860699558 |
| chr3:87226763-87228021 | -0.134872159 | 0.669423263 | 0.860699891 |
| chr6:32971541-32972405 | 0.13635691 | 0.669436235 | 0.860702126 |
| chr9:133133702-133134023 | -0.166155439 | 0.669481428 | 0.860744424 |
| chr17:59106037-59107536 | -0.132142257 | 0.669491601 | 0.860744424 |
| chr1:204313801-204314710 | -0.140301337 | 0.669506531 | 0.860749176 |

| | | | |
|---|---|---|---|
| chr3:102745905-102746167 | 0.185008931 | 0.669557624 | 0.860800421 |
| chr2:232432773-232433043 | -0.210529366 | 0.66960971 | 0.86085294 |
| chr2:53698193-53698422 | -0.205006806 | 0.669630702 | 0.860856872 |
| chr10:86194707-86194907 | 0.215865145 | 0.669635239 | 0.860856872 |
| chr17:32909973-32910287 | -0.183510416 | 0.669702214 | 0.860913319 |
| chr3:12830829-12831075 | 0.179143961 | 0.669725802 | 0.860913319 |
| chr2:239003002-239003396 | -0.173948779 | 0.669728215 | 0.860913319 |
| chr5:179908638-179908929 | -0.178633919 | 0.669732479 | 0.860913319 |
| chr5:163379637-163379888 | -0.188173368 | 0.669735327 | 0.860913319 |
| chr7:7062152-7062377 | -0.198679154 | 0.66974933 | 0.860916877 |
| chr5:94346592-94346792 | 0.220231461 | 0.669793266 | 0.860958909 |
| chr6:36724615-36724867 | 0.179654853 | 0.669817941 | 0.860976183 |
| chr5:86501012-86501274 | -0.189118351 | 0.669830965 | 0.86097848 |
| chr12:124904693-124905032 | -0.174153215 | 0.669868237 | 0.861000817 |
| chr13:81874247-81874629 | 0.163111939 | 0.669870817 | 0.861000817 |
| chr7:100015387-100015924 | -0.154816988 | 0.669887003 | 0.861007179 |
| chr22:29766372-29767674 | 0.137669899 | 0.669914567 | 0.861025271 |
| chr16:3820215-3820454 | 0.220164429 | 0.669927922 | 0.861025271 |
| chr11:64237630-64237946 | -0.157674482 | 0.669934791 | 0.861025271 |
| chr2:30146320-30148020 | -0.133968112 | 0.670026704 | 0.861128956 |
| chr3:7010410-7010610 | 0.211996323 | 0.670049054 | 0.861141916 |
| chr6:27472225-27473563 | -0.135791571 | 0.670059265 | 0.861141916 |
| chr12:12433330-12433937 | 0.156898169 | 0.670104482 | 0.861181054 |
| chr6:44456314-44456514 | 0.210642043 | 0.670112197 | 0.861181054 |
| chr6:92716016-92716479 | 0.168205415 | 0.670140627 | 0.861202031 |
| chr1:215181598-215181829 | -0.189621713 | 0.670151 | 0.861202031 |
| chr11:364452-364737 | 0.188526338 | 0.670193286 | 0.861235294 |
| chr6:43097583-43097789 | 0.226346914 | 0.670199363 | 0.861235294 |
| chr16:19717720-19718470 | 0.14420679 | 0.67022282 | 0.861246351 |
| chr11:118572186-118572919 | 0.145971147 | 0.670252465 | 0.861246351 |
| chr10:42952133-42952661 | 0.153381445 | 0.670255092 | 0.861246351 |
| chr9:131408537-131408746 | -0.205720561 | 0.670257549 | 0.861246351 |
| chr2:138275337-138275696 | -0.166975116 | 0.670264169 | 0.861246351 |
| chr3:127816699-127817088 | -0.164706023 | 0.670284252 | 0.861250385 |
| chr1:148038400-148039082 | -0.139186028 | 0.670291493 | 0.861250385 |
| chr22:46150242-46150482 | -0.193684255 | 0.670304146 | 0.861250385 |
| chr20:25057946-25058326 | 0.161609862 | 0.670312269 | 0.861250385 |
| chr1:56577321-56577600 | -0.185973446 | 0.670332287 | 0.861253054 |
| chr14:75258691-75259765 | -0.134650118 | 0.670336828 | 0.861253054 |
| chr9:35826036-35826539 | 0.156046031 | 0.670368329 | 0.861279086 |
| chr7:22500031-22500394 | 0.163551458 | 0.670389443 | 0.86129177 |
| chr7:155002680-155004140 | -0.135645262 | 0.670421482 | 0.861318491 |
| chr19:47600267-47600934 | 0.140867661 | 0.670438818 | 0.861326321 |
| chr1:150973893-150974314 | 0.145552056 | 0.670458996 | 0.861337803 |
| chr1:206809334-206809811 | -0.149983238 | 0.670486703 | 0.861358955 |
| chrX:40996942-40997491 | -0.161681785 | 0.670515115 | 0.861361857 |
| chrX:39975475-39975946 | -0.158333377 | 0.670546948 | 0.861361857 |
| chr19:49142902-49143216 | -0.154297774 | 0.670551667 | 0.861361857 |
| chr5:149344854-149345078 | 0.174121234 | 0.670562003 | 0.861361857 |
| chr22:41589195-41589575 | 0.16309594 | 0.670571675 | 0.861361857 |

| | | | |
|---|---|---|---|
| chr21:32612052-32613328 | 0.137937959 | 0.670581513 | 0.861361857 |
| chr6:135251461-135251820 | -0.158555355 | 0.670586651 | 0.861361857 |
| chr11:20110882-20111257 | -0.152501678 | 0.670589838 | 0.861361857 |
| chr9:33403428-33403785 | 0.16984334 | 0.670590137 | 0.861361857 |
| chr7:137953630-137954338 | 0.142747152 | 0.670622583 | 0.861389093 |
| chr17:75756050-75756283 | -0.167225113 | 0.670655433 | 0.861416847 |
| chr17:16521277-16521537 | -0.183361496 | 0.670670879 | 0.861422246 |
| chr12:110698279-110698685 | 0.156707436 | 0.670693263 | 0.861428926 |
| chr4:52832755-52833093 | -0.221255295 | 0.670698565 | 0.861428926 |
| chr6:20686206-20686502 | -0.193301168 | 0.670716151 | 0.861437073 |
| chr1:89822180-89822380 | 0.203601811 | 0.670759002 | 0.861461658 |
| chrX:35383191-35383455 | -0.199599107 | 0.6707636 | 0.861461658 |
| chr17:82647739-82648674 | 0.139413261 | 0.670772918 | 0.861461658 |
| chr20:52747822-52748022 | -0.213129491 | 0.670812384 | 0.861461658 |
| chr16:68539007-68540274 | 0.140356522 | 0.670815843 | 0.861461658 |
| chr7:99306756-99306986 | -0.219893505 | 0.670821484 | 0.861461658 |
| chr2:176652417-176652630 | 0.202628108 | 0.670822414 | 0.861461658 |
| chr19:18365126-18365326 | -0.21866594 | 0.670833069 | 0.861461658 |
| chr4:101981646-101982119 | -0.155817456 | 0.67083648 | 0.861461658 |
| chr19:10644707-10645031 | -0.177976804 | 0.670891865 | 0.861518343 |
| chr1:37437195-37437395 | -0.234931389 | 0.670933968 | 0.86155797 |
| chr12:67362611-67362955 | -0.197735848 | 0.671018704 | 0.861645441 |
| chr22:37563292-37563496 | 0.200688456 | 0.671037502 | 0.861645441 |
| chr13:34124738-34125381 | 0.150939269 | 0.671037832 | 0.861645441 |
| chr17:81119409-81120013 | -0.142290225 | 0.671052036 | 0.861645441 |
| chr9:130936604-130936906 | 0.179499564 | 0.671058313 | 0.861645441 |
| chr2:71067795-71069012 | -0.134973947 | 0.671079166 | 0.861657777 |
| chr16:48365632-48366347 | 0.146410128 | 0.671151635 | 0.861723321 |
| chr3:63417427-63417881 | 0.151258417 | 0.671161095 | 0.861723321 |
| chr1:43914399-43914645 | 0.214834018 | 0.67121204 | 0.861723321 |
| chr8:38768354-38768554 | -0.221328781 | 0.671219789 | 0.861723321 |
| chr11:84909779-84909979 | 0.252062772 | 0.671245458 | 0.861723321 |
| chr15:101294686-101295938 | 0.1348927 | 0.671271553 | 0.861723321 |
| chr2:174697224-174697551 | 0.157543201 | 0.671281584 | 0.861723321 |
| chr9:124771026-124771870 | 0.13849496 | 0.671286354 | 0.861723321 |
| chr3:150344906-150345489 | -0.143384313 | 0.671291545 | 0.861723321 |
| chr10:100342299-100342636 | -0.160207579 | 0.671299006 | 0.861723321 |
| chr3:8622939-8623261 | 0.153230635 | 0.671299692 | 0.861723321 |
| chr17:75557075-75558214 | -0.137023121 | 0.671302621 | 0.861723321 |
| chr2:216209662-216209968 | 0.199656413 | 0.671312618 | 0.861723321 |
| chr3:183884481-183885356 | 0.139036845 | 0.671318879 | 0.861723321 |
| chr18:23663024-23663224 | -0.216633925 | 0.671336493 | 0.861723321 |
| chr12:6323263-6323605 | 0.163888858 | 0.671351399 | 0.861723321 |
| chr15:45401715-45402020 | -0.165304245 | 0.671370451 | 0.861723321 |
| chr10:79240157-79240426 | -0.155426685 | 0.67140894 | 0.861723321 |
| chr1:198404772-198405128 | -0.195824797 | 0.671411613 | 0.861723321 |
| chrX:50580062-50580269 | -0.216590811 | 0.671413559 | 0.861723321 |
| chr1:145961896-145962160 | 0.152793588 | 0.671420142 | 0.861723321 |
| chr1:8026043-8026558 | 0.152450232 | 0.671428621 | 0.861723321 |
| chr22:21468130-21468746 | 0.15894406 | 0.67148835 | 0.861723321 |

| | | | |
|---|---|---|---|
| chr3:64627738-64628249 | -0.13677494 | 0.671494136 | 0.861723321 |
| chr12:102513907-102514122 | -0.234415253 | 0.67149768 | 0.861723321 |
| chr13:94601457-94602389 | 0.137480116 | 0.67149876 | 0.861723321 |
| chr4:147683327-147683540 | 0.174140388 | 0.671499246 | 0.861723321 |
| chr10:74115734-74115977 | 0.205315131 | 0.671517167 | 0.861723321 |
| chr9:129333743-129334822 | 0.138401875 | 0.671518887 | 0.861723321 |
| chr12:54888458-54888697 | -0.211180183 | 0.671525327 | 0.861723321 |
| chr9:110255848-110257185 | 0.134326263 | 0.671534057 | 0.861723321 |
| chr1:20640189-20640458 | -0.202767373 | 0.671534313 | 0.861723321 |
| chr9:136733528-136733732 | -0.213738708 | 0.671536716 | 0.861723321 |
| chr2:231642659-231643120 | 0.147648301 | 0.671550948 | 0.861723321 |
| chr8:119889307-119889507 | -0.249448381 | 0.671567539 | 0.861723321 |
| chrX:74274735-74274935 | -0.249448381 | 0.671567539 | 0.861723321 |
| chr1:188647941-188648326 | -0.177407813 | 0.671570489 | 0.861723321 |
| chr1:116372530-116374330 | 0.132487006 | 0.67157362 | 0.861723321 |
| chr5:142375704-142375904 | -0.238785475 | 0.671600762 | 0.861723321 |
| chr19:33778052-33778352 | 0.161456792 | 0.671607218 | 0.861723321 |
| chr19:1618247-1618447 | -0.203707886 | 0.671611722 | 0.861723321 |
| chr5:133399360-133399560 | -0.222113395 | 0.671615672 | 0.861723321 |
| chr2:85818368-85818778 | 0.159646671 | 0.671624657 | 0.861723321 |
| chr10:62112253-62112775 | 0.145820403 | 0.671634083 | 0.861723321 |
| chr5:81043763-81043995 | 0.199356863 | 0.671636302 | 0.861723321 |
| chr6:96182304-96182613 | 0.214452458 | 0.671670531 | 0.861752807 |
| chr2:90392541-90392741 | 0.231283218 | 0.671686173 | 0.861758447 |
| chr15:84164433-84164633 | 0.224016367 | 0.671707627 | 0.861759557 |
| chr22:24259352-24259758 | 0.188249683 | 0.671709532 | 0.861759557 |
| chr1:26620318-26621649 | -0.137778253 | 0.671732579 | 0.86177213 |
| chr1:178031820-178032185 | 0.151692409 | 0.671741953 | 0.86177213 |
| chr14:46872368-46872568 | 0.226511722 | 0.671753074 | 0.86177213 |
| chr14:63542348-63543959 | 0.13349817 | 0.671775298 | 0.861786212 |
| chrX:105273444-105273648 | 0.203090072 | 0.67179704 | 0.861799675 |
| chr4:112706260-112706839 | 0.156019202 | 0.671812038 | 0.861804487 |
| chr7:34747881-34748893 | -0.137733533 | 0.671836416 | 0.86180751 |
| chr4:153574443-153574645 | 0.21175651 | 0.67183689 | 0.86180751 |
| chr6:156564365-156564687 | -0.177415801 | 0.671885929 | 0.861816902 |
| chr1:202957633-202958619 | 0.134632118 | 0.671889615 | 0.861816902 |
| chr1:28517950-28518410 | 0.146332203 | 0.671889857 | 0.861816902 |
| chr16:35391376-35391652 | -0.197277212 | 0.671889942 | 0.861816902 |
| chr3:84987843-84988043 | 0.232068032 | 0.671911698 | 0.861816902 |
| chr4:105497022-105497222 | 0.232068032 | 0.671911698 | 0.861816902 |
| chr10:131890887-131891087 | -0.213429818 | 0.671945708 | 0.861846098 |
| chr12:110280683-110281971 | -0.137235852 | 0.671992801 | 0.861865804 |
| chr3:118685745-118685945 | 0.238367159 | 0.671992818 | 0.861865804 |
| chr14:90512395-90512639 | -0.16919279 | 0.671994817 | 0.861865804 |
| chr5:165227615-165227815 | -0.230610258 | 0.672034193 | 0.861901879 |
| chr14:80037453-80037695 | -0.197907317 | 0.672064502 | 0.861912763 |
| chr3:32557722-32557922 | 0.23613338 | 0.672067177 | 0.861912763 |
| chr9:123377337-123377872 | -0.155900452 | 0.67209188 | 0.861912763 |
| chr4:153507732-153507955 | 0.203768382 | 0.672103399 | 0.861912763 |
| chr15:64093650-64094413 | 0.137212047 | 0.672106922 | 0.861912763 |

| | | | |
|---|---|---|---|
| chr1:186212566-186212952 | 0.150334209 | 0.672110173 | 0.861912763 |
| chr16:85112324-85112583 | -0.178914422 | 0.672143179 | 0.861922844 |
| chr2:174700061-174700280 | -0.204966767 | 0.672143603 | 0.861922844 |
| chr7:41889357-41889607 | -0.22520877 | 0.672160741 | 0.861922844 |
| chr12:98490576-98491323 | -0.144649655 | 0.672191608 | 0.861922844 |
| chr10:133329752-133330081 | 0.160465565 | 0.672194108 | 0.861922844 |
| chr2:120463693-120463898 | 0.196609782 | 0.672201807 | 0.861922844 |
| chr3:133964670-133964870 | 0.196609782 | 0.672201807 | 0.861922844 |
| chr20:54167825-54168448 | 0.135247413 | 0.672208942 | 0.861922844 |
| chr2:144240632-144241023 | -0.175204869 | 0.672229305 | 0.861922844 |
| chr4:71607965-71608229 | 0.201022006 | 0.672230524 | 0.861922844 |
| chr1:47388749-47389122 | -0.158250832 | 0.672262159 | 0.861948981 |
| chr11:120013313-120013558 | -0.217808977 | 0.672286205 | 0.861960918 |
| chr14:89422748-89422948 | -0.196921839 | 0.67230015 | 0.861960918 |
| chr5:95697842-95698696 | -0.140817574 | 0.672308721 | 0.861960918 |
| chr10:124690948-124691343 | 0.1568798 | 0.672316466 | 0.861960918 |
| chr3:21484124-21484874 | 0.145663686 | 0.67234672 | 0.861984954 |
| chr14:70641206-70641736 | 0.155151524 | 0.672357714 | 0.861984954 |
| chr21:26389956-26390561 | -0.161124096 | 0.672381682 | 0.862001258 |
| chr5:170638668-170639055 | 0.152047248 | 0.672420439 | 0.86203238 |
| chr6:153312670-153312947 | -0.1810524 | 0.67243438 | 0.86203238 |
| chr2:84458531-84459889 | 0.135364808 | 0.672439709 | 0.86203238 |
| chr5:18465934-18466557 | 0.148931617 | 0.672466606 | 0.862052438 |
| chr17:45171846-45172126 | 0.179188367 | 0.672543981 | 0.862107476 |
| chr20:45436757-45437387 | 0.149764893 | 0.672565566 | 0.862107476 |
| chr17:36948458-36949249 | -0.142651067 | 0.672566841 | 0.862107476 |
| chr17:73091891-73093085 | -0.136229737 | 0.67257272 | 0.862107476 |
| chr3:141192220-141192706 | -0.144855262 | 0.672572783 | 0.862107476 |
| chr5:56951249-56952689 | -0.137655376 | 0.672577048 | 0.862107476 |
| chr5:181161174-181161449 | -0.169866253 | 0.672590009 | 0.862109667 |
| chr17:75855021-75856189 | -0.134755722 | 0.672640272 | 0.86215967 |
| chr6:99334402-99335010 | 0.154144817 | 0.672673734 | 0.862188137 |
| chr6:75266799-75267008 | 0.231343152 | 0.672693815 | 0.862199452 |
| chr6:140441340-140441668 | 0.168904215 | 0.672706954 | 0.86220187 |
| chr10:47165307-47165531 | -0.2128879 | 0.672733424 | 0.862206952 |
| chr11:62633317-62633546 | -0.2128879 | 0.672733424 | 0.862206952 |
| chr15:49045734-49046874 | -0.13691198 | 0.672752617 | 0.862209405 |
| chr13:30072364-30073131 | 0.151015754 | 0.672762407 | 0.862209405 |
| chr3:150303556-150304122 | 0.146805761 | 0.67277766 | 0.862209405 |
| chr8:99838081-99838375 | -0.175632399 | 0.672780349 | 0.862209405 |
| chr8:90991249-90991459 | -0.207344858 | 0.672812455 | 0.86221357 |
| chr3:115692859-115693274 | 0.155286116 | 0.672814484 | 0.86221357 |
| chr6:42044250-42044515 | 0.165140631 | 0.672817358 | 0.86221357 |
| chr2:32277511-32278564 | -0.137825669 | 0.672841804 | 0.862230478 |
| chr1:19341539-19341783 | -0.184588589 | 0.67285376 | 0.862231379 |
| chr15:78909868-78910082 | -0.186363239 | 0.672874458 | 0.862243481 |
| chr6:24581189-24581422 | 0.197876678 | 0.672912221 | 0.862262768 |
| chr7:44214320-44214678 | 0.156848863 | 0.672913148 | 0.862262768 |
| chr9:129584133-129584524 | -0.166149055 | 0.672923269 | 0.862262768 |
| chr8:108918421-108918928 | 0.152155924 | 0.6729412 | 0.862271324 |

| | | | |
|---|---|---|---|
| chr10:27360606-27360865 | -0.180509283 | 0.672955287 | 0.862272083 |
| chr17:8295279-8295875 | 0.143348172 | 0.672969375 | 0.862272083 |
| chr9:77648206-77649175 | -0.144036652 | 0.672975553 | 0.862272083 |
| chr20:56171119-56171453 | -0.153765518 | 0.672988677 | 0.86227448 |
| chr7:43905921-43906899 | -0.141881327 | 0.673026891 | 0.862296517 |
| chr20:37289469-37290046 | 0.148854542 | 0.673028385 | 0.862296517 |
| chr12:110937136-110937352 | 0.198531344 | 0.673068572 | 0.862333586 |
| chr14:80056141-80056557 | 0.158114032 | 0.67310726 | 0.862365292 |
| chr19:6504830-6505047 | -0.19331407 | 0.673115829 | 0.862365292 |
| chr9:132855092-132855292 | 0.223432681 | 0.673157978 | 0.862404872 |
| chr11:3807851-3808226 | 0.14959707 | 0.673195832 | 0.86242796 |
| chr5:148230346-148230589 | -0.219400099 | 0.673198511 | 0.86242796 |
| chr7:19754552-19754789 | 0.200233453 | 0.67322685 | 0.862449846 |
| chr1:49995800-49996128 | -0.174162644 | 0.67325804 | 0.862475381 |
| chr5:56909106-56910534 | -0.133208266 | 0.673301168 | 0.862504877 |
| chr2:112480894-112481796 | 0.140046828 | 0.673303577 | 0.862504877 |
| chr9:76665976-76666316 | -0.15891568 | 0.673321577 | 0.862513514 |
| chr2:113910429-113910660 | 0.213628121 | 0.673383885 | 0.862569038 |
| chr16:84084854-84085159 | -0.160596065 | 0.673387436 | 0.862569038 |
| chr8:88175778-88175978 | 0.223973701 | 0.673458165 | 0.862645216 |
| chr3:198080001-198081142 | -0.142673621 | 0.673470249 | 0.862646273 |
| chr20:41723216-41723520 | -0.166923525 | 0.673491169 | 0.862658649 |
| chr4:99405613-99405830 | -0.199500955 | 0.673513335 | 0.86267262 |
| chr15:45268179-45268511 | 0.169311627 | 0.673532475 | 0.862674847 |
| chr3:122894123-122894350 | -0.1929812 | 0.673537592 | 0.862674847 |
| chr10:86939411-86939818 | 0.147464534 | 0.67355949 | 0.862683697 |
| chr6:112043605-112044023 | -0.151461074 | 0.673567019 | 0.862683697 |
| chr17:59859953-59860250 | 0.138135113 | 0.673606287 | 0.862704702 |
| chr1:55277561-55277866 | 0.16000825 | 0.673607658 | 0.862704702 |
| chr2:88740066-88740345 | -0.177907436 | 0.673617586 | 0.862704702 |
| chr2:15097028-15097228 | -0.222323839 | 0.673628457 | 0.862704702 |
| chr16:3527318-3527552 | 0.198065974 | 0.673677034 | 0.862745206 |
| chr9:131530641-131530959 | -0.165837454 | 0.673682604 | 0.862745206 |
| chrX:56563234-56564015 | 0.144159472 | 0.67369807 | 0.862750594 |
| chr5:118678274-118678519 | 0.210447811 | 0.673710427 | 0.862751999 |
| chr16:24504800-24505142 | 0.17037155 | 0.673732288 | 0.862754882 |
| chr8:127865803-127866346 | -0.141605368 | 0.673735198 | 0.862754882 |
| chr8:8779712-8779915 | -0.224237495 | 0.673779004 | 0.862783325 |
| chr6:95974058-95974262 | -0.214529247 | 0.673784053 | 0.862783325 |
| chr9:90914949-90915257 | 0.169844256 | 0.673798156 | 0.862783325 |
| chr5:88243682-88243882 | -0.233606059 | 0.673807153 | 0.862783325 |
| chr16:1408037-1408769 | 0.142538868 | 0.673813711 | 0.862783325 |
| chr14:76977832-76978317 | -0.152175438 | 0.673834783 | 0.862795888 |
| chr1:32248126-32248944 | -0.142774143 | 0.673849496 | 0.862800309 |
| chr20:31968416-31968754 | 0.164675071 | 0.673942407 | 0.862904853 |
| chr3:8095864-8096064 | 0.205784827 | 0.6739645 | 0.862913524 |
| chr3:12046412-12046648 | 0.220147057 | 0.673971703 | 0.862913524 |
| chr1:160124878-160125078 | 0.19698837 | 0.674002885 | 0.862919963 |
| chr7:56196308-56196596 | 0.184461554 | 0.674005727 | 0.862919963 |
| chr17:51891241-51891545 | -0.196294333 | 0.674010519 | 0.862919963 |

| | | | |
|---|---|---|---|
| chr2:131092879-131094241 | -0.136089425 | 0.674070333 | 0.862982123 |
| chr17:73659643-73659843 | -0.214620831 | 0.674082831 | 0.862983703 |
| chr1:188650661-188650861 | 0.239808524 | 0.67410843 | 0.863002057 |
| chr17:50671022-50671351 | 0.162741116 | 0.674152144 | 0.863026752 |
| chr15:44426800-44428044 | 0.133553278 | 0.674152441 | 0.863026752 |
| chr6:129775146-129775467 | 0.168034862 | 0.67416151 | 0.863026752 |
| chr17:65953580-65953949 | -0.167573013 | 0.674195622 | 0.863043212 |
| chr6:89828980-89830244 | -0.132623523 | 0.674196895 | 0.863043212 |
| chr18:51110301-51110662 | -0.166998729 | 0.67423773 | 0.863081066 |
| chr8:111336173-111336373 | -0.223737161 | 0.674317562 | 0.863168836 |
| chr3:175731988-175732213 | -0.222563113 | 0.674344353 | 0.86318871 |
| chr6:68400557-68400768 | 0.219907804 | 0.674369285 | 0.863206204 |
| chr21:39445288-39446193 | 0.134487561 | 0.674394337 | 0.86320667 |
| chr17:18314594-18315498 | -0.13537595 | 0.674394799 | 0.86320667 |
| chr6:108541816-108542474 | -0.16119411 | 0.674403446 | 0.86320667 |
| chr16:21511928-21512155 | -0.191814237 | 0.67443388 | 0.863231203 |
| chr2:45348626-45348826 | -0.236586747 | 0.674505546 | 0.863289825 |
| chr8:133683994-133684235 | -0.206042364 | 0.674519386 | 0.863289825 |
| chr2:52976442-52976646 | -0.208131782 | 0.674540569 | 0.863289825 |
| chr1:32241035-32241720 | 0.148101491 | 0.674559183 | 0.863289825 |
| chr8:27361976-27362360 | -0.171232385 | 0.674561025 | 0.863289825 |
| chr1:58430368-58430629 | 0.212914116 | 0.674571949 | 0.863289825 |
| chr11:83115650-83115884 | -0.222990285 | 0.674573607 | 0.863289825 |
| chr2:102718026-102718247 | -0.222990285 | 0.674573607 | 0.863289825 |
| chr8:99923501-99923714 | 0.2013495 | 0.674581083 | 0.863289825 |
| chr18:9612268-9612563 | 0.171778573 | 0.674603061 | 0.863303533 |
| chr13:97141543-97141982 | -0.149222599 | 0.674650345 | 0.863336542 |
| chr7:65269154-65269781 | -0.145174026 | 0.674655076 | 0.863336542 |
| chr10:119542392-119543683 | 0.136165194 | 0.674671066 | 0.863336542 |
| chr6:72622086-72622558 | 0.151555753 | 0.67469806 | 0.863336542 |
| chr14:23528543-23528786 | 0.175053619 | 0.674708706 | 0.863336542 |
| chr4:121227606-121227815 | -0.210511807 | 0.674726293 | 0.863336542 |
| chr8:25577193-25577393 | -0.210511807 | 0.674726293 | 0.863336542 |
| chr20:41232391-41232956 | 0.144450084 | 0.67472852 | 0.863336542 |
| chr2:216012856-216013859 | -0.135531044 | 0.674730262 | 0.863336542 |
| chr1:156060872-156061448 | 0.139110962 | 0.67476368 | 0.863354853 |
| chr22:37607049-37607395 | -0.154890152 | 0.674771607 | 0.863354853 |
| chr4:86927971-86928418 | -0.177872689 | 0.674784116 | 0.863354853 |
| chr3:3636768-3637456 | -0.138422273 | 0.674803135 | 0.863354853 |
| chr3:123329888-123330099 | -0.198495164 | 0.674808256 | 0.863354853 |
| chr1:1399310-1399843 | -0.147143733 | 0.674812179 | 0.863354853 |
| chr17:82494216-82495134 | -0.134199375 | 0.674941848 | 0.863506333 |
| chr3:171566895-171567101 | -0.211448627 | 0.67495403 | 0.8635075 |
| chr9:108565783-108566082 | 0.176773523 | 0.674979105 | 0.863525161 |
| chr16:80613850-80614430 | -0.137179067 | 0.67499475 | 0.863530758 |
| chr12:75142617-75142817 | 0.231614865 | 0.675007429 | 0.863531834 |
| chr8:8454654-8454900 | 0.185740058 | 0.675020072 | 0.863531834 |
| chr9:12329401-12329689 | 0.168191692 | 0.675038527 | 0.863531834 |
| chr19:19366798-19367144 | -0.172557945 | 0.675063269 | 0.863531834 |
| chr9:111630772-111631737 | -0.134082194 | 0.675065025 | 0.863531834 |

| | | | |
|---|---|---|---|
| chr4:118823324-118823760 | -0.157542002 | 0.675081858 | 0.863531834 |
| chr15:76553210-76553410 | 0.226608968 | 0.675083921 | 0.863531834 |
| chr12:125330758-125331266 | -0.15436753 | 0.675086963 | 0.863531834 |
| chr18:79379011-79379211 | -0.201335513 | 0.675099351 | 0.863531834 |
| chr20:17431885-17432109 | 0.19894293 | 0.67510829 | 0.863531834 |
| chr2:182649343-182649590 | 0.192340435 | 0.675170271 | 0.863583186 |
| chr8:2025240-2025556 | -0.19279857 | 0.675177959 | 0.863583186 |
| chr3:18423418-18423621 | -0.20333731 | 0.675191354 | 0.863583186 |
| chr2:8539693-8540479 | -0.136214454 | 0.67519352 | 0.863583186 |
| chr1:240692055-240692401 | -0.167532346 | 0.675218259 | 0.863600411 |
| chr9:36995889-36996133 | -0.186082487 | 0.67524413 | 0.863619085 |
| chr2:122236547-122236846 | -0.169826135 | 0.675309403 | 0.863674105 |
| chr12:67161394-67161667 | 0.177398346 | 0.675309693 | 0.863674105 |
| chr3:27369070-27369708 | -0.137308775 | 0.675330625 | 0.863686459 |
| chr19:49157453-49158014 | 0.145270179 | 0.675355229 | 0.863703509 |
| chr5:57613797-57614155 | 0.191672431 | 0.675392454 | 0.8637367 |
| chr3:50602012-50602212 | 0.188062095 | 0.675411037 | 0.863746048 |
| chr8:142788270-142788801 | -0.146840796 | 0.67543058 | 0.863756624 |
| chr2:239157867-239158067 | 0.229683578 | 0.675443258 | 0.863758421 |
| chr17:62294425-62294738 | -0.2048856 | 0.675486993 | 0.863775626 |
| chr4:117835239-117835517 | -0.184760102 | 0.675497539 | 0.863775626 |
| chr9:125706645-125707675 | 0.139366636 | 0.675508936 | 0.863775626 |
| chr1:214550974-214552662 | -0.132454638 | 0.675522442 | 0.863775626 |
| chr14:63923159-63923439 | -0.174425868 | 0.67553153 | 0.863775626 |
| chr14:99635307-99635520 | -0.211968759 | 0.675532536 | 0.863775626 |
| chr17:17860214-17860513 | -0.164908489 | 0.675535625 | 0.863775626 |
| chr3:113092692-113092932 | 0.212924641 | 0.675550795 | 0.86378061 |
| chr2:15989666-15990080 | 0.155095987 | 0.675579916 | 0.863789864 |
| chr1:202274657-202274952 | -0.15155149 | 0.67558058 | 0.863789864 |
| chr1:148151360-148151805 | 0.152336575 | 0.675629221 | 0.86382549 |
| chr3:76222155-76222637 | -0.161506479 | 0.675630991 | 0.86382549 |
| chr2:138917141-138917346 | -0.227828917 | 0.675681783 | 0.863876015 |
| chr17:73826576-73826994 | -0.176204927 | 0.675701507 | 0.86388032 |
| chr6:30219255-30219532 | 0.204513048 | 0.675707699 | 0.86388032 |
| chr2:191646558-191646767 | 0.20037694 | 0.67584726 | 0.864044329 |
| chr16:86555037-86555668 | -0.140388941 | 0.675864547 | 0.864052012 |
| chr6:132538924-132539124 | -0.235435879 | 0.675948983 | 0.864117722 |
| chr9:19408674-19409635 | 0.136463556 | 0.675955622 | 0.864117722 |
| chr19:47428051-47428599 | 0.140923692 | 0.675959069 | 0.864117722 |
| chr3:149626751-149626960 | -0.200771625 | 0.675961056 | 0.864117722 |
| chr15:91856107-91856367 | -0.166627957 | 0.676017077 | 0.864170818 |
| chr1:178093139-178094734 | 0.132272046 | 0.676025147 | 0.864170818 |
| chr1:200310363-200310684 | -0.186533186 | 0.676042802 | 0.864178969 |
| chr11:123478366-123478650 | 0.183950428 | 0.676065243 | 0.864193237 |
| chr7:43729106-43729934 | -0.138339558 | 0.676100178 | 0.864202734 |
| chr11:72585692-72585906 | 0.181905151 | 0.67610371 | 0.864202734 |
| chr9:36273362-36273792 | 0.160745669 | 0.676106508 | 0.864202734 |
| chr6:73308987-73309606 | 0.146860594 | 0.676130441 | 0.864218908 |
| chr17:51165306-51166929 | -0.131393214 | 0.676159297 | 0.864231064 |
| chr6:4582696-4582943 | 0.205504109 | 0.67616251 | 0.864231064 |

| | | | |
|---|---|---|---|
| chr8:142603653-142603853 | 0.181129492 | 0.676197488 | 0.864261355 |
| chr17:65279523-65279772 | -0.187286448 | 0.676221396 | 0.864263979 |
| chr22:42437491-42437968 | 0.158548915 | 0.67624498 | 0.864263979 |
| chr3:171328728-171329017 | -0.186030527 | 0.676264149 | 0.864263979 |
| chr11:20387052-20388472 | -0.13208545 | 0.676272108 | 0.864263979 |
| chr11:114439206-114440015 | 0.140409225 | 0.676286016 | 0.864263979 |
| chr17:59899207-59899472 | 0.143616044 | 0.676296793 | 0.864263979 |
| chrX:100819797-100820923 | -0.133548517 | 0.676299097 | 0.864263979 |
| chr4:100494279-100494703 | -0.168237608 | 0.676301663 | 0.864263979 |
| chr11:34165722-34166442 | -0.139836633 | 0.676310197 | 0.864263979 |
| chr9:92706545-92706803 | 0.195709204 | 0.676312338 | 0.864263979 |
| chr19:48644837-48645332 | 0.149857856 | 0.676328103 | 0.864269711 |
| chr14:29585102-29585364 | -0.185741223 | 0.676434997 | 0.864365163 |
| chr19:28289631-28289969 | -0.166211593 | 0.67644536 | 0.864365163 |
| chr2:224449597-224449845 | 0.19817623 | 0.67645723 | 0.864365163 |
| chr17:75861347-75861673 | -0.168998293 | 0.676470279 | 0.864365163 |
| chr12:3076654-3077426 | 0.140322746 | 0.676472206 | 0.864365163 |
| chr6:8556752-8556957 | 0.217498403 | 0.676480098 | 0.864365163 |
| chr12:85278530-85278782 | -0.206806612 | 0.676481764 | 0.864365163 |
| chr20:14219700-14220951 | 0.141295088 | 0.676518951 | 0.864386133 |
| chr6:89352136-89353235 | 0.134781523 | 0.676536685 | 0.864386133 |
| chr20:60535239-60535566 | -0.173102098 | 0.676537491 | 0.864386133 |
| chr1:36542715-36543000 | -0.20234935 | 0.67654595 | 0.864386133 |
| chr1:144412057-144413431 | 0.131250011 | 0.676554582 | 0.864386133 |
| chr2:219611494-219611697 | -0.22886308 | 0.676596344 | 0.864408706 |
| chr4:77317958-77318192 | 0.198149722 | 0.676605457 | 0.864408706 |
| chr16:10520607-10520857 | -0.19100859 | 0.676606094 | 0.864408706 |
| chr7:32942092-32943436 | 0.131679419 | 0.676622381 | 0.8644151 |
| chr3:129690465-129690681 | 0.192298911 | 0.676635345 | 0.864417251 |
| chr11:66616113-66617410 | 0.13136031 | 0.676658408 | 0.864424118 |
| chr2:3959415-3959615 | -0.220194675 | 0.676669729 | 0.864424118 |
| chr3:75453289-75453553 | -0.207357343 | 0.676674566 | 0.864424118 |
| chr16:29595038-29595799 | 0.13990361 | 0.67670535 | 0.864439006 |
| chr13:66312792-66313076 | 0.186404419 | 0.676708784 | 0.864439006 |
| chr6:34536417-34536994 | -0.138423666 | 0.676768401 | 0.864498365 |
| chr5:180313449-180314090 | 0.137829856 | 0.676777817 | 0.864498365 |
| chr2:47121653-47121919 | -0.215044111 | 0.676801904 | 0.86451472 |
| chr1:16056059-16056410 | 0.155777426 | 0.676847649 | 0.86455874 |
| chr6:28124222-28125064 | -0.143544478 | 0.676885474 | 0.864592641 |
| chr18:10578591-10578814 | -0.215508557 | 0.676934156 | 0.864621438 |
| chr1:194102511-194102875 | -0.174800444 | 0.676935044 | 0.864621438 |
| chr1:94541412-94542296 | 0.147421339 | 0.676958551 | 0.864621438 |
| chr10:88879859-88881122 | 0.138242328 | 0.676966781 | 0.864621438 |
| chr5:55146558-55146892 | -0.161333366 | 0.676975115 | 0.864621438 |
| chr16:80466360-80466747 | 0.158301119 | 0.676979251 | 0.864621438 |
| chr4:184506747-184506947 | 0.205945047 | 0.676987009 | 0.864621438 |
| chr12:46072026-46072647 | -0.146083136 | 0.677007854 | 0.864622928 |
| chr7:9251398-9251598 | 0.211043773 | 0.677020074 | 0.864622928 |
| chr10:63712197-63712779 | 0.142211137 | 0.677022028 | 0.864622928 |
| chr3:177359864-177360326 | 0.139022288 | 0.677033395 | 0.864623034 |

| | | | |
|---|---|---|---|
| chr8:37694465-37694684 | 0.170405255 | 0.677048761 | 0.864628247 |
| chr1:168397043-168397522 | -0.159254886 | 0.677150338 | 0.864725517 |
| chr14:28652800-28653246 | 0.16092678 | 0.677154025 | 0.864725517 |
| chr3:124032690-124033418 | 0.139655969 | 0.677158785 | 0.864725517 |
| chr18:70348441-70348860 | 0.159124347 | 0.677231131 | 0.864803488 |
| chr10:93458925-93459237 | -0.184848079 | 0.67726437 | 0.86483152 |
| chr15:78506944-78508131 | 0.132960702 | 0.677279564 | 0.86483651 |
| chr9:83979526-83981279 | -0.130512028 | 0.677327834 | 0.864868805 |
| chr3:133805415-133806581 | 0.133152846 | 0.677338144 | 0.864868805 |
| chr22:20885467-20886113 | -0.143023137 | 0.677338718 | 0.864868805 |
| chr10:132751624-132751833 | -0.205812183 | 0.677364695 | 0.864887562 |
| chr6:7261086-7261871 | 0.137028327 | 0.677390798 | 0.864895232 |
| chr6:122744241-122744554 | -0.179465023 | 0.677393279 | 0.864895232 |
| chr5:17663089-17663349 | 0.191916522 | 0.677441114 | 0.864922349 |
| chr17:60602032-60602243 | 0.139285171 | 0.677451535 | 0.864922349 |
| chr6:2695530-2695749 | -0.221453914 | 0.677457343 | 0.864922349 |
| chr16:57757299-57757526 | -0.172575211 | 0.677459669 | 0.864922349 |
| chr3:124768873-124769145 | 0.165933177 | 0.677523125 | 0.864988951 |
| chr20:48184291-48184728 | 0.139032755 | 0.677549093 | 0.865007275 |
| chr1:45803357-45803760 | 0.148518641 | 0.677560056 | 0.865007275 |
| chr8:119354143-119354440 | -0.167155374 | 0.677574019 | 0.865010688 |
| chr15:78640906-78641436 | -0.154898383 | 0.677627476 | 0.865020841 |
| chr14:64800064-64800302 | 0.172149613 | 0.677636521 | 0.865020841 |
| chr19:43754427-43755385 | 0.133573686 | 0.677640589 | 0.865020841 |
| chr7:1359715-1360016 | 0.168155213 | 0.677655299 | 0.865020841 |
| chr2:39905465-39905666 | -0.216156094 | 0.67765759 | 0.865020841 |
| chr9:19379735-19379975 | 0.178787559 | 0.677667349 | 0.865020841 |
| chr2:79219442-79219680 | -0.196814402 | 0.67767691 | 0.865020841 |
| chr4:129344071-129344297 | -0.196814402 | 0.67767691 | 0.865020841 |
| chr2:160407489-160408052 | -0.15426763 | 0.677690505 | 0.865020841 |
| chr17:75456083-75457537 | -0.13204577 | 0.677697167 | 0.865020841 |
| chr3:149717514-149717752 | -0.179699395 | 0.677713013 | 0.865020841 |
| chr2:84870455-84870730 | -0.203791319 | 0.677717446 | 0.865020841 |
| chrX:115958107-115959304 | 0.139860137 | 0.677742684 | 0.865038645 |
| chr22:30890712-30890932 | 0.186655505 | 0.677802451 | 0.865100518 |
| chr4:125313511-125313772 | -0.161701452 | 0.677820539 | 0.865109194 |
| chr12:55619481-55620073 | 0.147208385 | 0.677837765 | 0.865115955 |
| chr7:43728683-43728883 | 0.198159507 | 0.677861284 | 0.865115955 |
| chr11:117941391-117941591 | 0.207694126 | 0.677863108 | 0.865115955 |
| chr16:89958037-89958298 | -0.176739099 | 0.677870999 | 0.865115955 |
| chr13:45464528-45465631 | -0.131970356 | 0.677886737 | 0.865119683 |
| chr1:57424583-57424941 | -0.169894764 | 0.677896612 | 0.865119683 |
| chr20:46253478-46253766 | -0.15945446 | 0.677907793 | 0.865119683 |
| chr21:10432137-10432670 | 0.161109358 | 0.677947633 | 0.86513791 |
| chr19:51926892-51927092 | 0.207753946 | 0.677962287 | 0.86513791 |
| chr14:59483726-59484862 | 0.131036229 | 0.677973407 | 0.86513791 |
| chr10:87017779-87018009 | -0.203500505 | 0.677978213 | 0.86513791 |
| chr20:54320074-54320369 | 0.142025096 | 0.677978871 | 0.86513791 |
| chr2:219497895-219498231 | -0.158256795 | 0.677989821 | 0.86513791 |
| chr14:90344376-90344578 | 0.199033252 | 0.678011186 | 0.865150764 |

| | | | |
|---|---|---|---|
| chr2:105853433-105854076 | -0.146303003 | 0.678044005 | 0.865178234 |
| chrX:34764758-34765076 | -0.180989468 | 0.678068797 | 0.86519546 |
| chr14:100502367-100502579 | -0.20876926 | 0.678104866 | 0.865227075 |
| chr15:58241984-58242184 | -0.206669945 | 0.678146763 | 0.865261109 |
| chr1:206612065-206612970 | -0.131413913 | 0.678154125 | 0.865261109 |
| chr14:65254973-65256160 | 0.132635939 | 0.678171514 | 0.865268887 |
| chr17:28902387-28903509 | 0.134567717 | 0.678195954 | 0.865285661 |
| chr2:219597344-219598382 | 0.132259135 | 0.678236743 | 0.865311401 |
| chr2:215988501-215989167 | 0.156395508 | 0.678238715 | 0.865311401 |
| chr11:66479860-66480906 | -0.132839655 | 0.678251269 | 0.865313009 |
| chr1:181423341-181423921 | 0.140033882 | 0.67826679 | 0.865318404 |
| chr20:49854022-49854229 | 0.192770475 | 0.678304707 | 0.865335955 |
| chr2:25672769-25673264 | -0.150807551 | 0.678309168 | 0.865335955 |
| chr3:72849052-72849252 | -0.203266253 | 0.678314428 | 0.865335955 |
| chr8:54145843-54146241 | -0.174828829 | 0.67835873 | 0.865378063 |
| chr1:101520384-101520584 | -0.223714188 | 0.678374034 | 0.865383178 |
| chr1:201113875-201114966 | 0.131026331 | 0.678427134 | 0.865420972 |
| chr10:28676926-28677758 | -0.136197559 | 0.678428766 | 0.865420972 |
| chr13:95189150-95189652 | -0.158146848 | 0.678446181 | 0.865420972 |
| chr11:77780026-77780249 | 0.216886337 | 0.678457484 | 0.865420972 |
| chr6:13574200-13575464 | -0.131342667 | 0.6784815 | 0.865420972 |
| chr4:75916039-75916239 | 0.212500338 | 0.678481738 | 0.865420972 |
| chr3:102098252-102098452 | 0.223385691 | 0.678482723 | 0.865420972 |
| chr6:7723066-7723382 | -0.167467564 | 0.678512556 | 0.865421861 |
| chr19:4915229-4915482 | -0.185042772 | 0.678514799 | 0.865421861 |
| chr1:185719094-185719411 | -0.175918303 | 0.678526715 | 0.865421861 |
| chr1:45686449-45687467 | -0.132155051 | 0.678528599 | 0.865421861 |
| chr2:121718702-121719050 | -0.16488663 | 0.678542016 | 0.865424568 |
| chr13:22650369-22650571 | 0.21431022 | 0.678558903 | 0.865431701 |
| chr12:113462479-113462699 | 0.195718467 | 0.67859025 | 0.86545715 |
| chr19:10010122-10010322 | -0.212578838 | 0.67860296 | 0.86545715 |
| chr10:101783000-101783200 | -0.178113109 | 0.678612743 | 0.86545715 |
| chr15:74460863-74461653 | -0.136604925 | 0.678626892 | 0.86546079 |
| chr11:64723511-64723770 | 0.169530499 | 0.678645582 | 0.865470221 |
| chr10:130045089-130045351 | -0.181585482 | 0.678686425 | 0.865507902 |
| chr15:67863409-67863963 | -0.149384863 | 0.678731494 | 0.865537682 |
| chr5:72366400-72366840 | -0.165464532 | 0.678739642 | 0.865537682 |
| chr2:104481450-104481841 | 0.155757867 | 0.67876185 | 0.865537682 |
| chr14:91173210-91173679 | 0.153605905 | 0.678764975 | 0.865537682 |
| chr8:123472149-123472420 | 0.164097209 | 0.678766259 | 0.865537682 |
| chrX:132169605-132170078 | -0.149032895 | 0.678809288 | 0.865558272 |
| chr11:44927453-44927696 | -0.205302044 | 0.678810446 | 0.865558272 |
| chr7:19663859-19664214 | 0.165817348 | 0.678816295 | 0.865558272 |
| chr12:122749913-122750376 | -0.142536966 | 0.678833852 | 0.865560754 |
| chr17:49557553-49557753 | -0.175265034 | 0.678840834 | 0.865560754 |
| chr18:24359690-24359933 | 0.19415215 | 0.678855828 | 0.865565468 |
| chr10:13708631-13709088 | -0.152204169 | 0.678880848 | 0.865582965 |
| chr1:22043258-22043542 | -0.190242119 | 0.67891759 | 0.865615408 |
| chr9:36772533-36772974 | -0.171139522 | 0.678970034 | 0.865651636 |
| chr6:96305056-96305419 | 0.17004154 | 0.678972867 | 0.865651636 |

| | | | |
|---|---|---|---|
| chr22:49724842-49725079 | 0.197384003 | 0.678979897 | 0.865651636 |
| chr1:233607726-233607994 | -0.19845149 | 0.679001044 | 0.865664194 |
| chr13:25114195-25114482 | 0.165291787 | 0.679022351 | 0.86566643 |
| chr14:105613560-105613820 | -0.202780837 | 0.679025395 | 0.86566643 |
| chr2:100177875-100178487 | -0.142411248 | 0.679044429 | 0.865668282 |
| chr4:168048682-168048885 | -0.208638102 | 0.679049443 | 0.865668282 |
| chr1:85297670-85298550 | -0.135041393 | 0.679086436 | 0.865701038 |
| chr7:1169368-1170424 | -0.134332425 | 0.679127038 | 0.865724871 |
| chr4:44448503-44449254 | 0.136853188 | 0.679132474 | 0.865724871 |
| chr20:31662200-31662454 | 0.164619587 | 0.679144691 | 0.865724871 |
| chr17:78491607-78491921 | 0.175840549 | 0.679150326 | 0.865724871 |
| chr6:134080254-134080454 | -0.214008096 | 0.679165441 | 0.865729736 |
| chr10:3955652-3955875 | -0.214694511 | 0.679184866 | 0.865740094 |
| chr5:72082134-72082372 | 0.180272025 | 0.67928941 | 0.86585895 |
| chr6:109164413-109164613 | -0.218255133 | 0.679318312 | 0.865869065 |
| chr2:209334501-209334804 | -0.169088216 | 0.679324182 | 0.865869065 |
| chr16:3182589-3183092 | 0.143573905 | 0.679341953 | 0.865869065 |
| chr11:81470525-81470782 | -0.19858714 | 0.679357463 | 0.865869065 |
| chr3:164143373-164143590 | -0.218111652 | 0.679381868 | 0.865869065 |
| chr16:679387-680433 | 0.134455325 | 0.679382335 | 0.865869065 |
| chr1:155648628-155648828 | 0.208992351 | 0.679387689 | 0.865869065 |
| chr1:74690186-74690474 | -0.210904272 | 0.679389257 | 0.865869065 |
| chr20:56893247-56893564 | -0.151370692 | 0.679408355 | 0.865869065 |
| chr5:2375079-2375333 | -0.206768105 | 0.679413522 | 0.865869065 |
| chr3:169813345-169813608 | -0.181851146 | 0.679421652 | 0.865869065 |
| chr4:149708839-149709039 | 0.222660061 | 0.679444418 | 0.865883677 |
| chr18:58202321-58202596 | -0.178062041 | 0.679491162 | 0.865928845 |
| chr16:61050724-61050938 | -0.213694614 | 0.679515476 | 0.865945428 |
| chr2:6917029-6917947 | -0.138453144 | 0.679540618 | 0.865963066 |
| chr1:156302126-156302326 | 0.237910142 | 0.679564154 | 0.865968328 |
| chr11:1389445-1389960 | 0.144087722 | 0.679583752 | 0.865968328 |
| chr6:3251511-3251974 | -0.151128365 | 0.679588293 | 0.865968328 |
| chrX:145405107-145405344 | -0.227320065 | 0.679589955 | 0.865968328 |
| chrX:64975539-64975964 | -0.169993229 | 0.679610212 | 0.865974881 |
| chr5:140174709-140175934 | -0.132713521 | 0.679617701 | 0.865974881 |
| chr17:28867693-28867893 | 0.221803519 | 0.679676622 | 0.866004506 |
| chr17:48024998-48025695 | 0.132428494 | 0.679677699 | 0.866004506 |
| chr2:23385421-23385641 | -0.170022365 | 0.679683815 | 0.866004506 |
| chr18:12038905-12039316 | -0.158968065 | 0.67968616 | 0.866004506 |
| chr3:47234934-47235266 | -0.20435857 | 0.679702273 | 0.866010634 |
| chr1:166893772-166893981 | -0.211878235 | 0.679720633 | 0.866015091 |
| chr17:10798398-10798731 | -0.161532504 | 0.679736104 | 0.866015091 |
| chr15:98444303-98444530 | -0.184448175 | 0.679739678 | 0.866015091 |
| chr1:26692870-26693377 | 0.143930165 | 0.679798083 | 0.866060864 |
| chr17:49569878-49570407 | 0.13868986 | 0.679798211 | 0.866060864 |
| chr1:171381283-171381483 | 0.205155938 | 0.679811659 | 0.866062675 |
| chr11:93655862-93656191 | -0.176466677 | 0.679829934 | 0.866062675 |
| chr16:57299551-57300868 | -0.129869555 | 0.679855978 | 0.866062675 |
| chr14:76346212-76346451 | -0.201856308 | 0.679862603 | 0.866062675 |
| chr10:114110240-114110927 | -0.149356443 | 0.679868638 | 0.866062675 |

| | | | |
|---|---|---|---|
| chr11:77046035-77046287 | -0.168545986 | 0.679869747 | 0.866062675 |
| chr15:70289326-70290000 | -0.143060104 | 0.679878754 | 0.866062675 |
| chr11:46392520-46392875 | -0.147018737 | 0.679905486 | 0.866082329 |
| chr1:35058470-35058728 | -0.200923925 | 0.679996202 | 0.86618074 |
| chr12:6937697-6938328 | -0.138181552 | 0.680005352 | 0.86618074 |
| chr16:74430031-74430438 | -0.152179676 | 0.680074813 | 0.866229681 |
| chr6:40536477-40536698 | -0.190416509 | 0.680087077 | 0.866229681 |
| chr5:10364247-10364645 | -0.156293799 | 0.68009789 | 0.866229681 |
| chr8:60714009-60715051 | -0.134004752 | 0.680103846 | 0.866229681 |
| chr20:53983515-53983922 | 0.13480287 | 0.680140935 | 0.866229681 |
| chr17:79009692-79010367 | -0.14479851 | 0.680158329 | 0.866229681 |
| chr6:36874472-36875242 | -0.139998402 | 0.680169236 | 0.866229681 |
| chr7:6483714-6484530 | -0.135560741 | 0.68017012 | 0.866229681 |
| chr18:32769822-32770419 | -0.15704431 | 0.680184566 | 0.866229681 |
| chr3:66372749-66373037 | 0.163239745 | 0.680192066 | 0.866229681 |
| chr1:244186623-244186942 | -0.195604702 | 0.680195154 | 0.866229681 |
| chr14:36647689-36648643 | -0.139802223 | 0.680219995 | 0.866229681 |
| chr4:60611910-60612110 | -0.212711232 | 0.680228213 | 0.866229681 |
| chr11:122628957-12262922б | 0.177665448 | 0.680238627 | 0.866229681 |
| chr7:51847797-51848042 | -0.17674588 | 0.680240346 | 0.866229681 |
| chr2:216666436-216666918 | 0.14852572 | 0.680270597 | 0.866229681 |
| chr5:34838620-34839608 | -0.132191108 | 0.680284395 | 0.866229681 |
| chr3:94069318-94069593 | 0.188312227 | 0.680297367 | 0.866229681 |
| chr9:89056734-89056934 | 0.230243833 | 0.680298881 | 0.866229681 |
| chr16:12228703-12229044 | 0.161530336 | 0.680304699 | 0.866229681 |
| chr3:126961186-126961558 | -0.173570367 | 0.680306234 | 0.866229681 |
| chr8:139837585-139837949 | -0.16854849 | 0.680310525 | 0.866229681 |
| chr6:43807545-43807745 | 0.201519944 | 0.680335972 | 0.866229681 |
| chrX:130299762-130300163 | 0.167339367 | 0.680337316 | 0.866229681 |
| chr11:118053423-11805363З | 0.183105033 | 0.680343066 | 0.866229681 |
| chr7:141074601-141074811 | 0.186212586 | 0.680347426 | 0.866229681 |
| chr11:60887553-60887908 | 0.158243305 | 0.680355873 | 0.866229681 |
| chr8:111100646-111100970 | 0.178614675 | 0.68036032 | 0.866229681 |
| chr9:89603037-89603294 | 0.165587909 | 0.680392458 | 0.866256204 |
| chr4:151362736-151363097 | -0.151329406 | 0.680408644 | 0.866262418 |
| chr1:113698919-113699237 | -0.168677415 | 0.680449497 | 0.866300036 |
| chr7:1608620-1608916 | 0.184838753 | 0.680462325 | 0.866301973 |
| chr3:153226915-153227172 | -0.176135902 | 0.680482908 | 0.866313784 |
| chr20:63748356-63748664 | 0.159922831 | 0.680545377 | 0.866375351 |
| chr6:98525889-98526099 | -0.209042579 | 0.680567116 | 0.866375351 |
| chr12:76896246-76896476 | -0.208608837 | 0.680574103 | 0.866375351 |
| chr15:57591536-57592273 | -0.14066109 | 0.680586497 | 0.866375351 |
| chr14:59254397-59255006 | -0.138614844 | 0.680587804 | 0.866375351 |
| chr3:120468978-120469400 | 0.166815983 | 0.68067004 | 0.866420771 |
| chr1:114524988-114525290 | 0.143088303 | 0.680677349 | 0.866420771 |
| chr4:64538356-64538686 | 0.203699735 | 0.680677617 | 0.866420771 |
| chr4:2419278-2419478 | 0.191937892 | 0.680692061 | 0.866420771 |
| chr1:8794678-8794900 | -0.239528855 | 0.680696713 | 0.866420771 |
| chr9:94919390-94920227 | -0.13578329 | 0.68071263 | 0.866420771 |
| chr8:129939050-129940474 | -0.129548529 | 0.680713062 | 0.866420771 |

| | | | |
|---|---|---|---|
| chr9:131502711-131504439 | -0.131674572 | 0.680713946 | 0.866420771 |
| chr17:47875068-47875350 | 0.166131139 | 0.680754535 | 0.866447562 |
| chr1:199441890-199442090 | -0.223611268 | 0.680766154 | 0.866447562 |
| chr15:71572818-71573082 | -0.182015941 | 0.680779944 | 0.866447562 |
| chr3:172516650-172517212 | -0.156732992 | 0.680780228 | 0.866447562 |
| chr12:80778166-80778472 | 0.169671359 | 0.680808207 | 0.86645226 |
| chr2:25877472-25879046 | -0.130641278 | 0.680821487 | 0.86645226 |
| chr21:33854324-33854524 | -0.209525522 | 0.680828514 | 0.86645226 |
| chr10:76086970-76087316 | -0.161359064 | 0.680829151 | 0.86645226 |
| chr2:55616556-55618060 | -0.12902427 | 0.680851599 | 0.866454808 |
| chr5:181184125-181184457 | 0.140016907 | 0.68087225 | 0.866454808 |
| chr9:92600810-92601019 | -0.215715366 | 0.680884801 | 0.866454808 |
| chr8:87763969-87764231 | 0.203271265 | 0.680926682 | 0.866454808 |
| chr2:174660992-174661474 | -0.167570941 | 0.680941253 | 0.866454808 |
| chr4:129900443-129900643 | 0.227609292 | 0.680946514 | 0.866454808 |
| chr3:111715985-111716251 | -0.166613487 | 0.680952695 | 0.866454808 |
| chr19:36849827-36851263 | 0.132763895 | 0.680954991 | 0.866454808 |
| chr3:123723588-123723936 | 0.155326949 | 0.680956387 | 0.866454808 |
| chr5:157744369-157744639 | 0.166909046 | 0.68096113 | 0.866454808 |
| chr19:41501389-41501776 | -0.175861832 | 0.680962392 | 0.866454808 |
| chr7:47195732-47195936 | -0.195643699 | 0.680978091 | 0.866454808 |
| chr3:10697112-10697349 | -0.201620334 | 0.680978159 | 0.866454808 |
| chr6:130933677-130934327 | -0.146356793 | 0.681012009 | 0.86646714 |
| chr1:159881648-159882282 | 0.141577707 | 0.68101718 | 0.86646714 |
| chr15:36883815-36884015 | 0.196255138 | 0.681025172 | 0.86646714 |
| chr2:45283828-45284115 | 0.177414514 | 0.681033085 | 0.86646714 |
| chr3:81987392-81987754 | -0.205619533 | 0.681097554 | 0.866534774 |
| chr10:88991615-88991845 | -0.182452735 | 0.681120597 | 0.866545588 |
| chr9:36855489-36855761 | -0.186326144 | 0.681128672 | 0.866545588 |
| chr2:236175630-236176158 | 0.160866089 | 0.681178923 | 0.866584396 |
| chr4:113556881-113557225 | -0.167812421 | 0.681181796 | 0.866584396 |
| chr1:6579688-6580214 | 0.140820733 | 0.681215698 | 0.866605578 |
| chr5:178099301-178099511 | -0.187492497 | 0.681257004 | 0.866605578 |
| chr1:34014036-34014281 | -0.210804595 | 0.681263485 | 0.866605578 |
| chr1:59976657-59977137 | -0.165357359 | 0.681264846 | 0.866605578 |
| chr6:29016528-29016971 | 0.153415871 | 0.681265468 | 0.866605578 |
| chr16:10743634-10744409 | 0.131233252 | 0.681292344 | 0.866605578 |
| chr14:49461798-49462149 | 0.154080388 | 0.681306718 | 0.866605578 |
| chr13:24922410-24923225 | 0.134681052 | 0.68131281 | 0.866605578 |
| chr12:56117858-56118810 | 0.130320256 | 0.681327507 | 0.866605578 |
| chr20:46681392-46681610 | 0.194840052 | 0.681327984 | 0.866605578 |
| chr2:135052083-135052846 | -0.137344649 | 0.681332735 | 0.866605578 |
| chr2:84474191-84474561 | 0.15228971 | 0.681337487 | 0.866605578 |
| chr4:105613327-105613640 | 0.17323293 | 0.681345479 | 0.866605578 |
| chr10:44316359-44316880 | -0.139132674 | 0.681370823 | 0.866622525 |
| chr20:51639215-51639730 | -0.13809563 | 0.681381423 | 0.866622525 |
| chr2:170967169-170967443 | -0.195055837 | 0.681426655 | 0.86665463 |
| chr3:133040829-133041065 | 0.187006549 | 0.681441385 | 0.86665463 |
| chr4:23334320-23334533 | 0.213202611 | 0.68144675 | 0.86665463 |
| chr14:23483366-23483612 | -0.172168414 | 0.681451909 | 0.86665463 |

| | | | |
|---|---|---|---|
| chr14:21723439-21723730 | -0.177806686 | 0.681544334 | 0.866751062 |
| chr1:36369041-36369357 | 0.166165897 | 0.681565033 | 0.866751062 |
| chr16:57186767-57187190 | 0.154957316 | 0.681574314 | 0.866751062 |
| chr10:11687301-11687728 | 0.143412183 | 0.681589702 | 0.866751062 |
| chr20:12012054-12012639 | -0.150387193 | 0.681593432 | 0.866751062 |
| chr6:26198987-26199874 | 0.134278594 | 0.681595606 | 0.866751062 |
| chr13:39415474-39415674 | -0.190081774 | 0.68165225 | 0.866808708 |
| chr3:51637888-51638265 | -0.161111304 | 0.6817125 | 0.866870937 |
| chr2:164115007-164115259 | 0.19139371 | 0.681741589 | 0.866873322 |
| chr2:25129244-25129548 | 0.156669266 | 0.681763686 | 0.866873322 |
| chr19:12793472-12794011 | -0.142831563 | 0.681774733 | 0.866873322 |
| chr3:119497897-119499041 | -0.129928235 | 0.681786947 | 0.866873322 |
| chr3:158081330-158081530 | -0.213398032 | 0.681791638 | 0.866873322 |
| chr5:84451160-84451373 | -0.213398032 | 0.681791638 | 0.866873322 |
| chr1:193121539-193122743 | -0.128545829 | 0.681793571 | 0.866873322 |
| chr9:69389514-69390011 | 0.154706058 | 0.681824797 | 0.866878048 |
| chr3:115922749-115922974 | -0.18315802 | 0.681825476 | 0.866878048 |
| chr4:141208697-141208897 | -0.236336687 | 0.681831229 | 0.866878048 |
| chr1:145996469-145997339 | 0.130990552 | 0.681871145 | 0.866892673 |
| chr16:31093178-31093390 | -0.176600124 | 0.681875064 | 0.866892673 |
| chr6:13538159-13538383 | 0.214659065 | 0.681882542 | 0.866892673 |
| chr9:77494944-77495169 | -0.217175242 | 0.681889657 | 0.866892673 |
| chr5:65803337-65803604 | 0.188678531 | 0.681899302 | 0.866892673 |
| chr3:127406374-127406574 | -0.192438438 | 0.681934238 | 0.866922703 |
| chr2:241595637-241595903 | -0.194011172 | 0.682022629 | 0.866994725 |
| chr5:20033124-20033533 | 0.174514096 | 0.682026295 | 0.866994725 |
| chr9:36729033-36729276 | -0.187759458 | 0.682043089 | 0.866994725 |
| chr2:42743445-42743699 | -0.182347836 | 0.68205024 | 0.866994725 |
| chr11:87000075-87000587 | -0.145088483 | 0.682054568 | 0.866994725 |
| chr1:220093856-220094852 | -0.131971472 | 0.682058783 | 0.866994725 |
| chr20:57074846-57075546 | 0.13250281 | 0.682076349 | 0.867002671 |
| chr2:12326467-12326681 | -0.21499382 | 0.68211035 | 0.867031507 |
| chr1:23590490-23590738 | -0.214938098 | 0.682124785 | 0.867035472 |
| chr8:90868783-90868983 | 0.23796489 | 0.682168678 | 0.867075845 |
| chr18:67360062-67360827 | -0.144982871 | 0.682199534 | 0.867075845 |
| chr5:31981424-31981862 | 0.15747355 | 0.682201315 | 0.867075845 |
| chr21:38505690-38506006 | -0.167627717 | 0.682201813 | 0.867075845 |
| chr7:11557659-11558041 | 0.157331426 | 0.682256823 | 0.867117664 |
| chr13:21670859-21671201 | 0.15574687 | 0.682266687 | 0.867117664 |
| chr4:121785603-121786067 | -0.169651175 | 0.682275034 | 0.867117664 |
| chr11:6282021-6282221 | 0.226152385 | 0.682279983 | 0.867117664 |
| chr12:1605130-1606428 | -0.135792042 | 0.682296953 | 0.867124849 |
| chr6:109504729-109505337 | 0.15767442 | 0.682312137 | 0.867129763 |
| chr6:10099031-10099567 | 0.148336695 | 0.682347773 | 0.867160669 |
| chr13:31836023-31836397 | 0.169791688 | 0.682399731 | 0.867199708 |
| chr5:38556339-38557536 | 0.133516587 | 0.682411368 | 0.867199708 |
| chr3:81339077-81339402 | -0.159198282 | 0.682412445 | 0.867199708 |
| chr1:32222019-32222724 | 0.138225399 | 0.682426099 | 0.867202532 |
| chr22:17075052-17075260 | 0.212545711 | 0.682473973 | 0.867202532 |
| chr1:186579278-186579481 | 0.208592515 | 0.682511302 | 0.867202532 |

| | | | |
|---|---|---|---|
| chrX:108399431-108399758 | -0.173646497 | 0.682528853 | 0.867202532 |
| chr1:145156902-145157322 | 0.159487076 | 0.682533919 | 0.867202532 |
| chr4:110050065-110050439 | 0.183691315 | 0.682545445 | 0.867202532 |
| chr8:48925495-48926119 | 0.146399791 | 0.682565901 | 0.867202532 |
| chr6:3258548-3259844 | 0.131662822 | 0.682573197 | 0.867202532 |
| chr3:28836258-28836497 | -0.196367092 | 0.682574559 | 0.867202532 |
| chr3:198167680-198167995 | -0.196367092 | 0.682574559 | 0.867202532 |
| chr15:40155208-40155630 | 0.154322415 | 0.682575512 | 0.867202532 |
| chr4:89753490-89753982 | 0.175927751 | 0.682577661 | 0.867202532 |
| chr5:35430427-35430715 | -0.187394476 | 0.682595229 | 0.867202532 |
| chr17:58022689-58023505 | -0.135285565 | 0.682600469 | 0.867202532 |
| chr2:54522748-54523022 | -0.165103747 | 0.682613741 | 0.867202532 |
| chr20:3203890-3204826 | 0.13512252 | 0.682618436 | 0.867202532 |
| chr12:8913015-8913250 | 0.169742576 | 0.682636684 | 0.867202532 |
| chr3:62852403-62852615 | 0.191469767 | 0.682649279 | 0.867202532 |
| chr10:17430807-17431038 | -0.187084574 | 0.682663421 | 0.867202532 |
| chr8:144427824-144428954 | 0.132369576 | 0.682665127 | 0.867202532 |
| chrX:3815545-3815745 | 0.172773601 | 0.682670172 | 0.867202532 |
| chr8:90119046-90119341 | -0.170424423 | 0.68267285 | 0.867202532 |
| chr6:76876815-76877304 | -0.141031557 | 0.68267498 | 0.867202532 |
| chr4:98261769-98261969 | 0.168944815 | 0.682710263 | 0.86722335 |
| chr2:173652778-173653063 | -0.174403957 | 0.682718934 | 0.86722335 |
| chr16:75199374-75199663 | -0.152477947 | 0.682725323 | 0.86722335 |
| chr14:96104619-96105221 | -0.140724441 | 0.682739194 | 0.867226593 |
| chr3:185585825-185586733 | 0.130601356 | 0.682756128 | 0.867233726 |
| chr12:130590160-130590438 | 0.200732408 | 0.68281337 | 0.867291378 |
| chr17:21126704-21127585 | 0.132569297 | 0.682835265 | 0.867291378 |
| chr1:65297582-65298218 | -0.138662131 | 0.68284555 | 0.867291378 |
| chr3:123066859-123068109 | 0.132737742 | 0.682846793 | 0.867291378 |
| chr8:131896478-131896828 | -0.176181175 | 0.682920727 | 0.867359321 |
| chr15:74473740-74473940 | -0.216769695 | 0.682922926 | 0.867359321 |
| chr8:80029565-80030539 | 0.129538659 | 0.682943029 | 0.867370475 |
| chr10:86230031-86230231 | -0.189273742 | 0.682994516 | 0.867421488 |
| chr5:24005431-24005663 | -0.195075841 | 0.683028053 | 0.867449703 |
| chr12:97943488-97943736 | -0.200088422 | 0.683072585 | 0.86749188 |
| chr21:25573903-25574145 | -0.190816919 | 0.683086718 | 0.867495451 |
| chr12:21675163-21675634 | 0.157573268 | 0.683124353 | 0.867528867 |
| chr3:197599034-197599825 | 0.139405009 | 0.683137301 | 0.867530931 |
| chr5:13688640-13688997 | -0.184757432 | 0.683196395 | 0.867578687 |
| chr1:6634713-6634928 | -0.201000782 | 0.683197552 | 0.867578687 |
| chrX:107118250-107119013 | -0.139851197 | 0.683215264 | 0.867586801 |
| chr15:31499395-31499678 | 0.187532985 | 0.683236464 | 0.867599342 |
| chr6:19332768-19333079 | 0.18571786 | 0.683267226 | 0.867624027 |
| chr15:40382664-40383397 | -0.136842659 | 0.68333831 | 0.86769991 |
| chr5:132962543-132962935 | -0.151803101 | 0.683353195 | 0.867704432 |
| chr1:154943718-154944005 | 0.171460947 | 0.683374994 | 0.867717732 |
| chr3:156668117-156668479 | -0.165343283 | 0.683431845 | 0.867767299 |
| chr1:204333579-204333851 | 0.166926711 | 0.683441629 | 0.867767299 |
| chr17:50841911-50842165 | 0.161203441 | 0.683452804 | 0.867767299 |
| chr10:35335739-35336856 | -0.135084366 | 0.683459332 | 0.867767299 |

| | | | |
|---|---|---|---|
| chr20:57514249-57514595 | -0.146126501 | 0.683480346 | 0.867779599 |
| chr12:120457234-120457473 | 0.189466266 | 0.683509398 | 0.8677863 |
| chrX:101624073-101624331 | 0.171001042 | 0.683518043 | 0.8677863 |
| chr14:89214095-89214872 | 0.136161256 | 0.683528713 | 0.8677863 |
| chr17:31489232-31489486 | -0.203548817 | 0.683530925 | 0.8677863 |
| chr15:57306477-57307149 | 0.144937124 | 0.683546373 | 0.867791533 |
| chr15:64355442-64356736 | 0.130018739 | 0.683570758 | 0.867792039 |
| chr8:56929046-56929626 | -0.150802327 | 0.683601777 | 0.867792039 |
| chr2:99559197-99559399 | 0.208876287 | 0.683619306 | 0.867792039 |
| chr2:31170126-31170423 | -0.17033081 | 0.683619549 | 0.867792039 |
| chr3:50316694-50317241 | -0.143127553 | 0.683620168 | 0.867792039 |
| chr1:9124483-9124797 | -0.160144208 | 0.683624263 | 0.867792039 |
| chr12:108612241-108612441 | -0.208019427 | 0.683626051 | 0.867792039 |
| chr1:185596056-185596425 | 0.152027301 | 0.683647107 | 0.867797189 |
| chr13:102845385-102846653 | -0.130843393 | 0.683652759 | 0.867797189 |
| chr21:42113970-42114454 | -0.142431421 | 0.683680604 | 0.867804555 |
| chr13:65926646-65927517 | -0.143971373 | 0.683706653 | 0.867804555 |
| chr11:18732068-18732355 | -0.169290975 | 0.683710322 | 0.867804555 |
| chr13:23980126-23980438 | 0.157294014 | 0.68371367 | 0.867804555 |
| chr8:86084759-86085000 | 0.171004294 | 0.68372429 | 0.867804555 |
| chr21:29386141-29386533 | 0.147012437 | 0.683726517 | 0.867804555 |
| chr9:32379352-32379610 | 0.165488599 | 0.683749896 | 0.867819853 |
| chr2:213150499-213150709 | -0.178099506 | 0.683797244 | 0.867865571 |
| chr1:92715380-92716032 | 0.141448501 | 0.683819009 | 0.86787882 |
| chr9:38400698-38400924 | -0.190508547 | 0.683841227 | 0.867891607 |
| chr9:121278723-121279224 | -0.148153861 | 0.683851738 | 0.867891607 |
| chr5:112608045-112608344 | -0.19113009 | 0.683871771 | 0.867902656 |
| chr2:86703709-86703979 | 0.186205475 | 0.683883987 | 0.867903783 |
| chr17:72720129-72720329 | -0.198425626 | 0.683897445 | 0.867906489 |
| chr16:81136770-81136970 | -0.189569766 | 0.683924552 | 0.867926514 |
| chr3:63753554-63754258 | 0.130579326 | 0.683944351 | 0.867937264 |
| chr15:92638251-92639133 | 0.132721387 | 0.683968485 | 0.867953516 |
| chr12:14207756-14208556 | 0.139378728 | 0.683982426 | 0.867955158 |
| chr9:93302721-93303017 | 0.166276627 | 0.683992435 | 0.867955158 |
| chr2:11930702-11930902 | 0.203787652 | 0.68403529 | 0.867995164 |
| chr14:60331203-60332081 | -0.134320148 | 0.684051895 | 0.86800186 |
| chr8:58318368-58318651 | -0.193743062 | 0.684079789 | 0.868004788 |
| chr19:18410481-18410736 | 0.190625782 | 0.684087585 | 0.868004788 |
| chr17:64236936-64237136 | 0.179602797 | 0.684088188 | 0.868004788 |
| chr8:93068311-93068834 | 0.140967291 | 0.684120338 | 0.8680051 |
| chr2:218269629-218270663 | 0.128996241 | 0.68412455 | 0.8680051 |
| chr6:148746634-148748117 | -0.131717899 | 0.684128409 | 0.8680051 |
| chr10:28735183-28735445 | 0.2097555 | 0.684133747 | 0.8680051 |
| chr5:171568979-171569223 | -0.230546777 | 0.684160592 | 0.868024786 |
| chr6:31272061-31272301 | -0.203060438 | 0.684184931 | 0.868027343 |
| chr11:84814651-84814856 | -0.241812625 | 0.684228527 | 0.868027343 |
| chr3:27680832-27681985 | 0.13283432 | 0.684233817 | 0.868027343 |
| chr8:68622919-68623296 | 0.168493002 | 0.684238039 | 0.868027343 |
| chr6:84284528-84284740 | -0.207595406 | 0.68424594 | 0.868027343 |
| chr2:195656708-195657496 | -0.128266633 | 0.684247933 | 0.868027343 |

| | | | |
|---|---|---|---|
| chr14:90235147-90235706 | 0.141645896 | 0.684260489 | 0.868027343 |
| chr8:130604322-130604603 | 0.174534956 | 0.68427336 | 0.868027343 |
| chr12:56007207-56007922 | 0.136566613 | 0.684277444 | 0.868027343 |
| chr10:14058492-14058692 | -0.22485977 | 0.684286502 | 0.868027343 |
| chr3:14690617-14690871 | 0.197871177 | 0.684313075 | 0.868027343 |
| chr7:50632473-50633253 | -0.134526537 | 0.684314431 | 0.868027343 |
| chr3:15630171-15630459 | 0.154888075 | 0.684332098 | 0.868027343 |
| chr5:36913966-36914212 | -0.204885286 | 0.684338005 | 0.868027343 |
| chr12:6752758-6753952 | -0.129566238 | 0.684350446 | 0.868027343 |
| chr6:127069931-127070149 | -0.194300349 | 0.684360096 | 0.868027343 |
| chr14:71319691-71321466 | 0.127955002 | 0.684364355 | 0.868027343 |
| chr2:153059762-153059990 | 0.19570626 | 0.684366524 | 0.868027343 |
| chr6:166997763-166998480 | -0.144575265 | 0.68438172 | 0.868032248 |
| chr4:75672540-75673814 | -0.130655533 | 0.684396971 | 0.868037223 |
| chr10:103554863-103555216 | 0.153994739 | 0.684422276 | 0.868042437 |
| chr3:150756446-150756684 | -0.171359261 | 0.684436723 | 0.868042437 |
| chr14:52684043-52684357 | 0.188047533 | 0.684446153 | 0.868042437 |
| chr3:75192180-75192387 | -0.19146329 | 0.684455802 | 0.868042437 |
| chr4:124610254-124610837 | 0.154591338 | 0.684467379 | 0.868042437 |
| chr15:83438439-83438654 | -0.210268069 | 0.684469056 | 0.868042437 |
| chr5:81838627-81838900 | -0.171322086 | 0.684481835 | 0.868044276 |
| chr6:83215193-83215499 | 0.154382455 | 0.684504911 | 0.868059175 |
| chr5:154712410-154713266 | -0.135720222 | 0.684529763 | 0.868074409 |
| chr8:101967693-101967900 | 0.192889911 | 0.684557988 | 0.868074409 |
| chr21:26845025-26845655 | 0.143171201 | 0.684568152 | 0.868074409 |
| chr8:86618003-86619322 | 0.129815281 | 0.684574387 | 0.868074409 |
| chr9:89318076-89319125 | 0.130294783 | 0.684582536 | 0.868074409 |
| chr7:157132784-157133134 | 0.162136995 | 0.684587865 | 0.868074409 |
| chr16:73821579-73821796 | 0.218624318 | 0.684605721 | 0.868074409 |
| chr6:31769635-31770160 | 0.144867256 | 0.684618844 | 0.868074409 |
| chr6:24720670-24721780 | -0.128691268 | 0.684618889 | 0.868074409 |
| chr1:151341388-151341589 | 0.172492842 | 0.684646199 | 0.868094673 |
| chr6:5746457-5746707 | 0.191973612 | 0.684721657 | 0.868155857 |
| chr2:98444492-98445495 | 0.137095399 | 0.684722211 | 0.868155857 |
| chr21:42593086-42593699 | -0.141935786 | 0.684728445 | 0.868155857 |
| chr5:170809548-170809909 | 0.149379779 | 0.684759332 | 0.868178269 |
| chr11:46936530-46937166 | 0.139086204 | 0.684768783 | 0.868178269 |
| chr17:28727970-28728250 | 0.14880221 | 0.684809026 | 0.868199676 |
| chr10:133318468-133319015 | 0.141591444 | 0.684826554 | 0.868199676 |
| chr3:129662870-129663550 | -0.130909636 | 0.684832082 | 0.868199676 |
| chr6:149717332-149718488 | 0.130024134 | 0.684851213 | 0.868199676 |
| chr19:42061539-42061837 | 0.184492101 | 0.684863384 | 0.868199676 |
| chr6:153947669-153948170 | -0.153379527 | 0.684871623 | 0.868199676 |
| chr12:94753429-94753629 | 0.209507357 | 0.684875313 | 0.868199676 |
| chr10:59563087-59563287 | -0.224344256 | 0.684876315 | 0.868199676 |
| chr17:49576383-49576667 | -0.156559454 | 0.684894365 | 0.868208194 |
| chr15:63857538-63857766 | 0.196389518 | 0.684923961 | 0.868231346 |
| chr13:45047374-45047688 | -0.15411527 | 0.68493942 | 0.868236578 |
| chr1:4656331-4656572 | 0.186957531 | 0.684957506 | 0.868239824 |
| chr11:95150399-95150870 | 0.146999259 | 0.684982214 | 0.868239824 |

| | | | |
|---|---|---|---|
| chr1:202836515-202837183 | -0.13359651 | 0.68499159 | 0.868239824 |
| chr2:70301212-70302351 | -0.130925465 | 0.685004038 | 0.868239824 |
| chr20:57881390-57881706 | 0.157730345 | 0.685005683 | 0.868239824 |
| chr8:127734008-127734339 | -0.136578063 | 0.685016347 | 0.868239824 |
| chr3:160719731-160719931 | 0.19451337 | 0.685021301 | 0.868239824 |
| chr17:65470804-65471326 | 0.149579396 | 0.685040927 | 0.868250337 |
| chr4:118685142-118685965 | -0.130921039 | 0.685061898 | 0.868255063 |
| chr3:63617674-63617874 | -0.170085694 | 0.685067319 | 0.868255063 |
| chr16:67561820-67563328 | 0.128251182 | 0.685083717 | 0.868261484 |
| chr5:11961784-11961984 | -0.201845876 | 0.68517742 | 0.868365774 |
| chrX:106415321-106415584 | -0.176232458 | 0.685188671 | 0.868365774 |
| chr15:76905064-76905773 | -0.137803825 | 0.685212902 | 0.86838212 |
| chr5:157668979-157669289 | -0.154462706 | 0.685245993 | 0.868392894 |
| chr14:67503204-67503611 | -0.163744841 | 0.685256342 | 0.868392894 |
| chr10:93039822-93040031 | 0.194188609 | 0.685300454 | 0.868392894 |
| chr15:83838009-83838243 | 0.194188609 | 0.685300454 | 0.868392894 |
| chr14:87745575-87745784 | 0.19319252 | 0.685309224 | 0.868392894 |
| chr3:63193704-63194226 | 0.134959334 | 0.685310618 | 0.868392894 |
| chr15:65517158-65518102 | -0.132353889 | 0.685312921 | 0.868392894 |
| chr19:46600431-46601443 | 0.130674895 | 0.685316571 | 0.868392894 |
| chr6:41072173-41073353 | 0.127604573 | 0.685323405 | 0.868392894 |
| chr4:53313493-53313811 | 0.168239677 | 0.685357078 | 0.868418436 |
| chr3:89314463-89314771 | -0.162472832 | 0.685372721 | 0.868418436 |
| chr2:176086115-176086521 | -0.161109132 | 0.685377564 | 0.868418436 |
| chr6:151961628-151961936 | -0.199824213 | 0.685468817 | 0.868506612 |
| chr7:114865637-114865837 | -0.220779975 | 0.685473966 | 0.868506612 |
| chr4:142230938-142231187 | -0.224534475 | 0.685493048 | 0.868506612 |
| chr9:85332340-85332817 | 0.160588396 | 0.685495262 | 0.868506612 |
| chr4:98658329-98659135 | -0.138776873 | 0.685503829 | 0.868506612 |
| chr5:170434946-170435365 | 0.155551508 | 0.685554166 | 0.868546989 |
| chr13:110094638-110094946 | 0.181057181 | 0.685568743 | 0.868546989 |
| chr6:7505286-7505729 | -0.149641218 | 0.685571612 | 0.868546989 |
| chr6:67067035-67067349 | -0.201644607 | 0.685611489 | 0.868546989 |
| chr5:149551236-149552054 | -0.129311481 | 0.685617935 | 0.868546989 |
| chr12:113477385-113477642 | -0.201666862 | 0.685618671 | 0.868546989 |
| chr11:19829165-19829375 | 0.192155205 | 0.685625137 | 0.868546989 |
| chr1:113928969-113930902 | -0.125343899 | 0.685626382 | 0.868546989 |
| chr6:131062329-131062614 | 0.158917147 | 0.685642483 | 0.868553026 |
| chr3:64685622-64686916 | -0.126045111 | 0.685737828 | 0.868659445 |
| chr4:142845343-142845548 | -0.183332211 | 0.685759608 | 0.86865951 |
| chr11:120172290-120172537 | -0.18536581 | 0.685760553 | 0.86865951 |
| chr7:32964540-32965048 | -0.148878933 | 0.685834039 | 0.868734351 |
| chr17:62895641-62895981 | -0.154284605 | 0.685842312 | 0.868734351 |
| chr9:35790148-35790610 | 0.159342353 | 0.685872916 | 0.868758755 |
| chr4:128060642-128062539 | 0.126614772 | 0.685908551 | 0.868789529 |
| chr10:8971559-8971954 | 0.157132917 | 0.685953529 | 0.868808319 |
| chr20:5610401-5610601 | 0.193779936 | 0.68595672 | 0.868808319 |
| chr3:50089169-50089604 | -0.169663914 | 0.685957402 | 0.868808319 |
| chr6:24774682-24775781 | -0.128615179 | 0.685997688 | 0.868844982 |
| chr1:200509358-200509708 | -0.162679696 | 0.686054697 | 0.868895367 |

| | | | |
|---|---|---|---|
| chr3:113695963-113696438 | -0.135040868 | 0.686073179 | 0.868895367 |
| chr6:70884391-70884668 | 0.165004416 | 0.68607516 | 0.868895367 |
| chr1:108121497-108121734 | -0.161194361 | 0.68608283 | 0.868895367 |
| chr1:101309336-101309569 | -0.190439663 | 0.68612593 | 0.868901399 |
| chr2:159788243-159788625 | -0.14823721 | 0.686152786 | 0.868901399 |
| chr8:86508052-86509020 | 0.127078417 | 0.686157286 | 0.868901399 |
| chr10:68153521-68154067 | -0.142397366 | 0.686199355 | 0.868901399 |
| chrX:77899159-77899596 | 0.138208323 | 0.686202492 | 0.868901399 |
| chr3:30682010-30682215 | -0.19885008 | 0.686205898 | 0.868901399 |
| chr19:344469-345182 | 0.144380898 | 0.686207459 | 0.868901399 |
| chr1:26233687-26234754 | 0.132076063 | 0.686207674 | 0.868901399 |
| chr8:104223041-104224357 | -0.128901467 | 0.686210199 | 0.868901399 |
| chr3:139677417-139678167 | 0.132289807 | 0.686232066 | 0.868901399 |
| chr2:241009637-241010006 | 0.159601198 | 0.686235229 | 0.868901399 |
| chr5:112245704-112245942 | -0.197925135 | 0.686236857 | 0.868901399 |
| chr6:40420030-40420414 | -0.187201701 | 0.686245054 | 0.868901399 |
| chr11:1554312-1554512 | 0.205798745 | 0.686246354 | 0.868901399 |
| chr3:164477771-164477991 | -0.203686218 | 0.686289062 | 0.86890808 |
| chr2:231621040-231621921 | 0.136927846 | 0.686290973 | 0.86890808 |
| chr11:76629846-76630081 | -0.196837279 | 0.686302468 | 0.86890808 |
| chr16:85777738-85778246 | 0.142253219 | 0.686326905 | 0.86890808 |
| chr22:38180995-38181380 | 0.156531827 | 0.68632888 | 0.86890808 |
| chr14:67571364-67571606 | 0.171957689 | 0.686336809 | 0.86890808 |
| chr4:105210960-105211348 | 0.181176333 | 0.686338528 | 0.86890808 |
| chr1:179036882-179037096 | -0.20345305 | 0.686371143 | 0.86890808 |
| chr20:17682142-17682926 | 0.129058962 | 0.686372189 | 0.86890808 |
| chr4:120049694-120050222 | 0.148133242 | 0.686372644 | 0.86890808 |
| chr15:40754773-40755018 | 0.169801588 | 0.686376373 | 0.86890808 |
| chr3:56682237-56683483 | -0.134261551 | 0.686445971 | 0.868960165 |
| chr10:130221698-130221898 | -0.227287224 | 0.686460077 | 0.868960165 |
| chr11:46295507-46295870 | -0.147756407 | 0.686476031 | 0.868960165 |
| chr8:81768110-81768355 | -0.195097357 | 0.686476795 | 0.868960165 |
| chr19:55461490-55462095 | 0.132634539 | 0.686482089 | 0.868960165 |
| chr2:231227346-231227574 | 0.204206769 | 0.686485562 | 0.868960165 |
| chr2:231365910-231366110 | 0.190781302 | 0.686524334 | 0.868994887 |
| chr5:171418466-171419752 | 0.1291621 | 0.686572486 | 0.869022282 |
| chrX:71181952-71182676 | -0.140185286 | 0.686579932 | 0.869022282 |
| chr4:71186370-71187328 | 0.141545116 | 0.686580002 | 0.869022282 |
| chr11:69278048-69278528 | -0.139254205 | 0.686618685 | 0.869056889 |
| chr6:117556035-117556262 | -0.192938836 | 0.686637706 | 0.869066608 |
| chr5:127885147-127885475 | 0.178749549 | 0.686661649 | 0.869082556 |
| chr10:97975786-97975986 | -0.209774243 | 0.686679375 | 0.869090635 |
| chr2:16490613-16490970 | -0.187422913 | 0.686701523 | 0.869104311 |
| chr11:124746551-124747249 | -0.134615354 | 0.686718267 | 0.869111146 |
| chr10:73110288-73110616 | 0.152022526 | 0.686766756 | 0.869158159 |
| chr8:102576904-102577237 | -0.147814975 | 0.686803927 | 0.86918138 |
| chr7:1137627-1137841 | -0.165660435 | 0.686809479 | 0.86918138 |
| chr15:34583963-34584315 | 0.160341211 | 0.686819136 | 0.86918138 |
| chr15:72230733-72231981 | 0.128293038 | 0.686862523 | 0.869212849 |
| chr20:54103378-54104117 | 0.127650827 | 0.686874862 | 0.869212849 |

| | | | |
|---|---|---|---|
| chr15:31372648-31372848 | -0.188416392 | 0.686878035 | 0.869212849 |
| chr9:127802453-127803883 | 0.130628196 | 0.686915043 | 0.869245324 |
| chr13:30355402-30355658 | -0.183368643 | 0.686968545 | 0.869284316 |
| chr14:52299783-52300014 | -0.183368643 | 0.686968545 | 0.869284316 |
| chr2:79935809-79936055 | 0.172370311 | 0.687000057 | 0.869299316 |
| chr3:133377504-133377709 | -0.186017143 | 0.68700309 | 0.869299316 |
| chr3:58842197-58842463 | -0.195522302 | 0.687037467 | 0.869310261 |
| chr14:88868650-88868919 | -0.196569321 | 0.687037542 | 0.869310261 |
| chr8:118000355-118000826 | 0.153741407 | 0.687045777 | 0.869310261 |
| chr9:130214446-130214667 | -0.191560917 | 0.687102838 | 0.869355316 |
| chr7:32604102-32604385 | -0.177840394 | 0.687104077 | 0.869355316 |
| chr1:200200909-200201118 | -0.18369087 | 0.687136612 | 0.869355921 |
| chr5:67342849-67343117 | -0.189765919 | 0.68714823 | 0.869355921 |
| chr22:19144329-19144977 | 0.134638978 | 0.687149298 | 0.869355921 |
| chr8:107240524-107240755 | -0.192616362 | 0.687160645 | 0.869355921 |
| chr5:168086354-168086729 | -0.145567313 | 0.687161286 | 0.869355921 |
| chrX:22081534-22081791 | 0.189546353 | 0.68719279 | 0.869367131 |
| chr17:7834887-7835087 | -0.165413314 | 0.687192839 | 0.869367131 |
| chr2:9607518-9608397 | 0.142214777 | 0.687235871 | 0.869387274 |
| chr1:183281497-183282076 | 0.140237512 | 0.687239905 | 0.869387274 |
| chr4:98928143-98929802 | -0.126185542 | 0.687258537 | 0.869387274 |
| chr8:126377186-126377402 | -0.19260714 | 0.687264045 | 0.869387274 |
| chr12:102618600-10261880C | -0.226729018 | 0.687269338 | 0.869387274 |
| chr3:49900412-49901085 | 0.144261633 | 0.687287568 | 0.869387274 |
| chr16:4650494-4650738 | 0.176043238 | 0.687292025 | 0.869387274 |
| chr12:62210374-62210657 | -0.175014073 | 0.687306899 | 0.869387274 |
| chr3:187181892-187182092 | -0.229362038 | 0.687310878 | 0.869387274 |
| chr6:33277000-33277408 | -0.148574002 | 0.687349406 | 0.869418316 |
| chrX:120741509-120741787 | -0.214232014 | 0.687358113 | 0.869418316 |
| chr16:78503583-78503783 | -0.21241027 | 0.687400304 | 0.86945733 |
| chr5:168269241-168269730 | 0.136590526 | 0.687508228 | 0.869579482 |
| chr4:53365175-53365465 | 0.165478238 | 0.687536197 | 0.869600503 |
| chr10:119000564-11900120S | -0.13898768 | 0.687556214 | 0.869611466 |
| chr16:19997915-19998115 | -0.208051554 | 0.687570239 | 0.86961485 |
| chr2:69375796-69376029 | 0.186115606 | 0.687586151 | 0.869620621 |
| chr10:73850955-73851472 | -0.151354601 | 0.687650566 | 0.869675462 |
| chr3:87398316-87398633 | -0.193062492 | 0.687652213 | 0.869675462 |
| chr20:25057125-25057717 | -0.139113448 | 0.68770385 | 0.869726412 |
| chr6:110814549-110816227 | -0.126355294 | 0.687740129 | 0.869749517 |
| chr18:832103-832549 | 0.153373134 | 0.687744822 | 0.869749517 |
| chr10:73664693-73664957 | 0.190724033 | 0.687814981 | 0.869814992 |
| chr5:172364861-172365106 | -0.199220946 | 0.6878193 | 0.869814992 |
| chr11:46308477-46308843 | -0.154638087 | 0.687837099 | 0.869823145 |
| chr5:8457291-8458458 | 0.131566777 | 0.687854015 | 0.869825807 |
| chr20:21235487-21235953 | 0.145710253 | 0.687861909 | 0.869825807 |
| chr9:84538509-84538709 | -0.22435433 | 0.687885053 | 0.869840719 |
| chr16:66696686-66697289 | 0.141570633 | 0.687904214 | 0.869850593 |
| chr20:50160253-50160466 | -0.188296403 | 0.687960348 | 0.869907218 |
| chr3:138180673-138180902 | 0.188153031 | 0.687982511 | 0.869920887 |
| chr3:77826015-77826215 | -0.215299826 | 0.687997641 | 0.869925662 |

| | | | |
|---|---|---|---|
| chr1:151008085-151009180 | -0.127343959 | 0.688034183 | 0.86995751 |
| chr1:147402279-147402627 | 0.152413864 | 0.688066822 | 0.8699624 |
| chr1:216306901-216307162 | -0.164763453 | 0.688070204 | 0.8699624 |
| chr1:49229674-49229879 | 0.210610018 | 0.688075844 | 0.8699624 |
| chr2:89849098-89849298 | 0.206662925 | 0.688083466 | 0.8699624 |
| chr8:139698420-139698833 | 0.149779028 | 0.688171417 | 0.870052971 |
| chr20:53973906-53974293 | 0.131310399 | 0.688177812 | 0.870052971 |
| chr16:68234070-68234673 | 0.130156468 | 0.688191436 | 0.87005584 |
| chr3:13461503-13461890 | 0.142953215 | 0.68824755 | 0.870112426 |
| chr2:211475513-211475782 | -0.189651136 | 0.688275795 | 0.870133777 |
| chr5:177496173-177496589 | 0.141121014 | 0.688319247 | 0.870166628 |
| chr3:98482485-98483564 | 0.130887844 | 0.688327534 | 0.870166628 |
| chr17:47721295-47721495 | -0.171056287 | 0.68833585 | 0.870166628 |
| chr11:78040057-78040284 | -0.176901249 | 0.688356536 | 0.870175685 |
| chr1:233394964-233395188 | 0.179477445 | 0.688365727 | 0.870175685 |
| chr1:44595215-44595444 | 0.180728919 | 0.688388968 | 0.870190707 |
| chr9:128321933-128323325 | 0.130114838 | 0.688413369 | 0.870207195 |
| chr9:76996751-76997048 | 0.166814817 | 0.688455112 | 0.870222699 |
| chr2:70132840-70133042 | -0.187363411 | 0.688466478 | 0.870222699 |
| chr2:1701686-1702124 | 0.160738995 | 0.688503273 | 0.870222699 |
| chr17:8120693-8121029 | -0.146132871 | 0.688504386 | 0.870222699 |
| chr6:158560514-158560794 | 0.169097626 | 0.68851612 | 0.870222699 |
| chr10:37754809-37755014 | 0.191065175 | 0.688525394 | 0.870222699 |
| chr6:26065976-26066228 | -0.175328871 | 0.688526465 | 0.870222699 |
| chr17:51193716-51193998 | -0.180990639 | 0.688543663 | 0.870222699 |
| chr1:31154647-31155729 | 0.130761739 | 0.688560978 | 0.870222699 |
| chr10:73096076-73097313 | -0.128221365 | 0.688561988 | 0.870222699 |
| chr7:46071638-46072340 | -0.14955436 | 0.688571476 | 0.870222699 |
| chr3:23938725-23938999 | 0.182582538 | 0.68857262 | 0.870222699 |
| chr12:45615434-45615634 | 0.221861436 | 0.68858742 | 0.870222699 |
| chr19:30317854-30318128 | 0.18699377 | 0.688589389 | 0.870222699 |
| chr20:57439412-57439753 | -0.141889819 | 0.688595994 | 0.870222699 |
| chr2:176489911-176490202 | -0.209265474 | 0.688630132 | 0.87024012 |
| chr5:146565809-146566111 | -0.159246776 | 0.688632494 | 0.87024012 |
| chr8:104351201-104351422 | -0.195110958 | 0.688653695 | 0.8702472 |
| chr1:150022443-150022722 | -0.16414609 | 0.688679752 | 0.8702472 |
| chr10:102503656-102504116 | 0.132664008 | 0.688690598 | 0.8702472 |
| chr17:78505117-78505332 | -0.184962587 | 0.688691271 | 0.8702472 |
| chr7:138303892-138304298 | 0.160641946 | 0.688694885 | 0.8702472 |
| chr1:115089386-115090233 | -0.131451693 | 0.688708539 | 0.870249427 |
| chr13:94595627-94595910 | 0.160968383 | 0.688719363 | 0.870249427 |
| chr3:182979801-182981277 | -0.129570701 | 0.688767505 | 0.870290378 |
| chr18:12287814-12288232 | 0.162618924 | 0.688774488 | 0.870290378 |
| chr9:35731899-35733492 | 0.127633533 | 0.688803041 | 0.870294484 |
| chr15:80977579-80977873 | -0.158639598 | 0.688811733 | 0.870294484 |
| chr20:34525807-34526382 | 0.138400898 | 0.688811813 | 0.870294484 |
| chr21:44485186-44485411 | 0.20516302 | 0.68884479 | 0.870321799 |
| chr4:147596472-147596672 | -0.21970619 | 0.688880081 | 0.870326818 |
| chr5:33470596-33470926 | -0.162053064 | 0.688904773 | 0.870326818 |
| chr10:80567330-80567590 | -0.167526079 | 0.688905255 | 0.870326818 |

| | | | |
|---|---|---|---|
| chr4:119212056-119213329 | -0.125470897 | 0.688910748 | 0.870326818 |
| chr1:203038432-203038708 | 0.149921211 | 0.688934584 | 0.870326818 |
| chr7:4711307-4711521 | 0.196714129 | 0.688936411 | 0.870326818 |
| chrX:144275534-144275845 | 0.176658696 | 0.688938726 | 0.870326818 |
| chr14:73713456-73714576 | 0.130386893 | 0.688939632 | 0.870326818 |
| chrX:103686469-103687631 | -0.125680524 | 0.688956109 | 0.870333284 |
| chr11:72793540-72794250 | 0.130883686 | 0.688997456 | 0.870351332 |
| chr22:21001519-21002336 | 0.131105662 | 0.68899975 | 0.870351332 |
| chr8:108659634-108660508 | 0.138817923 | 0.689013959 | 0.870351332 |
| chr2:39837894-39838343 | -0.150204425 | 0.689015832 | 0.870351332 |
| chr1:23403802-23404279 | 0.154721177 | 0.689049229 | 0.870379169 |
| chr13:95834176-95834455 | 0.181671302 | 0.689075852 | 0.870398449 |
| chr12:108129487-108129820 | -0.166784576 | 0.689091495 | 0.870400598 |
| chr10:73641754-73642016 | -0.180566356 | 0.68910391 | 0.870400598 |
| chr8:52089020-52089350 | -0.154558484 | 0.689111632 | 0.870400598 |
| chr14:82075696-82075948 | -0.188352066 | 0.689128383 | 0.870407407 |
| chr1:228903155-228903387 | 0.179732835 | 0.689142106 | 0.870410393 |
| chr7:36258376-36258902 | -0.141103381 | 0.68916868 | 0.870418746 |
| chr5:116343990-116344190 | -0.215983929 | 0.68917144 | 0.870418746 |
| chr16:74667395-74667648 | -0.162357123 | 0.689205704 | 0.870447673 |
| chr19:35931363-35931990 | 0.130388887 | 0.689233029 | 0.87044946 |
| chr13:53151796-53152142 | -0.178717912 | 0.689239257 | 0.87044946 |
| chr4:38236455-38236670 | -0.212336722 | 0.689242798 | 0.87044946 |
| chr16:56245949-56246749 | -0.142323055 | 0.68925256 | 0.87044946 |
| chr9:83431767-83432614 | -0.134360816 | 0.689282145 | 0.870463075 |
| chr1:47314768-47315066 | -0.148156274 | 0.689286062 | 0.870463075 |
| chr1:32400885-32401231 | 0.1555195 | 0.689313425 | 0.870467202 |
| chr12:131108340-131108540 | 0.206139269 | 0.689319723 | 0.870467202 |
| chr1:34866153-34866704 | 0.140226312 | 0.689323411 | 0.870467202 |
| chr2:20916701-20916912 | -0.173632101 | 0.689341541 | 0.87047575 |
| chr1:179809810-179810197 | 0.140678454 | 0.689379441 | 0.870509262 |
| chr13:20174717-20175246 | -0.143556483 | 0.68941025 | 0.870533819 |
| chr1:51313413-51314046 | -0.144711076 | 0.689450437 | 0.870553457 |
| chr1:150537789-150538007 | 0.178921866 | 0.68945433 | 0.870553457 |
| chr3:100634646-100635190 | 0.146483131 | 0.689469511 | 0.870553457 |
| chr3:42581606-42582552 | -0.130291664 | 0.689471248 | 0.870553457 |
| chr19:18203918-18204775 | 0.129904458 | 0.689500894 | 0.870576543 |
| chr6:12638071-12638321 | -0.187466184 | 0.689539124 | 0.870600719 |
| chr16:53099397-53100207 | 0.130905475 | 0.689543907 | 0.870600719 |
| chr6:87322333-87323191 | 0.131334824 | 0.689554129 | 0.870600719 |
| chr9:132240943-132241712 | -0.129200913 | 0.689600002 | 0.870617755 |
| chr6:157900713-157901016 | 0.180610558 | 0.689605784 | 0.870617755 |
| chr11:86167384-86167827 | 0.159149275 | 0.689607464 | 0.870617755 |
| chr15:58770635-58772227 | 0.126317862 | 0.689617498 | 0.870617755 |
| chr5:139905083-139905399 | -0.175454715 | 0.689624435 | 0.870617755 |
| chr3:133444087-133444403 | -0.154255127 | 0.689639187 | 0.870622034 |
| chr11:118180357-118180655 | 0.179873694 | 0.689680765 | 0.870652012 |
| chr22:17084623-17085641 | 0.130743655 | 0.68970026 | 0.870652012 |
| chr3:170614714-170614989 | -0.180289767 | 0.689712689 | 0.870652012 |
| chr6:379720-379947 | 0.179323319 | 0.68971691 | 0.870652012 |

| | | | |
|---|---|---|---|
| chr9:17268479-17268821 | 0.16422515 | 0.689727279 | 0.870652012 |
| chr6:36663024-36663264 | -0.197919264 | 0.68973111 | 0.870652012 |
| chr11:116691805-116692059 | 0.17940954 | 0.689749182 | 0.87066048 |
| chr12:44317624-44317934 | -0.188399035 | 0.689795731 | 0.870704895 |
| chr9:34665177-34665754 | -0.134578536 | 0.689859331 | 0.870756094 |
| chr11:43573336-43573551 | -0.202163763 | 0.689864205 | 0.870756094 |
| chr12:52278994-52279360 | 0.153880791 | 0.689870998 | 0.870756094 |
| chr18:6482486-6483275 | -0.142810288 | 0.689889399 | 0.870756094 |
| chr13:38582197-38582469 | -0.169773197 | 0.689901321 | 0.870756094 |
| chr10:86693884-86694155 | -0.15371925 | 0.689906836 | 0.870756094 |
| chr22:29232392-29233016 | 0.150198313 | 0.689915843 | 0.870756094 |
| chr4:175444579-175444874 | -0.182584529 | 0.689958944 | 0.870796149 |
| chr17:60372684-60373425 | 0.128557228 | 0.689976256 | 0.870803655 |
| chr9:128117688-128118115 | -0.14904819 | 0.690005498 | 0.870814111 |
| chrX:19382847-19383106 | 0.179016633 | 0.690031611 | 0.870814111 |
| chr11:64688418-64688690 | 0.161101873 | 0.690042011 | 0.870814111 |
| chr20:16574378-16575263 | 0.131799441 | 0.690052994 | 0.870814111 |
| chr2:200672632-200672832 | -0.215220883 | 0.690062544 | 0.870814111 |
| chr20:13181019-13181219 | -0.215220883 | 0.690062544 | 0.870814111 |
| chr7:109692662-109692906 | -0.185025305 | 0.690075461 | 0.870814111 |
| chr8:60619707-60620063 | -0.185025305 | 0.690075461 | 0.870814111 |
| chrX:95318866-95319246 | -0.157503517 | 0.690140675 | 0.870882062 |
| chr1:168603468-168603916 | 0.140784824 | 0.690194766 | 0.870935976 |
| chr14:92793628-92794580 | 0.12661307 | 0.690215152 | 0.870939725 |
| chr1:180555399-180555769 | 0.15442677 | 0.690220471 | 0.870939725 |
| chr5:173887552-173888482 | -0.127146806 | 0.690288943 | 0.871011781 |
| chr8:101375868-101376410 | 0.136357303 | 0.690319554 | 0.871036062 |
| chr1:180515876-180516129 | 0.167235408 | 0.690342624 | 0.871050827 |
| chr16:30354935-30356001 | 0.128699031 | 0.690358561 | 0.871051174 |
| chr7:106217059-106217314 | -0.185658606 | 0.690365635 | 0.871051174 |
| chr1:48777527-48777743 | 0.194835642 | 0.690387099 | 0.871063912 |
| chr6:139550448-139550702 | 0.174939308 | 0.690408798 | 0.871071417 |
| chr2:219160105-219161036 | -0.128112961 | 0.690422725 | 0.871071417 |
| chr1:46530797-46531095 | -0.15811385 | 0.690427152 | 0.871071417 |
| chr3:158097653-158098597 | -0.128220431 | 0.690453545 | 0.871090372 |
| chr9:70480763-70480983 | 0.180728471 | 0.690481893 | 0.871111792 |
| chr7:52355178-52355400 | -0.192366751 | 0.690526967 | 0.871145489 |
| chr11:71488144-71488517 | -0.156936435 | 0.690531341 | 0.871145489 |
| chr20:49592980-49593339 | 0.153544923 | 0.690578821 | 0.871183227 |
| chr1:113976645-113977186 | 0.130387495 | 0.690583994 | 0.871183227 |
| chr13:24160255-24161460 | 0.126982226 | 0.690623387 | 0.871218577 |
| chr5:151020285-151020710 | 0.139783628 | 0.690649528 | 0.87123721 |
| chr1:164589004-164589423 | -0.148410007 | 0.690709649 | 0.871298706 |
| chr21:44793347-44793591 | 0.183932926 | 0.690777337 | 0.871360114 |
| chr17:60352623-60352925 | 0.136205208 | 0.690788408 | 0.871360114 |
| chr6:136792072-136793364 | -0.12705432 | 0.690793647 | 0.871360114 |
| chr15:63436796-63437023 | 0.174169535 | 0.690803817 | 0.871360114 |
| chr19:51690351-51690652 | 0.146341651 | 0.690830979 | 0.87138003 |
| chr9:97013353-97013896 | -0.147766992 | 0.690874743 | 0.871414182 |
| chr15:84034885-84035165 | 0.164250887 | 0.690886355 | 0.871414182 |

| | | | |
|---|---|---|---|
| chr4:84239546-84239771 | 0.201319277 | 0.690892174 | 0.871414182 |
| chr2:174003609-174003964 | -0.152655893 | 0.69090766 | 0.871414343 |
| chr3:106812566-106812897 | 0.168444006 | 0.690931487 | 0.871414343 |
| chr12:120325488-120325778 | 0.159217484 | 0.690943264 | 0.871414343 |
| chr18:21740292-21741429 | 0.128071419 | 0.690946795 | 0.871414343 |
| chr6:99588527-99589676 | -0.130499193 | 0.690952673 | 0.871414343 |
| chr11:64850949-64851271 | -0.162378957 | 0.690970736 | 0.871414343 |
| chr6:75284295-75285314 | -0.125653838 | 0.690985403 | 0.871414343 |
| chr7:6453696-6454076 | 0.16053554 | 0.690993112 | 0.871414343 |
| chr1:106944739-106945030 | 0.143434568 | 0.690994657 | 0.871414343 |
| chr3:195907610-195908074 | -0.142120781 | 0.691032778 | 0.871448074 |
| chr5:154520937-154521427 | -0.140698939 | 0.69106609 | 0.871465179 |
| chr8:143573138-143573338 | 0.158903005 | 0.691070076 | 0.871465179 |
| chr8:121908332-121908618 | 0.18475787 | 0.691080463 | 0.871465179 |
| chr9:19493148-19493603 | -0.161967215 | 0.691134278 | 0.871518698 |
| chr20:60613881-60614086 | 0.203477422 | 0.691226749 | 0.871606615 |
| chr22:32823152-32823375 | 0.203477422 | 0.691226749 | 0.871606615 |
| chr1:109350734-109350934 | -0.223619378 | 0.691273564 | 0.871651302 |
| chr7:92832762-92834147 | 0.130305663 | 0.691300298 | 0.871670668 |
| chr15:32649358-32649558 | 0.199589147 | 0.691324964 | 0.871687424 |
| chr7:44581484-44582912 | -0.124853793 | 0.691352089 | 0.871707281 |
| chr6:30482096-30482403 | 0.166501894 | 0.691401787 | 0.871755598 |
| chr19:48501498-48502564 | 0.125307707 | 0.691421708 | 0.871766371 |
| chr14:74946144-74946653 | 0.138453942 | 0.691450762 | 0.871776112 |
| chr2:172461573-172461773 | -0.202161145 | 0.69145219 | 0.871776112 |
| chr16:67529101-67529312 | -0.181142516 | 0.69146393 | 0.87177657 |
| chr17:76079574-76079910 | 0.161861536 | 0.691477234 | 0.871777428 |
| chr7:74279617-74279963 | 0.167649598 | 0.691498951 | 0.871777428 |
| chr19:54489509-54489717 | -0.210742737 | 0.691510121 | 0.871777428 |
| chr1:190615894-190616094 | -0.210742737 | 0.691510121 | 0.871777428 |
| chr16:2155128-2155829 | 0.130112216 | 0.691528906 | 0.871786765 |
| chr5:55978232-55978447 | -0.192349891 | 0.691550656 | 0.871799842 |
| chr3:64104201-64105454 | -0.124370515 | 0.691580011 | 0.871822503 |
| chr10:29517391-29517650 | 0.181952806 | 0.69167694 | 0.871921251 |
| chr1:108512973-108513200 | 0.194945551 | 0.691689743 | 0.871921251 |
| chr9:132807129-132807329 | 0.188334071 | 0.691692482 | 0.871921251 |
| chr6:127121039-127121734 | -0.141211913 | 0.691710761 | 0.871929949 |
| chr13:47308683-47308883 | -0.20081391 | 0.691772342 | 0.871993229 |
| chr15:59741412-59741660 | 0.188299326 | 0.691808827 | 0.872024873 |
| chr1:155083219-155083500 | -0.144083674 | 0.691859476 | 0.872068098 |
| chr7:108091571-108091831 | 0.17393245 | 0.691870769 | 0.872068098 |
| chr17:2593209-2594268 | 0.131129761 | 0.691877263 | 0.872068098 |
| chr10:88975236-88975589 | 0.184827439 | 0.691919301 | 0.872104585 |
| chr6:43143520-43143941 | 0.142977093 | 0.691928975 | 0.872104585 |
| chr13:75695884-75696192 | -0.163283744 | 0.691947746 | 0.872113898 |
| chr14:50860577-50860838 | -0.169923367 | 0.691959178 | 0.872113961 |
| chr2:71299458-71299692 | 0.204598308 | 0.691974818 | 0.872114619 |
| chr3:172341552-172341866 | 0.158192123 | 0.691988734 | 0.872114619 |
| chr15:65422402-65422645 | -0.168567227 | 0.691997946 | 0.872114619 |
| chr12:121672312-121672921 | -0.135219758 | 0.692005573 | 0.872114619 |

| | | | |
|---|---|---|---|
| chr8:103076094-103076307 | -0.180927008 | 0.69201661 | 0.872114619 |
| chr11:28107479-28108683 | -0.125872169 | 0.692032753 | 0.872120618 |
| chr3:50292105-50292776 | -0.133986175 | 0.692049169 | 0.872126963 |
| chr7:148286220-148286482 | -0.16400115 | 0.692063978 | 0.872131281 |
| chr1:217660626-217661081 | -0.145350351 | 0.692080832 | 0.872138176 |
| chr19:45001221-45001566 | -0.149198134 | 0.692127002 | 0.872182014 |
| chr14:53432312-53432669 | 0.150628713 | 0.692147228 | 0.872193156 |
| chr3:158746895-158747171 | -0.163518529 | 0.692168903 | 0.872206126 |
| chr15:63157116-63158275 | 0.129592958 | 0.692182744 | 0.872209222 |
| chr13:32124702-32125023 | 0.182530197 | 0.692212993 | 0.872232995 |
| chr2:197247994-197248196 | -0.215833969 | 0.692250736 | 0.872246746 |
| chr8:23224426-23225495 | -0.131132548 | 0.692255565 | 0.872246746 |
| chr16:74618024-74618260 | -0.171475976 | 0.692258057 | 0.872246746 |
| chr11:61385641-61385873 | 0.175915121 | 0.692277467 | 0.872247463 |
| chr1:21195260-21195509 | 0.193574096 | 0.692294635 | 0.872247463 |
| chr3:186821470-186821709 | 0.183035268 | 0.692317028 | 0.872247463 |
| chr16:2135679-2136479 | 0.135921895 | 0.692324504 | 0.872247463 |
| chr6:71288204-71289489 | 0.128763266 | 0.692335805 | 0.872247463 |
| chr12:56448756-56449600 | 0.130792251 | 0.69233941 | 0.872247463 |
| chr7:29946862-29947170 | 0.163882829 | 0.692370667 | 0.872247463 |
| chr15:82895443-82895750 | 0.178436846 | 0.692372276 | 0.872247463 |
| chr11:61697096-61697424 | 0.152706016 | 0.692378038 | 0.872247463 |
| chr7:1113484-1113718 | 0.187969406 | 0.692391346 | 0.872247463 |
| chr5:81971612-81972661 | 0.12614408 | 0.692399425 | 0.872247463 |
| chr12:95336518-95336771 | 0.169739168 | 0.69240602 | 0.872247463 |
| chr19:3185432-3185992 | 0.140433836 | 0.692406616 | 0.872247463 |
| chr8:116765780-116767106 | -0.124551039 | 0.69245314 | 0.87229173 |
| chr3:75429613-75429963 | -0.141603858 | 0.692490724 | 0.872311252 |
| chr5:151020776-151021152 | 0.153241956 | 0.692502681 | 0.872311252 |
| chr14:72742324-72742613 | -0.192291724 | 0.692502791 | 0.872311252 |
| chr3:129892859-129894212 | -0.125508068 | 0.692547622 | 0.872353382 |
| chr8:89253269-89253859 | 0.133252046 | 0.692564344 | 0.872360104 |
| chr10:71384318-71384518 | -0.191761434 | 0.692597955 | 0.872383982 |
| chr2:39150932-39151263 | -0.173238868 | 0.692606071 | 0.872383982 |
| chr16:72848541-72849448 | 0.129448439 | 0.692623384 | 0.872391447 |
| chr2:11784831-11785294 | -0.146127526 | 0.692653204 | 0.872414666 |
| chr9:34651943-34652520 | -0.139927678 | 0.692673134 | 0.872425427 |
| chr6:158999113-159000406 | -0.125688086 | 0.692691115 | 0.872433733 |
| chr20:59527903-59528138 | -0.168503511 | 0.692718766 | 0.872454218 |
| chr8:118092665-118092865 | 0.195398271 | 0.692762686 | 0.872462597 |
| chr19:33269120-33269353 | 0.184256341 | 0.692774217 | 0.872462597 |
| chr2:3493700-3494036 | -0.155010614 | 0.692789236 | 0.872462597 |
| chr6:65776345-65776679 | 0.153260532 | 0.692811081 | 0.872462597 |
| chr1:38040561-38040785 | 0.184337734 | 0.692827933 | 0.872462597 |
| chr5:142717636-142717855 | -0.167671238 | 0.692829421 | 0.872462597 |
| chr19:37077782-37078818 | 0.133641919 | 0.692837486 | 0.872462597 |
| chr4:103204145-103204345 | 0.209739486 | 0.69283873 | 0.872462597 |
| chr12:123971781-123973601 | 0.124846643 | 0.69284514 | 0.872462597 |
| chr8:75849412-75849612 | 0.206642087 | 0.69286077 | 0.872462597 |
| chr8:126794161-126794361 | 0.206642087 | 0.69286077 | 0.872462597 |

| | | | |
|---|---|---|---|
| chr1:46488846-46489124 | -0.164432682 | 0.692862058 | 0.872462597 |
| chr7:2403030-2404247 | -0.126388912 | 0.69287615 | 0.872466005 |
| chr5:177113898-177114253 | 0.140905941 | 0.69291343 | 0.87249611 |
| chr17:77790159-77790397 | 0.181184347 | 0.692938659 | 0.87249611 |
| chr10:32414970-32415231 | -0.186389218 | 0.692949829 | 0.87249611 |
| chr9:133347489-133348447 | 0.129582739 | 0.692955562 | 0.87249611 |
| chr7:30977955-30978258 | 0.162528939 | 0.692956994 | 0.87249611 |
| chrX:140701555-140701844 | 0.164569872 | 0.692989619 | 0.872522851 |
| chrX:23742869-23743655 | -0.129813653 | 0.693014765 | 0.872540173 |
| chr7:43838577-43839209 | -0.137471527 | 0.693044533 | 0.872563315 |
| chr12:95473213-95473682 | -0.140346886 | 0.693078372 | 0.872591581 |
| chr5:148741301-148741655 | -0.165557977 | 0.693102488 | 0.872607605 |
| chr15:40125096-40125342 | 0.182536112 | 0.693115546 | 0.872609707 |
| chr20:64057789-64058092 | -0.145204468 | 0.693144467 | 0.872631779 |
| chr2:131754785-131755021 | -0.21355466 | 0.693162027 | 0.872639548 |
| chr8:42171668-42172054 | -0.156352794 | 0.693225354 | 0.872654604 |
| chr1:108327669-108328055 | 0.148727998 | 0.693231121 | 0.872654604 |
| chr2:84925550-84926370 | -0.127430473 | 0.693234072 | 0.872654604 |
| chr4:118224202-118224575 | -0.165773255 | 0.693234771 | 0.872654604 |
| chr3:158935616-158936006 | -0.182138845 | 0.693240747 | 0.872654604 |
| chr22:21713018-21713637 | 0.145612697 | 0.693242321 | 0.872654604 |
| chr17:66951579-66952623 | 0.125483801 | 0.693275218 | 0.872681678 |
| chr1:63149203-63149436 | -0.184962752 | 0.693288273 | 0.872683774 |
| chrX:116205061-116205263 | 0.19398059 | 0.693321952 | 0.872697209 |
| chr15:79596055-79596423 | 0.143187322 | 0.693325068 | 0.872697209 |
| chr3:15557584-15557966 | -0.141717533 | 0.693333115 | 0.872697209 |
| chrX:15674309-15674517 | -0.169562777 | 0.693399344 | 0.872724689 |
| chr21:39153438-39153725 | 0.148777115 | 0.693404235 | 0.872724689 |
| chr7:51455236-51455453 | -0.1852181 | 0.693420361 | 0.872724689 |
| chr11:69293312-69293512 | -0.16528717 | 0.693422592 | 0.872724689 |
| chr1:157112032-157112232 | 0.184792559 | 0.693445318 | 0.872724689 |
| chr2:5077474-5077674 | -0.208079013 | 0.693447654 | 0.872724689 |
| chr5:139064601-139064948 | 0.151284042 | 0.693451759 | 0.872724689 |
| chr5:142716989-142717273 | -0.150177673 | 0.693469184 | 0.872724689 |
| chr3:128776035-128776235 | -0.209316921 | 0.693472375 | 0.872724689 |
| chr17:58335938-58336175 | -0.163594658 | 0.693473684 | 0.872724689 |
| chr20:57305674-57306097 | -0.140289725 | 0.693480237 | 0.872724689 |
| chr3:4701203-4701450 | 0.201957406 | 0.693548522 | 0.872796289 |
| chr10:70815731-70816901 | 0.128317256 | 0.693607695 | 0.872856419 |
| chr15:77702662-77702879 | -0.171133889 | 0.693642332 | 0.872869348 |
| chr19:46784774-46785103 | -0.146166904 | 0.693662424 | 0.872869348 |
| chr8:101459828-101460066 | 0.177522555 | 0.693675222 | 0.872869348 |
| chr19:7652069-7652386 | -0.169298727 | 0.693677775 | 0.872869348 |
| chr10:118045984-118047549 | -0.126421633 | 0.693692452 | 0.872869348 |
| chr6:28383231-28383927 | 0.134754048 | 0.693693586 | 0.872869348 |
| chr3:155730547-155731364 | -0.128830095 | 0.693705886 | 0.872869348 |
| chrX:140391190-140391405 | -0.19425964 | 0.693713999 | 0.872869348 |
| chr10:21495627-21497413 | -0.12686462 | 0.693720496 | 0.872869348 |
| chr15:70355487-70355792 | 0.1644894 | 0.693745818 | 0.872886875 |
| chr7:67302103-67303164 | 0.124986424 | 0.6937643 | 0.872888696 |

| | | | |
|---|---|---|---|
| chr12:46493558-46494284 | 0.134960827 | 0.693781367 | 0.872888696 |
| chrX:106035167-106035370 | 0.191606173 | 0.693784553 | 0.872888696 |
| chr1:162056035-162056367 | 0.146728228 | 0.693792834 | 0.872888696 |
| chr11:65117491-65118140 | 0.129352271 | 0.693870043 | 0.872971502 |
| chrX:152926184-152926677 | 0.144078176 | 0.693894238 | 0.872987609 |
| chr3:155034090-155034290 | -0.21048789 | 0.693933092 | 0.873022156 |
| chr5:68406727-68407169 | -0.157903646 | 0.693956123 | 0.873036796 |
| chr8:38734486-38735159 | 0.131964635 | 0.694002816 | 0.873067629 |
| chrX:15789907-15791050 | -0.128597085 | 0.694010099 | 0.873067629 |
| chr19:8252992-8253300 | 0.176091296 | 0.694021716 | 0.873067629 |
| chr12:57546403-57546704 | 0.149598461 | 0.694029957 | 0.873067629 |
| chr6:78861599-78861799 | 0.199111386 | 0.694065618 | 0.873067629 |
| chr1:194295690-194296105 | -0.141106476 | 0.694075349 | 0.873067629 |
| chr19:2783193-2783635 | 0.137789242 | 0.69408989 | 0.873067629 |
| chr1:224565272-224565544 | -0.195802725 | 0.694103231 | 0.873067629 |
| chr14:87936003-87936215 | 0.191294222 | 0.694104133 | 0.873067629 |
| chr8:128259252-128259513 | -0.162603492 | 0.694108708 | 0.873067629 |
| chr1:175221082-175221290 | -0.171366841 | 0.694143601 | 0.873067629 |
| chr17:50178187-50178785 | -0.130875786 | 0.694151048 | 0.873067629 |
| chr18:12237027-12237601 | -0.14342698 | 0.694152196 | 0.873067629 |
| chr17:55295979-55296282 | 0.147790341 | 0.694157997 | 0.873067629 |
| chr12:65074925-65075262 | -0.153221565 | 0.694165975 | 0.873067629 |
| chr20:49482741-49482971 | 0.165369707 | 0.694169986 | 0.873067629 |
| chr12:94057803-94058003 | -0.192801974 | 0.694176355 | 0.873067629 |
| chr2:211352075-211352362 | -0.171917682 | 0.694185732 | 0.873067629 |
| chr18:158182-159001 | 0.127108462 | 0.694215514 | 0.873078211 |
| chr12:74269971-74270405 | -0.161206599 | 0.694216935 | 0.873078211 |
| chr1:151175719-151176207 | 0.138118524 | 0.694242657 | 0.873091464 |
| chr14:50003117-50003704 | -0.133545365 | 0.694255072 | 0.873091464 |
| chr15:45237110-45237378 | -0.20063277 | 0.694276745 | 0.873091464 |
| chr4:120066340-120067547 | -0.123882637 | 0.694277913 | 0.873091464 |
| chr2:1649514-1649762 | -0.206356853 | 0.694284447 | 0.873091464 |
| chr17:28371312-28371741 | -0.148070891 | 0.694338288 | 0.873124444 |
| chr6:188705-189111 | -0.146000822 | 0.694340049 | 0.873124444 |
| chr4:149740524-149740811 | -0.18795758 | 0.694344859 | 0.873124444 |
| chr4:107719617-107720991 | -0.123537067 | 0.694380946 | 0.873147843 |
| chr2:4654024-4654297 | 0.18099617 | 0.694386257 | 0.873147843 |
| chr19:48363632-48364569 | 0.125883674 | 0.694437942 | 0.873192195 |
| chr1:58576302-58577708 | -0.124463147 | 0.694444321 | 0.873192195 |
| chr17:44070567-44070932 | 0.141302375 | 0.69448368 | 0.873203474 |
| chr19:49363359-49364068 | 0.126604098 | 0.694498359 | 0.873203474 |
| chr22:25333571-25333773 | -0.207222742 | 0.694507062 | 0.873203474 |
| chr3:66661881-66662203 | -0.175928114 | 0.694521462 | 0.873203474 |
| chr12:6285098-6285442 | 0.161203925 | 0.694524507 | 0.873203474 |
| chr2:131822452-131822782 | 0.150509729 | 0.694543867 | 0.873203474 |
| chr2:13113099-13113400 | -0.172702097 | 0.694545486 | 0.873203474 |
| chr14:61761717-61762933 | -0.124804145 | 0.69455541 | 0.873203474 |
| chr2:218004249-218004603 | -0.160788358 | 0.694555858 | 0.873203474 |
| chr15:92400934-92401134 | -0.214007114 | 0.694574509 | 0.873212596 |
| chr5:73454433-73454794 | -0.148714222 | 0.694605567 | 0.873237312 |

| | | | |
|---|---|---|---|
| chr12:57905279-57905814 | -0.130008723 | 0.694639244 | 0.873253304 |
| chr11:69131590-69131831 | -0.16616294 | 0.694641081 | 0.873253304 |
| chr10:26557129-26557391 | 0.183721431 | 0.69472183 | 0.873290648 |
| chr12:42482834-42483270 | 0.149420863 | 0.694728934 | 0.873290648 |
| chr1:206854655-206854861 | -0.182688874 | 0.694731043 | 0.873290648 |
| chr7:56051133-56052219 | 0.126985279 | 0.694742309 | 0.873290648 |
| chr12:46992009-46992451 | 0.150463065 | 0.694750207 | 0.873290648 |
| chrX:57286422-57287220 | -0.131032825 | 0.694750547 | 0.873290648 |
| chr2:211745335-211745691 | -0.148907728 | 0.694750568 | 0.873290648 |
| chr1:68120182-68120504 | 0.153832554 | 0.69477669 | 0.873296066 |
| chr1:145448315-145448530 | 0.194906278 | 0.694786417 | 0.873296066 |
| chr9:83928129-83928329 | -0.228073775 | 0.694789071 | 0.873296066 |
| chr11:9314389-9315129 | 0.128517272 | 0.694828659 | 0.873323252 |
| chr12:54032634-54033070 | 0.139690402 | 0.694839761 | 0.873323252 |
| chr16:68731898-68732375 | 0.141218807 | 0.694844893 | 0.873323252 |
| chr14:21436936-21437755 | 0.126827887 | 0.694881793 | 0.873355304 |
| chr1:185681089-185681473 | -0.157627514 | 0.694902907 | 0.873367515 |
| chr12:71195585-71196001 | -0.169015892 | 0.694915499 | 0.873369015 |
| chr4:74623113-74623481 | -0.155360589 | 0.694930333 | 0.873373332 |
| chr20:32040200-32040483 | 0.173269322 | 0.695016827 | 0.873454924 |
| chr4:128809461-128810212 | 0.127067653 | 0.695018053 | 0.873454924 |
| chr3:64668235-64668540 | 0.133774656 | 0.695029566 | 0.873455067 |
| chr16:10521770-10521990 | 0.185977032 | 0.695076345 | 0.873470468 |
| chr16:82368038-82368328 | 0.16896827 | 0.695085397 | 0.873470468 |
| chr17:70704893-70705096 | 0.192768864 | 0.695096192 | 0.873470468 |
| chr15:101664384-101664641 | 0.162101936 | 0.695101055 | 0.873470468 |
| chr12:100554632-100554946 | -0.163569753 | 0.695102703 | 0.873470468 |
| chr3:165268104-165268609 | 0.162435349 | 0.695110219 | 0.873470468 |
| chr8:111416191-111416677 | -0.15088392 | 0.695122967 | 0.873472161 |
| chr17:41324164-41324494 | 0.164453399 | 0.695144355 | 0.873484712 |
| chr8:85219870-85220954 | 0.123563894 | 0.695223695 | 0.873556506 |
| chr2:95766465-95766683 | -0.216557053 | 0.695224293 | 0.873556506 |
| chr2:52613116-52613406 | -0.174367041 | 0.695253831 | 0.873579296 |
| chr6:12290232-12290514 | -0.154547419 | 0.695272562 | 0.873588506 |
| chr12:124566350-124567066 | 0.132242094 | 0.695303557 | 0.873599846 |
| chr5:125402840-125403193 | 0.189581524 | 0.695304391 | 0.873599846 |
| chr20:49888532-49888809 | 0.1426675 | 0.69533966 | 0.873629834 |
| chr6:56359311-56359542 | -0.192501173 | 0.695377284 | 0.87366278 |
| chr10:43137671-43138713 | -0.129898992 | 0.695393496 | 0.873668821 |
| chr6:82257392-82257720 | -0.183872214 | 0.695412701 | 0.873671711 |
| chr5:63986675-63986875 | 0.218608638 | 0.6954186 | 0.873671711 |
| chr12:75611086-75611344 | -0.183719481 | 0.695447448 | 0.873690526 |
| chr14:53929902-53930163 | 0.164983903 | 0.69546885 | 0.873690526 |
| chr13:41507782-41508070 | 0.194627818 | 0.695479089 | 0.873690526 |
| chr9:120366693-120367028 | -0.156261537 | 0.695479187 | 0.873690526 |
| chr2:85285480-85285804 | 0.14536461 | 0.695571046 | 0.873789424 |
| chr7:105760323-105760583 | -0.177228396 | 0.69558072 | 0.873789424 |
| chr3:4571780-4572002 | 0.165069002 | 0.695616758 | 0.873820369 |
| chr12:105092311-105092879 | -0.136669722 | 0.695641806 | 0.873837508 |
| chr1:153375637-153375864 | -0.18514208 | 0.695662139 | 0.873842018 |

| | | | |
|---|---|---|---|
| chr11:118689511-118690466 | -0.126666307 | 0.695668206 | 0.873842018 |
| chr19:56314457-56315609 | 0.124704044 | 0.695705073 | 0.873874002 |
| chr20:31557419-31557619 | -0.176565594 | 0.695728856 | 0.873878314 |
| chr2:27369727-27371047 | -0.124050392 | 0.695731316 | 0.873878314 |
| chr22:18078194-18078396 | -0.174590276 | 0.695758752 | 0.873898449 |
| chr19:58475428-58476532 | 0.124548603 | 0.695779121 | 0.873906342 |
| chr14:35260363-35260715 | 0.173808207 | 0.695787847 | 0.873906342 |
| chr14:95452485-95452772 | -0.151961126 | 0.695802484 | 0.873910401 |
| chr7:140564738-140564938 | -0.204988784 | 0.695829899 | 0.873922717 |
| chr3:161630824-161631124 | -0.157399261 | 0.695835101 | 0.873922717 |
| chr8:37694883-37695797 | 0.126675099 | 0.695855769 | 0.873928146 |
| chr8:232688-233014 | 0.176146567 | 0.695862235 | 0.873928146 |
| chr16:85821389-85821709 | 0.143829411 | 0.695885797 | 0.873943412 |
| chr1:183609228-183609619 | -0.148080871 | 0.695926249 | 0.873967195 |
| chr1:175191851-175193511 | -0.123219484 | 0.695927546 | 0.873967195 |
| chr2:71466638-71466886 | 0.163937342 | 0.695943741 | 0.873973208 |
| chr6:43806360-43806663 | -0.161518198 | 0.695981249 | 0.87399006 |
| chr5:21458987-21459905 | -0.12635855 | 0.69598311 | 0.87399006 |
| chr12:47808545-47808765 | -0.197599982 | 0.695992975 | 0.87399006 |
| chr18:9613559-9614247 | 0.142154826 | 0.696002786 | 0.87399006 |
| chr16:48353702-48353993 | -0.191478496 | 0.696080763 | 0.874073653 |
| chr8:124581054-124581254 | 0.168468484 | 0.696105433 | 0.874090306 |
| chr2:9170846-9171122 | 0.156596612 | 0.696166544 | 0.874152717 |
| chr9:77933102-77933412 | -0.170874578 | 0.696181188 | 0.87415678 |
| chr6:138517625-138518016 | -0.146743112 | 0.696208177 | 0.874162087 |
| chr8:102967107-102967350 | 0.17087383 | 0.696208232 | 0.874162087 |
| chr3:9554284-9554559 | 0.180747921 | 0.696280653 | 0.874215321 |
| chr6:24666319-24667395 | 0.126235364 | 0.696284153 | 0.874215321 |
| chr1:198079487-198079942 | -0.158370999 | 0.696293182 | 0.874215321 |
| chr6:3742437-3742681 | -0.189423266 | 0.696296267 | 0.874215321 |
| chr1:171684744-171685005 | 0.193020027 | 0.696392964 | 0.8743157 |
| chr7:136874496-136874696 | 0.207046701 | 0.696441403 | 0.8743157 |
| chr8:91056010-91056219 | 0.207046701 | 0.696441403 | 0.8743157 |
| chr12:9634133-9634505 | -0.188413328 | 0.69644585 | 0.8743157 |
| chr6:14210678-14211836 | -0.128543557 | 0.696452587 | 0.8743157 |
| chr6:125113077-125113310 | -0.18924245 | 0.696462516 | 0.8743157 |
| chr18:10578039-10578324 | -0.168107431 | 0.696464334 | 0.8743157 |
| chr2:263926-265309 | 0.122879961 | 0.696477323 | 0.8743157 |
| chr3:64488983-64490393 | -0.121641183 | 0.696504152 | 0.8743157 |
| chr10:3785083-3786104 | 0.123387692 | 0.696509599 | 0.8743157 |
| chr3:147393202-147393863 | -0.127332864 | 0.696531727 | 0.8743157 |
| chr15:78661194-78661394 | 0.210591474 | 0.696533334 | 0.8743157 |
| chr1:159936432-159936754 | 0.147748167 | 0.696535756 | 0.8743157 |
| chr9:132944309-132945255 | -0.128252201 | 0.696546248 | 0.8743157 |
| chr2:210248691-210249018 | 0.156422852 | 0.696547379 | 0.8743157 |
| chr3:159092002-159092225 | -0.20896293 | 0.696572599 | 0.874318519 |
| chr9:35161654-35162890 | -0.127070375 | 0.696578099 | 0.874318519 |
| chr19:38975920-38976400 | 0.149822982 | 0.696583857 | 0.874318519 |
| chr10:122981174-122981428 | 0.15985691 | 0.69668213 | 0.874427542 |
| chr2:30711596-30712073 | 0.133347729 | 0.696710782 | 0.87444918 |

| | | | |
|---|---|---|---|
| chr6:49462708-49463887 | -0.124682698 | 0.696757886 | 0.874488612 |
| chr15:92644191-92644901 | 0.130429239 | 0.696765025 | 0.874488612 |
| chr2:216777654-216778049 | 0.141112717 | 0.696813212 | 0.874534765 |
| chr4:168943063-168943308 | -0.177837419 | 0.696841371 | 0.874540047 |
| chr12:111841767-111843052 | 0.126295245 | 0.696852766 | 0.874540047 |
| chr6:26569160-26569375 | -0.168375681 | 0.696886714 | 0.874540047 |
| chr9:117411600-117411901 | 0.150031489 | 0.696893514 | 0.874540047 |
| chr3:67113803-67114295 | -0.14014653 | 0.696901882 | 0.874540047 |
| chr5:16508889-16509435 | -0.146880652 | 0.696917753 | 0.874540047 |
| chr7:35073522-35073793 | -0.167525957 | 0.696937477 | 0.874540047 |
| chr18:46842687-46842907 | -0.220111835 | 0.696943772 | 0.874540047 |
| chr6:16105970-16106218 | -0.220111835 | 0.696943772 | 0.874540047 |
| chr8:18209730-18210355 | 0.137488767 | 0.696959443 | 0.874540047 |
| chr20:40359478-40359751 | -0.187840167 | 0.696995046 | 0.874540047 |
| chrX:110045954-110046329 | 0.175485257 | 0.697019067 | 0.874540047 |
| chr16:71894822-71896315 | 0.123381309 | 0.697022493 | 0.874540047 |
| chr12:54427273-54427496 | 0.183122226 | 0.697027182 | 0.874540047 |
| chr2:38666589-38666913 | 0.15711106 | 0.697041129 | 0.874540047 |
| chr7:7595536-7595787 | 0.174184918 | 0.697049153 | 0.874540047 |
| chr7:74183662-74184045 | -0.182854907 | 0.697051796 | 0.874540047 |
| chr5:130199005-130199242 | -0.179015783 | 0.69705492 | 0.874540047 |
| chr3:159059011-159059211 | -0.205059879 | 0.697067008 | 0.874540047 |
| chr4:47032589-47033052 | -0.145092992 | 0.697070103 | 0.874540047 |
| chrX:132376011-132376229 | 0.225735114 | 0.697083658 | 0.874540047 |
| chr3:134002930-134003464 | 0.138951815 | 0.697099994 | 0.874540047 |
| chr15:41402086-41402680 | -0.135100927 | 0.697102171 | 0.874540047 |
| chr3:185983563-185984067 | 0.148492573 | 0.697116923 | 0.874540047 |
| chr1:234356606-234357163 | -0.14003945 | 0.697129525 | 0.874540047 |
| chr14:77203977-77204177 | 0.171794945 | 0.697179529 | 0.874540047 |
| chr14:92200078-92200279 | 0.203473168 | 0.697185773 | 0.874540047 |
| chr19:14450491-14450880 | -0.158969309 | 0.697190354 | 0.874540047 |
| chr6:89598058-89598412 | -0.149811179 | 0.697208654 | 0.874540047 |
| chr8:68432579-68432830 | -0.187745851 | 0.697214376 | 0.874540047 |
| chr5:141651238-141651570 | 0.139352696 | 0.697229342 | 0.874540047 |
| chr20:31475786-31476397 | 0.143959605 | 0.69723009 | 0.874540047 |
| chr13:24511300-24511755 | 0.142826768 | 0.697246998 | 0.874540047 |
| chr1:50968030-50968801 | 0.128056116 | 0.697248895 | 0.874540047 |
| chr22:41696951-41697582 | -0.15101615 | 0.697249202 | 0.874540047 |
| chr5:155139408-155139973 | -0.14427908 | 0.697249521 | 0.874540047 |
| chr6:75906960-75907171 | 0.211327156 | 0.697252962 | 0.874540047 |
| chrX:42195503-42195812 | -0.176967198 | 0.697268805 | 0.874540047 |
| chr10:104254462-104255574 | -0.129732668 | 0.697277716 | 0.874540047 |
| chr2:206981815-206982015 | -0.198490555 | 0.697288649 | 0.874540047 |
| chr1:59879662-59879898 | 0.173470227 | 0.697294767 | 0.874540047 |
| chr1:145990539-145990739 | 0.172485565 | 0.697296796 | 0.874540047 |
| chr12:48336634-48336834 | 0.20969703 | 0.697310409 | 0.874542805 |
| chr1:88891306-88892052 | -0.129641391 | 0.697323403 | 0.874544787 |
| chr11:79076615-79076890 | 0.160781017 | 0.697424136 | 0.87461482 |
| chrX:71664239-71664544 | 0.149035039 | 0.697438153 | 0.87461482 |
| chr6:51224652-51225147 | -0.150494049 | 0.697466294 | 0.87461482 |

| | | | |
|---|---|---|---|
| chr6:150046025-150046232 | -0.225979703 | 0.697476647 | 0.87461482 |
| chr1:157124940-157125225 | 0.168072587 | 0.697477508 | 0.87461482 |
| chr5:49599944-49600779 | 0.142386446 | 0.697491485 | 0.87461482 |
| chr15:85655083-85655409 | -0.192266925 | 0.697498999 | 0.87461482 |
| chr11:69437172-69437474 | 0.156985217 | 0.697502546 | 0.87461482 |
| chr3:172710202-172710530 | 0.139361185 | 0.697517986 | 0.87461482 |
| chr18:1521599-1521864 | -0.167455854 | 0.697522284 | 0.87461482 |
| chr15:68875712-68876295 | -0.14529183 | 0.697533649 | 0.87461482 |
| chr14:79363341-79363603 | 0.194625351 | 0.697536588 | 0.87461482 |
| chr6:111087183-111087869 | 0.135794388 | 0.697541593 | 0.87461482 |
| chr16:3003738-3004306 | 0.128934797 | 0.697558393 | 0.87461482 |
| chr3:81218095-81218295 | 0.190072918 | 0.697558419 | 0.87461482 |
| chr6:143410883-143411122 | -0.183944872 | 0.697562942 | 0.87461482 |
| chr22:41301069-41301316 | 0.196609417 | 0.697586871 | 0.87461482 |
| chr17:4555274-4555682 | 0.138356022 | 0.697597817 | 0.87461482 |
| chr14:53713847-53714076 | -0.206970603 | 0.697600785 | 0.87461482 |
| chr15:62391609-62392178 | 0.14062962 | 0.697607537 | 0.87461482 |
| chr13:29215679-29215892 | 0.221049184 | 0.697631526 | 0.874630585 |
| chr6:73223720-73223983 | 0.201899835 | 0.697647807 | 0.874636685 |
| chr6:144284722-144285543 | 0.131540483 | 0.697705409 | 0.874694588 |
| chr8:99415821-99416068 | -0.177351812 | 0.697730557 | 0.874711803 |
| chr1:162040707-162041061 | 0.160326961 | 0.697760334 | 0.874734821 |
| chr2:62281315-62281613 | 0.212974339 | 0.697776339 | 0.874740574 |
| chrX:71283094-71284173 | -0.127255596 | 0.697798858 | 0.874754492 |
| chr1:116504717-116504998 | 0.17152467 | 0.697821752 | 0.874768881 |
| chr6:31888776-31889074 | -0.164336272 | 0.697858001 | 0.874774992 |
| chr10:79084122-79084397 | -0.164358184 | 0.697859147 | 0.874774992 |
| chr14:26731305-26731568 | 0.159861424 | 0.697860877 | 0.874774992 |
| chr10:127102204-127102404 | -0.199191997 | 0.697883441 | 0.874788964 |
| chr11:32385999-32386353 | 0.146410215 | 0.697935682 | 0.874824436 |
| chr3:159813971-159814835 | -0.134423082 | 0.697951566 | 0.874824436 |
| chr20:13820002-13820691 | -0.137653819 | 0.697952979 | 0.874824436 |
| chr14:66932026-66932449 | 0.156760831 | 0.697957409 | 0.874824436 |
| chr13:82269314-82269601 | -0.177763191 | 0.697984594 | 0.874844199 |
| chr2:173159815-173160062 | -0.175642492 | 0.698030282 | 0.874854124 |
| chrX:103628594-103629225 | -0.131133833 | 0.698039112 | 0.874854124 |
| chr11:82971473-82971673 | 0.212690879 | 0.698063671 | 0.874854124 |
| chr17:1484663-1485107 | 0.142539665 | 0.698064519 | 0.874854124 |
| chr6:28848368-28849126 | 0.129451362 | 0.698066812 | 0.874854124 |
| chr5:91562116-91562399 | -0.212931272 | 0.698073678 | 0.874854124 |
| chr14:50201944-50202144 | -0.2256234 | 0.698092134 | 0.874854124 |
| chr2:121944149-121944432 | -0.166757882 | 0.698092974 | 0.874854124 |
| chr12:109097227-109098648 | 0.122153165 | 0.698115782 | 0.874854124 |
| chr3:153138637-153139063 | -0.149556875 | 0.698135595 | 0.874854124 |
| chr12:54419152-54419569 | 0.150449349 | 0.698141289 | 0.874854124 |
| chr20:63106185-63106385 | 0.173120436 | 0.698148154 | 0.874854124 |
| chr17:81295216-81295879 | -0.132570137 | 0.698151859 | 0.874854124 |
| chr20:45892164-45892394 | 0.156885379 | 0.698162791 | 0.874854124 |
| chrX:81121532-81121983 | -0.13439917 | 0.698180657 | 0.874854124 |
| chr1:224042825-224043108 | 0.184948772 | 0.698200469 | 0.874854124 |

| | | | |
|---|---|---|---|
| chr8:77244514-77244929 | 0.150639702 | 0.698206558 | 0.874854124 |
| chr1:32964513-32965188 | 0.131367363 | 0.698207494 | 0.874854124 |
| chr2:29806459-29806869 | -0.147931624 | 0.698209451 | 0.874854124 |
| chr3:62622592-62622909 | 0.132907084 | 0.69823284 | 0.874869124 |
| chr6:142414350-142414550 | 0.220690161 | 0.698284243 | 0.874919223 |
| chr2:120437314-120437541 | 0.18008599 | 0.69830707 | 0.874933518 |
| chr2:88711947-88712410 | 0.151342514 | 0.698333691 | 0.874950255 |
| chr2:126245013-126245290 | -0.168421358 | 0.698343267 | 0.874950255 |
| chr6:27892985-27893572 | -0.127930978 | 0.698373347 | 0.874973635 |
| chr3:150183977-150184860 | -0.130272364 | 0.698401149 | 0.874977782 |
| chr5:16465152-16466471 | 0.122581636 | 0.698412146 | 0.874977782 |
| chr11:72553789-72554001 | -0.191695289 | 0.698421758 | 0.874977782 |
| chr1:46132439-46133681 | 0.123238265 | 0.698431477 | 0.874977782 |
| chr13:99498409-99498889 | -0.127581033 | 0.698433754 | 0.874977782 |
| chr4:148770380-148770580 | 0.203146461 | 0.69844599 | 0.874978805 |
| chr17:19338123-19338499 | -0.146248561 | 0.698480861 | 0.875008184 |
| chr21:33947471-33948372 | 0.130651666 | 0.698536584 | 0.875051655 |
| chr8:132120386-132120631 | -0.178172976 | 0.698538403 | 0.875051655 |
| chr12:60100859-60101222 | 0.152895423 | 0.698564 | 0.875069414 |
| chr1:114510059-114511693 | -0.120133649 | 0.698615042 | 0.875106534 |
| chr4:46271122-46271373 | 0.177745272 | 0.698616475 | 0.875106534 |
| chr3:163328698-163328898 | -0.204164308 | 0.698684821 | 0.875177837 |
| chr16:1387903-1388103 | 0.208821015 | 0.698714671 | 0.875200921 |
| chr4:87033971-87034189 | -0.194122313 | 0.698748249 | 0.875228672 |
| chr6:136791436-136791645 | -0.164618274 | 0.698798194 | 0.875276923 |
| chr11:9015653-9015884 | 0.180480777 | 0.698816781 | 0.875281816 |
| chr1:204226221-204226696 | -0.138131853 | 0.698836456 | 0.875281816 |
| chr6:33409961-33411054 | 0.12511236 | 0.698839502 | 0.875281816 |
| chr12:43675080-43676069 | 0.129650097 | 0.698847793 | 0.875281816 |
| chr16:70686775-70687276 | -0.142304032 | 0.69889757 | 0.87532789 |
| chr1:30782933-30783223 | 0.150175414 | 0.698907428 | 0.87532789 |
| chr12:114662641-114662853 | 0.184389461 | 0.698941781 | 0.875339454 |
| chr2:114551423-114551676 | -0.168982056 | 0.698942376 | 0.875339454 |
| chr19:41904193-41904799 | 0.135377867 | 0.69896612 | 0.875339454 |
| chr10:91370022-91370324 | -0.168470172 | 0.698980609 | 0.875339454 |
| chr1:242525632-242525883 | 0.205130388 | 0.698983378 | 0.875339454 |
| chr19:39411865-39412980 | 0.122885024 | 0.698985206 | 0.875339454 |
| chr16:70739056-70739256 | 0.192327339 | 0.699036959 | 0.875389956 |
| chr3:63926361-63926620 | 0.135837118 | 0.699061921 | 0.875406909 |
| chr18:59783140-59783543 | -0.153185847 | 0.699120541 | 0.875459032 |
| chr22:44498176-44498651 | -0.142621585 | 0.699133574 | 0.875459032 |
| chr5:140690672-140691854 | -0.124709164 | 0.699137822 | 0.875459032 |
| chr3:126389117-126389619 | 0.139134202 | 0.699153922 | 0.875460012 |
| chr1:18666364-18667011 | -0.152045834 | 0.699172576 | 0.875460012 |
| chr17:44198527-44200891 | 0.121050796 | 0.699182318 | 0.875460012 |
| chr3:173993439-173993684 | -0.172638547 | 0.699184307 | 0.875460012 |
| chr9:133147573-133147880 | -0.140387848 | 0.699220377 | 0.875490868 |
| chr18:8610214-8610442 | 0.168446448 | 0.699262751 | 0.875505514 |
| chr13:25037371-25037571 | -0.186396245 | 0.699265293 | 0.875505514 |
| chr4:139889304-139889553 | -0.187476421 | 0.699276917 | 0.875505514 |

| | | | |
|---|---|---|---|
| chr4:169634992-169635363 | -0.144242548 | 0.699277779 | 0.875505514 |
| chr1:107782291-107782730 | 0.152264265 | 0.699382855 | 0.875603436 |
| chr11:85029803-85030394 | 0.145716135 | 0.699407328 | 0.875603436 |
| chr8:144754160-144754360 | 0.157358702 | 0.699411448 | 0.875603436 |
| chr2:115083259-115083593 | -0.154874509 | 0.699422446 | 0.875603436 |
| chr1:231691719-231691940 | 0.177239739 | 0.699422632 | 0.875603436 |
| chr17:72556971-72557275 | 0.143314186 | 0.699424556 | 0.875603436 |
| chr10:17412459-17412788 | 0.176574364 | 0.699445472 | 0.875615315 |
| chr1:190727744-190728046 | 0.201586029 | 0.699474444 | 0.875634588 |
| chr3:148309305-148309935 | -0.135396237 | 0.699509969 | 0.875634588 |
| chr13:42110710-42110913 | -0.174841435 | 0.699519992 | 0.875634588 |
| chr8:124781222-124781944 | 0.128561715 | 0.699538577 | 0.875634588 |
| chr3:81496495-81497336 | -0.131291555 | 0.699548978 | 0.875634588 |
| chr9:91948540-91949333 | -0.133891774 | 0.699549189 | 0.875634588 |
| chr3:9374694-9374909 | 0.191987642 | 0.699559294 | 0.875634588 |
| chr8:119963370-119963573 | -0.206314356 | 0.699560394 | 0.875634588 |
| chr8:89689236-89689468 | 0.152609225 | 0.699563719 | 0.875634588 |
| chr2:88967879-88968163 | -0.168612116 | 0.699605068 | 0.875672038 |
| chr16:85290337-85290537 | -0.179528877 | 0.699619101 | 0.875674336 |
| chr20:54726743-54727025 | 0.133296482 | 0.69962976 | 0.875674336 |
| chr1:157377109-157377564 | -0.150764978 | 0.699669759 | 0.875705961 |
| chr17:75679733-75680231 | -0.142113717 | 0.699686986 | 0.875705961 |
| chr12:60689147-60689411 | -0.181096029 | 0.699690331 | 0.875705961 |
| chr2:32772673-32772996 | 0.168291665 | 0.699708934 | 0.875705961 |
| chr20:3807994-3808354 | -0.14203481 | 0.699729489 | 0.875705961 |
| chr14:80987234-80987806 | -0.137967188 | 0.69975395 | 0.875705961 |
| chr5:177005816-177007229 | 0.124217196 | 0.699757699 | 0.875705961 |
| chr18:48712039-48712251 | -0.198367712 | 0.699800704 | 0.875705961 |
| chr3:156450087-156450301 | -0.183396436 | 0.699802772 | 0.875705961 |
| chr4:56387012-56387990 | 0.124570206 | 0.699815151 | 0.875705961 |
| chr22:30635285-30635909 | 0.132634834 | 0.699815283 | 0.875705961 |
| chr8:62701700-62702033 | -0.19725803 | 0.699821746 | 0.875705961 |
| chr3:26244342-26245350 | -0.12649918 | 0.699839398 | 0.875705961 |
| chr11:2532695-2533020 | 0.154955759 | 0.699844297 | 0.875705961 |
| chr11:76680926-76681180 | -0.155207924 | 0.699850585 | 0.875705961 |
| chr15:64093139-64093586 | 0.135577338 | 0.699852803 | 0.875705961 |
| chr12:98325232-98325468 | 0.18650179 | 0.69987208 | 0.875705961 |
| chrX:140056890-140057451 | 0.137774974 | 0.69988172 | 0.875705961 |
| chr3:7505264-7505480 | 0.193917697 | 0.699894729 | 0.875705961 |
| chr1:67832098-67832456 | -0.144122897 | 0.699896329 | 0.875705961 |
| chr10:5466044-5467133 | -0.123871255 | 0.699899481 | 0.875705961 |
| chr10:128011230-128011683 | 0.13843413 | 0.699916964 | 0.875705961 |
| chr17:60656542-60656948 | -0.129218349 | 0.699917892 | 0.875705961 |
| chr7:8072558-8072758 | -0.195327509 | 0.699956912 | 0.875739707 |
| chr12:6336120-6336915 | -0.124800615 | 0.699967723 | 0.875739707 |
| chr10:6416950-6417258 | 0.16490665 | 0.699980427 | 0.875741302 |
| chr3:154040332-154040657 | -0.149364285 | 0.700023464 | 0.875765556 |
| chr19:41959228-41959772 | 0.133321001 | 0.70002527 | 0.875765556 |
| chr5:179806706-179807154 | -0.134620449 | 0.700034102 | 0.875765556 |
| chrX:126165938-126166556 | -0.143468391 | 0.700071607 | 0.875798176 |

| | | | |
|---|---|---|---|
| chr7:47693056-47693728 | 0.132874234 | 0.700093392 | 0.875811129 |
| chr19:18006990-18007245 | -0.182350443 | 0.700118555 | 0.875828309 |
| chr4:132564884-132565564 | 0.132170239 | 0.700134459 | 0.875829736 |
| chr3:127683669-127683924 | 0.180783268 | 0.700142557 | 0.875829736 |
| chr7:6503384-6504714 | -0.120604727 | 0.70016533 | 0.875843924 |
| chr10:131246927-131247159 | 0.201345753 | 0.700205687 | 0.875880109 |
| chr1:193229298-193229636 | -0.165002232 | 0.700267848 | 0.875926943 |
| chr10:46256112-46256379 | 0.16660342 | 0.700271146 | 0.875926943 |
| chr3:179672950-179673281 | -0.151679244 | 0.700288159 | 0.875926943 |
| chr2:223957162-223957706 | 0.141020605 | 0.700288855 | 0.875926943 |
| chr10:71916011-71916231 | 0.169923493 | 0.700409926 | 0.876064078 |
| chr1:19311439-19312383 | 0.129112492 | 0.700433627 | 0.876079421 |
| chr2:216633057-216634066 | 0.135167531 | 0.700487195 | 0.876127301 |
| chr1:245516266-245516728 | -0.153814223 | 0.700500427 | 0.876127301 |
| chr8:125787736-125788093 | 0.147984358 | 0.700506211 | 0.876127301 |
| chr2:12717327-12717764 | -0.142407598 | 0.700550174 | 0.87616208 |
| chr6:108248059-108248281 | -0.18832296 | 0.700556888 | 0.87616208 |
| chr4:153152946-153153936 | 0.127417363 | 0.700584727 | 0.876178656 |
| chr15:92391656-92391968 | -0.16170049 | 0.700610749 | 0.876178656 |
| chr11:78852017-78852220 | -0.181204074 | 0.70061274 | 0.876178656 |
| chr11:12728417-12728681 | 0.159208564 | 0.700625365 | 0.876178656 |
| chr15:101008276-101009165 | -0.124563713 | 0.700629968 | 0.876178656 |
| chr6:42960282-42960881 | 0.135252837 | 0.700650977 | 0.876178656 |
| chr5:150263449-150263675 | 0.180890911 | 0.700660653 | 0.876178656 |
| chr3:113848115-113848388 | -0.169542588 | 0.700661622 | 0.876178656 |
| chr8:144137163-144137933 | 0.124937852 | 0.700675174 | 0.876181303 |
| chr11:116276383-116276914 | -0.143506725 | 0.70069097 | 0.876186756 |
| chr1:221478104-221478393 | 0.166795732 | 0.700709604 | 0.876195758 |
| chr1:200454723-200454923 | 0.181671071 | 0.700783547 | 0.87626784 |
| chr6:129500938-129501184 | 0.181612504 | 0.700800262 | 0.87626784 |
| chr7:5183278-5183478 | 0.181585166 | 0.700801558 | 0.87626784 |
| chr14:79011136-79011377 | 0.189922836 | 0.70082493 | 0.876281824 |
| chr22:39292529-39292963 | -0.14738818 | 0.700835615 | 0.876281824 |
| chr19:38300669-38301071 | 0.143233069 | 0.70084957 | 0.876284974 |
| chr19:3979673-3979908 | -0.176492961 | 0.700882622 | 0.876310847 |
| chr3:164490873-164491073 | -0.199779182 | 0.700893691 | 0.876310847 |
| chr3:197736095-197737334 | -0.123117895 | 0.700904574 | 0.876310847 |
| chr2:200525835-200526756 | -0.123431029 | 0.700957134 | 0.876333045 |
| chr5:102216878-102217109 | 0.186711557 | 0.700979438 | 0.876333045 |
| chr10:8670476-8670790 | 0.167949762 | 0.700980867 | 0.876333045 |
| chr15:40045108-40045432 | -0.144075826 | 0.70098717 | 0.876333045 |
| chr2:112795689-112796007 | 0.151664016 | 0.7010189 | 0.876333045 |
| chr1:112618793-112619293 | -0.135851644 | 0.701019903 | 0.876333045 |
| chr11:70695556-70695756 | 0.187706924 | 0.701028829 | 0.876333045 |
| chr14:20247739-20247948 | 0.187706924 | 0.701028829 | 0.876333045 |
| chr14:85611873-85612087 | 0.187706924 | 0.701028829 | 0.876333045 |
| chrX:70043039-70043283 | 0.186781259 | 0.7010367 | 0.876333045 |
| chr3:128584396-128584803 | -0.137972263 | 0.701061176 | 0.876349345 |
| chr13:22103083-22103315 | -0.196951712 | 0.701072735 | 0.876349496 |
| chr8:106755159-106755496 | -0.168301628 | 0.701109374 | 0.876380998 |

| | | | |
|---|---|---|---|
| chr2:73983412-73984361 | -0.123299416 | 0.701166421 | 0.876438008 |
| chr4:106369450-106369692 | -0.190113175 | 0.701178781 | 0.876439161 |
| chrX:75155948-75156520 | 0.137060751 | 0.701199064 | 0.876450216 |
| chrX:48696272-48697162 | 0.126579579 | 0.701248395 | 0.876497577 |
| chr21:29348611-29348816 | 0.191811908 | 0.701281465 | 0.876524613 |
| chr6:8131298-8131644 | 0.149130595 | 0.701301801 | 0.876525912 |
| chr13:97432850-97434168 | -0.120517268 | 0.701311283 | 0.876525912 |
| chr12:92899301-92899651 | -0.162754077 | 0.701316823 | 0.876525912 |
| chr9:135282061-135282581 | 0.144013431 | 0.701328962 | 0.876526787 |
| chr13:41011433-41011658 | -0.182346786 | 0.701340739 | 0.876527209 |
| chr1:115559163-115559363 | 0.198576386 | 0.701354233 | 0.876529776 |
| chr6:88780752-88781075 | -0.166191514 | 0.701425694 | 0.876594505 |
| chr9:128420585-128421643 | 0.126602444 | 0.70143772 | 0.876594505 |
| chr10:8407378-8408145 | -0.134616306 | 0.701440347 | 0.876594505 |
| chr6:2988321-2989351 | 0.122741517 | 0.701481264 | 0.876631341 |
| chr2:9475241-9475441 | -0.173761002 | 0.701512191 | 0.876655692 |
| chr22:49853362-49853869 | -0.132599328 | 0.701542075 | 0.876678739 |
| chr1:54354104-54354332 | -0.187358842 | 0.701566795 | 0.876695331 |
| chr11:27328701-27328908 | 0.218077814 | 0.70158899 | 0.876708769 |
| chr18:24076283-24076595 | 0.166535465 | 0.701684095 | 0.876796866 |
| chr1:200908525-200909160 | 0.131269687 | 0.701685594 | 0.876796866 |
| chr19:11434817-11435753 | -0.123920209 | 0.701702384 | 0.876796866 |
| chr3:161208971-161209180 | 0.178643824 | 0.701705263 | 0.876796866 |
| chr9:76587088-76587372 | 0.169063722 | 0.701751978 | 0.876840939 |
| chr6:93965342-93965542 | 0.217966502 | 0.70180136 | 0.876884675 |
| chr15:72628991-72629384 | -0.16445541 | 0.701809869 | 0.876884675 |
| chr15:22238404-22238845 | -0.164589771 | 0.701824041 | 0.876887562 |
| chr19:17576836-17577057 | 0.167663184 | 0.701836965 | 0.876887562 |
| chr4:19913714-19913959 | 0.177803504 | 0.701846513 | 0.876887562 |
| chr8:110832425-110832950 | 0.130920584 | 0.701913911 | 0.876957469 |
| chr22:21952519-21953119 | -0.134194887 | 0.701937622 | 0.876959464 |
| chr7:35520095-35520540 | 0.140835013 | 0.701944365 | 0.876959464 |
| chr2:159821854-159822443 | -0.15571237 | 0.701949843 | 0.876959464 |
| chr12:68584561-68585144 | 0.134617668 | 0.701967505 | 0.87696723 |
| chr7:876016-877185 | -0.122303646 | 0.702057187 | 0.87706497 |
| chr9:36407139-36407377 | -0.175987816 | 0.702084876 | 0.877085261 |
| chr12:96047658-96047872 | -0.204999787 | 0.702140518 | 0.877130989 |
| chr5:18523060-18523311 | 0.189474362 | 0.702144376 | 0.877130989 |
| chr2:96353096-96353333 | 0.191528406 | 0.702212502 | 0.877185807 |
| chr17:41417612-41417841 | 0.167796566 | 0.702215122 | 0.877185807 |
| chr4:109453192-109453482 | 0.160028735 | 0.702222602 | 0.877185807 |
| chr14:101964273-101964834 | -0.129697024 | 0.702267049 | 0.877192732 |
| chr10:92419995-92420360 | 0.158065597 | 0.702273678 | 0.877192732 |
| chr11:46382009-46382233 | -0.151934522 | 0.702277614 | 0.877192732 |
| chr14:70622213-70622417 | 0.18895621 | 0.702278847 | 0.877192732 |
| chr15:59917437-59917869 | -0.150584333 | 0.702294822 | 0.877192732 |
| chr7:34958468-34958668 | -0.208086924 | 0.702296836 | 0.877192732 |
| chr7:1368813-1369512 | 0.128150722 | 0.702350037 | 0.877244822 |
| chr6:1793022-1793222 | -0.204857295 | 0.702366008 | 0.877244822 |
| chr11:47578415-47579321 | 0.123213835 | 0.702372887 | 0.877244822 |

| | | | |
|---|---|---|---|
| chr1:24392257-24392551 | 0.176830338 | 0.702389126 | 0.877250805 |
| chr17:59892215-59893914 | -0.118203696 | 0.702409504 | 0.877253924 |
| chr19:13843554-13843859 | -0.152165744 | 0.702414521 | 0.877253924 |
| chr5:77697790-77698044 | -0.211880049 | 0.702440701 | 0.87727232 |
| chr4:15681267-15682308 | 0.121851523 | 0.702490292 | 0.877319955 |
| chr7:52154446-52154701 | -0.171794311 | 0.702515065 | 0.877336592 |
| chr7:22418386-22418626 | 0.185874235 | 0.70253036 | 0.877341394 |
| chr12:77969492-77969892 | 0.171224706 | 0.702583614 | 0.877381995 |
| chr5:77210394-77210594 | -0.220291385 | 0.702585772 | 0.877381995 |
| chr8:122549279-122549562 | -0.157186298 | 0.702618286 | 0.877408298 |
| chr11:69409708-69409938 | -0.199796404 | 0.702656605 | 0.877441849 |
| chr5:94043414-94043679 | -0.194847491 | 0.702675112 | 0.877450659 |
| chr6:28695474-28696517 | 0.127241326 | 0.702715207 | 0.877486426 |
| chr6:107973634-107973963 | -0.157364617 | 0.702743706 | 0.877502469 |
| chr5:180090423-180090832 | 0.147708313 | 0.702752124 | 0.877502469 |
| chr8:53851476-53851676 | 0.1982815 | 0.702772227 | 0.877502469 |
| chr1:207789918-207790135 | -0.196905973 | 0.702773864 | 0.877502469 |
| chr20:62220289-62220536 | 0.154145996 | 0.702790689 | 0.877508497 |
| chr8:143048067-143048524 | 0.142763521 | 0.702807801 | 0.877508497 |
| chr7:3909363-3909568 | 0.199577412 | 0.702813435 | 0.877508497 |
| chr8:122348187-122348660 | 0.159959117 | 0.702827342 | 0.877508497 |
| chr22:20507272-20507707 | 0.140064373 | 0.702835954 | 0.877508497 |
| chrX:101348200-101349221 | -0.122351865 | 0.702879665 | 0.877548772 |
| chr6:48916541-48916908 | 0.15539465 | 0.702901355 | 0.877561553 |
| chr14:60625183-60626054 | -0.129729491 | 0.70292174 | 0.877572084 |
| chrX:40575406-40575723 | 0.17184598 | 0.702932697 | 0.877572084 |
| chr20:10674751-10675048 | 0.13403302 | 0.702969657 | 0.877603927 |
| chr4:103948659-103948945 | 0.161873487 | 0.70298559 | 0.877609519 |
| chr2:61888138-61889184 | -0.123613107 | 0.703026331 | 0.877646081 |
| chr2:2110869-2111083 | -0.19116462 | 0.703046509 | 0.877656972 |
| chr8:129731134-129731486 | -0.141259794 | 0.703077228 | 0.87768102 |
| chr14:101809695-10181051[illegible] | -0.127344055 | 0.703097572 | 0.877692117 |
| chr16:660407-661214 | 0.128003684 | 0.70314805 | 0.877737655 |
| chr3:5144408-5145082 | -0.131345956 | 0.703158475 | 0.877737655 |
| chr10:50471646-50471898 | 0.160976695 | 0.703168418 | 0.877737655 |
| chr11:70227119-70227319 | 0.182585735 | 0.703208281 | 0.877773115 |
| chr5:110058188-110058458 | 0.172965059 | 0.703234722 | 0.877783648 |
| chr12:50088664-50088885 | 0.15857405 | 0.703241527 | 0.877783648 |
| chrX:47650172-47651228 | 0.120990557 | 0.703254369 | 0.877783648 |
| chr7:47721599-47721994 | 0.142900846 | 0.703262543 | 0.877783648 |
| chr1:29181127-29182376 | -0.122414605 | 0.703286542 | 0.877791625 |
| chr3:81249521-81250042 | -0.132383424 | 0.703300211 | 0.877791625 |
| chr3:166182595-166182838 | -0.185830167 | 0.703303302 | 0.877791625 |
| chr5:181260684-181262303 | 0.120372247 | 0.703358133 | 0.87784576 |
| chr1:84782475-84782814 | -0.144591229 | 0.703406933 | 0.877867866 |
| chr22:31253884-31254377 | 0.138894585 | 0.703414247 | 0.877867866 |
| chr10:48656238-48656655 | -0.136419603 | 0.70343806 | 0.877867866 |
| chr1:207091588-207091788 | 0.178245317 | 0.703459395 | 0.877867866 |
| chr16:3910307-3910507 | 0.212289531 | 0.703475375 | 0.877867866 |
| chr2:219277663-219278027 | 0.136793755 | 0.703477414 | 0.877867866 |

| | | | |
|---|---|---|---|
| chr3:47781324-47782534 | -0.121730764 | 0.703492582 | 0.877867866 |
| chr5:165630233-165630723 | -0.139236709 | 0.703499376 | 0.877867866 |
| chr19:44212235-44212945 | -0.13280565 | 0.703517212 | 0.877867866 |
| chr1:77978139-77978494 | -0.130140473 | 0.70352566 | 0.877867866 |
| chr4:121898211-121898474 | 0.165944508 | 0.703527727 | 0.877867866 |
| chr15:34337245-34338332 | -0.123264506 | 0.70354029 | 0.877867866 |
| chr5:59174354-59174568 | -0.200814343 | 0.703553005 | 0.877867866 |
| chr10:13847271-13847494 | -0.176108935 | 0.703554188 | 0.877867866 |
| chr12:133079917-133081173 | 0.120335573 | 0.703557382 | 0.877867866 |
| chr8:133202903-133203257 | -0.138342681 | 0.70355916 | 0.877867866 |
| chr2:106193783-106194714 | 0.122030552 | 0.703631968 | 0.877939333 |
| chr16:75565041-75565632 | -0.126461072 | 0.703651732 | 0.877939333 |
| chr3:130705518-130705813 | 0.159529841 | 0.703661304 | 0.877939333 |
| chr11:10293511-10294352 | 0.134860512 | 0.703671369 | 0.877939333 |
| chr6:151334755-151335122 | -0.159622263 | 0.703673726 | 0.877939333 |
| chr1:107964549-107965315 | 0.127904529 | 0.703687453 | 0.877942163 |
| chr2:95115399-95115792 | 0.147515514 | 0.703716507 | 0.877959155 |
| chr8:92475978-92476219 | -0.197334931 | 0.703723988 | 0.877959155 |
| chr20:22691735-22692137 | 0.153864261 | 0.703753851 | 0.877982115 |
| chr6:167750784-167751424 | -0.131940181 | 0.7037883 | 0.877998003 |
| chr8:79799503-79799982 | 0.138890183 | 0.703789504 | 0.877998003 |
| chr2:222884907-222885322 | 0.168605585 | 0.703816928 | 0.87801792 |
| chr6:35259079-35260234 | 0.121243931 | 0.7038661 | 0.878064966 |
| chr9:126582324-126582957 | -0.138263965 | 0.703887002 | 0.878076746 |
| chr14:20460961-20461259 | 0.133405916 | 0.703943955 | 0.878121606 |
| chr3:196342977-196343816 | 0.126358788 | 0.703945884 | 0.878121606 |
| chr4:38062696-38062896 | -0.200155352 | 0.704005258 | 0.878181374 |
| chr10:100516801-100517013 | -0.193876451 | 0.70402517 | 0.878191915 |
| chr10:114697089-114697683 | 0.144796125 | 0.704048414 | 0.878195511 |
| chr5:119470146-119470346 | 0.212121256 | 0.704050976 | 0.878195511 |
| chr1:223456137-223456461 | 0.179499098 | 0.704132406 | 0.878269942 |
| chr4:98642971-98643453 | -0.134013932 | 0.704134715 | 0.878269942 |
| chr17:66007689-66007898 | -0.206988666 | 0.704145034 | 0.878269942 |
| chr7:89545461-89545732 | -0.166088293 | 0.704159262 | 0.878273391 |
| chr3:132659366-132660678 | -0.121920966 | 0.704197269 | 0.878301728 |
| chr15:73992143-73992541 | 0.132641581 | 0.704204907 | 0.878301728 |
| chr11:82250151-82250388 | 0.178458688 | 0.704260932 | 0.878350695 |
| chr9:114397837-114398822 | 0.121145614 | 0.704267094 | 0.878350695 |
| chrX:40013019-40014606 | 0.121840147 | 0.704293218 | 0.878368979 |
| chr11:65491684-65491999 | 0.145204185 | 0.704331695 | 0.878379932 |
| chr11:42354063-42354263 | -0.20622636 | 0.704335298 | 0.878379932 |
| chr17:67717344-67718696 | -0.119214196 | 0.704361124 | 0.878379932 |
| chr17:60134833-60135947 | 0.118749059 | 0.704362919 | 0.878379932 |
| chr19:8307758-8308490 | 0.130999219 | 0.704363204 | 0.878379932 |
| chrX:86147606-86148628 | -0.124660881 | 0.704370784 | 0.878379932 |
| chr9:84404210-84404410 | 0.194237279 | 0.704385176 | 0.878383187 |
| chr2:35112833-35113075 | -0.173126957 | 0.704396321 | 0.878383187 |
| chr7:6965135-6965422 | -0.154942801 | 0.704436742 | 0.878419296 |
| chr11:93234663-93234949 | 0.158390261 | 0.704518307 | 0.878506709 |
| chr9:128771134-128771388 | -0.170067051 | 0.704533508 | 0.878511367 |

| | | | |
|---|---|---|---|
| chr17:68541319-68541537 | -0.162948372 | 0.704551388 | 0.878519037 |
| chr15:75758384-75758891 | -0.134953281 | 0.704575872 | 0.878519037 |
| chr6:75748577-75749845 | 0.120464148 | 0.704616725 | 0.878519037 |
| chr17:40168038-40168707 | 0.138217679 | 0.704618033 | 0.878519037 |
| chr19:2900431-2901074 | 0.130968138 | 0.704619801 | 0.878519037 |
| chr1:104132466-104132685 | -0.18639757 | 0.704625239 | 0.878519037 |
| chr7:25124887-25125955 | -0.119934493 | 0.704631206 | 0.878519037 |
| chr11:44979943-44980279 | -0.145775883 | 0.704632496 | 0.878519037 |
| chr14:94143369-94143586 | -0.200064437 | 0.70464285 | 0.878519037 |
| chr15:65982113-65982355 | -0.154228953 | 0.704696002 | 0.878547469 |
| chr15:66862717-66863203 | -0.135479765 | 0.704696194 | 0.878547469 |
| chr7:64758066-64758634 | -0.136015086 | 0.704725376 | 0.878547469 |
| chr10:48655802-48656094 | -0.161551632 | 0.704726906 | 0.878547469 |
| chr16:11172855-11173055 | -0.182017433 | 0.704731617 | 0.878547469 |
| chr3:75294160-75294550 | -0.139969648 | 0.704734451 | 0.878547469 |
| chr5:6823032-6823545 | 0.1426863 | 0.70474728 | 0.878549168 |
| chr4:76219451-76219745 | -0.177903945 | 0.704767767 | 0.878560414 |
| chr1:174652776-174652976 | 0.194316495 | 0.704832459 | 0.878626764 |
| chr2:200483701-200484220 | 0.136271651 | 0.704862373 | 0.87864976 |
| chr8:61714037-61715061 | -0.120267295 | 0.704893445 | 0.878666052 |
| chr9:90947103-90947358 | 0.171357786 | 0.704898378 | 0.878666052 |
| chr3:114056143-114057288 | -0.119324294 | 0.704911273 | 0.87866783 |
| chr10:90751449-90751664 | 0.178113648 | 0.704928896 | 0.878675504 |
| chr2:157825723-157826387 | -0.126842955 | 0.704950039 | 0.878685221 |
| chr22:40683093-40683355 | 0.176052177 | 0.704965967 | 0.878685221 |
| chr5:134176242-134177558 | -0.121855716 | 0.704979233 | 0.878685221 |
| chr4:48830721-48831635 | 0.120540949 | 0.704989869 | 0.878685221 |
| chr10:63217184-63217389 | -0.193279989 | 0.705002833 | 0.878685221 |
| chr3:75604449-75604861 | 0.140977694 | 0.705008508 | 0.878685221 |
| chr2:23375461-23376003 | -0.142492167 | 0.705023445 | 0.878685221 |
| chr20:42678761-42679228 | -0.137798181 | 0.705028434 | 0.878685221 |
| chr3:105997459-105997659 | 0.204282941 | 0.705041836 | 0.878687631 |
| chr3:101744728-101745051 | 0.166535492 | 0.705081995 | 0.878723388 |
| chr20:63706862-63707092 | -0.150889907 | 0.705116303 | 0.878751852 |
| chr2:120222906-120223971 | 0.121551197 | 0.705160294 | 0.878792382 |
| chr12:80186642-80187155 | -0.130652528 | 0.705193056 | 0.878818918 |
| chr1:205335624-205336037 | 0.129856323 | 0.705208901 | 0.87882437 |
| chr21:28992116-28992316 | 0.161095272 | 0.705224629 | 0.878826758 |
| chrX:94058661-94059074 | -0.151544961 | 0.705233756 | 0.878826758 |
| chr12:113161435-113161961 | 0.131238149 | 0.705246233 | 0.878828013 |
| chr12:6423588-6424097 | 0.14112499 | 0.705285372 | 0.878862492 |
| chr15:55775292-55775947 | 0.125015316 | 0.705305273 | 0.878872997 |
| chr9:129625983-129626614 | 0.127535587 | 0.705321915 | 0.878879442 |
| chr17:74872667-74873778 | 0.119292303 | 0.705371725 | 0.878927215 |
| chr2:45009195-45010480 | 0.120341348 | 0.705388197 | 0.878933447 |
| chr1:207329408-207329749 | 0.154359051 | 0.70540644 | 0.878941803 |
| chr11:110429710-110430867 | -0.125356321 | 0.705426854 | 0.878941803 |
| chr1:37940335-37940774 | 0.143938596 | 0.705440033 | 0.878941803 |
| chr14:80941187-80942087 | -0.120558394 | 0.705465949 | 0.878941803 |
| chr15:91224812-91225018 | 0.176151344 | 0.705471236 | 0.878941803 |

| | | | |
|---|---|---|---|
| chr9:126804766-126805343 | 0.127153217 | 0.705509715 | 0.878941803 |
| chr3:31704682-31705291 | -0.137552009 | 0.705520391 | 0.878941803 |
| chr6:27295276-27295616 | 0.14561258 | 0.705528594 | 0.878941803 |
| chr2:106582541-106582874 | 0.142191206 | 0.705530933 | 0.878941803 |
| chr3:62718709-62719297 | -0.130396864 | 0.705531807 | 0.878941803 |
| chr17:73998672-73998940 | 0.160856757 | 0.70554031 | 0.878941803 |
| chr11:3809135-3809423 | -0.142765576 | 0.705542315 | 0.878941803 |
| chr15:45705513-45705976 | 0.136739531 | 0.705561335 | 0.878941803 |
| chr2:238864265-238864609 | 0.147731097 | 0.705572124 | 0.878941803 |
| chr13:28100538-28100875 | -0.153650833 | 0.705573152 | 0.878941803 |
| chr19:3163419-3163619 | -0.190555491 | 0.70558805 | 0.878941803 |
| chr12:88425014-88425264 | -0.202326382 | 0.705589975 | 0.878941803 |
| chr1:153465784-153466058 | 0.17685497 | 0.705621693 | 0.878941803 |
| chr5:142982098-142983140 | -0.132721813 | 0.705634547 | 0.878941803 |
| chr9:83817148-83818367 | 0.124280325 | 0.705638585 | 0.878941803 |
| chr12:130716243-130716721 | 0.14462761 | 0.705663825 | 0.878941803 |
| chr3:125582778-125583082 | 0.161473604 | 0.705665794 | 0.878941803 |
| chr1:181937013-181937787 | -0.133160212 | 0.705690316 | 0.878941803 |
| chr8:140393810-140394117 | -0.147993458 | 0.705717658 | 0.878941803 |
| chr8:128887295-128887741 | -0.153279402 | 0.705728652 | 0.878941803 |
| chr1:112716266-112716611 | -0.129436483 | 0.705745638 | 0.878941803 |
| chr14:70567913-70568210 | 0.155552089 | 0.705749295 | 0.878941803 |
| chr1:111920650-111920866 | 0.146703126 | 0.705771074 | 0.878941803 |
| chr3:13861500-13861767 | -0.165553333 | 0.705784678 | 0.878941803 |
| chr1:161612328-161613300 | -0.125546957 | 0.70578841 | 0.878941803 |
| chr17:72540228-72540640 | -0.139110376 | 0.705788505 | 0.878941803 |
| chr13:53198135-53198397 | -0.173559722 | 0.705797677 | 0.878941803 |
| chr4:37793620-37793820 | -0.191352252 | 0.705802253 | 0.878941803 |
| chr7:8090491-8091005 | 0.129588126 | 0.705817873 | 0.878941803 |
| chr22:38985153-38985353 | -0.19756953 | 0.705820105 | 0.878941803 |
| chr21:45287730-45288408 | -0.124243882 | 0.70582637 | 0.878941803 |
| chr11:70202972-70203839 | -0.125548923 | 0.705827214 | 0.878941803 |
| chr18:79393778-79394095 | -0.154702173 | 0.705830807 | 0.878941803 |
| chr6:138287200-138287551 | -0.1425094 | 0.705868334 | 0.878963867 |
| chr11:63847376-63847695 | 0.151081506 | 0.70587669 | 0.878963867 |
| chr2:154493135-154493402 | -0.195077213 | 0.70588294 | 0.878963867 |
| chr12:131583327-131583527 | -0.203576483 | 0.705925489 | 0.879002565 |
| chr10:109956720-109957007 | 0.165021042 | 0.705948955 | 0.879005863 |
| chr5:157761985-157762279 | 0.151859939 | 0.705951082 | 0.879005863 |
| chr12:6535082-6535312 | 0.15463979 | 0.705962955 | 0.879006363 |
| chr14:98460823-98461035 | -0.205653437 | 0.705992162 | 0.879028445 |
| chr9:13023587-13023930 | 0.173817425 | 0.706004324 | 0.879029303 |
| chr20:52298008-52298209 | 0.190630317 | 0.70605469 | 0.879038765 |
| chr6:99330137-99330399 | -0.157234365 | 0.706060076 | 0.879038765 |
| chr4:41178425-41178625 | -0.205508396 | 0.706068615 | 0.879038765 |
| chr2:157782139-157782340 | 0.180686514 | 0.706092841 | 0.879038765 |
| chr2:127701133-127701630 | -0.126858126 | 0.706107929 | 0.879038765 |
| chr8:66712232-66713205 | 0.126875166 | 0.706120723 | 0.879038765 |
| chr2:152334919-152335827 | -0.124135734 | 0.706127501 | 0.879038765 |
| chr13:46206261-46206524 | -0.164754772 | 0.706135756 | 0.879038765 |

| | | | |
|---|---|---|---|
| chr5:119021934-119022173 | -0.171984415 | 0.706152807 | 0.879038765 |
| chr20:52192074-52192610 | 0.129852237 | 0.706154147 | 0.879038765 |
| chr6:170115381-170115793 | -0.15550202 | 0.706162258 | 0.879038765 |
| chr10:11874530-11875043 | -0.130690263 | 0.706181966 | 0.879038765 |
| chr2:106200678-106200939 | 0.193224807 | 0.706184101 | 0.879038765 |
| chr8:89535165-89535923 | 0.124672899 | 0.706191904 | 0.879038765 |
| chr2:212673897-212674186 | 0.17817682 | 0.706194034 | 0.879038765 |
| chr11:118263693-118264595 | 0.129516186 | 0.706195571 | 0.879038765 |
| chr3:194070590-194071447 | 0.12492765 | 0.706206954 | 0.879038765 |
| chr6:157322996-157324352 | -0.122319556 | 0.706255378 | 0.879076904 |
| chr9:7799226-7799976 | -0.129347314 | 0.706260541 | 0.879076904 |
| chr3:107525170-107525446 | -0.144714846 | 0.706273944 | 0.879079307 |
| chr11:18522516-18522716 | 0.162582042 | 0.70630857 | 0.879085748 |
| chr16:30534701-30535144 | -0.144679175 | 0.706312371 | 0.879085748 |
| chr1:11680600-11681749 | 0.127432519 | 0.706313538 | 0.879085748 |
| chr6:33794895-33795328 | 0.144962146 | 0.706358389 | 0.879098001 |
| chr12:104177447-104178545 | -0.123693078 | 0.706361817 | 0.879098001 |
| chr7:65037703-65038541 | -0.124878317 | 0.706364935 | 0.879098001 |
| chr9:103242115-103242315 | -0.200516222 | 0.70637994 | 0.879098001 |
| chr7:158587718-158588054 | -0.143103374 | 0.706380749 | 0.879098001 |
| chr6:111482166-111482956 | -0.127966609 | 0.706413843 | 0.879114946 |
| chr20:3796837-3797053 | 0.153040966 | 0.706422706 | 0.879114946 |
| chr3:130012249-130012533 | 0.184245169 | 0.706428785 | 0.879114946 |
| chr6:87589571-87590480 | -0.121934049 | 0.706457592 | 0.879136517 |
| chr3:120095513-120095892 | 0.124005898 | 0.706477163 | 0.879142524 |
| chr3:7487901-7488101 | 0.206202167 | 0.706485366 | 0.879142524 |
| chr17:63549666-63551242 | -0.118046007 | 0.706524417 | 0.879153285 |
| chr7:46764346-46764607 | -0.179180293 | 0.706533174 | 0.879153285 |
| chr1:53954483-53954994 | 0.138026403 | 0.706533281 | 0.879153285 |
| chrX:135032759-135033003 | 0.162030079 | 0.706548787 | 0.879153285 |
| chr12:119804309-119804651 | 0.147228963 | 0.706551384 | 0.879153285 |
| chr20:49431616-49431816 | -0.190342914 | 0.706596589 | 0.879179013 |
| chr19:16957127-16957432 | -0.161557896 | 0.706596741 | 0.879179013 |
| chr11:93648272-93648587 | 0.159322099 | 0.706606484 | 0.879179013 |
| chr21:38699123-38699324 | 0.170292691 | 0.70662144 | 0.879183346 |
| chr7:6546817-6547203 | -0.131567067 | 0.706646857 | 0.879200693 |
| chr20:56606299-56606921 | 0.126825868 | 0.706683968 | 0.87923259 |
| chr21:41650897-41651131 | -0.182190902 | 0.706763363 | 0.879317092 |
| chr3:63993588-63994195 | -0.119080042 | 0.706889814 | 0.879460135 |
| chr12:64221522-64222748 | -0.119742726 | 0.706918084 | 0.879469972 |
| chr6:136608214-136608716 | -0.139827914 | 0.706920677 | 0.879469972 |
| chrX:57591713-57593173 | 0.12480515 | 0.706946714 | 0.879479966 |
| chr15:67813755-67813955 | -0.199673443 | 0.706956475 | 0.879479966 |
| chr8:88415393-88415906 | 0.130143154 | 0.706963145 | 0.879479966 |
| chr6:54717868-54718163 | -0.15041066 | 0.706985078 | 0.879492973 |
| chr16:73878641-73878910 | -0.171250584 | 0.707006219 | 0.879494381 |
| chr19:45450659-45451229 | -0.129252115 | 0.70701117 | 0.879494381 |
| chr16:85554468-85554704 | -0.146839608 | 0.707037785 | 0.879494381 |
| chr1:36306084-36307442 | 0.120208982 | 0.707046539 | 0.879494381 |
| chr8:120444433-120445660 | -0.121405235 | 0.707052276 | 0.879494381 |

| | | | |
|---|---|---|---|
| chr3:197749347-197750255 | 0.123170308 | 0.707059714 | 0.879494381 |
| chr5:174935792-174936203 | 0.140346015 | 0.707078901 | 0.879494381 |
| chr19:925369-925843 | -0.134515179 | 0.707086571 | 0.879494381 |
| chr3:150497876-150498503 | 0.131545678 | 0.707093938 | 0.879494381 |
| chr17:63961746-63962148 | -0.131383757 | 0.707100994 | 0.879494381 |
| chr4:122587946-122588173 | 0.193828088 | 0.707162798 | 0.879519057 |
| chr4:5980066-5980274 | 0.193828088 | 0.707162798 | 0.879519057 |
| chr10:127524269-127524524 | 0.175243445 | 0.707167554 | 0.879519057 |
| chr20:31514177-31515278 | -0.121392029 | 0.707177796 | 0.879519057 |
| chr16:81214797-81215166 | -0.138542215 | 0.707178226 | 0.879519057 |
| chr18:2570697-2572293 | -0.120602887 | 0.707215919 | 0.879551659 |
| chr7:5440027-5440235 | 0.165925754 | 0.707228358 | 0.879552852 |
| chr1:213419346-213419609 | 0.194886747 | 0.707266713 | 0.879567277 |
| chr11:45105524-45105804 | -0.16836946 | 0.707279659 | 0.879567277 |
| chr9:76658576-76659103 | -0.140670855 | 0.70728489 | 0.879567277 |
| chr11:32893263-32894427 | -0.123457412 | 0.707285874 | 0.879567277 |
| chr1:109193759-109193959 | 0.169822542 | 0.707328027 | 0.879601095 |
| chr4:82900100-82901464 | -0.118219671 | 0.707339603 | 0.879601095 |
| chr17:62860678-62861891 | 0.120086547 | 0.707351532 | 0.879601095 |
| chr10:98916812-98917126 | -0.15541965 | 0.707360582 | 0.879601095 |
| chr9:128847164-128847487 | 0.164301473 | 0.7073792 | 0.879601095 |
| chr20:53773485-53774935 | 0.116736183 | 0.707385947 | 0.879601095 |
| chr1:151416319-151416915 | -0.131579791 | 0.707393426 | 0.879601095 |
| chr20:37178849-37179693 | 0.125287816 | 0.707413688 | 0.879612015 |
| chr5:174951211-174951523 | 0.152323139 | 0.707479229 | 0.879679234 |
| chr20:47809626-47810076 | 0.122144443 | 0.707553597 | 0.879731796 |
| chr2:129923946-129924182 | -0.166495239 | 0.707571707 | 0.879731796 |
| chr1:106285246-106285897 | -0.12877745 | 0.70757888 | 0.879731796 |
| chr2:31049344-31049550 | -0.201168287 | 0.707601064 | 0.879731796 |
| chr3:50345647-50345891 | 0.162742181 | 0.707603882 | 0.879731796 |
| chr11:134037400-134037600 | 0.167259181 | 0.707618284 | 0.879731796 |
| chr21:45292603-45292850 | 0.14793116 | 0.70762346 | 0.879731796 |
| chr1:222919305-222919701 | -0.171759999 | 0.707629861 | 0.879731796 |
| chr5:178247905-178248130 | 0.166557646 | 0.707644425 | 0.879731796 |
| chr14:69727434-69727634 | -0.187668932 | 0.707644665 | 0.879731796 |
| chr1:9942900-9943609 | 0.127645231 | 0.707647797 | 0.879731796 |
| chr6:99242071-99242347 | 0.170693383 | 0.707669779 | 0.879740903 |
| chr17:17168011-17168223 | -0.182758846 | 0.707689567 | 0.879740903 |
| chr17:61967850-61968059 | -0.182758846 | 0.707689567 | 0.879740903 |
| chr4:42401396-42401596 | 0.200672379 | 0.707709383 | 0.879751263 |
| chr17:28735295-28735495 | 0.171958799 | 0.707773944 | 0.879817245 |
| chr1:205977460-205977717 | 0.136524357 | 0.707816113 | 0.879855389 |
| chr4:2247965-2248167 | 0.189260758 | 0.707830524 | 0.879857417 |
| chr3:62873653-62874648 | 0.119235419 | 0.707840711 | 0.879857417 |
| chr3:66415150-66415356 | -0.185714051 | 0.707899695 | 0.879878372 |
| chr3:123781277-123781522 | -0.185714051 | 0.707899695 | 0.879878372 |
| chr17:4625929-4626163 | -0.183196413 | 0.707901715 | 0.879878372 |
| chr7:124139639-124139863 | -0.19243769 | 0.707912682 | 0.879878372 |
| chr17:69335086-69335372 | -0.142728701 | 0.70791982 | 0.879878372 |
| chr1:10694921-10695179 | 0.184026374 | 0.707942456 | 0.879878372 |

| | | | |
|---|---|---|---|
| chr17:80052044-80052244 | 0.186380584 | 0.707949436 | 0.879878372 |
| chr3:155791613-155791833 | 0.186380584 | 0.707949436 | 0.879878372 |
| chr5:159916180-159916769 | 0.140233239 | 0.708032079 | 0.879938917 |
| chr14:106883158-106883452 | 0.164309665 | 0.708032254 | 0.879938917 |
| chr20:53621400-53622357 | -0.11707664 | 0.708032602 | 0.879938917 |
| chr15:40274684-40275026 | 0.139611729 | 0.708052064 | 0.879948831 |
| chr15:80419012-80419212 | 0.197374937 | 0.708087244 | 0.879978279 |
| chr2:154494358-154494577 | -0.193668871 | 0.708137988 | 0.880020816 |
| chr1:27141632-27142142 | -0.139891758 | 0.708156053 | 0.880020816 |
| chr1:67148665-67149483 | 0.127636496 | 0.70816712 | 0.880020816 |
| chr16:68040589-68041116 | 0.137531898 | 0.708172673 | 0.880020816 |
| chr1:2199148-2199683 | 0.1314037 | 0.708178898 | 0.880020816 |
| chr8:104776832-104777112 | -0.14638203 | 0.708261565 | 0.880044161 |
| chr17:55420998-55422460 | 0.118479326 | 0.708270585 | 0.880044161 |
| chr17:72501463-72501724 | 0.169687785 | 0.708272864 | 0.880044161 |
| chr7:76997404-76997924 | 0.146103292 | 0.708281852 | 0.880044161 |
| chr4:62595320-62595593 | 0.184899667 | 0.708300089 | 0.880044161 |
| chr20:325218-325562 | 0.131915694 | 0.708302337 | 0.880044161 |
| chr12:76621466-76622015 | -0.134105915 | 0.708308446 | 0.880044161 |
| chr2:36597289-36598372 | -0.120166411 | 0.708315442 | 0.880044161 |
| chr8:31033378-31034318 | -0.122724476 | 0.708316667 | 0.880044161 |
| chrX:120260432-120260632 | 0.186044014 | 0.708323146 | 0.880044161 |
| chrX:96481960-96482382 | 0.152719331 | 0.708325547 | 0.880044161 |
| chr11:48116946-48117146 | -0.194657147 | 0.70833551 | 0.880044161 |
| chr15:97984290-97984492 | 0.191469031 | 0.708417734 | 0.880128379 |
| chr4:144021666-144021893 | 0.181948114 | 0.70842627 | 0.880128379 |
| chr12:12716845-12717685 | -0.123134725 | 0.708505609 | 0.880212676 |
| chr15:75104625-75105285 | 0.129271379 | 0.708540238 | 0.880241425 |
| chr3:128226837-128227099 | 0.157246303 | 0.708571323 | 0.880256605 |
| chr1:196309420-196309761 | 0.141729138 | 0.708575434 | 0.880256605 |
| chr3:49686740-49687001 | -0.164360706 | 0.708598371 | 0.880270828 |
| chr6:7615282-7615733 | 0.14888608 | 0.708621299 | 0.880285038 |
| chr8:101501449-101501696 | -0.147021841 | 0.708634579 | 0.880287263 |
| chr5:150700457-150701502 | 0.121158903 | 0.708685318 | 0.880336021 |
| chr11:67303968-67304223 | -0.138291756 | 0.708698159 | 0.880337699 |
| chr10:72354325-72355476 | -0.119196396 | 0.708723881 | 0.880354005 |
| chr10:6495725-6495990 | 0.178867022 | 0.708734265 | 0.880354005 |
| chr1:150259942-150260565 | 0.124408995 | 0.708766051 | 0.880370784 |
| chr1:48471680-48472723 | -0.119361531 | 0.708770752 | 0.880370784 |
| chr5:35598449-35599341 | -0.129430464 | 0.708791642 | 0.880372854 |
| chr11:12281833-12282114 | -0.15771183 | 0.708795398 | 0.880372854 |
| chr6:111057358-111057695 | 0.151646036 | 0.708954344 | 0.880552814 |
| chr2:98412690-98413172 | -0.144706053 | 0.70896327 | 0.880552814 |
| chr13:80340987-80341578 | 0.12575357 | 0.708975556 | 0.8805538 |
| chr17:75240479-75240985 | -0.134652993 | 0.709044404 | 0.880615603 |
| chr7:44885351-44886190 | -0.120026712 | 0.709076018 | 0.880615603 |
| chr3:7787937-7788182 | 0.177063939 | 0.709081188 | 0.880615603 |
| chr1:64736576-64736776 | -0.201260209 | 0.709082781 | 0.880615603 |
| chr1:91440938-91441144 | -0.201260209 | 0.709082781 | 0.880615603 |
| chr3:31210145-31210345 | 0.209992753 | 0.709111034 | 0.880634133 |

| | | | |
|---|---|---|---|
| chr1:116109868-116110685 | 0.130327495 | 0.709127101 | 0.880634133 |
| chr20:54123283-54124174 | -0.116283506 | 0.709139247 | 0.880634133 |
| chr7:44050041-44050382 | -0.158276363 | 0.709143675 | 0.880634133 |
| chr9:34989132-34989848 | -0.122538565 | 0.709158992 | 0.880638882 |
| chrX:54999716-55000621 | -0.121312166 | 0.709204139 | 0.880680672 |
| chr1:52054706-52055670 | 0.122364685 | 0.709275022 | 0.880744981 |
| chr17:49707660-49708526 | 0.119613904 | 0.709291864 | 0.880744981 |
| chr8:54632843-54633137 | -0.157065259 | 0.709310318 | 0.880744981 |
| chr22:47193190-47193433 | -0.164438742 | 0.709310919 | 0.880744981 |
| chr4:154044084-154044439 | 0.171218518 | 0.709323843 | 0.880744981 |
| chr4:169940558-169940794 | -0.163418844 | 0.709324894 | 0.880744981 |
| chr12:70558057-70558327 | -0.18788862 | 0.709341491 | 0.880751317 |
| chr2:176002654-176002904 | -0.150276714 | 0.709370718 | 0.880769546 |
| chr3:185943396-185943958 | -0.132707585 | 0.709379163 | 0.880769546 |
| chr5:16934148-16934384 | -0.144648731 | 0.709393657 | 0.880773269 |
| chr1:84077380-84078847 | -0.120230947 | 0.709459418 | 0.880830808 |
| chr8:143218481-143218828 | 0.145520367 | 0.709462992 | 0.880830808 |
| chr8:100914072-100914334 | 0.155316175 | 0.709503717 | 0.88085774 |
| chr6:131770347-131770564 | 0.208662433 | 0.709511518 | 0.88085774 |
| chrX:108020453-108020799 | 0.18124345 | 0.709523756 | 0.88085774 |
| chr6:12287269-12287702 | 0.137515472 | 0.709530668 | 0.88085774 |
| chr9:130187118-130187519 | -0.14336872 | 0.70957115 | 0.880880332 |
| chr20:42193121-42193389 | -0.136505135 | 0.70957468 | 0.880880332 |
| chr5:28808612-28809046 | 0.158532376 | 0.709599155 | 0.880880332 |
| chr10:329343-330142 | 0.126366513 | 0.709605947 | 0.880880332 |
| chr10:103396348-103396871 | 0.123610031 | 0.709616616 | 0.880880332 |
| chr6:33711151-33711985 | 0.123208681 | 0.709617844 | 0.880880332 |
| chr6:26521533-26522230 | 0.126411163 | 0.709630913 | 0.880882283 |
| chr2:172102141-172102357 | 0.14936965 | 0.709650071 | 0.880891793 |
| chr13:30286588-30286891 | 0.177369246 | 0.709690364 | 0.880922736 |
| chr12:54658706-54658940 | 0.178408339 | 0.709706119 | 0.880922736 |
| chr11:27698947-27699759 | -0.127467967 | 0.709713116 | 0.880922736 |
| chr5:137075203-137075525 | 0.147791956 | 0.709724928 | 0.880922736 |
| chr1:116151764-116152713 | 0.121385117 | 0.709741307 | 0.880922736 |
| chr15:95059212-95059918 | -0.130547564 | 0.70974398 | 0.880922736 |
| chr1:28232680-28233176 | 0.130817886 | 0.709761842 | 0.880929828 |
| chr10:102231886-10223208 | -0.179363986 | 0.70978418 | 0.880929828 |
| chr5:127878203-127878527 | -0.144213908 | 0.709784185 | 0.880929828 |
| chr12:55143345-55143573 | -0.195237119 | 0.709825456 | 0.880952511 |
| chr12:95241206-95241408 | -0.195237119 | 0.709825456 | 0.880952511 |
| chr14:97480572-97480772 | -0.203689589 | 0.70985635 | 0.880968847 |
| chr2:219294607-219295214 | 0.126130408 | 0.709861614 | 0.880968847 |
| chr19:29879083-29879291 | 0.175043817 | 0.709889019 | 0.880986259 |
| chr19:10936636-10937001 | -0.141310952 | 0.70989864 | 0.880986259 |
| chr8:108722903-108723223 | -0.130274582 | 0.709919749 | 0.880986494 |
| chr2:231612547-231612862 | 0.139839451 | 0.709921825 | 0.880986494 |
| chr20:49046003-49047217 | 0.11780742 | 0.709942004 | 0.880997266 |
| chr22:20957107-20957373 | -0.154035745 | 0.709969078 | 0.881016595 |
| chr2:74261615-74261889 | -0.150747783 | 0.710000876 | 0.881041785 |
| chr11:32312864-32313314 | -0.13296084 | 0.710012527 | 0.881041975 |

| | | | |
|---|---|---|---|
| chr16:67191937-67192489 | -0.130644162 | 0.710031741 | 0.881051548 |
| chr14:54722796-54723171 | 0.154524567 | 0.710066246 | 0.881080095 |
| chr8:30722837-30723505 | 0.124173368 | 0.71008162 | 0.881084904 |
| chr1:46394092-46394848 | 0.130594119 | 0.710113634 | 0.881110359 |
| chr2:207896977-207897267 | 0.156289703 | 0.710131409 | 0.88111749 |
| chr16:17340208-17340420 | 0.177126966 | 0.71014238 | 0.88111749 |
| chr3:64028857-64029077 | 0.128835291 | 0.710212991 | 0.881190831 |
| chr14:74531465-74531837 | -0.139434161 | 0.71026138 | 0.881221157 |
| chr6:89311779-89312054 | 0.161325145 | 0.710281678 | 0.881221157 |
| chr4:94452626-94452961 | -0.130464463 | 0.710288568 | 0.881221157 |
| chr16:87077324-87077594 | 0.162587666 | 0.710305803 | 0.881221157 |
| chr3:160564800-160566447 | -0.11642958 | 0.710326564 | 0.881221157 |
| chr16:68023095-68023765 | 0.131479755 | 0.71032849 | 0.881221157 |
| chr7:88102681-88102896 | -0.213857843 | 0.710334992 | 0.881221157 |
| chr1:156090695-156091106 | -0.135847944 | 0.710336639 | 0.881221157 |
| chr17:74775997-74776960 | -0.119875503 | 0.710363361 | 0.881221157 |
| chr8:80752849-80753636 | 0.12130531 | 0.710381599 | 0.881221157 |
| chr3:153980148-153980387 | 0.172393553 | 0.710404959 | 0.881221157 |
| chr12:76548488-76548903 | -0.161525714 | 0.71041281 | 0.881221157 |
| chr9:95316792-95317961 | -0.120073895 | 0.710420076 | 0.881221157 |
| chr3:187412557-187412757 | -0.207367927 | 0.710428836 | 0.881221157 |
| chr2:74009870-74010682 | -0.120762298 | 0.710433937 | 0.881221157 |
| chr7:151875965-151876360 | 0.141563655 | 0.710439562 | 0.881221157 |
| chr2:26764471-26764773 | 0.151151033 | 0.710456767 | 0.881221157 |
| chr4:20422450-20422798 | 0.151025561 | 0.710459881 | 0.881221157 |
| chr5:96823854-96824237 | 0.148403302 | 0.710466216 | 0.881221157 |
| chr9:131125095-131126154 | 0.121018861 | 0.710467451 | 0.881221157 |
| chr14:83816821-83817191 | 0.154654238 | 0.710505762 | 0.881254411 |
| chr14:23360616-23360885 | -0.160400612 | 0.710534552 | 0.881262933 |
| chr9:136399889-136400606 | -0.132168929 | 0.710546795 | 0.881262933 |
| chr16:83661751-83662038 | -0.189057552 | 0.710547137 | 0.881262933 |
| chr5:155812048-155812359 | 0.165644788 | 0.710565436 | 0.881271364 |
| chr1:228180355-228180565 | -0.178156765 | 0.710578791 | 0.881273662 |
| chr10:122911904-12291220[illegible] | 0.16227405 | 0.710629338 | 0.881322087 |
| chr2:219217895-219219191 | 0.119965875 | 0.710646526 | 0.881322313 |
| chr13:24660035-24660362 | -0.147922403 | 0.710657079 | 0.881322313 |
| chr2:91766464-91766709 | -0.161363998 | 0.710664028 | 0.881322313 |
| chr5:86566067-86566327 | 0.162175495 | 0.710682462 | 0.881324859 |
| chr13:21998995-21999308 | 0.167192129 | 0.710700486 | 0.881324859 |
| chr14:93873242-93873665 | 0.156089206 | 0.710700587 | 0.881324859 |
| chr10:101840695-101840984 | -0.138809268 | 0.710728246 | 0.881344894 |
| chr1:225652681-225654105 | -0.119134478 | 0.710757555 | 0.88134653 |
| chr5:180293171-180293371 | -0.158088251 | 0.71076 | 0.88134653 |
| chr16:68590182-68590428 | 0.178668826 | 0.710764073 | 0.88134653 |
| chr1:169712057-169712361 | 0.13631461 | 0.710778157 | 0.881349731 |
| chr2:197543729-197543929 | 0.211259022 | 0.710792935 | 0.881353793 |
| chr4:13953152-13953378 | 0.184325083 | 0.710827655 | 0.881382581 |
| chr2:149388460-149388736 | -0.151674055 | 0.710842421 | 0.881386626 |
| chr1:236456498-236456699 | -0.191736131 | 0.71086027 | 0.881394495 |
| chr1:42479668-42479994 | 0.15887515 | 0.710891296 | 0.88141244 |

| | | | |
|---|---|---|---|
| chr17:57077059-57077273 | 0.176925007 | 0.71089775 | 0.88141244 |
| chr10:69022998-69023330 | -0.180543661 | 0.710923312 | 0.881421953 |
| chr11:12110221-12111150 | -0.121502198 | 0.710935697 | 0.881421953 |
| chr16:71367431-71367666 | 0.157085676 | 0.710955131 | 0.881421953 |
| chr17:81309072-81309447 | 0.134917338 | 0.710958362 | 0.881421953 |
| chr4:142883518-142883828 | -0.15853712 | 0.710983824 | 0.881421953 |
| chr5:26382119-26382328 | -0.199631453 | 0.710994248 | 0.881421953 |
| chr5:68068173-68068403 | -0.199631453 | 0.710994248 | 0.881421953 |
| chr16:67816515-67816947 | -0.136747885 | 0.710997451 | 0.881421953 |
| chr19:44499945-44500918 | 0.120779281 | 0.711018735 | 0.881434079 |
| chr1:31182163-31182717 | 0.132685332 | 0.711049624 | 0.88145811 |
| chr20:34538760-34538973 | -0.189122667 | 0.711104931 | 0.881483286 |
| chr19:50783595-50783814 | -0.181606604 | 0.711109846 | 0.881483286 |
| chr3:160538554-160538792 | -0.201446387 | 0.711118 | 0.881483286 |
| chr17:8383080-8384012 | -0.121151425 | 0.711119067 | 0.881483286 |
| chr8:86564348-86564717 | -0.142253009 | 0.711127454 | 0.881483286 |
| chr16:8685946-8686629 | 0.133602264 | 0.711175077 | 0.881528056 |
| chr10:77889329-77889739 | -0.129750099 | 0.711241693 | 0.881595742 |
| chr1:53340435-53340678 | -0.18752803 | 0.711257375 | 0.881595742 |
| chr22:41688498-41689256 | -0.131678356 | 0.711264201 | 0.881595742 |
| chr10:51960501-51960720 | 0.187046041 | 0.71133949 | 0.881674798 |
| chr16:11871293-11871514 | 0.201173261 | 0.711363983 | 0.881677192 |
| chr20:33457226-33457456 | -0.172102282 | 0.711374846 | 0.881677192 |
| chr19:5293208-5293726 | 0.1351762 | 0.711375941 | 0.881677192 |
| chr18:32158053-32158521 | -0.139108165 | 0.711403916 | 0.881697602 |
| chr18:11980761-11981710 | 0.117182425 | 0.711421191 | 0.88170475 |
| chr10:92848059-92848996 | 0.120807895 | 0.711450126 | 0.881712661 |
| chr19:39824457-39825109 | 0.12645279 | 0.711450588 | 0.881712661 |
| chr6:116511393-116511793 | -0.141644444 | 0.711481844 | 0.881714612 |
| chr5:34914791-34916069 | -0.117578563 | 0.711519239 | 0.881714612 |
| chr3:9890216-9891097 | 0.120545491 | 0.711522618 | 0.881714612 |
| chr5:140357424-140357985 | 0.129667335 | 0.711533505 | 0.881714612 |
| chr14:68853633-68854265 | 0.123969373 | 0.711536019 | 0.881714612 |
| chr1:117674558-117674883 | 0.141635235 | 0.711538844 | 0.881714612 |
| chr16:4200065-4200787 | 0.13168197 | 0.711539751 | 0.881714612 |
| chr20:38790311-38790581 | 0.182799627 | 0.711550771 | 0.881714612 |
| chrX:48918901-48919422 | 0.123049085 | 0.711555729 | 0.881714612 |
| chr8:23246215-23247270 | 0.118774975 | 0.711613306 | 0.881753577 |
| chr15:47987985-47988214 | -0.165727549 | 0.711619825 | 0.881753577 |
| chr2:111120266-111121016 | -0.121914821 | 0.711621697 | 0.881753577 |
| chr2:173816313-173816513 | -0.197675883 | 0.711666714 | 0.881795096 |
| chr15:43105825-43106252 | -0.142765367 | 0.711705753 | 0.881807092 |
| chrX:47623614-47624064 | 0.134192604 | 0.711719897 | 0.881807092 |
| chr20:54468112-54468565 | 0.120342127 | 0.711737787 | 0.881807092 |
| chr9:16871581-16871915 | -0.17852934 | 0.711759158 | 0.881807092 |
| chr11:83007010-83007375 | -0.153265114 | 0.711762057 | 0.881807092 |
| chr11:48109371-48109675 | -0.146372229 | 0.711777089 | 0.881807092 |
| chr19:12722563-12722860 | -0.157159128 | 0.711781667 | 0.881807092 |
| chr17:1109019-1109319 | 0.156442414 | 0.711794845 | 0.881807092 |
| chr3:63662539-63662836 | -0.126603767 | 0.71179523 | 0.881807092 |

| | | | |
|---|---|---|---|
| chr20:42645981-42646181 | 0.175439942 | 0.71179726 | 0.881807092 |
| chrX:118823627-118823865 | -0.186298296 | 0.711822733 | 0.881807092 |
| chrX:7120602-7120876 | 0.161710418 | 0.7118235 | 0.881807092 |
| chr12:8032368-8033712 | -0.122528479 | 0.711826006 | 0.881807092 |
| chr1:6204983-6205529 | 0.131517307 | 0.711867154 | 0.881843808 |
| chr8:102654658-102655117 | -0.122857591 | 0.71192008 | 0.881895114 |
| chr14:54893235-54893472 | -0.164949354 | 0.711988488 | 0.881903979 |
| chr16:89722616-89722885 | 0.136284393 | 0.71198937 | 0.881903979 |
| chr3:61251029-61251689 | 0.123564107 | 0.711991533 | 0.881903979 |
| chr20:51735835-51736196 | 0.139185674 | 0.711997036 | 0.881903979 |
| chr7:74658121-74658791 | 0.124983209 | 0.712005955 | 0.881903979 |
| chr2:119240594-119240879 | -0.159447527 | 0.712016629 | 0.881903979 |
| chr17:49704040-49704242 | -0.149105053 | 0.712016855 | 0.881903979 |
| chr4:56129487-56129932 | 0.143784011 | 0.71202403 | 0.881903979 |
| chr6:158818096-158819717 | 0.118826901 | 0.712039333 | 0.881903979 |
| chr9:124711790-124712328 | 0.134319149 | 0.712042334 | 0.881903979 |
| chr3:148764001-148764300 | -0.16507861 | 0.712079424 | 0.88193565 |
| chr4:157076000-157076807 | -0.131598222 | 0.712102325 | 0.88193565 |
| chr2:218312216-218312557 | -0.147496219 | 0.712105165 | 0.88193565 |
| chr3:112783859-112784064 | -0.209976642 | 0.712134205 | 0.88193565 |
| chr6:52815488-52815840 | -0.143631041 | 0.712135742 | 0.88193565 |
| chr10:66244388-66244597 | 0.197180337 | 0.712136967 | 0.88193565 |
| chr9:134170743-134171094 | 0.141965027 | 0.712154124 | 0.881938732 |
| chr3:195894648-195894894 | -0.145590228 | 0.712169485 | 0.881938732 |
| chr5:139517748-139518534 | -0.124454864 | 0.712173986 | 0.881938732 |
| chr8:127725717-127726245 | -0.127814188 | 0.712210524 | 0.881956356 |
| chr6:10510581-10510869 | 0.148716136 | 0.712211238 | 0.881956356 |
| chr17:50639050-50639636 | 0.125128274 | 0.712245024 | 0.88198394 |
| chr12:106224210-106224439 | -0.157659005 | 0.712296311 | 0.882023643 |
| chr2:46286407-46287035 | 0.134700413 | 0.712310119 | 0.882023643 |
| chr18:58045482-58045828 | -0.151124828 | 0.712322938 | 0.882023643 |
| chr12:113319236-113319457 | 0.180672095 | 0.712323132 | 0.882023643 |
| chr15:100463528-100463925 | 0.130284244 | 0.712359091 | 0.882053915 |
| chr10:27635611-27635812 | 0.219823869 | 0.71244657 | 0.882146249 |
| chr9:100427536-100427929 | -0.137492921 | 0.712456687 | 0.882146249 |
| chr5:34929086-34930349 | 0.11611457 | 0.712509168 | 0.882196972 |
| chr5:41945461-41946217 | -0.12831353 | 0.71252716 | 0.882204994 |
| chr6:79949233-79949494 | 0.14296774 | 0.712554127 | 0.882224127 |
| chr9:112487526-112487726 | 0.169771797 | 0.712583609 | 0.882236019 |
| chr6:29942259-29942612 | -0.143196156 | 0.712586761 | 0.882236019 |
| chr3:172332555-172332840 | -0.141590659 | 0.712635539 | 0.882282155 |
| chr9:117413942-117414142 | -0.174554316 | 0.712670807 | 0.882311562 |
| chr10:128047420-128047664 | -0.165547819 | 0.712708623 | 0.882344123 |
| chr15:75138733-75139048 | -0.145115371 | 0.712772297 | 0.882398806 |
| chr12:64890742-64891210 | 0.132777244 | 0.712775826 | 0.882398806 |
| chr5:112963779-112964340 | -0.127950989 | 0.712791421 | 0.882403856 |
| chr12:2853684-2853925 | 0.158042671 | 0.712830354 | 0.882414817 |
| chr4:100372521-100372735 | -0.209608922 | 0.712831377 | 0.882414817 |
| chr3:184319804-184320379 | -0.128654328 | 0.712839726 | 0.882414817 |
| chr7:148557730-148557943 | 0.17633318 | 0.712846341 | 0.882414817 |

| | | | |
|---|---|---|---|
| chr3:192449716-192449926 | -0.213871374 | 0.712899671 | 0.88245901 |
| chr11:129279308-129279928 | 0.132191221 | 0.712908443 | 0.88245901 |
| chr8:38410710-38411133 | 0.163156946 | 0.712916593 | 0.88245901 |
| chr11:67239616-67240228 | -0.120771094 | 0.712943425 | 0.882477563 |
| chr6:102516759-102516960 | 0.189658047 | 0.712954616 | 0.882477563 |
| chr10:12875266-12875466 | -0.177414569 | 0.713007391 | 0.88252863 |
| chr20:7161763-7161965 | 0.195120604 | 0.713051855 | 0.882569408 |
| chr11:67118609-67119537 | 0.118192619 | 0.713073883 | 0.882580766 |
| chr20:63641770-63642468 | -0.121889546 | 0.713084069 | 0.882580766 |
| chr9:19127378-19127776 | 0.137175285 | 0.713114672 | 0.882604387 |
| chr12:76558809-76560245 | -0.11596948 | 0.713149376 | 0.882633083 |
| chr13:46934474-46935070 | -0.133515238 | 0.71318155 | 0.882657036 |
| chr3:98329158-98329852 | 0.133792804 | 0.713207356 | 0.882657036 |
| chr7:47607856-47608056 | 0.1802734 | 0.713208044 | 0.882657036 |
| chr12:12561563-12561865 | -0.154817837 | 0.713214808 | 0.882657036 |
| chr5:174312087-174312326 | 0.156816648 | 0.713246106 | 0.882681512 |
| chr6:169212978-169213515 | 0.139685094 | 0.713265801 | 0.882691629 |
| chr6:11601978-11602258 | -0.190835306 | 0.713292835 | 0.882693964 |
| chr6:148947654-148948463 | 0.126548271 | 0.713298697 | 0.882693964 |
| chr11:64876982-64877567 | 0.12751979 | 0.713302248 | 0.882693964 |
| chr15:87190345-87190554 | -0.192774694 | 0.713342443 | 0.882706777 |
| chr2:170928202-170929632 | 0.115701295 | 0.713342835 | 0.882706777 |
| chr13:20294483-20294873 | -0.145801956 | 0.713351941 | 0.882706777 |
| chr10:97572280-97572583 | -0.136544354 | 0.713366423 | 0.882706777 |
| chr22:41589650-41590314 | 0.120333925 | 0.713370204 | 0.882706777 |
| chr20:36773342-36774567 | 0.118363766 | 0.713408936 | 0.882740448 |
| chr20:49610648-49611576 | 0.119759203 | 0.713483216 | 0.882818103 |
| chr12:92129874-92130084 | -0.200149189 | 0.713496409 | 0.88282017 |
| chr8:29262994-29263304 | -0.145515702 | 0.713553953 | 0.882877113 |
| chr15:65029030-65029800 | 0.123407027 | 0.713579576 | 0.88288048 |
| chr17:44007587-44007787 | 0.159716223 | 0.713579719 | 0.88288048 |
| chr17:21511609-21511889 | -0.160858282 | 0.713614265 | 0.882899154 |
| chr2:219029717-219030007 | 0.140737677 | 0.713617858 | 0.882899154 |
| chr3:179652373-179653775 | -0.117146977 | 0.713634695 | 0.882905729 |
| chr19:48713720-48714687 | 0.117078086 | 0.713659863 | 0.882913909 |
| chr8:115987658-115988583 | -0.120209869 | 0.713664352 | 0.882913909 |
| chr6:79466326-79466552 | 0.175834344 | 0.713688612 | 0.882917434 |
| chr12:57693435-57694476 | 0.116586091 | 0.71369886 | 0.882917434 |
| chr1:67686367-67686934 | -0.124556829 | 0.713701771 | 0.882917434 |
| chr22:40950905-40951945 | -0.118244222 | 0.713720667 | 0.882926555 |
| chr6:170026075-170026867 | -0.120467435 | 0.71374101 | 0.882931841 |
| chr4:48767126-48767335 | 0.19176587 | 0.713747987 | 0.882931841 |
| chr2:74489830-74490093 | 0.171746363 | 0.713812572 | 0.88299739 |
| chr14:79093683-79093983 | 0.167158586 | 0.713824023 | 0.88299739 |
| chr3:13283586-13283823 | -0.153794497 | 0.713850589 | 0.883014787 |
| chr1:205175290-205175768 | -0.142264693 | 0.713861492 | 0.883014787 |
| chr5:171409809-171410009 | -0.187197676 | 0.71387266 | 0.883014787 |
| chr17:35312688-35313021 | -0.157577346 | 0.713887185 | 0.883018498 |
| chr12:70609274-70609526 | -0.17278358 | 0.71390539 | 0.883026761 |
| chr13:108218011-108218959 | -0.120643907 | 0.713961546 | 0.883050565 |

| | | | |
|---|---|---|---|
| chr1:226974050-226974281 | 0.161856743 | 0.713966323 | 0.883050565 |
| chr2:27217222-27218036 | 0.121991629 | 0.713968459 | 0.883050565 |
| chr3:194860758-194861119 | -0.155077597 | 0.713992992 | 0.883050565 |
| chr1:156053671-156055016 | 0.116306387 | 0.713993984 | 0.883050565 |
| chr2:101474263-101474568 | -0.170624634 | 0.714006028 | 0.883050565 |
| chr14:103391109-103391378 | -0.169104933 | 0.714034529 | 0.883050565 |
| chr8:108773983-108774386 | -0.144333417 | 0.714050182 | 0.883050565 |
| chr3:164478425-164478684 | 0.166955917 | 0.714051209 | 0.883050565 |
| chr3:168620895-168621248 | -0.13632163 | 0.714051368 | 0.883050565 |
| chr8:60136127-60136514 | -0.13857189 | 0.71406221 | 0.883050565 |
| chr19:14434015-14434399 | 0.160473995 | 0.71407397 | 0.883050565 |
| chr20:54173688-54174462 | 0.115850714 | 0.714078427 | 0.883050565 |
| chr16:8868161-8869372 | 0.116614512 | 0.714085981 | 0.883050565 |
| chr5:134737797-134738045 | -0.150464951 | 0.714109292 | 0.88306514 |
| chr4:83548521-83548765 | -0.161055878 | 0.714136162 | 0.883084115 |
| chrX:73662293-73662543 | 0.192890736 | 0.714180663 | 0.883124891 |
| chr12:130838256-130839750 | -0.119916961 | 0.714218201 | 0.883157056 |
| chr2:74334903-74335113 | -0.166444596 | 0.714245754 | 0.883168099 |
| chr13:51564774-51565052 | -0.193291586 | 0.714250184 | 0.883168099 |
| chr17:38730829-38731100 | 0.149665406 | 0.714273104 | 0.883182187 |
| chr22:47631265-47632068 | -0.126545334 | 0.714289364 | 0.88318804 |
| chrX:53093706-53094642 | 0.118812041 | 0.714341686 | 0.883226251 |
| chr7:2760578-2760892 | -0.140566455 | 0.714343322 | 0.883226251 |
| chr8:73633213-73633423 | -0.182466339 | 0.714366173 | 0.883240252 |
| chr6:91534690-91534908 | -0.206945657 | 0.714378179 | 0.883240845 |
| chr2:134119644-134120274 | -0.121170252 | 0.714460211 | 0.883328013 |
| chr7:25951911-25952316 | 0.151983556 | 0.714500088 | 0.883363062 |
| chr9:113222061-113222267 | -0.151496696 | 0.714514345 | 0.883366435 |
| chr17:69526076-69526304 | 0.156835533 | 0.714555777 | 0.883381074 |
| chr3:78268694-78269098 | -0.15047707 | 0.714570874 | 0.883381074 |
| chr2:205948111-205948570 | 0.140914853 | 0.714576957 | 0.883381074 |
| chr6:82589702-82590038 | -0.151722655 | 0.714579954 | 0.883381074 |
| chr17:54891206-54891448 | 0.158084223 | 0.714583832 | 0.883381074 |
| chr4:128068926-128069285 | -0.133393516 | 0.714599516 | 0.883386211 |
| chr10:80204270-80206018 | 0.116327863 | 0.714616046 | 0.883392393 |
| chr17:27198799-27199008 | -0.210166776 | 0.714643394 | 0.883404135 |
| chr1:183267964-183268421 | -0.153013612 | 0.714648604 | 0.883404135 |
| chr17:44218961-44219501 | 0.128625951 | 0.714663198 | 0.883407923 |
| chr6:167351451-167351657 | 0.188512733 | 0.714710295 | 0.883439671 |
| chr3:138329705-138330110 | 0.126088513 | 0.71471194 | 0.883439671 |
| chr10:119725434-119726576 | 0.116361965 | 0.714738969 | 0.883458828 |
| chr17:67377024-67379199 | 0.114413762 | 0.714822626 | 0.883547979 |
| chr20:54479634-54480488 | 0.115192039 | 0.714899154 | 0.883620084 |
| chr3:22429622-22429835 | -0.181963495 | 0.714911841 | 0.883620084 |
| chr9:98796424-98796957 | -0.129018707 | 0.714915558 | 0.883620084 |
| chr12:67222421-67222648 | -0.166640329 | 0.714956449 | 0.883647445 |
| chr9:131277994-131278345 | 0.136036595 | 0.71496076 | 0.883647445 |
| chr5:87171345-87171545 | -0.200914435 | 0.714991099 | 0.883670689 |
| chr16:84277283-84277557 | -0.152744213 | 0.71506169 | 0.883730309 |
| chr6:30516026-30516418 | -0.140517692 | 0.715062533 | 0.883730309 |

| | | | |
|---|---|---|---|
| chr10:5537338-5537538 | 0.190901892 | 0.715090247 | 0.883730309 |
| chr3:179147210-179149156 | -0.115155647 | 0.715094129 | 0.883730309 |
| chr8:128501240-128501482 | -0.168642149 | 0.715097007 | 0.883730309 |
| chr12:67413574-67413899 | -0.151038321 | 0.715133815 | 0.883754569 |
| chr22:41682336-41682557 | -0.179837449 | 0.715150343 | 0.883754569 |
| chr3:102112837-102113092 | 0.154936091 | 0.71515124 | 0.883754569 |
| chr11:14891498-14892619 | -0.115327432 | 0.715194595 | 0.883793892 |
| chr12:54690121-54690322 | 0.181432506 | 0.715299053 | 0.883894464 |
| chr12:85814213-85814421 | 0.181432506 | 0.715299053 | 0.883894464 |
| chr17:75323071-75323375 | -0.157168678 | 0.715330458 | 0.883919016 |
| chr19:45001763-45002345 | 0.119595113 | 0.715409552 | 0.884002494 |
| chrY:56849168-56849506 | 0.154923901 | 0.715431317 | 0.884015132 |
| chr5:132238274-132238630 | -0.141136262 | 0.715471837 | 0.884020764 |
| chr5:35044466-35044741 | 0.165282483 | 0.715476736 | 0.884020764 |
| chrX:48539279-48540317 | 0.116510313 | 0.71548201 | 0.884020764 |
| chr5:81657480-81657744 | 0.173099101 | 0.715487662 | 0.884020764 |
| chr1:188422034-188422234 | 0.215690943 | 0.715493561 | 0.884020764 |
| chr6:34136480-34136759 | 0.153717784 | 0.715509943 | 0.884023933 |
| chr8:118662446-118662731 | 0.154064504 | 0.715519201 | 0.884023933 |
| chr7:49777072-49777318 | 0.17079171 | 0.715531758 | 0.884024949 |
| chr3:48847038-48847714 | 0.133601484 | 0.715545681 | 0.884024949 |
| chr2:20180151-20180373 | -0.152200923 | 0.715554636 | 0.884024949 |
| chrX:109475024-109475719 | 0.128448811 | 0.715621131 | 0.884092844 |
| chr6:11011804-11012017 | -0.188480267 | 0.715650833 | 0.884100651 |
| chr17:68291051-68292722 | -0.113563478 | 0.715658572 | 0.884100651 |
| chr11:19244182-19244399 | -0.149577584 | 0.7156716 | 0.884100651 |
| chr5:94916701-94916953 | -0.170920708 | 0.715679102 | 0.884100651 |
| chr19:984080-984501 | -0.13184842 | 0.715694475 | 0.884100651 |
| chr1:37629108-37629379 | -0.185493956 | 0.715696681 | 0.884100651 |
| chr19:8005590-8005828 | 0.172871492 | 0.715725084 | 0.884121484 |
| chr3:43286129-43286860 | -0.127840999 | 0.715741104 | 0.884127019 |
| chr1:159780489-159781581 | -0.115294674 | 0.715787159 | 0.884169655 |
| chr1:160954245-160954501 | 0.157820679 | 0.715855324 | 0.8842396 |
| chr9:89738649-89739041 | 0.143901946 | 0.715938358 | 0.88431419 |
| chr8:96408580-96408822 | 0.169170366 | 0.715938793 | 0.88431419 |
| chrX:106481267-106481686 | -0.138171829 | 0.715954113 | 0.884318858 |
| chr3:3150623-3150850 | -0.187239966 | 0.715995758 | 0.884347216 |
| chr9:117783248-117783449 | -0.183487724 | 0.716000155 | 0.884347216 |
| chr4:150960861-150961165 | -0.144400375 | 0.716068273 | 0.884417094 |
| chr7:43891076-43891366 | -0.169134171 | 0.716137141 | 0.884487895 |
| chr1:44730460-44731594 | 0.116903675 | 0.716162089 | 0.88450445 |
| chr14:100478151-100478418 | -0.189154492 | 0.716195903 | 0.884525847 |
| chr11:72446923-72447305 | -0.1380111 | 0.716210188 | 0.884525847 |
| chr1:113714010-113714311 | 0.145415019 | 0.716214045 | 0.884525847 |
| chr12:64678161-64678600 | 0.136232254 | 0.716242461 | 0.884546683 |
| chr1:44819279-44820395 | 0.11750535 | 0.716259388 | 0.88455333 |
| chr20:59725649-59725889 | 0.147932518 | 0.71628649 | 0.884567636 |
| chr3:176916943-176917221 | 0.145965397 | 0.716299911 | 0.884567636 |
| chr5:172707242-172707813 | 0.128715985 | 0.716305606 | 0.884567636 |
| chr1:23619483-23620023 | 0.148528989 | 0.716329468 | 0.884570534 |

| | | | |
|---|---|---|---|
| chr5:120655846-120656157 | -0.150741045 | 0.716331042 | 0.884570534 |
| chr1:63773752-63774490 | 0.1255798 | 0.716389169 | 0.884628056 |
| chr10:97279426-97279626 | -0.186409692 | 0.716411942 | 0.884640716 |
| chr6:130238949-130239257 | -0.155023057 | 0.716422512 | 0.884640716 |
| chr7:143361857-143362057 | 0.179623872 | 0.716468338 | 0.884669856 |
| chr1:179742771-179743077 | 0.157396212 | 0.71647275 | 0.884669856 |
| chr1:187224040-187224288 | 0.185185039 | 0.716480748 | 0.884669856 |
| chr7:100483320-100483544 | -0.153384634 | 0.716531546 | 0.884702547 |
| chr1:15835584-15836218 | -0.122235957 | 0.716542918 | 0.884702547 |
| chr1:161565911-161566463 | -0.138207885 | 0.716549494 | 0.884702547 |
| chr16:66880047-66881023 | 0.119152932 | 0.71655341 | 0.884702547 |
| chr5:180464494-180464703 | 0.155448587 | 0.716590498 | 0.884707497 |
| chr1:58932903-58933299 | 0.148005807 | 0.716594312 | 0.884707497 |
| chr5:180454132-180454395 | 0.177225065 | 0.716599127 | 0.884707497 |
| chr10:4951286-4951491 | 0.195675949 | 0.716603605 | 0.884707497 |
| chr9:133022293-133022748 | 0.131567511 | 0.716620882 | 0.884714572 |
| chr11:72081929-72082132 | 0.152549618 | 0.71663815 | 0.884721636 |
| chr5:923563-923772 | 0.165999161 | 0.716660319 | 0.884721761 |
| chr11:8092718-8092938 | -0.164095531 | 0.716664431 | 0.884721761 |
| chr17:36152639-36152930 | 0.15087295 | 0.716676443 | 0.884721761 |
| chr2:176097251-176098109 | 0.119179913 | 0.716684437 | 0.884721761 |
| chr6:79431537-79431818 | -0.155972504 | 0.716696622 | 0.884722548 |
| chr5:171817974-171818252 | 0.167248681 | 0.716761332 | 0.884780255 |
| chr2:236824890-236825277 | 0.145530345 | 0.716766464 | 0.884780255 |
| chr18:79964347-79964734 | -0.134088053 | 0.716779627 | 0.88478225 |
| chr10:78328178-78328378 | -0.204301328 | 0.716813523 | 0.884804122 |
| chr8:134685595-134686131 | -0.123935474 | 0.716820442 | 0.884804122 |
| chr7:50308698-50308924 | -0.203294412 | 0.71685522 | 0.884815691 |
| chr17:58993349-58993749 | 0.121393511 | 0.71687943 | 0.884815691 |
| chr6:90215060-90215379 | 0.161621258 | 0.716897763 | 0.884815691 |
| chr11:71452607-71453541 | 0.116217084 | 0.716921489 | 0.884815691 |
| chr3:4290406-4290839 | -0.147500188 | 0.716934826 | 0.884815691 |
| chr2:138275935-138276234 | -0.14572033 | 0.716944947 | 0.884815691 |
| chr2:168480419-168480698 | -0.158930358 | 0.716951854 | 0.884815691 |
| chr7:26200615-26201885 | -0.113350735 | 0.716952011 | 0.884815691 |
| chr6:27332728-27333405 | 0.124170828 | 0.716952563 | 0.884815691 |
| chr14:90375956-90376161 | 0.180326027 | 0.716966054 | 0.884815691 |
| chr9:94374195-94374924 | 0.125280459 | 0.716973777 | 0.884815691 |
| chr6:2245222-2246513 | 0.11712401 | 0.716991477 | 0.884815691 |
| chr11:64204552-64204767 | -0.1684888 | 0.716999693 | 0.884815691 |
| chr13:52720966-52721195 | 0.171337772 | 0.71700019 | 0.884815691 |
| chr13:33004536-33004736 | 0.190772056 | 0.717003031 | 0.884815691 |
| chr6:43509890-43511129 | -0.119405343 | 0.717032378 | 0.884834408 |
| chr4:147731619-147733305 | 0.114593328 | 0.717041295 | 0.884834408 |
| chr5:59673572-59673777 | -0.200858236 | 0.717074509 | 0.884861145 |
| chr10:415938-416606 | 0.12858186 | 0.717098397 | 0.88487637 |
| chr14:24212942-24213729 | 0.115917083 | 0.71713317 | 0.884905028 |
| chr1:153544937-153545792 | -0.117849781 | 0.717155615 | 0.884918474 |
| chr3:99569077-99569344 | 0.161951724 | 0.717202091 | 0.884949338 |
| chr1:151485677-151485877 | -0.167732359 | 0.717203727 | 0.884949338 |

| | | | |
|---|---|---|---|
| chr1:222643649-222644770 | -0.115822983 | 0.717243043 | 0.884983598 |
| chr7:156949477-156950480 | -0.120913634 | 0.717281804 | 0.885004706 |
| chr14:51776266-51776562 | 0.142997974 | 0.717283251 | 0.885004706 |
| chr20:36154836-36155240 | 0.147565953 | 0.717320882 | 0.885018708 |
| chr6:43496488-43497004 | -0.127420857 | 0.71733105 | 0.885018708 |
| chr2:162890758-162891046 | -0.163439538 | 0.717350089 | 0.885018708 |
| chr17:58084471-58084952 | 0.121569819 | 0.717354651 | 0.885018708 |
| chr18:12092742-12093121 | 0.131874066 | 0.71736347 | 0.885018708 |
| chr1:114715951-114717250 | 0.113100831 | 0.717386586 | 0.885018708 |
| chr5:151115385-151115624 | 0.156519932 | 0.717390093 | 0.885018708 |
| chr17:9408980-9409427 | -0.143376899 | 0.717406757 | 0.885018708 |
| chr1:42334802-42335973 | -0.119813577 | 0.717413611 | 0.885018708 |
| chr1:205679661-205680888 | 0.11656789 | 0.717422165 | 0.885018708 |
| chr17:40423195-40423650 | 0.136576663 | 0.717440003 | 0.885018708 |
| chr19:46617142-46617437 | -0.158083263 | 0.717444028 | 0.885018708 |
| chr4:98467362-98468101 | -0.126568586 | 0.717444755 | 0.885018708 |
| chr6:126173912-126174112 | -0.195262854 | 0.717519382 | 0.885096516 |
| chr2:45670682-45670934 | -0.153761995 | 0.717589788 | 0.885157796 |
| chr7:140493159-140493431 | -0.166374857 | 0.717592164 | 0.885157796 |
| chr1:148951508-148952697 | 0.116970684 | 0.717652006 | 0.885215935 |
| chr8:122782934-122783310 | -0.120778002 | 0.717662404 | 0.885215935 |
| chr3:61655002-61655305 | -0.127567966 | 0.717703111 | 0.885251896 |
| chr15:85289703-85290063 | -0.144564776 | 0.71772786 | 0.885255107 |
| chr5:23527206-23527789 | 0.117994092 | 0.717728822 | 0.885255107 |
| chr17:11996746-11998054 | -0.115783628 | 0.717753006 | 0.885259863 |
| chr5:97231073-97231374 | 0.148560593 | 0.717767733 | 0.885259863 |
| chr14:106210744-106210944 | -0.183091558 | 0.717776734 | 0.885259863 |
| chr5:141848781-141850291 | 0.114803115 | 0.717778892 | 0.885259863 |
| chr3:50568958-50569804 | 0.120169547 | 0.717814063 | 0.88528899 |
| chr9:101957723-101958147 | 0.141196236 | 0.717859005 | 0.885330168 |
| chr2:223993683-223993942 | -0.173633899 | 0.717875774 | 0.885333028 |
| chr5:72391073-72391293 | -0.19683661 | 0.717889524 | 0.885333028 |
| chr6:28335354-28336469 | -0.116662176 | 0.717895987 | 0.885333028 |
| chr5:68962886-68963146 | 0.167833694 | 0.71792925 | 0.885338395 |
| chr16:12935021-12935616 | -0.149302128 | 0.71793025 | 0.885338395 |
| chr9:122904512-122905783 | 0.114650463 | 0.717935003 | 0.885338395 |
| chr10:45972270-45972708 | 0.139001274 | 0.717962472 | 0.885348542 |
| chr19:12669655-12669864 | -0.172397173 | 0.717966341 | 0.885348542 |
| chr9:33170917-33171117 | -0.163707969 | 0.71798773 | 0.885360669 |
| chr15:63610187-63610581 | 0.14002528 | 0.718012931 | 0.885366492 |
| chr10:102151811-102152844 | 0.115661701 | 0.718015562 | 0.885366492 |
| chr11:69644177-69644591 | 0.12415988 | 0.71805416 | 0.885399836 |
| chr22:23786776-23787682 | 0.119273861 | 0.718079917 | 0.885404843 |
| chr1:115641476-115643054 | -0.11421834 | 0.718081331 | 0.885404843 |
| chr14:78230285-78230543 | -0.184256172 | 0.718139726 | 0.88544277 |
| chr6:169701358-169702935 | -0.113723059 | 0.71814792 | 0.88544277 |
| chr12:202592-202807 | 0.17472123 | 0.718152938 | 0.88544277 |
| chr1:182029286-182030428 | 0.114189115 | 0.718158315 | 0.88544277 |
| chr7:1473905-1474350 | 0.127707404 | 0.718181996 | 0.88545772 |
| chr2:10924802-10925060 | -0.178903462 | 0.718211422 | 0.885474499 |

| | | | |
|---|---|---|---|
| chr18:15000155-15000668 | 0.14058124 | 0.718223496 | 0.885474499 |
| chr2:130336864-130337683 | -0.117550986 | 0.718233651 | 0.885474499 |
| chr4:76748784-76749191 | 0.161061 | 0.718246113 | 0.885474499 |
| chrX:48957538-48958716 | -0.115051305 | 0.718253387 | 0.885474499 |
| chr1:50348724-50349033 | -0.145878352 | 0.718299944 | 0.885515742 |
| chr8:107117029-107117229 | 0.201167555 | 0.718358427 | 0.885515742 |
| chr17:17919302-17919546 | -0.161052973 | 0.718362407 | 0.885515742 |
| chr12:93723220-93723778 | 0.133744664 | 0.71836847 | 0.885515742 |
| chr1:39183698-39184067 | -0.138782891 | 0.718369326 | 0.885515742 |
| chr1:77218948-77219624 | 0.122486845 | 0.718374181 | 0.885515742 |
| chr18:48031047-48031283 | -0.162106504 | 0.71837817 | 0.885515742 |
| chr7:38330960-38331165 | -0.17499443 | 0.718383281 | 0.885515742 |
| chr10:117084421-117084627 | -0.18180999 | 0.718390855 | 0.885515742 |
| chr1:201023601-201023919 | 0.12718137 | 0.718424587 | 0.885517198 |
| chr2:120598827-120599064 | 0.175440786 | 0.718425384 | 0.885517198 |
| chr2:209368883-209369449 | 0.119742295 | 0.718426707 | 0.885517198 |
| chr8:126394980-126395278 | 0.139053785 | 0.71845771 | 0.885541167 |
| chr6:108260208-108261704 | 0.115354532 | 0.718478092 | 0.885552044 |
| chr17:81976705-81978207 | 0.114812008 | 0.718498151 | 0.885562522 |
| chr6:26420641-26420907 | -0.158112153 | 0.71851565 | 0.885569845 |
| chr1:182143278-182143478 | 0.176131256 | 0.718530735 | 0.885574193 |
| chr2:71166206-71166535 | 0.164907222 | 0.718552596 | 0.88558689 |
| chr19:33831926-33832182 | -0.1686615 | 0.718564467 | 0.885587277 |
| chr6:56542418-56543304 | -0.117482514 | 0.718650489 | 0.885675469 |
| chr2:241063661-241064584 | 0.123847923 | 0.718665992 | 0.885675469 |
| chr11:77066302-77067224 | -0.116956508 | 0.718670703 | 0.885675469 |
| chr1:112620227-112620427 | 0.155503459 | 0.718736646 | 0.885742489 |
| chr17:50447922-50448122 | 0.192348719 | 0.718761077 | 0.885758352 |
| chr1:15799888-15800867 | 0.12113615 | 0.718827523 | 0.885799418 |
| chr10:72254934-72255393 | 0.136052732 | 0.718839497 | 0.885799418 |
| chr17:58565700-58566190 | 0.121028929 | 0.718865632 | 0.885799418 |
| chr1:220690073-220690720 | 0.119919074 | 0.718894638 | 0.885799418 |
| chr8:144785968-144786208 | -0.147284129 | 0.718899922 | 0.885799418 |
| chr7:2835204-2835440 | -0.164336808 | 0.718904611 | 0.885799418 |
| chr20:36199808-36200049 | 0.157596046 | 0.718910484 | 0.885799418 |
| chr15:75928551-75928914 | 0.163384249 | 0.718915315 | 0.885799418 |
| chr6:8590470-8590689 | -0.18143939 | 0.718915437 | 0.885799418 |
| chr7:47819405-47819633 | 0.161508856 | 0.718917524 | 0.885799418 |
| chr15:70302523-70302771 | -0.174230557 | 0.718921569 | 0.885799418 |
| chr10:79066425-79066658 | 0.141329381 | 0.718944277 | 0.885809508 |
| chr12:54945167-54945367 | -0.210956483 | 0.7189618 | 0.885809508 |
| chr19:28236157-28236434 | -0.183726104 | 0.71896444 | 0.885809508 |
| chr3:4978206-4979553 | -0.112238523 | 0.718990337 | 0.885827172 |
| chr3:63818332-63819057 | 0.115303533 | 0.719009165 | 0.885836124 |
| chr1:233172335-233172535 | -0.194162981 | 0.719031125 | 0.88583651 |
| chr5:150544629-150545090 | 0.129210193 | 0.719037048 | 0.88583651 |
| chr1:205003418-205004233 | -0.120861415 | 0.71909214 | 0.88583651 |
| chr2:85951708-85951978 | -0.147702326 | 0.719094235 | 0.88583651 |
| chr20:42161226-42161748 | 0.135530366 | 0.719098093 | 0.88583651 |
| chr11:15216806-15217006 | 0.175903358 | 0.71910462 | 0.88583651 |

| | | | |
|---|---|---|---|
| chr1:222634452-222634748 | -0.156252867 | 0.719108887 | 0.88583651 |
| chr1:155226982-155227209 | -0.142103688 | 0.719110047 | 0.88583651 |
| chr5:38854162-38854394 | 0.17989768 | 0.719113528 | 0.88583651 |
| chr2:10122198-10123021 | -0.115780564 | 0.719143678 | 0.885841747 |
| chr19:10095864-10096477 | 0.121819095 | 0.719148762 | 0.885841747 |
| chr6:30882764-30882964 | 0.148362436 | 0.719157788 | 0.885841747 |
| chr13:24339571-24339853 | 0.181081032 | 0.719174012 | 0.885841747 |
| chr17:18250374-18250630 | 0.147537358 | 0.71917765 | 0.885841747 |
| chrX:73618583-73618807 | -0.191443248 | 0.719193529 | 0.885841747 |
| chr14:93788235-93788969 | -0.125097625 | 0.719198708 | 0.885841747 |
| chr1:180501941-180503377 | 0.112831689 | 0.719271148 | 0.88589897 |
| chr11:123061973-123063122 | 0.113892634 | 0.719278557 | 0.88589897 |
| chr2:199458425-199459871 | -0.114319573 | 0.719279852 | 0.88589897 |
| chr11:122580568-122580825 | -0.153169351 | 0.719330283 | 0.885937138 |
| chr16:50680605-50680996 | 0.166208689 | 0.719333966 | 0.885937138 |
| chr5:126516949-126517155 | -0.174190597 | 0.719348478 | 0.88594077 |
| chr6:131782672-131783132 | 0.138740487 | 0.719403185 | 0.885975317 |
| chr2:95192963-95193184 | -0.171360547 | 0.719405264 | 0.885975317 |
| chr1:119886178-119886500 | 0.146378611 | 0.719422087 | 0.885975317 |
| chrX:106999528-107000546 | 0.116835066 | 0.71942278 | 0.885975317 |
| chr8:123166688-123167870 | -0.116246548 | 0.719452969 | 0.885998255 |
| chr5:40446815-40447311 | 0.133314133 | 0.719489282 | 0.886028733 |
| chr5:68287776-68288655 | -0.123960089 | 0.719521341 | 0.886041477 |
| chr6:33454123-33454830 | 0.120909617 | 0.719536692 | 0.886041477 |
| chr17:67992757-67994347 | -0.113290001 | 0.719551046 | 0.886041477 |
| chr17:72176333-72176842 | -0.128366866 | 0.719556188 | 0.886041477 |
| chr19:49451746-49452126 | 0.124442854 | 0.719557449 | 0.886041477 |
| chr16:2983664-2983890 | 0.156233732 | 0.719600918 | 0.886080763 |
| chr1:179228756-179229796 | -0.119636426 | 0.719661148 | 0.886127778 |
| chr15:66071145-66071396 | 0.159012372 | 0.71966223 | 0.886127778 |
| chr7:154927639-154928053 | -0.137470741 | 0.719685507 | 0.886134115 |
| chr14:49306208-49306518 | -0.151991824 | 0.719696144 | 0.886134115 |
| chr8:78854149-78854614 | 0.132563316 | 0.719702071 | 0.886134115 |
| chr22:37545157-37545475 | -0.147719896 | 0.719737439 | 0.886154899 |
| chr7:130322648-130322868 | 0.165879096 | 0.71976103 | 0.886154899 |
| chr5:1349320-1349585 | 0.142591936 | 0.719761354 | 0.886154899 |
| chr12:111331230-111331430 | 0.182898418 | 0.71976687 | 0.886154899 |
| chr3:139534619-139534952 | -0.156724613 | 0.719783217 | 0.886154899 |
| chr12:58341813-58342117 | -0.15001509 | 0.719793546 | 0.886154899 |
| chr2:64750583-64751853 | -0.112394756 | 0.719799909 | 0.886154899 |
| chr3:192597344-192598274 | 0.117783777 | 0.719887906 | 0.886248994 |
| chr11:69703109-69703344 | -0.171234771 | 0.719975645 | 0.886335467 |
| chr15:100042765-100043325 | -0.130424242 | 0.719987116 | 0.886335467 |
| chr16:84959025-84959432 | 0.133173007 | 0.719997566 | 0.886335467 |
| chr8:122252610-122252918 | 0.17102774 | 0.720004418 | 0.886335467 |
| chr16:52912701-52913016 | -0.163444743 | 0.720038521 | 0.886347075 |
| chr21:43880228-43880428 | -0.172915714 | 0.720040255 | 0.886347075 |
| chr2:60739703-60739911 | -0.184934054 | 0.72004855 | 0.886347075 |
| chr16:27313512-27313715 | -0.162256504 | 0.720099954 | 0.88637811 |
| chr15:33226721-33226992 | -0.168834332 | 0.72010539 | 0.88637811 |

| | | | |
|---|---|---|---|
| chr15:80943217-80943563 | 0.130799841 | 0.720108467 | 0.88637811 |
| chr5:180315746-180315993 | -0.139611837 | 0.720186008 | 0.886445075 |
| chr6:150080680-150080993 | -0.139611837 | 0.720186008 | 0.886445075 |
| chr8:63038458-63039736 | -0.114478042 | 0.720287791 | 0.886556112 |
| chr10:73601444-73601882 | -0.127063018 | 0.720304959 | 0.886557729 |
| chr6:32984812-32985043 | 0.163606368 | 0.720312245 | 0.886557729 |
| chr17:55939167-55939367 | 0.179892499 | 0.720331995 | 0.886567796 |
| chr3:62501048-62501271 | 0.143333292 | 0.720351328 | 0.886577349 |
| chr8:126066938-126067161 | 0.144881062 | 0.720378902 | 0.886597045 |
| chr8:90398658-90399637 | 0.114923847 | 0.720452512 | 0.886672218 |
| chr4:22607366-22607714 | -0.146989487 | 0.720463126 | 0.886672218 |
| chr2:240914426-240914653 | 0.153468023 | 0.72053812 | 0.886730681 |
| chr5:37927384-37927584 | -0.184457172 | 0.72053991 | 0.886730681 |
| chr6:20534272-20534633 | 0.137137596 | 0.720545348 | 0.886730681 |
| chr10:90148572-90148881 | -0.157197127 | 0.720595083 | 0.886777545 |
| chr7:65373604-65374310 | -0.121338007 | 0.720606576 | 0.886777545 |
| chr6:149045451-149045766 | -0.150051529 | 0.720650895 | 0.886817842 |
| chr7:78770455-78770655 | -0.198282619 | 0.720706767 | 0.886861802 |
| chr17:67170074-67170276 | 0.168264519 | 0.720709768 | 0.886861802 |
| chr13:67119557-67119924 | -0.138835111 | 0.720740315 | 0.886885149 |
| chr12:117025475-117025936 | 0.128298195 | 0.72075822 | 0.886892937 |
| chrX:48003454-48003720 | 0.175702797 | 0.720779958 | 0.88690287 |
| chr16:30609529-30610758 | 0.116237672 | 0.720789442 | 0.88690287 |
| chr13:27533095-27533651 | -0.128202767 | 0.720829194 | 0.88693754 |
| chr12:109899895-109901021 | -0.117907914 | 0.7209138 | 0.887016693 |
| chr5:43110858-43111308 | -0.128435822 | 0.720920008 | 0.887016693 |
| chr18:26468857-26469115 | 0.162142858 | 0.720928253 | 0.887016693 |
| chr5:71631682-71631993 | 0.153824468 | 0.720975082 | 0.887047767 |
| chr14:89859720-89859928 | 0.18447647 | 0.720976662 | 0.887047767 |
| chr1:23867663-23868533 | 0.126237851 | 0.721038855 | 0.887096703 |
| chr10:8084820-8085387 | -0.13078752 | 0.721039591 | 0.887096703 |
| chr8:98202381-98202621 | 0.17864093 | 0.721068561 | 0.887108669 |
| chr2:202032862-202034315 | 0.11412655 | 0.721092858 | 0.887108669 |
| chr20:51631261-51631663 | -0.138740918 | 0.721102538 | 0.887108669 |
| chr8:120642476-120642733 | 0.174558347 | 0.72110613 | 0.887108669 |
| chr4:169810342-169810641 | 0.148087353 | 0.721107206 | 0.887108669 |
| chr14:60723949-60724601 | -0.115527616 | 0.72112368 | 0.887114692 |
| chr10:101818061-101819173 | -0.113743053 | 0.721187953 | 0.887168133 |
| chr14:59918869-59919069 | -0.191284927 | 0.721190278 | 0.887168133 |
| chr17:55348840-55349194 | -0.138124991 | 0.721227462 | 0.88719963 |
| chr2:20040569-20040803 | 0.186089976 | 0.721249431 | 0.887210205 |
| chr3:122533884-122534432 | 0.125037735 | 0.721259217 | 0.887210205 |
| chr5:10695078-10695495 | 0.133948962 | 0.721294168 | 0.887224837 |
| chr11:59680562-59680846 | -0.146764503 | 0.72129427 | 0.887224837 |
| chr3:119240254-119241187 | 0.115662864 | 0.721380462 | 0.887316613 |
| chr15:100341569-100342375 | 0.115820085 | 0.721410406 | 0.887338881 |
| chr13:51826649-51826995 | -0.14981354 | 0.721421728 | 0.887338881 |
| chr11:85781790-85782012 | -0.156644837 | 0.721447279 | 0.887354184 |
| chr10:49418381-49418587 | 0.1635456 | 0.721465097 | 0.887354184 |
| chr3:145370591-145370828 | 0.189154652 | 0.721468912 | 0.887354184 |

| | | | |
|---|---|---|---|
| chr3:74789154-74789391 | -0.174483083 | 0.721499428 | 0.887377473 |
| chr21:31219214-31219512 | -0.161951513 | 0.72151887 | 0.887386882 |
| chr3:20056719-20056971 | 0.158241577 | 0.72153635 | 0.887386882 |
| chr13:98653381-98653740 | -0.13337587 | 0.721558475 | 0.887386882 |
| chrX:147602111-147602616 | -0.139805298 | 0.721565473 | 0.887386882 |
| chr4:111475055-111475287 | -0.171871591 | 0.72156902 | 0.887386882 |
| chr2:43018461-43018661 | -0.190466254 | 0.721576567 | 0.887386882 |
| chrX:24494268-24494481 | -0.158163103 | 0.721591466 | 0.887387642 |
| chr6:24341005-24341552 | 0.137430569 | 0.721600348 | 0.887387642 |
| chr6:93416577-93417063 | 0.123984064 | 0.721617387 | 0.887390635 |
| chr14:22929051-22929872 | 0.113089687 | 0.721625944 | 0.887390635 |
| chr15:96067725-96067925 | -0.181415148 | 0.721644832 | 0.887397315 |
| chr14:52043706-52044103 | -0.133577561 | 0.721659492 | 0.887397315 |
| chr13:46831642-46831926 | 0.137511723 | 0.721666121 | 0.887397315 |
| chr19:36014131-36014651 | 0.118054329 | 0.721700066 | 0.887412206 |
| chr8:135783663-135783864 | -0.196943921 | 0.721706845 | 0.887412206 |
| chr7:5840810-5841046 | -0.172768405 | 0.721712976 | 0.887412206 |
| chr17:50139896-50140723 | -0.12029811 | 0.72172795 | 0.887416378 |
| chr4:95250699-95251087 | 0.140405093 | 0.721815838 | 0.887497385 |
| chr5:151288264-151288510 | 0.151198343 | 0.721816998 | 0.887497385 |
| chr2:187453130-187453347 | -0.171730365 | 0.721835227 | 0.887505556 |
| chr8:107383642-107383882 | -0.190572016 | 0.721872165 | 0.88753673 |
| chr10:49538587-49539341 | -0.114472891 | 0.721885815 | 0.887539271 |
| chr5:113510936-113511136 | -0.158535424 | 0.721918261 | 0.887557124 |
| chr15:84928132-84928524 | -0.123615258 | 0.721923503 | 0.887557124 |
| chr17:74514057-74514281 | -0.170883806 | 0.721953177 | 0.887579365 |
| chr20:18507281-18508242 | 0.113359682 | 0.72201177 | 0.887624417 |
| chr2:85354025-85355352 | -0.113255746 | 0.722034734 | 0.887624417 |
| chr4:71732781-71733010 | -0.179989093 | 0.722036896 | 0.887624417 |
| chr1:1678709-1679047 | 0.163956041 | 0.722047067 | 0.887624417 |
| chr8:59394762-59395159 | -0.147325797 | 0.722047744 | 0.887624417 |
| chr1:85200429-85202195 | -0.112259432 | 0.722082238 | 0.887652579 |
| chr7:2241868-2243113 | -0.114574169 | 0.722130155 | 0.887697241 |
| chr3:184336778-184337112 | 0.132212553 | 0.722151571 | 0.887709325 |
| chr14:44912783-44912992 | -0.153110072 | 0.722172742 | 0.887721108 |
| chr4:155809875-155810148 | -0.155037575 | 0.722208527 | 0.887750855 |
| chr5:17747598-17747939 | 0.158401477 | 0.722311031 | 0.887838493 |
| chr17:57987241-57989289 | -0.111423431 | 0.722312829 | 0.887838493 |
| chr2:65553288-65553558 | 0.14207575 | 0.722314585 | 0.887838493 |
| chr17:68101047-68102135 | -0.112969016 | 0.722332939 | 0.88784681 |
| chr12:54279478-54281088 | 0.111290269 | 0.722354919 | 0.887852024 |
| chr9:37943590-37943840 | 0.161902906 | 0.722360356 | 0.887852024 |
| chr2:199471445-199471847 | -0.126451792 | 0.722377595 | 0.887858284 |
| chr8:76175654-76175921 | -0.158249986 | 0.722394532 | 0.887858284 |
| chr1:45230639-45230839 | 0.184383699 | 0.722408316 | 0.887858284 |
| chr2:10520385-10520603 | -0.160916593 | 0.722411799 | 0.887858284 |
| chr1:184025406-184025642 | -0.177816444 | 0.722426941 | 0.887862653 |
| chr15:69452393-69453416 | -0.115656475 | 0.722458034 | 0.887863614 |
| chr2:119544013-119544648 | 0.120331902 | 0.72246178 | 0.887863614 |
| chr5:100005016-100005252 | -0.167341093 | 0.722462486 | 0.887863614 |

| | | | |
|---|---|---|---|
| chr19:51408806-51409208 | 0.147532704 | 0.722531399 | 0.887934063 |
| chr5:153974217-153974625 | -0.148492063 | 0.722552466 | 0.887938488 |
| chr1:52921418-52921925 | 0.129006933 | 0.722560217 | 0.887938488 |
| chr1:47569881-47570122 | 0.144986553 | 0.722569766 | 0.887938488 |
| chr3:149386480-149387325 | 0.114347993 | 0.722591165 | 0.887945934 |
| chr8:107643687-107643908 | 0.167858755 | 0.722599002 | 0.887945934 |
| chr20:34875882-34876691 | 0.113451057 | 0.72261222 | 0.887947935 |
| chr2:219433948-219434150 | -0.164502818 | 0.722639063 | 0.887953608 |
| chr17:50151656-50152479 | -0.115203361 | 0.722645602 | 0.887953608 |
| chr4:150242907-150243154 | -0.163275292 | 0.722651603 | 0.887953608 |
| chr5:169218962-169219241 | -0.158047588 | 0.722680695 | 0.887960248 |
| chr13:41882687-41883101 | -0.162855705 | 0.722690738 | 0.887960248 |
| chr2:219010198-219010522 | -0.138599892 | 0.722691773 | 0.887960248 |
| chr18:3447899-3448516 | -0.115450799 | 0.722717523 | 0.887977647 |
| chrX:49190806-49191590 | 0.112899843 | 0.722733581 | 0.887983138 |
| chr8:123381661-123381935 | -0.145990897 | 0.722749399 | 0.887988334 |
| chr3:47004633-47004833 | -0.182021802 | 0.722787128 | 0.88802045 |
| chr3:141000086-141000499 | -0.131402144 | 0.722834944 | 0.888055831 |
| chr19:49019610-49019810 | -0.160524485 | 0.722839107 | 0.888055831 |
| chr3:12869031-12869695 | -0.120918736 | 0.72287204 | 0.8880814 |
| chr3:4867965-4869098 | -0.113451363 | 0.722892909 | 0.8880814 |
| chr11:126382582-126383143 | 0.120872637 | 0.72289469 | 0.8880814 |
| chr8:9556422-9556672 | 0.160863132 | 0.722911222 | 0.888085881 |
| chr3:63350875-63351087 | -0.146141944 | 0.722929337 | 0.888085881 |
| chr17:42322872-42323196 | 0.158610026 | 0.722933109 | 0.888085881 |
| chr4:41990072-41991080 | 0.114572762 | 0.722991102 | 0.888142884 |
| chr3:169813041-169813255 | 0.177209693 | 0.723023948 | 0.888168993 |
| chr11:108121286-108122055 | -0.119753791 | 0.723047974 | 0.888184268 |
| chr4:109702037-109702270 | -0.139602455 | 0.723072105 | 0.888194332 |
| chr11:74255849-74256051 | -0.173687282 | 0.723079351 | 0.888194332 |
| chr7:151459848-151460421 | 0.128415365 | 0.723092473 | 0.888196212 |
| chr7:8404129-8404457 | 0.136319003 | 0.723118869 | 0.888214395 |
| chr9:86099226-86099744 | 0.132530576 | 0.723141776 | 0.888228294 |
| chr4:73341456-73341666 | 0.173291779 | 0.7231671 | 0.888245159 |
| chr12:57243040-57243321 | -0.13730628 | 0.723192173 | 0.888261717 |
| chr10:95720774-95721009 | -0.155602762 | 0.723230567 | 0.888292253 |
| chr11:35392287-35392513 | -0.154378806 | 0.723264566 | 0.888292253 |
| chr1:159900916-159901171 | 0.141258598 | 0.723276852 | 0.888292253 |
| chr6:6748427-6748687 | -0.162821766 | 0.723297592 | 0.888292253 |
| chr10:71348006-71348279 | 0.145444825 | 0.72329965 | 0.888292253 |
| chr16:1300861-1301095 | 0.160603825 | 0.72330961 | 0.888292253 |
| chr21:45923147-45923621 | 0.132132305 | 0.723312541 | 0.888292253 |
| chr1:47710004-47710208 | 0.157296993 | 0.723321469 | 0.888292253 |
| chr14:77259733-77260153 | 0.14202013 | 0.723333578 | 0.888292253 |
| chr22:38701403-38701744 | -0.128777838 | 0.723342329 | 0.888292253 |
| chr9:98862219-98862423 | -0.19418681 | 0.72338561 | 0.888292253 |
| chr4:145808394-145808594 | -0.19418681 | 0.72338561 | 0.888292253 |
| chr2:75560406-75560606 | -0.19418681 | 0.72338561 | 0.888292253 |
| chr17:58531894-58532223 | 0.128264303 | 0.723388605 | 0.888292253 |
| chr4:48906146-48907257 | 0.116087524 | 0.723392941 | 0.888292253 |

| | | | |
|---|---|---|---|
| chr8:142145542-142145742 | 0.158447825 | 0.723406267 | 0.888292253 |
| chrX:117991837-117992121 | 0.165736167 | 0.72341478 | 0.888292253 |
| chr5:164475421-164475621 | 0.199689831 | 0.723428026 | 0.888292253 |
| chr11:120912590-120912790 | -0.194204664 | 0.723438421 | 0.888292253 |
| chr6:53926962-53927688 | -0.12099333 | 0.723461069 | 0.888292253 |
| chr5:1130617-1130942 | 0.135803912 | 0.723475345 | 0.888292253 |
| chr3:64111736-64112366 | 0.113426272 | 0.72348281 | 0.888292253 |
| chr5:176361055-176362379 | 0.11451473 | 0.723483677 | 0.888292253 |
| chr5:140557138-140558025 | -0.116073736 | 0.723497175 | 0.888294592 |
| chr11:76838797-76838997 | 0.163624472 | 0.723520014 | 0.888308398 |
| chr4:110472978-110473183 | 0.171138722 | 0.723554723 | 0.888327859 |
| chr6:26003177-26003631 | -0.151326551 | 0.723562772 | 0.888327859 |
| chr5:14685705-14685982 | 0.144758078 | 0.723610533 | 0.888327859 |
| chr15:72604887-72605087 | -0.178231389 | 0.723622714 | 0.888327859 |
| chr15:73156688-73156888 | -0.178231389 | 0.723622714 | 0.888327859 |
| chr11:1832551-1832899 | 0.160254572 | 0.723632476 | 0.888327859 |
| chrX:43654455-43655531 | -0.113465237 | 0.723632626 | 0.888327859 |
| chr16:28845241-28845692 | 0.125474479 | 0.723647928 | 0.888327859 |
| chr15:48112352-48112552 | 0.177705968 | 0.723651801 | 0.888327859 |
| chr17:71883020-71883256 | 0.177705968 | 0.723651801 | 0.888327859 |
| chr16:1492951-1493441 | -0.126603386 | 0.723697863 | 0.888359732 |
| chr2:201177378-201177650 | 0.149709407 | 0.723700954 | 0.888359732 |
| chr6:148746121-148746422 | 0.140939078 | 0.723748121 | 0.888403398 |
| chr9:100668027-100668244 | 0.185935571 | 0.723759779 | 0.888403477 |
| chr5:140113642-140114214 | 0.12478038 | 0.72377339 | 0.888405951 |
| chr17:1669414-1669683 | -0.167337903 | 0.723806184 | 0.888424468 |
| chr2:23330192-23330494 | -0.149930088 | 0.723813391 | 0.888424468 |
| chr14:27465991-27466191 | -0.192002387 | 0.72382326 | 0.888424468 |
| chr12:25655311-25656149 | -0.124203532 | 0.723848586 | 0.888426934 |
| chr20:8751548-8751777 | -0.16438087 | 0.723856296 | 0.888426934 |
| chr18:48949199-48949449 | 0.151342898 | 0.723862905 | 0.888426934 |
| chr4:107930964-107932015 | -0.115937459 | 0.723874001 | 0.888426934 |
| chr21:31888895-31889359 | -0.145603795 | 0.723888007 | 0.888426934 |
| chr7:100563961-100564280 | 0.138619353 | 0.723895592 | 0.888426934 |
| chr17:63389985-63390470 | 0.142544349 | 0.723920736 | 0.888426934 |
| chr4:148123011-148123395 | -0.150795863 | 0.723922115 | 0.888426934 |
| chr10:87862964-87863767 | -0.113205846 | 0.723929624 | 0.888426934 |
| chr5:167737308-167737508 | -0.1942729 | 0.723950104 | 0.888437838 |
| chr8:97452705-97452989 | 0.138478373 | 0.723969818 | 0.888447801 |
| chr8:102632615-102632851 | -0.146245056 | 0.724008541 | 0.888481093 |
| chr15:62935293-62935867 | -0.135731229 | 0.724024479 | 0.888486421 |
| chr9:96619758-96620045 | 0.144470719 | 0.724067357 | 0.888518857 |
| chr9:109319435-109319656 | -0.149867985 | 0.724095623 | 0.888518857 |
| chr2:860426-860788 | 0.140836556 | 0.724104042 | 0.888518857 |
| chr3:62559945-62560163 | 0.133306937 | 0.724108734 | 0.888518857 |
| chr7:157277286-157277520 | -0.146141049 | 0.724118552 | 0.888518857 |
| chr5:14427176-14427430 | -0.172027926 | 0.724123689 | 0.888518857 |
| chr8:97250921-97251186 | -0.179234389 | 0.724132084 | 0.888518857 |
| chr12:54324544-54325487 | 0.11322089 | 0.724160088 | 0.888531719 |
| chr3:167813216-167813562 | 0.146190964 | 0.724165759 | 0.888531719 |

| | | | |
|---|---|---|---|
| chr17:21687090-21687290 | 0.181000411 | 0.724181354 | 0.888536626 |
| chrX:148093073-148093335 | -0.154833656 | 0.724209733 | 0.888557217 |
| chr1:39258719-39258934 | 0.164601003 | 0.724249766 | 0.888592106 |
| chr5:15103806-15104006 | -0.183522424 | 0.724272772 | 0.888600858 |
| chr1:220544961-220545360 | 0.141372425 | 0.724280094 | 0.888600858 |
| chr14:97498743-97498969 | 0.175607003 | 0.724328062 | 0.888645479 |
| chr8:38996180-38997312 | -0.112701656 | 0.724352024 | 0.888660648 |
| chr11:76207544-76207844 | 0.130259325 | 0.724376456 | 0.888676393 |
| chr9:88782187-88782433 | 0.188610864 | 0.724400457 | 0.888691609 |
| chr3:157119505-157119748 | -0.166203768 | 0.724429968 | 0.888713584 |
| chr3:10115345-10116397 | 0.112468074 | 0.72449107 | 0.888770213 |
| chr6:150004480-150005717 | -0.113635699 | 0.724505653 | 0.888770213 |
| chr3:114290872-114291217 | -0.133307078 | 0.724521294 | 0.888770213 |
| chr19:40226498-40226847 | 0.146882377 | 0.724522527 | 0.888770213 |
| chr3:120094059-120095415 | -0.10979397 | 0.724544613 | 0.888783077 |
| chr16:79334277-79334507 | -0.169221304 | 0.724559363 | 0.888786603 |
| chr1:54405238-54405438 | -0.158962159 | 0.724571704 | 0.888786603 |
| chr15:64651590-64651880 | 0.162828083 | 0.724596909 | 0.888786603 |
| chr19:46660909-46661749 | -0.11314569 | 0.724608667 | 0.888786603 |
| chr16:35492276-35492627 | 0.141805454 | 0.724611956 | 0.888786603 |
| chr1:173791882-173792082 | -0.196831418 | 0.724629499 | 0.888786603 |
| chr14:64986619-64987521 | 0.113369922 | 0.724655892 | 0.888786603 |
| chr4:64623249-64623785 | -0.133578712 | 0.724658311 | 0.888786603 |
| chr17:63082829-63083152 | -0.144039006 | 0.724661753 | 0.888786603 |
| chr3:14124201-14126140 | 0.109917698 | 0.724671708 | 0.888786603 |
| chr3:42881043-42881318 | 0.148396396 | 0.724675084 | 0.888786603 |
| chr5:10432262-10432584 | 0.151887578 | 0.724686898 | 0.888786867 |
| chr4:40458830-40459030 | 0.191180803 | 0.724717227 | 0.888809836 |
| chr9:95048257-95049031 | 0.126613292 | 0.724748789 | 0.88882044 |
| chr6:158853040-158853638 | -0.122922538 | 0.724751138 | 0.88882044 |
| chr19:57813979-57814218 | 0.149039057 | 0.724760673 | 0.88882044 |
| chr7:33108625-33109892 | -0.11133876 | 0.724791454 | 0.888843962 |
| chr13:30464500-30465171 | -0.117010083 | 0.724817345 | 0.888858626 |
| chr2:1817543-1817847 | 0.132421424 | 0.724838345 | 0.888858626 |
| chr9:79140835-79141035 | 0.170327449 | 0.724842944 | 0.888858626 |
| chr19:40370151-40370351 | 0.185814259 | 0.724878537 | 0.888858626 |
| chr14:22579998-22580198 | -0.204402749 | 0.724883362 | 0.888858626 |
| chr2:102520160-102520398 | -0.179260975 | 0.724894995 | 0.888858626 |
| chr14:94598581-94598827 | 0.157506115 | 0.724923225 | 0.888858626 |
| chr2:14125586-14126019 | -0.13242934 | 0.724932278 | 0.888858626 |
| chr6:73579951-73580157 | -0.19718395 | 0.724936617 | 0.888858626 |
| chr5:178626066-178627499 | -0.113052289 | 0.724936644 | 0.888858626 |
| chr15:67637204-67637404 | 0.205280154 | 0.724938015 | 0.888858626 |
| chr9:129607478-129607978 | 0.123029142 | 0.724942618 | 0.888858626 |
| chr3:129225104-129225304 | -0.171121654 | 0.725022713 | 0.888942607 |
| chr12:53123129-53123449 | -0.142522944 | 0.725046745 | 0.888957847 |
| chr13:87124052-87124344 | 0.149236624 | 0.725105774 | 0.888977727 |
| chr15:62928268-62928488 | 0.172959898 | 0.725108985 | 0.888977727 |
| chr6:105810514-105810731 | -0.174054974 | 0.725127071 | 0.888977727 |
| chr6:26322038-26322409 | -0.126508179 | 0.725128467 | 0.888977727 |

| | | | |
|---|---|---|---|
| chr12:9135351-9135582 | -0.181525911 | 0.725163421 | 0.888977727 |
| chr3:135582842-135583146 | 0.163450297 | 0.72516423 | 0.888977727 |
| chr5:44538781-44539532 | -0.121645225 | 0.725170249 | 0.888977727 |
| chr13:43981195-43981395 | -0.179485584 | 0.725172778 | 0.888977727 |
| chr9:129524902-129525286 | -0.141765637 | 0.725182827 | 0.888977727 |
| chr16:74499378-74499578 | 0.184579767 | 0.725192309 | 0.888977727 |
| chr1:117591245-117591692 | 0.119703629 | 0.725201581 | 0.888977727 |
| chr13:108779388-108779591 | -0.186658351 | 0.725202185 | 0.888977727 |
| chr1:203172129-203172543 | -0.120305046 | 0.72522687 | 0.888993765 |
| chr3:75001454-75001654 | 0.19409409 | 0.725298163 | 0.889056743 |
| chr3:48558196-48558431 | -0.143832105 | 0.725301454 | 0.889056743 |
| chr6:111591241-111592097 | -0.115541015 | 0.725344741 | 0.889095581 |
| chr14:63493262-63493634 | -0.134998024 | 0.72538849 | 0.889127045 |
| chr8:143828851-143829850 | 0.112487525 | 0.725401543 | 0.889127045 |
| chr16:719846-720173 | -0.129725258 | 0.725405223 | 0.889127045 |
| chr19:495313-495941 | 0.133741754 | 0.72543159 | 0.889133949 |
| chr4:41258901-41259154 | -0.154373165 | 0.725434063 | 0.889133949 |
| chr10:97848958-97849897 | 0.116779423 | 0.725481838 | 0.889178281 |
| chr5:59127523-59128131 | -0.120459829 | 0.725522398 | 0.8892002 |
| chr16:89522741-89522941 | 0.174554684 | 0.725522932 | 0.8892002 |
| chr1:202624252-202624811 | -0.12196336 | 0.725536184 | 0.889202219 |
| chrX:9874031-9874313 | 0.159304194 | 0.725561771 | 0.889219354 |
| chr12:98503297-98504395 | -0.110288148 | 0.725591107 | 0.889228524 |
| chr22:41204904-41205806 | -0.120098069 | 0.725592464 | 0.889228524 |
| chr17:47970526-47971501 | 0.111387666 | 0.725617742 | 0.88924528 |
| chr17:60422795-60423060 | 0.119973217 | 0.725678879 | 0.88930598 |
| chr17:68200783-68201077 | -0.125316604 | 0.725716393 | 0.889314579 |
| chr3:163854708-163854908 | -0.199941577 | 0.725716556 | 0.889314579 |
| chr11:82864252-82864640 | -0.131726128 | 0.725720716 | 0.889314579 |
| chr4:25030137-25030378 | 0.168813191 | 0.725838296 | 0.889424945 |
| chrX:101097540-101098350 | -0.11284767 | 0.725840189 | 0.889424945 |
| chr8:104835841-104836121 | -0.148814225 | 0.725863223 | 0.889424945 |
| chr3:137105335-137105608 | 0.148505955 | 0.725887648 | 0.889424945 |
| chr7:3404499-3404826 | 0.14520558 | 0.725912536 | 0.889424945 |
| chr8:109745165-109745387 | -0.182892045 | 0.725913456 | 0.889424945 |
| chr12:76747761-76748143 | 0.138164128 | 0.725921969 | 0.889424945 |
| chr19:29812309-29812791 | 0.131012074 | 0.725931925 | 0.889424945 |
| chr14:67763432-67763753 | 0.131271938 | 0.725934319 | 0.889424945 |
| chr10:74176280-74177337 | -0.112086735 | 0.725934336 | 0.889424945 |
| chr11:12683901-12684158 | 0.171157367 | 0.725938467 | 0.889424945 |
| chr10:44347716-44347916 | -0.19964603 | 0.725961488 | 0.889429263 |
| chr8:80102066-80102413 | 0.149612762 | 0.725965208 | 0.889429263 |
| chr16:77063519-77064137 | -0.122489366 | 0.725988188 | 0.889443197 |
| chr20:61896309-61896588 | 0.137640468 | 0.72601881 | 0.889466491 |
| chr1:11463654-11463899 | 0.186177559 | 0.726047686 | 0.889470981 |
| chr8:97928997-97929549 | 0.135578177 | 0.726082973 | 0.889470981 |
| chr10:29535573-29535796 | 0.168735816 | 0.726083905 | 0.889470981 |
| chr11:765368-765568 | 0.168735816 | 0.726083905 | 0.889470981 |
| chr7:98421000-98421336 | -0.132635713 | 0.726096225 | 0.889470981 |
| chr16:84883366-84883645 | -0.148256342 | 0.726099047 | 0.889470981 |

| | | | |
|---|---|---|---|
| chr22:17952440-17952678 | 0.167702259 | 0.726103735 | 0.889470981 |
| chr1:239952710-239953354 | -0.121972007 | 0.726125497 | 0.889481948 |
| chr17:5237593-5237861 | 0.178514893 | 0.726135905 | 0.889481948 |
| chr12:51537236-51537455 | -0.166551349 | 0.726172241 | 0.889512236 |
| chr10:70887780-70888389 | 0.122776472 | 0.726211178 | 0.889545533 |
| chr1:175178512-175178849 | 0.13321234 | 0.726222642 | 0.889545533 |
| chr19:39031270-39032061 | 0.113146856 | 0.726234702 | 0.889546084 |
| chr14:41419053-41419261 | 0.18358468 | 0.726298879 | 0.889562805 |
| chr17:68344783-68344983 | -0.16224572 | 0.726305436 | 0.889562805 |
| chr17:69100041-69100241 | -0.177297134 | 0.726311151 | 0.889562805 |
| chr8:90082612-90083213 | -0.115996369 | 0.7263226 | 0.889562805 |
| chr1:220217496-220217697 | -0.172364611 | 0.726324919 | 0.889562805 |
| chr8:54021586-54023030 | -0.112347654 | 0.726328079 | 0.889562805 |
| chr2:85066822-85067428 | -0.121424898 | 0.72634912 | 0.889562805 |
| chr13:20950340-20951048 | -0.125256932 | 0.726358625 | 0.889562805 |
| chr15:65384869-65385333 | -0.122795712 | 0.726373555 | 0.889562805 |
| chr1:40850066-40850292 | 0.16328249 | 0.726375672 | 0.889562805 |
| chr3:48187782-48188003 | -0.15785119 | 0.726382936 | 0.889562805 |
| chr1:47951415-47951886 | 0.139390205 | 0.726398944 | 0.889562805 |
| chr2:191322128-191322362 | 0.181721997 | 0.726399279 | 0.889562805 |
| chr8:122781093-122781669 | -0.114983323 | 0.726424031 | 0.889578898 |
| chr19:2484286-2484519 | -0.175157762 | 0.726445848 | 0.889580894 |
| chr15:77926744-77927162 | -0.12851366 | 0.726480326 | 0.889580894 |
| chr20:36605355-36606263 | 0.114672002 | 0.726484253 | 0.889580894 |
| chr15:37101194-37102012 | 0.11638291 | 0.726486062 | 0.889580894 |
| chr3:129439435-129439635 | -0.147066193 | 0.726489713 | 0.889580894 |
| chr22:37696635-37697096 | -0.127937388 | 0.726516504 | 0.889580894 |
| chr1:188209333-188209621 | 0.16534634 | 0.726518077 | 0.889580894 |
| chr5:116382729-116382952 | 0.177144626 | 0.72651854 | 0.889580894 |
| chr3:62051456-62051954 | -0.116758341 | 0.726622716 | 0.889689992 |
| chr7:8536550-8536811 | 0.147681326 | 0.726635861 | 0.889689992 |
| chr20:13788299-13788499 | -0.153747831 | 0.726642475 | 0.889689992 |
| chr2:75157561-75157761 | -0.167039965 | 0.726655421 | 0.889691627 |
| chr3:195002015-195002278 | 0.148757008 | 0.72669134 | 0.889698207 |
| chr15:69159969-69160902 | 0.112557772 | 0.726700448 | 0.889698207 |
| chr11:747181-747670 | 0.129976132 | 0.726712481 | 0.889698207 |
| chr15:76139430-76139813 | 0.13313385 | 0.72671405 | 0.889698207 |
| chr5:73498048-73499082 | -0.113816927 | 0.726718854 | 0.889698207 |
| chr1:226122419-226123034 | 0.117736261 | 0.726749779 | 0.889721852 |
| chr1:30929293-30929749 | 0.134140339 | 0.726763425 | 0.889724342 |
| chr8:55886568-55886861 | -0.145563924 | 0.726848717 | 0.88977462 |
| chr6:51367402-51367610 | -0.193844448 | 0.726873527 | 0.88977462 |
| chr2:1707712-1708087 | 0.140151462 | 0.726877001 | 0.88977462 |
| chr11:67302156-67302432 | 0.131651125 | 0.726880635 | 0.88977462 |
| chr3:98602116-98602408 | 0.143742436 | 0.726880798 | 0.88977462 |
| chr8:125633841-125634041 | 0.156285335 | 0.726883101 | 0.88977462 |
| chr14:50961831-50962053 | 0.169068983 | 0.72690334 | 0.88977462 |
| chr17:2399464-2400251 | -0.114145373 | 0.726907239 | 0.88977462 |
| chr12:106247851-10624836 | -0.127556436 | 0.726928114 | 0.88977462 |
| chr9:76756320-76756520 | -0.185995786 | 0.72693127 | 0.88977462 |

| | | | |
|---|---|---|---|
| chr2:10043388-10044369 | 0.115510434 | 0.726934641 | 0.88977462 |
| chr3:11205993-11206247 | 0.157826282 | 0.726943844 | 0.88977462 |
| chr5:149129202-149129513 | -0.128751529 | 0.726969698 | 0.889792052 |
| chr7:87933712-87934622 | -0.113828536 | 0.726995809 | 0.88980653 |
| chr2:103789409-103789650 | -0.164350644 | 0.727010237 | 0.88980653 |
| chr7:148527590-148527970 | -0.125135827 | 0.727023192 | 0.88980653 |
| chr3:112813683-112813987 | -0.175215189 | 0.727037294 | 0.88980653 |
| chr6:107488962-107489238 | 0.147300706 | 0.727039592 | 0.88980653 |
| chr21:33266030-33266683 | 0.122774998 | 0.727077276 | 0.889829882 |
| chr19:38930120-38931113 | 0.111574706 | 0.727098757 | 0.889829882 |
| chr19:14366674-14367007 | 0.143904694 | 0.727100833 | 0.889829882 |
| chr19:12890547-12891490 | 0.115837668 | 0.727109761 | 0.889829882 |
| chr2:118982139-118982469 | 0.153043628 | 0.727116738 | 0.889829882 |
| chr3:49802571-49802984 | 0.134877887 | 0.727175002 | 0.889880124 |
| chr8:85176752-85178088 | 0.111564936 | 0.727181021 | 0.889880124 |
| chr17:62268413-62268613 | -0.184208597 | 0.727212174 | 0.889904035 |
| chr19:50476644-50477011 | 0.128408996 | 0.727267513 | 0.889952533 |
| chr14:45347661-45347909 | 0.194955428 | 0.727275036 | 0.889952533 |
| chr13:110705776-110706595 | -0.112907631 | 0.727305587 | 0.889964574 |
| chr20:35953710-35955211 | 0.111530121 | 0.727312144 | 0.889964574 |
| chr13:26972236-26972476 | 0.152103999 | 0.727319721 | 0.889964574 |
| chr18:26089854-26091393 | -0.110580533 | 0.727343114 | 0.889978985 |
| chr17:62652245-62653687 | -0.110674602 | 0.727370281 | 0.889998015 |
| chr11:70720419-70720736 | 0.157822071 | 0.727384273 | 0.890000922 |
| chr5:174119677-174119919 | -0.165698124 | 0.727398328 | 0.890003907 |
| chr19:50909256-50909978 | 0.119073096 | 0.727458712 | 0.890053185 |
| chr18:48540492-48540769 | 0.139150412 | 0.727461835 | 0.890053185 |
| chr2:238256382-238256862 | 0.127576327 | 0.727478802 | 0.890059732 |
| chr3:28348122-28349671 | -0.109057647 | 0.727509168 | 0.890080013 |
| chr2:200342366-200342644 | -0.166254184 | 0.727518986 | 0.890080013 |
| chr1:180917221-180917639 | -0.125895354 | 0.727551404 | 0.890080013 |
| chr6:90412797-90413284 | -0.149670588 | 0.72755259 | 0.890080013 |
| chr1:236124970-236125353 | -0.133303582 | 0.727555949 | 0.890080013 |
| chr1:109789144-109789517 | -0.127932586 | 0.727574397 | 0.890080013 |
| chr5:173122117-173122317 | 0.211994024 | 0.727576694 | 0.890080013 |
| chr11:18764328-18764587 | 0.150283898 | 0.727611842 | 0.8901088 |
| chr3:65732392-65732617 | 0.162464872 | 0.727626301 | 0.890111761 |
| chr14:50531909-50533385 | 0.110277509 | 0.727637497 | 0.890111761 |
| chr11:119334163-119334875 | -0.115049402 | 0.727681645 | 0.890151555 |
| chr3:63262281-63262824 | 0.121011604 | 0.727714036 | 0.890173638 |
| chr19:530957-531413 | -0.132215676 | 0.727722933 | 0.890173638 |
| chr17:18377394-18377594 | -0.183738853 | 0.727765129 | 0.890211042 |
| chr5:1601558-1601949 | 0.126622672 | 0.727798711 | 0.890237909 |
| chr10:125718903-125720116 | 0.110219865 | 0.727861037 | 0.890299932 |
| chr2:8882067-8882457 | -0.135577358 | 0.727891753 | 0.89032329 |
| chr9:129482440-129483189 | 0.116430602 | 0.727937104 | 0.890364548 |
| chr15:31392245-31392557 | 0.133775691 | 0.728026514 | 0.890452881 |
| chr7:36084411-36084919 | -0.119520018 | 0.728032565 | 0.890452881 |
| chr5:150001741-150002036 | -0.126166537 | 0.728053055 | 0.890463729 |
| chrX:11533116-11533487 | -0.132064934 | 0.728066175 | 0.890465562 |

| | | | |
|---|---|---|---|
| chr7:49874258-49874458 | -0.183571769 | 0.728089666 | 0.890480078 |
| chr14:105491594-105492002 | 0.114603237 | 0.728103392 | 0.890482651 |
| chr17:49914135-49914497 | -0.140915862 | 0.728146361 | 0.890520989 |
| chr13:25981065-25981388 | -0.157991017 | 0.728186692 | 0.89055412 |
| chr6:52724465-52725264 | -0.120751802 | 0.728204825 | 0.89055412 |
| chr20:51966316-51966755 | -0.11744502 | 0.728217143 | 0.89055412 |
| chr1:43389376-43390258 | 0.117510766 | 0.728227214 | 0.89055412 |
| chr2:16034079-16034451 | -0.139125329 | 0.728247005 | 0.89055412 |
| chrX:107360797-107361483 | -0.125357058 | 0.728253173 | 0.89055412 |
| chr2:85259065-85259638 | 0.123977932 | 0.72825645 | 0.89055412 |
| chrX:154527105-154527460 | 0.138827132 | 0.728266433 | 0.89055412 |
| chr1:70353823-70355164 | -0.110728379 | 0.728339059 | 0.890600108 |
| chr17:56244607-56244808 | -0.177271999 | 0.728341259 | 0.890600108 |
| chr18:71682478-71682950 | 0.135226502 | 0.728346307 | 0.890600108 |
| chr2:112054091-112054775 | 0.112331656 | 0.728350534 | 0.890600108 |
| chr2:57549571-57549938 | -0.146020359 | 0.728363014 | 0.890601156 |
| chr20:54995513-54995733 | -0.166747379 | 0.728389481 | 0.890609966 |
| chr10:1050627-1050827 | -0.139053153 | 0.728393465 | 0.890609966 |
| chr8:69556250-69556546 | -0.156829183 | 0.728431912 | 0.89063624 |
| chr11:45843125-45843630 | -0.117030985 | 0.728448247 | 0.89063624 |
| chr10:103485565-103486043 | 0.125908691 | 0.728449825 | 0.89063624 |
| chr1:201916718-201916937 | -0.160825482 | 0.728496813 | 0.890658404 |
| chr4:155787138-155787489 | 0.129285707 | 0.728501948 | 0.890658404 |
| chr6:106736643-106736843 | -0.186095673 | 0.72851781 | 0.890658404 |
| chr6:131923481-131923696 | -0.186095673 | 0.72851781 | 0.890658404 |
| chr9:134024864-134025567 | -0.114247486 | 0.728535787 | 0.890658404 |
| chr17:75764560-75765600 | -0.111651275 | 0.728553878 | 0.890658404 |
| chr4:8238135-8238458 | 0.144601594 | 0.728563939 | 0.890658404 |
| chr9:94065600-94065800 | 0.177997424 | 0.728568638 | 0.890658404 |
| chr10:31319345-31319545 | -0.16271469 | 0.728576708 | 0.890658404 |
| chr6:169723528-169723796 | -0.140332794 | 0.728584194 | 0.890658404 |
| chr11:9568825-9569186 | -0.125957342 | 0.728600616 | 0.890660555 |
| chr3:139195965-139196438 | -0.135006328 | 0.728609202 | 0.890660555 |
| chr8:116832936-116833337 | 0.132949772 | 0.728644191 | 0.890689117 |
| chr14:20653069-20653290 | 0.143032635 | 0.728670664 | 0.890694768 |
| chr8:89508064-89508306 | 0.175707539 | 0.728695351 | 0.890694768 |
| chr5:59800139-59800397 | 0.155095229 | 0.728718349 | 0.890694768 |
| chr7:72817663-72817944 | -0.175831457 | 0.728745903 | 0.890694768 |
| chr10:5493616-5494158 | -0.115462059 | 0.728765002 | 0.890694768 |
| chr13:34841262-34841486 | 0.193095612 | 0.7287739 | 0.890694768 |
| chrX:100368190-100368725 | 0.131516941 | 0.728781679 | 0.890694768 |
| chr14:88667422-88667799 | 0.133715105 | 0.728782994 | 0.890694768 |
| chr17:76840054-76840399 | -0.139110938 | 0.72878671 | 0.890694768 |
| chr17:51841082-51841282 | 0.181341561 | 0.728786873 | 0.890694768 |
| chr1:46340485-46341258 | -0.113506229 | 0.728789872 | 0.890694768 |
| chr2:94870372-94870572 | -0.159452245 | 0.728798172 | 0.890694768 |
| chr21:18165018-18165269 | 0.15523134 | 0.728799933 | 0.890694768 |
| chr4:111136582-111136783 | 0.167753309 | 0.7288488 | 0.89070838 |
| chr12:95574942-95575189 | -0.161295671 | 0.728851327 | 0.89070838 |
| chr15:75694542-75694945 | -0.127408447 | 0.728859911 | 0.89070838 |

| | | | |
|---|---|---|---|
| chr2:151849777-151850075 | -0.158815317 | 0.728865443 | 0.89070838 |
| chr5:139262669-139262946 | 0.13865676 | 0.728869195 | 0.89070838 |
| chr3:89503938-89504658 | -0.12721457 | 0.728885513 | 0.890714116 |
| chr1:196600801-196601181 | 0.142798348 | 0.728900715 | 0.890717031 |
| chr8:85854322-85854522 | 0.152090788 | 0.728911148 | 0.890717031 |
| chr11:69601399-69601599 | -0.17516034 | 0.728977696 | 0.890784144 |
| chr13:24148830-24149099 | 0.139447371 | 0.72900023 | 0.890797474 |
| chr17:82106073-82106721 | 0.119561663 | 0.729037461 | 0.890828762 |
| chr12:48851795-48852568 | 0.114607185 | 0.729061489 | 0.89083254 |
| chr12:72282800-72283031 | -0.166078276 | 0.729072781 | 0.89083254 |
| chr8:109778633-109778843 | -0.164719947 | 0.729075432 | 0.89083254 |
| chr8:110791848-110792048 | -0.196801733 | 0.729114933 | 0.890866598 |
| chr19:42088023-42088229 | 0.189493073 | 0.729201059 | 0.890957624 |
| chr12:30754170-30755393 | -0.111904394 | 0.729221508 | 0.890968402 |
| chr6:4806109-4806337 | -0.158540357 | 0.729234236 | 0.890969745 |
| chr20:59598505-59598968 | 0.121813301 | 0.729261716 | 0.890975234 |
| chr5:107672345-107673120 | 0.113424784 | 0.729272212 | 0.890975234 |
| chr7:116173085-116173337 | 0.173191889 | 0.729273613 | 0.890975234 |
| chr5:41708199-41708488 | -0.14385066 | 0.72932034 | 0.891018114 |
| chr20:50579999-50580355 | 0.121146976 | 0.729391591 | 0.891061041 |
| chr5:15426870-15427070 | -0.18786046 | 0.729392552 | 0.891061041 |
| chr10:11819913-11820132 | 0.15653378 | 0.729394655 | 0.891061041 |
| chr16:30615863-30616078 | 0.173943377 | 0.729401994 | 0.891061041 |
| chr3:67516593-67516879 | 0.152830944 | 0.72943596 | 0.891085052 |
| chr1:28328743-28329218 | -0.121805494 | 0.729444908 | 0.891085052 |
| chr1:198937286-198937725 | -0.111915102 | 0.729457057 | 0.891085686 |
| chr2:65265181-65265413 | -0.160859015 | 0.729478409 | 0.891088707 |
| chr4:161320436-161320740 | -0.1387077 | 0.729482789 | 0.891088707 |
| chr12:83826195-83826406 | -0.15333396 | 0.729512992 | 0.891111394 |
| chr1:31880963-31881219 | 0.170076714 | 0.729525384 | 0.891112326 |
| chr2:141455204-141455477 | 0.162492002 | 0.729550777 | 0.891129136 |
| chr12:112261700-112261900 | 0.174048764 | 0.729577687 | 0.891143349 |
| chr19:39397622-39398006 | 0.128527861 | 0.729589822 | 0.891143349 |
| chr14:69242642-69242882 | 0.153593877 | 0.72961265 | 0.891143349 |
| chr1:212588250-212588450 | -0.193099573 | 0.72961553 | 0.891143349 |
| chr3:184305685-184305948 | 0.147526433 | 0.729620564 | 0.891143349 |
| chr9:135907430-135908213 | 0.112728571 | 0.729644114 | 0.891149752 |
| chr3:66254048-66254311 | -0.160767105 | 0.729655283 | 0.891149752 |
| chr1:226149646-226150042 | 0.142147288 | 0.729664651 | 0.891149752 |
| chr2:204544831-204546269 | -0.1113944 | 0.729672628 | 0.891149752 |
| chr16:67199472-67199835 | 0.128839659 | 0.729683959 | 0.891149752 |
| chr17:50056882-50057112 | -0.140810571 | 0.729722583 | 0.891182718 |
| chr1:201153919-201154735 | 0.110463196 | 0.729759283 | 0.891213334 |
| chr7:92621026-92621272 | -0.172115463 | 0.729850629 | 0.891254974 |
| chr3:11774336-11774663 | -0.124501638 | 0.729855826 | 0.891254974 |
| chr7:149646903-149647118 | 0.168023275 | 0.72985989 | 0.891254974 |
| chr12:112932428-112932780 | 0.141067695 | 0.729861622 | 0.891254974 |
| chr17:9786612-9786815 | -0.194601101 | 0.729864567 | 0.891254974 |
| chr11:35662055-35663295 | -0.112339741 | 0.729874159 | 0.891254974 |
| chr1:65419761-65421003 | -0.108425067 | 0.729876719 | 0.891254974 |

| | | | |
|---|---|---|---|
| chr14:21070936-21071438 | -0.126277786 | 0.729896053 | 0.891254974 |
| chr11:79708349-79708675 | 0.142787556 | 0.729899348 | 0.891254974 |
| chr1:203342491-203342814 | -0.12226965 | 0.729909698 | 0.891254974 |
| chr1:202812442-202812858 | -0.116726668 | 0.729925488 | 0.891258043 |
| chr7:150322837-150323958 | -0.113187514 | 0.729946945 | 0.891258043 |
| chr7:37448744-37449137 | 0.123927057 | 0.729947107 | 0.891258043 |
| chr10:43384196-43384520 | -0.158256719 | 0.729975563 | 0.891273237 |
| chr3:171459602-171461325 | -0.109308564 | 0.729987657 | 0.891273237 |
| chr1:53837596-53837829 | 0.145271388 | 0.729999069 | 0.891273237 |
| chr9:35696598-35696886 | -0.141780514 | 0.730024004 | 0.891273237 |
| chr10:7475297-7475501 | 0.176020782 | 0.730028511 | 0.891273237 |
| chr14:58199613-58200679 | 0.109851115 | 0.730029344 | 0.891273237 |
| chr8:9074907-9075261 | 0.142824195 | 0.730042222 | 0.891274758 |
| chr7:38335540-38336117 | -0.114600681 | 0.73005763 | 0.891279368 |
| chr14:35808557-35809505 | 0.110759063 | 0.730089183 | 0.891296624 |
| chr17:32127263-32127932 | 0.123493025 | 0.730095029 | 0.891296624 |
| chr10:78682011-78682285 | 0.156524584 | 0.730168245 | 0.891371804 |
| chr19:42220035-42220661 | 0.114277998 | 0.730216837 | 0.891416921 |
| chrX:47233181-47233913 | 0.111413505 | 0.73024442 | 0.891429243 |
| chr22:31977149-31977602 | -0.127449165 | 0.730250199 | 0.891429243 |
| chr1:157155689-157156358 | 0.118249324 | 0.730264672 | 0.891430051 |
| chr17:58391445-58392031 | 0.11491503 | 0.730274129 | 0.891430051 |
| chr10:86334728-86335009 | 0.132299054 | 0.730297873 | 0.891434148 |
| chr15:98484934-98485254 | -0.153931041 | 0.73030147 | 0.891434148 |
| chr1:110338250-110339551 | -0.10770948 | 0.730320461 | 0.891434148 |
| chr2:73366542-73366764 | -0.179350961 | 0.730328132 | 0.891434148 |
| chr17:81034788-81035488 | -0.115040989 | 0.730335656 | 0.891434148 |
| chr20:56504491-56504716 | 0.151155087 | 0.730361312 | 0.891444319 |
| chr5:168346440-168346811 | -0.123495415 | 0.730367257 | 0.891444319 |
| chr2:228048772-228049125 | -0.147315918 | 0.730386825 | 0.891454001 |
| chr2:188299044-188299305 | -0.167907258 | 0.730409124 | 0.891467018 |
| chr5:176574027-176574351 | -0.131261226 | 0.730442021 | 0.891492968 |
| chr14:26984659-26984891 | 0.161943944 | 0.730479229 | 0.891524179 |
| chr20:1324961-1325532 | -0.121933477 | 0.730504667 | 0.891531902 |
| chr4:139470265-139470491 | 0.173350907 | 0.730508828 | 0.891531902 |
| chr2:52810532-52810759 | -0.167844255 | 0.730542343 | 0.891540619 |
| chr3:156673842-156675373 | -0.107588144 | 0.730549292 | 0.891540619 |
| chr1:240652510-240653366 | 0.117338121 | 0.730551264 | 0.891540619 |
| chr2:164826515-164826885 | -0.133577964 | 0.730582216 | 0.891540619 |
| chr3:71512099-71512360 | -0.14956603 | 0.730584781 | 0.891540619 |
| chr8:406466-407125 | 0.120846299 | 0.730585784 | 0.891540619 |
| chr17:7251471-7252677 | -0.109788658 | 0.730625307 | 0.89157465 |
| chr17:58713838-58714061 | 0.126168974 | 0.730680619 | 0.891604942 |
| chrX:37846902-37847885 | 0.115014664 | 0.730691215 | 0.891604942 |
| chr10:12665713-12666177 | 0.134275075 | 0.730692334 | 0.891604942 |
| chr12:15590139-15590886 | -0.120881982 | 0.730696676 | 0.891604942 |
| chr6:157379795-157380290 | -0.122001578 | 0.730788916 | 0.891685115 |
| chr7:32242926-32243158 | -0.163752196 | 0.730810769 | 0.891685115 |
| chr15:40894808-40895180 | -0.124515717 | 0.730812663 | 0.891685115 |
| chr17:15257350-15257610 | -0.141787266 | 0.730812846 | 0.891685115 |

| | | | |
|---|---|---|---|
| chr3:130892965-130894556 | -0.108730533 | 0.730820568 | 0.891685115 |
| chr11:46625335-46625715 | 0.150029447 | 0.730846433 | 0.891702475 |
| chr2:86442414-86442987 | 0.113795887 | 0.730867895 | 0.891714461 |
| chr19:46746344-46746681 | -0.130623261 | 0.730901507 | 0.89174127 |
| chr1:83998465-83999621 | -0.113300574 | 0.730929731 | 0.891752574 |
| chr8:142669611-142670246 | 0.115640213 | 0.730934049 | 0.891752574 |
| chr12:54017323-54018495 | -0.111153014 | 0.731014475 | 0.891822708 |
| chr4:29276246-29276496 | -0.174281549 | 0.731014813 | 0.891822708 |
| chr9:26975681-26976068 | -0.140875633 | 0.731048085 | 0.891849099 |
| chr3:25556590-25556790 | -0.182552465 | 0.731085132 | 0.891875247 |
| chr10:128117275-128117485 | 0.185524715 | 0.731094661 | 0.891875247 |
| chr20:9903078-9903339 | -0.165945329 | 0.731114307 | 0.891875247 |
| chr14:102591831-102592751 | 0.110195899 | 0.731116078 | 0.891875247 |
| chr4:8267637-8267872 | 0.178578771 | 0.73118686 | 0.891937274 |
| chr8:56787335-56787856 | -0.128324024 | 0.731190206 | 0.891937274 |
| chr8:29320420-29320691 | -0.153474994 | 0.731272029 | 0.892022884 |
| chr17:80036407-80036737 | 0.126492153 | 0.731295317 | 0.892037089 |
| chr5:171064010-171064246 | -0.176271766 | 0.731311649 | 0.89204281 |
| chr17:44138000-44138576 | 0.119688327 | 0.731337687 | 0.89206037 |
| chr14:21868518-21868810 | 0.14111962 | 0.73136236 | 0.892076264 |
| chr15:63173046-63173489 | -0.146012394 | 0.731381411 | 0.892079909 |
| chr17:57975220-57975420 | -0.152231721 | 0.731388633 | 0.892079909 |
| chr19:4400477-4400816 | 0.132423938 | 0.731404338 | 0.892084863 |
| chrX:12741386-12741899 | -0.123114056 | 0.731426686 | 0.89209792 |
| chr5:31809097-31809521 | -0.126160585 | 0.731445507 | 0.892106675 |
| chr1:100949601-100950043 | -0.146201969 | 0.731482091 | 0.892137094 |
| chr1:200296109-200296662 | 0.119515391 | 0.731515371 | 0.892155935 |
| chr7:48337023-48337372 | 0.131693643 | 0.731520827 | 0.892155935 |
| chr10:62804315-62805226 | -0.110985929 | 0.731545962 | 0.892169072 |
| chr5:171387393-171388689 | -0.107716422 | 0.731574801 | 0.892169072 |
| chr16:69796335-69796851 | 0.12918571 | 0.731584974 | 0.892169072 |
| chr10:100912547-100913506 | -0.111196413 | 0.73159777 | 0.892169072 |
| chr11:110098643-110098937 | -0.149477223 | 0.731601117 | 0.892169072 |
| chr5:78595025-78595480 | 0.127948445 | 0.731616134 | 0.892169072 |
| chr3:1275493-1275794 | 0.145702678 | 0.731617015 | 0.892169072 |
| chr12:12883351-12883898 | 0.123542535 | 0.731632447 | 0.892169072 |
| chr11:117817847-117818072 | -0.141329973 | 0.731636392 | 0.892169072 |
| chr5:170403530-170403882 | -0.151133202 | 0.731707967 | 0.892242152 |
| chrX:51893149-51893932 | -0.114186722 | 0.731798938 | 0.892320119 |
| chr11:47454249-47454526 | 0.13911097 | 0.731801296 | 0.892320119 |
| chr16:21525489-21525710 | 0.151314205 | 0.731806843 | 0.892320119 |
| chr11:67674406-67674998 | -0.114808454 | 0.731846836 | 0.892349368 |
| chr2:215454462-215454777 | 0.147769563 | 0.731858919 | 0.892349368 |
| chr3:190318808-190319008 | -0.185573458 | 0.731867987 | 0.892349368 |
| chr3:179015444-179015845 | -0.130545589 | 0.731879848 | 0.892349368 |
| chr12:2960504-2960984 | 0.124001017 | 0.731889062 | 0.892349368 |
| chr16:84013809-84014730 | -0.107143533 | 0.731908942 | 0.892359407 |
| chr1:199973403-199973625 | 0.151189539 | 0.731996603 | 0.892452085 |
| chr21:21508992-21509235 | -0.179177666 | 0.732043322 | 0.892494843 |
| chr22:50244975-50245199 | 0.151245057 | 0.732109656 | 0.892561514 |

| | | | |
|---|---|---|---|
| chr2:36545074-36545367 | 0.159757069 | 0.732123356 | 0.892564015 |
| chr16:67254929-67255375 | 0.13812679 | 0.732140273 | 0.892570437 |
| chr5:177561322-177561613 | 0.138288029 | 0.732164294 | 0.89258552 |
| chr20:63909695-63909931 | -0.140210521 | 0.732254921 | 0.892647706 |
| chr8:13819056-13819256 | 0.189210619 | 0.732258275 | 0.892647706 |
| chr1:204758628-204758976 | -0.132783005 | 0.73227348 | 0.892647706 |
| chr3:162022675-162022938 | -0.156584546 | 0.732300412 | 0.892647706 |
| chr7:25851683-25851947 | -0.135718939 | 0.732305914 | 0.892647706 |
| chr1:119782581-119783172 | -0.12070766 | 0.732311236 | 0.892647706 |
| chr16:24042120-24042324 | 0.163143805 | 0.732318128 | 0.892647706 |
| chr11:355792-356518 | -0.126801843 | 0.73232154 | 0.892647706 |
| chr3:11643732-11644103 | 0.125757237 | 0.732325401 | 0.892647706 |
| chr4:83535433-83536813 | -0.110322167 | 0.732336087 | 0.892647706 |
| chr5:9436864-9437074 | -0.187127779 | 0.732343454 | 0.892647706 |
| chr3:194871550-194871849 | -0.136290802 | 0.732378456 | 0.892661259 |
| chr11:59010631-59010831 | 0.179584153 | 0.732387195 | 0.892661259 |
| chr9:69173806-69174252 | -0.132235465 | 0.732389523 | 0.892661259 |
| chr14:69013997-69014600 | 0.119409947 | 0.732412501 | 0.892675066 |
| chr1:150010097-150011082 | 0.109876278 | 0.732429252 | 0.892681282 |
| chr8:123596396-123596986 | 0.120676988 | 0.732494749 | 0.892746909 |
| chr3:107240361-107241131 | -0.113941961 | 0.732531615 | 0.892777639 |
| chr13:102596154-102597240 | -0.108695305 | 0.73255237 | 0.892788734 |
| chr22:18528531-18529386 | 0.114921518 | 0.732588719 | 0.892818832 |
| chr12:78470710-78470920 | -0.169007334 | 0.732610602 | 0.892831301 |
| chr1:116834631-116834831 | 0.169329791 | 0.732668737 | 0.892887948 |
| chr6:27830710-27831203 | 0.126572786 | 0.732711555 | 0.892904785 |
| chr3:117407345-117407625 | 0.157921331 | 0.732716535 | 0.892904785 |
| chr7:87702328-87702584 | -0.183158911 | 0.732717513 | 0.892904785 |
| chr5:12691261-12691713 | -0.128010963 | 0.732801181 | 0.892968826 |
| chr15:67542547-67543352 | 0.121204453 | 0.732809946 | 0.892968826 |
| chr15:34583417-34583859 | 0.121994432 | 0.732820481 | 0.892968826 |
| chr1:179953960-179955576 | -0.107329109 | 0.732827096 | 0.892968826 |
| chr17:36731038-36731573 | -0.125300014 | 0.732828336 | 0.892968826 |
| chr1:74733714-74734053 | -0.120668827 | 0.732842134 | 0.892971439 |
| chr8:113980255-113980657 | -0.133343083 | 0.732886987 | 0.893011891 |
| chr6:110415441-110415717 | -0.156893981 | 0.732907199 | 0.893022318 |
| chrX:12702450-12702789 | -0.142856791 | 0.732983904 | 0.893088585 |
| chr1:94025882-94026166 | 0.153689441 | 0.733011996 | 0.893088585 |
| chr12:107093101-107093340 | -0.132059202 | 0.733019796 | 0.893088585 |
| chr3:28241091-28242055 | -0.108554464 | 0.73302306 | 0.893088585 |
| chr8:13033883-13034232 | 0.139200686 | 0.733026945 | 0.893088585 |
| chr2:232876312-232876896 | -0.120969092 | 0.733034679 | 0.893088585 |
| chr6:84422428-84422699 | -0.156762047 | 0.733043176 | 0.893088585 |
| chr10:43362031-43362799 | 0.110780037 | 0.733076558 | 0.89310909 |
| chr6:31619898-31621138 | 0.108400161 | 0.733083318 | 0.89310909 |
| chr3:19130455-19130863 | -0.148202255 | 0.733116176 | 0.893122891 |
| chr6:89117291-89117860 | 0.129064973 | 0.733117959 | 0.893122891 |
| chr8:116693684-116693981 | -0.14473053 | 0.733140429 | 0.893129973 |
| chrX:103687756-103688182 | 0.122252397 | 0.733166284 | 0.893129973 |
| chr1:203504483-203505027 | -0.114749836 | 0.73317228 | 0.893129973 |

| | | | |
|---|---|---|---|
| chr4:55395408-55396674 | 0.109317656 | 0.733172748 | 0.893129973 |
| chr6:24933936-24934149 | -0.176024197 | 0.733210641 | 0.893129973 |
| chr2:38602205-38603344 | 0.107883655 | 0.733213438 | 0.893129973 |
| chr11:30360794-30361139 | -0.16283988 | 0.733223502 | 0.893129973 |
| chr17:62381470-62382068 | -0.121722667 | 0.73325599 | 0.893129973 |
| chr6:117711484-117711684 | -0.174945297 | 0.733265357 | 0.893129973 |
| chr6:170584121-170585536 | -0.107960157 | 0.733266016 | 0.893129973 |
| chr10:78973146-78974332 | -0.10985477 | 0.733268896 | 0.893129973 |
| chr17:50865557-50865793 | 0.123640083 | 0.733293667 | 0.893129973 |
| chr12:123070023-123070252 | -0.158518086 | 0.733311634 | 0.893129973 |
| chr6:26123549-26124476 | -0.109785972 | 0.733312822 | 0.893129973 |
| chr20:63215998-63216641 | -0.111040328 | 0.733323758 | 0.893129973 |
| chr19:4717053-4717253 | -0.175794905 | 0.733325065 | 0.893129973 |
| chr5:90265278-90265497 | -0.17626637 | 0.733326953 | 0.893129973 |
| chr10:11285248-11285448 | -0.166714656 | 0.733339736 | 0.893129973 |
| chr10:100883402-100883708 | 0.145790701 | 0.733356565 | 0.893129973 |
| chr14:35413232-35413526 | -0.128187319 | 0.733356898 | 0.893129973 |
| chr17:42403124-42403432 | 0.154580948 | 0.733436488 | 0.893197861 |
| chr10:98134377-98135291 | -0.111485544 | 0.73344493 | 0.893197861 |
| chr16:72927567-72928378 | -0.112475333 | 0.733454345 | 0.893197861 |
| chr12:56636252-56636572 | -0.117669294 | 0.733459271 | 0.893197861 |
| chr19:51366286-51366656 | -0.118053977 | 0.733473271 | 0.893200715 |
| chr19:40777864-40778997 | 0.108303547 | 0.733487415 | 0.893203743 |
| chr18:78999972-79000281 | -0.14486779 | 0.733539582 | 0.89324362 |
| chr1:205376057-205376552 | 0.119312661 | 0.733543477 | 0.89324362 |
| chr3:125698323-125698591 | 0.152859688 | 0.733583595 | 0.893266485 |
| chr6:27824039-27824239 | -0.160410962 | 0.73360416 | 0.893266485 |
| chr1:244969540-244971478 | -0.106380878 | 0.73361518 | 0.893266485 |
| chrX:101759109-101759419 | -0.138475041 | 0.733630172 | 0.893266485 |
| chr1:115624931-115625458 | -0.125366446 | 0.733666241 | 0.893266485 |
| chr5:56814886-56815957 | 0.1100741 | 0.733672815 | 0.893266485 |
| chr1:205843229-205843436 | 0.15340143 | 0.733682076 | 0.893266485 |
| chr3:24583042-24583332 | -0.149551808 | 0.733684103 | 0.893266485 |
| chr11:6682804-6683765 | -0.109870454 | 0.733687266 | 0.893266485 |
| chr16:20729834-20730075 | -0.178527584 | 0.733689306 | 0.893266485 |
| chr1:162375626-162375957 | -0.151805933 | 0.733701359 | 0.893266485 |
| chr1:231725218-231725522 | 0.14079369 | 0.733702151 | 0.893266485 |
| chr1:48688490-48688690 | 0.171168014 | 0.733718768 | 0.893272522 |
| chr1:33127041-33127702 | 0.111818411 | 0.733738084 | 0.893281845 |
| chr5:179914069-179914397 | 0.134216799 | 0.733844006 | 0.893370458 |
| chr20:33992956-33993990 | 0.107149764 | 0.733844429 | 0.893370458 |
| chr13:100675387-100675601 | -0.146832228 | 0.733845849 | 0.893370458 |
| chr6:38639205-38640400 | 0.111272534 | 0.7339006 | 0.893412069 |
| chr3:2980653-2980996 | 0.131211056 | 0.733903349 | 0.893412069 |
| chr14:55438133-55438347 | 0.155923445 | 0.733953815 | 0.893442082 |
| chr3:126654571-126655313 | 0.114919011 | 0.733956095 | 0.893442082 |
| chr3:112990647-112991550 | 0.108562138 | 0.733971854 | 0.893442082 |
| chr12:13044019-13044919 | -0.115123813 | 0.733991415 | 0.893442082 |
| chr9:93451418-93451767 | 0.1300271 | 0.733996357 | 0.893442082 |
| chr5:178130437-178131487 | -0.108224034 | 0.734011733 | 0.893442082 |

| | | | |
|---|---|---|---|
| chr4:109814846-109816072 | -0.108816454 | 0.734016444 | 0.893442082 |
| chr5:18281939-18282240 | -0.136999546 | 0.734021287 | 0.893442082 |
| chr17:63612208-63612744 | 0.116531748 | 0.734038838 | 0.893449251 |
| chr19:47256369-47257807 | 0.106506897 | 0.734055172 | 0.89345494 |
| chr17:50707158-50708142 | -0.109959311 | 0.734101086 | 0.893486264 |
| chr6:37051472-37051702 | 0.152105678 | 0.734104229 | 0.893486264 |
| chr4:31370910-31371158 | 0.149381722 | 0.734146359 | 0.893519786 |
| chr2:176643851-176644051 | -0.19357293 | 0.734155094 | 0.893519786 |
| chr1:43649506-43650669 | 0.108064433 | 0.734201735 | 0.893562358 |
| chr10:86795464-86795711 | 0.143574959 | 0.734250898 | 0.893596879 |
| chr11:117143469-117143750 | -0.148741156 | 0.734262236 | 0.893596879 |
| chr5:134289575-134289865 | -0.144044327 | 0.734279383 | 0.893596879 |
| chr20:54680681-54681390 | 0.108267823 | 0.734287382 | 0.893596879 |
| chr3:122086752-122087015 | 0.15702332 | 0.734288412 | 0.893596879 |
| chr18:35355102-35355490 | -0.124832109 | 0.734351331 | 0.893659255 |
| chr1:194044801-194045019 | -0.167049376 | 0.73437518 | 0.893668741 |
| chr8:90174455-90174742 | 0.135774852 | 0.734382453 | 0.893668741 |
| chr16:96981-97290 | -0.129119664 | 0.734427708 | 0.893709619 |
| chr3:81643774-81644168 | 0.130984049 | 0.73445554 | 0.893724964 |
| chr16:3706778-3707025 | 0.160288662 | 0.734467457 | 0.893724964 |
| chr5:60860364-60860589 | 0.181384178 | 0.734475311 | 0.893724964 |
| chr15:65385448-65385971 | 0.116246881 | 0.73451503 | 0.893737907 |
| chr1:150560015-150560357 | 0.115658012 | 0.734515921 | 0.893737907 |
| chr17:43125066-43125724 | 0.113965892 | 0.734527377 | 0.893737907 |
| chr4:147683749-147684757 | 0.10799542 | 0.73454361 | 0.893737907 |
| chr21:42818355-42818558 | -0.173070822 | 0.734561332 | 0.893737907 |
| chr1:102897614-102897814 | -0.180104081 | 0.734562282 | 0.893737907 |
| chr8:98166761-98167063 | -0.130034515 | 0.734585633 | 0.893737907 |
| chr1:25232176-25232898 | 0.118356236 | 0.734586412 | 0.893737907 |
| chr11:69101214-69102006 | 0.111779618 | 0.734598146 | 0.893737907 |
| chr7:47876207-47876696 | -0.138159652 | 0.734604067 | 0.893737907 |
| chr3:87307821-87308110 | -0.142186221 | 0.734614254 | 0.893737907 |
| chr5:170631808-170632126 | 0.128895424 | 0.734658208 | 0.893777191 |
| chr6:138231236-138231888 | -0.116185712 | 0.734698993 | 0.893807719 |
| chr11:11411628-11411828 | -0.177341935 | 0.734706632 | 0.893807719 |
| chr11:130213152-130213496 | 0.139010765 | 0.734719285 | 0.893808921 |
| chr11:94768805-94769103 | 0.140871971 | 0.734746193 | 0.893811973 |
| chr7:2537944-2538375 | 0.126446283 | 0.734753186 | 0.893811973 |
| chr3:119463125-119464076 | 0.108250968 | 0.73475679 | 0.893811973 |
| chr5:113043625-113043825 | -0.15631801 | 0.734792928 | 0.893839136 |
| chr17:60614231-60614704 | -0.110557376 | 0.73480245 | 0.893839136 |
| chr8:66112895-66114350 | 0.106074927 | 0.734858071 | 0.893858138 |
| chr19:49135841-49136383 | 0.123543404 | 0.734863758 | 0.893858138 |
| chr22:35371923-35372322 | 0.134544788 | 0.734875749 | 0.893858138 |
| chr17:60704436-60704804 | 0.114525161 | 0.734879802 | 0.893858138 |
| chr8:143815661-143816180 | 0.116688393 | 0.734901522 | 0.893858138 |
| chr10:24503652-24503936 | -0.146282674 | 0.734912069 | 0.893858138 |
| chr10:50224371-50224678 | -0.140803757 | 0.734913697 | 0.893858138 |
| chr12:19165554-19165754 | 0.175725838 | 0.734923358 | 0.893858138 |
| chr4:30950802-30951164 | -0.149299144 | 0.734925002 | 0.893858138 |

| | | | |
|---|---|---|---|
| chr8:109333393-109334807 | -0.10550613 | 0.734935812 | 0.893858138 |
| chr2:171541164-171541547 | -0.129549676 | 0.734956937 | 0.893858138 |
| chr14:100284909-100286022 | -0.108878122 | 0.734961278 | 0.893858138 |
| chr5:5340372-5340589 | 0.16382255 | 0.734980409 | 0.893858138 |
| chr18:10687692-10687892 | 0.192325221 | 0.734981393 | 0.893858138 |
| chr1:60925438-60925790 | -0.139508523 | 0.73500151 | 0.893868417 |
| chr5:140261453-140261653 | 0.188611006 | 0.735014679 | 0.893870244 |
| chr17:59829368-59829921 | -0.111102147 | 0.735026871 | 0.893870884 |
| chr5:60295463-60295700 | 0.160641229 | 0.735095138 | 0.893898076 |
| chr6:7389274-7390272 | 0.108415735 | 0.735098685 | 0.893898076 |
| chr12:12788734-12789042 | -0.13780296 | 0.735115041 | 0.893898076 |
| chr4:70992809-70994208 | -0.107437464 | 0.735141543 | 0.893898076 |
| chr3:184234290-184234682 | 0.129035694 | 0.735141749 | 0.893898076 |
| chr8:48514045-48515245 | -0.1078282 | 0.735150253 | 0.893898076 |
| chr8:101607637-101608502 | 0.110670605 | 0.735169281 | 0.893898076 |
| chr18:3129332-3129595 | 0.145194361 | 0.735171329 | 0.893898076 |
| chr7:74455289-74455610 | 0.141079743 | 0.735186665 | 0.893898076 |
| chr19:14207217-14207665 | -0.128843424 | 0.735191986 | 0.893898076 |
| chr11:10285401-10285635 | -0.181458566 | 0.735197084 | 0.893898076 |
| chr16:66552349-66552742 | -0.120769802 | 0.735202898 | 0.893898076 |
| chr5:11890837-11891037 | 0.174588354 | 0.73521256 | 0.893898076 |
| chr6:91363645-91363845 | 0.174588354 | 0.73521256 | 0.893898076 |
| chr2:240851183-240851580 | -0.126768991 | 0.73524583 | 0.893906192 |
| chr5:150508176-150508517 | 0.134652155 | 0.735246562 | 0.893906192 |
| chr17:10286437-10286654 | -0.18879915 | 0.735254234 | 0.893906192 |
| chr10:124403534-124403829 | -0.146885421 | 0.735279815 | 0.893923109 |
| chr5:159975578-159975813 | 0.152234158 | 0.735332367 | 0.893971214 |
| chr11:119084364-119085711 | 0.108159209 | 0.735342718 | 0.893971214 |
| chr8:108222007-108222351 | 0.145880252 | 0.735359001 | 0.893976826 |
| chr1:163174920-163175385 | 0.137001488 | 0.735403924 | 0.894008609 |
| chr6:34888964-34889492 | -0.115703187 | 0.735408481 | 0.894008609 |
| chr2:70304002-70304239 | 0.141428653 | 0.735444518 | 0.894032512 |
| chr3:160720257-160720670 | 0.133085877 | 0.735452939 | 0.894032512 |
| chr20:63834939-63835365 | 0.115491696 | 0.735464247 | 0.894032512 |
| chr7:1536880-1537640 | 0.109507781 | 0.735516563 | 0.894032512 |
| chr3:119580186-119580411 | -0.152862797 | 0.735530977 | 0.894032512 |
| chr20:53652460-53652707 | 0.112956419 | 0.735535467 | 0.894032512 |
| chr4:67465705-67465958 | -0.163021018 | 0.735538343 | 0.894032512 |
| chr8:140944279-140944479 | 0.167825296 | 0.735570645 | 0.894032512 |
| chr19:48553708-48554115 | 0.124982681 | 0.735571515 | 0.894032512 |
| chr9:34178535-34179895 | -0.110122248 | 0.735583069 | 0.894032512 |
| chr5:10752737-10753113 | -0.136360505 | 0.735595422 | 0.894032512 |
| chr8:123517060-123517460 | -0.13568715 | 0.735599141 | 0.894032512 |
| chr15:72473604-72474626 | 0.108333365 | 0.735601605 | 0.894032512 |
| chr6:40714729-40715332 | -0.116781955 | 0.735605255 | 0.894032512 |
| chr2:188363861-188364503 | -0.119912154 | 0.735615934 | 0.894032512 |
| chr12:8697360-8698808 | -0.110062811 | 0.73562115 | 0.894032512 |
| chr3:112561171-112562653 | -0.105316295 | 0.735626501 | 0.894032512 |
| chr12:124939177-124940493 | 0.105610764 | 0.735643189 | 0.894038613 |
| chr3:58463538-58464089 | -0.123914924 | 0.735691406 | 0.894083031 |

| | | | |
|---|---|---|---|
| chr16:30092303-30092848 | -0.118482546 | 0.735704285 | 0.894084502 |
| chr22:30968877-30969631 | 0.111124928 | 0.735720457 | 0.894089974 |
| chr3:49104454-49105041 | 0.118831337 | 0.735743874 | 0.89410425 |
| chr10:78464493-78464812 | -0.157919825 | 0.73577138 | 0.894123496 |
| chr6:77457817-77458081 | -0.148422234 | 0.73581952 | 0.894149683 |
| chr8:37842895-37843295 | -0.13406613 | 0.735844798 | 0.894149683 |
| chr2:117814313-117815274 | -0.109246223 | 0.7358655 | 0.894149683 |
| chr4:152307720-152307998 | 0.140006317 | 0.735874147 | 0.894149683 |
| chr3:127227056-127227567 | 0.122711765 | 0.735879314 | 0.894149683 |
| chr1:198156771-198157845 | -0.106253781 | 0.735880029 | 0.894149683 |
| chr4:99946515-99946844 | 0.119934284 | 0.735891205 | 0.894149683 |
| chr1:109946812-109947030 | 0.165764856 | 0.735903406 | 0.894149683 |
| chr19:3055740-3056391 | 0.123471023 | 0.735911236 | 0.894149683 |
| chr8:109666041-109666712 | -0.117699165 | 0.73592509 | 0.894149683 |
| chr6:144266068-144266606 | -0.119127968 | 0.735925417 | 0.894149683 |
| chr16:75251518-75251996 | 0.112410601 | 0.735932965 | 0.894149683 |
| chr2:121402342-121402573 | 0.161674083 | 0.735984095 | 0.894197626 |
| chr15:89383624-89383864 | 0.131847041 | 0.736001922 | 0.894205107 |
| chr19:3863766-3863966 | -0.191919787 | 0.736027005 | 0.894218224 |
| chr15:39976290-39976955 | -0.117715285 | 0.736043598 | 0.894218224 |
| chr20:53685954-53686533 | 0.106799963 | 0.73604773 | 0.894218224 |
| chr16:84585903-84586156 | -0.142038478 | 0.73607545 | 0.894237722 |
| chr3:50455671-50456015 | -0.160092539 | 0.736100706 | 0.894254226 |
| chr10:60172370-60172714 | 0.123716471 | 0.736162566 | 0.89429741 |
| chr7:76437940-76438166 | 0.169385712 | 0.736169844 | 0.89429741 |
| chr8:2024077-2024290 | -0.16190246 | 0.736171267 | 0.89429741 |
| chr10:42552094-42553344 | -0.108861094 | 0.736199884 | 0.894311771 |
| chr10:35126376-35127768 | -0.106879997 | 0.736206432 | 0.894311771 |
| chr12:123150965-123151476 | -0.130062178 | 0.736226104 | 0.894321489 |
| chr10:72035595-72035795 | -0.167755288 | 0.736248552 | 0.894331934 |
| chr8:142615378-142615646 | -0.127642674 | 0.736258047 | 0.894331934 |
| chr2:148646080-148646368 | -0.119796484 | 0.736277747 | 0.894335665 |
| chr8:87409689-87409931 | -0.186026419 | 0.736284463 | 0.894335665 |
| chr7:38632443-38632741 | 0.126664417 | 0.736363014 | 0.894413793 |
| chr11:57778442-57778892 | -0.117996127 | 0.736374094 | 0.894413793 |
| chr2:151947259-151947506 | 0.137241136 | 0.736397906 | 0.894413793 |
| chr12:122774106-122775000 | -0.115103006 | 0.736399902 | 0.894413793 |
| chr2:241314986-241316242 | 0.106412047 | 0.736423949 | 0.894413793 |
| chr1:109546319-109546583 | 0.1276414 | 0.736439694 | 0.894413793 |
| chr22:29478591-29478791 | 0.178110214 | 0.736444679 | 0.894413793 |
| chr20:59708967-59709167 | 0.172487157 | 0.736478435 | 0.894413793 |
| chr1:244451090-244452427 | -0.105924691 | 0.736480637 | 0.894413793 |
| chr8:124030371-124030731 | -0.121189941 | 0.736493552 | 0.894413793 |
| chr13:31825059-31825377 | 0.146916802 | 0.736505544 | 0.894413793 |
| chr20:51801254-51801548 | -0.129193675 | 0.736515773 | 0.894413793 |
| chr19:35267610-35267843 | -0.139482801 | 0.73651924 | 0.894413793 |
| chr10:98375950-98376150 | -0.157126952 | 0.736539381 | 0.894413793 |
| chr16:49904246-49904501 | 0.174680896 | 0.736544948 | 0.894413793 |
| chr1:185317041-185318232 | -0.105727133 | 0.736546515 | 0.894413793 |
| chr6:53437550-53437897 | -0.131069791 | 0.736554432 | 0.894413793 |

| | | | |
|---|---|---|---|
| chr21:25608313-25608587 | 0.132841766 | 0.736558899 | 0.894413793 |
| chr13:95676136-95677792 | -0.106856781 | 0.736578257 | 0.894423126 |
| chr19:47048529-47049034 | -0.113051153 | 0.736624583 | 0.894459518 |
| chr16:74984470-74985518 | -0.106796076 | 0.736635417 | 0.894459518 |
| chr17:58218553-58219691 | -0.107087613 | 0.736643248 | 0.894459518 |
| chr8:144792161-144792700 | 0.110855462 | 0.73666468 | 0.894471367 |
| chr10:44306856-44307067 | -0.168369876 | 0.736688789 | 0.894486465 |
| chr10:70681432-70681632 | -0.169423417 | 0.736722443 | 0.894513153 |
| chr7:151856137-151856337 | -0.158492761 | 0.736738443 | 0.894514751 |
| chr1:27959402-27960460 | 0.111169211 | 0.736775826 | 0.894514751 |
| chr1:173431231-173431566 | 0.126320163 | 0.73677875 | 0.894514751 |
| chr11:36702504-36702961 | -0.127096473 | 0.736783309 | 0.894514751 |
| chr15:93041647-93041888 | -0.145976708 | 0.736787679 | 0.894514751 |
| chr8:22054106-22054324 | 0.147900192 | 0.736793859 | 0.894514751 |
| chr17:21275636-21276221 | 0.114765815 | 0.736817942 | 0.894514751 |
| chr17:12789280-12790287 | 0.112321083 | 0.736834607 | 0.894514751 |
| chr14:64619482-64619684 | 0.18475085 | 0.736842229 | 0.894514751 |
| chr5:137580194-137580394 | 0.197489477 | 0.736860338 | 0.894514751 |
| chr3:13352457-13352734 | 0.148087492 | 0.736872596 | 0.894514751 |
| chr6:32440048-32440248 | -0.204362904 | 0.736880353 | 0.894514751 |
| chr7:96916315-96916530 | -0.204362904 | 0.736880353 | 0.894514751 |
| chr20:57628994-57629248 | -0.148490567 | 0.7368872 | 0.894514751 |
| chr12:2959921-2960274 | 0.126263008 | 0.736915974 | 0.894535508 |
| chr19:46859807-46861061 | -0.106879047 | 0.736939175 | 0.894540172 |
| chr6:28606524-28607424 | -0.110667786 | 0.736943166 | 0.894540172 |
| chr9:134407362-134407674 | 0.126716377 | 0.736989879 | 0.894582702 |
| chr8:38230856-38232344 | 0.108710832 | 0.737016306 | 0.894600608 |
| chr9:133186117-133186381 | 0.15246051 | 0.737039383 | 0.894602137 |
| chr17:42997894-42998640 | 0.111026095 | 0.737045026 | 0.894602137 |
| chr11:71146166-71146366 | 0.173657292 | 0.737052592 | 0.894602137 |
| chr6:107957673-107959022 | 0.107014644 | 0.737104754 | 0.894628591 |
| chr12:72972380-72972694 | -0.173720502 | 0.737105599 | 0.894628591 |
| chr17:21042859-21043845 | -0.107299133 | 0.737120426 | 0.894628591 |
| chr6:11229751-11230075 | 0.130156778 | 0.737124918 | 0.894628591 |
| chr20:63682004-63682227 | -0.138093347 | 0.737140896 | 0.894628591 |
| chr3:149282901-149283170 | -0.147323599 | 0.737144443 | 0.894628591 |
| chr6:53347939-53349318 | -0.107632443 | 0.737185083 | 0.894663743 |
| chr15:44784039-44785019 | 0.116114013 | 0.737198558 | 0.894665926 |
| chr3:156382548-156382787 | -0.138367933 | 0.73723837 | 0.894674812 |
| chr8:143018326-143018643 | 0.123846103 | 0.737251098 | 0.894674812 |
| chr2:207149875-207150180 | -0.168437436 | 0.737259384 | 0.894674812 |
| chr1:201436492-201436791 | 0.123007842 | 0.737259415 | 0.894674812 |
| chr7:11181793-11182044 | 0.18383248 | 0.737264263 | 0.894674812 |
| chr18:23135016-23135522 | 0.118863895 | 0.737302031 | 0.894703068 |
| chr12:117236520-117236794 | -0.166974568 | 0.737335015 | 0.894703068 |
| chr13:113724830-113725404 | 0.125124134 | 0.737354954 | 0.894703068 |
| chr1:54575292-54575492 | 0.168747697 | 0.737356324 | 0.894703068 |
| chr12:78229585-78229785 | 0.187493021 | 0.737360413 | 0.894703068 |
| chr14:72575235-72575519 | -0.153472263 | 0.737373596 | 0.894703068 |
| chrX:140123479-140123733 | -0.156785207 | 0.737376906 | 0.894703068 |

| | | | |
|---|---|---|---|
| chr7:120383914-120384229 | -0.191083612 | 0.737380962 | 0.894703068 |
| chr19:15379190-15380334 | 0.109299676 | 0.737409282 | 0.894723263 |
| chr9:33742599-33742858 | 0.159914359 | 0.737455734 | 0.894765455 |
| chr12:14239340-14239822 | -0.120295632 | 0.737518781 | 0.894813292 |
| chr16:89972108-89972308 | 0.145570941 | 0.737530249 | 0.894813292 |
| chr10:24723793-24723993 | -0.144710926 | 0.737531674 | 0.894813292 |
| chr3:53845622-53845883 | 0.135897338 | 0.737541874 | 0.894813292 |
| chr16:67513954-67514225 | 0.1458948 | 0.737557949 | 0.894815831 |
| chr15:95559887-95560090 | 0.16547293 | 0.737577146 | 0.894815831 |
| chr7:2451360-2452155 | 0.111082775 | 0.737579001 | 0.894815831 |
| chr4:139176335-139178709 | 0.104277045 | 0.737591668 | 0.89481703 |
| chr21:42514235-42514825 | -0.114887105 | 0.737660771 | 0.894855855 |
| chr8:57491415-57491616 | -0.182576799 | 0.73766717 | 0.894855855 |
| chr10:104337850-104338833 | 0.112255976 | 0.737670784 | 0.894855855 |
| chr3:140863675-140863970 | -0.140711116 | 0.737676268 | 0.894855855 |
| chr6:142842896-142843144 | 0.149749567 | 0.737691368 | 0.894855855 |
| chr10:97531730-97531937 | -0.15930159 | 0.737703525 | 0.894855855 |
| chr3:66475402-66476307 | -0.114299433 | 0.737705423 | 0.894855855 |
| chr7:135707293-135707493 | 0.155552366 | 0.737750107 | 0.894895891 |
| chr3:121111943-121112176 | -0.157809484 | 0.737780242 | 0.894901166 |
| chr11:62123654-62124338 | 0.111189948 | 0.73779646 | 0.894901166 |
| chr16:89686358-89687450 | -0.107670828 | 0.737798677 | 0.894901166 |
| chr10:23686831-23687031 | 0.172861976 | 0.737801174 | 0.894901166 |
| chr19:48203700-48204418 | -0.110376409 | 0.737830037 | 0.894918468 |
| chr19:52171471-52171807 | 0.133195406 | 0.73784661 | 0.894918468 |
| chr2:20767048-20767248 | 0.177551435 | 0.737850478 | 0.894918468 |
| chr3:134145312-134145529 | 0.15731835 | 0.737933926 | 0.894953441 |
| chrX:74614426-74615085 | 0.110426046 | 0.737947542 | 0.894953441 |
| chr17:74379520-74379750 | -0.134867032 | 0.73794844 | 0.894953441 |
| chr10:88934865-88935133 | -0.133075315 | 0.737950134 | 0.894953441 |
| chr5:150847082-150847440 | 0.125484362 | 0.737969821 | 0.894953441 |
| chr15:64960034-64960234 | -0.173750162 | 0.737970958 | 0.894953441 |
| chr17:59440634-59441157 | 0.112684859 | 0.73798879 | 0.894953441 |
| chr4:81069716-81070142 | -0.121672391 | 0.738004287 | 0.894953441 |
| chr2:51102797-51102997 | -0.205816273 | 0.738027607 | 0.894953441 |
| chr17:49106667-49106891 | -0.153644886 | 0.738030258 | 0.894953441 |
| chr2:231672876-231673080 | -0.165770304 | 0.73804178 | 0.894953441 |
| chr9:15416923-15417201 | -0.145397693 | 0.738043758 | 0.894953441 |
| chrX:132737804-132738067 | -0.153835636 | 0.738055965 | 0.894953441 |
| chr2:198054375-198054625 | 0.142502961 | 0.738066375 | 0.894953441 |
| chr16:85611642-85612494 | -0.106623385 | 0.738075873 | 0.894953441 |
| chr17:61833544-61834061 | 0.107750682 | 0.738077267 | 0.894953441 |
| chr14:23094525-23095905 | 0.104993165 | 0.738077874 | 0.894953441 |
| chr5:180860532-180861902 | 0.105697703 | 0.738128481 | 0.894976335 |
| chr2:235551969-235552257 | 0.140872113 | 0.738164626 | 0.894976335 |
| chr5:159460356-159460585 | -0.178115555 | 0.738184038 | 0.894976335 |
| chr3:160999368-160999631 | -0.170734267 | 0.738184421 | 0.894976335 |
| chr20:49691374-49691604 | -0.164698681 | 0.738195063 | 0.894976335 |
| chr12:55659638-55660075 | 0.132826446 | 0.738195732 | 0.894976335 |
| chrX:129336987-129337776 | -0.114204542 | 0.738197677 | 0.894976335 |

| | | | |
|---|---|---|---|
| chr13:107680156-107680435 | -0.151216453 | 0.738201496 | 0.894976335 |
| chr11:69587494-69587703 | -0.179238982 | 0.738201879 | 0.894976335 |
| chr5:54296363-54296677 | -0.152092702 | 0.738257369 | 0.895029448 |
| chr12:72017067-72017406 | -0.122341312 | 0.738325288 | 0.895051917 |
| chr9:132354562-132355687 | -0.107666385 | 0.738337064 | 0.895051917 |
| chr22:42649223-42649697 | 0.12443204 | 0.738340292 | 0.895051917 |
| chr10:29990466-29990666 | -0.175952511 | 0.738347598 | 0.895051917 |
| chr10:26438665-26439097 | 0.137895437 | 0.738352 | 0.895051917 |
| chr2:32946244-32948188 | -0.105667609 | 0.738368091 | 0.895051917 |
| chr9:126513914-126514509 | 0.122482862 | 0.738369758 | 0.895051917 |
| chr3:128297714-128298095 | -0.134937027 | 0.738380126 | 0.895051917 |
| chr5:154732312-154732512 | 0.170053241 | 0.738400267 | 0.895051917 |
| chr8:29316763-29317026 | 0.146677032 | 0.738402831 | 0.895051917 |
| chr10:119029088-119030001 | 0.108645436 | 0.738404398 | 0.895051917 |
| chr9:133346929-133347234 | 0.122960873 | 0.738459315 | 0.895104324 |
| chr3:192916956-192918478 | 0.106259823 | 0.738492557 | 0.895106856 |
| chr2:84907171-84907812 | -0.114333911 | 0.738495893 | 0.895106856 |
| chr15:100464365-100464565 | 0.16423181 | 0.738505426 | 0.895106856 |
| chr15:77519467-77519741 | 0.16246685 | 0.738508132 | 0.895106856 |
| chr5:43514542-43515472 | 0.107085159 | 0.738534113 | 0.895124186 |
| chr5:173643953-173644213 | 0.143156617 | 0.738554225 | 0.895134404 |
| chr10:1845681-1845997 | 0.159954577 | 0.738582151 | 0.89515409 |
| chr15:86624651-86624976 | -0.132420814 | 0.738605465 | 0.89516252 |
| chrX:106168138-106168398 | -0.155465603 | 0.738612472 | 0.89516252 |
| chr3:46078899-46079120 | 0.153692075 | 0.738625281 | 0.895163885 |
| chr14:22982475-22983027 | 0.112241032 | 0.738686585 | 0.895224022 |
| chr6:28542446-28543370 | -0.107938759 | 0.738707516 | 0.895235228 |
| chr10:36889033-36889247 | -0.158427465 | 0.738720089 | 0.895236306 |
| chr3:136250148-136250869 | -0.107205322 | 0.738735469 | 0.895240785 |
| chr12:79373966-79374218 | -0.138866201 | 0.73879162 | 0.89526488 |
| chr14:101716933-101717336 | -0.140106037 | 0.738824789 | 0.89526488 |
| chr19:6459731-6460503 | 0.113105425 | 0.738844554 | 0.89526488 |
| chr5:154700932-154701590 | 0.113615691 | 0.738848617 | 0.89526488 |
| chr13:46552929-46553462 | 0.108534471 | 0.738862395 | 0.89526488 |
| chr14:44379651-44379936 | 0.161343313 | 0.738872246 | 0.89526488 |
| chr2:113977387-113977603 | 0.161343313 | 0.738872246 | 0.89526488 |
| chr3:42071882-42072141 | 0.161343313 | 0.738872246 | 0.89526488 |
| chr3:38521463-38521668 | 0.158958847 | 0.738875058 | 0.89526488 |
| chr5:142223730-142223977 | -0.150417642 | 0.738879152 | 0.89526488 |
| chr15:55435257-55435552 | 0.138111284 | 0.738886431 | 0.89526488 |
| chr19:50836077-50836299 | 0.156119756 | 0.738895562 | 0.89526488 |
| chr13:95380543-95380925 | 0.14144753 | 0.738933806 | 0.895281659 |
| chr1:47333461-47334825 | 0.106962966 | 0.738971218 | 0.895281659 |
| chr1:230581851-230582176 | 0.163037247 | 0.738981957 | 0.895281659 |
| chr2:215950036-215950328 | -0.133227954 | 0.738985897 | 0.895281659 |
| chr7:2151818-2152202 | -0.128042056 | 0.738992791 | 0.895281659 |
| chr6:6927849-6928366 | 0.119949956 | 0.738997379 | 0.895281659 |
| chr9:114155285-114155575 | 0.130350192 | 0.739005441 | 0.895281659 |
| chr8:142475502-142476148 | 0.111249201 | 0.739011538 | 0.895281659 |
| chr7:55181229-55181509 | -0.139692142 | 0.739014569 | 0.895281659 |

| | | | |
|---|---|---|---|
| chr6:109596160-109596360 | 0.170661741 | 0.739034882 | 0.895285025 |
| chrX:48675982-48677231 | 0.108130843 | 0.739040717 | 0.895285025 |
| chr7:17232686-17232943 | -0.163103564 | 0.739061767 | 0.895285449 |
| chr17:68011131-68011426 | -0.130606441 | 0.739064436 | 0.895285449 |
| chr3:18693663-18694437 | -0.118235213 | 0.739079445 | 0.895289476 |
| chr3:184149859-184150059 | 0.158132794 | 0.739093646 | 0.895292524 |
| chr3:123286288-123286673 | 0.11966758 | 0.739133152 | 0.895315158 |
| chr2:202871174-202872191 | 0.110519995 | 0.739135881 | 0.895315158 |
| chr4:53591301-53591907 | -0.108641784 | 0.739167814 | 0.895315158 |
| chr2:39093593-39093815 | 0.176341124 | 0.739186109 | 0.895315158 |
| chr10:111598474-111598746 | 0.139002231 | 0.739193503 | 0.895315158 |
| chr17:83200488-83200696 | -0.138374388 | 0.739200304 | 0.895315158 |
| chr2:214310350-214310651 | -0.138374388 | 0.739200304 | 0.895315158 |
| chr1:114316263-114316532 | 0.122786475 | 0.739213973 | 0.895315158 |
| chr1:204493551-204495304 | -0.104591128 | 0.739224507 | 0.895315158 |
| chr1:9589427-9589690 | -0.153591395 | 0.739229179 | 0.895315158 |
| chr16:74606388-74607590 | -0.106782074 | 0.739256621 | 0.895334242 |
| chr10:57657197-57657493 | -0.133483518 | 0.739331571 | 0.895410863 |
| chr14:76086486-76086686 | 0.179821423 | 0.739358512 | 0.895422282 |
| chr9:68574246-68574446 | -0.195017559 | 0.739364372 | 0.895422282 |
| chr11:64489875-64490115 | 0.152651393 | 0.739397618 | 0.895448392 |
| chr7:73830728-73831164 | -0.11452318 | 0.739431803 | 0.895475638 |
| chr17:16589068-16589703 | 0.114764785 | 0.739488706 | 0.895530395 |
| chr9:109705507-109705971 | 0.125997038 | 0.739534779 | 0.895559336 |
| chr18:3885267-3885488 | -0.164442045 | 0.73953598 | 0.895559336 |
| chrX:48071085-48072092 | 0.112717058 | 0.739576647 | 0.895594428 |
| chr22:37044070-37044442 | -0.131542912 | 0.739597419 | 0.895601185 |
| chr14:60443448-60443672 | -0.187856673 | 0.739610536 | 0.895601185 |
| chr11:17547055-17547454 | -0.12371565 | 0.739617293 | 0.895601185 |
| chr17:52171025-52171289 | 0.142402519 | 0.73964379 | 0.895614852 |
| chr11:62295509-62295762 | 0.161732018 | 0.739651956 | 0.895614852 |
| chr20:42302853-42303091 | -0.165303669 | 0.73968405 | 0.895629277 |
| chr19:11639334-11640172 | 0.111923801 | 0.739687247 | 0.895629277 |
| chr1:51331276-51331523 | 0.128937146 | 0.739719566 | 0.895649544 |
| chr1:151581913-151582369 | -0.113219395 | 0.739734471 | 0.895649544 |
| chr11:58579619-58580057 | 0.114543682 | 0.739746025 | 0.895649544 |
| chr3:189766417-189766723 | -0.12942682 | 0.739750743 | 0.895649544 |
| chr13:95426649-95426870 | 0.181242367 | 0.739762465 | 0.895649584 |
| chrX:53683767-53684629 | -0.106098692 | 0.739819075 | 0.89570397 |
| chr1:116498879-116499373 | -0.115585012 | 0.739831315 | 0.895704636 |
| chr16:53550046-53550258 | 0.168976336 | 0.739855903 | 0.895720251 |
| chr1:23344151-23344849 | 0.110707436 | 0.739907204 | 0.895768206 |
| chr9:126722136-126723207 | -0.108159489 | 0.739933759 | 0.895773256 |
| chr9:37422226-37423064 | -0.109800755 | 0.73995673 | 0.895773256 |
| chr20:63345263-63345464 | 0.156293878 | 0.739980887 | 0.895773256 |
| chr2:206323529-206323830 | -0.145495075 | 0.739989548 | 0.895773256 |
| chr6:120764440-120764759 | 0.140524565 | 0.740015166 | 0.895773256 |
| chr5:177402472-177403040 | -0.110855623 | 0.740019558 | 0.895773256 |
| chr12:85157066-85157276 | -0.168097365 | 0.740030048 | 0.895773256 |
| chr16:1932363-1932563 | 0.154092451 | 0.740033279 | 0.895773256 |

| | | | |
|---|---|---|---|
| chr15:90143177-90143722 | 0.119158047 | 0.740043586 | 0.895773256 |
| chr6:164005330-164005796 | -0.132442033 | 0.740062973 | 0.895773256 |
| chr3:58043987-58044242 | 0.145307461 | 0.740085682 | 0.895773256 |
| chr10:92823076-92823286 | 0.171298847 | 0.740094795 | 0.895773256 |
| chr4:83256824-83257141 | 0.136667585 | 0.740099288 | 0.895773256 |
| chr12:117587728-117588015 | 0.138663425 | 0.740105254 | 0.895773256 |
| chr13:79423920-79424393 | -0.142955117 | 0.740107527 | 0.895773256 |
| chr11:25180733-25180960 | 0.181470931 | 0.740110409 | 0.895773256 |
| chr10:102409258-102409611 | -0.13176614 | 0.740112142 | 0.895773256 |
| chr18:25351772-25352464 | 0.120519898 | 0.740121809 | 0.895773256 |
| chr17:47802392-47802920 | 0.115403322 | 0.740146253 | 0.895780842 |
| chr6:131199674-131200075 | 0.12408629 | 0.740155183 | 0.895780842 |
| chr14:24160342-24160542 | 0.139100685 | 0.74016315 | 0.895780842 |
| chr3:44337999-44339013 | -0.107516206 | 0.740197984 | 0.895799169 |
| chr7:7830204-7830418 | 0.145501635 | 0.740212214 | 0.895799169 |
| chr6:51618665-51618938 | 0.155216315 | 0.740217331 | 0.895799169 |
| chr17:28897170-28897885 | 0.115120971 | 0.740225058 | 0.895799169 |
| chr20:25537566-25537779 | 0.158141735 | 0.740257408 | 0.895824169 |
| chr2:127857758-127858859 | -0.107066805 | 0.74027934 | 0.895826028 |
| chr5:165457838-165458263 | -0.130720266 | 0.740282327 | 0.895826028 |
| chr6:49607117-49607429 | -0.13580598 | 0.740347193 | 0.89586977 |
| chr16:50719060-50719683 | -0.11823191 | 0.740359094 | 0.89586977 |
| chr1:23825162-23825599 | -0.131470595 | 0.740359846 | 0.89586977 |
| chr3:177196926-177198114 | -0.10782037 | 0.740365242 | 0.89586977 |
| chr22:39255273-39255473 | 0.183394094 | 0.740439068 | 0.895921123 |
| chr10:27656662-27656930 | -0.170027921 | 0.740446877 | 0.895921123 |
| chr7:74449833-74450229 | 0.132938527 | 0.740447478 | 0.895921123 |
| chr6:28489357-28489667 | 0.127004321 | 0.740461828 | 0.895921123 |
| chr17:68462796-68463311 | -0.11004451 | 0.740466145 | 0.895921123 |
| chr20:57029610-57029878 | -0.136175862 | 0.740481952 | 0.895926101 |
| chr19:10027630-10028093 | -0.120138069 | 0.740505682 | 0.895940665 |
| chrX:72181196-72182076 | 0.107888644 | 0.740541147 | 0.895969426 |
| chr3:66649518-66649718 | 0.188061758 | 0.740588628 | 0.89600339 |
| chr16:74480141-74480495 | -0.119728688 | 0.740592606 | 0.89600339 |
| chr2:65129964-65131019 | -0.104735128 | 0.740607782 | 0.896007602 |
| chr1:66503907-66504153 | -0.152013981 | 0.740637032 | 0.896028842 |
| chr10:60032779-60033095 | -0.135856435 | 0.740678682 | 0.896065082 |
| chrX:24426333-24426542 | 0.170737996 | 0.740710623 | 0.896072497 |
| chr5:151500717-151501113 | -0.118568464 | 0.740719381 | 0.896072497 |
| chr10:96147040-96147303 | -0.172128853 | 0.740726903 | 0.896072497 |
| chr12:92570385-92570669 | 0.138499337 | 0.74073159 | 0.896072497 |
| chr2:197804296-197804735 | 0.112868808 | 0.740753635 | 0.896082327 |
| chr1:151197127-151197522 | -0.110846232 | 0.740763105 | 0.896082327 |
| chrX:153495068-153495325 | 0.14187472 | 0.740807896 | 0.896116805 |
| chr8:134743738-134744294 | -0.11578525 | 0.740819544 | 0.896116805 |
| chr17:7239379-7240011 | -0.108906828 | 0.740836105 | 0.896116805 |
| chr14:81400128-81401138 | -0.112951541 | 0.740844185 | 0.896116805 |
| chr14:45140908-45141109 | -0.149691469 | 0.740851784 | 0.896116805 |
| chr6:93417340-93417630 | 0.128350499 | 0.740861779 | 0.896116805 |
| chr15:100035112-100035326 | -0.174535152 | 0.740901544 | 0.896150756 |

| | | | |
|---|---|---|---|
| chr13:110715253-110715918 | -0.106248459 | 0.740922139 | 0.896150782 |
| chr12:120194406-120195361 | 0.108254212 | 0.740924957 | 0.896150782 |
| chr19:40433701-40434549 | 0.110905161 | 0.740936977 | 0.896151174 |
| chr3:42600211-42601399 | -0.105917431 | 0.74104378 | 0.896266204 |
| chr10:148475-148804 | 0.153494019 | 0.741078774 | 0.89629438 |
| chr5:173573114-173573381 | 0.126003672 | 0.741129969 | 0.896327803 |
| chr14:103207030-103207365 | 0.142360122 | 0.74113978 | 0.896327803 |
| chr8:114503960-114504642 | -0.120608325 | 0.741144617 | 0.896327803 |
| chr1:15524174-15525252 | 0.10551012 | 0.741153201 | 0.896327803 |
| chr11:107916572-107916892 | -0.133357789 | 0.741167727 | 0.896329959 |
| chr22:42269670-42270776 | -0.114710109 | 0.741182764 | 0.896329959 |
| chr12:75977901-75978355 | -0.114397272 | 0.741216848 | 0.896329959 |
| chr11:47214239-47215529 | 0.104532367 | 0.741217664 | 0.896329959 |
| chr12:57111606-57112658 | 0.106881997 | 0.74122089 | 0.896329959 |
| chr2:45815452-45815704 | -0.163095007 | 0.741225173 | 0.896329959 |
| chr12:77324529-77325458 | -0.112577309 | 0.741256493 | 0.896346037 |
| chr2:37961977-37962651 | -0.122097806 | 0.741271004 | 0.896346037 |
| chr16:21159306-21159981 | -0.116381987 | 0.741290961 | 0.896346037 |
| chr1:7387522-7387759 | -0.163741432 | 0.741300546 | 0.896346037 |
| chrX:118014283-118014497 | -0.161008788 | 0.741307452 | 0.896346037 |
| chr19:48271181-48272089 | 0.113357851 | 0.741308658 | 0.896346037 |
| chr20:38460286-38460585 | -0.147436345 | 0.741339399 | 0.896352442 |
| chr5:163079664-163079875 | -0.166398447 | 0.741342587 | 0.896352442 |
| chr22:41832534-41833740 | -0.106664352 | 0.741349051 | 0.896352442 |
| chr14:81669518-81669718 | 0.165964165 | 0.741377768 | 0.896373019 |
| chr9:85425099-85425513 | -0.130127097 | 0.7414897 | 0.896494205 |
| chr14:20775861-20776070 | -0.173753846 | 0.741513453 | 0.896496881 |
| chr5:73448410-73448956 | 0.114455524 | 0.741515313 | 0.896496881 |
| chr2:178193554-178195535 | 0.103546751 | 0.741599356 | 0.896584342 |
| chr1:163807147-163807917 | -0.120546662 | 0.741669844 | 0.896655413 |
| chr2:207165978-207167104 | -0.10457702 | 0.741682821 | 0.896656954 |
| chr1:33095083-33095283 | 0.155548944 | 0.741708841 | 0.896674264 |
| chr1:152036242-152037186 | 0.105462569 | 0.741732579 | 0.896681104 |
| chr10:95907032-95908347 | -0.106128892 | 0.741758157 | 0.896681104 |
| chr5:109849378-109849578 | -0.179044756 | 0.741778788 | 0.896681104 |
| chr2:60189076-60189637 | -0.127998774 | 0.741779685 | 0.896681104 |
| chr11:68121288-68122108 | 0.108028912 | 0.741786332 | 0.896681104 |
| chr15:75450590-75451306 | 0.105250262 | 0.741786352 | 0.896681104 |
| chr5:35183343-35183564 | 0.14416258 | 0.741798402 | 0.896681104 |
| chr12:12872088-12873180 | -0.105729831 | 0.741815376 | 0.896681104 |
| chr10:24068931-24069141 | 0.172208297 | 0.741819824 | 0.896681104 |
| chr6:129252138-129252456 | -0.139810204 | 0.741874211 | 0.896732699 |
| chr17:63305205-63305503 | -0.132059898 | 0.741906253 | 0.896751031 |
| chr15:79768959-79769529 | -0.124152723 | 0.741927071 | 0.896751031 |
| chr1:150806071-150806301 | 0.157687588 | 0.741935096 | 0.896751031 |
| chr13:31846394-31847231 | 0.10979298 | 0.741936191 | 0.896751031 |
| chr14:60188165-60188439 | 0.145970176 | 0.741954294 | 0.896758766 |
| chr6:70566690-70567350 | -0.107516088 | 0.74198517 | 0.896767638 |
| chr16:57446819-57448024 | 0.111341633 | 0.741991279 | 0.896767638 |
| chr1:28505582-28506258 | 0.109369048 | 0.741996746 | 0.896767638 |

| | | | |
|---|---|---|---|
| chr16:85585955-85586290 | -0.134586186 | 0.742013726 | 0.896768568 |
| chr22:19122927-19123163 | 0.135362283 | 0.742020924 | 0.896768568 |
| chr1:94045507-94045958 | 0.123816682 | 0.742034914 | 0.896771331 |
| chrX:86369959-86370159 | -0.179358305 | 0.74206256 | 0.896777687 |
| chr1:179881873-179883324 | -0.103002225 | 0.742063581 | 0.896777687 |
| chr3:195270310-195271911 | -0.104230188 | 0.742100523 | 0.896808187 |
| chr6:90610853-90611671 | -0.11566039 | 0.742138901 | 0.896840421 |
| chr4:120977612-120977872 | 0.144097772 | 0.742178818 | 0.896874514 |
| chr1:159945474-159946488 | -0.104762422 | 0.742269865 | 0.896968952 |
| chr5:147869578-147869778 | -0.177485156 | 0.74228038 | 0.896968952 |
| chr22:20283150-20283352 | 0.168483357 | 0.742293784 | 0.896971003 |
| chr6:124683557-124683779 | -0.169665825 | 0.742339337 | 0.896978248 |
| chrX:136608597-136608923 | -0.15296924 | 0.742343691 | 0.896978248 |
| chr2:65430443-65431571 | 0.106002786 | 0.742345269 | 0.896978248 |
| chr11:36540979-36541363 | -0.125469804 | 0.742346606 | 0.896978248 |
| chr20:43192031-43192329 | 0.136627353 | 0.742392685 | 0.89701978 |
| chr13:40780688-40781083 | -0.119696906 | 0.742428187 | 0.897039954 |
| chr2:186951630-186951945 | -0.142824401 | 0.742434231 | 0.897039954 |
| chr3:126357326-126357673 | 0.117683707 | 0.742445401 | 0.897039954 |
| chr20:38143844-38144250 | -0.127200586 | 0.742456211 | 0.897039954 |
| chr5:139560682-139562057 | -0.104588733 | 0.742474206 | 0.89704755 |
| chr17:4789156-4789404 | -0.147756157 | 0.742502258 | 0.897059533 |
| chr2:192162462-192162751 | -0.144960845 | 0.742507539 | 0.897059533 |
| chr22:22195315-22195754 | -0.129931992 | 0.742566662 | 0.897116817 |
| chr16:31477895-31478270 | 0.119971998 | 0.742617017 | 0.897137405 |
| chr5:181188792-181189254 | -0.108167704 | 0.742621909 | 0.897137405 |
| chr5:50667420-50668224 | 0.113632105 | 0.742631533 | 0.897137405 |
| chr2:208545608-208545867 | -0.168136596 | 0.742661436 | 0.897137405 |
| chr6:57221345-57222604 | -0.105596856 | 0.742677231 | 0.897137405 |
| chr8:57065029-57065348 | -0.166695167 | 0.742695696 | 0.897137405 |
| chrX:49001498-49001734 | -0.149357045 | 0.742696273 | 0.897137405 |
| chr9:85248522-85248731 | 0.184704178 | 0.742710974 | 0.897137405 |
| chr5:4942151-4942561 | -0.134015167 | 0.742714717 | 0.897137405 |
| chr11:57311136-57311827 | -0.109226888 | 0.742720254 | 0.897137405 |
| chr15:84715446-84715807 | 0.122222649 | 0.742723105 | 0.897137405 |
| chr16:21301143-21301931 | 0.108387302 | 0.742725187 | 0.897137405 |
| chr3:167261933-167262133 | 0.192472713 | 0.742735915 | 0.897137405 |
| chr6:113654452-113654721 | 0.15659387 | 0.742775802 | 0.897155566 |
| chr6:85678531-85679511 | -0.104128303 | 0.74278366 | 0.897155566 |
| chr8:90888329-90888529 | -0.165533381 | 0.742786077 | 0.897155566 |
| chr11:65182988-65183256 | -0.140533814 | 0.742830487 | 0.897195062 |
| chr13:27620460-27622139 | 0.102731944 | 0.742865621 | 0.897223355 |
| chr1:212608244-212609015 | 0.105204134 | 0.742897417 | 0.897241364 |
| chr3:193743861-193744428 | -0.11496128 | 0.742903952 | 0.897241364 |
| chr4:126604372-126604673 | -0.142869355 | 0.742939634 | 0.897268741 |
| chr18:9073669-9074142 | -0.121093181 | 0.742950041 | 0.897268741 |
| chr2:274599-274799 | -0.15819252 | 0.742967474 | 0.897275653 |
| chr8:91155274-91155655 | -0.159434679 | 0.743019238 | 0.897324025 |
| chr22:29553335-29554042 | 0.108417948 | 0.743059745 | 0.897337708 |
| chr21:16587066-16587388 | 0.145267772 | 0.743060752 | 0.897337708 |

| | | | |
|---|---|---|---|
| chr5:154682380-154683298 | -0.104425353 | 0.743082106 | 0.897337708 |
| chr10:97407169-97407826 | -0.109046165 | 0.743093292 | 0.897337708 |
| chr15:92645074-92646010 | 0.106594824 | 0.743095297 | 0.897337708 |
| chr10:73772145-73773244 | 0.105715514 | 0.743100836 | 0.897337708 |
| chr6:154732976-154734529 | -0.102810557 | 0.74314831 | 0.897380893 |
| chr8:73293317-73294753 | 0.102839073 | 0.743198893 | 0.897406169 |
| chr8:143941928-143942418 | 0.109270131 | 0.743203324 | 0.897406169 |
| chr5:131775084-131775329 | 0.162635754 | 0.743204378 | 0.897406169 |
| chr4:106502096-106502459 | 0.126752394 | 0.743252702 | 0.897450377 |
| chr2:101120329-101120996 | -0.114466451 | 0.74326686 | 0.897453329 |
| chr2:168247351-168248131 | 0.110629663 | 0.743303398 | 0.897483303 |
| chr8:127851905-127852441 | -0.108344325 | 0.74332849 | 0.897499458 |
| chr12:65840860-65841060 | 0.183039544 | 0.743401816 | 0.897564459 |
| chr10:42646713-42647171 | 0.122049185 | 0.743441782 | 0.897564459 |
| chr10:73786046-73786416 | 0.118747459 | 0.743469354 | 0.897564459 |
| chr4:82500254-82500616 | -0.142956345 | 0.743478661 | 0.897564459 |
| chr4:35226828-35227233 | -0.132612273 | 0.743491656 | 0.897564459 |
| chr5:177124467-177124766 | -0.128905921 | 0.743491696 | 0.897564459 |
| chr1:181568141-181568404 | 0.157735074 | 0.743498823 | 0.897564459 |
| chr9:104394975-104395175 | -0.171717755 | 0.74349941 | 0.897564459 |
| chr2:204072460-204072741 | 0.141468247 | 0.743499923 | 0.897564459 |
| chr8:129962622-129963228 | -0.10913372 | 0.743501041 | 0.897564459 |
| chr8:78804286-78805262 | 0.106882516 | 0.743539365 | 0.897564459 |
| chr17:81328544-81328823 | -0.122327693 | 0.743562695 | 0.897564459 |
| chr1:24688681-24688881 | 0.167584336 | 0.743568279 | 0.897564459 |
| chr21:26846111-26846311 | 0.141446455 | 0.743574525 | 0.897564459 |
| chr12:6723733-6724748 | -0.105970502 | 0.743574942 | 0.897564459 |
| chrX:39705315-39705640 | -0.143043657 | 0.743589693 | 0.897564459 |
| chr14:61924998-61925237 | -0.151506945 | 0.743603369 | 0.897564459 |
| chr20:60132202-60132945 | -0.11613363 | 0.743621313 | 0.897564459 |
| chr16:57600330-57600726 | 0.135732827 | 0.743622022 | 0.897564459 |
| chr8:79880103-79880347 | -0.164879626 | 0.74362938 | 0.897564459 |
| chr22:31489374-31490515 | 0.108108454 | 0.743641793 | 0.897564459 |
| chr20:50958226-50959176 | 0.105547732 | 0.743654307 | 0.897564459 |
| chr4:159719272-159719517 | -0.173714208 | 0.743661101 | 0.897564459 |
| chr14:66507380-66508716 | 0.102998305 | 0.743663466 | 0.897564459 |
| chr15:85759589-85759833 | 0.150741688 | 0.743721834 | 0.897616651 |
| chr2:86486765-86486965 | 0.186670147 | 0.743730139 | 0.897616651 |
| chr10:87850808-87851008 | -0.163622192 | 0.743772965 | 0.897650208 |
| chr17:60107269-60108298 | -0.103179365 | 0.743793751 | 0.897650208 |
| chr8:91040321-91041243 | -0.106623789 | 0.743795855 | 0.897650208 |
| chr14:79413250-79413936 | -0.118144248 | 0.743804804 | 0.897650208 |
| chr3:158659774-158660058 | 0.13377634 | 0.743835906 | 0.897670771 |
| chr17:32876264-32877497 | 0.10673116 | 0.743845274 | 0.897670771 |
| chr13:34833630-34833889 | -0.140767242 | 0.743858074 | 0.897672079 |
| chr15:84894827-84895414 | -0.116870222 | 0.743894989 | 0.897702489 |
| chr9:133025428-133025628 | 0.159763255 | 0.743938238 | 0.897731331 |
| chr16:4511509-4511761 | 0.141697626 | 0.743972322 | 0.897731331 |
| chr4:147707957-147708471 | -0.117721281 | 0.743973118 | 0.897731331 |
| chr1:58784962-58785401 | -0.109807658 | 0.743979244 | 0.897731331 |

| | | | |
|---|---|---|---|
| chr16:88527446-88527711 | 0.125942916 | 0.743983296 | 0.897731331 |
| chr12:111791901-111792101 | -0.174642055 | 0.743989188 | 0.897731331 |
| chr17:79837190-79837617 | -0.116458712 | 0.74401013 | 0.897733873 |
| chr2:20917009-20917256 | -0.151955192 | 0.744023494 | 0.897733873 |
| chr1:154819601-154819947 | 0.126645969 | 0.744028989 | 0.897733873 |
| chr19:40529119-40529478 | 0.121635303 | 0.744064412 | 0.897733873 |
| chr10:27446149-27446612 | -0.143930788 | 0.744083255 | 0.897733873 |
| chr12:89708735-89710070 | -0.102263364 | 0.744086713 | 0.897733873 |
| chr1:199113244-199113625 | -0.115609823 | 0.744094852 | 0.897733873 |
| chr17:65027326-65027601 | 0.139091622 | 0.744096207 | 0.897733873 |
| chr21:28745135-28745565 | 0.13073813 | 0.744108287 | 0.897733873 |
| chr2:116094283-116094552 | -0.156748952 | 0.744120174 | 0.897733873 |
| chr4:86848801-86849110 | -0.156748952 | 0.744120174 | 0.897733873 |
| chr20:36573251-36573755 | 0.113279391 | 0.744139096 | 0.897742566 |
| chr10:89542822-89543100 | -0.172284783 | 0.744154219 | 0.897746675 |
| chr16:81095817-81097107 | -0.102686332 | 0.744270912 | 0.897859386 |
| chr8:106269695-106270775 | -0.103192985 | 0.744281909 | 0.897859386 |
| chr20:13784403-13785714 | 0.104447401 | 0.744292289 | 0.897859386 |
| chr14:75577681-75578075 | 0.114263798 | 0.744298427 | 0.897859386 |
| chr9:34072403-34072847 | -0.125070483 | 0.744306237 | 0.897859386 |
| chr21:39107288-39107525 | 0.146811724 | 0.744324137 | 0.897866844 |
| chrX:136577624-136577897 | -0.140242902 | 0.744345658 | 0.897878669 |
| chr9:82934963-82935291 | -0.137160489 | 0.74435809 | 0.897879529 |
| chr16:86505536-86505818 | -0.153589897 | 0.74441394 | 0.897903266 |
| chr17:69523294-69523515 | -0.153589897 | 0.74441394 | 0.897903266 |
| chr2:193695476-193695699 | -0.153589897 | 0.74441394 | 0.897903266 |
| chr19:58361945-58363152 | 0.104019442 | 0.744424642 | 0.897903266 |
| chr10:124801447-124801964 | 0.117697144 | 0.744441819 | 0.897909848 |
| chr5:139157586-139157850 | -0.145110316 | 0.744480923 | 0.897942879 |
| chr17:19714658-19714871 | 0.153451926 | 0.744501601 | 0.897953685 |
| chr20:31332836-31333243 | -0.148714416 | 0.744538771 | 0.897983541 |
| chr6:7618224-7618431 | -0.152240632 | 0.744549795 | 0.897983541 |
| chr5:171174614-171174892 | 0.146338106 | 0.744646782 | 0.898071433 |
| chr15:70072957-70073183 | 0.155202718 | 0.744653561 | 0.898071433 |
| chr16:58024630-58024857 | 0.160977886 | 0.74465995 | 0.898071433 |
| chr7:44884971-44885215 | 0.141438361 | 0.744669552 | 0.898071433 |
| chr6:31366896-31367363 | 0.126836439 | 0.744705274 | 0.898100377 |
| chr3:138114997-138115243 | 0.126616227 | 0.74474644 | 0.898121983 |
| chr3:26712442-26712738 | 0.137266297 | 0.744746632 | 0.898121983 |
| chr15:38453947-38454719 | 0.108257189 | 0.74478999 | 0.898160135 |
| chr6:150492510-150492839 | -0.141805757 | 0.744839562 | 0.89819682 |
| chr17:60315681-60316145 | -0.106029477 | 0.744855241 | 0.89819682 |
| chr20:8946053-8946278 | -0.171394979 | 0.744855579 | 0.89819682 |
| chr20:35561280-35561553 | 0.138309481 | 0.744923489 | 0.898264574 |
| chr2:216109017-216109777 | -0.105422185 | 0.7449894 | 0.898329915 |
| chr1:26787550-26788392 | -0.112459328 | 0.745021929 | 0.898355002 |
| chr17:66567222-66567592 | -0.117881741 | 0.745042187 | 0.898362718 |
| chr3:15858914-15859138 | 0.125694598 | 0.745057314 | 0.898362718 |
| chr7:151110533-151110767 | -0.13760751 | 0.745063502 | 0.898362718 |
| chr5:62661020-62661329 | -0.149521434 | 0.745089852 | 0.898365276 |

| | | | |
|---|---|---|---|
| chr2:68122423-68122720 | -0.13247408 | 0.745093892 | 0.898365276 |
| chr7:100101252-100102147 | -0.104970008 | 0.745104326 | 0.898365276 |
| chr1:94118114-94118505 | -0.12750183 | 0.745112522 | 0.898365276 |
| chr6:78436443-78436643 | 0.188536861 | 0.745134105 | 0.898375061 |
| chr2:101991788-101992035 | 0.154140613 | 0.745144087 | 0.898375061 |
| chr8:124232763-124232998 | 0.13979726 | 0.745157943 | 0.89837763 |
| chr1:201060614-201060943 | -0.118187008 | 0.74518736 | 0.898396852 |
| chr18:48133734-48134031 | -0.129993419 | 0.745197337 | 0.898396852 |
| chr1:28581360-28582253 | 0.107884085 | 0.745234295 | 0.898401109 |
| chr11:65134325-65135402 | 0.105095733 | 0.745235791 | 0.898401109 |
| chr17:38751773-38752087 | 0.124995861 | 0.745248101 | 0.898401109 |
| chr6:109296510-109296825 | 0.128663085 | 0.745264728 | 0.898401109 |
| chr8:72510072-72510400 | -0.13783347 | 0.745269895 | 0.898401109 |
| chr16:20900220-20901341 | 0.103905984 | 0.745274979 | 0.898401109 |
| chr13:30408274-30408790 | 0.10990919 | 0.745293261 | 0.898401109 |
| chr10:125201238-125201506 | -0.159957599 | 0.74530543 | 0.898401109 |
| chr13:19957368-19959103 | 0.101923283 | 0.745310522 | 0.898401109 |
| chrX:17046523-17046807 | 0.162516345 | 0.745318119 | 0.898401109 |
| chr6:139109645-139109856 | 0.167489867 | 0.745337904 | 0.898410824 |
| chr8:72155261-72155489 | -0.159237438 | 0.745360558 | 0.89841688 |
| chr8:81784265-81784603 | -0.138571554 | 0.745368701 | 0.89841688 |
| chr2:21083761-21083978 | -0.174166882 | 0.745384335 | 0.89841688 |
| chr2:6214557-6214775 | -0.159747474 | 0.745409339 | 0.89841688 |
| chr9:34484727-34485064 | -0.13591163 | 0.745424634 | 0.89841688 |
| chr17:64240256-64240566 | -0.136671596 | 0.745449548 | 0.89841688 |
| chr19:17226026-17226731 | -0.110892939 | 0.745464806 | 0.89841688 |
| chr11:118640872-118641072 | 0.156156098 | 0.745475729 | 0.89841688 |
| chr16:15022119-15022499 | 0.132940067 | 0.745482124 | 0.89841688 |
| chr10:110803931-110804261 | -0.127637233 | 0.745485046 | 0.89841688 |
| chr4:7434803-7435060 | 0.143713139 | 0.74548669 | 0.89841688 |
| chr3:72729107-72729388 | 0.138379134 | 0.745493091 | 0.89841688 |
| chr2:168346702-168347200 | -0.122403186 | 0.745495357 | 0.89841688 |
| chr12:81495578-81495778 | 0.193368891 | 0.74552446 | 0.898437822 |
| chr14:24431681-24432091 | 0.109496756 | 0.745537248 | 0.898439103 |
| chr11:93542819-93543061 | 0.161260509 | 0.745567914 | 0.898447796 |
| chr11:124863840-124864074 | 0.161260509 | 0.745567914 | 0.898447796 |
| chr4:139048597-139048938 | 0.147314689 | 0.745599297 | 0.898458235 |
| chr16:46947916-46948129 | -0.166500909 | 0.745600028 | 0.898458235 |
| chr14:52553134-52553600 | 0.110860353 | 0.745656991 | 0.898512745 |
| chr3:63652139-63652806 | 0.10819264 | 0.745688382 | 0.898513654 |
| chr15:40782584-40782987 | 0.121696197 | 0.745688957 | 0.898513654 |
| chr1:233299512-233299759 | 0.147621409 | 0.745698164 | 0.898513654 |
| chr1:150149483-150150245 | 0.10795016 | 0.745704651 | 0.898513654 |
| chr16:71351492-71351737 | 0.154613041 | 0.745731599 | 0.898523142 |
| chr6:7476014-7476303 | 0.158571971 | 0.745735979 | 0.898523142 |
| chr3:146334213-146334502 | -0.140767952 | 0.745853127 | 0.898650161 |
| chr3:11601828-11602059 | -0.168728761 | 0.745868254 | 0.898651308 |
| chr3:40524310-40525285 | -0.106260005 | 0.745877536 | 0.898651308 |
| chr5:27091947-27092147 | -0.1746555 | 0.745892565 | 0.89865214 |
| chr14:70045817-70046387 | -0.112935269 | 0.745909061 | 0.89865214 |

| | | | |
|---|---|---|---|
| chr18:31206807-31207319 | -0.114095393 | 0.745923932 | 0.89865214 |
| chr2:14779287-14779607 | -0.164120928 | 0.745926768 | 0.89865214 |
| chr19:3343959-3344512 | -0.119381725 | 0.745936869 | 0.89865214 |
| chr16:23073348-23073567 | 0.165183899 | 0.745997527 | 0.89869454 |
| chr10:93701909-93702269 | -0.127344483 | 0.745999185 | 0.89869454 |
| chr6:159789487-159790606 | -0.102773011 | 0.74600725 | 0.89869454 |
| chr19:1605371-1606201 | -0.109782855 | 0.74603097 | 0.898708985 |
| chr2:9212266-9212598 | 0.126149687 | 0.746072952 | 0.898739227 |
| chr9:133533176-133533376 | 0.14585501 | 0.746089932 | 0.898739227 |
| chr22:36659636-36660282 | 0.114628138 | 0.746091263 | 0.898739227 |
| chr2:135151720-135151920 | -0.170965519 | 0.746127011 | 0.898766207 |
| chr7:158432806-158433022 | -0.156959852 | 0.746143107 | 0.898766207 |
| chr15:60398453-60398972 | 0.112559963 | 0.746150366 | 0.898766207 |
| chr7:47571761-47572378 | 0.107843396 | 0.746171619 | 0.898766207 |
| chr2:162579178-162579453 | -0.145584202 | 0.74617231 | 0.898766207 |
| chr3:58471198-58471884 | 0.112893406 | 0.746225303 | 0.898815908 |
| chr6:115000791-115001035 | -0.163038023 | 0.74627072 | 0.898856482 |
| chrX:49282550-49283017 | 0.124332425 | 0.746340386 | 0.898915414 |
| chr11:59615584-59616572 | 0.102213674 | 0.746357948 | 0.898915414 |
| chr2:84946349-84946549 | -0.172860858 | 0.746367069 | 0.898915414 |
| chr11:118620852-118621436 | 0.113444347 | 0.746367149 | 0.898915414 |
| chr9:110255469-110255778 | -0.119346473 | 0.746400144 | 0.898915414 |
| chr2:208040848-208041048 | -0.155790546 | 0.74642236 | 0.898915414 |
| chr6:47168234-47168628 | -0.125945192 | 0.746436418 | 0.898915414 |
| chr20:20051932-20052970 | -0.104096397 | 0.746443002 | 0.898915414 |
| chr3:151327592-151327929 | -0.140440142 | 0.746445287 | 0.898915414 |
| chr21:45375337-45375603 | 0.154906312 | 0.746453368 | 0.898915414 |
| chr11:74036659-74036924 | -0.142469214 | 0.746463398 | 0.898915414 |
| chr3:100068825-100069153 | 0.126092843 | 0.74647457 | 0.898915414 |
| chr3:36992717-36993890 | 0.101426888 | 0.746477893 | 0.898915414 |
| chr21:42991993-42992257 | 0.172759061 | 0.746490306 | 0.898915414 |
| chr18:26103665-26103929 | -0.15425847 | 0.746495625 | 0.898915414 |
| chr12:102169973-102170490 | 0.123375869 | 0.746531639 | 0.898919791 |
| chr18:58175208-58175502 | 0.135400176 | 0.746534093 | 0.898919791 |
| chr5:119288678-119288978 | 0.152800677 | 0.746534456 | 0.898919791 |
| chr18:48037633-48038167 | 0.119385613 | 0.746570581 | 0.898946038 |
| chr4:48896920-48897198 | -0.150992138 | 0.746580121 | 0.898946038 |
| chr3:193289299-193289663 | -0.131497817 | 0.74659145 | 0.898946038 |
| chr13:31322134-31322334 | -0.168118346 | 0.746617304 | 0.898953257 |
| chr1:32520451-32520884 | 0.12023418 | 0.746622964 | 0.898953257 |
| chr1:32072500-32073651 | 0.103534971 | 0.746640636 | 0.898953257 |
| chr17:75412689-75412920 | -0.13876921 | 0.746644374 | 0.898953257 |
| chr6:27752005-27752849 | -0.109153827 | 0.746664638 | 0.898963528 |
| chr3:50336998-50337807 | -0.10623206 | 0.7466853 | 0.898966507 |
| chr12:93564169-93564551 | 0.135016306 | 0.74670357 | 0.898966507 |
| chr8:95112386-95113724 | -0.102149829 | 0.746717362 | 0.898966507 |
| chr20:50335136-50335510 | 0.112540749 | 0.746728082 | 0.898966507 |
| chr4:110480279-110480479 | 0.157037322 | 0.746739821 | 0.898966507 |
| chr1:10430223-10431114 | 0.114062871 | 0.746742608 | 0.898966507 |
| chr4:120948425-120948712 | -0.141943398 | 0.74674924 | 0.898966507 |

| | | | |
|---|---|---|---|
| chr3:172219017-172219336 | -0.133636337 | 0.746784032 | 0.898994267 |
| chr3:144323184-144323599 | -0.132276885 | 0.746841338 | 0.899033429 |
| chr9:16727049-16727272 | 0.149956312 | 0.746862233 | 0.899033429 |
| chr6:63571796-63572665 | -0.102955762 | 0.746869764 | 0.899033429 |
| chr22:35150496-35150778 | -0.16394321 | 0.746870708 | 0.899033429 |
| chr2:208192601-208192956 | 0.153914021 | 0.746876441 | 0.899033429 |
| chr17:59989895-59990129 | 0.148138124 | 0.746887502 | 0.899033429 |
| chr12:42744559-42745214 | -0.118421443 | 0.746898697 | 0.899033429 |
| chr14:96095546-96096299 | 0.104013612 | 0.746917768 | 0.899042261 |
| chr2:4269894-4270134 | -0.184005657 | 0.746958717 | 0.899077427 |
| chr2:103386839-103387233 | 0.120460823 | 0.746986364 | 0.899093294 |
| chr1:230797943-230798161 | -0.158617933 | 0.747006902 | 0.899093294 |
| chr5:147935075-147935391 | 0.137638423 | 0.747020935 | 0.899093294 |
| chr17:41792823-41793559 | -0.112683194 | 0.747024651 | 0.899093294 |
| chr9:35079404-35080279 | -0.108135131 | 0.747030571 | 0.899093294 |
| chr2:74472114-74472996 | 0.106235342 | 0.747053173 | 0.899106374 |
| chr2:65040657-65040985 | -0.118896601 | 0.747087991 | 0.899111928 |
| chr3:149044906-149045236 | -0.160640842 | 0.7471023 | 0.899111928 |
| chr17:57671889-57672261 | -0.112794477 | 0.747102968 | 0.899111928 |
| chr11:73787549-73788630 | 0.104612892 | 0.747104724 | 0.899111928 |
| chr1:182101418-182101691 | 0.139797225 | 0.747120637 | 0.899112431 |
| chr18:62325916-62326121 | -0.146055997 | 0.747129924 | 0.899112431 |
| chr11:126480709-126481064 | -0.126597553 | 0.747140346 | 0.899112431 |
| chr11:17991917-17992117 | -0.163325002 | 0.747181168 | 0.89913756 |
| chr17:17365770-17366075 | 0.146859941 | 0.747190408 | 0.89913756 |
| chr8:63167904-63169230 | -0.102039193 | 0.747196432 | 0.89913756 |
| chrX:107742285-107742485 | -0.162475515 | 0.747253885 | 0.899178135 |
| chr3:96595807-96596007 | -0.170120652 | 0.747275681 | 0.899178135 |
| chr2:169452122-169452442 | 0.125062016 | 0.747276503 | 0.899178135 |
| chr8:103225956-103226329 | 0.130627436 | 0.747277091 | 0.899178135 |
| chr17:66200690-66201012 | 0.118302989 | 0.747291224 | 0.899181019 |
| chr15:83734495-83734924 | 0.129000282 | 0.747319959 | 0.899189519 |
| chr19:55604931-55605682 | 0.104370016 | 0.747321758 | 0.899189519 |
| chr2:34096637-34096921 | 0.142744932 | 0.747342295 | 0.899200108 |
| chr4:138255953-138256338 | -0.14701606 | 0.747367555 | 0.899205376 |
| chr4:138978497-138979000 | 0.116296086 | 0.747370144 | 0.899205376 |
| chrX:100409716-100410646 | -0.105987347 | 0.747408138 | 0.899215164 |
| chr1:148285906-148286184 | 0.141298461 | 0.747413487 | 0.899215164 |
| chr17:67293371-67293604 | 0.141298461 | 0.747413487 | 0.899215164 |
| chr6:37102295-37102979 | -0.112131648 | 0.7474366 | 0.89921771 |
| chr5:110631662-110631892 | 0.162021559 | 0.747448344 | 0.89921771 |
| chr16:58499831-58500368 | 0.109411736 | 0.74745081 | 0.89921771 |
| chr17:76398084-76398301 | -0.13907633 | 0.74749635 | 0.899254851 |
| chr2:200735687-200736067 | 0.124316929 | 0.747505155 | 0.899254851 |
| chr13:113297532-113297885 | -0.1302331 | 0.747538093 | 0.899256606 |
| chr8:15561577-15562032 | -0.147399415 | 0.747558351 | 0.899256606 |
| chr20:7285958-7286332 | -0.124272195 | 0.747561689 | 0.899256606 |
| chr2:152016946-152017270 | -0.148487122 | 0.747589218 | 0.899256606 |
| chr1:90133262-90133588 | -0.149102715 | 0.747595929 | 0.899256606 |
| chr1:208036264-208036512 | -0.149102715 | 0.747595929 | 0.899256606 |

| | | | |
|---|---|---|---|
| chr1:244862893-244863808 | 0.102632492 | 0.747610849 | 0.899256606 |
| chr9:97745330-97745530 | 0.165858198 | 0.747616064 | 0.899256606 |
| chr20:49935805-49937041 | 0.102277733 | 0.747621368 | 0.899256606 |
| chr19:10380381-10381114 | 0.106560728 | 0.747623977 | 0.899256606 |
| chrX:71254035-71254845 | -0.10997713 | 0.747646197 | 0.899269216 |
| chr1:36299895-36300366 | 0.114282038 | 0.747660611 | 0.899272437 |
| chr20:44867664-44868058 | 0.128571211 | 0.747680528 | 0.899282277 |
| chr12:47397717-47397991 | 0.140505313 | 0.747700088 | 0.899291686 |
| chr11:128031623-128031823 | 0.160744462 | 0.747721305 | 0.899301301 |
| chr19:29873352-29873778 | 0.126942349 | 0.747731556 | 0.899301301 |
| chr1:116340833-116341061 | 0.140092635 | 0.747749332 | 0.899308565 |
| chrX:18425127-18425779 | -0.106207671 | 0.74778311 | 0.899320884 |
| chr1:68496577-68497694 | -0.102848392 | 0.747809239 | 0.899320884 |
| chr3:192188287-192188512 | 0.159826014 | 0.747815175 | 0.899320884 |
| chr2:124027212-124027482 | 0.13926618 | 0.747833816 | 0.899320884 |
| chr20:24931130-24931348 | 0.149859873 | 0.747854156 | 0.899320884 |
| chr12:3456650-3457219 | -0.128948149 | 0.747858669 | 0.899320884 |
| chr11:123145200-123145544 | -0.133926924 | 0.747868742 | 0.899320884 |
| chr1:20661236-20661981 | 0.106506449 | 0.747873783 | 0.899320884 |
| chr8:70271624-70271838 | 0.173110887 | 0.747876947 | 0.899320884 |
| chr6:167142293-167142493 | 0.173110887 | 0.747876947 | 0.899320884 |
| chr1:199422235-199422672 | 0.11640599 | 0.747908402 | 0.899333843 |
| chr8:92506311-92506655 | -0.154527926 | 0.747919751 | 0.899333843 |
| chr4:44425572-44425976 | 0.123777624 | 0.747928284 | 0.899333843 |
| chr2:112764048-112764248 | -0.147655193 | 0.747934672 | 0.899333843 |
| chr4:103018961-103019273 | -0.122532271 | 0.747954027 | 0.899343002 |
| chr2:128596095-128596332 | 0.165497356 | 0.747995873 | 0.899365091 |
| chr22:24968193-24968402 | 0.165497356 | 0.747995873 | 0.899365091 |
| chr6:47656285-47656621 | -0.120896213 | 0.748021366 | 0.89938163 |
| chr1:56409148-56409425 | 0.132389052 | 0.748083962 | 0.899438204 |
| chr12:74889928-74890550 | 0.109895379 | 0.748110504 | 0.899438204 |
| chr17:69609596-69609873 | -0.136526964 | 0.748124598 | 0.899438204 |
| chr4:110197729-110198163 | -0.111851588 | 0.74812561 | 0.899438204 |
| chr13:54316872-54317157 | -0.158892312 | 0.748127112 | 0.899438204 |
| chr12:14770802-14771314 | -0.107946443 | 0.748142398 | 0.899442468 |
| chr10:3208606-3208860 | 0.153282977 | 0.748193595 | 0.899483001 |
| chrX:64205026-64206503 | -0.101465283 | 0.748201188 | 0.899483001 |
| chr2:131529397-131529710 | -0.140664949 | 0.748212582 | 0.899483001 |
| chr5:74865378-74866362 | 0.109174645 | 0.74823227 | 0.899483001 |
| chr2:188607416-188607773 | -0.139040385 | 0.748243888 | 0.899483001 |
| chr2:133504483-133505585 | 0.105703315 | 0.748246548 | 0.899483001 |
| chr12:52694673-52694887 | 0.157923341 | 0.74828723 | 0.899511375 |
| chr1:178542083-178542400 | 0.116740277 | 0.748293631 | 0.899511375 |
| chr12:114770237-114770437 | -0.176027491 | 0.748338309 | 0.899539357 |
| chr17:43309820-43310137 | 0.130762011 | 0.748369124 | 0.899539357 |
| chr6:53064694-53065790 | -0.103259154 | 0.748369889 | 0.899539357 |
| chr5:57338107-57338457 | 0.121806944 | 0.748370727 | 0.899539357 |
| chr15:72460422-72460622 | -0.169331955 | 0.748384082 | 0.899539357 |
| chr17:82888642-82889368 | 0.107857438 | 0.748387348 | 0.899539357 |
| chr19:40810683-40811219 | 0.11538574 | 0.748414996 | 0.899558478 |

| | | | |
|---|---|---|---|
| chr16:79600159-79600528 | -0.126387719 | 0.748474121 | 0.899613881 |
| chr1:203812073-203812374 | 0.144076404 | 0.748491075 | 0.899613881 |
| chr14:29085657-29085913 | 0.138805695 | 0.748496313 | 0.899613881 |
| chr10:93445446-93445994 | -0.105255446 | 0.748536535 | 0.899634666 |
| chr6:85361360-85361631 | -0.149781146 | 0.748551264 | 0.899634666 |
| chr14:82347881-82348145 | 0.150656995 | 0.748554601 | 0.899634666 |
| chr1:53238214-53239079 | -0.105474105 | 0.748560572 | 0.899634666 |
| chr20:19262832-19263110 | -0.14175585 | 0.748584427 | 0.899647856 |
| chr1:160342883-160343851 | 0.105280529 | 0.748622565 | 0.899647856 |
| chr11:75161759-75162032 | 0.162096672 | 0.748627105 | 0.899647856 |
| chr1:201480308-201481244 | 0.101022488 | 0.748630749 | 0.899647856 |
| chr14:54366734-54367520 | -0.107343578 | 0.748635436 | 0.899647856 |
| chrX:16175380-16175580 | 0.200930333 | 0.748641995 | 0.899647856 |
| chr3:99984666-99984975 | 0.143031812 | 0.748654479 | 0.899648749 |
| chr17:30697852-30698213 | -0.128911065 | 0.748692309 | 0.899673706 |
| chr12:27142181-27142428 | -0.163589984 | 0.748699463 | 0.899673706 |
| chr14:24308173-24308439 | -0.125992951 | 0.748710552 | 0.899673706 |
| chr3:112646690-112646911 | 0.139363411 | 0.748722215 | 0.899673706 |
| chr22:40370488-40370840 | -0.130309857 | 0.748749814 | 0.89969276 |
| chr3:100509544-100510260 | 0.110855082 | 0.748783129 | 0.899718682 |
| chr17:55344468-55344811 | 0.117638707 | 0.748805252 | 0.899731155 |
| chr11:62678796-62679392 | 0.112182629 | 0.74888103 | 0.899808096 |
| chr21:33924015-33924297 | -0.138274386 | 0.748913302 | 0.899832762 |
| chr2:218687495-218687772 | 0.141452473 | 0.748985483 | 0.899880036 |
| chr12:17062236-17062709 | 0.130725726 | 0.748992561 | 0.899880036 |
| chr1:178655640-178656323 | 0.110863685 | 0.748999146 | 0.899880036 |
| chr1:147892181-147892459 | -0.1222957 | 0.748999626 | 0.899880036 |
| chr1:199948929-199949191 | -0.136065246 | 0.749024426 | 0.899894923 |
| chr2:74066729-74066929 | 0.16854444 | 0.749035505 | 0.899894923 |
| chr1:43974525-43975524 | 0.105633011 | 0.74906597 | 0.899898329 |
| chr16:27515933-27516163 | -0.14190035 | 0.749072215 | 0.899898329 |
| chr15:92969213-92969413 | -0.167458778 | 0.749073574 | 0.899898329 |
| chr1:230268807-230269214 | -0.127451298 | 0.749107512 | 0.89992499 |
| chr2:100267798-100268052 | -0.13933985 | 0.749162138 | 0.899976503 |
| chr13:95980608-95980808 | -0.17248396 | 0.749216076 | 0.899995037 |
| chr1:202008187-202008538 | 0.110257561 | 0.749216734 | 0.899995037 |
| chr10:6574424-6574664 | 0.15561108 | 0.749248476 | 0.899995037 |
| chrX:48834495-48834966 | -0.123632806 | 0.749256217 | 0.899995037 |
| chr1:65958980-65959180 | 0.167517136 | 0.749260624 | 0.899995037 |
| chr8:74850046-74850517 | -0.130506073 | 0.749268124 | 0.899995037 |
| chr17:59534634-59534925 | 0.130543785 | 0.749268375 | 0.899995037 |
| chr10:110785990-110786258 | -0.150368043 | 0.749271533 | 0.899995037 |
| chr15:80989387-80990137 | 0.104247261 | 0.749307744 | 0.900024423 |
| chr1:46198126-46198749 | 0.114957311 | 0.749333347 | 0.900038532 |
| chr3:164024261-164024461 | -0.167719195 | 0.749342984 | 0.900038532 |
| chrX:149510988-149511560 | -0.110383407 | 0.749391579 | 0.900052743 |
| chr21:38821903-38822103 | -0.154043783 | 0.749402733 | 0.900052743 |
| chr17:41688419-41689634 | 0.101401074 | 0.749416102 | 0.900052743 |
| chr10:49751835-49752137 | -0.151808765 | 0.749417568 | 0.900052743 |
| chr20:44187111-44187657 | 0.117103926 | 0.749422853 | 0.900052743 |

| | | | |
|---|---|---|---|
| chr19:2528710-2529144 | 0.120544962 | 0.749426621 | 0.900052743 |
| chr9:26946694-26947686 | -0.103800258 | 0.749437042 | 0.900052743 |
| chr20:13681375-13681593 | -0.170749025 | 0.749450164 | 0.900054395 |
| chr19:10478941-10479325 | 0.132866229 | 0.749527153 | 0.90011911 |
| chr12:6943493-6944942 | 0.101850753 | 0.749527545 | 0.90011911 |
| chr5:82225579-82225903 | -0.124682655 | 0.749539451 | 0.900119299 |
| chr13:100673880-100675277 | -0.102751042 | 0.749582688 | 0.900157115 |
| chr8:116874082-116875161 | -0.10053498 | 0.74959454 | 0.900157239 |
| chr17:75046495-75047299 | 0.10361354 | 0.749652987 | 0.900213318 |
| chr4:54322916-54323137 | 0.150738805 | 0.749688654 | 0.900234858 |
| chr1:183228056-183228294 | 0.133888128 | 0.749694422 | 0.900234858 |
| chr12:102301045-102301408 | -0.127940319 | 0.74971843 | 0.900236575 |
| chr14:50147019-50147315 | 0.15211799 | 0.749745537 | 0.900236575 |
| chr16:68396581-68396781 | 0.168208825 | 0.749746582 | 0.900236575 |
| chr2:5981739-5981939 | -0.171034671 | 0.749751461 | 0.900236575 |
| chr5:152659320-152659956 | -0.12204879 | 0.749754598 | 0.900236575 |
| chr6:4155859-4156203 | -0.131470563 | 0.74983879 | 0.900323556 |
| chr2:220533411-220533864 | -0.119747946 | 0.749876772 | 0.900355053 |
| chr14:76375615-76376394 | -0.112385741 | 0.74990247 | 0.900371798 |
| chrX:19990351-19991469 | 0.103803579 | 0.749937812 | 0.900390683 |
| chr5:58150945-58151170 | -0.169841706 | 0.749941701 | 0.900390683 |
| chr7:35537200-35537929 | -0.107909295 | 0.74997166 | 0.900402722 |
| chr20:33686279-33686845 | 0.109612128 | 0.749975231 | 0.900402722 |
| chr3:18837229-18837615 | -0.12621085 | 0.749994654 | 0.900411933 |
| chr11:47305413-47306130 | 0.113562242 | 0.750007294 | 0.900413 |
| chr14:77653444-77653644 | -0.149429235 | 0.750040908 | 0.900433118 |
| chr15:40296410-40297112 | 0.112072379 | 0.750047555 | 0.900433118 |
| chr8:89725808-89726225 | 0.10788932 | 0.750087931 | 0.900446044 |
| chr13:34347772-34348598 | 0.105864268 | 0.750089891 | 0.900446044 |
| chr15:33964161-33964361 | -0.174702001 | 0.750123063 | 0.900446044 |
| chr20:362333-362871 | 0.113067831 | 0.750126915 | 0.900446044 |
| chr1:106335790-106335990 | 0.139449797 | 0.750142356 | 0.900446044 |
| chr2:142863766-142863966 | -0.170769004 | 0.750144244 | 0.900446044 |
| chr6:131631543-131631916 | 0.116847062 | 0.750147901 | 0.900446044 |
| chr10:1240840-1241093 | -0.155146847 | 0.750155291 | 0.900446044 |
| chr2:73293507-73293967 | -0.11031289 | 0.750164089 | 0.900446044 |
| chr17:62477685-62479909 | 0.099975149 | 0.75018076 | 0.90045195 |
| chr1:162696331-162696881 | 0.113309138 | 0.750198749 | 0.900452957 |
| chr2:242064346-242064546 | -0.175808469 | 0.750222395 | 0.900452957 |
| chr2:231648586-231648976 | 0.117635476 | 0.750224545 | 0.900452957 |
| chr3:195895027-195895325 | 0.12069976 | 0.750228607 | 0.900452957 |
| chr11:94543746-94544527 | -0.108474341 | 0.750252745 | 0.900467823 |
| chr4:139484739-139485404 | -0.116243088 | 0.750277856 | 0.900472385 |
| chr5:159100273-159100491 | 0.147721987 | 0.75028005 | 0.900472385 |
| chr4:70652094-70652454 | -0.127073619 | 0.750292789 | 0.900473569 |
| chr2:154375337-154375537 | 0.162054523 | 0.750380669 | 0.900546981 |
| chr3:135965204-135966586 | -0.100394761 | 0.750387728 | 0.900546981 |
| chr2:236045386-236045897 | -0.127628196 | 0.750389217 | 0.900546981 |
| chr8:96333147-96333512 | 0.11308974 | 0.750434237 | 0.900556988 |
| chr19:4639166-4639605 | 0.113843516 | 0.750485051 | 0.900556988 |

| | | | |
|---|---|---|---|
| chr10:69723708-69723908 | 0.156471307 | 0.750490371 | 0.900556988 |
| chr22:31676712-31677040 | -0.127138194 | 0.750492881 | 0.900556988 |
| chr19:12669381-12669612 | 0.1432395 | 0.750505477 | 0.900556988 |
| chr2:127771051-127771313 | -0.151290707 | 0.750515766 | 0.900556988 |
| chr6:37423599-37424151 | -0.114913588 | 0.750521524 | 0.900556988 |
| chr3:161508896-161509238 | 0.115917035 | 0.750524731 | 0.900556988 |
| chr2:128715091-128715291 | -0.181126499 | 0.750528291 | 0.900556988 |
| chr8:25599566-25599818 | -0.169322516 | 0.750531414 | 0.900556988 |
| chr4:150722891-150723279 | 0.129736792 | 0.750538826 | 0.900556988 |
| chr8:27597168-27597467 | -0.139823273 | 0.750554326 | 0.900556988 |
| chr1:197801124-197801628 | 0.119366248 | 0.750556194 | 0.900556988 |
| chr20:49713112-49714299 | 0.107219027 | 0.7505621 | 0.900556988 |
| chr21:34510600-34510975 | -0.129735032 | 0.750599701 | 0.900588001 |
| chr3:127549775-127550043 | 0.125691863 | 0.750665796 | 0.900628131 |
| chr1:35172681-35172881 | -0.163057206 | 0.750700322 | 0.900628131 |
| chr10:21926536-21926783 | -0.163057206 | 0.750700322 | 0.900628131 |
| chr3:77038365-77038896 | -0.10856801 | 0.750714843 | 0.900628131 |
| chr5:10775654-10775942 | 0.130473252 | 0.750722207 | 0.900628131 |
| chr1:113505950-113506150 | -0.177489104 | 0.75073527 | 0.900628131 |
| chr11:62632624-62632880 | 0.145217396 | 0.750745894 | 0.900628131 |
| chr14:76773486-76773851 | 0.121878776 | 0.750778079 | 0.900628131 |
| chr1:159829239-159829577 | -0.119748363 | 0.750780912 | 0.900628131 |
| chr8:104130201-104130591 | 0.120565442 | 0.750783791 | 0.900628131 |
| chrX:56803410-56803680 | 0.150122445 | 0.750788646 | 0.900628131 |
| chr3:149214779-149215067 | -0.13292522 | 0.750794292 | 0.900628131 |
| chr1:116595537-116595745 | 0.149341273 | 0.750809959 | 0.900628131 |
| chr1:146036956-146037607 | 0.101530772 | 0.750814186 | 0.900628131 |
| chr2:216216054-216216281 | 0.152929134 | 0.750821215 | 0.900628131 |
| chr2:159557530-159557745 | 0.152929134 | 0.750821215 | 0.900628131 |
| chr17:50087416-50087904 | 0.112990425 | 0.750902576 | 0.900707025 |
| chr7:156650120-156650446 | 0.128764624 | 0.750910496 | 0.900707025 |
| chr1:223133522-223134363 | 0.10338325 | 0.750976576 | 0.900740231 |
| chr11:81988894-81989153 | -0.146411667 | 0.750986564 | 0.900740231 |
| chr9:36023513-36023735 | 0.167562271 | 0.750990832 | 0.900740231 |
| chr3:164451889-164452159 | -0.136327591 | 0.750996887 | 0.900740231 |
| chr14:96636320-96636520 | 0.164118605 | 0.750996958 | 0.900740231 |
| chr8:63300626-63301085 | -0.118738999 | 0.751060176 | 0.900801954 |
| chr10:79608840-79609068 | 0.151882813 | 0.751108571 | 0.900820906 |
| chr9:91732061-91732304 | -0.15156884 | 0.751113689 | 0.900820906 |
| chr14:21510758-21511704 | 0.101278459 | 0.751120195 | 0.900820906 |
| chr20:41017492-41018384 | 0.103077913 | 0.751123005 | 0.900820906 |
| chr16:75844434-75844709 | -0.165995134 | 0.751158835 | 0.900828345 |
| chr16:58487739-58488042 | 0.135669147 | 0.751162047 | 0.900828345 |
| chr2:113656047-113656867 | -0.111773711 | 0.751174234 | 0.900828345 |
| chr13:74133792-74134177 | -0.115880154 | 0.751177236 | 0.900828345 |
| chr9:113998834-113999147 | 0.126541782 | 0.751191606 | 0.900828345 |
| chr1:214865698-214865924 | 0.159976499 | 0.751202175 | 0.900828345 |
| chr3:68876204-68876409 | -0.164767808 | 0.751211505 | 0.900828345 |
| chr7:9031043-9031266 | -0.16287673 | 0.751224229 | 0.900829504 |
| chr1:201447948-201448384 | 0.109369636 | 0.751246168 | 0.900841714 |

| | | | |
|---|---|---|---|
| chr5:38845529-38846428 | 0.103478489 | 0.751277658 | 0.900862558 |
| chr8:123273590-123275416 | -0.098518119 | 0.751287064 | 0.900862558 |
| chr2:136378796-136379080 | 0.134025909 | 0.75141973 | 0.901007536 |
| chr1:37595517-37596252 | -0.105187353 | 0.751455225 | 0.901027333 |
| chr5:16568943-16569144 | -0.151323713 | 0.75146384 | 0.901027333 |
| chr5:134904380-134906016 | -0.100290851 | 0.751471518 | 0.901027333 |
| chr10:21364455-21364857 | 0.117428443 | 0.751503264 | 0.901041054 |
| chr2:32135452-32135652 | -0.16557179 | 0.751506481 | 0.901041054 |
| chr9:133121125-133121767 | 0.108411331 | 0.751523586 | 0.901047462 |
| chr2:75710362-75711202 | -0.102381278 | 0.751548522 | 0.90106326 |
| chr6:27837947-27839103 | 0.101854894 | 0.751580595 | 0.901081914 |
| chr12:131229485-131230034 | -0.112596292 | 0.751587601 | 0.901081914 |
| chr3:183099130-183099892 | -0.101339951 | 0.751606973 | 0.901091041 |
| chr2:227480126-227480379 | -0.147238762 | 0.751650799 | 0.901121354 |
| chr2:220111758-220112022 | -0.176399382 | 0.75166408 | 0.901121354 |
| chr1:229249132-229249444 | -0.145273269 | 0.751667539 | 0.901121354 |
| chr15:96330864-96331125 | -0.121724557 | 0.751689654 | 0.901133767 |
| chr11:74702432-74702715 | -0.122136112 | 0.751753555 | 0.901184336 |
| chr1:145129201-145129401 | -0.162305699 | 0.75175536 | 0.901184336 |
| chr22:23128444-23128735 | -0.128882456 | 0.751773238 | 0.901191668 |
| chr1:158203738-158204249 | -0.118047214 | 0.751810687 | 0.90122246 |
| chr9:62864528-62864865 | 0.137815979 | 0.751884031 | 0.90129628 |
| chr8:17246269-17247746 | 0.101009043 | 0.751940201 | 0.90134951 |
| chr17:51120278-51121307 | -0.100120087 | 0.751969308 | 0.901352241 |
| chr8:122236249-122236568 | -0.116991239 | 0.751976444 | 0.901352241 |
| chr19:55461123-55461371 | 0.13426635 | 0.751983628 | 0.901352241 |
| chr15:65832598-65833208 | 0.105910906 | 0.751989533 | 0.901352241 |
| chr10:96720809-96721033 | -0.139722186 | 0.752002389 | 0.90135355 |
| chr4:30789201-30789426 | -0.158266647 | 0.752046178 | 0.901391935 |
| chr13:31044428-31044723 | -0.122706432 | 0.752081891 | 0.901418115 |
| chr2:88099813-88100085 | -0.130197164 | 0.752091549 | 0.901418115 |
| chr20:59519406-59519762 | 0.110373781 | 0.752143351 | 0.90144122 |
| chr2:127857474-127857674 | -0.136817179 | 0.75214951 | 0.90144122 |
| chr6:157824087-157824287 | -0.164322211 | 0.752154827 | 0.90144122 |
| chr2:6526295-6526510 | -0.158835589 | 0.752157886 | 0.90144122 |
| chr6:105077993-105078213 | -0.15911503 | 0.752194278 | 0.901470734 |
| chr22:27933247-27933486 | 0.160598607 | 0.752218512 | 0.901485678 |
| chr17:74192901-74193544 | -0.109826196 | 0.752241798 | 0.901499485 |
| chr20:38893834-38894104 | -0.14094627 | 0.752301397 | 0.901528168 |
| chr10:85464266-85464543 | 0.148408852 | 0.752323243 | 0.901528168 |
| chr11:117327587-117328327 | 0.104199878 | 0.752332097 | 0.901528168 |
| chr12:116687920-116688213 | -0.137085617 | 0.752332144 | 0.901528168 |
| chr12:79183389-79183589 | 0.170623504 | 0.752335913 | 0.901528168 |
| chr12:108967877-108968120 | -0.139560324 | 0.752344957 | 0.901528168 |
| chr12:121203091-121203430 | -0.126554997 | 0.752359202 | 0.901528168 |
| chr20:59152855-59153066 | 0.139573566 | 0.75235986 | 0.901528168 |
| chr2:72917254-72917610 | 0.11538425 | 0.752395094 | 0.901556289 |
| chr1:178083795-178084030 | 0.165556858 | 0.752424655 | 0.90157541 |
| chr12:69359330-69360545 | -0.099376209 | 0.752434585 | 0.90157541 |
| chr14:64235537-64235771 | 0.144053484 | 0.752476591 | 0.901611643 |

| | | | |
|---|---|---|---|
| chr10:13347662-13348656 | -0.105670784 | 0.752587519 | 0.901669382 |
| chr6:70166150-70166658 | -0.11696344 | 0.752591254 | 0.901669382 |
| chr16:19543633-19543989 | -0.138587329 | 0.752617778 | 0.901669382 |
| chr6:17987728-17988345 | 0.112070589 | 0.752623801 | 0.901669382 |
| chr1:113548542-113548792 | -0.167416058 | 0.752631963 | 0.901669382 |
| chr1:97050540-97050750 | 0.162884355 | 0.752640864 | 0.901669382 |
| chr12:44455066-44455266 | -0.156748865 | 0.752650988 | 0.901669382 |
| chr22:45903265-45903587 | 0.141568049 | 0.752658706 | 0.901669382 |
| chr14:53306196-53306488 | 0.133492757 | 0.752670542 | 0.901669382 |
| chr9:63817395-63818156 | 0.115032462 | 0.752671025 | 0.901669382 |
| chr2:104607514-104607714 | -0.163697972 | 0.752672875 | 0.901669382 |
| chr2:204177963-204178479 | 0.120614092 | 0.752677465 | 0.901669382 |
| chr9:114082528-114082844 | 0.116411844 | 0.752677761 | 0.901669382 |
| chr8:126341512-126342235 | 0.107161816 | 0.75271197 | 0.901696266 |
| chr1:37989475-37990410 | -0.103165486 | 0.752746684 | 0.901723753 |
| chr6:28137172-28137515 | -0.126103003 | 0.752816703 | 0.901782597 |
| chr14:69726793-69727309 | -0.11271433 | 0.752838314 | 0.901782597 |
| chr3:113536047-113536255 | -0.127852507 | 0.752845566 | 0.901782597 |
| chr17:50478420-50479208 | 0.102820137 | 0.752851825 | 0.901782597 |
| chr8:130106605-130106961 | -0.131752971 | 0.752854652 | 0.901782597 |
| chr5:18394276-18394476 | 0.140751049 | 0.752908561 | 0.901815983 |
| chr1:47536793-47537008 | 0.162867763 | 0.752909825 | 0.901815983 |
| chr3:114383850-114384133 | -0.127082575 | 0.752917834 | 0.901815983 |
| chr9:92772305-92772616 | -0.155285037 | 0.752994882 | 0.90189417 |
| chr16:30634066-30634847 | 0.103586399 | 0.753012509 | 0.901901184 |
| chr4:104494761-104495451 | -0.107915186 | 0.753045604 | 0.901915003 |
| chr17:49556316-49556643 | -0.115818001 | 0.753047588 | 0.901915003 |
| chr5:43627025-43627381 | -0.118850792 | 0.753062035 | 0.901918207 |
| chr8:20196985-20197695 | -0.106943897 | 0.75310165 | 0.901945604 |
| chr4:87566330-87566649 | -0.135641121 | 0.753108453 | 0.901945604 |
| chr1:149927479-149928040 | -0.102723766 | 0.753129476 | 0.901954429 |
| chr2:161414131-161414963 | -0.105278875 | 0.753147391 | 0.901954429 |
| chr17:58249669-58249960 | 0.116172324 | 0.753151136 | 0.901954429 |
| chr10:77846988-77847188 | 0.162145407 | 0.753196482 | 0.901980539 |
| chr12:82909244-82909444 | 0.162145407 | 0.753196482 | 0.901980539 |
| chr12:101954697-101955045 | 0.123010378 | 0.753240385 | 0.902019017 |
| chr1:169601202-169601402 | 0.150624105 | 0.753252463 | 0.902019383 |
| chr1:74502445-74502717 | 0.15467903 | 0.753265978 | 0.902021469 |
| chrX:49884375-49884606 | 0.149792551 | 0.753310014 | 0.902060104 |
| chr1:178299966-178300223 | 0.135059674 | 0.75333948 | 0.902070855 |
| chrX:101051299-101052331 | -0.100571617 | 0.753349507 | 0.902070855 |
| chr8:48102668-48102906 | -0.1509521 | 0.753361029 | 0.902070855 |
| chr15:71114298-71114498 | -0.142737195 | 0.753385303 | 0.902070855 |
| chr22:43090273-43090569 | 0.136610411 | 0.753412514 | 0.902070855 |
| chr7:66352269-66352469 | -0.173928332 | 0.753413145 | 0.902070855 |
| chr7:149017-149932 | 0.099212642 | 0.753417112 | 0.902070855 |
| chr12:1831048-1831377 | -0.119396621 | 0.753418251 | 0.902070855 |
| chr15:97235367-97235666 | 0.124377704 | 0.753427371 | 0.902070855 |
| chr15:80906363-80906870 | -0.113577351 | 0.753436722 | 0.902070855 |
| chr12:78340316-78340538 | 0.154465938 | 0.753469587 | 0.902096107 |

| | | | |
|---|---|---|---|
| chr5:71757871-71758249 | -0.121732573 | 0.753485457 | 0.902101013 |
| chr15:89419419-89419637 | -0.131603683 | 0.753516251 | 0.902114215 |
| chr11:118400625-118400837 | 0.139091692 | 0.75353704 | 0.902114215 |
| chr5:151316055-151316538 | -0.118066456 | 0.753540835 | 0.902114215 |
| chr20:41925038-41925296 | -0.145386588 | 0.753555352 | 0.902114215 |
| chr3:106540831-106541065 | -0.145386588 | 0.753555352 | 0.902114215 |
| chr13:30132551-30132959 | -0.111426487 | 0.753592187 | 0.902144216 |
| chr1:223402891-223403116 | -0.171841541 | 0.753645811 | 0.902194315 |
| chr3:2714856-2715056 | -0.182409335 | 0.753659899 | 0.902197084 |
| chr3:194486616-194487276 | -0.104281962 | 0.753678951 | 0.902205796 |
| chr17:60174748-60175066 | -0.104747794 | 0.753692306 | 0.902207687 |
| chr4:149991876-149992188 | 0.147095826 | 0.753713171 | 0.902218569 |
| chr2:64153857-64154057 | -0.161017985 | 0.753747692 | 0.902245796 |
| chr10:119841713-119842079 | -0.139983488 | 0.753761898 | 0.902248706 |
| chr13:110500015-110500275 | -0.16226846 | 0.753828898 | 0.902279913 |
| chr2:172555401-172556746 | 0.100119127 | 0.753833867 | 0.902279913 |
| chr3:100819995-100820234 | 0.147466171 | 0.753840183 | 0.902279913 |
| chr17:56819487-56819997 | 0.110410485 | 0.753842775 | 0.902279913 |
| chr2:135797537-135797907 | 0.111837109 | 0.753846848 | 0.902279913 |
| chr1:109181150-109181458 | -0.133001321 | 0.753878954 | 0.902286973 |
| chr14:54805511-54806211 | 0.105801712 | 0.753879768 | 0.902286973 |
| chr13:113816978-113817279 | 0.132612437 | 0.753890861 | 0.902286973 |
| chr8:37328384-37328824 | -0.122808117 | 0.75389985 | 0.902286973 |
| chr6:30744266-30744721 | 0.111844827 | 0.753945038 | 0.902326962 |
| chr8:76678020-76678279 | -0.164381414 | 0.753975546 | 0.902343359 |
| chr4:15650003-15650206 | 0.154881333 | 0.753986978 | 0.902343359 |
| chr2:184597609-184598004 | 0.123285696 | 0.753994069 | 0.902343359 |
| chr18:9808789-9809256 | -0.125678322 | 0.754024 | 0.90235913 |
| chr6:82588158-82588358 | -0.159945318 | 0.7540308 | 0.90235913 |
| chr6:158704191-158704487 | 0.12868122 | 0.754052752 | 0.902371306 |
| chrX:74835681-74835891 | -0.159829476 | 0.75406555 | 0.902372528 |
| chr10:89793713-89793991 | 0.130922494 | 0.754086204 | 0.90238315 |
| chr16:4316986-4317223 | -0.133883084 | 0.754113453 | 0.902401665 |
| chr17:69026811-69027011 | -0.174934338 | 0.754151692 | 0.902433329 |
| chr15:92942762-92943060 | 0.13388342 | 0.754172532 | 0.902444174 |
| chr2:144170370-144170695 | -0.13387181 | 0.754187687 | 0.902448214 |
| chr3:175656466-175656780 | -0.132890306 | 0.754222435 | 0.902475699 |
| chr1:216183054-216183385 | 0.116385267 | 0.754250545 | 0.902488045 |
| chr14:54312214-54312552 | -0.123644284 | 0.754256309 | 0.902488045 |
| chr5:181246131-181247107 | -0.100647024 | 0.754282044 | 0.902491786 |
| chr8:108464382-108464867 | -0.114326977 | 0.75428711 | 0.902491786 |
| chr2:241415499-241415767 | -0.144904863 | 0.754294771 | 0.902491786 |
| chr9:122940638-122941522 | 0.099427752 | 0.754364477 | 0.902551225 |
| chr11:67805263-67806152 | -0.099244414 | 0.754370538 | 0.902551225 |
| chr16:89344302-89344528 | -0.152985829 | 0.754379788 | 0.902551225 |
| chr1:145845247-145846486 | 0.099967455 | 0.754398717 | 0.902559779 |
| chr15:91491184-91491451 | 0.155942665 | 0.754420735 | 0.902569705 |
| chr3:12194620-12194913 | -0.126949726 | 0.754440166 | 0.902569705 |
| chr16:89602557-89603178 | -0.112512571 | 0.754442352 | 0.902569705 |
| chr11:40307005-40307277 | 0.136592798 | 0.754511333 | 0.902630534 |

| | | | |
|---|---|---|---|
| chr6:43668232-43668608 | 0.112369352 | 0.754518447 | 0.902630534 |
| chr9:112705852-112706090 | -0.150146889 | 0.754540348 | 0.902630534 |
| chr8:126667618-126667893 | -0.129132247 | 0.754558494 | 0.902630534 |
| chr12:98532753-98533059 | 0.125951966 | 0.75457881 | 0.902630534 |
| chr1:40531128-40532046 | -0.104496959 | 0.75459072 | 0.902630534 |
| chr4:89140342-89140822 | 0.132621199 | 0.754590835 | 0.902630534 |
| chr8:128653345-128653660 | 0.118769756 | 0.754595866 | 0.902630534 |
| chr6:85498637-85499382 | -0.111078769 | 0.754604874 | 0.902630534 |
| chr16:88238792-88239088 | -0.121878732 | 0.754612504 | 0.902630534 |
| chr3:118501109-118501315 | -0.152167423 | 0.754623505 | 0.902630534 |
| chr16:88098774-88098987 | 0.151308538 | 0.754634562 | 0.902630534 |
| chr3:183824775-183825305 | -0.11230334 | 0.75464736 | 0.902631751 |
| chr20:14821253-14821492 | 0.154582894 | 0.754665044 | 0.902638812 |
| chr2:199733550-199733750 | -0.171412251 | 0.754686555 | 0.902650451 |
| chr13:42969594-42969908 | -0.120075017 | 0.754717578 | 0.902662372 |
| chr4:128479800-128480000 | 0.180155919 | 0.754730973 | 0.902662372 |
| chr13:112995555-112995854 | -0.15671177 | 0.75474486 | 0.902662372 |
| chr3:11335017-11335338 | 0.126420206 | 0.754749584 | 0.902662372 |
| chr4:37970732-37971155 | 0.12112649 | 0.754755425 | 0.902662372 |
| chr8:86320875-86321244 | 0.12735187 | 0.754826748 | 0.902716715 |
| chr13:109509546-109509815 | -0.140872453 | 0.754832694 | 0.902716715 |
| chr8:71547623-71547999 | 0.132716306 | 0.754851311 | 0.902716715 |
| chr16:67183450-67184350 | 0.100193513 | 0.754852976 | 0.902716715 |
| chr3:165545147-165545392 | 0.158183136 | 0.754867378 | 0.902716715 |
| chr12:67441819-67442123 | -0.123304942 | 0.754880201 | 0.902716715 |
| chr3:125921822-125922057 | -0.130023382 | 0.754883334 | 0.902716715 |
| chr1:241156477-241156913 | -0.115616652 | 0.754912503 | 0.902736478 |
| chr10:22086748-22087017 | 0.134357384 | 0.75493265 | 0.902736478 |
| chr14:65101701-65103379 | 0.097941498 | 0.754935205 | 0.902736478 |
| chr3:24496640-24496866 | -0.127176232 | 0.754975012 | 0.902753067 |
| chr8:61288130-61288407 | 0.136836665 | 0.754977736 | 0.902753067 |
| chr11:15327797-15328016 | 0.147989506 | 0.754984424 | 0.902753067 |
| chr5:23727404-23727604 | 0.161018296 | 0.755006321 | 0.902765162 |
| chr8:21326045-21326302 | -0.157313328 | 0.755046778 | 0.902799448 |
| chr17:79796741-79797807 | 0.099102202 | 0.755091232 | 0.902816412 |
| chr17:79778479-79778739 | -0.129375185 | 0.755123938 | 0.902816412 |
| chr3:192449089-192449311 | -0.156210668 | 0.755124473 | 0.902816412 |
| chrX:76454713-76454980 | 0.14313104 | 0.755133847 | 0.902816412 |
| chr22:43346414-43346682 | 0.140026519 | 0.755133873 | 0.902816412 |
| chr19:14073033-14073242 | 0.151004024 | 0.755139332 | 0.902816412 |
| chr8:104955493-104955706 | 0.138596738 | 0.755143445 | 0.902816412 |
| chr11:61361724-61362838 | -0.100490508 | 0.755172657 | 0.902837249 |
| chr16:85913382-85913705 | -0.13388347 | 0.755198255 | 0.902841866 |
| chr12:124940615-124940962 | 0.105939068 | 0.75520257 | 0.902841866 |
| chr8:56074190-56075121 | 0.101795847 | 0.755217756 | 0.902841866 |
| chr10:89466557-89467105 | 0.109536989 | 0.755223651 | 0.902841866 |
| chr4:166252862-166253140 | -0.139668128 | 0.755260901 | 0.902872311 |
| chr3:168040677-168040950 | -0.126115507 | 0.755282675 | 0.902884254 |
| chr7:54996020-54996220 | -0.159084601 | 0.755317109 | 0.902894165 |
| chr3:55568976-55569521 | 0.120283162 | 0.755335935 | 0.902894165 |

| | | | |
|---|---|---|---|
| chr19:48332684-48333521 | -0.100825117 | 0.755338886 | 0.902894165 |
| chr5:101663561-101663948 | -0.121344191 | 0.75534358 | 0.902894165 |
| chr1:154206228-154206538 | -0.137747955 | 0.755349885 | 0.902894165 |
| chr9:122498842-122499352 | 0.113412351 | 0.755390774 | 0.902900334 |
| chr5:113205865-113206130 | -0.131854015 | 0.755401641 | 0.902900334 |
| chr8:72287181-72287407 | -0.14558906 | 0.755410018 | 0.902900334 |
| chr2:35099327-35099645 | -0.134534685 | 0.755418197 | 0.902900334 |
| chr2:153195285-153195515 | -0.146784074 | 0.755419419 | 0.902900334 |
| chr8:58807992-58808243 | 0.132153928 | 0.755425749 | 0.902900334 |
| chr19:55063011-55063459 | -0.106671819 | 0.755448874 | 0.902913889 |
| chr16:75129995-75130309 | -0.128317006 | 0.755502002 | 0.90294774 |
| chr6:26056171-26056635 | -0.109667902 | 0.755508829 | 0.90294774 |
| chr17:74284703-74284970 | -0.12988437 | 0.755525354 | 0.90294774 |
| chr7:73521724-73522883 | 0.09967192 | 0.755561139 | 0.90294774 |
| chr19:55301782-55302332 | -0.110119393 | 0.755564033 | 0.90294774 |
| chr19:49533357-49534423 | -0.102902464 | 0.755567153 | 0.90294774 |
| chr2:11746387-11747387 | -0.099761369 | 0.755567605 | 0.90294774 |
| chr3:25962309-25962572 | -0.140015267 | 0.75557593 | 0.90294774 |
| chr4:127844135-127844354 | 0.154775451 | 0.755583256 | 0.90294774 |
| chr2:10812306-10813315 | -0.099956765 | 0.755595374 | 0.902948139 |
| chr1:198935444-198935908 | -0.1017874 | 0.755627315 | 0.902967087 |
| chr11:20128307-20128580 | -0.138102154 | 0.7556348 | 0.902967087 |
| chr19:7488890-7489262 | 0.123137628 | 0.75565315 | 0.902974932 |
| chr5:42057106-42057607 | 0.114531794 | 0.755759386 | 0.903081972 |
| chr1:110136936-110137195 | -0.136263123 | 0.755766298 | 0.903081972 |
| chr10:69629869-69630370 | -0.113170246 | 0.755783318 | 0.903088226 |
| chr5:38083656-38083969 | -0.155982514 | 0.755797935 | 0.903091609 |
| chr7:48559440-48559669 | -0.164144377 | 0.755830884 | 0.903091983 |
| chr4:88007238-88008571 | 0.10156531 | 0.755835562 | 0.903091983 |
| chr12:111692915-111693187 | 0.135714413 | 0.755836342 | 0.903091983 |
| chr7:16603646-16603888 | 0.143898701 | 0.755856918 | 0.903091983 |
| chr15:100671935-100672287 | 0.123008195 | 0.75585718 | 0.903091983 |
| chr17:73164272-73164931 | -0.110410603 | 0.75587908 | 0.903104066 |
| chr14:77990933-77991737 | 0.109443058 | 0.755910361 | 0.903122571 |
| chr18:45916362-45916660 | 0.135314206 | 0.755918141 | 0.903122571 |
| chr4:18882429-18882864 | -0.13359244 | 0.755946399 | 0.903131842 |
| chr6:30060558-30061965 | -0.10143863 | 0.755949475 | 0.903131842 |
| chr5:132776375-132776609 | 0.133582397 | 0.755975988 | 0.903134522 |
| chr12:124715456-124715890 | -0.118498357 | 0.75597759 | 0.903134522 |
| chr8:102961326-102961791 | -0.112108242 | 0.755987079 | 0.903134522 |
| chr2:134412093-134412327 | 0.137211434 | 0.756004909 | 0.903137117 |
| chr14:23115074-23115331 | 0.132931783 | 0.75602007 | 0.903137117 |
| chr9:36400207-36401443 | -0.099338169 | 0.756024612 | 0.903137117 |
| chr5:9898400-9898614 | 0.137263721 | 0.756045943 | 0.903139316 |
| chr9:84315317-84315588 | 0.138991296 | 0.756065213 | 0.903139316 |
| chr17:7436388-7436653 | -0.129927223 | 0.756078505 | 0.903139316 |
| chr6:111096274-111096474 | -0.156545985 | 0.75608166 | 0.903139316 |
| chr20:53865921-53866597 | 0.097293625 | 0.756085387 | 0.903139316 |
| chr14:20305256-20306201 | -0.103223084 | 0.756114762 | 0.903150317 |
| chr5:114305445-114305813 | 0.130625686 | 0.756129611 | 0.903150317 |

| | | | |
|---|---|---|---|
| chr1:84479113-84479907 | 0.106163604 | 0.75613483 | 0.903150317 |
| chr1:114389753-114389986 | 0.116072124 | 0.756148136 | 0.903150317 |
| chr4:39383760-39384062 | -0.133417277 | 0.756157448 | 0.903150317 |
| chr10:8672481-8672712 | -0.152867514 | 0.756172494 | 0.903150317 |
| chr8:86514175-86515287 | 0.097677785 | 0.756177107 | 0.903150317 |
| chr5:139422345-139422640 | 0.134288167 | 0.756205455 | 0.903170097 |
| chr2:36357670-36358100 | -0.110498914 | 0.756312759 | 0.903284174 |
| chr6:27132279-27133247 | -0.099289035 | 0.756342498 | 0.903293722 |
| chr20:54188170-54188463 | -0.10321308 | 0.756344331 | 0.903293722 |
| chr19:17555343-17555807 | 0.11117565 | 0.756378353 | 0.903311119 |
| chr13:114314158-114314757 | -0.108059865 | 0.756382909 | 0.903311119 |
| chr1:161165759-161167104 | -0.098550575 | 0.756394265 | 0.903311119 |
| chrX:107628156-107628705 | 0.101061042 | 0.756436964 | 0.903348032 |
| chr3:45656834-45657039 | -0.161307345 | 0.756480462 | 0.903385898 |
| chr4:76982999-76983238 | -0.160392422 | 0.756494779 | 0.903388915 |
| chr18:26342712-26343147 | 0.129685191 | 0.756534377 | 0.903393411 |
| chr1:50272043-50272280 | -0.144190355 | 0.756536794 | 0.903393411 |
| chr2:95207119-95208108 | 0.102906881 | 0.756541516 | 0.903393411 |
| chr22:39502029-39502730 | -0.105201265 | 0.756551407 | 0.903393411 |
| chrX:107537880-107538149 | 0.132953333 | 0.756557496 | 0.903393411 |
| chr20:21767540-21768102 | 0.12125832 | 0.756572142 | 0.903396822 |
| chr1:196808805-196809212 | -0.135441891 | 0.756598681 | 0.903414432 |
| chr8:64382142-64382418 | -0.129710448 | 0.756613436 | 0.903417973 |
| chr7:131556617-131556852 | -0.136214134 | 0.756690253 | 0.903488847 |
| chr12:113955226-113955507 | -0.127679299 | 0.756709027 | 0.903488847 |
| chr7:108477707-108477990 | 0.14274013 | 0.756730851 | 0.903488847 |
| chrX:38803642-38804320 | -0.104097392 | 0.756733538 | 0.903488847 |
| chr1:80160253-80160781 | -0.118913463 | 0.756735018 | 0.903488847 |
| chr4:99099355-99100132 | 0.110536648 | 0.756743542 | 0.903488847 |
| chr4:86855530-86856044 | -0.112961636 | 0.756755442 | 0.903488975 |
| chr13:75789344-75789747 | -0.147229478 | 0.756778777 | 0.903502757 |
| chr12:130924227-130924427 | -0.176446785 | 0.756808125 | 0.903523717 |
| chr1:241807009-241807209 | 0.162964338 | 0.756838173 | 0.903527064 |
| chr13:92752530-92753120 | 0.117971776 | 0.756840369 | 0.903527064 |
| chr3:101686119-101687455 | -0.097561235 | 0.756852743 | 0.903527064 |
| chr2:11738480-11739249 | -0.105811497 | 0.756869457 | 0.903527064 |
| chr2:74482330-74483635 | -0.098404065 | 0.756874889 | 0.903527064 |
| chr6:9875632-9876427 | -0.108583194 | 0.756881681 | 0.903527064 |
| chr1:166166673-166167505 | 0.103840196 | 0.756901144 | 0.903536222 |
| chr3:115217608-115218034 | -0.122287021 | 0.756926155 | 0.903539814 |
| chr8:29521229-29521528 | 0.139995066 | 0.756927738 | 0.903539814 |
| chr18:8563807-8564080 | -0.16175493 | 0.756958546 | 0.903545135 |
| chr14:92039319-92040565 | 0.099153628 | 0.7569835 | 0.903545135 |
| chr12:119020874-11902112€ | 0.160868003 | 0.756989531 | 0.903545135 |
| chr6:100944611-100944882 | -0.14915836 | 0.757016174 | 0.903545135 |
| chr3:124814699-124815263 | -0.107053722 | 0.757016653 | 0.903545135 |
| chr19:11528729-11529536 | -0.106756024 | 0.757025816 | 0.903545135 |
| chr12:98373067-98373328 | 0.125692976 | 0.757026299 | 0.903545135 |
| chr1:113958090-113958367 | 0.1174423 | 0.757026534 | 0.903545135 |
| chr11:3217120-3217590 | -0.107426961 | 0.757059873 | 0.903570852 |

| | | | |
|---|---|---|---|
| chr6:29552594-29552803 | -0.158792049 | 0.757084073 | 0.903585661 |
| chr6:125213976-125214311 | -0.132689459 | 0.75710566 | 0.90359735 |
| chrX:142495080-142495280 | -0.171465418 | 0.75714536 | 0.903630656 |
| chr22:19970597-19970833 | 0.131021482 | 0.757207431 | 0.903666166 |
| chr17:44132555-44132764 | 0.168477077 | 0.757211965 | 0.903666166 |
| chr11:65890386-65890779 | 0.105620329 | 0.757217654 | 0.903666166 |
| chr12:53825198-53825884 | 0.105529532 | 0.757235413 | 0.903666166 |
| chr19:10581047-10581349 | -0.127681564 | 0.75724613 | 0.903666166 |
| chr6:27556460-27556746 | 0.134405475 | 0.757253734 | 0.903666166 |
| chr20:30376608-30377801 | 0.100165511 | 0.75725767 | 0.903666166 |
| chr3:39406303-39407159 | 0.100677217 | 0.757275014 | 0.903672789 |
| chrX:103607398-103608944 | -0.099705832 | 0.757296152 | 0.903683939 |
| chr20:53829211-53829607 | 0.101263215 | 0.757377173 | 0.903752776 |
| chr4:81351733-81351986 | 0.141738447 | 0.757383854 | 0.903752776 |
| chr16:56372839-56373283 | 0.11667767 | 0.75739852 | 0.903752776 |
| chr6:27791946-27792571 | 0.108677968 | 0.757401018 | 0.903752776 |
| chr1:110240503-110240707 | 0.15266672 | 0.757441565 | 0.903777541 |
| chr11:61141668-61141868 | -0.150492179 | 0.757450658 | 0.903777541 |
| chr5:126544182-126544506 | 0.120527844 | 0.757457158 | 0.903777541 |
| chr1:145287239-145288138 | 0.098689008 | 0.757501157 | 0.90379537 |
| chr6:67009907-67010214 | -0.130048569 | 0.757504503 | 0.90379537 |
| chr1:176206601-176207905 | -0.09704942 | 0.757507487 | 0.90379537 |
| chr2:47323000-47323514 | -0.1095149 | 0.757528579 | 0.903800751 |
| chr4:140150681-140152446 | 0.09670524 | 0.757535588 | 0.903800751 |
| chr6:53544350-53545314 | -0.098712883 | 0.757571952 | 0.903804052 |
| chr18:35240409-35241339 | -0.099666142 | 0.757587126 | 0.903804052 |
| chr1:206416348-206416549 | 0.138074532 | 0.757589998 | 0.903804052 |
| chr20:50172430-50172644 | -0.143084496 | 0.757602745 | 0.903804052 |
| chr10:17388738-17388990 | -0.13733914 | 0.757606318 | 0.903804052 |
| chr1:108118271-108118740 | -0.109752254 | 0.757609129 | 0.903804052 |
| chr4:36243711-36244995 | 0.099278126 | 0.757629684 | 0.903814502 |
| chr5:138133408-138133717 | -0.131550884 | 0.757728053 | 0.903917778 |
| chr11:46367660-46367901 | 0.132439002 | 0.757770821 | 0.903953185 |
| chr6:26021061-26022001 | -0.098973089 | 0.757783778 | 0.903953185 |
| chr11:73950841-73951234 | -0.117588204 | 0.757796138 | 0.903953185 |
| chr5:93070962-93071162 | -0.161518059 | 0.757804924 | 0.903953185 |
| chr4:84596837-84597130 | -0.129945091 | 0.757836244 | 0.903976473 |
| chr16:80718356-80719240 | 0.100909527 | 0.75786318 | 0.903989949 |
| chr1:55378570-55378780 | 0.140959538 | 0.757871138 | 0.903989949 |
| chr3:143390020-143390220 | 0.185885711 | 0.757914436 | 0.904027521 |
| chr2:188305365-188305587 | 0.143261388 | 0.75793309 | 0.904035699 |
| chr1:11735965-11736266 | 0.126082585 | 0.757945643 | 0.904036598 |
| chr4:77156516-77157335 | 0.098118022 | 0.758001621 | 0.904089292 |
| chr5:74935042-74935251 | -0.163690668 | 0.758016776 | 0.904093294 |
| chr19:43668175-43669083 | 0.097395382 | 0.75803048 | 0.904095566 |
| chr12:7986786-7987051 | -0.146089121 | 0.758114619 | 0.90416047 |
| chr4:107037072-107037514 | 0.118971955 | 0.75811606 | 0.90416047 |
| chr16:74941743-74941943 | 0.148940984 | 0.758130012 | 0.90416047 |
| chr5:172462180-172462503 | -0.128870856 | 0.758132099 | 0.90416047 |
| chr1:93449160-93449418 | -0.134507596 | 0.758159915 | 0.904172394 |

| | | | |
|---|---|---|---|
| chr5:152349747-152350269 | 0.122486515 | 0.758165698 | 0.904172394 |
| chr10:102753014-102753427 | -0.136789886 | 0.758212061 | 0.904213611 |
| chr7:46886583-46887079 | 0.11981533 | 0.758240115 | 0.90421855 |
| chr16:88857769-88858036 | -0.111866375 | 0.758245121 | 0.90421855 |
| chr6:112180184-112180384 | -0.162203988 | 0.758251665 | 0.90421855 |
| chr7:66413371-66414078 | 0.102682563 | 0.758263407 | 0.90421855 |
| chr3:50542461-50542747 | 0.134291789 | 0.75828504 | 0.904230218 |
| chr2:27663284-27664440 | -0.100858689 | 0.758296794 | 0.904230218 |
| chr1:85240169-85240646 | -0.127077468 | 0.75832784 | 0.904253167 |
| chr9:122350566-122350954 | 0.112771944 | 0.758360748 | 0.904262059 |
| chr17:7652771-7652993 | -0.16217073 | 0.758369089 | 0.904262059 |
| chr9:15422071-15423352 | -0.099315263 | 0.758379522 | 0.904262059 |
| chr11:17978704-17978985 | 0.130970151 | 0.758382503 | 0.904262059 |
| chr17:28570850-28571307 | 0.110160388 | 0.758420145 | 0.904292869 |
| chr3:145776840-145777074 | -0.141310317 | 0.758473187 | 0.904332182 |
| chr22:34111305-34111586 | -0.137370383 | 0.758479849 | 0.904332182 |
| chr3:48188034-48188881 | -0.101756303 | 0.758490673 | 0.904332182 |
| chr8:80300000-80300288 | 0.126164357 | 0.758516113 | 0.904332182 |
| chr15:45402068-45402952 | 0.100083487 | 0.758517766 | 0.904332182 |
| chr1:185400931-185401131 | -0.171392383 | 0.758533203 | 0.904332182 |
| chr1:43549901-43550579 | -0.100367717 | 0.758535734 | 0.904332182 |
| chrX:41924343-41924699 | 0.118643897 | 0.75855378 | 0.904339626 |
| chr1:150235657-150236681 | -0.097596244 | 0.758605145 | 0.90438341 |
| chr3:152071977-152072319 | -0.121756341 | 0.758616938 | 0.90438341 |
| chr2:97645304-97646252 | -0.099337158 | 0.758631346 | 0.90438341 |
| chr8:56909676-56909933 | -0.154974079 | 0.758637719 | 0.90438341 |
| chr18:12658465-12658715 | 0.133558727 | 0.758650275 | 0.904384307 |
| chr5:10034157-10034357 | 0.141604946 | 0.758728201 | 0.90442215 |
| chr1:219839766-219839979 | -0.153730196 | 0.758736934 | 0.90442215 |
| chr9:108468397-108468682 | -0.135827502 | 0.758740055 | 0.90442215 |
| chr16:58515271-58516585 | -0.099423311 | 0.758754806 | 0.90442215 |
| chr8:131452249-131452567 | 0.132306017 | 0.758781525 | 0.90442215 |
| chr6:56378643-56379288 | 0.110477329 | 0.758782413 | 0.90442215 |
| chr4:56960356-56960681 | -0.129986193 | 0.758784855 | 0.90442215 |
| chr15:40310109-40310507 | 0.123575632 | 0.758788963 | 0.90442215 |
| chr8:42897240-42897466 | 0.126660772 | 0.758792004 | 0.90442215 |
| chr1:26112065-26112412 | 0.119084775 | 0.758800056 | 0.90442215 |
| chr15:75546718-75546918 | -0.168490444 | 0.758817367 | 0.904428713 |
| chr17:18857707-18858645 | -0.101062272 | 0.758838475 | 0.90443497 |
| chr11:12724346-12725009 | -0.104758002 | 0.758846223 | 0.90443497 |
| chr9:125200089-125201088 | -0.098922458 | 0.758860579 | 0.90443801 |
| chr1:150981191-150982497 | 0.097457004 | 0.758900638 | 0.904467166 |
| chr22:25057318-25057584 | 0.128463294 | 0.758917129 | 0.904467166 |
| chr15:86903664-86903974 | 0.132157597 | 0.758920454 | 0.904467166 |
| chr17:44218397-44218813 | 0.106812111 | 0.758950141 | 0.904488478 |
| chr19:5048735-5048935 | 0.163035904 | 0.758981338 | 0.904511588 |
| chr1:224205501-224205862 | -0.115531193 | 0.759002563 | 0.904520163 |
| chr8:89983707-89984888 | -0.096055526 | 0.759012143 | 0.904520163 |
| chr12:1836782-1837478 | 0.104513799 | 0.759095439 | 0.904602008 |
| chr5:172640712-172641720 | -0.100979002 | 0.759113874 | 0.904602008 |

| | | | |
|---|---|---|---|
| chr2:121648813-121650182 | -0.098040466 | 0.759116239 | 0.904602008 |
| chr5:126864040-126864287 | -0.136070055 | 0.759133631 | 0.904608664 |
| chr12:57741899-57742385 | -0.104083174 | 0.759182919 | 0.904653327 |
| chr10:78302749-78303904 | -0.099066944 | 0.75919626 | 0.904655156 |
| chr9:121326162-121326445 | -0.128516291 | 0.759245795 | 0.90465622 |
| chr9:127955152-127956033 | -0.100473974 | 0.759263685 | 0.90465622 |
| chr11:26284596-26285188 | 0.110091395 | 0.759263876 | 0.90465622 |
| chr2:219178536-219178742 | 0.126208224 | 0.759264989 | 0.90465622 |
| chr18:3216439-3216725 | 0.13983297 | 0.759266603 | 0.90465622 |
| chr18:32271796-32272020 | 0.146120599 | 0.759267994 | 0.90465622 |
| chr2:185738624-185739438 | 0.101740239 | 0.759289134 | 0.904661205 |
| chr4:148034595-148035165 | 0.108887544 | 0.759295791 | 0.904661205 |
| chr7:151085716-151086361 | 0.102373673 | 0.759342585 | 0.904702889 |
| chr2:130181253-130182487 | 0.098049914 | 0.759354888 | 0.90470348 |
| chr12:46509122-46509435 | 0.136663607 | 0.759370418 | 0.904707916 |
| chr8:134778222-134778452 | -0.14379116 | 0.759451349 | 0.904761398 |
| chr19:17488727-17489447 | -0.103001438 | 0.759458266 | 0.904761398 |
| chr15:39782318-39783023 | -0.100391762 | 0.759459133 | 0.904761398 |
| chr1:111045352-111045752 | -0.137193886 | 0.759478461 | 0.904761398 |
| chr17:75459499-75459769 | -0.116745023 | 0.759484239 | 0.904761398 |
| chr10:23439694-23440445 | -0.107908726 | 0.759486158 | 0.904761398 |
| chr1:75079967-75080230 | -0.13533181 | 0.759503051 | 0.904764337 |
| chr8:103900958-103901158 | -0.159208544 | 0.759512241 | 0.904764337 |
| chr17:45171323-45171702 | -0.114744235 | 0.759528686 | 0.904769861 |
| chr10:104824434-104824694 | -0.138025464 | 0.759569261 | 0.904804128 |
| chr17:57004977-57005204 | 0.126808986 | 0.759596799 | 0.904822865 |
| chr10:3882942-3883142 | -0.165095591 | 0.759617127 | 0.904831904 |
| chr6:19268217-19268450 | -0.147661158 | 0.759628006 | 0.904831904 |
| chr10:15567785-15568047 | -0.167877902 | 0.759686896 | 0.904868565 |
| chr8:38808060-38808356 | 0.153680966 | 0.759715412 | 0.904868565 |
| chr2:156332114-156333458 | -0.096524201 | 0.759741386 | 0.904868565 |
| chr5:1056814-1057191 | -0.121398908 | 0.759744231 | 0.904868565 |
| chr2:118269464-118269725 | -0.130681607 | 0.75975321 | 0.904868565 |
| chr16:31477494-31477793 | 0.116947521 | 0.759755852 | 0.904868565 |
| chr10:21078774-21078975 | 0.160518391 | 0.759760598 | 0.904868565 |
| chr3:75983517-75983717 | 0.176070336 | 0.759787114 | 0.904868565 |
| chr1:147402731-147403062 | -0.113824719 | 0.75978902 | 0.904868565 |
| chr12:120971809-120972009 | 0.154768381 | 0.759806254 | 0.904868565 |
| chr2:200626997-200627197 | -0.165011347 | 0.759809137 | 0.904868565 |
| chr13:74135250-74136009 | -0.101958804 | 0.759810555 | 0.904868565 |
| chr5:106658277-106658477 | -0.156905278 | 0.759812307 | 0.904868565 |
| chr5:159533278-159533498 | -0.156932418 | 0.759852955 | 0.904902909 |
| chr18:3593428-3594555 | -0.098767528 | 0.759907255 | 0.904953508 |
| chr9:35071764-35072459 | -0.103444729 | 0.759945281 | 0.904984727 |
| chr12:76472766-76473213 | -0.124715233 | 0.759964323 | 0.904993338 |
| chr19:15433333-15433533 | 0.146464891 | 0.760004366 | 0.90502563 |
| chr12:77877647-77878013 | -0.126104753 | 0.760015063 | 0.90502563 |
| chr19:29014598-29014798 | -0.151790887 | 0.760062691 | 0.905042639 |
| chr6:31806625-31807186 | 0.102647501 | 0.760073904 | 0.905042639 |
| chr1:12528361-12528642 | -0.143791116 | 0.760076177 | 0.905042639 |

| | | | |
|---|---|---|---|
| chr5:180915881-180916108 | -0.156725489 | 0.76008132 | 0.905042639 |
| chr1:198966074-198966284 | 0.151708827 | 0.760088405 | 0.905042639 |
| chr2:64653237-64654440 | -0.097620666 | 0.760108765 | 0.905051016 |
| chr3:124492887-124493133 | -0.126584785 | 0.760126722 | 0.905051016 |
| chr2:25951001-25951405 | -0.109788834 | 0.760130876 | 0.905051016 |
| chr17:76026552-76027758 | 0.099529656 | 0.760155553 | 0.905066333 |
| chr3:6035835-6036505 | -0.111454547 | 0.760177788 | 0.905073192 |
| chr14:76918911-76919119 | 0.123067364 | 0.760193823 | 0.905073192 |
| chr1:241533838-241534123 | -0.124855624 | 0.760208067 | 0.905073192 |
| chr16:88686021-88686927 | 0.100333864 | 0.76021649 | 0.905073192 |
| chr13:105580483-105580721 | 0.140587408 | 0.760220375 | 0.905073192 |
| chr9:127599227-127599689 | 0.116753749 | 0.760234757 | 0.905076252 |
| chr9:125408415-125408722 | 0.127465242 | 0.76025024 | 0.905080621 |
| chr3:62188762-62189580 | -0.09807897 | 0.76028354 | 0.905106202 |
| chrX:119189257-119189457 | -0.163481948 | 0.760302848 | 0.905108019 |
| chr14:77468879-77469102 | 0.144109179 | 0.760325307 | 0.905108019 |
| chr12:104741775-104742014 | 0.152018048 | 0.760333531 | 0.905108019 |
| chr13:27477150-27477569 | 0.122453389 | 0.760340518 | 0.905108019 |
| chr10:127344896-127345096 | 0.186193891 | 0.760344129 | 0.905108019 |
| chr2:213246781-213247012 | -0.150244489 | 0.76035979 | 0.905112599 |
| chr9:129473978-129474269 | -0.135807095 | 0.76038908 | 0.905133405 |
| chr10:81117043-81117313 | 0.125488712 | 0.760418097 | 0.905138979 |
| chr14:51750634-51750907 | 0.128644536 | 0.760428881 | 0.905138979 |
| chr5:7868687-7869840 | 0.098255203 | 0.760433875 | 0.905138979 |
| chr16:69083594-69083794 | 0.149876807 | 0.760441016 | 0.905138979 |
| chr10:93080342-93080579 | 0.129453308 | 0.760480279 | 0.905155479 |
| chr19:18880055-18880336 | -0.140030607 | 0.76050345 | 0.905155479 |
| chr6:72412036-72412575 | 0.110103287 | 0.76050906 | 0.905155479 |
| chr17:27654633-27654946 | 0.114427375 | 0.760510603 | 0.905155479 |
| chrX:130916952-130917185 | -0.141266381 | 0.760525517 | 0.905155479 |
| chr4:30380466-30380695 | -0.147090093 | 0.760532972 | 0.905155479 |
| chr16:78248958-78249455 | -0.109243543 | 0.760537571 | 0.905155479 |
| chrX:15349077-15349357 | 0.147496729 | 0.760575605 | 0.90516978 |
| chr1:42967083-42967394 | -0.126067097 | 0.760589717 | 0.90516978 |
| chr3:46558314-46558676 | 0.112211155 | 0.76059759 | 0.90516978 |
| chr18:9340895-9341114 | -0.150059498 | 0.760599974 | 0.90516978 |
| chr11:17386692-17386915 | 0.116925618 | 0.760617586 | 0.90516978 |
| chr5:51034574-51034846 | 0.147561939 | 0.760629913 | 0.90516978 |
| chr17:10797887-10798252 | -0.128932837 | 0.760632281 | 0.90516978 |
| chr2:212766889-212767416 | 0.112633898 | 0.760661195 | 0.905181475 |
| chrX:101906248-101906483 | 0.156166043 | 0.760705548 | 0.905181475 |
| chr18:3247073-3248167 | 0.09593592 | 0.760710545 | 0.905181475 |
| chr12:99619911-99620225 | 0.127901699 | 0.760711637 | 0.905181475 |
| chr19:55166456-55167301 | -0.098302335 | 0.760713717 | 0.905181475 |
| chr11:68030139-68030965 | -0.098937328 | 0.76071712 | 0.905181475 |
| chr8:54089673-54090471 | -0.105555178 | 0.760724803 | 0.905181475 |
| chr1:235327931-235329142 | 0.09558292 | 0.760755426 | 0.90519311 |
| chr1:33080863-33081608 | 0.102327757 | 0.76075821 | 0.90519311 |
| chr20:38057577-38057797 | 0.148134983 | 0.760827309 | 0.905261271 |
| chr1:49630234-49630436 | 0.15137546 | 0.760859978 | 0.905285062 |

| | | | |
|---|---|---|---|
| chr1:244439379-244439615 | -0.144680637 | 0.760875842 | 0.905285062 |
| chr22:24834936-24835179 | -0.143958523 | 0.760882749 | 0.905285062 |
| chr15:58555972-58556172 | 0.151222424 | 0.760913287 | 0.905307338 |
| chr19:38266120-38266325 | -0.137326125 | 0.760933093 | 0.905316846 |
| chr9:132507736-132507997 | -0.134518644 | 0.761000084 | 0.905370701 |
| chr3:119702584-119703306 | -0.099447074 | 0.761006767 | 0.905370701 |
| chr11:29449032-29449246 | 0.173260845 | 0.761025083 | 0.905370701 |
| chr8:63085921-63086256 | 0.120665512 | 0.761025624 | 0.905370701 |
| chr9:6645623-6645999 | 0.115618165 | 0.761044773 | 0.905371207 |
| chr3:185308144-185308467 | -0.133264456 | 0.761049682 | 0.905371207 |
| chr19:35748192-35748670 | 0.100899016 | 0.761085682 | 0.905380592 |
| chr12:10190894-10191110 | -0.151120424 | 0.761092319 | 0.905380592 |
| chr21:33641438-33642708 | 0.095947878 | 0.761093152 | 0.905380592 |
| chr20:58870275-58870659 | 0.105782873 | 0.761119928 | 0.905380592 |
| chr12:7933199-7933574 | -0.111096234 | 0.761128308 | 0.905380592 |
| chr18:50357259-50357512 | 0.13894153 | 0.761128468 | 0.905380592 |
| chr17:65774450-65774650 | 0.15110561 | 0.761146125 | 0.905385109 |
| chr20:42459548-42459825 | 0.13519238 | 0.761167254 | 0.905385109 |
| chr3:190884133-190884531 | -0.125824613 | 0.761167714 | 0.905385109 |
| chr15:72155073-72155525 | -0.114032291 | 0.76120334 | 0.905410121 |
| chr16:72627061-72627585 | 0.109450592 | 0.761230741 | 0.905410121 |
| chr18:59057296-59057535 | -0.149670943 | 0.761250912 | 0.905410121 |
| chr2:74528987-74530170 | -0.096638499 | 0.761256516 | 0.905410121 |
| chr8:97530408-97530671 | -0.129522642 | 0.761264054 | 0.905410121 |
| chr5:122252325-122252525 | -0.154050243 | 0.761269174 | 0.905410121 |
| chr8:63391368-63391747 | -0.120152059 | 0.761288998 | 0.905410121 |
| chr3:61959376-61959576 | 0.119097327 | 0.761300022 | 0.905410121 |
| chr2:121044491-121044730 | -0.136540299 | 0.761307387 | 0.905410121 |
| chr11:123103264-123103517 | 0.14777516 | 0.76130962 | 0.905410121 |
| chr12:112108423-112109228 | 0.100131168 | 0.761318724 | 0.905410121 |
| chr3:167721528-167721733 | -0.146575084 | 0.761349399 | 0.905425377 |
| chr6:125141469-125141705 | -0.151302034 | 0.761355185 | 0.905425377 |
| chr1:19799604-19799860 | -0.13955212 | 0.761390363 | 0.905453158 |
| chr3:139520113-139520383 | 0.132129645 | 0.761414181 | 0.905467428 |
| chr7:141533152-141533639 | -0.122401581 | 0.761443717 | 0.9054885 |
| chr1:155729021-155729444 | 0.113047251 | 0.761484541 | 0.905517645 |
| chr3:79314515-79314924 | 0.117386288 | 0.761501787 | 0.905517645 |
| chr3:15601271-15602396 | 0.095460453 | 0.76150368 | 0.905517645 |
| chr6:43070079-43070279 | -0.151782481 | 0.761529856 | 0.905534718 |
| chr6:52995079-52996008 | -0.096248427 | 0.761557086 | 0.905553044 |
| chr5:140372906-140373426 | -0.107814715 | 0.761604199 | 0.905577942 |
| chr7:33905494-33905896 | -0.114402919 | 0.76160874 | 0.905577942 |
| chr19:48999721-49000083 | 0.11862366 | 0.761613481 | 0.905577942 |
| chr11:102346724-102347601 | 0.101236333 | 0.761669922 | 0.90559418 |
| chr1:68205273-68205542 | -0.139846721 | 0.761719735 | 0.90559418 |
| chr3:185858686-185858886 | -0.169490181 | 0.761720687 | 0.90559418 |
| chrX:110317395-110317649 | 0.123572163 | 0.761728926 | 0.90559418 |
| chr8:94797192-94797444 | 0.140874986 | 0.761730264 | 0.90559418 |
| chr1:207259401-207259983 | 0.103619365 | 0.761736323 | 0.90559418 |
| chr2:16773195-16773409 | -0.137796522 | 0.761741755 | 0.90559418 |

| | | | |
|---|---|---|---|
| chr21:40643022-40643224 | -0.16277853 | 0.761743584 | 0.90559418 |
| chr20:30585169-30586564 | 0.09705676 | 0.7617477 | 0.90559418 |
| chr13:33514372-33514582 | -0.152505189 | 0.761750429 | 0.90559418 |
| chr6:150865339-150867096 | -0.097233853 | 0.761784719 | 0.90559418 |
| chr15:34318510-34318988 | -0.114623015 | 0.761784804 | 0.90559418 |
| chr7:492816-493052 | -0.130927167 | 0.7617881 | 0.90559418 |
| chr8:143691397-143691597 | -0.154197488 | 0.761792604 | 0.90559418 |
| chr9:32378563-32378811 | -0.131638951 | 0.761815868 | 0.905607785 |
| chr3:50157155-50157355 | -0.159052185 | 0.76183886 | 0.905620203 |
| chr7:151350315-151350575 | -0.126821347 | 0.761852863 | 0.905620203 |
| chr9:113376656-113377395 | 0.10110541 | 0.761861772 | 0.905620203 |
| chr19:45886446-45886992 | -0.104048136 | 0.761914864 | 0.905669264 |
| chr20:4869638-4869878 | 0.139549737 | 0.761941237 | 0.905686562 |
| chr12:62259353-62260915 | 0.095542751 | 0.761973857 | 0.905711285 |
| chr2:26033174-26034642 | -0.095377825 | 0.761986874 | 0.905712707 |
| chr6:32129976-32130588 | -0.102180837 | 0.761999385 | 0.905713528 |
| chr6:134156802-134157002 | 0.158324952 | 0.762082907 | 0.905774498 |
| chr13:57630850-57631050 | 0.149388397 | 0.762095307 | 0.905774498 |
| chr12:64363389-64363802 | 0.13890701 | 0.7620964 | 0.905774498 |
| chr21:40185598-40185808 | -0.143892372 | 0.762102664 | 0.905774498 |
| chr7:150988469-150988741 | 0.120325832 | 0.762128556 | 0.905774498 |
| chr17:56144305-56144567 | -0.139528409 | 0.76215697 | 0.905774498 |
| chr11:75456856-75457131 | 0.136573854 | 0.762168619 | 0.905774498 |
| chr1:109548376-109549250 | -0.098279935 | 0.762169055 | 0.905774498 |
| chr14:20681531-20681802 | -0.106770624 | 0.762178417 | 0.905774498 |
| chr8:91081700-91081928 | 0.142396535 | 0.76218387 | 0.905774498 |
| chr22:40636378-40637128 | 0.101346956 | 0.762188185 | 0.905774498 |
| chr11:34331873-34332719 | 0.107645595 | 0.762196327 | 0.905774498 |
| chr20:54619256-54619602 | 0.102644085 | 0.762204358 | 0.905774498 |
| chr11:12723809-12724272 | 0.103970439 | 0.762248055 | 0.905812377 |
| chr16:68022585-68022941 | -0.121702353 | 0.762274892 | 0.905822533 |
| chr3:64717454-64718068 | 0.098959683 | 0.762306231 | 0.905822533 |
| chr14:91535012-91535661 | -0.102293551 | 0.762321042 | 0.905822533 |
| chr15:85307057-85307319 | -0.150632832 | 0.762327887 | 0.905822533 |
| chr15:101651772-101653213 | 0.09572683 | 0.762329938 | 0.905822533 |
| chr8:87011171-87011407 | 0.133447079 | 0.762351461 | 0.905822533 |
| chr5:27682426-27683029 | -0.10758576 | 0.762355725 | 0.905822533 |
| chr22:24196715-24196936 | 0.144192201 | 0.76235704 | 0.905822533 |
| chr2:158024409-158024627 | 0.150046118 | 0.762374446 | 0.905822533 |
| chr14:71050272-71050472 | -0.168974774 | 0.762379435 | 0.905822533 |
| chr7:150946011-150946231 | -0.136475157 | 0.762389186 | 0.905822533 |
| chr1:66182576-66183124 | 0.117064942 | 0.762425526 | 0.905822533 |
| chr3:129383289-129383689 | 0.108695429 | 0.762427418 | 0.905822533 |
| chr10:13271543-13271886 | -0.125096275 | 0.762429613 | 0.905822533 |
| chr12:124993985-124994520 | -0.106656909 | 0.7624498 | 0.905822533 |
| chr4:129985498-129985745 | -0.162032482 | 0.762457574 | 0.905822533 |
| chr3:147842937-147843137 | -0.162032482 | 0.762457574 | 0.905822533 |
| chr17:48148405-48148605 | -0.148675695 | 0.762486195 | 0.90584249 |
| chr8:22945428-22945808 | -0.132007479 | 0.762500211 | 0.905845097 |
| chr10:34954162-34954624 | -0.104884059 | 0.762561501 | 0.905890385 |

| | | | |
|---|---|---|---|
| chrX:110173908-110174181 | 0.137719478 | 0.762561979 | 0.905890385 |
| chr2:176079714-176080199 | 0.116564032 | 0.762609796 | 0.905919857 |
| chr2:46296932-46297524 | -0.106821934 | 0.762610434 | 0.905919857 |
| chr12:52534540-52534740 | -0.172137206 | 0.762658354 | 0.905946777 |
| chr8:107416616-107416816 | 0.148477194 | 0.762673628 | 0.905946777 |
| chr11:63838578-63839386 | -0.098069691 | 0.762678509 | 0.905946777 |
| chr1:163321615-163322343 | 0.0973802 | 0.76268039 | 0.905946777 |
| chr17:58639339-58640182 | 0.095955551 | 0.762699492 | 0.905951497 |
| chr1:154925280-154925480 | -0.159387558 | 0.762708011 | 0.905951497 |
| chr18:54388799-54389059 | -0.144175358 | 0.762744075 | 0.905980289 |
| chr2:76748752-76748990 | -0.14484928 | 0.762768534 | 0.905982202 |
| chr1:41353706-41354242 | 0.107937359 | 0.762769334 | 0.905982202 |
| chr11:43680285-43681334 | 0.097041341 | 0.762795531 | 0.905999275 |
| chr1:223119466-223119732 | -0.133879411 | 0.762816201 | 0.906009781 |
| chr14:60734562-60735300 | -0.101124646 | 0.762842712 | 0.906027224 |
| chr1:182910210-182910478 | -0.129918433 | 0.762870293 | 0.906044972 |
| chr15:37134755-37134955 | -0.153555843 | 0.762881306 | 0.906044972 |
| chr10:112755744-112756083 | -0.119835996 | 0.762895767 | 0.906048104 |
| chr9:82332252-82332458 | -0.159105856 | 0.762914102 | 0.906055835 |
| chrX:3814422-3815409 | -0.096317782 | 0.762945668 | 0.906077268 |
| chrX:47190546-47191055 | -0.104193236 | 0.762955799 | 0.906077268 |
| chr3:37974697-37975038 | -0.123560878 | 0.763009641 | 0.906080006 |
| chr13:32984823-32985573 | 0.098543491 | 0.763023813 | 0.906080006 |
| chr7:29113653-29114099 | -0.122847196 | 0.763028436 | 0.906080006 |
| chr1:161530210-161530528 | 0.103765985 | 0.76302956 | 0.906080006 |
| chr16:35524090-35524290 | -0.162724091 | 0.763032106 | 0.906080006 |
| chr17:48338391-48338602 | -0.157413209 | 0.763034521 | 0.906080006 |
| chr2:88591141-88591349 | 0.151679633 | 0.763040882 | 0.906080006 |
| chr2:108719191-108720042 | 0.098098807 | 0.763102285 | 0.906132478 |
| chr10:67697717-67697956 | -0.136347009 | 0.763108722 | 0.906132478 |
| chr6:106339314-106339856 | 0.108314719 | 0.763126531 | 0.906139581 |
| chr8:113817505-113817717 | 0.148424704 | 0.76318506 | 0.906184369 |
| chrX:138026297-138026525 | -0.14207138 | 0.763199303 | 0.906184369 |
| chr11:34051222-34052474 | 0.096457011 | 0.763206576 | 0.906184369 |
| chr1:28087990-28088190 | 0.131804964 | 0.763211702 | 0.906184369 |
| chr20:25859662-25860120 | -0.108411408 | 0.763231744 | 0.906184369 |
| chrX:3712944-3714146 | -0.100678006 | 0.76323521 | 0.906184369 |
| chr3:18559379-18559728 | 0.113560152 | 0.763258159 | 0.906197575 |
| chr2:20577709-20578230 | 0.108777163 | 0.763270565 | 0.906198262 |
| chr6:152643001-152643510 | 0.112421635 | 0.763311671 | 0.90620119 |
| chr18:32372254-32373103 | -0.098538004 | 0.763324532 | 0.90620119 |
| chr3:140529056-140529478 | -0.12097834 | 0.763327656 | 0.90620119 |
| chr11:86461655-86461880 | -0.148319174 | 0.763352442 | 0.90620119 |
| chr11:33700437-33701262 | -0.099867844 | 0.763358071 | 0.90620119 |
| chr3:4999868-5000167 | 0.122590869 | 0.763364812 | 0.90620119 |
| chr4:87018502-87018702 | -0.172688771 | 0.763375823 | 0.90620119 |
| chr10:17454558-17454815 | 0.127094046 | 0.763377282 | 0.90620119 |
| chr22:45163456-45164123 | -0.104793974 | 0.763387136 | 0.90620119 |
| chr3:2446586-2446849 | 0.14098246 | 0.763391301 | 0.90620119 |
| chr5:132065425-132065640 | -0.127678564 | 0.763409179 | 0.906208374 |

| | | | |
|---|---|---|---|
| chr12:80936482-80938068 | -0.095351885 | 0.763441503 | 0.906230846 |
| chr2:144512878-144513175 | -0.149984588 | 0.763451764 | 0.906230846 |
| chr3:142994008-142994330 | -0.123526323 | 0.763506591 | 0.906270739 |
| chr3:149004721-149004942 | 0.145409057 | 0.763509028 | 0.906270739 |
| chr20:32277088-32277352 | -0.144886274 | 0.763522794 | 0.906272034 |
| chr20:23782267-23782953 | 0.111857652 | 0.763542838 | 0.906272034 |
| chr4:37802057-37802462 | -0.112075616 | 0.763545602 | 0.906272034 |
| chr1:75112246-75112949 | 0.109960401 | 0.763602447 | 0.906282556 |
| chr11:75532368-75532654 | -0.116816932 | 0.763606257 | 0.906282556 |
| chr1:30801799-30801999 | 0.143491719 | 0.763614787 | 0.906282556 |
| chr1:67376192-67376938 | 0.099046439 | 0.763642195 | 0.906282556 |
| chr17:81683435-81684205 | 0.097764458 | 0.763643359 | 0.906282556 |
| chr7:150340490-150340729 | -0.13873212 | 0.763644803 | 0.906282556 |
| chr6:168355865-168356188 | -0.136188729 | 0.763650764 | 0.906282556 |
| chr2:183891591-183891791 | -0.165073059 | 0.763652373 | 0.906282556 |
| chr4:40749671-40750877 | 0.098725004 | 0.763660919 | 0.906282556 |
| chr21:38356018-38356333 | -0.110545696 | 0.763693294 | 0.906292144 |
| chr16:3152375-3153101 | -0.097501569 | 0.763703197 | 0.906292144 |
| chr8:104349350-104349553 | 0.140757276 | 0.763704483 | 0.906292144 |
| chr14:69643421-69644022 | 0.102935919 | 0.763756676 | 0.906322936 |
| chrX:141945379-141945790 | 0.119081482 | 0.763758577 | 0.906322936 |
| chr2:38555208-38555497 | -0.123649621 | 0.763765915 | 0.906322936 |
| chr9:87502052-87502380 | 0.138599398 | 0.763787282 | 0.906334254 |
| chr11:79471100-79471474 | 0.119460215 | 0.763828187 | 0.906349517 |
| chr14:22319820-22320059 | 0.133921621 | 0.7638572 | 0.906349517 |
| chr17:56592524-56594467 | -0.094936409 | 0.763857851 | 0.906349517 |
| chr4:48588663-48588944 | -0.137498239 | 0.763875714 | 0.906349517 |
| chr1:110460293-110460633 | -0.117604397 | 0.76387663 | 0.906349517 |
| chr1:31957899-31958099 | 0.167838165 | 0.763880157 | 0.906349517 |
| chr13:60329694-60329911 | 0.141820748 | 0.763895887 | 0.906349517 |
| chr9:94867242-94867489 | 0.153927271 | 0.763906355 | 0.906349517 |
| chr15:33132721-33133023 | 0.119282954 | 0.763922131 | 0.906349517 |
| chr2:56861123-56861514 | -0.134982765 | 0.763929875 | 0.906349517 |
| chr11:33362465-33362743 | 0.141468076 | 0.763930262 | 0.906349517 |
| chr15:40847384-40847683 | 0.111554622 | 0.763975832 | 0.906377074 |
| chr1:145926800-145927834 | 0.095106752 | 0.763977147 | 0.906377074 |
| chr14:61188158-61188407 | -0.154018862 | 0.764042548 | 0.906440631 |
| chr6:137571957-137572157 | -0.164843401 | 0.764060318 | 0.906441636 |
| chr8:138767664-138768144 | 0.107097368 | 0.764070628 | 0.906441636 |
| chr8:11284011-11285290 | -0.097690953 | 0.764086792 | 0.906441636 |
| chr19:12163194-12163394 | -0.156133882 | 0.764090716 | 0.906441636 |
| chr17:78422050-78422350 | 0.108985218 | 0.76410256 | 0.906441653 |
| chr14:71753150-71753449 | -0.118184721 | 0.764143304 | 0.906475952 |
| chr1:182616111-182616518 | 0.11185895 | 0.764156865 | 0.906478005 |
| chr2:120066929-120067564 | 0.107287857 | 0.764192959 | 0.906506787 |
| chr17:57451955-57452300 | 0.111916453 | 0.764215356 | 0.906519321 |
| chr10:62374247-62374482 | -0.129961378 | 0.764231478 | 0.906521321 |
| chr6:90940908-90941144 | 0.134751377 | 0.764240705 | 0.906521321 |
| chr8:111086815-111087041 | 0.1437041 | 0.764300868 | 0.90657865 |
| chr2:146405467-146405882 | 0.123327315 | 0.764324313 | 0.906589419 |

| | | | |
|---|---|---|---|
| chr1:179306183-179306440 | 0.155673459 | 0.764377759 | 0.906589419 |
| chr17:45705430-45705819 | 0.118476439 | 0.76437904 | 0.906589419 |
| chr2:127438223-127438464 | 0.152480589 | 0.76439108 | 0.906589419 |
| chr2:132448807-132449142 | 0.131023828 | 0.764392989 | 0.906589419 |
| chr5:180556573-180556773 | -0.16137554 | 0.764399738 | 0.906589419 |
| chr12:69900617-69901006 | 0.123104796 | 0.76440423 | 0.906589419 |
| chr3:50436122-50436443 | -0.123748143 | 0.764404875 | 0.906589419 |
| chr13:29849745-29850959 | -0.095195419 | 0.764432509 | 0.906589419 |
| chr11:34438468-34439497 | 0.096882756 | 0.764448022 | 0.906589419 |
| chr6:87832298-87832546 | -0.151780874 | 0.764465528 | 0.906589419 |
| chr20:63980675-63981461 | -0.097888794 | 0.764474397 | 0.906589419 |
| chr12:65699275-65699520 | 0.14090159 | 0.764486248 | 0.906589419 |
| chrX:76148421-76148621 | 0.155670903 | 0.764487271 | 0.906589419 |
| chr2:87483137-87483427 | 0.137029062 | 0.764487425 | 0.906589419 |
| chr5:141747140-141747393 | -0.132895407 | 0.764510644 | 0.906602922 |
| chr12:22443415-22443749 | -0.112083315 | 0.764528465 | 0.906609293 |
| chr3:167209082-167209372 | 0.117283378 | 0.764541158 | 0.906609293 |
| chr17:80970470-80970698 | 0.129463883 | 0.764558818 | 0.906609293 |
| chr12:632107-632674 | 0.114310011 | 0.764563346 | 0.906609293 |
| chr3:130003503-130003709 | 0.146263077 | 0.764673449 | 0.90669683 |
| chr1:54229408-54229822 | -0.127617142 | 0.764676014 | 0.90669683 |
| chr19:1030102-1030408 | -0.132340774 | 0.7646777 | 0.90669683 |
| chr1:149937327-149937748 | 0.100538456 | 0.764690881 | 0.90669683 |
| chr5:177496781-177497945 | 0.095052382 | 0.764696334 | 0.90669683 |
| chr9:137200667-137201258 | -0.0992407 | 0.764722067 | 0.906700577 |
| chr17:43483541-43484250 | -0.099303409 | 0.76472734 | 0.906700577 |
| chr8:76924938-76925535 | -0.114513041 | 0.764734995 | 0.906700577 |
| chr17:50893319-50893579 | -0.137054234 | 0.764759002 | 0.90671501 |
| chr3:150371839-150372141 | -0.116386935 | 0.764774117 | 0.906718901 |
| chr2:70420673-70420958 | -0.142175156 | 0.764847073 | 0.90678053 |
| chr12:121686797-121687388 | 0.097447642 | 0.764849767 | 0.90678053 |
| chr1:67645602-67645821 | -0.137431082 | 0.764881727 | 0.906801987 |
| chr17:66302045-66303724 | 0.095375104 | 0.764891535 | 0.906801987 |
| chr8:102272542-102272790 | 0.132237479 | 0.764954995 | 0.90682809 |
| chr3:164208947-164209285 | -0.133148525 | 0.764975743 | 0.90682809 |
| chr6:108838156-108838363 | 0.150394279 | 0.764988585 | 0.90682809 |
| chr16:88662883-88663857 | 0.095495703 | 0.764998343 | 0.90682809 |
| chr5:14156488-14156925 | -0.112392954 | 0.764998704 | 0.90682809 |
| chrX:46912101-46912646 | 0.106674564 | 0.765011039 | 0.90682809 |
| chr1:26960119-26960836 | 0.105521511 | 0.765021676 | 0.90682809 |
| chr3:98434428-98434628 | 0.152216075 | 0.765025476 | 0.90682809 |
| chr16:80932109-80932668 | -0.110607801 | 0.765040775 | 0.90682809 |
| chr13:46905335-46905604 | -0.117503354 | 0.765042449 | 0.90682809 |
| chr20:50186302-50186539 | 0.109491811 | 0.765047837 | 0.90682809 |
| chr11:128956985-128957258 | -0.139568692 | 0.765055574 | 0.90682809 |
| chr11:119361015-119361792 | 0.097210949 | 0.765072339 | 0.906833933 |
| chr20:55266550-55266814 | 0.142500978 | 0.765111308 | 0.906859769 |
| chr1:225474923-225475515 | 0.107194156 | 0.765125367 | 0.906859769 |
| chr9:129465835-129466035 | 0.15766625 | 0.765129643 | 0.906859769 |
| chr19:1039761-1040217 | 0.103120964 | 0.765181718 | 0.906907462 |

| | | | |
|---|---|---|---|
| chr10:4512896-4513182 | -0.128355552 | 0.765199182 | 0.906914132 |
| chr16:21300500-21301049 | 0.099401164 | 0.765220808 | 0.906920685 |
| chr2:215311729-215312885 | -0.095296528 | 0.765229757 | 0.906920685 |
| chr1:65999926-66000443 | -0.124395286 | 0.765251035 | 0.906920685 |
| chr7:102286706-102286906 | -0.127934093 | 0.765252056 | 0.906920685 |
| chr8:127745259-127745549 | -0.113370345 | 0.765268547 | 0.906926201 |
| chr15:101440934-101441300 | -0.116867862 | 0.76530548 | 0.906946472 |
| chr4:56861535-56861769 | 0.154039225 | 0.765309325 | 0.906946472 |
| chr16:85182802-85183032 | 0.12915044 | 0.765327119 | 0.906953532 |
| chr7:2440605-2440816 | 0.125007075 | 0.765358824 | 0.906977076 |
| chr6:88952607-88952836 | 0.140543819 | 0.765383251 | 0.90697788 |
| chr2:54953232-54953572 | -0.121846622 | 0.765389272 | 0.90697788 |
| chr4:71004274-71004631 | 0.146075139 | 0.765395013 | 0.90697788 |
| chr19:23249834-23250648 | -0.109015959 | 0.765476857 | 0.907051904 |
| chr10:116138955-116139247 | -0.122648053 | 0.765492818 | 0.907051904 |
| chr1:52365701-52366681 | -0.094807874 | 0.765492995 | 0.907051904 |
| chr4:111583441-111583698 | -0.160433022 | 0.765506388 | 0.907053746 |
| chr3:171346960-171347187 | 0.166234498 | 0.765583422 | 0.907111643 |
| chr10:8471578-8471840 | 0.144353247 | 0.765583508 | 0.907111643 |
| chr1:197130756-197131228 | -0.114530705 | 0.765590767 | 0.907111643 |
| chr1:26771768-26771982 | -0.151890311 | 0.765646911 | 0.907159262 |
| chr15:75683759-75683960 | 0.131887477 | 0.765665186 | 0.907159262 |
| chr17:63842361-63843352 | -0.093599539 | 0.765666475 | 0.907159262 |
| chr2:240834660-240834860 | -0.163214008 | 0.7656833 | 0.90716517 |
| chr6:132919548-132919948 | 0.120444706 | 0.76572098 | 0.907195784 |
| chr9:99211790-99212072 | 0.13673666 | 0.76578862 | 0.907261892 |
| chr16:71626896-71627197 | -0.107964353 | 0.765825177 | 0.907284911 |
| chr8:122942731-122942931 | -0.128301703 | 0.765831731 | 0.907284911 |
| chr3:143847793-143847993 | 0.14262261 | 0.765853505 | 0.907289017 |
| chr2:9422344-9424023 | -0.093603167 | 0.765863773 | 0.907289017 |
| chr7:6284662-6285215 | -0.110946804 | 0.76587072 | 0.907289017 |
| chr1:150066826-150068016 | 0.094792505 | 0.765902645 | 0.907301658 |
| chr12:121146538-121146798 | -0.146376191 | 0.765905073 | 0.907301658 |
| chr8:60795794-60796017 | -0.149221678 | 0.765941444 | 0.90731768 |
| chr14:87934285-87934485 | 0.147953257 | 0.765942282 | 0.90731768 |
| chr7:66591895-66592782 | 0.099727314 | 0.765964959 | 0.907330516 |
| chr15:82040924-82041166 | -0.131256206 | 0.765979836 | 0.907334111 |
| chr8:125230740-125230989 | 0.117890039 | 0.766014126 | 0.907360702 |
| chrX:104156495-104157415 | 0.096170188 | 0.766071669 | 0.907414835 |
| chr8:80485260-80486957 | 0.093962547 | 0.766140967 | 0.907462541 |
| chr14:99812549-99812779 | -0.155456856 | 0.766146236 | 0.907462541 |
| chr3:107884976-107885411 | -0.118920472 | 0.766147474 | 0.907462541 |
| chr13:27242570-27242777 | 0.156733115 | 0.766172137 | 0.907470599 |
| chrX:78092220-78092618 | -0.11931813 | 0.766177964 | 0.907470599 |
| chr15:68277331-68278438 | -0.09587779 | 0.766220553 | 0.90748425 |
| chr12:15788712-15790206 | -0.094589582 | 0.766231832 | 0.90748425 |
| chr5:139177978-139178178 | 0.1528861 | 0.766237903 | 0.90748425 |
| chr1:67684855-67686142 | 0.093935093 | 0.766239787 | 0.90748425 |
| chr11:74311333-74311837 | 0.105908088 | 0.766255168 | 0.90748425 |
| chr10:75262760-75263052 | -0.125161235 | 0.766270366 | 0.90748425 |

| | | | |
|---|---|---|---|
| chr2:130797433-130798352 | 0.097385512 | 0.76627852 | 0.90748425 |
| chr2:27095671-27095974 | 0.118892284 | 0.76628424 | 0.90748425 |
| chr9:95603033-95603266 | -0.154197113 | 0.766311015 | 0.907491517 |
| chr15:89121504-89121733 | -0.137586787 | 0.766314063 | 0.907491517 |
| chr3:144530874-144531222 | -0.124050285 | 0.766331018 | 0.90749757 |
| chr3:133572956-133574073 | -0.093095065 | 0.766368003 | 0.907514178 |
| chr8:38286790-38286990 | -0.164634783 | 0.766368731 | 0.907514178 |
| chr8:61095404-61095647 | 0.133811387 | 0.766411723 | 0.907549313 |
| chr1:241985927-241986235 | 0.123499464 | 0.76642209 | 0.907549313 |
| chr2:126776421-126777274 | 0.098838474 | 0.766441545 | 0.907554333 |
| chr2:96407368-96407619 | -0.122645158 | 0.766450019 | 0.907554333 |
| chr1:202184610-202185058 | -0.098147878 | 0.766475341 | 0.907557328 |
| chr2:170873050-170873773 | -0.104423059 | 0.766476237 | 0.907557328 |
| chr14:62163019-62163303 | -0.123373272 | 0.766535628 | 0.907608331 |
| chr17:57917034-57917326 | 0.114717855 | 0.766554188 | 0.907608331 |
| chr2:70141092-70141642 | 0.095802847 | 0.766554848 | 0.907608331 |
| chr1:171142504-171142754 | 0.143008567 | 0.766632342 | 0.907686059 |
| chr16:89842698-89843285 | -0.104104606 | 0.766670141 | 0.90769297 |
| chr16:25055649-25055942 | 0.123648653 | 0.766671532 | 0.90769297 |
| chr1:64469841-64470653 | -0.100543218 | 0.766673815 | 0.90769297 |
| chr14:23944422-23944672 | 0.131398308 | 0.766690519 | 0.90769297 |
| chr20:35411873-35412363 | -0.104811185 | 0.766697411 | 0.90769297 |
| chr17:60720654-60721312 | 0.09519143 | 0.766713574 | 0.90769808 |
| chr17:9824770-9824985 | -0.15310396 | 0.766744462 | 0.907720623 |
| chr17:8630616-8631579 | -0.095623948 | 0.766777108 | 0.907731592 |
| chr10:100371160-10037151९ | 0.110568753 | 0.7667846 | 0.907731592 |
| chr3:170999836-171000116 | 0.12780711 | 0.766794933 | 0.907731592 |
| chr12:66169159-66170164 | -0.094221172 | 0.766807976 | 0.907731592 |
| chr17:58577620-58578314 | -0.095562054 | 0.766812961 | 0.907731592 |
| chr3:169619775-169619975 | -0.146106491 | 0.766860477 | 0.907773815 |
| chr11:24591979-24592291 | -0.129128848 | 0.76687485 | 0.907776804 |
| chr17:48107153-48108367 | -0.094368801 | 0.766902591 | 0.907786422 |
| chr5:180789990-180790319 | 0.113986915 | 0.766906671 | 0.907786422 |
| chr5:16744388-16744824 | -0.115006516 | 0.766926389 | 0.907789268 |
| chr19:17215200-17215855 | -0.103854785 | 0.766932769 | 0.907789268 |
| chr3:172076258-172076458 | -0.158747579 | 0.766999281 | 0.907842507 |
| chr3:72336815-72337297 | -0.116056457 | 0.767004338 | 0.907842507 |
| chr1:47527759-47528029 | -0.129391599 | 0.767013293 | 0.907842507 |
| chr11:57480080-57480658 | -0.106301163 | 0.767037015 | 0.907844032 |
| chr7:156504903-156505103 | -0.164974971 | 0.767038278 | 0.907844032 |
| chr10:110304398-11030550٤ | -0.094623397 | 0.767055893 | 0.907850857 |
| chr3:157499339-157500002 | 0.0971526 | 0.767074076 | 0.907854197 |
| chr21:14027130-14027774 | 0.102130068 | 0.767082412 | 0.907854197 |
| chr7:152463369-152464723 | -0.092780137 | 0.767104577 | 0.907866406 |
| chr8:104389979-104390201 | -0.129843659 | 0.767140915 | 0.907890727 |
| chr14:69357008-69358197 | 0.095390568 | 0.767148824 | 0.907890727 |
| chrX:70569924-70570163 | -0.163069631 | 0.767199767 | 0.907898733 |
| chr1:148522130-148523317 | 0.093478374 | 0.767200412 | 0.907898733 |
| chr12:109805128-10980558९ | -0.115907463 | 0.767202236 | 0.907898733 |
| chr19:33771630-33771850 | -0.135063522 | 0.767212552 | 0.907898733 |

| | | | |
|---|---|---|---|
| chr12:126096083-126096387 | 0.153783043 | 0.767214835 | 0.907898733 |
| chr15:69380455-69380658 | -0.142692644 | 0.767246507 | 0.907922191 |
| chr3:106360306-106360580 | -0.146358402 | 0.767259918 | 0.907924039 |
| chr18:54829704-54830199 | -0.115522418 | 0.767306738 | 0.907947723 |
| chr14:76267617-76268277 | -0.104842537 | 0.767320836 | 0.907947723 |
| chr6:28735308-28735632 | 0.11132895 | 0.767330516 | 0.907947723 |
| chr13:50403256-50403749 | 0.109731171 | 0.767357246 | 0.907947723 |
| chr10:23265068-23265277 | -0.159643493 | 0.767365614 | 0.907947723 |
| chr13:103676377-103676579 | -0.159643493 | 0.767365614 | 0.907947723 |
| chr20:16572947-16573863 | 0.096814596 | 0.767381017 | 0.907947723 |
| chr20:31869749-31870327 | 0.105064389 | 0.76738416 | 0.907947723 |
| chr3:84943425-84943630 | 0.152846841 | 0.76738658 | 0.907947723 |
| chr7:98797065-98797626 | -0.100652052 | 0.767404218 | 0.907954571 |
| chr11:113314312-113315022 | 0.098517169 | 0.767421901 | 0.907961473 |
| chr10:95296876-95297139 | -0.159194489 | 0.767533291 | 0.90807924 |
| chr1:196650369-196650569 | -0.154685613 | 0.767602108 | 0.908133805 |
| chr6:75893320-75893679 | -0.115696367 | 0.767603115 | 0.908133805 |
| chr14:83958463-83958676 | -0.1489476 | 0.767629302 | 0.908144838 |
| chr11:9565943-9566294 | 0.113916341 | 0.767636145 | 0.908144838 |
| chr16:75096716-75097087 | -0.10018608 | 0.767653265 | 0.908145639 |
| chr10:11345787-11346193 | -0.118109248 | 0.767660527 | 0.908145639 |
| chr3:62453614-62453950 | 0.108257666 | 0.767788282 | 0.908237243 |
| chr8:127552890-127553240 | -0.118628389 | 0.767792863 | 0.908237243 |
| chr6:130148229-130148438 | 0.157811858 | 0.767805676 | 0.908237243 |
| chr20:49763451-49763709 | -0.125903788 | 0.767810991 | 0.908237243 |
| chrX:112832642-112833117 | -0.108044803 | 0.767815163 | 0.908237243 |
| chr17:16653166-16654062 | -0.096576981 | 0.767827273 | 0.908237243 |
| chr6:144285804-144286526 | -0.10417288 | 0.76783988 | 0.908237243 |
| chr11:70827443-70827643 | -0.165006062 | 0.767848935 | 0.908237243 |
| chr1:243254363-243254688 | 0.106282897 | 0.767850714 | 0.908237243 |
| chr10:88835439-88835922 | 0.109218755 | 0.767856495 | 0.908237243 |
| chr8:61724780-61725087 | -0.120669472 | 0.767916555 | 0.908294262 |
| chr7:51811815-51812158 | -0.13872708 | 0.767947677 | 0.908310567 |
| chr14:103057103-103058271 | 0.095461622 | 0.767959275 | 0.908310567 |
| chr11:66983987-66984284 | -0.119972684 | 0.767965903 | 0.908310567 |
| chr6:149746435-149746693 | -0.111579993 | 0.76798812 | 0.908322822 |
| chr1:109425914-109427013 | -0.093210491 | 0.768011854 | 0.908336873 |
| chr11:10735543-10736113 | -0.102480568 | 0.768053759 | 0.908372412 |
| chr9:94553081-94553452 | 0.106030598 | 0.76808894 | 0.908391756 |
| chr5:149434428-149434760 | -0.112442854 | 0.768112393 | 0.908391756 |
| chr1:145765676-145765879 | 0.138216033 | 0.768115022 | 0.908391756 |
| chr5:13866863-13867139 | -0.134023898 | 0.768117536 | 0.908391756 |
| chr17:55526395-55526708 | -0.108190603 | 0.76813629 | 0.908399914 |
| chr16:12289051-12289580 | -0.107391487 | 0.768244664 | 0.908514055 |
| chr12:124988818-124989382 | -0.099169916 | 0.768303159 | 0.908569207 |
| chr20:54096487-54097451 | 0.092418184 | 0.768341089 | 0.908600039 |
| chr3:63539738-63539988 | 0.121267615 | 0.768359665 | 0.908607983 |
| chr6:73461335-73462444 | -0.096755847 | 0.768421482 | 0.908641401 |
| chr12:104609535-104609847 | -0.120844861 | 0.768441105 | 0.908641401 |
| chr4:82017836-82018206 | -0.161569586 | 0.768445206 | 0.908641401 |

| | | | |
|---|---|---|---|
| chr3:123115932-123116132 | -0.167747672 | 0.768455682 | 0.908641401 |
| chrX:131493172-131493372 | -0.164495025 | 0.768470458 | 0.908641401 |
| chr14:25406833-25407073 | 0.132568113 | 0.768471487 | 0.908641401 |
| chr3:16319374-16319668 | 0.12109468 | 0.768477805 | 0.908641401 |
| chr3:127629524-127630028 | -0.10850371 | 0.768482795 | 0.908641401 |
| chr1:39818261-39818461 | 0.154326395 | 0.768536652 | 0.908677036 |
| chr10:89223994-89224194 | 0.154326395 | 0.768536652 | 0.908677036 |
| chr14:61332062-61332356 | -0.13521522 | 0.768578082 | 0.908711998 |
| chr3:146422397-146422745 | -0.109990099 | 0.768635716 | 0.908762604 |
| chr14:20332721-20333519 | -0.095651388 | 0.768644604 | 0.908762604 |
| chr11:6637975-6638241 | -0.136224551 | 0.768660506 | 0.908767382 |
| chr10:122268092-122268374 | 0.122488598 | 0.768710301 | 0.908809386 |
| chr9:40951909-40952118 | -0.148235473 | 0.768726231 | 0.908809386 |
| chr1:53945378-53946655 | -0.092372348 | 0.768733609 | 0.908809386 |
| chr8:97375683-97375992 | 0.112968681 | 0.768743478 | 0.908809386 |
| chr10:103192467-103192876 | 0.114074905 | 0.76879902 | 0.908861026 |
| chr22:23168364-23168660 | 0.137821844 | 0.768837357 | 0.908892324 |
| chr3:18424789-18425189 | 0.112327231 | 0.768913506 | 0.908968321 |
| chr11:72080253-72081194 | 0.093593484 | 0.768960293 | 0.908997668 |
| chr20:54207471-54208890 | -0.091142553 | 0.768962058 | 0.908997668 |
| chr4:9153773-9153973 | 0.133166405 | 0.768974804 | 0.90899871 |
| chr9:133726404-133726650 | 0.137440402 | 0.769018308 | 0.909036112 |
| chr2:68943567-68944188 | 0.10983043 | 0.769045696 | 0.909054292 |
| chr6:96520259-96520482 | 0.119847059 | 0.769057416 | 0.909054292 |
| chr9:137223091-137223985 | 0.094821532 | 0.769074929 | 0.909060969 |
| chr13:45715452-45715652 | -0.141475723 | 0.76909643 | 0.90907236 |
| chr1:21607719-21608575 | 0.096924379 | 0.769140979 | 0.909104056 |
| chr2:27032348-27032548 | 0.141240855 | 0.769156037 | 0.909104056 |
| chr17:37489247-37490407 | -0.09665117 | 0.76915884 | 0.909104056 |
| chr9:132411185-132411472 | -0.115546127 | 0.769171318 | 0.909104781 |
| chr3:4688161-4688799 | 0.096676239 | 0.769203895 | 0.909125106 |
| chr9:37800430-37801768 | 0.095911135 | 0.769212245 | 0.909125106 |
| chr1:93346535-93347022 | -0.114107546 | 0.769225247 | 0.90912645 |
| chr18:54679607-54679863 | 0.144173936 | 0.769245156 | 0.909135956 |
| chr2:160694694-160695040 | -0.152588204 | 0.769321379 | 0.909212017 |
| chr21:42761255-42761535 | -0.129211606 | 0.769390615 | 0.909278443 |
| chr3:68932261-68932925 | 0.100929422 | 0.769401319 | 0.909278443 |
| chr22:37676859-37677059 | -0.131306924 | 0.769428925 | 0.909289198 |
| chr12:45130854-45131162 | 0.125590637 | 0.769439844 | 0.909289198 |
| chr15:64825163-64826262 | 0.094542696 | 0.769446021 | 0.909289198 |
| chr6:80003784-80005014 | -0.092725619 | 0.769466161 | 0.909298975 |
| chr15:48810392-48811510 | -0.092595035 | 0.7695076 | 0.909328383 |
| chr17:77834524-77834868 | -0.114700478 | 0.769522047 | 0.909328383 |
| chr7:36079193-36079454 | 0.135038391 | 0.769531672 | 0.909328383 |
| chr13:102580581-102580812 | -0.144270515 | 0.769543711 | 0.909328383 |
| chr9:4339442-4339642 | 0.142442079 | 0.769550386 | 0.909328383 |
| chr2:231677706-231677906 | 0.140469396 | 0.769595085 | 0.909367177 |
| chr3:108879926-108880269 | -0.114183254 | 0.769646074 | 0.909380749 |
| chr6:157198381-157198947 | -0.10158019 | 0.769654954 | 0.909380749 |
| chr14:69657893-69658359 | 0.105918183 | 0.769655046 | 0.909380749 |

| | | | |
|---|---|---|---|
| chr11:71241339-71241701 | -0.110413643 | 0.769664537 | 0.909380749 |
| chr7:6516345-6516764 | -0.101768287 | 0.769665913 | 0.909380749 |
| chr3:126475402-126476780 | 0.093235624 | 0.769695166 | 0.909397978 |
| chrX:110412974-110413460 | 0.108210785 | 0.769710371 | 0.909397978 |
| chr20:3807345-3807831 | 0.098237798 | 0.769722311 | 0.909397978 |
| chr6:117795248-117795592 | 0.119710624 | 0.769727969 | 0.909397978 |
| chr3:50088530-50089074 | -0.107894714 | 0.769752696 | 0.909399735 |
| chr2:31922556-31922845 | -0.141042013 | 0.769759722 | 0.909399735 |
| chr1:156616479-156616895 | 0.108302173 | 0.769771823 | 0.909399735 |
| chr8:89690335-89691042 | 0.101193722 | 0.769776971 | 0.909399735 |
| chr11:76444393-76446074 | 0.09203776 | 0.7697888 | 0.909399735 |
| chr7:32464218-32465159 | -0.099311873 | 0.769831357 | 0.909421666 |
| chr3:97821495-97822781 | 0.095586653 | 0.769842472 | 0.909421666 |
| chr8:143738753-143739830 | -0.093075262 | 0.769848823 | 0.909421666 |
| chr20:34112810-34113010 | 0.132742049 | 0.769860399 | 0.909421666 |
| chr11:71746405-71746630 | 0.14811008 | 0.769888952 | 0.909421666 |
| chr12:75289843-75290093 | 0.14811008 | 0.769888952 | 0.909421666 |
| chr20:53915167-53916493 | -0.090873212 | 0.769900946 | 0.909421666 |
| chr1:161073816-161074018 | 0.129115771 | 0.769914152 | 0.909421666 |
| chr16:56356437-56356767 | -0.115478908 | 0.769914184 | 0.909421666 |
| chr17:75589054-75589750 | 0.094961372 | 0.769960799 | 0.909462707 |
| chr10:11757932-11758245 | 0.109354666 | 0.769986125 | 0.909477706 |
| chr5:1167775-1168637 | 0.098314729 | 0.769997237 | 0.909477706 |
| chr10:14554099-14554299 | -0.150020547 | 0.770074474 | 0.909515838 |
| chr12:107318646-107319208 | -0.104772053 | 0.770080818 | 0.909515838 |
| chr1:16499149-16499415 | 0.128144665 | 0.770083891 | 0.909515838 |
| chr7:5344504-5345010 | -0.111553491 | 0.770087586 | 0.909515838 |
| chr6:149454458-149454682 | -0.131375894 | 0.770092187 | 0.909515838 |
| chr16:24517031-24517339 | 0.124016896 | 0.770102817 | 0.909515838 |
| chr11:121466339-121466885 | 0.099914807 | 0.770112611 | 0.909515838 |
| chr5:134649181-134649693 | -0.109192786 | 0.77016829 | 0.909567575 |
| chr19:16471845-16472401 | 0.102895394 | 0.770192803 | 0.909582505 |
| chr8:125432021-125432282 | 0.098557782 | 0.770288737 | 0.909649418 |
| chr16:89910356-89910587 | -0.121063066 | 0.770294436 | 0.909649418 |
| chr9:130451531-130451978 | 0.107936364 | 0.770298803 | 0.909649418 |
| chr12:34363100-34363324 | -0.127945542 | 0.770322163 | 0.909649418 |
| chr1:180533435-180533864 | -0.111315423 | 0.770326033 | 0.909649418 |
| chr17:55212934-55213134 | 0.156293206 | 0.770330273 | 0.909649418 |
| chr11:35618084-35618458 | 0.105724454 | 0.77035229 | 0.909649418 |
| chr13:46551563-46551845 | -0.101159882 | 0.770357859 | 0.909649418 |
| chr1:25523683-25523883 | 0.143557632 | 0.770358787 | 0.909649418 |
| chr1:169926103-169926322 | 0.139283791 | 0.770369717 | 0.909649418 |
| chr7:6589740-6591009 | -0.093180767 | 0.770405778 | 0.909649418 |
| chr14:76797585-76797785 | 0.158758212 | 0.770408969 | 0.909649418 |
| chr20:40664161-40664560 | 0.118592078 | 0.770409228 | 0.909649418 |
| chr8:37736930-37737332 | 0.11274587 | 0.770442417 | 0.909649418 |
| chr11:79117596-79117874 | -0.120040275 | 0.770458212 | 0.909649418 |
| chr3:161424048-161424248 | -0.125039846 | 0.770459848 | 0.909649418 |
| chr19:58572679-58573837 | 0.091721663 | 0.770463618 | 0.909649418 |
| chr20:58461899-58462140 | 0.124267586 | 0.770499354 | 0.909649418 |

| | | | |
|---|---|---|---|
| chr3:32901964-32902283 | -0.145148218 | 0.770513426 | 0.909649418 |
| chr1:151155978-151156346 | -0.10249969 | 0.770520424 | 0.909649418 |
| chr1:225911178-225911595 | -0.11759733 | 0.77052112 | 0.909649418 |
| chr1:148263080-148264212 | 0.091843463 | 0.770561813 | 0.909649418 |
| chr3:82643821-82644282 | -0.112196867 | 0.770566203 | 0.909649418 |
| chr11:82791732-82792065 | -0.119645443 | 0.77057607 | 0.909649418 |
| chr1:10761904-10762104 | 0.144803652 | 0.770595488 | 0.909649418 |
| chr17:42578495-42578745 | 0.119907392 | 0.770607763 | 0.909649418 |
| chr10:116654843-116655303 | 0.11025384 | 0.770620236 | 0.909649418 |
| chr3:151085150-151087291 | -0.091289356 | 0.770620487 | 0.909649418 |
| chr2:205563774-205563979 | -0.160051152 | 0.770634596 | 0.909649418 |
| chr6:137076496-137077020 | -0.106853973 | 0.770640767 | 0.909649418 |
| chr19:29610629-29610984 | 0.108125132 | 0.770665542 | 0.909649418 |
| chr3:121574425-121574805 | 0.106390504 | 0.770666021 | 0.909649418 |
| chr8:128403409-128403609 | 0.139182409 | 0.770668641 | 0.909649418 |
| chr8:111934536-111934736 | -0.143590846 | 0.770693183 | 0.909649418 |
| chr16:69530715-69530960 | 0.141619881 | 0.770698783 | 0.909649418 |
| chr17:72126588-72126907 | -0.114765296 | 0.770702121 | 0.909649418 |
| chr1:174603835-174604139 | -0.139075058 | 0.77071215 | 0.909649418 |
| chr1:40449783-40451105 | -0.095601248 | 0.770731401 | 0.909649418 |
| chr19:50804352-50805359 | -0.094846399 | 0.770733993 | 0.909649418 |
| chr3:119973317-119973692 | 0.114799015 | 0.770742942 | 0.909649418 |
| chr22:43896179-43896518 | -0.132822609 | 0.770744348 | 0.909649418 |
| chrX:47361702-47362462 | 0.095286308 | 0.770758113 | 0.909649418 |
| chr6:85642725-85644313 | -0.090924081 | 0.770759954 | 0.909649418 |
| chr4:178297049-178297359 | -0.133703598 | 0.770823155 | 0.909709996 |
| chr20:53805306-53806514 | 0.090540808 | 0.77083994 | 0.909715793 |
| chr15:90836451-90836776 | 0.132116666 | 0.770871017 | 0.909722942 |
| chr9:99807353-99807587 | 0.133686902 | 0.77088078 | 0.909722942 |
| chr20:62375011-62375217 | 0.118253539 | 0.770890164 | 0.909722942 |
| chr5:138491833-138492397 | 0.100454736 | 0.770893489 | 0.909722942 |
| chr22:49776674-49776874 | -0.164089711 | 0.770918721 | 0.90972439 |
| chr13:28563884-28564535 | -0.098092938 | 0.770922481 | 0.90972439 |
| chr22:19780364-19780693 | 0.117162744 | 0.770930335 | 0.90972439 |
| chr13:113153512-113153761 | -0.128724422 | 0.770954773 | 0.909734629 |
| chr11:107689655-107689877 | 0.1472851 | 0.770962757 | 0.909734629 |
| chr6:141326498-141326802 | 0.140150007 | 0.771010335 | 0.909776162 |
| chr11:36447813-36448516 | 0.100798454 | 0.771037768 | 0.909776162 |
| chr6:26520469-26521479 | 0.094255529 | 0.771043014 | 0.909776162 |
| chr17:68868628-68868944 | 0.131632034 | 0.771045449 | 0.909776162 |
| chr2:86194880-86195991 | 0.092925356 | 0.771068317 | 0.909789134 |
| chr11:6473394-6474463 | 0.093782865 | 0.771083438 | 0.909792965 |
| chr2:158967580-158968799 | -0.094128971 | 0.771098951 | 0.909797259 |
| chr1:247884981-247885318 | -0.130721868 | 0.771117954 | 0.90980567 |
| chr22:37063464-37063729 | -0.138863758 | 0.77114107 | 0.909817423 |
| chr1:109740643-109741115 | 0.102735479 | 0.771152297 | 0.909817423 |
| chr4:151014564-151016401 | -0.090928637 | 0.771168719 | 0.909817423 |
| chr8:88437409-88437913 | -0.111768947 | 0.771175411 | 0.909817423 |
| chr20:61100550-61100913 | 0.127714032 | 0.77122646 | 0.90986364 |
| chr12:58919863-58920780 | 0.095087366 | 0.771301921 | 0.909897688 |

| | | | |
|---|---|---|---|
| chrX:41293362-41293594 | -0.143302497 | 0.771307147 | 0.909897688 |
| chr1:147389772-147390314 | 0.101167998 | 0.771307618 | 0.909897688 |
| chr1:11736774-11736974 | -0.147929983 | 0.77131221 | 0.909897688 |
| chr3:61944849-61945167 | -0.107200274 | 0.771324753 | 0.909897688 |
| chr14:36942293-36942499 | 0.136166452 | 0.771326571 | 0.909897688 |
| chr8:34535765-34535965 | 0.155505354 | 0.771367909 | 0.909932443 |
| chr3:151168349-151168800 | -0.111648451 | 0.771397048 | 0.909952807 |
| chr9:37415220-37415557 | 0.109708951 | 0.771469988 | 0.910024839 |
| chr14:31229625-31230123 | 0.099398068 | 0.771525352 | 0.910076135 |
| chr2:86372895-86373169 | -0.132885287 | 0.771576635 | 0.910101219 |
| chrX:38326853-38327902 | -0.094504018 | 0.771584941 | 0.910101219 |
| chr11:130146295-130146832 | 0.102709376 | 0.771586929 | 0.910101219 |
| chr16:50374723-50375209 | 0.111834841 | 0.771622327 | 0.910101219 |
| chr9:78601986-78602271 | 0.138441687 | 0.77162313 | 0.910101219 |
| chr6:70604557-70604758 | -0.159044988 | 0.771636175 | 0.910101219 |
| chr1:181166480-181167152 | 0.097743488 | 0.771650824 | 0.910101219 |
| chr7:33062083-33063569 | 0.090737952 | 0.771655114 | 0.910101219 |
| chr12:90305502-90305752 | -0.138623873 | 0.771655506 | 0.910101219 |
| chr1:172041228-172041581 | -0.123768747 | 0.771665395 | 0.910101219 |
| chr17:49996625-49996900 | 0.113561787 | 0.771683557 | 0.910104314 |
| chr4:110198436-110198817 | 0.101194301 | 0.771691775 | 0.910104314 |
| chrX:103211045-103211342 | -0.126079267 | 0.771703914 | 0.910104622 |
| chr19:2599060-2599423 | -0.12673175 | 0.771751042 | 0.910146193 |
| chr5:83494417-83494617 | 0.144147116 | 0.771800913 | 0.910190998 |
| chr4:146636916-146637238 | -0.119762888 | 0.771844577 | 0.910228482 |
| chr8:37090933-37091170 | -0.142903865 | 0.77189816 | 0.910267438 |
| chr1:211577798-211578147 | -0.104690937 | 0.771912933 | 0.910267438 |
| chr17:65628047-65628711 | 0.097435558 | 0.77191325 | 0.910267438 |
| chr11:86955245-86956550 | 0.093108457 | 0.771977154 | 0.910328786 |
| chr3:64022075-64023969 | -0.089787398 | 0.772018064 | 0.910363017 |
| chr11:119196949-119197199 | 0.11950963 | 0.772042759 | 0.910378126 |
| chr6:108557886-108558766 | -0.093112064 | 0.772056734 | 0.910380595 |
| chr20:50165743-50166417 | 0.094267624 | 0.772112019 | 0.910418006 |
| chr5:157912686-157913328 | -0.104420679 | 0.772112225 | 0.910418006 |
| chr6:148464661-148465053 | -0.11407664 | 0.772131791 | 0.910423863 |
| chr5:34452052-34452590 | -0.107886587 | 0.772140955 | 0.910423863 |
| chr5:140205454-140205742 | -0.142236608 | 0.772208421 | 0.910484791 |
| chr11:114348321-114348535 | -0.153765587 | 0.772216395 | 0.910484791 |
| chr16:67431480-67431686 | 0.126258484 | 0.772270728 | 0.910529562 |
| chr17:55259565-55259847 | 0.127028068 | 0.772278134 | 0.910529562 |
| chr1:44327454-44328446 | 0.095014162 | 0.772326516 | 0.910558877 |
| chr19:45478422-45479308 | 0.094928272 | 0.772326765 | 0.910558877 |
| chr1:201702967-201703213 | -0.113291115 | 0.772354229 | 0.910570115 |
| chr4:128159032-128159297 | -0.145834664 | 0.772361186 | 0.910570115 |
| chr4:109804240-109804557 | 0.113869143 | 0.772371949 | 0.910570115 |
| chr20:19265715-19265915 | -0.14665918 | 0.772398799 | 0.910586699 |
| chr1:109620159-109621255 | 0.093365409 | 0.772414548 | 0.910586699 |
| chr17:60469784-60470467 | 0.093294554 | 0.772452839 | 0.910586699 |
| chr8:127573399-127573932 | 0.102321462 | 0.772465811 | 0.910586699 |
| chr6:158793461-158793718 | 0.134294438 | 0.77247354 | 0.910586699 |

| | | | |
|---|---|---|---|
| chr4:153750213-153750546 | -0.116340819 | 0.772480152 | 0.910586699 |
| chr1:155941463-155941829 | 0.115096536 | 0.772497466 | 0.910586699 |
| chr2:114442368-114442877 | -0.117572926 | 0.772511002 | 0.910586699 |
| chr2:159147401-159147692 | 0.149142271 | 0.772514143 | 0.910586699 |
| chr2:71995212-71995412 | 0.149142271 | 0.772514143 | 0.910586699 |
| chr9:134412240-134412741 | -0.098198118 | 0.772521628 | 0.910586699 |
| chrX:68828703-68829329 | -0.095874975 | 0.772535944 | 0.910586699 |
| chr10:93554083-93554508 | -0.1185817 | 0.772550227 | 0.910586699 |
| chr9:37453903-37454131 | -0.147772733 | 0.772561425 | 0.910586699 |
| chr10:101380457-101380970 | -0.116104324 | 0.772568468 | 0.910586699 |
| chr12:12837072-12837479 | -0.118814771 | 0.772576163 | 0.910586699 |
| chr5:62779007-62779240 | 0.129828157 | 0.772627208 | 0.910632856 |
| chr8:129682037-129682385 | -0.10566214 | 0.77264199 | 0.91063627 |
| chr12:104877194-104877394 | 0.161668523 | 0.772658474 | 0.91064169 |
| chr4:145840166-145840479 | -0.112520581 | 0.772673465 | 0.910645352 |
| chr1:166757214-166757414 | 0.178970473 | 0.772696722 | 0.910658754 |
| chr19:55338659-55339979 | 0.090788559 | 0.772762825 | 0.910714628 |
| chr6:123803591-123803791 | -0.143156067 | 0.772780134 | 0.910714628 |
| chr19:46731172-46731433 | 0.124761146 | 0.772789773 | 0.910714628 |
| chr5:134246816-134247203 | 0.114962764 | 0.772791674 | 0.910714628 |
| chr5:175687651-175687851 | -0.143641848 | 0.772830981 | 0.91073325 |
| chr3:129627203-129628203 | 0.094094969 | 0.772838823 | 0.91073325 |
| chr9:118595774-118596212 | 0.11562287 | 0.7728455 | 0.91073325 |
| chr2:109326640-109326900 | -0.148989996 | 0.77285502 | 0.91073325 |
| chr3:141769671-141769908 | -0.130749202 | 0.772874957 | 0.910742737 |
| chr17:48944484-48945164 | -0.096894782 | 0.77289517 | 0.910752548 |
| chr13:36526873-36527219 | -0.11533635 | 0.772963205 | 0.910809085 |
| chr1:36650438-36650770 | 0.11739599 | 0.772966923 | 0.910809085 |
| chr4:148214903-148215103 | -0.140564429 | 0.772981357 | 0.910812087 |
| chr14:50859404-50859892 | 0.106195709 | 0.773010812 | 0.910831977 |
| chr6:27591778-27591987 | -0.120636309 | 0.773022012 | 0.910831977 |
| chr17:4709746-4710867 | -0.092559154 | 0.773090582 | 0.910898763 |
| chr15:33067945-33068468 | -0.100121799 | 0.773106383 | 0.910903373 |
| chr15:60397467-60398293 | 0.093747753 | 0.773163059 | 0.910905072 |
| chr12:48119078-48119499 | -0.104622238 | 0.773170348 | 0.910905072 |
| chr11:6389814-6390945 | -0.092893479 | 0.773178628 | 0.910905072 |
| chr19:17074938-17075819 | 0.096313823 | 0.773181497 | 0.910905072 |
| chr3:146496504-146496713 | 0.141032496 | 0.77318679 | 0.910905072 |
| chr12:121873265-121873535 | 0.160938867 | 0.773198903 | 0.910905072 |
| chr19:5711138-5711338 | 0.154550855 | 0.773212031 | 0.910905072 |
| chr4:150668966-150669166 | 0.163595052 | 0.773213098 | 0.910905072 |
| chr16:27428528-27428813 | 0.12259152 | 0.773215399 | 0.910905072 |
| chr6:100116642-100116913 | 0.134403424 | 0.773232541 | 0.910905072 |
| chr8:88203406-88203702 | 0.117461146 | 0.773251967 | 0.910905072 |
| chr15:64334533-64334945 | 0.1086915 | 0.773260072 | 0.910905072 |
| chr10:103451158-103451797 | 0.098335427 | 0.773262373 | 0.910905072 |
| chr1:149811105-149811310 | -0.124030647 | 0.773294237 | 0.910915166 |
| chr16:81180984-81181691 | 0.096512006 | 0.773294718 | 0.910915166 |
| chr14:29906594-29906934 | -0.118914502 | 0.773324335 | 0.910935924 |
| chr5:171543588-171543788 | 0.146781476 | 0.773337658 | 0.910935924 |

| | | | |
|---|---|---|---|
| chr1:191831595-191831923 | -0.112874802 | 0.773348006 | 0.910935924 |
| chr17:40499521-40499789 | 0.123134593 | 0.773368206 | 0.910938162 |
| chr7:46558451-46558663 | -0.150677793 | 0.77337779 | 0.910938162 |
| chr19:41439422-41440187 | 0.095600475 | 0.773400388 | 0.910938162 |
| chr19:1354744-1354999 | 0.131517916 | 0.773404296 | 0.910938162 |
| chr16:80277275-80277521 | -0.134422448 | 0.773417963 | 0.910938162 |
| chr15:92383573-92383882 | -0.120989474 | 0.773423006 | 0.910938162 |
| chrX:70289597-70290750 | -0.094323586 | 0.773433127 | 0.910938162 |
| chr12:121839379-121840290 | 0.092455268 | 0.773461386 | 0.910957442 |
| chr7:41926118-41926385 | -0.137492806 | 0.773493633 | 0.910963362 |
| chr14:93893045-93893356 | 0.120013276 | 0.773494174 | 0.910963362 |
| chr12:123396196-123396438 | -0.155262301 | 0.773502079 | 0.910963362 |
| chr10:8868208-8868662 | 0.109974038 | 0.773537806 | 0.910976342 |
| chr8:97529809-97530285 | -0.108922665 | 0.773539911 | 0.910976342 |
| chr3:128123312-128124194 | -0.093817699 | 0.773548769 | 0.910976342 |
| chr12:57752078-57752858 | 0.091119732 | 0.77356411 | 0.910980408 |
| chr12:56172085-56172314 | -0.134124532 | 0.773582102 | 0.910987594 |
| chr2:238970543-238970798 | -0.134876586 | 0.773610033 | 0.911006485 |
| chr3:149479578-149479894 | -0.120620422 | 0.773625821 | 0.911011076 |
| chr11:73195242-73195637 | -0.10094147 | 0.773648399 | 0.911023661 |
| chr6:112372312-112372560 | -0.145727633 | 0.773738332 | 0.911089017 |
| chr4:118836428-118836825 | -0.098841683 | 0.773739349 | 0.911089017 |
| chr14:57714399-57714856 | 0.118259471 | 0.773739572 | 0.911089017 |
| chr15:81266416-81266740 | 0.113584177 | 0.773760617 | 0.911099797 |
| chr11:69617379-69617579 | -0.149253668 | 0.773807914 | 0.911116874 |
| chr7:98754065-98754275 | 0.14520084 | 0.773814384 | 0.911116874 |
| chr14:67067868-67068096 | -0.142936993 | 0.773817134 | 0.911116874 |
| chr4:1604971-1605220 | 0.124608104 | 0.773834575 | 0.911116874 |
| chr5:7261074-7261359 | 0.124608104 | 0.773834575 | 0.911116874 |
| chr6:15524278-15524486 | -0.1577204 | 0.773850188 | 0.911121256 |
| chr20:62652018-62652875 | 0.094082108 | 0.773908884 | 0.91116213 |
| chr8:17495113-17495426 | 0.114523681 | 0.773911607 | 0.91116213 |
| chr22:23767717-23768135 | -0.104339626 | 0.773922947 | 0.91116213 |
| chr1:72623285-72623771 | -0.107774546 | 0.77393247 | 0.91116213 |
| chr4:101646949-101647465 | -0.107011902 | 0.773956028 | 0.911175865 |
| chr12:69738305-69740031 | -0.090119595 | 0.773968978 | 0.911177111 |
| chr20:62937342-62938604 | 0.091504242 | 0.774014494 | 0.911216695 |
| chr19:49442735-49443334 | 0.098008523 | 0.774038818 | 0.911231329 |
| chr15:70512808-70513074 | 0.114574788 | 0.774068613 | 0.911249928 |
| chr2:163375518-163375833 | -0.11808859 | 0.774078402 | 0.911249928 |
| chrX:108165460-108165748 | -0.117380499 | 0.774121576 | 0.911271023 |
| chr1:46603811-46604570 | 0.09325675 | 0.774130513 | 0.911271023 |
| chr6:3162410-3163343 | 0.095918894 | 0.774132034 | 0.911271023 |
| chr12:97568427-97569078 | -0.093788825 | 0.774147215 | 0.911271023 |
| chr15:90400551-90401272 | 0.099346698 | 0.774191399 | 0.911271023 |
| chr20:57265710-57266952 | 0.089132801 | 0.774205935 | 0.911271023 |
| chr1:226195156-226195463 | -0.123547631 | 0.774207568 | 0.911271023 |
| chr5:64612426-64612678 | 0.132664981 | 0.774208096 | 0.911271023 |
| chr1:216371321-216371521 | -0.140304132 | 0.774215252 | 0.911271023 |
| chr12:79635746-79635958 | -0.140304132 | 0.774215252 | 0.911271023 |

| | | | |
|---|---|---|---|
| chr15:29925923-29926137 | -0.149995318 | 0.774339086 | 0.911389713 |
| chr11:57514923-57515123 | 0.145876697 | 0.77433988 | 0.911389713 |
| chr8:26046577-26047092 | -0.106361955 | 0.774364494 | 0.911404683 |
| chr17:18094006-18094245 | 0.134958544 | 0.774397078 | 0.911423096 |
| chr2:132669853-132671026 | -0.095527687 | 0.774403928 | 0.911423096 |
| chr20:55665858-55666058 | -0.140902815 | 0.774441237 | 0.911453005 |
| chr2:86622873-86624086 | 0.092601932 | 0.774480301 | 0.911472877 |
| chr11:81071710-81071920 | 0.136548536 | 0.774483609 | 0.911472877 |
| chr8:67459819-67460033 | 0.1401462 | 0.774493809 | 0.911472877 |
| chr10:32171079-32171563 | 0.103555771 | 0.774525759 | 0.911496306 |
| chr12:113809616-113809828 | -0.149050347 | 0.774543161 | 0.911496306 |
| chr7:18624563-18624763 | 0.158181199 | 0.774549405 | 0.911496306 |
| chr1:114551264-114551539 | 0.104763724 | 0.774578037 | 0.911502586 |
| chr8:8270470-8270789 | 0.121656602 | 0.774588142 | 0.911502586 |
| chr12:75720901-75721645 | -0.1005526 | 0.774591219 | 0.911502586 |
| chr10:17290294-17290517 | 0.127605778 | 0.774602326 | 0.911502586 |
| chr14:80996408-80996643 | 0.12209211 | 0.774647614 | 0.911541879 |
| chr2:177212095-177213457 | -0.089869308 | 0.774668418 | 0.91155236 |
| chr14:70192992-70193258 | -0.126906631 | 0.774717046 | 0.911595581 |
| chr7:54705500-54705715 | -0.159949144 | 0.774738105 | 0.911603778 |
| chr3:119098713-119098926 | 0.150793812 | 0.774747806 | 0.911603778 |
| chr16:1923638-1923855 | -0.133597838 | 0.774775396 | 0.911614928 |
| chr19:55279759-55280624 | 0.092256035 | 0.774781078 | 0.911614928 |
| chr15:45314976-45315345 | -0.108860034 | 0.77481365 | 0.911633528 |
| chrX:10254817-10255051 | 0.147973383 | 0.774820682 | 0.911633528 |
| chr7:41003117-41003322 | 0.14667485 | 0.77486896 | 0.911655781 |
| chr11:67464518-67464975 | -0.096422544 | 0.774885565 | 0.911655781 |
| chr4:107380375-107380601 | 0.136890612 | 0.774891489 | 0.911655781 |
| chr11:102950770-102951067 | -0.132952409 | 0.774895089 | 0.911655781 |
| chr2:54052807-54053007 | -0.148778979 | 0.774899085 | 0.911655781 |
| chr9:79129937-79130158 | -0.150996341 | 0.774951411 | 0.911703343 |
| chr6:25489888-25490438 | -0.106306141 | 0.775073879 | 0.911833422 |
| chr5:160199314-160199964 | -0.096596972 | 0.77516499 | 0.911926608 |
| chr17:62304274-62304516 | 0.146257827 | 0.775179093 | 0.911929197 |
| chr1:241265392-241265611 | 0.149201038 | 0.775211998 | 0.911932522 |
| chr12:14723796-14724128 | -0.120523049 | 0.77522096 | 0.911932522 |
| chr3:129650264-129650542 | -0.125495518 | 0.775224501 | 0.911932522 |
| chr2:169452692-169452892 | -0.138469171 | 0.775229526 | 0.911932522 |
| chr8:97259017-97259654 | 0.098691159 | 0.775262772 | 0.911957629 |
| chr10:99779840-99780162 | 0.115311653 | 0.775332306 | 0.912025422 |
| chr3:50616770-50617250 | 0.098976852 | 0.77536424 | 0.912041499 |
| chr5:912713-913049 | -0.114440867 | 0.775369779 | 0.912041499 |
| chr15:86743380-86743648 | 0.122324826 | 0.775395226 | 0.912057429 |
| chr2:84472516-84472716 | 0.131787155 | 0.77541038 | 0.912061253 |
| chr20:19235380-19235717 | -0.139679194 | 0.775435827 | 0.912067432 |
| chr14:75559875-75560357 | 0.103509431 | 0.775452388 | 0.912067432 |
| chr5:17838617-17838817 | -0.153973131 | 0.775484038 | 0.912067432 |
| chr6:96836657-96838004 | 0.091688935 | 0.775498148 | 0.912067432 |
| chr19:38214067-38214267 | -0.145574078 | 0.775506722 | 0.912067432 |
| chr1:227280935-227281140 | -0.151493562 | 0.77550909 | 0.912067432 |

| | | | |
|---|---|---|---|
| chr13:21925976-21926176 | -0.151493562 | 0.77550909 | 0.912067432 |
| chr2:30897602-30897908 | -0.13102948 | 0.775515598 | 0.912067432 |
| chr6:15248559-15249096 | -0.095587956 | 0.775522764 | 0.912067432 |
| chr4:87391164-87391814 | -0.099516593 | 0.775553825 | 0.912076185 |
| chr3:163328025-163328225 | -0.142834675 | 0.775554015 | 0.912076185 |
| chr11:23834354-23834581 | 0.134582581 | 0.775621672 | 0.912131772 |
| chr2:148644069-148645566 | -0.089537616 | 0.775651188 | 0.912131772 |
| chr5:6490731-6490957 | -0.140439055 | 0.775652088 | 0.912131772 |
| chr11:76371151-76371472 | -0.120042314 | 0.775661758 | 0.912131772 |
| chr3:107970494-107970694 | 0.144544783 | 0.775673051 | 0.912131772 |
| chr3:141729272-141729557 | -0.123812145 | 0.775674929 | 0.912131772 |
| chr2:96061518-96061718 | 0.147765139 | 0.775684611 | 0.912131772 |
| chrX:67780777-67781075 | 0.130143548 | 0.775719922 | 0.912159296 |
| chr8:143320821-143321170 | 0.121754416 | 0.775758497 | 0.912189821 |
| chr16:2751979-2753123 | 0.092364183 | 0.775778981 | 0.912189821 |
| chr13:48054583-48054808 | 0.150128147 | 0.775781597 | 0.912189821 |
| chr7:25856717-25857351 | 0.093507679 | 0.775824073 | 0.912208369 |
| chr13:97031154-97031354 | -0.161952223 | 0.775832855 | 0.912208369 |
| chr6:73520820-73521878 | -0.089589307 | 0.775833087 | 0.912208369 |
| chr8:144109482-144109828 | 0.10372619 | 0.775859769 | 0.912216778 |
| chr15:75843723-75844251 | 0.101587108 | 0.77586405 | 0.912216778 |
| chr1:183234614-183235296 | 0.099723106 | 0.77589924 | 0.912244155 |
| chr20:49982268-49983178 | 0.091252068 | 0.775939243 | 0.912254251 |
| chr17:42050515-42050747 | 0.129627029 | 0.775941295 | 0.912254251 |
| chr12:68830265-68830587 | -0.113450411 | 0.775943545 | 0.912254251 |
| chr15:93035806-93036284 | -0.099381513 | 0.776040433 | 0.91235416 |
| chr5:21802967-21803167 | -0.132246531 | 0.776073277 | 0.91237667 |
| chr14:64284950-64285155 | 0.151589554 | 0.776091473 | 0.91237667 |
| chr8:28386159-28386854 | -0.099019387 | 0.776095302 | 0.91237667 |
| chr1:55702091-55702366 | 0.125917992 | 0.776114046 | 0.912384707 |
| chr3:25324274-25324474 | -0.161618086 | 0.776151039 | 0.912411817 |
| chr14:24146499-24147613 | 0.089364027 | 0.776160923 | 0.912411817 |
| chr19:12937984-12938714 | -0.095103823 | 0.776199809 | 0.912443531 |
| chr1:39899035-39899341 | 0.109066435 | 0.776220902 | 0.912454327 |
| chr7:2707595-2707795 | -0.135049906 | 0.776254248 | 0.912479527 |
| chr18:50402045-50402261 | 0.138833678 | 0.776276591 | 0.912491792 |
| chr3:139564386-139564659 | -0.123102967 | 0.776329802 | 0.91254034 |
| chr9:126914510-126915505 | 0.093282738 | 0.77635011 | 0.912550213 |
| chr3:38346383-38347611 | 0.093724741 | 0.776402763 | 0.912593855 |
| chr17:7915624-7916107 | -0.104928178 | 0.776412053 | 0.912593855 |
| chr12:49784940-49785140 | -0.148648924 | 0.77642297 | 0.912593855 |
| chr9:26892221-26893012 | 0.093401845 | 0.776446589 | 0.912607616 |
| chr18:61334016-61334768 | -0.107044347 | 0.776466734 | 0.912617296 |
| chr17:68172431-68172929 | -0.097307349 | 0.776485413 | 0.91262525 |
| chr3:131444264-131444715 | -0.113143827 | 0.776521256 | 0.912653378 |
| chr9:77458041-77458322 | -0.115643935 | 0.776547185 | 0.912660604 |
| chr3:197943025-197943257 | 0.139345314 | 0.776551977 | 0.912660604 |
| chr21:43333303-43334088 | 0.093838186 | 0.776563138 | 0.912660604 |
| chr5:65926343-65926722 | 0.104211161 | 0.77657758 | 0.912663578 |
| chr1:41516572-41516858 | -0.117449596 | 0.776604847 | 0.912681625 |

| | | | |
|---|---|---|---|
| chr5:179763222-179763459 | -0.120580701 | 0.776700331 | 0.912775315 |
| chr16:69565503-69566922 | 0.091806346 | 0.776708394 | 0.912775315 |
| chr17:75773991-75774202 | 0.113618522 | 0.776743242 | 0.912802269 |
| chr10:49253817-49254039 | 0.156421867 | 0.776811029 | 0.91281337 |
| chr9:100098700-100099981 | -0.08977539 | 0.776829758 | 0.91281337 |
| chr2:119759338-119760471 | -0.090522809 | 0.776855283 | 0.91281337 |
| chr1:201042765-201043535 | -0.093054407 | 0.776856522 | 0.91281337 |
| chr1:146401331-146401946 | -0.107053341 | 0.776867311 | 0.91281337 |
| chr11:123561111-123561325 | -0.136688746 | 0.776867445 | 0.91281337 |
| chr8:123767812-123769136 | -0.088819071 | 0.776870157 | 0.91281337 |
| chr4:140426815-140427987 | -0.092805383 | 0.776872396 | 0.91281337 |
| chr11:22326994-22327194 | 0.161751837 | 0.776876938 | 0.91281337 |
| chr22:20319831-20320367 | -0.099839602 | 0.776887393 | 0.91281337 |
| chr11:7612131-7612571 | 0.107675092 | 0.776891628 | 0.91281337 |
| chr21:42120038-42120391 | 0.103033351 | 0.776895648 | 0.91281337 |
| chr6:50843904-50844202 | -0.117085447 | 0.776929929 | 0.912831674 |
| chr17:57612841-57613041 | 0.129550062 | 0.776945028 | 0.912831674 |
| chr9:136847829-136848031 | 0.115136966 | 0.776946966 | 0.912831674 |
| chr3:119241457-119241945 | 0.09601194 | 0.776968279 | 0.912837953 |
| chr1:39092423-39092670 | 0.128503806 | 0.776976138 | 0.912837953 |
| chr5:170861639-170862807 | 0.090440848 | 0.777057053 | 0.912910879 |
| chr10:116609532-116610229 | 0.092501068 | 0.777072177 | 0.912910879 |
| chrX:142804229-142804429 | -0.151661985 | 0.777089318 | 0.912910879 |
| chr2:74457466-74458589 | -0.089253168 | 0.777094089 | 0.912910879 |
| chr8:1695848-1696101 | -0.13169829 | 0.777097782 | 0.912910879 |
| chr20:60050184-60050608 | -0.095103105 | 0.777112017 | 0.912913605 |
| chr11:34263432-34263833 | 0.102170871 | 0.777171013 | 0.912956096 |
| chr14:105050600-105050847 | 0.11464769 | 0.777186287 | 0.912956096 |
| chr1:46900525-46900758 | 0.144041928 | 0.777201386 | 0.912956096 |
| chr6:108835261-108835633 | -0.109811985 | 0.777223845 | 0.912956096 |
| chr4:40673075-40673275 | -0.136903444 | 0.777227612 | 0.912956096 |
| chr8:2011166-2011501 | 0.111622143 | 0.777230212 | 0.912956096 |
| chr6:15245182-15246104 | -0.090887867 | 0.777231593 | 0.912956096 |
| chr22:50251974-50252322 | -0.120307556 | 0.777247698 | 0.912961018 |
| chr16:84951016-84951219 | 0.12504886 | 0.777302665 | 0.913011587 |
| chr5:2151343-2151549 | -0.128866411 | 0.777358478 | 0.9130519 |
| chr17:15684187-15684695 | -0.110871408 | 0.777360819 | 0.9130519 |
| chr9:34458215-34458943 | -0.093304297 | 0.777396191 | 0.913073313 |
| chr17:52383512-52383970 | -0.100135428 | 0.777402883 | 0.913073313 |
| chr10:81179837-81180174 | 0.118287313 | 0.777417868 | 0.913076916 |
| chr8:86199738-86200167 | 0.103636042 | 0.777456037 | 0.913094599 |
| chr5:74876109-74876375 | -0.137500546 | 0.777463287 | 0.913094599 |
| chr19:9847103-9847576 | 0.111158219 | 0.777498965 | 0.913094599 |
| chr6:139981800-139982130 | 0.114652431 | 0.777520465 | 0.913094599 |
| chr16:3189496-3189754 | 0.106886682 | 0.77753947 | 0.913094599 |
| chr22:46390681-46391189 | 0.111852216 | 0.777558479 | 0.913094599 |
| chr13:20943665-20943925 | 0.121372751 | 0.777562216 | 0.913094599 |
| chr8:127819436-127820319 | 0.090928018 | 0.777566522 | 0.913094599 |
| chr19:44552360-44552614 | 0.127247309 | 0.777574283 | 0.913094599 |
| chr2:36695846-36696065 | -0.14913825 | 0.777587451 | 0.913094599 |

| | | | |
|---|---|---|---|
| chr4:87839401-87839618 | -0.14913825 | 0.777587451 | 0.913094599 |
| chr6:111011755-111011956 | -0.14913825 | 0.777587451 | 0.913094599 |
| chr10:27742212-27742428 | -0.130262747 | 0.777591563 | 0.913094599 |
| chr11:124799377-124800176 | 0.099319295 | 0.777599759 | 0.913094599 |
| chr1:43358255-43359284 | 0.091052256 | 0.777616917 | 0.91309832 |
| chr4:107989344-107990379 | -0.091239433 | 0.777634694 | 0.91309832 |
| chr9:114906960-114907160 | 0.131189712 | 0.777651548 | 0.91309832 |
| chr14:30296039-30296268 | -0.137624864 | 0.777663923 | 0.91309832 |
| chr19:50376227-50377461 | -0.091187857 | 0.777672993 | 0.91309832 |
| chr14:62067347-62067724 | 0.130313257 | 0.77767443 | 0.91309832 |
| chr1:201610846-201611079 | 0.120047553 | 0.777702384 | 0.913105505 |
| chr9:127611916-127612649 | -0.094855469 | 0.777704383 | 0.913105505 |
| chr7:76048715-76048926 | 0.126056223 | 0.777735526 | 0.913128078 |
| chr10:125141970-125142170 | -0.146487957 | 0.777764976 | 0.913140517 |
| chr2:383256-383481 | 0.135749867 | 0.777771328 | 0.913140517 |
| chr1:150363342-150364837 | -0.088316466 | 0.777781874 | 0.913140517 |
| chr5:135210234-135210434 | -0.152326269 | 0.777831048 | 0.913184257 |
| chr8:60660452-60660860 | -0.102341264 | 0.777854845 | 0.913198204 |
| chr5:139341462-139342590 | 0.089785633 | 0.777867072 | 0.913198565 |
| chr5:108148694-108149138 | 0.110679637 | 0.777938295 | 0.913266562 |
| chr9:99385890-99386160 | -0.124770312 | 0.777960545 | 0.913266562 |
| chr4:112321852-112322074 | -0.141776588 | 0.77796402 | 0.913266562 |
| chr1:62435899-62437499 | -0.088437492 | 0.777972668 | 0.913266562 |
| chr7:38782094-38782294 | -0.154606225 | 0.778008255 | 0.913294345 |
| chr14:39218080-39218723 | -0.10360446 | 0.778040759 | 0.913318509 |
| chr7:121449414-121449614 | -0.135995975 | 0.778091179 | 0.913353376 |
| chr1:923626-923842 | -0.135294209 | 0.778094302 | 0.913353376 |
| chr5:159106657-159107110 | 0.109498378 | 0.77813754 | 0.913390137 |
| chr13:19633219-19634581 | 0.08978519 | 0.77823352 | 0.913468633 |
| chr12:31897793-31898198 | -0.113304162 | 0.778237888 | 0.913468633 |
| chr2:237165479-237165847 | 0.127332016 | 0.778242142 | 0.913468633 |
| chr10:130547426-130547626 | -0.144559653 | 0.778261697 | 0.913468633 |
| chr19:49338646-49339294 | 0.094207474 | 0.778264858 | 0.913468633 |
| chr6:150016991-150017264 | -0.127177365 | 0.778289883 | 0.913468633 |
| chr12:95714962-95715182 | -0.167974984 | 0.778309747 | 0.913468633 |
| chr10:11270600-11270918 | -0.110173026 | 0.778310901 | 0.913468633 |
| chr2:9984232-9984490 | -0.121566325 | 0.778311708 | 0.913468633 |
| chr21:29382030-29382349 | 0.107059593 | 0.778348571 | 0.913473712 |
| chr1:77533730-77533939 | -0.142617332 | 0.778355321 | 0.913473712 |
| chr3:155737960-155738160 | -0.137804408 | 0.778363835 | 0.913473712 |
| chr14:92944175-92944446 | 0.116188985 | 0.778371646 | 0.913473712 |
| chrX:74063404-74064058 | 0.104243832 | 0.778375645 | 0.913473712 |
| chr5:174394242-174394492 | -0.135075112 | 0.778432334 | 0.913512904 |
| chr1:150487119-150487713 | 0.091166045 | 0.778432885 | 0.913512904 |
| chr6:76876448-76876706 | -0.125295873 | 0.778455117 | 0.913525002 |
| chr6:73451196-73451582 | -0.111414538 | 0.778470568 | 0.913529143 |
| chr1:223712265-223712566 | 0.114246122 | 0.778530751 | 0.9135707 |
| chr6:149926171-149926490 | -0.104955941 | 0.778531514 | 0.9135707 |
| chr15:65650993-65651225 | 0.136030491 | 0.778541751 | 0.9135707 |
| chr10:76303817-76304063 | -0.131787694 | 0.778594688 | 0.913618828 |

| | | | |
|---|---|---|---|
| chr1:1068658-1069526 | -0.091617167 | 0.778622259 | 0.913626583 |
| chr12:113174461-113174747 | 0.116288795 | 0.778625145 | 0.913626583 |
| chr4:127605428-127605642 | 0.142217626 | 0.778650431 | 0.913642262 |
| chr1:84771945-84772217 | -0.12863337 | 0.778691137 | 0.913655284 |
| chr20:11890524-11891690 | -0.089789514 | 0.778694703 | 0.913655284 |
| chr20:20435582-20435782 | -0.15269365 | 0.778705012 | 0.913655284 |
| chr1:169940548-169940748 | 0.149648078 | 0.778709226 | 0.913655284 |
| chr2:216023077-216023381 | -0.110198574 | 0.77872997 | 0.913665632 |
| chr20:30724017-30724601 | -0.101534561 | 0.778763808 | 0.913674001 |
| chr6:160455938-160456228 | -0.126345228 | 0.778770517 | 0.913674001 |
| chr7:149496818-149498235 | 0.090895007 | 0.778772877 | 0.913674001 |
| chr1:108624126-108624370 | -0.122636717 | 0.778800371 | 0.913678761 |
| chr7:149697866-149698066 | -0.155564408 | 0.778800783 | 0.913678761 |
| chr13:60134441-60134675 | -0.134151083 | 0.778828644 | 0.913697458 |
| chr10:29449065-29449265 | -0.1421371 | 0.778846073 | 0.913703915 |
| chr14:105431629-105432042 | -0.11886451 | 0.778897322 | 0.913750048 |
| chr2:126886132-126887101 | -0.092353405 | 0.77892498 | 0.913767224 |
| chr6:111551562-111551914 | 0.108858221 | 0.778935951 | 0.913767224 |
| chr20:19583647-19584202 | 0.100190814 | 0.778956895 | 0.913767224 |
| chr5:177022479-177023795 | 0.091568589 | 0.778959832 | 0.913767224 |
| chr9:108535677-108535884 | -0.148053813 | 0.778971963 | 0.913767224 |
| chr15:78618408-78618686 | 0.111786937 | 0.778984792 | 0.913767224 |
| chr16:78796337-78796541 | 0.139393672 | 0.778995443 | 0.913767224 |
| chr6:40328307-40329002 | 0.097533643 | 0.77901089 | 0.913771354 |
| chr11:16956748-16957140 | -0.112476883 | 0.77904166 | 0.9137818 |
| chr11:33257973-33258782 | -0.091932749 | 0.779046196 | 0.9137818 |
| chr17:80454494-80455475 | 0.09166889 | 0.779055573 | 0.9137818 |
| chr15:41543825-41544526 | 0.097457011 | 0.779097399 | 0.913816871 |
| chr1:101357359-101357721 | -0.109821675 | 0.779126683 | 0.913835589 |
| chr6:73388558-73389392 | -0.094512052 | 0.77913721 | 0.913835589 |
| chr1:3491197-3491562 | 0.109817707 | 0.779219216 | 0.913892447 |
| chr15:42495407-42495828 | -0.098629603 | 0.779222565 | 0.913892447 |
| chr4:39044351-39045368 | 0.092584906 | 0.779224644 | 0.913892447 |
| chr16:15590878-15591128 | -0.128847095 | 0.779233397 | 0.913892447 |
| chr1:226602452-226602667 | -0.137145535 | 0.779276251 | 0.91390704 |
| chr10:44801915-44802329 | -0.107425404 | 0.779278539 | 0.91390704 |
| chr20:53753149-53754169 | -0.087624548 | 0.779282672 | 0.91390704 |
| chr8:115620757-115621014 | -0.123951595 | 0.779293549 | 0.91390704 |
| chr10:116877399-116877731 | 0.107660889 | 0.779327603 | 0.913932987 |
| chr3:158050958-158051245 | -0.140570452 | 0.779369391 | 0.913968004 |
| chr8:98358138-98358470 | -0.106919865 | 0.779435959 | 0.914025074 |
| chr11:73377136-73377411 | 0.104293085 | 0.779441914 | 0.914025074 |
| chr22:20777661-20778201 | 0.107995249 | 0.779467881 | 0.914041536 |
| chr16:88804487-88804878 | 0.098835356 | 0.779513424 | 0.914080952 |
| chr4:57163579-57164340 | 0.091921209 | 0.779535013 | 0.914092279 |
| chr2:234463983-234464342 | 0.135821483 | 0.779561301 | 0.914109115 |
| chr10:126504150-126504637 | 0.10514181 | 0.779594534 | 0.914134094 |
| chr14:76953346-76953970 | -0.093992234 | 0.779635087 | 0.914157835 |
| chr6:11537257-11538100 | 0.090785286 | 0.779649021 | 0.914157835 |
| chr14:75419415-75419659 | 0.129996929 | 0.779650573 | 0.914157835 |

| | | | |
|---|---|---|---|
| chr8:120917373-120917620 | 0.123484789 | 0.779663389 | 0.914158873 |
| chr8:127748842-127749042 | 0.113265399 | 0.779686793 | 0.914172325 |
| chr6:85551289-85551489 | -0.143375061 | 0.779726442 | 0.914197809 |
| chr13:47095551-47095785 | -0.120147282 | 0.77973239 | 0.914197809 |
| chr14:63437961-63438660 | 0.097607456 | 0.779771654 | 0.914229855 |
| chr5:150636566-150636843 | 0.125363742 | 0.779803478 | 0.914253177 |
| chr4:123102980-123103223 | -0.12837963 | 0.779874094 | 0.914319863 |
| chr1:173703476-173703805 | 0.12794032 | 0.779888653 | 0.914319863 |
| chr17:42676827-42677448 | 0.090698028 | 0.779913783 | 0.914319863 |
| chr20:46562323-46563002 | 0.096046924 | 0.779940209 | 0.914319863 |
| chr16:10942249-10942696 | 0.09582868 | 0.779940301 | 0.914319863 |
| chr5:69334818-69335070 | 0.111214627 | 0.77994801 | 0.914319863 |
| chr5:157842864-157843064 | -0.134072103 | 0.7799485 | 0.914319863 |
| chr18:3014026-3014240 | 0.123083734 | 0.77995582 | 0.914319863 |
| chr14:61089360-61089609 | -0.135098108 | 0.780022535 | 0.914384081 |
| chr14:51157135-51157593 | -0.111740591 | 0.780076014 | 0.914432782 |
| chr13:21139944-21141092 | 0.088968703 | 0.780116681 | 0.914459245 |
| chr18:14798443-14798886 | 0.109452066 | 0.780136537 | 0.914459245 |
| chr6:39884475-39884771 | 0.135578232 | 0.780146362 | 0.914459245 |
| chr5:109988003-109988243 | -0.132020208 | 0.780165249 | 0.914459245 |
| chr16:641722-642035 | 0.107245621 | 0.780177224 | 0.914459245 |
| chr2:211927414-211927730 | -0.124704273 | 0.780187976 | 0.914459245 |
| chr10:79184271-79184480 | -0.127266741 | 0.780190086 | 0.914459245 |
| chr16:4679589-4680336 | -0.098449671 | 0.780194067 | 0.914459245 |
| chr1:203702152-203702473 | -0.104212664 | 0.780228725 | 0.914468325 |
| chr18:31513590-31513842 | 0.132910413 | 0.780233116 | 0.914468325 |
| chr4:154807075-154807335 | -0.131948544 | 0.780244209 | 0.914468325 |
| chr3:168301495-168301702 | 0.134417992 | 0.78025963 | 0.914468325 |
| chr16:84356648-84356965 | -0.115931545 | 0.780268019 | 0.914468325 |
| chr7:151012918-151013141 | -0.114758109 | 0.780273422 | 0.914468325 |
| chr19:48954527-48955204 | -0.09501367 | 0.780309513 | 0.914491446 |
| chr9:114107846-114108280 | 0.100358307 | 0.78031702 | 0.914491446 |
| chr2:3198121-3198389 | 0.135468167 | 0.780334132 | 0.914495312 |
| chr2:84459980-84460188 | -0.122543168 | 0.780345042 | 0.914495312 |
| chr17:62690291-62690491 | -0.114231771 | 0.780356124 | 0.914495312 |
| chr8:2001092-2001498 | 0.112455483 | 0.780386528 | 0.914516954 |
| chr4:101348298-101348614 | -0.105955722 | 0.780419781 | 0.914534025 |
| chr3:52057026-52057733 | -0.093242397 | 0.780424966 | 0.914534025 |
| chr1:159915944-159916654 | 0.093218234 | 0.780460986 | 0.914546298 |
| chr16:19523697-19524195 | 0.09776179 | 0.780472736 | 0.914546298 |
| chr3:160608411-160608654 | 0.137824344 | 0.780478991 | 0.914546298 |
| chr21:18760241-18760534 | 0.11791371 | 0.780493727 | 0.914546298 |
| chr6:44237410-44238278 | 0.090985029 | 0.780495119 | 0.914546298 |
| chr10:85260025-85260248 | 0.130559554 | 0.780523766 | 0.914565879 |
| chr5:176553994-176554370 | 0.112161575 | 0.780542295 | 0.914573604 |
| chr1:206882602-206882950 | -0.111673095 | 0.780585174 | 0.91460986 |
| chr2:240560177-240560525 | 0.099896407 | 0.780614646 | 0.91462352 |
| chr19:15124762-15126031 | 0.090979948 | 0.7806298 | 0.91462352 |
| chr5:180718865-180719092 | 0.135480303 | 0.780657974 | 0.91462352 |
| chr10:99956428-99956726 | -0.137821875 | 0.780659871 | 0.91462352 |

| | | | |
|---|---|---|---|
| chr2:8044055-8044290 | 0.155958176 | 0.780683787 | 0.91462352 |
| chr7:73556919-73558160 | 0.089132037 | 0.780696147 | 0.91462352 |
| chr4:67160397-67160749 | -0.114066049 | 0.78070275 | 0.91462352 |
| chr17:11144454-11144757 | -0.129563627 | 0.780706979 | 0.91462352 |
| chr5:134114272-134114833 | -0.096892143 | 0.780710694 | 0.91462352 |
| chr6:83430186-83431520 | -0.089223835 | 0.780716201 | 0.91462352 |
| chr6:170439464-170439664 | 0.137308957 | 0.780793545 | 0.914700145 |
| chr14:56725013-56725265 | 0.132572695 | 0.780895045 | 0.914796204 |
| chr3:97914437-97914670 | -0.141810204 | 0.780907707 | 0.914796204 |
| chr14:97978382-97978724 | -0.108895971 | 0.780911358 | 0.914796204 |
| chr3:174973404-174974062 | 0.092447654 | 0.780960036 | 0.914824144 |
| chr13:20127305-20128443 | 0.090898855 | 0.780969118 | 0.914824144 |
| chr14:22653082-22653410 | -0.107461535 | 0.780971028 | 0.914824144 |
| chr17:27861482-27861703 | 0.156187347 | 0.780994859 | 0.914838074 |
| chrX:82087578-82088076 | -0.112723884 | 0.781053384 | 0.914878386 |
| chr7:72879047-72879538 | 0.100481623 | 0.78105763 | 0.914878386 |
| chr21:10719112-10719395 | 0.12459129 | 0.781073444 | 0.914878386 |
| chr15:41115578-41116934 | -0.088985138 | 0.781085213 | 0.914878386 |
| chr4:127781628-127783017 | -0.087574078 | 0.781094117 | 0.914878386 |
| chr12:42024168-42024510 | 0.112167356 | 0.781109802 | 0.914878386 |
| chr10:24206955-24207155 | -0.134602175 | 0.781112854 | 0.914878386 |
| chr15:100873107-10087337€ | 0.11299236 | 0.781137759 | 0.914881472 |
| chrX:57708927-57709152 | 0.127483548 | 0.781139495 | 0.914881472 |
| chr1:185045041-185046371 | -0.089093164 | 0.78115131 | 0.914881472 |
| chr6:107027956-107028918 | 0.089771442 | 0.7812107 | 0.914937045 |
| chr20:49768397-49769064 | 0.099135711 | 0.781243317 | 0.91494126 |
| chr1:195717752-195717980 | -0.11659482 | 0.781264269 | 0.91494126 |
| chr20:58372974-58373191 | 0.12218882 | 0.781266983 | 0.91494126 |
| chr6:16277320-16277657 | -0.105986784 | 0.781269251 | 0.91494126 |
| chr5:142782448-142782651 | -0.151382381 | 0.781275904 | 0.91494126 |
| chr3:182909277-182909550 | -0.126172227 | 0.781297421 | 0.91494126 |
| chr16:11312727-11313011 | -0.113275187 | 0.781304869 | 0.91494126 |
| chr21:20997355-20998734 | -0.090221529 | 0.781309827 | 0.91494126 |
| chr3:161220887-161221836 | -0.088515481 | 0.781355633 | 0.91496408 |
| chr5:64506908-64507175 | -0.127456641 | 0.781364988 | 0.91496408 |
| chr4:78938912-78939621 | -0.092473669 | 0.781365138 | 0.91496408 |
| chr1:165109069-165109387 | -0.125978683 | 0.781464401 | 0.91506633 |
| chr12:63779458-63780512 | 0.088426131 | 0.781480465 | 0.915071157 |
| chr7:34759535-34759996 | -0.098916395 | 0.781538207 | 0.915117965 |
| chr14:69014826-69015066 | 0.128315749 | 0.781544327 | 0.915117965 |
| chr3:64612528-64613596 | 0.087681843 | 0.781577309 | 0.9151426 |
| chr1:228004312-228004861 | -0.098276754 | 0.781640923 | 0.915191936 |
| chr20:13994998-13996237 | 0.08722504 | 0.781645045 | 0.915191936 |
| chr20:53943783-53944279 | 0.088458445 | 0.781655277 | 0.915191936 |
| chr17:81651793-81651995 | 0.130305012 | 0.781680957 | 0.915208018 |
| chr1:26275689-26276107 | -0.109890117 | 0.781713481 | 0.915219133 |
| chr6:33968020-33968227 | 0.139380828 | 0.781717855 | 0.915219133 |
| chr2:199955072-199956134 | -0.088893709 | 0.781728631 | 0.915219133 |
| chr13:63855620-63855962 | -0.11723296 | 0.781738229 | 0.915219133 |
| chr2:169767380-169767580 | 0.146976766 | 0.781776047 | 0.915225393 |

| | | | |
|---|---|---|---|
| chr15:25933802-25934121 | -0.132256541 | 0.781776783 | 0.915225393 |
| chr8:96056078-96056297 | 0.137250516 | 0.78177941 | 0.915225393 |
| chr11:19712535-19713071 | 0.096422086 | 0.781853467 | 0.915298106 |
| chr1:204431370-204431587 | -0.114095276 | 0.781882162 | 0.915317714 |
| chr9:69672084-69672576 | 0.104755028 | 0.781943259 | 0.915375253 |
| chr4:47490619-47490865 | 0.117797636 | 0.781986334 | 0.915377516 |
| chr14:52900675-52901100 | -0.09490894 | 0.781987155 | 0.915377516 |
| chr12:69091505-69091705 | 0.151422522 | 0.782004778 | 0.915377516 |
| chr1:158124989-158125379 | -0.095133681 | 0.782016714 | 0.915377516 |
| chr19:1940718-1941258 | 0.104710115 | 0.782020232 | 0.915377516 |
| chr16:80500583-80500919 | 0.116408863 | 0.782022233 | 0.915377516 |
| chr7:138212351-138212560 | -0.159363822 | 0.782028819 | 0.915377516 |
| chr2:142903551-142904183 | -0.09221505 | 0.782076297 | 0.915419105 |
| chr13:46617584-46617793 | -0.118784678 | 0.782092028 | 0.915423534 |
| chr7:5045529-5046689 | -0.08889685 | 0.782123546 | 0.915446441 |
| chr12:70242678-70243958 | -0.087705737 | 0.78217393 | 0.91548811 |
| chr5:21044179-21044392 | 0.135877405 | 0.782183043 | 0.91548811 |
| chr1:203204706-203204935 | -0.125409052 | 0.782215427 | 0.915492031 |
| chr9:4604604-4604919 | -0.114912792 | 0.782222584 | 0.915492031 |
| chr8:124525892-124526320 | 0.094266564 | 0.782243659 | 0.915492031 |
| chr1:93878861-93879613 | -0.096107476 | 0.782282126 | 0.915492031 |
| chr8:126557574-126558751 | -0.086267022 | 0.78229055 | 0.915492031 |
| chr9:4679269-4680198 | -0.089747296 | 0.782291838 | 0.915492031 |
| chr5:36066803-36067009 | 0.13818095 | 0.782297545 | 0.915492031 |
| chr6:37432815-37433423 | 0.097641204 | 0.782300317 | 0.915492031 |
| chr5:173056788-173057636 | -0.08820043 | 0.782331358 | 0.915492031 |
| chr15:78979686-78979981 | -0.107468603 | 0.782332112 | 0.915492031 |
| chr8:126383717-126384033 | -0.097331185 | 0.782370437 | 0.915492031 |
| chr1:39883139-39883689 | -0.098340506 | 0.782378157 | 0.915492031 |
| chr4:157805074-157805316 | -0.122539621 | 0.782382917 | 0.915492031 |
| chr7:129592428-129592662 | -0.122539621 | 0.782382917 | 0.915492031 |
| chr12:6851224-6851550 | -0.095509501 | 0.782383893 | 0.915492031 |
| chr1:200290354-200290755 | -0.098736226 | 0.782389208 | 0.915492031 |
| chr18:54158508-54158708 | -0.150200325 | 0.782389512 | 0.915492031 |
| chr17:47650401-47650868 | -0.093006047 | 0.782408242 | 0.915492406 |
| chr2:223101673-223102013 | -0.110399645 | 0.782413728 | 0.915492406 |
| chr15:101501718-101502293 | 0.095373394 | 0.782433866 | 0.915501988 |
| chr5:126366232-126366482 | -0.125018786 | 0.782446847 | 0.915503197 |
| chr6:474883-475083 | 0.149601715 | 0.782473004 | 0.915519821 |
| chr1:204073826-204074789 | -0.089026068 | 0.782502239 | 0.915528887 |
| chr1:181119217-181119787 | -0.088469762 | 0.78250465 | 0.915528887 |
| chr11:126383238-126383772 | -0.098695637 | 0.782520298 | 0.915533215 |
| chr11:94046371-94046769 | -0.130624931 | 0.782534886 | 0.915534979 |
| chr8:144900951-144902544 | -0.086768636 | 0.782552071 | 0.915534979 |
| chr21:39030531-39030767 | -0.129929366 | 0.782557651 | 0.915534979 |
| chr5:127975954-127976247 | 0.134700567 | 0.782594685 | 0.915564326 |
| chr3:28328656-28328900 | -0.134414246 | 0.782629719 | 0.91557988 |
| chr8:110449783-110449983 | -0.158811033 | 0.78263685 | 0.91557988 |
| chr7:42176464-42176673 | -0.14797041 | 0.782651162 | 0.91557988 |
| chr3:127455383-127455973 | 0.096639627 | 0.782663847 | 0.91557988 |

| | | | |
|---|---|---|---|
| chr5:171122269-171122469 | -0.1397052 | 0.782667726 | 0.91557988 |
| chr7:65872751-65873681 | -0.091198004 | 0.782697496 | 0.915587381 |
| chr5:118774470-118774720 | 0.135196864 | 0.782720692 | 0.915587381 |
| chr8:124538358-124539493 | 0.086919842 | 0.782748265 | 0.915587381 |
| chr11:6703507-6703812 | -0.115854368 | 0.782749188 | 0.915587381 |
| chr10:101213676-101214424 | 0.097173507 | 0.782752108 | 0.915587381 |
| chr6:24547328-24547548 | 0.143251327 | 0.782756805 | 0.915587381 |
| chr3:57397479-57397702 | 0.137907977 | 0.782757785 | 0.915587381 |
| chr3:184748669-184748874 | 0.13272612 | 0.782783591 | 0.915603589 |
| chr11:11454698-11455168 | -0.110134403 | 0.782819431 | 0.915631533 |
| chr2:95323390-95324061 | -0.095167423 | 0.782849261 | 0.915652445 |
| chr15:29821435-29821711 | -0.119397412 | 0.782864586 | 0.915655093 |
| chr11:85590903-85591224 | 0.113016606 | 0.782886176 | 0.915655093 |
| chr1:59296387-59297303 | 0.087475384 | 0.782902115 | 0.915655093 |
| chr17:18625512-18625972 | -0.104173899 | 0.782920656 | 0.915655093 |
| chr1:35882756-35883456 | 0.092734858 | 0.782922626 | 0.915655093 |
| chr8:126556785-126557163 | -0.093040689 | 0.782946335 | 0.915655093 |
| chr15:83066329-83066688 | 0.105696135 | 0.782956442 | 0.915655093 |
| chr2:171598760-171599081 | 0.104013845 | 0.783003841 | 0.915655093 |
| chr14:61477889-61478135 | -0.133905901 | 0.783017772 | 0.915655093 |
| chr1:149792508-149792800 | 0.10743328 | 0.783025279 | 0.915655093 |
| chr5:176536769-176538217 | -0.08672588 | 0.783027879 | 0.915655093 |
| chr20:21125228-21126561 | 0.087376587 | 0.783037098 | 0.915655093 |
| chr12:78119374-78119819 | 0.10471692 | 0.783039293 | 0.915655093 |
| chr1:195663989-195664317 | 0.115647572 | 0.783042239 | 0.915655093 |
| chr12:10867391-10867591 | -0.147317256 | 0.783042729 | 0.915655093 |
| chr3:189790797-189790997 | -0.147317256 | 0.783042729 | 0.915655093 |
| chr20:54740586-54740948 | 0.094387985 | 0.783155247 | 0.915762394 |
| chr14:99982549-99982755 | 0.129950949 | 0.783164705 | 0.915762394 |
| chr5:17402582-17402828 | 0.130069455 | 0.783170345 | 0.915762394 |
| chr15:78539876-78541181 | -0.089412943 | 0.783217413 | 0.915803455 |
| chr11:67303074-67303778 | -0.091025006 | 0.783240819 | 0.915816848 |
| chr1:151723675-151724502 | 0.0911744 | 0.783270094 | 0.915837102 |
| chr2:219737267-219737598 | 0.11087489 | 0.783310508 | 0.915844665 |
| chr4:139403436-139403740 | -0.12401654 | 0.783311827 | 0.915844665 |
| chr19:3852674-3852947 | 0.130962751 | 0.783312421 | 0.915844665 |
| chr2:114543474-114543674 | 0.135114253 | 0.783379812 | 0.915882462 |
| chr11:86940054-86940867 | -0.098231268 | 0.783389785 | 0.915882462 |
| chr17:8220740-8221039 | -0.101042776 | 0.783392994 | 0.915882462 |
| chr3:72887857-72888402 | 0.097642073 | 0.783439119 | 0.915882462 |
| chr2:152163655-152164081 | -0.101785407 | 0.783448496 | 0.915882462 |
| chr13:41276768-41276993 | 0.143745884 | 0.783448626 | 0.915882462 |
| chr1:152347204-152347595 | -0.105237819 | 0.78344957 | 0.915882462 |
| chr12:45216367-45216950 | 0.096128759 | 0.783451359 | 0.915882462 |
| chr1:64897706-64897906 | 0.13567475 | 0.783454589 | 0.915882462 |
| chr4:120719048-120719314 | -0.118030642 | 0.78346428 | 0.915882462 |
| chr14:92783845-92784175 | 0.102607838 | 0.783481251 | 0.915888327 |
| chr8:102738218-102738501 | -0.104634673 | 0.783520562 | 0.915920308 |
| chr7:39833251-39833945 | 0.09707243 | 0.783535311 | 0.915923575 |
| chr5:36744398-36745668 | -0.086950606 | 0.783548533 | 0.915925057 |

| | | | |
|---|---|---|---|
| chr17:43778110-43778843 | 0.092928598 | 0.783569411 | 0.91593549 |
| chr17:50915202-50915476 | 0.10294249 | 0.783598958 | 0.915956055 |
| chr3:4655977-4656317 | 0.100899886 | 0.783611469 | 0.915956705 |
| chr11:62672231-62672585 | 0.102115603 | 0.783626181 | 0.915959928 |
| chr1:239592947-239593173 | -0.143697313 | 0.78367647 | 0.91597456 |
| chr3:122001124-122001324 | -0.133717902 | 0.783679371 | 0.91597456 |
| chr15:62570105-62570417 | 0.115250049 | 0.783683958 | 0.91597456 |
| chr5:145981860-145982578 | 0.093740463 | 0.783686517 | 0.91597456 |
| chr11:113474288-113474678 | 0.105308332 | 0.783738998 | 0.916015095 |
| chr17:43006613-43006846 | -0.133709869 | 0.783745107 | 0.916015095 |
| chr8:48121182-48121453 | -0.118241583 | 0.783773919 | 0.916028938 |
| chr1:52033183-52034005 | -0.092947739 | 0.783789211 | 0.916028938 |
| chr3:49723814-49724214 | 0.100361148 | 0.783815559 | 0.916028938 |
| chr2:160918952-160919251 | -0.120549606 | 0.783816654 | 0.916028938 |
| chr15:59657045-59658152 | -0.089358669 | 0.783818705 | 0.916028938 |
| chr2:62042954-62043259 | -0.114465508 | 0.783828682 | 0.916028938 |
| chr14:38617763-38617967 | -0.13227865 | 0.783861076 | 0.916030416 |
| chr8:134623113-134623376 | 0.124421136 | 0.783862363 | 0.916030416 |
| chr9:136843761-136844019 | -0.111945691 | 0.783865813 | 0.916030416 |
| chr8:138953151-138953351 | 0.141632575 | 0.783886675 | 0.916040825 |
| chr5:75053507-75053940 | -0.10368962 | 0.783914854 | 0.916059784 |
| chr8:26576857-26578046 | 0.088367696 | 0.783955869 | 0.916073977 |
| chr11:8006409-8006717 | -0.111052571 | 0.783961412 | 0.916073977 |
| chr1:151326991-151328383 | 0.086672563 | 0.783962867 | 0.916073977 |
| chr8:57575373-57575573 | -0.140656947 | 0.784007418 | 0.91610448 |
| chr21:19529024-19529407 | 0.105671446 | 0.78402216 | 0.91610448 |
| chr17:4997242-4997997 | -0.091801101 | 0.78402484 | 0.91610448 |
| chr2:74464964-74465801 | 0.090439072 | 0.784053405 | 0.9161098 |
| chr14:100096526-100096998 | 0.105696378 | 0.784064543 | 0.9161098 |
| chr15:38872937-38873494 | -0.103876608 | 0.784065261 | 0.9161098 |
| chr20:50645420-50646070 | 0.090540814 | 0.784136338 | 0.916154197 |
| chr17:50427490-50427690 | 0.115141717 | 0.784141369 | 0.916154197 |
| chr21:33380513-33380920 | 0.09988476 | 0.784144142 | 0.916154197 |
| chr20:555599-555799 | -0.159080792 | 0.784157356 | 0.916154197 |
| chr14:81062851-81063185 | -0.107546818 | 0.784163043 | 0.916154197 |
| chr3:64530279-64530797 | -0.088993049 | 0.784178372 | 0.916158137 |
| chr17:63478745-63479277 | 0.089930444 | 0.784250937 | 0.916220077 |
| chr14:59869789-59871161 | 0.091031434 | 0.784255305 | 0.916220077 |
| chr12:79690155-79691777 | 0.085654417 | 0.784362188 | 0.916330974 |
| chr3:184469384-184469739 | 0.107308969 | 0.784396089 | 0.916355827 |
| chr16:56964575-56964902 | 0.113951517 | 0.784407381 | 0.916355827 |
| chr17:81500166-81500366 | -0.138609443 | 0.784423676 | 0.91635607 |
| chr1:235068259-235068594 | 0.105699905 | 0.784431508 | 0.91635607 |
| chr3:46488722-46489207 | -0.106331694 | 0.784460975 | 0.916376522 |
| chr10:114303976-114304258 | -0.105570372 | 0.78448026 | 0.916381903 |
| chr5:109946081-109946647 | -0.10310019 | 0.784490577 | 0.916381903 |
| chr16:30806214-30806414 | 0.127672043 | 0.784501461 | 0.916381903 |
| chr5:14142632-14143396 | 0.087684007 | 0.784515654 | 0.916384512 |
| chr7:4973749-4975065 | -0.088009774 | 0.784540297 | 0.916386667 |
| chr12:55973407-55974231 | 0.086659684 | 0.784541419 | 0.916386667 |

| | | | |
|---|---|---|---|
| chr15:93001717-93002188 | -0.108066453 | 0.784562068 | 0.916396817 |
| chr6:141552526-141552972 | 0.100887261 | 0.784578519 | 0.916402062 |
| chr1:189266721-189266921 | 0.147525546 | 0.784609661 | 0.916417723 |
| chr6:28719619-28719931 | 0.109926446 | 0.784618375 | 0.916417723 |
| chr3:139562114-139562464 | -0.108198186 | 0.784627808 | 0.916417723 |
| chr13:87680425-87680637 | -0.123221653 | 0.784662372 | 0.916444123 |
| chr11:90178841-90179096 | -0.124108175 | 0.784684757 | 0.916448099 |
| chr17:62790720-62790953 | -0.117310544 | 0.784724354 | 0.916448099 |
| chr15:89416314-89416864 | 0.092213346 | 0.784744628 | 0.916448099 |
| chr6:111068310-111068854 | -0.101845334 | 0.784751558 | 0.916448099 |
| chr10:100998138-100998561 | -0.097836753 | 0.784752326 | 0.916448099 |
| chr8:124474097-124475612 | -0.085022989 | 0.784753837 | 0.916448099 |
| chr7:143737585-143737785 | -0.151596733 | 0.784762642 | 0.916448099 |
| chr1:183431695-183431947 | -0.135759788 | 0.78477757 | 0.916448099 |
| chr1:53393349-53393656 | 0.112974544 | 0.784786846 | 0.916448099 |
| chr10:48690633-48690863 | -0.141792585 | 0.784795708 | 0.916448099 |
| chrX:53441547-53442082 | 0.093716754 | 0.784797343 | 0.916448099 |
| chr6:150068752-150069466 | -0.089697265 | 0.784812777 | 0.916451446 |
| chr7:66638359-66638559 | 0.151231644 | 0.784824131 | 0.916451446 |
| chr8:118861349-118861691 | 0.113412712 | 0.784861846 | 0.91648152 |
| chr3:150421882-150422086 | -0.128025798 | 0.784874552 | 0.916482389 |
| chr3:49786371-49787218 | -0.091070508 | 0.784898154 | 0.916490827 |
| chr1:41918797-41919073 | 0.125744309 | 0.7849057 | 0.916490827 |
| chr8:106624748-106625075 | 0.117879305 | 0.784958404 | 0.916535795 |
| chr7:44999506-45000345 | -0.086983411 | 0.784972935 | 0.916535795 |
| chr6:34465991-34466191 | 0.130119041 | 0.784980097 | 0.916535795 |
| chr19:50136611-50136835 | 0.132053081 | 0.785018146 | 0.916544093 |
| chr3:111691341-111691658 | -0.107569228 | 0.785021564 | 0.916544093 |
| chr12:68636366-68636673 | -0.112197913 | 0.78502309 | 0.916544093 |
| chr16:19400767-19400967 | -0.152535812 | 0.785059726 | 0.916572901 |
| chr11:20397261-20397580 | 0.113065736 | 0.785083743 | 0.916581031 |
| chr15:57612070-57612278 | -0.133532518 | 0.785090614 | 0.916581031 |
| chr19:46826625-46826909 | 0.118012572 | 0.785138943 | 0.916613424 |
| chr10:102854058-102854756 | 0.091558288 | 0.78516426 | 0.916613424 |
| chr2:158457295-158458349 | -0.088334327 | 0.785195098 | 0.916613424 |
| chr3:79724263-79724504 | -0.125760917 | 0.785195906 | 0.916613424 |
| chr1:179677813-179678040 | 0.132527779 | 0.785197391 | 0.916613424 |
| chr8:109325864-109326200 | 0.099076176 | 0.785199774 | 0.916613424 |
| chr2:138105486-138105850 | 0.131973483 | 0.7852021 | 0.916613424 |
| chr19:42606705-42606943 | -0.130774424 | 0.785287407 | 0.916697348 |
| chr1:115706845-115707096 | -0.114971915 | 0.785302433 | 0.916697348 |
| chr9:129737597-129738299 | 0.094756686 | 0.785312993 | 0.916697348 |
| chr3:186722487-186722768 | -0.121569261 | 0.785321848 | 0.916697348 |
| chr2:220619477-220620457 | -0.090704478 | 0.785335207 | 0.916698977 |
| chrX:55717427-55718499 | -0.088104191 | 0.785351696 | 0.916704259 |
| chr15:69491244-69491465 | 0.145622791 | 0.785385119 | 0.916720598 |
| chr4:15478996-15479955 | -0.089402052 | 0.785403741 | 0.916720598 |
| chr6:4677940-4678140 | 0.153656662 | 0.785410602 | 0.916720598 |
| chr5:101423587-101423792 | 0.148365316 | 0.785425093 | 0.916720598 |
| chr11:94623211-94623491 | -0.119340585 | 0.785425515 | 0.916720598 |

| | | | |
|---|---|---|---|
| chr9:41229448-41230076 | -0.099477558 | 0.785492872 | 0.916771572 |
| chr1:41754419-41754711 | -0.136888225 | 0.785493119 | 0.916771572 |
| chr6:129111087-129111318 | 0.143434152 | 0.785524606 | 0.916794357 |
| chr1:239616770-239617095 | -0.126116135 | 0.785537136 | 0.916795017 |
| chr1:28088276-28088933 | -0.089949792 | 0.785583913 | 0.916825776 |
| chr3:112230646-112230846 | -0.155088656 | 0.785587423 | 0.916825776 |
| chr6:97480843-97481259 | 0.119501993 | 0.785660914 | 0.916862677 |
| chr1:198088393-198088910 | -0.10156091 | 0.785670506 | 0.916862677 |
| chr8:144786686-144787642 | -0.08673106 | 0.785673769 | 0.916862677 |
| chrX:53686307-53686889 | -0.09256112 | 0.78568591 | 0.916862677 |
| chr1:203326902-203328241 | -0.085608325 | 0.785686798 | 0.916862677 |
| chr8:60396939-60397304 | -0.108703454 | 0.785690838 | 0.916862677 |
| chr5:150605650-150605850 | 0.132398145 | 0.78574494 | 0.916911847 |
| chr1:6513403-6513639 | 0.129630577 | 0.785784366 | 0.91694389 |
| chr12:114476081-114476478 | -0.118673863 | 0.785835933 | 0.916976471 |
| chr22:30356447-30357388 | 0.090108075 | 0.785863987 | 0.916976471 |
| chr1:108214762-108215020 | -0.121319461 | 0.785864917 | 0.916976471 |
| chr5:36606479-36607357 | -0.096337553 | 0.785866479 | 0.916976471 |
| chr6:88908481-88908902 | 0.117938603 | 0.785872124 | 0.916976471 |
| chr5:90381317-90381726 | 0.11217248 | 0.785911015 | 0.917005509 |
| chr10:71964801-71965036 | 0.124938622 | 0.785920947 | 0.917005509 |
| chr17:7035544-7036384 | 0.089325641 | 0.78593533 | 0.917008328 |
| chr14:90805637-90805929 | -0.124178522 | 0.786061938 | 0.917110855 |
| chr2:7962058-7962332 | 0.133477988 | 0.786068284 | 0.917110855 |
| chr8:127496476-127496710 | -0.11233534 | 0.78607326 | 0.917110855 |
| chr14:55448035-55448642 | -0.096245868 | 0.786078477 | 0.917110855 |
| chr2:72581651-72582089 | -0.107990539 | 0.786083049 | 0.917110855 |
| chr20:23601490-23601833 | 0.111342128 | 0.786106612 | 0.917115701 |
| chr19:41308558-41308813 | 0.114181558 | 0.786115058 | 0.917115701 |
| chr20:46445010-46445213 | -0.143806891 | 0.786123111 | 0.917115701 |
| chr1:97202602-97202941 | -0.116891735 | 0.786140422 | 0.917121933 |
| chr7:6373810-6375034 | -0.086005071 | 0.786159178 | 0.917129849 |
| chr8:90987721-90988056 | 0.109014704 | 0.786191779 | 0.91713227 |
| chr3:14990381-14990591 | -0.127791428 | 0.786192717 | 0.91713227 |
| chr5:80936119-80936355 | -0.130219315 | 0.786197162 | 0.91713227 |
| chr5:121274086-121274352 | 0.126573878 | 0.786211099 | 0.917134566 |
| chr14:103854737-103856018 | -0.088473934 | 0.78622313 | 0.917134637 |
| chr10:45000754-45001871 | 0.08967117 | 0.786290374 | 0.917184987 |
| chr5:33969736-33970152 | 0.103939213 | 0.786298889 | 0.917184987 |
| chr12:67873795-67873995 | 0.153959815 | 0.786302203 | 0.917184987 |
| chr14:90398396-90398681 | -0.106144748 | 0.786344598 | 0.917213925 |
| chr19:4246708-4247617 | 0.087453832 | 0.786350953 | 0.917213925 |
| chr4:130957696-130957896 | -0.147394466 | 0.78637995 | 0.917233637 |
| chr13:42039777-42041525 | -0.085396823 | 0.786391794 | 0.917233637 |
| chr1:112337037-112337490 | 0.102973068 | 0.786417376 | 0.917249511 |
| chr5:10521571-10521979 | 0.11613694 | 0.786462584 | 0.917287185 |
| chr11:18721713-18721917 | -0.13267435 | 0.786473619 | 0.917287185 |
| chr17:31494950-31495304 | -0.131340756 | 0.7865026 | 0.917291722 |
| chr16:48734540-48734962 | -0.104955856 | 0.78651632 | 0.917291722 |
| chr3:96618302-96618516 | 0.14983929 | 0.786522025 | 0.917291722 |

| | | | |
|---|---|---|---|
| chr17:54712835-54713228 | 0.117136514 | 0.786525396 | 0.917291722 |
| chr11:46700229-46701185 | 0.086523845 | 0.786540986 | 0.917295943 |
| chr16:9008793-9009113 | 0.103321194 | 0.786602936 | 0.917354228 |
| chr4:128810491-128810736 | 0.10936585 | 0.786658281 | 0.917391004 |
| chr10:11411988-11412211 | -0.133889966 | 0.786664497 | 0.917391004 |
| chr4:140332852-140333311 | 0.108636383 | 0.786678181 | 0.917391004 |
| chr3:24680556-24680897 | -0.107689282 | 0.786686229 | 0.917391004 |
| chrX:47144597-47145390 | -0.08863265 | 0.786694335 | 0.917391004 |
| chr4:145956282-145956613 | -0.117499969 | 0.786720753 | 0.917396052 |
| chr3:9715176-9715449 | 0.11184435 | 0.78672261 | 0.917396052 |
| chr12:1594561-1595021 | -0.103004127 | 0.786769625 | 0.917405514 |
| chr21:39838177-39838377 | -0.137735525 | 0.786770329 | 0.917405514 |
| chr1:109640244-109640558 | 0.097308805 | 0.7867786 | 0.917405514 |
| chr16:4351103-4351573 | 0.09249377 | 0.786778616 | 0.917405514 |
| chr6:30212911-30213892 | 0.087913677 | 0.786815115 | 0.917419852 |
| chr10:28375578-28376436 | -0.09278616 | 0.786823822 | 0.917419852 |
| chr3:24475836-24476309 | -0.1060194 | 0.786833346 | 0.917419852 |
| chr21:29904708-29905144 | -0.111877213 | 0.786838806 | 0.917419852 |
| chr12:91920252-91920700 | -0.110165349 | 0.786876075 | 0.917449346 |
| chr4:140851129-140851353 | -0.131666163 | 0.786909017 | 0.917462013 |
| chr20:5066017-5066306 | 0.13570308 | 0.786926214 | 0.917462013 |
| chr19:49127929-49128649 | 0.088542624 | 0.786932894 | 0.917462013 |
| chr11:46276315-46276947 | 0.095640356 | 0.786955212 | 0.917462013 |
| chr1:44784168-44784368 | -0.13293713 | 0.786958361 | 0.917462013 |
| chr20:20711994-20712206 | 0.112051468 | 0.786958783 | 0.917462013 |
| chr1:47313768-47314606 | 0.086396431 | 0.787011144 | 0.917494228 |
| chr14:77316000-77316279 | -0.101182412 | 0.787021597 | 0.917494228 |
| chr3:36700813-36701267 | -0.115138917 | 0.787026299 | 0.917494228 |
| chr13:101544937-101545304 | -0.11426191 | 0.787058608 | 0.917494228 |
| chr8:93127746-93128009 | -0.115784666 | 0.787061935 | 0.917494228 |
| chr11:126062603-126063589 | -0.086219451 | 0.787080746 | 0.917494228 |
| chr10:19178743-19179003 | -0.139433184 | 0.787082153 | 0.917494228 |
| chr1:61338008-61338700 | -0.095366683 | 0.78708524 | 0.917494228 |
| chr8:10417768-10418071 | 0.119063068 | 0.78711297 | 0.917494228 |
| chr11:63574048-63574314 | -0.143501295 | 0.787113973 | 0.917494228 |
| chr7:101882674-101883503 | -0.089317437 | 0.787127395 | 0.917494228 |
| chr3:33798142-33799304 | 0.08561502 | 0.787132111 | 0.917494228 |
| chr17:58729418-58729817 | 0.092551165 | 0.787142775 | 0.917494228 |
| chr9:15104878-15105102 | 0.14018383 | 0.787159414 | 0.917494228 |
| chr2:31137996-31138232 | -0.117579572 | 0.78716603 | 0.917494228 |
| chr1:16709750-16710112 | -0.102276649 | 0.787200362 | 0.917520287 |
| chr8:129786762-129787269 | -0.1124547 | 0.787215095 | 0.917522232 |
| chrX:51641537-51641751 | 0.132025649 | 0.787225981 | 0.917522232 |
| chr8:144479053-144479267 | 0.107570516 | 0.787270266 | 0.917549952 |
| chr11:65560072-65560407 | 0.107911295 | 0.787273714 | 0.917549952 |
| chr2:3805139-3805489 | -0.104429222 | 0.787309652 | 0.917577881 |
| chr6:116010875-116011075 | 0.135045532 | 0.787357397 | 0.917619568 |
| chr16:72099283-72099686 | 0.093348726 | 0.787413224 | 0.917670673 |
| chr4:76147724-76148980 | 0.086119932 | 0.787426884 | 0.917672635 |
| chr7:74312831-74313031 | -0.145494203 | 0.787469721 | 0.917708599 |

| | | | |
|---|---|---|---|
| chr1:236490005-236490310 | 0.109554666 | 0.787505927 | 0.917719228 |
| chr1:190381103-190381418 | -0.129894627 | 0.787524746 | 0.917719228 |
| chr7:6271022-6271294 | -0.102021852 | 0.78752961 | 0.917719228 |
| chr10:103686193-103686629 | -0.102210421 | 0.787537098 | 0.917719228 |
| chr16:72241892-72242190 | -0.107713688 | 0.787540532 | 0.917719228 |
| chr5:14581431-14581955 | 0.090728763 | 0.787550705 | 0.917719228 |
| chr10:100228707-100229771 | -0.088876856 | 0.787592888 | 0.917754426 |
| chr2:55010158-55010404 | -0.123274557 | 0.787624115 | 0.917772677 |
| chr3:179105128-179105328 | -0.133414136 | 0.787632507 | 0.917772677 |
| chr20:22728218-22728433 | 0.141446482 | 0.787657696 | 0.917788071 |
| chr21:33299477-33299677 | -0.140044772 | 0.787705959 | 0.917815099 |
| chr8:123220669-123220997 | -0.099120337 | 0.787715404 | 0.917815099 |
| chr9:127942826-127943078 | 0.110596086 | 0.787723922 | 0.917815099 |
| chr5:140351667-140351985 | 0.119576236 | 0.787739279 | 0.917815099 |
| chr2:83213985-83214206 | 0.144093988 | 0.787740784 | 0.917815099 |
| chr9:92133931-92134510 | 0.092902752 | 0.787794276 | 0.917863467 |
| chr6:167825738-167826054 | 0.101989384 | 0.787811118 | 0.917867219 |
| chr4:109729416-109730763 | 0.085477991 | 0.787821455 | 0.917867219 |
| chr21:29434099-29434299 | 0.140016322 | 0.787837194 | 0.917867754 |
| chr16:87316867-87317983 | 0.08647274 | 0.787850655 | 0.917867754 |
| chr7:34823743-34823971 | 0.117168472 | 0.787857852 | 0.917867754 |
| chr5:71011251-71011484 | 0.134064234 | 0.787875979 | 0.917874917 |
| chr16:23297033-23297271 | 0.122504376 | 0.787893478 | 0.917878796 |
| chr4:86451603-86451823 | 0.125135617 | 0.787914719 | 0.917878796 |
| chr9:134442556-134443189 | -0.094541212 | 0.787915247 | 0.917878796 |
| chr10:78657613-78657899 | 0.108714546 | 0.787955849 | 0.917912139 |
| chr3:15630708-15630919 | 0.113927863 | 0.787981438 | 0.917913503 |
| chr6:149922682-149923330 | 0.090714671 | 0.787994805 | 0.917913503 |
| chr6:43174285-43174894 | 0.088501895 | 0.788004612 | 0.917913503 |
| chr1:65209394-65209647 | -0.121665636 | 0.788023476 | 0.917913503 |
| chr3:171646431-171646631 | 0.133751896 | 0.788025832 | 0.917913503 |
| chr4:56978679-56979219 | -0.089859484 | 0.788028898 | 0.917913503 |
| chr1:161225528-161226244 | -0.086648799 | 0.78806887 | 0.917930035 |
| chr16:89170569-89170810 | -0.101725422 | 0.788069717 | 0.917930035 |
| chr3:105367141-105367414 | -0.098082746 | 0.788086784 | 0.917930035 |
| chr16:49491680-49492123 | -0.111446985 | 0.788094971 | 0.917930035 |
| chr19:50203238-50203749 | 0.091776142 | 0.788114317 | 0.917930035 |
| chr12:130748289-130748514 | 0.122821221 | 0.788123506 | 0.917930035 |
| chr3:54146666-54146903 | -0.136173039 | 0.788136699 | 0.917930035 |
| chr8:63137677-63137972 | 0.13864779 | 0.78813893 | 0.917930035 |
| chr16:67833575-67834111 | 0.094822935 | 0.788164081 | 0.917935891 |
| chrX:4675970-4676170 | -0.146319043 | 0.788178755 | 0.917935891 |
| chr8:56241978-56242270 | -0.106418514 | 0.788179899 | 0.917935891 |
| chr11:64723108-64723394 | -0.106252762 | 0.788205967 | 0.917944316 |
| chr4:139557136-139557474 | -0.096071403 | 0.788211094 | 0.917944316 |
| chr7:46081753-46081990 | -0.133825676 | 0.788226879 | 0.917948748 |
| chr6:116570842-116571852 | 0.084929889 | 0.788289277 | 0.918007463 |
| chr11:124739531-124740251 | 0.090821602 | 0.788308443 | 0.918015829 |
| chrX:69538063-69538610 | -0.09867508 | 0.788353128 | 0.918041416 |
| chr5:12175987-12176211 | 0.126836044 | 0.788357109 | 0.918041416 |

| | | | |
|---|---|---|---|
| chr14:58244314-58245373 | -0.0846148 | 0.788383558 | 0.918041416 |
| chr1:225777254-225778417 | 0.086008621 | 0.788390074 | 0.918041416 |
| chr1:75134917-75135117 | -0.142565422 | 0.788390321 | 0.918041416 |
| chr3:53110272-53110511 | -0.132826513 | 0.788429819 | 0.918066436 |
| chr4:38782819-38783165 | 0.107880094 | 0.788456612 | 0.918066436 |
| chr3:136195742-136197216 | -0.083808041 | 0.788466164 | 0.918066436 |
| chr2:174667246-174667657 | -0.111765174 | 0.788471033 | 0.918066436 |
| chrX:38219963-38220317 | -0.108138703 | 0.78848408 | 0.918066436 |
| chr2:24060959-24061358 | -0.109272311 | 0.788493477 | 0.918066436 |
| chr11:113875058-113876103 | 0.088983302 | 0.788495681 | 0.918066436 |
| chr14:36628061-36628342 | -0.129273352 | 0.78852025 | 0.918081092 |
| chr7:31744320-31744585 | -0.107923841 | 0.788540672 | 0.918090919 |
| chr6:36027444-36028583 | -0.087319984 | 0.788558945 | 0.918098243 |
| chr9:123081828-123082116 | 0.123060123 | 0.788571506 | 0.918098917 |
| chr10:18312827-18313083 | 0.119832553 | 0.788656066 | 0.918183414 |
| chr14:69191275-69192186 | 0.086642378 | 0.788762213 | 0.918293043 |
| chr1:162159494-162160182 | -0.091304838 | 0.788822297 | 0.918349039 |
| chr2:182763378-182763625 | -0.121693419 | 0.788860752 | 0.918366101 |
| chr9:109880510-109880837 | -0.114406977 | 0.788860923 | 0.918366101 |
| chr15:48331220-48332468 | -0.085608423 | 0.788921379 | 0.918422527 |
| chr5:25947204-25947431 | 0.137233765 | 0.788945363 | 0.918436494 |
| chr17:78040841-78041444 | -0.096555586 | 0.788960744 | 0.918440446 |
| chrX:19739729-19739993 | -0.123094778 | 0.788987929 | 0.918455903 |
| chr18:5295673-5297085 | -0.08496623 | 0.788997996 | 0.918455903 |
| chr4:76563377-76563994 | -0.096833019 | 0.789015111 | 0.918461872 |
| chr12:98502760-98503105 | -0.095149349 | 0.789037418 | 0.918473886 |
| chr1:82163051-82163381 | 0.127412762 | 0.789057171 | 0.918482925 |
| chr17:68456696-68458196 | -0.08351355 | 0.78907823 | 0.918493485 |
| chr14:75428203-75428750 | 0.09837275 | 0.789095216 | 0.91849585 |
| chr19:54189339-54191356 | 0.083205484 | 0.789113143 | 0.91849585 |
| chr19:42268139-42269342 | 0.083993191 | 0.7891226 | 0.91849585 |
| chr12:48971304-48971962 | 0.094634469 | 0.789130462 | 0.91849585 |
| chr6:104019226-104019640 | 0.120844181 | 0.789140199 | 0.91849585 |
| chr1:31195168-31195460 | 0.135922949 | 0.789170942 | 0.91851768 |
| chr20:32358831-32359061 | 0.11019621 | 0.789229957 | 0.91856441 |
| chr10:79467492-79467741 | 0.112001062 | 0.789252092 | 0.91856441 |
| chr11:32809287-32809712 | 0.109096896 | 0.789265813 | 0.91856441 |
| chr9:137564504-137564704 | -0.145354616 | 0.789267884 | 0.91856441 |
| chr14:24241941-24243278 | 0.084102174 | 0.789271032 | 0.91856441 |
| chr6:138852588-138852813 | -0.140240203 | 0.789300468 | 0.918572473 |
| chr20:25065532-25065774 | -0.13006994 | 0.789313143 | 0.918572473 |
| chr11:20031756-20032297 | 0.093953355 | 0.789333837 | 0.918572473 |
| chr17:58254816-58255175 | -0.095511141 | 0.789337263 | 0.918572473 |
| chr16:2719295-2719817 | 0.098594772 | 0.789337902 | 0.918572473 |
| chr6:18425289-18425576 | 0.124087565 | 0.789360994 | 0.918585393 |
| chr4:118351865-118352557 | -0.092964519 | 0.789374427 | 0.918587074 |
| chr13:100088582-100089375 | -0.088834151 | 0.789456409 | 0.918638465 |
| chr6:27605519-27605873 | 0.094984451 | 0.789465881 | 0.918638465 |
| chr5:118548897-118549108 | -0.141829528 | 0.789469732 | 0.918638465 |
| chr1:203172687-203172999 | -0.099312592 | 0.789477156 | 0.918638465 |

| | | | |
|---|---|---|---|
| chr6:52517678-52517878 | 0.137581961 | 0.789486101 | 0.918638465 |
| chr14:95525023-95525281 | 0.117535524 | 0.789490524 | 0.918638465 |
| chr6:98835264-98835509 | -0.111194022 | 0.789515535 | 0.918653617 |
| chr21:42145558-42145867 | -0.10314932 | 0.789588307 | 0.91872434 |
| chr4:141134117-141134331 | 0.124920498 | 0.789620538 | 0.918744265 |
| chr3:196739584-196740732 | 0.086387083 | 0.789629413 | 0.918744265 |
| chr16:83701597-83701891 | 0.110937216 | 0.789646372 | 0.918750046 |
| chr17:76736935-76738315 | 0.084090549 | 0.789660367 | 0.918752378 |
| chr8:58234518-58234742 | -0.132804393 | 0.789695468 | 0.918779267 |
| chr3:63475054-63475295 | -0.107875087 | 0.789725685 | 0.918800472 |
| chr1:84772432-84772681 | 0.113400284 | 0.789792562 | 0.918838551 |
| chr14:22903375-22903616 | 0.119512523 | 0.789813373 | 0.918838551 |
| chr1:223707235-223708039 | -0.092815467 | 0.789814055 | 0.918838551 |
| chr5:101995604-101995937 | -0.105578632 | 0.789816305 | 0.918838551 |
| chr22:26658632-26659221 | 0.09556425 | 0.789818375 | 0.918838551 |
| chr14:81269428-81269715 | 0.104521277 | 0.789845691 | 0.918856379 |
| chr1:114884639-114884922 | -0.105037521 | 0.789896951 | 0.91890206 |
| chr1:191297234-191297506 | -0.126600861 | 0.789920257 | 0.918915221 |
| chr8:124199980-124200761 | -0.087069051 | 0.789936478 | 0.918920139 |
| chr5:94086699-94086931 | 0.122263471 | 0.789976617 | 0.918932055 |
| chr10:78424798-78425016 | -0.123760729 | 0.78997911 | 0.918932055 |
| chr6:64778158-64778592 | -0.103585884 | 0.7899827 | 0.918932055 |
| chr1:205577008-205577256 | 0.106040683 | 0.790022856 | 0.918964814 |
| chr14:105528140-105528740 | 0.091990504 | 0.79004138 | 0.918972411 |
| chr21:44618621-44618845 | 0.119052862 | 0.790055837 | 0.918975276 |
| chr2:26758026-26758373 | -0.103009199 | 0.790169639 | 0.91908213 |
| chr1:10377449-10377706 | -0.126414685 | 0.790171691 | 0.91908213 |
| chrX:119015484-119015684 | 0.140194517 | 0.790195807 | 0.919096228 |
| chr6:149431928-149432357 | -0.095193535 | 0.790223018 | 0.919111313 |
| chr19:42450392-42450771 | -0.103333077 | 0.790245148 | 0.919111313 |
| chr5:10029861-10030061 | 0.126940733 | 0.790254801 | 0.919111313 |
| chr1:234723622-234725059 | 0.084019747 | 0.790258985 | 0.919111313 |
| chr5:134577809-134578042 | 0.111698774 | 0.790268754 | 0.919111313 |
| chr22:26574566-26574960 | 0.117648434 | 0.790281907 | 0.919112661 |
| chr15:75335781-75336399 | 0.092386365 | 0.790319081 | 0.919128065 |
| chr15:85543049-85543354 | -0.113897208 | 0.790319144 | 0.919128065 |
| chr12:66049052-66049409 | -0.112455057 | 0.790339806 | 0.919138144 |
| chr13:68745474-68745799 | -0.115786733 | 0.790413724 | 0.919210157 |
| chr15:40564528-40565344 | -0.086419395 | 0.790442962 | 0.919222487 |
| chr12:68487670-68487870 | 0.142428038 | 0.790448321 | 0.919222487 |
| chr20:2351881-2352153 | -0.120189296 | 0.79051219 | 0.919274667 |
| chrX:103310517-103311356 | -0.089131347 | 0.79052477 | 0.919274667 |
| chr7:98836915-98837178 | -0.115002762 | 0.790529183 | 0.919274667 |
| chr13:46143146-46143624 | -0.098945061 | 0.790544809 | 0.919278886 |
| chr12:162560-162778 | -0.140844652 | 0.790581667 | 0.919307794 |
| chr3:156004197-156004449 | -0.127926057 | 0.790616351 | 0.919334173 |
| chr6:108518983-108519368 | -0.107374193 | 0.790661165 | 0.919372331 |
| chrX:119181495-119181789 | 0.113215852 | 0.790700878 | 0.919386238 |
| chr6:73675525-73675797 | -0.113763953 | 0.790704348 | 0.919386238 |
| chr7:36389352-36390588 | -0.083275029 | 0.790709122 | 0.919386238 |

| | | | |
|---|---|---|---|
| chr6:24695602-24695931 | 0.106464488 | 0.790756255 | 0.919406771 |
| chr3:10653561-10653907 | -0.123375475 | 0.790758116 | 0.919406771 |
| chr17:78290680-78290890 | -0.133421592 | 0.790784324 | 0.919406771 |
| chr12:120228389-120229251 | 0.08649899 | 0.790789997 | 0.919406771 |
| chr12:103401529-103401750 | 0.125300286 | 0.790792387 | 0.919406771 |
| chr4:126627372-126627572 | 0.133489917 | 0.790798777 | 0.919406771 |
| chr5:152159995-152160326 | 0.110488833 | 0.790825006 | 0.919423315 |
| chr6:27145957-27146210 | 0.104300325 | 0.790872313 | 0.919434001 |
| chr14:106564135-106564530 | -0.111968724 | 0.790872953 | 0.919434001 |
| chr6:35297556-35298099 | 0.09305216 | 0.790881234 | 0.919434001 |
| chr2:191329750-191330290 | 0.090207237 | 0.790897183 | 0.919434001 |
| chr17:16380539-16381465 | 0.086514464 | 0.790899537 | 0.919434001 |
| chr2:218671432-218672565 | 0.086488 | 0.790912606 | 0.919434001 |
| chr9:135076926-135077151 | -0.129570217 | 0.79093012 | 0.919434001 |
| chr2:219684791-219685747 | 0.086830806 | 0.790930194 | 0.919434001 |
| chr6:89081293-89082143 | 0.086009051 | 0.790979815 | 0.919471092 |
| chr10:84054811-84055025 | 0.124146398 | 0.790986101 | 0.919471092 |
| chr3:16366653-16366925 | -0.107381954 | 0.791004536 | 0.919478573 |
| chr7:74657144-74658024 | -0.087975222 | 0.791023284 | 0.919486416 |
| chr16:81006336-81007859 | -0.082875846 | 0.791071732 | 0.919528782 |
| chr19:54096592-54096957 | 0.108575638 | 0.791110694 | 0.919540686 |
| chr17:68205063-68206395 | 0.08380658 | 0.791116576 | 0.919540686 |
| chr5:35202402-35202802 | -0.098087271 | 0.791117976 | 0.919540686 |
| chr9:134456517-134456765 | 0.12320323 | 0.791142516 | 0.919555261 |
| chr6:142698855-142699058 | 0.127551013 | 0.791223226 | 0.919635121 |
| chr5:60487943-60488437 | 0.095118899 | 0.791264572 | 0.919669226 |
| chr15:59893068-59893268 | -0.145253784 | 0.791321629 | 0.919684393 |
| chr3:83523538-83523834 | -0.108180513 | 0.791328111 | 0.919684393 |
| chrX:107628751-107629166 | -0.094637422 | 0.791329747 | 0.919684393 |
| chr6:42547430-42548059 | 0.094662644 | 0.791332716 | 0.919684393 |
| chr11:118241513-118241999 | 0.094203465 | 0.791337636 | 0.919684393 |
| chr8:60678227-60678779 | 0.092357695 | 0.791363753 | 0.919688426 |
| chr1:6246728-6247060 | 0.102548892 | 0.791365112 | 0.919688426 |
| chr15:40941851-40942096 | 0.110305355 | 0.791391861 | 0.919690621 |
| chr19:37932370-37932664 | -0.107738733 | 0.791407138 | 0.919690621 |
| chr6:17167370-17167916 | -0.097109486 | 0.791427887 | 0.919690621 |
| chr11:117703328-117703528 | 0.119939137 | 0.791445625 | 0.919690621 |
| chr7:100479049-100479584 | 0.08951908 | 0.791447874 | 0.919690621 |
| chr12:66237583-66238115 | 0.093822744 | 0.791459678 | 0.919690621 |
| chr7:157009509-157009856 | 0.104225628 | 0.791466534 | 0.919690621 |
| chr10:42637839-42638921 | -0.08394335 | 0.791468663 | 0.919690621 |
| chr5:143003053-143003335 | 0.116882736 | 0.791475028 | 0.919690621 |
| chr11:123144776-123144976 | 0.138927957 | 0.791523591 | 0.919725145 |
| chr3:60742695-60743157 | -0.107521704 | 0.791534019 | 0.919725145 |
| chr1:155910585-155911621 | 0.087208622 | 0.791568112 | 0.919725145 |
| chr14:36683103-36683681 | 0.096148564 | 0.791568602 | 0.919725145 |
| chr9:131383301-131383550 | -0.120460086 | 0.791570405 | 0.919725145 |
| chr3:107271800-107272172 | 0.108464822 | 0.791576759 | 0.919725145 |
| chr2:192940890-192941092 | -0.134928724 | 0.791620267 | 0.919761749 |
| chr14:74628271-74628507 | -0.12844626 | 0.791676752 | 0.9198015 |

| | | | |
|---|---|---|---|
| chr5:181115223-181115451 | -0.115688414 | 0.791678488 | 0.9198015 |
| chr10:48333469-48334233 | -0.093558583 | 0.791702009 | 0.919807307 |
| chr3:169907986-169908213 | -0.121743399 | 0.791718963 | 0.919807307 |
| chr6:6601026-6601226 | 0.131180994 | 0.7917195 | 0.919807307 |
| chr4:40719075-40719329 | -0.11173075 | 0.791734998 | 0.91980949 |
| chr1:154955710-154956562 | 0.087706722 | 0.791745388 | 0.91980949 |
| chr3:107064452-107064803 | -0.097703219 | 0.791770372 | 0.919824569 |
| chr8:106333837-106334037 | 0.151869588 | 0.791802217 | 0.919847617 |
| chr19:50556383-50557020 | 0.093884331 | 0.791837958 | 0.919875192 |
| chr5:10220518-10220762 | 0.11935317 | 0.791895516 | 0.919928109 |
| chr3:166510477-166510700 | 0.125244839 | 0.791913767 | 0.919935364 |
| chr19:13118017-13118217 | 0.130407165 | 0.791978801 | 0.919983323 |
| chr3:61706435-61706657 | -0.103834306 | 0.791987549 | 0.919983323 |
| chr17:44865237-44865487 | 0.125196138 | 0.791995881 | 0.919983323 |
| chr3:159787193-159787523 | 0.104013095 | 0.792003188 | 0.919983323 |
| chr1:12143747-12143991 | 0.12692419 | 0.792017937 | 0.919983323 |
| chr17:82698466-82699463 | -0.08463755 | 0.792027093 | 0.919983323 |
| chr15:84631000-84631942 | 0.087927438 | 0.792063158 | 0.919995487 |
| chrX:14872903-14873844 | -0.083931463 | 0.79207251 | 0.919995487 |
| chr12:78617335-78617551 | 0.141611292 | 0.792073586 | 0.919995487 |
| chr19:2096246-2096999 | 0.086940354 | 0.79209103 | 0.920001802 |
| chr2:31151635-31151857 | 0.129145053 | 0.792120713 | 0.920022333 |
| chr14:23078823-23079049 | -0.120628084 | 0.792184688 | 0.920082691 |
| chr1:221715341-221715832 | -0.098469237 | 0.792207908 | 0.920089816 |
| chr3:125211817-125212343 | -0.096905675 | 0.792221209 | 0.920089816 |
| chrX:54044128-54045266 | 0.084537925 | 0.792230813 | 0.920089816 |
| chr1:177945950-177946274 | -0.096450325 | 0.792246839 | 0.920089816 |
| chr18:72935833-72936077 | -0.133355719 | 0.792250864 | 0.920089816 |
| chr14:20801816-20802034 | 0.123277499 | 0.792293071 | 0.920113472 |
| chr5:35924936-35925328 | -0.098412859 | 0.792295251 | 0.920113472 |
| chr7:87220982-87221225 | -0.107279557 | 0.792313362 | 0.92012047 |
| chr1:116418228-116418989 | 0.08609978 | 0.792325293 | 0.92012047 |
| chr17:47843118-47843352 | 0.102590136 | 0.792339637 | 0.920123182 |
| chr7:149369554-149370072 | 0.09359827 | 0.792361576 | 0.920134714 |
| chr1:234154715-234154999 | 0.113488987 | 0.792408793 | 0.920166374 |
| chr2:104046967-104047251 | 0.117780145 | 0.79241843 | 0.920166374 |
| chr11:69173705-69173956 | 0.128553035 | 0.792450083 | 0.920166374 |
| chr8:129983938-129984658 | -0.087356888 | 0.792451759 | 0.920166374 |
| chr1:43979856-43980433 | 0.096579272 | 0.792453474 | 0.920166374 |
| chr11:118892516-118892738 | -0.117833449 | 0.792471661 | 0.920166374 |
| chr15:40104812-40105095 | 0.096184001 | 0.792478153 | 0.920166374 |
| chr2:232696950-232698044 | -0.085561677 | 0.792487881 | 0.920166374 |
| chr21:40660605-40660845 | -0.123872023 | 0.792496922 | 0.920166374 |
| chr6:68572493-68572791 | -0.113263883 | 0.792527652 | 0.92018811 |
| chr3:72847706-72848982 | 0.085047852 | 0.79254862 | 0.920198511 |
| chr3:138462962-138463195 | -0.112358605 | 0.792574881 | 0.920203009 |
| chr17:15999086-16000362 | -0.084195796 | 0.792576513 | 0.920203009 |
| chr1:111334267-111334467 | -0.140572694 | 0.792598981 | 0.920215151 |
| chr2:211055459-211055740 | 0.119795696 | 0.792623245 | 0.920225189 |
| chr12:115471043-115471291 | -0.133246052 | 0.792644996 | 0.920225189 |

| | | | |
|---|---|---|---|
| chr10:42885805-42886026 | -0.136533926 | 0.792665167 | 0.920225189 |
| chr18:70433849-70434257 | -0.097668514 | 0.79266639 | 0.920225189 |
| chr6:159104098-159104462 | -0.100787121 | 0.792667676 | 0.920225189 |
| chr11:61659827-61660094 | 0.126404575 | 0.792716248 | 0.920233678 |
| chr12:67377051-67377299 | 0.126404575 | 0.792716248 | 0.920233678 |
| chr17:18563519-18564020 | 0.102618824 | 0.792728698 | 0.920233678 |
| chr11:65524414-65524641 | 0.110848986 | 0.792729897 | 0.920233678 |
| chr12:62292083-62292314 | 0.122559333 | 0.792757588 | 0.920233678 |
| chrX:17356067-17356292 | 0.133899211 | 0.792763174 | 0.920233678 |
| chr1:211327311-211327980 | -0.090364294 | 0.792775934 | 0.920233678 |
| chr11:84398113-84398331 | 0.12163953 | 0.792776989 | 0.920233678 |
| chr4:163172429-163172763 | -0.123796366 | 0.792798374 | 0.920233678 |
| chr6:36340976-36341208 | -0.128643173 | 0.792803086 | 0.920233678 |
| chr7:21345045-21345270 | 0.122883359 | 0.792807099 | 0.920233678 |
| chr13:110914951-110915677 | -0.084204854 | 0.792824862 | 0.920240355 |
| chr15:64387505-64388106 | 0.087414849 | 0.792842597 | 0.920241483 |
| chr22:38353293-38353493 | -0.138390933 | 0.792849853 | 0.920241483 |
| chr19:55447807-55448106 | -0.107341428 | 0.792882282 | 0.920265182 |
| chr2:169733439-169734414 | 0.08951253 | 0.792905225 | 0.920277871 |
| chr3:24185839-24186439 | 0.092076682 | 0.792919981 | 0.920281057 |
| chr6:26659306-26660380 | -0.086747295 | 0.792966138 | 0.920304981 |
| chr17:7572329-7573209 | -0.08551462 | 0.792967517 | 0.920304981 |
| chrX:101407256-101407456 | -0.107116318 | 0.792987611 | 0.920304981 |
| chr16:85946819-85947076 | -0.115644974 | 0.792988639 | 0.920304981 |
| chr1:216119676-216119962 | -0.105218866 | 0.79303593 | 0.920345925 |
| chr11:86214802-86215124 | -0.106434175 | 0.793076804 | 0.92037942 |
| chr7:150777851-150778051 | -0.141363017 | 0.793090957 | 0.920381905 |
| chr10:99731797-99733009 | -0.083390661 | 0.793150308 | 0.920436841 |
| chr3:161293477-161293705 | 0.123142824 | 0.793235565 | 0.92050238 |
| chr11:22337584-22337951 | -0.101659083 | 0.793258673 | 0.92050238 |
| chr11:134045564-134046022 | -0.101659083 | 0.793258673 | 0.92050238 |
| chr3:18840548-18840822 | -0.108004532 | 0.793293096 | 0.92050238 |
| chr1:89085879-89086089 | -0.144693684 | 0.793296986 | 0.92050238 |
| chr17:60142543-60143444 | -0.082156853 | 0.793312392 | 0.92050238 |
| chr5:81301281-81302008 | -0.085887996 | 0.793320017 | 0.92050238 |
| chr4:115054906-115055181 | -0.113157857 | 0.793320082 | 0.92050238 |
| chr2:181613214-181613613 | -0.110123253 | 0.793322567 | 0.92050238 |
| chr17:74852468-74852767 | 0.108639668 | 0.793340887 | 0.92050238 |
| chr2:237858359-237858585 | 0.130231826 | 0.793345693 | 0.92050238 |
| chr19:58310828-58311028 | 0.136802152 | 0.793350946 | 0.92050238 |
| chr3:109098265-109098662 | -0.09754001 | 0.79336338 | 0.920502869 |
| chr4:14872126-14872440 | -0.111086319 | 0.793410647 | 0.92054377 |
| chr2:27322311-27323451 | -0.084847074 | 0.793425753 | 0.920547358 |
| chr3:164451123-164451323 | -0.131858021 | 0.793454078 | 0.920554022 |
| chr5:11962155-11962355 | 0.136346548 | 0.793464576 | 0.920554022 |
| chr10:119596318-119597421 | 0.08309987 | 0.793473337 | 0.920554022 |
| chr2:149204870-149205070 | 0.15913771 | 0.793479554 | 0.920554022 |
| chr14:21613178-21613378 | -0.143410365 | 0.79349453 | 0.920555383 |
| chr1:235503877-235505289 | 0.082046237 | 0.793504755 | 0.920555383 |
| chr10:70903379-70903636 | 0.106294486 | 0.793521731 | 0.920561139 |

| | | | |
|---|---|---|---|
| chr3:38453065-38453974 | 0.085785621 | 0.793551516 | 0.920578374 |
| chr8:66032144-66032651 | -0.094846061 | 0.793596582 | 0.920578374 |
| chr4:105473062-105474498 | 0.082776761 | 0.793609071 | 0.920578374 |
| chr2:43961964-43962348 | -0.11961694 | 0.793617038 | 0.920578374 |
| chr8:15378349-15378549 | -0.132435502 | 0.793621384 | 0.920578374 |
| chr20:44370726-44370932 | -0.133260272 | 0.793632064 | 0.920578374 |
| chr15:93030665-93031004 | -0.093136796 | 0.793639076 | 0.920578374 |
| chr7:6927360-6927560 | -0.138746197 | 0.793656938 | 0.920578374 |
| chr6:33577079-33577423 | -0.103926328 | 0.79366085 | 0.920578374 |
| chr17:49567839-49568559 | -0.087306783 | 0.79366381 | 0.920578374 |
| chr2:176106704-176106931 | -0.11675358 | 0.793677233 | 0.920578374 |
| chr7:1177052-1177316 | -0.10239587 | 0.793702983 | 0.920578374 |
| chr4:4327672-4327872 | 0.131451812 | 0.793704891 | 0.920578374 |
| chr20:51735009-51735231 | -0.104548845 | 0.793712576 | 0.920578374 |
| chrX:131747143-131747530 | -0.107995725 | 0.793716806 | 0.920578374 |
| chr12:51951143-51951740 | 0.090029629 | 0.79375938 | 0.920613817 |
| chr1:235601182-235601573 | 0.101535577 | 0.793786354 | 0.920631167 |
| chr8:121531550-121531750 | 0.128612478 | 0.793831279 | 0.920669334 |
| chr10:89643793-89645780 | -0.081979696 | 0.793870168 | 0.920676335 |
| chr2:220076799-220077182 | 0.108975497 | 0.793887508 | 0.920676335 |
| chr10:97759303-97759737 | 0.095424674 | 0.793913122 | 0.920676335 |
| chr18:13983516-13983857 | -0.112894673 | 0.793924677 | 0.920676335 |
| chr1:11967200-11967438 | 0.127030743 | 0.793925979 | 0.920676335 |
| chr22:38506181-38506948 | 0.086075795 | 0.793929091 | 0.920676335 |
| chr9:113580491-113581102 | -0.087453197 | 0.793934298 | 0.920676335 |
| chr8:78972766-78973078 | 0.105018704 | 0.793973654 | 0.920676335 |
| chr4:77157427-77158441 | -0.085435568 | 0.793975057 | 0.920676335 |
| chr7:73005461-73006389 | -0.086610952 | 0.79397727 | 0.920676335 |
| chr14:68941817-68942114 | -0.103067997 | 0.793993475 | 0.920676335 |
| chrX:45851498-45851983 | -0.09946045 | 0.794009547 | 0.920676335 |
| chr6:87796494-87797109 | -0.09187195 | 0.794013522 | 0.920676335 |
| chr3:93841413-93841989 | -0.092763223 | 0.794021482 | 0.920676335 |
| chr9:132259162-132259661 | 0.095250791 | 0.794028195 | 0.920676335 |
| chr7:158055164-158055440 | -0.124208809 | 0.794029568 | 0.920676335 |
| chr19:19063455-19064270 | -0.088525635 | 0.794048828 | 0.920684735 |
| chr1:216560776-216560988 | -0.13286935 | 0.79410157 | 0.920713014 |
| chr14:104108293-104108803 | 0.090728143 | 0.794142607 | 0.920713014 |
| chrX:54182400-54182651 | 0.113690594 | 0.794155054 | 0.920713014 |
| chr1:144418710-144419480 | 0.083992212 | 0.794160075 | 0.920713014 |
| chr13:38264633-38265059 | -0.110899302 | 0.794161222 | 0.920713014 |
| chr8:104418074-104418359 | 0.095094918 | 0.794161346 | 0.920713014 |
| chr9:113548572-113548818 | 0.125052931 | 0.794169312 | 0.920713014 |
| chr3:129625950-129626153 | 0.104166208 | 0.794193093 | 0.920713014 |
| chr17:73547079-73547288 | 0.109152168 | 0.794193298 | 0.920713014 |
| chr20:57495157-57495751 | 0.086516353 | 0.794193381 | 0.920713014 |
| chr12:54993156-54993528 | 0.101549727 | 0.794269312 | 0.920746499 |
| chr13:41919428-41919933 | -0.103404468 | 0.794280921 | 0.920746499 |
| chr12:18383274-18383487 | 0.132092289 | 0.794284318 | 0.920746499 |
| chr9:114505149-114505810 | -0.084999607 | 0.79429076 | 0.920746499 |
| chr12:56151701-56152684 | 0.084458083 | 0.794304033 | 0.920746499 |

| | | | |
|---|---|---|---|
| chr10:14959116-14960214 | -0.082459916 | 0.794304816 | 0.920746499 |
| chr5:142720595-142721267 | 0.091375918 | 0.794306382 | 0.920746499 |
| chr2:46202217-46202417 | 0.13760075 | 0.794350863 | 0.920784131 |
| chr2:190996860-190997185 | 0.116449031 | 0.794366207 | 0.920787987 |
| chr12:55068501-55068810 | -0.105515687 | 0.794450312 | 0.920829879 |
| chr4:120203612-120203812 | 0.142401037 | 0.794460692 | 0.920829879 |
| chr16:1826715-1827554 | 0.086700685 | 0.794486867 | 0.920829879 |
| chr5:125924484-125924710 | -0.126906316 | 0.794504657 | 0.920829879 |
| chr1:197976624-197976895 | -0.121107794 | 0.794522729 | 0.920829879 |
| chr2:127174916-127175232 | 0.101377089 | 0.794523586 | 0.920829879 |
| chr8:38175941-38177544 | 0.083790184 | 0.794545896 | 0.920829879 |
| chr1:41331322-41331541 | -0.1256013 | 0.794547708 | 0.920829879 |
| chr19:48835682-48836062 | 0.100832676 | 0.794549815 | 0.920829879 |
| chr15:78216728-78217091 | -0.101387303 | 0.794550594 | 0.920829879 |
| chr1:59523197-59523448 | -0.12972248 | 0.794557963 | 0.920829879 |
| chr1:85581262-85581504 | 0.102576398 | 0.79456276 | 0.920829879 |
| chr5:16552649-16553047 | -0.093699679 | 0.794565012 | 0.920829879 |
| chr8:119376382-119376743 | 0.117692197 | 0.794570973 | 0.920829879 |
| chr12:101876920-101878328 | -0.08305941 | 0.794591777 | 0.920829879 |
| chr12:106386208-106386493 | -0.121607028 | 0.794594633 | 0.920829879 |
| chr16:83480911-83481179 | 0.123660809 | 0.794621825 | 0.920847464 |
| chr20:54110384-54111216 | 0.082601822 | 0.794637591 | 0.920851807 |
| chrX:20265745-20265975 | 0.111877146 | 0.794698163 | 0.920901351 |
| chr12:96586688-96586969 | 0.141163317 | 0.794708323 | 0.920901351 |
| chr9:25964283-25964509 | -0.120436174 | 0.794733111 | 0.920901351 |
| chr12:12018518-12018718 | -0.145093408 | 0.79475097 | 0.920901351 |
| chr3:111714402-111714602 | -0.120961929 | 0.794780564 | 0.920901351 |
| chr17:17836019-17836728 | -0.084542062 | 0.794786423 | 0.920901351 |
| chr3:122921787-122922489 | 0.093161278 | 0.794787812 | 0.920901351 |
| chr3:44862425-44862751 | -0.117776412 | 0.794790615 | 0.920901351 |
| chr12:107105205-107105613 | 0.113121198 | 0.794809116 | 0.920901351 |
| chr2:1625195-1626019 | -0.085490526 | 0.794812189 | 0.920901351 |
| chr4:79089075-79089275 | -0.145601946 | 0.79481255 | 0.920901351 |
| chr1:202009594-202009874 | 0.09435551 | 0.794860825 | 0.920941952 |
| chr1:179599654-179600011 | 0.098994672 | 0.794871631 | 0.920941952 |
| chr20:10536531-10536829 | -0.122991289 | 0.794889655 | 0.920946806 |
| chr17:64479689-64479889 | 0.128504278 | 0.794904206 | 0.920946806 |
| chr8:107344544-107344876 | -0.126208059 | 0.794911878 | 0.920946806 |
| chr1:62716535-62716735 | -0.145292621 | 0.794936238 | 0.920961103 |
| chr5:35651095-35651514 | 0.107882381 | 0.794970158 | 0.920986475 |
| chr5:161676192-161676440 | -0.118913861 | 0.794997875 | 0.920987959 |
| chr7:152314137-152314411 | 0.128503732 | 0.795017087 | 0.920987959 |
| chr13:41992184-41992445 | -0.13047113 | 0.795025633 | 0.920987959 |
| chr17:38257208-38257597 | 0.100932684 | 0.79502855 | 0.920987959 |
| chr19:48510475-48511346 | -0.087663567 | 0.795036969 | 0.920987959 |
| chr3:194315223-194316196 | 0.08549423 | 0.795043559 | 0.920987959 |
| chr2:10303116-10303344 | -0.102268627 | 0.795075924 | 0.920996421 |
| chr11:46188091-46188402 | 0.108050982 | 0.795088717 | 0.920996421 |
| chr10:110137111-110137412 | -0.114422781 | 0.795112617 | 0.920996421 |
| chr4:158833640-158833914 | 0.133102975 | 0.795116487 | 0.920996421 |

| | | | |
|---|---|---|---|
| chr18:32289576-32289819 | 0.133102975 | 0.795116487 | 0.920996421 |
| chr8:28490106-28490868 | 0.085780325 | 0.795128235 | 0.920996421 |
| chr2:199940468-199940757 | 0.112287839 | 0.795135003 | 0.920996421 |
| chr11:62726997-62728563 | 0.081664715 | 0.795185129 | 0.92102952 |
| chr13:51815535-51816216 | -0.091382302 | 0.795187621 | 0.92102952 |
| chr15:45714541-45714816 | -0.112128903 | 0.795225352 | 0.921035778 |
| chr8:30727194-30728142 | -0.086838834 | 0.795233322 | 0.921035778 |
| chrX:45506871-45507293 | 0.104187792 | 0.795239702 | 0.921035778 |
| chr8:72778902-72779229 | 0.10944372 | 0.795241105 | 0.921035778 |
| chr2:216411978-216413073 | -0.082924448 | 0.795270833 | 0.921048987 |
| chr2:145594461-145594665 | 0.12370938 | 0.795276552 | 0.921048987 |
| chr5:178114494-178114763 | -0.096625385 | 0.795329538 | 0.92109125 |
| chr9:131393121-131393356 | -0.114343614 | 0.795353002 | 0.92109125 |
| chr8:109469068-109469334 | 0.129990078 | 0.795356399 | 0.92109125 |
| chr7:107995428-107995826 | -0.106108213 | 0.795370754 | 0.92109125 |
| chr8:135690227-135690427 | 0.124965811 | 0.795377411 | 0.92109125 |
| chrX:30126854-30127054 | -0.154421038 | 0.795393908 | 0.92109125 |
| chr12:94650541-94650757 | -0.132112871 | 0.795397192 | 0.92109125 |
| chr1:186155715-186155915 | 0.126091184 | 0.795420595 | 0.92110443 |
| chr1:109213476-109214655 | -0.082637055 | 0.795473846 | 0.921152174 |
| chr3:139923464-139923982 | 0.1104862 | 0.795507093 | 0.921169287 |
| chr17:40287500-40288287 | 0.086839746 | 0.795516702 | 0.921169287 |
| chr7:135520731-135521157 | -0.103967337 | 0.795524691 | 0.921169287 |
| chr6:20528803-20529372 | -0.102489273 | 0.795538017 | 0.921170796 |
| chr5:175706525-175706780 | 0.12044242 | 0.795555099 | 0.921176655 |
| chr6:29813047-29813340 | -0.120038921 | 0.795572071 | 0.921182386 |
| chr4:89024161-89024412 | 0.11545034 | 0.795598344 | 0.921198886 |
| chr7:26279484-26279735 | 0.118900995 | 0.795613937 | 0.921200183 |
| chr5:133384341-133384569 | -0.138835942 | 0.795623509 | 0.921200183 |
| chrX:24371426-24371750 | -0.107252865 | 0.795643903 | 0.921205903 |
| chr2:113889393-113890303 | -0.082899586 | 0.795652495 | 0.921205903 |
| chr4:75895745-75896058 | -0.109065542 | 0.795681815 | 0.92122593 |
| chr5:1737931-1738209 | -0.11847603 | 0.795724675 | 0.92124149 |
| chr8:17386833-17387221 | -0.102223737 | 0.795733797 | 0.92124149 |
| chr1:174217720-174217991 | -0.109239606 | 0.795740471 | 0.92124149 |
| chr16:81890404-81890698 | 0.099791623 | 0.795743347 | 0.92124149 |
| chr6:87154866-87156299 | -0.081518675 | 0.795761566 | 0.921248662 |
| chr8:143597626-143597899 | 0.093926068 | 0.795791131 | 0.92126897 |
| chr9:96418111-96419502 | 0.082420936 | 0.795829672 | 0.921289978 |
| chr1:43946238-43947077 | 0.084688264 | 0.795842627 | 0.921289978 |
| chr19:35123446-35123867 | 0.102740403 | 0.795854493 | 0.921289978 |
| chr6:11093286-11094394 | -0.083043471 | 0.795857373 | 0.921289978 |
| chr2:113136992-113137231 | -0.117477949 | 0.795888452 | 0.921312036 |
| chr15:101118580-101118793 | -0.123271034 | 0.795913265 | 0.921318595 |
| chr12:56157623-56158508 | 0.083181001 | 0.795918166 | 0.921318595 |
| chr10:86770437-86770751 | -0.102234982 | 0.795936985 | 0.921326459 |
| chr11:31107055-31107365 | -0.108842703 | 0.795969631 | 0.921345297 |
| chr12:113030678-113031222 | -0.09646457 | 0.795977308 | 0.921345297 |
| chr5:112605109-112605337 | -0.123535972 | 0.796033194 | 0.921396066 |
| chr4:64625264-64625707 | -0.110710004 | 0.796097019 | 0.921456023 |

| | | | |
|---|---|---|---|
| chr6:27374477-27375448 | -0.086133876 | 0.796123178 | 0.921472381 |
| chr6:28355757-28356079 | -0.10626138 | 0.796148732 | 0.921475445 |
| chr6:115643808-115644106 | -0.110966185 | 0.796149878 | 0.921475445 |
| chr17:57860915-57861807 | -0.084513489 | 0.796185061 | 0.921494349 |
| chr10:2839155-2839771 | -0.094895689 | 0.796199655 | 0.921494349 |
| chr1:6784468-6785566 | 0.082474556 | 0.796225683 | 0.921494349 |
| chr14:90004069-90004269 | -0.130897639 | 0.796235853 | 0.921494349 |
| chr14:23588968-23589433 | 0.09123877 | 0.796237581 | 0.921494349 |
| chr5:112227837-112228041 | 0.128369101 | 0.796246766 | 0.921494349 |
| chr1:183214444-183214778 | 0.111762182 | 0.79628073 | 0.921494349 |
| chr14:58941188-58941509 | -0.121462541 | 0.796281932 | 0.921494349 |
| chr6:26305159-26305656 | -0.092790095 | 0.796306824 | 0.921494349 |
| chr22:41003106-41003414 | 0.11432229 | 0.796312849 | 0.921494349 |
| chr5:116502612-116502812 | -0.126441833 | 0.796314092 | 0.921494349 |
| chr9:34540965-34541165 | -0.13520162 | 0.796321273 | 0.921494349 |
| chr16:69761990-69762924 | 0.087644839 | 0.796322556 | 0.921494349 |
| chr3:137013868-137014576 | 0.088711824 | 0.796357592 | 0.921506315 |
| chr9:97632833-97634109 | 0.081683324 | 0.796366264 | 0.921506315 |
| chr3:166052535-166052746 | 0.125985248 | 0.796368976 | 0.921506315 |
| chr14:76961749-76962285 | -0.088197566 | 0.796422786 | 0.921552021 |
| chr12:109999010-109999524 | -0.087842856 | 0.796443951 | 0.921552021 |
| chr6:26234717-26235152 | 0.093125788 | 0.796464008 | 0.921552021 |
| chr8:107632167-107632445 | 0.120865369 | 0.796474327 | 0.921552021 |
| chr8:124371620-124373362 | 0.080397034 | 0.796475956 | 0.921552021 |
| chr17:40936686-40936980 | -0.13170031 | 0.796484605 | 0.921552021 |
| chr1:109996483-109996824 | -0.116315889 | 0.79649888 | 0.921552021 |
| chr10:97599108-97599360 | 0.123437918 | 0.796504917 | 0.921552021 |
| chr3:127690924-127691232 | 0.09683753 | 0.796536535 | 0.921552021 |
| chr22:22559040-22559440 | -0.100261654 | 0.796538135 | 0.921552021 |
| chr2:96260389-96261056 | -0.087439729 | 0.796540775 | 0.921552021 |
| chr2:237085247-237085447 | -0.130705563 | 0.796585891 | 0.921578579 |
| chr2:36663946-36664450 | 0.096929637 | 0.796592884 | 0.921578579 |
| chr17:4639333-4639554 | 0.135287792 | 0.796611518 | 0.921578579 |
| chr11:57711591-57712824 | 0.081282985 | 0.796620967 | 0.921578579 |
| chrX:144219024-144219501 | 0.099067644 | 0.796626919 | 0.921578579 |
| chr14:99341194-99341440 | 0.135742024 | 0.796640135 | 0.921578579 |
| chr4:105632710-105632910 | -0.127616603 | 0.796652595 | 0.921578579 |
| chr1:1324548-1325161 | -0.089405678 | 0.796660641 | 0.921578579 |
| chr2:150387954-150388466 | -0.105808516 | 0.796671979 | 0.921578579 |
| chr14:64559135-64559335 | 0.130724547 | 0.796687162 | 0.921582229 |
| chr15:59492939-59493259 | 0.098056518 | 0.796730044 | 0.921617921 |
| chr3:150748238-150748551 | -0.101533378 | 0.796772517 | 0.921653137 |
| chr17:49968265-49968887 | 0.086266515 | 0.796795376 | 0.921654221 |
| chr12:8949282-8950312 | -0.082316675 | 0.796797512 | 0.921654221 |
| chr2:141911705-141912038 | -0.101361428 | 0.796826373 | 0.921673691 |
| chr11:13668008-13669491 | 0.081269326 | 0.796844889 | 0.921681195 |
| chr13:35283925-35284156 | 0.12219471 | 0.796877357 | 0.921702847 |
| chr20:2743119-2743319 | 0.123921556 | 0.796887667 | 0.921702847 |
| chr6:111523656-111523858 | -0.141310175 | 0.796950221 | 0.921761285 |
| chr19:41226475-41226705 | -0.124665391 | 0.797023807 | 0.92181951 |

| | | | |
|---|---|---|---|
| chr16:74296221-74297203 | -0.082697804 | 0.797024624 | 0.92181951 |
| chr8:6928690-6928914 | 0.122063107 | 0.797079911 | 0.921869539 |
| chr3:128787382-128787603 | 0.11611313 | 0.797121963 | 0.921904259 |
| chr19:44927062-44927433 | 0.096406681 | 0.797153169 | 0.92192258 |
| chr19:13794993-13795495 | -0.089872264 | 0.797161869 | 0.92192258 |
| chr11:67312260-67312460 | 0.110640646 | 0.797178005 | 0.921926778 |
| chr5:126680095-126680716 | -0.089689262 | 0.797190359 | 0.921926778 |
| chr3:168211371-168211744 | -0.096340076 | 0.797201595 | 0.921926778 |
| chr6:31660668-31660946 | 0.100315715 | 0.797228258 | 0.921943697 |
| chr19:46728784-46729035 | -0.108447393 | 0.797241942 | 0.921945607 |
| chr16:68085126-68085508 | -0.094266429 | 0.797274128 | 0.921956464 |
| chr14:61380969-61381351 | 0.094429691 | 0.797285976 | 0.921956464 |
| chr16:70065060-70065309 | 0.11108652 | 0.797287427 | 0.921956464 |
| chr11:1572176-1572468 | 0.102423425 | 0.797310835 | 0.92195911 |
| chr7:151262451-151262688 | 0.121149217 | 0.797326638 | 0.92195911 |
| chr6:22362504-22362877 | -0.125841491 | 0.797349458 | 0.92195911 |
| chr17:63826440-63826688 | 0.094324265 | 0.797349724 | 0.92195911 |
| chr3:135325918-135326167 | 0.117996469 | 0.797349878 | 0.92195911 |
| chr5:58850081-58850630 | -0.098003459 | 0.797376918 | 0.921976462 |
| chr22:36775947-36776648 | 0.084447802 | 0.79741419 | 0.921987784 |
| chr3:65854803-65855356 | -0.097047302 | 0.797426688 | 0.921987784 |
| chr8:48445587-48445894 | -0.122641875 | 0.797432425 | 0.921987784 |
| chr5:173332051-173332324 | -0.092250691 | 0.797434841 | 0.921987784 |
| chr17:40355050-40355365 | -0.103966726 | 0.797486697 | 0.922009182 |
| chr13:25648974-25650069 | 0.085827908 | 0.797498635 | 0.922009182 |
| chr1:1471452-1471887 | -0.094157885 | 0.79750171 | 0.922009182 |
| chr3:142828724-142829058 | 0.091176156 | 0.797509509 | 0.922009182 |
| chr1:7770703-7771605 | -0.089089272 | 0.797513515 | 0.922009182 |
| chr20:41249752-41250369 | 0.086444079 | 0.797543082 | 0.922029453 |
| chr14:23456964-23457204 | -0.116479742 | 0.797591035 | 0.922050133 |
| chr9:129483303-129483823 | -0.089033045 | 0.797598146 | 0.922050133 |
| chr8:81281093-81281689 | -0.087414814 | 0.797603857 | 0.922050133 |
| chr7:85186605-85187208 | -0.092113369 | 0.797616996 | 0.922050133 |
| chr3:189756407-189756966 | -0.095926249 | 0.797640547 | 0.922050133 |
| chr8:100907370-100907701 | 0.095682552 | 0.797645183 | 0.922050133 |
| chr19:41409406-41409608 | 0.121770292 | 0.797645206 | 0.922050133 |
| chr10:1715776-1716268 | -0.091794092 | 0.797658765 | 0.922051895 |
| chr19:5340713-5341111 | -0.101977481 | 0.797693594 | 0.922052558 |
| chr19:38698968-38699292 | 0.113322388 | 0.797698007 | 0.922052558 |
| chr20:53952444-53952802 | 0.083489182 | 0.797702234 | 0.922052558 |
| chr3:112767112-112767312 | 0.133185305 | 0.797707473 | 0.922052558 |
| chr1:220045980-220046893 | -0.082185161 | 0.797722341 | 0.922055833 |
| chr5:44002557-44003163 | 0.085963859 | 0.797746522 | 0.922063212 |
| chr19:39532315-39533309 | 0.084947503 | 0.797766095 | 0.922063212 |
| chr14:102046377-102046776 | 0.093167034 | 0.797777845 | 0.922063212 |
| chr11:19995023-19995550 | 0.088076097 | 0.797778629 | 0.922063212 |
| chr3:196711608-196713056 | -0.082792601 | 0.797788894 | 0.922063212 |
| chr1:206117619-206117916 | 0.099098522 | 0.797858065 | 0.922118407 |
| chr9:91954884-91955196 | -0.114756351 | 0.797860719 | 0.922118407 |
| chrX:47217672-47218813 | 0.08131775 | 0.797905182 | 0.922155885 |

| | | | |
|---|---|---|---|
| chr5:6632536-6633955 | 0.080733234 | 0.797942898 | 0.922185564 |
| chr11:61695064-61695422 | 0.097992619 | 0.797962732 | 0.922194576 |
| chr2:128465661-128465916 | 0.127042258 | 0.797987231 | 0.922199177 |
| chr5:40755400-40756343 | 0.082026043 | 0.797993353 | 0.922199177 |
| chr9:37650927-37651137 | -0.122939092 | 0.798002819 | 0.922199177 |
| chr10:80158929-80159359 | 0.102270099 | 0.798036022 | 0.922223638 |
| chr17:15502829-15503119 | -0.10441899 | 0.798075084 | 0.922248068 |
| chr8:73877937-73879363 | 0.080843707 | 0.798081235 | 0.922248068 |
| chr3:150364971-150365218 | -0.131970602 | 0.798161087 | 0.922326433 |
| chr6:31814544-31815704 | 0.082144964 | 0.798184646 | 0.922328883 |
| chr3:141402008-141403123 | -0.083565587 | 0.798187282 | 0.922328883 |
| chr7:111735723-111735994 | -0.106224438 | 0.798199847 | 0.922329493 |
| chr5:10059849-10060094 | -0.116511557 | 0.798247988 | 0.922361493 |
| chr15:23447250-23447512 | -0.133841551 | 0.798251616 | 0.922361493 |
| chr6:106997590-106997931 | 0.105222813 | 0.798314872 | 0.922414499 |
| chr3:185354138-185354566 | -0.091526977 | 0.798321567 | 0.922414499 |
| chr4:105497542-105497775 | 0.131673817 | 0.798338937 | 0.922420659 |
| chr14:88899272-88899513 | -0.111266002 | 0.798360391 | 0.922430251 |
| chr5:171871288-171871488 | -0.138684157 | 0.798371316 | 0.922430251 |
| chr5:134525121-134525510 | 0.08825537 | 0.798423213 | 0.922476302 |
| chr6:70667577-70668663 | -0.082769944 | 0.798449297 | 0.922492529 |
| chr17:76005752-76005952 | -0.11873328 | 0.798497265 | 0.922525282 |
| chr12:22410510-22410883 | -0.095242388 | 0.798512319 | 0.922525282 |
| chr10:14327283-14327717 | -0.110532296 | 0.798525245 | 0.922525282 |
| chr14:103889668-103890022 | -0.097852425 | 0.798525805 | 0.922525282 |
| chr8:104537163-104537654 | -0.09795408 | 0.798569729 | 0.922562115 |
| chr17:27456079-27456478 | 0.09834772 | 0.798614615 | 0.922587792 |
| chr17:17913939-17914402 | -0.095201381 | 0.798616036 | 0.922587792 |
| chr14:75278169-75279161 | 0.082194776 | 0.798629351 | 0.922589264 |
| chr7:38546048-38546258 | 0.13571724 | 0.798644562 | 0.922592926 |
| chr4:52312761-52313017 | -0.116801634 | 0.798684895 | 0.92260833 |
| chr5:151523451-151523658 | -0.118719175 | 0.798706637 | 0.92260833 |
| chr10:7260396-7260731 | -0.099685836 | 0.79870809 | 0.92260833 |
| chr13:62120505-62120705 | 0.14593394 | 0.798714696 | 0.92260833 |
| chr10:103053961-103054267 | 0.1086543 | 0.798729264 | 0.92260833 |
| chr9:98315537-98315943 | 0.095236373 | 0.798730142 | 0.92260833 |
| chr8:730726-731762 | -0.082512503 | 0.798763511 | 0.922632965 |
| chr20:25750726-25751704 | 0.082655293 | 0.798822863 | 0.922667263 |
| chr8:126012793-126013017 | -0.124287036 | 0.79882822 | 0.922667263 |
| chr6:25278813-25279549 | -0.083308207 | 0.798844759 | 0.922667263 |
| chr1:152870977-152871679 | -0.09099039 | 0.798851245 | 0.922667263 |
| chr4:108219357-108219557 | -0.143541118 | 0.798853413 | 0.922667263 |
| chrX:38006409-38006638 | -0.120726352 | 0.79886642 | 0.922668377 |
| chr18:27163252-27163603 | -0.102523082 | 0.798904305 | 0.9226875 |
| chr6:164106468-164106756 | -0.115728513 | 0.79891531 | 0.9226875 |
| chr2:27123283-27124330 | 0.085809403 | 0.798919103 | 0.9226875 |
| chr3:194632716-194634034 | 0.081949548 | 0.798959767 | 0.922716688 |
| chr1:183922228-183922540 | 0.104674894 | 0.798975306 | 0.922716688 |
| chr19:1122976-1123368 | 0.102753667 | 0.798983949 | 0.922716688 |
| chr7:50584968-50585649 | 0.094536856 | 0.798992546 | 0.922716688 |

| | | | |
|---|---|---|---|
| chr17:55718074-55718434 | 0.099009943 | 0.799008556 | 0.922721271 |
| chr7:73833334-73833731 | 0.095663667 | 0.799044327 | 0.922728414 |
| chr5:160923838-160924346 | 0.086481204 | 0.799046711 | 0.922728414 |
| chr6:39792242-39792451 | 0.122079601 | 0.79905087 | 0.922728414 |
| chr1:201450902-201451333 | 0.093046291 | 0.799092216 | 0.922751174 |
| chr5:168591923-168592492 | 0.087248793 | 0.799094665 | 0.922751174 |
| chr13:30796288-30796559 | 0.10441518 | 0.799123244 | 0.922769718 |
| chr17:7306785-7307717 | -0.081225021 | 0.799148673 | 0.922769718 |
| chr7:6385473-6386253 | 0.088063652 | 0.799160098 | 0.922769718 |
| chr4:36187779-36187979 | 0.140787748 | 0.799170939 | 0.922769718 |
| chr5:146685091-146685291 | 0.140787748 | 0.799170939 | 0.922769718 |
| chr8:73847204-73847481 | -0.09833313 | 0.799206835 | 0.922797259 |
| chr10:21278755-21279153 | 0.093035498 | 0.799232845 | 0.922813385 |
| chr11:2399895-2401038 | 0.081061672 | 0.799271502 | 0.922826515 |
| chr12:107929459-107929788 | -0.095950751 | 0.799277047 | 0.922826515 |
| chr8:87662352-87663066 | -0.087318388 | 0.799291472 | 0.922826515 |
| chr1:98048236-98048690 | -0.097486229 | 0.799292392 | 0.922826515 |
| chr4:146301231-146301431 | -0.139176801 | 0.799315072 | 0.922838795 |
| chr14:81868208-81868479 | 0.114377898 | 0.799349897 | 0.922858874 |
| chr8:121427533-121427742 | 0.110550275 | 0.79936873 | 0.922858874 |
| chr15:93321816-93322313 | 0.097699604 | 0.799387895 | 0.922858874 |
| chr6:36643193-36643566 | 0.110130291 | 0.799390147 | 0.922858874 |
| chr11:117888029-117888336 | -0.106755922 | 0.799417609 | 0.922858874 |
| chr11:19240558-19241554 | -0.081131561 | 0.799439717 | 0.922858874 |
| chr2:190880247-190882151 | -0.079945652 | 0.799448064 | 0.922858874 |
| chr10:99329680-99330293 | 0.091097007 | 0.799450452 | 0.922858874 |
| chr5:114519412-114519612 | -0.134966313 | 0.799470451 | 0.922858874 |
| chr2:151011475-151011778 | -0.110292236 | 0.799495642 | 0.922858874 |
| chr13:99606351-99607049 | -0.084911794 | 0.799515201 | 0.922858874 |
| chr4:51921542-51921873 | 0.11947788 | 0.799520422 | 0.922858874 |
| chr15:79157499-79158197 | 0.09303281 | 0.79952302 | 0.922858874 |
| chr1:156368847-156369374 | 0.088968857 | 0.79954072 | 0.922858874 |
| chr5:179807278-179807579 | -0.09892974 | 0.799545051 | 0.922858874 |
| chr19:17135524-17135813 | 0.110002007 | 0.799545711 | 0.922858874 |
| chr3:95899554-95899833 | 0.126150896 | 0.799555943 | 0.922858874 |
| chr6:48541818-48542029 | 0.126150896 | 0.799555943 | 0.922858874 |
| chr2:26840997-26841297 | -0.112218505 | 0.799580136 | 0.922858874 |
| chr8:107116624-107116843 | 0.124130513 | 0.799594097 | 0.922858874 |
| chr1:40691379-40692664 | 0.081034466 | 0.799598743 | 0.922858874 |
| chr2:61538021-61539085 | -0.079951191 | 0.799619311 | 0.922858874 |
| chr17:4903407-4903828 | 0.098243089 | 0.79962462 | 0.922858874 |
| chr11:62394888-62395141 | 0.10708503 | 0.799627431 | 0.922858874 |
| chrX:40646908-40648011 | -0.082723197 | 0.799641573 | 0.922858874 |
| chr1:24018608-24019275 | -0.091173241 | 0.799645616 | 0.922858874 |
| chr4:10006281-10006582 | 0.100954037 | 0.799663142 | 0.922865201 |
| chr15:68818114-68818331 | -0.107592017 | 0.799693598 | 0.922883848 |
| chr1:203281129-203281367 | 0.100300885 | 0.799703389 | 0.922883848 |
| chr10:11036335-11036841 | -0.09733987 | 0.799726326 | 0.922888163 |
| chr11:57666915-57668387 | 0.080043728 | 0.799731217 | 0.922888163 |
| chr17:56977678-56978336 | 0.082893527 | 0.799761022 | 0.922908657 |

| | | | |
|---|---|---|---|
| chr3:42701503-42701758 | -0.116571346 | 0.799829954 | 0.922974303 |
| chr13:38735662-38736170 | -0.097308585 | 0.799845018 | 0.922977785 |
| chr15:63834303-63834775 | 0.096922061 | 0.799883797 | 0.922980958 |
| chr9:128946944-128948260 | -0.081920308 | 0.799884815 | 0.922980958 |
| chr3:107543813-107544048 | -0.129143787 | 0.799896615 | 0.922980958 |
| chr14:69460181-69460454 | -0.104640934 | 0.79990025 | 0.922980958 |
| chr17:17694296-17694856 | -0.088468657 | 0.799907997 | 0.922980958 |
| chr6:99579816-99580170 | -0.102667432 | 0.799993655 | 0.923031655 |
| chr17:15699336-15699963 | -0.084475302 | 0.800001366 | 0.923031655 |
| chr17:43694442-43694766 | 0.101436553 | 0.800004798 | 0.923031655 |
| chr11:49177997-49178287 | 0.110369041 | 0.800018523 | 0.923031655 |
| chr15:60374399-60375124 | 0.084163896 | 0.800030049 | 0.923031655 |
| chr8:23353921-23354224 | 0.106473334 | 0.800044592 | 0.923031655 |
| chr8:95214124-95214371 | 0.106473334 | 0.800044592 | 0.923031655 |
| chr11:65615028-65615929 | 0.080056621 | 0.800048306 | 0.923031655 |
| chr5:31531548-31532627 | -0.080315603 | 0.800076467 | 0.923050246 |
| chr6:36441423-36441732 | -0.10072466 | 0.800119706 | 0.923086232 |
| chr7:57867917-57868153 | -0.122798405 | 0.800170949 | 0.923131451 |
| chr19:41706689-41707246 | -0.090474045 | 0.800212672 | 0.923136728 |
| chr6:109382012-109383463 | 0.079711525 | 0.800216485 | 0.923136728 |
| chr5:61290953-61291198 | -0.116207265 | 0.800248766 | 0.923136728 |
| chr2:230987012-230987573 | -0.096631122 | 0.800257041 | 0.923136728 |
| chr9:85258816-85259071 | -0.11261035 | 0.800261572 | 0.923136728 |
| chr14:63864373-63864706 | -0.099598683 | 0.800264007 | 0.923136728 |
| chr4:73258523-73259766 | 0.080016855 | 0.800265234 | 0.923136728 |
| chr2:240025050-240025755 | -0.090553136 | 0.800271906 | 0.923136728 |
| chr14:66109530-66109783 | -0.114850483 | 0.800293164 | 0.923138847 |
| chr16:50644910-50645110 | -0.143758525 | 0.800297839 | 0.923138847 |
| chr8:52407679-52407928 | 0.13060436 | 0.800317524 | 0.923147656 |
| chr5:150778264-150779030 | -0.086409771 | 0.800329866 | 0.923147994 |
| chr16:58644842-58645053 | 0.132037649 | 0.800362286 | 0.923154163 |
| chr4:179405934-179406134 | -0.145316844 | 0.80036838 | 0.923154163 |
| chr12:56126180-56126490 | 0.088346145 | 0.800371358 | 0.923154163 |
| chr19:38403523-38403723 | -0.11845758 | 0.800410883 | 0.923173959 |
| chr1:165686130-165686385 | -0.131542578 | 0.800418279 | 0.923173959 |
| chr8:124593442-124593673 | -0.10170524 | 0.800424666 | 0.923173959 |
| chr1:43783518-43783958 | 0.100972555 | 0.800448898 | 0.923188011 |
| chr22:22297614-22298347 | -0.089270197 | 0.800513373 | 0.923240193 |
| chr6:144278690-144278978 | 0.11271081 | 0.800522814 | 0.923240193 |
| chr15:35078583-35078808 | 0.119393604 | 0.800579771 | 0.923240193 |
| chr10:102869477-102869677 | -0.136318978 | 0.800581136 | 0.923240193 |
| chr13:76903670-76903870 | -0.136318978 | 0.800581136 | 0.923240193 |
| chr18:3045268-3045922 | 0.090168188 | 0.800582478 | 0.923240193 |
| chr11:46537912-46538194 | 0.10697597 | 0.800584945 | 0.923240193 |
| chr15:99058941-99059619 | 0.085839411 | 0.800590537 | 0.923240193 |
| chr15:46091737-46092147 | -0.113171221 | 0.800604523 | 0.923242426 |
| chr10:533712-533934 | 0.114514085 | 0.800634154 | 0.923247615 |
| chr3:62659696-62659992 | 0.09167728 | 0.800640342 | 0.923247615 |
| chr9:96339781-96340001 | 0.136500671 | 0.800645171 | 0.923247615 |
| chr3:195850857-195851266 | 0.087767944 | 0.800728573 | 0.923327644 |

| | | | |
|---|---|---|---|
| chr3:58532822-58533212 | 0.101498905 | 0.800738673 | 0.923327644 |
| chr3:131349875-131350233 | -0.103000068 | 0.800756943 | 0.923330372 |
| chr6:87723178-87723551 | 0.098846791 | 0.800788072 | 0.923330372 |
| chr5:112503462-112504410 | 0.084846488 | 0.800794848 | 0.923330372 |
| chr19:7294839-7295043 | 0.119793153 | 0.800797079 | 0.923330372 |
| chr5:76403074-76403343 | 0.108472648 | 0.800801292 | 0.923330372 |
| chr10:102713620-102715089 | 0.079580104 | 0.800847882 | 0.923370196 |
| chr2:38603449-38603760 | 0.087444428 | 0.800883506 | 0.923386238 |
| chr3:157618184-157618495 | -0.097687004 | 0.800889504 | 0.923386238 |
| chr11:15258532-15258817 | -0.108885872 | 0.800897948 | 0.923386238 |
| chr3:170593104-170593304 | 0.138243802 | 0.800932536 | 0.923412221 |
| chr2:47570820-47571606 | 0.084151586 | 0.800950685 | 0.923419251 |
| chr20:38446430-38447222 | -0.087176075 | 0.801008485 | 0.923464404 |
| chr3:111621658-111621886 | 0.117049113 | 0.801016356 | 0.923464404 |
| chr19:7535104-7535769 | -0.091890873 | 0.801030634 | 0.923464404 |
| chr16:74774017-74775424 | -0.080605877 | 0.801061442 | 0.923464404 |
| chr1:48878315-48878924 | 0.100196047 | 0.801077882 | 0.923464404 |
| chr11:70796582-70796914 | 0.111873311 | 0.801113747 | 0.923464404 |
| chr1:202969002-202969348 | -0.086303903 | 0.801118293 | 0.923464404 |
| chr9:37903417-37904544 | -0.080951703 | 0.801123655 | 0.923464404 |
| chr1:153415799-153416159 | -0.101157869 | 0.801150147 | 0.923464404 |
| chr17:39164595-39165069 | 0.103823398 | 0.801152863 | 0.923464404 |
| chr3:45593189-45593584 | 0.093516533 | 0.801165583 | 0.923464404 |
| chr2:14866925-14867574 | -0.09061973 | 0.801166771 | 0.923464404 |
| chr3:184523889-184524192 | -0.121293046 | 0.801180851 | 0.923464404 |
| chr1:231241765-231242098 | 0.10890905 | 0.801181399 | 0.923464404 |
| chr2:22375122-22375322 | -0.129864736 | 0.801192629 | 0.923464404 |
| chr17:69561132-69561390 | -0.11576648 | 0.801199812 | 0.923464404 |
| chr2:229407271-229407486 | 0.121924673 | 0.801200435 | 0.923464404 |
| chr5:15463306-15463572 | -0.116630186 | 0.801229879 | 0.923464404 |
| chr8:128217441-128217695 | 0.103416768 | 0.801231195 | 0.923464404 |
| chr1:39102621-39102821 | -0.131179796 | 0.801246593 | 0.923464404 |
| chr2:85888714-85889509 | -0.083670006 | 0.801250677 | 0.923464404 |
| chr4:68904725-68905261 | -0.09179673 | 0.801254999 | 0.923464404 |
| chr13:99496641-99497051 | -0.08582762 | 0.801282319 | 0.923471437 |
| chr7:156908896-156909296 | 0.095570758 | 0.801285206 | 0.923471437 |
| chr2:54206705-54206905 | 0.135463721 | 0.80130697 | 0.92348263 |
| chr4:111204556-111205106 | -0.103498793 | 0.801376665 | 0.92354906 |
| chr1:219141825-219142030 | -0.126608217 | 0.801403565 | 0.92356617 |
| chr9:122374871-122375071 | 0.123554229 | 0.801428741 | 0.923581293 |
| chr8:49301141-49301390 | -0.121244388 | 0.801499408 | 0.923596979 |
| chr21:28893585-28893972 | -0.120180102 | 0.801500563 | 0.923596979 |
| chr11:112270307-112270909 | -0.095820448 | 0.80152369 | 0.923596979 |
| chr3:63406000-63406227 | 0.098585902 | 0.801524461 | 0.923596979 |
| chr1:204151718-204152651 | -0.081238237 | 0.801529559 | 0.923596979 |
| chr10:80505336-80505879 | -0.086787853 | 0.801530029 | 0.923596979 |
| chr3:140922714-140922943 | 0.111463892 | 0.801540083 | 0.923596979 |
| chr3:9983016-9983282 | 0.099982551 | 0.801554103 | 0.923596979 |
| chr2:219021211-219021440 | -0.11466647 | 0.801567773 | 0.923596979 |
| chr18:57444274-57444513 | 0.110245091 | 0.801580312 | 0.923596979 |

| | | | |
|---|---|---|---|
| chr20:54267162-54268321 | -0.078376134 | 0.801590268 | 0.923596979 |
| chr6:139659270-139659517 | -0.11049007 | 0.801593046 | 0.923596979 |
| chr20:50306096-50306355 | -0.096104864 | 0.801599054 | 0.923596979 |
| chr9:137086300-137087441 | 0.080181593 | 0.801614207 | 0.92360055 |
| chr11:14380933-14381530 | 0.092676055 | 0.801654818 | 0.923633452 |
| chr10:72048580-72048902 | -0.095849567 | 0.801667572 | 0.923634258 |
| chr5:32099803-32100003 | 0.131401341 | 0.801704875 | 0.923650442 |
| chr15:68783747-68783956 | 0.112869116 | 0.801705728 | 0.923650442 |
| chr6:89895342-89895731 | 0.096209882 | 0.801720004 | 0.923653001 |
| chr2:131492270-131493239 | -0.08005488 | 0.801761808 | 0.923687274 |
| chr5:107930953-107931206 | -0.130409352 | 0.80180218 | 0.923695673 |
| chr1:146369596-146370437 | 0.081024771 | 0.801803613 | 0.923695673 |
| chr1:9005209-9005471 | -0.115771553 | 0.801805264 | 0.923695673 |
| chr16:2085755-2085955 | 0.119162817 | 0.801826757 | 0.923706546 |
| chr15:49155033-49156114 | -0.080703342 | 0.801847507 | 0.923716562 |
| chr3:104206956-104207659 | -0.089865596 | 0.8019258 | 0.923770012 |
| chr3:79878180-79878557 | -0.107731211 | 0.801928554 | 0.923770012 |
| chr1:165827361-165829220 | -0.079534598 | 0.801941431 | 0.923770012 |
| chr5:101136512-101136793 | 0.111063102 | 0.80194501 | 0.923770012 |
| chr2:43226338-43227930 | 0.079210108 | 0.801954186 | 0.923770012 |
| chr7:115593379-115593806 | -0.112756556 | 0.802000575 | 0.923793139 |
| chr2:28869981-28870955 | -0.081411531 | 0.802021096 | 0.923793139 |
| chr7:4479401-4479758 | 0.095005811 | 0.802031224 | 0.923793139 |
| chr13:30306702-30307862 | -0.080323969 | 0.802033095 | 0.923793139 |
| chr6:31474655-31474855 | 0.134342372 | 0.802038242 | 0.923793139 |
| chr11:65575635-65575909 | 0.097539688 | 0.802046603 | 0.923793139 |
| chr8:23474075-23474496 | -0.108014914 | 0.802064559 | 0.923799935 |
| chr17:75011055-75011358 | 0.099911249 | 0.802096981 | 0.923815184 |
| chr11:74454437-74454641 | 0.123992114 | 0.802101912 | 0.923815184 |
| chr2:70785372-70785608 | -0.114771543 | 0.802115772 | 0.92381726 |
| chr17:70552463-70552794 | -0.100436289 | 0.802176762 | 0.923823126 |
| chr19:10568737-10569336 | 0.08269741 | 0.802180789 | 0.923823126 |
| chr3:168511464-168511673 | 0.126087142 | 0.802185532 | 0.923823126 |
| chr15:90340397-90340686 | 0.098286843 | 0.802189708 | 0.923823126 |
| chr13:24539896-24540150 | 0.125278043 | 0.802190964 | 0.923823126 |
| chr2:229271287-229271487 | 0.130934583 | 0.802199627 | 0.923823126 |
| chr10:29239957-29240187 | -0.119041177 | 0.802205264 | 0.923823126 |
| chr14:45135672-45136244 | 0.081645797 | 0.80222564 | 0.923831028 |
| chr8:77745352-77745732 | 0.100731879 | 0.802236239 | 0.923831028 |
| chrX:38260302-38260521 | 0.12522795 | 0.802277027 | 0.923845792 |
| chr3:185363448-185363790 | -0.104944241 | 0.802298319 | 0.923845792 |
| chr9:127195342-127195664 | 0.105583154 | 0.802306375 | 0.923845792 |
| chr6:151446235-151446622 | -0.095444931 | 0.80233984 | 0.923845792 |
| chr3:23916494-23917782 | 0.078814761 | 0.802341475 | 0.923845792 |
| chr15:64380707-64381768 | 0.081877643 | 0.802343675 | 0.923845792 |
| chr3:50226134-50226419 | -0.099765837 | 0.802344178 | 0.923845792 |
| chr11:21355335-21355629 | 0.111598249 | 0.802355158 | 0.923845792 |
| chr2:239400876-239401987 | -0.080022003 | 0.802357574 | 0.923845792 |
| chr8:89785494-89785741 | 0.118780422 | 0.802408922 | 0.923888399 |
| chr3:114248584-114248883 | -0.126099357 | 0.802418694 | 0.923888399 |

| | | | |
|---|---|---|---|
| chr3:192753076-192753362 | -0.127269397 | 0.802459174 | 0.923920945 |
| chr11:9613161-9614515 | 0.08091943 | 0.802471674 | 0.923920945 |
| chr6:146985238-146985448 | 0.132680243 | 0.802483136 | 0.923920945 |
| chr16:84622166-84622420 | 0.097988506 | 0.802501701 | 0.923928437 |
| chr14:64337875-64339342 | 0.078554999 | 0.80253774 | 0.923930979 |
| chr11:72143887-72144311 | 0.100277368 | 0.802542469 | 0.923930979 |
| chr1:1364350-1364550 | 0.133569625 | 0.80255111 | 0.923930979 |
| chr3:67863984-67864210 | -0.111249668 | 0.802555796 | 0.923930979 |
| chr21:42878728-42879893 | -0.080220833 | 0.8025642 | 0.923930979 |
| chrX:151488692-151488989 | -0.111613827 | 0.80258024 | 0.923935563 |
| chr6:7561387-7562005 | -0.088852682 | 0.802600603 | 0.923939841 |
| chr3:23944887-23946745 | 0.077949104 | 0.802608073 | 0.923939841 |
| chr2:1104879-1105158 | 0.119819755 | 0.802629906 | 0.923951093 |
| chr17:19978543-19978749 | -0.117267396 | 0.80264794 | 0.923957971 |
| chr4:40265025-40265485 | 0.104457104 | 0.802673103 | 0.923960079 |
| chr17:41496743-41497162 | 0.090280265 | 0.802674929 | 0.923960079 |
| chr7:99919177-99920003 | 0.085054204 | 0.802686898 | 0.923960079 |
| chr2:200705717-200705986 | -0.115255309 | 0.802701884 | 0.923960079 |
| chr15:58749299-58750329 | 0.079499539 | 0.802710064 | 0.923960079 |
| chr19:11344446-11344785 | -0.096204525 | 0.802743039 | 0.923980634 |
| chr16:4271144-4271456 | -0.091629434 | 0.802786014 | 0.923980634 |
| chr4:153518092-153518362 | 0.127205052 | 0.802799366 | 0.923980634 |
| chr5:113474219-113474473 | -0.131839992 | 0.802799446 | 0.923980634 |
| chr4:23717641-23717868 | 0.121448418 | 0.802802562 | 0.923980634 |
| chr10:77857178-77857715 | 0.084865681 | 0.802805069 | 0.923980634 |
| chr5:74632203-74632410 | -0.116198282 | 0.802812335 | 0.923980634 |
| chr17:51322454-51322746 | -0.09240544 | 0.802830748 | 0.923984243 |
| chr15:101258173-101258504 | 0.098077865 | 0.802839588 | 0.923984243 |
| chr3:12788206-12788406 | 0.13357385 | 0.802864564 | 0.923999108 |
| chr8:127386347-127386547 | 0.116519746 | 0.802897342 | 0.924009102 |
| chr3:58526758-58526982 | -0.119587768 | 0.802897366 | 0.924009102 |
| chr17:60641753-60642217 | 0.080624559 | 0.802919387 | 0.924020566 |
| chr15:98295188-98295388 | 0.120806635 | 0.802972272 | 0.924067549 |
| chr6:142375245-142375571 | -0.108738475 | 0.803008286 | 0.924076489 |
| chr8:120124181-120124543 | -0.090487451 | 0.803012101 | 0.924076489 |
| chr12:47809077-47809324 | 0.11447543 | 0.803016935 | 0.924076489 |
| chr12:83955260-83955558 | 0.108848251 | 0.803028282 | 0.924076489 |
| chr15:64460078-64460670 | 0.08445719 | 0.803040585 | 0.924076769 |
| chr15:96353978-96354768 | 0.084822052 | 0.803053917 | 0.924078232 |
| chr1:201348140-201348450 | -0.090540467 | 0.803114367 | 0.924133913 |
| chr9:111897478-111897757 | -0.101399127 | 0.803147986 | 0.924158719 |
| chr16:15642628-15643407 | 0.082665087 | 0.803216837 | 0.924195802 |
| chr6:131459988-131460317 | 0.096248589 | 0.803217639 | 0.924195802 |
| chr9:90755635-90756064 | 0.100890329 | 0.803222404 | 0.924195802 |
| chr14:99475249-99475538 | -0.104940096 | 0.80322846 | 0.924195802 |
| chr7:66075404-66076493 | 0.081415929 | 0.803259357 | 0.92420249 |
| chr1:240888097-240888317 | -0.126577886 | 0.80325946 | 0.92420249 |
| chr13:112587331-112588349 | -0.082537478 | 0.803270458 | 0.92420249 |
| chr7:82443209-82444718 | 0.081255732 | 0.803286612 | 0.924207198 |
| chr15:34166028-34166323 | 0.107599898 | 0.803303304 | 0.924212525 |

| | | | |
|---|---|---|---|
| chr4:134201742-134202120 | 0.101985875 | 0.803320086 | 0.924217956 |
| chr6:72677121-72677542 | -0.117971698 | 0.803373241 | 0.924256006 |
| chr4:87848715-87848915 | 0.137524949 | 0.803380625 | 0.924256006 |
| chr11:62591450-62592355 | 0.080709408 | 0.803389346 | 0.924256006 |
| chr4:128298897-128299142 | -0.118921573 | 0.803411506 | 0.924267622 |
| chr1:59947397-59947711 | -0.097703554 | 0.803450808 | 0.924298958 |
| chr1:193031451-193031713 | 0.104029548 | 0.803510611 | 0.924346161 |
| chr3:63911629-63913500 | -0.07699264 | 0.803515967 | 0.924346161 |
| chr3:153704221-153704543 | -0.10473634 | 0.803573348 | 0.924387618 |
| chr1:157294404-157294694 | -0.1113503 | 0.803576133 | 0.924387618 |
| chr4:122914392-122914625 | -0.126959241 | 0.803595411 | 0.924391725 |
| chr4:118424098-118424408 | 0.098146157 | 0.803605216 | 0.924391725 |
| chr9:89999957-90000226 | 0.111704437 | 0.803617445 | 0.924391725 |
| chr8:61265693-61265893 | -0.134032095 | 0.803640525 | 0.924391725 |
| chr3:6392189-6392410 | 0.122226494 | 0.803643418 | 0.924391725 |
| chr3:183295218-183295804 | 0.083833031 | 0.803652089 | 0.924391725 |
| chr16:79200500-79200767 | -0.118603066 | 0.803664221 | 0.924391803 |
| chr3:164912671-164913029 | 0.107693387 | 0.803752649 | 0.92445335 |
| chr6:166862746-166863074 | 0.106088567 | 0.803762211 | 0.92445335 |
| chr3:96986291-96986584 | 0.10488854 | 0.803769744 | 0.92445335 |
| chr17:47530406-47531320 | -0.083006991 | 0.803774941 | 0.92445335 |
| chr1:223749189-223749414 | 0.119185052 | 0.803787784 | 0.92445335 |
| chr18:3441378-3441582 | -0.126647678 | 0.803806211 | 0.92445335 |
| chr15:31462205-31462415 | -0.127446103 | 0.803814346 | 0.92445335 |
| chr3:155434197-155434397 | 0.129029083 | 0.803822539 | 0.92445335 |
| chr3:100090075-100090829 | 0.088179534 | 0.803838967 | 0.92445335 |
| chr14:91317546-91317937 | -0.096107901 | 0.803859023 | 0.92445335 |
| chr16:67546895-67547208 | -0.096107901 | 0.803859023 | 0.92445335 |
| chr14:101760370-101761187 | -0.079178515 | 0.80387269 | 0.92445335 |
| chr12:98439716-98439945 | -0.131622477 | 0.803874576 | 0.92445335 |
| chr4:150209429-150209751 | -0.096101068 | 0.803912723 | 0.924483343 |
| chr6:109892483-109893389 | -0.082501733 | 0.803958233 | 0.924509994 |
| chr1:189859440-189859834 | 0.099469697 | 0.80396003 | 0.924509994 |
| chr19:29860540-29860865 | -0.09950811 | 0.804004962 | 0.924522915 |
| chr15:68252977-68253294 | -0.100871877 | 0.804009998 | 0.924522915 |
| chr16:89720876-89721737 | -0.08040331 | 0.804011649 | 0.924522915 |
| chrX:49922301-49923205 | 0.080428502 | 0.80401953 | 0.924522915 |
| chr3:149129187-149130404 | -0.07891623 | 0.804033833 | 0.924525487 |
| chr6:38176689-38177692 | 0.082930176 | 0.80405954 | 0.924534093 |
| chr1:23777963-23778748 | -0.08867337 | 0.80406545 | 0.924534093 |
| chr20:48572188-48572659 | 0.083163939 | 0.804089596 | 0.924544327 |
| chr12:7129862-7130494 | 0.085714715 | 0.804098482 | 0.924544327 |
| chr12:106903575-106903922 | 0.090324345 | 0.804150351 | 0.92459009 |
| chr12:109845109-109845641 | 0.084219001 | 0.804201994 | 0.924601634 |
| chr12:64818330-64818587 | 0.113772511 | 0.804205088 | 0.924601634 |
| chr14:33057578-33057859 | 0.113772511 | 0.804205088 | 0.924601634 |
| chr11:33971064-33971264 | 0.125507513 | 0.804215899 | 0.924601634 |
| chr7:1459320-1459978 | 0.078854042 | 0.804220727 | 0.924601634 |
| chr1:149103080-149103709 | -0.090802199 | 0.804257921 | 0.924608737 |
| chr11:119011468-119011740 | -0.117279795 | 0.804278293 | 0.924608737 |

| | | | |
|---|---|---|---|
| chr12:70589983-70590293 | -0.096017481 | 0.804280706 | 0.924608737 |
| chr16:85753125-85753376 | -0.097034483 | 0.804285847 | 0.924608737 |
| chr1:201709506-201709810 | 0.089150055 | 0.804287241 | 0.924608737 |
| chr7:37198909-37199188 | -0.097678328 | 0.804321907 | 0.924625803 |
| chr1:48916216-48916845 | -0.091159274 | 0.804336332 | 0.924625803 |
| chr2:50491430-50491859 | 0.093972927 | 0.804346245 | 0.924625803 |
| chr1:222711698-222713524 | -0.077875189 | 0.804350355 | 0.924625803 |
| chr2:196170820-196172006 | -0.07857397 | 0.804364823 | 0.924628563 |
| chr11:36258647-36259118 | -0.093230089 | 0.804379552 | 0.924629721 |
| chr3:62844895-62845143 | 0.108286815 | 0.804389966 | 0.924629721 |
| chr20:31879418-31880351 | 0.08008446 | 0.804409204 | 0.924637963 |
| chr7:47747151-47747501 | -0.096440737 | 0.804435775 | 0.924654634 |
| chr8:85245316-85246317 | 0.081080421 | 0.804462368 | 0.924664712 |
| chr3:193418750-193419172 | 0.102777998 | 0.804473209 | 0.924664712 |
| chr3:152056623-152057129 | 0.101275981 | 0.804486568 | 0.924664712 |
| chr12:122752414-122753394 | 0.080328721 | 0.804501919 | 0.924664712 |
| chr8:78615747-78616047 | -0.107504059 | 0.804504881 | 0.924664712 |
| chr11:19801129-19801369 | -0.116000034 | 0.80455653 | 0.924710203 |
| chr2:202918958-202919158 | 0.149512506 | 0.804611542 | 0.924759559 |
| chr2:222899877-222900121 | -0.119987688 | 0.804638314 | 0.924768152 |
| chr12:70733471-70733716 | 0.116337858 | 0.804643156 | 0.924768152 |
| chr15:44536368-44537633 | 0.078994679 | 0.804672162 | 0.924783797 |
| chr3:158795030-158795363 | 0.095503742 | 0.804690703 | 0.924783797 |
| chr1:119987052-119987384 | 0.118977225 | 0.804692977 | 0.924783797 |
| chr14:28755051-28755251 | -0.119780574 | 0.804711788 | 0.924791544 |
| chr2:239882468-239882757 | 0.112060965 | 0.804815736 | 0.924897131 |
| chr1:205228525-205228759 | 0.100631524 | 0.804859224 | 0.924917855 |
| chr1:155194196-155195207 | 0.080047572 | 0.804862244 | 0.924917855 |
| chr9:126690445-126690659 | 0.107203722 | 0.804924375 | 0.924917855 |
| chr8:122572350-122572630 | -0.09154821 | 0.804927722 | 0.924917855 |
| chr6:28810812-28811012 | 0.120097152 | 0.804930059 | 0.924917855 |
| chr12:76763621-76764832 | -0.077682646 | 0.804931387 | 0.924917855 |
| chr4:37826083-37827553 | -0.079232283 | 0.804941611 | 0.924917855 |
| chr21:42666087-42666342 | 0.100135399 | 0.804942254 | 0.924917855 |
| chr5:171436594-171436794 | -0.120862338 | 0.804951197 | 0.924917855 |
| chr1:156694209-156694529 | 0.089207558 | 0.804964514 | 0.924917855 |
| chr1:230752122-230752756 | -0.085450768 | 0.804966552 | 0.924917855 |
| chr2:62456891-62457616 | -0.084711804 | 0.804979157 | 0.924918468 |
| chr7:149685512-149685774 | -0.125237931 | 0.805019377 | 0.924936601 |
| chr17:21457492-21457722 | -0.116749025 | 0.805030474 | 0.924936601 |
| chr14:91331706-91332390 | -0.084618167 | 0.805031153 | 0.924936601 |
| chr15:26891406-26891731 | 0.103900494 | 0.805054183 | 0.924949192 |
| chr12:45728695-45729726 | 0.07926942 | 0.805104919 | 0.924993614 |
| chr1:32273063-32273371 | 0.107414217 | 0.805124414 | 0.924999156 |
| chr16:81585855-81586151 | 0.094919289 | 0.805133888 | 0.924999156 |
| chr21:39380076-39381335 | 0.078056873 | 0.805190713 | 0.925050571 |
| chr18:503994-504285 | 0.110652975 | 0.805219762 | 0.925066338 |
| chr6:161152970-161153170 | -0.139213549 | 0.805228583 | 0.925066338 |
| chr1:109918364-109918566 | -0.110759449 | 0.805249885 | 0.925076941 |
| chr5:56421991-56422428 | 0.096081419 | 0.805284046 | 0.925102314 |

| | | | |
|---|---|---|---|
| chr17:75632910-75634242 | -0.078299287 | 0.805299742 | 0.925103935 |
| chr8:86414455-86414670 | 0.116004952 | 0.805309603 | 0.925103935 |
| chr13:99426316-99426516 | -0.136004278 | 0.805345474 | 0.925110923 |
| chr1:114924557-114924763 | 0.106966574 | 0.80534654 | 0.925110923 |
| chr2:161223230-161223432 | -0.135389628 | 0.805351908 | 0.925110923 |
| chr22:37567417-37567617 | -0.118287201 | 0.805380715 | 0.925116189 |
| chrX:107849759-107850034 | -0.114582746 | 0.805395511 | 0.925116189 |
| chr5:124263124-124264226 | 0.08147141 | 0.805396066 | 0.925116189 |
| chr2:19863271-19863960 | 0.084042805 | 0.805405063 | 0.925116189 |
| chr21:17513344-17513555 | -0.111903979 | 0.805416861 | 0.925116189 |
| chr19:45178103-45179090 | 0.078061983 | 0.805462405 | 0.925154633 |
| chr11:14088953-14089153 | -0.125500424 | 0.805476449 | 0.925156895 |
| chr1:9539100-9539569 | 0.082736832 | 0.805498121 | 0.925162855 |
| chr20:3855121-3855442 | 0.096157061 | 0.805509983 | 0.925162855 |
| chr11:75388623-75389010 | 0.089222409 | 0.80552169 | 0.925162855 |
| chr11:71786461-71787813 | 0.077607008 | 0.805529935 | 0.925162855 |
| chr10:114173909-114174996 | -0.078790139 | 0.805571748 | 0.925197009 |
| chr20:10219489-10219836 | 0.090205685 | 0.805612622 | 0.925228954 |
| chr1:110963205-110963613 | -0.090492939 | 0.805651965 | 0.925228954 |
| chr2:189542171-189542467 | 0.103136292 | 0.805675571 | 0.925228954 |
| chr1:220089619-220090043 | -0.082105929 | 0.805683057 | 0.925228954 |
| chr9:130681548-130682152 | 0.090765977 | 0.805686161 | 0.925228954 |
| chr1:205210686-205211947 | -0.07804962 | 0.805687783 | 0.925228954 |
| chr3:18234908-18235239 | -0.100049233 | 0.805695 | 0.925228954 |
| chr9:100216234-100216553 | -0.103805576 | 0.805705545 | 0.925228954 |
| chr7:98470498-98470961 | 0.086305313 | 0.805740642 | 0.925228954 |
| chr10:92290301-92291566 | 0.078810642 | 0.805742471 | 0.925228954 |
| chr14:34496929-34497306 | 0.099598888 | 0.80574711 | 0.925228954 |
| chr3:150330947-150331176 | 0.124398191 | 0.805762139 | 0.925228954 |
| chr2:36775145-36775403 | -0.109822567 | 0.805763364 | 0.925228954 |
| chr2:141565607-141566000 | 0.091805328 | 0.805779972 | 0.925228954 |
| chr1:220697554-220697798 | -0.126194322 | 0.805780691 | 0.925228954 |
| chr8:56010637-56010905 | -0.105942219 | 0.805816484 | 0.925249094 |
| chr14:45551069-45551707 | 0.093330518 | 0.805822382 | 0.925249094 |
| chr3:65425920-65426120 | 0.124374814 | 0.805871208 | 0.925291291 |
| chr17:40317986-40318319 | 0.087799713 | 0.805887552 | 0.925296192 |
| chr12:33605817-33606113 | 0.110840411 | 0.805903935 | 0.925301136 |
| chr1:169466294-169466598 | -0.107278548 | 0.805917495 | 0.925302839 |
| chr6:25279698-25279898 | 0.109590847 | 0.805961925 | 0.925339985 |
| chr10:79445558-79446259 | 0.080543521 | 0.806015458 | 0.925387582 |
| chr3:193461701-193462065 | -0.100965118 | 0.806050549 | 0.92540682 |
| chr17:50630622-50630965 | 0.088593732 | 0.80605637 | 0.92540682 |
| chr2:63023484-63023812 | -0.094789219 | 0.806074779 | 0.925414089 |
| chr5:14136355-14136578 | 0.115127098 | 0.806118744 | 0.925425061 |
| chr4:55336663-55336980 | 0.099211236 | 0.806128965 | 0.925425061 |
| chr8:48083941-48084297 | -0.114875801 | 0.806171603 | 0.925425061 |
| chr1:204565674-204566183 | -0.082565825 | 0.806191252 | 0.925425061 |
| chr3:150545904-150547232 | 0.076860432 | 0.806192055 | 0.925425061 |
| chr11:84384724-84384940 | -0.116062593 | 0.806195204 | 0.925425061 |
| chr6:42096329-42096529 | -0.131654699 | 0.806198997 | 0.925425061 |

| | | | |
|---|---|---|---|
| chr5:140868781-140869647 | -0.084992622 | 0.806216689 | 0.925425061 |
| chr3:49803191-49803445 | 0.109796455 | 0.806228524 | 0.925425061 |
| chr1:202852335-202852629 | -0.089802245 | 0.806235292 | 0.925425061 |
| chr1:202966770-202967713 | -0.079200659 | 0.806236618 | 0.925425061 |
| chr19:12791261-12791596 | -0.091439101 | 0.806244419 | 0.925425061 |
| chr2:9142640-9143199 | -0.084376449 | 0.806255944 | 0.925425061 |
| chr17:64992433-64992633 | 0.108962132 | 0.806272259 | 0.925425061 |
| chr10:95896518-95896838 | -0.105579136 | 0.80629302 | 0.925425061 |
| chrX:53421908-53423164 | -0.078535355 | 0.806298092 | 0.925425061 |
| chr5:178107915-178108720 | -0.082320575 | 0.806298365 | 0.925425061 |
| chr11:87037189-87038215 | 0.079028733 | 0.806321211 | 0.925425061 |
| chr2:55163881-55164139 | 0.10512641 | 0.806327425 | 0.925425061 |
| chr9:34591750-34591950 | 0.113087626 | 0.806331719 | 0.925425061 |
| chr1:202860722-202862276 | -0.076801252 | 0.806338971 | 0.925425061 |
| chr1:227318059-227319477 | 0.078639643 | 0.806350047 | 0.925425061 |
| chr21:20971372-20971593 | -0.130435587 | 0.806398181 | 0.925460571 |
| chr3:168870899-168871212 | 0.111002667 | 0.806405145 | 0.925460571 |
| chr4:118765435-118765635 | -0.128254671 | 0.806418252 | 0.925461752 |
| chr1:220368553-220368876 | -0.097331285 | 0.806456446 | 0.925491722 |
| chr5:114361658-114362619 | 0.079395691 | 0.806495649 | 0.92552285 |
| chr17:1628128-1629546 | 0.079876085 | 0.80653347 | 0.92555239 |
| chr16:71537713-71537990 | 0.096260446 | 0.806546644 | 0.925553646 |
| chr3:63333865-63334507 | -0.084774545 | 0.806583501 | 0.92558208 |
| chr10:84139160-84140287 | 0.077746875 | 0.806619056 | 0.925609017 |
| chr21:26169629-26171243 | 0.078337215 | 0.806691212 | 0.925659128 |
| chr1:195886115-195886326 | 0.125344552 | 0.80670113 | 0.925659128 |
| chr4:76948710-76950335 | 0.07778034 | 0.806701601 | 0.925659128 |
| chr1:233707542-233707752 | 0.117702321 | 0.806711049 | 0.925659128 |
| chr10:5439796-5440036 | 0.119784413 | 0.806759464 | 0.925674991 |
| chr3:71582802-71583429 | -0.083732955 | 0.806759669 | 0.925674991 |
| chr1:160101031-160101360 | 0.086980395 | 0.806761116 | 0.925674991 |
| chr7:131606437-131606637 | -0.125641149 | 0.80678714 | 0.925689837 |
| chr3:192900404-192900604 | 0.12556569 | 0.806798217 | 0.925689837 |
| chr15:65270842-65271270 | 0.093061202 | 0.806834624 | 0.925717747 |
| chr10:31353696-31353923 | -0.123699602 | 0.806867701 | 0.925741835 |
| chr1:165697790-165698905 | -0.078415519 | 0.806890979 | 0.92575089 |
| chr12:121773241-121773560 | 0.090495167 | 0.806902722 | 0.92575089 |
| chr17:58388414-58388664 | 0.096082325 | 0.806912035 | 0.92575089 |
| chr17:78484294-78485218 | -0.080598862 | 0.806932067 | 0.92575089 |
| chr1:9300223-9300423 | 0.122858374 | 0.806942256 | 0.92575089 |
| chr17:17237060-17237843 | -0.081497438 | 0.806964764 | 0.92575089 |
| chr2:9799711-9799977 | 0.093353484 | 0.806988521 | 0.92575089 |
| chr11:46919976-46920185 | 0.110818145 | 0.807015181 | 0.92575089 |
| chr1:231604088-231604288 | 0.11953853 | 0.807015412 | 0.92575089 |
| chr12:122966185-122966980 | 0.079311258 | 0.807022823 | 0.92575089 |
| chr19:16572368-16572723 | -0.089608184 | 0.807029488 | 0.92575089 |
| chr3:186241851-186242146 | 0.108243598 | 0.807041403 | 0.92575089 |
| chr3:161371358-161373233 | -0.076116858 | 0.80705391 | 0.92575089 |
| chr16:19991602-19992017 | -0.103378213 | 0.807058749 | 0.92575089 |
| chr17:66255087-66255351 | 0.096191203 | 0.807078085 | 0.92575089 |

| | | | |
|---|---|---|---|
| chr19:2269203-2269599 | -0.089419951 | 0.807086547 | 0.92575089 |
| chr5:38404701-38404945 | -0.106069542 | 0.807104246 | 0.92575089 |
| chr15:84878312-84878867 | 0.086943893 | 0.807107023 | 0.92575089 |
| chr19:14571783-14572325 | 0.088792315 | 0.80710767 | 0.92575089 |
| chr7:44748066-44749693 | -0.077358078 | 0.807117234 | 0.92575089 |
| chr14:45665029-45665340 | 0.096352486 | 0.807144224 | 0.92575813 |
| chr4:42709729-42710140 | 0.092630189 | 0.807150468 | 0.92575813 |
| chr7:95596468-95597079 | -0.089627275 | 0.807176774 | 0.92575813 |
| chr1:33472162-33472965 | -0.080334179 | 0.807189602 | 0.92575813 |
| chr16:69385513-69386276 | 0.083672133 | 0.80720136 | 0.92575813 |
| chr19:42075636-42076515 | 0.078907208 | 0.807222101 | 0.92575813 |
| chr16:27179548-27179879 | -0.092408737 | 0.807261153 | 0.92575813 |
| chr17:75270087-75270962 | 0.07956263 | 0.807277453 | 0.92575813 |
| chr11:73289297-73289758 | 0.084781101 | 0.807280711 | 0.92575813 |
| chr8:143454633-143454898 | -0.118178283 | 0.807291721 | 0.92575813 |
| chr3:50616013-50616238 | -0.096600225 | 0.807320277 | 0.92575813 |
| chr19:51021108-51021313 | 0.110243586 | 0.807321244 | 0.92575813 |
| chr7:1131270-1131941 | -0.08770529 | 0.807322239 | 0.92575813 |
| chr2:171893842-171894693 | -0.082192773 | 0.807328325 | 0.92575813 |
| chr2:237490280-237490504 | 0.108299796 | 0.807342036 | 0.92575813 |
| chr1:162822378-162823035 | 0.087283131 | 0.807349723 | 0.92575813 |
| chr18:9951234-9951504 | -0.11762868 | 0.807361597 | 0.92575813 |
| chr1:145425039-145426042 | -0.079072128 | 0.807361715 | 0.92575813 |
| chr3:161420653-161421160 | 0.080864651 | 0.807368668 | 0.92575813 |
| chr17:58819568-58819944 | -0.085144563 | 0.807379284 | 0.92575813 |
| chr5:40999228-40999434 | 0.131181457 | 0.80738497 | 0.92575813 |
| chr10:58531102-58531415 | -0.098735944 | 0.807400719 | 0.92575813 |
| chr2:226799748-226800108 | -0.088639752 | 0.807408753 | 0.92575813 |
| chr18:10140987-10141313 | -0.099454993 | 0.807413517 | 0.92575813 |
| chr13:19984797-19984997 | -0.122058995 | 0.807447905 | 0.925774509 |
| chr8:123540872-123542194 | 0.077941972 | 0.807451968 | 0.925774509 |
| chr9:128684345-128685130 | -0.081730674 | 0.807477437 | 0.925789857 |
| chr1:16559727-16560074 | -0.099068423 | 0.807513682 | 0.925809728 |
| chr13:45511636-45511923 | 0.099826161 | 0.807518934 | 0.925809728 |
| chr7:152336887-152337301 | -0.101031981 | 0.807531769 | 0.925810591 |
| chrX:57156518-57156924 | -0.104140823 | 0.807566628 | 0.925836702 |
| chr1:155609614-155610305 | 0.080824238 | 0.807622194 | 0.925882539 |
| chr11:73393594-73393880 | 0.092309533 | 0.807650141 | 0.925882539 |
| chr5:149731379-149731809 | 0.086347769 | 0.807654341 | 0.925882539 |
| chr13:31928086-31928348 | 0.112053679 | 0.807654944 | 0.925882539 |
| chr5:141879120-141879539 | 0.084047241 | 0.807681596 | 0.925899239 |
| chr15:74545933-74546749 | 0.081237902 | 0.807707247 | 0.925907496 |
| chr8:129904623-129904823 | 0.139645202 | 0.807712967 | 0.925907496 |
| chr17:44898810-44900083 | 0.079379971 | 0.807739777 | 0.925924377 |
| chr10:103564650-103565490 | -0.086746582 | 0.807768787 | 0.925943779 |
| chr16:30400543-30401033 | 0.085805585 | 0.807812121 | 0.9259796 |
| chr11:16883092-16883519 | -0.095353422 | 0.807846605 | 0.92600477 |
| chr15:76469031-76469231 | 0.14071298 | 0.807865634 | 0.92600477 |
| chr8:115692041-115692258 | -0.116429692 | 0.807880128 | 0.92600477 |
| chr11:35404510-35404808 | 0.104823408 | 0.807882584 | 0.92600477 |

| | | | |
|---|---|---|---|
| chr6:101011222-101011506 | -0.108671835 | 0.807894506 | 0.92600477 |
| chr3:77348240-77348641 | -0.094926881 | 0.80791453 | 0.926012146 |
| chr2:176088328-176088672 | 0.091906921 | 0.807925112 | 0.926012146 |
| chr11:80197166-80197655 | -0.102366115 | 0.807971239 | 0.926030503 |
| chr6:26251495-26251905 | 0.090339596 | 0.807981511 | 0.926030503 |
| chr7:16668995-16669253 | -0.108257668 | 0.807997363 | 0.926030503 |
| chr20:54449189-54449472 | 0.081913983 | 0.808005619 | 0.926030503 |
| chr3:79872510-79872781 | 0.112626264 | 0.808013315 | 0.926030503 |
| chr3:134485373-134486752 | 0.076523138 | 0.808013643 | 0.926030503 |
| chr3:167813656-167813891 | -0.114423576 | 0.808036269 | 0.926030976 |
| chr1:20732756-20733422 | 0.088113232 | 0.808040215 | 0.926030976 |
| chr11:71412590-71412799 | -0.131042474 | 0.808050313 | 0.926030976 |
| chr5:117713682-117713921 | -0.128238452 | 0.808088899 | 0.926061345 |
| chr1:34589909-34590200 | 0.118535336 | 0.808109205 | 0.926069861 |
| chr6:90098029-90098259 | 0.134407468 | 0.808120502 | 0.926069861 |
| chr6:130697997-130698282 | -0.104589009 | 0.808179696 | 0.926094003 |
| chr11:1693248-1693514 | -0.103455985 | 0.808211467 | 0.926094003 |
| chr5:136192886-136193337 | 0.090615822 | 0.808211916 | 0.926094003 |
| chr20:23048547-23049018 | -0.087932733 | 0.808214322 | 0.926094003 |
| chr4:68864500-68864900 | 0.097035669 | 0.80822512 | 0.926094003 |
| chr15:59875167-59875532 | -0.111268821 | 0.808260015 | 0.926094003 |
| chr21:43711516-43711761 | -0.110293053 | 0.808274256 | 0.926094003 |
| chr2:151344948-151345234 | 0.103618196 | 0.808275541 | 0.926094003 |
| chr3:46895229-46895464 | 0.10549802 | 0.80830161 | 0.926094003 |
| chr20:32168581-32168786 | 0.124370409 | 0.808318655 | 0.926094003 |
| chr21:31459563-31459795 | 0.124370409 | 0.808318655 | 0.926094003 |
| chr1:112715128-112715931 | 0.078607155 | 0.808319891 | 0.926094003 |
| chr8:124450168-124451676 | -0.076288734 | 0.808329396 | 0.926094003 |
| chr2:74530431-74530846 | -0.08644446 | 0.808345903 | 0.926094003 |
| chr7:98241478-98241781 | 0.099597177 | 0.808358498 | 0.926094003 |
| chr1:38990784-38991394 | 0.081176917 | 0.808385497 | 0.926094003 |
| chr6:22828120-22828822 | -0.086335428 | 0.808386708 | 0.926094003 |
| chr22:19687796-19688092 | 0.108787811 | 0.808392741 | 0.926094003 |
| chr20:30723520-30723855 | -0.103337518 | 0.808417661 | 0.926094003 |
| chr2:110211838-110212599 | 0.08824606 | 0.808420293 | 0.926094003 |
| chr9:34665906-34666293 | 0.083401727 | 0.808431677 | 0.926094003 |
| chr11:91484121-91484330 | -0.130866704 | 0.808439895 | 0.926094003 |
| chr15:31382856-31383056 | -0.125564459 | 0.808463378 | 0.926094003 |
| chr8:112761940-112762167 | 0.121786339 | 0.808464154 | 0.926094003 |
| chr20:50190340-50191465 | 0.077010434 | 0.808485357 | 0.926094003 |
| chr5:10452784-10453253 | 0.085260641 | 0.808488321 | 0.926094003 |
| chr6:75714232-75714432 | 0.127036302 | 0.808488374 | 0.926094003 |
| chr7:5383030-5383230 | -0.115051367 | 0.808492001 | 0.926094003 |
| chr4:128265614-128265851 | 0.116163392 | 0.808492078 | 0.926094003 |
| chrX:51614499-51614755 | -0.123623345 | 0.808522081 | 0.926114525 |
| chr1:11645779-11646082 | -0.101256357 | 0.808543203 | 0.926124874 |
| chr4:89110603-89111938 | 0.076623913 | 0.808630154 | 0.926209767 |
| chr2:202870743-202871103 | -0.086167387 | 0.808645681 | 0.926209767 |
| chr1:79617809-79618027 | -0.124154178 | 0.808653821 | 0.926209767 |
| chr16:87461398-87461920 | -0.081752576 | 0.808685954 | 0.926209767 |

| | | | |
|---|---|---|---|
| chr1:62180665-62180995 | -0.096599943 | 0.808689168 | 0.926209767 |
| chr22:38705306-38706010 | 0.079144445 | 0.808689847 | 0.926209767 |
| chr3:115179311-115179511 | -0.124945363 | 0.808726826 | 0.926238275 |
| chr16:3645953-3646591 | 0.083670251 | 0.808755994 | 0.926257837 |
| chrX:81958018-81958225 | -0.131853195 | 0.8087891 | 0.926278669 |
| chr5:80487797-80488700 | -0.077497869 | 0.808798362 | 0.926278669 |
| chr20:61939664-61939933 | -0.105243971 | 0.808856667 | 0.926331598 |
| chr2:127885857-127886601 | -0.083513425 | 0.808899024 | 0.926338892 |
| chr11:83083114-83083349 | -0.11147607 | 0.808922599 | 0.926338892 |
| chrX:49754330-49754538 | -0.126893944 | 0.808931982 | 0.926338892 |
| chr19:3500126-3500369 | -0.118391862 | 0.808934384 | 0.926338892 |
| chr11:67379225-67379982 | -0.0823271 | 0.808938209 | 0.926338892 |
| chr7:38354259-38354587 | -0.097798601 | 0.808939013 | 0.926338892 |
| chr10:60870445-60870645 | 0.121560602 | 0.808947664 | 0.926338892 |
| chr20:50794458-50795655 | 0.075048855 | 0.80906647 | 0.926461092 |
| chr2:86440623-86441725 | -0.076672617 | 0.809087329 | 0.926471132 |
| chr18:12246903-12247484 | -0.093938054 | 0.809130359 | 0.926480592 |
| chr1:35930557-35931673 | 0.078276673 | 0.809145148 | 0.926480592 |
| chr18:998918-999443 | 0.093838738 | 0.809147332 | 0.926480592 |
| chr3:64533502-64534188 | 0.076550978 | 0.809150265 | 0.926480592 |
| chr7:1845033-1845411 | 0.095966912 | 0.809174775 | 0.926480592 |
| chr2:241559452-241559679 | 0.101796636 | 0.809177707 | 0.926480592 |
| chr19:49503141-49503412 | -0.10399182 | 0.809180231 | 0.926480592 |
| chr7:130695952-130696159 | 0.123826185 | 0.809208182 | 0.926498749 |
| chr10:22320598-22321598 | 0.079545302 | 0.80922701 | 0.926506462 |
| chr6:3848596-3848796 | 0.118493097 | 0.809252469 | 0.926521766 |
| chr20:62065859-62066832 | 0.079129278 | 0.809302295 | 0.926560028 |
| chr1:199909067-199909453 | -0.097056846 | 0.809310074 | 0.926560028 |
| chr5:26076187-26076387 | -0.1334366 | 0.809339227 | 0.92657956 |
| chr8:119504377-119504577 | -0.128286332 | 0.809372511 | 0.926603821 |
| chr9:136484343-136484839 | 0.083325803 | 0.809400434 | 0.926607163 |
| chr15:75150178-75150399 | -0.111280226 | 0.809414687 | 0.926607163 |
| chr3:58491640-58492391 | 0.081125543 | 0.809424622 | 0.926607163 |
| chr11:118400955-118402324 | -0.078775038 | 0.809429395 | 0.926607163 |
| chrX:108139411-108139683 | -0.104724068 | 0.809435897 | 0.926607163 |
| chr3:78645829-78646042 | -0.120659139 | 0.809455146 | 0.926613125 |
| chr3:13657700-13658366 | 0.08266797 | 0.809468974 | 0.926613125 |
| chr9:124940546-124941932 | 0.077213385 | 0.809477384 | 0.926613125 |
| chr17:22310701-22311098 | -0.091395747 | 0.809520347 | 0.926622903 |
| chr16:81137107-81137336 | -0.109164718 | 0.809526832 | 0.926622903 |
| chr6:57268000-57268414 | 0.095079681 | 0.809528271 | 0.926622903 |
| chr14:59570075-59570426 | -0.095440931 | 0.8095343 | 0.926622903 |
| chr15:44711110-44712245 | -0.077707224 | 0.809557683 | 0.926635826 |
| chr3:161339404-161339922 | -0.083102038 | 0.809608093 | 0.926679683 |
| chr17:76681284-76681579 | -0.093340259 | 0.809634956 | 0.926683399 |
| chr13:96299332-96299532 | -0.138498086 | 0.80963926 | 0.926683399 |
| chr5:82174758-82174976 | -0.127602333 | 0.809662044 | 0.926683399 |
| chr12:30440903-30441131 | -0.107858128 | 0.809663879 | 0.926683399 |
| chr1:8424140-8424421 | 0.10599772 | 0.809680005 | 0.926683399 |
| chr11:31870851-31871367 | -0.090089262 | 0.809683905 | 0.926683399 |

| | | | |
|---|---|---|---|
| chr5:136132544-136133460 | -0.079112161 | 0.809718128 | 0.926690278 |
| chr16:71894279-71894669 | 0.086836158 | 0.809724933 | 0.926690278 |
| chr1:25875906-25876106 | -0.126025648 | 0.80975435 | 0.926690278 |
| chr21:33491314-33492078 | 0.080084587 | 0.809778382 | 0.926690278 |
| chr8:86198838-86199063 | -0.110279289 | 0.809781448 | 0.926690278 |
| chr14:52650871-52651157 | 0.104752322 | 0.809785784 | 0.926690278 |
| chr9:78331690-78331890 | -0.129676725 | 0.809790254 | 0.926690278 |
| chr5:163467749-163467949 | -0.129918146 | 0.809820519 | 0.926690278 |
| chr1:40338739-40339046 | -0.099981878 | 0.809824759 | 0.926690278 |
| chr2:209208190-209208452 | 0.110087853 | 0.809828482 | 0.926690278 |
| chr3:124174552-124174752 | -0.120556412 | 0.809835352 | 0.926690278 |
| chr5:15003507-15003766 | 0.095690583 | 0.809841034 | 0.926690278 |
| chr2:88690963-88692201 | -0.075878293 | 0.809847142 | 0.926690278 |
| chr6:35920561-35921497 | -0.080303637 | 0.809893492 | 0.926729475 |
| chr6:111552069-111552744 | -0.08508552 | 0.809934205 | 0.926762222 |
| chr6:57316524-57317113 | -0.080923873 | 0.809947528 | 0.926763626 |
| chr3:190862590-190863318 | -0.080659495 | 0.810020676 | 0.926833483 |
| chr14:28653559-28653843 | 0.106470925 | 0.810041255 | 0.926843189 |
| chr11:72123571-72123863 | -0.119782505 | 0.810064131 | 0.926855524 |
| chr10:69392466-69392866 | -0.088374309 | 0.810076744 | 0.926856115 |
| chr20:33654176-33654446 | -0.094324794 | 0.810125642 | 0.926883807 |
| chr14:22644032-22644349 | 0.097647645 | 0.810131271 | 0.926883807 |
| chr11:9384400-9385561 | 0.078834052 | 0.810137238 | 0.926883807 |
| chr2:3001660-3001911 | 0.103884665 | 0.810219716 | 0.926961646 |
| chr6:46821690-46821890 | 0.128442653 | 0.810232724 | 0.926961646 |
| chr5:136905705-136905920 | -0.109442985 | 0.810241566 | 0.926961646 |
| chr5:10863808-10864097 | -0.096465502 | 0.810278145 | 0.926989654 |
| chr14:90382685-90383768 | 0.076212159 | 0.810297858 | 0.926998365 |
| chr9:73515329-73515550 | 0.115541502 | 0.810313233 | 0.927002114 |
| chr19:47729979-47730437 | -0.090708862 | 0.810358803 | 0.927009784 |
| chr17:58992691-58992952 | 0.083115118 | 0.81036036 | 0.927009784 |
| chr2:218710437-218711449 | 0.077463356 | 0.810364626 | 0.927009784 |
| chr5:154296553-154297303 | -0.079952593 | 0.810368332 | 0.927009784 |
| chr4:124429387-124429601 | -0.116959832 | 0.810414739 | 0.927048481 |
| chr8:13566661-13567014 | -0.09460508 | 0.810428957 | 0.927048481 |
| chr7:73667494-73667989 | -0.084969047 | 0.810441873 | 0.927048481 |
| chrX:16869794-16870951 | 0.077421049 | 0.810450556 | 0.927048481 |
| chr8:54344661-54344871 | -0.138212479 | 0.810467007 | 0.927051169 |
| chr1:114780132-114781114 | -0.075293267 | 0.810477104 | 0.927051169 |
| chr6:7001380-7001685 | 0.101316693 | 0.810511619 | 0.927065505 |
| chr17:20226824-20227024 | -0.131597302 | 0.810521557 | 0.927065505 |
| chr16:795589-796115 | 0.084951528 | 0.810525935 | 0.927065505 |
| chr15:81303776-81304090 | 0.094939205 | 0.810590845 | 0.927087057 |
| chr11:79160805-79161033 | -0.105277631 | 0.810602209 | 0.927087057 |
| chr5:174327362-174327697 | 0.091562548 | 0.810610359 | 0.927087057 |
| chr10:98170471-98170840 | 0.090163421 | 0.810621784 | 0.927087057 |
| chrX:108699438-108699686 | -0.106937568 | 0.810633045 | 0.927087057 |
| chr22:30206978-30207321 | -0.100661229 | 0.81063757 | 0.927087057 |
| chr10:8628239-8628780 | -0.086475144 | 0.81064245 | 0.927087057 |
| chr6:113526594-113526876 | -0.101853165 | 0.810661517 | 0.927087057 |

| | | | |
|---|---|---|---|
| chr2:127884719-127885110 | -0.088648125 | 0.810671748 | 0.927087057 |
| chr20:53547660-53548174 | -0.08505387 | 0.810677367 | 0.927087057 |
| chr9:25974717-25974917 | 0.135917389 | 0.810677872 | 0.927087057 |
| chr2:16757229-16757536 | 0.091459618 | 0.810736207 | 0.927139931 |
| chr3:194671560-194671856 | 0.099982595 | 0.810805539 | 0.927205379 |
| chr12:64832035-64832297 | -0.110408831 | 0.810851484 | 0.927239626 |
| chr2:100416895-100417790 | 0.08479104 | 0.810887333 | 0.927239626 |
| chr8:125479716-125479964 | -0.098409806 | 0.810890321 | 0.927239626 |
| chr3:113086903-113087319 | -0.101461587 | 0.810894175 | 0.927239626 |
| chr12:55682195-55682500 | 0.09373806 | 0.810906379 | 0.927239626 |
| chr3:197750583-197750785 | 0.104980353 | 0.810918591 | 0.927239626 |
| chr16:11528264-11528464 | 0.128582653 | 0.810920197 | 0.927239626 |
| chr17:39450907-39451496 | 0.088868851 | 0.810933679 | 0.927241205 |
| chr2:66425843-66426043 | 0.1194137 | 0.8109583 | 0.927248543 |
| chr19:4495639-4495914 | -0.109515869 | 0.810967171 | 0.927248543 |
| chr3:115085294-115085522 | 0.110609265 | 0.810976402 | 0.927248543 |
| chr6:142232087-142232878 | -0.083040476 | 0.81102062 | 0.927285263 |
| chr15:92200956-92201156 | -0.127827387 | 0.811074254 | 0.927332748 |
| chrX:101234072-101234327 | 0.117404995 | 0.811147911 | 0.927403125 |
| chr8:111081030-111081269 | 0.114037444 | 0.811195729 | 0.927408861 |
| chr17:28725599-28726345 | -0.078099606 | 0.811196474 | 0.927408861 |
| chr13:94548904-94550315 | 0.075532303 | 0.811197049 | 0.927408861 |
| chr2:19819120-19819320 | -0.115248765 | 0.811201343 | 0.927408861 |
| chr20:53336958-53337189 | -0.10702633 | 0.811219047 | 0.927409763 |
| chrX:15565257-15565522 | -0.1120008 | 0.811226339 | 0.927409763 |
| chr17:82064759-82065046 | 0.096186423 | 0.811254145 | 0.927427714 |
| chr19:51107931-51108950 | -0.077471056 | 0.811283087 | 0.927446963 |
| chr1:203722299-203722509 | 0.106425851 | 0.811308973 | 0.927462718 |
| chr10:122953913-122954588 | 0.078057931 | 0.811353048 | 0.927499265 |
| chr3:189897284-189897705 | -0.10726901 | 0.811399555 | 0.927538591 |
| chr16:72996578-72997549 | 0.075799721 | 0.811463098 | 0.92759739 |
| chr19:42290818-42291154 | 0.095139829 | 0.81150754 | 0.927634354 |
| chr18:3481809-3482331 | -0.092361771 | 0.811531937 | 0.927648402 |
| chr11:842716-842971 | -0.103735727 | 0.811652681 | 0.927750737 |
| chr1:156421586-156422155 | 0.081416916 | 0.811653769 | 0.927750737 |
| chr1:61424347-61424681 | 0.110251506 | 0.811657786 | 0.927750737 |
| chr10:80460742-80461373 | -0.077869723 | 0.81168486 | 0.927767842 |
| chr17:76121417-76121727 | -0.090429104 | 0.811705352 | 0.927777425 |
| chr2:231563419-231563701 | 0.101216858 | 0.811734969 | 0.927791362 |
| chr8:123819199-123819800 | 0.082866051 | 0.811749954 | 0.927791362 |
| chr20:10447074-10447356 | -0.107742497 | 0.811757366 | 0.927791362 |
| chr3:184284657-184285230 | -0.082998652 | 0.811767643 | 0.927791362 |
| chr1:18956571-18957037 | 0.081581406 | 0.811778089 | 0.927791362 |
| chr17:55549155-55549458 | 0.095184464 | 0.81181083 | 0.927804012 |
| chr15:73676926-73677397 | 0.091122492 | 0.811834839 | 0.927804012 |
| chr2:210188791-210189081 | -0.120131371 | 0.811851038 | 0.927804012 |
| chr3:25911158-25911358 | 0.125275458 | 0.811855462 | 0.927804012 |
| chr3:32177139-32177772 | 0.085450581 | 0.811857627 | 0.927804012 |
| chr3:184249386-184250594 | 0.075809287 | 0.81186181 | 0.927804012 |
| chr19:35090293-35090793 | -0.088071778 | 0.811881604 | 0.927806004 |

| | | | |
|---|---|---|---|
| chr7:146089963-146090356 | -0.094239922 | 0.811895305 | 0.927806004 |
| chr20:38033126-38034203 | 0.078151074 | 0.81189988 | 0.927806004 |
| chr11:60265711-60265965 | -0.125470119 | 0.811929673 | 0.927817667 |
| chr2:225665456-225665752 | -0.120963872 | 0.811934303 | 0.927817667 |
| chr15:98799336-98799776 | -0.089505355 | 0.811963098 | 0.927836734 |
| chr2:218152831-218153031 | -0.132922983 | 0.811997201 | 0.927861026 |
| chr5:36734556-36734765 | 0.113935934 | 0.812008576 | 0.927861026 |
| chr2:8583403-8584182 | 0.078073434 | 0.812136873 | 0.927968433 |
| chr21:42955546-42956028 | -0.088081545 | 0.812138114 | 0.927968433 |
| chr5:172162437-172162643 | -0.11062935 | 0.812138905 | 0.927968433 |
| chr6:30942526-30942966 | 0.088321171 | 0.812199023 | 0.928023286 |
| chr1:166902310-166902579 | -0.099528436 | 0.812273372 | 0.928094398 |
| chr1:202355858-202356163 | -0.099253966 | 0.812294355 | 0.928104533 |
| chr1:156746710-156747310 | 0.07987096 | 0.812313766 | 0.928112871 |
| chr12:68807336-68809101 | -0.074702408 | 0.812340859 | 0.928129987 |
| chr11:2859803-2860025 | -0.127836743 | 0.812366155 | 0.928131905 |
| chr17:1997274-1997805 | -0.08930729 | 0.812366764 | 0.928131905 |
| chr1:152909338-152909538 | 0.123374677 | 0.81242542 | 0.928182302 |
| chr17:20155597-20156622 | 0.077034521 | 0.812435103 | 0.928182302 |
| chr8:127509238-127509438 | -0.110290419 | 0.812464558 | 0.92819507 |
| chr5:180618754-180619197 | -0.086018092 | 0.812481558 | 0.92819507 |
| chr19:10702018-10702218 | 0.110635897 | 0.812482621 | 0.92819507 |
| chr6:3990848-3991241 | 0.089500941 | 0.81250154 | 0.928202845 |
| chr5:141826268-141826468 | -0.113190815 | 0.812531354 | 0.928210543 |
| chr1:207404987-207405217 | 0.102318798 | 0.812532507 | 0.928210543 |
| chr17:59027737-59028144 | 0.078652421 | 0.812578033 | 0.92821318 |
| chr12:93006490-93006690 | -0.131149838 | 0.812584619 | 0.92821318 |
| chr12:69470018-69471099 | -0.076654097 | 0.812586038 | 0.92821318 |
| chr15:67125043-67126007 | -0.081668044 | 0.812599699 | 0.92821318 |
| chr10:69332904-69333779 | -0.078243428 | 0.812600404 | 0.92821318 |
| chr8:90606056-90606342 | 0.100926216 | 0.812607501 | 0.92821318 |
| chr2:160970133-160970527 | 0.099685897 | 0.81262246 | 0.92821643 |
| chr8:144467605-144467918 | 0.083461664 | 0.812637141 | 0.928219362 |
| chrX:48521293-48522162 | 0.077082077 | 0.812682593 | 0.92825744 |
| chr15:63503730-63504903 | -0.075220034 | 0.812747687 | 0.928299543 |
| chr17:64938874-64939295 | -0.085283318 | 0.812761017 | 0.928299543 |
| chr9:136841063-136841701 | -0.081706113 | 0.812770177 | 0.928299543 |
| chr1:116537519-116537742 | -0.104969115 | 0.812777777 | 0.928299543 |
| chr17:59859494-59859765 | 0.078423406 | 0.81278003 | 0.928299543 |
| chr9:136109363-136109995 | 0.081611802 | 0.812805896 | 0.928301623 |
| chr5:11845243-11845443 | -0.138721825 | 0.812828227 | 0.928301623 |
| chr12:7188593-7188862 | -0.100125557 | 0.812852704 | 0.928301623 |
| chr4:153248748-153249800 | -0.076430093 | 0.812856843 | 0.928301623 |
| chr11:17479388-17479868 | -0.089861947 | 0.812862591 | 0.928301623 |
| chr5:38856240-38856440 | 0.123917784 | 0.812865984 | 0.928301623 |
| chr16:70439088-70439654 | 0.08147846 | 0.812866659 | 0.928301623 |
| chr10:92238872-92239193 | -0.093572765 | 0.812921805 | 0.928313139 |
| chr6:166774873-166775073 | -0.122235817 | 0.812932995 | 0.928313139 |
| chr6:41584789-41585309 | 0.085920932 | 0.812937952 | 0.928313139 |
| chr2:188280859-188281224 | 0.101154273 | 0.812955315 | 0.928313139 |

| | | | |
|---|---|---|---|
| chr3:125936280-125936841 | 0.087780649 | 0.812955625 | 0.928313139 |
| chr11:62975193-62975395 | 0.114497349 | 0.812959107 | 0.928313139 |
| chr13:113490561-113491333 | 0.079411058 | 0.812961551 | 0.928313139 |
| chr1:166910410-166910748 | -0.089525264 | 0.812981215 | 0.928316197 |
| chr22:36286535-36286816 | -0.105985691 | 0.812991003 | 0.928316197 |
| chr2:173963727-173964488 | 0.076398235 | 0.813000576 | 0.928316197 |
| chr17:36938224-36938435 | 0.112313022 | 0.8130324 | 0.928338701 |
| chr3:87006920-87007350 | 0.102517094 | 0.81311172 | 0.928399065 |
| chr8:98840807-98841179 | -0.096320876 | 0.813114167 | 0.928399065 |
| chrX:93874628-93874828 | 0.136074983 | 0.813121616 | 0.928399065 |
| chr4:123587862-123588500 | 0.083313749 | 0.813189505 | 0.928462744 |
| chr19:36666463-36667148 | 0.080008439 | 0.813206588 | 0.928465203 |
| chrX:40068118-40068398 | 0.107593089 | 0.813232397 | 0.928465203 |
| chr7:4245901-4246154 | 0.099087487 | 0.813242565 | 0.928465203 |
| chr6:55069190-55069457 | 0.10936054 | 0.81324413 | 0.928465203 |
| chr10:18226415-18226683 | 0.106324538 | 0.813252246 | 0.928465203 |
| chr10:110643893-110644280 | 0.100211532 | 0.813278564 | 0.928470102 |
| chr8:49911557-49911829 | 0.101030914 | 0.813308736 | 0.928470102 |
| chr7:39682112-39682312 | 0.121790352 | 0.813318943 | 0.928470102 |
| chr20:64255262-64256456 | -0.077033791 | 0.813321272 | 0.928470102 |
| chr11:85941257-85941852 | -0.083970023 | 0.813322442 | 0.928470102 |
| chr10:73653666-73654142 | 0.093832202 | 0.813329243 | 0.928470102 |
| chr18:74551971-74552287 | -0.093903582 | 0.813399072 | 0.928533105 |
| chr3:170907871-170908943 | -0.075192187 | 0.813408669 | 0.928533105 |
| chr14:51283372-51283710 | -0.092410367 | 0.813433715 | 0.928547862 |
| chr1:174942946-174943146 | 0.118013372 | 0.813489203 | 0.928594498 |
| chrX:13724869-13725234 | -0.102458664 | 0.813507569 | 0.928594498 |
| chr20:35741948-35742789 | -0.075377482 | 0.813510927 | 0.928594498 |
| chr9:2014710-2015512 | -0.076526994 | 0.813536977 | 0.928610398 |
| chr11:112024700-112025771 | 0.077127593 | 0.813574915 | 0.928639869 |
| chr9:35051435-35051676 | -0.106720308 | 0.813596122 | 0.928650241 |
| chr9:94630352-94630660 | -0.101600529 | 0.813649633 | 0.928697485 |
| chr2:118032448-118032657 | -0.11042545 | 0.813751895 | 0.928791645 |
| chr16:16973802-16974147 | -0.105550221 | 0.813757495 | 0.928791645 |
| chr19:46850171-46851195 | -0.07631778 | 0.813768493 | 0.928791645 |
| chr15:100512388-100512674 | -0.114277299 | 0.81380457 | 0.928818985 |
| chr11:46237204-46237475 | 0.104205913 | 0.813831047 | 0.928822851 |
| chr20:35454502-35455600 | 0.07607032 | 0.813838845 | 0.928822851 |
| chr6:149316795-149317940 | -0.076708593 | 0.813860103 | 0.928822851 |
| chr2:71871583-71871935 | -0.089836739 | 0.813879374 | 0.928822851 |
| chr2:109407763-109408259 | -0.08282368 | 0.813888433 | 0.928822851 |
| chr17:66964207-66964481 | 0.089002778 | 0.813890748 | 0.928822851 |
| chr16:89711566-89712323 | 0.078921791 | 0.813894826 | 0.928822851 |
| chr14:82271944-82272157 | -0.126654566 | 0.813904934 | 0.928822851 |
| chr10:103350284-103351321 | 0.080695137 | 0.81393207 | 0.928839647 |
| chr12:57088455-57089080 | -0.076536861 | 0.813943897 | 0.928839647 |
| chr10:5623090-5623483 | 0.082533137 | 0.813962855 | 0.928847447 |
| chr20:21505106-21505647 | 0.087287477 | 0.813975185 | 0.928847685 |
| chr15:80058877-80060355 | -0.074614397 | 0.814014494 | 0.928878707 |
| chr17:68976487-68976821 | 0.098308846 | 0.814032861 | 0.928884708 |

| | | | |
|---|---|---|---|
| chr16:20383464-20383664 | -0.119888963 | 0.81405301 | 0.928884708 |
| chr15:20356221-20356723 | -0.085416222 | 0.814056122 | 0.928884708 |
| chr1:161399123-161400221 | 0.075011832 | 0.814068698 | 0.928885225 |
| chr8:49650641-49650841 | 0.120639438 | 0.814087285 | 0.928892601 |
| chr12:56636672-56637284 | 0.076239291 | 0.814112429 | 0.928907458 |
| chr8:52467002-52467742 | -0.078360651 | 0.814169281 | 0.928949921 |
| chr2:28527200-28527435 | 0.116428364 | 0.814173892 | 0.928949921 |
| chr6:170465029-170465274 | 0.109519469 | 0.814207651 | 0.928974606 |
| chr1:63367180-63368154 | -0.075451789 | 0.814252782 | 0.929012265 |
| chr3:33717089-33717289 | 0.110519527 | 0.81428207 | 0.929022381 |
| chr5:139513600-139513870 | 0.093620865 | 0.814290588 | 0.929022381 |
| chr13:52707442-52707747 | -0.102584313 | 0.814298023 | 0.929022381 |
| chr11:125902783-125903369 | 0.084236201 | 0.814324123 | 0.929027224 |
| chr4:98260121-98261609 | -0.074632333 | 0.814333617 | 0.929027224 |
| chr14:98230928-98231128 | -0.12846377 | 0.814343057 | 0.929027224 |
| chr10:16853290-16853524 | 0.113719997 | 0.814350767 | 0.929027224 |
| chr17:72351798-72352425 | 0.080550138 | 0.814373218 | 0.929039004 |
| chr11:1548452-1548714 | -0.110622287 | 0.814399162 | 0.929054769 |
| chr11:93661381-93662173 | 0.077882879 | 0.814434025 | 0.929077079 |
| chr3:197554597-197556276 | -0.074687048 | 0.81444297 | 0.929077079 |
| chr2:120470290-120471002 | -0.080772211 | 0.814469731 | 0.929090169 |
| chr12:54216949-54217296 | -0.090688834 | 0.814478696 | 0.929090169 |
| chr12:42237419-42238926 | 0.074770149 | 0.814516532 | 0.92911308 |
| chr1:46489191-46489999 | 0.07954517 | 0.814523033 | 0.92911308 |
| chrX:13750223-13750528 | -0.105034593 | 0.814570446 | 0.929153331 |
| chr3:147667225-147667425 | 0.143036081 | 0.814607774 | 0.929182077 |
| chr20:54148833-54149489 | -0.073904772 | 0.814621871 | 0.929183022 |
| chr7:151080623-151081075 | 0.079499125 | 0.81463794 | 0.929183022 |
| chr6:28474299-28474801 | -0.079308876 | 0.814644983 | 0.929183022 |
| chr1:171313516-171314301 | -0.075645823 | 0.814694123 | 0.929225238 |
| chr7:105816544-105816814 | -0.109047052 | 0.814727501 | 0.929249477 |
| chr6:4358482-4359114 | 0.077624495 | 0.81479588 | 0.929309911 |
| chrX:53093055-53093475 | 0.08440995 | 0.814804744 | 0.929309911 |
| chr6:87472503-87473617 | -0.0752284 | 0.814829514 | 0.929313644 |
| chr21:26732516-26732940 | -0.090729254 | 0.814834681 | 0.929313644 |
| chr1:201170477-201171668 | 0.073466677 | 0.814844403 | 0.929313644 |
| chr9:124771962-124772241 | 0.091460875 | 0.814880208 | 0.929340646 |
| chr11:75791145-75791359 | 0.102094851 | 0.814920338 | 0.92937258 |
| chr4:113625526-113625736 | -0.11722537 | 0.814950362 | 0.929392987 |
| chr1:32362153-32362497 | 0.092888013 | 0.81498672 | 0.929404637 |
| chr8:101493025-101493520 | -0.077899524 | 0.814996199 | 0.929404637 |
| chr17:50150934-50151300 | -0.080791496 | 0.814996966 | 0.929404637 |
| chrX:82813530-82814175 | 0.077979893 | 0.815018862 | 0.929415774 |
| chr7:32433741-32433941 | -0.114557694 | 0.815078446 | 0.929469888 |
| chr17:56469300-56469593 | -0.101483461 | 0.81510106 | 0.929479726 |
| chr1:196143950-196144300 | 0.093453225 | 0.815111335 | 0.929479726 |
| chr6:47413999-47414333 | -0.092056531 | 0.815142661 | 0.929501614 |
| chr20:21529925-21530139 | 0.121455851 | 0.815160992 | 0.929508684 |
| chr20:57243630-57245119 | 0.072542253 | 0.815182044 | 0.929518857 |
| chr17:81967746-81968232 | -0.085286037 | 0.815198993 | 0.929522951 |

| | | | |
|---|---|---|---|
| chr11:123083485-123083922 | -0.089443129 | 0.815265294 | 0.929522951 |
| chr19:47296178-47296468 | 0.110934205 | 0.815266508 | 0.929522951 |
| chr4:100780102-100780461 | 0.094322547 | 0.815282985 | 0.929522951 |
| chr1:97525443-97525972 | 0.084923509 | 0.815299582 | 0.929522951 |
| chr16:23452648-23453570 | 0.078699073 | 0.815305562 | 0.929522951 |
| chr8:139646947-139647155 | -0.099821963 | 0.815308508 | 0.929522951 |
| chr5:122516170-122516627 | 0.090017068 | 0.815322019 | 0.929522951 |
| chr6:20182935-20183508 | 0.080950721 | 0.815327625 | 0.929522951 |
| chr7:538880-539088 | -0.092795749 | 0.815328927 | 0.929522951 |
| chr4:43132103-43132337 | -0.110514722 | 0.815331297 | 0.929522951 |
| chr4:169492557-169492757 | -0.118322472 | 0.815361341 | 0.929522951 |
| chr3:19910659-19910859 | -0.126027412 | 0.815363285 | 0.929522951 |
| chr16:67718935-67719816 | -0.076808454 | 0.815366007 | 0.929522951 |
| chr17:39958986-39959226 | 0.111945412 | 0.815387149 | 0.929522951 |
| chr15:74835332-74835690 | 0.082851141 | 0.815404508 | 0.929522951 |
| chr3:48187538-48187776 | 0.103002824 | 0.815427615 | 0.929522951 |
| chr19:13905914-13906337 | 0.090471037 | 0.815429782 | 0.929522951 |
| chr10:131900381-131901290 | 0.075161755 | 0.815436214 | 0.929522951 |
| chr2:106839890-106840203 | 0.092481838 | 0.815448975 | 0.929522951 |
| chr8:109652461-109652764 | 0.091279723 | 0.815449997 | 0.929522951 |
| chr7:100725257-100725977 | 0.084027054 | 0.815452523 | 0.929522951 |
| chr19:4800821-4801164 | 0.08798139 | 0.815476197 | 0.929531515 |
| chr8:66612755-66613843 | 0.07413304 | 0.815484299 | 0.929531515 |
| chr10:110497325-110498033 | -0.07629187 | 0.815521019 | 0.929559542 |
| chr10:69901664-69901914 | -0.100668266 | 0.815596257 | 0.929631471 |
| chr17:66625329-66625825 | 0.082821195 | 0.815612323 | 0.929635954 |
| chr16:81277470-81278102 | -0.078095318 | 0.815638737 | 0.929652232 |
| chr17:67285450-67285664 | 0.103688459 | 0.815659449 | 0.929662011 |
| chr3:125150572-125150828 | 0.096525424 | 0.815679692 | 0.929671254 |
| chr5:94778549-94778793 | 0.115299936 | 0.815725596 | 0.929704436 |
| chr10:8908134-8908352 | 0.110751565 | 0.815733073 | 0.929704436 |
| chr14:62377964-62378470 | -0.085241137 | 0.81575832 | 0.929717215 |
| chrX:131366489-131366735 | 0.11462578 | 0.815775187 | 0.929717215 |
| chr12:6607375-6607675 | -0.08918262 | 0.815784244 | 0.929717215 |
| chr14:88632588-88632788 | 0.116437246 | 0.815792821 | 0.929717215 |
| chr8:98426780-98427135 | 0.08523164 | 0.815815405 | 0.929729125 |
| chr2:220165485-220165754 | 0.105716908 | 0.815910795 | 0.929814302 |
| chrX:120213555-120213959 | 0.094607033 | 0.815914416 | 0.929814302 |
| chr14:101381415-101381800 | 0.100107985 | 0.815937212 | 0.929826451 |
| chr22:41597307-41597688 | -0.090143647 | 0.815949948 | 0.929827135 |
| chr1:246497384-246497688 | -0.090507676 | 0.81596764 | 0.929833468 |
| chr10:103931222-103931655 | -0.091711405 | 0.816028127 | 0.929888565 |
| chr2:55050194-55051004 | -0.073938418 | 0.816043472 | 0.929892222 |
| chr19:10113060-10113265 | 0.099570282 | 0.816068834 | 0.929907293 |
| chr1:159899899-159900785 | 0.07442609 | 0.816100698 | 0.929929772 |
| chr5:115098169-115098408 | -0.103132922 | 0.81616713 | 0.929984789 |
| chr6:118893432-118895347 | -0.072985907 | 0.816201113 | 0.929984789 |
| chr3:71753720-71754607 | 0.079369886 | 0.816207753 | 0.929984789 |
| chr8:118711923-118712123 | 0.12421487 | 0.816211574 | 0.929984789 |
| chr2:40451514-40452333 | 0.081626356 | 0.816220968 | 0.929984789 |

| | | | |
|---|---|---|---|
| chr4:119300068-119301147 | -0.074022369 | 0.816226026 | 0.929984789 |
| chr10:74889684-74889884 | 0.116240707 | 0.816250058 | 0.929984789 |
| chr8:127894703-127894905 | 0.091641346 | 0.816255846 | 0.929984789 |
| chr4:81052489-81052795 | -0.094304454 | 0.816258216 | 0.929984789 |
| chr13:104993816-104994026 | -0.127683987 | 0.816336886 | 0.93006059 |
| chr4:95549689-95549897 | -0.103301425 | 0.816384689 | 0.930101223 |
| chr7:1086706-1087391 | -0.078535739 | 0.8164061 | 0.930111786 |
| chrX:82218483-82218683 | 0.116869739 | 0.816461532 | 0.930128552 |
| chr2:204360267-204360467 | 0.122194625 | 0.816468705 | 0.930128552 |
| chr19:38899328-38900293 | 0.074709987 | 0.816495546 | 0.930128552 |
| chr7:46352427-46352644 | 0.108256315 | 0.81650817 | 0.930128552 |
| chr18:712284-713286 | -0.07476977 | 0.816512026 | 0.930128552 |
| chr6:144931511-144931788 | -0.10161569 | 0.816535184 | 0.930128552 |
| chr19:51702810-51703187 | -0.089133642 | 0.816547681 | 0.930128552 |
| chr20:50460557-50461069 | -0.080380275 | 0.81656774 | 0.930128552 |
| chr3:38662632-38662874 | 0.107217177 | 0.816567957 | 0.930128552 |
| chr11:130609085-130609318 | 0.108235626 | 0.816575624 | 0.930128552 |
| chr12:98593075-98594403 | -0.073158872 | 0.816577082 | 0.930128552 |
| chr1:161209810-161210218 | -0.085095028 | 0.816613918 | 0.930128552 |
| chr1:161132237-161133355 | 0.074442532 | 0.816617204 | 0.930128552 |
| chr9:42743507-42743758 | -0.107980807 | 0.816617651 | 0.930128552 |
| chr5:59039127-59039565 | 0.09410972 | 0.816625197 | 0.930128552 |
| chr20:40903412-40903770 | 0.089032906 | 0.816626969 | 0.930128552 |
| chr15:99250862-99251853 | 0.073593576 | 0.816627182 | 0.930128552 |
| chr17:44911127-44911459 | 0.09331817 | 0.81666892 | 0.930131314 |
| chr4:124939758-124940280 | -0.082745433 | 0.816676937 | 0.930131314 |
| chr3:19653255-19653455 | 0.124205044 | 0.816694571 | 0.930131314 |
| chr4:88104877-88105238 | 0.090473578 | 0.8167234 | 0.930131314 |
| chr1:182186388-182186605 | -0.121319552 | 0.816733151 | 0.930131314 |
| chr16:89192244-89192648 | 0.087416822 | 0.816735158 | 0.930131314 |
| chr6:122726699-122726940 | -0.113894102 | 0.816737456 | 0.930131314 |
| chr12:124862950-124864319 | 0.074948579 | 0.816742758 | 0.930131314 |
| chr11:1435824-1436236 | 0.103752542 | 0.816750913 | 0.930131314 |
| chr3:31531890-31533103 | 0.072700031 | 0.816750999 | 0.930131314 |
| chr5:76083211-76083419 | 0.109399734 | 0.816810689 | 0.930179727 |
| chr7:33953494-33953715 | 0.123033102 | 0.816825013 | 0.930179727 |
| chr21:33362073-33362526 | -0.086359694 | 0.816829931 | 0.930179727 |
| chr10:44301002-44301676 | 0.078026729 | 0.816883595 | 0.930227013 |
| chr22:46399199-46399399 | -0.122999296 | 0.816897122 | 0.930228592 |
| chr6:10412131-10413499 | -0.073424885 | 0.816909882 | 0.930229298 |
| chr8:86892980-86893614 | 0.083003026 | 0.816937394 | 0.930246802 |
| chr12:78473216-78473430 | -0.115740898 | 0.816958301 | 0.930253999 |
| chr13:74134273-74134664 | 0.080462914 | 0.816983443 | 0.930253999 |
| chr6:12893932-12894132 | 0.122692259 | 0.816983847 | 0.930253999 |
| chr14:68468689-68469274 | 0.083362191 | 0.816992278 | 0.930253999 |
| chr16:3879886-3880148 | 0.098877537 | 0.817010514 | 0.930257196 |
| chr1:206685272-206686639 | 0.073547227 | 0.817030266 | 0.930257196 |
| chr5:103099087-103099292 | -0.123765536 | 0.817031509 | 0.930257196 |
| chr4:55386113-55386412 | -0.096350068 | 0.817058809 | 0.930274456 |
| chr11:34174158-34175109 | -0.079406649 | 0.817084815 | 0.930277525 |

| | | | |
|---|---|---|---|
| chr4:74308188-74308446 | -0.10937759 | 0.817085787 | 0.930277525 |
| chr14:49950959-49951228 | 0.09690236 | 0.817122647 | 0.930285902 |
| chr8:57823748-57824173 | -0.091454499 | 0.817132721 | 0.930285902 |
| chr17:31549427-31550353 | 0.077807099 | 0.817144769 | 0.930285902 |
| chr10:126935392-126935638 | 0.11197721 | 0.81715604 | 0.930285902 |
| chr16:80991126-80991368 | 0.110851894 | 0.817165168 | 0.930285902 |
| chr3:120169496-120169710 | 0.107042054 | 0.817165992 | 0.930285902 |
| chr20:62237951-62238704 | 0.073607819 | 0.817186749 | 0.93029571 |
| chr2:122415349-122415783 | -0.106729047 | 0.817214738 | 0.930307361 |
| chr5:142866879-142867412 | -0.084115597 | 0.8172272 | 0.930307361 |
| chr1:151197733-151199076 | 0.072616296 | 0.817233407 | 0.930307361 |
| chr6:100948547-100948997 | -0.097552301 | 0.817301732 | 0.930371316 |
| chrX:129677837-129678105 | -0.100646604 | 0.817361101 | 0.930425076 |
| chr3:9704336-9704707 | 0.090708515 | 0.817388699 | 0.930442669 |
| chr5:135830148-135830348 | -0.104879345 | 0.817464796 | 0.930456252 |
| chr3:187063786-187064153 | 0.086034163 | 0.817468928 | 0.930456252 |
| chr12:64273740-64274403 | -0.079970689 | 0.817475253 | 0.930456252 |
| chr17:28728433-28729248 | -0.074126898 | 0.817481257 | 0.930456252 |
| chr10:92573465-92574872 | 0.073273849 | 0.817481912 | 0.930456252 |
| chr3:42590267-42591044 | -0.075027714 | 0.817492062 | 0.930456252 |
| chr20:45373690-45374194 | -0.081109686 | 0.817492568 | 0.930456252 |
| chr12:61654782-61655077 | -0.109157064 | 0.81749778 | 0.930456252 |
| chr9:129884956-129885378 | -0.086351446 | 0.817554861 | 0.930507399 |
| chr15:99792129-99792522 | 0.087624069 | 0.817584692 | 0.930514632 |
| chr1:66446471-66446719 | -0.110028376 | 0.817601822 | 0.930514632 |
| chr1:212477071-212477482 | -0.085206266 | 0.817603487 | 0.930514632 |
| chr14:75132458-75132668 | 0.114451731 | 0.817609794 | 0.930514632 |
| chr4:128723518-128723771 | 0.104226085 | 0.817630249 | 0.930518214 |
| chr10:118341729-118342703 | -0.076383625 | 0.817645438 | 0.930518214 |
| chr2:96335545-96336611 | 0.075167961 | 0.817664065 | 0.930518214 |
| chr15:80152252-80153305 | -0.073648209 | 0.817668366 | 0.930518214 |
| chr6:76020368-76020638 | -0.126176963 | 0.817673662 | 0.930518214 |
| chr2:193240823-193241103 | -0.109072761 | 0.817708415 | 0.930543943 |
| chr6:140230309-140230880 | -0.087950573 | 0.817733106 | 0.93055822 |
| chr20:50499467-50499677 | 0.103194071 | 0.817746905 | 0.930560102 |
| chr2:135916299-135916499 | -0.126637259 | 0.817765836 | 0.930567824 |
| chr3:149224792-149225420 | -0.083736626 | 0.817786465 | 0.930571109 |
| chr10:103931705-103932033 | -0.094001765 | 0.817802718 | 0.930571109 |
| chr17:60170600-60170908 | 0.076991263 | 0.817805158 | 0.930571109 |
| chr3:143973145-143973383 | -0.083238271 | 0.817820115 | 0.930574309 |
| chr1:198276714-198276959 | -0.108845313 | 0.817841793 | 0.930585156 |
| chr3:70550787-70551048 | 0.103770272 | 0.817883993 | 0.930606806 |
| chr18:7221460-7221896 | -0.094484891 | 0.817904675 | 0.930606806 |
| chr2:153163208-153163417 | 0.120271761 | 0.817906444 | 0.930606806 |
| chr11:76235049-76236503 | -0.072221939 | 0.817909401 | 0.930606806 |
| chr5:151427320-151427520 | 0.120699883 | 0.817941337 | 0.930619641 |
| chr3:36843671-36843990 | 0.090009766 | 0.817944973 | 0.930619641 |
| chr1:204515876-204517004 | -0.072962871 | 0.817966142 | 0.930629907 |
| chr5:10249360-10251122 | -0.072058187 | 0.817979328 | 0.93063109 |
| chr9:131366937-131367183 | 0.109905319 | 0.818010236 | 0.930640819 |

| | | | |
|---|---|---|---|
| chr7:49852678-49853140 | -0.083705936 | 0.818017909 | 0.930640819 |
| chr5:18390290-18390527 | -0.103027541 | 0.818024316 | 0.930640819 |
| chr8:66954502-66955219 | -0.080860291 | 0.818128729 | 0.930742528 |
| chr17:56425611-56425827 | -0.107406748 | 0.818138012 | 0.930742528 |
| chr17:65432660-65432860 | -0.113668819 | 0.818182055 | 0.93077375 |
| chr15:72599660-72600002 | -0.106700053 | 0.818195863 | 0.93077375 |
| chr3:147405806-147406006 | -0.112456336 | 0.818201899 | 0.93077375 |
| chr3:17515600-17515802 | 0.125820941 | 0.818234195 | 0.930775368 |
| chr5:133496889-133497127 | -0.114755114 | 0.818234257 | 0.930775368 |
| chr1:145691257-145691543 | 0.094338373 | 0.818239764 | 0.930775368 |
| chr2:191050063-191050368 | -0.092754539 | 0.81828341 | 0.930811198 |
| chr4:18418824-18419024 | -0.11504115 | 0.818381011 | 0.930902858 |
| chr17:45247348-45248143 | 0.075739865 | 0.818394204 | 0.930902858 |
| chr3:183122392-183122633 | -0.1114706 | 0.818420621 | 0.930902858 |
| chr22:19751192-19751495 | 0.102838755 | 0.818421029 | 0.930902858 |
| chr10:93296430-93296684 | -0.11627939 | 0.81842661 | 0.930902858 |
| chr5:98990039-98990319 | -0.112904615 | 0.818441092 | 0.930902858 |
| chr19:48264434-48264634 | -0.134463831 | 0.818449034 | 0.930902858 |
| chr10:5413151-5413351 | 0.10532624 | 0.818509379 | 0.930937692 |
| chr3:141192804-141193011 | -0.114684139 | 0.818530001 | 0.930937692 |
| chr14:72714987-72715256 | 0.098827152 | 0.818541978 | 0.930937692 |
| chr8:133696726-133696926 | -0.114220461 | 0.818552052 | 0.930937692 |
| chr7:51620825-51621025 | -0.116343762 | 0.818574633 | 0.930937692 |
| chr6:118934336-118935397 | -0.073114389 | 0.818578272 | 0.930937692 |
| chr12:77724440-77724725 | -0.095295578 | 0.818594516 | 0.930937692 |
| chr2:30026294-30026724 | -0.087625141 | 0.818596649 | 0.930937692 |
| chr11:75204934-75205260 | -0.086526257 | 0.818610254 | 0.930937692 |
| chr1:211542059-211542325 | -0.096106396 | 0.818619371 | 0.930937692 |
| chr1:31939797-31940193 | -0.082266211 | 0.818625327 | 0.930937692 |
| chr16:10688416-10688642 | 0.11461998 | 0.818625458 | 0.930937692 |
| chr6:137717512-137717712 | 0.129295732 | 0.818670995 | 0.93097566 |
| chr3:129278381-129279946 | 0.072627906 | 0.81869556 | 0.930986323 |
| chr16:79836125-79836526 | 0.099353672 | 0.818704673 | 0.930986323 |
| chr7:43108409-43108718 | -0.088815942 | 0.818718305 | 0.930988008 |
| chr10:122518448-122518648 | 0.135650385 | 0.818802953 | 0.931034803 |
| chr3:130276220-130276502 | 0.091969078 | 0.81881111 | 0.931034803 |
| chr5:28710459-28710659 | -0.124525471 | 0.818821923 | 0.931034803 |
| chr5:61298296-61298496 | -0.124525471 | 0.818821923 | 0.931034803 |
| chr5:140107337-140108357 | 0.074358125 | 0.818826478 | 0.931034803 |
| chr3:39260992-39261252 | -0.115406833 | 0.818840633 | 0.931034803 |
| chr6:112010452-112010721 | -0.102576171 | 0.818844514 | 0.931034803 |
| chr7:73725745-73726420 | 0.079910787 | 0.818891999 | 0.931056998 |
| chr20:13008786-13009421 | 0.08155839 | 0.818898455 | 0.931056998 |
| chr1:203305101-203305929 | 0.073577494 | 0.818917971 | 0.931056998 |
| chr5:40834847-40835733 | 0.073478172 | 0.818918798 | 0.931056998 |
| chr5:12294233-12294496 | -0.100038762 | 0.818948833 | 0.931056998 |
| chr17:75288763-75289887 | 0.072174699 | 0.818953222 | 0.931056998 |
| chr5:144247203-144247453 | 0.107204564 | 0.81895772 | 0.931056998 |
| chr4:162407075-162407285 | -0.11502747 | 0.818973204 | 0.931056998 |
| chr9:33415290-33416129 | -0.075023556 | 0.818975688 | 0.931056998 |

| | | | |
|---|---|---|---|
| chr9:37034232-37035101 | -0.079181559 | 0.819001336 | 0.931056998 |
| chr10:96626999-96627199 | 0.127365361 | 0.81902979 | 0.931056998 |
| chr16:85253582-85253853 | 0.090432355 | 0.819034432 | 0.931056998 |
| chr10:86100532-86100804 | -0.091293952 | 0.819034468 | 0.931056998 |
| chr17:2347673-2348276 | -0.076727302 | 0.819042324 | 0.931056998 |
| chr16:3654390-3654590 | -0.105487286 | 0.819046304 | 0.931056998 |
| chr13:25371617-25371869 | -0.105945666 | 0.819062008 | 0.931061037 |
| chr2:60642045-60642245 | -0.119018402 | 0.819082433 | 0.931070442 |
| chr13:34488539-34488787 | 0.115716135 | 0.819100776 | 0.93107748 |
| chr17:58692222-58693179 | -0.071507166 | 0.819174581 | 0.931133403 |
| chr11:36509637-36510728 | -0.074414938 | 0.819185925 | 0.931133403 |
| chr9:129726524-129727058 | -0.082084823 | 0.819189844 | 0.931133403 |
| chr5:147822732-147822988 | -0.103795668 | 0.819216808 | 0.931133403 |
| chr6:7542901-7543316 | 0.083322 | 0.819217543 | 0.931133403 |
| chr1:197854930-197855166 | 0.104836961 | 0.819223302 | 0.931133403 |
| chr3:14431891-14432666 | 0.075053377 | 0.819251937 | 0.931133403 |
| chr17:7456283-7456681 | 0.091068895 | 0.819255812 | 0.931133403 |
| chr2:241492106-241492364 | -0.100712343 | 0.819259345 | 0.931133403 |
| chr16:10988617-10988817 | 0.117089211 | 0.819290288 | 0.93115476 |
| chr12:93243517-93244046 | -0.086069252 | 0.819333142 | 0.931175648 |
| chr8:82023196-82023650 | -0.092212334 | 0.819341711 | 0.931175648 |
| chr21:43711854-43712178 | 0.09587648 | 0.819345125 | 0.931175648 |
| chr1:201254382-201255084 | 0.076592591 | 0.81938351 | 0.931178155 |
| chr17:9972079-9972332 | -0.108101137 | 0.819388815 | 0.931178155 |
| chr17:72487993-72488193 | -0.114385439 | 0.819403424 | 0.931178155 |
| chr3:61953048-61953285 | 0.087937728 | 0.819407879 | 0.931178155 |
| chr19:7748853-7749121 | 0.104124715 | 0.819417234 | 0.931178155 |
| chr2:42663125-42663545 | -0.086206176 | 0.819436072 | 0.931178155 |
| chr15:75023335-75023978 | 0.076555436 | 0.819440301 | 0.931178155 |
| chr1:109151194-109151561 | 0.083096796 | 0.819455725 | 0.931178155 |
| chr17:76557261-76558079 | -0.075497397 | 0.819456707 | 0.931178155 |
| chr5:140346362-140346981 | -0.079204235 | 0.819498722 | 0.931212087 |
| chr11:111423029-111423323 | 0.094051869 | 0.819512101 | 0.93121348 |
| chr22:17339144-17339589 | -0.097468193 | 0.819554616 | 0.93123828 |
| chr6:36986031-36986730 | 0.076342429 | 0.819558233 | 0.93123828 |
| chr9:111597952-111598707 | 0.078552589 | 0.819588635 | 0.931258895 |
| chr8:89900830-89901288 | 0.075407085 | 0.819600684 | 0.931258895 |
| chr22:17988271-17988569 | 0.095638855 | 0.81961329 | 0.931259409 |
| chr1:157691364-157691674 | 0.098667038 | 0.819635467 | 0.931270797 |
| chr17:56435195-56435404 | 0.104595642 | 0.819665251 | 0.931288193 |
| chr3:79018008-79018208 | 0.094084539 | 0.819707351 | 0.931288193 |
| chr2:200626440-200626640 | 0.119262547 | 0.819710414 | 0.931288193 |
| chr20:44844214-44844414 | 0.119262547 | 0.819710414 | 0.931288193 |
| chr6:32195444-32196645 | 0.0726871 | 0.819711549 | 0.931288193 |
| chr14:55579564-55580528 | 0.07454397 | 0.819809059 | 0.931374252 |
| chr10:86139112-86139440 | 0.091684921 | 0.819811608 | 0.931374252 |
| chr7:36967332-36967634 | 0.092123362 | 0.819841515 | 0.931394419 |
| chr2:43168793-43168993 | -0.106398862 | 0.819889584 | 0.931435217 |
| chr3:111904734-111904993 | -0.110251515 | 0.819929879 | 0.931467185 |
| chr19:35845410-35845641 | 0.100261311 | 0.819973692 | 0.931503147 |

| | | | |
|---|---|---|---|
| chr11:2699285-2699485 | -0.108142777 | 0.820019582 | 0.931535658 |
| chr1:106258709-106258946 | 0.088318044 | 0.820026626 | 0.931535658 |
| chr17:5419553-5420446 | -0.074636388 | 0.820046053 | 0.931543916 |
| chr14:106334890-106335214 | 0.095756591 | 0.820105318 | 0.931597427 |
| chr11:66592648-66593670 | 0.073825335 | 0.820124955 | 0.931603203 |
| chr20:5750212-5750586 | -0.094880634 | 0.820144045 | 0.931603203 |
| chr1:173714702-173715632 | -0.072597625 | 0.820146877 | 0.931603203 |
| chr5:155504763-155504963 | -0.122689197 | 0.820209499 | 0.931624089 |
| chr22:26364293-26364829 | -0.085831412 | 0.820213275 | 0.931624089 |
| chr16:74309902-74310278 | -0.091257249 | 0.820233263 | 0.931624089 |
| chr9:79026331-79026578 | 0.111795033 | 0.820246035 | 0.931624089 |
| chr11:59560262-59560584 | -0.079206916 | 0.82025432 | 0.931624089 |
| chr4:119628359-119629423 | 0.073093746 | 0.82026384 | 0.931624089 |
| chr12:101894430-101895175 | -0.074258858 | 0.820264689 | 0.931624089 |
| chr11:118295922-118296126 | -0.109012999 | 0.820269018 | 0.931624089 |
| chr2:229921190-229923081 | -0.072335977 | 0.820274693 | 0.931624089 |
| chr16:27203332-27204335 | 0.07190917 | 0.820345735 | 0.931684987 |
| chr22:42688423-42688623 | -0.114963753 | 0.820352631 | 0.931684987 |
| chr4:107867350-107867774 | -0.090026951 | 0.82039037 | 0.931714037 |
| chr4:48779995-48780765 | -0.072337837 | 0.820422315 | 0.931732908 |
| chr20:10240016-10240329 | 0.085958047 | 0.820431307 | 0.931732908 |
| chr1:67561718-67562111 | -0.085698417 | 0.820455688 | 0.931741197 |
| chr21:14382688-14383739 | 0.072728554 | 0.820462926 | 0.931741197 |
| chr13:35978602-35978917 | -0.096723685 | 0.820485967 | 0.931753554 |
| chr10:96513485-96514609 | -0.073406451 | 0.820572099 | 0.931816811 |
| chr1:244548682-244548887 | 0.113466039 | 0.820576254 | 0.931816811 |
| chr11:14908585-14908826 | 0.093852015 | 0.820578154 | 0.931816811 |
| chr10:110530033-110530748 | 0.077305805 | 0.820593737 | 0.931820385 |
| chr8:30095000-30095835 | -0.080395086 | 0.820640844 | 0.931820385 |
| chr2:111413987-111414219 | -0.115637068 | 0.820649009 | 0.931820385 |
| chr20:49216939-49217417 | 0.079891131 | 0.820657866 | 0.931820385 |
| chr8:122156197-122156406 | -0.107814819 | 0.820681821 | 0.931820385 |
| chr8:112026739-112027054 | 0.096719391 | 0.820700824 | 0.931820385 |
| chr2:219403005-219403215 | 0.09952301 | 0.820701131 | 0.931820385 |
| chr10:110557078-110557349 | -0.100229758 | 0.820703304 | 0.931820385 |
| chr12:56687735-56689050 | -0.072153323 | 0.820711843 | 0.931820385 |
| chr3:197456307-197456834 | -0.083479546 | 0.820714553 | 0.931820385 |
| chr3:124688914-124689174 | 0.098354676 | 0.820715076 | 0.931820385 |
| chr1:1375067-1375679 | -0.076559833 | 0.820738376 | 0.931833032 |
| chr10:120942852-120943422 | 0.08139301 | 0.820778118 | 0.931864346 |
| chr17:76466006-76466351 | 0.089482048 | 0.820798867 | 0.931874095 |
| chr11:35821802-35822435 | -0.077894575 | 0.820814706 | 0.931878269 |
| chr5:1448431-1448631 | -0.116844789 | 0.82082905 | 0.931880747 |
| chr9:88915721-88916052 | -0.100499125 | 0.820843838 | 0.931883728 |
| chr17:9628604-9628926 | 0.096939932 | 0.820857273 | 0.931885173 |
| chr2:9003417-9004599 | -0.073073416 | 0.820929111 | 0.931952919 |
| chr14:64539879-64540903 | 0.071283336 | 0.820973706 | 0.93197003 |
| chr11:62208855-62209065 | 0.117816834 | 0.820985344 | 0.93197003 |
| chr2:60485127-60485403 | -0.120790684 | 0.820991402 | 0.93197003 |
| chrX:18354608-18354995 | 0.094332326 | 0.820992836 | 0.93197003 |

| | | | |
|---|---|---|---|
| chr11:69718254-69718469 | -0.108000183 | 0.821022538 | 0.931984808 |
| chr16:84851182-84851829 | 0.07615468 | 0.821061179 | 0.931984808 |
| chr8:67380975-67381366 | -0.081704496 | 0.821061496 | 0.931984808 |
| chr5:149915709-149916036 | -0.098813461 | 0.821066466 | 0.931984808 |
| chr6:89819024-89820410 | 0.071891059 | 0.821066671 | 0.931984808 |
| chr6:34223576-34224539 | -0.074734849 | 0.821089773 | 0.931991925 |
| chr12:47943767-47943967 | -0.104660624 | 0.821097268 | 0.931991925 |
| chr3:66972247-66972523 | 0.102961317 | 0.821121629 | 0.932005769 |
| chr19:47745293-47745844 | 0.080190798 | 0.821162216 | 0.932034677 |
| chr3:53045920-53047140 | -0.076762899 | 0.821173501 | 0.932034677 |
| chr6:28865971-28866919 | 0.073418678 | 0.821215217 | 0.932034677 |
| chr11:36034805-36035166 | 0.090852284 | 0.821223567 | 0.932034677 |
| chr19:16896851-16897579 | 0.075837615 | 0.821234696 | 0.932034677 |
| chr19:35879852-35880060 | 0.101626364 | 0.821238454 | 0.932034677 |
| chr21:36113687-36113955 | 0.095568469 | 0.821238656 | 0.932034677 |
| chr1:107153103-107153390 | -0.078366017 | 0.821264782 | 0.932034677 |
| chr8:101460190-101460473 | -0.090577558 | 0.82127185 | 0.932034677 |
| chr4:82271871-82272286 | -0.089715064 | 0.821277283 | 0.932034677 |
| chr1:190321015-190321249 | 0.098770556 | 0.821306115 | 0.932034677 |
| chr3:197959812-197960857 | -0.072859821 | 0.821314182 | 0.932034677 |
| chr12:19431679-19431885 | -0.113556784 | 0.821314302 | 0.932034677 |
| chr2:150309270-150309595 | 0.104169292 | 0.821324359 | 0.932034677 |
| chr17:66191600-66192626 | 0.074177861 | 0.821329559 | 0.932034677 |
| chr11:2991988-2993099 | 0.072362992 | 0.821367526 | 0.932057282 |
| chr17:57264842-57265151 | 0.090874529 | 0.821373807 | 0.932057282 |
| chr9:127561486-127562248 | -0.076730996 | 0.821390834 | 0.9320628 |
| chr8:22622961-22623161 | 0.106136918 | 0.821418127 | 0.932069077 |
| chr11:81989203-81989403 | -0.114275571 | 0.821420695 | 0.932069077 |
| chr1:168586047-168586328 | -0.100294779 | 0.821435619 | 0.932072208 |
| chr6:31442510-31442953 | -0.090394576 | 0.821488131 | 0.932109589 |
| chr1:42657955-42659228 | -0.073796754 | 0.821507366 | 0.932109589 |
| chr1:47852998-47853206 | -0.110376636 | 0.821527447 | 0.932109589 |
| chr8:121240453-121240833 | 0.089279904 | 0.82154349 | 0.932109589 |
| chr1:112594316-112594869 | -0.082821044 | 0.821550979 | 0.932109589 |
| chr20:5950291-5951131 | -0.074551305 | 0.821552793 | 0.932109589 |
| chr7:76089878-76090189 | -0.090005529 | 0.821580838 | 0.932109589 |
| chr16:69726277-69727120 | 0.071939587 | 0.821593475 | 0.932109589 |
| chr9:87386522-87386722 | -0.12375175 | 0.821599252 | 0.932109589 |
| chr17:14779235-14779516 | 0.10152058 | 0.82161602 | 0.932109589 |
| chr5:25083375-25083688 | -0.09632393 | 0.821619015 | 0.932109589 |
| chr3:5426334-5426722 | -0.080303654 | 0.821624109 | 0.932109589 |
| chr20:47321124-47321853 | -0.070765608 | 0.821627258 | 0.932109589 |
| chr16:73573223-73573423 | -0.116986638 | 0.821638873 | 0.932109589 |
| chr17:19063431-19064084 | -0.077673906 | 0.821695743 | 0.932137443 |
| chr10:21173885-21174836 | 0.072604044 | 0.821714224 | 0.932137443 |
| chr6:140106025-140106225 | -0.1246564 | 0.821727534 | 0.932137443 |
| chr8:76999867-77001046 | 0.07096293 | 0.821728031 | 0.932137443 |
| chr19:45792665-45793322 | 0.075166432 | 0.821739791 | 0.932137443 |
| chr1:146037661-146038403 | 0.071696561 | 0.821742169 | 0.932137443 |
| chr16:48526584-48526851 | -0.095553353 | 0.821748584 | 0.932137443 |

| | | | |
|---|---|---|---|
| chr8:66924607-66926118 | 0.072355403 | 0.821797946 | 0.932153994 |
| chr6:44593510-44593776 | -0.098902698 | 0.821803168 | 0.932153994 |
| chr4:112285035-112285323 | -0.087050447 | 0.821808159 | 0.932153994 |
| chr1:214216704-214217152 | -0.081073804 | 0.821829576 | 0.932153994 |
| chr16:67236520-67236909 | -0.081471371 | 0.821833053 | 0.932153994 |
| chr10:112471905-112472137 | -0.111314271 | 0.821836684 | 0.932153994 |
| chr2:26244305-26245538 | -0.072799134 | 0.821854988 | 0.932153994 |
| chr11:125111433-125112099 | -0.072504998 | 0.821861333 | 0.932153994 |
| chr5:180471033-180471281 | -0.093149541 | 0.821872665 | 0.932153994 |
| chr6:18043187-18043515 | -0.093793652 | 0.821899448 | 0.932159475 |
| chr3:71688235-71688572 | -0.094580791 | 0.82190183 | 0.932159475 |
| chr11:22130723-22131025 | 0.094391285 | 0.8219192 | 0.932165378 |
| chr17:50345797-50346434 | 0.075286966 | 0.821957426 | 0.932194273 |
| chr4:37541763-37542064 | 0.110589202 | 0.821973456 | 0.932194273 |
| chr3:186277564-186277764 | 0.11663236 | 0.821983191 | 0.932194273 |
| chr6:32113970-32114387 | 0.087500571 | 0.821993342 | 0.932194273 |
| chr3:112734702-112735248 | -0.077638301 | 0.822009394 | 0.93219868 |
| chr19:53979911-53980774 | 0.073687808 | 0.822045677 | 0.932204258 |
| chr10:26227712-26227955 | -0.118484868 | 0.822046105 | 0.932204258 |
| chr4:68682842-68683347 | -0.086097449 | 0.822058654 | 0.932204258 |
| chr19:54399283-54399487 | -0.114527652 | 0.822089017 | 0.932204258 |
| chr11:65315279-65315818 | 0.079419048 | 0.822094418 | 0.932204258 |
| chr7:5983990-5984247 | -0.099967287 | 0.822105713 | 0.932204258 |
| chr2:233443600-233443860 | -0.102038921 | 0.82214264 | 0.932204258 |
| chr2:31467578-31467787 | -0.126555074 | 0.822162923 | 0.932204258 |
| chr22:31399140-31400068 | 0.073884368 | 0.822165071 | 0.932204258 |
| chr3:109050649-109050903 | -0.110090017 | 0.82217152 | 0.932204258 |
| chr12:79119910-79120110 | 0.115062316 | 0.822171874 | 0.932204258 |
| chr1:23958996-23960127 | 0.074246757 | 0.822174085 | 0.932204258 |
| chr3:28916750-28916952 | 0.108187237 | 0.822184641 | 0.932204258 |
| chr3:69920103-69920348 | 0.108187237 | 0.822184641 | 0.932204258 |
| chr3:63931417-63931628 | 0.082385104 | 0.822202054 | 0.932210207 |
| chr3:184711629-184712631 | 0.075099128 | 0.822214417 | 0.932210429 |
| chr5:26121468-26121687 | -0.117977508 | 0.822232394 | 0.932217018 |
| chr13:30103092-30103399 | -0.098415459 | 0.822279493 | 0.932252707 |
| chr6:33000107-33000468 | -0.094782937 | 0.822295036 | 0.932252707 |
| chr15:58932977-58934320 | 0.071743699 | 0.822311826 | 0.932252707 |
| chr13:33682559-33682782 | -0.110671313 | 0.822318153 | 0.932252707 |
| chr20:54980209-54980660 | -0.084794288 | 0.82233994 | 0.932252707 |
| chr11:60889994-60890289 | 0.095549599 | 0.822346921 | 0.932252707 |
| chr2:27016916-27017117 | -0.104783311 | 0.822355895 | 0.932252707 |
| chr11:65873449-65873979 | -0.075733034 | 0.822363673 | 0.932252707 |
| chr2:158179498-158179941 | -0.094328262 | 0.822377473 | 0.932252707 |
| chr6:107115411-107115622 | 0.09629023 | 0.822396649 | 0.932252707 |
| chr2:25166105-25166321 | 0.100856648 | 0.822397709 | 0.932252707 |
| chr14:91303244-91303713 | 0.077016853 | 0.822419044 | 0.932263101 |
| chr8:121248845-121249109 | 0.087423007 | 0.822449496 | 0.932283827 |
| chrX:103831321-103832633 | 0.072818488 | 0.822464649 | 0.932287211 |
| chr4:73550665-73551163 | -0.089809835 | 0.822494429 | 0.932307175 |
| chr3:63661278-63661563 | -0.077967299 | 0.822545799 | 0.932344294 |

| | | | |
|---|---|---|---|
| chr15:71337176-71337758 | 0.080676782 | 0.822573616 | 0.932344294 |
| chr1:173824038-173825122 | -0.070688423 | 0.822582982 | 0.932344294 |
| chr1:64505207-64506656 | -0.073430286 | 0.822595332 | 0.932344294 |
| chr11:66192343-66192543 | 0.109629669 | 0.822638621 | 0.932344294 |
| chr6:130313271-130313530 | 0.104517777 | 0.822643964 | 0.932344294 |
| chr1:247183244-247183477 | -0.105453162 | 0.822651501 | 0.932344294 |
| chr2:125761349-125761560 | -0.115687147 | 0.822668871 | 0.932344294 |
| chr17:79019645-79019847 | 0.111473593 | 0.822674255 | 0.932344294 |
| chrX:13652875-13653728 | -0.072013993 | 0.822677226 | 0.932344294 |
| chrX:10217336-10217600 | -0.101824434 | 0.822682838 | 0.932344294 |
| chr11:66438116-66439496 | 0.072329969 | 0.822683654 | 0.932344294 |
| chr20:51744546-51744907 | 0.077999846 | 0.822691719 | 0.932344294 |
| chr7:128105445-128105645 | 0.116850459 | 0.822705087 | 0.932344294 |
| chr5:110338259-110338662 | -0.088730455 | 0.822721631 | 0.932344294 |
| chr7:31056887-31057121 | 0.100170028 | 0.822728639 | 0.932344294 |
| chr2:95402452-95403505 | 0.073622844 | 0.82273525 | 0.932344294 |
| chr5:176803989-176804254 | -0.099351808 | 0.822746201 | 0.932344294 |
| chr21:41761278-41761554 | 0.092075628 | 0.822777187 | 0.932365618 |
| chr6:44166612-44166898 | -0.104517212 | 0.822803725 | 0.932381902 |
| chr12:52971602-52972299 | -0.074272781 | 0.822875071 | 0.932448959 |
| chr20:13324984-13325249 | 0.098219057 | 0.822913145 | 0.932478312 |
| chr12:13404790-13405198 | -0.095840074 | 0.822928602 | 0.932482037 |
| chr14:81164136-81164606 | -0.076352429 | 0.823046744 | 0.932600598 |
| chr3:135744539-135744739 | -0.108304326 | 0.823067339 | 0.932600598 |
| chr20:13458916-13459250 | 0.088590569 | 0.823069748 | 0.932600598 |
| chr6:26232773-26233367 | 0.07839803 | 0.823084734 | 0.932603787 |
| chr7:138865984-138866256 | -0.104409333 | 0.823105582 | 0.932613618 |
| chr3:20041131-20041842 | 0.073316768 | 0.823201917 | 0.932688191 |
| chr15:62391160-62391474 | 0.094101737 | 0.823213304 | 0.932688191 |
| chr1:164673733-164673973 | -0.105186841 | 0.82321956 | 0.932688191 |
| chr1:86780920-86781231 | -0.09361085 | 0.823220089 | 0.932688191 |
| chr1:183804383-183805661 | -0.072707284 | 0.823308286 | 0.932772567 |
| chr3:51390750-51391347 | -0.076852066 | 0.823318909 | 0.932772567 |
| chr12:58176350-58176554 | 0.110296667 | 0.823356904 | 0.932801821 |
| chr2:102976074-102976327 | 0.100734909 | 0.823399921 | 0.932836762 |
| chr8:132057816-132058045 | -0.095638222 | 0.82342112 | 0.932843607 |
| chr21:43344101-43344325 | 0.094519675 | 0.823432407 | 0.932843607 |
| chr8:130094688-130095446 | 0.07354206 | 0.82344506 | 0.932843607 |
| chr2:218567902-218569232 | -0.070574443 | 0.823465805 | 0.932843607 |
| chr15:89377888-89378850 | 0.071483207 | 0.823500219 | 0.932843607 |
| chr21:39387503-39387932 | -0.073961271 | 0.823506918 | 0.932843607 |
| chr19:4791176-4792092 | 0.073014868 | 0.823531446 | 0.932843607 |
| chr6:24991928-24992238 | 0.093273875 | 0.82355977 | 0.932843607 |
| chr11:46176426-46176786 | 0.087371527 | 0.823573502 | 0.932843607 |
| chr22:31272877-31273193 | -0.095009889 | 0.823577864 | 0.932843607 |
| chr3:72487452-72487746 | 0.096098128 | 0.823580931 | 0.932843607 |
| chr5:131019661-131020022 | -0.105486184 | 0.823597399 | 0.932843607 |
| chr10:12129042-12130110 | -0.071295354 | 0.823617834 | 0.932843607 |
| chr14:74860969-74861247 | 0.093189485 | 0.823620421 | 0.932843607 |
| chr6:47047481-47047904 | -0.080585392 | 0.8236249 | 0.932843607 |

| | | | |
|---|---|---|---|
| chr12:75908977-75909391 | -0.090918775 | 0.823653764 | 0.932843607 |
| chr5:16134132-16134435 | -0.109387126 | 0.82366104 | 0.932843607 |
| chr1:117758494-117759445 | -0.07252838 | 0.823667664 | 0.932843607 |
| chr3:157397839-157398075 | 0.119294416 | 0.823668343 | 0.932843607 |
| chr16:30839080-30839350 | -0.099101409 | 0.823669637 | 0.932843607 |
| chr6:6599371-6599755 | -0.088914665 | 0.823681095 | 0.932843607 |
| chr14:26255307-26255517 | 0.103916013 | 0.823697405 | 0.932843607 |
| chr15:38817657-38818146 | 0.079581225 | 0.823699664 | 0.932843607 |
| chr8:10054035-10055239 | -0.073064547 | 0.82370664 | 0.932843607 |
| chr5:136769951-136770374 | -0.084504327 | 0.823715095 | 0.932843607 |
| chr9:94258780-94259710 | -0.074618964 | 0.823761957 | 0.932843607 |
| chr11:45847780-45848330 | -0.074604111 | 0.823763386 | 0.932843607 |
| chr1:143735779-143736163 | 0.090456606 | 0.823784439 | 0.932843607 |
| chr8:118743471-118743923 | -0.085173513 | 0.823789484 | 0.932843607 |
| chr11:66222702-66223056 | 0.080126304 | 0.823832598 | 0.932843607 |
| chr18:48453348-48453750 | -0.0857136 | 0.823844247 | 0.932843607 |
| chr1:191603440-191603640 | 0.134135166 | 0.82384673 | 0.932843607 |
| chr8:104325453-104326375 | -0.071364566 | 0.823869879 | 0.932843607 |
| chr1:61050581-61050792 | -0.102597886 | 0.823877343 | 0.932843607 |
| chr2:26720286-26720634 | -0.089980293 | 0.823882149 | 0.932843607 |
| chr17:3754480-3755293 | -0.071492727 | 0.823888925 | 0.932843607 |
| chr17:82497186-82497428 | 0.085103829 | 0.823901832 | 0.932843607 |
| chr19:13817320-13817715 | 0.085963188 | 0.823904495 | 0.932843607 |
| chr11:35373398-35373716 | -0.110396303 | 0.82390579 | 0.932843607 |
| chr14:39432011-39433333 | 0.069415925 | 0.823911404 | 0.932843607 |
| chr3:113018837-113019966 | -0.070592334 | 0.823932518 | 0.932843607 |
| chr15:57866232-57866432 | 0.115584329 | 0.82395973 | 0.932843607 |
| chr12:57005981-57006305 | 0.093441697 | 0.8239634 | 0.932843607 |
| chr17:57130735-57131380 | -0.074453169 | 0.823993374 | 0.932843607 |
| chr15:90034538-90034862 | -0.088339565 | 0.823995753 | 0.932843607 |
| chr4:56977415-56977885 | -0.073450174 | 0.823997394 | 0.932843607 |
| chr8:52629431-52629760 | -0.09395226 | 0.824011446 | 0.932843607 |
| chr1:110615711-110616034 | -0.089300966 | 0.824013879 | 0.932843607 |
| chr11:96096078-96096311 | 0.107706796 | 0.824040645 | 0.932843607 |
| chr22:42500185-42500744 | 0.081692477 | 0.824057342 | 0.932843607 |
| chr16:22627000-22627282 | 0.098037076 | 0.824067339 | 0.932843607 |
| chr21:46668128-46668367 | -0.09857323 | 0.824073161 | 0.932843607 |
| chr7:18570928-18571202 | -0.096806654 | 0.824074046 | 0.932843607 |
| chr7:151626471-151626684 | 0.10319879 | 0.82408127 | 0.932843607 |
| chr5:43631081-43631646 | -0.078240119 | 0.824081633 | 0.932843607 |
| chr1:20634564-20634764 | -0.111766714 | 0.824091551 | 0.932843607 |
| chr17:63406726-63407085 | 0.077222448 | 0.824099916 | 0.932843607 |
| chr7:32800508-32800938 | 0.085083137 | 0.824154478 | 0.932871004 |
| chr10:106571665-106572110 | -0.09953586 | 0.82415653 | 0.932871004 |
| chr18:11851855-11852113 | 0.091922656 | 0.824160644 | 0.932871004 |
| chr1:150464427-150464679 | -0.093983791 | 0.824176531 | 0.932875206 |
| chr20:50112580-50113349 | -0.072085729 | 0.824214647 | 0.932904568 |
| chr9:83334309-83334509 | -0.107094348 | 0.824243326 | 0.932923247 |
| chr5:159444424-159444758 | -0.092274611 | 0.824260973 | 0.93292944 |
| chr20:18363754-18364047 | -0.105438413 | 0.824282577 | 0.93294011 |

| | | | |
|---|---|---|---|
| chr17:76383382-76384004 | -0.07571098 | 0.824335428 | 0.93297397 |
| chr16:2428885-2429541 | -0.073792388 | 0.824348428 | 0.93297397 |
| chr18:63994812-63995094 | -0.096121067 | 0.82435616 | 0.93297397 |
| chr5:43483333-43484744 | 0.070550108 | 0.824361198 | 0.93297397 |
| chr12:68331603-68331814 | 0.098641937 | 0.824383734 | 0.932985694 |
| chr11:37141693-37142062 | -0.088657399 | 0.824414585 | 0.933006828 |
| chr5:20545329-20545557 | 0.116847825 | 0.824438293 | 0.933007096 |
| chr19:46745168-46745434 | 0.093173506 | 0.824439175 | 0.933007096 |
| chr7:6901672-6902000 | -0.113167102 | 0.824519344 | 0.933077113 |
| chr1:147515377-147515860 | -0.079124802 | 0.8245343 | 0.933077113 |
| chr8:139706719-139706919 | -0.098958137 | 0.824575962 | 0.933077113 |
| chr6:72247056-72247382 | -0.093773627 | 0.82457821 | 0.933077113 |
| chr1:16111835-16112035 | -0.116775484 | 0.824606363 | 0.933077113 |
| chr22:20495723-20496908 | 0.072188046 | 0.824606594 | 0.933077113 |
| chr17:74987412-74987765 | -0.077775139 | 0.824608035 | 0.933077113 |
| chr14:92040710-92040981 | -0.082258657 | 0.824628886 | 0.933077113 |
| chr11:27158739-27159176 | -0.083342383 | 0.824631971 | 0.933077113 |
| chr22:42090122-42091399 | -0.071790733 | 0.824633084 | 0.933077113 |
| chr11:66466338-66467504 | 0.071025918 | 0.824634999 | 0.933077113 |
| chr8:144409022-144409918 | 0.072413909 | 0.824690313 | 0.933098676 |
| chr8:26265947-26266338 | 0.087910479 | 0.82469422 | 0.933098676 |
| chr1:167220001-167221026 | 0.069935875 | 0.824697296 | 0.933098676 |
| chr8:143840668-143841520 | 0.071106308 | 0.824702767 | 0.933098676 |
| chr4:53644523-53645063 | -0.074087171 | 0.824800169 | 0.9331951 |
| chr1:50959603-50960869 | -0.07055947 | 0.824845992 | 0.933199805 |
| chr20:56821058-56821277 | 0.093033947 | 0.824863635 | 0.933199805 |
| chrX:146616103-146616336 | 0.111558807 | 0.824873187 | 0.933199805 |
| chr9:97399098-97399575 | -0.083578352 | 0.824894808 | 0.933199805 |
| chr1:168235268-168235606 | -0.083476994 | 0.824903535 | 0.933199805 |
| chr16:89094347-89094565 | -0.097809563 | 0.824905236 | 0.933199805 |
| chr8:103373209-103373516 | 0.079462838 | 0.824905299 | 0.933199805 |
| chr2:173075653-173076834 | 0.072166489 | 0.824911935 | 0.933199805 |
| chr5:10352974-10354451 | 0.069330665 | 0.824913941 | 0.933199805 |
| chr3:113745167-113747673 | -0.068820051 | 0.82493555 | 0.933210472 |
| chr10:86178457-86178714 | -0.090257195 | 0.824961232 | 0.933215771 |
| chr10:79502541-79502961 | -0.091904288 | 0.824969416 | 0.933215771 |
| chr14:92973259-92974573 | 0.070070651 | 0.824978704 | 0.933215771 |
| chr4:90314908-90315407 | 0.082647277 | 0.824988951 | 0.933215771 |
| chr1:98794296-98794612 | -0.089383402 | 0.825047827 | 0.933268591 |
| chr3:101848899-101850144 | 0.069385112 | 0.825079253 | 0.933283569 |
| chr12:93345060-93345290 | 0.109392895 | 0.825101962 | 0.933283569 |
| chr2:54176951-54177233 | -0.117403991 | 0.825119792 | 0.933283569 |
| chr1:117928840-117930141 | 0.070732059 | 0.825122116 | 0.933283569 |
| chr17:50397220-50398032 | 0.073363344 | 0.8251222 | 0.933283569 |
| chr3:170085641-170085841 | 0.105530337 | 0.82513415 | 0.933283569 |
| chr1:104933949-104934188 | 0.111083987 | 0.825179123 | 0.93332066 |
| chr10:95963286-95963551 | 0.10565737 | 0.825211835 | 0.933343881 |
| chr6:26216313-26217390 | 0.070933014 | 0.825232373 | 0.933344255 |
| chr3:134381700-134382287 | -0.073176355 | 0.825236528 | 0.933344255 |
| chr11:47161633-47161833 | 0.096194253 | 0.825262633 | 0.933357918 |

| | | | |
|---|---|---|---|
| chr5:32443617-32445054 | 0.070046768 | 0.825272971 | 0.933357918 |
| chr11:75814430-75815374 | -0.069610411 | 0.825327582 | 0.933380782 |
| chr5:57209291-57209620 | 0.095943861 | 0.82533209 | 0.933380782 |
| chr2:101110448-101110850 | 0.084252126 | 0.825338937 | 0.933380782 |
| chr17:78272613-78273019 | -0.08456437 | 0.825344986 | 0.933380782 |
| chr17:58587654-58588200 | 0.070130169 | 0.825354096 | 0.933380782 |
| chr3:187678942-187679189 | -0.103893875 | 0.825379133 | 0.93339532 |
| chr10:13161089-13161791 | -0.072344372 | 0.825429616 | 0.933438634 |
| chr9:35748881-35749795 | 0.074184877 | 0.825487601 | 0.933469342 |
| chr4:118981800-118982143 | 0.084934944 | 0.825509585 | 0.933469342 |
| chr20:53790838-53791560 | -0.069308293 | 0.825514387 | 0.933469342 |
| chr12:99390706-99391117 | 0.089413376 | 0.825531527 | 0.933469342 |
| chr15:65868962-65870016 | 0.070211779 | 0.82553731 | 0.933469342 |
| chr3:27263267-27264051 | -0.078141555 | 0.825568925 | 0.933469342 |
| chr1:234676091-234676512 | 0.08561455 | 0.825570753 | 0.933469342 |
| chrX:70263028-70263341 | -0.09607528 | 0.825574574 | 0.933469342 |
| chr1:184973855-184974459 | -0.082986578 | 0.825588585 | 0.933469342 |
| chr10:109866927-109867146 | -0.112808088 | 0.825591543 | 0.933469342 |
| chr3:113317801-113318014 | -0.109018136 | 0.82559533 | 0.933469342 |
| chr10:86688761-86688961 | 0.089406512 | 0.825622199 | 0.933469342 |
| chr20:19530926-19531165 | -0.114249947 | 0.825627536 | 0.933469342 |
| chr6:169724329-169725533 | -0.07217072 | 0.825641111 | 0.933469342 |
| chr12:16606196-16606743 | -0.086365098 | 0.825644993 | 0.933469342 |
| chr9:34380736-34380941 | -0.09321461 | 0.825651696 | 0.933469342 |
| chr7:152436204-152437007 | -0.072200227 | 0.825708847 | 0.933520181 |
| chr1:108199180-108200758 | 0.069147093 | 0.825730886 | 0.933523318 |
| chr15:40357808-40358420 | 0.075535727 | 0.825735988 | 0.933523318 |
| chr14:35480402-35480603 | 0.111505362 | 0.825757115 | 0.933533428 |
| chr6:12411249-12411449 | 0.126205009 | 0.825769864 | 0.933533824 |
| chr6:46207384-46207643 | -0.118958138 | 0.82579324 | 0.933533824 |
| chr19:38849870-38850949 | 0.069350688 | 0.825804077 | 0.933533824 |
| chr2:70419991-70420409 | -0.086136508 | 0.825817087 | 0.933533824 |
| chrX:111568503-111568714 | 0.10442545 | 0.825827531 | 0.933533824 |
| chr11:66268246-66268957 | 0.071673029 | 0.825830566 | 0.933533824 |
| chr3:20668427-20668663 | -0.110173562 | 0.825846871 | 0.933538482 |
| chr13:71864651-71864851 | 0.099467479 | 0.82590312 | 0.933585732 |
| chr20:54199630-54200004 | 0.070721293 | 0.825916513 | 0.933585732 |
| chr9:93223274-93223769 | 0.082895363 | 0.825925223 | 0.933585732 |
| chr8:139772216-139772468 | 0.100328086 | 0.825954352 | 0.933594104 |
| chr14:102592907-102593234 | 0.083649249 | 0.825962527 | 0.933594104 |
| chr5:35003207-35003719 | -0.083284106 | 0.82597586 | 0.933594104 |
| chr4:82654985-82655185 | 0.129527812 | 0.825981367 | 0.933594104 |
| chrX:39613573-39613963 | -0.097439988 | 0.826077165 | 0.933658622 |
| chr8:132491916-132492116 | 0.118468591 | 0.826078783 | 0.933658622 |
| chr6:101393293-101393919 | -0.077046239 | 0.826090699 | 0.933658622 |
| chr17:49952508-49952837 | 0.079729669 | 0.826104539 | 0.933658622 |
| chr6:106360221-106361661 | 0.069832567 | 0.826106475 | 0.933658622 |
| chr4:75573353-75574001 | 0.074057818 | 0.826118183 | 0.933658622 |
| chr15:29103602-29104142 | 0.081594977 | 0.826124712 | 0.933658622 |
| chr13:86758231-86758443 | 0.111744307 | 0.82613593 | 0.933658622 |

| | | | |
|---|---|---|---|
| chr3:135118514-135118714 | -0.109878879 | 0.82619142 | 0.933707562 |
| chr13:34336487-34336805 | 0.100383675 | 0.82623089 | 0.933732051 |
| chr3:193970247-193970468 | 0.100391895 | 0.826260813 | 0.933732051 |
| chr9:133406845-133407153 | -0.084152833 | 0.826263785 | 0.933732051 |
| chr10:91908255-91909555 | -0.070978921 | 0.826274249 | 0.933732051 |
| chr16:81886100-81886300 | 0.097383048 | 0.826282622 | 0.933732051 |
| chrX:101799386-101799817 | 0.077477833 | 0.826302809 | 0.933732051 |
| chr2:61196456-61196679 | -0.115546638 | 0.82630944 | 0.933732051 |
| chr22:42508469-42508749 | 0.110515607 | 0.826310579 | 0.933732051 |
| chr6:28789674-28790178 | -0.077018087 | 0.826342267 | 0.933740646 |
| chr1:161998873-161999082 | 0.10731086 | 0.826342558 | 0.933740646 |
| chr20:3846144-3847400 | 0.070615232 | 0.826355743 | 0.933741775 |
| chr6:149863600-149865203 | -0.069514515 | 0.826406079 | 0.933784881 |
| chr3:96573124-96573362 | 0.114322901 | 0.826464369 | 0.933827501 |
| chr15:74100099-74100404 | -0.088792069 | 0.826468173 | 0.933827501 |
| chr4:113684801-113685001 | 0.11100537 | 0.826482102 | 0.933829469 |
| chr12:64018642-64019045 | -0.085902273 | 0.826517484 | 0.933843885 |
| chr8:22599269-22599690 | -0.080824395 | 0.826519236 | 0.933843885 |
| chr6:153790124-153790389 | -0.092336698 | 0.826535631 | 0.933848639 |
| chr3:186570162-186571042 | -0.069665881 | 0.826557071 | 0.933859092 |
| chr9:89999267-89999542 | 0.099912438 | 0.826589094 | 0.933881501 |
| chr3:47764440-47764661 | -0.110986479 | 0.826636322 | 0.933907319 |
| chr3:160134614-160134846 | -0.110986479 | 0.826636322 | 0.933907319 |
| chr14:69580112-69580518 | 0.079469892 | 0.826658135 | 0.933917052 |
| chr11:119205735-119206505 | 0.072241804 | 0.826669315 | 0.933917052 |
| chr11:111936814-111937691 | -0.075082923 | 0.826683613 | 0.933919435 |
| chr1:237124964-237125238 | 0.096113283 | 0.826758055 | 0.933979818 |
| chr3:98457609-98458075 | 0.094089295 | 0.826800939 | 0.933979818 |
| chr1:50369898-50370161 | 0.093292191 | 0.826827438 | 0.933979818 |
| chr9:126705293-126705619 | -0.083966946 | 0.826834224 | 0.933979818 |
| chr6:104325697-104325911 | -0.105485956 | 0.826834624 | 0.933979818 |
| chr11:43556943-43557182 | -0.11177303 | 0.826836988 | 0.933979818 |
| chr16:1929598-1929980 | 0.080035199 | 0.826844506 | 0.933979818 |
| chr2:20223794-20224359 | -0.075373584 | 0.82685457 | 0.933979818 |
| chr20:37945319-37945519 | 0.097630353 | 0.826857786 | 0.933979818 |
| chr6:39985116-39985316 | -0.124045101 | 0.826891631 | 0.933979818 |
| chr18:6219165-6219432 | -0.098660453 | 0.826897762 | 0.933979818 |
| chr7:75934661-75934861 | -0.12098131 | 0.826911543 | 0.933979818 |
| chr17:28552543-28552811 | 0.080303045 | 0.826916667 | 0.933979818 |
| chr1:199463349-199463877 | -0.074991856 | 0.826944705 | 0.933979818 |
| chr12:642621-643240 | 0.079781778 | 0.826949263 | 0.933979818 |
| chr16:49921912-49922139 | 0.102259294 | 0.826949988 | 0.933979818 |
| chr15:70349209-70349688 | -0.081428581 | 0.826951285 | 0.933979818 |
| chr21:46325266-46325532 | 0.082355915 | 0.826956472 | 0.933979818 |
| chr11:73307252-73308344 | -0.068981284 | 0.827030028 | 0.934045795 |
| chr2:156399596-156399796 | -0.082764814 | 0.82703927 | 0.934045795 |
| chr3:58237025-58238087 | 0.073617285 | 0.827101287 | 0.934091247 |
| chr1:163796344-163796889 | -0.087588041 | 0.827106059 | 0.934091247 |
| chr6:39137333-39137554 | -0.105336357 | 0.827124917 | 0.934091247 |
| chr7:45574594-45575308 | 0.080239049 | 0.827128278 | 0.934091247 |

| | | | |
|---|---|---|---|
| chr3:47788629-47789510 | -0.072416957 | 0.827155627 | 0.934108365 |
| chr2:33291137-33291912 | 0.078368109 | 0.827185946 | 0.934128836 |
| chr16:53131442-53131738 | -0.092380582 | 0.827227041 | 0.934161476 |
| chr8:60708223-60708652 | 0.08464512 | 0.827262361 | 0.934181233 |
| chr19:37931997-37932310 | 0.090860183 | 0.827279276 | 0.934181233 |
| chr4:155666700-155668275 | -0.06976608 | 0.827310336 | 0.934181233 |
| chr2:103823875-103824182 | 0.08955273 | 0.827331858 | 0.934181233 |
| chr19:18437163-18437504 | 0.088808276 | 0.827342161 | 0.934181233 |
| chr13:91375146-91375437 | 0.095030307 | 0.827347466 | 0.934181233 |
| chr8:101063593-101064301 | -0.075148669 | 0.827351306 | 0.934181233 |
| chr17:57695869-57696261 | 0.084677418 | 0.827356861 | 0.934181233 |
| chr5:23820733-23821273 | 0.078553599 | 0.827366093 | 0.934181233 |
| chr11:67698339-67698622 | 0.091318253 | 0.827369445 | 0.934181233 |
| chr6:138263162-138263374 | 0.115611429 | 0.827378649 | 0.934181233 |
| chr11:35163834-35164058 | -0.112508037 | 0.827418633 | 0.934212612 |
| chr3:127651569-127651781 | -0.106837573 | 0.827446963 | 0.934230832 |
| chr3:119289741-119289963 | 0.103434938 | 0.827460418 | 0.934232258 |
| chr14:94681403-94681713 | 0.088009403 | 0.827495066 | 0.934237402 |
| chr11:61567606-61567923 | -0.086130984 | 0.827498058 | 0.934237402 |
| chr9:123385183-123385684 | -0.076353785 | 0.827502369 | 0.934237402 |
| chr6:118618798-118619222 | -0.079696936 | 0.827548516 | 0.934237402 |
| chr6:32128130-32128569 | 0.073203913 | 0.827559626 | 0.934237402 |
| chr3:134261035-134261666 | 0.086073543 | 0.827567434 | 0.934237402 |
| chr11:73242353-73242959 | -0.07499983 | 0.827601688 | 0.934237402 |
| chrX:66750742-66751724 | -0.073125587 | 0.827615963 | 0.934237402 |
| chr7:3300250-3301360 | -0.070268013 | 0.827615977 | 0.934237402 |
| chr16:79261534-79261792 | -0.093818679 | 0.827618811 | 0.934237402 |
| chr3:122169240-122169618 | -0.097846889 | 0.827621166 | 0.934237402 |
| chr2:138090455-138090687 | -0.09084524 | 0.827625383 | 0.934237402 |
| chr4:13798841-13799219 | -0.08997055 | 0.82763263 | 0.934237402 |
| chr15:40085389-40085626 | 0.097062458 | 0.827635674 | 0.934237402 |
| chr2:109062640-109062916 | 0.106202454 | 0.827649719 | 0.934239492 |
| chr8:94311661-94311911 | -0.0945452 | 0.827690693 | 0.93427198 |
| chr4:136687182-136687439 | 0.104648428 | 0.827789275 | 0.934351852 |
| chr4:185122639-185122867 | -0.116712791 | 0.827789656 | 0.934351852 |
| chr14:105343120-105343320 | 0.110688354 | 0.827798036 | 0.934351852 |
| chr10:31961850-31962054 | 0.100815113 | 0.827853529 | 0.934390359 |
| chr18:22563173-22563589 | -0.086354637 | 0.827859596 | 0.934390359 |
| chr7:522420-522647 | -0.089407495 | 0.827868736 | 0.934390359 |
| chr17:68134051-68134375 | -0.087310568 | 0.827917285 | 0.93442701 |
| chr7:148856329-148856602 | 0.091268672 | 0.827941929 | 0.93442701 |
| chr1:44799673-44800606 | 0.069791451 | 0.827963715 | 0.93442701 |
| chr8:85843425-85843625 | -0.097113465 | 0.827973709 | 0.93442701 |
| chr20:63894301-63895236 | 0.069244713 | 0.827989397 | 0.93442701 |
| chr18:38554035-38554235 | -0.133544035 | 0.827996926 | 0.93442701 |
| chr9:87834419-87834771 | -0.087732551 | 0.8279976 | 0.93442701 |
| chr2:206085012-206086339 | -0.069325352 | 0.828009585 | 0.93442701 |
| chr2:71393675-71393932 | 0.112293445 | 0.828022683 | 0.93442701 |
| chr8:141988551-141988830 | 0.09049896 | 0.828033264 | 0.93442701 |
| chr13:26186210-26186930 | 0.072718955 | 0.828035357 | 0.93442701 |

| | | | |
|---|---|---|---|
| chr12:5945110-5945545 | -0.087037021 | 0.828084682 | 0.934449473 |
| chr8:97277442-97278402 | 0.069849999 | 0.828088729 | 0.934449473 |
| chr5:150286825-150287172 | -0.078031972 | 0.828119005 | 0.934449473 |
| chr17:66301741-66302003 | 0.090112953 | 0.828122877 | 0.934449473 |
| chr10:129658788-129659000 | 0.100234384 | 0.82812579 | 0.934449473 |
| chr2:158968804-158969434 | 0.075332029 | 0.828128463 | 0.934449473 |
| chrX:43958411-43958737 | 0.100485796 | 0.828154396 | 0.934449473 |
| chr5:140841450-140842050 | -0.079465047 | 0.828182351 | 0.934449473 |
| chr5:65093865-65094107 | -0.10992234 | 0.828183182 | 0.934449473 |
| chr10:52908786-52909021 | -0.116862554 | 0.828188676 | 0.934449473 |
| chr6:89411795-89412827 | -0.071738931 | 0.82820003 | 0.934449473 |
| chr17:64935092-64935347 | -0.090518941 | 0.828218105 | 0.934449473 |
| chr20:56843497-56843698 | 0.093083477 | 0.828220239 | 0.934449473 |
| chr1:183089050-183089361 | 0.097172692 | 0.828239485 | 0.934449473 |
| chr6:2749977-2750232 | 0.102986509 | 0.828256021 | 0.934449473 |
| chr6:160992824-160993052 | 0.105275292 | 0.828290775 | 0.934449473 |
| chr7:76302245-76302796 | -0.074143843 | 0.828302703 | 0.934449473 |
| chr1:156890176-156890384 | -0.095266002 | 0.828316707 | 0.934449473 |
| chrX:63350354-63351698 | -0.070565995 | 0.828343005 | 0.934449473 |
| chr1:113579666-113579948 | 0.087736589 | 0.828368376 | 0.934449473 |
| chr19:48398711-48399011 | 0.084377317 | 0.828369138 | 0.934449473 |
| chr11:73625699-73626079 | -0.08074539 | 0.828370515 | 0.934449473 |
| chr20:3888656-3889504 | 0.072025335 | 0.82838574 | 0.934449473 |
| chr3:191328961-191330272 | -0.070474622 | 0.82838767 | 0.934449473 |
| chr10:72691862-72692857 | 0.070373774 | 0.828392581 | 0.934449473 |
| chr5:149134263-149134463 | -0.112892514 | 0.828419357 | 0.934449473 |
| chr10:69451236-69452028 | 0.071833304 | 0.828433123 | 0.934449473 |
| chr5:180831086-180831966 | 0.071733758 | 0.828443566 | 0.934449473 |
| chr2:47755491-47755733 | -0.104518295 | 0.828452782 | 0.934449473 |
| chr1:236794842-236795998 | -0.068255043 | 0.828461363 | 0.934449473 |
| chr5:170745531-170745931 | -0.080415587 | 0.828473574 | 0.934449473 |
| chr5:11048863-11049242 | -0.087476764 | 0.828474799 | 0.934449473 |
| chr11:114262755-114262958 | -0.109761532 | 0.828477706 | 0.934449473 |
| chr3:32019130-32019429 | -0.09186477 | 0.828483548 | 0.934449473 |
| chr7:100604085-100604940 | 0.07351515 | 0.828514387 | 0.934449473 |
| chr20:59495925-59496222 | -0.082750544 | 0.828517075 | 0.934449473 |
| chr1:246003190-246003419 | -0.09940334 | 0.828517556 | 0.934449473 |
| chr6:108294599-108295634 | -0.070323083 | 0.828518694 | 0.934449473 |
| chr1:229674251-229674460 | -0.111470618 | 0.828540869 | 0.934460728 |
| chr11:123580558-123580888 | 0.086394248 | 0.828580735 | 0.934491935 |
| chr9:97387065-97387292 | -0.093735709 | 0.82859753 | 0.93449368 |
| chr6:1336308-1336578 | -0.092990242 | 0.828606675 | 0.93449368 |
| chr3:150470081-150470768 | -0.073082365 | 0.828640663 | 0.934518257 |
| chr11:111870792-111871779 | -0.073096436 | 0.828673984 | 0.934541128 |
| chr2:68925406-68925624 | 0.107946054 | 0.828685336 | 0.934541128 |
| chr10:86585420-86585838 | -0.075288608 | 0.828747286 | 0.934597235 |
| chr3:31132229-31132431 | 0.105061263 | 0.828849005 | 0.934698189 |
| chr6:23952323-23952523 | -0.108890653 | 0.828870444 | 0.934708609 |
| chr8:141000960-141002177 | 0.068999239 | 0.828940464 | 0.934770878 |
| chr17:56833110-56835155 | 0.067262645 | 0.828958038 | 0.934770878 |

| | | | |
|---|---|---|---|
| chr17:46033777-46034031 | 0.114130195 | 0.828964937 | 0.934770878 |
| chr14:26448458-26448843 | -0.087310934 | 0.828974461 | 0.934770878 |
| chr3:188153533-188154633 | -0.069523793 | 0.829006603 | 0.934793365 |
| chr6:138590976-138591228 | 0.089924321 | 0.829045032 | 0.934812221 |
| chr9:125536181-125536470 | 0.089894735 | 0.829047726 | 0.934812221 |
| chr6:131823089-131823825 | 0.075048769 | 0.829068476 | 0.934821861 |
| chr2:207474951-207475214 | -0.103905028 | 0.829107076 | 0.934851628 |
| chr7:44531103-44531503 | -0.088564695 | 0.829135437 | 0.934852362 |
| chr3:184777671-184778047 | 0.087787624 | 0.829151066 | 0.934852362 |
| chr18:47406066-47406333 | -0.103565017 | 0.829154097 | 0.934852362 |
| chr2:65719482-65719758 | -0.106871854 | 0.829160245 | 0.934852362 |
| chr11:97268721-97269000 | -0.104161941 | 0.82916923 | 0.934852362 |
| chr3:34106810-34107125 | 0.097470917 | 0.829188887 | 0.934852362 |
| chr9:127764247-127764447 | -0.10908283 | 0.829211417 | 0.934852362 |
| chr1:53182142-53182406 | 0.096400236 | 0.829219281 | 0.934852362 |
| chr3:69385679-69386812 | -0.072167178 | 0.829232192 | 0.934852362 |
| chr12:89919360-89919813 | -0.075123271 | 0.829238765 | 0.934852362 |
| chr2:218607492-218608017 | 0.084087762 | 0.829241936 | 0.934852362 |
| chr5:51378855-51379055 | -0.099794145 | 0.829294399 | 0.934878125 |
| chr4:139453068-139454340 | 0.06767762 | 0.829299599 | 0.934878125 |
| chr12:69779886-69780224 | -0.082525599 | 0.829301392 | 0.934878125 |
| chr1:175125346-175125600 | -0.087444489 | 0.829365913 | 0.934921424 |
| chr8:66741801-66742114 | 0.090725187 | 0.829373843 | 0.934921424 |
| chr12:132804308-132804856 | 0.080555216 | 0.829376406 | 0.934921424 |
| chr3:38004402-38005067 | 0.07116269 | 0.829389694 | 0.934922648 |
| chr4:6673258-6673570 | 0.086069161 | 0.829407742 | 0.934929238 |
| chrX:41084198-41085447 | -0.068122186 | 0.82943444 | 0.934940298 |
| chr17:54678188-54678428 | 0.109173295 | 0.829441958 | 0.934940298 |
| chr15:89917651-89918026 | -0.087075554 | 0.829489362 | 0.934979977 |
| chr3:97264246-97264556 | -0.093750452 | 0.829501827 | 0.934980273 |
| chr22:19941607-19942036 | 0.083629694 | 0.829520724 | 0.934987819 |
| chr5:116299898-116300350 | 0.084635601 | 0.829538861 | 0.934994507 |
| chr13:112690349-112690573 | 0.098082243 | 0.829555087 | 0.934999043 |
| chr1:171424490-171424914 | 0.073244277 | 0.829588757 | 0.935021148 |
| chr1:26692424-26692713 | -0.086109175 | 0.829599106 | 0.935021148 |
| chr20:19935899-19936208 | 0.085652019 | 0.829631499 | 0.935043903 |
| chr5:39376269-39376707 | -0.082730607 | 0.829676726 | 0.935078237 |
| chr22:41512799-41513080 | 0.085868529 | 0.82968639 | 0.935078237 |
| chr17:64540109-64540380 | 0.087918891 | 0.829698573 | 0.935078237 |
| chr9:41074616-41074921 | 0.104043483 | 0.829762699 | 0.935102689 |
| chr11:93543354-93543554 | -0.107545109 | 0.829772376 | 0.935102689 |
| chr11:406894-407486 | -0.077587257 | 0.829772985 | 0.935102689 |
| chr7:100158254-100158912 | -0.074626923 | 0.829777432 | 0.935102689 |
| chr21:37565187-37565387 | -0.095623298 | 0.82978129 | 0.935102689 |
| chr3:129120590-129120839 | -0.079630462 | 0.82986979 | 0.935163987 |
| chr5:139452027-139452243 | 0.100281256 | 0.829875479 | 0.935163987 |
| chr8:139246143-139246368 | 0.093940994 | 0.829887696 | 0.935163987 |
| chr8:55772604-55773788 | 0.068310464 | 0.829895563 | 0.935163987 |
| chr5:565787-565988 | -0.116714915 | 0.829896709 | 0.935163987 |
| chr8:67342986-67344211 | 0.069996833 | 0.829939109 | 0.93518095 |

| | | | |
|---|---|---|---|
| chr12:12704629-12704961 | -0.083735015 | 0.829943007 | 0.93518095 |
| chr1:92784718-92785614 | -0.070316174 | 0.829953331 | 0.93518095 |
| chr7:95477067-95477294 | 0.106168866 | 0.829967841 | 0.93518095 |
| chr5:154610343-154610698 | 0.076591619 | 0.829972788 | 0.93518095 |
| chr17:75666996-75667836 | -0.06964753 | 0.83003901 | 0.93523009 |
| chr2:64516512-64517248 | 0.073654884 | 0.830040812 | 0.93523009 |
| chr3:27533781-27534297 | 0.074703332 | 0.830058244 | 0.935235979 |
| chr8:119919769-119920170 | -0.084079794 | 0.830082803 | 0.935249897 |
| chr3:161403098-161403807 | -0.073407968 | 0.83009525 | 0.935250169 |
| chr9:84954079-84954470 | -0.077657669 | 0.830135552 | 0.935281824 |
| chr11:65112562-65112762 | 0.090663987 | 0.830191014 | 0.935330558 |
| chrX:119411983-119412290 | -0.091598585 | 0.830225447 | 0.935349449 |
| chr8:48819108-48819451 | -0.094107058 | 0.830232197 | 0.935349449 |
| chr14:37142840-37143040 | 0.107157408 | 0.83027017 | 0.935378477 |
| chr2:10409116-10409467 | 0.073196751 | 0.830285776 | 0.935382305 |
| chr8:72374767-72375268 | 0.093720601 | 0.830333924 | 0.935422794 |
| chr5:138609274-138610231 | -0.070012038 | 0.830351911 | 0.935429304 |
| chr15:35751617-35751934 | -0.094337193 | 0.830373009 | 0.935439319 |
| chr19:41697178-41697446 | 0.101533611 | 0.830423506 | 0.935460785 |
| chr8:129728579-129728895 | -0.078181235 | 0.830446758 | 0.935460785 |
| chr18:35971797-35972996 | -0.069434421 | 0.830464482 | 0.935460785 |
| chr11:68751040-68751807 | -0.070246035 | 0.830484749 | 0.935460785 |
| chr15:94327434-94327634 | 0.120367517 | 0.830487695 | 0.935460785 |
| chr14:37566795-37567282 | -0.077823357 | 0.830498005 | 0.935460785 |
| chr19:33047380-33047726 | -0.087842925 | 0.830501198 | 0.935460785 |
| chr6:35332114-35332689 | -0.075285535 | 0.8305018 | 0.935460785 |
| chr2:19131908-19132342 | -0.08090887 | 0.830501944 | 0.935460785 |
| chr14:89619187-89619572 | 0.095683035 | 0.830532304 | 0.935479363 |
| chr11:14540078-14540354 | 0.110559624 | 0.830542855 | 0.935479363 |
| chr6:143536421-143537720 | 0.069276213 | 0.830626926 | 0.935560303 |
| chr9:26613868-26614068 | -0.109847265 | 0.830663593 | 0.93558785 |
| chr22:37019660-37020023 | -0.089207952 | 0.830689473 | 0.935593161 |
| chr2:190319063-190320203 | -0.067988384 | 0.830702758 | 0.935593161 |
| chr13:100606888-100607246 | -0.100922959 | 0.83070494 | 0.935593161 |
| chr3:117355663-117355863 | -0.116409842 | 0.830752244 | 0.935620767 |
| chr3:189652116-189652354 | 0.099629645 | 0.830753873 | 0.935620767 |
| chr10:30494121-30494325 | 0.109649635 | 0.830780767 | 0.935637304 |
| chr12:104065203-104065409 | -0.093620595 | 0.830811408 | 0.935645253 |
| chr6:145634057-145634313 | 0.097490651 | 0.830814315 | 0.935645253 |
| chr1:201184731-201185475 | -0.069876814 | 0.830835107 | 0.935645253 |
| chr11:61626541-61627261 | 0.071805079 | 0.83083667 | 0.935645253 |
| chr5:179819488-179819837 | 0.076929103 | 0.830868081 | 0.935661023 |
| chr11:67572737-67572996 | -0.089852279 | 0.830885081 | 0.935661023 |
| chr5:25181191-25181415 | 0.110823304 | 0.830887308 | 0.935661023 |
| chr12:122981403-122981616 | 0.094372418 | 0.830913964 | 0.935677289 |
| chr4:15956264-15956640 | 0.080359356 | 0.830934953 | 0.935687173 |
| chr1:202122869-202123069 | 0.087974625 | 0.830955665 | 0.935696745 |
| chr8:67064012-67064683 | 0.069493644 | 0.830980669 | 0.935711149 |
| chr8:86871926-86872206 | -0.093741832 | 0.831043767 | 0.935760546 |
| chr14:101761346-101761657 | 0.075931737 | 0.831048963 | 0.935760546 |

| | | | |
|---|---|---|---|
| chr6:28587126-28587679 | 0.073299284 | 0.831068235 | 0.935767572 |
| chr11:62771110-62771866 | 0.068968389 | 0.831088463 | 0.935767572 |
| chr8:94808645-94808886 | 0.091258425 | 0.831100569 | 0.935767572 |
| chr19:18152848-18153304 | 0.074789787 | 0.831107431 | 0.935767572 |
| chr1:225426915-225428832 | -0.067921837 | 0.831116267 | 0.935767572 |
| chr1:9689062-9689734 | -0.080448662 | 0.831140103 | 0.93577551 |
| chr9:129567839-129568338 | 0.07528643 | 0.831158643 | 0.93577551 |
| chr3:48472730-48473638 | -0.073247631 | 0.831170808 | 0.93577551 |
| chr2:177254396-177254805 | -0.080512463 | 0.831180059 | 0.93577551 |
| chr8:65644244-65645431 | -0.067336051 | 0.831184382 | 0.93577551 |
| chr15:99014559-99015903 | 0.068473937 | 0.831202509 | 0.935782169 |
| chr6:18749427-18749646 | -0.1071328 | 0.831230264 | 0.935794151 |
| chr17:42980385-42980664 | -0.083779968 | 0.831237578 | 0.935794151 |
| chr6:109008905-109009967 | -0.07038332 | 0.831272206 | 0.935813085 |
| chr17:4074883-4075150 | 0.115774939 | 0.831294148 | 0.935813085 |
| chr19:37654810-37655221 | 0.086875858 | 0.831298224 | 0.935813085 |
| chr19:12440919-12441270 | 0.082772376 | 0.831303481 | 0.935813085 |
| chr1:236604223-236605231 | -0.068709688 | 0.831315464 | 0.935813085 |
| chr9:134217904-134218144 | -0.093046838 | 0.831331212 | 0.935817065 |
| chr19:10543501-10544228 | -0.072390817 | 0.831352506 | 0.935820149 |
| chr10:24945651-24946018 | 0.103625785 | 0.831358379 | 0.935820149 |
| chr13:30616942-30618434 | -0.06677912 | 0.83142565 | 0.935882124 |
| chr10:94145106-94145434 | -0.103898818 | 0.831471474 | 0.935919955 |
| chr15:59313905-59314105 | -0.11463199 | 0.831488145 | 0.935924971 |
| chr2:111205779-111206133 | -0.097689884 | 0.831520368 | 0.935947492 |
| chr5:145936391-145936922 | -0.079207444 | 0.831554654 | 0.935972334 |
| chr11:37946224-37946504 | -0.091314646 | 0.831614807 | 0.936026291 |
| chr2:200962992-200964161 | -0.067371563 | 0.831640657 | 0.936038727 |
| chr5:115799693-115799893 | 0.102271089 | 0.831651799 | 0.936038727 |
| chr2:105328953-105330356 | 0.069909894 | 0.831673554 | 0.936038727 |
| chr14:70817690-70818040 | -0.076916125 | 0.831683341 | 0.936038727 |
| chr10:114042861-114043380 | -0.078816968 | 0.831692451 | 0.936038727 |
| chr1:51769029-51769240 | -0.103465992 | 0.831699154 | 0.936038727 |
| chr17:43170915-43171332 | 0.07320288 | 0.83171437 | 0.936042103 |
| chr11:116605639-116605895 | -0.099590561 | 0.831735135 | 0.936051724 |
| chrX:17608946-17609244 | 0.109983044 | 0.831780175 | 0.936080291 |
| chr11:74739844-74740136 | 0.086606151 | 0.831802461 | 0.936080291 |
| chr10:88303290-88303532 | 0.107785078 | 0.831802985 | 0.936080291 |
| chr8:77288271-77288524 | 0.107616468 | 0.831809839 | 0.936080291 |
| chr11:12377236-12378385 | -0.069938276 | 0.831821603 | 0.936080291 |
| chr9:97205523-97205727 | -0.113892041 | 0.831842288 | 0.936089821 |
| chr14:37482495-37482769 | 0.090928427 | 0.831857403 | 0.936092954 |
| chr20:51766847-51767976 | -0.066960409 | 0.831871207 | 0.936092954 |
| chr20:51843092-51843658 | 0.075280699 | 0.831881723 | 0.936092954 |
| chr15:72670612-72671471 | 0.072260753 | 0.831905373 | 0.936093819 |
| chr14:103281479-103281811 | 0.084254399 | 0.831906927 | 0.936093819 |
| chr1:245107567-245107767 | -0.108124371 | 0.831964618 | 0.936126797 |
| chr9:137168227-137168722 | 0.076381295 | 0.831970111 | 0.936126797 |
| chr20:58981895-58982130 | 0.0784032 | 0.83198211 | 0.936126797 |
| chr8:102585197-102585530 | 0.077972644 | 0.832001763 | 0.936126797 |

| | | | |
|---|---|---|---|
| chr17:77887987-77888189 | 0.099813189 | 0.832031196 | 0.936126797 |
| chr1:204127702-204127921 | -0.078052522 | 0.832034783 | 0.936126797 |
| chr3:46976369-46977237 | 0.069854922 | 0.832040577 | 0.936126797 |
| chr8:123486535-123486916 | -0.077865398 | 0.832041197 | 0.936126797 |
| chr1:114864319-114864524 | 0.102701126 | 0.832054472 | 0.936126797 |
| chr2:42770538-42770873 | -0.079255128 | 0.832058408 | 0.936126797 |
| chr3:48507382-48508005 | -0.072764356 | 0.832125947 | 0.936189036 |
| chr4:112854464-112854670 | -0.114097023 | 0.832143896 | 0.936192306 |
| chr2:194029620-194029890 | -0.107293982 | 0.832168418 | 0.936192306 |
| chr9:6829050-6829453 | -0.081307057 | 0.832179784 | 0.936192306 |
| chr10:29496504-29496705 | 0.102919875 | 0.832190075 | 0.936192306 |
| chr15:32651286-32651595 | -0.083909711 | 0.832191894 | 0.936192306 |
| chr6:7813766-7814023 | -0.106838358 | 0.832203264 | 0.936192306 |
| chr11:60941811-60942084 | 0.088658672 | 0.832214382 | 0.936192306 |
| chr12:112005964-11200621€ | -0.092797075 | 0.832235943 | 0.936202816 |
| chr3:163304391-163304762 | -0.095813415 | 0.832316263 | 0.936279424 |
| chr1:94325706-94325906 | -0.110699384 | 0.832353943 | 0.936292414 |
| chr16:81809252-81809452 | -0.098905679 | 0.832382028 | 0.936292414 |
| chr7:34927993-34928303 | -0.081597393 | 0.832402848 | 0.936292414 |
| chr12:7374551-7374878 | 0.09199505 | 0.83241544 | 0.936292414 |
| chr1:244703312-244703529 | -0.116905398 | 0.832421248 | 0.936292414 |
| chr1:195046194-195046479 | 0.095450542 | 0.832421784 | 0.936292414 |
| chr5:38781055-38781265 | -0.100963424 | 0.832424047 | 0.936292414 |
| chrX:71255207-71255451 | -0.086370266 | 0.832425567 | 0.936292414 |
| chr11:93783829-93785057 | -0.068232613 | 0.832442277 | 0.936297465 |
| chr3:111071083-111072634 | 0.068007524 | 0.832470094 | 0.936301285 |
| chr14:89261070-89261330 | 0.10596794 | 0.832470113 | 0.936301285 |
| chr5:89995337-89995629 | -0.113945868 | 0.832512248 | 0.936302004 |
| chr17:45172583-45172983 | 0.081126768 | 0.83254935 | 0.936302004 |
| chr2:108069615-108069824 | -0.106842829 | 0.832565284 | 0.936302004 |
| chr5:142769695-142771124 | -0.067526343 | 0.832571792 | 0.936302004 |
| chr4:52711893-52713113 | 0.067054463 | 0.832578436 | 0.936302004 |
| chr10:2322521-2322757 | -0.114248693 | 0.832588924 | 0.936302004 |
| chr17:75927377-75927695 | 0.085246776 | 0.832596477 | 0.936302004 |
| chr3:143594249-143594635 | -0.086436549 | 0.832600144 | 0.936302004 |
| chr3:86050446-86050781 | 0.09738564 | 0.832601253 | 0.936302004 |
| chr8:136714125-136714404 | 0.088421311 | 0.832604911 | 0.936302004 |
| chr1:6464508-6464778 | -0.092817034 | 0.832606858 | 0.936302004 |
| chr10:114306724-11430692€ | -0.094975054 | 0.83262961 | 0.936302004 |
| chr12:40693168-40693467 | -0.094975054 | 0.83262961 | 0.936302004 |
| chrX:141405532-141406295 | 0.076374593 | 0.832654307 | 0.936302943 |
| chr5:141199380-141199923 | -0.078433245 | 0.832654884 | 0.936302943 |
| chr3:192772182-192772837 | -0.0768254 | 0.832679295 | 0.936315021 |
| chr20:62972390-62972626 | 0.083150708 | 0.832690065 | 0.936315021 |
| chrX:130040490-130040944 | 0.080628029 | 0.832739413 | 0.936352772 |
| chr2:116819442-116819642 | 0.121607411 | 0.832767159 | 0.936352772 |
| chr5:176609443-176609950 | 0.075160078 | 0.832769523 | 0.936352772 |
| chr5:178223156-178223456 | 0.085233753 | 0.83277252 | 0.936352772 |
| chr8:123462875-123463127 | -0.084796914 | 0.832801335 | 0.936371431 |
| chr20:63291178-63291531 | 0.076641779 | 0.832818073 | 0.936376509 |

| | | | |
|---|---|---|---|
| chr1:197908780-197909250 | -0.076477195 | 0.832868264 | 0.936393384 |
| chr22:31616354-31616554 | 0.096603914 | 0.832871505 | 0.936393384 |
| chr7:74890661-74890953 | -0.086946507 | 0.832871659 | 0.936393384 |
| chr17:68102276-68102704 | -0.072644721 | 0.832881965 | 0.936393384 |
| chr2:197012137-197012363 | -0.102055243 | 0.832916865 | 0.936405343 |
| chr14:105473408-105474854 | 0.066513705 | 0.832917044 | 0.936405343 |
| chr16:85458844-85459364 | -0.072027442 | 0.832949758 | 0.936428382 |
| chr1:2684039-2684270 | 0.09635642 | 0.832988632 | 0.936431725 |
| chr6:17389368-17389796 | 0.0756436 | 0.832993239 | 0.936431725 |
| chr9:109440734-109441117 | -0.07571674 | 0.832996416 | 0.936431725 |
| chr16:66315217-66315538 | 0.086184808 | 0.833017143 | 0.936431725 |
| chr15:78726800-78727259 | -0.072267085 | 0.833038718 | 0.936431725 |
| chr11:83312100-83312371 | -0.101184554 | 0.833040026 | 0.936431725 |
| chr21:38267532-38267847 | 0.104919297 | 0.833042634 | 0.936431725 |
| chrX:41333050-41334777 | 0.066417287 | 0.833061806 | 0.936431725 |
| chr12:98644119-98645931 | -0.066216482 | 0.833062725 | 0.936431725 |
| chr1:202143082-202144954 | 0.066233695 | 0.833114655 | 0.936435284 |
| chr2:178450439-178451497 | -0.066271865 | 0.833116253 | 0.936435284 |
| chr5:150350510-150350912 | 0.076269597 | 0.833119318 | 0.936435284 |
| chrX:48573845-48575067 | -0.067728243 | 0.833126781 | 0.936435284 |
| chr2:37232261-37232486 | -0.086625068 | 0.833126998 | 0.936435284 |
| chr4:82373119-82373527 | -0.071972897 | 0.833151683 | 0.936448784 |
| chrX:68018551-68019185 | 0.076367816 | 0.833171434 | 0.936448784 |
| chr14:20428678-20429111 | 0.080820029 | 0.833175674 | 0.936448784 |
| chr6:152124396-152124760 | -0.089363511 | 0.833230047 | 0.936496159 |
| chr5:139775102-139775455 | -0.078695426 | 0.833261181 | 0.936509207 |
| chr7:33185760-33186043 | 0.082358467 | 0.833266101 | 0.936509207 |
| chr20:3786744-3786954 | 0.088509316 | 0.833297605 | 0.936509338 |
| chr17:60142082-60142446 | -0.067908648 | 0.83329826 | 0.936509338 |
| chr18:31241829-31242293 | 0.073562116 | 0.833302885 | 0.936509338 |
| chr16:73949542-73949851 | 0.092791919 | 0.833320577 | 0.936515485 |
| chr6:6803253-6803453 | -0.105261828 | 0.833352481 | 0.936537603 |
| chr15:98746040-98746617 | -0.076590308 | 0.833368853 | 0.936542266 |
| chr9:92808599-92809410 | -0.074432462 | 0.83339261 | 0.936546535 |
| chr2:172299861-172300129 | -0.098213173 | 0.83340466 | 0.936546535 |
| chr1:181669337-181669537 | 0.114260155 | 0.833426541 | 0.936546535 |
| chr2:59436354-59436554 | -0.126096218 | 0.833429532 | 0.936546535 |
| chr14:103103520-103103841 | -0.08431918 | 0.833445327 | 0.936546535 |
| chr6:87421589-87421963 | 0.077631694 | 0.833445989 | 0.936546535 |
| chr1:23727379-23727765 | -0.095133385 | 0.833481085 | 0.936551553 |
| chr8:144852545-144853628 | -0.069095416 | 0.833483999 | 0.936551553 |
| chr20:8534198-8534686 | 0.084984267 | 0.833487124 | 0.936551553 |
| chr12:24836384-24836614 | -0.118553476 | 0.833526547 | 0.936582116 |
| chr20:23606538-23606919 | -0.07691591 | 0.833616475 | 0.936652373 |
| chr16:48630482-48630766 | 0.093217388 | 0.833626147 | 0.936652373 |
| chr2:70994729-70995215 | -0.071395004 | 0.833638243 | 0.936652373 |
| chr9:34637336-34638122 | -0.072903226 | 0.833639698 | 0.936652373 |
| chr3:170961635-170961860 | 0.105021823 | 0.833650195 | 0.936652373 |
| chr2:135004268-135004611 | 0.082938205 | 0.833684806 | 0.936670415 |
| chr6:7553696-7554263 | -0.081626217 | 0.833701863 | 0.936670415 |

| | | | |
|---|---|---|---|
| chr7:8211181-8211444 | 0.093299334 | 0.833704418 | 0.936670415 |
| chr19:1095307-1095575 | -0.089926268 | 0.833715151 | 0.936670415 |
| chr10:24494218-24494523 | -0.093981977 | 0.833732278 | 0.936675628 |
| chr1:62687410-62688729 | -0.067969172 | 0.833744241 | 0.936675628 |
| chr8:144477727-144478509 | 0.068063396 | 0.833794787 | 0.936718681 |
| chr14:103111233-103111778 | -0.076392321 | 0.833816655 | 0.936729514 |
| chr17:78646858-78647058 | 0.112481617 | 0.833831477 | 0.93673243 |
| chr16:87950605-87951701 | 0.066087791 | 0.833856684 | 0.936747014 |
| chr20:51109241-51109464 | 0.093483747 | 0.833897181 | 0.936778774 |
| chr17:37406509-37407381 | 0.072553843 | 0.833917687 | 0.93678618 |
| chr14:103075057-103075263 | -0.090499866 | 0.833928226 | 0.93678618 |
| chr2:182101400-182101600 | 0.112222642 | 0.83397414 | 0.936799613 |
| chr20:8156738-8156955 | -0.109786107 | 0.833974719 | 0.936799613 |
| chr22:49852916-49853231 | -0.083728932 | 0.833976863 | 0.936799613 |
| chr10:29099445-29099744 | -0.090943668 | 0.834051557 | 0.936841272 |
| chr12:93569708-93569908 | -0.090943668 | 0.834051557 | 0.936841272 |
| chr2:214069270-214069579 | -0.087220064 | 0.834058946 | 0.936841272 |
| chr1:629810-630112 | -0.067630759 | 0.834073918 | 0.936841272 |
| chr3:169052991-169053191 | 0.112699038 | 0.834075084 | 0.936841272 |
| chr16:30075279-30076074 | 0.071353215 | 0.834117568 | 0.936875257 |
| chr1:195030985-195031260 | -0.093322633 | 0.834194596 | 0.936921995 |
| chr5:96614072-96614365 | -0.088143123 | 0.834207776 | 0.936921995 |
| chr2:234135730-234136001 | 0.096954001 | 0.834214011 | 0.936921995 |
| chr14:89490963-89491163 | 0.118777752 | 0.834220247 | 0.936921995 |
| chr3:193605888-193606243 | -0.080599582 | 0.834220318 | 0.936921995 |
| chr5:9482582-9482807 | 0.107171569 | 0.834244396 | 0.936935303 |
| chr18:10131950-10132175 | -0.100401328 | 0.834319499 | 0.936987553 |
| chr12:57744481-57745666 | -0.066360403 | 0.834323459 | 0.936987553 |
| chr1:224106588-224107031 | 0.077349178 | 0.834327606 | 0.936987553 |
| chr17:2025276-2025591 | 0.078582294 | 0.834340275 | 0.936988048 |
| chr11:68618077-68618277 | 0.108663092 | 0.83439117 | 0.937031471 |
| chrX:24002854-24003085 | -0.101721197 | 0.834450606 | 0.937082886 |
| chr6:8167784-8168043 | -0.117948282 | 0.834466366 | 0.937082886 |
| chr16:2968849-2969381 | -0.070006934 | 0.834473658 | 0.937082886 |
| chr2:211035634-211036084 | -0.087000139 | 0.834514468 | 0.937082886 |
| chr10:68526938-68527970 | 0.066867639 | 0.83452597 | 0.937082886 |
| chr9:99819337-99820423 | 0.06624514 | 0.834545527 | 0.937082886 |
| chr14:77980519-77981344 | -0.068386344 | 0.834547938 | 0.937082886 |
| chr11:48090710-48091518 | 0.071570259 | 0.834554261 | 0.937082886 |
| chr3:113495103-113495306 | -0.098401831 | 0.834555167 | 0.937082886 |
| chr3:135412554-135412776 | -0.10472761 | 0.834578092 | 0.937082886 |
| chr7:191442-191930 | 0.074929816 | 0.834582294 | 0.937082886 |
| chr7:45136104-45136421 | 0.084204113 | 0.834598457 | 0.937082886 |
| chr12:132851487-132851718 | -0.110208384 | 0.834617432 | 0.937082886 |
| chr2:235836631-235836967 | 0.095153053 | 0.83461799 | 0.937082886 |
| chr6:27177569-27178573 | -0.068957413 | 0.834620402 | 0.937082886 |
| chr8:143770897-143771932 | 0.068903836 | 0.834652433 | 0.937088903 |
| chr17:21213722-21215012 | -0.067262422 | 0.834672018 | 0.937088903 |
| chr19:38869759-38870233 | 0.075287411 | 0.834672101 | 0.937088903 |
| chr3:149401236-149401984 | -0.069441554 | 0.834694406 | 0.937088903 |

| | | | |
|---|---|---|---|
| chr20:35884305-35884547 | 0.095955659 | 0.834698689 | 0.937088903 |
| chr7:6300438-6300638 | 0.10614564 | 0.834699142 | 0.937088903 |
| chr3:61628794-61629037 | 0.078954064 | 0.834756301 | 0.937139259 |
| chr2:160252419-160252619 | -0.10635419 | 0.83477923 | 0.937139259 |
| chr2:238263482-238263805 | 0.09053396 | 0.834782301 | 0.937139259 |
| chr6:30481624-30481824 | -0.105644032 | 0.834792918 | 0.937139259 |
| chr11:71072091-71072291 | -0.10313606 | 0.834819061 | 0.937145739 |
| chr19:51927264-51927507 | -0.108396074 | 0.834824071 | 0.937145739 |
| chr1:231741526-231741802 | -0.101373387 | 0.834835383 | 0.937145739 |
| chrX:15336040-15336304 | -0.083735836 | 0.834868712 | 0.937166042 |
| chr10:43689485-43689863 | -0.082772032 | 0.834877931 | 0.937166042 |
| chr18:75896869-75897264 | -0.082285161 | 0.834911914 | 0.937190458 |
| chr1:58783792-58784782 | -0.065638279 | 0.834936061 | 0.937203834 |
| chr11:75515131-75515361 | -0.095324348 | 0.834965375 | 0.937215001 |
| chr8:66438881-66439841 | 0.069944288 | 0.83498002 | 0.937215001 |
| chr3:136861554-136862598 | 0.066083512 | 0.834991008 | 0.937215001 |
| chr6:127640550-127640750 | 0.114499104 | 0.834994936 | 0.937215001 |
| chr21:44773506-44773966 | 0.075918862 | 0.835056751 | 0.937270653 |
| chr14:24441914-24443099 | -0.065279824 | 0.835079844 | 0.937282492 |
| chr10:114166680-114166880 | -0.117173901 | 0.835091764 | 0.937282492 |
| chr7:56092949-56093298 | 0.078930655 | 0.835106577 | 0.937285388 |
| chr1:226182083-226182290 | -0.097837449 | 0.835145294 | 0.937315112 |
| chr1:27663163-27663418 | 0.085963564 | 0.835163805 | 0.937322159 |
| chr1:153513172-153513372 | -0.091643777 | 0.835194957 | 0.937342524 |
| chr2:230160339-230160807 | 0.084465424 | 0.835206418 | 0.937342524 |
| chr8:38089944-38090238 | 0.092503802 | 0.835246523 | 0.937373804 |
| chr14:22047150-22047399 | -0.102322786 | 0.835283976 | 0.937402107 |
| chr20:52191200-52192015 | -0.068137789 | 0.835312149 | 0.937412765 |
| chr7:974919-975758 | -0.068031466 | 0.835317942 | 0.937412765 |
| chr3:186259572-186259934 | -0.076707406 | 0.835359916 | 0.937446139 |
| chr5:172187364-172187749 | 0.073465984 | 0.835391333 | 0.937453253 |
| chr3:167406893-167407213 | 0.078896637 | 0.835399192 | 0.937453253 |
| chr2:86721093-86721293 | 0.111052521 | 0.83541892 | 0.937453253 |
| chr12:2890343-2891560 | -0.067146143 | 0.835435269 | 0.937453253 |
| chr8:70205543-70205743 | 0.11131882 | 0.835435733 | 0.937453253 |
| chr1:233326968-233328569 | 0.065686191 | 0.835454813 | 0.937453253 |
| chr17:49910678-49910896 | 0.090650484 | 0.835464016 | 0.937453253 |
| chr20:53869756-53870103 | 0.066449104 | 0.835474462 | 0.937453253 |
| chr16:84806366-84807085 | -0.072526853 | 0.835476368 | 0.937453253 |
| chr4:121696231-121697518 | -0.066034095 | 0.835551885 | 0.937524258 |
| chr8:74322301-74322503 | -0.090398654 | 0.835588035 | 0.937547567 |
| chr5:37847938-37848201 | 0.097618119 | 0.835597131 | 0.937547567 |
| chr6:87772937-87773484 | 0.073760155 | 0.835633914 | 0.937564773 |
| chr1:222738802-222739093 | -0.095666545 | 0.835639454 | 0.937564773 |
| chr11:71240880-71241291 | -0.073262174 | 0.835649175 | 0.937564773 |
| chr20:50152948-50153350 | 0.072228062 | 0.835712982 | 0.937622634 |
| chr20:63864814-63866026 | 0.065341196 | 0.835735139 | 0.937633763 |
| chr8:106159219-106159419 | -0.105802184 | 0.835768075 | 0.937641434 |
| chr3:179260830-179261840 | 0.069489191 | 0.835776436 | 0.937641434 |
| chr5:17046037-17046240 | -0.105520985 | 0.835798502 | 0.937641434 |

| | | | |
|---|---|---|---|
| chr3:194404496-194404915 | 0.092341346 | 0.835803146 | 0.937641434 |
| chr1:167820136-167820480 | 0.0793015 | 0.835803163 | 0.937641434 |
| chr4:148008219-148008419 | 0.107813188 | 0.835874793 | 0.937708063 |
| chr22:24156574-24157224 | 0.071252779 | 0.835890587 | 0.937712051 |
| chr3:150503479-150503716 | 0.087791251 | 0.835906205 | 0.937715843 |
| chr17:82292291-82293080 | 0.067960035 | 0.835920879 | 0.937718576 |
| chr12:47370432-47370746 | 0.084984297 | 0.835937477 | 0.937723466 |
| chr11:72041405-72041978 | 0.071276192 | 0.835981112 | 0.937758685 |
| chr3:149273152-149273497 | -0.076473609 | 0.83600882 | 0.937764736 |
| chr8:94406074-94406334 | -0.085845297 | 0.83601485 | 0.937764736 |
| chr1:9588302-9588830 | 0.075368098 | 0.836023222 | 0.937764736 |
| chr3:107370989-107371217 | -0.1055299 | 0.83607108 | 0.937804689 |
| chr2:139645454-139646037 | 0.078386896 | 0.83611055 | 0.937835233 |
| chr16:3158541-3159345 | 0.066581917 | 0.836125864 | 0.937838681 |
| chr1:3907261-3907491 | -0.101206755 | 0.836213798 | 0.937923384 |
| chr3:140216265-140216526 | 0.091155034 | 0.836225863 | 0.937923384 |
| chr1:155093999-155094636 | -0.074383262 | 0.836282993 | 0.937966751 |
| chr3:101803225-101803583 | -0.07779638 | 0.836297771 | 0.937966751 |
| chrX:52577072-52577363 | 0.090713225 | 0.836301252 | 0.937966751 |
| chr15:76313842-76314213 | -0.074961202 | 0.836442319 | 0.938083473 |
| chr1:157045481-157046653 | -0.067039946 | 0.836449046 | 0.938083473 |
| chr5:145681641-145681962 | 0.078710721 | 0.836452197 | 0.938083473 |
| chr13:20118405-20119483 | 0.066645232 | 0.836467326 | 0.938083473 |
| chr6:158660319-158660531 | 0.096635121 | 0.836498092 | 0.938083473 |
| chr12:52976258-52976550 | 0.076645924 | 0.836498189 | 0.938083473 |
| chr3:170811993-170812279 | 0.076469619 | 0.836509329 | 0.938083473 |
| chr7:44587950-44588550 | 0.077388386 | 0.836512715 | 0.938083473 |
| chr5:151080516-151081504 | 0.066693476 | 0.836515509 | 0.938083473 |
| chr14:93033162-93033371 | -0.08664669 | 0.836544377 | 0.938089112 |
| chr13:44879459-44879664 | 0.097048673 | 0.836571369 | 0.938089112 |
| chr8:101288075-101288342 | 0.083738751 | 0.836581017 | 0.938089112 |
| chr5:155884098-155884444 | -0.089317169 | 0.836584377 | 0.938089112 |
| chr17:44161762-44161983 | 0.1078761 | 0.836601449 | 0.938089112 |
| chr9:98808415-98808799 | 0.084152703 | 0.836608488 | 0.938089112 |
| chr1:161314157-161314986 | 0.068420485 | 0.836621498 | 0.938089112 |
| chr9:111728656-111729277 | -0.080103054 | 0.836632563 | 0.938089112 |
| chr8:25458198-25459339 | -0.064751994 | 0.83664186 | 0.938089112 |
| chr2:70919812-70920172 | 0.079965948 | 0.836661936 | 0.938089112 |
| chr3:149879894-149880178 | -0.082122063 | 0.836673175 | 0.938089112 |
| chr17:10697035-10698017 | -0.067246261 | 0.836679518 | 0.938089112 |
| chr3:85360739-85361099 | -0.075632474 | 0.836710735 | 0.938089112 |
| chr3:187062539-187062751 | -0.095013414 | 0.836713847 | 0.938089112 |
| chr17:63826930-63827825 | -0.065264753 | 0.836719703 | 0.938089112 |
| chr12:101323777-101323977 | -0.100793287 | 0.836728051 | 0.938089112 |
| chr11:75384491-75384801 | 0.078386087 | 0.83672867 | 0.938089112 |
| chr3:64244370-64244888 | -0.066511345 | 0.836750319 | 0.938097112 |
| chr17:29293509-29294508 | -0.0670847 | 0.836783786 | 0.938097112 |
| chr3:3015553-3015818 | 0.080944374 | 0.836784873 | 0.938097112 |
| chr5:151729698-151729898 | 0.098686192 | 0.836803111 | 0.938097112 |
| chr10:20290515-20290792 | 0.105924963 | 0.836822364 | 0.938097112 |

| | | | |
|---|---|---|---|
| chr2:111049744-111050167 | -0.078847131 | 0.836830752 | 0.938097112 |
| chr1:49765166-49765370 | -0.101082722 | 0.83683179 | 0.938097112 |
| chr11:61949588-61949807 | 0.092390118 | 0.836833751 | 0.938097112 |
| chr10:34666241-34666654 | -0.078834854 | 0.836885764 | 0.938125327 |
| chr3:100737433-100737633 | 0.100696332 | 0.836888039 | 0.938125327 |
| chr10:21101284-21101621 | -0.101350586 | 0.836895651 | 0.938125327 |
| chrX:9379109-9379357 | -0.106769714 | 0.836990524 | 0.938217949 |
| chr17:7435836-7436164 | -0.081098986 | 0.837018179 | 0.938235223 |
| chr3:63607688-63608011 | -0.077892827 | 0.837102718 | 0.938279215 |
| chr6:30516535-30516784 | 0.094706706 | 0.837102972 | 0.938279215 |
| chr15:45187984-45188338 | -0.083866789 | 0.837114969 | 0.938279215 |
| chr17:45640382-45640618 | 0.098636008 | 0.837116572 | 0.938279215 |
| chr5:29697803-29698293 | -0.073471904 | 0.837131666 | 0.938279215 |
| chr19:45558415-45558729 | 0.082704904 | 0.83713786 | 0.938279215 |
| chr12:71747043-71747415 | -0.075872736 | 0.837151364 | 0.938279215 |
| chr4:105596745-105596954 | -0.105456839 | 0.837163632 | 0.938279215 |
| chr19:49766256-49767043 | 0.065785043 | 0.837167635 | 0.938279215 |
| chr17:67334863-67335207 | -0.078645328 | 0.837198745 | 0.938292101 |
| chr5:14268285-14268641 | 0.076764134 | 0.837203624 | 0.938292101 |
| chr11:72573399-72573686 | 0.078831019 | 0.837276241 | 0.938337543 |
| chr1:6482951-6483389 | -0.079171911 | 0.837290652 | 0.938337543 |
| chr3:18684300-18684500 | 0.11550258 | 0.837308697 | 0.938337543 |
| chr19:58543544-58544778 | 0.065052287 | 0.837318283 | 0.938337543 |
| chr10:111102417-111102820 | 0.085206269 | 0.837339262 | 0.938337543 |
| chr17:51669248-51669450 | 0.101682843 | 0.837343763 | 0.938337543 |
| chr6:134063876-134064396 | -0.073708551 | 0.83736325 | 0.938337543 |
| chr8:26449008-26449968 | 0.066256388 | 0.837365477 | 0.938337543 |
| chr1:45205721-45205997 | -0.101257843 | 0.837365898 | 0.938337543 |
| chrX:40105308-40105541 | 0.086436452 | 0.837367568 | 0.938337543 |
| chr11:62646218-62646934 | 0.067254063 | 0.83737888 | 0.938337543 |
| chr22:43812251-43812573 | 0.082910718 | 0.837395366 | 0.938340001 |
| chr12:123364573-123366020 | -0.064805055 | 0.837405566 | 0.938340001 |
| chr3:150922663-150922938 | 0.078373162 | 0.837453279 | 0.938367761 |
| chr5:112823285-112823990 | -0.07054336 | 0.837454833 | 0.938367761 |
| chr2:20224586-20225617 | 0.066802938 | 0.837512722 | 0.938418902 |
| chr1:111619536-111620588 | 0.066828957 | 0.837533741 | 0.93842873 |
| chr3:155382466-155382923 | 0.076540812 | 0.837570056 | 0.938430835 |
| chr19:11355684-11355905 | 0.103776675 | 0.837572362 | 0.938430835 |
| chr18:5868296-5868498 | 0.103776675 | 0.837572362 | 0.938430835 |
| chr19:18224433-18224925 | 0.070882834 | 0.837691776 | 0.938541323 |
| chr1:187817778-187818121 | -0.083756143 | 0.837696782 | 0.938541323 |
| chr18:589508-589708 | -0.107851977 | 0.837707722 | 0.938541323 |
| chr15:48250614-48250867 | -0.092635735 | 0.837743812 | 0.938557338 |
| chr20:50068291-50068536 | -0.083399947 | 0.837746515 | 0.938557338 |
| chr2:47664880-47665106 | 0.100717514 | 0.83778186 | 0.938579567 |
| chr8:129023816-129024245 | -0.085541016 | 0.837790855 | 0.938579567 |
| chr10:73888420-73889257 | 0.067869915 | 0.837845038 | 0.93859838 |
| chr2:222939271-222939551 | 0.086057594 | 0.83785099 | 0.93859838 |
| chr14:53963542-53963994 | 0.07290227 | 0.837854108 | 0.93859838 |
| chr19:58575412-58575978 | -0.067908411 | 0.837856647 | 0.93859838 |

| | | | |
|---|---|---|---|
| chr1:154391803-154392068 | -0.115431461 | 0.83787181 | 0.938598459 |
| chr6:72252655-72252901 | 0.110593209 | 0.837883947 | 0.938598459 |
| chr22:36387912-36388637 | 0.06646964 | 0.837893466 | 0.938598459 |
| chr7:151992353-151992689 | -0.078962141 | 0.837934941 | 0.938631195 |
| chr11:125591976-125593464 | -0.06471341 | 0.83798213 | 0.938651426 |
| chr19:38849278-38849677 | 0.069576333 | 0.837997429 | 0.938651426 |
| chr14:72666745-72667094 | -0.093976089 | 0.837997456 | 0.938651426 |
| chr6:110179483-110180721 | -0.065096808 | 0.838004197 | 0.938651426 |
| chr12:31323883-31324899 | -0.068030416 | 0.83803232 | 0.938651426 |
| chr12:72466378-72466578 | -0.107743372 | 0.838037807 | 0.938651426 |
| chr9:124840678-124840904 | -0.109405174 | 0.838038754 | 0.938651426 |
| chr5:154610871-154611071 | -0.095447712 | 0.838061317 | 0.938662977 |
| chr8:73971607-73972925 | -0.064249148 | 0.838087469 | 0.938667846 |
| chr3:194212904-194213472 | -0.074443961 | 0.838090166 | 0.938667846 |
| chr2:219279682-219280141 | -0.073202575 | 0.838118871 | 0.938686275 |
| chr3:181700483-181700781 | -0.079346728 | 0.838158688 | 0.938702371 |
| chr12:56487673-56488688 | -0.068670175 | 0.838169371 | 0.938702371 |
| chr6:146700117-146700373 | 0.090702646 | 0.838181704 | 0.938702371 |
| chr17:18217487-18217687 | 0.097842798 | 0.838194557 | 0.938702371 |
| chr11:1749487-1749711 | -0.094202874 | 0.838196965 | 0.938702371 |
| chr6:30214074-30214387 | 0.077707301 | 0.838206749 | 0.938702371 |
| chr6:99152072-99152338 | 0.085308674 | 0.838240622 | 0.938726585 |
| chr6:27146559-27147660 | -0.065253389 | 0.838269291 | 0.938744971 |
| chrX:138775612-138775812 | 0.118760826 | 0.838284447 | 0.938748223 |
| chr11:910720-911327 | -0.072035338 | 0.838301534 | 0.938753638 |
| chr6:87210658-87210871 | -0.112912115 | 0.838367309 | 0.93877401 |
| chr6:46848888-46849345 | -0.07913374 | 0.838367602 | 0.93877401 |
| chr6:137358381-137358581 | 0.119885329 | 0.838368258 | 0.93877401 |
| chr18:50194688-50195591 | -0.068802643 | 0.838368734 | 0.93877401 |
| chr11:35391014-35391650 | -0.070871709 | 0.83838142 | 0.938774495 |
| chr5:95759712-95759969 | -0.083827946 | 0.838422055 | 0.938796368 |
| chr1:43367555-43368584 | 0.066283156 | 0.838425458 | 0.938796368 |
| chr10:84762297-84762497 | -0.100851696 | 0.838439408 | 0.938798268 |
| chr6:27541651-27541966 | 0.083159965 | 0.838473363 | 0.938814066 |
| chr13:19905525-19905741 | 0.097712389 | 0.838486662 | 0.938814066 |
| chr22:42444075-42444275 | 0.108180923 | 0.838492192 | 0.938814066 |
| chr9:6006912-6007558 | 0.069007217 | 0.838514587 | 0.938814066 |
| chr3:172467060-172467618 | -0.074632699 | 0.838533744 | 0.938814066 |
| chr5:122998570-122998807 | -0.107963371 | 0.838537187 | 0.938814066 |
| chr2:118185683-118186070 | -0.081989309 | 0.838539554 | 0.938814066 |
| chr8:122008626-122009045 | 0.082563282 | 0.838551537 | 0.938814066 |
| chr1:109725438-109726049 | -0.070033498 | 0.838571896 | 0.938823142 |
| chr20:6767578-6768158 | -0.075985835 | 0.838633023 | 0.938861448 |
| chr13:29963808-29964022 | -0.099489782 | 0.838641066 | 0.938861448 |
| chr5:181221759-181222809 | -0.063857106 | 0.838647659 | 0.938861448 |
| chr9:112750207-112751176 | 0.067484591 | 0.838659976 | 0.938861448 |
| chr1:41360884-41361466 | 0.066132731 | 0.838674207 | 0.938861448 |
| chr6:117840798-117841031 | 0.110980875 | 0.838688244 | 0.938861448 |
| chr2:113370544-113370864 | 0.083670706 | 0.838691884 | 0.938861448 |
| chr14:75293926-75294505 | 0.066500285 | 0.838718748 | 0.938866369 |

| | | | |
|---|---|---|---|
| chr4:105586796-105587041 | 0.095746741 | 0.838720787 | 0.938866369 |
| chr5:36876177-36876377 | 0.086694748 | 0.838740648 | 0.938874885 |
| chr2:65505219-65505745 | -0.077059894 | 0.838793063 | 0.938892782 |
| chr3:149454841-149455074 | 0.097534808 | 0.838793084 | 0.938892782 |
| chr16:4585294-4585810 | -0.073518223 | 0.838795239 | 0.938892782 |
| chr5:32193559-32194157 | -0.068959257 | 0.83880565 | 0.938892782 |
| chr17:49394847-49395309 | 0.07361795 | 0.838818043 | 0.938892939 |
| chr6:47097442-47097687 | -0.096194483 | 0.838843465 | 0.938903125 |
| chr11:44538499-44538768 | 0.092001762 | 0.838851651 | 0.938903125 |
| chr3:37862377-37862877 | -0.078685209 | 0.838878361 | 0.938917301 |
| chr6:31738442-31738709 | -0.08088115 | 0.838888824 | 0.938917301 |
| chrX:74292135-74293061 | 0.065960738 | 0.83891544 | 0.938933375 |
| chr11:67428037-67428618 | 0.068551652 | 0.838954624 | 0.938953939 |
| chr4:39324642-39324980 | 0.105509427 | 0.838958323 | 0.938953939 |
| chr3:190591137-190591500 | 0.073800914 | 0.83897297 | 0.938956617 |
| chr11:59144566-59145298 | 0.071079981 | 0.839016942 | 0.938962586 |
| chr17:79075354-79075575 | 0.087317476 | 0.839060427 | 0.938962586 |
| chr1:107140901-107142579 | -0.063331723 | 0.839063298 | 0.938962586 |
| chr14:22994673-22995019 | -0.082392346 | 0.839078838 | 0.938962586 |
| chr18:31819674-31819945 | 0.086169989 | 0.839096595 | 0.938962586 |
| chr2:121829248-121829523 | -0.111523053 | 0.839099749 | 0.938962586 |
| chr11:100861296-10086174( | -0.085130353 | 0.839106109 | 0.938962586 |
| chr5:138036008-138036415 | 0.072841409 | 0.839115036 | 0.938962586 |
| chr6:79525887-79526122 | -0.096993955 | 0.839121515 | 0.938962586 |
| chr1:153441271-153441471 | 0.102745532 | 0.839129219 | 0.938962586 |
| chr1:242139462-242139836 | 0.082819858 | 0.839135911 | 0.938962586 |
| chr8:133189536-133189736 | -0.099858907 | 0.839136802 | 0.938962586 |
| chr16:10652844-10653380 | 0.074568721 | 0.839142771 | 0.938962586 |
| chr3:115758136-115758361 | -0.099737133 | 0.839149866 | 0.938962586 |
| chr14:43539263-43539506 | -0.085051356 | 0.839162545 | 0.938963061 |
| chr22:17367035-17367263 | -0.0964647 | 0.839202093 | 0.938991426 |
| chr21:42337805-42338659 | 0.067513316 | 0.839215848 | 0.938991426 |
| chr17:49987873-49988210 | 0.079259882 | 0.83922466 | 0.938991426 |
| chr1:52552948-52553201 | -0.092033442 | 0.839260631 | 0.939009238 |
| chr2:211157920-211158177 | 0.093703816 | 0.839265089 | 0.939009238 |
| chr1:61641727-61642075 | 0.074806845 | 0.839299881 | 0.939034453 |
| chr2:26177849-26178090 | -0.102633625 | 0.839333118 | 0.939053683 |
| chr6:88989362-88989600 | -0.108496526 | 0.83934158 | 0.939053683 |
| chr1:169609433-169609651 | 0.099662057 | 0.839377334 | 0.939076952 |
| chr1:110780998-110781233 | 0.09626973 | 0.83938689 | 0.939076952 |
| chr1:191432803-191433241 | -0.075932153 | 0.839437329 | 0.939113643 |
| chr1:15153285-15153852 | -0.072758558 | 0.839455721 | 0.939113643 |
| chr22:45413491-45413741 | -0.103493212 | 0.839465665 | 0.939113643 |
| chr6:100925658-100926137 | 0.081394131 | 0.839478225 | 0.939113643 |
| chr12:88480983-88481362 | -0.089553511 | 0.839489161 | 0.939113643 |
| chr3:52897311-52897781 | -0.079934073 | 0.839509534 | 0.939113643 |
| chr3:138915063-138916466 | 0.064535455 | 0.83952368 | 0.939113643 |
| chr3:63999476-63999885 | 0.067127637 | 0.839529044 | 0.939113643 |
| chr10:93115795-93116140 | 0.080384508 | 0.839529994 | 0.939113643 |
| chr8:72536313-72537476 | -0.066957455 | 0.839588894 | 0.93911609 |

| | | | |
|---|---|---|---|
| chr3:126198326-126198623 | 0.08481089 | 0.839589588 | 0.93911609 |
| chr2:199011744-199011944 | -0.106712372 | 0.839599323 | 0.93911609 |
| chr3:74408947-74409414 | -0.075072787 | 0.839600744 | 0.93911609 |
| chr9:136245232-136246082 | 0.067971819 | 0.839610233 | 0.93911609 |
| chr10:70246166-70246366 | -0.107165715 | 0.839620659 | 0.93911609 |
| chrX:92221397-92221659 | 0.097247602 | 0.839625256 | 0.93911609 |
| chr6:148972330-148972807 | -0.076995326 | 0.839630233 | 0.93911609 |
| chr3:138174089-138175041 | 0.066234039 | 0.839676912 | 0.939138776 |
| chrX:63519170-63519773 | 0.073040561 | 0.839699378 | 0.939138776 |
| chr8:102806179-102807814 | 0.063062926 | 0.83971379 | 0.939138776 |
| chr4:25993723-25994049 | -0.084762432 | 0.83971987 | 0.939138776 |
| chr4:160027994-160028194 | 0.106628472 | 0.839724142 | 0.939138776 |
| chr2:101205588-101205989 | -0.079475607 | 0.839736518 | 0.939138776 |
| chr17:41836137-41836388 | 0.089233501 | 0.839753909 | 0.939138776 |
| chrX:136767168-136767452 | 0.081538332 | 0.839768729 | 0.939138776 |
| chr2:159807443-159807740 | -0.086422707 | 0.839769384 | 0.939138776 |
| chr12:131133573-131133773 | -0.096962552 | 0.839773084 | 0.939138776 |
| chr3:52685464-52686343 | 0.068735037 | 0.839799208 | 0.939154284 |
| chr17:64420184-64420682 | -0.070738381 | 0.839837866 | 0.939183808 |
| chr18:3330518-3330976 | -0.077276302 | 0.839863304 | 0.939198548 |
| chr1:114428154-114428627 | 0.068531767 | 0.839896981 | 0.939209327 |
| chr14:98561628-98561828 | -0.107151315 | 0.839897458 | 0.939209327 |
| chr6:85958331-85958778 | -0.075501844 | 0.839929826 | 0.939231814 |
| chr11:7597412-7598059 | -0.067867223 | 0.839986569 | 0.939275455 |
| chr4:72637022-72637413 | -0.086823549 | 0.83999337 | 0.939275455 |
| chr17:83050929-83052363 | -0.06319367 | 0.8400253 | 0.939286015 |
| chr15:75842783-75843650 | 0.068881489 | 0.8400275 | 0.939286015 |
| chr8:127743068-127743269 | 0.08124083 | 0.840042315 | 0.939286015 |
| chr1:70034509-70034749 | -0.097647904 | 0.840051849 | 0.939286015 |
| chr2:160157940-160158472 | -0.075708127 | 0.840088905 | 0.939312372 |
| chr14:73723877-73724380 | -0.070199746 | 0.840099939 | 0.939312372 |
| chr13:49401163-49401688 | -0.073021821 | 0.84012871 | 0.939330834 |
| chr17:75326417-75326728 | 0.085810276 | 0.840149753 | 0.939340654 |
| chr19:18157219-18157435 | 0.085221133 | 0.840171123 | 0.939342697 |
| chr6:44735557-44735757 | -0.105646895 | 0.840176099 | 0.939342697 |
| chr17:49515484-49515690 | -0.088016264 | 0.840212091 | 0.939369231 |
| chr2:98837155-98837404 | -0.09855454 | 0.840246161 | 0.939383818 |
| chr1:244834335-244835995 | -0.063256308 | 0.840249659 | 0.939383818 |
| chr2:172095345-172096057 | 0.069479771 | 0.840273091 | 0.939384253 |
| chr1:175757428-175757810 | -0.077598013 | 0.840274568 | 0.939384253 |
| chr13:112931269-112931916 | 0.070882866 | 0.84030879 | 0.939402095 |
| chr2:201993091-201993341 | 0.108406454 | 0.840321035 | 0.939402095 |
| chr2:156903425-156903683 | 0.111594197 | 0.840327308 | 0.939402095 |
| chr14:20683171-20684376 | -0.063605025 | 0.840350969 | 0.939413605 |
| chr14:63748873-63749532 | 0.070460266 | 0.840369814 | 0.939413605 |
| chr14:93106230-93106430 | -0.091118722 | 0.840374385 | 0.939413605 |
| chr1:67535442-67535765 | -0.07831591 | 0.840400505 | 0.939429097 |
| chr2:71930050-71930274 | 0.087601896 | 0.840428784 | 0.939433241 |
| chr2:153232534-153232892 | 0.074897699 | 0.840446133 | 0.939433241 |
| chr6:151675736-151676277 | 0.068483539 | 0.840450323 | 0.939433241 |

| | | | |
|---|---|---|---|
| chrX:118016536-118016866 | 0.095709774 | 0.840453254 | 0.939433241 |
| chr16:82011111-82011760 | 0.070685266 | 0.840493961 | 0.939454357 |
| chr8:127849303-127849596 | -0.071176339 | 0.840498923 | 0.939454357 |
| chr11:74487919-74488119 | 0.093032652 | 0.840508928 | 0.939454357 |
| chr12:121038692-121039450 | 0.070240775 | 0.840542424 | 0.939478092 |
| chr6:89080307-89081080 | -0.065068664 | 0.840573788 | 0.939496702 |
| chr13:43905456-43905840 | -0.080685202 | 0.840594484 | 0.939496702 |
| chr5:169512459-169512659 | 0.101879823 | 0.840595859 | 0.939496702 |
| chr17:55177212-55177433 | -0.09730165 | 0.840617471 | 0.939498071 |
| chr1:181479197-181479660 | 0.071552549 | 0.840642201 | 0.939498071 |
| chr20:38004409-38004773 | 0.08431473 | 0.840643842 | 0.939498071 |
| chr1:198510562-198510819 | 0.095471998 | 0.84064613 | 0.939498071 |
| chr1:41417215-41417415 | 0.092045511 | 0.840673681 | 0.93950817 |
| chr2:168759282-168759537 | 0.089047304 | 0.840679689 | 0.93950817 |
| chr5:61621087-61621341 | -0.087245172 | 0.840722419 | 0.939512797 |
| chr3:139480673-139480928 | 0.101794085 | 0.840727592 | 0.939512797 |
| chr9:120985805-120986102 | -0.090303855 | 0.840744044 | 0.939512797 |
| chr5:126500088-126500346 | 0.092431244 | 0.840746414 | 0.939512797 |
| chr6:139343106-139343320 | 0.09938188 | 0.840756974 | 0.939512797 |
| chr11:66709828-66710056 | 0.087358978 | 0.840757399 | 0.939512797 |
| chr1:19659750-19660075 | 0.081290821 | 0.840878 | 0.939628162 |
| chr8:122694139-122694497 | -0.078074618 | 0.840885164 | 0.939628162 |
| chr20:62900094-62901098 | 0.065576504 | 0.840898379 | 0.939629225 |
| chr3:149969969-149971842 | -0.063012875 | 0.840933006 | 0.939645424 |
| chr6:34236643-34237080 | 0.072443441 | 0.840937548 | 0.939645424 |
| chr20:18019192-18019475 | -0.083733601 | 0.840954103 | 0.939645424 |
| chr9:128082282-128082563 | -0.084478754 | 0.84096193 | 0.939645424 |
| chr4:85747971-85748171 | -0.102324575 | 0.840979419 | 0.939651263 |
| chr8:23410110-23410704 | -0.074278922 | 0.841024787 | 0.939688252 |
| chr6:87436455-87436835 | 0.075966242 | 0.84104841 | 0.939700943 |
| chr19:4084520-4084725 | 0.102926201 | 0.84110819 | 0.939726916 |
| chr15:64162619-64163924 | 0.063826862 | 0.841109715 | 0.939726916 |
| chr2:241762338-241762846 | -0.078472866 | 0.84113275 | 0.939726916 |
| chr1:204837547-204837749 | 0.084005901 | 0.841136051 | 0.939726916 |
| chr11:124069388-124069699 | -0.084982906 | 0.841140023 | 0.939726916 |
| chr20:6708515-6708805 | 0.090560131 | 0.841154344 | 0.939726916 |
| chr3:115819796-115820104 | 0.075694149 | 0.841160266 | 0.939726916 |
| chr10:81138685-81139139 | 0.090942503 | 0.841169772 | 0.939726916 |
| chr3:16884883-16885307 | 0.08119642 | 0.84119979 | 0.93973309 |
| chr19:50200107-50200323 | -0.085993827 | 0.841211215 | 0.93973309 |
| chr10:29874036-29874278 | -0.107247847 | 0.841212156 | 0.93973309 |
| chr6:11575413-11575870 | 0.079536663 | 0.841225442 | 0.93973309 |
| chr3:134299659-134299964 | 0.088550942 | 0.841248288 | 0.93973309 |
| chr17:56629853-56630133 | 0.077804944 | 0.841259134 | 0.93973309 |
| chr20:47985527-47985757 | -0.069284709 | 0.84126115 | 0.93973309 |
| chr15:40340751-40341114 | -0.075144279 | 0.841335921 | 0.939787751 |
| chr14:22588915-22590049 | -0.063195251 | 0.841342084 | 0.939787751 |
| chr8:22036775-22037030 | 0.090895018 | 0.841357429 | 0.939787751 |
| chr1:172429289-172429690 | 0.072713597 | 0.841359145 | 0.939787751 |
| chr5:15105835-15106138 | 0.082680735 | 0.841407011 | 0.939817595 |

| | | | |
|---|---|---|---|
| chr16:78016827-78017133 | 0.09172065 | 0.841417362 | 0.939817595 |
| chr9:127899501-127899749 | 0.084118509 | 0.841422659 | 0.939817595 |
| chr15:74616362-74616667 | 0.075390344 | 0.841450133 | 0.939832385 |
| chr20:50190052-50190257 | 0.077558221 | 0.841470795 | 0.939832385 |
| chr17:17836895-17837172 | -0.074562692 | 0.841484584 | 0.939832385 |
| chr22:30298286-30298788 | 0.067664591 | 0.841484965 | 0.939832385 |
| chr13:28376690-28377120 | 0.079161954 | 0.841514126 | 0.939835349 |
| chr6:49386984-49387507 | -0.080855457 | 0.841530195 | 0.939835349 |
| chr1:218000228-218000437 | -0.096839078 | 0.841536106 | 0.939835349 |
| chr14:103871912-103872112 | 0.092426608 | 0.841536682 | 0.939835349 |
| chr22:25072362-25072759 | -0.093183806 | 0.841573998 | 0.939863326 |
| chr18:62957879-62958211 | -0.08700086 | 0.841608181 | 0.939882892 |
| chr22:39320176-39320451 | 0.082736309 | 0.841616052 | 0.939882892 |
| chr1:204446301-204446673 | 0.070855992 | 0.841678102 | 0.939938487 |
| chr1:202315626-202315917 | 0.0725018 | 0.841761316 | 0.939992237 |
| chr13:30972826-30973026 | 0.093069005 | 0.841762221 | 0.939992237 |
| chr14:55430001-55430460 | 0.078115579 | 0.841763036 | 0.939992237 |
| chr3:145249634-145249994 | 0.089043934 | 0.841808191 | 0.940003476 |
| chr12:107104725-107105012 | -0.082904585 | 0.841808567 | 0.940003476 |
| chr4:61922785-61923068 | 0.092281035 | 0.841820252 | 0.940003476 |
| chr14:43357093-43357486 | 0.086402452 | 0.841822173 | 0.940003476 |
| chr17:55585812-55586012 | 0.098047525 | 0.841920812 | 0.940099919 |
| chr9:120252344-120252553 | -0.115787303 | 0.841951996 | 0.94010929 |
| chr3:74883713-74884032 | 0.090768071 | 0.841953743 | 0.94010929 |
| chr3:58021573-58021773 | -0.101491236 | 0.841998717 | 0.940129297 |
| chr19:12832751-12832951 | 0.112039537 | 0.84200499 | 0.940129297 |
| chr11:68146954-68147351 | 0.075387671 | 0.84200847 | 0.940129297 |
| chr6:82415986-82416219 | -0.104499832 | 0.842043738 | 0.940154975 |
| chr3:192444565-192444800 | -0.087752593 | 0.842090998 | 0.940194041 |
| chr9:19229784-19231397 | -0.063306672 | 0.842114405 | 0.940201802 |
| chr11:20671920-20672373 | 0.079838479 | 0.842122491 | 0.940201802 |
| chr22:50185710-50185925 | 0.095102819 | 0.842137515 | 0.940204877 |
| chr12:101459868-101460292 | 0.07588229 | 0.842199586 | 0.940255169 |
| chr3:158791397-158791610 | -0.100408915 | 0.842207105 | 0.940255169 |
| chr2:128027182-128028626 | 0.062611853 | 0.842220033 | 0.940255902 |
| chr10:52488709-52489272 | -0.077580234 | 0.84225814 | 0.940284745 |
| chr21:39125807-39126306 | -0.071521955 | 0.842274587 | 0.940289407 |
| chr19:48333880-48334208 | -0.071684578 | 0.842329166 | 0.940329398 |
| chr6:158002249-158002824 | -0.071344313 | 0.842338831 | 0.940329398 |
| chr15:44663032-44664167 | 0.064233809 | 0.84234996 | 0.940329398 |
| chr6:154928454-154928684 | -0.089064941 | 0.842359499 | 0.940329398 |
| chr1:215482087-215482451 | -0.069935425 | 0.842394502 | 0.940354771 |
| chr2:7394285-7394548 | 0.097385867 | 0.842418283 | 0.940367618 |
| chr2:74379859-74380621 | 0.068471543 | 0.842448487 | 0.940367721 |
| chr11:117426342-117426544 | -0.097229049 | 0.842454445 | 0.940367721 |
| chr17:59619103-59620347 | 0.062073423 | 0.842456988 | 0.940367721 |
| chr5:35198177-35198452 | -0.078675767 | 0.842471348 | 0.940367721 |
| chr6:79868013-79868494 | -0.077075713 | 0.842488216 | 0.940367721 |
| chr3:196081394-196082532 | 0.063926243 | 0.842492012 | 0.940367721 |
| chr1:215923361-215923797 | 0.075725483 | 0.842552957 | 0.940409237 |

| | | | |
|---|---|---|---|
| chr11:32884870-32885070 | 0.103119489 | 0.842553754 | 0.940409237 |
| chr12:51249306-51249506 | 0.102509281 | 0.842597175 | 0.940432118 |
| chr11:13462558-13463874 | -0.063418406 | 0.842598801 | 0.940432118 |
| chr9:132877990-132878461 | 0.075050923 | 0.842613795 | 0.940435154 |
| chr13:45281050-45281418 | -0.074132171 | 0.842628221 | 0.940437557 |
| chr7:1432533-1432891 | -0.072948889 | 0.842645638 | 0.940443296 |
| chr9:77879412-77879968 | 0.073140131 | 0.842660685 | 0.940446392 |
| chr4:123396612-123396945 | -0.070350993 | 0.842673457 | 0.940446947 |
| chr3:139111688-139112220 | -0.071669414 | 0.842693305 | 0.940450015 |
| chr11:13487061-13487540 | 0.069323089 | 0.842700753 | 0.940450015 |
| chr13:98416256-98416843 | 0.067234477 | 0.842715531 | 0.940452809 |
| chr9:129178130-129178517 | 0.067260646 | 0.842731563 | 0.940457003 |
| chr17:76385817-76386046 | 0.077824523 | 0.842755766 | 0.940465448 |
| chr6:6584737-6585084 | 0.092510358 | 0.842763679 | 0.940465448 |
| chr11:130448357-130449074 | 0.06895789 | 0.842786606 | 0.940477336 |
| chr17:79239078-79239296 | -0.104431811 | 0.842856324 | 0.940541438 |
| chr2:71933998-71934203 | 0.09091037 | 0.842909615 | 0.940587206 |
| chr17:46192661-46193123 | -0.073037025 | 0.843011366 | 0.940685373 |
| chr10:127243598-127243824 | -0.091651135 | 0.843024309 | 0.940685373 |
| chr4:151099104-151100253 | -0.062946054 | 0.843037224 | 0.940685373 |
| chr2:78672994-78673542 | 0.069618328 | 0.843058374 | 0.940685373 |
| chr5:141618119-141619435 | -0.06303774 | 0.843058972 | 0.940685373 |
| chr7:99465596-99466376 | -0.066933644 | 0.843079698 | 0.9406948 |
| chr21:25801346-25801910 | 0.07121946 | 0.843104103 | 0.940708332 |
| chr1:218402739-218403007 | -0.081845682 | 0.843130148 | 0.940713768 |
| chr7:36291439-36291863 | 0.069537257 | 0.84313353 | 0.940713768 |
| chr10:104268538-104269385 | -0.06387811 | 0.843149456 | 0.940717839 |
| chr21:46153343-46153582 | 0.084748584 | 0.843168512 | 0.94072366 |
| chr6:7312335-7313524 | -0.065113729 | 0.84317936 | 0.94072366 |
| chr1:26472642-26473231 | -0.070930172 | 0.843191506 | 0.94072366 |
| chr12:48119625-48119839 | 0.095733914 | 0.84320718 | 0.94072745 |
| chr17:10119619-10120247 | 0.06620758 | 0.843225542 | 0.940734238 |
| chr2:11561594-11561850 | -0.084682076 | 0.843246569 | 0.940743999 |
| chr13:70353126-70353326 | 0.096569248 | 0.843337083 | 0.940828856 |
| chr5:20109258-20109719 | -0.068156652 | 0.843347189 | 0.940828856 |
| chr6:87682259-87682533 | 0.096652195 | 0.843371487 | 0.940832672 |
| chr12:43338951-43339185 | -0.096655313 | 0.843375168 | 0.940832672 |
| chr14:49585899-49587179 | 0.061857506 | 0.843401896 | 0.940848791 |
| chr5:38528151-38528452 | -0.094828301 | 0.843424504 | 0.940860314 |
| chr14:41679596-41679856 | 0.095025721 | 0.84355206 | 0.940988905 |
| chr12:71217014-71217322 | -0.0792535 | 0.843569837 | 0.940995036 |
| chr1:61842899-61843560 | -0.075604668 | 0.843592251 | 0.94100634 |
| chr20:63664825-63665711 | 0.06214434 | 0.843626459 | 0.941015109 |
| chr5:179732397-179733507 | -0.062655038 | 0.843641776 | 0.941015109 |
| chr6:49523014-49523464 | 0.079225027 | 0.843654897 | 0.941015109 |
| chr21:42513448-42513648 | 0.090572091 | 0.843656048 | 0.941015109 |
| chr12:68331965-68332756 | 0.063051878 | 0.843661653 | 0.941015109 |
| chr22:30255948-30256238 | -0.080140356 | 0.843699359 | 0.941015109 |
| chr9:122912900-122914160 | -0.064752095 | 0.843722498 | 0.941015109 |
| chr16:89443083-89443823 | 0.067151209 | 0.843724371 | 0.941015109 |

| | | | |
|---|---|---|---|
| chr15:78944730-78945476 | -0.066413164 | 0.843724824 | 0.941015109 |
| chr10:55668351-55668629 | -0.092708249 | 0.843734137 | 0.941015109 |
| chr2:218406001-218407139 | 0.062851733 | 0.843744631 | 0.941015109 |
| chr5:17132637-17132843 | 0.094439184 | 0.843755971 | 0.941015109 |
| chr12:6873219-6873993 | -0.06495436 | 0.843759769 | 0.941015109 |
| chr10:32445723-32446638 | -0.065607344 | 0.843776371 | 0.941017862 |
| chr11:44096223-44096580 | 0.092397628 | 0.843799775 | 0.941017862 |
| chr17:50783138-50783476 | 0.070994355 | 0.843801789 | 0.941017862 |
| chr6:53956064-53956669 | -0.067371288 | 0.843824874 | 0.941017862 |
| chrX:151562139-151562717 | 0.078721415 | 0.843845048 | 0.941017862 |
| chr21:28781647-28781853 | 0.091887032 | 0.8438487 | 0.941017862 |
| chr7:31134469-31134788 | 0.080346391 | 0.843852642 | 0.941017862 |
| chr10:71889490-71889764 | 0.085506512 | 0.843860488 | 0.941017862 |
| chr12:60210060-60210262 | 0.097847814 | 0.84387297 | 0.941018086 |
| chr15:76329947-76330147 | -0.097868317 | 0.843938974 | 0.941075874 |
| chr3:108626894-108627094 | -0.109755932 | 0.843958941 | 0.941075874 |
| chr1:182720561-182720777 | 0.091835086 | 0.843961639 | 0.941075874 |
| chr14:52370382-52371071 | -0.068760343 | 0.844006623 | 0.941112339 |
| chr1:223142728-223143483 | -0.064527448 | 0.844033892 | 0.94111826 |
| chr20:23049260-23050008 | 0.067055641 | 0.844036499 | 0.94111826 |
| chr20:53898006-53898828 | 0.061989221 | 0.844069663 | 0.941141543 |
| chr1:92858896-92859667 | 0.073170507 | 0.844087203 | 0.941147405 |
| chr11:73193843-73194404 | -0.065544495 | 0.844155051 | 0.941202214 |
| chr15:65323159-65323434 | 0.082802569 | 0.84416694 | 0.941202214 |
| chr1:110254652-110254909 | 0.083626761 | 0.84418316 | 0.941202214 |
| chr1:21251716-21252497 | -0.066649416 | 0.844199228 | 0.941202214 |
| chrX:107542619-107542819 | -0.102218355 | 0.844204155 | 0.941202214 |
| chr2:130156045-130156439 | 0.074550826 | 0.844229249 | 0.941202214 |
| chr18:34222404-34222770 | 0.082577613 | 0.844229478 | 0.941202214 |
| chr2:77522193-77522476 | 0.093177138 | 0.844234629 | 0.941202214 |
| chr10:101783302-101783604 | -0.074231162 | 0.844295228 | 0.941255668 |
| chr16:89507745-89509214 | 0.062816338 | 0.844307145 | 0.941255668 |
| chr8:115485120-115485478 | 0.090941008 | 0.844361042 | 0.941256125 |
| chr17:57849572-57850770 | -0.061920594 | 0.844361556 | 0.941256125 |
| chr14:75277037-75277507 | -0.067529672 | 0.844365978 | 0.941256125 |
| chr12:74111685-74112255 | -0.071242517 | 0.844375499 | 0.941256125 |
| chr11:118415684-118416060 | 0.07731717 | 0.844379388 | 0.941256125 |
| chr20:56550646-56551085 | 0.067601205 | 0.844381261 | 0.941256125 |
| chr15:94844562-94845486 | -0.066945064 | 0.844435976 | 0.941301754 |
| chr2:218852937-218853181 | -0.081153597 | 0.844446764 | 0.941301754 |
| chr20:54155357-54155583 | -0.068262135 | 0.844476065 | 0.941311794 |
| chr2:96927855-96928269 | 0.071456281 | 0.844515333 | 0.941311794 |
| chr5:177455207-177455899 | -0.064127869 | 0.84452768 | 0.941311794 |
| chr3:142184768-142185023 | 0.110902705 | 0.844530541 | 0.941311794 |
| chr8:132774902-132776024 | 0.062776649 | 0.844532244 | 0.941311794 |
| chr12:103677462-103677662 | 0.110572857 | 0.8445325 | 0.941311794 |
| chr9:34675038-34675391 | 0.076301098 | 0.844553601 | 0.941311794 |
| chr10:101569843-101570547 | -0.067783671 | 0.844559859 | 0.941311794 |
| chr11:40329189-40329531 | 0.079169955 | 0.844573365 | 0.941311794 |
| chr5:168697195-168697628 | -0.071205358 | 0.844594595 | 0.941311794 |

| | | | |
|---|---|---|---|
| chr1:114345919-114347103 | -0.06131621 | 0.844596614 | 0.941311794 |
| chr4:68905799-68906760 | -0.066376831 | 0.844605723 | 0.941311794 |
| chr8:130302978-130303217 | -0.090063633 | 0.844615478 | 0.941311794 |
| chr8:140834896-140835145 | -0.084351026 | 0.844645697 | 0.941331781 |
| chr2:73908044-73908267 | 0.091377745 | 0.844706292 | 0.941379542 |
| chr17:38585092-38585524 | 0.071083303 | 0.844724081 | 0.941379542 |
| chr11:72223043-72224177 | -0.063073903 | 0.84472541 | 0.941379542 |
| chr15:41332050-41333214 | -0.062581754 | 0.844759308 | 0.941394485 |
| chr16:20675764-20676314 | 0.072432146 | 0.844770585 | 0.941394485 |
| chr3:47163956-47164539 | -0.069844481 | 0.844775678 | 0.941394485 |
| chr5:13715929-13716129 | -0.095734612 | 0.84481431 | 0.941394998 |
| chr17:48184774-48184974 | -0.096232686 | 0.844819446 | 0.941394998 |
| chr1:188071602-188072201 | -0.077951099 | 0.844821731 | 0.941394998 |
| chr19:3970707-3971694 | 0.064195002 | 0.844839755 | 0.941394998 |
| chr8:101041735-101041985 | 0.085610516 | 0.844843789 | 0.941394998 |
| chr17:50950299-50951028 | 0.06431487 | 0.844860874 | 0.941394998 |
| chr6:5214788-5215004 | 0.095353668 | 0.844862142 | 0.941394998 |
| chr1:28247130-28247513 | -0.071173259 | 0.84490983 | 0.941434444 |
| chr16:84819618-84820212 | -0.064651383 | 0.844923269 | 0.941435728 |
| chr10:102347316-102347605 | -0.097420055 | 0.844942052 | 0.941442966 |
| chr2:201440209-201440477 | 0.086528963 | 0.845018615 | 0.941496056 |
| chr3:23112557-23112866 | 0.084185741 | 0.845032457 | 0.941496056 |
| chr1:169849769-169850124 | -0.080042161 | 0.845034018 | 0.941496056 |
| chr5:148558930-148559246 | 0.081198493 | 0.84503885 | 0.941496056 |
| chr16:89501511-89501711 | 0.102537565 | 0.84507711 | 0.941524993 |
| chr10:15419405-15419632 | 0.09114815 | 0.845117935 | 0.94154804 |
| chr8:116447958-116448356 | -0.078220826 | 0.845122373 | 0.94154804 |
| chr1:244652551-244653495 | 0.061561354 | 0.845171021 | 0.941564811 |
| chr5:19885348-19885744 | -0.08221012 | 0.845172766 | 0.941564811 |
| chr3:20108675-20109488 | -0.06715672 | 0.845174292 | 0.941564811 |
| chr2:85814226-85814531 | 0.075288849 | 0.845214498 | 0.941595912 |
| chr2:29295901-29296101 | -0.095104679 | 0.845229809 | 0.941599279 |
| chr14:88554332-88555246 | 0.066794411 | 0.845284093 | 0.941632716 |
| chr8:108266126-108266752 | 0.06949512 | 0.845284403 | 0.941632716 |
| chr1:67451374-67451708 | -0.083994113 | 0.84530106 | 0.941637582 |
| chr8:37849517-37850151 | 0.067215292 | 0.845341238 | 0.941668649 |
| chr12:62192377-62193168 | -0.067579063 | 0.845360479 | 0.941668785 |
| chr3:195441904-195442185 | 0.07686364 | 0.84536594 | 0.941668785 |
| chr10:93763042-93763435 | -0.073400193 | 0.845409409 | 0.941703515 |
| chr8:140464022-140465279 | 0.061032575 | 0.845423584 | 0.941705615 |
| chr12:113196951-113197250 | 0.076739587 | 0.84548306 | 0.941758174 |
| chr19:39402463-39403076 | 0.067266887 | 0.845512279 | 0.941762703 |
| chr8:108442881-108444213 | -0.06092248 | 0.845545285 | 0.941762703 |
| chr13:37099099-37099364 | -0.086635655 | 0.845546206 | 0.941762703 |
| chr3:142608696-142608928 | -0.096990087 | 0.845554327 | 0.941762703 |
| chr10:92839215-92839625 | 0.081659865 | 0.845567317 | 0.941762703 |
| chr20:57089646-57090192 | 0.063885163 | 0.845576318 | 0.941762703 |
| chr17:49260454-49261264 | 0.066247508 | 0.84559957 | 0.941762703 |
| chr1:64245533-64245761 | 0.093395374 | 0.84561426 | 0.941762703 |
| chr20:31723223-31724122 | 0.062485249 | 0.845614394 | 0.941762703 |

| | | | |
|---|---|---|---|
| chr7:74206513-74207083 | -0.070643752 | 0.845637754 | 0.941762703 |
| chr19:8343189-8343508 | 0.078116371 | 0.845647031 | 0.941762703 |
| chr19:55587215-55587685 | -0.068539642 | 0.845648838 | 0.941762703 |
| chr5:10267380-10267878 | -0.071122859 | 0.84565067 | 0.941762703 |
| chr11:66022410-66022663 | -0.082512987 | 0.8456592 | 0.941762703 |
| chr12:118016400-118017025 | -0.0646748 | 0.845677018 | 0.941768858 |
| chr5:892107-892359 | 0.081118072 | 0.845700212 | 0.941781 |
| chr19:46788215-46788897 | -0.063718584 | 0.845749871 | 0.941822612 |
| chr14:49505371-49505829 | 0.067241962 | 0.845765101 | 0.941825884 |
| chr5:34188297-34188621 | 0.074285426 | 0.845799582 | 0.94184993 |
| chr3:100697320-100697520 | 0.099630754 | 0.845811279 | 0.94184993 |
| chr20:39081211-39081457 | 0.087229773 | 0.84584002 | 0.941865344 |
| chr1:32703174-32703387 | 0.08926235 | 0.845849706 | 0.941865344 |
| chr3:159766645-159766871 | 0.08180181 | 0.845896669 | 0.94187343 |
| chr17:21042249-21042592 | 0.069004267 | 0.845899122 | 0.94187343 |
| chr14:74988186-74988451 | 0.094632473 | 0.845908525 | 0.94187343 |
| chr8:26291187-26292143 | 0.065237008 | 0.845913142 | 0.94187343 |
| chr19:2613910-2614110 | 0.09612651 | 0.845918431 | 0.94187343 |
| chr1:42340160-42340486 | -0.076019812 | 0.846023563 | 0.941954821 |
| chrX:68462326-68462567 | 0.095957264 | 0.846029028 | 0.941954821 |
| chr9:71768350-71769777 | -0.061588547 | 0.8460376 | 0.941954821 |
| chr4:132871455-132871663 | 0.091859764 | 0.846052624 | 0.941954821 |
| chr5:53777985-53778202 | 0.091783336 | 0.846061445 | 0.941954821 |
| chr5:175288009-175288209 | -0.098851841 | 0.84606529 | 0.941954821 |
| chr11:94116477-94116738 | -0.093122405 | 0.846088108 | 0.941956805 |
| chr7:73647138-73647430 | -0.094392848 | 0.846100829 | 0.941956805 |
| chr1:30302844-30303282 | -0.080926923 | 0.846103953 | 0.941956805 |
| chr2:46542279-46542796 | -0.06649352 | 0.846131096 | 0.941973336 |
| chr1:51228511-51228879 | 0.077004582 | 0.846152517 | 0.941983497 |
| chr12:78369893-78370093 | 0.107531773 | 0.846207111 | 0.942028224 |
| chr5:55711434-55712784 | -0.062678374 | 0.846217283 | 0.942028224 |
| chr5:177599635-177600582 | 0.064628595 | 0.846235601 | 0.94203493 |
| chr20:50151054-50151302 | 0.072397075 | 0.846288702 | 0.942080355 |
| chrX:105503356-105503643 | 0.086569639 | 0.846362097 | 0.942135261 |
| chr1:36449953-36450952 | -0.064104089 | 0.846362617 | 0.942135261 |
| chr15:90046812-90047202 | 0.068446966 | 0.846401557 | 0.942152716 |
| chr8:96068523-96068771 | 0.097531479 | 0.84640289 | 0.942152716 |
| chr10:86521257-86522297 | 0.062194224 | 0.84643689 | 0.942171719 |
| chr13:39047608-39047925 | 0.074642051 | 0.846455048 | 0.942171719 |
| chr2:64049704-64050114 | -0.080282941 | 0.84645685 | 0.942171719 |
| chr16:29678799-29679675 | -0.066358043 | 0.846490417 | 0.942187221 |
| chr21:8219409-8220407 | -0.060685577 | 0.846495371 | 0.942187221 |
| chr20:35995007-35995397 | -0.076647035 | 0.846508968 | 0.942188669 |
| chr6:136288831-136290558 | -0.060634007 | 0.846573334 | 0.942246623 |
| chr8:128197232-128198006 | -0.063218408 | 0.846590593 | 0.942252145 |
| chr6:3068194-3069194 | 0.06280388 | 0.84662987 | 0.942282173 |
| chr3:151818464-151818752 | 0.079391022 | 0.846647046 | 0.942287602 |
| chr1:52169097-52169369 | -0.111342456 | 0.846708135 | 0.942336957 |
| chr1:46333574-46333891 | 0.078553227 | 0.846738984 | 0.942336957 |
| chr8:78515957-78516743 | -0.065376674 | 0.846739127 | 0.942336957 |

| | | | |
|---|---|---|---|
| chr15:78640534-78640834 | 0.089672099 | 0.846740586 | 0.942336957 |
| chr8:141059255-141059462 | -0.081522286 | 0.84677088 | 0.94235153 |
| chr16:78297056-78297376 | -0.08149768 | 0.846778278 | 0.94235153 |
| chr17:28234601-28234807 | -0.088278768 | 0.846829658 | 0.942395022 |
| chr15:58866150-58866630 | -0.06625446 | 0.846864042 | 0.942415642 |
| chr6:130995501-130996272 | 0.069508164 | 0.846891394 | 0.942415642 |
| chr15:93900559-93900888 | -0.083567698 | 0.846907423 | 0.942415642 |
| chr3:48439807-48440730 | 0.065056234 | 0.84690892 | 0.942415642 |
| chr7:148865337-148865852 | -0.07210741 | 0.846909685 | 0.942415642 |
| chr1:84224176-84224376 | 0.10853995 | 0.846941414 | 0.942437263 |
| chr20:62925801-62926771 | 0.062974819 | 0.846967599 | 0.942446675 |
| chr1:92960890-92961731 | 0.067294366 | 0.846983686 | 0.942446675 |
| chr20:15818674-15819015 | -0.090841324 | 0.846988882 | 0.942446675 |
| chr1:164440224-164440571 | -0.076738931 | 0.846999073 | 0.942446675 |
| chr17:12020586-12021649 | -0.062577824 | 0.847072208 | 0.942514365 |
| chr9:130693485-130694169 | -0.064212288 | 0.847095764 | 0.942526888 |
| chr9:99293152-99293352 | 0.11121412 | 0.847134939 | 0.942554097 |
| chr11:76810253-76810502 | -0.080958162 | 0.847161405 | 0.942554097 |
| chr19:35291477-35291709 | -0.093847048 | 0.84717809 | 0.942554097 |
| chr1:53715542-53715742 | 0.098803621 | 0.847210253 | 0.942554097 |
| chr9:38023062-38023350 | -0.078572982 | 0.847218172 | 0.942554097 |
| chr20:16294639-16294839 | 0.109904735 | 0.847277834 | 0.942554097 |
| chr5:35444050-35444426 | -0.081445521 | 0.847279133 | 0.942554097 |
| chr17:16427862-16428445 | 0.069196793 | 0.847282054 | 0.942554097 |
| chr15:96330178-96330835 | 0.062891584 | 0.847289739 | 0.942554097 |
| chr17:17729729-17729982 | -0.08064545 | 0.847308617 | 0.942554097 |
| chr5:177425846-177427149 | -0.060504096 | 0.84731524 | 0.942554097 |
| chr7:150286206-150286412 | -0.092379231 | 0.847318632 | 0.942554097 |
| chr5:177454398-177454859 | -0.063638668 | 0.847333775 | 0.942554097 |
| chr19:56091365-56091565 | 0.093484622 | 0.847338808 | 0.942554097 |
| chr10:116797030-116797251 | 0.087669576 | 0.84735463 | 0.942554097 |
| chr17:17146815-17147031 | -0.086472545 | 0.847360561 | 0.942554097 |
| chr1:182786766-182787003 | -0.081574969 | 0.847360609 | 0.942554097 |
| chr9:19379997-19380729 | -0.062617908 | 0.847371667 | 0.942554097 |
| chr20:3768315-3768994 | 0.063679962 | 0.847375133 | 0.942554097 |
| chr4:55635789-55636512 | 0.069251303 | 0.847381341 | 0.942554097 |
| chr18:12046975-12047251 | -0.082631891 | 0.847385038 | 0.942554097 |
| chr21:39852374-39852693 | 0.082927435 | 0.847396447 | 0.942554097 |
| chr17:81821759-81822248 | 0.069033567 | 0.847403161 | 0.942554097 |
| chr7:55079158-55079385 | -0.099216423 | 0.847415451 | 0.942554097 |
| chr11:73498466-73498765 | -0.086814507 | 0.84746081 | 0.942590866 |
| chr8:88072916-88073260 | 0.080746265 | 0.847540563 | 0.942648224 |
| chr4:150976746-150977099 | -0.076702295 | 0.84756422 | 0.942648224 |
| chr10:70283810-70284183 | -0.081257468 | 0.847566855 | 0.942648224 |
| chr2:27071269-27072247 | 0.062706448 | 0.847571097 | 0.942648224 |
| chr16:68328133-68328425 | 0.079539999 | 0.847573892 | 0.942648224 |
| chr4:77075288-77075791 | 0.065746408 | 0.847602738 | 0.942666623 |
| chr1:203995986-203996400 | -0.07038108 | 0.847663015 | 0.942715178 |
| chr2:208407748-208408009 | 0.08617975 | 0.847677359 | 0.942715178 |
| chr12:109864925-109865248 | -0.076000814 | 0.847683307 | 0.942715178 |

| | | | |
|---|---|---|---|
| chr7:7639990-7641484 | 0.059834506 | 0.847711529 | 0.942722572 |
| chr3:105868364-105869704 | -0.060297209 | 0.847724082 | 0.942722572 |
| chr1:85830306-85830613 | -0.08456176 | 0.847751382 | 0.942722572 |
| chr3:159051917-159052117 | 0.113729014 | 0.847751473 | 0.942722572 |
| chr5:30495598-30495798 | 0.113729014 | 0.847751473 | 0.942722572 |
| chr2:74002523-74002834 | -0.070952671 | 0.847798929 | 0.942755348 |
| chr3:145940052-145940252 | -0.104303998 | 0.847805555 | 0.942755348 |
| chr4:118591396-118592142 | 0.063045005 | 0.847847208 | 0.942787984 |
| chr17:65231746-65232065 | 0.068407617 | 0.847866107 | 0.942789626 |
| chr6:170480375-170480575 | 0.0959387 | 0.847884162 | 0.942789626 |
| chr5:112630249-112630649 | -0.070926293 | 0.847885599 | 0.942789626 |
| chr1:245434400-245434664 | 0.085902701 | 0.84791668 | 0.942804692 |
| chr3:49021299-49022210 | 0.062191482 | 0.847923757 | 0.942804692 |
| chr2:7111705-7111976 | 0.08336721 | 0.847979623 | 0.942836827 |
| chr12:82473834-82474034 | -0.10253652 | 0.848000317 | 0.942836827 |
| chr1:179016929-179017421 | 0.067775887 | 0.848025416 | 0.942836827 |
| chr10:86711014-86711871 | 0.061554489 | 0.848034885 | 0.942836827 |
| chr8:94990833-94991502 | 0.067700756 | 0.848035807 | 0.942836827 |
| chr9:129401397-129401597 | -0.0964153 | 0.84803737 | 0.942836827 |
| chr19:12400869-12401507 | 0.06805202 | 0.848069064 | 0.942836827 |
| chr5:9498580-9499244 | -0.068476705 | 0.848069819 | 0.942836827 |
| chr11:2137217-2137489 | 0.087188689 | 0.848073687 | 0.942836827 |
| chr13:96994627-96994939 | -0.094416554 | 0.848075709 | 0.942836827 |
| chr10:100146873-100147144 | 0.091738219 | 0.848134469 | 0.942887096 |
| chr6:96897314-96898296 | -0.06121019 | 0.848237823 | 0.942887096 |
| chr22:10708191-10708591 | -0.092112029 | 0.848242568 | 0.942887096 |
| chr1:205791623-205791907 | 0.069917259 | 0.848247879 | 0.942887096 |
| chr4:119659624-119659903 | -0.090183757 | 0.848267122 | 0.942887096 |
| chr13:40504474-40504726 | -0.089715133 | 0.848267494 | 0.942887096 |
| chr3:154430051-154430323 | 0.07890541 | 0.848277831 | 0.942887096 |
| chr17:81917854-81918380 | -0.064962158 | 0.848282733 | 0.942887096 |
| chr17:79892460-79893075 | 0.066683191 | 0.848292683 | 0.942887096 |
| chr3:76986994-76987450 | 0.07587817 | 0.84829879 | 0.942887096 |
| chr2:36355111-36356007 | -0.062452039 | 0.848309437 | 0.942887096 |
| chr6:38671959-38672159 | 0.112875798 | 0.848311354 | 0.942887096 |
| chr6:41427898-41428108 | -0.094171042 | 0.848316725 | 0.942887096 |
| chr5:1344390-1344619 | -0.081511574 | 0.848319531 | 0.942887096 |
| chr12:77667518-77667718 | 0.09566276 | 0.848322539 | 0.942887096 |
| chr3:37998323-37998523 | 0.082949094 | 0.848327215 | 0.942887096 |
| chr8:140637433-140637716 | -0.068461769 | 0.848330123 | 0.942887096 |
| chr17:24421345-24421611 | 0.086405882 | 0.848353659 | 0.942899579 |
| chr17:28040993-28042062 | -0.060872312 | 0.848395863 | 0.942932809 |
| chr12:120646964-120647356 | 0.071607919 | 0.848442051 | 0.942968024 |
| chr2:134585010-134585290 | 0.106060783 | 0.848453051 | 0.942968024 |
| chr16:4371675-4371924 | 0.074224271 | 0.848464468 | 0.942968024 |
| chr22:39598208-39598490 | -0.080213365 | 0.848479579 | 0.942971141 |
| chr12:57456183-57456410 | 0.07591653 | 0.848497531 | 0.942975241 |
| chr11:126113569-126113945 | -0.070882487 | 0.848519993 | 0.942975241 |
| chr5:95861496-95861826 | 0.076665966 | 0.84852019 | 0.942975241 |
| chr4:86934546-86936279 | -0.060350685 | 0.848540777 | 0.942977053 |

| | | | |
|---|---|---|---|
| chr16:81599109-81599653 | 0.067443825 | 0.848546433 | 0.942977053 |
| chr14:48580798-48581026 | 0.094876812 | 0.848567663 | 0.942986265 |
| chr5:6360153-6360540 | -0.082736322 | 0.848579337 | 0.942986265 |
| chr17:66462942-66463245 | -0.073485808 | 0.848611198 | 0.942988708 |
| chr8:130749645-130750019 | 0.069828194 | 0.84863214 | 0.942988708 |
| chr6:40268203-40268403 | -0.102186997 | 0.848638852 | 0.942988708 |
| chr1:68525858-68526229 | -0.071591404 | 0.84864521 | 0.942988708 |
| chr19:36919621-36919821 | 0.104068955 | 0.848648827 | 0.942988708 |
| chr1:207422406-207422628 | -0.084857117 | 0.848724201 | 0.942988708 |
| chr8:58513277-58513788 | 0.069133507 | 0.848727015 | 0.942988708 |
| chr1:157963695-157963965 | 0.079913529 | 0.848728461 | 0.942988708 |
| chr9:111661229-111662360 | -0.063397765 | 0.848736244 | 0.942988708 |
| chr14:90674192-90674392 | -0.098471047 | 0.848758073 | 0.942988708 |
| chr12:132494287-132494531 | -0.072753137 | 0.848771002 | 0.942988708 |
| chr17:39461220-39461834 | -0.067518626 | 0.848786718 | 0.942988708 |
| chr18:4017556-4017820 | -0.08277879 | 0.848792339 | 0.942988708 |
| chr1:169081595-169081926 | 0.080133263 | 0.848794928 | 0.942988708 |
| chr20:36064227-36065134 | -0.064356444 | 0.848801464 | 0.942988708 |
| chr17:67244654-67246449 | 0.060147965 | 0.84880323 | 0.942988708 |
| chr20:10341817-10342050 | -0.101927319 | 0.848807167 | 0.942988708 |
| chr9:19221317-19221729 | -0.082753631 | 0.848815683 | 0.942988708 |
| chr1:173527199-173527444 | 0.081274401 | 0.848832458 | 0.942988708 |
| chr2:88307904-88308209 | -0.08044709 | 0.84884121 | 0.942988708 |
| chr5:11797194-11797598 | -0.07521156 | 0.848844465 | 0.942988708 |
| chr1:178361867-178362067 | -0.105419607 | 0.84885229 | 0.942988708 |
| chr16:74666375-74667199 | -0.061133677 | 0.848884453 | 0.94300487 |
| chr8:102238482-102239858 | 0.059270468 | 0.848894684 | 0.94300487 |
| chr3:37223129-37223439 | 0.095779277 | 0.848927639 | 0.94300487 |
| chr3:108412415-108412682 | -0.078655623 | 0.848927749 | 0.94300487 |
| chr11:66035731-66035962 | 0.088575569 | 0.848936256 | 0.94300487 |
| chrX:49106985-49107278 | 0.088291304 | 0.848940682 | 0.94300487 |
| chr7:72494133-72494342 | 0.093123836 | 0.849005012 | 0.943056178 |
| chr1:175178134-175178406 | 0.079117976 | 0.849014022 | 0.943056178 |
| chr1:114764753-114765078 | 0.066030692 | 0.849023796 | 0.943056178 |
| chr13:112874596-112874862 | -0.085168353 | 0.849099553 | 0.943070709 |
| chr10:77907885-77908556 | -0.066928211 | 0.849122828 | 0.943070709 |
| chr9:68704655-68705598 | 0.065080153 | 0.849129085 | 0.943070709 |
| chr7:15168818-15169018 | 0.096825713 | 0.849137972 | 0.943070709 |
| chr19:1863622-1864582 | 0.062134 | 0.84915256 | 0.943070709 |
| chr9:109355287-109355487 | 0.104268098 | 0.84916238 | 0.943070709 |
| chr11:74050897-74051113 | -0.095412645 | 0.849181604 | 0.943070709 |
| chr10:129031630-129031841 | 0.091176945 | 0.849192982 | 0.943070709 |
| chr20:32731907-32732137 | 0.09106303 | 0.849196219 | 0.943070709 |
| chr3:187139099-187140029 | -0.062514428 | 0.849219114 | 0.943070709 |
| chr6:109527057-109527536 | -0.07674395 | 0.849233046 | 0.943070709 |
| chr20:56827153-56827376 | -0.078954792 | 0.849248932 | 0.943070709 |
| chr8:33472575-33473535 | -0.062233114 | 0.849248995 | 0.943070709 |
| chr11:65204785-65204985 | -0.092012815 | 0.849252595 | 0.943070709 |
| chr13:42187640-42187908 | 0.096882791 | 0.849259541 | 0.943070709 |
| chr12:58443676-58443891 | -0.10098572 | 0.849263318 | 0.943070709 |

| | | | |
|---|---|---|---|
| chr21:44479713-44479952 | 0.088310114 | 0.849269782 | 0.943070709 |
| chr15:90062825-90063025 | 0.085707188 | 0.849281873 | 0.943070709 |
| chr9:113112767-113113044 | -0.087267717 | 0.849292012 | 0.943070709 |
| chr8:28657289-28657503 | -0.097638641 | 0.849296916 | 0.943070709 |
| chr17:63834898-63835190 | 0.066078978 | 0.849322752 | 0.943070709 |
| chr1:157163362-157164011 | -0.066139798 | 0.849330137 | 0.943070709 |
| chr4:129498313-129498513 | 0.110063061 | 0.849345229 | 0.943070709 |
| chr12:120580994-120581856 | 0.062483774 | 0.849348391 | 0.943070709 |
| chr5:32312187-32313769 | -0.059715445 | 0.84935211 | 0.943070709 |
| chr1:3353120-3353320 | -0.105027801 | 0.849365297 | 0.943070709 |
| chr3:129316502-129316901 | -0.067100731 | 0.849369197 | 0.943070709 |
| chr3:152105269-152105617 | 0.087858172 | 0.849406689 | 0.943085402 |
| chr6:116532105-116532438 | 0.081633681 | 0.849418634 | 0.943085402 |
| chr5:2634540-2634800 | 0.093425027 | 0.849419354 | 0.943085402 |
| chr8:106326501-106326857 | -0.075430835 | 0.84943514 | 0.943089263 |
| chr20:25076648-25077257 | -0.067351532 | 0.84945894 | 0.94309793 |
| chr17:73278573-73279301 | 0.064750674 | 0.849502609 | 0.94309793 |
| chr2:54107013-54107226 | -0.08385421 | 0.849509346 | 0.94309793 |
| chr18:3246065-3246355 | -0.065782341 | 0.849510967 | 0.94309793 |
| chr8:115076722-115077041 | -0.069372157 | 0.84951245 | 0.94309793 |
| chr2:119352348-119352587 | 0.089509389 | 0.849519936 | 0.94309793 |
| chr11:95138515-95138772 | -0.099300133 | 0.849535777 | 0.94309793 |
| chr5:177554447-177554871 | -0.066098086 | 0.849552101 | 0.94309793 |
| chr7:151310636-151310857 | -0.091371964 | 0.849554678 | 0.94309793 |
| chr15:71730631-71730849 | -0.088079715 | 0.849580832 | 0.94309793 |
| chr13:90427666-90427894 | -0.100422297 | 0.849584022 | 0.94309793 |
| chr4:151489379-151490015 | -0.06689695 | 0.849590648 | 0.94309793 |
| chr11:65711742-65712825 | 0.059849331 | 0.849633683 | 0.943132038 |
| chr18:51051183-51051419 | 0.083977278 | 0.849654388 | 0.943136253 |
| chr1:113178598-113178825 | -0.090484228 | 0.849662098 | 0.943136253 |
| chr1:165924685-165925063 | 0.070686532 | 0.849674501 | 0.943136358 |
| chr1:246968298-246968554 | -0.079381183 | 0.849709158 | 0.943161163 |
| chr2:38536057-38536426 | -0.069782693 | 0.849801468 | 0.943237362 |
| chr15:78660516-78660716 | 0.093829574 | 0.849802427 | 0.943237362 |
| chr7:2249474-2249728 | -0.085651173 | 0.849842516 | 0.943268194 |
| chr1:203411238-203411476 | -0.087864602 | 0.849884505 | 0.943301135 |
| chr20:36236333-36236534 | 0.095927477 | 0.849910503 | 0.943316326 |
| chr12:95449715-95450034 | 0.079725422 | 0.849925633 | 0.943319454 |
| chr1:43073722-43073925 | 0.089770078 | 0.849996457 | 0.943379248 |
| chr3:51973936-51974233 | 0.072861472 | 0.850004131 | 0.943379248 |
| chr1:171715870-171716277 | -0.073511522 | 0.850043818 | 0.943401386 |
| chr2:7574996-7575196 | 0.104345858 | 0.850081944 | 0.943401386 |
| chr4:39469404-39469759 | 0.074963514 | 0.850085678 | 0.943401386 |
| chr19:51315346-51315546 | -0.098885392 | 0.850090852 | 0.943401386 |
| chr3:27712077-27712756 | -0.06515327 | 0.850096605 | 0.943401386 |
| chr20:63642576-63643002 | 0.069938949 | 0.850097953 | 0.943401386 |
| chr14:87014881-87015286 | -0.076261411 | 0.850125012 | 0.943417752 |
| chr15:32670284-32670528 | -0.092458167 | 0.850159194 | 0.94344202 |
| chr2:75789017-75789235 | -0.096932063 | 0.850186472 | 0.943446921 |
| chr2:159797851-159798347 | -0.072822355 | 0.850188236 | 0.943446921 |

| | | | |
|---|---|---|---|
| chr9:33473591-33474251 | -0.062408262 | 0.850289525 | 0.943541702 |
| chr15:100663127-100663427 | -0.080601593 | 0.850306107 | 0.943541702 |
| chr5:42382678-42382939 | 0.091837198 | 0.850335901 | 0.943541702 |
| chr3:87359788-87360181 | 0.067896727 | 0.850342613 | 0.943541702 |
| chr17:58969109-58970006 | -0.060369736 | 0.850344415 | 0.943541702 |
| chr7:104871080-104871301 | 0.101609412 | 0.850376518 | 0.943541702 |
| chr7:144437653-144437918 | 0.081255015 | 0.850380328 | 0.943541702 |
| chr8:123072302-123073524 | 0.059474083 | 0.850383217 | 0.943541702 |
| chrX:18652657-18652898 | 0.092181381 | 0.850384476 | 0.943541702 |
| chr20:63626741-63626941 | 0.080324417 | 0.850416232 | 0.943555142 |
| chr16:20049880-20050239 | -0.075231192 | 0.850446884 | 0.943555142 |
| chr11:62112473-62112767 | 0.075004236 | 0.85045017 | 0.943555142 |
| chr1:2139749-2139949 | -0.105171067 | 0.850458161 | 0.943555142 |
| chr11:1685535-1685758 | -0.105171067 | 0.850458161 | 0.943555142 |
| chr3:42544875-42545147 | 0.090370607 | 0.850496972 | 0.943584539 |
| chr4:104966681-104966940 | 0.073643877 | 0.850530601 | 0.943608185 |
| chr19:36528011-36529137 | 0.061537601 | 0.850553241 | 0.943616186 |
| chr3:3962941-3963225 | 0.08089209 | 0.850567976 | 0.943616186 |
| chr11:5600348-5600798 | -0.075135946 | 0.850574758 | 0.943616186 |
| chr12:69585219-69586056 | 0.060054185 | 0.850608623 | 0.943640093 |
| chr12:74365425-74365907 | 0.077416878 | 0.850625985 | 0.943645691 |
| chr14:90676454-90676682 | 0.092111492 | 0.850640243 | 0.943647846 |
| chr8:85540240-85540449 | 0.086358624 | 0.85067737 | 0.943675369 |
| chr16:56702046-56702395 | -0.075196581 | 0.850712423 | 0.943700592 |
| chr17:58914049-58914675 | 0.062003006 | 0.85076458 | 0.943707079 |
| chr3:165086820-165087089 | 0.096460259 | 0.85077722 | 0.943707079 |
| chr6:116042097-116042343 | 0.096460259 | 0.85077722 | 0.943707079 |
| chr1:190119735-190119987 | -0.088565906 | 0.850779429 | 0.943707079 |
| chr19:5086681-5087084 | 0.074766594 | 0.850779853 | 0.943707079 |
| chr14:39638723-39639750 | -0.062770216 | 0.850814078 | 0.943714554 |
| chr1:11805362-11806443 | 0.060759466 | 0.850817848 | 0.943714554 |
| chr1:200890894-200891790 | -0.060874978 | 0.850840744 | 0.943714554 |
| chr3:160514931-160515138 | -0.097949814 | 0.850849656 | 0.943714554 |
| chr8:133481217-133481511 | 0.075552914 | 0.850849795 | 0.943714554 |
| chr3:111012693-111012994 | -0.074022598 | 0.850868458 | 0.943714554 |
| chr1:156920211-156920756 | -0.067804521 | 0.850872807 | 0.943714554 |
| chr7:76548921-76549910 | -0.061772169 | 0.850924682 | 0.943751508 |
| chr20:45469371-45469961 | 0.068703993 | 0.850949035 | 0.943751508 |
| chr11:59823063-59823281 | 0.099790874 | 0.850952556 | 0.943751508 |
| chr6:82783899-82784192 | -0.076224604 | 0.850978971 | 0.943751508 |
| chr17:53648058-53648258 | -0.102757378 | 0.850985296 | 0.943751508 |
| chr5:177109913-177110162 | -0.075050149 | 0.851000294 | 0.943751508 |
| chr22:29072596-29073135 | -0.063488212 | 0.851000304 | 0.943751508 |
| chr19:44093991-44094709 | 0.066116262 | 0.851004662 | 0.943751508 |
| chr4:25988210-25988470 | 0.082956981 | 0.851018511 | 0.943753207 |
| chr5:143086910-143087590 | -0.066110253 | 0.851031576 | 0.943754037 |
| chr4:113645838-113646203 | -0.070845245 | 0.85109097 | 0.943806242 |
| chr14:89177465-89178001 | -0.067320327 | 0.851131567 | 0.943810785 |
| chr5:173010582-173011011 | 0.072542126 | 0.85115057 | 0.943810785 |
| chr12:33439397-33440099 | -0.06791466 | 0.851160743 | 0.943810785 |

| | | | |
|---|---|---|---|
| chr12:3214516-3214787 | 0.07851111 | 0.851165327 | 0.943810785 |
| chr13:24127041-24127625 | 0.062151645 | 0.851172547 | 0.943810785 |
| chrX:107764466-107764666 | 0.088777561 | 0.851180183 | 0.943810785 |
| chr15:76150996-76151269 | 0.075559478 | 0.851190003 | 0.943810785 |
| chr5:72940832-72941032 | 0.090178833 | 0.851193608 | 0.943810785 |
| chr6:41022396-41022762 | 0.076252519 | 0.851223057 | 0.94382978 |
| chr11:119126798-119127245 | -0.072746199 | 0.851242556 | 0.943831872 |
| chr12:100168718-100169040 | -0.076783217 | 0.851249581 | 0.943831872 |
| chr15:78733988-78734464 | 0.077174014 | 0.85127567 | 0.943844583 |
| chr1:211331313-211331702 | 0.067765323 | 0.851287969 | 0.943844583 |
| chr2:94586105-94586393 | -0.076906679 | 0.851297999 | 0.943844583 |
| chr2:162153652-162153962 | 0.07722922 | 0.851314975 | 0.943849747 |
| chr12:118035296-118035496 | -0.09162704 | 0.851338157 | 0.94386078 |
| chr8:73008525-73009319 | -0.06048879 | 0.851356415 | 0.94386078 |
| chr17:69762088-69762323 | -0.087047911 | 0.851363647 | 0.94386078 |
| chr6:83536113-83536514 | 0.075837254 | 0.851378128 | 0.94386078 |
| chr8:70668394-70668703 | 0.067385274 | 0.851386518 | 0.94386078 |
| chr17:40314577-40314926 | -0.070002176 | 0.851416296 | 0.943874431 |
| chr11:102476987-102477187 | -0.098773477 | 0.851423469 | 0.943874431 |
| chr8:25201426-25201626 | -0.103332149 | 0.851452032 | 0.943880361 |
| chr18:4477279-4477845 | 0.073582278 | 0.851473205 | 0.943880361 |
| chr13:19782009-19783227 | -0.060363102 | 0.851481667 | 0.943880361 |
| chr4:152778931-152779145 | -0.074696056 | 0.851487274 | 0.943880361 |
| chr12:117628481-117629245 | -0.066213783 | 0.851490412 | 0.943880361 |
| chr1:155950380-155950580 | 0.089687315 | 0.851548163 | 0.943930723 |
| chr1:27321909-27322595 | 0.063841955 | 0.851570848 | 0.943939408 |
| chr8:127884790-127885046 | 0.077714188 | 0.851580637 | 0.943939408 |
| chr9:128723646-128725146 | -0.059270966 | 0.851617039 | 0.943955383 |
| chr12:68800112-68800619 | -0.067067091 | 0.851619689 | 0.943955383 |
| chr9:126860077-126861377 | 0.063010159 | 0.851650334 | 0.943968289 |
| chrX:49100857-49102013 | -0.06021746 | 0.851677738 | 0.943968289 |
| chr7:99407754-99408194 | 0.06800414 | 0.851691875 | 0.943968289 |
| chr4:121464901-121465240 | -0.078365528 | 0.851692505 | 0.943968289 |
| chr16:5071636-5072326 | 0.063519594 | 0.851705962 | 0.943968289 |
| chr2:130144959-130145284 | -0.081398472 | 0.851716562 | 0.943968289 |
| chr8:91069712-91070841 | 0.058772532 | 0.851717571 | 0.943968289 |
| chr2:106805777-106805977 | 0.103104683 | 0.851803181 | 0.944037787 |
| chr9:122827189-122827428 | 0.078895594 | 0.851805916 | 0.944037787 |
| chr1:22416638-22416863 | 0.08642908 | 0.851836173 | 0.944037787 |
| chr2:195026125-195026441 | -0.086850236 | 0.851841956 | 0.944037787 |
| chr7:152955992-152956334 | 0.084108582 | 0.851851077 | 0.944037787 |
| chrX:93932535-93933071 | 0.071552298 | 0.851854201 | 0.944037787 |
| chr10:60140273-60140515 | -0.084790195 | 0.851877529 | 0.944049985 |
| chr10:43803873-43804124 | 0.097882138 | 0.851915161 | 0.94407545 |
| chr6:34887821-34888313 | -0.063390475 | 0.851939783 | 0.94407545 |
| chr1:174490637-174490965 | 0.083382431 | 0.851953824 | 0.94407545 |
| chr15:74956221-74956530 | 0.071896908 | 0.851965444 | 0.94407545 |
| chr5:43556202-43557696 | 0.058960446 | 0.851966369 | 0.94407545 |
| chr7:101173439-101173795 | 0.068144688 | 0.851974435 | 0.94407545 |
| chr8:106657278-106658617 | -0.058528387 | 0.852007699 | 0.944098657 |

| | | | |
|---|---|---|---|
| chr17:77088512-77089003 | 0.064440601 | 0.85202998 | 0.944100942 |
| chr16:85524543-85524752 | -0.074056522 | 0.852034404 | 0.944100942 |
| chr3:121545114-121546264 | -0.060371109 | 0.852069545 | 0.944126226 |
| chr15:59587065-59587395 | 0.079674416 | 0.852089783 | 0.94413198 |
| chr8:66476609-66476888 | -0.088576641 | 0.852105969 | 0.94413198 |
| chr6:36885591-36886244 | 0.062359368 | 0.852115143 | 0.94413198 |
| chr14:24505485-24505714 | -0.092871737 | 0.852124818 | 0.94413198 |
| chr14:105899294-105899591 | -0.084693564 | 0.852141408 | 0.94413198 |
| chr2:96815825-96817003 | -0.05962831 | 0.852148669 | 0.94413198 |
| chr20:31738720-31739607 | 0.060310914 | 0.852214947 | 0.944179796 |
| chr16:71643860-71644060 | -0.084981695 | 0.852223305 | 0.944179796 |
| chr5:134225954-134226838 | -0.059044518 | 0.852239927 | 0.944179796 |
| chr6:143427255-143427580 | 0.072596591 | 0.852247038 | 0.944179796 |
| chr3:173221092-173221350 | 0.080387855 | 0.85225344 | 0.944179796 |
| chr3:196287068-196288408 | -0.059604317 | 0.852279287 | 0.944194779 |
| chr13:44483694-44483998 | 0.079019836 | 0.852308312 | 0.944209678 |
| chr5:9545823-9546333 | -0.065441866 | 0.852317505 | 0.944209678 |
| chr3:23518543-23518743 | -0.093771134 | 0.852329704 | 0.944209678 |
| chr5:178436251-178436451 | -0.090076364 | 0.852369063 | 0.944239628 |
| chr17:75092857-75093124 | 0.072935583 | 0.852382497 | 0.944240859 |
| chr7:117828261-117828477 | -0.094948132 | 0.852419087 | 0.94426264 |
| chr8:53317160-53317381 | 0.088792436 | 0.852426807 | 0.94426264 |
| chr14:84442645-84442948 | 0.079417609 | 0.852458533 | 0.944272329 |
| chr5:126234621-126234880 | 0.089091475 | 0.852463816 | 0.944272329 |
| chr12:109879949-109880837 | -0.06169303 | 0.852472525 | 0.944272329 |
| chr1:225882278-225883347 | 0.060630874 | 0.852487172 | 0.944274902 |
| chr1:26321542-26321831 | -0.077375105 | 0.852508675 | 0.944285069 |
| chr16:72008392-72009234 | -0.059062229 | 0.852565147 | 0.944321702 |
| chr11:61971774-61972238 | 0.063363902 | 0.85257041 | 0.944321702 |
| chr17:77287769-77287979 | 0.072124267 | 0.852578829 | 0.944321702 |
| chr1:48284818-48285110 | 0.089401583 | 0.852591045 | 0.944321702 |
| chr11:7442723-7442927 | -0.094719507 | 0.852605454 | 0.944324011 |
| chr1:107196851-107197351 | -0.061688305 | 0.852662767 | 0.944373839 |
| chr6:109690438-109690681 | -0.077498485 | 0.852681467 | 0.944380899 |
| chr9:97982691-97983444 | -0.061856939 | 0.852737897 | 0.944429747 |
| chr1:224079545-224079896 | 0.081631144 | 0.852770325 | 0.944432766 |
| chr3:155462718-155463018 | 0.077408413 | 0.852774722 | 0.944432766 |
| chr11:71396819-71397019 | -0.089911988 | 0.852786429 | 0.944432766 |
| chr1:145730978-145731258 | 0.072814332 | 0.852792537 | 0.944432766 |
| chr11:48307668-48308135 | 0.073360254 | 0.852802252 | 0.944432766 |
| chr14:60862303-60862690 | -0.067399104 | 0.852825338 | 0.944444681 |
| chr2:43609863-43610170 | -0.082445684 | 0.852897013 | 0.944489247 |
| chr1:113631142-113631374 | 0.088095081 | 0.852909125 | 0.944489247 |
| chr1:28785166-28785429 | 0.077695991 | 0.852924209 | 0.944489247 |
| chr3:100710623-100710883 | 0.068057764 | 0.852965071 | 0.944489247 |
| chr6:28079930-28081405 | -0.058868106 | 0.852968034 | 0.944489247 |
| chr17:40121185-40122336 | -0.061297854 | 0.852998869 | 0.944489247 |
| chr3:149512278-149512568 | 0.072434857 | 0.853000931 | 0.944489247 |
| chr8:47737445-47738672 | 0.059586376 | 0.853009447 | 0.944489247 |
| chr1:116134187-116134452 | -0.074905585 | 0.853010046 | 0.944489247 |

| | | | |
|---|---|---|---|
| chr11:3840654-3841892 | -0.059744506 | 0.853018301 | 0.944489247 |
| chr7:6953781-6954128 | -0.073355733 | 0.853039745 | 0.944489247 |
| chr11:77548336-77548553 | 0.086360723 | 0.853055886 | 0.944489247 |
| chr9:36008790-36009150 | 0.074387292 | 0.853056918 | 0.944489247 |
| chr20:17504672-17505122 | -0.071941711 | 0.853059363 | 0.944489247 |
| chr8:92964931-92966393 | 0.05784449 | 0.853060497 | 0.944489247 |
| chr19:9055443-9055765 | 0.07301466 | 0.853067099 | 0.944489247 |
| chr11:77820632-77821802 | 0.058506864 | 0.853106751 | 0.944489247 |
| chr1:214753602-214753834 | -0.085369633 | 0.853110402 | 0.944489247 |
| chr8:88640502-88640702 | 0.102447978 | 0.853121195 | 0.944489247 |
| chr12:107317775-107318553 | -0.061735526 | 0.853122385 | 0.944489247 |
| chr11:77789818-77790098 | -0.090182255 | 0.85312444 | 0.944489247 |
| chr3:166243221-166243421 | 0.104936868 | 0.853164799 | 0.944507498 |
| chr17:49315528-49315805 | 0.075696147 | 0.853200923 | 0.944507498 |
| chr11:73530275-73530500 | -0.089512457 | 0.853204695 | 0.944507498 |
| chr8:116703647-116703847 | 0.088377742 | 0.853219949 | 0.944507498 |
| chr21:46538756-46539121 | 0.084350592 | 0.853228457 | 0.944507498 |
| chr11:71120472-71120894 | -0.072116244 | 0.853237897 | 0.944507498 |
| chr12:131834841-131835042 | 0.078149961 | 0.853239014 | 0.944507498 |
| chr1:202457150-202457431 | -0.089912018 | 0.85323954 | 0.944507498 |
| chr8:144326485-144327322 | 0.06010284 | 0.853280375 | 0.944529134 |
| chr19:6753331-6753557 | -0.100670888 | 0.853303957 | 0.944529134 |
| chr6:31157982-31158783 | -0.059931188 | 0.853338699 | 0.944529134 |
| chr2:113756066-113757052 | 0.059436949 | 0.853340677 | 0.944529134 |
| chr8:27602136-27602451 | -0.078794165 | 0.853344749 | 0.944529134 |
| chr2:152097870-152098573 | -0.066746011 | 0.853361153 | 0.944529134 |
| chr18:657107-658767 | -0.058858083 | 0.853376847 | 0.944529134 |
| chr8:69663757-69664086 | 0.071487904 | 0.85339523 | 0.944529134 |
| chr2:215166390-215166618 | 0.090295068 | 0.853420325 | 0.944529134 |
| chr1:220786594-220787873 | -0.059165318 | 0.853425462 | 0.944529134 |
| chr1:114511769-114512395 | 0.058794827 | 0.853434711 | 0.944529134 |
| chr15:69073947-69074778 | -0.062123671 | 0.853451099 | 0.944529134 |
| chr13:113296775-113297399 | -0.063010806 | 0.853466579 | 0.944529134 |
| chr12:5965935-5966205 | 0.082347694 | 0.853482273 | 0.944529134 |
| chr6:170231729-170231929 | 0.088002987 | 0.853483265 | 0.944529134 |
| chr4:153571117-153571498 | 0.074455588 | 0.853485409 | 0.944529134 |
| chr2:171159696-171161395 | 0.059317365 | 0.853496572 | 0.944529134 |
| chr8:91680732-91680963 | 0.084343709 | 0.853512219 | 0.944529134 |
| chr1:180252617-180252916 | -0.068927718 | 0.853513132 | 0.944529134 |
| chr10:95430439-95430707 | -0.076033374 | 0.853529656 | 0.944529134 |
| chr9:134449584-134449938 | 0.06705796 | 0.853535014 | 0.944529134 |
| chr13:23142197-23142911 | -0.070839118 | 0.853540497 | 0.944529134 |
| chr2:182186215-182186435 | -0.086189086 | 0.853542609 | 0.944529134 |
| chr6:154786018-154786496 | -0.069630766 | 0.853556299 | 0.944530643 |
| chr13:99198766-99198989 | -0.075138958 | 0.853633057 | 0.944601939 |
| chr21:32412053-32412294 | 0.080068983 | 0.853646702 | 0.944603397 |
| chr14:93219203-93219463 | 0.07911488 | 0.853666036 | 0.944611149 |
| chr20:36085839-36086039 | 0.097144704 | 0.853678967 | 0.944611816 |
| chr3:183697578-183698258 | -0.064085238 | 0.853721737 | 0.94462351 |
| chr16:80900212-80900625 | -0.068013853 | 0.853725036 | 0.94462351 |

| | | | |
|---|---|---|---|
| chr5:170211617-170211817 | 0.087176777 | 0.853726521 | 0.94462351 |
| chr11:69088667-69089018 | 0.068797225 | 0.853797371 | 0.944688262 |
| chr18:24138824-24139811 | 0.061227458 | 0.853835238 | 0.944713338 |
| chr1:217424383-217425377 | -0.061556107 | 0.853853721 | 0.944713338 |
| chr9:21095643-21095898 | 0.087317212 | 0.853857023 | 0.944713338 |
| chr1:205569491-205569863 | -0.070872764 | 0.853893021 | 0.944739525 |
| chr10:125934304-125934606 | -0.074435338 | 0.853917636 | 0.944753117 |
| chr3:194333332-194334085 | 0.062938164 | 0.853941691 | 0.944758872 |
| chr8:142993900-142994244 | 0.079629633 | 0.853951165 | 0.944758872 |
| chr1:37931166-37932108 | -0.060473031 | 0.853971615 | 0.944758872 |
| chr11:41459990-41460443 | -0.07305895 | 0.85398522 | 0.944758872 |
| chr2:240566384-240566835 | -0.064931886 | 0.853996216 | 0.944758872 |
| chr1:223363897-223364957 | 0.060122522 | 0.853996819 | 0.944758872 |
| chr19:40285273-40285756 | -0.060975622 | 0.854074052 | 0.944830672 |
| chr3:62948681-62948881 | 0.072920823 | 0.854095575 | 0.944840526 |
| chr5:133612443-133612862 | -0.073379046 | 0.854107622 | 0.944840526 |
| chr16:27181132-27181389 | -0.07700696 | 0.854169286 | 0.944893384 |
| chr20:37534532-37534781 | 0.079887448 | 0.854180068 | 0.944893384 |
| chr8:142988190-142988398 | 0.083964138 | 0.854264155 | 0.944972758 |
| chr11:121479439-121479688 | 0.08354598 | 0.854293624 | 0.944991714 |
| chr8:26382616-26383714 | -0.058774299 | 0.854332626 | 0.944999048 |
| chr1:96234396-96234608 | -0.087990838 | 0.854334425 | 0.944999048 |
| chr20:18287842-18288959 | 0.058806551 | 0.854337254 | 0.944999048 |
| chr2:61671096-61671340 | -0.105761889 | 0.854355923 | 0.945001685 |
| chr19:35060706-35061094 | -0.079062024 | 0.85436689 | 0.945001685 |
| chr17:44708403-44708722 | -0.07740758 | 0.854383731 | 0.945001685 |
| chr1:27719617-27719817 | -0.09349576 | 0.854388971 | 0.945001685 |
| chr5:173530602-173530940 | 0.070549892 | 0.854463921 | 0.945069711 |
| chr22:31148901-31149575 | 0.067855175 | 0.854475142 | 0.945069711 |
| chr1:199093108-199093336 | -0.083179274 | 0.854523194 | 0.945109215 |
| chr11:45655360-45655648 | -0.085552858 | 0.854573575 | 0.945141368 |
| chr16:47037486-47038368 | -0.064143629 | 0.854576936 | 0.945141368 |
| chr2:219627502-219628280 | 0.063907139 | 0.854660695 | 0.94522036 |
| chr12:131928708-131929511 | 0.060217257 | 0.85469944 | 0.945249566 |
| chr12:75605088-75605468 | 0.079779682 | 0.854724568 | 0.945255004 |
| chr9:5843764-5844224 | -0.066309724 | 0.854729029 | 0.945255004 |
| chr3:23202330-23203824 | -0.058086951 | 0.854774108 | 0.945273084 |
| chr12:68912940-68913333 | 0.074057128 | 0.854794883 | 0.945273084 |
| chr17:47323623-47324843 | -0.059578565 | 0.854800567 | 0.945273084 |
| chr2:175745649-175745871 | -0.092436867 | 0.854802087 | 0.945273084 |
| chr1:160681500-160681917 | -0.072114658 | 0.854810515 | 0.945273084 |
| chr5:178164189-178164567 | 0.06588906 | 0.854824297 | 0.945273084 |
| chr1:108746318-108747580 | -0.057543578 | 0.854840001 | 0.945273084 |
| chr1:18902433-18903281 | 0.061184074 | 0.854844073 | 0.945273084 |
| chr9:113275070-113276049 | 0.058334029 | 0.854862888 | 0.94527311 |
| chr16:2035990-2036212 | 0.079014052 | 0.85486877 | 0.94527311 |
| chr2:102235058-102235374 | 0.073907634 | 0.854885025 | 0.945277443 |
| chr16:87853663-87853982 | 0.06645523 | 0.854950956 | 0.945323609 |
| chr7:7114808-7115329 | -0.066328806 | 0.854951478 | 0.945323609 |
| chr5:111158152-111158398 | -0.079593439 | 0.854963789 | 0.945323609 |

| | | | |
|---|---|---|---|
| chr15:99703732-99703956 | 0.08022944 | 0.85498004 | 0.945327936 |
| chr3:38203724-38203961 | -0.092555218 | 0.855005338 | 0.945342266 |
| chr4:26055967-26056167 | -0.097039295 | 0.855058975 | 0.945387928 |
| chr14:106349015-106349374 | 0.067873652 | 0.855104552 | 0.945424677 |
| chr1:150696968-150697819 | -0.058076706 | 0.855144239 | 0.945454914 |
| chr3:123201429-123202489 | 0.058452772 | 0.855174276 | 0.945474481 |
| chr1:80083854-80084158 | 0.077736877 | 0.855193884 | 0.945482516 |
| chr15:66356262-66357156 | -0.060087357 | 0.855221882 | 0.945490854 |
| chr3:187112827-187113113 | -0.096895972 | 0.855226105 | 0.945490854 |
| chr12:50747916-50748162 | 0.085059869 | 0.855251041 | 0.945493055 |
| chr3:48548789-48548989 | 0.094815697 | 0.855263019 | 0.945493055 |
| chr8:81253933-81254206 | 0.079458833 | 0.855271794 | 0.945493055 |
| chr5:176660787-176661138 | -0.066770648 | 0.855286991 | 0.945493055 |
| chr2:195657993-195658306 | 0.065778902 | 0.855294538 | 0.945493055 |
| chr17:63113258-63113458 | 0.086988483 | 0.855302134 | 0.945493055 |
| chr8:128014745-128015009 | 0.070593798 | 0.855324699 | 0.945504358 |
| chr2:111946616-111947095 | 0.064308327 | 0.855354073 | 0.945511424 |
| chr2:14634372-14635783 | -0.058360944 | 0.855355771 | 0.945511424 |
| chr11:74748428-74749001 | 0.060423085 | 0.855393722 | 0.945539734 |
| chr2:24792776-24794011 | -0.05752476 | 0.855435022 | 0.945571745 |
| chr4:102165765-102165971 | -0.093529641 | 0.855486648 | 0.945615169 |
| chr17:47957448-47958441 | 0.058479719 | 0.855540815 | 0.945661401 |
| chr8:86482596-86482865 | 0.074738797 | 0.855652719 | 0.945718669 |
| chr19:11131012-11131222 | 0.081978485 | 0.855659439 | 0.945718669 |
| chr3:77815514-77815819 | -0.074348805 | 0.855667114 | 0.945718669 |
| chr1:236394994-236395351 | 0.076096576 | 0.855672318 | 0.945718669 |
| chr16:89599440-89599667 | -0.076388925 | 0.855682327 | 0.945718669 |
| chr12:6470407-6471263 | -0.058971491 | 0.855689057 | 0.945718669 |
| chr10:50367812-50368019 | -0.087720299 | 0.855690237 | 0.945718669 |
| chr3:141875529-141877430 | -0.057195992 | 0.855697881 | 0.945718669 |
| chr7:6104197-6105053 | -0.058356499 | 0.855703709 | 0.945718669 |
| chr17:7614765-7615648 | -0.058845983 | 0.855722504 | 0.9457258 |
| chr2:39379777-39379990 | 0.087605117 | 0.855759519 | 0.945735535 |
| chr3:135762516-135762761 | -0.088947098 | 0.855768341 | 0.945735535 |
| chr3:144853061-144853368 | -0.088947098 | 0.855768341 | 0.945735535 |
| chr19:29858057-29858368 | -0.070574056 | 0.855807297 | 0.945764945 |
| chr16:67857796-67858228 | -0.066365818 | 0.855852047 | 0.945790709 |
| chr14:102096567-102096872 | 0.069423804 | 0.855855297 | 0.945790709 |
| chr2:48600282-48600507 | 0.080808808 | 0.855871124 | 0.945794558 |
| chr5:79774556-79774877 | 0.070355635 | 0.855932609 | 0.945848862 |
| chr6:42979057-42979381 | 0.069543114 | 0.855980093 | 0.945885934 |
| chr16:68843344-68844108 | -0.062324881 | 0.855994601 | 0.945885934 |
| chr7:30771152-30771873 | 0.064769127 | 0.856007881 | 0.945885934 |
| chr1:28642605-28643550 | 0.059273563 | 0.856033664 | 0.945885934 |
| chr2:241470441-241470662 | -0.091808345 | 0.856033784 | 0.945885934 |
| chr2:24122929-24124021 | -0.05935722 | 0.856040226 | 0.945885934 |
| chr9:112890830-112891072 | -0.086667749 | 0.856069438 | 0.945904572 |
| chr21:44742472-44742729 | -0.091419554 | 0.856092582 | 0.945907494 |
| chr6:45445757-45446156 | 0.067044311 | 0.856097268 | 0.945907494 |
| chr16:72811538-72811871 | -0.086578932 | 0.856109119 | 0.945907494 |

| | | | |
|---|---|---|---|
| chr2:212979840-212980281 | 0.076239793 | 0.856149469 | 0.945913937 |
| chr17:48678507-48678905 | 0.073612424 | 0.856149648 | 0.945913937 |
| chr1:248788515-248788786 | 0.079501756 | 0.856151986 | 0.945913937 |
| chr17:64025261-64025648 | 0.066776212 | 0.856174795 | 0.945925498 |
| chr14:102517381-102517643 | 0.074056767 | 0.856187367 | 0.945925749 |
| chr2:202060361-202060728 | -0.090913427 | 0.856227525 | 0.945946063 |
| chr7:38731929-38732433 | -0.074646572 | 0.856230446 | 0.945946063 |
| chr10:119206581-119207711 | -0.059021014 | 0.856246133 | 0.945949755 |
| chr12:56257838-56258858 | 0.059466788 | 0.856278714 | 0.94597211 |
| chr1:110089101-110089320 | 0.088681839 | 0.856309297 | 0.945992256 |
| chr3:133393190-133393571 | 0.0681575 | 0.856342402 | 0.945995348 |
| chr5:25515590-25515817 | 0.089454906 | 0.856343546 | 0.945995348 |
| chr2:199488866-199489123 | -0.075350473 | 0.856349134 | 0.945995348 |
| chr5:137470345-137470558 | -0.095353537 | 0.856403518 | 0.946019648 |
| chr19:18488390-18488590 | -0.089452732 | 0.856432448 | 0.946019648 |
| chr6:107490731-107491023 | -0.080703086 | 0.856446115 | 0.946019648 |
| chr3:27677571-27677935 | 0.071766914 | 0.856453346 | 0.946019648 |
| chr4:92163459-92163666 | -0.090042116 | 0.856464753 | 0.946019648 |
| chr15:84570664-84571322 | 0.060727999 | 0.856470791 | 0.946019648 |
| chr4:102474644-102475466 | 0.064297458 | 0.856481353 | 0.946019648 |
| chr1:218784031-218784231 | -0.088920875 | 0.856481937 | 0.946019648 |
| chr3:136740652-136740908 | 0.086609686 | 0.856483429 | 0.946019648 |
| chrX:63488473-63488673 | 0.096105053 | 0.856494597 | 0.946019648 |
| chr1:187476730-187476930 | 0.09259565 | 0.856560501 | 0.946077682 |
| chr8:16826009-16826298 | 0.083137067 | 0.856575034 | 0.946077682 |
| chr19:18340511-18340841 | -0.083378213 | 0.856584181 | 0.946077682 |
| chrX:103157945-103158419 | 0.069127267 | 0.856612453 | 0.946093244 |
| chr8:70607438-70608761 | -0.057622179 | 0.856633758 | 0.946093244 |
| chr19:39858509-39858998 | -0.063110223 | 0.856642488 | 0.946093244 |
| chr3:51982481-51983582 | 0.060162396 | 0.856655474 | 0.946093244 |
| chr14:95581185-95581545 | 0.080887724 | 0.856660009 | 0.946093244 |
| chr17:59753493-59753764 | 0.066490964 | 0.856674142 | 0.946095217 |
| chr3:129587620-129587985 | -0.070415276 | 0.856704343 | 0.946114488 |
| chr18:32624303-32624608 | -0.075828608 | 0.856725451 | 0.946114488 |
| chr1:112707027-112707802 | -0.058565363 | 0.856728636 | 0.946114488 |
| chrX:147912197-147912435 | 0.076344115 | 0.856752481 | 0.946127185 |
| chr3:122415793-122416636 | 0.057503193 | 0.856795509 | 0.946161065 |
| chr2:206392183-206392507 | 0.077254631 | 0.856826284 | 0.946181414 |
| chr8:134831654-134833047 | -0.056581086 | 0.856850966 | 0.946195033 |
| chr3:124592680-124593169 | -0.064059374 | 0.856867562 | 0.946199723 |
| chrX:18119919-18120240 | -0.072405638 | 0.856939893 | 0.946264309 |
| chrX:71622785-71623065 | 0.081675358 | 0.856953911 | 0.946264309 |
| chr5:128846386-128846796 | -0.076063059 | 0.856983586 | 0.946264309 |
| chr12:115468231-115468431 | 0.095935645 | 0.856990625 | 0.946264309 |
| chr8:108555874-108556105 | -0.079891895 | 0.857016071 | 0.946264309 |
| chr12:1804564-1804783 | 0.077776109 | 0.85701721 | 0.946264309 |
| chr11:32892252-32893160 | 0.060120009 | 0.857024633 | 0.946264309 |
| chr3:40913808-40914008 | -0.102454473 | 0.857024966 | 0.946264309 |
| chr1:52904934-52905168 | 0.080848546 | 0.857037199 | 0.946264309 |
| chr8:103279118-103279343 | -0.080654779 | 0.85706799 | 0.94628467 |

| | | | |
|---|---|---|---|
| chrX:103533501-103534085 | -0.069213186 | 0.857130406 | 0.946339947 |
| chr4:145003479-145004382 | -0.060353233 | 0.857178564 | 0.946350228 |
| chr2:120487194-120487432 | -0.084755279 | 0.85718577 | 0.946350228 |
| chr4:48290421-48290724 | 0.070728822 | 0.857189359 | 0.946350228 |
| chr22:50582396-50583426 | -0.058903049 | 0.857197015 | 0.946350228 |
| chr3:149771699-149771962 | 0.075963294 | 0.857204255 | 0.946350228 |
| chr3:120742104-120743251 | -0.057018943 | 0.857213824 | 0.946350228 |
| chr12:57831180-57831751 | -0.065672423 | 0.857254783 | 0.946355065 |
| chr8:90627888-90628120 | 0.090340883 | 0.857259157 | 0.946355065 |
| chr5:42772317-42772626 | 0.080379064 | 0.857266684 | 0.946355065 |
| chr3:138118750-138119230 | -0.062766242 | 0.857267609 | 0.946355065 |
| chr7:423857-424086 | 0.081347193 | 0.857283939 | 0.946359458 |
| chr9:34701790-34702125 | 0.076779277 | 0.85730021 | 0.946363785 |
| chr2:136932745-136933012 | 0.081670551 | 0.857317451 | 0.946369183 |
| chr4:40916931-40917212 | -0.077617549 | 0.857388416 | 0.946385629 |
| chr10:9258976-9259176 | -0.097578477 | 0.857390639 | 0.946385629 |
| chr1:168578715-168578972 | 0.084522629 | 0.857401549 | 0.946385629 |
| chr2:204325457-204325823 | -0.065142248 | 0.85740576 | 0.946385629 |
| chr4:77861985-77862196 | -0.072398216 | 0.857409882 | 0.946385629 |
| chr9:137311093-137311845 | -0.060675944 | 0.857421895 | 0.946385629 |
| chr11:79461078-79461377 | -0.073021644 | 0.857424715 | 0.946385629 |
| chr1:179369889-179370089 | 0.083085264 | 0.85743116 | 0.946385629 |
| chr4:156970614-156972307 | 0.057120397 | 0.85748443 | 0.946426128 |
| chr8:105773009-105773304 | -0.072617516 | 0.857499752 | 0.946426128 |
| chr11:66447236-66447436 | 0.095448962 | 0.857512203 | 0.946426128 |
| chr3:142007724-142008056 | 0.079955804 | 0.857520374 | 0.946426128 |
| chr1:195897168-195897404 | -0.078176113 | 0.857529612 | 0.946426128 |
| chr11:27395725-27395925 | -0.089782107 | 0.857564874 | 0.946451413 |
| chr5:146966097-146966429 | -0.071666796 | 0.857658214 | 0.946538739 |
| chr5:69559919-69560138 | 0.09210736 | 0.85770053 | 0.946538739 |
| chr2:108417096-108417599 | -0.067751867 | 0.85770839 | 0.946538739 |
| chr11:125300906-125301106 | 0.098243349 | 0.857720284 | 0.946538739 |
| chr7:129496051-129496251 | 0.094962459 | 0.857727433 | 0.946538739 |
| chr11:125165285-125165587 | 0.084252626 | 0.857729525 | 0.946538739 |
| chr7:151782605-151783239 | -0.063007638 | 0.857735013 | 0.946538739 |
| chr1:204989352-204989635 | 0.064919482 | 0.857760321 | 0.946538739 |
| chr14:92684857-92685108 | 0.074459197 | 0.857766503 | 0.946538739 |
| chr4:55545744-55547311 | 0.05704812 | 0.857771435 | 0.946538739 |
| chr22:45645623-45645877 | -0.084310829 | 0.857779886 | 0.946538739 |
| chr1:180229535-180229764 | 0.071774705 | 0.857811926 | 0.946554608 |
| chr3:138197080-138197389 | 0.079234556 | 0.85782295 | 0.946554608 |
| chr1:53837946-53838803 | 0.05828926 | 0.857831328 | 0.946554608 |
| chrX:109645074-109645276 | 0.088098431 | 0.857875153 | 0.946578244 |
| chr16:73219407-73219607 | -0.083619385 | 0.857877456 | 0.946578244 |
| chr14:76546879-76547230 | 0.071577782 | 0.857897734 | 0.946586988 |
| chr1:99180337-99180578 | -0.093093455 | 0.857928266 | 0.946607044 |
| chr15:98339126-98339326 | 0.097813864 | 0.857941001 | 0.946607464 |
| chr19:38343277-38343477 | 0.087100085 | 0.857972377 | 0.946628452 |
| chr20:16761269-16761586 | 0.069153366 | 0.85807988 | 0.946725536 |
| chr2:181430461-181430915 | -0.074436003 | 0.858091638 | 0.946725536 |

| | | | |
|---|---|---|---|
| chr2:91659942-91660187 | -0.091711754 | 0.858097437 | 0.946725536 |
| chr19:44213057-44213257 | -0.092846545 | 0.858120658 | 0.946732435 |
| chr14:57349394-57349622 | -0.09115087 | 0.858128401 | 0.946732435 |
| chr13:39373072-39373272 | 0.092301927 | 0.858149531 | 0.946742115 |
| chr14:63852538-63854190 | 0.05668692 | 0.858163443 | 0.946743832 |
| chr5:174600025-174600573 | 0.066300094 | 0.858204624 | 0.946775632 |
| chr15:44476610-44476854 | 0.105077212 | 0.858226022 | 0.946785606 |
| chr2:121755284-121756081 | 0.057888734 | 0.858238912 | 0.946786194 |
| chr12:81999850-82000050 | 0.091599361 | 0.858297847 | 0.946811635 |
| chr7:945447-946245 | 0.061317956 | 0.858305455 | 0.946811635 |
| chr6:68813085-68813285 | 0.102430388 | 0.858309839 | 0.946811635 |
| chr17:60627545-60627745 | 0.066586822 | 0.858311401 | 0.946811635 |
| chr1:187582997-187583231 | 0.093885699 | 0.858326706 | 0.946814887 |
| chr7:66844521-66845343 | -0.058237495 | 0.858369468 | 0.946847175 |
| chr1:201649291-201649630 | -0.067480271 | 0.858380691 | 0.946847175 |
| chr19:18328422-18328813 | -0.068047488 | 0.858437912 | 0.946896662 |
| chr15:99123483-99124442 | 0.060235993 | 0.858474491 | 0.946897164 |
| chr8:42391426-42392539 | 0.05791097 | 0.858483899 | 0.946897164 |
| chr6:166400878-166401129 | 0.085036369 | 0.858488516 | 0.946897164 |
| chr1:39988727-39988996 | -0.083930319 | 0.858497287 | 0.946897164 |
| chr1:119475900-119476334 | -0.071252581 | 0.85850621 | 0.946897164 |
| chr3:151304716-151305750 | -0.056922686 | 0.858512516 | 0.946897164 |
| chr10:22316893-22317194 | -0.072971522 | 0.858551846 | 0.946905286 |
| chr12:8180132-8180546 | -0.072713587 | 0.858560891 | 0.946905286 |
| chr19:42206996-42207380 | 0.071685634 | 0.858566269 | 0.946905286 |
| chr5:72269150-72269439 | 0.077419081 | 0.85857351 | 0.946905286 |
| chr8:11808669-11809182 | 0.064899597 | 0.85858167 | 0.946905286 |
| chr20:53503511-53504470 | -0.060285324 | 0.858615266 | 0.946925869 |
| chr5:141637898-141638374 | -0.062606869 | 0.858634508 | 0.946925869 |
| chr8:90019353-90019735 | -0.073840403 | 0.858637409 | 0.946925869 |
| chr20:19047571-19047948 | 0.067874108 | 0.858662526 | 0.94693994 |
| chr1:152168238-152168439 | 0.083615863 | 0.858691326 | 0.946958071 |
| chr1:169674893-169675106 | -0.091176274 | 0.858713941 | 0.946961805 |
| chr14:66197340-66197556 | -0.097377696 | 0.858722388 | 0.946961805 |
| chr14:26401470-26401940 | -0.068057794 | 0.858731789 | 0.946961805 |
| chr10:97291864-97292963 | -0.056889157 | 0.858750627 | 0.94696895 |
| chr10:97413994-97414194 | 0.089974618 | 0.858800924 | 0.947010786 |
| chr17:65339341-65339541 | 0.090183346 | 0.858857423 | 0.947048279 |
| chr3:50359331-50359876 | 0.061329033 | 0.858864576 | 0.947048279 |
| chr5:149799011-149799226 | -0.088522312 | 0.858872005 | 0.947048279 |
| chr3:157412648-157412950 | -0.094848625 | 0.858896498 | 0.947061657 |
| chr20:53656270-53656635 | 0.057960444 | 0.858921888 | 0.947063129 |
| chr6:34163606-34164074 | 0.069181156 | 0.858922553 | 0.947063129 |
| chr1:107738685-107739221 | -0.067354867 | 0.858939411 | 0.947068087 |
| chr20:33096001-33096231 | -0.081883103 | 0.858976929 | 0.947089512 |
| chr4:95205001-95205201 | 0.093893924 | 0.858983563 | 0.947089512 |
| chr3:113696507-113697551 | -0.056276737 | 0.858996071 | 0.947089675 |
| chr9:113853390-113853622 | 0.086448253 | 0.859025046 | 0.947107993 |
| chr2:215979715-215979915 | -0.077304077 | 0.859043483 | 0.947114692 |
| chr14:25431358-25431962 | 0.065519632 | 0.859061866 | 0.947121331 |

| | | | |
|---|---|---|---|
| chr16:4374917-4375371 | -0.06133832 | 0.859116908 | 0.947168387 |
| chr2:221372561-221372817 | -0.090885995 | 0.859144171 | 0.947176204 |
| chr21:29386697-29387207 | -0.063712348 | 0.859149835 | 0.947176204 |
| chr10:113923006-113923481 | -0.071103516 | 0.859175607 | 0.947176204 |
| chr2:84486563-84486763 | 0.087538429 | 0.85918435 | 0.947176204 |
| chr9:127569019-127569797 | -0.058694139 | 0.859190683 | 0.947176204 |
| chr4:86593715-86595249 | -0.056093919 | 0.859208332 | 0.947176204 |
| chr5:152405242-152405469 | -0.084297798 | 0.859218879 | 0.947176204 |
| chr4:152063622-152064381 | -0.061700996 | 0.859222892 | 0.947176204 |
| chr14:101505572-101505827 | -0.076337697 | 0.859236306 | 0.947177365 |
| chr3:42744371-42744681 | -0.077177049 | 0.859256866 | 0.947186401 |
| chr21:34846580-34846805 | -0.087697596 | 0.859286024 | 0.947194531 |
| chr16:74843701-74843901 | 0.081276397 | 0.859288965 | 0.947194531 |
| chr5:66209393-66209846 | -0.066359374 | 0.859321065 | 0.947212569 |
| chr6:125924216-125924509 | 0.073040716 | 0.859330053 | 0.947212569 |
| chr3:187713571-187713771 | 0.086105652 | 0.859346394 | 0.947216954 |
| chr22:41091222-41092091 | -0.058045048 | 0.859372566 | 0.947225067 |
| chr12:70499059-70499319 | 0.084027849 | 0.859381632 | 0.947225067 |
| chr2:35343255-35343676 | -0.0721417 | 0.859400143 | 0.947225067 |
| chr22:37032645-37033181 | 0.065797968 | 0.859403203 | 0.947225067 |
| chr11:57741132-57742003 | -0.059057778 | 0.859426401 | 0.94723701 |
| chr2:7767938-7768138 | 0.094581759 | 0.859457664 | 0.947257841 |
| chr11:86274533-86274952 | -0.071503292 | 0.859552795 | 0.94733696 |
| chr10:116682388-116682841 | -0.06588254 | 0.859558278 | 0.94733696 |
| chr15:84826193-84826970 | -0.06072347 | 0.859570038 | 0.94733696 |
| chr15:45179243-45179501 | -0.077942327 | 0.859578905 | 0.94733696 |
| chr3:15470264-15470653 | 0.067610302 | 0.859620965 | 0.947365399 |
| chr1:165630388-165631387 | 0.056366359 | 0.859629437 | 0.947365399 |
| chr3:132262032-132262232 | -0.097062507 | 0.859654754 | 0.947379673 |
| chr17:78781878-78782822 | 0.05817918 | 0.859690878 | 0.947392726 |
| chr16:84103916-84104116 | 0.080480198 | 0.859691327 | 0.947392726 |
| chr11:123023159-123023454 | 0.08496155 | 0.859739194 | 0.947422445 |
| chr1:21022303-21022560 | 0.086457186 | 0.85974633 | 0.947422445 |
| chr17:50094306-50094778 | 0.060903343 | 0.85975539 | 0.947422445 |
| chr8:90692757-90693044 | -0.078858689 | 0.859796231 | 0.947429029 |
| chr7:5423710-5424368 | -0.057380037 | 0.859799706 | 0.947429029 |
| chr2:73921248-73921508 | -0.075257923 | 0.85980004 | 0.947429029 |
| chr1:54125736-54125946 | -0.079626103 | 0.859810824 | 0.947429029 |
| chr5:81165576-81165874 | 0.068870794 | 0.859878532 | 0.947458235 |
| chr16:9123885-9124163 | -0.080229715 | 0.859879409 | 0.947458235 |
| chr4:128782399-128782687 | -0.083404955 | 0.859885849 | 0.947458235 |
| chr8:116788322-116788715 | -0.065579218 | 0.859886791 | 0.947458235 |
| chr8:101097934-101098184 | 0.077842894 | 0.859930698 | 0.947492989 |
| chr3:125519501-125520550 | 0.057890557 | 0.859943939 | 0.947493953 |
| chr20:19205951-19206269 | -0.075757672 | 0.859964147 | 0.947502593 |
| chr17:59094855-59095238 | -0.058109665 | 0.860033466 | 0.947542205 |
| chr2:42545729-42546256 | 0.065057512 | 0.860034386 | 0.947542205 |
| chr6:26571679-26572257 | -0.060130904 | 0.860037198 | 0.947542205 |
| chr1:201398647-201399037 | -0.062210567 | 0.860066695 | 0.947557543 |
| chr4:95045635-95045868 | 0.087622641 | 0.860075853 | 0.947557543 |

| | | | |
|---|---|---|---|
| chr5:178699857-178700221 | 0.070217761 | 0.860090727 | 0.947559078 |
| chr10:97334491-97335182 | -0.05733239 | 0.860101979 | 0.947559078 |
| chr2:213251647-213251853 | -0.095200907 | 0.860135188 | 0.947582038 |
| chr22:21641440-21641814 | 0.067055267 | 0.860191793 | 0.947604078 |
| chr22:37601555-37602091 | -0.060762279 | 0.860195117 | 0.947604078 |
| chr2:64453268-64453474 | 0.076564574 | 0.860195473 | 0.947604078 |
| chr1:166875655-166876800 | -0.055489273 | 0.860204662 | 0.947604078 |
| chr17:47492105-47492889 | -0.058527666 | 0.860235271 | 0.947624172 |
| chr17:70447572-70447895 | 0.070415252 | 0.860280158 | 0.947659792 |
| chr4:78056269-78057114 | 0.057628112 | 0.860292342 | 0.947659792 |
| chr13:46553757-46554875 | 0.055683862 | 0.860321774 | 0.94766574 |
| chr12:99043399-99043630 | 0.083461164 | 0.860322477 | 0.94766574 |
| chr12:6843200-6843524 | -0.075325236 | 0.86035541 | 0.947684918 |
| chr3:170258414-170258614 | 0.093968964 | 0.860375001 | 0.947684918 |
| chr6:33744192-33744703 | -0.06499938 | 0.860376993 | 0.947684918 |
| chr8:133102856-133103158 | -0.073106301 | 0.860448859 | 0.947744069 |
| chr8:143943061-143943653 | 0.057535968 | 0.860455432 | 0.947744069 |
| chr1:110106086-110106456 | -0.06563384 | 0.860476784 | 0.947753962 |
| chr21:38170725-38170925 | 0.076850032 | 0.860500685 | 0.947765322 |
| chr17:56870449-56870970 | 0.065159386 | 0.860523359 | 0.947765322 |
| chr1:107965457-107966060 | 0.05855321 | 0.860524205 | 0.947765322 |
| chr4:23036537-23036904 | 0.075570724 | 0.860563941 | 0.947795462 |
| chr8:135054632-135054934 | 0.084296486 | 0.860623628 | 0.947847575 |
| chr6:83192858-83194417 | 0.055595887 | 0.860674214 | 0.947889663 |
| chr17:1643068-1643540 | -0.06376478 | 0.860688075 | 0.947891304 |
| chr1:201222848-201223232 | -0.061038168 | 0.860707785 | 0.947898476 |
| chr14:30791366-30791709 | 0.079013178 | 0.86071933 | 0.947898476 |
| chr9:14442123-14442637 | -0.06595358 | 0.860741098 | 0.947908825 |
| chr5:172983364-172984671 | -0.055869352 | 0.860788711 | 0.947926583 |
| chr3:124559322-124559559 | -0.077785097 | 0.860791565 | 0.947926583 |
| chr3:58414642-58414903 | -0.079881242 | 0.860799581 | 0.947926583 |
| chr13:101710067-101710366 | -0.091424822 | 0.860806709 | 0.947926583 |
| chr7:92763625-92764021 | -0.071166861 | 0.860836298 | 0.947933123 |
| chr4:135371156-135371454 | -0.076588547 | 0.860857922 | 0.947933123 |
| chr19:49593420-49593657 | 0.06634031 | 0.860867065 | 0.947933123 |
| chr2:42386763-42386994 | 0.079948793 | 0.860869085 | 0.947933123 |
| chr2:210503046-210503322 | 0.077106981 | 0.860887737 | 0.947933123 |
| chr17:8249135-8249470 | -0.064678614 | 0.860888359 | 0.947933123 |
| chr3:100537689-100537889 | -0.090470194 | 0.860901827 | 0.947933123 |
| chr17:20908389-20908926 | -0.061076229 | 0.860912267 | 0.947933123 |
| chr6:151389965-151391214 | -0.055095809 | 0.860940969 | 0.947933123 |
| chr12:107892094-107892314 | -0.084754012 | 0.860944624 | 0.947933123 |
| chr9:105162344-105162544 | -0.087247849 | 0.860957861 | 0.947933123 |
| chr11:66311628-66313118 | 0.055754181 | 0.860961106 | 0.947933123 |
| chr2:65129557-65129757 | 0.06913886 | 0.861012573 | 0.947976167 |
| chr12:2583094-2583512 | -0.072127568 | 0.861033868 | 0.947977906 |
| chr3:119257233-119257986 | -0.05948555 | 0.861043851 | 0.947977906 |
| chr5:132007858-132008193 | 0.063672184 | 0.861060176 | 0.947977906 |
| chr8:71361554-71362112 | 0.062628914 | 0.861063642 | 0.947977906 |
| chr3:22089268-22089663 | -0.064083849 | 0.861097203 | 0.948001234 |

| | | | |
|---|---|---|---|
| chr3:134092134-134092334 | 0.094221466 | 0.861143095 | 0.948024514 |
| chr3:159856130-159856332 | 0.094221466 | 0.861143095 | 0.948024514 |
| chr14:52446288-52446713 | -0.070228492 | 0.861178003 | 0.948049323 |
| chr7:151351145-151351349 | 0.085469142 | 0.861194861 | 0.94805426 |
| chr4:83334769-83335304 | 0.061649986 | 0.861214765 | 0.948062551 |
| chr5:86118578-86118853 | -0.082238305 | 0.86125807 | 0.948089352 |
| chr4:147526408-147526672 | 0.076637841 | 0.861275212 | 0.948089352 |
| chr2:130693547-130693791 | -0.090197932 | 0.861276232 | 0.948089352 |
| chr12:12582779-12583446 | 0.062130285 | 0.861300998 | 0.948102993 |
| chr11:15819700-15820052 | -0.067329948 | 0.861338939 | 0.948131137 |
| chr17:6082174-6082454 | 0.088820943 | 0.86138031 | 0.948158772 |
| chr19:11089841-11090785 | 0.05774148 | 0.861388793 | 0.948158772 |
| chr1:202600173-202600664 | -0.059482322 | 0.861430488 | 0.948179588 |
| chr5:159621264-159621478 | -0.086366915 | 0.861440709 | 0.948179588 |
| chr3:49007072-49008168 | 0.056407526 | 0.861444829 | 0.948179588 |
| chr12:96770247-96770642 | 0.076611401 | 0.861465715 | 0.948188956 |
| chr9:86276110-86276310 | -0.091449061 | 0.861537083 | 0.948211196 |
| chr1:89820567-89822014 | 0.056614379 | 0.861549241 | 0.948211196 |
| chr12:113190771-113191127 | -0.06947235 | 0.861553163 | 0.948211196 |
| chr10:124387458-124387738 | 0.075860833 | 0.861562982 | 0.948211196 |
| chr17:48307877-48308089 | 0.091259162 | 0.8615727 | 0.948211196 |
| chr10:110168502-110168859 | -0.074650304 | 0.861574343 | 0.948211196 |
| chr4:80202131-80202543 | 0.067542223 | 0.861586207 | 0.948211196 |
| chr20:60851775-60852127 | 0.069499705 | 0.861589634 | 0.948211196 |
| chr1:179081549-179083091 | -0.054815517 | 0.861597297 | 0.948211196 |
| chr5:108697440-108697640 | 0.096632201 | 0.861632813 | 0.948213336 |
| chr5:135821477-135821818 | -0.074798383 | 0.861660542 | 0.948213336 |
| chr18:50644893-50645159 | -0.076747548 | 0.861674564 | 0.948213336 |
| chr17:49580133-49580461 | 0.066131589 | 0.861676313 | 0.948213336 |
| chr7:100177156-100177864 | -0.058363579 | 0.861677776 | 0.948213336 |
| chr4:112231257-112232611 | -0.054731021 | 0.86167783 | 0.948213336 |
| chr12:46160631-46161034 | 0.072232699 | 0.861685868 | 0.948213336 |
| chr1:38028572-38028813 | 0.082030296 | 0.861711588 | 0.94821375 |
| chr1:54764087-54764965 | -0.055712173 | 0.861722805 | 0.94821375 |
| chr8:118004526-118004870 | -0.072993617 | 0.86172337 | 0.94821375 |
| chr7:32062141-32062492 | -0.065356259 | 0.861736939 | 0.948215063 |
| chr20:62395624-62395900 | -0.066002668 | 0.86178201 | 0.94825104 |
| chr6:152057942-152058188 | -0.088641909 | 0.861891398 | 0.948319457 |
| chr8:85106594-85108149 | -0.055693242 | 0.861892297 | 0.948319457 |
| chr1:168321922-168322187 | -0.076368656 | 0.861893124 | 0.948319457 |
| chr2:37671181-37671381 | -0.077556952 | 0.861902225 | 0.948319457 |
| chr5:141524621-141525306 | 0.059017305 | 0.861906072 | 0.948319457 |
| chr1:61547538-61547829 | -0.089598794 | 0.861926299 | 0.948328095 |
| chr5:168211286-168211514 | -0.083781493 | 0.861947832 | 0.948328433 |
| chr1:48202187-48202387 | -0.08749124 | 0.86195667 | 0.948328433 |
| chr1:160061442-160061714 | -0.071895991 | 0.861963736 | 0.948328433 |
| chr10:99682781-99683081 | -0.076477446 | 0.861981882 | 0.948332194 |
| chr11:393644-394329 | -0.05887513 | 0.862012841 | 0.948332194 |
| chr6:122491986-122492223 | 0.071407816 | 0.86202586 | 0.948332194 |
| chr11:111230377-111230766 | 0.067494601 | 0.862030969 | 0.948332194 |

| | | | |
|---|---|---|---|
| chr8:123086378-123086587 | -0.082496776 | 0.862072416 | 0.948332194 |
| chr12:30844596-30844796 | -0.088551011 | 0.862074881 | 0.948332194 |
| chr1:198228918-198229462 | 0.059131011 | 0.862077159 | 0.948332194 |
| chr6:39169667-39169894 | 0.083210425 | 0.862129302 | 0.948332194 |
| chr6:154852513-154853219 | 0.062062833 | 0.862132894 | 0.948332194 |
| chr1:45508440-45508690 | 0.079005993 | 0.862136145 | 0.948332194 |
| chr2:134189142-134189484 | -0.080397822 | 0.862140423 | 0.948332194 |
| chr17:17205369-17205653 | 0.068508655 | 0.862146453 | 0.948332194 |
| chr5:63401468-63401759 | -0.074990239 | 0.862149387 | 0.948332194 |
| chr8:141160986-141161290 | 0.072749851 | 0.862155734 | 0.948332194 |
| chr7:117827956-117828249 | 0.078128743 | 0.862163486 | 0.948332194 |
| chr11:73478813-73479110 | 0.082624154 | 0.8621693 | 0.948332194 |
| chr20:5127677-5127877 | -0.07922914 | 0.862177559 | 0.948332194 |
| chr14:103865090-103865290 | 0.092104779 | 0.862349065 | 0.948507222 |
| chr6:10465713-10465913 | -0.097707815 | 0.86239687 | 0.948546186 |
| chr6:75243267-75244220 | -0.055416143 | 0.862447839 | 0.948572747 |
| chr7:901178-902093 | -0.0554426 | 0.862456746 | 0.948572747 |
| chr1:43853183-43853397 | -0.094216702 | 0.862458158 | 0.948572747 |
| chr17:78852720-78853012 | -0.073328003 | 0.862528202 | 0.948617971 |
| chr1:117842721-117843172 | 0.063987192 | 0.862550222 | 0.948617971 |
| chr9:133355944-133356228 | -0.067488476 | 0.862560494 | 0.948617971 |
| chr9:15552667-15553754 | -0.055678752 | 0.8625609 | 0.948617971 |
| chr2:164538297-164538597 | -0.068584434 | 0.862564184 | 0.948617971 |
| chr2:74019551-74019751 | 0.090369107 | 0.862581037 | 0.948617971 |
| chr20:18198625-18198845 | 0.094888305 | 0.86258594 | 0.948617971 |
| chr13:99970192-99970407 | 0.083971768 | 0.862621877 | 0.948643876 |
| chr5:160315976-160316267 | -0.069023637 | 0.862636085 | 0.948645886 |
| chr9:119868697-119868950 | 0.086644728 | 0.862688935 | 0.94869039 |
| chr11:12109948-12110148 | -0.077111931 | 0.862739897 | 0.948732815 |
| chr19:38122723-38123065 | -0.078509911 | 0.862758598 | 0.948738107 |
| chr17:75033669-75034700 | 0.054738014 | 0.862805119 | 0.948738107 |
| chr20:19903280-19903480 | 0.092615707 | 0.862808241 | 0.948738107 |
| chr4:134935405-134935605 | 0.092615707 | 0.862808241 | 0.948738107 |
| chr3:54891451-54892027 | -0.060659231 | 0.86281217 | 0.948738107 |
| chr10:95264200-95264409 | 0.084319364 | 0.862820824 | 0.948738107 |
| chr3:138115352-138115862 | 0.061473315 | 0.862831383 | 0.948738107 |
| chr10:81286379-81286773 | 0.074838456 | 0.862925487 | 0.948815178 |
| chr22:50498181-50498879 | 0.065223148 | 0.862926242 | 0.948815178 |
| chr17:75524836-75526639 | 0.053962247 | 0.862994347 | 0.948859643 |
| chr1:224113744-224114822 | 0.055409175 | 0.863000947 | 0.948859643 |
| chr17:74888498-74888974 | 0.064796865 | 0.863003832 | 0.948859643 |
| chr13:106284108-106284789 | 0.062745736 | 0.863108676 | 0.948933893 |
| chr5:42843613-42843813 | 0.097121082 | 0.8631222 | 0.948933893 |
| chr14:74111654-74111871 | 0.088765204 | 0.863159045 | 0.948933893 |
| chr3:162887158-162887358 | 0.097863454 | 0.863160215 | 0.948933893 |
| chr1:202464178-202464379 | 0.080300111 | 0.863165097 | 0.948933893 |
| chr2:168890230-168890711 | -0.058663852 | 0.863165109 | 0.948933893 |
| chr7:102272219-102272514 | -0.070609938 | 0.863168947 | 0.948933893 |
| chr17:82636446-82637066 | -0.062402821 | 0.863219878 | 0.948933893 |
| chr20:45155755-45156050 | 0.08401071 | 0.863222407 | 0.948933893 |

| | | | |
|---|---|---|---|
| chr7:32494924-32496341 | -0.053914439 | 0.863222521 | 0.948933893 |
| chr2:106587439-106587716 | 0.072750594 | 0.863224458 | 0.948933893 |
| chr6:28747265-28748161 | -0.057174034 | 0.863246846 | 0.948933893 |
| chr4:141900813-141901033 | 0.080758352 | 0.863271621 | 0.948933893 |
| chr2:202265133-202266210 | -0.055229619 | 0.863275971 | 0.948933893 |
| chr7:66681719-66682701 | -0.0563137 | 0.863281035 | 0.948933893 |
| chr12:64824304-64825951 | 0.054453725 | 0.863285714 | 0.948933893 |
| chr15:97960159-97961493 | -0.053967107 | 0.863290689 | 0.948933893 |
| chr2:203815262-203815637 | -0.063745167 | 0.863305092 | 0.948933893 |
| chr8:87283638-87283858 | 0.08143712 | 0.863306671 | 0.948933893 |
| chr19:15563445-15563725 | -0.07655049 | 0.863325803 | 0.948941309 |
| chr3:39052258-39052572 | -0.066972614 | 0.863437456 | 0.94905042 |
| chr11:72617176-72617417 | -0.077714907 | 0.863484747 | 0.949076046 |
| chr11:75280210-75280504 | 0.065103958 | 0.86349423 | 0.949076046 |
| chr11:124622268-124623244 | 0.058038454 | 0.863501827 | 0.949076046 |
| chr12:75236023-75236697 | -0.061452908 | 0.863510317 | 0.949076046 |
| chr14:89848268-89848469 | 0.088703383 | 0.86353601 | 0.94908879 |
| chr19:47318392-47318634 | -0.096898603 | 0.86356036 | 0.94908879 |
| chr16:75471738-75472120 | -0.062201802 | 0.863561201 | 0.94908879 |
| chr6:43723835-43724179 | -0.06392563 | 0.863571458 | 0.94908879 |
| chr15:42858684-42858956 | 0.075644175 | 0.863628469 | 0.949137832 |
| chr13:19794516-19795145 | -0.057223363 | 0.863663361 | 0.949162566 |
| chr2:176817352-176817552 | -0.081198245 | 0.863685412 | 0.949173186 |
| chrX:132413479-132413886 | -0.067771076 | 0.863746398 | 0.949226592 |
| chr19:29845610-29845879 | 0.063768155 | 0.863780896 | 0.949244029 |
| chr7:151519634-151520828 | 0.054282069 | 0.863813225 | 0.949244029 |
| chr19:10184569-10185202 | -0.05930199 | 0.863814429 | 0.949244029 |
| chr17:58870869-58871243 | 0.059492994 | 0.863844808 | 0.949244029 |
| chr16:88562025-88562301 | -0.069443273 | 0.863855002 | 0.949244029 |
| chr5:54671150-54671697 | -0.068297245 | 0.86388677 | 0.949244029 |
| chr16:69175927-69176173 | -0.094566091 | 0.86388751 | 0.949244029 |
| chr5:177607406-177607814 | 0.060498266 | 0.863894906 | 0.949244029 |
| chr11:61587660-61588579 | -0.056815867 | 0.863901255 | 0.949244029 |
| chr8:90203462-90203670 | -0.083878459 | 0.863913438 | 0.949244029 |
| chr4:139300806-139302593 | -0.053934734 | 0.863922094 | 0.949244029 |
| chr6:73840756-73840991 | 0.090152579 | 0.863923316 | 0.949244029 |
| chr7:3495797-3496006 | 0.090152579 | 0.863923316 | 0.949244029 |
| chrX:107672934-107673299 | -0.059585554 | 0.863981478 | 0.949293944 |
| chr7:151368107-151368522 | -0.061060147 | 0.86400458 | 0.949293944 |
| chr17:60435554-60436481 | -0.053853594 | 0.864030038 | 0.949293944 |
| chr17:28357019-28357988 | 0.055970235 | 0.864055208 | 0.949293944 |
| chr2:131775973-131776188 | -0.089078898 | 0.86406099 | 0.949293944 |
| chr19:737872-738096 | 0.090787076 | 0.86406374 | 0.949293944 |
| chr5:69024273-69024487 | 0.090787076 | 0.86406374 | 0.949293944 |
| chr19:44906229-44906778 | 0.060675241 | 0.864067859 | 0.949293944 |
| chr6:130293844-130294077 | 0.075663186 | 0.864143337 | 0.949344822 |
| chr2:15730414-15730969 | 0.06083245 | 0.864149647 | 0.949344822 |
| chr1:112674261-112675473 | 0.055637617 | 0.86415134 | 0.949344822 |
| chr2:105369831-105370212 | 0.062903376 | 0.864171129 | 0.949352951 |
| chr3:111978587-111979538 | -0.054343115 | 0.864200095 | 0.949371161 |

| | | | |
|---|---|---|---|
| chr1:28104820-28105632 | -0.058266232 | 0.864240199 | 0.949396481 |
| chr9:110183298-110183562 | 0.074544922 | 0.86426361 | 0.949396481 |
| chr11:60934063-60934263 | -0.08860558 | 0.86426824 | 0.949396481 |
| chr3:129604599-129604886 | -0.06852392 | 0.864285038 | 0.949396481 |
| chr15:91006233-91006447 | 0.080282531 | 0.864285097 | 0.949396481 |
| chr5:68450204-68450865 | 0.062291685 | 0.864313879 | 0.949414486 |
| chr15:63487344-63487615 | -0.069157399 | 0.864343896 | 0.949433847 |
| chr17:27353497-27354210 | -0.056308296 | 0.86439292 | 0.949471793 |
| chr19:45370068-45370992 | 0.056433482 | 0.864406818 | 0.949471793 |
| chrX:107825652-107827182 | 0.055218412 | 0.864416429 | 0.949471793 |
| chrX:71095681-71096585 | -0.058841143 | 0.864446184 | 0.949471793 |
| chr16:82819525-82819751 | 0.087026688 | 0.864461751 | 0.949471793 |
| chr19:17511914-17512404 | 0.065525054 | 0.864476698 | 0.949471793 |
| chr19:9674799-9675480 | 0.062350019 | 0.864478968 | 0.949471793 |
| chr1:174238935-174239845 | 0.058419373 | 0.864484916 | 0.949471793 |
| chr4:127538798-127539091 | -0.075423596 | 0.864489966 | 0.949471793 |
| chr7:50812884-50813202 | -0.066801386 | 0.86450365 | 0.949473212 |
| chr22:20957744-20958135 | 0.065649564 | 0.864541864 | 0.949487376 |
| chr16:8679242-8679536 | -0.066992288 | 0.864558819 | 0.949487376 |
| chr11:86244296-86245633 | -0.053759821 | 0.864564026 | 0.949487376 |
| chr17:66829434-66830077 | 0.058877465 | 0.864566114 | 0.949487376 |
| chr4:121277597-121277938 | 0.064188673 | 0.864603723 | 0.949500986 |
| chr17:20381354-20381565 | 0.08134291 | 0.864619044 | 0.949500986 |
| chr14:102271838-102272038 | 0.090652404 | 0.864628074 | 0.949500986 |
| chr20:56788307-56788645 | 0.060547974 | 0.864638441 | 0.949500986 |
| chr6:166097571-166098232 | 0.063083922 | 0.864647378 | 0.949500986 |
| chr20:24303489-24303789 | 0.0787748 | 0.864666071 | 0.949500986 |
| chr11:67442957-67443869 | 0.056362004 | 0.864672726 | 0.949500986 |
| chr17:7004812-7005012 | 0.080623863 | 0.864677643 | 0.949500986 |
| chr16:84699402-84700540 | -0.054864552 | 0.864740433 | 0.94954738 |
| chr2:62409442-62409966 | -0.065194877 | 0.864744677 | 0.94954738 |
| chr5:173166350-173166581 | 0.074310354 | 0.864761528 | 0.949549835 |
| chr5:105943271-105943555 | -0.085301451 | 0.864771698 | 0.949549835 |
| chr2:126634434-126634761 | -0.079878479 | 0.864784601 | 0.949550396 |
| chr19:13952023-13952820 | 0.056471787 | 0.864823911 | 0.949558324 |
| chr8:123740935-123741135 | 0.08497264 | 0.864840029 | 0.949558324 |
| chr5:4862569-4862848 | 0.078690188 | 0.864840684 | 0.949558324 |
| chr1:109776859-109777212 | 0.060820965 | 0.864852384 | 0.949558324 |
| chr12:43805404-43806717 | -0.055193499 | 0.864861584 | 0.949558324 |
| chr11:63509000-63509238 | -0.089006959 | 0.86489911 | 0.949558324 |
| chr12:18224786-18225008 | -0.089006959 | 0.86489911 | 0.949558324 |
| chr12:85403553-85403834 | -0.089006959 | 0.86489911 | 0.949558324 |
| chr1:121016669-121017081 | 0.060159668 | 0.864903357 | 0.949558324 |
| chr17:42388165-42389015 | -0.054726869 | 0.864916795 | 0.949559472 |
| chr21:6565307-6565545 | -0.086638085 | 0.864984841 | 0.949620571 |
| chr14:70873609-70873900 | -0.077456548 | 0.865041425 | 0.949624303 |
| chr16:30927959-30928475 | 0.059402607 | 0.865044164 | 0.949624303 |
| chr1:212721486-212721940 | -0.06535514 | 0.865044477 | 0.949624303 |
| chr17:76000463-76000678 | -0.074417699 | 0.865051155 | 0.949624303 |
| chr2:224369096-224369296 | 0.096085045 | 0.865052246 | 0.949624303 |

| | | | |
|---|---|---|---|
| chr13:106349112-106349629 | 0.066055914 | 0.865065495 | 0.949624303 |
| chr2:211835765-211835980 | -0.08651785 | 0.865087389 | 0.949624303 |
| chr20:16279370-16279570 | -0.08651785 | 0.865087389 | 0.949624303 |
| chr3:38031167-38031586 | 0.067961647 | 0.865101327 | 0.949625998 |
| chr19:48468788-48469491 | -0.054548325 | 0.865136676 | 0.949649505 |
| chr10:69237600-69237840 | -0.073463803 | 0.865156635 | 0.949649505 |
| chr8:133062701-133062901 | -0.084774946 | 0.865159924 | 0.949649505 |
| chr5:122492735-122493027 | -0.072941094 | 0.865179933 | 0.949657863 |
| chrX:47129049-47129836 | -0.056047264 | 0.865356703 | 0.949835682 |
| chr9:83965069-83965269 | -0.083569171 | 0.865366726 | 0.949835682 |
| chr3:25033403-25033786 | -0.066416235 | 0.865403125 | 0.949849221 |
| chr4:82721894-82722159 | -0.073939746 | 0.865403855 | 0.949849221 |
| chrX:149493734-149493975 | 0.071329623 | 0.865429438 | 0.949863695 |
| chr14:90454093-90454797 | -0.055352775 | 0.865446333 | 0.949868632 |
| chr7:151057833-151058408 | 0.05473391 | 0.865466218 | 0.94987685 |
| chr3:145197088-145197363 | 0.080059774 | 0.865496529 | 0.949887665 |
| chr15:55064028-55064281 | 0.081129604 | 0.865500865 | 0.949887665 |
| chr4:106935520-106935744 | -0.088744798 | 0.865553419 | 0.949918979 |
| chr17:59004717-59005616 | 0.053857958 | 0.865554193 | 0.949918979 |
| chr10:128263945-128264166 | -0.081595545 | 0.865567364 | 0.949919126 |
| chr7:35916984-35917304 | -0.064060031 | 0.865587341 | 0.949919126 |
| chr20:25065855-25066082 | -0.075751456 | 0.865591519 | 0.949919126 |
| chr12:93299581-93299839 | -0.084591424 | 0.865656838 | 0.949977202 |
| chr4:139555188-139556995 | 0.053310836 | 0.865692204 | 0.949999036 |
| chr4:48341841-48342229 | 0.06010255 | 0.86570153 | 0.949999036 |
| chr20:23421094-23421945 | 0.056037241 | 0.865744841 | 0.950031367 |
| chr13:34162152-34162491 | 0.070095535 | 0.86575579 | 0.950031367 |
| chr22:41444380-41444683 | -0.061197423 | 0.865773249 | 0.950036918 |
| chr14:90491485-90491807 | -0.068960479 | 0.865790822 | 0.950042596 |
| chr7:149261651-149262703 | -0.054977061 | 0.865818548 | 0.950046755 |
| chr22:37791829-37792185 | 0.07004937 | 0.86581941 | 0.950046755 |
| chr1:24562687-24562988 | 0.075769131 | 0.865845108 | 0.950055753 |
| chr9:127869919-127870229 | 0.068017786 | 0.865852409 | 0.950055753 |
| chr1:145885720-145886321 | -0.055248526 | 0.865868078 | 0.950056797 |
| chr1:66995985-66996394 | -0.0653933 | 0.865878159 | 0.950056797 |
| chr7:151695009-151695615 | -0.061615366 | 0.865918003 | 0.950086909 |
| chr17:32319471-32319760 | -0.071967445 | 0.865932071 | 0.95008874 |
| chr7:38748056-38748304 | 0.07270866 | 0.865964106 | 0.950110283 |
| chr14:64234902-64235242 | -0.068503627 | 0.865983536 | 0.950117997 |
| chr6:50952148-50952402 | -0.07643754 | 0.866023454 | 0.95013528 |
| chr12:108691440-108692100 | -0.055059975 | 0.86602409 | 0.95013528 |
| chr6:107218350-107218550 | 0.090100645 | 0.866037149 | 0.950136003 |
| chr18:11136351-11136583 | -0.078483565 | 0.866091522 | 0.950171668 |
| chr10:33801661-33801995 | 0.07069109 | 0.866102871 | 0.950171668 |
| chr1:236281358-236282658 | -0.053963193 | 0.866116529 | 0.950171668 |
| chr1:41661561-41661823 | -0.072681373 | 0.866120841 | 0.950171668 |
| chr1:236388841-236389822 | 0.055160946 | 0.866138782 | 0.950171668 |
| chr14:92852801-92853001 | -0.087050072 | 0.866159565 | 0.950171668 |
| chr14:90301264-90301500 | -0.083352995 | 0.866169576 | 0.950171668 |
| chr2:8319692-8319894 | 0.089476959 | 0.866179563 | 0.950171668 |

| | | | |
|---|---|---|---|
| chr10:132155807-132156134 | 0.077380858 | 0.866181264 | 0.950171668 |
| chr1:198936903-198937103 | 0.065405163 | 0.866197896 | 0.950176309 |
| chr17:31936544-31937377 | 0.059928228 | 0.866213981 | 0.95018035 |
| chr21:21163958-21164224 | -0.075421743 | 0.866227252 | 0.950181305 |
| chr14:102472839-102473200 | 0.068439296 | 0.86628556 | 0.95023166 |
| chr17:81512681-81513150 | -0.057364982 | 0.866334801 | 0.950260554 |
| chr5:39283221-39283632 | 0.064602687 | 0.866344462 | 0.950260554 |
| chr9:4924346-4924559 | -0.095730079 | 0.866376025 | 0.950260554 |
| chr5:112186446-112186659 | -0.080771098 | 0.866377944 | 0.950260554 |
| chr3:128052852-128053068 | 0.075065528 | 0.866384878 | 0.950260554 |
| chr21:17105991-17106350 | 0.064643168 | 0.866386312 | 0.950260554 |
| chr3:184174470-184175565 | 0.054147898 | 0.866403803 | 0.95026274 |
| chr3:156082281-156082614 | 0.071260577 | 0.866422092 | 0.95026274 |
| chr16:82112068-82112508 | 0.071629905 | 0.866426179 | 0.95026274 |
| chr2:55779475-55780234 | 0.057696503 | 0.866437913 | 0.95026274 |
| chr1:110214623-110214827 | -0.074747078 | 0.866474911 | 0.950282884 |
| chr17:47895531-47896747 | -0.052726234 | 0.866481085 | 0.950282884 |
| chr9:38371600-38371800 | -0.085041073 | 0.866518083 | 0.950307563 |
| chr9:134135039-134136190 | 0.055648494 | 0.86653956 | 0.950307563 |
| chr8:1892492-1892759 | 0.071790514 | 0.866547078 | 0.950307563 |
| chr3:64575354-64575855 | 0.054313612 | 0.866560134 | 0.950307563 |
| chr17:7351715-7352289 | -0.058187638 | 0.866565601 | 0.950307563 |
| chrX:71365761-71366711 | 0.05410163 | 0.866587156 | 0.950317601 |
| chr5:178213845-178214045 | -0.069636521 | 0.866608174 | 0.950327048 |
| chr17:7102367-7102598 | -0.089597172 | 0.86662763 | 0.950331674 |
| chrX:69133835-69134319 | 0.06143039 | 0.866637198 | 0.950331674 |
| chr7:2757920-2758545 | 0.058203683 | 0.866693912 | 0.950346808 |
| chr1:10796369-10796897 | 0.060452607 | 0.866719375 | 0.950346808 |
| chr15:89893619-89894544 | 0.054157572 | 0.866731279 | 0.950346808 |
| chr20:44228320-44229005 | -0.059573272 | 0.866732016 | 0.950346808 |
| chr12:11555841-11556042 | 0.08783075 | 0.866740977 | 0.950346808 |
| chr13:26719639-26719839 | 0.08783075 | 0.866740977 | 0.950346808 |
| chr2:48806360-48806628 | -0.090684594 | 0.866741535 | 0.950346808 |
| chr20:45803837-45804037 | 0.08974462 | 0.866757028 | 0.950346808 |
| chr22:41946310-41947282 | -0.055618361 | 0.866778531 | 0.950346808 |
| chr17:3965074-3965301 | 0.073703136 | 0.866779517 | 0.950346808 |
| chr4:122716909-122717218 | 0.067165719 | 0.866787433 | 0.950346808 |
| chr1:108963016-108963885 | -0.054638736 | 0.866828454 | 0.950378185 |
| chr6:41895190-41895801 | 0.057028306 | 0.86688335 | 0.950396056 |
| chr20:25187602-25187980 | 0.072158493 | 0.866907016 | 0.950396056 |
| chr8:90676232-90676524 | 0.06980663 | 0.866908336 | 0.950396056 |
| chr9:100411354-100412235 | 0.054643835 | 0.866916196 | 0.950396056 |
| chr20:63381857-63382194 | 0.065140476 | 0.866921972 | 0.950396056 |
| chr17:18033469-18033754 | 0.070114451 | 0.866922175 | 0.950396056 |
| chr15:75139127-75139434 | 0.068222869 | 0.86693158 | 0.950396056 |
| chr8:126062704-126062921 | -0.074753136 | 0.866965908 | 0.95042009 |
| chr12:78863450-78864854 | -0.0537326 | 0.86699174 | 0.950434811 |
| chr6:46035382-46035632 | -0.082752059 | 0.867058586 | 0.950473233 |
| chr7:36299993-36300340 | -0.062344362 | 0.867064735 | 0.950473233 |
| chr5:181206672-181208049 | -0.052555712 | 0.867094799 | 0.950473233 |

| | | | |
|---|---|---|---|
| chr9:113400910-113401717 | 0.053662747 | 0.867102456 | 0.950473233 |
| chr4:55361044-55361298 | -0.076263921 | 0.86711304 | 0.950473233 |
| chr4:139619697-139619926 | -0.076263921 | 0.86711304 | 0.950473233 |
| chr1:73770218-73770461 | -0.087413289 | 0.867113622 | 0.950473233 |
| chr11:64742834-64743143 | 0.064058823 | 0.867169827 | 0.950521243 |
| chr9:127586473-127586825 | -0.065678098 | 0.867196668 | 0.950537067 |
| chr5:115309821-115310073 | 0.084467952 | 0.867255748 | 0.950587549 |
| chr2:112275227-112276447 | 0.053481004 | 0.867282311 | 0.950587549 |
| chr2:70900043-70900713 | 0.055663479 | 0.867286867 | 0.950587549 |
| chr17:7478797-7479262 | -0.062091989 | 0.867301872 | 0.950587549 |
| chr4:143348253-143348490 | 0.077428597 | 0.867304756 | 0.950587549 |
| chr5:178526206-178526406 | -0.077294928 | 0.86732454 | 0.950592694 |
| chr1:193283714-193284024 | -0.070657439 | 0.867336688 | 0.950592694 |
| chr3:34283846-34284101 | -0.080898376 | 0.867346669 | 0.950592694 |
| chr7:149598959-149599272 | -0.063675669 | 0.867447342 | 0.950689432 |
| chr17:18087633-18088188 | -0.059130505 | 0.867502653 | 0.950717998 |
| chr8:125717446-125717646 | 0.077521441 | 0.867505169 | 0.950717998 |
| chr1:35883721-35884179 | -0.062683208 | 0.867512123 | 0.950717998 |
| chr20:1423754-1424155 | -0.06406483 | 0.867524283 | 0.950717998 |
| chr22:19593064-19593295 | 0.085142973 | 0.867541657 | 0.950717998 |
| chr16:72983906-72984219 | 0.066367226 | 0.867547854 | 0.950717998 |
| chr9:130681108-130681516 | 0.063524291 | 0.867622904 | 0.950780068 |
| chr8:28622211-28623225 | 0.054072169 | 0.867637403 | 0.950780068 |
| chr1:162330031-162330485 | 0.075825119 | 0.867645401 | 0.950780068 |
| chr12:116298999-116299455 | -0.062773207 | 0.867654678 | 0.950780068 |
| chr5:38441750-38442111 | 0.059413622 | 0.867666538 | 0.950780068 |
| chr22:41621580-41621898 | 0.063856652 | 0.86769285 | 0.950795303 |
| chr14:54577116-54577317 | 0.082951463 | 0.867736438 | 0.950825706 |
| chr22:30634461-30635161 | 0.056786871 | 0.867745414 | 0.950825706 |
| chr1:23484172-23484886 | -0.056114078 | 0.867862272 | 0.950932509 |
| chr12:68358698-68359168 | 0.062854503 | 0.86789324 | 0.950932509 |
| chr12:90024075-90024671 | 0.060631984 | 0.867902659 | 0.950932509 |
| chr5:129440602-129440832 | 0.080494558 | 0.867932987 | 0.950932509 |
| chr12:116520258-116520512 | -0.080361938 | 0.867957889 | 0.950932509 |
| chr20:50744149-50744949 | 0.052038178 | 0.867971163 | 0.950932509 |
| chr5:151570133-151570853 | -0.058104463 | 0.867971856 | 0.950932509 |
| chr1:54672049-54672324 | 0.06843537 | 0.867979677 | 0.950932509 |
| chr3:196754429-196755060 | 0.061551272 | 0.867990194 | 0.950932509 |
| chr3:184298668-184299781 | 0.05306816 | 0.86800981 | 0.950932509 |
| chr10:101161699-101162326 | -0.060694038 | 0.868011514 | 0.950932509 |
| chr2:226801059-226801587 | 0.056725978 | 0.868015942 | 0.950932509 |
| chr7:99551974-99552286 | -0.068721587 | 0.86801723 | 0.950932509 |
| chr1:84775106-84775392 | -0.07119118 | 0.868025544 | 0.950932509 |
| chr12:117856615-117856903 | 0.072115324 | 0.868029046 | 0.950932509 |
| chr10:106705636-106705838 | -0.091810843 | 0.868051689 | 0.950943719 |
| chr11:73585968-73586211 | -0.08145896 | 0.868074573 | 0.950955192 |
| chr4:142282936-142283339 | -0.071129525 | 0.868096385 | 0.950965491 |
| chr10:22218532-22218753 | -0.082571257 | 0.868113796 | 0.950970967 |
| chr10:120221979-120222179 | -0.085754022 | 0.868142873 | 0.950989224 |
| chr3:120108598-120108798 | -0.093507876 | 0.8681615 | 0.950994904 |

| | | | |
|---|---|---|---|
| chr2:118003340-118003718 | -0.059738554 | 0.868172882 | 0.950994904 |
| chr6:151334438-151334666 | 0.087065347 | 0.868197813 | 0.951008618 |
| chr9:134251602-134251844 | 0.088064627 | 0.868227978 | 0.951028064 |
| chrX:7191621-7192138 | -0.064941148 | 0.868273857 | 0.951064723 |
| chr1:80130892-80131211 | -0.068574517 | 0.868312025 | 0.951079729 |
| chrX:149938331-149940804 | 0.051411075 | 0.868317764 | 0.951079729 |
| chr6:15908094-15908646 | 0.059025899 | 0.868324795 | 0.951079729 |
| chr14:51087037-51087237 | 0.086711459 | 0.868363585 | 0.951093487 |
| chr1:22052407-22052896 | 0.058480277 | 0.868377284 | 0.951093487 |
| chr1:154714604-154714864 | -0.074965006 | 0.868381059 | 0.951093487 |
| chr3:57555540-57556536 | 0.055307437 | 0.868401805 | 0.951093487 |
| chr11:46413808-46414051 | -0.069098242 | 0.868404321 | 0.951093487 |
| chr3:150865049-150865667 | 0.056171156 | 0.868419979 | 0.951093487 |
| chr6:29005338-29005775 | 0.070978005 | 0.868424246 | 0.951093487 |
| chr17:2030432-2030924 | -0.058732608 | 0.868441919 | 0.951099248 |
| chr1:236169080-236169396 | 0.065724694 | 0.86852485 | 0.951176477 |
| chr1:23367846-23368602 | -0.055413372 | 0.868571457 | 0.951213924 |
| chr17:3846115-3846606 | 0.05781441 | 0.868594205 | 0.951225241 |
| chr17:41624648-41624877 | 0.08310251 | 0.868607744 | 0.951226472 |
| chr16:23460183-23460383 | -0.093069978 | 0.868702494 | 0.951316638 |
| chr15:68651203-68651412 | 0.076817829 | 0.868751669 | 0.951353018 |
| chr1:156961753-156962061 | 0.067118928 | 0.868760547 | 0.951353018 |
| chr11:119741308-119741540 | 0.079001294 | 0.868779629 | 0.951355091 |
| chr10:81082007-81082219 | -0.079206534 | 0.868787273 | 0.951355091 |
| chr5:162683365-162683620 | 0.076808036 | 0.868808897 | 0.951365174 |
| chr15:41559603-41559817 | 0.080769327 | 0.868828686 | 0.951373247 |
| chr1:203440969-203441376 | 0.059608992 | 0.868857015 | 0.951390671 |
| chr13:27339404-27339648 | 0.066529878 | 0.86890581 | 0.951430504 |
| chr2:239824901-239825108 | -0.089528721 | 0.86900895 | 0.951523644 |
| chr2:177906105-177906440 | 0.060987537 | 0.869042861 | 0.951523644 |
| chr11:72722447-72722742 | 0.064179831 | 0.869055192 | 0.951523644 |
| chr15:73225678-73225878 | -0.093640144 | 0.86906037 | 0.951523644 |
| chr7:28921800-28922088 | 0.072678241 | 0.869062101 | 0.951523644 |
| chr6:14373826-14374109 | 0.093333636 | 0.869065381 | 0.951523644 |
| chr2:11154695-11155822 | 0.052504968 | 0.869107357 | 0.95154705 |
| chr14:36105463-36105676 | -0.077484115 | 0.869111596 | 0.95154705 |
| chr9:108933443-108934955 | 0.052338841 | 0.869132146 | 0.951555952 |
| chr16:28277647-28278403 | 0.057013785 | 0.86916665 | 0.951580131 |
| chr1:247750129-247750364 | 0.078261097 | 0.869199079 | 0.951587133 |
| chr2:174157818-174158387 | -0.061363977 | 0.869209843 | 0.951587133 |
| chr16:85645020-85645576 | -0.057454139 | 0.869210303 | 0.951587133 |
| chr5:143418005-143418350 | 0.068183804 | 0.869233042 | 0.951598431 |
| chr6:692319-693671 | -0.053667388 | 0.869268754 | 0.95162393 |
| chr17:38598595-38598937 | 0.062237331 | 0.869293517 | 0.951636642 |
| chr19:10625261-10625619 | 0.062047717 | 0.869305206 | 0.951636642 |
| chr1:169107031-169107264 | -0.061312883 | 0.869342235 | 0.951657763 |
| chr17:59418781-59419032 | 0.065282868 | 0.869366283 | 0.951657763 |
| chr6:7532291-7532570 | 0.068786104 | 0.869371131 | 0.951657763 |
| chr4:82890690-82891623 | -0.053010805 | 0.869386201 | 0.951657763 |
| chr3:154683946-154684505 | -0.059986679 | 0.869397492 | 0.951657763 |

| | | | |
|---|---|---|---|
| chr1:18875045-18875528 | -0.064707745 | 0.869399021 | 0.951657763 |
| chr15:92222593-92222802 | -0.09245329 | 0.869425151 | 0.951665104 |
| chr8:38785117-38785487 | -0.072685365 | 0.869430568 | 0.951665104 |
| chr1:42757050-42757300 | 0.071613191 | 0.869446219 | 0.951668641 |
| chr5:108417283-108417646 | 0.070668223 | 0.869523173 | 0.951739276 |
| chr2:39486180-39486463 | -0.08872715 | 0.86956351 | 0.951769831 |
| chr12:57782509-57783397 | -0.05403824 | 0.869590199 | 0.951785447 |
| chr1:195209344-195209666 | -0.080341376 | 0.869606488 | 0.951785745 |
| chr7:64665806-64666606 | -0.055097949 | 0.869621787 | 0.951785745 |
| chr13:65868960-65869182 | 0.07831169 | 0.869635059 | 0.951785745 |
| chr6:43629141-43630368 | -0.051978432 | 0.869640158 | 0.951785745 |
| chr17:36591753-36592702 | -0.059335504 | 0.869665821 | 0.951793626 |
| chr7:47534759-47535023 | -0.074486321 | 0.869692138 | 0.951793626 |
| chr9:128504470-128505485 | 0.054180269 | 0.86969541 | 0.951793626 |
| chr7:1183563-1183805 | -0.069555404 | 0.869697047 | 0.951793626 |
| chr20:57809591-57810080 | 0.055750881 | 0.869728813 | 0.951814796 |
| chr4:94451647-94452510 | 0.052851222 | 0.869769679 | 0.951830735 |
| chr17:64320172-64321629 | 0.051130737 | 0.869775258 | 0.951830735 |
| chr1:197807272-197807486 | 0.075388965 | 0.869780645 | 0.951830735 |
| chr4:54232913-54233154 | -0.07175892 | 0.869798411 | 0.951836582 |
| chr12:11176935-11177180 | 0.08707009 | 0.869830874 | 0.951858513 |
| chr14:72864728-72864965 | 0.081139023 | 0.869880449 | 0.951889383 |
| chr8:134504735-134505003 | -0.074108581 | 0.86988393 | 0.951889383 |
| chr5:159316814-159317099 | 0.070085055 | 0.869928254 | 0.95191792 |
| chr17:59220231-59221333 | -0.051687461 | 0.869942036 | 0.95191792 |
| chr22:42581571-42581775 | 0.088022894 | 0.869947279 | 0.95191792 |
| chr16:10181653-10181891 | 0.071609513 | 0.869962017 | 0.951920452 |
| chr16:85717925-85718156 | 0.067898938 | 0.870020703 | 0.951954499 |
| chr2:197107293-197107514 | -0.075170263 | 0.870027955 | 0.951954499 |
| chr10:111076123-111077351 | -0.053880344 | 0.870030404 | 0.951954499 |
| chr9:104868799-104869275 | -0.061411953 | 0.870062717 | 0.95197626 |
| chr9:33524028-33524333 | -0.072728294 | 0.87011729 | 0.952022376 |
| chr22:42581916-42582148 | -0.079190794 | 0.870131794 | 0.952024651 |
| chr8:42152542-42153735 | -0.052366148 | 0.870173629 | 0.952054635 |
| chr8:27274324-27274965 | 0.056551078 | 0.870222827 | 0.952054635 |
| chr1:52842534-52843447 | -0.055118321 | 0.870231862 | 0.952054635 |
| chr1:38527306-38527762 | 0.065722177 | 0.87025059 | 0.952054635 |
| chr14:92987798-92988108 | 0.065111401 | 0.870252575 | 0.952054635 |
| chr3:194112530-194112919 | 0.065996967 | 0.870257352 | 0.952054635 |
| chr3:68972412-68972713 | -0.063961699 | 0.870258032 | 0.952054635 |
| chr10:103368347-103368976 | 0.053941171 | 0.870258602 | 0.952054635 |
| chr7:65750338-65751449 | 0.053505311 | 0.870284147 | 0.952067364 |
| chr6:27772265-27772846 | 0.056004808 | 0.870295088 | 0.952067364 |
| chr1:85024828-85025100 | -0.08036011 | 0.87033196 | 0.952092141 |
| chr6:31777026-31777430 | -0.061686727 | 0.870342589 | 0.952092141 |
| chr1:16612959-16614012 | 0.052904424 | 0.870391578 | 0.952132138 |
| chr2:119826452-119826864 | -0.070470512 | 0.870422655 | 0.95215254 |
| chr7:148929199-148929488 | 0.076970474 | 0.870444391 | 0.952162724 |
| chr8:28092738-28093566 | -0.055967369 | 0.870468762 | 0.952170724 |
| chr8:2011575-2012096 | -0.064191334 | 0.870476559 | 0.952170724 |

| | | | |
|---|---|---|---|
| chr3:44761210-44762380 | -0.053990969 | 0.870500896 | 0.952183752 |
| chr3:189304379-189305309 | 0.052294874 | 0.870542998 | 0.952187345 |
| chr15:74621133-74621472 | 0.059914177 | 0.870543518 | 0.952187345 |
| chr6:107489739-107490124 | 0.065252827 | 0.870545059 | 0.952187345 |
| chr17:9928883-9929134 | -0.067876012 | 0.870554288 | 0.952187345 |
| chr13:45813104-45813319 | 0.077745099 | 0.87057326 | 0.952187345 |
| chr2:198775751-198775976 | -0.090216598 | 0.870578743 | 0.952187345 |
| chr14:65940115-65940356 | 0.072456284 | 0.870591917 | 0.952188161 |
| chr1:160997296-160997673 | -0.063229669 | 0.870655177 | 0.952243758 |
| chr2:28286443-28286696 | 0.080611901 | 0.870690354 | 0.952268638 |
| chr13:60618177-60618458 | 0.069880565 | 0.870740994 | 0.95231043 |
| chr19:33905732-33906503 | -0.05633468 | 0.870767513 | 0.952318926 |
| chr17:58959992-58960667 | -0.051963549 | 0.87077362 | 0.952318926 |
| chr3:11545103-11545408 | 0.070775546 | 0.870793987 | 0.952320072 |
| chr16:11192315-11192515 | -0.083232275 | 0.870799525 | 0.952320072 |
| chr3:184345676-184346086 | 0.056543707 | 0.870835611 | 0.952345943 |
| chr1:115138759-115139407 | 0.054893194 | 0.870893086 | 0.952395205 |
| chr1:106401060-106401276 | 0.064720241 | 0.870928949 | 0.952411433 |
| chr4:54971429-54971934 | -0.064088598 | 0.870932785 | 0.952411433 |
| chr15:100959209-100959432 | 0.079025246 | 0.871027481 | 0.952501394 |
| chr17:29639363-29639825 | 0.063198787 | 0.871048257 | 0.952503492 |
| chr2:24920094-24920861 | -0.055272946 | 0.871054262 | 0.952503492 |
| chr12:48716415-48717338 | 0.052334504 | 0.871099581 | 0.952514936 |
| chr4:118746370-118746822 | -0.065563874 | 0.87113325 | 0.952514936 |
| chr15:45802113-45802318 | 0.085001297 | 0.871136975 | 0.952514936 |
| chrX:107442883-107443363 | -0.057538656 | 0.871138567 | 0.952514936 |
| chr3:195747745-195748218 | -0.058734318 | 0.871142932 | 0.952514936 |
| chr4:75723960-75725377 | -0.051463895 | 0.871143695 | 0.952514936 |
| chr2:170815740-170816052 | -0.06152312 | 0.871167782 | 0.952514936 |
| chr11:102268300-102269041 | -0.053543246 | 0.871173253 | 0.952514936 |
| chr3:126973043-126973287 | 0.075639988 | 0.87117661 | 0.952514936 |
| chr2:27371283-27371483 | 0.067545041 | 0.871191523 | 0.95251765 |
| chr17:58351718-58352817 | -0.051102565 | 0.871224421 | 0.952533226 |
| chr4:99088187-99089621 | -0.052041061 | 0.871230633 | 0.952533226 |
| chr13:50320319-50320539 | 0.075998394 | 0.871253661 | 0.952540908 |
| chr19:18986838-18987117 | 0.079316379 | 0.871262523 | 0.952540908 |
| chr1:205894138-205894745 | 0.056473605 | 0.871308707 | 0.952559919 |
| chrX:38654324-38654633 | -0.077529133 | 0.871321538 | 0.952559919 |
| chr3:129439754-129440719 | 0.051261302 | 0.871355454 | 0.952559919 |
| chr9:112867158-112867883 | 0.055998865 | 0.871358246 | 0.952559919 |
| chrX:107449745-107449954 | 0.073250425 | 0.871360241 | 0.952559919 |
| chrX:135985224-135985855 | 0.058814038 | 0.871361201 | 0.952559919 |
| chr14:55191025-55192149 | 0.051560587 | 0.871381711 | 0.952559919 |
| chr16:88634053-88634674 | -0.054607912 | 0.871387867 | 0.952559919 |
| chr22:33000844-33001053 | -0.082049704 | 0.871403867 | 0.952559919 |
| chr12:98698701-98699185 | 0.060839015 | 0.871422875 | 0.952559919 |
| chr3:165210218-165210447 | -0.072912999 | 0.87142744 | 0.952559919 |
| chr9:33328840-33329599 | 0.055936604 | 0.871429095 | 0.952559919 |
| chr1:116020766-116021022 | -0.067172826 | 0.871469687 | 0.952566054 |
| chr22:45435521-45435975 | 0.067454714 | 0.871471155 | 0.952566054 |

| | | | |
|---|---|---|---|
| chr15:33547962-33548169 | 0.084692476 | 0.871472132 | 0.952566054 |
| chr16:85365232-85365469 | -0.069982732 | 0.871496602 | 0.952566054 |
| chr19:48321198-48322241 | 0.051854141 | 0.871496868 | 0.952566054 |
| chr11:40188936-40189139 | 0.079322091 | 0.871572268 | 0.95262944 |
| chr8:10425175-10425463 | -0.07129404 | 0.871579725 | 0.95262944 |
| chr17:2353147-2353401 | 0.070904266 | 0.871624393 | 0.952663811 |
| chr3:188061036-188061546 | 0.057421897 | 0.871636037 | 0.952663811 |
| chr4:41734396-41734657 | -0.084031103 | 0.871649954 | 0.952665432 |
| chr8:126061525-126062092 | 0.058853195 | 0.871684172 | 0.952689241 |
| chr2:142974235-142974722 | -0.060905753 | 0.871749351 | 0.952717601 |
| chr1:151501497-151501721 | -0.068088762 | 0.871783532 | 0.952717601 |
| chr14:49862045-49863342 | 0.050568899 | 0.871791749 | 0.952717601 |
| chr5:167245983-167246183 | -0.083975495 | 0.871808692 | 0.952717601 |
| chrX:129347216-129347490 | 0.083728977 | 0.87181852 | 0.952717601 |
| chr20:38730948-38731823 | 0.055249752 | 0.871826151 | 0.952717601 |
| chr18:36187074-36187636 | 0.058465029 | 0.871826519 | 0.952717601 |
| chr10:114304606-114305111 | 0.057859709 | 0.871846622 | 0.952717601 |
| chr22:22692776-22693070 | -0.070800384 | 0.871875819 | 0.952717601 |
| chr1:156445827-156446167 | -0.059372241 | 0.871892306 | 0.952717601 |
| chr11:75597755-75597967 | -0.071773669 | 0.871899582 | 0.952717601 |
| chr3:156337459-156337725 | -0.078826429 | 0.871912683 | 0.952717601 |
| chr1:149812061-149812793 | -0.052681299 | 0.871925392 | 0.952717601 |
| chr22:21629852-21630295 | 0.055848266 | 0.871929405 | 0.952717601 |
| chr1:40161063-40162048 | 0.052999988 | 0.871930581 | 0.952717601 |
| chr7:45511796-45512041 | -0.076905506 | 0.871952557 | 0.952717601 |
| chr9:99104315-99105251 | -0.052983847 | 0.871955679 | 0.952717601 |
| chr7:2873695-2874507 | -0.054482675 | 0.871956263 | 0.952717601 |
| chr2:73163763-73164116 | -0.060686808 | 0.871964784 | 0.952717601 |
| chr16:87383665-87384570 | 0.051361314 | 0.871992053 | 0.952717601 |
| chr12:125080019-125080219 | -0.081079928 | 0.872003865 | 0.952717601 |
| chr16:66830388-66831197 | 0.055519584 | 0.872043936 | 0.952717601 |
| chr1:198188325-198188574 | 0.074132625 | 0.872048429 | 0.952717601 |
| chr1:37514638-37515304 | 0.054926994 | 0.872052784 | 0.952717601 |
| chr6:44127234-44127863 | 0.053875975 | 0.872059682 | 0.952717601 |
| chr3:122383873-122384682 | 0.051438129 | 0.872068472 | 0.952717601 |
| chr3:190321856-190322647 | -0.055680674 | 0.87207322 | 0.952717601 |
| chr1:24415334-24415841 | -0.055550455 | 0.872101769 | 0.952717601 |
| chr1:44673967-44674940 | -0.051164638 | 0.872104978 | 0.952717601 |
| chr10:88118292-88118539 | -0.075107045 | 0.8721082 | 0.952717601 |
| chr15:75451544-75452340 | 0.050784414 | 0.872133041 | 0.952717601 |
| chr13:97849350-97849557 | 0.077555027 | 0.872141627 | 0.952717601 |
| chr14:64812471-64812764 | 0.061862353 | 0.872166719 | 0.952717601 |
| chr16:88332580-88332929 | -0.06510933 | 0.872169147 | 0.952717601 |
| chr17:19533447-19533944 | 0.065207801 | 0.87217462 | 0.952717601 |
| chr1:42846197-42846800 | -0.054888149 | 0.872178966 | 0.952717601 |
| chr20:129886-130285 | 0.070205573 | 0.87218115 | 0.952717601 |
| chr3:148698072-148698272 | 0.078579276 | 0.872192432 | 0.952717601 |
| chr1:214628212-214628412 | -0.087506376 | 0.872210766 | 0.952717601 |
| chr13:67061866-67062097 | -0.087506376 | 0.872210766 | 0.952717601 |
| chr15:34987865-34988900 | 0.052082345 | 0.872219914 | 0.952717601 |

| | | | |
|---|---|---|---|
| chr1:218469932-218470197 | -0.070262424 | 0.87227779 | 0.952763012 |
| chr3:129343689-129344577 | -0.05354268 | 0.872297675 | 0.952763012 |
| chrX:49268863-49270312 | -0.050620446 | 0.872308199 | 0.952763012 |
| chr7:5407410-5407610 | -0.071085217 | 0.872311227 | 0.952763012 |
| chr1:155645173-155645373 | 0.08723119 | 0.87234733 | 0.952775282 |
| chr11:48183406-48183606 | 0.08723119 | 0.87234733 | 0.952775282 |
| chr17:352709-352965 | 0.074202852 | 0.872400143 | 0.952810031 |
| chr1:151473227-151473729 | -0.056949799 | 0.872417217 | 0.952810031 |
| chr5:19987732-19988827 | -0.052973341 | 0.872425783 | 0.952810031 |
| chr13:113731300-113731500 | 0.080620819 | 0.872440303 | 0.952810031 |
| chr1:76687212-76687816 | -0.055462646 | 0.872441322 | 0.952810031 |
| chr3:111895168-111895404 | 0.075487828 | 0.872458842 | 0.952815584 |
| chr10:60473649-60473869 | 0.090570404 | 0.872497412 | 0.952839942 |
| chr7:75993810-75994096 | -0.062736827 | 0.872508437 | 0.952839942 |
| chr10:51041680-51041880 | 0.094664568 | 0.872519668 | 0.952839942 |
| chr1:36960966-36961236 | 0.080101083 | 0.872530888 | 0.952839942 |
| chr4:31674297-31674680 | 0.06415449 | 0.872560891 | 0.952859126 |
| chr5:140391769-140392038 | -0.071270116 | 0.872655002 | 0.952941857 |
| chr4:124863151-124863356 | -0.079749514 | 0.872661524 | 0.952941857 |
| chr20:38290948-38291555 | 0.058264006 | 0.872704146 | 0.952974819 |
| chr6:86958919-86959119 | 0.091234254 | 0.872748346 | 0.953009503 |
| chr9:129697056-129697502 | 0.063991533 | 0.872825692 | 0.953077441 |
| chr1:106258253-106258635 | -0.057544388 | 0.87283544 | 0.953077441 |
| chr7:35694252-35695795 | 0.050421042 | 0.872855794 | 0.953086083 |
| chr6:102114453-102114789 | -0.066057678 | 0.872876958 | 0.953095611 |
| chr13:49669815-49670035 | 0.074779761 | 0.8728941 | 0.953100746 |
| chr1:145558913-145559182 | 0.081961914 | 0.872932486 | 0.953106709 |
| chr20:62794023-62794661 | -0.051765656 | 0.872939983 | 0.953106709 |
| chr8:88110865-88111491 | -0.056962632 | 0.872952677 | 0.953106709 |
| chr12:132159431-132159728 | -0.068500253 | 0.872959724 | 0.953106709 |
| chr5:76137996-76138196 | 0.082529061 | 0.872961756 | 0.953106709 |
| chr17:62065614-62066456 | -0.050778739 | 0.872975269 | 0.95310788 |
| chr5:95822225-95822505 | -0.063677491 | 0.873002276 | 0.953123786 |
| chr8:133360034-133360345 | 0.061279476 | 0.873016731 | 0.953125987 |
| chr4:169941096-169941552 | 0.057113264 | 0.873030879 | 0.953127851 |
| chr12:109264162-109264519 | 0.060586549 | 0.873064942 | 0.953145094 |
| chr22:40732686-40733054 | -0.064358997 | 0.873083911 | 0.953145094 |
| chr11:134139856-134140140 | -0.067473888 | 0.873083991 | 0.953145094 |
| chrX:95903650-95904036 | -0.06913335 | 0.873146598 | 0.953199861 |
| chr6:131534175-131534471 | -0.066700416 | 0.873169904 | 0.953200312 |
| chr2:121529921-121531841 | 0.050122719 | 0.873171892 | 0.953200312 |
| chr2:219245088-219245620 | -0.055656572 | 0.873189511 | 0.953205966 |
| chr5:5679757-5680004 | 0.081101772 | 0.87321123 | 0.953216094 |
| chr14:90554398-90554668 | 0.065771026 | 0.873252847 | 0.953237589 |
| chr18:79046893-79047113 | 0.0884569 | 0.873255802 | 0.953237589 |
| chr13:45120105-45121200 | -0.052072597 | 0.873281861 | 0.953249542 |
| chr18:70490475-70490699 | 0.08074098 | 0.873309976 | 0.953249542 |
| chr3:5806306-5806539 | 0.08074098 | 0.873309976 | 0.953249542 |
| chr20:62860414-62861024 | -0.055537846 | 0.873327696 | 0.953249542 |
| chr6:8983197-8983416 | 0.072238411 | 0.873328957 | 0.953249542 |

| | | | |
|---|---|---|---|
| chr12:132084818-132085029 | 0.068070997 | 0.873368866 | 0.953270175 |
| chr2:120279105-120279339 | 0.074699974 | 0.873374 | 0.953270175 |
| chr5:146049488-146049970 | -0.058880317 | 0.873396055 | 0.953270175 |
| chr13:99981160-99982057 | -0.054183709 | 0.873397625 | 0.953270175 |
| chr17:66673910-66674365 | 0.055643569 | 0.873421257 | 0.953282389 |
| chr8:122925604-122925877 | -0.064602586 | 0.873460485 | 0.953305678 |
| chr15:41018470-41018917 | 0.057134479 | 0.873467478 | 0.953305678 |
| chr22:31749726-31750410 | 0.05276696 | 0.873487305 | 0.953305985 |
| chr11:66842983-66843915 | -0.051507791 | 0.873492643 | 0.953305985 |
| chrX:54826085-54826364 | 0.072562349 | 0.873528637 | 0.953331689 |
| chr14:94670795-94671039 | 0.074414231 | 0.873567456 | 0.953347408 |
| chr2:169584166-169585114 | -0.052259734 | 0.873567924 | 0.953347408 |
| chr3:163737778-163738103 | -0.070369554 | 0.873616894 | 0.953379281 |
| chr8:107908382-107908671 | -0.066668818 | 0.873622016 | 0.953379281 |
| chr10:104297889-104298105 | -0.076942203 | 0.873655772 | 0.953402541 |
| chr8:117474332-117474592 | 0.073683943 | 0.873706649 | 0.953444482 |
| chr9:93304855-93305124 | 0.068425475 | 0.873719478 | 0.953444903 |
| chr5:114360993-114361504 | -0.055704231 | 0.873759817 | 0.953466963 |
| chr6:170208918-170209157 | 0.07298925 | 0.873764582 | 0.953466963 |
| chr15:40232290-40232551 | -0.08330608 | 0.873873182 | 0.95356222 |
| chr18:65036982-65037402 | 0.070429762 | 0.873876765 | 0.95356222 |
| chr17:1650031-1650231 | 0.071184384 | 0.873933631 | 0.953584356 |
| chr1:226828082-226828455 | 0.060694836 | 0.873938341 | 0.953584356 |
| chr19:41570676-41570964 | -0.066058508 | 0.873941719 | 0.953584356 |
| chr1:19241392-19241643 | 0.072795101 | 0.873969947 | 0.953584356 |
| chr14:103693532-103693799 | -0.06334923 | 0.873985455 | 0.953584356 |
| chr8:30339452-30340344 | -0.05359437 | 0.873985473 | 0.953584356 |
| chrX:9464482-9465183 | -0.054340677 | 0.874002396 | 0.953584356 |
| chr7:34463868-34464211 | 0.063006721 | 0.874021051 | 0.953584356 |
| chr6:85491277-85491477 | 0.082051595 | 0.874030292 | 0.953584356 |
| chr1:199242493-199242952 | -0.061133239 | 0.874031231 | 0.953584356 |
| chr18:9643637-9643946 | -0.079811581 | 0.8740464 | 0.953584356 |
| chr8:42385456-42385690 | 0.085672375 | 0.874070533 | 0.953584356 |
| chr6:150264467-150264679 | 0.083883842 | 0.874089023 | 0.953584356 |
| chr17:47531595-47532221 | -0.057401074 | 0.874092017 | 0.953584356 |
| chr17:74332574-74332774 | -0.080777913 | 0.874096011 | 0.953584356 |
| chr16:19086399-19087004 | 0.053582028 | 0.874096177 | 0.953584356 |
| chr1:107750432-107750662 | -0.073295591 | 0.874144106 | 0.953592581 |
| chr1:39799196-39800182 | -0.052911983 | 0.874162518 | 0.953592581 |
| chr2:108580270-108580470 | -0.085858382 | 0.874170448 | 0.953592581 |
| chr17:72421203-72421562 | 0.062829153 | 0.87417428 | 0.953592581 |
| chr19:47079423-47079823 | 0.058561421 | 0.87417699 | 0.953592581 |
| chr15:92226111-92226352 | 0.070130609 | 0.874178389 | 0.953592581 |
| chr3:95154698-95155215 | -0.055229014 | 0.874220553 | 0.953604561 |
| chr15:78075885-78076141 | -0.062567421 | 0.874223307 | 0.953604561 |
| chr4:119626977-119627507 | -0.055308638 | 0.874226707 | 0.953604561 |
| chr11:4186982-4187477 | 0.057564365 | 0.874246228 | 0.953605718 |
| chr20:34302655-34303604 | 0.052082314 | 0.87428372 | 0.953605718 |
| chr8:144684098-144684661 | 0.053349457 | 0.874291543 | 0.953605718 |
| chr16:27779957-27780233 | 0.069699272 | 0.874307943 | 0.953605718 |

| | | | |
|---|---|---|---|
| chr3:164634774-164635110 | -0.07417115 | 0.874315054 | 0.953605718 |
| chr14:37529714-37529937 | 0.074776091 | 0.874319435 | 0.953605718 |
| chr9:88578528-88578965 | -0.060284566 | 0.874324598 | 0.953605718 |
| chr2:165804802-165805087 | -0.066000947 | 0.874340593 | 0.953605718 |
| chr3:112111820-112112139 | 0.057965315 | 0.874341138 | 0.953605718 |
| chr19:9054690-9055238 | -0.05637873 | 0.874363415 | 0.953605718 |
| chr4:110198930-110199201 | -0.063464499 | 0.87437393 | 0.953605718 |
| chr3:162863587-162863889 | -0.063017368 | 0.874380408 | 0.953605718 |
| chr4:88996091-88996417 | 0.075395675 | 0.87438956 | 0.953605718 |
| chr6:161238564-161238764 | -0.081117322 | 0.874446987 | 0.953654773 |
| chr17:57175953-57176154 | -0.078605912 | 0.874479136 | 0.953668535 |
| chr1:109960391-109960826 | 0.056237848 | 0.874493458 | 0.953668535 |
| chr1:56389874-56390230 | -0.062163038 | 0.874496945 | 0.953668535 |
| chr5:178203975-178204839 | -0.051047887 | 0.874537793 | 0.953699507 |
| chr17:77430117-77430331 | 0.070343312 | 0.874562973 | 0.953701575 |
| chr6:146543134-146544566 | 0.050912846 | 0.874573519 | 0.953701575 |
| chr10:18406216-18406479 | 0.072555104 | 0.874593541 | 0.953701575 |
| chr20:46939067-46939313 | -0.065985448 | 0.874625295 | 0.953701575 |
| chr6:166998924-166999830 | -0.051817393 | 0.874629465 | 0.953701575 |
| chr8:102738602-102739107 | 0.053586562 | 0.874630001 | 0.953701575 |
| chr15:101567520-101567720 | 0.074110568 | 0.874632821 | 0.953701575 |
| chr8:123381240-123381546 | -0.067746012 | 0.874639264 | 0.953701575 |
| chr15:75024220-75024440 | -0.068106763 | 0.874675075 | 0.953724087 |
| chr19:49676357-49677373 | 0.050249709 | 0.874707176 | 0.953724087 |
| chr13:27956892-27957247 | 0.06414061 | 0.874713771 | 0.953724087 |
| chr14:100383737-100384173 | -0.057967788 | 0.874719081 | 0.953724087 |
| chr4:140154066-140154577 | -0.053425469 | 0.874734476 | 0.953724087 |
| chr11:71447891-71448132 | 0.062170454 | 0.874743924 | 0.953724087 |
| chr8:127270889-127271219 | 0.058548575 | 0.87474704 | 0.953724087 |
| chr13:37636916-37637157 | 0.081174717 | 0.874772392 | 0.953738158 |
| chr4:26861060-26861288 | -0.061942252 | 0.87480186 | 0.953748047 |
| chr10:23158079-23158290 | 0.071752404 | 0.874806358 | 0.953748047 |
| chr11:79100094-79100411 | 0.060741399 | 0.874856019 | 0.953788618 |
| chr8:102616922-102617473 | -0.054649755 | 0.874905137 | 0.953828596 |
| chr6:27177158-27177389 | -0.067858266 | 0.87496049 | 0.95387537 |
| chr8:140322240-140322453 | 0.081692083 | 0.875043555 | 0.953952355 |
| chr7:33982441-33982641 | -0.075466067 | 0.875089914 | 0.953962852 |
| chr20:16821524-16821755 | -0.07398647 | 0.87509477 | 0.953962852 |
| chr9:14353442-14353742 | 0.076528841 | 0.875099767 | 0.953962852 |
| chr2:38446460-38447005 | 0.058450771 | 0.875102985 | 0.953962852 |
| chr20:59221118-59221332 | -0.066722676 | 0.875163204 | 0.954014925 |
| chr19:15195401-15196372 | 0.052021249 | 0.87520431 | 0.954027533 |
| chr2:188130390-188130592 | -0.075213223 | 0.875223601 | 0.954027533 |
| chr17:64927480-64927864 | 0.060433538 | 0.875234237 | 0.954027533 |
| chr10:33334771-33335382 | 0.054744096 | 0.87525189 | 0.954027533 |
| chr20:17617532-17618063 | -0.055457406 | 0.875259647 | 0.954027533 |
| chr17:76353257-76354480 | 0.050549993 | 0.875276278 | 0.954027533 |
| chr3:95509957-95510205 | 0.07931863 | 0.875279334 | 0.954027533 |
| chr18:48410262-48410889 | -0.05358412 | 0.875284755 | 0.954027533 |
| chr16:20107141-20107409 | 0.07562744 | 0.875305129 | 0.954027533 |

| | | | |
|---|---|---|---|
| chr3:58177730-58178100 | 0.060790586 | 0.87531845 | 0.954027533 |
| chr17:53931881-53932115 | -0.085143802 | 0.875337966 | 0.954027533 |
| chr4:112523634-112524468 | -0.052343737 | 0.875342737 | 0.954027533 |
| chr1:224357767-224357987 | -0.068891308 | 0.875358278 | 0.954027533 |
| chr2:191299118-191299332 | 0.07844107 | 0.875382305 | 0.954027533 |
| chr8:120862360-120862569 | -0.071810338 | 0.875386551 | 0.954027533 |
| chr21:45910237-45910449 | 0.069690346 | 0.87540066 | 0.954027533 |
| chr11:118546154-118546354 | -0.085101421 | 0.875424676 | 0.954027533 |
| chr11:58900709-58901194 | 0.05842804 | 0.875425557 | 0.954027533 |
| chr2:101253156-101253562 | -0.055676058 | 0.875428919 | 0.954027533 |
| chr21:29073020-29074200 | -0.05000662 | 0.8754413 | 0.954027533 |
| chr15:98659269-98659570 | 0.057900572 | 0.875455838 | 0.954027533 |
| chr14:81453027-81453504 | 0.058051572 | 0.875456329 | 0.954027533 |
| chr19:11143119-11143583 | -0.059371376 | 0.875462286 | 0.954027533 |
| chr8:28401155-28402886 | 0.049956805 | 0.875489787 | 0.954027533 |
| chr9:131421010-131421292 | 0.065113845 | 0.875496041 | 0.954027533 |
| chr22:30423554-30424402 | 0.050483578 | 0.875499813 | 0.954027533 |
| chr4:88523234-88524165 | 0.050695928 | 0.87551095 | 0.954027533 |
| chr5:134004078-134005494 | -0.0502546 | 0.875528585 | 0.954028404 |
| chr5:41903814-41904670 | 0.0493525 | 0.875543533 | 0.954028404 |
| chr1:54996418-54996744 | 0.061653053 | 0.875553287 | 0.954028404 |
| chr12:101773635-101773999 | -0.058615774 | 0.875561554 | 0.954028404 |
| chr12:108560971-108561535 | -0.053399418 | 0.875598376 | 0.954054959 |
| chr8:47765573-47765913 | 0.067337919 | 0.875617086 | 0.954061778 |
| chr9:134099142-134099534 | -0.058762227 | 0.875637667 | 0.954062299 |
| chr11:65550938-65551226 | -0.057195176 | 0.875644838 | 0.954062299 |
| chr8:144528501-144529251 | 0.051541997 | 0.875654918 | 0.954062299 |
| chr10:29022576-29022924 | 0.062115397 | 0.875744332 | 0.954139063 |
| chr17:60599350-60601488 | -0.048301112 | 0.875764806 | 0.954139063 |
| chr14:60649100-60649901 | 0.051773152 | 0.875775141 | 0.954139063 |
| chr7:2670991-2671269 | 0.06126425 | 0.875775184 | 0.954139063 |
| chr14:73526108-73526344 | 0.07015633 | 0.875810278 | 0.954151504 |
| chr22:26589812-26590418 | 0.053992241 | 0.875813679 | 0.954151504 |
| chr1:201149510-201149797 | 0.056865895 | 0.875823961 | 0.954151504 |
| chr12:108514721-108515892 | -0.049888917 | 0.875853236 | 0.95416983 |
| chrX:46836627-46837525 | 0.051465229 | 0.875899634 | 0.954172969 |
| chr16:70460988-70461214 | 0.071560944 | 0.875899792 | 0.954172969 |
| chr19:12140256-12140591 | -0.061170556 | 0.87591314 | 0.954172969 |
| chr12:40011189-40011476 | -0.065255486 | 0.875914156 | 0.954172969 |
| chr11:3422237-3423438 | 0.04919392 | 0.875918383 | 0.954172969 |
| chr14:68987722-68988102 | 0.057721528 | 0.875944336 | 0.954184228 |
| chr10:23095152-23095907 | -0.050900353 | 0.875953624 | 0.954184228 |
| chr8:142984169-142984632 | -0.059238571 | 0.875980163 | 0.954199046 |
| chr7:44153557-44153790 | -0.066167112 | 0.875999549 | 0.954199046 |
| chr2:219387275-219388750 | -0.049036188 | 0.876004587 | 0.954199046 |
| chr16:84617283-84618398 | -0.050313755 | 0.876027616 | 0.954210152 |
| chr11:126303629-126304638 | 0.050766022 | 0.87603969 | 0.954210152 |
| chr6:150600786-150601268 | 0.061545317 | 0.876058691 | 0.954217284 |
| chr12:103504001-103504240 | -0.069987889 | 0.876102875 | 0.954251844 |
| chr5:37630764-37630976 | 0.086759838 | 0.876148939 | 0.954288452 |

| | | | |
|---|---|---|---|
| chr6:127464931-127465220 | 0.063240803 | 0.87619223 | 0.954304089 |
| chr2:98730166-98730633 | -0.058345621 | 0.876200498 | 0.954304089 |
| chr17:75130755-75131964 | 0.050446164 | 0.87620066 | 0.954304089 |
| chr6:161695026-161695314 | 0.07057491 | 0.876215009 | 0.954306152 |
| chr19:33795299-33796552 | 0.050577396 | 0.876264039 | 0.954339396 |
| chr8:22382923-22383212 | -0.082193622 | 0.876270443 | 0.954339396 |
| chr6:106223384-106223616 | -0.082367144 | 0.876303503 | 0.954361837 |
| chr7:50854857-50855263 | 0.056620993 | 0.876323207 | 0.954364684 |
| chr12:120394642-120395112 | 0.054861379 | 0.876341619 | 0.954364684 |
| chr13:112885673-112886157 | 0.057424769 | 0.876343484 | 0.954364684 |
| chr2:112259504-112259711 | -0.066604472 | 0.876409854 | 0.954377029 |
| chr4:55184342-55184565 | 0.075012422 | 0.876410408 | 0.954377029 |
| chr17:68424741-68425446 | -0.051101597 | 0.87641582 | 0.954377029 |
| chr2:9292478-9292678 | -0.077446193 | 0.876429161 | 0.954377029 |
| chr17:62892913-62893317 | -0.055676987 | 0.876429366 | 0.954377029 |
| chr8:86871086-86871406 | 0.056472178 | 0.876443205 | 0.954377029 |
| chr9:93682255-93682522 | -0.062005001 | 0.876481203 | 0.954377029 |
| chr5:40776108-40776546 | -0.069969846 | 0.876486761 | 0.954377029 |
| chr20:24039638-24039879 | 0.078274498 | 0.876493249 | 0.954377029 |
| chr17:6443689-6444496 | -0.052117238 | 0.876497377 | 0.954377029 |
| chr5:113330682-113331063 | -0.067929355 | 0.876502632 | 0.954377029 |
| chr5:93612541-93612741 | -0.072452562 | 0.876505689 | 0.954377029 |
| chr12:45276267-45276472 | -0.080744026 | 0.876516744 | 0.954377029 |
| chr17:59209327-59210730 | -0.049336548 | 0.87653819 | 0.954386818 |
| chr6:154474773-154475501 | -0.052636877 | 0.876590302 | 0.954415867 |
| chr14:70600518-70601175 | -0.052583299 | 0.876594719 | 0.954415867 |
| chr5:134125279-134125616 | -0.061248179 | 0.876602237 | 0.954415867 |
| chr17:60036898-60037861 | -0.048697201 | 0.8766413 | 0.954435852 |
| chr13:80979076-80979364 | -0.067251154 | 0.876681906 | 0.954435852 |
| chr2:200481455-200481754 | -0.06217675 | 0.876729897 | 0.954435852 |
| chr7:726179-726818 | 0.052235423 | 0.876736568 | 0.954435852 |
| chr17:10177299-10177544 | 0.071404208 | 0.876750795 | 0.954435852 |
| chr3:83602264-83602511 | -0.069024663 | 0.876752996 | 0.954435852 |
| chr17:60372281-60372536 | 0.054018826 | 0.876756958 | 0.954435852 |
| chr5:24232949-24233369 | -0.062389826 | 0.876757078 | 0.954435852 |
| chr1:178629551-178629779 | 0.077077331 | 0.876781817 | 0.954435852 |
| chr3:66220584-66221419 | -0.049798278 | 0.876788219 | 0.954435852 |
| chr6:75297206-75297526 | 0.06440009 | 0.876793684 | 0.954435852 |
| chr12:85230760-85231071 | -0.065364962 | 0.876809472 | 0.954435852 |
| chr22:43151338-43151594 | 0.076214736 | 0.876815127 | 0.954435852 |
| chr1:223556893-223557165 | -0.069630902 | 0.876832364 | 0.954435852 |
| chr5:139633339-139633577 | 0.061254623 | 0.876838339 | 0.954435852 |
| chr4:87073459-87074071 | 0.058866906 | 0.8768431 | 0.954435852 |
| chr1:146038547-146038836 | -0.053264821 | 0.876847548 | 0.954435852 |
| chr1:237756498-237756860 | -0.069854945 | 0.876853283 | 0.954435852 |
| chr17:57868238-57868575 | 0.058904632 | 0.876857265 | 0.954435852 |
| chr2:95526183-95527066 | 0.052558403 | 0.876888017 | 0.954450567 |
| chr1:159161056-159161408 | 0.061165933 | 0.876905308 | 0.954450567 |
| chr17:4420633-4420848 | -0.078962487 | 0.876915485 | 0.954450567 |
| chr20:51624269-51624701 | -0.053274092 | 0.876935859 | 0.954450567 |

| | | | |
|---|---|---|---|
| chr7:2680690-2681249 | 0.054114881 | 0.876938295 | 0.954450567 |
| chr8:38386021-38386785 | 0.051396543 | 0.876949783 | 0.954450567 |
| chr1:208528946-208529152 | -0.079973063 | 0.876966034 | 0.954450567 |
| chr5:43064332-43065217 | 0.049727164 | 0.876970437 | 0.954450567 |
| chr11:17077364-17078437 | -0.049278521 | 0.876999653 | 0.954454854 |
| chr8:142775943-142776176 | 0.075665842 | 0.877005018 | 0.954454854 |
| chr10:63703820-63704248 | 0.060822457 | 0.877011746 | 0.954454854 |
| chr14:75084488-75084909 | 0.057662605 | 0.877066361 | 0.954493831 |
| chr3:59857767-59858068 | -0.06589792 | 0.877102972 | 0.954493831 |
| chr6:111985186-111985602 | 0.059839149 | 0.877129751 | 0.954493831 |
| chr1:109532859-109533301 | 0.053687892 | 0.877130839 | 0.954493831 |
| chr10:6786536-6786900 | 0.071258039 | 0.877134259 | 0.954493831 |
| chr11:118145307-118145921 | -0.051741375 | 0.877162695 | 0.954493831 |
| chr6:97186479-97186685 | 0.073368992 | 0.877165803 | 0.954493831 |
| chr17:16568707-16569547 | 0.05278735 | 0.87716953 | 0.954493831 |
| chr2:158998458-158998725 | -0.066050373 | 0.877183052 | 0.954493831 |
| chr10:7519488-7519772 | 0.063309885 | 0.87718714 | 0.954493831 |
| chr16:46900448-46901103 | -0.054080705 | 0.877204019 | 0.954493831 |
| chr10:101353637-101354847 | -0.049821108 | 0.877215704 | 0.954493831 |
| chr16:67841774-67842473 | 0.050663594 | 0.877226201 | 0.954493831 |
| chr12:9647720-9648932 | -0.050084252 | 0.877243918 | 0.954493831 |
| chr8:78600806-78601159 | 0.066428678 | 0.877248912 | 0.954493831 |
| chr16:3175338-3175889 | 0.053136878 | 0.877254305 | 0.954493831 |
| chr15:32641681-32641881 | 0.083220335 | 0.877265058 | 0.954493831 |
| chr6:133891286-133891672 | -0.065609029 | 0.877303784 | 0.954493831 |
| chr4:146521528-146521744 | -0.066798787 | 0.877324347 | 0.954493831 |
| chr1:204457182-204457401 | 0.070106314 | 0.877339895 | 0.954493831 |
| chr14:103607130-103607348 | -0.071638379 | 0.87734495 | 0.954493831 |
| chr9:69412658-69412986 | 0.062881625 | 0.877350095 | 0.954493831 |
| chr19:34501498-34501910 | 0.05597648 | 0.877353254 | 0.954493831 |
| chr8:81708665-81708865 | -0.084446312 | 0.877372932 | 0.954493831 |
| chr3:90091334-90091942 | 0.056484099 | 0.877385863 | 0.954493831 |
| chr6:169800832-169801032 | 0.073205943 | 0.877387371 | 0.954493831 |
| chr8:96235033-96235868 | -0.049309431 | 0.87739711 | 0.954493831 |
| chr3:141952711-141952936 | -0.070330523 | 0.877410012 | 0.954493831 |
| chr14:31045404-31045604 | -0.082573145 | 0.877414324 | 0.954493831 |
| chr5:142324442-142325484 | 0.050122112 | 0.877421276 | 0.954493831 |
| chr1:29282968-29283734 | 0.051107811 | 0.877438304 | 0.954498804 |
| chr14:90827780-90828516 | -0.051202554 | 0.877474917 | 0.954517493 |
| chr7:143836557-143836861 | 0.061599799 | 0.877491083 | 0.954517493 |
| chr22:27373225-27373727 | -0.060647193 | 0.877495891 | 0.954517493 |
| chr8:53815401-53815638 | 0.079463732 | 0.877505314 | 0.954517493 |
| chr4:139633747-139634073 | -0.060601688 | 0.877603775 | 0.954611042 |
| chr12:124683207-124683440 | -0.072934909 | 0.87763269 | 0.954620372 |
| chr3:138068452-138068971 | -0.054495805 | 0.87763727 | 0.954620372 |
| chr3:101188068-101188304 | -0.086190972 | 0.877668448 | 0.954632221 |
| chr3:185824013-185824240 | -0.072680629 | 0.877682997 | 0.954632221 |
| chr2:100322178-100322718 | 0.054736178 | 0.87768554 | 0.954632221 |
| chr1:183635111-183636440 | 0.048547575 | 0.877712829 | 0.954634663 |
| chr9:92669766-92670714 | 0.04983673 | 0.877742422 | 0.954634663 |

| | | | |
|---|---|---|---|
| chr1:38138648-38139257 | -0.056268636 | 0.877753264 | 0.954634663 |
| chr7:35249238-35249833 | 0.05610303 | 0.87776471 | 0.954634663 |
| chr8:78924804-78925112 | -0.06281068 | 0.877770819 | 0.954634663 |
| chr14:34375456-34375667 | 0.075929823 | 0.877774998 | 0.954634663 |
| chr15:60520887-60521089 | 0.075929823 | 0.877774998 | 0.954634663 |
| chr7:1579475-1579864 | 0.053961492 | 0.877830843 | 0.954681847 |
| chr20:60412332-60412555 | 0.075558385 | 0.877870958 | 0.954701869 |
| chr15:82942431-82943157 | 0.050045888 | 0.877880176 | 0.954701869 |
| chr2:152097450-152097828 | -0.06212287 | 0.877886633 | 0.954701869 |
| chr14:58873865-58874065 | 0.081612187 | 0.877955514 | 0.954763226 |
| chrX:104358748-104359078 | 0.071916609 | 0.877971434 | 0.954766988 |
| chr19:58466533-58467490 | -0.04965984 | 0.877986584 | 0.954769913 |
| chr16:54285640-54285840 | 0.067664251 | 0.878044616 | 0.954813983 |
| chr1:188500408-188500652 | -0.083412933 | 0.878066028 | 0.954813983 |
| chr1:50450415-50450646 | -0.080872973 | 0.878072393 | 0.954813983 |
| chr7:135847246-135847557 | -0.071075655 | 0.878076956 | 0.954813983 |
| chr5:71893776-71893976 | 0.081039393 | 0.878107653 | 0.954832567 |
| chr19:57487556-57487938 | -0.058426884 | 0.878137529 | 0.954832567 |
| chr1:177170683-177171346 | 0.050793494 | 0.878139688 | 0.954832567 |
| chr2:69013462-69013943 | -0.060315468 | 0.878172685 | 0.954832567 |
| chr19:40425751-40426579 | 0.049777805 | 0.878176567 | 0.954832567 |
| chr10:132918610-132918862 | 0.077004593 | 0.878190405 | 0.954832567 |
| chr2:162074338-162074683 | -0.069807031 | 0.878194824 | 0.954832567 |
| chr1:227474161-227474397 | 0.074443764 | 0.878205134 | 0.954832567 |
| chr11:4094233-4095330 | 0.050116594 | 0.878206201 | 0.954832567 |
| chr8:37996321-37996537 | -0.073047306 | 0.878221866 | 0.95483605 |
| chr2:68404255-68404455 | 0.085250572 | 0.878265077 | 0.954868163 |
| chr19:55484601-55484838 | 0.065128812 | 0.878276326 | 0.954868163 |
| chr14:73026722-73027533 | 0.049239317 | 0.878298992 | 0.954879257 |
| chr2:66391545-66391885 | -0.073811703 | 0.87834651 | 0.954902229 |
| chr2:60522589-60522795 | 0.075647976 | 0.878361008 | 0.954902229 |
| chr3:57006680-57007435 | 0.055762086 | 0.878366333 | 0.954902229 |
| chr5:149271982-149272608 | -0.054478201 | 0.878369972 | 0.954902229 |
| chr1:111649255-111649541 | -0.071104224 | 0.878441441 | 0.954950687 |
| chr22:46071539-46071742 | 0.069889124 | 0.878443774 | 0.954950687 |
| chr12:122500212-122501498 | 0.049021247 | 0.878455958 | 0.954950687 |
| chr9:81413438-81413686 | 0.080042736 | 0.878476862 | 0.954950687 |
| chr5:154070052-154070252 | 0.080042736 | 0.878476862 | 0.954950687 |
| chr17:59565214-59566335 | 0.04814641 | 0.878490844 | 0.954952338 |
| chr5:148107663-148107889 | 0.07125859 | 0.878529138 | 0.954980417 |
| chr10:30231286-30231548 | -0.07036413 | 0.878607645 | 0.955039611 |
| chr18:12741282-12741720 | 0.058320445 | 0.878620487 | 0.955039611 |
| chr20:13915449-13915685 | 0.072211163 | 0.878631482 | 0.955039611 |
| chr9:85940717-85941735 | 0.048580681 | 0.878637815 | 0.955039611 |
| chr5:147234772-147235092 | 0.073561403 | 0.878645915 | 0.955039611 |
| chr17:4980638-4980843 | -0.073158084 | 0.878765659 | 0.955145499 |
| chr15:40806957-40807945 | -0.049506647 | 0.878768264 | 0.955145499 |
| chr20:51345534-51346117 | -0.054370525 | 0.87878921 | 0.955154716 |
| chr1:169849218-169849684 | 0.056368255 | 0.878826943 | 0.955182178 |
| chr3:56468290-56468620 | -0.064672673 | 0.878853728 | 0.955197741 |

| | | | |
|---|---|---|---|
| chr9:83956375-83957178 | -0.050628599 | 0.878885873 | 0.955219055 |
| chr8:132936804-132937006 | 0.076665129 | 0.878898272 | 0.955219055 |
| chr10:129710172-129710723 | -0.060885405 | 0.878946321 | 0.955257727 |
| chrX:72572295-72573238 | -0.05158468 | 0.878970426 | 0.955270375 |
| chr12:68932483-68933614 | 0.050307535 | 0.879012315 | 0.955289206 |
| chr6:81753069-81753605 | 0.050349668 | 0.879014023 | 0.955289206 |
| chr16:27368173-27368414 | -0.064519481 | 0.879036298 | 0.955289206 |
| chr4:113952480-113952777 | -0.060928224 | 0.879037623 | 0.955289206 |
| chr19:46757319-46757675 | 0.059354618 | 0.879054477 | 0.955293972 |
| chr16:5490416-5490780 | -0.067661093 | 0.879086939 | 0.9553157 |
| chr2:42238616-42239072 | -0.058716519 | 0.879117046 | 0.955334869 |
| chr20:53785323-53787270 | 0.047049353 | 0.879147951 | 0.955342187 |
| chr6:73842483-73842767 | -0.079644678 | 0.879148717 | 0.955342187 |
| chr5:173293774-173294415 | 0.052742632 | 0.879175342 | 0.955353972 |
| chr2:157875780-157877056 | 0.048141223 | 0.879184498 | 0.955353972 |
| chr7:39458608-39458827 | 0.073191724 | 0.879222989 | 0.955380371 |
| chr6:102516364-102516677 | 0.062994523 | 0.87923373 | 0.955380371 |
| chr11:84463327-84463527 | -0.085467301 | 0.879260697 | 0.955389612 |
| chr11:57460328-57460598 | -0.065586479 | 0.879267172 | 0.955389612 |
| chr4:77819333-77819533 | -0.063205364 | 0.879285857 | 0.955396366 |
| chr19:1260204-1260643 | -0.055576325 | 0.879317132 | 0.955403857 |
| chr15:42273033-42273877 | -0.049750718 | 0.879322412 | 0.955403857 |
| chr3:44914995-44915215 | 0.075038154 | 0.879337531 | 0.955403857 |
| chr3:77039781-77040184 | 0.05651641 | 0.879342627 | 0.955403857 |
| chr16:75148250-75149554 | -0.04778485 | 0.87935718 | 0.95540612 |
| chr6:131694900-131695156 | -0.064950055 | 0.879426161 | 0.955455142 |
| chr20:50464337-50464657 | 0.05833966 | 0.879433564 | 0.955455142 |
| chr5:21494089-21494360 | 0.068270537 | 0.879439709 | 0.955455142 |
| chr6:14799891-14800221 | -0.061389287 | 0.879513226 | 0.955514823 |
| chr3:30548622-30548898 | 0.06730318 | 0.879519583 | 0.955514823 |
| chrX:74550315-74550694 | -0.05924684 | 0.879554383 | 0.955539082 |
| chr10:8332319-8332790 | -0.056271229 | 0.879606925 | 0.955564396 |
| chr13:27424167-27425028 | 0.049284352 | 0.879623619 | 0.955564396 |
| chr19:12734597-12734930 | 0.058611126 | 0.879624693 | 0.955564396 |
| chr17:48602625-48602905 | 0.060003467 | 0.879631691 | 0.955564396 |
| chr19:48372948-48373157 | -0.07404105 | 0.879640039 | 0.955564396 |
| chr2:232611436-232611931 | -0.054551777 | 0.879676098 | 0.955579362 |
| chr3:8443978-8444831 | 0.048516613 | 0.879678759 | 0.955579362 |
| chr15:69413909-69414589 | -0.050596766 | 0.879721706 | 0.955612466 |
| chr5:57893333-57893578 | 0.080078082 | 0.87974081 | 0.955619671 |
| chr11:126079734-126080014 | -0.061162474 | 0.879757716 | 0.955621622 |
| chr5:147062546-147062777 | -0.070781487 | 0.87978333 | 0.955621622 |
| chr17:54172660-54172973 | 0.064560579 | 0.879796697 | 0.955621622 |
| chr11:86455935-86456421 | -0.054337639 | 0.879801458 | 0.955621622 |
| chr5:143569008-143569208 | 0.077622105 | 0.879804966 | 0.955621622 |
| chr20:50761240-50762809 | 0.046827671 | 0.879838351 | 0.955642243 |
| chr17:51940706-51940906 | -0.085020769 | 0.879848895 | 0.955642243 |
| chr17:74759988-74760385 | -0.052343345 | 0.879872696 | 0.955646208 |
| chr3:128497583-128498175 | 0.053188758 | 0.87988957 | 0.955646208 |
| chr13:45535299-45535576 | -0.07366861 | 0.879891744 | 0.955646208 |

| | | | |
|---|---|---|---|
| chr15:100849263-100850405 | 0.049179299 | 0.879902435 | 0.955646208 |
| chr15:22224229-22224479 | -0.071626852 | 0.879920492 | 0.955652274 |
| chr11:518501-519176 | -0.051851907 | 0.879980015 | 0.955697935 |
| chr1:111448528-111449816 | -0.04824879 | 0.87998748 | 0.955697935 |
| chr4:139657032-139657376 | -0.056992402 | 0.880006806 | 0.955705378 |
| chr11:71153048-71153253 | 0.072030292 | 0.880040444 | 0.955720356 |
| chr3:48663328-48664412 | -0.048150348 | 0.880045544 | 0.955720356 |
| chrX:124228916-124229158 | 0.071725595 | 0.880078063 | 0.955742125 |
| chr7:1159888-1160579 | -0.049099384 | 0.880104301 | 0.955746689 |
| chr17:57927368-57927661 | -0.058368695 | 0.880119526 | 0.955746689 |
| chr14:104687835-104688398 | 0.051398161 | 0.880132191 | 0.955746689 |
| chr13:39578998-39579198 | -0.084515944 | 0.880135235 | 0.955746689 |
| chr17:65394730-65394954 | 0.066604568 | 0.880144634 | 0.955746689 |
| chr17:57255691-57256724 | -0.047948418 | 0.880170529 | 0.955757037 |
| chr3:190230077-190230352 | 0.07153277 | 0.880179111 | 0.955757037 |
| chr17:47649076-47650100 | -0.047573472 | 0.88019897 | 0.955757846 |
| chr1:156927817-156928039 | 0.063655928 | 0.880204803 | 0.955757846 |
| chr21:45515750-45515950 | 0.078258415 | 0.880227792 | 0.955760411 |
| chr3:33095538-33095738 | -0.066557681 | 0.880232113 | 0.955760411 |
| chr6:6546685-6547379 | -0.054722384 | 0.880301971 | 0.955822719 |
| chr10:22252406-22252806 | -0.06737854 | 0.880322071 | 0.955830999 |
| chr15:90931688-90932353 | -0.048601586 | 0.880404553 | 0.955891111 |
| chr2:51710221-51710604 | 0.064409632 | 0.880415356 | 0.955891111 |
| chr19:32569943-32570664 | -0.052850571 | 0.880418957 | 0.955891111 |
| chr17:56478882-56479181 | -0.061954408 | 0.88043442 | 0.955891111 |
| chr1:193275813-193276192 | -0.060730436 | 0.880439812 | 0.955891111 |
| chr5:172123205-172123606 | -0.055796526 | 0.880473753 | 0.955914416 |
| chr1:165482147-165482418 | -0.069572275 | 0.880486395 | 0.955914596 |
| chr15:83066862-83067710 | 0.049632256 | 0.880577285 | 0.955973839 |
| chr12:122895806-122896431 | -0.053644735 | 0.880577733 | 0.955973839 |
| chr3:124834696-124834899 | -0.067943046 | 0.880591419 | 0.955973839 |
| chr4:25441617-25441893 | 0.065935201 | 0.880598978 | 0.955973839 |
| chr5:20647229-20647485 | 0.072383256 | 0.880603345 | 0.955973839 |
| chr11:65546329-65546957 | 0.049650854 | 0.880644402 | 0.956004865 |
| chr6:2853696-2854472 | -0.054290287 | 0.880726171 | 0.956080086 |
| chr1:43934258-43934525 | 0.058085148 | 0.880797741 | 0.956144232 |
| chr17:72683387-72684815 | 0.048175738 | 0.880814869 | 0.95614928 |
| chr15:75453892-75454458 | 0.048197852 | 0.880830543 | 0.956152748 |
| chr6:3819235-3819707 | 0.061986028 | 0.880857634 | 0.956160509 |
| chr9:70601066-70601343 | -0.063617856 | 0.88086522 | 0.956160509 |
| chr1:106435419-106435704 | -0.055813489 | 0.880877338 | 0.956160509 |
| chr17:45051252-45051751 | -0.052898955 | 0.880887608 | 0.956160509 |
| chr5:138133109-138133325 | -0.076355363 | 0.880954296 | 0.956207356 |
| chr17:30477198-30477739 | -0.055799313 | 0.880955727 | 0.956207356 |
| chr14:66858734-66858966 | -0.074036818 | 0.880974984 | 0.956214713 |
| chr17:47423424-47423692 | -0.077399657 | 0.88106018 | 0.956293639 |
| chr14:74979575-74979834 | -0.063543166 | 0.881114089 | 0.956338604 |
| chr21:43956451-43956720 | 0.068861003 | 0.88115625 | 0.956370817 |
| chr2:181891465-181892897 | 0.047245963 | 0.881222945 | 0.956416462 |
| chr2:25130190-25130390 | -0.069755558 | 0.881240141 | 0.956416462 |

| | | | |
|---|---|---|---|
| chr9:91361191-91362300 | 0.049966982 | 0.881249471 | 0.956416462 |
| chr5:68315740-68316566 | 0.055412712 | 0.881273228 | 0.956416462 |
| chr10:97566481-97566741 | 0.064540035 | 0.881276347 | 0.956416462 |
| chr3:113472114-113472392 | 0.065675042 | 0.881280361 | 0.956416462 |
| chr8:31034663-31034915 | -0.06492493 | 0.881285681 | 0.956416462 |
| chr6:149723900-149724171 | -0.061385087 | 0.8813335 | 0.95645481 |
| chr10:101291625-101291995 | 0.060683926 | 0.881388311 | 0.95649544 |
| chr11:18321945-18322729 | 0.047425372 | 0.881397597 | 0.95649544 |
| chr10:11986683-11986982 | -0.065129819 | 0.881412792 | 0.95649544 |
| chr5:21485570-21486040 | -0.049582138 | 0.881421892 | 0.95649544 |
| chr19:35693819-35694164 | 0.055125614 | 0.881433355 | 0.95649544 |
| chrX:119655161-119655370 | 0.07567454 | 0.881506624 | 0.956545478 |
| chr17:62423308-62424535 | -0.046966587 | 0.881537397 | 0.956545478 |
| chr14:52553861-52554123 | 0.053643435 | 0.881568038 | 0.956545478 |
| chr17:75182503-75183403 | -0.048655604 | 0.881586459 | 0.956545478 |
| chr17:61393531-61394542 | -0.04637218 | 0.881594023 | 0.956545478 |
| chr8:27310747-27311805 | 0.047347654 | 0.8815961 | 0.956545478 |
| chr8:94405300-94405723 | -0.052175856 | 0.881615918 | 0.956545478 |
| chr14:54338588-54339029 | -0.060721322 | 0.881616163 | 0.956545478 |
| chr5:42636452-42636736 | -0.063567513 | 0.881624 | 0.956545478 |
| chr11:23540042-23540559 | -0.055322461 | 0.88162697 | 0.956545478 |
| chr6:4942143-4942410 | -0.064156145 | 0.881628744 | 0.956545478 |
| chr13:99614573-99614866 | 0.063120891 | 0.881629273 | 0.956545478 |
| chr22:35627650-35627908 | 0.066322948 | 0.881663559 | 0.956561015 |
| chr7:92721072-92721359 | 0.074770015 | 0.881668562 | 0.956561015 |
| chr3:129626343-129627187 | 0.048308249 | 0.881701965 | 0.95658371 |
| chr1:45688064-45688441 | 0.054182555 | 0.881725102 | 0.956595267 |
| chr11:57370720-57370923 | -0.082281169 | 0.881762641 | 0.956622449 |
| chr11:13276952-13277948 | -0.047470421 | 0.881803947 | 0.956649811 |
| chr19:52269294-52269997 | -0.052337509 | 0.881823667 | 0.956649811 |
| chr5:127492796-127493336 | 0.057561047 | 0.881825318 | 0.956649811 |
| chr22:28802275-28802499 | -0.064647659 | 0.881851653 | 0.956664836 |
| chr2:217082354-217082554 | -0.0826478 | 0.881878561 | 0.956669682 |
| chr6:2848827-2849088 | -0.072524459 | 0.881881092 | 0.956669682 |
| chr16:2720978-2721723 | -0.05025084 | 0.881918557 | 0.956692504 |
| chr2:78106753-78106953 | -0.080408585 | 0.881927554 | 0.956692504 |
| chr4:27692136-27692485 | 0.061313966 | 0.881939586 | 0.956692504 |
| chr4:84583482-84583900 | -0.052064207 | 0.881965475 | 0.956707042 |
| chr15:101687705-101687965 | 0.072374988 | 0.882001065 | 0.956722005 |
| chr22:44759523-44759889 | 0.062066064 | 0.882012543 | 0.956722005 |
| chr4:20395325-20395525 | -0.073031456 | 0.882016727 | 0.956722005 |
| chr15:40405367-40406222 | 0.048868646 | 0.882121426 | 0.95679676 |
| chr1:116168268-116168698 | -0.054679649 | 0.882122322 | 0.95679676 |
| chr17:45148078-45148806 | 0.047850437 | 0.882133318 | 0.95679676 |
| chr14:75560934-75561333 | 0.051539322 | 0.882152042 | 0.95679676 |
| chr4:67087982-67088222 | -0.080288527 | 0.882154133 | 0.95679676 |
| chr1:112704859-112705543 | 0.048067777 | 0.882168691 | 0.95679676 |
| chr14:37124389-37125060 | -0.054143075 | 0.882175704 | 0.95679676 |
| chr9:25677402-25678765 | 0.04724168 | 0.882185543 | 0.95679676 |
| chr5:176646954-176648248 | 0.046811673 | 0.882199078 | 0.956797896 |

| | | | |
|---|---|---|---|
| chr6:125546836-125547193 | 0.055110904 | 0.88224715 | 0.956826662 |
| chr14:22983351-22983560 | 0.059081962 | 0.882256532 | 0.956826662 |
| chr9:125407994-125408295 | 0.058570911 | 0.882265461 | 0.956826662 |
| chr8:127178530-127179062 | 0.053522294 | 0.882275551 | 0.956826662 |
| chr17:49764034-49764608 | 0.047583808 | 0.882295993 | 0.956835287 |
| chr4:190111204-190112025 | -0.048638993 | 0.882313719 | 0.956840969 |
| chr15:80615898-80616396 | 0.054964441 | 0.882355391 | 0.956872617 |
| chr14:79324883-79325520 | 0.052725504 | 0.882369294 | 0.956874151 |
| chr1:148159733-148159987 | 0.06404081 | 0.882391589 | 0.956875479 |
| chr2:5781973-5782459 | 0.05585153 | 0.88241285 | 0.956875479 |
| chr9:81529731-81530146 | -0.071426366 | 0.882417237 | 0.956875479 |
| chr20:51700847-51701135 | -0.056523981 | 0.882420472 | 0.956875479 |
| chr17:76547256-76547554 | 0.061782428 | 0.882444518 | 0.956888013 |
| chr4:82373701-82374655 | 0.046666548 | 0.882468881 | 0.956900889 |
| chr12:9378646-9378846 | 0.078972229 | 0.882529126 | 0.956949193 |
| chr8:123018957-123019216 | -0.058148738 | 0.882538406 | 0.956949193 |
| chr5:153756624-153756824 | 0.074769825 | 0.882565563 | 0.956964039 |
| chr3:145639882-145640197 | -0.074092204 | 0.882577076 | 0.956964039 |
| chr2:63861252-63861581 | -0.061306574 | 0.882615692 | 0.956991727 |
| chr14:23567081-23567987 | -0.048044994 | 0.882637326 | 0.956991727 |
| chr14:79278774-79280069 | -0.046753903 | 0.882640082 | 0.956991727 |
| chr2:84970591-84972257 | 0.047230453 | 0.882654451 | 0.956993765 |
| chr22:24142488-24142702 | 0.072564424 | 0.882801226 | 0.957102309 |
| chr12:7960524-7961010 | -0.057635007 | 0.882815041 | 0.957102309 |
| chr3:174090807-174091099 | -0.078182677 | 0.882817886 | 0.957102309 |
| chrX:123432425-123432722 | -0.060278497 | 0.882834045 | 0.957102309 |
| chr4:151604286-151604565 | -0.062626309 | 0.882853271 | 0.957102309 |
| chr17:75639794-75640493 | 0.048333255 | 0.882864899 | 0.957102309 |
| chr16:25257390-25258724 | 0.047090731 | 0.882871331 | 0.957102309 |
| chr1:9267151-9267385 | 0.066376806 | 0.882880325 | 0.957102309 |
| chr11:108593052-108593310 | 0.062492621 | 0.882892585 | 0.957102309 |
| chr3:176767342-176767817 | -0.053190878 | 0.882897091 | 0.957102309 |
| chr9:128083892-128084134 | 0.065309852 | 0.882898598 | 0.957102309 |
| chr1:157490434-157490851 | 0.05859292 | 0.882916535 | 0.957102309 |
| chr5:157917988-157918296 | 0.05663682 | 0.882916949 | 0.957102309 |
| chr8:89757108-89758757 | -0.046007691 | 0.88292989 | 0.957102797 |
| chr3:119274955-119275330 | 0.058678519 | 0.88294505 | 0.957105689 |
| chr3:143484752-143484952 | 0.078963026 | 0.882979162 | 0.957129125 |
| chr10:75099981-75100428 | 0.054128483 | 0.883028228 | 0.957160862 |
| chr8:123240579-123241647 | -0.046926138 | 0.883033424 | 0.957160862 |
| chr4:73434450-73434657 | 0.073570848 | 0.8830581 | 0.957174069 |
| chr12:53230509-53230945 | -0.049875796 | 0.883093121 | 0.957198488 |
| chr22:17009569-17009963 | 0.065999361 | 0.883126778 | 0.957221428 |
| chr4:71608292-71608663 | -0.056789141 | 0.883149942 | 0.957222277 |
| chr17:46266815-46267566 | 0.05243193 | 0.883156676 | 0.957222277 |
| chr19:35141104-35141718 | 0.050601931 | 0.883172599 | 0.957222277 |
| chr12:49092179-49092379 | -0.065517705 | 0.883177532 | 0.957222277 |
| chr1:182096205-182096405 | -0.070156256 | 0.883206748 | 0.95723049 |
| chr20:63628346-63628575 | 0.05758 | 0.883217957 | 0.95723049 |
| chr3:153988905-153989124 | 0.074830964 | 0.883222589 | 0.95723049 |

| | | | |
|---|---|---|---|
| chr17:51446216-51446557 | 0.055794312 | 0.883259247 | 0.95725668 |
| chr20:51015167-51015534 | -0.054765498 | 0.883307029 | 0.957289758 |
| chr3:58167431-58167655 | -0.06963985 | 0.883314755 | 0.957289758 |
| chr2:25196424-25196804 | -0.056089034 | 0.883361352 | 0.957326716 |
| chr11:134207537-134207742 | 0.082003783 | 0.883386461 | 0.957340387 |
| chr10:77405755-77406180 | -0.055088172 | 0.883456915 | 0.95739001 |
| chr6:33312577-33313018 | -0.053807948 | 0.88345724 | 0.95739001 |
| chr8:102577716-102578139 | -0.052820092 | 0.883471721 | 0.957392162 |
| chr1:160046556-160047053 | -0.054318117 | 0.883505187 | 0.957414887 |
| chr5:151040134-151040334 | 0.079368225 | 0.883538146 | 0.957437063 |
| chr1:1908789-1909500 | -0.049867271 | 0.883575356 | 0.957463844 |
| chr19:8832224-8833175 | 0.047496537 | 0.883597281 | 0.957474061 |
| chr10:91758680-91758880 | -0.076066735 | 0.883651824 | 0.957508343 |
| chrX:108735098-108735426 | -0.054364028 | 0.883657659 | 0.957508343 |
| chr1:205325415-205325723 | 0.052164648 | 0.883666407 | 0.957508343 |
| chr1:40665604-40666121 | 0.052725561 | 0.883707308 | 0.957517722 |
| chrX:17496489-17496713 | 0.07685072 | 0.883712248 | 0.957517722 |
| chr20:59003860-59004170 | -0.054235194 | 0.883712553 | 0.957517722 |
| chr7:17503903-17504152 | -0.0680068 | 0.883751259 | 0.957531372 |
| chr20:34112053-34112733 | 0.048060628 | 0.883753217 | 0.957531372 |
| chr21:39348360-39350017 | -0.045722452 | 0.883764184 | 0.957531372 |
| chr6:69601013-69601395 | 0.06358287 | 0.883787057 | 0.957531372 |
| chr6:27157999-27158420 | 0.051560613 | 0.883795126 | 0.957531372 |
| chr11:34654860-34655299 | 0.052687326 | 0.88380119 | 0.957531372 |
| chr4:38673145-38673492 | 0.058560306 | 0.883812628 | 0.957531372 |
| chr3:5305393-5305853 | -0.054513225 | 0.883903696 | 0.957609354 |
| chr3:131381111-131382295 | -0.046195416 | 0.883911499 | 0.957609354 |
| chr16:71722782-71724369 | 0.046185668 | 0.8839221 | 0.957609354 |
| chr9:20508301-20508517 | -0.080008984 | 0.883949359 | 0.957613928 |
| chr5:97764763-97765018 | -0.069957857 | 0.883951318 | 0.957613928 |
| chr2:220077571-220077938 | 0.067482802 | 0.884014908 | 0.957667886 |
| chr3:62007073-62007905 | -0.047542414 | 0.884026122 | 0.957667886 |
| chr15:41416454-41417938 | 0.047068555 | 0.884091444 | 0.957702458 |
| chr20:62776793-62777236 | -0.051014687 | 0.884100287 | 0.957702458 |
| chr16:27690638-27690848 | 0.070270736 | 0.88410202 | 0.957702458 |
| chr20:48051548-48052775 | 0.045937006 | 0.884108033 | 0.957702458 |
| chr2:127649719-127650060 | -0.055161084 | 0.884169703 | 0.957755722 |
| chr12:42483690-42483996 | 0.067078613 | 0.884194157 | 0.957762081 |
| chr14:22886521-22887197 | -0.048806632 | 0.884217469 | 0.957762081 |
| chr6:28266955-28267465 | 0.053825414 | 0.884220753 | 0.957762081 |
| chr12:26101402-26101706 | -0.059276716 | 0.884234221 | 0.957762081 |
| chr15:95846195-95846395 | -0.080645865 | 0.884245983 | 0.957762081 |
| chr16:67017924-67018309 | -0.053708178 | 0.884250572 | 0.957762081 |
| chr14:63134884-63135084 | 0.080244873 | 0.884298675 | 0.957785741 |
| chr9:120868225-120869416 | -0.045869111 | 0.884302142 | 0.957785741 |
| chr1:219432816-219433086 | -0.06465616 | 0.884314981 | 0.957785741 |
| chr16:81996988-81997499 | -0.052865147 | 0.884322418 | 0.957785741 |
| chr1:231528041-231529011 | 0.047181922 | 0.884358132 | 0.957787182 |
| chr2:159208451-159208653 | -0.069010659 | 0.884364067 | 0.957787182 |
| chr6:110484328-110484592 | -0.065133345 | 0.884386211 | 0.957787182 |

| | | | |
|---|---|---|---|
| chrX:140439400-140439826 | -0.062404274 | 0.884396136 | 0.957787182 |
| chr14:67765549-67765815 | -0.061240932 | 0.88439738 | 0.957787182 |
| chr19:39237800-39238160 | -0.059969986 | 0.884398749 | 0.957787182 |
| chr6:27809870-27810106 | -0.056810382 | 0.884435561 | 0.957808954 |
| chr8:75177459-75177659 | -0.081353624 | 0.884472639 | 0.957808954 |
| chr5:84502926-84503126 | -0.079499396 | 0.884476217 | 0.957808954 |
| chr19:46077615-46077858 | -0.067246771 | 0.884489971 | 0.957808954 |
| chr10:50275745-50275978 | -0.067525948 | 0.884490133 | 0.957808954 |
| chr9:126940628-126940872 | -0.065900697 | 0.884493855 | 0.957808954 |
| chr5:178437576-178437776 | 0.070238267 | 0.884516592 | 0.957812648 |
| chr12:6866889-6867176 | -0.05849227 | 0.884522531 | 0.957812648 |
| chr18:58850341-58850542 | 0.073732529 | 0.884541406 | 0.957812648 |
| chrX:70134114-70134428 | 0.059705672 | 0.884547268 | 0.957812648 |
| chr6:126033661-126033909 | 0.06377571 | 0.884599595 | 0.957855773 |
| chr11:33774082-33774938 | 0.048549746 | 0.884640478 | 0.957886504 |
| chr6:35522887-35523156 | -0.060338205 | 0.884671508 | 0.957906048 |
| chr10:97791931-97792310 | 0.052563145 | 0.88468353 | 0.957906048 |
| chr16:84228850-84229050 | 0.069940944 | 0.884752586 | 0.957963854 |
| chr2:84697977-84698244 | 0.075257556 | 0.884785051 | 0.957963854 |
| chr1:89358326-89358610 | -0.062057041 | 0.884785747 | 0.957963854 |
| chr8:125990533-125991057 | -0.051083493 | 0.884786928 | 0.957963854 |
| chr1:33633122-33633374 | 0.061584445 | 0.884853787 | 0.958022706 |
| chr3:156541087-156541396 | -0.056485204 | 0.884867743 | 0.958024278 |
| chr12:63985074-63985325 | 0.070435975 | 0.884880604 | 0.958024666 |
| chr1:63575354-63575610 | -0.064511339 | 0.884905316 | 0.958032356 |
| chr19:35745330-35745721 | 0.048440909 | 0.884912714 | 0.958032356 |
| chr1:151281148-151283115 | -0.044876563 | 0.884934209 | 0.958042091 |
| chr21:45505868-45506167 | 0.061216958 | 0.884964244 | 0.95806107 |
| chr1:22662529-22662752 | 0.066223105 | 0.884991364 | 0.958076894 |
| chr1:34597972-34598179 | -0.069590319 | 0.88503256 | 0.958107955 |
| chr8:109281564-109282498 | 0.046403214 | 0.885083006 | 0.958127948 |
| chr6:128586208-128586500 | -0.074238562 | 0.885085236 | 0.958127948 |
| chr17:79919275-79919496 | 0.063401013 | 0.885088543 | 0.958127948 |
| chr18:32930729-32931035 | -0.05872299 | 0.885147033 | 0.958176837 |
| chr4:24809443-24809695 | 0.070624 | 0.885158715 | 0.958176837 |
| chr10:28684951-28685559 | 0.050043663 | 0.885238645 | 0.958244451 |
| chr4:119454223-119455361 | -0.045870213 | 0.885251062 | 0.958244451 |
| chr5:87885867-87886067 | -0.077281203 | 0.885258695 | 0.958244451 |
| chr11:61733536-61733736 | 0.065415519 | 0.885298544 | 0.958261414 |
| chr6:152208238-152208535 | -0.072194629 | 0.885299379 | 0.958261414 |
| chr2:42975111-42975439 | -0.055882309 | 0.885329641 | 0.958280633 |
| chr16:1611922-1612497 | 0.047441116 | 0.885354856 | 0.958294388 |
| chr1:160098367-160099224 | -0.046212031 | 0.885371558 | 0.958298929 |
| chr6:10383277-10384005 | 0.051750602 | 0.885420869 | 0.958338765 |
| chr2:219229020-219230016 | -0.046309273 | 0.8854613 | 0.958358985 |
| chr2:20326216-20326523 | -0.057746251 | 0.885481082 | 0.958358985 |
| chr3:39383156-39384554 | -0.045757434 | 0.885486318 | 0.958358985 |
| chr4:37685786-37687097 | 0.045134485 | 0.885494879 | 0.958358985 |
| chrX:124719108-124719439 | 0.05970727 | 0.885502089 | 0.958358985 |
| chr5:168268940-168269148 | -0.061324345 | 0.885531748 | 0.958377548 |

| | | | |
|---|---|---|---|
| chr1:205648817-205649139 | -0.058794194 | 0.885596327 | 0.958433902 |
| chr6:81440991-81441418 | 0.056782675 | 0.885617192 | 0.958442944 |
| chr10:90911507-90911913 | 0.055902008 | 0.8856457 | 0.958460259 |
| chr12:57520370-57521051 | -0.046252751 | 0.885659938 | 0.95846213 |
| chr9:125943825-125944025 | 0.077559133 | 0.88572562 | 0.958519674 |
| chr6:89638303-89639180 | -0.04604641 | 0.88580733 | 0.958587342 |
| chr2:205944424-205944907 | -0.056230227 | 0.885823808 | 0.958587342 |
| chr3:1082076-1082494 | -0.052519793 | 0.885826798 | 0.958587342 |
| chr1:175146563-175146881 | -0.052003763 | 0.885838192 | 0.958587342 |
| chr1:168888240-168888458 | -0.072184461 | 0.885882338 | 0.958605781 |
| chr5:61805096-61805480 | -0.055897155 | 0.885885041 | 0.958605781 |
| chr4:176792219-176793046 | 0.049647709 | 0.885892764 | 0.958605781 |
| chr11:22625047-22626139 | -0.045566897 | 0.885914242 | 0.958607548 |
| chr17:2711501-2712118 | 0.049757899 | 0.885923888 | 0.958607548 |
| chr4:137611089-137611396 | -0.066384963 | 0.88593193 | 0.958607548 |
| chr3:134630531-134630785 | 0.066656441 | 0.885954608 | 0.958611302 |
| chr1:246583013-246583904 | -0.046721676 | 0.885966131 | 0.958611302 |
| chr3:169325061-169325285 | -0.067426425 | 0.885987062 | 0.958611302 |
| chr1:65467474-65467944 | 0.055059919 | 0.885989075 | 0.958611302 |
| chr8:28889775-28890688 | -0.046182743 | 0.886009152 | 0.958611302 |
| chr6:104228947-104229483 | -0.054541866 | 0.886012142 | 0.958611302 |
| chr17:18960916-18961135 | -0.062511241 | 0.886022975 | 0.958611302 |
| chr15:77753358-77753558 | 0.063245459 | 0.88604544 | 0.95861456 |
| chr14:103847066-103848305 | -0.046017557 | 0.886051009 | 0.95861456 |
| chr5:43105157-43106100 | 0.045988238 | 0.886079894 | 0.958630482 |
| chr13:25447431-25447766 | 0.059397641 | 0.886094666 | 0.958630482 |
| chr17:47889958-47890269 | 0.052711455 | 0.886103259 | 0.958630482 |
| chr20:59515645-59515985 | 0.05024804 | 0.886132981 | 0.958649102 |
| chr16:71383813-71384260 | -0.058302095 | 0.886161379 | 0.958654422 |
| chr15:82709795-82710346 | -0.049069239 | 0.886162922 | 0.958654422 |
| chr1:181276192-181276709 | -0.052194073 | 0.886218864 | 0.958701404 |
| chr3:86687555-86687805 | 0.071626026 | 0.886282895 | 0.958757137 |
| chr3:128348954-128349244 | -0.059627089 | 0.886325317 | 0.958779351 |
| chr6:28790267-28790567 | -0.057134662 | 0.886328457 | 0.958779351 |
| chr14:95255523-95255865 | 0.053416262 | 0.886347914 | 0.958786863 |
| chr16:81798252-81798484 | 0.064768301 | 0.886382722 | 0.958790754 |
| chr14:43665158-43665569 | -0.056631294 | 0.886398279 | 0.958790754 |
| chr6:63245995-63246238 | 0.063149905 | 0.886405725 | 0.958790754 |
| chr3:25592518-25592804 | -0.055659261 | 0.886411342 | 0.958790754 |
| chr3:124429082-124429282 | 0.072226282 | 0.886417989 | 0.958790754 |
| chr7:35975830-35976216 | 0.054704906 | 0.886435417 | 0.958790754 |
| chr15:100917963-100918534 | -0.049870323 | 0.886466323 | 0.958790754 |
| chr4:109652607-109653072 | -0.05519911 | 0.886475947 | 0.958790754 |
| chr20:35615905-35616695 | 0.045762513 | 0.886477078 | 0.958790754 |
| chr17:44006646-44007466 | 0.046930921 | 0.886482508 | 0.958790754 |
| chr20:24100745-24100957 | 0.064346514 | 0.886489157 | 0.958790754 |
| chr1:39605889-39606685 | -0.051297871 | 0.886559033 | 0.95882078 |
| chr7:151341137-151342115 | 0.046639958 | 0.886559784 | 0.95882078 |
| chr15:31391729-31391942 | 0.066240748 | 0.886565251 | 0.95882078 |
| chr8:6800749-6801085 | -0.055141485 | 0.886566973 | 0.95882078 |

| | | | |
|---|---|---|---|
| chr10:101214747-101214956 | 0.064717242 | 0.886625456 | 0.95882192 |
| chr3:46629769-46630204 | -0.058857817 | 0.886634251 | 0.95882192 |
| chr7:149644670-149645138 | -0.051997022 | 0.88663628 | 0.95882192 |
| chr3:30077338-30077539 | 0.073476054 | 0.886661167 | 0.95882192 |
| chr6:149863160-149863557 | -0.050157174 | 0.886663119 | 0.95882192 |
| chr7:52719652-52719987 | -0.061219578 | 0.886683768 | 0.95882192 |
| chr16:85718632-85719039 | -0.053011796 | 0.88668618 | 0.95882192 |
| chr2:86719837-86720486 | -0.051574336 | 0.886695394 | 0.95882192 |
| chr8:96261278-96263054 | 0.044599355 | 0.88670825 | 0.95882192 |
| chr10:42827315-42827515 | -0.077389554 | 0.886710062 | 0.95882192 |
| chr10:77711434-77711957 | -0.052456199 | 0.886714593 | 0.95882192 |
| chr7:40899070-40899270 | -0.074568222 | 0.886727134 | 0.95882192 |
| chr16:83199897-83200227 | -0.066981616 | 0.886739477 | 0.95882192 |
| chr1:59971403-59971754 | 0.054067485 | 0.886752848 | 0.95882192 |
| chr10:119686573-119686836 | 0.058289544 | 0.886759561 | 0.95882192 |
| chr12:124927013-124928163 | 0.045193242 | 0.886788026 | 0.95882192 |
| chr5:35136353-35136567 | -0.069837655 | 0.886796061 | 0.95882192 |
| chr18:45880003-45880320 | -0.062495492 | 0.886799281 | 0.95882192 |
| chr12:47080050-47080765 | -0.05104776 | 0.886805787 | 0.95882192 |
| chr22:19849822-19850044 | -0.060179923 | 0.886844673 | 0.958837132 |
| chr22:49632542-49632762 | 0.080316804 | 0.886844884 | 0.958837132 |
| chrX:13688895-13689708 | 0.04726179 | 0.886868223 | 0.958848836 |
| chrX:66815401-66815605 | -0.07349926 | 0.886925104 | 0.958879003 |
| chr4:25718148-25718372 | -0.078179245 | 0.886934682 | 0.958879003 |
| chr1:229203485-229203714 | 0.062482514 | 0.886945095 | 0.958879003 |
| chr14:98819423-98819973 | 0.049162915 | 0.886955869 | 0.958879003 |
| chr20:25017304-25017748 | -0.05015963 | 0.886958698 | 0.958879003 |
| chr6:73309702-73310567 | -0.04704707 | 0.886984615 | 0.958893243 |
| chr5:71475840-71476081 | -0.064686125 | 0.886999332 | 0.958893243 |
| chr11:86790389-86790744 | -0.061560468 | 0.887013888 | 0.958893243 |
| chr12:122131081-122131345 | 0.062695145 | 0.887032909 | 0.958893243 |
| chr3:135185438-135185738 | -0.072141891 | 0.887034443 | 0.958893243 |
| chr1:230632228-230632613 | 0.057236011 | 0.887060805 | 0.958899221 |
| chr15:90710534-90710734 | -0.077117937 | 0.887066356 | 0.958899221 |
| chr4:38621073-38621408 | 0.057441115 | 0.887078087 | 0.958899221 |
| chr6:119077590-119079314 | 0.045319263 | 0.887090031 | 0.958899221 |
| chr2:73700219-73700433 | 0.067397351 | 0.887109879 | 0.958907148 |
| chr9:136482985-136484222 | 0.045017446 | 0.887124419 | 0.958909336 |
| chr1:51517961-51519567 | -0.045733296 | 0.887157067 | 0.958921801 |
| chr14:54827512-54827712 | 0.077089957 | 0.887166285 | 0.958921801 |
| chr10:100185311-100186662 | -0.046209405 | 0.887173495 | 0.958921801 |
| chr10:27998592-27999056 | -0.058912306 | 0.88722414 | 0.958951942 |
| chr22:50627546-50628457 | 0.048637549 | 0.887237699 | 0.958951942 |
| chr15:100807553-100807764 | -0.073011827 | 0.887250752 | 0.958951942 |
| chr10:97731800-97732093 | -0.053688964 | 0.887251443 | 0.958951942 |
| chrX:134549151-134550200 | 0.046959653 | 0.887412015 | 0.95909956 |
| chr5:12934903-12935277 | -0.06312435 | 0.887416992 | 0.95909956 |
| chr18:35144009-35144292 | -0.062071255 | 0.887425575 | 0.95909956 |
| chr6:104186740-104187387 | -0.052340015 | 0.887464407 | 0.959127999 |
| chr5:142600667-142600867 | 0.068649966 | 0.887486795 | 0.959128818 |

| | | | |
|---|---|---|---|
| chr5:124596808-124597217 | -0.057202811 | 0.887495343 | 0.959128818 |
| chr3:129885629-129885829 | -0.07852959 | 0.887502717 | 0.959128818 |
| chr11:77589231-77590758 | -0.044960867 | 0.887542048 | 0.959138421 |
| chr2:233566023-233566334 | -0.06103688 | 0.88755625 | 0.959138421 |
| chr3:165522219-165522461 | -0.073201602 | 0.887561559 | 0.959138421 |
| chr3:85342936-85343305 | 0.055284904 | 0.887575282 | 0.959138421 |
| chr7:9221554-9221853 | -0.067098737 | 0.887578149 | 0.959138421 |
| chr1:8182394-8183082 | 0.052322255 | 0.88758671 | 0.959138421 |
| chr8:101252384-101252908 | 0.050333049 | 0.887614789 | 0.959150139 |
| chr17:14098315-14098580 | -0.059590264 | 0.88762259 | 0.959150139 |
| chr7:484784-485240 | 0.054231362 | 0.887651767 | 0.959161215 |
| chr1:181117532-181118187 | 0.044870428 | 0.887667658 | 0.959161215 |
| chr2:309538-309787 | -0.069353622 | 0.887670394 | 0.959161215 |
| chr12:87953361-87953561 | 0.072821428 | 0.887686123 | 0.959163161 |
| chr2:46063043-46063248 | 0.072301953 | 0.887705292 | 0.959163161 |
| chr4:78081587-78081836 | 0.066637398 | 0.88770975 | 0.959163161 |
| chr17:16214842-16215809 | 0.046293519 | 0.887725072 | 0.959166192 |
| chr5:32038628-32038828 | 0.069407489 | 0.887749318 | 0.959169908 |
| chr20:44264507-44264815 | 0.063613068 | 0.887753548 | 0.959169908 |
| chr8:128224698-128225045 | -0.051962086 | 0.887807984 | 0.959210933 |
| chr9:88076103-88077360 | 0.045608513 | 0.887816556 | 0.959210933 |
| chr20:10433614-10434810 | -0.044604325 | 0.887846031 | 0.959224556 |
| chr8:108343088-108343345 | 0.063407024 | 0.887854203 | 0.959224556 |
| chr4:153465575-153466923 | -0.045223493 | 0.887894476 | 0.959254541 |
| chr15:43185829-43186209 | 0.059378609 | 0.88791721 | 0.959256122 |
| chr2:200863785-200865122 | 0.045008177 | 0.887925745 | 0.959256122 |
| chr11:44066718-44067149 | 0.055950751 | 0.887933498 | 0.959256122 |
| chr12:3315075-3315724 | 0.051192554 | 0.887963314 | 0.959274808 |
| chr14:102086831-102087812 | 0.044950152 | 0.888019641 | 0.959320402 |
| chr3:128011715-128011948 | -0.064254665 | 0.888031949 | 0.959320402 |
| chr4:127732658-127733017 | 0.053677998 | 0.888043079 | 0.959320402 |
| chr3:130143737-130144002 | 0.057966016 | 0.888060962 | 0.959326195 |
| chr3:192423067-192423308 | 0.077332436 | 0.888092749 | 0.959347008 |
| chr4:140150284-140150487 | 0.055525768 | 0.888194099 | 0.959442963 |
| chr4:11011156-11011587 | -0.056063641 | 0.888239376 | 0.959478346 |
| chr11:79885179-79885379 | -0.080345591 | 0.888261883 | 0.95948913 |
| chr3:4695843-4696224 | 0.056775577 | 0.888295167 | 0.959499824 |
| chr2:236168320-236168654 | 0.055112288 | 0.888296828 | 0.959499824 |
| chr6:101393980-101394194 | -0.064583422 | 0.888329209 | 0.959521274 |
| chr4:88082256-88082512 | 0.080406627 | 0.888365573 | 0.959547026 |
| chr11:65510834-65511080 | 0.055504659 | 0.888408673 | 0.959569593 |
| chr3:17741974-17743315 | -0.043886549 | 0.8884212 | 0.959569593 |
| chr18:79680045-79680359 | -0.05268627 | 0.888427301 | 0.959569593 |
| chr14:64938498-64938910 | 0.052252817 | 0.88843656 | 0.959569593 |
| chr2:174771564-174771903 | 0.052244371 | 0.888461484 | 0.959577387 |
| chr1:175728363-175728563 | -0.074586526 | 0.888468823 | 0.959577387 |
| chr10:69319247-69319468 | 0.056949456 | 0.888492598 | 0.959580919 |
| chr12:74043246-74043764 | 0.047362039 | 0.888508107 | 0.959580919 |
| chrX:108199739-108200003 | -0.059596077 | 0.888509664 | 0.959580919 |
| chr17:28318446-28319501 | -0.045422915 | 0.888539547 | 0.959592961 |

| | | | |
|---|---|---|---|
| chr6:16206561-16206857 | 0.06017131 | 0.888570263 | 0.959592961 |
| chr3:184017281-184018412 | 0.04496853 | 0.888570445 | 0.959592961 |
| chrX:45850782-45851228 | -0.05635211 | 0.888574956 | 0.959592961 |
| chr10:69123321-69124677 | -0.044092264 | 0.888599719 | 0.959592961 |
| chr8:104759731-104760045 | -0.051316366 | 0.888602891 | 0.959592961 |
| chr9:121599969-121600186 | -0.06210717 | 0.88860848 | 0.959592961 |
| chr10:89235173-89235411 | -0.066918955 | 0.888650431 | 0.959619267 |
| chr10:12130273-12130600 | -0.04967622 | 0.888657889 | 0.959619267 |
| chr3:30319786-30320117 | -0.056569278 | 0.888670962 | 0.959619861 |
| chr2:216783435-216783656 | -0.073438722 | 0.888712 | 0.95965065 |
| chr7:150986151-150986382 | -0.059924543 | 0.888725773 | 0.959651998 |
| chr14:52014070-52014409 | -0.056139606 | 0.888758937 | 0.959674285 |
| chr2:131516609-131516893 | 0.056956743 | 0.888790661 | 0.959682274 |
| chr2:141942457-141942668 | 0.076661154 | 0.888808844 | 0.959682274 |
| chr6:138308406-138308922 | -0.053506402 | 0.888826476 | 0.959682274 |
| chr13:19439116-19439408 | 0.059726324 | 0.888828279 | 0.959682274 |
| chr19:18678118-18678348 | 0.066109302 | 0.888833937 | 0.959682274 |
| chr6:30396866-30397277 | -0.054400835 | 0.888841485 | 0.959682274 |
| chr1:228721410-228721781 | 0.052734029 | 0.888871367 | 0.959686983 |
| chr10:127906641-127907138 | -0.049196129 | 0.888891337 | 0.959686983 |
| chr2:86025278-86025519 | -0.066041323 | 0.888897705 | 0.959686983 |
| chr5:177403462-177404706 | 0.044933294 | 0.888910161 | 0.959686983 |
| chrX:46545089-46546021 | -0.046179927 | 0.888912193 | 0.959686983 |
| chr16:89720263-89720615 | -0.050780574 | 0.888920996 | 0.959686983 |
| chr1:175029662-175030156 | -0.046592237 | 0.888948034 | 0.959696736 |
| chr3:52239064-52239362 | 0.055038256 | 0.88895508 | 0.959696736 |
| chr9:83920676-83921763 | -0.044395108 | 0.888973929 | 0.959703563 |
| chr20:57622478-57622834 | 0.048630884 | 0.88900228 | 0.95971857 |
| chr22:40044099-40044867 | -0.046798973 | 0.889012881 | 0.95971857 |
| chr22:32985867-32986364 | 0.052750763 | 0.889027367 | 0.959720687 |
| chr2:209423461-209424581 | 0.04468315 | 0.889066545 | 0.959749458 |
| chr3:66989882-66990754 | -0.050866066 | 0.88912186 | 0.959795649 |
| chr19:48135466-48135755 | 0.058860812 | 0.889189946 | 0.959855624 |
| chr10:114703574-114703935 | 0.06182079 | 0.889234736 | 0.95989045 |
| chr7:32521736-32521977 | 0.061388099 | 0.889293036 | 0.959913033 |
| chr6:13747704-13747904 | 0.068438562 | 0.889311957 | 0.959913033 |
| chr2:42623431-42623651 | -0.066333611 | 0.889315563 | 0.959913033 |
| chr19:38336194-38337337 | 0.04419733 | 0.889318801 | 0.959913033 |
| chr16:71311680-71311880 | 0.069221264 | 0.889322274 | 0.959913033 |
| chr20:37693608-37694639 | 0.045277372 | 0.889330824 | 0.959913033 |
| chr3:157239111-157239472 | -0.049280063 | 0.889355418 | 0.959926057 |
| chr6:170030938-170031412 | 0.049511868 | 0.889391383 | 0.959929687 |
| chr2:109136536-109136778 | -0.073058961 | 0.889396365 | 0.959929687 |
| chr3:150968726-150968975 | -0.073058961 | 0.889396365 | 0.959929687 |
| chr2:79041065-79041529 | -0.05728687 | 0.889409584 | 0.959930432 |
| chr15:101489362-101489817 | 0.054004562 | 0.889431913 | 0.959941009 |
| chr3:136650642-136650872 | 0.069731445 | 0.889457231 | 0.959954813 |
| chr4:152535946-152536953 | -0.0443995 | 0.889476388 | 0.959959772 |
| chr8:130682800-130683092 | -0.063106043 | 0.88950064 | 0.959959772 |
| chr1:67832684-67833684 | 0.045099096 | 0.889504488 | 0.959959772 |

| | | | |
|---|---|---|---|
| chr17:80861874-80862077 | -0.064544398 | 0.889522548 | 0.959959772 |
| chr7:99144015-99144477 | -0.049721355 | 0.889524468 | 0.959959772 |
| chr1:205591748-205592265 | -0.04646911 | 0.889580217 | 0.960006414 |
| chr19:56172043-56172243 | -0.06909906 | 0.889594319 | 0.960008112 |
| chr15:76185718-76186035 | 0.052907977 | 0.889622665 | 0.960012208 |
| chr10:100373188-100374227 | -0.044636924 | 0.889623174 | 0.960012208 |
| chr10:120077897-120078171 | -0.066860403 | 0.889663883 | 0.960042618 |
| chr3:179071270-179072448 | 0.045519589 | 0.88968388 | 0.960046436 |
| chr1:109643282-109643832 | -0.047905127 | 0.889692481 | 0.960046436 |
| chr5:101977795-101977995 | 0.080148549 | 0.889733161 | 0.960071508 |
| chr8:104787670-104788172 | -0.050386165 | 0.889740776 | 0.960071508 |
| chr3:153034903-153035103 | -0.078094091 | 0.889800715 | 0.960082906 |
| chr7:4753721-4754223 | -0.0482285 | 0.889836809 | 0.960082906 |
| chr6:5260374-5261609 | 0.044800136 | 0.889843071 | 0.960082906 |
| chr10:100562844-100563382 | 0.0507169 | 0.889857307 | 0.960082906 |
| chr8:73356322-73356885 | 0.047240645 | 0.889859461 | 0.960082906 |
| chr1:39653546-39653804 | -0.061937364 | 0.889863308 | 0.960082906 |
| chr3:54243562-54243893 | 0.059913203 | 0.889867737 | 0.960082906 |
| chr2:223815215-223815441 | 0.068375376 | 0.889868916 | 0.960082906 |
| chr2:203650859-203651064 | -0.067405456 | 0.88987372 | 0.960082906 |
| chr12:120302118-120302637 | 0.049947267 | 0.88987664 | 0.960082906 |
| chr12:48853230-48853507 | 0.055467671 | 0.889931858 | 0.960123433 |
| chr12:109783865-109784304 | -0.049425358 | 0.889939264 | 0.960123433 |
| chr16:78257328-78257655 | 0.056940587 | 0.889984428 | 0.960158639 |
| chr16:88755379-88755767 | 0.048673012 | 0.890015128 | 0.96017824 |
| chr14:50018373-50018662 | 0.058654048 | 0.890040449 | 0.960192038 |
| chr2:30234116-30234582 | 0.052865671 | 0.890111716 | 0.96024497 |
| chr6:139219207-139219767 | -0.050235274 | 0.89011487 | 0.96024497 |
| chr5:126353982-126354195 | -0.069573697 | 0.890127111 | 0.96024497 |
| chr8:109551990-109552500 | 0.050579854 | 0.890162337 | 0.960263026 |
| chr18:14928470-14928775 | -0.055935835 | 0.890174498 | 0.960263026 |
| chr3:101512141-101512341 | 0.061080558 | 0.890181446 | 0.960263026 |
| chr8:37696526-37696756 | 0.054898676 | 0.89020431 | 0.960274171 |
| chr16:27457174-27457536 | 0.053791296 | 0.890226393 | 0.960284473 |
| chr3:114827581-114827925 | 0.057576261 | 0.890287188 | 0.960336533 |
| chr22:50783477-50783815 | 0.06232468 | 0.890357577 | 0.960393856 |
| chr16:24834480-24834784 | 0.057656639 | 0.890367548 | 0.960393856 |
| chr8:102682560-102682880 | 0.052586197 | 0.890377933 | 0.960393856 |
| chr1:168098330-168099071 | -0.046589398 | 0.890414845 | 0.960420151 |
| chr11:18012278-18013733 | 0.044239999 | 0.890427394 | 0.960420167 |
| chr17:41648662-41648884 | 0.060585001 | 0.890455359 | 0.96043681 |
| chr21:42437977-42438177 | -0.060281807 | 0.890516449 | 0.960469526 |
| chr16:4944661-4945229 | -0.046306091 | 0.890518689 | 0.960469526 |
| chr2:176653956-176654221 | 0.059913854 | 0.890533427 | 0.960469526 |
| chrX:15127113-15127314 | 0.076967933 | 0.890535833 | 0.960469526 |
| chr2:157971447-157972070 | 0.05207209 | 0.890602808 | 0.960528241 |
| chr11:407804-408022 | 0.065286701 | 0.890624156 | 0.960537744 |
| chr17:32912356-32912607 | -0.064520702 | 0.890705688 | 0.960579639 |
| chr13:44692204-44692729 | -0.051041476 | 0.890715397 | 0.960579639 |
| chr17:48018315-48018619 | 0.050838121 | 0.890718893 | 0.960579639 |

| | | | |
|---|---|---|---|
| chr21:39806044-39806290 | 0.061115229 | 0.890727704 | 0.960579639 |
| chr1:201025804-201026066 | 0.050982259 | 0.890736093 | 0.960579639 |
| chr14:102362615-102363751 | -0.044968708 | 0.890738221 | 0.960579639 |
| chr1:81801918-81802299 | 0.0525171 | 0.890754148 | 0.960583295 |
| chr20:57997123-57997587 | -0.049293432 | 0.89076684 | 0.960583463 |
| chr20:23087304-23087535 | -0.069495002 | 0.89080578 | 0.960611936 |
| chr8:56195843-56196110 | 0.061238746 | 0.890837311 | 0.960619188 |
| chr17:50346571-50346915 | -0.050651832 | 0.890840325 | 0.960619188 |
| chr8:108073144-108073517 | 0.050972341 | 0.890850117 | 0.960619188 |
| chr8:83693707-83693915 | -0.070816672 | 0.890887726 | 0.960628266 |
| chr1:30345508-30346113 | 0.054359402 | 0.890935035 | 0.960628266 |
| chr10:73413191-73414521 | -0.044057952 | 0.890939354 | 0.960628266 |
| chr4:104971576-104971834 | 0.055132637 | 0.890946167 | 0.960628266 |
| chr5:6435398-6435614 | 0.063918442 | 0.890946253 | 0.960628266 |
| chr1:111139322-111140827 | -0.043854362 | 0.890946795 | 0.960628266 |
| chr11:72091306-72091781 | 0.050751276 | 0.89096062 | 0.960628266 |
| chr12:80838223-80838547 | -0.059601297 | 0.89096588 | 0.960628266 |
| chr2:154855214-154855466 | -0.065495895 | 0.89098388 | 0.960628266 |
| chr5:123397683-123397966 | 0.06476704 | 0.890995364 | 0.960628266 |
| chr19:33041244-33041641 | 0.052817148 | 0.890997758 | 0.960628266 |
| chr14:61106879-61107129 | -0.057148244 | 0.89102152 | 0.960628266 |
| chr15:73244388-73244683 | -0.057148244 | 0.89102152 | 0.960628266 |
| chr8:122909449-122910054 | -0.044853646 | 0.891070879 | 0.96065396 |
| chr7:52260139-52260405 | 0.062966115 | 0.891081882 | 0.96065396 |
| chr5:140043174-140043493 | 0.05493823 | 0.891093164 | 0.96065396 |
| chr15:74781883-74782231 | -0.052905535 | 0.891095502 | 0.96065396 |
| chr4:84582227-84583398 | 0.043088574 | 0.891162383 | 0.960695551 |
| chr3:37967370-37967621 | -0.058271999 | 0.891169285 | 0.960695551 |
| chr5:169779686-169779893 | 0.071590988 | 0.891171696 | 0.960695551 |
| chr13:28021121-28021321 | 0.071423407 | 0.891213691 | 0.960727305 |
| chr6:2903129-2903554 | -0.052327303 | 0.891292127 | 0.960789494 |
| chr19:32971824-32972429 | -0.044572173 | 0.891296459 | 0.960789494 |
| chr5:140408894-140409165 | 0.066064968 | 0.891350773 | 0.960818235 |
| chr11:59776502-59776774 | 0.059983981 | 0.891355542 | 0.960818235 |
| chr8:73294887-73295186 | 0.049011017 | 0.89136074 | 0.960818235 |
| chr3:150762449-150764177 | -0.042633202 | 0.891408133 | 0.960830056 |
| chr15:60359742-60359964 | -0.064556243 | 0.891412066 | 0.960830056 |
| chr7:80942214-80943205 | 0.045329366 | 0.891420475 | 0.960830056 |
| chr10:24394648-24394880 | -0.071464304 | 0.891421866 | 0.960830056 |
| chr1:147529298-147529571 | -0.058923088 | 0.891446913 | 0.960843537 |
| chr4:118954325-118954575 | -0.062380021 | 0.89146263 | 0.960846961 |
| chr5:16288084-16288363 | 0.07103954 | 0.891486184 | 0.960851043 |
| chr1:22275085-22275675 | -0.052636794 | 0.891491497 | 0.960851043 |
| chr3:101724286-101725395 | -0.043164346 | 0.891562425 | 0.960903282 |
| chr8:94262006-94262569 | 0.045520863 | 0.891565047 | 0.960903282 |
| chr19:50431606-50432488 | -0.046342491 | 0.891593536 | 0.960915876 |
| chr3:72597809-72598029 | 0.06798915 | 0.891607536 | 0.960915876 |
| chr5:119037544-119037897 | 0.053394634 | 0.891614356 | 0.960915876 |
| chr20:57263838-57264852 | 0.042452663 | 0.891631938 | 0.96092131 |
| chr6:75504865-75505076 | -0.073099103 | 0.891649822 | 0.960927068 |

| | | | |
|---|---|---|---|
| chr17:59331249-59332520 | 0.043009197 | 0.89166585 | 0.960930826 |
| chr13:110455922-110456335 | 0.051731628 | 0.891688126 | 0.960941316 |
| chr21:17925055-17925327 | -0.061571468 | 0.891715635 | 0.960944425 |
| chr17:60345186-60345894 | 0.043523118 | 0.891716093 | 0.960944425 |
| chrX:71067705-71068811 | -0.04401271 | 0.891735479 | 0.960951801 |
| chr22:19959289-19959638 | 0.048731727 | 0.891811186 | 0.960985389 |
| chr2:11955518-11955733 | 0.072339468 | 0.891814381 | 0.960985389 |
| chr14:85028099-85028299 | -0.070218093 | 0.891825713 | 0.960985389 |
| chr19:49084840-49085776 | 0.043808234 | 0.891827132 | 0.960985389 |
| chr19:17859429-17860061 | 0.046165632 | 0.891838567 | 0.960985389 |
| chr17:75271607-75271921 | 0.052977527 | 0.891855429 | 0.960985389 |
| chr18:14132397-14132748 | 0.049861348 | 0.891863557 | 0.960985389 |
| chr1:207316127-207317076 | 0.043098635 | 0.891866984 | 0.960985389 |
| chr9:86176226-86176497 | -0.067989102 | 0.891976049 | 0.961057965 |
| chr6:63597819-63599072 | -0.044370596 | 0.891984514 | 0.961057965 |
| chr3:39288328-39288688 | 0.050107038 | 0.891985435 | 0.961057965 |
| chr15:65906466-65906687 | -0.061019329 | 0.892000882 | 0.961057965 |
| chr22:17570919-17571119 | -0.066743934 | 0.89200532 | 0.961057965 |
| chr20:8853827-8854085 | -0.057078446 | 0.892009596 | 0.961057965 |
| chr9:82740149-82740401 | 0.069254632 | 0.892064201 | 0.961103282 |
| chr17:63621510-63622571 | -0.042421048 | 0.892088403 | 0.961108228 |
| chr19:38361647-38362811 | 0.043902271 | 0.892093878 | 0.961108228 |
| chr12:95117176-95117481 | -0.057332467 | 0.892178304 | 0.96117875 |
| chr3:100254245-100254574 | 0.052169213 | 0.892184426 | 0.96117875 |
| chr6:146864686-146865000 | 0.070854724 | 0.89219778 | 0.961179622 |
| chr19:49443410-49444039 | -0.04566269 | 0.892259052 | 0.961222011 |
| chrX:139966200-139966409 | 0.069499538 | 0.892262216 | 0.961222011 |
| chr20:3209544-3210029 | 0.052489946 | 0.892302205 | 0.961231353 |
| chr15:87121336-87121712 | 0.051285352 | 0.892322874 | 0.961231353 |
| chr2:204012026-204012612 | 0.047360213 | 0.892334154 | 0.961231353 |
| chr12:1795857-1796475 | -0.04721477 | 0.892338462 | 0.961231353 |
| chr10:133324478-133324805 | -0.052215892 | 0.892352658 | 0.961231353 |
| chr1:107176820-107177101 | 0.049785483 | 0.892356494 | 0.961231353 |
| chr17:49575553-49575934 | -0.048532004 | 0.892391333 | 0.961231353 |
| chr1:248387688-248387986 | 0.065911168 | 0.892391388 | 0.961231353 |
| chr8:73665532-73665823 | 0.057546722 | 0.892395035 | 0.961231353 |
| chr6:100176052-100176297 | -0.064934744 | 0.892396339 | 0.961231353 |
| chr1:47842287-47842561 | -0.064430242 | 0.892420925 | 0.961244322 |
| chr16:29863001-29863785 | 0.045007298 | 0.892438615 | 0.961249864 |
| chr7:148883942-148885259 | -0.043024468 | 0.89249823 | 0.961273383 |
| chr12:67068325-67070044 | -0.043838587 | 0.892503605 | 0.961273383 |
| chr9:134164413-134165529 | 0.043204615 | 0.892506141 | 0.961273383 |
| chrX:115898697-115898969 | 0.063718804 | 0.892517685 | 0.961273383 |
| chr6:143744973-143745249 | 0.058414338 | 0.892523179 | 0.961273383 |
| chr14:24113939-24115437 | 0.042254621 | 0.892551681 | 0.961290568 |
| chr20:21810178-21810378 | 0.069639002 | 0.892602781 | 0.961332091 |
| chr13:33743560-33743984 | -0.051552392 | 0.892652378 | 0.961359574 |
| chr8:79783362-79784445 | 0.043746714 | 0.892653393 | 0.961359574 |
| chr5:170774899-170775168 | -0.06157273 | 0.892680538 | 0.961375296 |
| chr5:151619690-151620125 | 0.049842802 | 0.892704174 | 0.961387238 |

| | | | |
|---|---|---|---|
| chr5:10111604-10111889 | -0.052106038 | 0.892735269 | 0.961407213 |
| chr13:30486262-30486976 | 0.045509265 | 0.89276178 | 0.96142225 |
| chr19:5229183-5229811 | 0.051218628 | 0.892818251 | 0.961469551 |
| chr13:51674298-51674833 | -0.046702042 | 0.892847879 | 0.961476805 |
| chr6:19472140-19472520 | -0.054309323 | 0.892857934 | 0.961476805 |
| chr19:17309168-17309670 | -0.046140965 | 0.892862632 | 0.961476805 |
| chr13:44565150-44565350 | -0.079486308 | 0.892890245 | 0.961481402 |
| chr3:96984489-96984689 | 0.067262702 | 0.892891997 | 0.961481402 |
| chr4:165872663-165873000 | 0.056139265 | 0.892944456 | 0.96151359 |
| chr17:20116757-20117031 | -0.061401759 | 0.892946987 | 0.96151359 |
| chr14:39956476-39956676 | -0.069019783 | 0.89296115 | 0.961515329 |
| chr6:73653188-73654519 | -0.04254242 | 0.892991483 | 0.961528649 |
| chr5:35213608-35214472 | 0.043913721 | 0.892998619 | 0.961528649 |
| chr4:150875441-150875641 | -0.068505667 | 0.893017552 | 0.961530025 |
| chr14:59897790-59898065 | 0.056227955 | 0.893044884 | 0.961530025 |
| chr8:103414501-103415769 | 0.042010755 | 0.893058787 | 0.961530025 |
| chr13:40885249-40885477 | -0.067213417 | 0.893068522 | 0.961530025 |
| chr17:55750626-55751681 | -0.042909742 | 0.893085604 | 0.961530025 |
| chr12:77077548-77077763 | 0.072467847 | 0.89308872 | 0.961530025 |
| chr16:15147356-15147556 | 0.072467847 | 0.89308872 | 0.961530025 |
| chr1:28882069-28882452 | -0.048125153 | 0.893100289 | 0.961530025 |
| chr20:47704685-47705673 | 0.042029653 | 0.893123824 | 0.961532875 |
| chr11:65651586-65651874 | 0.052077281 | 0.893128035 | 0.961532875 |
| chr5:14793677-14794300 | -0.046142232 | 0.893160971 | 0.961552645 |
| chr1:215490093-215490350 | 0.056737065 | 0.893171496 | 0.961552645 |
| chrX:15493097-15493977 | -0.043860414 | 0.893190761 | 0.961559874 |
| chrX:104432304-104432787 | -0.052488697 | 0.893240197 | 0.961599584 |
| chr5:61822862-61823062 | -0.071384944 | 0.893255796 | 0.961602866 |
| chr3:124108091-124108355 | -0.064025228 | 0.893286638 | 0.961622557 |
| chr3:159872410-159872610 | -0.077257031 | 0.89331152 | 0.961635833 |
| chr12:132493629-132493829 | -0.055321723 | 0.893364138 | 0.961678964 |
| chr5:123035377-123037338 | -0.042356626 | 0.893403046 | 0.961701907 |
| chr6:44338396-44338866 | -0.049439739 | 0.893410554 | 0.961701907 |
| chr6:138817766-138818563 | 0.04368828 | 0.893473544 | 0.961744692 |
| chr3:65509156-65509851 | -0.04891417 | 0.893485293 | 0.961744692 |
| chr5:90547121-90547502 | 0.055282116 | 0.893516645 | 0.961744692 |
| chr4:167506541-167506831 | -0.067871249 | 0.893543122 | 0.961744692 |
| chr17:69005009-69005382 | -0.055937426 | 0.893548754 | 0.961744692 |
| chr4:147542358-147542777 | 0.05689096 | 0.893557551 | 0.961744692 |
| chr4:54217636-54218217 | 0.048805745 | 0.893574891 | 0.961744692 |
| chr2:98479980-98480551 | 0.048429263 | 0.893575464 | 0.961744692 |
| chr7:149634543-149634829 | -0.05856435 | 0.893576573 | 0.961744692 |
| chr2:187519342-187520123 | -0.046352286 | 0.893576808 | 0.961744692 |
| chr16:56691880-56692301 | -0.049900669 | 0.893588371 | 0.961744692 |
| chr5:95283725-95285293 | -0.042725006 | 0.893613575 | 0.961754207 |
| chr12:132226304-132226504 | 0.065485626 | 0.893627602 | 0.961754207 |
| chr19:58554911-58555253 | 0.048439696 | 0.893641724 | 0.961754207 |
| chr8:89273208-89273456 | 0.069407584 | 0.893647418 | 0.961754207 |
| chr17:61398883-61400741 | 0.041327685 | 0.89367 | 0.961765 |
| chr6:111105579-111105906 | 0.052547123 | 0.893702257 | 0.961786207 |

| | | | |
|---|---|---|---|
| chr11:86818470-86818884 | 0.053823516 | 0.893718193 | 0.961789848 |
| chr22:17876454-17876654 | -0.067184779 | 0.893740588 | 0.96180044 |
| chr16:69186731-69187793 | 0.044974369 | 0.89376247 | 0.96181048 |
| chr5:139651886-139652210 | 0.048291474 | 0.893779326 | 0.961815111 |
| chr1:110412530-110412868 | 0.051344959 | 0.893808498 | 0.961832994 |
| chr6:27162187-27162590 | -0.049682013 | 0.893921777 | 0.961850611 |
| chr1:9843514-9843883 | 0.054230192 | 0.893928054 | 0.961850611 |
| chr1:22024902-22025665 | -0.045600605 | 0.893928719 | 0.961850611 |
| chr13:26985346-26985589 | -0.056349073 | 0.893928963 | 0.961850611 |
| chr2:46276745-46276963 | -0.063986446 | 0.893929603 | 0.961850611 |
| chr20:47927895-47928288 | 0.04548622 | 0.893940159 | 0.961850611 |
| chr5:14364666-14364956 | -0.053002221 | 0.893942942 | 0.961850611 |
| chr3:107690612-107690830 | -0.065309402 | 0.893945906 | 0.961850611 |
| chr5:29114284-29114609 | 0.059884469 | 0.893956138 | 0.961850611 |
| chrX:35202204-35202426 | -0.073865285 | 0.893959163 | 0.961850611 |
| chr5:35954140-35954565 | -0.054770976 | 0.893962953 | 0.961850611 |
| chr9:113411252-113411504 | 0.055153121 | 0.894005388 | 0.961882761 |
| chr13:26221457-26222683 | -0.042648647 | 0.894038552 | 0.961894836 |
| chr17:76264802-76265418 | -0.043964949 | 0.894041718 | 0.961894836 |
| chrX:139776628-139776832 | -0.0700787 | 0.89406868 | 0.961910337 |
| chr4:153344205-153345514 | 0.04271992 | 0.894120312 | 0.96195238 |
| chr3:129245766-129246102 | -0.046260339 | 0.894140774 | 0.961960887 |
| chr1:206979417-206979642 | 0.056190288 | 0.894158299 | 0.961961455 |
| chr1:67853669-67853965 | 0.053326037 | 0.894177043 | 0.961961455 |
| chr10:123227850-123228059 | -0.07226934 | 0.894178966 | 0.961961455 |
| chr7:151226956-151227703 | 0.043249726 | 0.894262984 | 0.962038334 |
| chr6:29632795-29633553 | -0.044191935 | 0.894289976 | 0.962053865 |
| chr17:29370178-29370378 | -0.063405827 | 0.89430369 | 0.962055111 |
| chr12:104680114-104680345 | 0.054887329 | 0.894340666 | 0.962070463 |
| chr15:45320711-45320972 | 0.062757327 | 0.894348298 | 0.962070463 |
| chr9:70111026-70111236 | -0.066140575 | 0.894355629 | 0.962070463 |
| chr2:43293865-43294099 | -0.060894472 | 0.894385544 | 0.962074289 |
| chr1:39777038-39777238 | -0.071017167 | 0.894396854 | 0.962074289 |
| chr1:190436722-190436922 | -0.071017167 | 0.894396854 | 0.962074289 |
| chr19:45098302-45098514 | 0.064965586 | 0.894415156 | 0.962080469 |
| chr1:217868946-217869146 | -0.062736173 | 0.894439858 | 0.962093534 |
| chr4:57426821-57427098 | 0.060791565 | 0.894478437 | 0.962121525 |
| chr21:15483221-15483536 | -0.053708303 | 0.894513399 | 0.962144511 |
| chr4:155207598-155207803 | 0.062177791 | 0.894524922 | 0.962144511 |
| chr10:88131727-88132204 | -0.048648855 | 0.894547777 | 0.962155588 |
| chr22:31140341-31140756 | -0.049726926 | 0.894592734 | 0.962169321 |
| chr3:124300346-124300549 | 0.06451979 | 0.894598782 | 0.962169321 |
| chr5:13632757-13633037 | 0.062277932 | 0.894599567 | 0.962169321 |
| chr17:67603013-67603268 | 0.054703896 | 0.894610774 | 0.962169321 |
| chr10:43690098-43690391 | 0.050263116 | 0.89466882 | 0.962218243 |
| chrX:140503850-140504242 | 0.049643405 | 0.894710027 | 0.962242941 |
| chr6:87391375-87391637 | 0.056804074 | 0.8947169 | 0.962242941 |
| chr5:170420867-170421192 | -0.051066755 | 0.894738309 | 0.962252459 |
| chr10:119511833-119512134 | 0.054210292 | 0.894761139 | 0.962263358 |
| chr8:80052521-80052861 | 0.054073804 | 0.894774741 | 0.962263358 |

| | | | |
|---|---|---|---|
| chrX:139559690-139559945 | -0.060922861 | 0.894807771 | 0.962263358 |
| chr8:19816990-19818386 | -0.042247255 | 0.894809405 | 0.962263358 |
| chr14:90109756-90110289 | -0.046918075 | 0.894811236 | 0.962263358 |
| chr21:45910647-45910903 | -0.062159412 | 0.894867278 | 0.962305093 |
| chr2:172462109-172462333 | 0.057168695 | 0.894895085 | 0.962305093 |
| chr11:100575927-100576158 | -0.066181974 | 0.894903455 | 0.962305093 |
| chr16:71288877-71289850 | -0.042112804 | 0.894912164 | 0.962305093 |
| chr6:1311023-1312223 | 0.044145302 | 0.89491408 | 0.962305093 |
| chr2:10286270-10287067 | 0.044144177 | 0.894930537 | 0.962305093 |
| chr10:77140488-77140688 | 0.072313512 | 0.894937959 | 0.962305093 |
| chr15:72834145-72834345 | -0.072996537 | 0.894955868 | 0.962310845 |
| chr3:160413692-160413952 | 0.053986155 | 0.894984132 | 0.962326079 |
| chr9:114483523-114483910 | 0.046539389 | 0.894995154 | 0.962326079 |
| chr11:8309807-8310177 | -0.052079996 | 0.895032 | 0.962352192 |
| chr7:38660510-38660832 | 0.052947927 | 0.895077387 | 0.962387488 |
| chrX:130339400-130340339 | -0.04314844 | 0.895137402 | 0.962407524 |
| chr10:86966615-86966904 | 0.047274907 | 0.895144433 | 0.962407524 |
| chr2:37924876-37925841 | -0.042595606 | 0.895148628 | 0.962407524 |
| chr5:85247158-85247358 | 0.082934694 | 0.895150745 | 0.962407524 |
| chr9:117414262-117414811 | -0.048010664 | 0.895158824 | 0.962407524 |
| chr22:30667448-30668322 | -0.043195146 | 0.895176853 | 0.962413403 |
| chr2:60756021-60757061 | -0.042143483 | 0.895207614 | 0.9624256 |
| chr12:31682213-31682431 | -0.069999479 | 0.89522588 | 0.9624256 |
| chr13:23553993-23554221 | -0.069999479 | 0.89522588 | 0.9624256 |
| chr11:131694845-131695152 | -0.060380951 | 0.895288002 | 0.962470698 |
| chr13:35697544-35697948 | 0.05305095 | 0.895292951 | 0.962470698 |
| chr5:33934766-33935020 | -0.055955183 | 0.895306523 | 0.962471783 |
| chr20:1875779-1875979 | -0.071626425 | 0.895357516 | 0.962499195 |
| chr5:62143761-62143961 | -0.071626425 | 0.895357516 | 0.962499195 |
| chr3:142447142-142448336 | -0.042485197 | 0.895380782 | 0.962499195 |
| chr5:102226278-102226657 | 0.053830876 | 0.895382268 | 0.962499195 |
| chr17:49230728-49231223 | 0.046243705 | 0.895444037 | 0.962552091 |
| chr16:85420823-85421028 | -0.054194277 | 0.895461467 | 0.962557323 |
| chr1:62318135-62318490 | -0.058970599 | 0.895497482 | 0.962571284 |
| chrX:110408143-110408467 | 0.055468937 | 0.895502512 | 0.962571284 |
| chr4:10006957-10007211 | 0.058672081 | 0.895512143 | 0.962571284 |
| chr12:71636501-71636931 | -0.052888697 | 0.895548937 | 0.962590445 |
| chr1:88992447-88993356 | -0.041481737 | 0.895555094 | 0.962590445 |
| chr8:94623411-94623709 | -0.050377044 | 0.895640613 | 0.962598967 |
| chr14:55749502-55749883 | 0.053640279 | 0.895640647 | 0.962598967 |
| chr11:6251150-6251414 | -0.065292852 | 0.895642419 | 0.962598967 |
| chr8:116457186-116457402 | 0.066924058 | 0.895645227 | 0.962598967 |
| chr6:35449427-35449732 | 0.050515473 | 0.895646314 | 0.962598967 |
| chr5:146564089-146564313 | 0.064510481 | 0.895651385 | 0.962598967 |
| chr1:31358492-31358798 | 0.061995013 | 0.89565203 | 0.962598967 |
| chr6:138104754-138104999 | -0.060769905 | 0.895663527 | 0.962598967 |
| chr2:99489785-99490763 | 0.042059406 | 0.895699494 | 0.96262412 |
| chr12:104294295-104294537 | -0.059350118 | 0.89571242 | 0.96262451 |
| chr8:87143141-87143341 | 0.07323746 | 0.895751865 | 0.962628361 |
| chr8:121789247-121789447 | 0.07323746 | 0.895751865 | 0.962628361 |

| | | | |
|---|---|---|---|
| chr13:21212521-21212782 | -0.067375708 | 0.895753694 | 0.962628361 |
| chr8:27340595-27340957 | -0.051681959 | 0.895767202 | 0.962629377 |
| chr5:176442152-176442427 | 0.059110837 | 0.895788282 | 0.962629397 |
| chr3:13636122-13636411 | 0.05749037 | 0.895792348 | 0.962629397 |
| chr9:35489426-35490385 | 0.044237518 | 0.89586406 | 0.962681395 |
| chr13:31614679-31615002 | 0.062143905 | 0.895876622 | 0.962681395 |
| chr2:71747195-71748102 | -0.043840097 | 0.895878428 | 0.962681395 |
| chr21:46206698-46206898 | -0.064294975 | 0.895893906 | 0.962684527 |
| chr22:36301326-36301535 | 0.065079107 | 0.895930872 | 0.962705461 |
| chr5:149632402-149632602 | -0.06544981 | 0.895948335 | 0.962705461 |
| chr1:11053617-11053976 | 0.056879453 | 0.895951081 | 0.962705461 |
| chr7:141339658-141339888 | -0.062088951 | 0.895988896 | 0.96272414 |
| chr4:77121092-77121444 | -0.058154631 | 0.895993594 | 0.96272414 |
| chr20:10511568-10511860 | 0.051142569 | 0.896037879 | 0.962733708 |
| chr21:42183566-42183783 | -0.0577433 | 0.896039483 | 0.962733708 |
| chrX:32261956-32262400 | 0.057726023 | 0.896040192 | 0.962733708 |
| chr22:41301411-41301634 | -0.06176968 | 0.896073782 | 0.962737766 |
| chr1:222618366-222618744 | -0.051479853 | 0.896076164 | 0.962737766 |
| chr4:89598644-89598877 | -0.062942855 | 0.896108177 | 0.962737766 |
| chr17:66040499-66040717 | -0.062669649 | 0.896109099 | 0.962737766 |
| chr7:74566823-74567030 | -0.064055679 | 0.896117641 | 0.962737766 |
| chr14:78598053-78598408 | -0.048096757 | 0.896119359 | 0.962737766 |
| chr3:164221857-164222234 | -0.0606024 | 0.896133242 | 0.962739183 |
| chr15:91742778-91743012 | 0.062997946 | 0.896187139 | 0.962783587 |
| chr4:159427297-159427576 | 0.066369518 | 0.896228883 | 0.962814934 |
| chr15:40160710-40161410 | -0.043465163 | 0.896256304 | 0.962829064 |
| chr10:60100251-60100461 | -0.070637189 | 0.896267168 | 0.962829064 |
| chr11:72238156-72238510 | 0.049282145 | 0.896293123 | 0.962843446 |
| chr3:15631105-15631333 | 0.053321252 | 0.896319344 | 0.962845191 |
| chrX:9342466-9342840 | 0.047533599 | 0.89633744 | 0.962845191 |
| chr1:223041502-223041775 | 0.064475386 | 0.896341337 | 0.962845191 |
| chr4:84028357-84028677 | -0.055785087 | 0.896345012 | 0.962845191 |
| chr1:75797104-75797304 | -0.066518154 | 0.896374109 | 0.962854868 |
| chr12:105934374-105934636 | -0.059749824 | 0.896379153 | 0.962854868 |
| chr2:214447396-214447596 | 0.072998627 | 0.896418829 | 0.962875624 |
| chr12:123633476-123633971 | 0.043707565 | 0.896423608 | 0.962875624 |
| chr3:196317519-196318537 | 0.042259032 | 0.896460586 | 0.962897145 |
| chr4:153486273-153487180 | -0.042369886 | 0.896468779 | 0.962897145 |
| chr8:101155427-101155627 | 0.057345895 | 0.896496251 | 0.962899918 |
| chr19:39890670-39891007 | -0.056302123 | 0.896496494 | 0.962899918 |
| chr1:204319522-204319976 | -0.044940043 | 0.89651085 | 0.96290184 |
| chr3:154096946-154097207 | 0.069013874 | 0.896529461 | 0.962903058 |
| chr5:28215107-28215855 | 0.045118289 | 0.896537118 | 0.962903058 |
| chr3:89114088-89115161 | -0.042340012 | 0.896589721 | 0.962914499 |
| chr10:48568140-48568491 | 0.058138924 | 0.896593415 | 0.962914499 |
| chr12:120929038-120929659 | -0.045783676 | 0.896594586 | 0.962914499 |
| chr1:67429648-67429851 | 0.052317615 | 0.896598039 | 0.962914499 |
| chr20:54194622-54195133 | -0.041463458 | 0.896656762 | 0.962946113 |
| chr17:44186442-44187331 | 0.041707205 | 0.896661209 | 0.962946113 |
| chr4:26860226-26860887 | 0.042150788 | 0.896667843 | 0.962946113 |

| | | | |
|---|---|---|---|
| chr14:22766476-22767426 | -0.042627169 | 0.896687689 | 0.962946113 |
| chr1:216485219-216485474 | -0.067971111 | 0.896720709 | 0.962946113 |
| chrX:152861125-152861390 | -0.058892679 | 0.896723772 | 0.962946113 |
| chr1:224774443-224774657 | -0.069551883 | 0.896727202 | 0.962946113 |
| chr3:172490200-172490469 | -0.052562013 | 0.896728015 | 0.962946113 |
| chr10:70078264-70078466 | 0.06777737 | 0.896763954 | 0.96296089 |
| chr1:88736720-88737097 | 0.054786325 | 0.896807626 | 0.96296089 |
| chr19:48645654-48646864 | -0.042928291 | 0.896827313 | 0.96296089 |
| chr22:35921874-35922097 | 0.062712493 | 0.896828792 | 0.96296089 |
| chr22:37166555-37166826 | 0.062712493 | 0.896828792 | 0.96296089 |
| chr7:1087929-1088454 | 0.045193212 | 0.896830639 | 0.96296089 |
| chr22:24835276-24835492 | 0.060871223 | 0.896834358 | 0.96296089 |
| chr9:2087818-2088018 | 0.066981153 | 0.896842318 | 0.96296089 |
| chr6:87701267-87701670 | -0.047466381 | 0.896903427 | 0.963004424 |
| chr1:151006408-151006643 | 0.047183249 | 0.896909387 | 0.963004424 |
| chr11:118910427-118911540 | 0.041491861 | 0.896920568 | 0.963004424 |
| chr2:76096061-76096423 | -0.05324844 | 0.896946455 | 0.963018725 |
| chr1:181021972-181023351 | -0.040678231 | 0.896987446 | 0.963037105 |
| chr3:42547540-42547786 | -0.064794048 | 0.896988712 | 0.963037105 |
| chr21:36210090-36210379 | -0.057104293 | 0.897046445 | 0.963072548 |
| chr8:42139692-42140961 | -0.043697884 | 0.897046863 | 0.963072548 |
| chr6:30555834-30557383 | 0.040911461 | 0.89706125 | 0.963074501 |
| chr3:62531577-62531811 | 0.048798854 | 0.897078459 | 0.963079482 |
| chr20:62831485-62831722 | 0.052858254 | 0.897114417 | 0.963104591 |
| chr12:100348735-100348988 | -0.058294431 | 0.897142869 | 0.963121642 |
| chr2:100819958-100820238 | 0.055228816 | 0.897184934 | 0.963153305 |
| chr1:241029562-241030251 | -0.048709824 | 0.897224996 | 0.963154206 |
| chr8:58067480-58067846 | -0.053997684 | 0.897246649 | 0.963154206 |
| chr1:38489859-38490062 | -0.061384313 | 0.897262033 | 0.963154206 |
| chr12:2795777-2796217 | -0.049215586 | 0.897266144 | 0.963154206 |
| chr4:149980954-149981154 | -0.067960839 | 0.897274906 | 0.963154206 |
| chr11:102178291-102178643 | -0.052375489 | 0.897280718 | 0.963154206 |
| chr17:48892038-48893251 | -0.040512807 | 0.897291668 | 0.963154206 |
| chr10:99328986-99329677 | 0.044252531 | 0.897308815 | 0.963154206 |
| chr6:82040343-82040855 | -0.051081036 | 0.897323241 | 0.963154206 |
| chr12:57459151-57459468 | -0.04619796 | 0.897325374 | 0.963154206 |
| chr5:1948606-1948860 | 0.058604287 | 0.897345306 | 0.963154206 |
| chr11:118076730-118077329 | -0.045241449 | 0.897350303 | 0.963154206 |
| chr3:182739053-182739590 | 0.046737956 | 0.897358597 | 0.963154206 |
| chr17:67257948-67258232 | 0.054531066 | 0.897361756 | 0.963154206 |
| chr12:42144301-42145166 | -0.041978552 | 0.897401407 | 0.963167008 |
| chrX:52995137-52996092 | 0.041746394 | 0.897407656 | 0.963167008 |
| chr13:113028649-113029013 | -0.056934634 | 0.897411394 | 0.963167008 |
| chr1:155989142-155989694 | 0.045030399 | 0.89744571 | 0.963190346 |
| chr20:63115032-63115377 | 0.052911379 | 0.897473555 | 0.963206739 |
| chr7:33904955-33905186 | -0.053843328 | 0.89751108 | 0.963233521 |
| chr2:118013225-118014850 | 0.041057133 | 0.897539462 | 0.963250489 |
| chr1:201289880-201290159 | 0.050034797 | 0.89757234 | 0.963272282 |
| chr16:2964006-2964386 | -0.044978602 | 0.897621661 | 0.963311381 |
| chr17:17018672-17019013 | 0.049023271 | 0.897649237 | 0.963311381 |

| | | | |
|---|---|---|---|
| chr2:62411515-62411828 | -0.054944247 | 0.897653087 | 0.963311381 |
| chr10:110417424-110417631 | -0.061643644 | 0.897664877 | 0.963311381 |
| chr3:72724188-72724416 | 0.05671905 | 0.897677806 | 0.963311381 |
| chrX:18674460-18675412 | -0.041797685 | 0.897684206 | 0.963311381 |
| chr20:652095-652354 | -0.055157585 | 0.897724453 | 0.963340015 |
| chr9:113602687-113602950 | -0.054126168 | 0.897736035 | 0.963340015 |
| chr22:19947633-19948040 | 0.053115535 | 0.897765995 | 0.963341561 |
| chr2:43877945-43878171 | 0.049752044 | 0.897781744 | 0.963341561 |
| chr2:176814761-176815075 | 0.052885453 | 0.897784946 | 0.963341561 |
| chr8:80258558-80258763 | -0.062263133 | 0.897799981 | 0.963341561 |
| chr3:128879153-128879979 | 0.041259777 | 0.897800875 | 0.963341561 |
| chr20:34955370-34956172 | 0.042631778 | 0.897820546 | 0.963341561 |
| chr2:38741016-38741440 | -0.048442163 | 0.897825484 | 0.963341561 |
| chr13:113448680-113449740 | -0.04130361 | 0.897867422 | 0.963363127 |
| chr2:94207917-94208539 | 0.044898405 | 0.897877016 | 0.963363127 |
| chrX:17944960-17945407 | 0.059431943 | 0.897883302 | 0.963363127 |
| chr1:205955120-205955322 | 0.052739179 | 0.897930702 | 0.963400494 |
| chr1:175426269-175426469 | 0.066342325 | 0.89796997 | 0.963422219 |
| chr2:160210675-160211148 | 0.047906334 | 0.897976098 | 0.963422219 |
| chr4:105147799-105148044 | 0.053712956 | 0.898006951 | 0.963430185 |
| chr3:188037199-188037536 | 0.049053221 | 0.898008671 | 0.963430185 |
| chr12:109476905-109477959 | -0.042117403 | 0.898072224 | 0.963484878 |
| chr12:55721054-55721436 | 0.045442828 | 0.898099767 | 0.963500936 |
| chr10:102065081-102065879 | -0.042499273 | 0.898115409 | 0.963504226 |
| chr7:44776219-44776525 | -0.052528891 | 0.898130856 | 0.963507308 |
| chr1:202796185-202796832 | -0.044576959 | 0.898191076 | 0.963551081 |
| chr13:51120935-51121199 | -0.055433141 | 0.89819681 | 0.963551081 |
| chr1:156066425-156066627 | 0.058637365 | 0.898223868 | 0.963555584 |
| chrX:57120959-57121884 | -0.042189406 | 0.898228586 | 0.963555584 |
| chr5:26231400-26231600 | 0.06650769 | 0.898238734 | 0.963555584 |
| chr15:99565171-99566608 | 0.040787365 | 0.898321121 | 0.963615334 |
| chr12:124921672-124921928 | -0.045796891 | 0.89832659 | 0.963615334 |
| chr6:57089489-57090505 | -0.041529925 | 0.898332162 | 0.963615334 |
| chr20:9068291-9069647 | -0.040892347 | 0.898348401 | 0.963619263 |
| chr14:24263448-24264446 | 0.041446206 | 0.898378114 | 0.963628371 |
| chr10:14536967-14537299 | 0.049131915 | 0.898398667 | 0.963628371 |
| chr9:133375626-133376519 | -0.041671758 | 0.898400754 | 0.963628371 |
| chr20:48298814-48299424 | 0.041579101 | 0.898407198 | 0.963628371 |
| chr5:133051538-133052480 | -0.040866346 | 0.898529098 | 0.963739573 |
| chr1:174153515-174153809 | -0.057679493 | 0.898536029 | 0.963739573 |
| chr11:3037325-3037785 | 0.051685675 | 0.898582831 | 0.963776281 |
| chr17:9693383-9693583 | 0.074478606 | 0.898606476 | 0.96378815 |
| chr8:143966455-143966707 | 0.052890915 | 0.898669928 | 0.963842712 |
| chr22:35622124-35623024 | 0.042501169 | 0.898693267 | 0.963847037 |
| chr1:174883194-174883805 | -0.045049054 | 0.898699119 | 0.963847037 |
| chr3:88321678-88321981 | -0.054580174 | 0.898711732 | 0.963847073 |
| chr20:7859347-7859598 | 0.062526102 | 0.898730826 | 0.96384821 |
| chr4:147786574-147787171 | -0.04626689 | 0.89873795 | 0.96384821 |
| chr5:73591195-73591395 | 0.066287484 | 0.898791028 | 0.963891642 |
| chr13:45872791-45873094 | 0.058872941 | 0.898851789 | 0.963918914 |

| | | | |
|---|---|---|---|
| chr14:20657346-20658191 | 0.04083616 | 0.898879301 | 0.963918914 |
| chr1:170191883-170192083 | 0.066556614 | 0.898882359 | 0.963918914 |
| chr9:94726324-94727220 | 0.041445987 | 0.898884537 | 0.963918914 |
| chr12:89918570-89918928 | 0.050694087 | 0.898890257 | 0.963918914 |
| chr6:19269387-19270040 | 0.042986433 | 0.898899864 | 0.963918914 |
| chr17:81735687-81736142 | -0.04515509 | 0.89890452 | 0.963918914 |
| chr1:2195285-2195554 | 0.050406927 | 0.898939642 | 0.963940787 |
| chr8:86597424-86597645 | -0.06187327 | 0.898950078 | 0.963940787 |
| chr3:169657484-169657760 | 0.047206419 | 0.898968881 | 0.96394746 |
| chr3:147245443-147245643 | -0.061192817 | 0.898992401 | 0.963959189 |
| chr2:178480068-178481240 | 0.04035129 | 0.89902556 | 0.963981254 |
| chr12:48105673-48106640 | -0.042159481 | 0.899039379 | 0.963982582 |
| chr1:246608524-246609374 | -0.040374644 | 0.899081416 | 0.964011354 |
| chr11:125625442-125626647 | 0.040746826 | 0.89911269 | 0.964011354 |
| chr20:32652782-32653208 | -0.047621826 | 0.899116232 | 0.964011354 |
| chr4:41232594-41232871 | -0.061212339 | 0.899116539 | 0.964011354 |
| chr4:48275326-48275604 | -0.053995794 | 0.899134981 | 0.964017639 |
| chr1:246639368-246639644 | -0.054184511 | 0.899163745 | 0.964022145 |
| chr3:20340351-20340858 | -0.046938988 | 0.899164348 | 0.964022145 |
| chr10:69408475-69409142 | -0.042668554 | 0.899211698 | 0.964043728 |
| chr1:161421841-161422561 | -0.042126627 | 0.899219367 | 0.964043728 |
| chr22:41381021-41382396 | -0.039976136 | 0.899222224 | 0.964043728 |
| chr3:99753356-99753709 | -0.04921986 | 0.899239843 | 0.964049128 |
| chr4:169933623-169933885 | 0.057882171 | 0.899267553 | 0.96406003 |
| chr9:33233862-33234706 | -0.04176082 | 0.899275176 | 0.96406003 |
| chr1:150925594-150926788 | 0.040036694 | 0.899304117 | 0.964077568 |
| chr8:17155766-17156336 | -0.046581588 | 0.899330546 | 0.964083599 |
| chr5:178729885-178730241 | 0.05052014 | 0.899334908 | 0.964083599 |
| chr6:158536144-158537432 | 0.042408645 | 0.899385886 | 0.964124758 |
| chr3:4466495-4467559 | -0.040735351 | 0.89942309 | 0.964151151 |
| chr8:116755151-116756242 | -0.040845351 | 0.899437082 | 0.964152662 |
| chr22:39519628-39520825 | -0.041050117 | 0.899461344 | 0.964156243 |
| chr9:127396465-127397399 | 0.040182506 | 0.89946559 | 0.964156243 |
| chr1:181142364-181142811 | 0.043510623 | 0.899503642 | 0.964173505 |
| chr16:11493061-11493270 | -0.065123388 | 0.89950686 | 0.964173505 |
| chr2:230066975-230067329 | 0.047570586 | 0.899543285 | 0.964190516 |
| chr14:34368498-34368779 | -0.054362141 | 0.899547898 | 0.964190516 |
| chr2:230691114-230691376 | 0.055721811 | 0.89956902 | 0.964199668 |
| chr11:118911780-118912747 | 0.040248637 | 0.89961144 | 0.964226869 |
| chr22:41448219-41448812 | 0.040785401 | 0.899619566 | 0.964226869 |
| chr2:120540384-120540778 | 0.045349143 | 0.899667645 | 0.964264912 |
| chr17:10295820-10296083 | 0.053920511 | 0.899704363 | 0.964282384 |
| chr11:8721655-8721856 | 0.060116093 | 0.899709116 | 0.964282384 |
| chr17:63023014-63023464 | -0.047138713 | 0.899732686 | 0.964294157 |
| chr4:139167782-139168189 | -0.04747272 | 0.899789241 | 0.964341281 |
| chr3:194769027-194769706 | -0.040993014 | 0.899806146 | 0.96434591 |
| chr16:84746946-84747147 | 0.063386 | 0.899835876 | 0.964364284 |
| chr3:170499649-170499878 | -0.052381123 | 0.899859103 | 0.964367688 |
| chr8:100150403-100150899 | 0.041612514 | 0.899864224 | 0.964367688 |
| chr1:198724987-198725223 | 0.064948068 | 0.89988573 | 0.964376582 |

| | | | |
|---|---|---|---|
| chr4:53804904-53805109 | -0.064932179 | 0.8999065 | 0.964376582 |
| chr10:30671066-30671346 | -0.062220573 | 0.899919815 | 0.964376582 |
| chr3:53347134-53348108 | -0.041490025 | 0.899942626 | 0.964376582 |
| chr10:83806737-83807016 | 0.058505309 | 0.899953144 | 0.964376582 |
| chr17:57157496-57157733 | 0.058505309 | 0.899953144 | 0.964376582 |
| chr10:119602528-119602888 | 0.047567855 | 0.89997306 | 0.964376582 |
| chr17:10105841-10106129 | -0.059605787 | 0.899997923 | 0.964376582 |
| chrX:130092245-130092571 | -0.05703292 | 0.900000995 | 0.964376582 |
| chr14:56422652-56422939 | 0.050654997 | 0.900009996 | 0.964376582 |
| chr1:146039015-146039620 | 0.040232948 | 0.900010972 | 0.964376582 |
| chr16:68590474-68590851 | 0.048914593 | 0.900033717 | 0.964387468 |
| chr8:123173129-123173468 | 0.049842284 | 0.900057211 | 0.964399155 |
| chr14:24412169-24412415 | -0.050245186 | 0.900116198 | 0.96444249 |
| chr14:106802500-106802783 | 0.05201664 | 0.900136049 | 0.96444249 |
| chr20:54657481-54657896 | -0.041076375 | 0.90016064 | 0.96444249 |
| chr21:45128673-45128907 | -0.05509646 | 0.90016668 | 0.96444249 |
| chr10:49606963-49607273 | -0.048316761 | 0.900191026 | 0.96444249 |
| chr8:46215679-46215943 | 0.060867785 | 0.900192826 | 0.96444249 |
| chr1:109113608-109114527 | -0.041061503 | 0.900197494 | 0.96444249 |
| chr6:26205305-26205518 | 0.051991867 | 0.900252141 | 0.96444249 |
| chr1:145957766-145958307 | 0.040756845 | 0.900303755 | 0.96444249 |
| chr3:120469548-120469866 | -0.047458242 | 0.900310509 | 0.96444249 |
| chr8:128846873-128847121 | -0.061864415 | 0.900315141 | 0.96444249 |
| chr20:3173140-3173754 | 0.045710822 | 0.900338468 | 0.96444249 |
| chr5:7657461-7657688 | -0.063603408 | 0.900355015 | 0.96444249 |
| chr15:82849651-82850135 | 0.046530853 | 0.900358084 | 0.96444249 |
| chr3:95567462-95567760 | 0.058649925 | 0.90037491 | 0.96444249 |
| chr20:3094825-3095210 | 0.053635246 | 0.900376154 | 0.96444249 |
| chr14:66495320-66495812 | 0.046598037 | 0.900387019 | 0.96444249 |
| chr3:186269139-186269342 | 0.059298082 | 0.900390953 | 0.96444249 |
| chr9:129258325-129258778 | -0.043718387 | 0.900396177 | 0.96444249 |
| chr5:111671138-111671345 | 0.063994634 | 0.900401085 | 0.96444249 |
| chr1:158653276-158653477 | 0.072925735 | 0.900406423 | 0.96444249 |
| chr8:61614044-61614244 | -0.063508145 | 0.90040925 | 0.96444249 |
| chrX:18739750-18739950 | -0.063508145 | 0.90040925 | 0.96444249 |
| chr5:96483967-96484167 | -0.066894154 | 0.9004213 | 0.96444249 |
| chr22:36352935-36353152 | 0.066439025 | 0.900431138 | 0.96444249 |
| chr12:107450683-107450883 | 0.062157479 | 0.900444735 | 0.96444249 |
| chr1:67532459-67532748 | -0.0524853 | 0.900455534 | 0.96444249 |
| chr17:13017329-13018606 | -0.039659513 | 0.900465607 | 0.96444249 |
| chr1:108511761-108512194 | -0.046571053 | 0.900473118 | 0.96444249 |
| chr2:23926009-23927737 | 0.039468176 | 0.900475265 | 0.96444249 |
| chr16:19521454-19522308 | 0.04181733 | 0.900524897 | 0.964462084 |
| chr6:27824537-27824737 | -0.059907409 | 0.900530903 | 0.964462084 |
| chr3:50150834-50151446 | 0.042763167 | 0.900531321 | 0.964462084 |
| chr18:24413219-24413428 | -0.065603749 | 0.900580225 | 0.964493484 |
| chr2:11744045-11744461 | -0.041732874 | 0.900591826 | 0.964493484 |
| chr6:26285130-26285989 | 0.04051408 | 0.900620224 | 0.964493484 |
| chr9:81370157-81370453 | -0.050824445 | 0.900625434 | 0.964493484 |
| chr9:122838569-122838782 | -0.067323465 | 0.900626717 | 0.964493484 |

| | | | |
|---|---|---|---|
| chr9:109498217-109498493 | 0.051210019 | 0.900636165 | 0.964493484 |
| chr4:122450794-122451224 | -0.044669245 | 0.900651133 | 0.964496032 |
| chr14:101083673-101084118 | -0.046872509 | 0.900735829 | 0.964573251 |
| chr11:88183275-88183525 | 0.05955288 | 0.90078024 | 0.964591735 |
| chr1:111755017-111756425 | 0.039758268 | 0.900782947 | 0.964591735 |
| chr8:64457912-64458112 | 0.068488559 | 0.900809852 | 0.964591735 |
| chr8:133369395-133369627 | -0.059270477 | 0.900814377 | 0.964591735 |
| chr11:66288974-66289859 | 0.039757945 | 0.900827005 | 0.964591735 |
| chr3:129160724-129161882 | 0.039627929 | 0.900828624 | 0.964591735 |
| chr14:23010445-23010763 | 0.04688152 | 0.900848893 | 0.964599959 |
| chr5:80493486-80493902 | 0.045491453 | 0.900868511 | 0.964607485 |
| chr1:234531987-234532198 | 0.057866839 | 0.900959507 | 0.96467967 |
| chr8:106594699-106594956 | 0.053511615 | 0.900971126 | 0.96467967 |
| chr19:3990276-3990483 | -0.056445706 | 0.900997182 | 0.96467967 |
| chr22:39496482-39496682 | 0.065062728 | 0.900998877 | 0.96467967 |
| chr6:57536048-57536255 | 0.065062728 | 0.900998877 | 0.96467967 |
| chr5:95858721-95859177 | 0.044252846 | 0.901028994 | 0.964698437 |
| chr15:73964749-73965254 | -0.04558589 | 0.901130876 | 0.964794036 |
| chr11:72112073-72113020 | -0.039425228 | 0.901153625 | 0.964797752 |
| chr5:169265987-169266259 | 0.053735265 | 0.901177358 | 0.964797752 |
| chr4:116358581-116359225 | 0.043035846 | 0.90117759 | 0.964797752 |
| chr5:134738114-134738857 | -0.040450721 | 0.901194826 | 0.964797752 |
| chr1:172542330-172542559 | 0.068409105 | 0.901227316 | 0.964797752 |
| chr14:57871625-57871841 | -0.058927758 | 0.901234948 | 0.964797752 |
| chr15:74873698-74874051 | -0.047226868 | 0.90124829 | 0.964797752 |
| chr2:85741903-85742215 | -0.048830349 | 0.901252916 | 0.964797752 |
| chr14:58620233-58620459 | 0.060018717 | 0.901260283 | 0.964797752 |
| chr20:10383402-10383602 | -0.067547166 | 0.901273458 | 0.964797752 |
| chr11:124867726-124867926 | -0.06008244 | 0.901285447 | 0.964797752 |
| chr15:83372619-83372860 | -0.06008244 | 0.901285447 | 0.964797752 |
| chr4:88612336-88612536 | -0.068485448 | 0.901325232 | 0.964820008 |
| chrX:102976848-102977086 | -0.06317768 | 0.901331422 | 0.964820008 |
| chr10:23053134-23053420 | -0.055497391 | 0.901398868 | 0.964860198 |
| chr17:56780044-56780894 | 0.04036158 | 0.901407498 | 0.964860198 |
| chr15:79130634-79130834 | -0.067371241 | 0.901409626 | 0.964860198 |
| chr8:105054147-105054361 | -0.068636052 | 0.901419337 | 0.964860198 |
| chr8:108511837-108512298 | 0.049506253 | 0.90144924 | 0.964869924 |
| chr1:223665640-223665891 | 0.055391458 | 0.901463413 | 0.964869924 |
| chr17:63946839-63947065 | -0.046919371 | 0.901466201 | 0.964869924 |
| chr22:33852081-33852283 | -0.059340329 | 0.901502318 | 0.964895103 |
| chr7:73683113-73684124 | -0.041002677 | 0.901525745 | 0.964898421 |
| chr5:188610-188949 | 0.043842592 | 0.901530605 | 0.964898421 |
| chr3:100462779-100463249 | -0.047976373 | 0.901574065 | 0.964924818 |
| chr14:102222214-102222414 | -0.063459355 | 0.901581172 | 0.964924818 |
| chr5:19314867-19315374 | -0.045000688 | 0.901596991 | 0.964924818 |
| chrX:109890933-109891259 | 0.047537317 | 0.901605641 | 0.964924818 |
| chr17:7380868-7381135 | -0.052622972 | 0.901632825 | 0.964940433 |
| chr2:189466735-189467190 | -0.04921319 | 0.901662827 | 0.964959064 |
| chr14:67513892-67514494 | 0.043596428 | 0.901710556 | 0.964996665 |
| chr15:68730445-68730673 | 0.055177688 | 0.901736206 | 0.965010637 |

| | | | |
|---|---|---|---|
| chr11:72471125-72471644 | -0.04633898 | 0.901761788 | 0.965011506 |
| chr17:20009815-20010394 | -0.045330284 | 0.901777536 | 0.965011506 |
| chrX:53225125-53225779 | 0.042158387 | 0.901780996 | 0.965011506 |
| chr7:17696402-17697017 | -0.041503657 | 0.901787395 | 0.965011506 |
| chr14:70808312-70808768 | 0.044370048 | 0.901834311 | 0.965032472 |
| chr19:38618609-38619368 | 0.04070411 | 0.901836012 | 0.965032472 |
| chr12:51424594-51424913 | 0.045448281 | 0.901857213 | 0.965032472 |
| chr1:203267322-203267777 | -0.043389263 | 0.901857367 | 0.965032472 |
| chr1:173503487-173503689 | 0.060440421 | 0.901887736 | 0.965042505 |
| chrX:13734259-13735349 | 0.039455705 | 0.901916536 | 0.965042505 |
| chr7:26278210-26278438 | 0.062473369 | 0.901926321 | 0.965042505 |
| chr14:33781339-33781638 | 0.048519431 | 0.90193837 | 0.965042505 |
| chr13:49607031-49607402 | -0.04703247 | 0.901954668 | 0.965042505 |
| chr2:133674753-133675349 | 0.046093617 | 0.901985637 | 0.965042505 |
| chr5:72847961-72849055 | -0.040345919 | 0.901998553 | 0.965042505 |
| chr6:109236258-109236678 | 0.047414843 | 0.902017224 | 0.965042505 |
| chr3:134145862-134146073 | 0.061681593 | 0.902017478 | 0.965042505 |
| chr3:23183701-23184107 | -0.045058909 | 0.902023557 | 0.965042505 |
| chr6:105179361-105180528 | 0.041702968 | 0.902025077 | 0.965042505 |
| chr14:59969375-59969820 | -0.049659933 | 0.902032738 | 0.965042505 |
| chr9:107504507-107505158 | -0.044131217 | 0.902062471 | 0.965042505 |
| chr9:122093221-122093723 | 0.043126491 | 0.902076209 | 0.965042505 |
| chr15:97909355-97909649 | -0.049647541 | 0.90207673 | 0.965042505 |
| chr3:119185279-119185522 | -0.059675892 | 0.902079327 | 0.965042505 |
| chr5:151223513-151224294 | -0.041322568 | 0.902080856 | 0.965042505 |
| chr5:31395497-31396133 | 0.041744726 | 0.902107975 | 0.965049099 |
| chr6:147081081-147081582 | -0.04571748 | 0.902120754 | 0.965049099 |
| chr2:170530238-170530586 | 0.051121065 | 0.902133365 | 0.965049099 |
| chr1:48530764-48531281 | -0.04624307 | 0.902147617 | 0.965049099 |
| chr12:87120371-87120879 | -0.042862571 | 0.90215524 | 0.965049099 |
| chr11:73141859-73143067 | -0.039428226 | 0.90216259 | 0.965049099 |
| chr13:19587454-19588027 | -0.043196308 | 0.902188042 | 0.965055636 |
| chr12:116492898-116493183 | 0.051305558 | 0.902215799 | 0.965055636 |
| chr2:7429819-7430019 | -0.060411993 | 0.902219958 | 0.965055636 |
| chr10:124916227-124917255 | 0.039926969 | 0.902240361 | 0.965055636 |
| chr6:28597173-28597875 | -0.042827477 | 0.902250043 | 0.965055636 |
| chr17:8209964-8211007 | 0.04120305 | 0.902250231 | 0.965055636 |
| chr3:40179048-40179334 | -0.055679335 | 0.90226712 | 0.965055636 |
| chr4:94788697-94788984 | 0.056015586 | 0.90226946 | 0.965055636 |
| chr3:106609764-106610001 | -0.053261699 | 0.90230141 | 0.965075605 |
| chr12:108587979-108588179 | 0.061716128 | 0.902313321 | 0.965075605 |
| chr11:100687132-100688127 | 0.041398947 | 0.90233849 | 0.965079593 |
| chr1:49298397-49298639 | -0.061662865 | 0.902342241 | 0.965079593 |
| chr17:48047577-48049172 | 0.038398544 | 0.902357437 | 0.965082376 |
| chr3:107522668-107523648 | 0.039225904 | 0.90241068 | 0.965120999 |
| chr1:25430106-25431046 | -0.040923742 | 0.902418742 | 0.965120999 |
| chr3:120650304-120650888 | 0.044391788 | 0.90243625 | 0.965126253 |
| chr6:169722945-169723153 | -0.05244973 | 0.902493512 | 0.96517402 |
| chr11:57657281-57658031 | -0.040055401 | 0.902524677 | 0.965187628 |
| chr6:72621427-72621627 | -0.056823316 | 0.902556368 | 0.965187628 |

| | | | |
|---|---|---|---|
| chr16:2244180-2244488 | -0.054941125 | 0.902558762 | 0.965187628 |
| chr14:44189995-44190220 | -0.066147874 | 0.902565017 | 0.965187628 |
| chr3:13513559-13513931 | -0.048318965 | 0.902569219 | 0.965187628 |
| chr1:5913453-5913842 | 0.054097792 | 0.902599209 | 0.965206227 |
| chr7:3154974-3155257 | 0.051254041 | 0.902622137 | 0.965207819 |
| chr10:133373078-13337377‍8 | -0.04143719 | 0.902634994 | 0.965207819 |
| chr16:48165992-48166387 | -0.050304199 | 0.902638529 | 0.965207819 |
| chr6:38973118-38973461 | 0.054904591 | 0.902658595 | 0.965207819 |
| chr11:12653728-12653946 | 0.060953016 | 0.90267628 | 0.965207819 |
| chr14:53041057-53041333 | 0.060953016 | 0.90267628 | 0.965207819 |
| chr8:86592117-86592484 | 0.050475318 | 0.902697682 | 0.965217234 |
| chr13:21625263-21625602 | 0.056310233 | 0.902739637 | 0.965247932 |
| chr3:73070035-73070235 | 0.065202582 | 0.902751586 | 0.965247932 |
| chr1:201597443-201598309 | 0.039131706 | 0.902779427 | 0.96526423 |
| chr13:26872953-26873187 | -0.058252377 | 0.902821255 | 0.965293741 |
| chr2:191150376-191151496 | 0.040990296 | 0.90283329 | 0.965293741 |
| chr21:43788963-43789838 | -0.039866988 | 0.902864445 | 0.965293741 |
| chr17:44071051-44071310 | -0.051201201 | 0.902866397 | 0.965293741 |
| chr8:11429295-11429635 | 0.050852401 | 0.90287729 | 0.965293741 |
| chr3:99173082-99173351 | -0.054023821 | 0.902882617 | 0.965293741 |
| chr6:169737731-169737931 | -0.065374107 | 0.902939596 | 0.965309324 |
| chr8:62984125-62984334 | -0.065374107 | 0.902939596 | 0.965309324 |
| chr4:107050545-107050915 | 0.050163445 | 0.90294938 | 0.965309324 |
| chr16:49283124-49283324 | 0.066104581 | 0.902958953 | 0.965309324 |
| chr2:232439485-232439685 | 0.062493733 | 0.90296124 | 0.965309324 |
| chr1:146035453-146035864 | 0.039412711 | 0.903000726 | 0.965309324 |
| chr21:25607010-25608106 | 0.039829878 | 0.903011801 | 0.965309324 |
| chr7:7681204-7681803 | 0.042396891 | 0.90301634 | 0.965309324 |
| chr17:66915115-66915503 | -0.042356978 | 0.903026111 | 0.965309324 |
| chr22:39310956-39311321 | -0.049332946 | 0.903031465 | 0.965309324 |
| chr5:180353052-180353674 | 0.041947445 | 0.903035773 | 0.965309324 |
| chr3:138082517-138082984 | 0.044500355 | 0.903054912 | 0.965316315 |
| chrX:105454727-105454963 | -0.062504582 | 0.90307719 | 0.965326662 |
| chr5:14270166-14270521 | -0.046948957 | 0.903130618 | 0.965361185 |
| chr15:74768881-74769180 | 0.047983052 | 0.903134685 | 0.965361185 |
| chr22:36558324-36558524 | 0.06649939 | 0.903185977 | 0.965402543 |
| chr12:6765871-6766400 | 0.041426571 | 0.903239511 | 0.965437471 |
| chr5:41722287-41722650 | -0.055728409 | 0.903258586 | 0.965437471 |
| chr17:7220233-7220851 | -0.042665484 | 0.90326809 | 0.965437471 |
| chr6:24583462-24583711 | 0.053559922 | 0.903269054 | 0.965437471 |
| chr5:26538377-26538658 | 0.055612782 | 0.903291669 | 0.965448176 |
| chr12:14284041-14284391 | 0.051304505 | 0.90331869 | 0.965463589 |
| chr17:39703389-39703706 | 0.049993026 | 0.903348638 | 0.96548213 |
| chr1:245197867-245198258 | -0.06163125 | 0.903427152 | 0.965546523 |
| chr10:70221568-70221801 | 0.061247817 | 0.903447242 | 0.965546523 |
| chr6:28940696-28941816 | -0.03852441 | 0.903462805 | 0.965546523 |
| chr5:149654142-149654427 | -0.060612422 | 0.903483395 | 0.965546523 |
| chr3:161072181-161072414 | -0.054473797 | 0.90351344 | 0.965546523 |
| chr19:38233852-38234152 | 0.052980922 | 0.9035182 | 0.965546523 |
| chr11:129763953-12976457‍9 | 0.044446388 | 0.903525962 | 0.965546523 |

| | | | |
|---|---|---|---|
| chr15:73636436-73636763 | -0.04294176 | 0.903531033 | 0.965546523 |
| chr12:56941289-56941954 | -0.044615473 | 0.903534796 | 0.965546523 |
| chr20:38747975-38749007 | -0.039482129 | 0.903541695 | 0.965546523 |
| chr2:25664554-25665088 | -0.044145081 | 0.903548666 | 0.965546523 |
| chr16:23727782-23728229 | 0.045861021 | 0.903560104 | 0.965546523 |
| chr2:191646145-191646403 | -0.052533266 | 0.903590346 | 0.965564842 |
| chr12:12999770-13000073 | -0.048759217 | 0.903602451 | 0.965564842 |
| chr12:120903437-120903728 | -0.050710413 | 0.903654 | 0.965579762 |
| chr17:56960558-56961289 | -0.038743791 | 0.90366314 | 0.965579762 |
| chr22:22357247-22357660 | 0.047427039 | 0.903666813 | 0.965579762 |
| chr3:10783465-10783678 | -0.057390969 | 0.903666821 | 0.965579762 |
| chr10:116690878-116691078 | 0.065210563 | 0.903713591 | 0.965602746 |
| chr19:18223885-18224344 | 0.043925744 | 0.90373278 | 0.965602746 |
| chr7:73308673-73308993 | 0.049033336 | 0.903733018 | 0.965602746 |
| chr19:50329053-50329741 | 0.038785666 | 0.90373874 | 0.965602746 |
| chr2:208148873-208149083 | -0.067610738 | 0.903778428 | 0.965628986 |
| chr8:74090506-74090760 | 0.050333088 | 0.903800689 | 0.965628986 |
| chr1:204351636-204351916 | -0.047385396 | 0.903801106 | 0.965628986 |
| chr6:136644130-136644533 | 0.044404422 | 0.903815682 | 0.965631094 |
| chr10:93482130-93482845 | 0.041573549 | 0.903870849 | 0.965665513 |
| chr19:40603800-40604510 | 0.039209253 | 0.903873103 | 0.965665513 |
| chr12:100951791-100952085 | -0.051429466 | 0.903889924 | 0.965670019 |
| chr12:98566863-98567426 | -0.041710068 | 0.903905041 | 0.965672705 |
| chr19:7680753-7681240 | 0.046782128 | 0.903929045 | 0.965675203 |
| chr11:57335287-57336130 | -0.040170725 | 0.903932586 | 0.965675203 |
| chr9:128733200-128733737 | -0.044150713 | 0.903987503 | 0.965711544 |
| chr19:4338580-4338974 | -0.045052418 | 0.904011088 | 0.965711544 |
| chr14:59632386-59632676 | 0.049107024 | 0.904014943 | 0.965711544 |
| chr20:33719992-33720723 | -0.040011518 | 0.904046096 | 0.965711544 |
| chr12:40692666-40692928 | -0.05581295 | 0.904076113 | 0.965711544 |
| chr10:118300388-118300732 | 0.053247843 | 0.904079228 | 0.965711544 |
| chr19:39435700-39436290 | 0.040520302 | 0.904089145 | 0.965711544 |
| chr15:80633885-80634085 | -0.064412712 | 0.904100045 | 0.965711544 |
| chr2:196292256-196292844 | -0.042270442 | 0.904100229 | 0.965711544 |
| chr9:27535870-27536071 | -0.058765851 | 0.904129032 | 0.965711544 |
| chr16:89357044-89357279 | 0.052959563 | 0.904133185 | 0.965711544 |
| chr5:152207290-152207568 | -0.055443919 | 0.904137618 | 0.965711544 |
| chr7:44513841-44514107 | -0.055443919 | 0.904137618 | 0.965711544 |
| chr16:67544325-67544601 | 0.050735071 | 0.904143054 | 0.965711544 |
| chr11:122581001-122581438 | 0.044132793 | 0.904158665 | 0.965713115 |
| chr5:14352090-14352433 | 0.043537126 | 0.904192423 | 0.965713115 |
| chr3:77319772-77320033 | -0.053521321 | 0.904195233 | 0.965713115 |
| chr7:49438251-49438491 | 0.061572965 | 0.904207802 | 0.965713115 |
| chr4:147759984-147761045 | 0.038687404 | 0.904208029 | 0.965713115 |
| chr12:84497913-84498113 | -0.065446694 | 0.90422583 | 0.965713115 |
| chr5:135514149-135514423 | -0.061317239 | 0.904232749 | 0.965713115 |
| chr6:107418222-107418422 | 0.061336154 | 0.904259029 | 0.965717112 |
| chr12:57725819-57726900 | -0.038599119 | 0.904261699 | 0.965717112 |
| chrX:90601644-90602359 | -0.044635804 | 0.904415905 | 0.965868335 |
| chr2:28175052-28175439 | -0.04752646 | 0.904448291 | 0.965879244 |

| | | | |
|---|---|---|---|
| chr19:17327661-17328286 | 0.040967891 | 0.904451331 | 0.965879244 |
| chr1:246696517-246696717 | -0.050621619 | 0.904463969 | 0.965879278 |
| chr19:12847584-12847897 | 0.052766484 | 0.904511787 | 0.965909041 |
| chr19:15555784-15556177 | 0.04399397 | 0.90453209 | 0.965909041 |
| chr3:43690232-43691671 | -0.038691826 | 0.904566025 | 0.965909041 |
| chr3:129398894-129399923 | -0.03899886 | 0.904573172 | 0.965909041 |
| chr3:83233815-83234188 | -0.053393615 | 0.904616556 | 0.965909041 |
| chr2:65071596-65071832 | 0.05130678 | 0.904619073 | 0.965909041 |
| chr1:242129002-242129471 | -0.045605842 | 0.904630519 | 0.965909041 |
| chr1:32003603-32003848 | 0.058950131 | 0.904642218 | 0.965909041 |
| chr3:98593016-98594323 | -0.0382745 | 0.904644322 | 0.965909041 |
| chr5:102153264-102153667 | 0.046496699 | 0.904650634 | 0.965909041 |
| chr8:88244078-88244305 | 0.053149379 | 0.904652463 | 0.965909041 |
| chr2:85327494-85328596 | -0.038838887 | 0.904674568 | 0.965909041 |
| chr22:49960045-49960798 | 0.039550417 | 0.904683139 | 0.965909041 |
| chr16:85316368-85316687 | -0.044140062 | 0.90470856 | 0.965909041 |
| chr8:94008929-94009783 | -0.040728728 | 0.904742713 | 0.965909041 |
| chrX:68490652-68490997 | 0.047773709 | 0.904750527 | 0.965909041 |
| chr5:69341589-69341789 | -0.067609945 | 0.904755919 | 0.965909041 |
| chr5:159939613-159939813 | -0.067609945 | 0.904755919 | 0.965909041 |
| chr8:119054415-119054615 | -0.067609945 | 0.904755919 | 0.965909041 |
| chr1:234309420-234310328 | 0.040343568 | 0.904756755 | 0.965909041 |
| chr6:10747062-10748150 | -0.038178913 | 0.904759695 | 0.965909041 |
| chr3:43105645-43106261 | 0.043189014 | 0.9047776 | 0.965909041 |
| chr20:32256645-32256913 | -0.05769915 | 0.904781781 | 0.965909041 |
| chr10:123967665-123967935 | -0.051850185 | 0.904819047 | 0.965923209 |
| chr15:31214671-31214871 | 0.058949071 | 0.904824893 | 0.965923209 |
| chr3:61936372-61936703 | 0.04053678 | 0.904857496 | 0.965923209 |
| chr14:43898701-43899036 | 0.049057852 | 0.904860369 | 0.965923209 |
| chr16:3067563-3067785 | 0.058269929 | 0.904862816 | 0.965923209 |
| chr8:12665514-12665717 | -0.065548677 | 0.90487069 | 0.965923209 |
| chr5:179678079-179678826 | 0.039367897 | 0.9048965 | 0.965937303 |
| chr5:37413020-37413359 | -0.045010005 | 0.904987239 | 0.966020705 |
| chr2:24613122-24613637 | -0.044138912 | 0.905016798 | 0.966038799 |
| chrX:100641993-100642229 | 0.060140434 | 0.905051514 | 0.966062397 |
| chr2:112764327-112765162 | 0.039417952 | 0.905087172 | 0.966069754 |
| chr1:8259281-8259522 | -0.061576899 | 0.905091611 | 0.966069754 |
| chr10:49282862-49283062 | -0.060018281 | 0.90509623 | 0.966069754 |
| chr15:83930766-83931240 | 0.043599627 | 0.905125767 | 0.966075343 |
| chr22:50327005-50327475 | 0.047503422 | 0.905126683 | 0.966075343 |
| chr3:17894512-17894967 | -0.044500164 | 0.90522189 | 0.966150854 |
| chr1:202999342-202999794 | -0.042683767 | 0.905222649 | 0.966150854 |
| chr19:15556368-15556737 | -0.047086773 | 0.905279762 | 0.966179149 |
| chr13:36395362-36395562 | 0.059444922 | 0.905302618 | 0.966179149 |
| chr4:55345984-55347160 | -0.038636987 | 0.905311734 | 0.966179149 |
| chr8:120089360-120089597 | 0.051554485 | 0.905312979 | 0.966179149 |
| chr5:79858202-79858481 | -0.050014263 | 0.905315545 | 0.966179149 |
| chr1:31956075-31956356 | -0.050863333 | 0.905332609 | 0.966179149 |
| chr14:95314725-95315060 | 0.046566543 | 0.905340057 | 0.966179149 |
| chr22:41092383-41092685 | -0.047316162 | 0.905350037 | 0.966179149 |

| | | | |
|---|---|---|---|
| chr17:62170351-62170769 | -0.043906717 | 0.905420129 | 0.966220366 |
| chr11:16604942-16605274 | 0.050183901 | 0.905425002 | 0.966220366 |
| chr12:6533249-6534875 | -0.037819697 | 0.905426489 | 0.966220366 |
| chr9:95127206-95127936 | -0.043086144 | 0.905522214 | 0.966308532 |
| chr2:219021647-219021936 | -0.049463629 | 0.905534331 | 0.966308532 |
| chr16:22195652-22196208 | 0.03941054 | 0.905567724 | 0.966330708 |
| chr8:97569627-97570016 | -0.049105903 | 0.905625382 | 0.966362521 |
| chr19:48637209-48637912 | -0.038499139 | 0.905637042 | 0.966362521 |
| chr2:105235342-105235946 | -0.041756272 | 0.905637434 | 0.966362521 |
| chr18:79246228-79246895 | 0.039120326 | 0.905647985 | 0.966362521 |
| chr10:35218341-35218840 | 0.043705667 | 0.90569107 | 0.966388718 |
| chr15:42084486-42084691 | 0.054979616 | 0.905697761 | 0.966388718 |
| chr6:136549439-136551087 | 0.037513862 | 0.905761079 | 0.966435181 |
| chr5:43000137-43001294 | 0.038161773 | 0.905777195 | 0.966435181 |
| chr7:150276201-150276401 | 0.059517491 | 0.905779145 | 0.966435181 |
| chr2:161244534-161244941 | 0.04417906 | 0.905801109 | 0.966445158 |
| chr4:109559710-109561230 | 0.037791518 | 0.905853541 | 0.966477744 |
| chr14:50001040-50001945 | 0.037974711 | 0.905866838 | 0.966477744 |
| chr11:48210120-48210330 | -0.061927051 | 0.905894117 | 0.966477744 |
| chr1:67716903-67717103 | -0.060254535 | 0.905898937 | 0.966477744 |
| chr17:41777534-41777792 | -0.050033192 | 0.905910278 | 0.966477744 |
| chr16:68364715-68364973 | 0.050890035 | 0.905915161 | 0.966477744 |
| chr9:4297130-4298280 | 0.039311564 | 0.905919945 | 0.966477744 |
| chr3:183635369-183636993 | 0.03742947 | 0.905947394 | 0.966482038 |
| chr18:5057812-5058060 | 0.0619421 | 0.905959767 | 0.966482038 |
| chr19:50312211-50312691 | 0.041911386 | 0.905961812 | 0.966482038 |
| chr17:61927028-61928410 | -0.03702932 | 0.906032707 | 0.966502385 |
| chr19:5689989-5690559 | -0.042124258 | 0.906040058 | 0.966502385 |
| chr1:108034453-108034903 | 0.044191058 | 0.906046367 | 0.966502385 |
| chr8:63405909-63406186 | -0.055618319 | 0.906056938 | 0.966502385 |
| chr13:51911955-51912490 | 0.043200297 | 0.906057245 | 0.966502385 |
| chr6:41071836-41072065 | -0.045201242 | 0.906073785 | 0.966502385 |
| chr17:75572783-75573010 | -0.048100191 | 0.906080059 | 0.966502385 |
| chr9:124794950-124795187 | -0.05392893 | 0.906081795 | 0.966502385 |
| chr1:192457869-192458090 | -0.062095815 | 0.90611063 | 0.966519687 |
| chr3:140941369-140942734 | -0.037203156 | 0.906132488 | 0.966529547 |
| chr8:88088543-88088779 | 0.057162697 | 0.906160754 | 0.966546242 |
| chr8:25175252-25175578 | -0.048354913 | 0.906208258 | 0.966577033 |
| chrX:103698808-103699172 | 0.047042674 | 0.906214851 | 0.966577033 |
| chr5:38994423-38994623 | -0.064075286 | 0.906238981 | 0.966589315 |
| chr6:142566801-142567115 | -0.044925434 | 0.906259973 | 0.966598249 |
| chr1:182515890-182516262 | -0.053350873 | 0.906293557 | 0.966620614 |
| chr11:59171463-59171815 | 0.041866362 | 0.906309399 | 0.966624056 |
| chrX:70979494-70979739 | -0.055979678 | 0.906344122 | 0.966639765 |
| chr8:144291130-144291836 | 0.038913821 | 0.90634936 | 0.966639765 |
| chr4:113278372-113278671 | -0.047095547 | 0.906383673 | 0.966662906 |
| chr1:179193781-179194116 | 0.050700577 | 0.906402186 | 0.966669195 |
| chr3:130242884-130243280 | -0.044100247 | 0.906426674 | 0.966681856 |
| chr17:2392659-2393435 | -0.038619046 | 0.906460807 | 0.966704803 |
| chr1:200185799-200186039 | -0.046925439 | 0.906493271 | 0.966710352 |

| | | | |
|---|---|---|---|
| chr11:110377060-110377363 | -0.048378208 | 0.906493418 | 0.966710352 |
| chr6:70955830-70956540 | -0.040347099 | 0.906527703 | 0.966710352 |
| chr10:103879387-103879728 | 0.051920528 | 0.906533071 | 0.966710352 |
| chr13:75133265-75133670 | 0.054220746 | 0.906540044 | 0.966710352 |
| chr2:174341537-174341924 | 0.04463503 | 0.90654171 | 0.966710352 |
| chr9:100602413-100602671 | -0.048280304 | 0.906622192 | 0.966763657 |
| chr3:120678626-120678826 | 0.054952784 | 0.906642718 | 0.966763657 |
| chr13:37969633-37970055 | -0.045688004 | 0.9066443 | 0.966763657 |
| chr11:3057133-3058017 | -0.038414444 | 0.906664583 | 0.966763657 |
| chr2:199985659-199985993 | -0.046771693 | 0.906665481 | 0.966763657 |
| chr15:79432452-79432696 | 0.051454713 | 0.906667401 | 0.966763657 |
| chr14:94536914-94537147 | -0.052164621 | 0.906683489 | 0.966765903 |
| chr8:139720018-139720590 | 0.040659746 | 0.906694742 | 0.966765903 |
| chr9:97046199-97046598 | -0.043947131 | 0.906717049 | 0.966775787 |
| chr19:38351917-38352157 | 0.056259286 | 0.906729247 | 0.966775787 |
| chr1:37514199-37514421 | -0.051367697 | 0.906792743 | 0.966819476 |
| chr17:65794578-65794992 | 0.04511087 | 0.906828624 | 0.966819476 |
| chr3:123825815-123826091 | 0.054821982 | 0.906834962 | 0.966819476 |
| chr6:24257374-24257574 | 0.062134502 | 0.906836112 | 0.966819476 |
| chr6:108526788-108527196 | -0.049845091 | 0.906838848 | 0.966819476 |
| chr1:205528976-205529292 | -0.040967743 | 0.906857912 | 0.966819476 |
| chr3:185362846-185363255 | -0.040839359 | 0.906872987 | 0.966819476 |
| chr3:136186415-136186666 | 0.051552677 | 0.906878015 | 0.966819476 |
| chrX:129601110-129601361 | -0.050286901 | 0.906895451 | 0.966819476 |
| chr16:85491853-85492372 | 0.040398628 | 0.90693148 | 0.966819476 |
| chr1:51235800-51236545 | 0.038931037 | 0.90695334 | 0.966819476 |
| chr6:125311079-125311409 | 0.043872864 | 0.906964969 | 0.966819476 |
| chr8:144826711-144827875 | -0.037026055 | 0.906966472 | 0.966819476 |
| chr3:153768716-153769276 | 0.039770983 | 0.906970938 | 0.966819476 |
| chr10:22316103-22316785 | 0.039328239 | 0.906971112 | 0.966819476 |
| chr2:230712449-230713163 | 0.040801388 | 0.90698349 | 0.966819476 |
| chr10:72096142-72096482 | 0.048068219 | 0.907003494 | 0.966819476 |
| chr17:67047338-67047981 | 0.03903253 | 0.907013425 | 0.966819476 |
| chr8:96979420-96979620 | 0.063709391 | 0.9070305 | 0.966819476 |
| chr1:28668224-28669067 | -0.041550211 | 0.9070343 | 0.966819476 |
| chr8:120808652-120808926 | -0.042806464 | 0.907058409 | 0.966819476 |
| chr10:97497707-97497996 | 0.041354512 | 0.907060044 | 0.966819476 |
| chr8:121550558-121550758 | -0.061167788 | 0.907061365 | 0.966819476 |
| chr2:45992251-45992774 | 0.042816814 | 0.907100419 | 0.966819476 |
| chr15:59455977-59456695 | 0.040805226 | 0.907106868 | 0.966819476 |
| chr17:19111653-19112485 | -0.03997406 | 0.907114522 | 0.966819476 |
| chr17:47315790-47315990 | -0.057299952 | 0.907122069 | 0.966819476 |
| chrX:130330222-130330558 | -0.04458113 | 0.907133568 | 0.966819476 |
| chr16:1645939-1646335 | 0.043991085 | 0.907136146 | 0.966819476 |
| chrX:110418850-110419242 | -0.044391779 | 0.907173096 | 0.966828458 |
| chr8:8963234-8964016 | -0.04332297 | 0.907176258 | 0.966828458 |
| chr9:82811116-82811337 | 0.057776028 | 0.907182428 | 0.966828458 |
| chr10:11725278-11725506 | -0.050831527 | 0.907233564 | 0.96685795 |
| chr5:55324875-55325076 | 0.062039888 | 0.907247597 | 0.96685795 |
| chr1:207336219-207336602 | 0.052262863 | 0.907247956 | 0.96685795 |

| | | | |
|---|---|---|---|
| chr20:51564003-51564265 | 0.044558126 | 0.907279756 | 0.966868183 |
| chr13:113372986-113373252 | -0.057923558 | 0.907282795 | 0.966868183 |
| chr17:49821947-49822456 | -0.045409719 | 0.907298787 | 0.966871777 |
| chr2:27209780-27210339 | -0.040308725 | 0.907325917 | 0.966887242 |
| chr15:83283394-83283620 | 0.054079819 | 0.907351536 | 0.966901094 |
| chr4:109433213-109434211 | -0.037439736 | 0.90738319 | 0.966921378 |
| chrX:95762791-95763010 | 0.05334982 | 0.907483266 | 0.966997514 |
| chr2:33451047-33451540 | -0.049617483 | 0.907495891 | 0.966997514 |
| chr17:45371654-45372173 | 0.043984814 | 0.907501625 | 0.966997514 |
| chr2:236856144-236856875 | -0.040685606 | 0.907517389 | 0.966997514 |
| chr16:69124038-69124279 | 0.058253791 | 0.907518663 | 0.966997514 |
| chr15:85878927-85879191 | -0.049852598 | 0.907530359 | 0.966997514 |
| chr17:32349914-32350712 | -0.039788871 | 0.907544874 | 0.966999532 |
| chr2:162989554-162989849 | -0.046178799 | 0.907561296 | 0.966999622 |
| chr11:124800284-124801039 | 0.039936997 | 0.907581796 | 0.966999622 |
| chr12:45235718-45236037 | -0.047781308 | 0.907582819 | 0.966999622 |
| chr2:159185326-159185622 | -0.048679412 | 0.907629232 | 0.967023803 |
| chr17:76384311-76384993 | -0.038560999 | 0.907630756 | 0.967023803 |
| chr15:36161994-36162242 | 0.059164671 | 0.907675882 | 0.967058435 |
| chr5:181037469-181037718 | -0.051063124 | 0.907714884 | 0.967085967 |
| chr1:79489348-79489690 | 0.05657741 | 0.90773698 | 0.967085967 |
| chr6:106510471-106511027 | 0.04228044 | 0.907742767 | 0.967085967 |
| chr1:235866143-235866420 | -0.044698791 | 0.907764732 | 0.967085967 |
| chr13:29362986-29363324 | 0.043513593 | 0.907764831 | 0.967085967 |
| chr3:42750085-42750285 | 0.060172611 | 0.907792322 | 0.967101808 |
| chr7:43737548-43737877 | -0.052925003 | 0.907857596 | 0.967139224 |
| chr3:122101653-122101903 | -0.0482726 | 0.907860879 | 0.967139224 |
| chr3:63279255-63279972 | -0.038518646 | 0.907870914 | 0.967139224 |
| chr5:134253839-134254039 | 0.058189501 | 0.907878747 | 0.967139224 |
| chr6:96015222-96016787 | -0.036516037 | 0.907890555 | 0.967139224 |
| chr1:116538736-116538936 | -0.055159047 | 0.907937598 | 0.967175891 |
| chr18:79951375-79951922 | 0.039881738 | 0.907962459 | 0.967188927 |
| chr11:34570346-34571296 | 0.037656565 | 0.908014542 | 0.967226492 |
| chr20:53586449-53587026 | 0.037044998 | 0.90802297 | 0.967226492 |
| chr6:30617136-30618163 | -0.036848789 | 0.908104138 | 0.967299505 |
| chr3:129316015-129316487 | 0.038776523 | 0.908139363 | 0.967323578 |
| chr1:150321084-150321976 | 0.036992011 | 0.908154334 | 0.967324608 |
| chr3:45688825-45689621 | -0.037573725 | 0.908166172 | 0.967324608 |
| chr2:147082889-147083228 | -0.047773824 | 0.908178204 | 0.967324608 |
| chr20:10559382-10559582 | -0.060409826 | 0.908191075 | 0.96732487 |
| chr1:99264356-99265020 | -0.041975981 | 0.908229283 | 0.967330139 |
| chr5:37684998-37685237 | -0.055930304 | 0.908231416 | 0.967330139 |
| chr20:60206249-60206475 | -0.047094598 | 0.908233895 | 0.967330139 |
| chr14:99971190-99971879 | 0.038341546 | 0.908247166 | 0.967330826 |
| chr3:141782860-141783060 | -0.057886157 | 0.908287657 | 0.967333936 |
| chr6:101226402-101227286 | -0.0383734 | 0.908290886 | 0.967333936 |
| chr5:180885927-180886355 | 0.043892338 | 0.908307171 | 0.967333936 |
| chr1:151563372-151563750 | -0.040865846 | 0.908311384 | 0.967333936 |
| chr10:122846033-122846292 | -0.062226194 | 0.908313209 | 0.967333936 |
| chr10:101142232-101142546 | 0.044598929 | 0.908349839 | 0.967339583 |

| | | | |
|---|---|---|---|
| chr12:6872451-6872651 | -0.048489207 | 0.908353891 | 0.967339583 |
| chr8:69833323-69833630 | -0.047022826 | 0.908356386 | 0.967339583 |
| chr10:70775055-70775361 | 0.044524931 | 0.908389156 | 0.967350813 |
| chr1:230185940-230186490 | -0.041755046 | 0.908392181 | 0.967350813 |
| chr10:79068705-79069260 | -0.039334 | 0.908436838 | 0.967373143 |
| chr1:30824013-30824764 | -0.037485494 | 0.908450317 | 0.967373143 |
| chr22:20990994-20991253 | -0.051427757 | 0.908451026 | 0.967373143 |
| chr4:141254412-141254626 | -0.053821093 | 0.908485726 | 0.967396649 |
| chr17:78840280-78841483 | -0.036487402 | 0.908498565 | 0.967396876 |
| chr16:90019460-90020251 | 0.038623421 | 0.908536095 | 0.967423395 |
| chr2:73112870-73113653 | 0.039174875 | 0.908629265 | 0.967477967 |
| chr11:47714535-47715557 | 0.038257237 | 0.908665346 | 0.967477967 |
| chr3:100708934-100709839 | -0.036444741 | 0.908669854 | 0.967477967 |
| chr16:88784135-88785649 | -0.036737914 | 0.908672666 | 0.967477967 |
| chr18:48430374-48430574 | 0.067151742 | 0.908673452 | 0.967477967 |
| chr8:11802123-11803339 | 0.036965419 | 0.908703635 | 0.967477967 |
| chr20:20772585-20773084 | -0.043558231 | 0.908709282 | 0.967477967 |
| chrX:15838125-15838389 | -0.055967135 | 0.90871126 | 0.967477967 |
| chr2:6352972-6353228 | -0.051518743 | 0.908714848 | 0.967477967 |
| chr8:100736345-100736699 | -0.047153651 | 0.908740622 | 0.967477967 |
| chr14:50396474-50397325 | -0.038603925 | 0.908744975 | 0.967477967 |
| chr21:41436119-41436319 | -0.057682071 | 0.908753652 | 0.967477967 |
| chr7:5240186-5240960 | -0.038140833 | 0.908756317 | 0.967477967 |
| chr12:67647421-67647637 | -0.048310466 | 0.908799966 | 0.967477967 |
| chr2:222087881-222088310 | 0.046466125 | 0.908810189 | 0.967477967 |
| chr7:151933042-151933617 | 0.040594192 | 0.908810401 | 0.967477967 |
| chr6:33279972-33280464 | -0.039403127 | 0.908811625 | 0.967477967 |
| chr17:27293832-27295095 | -0.0369445 | 0.908814625 | 0.967477967 |
| chr21:29024459-29025291 | -0.036683443 | 0.908861914 | 0.967514867 |
| chr20:43692054-43692280 | 0.053485728 | 0.908888154 | 0.967529358 |
| chr17:67322164-67322431 | -0.046133818 | 0.908997763 | 0.967623202 |
| chr7:124709135-124709335 | 0.067912667 | 0.909001567 | 0.967623202 |
| chr9:83744282-83744488 | -0.059033421 | 0.909028183 | 0.967638091 |
| chr9:122810376-122810676 | 0.047041545 | 0.909050758 | 0.967648678 |
| chr7:45722055-45722288 | 0.051043104 | 0.909078578 | 0.967654764 |
| chr10:29392464-29392704 | -0.053032006 | 0.909081733 | 0.967654764 |
| chr20:47464121-47465285 | 0.035487414 | 0.909096471 | 0.96765701 |
| chr1:109499913-109500176 | 0.04417958 | 0.909133935 | 0.967679536 |
| chr2:119415658-119416206 | -0.042714632 | 0.909142893 | 0.967679536 |
| chr20:60352138-60352347 | 0.051787159 | 0.909163718 | 0.967684925 |
| chr1:159214244-159214444 | 0.056540125 | 0.909200132 | 0.967684925 |
| chr20:10172124-10172941 | 0.039973272 | 0.909212551 | 0.967684925 |
| chr10:121570968-121571168 | 0.066195017 | 0.90923347 | 0.967684925 |
| chr17:74260026-74260247 | 0.052947757 | 0.909260158 | 0.967684925 |
| chr14:27587218-27587474 | -0.05960857 | 0.909260311 | 0.967684925 |
| chr2:101883916-101884116 | 0.058793486 | 0.909266124 | 0.967684925 |
| chr2:127050216-127050507 | 0.043695625 | 0.9092792 | 0.967684925 |
| chr15:63068774-63068974 | 0.062608075 | 0.909294987 | 0.967684925 |
| chr11:119341862-119342062 | -0.05306694 | 0.909297586 | 0.967684925 |
| chr19:43765186-43765547 | -0.051077648 | 0.909298946 | 0.967684925 |

| | | | |
|---|---|---|---|
| chr1:32204930-32206029 | 0.037034019 | 0.909307648 | 0.967684925 |
| chr16:88706102-88706810 | 0.038496738 | 0.909319164 | 0.967684925 |
| chr17:64661503-64662947 | -0.036098456 | 0.909324767 | 0.967684925 |
| chr1:65147439-65149229 | -0.036244967 | 0.909378259 | 0.967702451 |
| chr16:83898449-83899962 | -0.036280195 | 0.909389943 | 0.967702451 |
| chr14:67359653-67360708 | 0.035898612 | 0.909392938 | 0.967702451 |
| chr10:81920111-81920521 | -0.048398595 | 0.90940049 | 0.967702451 |
| chr11:111727516-111727843 | -0.055719959 | 0.90941968 | 0.967702451 |
| chr15:64024806-64025028 | 0.053719521 | 0.909422207 | 0.967702451 |
| chr10:114767695-114767921 | -0.053343617 | 0.909442272 | 0.967702451 |
| chr17:6473240-6473475 | -0.053343617 | 0.909442272 | 0.967702451 |
| chr6:159761729-159762583 | 0.036351904 | 0.909458905 | 0.967704928 |
| chr2:194166873-194167073 | 0.06571013 | 0.909475442 | 0.967704928 |
| chr2:196535805-196536051 | -0.057225226 | 0.909494073 | 0.967704928 |
| chr13:32585628-32585834 | -0.048726567 | 0.909503982 | 0.967704928 |
| chr2:37813452-37813652 | -0.066739846 | 0.909515143 | 0.967704928 |
| chr1:85047969-85048978 | -0.037873742 | 0.909520378 | 0.967704928 |
| chr6:7899028-7899228 | 0.059833569 | 0.909580082 | 0.967742323 |
| chr2:64330912-64331179 | 0.050113217 | 0.909580784 | 0.967742323 |
| chr2:73176727-73178150 | -0.036484503 | 0.909597962 | 0.967747161 |
| chr12:49188230-49189392 | -0.037243867 | 0.90962219 | 0.9677595 |
| chr11:19954034-19954234 | 0.054685489 | 0.909665916 | 0.967780378 |
| chr1:156815623-156815846 | -0.056298735 | 0.90967526 | 0.967780378 |
| chr21:38170195-38170624 | -0.041133122 | 0.909680926 | 0.967780378 |
| chr18:36129012-36130296 | -0.035830107 | 0.90969418 | 0.967780378 |
| chr12:31325007-31325506 | -0.041008763 | 0.909713303 | 0.967780378 |
| chr9:75002212-75002500 | 0.049789579 | 0.909717597 | 0.967780378 |
| chr6:291891-292849 | 0.03662383 | 0.909747828 | 0.967799101 |
| chr3:149377547-149378244 | -0.039970847 | 0.909801882 | 0.967818385 |
| chr19:35658042-35658242 | -0.058082963 | 0.909827538 | 0.967818385 |
| chr2:178149132-178149386 | -0.052316511 | 0.909836341 | 0.967818385 |
| chr1:74504156-74504452 | 0.045401177 | 0.909838191 | 0.967818385 |
| chr3:194573841-194574213 | 0.042047228 | 0.90984643 | 0.967818385 |
| chr10:17742971-17743171 | 0.059502375 | 0.909853086 | 0.967818385 |
| chr6:74251659-74251873 | 0.056350544 | 0.909854373 | 0.967818385 |
| chr1:2548023-2548248 | 0.049096354 | 0.909868304 | 0.967819768 |
| chr11:134331593-134332712 | 0.037711827 | 0.909898736 | 0.967823244 |
| chr10:60175379-60175596 | 0.05174036 | 0.909911447 | 0.967823244 |
| chr10:86355484-86355694 | -0.05273418 | 0.909935629 | 0.967823244 |
| chr19:40623554-40624332 | 0.038407207 | 0.909940902 | 0.967823244 |
| chr4:161201507-161201774 | 0.055602743 | 0.909950131 | 0.967823244 |
| chr6:168895102-168895418 | -0.046764783 | 0.909959034 | 0.967823244 |
| chr14:97331626-97331995 | 0.046349028 | 0.909959989 | 0.967823244 |
| chr3:157437744-157438304 | -0.041041981 | 0.910000175 | 0.967849891 |
| chr17:80621658-80621921 | -0.056147372 | 0.910013351 | 0.967849891 |
| chr20:37192232-37192581 | -0.044134777 | 0.910027896 | 0.967849891 |
| chr1:116261363-116261720 | -0.04311304 | 0.910035569 | 0.967849891 |
| chr3:5136721-5136952 | 0.048850702 | 0.910058785 | 0.967861147 |
| chrX:41275777-41276540 | 0.039848454 | 0.910145472 | 0.96793901 |
| chr2:12400706-12400906 | 0.06178515 | 0.910157263 | 0.96793901 |

| | | | |
|---|---|---|---|
| chr3:40452712-40453578 | -0.036262371 | 0.910173221 | 0.967942547 |
| chr11:74634522-74634939 | -0.041317474 | 0.910193143 | 0.967950299 |
| chr14:21300961-21301376 | -0.044970898 | 0.910207506 | 0.967952139 |
| chr13:45850659-45851176 | -0.042197616 | 0.910235113 | 0.967955637 |
| chr8:47960546-47961074 | -0.03694568 | 0.910236062 | 0.967955637 |
| chr15:41827851-41828235 | 0.038539612 | 0.910257145 | 0.967964623 |
| chr15:56464879-56465650 | 0.036697802 | 0.910292368 | 0.967977813 |
| chr6:54873444-54873915 | -0.045567188 | 0.910294815 | 0.967977813 |
| chr8:124244973-124245543 | 0.041557446 | 0.91033336 | 0.967980484 |
| chr1:22428723-22428923 | 0.05964164 | 0.910346745 | 0.967980484 |
| chr4:100690893-100691098 | 0.056628322 | 0.910347859 | 0.967980484 |
| chr8:54278314-54278514 | 0.056628322 | 0.910347859 | 0.967980484 |
| chr11:1749989-1750264 | 0.046496926 | 0.910365123 | 0.967985407 |
| chr17:73369903-73370405 | 0.040682629 | 0.91039971 | 0.96800798 |
| chr10:33335657-33335879 | -0.049096133 | 0.910419402 | 0.96800798 |
| chrX:116240526-116240757 | -0.050536765 | 0.910437486 | 0.96800798 |
| chr11:75768049-75769018 | 0.037426738 | 0.910487154 | 0.96800798 |
| chr2:216588757-216588983 | -0.055730137 | 0.91050793 | 0.96800798 |
| chr1:202625420-202625693 | 0.042508568 | 0.910522215 | 0.96800798 |
| chr2:241355869-241356671 | 0.036390623 | 0.910552264 | 0.96800798 |
| chr3:111252561-111253135 | -0.039147953 | 0.910556428 | 0.96800798 |
| chr17:50559882-50560201 | 0.042370718 | 0.910563097 | 0.96800798 |
| chr19:14418976-14419719 | 0.038167014 | 0.910565975 | 0.96800798 |
| chr17:45584410-45584763 | 0.047637815 | 0.910569741 | 0.96800798 |
| chr2:208175342-208175664 | -0.041289436 | 0.91057069 | 0.96800798 |
| chr1:5514627-5514827 | -0.059212686 | 0.91057478 | 0.96800798 |
| chr5:52007343-52007681 | -0.04955772 | 0.910575694 | 0.96800798 |
| chr21:44873412-44874274 | -0.036445082 | 0.910575855 | 0.96800798 |
| chr11:32583436-32584176 | -0.036714059 | 0.910593485 | 0.968013291 |
| chr8:27872518-27873054 | -0.040518522 | 0.910647969 | 0.96805778 |
| chr20:34824894-34825915 | 0.036914304 | 0.910671246 | 0.968069094 |
| chr6:16760923-16761159 | 0.045572067 | 0.91076367 | 0.968145172 |
| chr2:26503131-26503946 | 0.037102231 | 0.910769519 | 0.968145172 |
| chr4:37132430-37132723 | -0.055824675 | 0.910780719 | 0.968145172 |
| chr2:168889687-168890123 | 0.043846138 | 0.910823863 | 0.968167155 |
| chr8:121642207-121642575 | -0.038460689 | 0.910826671 | 0.968167155 |
| chr4:167218238-167218574 | 0.045470323 | 0.910868559 | 0.968168702 |
| chr20:32359301-32359501 | -0.048797945 | 0.910892515 | 0.968168702 |
| chr3:141228464-141228712 | 0.045952383 | 0.910908301 | 0.968168702 |
| chr14:65594142-65594650 | 0.041598552 | 0.910918708 | 0.968168702 |
| chr14:93174820-93175158 | 0.045868408 | 0.910943399 | 0.968168702 |
| chr7:44241003-44241601 | -0.037608243 | 0.910946839 | 0.968168702 |
| chr3:157354789-157355044 | 0.048503844 | 0.910961837 | 0.968168702 |
| chr20:49908220-49908426 | 0.046528929 | 0.910962485 | 0.968168702 |
| chr20:24929945-24930860 | -0.036718019 | 0.91096926 | 0.968168702 |
| chr1:107276978-107277469 | -0.036166868 | 0.911002512 | 0.968168702 |
| chr9:36798354-36798554 | -0.05606835 | 0.911003327 | 0.968168702 |
| chr7:33334349-33334672 | 0.045756562 | 0.911005751 | 0.968168702 |
| chrX:17347867-17348276 | 0.043803748 | 0.911007272 | 0.968168702 |
| chr1:145607746-145608372 | -0.036309563 | 0.911009353 | 0.968168702 |

| | | | |
|---|---|---|---|
| chr12:67542941-67543141 | 0.061650733 | 0.911024815 | 0.968168702 |
| chr11:73965450-73965677 | -0.048247372 | 0.911030297 | 0.968168702 |
| chr1:43227267-43227513 | 0.047407012 | 0.911052917 | 0.968179313 |
| chr8:142738559-142739871 | -0.035719343 | 0.911094932 | 0.96820912 |
| chr17:28426174-28426446 | -0.049365438 | 0.911126956 | 0.96820912 |
| chr17:49548549-49548758 | 0.047105381 | 0.911128365 | 0.96820912 |
| chr20:19995390-19995617 | 0.050927083 | 0.911140642 | 0.96820912 |
| chr16:3191305-3192182 | -0.035986004 | 0.911146625 | 0.96820912 |
| chr1:16861777-16862046 | 0.054330217 | 0.911156783 | 0.96820912 |
| chr3:42819876-42820351 | 0.043372127 | 0.911170125 | 0.96820987 |
| chr3:141441429-141441719 | -0.042670886 | 0.911195201 | 0.968223089 |
| chr1:109532299-109532631 | 0.040029911 | 0.91122535 | 0.968226655 |
| chr2:27766084-27766284 | -0.059589691 | 0.911237143 | 0.968226655 |
| chr3:14017031-14017514 | -0.044286311 | 0.911244377 | 0.968226655 |
| chr3:19946546-19948086 | -0.034829242 | 0.911253264 | 0.968226655 |
| chr13:60379954-60380167 | -0.053765946 | 0.91126174 | 0.968226655 |
| chr22:19953618-19954012 | -0.042196591 | 0.91128518 | 0.968238135 |
| chr4:53119689-53119990 | -0.049486207 | 0.911316261 | 0.968247254 |
| chr19:2785092-2785571 | -0.039177642 | 0.911319036 | 0.968247254 |
| chr15:62392360-62392581 | -0.05019612 | 0.911349997 | 0.968257877 |
| chr2:46541761-46542136 | -0.039322764 | 0.911354309 | 0.968257877 |
| chr11:46347953-46348219 | 0.042368752 | 0.911372485 | 0.968263762 |
| chr8:123158642-123158842 | 0.04216997 | 0.911423752 | 0.968304804 |
| chr6:35688370-35689409 | 0.036841234 | 0.911479409 | 0.968350507 |
| chr6:32968409-32969430 | 0.035312235 | 0.911492637 | 0.968351134 |
| chr14:53539545-53539818 | -0.049259247 | 0.911534342 | 0.968382013 |
| chr19:39878464-39878694 | 0.055278251 | 0.911562044 | 0.968391075 |
| chrX:51487309-51487986 | -0.043691728 | 0.911568149 | 0.968391075 |
| chr5:22070313-22070701 | 0.043488044 | 0.911587206 | 0.968397894 |
| chr10:11611040-11611921 | -0.037120822 | 0.911635388 | 0.968435652 |
| chr9:137205356-137205759 | 0.042844308 | 0.911649533 | 0.968437251 |
| chr12:124922114-124922321 | 0.044299528 | 0.911662792 | 0.96843791 |
| chr3:123489479-123489722 | 0.050573204 | 0.911738482 | 0.968504886 |
| chr17:64836314-64837439 | -0.036144124 | 0.91177809 | 0.968533533 |
| chr11:47766654-47768028 | -0.034899819 | 0.911791961 | 0.968534841 |
| chr19:34364789-34365346 | -0.037834429 | 0.911851994 | 0.968573251 |
| chr4:77418431-77418631 | 0.055006279 | 0.911862043 | 0.968573251 |
| chr19:49817372-49817681 | -0.042503842 | 0.911866044 | 0.968573251 |
| chr11:86790822-86791073 | 0.054558272 | 0.911897711 | 0.968590456 |
| chr7:65980936-65981169 | 0.04973861 | 0.911907524 | 0.968590456 |
| chr17:40443237-40443526 | 0.04363174 | 0.911959921 | 0.968620894 |
| chr10:79367817-79368067 | -0.055290344 | 0.911961464 | 0.968620894 |
| chr19:50214680-50215714 | 0.036010353 | 0.912000704 | 0.968649145 |
| chr22:38733255-38733477 | -0.051455322 | 0.912042181 | 0.968668554 |
| chr17:3845609-3845833 | 0.04860345 | 0.912044263 | 0.968668554 |
| chr1:52697353-52698698 | -0.035681008 | 0.912102981 | 0.968706716 |
| chr17:57054303-57054638 | 0.043680028 | 0.912111053 | 0.968706716 |
| chr12:68100292-68100630 | -0.045538632 | 0.912118181 | 0.968706716 |
| chr19:48933408-48933998 | -0.04034583 | 0.912130764 | 0.968706716 |
| chr1:179447516-179447826 | -0.0505699 | 0.912154816 | 0.96871658 |

| | | | |
|---|---|---|---|
| chr4:48059224-48059797 | 0.043659944 | 0.912182441 | 0.96871658 |
| chr6:116279199-116280240 | -0.035083492 | 0.912189474 | 0.96871658 |
| chr5:112529574-112529847 | -0.047746837 | 0.912190623 | 0.96871658 |
| chr10:71955778-71956000 | 0.053682822 | 0.912215799 | 0.96871718 |
| chrX:96546346-96546546 | 0.056143537 | 0.912216474 | 0.96871718 |
| chr2:45938696-45939172 | 0.038035727 | 0.912246593 | 0.968735738 |
| chrX:93422761-93423072 | 0.045198427 | 0.91226449 | 0.968740889 |
| chrX:101617214-101618200 | 0.035744288 | 0.912282233 | 0.968740889 |
| chr17:78289788-78290110 | -0.040539258 | 0.912289373 | 0.968740889 |
| chr2:157492308-157492741 | -0.043134942 | 0.912353172 | 0.968787167 |
| chr8:144326181-144326413 | 0.044079818 | 0.912369211 | 0.968787167 |
| chr11:67262633-67262971 | 0.041262474 | 0.912370885 | 0.968787167 |
| chr2:96763737-96764638 | -0.036018934 | 0.912387865 | 0.968791771 |
| chr7:6348367-6349577 | -0.034692724 | 0.912414357 | 0.96880022 |
| chr5:73448140-73448348 | -0.051770552 | 0.912430686 | 0.96880022 |
| chr17:51152836-51154058 | 0.034920087 | 0.912433754 | 0.96880022 |
| chr9:122930978-122932044 | -0.035237183 | 0.912463741 | 0.968816635 |
| chr11:66267930-66268156 | 0.04300943 | 0.912474609 | 0.968816635 |
| chr8:144082425-144082939 | -0.037493616 | 0.91249849 | 0.968816635 |
| chr2:85679003-85679523 | 0.039217266 | 0.912506466 | 0.968816635 |
| chr10:111344923-111345160 | -0.056646861 | 0.912523095 | 0.968816635 |
| chr6:82247106-82248423 | -0.0348195 | 0.912525078 | 0.968816635 |
| chr16:67109441-67110004 | -0.038720265 | 0.9125524 | 0.968832218 |
| chrX:68710920-68711155 | 0.049070164 | 0.912595393 | 0.968864438 |
| chr2:72825287-72826238 | -0.036266598 | 0.912614363 | 0.968871153 |
| chr1:1890966-1891274 | 0.046120381 | 0.912647715 | 0.968893136 |
| chr5:170225899-170226099 | -0.052526051 | 0.912663493 | 0.968896462 |
| chr8:88434799-88435077 | 0.046487624 | 0.912709132 | 0.968924969 |
| chr11:65524775-65525174 | 0.038643933 | 0.912715637 | 0.968924969 |
| chr1:179025567-179026787 | -0.034649863 | 0.912730763 | 0.968927603 |
| chr8:101027019-101027641 | -0.039471526 | 0.912824273 | 0.969013445 |
| chr8:129816471-129816671 | -0.059247811 | 0.912867481 | 0.969045887 |
| chr5:42811732-42812272 | -0.038051263 | 0.912932224 | 0.969101188 |
| chr15:64095719-64096538 | 0.03565528 | 0.912975908 | 0.969134133 |
| chr17:18856072-18856460 | 0.038300578 | 0.913001762 | 0.969148151 |
| chr3:79886245-79886520 | -0.048021058 | 0.913029411 | 0.969159043 |
| chr18:22885510-22886185 | 0.03959872 | 0.913041364 | 0.969159043 |
| chr8:108051277-108051477 | 0.052058285 | 0.913049968 | 0.969159043 |
| chr10:15461342-15461623 | -0.049609041 | 0.91309541 | 0.969192103 |
| chr1:14731897-14732209 | -0.04606328 | 0.913106413 | 0.969192103 |
| chr10:46023132-46023750 | 0.03685691 | 0.913126113 | 0.969199588 |
| chr1:181044538-181044885 | 0.042179086 | 0.913140584 | 0.969201071 |
| chr14:93953725-93953932 | -0.052233992 | 0.913161067 | 0.969201071 |
| chr1:93447889-93448348 | -0.041201793 | 0.913165458 | 0.969201071 |
| chr2:120680087-120680588 | -0.042457053 | 0.913203829 | 0.969217698 |
| chr3:170751548-170751933 | -0.04349999 | 0.913206422 | 0.969217698 |
| chr2:116435601-116435858 | -0.056739963 | 0.913246029 | 0.969246308 |
| chr18:31956964-31957344 | -0.044198699 | 0.91330298 | 0.969284891 |
| chr10:116141214-116141467 | 0.047779874 | 0.913307683 | 0.969284891 |
| chr5:96968179-96968379 | -0.058270924 | 0.913329168 | 0.969293828 |

| | | | |
|---|---|---|---|
| chr1:242189677-242189877 | 0.057331219 | 0.913365885 | 0.969293828 |
| chr22:40345956-40346931 | 0.03719602 | 0.91336789 | 0.969293828 |
| chr6:150894928-150895408 | -0.044348062 | 0.913396382 | 0.969293828 |
| chr1:115676661-115676892 | -0.046914863 | 0.913399898 | 0.969293828 |
| chr10:96470136-96470495 | -0.041567308 | 0.913403073 | 0.969293828 |
| chr15:70375088-70375510 | 0.040984457 | 0.913433296 | 0.969293828 |
| chr7:97179611-97179821 | -0.057125551 | 0.913464587 | 0.969293828 |
| chr13:111153232-111154310 | 0.035822928 | 0.913507628 | 0.969293828 |
| chr17:63304703-63304920 | -0.048754836 | 0.913515098 | 0.969293828 |
| chr17:68398989-68399366 | 0.037625732 | 0.913520998 | 0.969293828 |
| chr7:36609418-36609618 | 0.058589705 | 0.913521664 | 0.969293828 |
| chr6:155186138-155186341 | 0.058589705 | 0.913521664 | 0.969293828 |
| chr20:62818700-62818918 | -0.046298055 | 0.913524026 | 0.969293828 |
| chr13:41284834-41285189 | -0.041861976 | 0.913541694 | 0.969293828 |
| chr11:59171900-59172266 | 0.037307711 | 0.913541753 | 0.969293828 |
| chr9:107422230-107422545 | 0.041544156 | 0.913542241 | 0.969293828 |
| chr12:133036902-133038038 | 0.035128756 | 0.913546263 | 0.969293828 |
| chr10:78974813-78975049 | -0.042446569 | 0.913556776 | 0.969293828 |
| chr3:196196661-196196914 | -0.044538506 | 0.91358019 | 0.969293828 |
| chr1:205141418-205142001 | -0.037354176 | 0.91358176 | 0.969293828 |
| chr10:86769929-86770345 | 0.043599388 | 0.913598325 | 0.96929798 |
| chr2:49068521-49068753 | 0.056441574 | 0.913637546 | 0.969314383 |
| chr17:7856890-7858284 | 0.034814752 | 0.913639086 | 0.969314383 |
| chr9:33076485-33077051 | 0.035774653 | 0.913657216 | 0.969320196 |
| chr14:94641865-94642078 | 0.058011294 | 0.913689223 | 0.969335747 |
| chr14:20624699-20625038 | 0.038326474 | 0.913697175 | 0.969335747 |
| chr5:171370957-171371218 | -0.045615803 | 0.913719712 | 0.969346235 |
| chr4:40309262-40309539 | -0.047524045 | 0.913792034 | 0.969409537 |
| chr5:112848107-112848583 | 0.039233323 | 0.913826247 | 0.96943241 |
| chr5:85452922-85453176 | -0.060380752 | 0.913850649 | 0.969444875 |
| chr20:50730658-50732292 | -0.033436969 | 0.913869419 | 0.969450157 |
| chrX:56052644-56052891 | -0.049428543 | 0.913880933 | 0.969450157 |
| chr12:80946716-80946944 | -0.051803781 | 0.913923806 | 0.969480158 |
| chr11:114280610-114280825 | 0.053115318 | 0.91393452 | 0.969480158 |
| chr17:42101592-42101827 | 0.048483124 | 0.913952535 | 0.969481771 |
| chr1:46747703-46748042 | -0.041857654 | 0.913972206 | 0.969481771 |
| chr2:125745563-125745882 | -0.043969344 | 0.913981684 | 0.969481771 |
| chr14:63155912-63156122 | -0.059631697 | 0.913988636 | 0.969481771 |
| chr12:79022928-79023128 | 0.062566968 | 0.913999304 | 0.969481771 |
| chr6:152636694-152637214 | 0.040150616 | 0.914046926 | 0.969513737 |
| chr11:44095495-44095762 | -0.045879398 | 0.914054748 | 0.969513737 |
| chr8:106314084-106314284 | 0.056771735 | 0.914128619 | 0.969545968 |
| chr1:61139351-61139760 | -0.041746238 | 0.914138635 | 0.969545968 |
| chr8:140192878-140193078 | -0.052227948 | 0.914161057 | 0.969545968 |
| chr3:184811930-184812923 | -0.035090565 | 0.914176105 | 0.969545968 |
| chr5:103425893-103426427 | -0.042109862 | 0.914178543 | 0.969545968 |
| chr16:29877459-29877906 | 0.042004692 | 0.914189541 | 0.969545968 |
| chr17:2041540-2042399 | -0.035794536 | 0.914196424 | 0.969545968 |
| chr19:1240293-1240561 | 0.045983531 | 0.914204804 | 0.969545968 |
| chr19:7925299-7925605 | 0.042670107 | 0.914218643 | 0.969545968 |

| | | | |
|---|---|---|---|
| chr17:31488511-31488847 | 0.044739585 | 0.914231689 | 0.969545968 |
| chr22:44023892-44024118 | 0.058140918 | 0.914232027 | 0.969545968 |
| chr2:222859955-222860173 | 0.049921465 | 0.914236978 | 0.969545968 |
| chr7:50614924-50615179 | 0.046292473 | 0.91431652 | 0.969590619 |
| chr3:118628746-118629213 | -0.039259395 | 0.914338862 | 0.969590619 |
| chr8:92612140-92612438 | 0.05385192 | 0.914347893 | 0.969590619 |
| chr1:58689023-58689459 | -0.041761949 | 0.914353364 | 0.969590619 |
| chr9:115118031-115118240 | 0.050380765 | 0.914354228 | 0.969590619 |
| chr18:13726090-13727313 | 0.034936437 | 0.914355007 | 0.969590619 |
| chr17:16217035-16217712 | 0.036754664 | 0.914393527 | 0.969594974 |
| chr10:91299051-91299252 | -0.050216949 | 0.914410795 | 0.969594974 |
| chr12:91811800-91812035 | 0.049543976 | 0.914412858 | 0.969594974 |
| chr20:7936263-7936838 | 0.040610505 | 0.914414037 | 0.969594974 |
| chr12:25524329-25524981 | -0.038213997 | 0.914422894 | 0.969594974 |
| chr2:119290876-119291193 | 0.041584225 | 0.914453103 | 0.969594974 |
| chr22:30822452-30822652 | 0.052314133 | 0.914468251 | 0.969594974 |
| chr20:462235-462768 | 0.038335988 | 0.914472351 | 0.969594974 |
| chr20:58179049-58180058 | -0.034935956 | 0.914473002 | 0.969594974 |
| chr3:169147307-169147880 | 0.03779132 | 0.914539578 | 0.969652145 |
| chr9:112903193-112903510 | -0.04294292 | 0.914582469 | 0.969674602 |
| chr11:110376704-110376991 | 0.045154187 | 0.914596633 | 0.969674602 |
| chr5:15499923-15500774 | 0.036136018 | 0.914598724 | 0.969674602 |
| chr5:136374056-136374330 | -0.049874963 | 0.914615116 | 0.969678563 |
| chr1:240628887-240629087 | 0.058121638 | 0.914652967 | 0.969705275 |
| chr1:225319021-225319275 | -0.046425537 | 0.914681191 | 0.96972178 |
| chr8:1823254-1824174 | -0.036318996 | 0.914724625 | 0.969728825 |
| chr3:49411516-49412716 | -0.034386899 | 0.914726316 | 0.969728825 |
| chr5:172958413-172959757 | -0.033887696 | 0.914726947 | 0.969728825 |
| chr14:100586241-100587401 | 0.034898199 | 0.914741654 | 0.969728825 |
| chr16:75188720-75189456 | 0.035521974 | 0.914751116 | 0.969728825 |
| chr8:123828029-123828230 | 0.049987066 | 0.914765448 | 0.969729206 |
| chr2:85524553-85524950 | 0.039397202 | 0.914799191 | 0.969729206 |
| chr13:51118888-51119088 | -0.051388863 | 0.914799686 | 0.969729206 |
| chr19:36488921-36490111 | -0.034808796 | 0.9148021 | 0.969729206 |
| chr12:80195227-80195437 | -0.056051128 | 0.914828217 | 0.969740068 |
| chr3:190811804-190812100 | 0.044279927 | 0.91485729 | 0.969740068 |
| chr6:146410878-146411241 | 0.043579724 | 0.914862898 | 0.969740068 |
| chrX:73214478-73215153 | 0.038182492 | 0.914862971 | 0.969740068 |
| chr3:187101148-187101369 | -0.053199988 | 0.914891817 | 0.969748168 |
| chr2:70341210-70341674 | -0.040273167 | 0.914895925 | 0.969748168 |
| chr1:37255477-37255726 | -0.049369813 | 0.914924396 | 0.969760825 |
| chr2:233566423-233566978 | 0.038633828 | 0.914933179 | 0.969760825 |
| chr16:84144585-84145618 | -0.034511538 | 0.914968562 | 0.969773909 |
| chr17:76624020-76624220 | -0.056150886 | 0.914970837 | 0.969773909 |
| chr4:153257159-153257359 | 0.055077337 | 0.915011063 | 0.969795167 |
| chr1:116138828-116139671 | 0.035245313 | 0.915027414 | 0.969795167 |
| chr6:134453884-134454138 | 0.048550391 | 0.915036192 | 0.969795167 |
| chr21:18951708-18952006 | -0.044794623 | 0.915045723 | 0.969795167 |
| chr11:18394256-18394717 | -0.035577429 | 0.915054178 | 0.969795167 |
| chr17:43530473-43530705 | 0.054116092 | 0.915075593 | 0.96980445 |

| | | | |
|---|---|---|---|
| chr20:17594491-17594841 | 0.038640273 | 0.915124002 | 0.969842339 |
| chr1:175743582-175743883 | -0.044268249 | 0.915138867 | 0.969844678 |
| chr9:136949613-136950627 | 0.034503459 | 0.915155506 | 0.969848898 |
| chr1:110495982-110496331 | -0.039944228 | 0.915198013 | 0.969880531 |
| chr12:54188617-54189286 | -0.035721701 | 0.915212246 | 0.9698822 |
| chr4:156983424-156983676 | -0.050958652 | 0.91526393 | 0.969923556 |
| chrX:136209353-136209649 | -0.04277695 | 0.915279467 | 0.969926607 |
| chr11:74170676-74171660 | 0.033761376 | 0.915301462 | 0.969936501 |
| chr19:44657248-44657573 | -0.042673452 | 0.915349578 | 0.969974074 |
| chr6:4134951-4136295 | 0.034094081 | 0.915380803 | 0.969993748 |
| chr1:7669539-7669909 | -0.043623775 | 0.915411834 | 0.969996658 |
| chrX:89871770-89872024 | 0.049127856 | 0.915418701 | 0.969996658 |
| chr9:34376801-34377383 | -0.035043133 | 0.915428101 | 0.969996658 |
| chr5:87169325-87169606 | -0.051317917 | 0.915434188 | 0.969996658 |
| chr19:6199398-6199962 | 0.035543262 | 0.915487287 | 0.970029517 |
| chr4:128227951-128228568 | 0.036624601 | 0.915508194 | 0.970029517 |
| chr1:45550394-45551357 | 0.035019337 | 0.91551283 | 0.970029517 |
| chr1:183881695-183882101 | 0.039604689 | 0.91552182 | 0.970029517 |
| chr5:139481741-139481950 | -0.051627038 | 0.915528498 | 0.970029517 |
| chr12:33267848-33268060 | -0.056779219 | 0.915543026 | 0.970031496 |
| chr2:134599340-134599547 | 0.059218613 | 0.915612211 | 0.97007992 |
| chr10:98309249-98309908 | 0.035967625 | 0.915619822 | 0.97007992 |
| chr2:119702843-119703194 | 0.039758417 | 0.915626711 | 0.97007992 |
| chr4:155972184-155972431 | -0.04694544 | 0.915670444 | 0.97011284 |
| chr6:154534502-154534702 | -0.060860622 | 0.91571703 | 0.970148372 |
| chr3:127254746-127255063 | -0.04704818 | 0.915736249 | 0.970148372 |
| chr1:236868883-236869143 | -0.046555526 | 0.915741968 | 0.970148372 |
| chr5:139649048-139649248 | -0.045430156 | 0.91579267 | 0.970188673 |
| chrX:56764413-56764667 | -0.048517812 | 0.915835713 | 0.970193973 |
| chr4:20015888-20016098 | -0.049065466 | 0.915844265 | 0.970193973 |
| chr6:16761378-16762387 | -0.034316289 | 0.915845361 | 0.970193973 |
| chr3:156305697-156306066 | 0.048047451 | 0.915859632 | 0.970193973 |
| chr5:112958168-112959044 | -0.03589327 | 0.915861293 | 0.970193973 |
| chr9:85741293-85742583 | 0.033762137 | 0.915873645 | 0.970193973 |
| chr14:76133138-76133412 | -0.049535525 | 0.915888652 | 0.970196457 |
| chrX:72238608-72239461 | -0.035212797 | 0.915908781 | 0.970198839 |
| chr1:211134007-211134517 | 0.038232766 | 0.915919992 | 0.970198839 |
| chr2:200811119-200812739 | -0.033080133 | 0.915928887 | 0.970198839 |
| chr6:26223846-26224493 | 0.035693425 | 0.915971302 | 0.970215086 |
| chr20:41488496-41488711 | 0.050357426 | 0.915978937 | 0.970215086 |
| chr1:217312892-217313214 | -0.041478209 | 0.915990275 | 0.970215086 |
| chr8:123395279-123396923 | 0.032960477 | 0.916013062 | 0.970215086 |
| chr14:76651567-76652011 | -0.038921052 | 0.916013434 | 0.970215086 |
| chr1:113389881-113390593 | -0.033801042 | 0.916027555 | 0.970215086 |
| chr12:45232993-45233193 | -0.051226049 | 0.916052399 | 0.970215086 |
| chr14:30214388-30214716 | 0.04564618 | 0.916057408 | 0.970215086 |
| chr18:46250386-46250791 | -0.041049242 | 0.916058187 | 0.970215086 |
| chr8:107495852-107496155 | 0.038880726 | 0.916076753 | 0.970221339 |
| chr5:149730666-149731271 | -0.036003298 | 0.916090114 | 0.970222078 |
| chr19:42283105-42284441 | 0.033908403 | 0.916105191 | 0.970224636 |

| | | | |
|---|---|---|---|
| chr7:151111092-151111382 | 0.041043249 | 0.916124095 | 0.970231246 |
| chr18:23662394-23662689 | -0.050279715 | 0.916187286 | 0.970284758 |
| chr1:11262320-11262978 | -0.035037763 | 0.916241374 | 0.970328628 |
| chr3:133574352-133574559 | -0.04014106 | 0.916260942 | 0.97033594 |
| chr17:7888351-7888646 | 0.03888521 | 0.916292013 | 0.97035036 |
| chr11:66497318-66497602 | -0.048696735 | 0.916305499 | 0.97035036 |
| chr10:122375451-122375665 | -0.0475057 | 0.916312551 | 0.97035036 |
| chr10:133324168-133324372 | 0.048767933 | 0.916327861 | 0.970353162 |
| chr5:127958594-127958909 | 0.050135957 | 0.916371805 | 0.970377123 |
| chr20:58623925-58624131 | 0.042657955 | 0.916375817 | 0.970377123 |
| chr1:19597235-19597806 | 0.039976635 | 0.916397271 | 0.97038643 |
| chr6:159074399-159074682 | -0.0524626 | 0.916441736 | 0.970420104 |
| chr12:61539074-61539274 | 0.062149922 | 0.916472502 | 0.97043927 |
| chr14:59444283-59444562 | -0.042665426 | 0.916488769 | 0.970443084 |
| chr2:130342032-130343259 | 0.033079353 | 0.91658159 | 0.970486227 |
| chr22:43187401-43188040 | -0.036541542 | 0.916612614 | 0.970486227 |
| chr2:137168561-137168975 | -0.042750431 | 0.916613027 | 0.970486227 |
| chr8:53621961-53622434 | -0.042522961 | 0.916617132 | 0.970486227 |
| chr1:32013127-32013999 | 0.033737007 | 0.916635941 | 0.970486227 |
| chr21:6369562-6371549 | -0.032376899 | 0.91663599 | 0.970486227 |
| chr9:97038857-97039784 | 0.034952984 | 0.916642149 | 0.970486227 |
| chr2:161624086-161625072 | 0.035908988 | 0.916643376 | 0.970486227 |
| chr15:35545440-35546719 | -0.033414972 | 0.916643506 | 0.970486227 |
| chr12:12695922-12696403 | -0.03775024 | 0.916707346 | 0.970530427 |
| chr1:12208926-12209169 | -0.049430946 | 0.916710587 | 0.970530427 |
| chr11:62831162-62832560 | -0.033086173 | 0.916744301 | 0.970534321 |
| chr10:99612281-99612783 | 0.038641936 | 0.916746597 | 0.970534321 |
| chr1:16963547-16963818 | 0.048410891 | 0.91675285 | 0.970534321 |
| chr6:42890312-42891302 | 0.034122367 | 0.916764931 | 0.970534321 |
| chr12:56590296-56590570 | -0.042638196 | 0.916828297 | 0.970570579 |
| chr20:51800108-51801151 | -0.033487581 | 0.916834677 | 0.970570579 |
| chr22:17715571-17715796 | -0.056530457 | 0.916874764 | 0.970570579 |
| chr4:137070884-137071309 | 0.045666933 | 0.916876558 | 0.970570579 |
| chr6:73394512-73395530 | -0.033140392 | 0.91687775 | 0.970570579 |
| chr16:12907740-12907949 | -0.05117263 | 0.916888696 | 0.970570579 |
| chr6:149565588-149567260 | 0.032741612 | 0.916908462 | 0.970570579 |
| chr5:134526552-134527867 | 0.033035533 | 0.916911451 | 0.970570579 |
| chr1:176843730-176844053 | -0.043321387 | 0.916913183 | 0.970570579 |
| chr4:10018808-10019169 | -0.040931222 | 0.916963569 | 0.970610505 |
| chr4:56884412-56885246 | -0.036295223 | 0.916990549 | 0.970625077 |
| chr19:33143879-33144256 | -0.040882397 | 0.917010496 | 0.970625077 |
| chr14:21383456-21384709 | 0.033127088 | 0.917021414 | 0.970625077 |
| chr15:43370685-43371785 | 0.034598133 | 0.917038176 | 0.970625077 |
| chr8:125090915-125092164 | -0.032882649 | 0.917040675 | 0.970625077 |
| chr6:4647462-4647906 | -0.041872109 | 0.917090294 | 0.970658094 |
| chr2:33441305-33441581 | 0.04448779 | 0.917109549 | 0.970658094 |
| chr17:76446958-76447162 | 0.050969734 | 0.91711332 | 0.970658094 |
| chr3:121546789-121547014 | 0.043867538 | 0.917122541 | 0.970658094 |
| chr1:187054974-187055215 | 0.047277997 | 0.917148405 | 0.970672059 |
| chr9:127503973-127504236 | -0.043231978 | 0.917187323 | 0.970679347 |

| | | | |
|---|---|---|---|
| chr4:130258700-130259070 | 0.043830908 | 0.91719777 | 0.970679347 |
| chr12:47837660-47837864 | 0.048658833 | 0.917224848 | 0.970679347 |
| chr1:167637257-167637628 | -0.046257084 | 0.917227192 | 0.970679347 |
| chr9:104910133-104910333 | -0.05905719 | 0.917227203 | 0.970679347 |
| chr6:41367701-41368020 | 0.041080648 | 0.917231301 | 0.970679347 |
| chr9:98455396-98455596 | -0.052682811 | 0.917269591 | 0.970699391 |
| chr3:196969111-196969563 | 0.038107367 | 0.917275579 | 0.970699391 |
| chr12:57846006-57847368 | 0.032777477 | 0.917296196 | 0.970707802 |
| chr6:73143444-73143644 | 0.061215323 | 0.917334602 | 0.970725111 |
| chr7:35800236-35802025 | -0.032424604 | 0.917346855 | 0.970725111 |
| chr11:120985750-120986403 | 0.039589927 | 0.917364571 | 0.970725111 |
| chr22:45896485-45896870 | 0.041200351 | 0.917377815 | 0.970725111 |
| chr12:108664872-108665112 | -0.044895902 | 0.9173843 | 0.970725111 |
| chr6:132780855-132781080 | -0.056762526 | 0.917396374 | 0.970725111 |
| chr20:62156765-62157247 | 0.037063634 | 0.917401235 | 0.970725111 |
| chr5:57567419-57567619 | 0.057955207 | 0.91743863 | 0.970743567 |
| chr1:115833913-115834169 | -0.044056107 | 0.917444016 | 0.970743567 |
| chr3:184185636-184186637 | -0.03321728 | 0.917475922 | 0.970763921 |
| chr3:86694598-86694922 | -0.050378762 | 0.917534447 | 0.97078988 |
| chr8:100157634-100158745 | 0.033364878 | 0.917547428 | 0.97078988 |
| chr20:25310430-25310667 | -0.044347492 | 0.917549219 | 0.97078988 |
| chr8:67595364-67595740 | 0.043039394 | 0.917553785 | 0.97078988 |
| chr1:153616215-153616605 | -0.037343864 | 0.917563806 | 0.97078988 |
| chr1:166433745-166434018 | 0.044605415 | 0.917606021 | 0.970806504 |
| chr7:77498681-77499358 | -0.038715289 | 0.917618856 | 0.970806504 |
| chrX:101462822-101463402 | 0.038937794 | 0.917633269 | 0.970806504 |
| chr8:90593591-90594019 | 0.037015243 | 0.917648526 | 0.970806504 |
| chr8:88424920-88425930 | 0.034569457 | 0.917654013 | 0.970806504 |
| chr17:40362494-40362885 | -0.038664125 | 0.91767436 | 0.970806504 |
| chr19:41859404-41860364 | 0.033053094 | 0.917681696 | 0.970806504 |
| chr6:40308717-40308989 | -0.043274074 | 0.917686196 | 0.970806504 |
| chr3:153927180-153927454 | -0.045370515 | 0.917693548 | 0.970806504 |
| chr13:82436176-82436462 | -0.04838532 | 0.917843732 | 0.970949597 |
| chr1:83583652-83584022 | 0.042083898 | 0.917892517 | 0.970949597 |
| chr3:159586556-159586895 | 0.041967491 | 0.917902694 | 0.970949597 |
| chr7:67625198-67625586 | -0.039350229 | 0.917903912 | 0.970949597 |
| chr3:20672138-20672338 | -0.056438067 | 0.917912395 | 0.970949597 |
| chr10:127636563-127636822 | 0.052017879 | 0.917935891 | 0.970949597 |
| chr8:134712325-134713612 | 0.032539381 | 0.91794062 | 0.970949597 |
| chr6:112042979-112043345 | -0.038756885 | 0.917941553 | 0.970949597 |
| chr5:163696305-163696713 | -0.039391956 | 0.917942861 | 0.970949597 |
| chr15:43168415-43168615 | -0.055532337 | 0.917988998 | 0.970984995 |
| chr11:45964864-45965393 | -0.037805456 | 0.918030669 | 0.971008787 |
| chr1:161487110-161487310 | -0.050867512 | 0.918063566 | 0.971008787 |
| chr11:393276-393512 | 0.041876407 | 0.918064886 | 0.971008787 |
| chrX:8743203-8743403 | 0.05840452 | 0.918072811 | 0.971008787 |
| chr17:60231197-60231707 | -0.033669142 | 0.918086951 | 0.971008787 |
| chr13:113393434-113393637 | -0.050552698 | 0.918087528 | 0.971008787 |
| chr22:27886455-27886818 | -0.048390026 | 0.918130858 | 0.971036281 |
| chr14:74257438-74257813 | 0.03900711 | 0.91814457 | 0.971036281 |

| | | | |
|---|---|---|---|
| chr13:36086186-36086386 | -0.051785007 | 0.918161428 | 0.971036281 |
| chr4:165710149-165710410 | -0.046283005 | 0.918166502 | 0.971036281 |
| chr6:63573112-63574016 | -0.033271238 | 0.918176889 | 0.971036281 |
| chr1:45326599-45326931 | 0.041253748 | 0.918204628 | 0.971052214 |
| chr11:63762949-63763277 | -0.043239859 | 0.918239376 | 0.971054387 |
| chr5:175432515-175432743 | 0.052038662 | 0.918243128 | 0.971054387 |
| chr9:88991258-88991955 | 0.037464147 | 0.918244703 | 0.971054387 |
| chr5:2208003-2208232 | 0.044293157 | 0.918264612 | 0.971062039 |
| chr3:66407789-66408148 | 0.038957971 | 0.918292921 | 0.971078574 |
| chr5:175718776-175718976 | 0.049348856 | 0.918370921 | 0.971147654 |
| chr1:26336139-26336894 | 0.035073883 | 0.918419096 | 0.971185194 |
| chr1:144460789-144461817 | -0.033746635 | 0.918463322 | 0.971214191 |
| chr12:43835427-43836591 | -0.033726034 | 0.918491343 | 0.971214191 |
| chr1:108990836-108991248 | -0.046823805 | 0.91849692 | 0.971214191 |
| chr4:156998975-156999539 | -0.036946162 | 0.918504552 | 0.971214191 |
| chr19:15112440-15112659 | -0.057708871 | 0.918519848 | 0.971214191 |
| chr3:141486579-141488044 | -0.032395144 | 0.918537414 | 0.971214191 |
| chr5:38551258-38551756 | 0.037169435 | 0.918543773 | 0.971214191 |
| chr7:140541479-140541974 | -0.040627871 | 0.91854792 | 0.971214191 |
| chr3:100492065-100493026 | -0.033170266 | 0.918596236 | 0.971231592 |
| chr3:120682377-120682940 | -0.035044298 | 0.918610218 | 0.971231592 |
| chr3:127831852-127832120 | 0.046626466 | 0.91862048 | 0.971231592 |
| chr6:125301424-125302322 | -0.033718957 | 0.918656446 | 0.971231592 |
| chr4:30714486-30714717 | -0.042431523 | 0.918656752 | 0.971231592 |
| chr1:48443284-48443558 | -0.0482658 | 0.91867354 | 0.971231592 |
| chr15:75704612-75704835 | -0.048094639 | 0.918683847 | 0.971231592 |
| chr2:229914865-229915155 | 0.053698313 | 0.918685374 | 0.971231592 |
| chr19:38719957-38720199 | 0.051336733 | 0.918688933 | 0.971231592 |
| chr14:78904064-78904435 | -0.042061739 | 0.918698737 | 0.971231592 |
| chr3:64463298-64463888 | -0.033079928 | 0.918715968 | 0.971231592 |
| chr11:7665545-7666069 | 0.03859473 | 0.918739644 | 0.971231592 |
| chr13:110801170-110801382 | -0.047805158 | 0.918741836 | 0.971231592 |
| chr16:1651513-1651776 | -0.047805158 | 0.918741836 | 0.971231592 |
| chr4:70838819-70840365 | -0.032045046 | 0.918779823 | 0.971258348 |
| chr15:97947593-97947793 | -0.053764871 | 0.918807251 | 0.971273942 |
| chr3:193572443-193572781 | 0.039870905 | 0.918824837 | 0.971279132 |
| chr6:78612008-78612286 | -0.04669203 | 0.918883823 | 0.971322897 |
| chr9:99563466-99563680 | -0.054991238 | 0.918896801 | 0.971322897 |
| chr1:98468219-98468419 | -0.055835933 | 0.918914768 | 0.971322897 |
| chr10:87842410-87842725 | -0.041865015 | 0.918916945 | 0.971322897 |
| chr2:70142014-70143039 | -0.032123348 | 0.918990229 | 0.97138696 |
| chr17:74037277-74037760 | 0.040016613 | 0.919031354 | 0.971416504 |
| chr3:187922857-187923124 | 0.052269498 | 0.919044463 | 0.971416504 |
| chr13:46479340-46479608 | 0.041900168 | 0.919056214 | 0.971416504 |
| chr10:42836688-42837085 | -0.040165517 | 0.919068985 | 0.971416602 |
| chr8:57052362-57052615 | -0.046603271 | 0.919095217 | 0.971419211 |
| chr3:85270375-85270900 | 0.035532609 | 0.919096809 | 0.971419211 |
| chr11:73288933-73289139 | -0.042817399 | 0.9191633 | 0.971476087 |
| chr9:86266688-86266949 | 0.047838786 | 0.919228194 | 0.971520411 |
| chr6:157184709-157184931 | -0.044545515 | 0.919232244 | 0.971520411 |

| | | | |
|---|---|---|---|
| chr2:149174968-149175171 | -0.056119506 | 0.919259631 | 0.971520411 |
| chr17:7844681-7845113 | 0.034957896 | 0.919274224 | 0.971520411 |
| chr11:71251814-71252033 | 0.041574908 | 0.919278818 | 0.971520411 |
| chr2:197145703-197146146 | 0.036926972 | 0.919290377 | 0.971520411 |
| chr7:51369874-51370131 | -0.046537453 | 0.919295299 | 0.971520411 |
| chr9:100323038-100323321 | -0.04859881 | 0.919313801 | 0.971520411 |
| chr14:59507170-59507680 | -0.035781943 | 0.919319352 | 0.971520411 |
| chr17:39002298-39002545 | 0.049598211 | 0.919346509 | 0.971535711 |
| chr22:17638160-17639249 | -0.033288768 | 0.919361882 | 0.971538557 |
| chr2:216498462-216499207 | 0.033541055 | 0.919407566 | 0.971570651 |
| chr8:100073514-100073786 | -0.044909993 | 0.919417613 | 0.971570651 |
| chr8:127668081-127668317 | -0.039176582 | 0.919437256 | 0.971578009 |
| chr17:57662669-57663354 | -0.032836311 | 0.919455107 | 0.971583473 |
| chr22:41285495-41286685 | 0.033039378 | 0.919566104 | 0.971671484 |
| chr12:102992403-102992728 | -0.046358634 | 0.919573074 | 0.971671484 |
| chr20:24699946-24700162 | 0.05028265 | 0.91957644 | 0.971671484 |
| chr5:44332490-44332690 | 0.052109376 | 0.919592456 | 0.971675008 |
| chr15:60255443-60255843 | -0.042365965 | 0.919644946 | 0.971713035 |
| chr6:26596649-26597321 | -0.033729691 | 0.919653809 | 0.971713035 |
| chr11:67507744-67508897 | -0.031920627 | 0.919688807 | 0.971736614 |
| chr11:46490051-46490313 | 0.044227964 | 0.919717684 | 0.971753725 |
| chr1:29556988-29557394 | 0.042527266 | 0.919752147 | 0.971776738 |
| chr1:68231994-68232194 | -0.048329915 | 0.919765082 | 0.971777004 |
| chr13:29418119-29418329 | -0.052503693 | 0.919796093 | 0.971796368 |
| chr3:189049456-189049732 | -0.049590156 | 0.919846997 | 0.971797513 |
| chr16:89453760-89453962 | 0.047464205 | 0.919852951 | 0.971797513 |
| chr2:229780810-229781054 | 0.047464205 | 0.919852951 | 0.971797513 |
| chr1:52056009-52056670 | -0.033504968 | 0.919855763 | 0.971797513 |
| chr18:8478543-8478988 | 0.039185572 | 0.919866539 | 0.971797513 |
| chr15:45198340-45199184 | 0.032089663 | 0.919873275 | 0.971797513 |
| chr22:30591739-30592429 | 0.033182032 | 0.920032042 | 0.971914968 |
| chr4:183636561-183637032 | -0.039290889 | 0.920034297 | 0.971914968 |
| chr5:133610597-133610797 | 0.050600223 | 0.920042791 | 0.971914968 |
| chr3:187679703-187679987 | -0.040869292 | 0.920042828 | 0.971914968 |
| chr11:74570582-74570821 | -0.045773237 | 0.920065918 | 0.971914968 |
| chr1:37634724-37634981 | -0.046805215 | 0.920066928 | 0.971914968 |
| chr2:88600205-88600440 | 0.044550338 | 0.920086397 | 0.971914968 |
| chr6:1609725-1610211 | -0.037434644 | 0.920096576 | 0.971914968 |
| chr2:217987953-217988823 | -0.032459376 | 0.920098614 | 0.971914968 |
| chr16:80676370-80676825 | -0.03955864 | 0.920119571 | 0.971916411 |
| chr9:98056254-98057202 | -0.032010856 | 0.92012535 | 0.971916411 |
| chr3:157532605-157532942 | -0.037243815 | 0.920167021 | 0.971941615 |
| chr7:66920684-66921298 | 0.034648046 | 0.920174581 | 0.971941615 |
| chr22:24189735-24190008 | 0.039205778 | 0.920226906 | 0.971983486 |
| chr20:38472353-38473414 | -0.032247885 | 0.92029433 | 0.972041302 |
| chr5:173757295-173757507 | -0.044110396 | 0.920333742 | 0.972064199 |
| chr20:10861603-10861959 | 0.040518121 | 0.920360611 | 0.972064199 |
| chr1:245986945-245987145 | 0.050400478 | 0.920365402 | 0.972064199 |
| chr20:56949313-56949696 | -0.034893496 | 0.920366754 | 0.972064199 |
| chr12:125285090-125285290 | -0.051838799 | 0.920435525 | 0.972123433 |

| | | | |
|---|---|---|---|
| chr12:110403969-110404214 | 0.039250156 | 0.920454606 | 0.97212503 |
| chr3:130144141-130144936 | -0.034072028 | 0.920474017 | 0.97212503 |
| chr22:50244353-50244655 | 0.040652412 | 0.920475099 | 0.97212503 |
| chr1:121184333-121185114 | 0.032826839 | 0.92050526 | 0.972130661 |
| chr1:168554177-168554393 | -0.05344893 | 0.920505806 | 0.972130661 |
| chr15:44812113-44812379 | 0.045601903 | 0.920524778 | 0.972137298 |
| chr15:89430729-89431080 | -0.036342491 | 0.920562389 | 0.97215041 |
| chr17:1203244-1204190 | 0.034435909 | 0.920564099 | 0.97215041 |
| chr17:55386819-55387223 | -0.044224608 | 0.920578267 | 0.97215041 |
| chr11:6638453-6638765 | 0.047429458 | 0.920587944 | 0.97215041 |
| chr2:45778449-45778878 | -0.036821399 | 0.920628073 | 0.97216576 |
| chr7:140512435-140512909 | 0.037256086 | 0.920635266 | 0.97216576 |
| chr10:5853324-5853954 | 0.03270382 | 0.920640543 | 0.97216576 |
| chr18:23502974-23503882 | -0.032603038 | 0.920688643 | 0.972197686 |
| chr1:119648004-119648652 | 0.033380679 | 0.920696153 | 0.972197686 |
| chr14:68539243-68539448 | 0.051590482 | 0.920727157 | 0.972208991 |
| chr8:116574878-116575138 | 0.050164371 | 0.920737923 | 0.972208991 |
| chr18:11655298-11655610 | -0.042446842 | 0.920744925 | 0.972208991 |
| chr1:109984434-109985108 | -0.03216922 | 0.9207638 | 0.972215523 |
| chr3:81714572-81714772 | -0.05684351 | 0.92084189 | 0.972237712 |
| chr3:51941821-51942602 | -0.032649238 | 0.920843306 | 0.972237712 |
| chr1:182603860-182604659 | -0.032520093 | 0.920845089 | 0.972237712 |
| chr6:64012291-64012688 | 0.038113002 | 0.920855693 | 0.972237712 |
| chr16:80803194-80803775 | -0.03492438 | 0.920855714 | 0.972237712 |
| chr4:88283599-88285135 | 0.031350475 | 0.920876545 | 0.972237712 |
| chr2:129862745-129862956 | -0.051034321 | 0.920878047 | 0.972237712 |
| chr8:127859500-127860454 | -0.031233859 | 0.920889457 | 0.972237712 |
| chrX:104112751-104113008 | -0.043826255 | 0.920899013 | 0.972237712 |
| chr14:42806521-42807154 | -0.036136662 | 0.920934959 | 0.972262266 |
| chr17:57857960-57858207 | 0.036883092 | 0.920967467 | 0.972265876 |
| chr18:23803197-23803467 | -0.052803343 | 0.920973373 | 0.972265876 |
| chr2:231604248-231605172 | -0.032628273 | 0.920986089 | 0.972265876 |
| chr8:81731828-81733067 | 0.031566337 | 0.921008323 | 0.972265876 |
| chr9:89179896-89180715 | -0.033229329 | 0.921017392 | 0.972265876 |
| chr5:62815107-62815379 | 0.04642509 | 0.921025927 | 0.972265876 |
| chr4:94256881-94257143 | -0.049090561 | 0.921027203 | 0.972265876 |
| chr3:151379443-151379736 | -0.041108093 | 0.921058439 | 0.972276458 |
| chr11:66868738-66868997 | 0.037797484 | 0.921071445 | 0.972276458 |
| chr2:112749688-112749971 | 0.041218326 | 0.921076383 | 0.972276458 |
| chr9:128203212-128203439 | 0.042986344 | 0.921096594 | 0.972276458 |
| chr16:53703550-53704346 | 0.032410764 | 0.921100674 | 0.972276458 |
| chr20:57091952-57092582 | 0.032426567 | 0.92118616 | 0.972353299 |
| chr5:56816037-56816300 | 0.039723193 | 0.921248426 | 0.972376003 |
| chr2:69741230-69742510 | -0.031990884 | 0.921255571 | 0.972376003 |
| chr2:48558332-48558607 | 0.046212381 | 0.921262797 | 0.972376003 |
| chr2:26970229-26971167 | -0.032582284 | 0.921291245 | 0.972376003 |
| chr6:30810784-30811035 | -0.042791725 | 0.921326043 | 0.972376003 |
| chr20:856545-856843 | -0.045037091 | 0.921352908 | 0.972376003 |
| chr7:99499817-99500113 | 0.039801501 | 0.92135665 | 0.972376003 |
| chr5:69493377-69493940 | 0.034501549 | 0.921370114 | 0.972376003 |

| | | | |
|---|---|---|---|
| chr3:183904512-183904751 | 0.045012848 | 0.921371289 | 0.972376003 |
| chr6:28795826-28796369 | 0.033625755 | 0.921379942 | 0.972376003 |
| chr10:69061485-69062218 | -0.035207431 | 0.921380313 | 0.972376003 |
| chr16:2520196-2520481 | -0.038022875 | 0.921403487 | 0.972376003 |
| chr11:77091063-77091564 | -0.033353653 | 0.921424578 | 0.972376003 |
| chr12:111341926-111342126 | 0.04962829 | 0.921426916 | 0.972376003 |
| chr3:5246613-5247008 | 0.038796118 | 0.921452688 | 0.972376003 |
| chr7:151303321-151303968 | 0.035777088 | 0.921457915 | 0.972376003 |
| chr11:43283063-43283353 | 0.044450271 | 0.921463165 | 0.972376003 |
| chr13:100573091-100573372 | 0.041426896 | 0.921472769 | 0.972376003 |
| chr3:30606256-30607427 | -0.032705459 | 0.921473431 | 0.972376003 |
| chr4:121252287-121252556 | -0.042214414 | 0.921474461 | 0.972376003 |
| chr5:140042612-140043014 | 0.039907083 | 0.921508811 | 0.972376003 |
| chr7:51738599-51738799 | -0.058560946 | 0.92150905 | 0.972376003 |
| chr2:27461652-27461893 | -0.046299106 | 0.92151212 | 0.972376003 |
| chr16:65813920-65814129 | 0.051094455 | 0.921512243 | 0.972376003 |
| chr2:25042607-25042842 | 0.0392608 | 0.921584664 | 0.972400004 |
| chr6:133040138-133040338 | -0.049573985 | 0.921588685 | 0.972400004 |
| chr8:123001490-123001690 | -0.044450048 | 0.921588786 | 0.972400004 |
| chr16:75733021-75733221 | 0.051190714 | 0.921595831 | 0.972400004 |
| chr11:62664550-62665628 | -0.031726165 | 0.921598443 | 0.972400004 |
| chr16:10382308-10382508 | -0.048653012 | 0.921627755 | 0.972417541 |
| chr5:6712123-6713621 | -0.030902424 | 0.92165349 | 0.972424089 |
| chr7:55593803-55594108 | 0.04387636 | 0.921659344 | 0.972424089 |
| chr3:103220376-103220694 | 0.043857212 | 0.921676503 | 0.972428804 |
| chr4:63316825-63317186 | -0.045297609 | 0.921692352 | 0.972432135 |
| chr12:50786774-50787216 | -0.036596879 | 0.921731028 | 0.97245955 |
| chr1:205798635-205798955 | -0.035862155 | 0.921754168 | 0.97246566 |
| chr17:75393508-75394476 | -0.031392052 | 0.921762203 | 0.97246566 |
| chr10:1612993-1613237 | 0.050134791 | 0.921785777 | 0.972477141 |
| chr19:4815675-4815875 | 0.047727396 | 0.921836958 | 0.972503639 |
| chr17:1056102-1056302 | 0.050417261 | 0.921843643 | 0.972503639 |
| chr17:44689217-44690283 | 0.032485607 | 0.921876465 | 0.972503639 |
| chr3:188185953-188186312 | 0.041059788 | 0.921881218 | 0.972503639 |
| chr12:102155262-102155462 | 0.053650558 | 0.921883166 | 0.972503639 |
| chr4:102431118-102431676 | 0.035679269 | 0.921887047 | 0.972503639 |
| chr11:22921034-22921286 | 0.042816031 | 0.921920894 | 0.972525955 |
| chr6:125889157-125889634 | -0.0394966 | 0.921978782 | 0.972560761 |
| chr11:118611040-118611293 | 0.040015803 | 0.922006674 | 0.972560761 |
| chr13:27984266-27984615 | -0.037168738 | 0.922009348 | 0.972560761 |
| chr1:160030848-160032431 | -0.031231339 | 0.922022138 | 0.972560761 |
| chr11:65644116-65644316 | -0.044201697 | 0.922039831 | 0.972560761 |
| chr11:129375318-129376571 | 0.032613332 | 0.922040827 | 0.972560761 |
| chr10:101066131-101066866 | 0.032354071 | 0.922054298 | 0.972560761 |
| chr16:87264723-87265005 | -0.046930006 | 0.922078359 | 0.972560761 |
| chr3:175005925-175006127 | -0.053237217 | 0.922095967 | 0.972560761 |
| chr13:27043108-27043330 | -0.043867899 | 0.922100463 | 0.972560761 |
| chr10:91185573-91185784 | 0.051289554 | 0.922123367 | 0.972560761 |
| chr16:29192327-29192547 | 0.051289554 | 0.922123367 | 0.972560761 |
| chr12:99261199-99261688 | 0.03927743 | 0.922131131 | 0.972560761 |

| | | | |
|---|---|---|---|
| chr19:50368598-50369049 | -0.036146395 | 0.922131591 | 0.972560761 |
| chr5:41924815-41925977 | 0.031398197 | 0.922159488 | 0.972575614 |
| chr3:12129321-12129521 | -0.056302222 | 0.92217106 | 0.972575614 |
| chr1:42682061-42682655 | -0.032333515 | 0.922220957 | 0.972608197 |
| chr1:191608240-191608583 | 0.042141542 | 0.922227341 | 0.972608197 |
| chr6:13336573-13336789 | -0.051439365 | 0.922314744 | 0.972686986 |
| chr1:111481649-111482106 | 0.036617701 | 0.922385772 | 0.972699552 |
| chr8:120060655-120060962 | 0.038757703 | 0.922404607 | 0.972699552 |
| chr12:123390048-123390378 | 0.040115415 | 0.922426806 | 0.972699552 |
| chr20:58342740-58343111 | 0.038703759 | 0.922430087 | 0.972699552 |
| chrX:100731495-100731888 | -0.040274071 | 0.922435764 | 0.972699552 |
| chr6:166707095-166707328 | -0.048451431 | 0.922441697 | 0.972699552 |
| chr1:176746053-176746351 | -0.040078697 | 0.922450169 | 0.972699552 |
| chr2:199704644-199705004 | -0.038953756 | 0.922451308 | 0.972699552 |
| chr11:65083833-65084597 | 0.03302683 | 0.922456161 | 0.972699552 |
| chr1:191264-191640 | 0.041606286 | 0.922468149 | 0.972699552 |
| chr17:59607851-59608116 | -0.040017245 | 0.922501279 | 0.972699552 |
| chr5:35436544-35436772 | 0.050096571 | 0.922517081 | 0.972699552 |
| chr4:177380979-177381179 | 0.050096571 | 0.922517081 | 0.972699552 |
| chr6:139344085-139344335 | 0.050096571 | 0.922517081 | 0.972699552 |
| chr6:159311703-159311918 | 0.050096571 | 0.922517081 | 0.972699552 |
| chr2:61470426-61471874 | 0.030725313 | 0.922589636 | 0.972762668 |
| chr12:8242847-8243872 | 0.030699503 | 0.922632183 | 0.972794143 |
| chr1:114756937-114758356 | -0.030125246 | 0.922686397 | 0.972830866 |
| chr15:40108769-40109546 | 0.031466115 | 0.92270201 | 0.972830866 |
| chrX:109777844-109778080 | 0.049327254 | 0.92272749 | 0.972830866 |
| chr9:137068241-137068481 | 0.038357806 | 0.922728368 | 0.972830866 |
| chrX:82692136-82692581 | -0.036767112 | 0.922730496 | 0.972830866 |
| chr6:17472547-17473074 | 0.040674333 | 0.922789464 | 0.97287965 |
| chr1:199819278-199819478 | 0.055740411 | 0.922901373 | 0.972984246 |
| chr16:56932763-56933267 | 0.036648872 | 0.92292253 | 0.972993163 |
| chr14:55707365-55707620 | 0.04233809 | 0.922965487 | 0.973025062 |
| chr22:46116374-46116804 | 0.037232268 | 0.923009697 | 0.973037708 |
| chr8:126410292-126410542 | -0.044069868 | 0.923042101 | 0.973037708 |
| chr20:62822736-62823080 | 0.033815291 | 0.923049754 | 0.973037708 |
| chr5:149125350-149125955 | 0.032614972 | 0.92306043 | 0.973037708 |
| chr2:102753676-102753923 | 0.047417891 | 0.923066912 | 0.973037708 |
| chr10:43420331-43421314 | 0.031979097 | 0.923067737 | 0.973037708 |
| chr12:6188456-6188682 | -0.051553456 | 0.923079075 | 0.973037708 |
| chr14:78927874-78928074 | -0.051553456 | 0.923079075 | 0.973037708 |
| chr17:58514540-58514805 | 0.033210878 | 0.923101158 | 0.973047599 |
| chr6:159726652-159727946 | -0.03093359 | 0.923135495 | 0.973062641 |
| chr11:15159095-15159388 | 0.04152337 | 0.923140827 | 0.973062641 |
| chr2:170930290-170930554 | -0.035384706 | 0.923167719 | 0.973077601 |
| chr9:128226098-128226298 | -0.051322055 | 0.923187545 | 0.973085112 |
| chr1:246492615-246492953 | 0.034535494 | 0.923234428 | 0.973121142 |
| chr5:2200636-2200861 | 0.043773321 | 0.923293687 | 0.973145678 |
| chr5:16585368-16585661 | -0.03916857 | 0.923295781 | 0.973145678 |
| chr4:87120769-87121193 | 0.037154252 | 0.923295807 | 0.973145678 |
| chr6:158605328-158605562 | 0.045477961 | 0.923314659 | 0.973152161 |

| | | | |
|---|---|---|---|
| chr20:16434157-16434440 | -0.0406526 | 0.923363862 | 0.973175111 |
| chr20:8131701-8132913 | 0.031090474 | 0.923365293 | 0.973175111 |
| chr16:68704301-68704707 | -0.034637959 | 0.923374536 | 0.973175111 |
| chr12:48789504-48789716 | 0.046311088 | 0.923402932 | 0.973191652 |
| chr12:45260165-45260626 | -0.03786691 | 0.923427501 | 0.973204159 |
| chr6:149834596-149834810 | 0.048676402 | 0.923446327 | 0.973210614 |
| chr11:59919081-59919285 | -0.051239104 | 0.923526571 | 0.973276253 |
| chr8:79765222-79765683 | 0.033605998 | 0.923547451 | 0.973276253 |
| chr20:49534292-49534579 | 0.037237482 | 0.923547619 | 0.973276253 |
| chr15:84653800-84654513 | 0.03132566 | 0.923560344 | 0.973276253 |
| chr9:97411712-97412812 | -0.031685699 | 0.923572121 | 0.973276253 |
| chr9:120714021-120714221 | 0.040950142 | 0.923598028 | 0.973290168 |
| chr16:4957684-4958654 | -0.030598551 | 0.923613887 | 0.973293494 |
| chr7:42186205-42186442 | -0.045405816 | 0.923634315 | 0.973301635 |
| chr1:185393868-185394157 | -0.041506548 | 0.923652265 | 0.973307164 |
| chrX:17121546-17121865 | -0.041775764 | 0.923665192 | 0.973307401 |
| chrX:130097478-130098392 | -0.032166817 | 0.923713562 | 0.973321151 |
| chr17:63882267-63882554 | 0.036001242 | 0.923714027 | 0.973321151 |
| chr19:33176430-33177826 | -0.030036822 | 0.923716349 | 0.973321151 |
| chr22:19962306-19962874 | -0.032739903 | 0.923744298 | 0.973337215 |
| chr7:63925598-63927072 | -0.030444719 | 0.923768048 | 0.973343587 |
| chr14:67816240-67817092 | -0.030953318 | 0.92378833 | 0.973343587 |
| chr9:26723517-26723822 | -0.039093536 | 0.923788455 | 0.973343587 |
| chr4:138919638-138919919 | -0.042408693 | 0.923825969 | 0.973363089 |
| chr5:32585355-32586284 | 0.030863318 | 0.923832371 | 0.973363089 |
| chr2:130920464-130920746 | -0.040731949 | 0.923893478 | 0.973402328 |
| chr17:16398454-16399051 | 0.035253871 | 0.923895021 | 0.973402328 |
| chr19:1253365-1253568 | 0.038336476 | 0.923919021 | 0.973414229 |
| chr17:65901502-65901843 | -0.043220166 | 0.923958691 | 0.97344264 |
| chr4:76090317-76090583 | -0.045017355 | 0.923994361 | 0.973445133 |
| chr14:73792279-73792510 | -0.04470742 | 0.9240082 | 0.973445133 |
| chr5:112390132-112391025 | 0.0323885 | 0.924011025 | 0.973445133 |
| chr20:24746328-24746688 | 0.040955995 | 0.924019941 | 0.973445133 |
| chr10:6050704-6050965 | 0.042433958 | 0.924037286 | 0.973445133 |
| chr11:44616887-44617131 | 0.042433958 | 0.924037286 | 0.973445133 |
| chr10:118754689-118755601 | 0.030431687 | 0.924055022 | 0.973446594 |
| chr3:181839173-181839373 | -0.047456937 | 0.924064081 | 0.973446594 |
| chr6:26097365-26097662 | 0.042399355 | 0.92409574 | 0.973466561 |
| chr14:30270251-30271138 | 0.031255391 | 0.924154426 | 0.973512175 |
| chr12:120200728-120201824 | 0.030546659 | 0.924165467 | 0.973512175 |
| chr11:58577905-58579297 | 0.029921814 | 0.924185682 | 0.973512175 |
| chr5:178839066-178839719 | 0.032885692 | 0.924189862 | 0.973512175 |
| chr1:181104509-181105874 | 0.029938064 | 0.924215322 | 0.973513052 |
| chr7:32775002-32775240 | 0.050081694 | 0.924217767 | 0.973513052 |
| chr17:42823777-42824590 | -0.03212829 | 0.924229528 | 0.973513052 |
| chr9:93453386-93453834 | -0.033731331 | 0.924241516 | 0.973513052 |
| chr3:120348572-120349936 | -0.03001951 | 0.924291299 | 0.973542347 |
| chr12:67545358-67545666 | -0.043349982 | 0.92430322 | 0.973542347 |
| chr12:75087663-75088044 | 0.04072899 | 0.924307446 | 0.973542347 |
| chr14:24366604-24367599 | -0.030148563 | 0.924339542 | 0.97356277 |

| | | | |
|---|---|---|---|
| chr5:93611911-93612339 | 0.036877164 | 0.924355177 | 0.973564695 |
| chr2:119234644-119234916 | 0.039782538 | 0.924366782 | 0.973564695 |
| chr8:22141756-22142097 | -0.033948383 | 0.924413437 | 0.973591146 |
| chr20:51707659-51707886 | -0.041170681 | 0.924435166 | 0.973591146 |
| chr11:568289-568866 | 0.032786505 | 0.924441798 | 0.973591146 |
| chr8:70403331-70403531 | 0.037767144 | 0.924442722 | 0.973591146 |
| chr12:89262160-89262360 | 0.049642127 | 0.924484938 | 0.973593309 |
| chr3:112649066-112649301 | -0.048820299 | 0.924491486 | 0.973593309 |
| chr8:94919547-94919820 | -0.038310695 | 0.924492309 | 0.973593309 |
| chr8:93991124-93992102 | 0.030518267 | 0.924510684 | 0.973593309 |
| chr2:218688540-218688740 | -0.052927801 | 0.924519075 | 0.973593309 |
| chr3:151918335-151918581 | 0.044762647 | 0.924521014 | 0.973593309 |
| chr17:63990339-63990691 | 0.034504686 | 0.924538223 | 0.973595787 |
| chr3:171967212-171967495 | 0.041651867 | 0.92456071 | 0.973595787 |
| chr17:56542447-56542710 | 0.043781098 | 0.924563489 | 0.973595787 |
| chr8:123705388-123705656 | 0.039917101 | 0.924574194 | 0.973595787 |
| chr7:34320875-34321704 | -0.031138745 | 0.924629869 | 0.973631377 |
| chr6:30364661-30364939 | 0.041679341 | 0.924636671 | 0.973631377 |
| chr16:67961411-67961941 | -0.034763908 | 0.924655863 | 0.973631377 |
| chr20:5529767-5530010 | 0.042889081 | 0.924658819 | 0.973631377 |
| chr14:105845712-105846098 | 0.038578773 | 0.924693661 | 0.973654684 |
| chr17:71085268-71085496 | 0.045680133 | 0.924714176 | 0.973662905 |
| chr1:45644061-45644315 | -0.044561211 | 0.924806134 | 0.973746349 |
| chr1:11893925-11894125 | 0.047655692 | 0.924838069 | 0.973755835 |
| chr6:133553617-133553933 | 0.046174436 | 0.92484056 | 0.973755835 |
| chr15:63601475-63601716 | -0.038320854 | 0.924888165 | 0.973792577 |
| chr4:119136620-119136863 | -0.043714765 | 0.925054748 | 0.973943242 |
| chr2:27280010-27280211 | 0.045283386 | 0.92508292 | 0.973943242 |
| chrX:99242047-99242271 | -0.049302603 | 0.92508585 | 0.973943242 |
| chr4:44725808-44727151 | 0.030779203 | 0.925085895 | 0.973943242 |
| chr8:94628681-94628883 | 0.044771641 | 0.925098649 | 0.973943242 |
| chr12:98296729-98296930 | 0.04859217 | 0.925112912 | 0.973943242 |
| chr4:2793840-2794073 | 0.045566951 | 0.925120241 | 0.973943242 |
| chr19:4670221-4670748 | 0.033627659 | 0.925170165 | 0.973968458 |
| chr7:151232261-151232656 | -0.03313753 | 0.925171846 | 0.973968458 |
| chr11:107564916-107565904 | -0.031736662 | 0.925182327 | 0.973968458 |
| chr14:66284455-66284726 | 0.044570663 | 0.925197403 | 0.973970948 |
| chr1:54715424-54716171 | -0.032093196 | 0.925229053 | 0.973989751 |
| chr15:40115777-40116062 | -0.040626741 | 0.925240689 | 0.973989751 |
| chr1:93908700-93910000 | -0.03084143 | 0.925254485 | 0.973990893 |
| chr6:18386981-18387697 | 0.031586103 | 0.925283579 | 0.974000583 |
| chr11:79481003-79481203 | 0.051046323 | 0.925289113 | 0.974000583 |
| chr16:55500012-55500220 | -0.048472597 | 0.925351656 | 0.974053036 |
| chr3:106121633-106122222 | -0.034872919 | 0.925413095 | 0.974104327 |
| chr6:31958803-31959372 | -0.031578423 | 0.925518056 | 0.974197335 |
| chr12:34133243-34133566 | -0.042274327 | 0.925526883 | 0.974197335 |
| chr2:62164149-62164349 | -0.054008744 | 0.925545469 | 0.974203516 |
| chr1:246827327-246827843 | 0.031848541 | 0.925596021 | 0.974243342 |
| chr18:59308551-59308991 | 0.036128307 | 0.925647999 | 0.974266851 |
| chr2:177753928-177754254 | -0.041227398 | 0.925650352 | 0.974266851 |

| | | | |
|---|---|---|---|
| chr8:129303391-129304032 | -0.0330479 | 0.925656502 | 0.974266851 |
| chr2:195561087-195561389 | 0.037195087 | 0.925708474 | 0.974296428 |
| chr14:20342690-20343803 | 0.029208484 | 0.925720741 | 0.974296428 |
| chr4:40897592-40898139 | -0.034148032 | 0.92572275 | 0.974296428 |
| chr15:22838006-22838942 | 0.030429601 | 0.925748792 | 0.974304335 |
| chr5:160021856-160022369 | 0.034150024 | 0.92576664 | 0.974304335 |
| chr3:170815293-170815788 | -0.03236127 | 0.92576841 | 0.974304335 |
| chr9:134068484-134068878 | 0.034754045 | 0.925795274 | 0.974316673 |
| chr6:121306077-121306378 | -0.038983866 | 0.925805565 | 0.974316673 |
| chr5:134637088-134637450 | -0.034241835 | 0.925828337 | 0.974327256 |
| chr20:23050754-23051182 | 0.033095922 | 0.925847382 | 0.974328435 |
| chr19:572108-572703 | 0.031715933 | 0.925862171 | 0.974328435 |
| chr1:109186091-109186334 | -0.041833341 | 0.925872403 | 0.974328435 |
| chr14:100708391-100708668 | 0.03673146 | 0.925891169 | 0.974328435 |
| chr14:59511998-59512198 | -0.045693513 | 0.925893038 | 0.974328435 |
| chr11:65355013-65355392 | 0.036015553 | 0.925920879 | 0.974344352 |
| chr2:218230912-218231149 | 0.041210454 | 0.925952423 | 0.974364164 |
| chr22:38755906-38756202 | -0.039027523 | 0.925987862 | 0.974387621 |
| chr20:50313290-50313534 | 0.036568888 | 0.926000149 | 0.974387621 |
| chr3:141089443-141089652 | 0.042231222 | 0.926091577 | 0.974470444 |
| chrX:119094383-119094764 | -0.036714731 | 0.926131952 | 0.974495652 |
| chr3:64030314-64031080 | -0.02920696 | 0.926151019 | 0.974495652 |
| chr17:19377593-19378693 | 0.029357469 | 0.926153688 | 0.974495652 |
| chr22:35350660-35351192 | 0.032554709 | 0.92616765 | 0.974496961 |
| chr6:32177597-32178733 | -0.029433668 | 0.926211092 | 0.97450103 |
| chr17:4806962-4807328 | -0.034878144 | 0.926215718 | 0.97450103 |
| chr17:17752086-17753012 | 0.030131975 | 0.926220849 | 0.97450103 |
| chr22:23290568-23290818 | -0.040431593 | 0.92622239 | 0.97450103 |
| chr16:89333671-89333982 | 0.037752914 | 0.926287584 | 0.97453967 |
| chr1:153483895-153484110 | -0.046640206 | 0.926296627 | 0.97453967 |
| chr11:66510438-66511093 | 0.032312518 | 0.926297273 | 0.97453967 |
| chr3:33658984-33659208 | -0.043703121 | 0.92634587 | 0.974567103 |
| chr9:128844970-128845753 | 0.031014755 | 0.926369298 | 0.974567103 |
| chr8:86341639-86343339 | 0.028810039 | 0.926372704 | 0.974567103 |
| chr21:43864182-43864623 | -0.034270668 | 0.926374497 | 0.974567103 |
| chr19:55545451-55545651 | -0.045799446 | 0.926386943 | 0.974567103 |
| chr1:224929280-224930186 | -0.030518642 | 0.926403487 | 0.974571126 |
| chr4:22515123-22516657 | 0.029694374 | 0.926458025 | 0.974615119 |
| chr17:60971624-60972206 | 0.031614357 | 0.926495939 | 0.974641622 |
| chr7:131109966-131110194 | -0.037543846 | 0.92653587 | 0.974662662 |
| chr12:71014253-71014532 | 0.037794185 | 0.926547496 | 0.974662662 |
| chr19:39212439-39212639 | -0.052068061 | 0.926555563 | 0.974662662 |
| chr5:17670515-17670800 | -0.04270146 | 0.926570449 | 0.974662662 |
| chr15:92808585-92809730 | 0.029096254 | 0.926585062 | 0.974662662 |
| chr8:127870266-127871274 | -0.029253349 | 0.92661976 | 0.974662662 |
| chr11:82680293-82680555 | 0.040013545 | 0.926634909 | 0.974662662 |
| chr8:80325152-80325550 | 0.039400116 | 0.926646454 | 0.974662662 |
| chr12:67992681-67992979 | 0.043503993 | 0.926650448 | 0.974662662 |
| chr6:147640729-147640975 | 0.043437111 | 0.926672165 | 0.974662662 |
| chr8:113246886-113247369 | -0.033805675 | 0.926677614 | 0.974662662 |

| | | | |
|---|---|---|---|
| chr7:16688233-16688587 | 0.034903869 | 0.926681636 | 0.974662662 |
| chr1:242936787-242937039 | 0.039182542 | 0.926702798 | 0.974662662 |
| chr8:56211174-56212391 | -0.029219743 | 0.926730634 | 0.974662662 |
| chr19:3987733-3987994 | -0.036533216 | 0.926745091 | 0.974662662 |
| chr16:84759956-84760256 | -0.041893219 | 0.92675135 | 0.974662662 |
| chr19:48400515-48400793 | 0.036064842 | 0.926756366 | 0.974662662 |
| chrX:97328872-97329203 | 0.037119468 | 0.926795601 | 0.974662662 |
| chr7:5282767-5283014 | 0.038999065 | 0.926797699 | 0.974662662 |
| chr5:36041037-36041509 | -0.035432316 | 0.926800013 | 0.974662662 |
| chr14:94433982-94434253 | 0.038008823 | 0.926801775 | 0.974662662 |
| chr7:42236276-42236660 | -0.034059384 | 0.926810256 | 0.974662662 |
| chr3:48556612-48557444 | -0.029972263 | 0.926813559 | 0.974662662 |
| chr20:40968776-40969192 | 0.033693099 | 0.926821229 | 0.974662662 |
| chr3:125915637-125916654 | -0.0287954 | 0.926873347 | 0.974704092 |
| chr3:195990016-195990799 | 0.032314048 | 0.926893278 | 0.974711674 |
| chr19:46579734-46579934 | 0.047910927 | 0.926943119 | 0.97474332 |
| chrX:14529417-14529632 | -0.045638777 | 0.926948814 | 0.97474332 |
| chr18:12422946-12423277 | 0.034354604 | 0.92696547 | 0.974747457 |
| chr10:81899438-81899738 | -0.041902603 | 0.926982525 | 0.974747688 |
| chr5:28411972-28412172 | -0.047484002 | 0.926991132 | 0.974747688 |
| chr11:47268867-47270373 | -0.028868028 | 0.927098295 | 0.974846994 |
| chrX:12974539-12975766 | 0.029749172 | 0.927141031 | 0.974878411 |
| chr3:150151343-150151578 | -0.048544253 | 0.927154396 | 0.974878411 |
| chr17:48026525-48026796 | 0.033149228 | 0.927171266 | 0.974878411 |
| chr12:32106515-32107043 | 0.033897835 | 0.927201235 | 0.974878411 |
| chr3:58518439-58518639 | -0.048753038 | 0.927204227 | 0.974878411 |
| chr9:114908013-114908598 | 0.030131034 | 0.927215433 | 0.974878411 |
| chr14:99224104-99224304 | -0.044929312 | 0.927221761 | 0.974878411 |
| chr12:46661359-46661686 | 0.03755568 | 0.92722996 | 0.974878411 |
| chr14:87615881-87616452 | -0.032872945 | 0.927297817 | 0.974930486 |
| chr17:74791495-74791811 | 0.035500297 | 0.927312589 | 0.974930486 |
| chr3:19688930-19689137 | -0.044947421 | 0.927333617 | 0.974930486 |
| chr6:28141478-28142198 | 0.031044994 | 0.927342935 | 0.974930486 |
| chr17:8247495-8248260 | 0.031201757 | 0.927358209 | 0.974930486 |
| chr2:179803976-179804489 | -0.04099077 | 0.927373994 | 0.974930486 |
| chr2:127859714-127860051 | 0.032430481 | 0.927387365 | 0.974930486 |
| chr19:48965660-48966153 | -0.032464106 | 0.927389824 | 0.974930486 |
| chr17:73595633-73595895 | -0.042705087 | 0.927415956 | 0.974930486 |
| chr6:5829681-5829881 | 0.047119341 | 0.927458506 | 0.974930486 |
| chr20:46349065-46349493 | 0.034486726 | 0.927458509 | 0.974930486 |
| chr5:35306552-35307356 | -0.031015193 | 0.92746096 | 0.974930486 |
| chr8:133695305-133695633 | 0.03569706 | 0.927461428 | 0.974930486 |
| chr1:154760521-154760721 | 0.046572497 | 0.927479776 | 0.974930486 |
| chr9:122229065-122229295 | 0.043762968 | 0.927493363 | 0.974930486 |
| chr8:120123493-120124071 | 0.031743906 | 0.927497928 | 0.974930486 |
| chr10:110567144-110568507 | 0.028914873 | 0.927507883 | 0.974930486 |
| chr16:19121866-19122255 | -0.037215198 | 0.927511349 | 0.974930486 |
| chr7:152675410-152676432 | -0.02918378 | 0.927525929 | 0.974930486 |
| chr2:9102802-9103745 | -0.030632409 | 0.927533967 | 0.974930486 |
| chr11:77994436-77995299 | -0.02881564 | 0.927566631 | 0.974951445 |

| | | | |
|---|---|---|---|
| chr7:32479618-32479952 | 0.033368713 | 0.927595691 | 0.974968614 |
| chr6:10850786-10851033 | 0.041846789 | 0.927620611 | 0.974978478 |
| chr22:42070358-42071357 | -0.030392192 | 0.927630525 | 0.974978478 |
| chr5:37837450-37837793 | -0.036911731 | 0.927659675 | 0.97499357 |
| chr4:113761033-113763128 | 0.028429901 | 0.927670332 | 0.97499357 |
| chr3:96364379-96364579 | 0.047168532 | 0.927689852 | 0.975000711 |
| chr3:14319629-14319946 | 0.034978888 | 0.927703402 | 0.975000881 |
| chr21:39197635-39197868 | -0.04248374 | 0.927737412 | 0.975000881 |
| chr6:28953571-28954172 | 0.031731996 | 0.927742211 | 0.975000881 |
| chr14:20469039-20470227 | -0.02886465 | 0.927751371 | 0.975000881 |
| chr17:58172462-58172756 | -0.034875009 | 0.927753638 | 0.975000881 |
| chr13:99086715-99088331 | -0.029171051 | 0.927779575 | 0.975014766 |
| chr12:75784592-75785005 | 0.033228225 | 0.927820195 | 0.975032059 |
| chr14:99604836-99605154 | 0.032148675 | 0.927821481 | 0.975032059 |
| chr1:112369635-112370067 | -0.031650049 | 0.927844728 | 0.975043117 |
| chr1:186605765-186606097 | -0.035679892 | 0.92787075 | 0.975050201 |
| chr19:7943561-7944126 | -0.03027396 | 0.927876921 | 0.975050201 |
| chr4:87353015-87353252 | -0.044074764 | 0.927897632 | 0.975058593 |
| chr4:24793632-24793950 | 0.037668813 | 0.927928055 | 0.975077189 |
| chr1:161291543-161291793 | -0.040082569 | 0.927977938 | 0.975116234 |
| chr4:74970172-74970460 | -0.040334152 | 0.928044619 | 0.975172928 |
| chr10:26846794-26846994 | 0.05222331 | 0.928065424 | 0.975181417 |
| chr5:33588761-33589290 | 0.033912295 | 0.928120227 | 0.975196471 |
| chr3:167929977-167930244 | -0.041965851 | 0.928161459 | 0.975196471 |
| chr1:202010219-202010897 | 0.029131916 | 0.928167907 | 0.975196471 |
| chr1:196377871-196378071 | -0.042853 | 0.928186516 | 0.975196471 |
| chr4:48779328-48779917 | -0.030181788 | 0.928188652 | 0.975196471 |
| chrX:48704009-48704454 | 0.03253924 | 0.928200424 | 0.975196471 |
| chr2:96771794-96772367 | -0.029949455 | 0.928204326 | 0.975196471 |
| chr8:26457837-26458131 | -0.0454667 | 0.928225963 | 0.975196471 |
| chr6:6686719-6687692 | 0.028699142 | 0.928227037 | 0.975196471 |
| chr7:39791071-39791303 | -0.047129859 | 0.92822943 | 0.975196471 |
| chr4:21512476-21512681 | -0.047129859 | 0.92822943 | 0.975196471 |
| chr7:6358995-6359254 | -0.041187366 | 0.92823248 | 0.975196471 |
| chr5:153964967-153965447 | -0.033174512 | 0.928252612 | 0.975204251 |
| chr8:28700855-28701118 | -0.03949171 | 0.928273675 | 0.97520982 |
| chr15:41585310-41586061 | -0.032045731 | 0.92829074 | 0.97520982 |
| chr12:47249873-47250233 | 0.037309848 | 0.928308157 | 0.97520982 |
| chr17:51434494-51434704 | -0.040530896 | 0.928308823 | 0.97520982 |
| chr20:62861695-62862256 | 0.029773029 | 0.92832264 | 0.975210964 |
| chrX:32225177-32225464 | -0.04622661 | 0.928364298 | 0.975241356 |
| chr8:124997640-124999046 | -0.028071532 | 0.928466603 | 0.975335455 |
| chr6:9250855-9251090 | 0.042837084 | 0.928492072 | 0.975348838 |
| chr17:59121864-59122465 | -0.029645809 | 0.928555991 | 0.975399287 |
| chr14:54472017-54472217 | -0.046911867 | 0.928576683 | 0.975399287 |
| chr2:121974845-121975583 | -0.031398519 | 0.928578768 | 0.975399287 |
| chr12:65278528-65279289 | 0.030855951 | 0.928591018 | 0.975399287 |
| chr15:81297199-81297399 | -0.041967332 | 0.928650217 | 0.975404846 |
| chr4:7070792-7071091 | 0.03372545 | 0.928674826 | 0.975404846 |
| chr5:10760613-10761882 | 0.028467884 | 0.928678036 | 0.975404846 |

| | | | |
|---|---|---|---|
| chr1:37388006-37388295 | -0.043072685 | 0.928683025 | 0.975404846 |
| chr10:117262277-117262765 | -0.034283778 | 0.928694044 | 0.975404846 |
| chrX:8190981-8191493 | -0.030893878 | 0.9286958 | 0.975404846 |
| chr1:1511006-1511206 | 0.042367665 | 0.928696209 | 0.975404846 |
| chr2:161844822-161845233 | -0.035742752 | 0.928699077 | 0.975404846 |
| chr1:176897866-176898193 | -0.034975937 | 0.928729071 | 0.975404846 |
| chr4:139740560-139741098 | 0.032790158 | 0.928729843 | 0.975404846 |
| chr5:16550383-16550666 | -0.034722679 | 0.928736341 | 0.975404846 |
| chr4:40476422-40477136 | -0.031515408 | 0.92878195 | 0.975439376 |
| chr11:45922073-45922410 | -0.031719845 | 0.928836413 | 0.975477485 |
| chr19:41397203-41397928 | 0.02951601 | 0.928843698 | 0.975477485 |
| chr2:42423161-42423397 | -0.047521753 | 0.92886494 | 0.975486423 |
| chr7:107387678-107387980 | -0.039933577 | 0.928981326 | 0.975562927 |
| chr16:88442985-88443608 | 0.030174793 | 0.928985553 | 0.975562927 |
| chr16:57084301-57084583 | -0.036422218 | 0.928986467 | 0.975562927 |
| chr9:34374992-34375282 | -0.035120063 | 0.928988716 | 0.975562927 |
| chr2:102505316-102505779 | 0.032870824 | 0.929016623 | 0.975575488 |
| chr16:11538994-11539194 | -0.048325297 | 0.929027381 | 0.975575488 |
| chr9:98034585-98035100 | 0.033214999 | 0.929038875 | 0.975575488 |
| chr1:147224956-147225689 | -0.029800175 | 0.929106651 | 0.975615242 |
| chr8:67456850-67457106 | -0.043875288 | 0.929131169 | 0.975615242 |
| chr20:19371978-19372178 | -0.049234469 | 0.929150288 | 0.975615242 |
| chr6:137443912-137444208 | 0.045637691 | 0.929152408 | 0.975615242 |
| chr1:84956719-84956919 | -0.052900683 | 0.929162727 | 0.975615242 |
| chr1:199202545-199202753 | 0.045385719 | 0.929189913 | 0.975615242 |
| chr15:78971526-78971769 | 0.042508658 | 0.929191792 | 0.975615242 |
| chr21:9065726-9065926 | 0.04463219 | 0.929206563 | 0.975615242 |
| chr2:157093751-157094016 | 0.04463219 | 0.929206563 | 0.975615242 |
| chr2:160278861-160279083 | 0.04463219 | 0.929206563 | 0.975615242 |
| chr20:53593059-53594837 | -0.027448761 | 0.929229843 | 0.975615242 |
| chr15:33194440-33195394 | 0.029058609 | 0.929240453 | 0.975615242 |
| chr5:98750538-98751041 | 0.03385903 | 0.929242259 | 0.975615242 |
| chr13:76473594-76473874 | 0.038064307 | 0.929256193 | 0.975616503 |
| chrX:48801537-48802731 | -0.028030113 | 0.929288266 | 0.975636808 |
| chr16:72847248-72847450 | -0.037578947 | 0.929361942 | 0.975697564 |
| chr14:76644971-76645221 | -0.03780588 | 0.929371603 | 0.975697564 |
| chr11:69044088-69044509 | 0.032303251 | 0.929420266 | 0.975735283 |
| chr1:196738777-196738977 | 0.04417001 | 0.929474317 | 0.975768542 |
| chr1:206490053-206490397 | -0.030681297 | 0.929477416 | 0.975768542 |
| chr1:203650208-203650784 | -0.030735155 | 0.929514987 | 0.97578484 |
| chr8:57537059-57537437 | 0.03708978 | 0.929527496 | 0.97578484 |
| chr1:195581123-195581342 | 0.041138425 | 0.929531146 | 0.97578484 |
| chr9:101458912-101459535 | -0.03070673 | 0.929562183 | 0.975798605 |
| chr2:46940687-46942384 | -0.028402874 | 0.929569729 | 0.975798605 |
| chr10:77846715-77846915 | 0.039711379 | 0.929597756 | 0.975803336 |
| chr7:149623985-149625176 | -0.028479092 | 0.929599706 | 0.975803336 |
| chr6:26382940-26383536 | -0.030196715 | 0.929616567 | 0.975807667 |
| chr11:33857912-33858120 | 0.044096422 | 0.929636963 | 0.975815708 |
| chr11:9569349-9569607 | -0.034737336 | 0.929663924 | 0.97583064 |
| chr8:108287144-108287430 | 0.035832417 | 0.929714585 | 0.975853897 |

| | | | |
|---|---|---|---|
| chr2:100262682-100263177 | 0.032071571 | 0.929720177 | 0.975853897 |
| chr5:39948942-39949194 | -0.040386494 | 0.92972429 | 0.975853897 |
| chr20:57381752-57382084 | 0.030937706 | 0.929747398 | 0.975864785 |
| chr8:123137781-123137981 | -0.040630724 | 0.929768262 | 0.975873315 |
| chr3:69739162-69739989 | 0.028732703 | 0.929781701 | 0.975874054 |
| chr2:112541749-112542847 | -0.02840521 | 0.929824029 | 0.975905111 |
| chr10:80829584-80829995 | -0.030610944 | 0.929881903 | 0.975910732 |
| chr19:48872102-48872549 | 0.029991945 | 0.929883718 | 0.975910732 |
| chr14:48792290-48792502 | 0.044189929 | 0.929887032 | 0.975910732 |
| chr6:41779809-41780475 | 0.029582888 | 0.929891422 | 0.975910732 |
| chr12:121802917-121803375 | 0.030695262 | 0.929893067 | 0.975910732 |
| chr17:48011399-48011644 | 0.035885481 | 0.929924159 | 0.975922861 |
| chr6:10185785-10186150 | -0.043172504 | 0.929936533 | 0.975922861 |
| chr4:56434877-56436537 | -0.027475406 | 0.929943611 | 0.975922861 |
| chr2:156110801-156111150 | 0.037032558 | 0.92996953 | 0.975922861 |
| chr3:49703496-49703698 | 0.041268147 | 0.929983107 | 0.975922861 |
| chr5:12320755-12320955 | -0.045599061 | 0.930002696 | 0.975922861 |
| chr1:39658079-39658304 | -0.040755154 | 0.930004609 | 0.975922861 |
| chr20:36792514-36792797 | 0.037325307 | 0.93000652 | 0.975922861 |
| chr15:43776127-43776328 | 0.0368497 | 0.930052812 | 0.975952434 |
| chr6:32190075-32190464 | -0.031550103 | 0.930067288 | 0.975952434 |
| chr19:1266394-1266781 | 0.033286791 | 0.930072913 | 0.975952434 |
| chr17:77274155-77274483 | 0.035639779 | 0.93008644 | 0.975953263 |
| chr8:105148391-105148591 | 0.048882622 | 0.930111273 | 0.975955738 |
| chr8:79652771-79653078 | -0.037893258 | 0.930114274 | 0.975955738 |
| chr7:39746220-39746467 | -0.034976005 | 0.930194058 | 0.97600096 |
| chr10:80078304-80079504 | -0.027790323 | 0.930207788 | 0.97600096 |
| chr1:109089290-109089725 | 0.030864864 | 0.930238777 | 0.97600096 |
| chr10:89701065-89701935 | 0.027779252 | 0.930241015 | 0.97600096 |
| chr21:23942968-23943202 | 0.045908492 | 0.930247187 | 0.97600096 |
| chr14:21269208-21270034 | -0.027767451 | 0.930247988 | 0.97600096 |
| chr9:136363025-136363959 | 0.028245897 | 0.930252076 | 0.97600096 |
| chr1:147593326-147593531 | -0.044497067 | 0.930262863 | 0.97600096 |
| chr2:101251856-101252157 | 0.031677972 | 0.930272012 | 0.97600096 |
| chr12:123670827-123671430 | 0.030392981 | 0.93033377 | 0.976052389 |
| chr2:24409976-24410191 | 0.042372044 | 0.930390286 | 0.976082513 |
| chrX:101287516-101287748 | 0.040467609 | 0.930391812 | 0.976082513 |
| chr8:41767291-41767541 | -0.043568697 | 0.930418016 | 0.976082513 |
| chr2:46464530-46464760 | -0.036904158 | 0.930422288 | 0.976082513 |
| chr8:25765744-25766479 | 0.031936187 | 0.930428444 | 0.976082513 |
| chr1:157580289-157580534 | -0.039418334 | 0.93045202 | 0.976082513 |
| chr10:61981463-61981882 | -0.037518647 | 0.930469678 | 0.976082513 |
| chr7:6847688-6848081 | 0.036540384 | 0.930484494 | 0.976082513 |
| chr1:67749518-67750032 | -0.031817332 | 0.930492042 | 0.976082513 |
| chr2:15383755-15384156 | -0.036520005 | 0.930497666 | 0.976082513 |
| chr10:68837705-68837942 | 0.038255893 | 0.930502612 | 0.976082513 |
| chr19:11137713-11138264 | -0.030293761 | 0.930523792 | 0.976091369 |
| chr11:62761172-62762087 | 0.028579224 | 0.930539864 | 0.976094864 |
| chr18:24727799-24728019 | -0.042737667 | 0.930554036 | 0.976096368 |
| chr4:54234314-54234540 | 0.039505251 | 0.93060336 | 0.976134742 |

| | | | |
|---|---|---|---|
| chr10:119892242-119892898 | 0.028412171 | 0.930663621 | 0.976184588 |
| chr19:17579646-17580271 | 0.030440201 | 0.930686493 | 0.976195215 |
| chr17:10646402-10646718 | -0.036971219 | 0.930716778 | 0.976213618 |
| chr2:60849616-60849816 | -0.043041203 | 0.930749551 | 0.976216123 |
| chr3:148991225-148992499 | -0.027699578 | 0.930750392 | 0.976216123 |
| chr20:31618838-31619097 | 0.041268686 | 0.930757389 | 0.976216123 |
| chr11:87097570-87097841 | 0.037140193 | 0.930785743 | 0.976232499 |
| chr2:208571293-208571737 | 0.034490232 | 0.930877182 | 0.976314263 |
| chr1:226436877-226437206 | 0.038984009 | 0.930889184 | 0.976314263 |
| chr15:70111345-70111597 | 0.037427283 | 0.930916987 | 0.976330058 |
| chr14:100214000-100214916 | 0.028052145 | 0.930956922 | 0.976358576 |
| chr11:13289153-13289398 | 0.040500108 | 0.930980361 | 0.976369795 |
| chr2:113080125-113080326 | 0.04111229 | 0.931013392 | 0.976387296 |
| chr4:126582756-126583115 | 0.032147061 | 0.931030415 | 0.976387296 |
| chr20:50120895-50121378 | 0.02935157 | 0.931052229 | 0.976387296 |
| chr3:152402356-152402591 | -0.044486454 | 0.931060183 | 0.976387296 |
| chr11:14537115-14537431 | 0.039379831 | 0.931060763 | 0.976387296 |
| chr2:194131350-194131648 | -0.03997295 | 0.931087014 | 0.976401461 |
| chr5:94021248-94021578 | -0.03643088 | 0.93113362 | 0.976424183 |
| chr9:100098141-100098482 | -0.030368389 | 0.931143996 | 0.976424183 |
| chr18:26549602-26550000 | -0.03153638 | 0.931161187 | 0.976424183 |
| chr8:95677387-95677631 | 0.042159237 | 0.931181956 | 0.976424183 |
| chr1:206744822-206745053 | 0.035725057 | 0.931191915 | 0.976424183 |
| chr5:152164908-152165154 | 0.042028221 | 0.93119423 | 0.976424183 |
| chr5:139293415-139294804 | 0.027756505 | 0.931197885 | 0.976424183 |
| chr6:97282597-97283761 | -0.027686257 | 0.931218484 | 0.976432421 |
| chr3:25897062-25897273 | -0.041914466 | 0.931268977 | 0.97647157 |
| chr9:128683698-128683914 | 0.035807014 | 0.931281309 | 0.97647157 |
| chr21:42809579-42809868 | -0.036962505 | 0.931299374 | 0.976477149 |
| chr12:104219435-104220636 | 0.027928808 | 0.931316833 | 0.976482093 |
| chr17:67800415-67800817 | 0.029790758 | 0.931387125 | 0.976542431 |
| chr3:194972039-194972268 | -0.041776254 | 0.931406968 | 0.976549872 |
| chr8:26133754-26134350 | -0.031172651 | 0.931449608 | 0.976577396 |
| chr19:9309455-9310319 | 0.028833364 | 0.931462312 | 0.976577396 |
| chr7:51029413-51029650 | -0.045716735 | 0.931471455 | 0.976577396 |
| chr11:34104996-34106366 | 0.027391736 | 0.931490692 | 0.976584202 |
| chr6:82006218-82006418 | 0.045682392 | 0.931529798 | 0.976589818 |
| chr6:145189642-145190241 | 0.030107446 | 0.931549238 | 0.976589818 |
| chr7:140913193-140913435 | -0.04237511 | 0.931549303 | 0.976589818 |
| chr2:101038712-101038980 | -0.036134869 | 0.931549509 | 0.976589818 |
| chr13:20128514-20128910 | 0.031263408 | 0.931583757 | 0.976589818 |
| chr5:2732980-2733180 | -0.041250047 | 0.93159793 | 0.976589818 |
| chr2:224038541-224038990 | 0.034999053 | 0.931606152 | 0.976589818 |
| chr13:20408175-20408500 | 0.036467784 | 0.93161617 | 0.976589818 |
| chr10:79251637-79251964 | -0.035600973 | 0.931619535 | 0.976589818 |
| chr21:41501110-41501342 | 0.038864752 | 0.931628842 | 0.976589818 |
| chr17:21366969-21367412 | -0.035213967 | 0.93164052 | 0.976589818 |
| chr9:81737945-81738145 | -0.042707279 | 0.931648995 | 0.976589818 |
| chr15:90994352-90995101 | 0.027519966 | 0.931672281 | 0.97659676 |
| chr8:132567999-132568380 | 0.036135894 | 0.931681109 | 0.97659676 |

| | | | |
|---|---|---|---|
| chr12:8237504-8237755 | -0.03836766 | 0.931771313 | 0.976677952 |
| chr20:44572627-44573041 | 0.035720757 | 0.931812299 | 0.976680013 |
| chr14:25833822-25834043 | -0.03854679 | 0.931820232 | 0.976680013 |
| chr6:31651850-31653169 | 0.027359189 | 0.931828368 | 0.976680013 |
| chr20:47493068-47494400 | 0.026578988 | 0.931831039 | 0.976680013 |
| chr7:66995401-66995854 | 0.029085706 | 0.931837014 | 0.976680013 |
| chr2:11124543-11124777 | 0.034920254 | 0.931887731 | 0.97671981 |
| chr3:71875423-71875705 | -0.03477537 | 0.931935444 | 0.976756458 |
| chr16:69143218-69143492 | -0.037684217 | 0.931985547 | 0.976795609 |
| chr4:82284763-82285463 | -0.027095071 | 0.931999014 | 0.976796363 |
| chr15:62390846-62391135 | -0.034303699 | 0.932050791 | 0.97683305 |
| chr7:41491396-41491748 | 0.036318757 | 0.932067181 | 0.97683305 |
| chr6:142372416-142372685 | 0.038685052 | 0.932072266 | 0.97683305 |
| chr10:130371890-130372201 | 0.040094582 | 0.932091862 | 0.976840227 |
| chr1:54950325-54951127 | -0.02939509 | 0.932135352 | 0.976872443 |
| chr17:70767282-70767571 | 0.041357466 | 0.932160034 | 0.976884949 |
| chr17:63911706-63911980 | -0.032514609 | 0.932225277 | 0.97693996 |
| chr17:75515368-75516601 | -0.026697779 | 0.932275809 | 0.976957563 |
| chr17:62335380-62335634 | 0.038686141 | 0.932292493 | 0.976957563 |
| chr1:172639394-172639860 | -0.030510417 | 0.932295675 | 0.976957563 |
| chr2:64644334-64644933 | 0.029375629 | 0.932300664 | 0.976957563 |
| chrX:20141199-20142398 | -0.027195818 | 0.932340165 | 0.976957563 |
| chr3:196431744-196432947 | 0.027456943 | 0.932345049 | 0.976957563 |
| chr5:134225386-134225712 | 0.030774499 | 0.932366006 | 0.976957563 |
| chr20:32457065-32457308 | 0.037133998 | 0.932371849 | 0.976957563 |
| chr20:10553413-10553632 | -0.03912194 | 0.932373245 | 0.976957563 |
| chr10:3172415-3173622 | 0.026462827 | 0.932388588 | 0.976957563 |
| chr1:200896891-200897132 | -0.033405683 | 0.932394379 | 0.976957563 |
| chr9:95736418-95736631 | 0.041091364 | 0.932395079 | 0.976957563 |
| chr18:11387927-11388230 | 0.036560456 | 0.932428019 | 0.976973277 |
| chr1:157403202-157403406 | 0.04367053 | 0.932444539 | 0.976973277 |
| chr20:58248001-58248265 | 0.035429745 | 0.93245961 | 0.976973277 |
| chr15:92714048-92715133 | -0.027973738 | 0.932464735 | 0.976973277 |
| chr3:94935657-94936614 | -0.028058756 | 0.932473828 | 0.976973277 |
| chr14:30521925-30522208 | -0.041388899 | 0.932557762 | 0.977047856 |
| chr1:215312779-215312979 | -0.045912002 | 0.932591249 | 0.97706958 |
| chr12:6867326-6867727 | 0.029716675 | 0.932655946 | 0.977124003 |
| chr1:150234634-150234860 | -0.031670765 | 0.932762534 | 0.977221818 |
| chr11:124481469-124482208 | 0.030738582 | 0.932791429 | 0.977221818 |
| chr17:49365732-49365984 | 0.03293223 | 0.932797108 | 0.977221818 |
| chr19:48415384-48415731 | 0.033080018 | 0.932801519 | 0.977221818 |
| chr10:35195914-35196347 | -0.030504221 | 0.932851706 | 0.977221818 |
| chr1:239765206-239765492 | -0.039777052 | 0.932859295 | 0.977221818 |
| chr17:66930945-66931347 | 0.029077734 | 0.932863158 | 0.977221818 |
| chr8:144462834-144463075 | -0.03081513 | 0.932863304 | 0.977221818 |
| chr10:100020586-100020825 | -0.040949365 | 0.932864095 | 0.977221818 |
| chr2:99140844-99142035 | 0.026597229 | 0.93288484 | 0.977225679 |
| chr18:34631616-34631903 | -0.037821578 | 0.932893288 | 0.977225679 |
| chr1:202207347-202207633 | 0.031126164 | 0.933006808 | 0.977324913 |
| chr1:165152783-165153025 | -0.039282187 | 0.93301353 | 0.977324913 |

| | | | |
|---|---|---|---|
| chrX:97175214-97175427 | 0.045020775 | 0.933060121 | 0.977360355 |
| chr17:35242876-35243133 | 0.036226349 | 0.933158051 | 0.977449572 |
| chr20:63082405-63082613 | 0.038501691 | 0.933256472 | 0.977539301 |
| chr1:51329418-51330456 | 0.027357022 | 0.933288638 | 0.977556324 |
| chr3:146317427-146317763 | 0.039770802 | 0.933319864 | 0.977556324 |
| chr16:71625780-71626811 | -0.026906579 | 0.933322444 | 0.977556324 |
| chr3:111321883-111322214 | 0.039153946 | 0.933323757 | 0.977556324 |
| chr3:166090894-166091348 | -0.035625492 | 0.933340154 | 0.977556697 |
| chr6:30752206-30752971 | -0.027746512 | 0.933350728 | 0.977556697 |
| chr15:67457004-67457296 | 0.034791003 | 0.933369185 | 0.977556697 |
| chr5:13987913-13988137 | 0.035444352 | 0.933375146 | 0.977556697 |
| chr2:201642329-201643652 | -0.026992675 | 0.93342784 | 0.977574424 |
| chr1:223845296-223846594 | 0.026815131 | 0.933429831 | 0.977574424 |
| chr7:31782421-31782858 | 0.028992656 | 0.933430347 | 0.977574424 |
| chr12:9813694-9814015 | 0.041816988 | 0.933445693 | 0.977577135 |
| chrX:129522263-129523248 | 0.028273582 | 0.933469638 | 0.977586986 |
| chr8:93456054-93456425 | -0.030776174 | 0.933480617 | 0.977586986 |
| chr13:112617046-112617302 | 0.040541436 | 0.933498051 | 0.977591882 |
| chr20:13221525-13222282 | 0.029102548 | 0.933531569 | 0.977601246 |
| chr15:81753044-81753352 | 0.038976305 | 0.93353251 | 0.977601246 |
| chr8:87750783-87751109 | -0.039944727 | 0.93356202 | 0.977609197 |
| chr5:154445282-154446261 | -0.026756504 | 0.933576891 | 0.977609197 |
| chr7:32661936-32662271 | -0.036155881 | 0.933578596 | 0.977609197 |
| chr19:3573726-3574421 | -0.027546157 | 0.933591138 | 0.977609197 |
| chr2:160632531-160632935 | 0.035361445 | 0.933616145 | 0.977622022 |
| chr11:86830894-86831094 | 0.044561787 | 0.933642219 | 0.977623058 |
| chr10:129888999-129889673 | 0.030541258 | 0.93365381 | 0.977623058 |
| chr9:92764780-92765643 | 0.026854673 | 0.933655411 | 0.977623058 |
| chr19:2819626-2820402 | -0.029067634 | 0.933729349 | 0.977687118 |
| chr6:7339172-7339424 | -0.037563592 | 0.933742516 | 0.977687544 |
| chr1:67760361-67760872 | 0.031639721 | 0.933771208 | 0.977704226 |
| chr15:30903446-30904262 | 0.027795402 | 0.93381961 | 0.977725635 |
| chr10:25081003-25081325 | -0.033762234 | 0.933820644 | 0.977725635 |
| chr22:50532105-50532452 | 0.037078114 | 0.933829937 | 0.977725635 |
| chr17:7435153-7435634 | -0.028436421 | 0.933851953 | 0.977730967 |
| chr17:68290395-68290763 | 0.028479028 | 0.933863151 | 0.977730967 |
| chr3:79766568-79766786 | -0.040258007 | 0.933897319 | 0.977730967 |
| chr5:140157138-140157783 | -0.028185045 | 0.933902541 | 0.977730967 |
| chr17:63071328-63071700 | 0.034436636 | 0.933907211 | 0.977730967 |
| chr7:8433521-8434429 | -0.027062236 | 0.933919868 | 0.977730967 |
| chr19:49869353-49870059 | -0.027004115 | 0.933937983 | 0.977730967 |
| chr13:38045741-38046069 | 0.03254849 | 0.933951135 | 0.977730967 |
| chr1:25338212-25338761 | 0.029534962 | 0.933961537 | 0.977730967 |
| chr13:110138188-110138973 | -0.028680624 | 0.933962634 | 0.977730967 |
| chr4:128287648-128288683 | 0.025996607 | 0.933990077 | 0.977739264 |
| chr20:15061950-15062295 | -0.033593945 | 0.934017741 | 0.977739264 |
| chr19:15640571-15640910 | 0.033534904 | 0.934029892 | 0.977739264 |
| chr3:111668946-111669221 | 0.034971852 | 0.934031556 | 0.977739264 |
| chr1:228486564-228487562 | 0.027073839 | 0.934042219 | 0.977739264 |
| chr20:32325835-32326247 | 0.031311725 | 0.934047123 | 0.977739264 |

| | | | |
|---|---|---|---|
| chr3:151381280-151381534 | 0.040615468 | 0.934083637 | 0.977764129 |
| chr16:50627663-50627894 | -0.041376801 | 0.934138098 | 0.977795765 |
| chr16:2531842-2532876 | -0.026737565 | 0.934139383 | 0.977795765 |
| chr8:61805625-61805904 | -0.035054549 | 0.934205431 | 0.977827823 |
| chr10:103666217-103666430 | 0.038943441 | 0.934217942 | 0.977827823 |
| chr10:93340866-93341091 | -0.043321148 | 0.934218241 | 0.977827823 |
| chr4:127581009-127581213 | 0.03970475 | 0.934229176 | 0.977827823 |
| chr19:38335584-38336110 | -0.027688759 | 0.934233818 | 0.977827823 |
| chr8:122703895-122704147 | 0.036664808 | 0.934281876 | 0.977864766 |
| chr2:138602180-138602432 | 0.037572428 | 0.934299487 | 0.97786984 |
| chr9:136547053-136547322 | -0.032765092 | 0.934381694 | 0.977942523 |
| chr1:91885790-91886428 | 0.028558254 | 0.934465288 | 0.977982517 |
| chr15:40679070-40679540 | -0.029940137 | 0.934490359 | 0.977982517 |
| chr1:240189485-240189689 | -0.042727727 | 0.934492298 | 0.977982517 |
| chr2:98385170-98385407 | 0.041267483 | 0.934522841 | 0.977982517 |
| chr22:30326824-30327307 | -0.028217251 | 0.934526509 | 0.977982517 |
| chrX:124929352-124929659 | 0.037790314 | 0.934537031 | 0.977982517 |
| chr3:23750696-23750896 | -0.041822907 | 0.934538845 | 0.977982517 |
| chr3:38628036-38628238 | -0.041822907 | 0.934538845 | 0.977982517 |
| chr3:179281334-179281575 | -0.041822907 | 0.934538845 | 0.977982517 |
| chr3:12479958-12480206 | 0.032534857 | 0.934551622 | 0.977982517 |
| chr13:46728859-46729162 | -0.03026773 | 0.934566278 | 0.977982517 |
| chr11:57567660-57568669 | 0.027744234 | 0.934573072 | 0.977982517 |
| chr17:28747383-28747583 | 0.032397963 | 0.934594619 | 0.977991709 |
| chr10:25367802-25368140 | 0.038955686 | 0.934703179 | 0.978091952 |
| chr14:98937947-98938147 | 0.043214333 | 0.934739163 | 0.978115868 |
| chr19:56140440-56141411 | -0.025994853 | 0.934751566 | 0.978115868 |
| chr7:12137099-12137790 | 0.029197381 | 0.934839959 | 0.978192255 |
| chr12:57815239-57815521 | -0.034521141 | 0.934850099 | 0.978192255 |
| chr12:132955932-132956729 | 0.028113122 | 0.934885843 | 0.978216298 |
| chr20:51542640-51542916 | 0.032035931 | 0.934910232 | 0.978228458 |
| chr13:25980357-25980588 | 0.038860304 | 0.934965085 | 0.978272494 |
| chr6:97282342-97282542 | -0.036844235 | 0.935025005 | 0.978316969 |
| chr1:109619643-109619952 | 0.030235275 | 0.935041555 | 0.978316969 |
| chr1:165914234-165914525 | -0.032690059 | 0.935045895 | 0.978316969 |
| chr19:47514882-47515883 | 0.025853666 | 0.935114976 | 0.978361557 |
| chr13:42414576-42414814 | 0.040057707 | 0.935115367 | 0.978361557 |
| chr2:9554940-9556522 | 0.0256781 | 0.935126817 | 0.978361557 |
| chr7:151039055-151039560 | 0.030210787 | 0.935149133 | 0.978371545 |
| chr17:60569621-60570119 | -0.026320826 | 0.935173566 | 0.978383749 |
| chr3:65669722-65669922 | 0.052113771 | 0.935219981 | 0.978418948 |
| chr5:155587560-155587760 | -0.044618222 | 0.935245165 | 0.978429769 |
| chr12:56266647-56267548 | 0.026900165 | 0.935280911 | 0.978429769 |
| chr15:101578279-101579145 | 0.027202794 | 0.935285364 | 0.978429769 |
| chr7:4610844-4611149 | -0.030705085 | 0.935290678 | 0.978429769 |
| chr11:74449398-74449664 | 0.033590731 | 0.935300992 | 0.978429769 |
| chr3:150504008-150504245 | 0.037625504 | 0.935306941 | 0.978429769 |
| chr11:79439799-79441222 | 0.026013879 | 0.935325891 | 0.978436235 |
| chr11:67428791-67428999 | -0.033499556 | 0.935355132 | 0.978450595 |
| chr12:51955035-51955235 | -0.040051132 | 0.935365159 | 0.978450595 |

| | | | |
|---|---|---|---|
| chr19:6516841-6517398 | 0.029284416 | 0.935422853 | 0.978490188 |
| chr1:190478101-190479010 | 0.026520923 | 0.935428549 | 0.978490188 |
| chr3:121152701-121152901 | 0.039759097 | 0.93545409 | 0.978503546 |
| chr8:98940960-98941536 | 0.027023779 | 0.935477534 | 0.978514711 |
| chr7:47734563-47734763 | -0.04292111 | 0.93550114 | 0.978523702 |
| chr4:1010482-1011647 | 0.026673023 | 0.935511671 | 0.978523702 |
| chr2:120312914-120313115 | -0.039182989 | 0.93553022 | 0.978529746 |
| chr13:36101244-36101495 | -0.037582211 | 0.935559722 | 0.978547246 |
| chr6:69894574-69894781 | 0.041105131 | 0.935587248 | 0.978553174 |
| chr8:116919772-116920025 | 0.03771295 | 0.935603062 | 0.978553174 |
| chr15:66683749-66683949 | -0.039540548 | 0.935612067 | 0.978553174 |
| chr5:173230591-173230895 | 0.035284035 | 0.935624748 | 0.978553174 |
| chr14:81219213-81219657 | -0.026272777 | 0.935631536 | 0.978553174 |
| chr5:177516618-177517582 | -0.025824287 | 0.935653025 | 0.978553174 |
| chr22:19193335-19193657 | 0.037723325 | 0.935654788 | 0.978553174 |
| chr3:183844629-183844901 | -0.035004058 | 0.935692821 | 0.978579593 |
| chr3:63343733-63344058 | 0.029037139 | 0.935719989 | 0.978592202 |
| chr7:72830062-72830312 | -0.03207306 | 0.93574848 | 0.978592202 |
| chr8:11970511-11970844 | -0.033004265 | 0.935762789 | 0.978592202 |
| chr9:23492175-23492394 | 0.037937694 | 0.935764852 | 0.978592202 |
| chr1:44905019-44905307 | 0.041243086 | 0.935768736 | 0.978592202 |
| chr19:3762542-3763188 | -0.026991866 | 0.935786527 | 0.978597452 |
| chr1:217921521-217921983 | 0.02900021 | 0.935803281 | 0.978600669 |
| chr20:62875549-62875940 | 0.02995205 | 0.935815147 | 0.978600669 |
| chr8:109975274-109976043 | 0.025829546 | 0.935838313 | 0.978603626 |
| chr9:34437446-34437650 | 0.038302425 | 0.935843519 | 0.978603626 |
| chrX:47572890-47573106 | -0.041633093 | 0.935862226 | 0.978609832 |
| chr2:31428114-31428411 | -0.036440512 | 0.935987928 | 0.978727919 |
| chr17:63361942-63362238 | 0.032242223 | 0.936004865 | 0.978732273 |
| chr8:122733547-122734103 | -0.027157256 | 0.936025155 | 0.978735494 |
| chr5:142012136-142013422 | -0.026353491 | 0.936046014 | 0.978735494 |
| chr5:90909938-90910323 | -0.03671749 | 0.936049618 | 0.978735494 |
| chr3:142888163-142889275 | -0.026368944 | 0.93605904 | 0.978735494 |
| chr3:3666350-3666566 | 0.040341251 | 0.936133337 | 0.978798948 |
| chr2:55049482-55049910 | 0.027533036 | 0.936153614 | 0.978798948 |
| chr9:100473332-100474101 | -0.02801186 | 0.93615805 | 0.978798948 |
| chr9:95168083-95168283 | 0.041091701 | 0.936199159 | 0.978828573 |
| chr22:30694830-30695167 | -0.03693291 | 0.936229335 | 0.978846767 |
| chr1:107835468-107835752 | 0.036229417 | 0.936244325 | 0.978847003 |
| chr6:9532526-9532829 | -0.039116125 | 0.936255112 | 0.978847003 |
| chr1:105199336-105199559 | -0.039089987 | 0.936355745 | 0.978908403 |
| chr17:68879426-68879690 | -0.037667934 | 0.936366672 | 0.978908403 |
| chrX:8988545-8988745 | -0.043082585 | 0.936367261 | 0.978908403 |
| chr5:177958314-177958566 | 0.036291769 | 0.936384351 | 0.978908403 |
| chr1:633858-634201 | -0.025362793 | 0.936389649 | 0.978908403 |
| chr12:64697916-64698346 | -0.029050308 | 0.936405192 | 0.978908403 |
| chr1:40373974-40374951 | 0.026666855 | 0.936408316 | 0.978908403 |
| chr5:65488091-65488351 | -0.036125404 | 0.936416047 | 0.978908403 |
| chrX:129037204-129037765 | 0.029740842 | 0.936438245 | 0.978918254 |
| chr3:169575714-169575964 | 0.03811776 | 0.936456079 | 0.978919369 |

| | | | |
|---|---|---|---|
| chr14:64193921-64194189 | -0.035665752 | 0.936487384 | 0.978919369 |
| chr2:99154585-99155469 | -0.026286375 | 0.936487547 | 0.978919369 |
| chr10:119080009-119081463 | 0.025260989 | 0.936493048 | 0.978919369 |
| chr6:126023437-126023875 | -0.033253691 | 0.936517998 | 0.978919369 |
| chr5:137156334-137156551 | 0.040809628 | 0.936527937 | 0.978919369 |
| chr5:125043783-125044486 | -0.027018178 | 0.936528744 | 0.978919369 |
| chr3:44975178-44976409 | -0.026970772 | 0.936616572 | 0.978952661 |
| chr15:62742467-62742925 | 0.031402787 | 0.936640958 | 0.978952661 |
| chrX:16123268-16123595 | 0.031842814 | 0.93664894 | 0.978952661 |
| chr14:100059407-100059985 | -0.026962838 | 0.93665602 | 0.978952661 |
| chr12:57498226-57498525 | 0.034605401 | 0.936659089 | 0.978952661 |
| chr14:88045002-88045327 | 0.037147903 | 0.936659515 | 0.978952661 |
| chr19:56309602-56309859 | -0.032437506 | 0.936660237 | 0.978952661 |
| chr2:12317837-12318037 | 0.042072283 | 0.936662806 | 0.978952661 |
| chr5:149760644-149760945 | -0.030970417 | 0.936678903 | 0.978956132 |
| chr11:1040267-1040467 | 0.047035041 | 0.936713878 | 0.978979332 |
| chr3:26280608-26280840 | 0.037454038 | 0.936754572 | 0.978998828 |
| chr16:71461585-71462470 | 0.026329332 | 0.936758086 | 0.978998828 |
| chr1:223860181-223860656 | 0.030331483 | 0.936784433 | 0.979013009 |
| chr17:81871919-81872128 | 0.031432267 | 0.936813691 | 0.979029447 |
| chr1:188061893-188062279 | -0.033301502 | 0.93682749 | 0.979029447 |
| chr19:49017784-49018190 | 0.029500422 | 0.936859864 | 0.979029447 |
| chrX:68822497-68822972 | -0.030950055 | 0.936885342 | 0.979029447 |
| chr19:35994931-35995295 | -0.029311562 | 0.9369091 | 0.979029447 |
| chr6:152338623-152338846 | -0.03663404 | 0.936912814 | 0.979029447 |
| chr2:218731607-218731876 | -0.038941591 | 0.936913952 | 0.979029447 |
| chr2:86298291-86298543 | -0.038342652 | 0.936915846 | 0.979029447 |
| chr13:36130439-36131699 | 0.025569672 | 0.936918728 | 0.979029447 |
| chr17:75940380-75940580 | 0.036104657 | 0.936927935 | 0.979029447 |
| chr1:241847683-241848875 | 0.025103731 | 0.93696051 | 0.979035755 |
| chr14:102524332-102525351 | 0.025484003 | 0.936963491 | 0.979035755 |
| chr3:181712859-181713063 | 0.035040531 | 0.936972304 | 0.979035755 |
| chr5:31825505-31825834 | -0.030351255 | 0.937022431 | 0.97907478 |
| chr12:69415681-69416188 | 0.029062614 | 0.937040434 | 0.97908024 |
| chr1:194818802-194819158 | -0.036541261 | 0.937074844 | 0.979100778 |
| chr17:73255687-73256011 | 0.030242812 | 0.937085647 | 0.979100778 |
| chr7:3215861-3216489 | -0.027119928 | 0.93710562 | 0.979108296 |
| chr11:17504599-17504876 | 0.031372631 | 0.937139649 | 0.979130498 |
| chr2:137838508-137838770 | -0.033736136 | 0.937174416 | 0.979153471 |
| chr17:47830952-47832001 | -0.025255444 | 0.937198685 | 0.979161152 |
| chr9:99079439-99079780 | 0.036302996 | 0.937210483 | 0.979161152 |
| chr19:17873772-17874446 | 0.02784347 | 0.937220105 | 0.979161152 |
| chr8:122902793-122903089 | -0.03260213 | 0.937242972 | 0.979170908 |
| chr14:24006932-24007648 | 0.028481064 | 0.937255002 | 0.979170908 |
| chr14:100417422-100417770 | -0.029621924 | 0.937283282 | 0.979173677 |
| chr7:149005855-149006069 | 0.034380601 | 0.937287774 | 0.979173677 |
| chr12:77010969-77011267 | 0.031077283 | 0.93729599 | 0.979173677 |
| chr17:43846273-43846520 | 0.034615239 | 0.937326429 | 0.979192126 |
| chr1:167564647-167564937 | 0.035029671 | 0.937362968 | 0.979193391 |
| chr10:436104-437577 | 0.024460971 | 0.937363922 | 0.979193391 |

| | | | |
|---|---|---|---|
| chr5:38816143-38816877 | -0.02838528 | 0.937367787 | 0.979193391 |
| chr9:33458387-33458714 | -0.038095739 | 0.937387265 | 0.979193391 |
| chr1:200680800-200681017 | 0.040192263 | 0.937398726 | 0.979193391 |
| chr11:67468884-67469528 | -0.026277375 | 0.937405094 | 0.979193391 |
| chr2:14297448-14297690 | 0.03395267 | 0.937423146 | 0.979193391 |
| chr13:41346393-41346616 | -0.037699266 | 0.937429876 | 0.979193391 |
| chr17:42407549-42407763 | 0.036655509 | 0.937501806 | 0.979241826 |
| chr19:2495359-2495602 | 0.036655509 | 0.937501806 | 0.979241826 |
| chr22:50342423-50342630 | 0.036338125 | 0.937526949 | 0.979242002 |
| chr8:133391018-133391219 | -0.038516695 | 0.937527535 | 0.979242002 |
| chr2:34570475-34570919 | -0.028457472 | 0.937573792 | 0.979267589 |
| chr6:55108841-55109175 | -0.032534456 | 0.937577869 | 0.979267589 |
| chr3:130491440-130491843 | 0.03273818 | 0.937590373 | 0.979267589 |
| chr17:7685968-7686478 | -0.028549847 | 0.937621701 | 0.979274545 |
| chr6:36197058-36197686 | 0.026403131 | 0.937627687 | 0.979274545 |
| chr8:73340855-73341084 | -0.038641337 | 0.937635375 | 0.979274545 |
| chr13:44336945-44337150 | -0.038434578 | 0.937667781 | 0.979295042 |
| chr2:238288695-238289341 | -0.025964719 | 0.937684648 | 0.979298146 |
| chr8:102317061-102317312 | 0.036889366 | 0.937696315 | 0.979298146 |
| chr12:125974321-125975116 | 0.027090622 | 0.937709803 | 0.979298884 |
| chr10:13099157-13099552 | -0.029489791 | 0.937739916 | 0.979310328 |
| chr1:231248729-231249116 | -0.03031847 | 0.937771449 | 0.979310328 |
| chr20:62623406-62624652 | -0.024784457 | 0.937781014 | 0.979310328 |
| chr3:8450724-8450977 | -0.034869797 | 0.937787718 | 0.979310328 |
| chr15:78872496-78873907 | 0.025006094 | 0.937809871 | 0.979310328 |
| chr2:216698859-216699177 | -0.0307069 | 0.937809944 | 0.979310328 |
| chr10:51962633-51962845 | 0.043722188 | 0.937810229 | 0.979310328 |
| chr21:41168399-41168606 | 0.034237469 | 0.937830506 | 0.979314136 |
| chr22:43089239-43089647 | -0.031058601 | 0.937852904 | 0.979314136 |
| chr12:112125250-112126142 | 0.025918837 | 0.937856453 | 0.979314136 |
| chr1:78114217-78114766 | 0.027413838 | 0.937885512 | 0.979314136 |
| chr14:74349357-74349557 | -0.037878877 | 0.93790006 | 0.979314136 |
| chr14:68459514-68459855 | 0.033542983 | 0.937905007 | 0.979314136 |
| chr6:113642834-113643356 | 0.027240113 | 0.937910541 | 0.979314136 |
| chr12:53286717-53286921 | 0.038036853 | 0.937916124 | 0.979314136 |
| chr11:116808141-116808702 | -0.029339207 | 0.937957729 | 0.979344232 |
| chr2:10914006-10914206 | -0.034443889 | 0.937979886 | 0.979344614 |
| chr9:87950879-87951118 | 0.036024197 | 0.937983658 | 0.979344614 |
| chr8:123060792-123061134 | -0.027996011 | 0.93801564 | 0.979364661 |
| chr7:50560908-50561482 | -0.027658318 | 0.938037172 | 0.979373797 |
| chr20:62628642-62628933 | 0.029935426 | 0.938051137 | 0.979375032 |
| chr1:170929811-170930236 | 0.029071025 | 0.938069895 | 0.979381272 |
| chr11:64185242-64186378 | -0.024676997 | 0.938089335 | 0.979388222 |
| chr15:65530089-65531183 | -0.025447675 | 0.938129957 | 0.979417287 |
| chrX:104565800-104566294 | -0.027800658 | 0.938143718 | 0.979418309 |
| chr11:78079173-78080225 | -0.025662717 | 0.938169393 | 0.979420447 |
| chr10:123351380-123351601 | 0.041622857 | 0.938173556 | 0.979420447 |
| chr9:114027503-114027796 | 0.034141411 | 0.938184113 | 0.979420447 |
| chr1:149740073-149740801 | 0.025598889 | 0.938228284 | 0.979444475 |
| chr3:13507340-13508165 | -0.026438678 | 0.938232695 | 0.979444475 |

| | | | |
|---|---|---|---|
| chr11:118057870-118058117 | 0.032828343 | 0.938293205 | 0.979478781 |
| chr11:59668236-59669876 | 0.024611885 | 0.938311879 | 0.979478781 |
| chr16:46620720-46621581 | -0.027186615 | 0.938315751 | 0.979478781 |
| chr1:78092717-78093030 | 0.032585877 | 0.938316691 | 0.979478781 |
| chr8:130357939-130358873 | 0.025118388 | 0.938382427 | 0.979521907 |
| chr4:681533-682042 | -0.027706227 | 0.938385521 | 0.979521907 |
| chr12:5870219-5870572 | -0.036636516 | 0.938406419 | 0.979521907 |
| chr17:18038979-18039991 | -0.024911712 | 0.93840914 | 0.979521907 |
| chr14:20998739-20998970 | -0.034808688 | 0.938440838 | 0.979526485 |
| chr3:169822378-169822978 | -0.028298446 | 0.93844395 | 0.979526485 |
| chr8:80998333-80998579 | 0.034685537 | 0.938451877 | 0.979526485 |
| chr1:183185787-183186558 | -0.025870743 | 0.938506302 | 0.979536542 |
| chr2:128304820-128305598 | -0.025297023 | 0.938527642 | 0.979536542 |
| chr8:51898335-51899773 | -0.024343269 | 0.938537585 | 0.979536542 |
| chr2:60674593-60675119 | -0.027281873 | 0.938540667 | 0.979536542 |
| chr2:203014416-203015565 | -0.02439909 | 0.938547777 | 0.979536542 |
| chr17:51121706-51122155 | 0.025259796 | 0.938556113 | 0.979536542 |
| chr17:8120104-8120525 | -0.02632813 | 0.938556233 | 0.979536542 |
| chr11:75428249-75428612 | -0.027603488 | 0.938563784 | 0.979536542 |
| chr5:73536849-73537049 | 0.036760897 | 0.938597343 | 0.979558223 |
| chr6:37624210-37624519 | -0.033296094 | 0.938653559 | 0.97960355 |
| chr6:108559278-108559893 | -0.025278716 | 0.93871658 | 0.979647538 |
| chr4:48340823-48341731 | 0.02442135 | 0.93872426 | 0.979647538 |
| chr17:9987929-9988129 | -0.038672361 | 0.938734065 | 0.979647538 |
| chr14:75417946-75418148 | 0.043627871 | 0.938814655 | 0.979718297 |
| chr10:94439313-94439599 | -0.037550027 | 0.938870007 | 0.979761884 |
| chr4:48930836-48931036 | 0.041299096 | 0.938893314 | 0.979761884 |
| chr22:45101975-45102175 | -0.039711904 | 0.938896607 | 0.979761884 |
| chr19:4015676-4015937 | -0.035429995 | 0.93890757 | 0.979761884 |
| chr1:46246967-46248563 | -0.024367961 | 0.93892128 | 0.979762847 |
| chr2:210249178-210249609 | 0.031932153 | 0.938960473 | 0.979786877 |
| chr6:125789913-125791932 | 0.024263456 | 0.938969883 | 0.979786877 |
| chr4:2468134-2468680 | 0.029149553 | 0.939014128 | 0.979814058 |
| chr7:147245344-147245726 | -0.031924859 | 0.939021507 | 0.979814058 |
| chr3:124559679-124560118 | 0.029049551 | 0.939039078 | 0.97981905 |
| chr6:110457639-110457987 | -0.031622628 | 0.939096941 | 0.979849373 |
| chr3:99843327-99843832 | 0.027793413 | 0.939114484 | 0.979849373 |
| chr12:32657611-32657940 | 0.031633974 | 0.939132669 | 0.979849373 |
| chr16:12215380-12215580 | -0.040850956 | 0.939133961 | 0.979849373 |
| chr5:169548400-169548600 | -0.043158398 | 0.939153701 | 0.979849373 |
| chr16:14097178-14097559 | 0.028506392 | 0.939158988 | 0.979849373 |
| chr9:86637575-86637781 | -0.039191043 | 0.93917339 | 0.979849373 |
| chr1:223670528-223670788 | 0.032474073 | 0.939174824 | 0.979849373 |
| chr5:176578382-176578683 | 0.028190828 | 0.939183232 | 0.979849373 |
| chr12:59736808-59737239 | 0.027203564 | 0.939221807 | 0.979876275 |
| chr17:80035486-80036339 | -0.025526551 | 0.939281844 | 0.979903243 |
| chr11:78187857-78189152 | -0.024090398 | 0.93930156 | 0.979903243 |
| chr1:201361652-201361980 | 0.02677012 | 0.939303122 | 0.979903243 |
| chr12:123445745-123446289 | -0.028069179 | 0.939309447 | 0.979903243 |
| chr1:157171957-157172157 | 0.038844963 | 0.9393116 | 0.979903243 |

| | | | |
|---|---|---|---|
| chr10:92771767-92772209 | 0.033982239 | 0.939353463 | 0.979933574 |
| chr2:96913516-96913783 | 0.033433023 | 0.939383531 | 0.979951599 |
| chr11:73194592-73195126 | 0.025404965 | 0.939397695 | 0.979953032 |
| chr17:81164884-81165533 | 0.026523918 | 0.939438318 | 0.979982067 |
| chr1:218475384-218475584 | 0.034206254 | 0.939473448 | 0.980005198 |
| chr17:58199965-58200379 | -0.025961058 | 0.939486072 | 0.980005198 |
| chr13:25301194-25302183 | -0.024626398 | 0.93952202 | 0.980029354 |
| chr5:179698116-179699430 | -0.023919842 | 0.939551519 | 0.980034285 |
| chrX:22192496-22192976 | -0.029666497 | 0.939552328 | 0.980034285 |
| chr8:30657571-30658689 | 0.024836771 | 0.939611832 | 0.98008301 |
| chr1:235598169-235598407 | 0.032488342 | 0.939632418 | 0.980091141 |
| chr2:95300457-95300986 | 0.027566573 | 0.939647244 | 0.980093264 |
| chr6:35727513-35727858 | 0.027973821 | 0.939726961 | 0.980163069 |
| chr12:69239093-69240346 | 0.024104135 | 0.939762904 | 0.980187216 |
| chr17:46044341-46044548 | 0.039047341 | 0.939800542 | 0.980212739 |
| chr9:97950861-97951211 | -0.029085373 | 0.939823359 | 0.980212739 |
| chr20:35092539-35093329 | 0.026166773 | 0.939837619 | 0.980212739 |
| chr17:59930777-59931185 | 0.025480301 | 0.939848331 | 0.980212739 |
| chr6:141496603-141496835 | -0.038950075 | 0.939851339 | 0.980212739 |
| chr22:11917827-11918095 | -0.035589311 | 0.939873248 | 0.980222247 |
| chr3:122392993-122393193 | 0.040794873 | 0.939897461 | 0.980234156 |
| chr16:1255271-1255471 | 0.041291013 | 0.939928646 | 0.980247171 |
| chr15:42155967-42156209 | -0.032968867 | 0.939965027 | 0.980247171 |
| chr5:172005744-172007408 | 0.023696045 | 0.939969876 | 0.980247171 |
| chr1:36553869-36554188 | -0.030534225 | 0.939992623 | 0.980247171 |
| chr3:184543822-184544362 | -0.026994897 | 0.940011576 | 0.980247171 |
| chr3:436740-436954 | 0.039642878 | 0.940012278 | 0.980247171 |
| chr1:214708383-214708838 | 0.028471385 | 0.940017578 | 0.980247171 |
| chr11:75103675-75103894 | 0.038019642 | 0.940041239 | 0.980247171 |
| chr14:23694165-23695396 | -0.02411977 | 0.940047056 | 0.980247171 |
| chr5:154386320-154386563 | -0.037249457 | 0.940048945 | 0.980247171 |
| chr12:59164921-59165398 | 0.027735003 | 0.940050666 | 0.980247171 |
| chr16:81175330-81175724 | 0.028891837 | 0.940076783 | 0.980252896 |
| chr5:66864642-66865079 | 0.030339316 | 0.940098654 | 0.980252896 |
| chr5:178222504-178222936 | -0.028462825 | 0.940104171 | 0.980252896 |
| chr1:184905424-184906258 | -0.026269222 | 0.94010733 | 0.980252896 |
| chr17:67825202-67826453 | -0.023749558 | 0.94012939 | 0.980262558 |
| chr14:95553067-95553459 | -0.032527261 | 0.940145826 | 0.980266357 |
| chr8:124972354-124973857 | -0.023516941 | 0.940193771 | 0.980303007 |
| chr2:218860965-218861302 | 0.029951781 | 0.940222329 | 0.980319444 |
| chr16:3596421-3596655 | -0.037667143 | 0.940239353 | 0.980323854 |
| chr6:157273719-157273993 | 0.028693827 | 0.940259702 | 0.980325229 |
| chr16:24729245-24729854 | 0.025933933 | 0.940277641 | 0.980325229 |
| chr12:6688823-6690069 | 0.023808591 | 0.940284043 | 0.980325229 |
| chr11:93740846-93742223 | -0.023894941 | 0.940291849 | 0.980325229 |
| chr16:71883233-71884755 | -0.023994016 | 0.940334497 | 0.980356353 |
| chr5:176388070-176388356 | 0.0284115 | 0.940367922 | 0.980377862 |
| chr1:197335833-197336080 | -0.032459706 | 0.940407543 | 0.980395123 |
| chr6:111483044-111484297 | -0.023654076 | 0.940410069 | 0.980395123 |
| chr2:121748257-121748510 | -0.035359379 | 0.940425571 | 0.980397944 |

| | | | |
|---|---|---|---|
| chr17:83096772-83097381 | 0.024654632 | 0.940470255 | 0.980429263 |
| chr19:40714957-40715249 | 0.026944478 | 0.940481204 | 0.980429263 |
| chr11:95089380-95090601 | 0.024327601 | 0.940533258 | 0.980438052 |
| chr12:96094156-96094582 | -0.027176928 | 0.940535348 | 0.980438052 |
| chr19:18322811-18323425 | -0.026086145 | 0.940541318 | 0.980438052 |
| chr3:632600-632906 | -0.030278794 | 0.9405545 | 0.980438052 |
| chr2:214543711-214543922 | 0.037027266 | 0.940561496 | 0.980438052 |
| chr10:74058156-74058448 | -0.033604897 | 0.940577678 | 0.980438052 |
| chr2:130129275-130129477 | -0.037395058 | 0.940579205 | 0.980438052 |
| chr3:100334326-100335347 | -0.024017687 | 0.940614873 | 0.980449399 |
| chr17:60356926-60357312 | -0.024869036 | 0.94063858 | 0.980449399 |
| chrX:138128085-138128602 | 0.028294696 | 0.940648372 | 0.980449399 |
| chr8:68257902-68258172 | -0.032128601 | 0.940663206 | 0.980449399 |
| chr2:69989447-69989691 | -0.034552001 | 0.940665486 | 0.980449399 |
| chr3:127738274-127738482 | 0.038780973 | 0.94067894 | 0.980449399 |
| chr9:129495735-129496582 | -0.02427868 | 0.940694353 | 0.980449399 |
| chr14:103901090-103901351 | -0.031478121 | 0.940697535 | 0.980449399 |
| chr16:71195-72111 | 0.024153332 | 0.940715017 | 0.980449399 |
| chr10:84783749-84784197 | 0.030267679 | 0.940729528 | 0.980449399 |
| chr1:62917732-62918006 | -0.038966709 | 0.940733745 | 0.980449399 |
| chr2:118088010-118088963 | 0.023808965 | 0.940743642 | 0.980449399 |
| chr17:66975607-66976240 | -0.025232627 | 0.940807956 | 0.98050309 |
| chr8:102700529-102700909 | -0.029570343 | 0.940853575 | 0.980537297 |
| chr19:38636801-38637241 | 0.027528607 | 0.940882534 | 0.980551616 |
| chr3:150798087-150798630 | 0.027070138 | 0.940904697 | 0.980551616 |
| chr1:186325351-186325557 | -0.035559535 | 0.940905706 | 0.980551616 |
| chr5:150560366-150560666 | -0.029632856 | 0.941004732 | 0.980621031 |
| chr2:66587311-66587512 | 0.034677842 | 0.941007215 | 0.980621031 |
| chr5:671443-671917 | -0.025825859 | 0.941018762 | 0.980621031 |
| chr5:127988152-127988395 | 0.037636396 | 0.941026064 | 0.980621031 |
| chr20:47501193-47502585 | 0.022925819 | 0.941036305 | 0.980621031 |
| chr1:241639695-241640844 | 0.023532011 | 0.941077595 | 0.980650721 |
| chr5:32342997-32343254 | 0.039074367 | 0.941098571 | 0.980659242 |
| chr2:207375174-207375864 | 0.026813807 | 0.941182446 | 0.98071596 |
| chr5:43469682-43470294 | -0.02653585 | 0.941188089 | 0.98071596 |
| chr1:77092587-77092823 | 0.042154011 | 0.941199015 | 0.98071596 |
| chr17:63362564-63362764 | 0.033774113 | 0.941233173 | 0.98071596 |
| chr17:41799254-41799507 | 0.03074415 | 0.941235474 | 0.98071596 |
| chr12:83929812-83930266 | 0.031601278 | 0.941250067 | 0.98071596 |
| chr16:9092139-9092339 | -0.030337439 | 0.941256035 | 0.98071596 |
| chr5:132813176-132813404 | 0.034164594 | 0.941265748 | 0.98071596 |
| chr11:71128204-71128462 | 0.038191325 | 0.941268196 | 0.98071596 |
| chr12:70358291-70358559 | -0.037499884 | 0.941324183 | 0.980760957 |
| chr3:136989892-136990123 | 0.031999445 | 0.941348886 | 0.980773359 |
| chr6:127737028-127737475 | 0.028332013 | 0.94137616 | 0.980775156 |
| chr3:155001155-155001634 | -0.027623853 | 0.941376212 | 0.980775156 |
| chr9:83396393-83396593 | 0.037495534 | 0.941410399 | 0.980791949 |
| chr12:95826396-95826625 | -0.035643774 | 0.941438425 | 0.980791949 |
| chr16:21584059-21584261 | -0.035643774 | 0.941438425 | 0.980791949 |
| chr1:31788066-31789551 | -0.023343051 | 0.941443531 | 0.980791949 |

| | | | |
|---|---|---|---|
| chr20:833386-834084 | -0.024870327 | 0.941482263 | 0.980797292 |
| chr2:73078016-73078240 | 0.033025919 | 0.941485234 | 0.980797292 |
| chr1:93814001-93814329 | 0.030343213 | 0.94152914 | 0.980797292 |
| chr11:111926554-111927571 | -0.023670855 | 0.941534201 | 0.980797292 |
| chr16:81053797-81054004 | -0.035969271 | 0.941536014 | 0.980797292 |
| chr12:92879838-92880261 | 0.030175417 | 0.941562687 | 0.980797292 |
| chr19:50173157-50173457 | -0.033896502 | 0.941565968 | 0.980797292 |
| chr10:98244314-98244514 | 0.042942858 | 0.94157434 | 0.980797292 |
| chr10:104349771-104350007 | -0.03071381 | 0.941582129 | 0.980797292 |
| chr7:48443170-48443485 | -0.034716752 | 0.941582956 | 0.980797292 |
| chr11:120284310-120284640 | 0.029530793 | 0.941589466 | 0.980797292 |
| chr6:12492084-12492715 | 0.025920919 | 0.941629044 | 0.980803443 |
| chr2:98736511-98736776 | 0.035134277 | 0.941629159 | 0.980803443 |
| chr2:120521111-120521560 | -0.027968029 | 0.941650112 | 0.980803443 |
| chr1:20507809-20508336 | -0.027477125 | 0.94166321 | 0.980803443 |
| chr6:119240984-119241376 | 0.029602335 | 0.94167122 | 0.980803443 |
| chr15:65086388-65086729 | -0.026476883 | 0.941672174 | 0.980803443 |
| chr5:74199550-74199925 | 0.03029345 | 0.941692805 | 0.980811599 |
| chr17:82450418-82451057 | 0.024537989 | 0.941752359 | 0.980842144 |
| chr2:10865003-10865234 | -0.034064226 | 0.941758657 | 0.980842144 |
| chr14:28609308-28609526 | 0.035214625 | 0.941760535 | 0.980842144 |
| chr15:75921753-75921998 | 0.033383479 | 0.941807423 | 0.980866443 |
| chr9:113710028-113710234 | -0.037917487 | 0.941809468 | 0.980866443 |
| chr11:64670365-64670673 | -0.033826053 | 0.941905467 | 0.980953089 |
| chr2:82993572-82994127 | 0.025733801 | 0.941933531 | 0.980968984 |
| chr2:97005435-97005648 | 0.03923739 | 0.941965928 | 0.980969139 |
| chr5:50440410-50441667 | 0.02336146 | 0.941981187 | 0.980969139 |
| chr6:54232850-54233050 | 0.036146132 | 0.941994713 | 0.980969139 |
| chr5:122845183-122846132 | -0.023556439 | 0.941996138 | 0.980969139 |
| chr3:132721881-132723362 | 0.022859307 | 0.941997694 | 0.980969139 |
| chr11:124953783-124954474 | -0.024024643 | 0.942042654 | 0.980999049 |
| chr19:9785400-9786308 | 0.024145971 | 0.942066783 | 0.980999049 |
| chr10:80444680-80445024 | -0.025922826 | 0.942078101 | 0.980999049 |
| chr5:106875813-106876118 | 0.030051484 | 0.942107782 | 0.980999049 |
| chr1:201408224-201408769 | -0.024520738 | 0.942112373 | 0.980999049 |
| chr18:63518498-63518823 | -0.029911596 | 0.942120691 | 0.980999049 |
| chr5:151658957-151659280 | 0.028240246 | 0.9421222 | 0.980999049 |
| chr1:52141690-52143017 | -0.023851632 | 0.942128841 | 0.980999049 |
| chr6:47787935-47788418 | 0.02755867 | 0.942192527 | 0.981052031 |
| chr1:151458055-151459670 | -0.022931862 | 0.942222935 | 0.981070361 |
| chr5:11849316-11849557 | 0.035183927 | 0.942243348 | 0.981076312 |
| chr6:167826289-167826914 | -0.026611349 | 0.942260845 | 0.981076312 |
| chr4:127783194-127783434 | 0.02750467 | 0.942275372 | 0.981076312 |
| chr17:67158138-67158370 | -0.032485626 | 0.942279867 | 0.981076312 |
| chr21:42315415-42315805 | -0.027410629 | 0.942319215 | 0.981103949 |
| chr14:94748870-94749125 | 0.030279776 | 0.942376977 | 0.981146435 |
| chr8:125644363-125644883 | 0.023870268 | 0.942407545 | 0.981146435 |
| chr1:165367432-165367640 | -0.034658285 | 0.942407966 | 0.981146435 |
| chr1:197156787-197156987 | -0.036517942 | 0.942434203 | 0.981146435 |
| chr7:40329604-40329819 | -0.035724039 | 0.942436775 | 0.981146435 |

| | | | |
|---|---|---|---|
| chr16:3488116-3488316 | 0.037225045 | 0.942436852 | 0.981146435 |
| chr15:59412638-59412840 | 0.035075515 | 0.942458666 | 0.981151251 |
| chr4:86891739-86892803 | -0.023170723 | 0.94247468 | 0.981151251 |
| chr15:31440758-31441319 | -0.026297816 | 0.942481117 | 0.981151251 |
| chr15:22223105-22223475 | -0.028127312 | 0.942492698 | 0.981151251 |
| chr7:92953633-92954012 | 0.028390182 | 0.942530279 | 0.981177042 |
| chr2:65406662-65406994 | 0.028546248 | 0.942550728 | 0.981178402 |
| chr4:56281174-56281375 | -0.039015807 | 0.942573843 | 0.981178402 |
| chr1:149849889-149850301 | 0.026052675 | 0.942580056 | 0.981178402 |
| chr10:104353284-104353977 | -0.023421449 | 0.942582807 | 0.981178402 |
| chr6:148578242-148578491 | -0.030744592 | 0.942599448 | 0.981182395 |
| chr1:182838822-182840191 | -0.022643972 | 0.942616683 | 0.981187005 |
| chr7:34840332-34840631 | 0.028771178 | 0.942665444 | 0.981224432 |
| chr20:51470375-51470586 | 0.033410191 | 0.942698009 | 0.981224702 |
| chr1:173056367-173056709 | 0.028851744 | 0.942699164 | 0.981224702 |
| chr3:113514544-113515648 | -0.022767618 | 0.942704122 | 0.981224702 |
| chr1:149886282-149887245 | -0.022682775 | 0.942725955 | 0.981234099 |
| chr12:131437070-131437849 | 0.025743462 | 0.942752038 | 0.981239904 |
| chr8:48921229-48922097 | -0.025675461 | 0.942757146 | 0.981239904 |
| chr11:112073893-112074679 | 0.02386434 | 0.942779493 | 0.981249834 |
| chr9:127784996-127786196 | 0.023343495 | 0.942793839 | 0.981251437 |
| chr4:127964587-127966172 | -0.022394516 | 0.942830394 | 0.981267735 |
| chr3:183085756-183085956 | 0.038735456 | 0.942835112 | 0.981267735 |
| chr11:11841288-11842085 | -0.024817322 | 0.94287493 | 0.981295847 |
| chr3:174440745-174441736 | -0.0228024 | 0.942888434 | 0.981296573 |
| chr13:32506415-32506625 | 0.039437515 | 0.942901628 | 0.981296976 |
| chr5:36151378-36152249 | -0.023047813 | 0.942918694 | 0.981301408 |
| chr17:981067-981267 | 0.034430504 | 0.942952958 | 0.981323739 |
| chr19:54200312-54201275 | -0.02308128 | 0.943015985 | 0.981357086 |
| chr3:98522509-98523253 | 0.022707909 | 0.943025558 | 0.981357086 |
| chr2:203340679-203341224 | 0.026194274 | 0.943042078 | 0.981357086 |
| chr3:139389268-139390340 | 0.022568415 | 0.9430621 | 0.981357086 |
| chr3:62366978-62367178 | -0.030102103 | 0.943067796 | 0.981357086 |
| chr3:149812432-149813921 | -0.022377041 | 0.943073984 | 0.981357086 |
| chr7:38425124-38425355 | -0.038948978 | 0.943074656 | 0.981357086 |
| chr5:80481089-80481589 | 0.0276071 | 0.943100606 | 0.981370761 |
| chr3:121749118-121750172 | -0.022511196 | 0.943121795 | 0.981379482 |
| chr3:13514167-13514394 | -0.029336641 | 0.94316537 | 0.981389242 |
| chr1:243748747-243749034 | 0.03174861 | 0.943171078 | 0.981389242 |
| chr11:45872753-45873097 | 0.027599875 | 0.94318708 | 0.981389242 |
| chr3:5767387-5767643 | 0.036029284 | 0.943222289 | 0.981389242 |
| chr8:63633704-63634023 | -0.037501777 | 0.943230131 | 0.981389242 |
| chr3:136751757-136753215 | 0.022353809 | 0.9432688 | 0.981389242 |
| chr3:113577848-113578203 | 0.026196544 | 0.943269773 | 0.981389242 |
| chr12:124567653-124568249 | 0.025198714 | 0.943303358 | 0.981389242 |
| chr10:70403869-70404292 | 0.024367712 | 0.943305129 | 0.981389242 |
| chr5:171630070-171630455 | 0.02591817 | 0.943315509 | 0.981389242 |
| chr6:26175469-26175669 | 0.034903154 | 0.943319906 | 0.981389242 |
| chr2:158650514-158650945 | -0.029388029 | 0.943325092 | 0.981389242 |
| chr1:207811896-207812386 | -0.025638316 | 0.943327376 | 0.981389242 |

| | | | |
|---|---|---|---|
| chr3:160006626-160006882 | -0.029412573 | 0.943330067 | 0.981389242 |
| chr15:82305438-82305651 | 0.034273472 | 0.943334536 | 0.981389242 |
| chr17:44315397-44315647 | 0.029350432 | 0.943348815 | 0.981389242 |
| chr22:20945650-20945906 | 0.036742632 | 0.943348913 | 0.981389242 |
| chr10:56361699-56362021 | 0.027180979 | 0.943375861 | 0.981398664 |
| chr21:44400935-44401267 | -0.030735514 | 0.943383586 | 0.981398664 |
| chr12:116708804-116709400 | 0.025842207 | 0.94343172 | 0.981423811 |
| chr19:6373000-6373676 | 0.023405295 | 0.943437375 | 0.981423811 |
| chr14:75050994-75051686 | 0.023908906 | 0.943446186 | 0.981423811 |
| chr16:88569545-88570550 | -0.022927054 | 0.943474224 | 0.981439653 |
| chr1:206808693-206809017 | -0.025434102 | 0.943499128 | 0.981452235 |
| chr9:96668172-96668394 | 0.035088939 | 0.943522741 | 0.981463474 |
| chr17:10106262-10106462 | 0.038315665 | 0.943535855 | 0.98146379 |
| chr2:3574799-3576218 | 0.022537551 | 0.943560203 | 0.981475793 |
| chr10:5997767-5997969 | -0.043545531 | 0.943576273 | 0.981479185 |
| chr4:41318810-41320076 | -0.022963146 | 0.943602708 | 0.981493357 |
| chr10:90937331-90937531 | -0.038779464 | 0.943641284 | 0.981501684 |
| chr4:165537083-165537292 | -0.035334603 | 0.943646659 | 0.981501684 |
| chr15:39920623-39921230 | 0.02474145 | 0.943654364 | 0.981501684 |
| chr14:81382454-81383387 | 0.02424914 | 0.943661952 | 0.981501684 |
| chr15:80502691-80502891 | -0.037911619 | 0.943688955 | 0.981516447 |
| chr16:89657377-89658166 | 0.022959108 | 0.943722258 | 0.981529996 |
| chr14:61322514-61322996 | -0.024627167 | 0.943727602 | 0.981529996 |
| chr2:27032695-27033245 | -0.024400674 | 0.943752009 | 0.981533307 |
| chr2:127387169-127388684 | 0.022345419 | 0.943756406 | 0.981533307 |
| chr6:159174269-159174516 | 0.03729549 | 0.943771432 | 0.981535612 |
| chr15:70362459-70362746 | -0.029923064 | 0.943799454 | 0.981551432 |
| chr9:32251601-32251847 | 0.034988175 | 0.943821167 | 0.981560691 |
| chr5:171336596-171337287 | 0.02461581 | 0.943835241 | 0.981562005 |
| chr8:23019243-23019575 | -0.028117615 | 0.943873561 | 0.981577806 |
| chrX:36867214-36867450 | 0.035764965 | 0.943877745 | 0.981577806 |
| chr12:6697853-6698062 | 0.033746267 | 0.943888867 | 0.981577806 |
| chr10:8079677-8079990 | 0.028098237 | 0.943907742 | 0.981584113 |
| chr10:26860163-26860402 | -0.026575599 | 0.943928826 | 0.981592717 |
| chr3:95027189-95027389 | 0.038680398 | 0.943972248 | 0.981609335 |
| chr17:67584949-67585332 | -0.027151198 | 0.943998982 | 0.981609335 |
| chr3:25664003-25665395 | 0.022165274 | 0.944000973 | 0.981609335 |
| chr9:5600507-5600779 | -0.030942789 | 0.944008269 | 0.981609335 |
| chr7:64109-64345 | 0.029148512 | 0.944008863 | 0.981609335 |
| chr2:199602267-199602606 | -0.027635093 | 0.94402555 | 0.981613365 |
| chr4:139082773-139084673 | 0.021979731 | 0.944050646 | 0.981625923 |
| chr3:1439399-1439751 | -0.028373598 | 0.944063968 | 0.981625923 |
| chr11:112219768-112220093 | -0.029007448 | 0.94407606 | 0.981625923 |
| chr19:10501988-10503259 | -0.022671242 | 0.944145622 | 0.981673978 |
| chr12:66145182-66145517 | 0.027596012 | 0.944160289 | 0.981673978 |
| chr2:122168842-122169102 | 0.031194217 | 0.944160713 | 0.981673978 |
| chr15:73781098-73781553 | 0.026621897 | 0.944231005 | 0.981732709 |
| chr7:8277398-8277763 | -0.027153603 | 0.944250811 | 0.981732709 |
| chr1:219837773-219837979 | -0.033925877 | 0.944257518 | 0.981732709 |
| chr14:54870286-54870585 | 0.029566002 | 0.94426845 | 0.981732709 |

| | | | |
|---|---|---|---|
| chr5:155783937-155784170 | 0.032606145 | 0.944314297 | 0.981752358 |
| chr9:129406369-129406689 | 0.029061822 | 0.944315684 | 0.981752358 |
| chr2:211248899-211249425 | 0.026365366 | 0.944330739 | 0.981752358 |
| chr5:159773173-159773749 | 0.025798685 | 0.944338601 | 0.981752358 |
| chr17:63600276-63601153 | 0.022622541 | 0.944364603 | 0.981756894 |
| chr9:121267691-121268268 | 0.024376464 | 0.94438599 | 0.981756894 |
| chr1:197800675-197800875 | -0.031603438 | 0.944393705 | 0.981756894 |
| chr19:49441492-49441718 | -0.030593365 | 0.944406858 | 0.981756894 |
| chr5:24368166-24368442 | 0.030903346 | 0.944425959 | 0.981756894 |
| chr4:88054886-88055251 | -0.028221139 | 0.944436549 | 0.981756894 |
| chr1:175887817-175888329 | -0.024228001 | 0.944446889 | 0.981756894 |
| chr17:65066915-65067252 | -0.027532008 | 0.944448835 | 0.981756894 |
| chr3:123448076-123448276 | 0.033769992 | 0.944468451 | 0.981756894 |
| chr11:42524657-42524870 | 0.033849598 | 0.944471094 | 0.981756894 |
| chr12:56041470-56042320 | 0.022463827 | 0.944495247 | 0.981768681 |
| chr19:45817128-45817428 | 0.026116185 | 0.944534155 | 0.981787037 |
| chr4:158410267-158410649 | 0.027935512 | 0.944551653 | 0.981787037 |
| chr2:134121276-134121572 | -0.026962163 | 0.944554642 | 0.981787037 |
| chr8:92167927-92168135 | -0.038632269 | 0.944564159 | 0.981787037 |
| chr15:74127559-74127837 | 0.028634655 | 0.944594738 | 0.981805502 |
| chr6:85354985-85355345 | 0.026507625 | 0.944625317 | 0.981819855 |
| chr2:36356076-36356577 | -0.024899874 | 0.944643951 | 0.981819855 |
| chr20:17525685-17525974 | -0.028095463 | 0.944646988 | 0.981819855 |
| chr17:17078329-17078783 | -0.025214697 | 0.944717603 | 0.98186697 |
| chr12:132001584-13200188€ | -0.027676445 | 0.944718782 | 0.98186697 |
| chr12:131132543-131132834 | 0.030877215 | 0.944730762 | 0.98186697 |
| chr22:20586515-20586715 | 0.033300676 | 0.944789251 | 0.981893017 |
| chr22:46267839-46268093 | 0.033300676 | 0.944789251 | 0.981893017 |
| chr5:179907014-179907285 | 0.027122512 | 0.94481636 | 0.981893017 |
| chr1:51236777-51236990 | 0.030293457 | 0.94481683 | 0.981893017 |
| chr5:180293486-180293848 | 0.023781079 | 0.944819899 | 0.981893017 |
| chr2:231706873-231708600 | -0.021848513 | 0.944879366 | 0.981940196 |
| chr16:81314607-81315874 | -0.021966406 | 0.944890927 | 0.981940196 |
| chr2:95165200-95166211 | 0.022385018 | 0.944918013 | 0.981952919 |
| chr5:174793722-174794166 | 0.026002697 | 0.944936887 | 0.981952919 |
| chrX:136908966-136910587 | 0.021641072 | 0.944945858 | 0.981952919 |
| chr22:42856882-42857537 | -0.024614795 | 0.944954432 | 0.981952919 |
| chr1:107127579-107127800 | -0.025957019 | 0.944984179 | 0.981955459 |
| chr15:69628670-69628887 | -0.031893459 | 0.944984299 | 0.981955459 |
| chr1:77682087-77683847 | 0.021671507 | 0.945019254 | 0.981955459 |
| chr14:61909502-61909703 | 0.036260606 | 0.945050373 | 0.981955459 |
| chr2:53473311-53473594 | -0.029818966 | 0.94505126 | 0.981955459 |
| chrX:103343135-103343410 | 0.02834178 | 0.945068909 | 0.981955459 |
| chr8:118307991-118308268 | -0.031424107 | 0.945079658 | 0.981955459 |
| chr9:126673639-126674089 | 0.026976391 | 0.945091236 | 0.981955459 |
| chr1:43529224-43529515 | 0.027571303 | 0.945103814 | 0.981955459 |
| chr22:36528887-36530196 | 0.022366541 | 0.945105143 | 0.981955459 |
| chr2:64917700-64918033 | -0.025703637 | 0.945121769 | 0.981955459 |
| chr9:134136806-134137099 | 0.028986634 | 0.945126995 | 0.981955459 |
| chr3:42191159-42191641 | -0.027697958 | 0.945138308 | 0.981955459 |

| | | | |
|---|---|---|---|
| chr17:63866093-63866418 | -0.025817795 | 0.945149857 | 0.981955459 |
| chr19:46634137-46634604 | 0.022946774 | 0.945155398 | 0.981955459 |
| chr7:123666075-123666305 | 0.03785355 | 0.945161926 | 0.981955459 |
| chr17:64129215-64129553 | -0.024058229 | 0.945183146 | 0.98196164 |
| chr12:121552757-121552957 | -0.03333232 | 0.945193506 | 0.98196164 |
| chr6:111727858-111728105 | -0.031521407 | 0.945235668 | 0.981980189 |
| chr13:30420896-30421990 | 0.022477762 | 0.945242852 | 0.981980189 |
| chr18:36107147-36107358 | 0.037333982 | 0.945254211 | 0.981980189 |
| chr10:7989192-7989424 | 0.033430384 | 0.945262624 | 0.981980189 |
| chr14:59028663-59028867 | 0.031634751 | 0.945297014 | 0.981991204 |
| chr1:41665298-41665687 | -0.02668913 | 0.945298859 | 0.981991204 |
| chr7:55774810-55775022 | 0.033567961 | 0.9453667 | 0.982048364 |
| chr1:146938176-146939301 | -0.022181395 | 0.945400387 | 0.982070043 |
| chr11:64284529-64284986 | -0.023291777 | 0.945453182 | 0.982111571 |
| chr1:114581077-114582274 | 0.021338882 | 0.945502959 | 0.982149963 |
| chr10:60757380-60757867 | -0.026717901 | 0.945527755 | 0.9821521 |
| chr8:73869717-73869917 | -0.037042709 | 0.945530652 | 0.9821521 |
| chr14:54440833-54441913 | 0.0217474 | 0.945597177 | 0.982207886 |
| chr3:79086476-79086821 | 0.026750523 | 0.945619134 | 0.982217378 |
| chr10:127198374-127198591 | -0.033241037 | 0.945647629 | 0.98223366 |
| chr14:96998357-96998600 | -0.036023303 | 0.945665744 | 0.982238194 |
| chrX:127256262-127256619 | 0.025686602 | 0.945677633 | 0.982238194 |
| chr6:134174652-134175483 | -0.022426785 | 0.945712218 | 0.982260801 |
| chr10:71912508-71912787 | 0.031812516 | 0.945828624 | 0.98236839 |
| chr1:179591442-179592524 | 0.021525577 | 0.945852273 | 0.982376629 |
| chr21:41572196-41572440 | 0.030435945 | 0.945862199 | 0.982376629 |
| chr4:144143011-144143211 | 0.037186299 | 0.945901056 | 0.982403669 |
| chr13:27170644-27171137 | -0.024607333 | 0.945945461 | 0.982420866 |
| chr14:93925669-93925960 | 0.029669843 | 0.945949576 | 0.982420866 |
| chr11:33310780-33311057 | 0.031642865 | 0.945963769 | 0.982420866 |
| chr22:26756741-26757138 | -0.027321485 | 0.9459689 | 0.982420866 |
| chr2:20353839-20354074 | 0.025711824 | 0.945991765 | 0.982431296 |
| chr5:39599814-39600112 | -0.031719224 | 0.946009573 | 0.982436474 |
| chr6:27388538-27389229 | 0.022923574 | 0.946039491 | 0.982446238 |
| chr20:46708142-46708371 | -0.031822222 | 0.946044619 | 0.982446238 |
| chr10:32957521-32957798 | -0.027446605 | 0.946070775 | 0.982453647 |
| chr2:33033561-33033854 | -0.031934342 | 0.946097766 | 0.982453647 |
| chr4:127880437-127881762 | 0.021155089 | 0.946119079 | 0.982453647 |
| chr9:37079359-37080323 | -0.022038566 | 0.946128634 | 0.982453647 |
| chr7:2232437-2233314 | 0.022078616 | 0.946137976 | 0.982453647 |
| chr17:76458627-76458894 | 0.027714956 | 0.946140464 | 0.982453647 |
| chr2:43024508-43024814 | 0.02655138 | 0.946141508 | 0.982453647 |
| chr15:29438724-29439001 | -0.028963043 | 0.946211434 | 0.982473631 |
| chr6:28281380-28281701 | 0.027210998 | 0.946218471 | 0.982473631 |
| chr11:65857145-65857400 | -0.026021683 | 0.94623828 | 0.982473631 |
| chr10:80256906-80257140 | -0.028258331 | 0.946248255 | 0.982473631 |
| chr17:67468089-67468318 | 0.030206847 | 0.946270346 | 0.982473631 |
| chr18:32091726-32093450 | 0.021119815 | 0.946302989 | 0.982473631 |
| chr11:108664724-108665209 | -0.022327482 | 0.946321013 | 0.982473631 |
| chr3:44477315-44477893 | 0.025522318 | 0.946322861 | 0.982473631 |

| | | | |
|---|---|---|---|
| chr2:10021731-10022086 | 0.026576115 | 0.94633661 | 0.982473631 |
| chr2:101387054-101387610 | 0.025396285 | 0.946342869 | 0.982473631 |
| chr10:97498137-97499032 | 0.021244039 | 0.946353493 | 0.982473631 |
| chr1:175319620-175319954 | 0.025138702 | 0.946365984 | 0.982473631 |
| chr1:157991938-157992339 | 0.026170248 | 0.946366831 | 0.982473631 |
| chr1:151165668-151166662 | -0.021538241 | 0.946374086 | 0.982473631 |
| chr20:13546546-13546787 | 0.030955417 | 0.946374211 | 0.982473631 |
| chr5:68288802-68289102 | 0.02952463 | 0.946391471 | 0.982473631 |
| chr6:113375570-113375804 | 0.02952463 | 0.946391471 | 0.982473631 |
| chr18:26548410-26549519 | 0.021852563 | 0.946391555 | 0.982473631 |
| chr2:70834860-70835174 | -0.030235946 | 0.946418189 | 0.982487969 |
| chr3:187750049-187750872 | 0.021845928 | 0.946500485 | 0.982528868 |
| chr10:46228587-46228914 | -0.024404496 | 0.946504317 | 0.982528868 |
| chr6:108127882-108128163 | -0.028762567 | 0.946508586 | 0.982528868 |
| chr1:2381679-2382273 | -0.02530093 | 0.946508879 | 0.982528868 |
| chr2:176816953-176817256 | -0.028450932 | 0.946537465 | 0.982545231 |
| chr17:8753916-8754152 | 0.032833038 | 0.946575649 | 0.982561444 |
| chr13:41075223-41075423 | -0.032856817 | 0.946589523 | 0.982561444 |
| chr3:79490203-79490460 | -0.029334988 | 0.946603902 | 0.982561444 |
| chr5:83077056-83078118 | -0.021317216 | 0.946604378 | 0.982561444 |
| chr20:50153390-50154037 | 0.022012543 | 0.946649891 | 0.982564065 |
| chr19:46373492-46374428 | 0.021655156 | 0.946650758 | 0.982564065 |
| chr9:120774659-120774960 | 0.030726463 | 0.946657154 | 0.982564065 |
| chr1:68111630-68112219 | 0.023856859 | 0.946658197 | 0.982564065 |
| chr8:93565513-93565797 | -0.027628858 | 0.946689198 | 0.982575556 |
| chr17:8125633-8125858 | -0.027583736 | 0.946698682 | 0.982575556 |
| chr8:137890977-137891208 | -0.028972872 | 0.946714193 | 0.982575556 |
| chr3:125374947-125376087 | -0.021718138 | 0.946720563 | 0.982575556 |
| chr2:111288227-111288593 | -0.028854807 | 0.946756571 | 0.982599618 |
| chr6:122033627-122033913 | -0.027357734 | 0.946778412 | 0.982608976 |
| chr5:431023-431290 | -0.025189173 | 0.946853641 | 0.982673741 |
| chr4:94083770-94083970 | 0.036297261 | 0.946868772 | 0.982676135 |
| chr8:115019079-115019407 | 0.029963248 | 0.946903296 | 0.982698654 |
| chr4:77203571-77203956 | -0.026606979 | 0.946919312 | 0.982698926 |
| chr3:63043710-63043920 | -0.026065271 | 0.946929209 | 0.982698926 |
| chr7:48321321-48321724 | -0.030072285 | 0.946949942 | 0.982707133 |
| chr16:2673057-2673877 | -0.021948275 | 0.946974424 | 0.982718579 |
| chr6:132401732-132401932 | 0.036940636 | 0.946986622 | 0.982718579 |
| chr5:114063554-114063825 | 0.031360693 | 0.947024153 | 0.982744216 |
| chr3:23079930-23080148 | -0.031429157 | 0.947168532 | 0.982880729 |
| chr3:38124363-38124563 | 0.036119031 | 0.947190276 | 0.982889982 |
| chr11:57800517-57800989 | -0.023281054 | 0.947214384 | 0.982892614 |
| chr13:84896001-84896379 | -0.030830354 | 0.947218469 | 0.982892614 |
| chr1:48109349-48109667 | -0.03064408 | 0.947255469 | 0.982911828 |
| chr11:83007930-83008130 | -0.031236514 | 0.947262641 | 0.982911828 |
| chr16:3673107-3673374 | 0.032917721 | 0.947290314 | 0.982927231 |
| chrX:18395102-18395304 | 0.035174972 | 0.947314703 | 0.982932843 |
| chr6:125212533-125212733 | -0.034704414 | 0.947331233 | 0.982932843 |
| chr15:32030856-32031211 | -0.026312046 | 0.947334207 | 0.982932843 |
| chr12:41189534-41190286 | 0.024005643 | 0.947369166 | 0.982955805 |

| | | | |
|---|---|---|---|
| chr3:151705136-151705714 | 0.024683341 | 0.947415601 | 0.982986505 |
| chr4:64539313-64539552 | 0.030848275 | 0.947439757 | 0.982986505 |
| chr15:91852689-91853904 | 0.021373044 | 0.94747688 | 0.982986505 |
| chr8:116839877-116840161 | -0.027086215 | 0.947488314 | 0.982986505 |
| chr8:66666864-66667833 | -0.02082211 | 0.947491434 | 0.982986505 |
| chr14:101175103-101175405 | 0.028572079 | 0.947493355 | 0.982986505 |
| chr15:94187019-94187252 | 0.031423282 | 0.94753076 | 0.982986505 |
| chr6:33384567-33384869 | -0.028079259 | 0.947541681 | 0.982986505 |
| chr10:93495650-93496925 | 0.021107414 | 0.947570625 | 0.982986505 |
| chr4:99561207-99561412 | 0.02742359 | 0.947580354 | 0.982986505 |
| chr15:57377466-57377755 | -0.025487708 | 0.9475862 | 0.982986505 |
| chr6:33208381-33208633 | -0.025625715 | 0.947593127 | 0.982986505 |
| chr19:56252668-56253131 | -0.027675176 | 0.947614976 | 0.982986505 |
| chr19:4323121-4323321 | -0.037184411 | 0.947616069 | 0.982986505 |
| chr2:115165596-115165796 | -0.037184411 | 0.947616069 | 0.982986505 |
| chr6:7267384-7268066 | 0.022252867 | 0.947621233 | 0.982986505 |
| chr6:3978309-3978621 | 0.028311724 | 0.947625598 | 0.982986505 |
| chr8:97775009-97775330 | 0.023233458 | 0.947639641 | 0.982986505 |
| chr6:85126888-85127409 | 0.023707845 | 0.947642507 | 0.982986505 |
| chr1:161764889-161765539 | -0.022654631 | 0.947756648 | 0.983091594 |
| chr11:119195709-119196103 | 0.023630014 | 0.947801766 | 0.983116831 |
| chr2:131153854-131154493 | 0.022849566 | 0.947834232 | 0.983116831 |
| chr1:152533393-152533623 | -0.029888395 | 0.947854299 | 0.983116831 |
| chrX:69615704-69616885 | 0.021480809 | 0.947856362 | 0.983116831 |
| chr4:156109841-156110487 | -0.022695761 | 0.947856968 | 0.983116831 |
| chr8:18564971-18565184 | 0.031700176 | 0.947869381 | 0.983116831 |
| chr21:39388055-39388986 | 0.02093983 | 0.947879231 | 0.983116831 |
| chr6:104527810-104528134 | -0.026751998 | 0.947895749 | 0.983116831 |
| chr4:124111933-124112162 | -0.031148589 | 0.947896454 | 0.983116831 |
| chr16:30875847-30876129 | -0.030551336 | 0.947919906 | 0.983127846 |
| chrX:104165755-104167057 | 0.020680913 | 0.947941811 | 0.983137257 |
| chr3:39808809-39809687 | 0.02210992 | 0.947968323 | 0.983151446 |
| chr8:114614566-114614834 | 0.029012417 | 0.948046381 | 0.983219093 |
| chr11:85150240-85150519 | 0.034338862 | 0.948060593 | 0.983220524 |
| chr1:63316772-63317674 | 0.021575425 | 0.948075498 | 0.983222674 |
| chr5:7078989-7079228 | 0.030396335 | 0.948106856 | 0.983241886 |
| chr6:141989322-141989626 | 0.026505819 | 0.94812462 | 0.983247001 |
| chr16:81444213-81445120 | -0.021113701 | 0.948183114 | 0.983288685 |
| chr1:1347943-1348190 | 0.029684846 | 0.948190481 | 0.983288685 |
| chr16:71645397-71645825 | -0.023835137 | 0.948218373 | 0.983288703 |
| chr2:219308141-219308440 | 0.028455909 | 0.948227989 | 0.983288703 |
| chr1:202958761-202958974 | 0.025630403 | 0.948235336 | 0.983288703 |
| chr8:55957899-55958177 | 0.028134766 | 0.948241831 | 0.983288703 |
| chr4:76067647-76068030 | 0.025561765 | 0.948260119 | 0.983294359 |
| chr1:46718614-46719536 | -0.021082728 | 0.948293332 | 0.983313776 |
| chrX:51496474-51497238 | 0.023397872 | 0.94830451 | 0.983313776 |
| chr17:64129743-64130641 | -0.020682571 | 0.948345299 | 0.983342763 |
| chr17:72121220-72121899 | -0.021771031 | 0.948382979 | 0.983368526 |
| chr17:80101964-80102242 | -0.02585073 | 0.948418491 | 0.983374285 |
| chr1:147890183-147890550 | -0.023421802 | 0.948432435 | 0.983374285 |

| | | | |
|---|---|---|---|
| chr12:70244104-70244314 | 0.026157778 | 0.948463201 | 0.983374285 |
| chr20:56202576-56202792 | 0.026550818 | 0.948498392 | 0.983374285 |
| chr5:141208597-141208806 | -0.034044063 | 0.948498394 | 0.983374285 |
| chr6:51959827-51960027 | -0.034044063 | 0.948498394 | 0.983374285 |
| chr20:53623225-53624265 | 0.020141991 | 0.948519568 | 0.983374285 |
| chr11:72328760-72328970 | 0.036224043 | 0.948538496 | 0.983374285 |
| chr7:2540699-2540917 | 0.028621919 | 0.948545371 | 0.983374285 |
| chr22:22370975-22371240 | -0.030348774 | 0.948546016 | 0.983374285 |
| chr3:89107392-89107990 | -0.022207785 | 0.948590202 | 0.983374285 |
| chr2:175536773-175537054 | -0.029572411 | 0.948593801 | 0.983374285 |
| chr3:13495106-13495306 | -0.02908357 | 0.948595993 | 0.983374285 |
| chr20:50478260-50478657 | -0.023172786 | 0.948604247 | 0.983374285 |
| chr10:131431672-131431886 | -0.033029291 | 0.948612471 | 0.983374285 |
| chr17:15260359-15261039 | -0.022220199 | 0.948615145 | 0.983374285 |
| chr2:202858687-202859072 | 0.02626957 | 0.948615368 | 0.983374285 |
| chr15:40756989-40757282 | -0.027220662 | 0.948619547 | 0.983374285 |
| chr3:107976299-107976663 | 0.027181434 | 0.948639424 | 0.983379552 |
| chr20:47784503-47787014 | 0.019933019 | 0.948650296 | 0.983379552 |
| chr22:17773847-17774230 | 0.024323487 | 0.94873452 | 0.983441864 |
| chr15:31632673-31632908 | 0.033053131 | 0.948753955 | 0.983441864 |
| chr1:26471525-26471859 | 0.026502156 | 0.94875504 | 0.983441864 |
| chr6:116941830-116942034 | -0.036658081 | 0.948781306 | 0.983441864 |
| chr8:90679356-90679577 | -0.030927296 | 0.948789901 | 0.983441864 |
| chr16:57475895-57476225 | 0.025772357 | 0.948804944 | 0.983441864 |
| chr3:113169716-113169962 | 0.03207184 | 0.948814805 | 0.983441864 |
| chr4:128525424-128526334 | 0.021318158 | 0.948836796 | 0.983441864 |
| chr3:132902528-132902908 | -0.028536687 | 0.948861374 | 0.983441864 |
| chr11:126268717-126269497 | 0.020768164 | 0.948865449 | 0.983441864 |
| chr7:41977199-41977408 | -0.030968239 | 0.948880914 | 0.983441864 |
| chr5:139087955-139088284 | 0.030290243 | 0.948882541 | 0.983441864 |
| chr5:32989919-32990119 | 0.03458033 | 0.948930489 | 0.983441864 |
| chr15:81323842-81324892 | 0.020588377 | 0.948957282 | 0.983441864 |
| chr22:25220264-25220464 | -0.033379728 | 0.948961151 | 0.983441864 |
| chr10:11742153-11742514 | -0.025295563 | 0.949000913 | 0.983441864 |
| chr11:18105761-18106305 | -0.022420152 | 0.949006825 | 0.983441864 |
| chr2:9333616-9333951 | 0.024881922 | 0.949013389 | 0.983441864 |
| chr2:51795509-51795844 | -0.028010994 | 0.949016448 | 0.983441864 |
| chr10:102501978-102503129 | 0.020192971 | 0.949031402 | 0.983441864 |
| chr2:64609335-64609767 | -0.023586214 | 0.949056275 | 0.983441864 |
| chr8:40355583-40355938 | 0.028811356 | 0.949076517 | 0.983441864 |
| chr17:28726848-28727479 | 0.020914918 | 0.949090246 | 0.983441864 |
| chrX:57909428-57909967 | -0.024024436 | 0.949103313 | 0.983441864 |
| chr6:81752112-81752974 | 0.020594725 | 0.949107272 | 0.983441864 |
| chr7:538373-538573 | -0.02487021 | 0.94913172 | 0.983441864 |
| chr1:181112436-181112836 | -0.021781114 | 0.949131949 | 0.983441864 |
| chr3:113211402-113212655 | 0.021010274 | 0.949135894 | 0.983441864 |
| chr5:129903760-129905007 | 0.020962321 | 0.949137323 | 0.983441864 |
| chr1:172470177-172470487 | 0.025518409 | 0.949143541 | 0.983441864 |
| chr10:102743550-102744552 | 0.020615783 | 0.949149125 | 0.983441864 |
| chr7:48701648-48701848 | -0.032834292 | 0.949155288 | 0.983441864 |

| | | | |
|---|---|---|---|
| chr2:241010917-241011117 | -0.033836562 | 0.949155443 | 0.983441864 |
| chr3:124175058-124175258 | -0.033836562 | 0.949155443 | 0.983441864 |
| chr14:82710293-82710606 | -0.024759576 | 0.949159631 | 0.983441864 |
| chr10:13056497-13056745 | 0.03053254 | 0.949224637 | 0.983446061 |
| chr8:76677642-76677842 | 0.034194267 | 0.949234185 | 0.983446061 |
| chr8:119669937-119670137 | -0.036358676 | 0.949262448 | 0.983446061 |
| chr3:19146584-19147646 | -0.020133788 | 0.949263747 | 0.983446061 |
| chr6:128376039-128376239 | 0.033351412 | 0.949289417 | 0.983446061 |
| chr3:156535194-156535542 | 0.023528224 | 0.949298887 | 0.983446061 |
| chr16:80924376-80924920 | 0.023286192 | 0.949300175 | 0.983446061 |
| chr1:225438108-225438308 | 0.032355851 | 0.949305953 | 0.983446061 |
| chr17:71460966-71461190 | 0.031319588 | 0.949308676 | 0.983446061 |
| chr17:17720244-17720459 | -0.02846651 | 0.949324576 | 0.983446061 |
| chr2:106792801-106793037 | -0.031144808 | 0.94932637 | 0.983446061 |
| chr4:140736682-140736895 | 0.029998465 | 0.949331676 | 0.983446061 |
| chr12:14454138-14454479 | -0.02388926 | 0.949342509 | 0.983446061 |
| chr1:164772688-164773066 | -0.023696757 | 0.949343373 | 0.983446061 |
| chr21:46635013-46636257 | -0.02009518 | 0.949384764 | 0.983450322 |
| chr2:133421509-133421966 | -0.02524524 | 0.949386153 | 0.983450322 |
| chr17:65556375-65556756 | 0.021910817 | 0.949399324 | 0.983450322 |
| chr20:8361026-8361262 | 0.033310752 | 0.949402545 | 0.983450322 |
| chr6:150891927-150892190 | 0.02791978 | 0.949415499 | 0.983450322 |
| chr4:56530251-56531044 | 0.02073869 | 0.94942967 | 0.983450322 |
| chr11:86802646-86802920 | -0.024640139 | 0.949437331 | 0.983450322 |
| chr16:25126809-25127129 | -0.026607909 | 0.949465321 | 0.983451459 |
| chr5:179059984-179060649 | 0.022318955 | 0.949472126 | 0.983451459 |
| chr17:67637867-67638617 | -0.020644256 | 0.949493624 | 0.983451459 |
| chr4:105585825-105586507 | 0.022146714 | 0.949508386 | 0.983451459 |
| chr14:94689509-94689895 | -0.024688751 | 0.949531601 | 0.983451459 |
| chr7:66996448-66997194 | -0.020467967 | 0.949539727 | 0.983451459 |
| chr6:57172012-57172969 | 0.020967524 | 0.949544708 | 0.983451459 |
| chr3:170974962-170975243 | -0.028281845 | 0.949559211 | 0.983451459 |
| chr16:87297966-87298256 | -0.025726906 | 0.949565548 | 0.983451459 |
| chr11:71813605-71814050 | -0.024807942 | 0.94956678 | 0.983451459 |
| chr10:56352057-56352257 | 0.03335133 | 0.949581417 | 0.983453325 |
| chr2:170679608-170679809 | -0.030654005 | 0.949611085 | 0.983458091 |
| chr5:155746442-155746924 | -0.0270428 | 0.949617316 | 0.983458091 |
| chr11:117734750-117734958 | 0.03108392 | 0.949633822 | 0.983458091 |
| chr6:121400817-121401312 | 0.023907419 | 0.94963736 | 0.983458091 |
| chr21:33894766-33895131 | 0.024236874 | 0.949675599 | 0.9834844 |
| chr14:56294173-56294489 | 0.026834045 | 0.949709563 | 0.98350628 |
| chr6:36154124-36154775 | 0.023179404 | 0.949736389 | 0.983520768 |
| chr11:18536020-18536931 | -0.021264075 | 0.949771198 | 0.983527531 |
| chr8:64102165-64102478 | -0.028629679 | 0.949772664 | 0.983527531 |
| chrX:55451765-55452314 | -0.023610495 | 0.94980174 | 0.983527531 |
| chr6:160391359-160391624 | 0.0292104 | 0.949805172 | 0.983527531 |
| chr22:18076098-18076298 | -0.028346008 | 0.949810957 | 0.983527531 |
| chr21:39312329-39312706 | 0.020826875 | 0.949819937 | 0.983527531 |
| chr5:142740484-142740863 | 0.023093261 | 0.949882468 | 0.983537653 |
| chr13:30049783-30050162 | -0.025055003 | 0.949894917 | 0.983537653 |

| | | | |
|---|---|---|---|
| chr5:112636124-112636771 | -0.021926851 | 0.949902911 | 0.983537653 |
| chr17:80101102-80101799 | -0.021288669 | 0.949906086 | 0.983537653 |
| chr3:101778857-101779963 | -0.019910351 | 0.949911906 | 0.983537653 |
| chr6:169751167-169752247 | 0.019973839 | 0.949913719 | 0.983537653 |
| chr8:28419496-28419770 | 0.028430068 | 0.949925311 | 0.983537653 |
| chr11:66871422-66871688 | -0.02548517 | 0.949932402 | 0.983537653 |
| chr14:23825654-23825887 | 0.03012361 | 0.949956838 | 0.983549664 |
| chr3:166444843-166445093 | -0.029949732 | 0.94999888 | 0.983579902 |
| chr3:138935740-138936586 | 0.020719518 | 0.950042041 | 0.983611297 |
| chr6:149200345-149200734 | -0.026651014 | 0.950071033 | 0.98361744 |
| chr10:97445375-97446559 | -0.020149698 | 0.950073649 | 0.98361744 |
| chr9:120109281-120109634 | -0.025165509 | 0.950143926 | 0.983676907 |
| chr3:190339232-190339463 | -0.029077325 | 0.950174473 | 0.98368386 |
| chr8:143131088-143131397 | 0.022950264 | 0.950188817 | 0.98368386 |
| chr5:140664170-140665218 | -0.020108366 | 0.950189156 | 0.98368386 |
| chr9:14251081-14251861 | 0.021077047 | 0.95022671 | 0.983709447 |
| chr12:6341972-6342215 | -0.026946298 | 0.950246714 | 0.983712288 |
| chr7:34900907-34901171 | 0.02578701 | 0.950255132 | 0.983712288 |
| chr10:29784411-29784745 | 0.024387787 | 0.950328064 | 0.983774497 |
| chr5:175847589-175847790 | -0.029241058 | 0.950348451 | 0.98378192 |
| chr3:43621706-43622502 | -0.020200052 | 0.950360914 | 0.98378192 |
| chr18:46173157-46174248 | -0.020514085 | 0.950385236 | 0.98378635 |
| chr11:111841513-111841713 | 0.033690899 | 0.950420529 | 0.98378635 |
| chr6:5030685-5031028 | 0.02281722 | 0.950435574 | 0.98378635 |
| chr2:38784634-38784892 | 0.029222179 | 0.950448091 | 0.98378635 |
| chr2:1489123-1489328 | 0.032466363 | 0.950467398 | 0.98378635 |
| chr17:82458037-82458966 | 0.019799838 | 0.950471323 | 0.98378635 |
| chr2:85420639-85421012 | -0.021320846 | 0.950473784 | 0.98378635 |
| chr18:34978315-34978515 | 0.031909679 | 0.950475206 | 0.98378635 |
| chr12:45989835-45991088 | 0.01958589 | 0.950480749 | 0.98378635 |
| chr7:102286248-102286488 | 0.025301062 | 0.950515661 | 0.983809196 |
| chr16:88468616-88469179 | 0.021675034 | 0.950552556 | 0.983828028 |
| chr13:38900220-38900550 | -0.024859288 | 0.950573629 | 0.983828028 |
| chr3:183178994-183179245 | -0.029686727 | 0.950579186 | 0.983828028 |
| chrX:124582815-124583123 | 0.027426461 | 0.950585215 | 0.983828028 |
| chr3:71064575-71065240 | 0.022008881 | 0.950615248 | 0.983843419 |
| chr8:128322749-128322949 | -0.028203889 | 0.950625767 | 0.983843419 |
| chrX:40735338-40736270 | -0.020248545 | 0.950645907 | 0.983850974 |
| chr1:59814301-59815501 | -0.019549719 | 0.950664483 | 0.983855878 |
| chr4:123479955-123480302 | -0.024831111 | 0.950676327 | 0.983855878 |
| chr5:177390503-177390805 | 0.023103939 | 0.95069639 | 0.983863353 |
| chr5:132490096-132491196 | -0.019881792 | 0.950716284 | 0.983870652 |
| chr4:54063763-54065040 | -0.020257881 | 0.950736078 | 0.983873645 |
| chr2:132287161-132287361 | 0.034654903 | 0.95077695 | 0.983873645 |
| chr20:20712301-20713115 | 0.01993076 | 0.950782245 | 0.983873645 |
| chr5:8938853-8939053 | 0.035905156 | 0.95080041 | 0.983873645 |
| chr10:92591518-92591928 | 0.021743819 | 0.950802542 | 0.983873645 |
| chr4:77308907-77309131 | 0.028279581 | 0.950829572 | 0.983873645 |
| chr17:75723116-75723723 | 0.020375468 | 0.950836177 | 0.983873645 |
| chr1:155244374-155245273 | -0.019892134 | 0.950844264 | 0.983873645 |

| | | | |
|---|---|---|---|
| chr6:106688253-106688504 | 0.0284275 | 0.950855434 | 0.983873645 |
| chr19:35900503-35900728 | 0.025354845 | 0.950855637 | 0.983873645 |
| chr17:82238804-82239156 | -0.021752806 | 0.950871309 | 0.983873645 |
| chr10:95243100-95243311 | -0.029358367 | 0.95090266 | 0.983873645 |
| chr3:30013063-30013263 | -0.031852253 | 0.950908171 | 0.983873645 |
| chr3:23810257-23811492 | -0.019764022 | 0.950926104 | 0.983873645 |
| chr3:169755277-169755477 | 0.029609339 | 0.950926606 | 0.983873645 |
| chr1:205939713-205940125 | -0.021405511 | 0.95094105 | 0.983873645 |
| chr8:119707473-119707773 | 0.026837269 | 0.950944477 | 0.983873645 |
| chr17:59093249-59094218 | -0.019425295 | 0.950950307 | 0.983873645 |
| chr18:80200251-80200576 | 0.025226452 | 0.950981934 | 0.983893082 |
| chr2:178112101-178113237 | -0.019657223 | 0.951026323 | 0.983925721 |
| chr8:92266524-92266724 | -0.030507451 | 0.951059906 | 0.98394718 |
| chr4:125316233-125316695 | -0.021560816 | 0.951082449 | 0.983957217 |
| chr1:13807295-13807495 | 0.035468459 | 0.95110552 | 0.9839678 |
| chr5:24642038-24642334 | -0.024505296 | 0.951121738 | 0.983971293 |
| chr11:125452743-125452989 | -0.030030663 | 0.951153174 | 0.983985085 |
| chr20:37605777-37606051 | 0.030519236 | 0.951160754 | 0.983985085 |
| chr6:48498269-48498469 | 0.034665718 | 0.951178667 | 0.983990331 |
| chr15:100049507-100049738 | -0.029885527 | 0.951359801 | 0.9841644 |
| chr12:64608888-64609396 | -0.020797755 | 0.951372621 | 0.9841644 |
| chr10:97632898-97634309 | 0.019443703 | 0.951484404 | 0.984262425 |
| chr1:150187340-150187831 | 0.021544618 | 0.951493071 | 0.984262425 |
| chr6:87407617-87408970 | 0.019443948 | 0.951514627 | 0.984268283 |
| chr12:71839436-71840768 | 0.019353557 | 0.951524425 | 0.984268283 |
| chr6:85593159-85594755 | -0.019082146 | 0.951577877 | 0.984277641 |
| chr9:137240328-137241393 | 0.019365494 | 0.951582279 | 0.984277641 |
| chr6:77124460-77125042 | -0.024228335 | 0.951605169 | 0.984277641 |
| chr3:163490129-163490596 | -0.022077328 | 0.951605681 | 0.984277641 |
| chr2:120262434-120262660 | 0.028103372 | 0.951621906 | 0.984277641 |
| chr10:113706042-113706326 | 0.030195611 | 0.951625982 | 0.984277641 |
| chr3:139353972-139354324 | 0.022168478 | 0.951633269 | 0.984277641 |
| chr8:54736425-54736764 | 0.024454891 | 0.951640863 | 0.984277641 |
| chr6:109690805-109691747 | -0.019852774 | 0.951662695 | 0.984277641 |
| chr2:138787145-138787355 | -0.028322228 | 0.951663239 | 0.984277641 |
| chr2:134989091-134989291 | -0.031045462 | 0.951699802 | 0.984277641 |
| chr1:149887461-149888686 | 0.019026494 | 0.951706359 | 0.984277641 |
| chr14:36771316-36771671 | -0.023056381 | 0.951711967 | 0.984277641 |
| chr10:91409384-91410985 | 0.019194177 | 0.951713315 | 0.984277641 |
| chr2:205776835-205777035 | -0.029668886 | 0.951748463 | 0.984287421 |
| chr20:36092970-36093171 | -0.029668886 | 0.951748463 | 0.984287421 |
| chr6:20487739-20487983 | -0.02943598 | 0.951777419 | 0.98430013 |
| chr7:149205940-149206476 | 0.020910407 | 0.95179509 | 0.98430013 |
| chr6:18263779-18265001 | -0.019529443 | 0.951804954 | 0.98430013 |
| chr11:71256128-71256378 | -0.025336697 | 0.951814212 | 0.98430013 |
| chr19:58154717-58155374 | -0.020527515 | 0.951831837 | 0.98430013 |
| chr7:6577256-6577860 | 0.019808813 | 0.951846141 | 0.98430013 |
| chr3:15292801-15293470 | -0.020496957 | 0.951864236 | 0.98430013 |
| chr10:8350530-8350773 | -0.030073368 | 0.951868027 | 0.98430013 |
| chr6:56370103-56370609 | -0.022811145 | 0.951876368 | 0.98430013 |

| | | | |
|---|---|---|---|
| chr11:18651115-18651536 | 0.022745335 | 0.951928107 | 0.984340348 |
| chr6:99746325-99746635 | -0.024722817 | 0.951961008 | 0.984349588 |
| chr16:87388123-87388766 | -0.019479688 | 0.951962737 | 0.984349588 |
| chr14:78700337-78700537 | 0.031788747 | 0.951982063 | 0.984356287 |
| chr2:80256530-80256763 | -0.031374127 | 0.951997173 | 0.984358628 |
| chr10:44932785-44933227 | -0.022657453 | 0.952027813 | 0.984364631 |
| chr3:62907310-62907616 | -0.02175397 | 0.952028673 | 0.984364631 |
| chr1:44986278-44986905 | 0.020353683 | 0.952052994 | 0.984366087 |
| chr5:141866032-141866507 | -0.020858562 | 0.952055826 | 0.984366087 |
| chr17:50661867-50662192 | -0.022353669 | 0.952075916 | 0.984366087 |
| chr12:55966179-55967310 | -0.01960658 | 0.952081469 | 0.984366087 |
| chr13:51591030-51591551 | 0.023568299 | 0.952129915 | 0.984385727 |
| chr4:155822571-155822900 | -0.027969508 | 0.952133265 | 0.984385727 |
| chr1:147680114-147680358 | 0.028204499 | 0.952143731 | 0.984385727 |
| chr10:127694579-127694841 | 0.027356537 | 0.952163578 | 0.984385727 |
| chr1:226637738-226638130 | 0.026423739 | 0.952169519 | 0.984385727 |
| chr11:101346618-101346966 | -0.025529348 | 0.952191934 | 0.984385727 |
| chr3:49539615-49540458 | -0.020368747 | 0.952217304 | 0.984385727 |
| chr9:120842452-120843599 | 0.019071299 | 0.952219191 | 0.984385727 |
| chr16:23275032-23275285 | 0.026994721 | 0.952253947 | 0.984385727 |
| chr13:26276951-26277279 | -0.032493481 | 0.952260631 | 0.984385727 |
| chr17:68035241-68036396 | -0.018929656 | 0.952262014 | 0.984385727 |
| chrX:123962950-123963363 | 0.020305474 | 0.952264485 | 0.984385727 |
| chr19:44112872-44113530 | 0.020491223 | 0.952281006 | 0.984385727 |
| chr16:28596947-28597285 | -0.025354501 | 0.952324098 | 0.984385727 |
| chr6:27096893-27097592 | -0.020415319 | 0.952326013 | 0.984385727 |
| chr7:54767291-54767491 | -0.032116334 | 0.952330951 | 0.984385727 |
| chr7:105141201-105141401 | -0.032116334 | 0.952330951 | 0.984385727 |
| chr13:34077268-34077468 | 0.02899523 | 0.952339379 | 0.984385727 |
| chr3:79018374-79019885 | -0.018820581 | 0.952354544 | 0.984385727 |
| chr16:89120103-89120315 | 0.028901398 | 0.952365086 | 0.984385727 |
| chr1:26993459-26993827 | 0.024321212 | 0.952370259 | 0.984385727 |
| chr12:10750110-10750504 | 0.023093126 | 0.952424622 | 0.984399639 |
| chr3:191416787-191417349 | -0.020944635 | 0.952431304 | 0.984399639 |
| chr16:87278674-87279042 | 0.022291072 | 0.952446892 | 0.984399639 |
| chr3:129328735-129328998 | -0.023257442 | 0.952460176 | 0.984399639 |
| chr11:80720754-80721071 | -0.024313444 | 0.952460708 | 0.984399639 |
| chr8:60948360-60948584 | -0.029237021 | 0.952460803 | 0.984399639 |
| chr11:207279-207961 | 0.01998041 | 0.952478111 | 0.984404249 |
| chr8:94080289-94080595 | 0.023806277 | 0.952534189 | 0.984448928 |
| chr5:173823945-173824561 | 0.020340481 | 0.952557164 | 0.984459394 |
| chr10:107728957-107729237 | 0.026426203 | 0.952571813 | 0.984461255 |
| chr12:121033362-121033652 | 0.030587464 | 0.952595709 | 0.984464341 |
| chr5:61801273-61801473 | -0.032321801 | 0.952629587 | 0.984464341 |
| chr5:31518082-31518282 | -0.032321801 | 0.952629587 | 0.984464341 |
| chr5:126718332-126718558 | -0.032321801 | 0.952629587 | 0.984464341 |
| chr2:9284634-9284839 | -0.032338108 | 0.95263904 | 0.984464341 |
| chr19:33684716-33685009 | -0.026700528 | 0.952709878 | 0.984524267 |
| chr3:188664857-188665081 | 0.029713352 | 0.95273574 | 0.984537714 |
| chr10:80443425-80443974 | -0.020677161 | 0.952763259 | 0.984540622 |

| | | | |
|---|---|---|---|
| chr6:3279701-3279924 | 0.029036448 | 0.952764252 | 0.984540622 |
| chr11:77598287-77598515 | -0.025712536 | 0.952837327 | 0.984600446 |
| chr15:75584310-75584510 | -0.0299984 | 0.952868969 | 0.984600446 |
| chr11:78646627-78647062 | -0.022099189 | 0.952870881 | 0.984600446 |
| chr1:31548929-31549733 | -0.020000702 | 0.952873546 | 0.984600446 |
| chr15:42704489-42704769 | -0.025462507 | 0.95292055 | 0.984635736 |
| chr22:41501342-41501542 | -0.029513359 | 0.95294539 | 0.984635801 |
| chr7:19116195-19116758 | 0.021792148 | 0.952948273 | 0.984635801 |
| chr9:115326673-115326878 | -0.02934214 | 0.952968198 | 0.984635801 |
| chr4:143335642-143337640 | -0.018565224 | 0.952972015 | 0.984635801 |
| chr17:7560996-7561383 | -0.021130752 | 0.953020333 | 0.984660735 |
| chrX:68808837-68809072 | -0.029431336 | 0.953038736 | 0.984660735 |
| chr19:51158819-51159019 | 0.034370625 | 0.953039759 | 0.984660735 |
| chr18:61155683-61156149 | 0.023362537 | 0.95304755 | 0.984660735 |
| chr2:27086355-27086822 | -0.019784573 | 0.95311612 | 0.984701606 |
| chr1:217747280-217747508 | -0.02844673 | 0.953116127 | 0.984701606 |
| chr12:9064202-9065435 | -0.018861352 | 0.953125663 | 0.984701606 |
| chr2:55175661-55175972 | -0.024615161 | 0.953200133 | 0.984765264 |
| chr1:110467726-110467983 | -0.025354338 | 0.953234474 | 0.984787465 |
| chr3:71641687-71641959 | -0.030255635 | 0.953275453 | 0.984816522 |
| chr5:177362885-177363480 | 0.019906968 | 0.953289518 | 0.984817774 |
| chr8:109539204-109540636 | -0.018249646 | 0.953311837 | 0.984820544 |
| chr11:17207438-17208605 | -0.018646454 | 0.953317905 | 0.984820544 |
| chr3:175057384-175057662 | 0.029260103 | 0.953348101 | 0.98483272 |
| chr4:143375654-143376052 | -0.022434494 | 0.953355398 | 0.98483272 |
| chr15:96317294-96317556 | -0.027143275 | 0.953394733 | 0.984860076 |
| chr2:47069500-47069965 | -0.019378247 | 0.953418458 | 0.984871306 |
| chr2:127700516-127700930 | -0.019716794 | 0.953443498 | 0.984883894 |
| chr11:65706327-65706601 | -0.024800938 | 0.953464754 | 0.984892573 |
| chr3:43769403-43770304 | 0.019444472 | 0.953512025 | 0.984928124 |
| chr1:170265164-170265364 | -0.031654799 | 0.953610845 | 0.984994084 |
| chr1:145730646-145730909 | 0.024620613 | 0.95362039 | 0.984994084 |
| chr12:46675902-46676115 | -0.027233617 | 0.953626286 | 0.984994084 |
| chr3:81760890-81761968 | -0.018373905 | 0.953635215 | 0.984994084 |
| chr11:129324118-129324424 | -0.023907468 | 0.953660499 | 0.984994084 |
| chr16:75000474-75000690 | 0.022542176 | 0.953667325 | 0.984994084 |
| chr22:36873685-36873920 | -0.026983912 | 0.95367466 | 0.984994084 |
| chr6:131085130-131085330 | 0.03083136 | 0.953687836 | 0.984994084 |
| chr17:9575692-9576054 | -0.022411637 | 0.953691578 | 0.984994084 |
| chr2:232890701-232891222 | 0.021620128 | 0.953721676 | 0.985011892 |
| chr8:128342732-128343149 | -0.021417969 | 0.953755619 | 0.985024711 |
| chr16:11024409-11024710 | -0.028318127 | 0.953799036 | 0.985024711 |
| chr6:33204434-33204796 | 0.020406244 | 0.953815814 | 0.985024711 |
| chr11:72040667-72041333 | 0.019474548 | 0.953823312 | 0.985024711 |
| chr8:30812289-30813086 | -0.019749919 | 0.953845667 | 0.985024711 |
| chr1:44500573-44500874 | 0.025570995 | 0.953845865 | 0.985024711 |
| chr2:42101147-42101560 | 0.021806315 | 0.953859867 | 0.985024711 |
| chr1:31814614-31814883 | -0.025660658 | 0.953873233 | 0.985024711 |
| chr1:96721330-96722566 | 0.018709977 | 0.953875101 | 0.985024711 |
| chr3:123584439-123585864 | 0.018270872 | 0.953897791 | 0.985024711 |

| | | | |
|---|---|---|---|
| chr14:48975112-48975362 | 0.026943539 | 0.953912016 | 0.985024711 |
| chr10:58086861-58087061 | 0.030497979 | 0.953919336 | 0.985024711 |
| chr13:34001037-34001245 | 0.030497979 | 0.953919336 | 0.985024711 |
| chr12:76347948-76349029 | 0.018623225 | 0.953923891 | 0.985024711 |
| chr3:39147093-39147392 | -0.022082882 | 0.953932207 | 0.985024711 |
| chr9:73470867-73471183 | -0.026990038 | 0.95398329 | 0.985024711 |
| chr3:182792847-182794511 | -0.018471073 | 0.9539894 | 0.985024711 |
| chr3:102132373-102132573 | 0.032502629 | 0.953990128 | 0.985024711 |
| chr14:93105095-93105295 | 0.026518701 | 0.95399425 | 0.985024711 |
| chr3:4906218-4906643 | -0.022632543 | 0.953997474 | 0.985024711 |
| chr3:195907156-195907462 | -0.021331847 | 0.954004056 | 0.985024711 |
| chr14:59190013-59190219 | 0.021912513 | 0.954018255 | 0.985026098 |
| chr6:2989453-2990112 | -0.019363698 | 0.954092724 | 0.985072026 |
| chr1:153515557-153516406 | -0.019806984 | 0.954108228 | 0.985072026 |
| chr3:181176694-181177158 | -0.023804322 | 0.954149617 | 0.985072026 |
| chr16:53080785-53080985 | -0.029362752 | 0.954149967 | 0.985072026 |
| chr2:157453898-157454100 | -0.029362752 | 0.954149967 | 0.985072026 |
| chr9:36572573-36573334 | -0.019974015 | 0.954156864 | 0.985072026 |
| chr3:122459068-122459268 | -0.028750863 | 0.954163923 | 0.985072026 |
| chr4:139651358-139651940 | -0.020320551 | 0.954165587 | 0.985072026 |
| chrX:129605445-129606053 | -0.019594725 | 0.954195045 | 0.985089165 |
| chr8:70401953-70403094 | 0.018097272 | 0.954212656 | 0.985094074 |
| chr16:67806436-67807392 | -0.018243719 | 0.954271936 | 0.985113615 |
| chrX:110317786-110318871 | -0.018772822 | 0.954276359 | 0.985113615 |
| chr14:22918593-22919815 | 0.018116962 | 0.954282224 | 0.985113615 |
| chr5:76372614-76372814 | -0.028426288 | 0.954287332 | 0.985113615 |
| chr3:63872266-63872918 | -0.018693546 | 0.954298883 | 0.985113615 |
| chr17:50467418-50467678 | -0.025259871 | 0.954308726 | 0.985113615 |
| chr10:96335029-96335229 | 0.030411164 | 0.954328843 | 0.985121109 |
| chr16:401495-402061 | 0.019462361 | 0.954374542 | 0.98515303 |
| chr6:53811178-53811480 | -0.0228346 | 0.954403833 | 0.98515303 |
| chr4:42568692-42568923 | 0.02607163 | 0.954433582 | 0.98515303 |
| chr3:168467332-168467771 | 0.022645516 | 0.954434556 | 0.98515303 |
| chr9:114662150-114662425 | 0.022230574 | 0.954464451 | 0.98515303 |
| chr6:127459326-127459545 | -0.026363895 | 0.954477289 | 0.98515303 |
| chr5:34474340-34474791 | 0.023367584 | 0.954491576 | 0.98515303 |
| chr4:3780318-3781051 | 0.018772818 | 0.954492583 | 0.98515303 |
| chr14:49962560-49962985 | -0.02033024 | 0.954494665 | 0.98515303 |
| chr2:148874921-148875567 | -0.018664422 | 0.954497519 | 0.98515303 |
| chr18:26550028-26550371 | -0.022172439 | 0.954505112 | 0.98515303 |
| chr4:40055796-40058155 | 0.017954262 | 0.954520471 | 0.98515303 |
| chr19:29476641-29476841 | -0.032034159 | 0.954538125 | 0.98515303 |
| chr1:174281282-174281503 | -0.027662375 | 0.954540955 | 0.98515303 |
| chr19:44643624-44644416 | -0.019793804 | 0.95456094 | 0.98515303 |
| chr3:140848778-140848978 | 0.027781912 | 0.954565483 | 0.98515303 |
| chr17:68614753-68615071 | -0.021187273 | 0.954580184 | 0.985154933 |
| chr6:99514854-99516174 | -0.018031062 | 0.954650247 | 0.98521397 |
| chr18:80257818-80258119 | 0.023173126 | 0.954675436 | 0.985226695 |
| chrX:74536211-74537121 | -0.018744691 | 0.954713149 | 0.985229033 |
| chr1:107936490-107937094 | -0.019608971 | 0.95472747 | 0.985229033 |

| | | | |
|---|---|---|---|
| chr17:49788315-49789445 | 0.017934553 | 0.954727814 | 0.985229033 |
| chr3:194134397-194135304 | 0.01799523 | 0.954729134 | 0.985229033 |
| chr6:31102398-31102674 | -0.026034517 | 0.954762573 | 0.985229218 |
| chr3:63918040-63918251 | -0.020285355 | 0.95476593 | 0.985229218 |
| chr1:94101171-94102028 | 0.019288808 | 0.954767888 | 0.985229218 |
| chr19:2385095-2385624 | 0.020573499 | 0.954797726 | 0.985246739 |
| chr21:42513769-42514035 | -0.022445246 | 0.954843366 | 0.985264671 |
| chr1:1273518-1274061 | 0.021182555 | 0.95485359 | 0.985264671 |
| chr12:52417785-52417985 | 0.029947384 | 0.954859104 | 0.985264671 |
| chr12:117155180-117155764 | 0.018948267 | 0.954866539 | 0.985264671 |
| chr3:183386019-183386280 | 0.024683076 | 0.954885181 | 0.985270638 |
| chr2:197168638-197169019 | 0.021681495 | 0.954900574 | 0.985273253 |
| chr2:126971775-126972503 | -0.019312083 | 0.954938073 | 0.985292431 |
| chr3:128011090-128011547 | -0.019918779 | 0.954956407 | 0.985292431 |
| chr22:35552541-35552764 | -0.026113899 | 0.954957738 | 0.985292431 |
| chr2:237966448-237967187 | -0.019138224 | 0.954996992 | 0.985310069 |
| chr14:60837862-60838116 | 0.028057368 | 0.955000552 | 0.985310069 |
| chr19:38109977-38110177 | -0.030071286 | 0.955053528 | 0.985317174 |
| chr3:113536545-113537761 | -0.017874313 | 0.955057349 | 0.985317174 |
| chr15:67833950-67834286 | 0.023423296 | 0.955064013 | 0.985317174 |
| chr3:13016489-13016689 | 0.024667648 | 0.955090812 | 0.985317174 |
| chr11:79212967-79213298 | 0.02451948 | 0.955092351 | 0.985317174 |
| chr17:4703754-4704431 | -0.019461754 | 0.955095287 | 0.985317174 |
| chr7:6729532-6730503 | 0.018344649 | 0.955097455 | 0.985317174 |
| chr4:88318793-88319308 | -0.019524426 | 0.955116745 | 0.985323809 |
| chr6:53651484-53652392 | -0.018423268 | 0.955130445 | 0.985324676 |
| chr2:12088017-12088292 | -0.028308584 | 0.955166253 | 0.985341227 |
| chr5:58181861-58182079 | -0.029957051 | 0.955172209 | 0.985341227 |
| chr11:67391706-67392317 | 0.018834364 | 0.955238665 | 0.98537667 |
| chr3:88093350-88093681 | -0.023021662 | 0.955249089 | 0.98537667 |
| chr6:157985709-157986479 | 0.018890246 | 0.955251186 | 0.98537667 |
| chr11:46346996-46347606 | 0.018671596 | 0.955259947 | 0.98537667 |
| chr5:19029676-19030413 | -0.020852951 | 0.955281063 | 0.98537667 |
| chr2:47076685-47077440 | 0.019308843 | 0.955295279 | 0.98537667 |
| chr19:45691735-45692773 | 0.017829042 | 0.955312559 | 0.98537667 |
| chr6:53151592-53151951 | 0.024643557 | 0.955324707 | 0.98537667 |
| chr3:49429208-49429858 | 0.020041147 | 0.955334504 | 0.98537667 |
| chr17:75426136-75426821 | -0.018982544 | 0.955346734 | 0.98537667 |
| chr11:10857664-10858872 | -0.018272375 | 0.955364576 | 0.98537667 |
| chr19:33241336-33241858 | -0.020007459 | 0.955384699 | 0.98537667 |
| chr8:138727200-138727765 | -0.019656457 | 0.955418965 | 0.98537667 |
| chr7:84529415-84529738 | -0.023065207 | 0.955424121 | 0.98537667 |
| chr2:15061890-15062112 | 0.028478764 | 0.955426027 | 0.98537667 |
| chr12:54307073-54307464 | 0.020473602 | 0.955431261 | 0.98537667 |
| chr2:27356087-27357397 | 0.017745244 | 0.955434656 | 0.98537667 |
| chr5:17343719-17344073 | -0.021315304 | 0.955442503 | 0.98537667 |
| chr17:47840714-47841535 | 0.018068113 | 0.955475581 | 0.98537667 |
| chr9:95563513-95563786 | -0.030286924 | 0.955518844 | 0.98537667 |
| chr10:102917485-102918639 | 0.017709003 | 0.955531288 | 0.98537667 |
| chr2:42827601-42828131 | 0.019605111 | 0.955546514 | 0.98537667 |

| | | | |
|---|---|---|---|
| chr2:189481132-189481427 | -0.021914007 | 0.955550579 | 0.98537667 |
| chr5:110219671-110219871 | -0.031338377 | 0.955552829 | 0.98537667 |
| chr3:141311425-141312293 | -0.017854265 | 0.955554926 | 0.98537667 |
| chr7:22442841-22443182 | 0.021690408 | 0.955563549 | 0.98537667 |
| chr1:159891604-159891886 | 0.021512789 | 0.955564093 | 0.98537667 |
| chr7:74606133-74606812 | -0.019937427 | 0.955577983 | 0.98537667 |
| chr4:77564290-77564638 | -0.025822224 | 0.955591135 | 0.98537667 |
| chr2:48856216-48856536 | 0.021857664 | 0.955613174 | 0.98537667 |
| chr21:38832828-38833199 | -0.02242382 | 0.955613362 | 0.98537667 |
| chr10:96586048-96586772 | -0.018796526 | 0.955618094 | 0.98537667 |
| chr8:85297406-85297969 | 0.020720256 | 0.95565387 | 0.985387066 |
| chrX:11757653-11758886 | -0.01795516 | 0.955654272 | 0.985387066 |
| chr5:162473388-162473595 | -0.029079464 | 0.955666757 | 0.985387066 |
| chr10:97333825-97334025 | -0.022842515 | 0.955687726 | 0.985395427 |
| chr14:33678161-33678425 | -0.025578836 | 0.955704901 | 0.985399771 |
| chr13:34681962-34682269 | 0.027669758 | 0.955717661 | 0.985399771 |
| chr2:19851151-19851530 | -0.022813975 | 0.955782455 | 0.985445931 |
| chr6:30259165-30260083 | 0.018760353 | 0.955788152 | 0.985445931 |
| chr2:10832350-10832684 | 0.020255576 | 0.955816805 | 0.985462212 |
| chr9:95448115-95448339 | 0.029892226 | 0.955848465 | 0.985481594 |
| chr1:161176830-161177922 | 0.018002991 | 0.955886839 | 0.985495293 |
| chr14:77086383-77086615 | -0.023899754 | 0.955887476 | 0.985495293 |
| chr17:49677394-49678445 | 0.017428218 | 0.955940718 | 0.985536924 |
| chr8:77310409-77310799 | 0.020795643 | 0.955970385 | 0.985549725 |
| chr2:1769786-1770095 | -0.021501191 | 0.955986126 | 0.985549725 |
| chr20:17226412-17227073 | -0.018888074 | 0.955991723 | 0.985549725 |
| chr6:44234724-44234996 | 0.022029447 | 0.956066504 | 0.985613557 |
| chr9:111857542-111857973 | 0.019664592 | 0.956079542 | 0.985613737 |
| chr1:65431194-65431416 | -0.02578126 | 0.956108569 | 0.985620781 |
| chr15:45200130-45201348 | 0.017421713 | 0.956112101 | 0.985620781 |
| chr1:77313478-77313955 | 0.018741493 | 0.956191149 | 0.985663395 |
| chr10:14837682-14839174 | -0.017424785 | 0.956191801 | 0.985663395 |
| chr11:1796309-1797173 | -0.018717395 | 0.956192031 | 0.985663395 |
| chr1:38208649-38208882 | -0.025640799 | 0.956218227 | 0.985675271 |
| chr13:27450936-27451323 | -0.0198391 | 0.956240018 | 0.985675271 |
| chr10:92074157-92074447 | -0.026917558 | 0.956242145 | 0.985675271 |
| chr16:84062794-84063123 | 0.020831335 | 0.956280865 | 0.985701923 |
| chr6:79536800-79537736 | 0.017708246 | 0.956320442 | 0.985729457 |
| chr4:47668952-47669332 | 0.023857934 | 0.956389064 | 0.985786928 |
| chr7:33459267-33459837 | 0.01866669 | 0.956429498 | 0.985815343 |
| chr11:128890494-12889071 | 0.028440864 | 0.956453505 | 0.98581699 |
| chr9:94165800-94167111 | -0.017166955 | 0.956456827 | 0.98581699 |
| chr2:155336214-155336462 | -0.028543068 | 0.956499635 | 0.98584785 |
| chr11:65719853-65720148 | -0.021788169 | 0.956527938 | 0.985851435 |
| chr3:141231206-141232268 | -0.017669534 | 0.956528846 | 0.985851435 |
| chr22:17423320-17423635 | -0.027072583 | 0.956586552 | 0.985868868 |
| chr15:42046469-42046990 | 0.01943079 | 0.956600999 | 0.985868868 |
| chr7:98433361-98433597 | 0.026066385 | 0.95660376 | 0.985868868 |
| chr22:33579520-33579824 | -0.022408178 | 0.9566213 | 0.985868868 |
| chr9:99906425-99907304 | -0.017591412 | 0.956628473 | 0.985868868 |

| | | | |
|---|---|---|---|
| chr19:51657072-51657272 | -0.027717742 | 0.956641866 | 0.985868868 |
| chr2:120552897-120553520 | 0.018424478 | 0.95664314 | 0.985868868 |
| chr8:97553405-97554000 | -0.018703819 | 0.956648693 | 0.985868868 |
| chr13:40993884-40994613 | -0.018000741 | 0.956711969 | 0.985911311 |
| chr1:155192807-155194161 | 0.017338211 | 0.956727943 | 0.985911311 |
| chr5:138575032-138576054 | 0.017289959 | 0.956740728 | 0.985911311 |
| chrX:23742353-23742632 | -0.021492965 | 0.956741348 | 0.985911311 |
| chr1:228739377-228739830 | 0.019985419 | 0.956766135 | 0.985914462 |
| chr1:43969664-43970250 | -0.018042933 | 0.956770139 | 0.985914462 |
| chr6:20127265-20127823 | -0.019778122 | 0.956867363 | 0.985998458 |
| chr2:218658641-218659942 | -0.017348332 | 0.95687739 | 0.985998458 |
| chr2:10302342-10303049 | 0.017667561 | 0.956906788 | 0.986015491 |
| chr19:34255096-34255296 | 0.02549105 | 0.956964124 | 0.98606131 |
| chr16:1303354-1303623 | -0.024140901 | 0.956988096 | 0.986071018 |
| chr11:68950326-68950616 | -0.023272471 | 0.957001576 | 0.986071018 |
| chr8:53863379-53863595 | 0.026649723 | 0.957012153 | 0.986071018 |
| chr8:126556435-126556640 | -0.019307553 | 0.957118021 | 0.986130825 |
| chr17:49585640-49585982 | -0.01917585 | 0.957118256 | 0.986130825 |
| chr1:241519030-241520124 | 0.017082118 | 0.957123075 | 0.986130825 |
| chr13:110712340-110713712 | -0.016912149 | 0.957128066 | 0.986130825 |
| chr8:98944112-98944860 | 0.01737178 | 0.957135607 | 0.986130825 |
| chr5:181217611-181218556 | 0.0170454 | 0.957160238 | 0.986130825 |
| chr8:132924456-132924673 | -0.027438101 | 0.957164698 | 0.986130825 |
| chr19:50325694-50325905 | -0.021472031 | 0.957173158 | 0.986130825 |
| chr3:161559429-161559742 | 0.020708709 | 0.957217882 | 0.986145657 |
| chr10:7219087-7219326 | 0.026357368 | 0.957223913 | 0.986145657 |
| chr12:116390020-116390234 | -0.022377219 | 0.957226166 | 0.986145657 |
| chr5:9523387-9523830 | 0.02141613 | 0.957254577 | 0.98615071 |
| chr17:49092536-49093459 | -0.018463928 | 0.957256811 | 0.98615071 |
| chr2:2464142-2464435 | 0.022641898 | 0.957289866 | 0.986153207 |
| chr2:159711880-159713045 | 0.017128976 | 0.957297154 | 0.986153207 |
| chr11:76860534-76861782 | -0.017171464 | 0.957316922 | 0.986153207 |
| chrX:20530392-20530637 | 0.027374896 | 0.957317395 | 0.986153207 |
| chr10:114471409-114471647 | 0.028265253 | 0.957340518 | 0.986153207 |
| chrX:48003858-48004379 | 0.018491368 | 0.957361477 | 0.986153207 |
| chr1:52403741-52405059 | -0.017101466 | 0.957385242 | 0.986153207 |
| chr1:160416397-160416738 | 0.020101618 | 0.957386159 | 0.986153207 |
| chr1:106961674-106961918 | 0.020206374 | 0.95740216 | 0.986153207 |
| chr10:99443648-99443849 | 0.027855463 | 0.957411477 | 0.986153207 |
| chr17:49035586-49036325 | 0.017953216 | 0.957414577 | 0.986153207 |
| chr2:88627203-88628325 | 0.016808108 | 0.957414606 | 0.986153207 |
| chr17:42798392-42798929 | 0.019322201 | 0.957436119 | 0.986153207 |
| chr22:20420875-20421108 | -0.023591841 | 0.957443825 | 0.986153207 |
| chr22:21681755-21682257 | 0.020390576 | 0.957459846 | 0.986153207 |
| chr18:31942267-31943913 | 0.016849219 | 0.957465161 | 0.986153207 |
| chr9:111980558-111980791 | 0.026739522 | 0.957513161 | 0.986169714 |
| chr14:60980714-60981787 | 0.016890941 | 0.957534823 | 0.986169714 |
| chrX:46447047-46447594 | -0.018899837 | 0.957538559 | 0.986169714 |
| chr5:116557742-116557985 | -0.025367482 | 0.957603607 | 0.986169714 |
| chr5:135403938-135404138 | -0.025367482 | 0.957603607 | 0.986169714 |

| | | | |
|---|---|---|---|
| chr11:126310542-126310810 | -0.022240941 | 0.957615825 | 0.986169714 |
| chr1:240608778-240609046 | 0.023488348 | 0.957644797 | 0.986169714 |
| chr10:92087551-92088086 | -0.019436342 | 0.957647407 | 0.986169714 |
| chr1:75265774-75265993 | 0.025864909 | 0.957655923 | 0.986169714 |
| chr4:151046011-151046270 | 0.02244968 | 0.957692567 | 0.986169714 |
| chr11:7892772-7893055 | -0.024852105 | 0.957693796 | 0.986169714 |
| chr1:1511517-1512274 | -0.018260939 | 0.957738258 | 0.986169714 |
| chr1:206606911-206607186 | 0.019948079 | 0.957738484 | 0.986169714 |
| chr3:141279236-141279662 | -0.019121408 | 0.957751012 | 0.986169714 |
| chr5:179368765-179368965 | -0.02509396 | 0.957766357 | 0.986169714 |
| chr19:1885343-1885666 | 0.02142436 | 0.957767594 | 0.986169714 |
| chr7:149195230-149196185 | -0.017130243 | 0.957780233 | 0.986169714 |
| chr16:67520845-67521827 | -0.017208681 | 0.957788106 | 0.986169714 |
| chr2:28710645-28710845 | -0.02742456 | 0.957788779 | 0.986169714 |
| chr19:16778905-16779118 | -0.02742456 | 0.957788779 | 0.986169714 |
| chr2:48634860-48635073 | -0.02742456 | 0.957788779 | 0.986169714 |
| chr3:5799521-5799721 | -0.02742456 | 0.957788779 | 0.986169714 |
| chr12:53482101-53482303 | -0.025603146 | 0.957791239 | 0.986169714 |
| chr22:25405083-25405283 | 0.025756267 | 0.957791517 | 0.986169714 |
| chr3:71926198-71926464 | -0.022553421 | 0.957802952 | 0.986169714 |
| chr11:73228332-73228591 | -0.022396859 | 0.957817529 | 0.986171471 |
| chr20:49846204-49846481 | -0.024315321 | 0.957838429 | 0.98617262 |
| chr4:85165884-85166226 | -0.022703163 | 0.957844386 | 0.98617262 |
| chr5:149030459-149030739 | 0.023404477 | 0.957888852 | 0.986186036 |
| chr9:89564450-89564785 | -0.02110482 | 0.957894253 | 0.986186036 |
| chr6:16128716-16129840 | -0.016979435 | 0.957896029 | 0.986186036 |
| chr12:131137819-131138079 | 0.022310954 | 0.957934323 | 0.986212209 |
| chr11:3854682-3855758 | -0.017151316 | 0.958019733 | 0.986278979 |
| chr2:173964560-173965007 | -0.017572468 | 0.958024922 | 0.986278979 |
| chr12:120102181-120102409 | -0.027392894 | 0.958063215 | 0.986296562 |
| chr9:77014749-77015118 | 0.021680594 | 0.958080752 | 0.986296562 |
| chr9:32217799-32217999 | 0.02779629 | 0.958122497 | 0.986296562 |
| chr17:39153478-39153958 | 0.019056814 | 0.9581463 | 0.986296562 |
| chr4:102760236-102761579 | -0.016832177 | 0.958158199 | 0.986296562 |
| chr13:99201461-99201715 | 0.020303541 | 0.958160633 | 0.986296562 |
| chr12:122946787-122947058 | 0.021835424 | 0.958164207 | 0.986296562 |
| chr12:105705793-105706061 | 0.02425419 | 0.958165368 | 0.986296562 |
| chr6:35488275-35488475 | 0.022631455 | 0.958166804 | 0.986296562 |
| chr8:144508293-144509157 | 0.016948313 | 0.958170723 | 0.986296562 |
| chr2:187554348-187554572 | -0.024619337 | 0.958204606 | 0.986316693 |
| chr3:113736470-113736873 | 0.0195059 | 0.95822671 | 0.986316693 |
| chr8:61792120-61792320 | 0.025897135 | 0.958228898 | 0.986316693 |
| chr13:24495358-24495746 | -0.020973311 | 0.958262908 | 0.986338449 |
| chr9:136848886-136849820 | -0.01690037 | 0.958282424 | 0.986345287 |
| chr6:160534016-160534535 | 0.019850566 | 0.958304489 | 0.986354749 |
| chr13:50125541-50125817 | 0.019121045 | 0.958321641 | 0.986358233 |
| chr3:49171071-49171281 | -0.024264775 | 0.958333621 | 0.986358233 |
| chr8:139633680-139634001 | 0.021965349 | 0.958383314 | 0.986393149 |
| chr1:169105620-169106837 | -0.016435556 | 0.958395546 | 0.986393149 |
| chr3:112043639-112043980 | 0.019168935 | 0.95842263 | 0.986393149 |

| | | | |
|---|---|---|---|
| chr18:11851151-11851606 | 0.018113066 | 0.958428325 | 0.986393149 |
| chr13:40665548-40665824 | 0.020226895 | 0.958431912 | 0.986393149 |
| chr11:100828348-10082864 | 0.021496243 | 0.958483978 | 0.986433484 |
| chr3:192409294-192410502 | -0.017261992 | 0.958496979 | 0.986433615 |
| chr4:142498568-142499205 | -0.02010818 | 0.958618658 | 0.986537381 |
| chr17:80316952-80317152 | -0.02780304 | 0.95862451 | 0.986537381 |
| chr20:51144383-51144686 | 0.023628992 | 0.958639233 | 0.986537381 |
| chr18:32919834-32920191 | 0.019862948 | 0.958649307 | 0.986537381 |
| chr1:85061857-85062381 | -0.018309515 | 0.958690623 | 0.986546725 |
| chr1:108898167-108898434 | -0.02149279 | 0.958699063 | 0.986546725 |
| chr17:9206939-9207222 | 0.023510082 | 0.95872124 | 0.986546725 |
| chr6:16294527-16294727 | 0.0299417 | 0.958739474 | 0.986546725 |
| chr5:90548244-90548496 | -0.022564249 | 0.958752485 | 0.986546725 |
| chr2:121297513-121297764 | -0.022320866 | 0.958755362 | 0.986546725 |
| chrX:48972392-48972597 | 0.025362406 | 0.958772617 | 0.986546725 |
| chr1:197435506-197435770 | -0.023715144 | 0.958773373 | 0.986546725 |
| chr3:21665444-21665757 | -0.020994182 | 0.958774267 | 0.986546725 |
| chr10:99548054-99548283 | 0.022621013 | 0.958797227 | 0.986557102 |
| chr17:27430958-27431210 | 0.022262047 | 0.958860713 | 0.986602367 |
| chr22:37844193-37844721 | -0.017655596 | 0.958866971 | 0.986602367 |
| chr18:46281220-46281692 | -0.019620134 | 0.95888929 | 0.986611224 |
| chr12:25387553-25387753 | -0.022579126 | 0.958901332 | 0.986611224 |
| chr1:195141365-195141578 | 0.024757785 | 0.958917148 | 0.986614248 |
| chr2:56191658-56191977 | 0.019863971 | 0.959066927 | 0.986747426 |
| chr1:147928221-147928898 | -0.01682997 | 0.959072343 | 0.986747426 |
| chr10:124125509-12412604 | -0.019363341 | 0.959134547 | 0.986798174 |
| chr3:134600874-134601107 | 0.021293114 | 0.959214012 | 0.986846758 |
| chr2:85552153-85552400 | -0.020804501 | 0.959221212 | 0.986846758 |
| chr5:148290666-148290923 | 0.023595268 | 0.959229616 | 0.986846758 |
| chr19:5805819-5806019 | -0.025888642 | 0.959233286 | 0.986846758 |
| chr9:130240664-130240929 | 0.021379156 | 0.959255417 | 0.986851503 |
| chr18:22223656-22224057 | -0.01888865 | 0.95927341 | 0.986851503 |
| chr16:46929763-46930012 | -0.023400721 | 0.959277272 | 0.986851503 |
| chr1:201381004-201381695 | 0.01662131 | 0.959300366 | 0.986851503 |
| chr20:16747489-16747879 | 0.020831387 | 0.959302296 | 0.986851503 |
| chr3:47380308-47381729 | 0.01631586 | 0.959322076 | 0.986852285 |
| chr6:107458890-107460275 | 0.016251638 | 0.959342347 | 0.986852285 |
| chr13:67175878-67176282 | -0.021433772 | 0.959348088 | 0.986852285 |
| chr20:60211056-60211395 | -0.018982173 | 0.959354573 | 0.986852285 |
| chr7:32846594-32847031 | 0.017210881 | 0.959384685 | 0.986870011 |
| chr1:205988754-205988985 | -0.020762187 | 0.959409391 | 0.986882175 |
| chr6:36546932-36547693 | -0.01725356 | 0.95942335 | 0.986883285 |
| chr17:41579550-41579924 | -0.01895858 | 0.959485869 | 0.986927346 |
| chr10:124106476-12410669 | 0.024303594 | 0.959491946 | 0.986927346 |
| chr8:23859382-23859608 | 0.024584134 | 0.959525404 | 0.98693288 |
| chr3:35638990-35639812 | -0.018171764 | 0.959541698 | 0.98693288 |
| chr15:42056966-42057817 | -0.016933215 | 0.959542026 | 0.98693288 |
| chr2:211025884-211026551 | 0.018250234 | 0.959575506 | 0.98693288 |
| chr12:90253892-90254095 | 0.025698385 | 0.959578519 | 0.98693288 |
| chr17:81222844-81223625 | 0.016345346 | 0.959584523 | 0.98693288 |

| | | | |
|---|---|---|---|
| chr3:128133581-128133781 | 0.027507907 | 0.959607911 | 0.98693288 |
| chr6:99467597-99467822 | -0.024844443 | 0.959626408 | 0.98693288 |
| chr19:6463999-6464374 | 0.018392972 | 0.959642481 | 0.98693288 |
| chr1:27365451-27366737 | 0.016219859 | 0.959660868 | 0.98693288 |
| chr2:196592338-196593923 | 0.016274508 | 0.959665619 | 0.98693288 |
| chr1:213931829-213932052 | -0.026676611 | 0.959671477 | 0.98693288 |
| chr2:214388828-214389054 | 0.026672118 | 0.959682547 | 0.98693288 |
| chr11:15770871-15771071 | -0.027229086 | 0.959684972 | 0.98693288 |
| chr2:71007614-71007820 | 0.023241967 | 0.959690534 | 0.98693288 |
| chr10:21327623-21327880 | -0.023378542 | 0.959717888 | 0.986936971 |
| chr7:142901449-142901874 | 0.019181351 | 0.959720273 | 0.986936971 |
| chr8:144575459-144575705 | -0.024061586 | 0.959739279 | 0.986942958 |
| chr11:62545149-62545353 | 0.020759605 | 0.959751857 | 0.986942958 |
| chr20:47972188-47972496 | 0.016899387 | 0.959808468 | 0.986987927 |
| chr3:139541989-139542198 | 0.025301136 | 0.959921166 | 0.987090569 |
| chr12:70049678-70049953 | 0.02214049 | 0.959987159 | 0.987145182 |
| chr1:212665234-212665987 | 0.017088959 | 0.960013717 | 0.987159244 |
| chr8:21683827-21684327 | 0.019822152 | 0.960036696 | 0.987169624 |
| chr19:45717156-45718003 | 0.016308258 | 0.960088236 | 0.987209372 |
| chr11:63606480-63606779 | -0.023234809 | 0.960108887 | 0.987214458 |
| chr7:152108489-152108689 | 0.027564131 | 0.96011895 | 0.987214458 |
| chr4:139875575-139875845 | -0.023062597 | 0.960143449 | 0.9872264 |
| chr1:151606409-151606863 | -0.017968742 | 0.960179487 | 0.987231889 |
| chr2:170715660-170715913 | -0.021837086 | 0.960195314 | 0.987231889 |
| chrX:8153831-8154352 | -0.019354118 | 0.960195398 | 0.987231889 |
| chr22:28873652-28873852 | -0.025849056 | 0.960216492 | 0.987231889 |
| chr1:200983363-200983594 | 0.02099551 | 0.960221824 | 0.987231889 |
| chr8:22089400-22090096 | 0.017907105 | 0.960269651 | 0.987231889 |
| chr5:31854578-31855611 | 0.016488713 | 0.960273924 | 0.987231889 |
| chr11:34633778-34633983 | 0.02325043 | 0.960275761 | 0.987231889 |
| chr11:123054377-123054589 | 0.02325043 | 0.960275761 | 0.987231889 |
| chr16:75254374-75254852 | -0.016350176 | 0.960292219 | 0.987231889 |
| chr6:90497511-90497756 | 0.025257033 | 0.96030051 | 0.987231889 |
| chr8:123154809-123155039 | 0.017933461 | 0.9603034 | 0.987231889 |
| chr7:148575645-148575992 | -0.019277488 | 0.960325743 | 0.987241612 |
| chr5:7754995-7755420 | -0.018458613 | 0.960348488 | 0.987249732 |
| chr3:155023932-155024390 | -0.017580662 | 0.96036192 | 0.987249732 |
| chr9:4662594-4663203 | 0.017650229 | 0.960372295 | 0.987249732 |
| chr1:111989540-111991737 | -0.015474282 | 0.960409031 | 0.987268837 |
| chr3:127574947-127575428 | -0.016900182 | 0.960419582 | 0.987268837 |
| chr18:24119044-24119280 | 0.022445382 | 0.960429535 | 0.987268837 |
| chrX:104566387-104566922 | 0.018392138 | 0.960473838 | 0.987301132 |
| chr13:105904143-105904832 | 0.017532789 | 0.960502172 | 0.987317013 |
| chr3:57165212-57165588 | 0.018500402 | 0.960556174 | 0.987359276 |
| chrX:119981741-119982004 | 0.023527752 | 0.960578417 | 0.987368894 |
| chr14:70359244-70360065 | -0.016444272 | 0.960616013 | 0.987394293 |
| chr8:125556433-125556727 | 0.019901513 | 0.960640133 | 0.987402995 |
| chr14:103980673-103980978 | -0.022131421 | 0.960650862 | 0.987402995 |
| chr8:232184-232565 | 0.018290038 | 0.960664106 | 0.987402995 |
| chr10:97425772-97426752 | -0.015773753 | 0.960698917 | 0.987402995 |

| | | | |
|---|---|---|---|
| chr20:48682601-48683073 | 0.016126506 | 0.960706141 | 0.987402995 |
| chrX:103941641-103941896 | -0.026100263 | 0.960709636 | 0.987402995 |
| chr1:26695958-26696158 | -0.021155211 | 0.960735768 | 0.987402995 |
| chr7:50595534-50595858 | -0.019235152 | 0.960742033 | 0.987402995 |
| chr17:67046560-67046905 | 0.018168865 | 0.960760802 | 0.987402995 |
| chr17:14069045-14070034 | 0.016098943 | 0.960763772 | 0.987402995 |
| chr3:150602864-150604182 | 0.015483296 | 0.960787788 | 0.987402995 |
| chr3:15859364-15860896 | -0.015514263 | 0.960789074 | 0.987402995 |
| chr14:55094645-55094903 | 0.0233056 | 0.960792006 | 0.987402995 |
| chr22:43014966-43015478 | 0.016945469 | 0.96091973 | 0.987503898 |
| chr4:84890380-84890668 | -0.021032082 | 0.9609296 | 0.987503898 |
| chr4:13548986-13549264 | 0.019550088 | 0.960941195 | 0.987503898 |
| chr3:141894656-141894891 | 0.023654578 | 0.960941742 | 0.987503898 |
| chr7:46432562-46433148 | -0.017674459 | 0.960964965 | 0.987514519 |
| chr21:38628488-38628688 | -0.025067226 | 0.961025882 | 0.987563873 |
| chr6:7109019-7109219 | 0.021041923 | 0.961067042 | 0.987592925 |
| chr2:143336857-143337394 | -0.018057149 | 0.961088408 | 0.987595668 |
| chr1:28775150-28775866 | 0.016799408 | 0.961098732 | 0.987595668 |
| chr10:3160704-3160954 | 0.023960513 | 0.961129143 | 0.987595668 |
| chr6:52909942-52910411 | 0.021684187 | 0.961133941 | 0.987595668 |
| chr6:10403639-10404754 | 0.015429302 | 0.961139938 | 0.987595668 |
| chr13:42498536-42498813 | -0.023022963 | 0.961167792 | 0.987595668 |
| chr19:506731-507634 | 0.015790389 | 0.961173675 | 0.987595668 |
| chr15:59104682-59105801 | 0.0157868 | 0.961187117 | 0.987595668 |
| chr8:109450901-109451440 | -0.016454197 | 0.961187985 | 0.987595668 |
| chr6:15455690-15456216 | 0.017572719 | 0.961200177 | 0.987595668 |
| chr10:34814383-34815572 | 0.015962314 | 0.961214539 | 0.987595668 |
| chr3:187220225-187220480 | 0.020979493 | 0.961224381 | 0.987595668 |
| chr7:144419042-144419251 | 0.023244221 | 0.96131185 | 0.987662815 |
| chr8:90859089-90859390 | -0.020965055 | 0.961315516 | 0.987662815 |
| chr10:101843591-101843850 | 0.019892135 | 0.961341994 | 0.987676776 |
| chr10:132711296-132711527 | -0.024613584 | 0.961378332 | 0.987700865 |
| chr15:84587823-84588434 | -0.016392435 | 0.961499371 | 0.987789834 |
| chr18:23281755-23281955 | 0.022735882 | 0.961502649 | 0.987789834 |
| chr14:58166424-58166728 | 0.017971127 | 0.96151587 | 0.987789834 |
| chr8:138102786-138103209 | -0.018293421 | 0.961532194 | 0.987789834 |
| chr5:75775073-75775273 | -0.025161377 | 0.961536786 | 0.987789834 |
| chr12:109311876-109312239 | -0.01895508 | 0.96154228 | 0.987789834 |
| chr1:201714691-201714891 | -0.01953425 | 0.961578154 | 0.987795433 |
| chr5:101103944-101104278 | -0.020204533 | 0.961595342 | 0.987795433 |
| chr19:16829176-16829514 | -0.020524979 | 0.961595813 | 0.987795433 |
| chr6:41920642-41921620 | -0.015675679 | 0.9616118 | 0.987795433 |
| chr11:27499174-27499470 | 0.0205768 | 0.961612189 | 0.987795433 |
| chr8:143231569-143231891 | 0.019610183 | 0.961638183 | 0.987808891 |
| chr20:49219090-49220073 | 0.015408081 | 0.961683974 | 0.987813206 |
| chr9:120439144-120439488 | -0.019102701 | 0.961693403 | 0.987813206 |
| chr5:57299019-57299306 | -0.021783091 | 0.961720622 | 0.987813206 |
| chr21:45941523-45941906 | -0.019342535 | 0.961722358 | 0.987813206 |
| chr12:62983943-62984143 | 0.024144349 | 0.961725421 | 0.987813206 |
| chr10:103800671-103801122 | -0.019491674 | 0.961737902 | 0.987813206 |

| | | | |
|---|---|---|---|
| chr16:15749498-15749841 | -0.02001738 | 0.961738823 | 0.987813206 |
| chr11:32870723-32870971 | 0.02306233 | 0.961745519 | 0.987813206 |
| chr10:96632934-96633394 | -0.016811866 | 0.961792303 | 0.987830976 |
| chr10:122980182-122981059 | -0.015562254 | 0.961802196 | 0.987830976 |
| chr22:41022474-41022902 | -0.017774394 | 0.961805067 | 0.987830976 |
| chr17:67365695-67366929 | 0.015222353 | 0.961814389 | 0.987830976 |
| chr6:10722547-10723258 | -0.015489257 | 0.961858583 | 0.987863124 |
| chr2:112583962-112585005 | -0.015339607 | 0.961915783 | 0.987908628 |
| chr1:44714944-44715207 | 0.021077593 | 0.961954229 | 0.98792413 |
| chr5:57953752-57954072 | 0.020305837 | 0.961956664 | 0.98792413 |
| chr2:241270036-241270441 | -0.018196942 | 0.962028439 | 0.9879846 |
| chrX:9885175-9885438 | 0.021337166 | 0.962085383 | 0.988029838 |
| chr4:86878634-86878931 | -0.020835814 | 0.962175737 | 0.988107118 |
| chr3:184361990-184363783 | -0.014855123 | 0.962186426 | 0.988107118 |
| chr10:84205639-84205930 | 0.017333137 | 0.962221455 | 0.988108979 |
| chr14:96139753-96140100 | 0.020900368 | 0.962240223 | 0.988108979 |
| chrX:48911229-48911429 | 0.021067889 | 0.962243143 | 0.988108979 |
| chr5:50668289-50668494 | 0.021742556 | 0.962248578 | 0.988108979 |
| chr14:54329155-54329470 | 0.018615502 | 0.96226832 | 0.988108979 |
| chr1:229715047-229715310 | 0.022310385 | 0.962286427 | 0.988108979 |
| chr17:80670929-80671251 | 0.022310385 | 0.962286427 | 0.988108979 |
| chr15:84035428-84035628 | -0.023198828 | 0.962323422 | 0.988108979 |
| chr8:102996712-102996924 | 0.023307543 | 0.962324969 | 0.988108979 |
| chr6:149749492-149750693 | -0.015044993 | 0.962331649 | 0.988108979 |
| chr17:34892943-34893173 | -0.025868404 | 0.96234114 | 0.988108979 |
| chr1:77448515-77448762 | 0.021355456 | 0.962344628 | 0.988108979 |
| chr11:17528167-17528468 | 0.018806224 | 0.962355884 | 0.988108979 |
| chr11:82174673-82174945 | -0.0229192 | 0.962376425 | 0.988116828 |
| chr10:51389339-51389698 | 0.019225806 | 0.962424349 | 0.988152792 |
| chr13:20495148-20495403 | 0.019262855 | 0.962492742 | 0.988209772 |
| chr8:121650889-121651089 | 0.025406221 | 0.962533605 | 0.988238484 |
| chr1:44294834-44295129 | -0.021004114 | 0.96255847 | 0.988250772 |
| chr19:54159387-54160244 | 0.01503524 | 0.962607833 | 0.988285544 |
| chr14:81533164-81534319 | -0.014743931 | 0.962621818 | 0.988285544 |
| chr6:26286334-26286632 | 0.017674383 | 0.96263664 | 0.988285544 |
| chr5:178361333-178361604 | 0.021516119 | 0.962686627 | 0.988285544 |
| chr12:59027245-59027520 | -0.021238797 | 0.962687575 | 0.988285544 |
| chr12:69676402-69676654 | 0.021514562 | 0.962697772 | 0.988285544 |
| chr3:170506636-170506870 | 0.022141128 | 0.962699309 | 0.988285544 |
| chr2:213175653-213176216 | -0.017701205 | 0.96271305 | 0.988285544 |
| chr13:22967797-22967997 | 0.022141338 | 0.962737677 | 0.988285544 |
| chr4:105520930-105521149 | 0.021734304 | 0.962738208 | 0.988285544 |
| chr2:100320774-100321793 | 0.015893485 | 0.962765827 | 0.988285544 |
| chr11:31958165-31958449 | -0.019751639 | 0.96276729 | 0.988285544 |
| chr6:151847073-151847305 | -0.025012679 | 0.962770803 | 0.988285544 |
| chr12:7638789-7639001 | 0.022034032 | 0.962805117 | 0.988285544 |
| chr10:24196522-24196760 | -0.021705874 | 0.962820077 | 0.988285544 |
| chr6:106608104-106608440 | -0.019801809 | 0.962839761 | 0.988285544 |
| chr3:38471365-38471614 | -0.024930492 | 0.962852619 | 0.988285544 |
| chr2:49687914-49688178 | 0.023488428 | 0.962852987 | 0.988285544 |

| | | | |
|---|---|---|---|
| chr15:82106938-82107178 | 0.022657537 | 0.962853549 | 0.988285544 |
| chr3:8530557-8530803 | -0.022720954 | 0.962859032 | 0.988285544 |
| chr1:59903248-59903448 | -0.022546828 | 0.962863201 | 0.988285544 |
| chr14:49324367-49324640 | 0.022217472 | 0.962890701 | 0.988295433 |
| chr19:2315938-2316191 | 0.022549821 | 0.962913701 | 0.988295433 |
| chr20:44531716-44532124 | -0.017737524 | 0.962919782 | 0.988295433 |
| chrX:123207837-123208095 | -0.020650354 | 0.962924429 | 0.988295433 |
| chr6:111957956-111958186 | -0.024260396 | 0.962939547 | 0.988297712 |
| chr17:28227363-28228110 | 0.015254241 | 0.962964153 | 0.988303387 |
| chr9:109151262-109151491 | 0.022964664 | 0.962970873 | 0.988303387 |
| chr19:17766617-17766890 | -0.021426181 | 0.962998335 | 0.988318333 |
| chr5:168076779-168077072 | -0.020104515 | 0.963038083 | 0.988345888 |
| chr15:76346699-76347043 | 0.020292212 | 0.963053547 | 0.98834852 |
| chr3:167032342-167032696 | 0.017860166 | 0.963112158 | 0.988395433 |
| chr3:63847573-63848292 | 0.014873169 | 0.963186841 | 0.988438267 |
| chr19:55407218-55408525 | 0.014774812 | 0.963189616 | 0.988438267 |
| chr5:143404765-143405743 | 0.014985186 | 0.963198967 | 0.988438267 |
| chr4:88823080-88823915 | 0.015333871 | 0.963205498 | 0.988438267 |
| chr9:21559835-21560044 | 0.023070858 | 0.963227318 | 0.98844742 |
| chr2:172455122-172455422 | 0.01901881 | 0.963270864 | 0.988478869 |
| chr10:132571946-132572545 | 0.016250457 | 0.963286689 | 0.988479479 |
| chr1:73401780-73401988 | 0.025874208 | 0.963297261 | 0.988479479 |
| chr1:146376362-146377632 | 0.014528748 | 0.963335043 | 0.988505011 |
| chr1:26190345-26190545 | -0.024146635 | 0.963429537 | 0.988588735 |
| chr12:106736493-106736748 | 0.020465197 | 0.963480326 | 0.98862523 |
| chr1:224142676-224143421 | 0.016541596 | 0.963490909 | 0.98862523 |
| chr21:41568600-41568800 | -0.02479716 | 0.963532691 | 0.988636414 |
| chr6:128327957-128328355 | -0.016922437 | 0.963556521 | 0.988636414 |
| chr5:179820553-179821985 | 0.014479825 | 0.963557261 | 0.988636414 |
| chr3:149530727-149530965 | -0.019561075 | 0.963564264 | 0.988636414 |
| chr2:218301326-218301672 | 0.017895195 | 0.963566323 | 0.988636414 |
| chr7:7004623-7004885 | -0.022271002 | 0.963620149 | 0.98866332 |
| chr1:156572370-156572823 | 0.015846822 | 0.963627251 | 0.98866332 |
| chr22:31753545-31754212 | 0.015170234 | 0.963635008 | 0.98866332 |
| chr11:74988474-74989410 | 0.014886028 | 0.963650178 | 0.98866332 |
| chr12:120335470-120335813 | 0.017265572 | 0.963657062 | 0.98866332 |
| chr10:97736694-97737992 | 0.014507188 | 0.963682436 | 0.988669691 |
| chr3:123365752-123366057 | 0.017621592 | 0.963689078 | 0.988669691 |
| chr6:42879307-42880253 | 0.015021929 | 0.963749823 | 0.988718772 |
| chr1:91404145-91404905 | 0.01639381 | 0.963773684 | 0.988730013 |
| chr5:33338789-33339014 | -0.023496838 | 0.963815126 | 0.988746747 |
| chr15:82985686-82986588 | -0.014952817 | 0.963835641 | 0.988746747 |
| chr14:47845837-47846287 | -0.019631814 | 0.963835858 | 0.988746747 |
| chr15:32615286-32616097 | -0.016150368 | 0.963866955 | 0.988746747 |
| chr15:78076863-78077401 | 0.016534441 | 0.963867822 | 0.988746747 |
| chr1:246503298-246503816 | -0.015007516 | 0.963871251 | 0.988746747 |
| chr5:178075891-178076475 | -0.014992045 | 0.963881035 | 0.988746747 |
| chr6:170305990-170307189 | 0.014415047 | 0.96392411 | 0.988746747 |
| chr13:65553130-65553401 | -0.023078692 | 0.963941288 | 0.988746747 |
| chr16:23359523-23359786 | -0.023078692 | 0.963941288 | 0.988746747 |

| | | | |
|---|---|---|---|
| chr10:128078546-128078761 | -0.020110661 | 0.963942448 | 0.988746747 |
| chr22:28679739-28680159 | 0.017638189 | 0.963944846 | 0.988746747 |
| chr2:27096321-27096528 | 0.019514519 | 0.963976384 | 0.988750393 |
| chr20:51680937-51681206 | 0.01833089 | 0.964029006 | 0.988750393 |
| chr2:191245080-191246399 | 0.014391518 | 0.964082793 | 0.988750393 |
| chrX:74382735-74383022 | -0.019016565 | 0.96410475 | 0.988750393 |
| chr1:214910176-214910439 | -0.019480464 | 0.964112577 | 0.988750393 |
| chr3:98522156-98522450 | -0.015999924 | 0.964118367 | 0.988750393 |
| chr22:22322079-22322283 | 0.02233976 | 0.964126399 | 0.988750393 |
| chr15:48963519-48963799 | -0.018033007 | 0.964139434 | 0.988750393 |
| chr8:64428907-64429172 | -0.021194069 | 0.964142559 | 0.988750393 |
| chr7:41696457-41696996 | -0.016841214 | 0.96415212 | 0.988750393 |
| chr12:132887055-132888441 | -0.01443179 | 0.964166397 | 0.988750393 |
| chr1:112358790-112359183 | -0.016778972 | 0.964172373 | 0.988750393 |
| chr3:34012037-34012453 | 0.016713758 | 0.964177962 | 0.988750393 |
| chr16:82091510-82091923 | 0.019464981 | 0.964186241 | 0.988750393 |
| chr4:38318792-38319289 | 0.015559584 | 0.96419061 | 0.988750393 |
| chr16:5025357-5025698 | -0.01794917 | 0.964190643 | 0.988750393 |
| chr6:57970627-57970868 | -0.021233161 | 0.964193018 | 0.988750393 |
| chr12:86668947-86669537 | -0.016125329 | 0.964194809 | 0.988750393 |
| chr14:64879871-64880887 | 0.015115875 | 0.964204561 | 0.988750393 |
| chr19:14363704-14364000 | -0.019519863 | 0.964206486 | 0.988750393 |
| chr13:110610442-110610642 | 0.023038759 | 0.964227955 | 0.988753593 |
| chr7:34165995-34166308 | 0.017250843 | 0.96424732 | 0.988753593 |
| chr8:141346597-141347267 | 0.01527168 | 0.96424832 | 0.988753593 |
| chr11:69554088-69554288 | -0.023170811 | 0.964321068 | 0.988814958 |
| chr12:98424814-98425138 | 0.016862811 | 0.964359377 | 0.988841006 |
| chrX:120244660-120245524 | 0.014979019 | 0.964425325 | 0.988892922 |
| chr16:89037631-89037831 | -0.019865343 | 0.964450745 | 0.988892922 |
| chr1:145867790-145868116 | 0.017979701 | 0.964464389 | 0.988892922 |
| chr1:206202849-206203635 | -0.014175614 | 0.96448265 | 0.988892922 |
| chr5:140564125-140565014 | -0.014221621 | 0.96450686 | 0.988892922 |
| chr2:119678718-119679488 | 0.01493025 | 0.96450868 | 0.988892922 |
| chr14:78767420-78767689 | 0.021194654 | 0.964552496 | 0.988892922 |
| chr3:98901057-98902184 | 0.013980128 | 0.964562984 | 0.988892922 |
| chr2:10724166-10724366 | 0.02418469 | 0.964569113 | 0.988892922 |
| chr6:7590030-7591241 | 0.014538043 | 0.964579424 | 0.988892922 |
| chr22:41620240-41621438 | 0.014255954 | 0.964620716 | 0.988892922 |
| chr16:67892869-67893570 | 0.015171337 | 0.964624169 | 0.988892922 |
| chr8:96313168-96313388 | 0.018475186 | 0.964630861 | 0.988892922 |
| chr5:16475289-16475489 | -0.024077966 | 0.964656648 | 0.988892922 |
| chr7:5710553-5710876 | 0.020269137 | 0.964660699 | 0.988892922 |
| chr2:167845062-167845371 | -0.019862461 | 0.964665581 | 0.988892922 |
| chr3:139329289-139329805 | -0.015384749 | 0.96468005 | 0.988892922 |
| chr1:205701069-205701440 | -0.017102592 | 0.964687158 | 0.988892922 |
| chr7:5487790-5488166 | 0.015853368 | 0.964687842 | 0.988892922 |
| chr19:48330150-48330410 | -0.016453913 | 0.964690003 | 0.988892922 |
| chrX:49171637-49172789 | 0.014452875 | 0.964701686 | 0.988892922 |
| chr21:33587967-33589327 | 0.014347744 | 0.964704577 | 0.988892922 |
| chr14:83071981-83072221 | -0.019982992 | 0.964730691 | 0.988892922 |

| | | | |
|---|---|---|---|
| chr20:49625511-49626206 | -0.015547317 | 0.964758796 | 0.988892922 |
| chr20:54668227-54669295 | 0.013857697 | 0.964761362 | 0.988892922 |
| chr11:67421045-67421699 | 0.01454521 | 0.964775775 | 0.988892922 |
| chr6:26272652-26273295 | -0.014609172 | 0.964783475 | 0.988892922 |
| chr14:30752063-30752272 | 0.021320884 | 0.964816201 | 0.988892922 |
| chr18:3585856-3586229 | 0.016546593 | 0.96481767 | 0.988892922 |
| chr1:111976435-111977242 | -0.014411935 | 0.964818891 | 0.988892922 |
| chr16:58129114-58129843 | -0.014446916 | 0.964820527 | 0.988892922 |
| chr6:4664421-4664621 | 0.025646286 | 0.964823003 | 0.988892922 |
| chr2:74690660-74690888 | 0.021626415 | 0.964868541 | 0.988918647 |
| chr20:2508515-2509540 | 0.015421672 | 0.964873915 | 0.988918647 |
| chr8:121145287-121145979 | -0.014551319 | 0.964901252 | 0.988920653 |
| chr11:93236848-93237065 | -0.022574705 | 0.96492123 | 0.988920653 |
| chr6:144415052-144415386 | 0.016496541 | 0.964928176 | 0.988920653 |
| chr10:132223130-132223485 | 0.016141733 | 0.964955598 | 0.988920653 |
| chr1:26889813-26890684 | 0.015692479 | 0.964967454 | 0.988920653 |
| chrX:101602975-101603276 | -0.023511393 | 0.964972684 | 0.988920653 |
| chr6:27753062-27753620 | 0.015094752 | 0.96497846 | 0.988920653 |
| chr3:173214636-173214986 | -0.017964906 | 0.964979124 | 0.988920653 |
| chr16:80974694-80975098 | -0.018175663 | 0.965042847 | 0.988972316 |
| chr8:67028215-67029020 | 0.014643891 | 0.965065315 | 0.988972316 |
| chr4:147786229-147786429 | -0.021254339 | 0.965068258 | 0.988972316 |
| chr15:80235295-80235512 | 0.022547726 | 0.965102879 | 0.988994567 |
| chr9:117556324-117556570 | -0.019451276 | 0.965136958 | 0.989000305 |
| chr12:6612899-6614507 | -0.013822898 | 0.965139055 | 0.989000305 |
| chr11:76598606-76598807 | -0.019669175 | 0.965149553 | 0.989000305 |
| chr14:77965931-77966192 | -0.019619264 | 0.965160108 | 0.989000305 |
| chr1:110034344-110035544 | -0.014180824 | 0.965199391 | 0.989027332 |
| chr12:42588859-42589259 | 0.017293688 | 0.965255809 | 0.989057374 |
| chr8:123178971-123179176 | 0.017364341 | 0.965279671 | 0.989057374 |
| chr3:62755432-62755694 | -0.01701078 | 0.965290378 | 0.989057374 |
| chr14:100440246-100440680 | -0.017369305 | 0.965296508 | 0.989057374 |
| chr6:30066607-30067672 | -0.013908556 | 0.965304819 | 0.989057374 |
| chr9:38392423-38393393 | -0.014588358 | 0.965306159 | 0.989057374 |
| chrX:57528720-57529182 | 0.017985418 | 0.965350482 | 0.989089561 |
| chr3:137032433-137032804 | -0.015695729 | 0.965409433 | 0.98912775 |
| chr3:146928692-146928892 | 0.021263966 | 0.96541495 | 0.98912775 |
| chr2:47520924-47521756 | 0.014687257 | 0.965442421 | 0.98912775 |
| chr3:192303357-192303569 | 0.021405821 | 0.965496807 | 0.98912775 |
| chr8:134691101-134691638 | 0.014847294 | 0.965515713 | 0.98912775 |
| chr8:73400415-73400671 | 0.022476475 | 0.965516787 | 0.98912775 |
| chr15:66588605-66589479 | -0.01483364 | 0.965516955 | 0.98912775 |
| chr6:161393855-161394168 | -0.0188676 | 0.965533215 | 0.98912775 |
| chr12:109432732-109433102 | -0.01605782 | 0.965557195 | 0.98912775 |
| chr11:111454691-111454943 | 0.019814788 | 0.965557948 | 0.98912775 |
| chr17:47949298-47949528 | -0.016608264 | 0.965559678 | 0.98912775 |
| chr10:1074975-1075456 | 0.017031567 | 0.965562942 | 0.98912775 |
| chr4:118798747-118798963 | 0.021883253 | 0.96556489 | 0.98912775 |
| chr2:175329053-175329447 | -0.016507616 | 0.96556853 | 0.98912775 |
| chr15:92694083-92694380 | 0.017540882 | 0.965581392 | 0.98912775 |

| | | | |
|---|---|---|---|
| chr1:78237594-78237919 | 0.016578768 | 0.965639044 | 0.989162122 |
| chr5:171365374-171365663 | 0.017749472 | 0.965648244 | 0.989162122 |
| chr4:162131408-162131662 | 0.020649589 | 0.965653675 | 0.989162122 |
| chr5:151646433-151646732 | 0.01765141 | 0.965710545 | 0.989187537 |
| chr8:97560067-97560503 | -0.015609262 | 0.965714406 | 0.989187537 |
| chr17:45133202-45133426 | 0.017464758 | 0.965733337 | 0.989187537 |
| chr10:70232405-70232752 | -0.017157169 | 0.965748739 | 0.989187537 |
| chr10:87635603-87636038 | -0.017338222 | 0.965769316 | 0.989187537 |
| chr19:39083956-39084694 | -0.015636099 | 0.965773711 | 0.989187537 |
| chr18:31497544-31498677 | 0.013940327 | 0.965775647 | 0.989187537 |
| chr12:48956461-48957927 | -0.013777786 | 0.965788294 | 0.989187537 |
| chr1:198830062-198830308 | 0.018974817 | 0.965827608 | 0.989187537 |
| chr11:69444524-69444808 | 0.017548089 | 0.965831039 | 0.989187537 |
| chr2:169715334-169715977 | -0.01540083 | 0.965831106 | 0.989187537 |
| chr5:58188214-58188609 | -0.016950276 | 0.96583426 | 0.989187537 |
| chr8:141115145-141115496 | 0.016128518 | 0.965846315 | 0.989187537 |
| chr19:32196944-32197186 | -0.022292035 | 0.965882539 | 0.989211413 |
| chr15:85837217-85837460 | -0.019314987 | 0.96591602 | 0.989223661 |
| chr13:42041633-42042045 | 0.014935279 | 0.965928764 | 0.989223661 |
| chr1:114443930-114444409 | 0.014628442 | 0.965933229 | 0.989223661 |
| chr1:16951649-16951959 | 0.017622848 | 0.965966104 | 0.989232444 |
| chr17:29580342-29580658 | 0.015701378 | 0.96596925 | 0.989232444 |
| chr16:82169765-82170787 | -0.013737442 | 0.965990315 | 0.989232444 |
| chr13:100594791-100595077 | 0.020982952 | 0.966002716 | 0.989232444 |
| chr6:73970087-73970287 | 0.02110905 | 0.966014918 | 0.989232444 |
| chr7:74084632-74085021 | 0.017397141 | 0.966019268 | 0.989232444 |
| chr19:48782493-48782762 | 0.017496957 | 0.966068309 | 0.989260085 |
| chr19:7605479-7605922 | -0.016515663 | 0.966087044 | 0.989260085 |
| chr1:31296454-31297315 | -0.014034416 | 0.966094698 | 0.989260085 |
| chr5:162128314-162128539 | -0.020033508 | 0.966097905 | 0.989260085 |
| chr1:144532122-144532612 | -0.016924136 | 0.966164281 | 0.989297783 |
| chr2:213923479-213923858 | 0.018948502 | 0.966168894 | 0.989297783 |
| chr12:104957703-104957918 | 0.018542543 | 0.966173454 | 0.989297783 |
| chr19:1074577-1075311 | -0.015133048 | 0.966196745 | 0.989305601 |
| chr16:74999730-75000362 | -0.014126444 | 0.966211586 | 0.989305601 |
| chr6:27795807-27796155 | 0.014805221 | 0.966223899 | 0.989305601 |
| chr1:2554929-2555129 | 0.023721449 | 0.966232735 | 0.989305601 |
| chr1:110116526-110116805 | -0.016266936 | 0.966260688 | 0.989321001 |
| chr9:113221022-113221914 | -0.013619322 | 0.966288967 | 0.989335782 |
| chr6:167791386-167791876 | -0.016609306 | 0.966300948 | 0.989335782 |
| chr8:41528742-41529524 | -0.013796439 | 0.966314712 | 0.989336654 |
| chr9:93133612-93133864 | 0.017047554 | 0.966377643 | 0.989374297 |
| chr6:139173332-139173532 | -0.021932693 | 0.966385961 | 0.989374297 |
| chr8:38067166-38067366 | 0.021175066 | 0.966401548 | 0.989374297 |
| chr5:113379010-113379262 | 0.017968736 | 0.966403128 | 0.989374297 |
| chr5:139648144-139648424 | -0.015407661 | 0.966440168 | 0.989398998 |
| chr8:126809938-126810234 | 0.017610864 | 0.966454351 | 0.989400298 |
| chr15:26082058-26082871 | 0.013839809 | 0.966475223 | 0.989400331 |
| chr3:10362183-10362383 | -0.020450956 | 0.966484506 | 0.989400331 |
| chr3:20459632-20459832 | 0.024173104 | 0.966493122 | 0.989400331 |

| | | | |
|---|---|---|---|
| chr14:20609290-20609589 | 0.014331876 | 0.966526258 | 0.989421033 |
| chr2:193960946-193961154 | 0.021558584 | 0.966548581 | 0.989424632 |
| chr1:31902206-31902433 | 0.021548391 | 0.966564142 | 0.989424632 |
| chr18:79950767-79951267 | 0.015185692 | 0.966601312 | 0.989424632 |
| chr1:1115851-1116619 | -0.014131561 | 0.96661143 | 0.989424632 |
| chr16:35432202-35432421 | 0.022469573 | 0.966615255 | 0.989424632 |
| chr4:88706570-88706858 | 0.018000976 | 0.966617397 | 0.989424632 |
| chr17:72592337-72593031 | 0.014010424 | 0.966620165 | 0.989424632 |
| chr1:203519468-203519735 | -0.016808519 | 0.966704692 | 0.989465792 |
| chr3:159331184-159331436 | 0.021903517 | 0.966705726 | 0.989465792 |
| chr1:155172716-155173009 | 0.015279127 | 0.966712457 | 0.989465792 |
| chr15:99733072-99733933 | 0.013384713 | 0.96671545 | 0.989465792 |
| chr3:162769501-162769732 | 0.022675161 | 0.966729705 | 0.989465792 |
| chr1:93585098-93585368 | 0.018025164 | 0.966750619 | 0.989465792 |
| chrX:10062903-10063278 | -0.017256794 | 0.966753861 | 0.989465792 |
| chr17:16438319-16439132 | 0.013354314 | 0.966768263 | 0.989465792 |
| chr2:108350149-108350367 | -0.01927583 | 0.966776598 | 0.989465792 |
| chr12:69966930-69967134 | 0.02092141 | 0.966796089 | 0.989472523 |
| chr7:66740295-66741582 | 0.013345889 | 0.966817179 | 0.989480891 |
| chr1:244719826-244720055 | 0.020207596 | 0.96685071 | 0.989501991 |
| chr17:7437001-7437318 | 0.015939048 | 0.966882888 | 0.989521707 |
| chr16:66324849-66325118 | 0.018126174 | 0.966941608 | 0.989541484 |
| chr11:9314085-9314285 | 0.019073938 | 0.966950754 | 0.989541484 |
| chr6:33313778-33314340 | -0.014432595 | 0.966958498 | 0.989541484 |
| chr7:4236618-4236900 | -0.018797446 | 0.96696339 | 0.989541484 |
| chr8:8386083-8387372 | -0.01371218 | 0.966966786 | 0.989541484 |
| chr6:144036304-144036639 | 0.018882864 | 0.967003544 | 0.989565884 |
| chr14:95524112-95524554 | -0.014939983 | 0.967018189 | 0.989567654 |
| chr6:169452186-169452689 | 0.016073765 | 0.967111432 | 0.989635324 |
| chr12:131784023-131784418 | -0.017264901 | 0.967142252 | 0.989635324 |
| chr3:48240899-48241541 | 0.014294154 | 0.967207793 | 0.989635324 |
| chr1:197222252-197222766 | -0.014110995 | 0.967214368 | 0.989635324 |
| chr1:205610090-205610393 | 0.018913068 | 0.967221164 | 0.989635324 |
| chr7:36440393-36440620 | -0.019908286 | 0.967228333 | 0.989635324 |
| chr7:34451447-34451719 | -0.020310295 | 0.967233351 | 0.989635324 |
| chr10:73874359-73874861 | -0.014411612 | 0.967233369 | 0.989635324 |
| chr3:151468319-151468654 | 0.015396068 | 0.967242454 | 0.989635324 |
| chr7:113742855-113743489 | -0.015520938 | 0.967243199 | 0.989635324 |
| chr6:28136707-28137143 | 0.015088915 | 0.967245202 | 0.989635324 |
| chr1:185799633-185799833 | -0.020803101 | 0.967262441 | 0.989635324 |
| chr5:749365-749608 | 0.021531627 | 0.967264786 | 0.989635324 |
| chr14:81220713-81221638 | 0.01291466 | 0.967265138 | 0.989635324 |
| chr5:178875592-178876145 | -0.016267734 | 0.967327771 | 0.989672979 |
| chr17:82417982-82419018 | 0.013342769 | 0.967335762 | 0.989672979 |
| chr5:13621244-13621478 | 0.020970523 | 0.967340691 | 0.989672979 |
| chr21:44338229-44338617 | -0.014619704 | 0.96740037 | 0.989697781 |
| chr4:986870-987107 | 0.017059068 | 0.967406247 | 0.989697781 |
| chr2:35391554-35391796 | -0.020967861 | 0.967410143 | 0.989697781 |
| chr6:136610331-136610531 | 0.019908551 | 0.967424669 | 0.989697781 |
| chr1:160085645-160085914 | 0.017003955 | 0.967434963 | 0.989697781 |

| | | | |
|---|---|---|---|
| chr1:200935756-200936148 | -0.015867848 | 0.967454115 | 0.989697781 |
| chr1:44213248-44214085 | -0.014349362 | 0.967481499 | 0.989697781 |
| chr5:97937888-97938284 | 0.016200509 | 0.967504426 | 0.989697781 |
| chr7:76282444-76282963 | 0.016179134 | 0.96751465 | 0.989697781 |
| chrX:81122087-81122311 | -0.018430803 | 0.967518375 | 0.989697781 |
| chr5:54904241-54904644 | 0.016273739 | 0.967526612 | 0.989697781 |
| chr6:36139347-36139652 | 0.017615045 | 0.967541691 | 0.989697781 |
| chr1:232387129-232387379 | -0.021283115 | 0.967548953 | 0.989697781 |
| chr20:31572549-31573727 | -0.012866477 | 0.967556218 | 0.989697781 |
| chr10:128541430-128541719 | -0.01916736 | 0.967558683 | 0.989697781 |
| chr2:111231664-111231905 | -0.020755812 | 0.967585586 | 0.98969823 |
| chr17:21295209-21295610 | -0.015372152 | 0.967629558 | 0.98969823 |
| chr4:125315656-125315856 | -0.016969486 | 0.967633684 | 0.98969823 |
| chr16:87203175-87203594 | -0.015377915 | 0.967634123 | 0.98969823 |
| chr1:229625009-229625209 | 0.018628047 | 0.967634747 | 0.98969823 |
| chr2:196232917-196233371 | -0.013912893 | 0.967636621 | 0.98969823 |
| chr9:122463814-122464761 | 0.014008868 | 0.967651196 | 0.989699926 |
| chr8:93250598-93250839 | 0.020149694 | 0.967697087 | 0.989722331 |
| chrX:7170997-7171197 | 0.022217937 | 0.967715862 | 0.989722331 |
| chr5:166341759-166341998 | -0.019811289 | 0.967724755 | 0.989722331 |
| chr8:30743829-30744558 | 0.013810934 | 0.967730486 | 0.989722331 |
| chr15:73053003-73053443 | 0.01460194 | 0.967737687 | 0.989722331 |
| chr8:106591873-106592108 | 0.017338129 | 0.967758241 | 0.989730142 |
| chr9:136790687-136791564 | 0.012834879 | 0.967791726 | 0.989744654 |
| chr12:75505874-75506074 | 0.022199801 | 0.96781407 | 0.989744654 |
| chr3:42831373-42831612 | -0.018989907 | 0.96782758 | 0.989744654 |
| chr15:37289157-37289683 | -0.016165298 | 0.967843183 | 0.989744654 |
| chr6:145028060-145028555 | 0.016685651 | 0.967847304 | 0.989744654 |
| chrX:15335012-15335917 | 0.013009138 | 0.967853899 | 0.989744654 |
| chr17:30708152-30709107 | 0.013607922 | 0.96787081 | 0.989744654 |
| chr13:19863177-19864978 | 0.01260421 | 0.967875769 | 0.989744654 |
| chr10:91206545-91206745 | 0.020654317 | 0.967927923 | 0.989784776 |
| chr10:119103787-119104975 | 0.012933331 | 0.967941341 | 0.989785288 |
| chr13:30216192-30216392 | 0.02113761 | 0.968053005 | 0.989882051 |
| chr12:123711917-123712931 | 0.013212967 | 0.968065105 | 0.989882051 |
| chr15:81281928-81282539 | -0.013788472 | 0.968082082 | 0.989882051 |
| chr17:59853143-59853754 | -0.012628475 | 0.968108428 | 0.989882051 |
| chr4:52751315-52751898 | -0.014054689 | 0.968115728 | 0.989882051 |
| chr17:22273841-22274041 | -0.020726257 | 0.968124682 | 0.989882051 |
| chr2:208820272-208820586 | -0.016666335 | 0.968126402 | 0.989882051 |
| chr1:77759006-77760106 | 0.01288601 | 0.968186153 | 0.989899803 |
| chr17:29615346-29615878 | 0.014026595 | 0.968200426 | 0.989899803 |
| chr8:11409500-11409713 | -0.020328867 | 0.968248257 | 0.989899803 |
| chr5:138503639-138504183 | 0.014744471 | 0.968253067 | 0.989899803 |
| chr3:128152952-128154011 | 0.013098018 | 0.968272301 | 0.989899803 |
| chr12:64672880-64673179 | -0.015714637 | 0.968301001 | 0.989899803 |
| chr11:27700898-27701098 | -0.018929862 | 0.96831309 | 0.989899803 |
| chr6:11144420-11144637 | -0.019129046 | 0.968322657 | 0.989899803 |
| chr6:112780536-112780769 | -0.019129046 | 0.968322657 | 0.989899803 |
| chr8:108162049-108162249 | -0.021799896 | 0.968354734 | 0.989899803 |

| | | | |
|---|---|---|---|
| chr13:30755699-30755935 | -0.017746566 | 0.968359029 | 0.989899803 |
| chr7:131844532-131844786 | 0.019396439 | 0.968367087 | 0.989899803 |
| chr20:62525453-62525725 | -0.016559281 | 0.968381602 | 0.989899803 |
| chr6:39114609-39115568 | 0.012942989 | 0.968383923 | 0.989899803 |
| chr12:122399942-122400187 | 0.01578692 | 0.968395696 | 0.989899803 |
| chr9:99221428-99222786 | -0.012590092 | 0.968404329 | 0.989899803 |
| chrX:71824836-71825051 | -0.022100606 | 0.96842279 | 0.989899803 |
| chr6:36758605-36758837 | 0.017841619 | 0.968424973 | 0.989899803 |
| chr4:104967012-104967519 | -0.013032334 | 0.968435778 | 0.989899803 |
| chr17:50479428-50479795 | -0.013895765 | 0.968439323 | 0.989899803 |
| chr8:87445578-87445821 | -0.019047068 | 0.968454357 | 0.989899803 |
| chr1:184270585-184270883 | -0.018599238 | 0.968467603 | 0.989899803 |
| chr5:150175621-150175868 | -0.01730065 | 0.968472991 | 0.989899803 |
| chr12:109387413-109388027 | 0.015124452 | 0.968486183 | 0.989899803 |
| chr15:92097569-92097874 | 0.015463595 | 0.968495737 | 0.989899803 |
| chr11:2305441-2305695 | -0.017029895 | 0.968514022 | 0.989899803 |
| chr1:162357747-162358004 | -0.01919986 | 0.968514339 | 0.989899803 |
| chr19:13150268-13150515 | -0.015731504 | 0.968525499 | 0.989899803 |
| chr6:36977121-36977384 | -0.019350881 | 0.968529914 | 0.989899803 |
| chr2:33583355-33583555 | 0.021017471 | 0.968548029 | 0.989899803 |
| chr2:86807383-86808709 | -0.012557119 | 0.968554132 | 0.989899803 |
| chr1:28736378-28737090 | 0.013258844 | 0.96855718 | 0.989899803 |
| chr11:47354988-47355887 | 0.01269662 | 0.968606345 | 0.989936847 |
| chr9:108272047-108272283 | -0.019289869 | 0.968635236 | 0.989952422 |
| chr15:94576978-94577253 | 0.015946831 | 0.968647424 | 0.989952422 |
| chr2:147045194-147045404 | -0.019999009 | 0.968672817 | 0.989955068 |
| chr13:20163403-20163701 | 0.015115226 | 0.968682054 | 0.989955068 |
| chr1:204241143-204241518 | 0.014553446 | 0.968726551 | 0.989955068 |
| chr4:100848917-100849227 | -0.016072846 | 0.968728757 | 0.989955068 |
| chr19:44094807-44095007 | -0.019930792 | 0.968745506 | 0.989955068 |
| chr3:125437904-125438109 | -0.019930792 | 0.968745506 | 0.989955068 |
| chr19:49890234-49890992 | 0.013591788 | 0.968752946 | 0.989955068 |
| chr5:130555857-130556118 | 0.018570137 | 0.968773422 | 0.989955068 |
| chr8:101515376-101515897 | 0.013750897 | 0.968783489 | 0.989955068 |
| chr8:70839699-70840124 | -0.015260483 | 0.968785757 | 0.989955068 |
| chr4:77075868-77076745 | 0.012602246 | 0.968792133 | 0.989955068 |
| chr21:42120406-42120898 | -0.013766645 | 0.968809188 | 0.98995728 |
| chr3:16264291-16265540 | 0.012266148 | 0.968825218 | 0.98995728 |
| chr10:80699530-80699992 | -0.013674698 | 0.968833058 | 0.98995728 |
| chr1:198549799-198550170 | -0.016978416 | 0.968849323 | 0.989960698 |
| chr3:69309829-69310116 | 0.016572773 | 0.968862971 | 0.989961442 |
| chr7:107725234-107725474 | -0.017852894 | 0.968897777 | 0.989983804 |
| chr2:45729721-45730020 | -0.017259385 | 0.968930347 | 0.989997594 |
| chr7:41940786-41941207 | -0.014060722 | 0.96897541 | 0.989997594 |
| chr11:36289252-36289885 | 0.013020097 | 0.968985564 | 0.989997594 |
| chr4:88967572-88968251 | 0.014126852 | 0.968988341 | 0.989997594 |
| chr5:16528444-16528929 | -0.014778844 | 0.969006664 | 0.989997594 |
| chr20:57596826-57597470 | -0.012925691 | 0.969022882 | 0.989997594 |
| chr12:69008505-69008728 | 0.018699642 | 0.969024467 | 0.989997594 |
| chr12:59973583-59973911 | -0.018565834 | 0.969035071 | 0.989997594 |

| | | | |
|---|---|---|---|
| chr12:6278493-6279015 | 0.013653496 | 0.969042369 | 0.989997594 |
| chr9:112332867-112334221 | 0.012284817 | 0.969063397 | 0.989997594 |
| chr12:57736045-57736454 | -0.0141762 | 0.969076435 | 0.989997594 |
| chr11:35305837-35306113 | -0.017254324 | 0.96908569 | 0.989997594 |
| chr17:16267839-16268039 | 0.021150387 | 0.96908992 | 0.989997594 |
| chr3:172321100-172321300 | -0.018971869 | 0.969097764 | 0.989997594 |
| chr5:56900796-56901042 | -0.017201111 | 0.969105081 | 0.989997594 |
| chr2:587177-587500 | -0.019026031 | 0.969146352 | 0.990022997 |
| chr1:85576518-85577252 | -0.012978632 | 0.969180586 | 0.990022997 |
| chr8:141925730-141925930 | 0.019337191 | 0.969198111 | 0.990022997 |
| chr6:4964819-4965030 | 0.020423563 | 0.969206365 | 0.990022997 |
| chr1:235093821-235094147 | -0.014648988 | 0.969206869 | 0.990022997 |
| chr1:215644745-215645049 | 0.016296894 | 0.969207473 | 0.990022997 |
| chr2:116552728-116553688 | 0.013109054 | 0.969233884 | 0.99003376 |
| chr7:6683989-6684238 | -0.018830994 | 0.969243852 | 0.99003376 |
| chr4:1746228-1746434 | 0.01712407 | 0.969281045 | 0.990045943 |
| chr17:27341042-27341251 | -0.019417466 | 0.969294115 | 0.990045943 |
| chr21:45270175-45270526 | -0.01659229 | 0.969315428 | 0.990045943 |
| chr15:66923444-66923699 | 0.016409733 | 0.969319339 | 0.990045943 |
| chr1:6475576-6475812 | -0.018112737 | 0.969320386 | 0.990045943 |
| chr3:98705728-98706014 | -0.016962874 | 0.969341596 | 0.99005441 |
| chr6:95203197-95203555 | 0.01469093 | 0.969368291 | 0.99005703 |
| chr12:132144179-132144594 | 0.013390997 | 0.969370004 | 0.99005703 |
| chr11:65057347-65057547 | 0.020558661 | 0.969420323 | 0.990091211 |
| chr3:113354511-113354711 | 0.019140671 | 0.969454287 | 0.990091211 |
| chr5:179506051-179506255 | 0.019140671 | 0.969454287 | 0.990091211 |
| chr19:39833153-39834416 | 0.01220272 | 0.969455157 | 0.990091211 |
| chr2:43950464-43950739 | -0.01680525 | 0.969488002 | 0.990111558 |
| chr17:75779245-75780200 | -0.012110457 | 0.969546911 | 0.990146036 |
| chr6:152118763-152118963 | 0.018028385 | 0.969547607 | 0.990146036 |
| chr1:160529558-160529758 | -0.021015173 | 0.969582099 | 0.990156664 |
| chr1:232600663-232600878 | -0.018006508 | 0.96958386 | 0.990156664 |
| chr12:74538558-74538788 | -0.014689631 | 0.969603218 | 0.990163236 |
| chr1:1960096-1960371 | -0.016613568 | 0.969649195 | 0.990196292 |
| chr20:22735074-22735317 | -0.017040904 | 0.969661434 | 0.990196292 |
| chr7:33310522-33310980 | 0.01399475 | 0.969679756 | 0.990201805 |
| chr1:31065285-31066187 | -0.01236695 | 0.969700588 | 0.990201832 |
| chr8:88374448-88374789 | -0.017420937 | 0.969708975 | 0.990201832 |
| chrX:24494721-24495030 | -0.014959025 | 0.969743997 | 0.990201832 |
| chr2:108786251-108787223 | 0.012399167 | 0.969748461 | 0.990201832 |
| chr9:100690380-100690829 | 0.013683374 | 0.969768393 | 0.990201832 |
| chr16:67285070-67285278 | 0.016901497 | 0.969778951 | 0.990201832 |
| chr14:95196312-95196618 | -0.015029973 | 0.969780414 | 0.990201832 |
| chr5:102421435-102422027 | 0.014082569 | 0.969800025 | 0.990201832 |
| chr17:52297303-52297617 | 0.014949513 | 0.969813971 | 0.990201832 |
| chr20:21302946-21303980 | -0.012192898 | 0.969837513 | 0.990201832 |
| chr2:85602198-85603199 | -0.012222632 | 0.96983857 | 0.990201832 |
| chr2:28751376-28752269 | 0.012192475 | 0.969842382 | 0.990201832 |
| chr19:14006212-14006993 | 0.012496818 | 0.969847784 | 0.990201832 |
| chr15:45457979-45458350 | 0.015203902 | 0.969881664 | 0.990212192 |

| | | | |
|---|---|---|---|
| chr20:50256685-50256923 | 0.01547636 | 0.969883778 | 0.990212192 |
| chr7:67620601-67620876 | 0.016351464 | 0.969928568 | 0.990243524 |
| chr13:42258915-42259186 | -0.018670486 | 0.969940313 | 0.990243524 |
| chr6:15691281-15691626 | 0.014813043 | 0.969998091 | 0.99027989 |
| chr8:19423995-19424204 | 0.020489318 | 0.970017438 | 0.99027989 |
| chr14:62117276-62117659 | -0.014755725 | 0.970027641 | 0.99027989 |
| chr20:58955908-58956653 | -0.01251497 | 0.970028947 | 0.99027989 |
| chr3:62011139-62011431 | 0.013332334 | 0.970054202 | 0.99027989 |
| chr12:49234474-49234715 | 0.017042095 | 0.970066227 | 0.99027989 |
| chr1:229963025-229963266 | 0.020126597 | 0.97009066 | 0.99027989 |
| chr12:112381545-112382775 | 0.012341076 | 0.970097366 | 0.99027989 |
| chr2:25700251-25700481 | -0.016225221 | 0.970120325 | 0.99027989 |
| chr3:169686956-169687343 | -0.015721425 | 0.970132105 | 0.99027989 |
| chr1:39191701-39191945 | -0.01681645 | 0.970145603 | 0.99027989 |
| chr5:36875859-36876107 | 0.014978734 | 0.970157281 | 0.99027989 |
| chr5:7816048-7816731 | -0.012786043 | 0.970165935 | 0.99027989 |
| chr3:109532304-109532751 | 0.014157247 | 0.970166302 | 0.99027989 |
| chr2:85181291-85181491 | 0.017524726 | 0.970175079 | 0.99027989 |
| chr17:27381025-27381343 | 0.015450625 | 0.970219751 | 0.99027989 |
| chr5:143507411-143507797 | 0.015846819 | 0.970220359 | 0.99027989 |
| chr3:42803820-42804795 | -0.012470032 | 0.970226791 | 0.99027989 |
| chr12:56299708-56300869 | 0.011958243 | 0.970229591 | 0.99027989 |
| chr15:100350716-100351006 | 0.013508169 | 0.970234419 | 0.99027989 |
| chr3:32845045-32845275 | -0.019956963 | 0.970262294 | 0.990289803 |
| chr1:32014195-32014712 | 0.013797015 | 0.970269979 | 0.990289803 |
| chr17:69506412-69506860 | -0.014894402 | 0.970287 | 0.990293984 |
| chr10:34655410-34655736 | 0.016121024 | 0.970347013 | 0.99030665 |
| chr15:72685777-72686766 | -0.012025587 | 0.97034731 | 0.99030665 |
| chr10:102963263-102963475 | 0.019618368 | 0.970389024 | 0.99030665 |
| chr6:40186219-40186492 | -0.019791034 | 0.970398397 | 0.99030665 |
| chr5:63962369-63962589 | 0.019319869 | 0.970400574 | 0.99030665 |
| chr8:29262534-29262743 | 0.019319869 | 0.970400574 | 0.99030665 |
| chr17:19718867-19719067 | -0.018004354 | 0.970409532 | 0.99030665 |
| chr3:159929195-159929888 | 0.013582404 | 0.97040969 | 0.99030665 |
| chr1:204258307-204258723 | -0.012729503 | 0.970424411 | 0.99030665 |
| chr8:133385494-133385724 | -0.017035393 | 0.970428657 | 0.99030665 |
| chr1:241957984-241958184 | -0.018326758 | 0.970472311 | 0.990311003 |
| chr8:141084310-141084838 | -0.012872403 | 0.970478551 | 0.990311003 |
| chr14:44767725-44767927 | 0.01693027 | 0.970485844 | 0.990311003 |
| chr17:40903694-40903894 | 0.018312752 | 0.970495441 | 0.990311003 |
| chr12:57078303-57079221 | 0.011863436 | 0.970497545 | 0.990311003 |
| chr3:197790748-197792026 | -0.012082773 | 0.970523 | 0.990323789 |
| chr9:101028316-101029424 | -0.011990064 | 0.97060745 | 0.990396772 |
| chr16:66895229-66895492 | -0.016108792 | 0.970633867 | 0.99040241 |
| chr7:25098512-25098712 | 0.01868748 | 0.970644345 | 0.99040241 |
| chr12:121663885-121664199 | 0.014050533 | 0.970651753 | 0.99040241 |
| chr4:107689148-107689348 | 0.020757668 | 0.970678422 | 0.990416433 |
| chr4:70505233-70505475 | -0.016342719 | 0.970692708 | 0.990417821 |
| chr5:170596983-170597183 | 0.016762817 | 0.970767655 | 0.990481101 |
| chr1:180155109-180155476 | -0.013487917 | 0.970792915 | 0.990485982 |

| | | | |
|---|---|---|---|
| chr15:79371647-79371847 | -0.019679608 | 0.970826314 | 0.990485982 |
| chr6:10084557-10084938 | -0.014763334 | 0.970839334 | 0.990485982 |
| chr2:46915327-46916776 | -0.011642902 | 0.970851654 | 0.990485982 |
| chr8:12753826-12754863 | -0.011994078 | 0.970863693 | 0.990485982 |
| chr3:88219097-88219811 | -0.01281301 | 0.970885738 | 0.990485982 |
| chr17:43791659-43791897 | 0.017674296 | 0.970894861 | 0.990485982 |
| chr2:189311221-189311699 | 0.013025679 | 0.970910378 | 0.990485982 |
| chr11:64230275-64230853 | 0.012616211 | 0.970918314 | 0.990485982 |
| chr8:119769019-119769345 | -0.013431737 | 0.970937395 | 0.990485982 |
| chr6:138088259-138088641 | 0.013898986 | 0.970944475 | 0.990485982 |
| chr20:36460787-36461838 | -0.011903779 | 0.970976823 | 0.990485982 |
| chr1:113904354-113905901 | -0.011269929 | 0.970977479 | 0.990485982 |
| chr9:131755580-131755898 | 0.015101589 | 0.970983261 | 0.990485982 |
| chrX:110421224-110421425 | 0.016221597 | 0.970988149 | 0.990485982 |
| chr5:43039444-43040143 | 0.011910797 | 0.970989124 | 0.990485982 |
| chr5:179795115-179796750 | 0.011520213 | 0.970992196 | 0.990485982 |
| chrX:130079849-130080142 | -0.014549304 | 0.971017586 | 0.990494985 |
| chr6:122530491-122530813 | -0.014240322 | 0.971026876 | 0.990494985 |
| chr1:156020278-156021133 | -0.012078025 | 0.971048403 | 0.990503757 |
| chr20:12570396-12570715 | 0.01533458 | 0.971113199 | 0.990556665 |
| chr11:26284106-26284330 | 0.016557915 | 0.971129366 | 0.990559969 |
| chr9:37964015-37964360 | 0.015421688 | 0.971259429 | 0.990675896 |
| chr1:203615260-203615536 | 0.014915403 | 0.971268879 | 0.990675896 |
| chr3:89478207-89478489 | 0.018212246 | 0.971297186 | 0.990691582 |
| chr20:31951548-31952301 | -0.011788304 | 0.971353548 | 0.99073588 |
| chr17:43034067-43034301 | -0.018574713 | 0.971378877 | 0.990740797 |
| chr21:38180021-38180402 | 0.015379412 | 0.97140354 | 0.990740797 |
| chr8:58074289-58074647 | 0.015372206 | 0.971416939 | 0.990740797 |
| chr9:126500799-126501201 | 0.0134386 | 0.971419307 | 0.990740797 |
| chr5:14871233-14871691 | -0.013468807 | 0.971431238 | 0.990740797 |
| chr3:188954643-188954928 | 0.015978428 | 0.97143595 | 0.990740797 |
| chr17:13114564-13114863 | -0.014087984 | 0.971456447 | 0.990747438 |
| chr8:95190628-95190948 | 0.013841311 | 0.971468322 | 0.990747438 |
| chr11:46616837-46618126 | 0.011445112 | 0.971510104 | 0.990772504 |
| chr11:20634090-20634298 | -0.019389113 | 0.971537302 | 0.990772504 |
| chr12:90023583-90023783 | 0.018448438 | 0.971541797 | 0.990772504 |
| chr12:32399081-32400223 | 0.011600413 | 0.971552931 | 0.990772504 |
| chr17:43994727-43995209 | 0.013739727 | 0.971586778 | 0.990772504 |
| chr20:51693583-51693900 | 0.01375025 | 0.971592111 | 0.990772504 |
| chr2:222052894-222053129 | 0.016228176 | 0.971594395 | 0.990772504 |
| chr22:28662941-28663146 | 0.017515394 | 0.971596705 | 0.990772504 |
| chr6:45714762-45715067 | 0.014365762 | 0.971615786 | 0.990772504 |
| chr2:190864037-190864237 | 0.020686922 | 0.971627296 | 0.990772504 |
| chr15:78638781-78638986 | 0.017215543 | 0.971635137 | 0.990772504 |
| chr5:135488811-135489264 | 0.013690488 | 0.971649598 | 0.990774064 |
| chr1:77978736-77979672 | -0.011303863 | 0.971662938 | 0.990774481 |
| chr6:117906787-117907605 | 0.012733107 | 0.971687662 | 0.990775735 |
| chr8:35595564-35595825 | -0.017439216 | 0.971703927 | 0.990775735 |
| chr17:73191524-73191748 | 0.014281206 | 0.971712668 | 0.990775735 |
| chr1:180609833-180610033 | 0.0178673 | 0.971721958 | 0.990775735 |

| | | | |
|---|---|---|---|
| chr1:8423435-8423714 | 0.017270722 | 0.971728821 | 0.990775735 |
| chr3:25272288-25272616 | -0.017189103 | 0.971755163 | 0.990789409 |
| chr3:63077851-63078088 | 0.014015072 | 0.971769548 | 0.990790892 |
| chr9:38363078-38363278 | 0.017423643 | 0.97180373 | 0.990812438 |
| chr6:25223543-25223749 | -0.018655206 | 0.971842406 | 0.990812438 |
| chr14:44649367-44649567 | -0.019074186 | 0.971856504 | 0.990812438 |
| chr8:102719568-102719785 | -0.016087827 | 0.971856551 | 0.990812438 |
| chr14:67445977-67446210 | 0.016317861 | 0.971880758 | 0.990812438 |
| chr6:42451273-42452044 | 0.011953884 | 0.971890631 | 0.990812438 |
| chr3:140737235-140737439 | -0.016445451 | 0.971900825 | 0.990812438 |
| chr9:111482712-111482985 | -0.013487403 | 0.971902557 | 0.990812438 |
| chr6:30086008-30086208 | 0.019906122 | 0.971921731 | 0.990812438 |
| chr17:7012267-7012725 | 0.012607291 | 0.97193218 | 0.990812438 |
| chr5:27021038-27021295 | 0.016815353 | 0.971932923 | 0.990812438 |
| chr3:46948043-46948679 | 0.013040076 | 0.971984247 | 0.990851576 |
| chr13:25658796-25659609 | -0.011948978 | 0.972063452 | 0.990887254 |
| chr6:158869090-158870498 | -0.011140309 | 0.972064743 | 0.990887254 |
| chr3:83426409-83426765 | -0.015466454 | 0.97207058 | 0.990887254 |
| chr2:187924316-187924609 | -0.016429027 | 0.972076751 | 0.990887254 |
| chr11:41659992-41660915 | 0.011746947 | 0.972095308 | 0.990887254 |
| chr8:96654448-96654648 | -0.020841096 | 0.972096839 | 0.990887254 |
| chr1:55071457-55071787 | 0.014350627 | 0.972149206 | 0.990911406 |
| chr7:13017391-13017911 | -0.013167063 | 0.97216966 | 0.990911406 |
| chr11:65314625-65315184 | -0.011886182 | 0.972201998 | 0.990911406 |
| chr2:18149722-18149947 | -0.01618789 | 0.972223592 | 0.990911406 |
| chr12:13864216-13864416 | -0.018253147 | 0.972227925 | 0.990911406 |
| chr10:111281380-111281821 | -0.014487125 | 0.972228606 | 0.990911406 |
| chr7:66114677-66115620 | 0.011581415 | 0.972237475 | 0.990911406 |
| chr9:4984227-4985671 | -0.011117341 | 0.972253441 | 0.990911406 |
| chr3:190701037-190701407 | -0.013315307 | 0.972259963 | 0.990911406 |
| chr20:64222403-64222639 | 0.014789332 | 0.972265961 | 0.990911406 |
| chr12:49130192-49131763 | 0.011046044 | 0.972274566 | 0.990911406 |
| chr17:60391253-60391467 | 0.011579189 | 0.972278735 | 0.990911406 |
| chr2:170361404-170361653 | 0.015250995 | 0.972294247 | 0.990911406 |
| chr5:139690153-139690353 | 0.014790414 | 0.972302756 | 0.990911406 |
| chr6:108947361-108947907 | 0.011683566 | 0.97231452 | 0.990911406 |
| chr8:144685569-144686208 | 0.011527514 | 0.972333583 | 0.990917654 |
| chr10:91887186-91887406 | -0.01933466 | 0.972370893 | 0.99093183 |
| chr14:80448357-80448617 | 0.017349767 | 0.972403926 | 0.99093183 |
| chr12:122111037-122111291 | -0.016584911 | 0.972406468 | 0.99093183 |
| chr5:36696108-36696415 | -0.014086549 | 0.972441348 | 0.99093183 |
| chr3:5972578-5973136 | -0.012354852 | 0.972451391 | 0.99093183 |
| chr14:87233290-87233576 | -0.014734422 | 0.972478671 | 0.99093183 |
| chr2:215796192-215796478 | -0.015298804 | 0.972479685 | 0.99093183 |
| chr9:37953680-37954536 | 0.011932065 | 0.972484437 | 0.99093183 |
| chr12:48157299-48158046 | 0.01136094 | 0.972501933 | 0.99093183 |
| chr6:53484705-53485000 | 0.015049655 | 0.972513163 | 0.99093183 |
| chr10:124736538-124736738 | 0.017702054 | 0.972519417 | 0.99093183 |
| chr11:101321436-101321636 | -0.018946944 | 0.972521154 | 0.99093183 |
| chr8:81685790-81686556 | -0.011330133 | 0.97253566 | 0.99093183 |

| | | | |
|---|---|---|---|
| chr17:41527482-41527704 | 0.014558651 | 0.972536755 | 0.99093183 |
| chr11:70183551-70184034 | -0.012943399 | 0.97254895 | 0.99093183 |
| chr19:45607011-45607919 | 0.011898089 | 0.972566018 | 0.99093183 |
| chr21:42973917-42974829 | 0.010973993 | 0.972569833 | 0.99093183 |
| chrX:139932039-139933806 | -0.010823113 | 0.972588926 | 0.99093183 |
| chr10:6086420-6086760 | -0.012523006 | 0.972609771 | 0.99093183 |
| chr5:94830545-94830824 | 0.015587307 | 0.972616018 | 0.99093183 |
| chr7:101542851-101543075 | -0.014515989 | 0.972619081 | 0.99093183 |
| chr19:45902506-45903353 | 0.011090314 | 0.972634692 | 0.990934559 |
| chr11:46243075-46243867 | 0.011318699 | 0.972682285 | 0.990969871 |
| chr5:175478820-175479089 | 0.013337837 | 0.972724142 | 0.990987255 |
| chr8:117037961-117038186 | -0.01650056 | 0.972725215 | 0.990987255 |
| chr12:124914408-124915089 | -0.011270407 | 0.972741145 | 0.990989515 |
| chr6:39934203-39934671 | 0.013081775 | 0.972810569 | 0.990989515 |
| chr9:113682272-113682773 | 0.011995236 | 0.972829595 | 0.990989515 |
| chr8:128316640-128316868 | 0.014167062 | 0.972835187 | 0.990989515 |
| chr9:131738843-131739134 | 0.013483083 | 0.972854041 | 0.990989515 |
| chr15:66619205-66619641 | -0.012774028 | 0.972857178 | 0.990989515 |
| chr10:60238455-60238695 | -0.016333234 | 0.972865785 | 0.990989515 |
| chr1:184388-184588 | 0.019066157 | 0.972890634 | 0.990989515 |
| chr4:121307658-121308103 | 0.014476205 | 0.972894774 | 0.990989515 |
| chr14:99995743-99996069 | -0.014473931 | 0.972898149 | 0.990989515 |
| chr19:49324711-49324911 | -0.018608344 | 0.972898994 | 0.990989515 |
| chr19:7595346-7595940 | -0.012679552 | 0.972906801 | 0.990989515 |
| chr2:135074037-135074800 | 0.011520785 | 0.972922292 | 0.990989515 |
| chr1:42766745-42767774 | 0.010994909 | 0.972923375 | 0.990989515 |
| chr4:83283967-83285026 | -0.010883205 | 0.972940349 | 0.990989515 |
| chr3:159941714-159942043 | -0.013882329 | 0.972941598 | 0.990989515 |
| chr8:96625078-96625327 | -0.016122431 | 0.972955377 | 0.990989515 |
| chr2:88170141-88170641 | 0.012712044 | 0.972960237 | 0.990989515 |
| chr1:200053729-200053934 | 0.016948981 | 0.973003644 | 0.991020554 |
| chr11:86314807-86315010 | 0.017964737 | 0.973030369 | 0.9910346 |
| chr16:53054398-53054598 | -0.016069078 | 0.973055317 | 0.991039132 |
| chr16:24944260-24944530 | -0.014619035 | 0.973065422 | 0.991039132 |
| chr17:75758783-75759011 | -0.01341543 | 0.973073621 | 0.991039132 |
| chr12:7970611-7971287 | 0.011865899 | 0.973115383 | 0.99106375 |
| chr20:24992451-24993197 | -0.011110264 | 0.973123662 | 0.99106375 |
| chr3:76186075-76186275 | -0.019412688 | 0.973146498 | 0.991070838 |
| chr1:40923044-40923489 | -0.012940077 | 0.97316106 | 0.991070838 |
| chr1:94247277-94247893 | -0.012874524 | 0.973173929 | 0.991070838 |
| chr19:38058062-38058754 | -0.011438093 | 0.973191487 | 0.991070838 |
| chr15:83000291-83000698 | -0.013549208 | 0.973205753 | 0.991070838 |
| chr4:103996073-103996372 | 0.01444556 | 0.973223602 | 0.991070838 |
| chr1:201374721-201375481 | -0.011247539 | 0.973252358 | 0.991070838 |
| chr1:7667621-7667892 | 0.014714596 | 0.973260305 | 0.991070838 |
| chr3:2156133-2156834 | 0.011464291 | 0.97327429 | 0.991070838 |
| chr1:55214408-55215965 | 0.010807105 | 0.973281157 | 0.991070838 |
| chr11:75563104-75563318 | -0.013268201 | 0.973285918 | 0.991070838 |
| chr1:79255416-79255894 | -0.011794601 | 0.973292632 | 0.991070838 |
| chr3:127233444-127233651 | 0.015911703 | 0.973300581 | 0.991070838 |

| | | | |
|---|---|---|---|
| chr2:218216794-218218134 | -0.010870964 | 0.973311705 | 0.991070838 |
| chr9:77624205-77624518 | 0.017756697 | 0.973343079 | 0.991089613 |
| chr6:4313031-4313659 | 0.012422717 | 0.973367026 | 0.991090877 |
| chr11:66371061-66372318 | -0.010565746 | 0.973390819 | 0.991090877 |
| chr20:738965-739395 | 0.013155652 | 0.973409528 | 0.991090877 |
| chr11:61392230-61392920 | 0.01163143 | 0.97345172 | 0.991090877 |
| chr11:75430574-75430831 | 0.01364398 | 0.973455087 | 0.991090877 |
| chr6:43053106-43054078 | 0.011062502 | 0.9734645 | 0.991090877 |
| chr2:152004548-152005039 | -0.012449663 | 0.973493245 | 0.991090877 |
| chr5:134184191-134184650 | -0.0119842 | 0.973512142 | 0.991090877 |
| chr11:134132784-134133049 | 0.016271692 | 0.973519665 | 0.991090877 |
| chr10:99430318-99431490 | 0.010533219 | 0.973523443 | 0.991090877 |
| chr12:54497607-54497807 | 0.018274438 | 0.973523634 | 0.991090877 |
| chr8:65841359-65842779 | 0.010496581 | 0.973548315 | 0.991090877 |
| chr3:26484160-26484422 | -0.013822923 | 0.973550106 | 0.991090877 |
| chr6:85483117-85483458 | -0.012488686 | 0.973553999 | 0.991090877 |
| chr4:161799798-161800006 | -0.017071534 | 0.973561103 | 0.991090877 |
| chr6:69769962-69770239 | -0.017071534 | 0.973561103 | 0.991090877 |
| chr5:24823606-24824073 | -0.011769141 | 0.973578297 | 0.991090877 |
| chr22:30445102-30445456 | 0.013503862 | 0.973583492 | 0.991090877 |
| chr3:171453347-171453562 | 0.017254685 | 0.973590081 | 0.991090877 |
| chr8:127751541-127752193 | 0.011081001 | 0.973619756 | 0.991107918 |
| chr4:180455273-180455576 | -0.014008034 | 0.973633761 | 0.991109006 |
| chr5:177351354-177352239 | 0.010828286 | 0.973666617 | 0.991129284 |
| chr9:111483007-111484607 | -0.010518969 | 0.973679781 | 0.991129518 |
| chr2:212686407-212686631 | 0.016140508 | 0.973719632 | 0.991148991 |
| chr6:131651680-131652204 | -0.011939433 | 0.973730021 | 0.991148991 |
| chr8:139377304-139377519 | -0.016688457 | 0.973737838 | 0.991148991 |
| chr4:41611897-41612128 | 0.015220806 | 0.973776087 | 0.991148991 |
| chr7:51389678-51389900 | -0.015768641 | 0.973777418 | 0.991148991 |
| chr21:40099933-40100133 | -0.017028253 | 0.973779881 | 0.991148991 |
| chr11:84270456-84270740 | 0.015932633 | 0.97378946 | 0.991148991 |
| chr9:69820522-69820961 | -0.01260646 | 0.973803655 | 0.991150273 |
| chr3:81302049-81302331 | 0.012964333 | 0.973840809 | 0.991174921 |
| chr11:57515623-57515972 | -0.013574709 | 0.973862375 | 0.991183706 |
| chr14:69531674-69532067 | -0.012189993 | 0.973901739 | 0.991210603 |
| chr3:66822239-66822646 | 0.012070003 | 0.97395173 | 0.991248316 |
| chr11:29159575-29159905 | 0.013357903 | 0.973972624 | 0.991256414 |
| chr19:56091751-56092140 | -0.013020174 | 0.973992321 | 0.991263294 |
| chr2:159118621-159119059 | 0.01352646 | 0.974043881 | 0.991292929 |
| chr14:31561150-31561795 | -0.010989337 | 0.974047314 | 0.991292929 |
| chr3:149494238-149494484 | -0.014019073 | 0.974097359 | 0.99129666 |
| chr6:125512714-125512949 | 0.015295812 | 0.974100079 | 0.99129666 |
| chr22:29617915-29618286 | 0.012892563 | 0.974108822 | 0.99129666 |
| chr5:178043139-178043339 | -0.016680304 | 0.974117706 | 0.99129666 |
| chr8:107781614-107782031 | -0.011987817 | 0.97412251 | 0.99129666 |
| chr1:180631696-180632783 | 0.01023292 | 0.974128605 | 0.99129666 |
| chr4:95548848-95549556 | -0.011087309 | 0.974201692 | 0.991345122 |
| chr1:150628980-150630126 | -0.010128462 | 0.974202104 | 0.991345122 |
| chr8:108248182-108249194 | -0.01022852 | 0.97424166 | 0.991367621 |

| | | | |
|---|---|---|---|
| chr1:117605122-117606699 | 0.010157723 | 0.974251263 | 0.991367621 |
| chr1:108512396-108512774 | 0.013907712 | 0.974270347 | 0.991367621 |
| chr8:143902516-143902848 | 0.012179735 | 0.974290802 | 0.991367621 |
| chr3:137474997-137475197 | 0.017572041 | 0.974291023 | 0.991367621 |
| chr8:95503259-95503683 | -0.012765746 | 0.974301845 | 0.991367621 |
| chr17:7330359-7330559 | -0.013648396 | 0.974368851 | 0.991417503 |
| chr5:36690179-36690946 | -0.010759025 | 0.974392379 | 0.991417503 |
| chr4:71095664-71095864 | -0.017909354 | 0.97439583 | 0.991417503 |
| chr13:43942221-43942421 | 0.016588434 | 0.974406388 | 0.991417503 |
| chr1:160375118-160375318 | -0.016489534 | 0.974415563 | 0.991417503 |
| chr2:112254507-112255292 | 0.010484531 | 0.974466833 | 0.991434079 |
| chr9:97922054-97923040 | -0.010310796 | 0.974470735 | 0.991434079 |
| chr3:52220573-52220804 | -0.0148173 | 0.974477937 | 0.991434079 |
| chr1:109041293-109042607 | -0.01018138 | 0.974487271 | 0.991434079 |
| chr8:125495325-125495539 | -0.014175658 | 0.974496552 | 0.991434079 |
| chr8:90501421-90501745 | 0.013287984 | 0.974553554 | 0.991478907 |
| chr12:64796207-64796599 | 0.01124383 | 0.974585213 | 0.991497952 |
| chr8:65636588-65637083 | -0.010589482 | 0.974616599 | 0.991507488 |
| chr13:32639588-32639813 | -0.015441976 | 0.974620468 | 0.991507488 |
| chr8:115256483-115256727 | -0.015148586 | 0.974717969 | 0.991573071 |
| chr6:20058989-20059272 | 0.015863805 | 0.974730404 | 0.991573071 |
| chr8:72018591-72018877 | -0.013563595 | 0.974736099 | 0.991573071 |
| chr20:63863849-63864560 | -0.010341748 | 0.974736698 | 0.991573071 |
| chr1:54542070-54542702 | -0.011232639 | 0.974797908 | 0.991622173 |
| chr3:196253440-196253955 | -0.011905498 | 0.974816586 | 0.991628008 |
| chr3:126782654-126782886 | -0.015730959 | 0.974856048 | 0.991653129 |
| chr1:190744598-190744798 | 0.017333305 | 0.974867166 | 0.991653129 |
| chr12:47651705-47651970 | 0.013876608 | 0.974884746 | 0.991657847 |
| chr1:168690156-168690371 | -0.016398768 | 0.974911238 | 0.991670649 |
| chr6:19851903-19852164 | -0.013811334 | 0.97492399 | 0.991670649 |
| chr3:143881144-143881389 | -0.017045328 | 0.974936158 | 0.991670649 |
| chr3:66038508-66039862 | -0.009942579 | 0.975013307 | 0.991735957 |
| chr20:63658517-63659000 | 0.010908609 | 0.975078644 | 0.99175045 |
| chr5:141951310-141951510 | -0.015484911 | 0.975084119 | 0.99175045 |
| chr14:105974818-105975198 | -0.012996519 | 0.975087833 | 0.99175045 |
| chr15:70526158-70526754 | -0.011383525 | 0.975090767 | 0.99175045 |
| chr6:118580621-118581132 | 0.011218291 | 0.975105876 | 0.99175045 |
| chr20:63740486-63741524 | -0.010022498 | 0.975107137 | 0.99175045 |
| chr8:18197342-18197666 | 0.012611352 | 0.97511816 | 0.99175045 |
| chr9:120792819-120793908 | -0.01012207 | 0.975143299 | 0.991754232 |
| chr1:155934084-155934786 | 0.010254459 | 0.97515206 | 0.991754232 |
| chr8:9338225-9338561 | -0.012573838 | 0.975173964 | 0.991754232 |
| chr20:16729633-16730502 | 0.010225801 | 0.975189174 | 0.991754232 |
| chr14:103628687-103629950 | 0.01005952 | 0.975196796 | 0.991754232 |
| chr15:62759583-62759783 | 0.016793405 | 0.975199539 | 0.991754232 |
| chr21:8704698-8704898 | 0.017879201 | 0.975240016 | 0.991782232 |
| chr5:6328326-6328528 | 0.015031976 | 0.975263991 | 0.99179345 |
| chr6:33768643-33768862 | -0.013410383 | 0.975288951 | 0.99180567 |
| chr3:168022719-168022925 | 0.016498717 | 0.975306064 | 0.991809909 |
| chr5:142108419-142109419 | 0.01026738 | 0.97532205 | 0.991813002 |

| | | | |
|---|---|---|---|
| chr15:88620876-88621636 | 0.010182195 | 0.975359579 | 0.991828147 |
| chr4:2363973-2364306 | 0.014196782 | 0.975366569 | 0.991828147 |
| chr1:54489384-54489727 | -0.011789229 | 0.975384983 | 0.991828147 |
| chr4:72705411-72705611 | -0.017631153 | 0.97538872 | 0.991828147 |
| chr5:54517204-54518469 | -0.010048024 | 0.975435026 | 0.99186207 |
| chr6:148251397-148251693 | -0.013551249 | 0.975458326 | 0.9918726 |
| chr1:152034371-152035420 | 0.0097525 | 0.975486869 | 0.99188846 |
| chr19:8311460-8311756 | -0.013524855 | 0.975512417 | 0.991901274 |
| chr6:33271320-33272244 | -0.010004331 | 0.975528436 | 0.9919044 |
| chr8:127553647-127553869 | -0.014185937 | 0.975578256 | 0.991927276 |
| chr11:71922668-71922876 | 0.015362411 | 0.975584486 | 0.991927276 |
| chr17:18157943-18158583 | 0.010515931 | 0.975589772 | 0.991927276 |
| chr9:131531141-131531661 | 0.01076586 | 0.975647909 | 0.991973223 |
| chr1:46179573-46179788 | 0.012681521 | 0.975665727 | 0.991978177 |
| chr3:194141234-194142172 | -0.009863499 | 0.97568879 | 0.991983899 |
| chr22:19291666-19292451 | 0.01061456 | 0.975697249 | 0.991983899 |
| chr9:6680809-6682212 | 0.009620086 | 0.975730053 | 0.991991217 |
| chr22:18062370-18062570 | 0.015538049 | 0.975730339 | 0.991991217 |
| chr18:11553400-11553671 | 0.015155718 | 0.975749825 | 0.991997865 |
| chr3:195994417-195994658 | -0.013672303 | 0.97577454 | 0.992009829 |
| chr20:37269691-37270060 | -0.012184219 | 0.975799564 | 0.992016573 |
| chr15:59736224-59736424 | -0.016544241 | 0.975807067 | 0.992016573 |
| chr14:106045186-106045682 | -0.011343094 | 0.975834185 | 0.992021115 |
| chr20:18582036-18582236 | -0.016311094 | 0.975837429 | 0.992021115 |
| chr17:47067214-47067677 | -0.011134265 | 0.975876051 | 0.992047216 |
| chr6:118876255-118876518 | 0.01449325 | 0.975927139 | 0.992076638 |
| chr8:128117579-128117858 | -0.011050548 | 0.975935133 | 0.992076638 |
| chr3:61904012-61904217 | 0.011942748 | 0.975943836 | 0.992076638 |
| chr2:96739761-96740360 | -0.010136218 | 0.975980619 | 0.992090833 |
| chr11:83219085-83219583 | 0.012157622 | 0.97599297 | 0.992090833 |
| chr11:69588498-69588698 | -0.015421488 | 0.976019253 | 0.992090833 |
| chr4:151408590-151409985 | 0.009487038 | 0.976036192 | 0.992090833 |
| chr7:149196281-149197018 | 0.010215202 | 0.976036631 | 0.992090833 |
| chr18:1522148-1522503 | 0.013533645 | 0.976050132 | 0.992090833 |
| chr15:45522020-45522542 | 0.011235797 | 0.976053432 | 0.992090833 |
| chr15:58839581-58839782 | -0.015422951 | 0.97607275 | 0.992090833 |
| chr1:220803502-220803963 | -0.011487539 | 0.976077539 | 0.992090833 |
| chr4:119490462-119490662 | 0.014836801 | 0.976088711 | 0.992090833 |
| chr11:94737387-94737612 | 0.014529338 | 0.976100228 | 0.992090833 |
| chr2:73828720-73829471 | -0.009859129 | 0.976158549 | 0.992121698 |
| chr2:85410400-85410600 | 0.014151431 | 0.976161986 | 0.992121698 |
| chr11:34668044-34668630 | -0.010254732 | 0.97616944 | 0.992121698 |
| chr15:76311045-76311789 | -0.009855813 | 0.976234379 | 0.992174538 |
| chr1:206982176-206982459 | 0.011115042 | 0.976302059 | 0.992177287 |
| chr6:1610457-1610748 | 0.012474379 | 0.976312702 | 0.992177287 |
| chr18:79644063-79644386 | 0.012284594 | 0.976319228 | 0.992177287 |
| chr20:33862769-33863163 | 0.012284594 | 0.976319228 | 0.992177287 |
| chr17:45060664-45061723 | 0.009519139 | 0.97632495 | 0.992177287 |
| chr18:32018411-32019103 | 0.009873888 | 0.976346206 | 0.992177287 |
| chr17:65595349-65595554 | -0.013850976 | 0.976346655 | 0.992177287 |

| | | | |
|---|---|---|---|
| chr12:51217082-51217407 | -0.012266475 | 0.976371209 | 0.992177287 |
| chr17:17280567-17281508 | 0.009557565 | 0.976372738 | 0.992177287 |
| chr7:55589520-55589720 | 0.016228217 | 0.976383644 | 0.992177287 |
| chr5:7530907-7531298 | -0.013010659 | 0.976394202 | 0.992177287 |
| chr8:40443000-40443353 | 0.01227471 | 0.976403463 | 0.992177287 |
| chr4:82225712-82226018 | 0.016572781 | 0.97640736 | 0.992177287 |
| chr8:68699181-68699741 | 0.010453756 | 0.976427003 | 0.992177287 |
| chr12:82686246-82687435 | 0.009250452 | 0.976431319 | 0.992177287 |
| chr13:70388988-70389331 | -0.012465698 | 0.976548562 | 0.992283262 |
| chrX:107566102-107566606 | 0.01057905 | 0.976592106 | 0.992306437 |
| chr14:32583467-32583676 | 0.014542011 | 0.976597271 | 0.992306437 |
| chr12:49253082-49253483 | 0.011491934 | 0.976617834 | 0.992314171 |
| chr7:35580722-35580922 | 0.016619668 | 0.976637769 | 0.992321268 |
| chr2:65382755-65383377 | 0.009703408 | 0.976675518 | 0.992337657 |
| chr8:60651600-60652785 | 0.009222788 | 0.976679801 | 0.992337657 |
| chr10:49161692-49162057 | 0.012192527 | 0.976700796 | 0.99234583 |
| chr9:87324762-87324962 | 0.015791004 | 0.976742141 | 0.992374678 |
| chr16:85471771-85472509 | 0.00953976 | 0.976800317 | 0.992403206 |
| chrX:103629314-103630274 | -0.009424916 | 0.976825567 | 0.992403206 |
| chr1:228191722-228192187 | -0.011209702 | 0.976825789 | 0.992403206 |
| chr6:109336525-109337017 | 0.010507828 | 0.976837239 | 0.992403206 |
| chr20:49979038-49979535 | -0.009899762 | 0.976860212 | 0.992403206 |
| chr15:31488677-31489248 | 0.010434041 | 0.976861689 | 0.992403206 |
| chr2:206084410-206084731 | 0.010765313 | 0.976866385 | 0.992403206 |
| chr18:58322049-58322322 | 0.013342532 | 0.976873835 | 0.992403206 |
| chr6:17804271-17804516 | 0.013650268 | 0.976909779 | 0.992417521 |
| chr11:102341839-10234209 | 0.012864986 | 0.97691383 | 0.992417521 |
| chr5:90391202-90391402 | 0.01653106 | 0.976929129 | 0.992417873 |
| chr11:86155288-86155488 | -0.013264169 | 0.976940081 | 0.992417873 |
| chr3:42490301-42490695 | -0.010930876 | 0.976968121 | 0.992433199 |
| chr3:53073077-53073309 | -0.012672584 | 0.97699162 | 0.992443913 |
| chr8:144478696-144478896 | 0.011967358 | 0.977018454 | 0.992458014 |
| chr9:124868534-124869300 | 0.009539019 | 0.977038745 | 0.992465468 |
| chr7:5995503-5996058 | -0.009877659 | 0.977107601 | 0.992505827 |
| chr11:59158463-59158802 | 0.011666072 | 0.977107949 | 0.992505827 |
| chr9:121369740-121370754 | -0.009287194 | 0.977126052 | 0.992505827 |
| chr17:29931371-29931590 | 0.012796212 | 0.97713029 | 0.992505827 |
| chr1:37553981-37554555 | -0.010009413 | 0.977160076 | 0.992522924 |
| chr8:75149052-75149284 | 0.013084109 | 0.977192564 | 0.992542766 |
| chr2:27428178-27429517 | 0.009126037 | 0.977217797 | 0.992555238 |
| chr6:47414580-47414858 | 0.013468517 | 0.977310616 | 0.992636355 |
| chr15:86893096-86893296 | 0.016116981 | 0.977326222 | 0.992639048 |
| chrX:20253012-20253263 | 0.013168871 | 0.977346025 | 0.99264219 |
| chr11:44927211-44927421 | -0.012988069 | 0.977355226 | 0.99264219 |
| chr6:27213772-27214009 | -0.011812421 | 0.977395413 | 0.992652845 |
| chr7:7352939-7353258 | 0.010723108 | 0.977396984 | 0.992652845 |
| chr22:46053551-46054371 | 0.010182718 | 0.977411846 | 0.992652845 |
| chr20:33299158-33299491 | 0.01185219 | 0.977426456 | 0.992652845 |
| chr4:74874836-74875115 | 0.01240933 | 0.977432148 | 0.992652845 |
| chr19:10520852-10521187 | 0.011288313 | 0.977443448 | 0.992652845 |

| | | | |
|---|---|---|---|
| chr9:107044995-107045224 | -0.014251852 | 0.977497816 | 0.992692417 |
| chr17:65561040-65561409 | 0.009998502 | 0.977527859 | 0.992692417 |
| chr6:81895645-81896150 | 0.011573533 | 0.977534119 | 0.992692417 |
| chrX:3144087-3144287 | -0.01643964 | 0.977534236 | 0.992692417 |
| chr18:57712359-57712920 | -0.010761035 | 0.977552 | 0.9926973 |
| chr8:64581585-64581947 | 0.011480967 | 0.977567345 | 0.992699726 |
| chr16:88821592-88821894 | 0.011442138 | 0.977593199 | 0.99270732 |
| chr2:176637551-176638488 | 0.009597792 | 0.977600735 | 0.99270732 |
| chr4:30717237-30718375 | -0.00907952 | 0.977622845 | 0.992716615 |
| chr10:32036992-32037254 | -0.012829002 | 0.97772948 | 0.992795495 |
| chr17:29117138-29117649 | 0.010278814 | 0.977729785 | 0.992795495 |
| chr8:144465408-144466339 | -0.008941005 | 0.97775335 | 0.992795495 |
| chr7:27135938-27136220 | -0.012673598 | 0.977758423 | 0.992795495 |
| chr1:30854187-30854396 | 0.012913445 | 0.977769808 | 0.992795495 |
| chrX:72713614-72715076 | 0.009142521 | 0.977778267 | 0.992795495 |
| chr4:146241384-146241787 | 0.01041127 | 0.97781038 | 0.992798178 |
| chr21:41751309-41751853 | 0.009818984 | 0.977813429 | 0.992798178 |
| chr14:76208480-76208910 | -0.011925514 | 0.977819781 | 0.992798178 |
| chr5:132012568-132012786 | 0.012097246 | 0.977843682 | 0.992803154 |
| chr1:20786549-20787453 | -0.009159594 | 0.977850597 | 0.992803154 |
| chr13:114281061-114282298 | -0.008970047 | 0.977948026 | 0.992843075 |
| chr6:113306637-113306904 | -0.012077995 | 0.977970316 | 0.992843075 |
| chr3:158091548-158091966 | -0.009988474 | 0.977976844 | 0.992843075 |
| chr21:19507908-19508278 | -0.012379151 | 0.977979973 | 0.992843075 |
| chr2:42828704-42829137 | -0.010300649 | 0.977982785 | 0.992843075 |
| chr3:112551019-112551220 | -0.013457655 | 0.977985765 | 0.992843075 |
| chr8:34108510-34108710 | -0.014845948 | 0.978002005 | 0.992843075 |
| chr5:71631234-71631586 | -0.011499571 | 0.978002742 | 0.992843075 |
| chr3:150830176-150830649 | -0.010016636 | 0.978013703 | 0.992843075 |
| chr8:121103791-121103997 | 0.013733626 | 0.978019493 | 0.992843075 |
| chr10:108784026-108784274 | -0.012603163 | 0.978053767 | 0.992864714 |
| chr11:73982629-73983728 | 0.008722886 | 0.978092613 | 0.992890993 |
| chr2:14632299-14632978 | 0.009102229 | 0.978111784 | 0.9928973 |
| chr7:148436972-148437198 | -0.012927261 | 0.978125755 | 0.992898329 |
| chr5:32232439-32232910 | 0.009676564 | 0.978160007 | 0.992919943 |
| chr10:128069383-128069849 | 0.010039613 | 0.978184636 | 0.992924638 |
| chr5:44022379-44022854 | -0.011114176 | 0.978196489 | 0.992924638 |
| chr6:167045914-167046114 | 0.013876566 | 0.978203508 | 0.992924638 |
| chr19:47778088-47778814 | 0.009333682 | 0.978249164 | 0.992957827 |
| chr20:55095858-55096058 | -0.012658835 | 0.978285068 | 0.992981116 |
| chr6:137166108-137166328 | -0.013913484 | 0.978304867 | 0.992982916 |
| chr6:5003337-5004473 | -0.008610379 | 0.978323427 | 0.992982916 |
| chr11:67317165-67318418 | -0.008661019 | 0.978328546 | 0.992982916 |
| chr22:24525499-24525788 | 0.010878129 | 0.97833868 | 0.992982916 |
| chr8:144524500-144524904 | -0.009653856 | 0.978370248 | 0.992987859 |
| chr12:100763021-100763307 | -0.013104859 | 0.978386198 | 0.992987859 |
| chr3:113966204-113966587 | -0.010036503 | 0.978393431 | 0.992987859 |
| chr19:45935411-45935672 | -0.012537401 | 0.978395388 | 0.992987859 |
| chr1:231211191-231211886 | 0.009376104 | 0.978414326 | 0.992993926 |
| chr6:10419110-10420171 | -0.008784069 | 0.978439304 | 0.993001424 |

| | | | |
|---|---|---|---|
| chr8:47659424-47659744 | 0.011762199 | 0.978459795 | 0.993001424 |
| chr12:88425351-88425667 | 0.011532084 | 0.97846391 | 0.993001424 |
| chr14:102314232-102314680 | -0.009654198 | 0.978479533 | 0.993001424 |
| chr12:49088152-49088788 | -0.009385994 | 0.978486512 | 0.993001424 |
| chr1:178542529-178543570 | 0.008721421 | 0.978536455 | 0.993029619 |
| chr15:88004037-88004237 | 0.015014537 | 0.978555267 | 0.993029619 |
| chr10:88582650-88583532 | -0.008815323 | 0.978565179 | 0.993029619 |
| chr20:53939521-53940274 | 0.008423669 | 0.978566135 | 0.993029619 |
| chr17:38296835-38297300 | -0.009954184 | 0.978587355 | 0.9930351 |
| chr6:108869908-108870298 | -0.01123221 | 0.978597457 | 0.9930351 |
| chr7:2488371-2488571 | 0.013002757 | 0.978619475 | 0.993043741 |
| chr15:90576394-90576594 | -0.01517265 | 0.978631892 | 0.993043741 |
| chr9:135961093-135961636 | 0.009172064 | 0.978645473 | 0.99304437 |
| chr16:49407906-49408243 | -0.01090883 | 0.978685134 | 0.993050178 |
| chr2:228749098-228749386 | 0.011971918 | 0.978689687 | 0.993050178 |
| chr17:65418691-65418950 | -0.010681814 | 0.978690078 | 0.993050178 |
| chr1:117895539-117896021 | 0.010662408 | 0.978763055 | 0.993100797 |
| chr5:149547017-149547343 | -0.009444151 | 0.978765887 | 0.993100797 |
| chr5:150000086-150001315 | -0.008444304 | 0.978933778 | 0.993257994 |
| chr2:89851045-89851416 | -0.011216791 | 0.978961339 | 0.993260485 |
| chr4:122066983-122067212 | 0.014986063 | 0.978962159 | 0.993260485 |
| chr14:20970953-20971928 | -0.008587348 | 0.979033058 | 0.993300109 |
| chr1:15154072-15154285 | -0.012073923 | 0.979040447 | 0.993300109 |
| chr12:70366613-70367378 | 0.008681844 | 0.979048042 | 0.993300109 |
| chr12:71439439-71440401 | -0.008889945 | 0.979062598 | 0.993300109 |
| chr11:76946343-76946543 | -0.013385664 | 0.979072911 | 0.993300109 |
| chr3:149436157-149436373 | -0.014229512 | 0.979078995 | 0.993300109 |
| chr6:157380799-157381586 | 0.008890309 | 0.979198303 | 0.993357784 |
| chr16:85759784-85760180 | -0.010056868 | 0.979209492 | 0.993357784 |
| chr7:55420640-55420993 | 0.010582917 | 0.979215773 | 0.993357784 |
| chr3:109803233-109803629 | 0.010963859 | 0.979226289 | 0.993357784 |
| chr3:96331374-96331886 | -0.01004014 | 0.979228345 | 0.993357784 |
| chr11:130062609-130062813 | 0.015234498 | 0.979230475 | 0.993357784 |
| chr2:113234267-113234467 | -0.011997486 | 0.979232093 | 0.993357784 |
| chr4:16133719-16134114 | 0.01065219 | 0.979239559 | 0.993357784 |
| chrX:2812439-2812775 | 0.012579313 | 0.97929497 | 0.993385862 |
| chr1:152398748-152399736 | 0.008558829 | 0.97930375 | 0.993385862 |
| chr18:62522252-62523469 | 0.008230659 | 0.979306132 | 0.993385862 |
| chr2:187544808-187545112 | -0.011365712 | 0.979399384 | 0.993445376 |
| chr2:152738471-152738756 | 0.012621031 | 0.979402269 | 0.993445376 |
| chr5:139492831-139493098 | 0.012407971 | 0.9794037 | 0.993445376 |
| chr6:119720212-119720481 | -0.012732237 | 0.979429816 | 0.993458715 |
| chr2:175864163-175864476 | 0.011239432 | 0.979477901 | 0.993494337 |
| chr6:30447264-30447585 | -0.010336434 | 0.979502573 | 0.99350621 |
| chr2:37535871-37536520 | 0.008812942 | 0.979574886 | 0.993513889 |
| chr3:99114613-99115107 | -0.009596218 | 0.979594636 | 0.993513889 |
| chr5:51960966-51961166 | 0.012854802 | 0.979609564 | 0.993513889 |
| chr7:158353970-158354190 | 0.012854802 | 0.979609564 | 0.993513889 |
| chr6:65638099-65638400 | -0.010075111 | 0.979613477 | 0.993513889 |
| chr3:124475495-124475817 | 0.013140779 | 0.979619719 | 0.993513889 |

| | | | |
|---|---|---|---|
| chr3:80899962-80900375 | -0.010565365 | 0.979620336 | 0.993513889 |
| chr6:170186512-170187015 | 0.009611089 | 0.979621951 | 0.993513889 |
| chr19:34905753-34905953 | -0.013066493 | 0.979626841 | 0.993513889 |
| chr5:112372591-112372829 | 0.011534711 | 0.979653698 | 0.993517573 |
| chr3:97793441-97793710 | -0.011858534 | 0.979658068 | 0.993517573 |
| chr21:29018993-29019514 | 0.009240251 | 0.979694913 | 0.993517573 |
| chr17:14203505-14203749 | 0.011557781 | 0.979703476 | 0.993517573 |
| chr4:99893832-99895019 | 0.008169509 | 0.979706639 | 0.993517573 |
| chr4:103896140-103896361 | 0.012105927 | 0.979708273 | 0.993517573 |
| chr19:42693837-42694041 | -0.013559777 | 0.979737597 | 0.993534161 |
| chr12:8026284-8027360 | 0.008142548 | 0.979778219 | 0.993562205 |
| chr8:121588485-121588721 | -0.010904766 | 0.979829577 | 0.993597088 |
| chr4:143184315-143185887 | -0.007930901 | 0.979850334 | 0.993597088 |
| chr3:50350642-50351321 | -0.008490948 | 0.979854833 | 0.993597088 |
| chr2:217441296-217441654 | 0.010032258 | 0.979874246 | 0.993597088 |
| chr3:132029206-132029642 | 0.008879013 | 0.979877455 | 0.993597088 |
| chr10:14879509-14879709 | -0.010582771 | 0.979917751 | 0.993622555 |
| chr17:49880498-49880855 | 0.009155693 | 0.979942517 | 0.993622555 |
| chr8:89901706-89903085 | -0.007848018 | 0.979953365 | 0.993622555 |
| chr14:63641345-63642587 | 0.007867911 | 0.979963669 | 0.993622555 |
| chr1:113247434-113247727 | -0.010455304 | 0.979967409 | 0.993622555 |
| chr16:82456640-82457089 | -0.008977889 | 0.979983641 | 0.993625864 |
| chr12:85468695-85469154 | 0.00931686 | 0.980022484 | 0.993652099 |
| chr13:34334741-34335013 | 0.012211579 | 0.980046819 | 0.993655858 |
| chr4:91746656-91746950 | -0.012616377 | 0.980052128 | 0.993655858 |
| chr15:43376138-43376405 | -0.011270029 | 0.980132087 | 0.993714956 |
| chr14:96037805-96038174 | -0.008614024 | 0.98014388 | 0.993714956 |
| chr2:98649446-98649761 | -0.010406379 | 0.980164357 | 0.993714956 |
| chr1:148388057-148389172 | -0.007946377 | 0.980176147 | 0.993714956 |
| chr10:94402105-94403960 | -0.007850074 | 0.980181997 | 0.993714956 |
| chr9:77433532-77434214 | -0.009030337 | 0.980211337 | 0.993714956 |
| chr19:31249807-31250062 | -0.013448043 | 0.98021678 | 0.993714956 |
| chr21:32771221-32772331 | 0.007937352 | 0.980229952 | 0.993714956 |
| chr15:32615063-32615263 | -0.012836448 | 0.980234763 | 0.993714956 |
| chr22:28813548-28814256 | 0.008221033 | 0.980245145 | 0.993714956 |
| chr17:64779548-64779921 | 0.00849594 | 0.980258284 | 0.993714956 |
| chr5:9901177-9901424 | 0.011433097 | 0.980266045 | 0.993714956 |
| chr1:188648436-188648947 | -0.010019365 | 0.980302077 | 0.993721855 |
| chr10:80371372-80371616 | -0.011374156 | 0.980327405 | 0.993721855 |
| chr19:29587410-29587845 | 0.009426273 | 0.980335176 | 0.993721855 |
| chr3:174955440-174956235 | 0.008196759 | 0.980337403 | 0.993721855 |
| chr15:40569103-40569417 | -0.008914215 | 0.980337697 | 0.993721855 |
| chr14:59188112-59189681 | -0.007695609 | 0.980387481 | 0.993759172 |
| chr8:110894008-110894872 | -0.008780876 | 0.980408412 | 0.993766746 |
| chr4:54658541-54659044 | 0.008946573 | 0.980428722 | 0.993766746 |
| chr10:23081184-23081724 | -0.009521605 | 0.980433862 | 0.993766746 |
| chr7:5116743-5117123 | -0.009892779 | 0.980481418 | 0.993789262 |
| chr8:127899530-127899878 | 0.009357002 | 0.980488901 | 0.993789262 |
| chr3:161274719-161275097 | -0.008643955 | 0.980494987 | 0.993789262 |
| chr4:157389160-157389414 | 0.011625204 | 0.980522988 | 0.993796973 |

| | | | |
|---|---|---|---|
| chr5:172478974-172479285 | 0.009121035 | 0.980528535 | 0.993796973 |
| chr22:26429099-26429369 | 0.010906336 | 0.980634411 | 0.993891135 |
| chr1:176559849-176560144 | 0.009229543 | 0.980695627 | 0.993902813 |
| chr12:113422064-11342260[illegible] | -0.008470186 | 0.980701528 | 0.993902813 |
| chr17:21284327-21284984 | 0.00789822 | 0.980703047 | 0.993902813 |
| chr8:53734351-53735201 | -0.007944859 | 0.980704329 | 0.993902813 |
| chr2:143322049-143322295 | -0.010401139 | 0.980714424 | 0.993902813 |
| chr5:177837464-177837792 | -0.009952846 | 0.980735436 | 0.993902813 |
| chr17:22521268-22521514 | -0.008995955 | 0.980743642 | 0.993902813 |
| chr1:220565722-220565996 | 0.011992969 | 0.980765116 | 0.993902813 |
| chr9:128656398-128657300 | 0.008037936 | 0.980783648 | 0.993902813 |
| chr17:80858121-80858377 | 0.010529328 | 0.980799174 | 0.993902813 |
| chr14:32993868-32994230 | -0.010183978 | 0.980809274 | 0.993902813 |
| chr8:94961861-94962205 | 0.009431261 | 0.980811717 | 0.993902813 |
| chr7:36152505-36153811 | 0.007522692 | 0.980814563 | 0.993902813 |
| chr1:234373014-234374045 | 0.007733116 | 0.980847628 | 0.993923175 |
| chr18:79068904-79069588 | 0.007866252 | 0.980869737 | 0.993924846 |
| chr2:16508065-16508589 | -0.008878412 | 0.980883378 | 0.993924846 |
| chr19:38506887-38507638 | -0.007656076 | 0.980896932 | 0.993924846 |
| chr6:2875264-2875775 | -0.008735678 | 0.980901164 | 0.993924846 |
| chr2:96174527-96174830 | 0.009564318 | 0.980970349 | 0.993981804 |
| chr6:138520527-138520727 | -0.011477346 | 0.981007207 | 0.993996129 |
| chr17:1908854-1909138 | -0.012158713 | 0.981012998 | 0.993996129 |
| chr1:246117237-246117446 | 0.012508576 | 0.981023404 | 0.993996129 |
| chr7:786465-786860 | 0.009216591 | 0.981071982 | 0.994008294 |
| chr17:4966882-4968070 | 0.007554775 | 0.981076438 | 0.994008294 |
| chr5:81750799-81751951 | -0.007469562 | 0.981084887 | 0.994008294 |
| chr10:72319635-72320359 | 0.008197294 | 0.981091882 | 0.994008294 |
| chr7:115724357-115724582 | -0.013028893 | 0.981100275 | 0.994008294 |
| chr8:37899059-37899827 | 0.007891382 | 0.981114258 | 0.994009318 |
| chr1:153658450-153659415 | -0.007730195 | 0.981134864 | 0.994009897 |
| chr7:33128908-33130065 | 0.007419625 | 0.981140775 | 0.994009897 |
| chr1:84634514-84635295 | -0.007830656 | 0.981159358 | 0.994011766 |
| chr8:114845276-114845499 | -0.011753614 | 0.981168783 | 0.994011766 |
| chr8:100932882-100933141 | 0.010119859 | 0.981192267 | 0.994011766 |
| chr12:92987336-92987660 | 0.010189489 | 0.981194512 | 0.994011766 |
| chr9:128627915-128628184 | 0.010749718 | 0.981236886 | 0.994041551 |
| chr19:58278267-58279481 | -0.007529204 | 0.981278393 | 0.994065998 |
| chr19:39341613-39342576 | 0.007580434 | 0.981295541 | 0.994065998 |
| chr4:124117441-124118147 | 0.008553618 | 0.98129994 | 0.994065998 |
| chr20:14764606-14765403 | -0.008181492 | 0.981367142 | 0.994120932 |
| chr14:67391495-67391872 | 0.00896752 | 0.98144293 | 0.994184561 |
| chr8:40253121-40253338 | 0.011816463 | 0.981494841 | 0.994224001 |
| chr3:150081934-150082677 | 0.008240271 | 0.98154058 | 0.994240137 |
| chr6:132180066-132180277 | 0.011054748 | 0.98154106 | 0.994240137 |
| chr18:48675285-48675739 | 0.008581476 | 0.981552439 | 0.994240137 |
| chr17:58417399-58417965 | -0.007402247 | 0.981562674 | 0.994240137 |
| chr14:49633610-49635456 | 0.007221996 | 0.981581845 | 0.994246413 |
| chr5:15009492-15009693 | 0.010615307 | 0.981631821 | 0.994273812 |
| chr1:40898601-40899029 | 0.008880841 | 0.981642521 | 0.994273812 |

| | | | |
|---|---|---|---|
| chr3:66593973-66594287 | -0.009823843 | 0.981647824 | 0.994273812 |
| chr3:63217504-63218480 | 0.007618858 | 0.981674848 | 0.99428804 |
| chr8:142718133-142718434 | -0.009736696 | 0.981718356 | 0.994318963 |
| chr17:49032647-49032895 | 0.008705908 | 0.981744888 | 0.994321678 |
| chr12:82479685-82480016 | 0.010371657 | 0.981772951 | 0.994321678 |
| chr1:204935628-204935992 | 0.008929144 | 0.981775615 | 0.994321678 |
| chr3:31947436-31948124 | 0.008221943 | 0.981794687 | 0.994321678 |
| chr8:65633791-65634560 | -0.007272121 | 0.981798826 | 0.994321678 |
| chr19:32345411-32346109 | 0.007790792 | 0.981808421 | 0.994321678 |
| chr1:119686706-119687047 | 0.009269897 | 0.9818303 | 0.994321678 |
| chr14:61269432-61270230 | 0.007737039 | 0.981838581 | 0.994321678 |
| chr2:134450786-134451176 | -0.008352144 | 0.981845015 | 0.994321678 |
| chr1:202661210-202661512 | -0.008329009 | 0.981855796 | 0.994321678 |
| chr19:50510591-50510880 | 0.009167445 | 0.981867703 | 0.994321678 |
| chr10:124418176-124419427 | -0.007288096 | 0.98187676 | 0.994321678 |
| chr17:80414649-80415812 | 0.007302684 | 0.982001843 | 0.994417361 |
| chr5:52786902-52787255 | 0.008612104 | 0.982002864 | 0.994417361 |
| chr1:181088258-181090064 | 0.007046481 | 0.982010181 | 0.994417361 |
| chr12:74897891-74898135 | -0.010343319 | 0.982045303 | 0.994433675 |
| chr5:60710377-60710577 | 0.011299421 | 0.982052248 | 0.994433675 |
| chrX:57335742-57335982 | -0.010787784 | 0.982088196 | 0.994456935 |
| chr5:32523747-32523955 | -0.011659256 | 0.982155112 | 0.994479217 |
| chr19:1021075-1021858 | 0.007287714 | 0.982161214 | 0.994479217 |
| chr3:122794693-122795450 | -0.007466024 | 0.982174438 | 0.994479217 |
| chr1:17881846-17882059 | -0.012514909 | 0.982178131 | 0.994479217 |
| chr3:194852956-194853608 | -0.00813414 | 0.982183825 | 0.994479217 |
| chr1:183159929-183160551 | -0.007309982 | 0.982188076 | 0.994479217 |
| chr8:118120629-118120873 | 0.009926474 | 0.982223657 | 0.994486966 |
| chr1:86515645-86515895 | 0.011105713 | 0.982247292 | 0.994486966 |
| chr11:76616027-76616309 | -0.009782456 | 0.982253998 | 0.994486966 |
| chr5:124661476-124661676 | 0.011979407 | 0.982254774 | 0.994486966 |
| chr11:76733940-76734257 | 0.008990545 | 0.982260625 | 0.994486966 |
| chr21:30194552-30194973 | -0.00960487 | 0.982319394 | 0.994500813 |
| chrX:134537264-134537733 | -0.009427462 | 0.982325281 | 0.994500813 |
| chr13:113585346-113585599 | 0.010074802 | 0.982339252 | 0.994500813 |
| chr20:30579656-30580535 | 0.007337532 | 0.982362262 | 0.994500813 |
| chr2:19362901-19363309 | -0.008026835 | 0.982377102 | 0.994500813 |
| chr3:144770908-144771108 | -0.012195662 | 0.982381283 | 0.994500813 |
| chr7:9020479-9020687 | -0.011468851 | 0.982396194 | 0.994500813 |
| chr14:70013359-70013571 | -0.010401208 | 0.982397085 | 0.994500813 |
| chr5:7851649-7852141 | 0.007758725 | 0.982399393 | 0.994500813 |
| chr10:45373687-45373937 | -0.010960316 | 0.982407116 | 0.994500813 |
| chr4:140737300-140737500 | -0.010507203 | 0.982417074 | 0.994500813 |
| chr17:49414559-49415468 | 0.007028169 | 0.982454734 | 0.994509579 |
| chr20:57718403-57719079 | 0.0070587 | 0.982475323 | 0.994509579 |
| chr5:33440125-33441458 | -0.006923593 | 0.982476074 | 0.994509579 |
| chr22:38570089-38570717 | 0.007542296 | 0.982477651 | 0.994509579 |
| chr13:23029027-23029227 | 0.01110785 | 0.982516972 | 0.994523502 |
| chr19:45514629-45514932 | -0.010133373 | 0.982517365 | 0.994523502 |
| chr1:246482301-246482519 | 0.010938089 | 0.982574996 | 0.994546196 |

| | | | |
|---|---|---|---|
| chr1:148264687-148265293 | 0.006986791 | 0.982586869 | 0.994546196 |
| chr1:180167885-180168491 | -0.007125022 | 0.982588417 | 0.994546196 |
| chr17:27946342-27946550 | -0.012326709 | 0.982591704 | 0.994546196 |
| chr6:12603452-12603657 | -0.010821124 | 0.982657806 | 0.994591859 |
| chr1:179293335-179294241 | -0.0069939 | 0.982662779 | 0.994591859 |
| chr10:21356780-21356980 | -0.011781428 | 0.982681957 | 0.994598131 |
| chr6:160930585-160931435 | -0.007529889 | 0.982724922 | 0.994628478 |
| chr11:20873138-20873400 | -0.009815496 | 0.982798406 | 0.994677236 |
| chr15:87477760-87478578 | 0.007390339 | 0.98279906 | 0.994677236 |
| chr2:42897446-42897817 | -0.009753875 | 0.982815227 | 0.994678566 |
| chr3:100400800-100401870 | 0.006814631 | 0.982826336 | 0.994678566 |
| chr8:109298981-109300116 | 0.007048527 | 0.982860046 | 0.994683245 |
| chr4:75990564-75991223 | -0.007275874 | 0.982865455 | 0.994683245 |
| chr6:52670658-52671479 | -0.007046787 | 0.982874048 | 0.994683245 |
| chr3:195687409-195687823 | -0.007962953 | 0.982890132 | 0.994683245 |
| chr17:55770287-55770532 | 0.008843937 | 0.982906278 | 0.994683245 |
| chr12:48618453-48618798 | 0.008992947 | 0.982932099 | 0.994683245 |
| chrX:48897618-48898439 | -0.006892675 | 0.982957165 | 0.994683245 |
| chr17:28951380-28952124 | 0.006839767 | 0.98295826 | 0.994683245 |
| chr10:125822791-125823669 | -0.006931377 | 0.98295937 | 0.994683245 |
| chr12:6089176-6089376 | -0.010500547 | 0.982960777 | 0.994683245 |
| chr3:138863029-138863663 | -0.007625562 | 0.982991947 | 0.994689625 |
| chr13:44364357-44364557 | 0.010621667 | 0.982993045 | 0.994689625 |
| chrX:24561201-24561489 | 0.008912031 | 0.983018851 | 0.994702602 |
| chr2:113603347-113604503 | -0.006750856 | 0.983045474 | 0.994716404 |
| chr10:91163160-91164240 | -0.007118764 | 0.983079025 | 0.994718474 |
| chr17:78257116-78257389 | -0.009349079 | 0.983109055 | 0.994718474 |
| chr8:80300443-80300952 | 0.007981562 | 0.983113793 | 0.994718474 |
| chr5:90337988-90338275 | 0.012053361 | 0.98312234 | 0.994718474 |
| chr4:65089665-65090203 | -0.007931112 | 0.983127901 | 0.994718474 |
| chr10:60139806-60140071 | 0.008624946 | 0.983130366 | 0.994718474 |
| chr20:41393292-41393632 | -0.007896049 | 0.983140803 | 0.994718474 |
| chr8:86834168-86834376 | 0.009967382 | 0.983151377 | 0.994718474 |
| chr10:123144915-123145239 | 0.008527533 | 0.983171358 | 0.994725555 |
| chr11:72684547-72684858 | -0.007721319 | 0.983189647 | 0.994728923 |
| chr1:200738500-200739881 | 0.006633333 | 0.9832021 | 0.994728923 |
| chr9:94681552-94681772 | 0.009496388 | 0.983234689 | 0.994728923 |
| chr2:232355124-232355329 | -0.0117 | 0.983239198 | 0.994728923 |
| chr11:64998041-64998323 | 0.010501805 | 0.983239598 | 0.994728923 |
| chr11:16785567-16785829 | 0.008429904 | 0.983279614 | 0.994738915 |
| chr6:107114102-107114819 | 0.007345822 | 0.983282039 | 0.994738915 |
| chr8:116920702-116921095 | 0.009092959 | 0.983292164 | 0.994738915 |
| chr6:20521360-20521600 | -0.010839228 | 0.983301404 | 0.994738915 |
| chr7:43799689-43799926 | 0.010567451 | 0.983340022 | 0.994754032 |
| chr4:8470603-8470812 | -0.011433597 | 0.983342313 | 0.994754032 |
| chr1:206734598-206734803 | 0.008164017 | 0.983371631 | 0.994770558 |
| chr1:156082807-156083011 | 0.008819821 | 0.983423037 | 0.994796183 |
| chr2:231906135-231906354 | -0.010240003 | 0.983427829 | 0.994796183 |
| chr20:54456376-54457303 | -0.006542591 | 0.983435912 | 0.994796183 |
| chr10:83166386-83166722 | 0.008466996 | 0.98345091 | 0.994798221 |

| | | | |
|---|---|---|---|
| chr2:176554119-176554782 | -0.007331992 | 0.983486953 | 0.994816364 |
| chr1:28170033-28170388 | 0.008919116 | 0.983502044 | 0.994816364 |
| chr8:88732978-88733207 | -0.009900332 | 0.983507797 | 0.994816364 |
| chr9:104977612-104977948 | 0.008460579 | 0.983530766 | 0.994826465 |
| chr9:109012302-109013843 | -0.006560562 | 0.983572933 | 0.9948553 |
| chr2:25544088-25544400 | 0.009927068 | 0.983585242 | 0.9948553 |
| chr2:111048958-111049572 | 0.007467399 | 0.983631961 | 0.994889422 |
| chr20:62977700-62978043 | -0.007401858 | 0.983694758 | 0.994939803 |
| chr1:175598654-175598854 | -0.011207628 | 0.983716898 | 0.994949063 |
| chrX:102338994-102339657 | 0.007643732 | 0.983766355 | 0.994964168 |
| chr20:46230779-46231205 | -0.00795334 | 0.983770083 | 0.994964168 |
| chr12:110543653-110543853 | 0.012601306 | 0.983812657 | 0.994964168 |
| chrX:74613987-74614372 | 0.007730734 | 0.983814143 | 0.994964168 |
| chr5:144203808-144204055 | 0.008881167 | 0.983822283 | 0.994964168 |
| chr6:30131062-30131281 | -0.009573467 | 0.983826841 | 0.994964168 |
| chr17:45385646-45385846 | -0.009336352 | 0.983846769 | 0.994964168 |
| chr20:58685787-58686203 | 0.007574241 | 0.983857847 | 0.994964168 |
| chr10:26890869-26891283 | -0.00760743 | 0.983876373 | 0.994964168 |
| chr5:79578884-79579184 | 0.00832959 | 0.983912501 | 0.994964168 |
| chr10:97327901-97328278 | 0.007367379 | 0.983920715 | 0.994964168 |
| chr5:152142935-152143172 | -0.008630428 | 0.983942759 | 0.994964168 |
| chr8:66176809-66177777 | -0.0065956 | 0.983944309 | 0.994964168 |
| chr19:29787364-29787580 | -0.00999378 | 0.983947786 | 0.994964168 |
| chr3:104102395-104102683 | 0.007771087 | 0.983984225 | 0.994964168 |
| chr5:43245713-43245970 | 0.009622772 | 0.984005182 | 0.994964168 |
| chr1:196503880-196504088 | -0.010302247 | 0.984011025 | 0.994964168 |
| chr12:117685645-117685875 | 0.009812671 | 0.984013652 | 0.994964168 |
| chr10:23438555-23439592 | 0.006440082 | 0.984024206 | 0.994964168 |
| chr15:41928174-41928435 | -0.00911623 | 0.984029633 | 0.994964168 |
| chr14:65373117-65373423 | -0.009068546 | 0.984053708 | 0.994964168 |
| chr3:14441713-14441946 | 0.008738291 | 0.984060703 | 0.994964168 |
| chr16:81368708-81368908 | 0.010843768 | 0.984076022 | 0.994964168 |
| chr6:158167741-158168883 | -0.006536336 | 0.984080462 | 0.994964168 |
| chr15:62344974-62345321 | 0.007409429 | 0.984094488 | 0.994964168 |
| chr5:93584273-93584473 | -0.008534737 | 0.984097058 | 0.994964168 |
| chr2:37396039-37396302 | -0.009657807 | 0.984128206 | 0.994964168 |
| chr1:220218657-220218947 | 0.008487885 | 0.984129326 | 0.994964168 |
| chr10:46545084-46545325 | 0.008487885 | 0.984129326 | 0.994964168 |
| chr1:66452368-66452636 | -0.009314017 | 0.984134402 | 0.994964168 |
| chr6:1381731-1382220 | -0.00779731 | 0.984156938 | 0.994964168 |
| chr2:172710576-172710999 | -0.007716512 | 0.984167782 | 0.994964168 |
| chr11:68974781-68975188 | -0.007970986 | 0.984169394 | 0.994964168 |
| chr4:54409057-54409570 | -0.008193231 | 0.984182971 | 0.994964168 |
| chrX:56729079-56729558 | -0.007534273 | 0.984196815 | 0.994964168 |
| chr3:138101399-138102063 | 0.007199829 | 0.98420604 | 0.994964168 |
| chr5:38033413-38033619 | -0.010061283 | 0.984212291 | 0.994964168 |
| chr20:45415918-45416562 | 0.006712265 | 0.984274202 | 0.994981774 |
| chr1:218171432-218171632 | 0.009755565 | 0.984284841 | 0.994981774 |
| chr3:66500344-66501621 | 0.006232308 | 0.984301301 | 0.994981774 |
| chr8:106870154-106870354 | -0.009901775 | 0.984308528 | 0.994981774 |

| | | | |
|---|---|---|---|
| chr7:100469264-100469547 | 0.008366656 | 0.984310544 | 0.994981774 |
| chr9:34702200-34702443 | -0.008893373 | 0.984312076 | 0.994981774 |
| chr14:24631576-24631818 | 0.009786149 | 0.984320605 | 0.994981774 |
| chr20:57326459-57327205 | -0.006321845 | 0.984343345 | 0.994982939 |
| chr9:93133988-93134544 | 0.006903774 | 0.98435594 | 0.994982939 |
| chr1:40783590-40783823 | 0.009254215 | 0.984375358 | 0.994982939 |
| chr19:46662049-46662306 | 0.007460687 | 0.984377143 | 0.994982939 |
| chr19:48899948-48900654 | 0.006572658 | 0.984404659 | 0.994982939 |
| chr7:20199405-20199669 | 0.00817722 | 0.984411216 | 0.994982939 |
| chr3:183825396-183826154 | 0.00662577 | 0.984450284 | 0.994982939 |
| chr7:35058735-35058935 | -0.009121972 | 0.984450445 | 0.994982939 |
| chr20:25233507-25233753 | -0.007997405 | 0.984462086 | 0.994982939 |
| chrX:115643089-115643289 | 0.010804221 | 0.98446336 | 0.994982939 |
| chr4:139329626-139330098 | 0.007560548 | 0.9844646 | 0.994982939 |
| chr1:187006765-187007032 | 0.009507584 | 0.984490672 | 0.994996165 |
| chrX:110847144-110847555 | -0.008305626 | 0.984622561 | 0.995075598 |
| chr11:111299137-11129940 | 0.008403117 | 0.984631606 | 0.995075598 |
| chr3:115325614-115325895 | -0.008303983 | 0.98464284 | 0.995075598 |
| chrX:55907951-55908368 | 0.007024961 | 0.984661648 | 0.995075598 |
| chr13:40537127-40537392 | 0.008647359 | 0.984677441 | 0.995075598 |
| chr10:53439050-53439350 | 0.00858868 | 0.9846782 | 0.995075598 |
| chr2:28392155-28393145 | -0.00612971 | 0.98467986 | 0.995075598 |
| chr2:165779105-165779368 | -0.00761361 | 0.984682712 | 0.995075598 |
| chr7:155724604-155724835 | 0.010229307 | 0.984686148 | 0.995075598 |
| chr12:82527275-82527490 | 0.009278909 | 0.984721052 | 0.995093697 |
| chrX:90854697-90855308 | -0.006724832 | 0.984742888 | 0.995093697 |
| chr3:4780822-4781056 | 0.010213081 | 0.984762218 | 0.995093697 |
| chr2:164697740-164697973 | -0.009304018 | 0.984777171 | 0.995093697 |
| chr7:47988889-47989132 | 0.008141745 | 0.984782642 | 0.995093697 |
| chr2:9994828-9995030 | 0.010899338 | 0.984783312 | 0.995093697 |
| chr5:166933320-166933520 | 0.010175329 | 0.98479564 | 0.995093697 |
| chr14:20934000-20934493 | 0.007876726 | 0.984813835 | 0.995093697 |
| chr11:43311560-43312564 | -0.006131101 | 0.984820941 | 0.995093697 |
| chr17:73068314-73068755 | -0.007460817 | 0.984836967 | 0.995096767 |
| chr1:29332240-29332480 | 0.009557743 | 0.984869337 | 0.995116352 |
| chr3:196073335-196073535 | 0.009806303 | 0.984896781 | 0.995130959 |
| chr9:109192805-109193247 | 0.007108466 | 0.984914728 | 0.99513597 |
| chr17:78708644-78708871 | -0.00854786 | 0.984946919 | 0.99514306 |
| chr12:121893904-12189436 | 0.007274386 | 0.984947721 | 0.99514306 |
| chr1:153538393-153538597 | 0.008423306 | 0.984983623 | 0.995156279 |
| chr20:32357566-32358610 | -0.006123861 | 0.984986779 | 0.995156279 |
| chr14:89012869-89013504 | 0.006269934 | 0.9850028 | 0.995158729 |
| chr8:140510656-140512367 | -0.005865917 | 0.985015181 | 0.995158729 |
| chr22:48721341-48721559 | -0.009336995 | 0.985031995 | 0.995162595 |
| chr7:67129264-67129515 | -0.00819655 | 0.985067612 | 0.995185456 |
| chr22:24521653-24522097 | -0.00703334 | 0.98509528 | 0.995200287 |
| chr13:70870001-70870201 | -0.010315742 | 0.985135586 | 0.995220369 |
| chr6:153565027-153565266 | 0.009279642 | 0.985141135 | 0.995220369 |
| chr1:234069859-234070145 | 0.008132854 | 0.985172513 | 0.995223361 |
| chr17:28576773-28577386 | 0.006239156 | 0.985215918 | 0.995223361 |

| | | | |
|---|---|---|---|
| chrX:81241753-81242114 | 0.008804397 | 0.985239409 | 0.995223361 |
| chr17:58925725-58926080 | 0.006451587 | 0.985247407 | 0.995223361 |
| chr14:75518774-75519046 | -0.007537743 | 0.985251239 | 0.995223361 |
| chr1:211326521-211326936 | -0.006932956 | 0.98525446 | 0.995223361 |
| chr22:41468327-41469367 | -0.005971226 | 0.985276648 | 0.995223361 |
| chr6:116876663-116876927 | -0.008391798 | 0.985278856 | 0.995223361 |
| chr3:165837288-165837540 | -0.009287076 | 0.985288297 | 0.995223361 |
| chr3:131729034-131729266 | 0.00875155 | 0.985301603 | 0.995223361 |
| chr20:49915373-49916153 | 0.00585673 | 0.985306699 | 0.995223361 |
| chr19:10899145-10899713 | 0.006306949 | 0.985324215 | 0.995223361 |
| chrX:120468422-120469528 | 0.005925214 | 0.985332765 | 0.995223361 |
| chr19:38384871-38385090 | 0.007797963 | 0.985342554 | 0.995223361 |
| chr2:117883172-117883372 | 0.010248518 | 0.985366963 | 0.995223361 |
| chr3:185937140-185938339 | -0.005836269 | 0.985380848 | 0.995223361 |
| chr9:6756993-6758715 | 0.005773115 | 0.985390775 | 0.995223361 |
| chr3:170037518-170039126 | -0.005753586 | 0.985404951 | 0.995223361 |
| chr9:129568557-129568864 | -0.00726603 | 0.985412079 | 0.995223361 |
| chr9:129823040-129823277 | 0.008160964 | 0.985414339 | 0.995223361 |
| chr3:58585383-58585662 | -0.007450261 | 0.985416861 | 0.995223361 |
| chr20:63979544-63980359 | -0.006033024 | 0.985463061 | 0.995243957 |
| chr14:76056273-76056639 | 0.007593036 | 0.985467736 | 0.995243957 |
| chr11:95836015-95836296 | 0.007761774 | 0.985485679 | 0.995243957 |
| chr20:58367829-58368044 | -0.008043324 | 0.985513688 | 0.995243957 |
| chr17:5031533-5031834 | -0.007767532 | 0.985528684 | 0.995243957 |
| chr3:157310725-157310925 | 0.009536188 | 0.98552909 | 0.995243957 |
| chrX:49184112-49185219 | 0.005808334 | 0.985535023 | 0.995243957 |
| chr17:43170040-43170641 | -0.00617304 | 0.985553858 | 0.995243957 |
| chr12:26195973-26196173 | 0.009104061 | 0.985572148 | 0.995243957 |
| chr12:54725551-54725798 | 0.009104061 | 0.985572148 | 0.995243957 |
| chr11:118917515-118917782 | 0.006598847 | 0.985580133 | 0.995243957 |
| chr3:13528270-13528845 | 0.006448521 | 0.985626831 | 0.995261006 |
| chr12:47900541-47901038 | 0.006279373 | 0.985630703 | 0.995261006 |
| chr14:76924685-76924958 | 0.007642229 | 0.985635984 | 0.995261006 |
| chrX:134172611-134172999 | -0.007563396 | 0.98569299 | 0.995296031 |
| chr19:46021535-46021774 | 0.007894174 | 0.98569665 | 0.995296031 |
| chr4:41934956-41935974 | -0.005836082 | 0.985733488 | 0.995307892 |
| chr3:17401978-17402311 | 0.008617075 | 0.985734376 | 0.995307892 |
| chr14:102747454-102747690 | 0.008471858 | 0.985766357 | 0.995327067 |
| chr20:53859739-53860101 | 0.005786323 | 0.985790997 | 0.99533883 |
| chr6:35485625-35485825 | -0.0079122 | 0.985815968 | 0.995350926 |
| chr17:41665874-41666228 | 0.006524746 | 0.985874724 | 0.995397134 |
| chr8:54551717-54552190 | 0.006666485 | 0.985900249 | 0.995400725 |
| chr2:105545781-105546014 | -0.008402849 | 0.985912608 | 0.995400725 |
| chr12:88816315-88816747 | -0.006727062 | 0.985933025 | 0.995400725 |
| chr8:85987286-85987555 | 0.006812974 | 0.985943498 | 0.995400725 |
| chr7:4261491-4261691 | 0.008861219 | 0.985954725 | 0.995400725 |
| chr19:10106084-10106652 | 0.006041538 | 0.985956227 | 0.995400725 |
| chr12:123232592-123233930 | -0.005575282 | 0.986020259 | 0.995449566 |
| chr17:73267863-73268176 | -0.007207984 | 0.986031453 | 0.995449566 |
| chr6:136927477-136927677 | -0.008330128 | 0.986045221 | 0.995449566 |

| | | | |
|---|---|---|---|
| chr21:39676014-39676477 | -0.006464744 | 0.986065755 | 0.995449566 |
| chr3:79937586-79937958 | -0.006871115 | 0.986080012 | 0.995449566 |
| chr2:171574129-171574401 | 0.007487713 | 0.986082554 | 0.995449566 |
| chr6:166894586-166895172 | -0.006558745 | 0.986175346 | 0.995502835 |
| chr10:43348500-43348830 | -0.006462514 | 0.986182503 | 0.995502835 |
| chr20:46013848-46014127 | -0.006934293 | 0.986211911 | 0.995502835 |
| chr11:125923109-125923321 | 0.008734958 | 0.986215088 | 0.995502835 |
| chr12:9319098-9319423 | -0.008094274 | 0.986222432 | 0.995502835 |
| chr16:75593969-75594169 | 0.00832509 | 0.986225715 | 0.995502835 |
| chr3:27134218-27134548 | -0.00691033 | 0.986242485 | 0.995502835 |
| chr11:8682257-8683138 | 0.005646618 | 0.986242875 | 0.995502835 |
| chr3:135055290-135055981 | 0.006704631 | 0.986292519 | 0.995502835 |
| chr17:29786038-29786448 | 0.006584196 | 0.986304806 | 0.995502835 |
| chr14:91635652-91635852 | -0.009311651 | 0.986313078 | 0.995502835 |
| chr11:43738918-43739298 | -0.007027435 | 0.986322457 | 0.995502835 |
| chr1:241579564-241579764 | 0.008620526 | 0.98632648 | 0.995502835 |
| chr14:95683797-95684086 | 0.007663037 | 0.986339337 | 0.995502835 |
| chr19:2637054-2637276 | 0.008248725 | 0.986344086 | 0.995502835 |
| chr16:28326962-28327238 | -0.007533581 | 0.986354033 | 0.995502835 |
| chr17:61411392-61412817 | 0.005297128 | 0.986356194 | 0.995502835 |
| chr13:46721251-46721451 | -0.007771557 | 0.986403625 | 0.995537594 |
| chr2:96868513-96868864 | -0.005947098 | 0.986436744 | 0.995557906 |
| chr1:233735974-233736210 | -0.007910562 | 0.986475823 | 0.995558077 |
| chr12:127677608-127677840 | -0.008252077 | 0.986491943 | 0.995558077 |
| chr10:38527244-38527711 | -0.00664788 | 0.986499069 | 0.995558077 |
| chr13:44048384-44048616 | 0.007823459 | 0.98651501 | 0.995558077 |
| chr4:146521805-146523068 | -0.00543999 | 0.986523819 | 0.995558077 |
| chr19:50024912-50026114 | 0.00538121 | 0.98653692 | 0.995558077 |
| chr1:222464852-222466031 | 0.005391338 | 0.986538079 | 0.995558077 |
| chr17:64145468-64146329 | 0.005346722 | 0.986549808 | 0.995558077 |
| chr6:73520243-73520558 | 0.005971023 | 0.986553852 | 0.995558077 |
| chr4:38958560-38958770 | -0.008562725 | 0.986592909 | 0.995584379 |
| chr8:27298265-27298525 | 0.007834373 | 0.986614927 | 0.995593485 |
| chr10:93741434-93741665 | -0.008342644 | 0.986657701 | 0.995599744 |
| chr7:76108884-76109110 | -0.008163766 | 0.986659459 | 0.995599744 |
| chr2:159921647-159922192 | 0.006065687 | 0.986660111 | 0.995599744 |
| chr1:20531025-20531376 | -0.007102743 | 0.986679978 | 0.995603242 |
| chr1:177087377-177087583 | -0.008761791 | 0.986700488 | 0.995603242 |
| chr1:157773775-157774091 | 0.006734351 | 0.986702558 | 0.995603242 |
| chr10:124461671-124462250 | -0.005710675 | 0.986732091 | 0.99561993 |
| chr11:18634147-18635013 | 0.005296721 | 0.986778162 | 0.995653305 |
| chr2:59425194-59425640 | 0.006661706 | 0.98681405 | 0.995664314 |
| chr9:100637777-100638337 | -0.006002956 | 0.986816585 | 0.995664314 |
| chr7:6483012-6483632 | -0.005510215 | 0.986828057 | 0.995664314 |
| chr2:7031499-7032246 | 0.005329788 | 0.986850966 | 0.995674318 |
| chr6:170553811-170554751 | 0.005225759 | 0.986874573 | 0.995685025 |
| chr19:4677169-4677456 | 0.007408799 | 0.986894581 | 0.995692101 |
| chr8:145002524-145003616 | -0.005242543 | 0.98694246 | 0.99572213 |
| chr8:26045297-26045927 | 0.005672382 | 0.987002339 | 0.99572213 |
| chr8:139742852-139743073 | 0.00715211 | 0.987009953 | 0.99572213 |

| | | | |
|---|---|---|---|
| chr14:106844212-106844491 | 0.006795635 | 0.987028363 | 0.99572213 |
| chr6:26532772-26533357 | 0.005437996 | 0.987060509 | 0.99572213 |
| chr16:14634046-14634317 | 0.005953698 | 0.987070416 | 0.99572213 |
| chr9:136545869-136546715 | 0.005385991 | 0.987070443 | 0.99572213 |
| chr10:101118659-101118859 | 0.008674986 | 0.987080904 | 0.99572213 |
| chr12:68185674-68185874 | 0.008674986 | 0.987080904 | 0.99572213 |
| chr18:112184-112424 | 0.007727089 | 0.987095779 | 0.99572213 |
| chr12:108562252-108563165 | 0.005211304 | 0.987100195 | 0.99572213 |
| chr19:37217676-37218471 | -0.005475346 | 0.987118518 | 0.99572213 |
| chr1:241485755-241486072 | -0.007111692 | 0.987119091 | 0.99572213 |
| chr3:130538973-130539321 | 0.006574113 | 0.987121689 | 0.99572213 |
| chr1:115838963-115839228 | -0.007188951 | 0.987146297 | 0.99572213 |
| chr6:101238733-101238936 | 0.007869906 | 0.987177605 | 0.99572213 |
| chr11:78715744-78716097 | 0.006187333 | 0.987182614 | 0.99572213 |
| chr4:149999615-149999817 | -0.007602048 | 0.987183994 | 0.99572213 |
| chr2:222971738-222972097 | -0.006364673 | 0.987184872 | 0.99572213 |
| chr19:16197623-16197976 | -0.006094495 | 0.987193715 | 0.99572213 |
| chr20:35992767-35992967 | -0.008373159 | 0.987197245 | 0.99572213 |
| chr21:32278682-32279542 | -0.005294819 | 0.98723353 | 0.995737086 |
| chr8:126510270-126510697 | -0.005277025 | 0.987249573 | 0.995737086 |
| chr17:49201162-49201527 | 0.006454104 | 0.987270185 | 0.995737086 |
| chr19:55216434-55217272 | 0.005270858 | 0.987277862 | 0.995737086 |
| chr2:220682603-220682991 | 0.006041086 | 0.987281045 | 0.995737086 |
| chr6:81534998-81535433 | -0.006845145 | 0.987290046 | 0.995737086 |
| chr15:69713036-69713280 | 0.007597877 | 0.987308905 | 0.995737558 |
| chr1:207401667-207401867 | 0.007825108 | 0.987324739 | 0.995737558 |
| chr20:327048-327620 | -0.005283033 | 0.9873295 | 0.995737558 |
| chr15:79993434-79993715 | -0.006876765 | 0.987367508 | 0.995762782 |
| chr12:88634255-88634455 | -0.007798852 | 0.987400203 | 0.995771644 |
| chr8:25169990-25170431 | -0.006402518 | 0.987402286 | 0.995771644 |
| chr20:36585509-36585956 | 0.005833684 | 0.987445844 | 0.995798704 |
| chr4:38319847-38320107 | 0.00638909 | 0.987455111 | 0.995798704 |
| chr12:94615453-94615709 | -0.006719484 | 0.987500795 | 0.995831668 |
| chr14:103715335-103715956 | 0.005041998 | 0.987571341 | 0.995884235 |
| chr15:67839567-67840724 | 0.005022171 | 0.987578924 | 0.995884235 |
| chr13:30963903-30964114 | 0.007113847 | 0.987591915 | 0.995884235 |
| chr1:167935485-167937109 | -0.004879393 | 0.987612266 | 0.99589165 |
| chr11:99020394-99021342 | 0.005178954 | 0.987632157 | 0.995898601 |
| chr2:119730603-119730803 | -0.008025981 | 0.98764873 | 0.995902207 |
| chr15:62482876-62483076 | -0.007933863 | 0.987679825 | 0.995920455 |
| chr9:33166849-33167737 | -0.005012826 | 0.987711782 | 0.995934247 |
| chr1:59759272-59759845 | -0.005576782 | 0.987722092 | 0.995934247 |
| chr10:91788238-91788902 | 0.005313652 | 0.987745794 | 0.995934247 |
| chr14:81805723-81806018 | 0.006422888 | 0.987757115 | 0.995934247 |
| chr6:112163387-112163727 | -0.006131161 | 0.987762895 | 0.995934247 |
| chr1:26864961-26865672 | -0.005156414 | 0.987771491 | 0.995934247 |
| chr19:48795110-48795767 | -0.005178513 | 0.987818767 | 0.995967285 |
| chr12:123983960-123984214 | 0.006921968 | 0.987835006 | 0.995967285 |
| chr3:33277186-33277915 | -0.005303787 | 0.987843253 | 0.995967285 |
| chr2:114068536-114068740 | -0.007863549 | 0.987890221 | 0.996001534 |

| | | | |
|---|---|---|---|
| chr1:42035178-42036276 | 0.004942136 | 0.98794845 | 0.996032064 |
| chr1:155226565-155226820 | -0.005924581 | 0.987965719 | 0.996032064 |
| chr17:59706834-59708208 | 0.004707307 | 0.987966849 | 0.996032064 |
| chr8:143540952-143541861 | -0.004909864 | 0.9879725 | 0.996032064 |
| chr12:110512514-110512714 | 0.008294608 | 0.988048933 | 0.99608658 |
| chr13:30743413-30743951 | -0.005371416 | 0.988072109 | 0.99608658 |
| chr11:63872933-63873201 | 0.007008496 | 0.988075618 | 0.99608658 |
| chr3:101840742-101840942 | -0.007572323 | 0.988078575 | 0.99608658 |
| chr6:91590343-91590654 | 0.006522952 | 0.988179985 | 0.996175705 |
| chr9:86052894-86053364 | -0.006116369 | 0.98819649 | 0.996179237 |
| chr4:139656671-139656871 | -0.006619553 | 0.988217322 | 0.996187131 |
| chr11:68459910-68461044 | -0.004636108 | 0.988289891 | 0.996187508 |
| chr14:100425158-100425358 | -0.006703576 | 0.988299362 | 0.996187508 |
| chr12:125032589-125033366 | 0.005257232 | 0.988331835 | 0.996187508 |
| chr5:163350089-163350303 | -0.006901862 | 0.988332072 | 0.996187508 |
| chrX:107894114-107894482 | -0.005897268 | 0.98834571 | 0.996187508 |
| chr6:27721019-27721560 | -0.005319024 | 0.988352846 | 0.996187508 |
| chr5:132963187-132964161 | -0.004643771 | 0.988357207 | 0.996187508 |
| chr17:41966510-41966931 | -0.005091874 | 0.988387653 | 0.996187508 |
| chr2:241659293-241659638 | 0.006305123 | 0.988398857 | 0.996187508 |
| chr2:113149833-113150078 | 0.007090775 | 0.98840231 | 0.996187508 |
| chr20:3970095-3970342 | -0.007277432 | 0.988404332 | 0.996187508 |
| chr5:108231441-108231860 | 0.006121875 | 0.988410788 | 0.996187508 |
| chr3:125632559-125632759 | 0.008489553 | 0.988418989 | 0.996187508 |
| chr17:73985445-73985681 | -0.006947135 | 0.988433186 | 0.996187508 |
| chr7:70829177-70829522 | 0.006250748 | 0.988437804 | 0.996187508 |
| chr10:48821604-48821880 | -0.007188807 | 0.988458317 | 0.996187508 |
| chr3:112378633-112378833 | -0.007719291 | 0.988460313 | 0.996187508 |
| chr8:108083539-108083775 | 0.005697469 | 0.988464264 | 0.996187508 |
| chr11:66727318-66727747 | -0.005512233 | 0.988476588 | 0.996187508 |
| chr15:64006777-64007081 | -0.006635202 | 0.988477723 | 0.996187508 |
| chr16:85114933-85115262 | -0.005224105 | 0.988515678 | 0.996212656 |
| chr6:42075445-42075799 | 0.005860771 | 0.988546523 | 0.996230638 |
| chr8:117862111-117862361 | 0.006241582 | 0.988570439 | 0.996241638 |
| chr8:30155930-30156797 | 0.004751791 | 0.988586276 | 0.996244494 |
| chr2:37982083-37982471 | -0.005490784 | 0.988612345 | 0.996244774 |
| chr3:82947130-82947449 | -0.006050189 | 0.988612558 | 0.996244774 |
| chr8:135335826-135336027 | -0.007015298 | 0.988706876 | 0.996324077 |
| chr15:58987425-58988560 | -0.004655675 | 0.988718643 | 0.996324077 |
| chr6:35342214-35342974 | 0.004740972 | 0.988730262 | 0.996324077 |
| chr15:98872959-98873182 | -0.005689795 | 0.988763385 | 0.996344351 |
| chr8:117761133-117761395 | -0.005921851 | 0.988815005 | 0.996347762 |
| chr5:69492262-69493352 | 0.00448139 | 0.988825989 | 0.996347762 |
| chr11:77035651-77035869 | 0.006331212 | 0.988831551 | 0.996347762 |
| chr1:113811481-113812766 | -0.004347165 | 0.988839351 | 0.996347762 |
| chr2:24708198-24708564 | -0.007262594 | 0.988841721 | 0.996347762 |
| chr15:100601879-100603370 | 0.004368743 | 0.988881262 | 0.996347762 |
| chr7:30938613-30939105 | 0.005117041 | 0.9889095 | 0.996347762 |
| chr2:68031605-68031825 | 0.007208256 | 0.988932373 | 0.996347762 |
| chr15:71304118-71304382 | -0.007149334 | 0.988936377 | 0.996347762 |

| | | | |
|---|---|---|---|
| chr21:45074130-45074548 | -0.005221327 | 0.988945513 | 0.996347762 |
| chr2:1887840-1888161 | 0.006005459 | 0.988950511 | 0.996347762 |
| chrX:105666488-105667005 | 0.005233779 | 0.98895077 | 0.996347762 |
| chr2:234496996-234497441 | -0.005698885 | 0.988970664 | 0.996347762 |
| chr3:155491718-155491998 | -0.0058723 | 0.988972112 | 0.996347762 |
| chr11:64342402-64343086 | -0.004639585 | 0.988991912 | 0.996347762 |
| chr9:86639889-86640336 | -0.005589161 | 0.9889963 | 0.996347762 |
| chr5:69032126-69032326 | 0.007087835 | 0.988999028 | 0.996347762 |
| chr17:4533928-4534289 | -0.005336569 | 0.989000832 | 0.996347762 |
| chr8:121097342-121097773 | -0.004794118 | 0.98904117 | 0.996375299 |
| chr6:88963227-88964163 | -0.004384383 | 0.989055016 | 0.996376148 |
| chr4:41692729-41693184 | 0.005576505 | 0.989118747 | 0.99642725 |
| chr2:63680954-63681206 | 0.007015252 | 0.989134396 | 0.996429914 |
| chr1:1073689-1074512 | 0.004479345 | 0.989151045 | 0.996433585 |
| chr16:77981-78333 | 0.004972989 | 0.989179393 | 0.996447072 |
| chr17:52160886-52161093 | 0.006673999 | 0.989190442 | 0.996447072 |
| chr10:117226880-11722712C | -0.005886397 | 0.98920846 | 0.996452121 |
| chr14:20461330-20462094 | -0.004226404 | 0.989310718 | 0.996527929 |
| chr5:80255374-80256559 | 0.004261361 | 0.989349559 | 0.996527929 |
| chr10:100999189-100999438 | 0.00517399 | 0.989356826 | 0.996527929 |
| chr1:235104782-235104983 | 0.006051269 | 0.989362149 | 0.996527929 |
| chr19:40444130-40444938 | -0.004314542 | 0.989368215 | 0.996527929 |
| chr17:16492099-16492338 | 0.006347875 | 0.989387256 | 0.996527929 |
| chr3:156107076-156107366 | 0.005673354 | 0.989388427 | 0.996527929 |
| chr8:140898252-140898527 | 0.005722773 | 0.989425671 | 0.996527929 |
| chr22:17381587-17381942 | 0.005332614 | 0.989454112 | 0.996527929 |
| chr1:63674463-63674714 | -0.005659112 | 0.989454753 | 0.996527929 |
| chr2:10689652-10690783 | -0.004379358 | 0.989468627 | 0.996527929 |
| chr4:32775000-32775376 | 0.005049786 | 0.989487055 | 0.996527929 |
| chr3:113035372-113035771 | -0.004755597 | 0.989503974 | 0.996527929 |
| chr11:48011146-48011403 | -0.005372275 | 0.989509409 | 0.996527929 |
| chr10:28991788-28992164 | -0.005157509 | 0.989518252 | 0.996527929 |
| chr4:38110005-38110205 | 0.007264116 | 0.989521123 | 0.996527929 |
| chr9:135023387-135023704 | -0.005979889 | 0.989525921 | 0.996527929 |
| chr4:121736823-121737073 | 0.005587955 | 0.989530534 | 0.996527929 |
| chrX:48761764-48762348 | -0.004629809 | 0.989535893 | 0.996527929 |
| chr8:92674536-92674736 | 0.007818524 | 0.98956406 | 0.996527929 |
| chr16:30893584-30893881 | -0.005184215 | 0.989567351 | 0.996527929 |
| chr9:74951809-74952042 | 0.005579578 | 0.989581432 | 0.996527929 |
| chr17:8119597-8119907 | 0.004666997 | 0.98958285 | 0.996527929 |
| chr14:77623257-77623478 | -0.005990015 | 0.989603264 | 0.99653539 |
| chr2:30324501-30324701 | -0.006880548 | 0.989647592 | 0.996536108 |
| chr2:80658321-80658521 | -0.006880548 | 0.989647592 | 0.996536108 |
| chr4:103995037-103995320 | -0.005853758 | 0.989672099 | 0.996536108 |
| chr20:62301847-62303139 | 0.004076285 | 0.989673361 | 0.996536108 |
| chr5:60007726-60008006 | -0.006070265 | 0.989676664 | 0.996536108 |
| chr3:61560650-61561997 | 0.004204161 | 0.989684105 | 0.996536108 |
| chr17:40443716-40444063 | 0.004898169 | 0.989707414 | 0.996536108 |
| chr11:65382290-65383015 | -0.00431241 | 0.989708024 | 0.996536108 |
| chr1:204110902-204111540 | -0.00437756 | 0.989738544 | 0.996549922 |

| | | | |
|---|---|---|---|
| chr1:52926995-52927991 | 0.004370375 | 0.989747756 | 0.996549922 |
| chr1:244327664-244327864 | -0.006995956 | 0.989791171 | 0.996569029 |
| chr15:98860219-98860627 | 0.004664718 | 0.989801622 | 0.996569029 |
| chr16:1200755-1200970 | -0.006177082 | 0.989814977 | 0.996569029 |
| chr19:40190858-40191547 | 0.004664129 | 0.989831609 | 0.996569029 |
| chr3:122483297-122483502 | -0.007049656 | 0.989831763 | 0.996569029 |
| chr9:69936485-69936740 | 0.006234751 | 0.989845295 | 0.996569558 |
| chr12:51993083-51993475 | 0.004723378 | 0.989859961 | 0.996571228 |
| chr14:22930359-22930889 | 0.004319627 | 0.98988926 | 0.996576665 |
| chr19:39996388-39997359 | -0.00431768 | 0.989900207 | 0.996576665 |
| chr4:143178890-143179307 | 0.00449989 | 0.98990438 | 0.996576665 |
| chr8:129826844-129827091 | 0.005578628 | 0.989961068 | 0.996620641 |
| chr14:76776771-76776978 | 0.005375592 | 0.989975833 | 0.99662241 |
| chr7:44036391-44036771 | 0.004176309 | 0.990009969 | 0.996631752 |
| chrX:63506384-63506593 | 0.006148576 | 0.990011126 | 0.996631752 |
| chr8:74965803-74966277 | -0.00471874 | 0.990113683 | 0.996705186 |
| chr6:156380373-156380801 | -0.005233732 | 0.990121539 | 0.996705186 |
| chr8:132478649-132478943 | 0.004644488 | 0.990123097 | 0.996705186 |
| chr11:34840639-34841064 | -0.005222102 | 0.990143492 | 0.996712622 |
| chr10:80203561-80204020 | -0.004454643 | 0.99021027 | 0.996765806 |
| chr7:2260879-2261215 | 0.004486444 | 0.990233547 | 0.996765806 |
| chr7:32765532-32765807 | -0.00542869 | 0.990237527 | 0.996765806 |
| chr5:142748628-142748855 | -0.005910489 | 0.990262089 | 0.996765806 |
| chr17:50708633-50708958 | -0.004771862 | 0.990286479 | 0.996765806 |
| chr12:122072273-122072874 | 0.004470139 | 0.99030743 | 0.996765806 |
| chr11:74443155-74443413 | 0.005308245 | 0.990312127 | 0.996765806 |
| chr22:50525200-50526277 | 0.004003319 | 0.99032023 | 0.996765806 |
| chr20:3659950-3660192 | -0.005921637 | 0.990324123 | 0.996765806 |
| chr7:5562027-5563905 | 0.003813682 | 0.990338995 | 0.996765806 |
| chr10:116684284-116684504 | -0.00578879 | 0.990344535 | 0.996765806 |
| chr15:72783330-72784640 | -0.00387979 | 0.990352432 | 0.996765806 |
| chr1:58620113-58620672 | 0.004058589 | 0.990379261 | 0.996779715 |
| chrX:8845109-8845408 | 0.006040068 | 0.990393258 | 0.99678071 |
| chr9:106953259-106953500 | 0.005973178 | 0.990434018 | 0.996808639 |
| chr3:128335351-128336182 | -0.003957105 | 0.990494586 | 0.9968447 |
| chr12:70013119-70013553 | -0.004839069 | 0.990495868 | 0.9968447 |
| chr5:25544696-25545048 | 0.004506718 | 0.990511332 | 0.99684717 |
| chr20:816064-816264 | 0.00567282 | 0.990530675 | 0.996853543 |
| chrX:15737826-15738682 | 0.003749683 | 0.990613723 | 0.996924028 |
| chr6:119129243-119129702 | 0.0043691 | 0.990694145 | 0.996987956 |
| chr12:120446020-120446927 | -0.003891045 | 0.990703271 | 0.996987956 |
| chr2:208101823-208102260 | 0.004727645 | 0.990749041 | 0.996998592 |
| chr2:124323319-124323586 | -0.005212026 | 0.990768691 | 0.996998592 |
| chr6:44001443-44001683 | -0.005069644 | 0.990780249 | 0.996998592 |
| chr4:140715091-140715434 | -0.005099246 | 0.990780705 | 0.996998592 |
| chr6:102629110-102629310 | 0.005870569 | 0.990784415 | 0.996998592 |
| chr20:35523342-35523542 | -0.005512746 | 0.990798904 | 0.996998592 |
| chr11:45804026-45805654 | 0.003651994 | 0.990814239 | 0.996998592 |
| chr3:67357828-67358368 | -0.004329431 | 0.990818102 | 0.996998592 |
| chr6:132568931-132569140 | -0.006438274 | 0.990833761 | 0.996998592 |

| | | | |
|---|---|---|---|
| chrX:37745473-37745673 | -0.005740734 | 0.990843958 | 0.996998592 |
| chr19:32357794-32358009 | 0.006090261 | 0.990879679 | 0.997010323 |
| chr1:34852686-34853293 | -0.00393336 | 0.990885872 | 0.997010323 |
| chr1:227779679-227779879 | 0.006056695 | 0.990894653 | 0.997010323 |
| chr3:194682536-194683186 | 0.003899792 | 0.990919141 | 0.997021869 |
| chr3:126083512-126084522 | -0.003728417 | 0.990936715 | 0.99702646 |
| chr3:69161677-69162088 | -0.004554316 | 0.990974085 | 0.997027227 |
| chr4:122151915-122153061 | 0.003647214 | 0.990975489 | 0.997027227 |
| chr3:165712377-165712577 | -0.005711846 | 0.990976515 | 0.997027227 |
| chr4:71540945-71541145 | 0.00554421 | 0.991020656 | 0.997058545 |
| chr10:43093373-43093743 | -0.004188167 | 0.991101977 | 0.997127269 |
| chr21:44964613-44964813 | -0.005625075 | 0.991149957 | 0.997162447 |
| chr20:3408659-3409021 | 0.004028266 | 0.991188984 | 0.997188618 |
| chr9:96441835-96442180 | 0.004495613 | 0.991212822 | 0.997189491 |
| chr1:114956161-114956904 | 0.003665334 | 0.991220228 | 0.997189491 |
| chr3:140128666-140128955 | -0.004402992 | 0.991228895 | 0.997189491 |
| chr7:36487850-36488072 | -0.005479515 | 0.99124988 | 0.99719751 |
| chr6:100904181-100904381 | 0.00685722 | 0.991274332 | 0.997209015 |
| chr2:222690616-222690921 | -0.004319455 | 0.991312841 | 0.997234662 |
| chr8:100458476-100458784 | 0.005518211 | 0.991352032 | 0.997260994 |
| chr11:6439104-6439304 | 0.005264344 | 0.991404058 | 0.997300236 |
| chr5:74212347-74212708 | -0.004523542 | 0.991466494 | 0.997329476 |
| chr6:3117996-3119272 | 0.003399804 | 0.991473468 | 0.997329476 |
| chr14:24108617-24109101 | -0.003881161 | 0.991477878 | 0.997329476 |
| chr21:28696935-28697144 | -0.005193173 | 0.991505288 | 0.997329476 |
| chr1:201530787-201531115 | -0.003827883 | 0.99151042 | 0.997329476 |
| chr16:10580444-10580939 | -0.003632674 | 0.99154132 | 0.997329476 |
| chr2:238234335-238234816 | 0.004227038 | 0.991551499 | 0.997329476 |
| chr1:113677941-113678151 | -0.004827867 | 0.99155183 | 0.997329476 |
| chr12:57583353-57583688 | 0.0040903 | 0.99156656 | 0.997329476 |
| chr14:51564436-51564752 | -0.00445303 | 0.991571897 | 0.997329476 |
| chr19:4140401-4140656 | -0.004573594 | 0.991576303 | 0.997329476 |
| chr17:78132350-78132574 | -0.004273744 | 0.991626691 | 0.99734611 |
| chr1:206933405-206933605 | 0.00429382 | 0.991641349 | 0.99734611 |
| chr11:72102786-72103814 | -0.003350049 | 0.991643341 | 0.99734611 |
| chr1:224930277-224930840 | 0.003682065 | 0.991650942 | 0.99734611 |
| chr4:117700567-117700911 | 0.004425221 | 0.991696066 | 0.99734611 |
| chr14:87888212-87888422 | 0.004795494 | 0.991699622 | 0.99734611 |
| chr12:32365550-32365753 | 0.00524011 | 0.991705859 | 0.99734611 |
| chr1:85258996-85260098 | -0.00334628 | 0.991710308 | 0.99734611 |
| chr8:53842554-53843630 | -0.003334732 | 0.991714377 | 0.99734611 |
| chr1:161037959-161039166 | -0.003340577 | 0.991723006 | 0.99734611 |
| chr18:4068702-4068908 | 0.005105935 | 0.991755893 | 0.997366094 |
| chr3:109300075-109300309 | -0.005485355 | 0.991801766 | 0.997374825 |
| chr16:85010849-85011957 | -0.003335519 | 0.991808116 | 0.997374825 |
| chr16:13426418-13426618 | -0.005665893 | 0.991808322 | 0.997374825 |
| chr5:119470865-119471170 | 0.004733278 | 0.991816643 | 0.997374825 |
| chrX:30756419-30756855 | 0.004881804 | 0.991879896 | 0.997425343 |
| chr11:87421407-87421660 | -0.004685894 | 0.991908544 | 0.997438025 |
| chr19:31596569-31597005 | -0.004471387 | 0.991921294 | 0.997438025 |

| | | | |
|---|---|---|---|
| chr6:125995972-125996431 | -0.003653076 | 0.991931561 | 0.997438025 |
| chr2:88055109-88056046 | -0.003322398 | 0.991944638 | 0.997438085 |
| chr9:33623982-33624309 | -0.004174061 | 0.991994882 | 0.997468566 |
| chr1:234867324-234867628 | 0.004335964 | 0.992001045 | 0.997468566 |
| chr15:43122793-43123484 | 0.003695843 | 0.992018607 | 0.997468566 |
| chr2:173680705-173680994 | -0.004532995 | 0.992027023 | 0.997468566 |
| chr5:27903554-27903885 | 0.005077332 | 0.992071286 | 0.997499981 |
| chr6:26956199-26957144 | 0.003353103 | 0.992104022 | 0.997519806 |
| chrX:13110830-13111254 | 0.004193007 | 0.992151755 | 0.997543779 |
| chr1:48236878-48237221 | -0.003836113 | 0.992153902 | 0.997543779 |
| chr6:110874305-110874922 | 0.003206884 | 0.992185536 | 0.997557642 |
| chr6:3433084-3433327 | -0.00450822 | 0.992193729 | 0.997557642 |
| chr13:48318106-48318346 | -0.00546629 | 0.992226698 | 0.997577699 |
| chr11:61680132-61680448 | 0.003911761 | 0.992253829 | 0.997586347 |
| chr1:203795371-203796544 | 0.003088257 | 0.99227753 | 0.997586347 |
| chr18:58768434-58768983 | 0.003565782 | 0.992288815 | 0.997586347 |
| chr1:7561541-7561767 | -0.004566057 | 0.992306278 | 0.997586347 |
| chr11:108008646-108009465 | -0.003177613 | 0.992309511 | 0.997586347 |
| chr3:167178574-167179069 | 0.003506483 | 0.992317166 | 0.997586347 |
| chr2:153197317-153197519 | -0.004856396 | 0.992326436 | 0.997586347 |
| chr3:107871325-107871663 | -0.003753832 | 0.9923706 | 0.997617656 |
| chr5:14582033-14582817 | 0.003150401 | 0.992394725 | 0.99762882 |
| chr20:9542720-9542956 | -0.00488193 | 0.992432366 | 0.99765357 |
| chr3:171543166-171543366 | 0.004630713 | 0.992463212 | 0.997671489 |
| chr14:60247969-60250012 | -0.002943966 | 0.99247925 | 0.997674522 |
| chr1:207453944-207454965 | -0.003134225 | 0.992518609 | 0.997700997 |
| chr8:125897481-125897710 | 0.004655146 | 0.99254194 | 0.997706348 |
| chr20:18466684-18467808 | 0.00299921 | 0.992549974 | 0.997706348 |
| chr5:61331401-61332379 | 0.002941427 | 0.992592605 | 0.997736111 |
| chr13:110944450-110944677 | -0.004151192 | 0.992639202 | 0.997769861 |
| chr9:117644644-117645210 | -0.003386565 | 0.992659673 | 0.997776136 |
| chr2:8836956-8837837 | -0.002990972 | 0.992674187 | 0.997776136 |
| chr1:174189767-174189967 | 0.00467828 | 0.992684511 | 0.997776136 |
| chr4:57190902-57191285 | -0.003569359 | 0.992720024 | 0.997792989 |
| chr1:110203105-110203315 | -0.003971609 | 0.992727323 | 0.997792989 |
| chr20:36835251-36836123 | -0.003072491 | 0.992801886 | 0.997854843 |
| chr6:30690647-30691238 | 0.003051822 | 0.992847646 | 0.997877169 |
| chr2:201116057-201116716 | 0.003038253 | 0.992850146 | 0.997877169 |
| chr4:44677990-44679052 | -0.00285185 | 0.992870468 | 0.997878513 |
| chr10:96271454-96271877 | -0.003322505 | 0.992877853 | 0.997878513 |
| chr7:128167310-128167682 | 0.003630042 | 0.992891952 | 0.997878513 |
| chr14:91255111-91255404 | 0.00346095 | 0.992906219 | 0.997878513 |
| chr1:110383398-110383693 | 0.003626999 | 0.9929166 | 0.997878513 |
| chr19:39970533-39971343 | 0.002872498 | 0.992948154 | 0.997897136 |
| chr9:83649170-83649512 | 0.003569189 | 0.992967514 | 0.997898821 |
| chr3:118636910-118637162 | -0.003887847 | 0.993003647 | 0.997898821 |
| chr17:14309094-14309707 | 0.002876498 | 0.993006735 | 0.997898821 |
| chr14:37145357-37145719 | 0.003927534 | 0.99300986 | 0.997898821 |
| chr11:78564238-78564712 | 0.00352536 | 0.993014949 | 0.997898821 |
| chr3:39177410-39177686 | 0.003429337 | 0.993033704 | 0.997904581 |

| | | | |
|---|---|---|---|
| chr17:5260696-5260926 | -0.004495831 | 0.993091007 | 0.997918405 |
| chr1:201119281-201119524 | 0.003230275 | 0.993098466 | 0.997918405 |
| chr16:84970578-84970938 | 0.003207398 | 0.993111514 | 0.997918405 |
| chr3:103690234-103690948 | -0.002982076 | 0.993113534 | 0.997918405 |
| chr18:449347-449862 | -0.003289557 | 0.993132636 | 0.997918405 |
| chr1:229162291-229162625 | -0.003678248 | 0.993146042 | 0.997918405 |
| chr8:20296313-20296513 | 0.005620753 | 0.993150786 | 0.997918405 |
| chrX:54639318-54640156 | -0.00277105 | 0.993151652 | 0.997918405 |
| chr9:96077222-96077422 | -0.004053743 | 0.993191295 | 0.997932931 |
| chr7:34184504-34184707 | -0.003897033 | 0.993192157 | 0.997932931 |
| chr15:68433867-68434108 | 0.004618556 | 0.993231482 | 0.997948768 |
| chr14:67532805-67533999 | -0.00275206 | 0.993233967 | 0.997948768 |
| chr1:224434675-224435560 | 0.00276918 | 0.993273542 | 0.997974164 |
| chr1:25905987-25907054 | -0.002786585 | 0.993296531 | 0.997974164 |
| chr4:103303130-103303337 | 0.004338096 | 0.993308897 | 0.997974164 |
| chr5:10509083-10509283 | -0.004001064 | 0.993319261 | 0.997974164 |
| chr9:125240907-125242161 | 0.002671703 | 0.993324367 | 0.997974164 |
| chr11:32006885-32007095 | 0.004656231 | 0.993401939 | 0.998003125 |
| chr1:167059032-167059675 | -0.00285125 | 0.993405794 | 0.998003125 |
| chrX:103538975-103539186 | -0.003927792 | 0.993422042 | 0.998003125 |
| chr4:174120478-174120814 | 0.003275068 | 0.993429445 | 0.998003125 |
| chr2:241149234-241150032 | 0.002691359 | 0.993429777 | 0.998003125 |
| chr22:31969968-31971176 | -0.002677077 | 0.993431343 | 0.998003125 |
| chr13:95404295-95404515 | -0.003916841 | 0.993459215 | 0.998018041 |
| chr8:81651406-81651606 | -0.004147543 | 0.993531051 | 0.99807712 |
| chr3:184541076-184541424 | 0.003232859 | 0.993563205 | 0.998096335 |
| chr1:153687017-153687496 | 0.003123242 | 0.993576686 | 0.998096757 |
| chr7:99500124-99500495 | -0.002866938 | 0.993598473 | 0.998096757 |
| chr4:147831620-147832338 | -0.00269612 | 0.993602703 | 0.998096757 |
| chr19:5804001-5804315 | -0.003392981 | 0.993634267 | 0.99810235 |
| chr3:62562586-62562788 | -0.00319462 | 0.993641442 | 0.99810235 |
| chr12:104677406-104678103 | 0.002709902 | 0.99364735 | 0.99810235 |
| chr1:77509260-77509518 | 0.003979633 | 0.993694591 | 0.998126857 |
| chr4:102500649-102502531 | 0.002516062 | 0.993697801 | 0.998126857 |
| chr8:144754583-144755907 | -0.002504894 | 0.993726451 | 0.998136418 |
| chr15:66025283-66025550 | 0.003663908 | 0.993735388 | 0.998136418 |
| chr19:19627650-19627850 | -0.003647656 | 0.9937464 | 0.998136418 |
| chr5:62434227-62434453 | -0.004307925 | 0.993765302 | 0.998142319 |
| chr17:39927262-39927858 | 0.00303066 | 0.993830464 | 0.998194683 |
| chr5:59361520-59361780 | 0.00349153 | 0.993897612 | 0.998233602 |
| chr9:134502489-134502843 | -0.003482161 | 0.993930646 | 0.998233602 |
| chr1:113146761-113147214 | 0.002894082 | 0.993930723 | 0.998233602 |
| chr8:125664258-125664879 | 0.00258429 | 0.993934002 | 0.998233602 |
| chr13:113951099-113951509 | 0.003016013 | 0.9939416 | 0.998233602 |
| chr8:88843550-88843907 | -0.003018951 | 0.99394783 | 0.998233602 |
| chr1:196148476-196148676 | 0.003549433 | 0.993960408 | 0.998233602 |
| chr1:23790836-23791441 | 0.002734894 | 0.994013957 | 0.998265338 |
| chr1:179814555-179815608 | 0.002385987 | 0.994054219 | 0.998265338 |
| chr3:165347828-165348052 | 0.003876105 | 0.994054929 | 0.998265338 |
| chr12:75488616-75488881 | 0.002992737 | 0.994074576 | 0.998265338 |

| | | | |
|---|---|---|---|
| chr2:10490800-10491098 | -0.00306846 | 0.994088624 | 0.998265338 |
| chr7:151745195-151745650 | -0.002815212 | 0.994091102 | 0.998265338 |
| chr10:69994322-69994566 | -0.003291698 | 0.994096466 | 0.998265338 |
| chr10:99246843-99247057 | 0.003936257 | 0.99409725 | 0.998265338 |
| chr17:61978459-61978659 | 0.003335973 | 0.994110351 | 0.998265338 |
| chr13:45807556-45807898 | -0.002936734 | 0.994122293 | 0.998265338 |
| chr1:217078203-217078411 | 0.003142283 | 0.994163507 | 0.998282901 |
| chr8:91197540-91197925 | -0.003052219 | 0.994171548 | 0.998282901 |
| chr13:70170760-70171198 | -0.002916232 | 0.994191778 | 0.998282901 |
| chr3:181711311-181712042 | 0.002448005 | 0.994192929 | 0.998282901 |
| chr15:57619895-57620125 | 0.003505521 | 0.994204927 | 0.998282901 |
| chr2:95991801-95992001 | 0.003481043 | 0.994274386 | 0.998339562 |
| chr6:1552562-1552768 | 0.003825938 | 0.994296809 | 0.998346748 |
| chr4:75468787-75469080 | -0.00316804 | 0.994307602 | 0.998346748 |
| chr14:24430433-24430738 | -0.002537802 | 0.994353495 | 0.998379745 |
| chr2:69894487-69894687 | 0.003419876 | 0.994393656 | 0.998406986 |
| chr7:96800800-96801102 | 0.003256341 | 0.9944264 | 0.998418093 |
| chr3:133578581-133578921 | 0.002497796 | 0.994450246 | 0.998418093 |
| chr5:112175000-112175309 | 0.002834605 | 0.99445401 | 0.998418093 |
| chr10:91632986-91633561 | 0.002377508 | 0.994466087 | 0.998418093 |
| chr2:143034328-143034723 | -0.002485929 | 0.994469871 | 0.998418093 |
| chr14:100546369-100547148 | 0.002271179 | 0.994514959 | 0.998443506 |
| chr1:54081663-54082031 | -0.00268417 | 0.994523415 | 0.998443506 |
| chr8:140481166-140481416 | -0.003441475 | 0.994534275 | 0.998443506 |
| chr15:90149737-90149938 | 0.003335261 | 0.994549979 | 0.998444465 |
| chr3:29453394-29453765 | -0.00295833 | 0.994584128 | 0.998444465 |
| chr2:11656597-11657374 | -0.002262554 | 0.994612101 | 0.998444465 |
| chr1:37473753-37474833 | 0.002165429 | 0.994620155 | 0.998444465 |
| chrX:133444890-133445764 | 0.002377782 | 0.994643385 | 0.998444465 |
| chr8:130675548-130675777 | -0.003062567 | 0.994648325 | 0.998444465 |
| chr15:98609626-98609849 | 0.00342975 | 0.994659043 | 0.998444465 |
| chr4:139014592-139016372 | -0.002104569 | 0.994661256 | 0.998444465 |
| chr13:31982597-31982901 | -0.002807219 | 0.994665541 | 0.998444465 |
| chr19:58386558-58387406 | 0.002149716 | 0.994666922 | 0.998444465 |
| chr10:97574528-97575077 | -0.00228487 | 0.99467857 | 0.998444465 |
| chr18:3512154-3512417 | 0.003164926 | 0.994709156 | 0.998462087 |
| chr2:190786167-190787204 | 0.002196121 | 0.994733088 | 0.998465604 |
| chr15:67915627-67916064 | 0.002544533 | 0.994745671 | 0.998465604 |
| chr5:172772009-172772326 | 0.00232697 | 0.994751753 | 0.998465604 |
| chr19:54354035-54354264 | 0.003260658 | 0.994776149 | 0.998477012 |
| chr3:149016320-149016520 | 0.002941964 | 0.994789313 | 0.998477145 |
| chr7:47266316-47266706 | -0.002598945 | 0.994805015 | 0.998479825 |
| chr13:41054324-41054571 | -0.002671944 | 0.994830922 | 0.99847994 |
| chr17:63857144-63857364 | -0.002697171 | 0.99483397 | 0.99847994 |
| chr22:42369073-42369709 | -0.002261594 | 0.994844223 | 0.99847994 |
| chr9:34701050-34701763 | -0.002269477 | 0.994886384 | 0.99850471 |
| chr20:31502953-31503683 | -0.002153671 | 0.994899347 | 0.99850471 |
| chr14:94306433-94306651 | 0.003252191 | 0.994917231 | 0.99850471 |
| chr6:18042858-18043059 | -0.003465473 | 0.994921029 | 0.99850471 |
| chr1:203039292-203039666 | -0.002163141 | 0.994957881 | 0.998507997 |

| | | | |
|---|---|---|---|
| chr2:61536606-61536869 | 0.002258765 | 0.994977055 | 0.998507997 |
| chr1:54406944-54407209 | -0.002541049 | 0.994978757 | 0.998507997 |
| chr6:4803414-4803635 | 0.003122777 | 0.994997064 | 0.998507997 |
| chr8:115335815-115336015 | -0.00324135 | 0.994999511 | 0.998507997 |
| chr5:150289717-150289941 | 0.002707893 | 0.995002494 | 0.998507997 |
| chr1:61049844-61050414 | -0.002159072 | 0.995021645 | 0.998514138 |
| chr11:18722049-18722280 | -0.002700617 | 0.99505894 | 0.998520253 |
| chr3:119042209-119042728 | 0.00230191 | 0.995059923 | 0.998520253 |
| chr5:148808915-148809115 | -0.003027836 | 0.995067892 | 0.998520253 |
| chr5:135778201-135778461 | 0.00331971 | 0.995081866 | 0.998520253 |
| chr6:36245784-36246069 | -0.002717547 | 0.995093379 | 0.998520253 |
| chr12:67924861-67925371 | 0.002486556 | 0.99510593 | 0.998520253 |
| chr17:74045693-74046066 | -0.002507914 | 0.995135916 | 0.998537266 |
| chr22:30451138-30451451 | 0.002923315 | 0.9952385 | 0.998625993 |
| chr5:1659830-1660030 | -0.002861079 | 0.99528093 | 0.998625993 |
| chr1:16612551-16612779 | 0.002337411 | 0.995292134 | 0.998625993 |
| chr10:96831855-96833231 | -0.001875902 | 0.995294251 | 0.998625993 |
| chrX:11100806-11101045 | 0.002548149 | 0.995318699 | 0.998625993 |
| chr1:231315773-231316119 | -0.002358016 | 0.995319985 | 0.998625993 |
| chr19:55075630-55075988 | 0.002217426 | 0.995328457 | 0.998625993 |
| chrX:8204180-8204487 | -0.002588428 | 0.995328606 | 0.998625993 |
| chr1:236531726-236532111 | 0.002085186 | 0.995348274 | 0.998627346 |
| chr14:34788918-34789159 | -0.003036332 | 0.995356021 | 0.998627346 |
| chr2:169329781-169330618 | 0.002023358 | 0.995369623 | 0.998627916 |
| chr11:3797196-3798207 | -0.001829246 | 0.995395222 | 0.998634339 |
| chr8:74984246-74985407 | -0.001928456 | 0.995402091 | 0.998634339 |
| chr11:20035688-20036259 | 0.002025939 | 0.995415549 | 0.998634765 |
| chrX:147550296-147550521 | -0.002887375 | 0.99543885 | 0.998636889 |
| chr11:1559411-1559846 | 0.00210776 | 0.995443733 | 0.998636889 |
| chr6:33391237-33392158 | 0.001812484 | 0.995485558 | 0.998652585 |
| chr17:82351766-82352422 | -0.001946585 | 0.995520094 | 0.998652585 |
| chr11:120301878-120302465 | -0.001857894 | 0.99552178 | 0.998652585 |
| chr3:87333230-87333921 | 0.001857323 | 0.995532887 | 0.998652585 |
| chr7:7779790-7780209 | -0.002172615 | 0.995540097 | 0.998652585 |
| chr1:19974873-19975073 | 0.002746579 | 0.995572262 | 0.998652585 |
| chr8:129535777-129536110 | 0.002123686 | 0.995593626 | 0.998652585 |
| chr3:99831994-99832451 | 0.001969498 | 0.995608486 | 0.998652585 |
| chr3:77255274-77255596 | 0.002110021 | 0.995614696 | 0.998652585 |
| chr3:61250795-61251017 | 0.002398506 | 0.99564255 | 0.998652585 |
| chr1:155140216-155140633 | 0.001957372 | 0.995644325 | 0.998652585 |
| chr18:76426772-76427016 | -0.002261484 | 0.995645291 | 0.998652585 |
| chr20:43681403-43681846 | 0.002175875 | 0.99565119 | 0.998652585 |
| chr11:60710550-60710798 | -0.002677568 | 0.995662486 | 0.998652585 |
| chr15:100145404-100145604 | 0.003168602 | 0.995665618 | 0.998652585 |
| chr15:69410326-69410621 | 0.002052312 | 0.995667914 | 0.998652585 |
| chr10:22713595-22713795 | -0.002537967 | 0.995686107 | 0.99865776 |
| chr9:114616627-114616898 | 0.002114547 | 0.995723276 | 0.998664155 |
| chr2:67214197-67214397 | 0.002657567 | 0.995727915 | 0.998664155 |
| chr14:52841353-52841696 | -0.00213437 | 0.995731585 | 0.998664155 |
| chr1:149899202-149900285 | 0.001689918 | 0.995758649 | 0.998678227 |

| | | | |
|---|---|---|---|
| chr6:111642451-111642895 | 0.0019888 | 0.995812154 | 0.998718816 |
| chr20:32445947-32446179 | 0.002495934 | 0.995865496 | 0.998756061 |
| chr17:56162808-56163575 | 0.001827982 | 0.99587536 | 0.998756061 |
| chr7:101059403-101059658 | -0.0020769 | 0.995910185 | 0.998768475 |
| chr3:182683615-182683905 | -0.002005553 | 0.995913808 | 0.998768475 |
| chr14:56213351-56213744 | 0.002416981 | 0.995936275 | 0.998777934 |
| chr19:44025184-44025400 | 0.00236286 | 0.995972568 | 0.998786588 |
| chr10:42690299-42690883 | -0.001898872 | 0.995983848 | 0.998786588 |
| chr3:99808163-99808613 | -0.001940315 | 0.995984011 | 0.998786588 |
| chr5:17203300-17203824 | 0.001746708 | 0.99600602 | 0.998795588 |
| chr11:2912280-2912968 | -0.001632341 | 0.996034459 | 0.998811034 |
| chrX:34918831-34919237 | -0.001886675 | 0.996051775 | 0.998812236 |
| chr7:347641-347841 | 0.00259795 | 0.99606583 | 0.998812236 |
| chr1:174114855-174115230 | 0.001764383 | 0.996085552 | 0.998812236 |
| chr16:81275790-81276046 | 0.002005383 | 0.996091169 | 0.998812236 |
| chr22:20393860-20394408 | -0.001731495 | 0.996100835 | 0.998812236 |
| chr16:1308806-1309699 | 0.001587956 | 0.996144144 | 0.998842591 |
| chr2:20122061-20122531 | -0.001833419 | 0.996178441 | 0.998858675 |
| chr1:93345640-93346020 | -0.001832005 | 0.996186256 | 0.998858675 |
| chr9:112555140-112555343 | 0.002442757 | 0.996274455 | 0.998934038 |
| chr17:18564112-18564391 | -0.00209019 | 0.996333722 | 0.998977122 |
| chr12:77667059-77667431 | -0.001967698 | 0.996354274 | 0.998977122 |
| chr12:103548709-103549091 | 0.001961604 | 0.996380865 | 0.998977122 |
| chr8:74366141-74366461 | 0.002155397 | 0.996382145 | 0.998977122 |
| chr1:205907842-205908221 | -0.001682913 | 0.996382613 | 0.998977122 |
| chr14:26394658-26394875 | 0.002150435 | 0.996404275 | 0.998980828 |
| chr16:85707646-85708145 | 0.001560476 | 0.996416705 | 0.998980828 |
| chr5:16721951-16722151 | -0.002234546 | 0.996425423 | 0.998980828 |
| chr2:149069266-149069869 | -0.001573824 | 0.996439388 | 0.998981757 |
| chr17:5438675-5439580 | 0.0015344 | 0.996499253 | 0.999003922 |
| chr4:74932785-74933149 | 0.001584701 | 0.996501958 | 0.999003922 |
| chr19:40897240-40897515 | 0.002069679 | 0.996507765 | 0.999003922 |
| chr4:94527575-94528041 | -0.001726011 | 0.996513649 | 0.999003922 |
| chr4:108118437-108118834 | -0.001584931 | 0.996544123 | 0.999021402 |
| chr8:125118129-125118840 | -0.001449894 | 0.996565636 | 0.999028089 |
| chr11:83533578-83533778 | 0.002559995 | 0.996579931 | 0.999028089 |
| chr10:100301233-100301463 | 0.002181414 | 0.996589908 | 0.999028089 |
| chr17:58710845-58711284 | -0.001411954 | 0.996615632 | 0.999040805 |
| chr14:92853049-92853556 | 0.001558015 | 0.996639948 | 0.99905211 |
| chr16:84504170-84505255 | -0.001317908 | 0.996684769 | 0.999071528 |
| chr14:37232938-37233228 | -0.001708777 | 0.996699832 | 0.999071528 |
| chr15:66671077-66671469 | -0.001450072 | 0.9967031 | 0.999071528 |
| chr1:44631543-44632562 | 0.001369019 | 0.996722331 | 0.999071528 |
| chr21:28992445-28993309 | -0.001316778 | 0.996724515 | 0.999071528 |
| chr8:26329880-26330102 | 0.002061895 | 0.996775163 | 0.999080852 |
| chrX:111096203-111096423 | 0.001987119 | 0.996780751 | 0.999080852 |
| chr3:184248751-184249166 | 0.001400888 | 0.996782641 | 0.999080852 |
| chr2:129877287-129877779 | -0.001428275 | 0.996785973 | 0.999080852 |
| chr16:9047700-9048160 | -0.001434646 | 0.99682563 | 0.999107531 |
| chr5:24023286-24023538 | 0.002058003 | 0.996871786 | 0.999123578 |

| | | | |
|---|---|---|---|
| chr5:5279044-5279244 | -0.001959285 | 0.996874388 | 0.999123578 |
| chr12:79515714-79515960 | 0.001707491 | 0.996893197 | 0.999123578 |
| chr11:126607317-126607590 | -0.00151093 | 0.996893799 | 0.999123578 |
| chr3:162731857-162732057 | -0.002131669 | 0.996912715 | 0.999129467 |
| chr2:187577397-187577677 | -0.001557891 | 0.996952594 | 0.999156366 |
| chr11:95068527-95068943 | -0.001323775 | 0.997044956 | 0.999211574 |
| chr5:153711788-153711998 | 0.001681681 | 0.997057632 | 0.999211574 |
| chr8:47959936-47960457 | 0.001235141 | 0.997065019 | 0.999211574 |
| chrX:77910356-77911055 | 0.001210529 | 0.997107466 | 0.999211574 |
| chr8:30137300-30137569 | -0.001910254 | 0.997150515 | 0.999211574 |
| chr3:72637111-72637311 | -0.001646238 | 0.997171462 | 0.999211574 |
| chr1:223993706-223993972 | 0.001631275 | 0.997176311 | 0.999211574 |
| chr9:135718715-135718915 | -0.001807558 | 0.997192733 | 0.999211574 |
| chr1:219812591-219812953 | 0.001324521 | 0.997197127 | 0.999211574 |
| chrX:103940000-103940273 | -0.001657701 | 0.997199947 | 0.999211574 |
| chr2:84798381-84798656 | -0.001401974 | 0.997204435 | 0.999211574 |
| chr2:17901265-17901465 | -0.002016029 | 0.997229263 | 0.999211574 |
| chr3:154053049-154053258 | -0.002016029 | 0.997229263 | 0.999211574 |
| chr1:177513028-177513298 | 0.00175751 | 0.997234596 | 0.999211574 |
| chr14:103910061-103910822 | -0.001161702 | 0.997239534 | 0.999211574 |
| chr5:429833-430083 | 0.001534632 | 0.997241732 | 0.999211574 |
| chr9:137459259-137459615 | -0.001222778 | 0.997245412 | 0.999211574 |
| chr8:60653499-60653816 | -0.001264359 | 0.99724977 | 0.999211574 |
| chr4:105666806-105667009 | 0.001902712 | 0.997271476 | 0.999211574 |
| chr4:162169056-162169338 | -0.001392547 | 0.99731885 | 0.999211574 |
| chr6:113994823-113995232 | -0.001225915 | 0.997346177 | 0.999211574 |
| chr13:36850847-36851234 | 0.001407572 | 0.997362367 | 0.999211574 |
| chr8:78530234-78530852 | -0.001085075 | 0.997374389 | 0.999211574 |
| chr2:235595473-235595694 | -0.001544044 | 0.997380276 | 0.999211574 |
| chr6:47308830-47309231 | 0.001242283 | 0.997383609 | 0.999211574 |
| chr10:24262892-24263442 | 0.00121929 | 0.997395697 | 0.999211574 |
| chr3:86759973-86760173 | -0.001457679 | 0.997395733 | 0.999211574 |
| chr3:128784324-128784579 | -0.00134566 | 0.9974053 | 0.999211574 |
| chr19:17865806-17866032 | -0.001529302 | 0.997410223 | 0.999211574 |
| chr12:64181484-64182009 | 0.001151406 | 0.997437729 | 0.999211574 |
| chr6:16511136-16511336 | -0.001643661 | 0.997441453 | 0.999211574 |
| chr6:33665791-33666039 | 0.001710032 | 0.997442931 | 0.999211574 |
| chr20:381954-382213 | -0.00132467 | 0.997459964 | 0.999211574 |
| chr17:9863396-9863596 | 0.002388031 | 0.997461427 | 0.999211574 |
| chr5:141192254-141192524 | 0.001631651 | 0.997464108 | 0.999211574 |
| chr3:142148844-142149776 | 0.001022586 | 0.997492371 | 0.999212113 |
| chr1:53616857-53617137 | 0.001309994 | 0.997499489 | 0.999212113 |
| chr3:157008064-157008388 | -0.00129265 | 0.997503768 | 0.999212113 |
| chr7:137093221-137093421 | -0.001930206 | 0.997517512 | 0.999212817 |
| chr1:177355567-177355992 | -0.00112366 | 0.997583346 | 0.999265699 |
| chr12:109713414-109714020 | 0.000995777 | 0.997604134 | 0.999270143 |
| chr1:218789400-218789769 | 0.001224658 | 0.997618836 | 0.999270143 |
| chr1:41500446-41500871 | -0.001040621 | 0.997626907 | 0.999270143 |
| chr9:5886125-5886520 | -0.001057155 | 0.997650993 | 0.999281206 |
| chr17:76905771-76906051 | -0.001281653 | 0.997681889 | 0.999293445 |

| | | | |
|---|---|---|---|
| chr10:93535438-93535638 | 0.001788146 | 0.997702338 | 0.999293445 |
| chr12:31486418-31486618 | 0.001788146 | 0.997702338 | 0.999293445 |
| chr21:32727276-32728778 | -0.000910355 | 0.997725417 | 0.999303498 |
| chr5:113663926-113664425 | -0.001151118 | 0.997744113 | 0.999309161 |
| chr8:1161599-1161799 | -0.001526052 | 0.997763718 | 0.999315734 |
| chr20:47360131-47361151 | -0.000840138 | 0.99784527 | 0.99938435 |
| chr2:164840535-164842637 | 0.000843811 | 0.997861806 | 0.999387848 |
| chr18:39614022-39614238 | 0.001417304 | 0.99790274 | 0.999409487 |
| chr12:96132935-96133174 | 0.001249142 | 0.997909499 | 0.999409487 |
| chrX:122271663-122271943 | 0.00128594 | 0.997937618 | 0.999411589 |
| chr19:4204717-4204917 | -0.001092831 | 0.997950495 | 0.999411589 |
| chr1:156910348-156910705 | 0.000940313 | 0.997963327 | 0.999411589 |
| chr6:131063010-131063904 | 0.000829732 | 0.997967446 | 0.999411589 |
| chr16:88570632-88570999 | -0.000932317 | 0.997987579 | 0.999411589 |
| chr2:156065975-156066191 | 0.00146398 | 0.997989858 | 0.999411589 |
| chr6:34190728-34191022 | 0.001014024 | 0.998014781 | 0.999422003 |
| chr8:15537852-15538212 | -0.001126927 | 0.998026344 | 0.999422003 |
| chr8:73305564-73305806 | -0.001244046 | 0.998052477 | 0.999428354 |
| chr3:133821849-133822233 | -0.000879399 | 0.998058774 | 0.999428354 |
| chr3:16767086-16767286 | -0.001259488 | 0.998094942 | 0.99943254 |
| chr22:22576429-22576629 | -0.001251542 | 0.998107069 | 0.99943254 |
| chr1:36463683-36463900 | -0.001029778 | 0.998113608 | 0.99943254 |
| chr5:139904386-139904586 | 0.001162865 | 0.998115129 | 0.99943254 |
| chr16:22218013-22218286 | 0.00090097 | 0.998194453 | 0.999457361 |
| chr4:150784490-150784759 | -0.000931267 | 0.998198558 | 0.999457361 |
| chr2:64645027-64645335 | -0.000862748 | 0.998206334 | 0.999457361 |
| chr19:14505478-14505678 | -0.001188181 | 0.998208707 | 0.999457361 |
| chr8:54569618-54569918 | 0.000855542 | 0.998217542 | 0.999457361 |
| chr5:16845052-16845256 | -0.001024223 | 0.998221817 | 0.999457361 |
| chr6:30981910-30982145 | 0.001063538 | 0.998237435 | 0.999457361 |
| chr6:18503624-18504292 | 0.000769101 | 0.998251512 | 0.999457361 |
| chr5:14246641-14246966 | -0.000929856 | 0.998263129 | 0.999457361 |
| chrX:110181571-110181995 | -0.000935232 | 0.998308463 | 0.999457361 |
| chr3:142224910-142225994 | -0.000679152 | 0.998309 | 0.999457361 |
| chr20:40268686-40268896 | -0.001074831 | 0.998313943 | 0.999457361 |
| chr4:139184198-139184417 | -0.00100003 | 0.998341847 | 0.999457361 |
| chr3:3170946-3171272 | 0.000907848 | 0.998343996 | 0.999457361 |
| chr3:152988312-152988545 | -0.001129145 | 0.99835411 | 0.999457361 |
| chr3:143152605-143152915 | -0.001056941 | 0.998355807 | 0.999457361 |
| chr9:127930361-127931019 | -0.000684562 | 0.998361665 | 0.999457361 |
| chr17:56913334-56914552 | -0.000626342 | 0.998420025 | 0.999502726 |
| chr19:35774700-35775621 | -0.000636584 | 0.998458435 | 0.999519575 |
| chr8:127793840-127795342 | -0.000606264 | 0.998462945 | 0.999519575 |
| chr3:79361896-79362138 | -0.000929868 | 0.998519291 | 0.999562921 |
| chr3:15667504-15667774 | 0.001581186 | 0.998552584 | 0.999568853 |
| chr17:63099110-63099310 | 0.001095869 | 0.998574046 | 0.999568853 |
| chr3:118842350-118842559 | -0.000875872 | 0.998587863 | 0.999568853 |
| chr12:46292793-46293132 | 0.000679854 | 0.998602731 | 0.999568853 |
| chr19:45615890-45616390 | -0.00059394 | 0.998604782 | 0.999568853 |
| chr7:70694347-70694835 | -0.000644674 | 0.998624359 | 0.999568853 |

| | | | |
|---|---|---|---|
| chr22:29581837-29582178 | -0.000641591 | 0.998626501 | 0.999568853 |
| chr7:151632127-151632710 | -0.000618769 | 0.99862958 | 0.999568853 |
| chr7:17689450-17690486 | -0.000576006 | 0.9986433 | 0.999569528 |
| chr13:32811346-32811848 | -0.000632179 | 0.998659343 | 0.999570943 |
| chr11:83103995-83104265 | 0.000880801 | 0.998670804 | 0.999570943 |
| chr14:50260625-50260914 | -0.000545703 | 0.998760825 | 0.999643798 |
| chr11:12888685-12888977 | 0.000620431 | 0.998769687 | 0.999643798 |
| chr1:46528084-46528306 | -0.000602329 | 0.99880545 | 0.999649091 |
| chr16:74813078-74813612 | 0.000496827 | 0.998828448 | 0.999649091 |
| chr10:25523031-25523291 | 0.000592738 | 0.998830238 | 0.999649091 |
| chr1:151070305-151071368 | 0.000466351 | 0.99883148 | 0.999649091 |
| chr17:44141611-44142134 | 0.000492391 | 0.998849728 | 0.999649091 |
| chr16:5024679-5024958 | -0.000611039 | 0.998856733 | 0.999649091 |
| chr20:3820070-3821015 | 0.000459725 | 0.998866301 | 0.999649091 |
| chr7:44375381-44375719 | -0.000497688 | 0.998938379 | 0.999706146 |
| chr21:41396510-41397106 | -0.000465213 | 0.998949406 | 0.999706146 |
| chr5:169583284-169584251 | -0.000399581 | 0.999010985 | 0.999723115 |
| chr19:58401113-58401812 | 0.000397594 | 0.999014925 | 0.999723115 |
| chr7:817226-817647 | 0.000469522 | 0.999025609 | 0.999723115 |
| chr15:101251867-101252563 | 0.000393514 | 0.999069642 | 0.999723115 |
| chr5:173328175-173329346 | 0.0003613 | 0.999091913 | 0.999723115 |
| chr5:31664667-31665028 | 0.000421776 | 0.999091936 | 0.999723115 |
| chr6:53061496-53062068 | -0.000374639 | 0.999100144 | 0.999723115 |
| chr22:45164282-45164690 | -0.000438723 | 0.999101803 | 0.999723115 |
| chr7:1044122-1045058 | 0.000360177 | 0.999108978 | 0.999723115 |
| chr1:187688878-187689078 | 0.00117536 | 0.999109163 | 0.999723115 |
| chr11:69048607-69049253 | -0.000369479 | 0.999109884 | 0.999723115 |
| chr2:215442566-215442781 | 0.000431918 | 0.999160378 | 0.999738272 |
| chr5:685490-685709 | -0.000485638 | 0.999186272 | 0.999738272 |
| chr4:67700583-67701520 | 0.000323827 | 0.999187882 | 0.999738272 |
| chr17:82450106-82450325 | 0.000576467 | 0.999188192 | 0.999738272 |
| chr7:113171493-113171798 | -0.000257473 | 0.99919735 | 0.999738272 |
| chr2:182030973-182031317 | 0.000393214 | 0.999203318 | 0.999738272 |
| chr4:76689092-76689813 | -0.000321628 | 0.99924818 | 0.999750018 |
| chr8:106397905-106398194 | -0.000372708 | 0.999253573 | 0.999750018 |
| chr4:77104905-77105359 | 0.000319068 | 0.999254201 | 0.999750018 |
| chr11:110295728-110295986 | -0.000397742 | 0.999286859 | 0.999756611 |
| chr13:23888644-23889882 | 0.000283395 | 0.999294979 | 0.999756611 |
| chr3:105930788-105930988 | 0.000338513 | 0.999299934 | 0.999756611 |
| chr6:106325083-106326015 | 0.000271892 | 0.999317858 | 0.999761489 |
| chr9:138061706-138061906 | 0.000318645 | 0.999342449 | 0.999773037 |
| chr4:37603052-37603534 | -0.000282615 | 0.999394297 | 0.999811853 |
| chr1:203661374-203661665 | -0.000304255 | 0.999421055 | 0.99982333 |
| chr6:113792941-113793235 | 0.000419767 | 0.999431867 | 0.99982333 |
| chr12:132986060-132986774 | 0.000227283 | 0.999457518 | 0.999826455 |
| chr1:205066222-205066447 | -0.000234888 | 0.999482267 | 0.999826455 |
| chr1:37275365-37275643 | -0.000287133 | 0.999485025 | 0.999826455 |
| chr4:146335857-146336707 | -0.000219773 | 0.999487186 | 0.999826455 |
| chr19:10288649-10288986 | 0.000205573 | 0.999526519 | 0.99985004 |
| chr16:85931335-85931561 | -0.000274892 | 0.999536861 | 0.99985004 |

| | | | |
|---|---|---|---|
| chr5:175494298-175494588 | -0.000222203 | 0.999601283 | 0.999897332 |
| chr1:149708343-149708922 | -0.000157063 | 0.999610237 | 0.999897332 |
| chr2:240981997-240982243 | 0.000289432 | 0.99964087 | 0.999914919 |
| chr10:60549776-60549999 | 0.000250575 | 0.999666204 | 0.99991595 |
| chr20:30513281-30513503 | -8.87E-05 | 0.999668 | 0.99991595 |
| chr8:142675894-142676552 | -0.000130301 | 0.999710324 | 0.9999392 |
| chr9:101387863-101388117 | -6.45E-05 | 0.999723249 | 0.9999392 |
| chr19:48543263-48543826 | -0.000112352 | 0.999731892 | 0.9999392 |
| chr19:36114685-36115871 | 0.000101974 | 0.999743445 | 0.9999392 |
| chr3:72996403-72997388 | 8.14E-05 | 0.999797482 | 0.999980194 |
| chr3:64329098-64329373 | 6.96E-05 | 0.999826976 | 0.999987665 |
| chr1:202192851-202193184 | 7.54E-05 | 0.999831054 | 0.999987665 |
| chr2:128421983-128422183 | 2.61E-05 | 0.999858395 | 0.999996576 |
| chr3:184314334-184315571 | 3.65E-05 | 0.999910604 | 0.999996576 |
| chr1:145978979-145979835 | 3.00E-05 | 0.999923474 | 0.999996576 |
| chr15:40038687-40039746 | 2.97E-05 | 0.999928132 | 0.999996576 |
| chr4:138919149-138919461 | 5.95E-06 | 0.999930958 | 0.999996576 |
| chr2:74834802-74835618 | -2.45E-05 | 0.999933959 | 0.999996576 |
| chr5:5436406-5436662 | -0.000120639 | 0.999934559 | 0.999996576 |
| chr2:197615086-197615360 | -8.86E-05 | 0.999960214 | 0.999996576 |
| chr3:162316786-162316986 | 0.000290934 | 0.999973316 | 0.999996576 |
| chr6:139878085-139878287 | 0.00027689 | 0.999979266 | 0.999996576 |
| chr2:10395569-10396037 | 1.42E-05 | 0.999983525 | 0.999996576 |
| chr7:85041631-85041935 | -4.60E-07 | 0.999998502 | 0.999998502 |
